# Dual domestication, diversity, and differential introgression in Old World cotton diploids

**DOI:** 10.1101/2021.10.20.465142

**Authors:** Corrinne E. Grover, Mark A. Arick, Adam Thrash, Joel Sharbrough, Guanjing Hu, Daojun Yuan, Emma R. Miller, Thiruvarangan Ramaraj, Daniel G. Peterson, Joshua A. Udall, Jonathan F. Wendel

## Abstract

Domestication in the cotton genus is remarkable in that it has occurred independently four different times at two different ploidy levels. Relatively little is known about genome evolution and domestication in the cultivated diploid species *Gossypium herbaceum* and *G. arboreum*, because of the absence of wild representatives for the latter species, their ancient domestication, and their joint history of human-mediated dispersal and interspecific gene flow. Using in-depth resequencing of a broad sampling from both species, we confirm their independent domestication, as opposed to a progenitor-derivative relationship, showing that diversity (mean π = 2.3×10^-3^) within species is similar, and that divergence between species is modest (weighted F_ST_=0.4430). Individual accessions were homozygous for ancestral SNPs at over half of variable sites, while fixed, derived sites were at modest frequencies. Notably, two chromosomes with a paucity of fixed, derived sites (*i.e*., chromosomes 7 and 10) were also strongly implicated in introgression analyses. Collectively, these data demonstrate variable permeability to introgression among chromosomes, which we propose is due to divergent selection under domestication and/or the phenomenon of F_2_ breakdown in interspecific crosses. Our analyses provide insight into the evolutionary forces influencing diversity and divergence in the diploid cultivated species, and establish a foundation for understanding the contribution of introgression and/or strong parallel selection to the extensive morphological similarities shared between species.

**Significance statement:** The cotton genus (*Gossypium*) contains four different species that were independently domesticated at least 4,000 years ago. Relatively little is understood about diversity and evolution in the two diploid African-Asian sister-species *G. herbaceum* and *G. arboreum*, despite their historical importance in the region and contemporary cultivation, largely in the Indian subcontinent. Here we address questions regarding the relationship between the two species, their contemporary levels of diversity, and their patterns of interspecific gene flow accompanying their several millennia history of human-mediated dispersal and contact. We validate independent domestication of the two species and document the genomic distribution of interspecific genetic exchange.

## Introduction

Domestication is an important directional and in many cases diversifying evolutionary process that transformed wild plants and animals into their modern domesticated forms. Intentional selection applied to wild populations differentiates domesticates from their progenitors on both the phenotypic and genetic levels, a process usually accompanied by an overall reduction in genetic diversity in the domesticate relative to its ancestral gene pool. In some crops, domestication has occurred independently more than once (*e.g.*, rice and common bean; (Wang et al. 2014; Sang & Ge 2007; Bellucci et al. 2014)), resulting in convergent phenotypes with potentially divergent genetic underpinnings.

The cotton genus (*Gossypium*) provides an example of a crop having multiple, independent domestications that span both continents and ploidy levels. While the two cultivated polyploid species (*i.e., G. hirsutum and G. barbadense*) dominate contemporary worldwide commerce, cotton also has been domesticated twice at the diploid level. Colloquially known as the “A-genome cottons”, *G. arboreum* and *G. herbaceum* were both domesticated during the same approximate timeframe as the polyploid species (4,000 - 8,000 years ago), albeit in southwestern Asia and Africa (versus the American tropics for the polyploid species; reviewed in (Wendel & Grover 2015; Hu et al. 2021)). Although fiber quality from both A-genome cotton species is inferior to that of the tetraploids, they possess spinnable fiber and are the closest living relatives to the maternal progenitor of the polyploid species (including *G. hirsutum* and *G. barbadense*; reviewed in (Wendel & Grover 2015; Hu et al. 2021)).

Given their historical and modern importance as crops in parts of Africa-Asia, it is surprising that so little is known regarding their origin, domestication, and modern patterns of diversity. Although *G. herbaceum* is native to the savannahs of Southern Africa (Vollesen 1987; Wendel et al. 1989; Khadi et al. 2010), the center of early diversification was likely in Northern Africa or the Near East (Fryxell 1979). *Gossypium herbaceum* expanded bidirectionally (east-west) through the Persian Gulf States and Indian subcontinent (Kulkarni et al. 2009; Kranthi 2018). The natural and human histories of *G. arboreum* are less clear, as no true wild forms have been identified. Some have suggested that *G. arboreum* may be the derivative of an early landrace of *G. herbaceum* that became isolated due to a reciprocal translocation (Gerstel 1953; Hutchinson 1954a; Gulati & Turner 1929; Gennur et al. 1986), although recent research indicates that the two sister species separated long prior to domestication and perhaps prior to hominin (*i.e*., modern and extinct human species) evolution (Wendel et al. 1989; Renny-Byfield et al. 2016; Huang et al. 2020; Du et al. 2018). While little is known about the history of *G. arboreum* prior to domestication, archaeological evidence and genetic diversity analyses suggest the Indus Valley as a candidate for the origin of *G. arboreum* (Wendel et al. 2010; Gulati & Turner 1928), although this may instead represent a secondary center of diversity following initial domestication elsewhere (Hutchinson 1954b; Wendel et al. 2010).

Analyses of the A-genome diploids suggest that diversity within species is low (Du et al. 2018; Page et al. 2013; Wendel et al. 1989; Fang, Gong, et al. 2017; Jena et al. 2011). Recent resequencing among predominantly Chinese accessions of *G. arboreum* (Du et al. 2018) suggests that, while diversity is low among those regionally restricted domesticated accessions (*π*=0.002), it is similar to that recently reported (Yuan et al. 2021) for wild accessions of the domesticated polyploid species *G. hirsutum* and *G. barbadense* (*π*=0.003 in both). This observation is similar to previous reports that diversity in the diploid species is roughly equivalent to that found in the tetraploids (Wendel et al. 1989; Stanton et al. 1994). Relative diversity between the two diploid species is unclear, with the few direct comparisons reporting conflicting results (Wendel et al. 1989; Jena et al. 2011) perhaps due to differences in germplasm evaluated and/or the markers used for diversity analysis (*i.e*., allozymes versus AFLP markers, respectively).

Throughout their pre-colonial history, cultivation of the A-genome diploids has been limited to Asia (Khadi et al. 2010; Basu 1996; Guo et al. 2006; Wendel et al. 1989), and their derivatives are still grown in many Asian regions (*e.g.*, India, Myanmar, and Thailand) (Kranthi 2018), where pests and growing conditions make these species more competitive than the polyploid cultivars. In addition, A-genome diploid cottons are also used as genetic resources for introducing stress tolerance and/or disease resistance into the commercially more important polyploid cultivars (Kulkarni et al. 2009). Finally, the A-genome diploids also are of interest in that they provide a parallel to the dual domestication of cotton at the polyploid level (Yuan et al. 2021). In an effort to clarify the species history and dual domestication of these sister taxa, we employed high throughput DNA sequencing and computational biology techniques to analyze a diverse assemblage of accessions of both species. We use this whole genome approach to improve our understanding of the modern gene pools of these species and their interrelationships to each other.

## Results

### Sample selection and verification

We resequenced 80 *G. herbaceum* and *G. arboreum* accessions, selected to represent the diversity of the A-genome clade (Supplementary Table 1). These newly sequenced accessions averaged 38✕ genome equivalent coverage (18✕ - 64✕; median = 35✕) of the ∼1700 Mbp genomes (Hendrix & Stewart 2005), a depth suitable for accurate SNP detection and diversity analysis. In addition, we included representatives of existing resequencing datasets from both A-genome species (Du et al. 2018; Huang et al. 2020; Page et al. 2013), evaluating an additional 292 accessions whose average coverage was approximately one-third of the resequencing depth (median = 9.9X) of accessions sequenced specifically for this study. Of the 372 total accessions, 154 were excluded due to low coverage (i.e., <10✕ coverage; all samples were from (Du et al. 2018)). Phylogenetic and PCA analysis of the remaining 218 samples (Supplementary Figure 1) led to the exclusion of seven samples due to incorrect species assignment, suggesting sample and/or germplasm (source) misidentification, and a further four were excluded as putative hybrid and/or contaminated samples (Supplementary Table 1). Notably, the remaining samples originating from (Du et al. 2018) were distinct on both the whole-genome and genic-only PCAs (Supplementary Figure 2); these were consequently excluded (Supplementary Table 1) for possible batch effects due to PCR selection (Tom et al. 2017; Buckley et al. 2017; Aird et al. 2011; Jones et al. 2015). All other samples were retained for further analyses, resulting in a dataset composed of 21 *G. herbaceum* and 99 *G. arboreum* accessions (17 and 54 newly sequenced, respectively).

### Diversity and divergence within and among A-genome species

Single nucleotide polymorphisms (SNPs) within and between A-genome species were identified using the outgroup *G. longicalyx* as the reference sequence. Derived SNPs (relative to the ancestor, *G. longicalyx*) that were shared by all accessions of both species were excluded as uninformative. In total, 12.1 million (M) variant sites (non-ancestral) were detected and distributed evenly across the *G. longicalyx* reference (Supplementary Table 2), representing <1% of the genome. In general, individual accessions were homozygous for the ancestral (*G. longicalyx*) SNP at 50 - 65% of variable sites, ranging from 5.6 - 7.9M sites per sample (Supplementary Table 3). The number of sites fixed for the derived allele (*i.e.*, homozygous derived) varied narrowly among samples, from 1.6 - 2.1 M sites per sample, while heterozygous sites varied more broadly, from 1.7 - 4.8 M sites per sample. While the number of homozygous reference and heterozygous sites per sample is similar between *G. herbaceum* and *G. arboreum*, the number of homozygous derived sites was generally lower for *G. herbaceum* (Mann–Whitney *U*, *p* =6.157e-06). Fixed differences between species are relatively rare (<3% of sites), and evenly distributed across most chromosomes. Notably, chromosomes 7 and 10 from *G. herbaceum* had an order of magnitude fewer fixed, derived sites than the other chromosomes (2,385 and 2,074 versus 23,243 - 38,434 for other chromosomes; Supplementary Table 4); *G. arboreum* also shared the lack of fixed sites for chromosome 7 (619 versus 5,824 - 10,693). In total, *G. arboreum* had approximately threefold fewer fixed, derived sites, likely due to the greater sampling in that species. Interestingly, *G. arboreum* had nearly twice the number of variant sites as *G. herbaceum* (7.6 M versus 4.0 M) when including sites where more than 10% of the population is either fixed or heterozygous for the derived allele (allele is absent in the other species).

Most SNPs occurred in the intergenic space (Table 1; 93%, or 11.3M SNPs), a third of which were located proximal to genes (∼31%, or 3.2 M within 5 kb of genes). Only 7% of SNPs (0.9 M) were located in genic regions, with slightly fewer occurring in exons versus introns (0.38M versus 0.44M, respectively). Most exonic SNPs resulted in missense mutations (60.5%, or 228,822 SNPs) or silent changes (36.9%, 139,515 SNPs); less than 3% of SNPs resulted in a nonsense mutation. Despite the significant difference in the number of accessions represented by each species (21 *G. herbaceum* versus 99 *G. arboreum*), a similar number and pattern of SNP distribution within species was observed, which was also consistent with the overall pattern of similarity between the two species (Table 1).

**Table 1.** Distribution of SNPs among genomic regions. Subcategories are listed under each main category. Percentages are relative to category or subcategory

|  |  |  | among *G. herbaceum* |  | among *G. arboreum* |  |
| --- | --- | --- | --- | --- | --- | --- |
|  | Number of SNPs | Relative percent | Number of SNPs | Relative percent | Number of SNPs | Relative percent |
| **Intergenic** | **11,315,283** | **92.90%** | **10,745,009** | **93.14%** | **10,689,194** | **93.38%** |
| Downstream | 1,722,429 | 15.22% | 1,565,393 | 14.57% | 1,529,060 | 14.30% |
| Upstream | 1,781,384 | 15.74% | 1,618,137 | 15.06% | 1,585,525 | 14.83% |
| **Genic** | **867,344** | **7.10%** | **791,102** | **6.86%** | **758,164** | **6.62%** |
| Exon | 375,895 | 43.34% | 350,339 | 44.28% | 338,689 | 44.67% |
| Intron | 437,952 | 50.49% | 393,628 | 49.76% | 375,592 | 49.54% |
| 5'UTR | 22,989 | 2.65% | 20,386 | 2.58% | 18,935 | 2.50% |
| 3'UTR | 30,508 | 3.52% | 26,749 | 3.38% | 24,948 | 3.29% |
| **Gene effects** | **Number of SNPs** | **Relative percent** | **Number of SNPs** | **Relative percent** | **Number of SNPs** | **Relative percent** |
|  | **378,133** |  | **352,389** |  | **340,633** |  |
| silent | 139,515 | 36.90% | 128,775 | 36.54% | 124,172 | 36.45% |
| missense | 228,822 | 60.51% | 214,193 | 60.78% | 207,202 | 60.83% |
| nonsense | 9,796 | 2.59% | 9,421 | 2.67% | 9,259 | 2.72% |

Indel polymorphisms within and between species were also characterized, using the outgroup *G. longicalyx* to polarize each as either an insertion or deletion. Indels that occurred prior to species divergence, and hence were shared between *G. arboreum* and *G. herbaceum*, were discarded. Deletions generally outweighed insertions by about 50-60% within and among species (2.1 M insertions versus 3.3M deletions; Supplementary Table 5), although the average size (4.4 and 4.8, respectively) and size distribution of each was similar (Supplementary Figure 2). As expected, most indels (85-90%) were located in intergenic regions, and over half of genic indels (444,663 out of 867,344) were located within introns. Indels located within exons frequently resulted in frameshift mutations in gene models (75,086 indels out of 101,475; 74%), affecting just over half (20,136) of the 38,378 total genes. As with SNPs, the indel profiles of *G. herbaceum* and *G. arboreum* were similar (Supplementary Table 5), including the difference in number of fixed indels, which was approximately three times greater in *G. herbaceum*. Notably, because the insertion and deletion rates are similar between these species (relative to the outgroup), each accession has, on average, gained ∼7 Mbp of sequence and lost ∼11 Mbp, leading to a net reduction in genome size due to small indels and further contributing to the divergence between species.

Nucleotide diversity (*π*) within and among samples was similar (Figure 1; Supplementary Table 6) between the two A-genome species, although it was slightly higher in the more abundantly sampled *G. arboreum* (mean π = 2.4×10^-3^, versus 2.2×10^-3^ in *G. herbaceum*). Diversity on individual chromosomes also followed a general pattern of higher diversity in *G. arboreum*, although the maximum *π* for chromosomes F03, F06, F08, and F12 was slightly higher in *G. herbaceum* (Table 2). In general, diversity was chromosomally similar (Figure 1) and correlated (Supplementary Figure 4) between *G. herbaceum* and *G. arboreum*. As expected, diversity in intergenic regions is much higher than in genic regions (by an order of magnitude); however, diversity in exon and intron regions was highly similar (Figure 1; Supplementary Table 7). Again, diversity in intergenic regions was slightly higher in *G. arboreum* than in *G. herbaceum*. Between sample divergence, as measured by the Weir and Cockerham weighted *F_ST_* implemented in vcftools, was modest (weighted *F_ST_*=0.4430). Mean *F_ST_* per chromosome (1 Mb window, 100 kb step; Figure 1C) varied from 0.3855 on Chromosome F10 to 0.4842 on Chromosome F05, but was highly variable along the chromosome (minimum *F_ST_*=0.0497, maximum *F_ST_*=0.8788).

**Figure 1.**
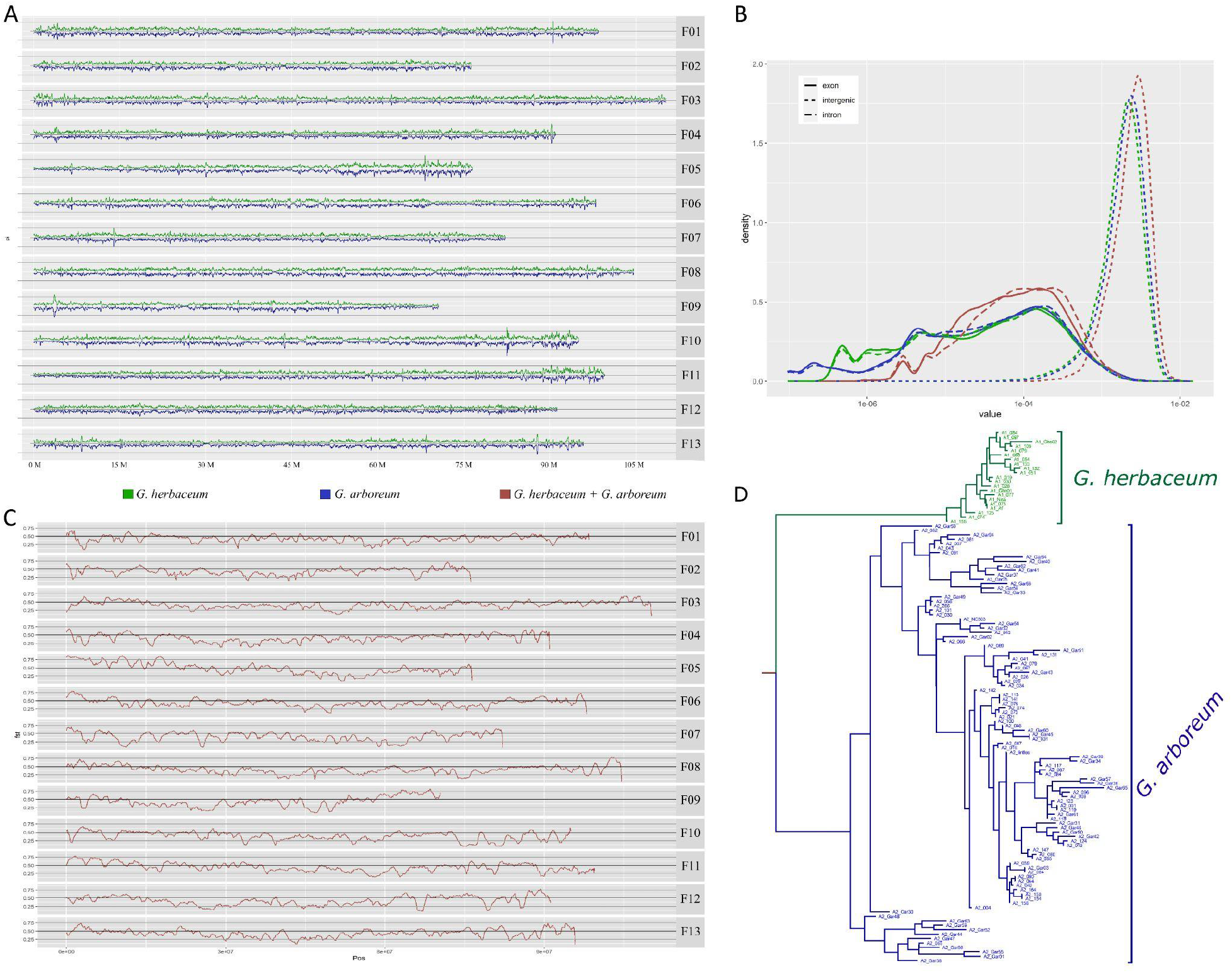
Diversity and divergence in *G. herbaceum* and *G. arboreum*. **A**. Nucleotide diversity (*π*) with *G. herbaceum* (top line, green) and *G. arboreum* (bottom line, blue) accessions partitioned by chromosome. **B**. Nucleotide diversity (*π*, x-axis) partitioned by intron (long dash), exon (solid line), and intergenic (small dash) regions displayed as a density plot. Within species divergence is shown for both *G. herbaceum* (green) and *G. arboreum* (blue), and the overall divergence for the A-genome (*G. herbaceum* + *G. arboreum*) is shown in red. **C**. Weighted *F_ST_* between *G. herbaceum* and *G. arboreum*, partitioned by chromosome. **D**. Phylogenetics of accessions passing quality filters. The *G. herbaceum* clade is shown in green and the *G. arboreum* clade is shown in blue. High-resolution versions of these images are found at https://github.com/Wendellab/A1A2resequencing. Phylogenetic reconstruction using genic SNPs from only these samples recovered two distinct clades, one for each species (Figure 1D).

**Table 2.** Nucleotide diversity within and between species. Combined diversity is given (both), as well as within species divergence for *G. herbaceum* (A1) and *G. arboreum* (A2).

|  | All chromosomes |  |  | Chromosome 1 |  |  | Chromosome 2 |  |  | Chromosome 3 |  |  | Chromosome 4 |  |  | Chromosome 5 |  |  | Chromosome 6 |  |  |
| --- | --- | --- | --- | --- | --- | --- | --- | --- | --- | --- | --- | --- | --- | --- | --- | --- | --- | --- | --- | --- | --- |
|  | both | A1 | A2 | both | A1 | A2 | both | A1 | A2 | both | A1 | A2 | both | A1 | A2 | both | A1 | A2 | both | A1 | A2 |
| **Mean** | 0.0029 | 0.0022 | 0.0024 | 0.0027 | 0.0020 | 0.0022 | 0.0031 | 0.0024 | 0.0025 | 0.0029 | 0.0022 | 0.0024 | 0.0027 | 0.0020 | 0.0022 | 0.0028 | 0.0021 | 0.0022 | 0.0029 | 0.0022 | 0.0024 |
| **Stdev** | 0.0013 | 0.0012 | 0.0012 | 0.0013 | 0.0011 | 0.0011 | 0.0013 | 0.0011 | 0.0011 | 0.0012 | 0.0011 | 0.0011 | 0.0013 | 0.0011 | 0.0011 | 0.0016 | 0.0015 | 0.0016 | 0.0013 | 0.0011 | 0.0012 |
| **Median** | 0.0027 | 0.0020 | 0.0022 | 0.0026 | 0.0019 | 0.0021 | 0.0030 | 0.0022 | 0.0024 | 0.0027 | 0.0020 | 0.0022 | 0.0026 | 0.0018 | 0.0020 | 0.0024 | 0.0019 | 0.0019 | 0.0028 | 0.0021 | 0.0022 |
| **Minimum** | 0.0000 | 0.0000 | 0.0000 | 0.0000 | 0.0000 | 0.0000 | 0.0001 | 0.0001 | 0.0001 | 0.0001 | 0.0000 | 0.0001 | 0.0000 | 0.0000 | 0.0000 | 0.0000 | 0.0000 | 0.0000 | 0.0001 | 0.0001 | 0.0001 |
| **Maximum** | 0.0170 | 0.0147 | 0.0169 | 0.0138 | 0.0113 | 0.0135 | 0.0082 | 0.0073 | 0.0072 | 0.0088 | 0.0078 | 0.0081 | 0.0119 | 0.0114 | 0.0100 | 0.0147 | 0.0144 | 0.0136 | 0.0088 | 0.0077 | 0.0079 |
| **Quartile 1** | 0.0019 | 0.0014 | 0.0015 | 0.0018 | 0.0013 | 0.0015 | 0.0021 | 0.0016 | 0.0017 | 0.0020 | 0.0014 | 0.0016 | 0.0018 | 0.0012 | 0.0014 | 0.0016 | 0.0010 | 0.0011 | 0.0019 | 0.0014 | 0.0015 |
| **Quartile 3** | 0.0036 | 0.0028 | 0.0030 | 0.0035 | 0.0026 | 0.0029 | 0.0039 | 0.0030 | 0.0032 | 0.0037 | 0.0028 | 0.0031 | 0.0034 | 0.0025 | 0.0028 | 0.0036 | 0.0028 | 0.0030 | 0.0037 | 0.0028 | 0.0030 |

|  | Chromosome 7 |  |  | Chromosome 8 |  |  | Chromosome 9 |  |  | Chromosome 10 |  |  | Chromosome 11 |  |  | Chromosome 12 |  |  | Chromosome 13 |  |  |
| --- | --- | --- | --- | --- | --- | --- | --- | --- | --- | --- | --- | --- | --- | --- | --- | --- | --- | --- | --- | --- | --- |
|  | both | A1 | A2 | both | A1 | A2 | both | A1 | A2 | both | A1 | A2 | both | A1 | A2 | both | A1 | A2 | both | A1 | A2 |
| **Mean** | 0.0028 | 0.0023 | 0.0023 | 0.0028 | 0.0023 | 0.0023 | 0.0028 | 0.0021 | 0.0023 | 0.0032 | 0.0025 | 0.0027 | 0.0030 | 0.0023 | 0.0024 | 0.0028 | 0.0021 | 0.0023 | 0.0029 | 0.0021 | 0.0023 |
| **Stdev** | 0.0012 | 0.0011 | 0.0011 | 0.0012 | 0.0011 | 0.0011 | 0.0013 | 0.0013 | 0.0013 | 0.0015 | 0.0013 | 0.0014 | 0.0013 | 0.0013 | 0.0013 | 0.0011 | 0.0010 | 0.0011 | 0.0013 | 0.0011 | 0.0012 |
| **Median** | 0.0026 | 0.0022 | 0.0022 | 0.0027 | 0.0022 | 0.0023 | 0.0026 | 0.0019 | 0.0021 | 0.0030 | 0.0023 | 0.0025 | 0.0028 | 0.0021 | 0.0023 | 0.0027 | 0.0019 | 0.0022 | 0.0027 | 0.0020 | 0.0022 |
| **Minimum** | 0.0000 | 0.0000 | 0.0000 | 0.0001 | 0.0001 | 0.0001 | 0.0002 | 0.0001 | 0.0001 | 0.0000 | 0.0000 | 0.0000 | 0.0002 | 0.0001 | 0.0002 | 0.0002 | 0.0001 | 0.0001 | 0.0001 | 0.0001 | 0.0001 |
| **Maximum** | 0.0103 | 0.0102 | 0.0098 | 0.0089 | 0.0081 | 0.0082 | 0.0125 | 0.0122 | 0.0122 | 0.0170 | 0.0147 | 0.0169 | 0.0106 | 0.0109 | 0.0103 | 0.0087 | 0.0072 | 0.0074 | 0.0119 | 0.0102 | 0.0113 |
| **Quartile 1** | 0.0019 | 0.0015 | 0.0015 | 0.0020 | 0.0015 | 0.0016 | 0.0018 | 0.0012 | 0.0014 | 0.0022 | 0.0017 | 0.0018 | 0.0020 | 0.0014 | 0.0015 | 0.0019 | 0.0013 | 0.0015 | 0.0019 | 0.0013 | 0.0016 |
| **Quartile 3** | 0.0035 | 0.0029 | 0.0029 | 0.0036 | 0.0029 | 0.0030 | 0.0035 | 0.0028 | 0.0029 | 0.0039 | 0.0031 | 0.0034 | 0.0037 | 0.0029 | 0.0031 | 0.0035 | 0.0027 | 0.0030 | 0.0036 | 0.0027 | 0.0030 |

Phylogenetic substructure was more prominent in *G. arboreum*, given the increased sampling relative to *G. herbaceum*. Most of the early diverging *G. herbaceum* lineages were collected on the African continent (Supplementary Table 1), with the exceptions of A1_Af (PI 630014) and A1_125 (PI 529698), which lacked collection information (both) and were from seed collections from Uzbekistan (A1_125). The latter (A1_125) may appear to be in conflict with the African origin of *G. herbaceum*; however, locality information in the U.S. National Plant Germplasm System (GRIN) can reflect secondary acquisition from another collection repository. Phylogenetics and geographic conflict in *G. arboreum* similarly reflect trade and secondary acquisitions in this domesticate-only species. For example, while accessions A2_073 and A2_074 have Texas, USA listed for location, the cultivar names (“Chinese naked” and “Chinese pale”, respectively) indicate they may have originated in China. As expected, many of the accessions trace to the Indian subcontinent (Supplementary Table 1), which encompasses some of the major locations for cultivation of *G. arboreum*.

When partitioned into genomic windows composed of 50 sequential genes, phylogenetic reconstruction generally recapitulates the results of the whole genome phylogeny vis-a-vis the distinction between *G. herbaceum* and *G. arboreum*. While 478 out of the 562 windows surveyed (85%) exhibit a tree topology that is consistent with complete isolation of *G. herbaceum* and *G. arboreum,* 69 gene windows (12%; Supplementary Table 8) exhibited at least one accession nested within the alternate species, potentially signaling introgression; fifteen genomic windows (3%) were excluded due to poor resolution and limited structure. Exemplar tree topologies of concordant and discordant windows are depicted in Supplementary Figure 5. Most of the affected *G. arboreum* accessions contain only a single window exhibiting possible introgression (19 out of 21 accessions); however, six of the nine *G. herbaceum* accessions (i.e., 67%) contain multiple regions exhibiting signs of introgression (median=9.5; range=1-37).

*Gossypium herbaceum* accession A1_155 is most notable in that it is nested within *G. arboreum* in 37 regions, comprising 6.6% of the windows. While the overall phylogenetic placement for *G. herbaceum* accession A1_155 (PI 630024) is reasonable considering it is reportedly an *africanum* (hence, wild accession), the number of regions nested within its sister species may suggest it is affected by lineage sorting, introgression, or both. Four of the other *G. herbaceum* accessions with unusually large numbers of phylogenetically discordant windows (*i.e.*, accessions A1_051, A1_054, A1_132, and A1_133; median = 11.5) form a clade, suggesting that there may have been some introgression in the ancestor to these four lineages. Notably, these regions generally appear concentrated in the gene-rich distal regions of the chromosomes (Figure 2).

**Figure 2.**
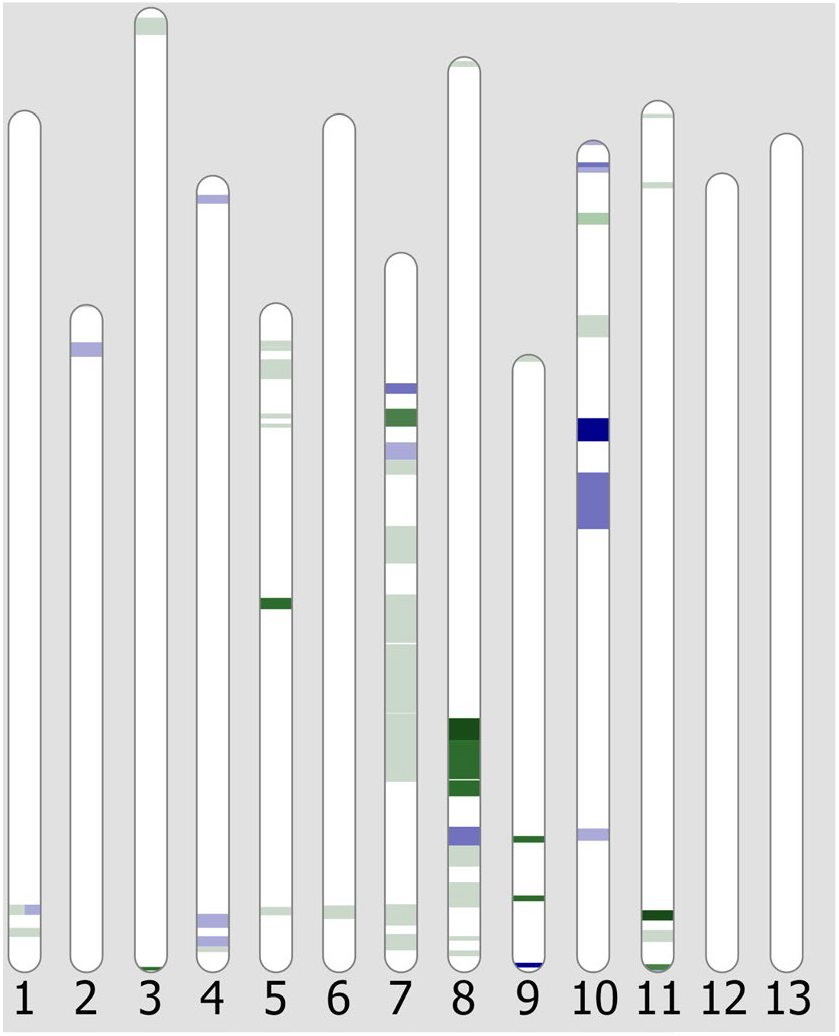
Ideogram displaying the distribution of introgressed windows in either species. Regions where at least one *G. herbaceum* accession exhibited introgression from *G. arboreum* are marked in green, and the converse pattern in which *G. arboreum* accessions exhibit signatures of introgression from *G. herbaceum* are marked in blue. The depth of the color indicates the number of accessions involved (maximum=5), with darker colors indicating the introgression was shared among more accessions. Chromosomes 12 and 13 do not exhibit introgression, but are included here for completeness.

These phylogenetically discordant windows were unevenly distributed, with <10% of windows affected on some chromosomes (*i.e.*, F01, F02, F06, F12, and F13) and others with >20% of windows exhibiting discordance (*i.e.*, F07, F08, F10). Chromosome F13 was the only chromosome where all phylogenetic windows exhibited a strict division between the two species (no discordance). Conversely, chromosome F07 exhibited the greatest number of discordant windows (11 of the 41 windows, or 26.8%), followed by chromosomes F08 (25.6%) and F10 (20.0% of windows). Notably, F07 and F10 also exhibit a paucity of fixed, derived sites, potentially indicating that these two are more permeable to introgression than are the other chromosomes, although we cannot disentangle the absence of gene flow from strong parallel selection.

### Transposable element divergence within and among A-genome species

Transposable element (TE) diversity within and between these two species was characterized by clustering reads representing ∼1% of each genome (based on published species-specific genome sizes (Hendrix & Stewart 2005)). As previously reported for many angiosperms, including cotton, *Ty3-Gypsy* elements comprise the majority of the repetitive sequence in both species (Figure 3), with slightly greater *Gypsy* representation in *G. arboreum* (976 Mbp, on average, versus 910 Mbp in *G. herbaceum*), consistent with the slightly greater average genome size (1710 Mbp, versus 1667 Mbp in *G. herbaceum*). Notably, *Gypsy* elements occupy slightly more of the *G. arboreum* genome (59%) versus *G. herbaceum* (55%). Comparisons of individual cluster abundance (Figure 3) also suggest a general overabundance of TEs in *G. arboreum*, with many of the clusters with the greatest average divergence between species annotated as *Gypsy*. In total, the *G. arboreum* accessions average ∼58 Mb more repetitive sequence than the *G. herbaceum* accessions, congruent with their ∼43 Mb difference in genome size.

**Figure 3.**
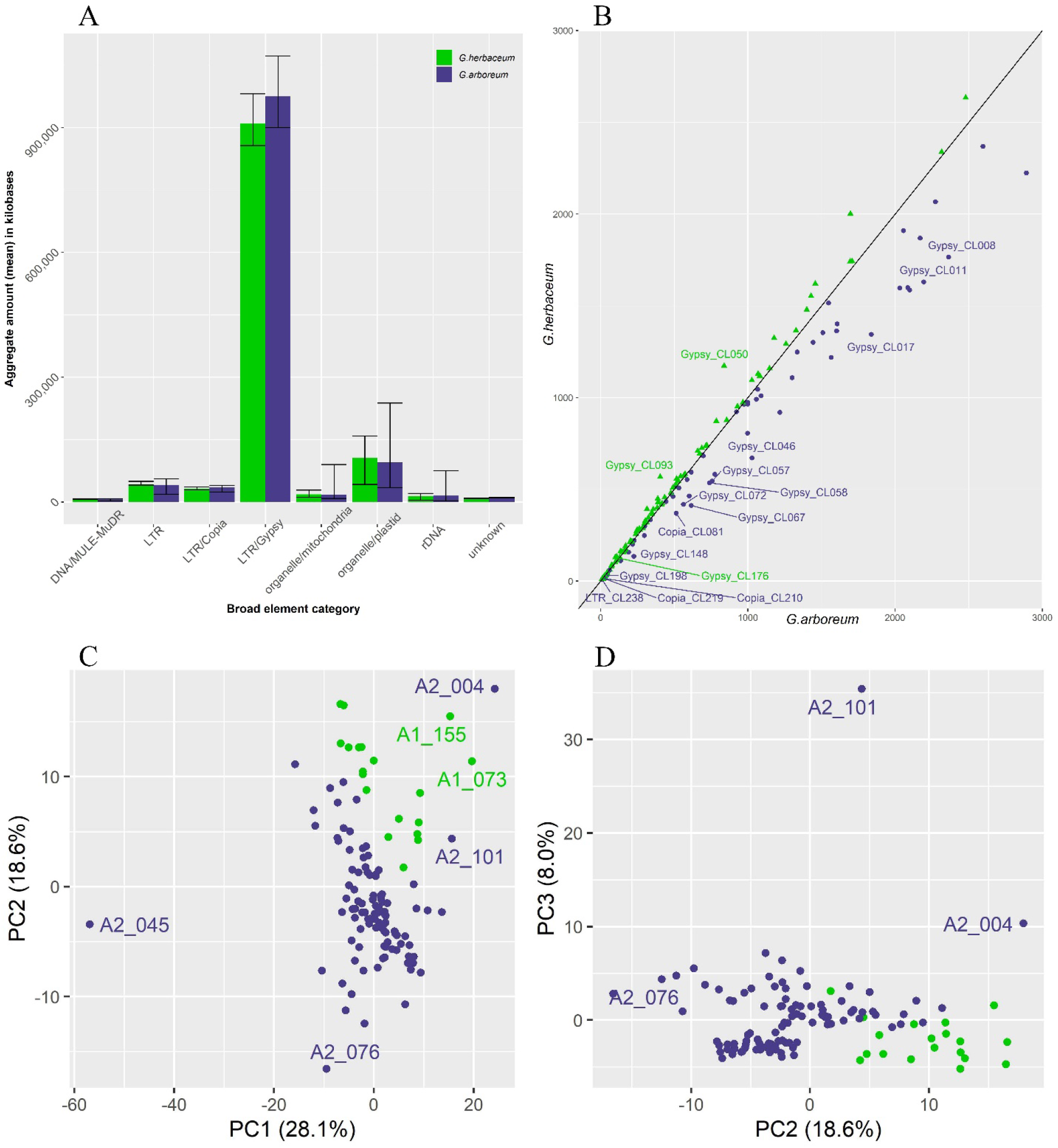
**A.** Read classification in *G. herbaceum* (19 accessions) and *G. arboreum* (99 accessions) by annotation. Transposable elements and/or other high copy sequence categories are shown on the x-axis with their corresponding total genomic amount (mean per sample) on the y-axis (in kilobases). Error bars indicate the minimum and maximum values for each category and species. **B.** Comparison of average cluster abundance between *G. herbaceum* and *G. arboreum*. Points indicate the difference in cluster abundance between *G. herbaceum* and *G. arboreum*, where green triangles (above the line) indicate an overabundance in *G. herbaceum* and blue circles (below the line) indicate an overabundance in *G. arboreum*. The black line indicates a 1:1 ratio for cluster abundance. Labeled clusters indicate those that deviate from 1:1 by more than 20% of the total for that cluster. **C-D**. First three principal component axes for cluster abundance in *G. herbaceum* (green) and *G. arboreum* (blue) accessions. In each panel, the proportion of variance explained is listed along the axis. Accessions whose profiles are distinct from the remaining samples are individually labeled (see Supplementary Table 1 for accession details).

PCA of cluster abundance distinguishes *G. herbaceum* accessions from *G. arboreum* along the first three axes (Figure 3), which collectively account for 55% of the variation. These axes exhibit overlap among accessions of the two species, suggesting that, while their repetitive profiles are somewhat distinct, they are not species-specific. These results remain consistent when removing *G. arboreum* accessions that appear somewhat distant from the others on the initial PCA (*i.e*., A2_004, A2_101, A2_045). Notably, many clusters appear correlated (Supplementary Figure 6), possibly due to cross-mobilization of similar elements, shared ancestry, or both. On an individual cluster basis, 64 clusters distinguish accessions of *G. herbaceum* from *G. arboreum* (t-test p < 0.01 with Benjamini-Hochberg correction), including seven of the top 10 most abundant clusters (Supplementary Table 9). Many of these clusters exhibit intercorrelation (Supplementary Figure 6; Supplementary Table 10), and the PCA loadings from several (*i.e.*, clusters 4, 8, 13, 15, and 21) are largely responsible for cluster separation along the second axis (Supplementary Figure 7). As expected, a majority (84.3%) of these clusters are annotated as *Gypsy*, similar to the overall proportion of *Gypsy* elements evaluated (80.1%).

### Synonymous substitution rates and population structure suggest little interspecific contact

Population structure analysis reveals two to three populations (Figure 4), one solely containing *G. herbaceum* accessions, and 1-2 populations comprising *G. arboreum,* depending on method (*i.e.*, STRUCTURE versus LEA, see methods). Congruence between the two methods is high, with the major difference being the presence of substructure in the *G. arboreum* population using LEA. Congruent with the PCA, this substructure distinguishes the previously sequenced (and primarily Chinese) *G. arboreum* samples from those sequenced here. Notably, LEA also detects a higher proportion of admixture between *G. herbaceum* and *G. arboreum* accessions than STRUCTURE, which may reflect phenomena such as lineage sorting or introgression. Notably, *G. herbaceum* accession A1_155, which had the greatest number of windows indicating possible introgression, is highlighted by both STRUCTURE and LEA as containing *G. arboreum* sequence, as is *G. herbaceum* accession A1_132, albeit to a lesser degree. STRUCTURE analysis including all samples (Supplementary Figure 8) confirms the species misidentifications suggested by PCA, as well as the distinctiveness of the *G. arboreum* accessions sequenced in Du et al (2018).

**Figure 4.**
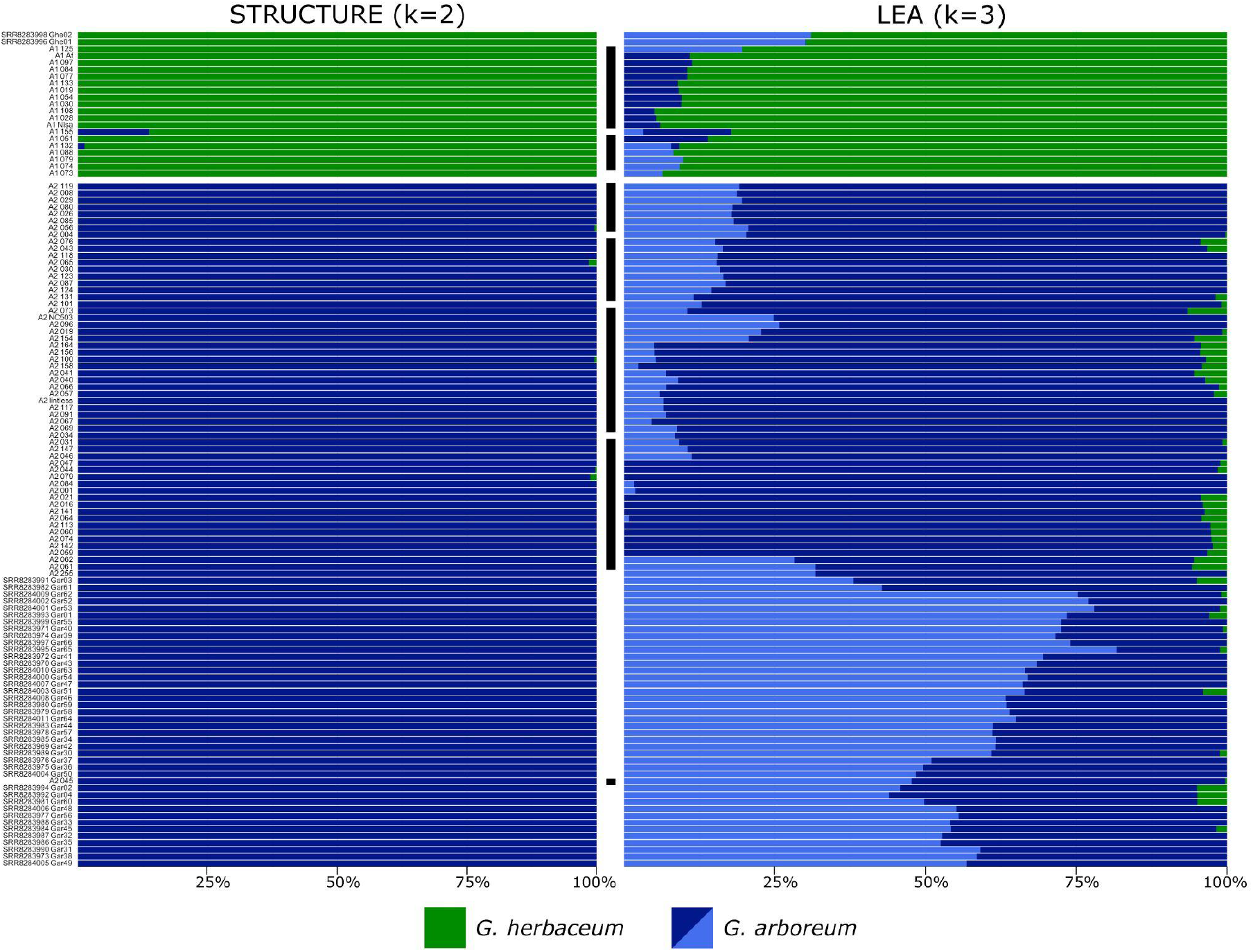
STRUCTURE (left) and LEA (right) analysis of *G. herbaceum* (green) and *G. arboreum* (blues). Newly sequenced accessions are noted with a black bar. Population optimization (see methods) for STRUCTURE recovered only two populations (k=2) split along species lines, whereas LEA recovered three populations (k=3): one *G. herbaceum* population (green) and two *G. arboreum* populations (blue). While both STRUCTURE and LEA are based on the same underlying algorithms, LEA appears more sensitive to lineage sorting and/or introgression. A high-resolution version of this image is available at https://github.com/Wendellab/A1A2resequencing and accession details are found in Supplementary Table 1.

Genome-wide synonymous substitution rates (*d_S_*) between *G. herbaceum* and *G. arboreum* were estimated for 562 genome windows each containing 50 orthologous genes (Supplementary Table 11; Supplementary Figure 9) for all samples. Two haplotypes (see methods) per each accession/species were extracted from the mapped reads for each gene present in the *G. longicalyx* reference annotation; however, those with ≥70% ambiguity (*i.e*., “N”) were removed from the analysis. More stringent filters were also tested and gave similar results, albeit with a lower estimated *d_S_* (Supplementary Table 11). Notably, accessions that were considered putative hybrid and/or contaminated samples were easily spotted due to excessively high or low *d_S_* values (Supplementary Figures 10 and 11). While the low *d_S_* values are consistent with mislabeled species and/or introgression from the sister taxon, those samples with excessive *d_S_* values (*i.e.*, *G. herbaceum* accessions A1_037 and A1_148 and *G. arboreum* accession A2_038) are likely introgressed and/or otherwise contaminated with germplasm from species other than *G. herbaceum* or *G. arboreum*. Excluding these samples and those not passing quality filters (QC accessions, filtered as per methods and noted in Supplementary Table 1), the overall mean *d_S_* between *G. herbaceum* or *G. arboreum* (Table 3) was smaller than previously estimated from ∼7,000 individual genes (*d_S_*= 0.0088 versus 0.0132 from (Renny-Byfield et al. 2016)). Notably, the 95% confidence interval (CI) was also broader than previously reported (Renny-Byfield et al. 2016), ranging from *d_S_*= 0.0031 - 0.0198 (versus *d_S_*=0.0127 - 0.0137), which may reflect the substantially higher sampling in the present analysis (21 *G. herbaceum* and 99 *G. arboreum* accessions, versus two accessions each in (Renny-Byfield et al. 2016)). Mean *d_S_* for each chromosomal window was generally close to the genome-wide mean (*i.e.*, within the 95% confidence interval; Figure 5A), although 13 windows of excess *d_S_* were observed and a single window with reduced *d_S_* (Supplementary Table 12). These windows are represented on approximately half of the chromosomes (6 of 13). Two of these windows are less notable, in that they slightly exceed the 95% CI (limit=0.0197), *i.e.*, position 93.1 Mb on F13 (*d_S_*=0.0201) and position 0.98 Mb on F06 (*d_S_*=0.0207). The other three windows, however, exceed the 95% CI by a larger margin: *d_S_*=0.0244 for the window on F07 at 48.6 Mb, *d_S_*=0.0269 for F09 at 16.7 Mb, and 0.0315 for F11 at 64.3 Mb. Notably, the window on F11 with excess *d_S_* (0.0315) is bordered by windows with far lower *d_S_* (0.0054 at position 56.3 Mb and 0.0105 at 69.8 Mb), which explain why no peak is observed in Figure 5.

**Figure 5.**
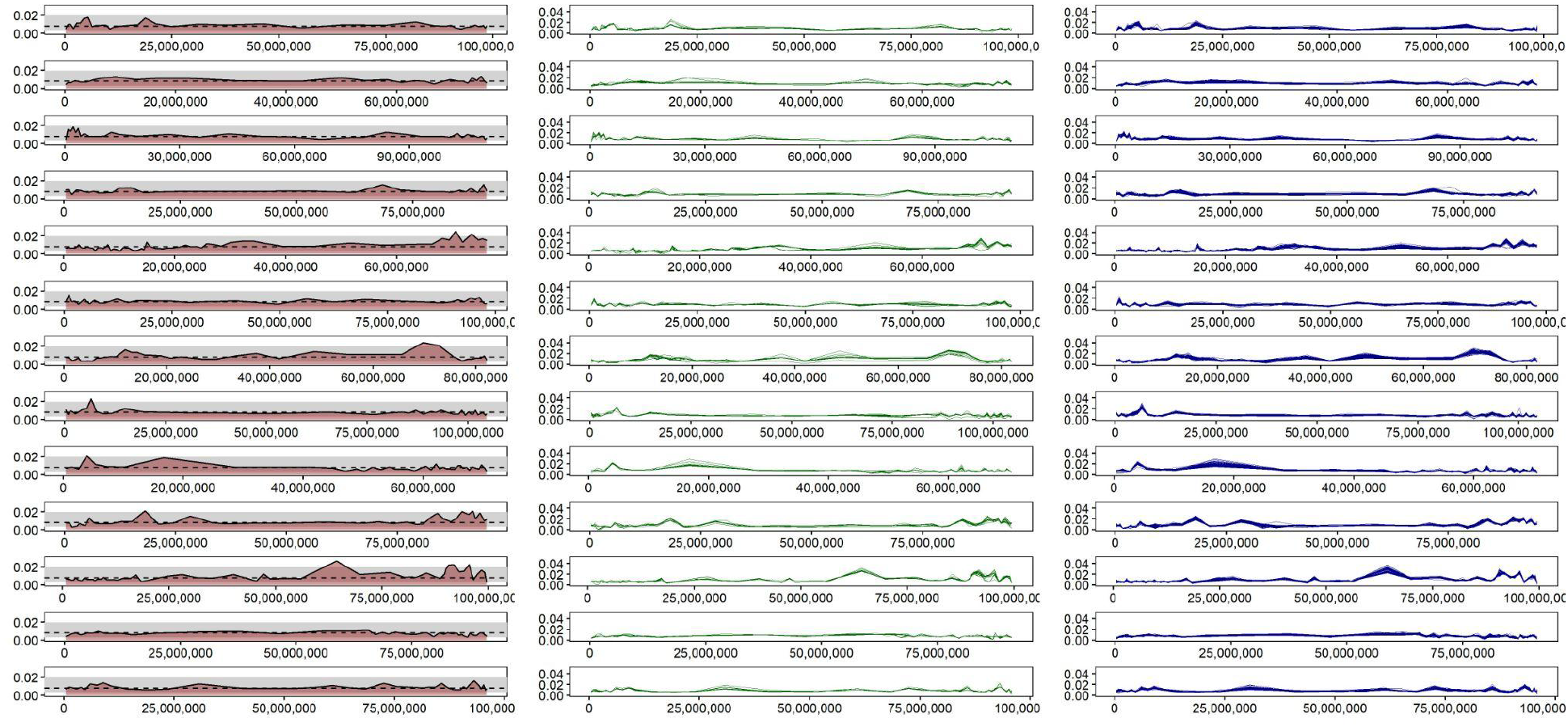
Synonymous substitution (*d_S_*) rates for 50-gene windows are shown by chromosome. The left panel depicts the overall estimate of *d_S_* for *G. herbaceum* versus *G. arboreum*. The middle and right panels depict the individual *d_S_* estimates for *G. herbaceum* (green) and *G. arboreum* (blue) accessions, respectively. These panels show the average *d_S_* of each accession relative to the other species.

**Table 3.** Summary statistics for all estimates of Ks between *G. herbaceum* (A1) and *G. arboreum* (A2) among pairwise permutations of 120 A1 and A2 individuals passing quality filters and using *G. longicalyx* (F1) as an outgroup.

| Filter Level* | Mean | Median | Mode | Range | Standard Deviation | 95% CIs | Divergence** | Divergence CI |
| --- | --- | --- | --- | --- | --- | --- | --- | --- |
| Low Filter | 0.0088 | 0.0080 | 0.0070 | 0.0002 - 0.0562 | 0.0041 | 0.0031 - 0.0198 | 964,912 | 340,000 - 2,171,000 |
| Medium Filter | 0.0064 | 0.0062 | 0.0063 | 0.0002 - 0.0421 | 0.0027 | 0.0022 - 0.0126 | 701,754 | 241,000 - 1,382,000 |
| High Filter | 0.0063 | 0.0060 | 0.0058 | 0.0002 - 0.0421 | 0.0025 | 0.0022 - 0.0124 | 690,789 | 220,000 - 1,360,000 |
| No Filter | 0.0088 | 0.0080 | 0.0074 | 0.0002 - 0.0708 | 0.0042 | 0.0029 - 0.0200 | 964,912 | 318,000 - 2,193,000 |

\* No filter includes all genes, low filter removes genes if $\geq 70\%$ of the nucleotides in the alignment were ambiguous, medium filter removes genes if $\geq 70\%$ of the nucleotides in the alignment were ambiguous and/or $>4$ sequences contained stop codons, and high filter removes genes if either $\geq 70\%$ of the nucleotides in the alignment were ambiguous and/or any sequences contained stop codons.
\*\* Divergence time is calculated via $T=dS/2r$ , where $dS$ is represented by the mean $dS$ or CI for each filter level. Time is given in years before present.

Individual genome-wide interspecific *d_S_* estimates largely mirror the overall mean, ranging from 0.0006 - 0.0370 among windows and accessions for QC accessions (0.0002 to 0.3206 including all accessions). While all accessions appear to follow similar patterns of *d_S_* variation across the genome (Figure 5B,C; Supplementary Figure 9), ∼22% of genomic windows (125 out of 562) have at least one QC accession whose average *d_S_* is outside the 95% CI for the genome (Supplementary Table 13). All accessions fall outside of the CI in at least 13 of these genomic windows (range: 13 - 44), and each of the 125 windows has a median of 2 accessions outside of the CI (range: 1 - 120 accessions). While over 67% of windows (84 out of 125) have at least one accession whose *d_S_* is lower than expected by the CI, only 33% had at least one accession exceed the *d_S_* CI (41 out of 125); however, windows exceeding the *d_S_* CI were generally represented by more accessions (median = 5, versus median = 2 for low *d_S_* windows). Notably, while the *d_S_* CI is exceeded by more than 75% of accessions for 10 genomic windows, only a single window has more than 75% of accessions below the *d_S_* CI (Supplementary Table 13). Also, while all accessions exceed the CI for 9 to 18 windows, only 116 of the 120 QC accessions were under the CI in at least one genomic window. That is, four of the most wild accessions (*i.e.*, A1_074, A1_073, A1_Af, and A1_Nisa; Supplementary Table 13) never exhibited excessively low *d_S_*, perhaps indicating a lack of post-speciation interspecific contact.

Estimated divergence time between species is similar to a previously described estimate that also used synonymous substitutions rates (Renny-Byfield et al. 2016); however, the range in divergence time calculated here is wider than previously estimated. Using our genome-wide assessment of *d_S_* and a Malvaceae-specific synonymous substitution rate (4.56E-09; see methods), we estimate divergence between *G. herbaceum* and *G. arboreum* at 0.97 million years ago (MYA; range 0.34 to 2.20 MYA; Table 3), over 1.20 million years more recent than the estimate by (Renny-Byfield et al. 2016) using a similar mutation rate (2.5 mya using 2.6E-09).

Notably, our estimates are more similar to other recent estimates (Huang et al. 2020), which report a peak *Ks*=0.0056 and a divergence time estimate of 0.70 MYA (range = 0.40 - 1.40 MYA) using coalescent simulations.

## Discussion

The two extant species of subgenus *Gossypium* (colloquially, the A-genome cottons) have been of great interest both because they have historically been important as sources of textile fiber and because their status as the closest relatives of the extinct A-genome donor of polyploid cottons. Disentangling the history of the A-genome cottons, including their species delimitation and infraspecific relationships, has historically been challenging due to their complex, overlapping morphologies (Wendel et al. 1989; Fryxell 1979) and putative history of introgression (Wendel et al. 1989). These challenges have led to germplasm misidentification (Wendel et al. 1989), which evidenced here in the form of five samples misidentified as *G. herbaceum* and two misidentified as *G. arboreum* (noted in Supplementary Table 1). Genetically and cytogenetically, however, these species are distinct and exhibit evidence of interspecific F_2_ breakdown (Gerstel 1953; Menzel & Brown 1954; Phillips 1961; Silow 1944; Stephens 1950), due in part to a putatively isolating chromosomal translocation. Because wild forms of *G. arboreum* are unknown and because wild forms of *G. herbaceum* are geographically disjunct from regions of cultivation (Fryxell 1979; Saunders & Others 1961; Vollesen 1987), this cytogenetic difference has historically caused some to question the independent evolution of these species (Hutchinson 1954b, 1959), rather suggesting that *G. herbaceum* subspecies *africanum* represents the ancestor to both modern *G. herbaceum* and all of *G. arboreum*. These arguments have been refuted based on observations that the reconstructed divergence time between *G. herbaceum* and *G. arboreum* (Wendel et al. 1989; Renny-Byfield et al. 2016; Huang et al. 2020; Page et al. 2013) predates human agronomic innovation, typically by more than two orders of magnitude. Indeed, our estimates are similar to those previously reported, suggesting that these species diverged approximately 960,000 years before present (ybp), well before domestication (circa 5,000 ybp) and comparable to previous estimates, including allozymes (1.4 million years; (Wendel et al. 1989)), cpDNA (715,000 years; (Chen et al. 2016)) and resequencing (400,000 - 2.5 million years; (Huang et al. 2020; Renny-Byfield et al. 2016; Page et al. 2013). Even the most recent date reconstructed here, which relies on the most stringently filtered data and encompasses the lowest end of the confidence interval, suggests that divergence between these species was well beyond an order of magnitude earlier than domestication (200,000 versus 5,000 ybp). Furthermore, phylogenetic reconstruction of both species using the outgroup *G. longicalyx* recovers a topology that clearly delineates all *G. herbaceum* accessions from *G. arboreum* and does not suggest a progenitor-derivative relationship between wild *G. herbaceum* and *G. arboreum*.

Independent evolution of *G. herbaceum* and *G. arboreum* is also supported by the prevalence of fixed homozygous derived sites in both species, 1.6 - 2.1 M in each, with the mean number of fixed, derived sites in *G. arboreum* slightly exceeding that in *G. herbaceum*. Fixed indels that differentiate species are likewise prevalent, with ∼37,000 indels fixed in *G. herbaceum* and 6,800 different indels fixed in *G. arboreum*. While *G. arboreum* has a much lower indel fixation rate than *G. herbaceum* in the present analysis, we note that the sampling of *G. arboreum* was approximately five times greater than *G. herbaceum* and therefore the threshold to achieve fixation was greater. Supporting this is the observation that the number of differentiating indels (disregarding fixation status) is similar between species, with *G. arboreum* having slightly more indels than *G. herbaceum* (4.5 M vs. 4.4 M, respectively). Notably, nucleotide diversity was similar between the two species, *i.e.* 0.0022 and 0.0024 for *G. herbaceum* and *G. arboreum*, respectively, which also does not support a founder effect of *G*.

*arboreum* being derived from *G. herbaceum*. The diversity within both species is similar to that found in among wild or semi-wild and domesticated accessions of *G. barbadense* (0.0021), and greater than the diversity found within the wild-to-domesticated continuum surveyed in *G. hirsutum* (0.0017; (Yuan et al. 2021)). Interspecific divergence between the two species was modest, giving a weighted *F_ST_* between *G. herbaceum* and *G. arboreum* (0.4430) similar to that between the species *G. mustelinum* and *G. ekmanianum* (0.4900; (Yuan et al. 2021)), polyploid species whose evolutionary independence is clear. Concordant with a previous analysis of limited sampling (Renny-Byfield et al. 2016), multidimensional-representation of transposable element abundances (*i.e.*, PCA; Figure 3) also distinguishes these species along the first three axes, with 64 of the top 239 clusters exhibiting species-specific abundances. Together, these analyses represent the first direct comparison of diversity and divergence using modern techniques and diverse accessions of both species, collectively indicating recent divergence of *G. herbaceum* and *G. arboreum* followed by independent domestication.

Although we find substantial evidence for independent evolution and domestication, we also find evidence for post-speciation bidirectional interspecific contact (*i.e.*, introgression) in both species. While phylogenetic reconstruction based on 50-gene windows typically results in a clear division between species, ∼12% of windows contain topologies consistent with introgression (*i.e*., the inclusion of one or few accessions with the alternate species).

Interestingly, we observe both species-specific differences and chromosomal differences in the evidence for introgression. In general, *G. herbaceum* retains more introgression than *G. arboreum* (median=9.5 windows, versus median = 1 in *G. arboreum*), despite the greater sampling in the latter. Furthermore, introgression has been differentially retained among chromosomes, with some chromosomes (*e.g.*, F13) exhibiting no lingering evidence of introgression while other chromosomes (*e.g.*, F07) retain evidence of introgression in over a quarter of the windows surveyed. Notably, two of the three chromosomes with the highest proportion of retained introgression (F07 and F10) were also exceptional in their dearth of species-specific fixed, derived sites. Studies from nearly a century ago provide potential insight into these observed differences in introgression permeability. Early research on crossing behavior in *G. herbaceum* and *G. arboreum* (Stephens 1949, 1950; Stebbins 1945; Skovsted 1933) noted F_2_ breakdown in hybrids between these species consistent with underlying genetic differentiation leading to a reduction in fertility. Stephens suggested “small scale structural differentiations” which, when combined with the low crossover rate in *Gossypium*, led generally to either gametes with near-parental structure or those which “carry deficiencies and their reciprocal duplications” (Stephens 1950). This predicts that subsequent generations would favor progeny which maximize the parental state. That is, “later generations would tend to eliminate the F_1_ type and to increase the number of parental type segregates” (Stephens 1950), a consequence which Stephens notes has been generally observed with interfertile species of cotton grown in mixed cultivation. Together, these observations may highlight chromosomes and/or regions that contain factors involved in F_2_ breakdown between *G. herbaceum* and *G. arboreum*, as well as indicating those chromosomes/regions that do not operate in reducing interspecific fertility and are therefore permeable to introgression. Alternatively, regions of fixed differences may indicate differential targets of selection leading either to a reduction in diversity in parts of the genome. Given the general interest in speciation genetics and islands of fertility, disentangling these two avenues would be a fruitful path for future investigation.

Cotton is an interesting model for domestication in that four species were domesticated in parallel at two different ploidy levels, providing a naturally replicated experiment for understanding convergent paths of crop evolution. Research into the evolution and domestication of the polyploid cultivars has been extensive and has yielded valuable insights in this regard (Applequist et al. 2001; Yuan et al. 2021; Grover, Yoo, et al. 2020; Said et al. 2013; Fang, Wang, et al. 2017; Fang, Gong, et al. 2017; Rapp et al. 2010; Chaudhary et al. 2008; Hovav et al. 2008; Chen et al. 2020; Gallagher et al. 2020; Hu et al. 2014, 2019; Fang, Guan, et al. 2017; Li et al. 2021). Understanding the evolution and domestication of the diploid species, however, is complicated by the lack of wild representatives for *G. arboreum*. Notwithstanding this limitation, most studies have focused on *G. arboreum*, for which many more accessions are available and sometimes with regional biases (Du et al. 2018), or have been limited in sampling (Renny-Byfield et al. 2016) or power of the genetic markers employed (Wendel et al. 1989). The analyses presented here combined resequencing of newly acquired accessions with existing resequencing to provide a global evaluation of diversity and domestication in the A-genome species, with special consideration for evidence that supports or refutes independent evolution of these sister taxa. From these analyses, we draw the conclusion that these species evolved independently with limited interspecific contact post-speciation. Subsequently, each species acquired a level of diversification and divergence that is similar to each other and to the two domesticated allopolyploids, *G. barbadense* and *G. hirsutum* (Yuan et al. 2021). While extensive morphological similarities exist between the two A-genome diploids (Stanton et al. 1994; Wendel et al. 1989), these reflect a shared history combined with a degree of phenotypic convergence and human-mediated introgression (Wendel et al. 1989; Hutchinson 1954b; Silow 1944), with chromosomal and regional barriers to the latter highlighted by the uneven distribution of introgression observed here.

## Methods

### Germplasm selection and sequencing

Based on previous assessments of diversity and with the goal of capturing as much of the A-genome gene pool as possible, we selected 25 previously unsequenced accessions from *G. herbaceum* and 56 from *G. arboreum* (Supplementary Table 1). All accessions were grown in either the greenhouse or field at Brigham Young University (BYU; Provo, Utah) or the Pohl Conservatory at Iowa State University (ISU; Ames, Iowa). Young leaves were collected and high-quality DNA was extracted at BYU using the Cetyl Trimethyl Ammonium Bromide (CTAB) method (Allen et al. 2006). PCR-free libraries were constructed and sequenced using Illumina instruments (PE150) at the Beijing Genomics Institute (BGI) or the DNA Sequencing Center (DNASC) at BYU. An average coverage of 38✕ genome equivalents was generated for each accession.

Existing sequencing data from these two species (Page et al. 2013; Du et al. 2018; Huang et al. 2020) were downloaded (Supplementary Table 1) from the Short Read Archive (SRA) hosted by the National Center for Biotechnology Information (NCBI). In total, 19 accessions of *G. herbaceum* and 273 accessions of *G. arboreum* were downloaded, most with relatively low (<10✕ average genome equivalent) coverage (Du et al. 2018).

### Read mapping and SNP inference

Raw reads were mapped to the phylogenetic outgroup *G. longicalyx* genome (Grover, Pan, et al. 2020) using bwa v0.7.17-rgxh5dw (Li & Durbin 2009) from Spack (Gamblin et al. 2015).

Single-nucleotide polymorphisms (SNPs) were called using the software suite provided by Sentieon (Kendig et al. 2019) (Spack version sentieon-genomics/201808.01-opfuvzr) and following the DNAseq guidelines. This pipeline is an optimization of existing methods, such as GATK (McKenna et al. 2010), and includes read deduplication, indel realignment, haplotyping, and joint genotyping. Parameters for mapping and SNP calling follow standard practices, and are available in detail at https://github.com/Wendellab/A1A2resequencing.

Previous results (Yuan et al. 2021) suggest that lower coverage datasets lack robustness and reproducibility. Therefore, SNP coverage for each accession was calculated by vcftools (Spack version 0.1.14-v5mvhea) (Danecek et al. 2011), and samples with insufficient depth (*i.e.*, < 10✕ average coverage for SNP sites present in >90% of samples) were removed from further analyses. SNP sites with more than two alternative nucleotides were excluded, and a minimum average read depth of 10, a maximum average read depth of 150, and a minor allele frequency of 5% were required for a site to be retained. For the purposes of principal component analysis (PCA) and phylogenetics (see below), all sites with indels or missing data were excluded. The outgroup (*G. longicalyx*) was removed from the VCF for PCA, and all sites monomorphic among the A-genome diploids were removed as uninformative. All filtering was completed in vcftools (Danecek et al. 2011), and specific parameters are available at https://github.com/Wendellab/A1A2resequencing.

### SNP and indel analyses

Gene-associated SNPs were evaluated by intersecting the filtered VCF with the relevant feature (*e.g.*, exon, intron, etc.) from the *G. longicalyx* annotation (Grover, Pan, et al. 2020) hosted by CottonGen (Yu et al. 2014). In each case, the Unix command grep was used to recover only the targeted feature(s), and intersectBed from bedtools2 (Spack version 2.27.1-s2mtpsu) (Quinlan 2014) was used to recover only SNP sites contained within those regions. Putative effects of each SNP (relative to the outgroup, *G. longicalyx*) were calculated by passing the entire filtered VCF to SNPEff (Cingolani, Platts, et al. 2012), which returned summary statistics as html. The SNPEff config file and parameters are available at https://github.com/Wendellab/A1A2resequencing.

Indels were placed in a separate VCF file using vcftools (Danecek et al. 2011) with the ‘--keep-only-indels’ flag. Samples that did not pass the SNP filtering were removed from the indel set. Because indels were mapped against the outgroup sequence, *G. longicalyx*, the reference state was considered ancestral, allowing the alternate state to be characterized specifically as an insertion or deletion; this was completed using ‘varTypè from SnpSift (Cingolani, Patel, et al. 2012). Indel effects were characterized using SNPEff, as above.

Nucleotide diversity (*π*) was calculated in 100 kb windows (sliding 20 kb) using the ‘-- window-pì function in vcftools. Diversity in genic regions was assessed by using the gene/feature specific VCF generated above (i.e., intersections between the full VCF and feature coordinates found in the annotation file for the *G. longicalyx* genome) prior to assessing diversity in vcftools. *F_ST_* between populations was similarly calculated in 100 kb windows (sliding 20 kb) and specifying the population of origin (Supplementary Table 1). Nucleotide diversity and *F_ST_* were only calculated for samples/sites passing the above filters.

### Synonymous substitution rates

Genome-wide synonymous substitution rates were calculated for windows of 50 genes each, with the last window along each chromosome containing slightly fewer genes. Two haplotypes for each accession were reconstructed (relative to the *G. longicalyx* reference) from the mapped reads using *bam2consensus* from BamBam v. 1.3 (Page et al. 2014) and requiring a minimum of 5 mapped reads. In constructing windows, we only used genes that had <70% missing data, to prevent short and/or phylogenetically uninformative genes from overly influencing divergence estimates. This resulted in 563 non-overlapping windows, with a mean of 42.23 windows per chromosome (range = 31 to 65 windows on chromosomes F02 and F05, respectively). The synonymous substitution rate (*d_S_*) between *G. herbaceum* and *G. arboreum* was then estimated for each window by permuting all combinations of haplotypes from G. *herbaceum* and *G. arboreum* with both haplotypes from the outgroup *G. longicalyx.* This resulted in eight separate haplotype permutations for each *G. herbaceum*-*G. arboreum* accession pair per genomic window for a total of 112,832 permutations of each genomic window using all accessions. The total synonymous distance between *G. herbaceum* and *G. arboreum* (outgroup=*G. longicalyx*) was estimated for each permutation of each window by employing model 0 (single ω estimated for the unrooted three-taxon tree) from codeml inside Phylogenetic Analysis by Maximum Likelihood (PAML) v. 4.9j (Yang 2007). Synonymous substitution rates inferred by codeml were extracted from codeml output using a custom script (dSPermutations.py), and visualized using ggplot2 (Wickham 2016) in R v 4.05 (R Core Team 2020). R code and PAML parsing scripts are available at https://github.com/Wendellab/A1A2resequencing. Because non-functional genes can inflate estimates of *d_S_*, we repeated the analysis using a series of filters with increasing stringency to iteratively remove genes based on the number of stop codons (*i.e.*, no limit, <4, and 0 for low, medium, and high stringency, respectively); all stringency filters removed genes with >70% ambiguity. Overall, these filters did not alter the conclusions drawn from these data, but their values are shown in Supplementary Table 11.

Divergence time between *G. herbaceum* and *G. arboreum* was estimated using a previously calculated rate of synonymous substitutions for the Malvaceae (4.56E-09 substitutions/year), which includes *Gossypium* (De La Torre et al. 2017)). We estimated divergence between *G. herbaceum* and *G. arboreum* using the equation *T* = *d_S_*/(2*r*), where *d_S_* is represented by the mean *d_S_* between species (excluding outliers) and *r* is the Malvaceae-specific synonymous substitution rate. The range in divergence time was calculated using the 95% confidence interval for each filter level.

### Phylogenetics and Principal Component Analysis (PCA)

For samples with a minimum 10✕ average read coverage per SNP, we generated a neighbor-joining tree using VCF-kit commit 25c7c03 (Cook & Andersen 2017) with default parameters. After pruning samples with incorrect or questionable identity, a new phylogeny was generated. We also inferred phylogenetic trees for the 50-gene windows used for *d_S_* analyses (“low filter” only, which removes sequences with >70% ambiguity; Supplementary Table 11) in RAxML v8.2.12. RAxML was run using the rapid bootstrapping algorithm (100 bootstrap replicates) assuming a GTRGAMMAIX model of molecular evolution, and *G. longicalyx* was specified as the outgroup to *G. herbaceum* and *G. arboreum*. Bifurcations with low bootstrap support (i.e., ≤ 60 bootstrap support) were collapsed into polytomies using a custom Python script (collapseLowSupportBranches.py) available at https://github.com/Wendellab/A1A2resequencing. Putative introgression was evaluated by screening for tree topologies that contain highly supported clades composed entirely of *G. herbaceum* or *G. arboreum* and which also include every representative accession for that species.

PCA was initially conducted for all samples passing the filters described above using the R v4.0.2 (R Core Team 2020) package SNPRelate v 1.22.0 (Zheng et al. 2012). Subsequently, misidentified or putative hybrid samples were removed to compute an exon-only PCA, using the VCF generated above. Data were visualized using ggplot2 (Wickham 2016).

### Population structure

Population structure was predicted using two datasets, one containing all samples (except for the outgroup, *G. longicalyx*), including those considered mislabeled by PCA, and the other containing only samples passing quality/identity filters (see above). The larger dataset containing all samples was thinned with vcftools to 1 SNP per 10 kb, and then both were filtered with vcftools to remove loci with more than 10% missing data and individuals with more than 95% missing data. Due to capacity limitations in STRUCTURE, a subset of 10,000 loci were randomly selected from each of the filtered VCFs (Burgos et al. 2014) and subsequently converted to STRUCTURE format via plink v1.9 (Purcell et al. 2007). Population information was added to each of these STRUCTURE input files using a custom python script available from https://github.com/Wendellab/A1A2resequencing. A third STRUCTURE dataset was created to further evaluate population structure in *G. arboreum* by removing *G. herbaceum* accessions prior to STRUCTURE conversion. Custom scripts and detailed parameters are available at https://github.com/Wendellab/A1A2resequencing.

STRUCTURE v2.3.4 (Pritchard et al. 2000; Hubisz et al. 2009; Falush et al. 2007, 2003) was run on each VCF using the range K=1 to K=5. Each individual K was run 16 times per file (for *G. herbaceum* and *G. arboreum*, together) or 8 times (*G. arboreum* only). STRUCTURE results were compressed into ZIP archives and uploaded to STRUCTURE Harvester (Earl & vonHoldt 2012), which uses the Evanno method (Gilbert 2016; Evanno et al. 2005) to determine the optimal K. The best K for each set of results was visualized using ggplot2 in R v4.0 to show membership proportion for individuals and to show which individuals had the most similar membership proportions.

A second evaluation of population structure was completed for *G. herbaceum* and *G. arboreum* using LEA (Frichot & François 2015), which implements a STRUCTURE-like admixture analysis in the R environment (here, in R v4.0). The original and filtered VCFs were thinned via plink to include a subset of markers in approximate linkage equilibrium using ‘-- indep-pairwisè to remove any pair of SNPs within a 50 SNP window (sliding 10 SNPs) with an allele count correlation (*r^2^*) value greater than 0.1 (Liu et al. 2020). A subset containing only *G. arboreum* accessions was created by filtering missing data (*i.e.*, keeping sites with <10% missing data and individuals with <95% missing data, as described above) and removing *G. herbaceum* accessions via vcftools. LEA was run 10 times per K (K = 1 to K = 10) for each dataset. The cross-entropy criterion was plotted against the number of inferred ancestral populations for each analysis, retaining results for the K-value with the minimum cross-entropy (*i.e.*, the lowest point on the curve). As with STRUCTURE, the best K for each set of results was visualized with ggplot2 in R v4.0 to show membership proportion for individuals and to show which individuals had the most similar membership proportions.

### Repeat analysis

Repetitive content was evaluated for each genome using the RepeatExplorer v2 pipeline (Novák et al. 2010). Forward reads from each library were filtered for quality and trimmed to a uniform 90 nt using Trimmomatic version 0.36-lkktrba (Bolger et al. 2014) from Spack and then randomly subsampled to represent a 1% genome size equivalent, using the average genome size for each species (Hendrix & Stewart 2005). Reads from each species were combined and used as input into the RepeatExplorer pipeline, and resulting clusters were annotated using a custom repeat library consisting of Repbase version 21.08 (Bao et al. 2015) and previously annotated cotton repeats (Grover et al. 2004, 2007, 2008; Hawkins et al. 2006; Paterson et al. 2012).

Clusters were filtered to include only those where either species, *i.e.*, *G. herbaceum* or *G. arboreum*, averaged 10 or more reads across accessions. The contribution of each cluster to the overall genome was calculated in R version 4.0.3 (R Core Team 2020) based on the genome sampling rate (1% of the total genome size) and the input read length (*i.e.*, 90 nt). PCA of accessions using cluster abundance was conducted using both prcomp from the R packages stats and PCAtools (Blighe & Lun 2020); in both cases, data were scaled. Clusters that differentiate *G. herbaceum* and *G. arboreum* were determined via t-test, and p-values were adjusted using the Benjamini & Hochberg correction (Benjamini & Hochberg 1995). All images were generated using ggplot2 (Wickham 2016). Code for all R analyses is available from https://github.com/Wendellab/A1A2resequencing.

## Supplementary Material

Supplementary data are available online.

## Acknowledgements

We thank the ResearchIT unit at Iowa State University for computational support. This work was supported by the National Science Foundation (to JFW and JAU) and the New Mexico Institute of Mining and Technology. We also used resources from the University of Colorado Boulder Research Computing Group, which is supported by the National Science Foundation (awards ACI-1532235 and ACI-1532236), the University of Colorado Boulder, and Colorado State University.

## Data Availability Statement

The data used in this article are available from the Short Read Archive (under PRJNA539957) at https://www.ncbi.nlm.nih.gov/sra for sequencing data and from Github (https://github.com/Wendellab/A1A2resequencing) for code and analyses.

