## Supplemental Files for "Dual domestication, diversity, and differential introgression in Old World cotton diploids"

**Supplementary Figure 1.** PCA and phylogenetic tree of all samples passing minimum read depth filters. Putatively mislabeled germplasm is marked in the PCA with red arrows and in the phylogenetic tree with red dots; possibly hybrid accessions are similarly marked in orange. Notably, there exists a clear division among samples that is associated with sequencing (further evaluated in Supplementary Figure 2).

**Supplementary Figure 2.** PCA labeled by sequencing origin. The PCA on the left represents all SNPs passing quality filters and the right PCA represents only genic SNPs passing quality filters. All accessions arising from Du et al (2018) are separated on PCA1 from the total SNP PCA (left) and on PCA2 from genic SNPs only. Conversely, the resequencing generated here and the Huang et al (2020) resequencing were generally placed on a continuous spectrum (with the exception of putative hybrids). Therefore, all Du et al (2018) samples were excluded due to possible artificial differences arising during library preparation and/or sequencing.

**Supplementary Figure 3.** Overall distribution of indel sizes in the two species, parsed as insertions (yellow) and deletions (grey).

**Supplementary Figure 4.** Correlation in diversity between *G. herbaceum* (x-axis) and *G. arboreum* (y-axis), partitioned by chromosome.

**Supplementary Figure 5.** Exemplar phylogenetic tree topologies of regions introgressed vs. non-introgressed regions. Cladograms from selected windows including both haplotypes from filtered individuals are depicted with low-support splits (i.e., bootstrap values <60) collapsed into polytomies. *G. longicalyx* (outgroup) is represented by black circles; *G. herbaceum* [A1] is represented in green; and *G. arboreum* [A2] is represented in blue. Panels A-D show topologies of several windows in which we failed to detect introgression: (A) chromosome F01, window 1, strong bootstrap support for *G. herbaceum* clade, (B) chromosome F01, window 3, strong bootstrap support for *G. arboreum* clade, (C) chromosome F02, window 6, strong bootstrap support for *G. arboreum* clade and *G. herbaceum* clade, and (D) chromosome F02, window 11, strong bootstrap support for *G. arboreum* clade and *G. herbaceum* clade. Panel E is an exemplar topology marked as "complex" due to low resolution and abundant discordance that cannot disentangle introgression from lineage sorting and/or other evolutionary phenomena. (E) chromosome F04, window 29, generally poor resolution with thirteen *G. herbaceum* individuals (Ghe02, A1_019, A1_028, A1_030, A1_051, A1_054, A1_077, A1_079, A1_084, A1_088, A1_097, A1_108, A1_125) intercalated within *G. arboreum* haplotypes. Panels F-H provide exemplar topologies of various patterns consistent with introgression: (F) chromosome F05, window 49, four *G. herbaceum* individuals (A1_051, A1_054, A1_132, A1_133) exhibited *G. arboreum* haplotypes, (G) chromosome F07, window 27, one *G. herbaceum* individual (A1_155) exhibited *G. arboreum* haplotypes, (H) chromosome F10, window 05, two *G. arboreum* individuals (Gar04, A2_041) exhibited *G. herbaceum* haplotypes.

**Supplementary Figure 6**. Correlation in cluster abundance for the top 100 most abundant clusters. Correlations are shown both as networks (lower left) and as a heatmap (upper right). For the heatmap, dark red coloration suggests a strong negative association between those elements whereas dark blue coloration suggests a strong positive association. The correlation network demonstrates the degree of interconnectedness among these clusters.

**Supplementary Figure 7**. PCA of repetitive clusters with directional loading shown for significant clusters. *G. herbaceum* is represented in green and *G. arboreum* is represented in blue.

**Supplementary Figure 8.** STRUCTURE analysis including all samples (including hybrid and misidentified germplasm) using *k=3*.

**Supplementary Figure 9**. Genome-wide ds, partitioned by chromosomes (in descending order, F01 - F13). The left panel (red) summarizes mean ds between *G. herbaceum* and *G. arboreum*, the middle panel (green) depicts all *G. herbaceum* accessions versus the *G. arboreum* mean, and the right panel (blue) depicts all *G. arboreum* accessions versus the *G. herbaceum* mean.

**Supplementary Figure 10.** The range in ds for each *G. herbaceum* accession relative to all *G. arboreum* samples. Outliers are highlighted.

**Supplementary Figure 10.** The range in ds for each *G. arboreum* accession relative to all *G. herbaceum* samples. Outliers are highlighted.

Figure S1

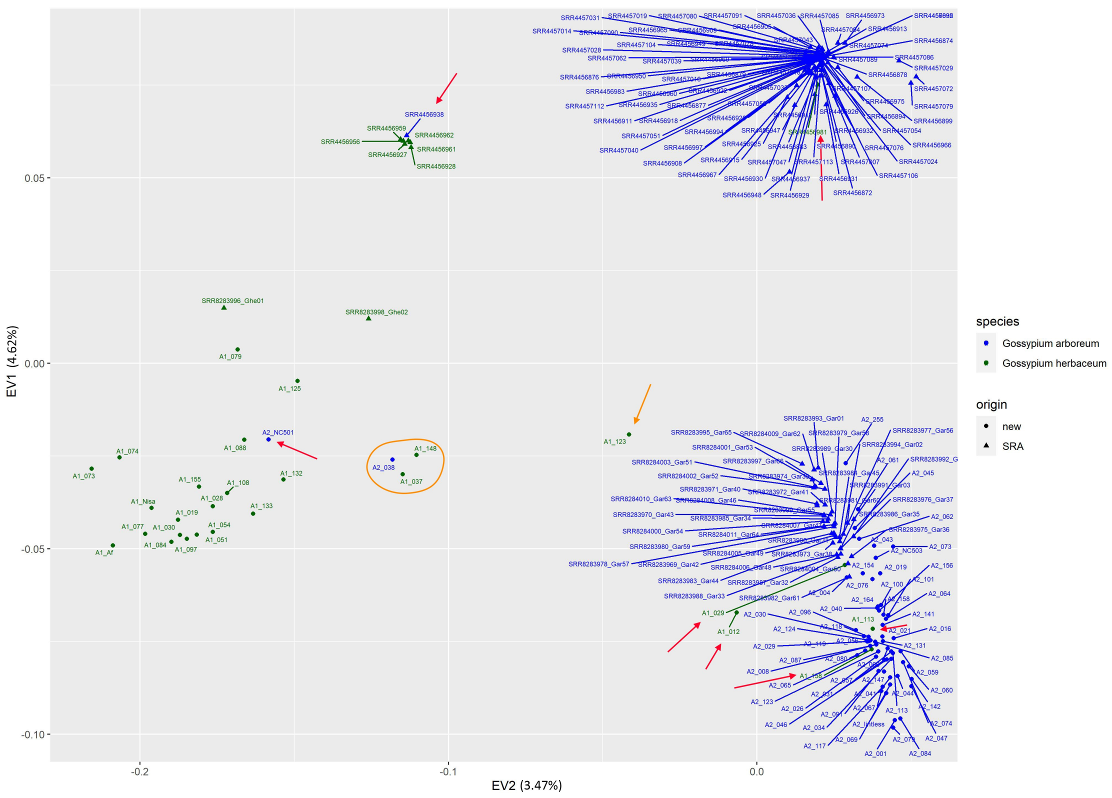
EV1 (4.62%)
EV2 (3.47%)
species
Gossypium arboreum
Gossypium herbaceum
origin
new
SRA

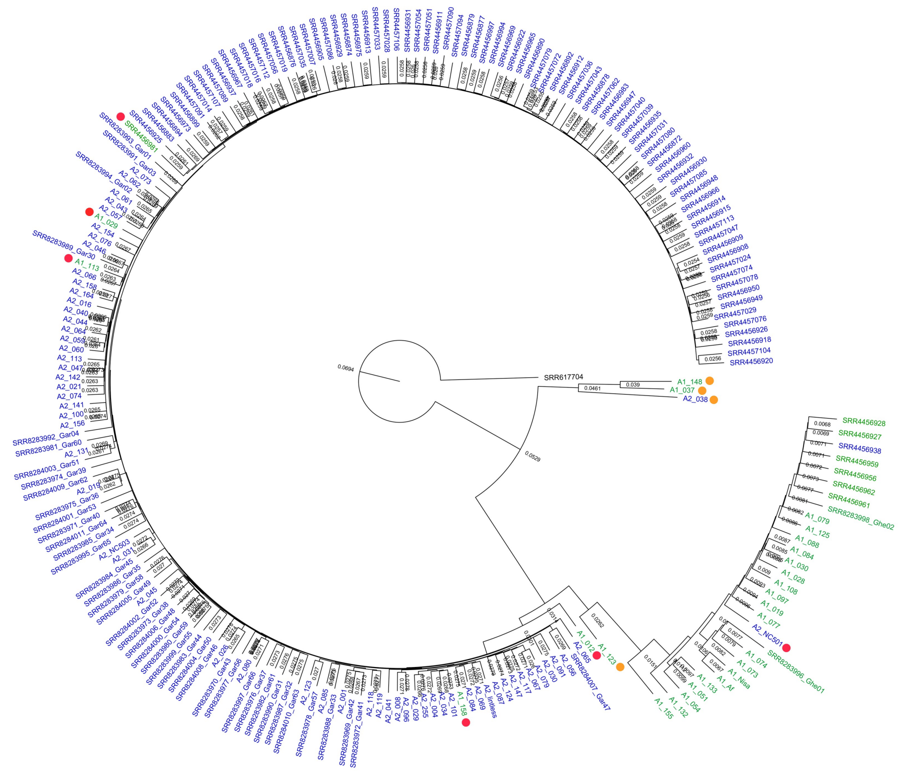

Figure S2

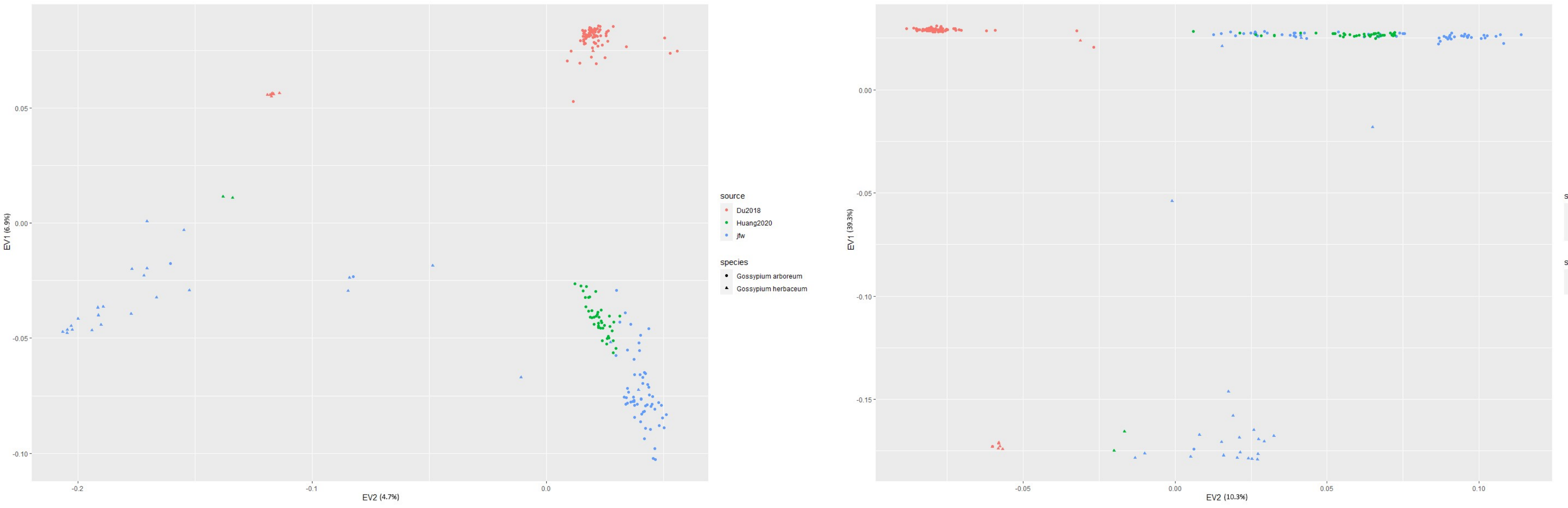

EV1 (6.9%)
EV2 (4.7%)
source
Du2018
Huang2020
jfw
species
Gossypium arboreum
Gossypium herbaceum
EV1 (39.3%)
EV2 (10.3%)

Figure S3

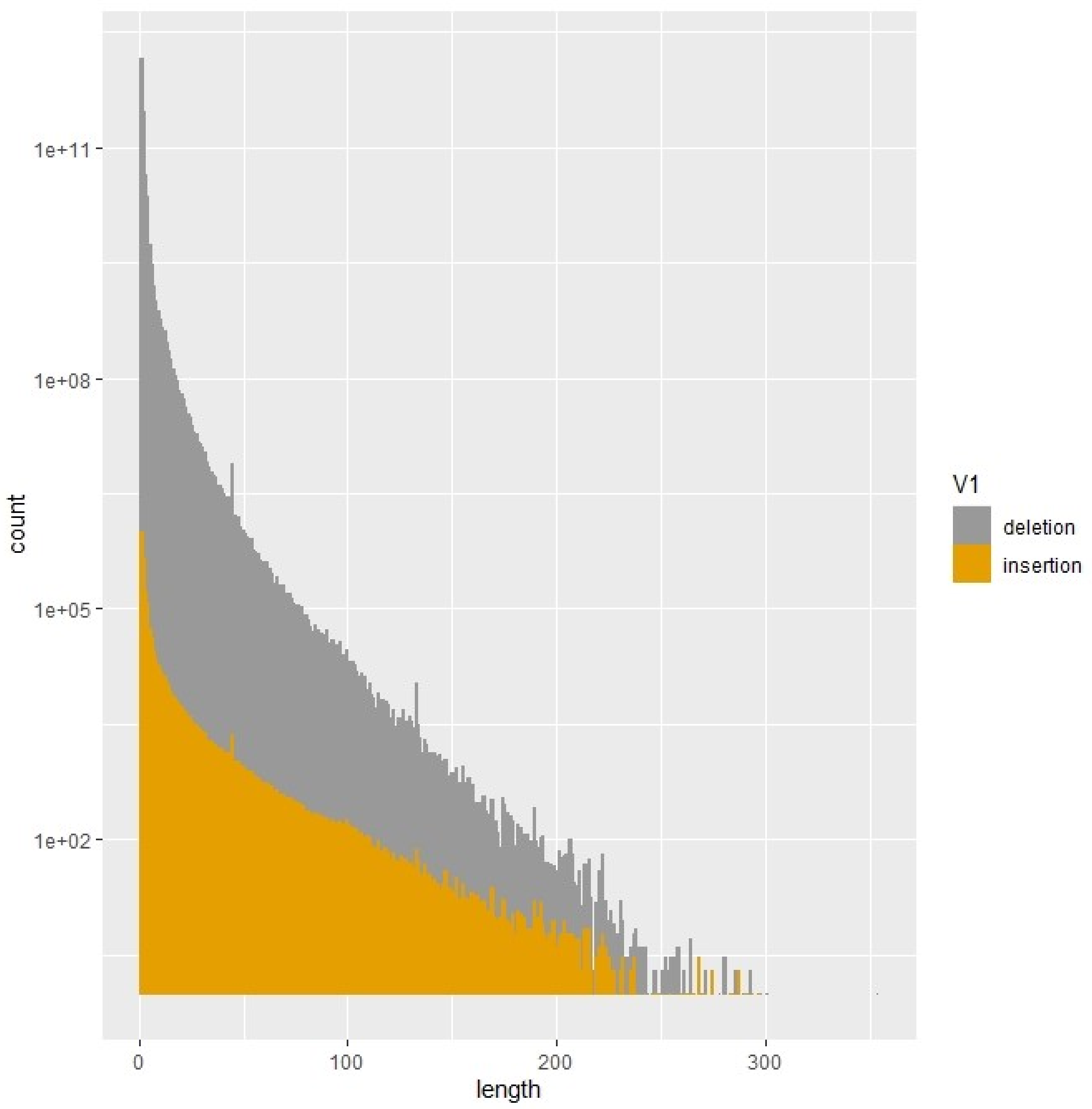

1e+11
1e+08
1e+05
1e+02
count
0
100
200
300
length
V1
deletion
insertion

Figure S4

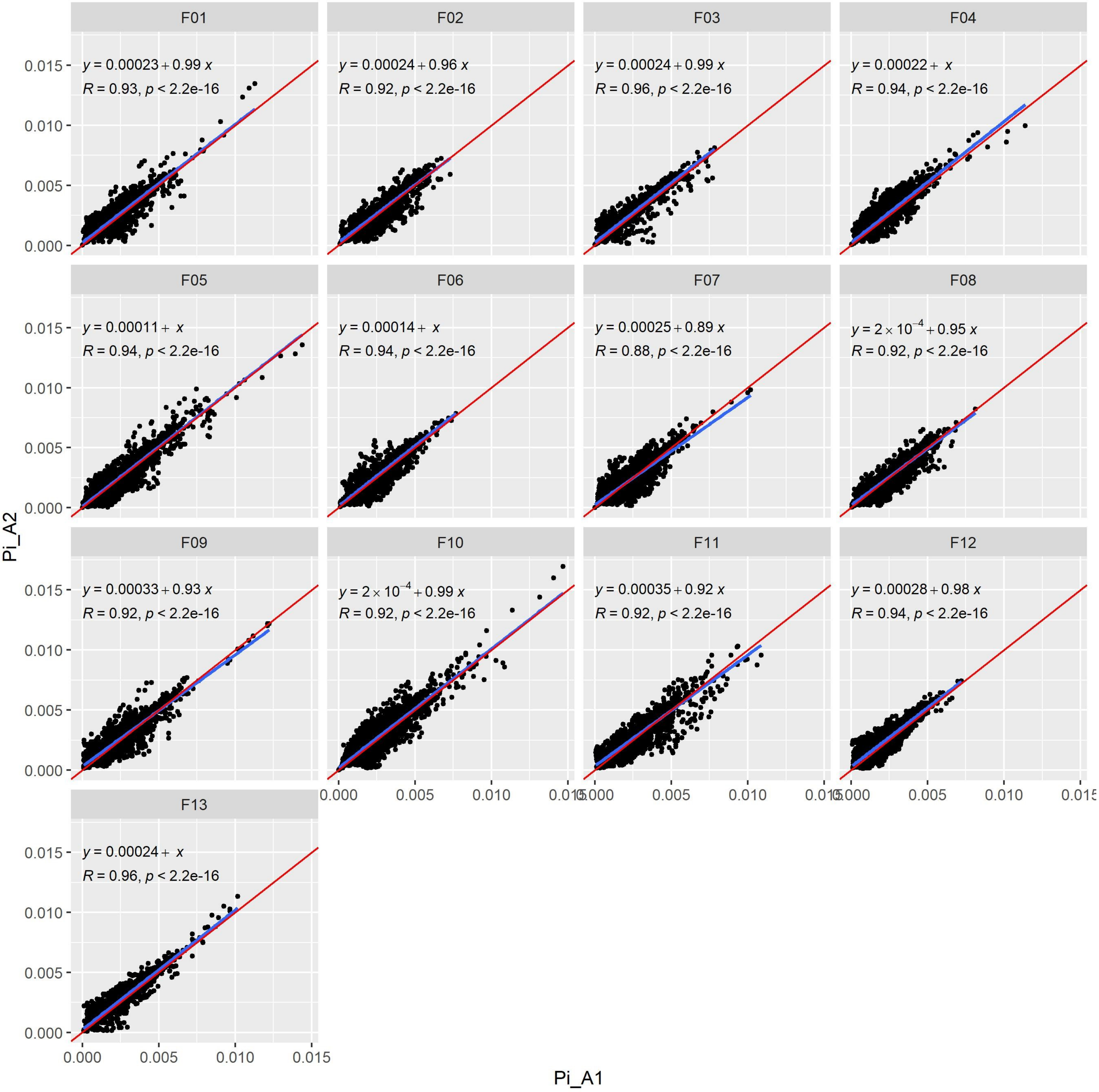

F01
y = 0.00023 + 0.99 x
R = 0.93, p < 2.2e-16
F02
y = 0.00024 + 0.96 x
R = 0.92, p < 2.2e-16
F03
y = 0.00024 + 0.99 x
R = 0.96, p < 2.2e-16
F04
y = 0.00022 + x
R = 0.94, p < 2.2e-16
F05
y = 0.00011 + x
R = 0.94, p < 2.2e-16
F06
y = 0.00014 + x
R = 0.94, p < 2.2e-16
F07
y = 0.00025 + 0.89 x
R = 0.88, p < 2.2e-16
F08
y = 2 × 10^-4 + 0.95 x
R = 0.92, p < 2.2e-16
F09
y = 0.00033 + 0.93 x
R = 0.92, p < 2.2e-16
F10
y = 2 × 10^-4 + 0.99 x
R = 0.92, p < 2.2e-16
F11
y = 0.00035 + 0.92 x
R = 0.92, p < 2.2e-16
F12
y = 0.00028 + 0.98 x
R = 0.94, p < 2.2e-16
F13
y = 0.00024 + x
R = 0.96, p < 2.2e-16
Pi_A2
Pi_A1
0.000
0.005
0.010
0.015

Figure S5

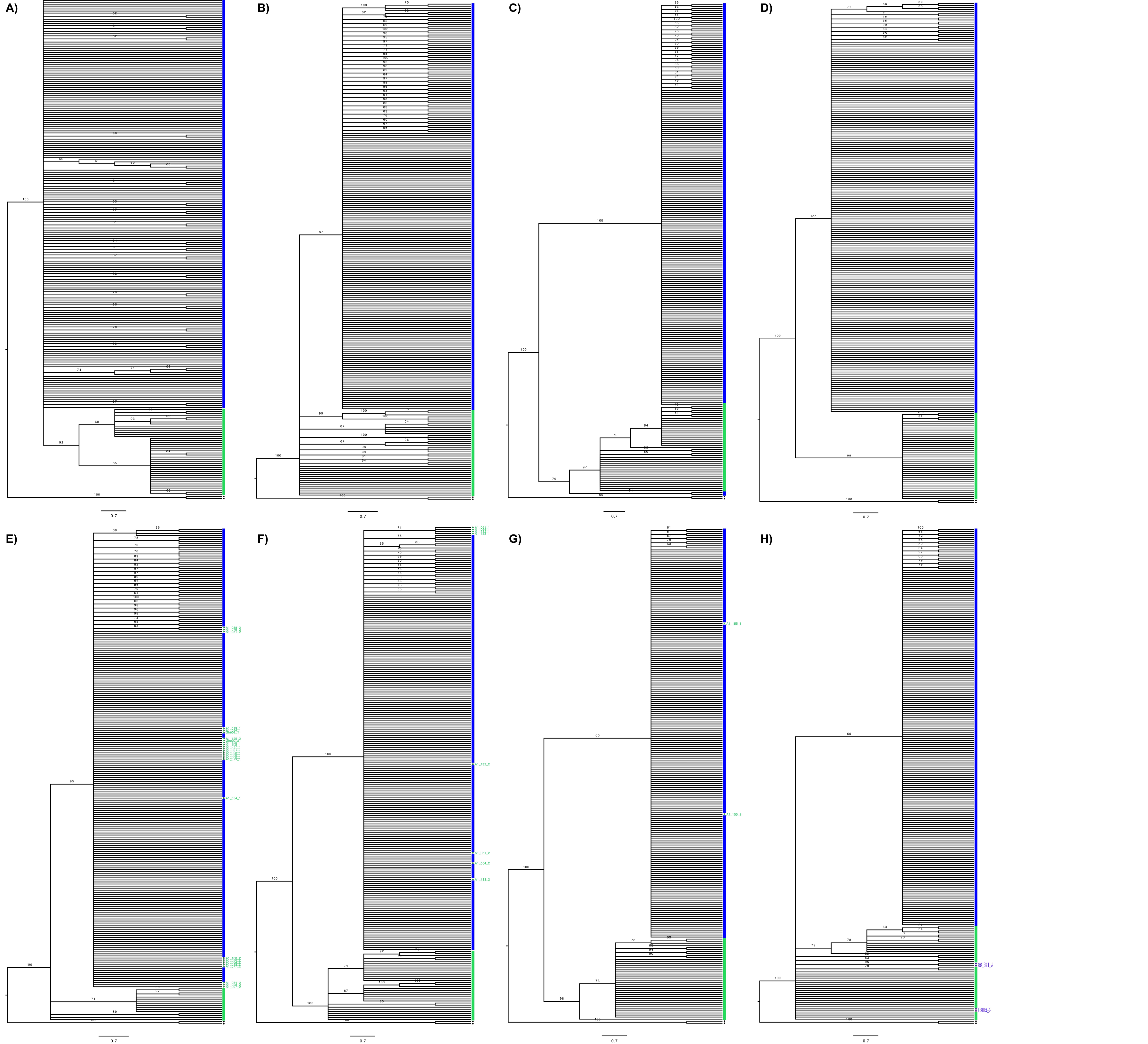

Figure S6

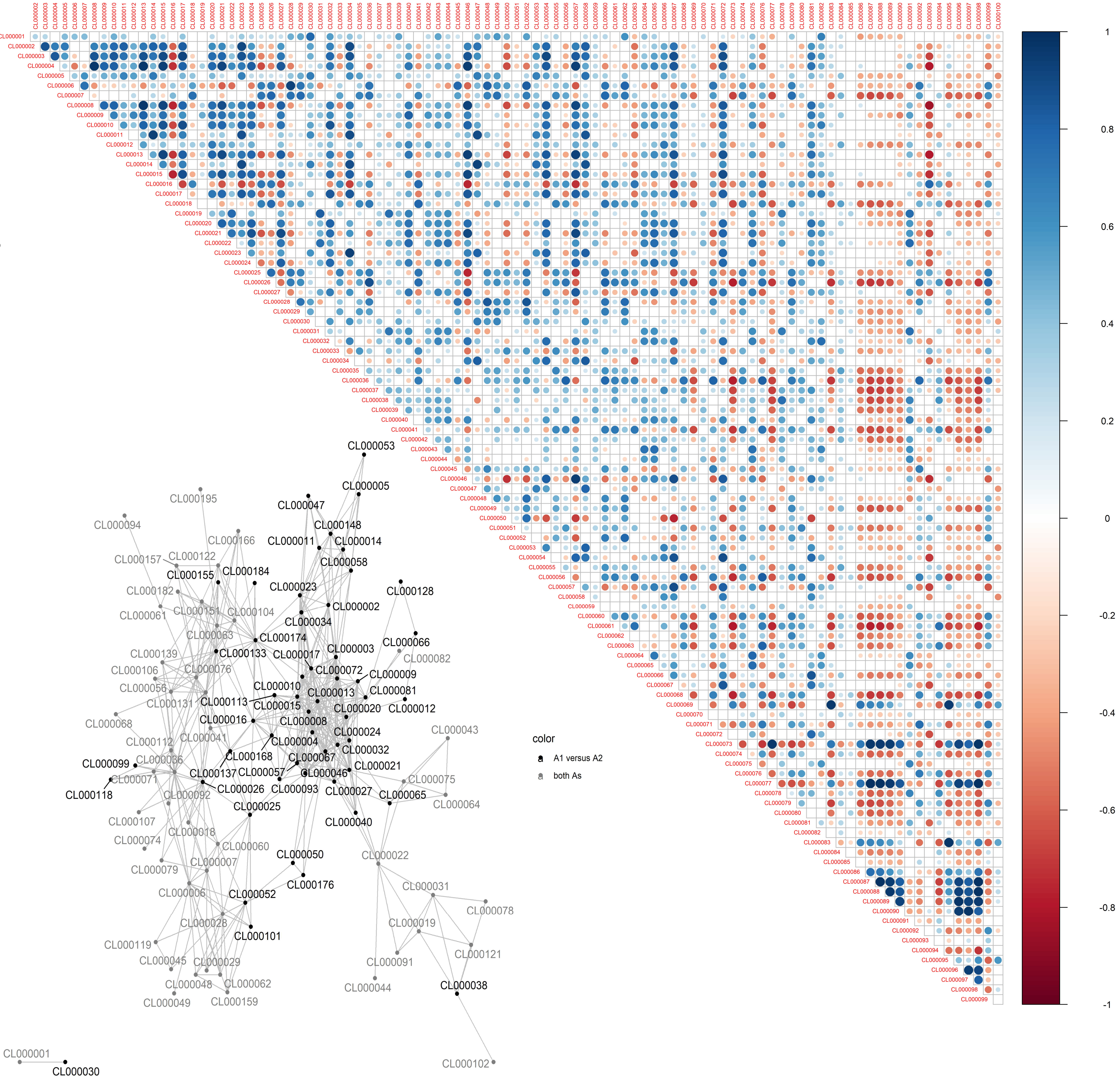

Figure S7

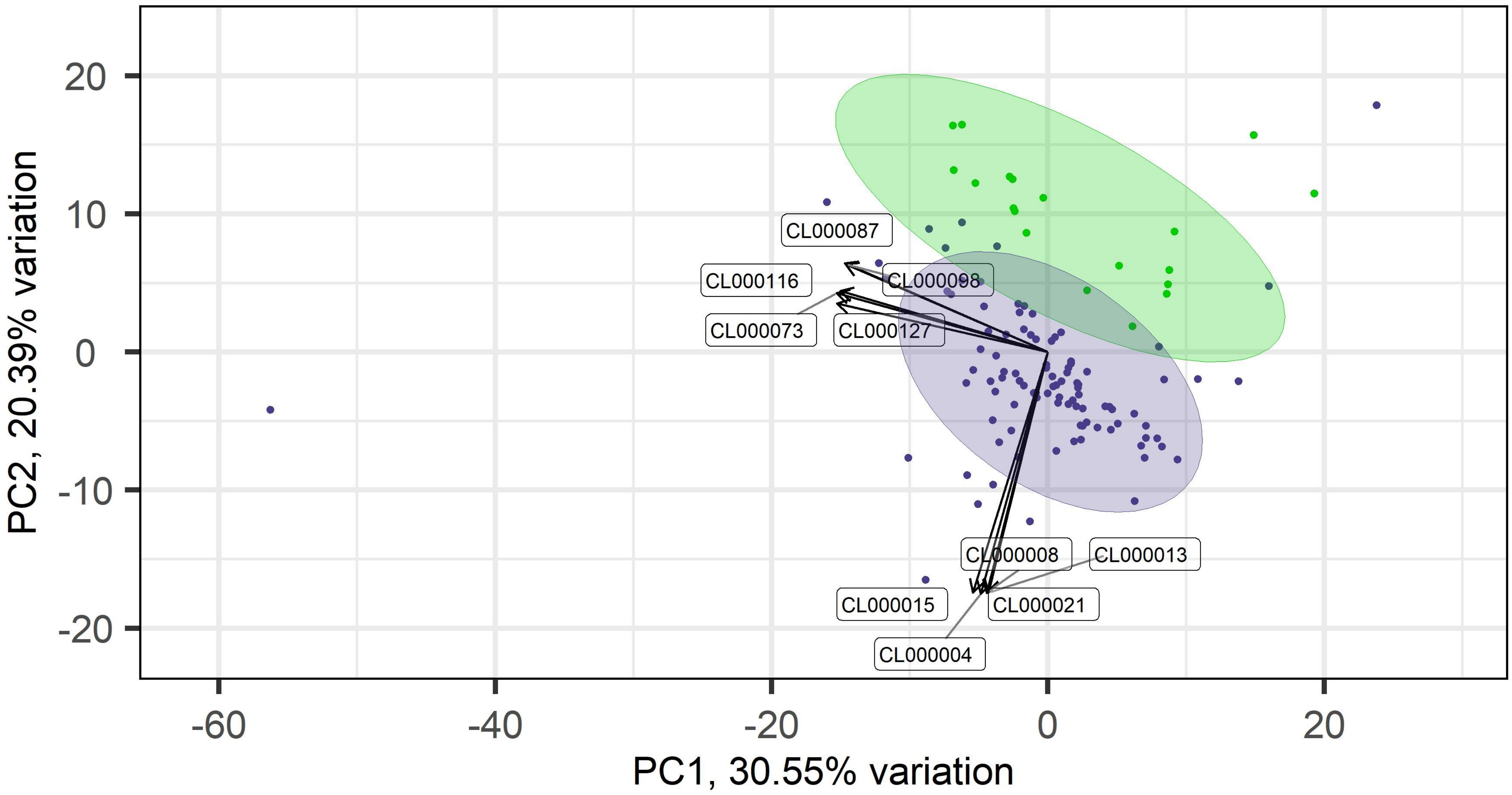

PC2, 20.39% variation
PC1, 30.55% variation
20
10
0
-10
-20
-60
-40
-20
0
20
CL000087
CL000116
CL000098
CL000073
CL000127
CL000008
CL000013
CL000015
CL000021
CL000004

FigureS8

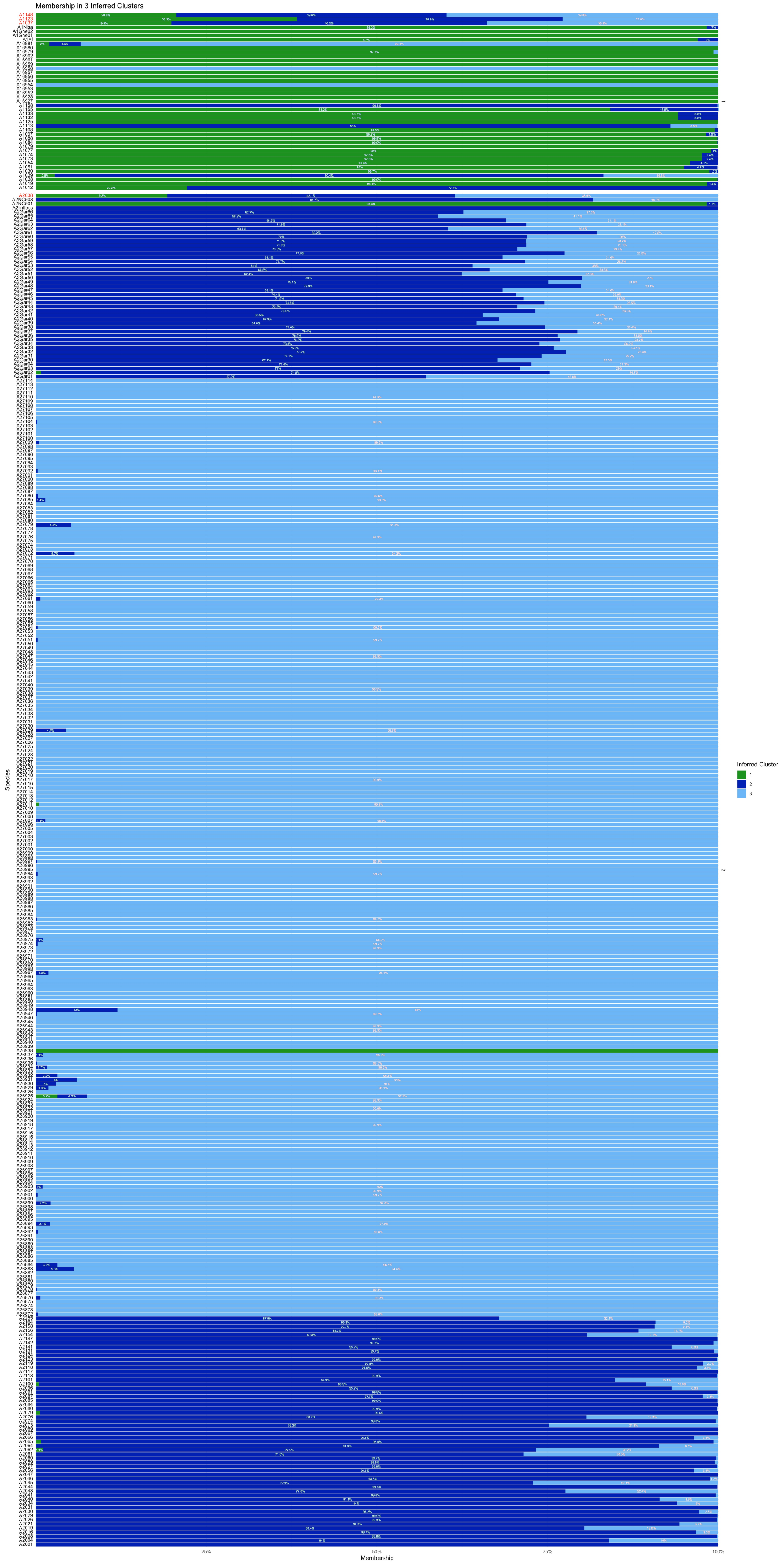
Membership in 3 Inferred Clusters
Species
Membership
25%
50%
75%
100%
Inferred Cluster
1
2
3

Figure S9

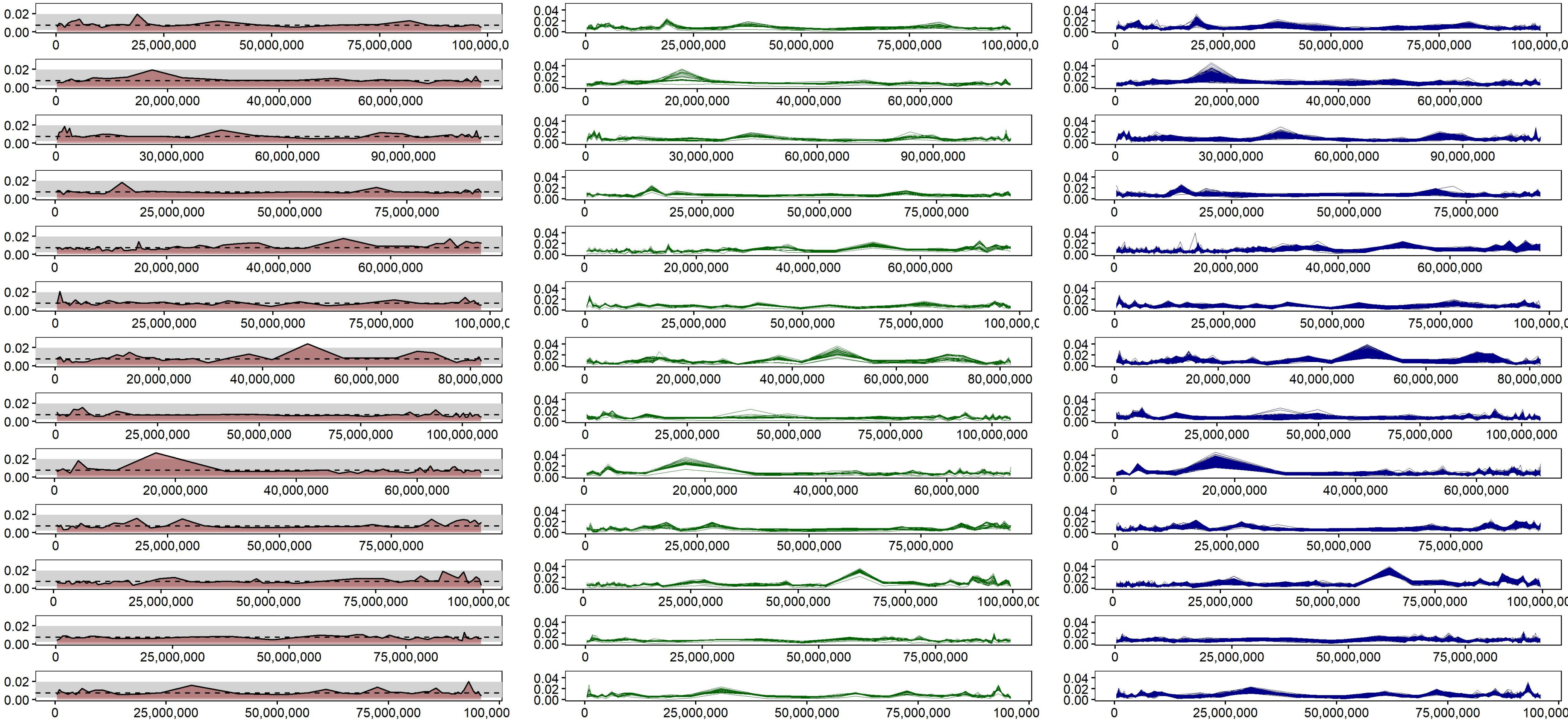
0.02
0.00
0.04
0
25,000,000
50,000,000
75,000,000
100,000,000
20,000,000
40,000,000
60,000,000
80,000,000
30,000,000
60,000,000
90,000,000

Figure S10

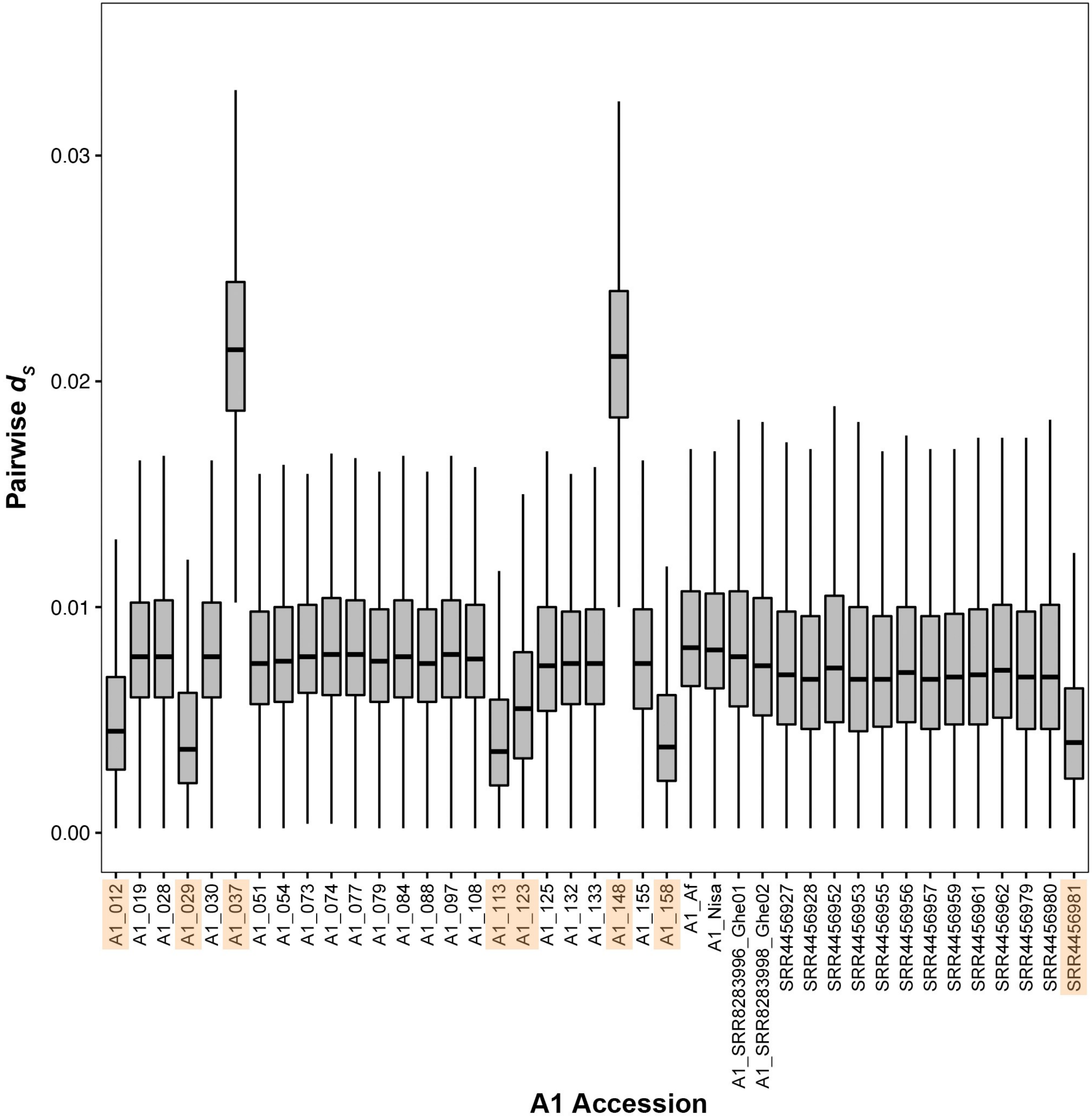
Pairwise $d_S$
0.03
0.02
0.01
0.00
A1_012
A1_019
A1_028
A1_029
A1_030
A1_037
A1_051
A1_054
A1_073
A1_074
A1_077
A1_079
A1_084
A1_088
A1_097
A1_108
A1_113
A1_123
A1_125
A1_132
A1_133
A1_148
A1_155
A1_158
A1_Af
A1_Nisa
A1_SRR8283996_Ghe01
A1_SRR8283998_Ghe02
SRR4456927
SRR4456928
SRR4456952
SRR4456953
SRR4456955
SRR4456956
SRR4456957
SRR4456959
SRR4456961
SRR4456962
SRR4456979
SRR4456980
SRR4456981
A1 Accession

FigureS11

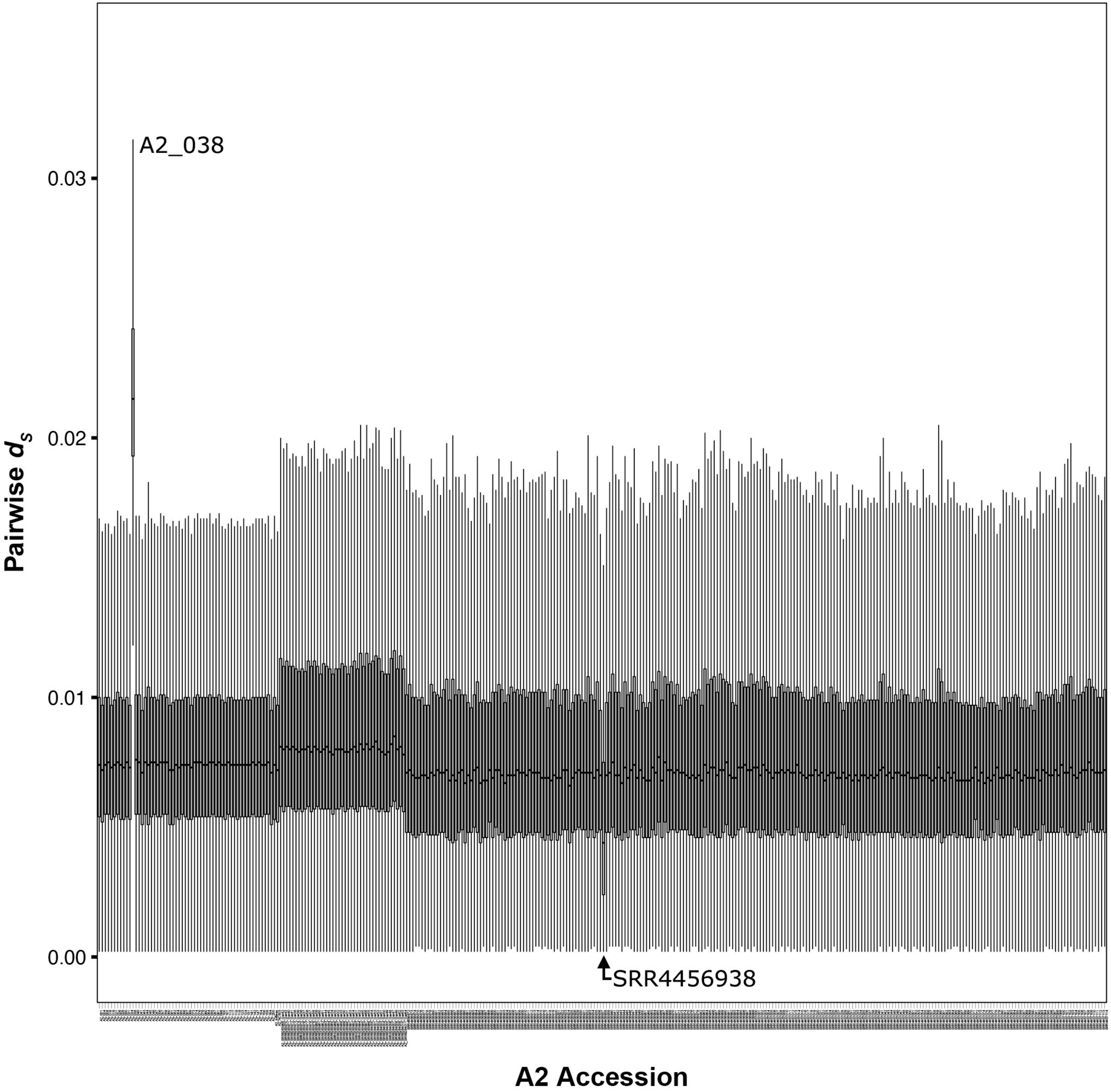

**Supplementary Table 1. A descriptive list of the accessions analyzed in this study. Locations collected are from GRIN or the reference listed for each and may reflect the source of the donation and not the original collection location.**

| Species | Accession | Alternate Name | Sample Name | GRIN (PI) | SRA ID | Location | Sequenced | Read Coverage | Sufficient Coverage | Pass Filter | In QC set |
|---|---|---|---|---|---|---|---|---|---|---|---|
| *Gossypium herbaceum* | GC0006 | | SRR4456958 | | SRR4456958 | China | Du 2018<br>10.1038/s41588-018-0116-x | 9.23 | no | no | no |
| *Gossypium herbaceum* | GC0010 | | SRR4456954 | | SRR4456954 | China | Du 2018<br>10.1038/s41588-018-0116-x | 9.07 | no | no | no |
| *Gossypium herbaceum* | GC0012 | HongXingCaoMian | SRR4456953 | | SRR4456953 | China | Du 2018<br>10.1038/s41588-018-0116-x | 8.23 | no | no | no |
| *Gossypium arboreum* | GA0144 | ChangRongZhongMian | SRR4457077 | | SRR4457077 | China | Du 2018<br>10.1038/s41588-018-0116-x | 9.34 | no | no | no |
| *Gossypium herbaceum* | GC0002 | GaoTaiCaoMian | SRR4456928 | | SRR4456928 | Gansu, China | Du 2018<br>10.1038/s41588-018-0116-x | 9.88 | no | no | no |
| *Gossypium herbaceum* | GC0007 | ZiHuaCaoMian | SRR4456957 | | SRR4456957 | Gansu, China | Du 2018<br>10.1038/s41588-018-0116-x | 9.63 | no | no | no |
| *Gossypium herbaceum* | GC0009 | LinZeCaoMian | SRR4456955 | | SRR4456955 | Gansu, China | Du 2018<br>10.1038/s41588-018-0116-x | 9.17 | no | no | no |
| *Gossypium herbaceum* | GC0013 | YuMenCaoMian | SRR4456952 | | SRR4456952 | Gansu, China | Du 2018<br>10.1038/s41588-018-0116-x | 7.68 | no | no | no |
| *Gossypium arboreum* | GA0030 | IndiaN.V50-70 | SRR4456964 | | SRR4456964 | India | Du 2018<br>10.1038/s41588-018-0116-x | 8.24 | no | no | no |
| *Gossypium arboreum* | GA0220 | Dharwar cotton | SRR4456884 | | SRR4456884 | India | Du 2018<br>10.1038/s41588-018-0116-x | 9.55 | no | no | no |
| *Gossypium arboreum* | GA0228 | NongLin-4 | SRR4456897 | | SRR4456897 | Japan | Du 2018<br>10.1038/s41588-018-0116-x | 8.63 | no | no | no |
| *Gossypium arboreum* | GA0229 | NongLin-5 | SRR4456898 | | SRR4456898 | Japan | Du 2018<br>10.1038/s41588-018-0116-x | 8.14 | no | no | no |
| *Gossypium arboreum* | GA0034 | 119S | SRR4456990 | | SRR4456990 | Pakistan | Du 2018<br>10.1038/s41588-018-0116-x | 7.88 | no | no | no |
| *Gossypium herbaceum* | GC0014 | Gusa | SRR4456962 | | SRR4456962 | Russia | Du 2018<br>10.1038/s41588-018-0116-x | 9.77 | no | no | no |
| *Gossypium herbaceum* | WC0001 | var. africanum | SRR4456979 | | SRR4456979 | South Africa | Du 2018<br>10.1038/s41588-018-0116-x | 8.76 | no | no | no |
| *Gossypium arboreum* | GA0198 | FuJianShenShanXiangZiHuaZhongMian | SRR4456904 | | SRR4456904 | South China<br>FuJian, China | Du 2018<br>10.1038/s41588-018-0116-x | 9.27 | no | no | no |
| *Gossypium arboreum* | GA0043 | GeBeiMian | SRR4457084 | | SRR4457084 | South China<br>GuangDong, China | Du 2018<br>10.1038/s41588-018-0116-x | 9.28 | no | no | no |
| *Gossypium arboreum* | GA0110 | GuangXiTianBaoXianZhongMian | SRR4457083 | | SRR4457083 | South China<br>GuangXi, China | Du 2018<br>10.1038/s41588-018-0116-x | 9.66 | no | no | no |
| *Gossypium arboreum* | GA0136 | FuChuanJiangTangZhongMian | SRR4456946 | | SRR4456946 | South China<br>GuangXi, China | Du 2018<br>10.1038/s41588-018-0116-x | 8.67 | no | no | no |
| *Gossypium arboreum* | GA0138 | SanJiangBaJiangZongXuZhongMian | SRR4456948 | | SRR4456948 | South China<br>GuangXi, China | Du 2018<br>10.1038/s41588-018-0116-x | 9.78 | no | no | no |
| *Gossypium arboreum* | GA0153 | QiangWeiNaLiangZhongMian-1 | SRR4457050 | | SRR4457050 | South China<br>GuangXi, China | Du 2018<br>10.1038/s41588-018-0116-x | 9.05 | no | no | no |
| *Gossypium arboreum* | GA0156 | BaMaNaTaoZhongMian | SRR4457057 | | SRR4457057 | South China<br>GuangXi, China | Du 2018<br>10.1038/s41588-018-0116-x | 9.57 | no | no | no |
| *Gossypium arboreum* | GA0160 | SanJiangBaJiangZhongMian | SRR4457048 | | SRR4457048 | South China<br>GuangXi, China | Du 2018<br>10.1038/s41588-018-0116-x | 8.88 | no | no | no |
| *Gossypium arboreum* | GA0187 | LeYeLuoXiZhongMian1-3 | SRR4456934 | | SRR4456934 | South China<br>GuangXi, China | Du 2018<br>10.1038/s41588-018-0116-x | 9.24 | no | no | no |
| *Gossypium arboreum* | GA0215 | DeBaoBaTouZhongMian | SRR4456889 | | SRR4456889 | South China<br>GuangXi, China | Du 2018<br>10.1038/s41588-018-0116-x | 9.53 | no | no | no |
| *Gossypium arboreum* | GA0107 | GuiZhouJianHeZhongMian | SRR4456919 | | SRR4456919 | South China<br>GuiZhou, China | Du 2018<br>10.1038/s41588-018-0116-x | 8.60 | no | no | no |
| *Gossypium arboreum* | GA0131 | MoYangXiaoHua | SRR4456941 | | SRR4456941 | South China<br>GuiZhou, China | Du 2018<br>10.1038/s41588-018-0116-x | 7.92 | no | no | no |
| *Gossypium arboreum* | GA0132 | LuoKunQuXiaoHua | SRR4456942 | | SRR4456942 | South China<br>GuiZhou, China | Du 2018<br>10.1038/s41588-018-0116-x | 8.15 | no | no | no |
| *Gossypium arboreum* | GA0133 | NaYeXiaoHua | SRR4456943 | | SRR4456943 | South China<br>GuiZhou, China | Du 2018<br>10.1038/s41588-018-0116-x | 9.25 | no | no | no |
| *Gossypium arboreum* | GA0134 | PingLeXiaoHua | SRR4456944 | | SRR4456944 | South China<br>GuiZhou, China | Du 2018<br>10.1038/s41588-018-0116-x | 8.68 | no | no | no |
| *Gossypium arboreum* | GA0135 | XiaZhaiXiaoHua | SRR4456945 | | SRR4456945 | South China<br>GuiZhou, China | Du 2018<br>10.1038/s41588-018-0116-x | 7.49 | no | no | no |
| *Gossypium arboreum* | GA0142 | AnLongXiaoHua | SRR4457075 | | SRR4457075 | South China<br>GuiZhou, China | Du 2018<br>10.1038/s41588-018-0116-x | 8.54 | no | no | no |
| *Gossypium arboreum* | GA0143 | ZhenFengTuMian | SRR4457078 | | SRR4457078 | South China<br>GuiZhou, China | Du 2018<br>10.1038/s41588-018-0116-x | 9.82 | no | no | no |
| *Gossypium arboreum* | GA0154 | WangMoSangLangDaMianHua | SRR4457051 | | SRR4457051 | South China<br>GuiZhou, China | Du 2018<br>10.1038/s41588-018-0116-x | 9.98 | no | no | no |
| *Gossypium arboreum* | GA0173 | TuQiaoXiaoHua | SRR4457111 | | SRR4457111 | South China<br>GuiZhou, China | Du 2018<br>10.1038/s41588-018-0116-x | 9.19 | no | no | no |
| *Gossypium arboreum* | GA0176 | NaShangQuXiaoHua | SRR4457106 | | SRR4457106 | South China<br>GuiZhou, China | Du 2018<br>10.1038/s41588-018-0116-x | 9.89 | no | no | no |
| *Gossypium arboreum* | GA0207 | ZiYunXiaoHua | SRR4457087 | | SRR4457087 | South China<br>GuiZhou, China | Du 2018<br>10.1038/s41588-018-0116-x | 9.70 | no | no | no |
| *Gossypium arboreum* | GA0208 | DaGuanXiaoHua | SRR4457088 | | SRR4457088 | South China<br>GuiZhou, China | Du 2018<br>10.1038/s41588-018-0116-x | 9.46 | no | no | no |

| Species | Accession | Alternate Name | Sample Name | GRIN (PI) | SRA ID | Location | Sequenced | Read Coverage | Sufficient Coverage | Pass Filter | In QC set |
|---|---|---|---|---|---|---|---|---|---|---|---|
| *Gossypium arboreum* | GA0204 | XinGanZhongMian | SRR4457092 | | SRR4457092 | South China JiangXi, China | Du 2018 10.1038/s41588-018-0116-x | 7.97 | no | no | no |
| *Gossypium arboreum* | GA0035 | PingZhanZhaiZhongMian | SRR4456987 | | SRR4456987 | South China YunNan, China | Du 2018 10.1038/s41588-018-0116-x | 7.95 | no | no | no |
| *Gossypium arboreum* | GA0038 | BaoShanTuMian | SRR4456986 | | SRR4456986 | South China YunNan, China | Du 2018 10.1038/s41588-018-0116-x | 7.97 | no | no | no |
| *Gossypium arboreum* | GA0040 | YunNanHuangRongTumian | SRR4456984 | | SRR4456984 | South China YunNan, China | Du 2018 10.1038/s41588-018-0116-x | 6.58 | no | no | no |
| *Gossypium arboreum* | GA0046 | KaiYuanTuMian | SRR4457049 | | SRR4457049 | South China YunNan, China | Du 2018 10.1038/s41588-018-0116-x | 8.53 | no | no | no |
| *Gossypium arboreum* | GA0191 | MengLaZhongMian | SRR4456974 | | SRR4456974 | South China YunNan, China | Du 2018 10.1038/s41588-018-0116-x | 8.82 | no | no | no |
| *Gossypium arboreum* | GA0196 | MoJiangNaHaZhongMian | SRR4456963 | | SRR4456963 | South China YunNan, China | Du 2018 10.1038/s41588-018-0116-x | 9.33 | no | no | no |
| *Gossypium arboreum* | GA0119 | MeiZhongMian | SRR4456881 | | SRR4456881 | USA | Du 2018 10.1038/s41588-018-0116-x | 9.51 | no | no | no |
| *Gossypium arboreum* | GA0007 | ChangZi-1 | SRR4456908 | | SRR4456908 | Yangtze River HuNan, China | Du 2018 10.1038/s41588-018-0116-x | 9.82 | no | no | no |
| *Gossypium arboreum* | GA0012 | TongNong-1 | SRR4456976 | | SRR4456976 | Yangtze River AnHui, China | Du 2018 10.1038/s41588-018-0116-x | 9.22 | no | no | no |
| *Gossypium arboreum* | GA0015 | WanGuiChiWuZi | SRR4457037 | | SRR4457037 | Yangtze River AnHui, China | Du 2018 10.1038/s41588-018-0116-x | 8.64 | no | no | no |
| *Gossypium arboreum* | GA0058 | TangShanBaiZiZhongMian | SRR4457059 | | SRR4457059 | Yangtze River AnHui, China | Du 2018 10.1038/s41588-018-0116-x | 9.44 | no | no | no |
| *Gossypium arboreum* | GA0079 | AnHuiShuChengZhongMian | SRR4457103 | | SRR4457103 | Yangtze River AnHui, China | Du 2018 10.1038/s41588-018-0116-x | 8.53 | no | no | no |
| *Gossypium arboreum* | GA0081 | AnHuiFuYangZiSeXiaoHua | SRR4456999 | | SRR4456999 | Yangtze River AnHui, China | Du 2018 10.1038/s41588-018-0116-x | 7.65 | no | no | no |
| *Gossypium arboreum* | GA0082 | AnHuiChuXianXiaoZiMian | SRR4456998 | | SRR4456998 | Yangtze River AnHui, China | Du 2018 10.1038/s41588-018-0116-x | 9.21 | no | no | no |
| *Gossypium arboreum* | GA0084 | AnHuiQuanJiaoXiaoZiMianMaoZi | SRR4456996 | | SRR4456996 | Yangtze River AnHui, China | Du 2018 10.1038/s41588-018-0116-x | 8.18 | no | no | no |
| *Gossypium arboreum* | GA0085 | AnHuiQuanJiaoXiaoZiMianGuangZi | SRR4457003 | | SRR4457003 | Yangtze River AnHui, China | Du 2018 10.1038/s41588-018-0116-x | 7.56 | no | no | no |
| *Gossypium arboreum* | GA0086 | AnHuiDingYuanZiHuaMian | SRR4457002 | | SRR4457002 | Yangtze River AnHui, China | Du 2018 10.1038/s41588-018-0116-x | 8.52 | no | no | no |
| *Gossypium arboreum* | GA0087 | FengYangZhongMian | SRR4457001 | | SRR4457001 | Yangtze River AnHui, China | Du 2018 10.1038/s41588-018-0116-x | 8.95 | no | no | no |
| *Gossypium arboreum* | GA0089 | TongChengZhongMianHuiZi | SRR4457005 | | SRR4457005 | Yangtze River AnHui, China | Du 2018 10.1038/s41588-018-0116-x | 8.10 | no | no | no |
| *Gossypium arboreum* | GA0090 | AnHuiGuiChiZhongMian | SRR4457004 | | SRR4457004 | Yangtze River AnHui, China | Du 2018 10.1038/s41588-018-0116-x | 8.45 | no | no | no |
| *Gossypium arboreum* | GA0091 | AnHuiGuiChiXiaoZiMianBaiZi | SRR4457027 | | SRR4457027 | Yangtze River AnHui, China | Du 2018 10.1038/s41588-018-0116-x | 8.31 | no | no | no |
| *Gossypium arboreum* | GA0094 | AnHuiTongLingZhongMian | SRR4457030 | | SRR4457030 | Yangtze River AnHui, China | Du 2018 10.1038/s41588-018-0116-x | 9.07 | no | no | no |
| *Gossypium arboreum* | GA0161 | AnHuiXuanChengZhongMianGuangZi | SRR4456903 | | SRR4456903 | Yangtze River AnHui, China | Du 2018 10.1038/s41588-018-0116-x | 9.27 | no | no | no |
| *Gossypium arboreum* | GA0162 | AnHuiLiuAnDaHuaMaoZi | SRR4456902 | | SRR4456902 | Yangtze River AnHui, China | Du 2018 10.1038/s41588-018-0116-x | 7.27 | no | no | no |
| *Gossypium arboreum* | GA0163 | AnHuiFeiXiXiaoHeiZi | SRR4456901 | | SRR4456901 | Yangtze River AnHui, China | Du 2018 10.1038/s41588-018-0116-x | 8.57 | no | no | no |
| *Gossypium arboreum* | GA0178 | AnHuiHuaiNingXianHeiZi | SRR4457108 | | SRR4457108 | Yangtze River AnHui, China | Du 2018 10.1038/s41588-018-0116-x | 9.52 | no | no | no |
| *Gossypium arboreum* | GA0197 | AnHuiTongChengZhongMianGuangZi | SRR4457034 | | SRR4457034 | Yangtze River AnHui, China | Du 2018 10.1038/s41588-018-0116-x | 8.90 | no | no | no |
| *Gossypium arboreum* | GA0213 | AnHuiTaiHeShuangFuQuXinGouXiang | SRR4456895 | | SRR4456895 | Yangtze River AnHui, China | Du 2018 10.1038/s41588-018-0116-x | 8.74 | no | no | no |
| *Gossypium arboreum* | GA0011 | HuangZhou | SRR4457041 | | SRR4457041 | Yangtze River HuBei, China | Du 2018 10.1038/s41588-018-0116-x | 8.51 | no | no | no |
| *Gossypium arboreum* | GA0025 | XiaoGanTieZiXiaoYiHua | SRR4456978 | | SRR4456978 | Yangtze River HuBei, China | Du 2018 10.1038/s41588-018-0116-x | 8.93 | no | no | no |
| *Gossypium arboreum* | GA0039 | SuiXianBaiZi | SRR4457042 | | SRR4457042 | Yangtze River HuBei, China | Du 2018 10.1038/s41588-018-0116-x | 7.36 | no | no | no |
| *Gossypium arboreum* | GA0044 | SuBianTieZi | SRR4457070 | | SRR4457070 | Yangtze River HuBei, China | Du 2018 10.1038/s41588-018-0116-x | 9.45 | no | no | no |
| *Gossypium arboreum* | GA0047 | XiaoGanTieZiDaYiHua | SRR4457032 | | SRR4457032 | Yangtze River HuBei, China | Du 2018 10.1038/s41588-018-0116-x | 8.72 | no | no | no |
| *Gossypium arboreum* | GA0048 | HuangGangAiJiaoCha | SRR4456896 | | SRR4456896 | Yangtze River HuBei, China | Du 2018 10.1038/s41588-018-0116-x | 8.31 | no | no | no |
| *Gossypium arboreum* | GA0050 | DaYeXiRongJian | SRR4456993 | | SRR4456993 | Yangtze River HuBei, China | Du 2018 10.1038/s41588-018-0116-x | 8.72 | no | no | no |
| *Gossypium arboreum* | GA0088 | HuaZhongZiGanZhongMian | SRR4457000 | | SRR4457000 | Yangtze River HuBei, China | Du 2018 10.1038/s41588-018-0116-x | 8.90 | no | no | no |
| *Gossypium arboreum* | GA0095 | XiaoGanQingPiSuZhongMianBaiZi(DaYiFen) | SRR4457023 | | SRR4457023 | Yangtze River HuBei, China | Du 2018 10.1038/s41588-018-0116-x | 8.95 | no | no | no |
| *Gossypium arboreum* | GA0097 | JiangLingZhongMian | SRR4457025 | | SRR4457025 | Yangtze River HuBei, China | Du 2018 10.1038/s41588-018-0116-x | 8.44 | no | no | no |

| Species | Accession | Alternate Name | Sample Name | GRIN (PI) | SRA ID | Location | Sequenced | Read Coverage | Sufficient Coverage | Pass Filter | In QC set |
|---|---|---|---|---|---|---|---|---|---|---|---|
| *Gossypium arboreum* | GA0182 | DaYeDaRong | SRR4456935 | | SRR4456935 | Yangtze River<br>HuBei, China | Du 2018<br>10.1038/s41588-018-0116-x | 9.83 | no | no | no |
| *Gossypium arboreum* | GA0017 | HuNaGuiYangTuMian | SRR4457039 | | SRR4457039 | Yangtze River<br>HuNan, China | Du 2018<br>10.1038/s41588-018-0116-x | 9.89 | no | no | no |
| *Gossypium arboreum* | GA0098 | HuNaMaYangTieZiMian | SRR4457026 | | SRR4457026 | Yangtze River<br>HuNan, China | Du 2018<br>10.1038/s41588-018-0116-x | 8.91 | no | no | no |
| *Gossypium arboreum* | GA0099 | HuNaChangSuiTieZiMian | SRR4457021 | | SRR4457021 | Yangtze River<br>HuNan, China | Du 2018<br>10.1038/s41588-018-0116-x | 8.72 | no | no | no |
| *Gossypium arboreum* | GA0014 | ZhongYinZaJiaoMian-5 | SRR4457044 | | SRR4457044 | Yangtze River<br>JiangSu, China | Du 2018<br>10.1038/s41588-018-0116-x | 9.56 | no | no | no |
| *Gossypium arboreum* | GA0028 | XiaoBaiHua(HaiMen) | SRR4456968 | | SRR4456968 | Yangtze River<br>JiangSu, China | Du 2018<br>10.1038/s41588-018-0116-x | 9.22 | no | no | no |
| *Gossypium arboreum* | GA0029 | JiangSuHongJingJiJiao | SRR4456965 | | SRR4456965 | Yangtze River<br>JiangSu, China | Du 2018<br>10.1038/s41588-018-0116-x | 9.81 | no | no | no |
| *Gossypium arboreum* | GA0031 | NaTongQingJingJiJiaoYaLingGuoHuangHua | SRR4456991 | | SRR4456991 | Yangtze River<br>JiangSu, China | Du 2018<br>10.1038/s41588-018-0116-x | 9.64 | no | no | no |
| *Gossypium arboreum* | GA0032 | NaTongZiJingJiJiaoBaiHuaHongXin | SRR4456992 | | SRR4456992 | Yangtze River<br>JiangSu, China | Du 2018<br>10.1038/s41588-018-0116-x | 8.42 | no | no | no |
| *Gossypium arboreum* | GA0033 | JiangSuChangShuJiJiaoMian | SRR4456989 | | SRR4456989 | Yangtze River<br>JiangSu, China | Du 2018<br>10.1038/s41588-018-0116-x | 7.15 | no | no | no |
| *Gossypium arboreum* | GA0037 | JiangYinBaiZi | SRR4456985 | | SRR4456985 | Yangtze River<br>JiangSu, China | Du 2018<br>10.1038/s41588-018-0116-x | 8.57 | no | no | no |
| *Gossypium arboreum* | GA0051 | NaTongQingJingTongMian | SRR4456977 | | SRR4456977 | Yangtze River<br>JiangSu, China | Du 2018<br>10.1038/s41588-018-0116-x | 8.88 | no | no | no |
| *Gossypium arboreum* | GA0052 | HaiMenLuHeZi | SRR4456983 | | SRR4456983 | Yangtze River<br>JiangSu, China | Du 2018<br>10.1038/s41588-018-0116-x | 9.76 | no | no | no |
| *Gossypium arboreum* | GA0053 | HaiMenZiMianHua | SRR4456994 | | SRR4456994 | Yangtze River<br>JiangSu, China | Du 2018<br>10.1038/s41588-018-0116-x | 9.85 | no | no | no |
| *Gossypium arboreum* | GA0054 | HaiMenJiJiaoMian | SRR4457008 | | SRR4457008 | Yangtze River<br>JiangSu, China | Du 2018<br>10.1038/s41588-018-0116-x | 8.59 | no | no | no |
| *Gossypium arboreum* | GA0055 | HaiMenHuangHuaQingJingJiJiaoMian | SRR4457020 | | SRR4457020 | Yangtze River<br>JiangSu, China | Du 2018<br>10.1038/s41588-018-0116-x | 9.21 | no | no | no |
| *Gossypium arboreum* | GA0059 | PiaoYangTieZi | SRR4457006 | | SRR4457006 | Yangtze River<br>JiangSu, China | Du 2018<br>10.1038/s41588-018-0116-x | 8.41 | no | no | no |
| *Gossypium arboreum* | GA0060 | BaiWanMian83-4 | SRR4457073 | | SRR4457073 | Yangtze River<br>JiangSu, China | Du 2018<br>10.1038/s41588-018-0116-x | 9.29 | no | no | no |
| *Gossypium arboreum* | GA0061 | ChangShuChangYinBaiHua | SRR4457061 | | SRR4457061 | Yangtze River<br>JiangSu, China | Du 2018<br>10.1038/s41588-018-0116-x | 6.36 | no | no | no |
| *Gossypium arboreum* | GA0062 | JiangSuChangShuLongZhong-1 | SRR4457060 | | SRR4457060 | Yangtze River<br>JiangSu, China | Du 2018<br>10.1038/s41588-018-0116-x | 7.68 | no | no | no |
| *Gossypium arboreum* | GA0063 | JiangSuChangShuYangMaoBai | SRR4457063 | | SRR4457063 | Yangtze River<br>JiangSu, China | Du 2018<br>10.1038/s41588-018-0116-x | 7.81 | no | no | no |
| *Gossypium arboreum* | GA0064 | JiangYinBaiZi111 | SRR4457062 | | SRR4457062 | Yangtze River<br>JiangSu, China | Du 2018<br>10.1038/s41588-018-0116-x | 9.70 | no | no | no |
| *Gossypium arboreum* | GA0065 | JiangYinJiJiaoMianGuangZi | SRR4457065 | | SRR4457065 | Yangtze River<br>JiangSu, China | Du 2018<br>10.1038/s41588-018-0116-x | 8.77 | no | no | no |
| *Gossypium arboreum* | GA0129 | ZhongYinZaJiaoZhong | SRR4457011 | | SRR4457011 | Yangtze River<br>JiangSu, China | Du 2018<br>10.1038/s41588-018-0116-x | 8.13 | no | no | no |
| *Gossypium arboreum* | GA0150 | RuDongJiJiaoYaGuo | SRR4457071 | | SRR4457071 | Yangtze River<br>JiangSu, China | Du 2018<br>10.1038/s41588-018-0116-x | 9.67 | no | no | no |
| *Gossypium arboreum* | GA0151 | ChangShuXiaoBaiZi | SRR4457052 | | SRR4457052 | Yangtze River<br>JiangSu, China | Du 2018<br>10.1038/s41588-018-0116-x | 9.01 | no | no | no |
| *Gossypium arboreum* | GA0170 | TaiCangBaiZi(Liu) | SRR4456910 | | SRR4456910 | Yangtze River<br>JiangSu, China | Du 2018<br>10.1038/s41588-018-0116-x | 8.48 | no | no | no |
| *Gossypium arboreum* | GA0175 | JiangYinBaiZi102 | SRR4457105 | | SRR4457105 | Yangtze River<br>JiangSu, China | Du 2018<br>10.1038/s41588-018-0116-x | 8.99 | no | no | no |
| *Gossypium arboreum* | GA0217 | JiangSuNanTongZhongMian | SRR4456891 | | SRR4456891 | Yangtze River<br>JiangSu, China | Du 2018<br>10.1038/s41588-018-0116-x | 8.40 | no | no | no |
| *Gossypium arboreum* | GA0180 | LianHuaZhongMian | SRR4457114 | | SRR4457114 | Yangtze River<br>JiangXi, China | Du 2018<br>10.1038/s41588-018-0116-x | 9.25 | no | no | no |
| *Gossypium arboreum* | GA0020 | JiaDingZhongMian | SRR4457046 | | SRR4457046 | Yangtze River<br>ShangHai, China | Du 2018<br>10.1038/s41588-018-0116-x | 9.08 | no | no | no |
| *Gossypium arboreum* | GA0066 | ShangHaiDaZiMian | SRR4457064 | | SRR4457064 | Yangtze River<br>ShangHai, China | Du 2018<br>10.1038/s41588-018-0116-x | 9.19 | no | no | no |
| *Gossypium arboreum* | GA0067 | ShangHaiJinJiaHuaZi | SRR4457067 | | SRR4457067 | Yangtze River<br>ShangHai, China | Du 2018<br>10.1038/s41588-018-0116-x | 9.04 | no | no | no |
| *Gossypium arboreum* | GA0068 | JiaDingBaiZi | SRR4457066 | | SRR4457066 | Yangtze River<br>ShangHai, China | Du 2018<br>10.1038/s41588-018-0116-x | 8.89 | no | no | no |
| *Gossypium arboreum* | GA0069 | PuDongHuoJiMian | SRR4457069 | | SRR4457069 | Yangtze River<br>ShangHai, China | Du 2018<br>10.1038/s41588-018-0116-x | 8.00 | no | no | no |
| *Gossypium arboreum* | GA0070 | JiangSuChuanShaQingJingZhongMian | SRR4457068 | | SRR4457068 | Yangtze River<br>ShangHai, China | Du 2018<br>10.1038/s41588-018-0116-x | 9.10 | no | no | no |
| *Gossypium arboreum* | GA0071 | JiangSuFengXianZhongMian | SRR4457097 | | SRR4457097 | Yangtze River<br>ShangHai, China | Du 2018<br>10.1038/s41588-018-0116-x | 9.57 | no | no | no |
| *Gossypium arboreum* | GA0072 | QingPuXiaoZiZhongMian | SRR4457098 | | SRR4457098 | Yangtze River<br>ShangHai, China | Du 2018<br>10.1038/s41588-018-0116-x | 9.10 | no | no | no |
| *Gossypium arboreum* | GA0073 | BaoShanZiHua | SRR4457095 | | SRR4457095 | Yangtze River<br>ShangHai, China | Du 2018<br>10.1038/s41588-018-0116-x | 8.98 | no | no | no |

| Species | Accession | Alternate Name | Sample Name | GRIN (PI) | SRA ID | Location | Sequenced | Read Coverage | Sufficient Coverage | Pass Filter | In QC set |
|---|---|---|---|---|---|---|---|---|---|---|---|
| *Gossypium arboreum* | GA0164 | SongJiangBaiZi | SRR4456900 | | SRR4456900 | Yangtze River ShangHai, China | Du 2018 10.1038/s41588-018-0116-x | 8.70 | no | no | no |
| *Gossypium arboreum* | GA0216 | ShangHaiZhongMian | SRR4456888 | | SRR4456888 | Yangtze River ShangHai, China | Du 2018 10.1038/s41588-018-0116-x | 8.94 | no | no | no |
| *Gossypium arboreum* | GA0100 | SiChuanShuangLiuZhongMian | SRR4457022 | | SRR4457022 | Yangtze River SiChuan, China | Du 2018 10.1038/s41588-018-0116-x | 9.16 | no | no | no |
| *Gossypium arboreum* | GA0101 | SiChuanMianYangZhongMian | SRR4456921 | | SRR4456921 | Yangtze River SiChuan, China | Du 2018 10.1038/s41588-018-0116-x | 7.12 | no | no | no |
| *Gossypium arboreum* | GA0102 | SiChuanSuiNingZhongMianHuiZi | SRR4456920 | | SRR4456920 | Yangtze River SiChuan, China | Du 2018 10.1038/s41588-018-0116-x | 9.77 | no | no | no |
| *Gossypium arboreum* | GA0103 | SiChuanSuiNingZhongMianBaiZi | SRR4456923 | | SRR4456923 | Yangtze River SiChuan, China | Du 2018 10.1038/s41588-018-0116-x | 9.15 | no | no | no |
| *Gossypium arboreum* | GA0105 | SiChuanZiZhongZhongMian | SRR4456917 | | SRR4456917 | Yangtze River SiChuan, China | Du 2018 10.1038/s41588-018-0116-x | 8.67 | no | no | no |
| *Gossypium arboreum* | GA0106 | SiChuanFuShunZhongMian | SRR4456916 | | SRR4456916 | Yangtze River SiChuan, China | Du 2018 10.1038/s41588-018-0116-x | 8.82 | no | no | no |
| *Gossypium arboreum* | GA0021 | ZheJiangYuYaoZiXueMianHua | SRR4456971 | | SRR4456971 | Yangtze River ZheJiang, China | Du 2018 10.1038/s41588-018-0116-x | 8.88 | no | no | no |
| *Gossypium arboreum* | GA0022 | ZheJiangShangYuDaKeLuShu | SRR4456970 | | SRR4456970 | Yangtze River ZheJiang, China | Du 2018 10.1038/s41588-018-0116-x | 9.37 | no | no | no |
| *Gossypium arboreum* | GA0074 | ZheJiangXiaoShanZiHongGengDaKeMianBaiZi | SRR4457096 | | SRR4457096 | Yangtze River ZheJiang, China | Du 2018 10.1038/s41588-018-0116-x | 9.42 | no | no | no |
| *Gossypium arboreum* | GA0075 | ZheJiangXiaoShanZiHongGengDaKeMianGuangZi | SRR4457101 | | SRR4457101 | Yangtze River ZheJiang, China | Du 2018 10.1038/s41588-018-0116-x | 9.21 | no | no | no |
| *Gossypium arboreum* | GA0076 | PingHuXiaoZhongHuiZi | SRR4457102 | | SRR4457102 | Yangtze River ZheJiang, China | Du 2018 10.1038/s41588-018-0116-x | 8.34 | no | no | no |
| *Gossypium arboreum* | GA0077 | ZheJiangHaiYanLuShu | SRR4456995 | | SRR4456995 | Yangtze River ZheJiang, China | Du 2018 10.1038/s41588-018-0116-x | 8.39 | no | no | no |
| *Gossypium arboreum* | GA0078 | ZheJiangDingHaiZhongMian | SRR4457100 | | SRR4457100 | Yangtze River ZheJiang, China | Du 2018 10.1038/s41588-018-0116-x | 9.35 | no | no | no |
| *Gossypium arboreum* | GA0130 | YuYaoZiBaiYi | SRR4457010 | | SRR4457010 | Yangtze River ZheJiang, China | Du 2018 10.1038/s41588-018-0116-x | 8.91 | no | no | no |
| *Gossypium arboreum* | GA0152 | ZheJiangZhenHaiHouShiHongJingZhongMian | SRR4457053 | | SRR4457053 | Yangtze River ZheJiang, China | Du 2018 10.1038/s41588-018-0116-x | 8.15 | no | no | no |
| *Gossypium arboreum* | GA0172 | PingHuXiaoZhongGuangZi | SRR4457110 | | SRR4457110 | Yangtze River ZheJiang, China | Du 2018 10.1038/s41588-018-0116-x | 8.89 | no | no | no |
| *Gossypium arboreum* | GA0188 | ZheJiangJinHuaZhongMian | SRR4457099 | | SRR4457099 | Yangtze River ZheJiang, China | Du 2018 10.1038/s41588-018-0116-x | 9.12 | no | no | no |
| *Gossypium arboreum* | GA0181 | LinZeZhongMian | SRR4456936 | | SRR4456936 | Yellow River GanSu, China | Du 2018 10.1038/s41588-018-0116-x | 8.74 | no | no | no |
| *Gossypium arboreum* | GA0200 | GanSuWuDuZhongMian | SRR4456887 | | SRR4456887 | Yellow River GanSu, China | Du 2018 10.1038/s41588-018-0116-x | 6.49 | no | no | no |
| *Gossypium arboreum* | GA0201 | GaoTaiZhongMian | SRR4457093 | | SRR4457093 | Yellow River GanSu, China | Du 2018 10.1038/s41588-018-0116-x | 7.95 | no | no | no |
| *Gossypium arboreum* | GA0023 | AnGuoLiuShuangShuDaQingJie | SRR4456940 | | SRR4456940 | Yellow River HeBei, China | Du 2018 10.1038/s41588-018-0116-x | 9.19 | no | no | no |
| *Gossypium arboreum* | GA0024 | ShiXiYa-1 | SRR4456972 | | SRR4456972 | Yellow River HeBei, China | Du 2018 10.1038/s41588-018-0116-x | 8.91 | no | no | no |
| *Gossypium arboreum* | GA0049 | WanZiZhongMian | SRR4456951 | | SRR4456951 | Yellow River HeBei, China | Du 2018 10.1038/s41588-018-0116-x | 8.68 | no | no | no |
| *Gossypium arboreum* | GA0121 | XiaoZiJieMian | SRR4457019 | | SRR4457019 | Yellow River HeBei, China | Du 2018 10.1038/s41588-018-0116-x | 9.85 | no | no | no |
| *Gossypium arboreum* | GA0123 | LeTingXiaoHeiZi | SRR4457017 | | SRR4457017 | Yellow River HeBei, China | Du 2018 10.1038/s41588-018-0116-x | 9.14 | no | no | no |
| *Gossypium arboreum* | GA0124 | AKeXiaoYangHua | SRR4457016 | | SRR4457016 | Yellow River HeBei, China | Du 2018 10.1038/s41588-018-0116-x | 9.79 | no | no | no |
| *Gossypium arboreum* | GA0125 | ZiJieZiHeZhongMian | SRR4457015 | | SRR4457015 | Yellow River HeBei, China | Du 2018 10.1038/s41588-018-0116-x | 9.04 | no | no | no |
| *Gossypium arboreum* | GA0147 | ShuLuBaiHuaBai | SRR4457082 | | SRR4457082 | Yellow River HeBei, China | Du 2018 10.1038/s41588-018-0116-x | 8.69 | no | no | no |
| *Gossypium arboreum* | GA0148 | WanXianZiJieXiaoBaiHua | SRR4457081 | | SRR4457081 | Yellow River HeBei, China | Du 2018 10.1038/s41588-018-0116-x | 9.18 | no | no | no |
| *Gossypium arboreum* | GA0155 | ShiJiaZhuangJianMian | SRR4457056 | | SRR4457056 | Yellow River HeBei, China | Du 2018 10.1038/s41588-018-0116-x | 9.72 | no | no | no |
| *Gossypium arboreum* | GA0158 | AnGuoBaiJieBaiHua | SRR4457055 | | SRR4457055 | Yellow River HeBei, China | Du 2018 10.1038/s41588-018-0116-x | 8.98 | no | no | no |
| *Gossypium arboreum* | GA0167 | ZiHeZhongMian | SRR4456905 | | SRR4456905 | Yellow River HeBei, China | Du 2018 10.1038/s41588-018-0116-x | 9.69 | no | no | no |
| *Gossypium arboreum* | GA0168 | BaiJieMian | SRR4457013 | | SRR4457013 | Yellow River HeBei, China | Du 2018 10.1038/s41588-018-0116-x | 8.95 | no | no | no |
| *Gossypium arboreum* | GA0171 | ZhaoXianHongJieMian | SRR4457109 | | SRR4457109 | Yellow River HeBei, China | Du 2018 10.1038/s41588-018-0116-x | 7.15 | no | no | no |
| *Gossypium arboreum* | GA0199 | DingXianXiaoBaiJieMian | SRR4456886 | | SRR4456886 | Yellow River HeBei, China | Du 2018 10.1038/s41588-018-0116-x | 9.05 | no | no | no |
| *Gossypium arboreum* | GA0219 | QianXiXiaoHeiZi | SRR4456885 | | SRR4456885 | Yellow River HeBei, China | Du 2018 10.1038/s41588-018-0116-x | 8.34 | no | no | no |
| *Gossypium arboreum* | GA0227 | ShiXiYaBaiZi | SRR4456933 | | SRR4456933 | Yellow River HeBei, China | Du 2018 10.1038/s41588-018-0116-x | 6.34 | no | no | no |

| Species | Accession | Alternate Name | Sample Name | GRIN (PI) | SRA ID | Location | Sequenced | Read Coverage | Sufficient Coverage | Pass Filter | In QC set |
|---|---|---|---|---|---|---|---|---|---|---|---|
| *Gossypium arboreum* | GA0009 | SuiXianZiHuaMian | SRR4456924 | | SRR4456924 | Yellow River<br>HeNan, China | Du 2018<br>10.1038/s41588-018-0116-x | 8.83 | no | no | no |
| *Gossypium arboreum* | GA0113 | HeNaShangQiuHuiZi | SRR4456876 | | SRR4456876 | Yellow River<br>HeNan, China | Du 2018<br>10.1038/s41588-018-0116-x | 9.69 | no | no | no |
| *Gossypium arboreum* | GA0114 | HeNaHuaiYangTuMianBaiZi | SRR4456877 | | SRR4456877 | Yellow River<br>HeNan, China | Du 2018<br>10.1038/s41588-018-0116-x | 9.67 | no | no | no |
| *Gossypium arboreum* | GA0116 | HeNaShangCaiZhongMian | SRR4456875 | | SRR4456875 | Yellow River<br>HeNan, China | Du 2018<br>10.1038/s41588-018-0116-x | 8.60 | no | no | no |
| *Gossypium arboreum* | GA0166 | HeNanShangQiuHeiZi | SRR4456906 | | SRR4456906 | Yellow River<br>HeNan, China | Du 2018<br>10.1038/s41588-018-0116-x | 8.85 | no | no | no |
| *Gossypium arboreum* | GA0019 | LiaoYang-1 | SRR4457045 | | SRR4457045 | Yellow River<br>LiaoNing, China | Du 2018<br>10.1038/s41588-018-0116-x | 8.94 | no | no | no |
| *Gossypium arboreum* | GA0036 | LiaoZhongHong | SRR4456988 | | SRR4456988 | Yellow River<br>LiaoNing, China | Du 2018<br>10.1038/s41588-018-0116-x | 8.46 | no | no | no |
| *Gossypium arboreum* | GA0042 | TieMuChi | SRR4457009 | | SRR4457009 | Yellow River<br>LiaoNing, China | Du 2018<br>10.1038/s41588-018-0116-x | 8.67 | no | no | no |
| *Gossypium arboreum* | GA0045 | LiaoNingZhongMian | SRR4457058 | | SRR4457058 | Yellow River<br>LiaoNing, China | Du 2018<br>10.1038/s41588-018-0116-x | 9.16 | no | no | no |
| *Gossypium arboreum* | GA0128 | ChiMuBaiZhong | SRR4457012 | | SRR4457012 | Yellow River<br>LiaoNing, China | Du 2018<br>10.1038/s41588-018-0116-x | 8.84 | no | no | no |
| *Gossypium arboreum* | GA0165 | FaKuBaiZi | SRR4456907 | | SRR4456907 | Yellow River<br>LiaoNing, China | Du 2018<br>10.1038/s41588-018-0116-x | 9.56 | no | no | no |
| *Gossypium arboreum* | GA0183 | ChiMuHeiZhong | SRR4456939 | | SRR4456939 | Yellow River<br>LiaoNing, China | Du 2018<br>10.1038/s41588-018-0116-x | 9.50 | no | no | no |
| *Gossypium arboreum* | GA0010 | DanXianXiaoZiHua | SRR4456982 | | SRR4456982 | Yellow River<br>ShanDong, China | Du 2018<br>10.1038/s41588-018-0116-x | 8.53 | no | no | no |
| *Gossypium arboreum* | GA0016 | ShanDongXiaoGuangXu | SRR4457038 | | SRR4457038 | Yellow River<br>ShanDong, China | Du 2018<br>10.1038/s41588-018-0116-x | 9.53 | no | no | no |
| *Gossypium arboreum* | GA0041 | XiWang-1 | SRR4456880 | | SRR4456880 | Yellow River<br>ShanDong, China | Du 2018<br>10.1038/s41588-018-0116-x | 8.81 | no | no | no |
| *Gossypium arboreum* | GA0112 | ShanDongJiYangXiaoMian200 | SRR4456879 | | SRR4456879 | Yellow River<br>ShanDong, China | Du 2018<br>10.1038/s41588-018-0116-x | 9.77 | no | no | no |
| *Gossypium arboreum* | GA0120 | ShanDongHuiMinYiWoHou | SRR4456882 | | SRR4456882 | Yellow River<br>ShanDong, China | Du 2018<br>10.1038/s41588-018-0116-x | 9.18 | no | no | no |
| *Gossypium arboreum* | GA0211 | DuiHua | SRR4456893 | | SRR4456893 | Yellow River<br>ShanDong, China | Du 2018<br>10.1038/s41588-018-0116-x | 9.44 | no | no | no |
| *Gossypium arboreum* | GA0118 | ShanXiNaZhengZhongMian | SRR4456873 | | SRR4456873 | Yellow River<br>ShanXi, China | Du 2018<br>10.1038/s41588-018-0116-x | 8.93 | no | no | no |
| *Gossypium herbaceum* | GC0016 | Local cotton | SRR4456980 | | SRR4456980 | | Du 2018<br>10.1038/s41588-018-0116-x | 9.50 | no | no | no |
| *Gossypium arboreum* | A2_038 | | A2_038 | PI 183188 | SRR8979981 | India | in this study; BGI | 25.67 | yes | no<br>hybrid/contaminated | no |
| *Gossypium herbaceum* | A1_148 | 7255 | A1_148 | PI 532172 | SRR8979970 | Oman | in this study; BGI | 27.69 | yes | no<br>hybrid/contaminated | no |
| *Gossypium herbaceum* | A1_037 | | A1_037 | PI 408795 | SRR8979987 | Uzbekistan | in this study; BGI | 36.59 | yes | no<br>hybrid/contaminated | no |
| *Gossypium arboreum* | GA0232 | | SRR4456938 | | SRR4456938 | | Du 2018<br>10.1038/s41588-018-0116-x | 10.31 | yes | no<br>incorrect species | no |
| *Gossypium arboreum* | A2_NC501 | | A2_NC501 | | SRR8979995 | | in this study | 20.09 | yes | no<br>incorrect species | no |
| *Gossypium herbaceum* | A1_012 | | A1_012 | PI 190542 | SRR8979930 | Iraq | in this study; BGI | 33.76 | yes | no<br>incorrect species | no |
| *Gossypium herbaceum* | A1_158 | DEZI KHAKI | A1_158 | PI 630027 | SRR8979968 | Pakistan* | in this study; BGI | 30.59 | yes | no<br>incorrect species | no |
| *Gossypium herbaceum* | A1_029 | | A1_029 | PI 408787 | SRR8979989 | Turkey | in this study; BGI | 23.73 | yes | no<br>incorrect species | no |
| *Gossypium herbaceum* | A1_113 | LINE 4241 | A1_113 | PI 529686 | SRR8979908 | Uzbekistan | in this study; BGI | 34.08 | yes | no<br>incorrect species | no |
| *Gossypium herbaceum* | GC0005 | CaoMian | SRR4456959 | | SRR4456959 | Gansu, China | Du 2018<br>10.1038/s41588-018-0116-x | 10.51 | yes | no<br>possible PCR bias | no |
| *Gossypium arboreum* | GA0214 | BaiMiLaHuangHuaHeiZi | SRR4456894 | | SRR4456894 | Japan | Du 2018<br>10.1038/s41588-018-0116-x | 10.93 | yes | no<br>possible PCR bias | no |
| *Gossypium arboreum* | GA0230 | NongLin-6 | SRR4456899 | | SRR4456899 | Japan | Du 2018<br>10.1038/s41588-018-0116-x | 15.33 | yes | no<br>possible PCR bias | no |
| *Gossypium arboreum* | GA0231 | Akguza | SRR4456925 | | SRR4456925 | Pakistan | Du 2018<br>10.1038/s41588-018-0116-x | 10.06 | yes | no<br>possible PCR bias | no |
| *Gossypium herbaceum* | GC0015 | L-02066-1-3 | SRR4456961 | | SRR4456961 | Russia | Du 2018<br>10.1038/s41588-018-0116-x | 10.02 | yes | no<br>possible PCR bias | no |
| *Gossypium arboreum* | GA0109 | GuangXiZuoXianZhongMian | SRR4456926 | | SRR4456926 | South China<br>GuangXi, China | Du 2018<br>10.1038/s41588-018-0116-x | 10.92 | yes | no<br>possible PCR bias | no |
| *Gossypium arboreum* | GA0137 | PingGuoJiuPingZhongMian | SRR4456947 | | SRR4456947 | South China<br>GuangXi, China | Du 2018<br>10.1038/s41588-018-0116-x | 10.45 | yes | no<br>possible PCR bias | no |
| *Gossypium arboreum* | GA0169 | DongLanChangJingZhongMian | SRR4456911 | | SRR4456911 | South China<br>GuangXi, China | Du 2018<br>10.1038/s41588-018-0116-x | 10.10 | yes | no<br>possible PCR bias | no |
| *Gossypium arboreum* | GA0179 | GuangXiFuZhongZhongMian | SRR4457113 | | SRR4457113 | South China<br>GuangXi, China | Du 2018<br>10.1038/s41588-018-0116-x | 10.62 | yes | no<br>possible PCR bias | no |
| *Gossypium arboreum* | GA0186 | JingXiSuiHuaiZhongMian | SRR4456931 | | SRR4456931 | South China<br>GuangXi, China | Du 2018<br>10.1038/s41588-018-0116-x | 11.35 | yes | no<br>possible PCR bias | no |

| Species | Accession | Alternate Name | Sample Name | GRIN (PI) | SRA ID | Location | Sequenced | Read Coverage | Sufficient Coverage | Pass Filter | In QC set |
|---|---|---|---|---|---|---|---|---|---|---|---|
| *Gossypium arboreum* | GA0206 | NanDanBaDiZhongMian | SRR4457090 | | SRR4457090 | South China<br>GuangXi, China | Du 2018<br>10.1038/s41588-018-0116-x | 10.36 | yes | no<br>possible PCR bias | no |
| *Gossypium arboreum* | GA0209 | NanDanHuaiLiZhongMian | SRR4457085 | | SRR4457085 | South China<br>GuangXi, China | Du 2018<br>10.1038/s41588-018-0116-x | 11.07 | yes | no<br>possible PCR bias | no |
| *Gossypium arboreum* | GA0108 | GuiZhouJinPingZhongMian | SRR4456918 | | SRR4456918 | South China<br>GuiZhou, China | Du 2018<br>10.1038/s41588-018-0116-x | 10.05 | yes | no<br>possible PCR bias | no |
| *Gossypium arboreum* | GA0139 | ShiQianZhongMian | SRR4456949 | | SRR4456949 | South China<br>GuiZhou, China | Du 2018<br>10.1038/s41588-018-0116-x | 10.41 | yes | no<br>possible PCR bias | no |
| *Gossypium arboreum* | GA0140 | ChangShunDaiHuaXiaoHua | SRR4456950 | | SRR4456950 | South China<br>GuiZhou, China | Du 2018<br>10.1038/s41588-018-0116-x | 11.01 | yes | no<br>possible PCR bias | no |
| *Gossypium arboreum* | GA0141 | CongJiangZhongMian | SRR4457076 | | SRR4457076 | South China<br>GuiZhou, China | Du 2018<br>10.1038/s41588-018-0116-x | 10.29 | yes | no<br>possible PCR bias | no |
| *Gossypium arboreum* | GA0157 | PingDongXiaoHua | SRR4457054 | | SRR4457054 | South China<br>GuiZhou, China | Du 2018<br>10.1038/s41588-018-0116-x | 10.68 | yes | no<br>possible PCR bias | no |
| *Gossypium arboreum* | GA0185 | XinPuXiaoHua | SRR4456932 | | SRR4456932 | South China<br>GuiZhou, China | Du 2018<br>10.1038/s41588-018-0116-x | 12.12 | yes | no<br>possible PCR bias | no |
| *Gossypium arboreum* | GA0193 | XueGuanXiaoHua | SRR4456892 | | SRR4456892 | South China<br>GuiZhou, China | Du 2018<br>10.1038/s41588-018-0116-x | 11.35 | yes | no<br>possible PCR bias | no |
| *Gossypium arboreum* | GA0210 | ZiJingZhongMian | SRR4457086 | | SRR4457086 | South China<br>GuiZhou, China | Du 2018<br>10.1038/s41588-018-0116-x | 11.04 | yes | no<br>possible PCR bias | no |
| *Gossypium arboreum* | GA0008 | NanLingZhongMian | SRR4456909 | | SRR4456909 | South China<br>YunNan, China | Du 2018<br>10.1038/s41588-018-0116-x | 10.01 | yes | no<br>possible PCR bias | no |
| *Gossypium arboreum* | GA0190 | YunNaFuNingZhongMian | SRR4456929 | | SRR4456929 | South China<br>YunNan, China | Du 2018<br>10.1038/s41588-018-0116-x | 10.51 | yes | no<br>possible PCR bias | no |
| *Gossypium arboreum* | GA0202 | XinPingTuMian | SRR4457094 | | SRR4457094 | South China<br>YunNan, China | Du 2018<br>10.1038/s41588-018-0116-x | 11.38 | yes | no<br>possible PCR bias | no |
| *Gossypium arboreum* | GA0146 | MeiZhongMian971(MaoZi) | SRR4457079 | | SRR4457079 | USA | Du 2018<br>10.1038/s41588-018-0116-x | 33.73 | yes | no<br>possible PCR bias | no |
| *Gossypium arboreum* | GA0149 | MeiZhongMian971(GuangZi) | SRR4457072 | | SRR4457072 | USA | Du 2018<br>10.1038/s41588-018-0116-x | 30.90 | yes | no<br>possible PCR bias | no |
| *Gossypium arboreum* | GA0223 | Nha92 | SRR4456883 | | SRR4456883 | Vietnam | Du 2018<br>10.1038/s41588-018-0116-x | 10.89 | yes | no<br>possible PCR bias | no |
| *Gossypium herbaceum* | GC0008 | HongKeCaoMian | SRR4456956 | | SRR4456956 | Xinjiang, China | Du 2018<br>10.1038/s41588-018-0116-x | 10.28 | yes | no<br>possible PCR bias | no |
| *Gossypium arboreum* | GA0003 | NaTongGuanYinHua | SRR4456912 | | SRR4456912 | Yangtze River<br>JiangSu, China | Du 2018<br>10.1038/s41588-018-0116-x | 10.56 | yes | no<br>possible PCR bias | no |
| *Gossypium arboreum* | GA0004 | ShangHaiYingHuMian | SRR4456913 | | SRR4456913 | Yangtze River<br>JiangSu, China | Du 2018<br>10.1038/s41588-018-0116-x | 10.39 | yes | no<br>possible PCR bias | no |
| *Gossypium arboreum* | GA0001 | ZheJiangYuYaoHuangDiDaBu | SRR4456914 | | SRR4456914 | Yangtze River<br>ZheJiang, China | Du 2018<br>10.1038/s41588-018-0116-x | 10.01 | yes | no<br>possible PCR bias | no |
| *Gossypium arboreum* | GA0002 | ZheJiangShangYuQingShanHongGeng | SRR4456915 | | SRR4456915 | Yangtze River<br>ZheJiang, China | Du 2018<br>10.1038/s41588-018-0116-x | 10.32 | yes | no<br>possible PCR bias | no |
| *Gossypium arboreum* | GA0080 | AnHuiFuYangEYangHua | SRR4457104 | | SRR4457104 | Yangtze River<br>AnHui, China | Du 2018<br>10.1038/s41588-018-0116-x | 10.28 | yes | no<br>possible PCR bias | no |
| *Gossypium arboreum* | GA0083 | AnHuiQuanJiaoZhongMian | SRR4456997 | | SRR4456997 | Yangtze River<br>AnHui, China | Du 2018<br>10.1038/s41588-018-0116-x | 11.26 | yes | no<br>possible PCR bias | no |
| *Gossypium arboreum* | GA0092 | AnHuiGuiChiXiaoZiMianHuiZi | SRR4457028 | | SRR4457028 | Yangtze River<br>AnHui, China | Du 2018<br>10.1038/s41588-018-0116-x | 10.31 | yes | no<br>possible PCR bias | no |
| *Gossypium arboreum* | GA0195 | AnHuiXuanChengZhongZiMianBaiZi | SRR4456960 | | SRR4456960 | Yangtze River<br>AnHui, China | Du 2018<br>10.1038/s41588-018-0116-x | 10.80 | yes | no<br>possible PCR bias | no |
| *Gossypium arboreum* | GA0096 | EChengXianZhongMian | SRR4457024 | | SRR4457024 | Yangtze River<br>HuBei, China | Du 2018<br>10.1038/s41588-018-0116-x | 10.10 | yes | no<br>possible PCR bias | no |
| *Gossypium arboreum* | GA0013 | ZhongYinZaJiaoMian-4 | SRR4457043 | | SRR4457043 | Yangtze River<br>JiangSu, China | Du 2018<br>10.1038/s41588-018-0116-x | 10.22 | yes | no<br>possible PCR bias | no |
| *Gossypium arboreum* | GA0027 | JiangSuChangShuHeiZi | SRR4456969 | | SRR4456969 | Yangtze River<br>JiangSu, China | Du 2018<br>10.1038/s41588-018-0116-x | 10.01 | yes | no<br>possible PCR bias | no |
| *Gossypium arboreum* | GA0056 | SiYangCangJiQingJingZhongMian | SRR4457036 | | SRR4457036 | Yangtze River<br>JiangSu, China | Du 2018<br>10.1038/s41588-018-0116-x | 10.45 | yes | no<br>possible PCR bias | no |
| *Gossypium arboreum* | GA0057 | JiangSuXuZhouZhongMian | SRR4457031 | | SRR4457031 | Yangtze River<br>JiangSu, China | Du 2018<br>10.1038/s41588-018-0116-x | 10.29 | yes | no<br>possible PCR bias | no |
| *Gossypium arboreum* | GA0218 | JiangSuTaiCangYangMaoGuangZi | SRR4456890 | | SRR4456890 | Yangtze River<br>JiangSu, China | Du 2018<br>10.1038/s41588-018-0116-x | 11.18 | yes | no<br>possible PCR bias | no |
| *Gossypium arboreum* | GA0093 | JiangXiXiuShuiZhongMianMaoZi | SRR4457029 | | SRR4457029 | Yangtze River<br>JiangXi, China | Du 2018<br>10.1038/s41588-018-0116-x | 20.56 | yes | no<br>possible PCR bias | no |
| *Gossypium arboreum* | GA0145 | XiuShuiZhongMianGuangZi | SRR4457080 | | SRR4457080 | Yangtze River<br>JiangXi, China | Du 2018<br>10.1038/s41588-018-0116-x | 10.17 | yes | no<br>possible PCR bias | no |
| *Gossypium arboreum* | GA0184 | HengFengTieZi | SRR4456937 | | SRR4456937 | Yangtze River<br>JiangXi, China | Du 2018<br>10.1038/s41588-018-0116-x | 12.66 | yes | no<br>possible PCR bias | no |
| *Gossypium arboreum* | GA0194 | DongXiangTieZi | SRR4456973 | | SRR4456973 | Yangtze River<br>JiangXi, China | Du 2018<br>10.1038/s41588-018-0116-x | 11.86 | yes | no<br>possible PCR bias | no |
| *Gossypium arboreum* | GA0104 | SiChuanSuiNingZhongMianGuangZi | SRR4456922 | | SRR4456922 | Yangtze River<br>SiChuan, China | Du 2018<br>10.1038/s41588-018-0116-x | 10.83 | yes | no<br>possible PCR bias | no |
| *Gossypium arboreum* | GA0159 | ZheJiangHaiYanHongGeng | SRR4457047 | | SRR4457047 | Yangtze River<br>ZheJiang, China | Du 2018<br>10.1038/s41588-018-0116-x | 10.20 | yes | no<br>possible PCR bias | no |
| *Gossypium arboreum* | GA0005 | ShanDongXiaoYuHua | SRR4457035 | | SRR4457035 | Yellow River<br>ShanDong, China | Du 2018<br>10.1038/s41588-018-0116-x | 11.25 | yes | no<br>possible PCR bias | no |
| *Gossypium arboreum* | GA0006 | DingTaoZhongMian | SRR4457007 | | SRR4457007 | Yellow River<br>ShanDong, China | Du 2018<br>10.1038/s41588-018-0116-x | 11.99 | yes | no<br>possible PCR bias | no |

| Species | Accession | Alternate Name | Sample Name | GRIN (PI) | SRA ID | Location | Sequenced | Read Coverage | Sufficient Coverage | Pass Filter | In QC set |
|---|---|---|---|---|---|---|---|---|---|---|---|
| *Gossypium arboreum* | GA0018 | YuanShiZhongMian | SRR4457040 | | SRR4457040 | Yellow River HeBei, China | Du 2018 10.1038/s41588-018-0116-x | 10.03 | yes | no possible PCR bias | no |
| *Gossypium arboreum* | GA0122 | SiHuaZhongMian | SRR4457018 | | SRR4457018 | Yellow River HeBei, China | Du 2018 10.1038/s41588-018-0116-x | 10.57 | yes | no possible PCR bias | no |
| *Gossypium arboreum* | GA0126 | YuXi3-3 | SRR4457014 | | SRR4457014 | Yellow River HeBei, China | Du 2018 10.1038/s41588-018-0116-x | 10.13 | yes | no possible PCR bias | no |
| *Gossypium arboreum* | GA0177 | ZhaoXianDaQingJie | SRR4457107 | | SRR4457107 | Yellow River HeBei, China | Du 2018 10.1038/s41588-018-0116-x | 10.54 | yes | no possible PCR bias | no |
| *Gossypium arboreum* | GA0189 | XiaoMaoZi | SRR4456930 | | SRR4456930 | Yellow River HeBei, China | Du 2018 10.1038/s41588-018-0116-x | 10.27 | yes | no possible PCR bias | no |
| *Gossypium arboreum* | GA0203 | DaZiJieMian | SRR4457091 | | SRR4457091 | Yellow River HeBei, China | Du 2018 10.1038/s41588-018-0116-x | 10.68 | yes | no possible PCR bias | no |
| *Gossypium arboreum* | GA0205 | XiaoBaiHuaZhongMian | SRR4457089 | | SRR4457089 | Yellow River HeBei, China | Du 2018 10.1038/s41588-018-0116-x | 10.65 | yes | no possible PCR bias | no |
| *Gossypium arboreum* | GA0212 | JinXianZhongMian | SRR4456967 | | SRR4456967 | Yellow River HeBei, China | Du 2018 10.1038/s41588-018-0116-x | 10.45 | yes | no possible PCR bias | no |
| *Gossypium arboreum* | GA0226 | ShiXiYaHeiZi | SRR4457074 | | SRR4457074 | Yellow River HeBei, China | Du 2018 10.1038/s41588-018-0116-x | 10.99 | yes | no possible PCR bias | no |
| *Gossypium arboreum* | GA0026 | ShangQiuXiaoBaiHua | SRR4456966 | | SRR4456966 | Yellow River HeNan, China | Du 2018 10.1038/s41588-018-0116-x | 10.68 | yes | no possible PCR bias | no |
| *Gossypium arboreum* | GA0115 | HeNaHuaiYangTuMianGuangZi | SRR4456874 | | SRR4456874 | Yellow River HeNan, China | Du 2018 10.1038/s41588-018-0116-x | 10.46 | yes | no possible PCR bias | no |
| *Gossypium arboreum* | GA0127 | ZhongMian-6 | SRR4457033 | | SRR4457033 | Yellow River LiaoNing, China | Du 2018 10.1038/s41588-018-0116-x | 11.20 | yes | no possible PCR bias | no |
| *Gossypium arboreum* | GA0111 | ShanDongHuiMinLunShanXiaoMian | SRR4456878 | | SRR4456878 | Yellow River ShanDong, China | Du 2018 10.1038/s41588-018-0116-x | 11.25 | yes | no possible PCR bias | no |
| *Gossypium arboreum* | GA0192 | ZongXuXiaoBenHua | SRR4456975 | | SRR4456975 | Yellow River ShanDong, China | Du 2018 10.1038/s41588-018-0116-x | 12.22 | yes | no possible PCR bias | no |
| *Gossypium arboreum* | GA0117 | ShanXiLinFenDongMaCeZhongMian | SRR4456872 | | SRR4456872 | Yellow River ShanXi, China | Du 2018 10.1038/s41588-018-0116-x | 10.52 | yes | no possible PCR bias | no |
| *Gossypium herbaceum* | GC0017 | | SRR4456981 | | SRR4456981 | China | Du 2018 10.1038/s41588-018-0116-x | 11.02 | yes | no incorrect species | no |
| *Gossypium arboreum* | GA0174 | JiJiaoZhongMian | SRR4457112 | | SRR4457112 | China | Du 2018 10.1038/s41588-018-0116-x | 10.05 | yes | no possible PCR bias | no |
| *Gossypium arboreum* | Gar36 | var. burmanicum | SRR8283975_Gar36 | | SRR8283975 | Burma | Huang 2020 10.1038/s41588-020-0607-4 | 17.34 | yes | yes | yes |
| *Gossypium arboreum* | Gar40 | var. indicum | SRR8283971_Gar40 | | SRR8283971 | India | Huang 2020 10.1038/s41588-020-0607-4 | 12.74 | yes | yes | yes |
| *Gossypium arboreum* | Gar44 | Desi Rui | SRR8283983_Gar44 | | SRR8283983 | India | Huang 2020 10.1038/s41588-020-0607-4 | 15.47 | yes | yes | yes |
| *Gossypium arboreum* | Gar54 | var. indicum | SRR8284000_Gar54 | | SRR8284000 | India | Huang 2020 10.1038/s41588-020-0607-4 | 14.68 | yes | yes | yes |
| *Gossypium arboreum* | Gar57 | var. indicum | SRR8283978_Gar57 | | SRR8283978 | India | Huang 2020 10.1038/s41588-020-0607-4 | 14.80 | yes | yes | yes |
| *Gossypium arboreum* | Gar60 | var. cernuum | SRR8283981_Gar60 | | SRR8283981 | India | Huang 2020 10.1038/s41588-020-0607-4 | 15.97 | yes | yes | yes |
| *Gossypium arboreum* | Gar64 | var. bengalense | SRR8284011_Gar64 | | SRR8284011 | India | Huang 2020 10.1038/s41588-020-0607-4 | 14.00 | yes | yes | yes |
| *Gossypium arboreum* | Gar66 | var. indicum | SRR8283997_Gar66 | | SRR8283997 | India | Huang 2020 10.1038/s41588-020-0607-4 | 13.93 | yes | yes | yes |
| *Gossypium arboreum* | Gar02 | Koresu | SRR8283994_Gar02 | | SRR8283994 | Iran via USSR | Huang 2020 10.1038/s41588-020-0607-4 | 16.51 | yes | yes | yes |
| *Gossypium herbaceum* | Ghe01 | var. africanum (Mutema) | SRR8283996_Ghe01 | | SRR8283996 | Kodoma Zimbabwe | Huang 2020 10.1038/s41588-020-0607-4 | 13.47 | yes | yes | yes |
| *Gossypium arboreum* | Gar30 | DC00092 | SRR8283989_Gar30 | | SRR8283989 | Lyallpur, Pakistan | Huang 2020 10.1038/s41588-020-0607-4 | 14.37 | yes | yes | yes |
| *Gossypium arboreum* | Gar01 | Malvi 9 | SRR8283993_Gar01 | | SRR8283993 | Madhya Pradesh India | Huang 2020 10.1038/s41588-020-0607-4 | 11.56 | yes | yes | yes |
| *Gossypium arboreum* | Gar41 | Verum | SRR8283972_Gar41 | | SRR8283972 | Madhya Pradesh India | Huang 2020 10.1038/s41588-020-0607-4 | 14.12 | yes | yes | yes |
| *Gossypium arboreum* | Gar31 | Indicum 00038 | SRR8283990_Gar31 | | SRR8283990 | Maharashtra India | Huang 2020 10.1038/s41588-020-0607-4 | 15.40 | yes | yes | yes |
| *Gossypium arboreum* | Gar32 | Dhulia 215 | SRR8283987_Gar32 | | SRR8283987 | Maharashtra India | Huang 2020 10.1038/s41588-020-0607-4 | 16.62 | yes | yes | yes |
| *Gossypium arboreum* | Gar33 | Burma Silk | SRR8283988_Gar33 | | SRR8283988 | Maharashtra India | Huang 2020 10.1038/s41588-020-0607-4 | 16.57 | yes | yes | yes |
| *Gossypium arboreum* | Gar34 | Kokati Khaki | SRR8283985_Gar34 | | SRR8283985 | Maharashtra India | Huang 2020 10.1038/s41588-020-0607-4 | 15.10 | yes | yes | yes |
| *Gossypium arboreum* | Gar35 | Indicum 00010 | SRR8283986_Gar35 | | SRR8283986 | Maharashtra India | Huang 2020 10.1038/s41588-020-0607-4 | 16.93 | yes | yes | yes |
| *Gossypium arboreum* | Gar37 | Jarilla | SRR8283976_Gar37 | | SRR8283976 | New Delhi, India | Huang 2020 10.1038/s41588-020-0607-4 | 17.38 | yes | yes | yes |
| *Gossypium arboreum* | Gar45 | An. B.B. 1000P7 | SRR8283984_Gar45 | | SRR8283984 | Pakistan | Huang 2020 10.1038/s41588-020-0607-4 | 15.78 | yes | yes | yes |
| *Gossypium arboreum* | Gar38 | White Flower | SRR8283973_Gar38 | | SRR8283973 | Pakistan via USSR | Huang 2020 10.1038/s41588-020-0607-4 | 16.18 | yes | yes | yes |
| *Gossypium arboreum* | Gar47 | var. roseum | SRR8284007_Gar47 | | SRR8284007 | Pakistan via USSR | Huang 2020 10.1038/s41588-020-0607-4 | 14.04 | yes | yes | yes |

| Species | Accession | Alternate Name | Sample Name | GRIN (PI) | SRA ID | Location | Sequenced | Read Coverage | Sufficient Coverage | Pass Filter | In QC set |
|---|---|---|---|---|---|---|---|---|---|---|---|
| *Gossypium arboreum* | Gar61 | Red Flower | SRR8283982_Gar61 | | SRR8283982 | Pakistan via USSR | Huang 2020 10.1038/s41588-020-0607-4 | 19.25 | yes | yes | yes |
| *Gossypium arboreum* | Gar43 | Kapas | SRR8283970_Gar43 | | SRR8283970 | Rajastan India | Huang 2020 10.1038/s41588-020-0607-4 | 14.14 | yes | yes | yes |
| *Gossypium arboreum* | Gar46 | Kapas | SRR8284008_Gar46 | | SRR8284008 | Rajastan India | Huang 2020 10.1038/s41588-020-0607-4 | 15.02 | yes | yes | yes |
| *Gossypium herbaceum* | Ghe02 | Russian No. 2531 | SRR8283998_Ghe02 | | SRR8283998 | Russian | Huang 2020 10.1038/s41588-020-0607-4 | 12.18 | yes | yes | yes |
| *Gossypium arboreum* | Gar42 | Nanking | SRR8283969_Gar42 | | SRR8283969 | Tashkent Uzbekistan | Huang 2020 10.1038/s41588-020-0607-4 | 15.38 | yes | yes | yes |
| *Gossypium arboreum* | Gar03 | Line O000824 II000I | SRR8283991_Gar03 | | SRR8283991 | Uzbekistan | Huang 2020 10.1038/s41588-020-0607-4 | 18.15 | yes | yes | yes |
| *Gossypium arboreum* | Gar04 | var. soudanense | SRR8283992_Gar04 | | SRR8283992 | West Iran via CSSR | Huang 2020 10.1038/s41588-020-0607-4 | 16.70 | yes | yes | yes |
| *Gossypium arboreum* | Gar65 | var. soudanense | SRR8283995_Gar65 | | SRR8283995 | West Iran via India | Huang 2020 10.1038/s41588-020-0607-4 | 11.58 | yes | yes | yes |
| *Gossypium arboreum* | Gar39 | var. cernuum | SRR8283974_Gar39 | | SRR8283974 | | Huang 2020 10.1038/s41588-020-0607-4 | 13.22 | yes | yes | yes |
| *Gossypium arboreum* | Gar48 | var. bengalense | SRR8284006_Gar48 | | SRR8284006 | | Huang 2020 10.1038/s41588-020-0607-4 | 17.28 | yes | yes | yes |
| *Gossypium arboreum* | Gar49 | var. bengalense | SRR8284005_Gar49 | | SRR8284005 | | Huang 2020 10.1038/s41588-020-0607-4 | 17.28 | yes | yes | yes |
| *Gossypium arboreum* | Gar50 | var. bengalense | SRR8284004_Gar50 | | SRR8284004 | | Huang 2020 10.1038/s41588-020-0607-4 | 18.86 | yes | yes | yes |
| *Gossypium arboreum* | Gar51 | var. carnuum | SRR8284003_Gar51 | | SRR8284003 | | Huang 2020 10.1038/s41588-020-0607-4 | 13.21 | yes | yes | yes |
| *Gossypium arboreum* | Gar52 | var. bengalense | SRR8284002_Gar52 | | SRR8284002 | | Huang 2020 10.1038/s41588-020-0607-4 | 12.88 | yes | yes | yes |
| *Gossypium arboreum* | Gar53 | var. carnuum | SRR8284001_Gar53 | | SRR8284001 | | Huang 2020 10.1038/s41588-020-0607-4 | 12.20 | yes | yes | yes |
| *Gossypium arboreum* | Gar55 | var. bengalense | SRR8283999_Gar55 | | SRR8283999 | | Huang 2020 10.1038/s41588-020-0607-4 | 13.09 | yes | yes | yes |
| *Gossypium arboreum* | Gar56 | var. bengalense | SRR8283977_Gar56 | | SRR8283977 | | Huang 2020 10.1038/s41588-020-0607-4 | 16.39 | yes | yes | yes |
| *Gossypium arboreum* | Gar58 | var. indicum | SRR8283979_Gar58 | | SRR8283979 | | Huang 2020 10.1038/s41588-020-0607-4 | 14.79 | yes | yes | yes |
| *Gossypium arboreum* | Gar59 | var. bengalense | SRR8283980_Gar59 | | SRR8283980 | | Huang 2020 10.1038/s41588-020-0607-4 | 15.43 | yes | yes | yes |
| *Gossypium arboreum* | Gar62 | var. burmanicum | SRR8284009_Gar62 | | SRR8284009 | | Huang 2020 10.1038/s41588-020-0607-4 | 13.04 | yes | yes | yes |
| *Gossypium arboreum* | Gar63 | var. bengalense | SRR8284010_Gar63 | | SRR8284010 | | Huang 2020 10.1038/s41588-020-0607-4 | 13.74 | yes | yes | yes |
| *Gossypium arboreum* | A2_164 | SELECTION OF SHIH | A2_164 | PI 529781 | SRR8979998 | China | in this study | 30.77 | yes | yes | yes |
| *Gossypium herbaceum* | A1_079 | VAR. KULJIANUM | A1_079 | PI 529655 | SRR8979915 | Zimbabwe | in this study | 52.41 | yes | yes | yes |
| *Gossypium arboreum* | A2_lintless | | A2_lintless | | SRR8980000 | | in this study | 42.63 | yes | yes | yes |
| *Gossypium arboreum* | A2_NC503 | | A2_NC503 | | SRR8979994 | | in this study | 33.04 | yes | yes | yes |
| *Gossypium herbaceum* | A1_019 | | A1_019 | PI 408777 | SRR8979931 | Afghanistan | in this study; BGI | 51.41 | yes | yes | yes |
| *Gossypium herbaceum* | A1_084 | Tzuyung | A1_084 | PI 441982 | SRR8979912 | China | in this study; BGI | 53.99 | yes | yes | yes |
| *Gossypium arboreum* | A2_154 | Liaochung 6 | A2_154 | PI 441979 | SRR8979932 | China | in this study; BGI | 23.27 | yes | yes | yes |
| *Gossypium arboreum* | A2_156 | JIANG YIN | A2_156 | PI 529773 | SRR8979933 | China | in this study; BGI | 29.79 | yes | yes | yes |
| *Gossypium arboreum* | A2_158 | SHIXIYA 1 | A2_158 | PI 529775 | SRR8979999 | China | in this study; BGI | 33.77 | yes | yes | yes |
| *Gossypium arboreum* | A2_141 | Chang Zi No. 1 | A2_141 | PI 452093 | SRR8979935 | China* | in this study; BGI | 34.01 | yes | yes | yes |
| *Gossypium arboreum* | A2_142 | Liaoyang No. 1 | A2_142 | PI 452094 | SRR8979936 | China* | in this study; BGI | 41.77 | yes | yes | yes |
| *Gossypium arboreum* | A2_021 | NANKING | A2_021 | PI 152088 | SRR8979975 | Former Soviet Union | in this study; BGI | 40.44 | yes | yes | yes |
| *Gossypium arboreum* | A2_079 | MUDHOL-2680 | A2_079 | PI 417887 | SRR8979953 | Gujarat, India* | in this study; BGI | 61.30 | yes | yes | yes |
| *Gossypium arboreum* | A2_080 | GAORANI-46 | A2_080 | PI 417888 | SRR8979921 | Gujarat, India* | in this study; BGI | 29.85 | yes | yes | yes |
| *Gossypium arboreum* | A2_084 | SANGUIENUM MINOR | A2_084 | PI 417892 | SRR8979920 | Gujarat, India* | in this study; BGI | 52.33 | yes | yes | yes |
| *Gossypium arboreum* | A2_085 | CHANG FEND | A2_085 | PI 417893 | SRR8979919 | Gujarat, India* | in this study; BGI | 31.51 | yes | yes | yes |
| *Gossypium arboreum* | A2_087 | NORTHERN-14 | A2_087 | PI 417895 | SRR8979918 | Gujarat, India* | in this study; BGI | 31.79 | yes | yes | yes |
| *Gossypium arboreum* | A2_091 | NANKING KHAKHI | A2_091 | PI 417899 | SRR8979925 | Gujarat, India* | in this study; BGI | 49.90 | yes | yes | yes |
| *Gossypium herbaceum* | A1_054 | WIGHTIANUM | A1_054 | PI 529646 | SRR8979985 | India | in this study; BGI | 38.37 | yes | yes | yes |
| *Gossypium arboreum* | A2_031 | AN. B.B. 1-P7 | A2_031 | PI 180365 | SRR8979979 | India | in this study; BGI | 33.88 | yes | yes | yes |
| *Gossypium arboreum* | A2_040 | | A2_040 | PI 183202 | SRR8979972 | India | in this study; BGI | 26.94 | yes | yes | yes |
| *Gossypium arboreum* | A2_057 | COMPANY NO. 3 | A2_057 | PI 408754 | SRR8979951 | India | in this study; BGI | 30.74 | yes | yes | yes |
| *Gossypium herbaceum* | A1_051 | G. Cot-11 | A1_051 | PI 442921 | SRR8979984 | India* | in this study; BGI | 58.92 | yes | yes | yes |
| *Gossypium herbaceum* | A1_133 | DIGVIJAY | A1_133 | PI 630008 | SRR8979971 | India* | in this study; BGI | 33.84 | yes | yes | yes |
| *Gossypium arboreum* | A2_026 | KAPAS | A2_026 | PI 179607 | SRR8979976 | India* | in this study; BGI | 28.68 | yes | yes | yes |
| *Gossypium arboreum* | A2_029 | KAPAS | A2_029 | PI 180245 | SRR8979977 | India* | in this study; BGI | 29.15 | yes | yes | yes |
| *Gossypium arboreum* | A2_030 | KAPAS | A2_030 | PI 180246 | SRR8979978 | India* | in this study; BGI | 36.83 | yes | yes | yes |
| *Gossypium herbaceum* | A1_030 | | A1_030 | PI 408788 | SRR8979986 | Indonesia | in this study; BGI | 46.34 | yes | yes | yes |
| *Gossypium arboreum* | A2_044 | YONG-GANG #103 | A2_044 | PI 185788 | SRR8979944 | Korea* | in this study; BGI | 52.03 | yes | yes | yes |
| *Gossypium arboreum* | A2_041 | K-1 | A2_041 | PI 183887 | SRR8979973 | Maharashtra India* | in this study; BGI | 34.68 | yes | yes | yes |
| *Gossypium arboreum* | A2_046 | DC-92 | A2_046 | PI 200422 | SRR8979946 | Pakistan* | in this study; BGI | 32.34 | yes | yes | yes |
| *Gossypium arboreum* | A2_047 | GAORANI | A2_047 | PI 213373 | SRR8979949 | Sudan* | in this study; BGI | 55.53 | yes | yes | yes |
| *Gossypium arboreum* | A2_001 | #1 | A2_001 | PI 529704 | SRR8979965 | Texas United States* | in this study; BGI | 58.49 | yes | yes | yes |

| Species | Accession | Alternate Name | Sample Name | GRIN (PI) | SRA ID | Location | Sequenced | Read Coverage | Sufficient Coverage | Pass Filter | In QC set |
|---|---|---|---|---|---|---|---|---|---|---|---|
| *Gossypium arboreum* | A2_008 | #43 | A2_008 | PI 529711 | SRR8979963 | Texas<br>United States* | in this study; BGI | 29.33 | yes | yes | yes |
| *Gossypium arboreum* | A2_016 | SMA-2 | A2_016 | PI 529719 | SRR8979962 | Texas<br>United States* | in this study; BGI | 37.01 | yes | yes | yes |
| *Gossypium arboreum* | A2_096 | INDIA 15 | A2_096 | PI 529724 | SRR8979924 | Texas<br>United States* | in this study; BGI | 27.42 | yes | yes | yes |
| *Gossypium arboreum* | A2_100 | NANKING | A2_100 | PI 529728 | SRR8979923 | Texas<br>United States* | in this study; BGI | 31.20 | yes | yes | yes |
| *Gossypium arboreum* | A2_113 | SMA4 | A2_113 | PI 529740 | SRR8979927 | Texas<br>United States* | in this study; BGI | 42.33 | yes | yes | yes |
| *Gossypium arboreum* | A2_117 | A2 117 | A2_117 | PI 529744 | SRR8979926 | Texas<br>United States* | in this study; BGI | 40.13 | yes | yes | yes |
| *Gossypium arboreum* | A2_118 | AKH 4 | A2_118 | PI 529745 | SRR8979938 | Texas<br>United States* | in this study; BGI | 32.83 | yes | yes | yes |
| *Gossypium arboreum* | A2_119 | AKH 235 | A2_119 | PI 529746 | SRR8979939 | Texas<br>United States* | in this study; BGI | 30.15 | yes | yes | yes |
| *Gossypium arboreum* | A2_123 | CJ 73 | A2_123 | PI 529750 | SRR8979940 | Texas<br>United States* | in this study; BGI | 33.47 | yes | yes | yes |
| *Gossypium arboreum* | A2_124 | DAVDAT | A2_124 | PI 529751 | SRR8979941 | Texas<br>United States* | in this study; BGI | 36.57 | yes | yes | yes |
| *Gossypium arboreum* | A2_131 | LBM | A2_131 | PI 529758 | SRR8979934 | Texas<br>United States* | in this study; BGI | 32.27 | yes | yes | yes |
| *Gossypium arboreum* | A2_147 | ACH 344 INDIA | A2_147 | PI 529767 | SRR8979937 | Texas<br>United States* | in this study; BGI | 36.04 | yes | yes | yes |
| *Gossypium herbaceum* | A1_028 | | A1_028 | PI 408786 | SRR8979988 | Turkey | in this study; BGI | 31.49 | yes | yes | yes |
| *Gossypium herbaceum* | A1_097 | A1 97 | A1_097 | PI 529670 | SRR8979910 | Turkey | in this study; BGI | 52.64 | yes | yes | yes |
| *Gossypium arboreum* | A2_056 | ROSEUM | A2_056 | PI 408753 | SRR8979948 | United States | in this study; BGI | 28.34 | yes | yes | yes |
| *Gossypium arboreum* | A2_059 | MAYOR N. | A2_059 | PI 408756 | SRR8979950 | United States | in this study; BGI | 36.75 | yes | yes | yes |
| *Gossypium arboreum* | A2_060 | NO. 78 | A2_060 | PI 408757 | SRR8979943 | United States | in this study; BGI | 40.07 | yes | yes | yes |
| *Gossypium arboreum* | A2_064 | TYPE 9 | A2_064 | PI 408761 | SRR8979957 | United States | in this study; BGI | 35.00 | yes | yes | yes |
| *Gossypium arboreum* | A2_065 | | A2_065 | PI 408762 | SRR8979954 | United States | in this study; BGI | 31.79 | yes | yes | yes |
| *Gossypium arboreum* | A2_066 | SMALL SPORT | A2_066 | PI 408763 | SRR8979955 | United States | in this study; BGI | 35.39 | yes | yes | yes |
| *Gossypium arboreum* | A2_067 | KOKATI NARROW | A2_067 | PI 408764 | SRR8979960 | United States | in this study; BGI | 44.18 | yes | yes | yes |
| *Gossypium arboreum* | A2_069 | ALIGARH A19 | A2_069 | PI 408766 | SRR8979961 | United States | in this study; BGI | 36.95 | yes | yes | yes |
| *Gossypium arboreum* | A2_074 | CHINESE PALE | A2_074 | PI 408771 | SRR8979959 | United States | in this study; BGI | 38.05 | yes | yes | yes |
| *Gossypium arboreum* | A2_076 | | A2_076 | PI 408773 | SRR8979952 | United States | in this study; BGI | 27.62 | yes | yes | yes |
| *Gossypium herbaceum* | A1_132 | | A1_132 | PI 358402 | SRR8979917 | Unknown | in this study; BGI | 27.03 | yes | yes | yes |
| *Gossypium herbaceum* | A1_Af | var. africanum | A1_Af | PI 630014 | SRR8979967 | Unknown* | in this study; BGI | 58.80 | yes | yes | yes |
| *Gossypium herbaceum* | A1_108 | LINE 3856 | A1_108 | PI 529681 | SRR8979911 | Uzbekistan | in this study; BGI | 34.38 | yes | yes | yes |
| *Gossypium herbaceum* | A1_077 | BOUMI RED | A1_077 | PI 529653 | SRR8979914 | Zimbabwe | in this study; BGI | 59.35 | yes | yes | yes |
| *Gossypium herbaceum* | A1_Nisa | | A1_Nisa | | SRR8979966 | | in this study; BGI | 39.45 | yes | yes | yes |
| *Gossypium herbaceum* | A1_088 | KRASNYJ | A1_088 | PI 529661 | SRR8979913 | Afghanistan | in this study; BYUSC | 27.98 | yes | yes | yes |
| *Gossypium arboreum* | A2_061 | | A2_061 | PI 408758 | SRR8979942 | Japan | in this study; BYUSC | 24.00 | yes | yes | yes |
| *Gossypium arboreum* | A2_043 | SAMJU | A2_043 | PI 185787 | SRR8979945 | Korea* | in this study; BYUSC | 33.66 | yes | yes | yes |
| *Gossypium arboreum* | A2_019 | VERUM | A2_019 | PI 129723 | SRR8979974 | Madhya Pradesh<br>India* | in this study; BYUSC | 34.86 | yes | yes | yes |
| *Gossypium arboreum* | A2_045 | 39 MOLLISANI | A2_045 | PI 200421 | SRR8979947 | Pakistan* | in this study; BYUSC | 27.49 | yes | yes | yes |
| *Gossypium herbaceum* | A1_074 | A1 57 | A1_074 | PI 529649 | SRR8979983 | Texas<br>United States* | in this study; BYUSC | 25.95 | yes | yes | yes |
| *Gossypium arboreum* | A2_062 | KORESU | A2_062 | PI 408759 | SRR8979956 | United States | in this study; BYUSC | 24.97 | yes | yes | yes |
| *Gossypium arboreum* | A2_073 | CHINESE NAKED | A2_073 | PI 408770 | SRR8979958 | United States | in this study; BYUSC | 35.27 | yes | yes | yes |
| *Gossypium herbaceum* | A1_125 | LINE 6969 | A1_125 | PI 529698 | SRR8979916 | Uzbekistan | in this study; BYUSC | 17.31 | yes | yes | yes |
| *Gossypium arboreum* | A2_255 | | A2_255 | PI 615756 | SRR8980001 | | Page 2013<br>10.1534/g3.113.007229 | 57.56 | yes | yes | yes |
| *Gossypium herbaceum* | A1_073 | M-881.8 | A1_073 | PI 485587 | SRR8979982 | Botswana | Page 2013<br>10.1534/g3.113.007229 | 41.94 | yes | yes | yes |
| *Gossypium arboreum* | A2_034 | NEGLECTUM | A2_034 | PI 183160 | SRR8979980 | India | Page 2013<br>10.1534/g3.113.007229 | 73.63 | yes | yes | yes |
| *Gossypium herbaceum* | A1_155 | AFRICANUM | A1_155 | PI 630024 | SRR8979969 | Sudan* | Page 2013<br>10.1534/g3.113.007229 | 49.22 | yes | yes | yes |
| *Gossypium arboreum* | A2_004 | #5 | A2_004 | PI 529707 | SRR8979964 | Texas<br>United States* | Page 2013<br>10.1534/g3.113.007229 | 57.56 | yes | yes | yes |
| *Gossypium arboreum* | A2_101 | GARO HILL | A2_101 | PI 529729 | SRR8979922 | Texas<br>United States* | Page 2013<br>10.1534/g3.113.007229 | 76.02 | yes | yes | yes |
| *Gossypium longicalyx* | F1-1 | | SRR617704 | | SRR617704 | | PRJNA178971 | 51.78 | yes | yes | yes |

Supplementary Table 2. Distribution of variant detection for all A-genome accessions among the 13 chromosomes.

| Chromosome | Length | Variants | Variants rate |
|---|---|---|---|
| F01 | 98,464,577 | 961,938 | 102 |
| F02 | 76,289,367 | 847,715 | 89 |
| F03 | 110,231,091 | 1,138,935 | 96 |
| F04 | 91,022,893 | 861,441 | 105 |
| F05 | 76,482,865 | 744,850 | 102 |
| F06 | 98,089,871 | 1,022,627 | 95 |
| F07 | 82,226,542 | 818,582 | 100 |
| F08 | 104,627,231 | 1,068,403 | 97 |
| F09 | 70,606,549 | 695,530 | 101 |
| F10 | 95,083,790 | 1,062,461 | 89 |
| F11 | 99,601,080 | 1,049,001 | 94 |
| F12 | 91,301,993 | 903,195 | 101 |
| F13 | 95,878,146 | 973,375 | 98 |
| **Total** | **1,189,905,995** | **12,148,053** | **97** |

Supplementary Table 3. Number of SNP sites (summarized and per sample) that are homozygous for the reference allele, heterozygous, or homozygous for the derived allele.

| *both* | Min. SNPs/sample | Max. SNPs/sample | Mean SNPs/sample | Median SNPs/sample |
|---|---|---|---|---|
| Reference | 5,577,272 | 7,910,974 | 6,630,940 | 6,521,348 |
| Heterozygous | 1,652,225 | 4,840,734 | 3,504,973 | 3,707,671 |
| Homozygous derived | 1,560,246 | 2,128,889 | 1,892,430 | 1,853,971 |
| Missing | 11,397 | 740,170 | 119,710 | 56,609 |

| *G. herbaceum* | Minimum number of SNPs/sample | Maximum number of SNPs/sample | Mean number of SNPs/sample | Median number of SNPs/sample |
|---|---|---|---|---|
| Reference | 6,296,046 | 7,910,974 | 6,816,084 | 6,712,165 |
| Heterozygous | 1,652,225 | 4,107,950 | 3,420,629 | 3,603,672 |
| Homozygous derived | 1,560,246 | 2,128,889 | 1,786,056 | 1,761,334 |
| Missing | 33,940 | 740,170 | 125,284 | 64,789 |

| *G. arboreum* | Minimum number of SNPs/sample | Maximum number of SNPs/sample | Mean number of SNPs/sample | Median number of SNPs/sample |
|---|---|---|---|---|
| Reference | 5,577,272 | 7,475,944 | 6,591,667 | 6,450,274 |
| Heterozygous | 2,136,325 | 4,840,734 | 3,522,864 | 3,824,722 |
| Homozygous derived | 1,645,667 | 2,122,040 | 1,914,994 | 1,865,170 |
| Missing | 11,397 | 463,623 | 118,528 | 45,339 |

| | A1_019 | A1_028 | A1_030 | A1_051 | A1_054 | A1_073 | A1_074 | A1_077 | A1_079 | A1_084 | A1_088 | A1_097 |
|---|---|---|---|---|---|---|---|---|---|---|---|---|
| Reference | 6,452,063 | 6,670,509 | 6,400,406 | 6,352,571 | 6,523,138 | 7,106,228 | 7,181,807 | 6,378,072 | 7,093,011 | 6,348,038 | 7,039,755 | 6,296,046 |
| Heterozygous | 3,902,637 | 3,598,850 | 3,966,282 | 3,999,813 | 3,811,429 | 3,285,823 | 3,140,183 | 4,001,517 | 3,047,275 | 4,025,200 | 3,211,928 | 4,107,950 |
| Homozygous derived | 1,752,812 | 1,816,897 | 1,740,525 | 1,761,334 | 1,764,663 | 1,636,978 | 1,715,538 | 1,732,943 | 1,919,422 | 1,739,492 | 1,803,517 | 1,710,117 |
| Missing | 40,541 | 61,797 | 40,840 | 34,335 | 48,823 | 119,024 | 110,525 | 35,521 | 88,345 | 35,323 | 92,853 | 33,940 |

| | A1_108 | A1_125 | A1_132 | A1_133 | A1_155 | A1_Af | A1_Nisa | SRR8283996 Ghe01 | SRR8283998 Ghe02 |
|---|---|---|---|---|---|---|---|---|---|
| Reference | 6,712,165 | 7,512,742 | 6,925,982 | 6,677,943 | 6,790,120 | 6,409,324 | 6,730,105 | 7,910,974 | 7,626,769 |
| Heterozygous | 3,567,470 | 2,434,590 | 3,271,381 | 3,603,672 | 3,719,846 | 4,080,814 | 3,663,523 | 1,740,796 | 1,652,225 |
| Homozygous derived | 1,802,981 | 1,957,081 | 1,818,344 | 1,801,649 | 1,560,246 | 1,616,843 | 1,694,481 | 2,032,415 | 2,128,889 |
| Missing | 65,437 | 243,640 | 132,346 | 64,789 | 77,841 | 41,072 | 59,944 | 463,868 | 740,170 |

| | A2_001 | A2_004 | A2_008 | A2_016 | A2_019 | A2_021 | A2_026 | A2_029 | A2_030 | A2_031 | A2_034 | A2_040 |
|---|---|---|---|---|---|---|---|---|---|---|---|---|
| Reference | 5,667,466 | 6,318,663 | 6,319,909 | 6,113,281 | 6,426,653 | 6,081,501 | 6,258,622 | 6,296,678 | 6,261,414 | 6,129,874 | 5,905,113 | 6,391,260 |
| Heterozygous | 4,693,279 | 4,040,486 | 3,932,536 | 4,173,518 | 3,905,478 | 4,235,748 | 4,004,900 | 3,942,868 | 4,001,783 | 4,154,330 | 4,538,757 | 3,821,740 |
| Homozygous derived | 1,775,911 | 1,751,103 | 1,865,170 | 1,833,406 | 1,785,326 | 1,805,161 | 1,853,930 | 1,876,026 | 1,858,374 | 1,837,799 | 1,679,541 | 1,893,741 |
| Missing | 11,397 | 37,801 | 30,438 | 27,848 | 30,596 | 25,643 | 30,601 | 32,481 | 26,482 | 26,050 | 24,642 | 41,312 |

| | A2_041 | A2_043 | A2_044 | A2_045 | A2_046 | A2_047 | A2_056 | A2_057 | A2_059 | A2_060 | A2_061 | A2_062 |
|---|---|---|---|---|---|---|---|---|---|---|---|---|
| Reference | 6,154,928 | 6,450,274 | 5,771,300 | 6,868,379 | 6,115,620 | 5,686,696 | 6,351,133 | 6,131,774 | 6,034,492 | 5,919,573 | 6,828,002 | 6,722,606 |
| Heterozygous | 4,142,062 | 3,842,087 | 4,584,228 | 3,129,686 | 4,181,017 | 4,684,587 | 3,919,228 | 4,162,221 | 4,272,866 | 4,414,818 | 3,352,717 | 3,532,309 |
| Homozygous derived | 1,818,880 | 1,824,359 | 1,778,164 | 1,852,142 | 1,824,522 | 1,764,109 | 1,839,068 | 1,822,290 | 1,816,618 | 1,793,835 | 1,897,150 | 1,822,362 |
| Missing | 32,183 | 31,333 | 14,361 | 297,846 | 26,894 | 12,661 | 38,624 | 31,768 | 24,077 | 19,827 | 70,184 | 70,776 |

| | A2_064 | A2_065 | A2_066 | A2_067 | A2_069 | A2_073 | A2_074 | A2_076 | A2_079 | A2_080 | A2_084 | A2_085 |
|---|---|---|---|---|---|---|---|---|---|---|---|---|
| Reference | 6,217,382 | 6,255,608 | 6,165,028 | 5,884,372 | 6,005,154 | 6,423,925 | 5,962,363 | 6,519,558 | 5,577,272 | 6,280,246 | 5,723,237 | 6,320,176 |
| Heterozygous | 4,047,963 | 4,017,063 | 4,109,692 | 4,440,350 | 4,308,961 | 3,919,523 | 4,358,805 | 3,615,675 | 4,840,734 | 3,976,733 | 4,635,977 | 3,929,428 |
| Homozygous derived | 1,853,277 | 1,830,881 | 1,841,203 | 1,810,385 | 1,815,088 | 1,769,602 | 1,806,277 | 1,842,208 | 1,717,217 | 1,859,948 | 1,776,640 | 1,860,548 |
| Missing | 29,431 | 44,501 | 32,130 | 12,946 | 18,850 | 35,003 | 20,608 | 170,612 | 12,830 | 31,126 | 12,199 | 37,901 |

| | A2_087 | A2_091 | A2_096 | A2_100 | A2_101 | A2_113 | A2_117 | A2_118 | A2_119 | A2_123 | A2_124 | A2_131 |
|---|---|---|---|---|---|---|---|---|---|---|---|---|
| Reference | 6,297,266 | 5,963,917 | 6,489,522 | 6,302,750 | 6,086,964 | 5,978,974 | 5,967,847 | 6,193,826 | 6,289,423 | 6,234,749 | 6,207,715 | 6,266,542 |
| Heterozygous | 3,950,580 | 4,335,453 | 3,667,263 | 3,984,216 | 4,371,700 | 4,315,871 | 4,349,069 | 4,139,818 | 3,974,165 | 4,015,678 | 4,048,788 | 3,984,668 |
| Homozygous derived | 1,863,259 | 1,833,178 | 1,888,440 | 1,807,813 | 1,645,667 | 1,837,892 | 1,810,753 | 1,788,524 | 1,854,012 | 1,852,287 | 1,863,356 | 1,865,338 |
| Missing | 36,948 | 15,505 | 102,828 | 53,274 | 43,722 | 15,316 | 20,384 | 25,885 | 30,453 | 45,339 | 28,194 | 31,505 |

Supplementary Table 4. Number of sites (per chromosome) fixed for the derived allele in that species. The allele may either be absent or segregating in the opposite species.

| Chromosome | ***Gossypium herbaceum*** | ***Gossypium arboreum*** |
|---|---:|---:|
| F01 | 35,144 | 9,345 |
| F02 | 32,148 | 8,304 |
| F03 | 42,381 | 10,693 |
| F04 | 25,375 | 7,263 |
| F05 | 23,880 | 5,824 |
| F06 | 32,961 | 8,743 |
| F07 | 2,385 | 619 |
| F08 | 27,399 | 7,452 |
| F09 | 23,243 | 5,949 |
| F10 | 2,074 | 7,528 |
| F11 | 33,932 | 9,534 |
| F12 | 33,335 | 8,174 |
| F13 | 38,434 | 9,833 |

Supplementary Table 5. Distribution of indels among genomic regions. Subcategories are listed under each main category. Percentages are relative to category or subcategory.

| | | | among *G. herbaceum* | | among *G. arboreum* | |
|---|---|---|---|---|---|---|
| | **Number of indels** | | **Number of indels** | **Fixed indels** | **Number of indels** | **Fixed Indels** |
| **Insertions** | 2,111,173 | | 1,629,250 | 13,754 | 1,673,249 | 2,175 |
| **Deletions** | 3,252,844 | | 2,727,010 | 23,506 | 2,778,580 | 4,668 |
| | **Number of indels** | **Relative percent** | **Number of indels** | **Relative percent** | **Number of indels** | **Relative percent** |
| **Intergenic** | **4,786,137** | **84.66%** | **3,953,422** | **90.30%** | **4,041,832** | **90.34%** |
| Downstream | 1,426,053 | 29.80% | 1,037,303 | 26.24% | 1,070,129 | 26.48% |
| Upstream | 1,468,219 | 30.68% | 1,084,755 | 27.44% | 1,121,612 | 27.75% |
| **Genic** | **867,344** | **15.34%** | **424,898** | **9.70%** | **432,340** | **9.66%** |
| Exon | 101,029 | 11.65% | 82,456 | 19.41% | 83,363 | 19.28% |
| Intron | 444,663 | 51.27% | 299,239 | 70.43% | 305,146 | 70.58% |
| 5'UTR | 24,689 | 2.85% | 16,315 | 3.84% | 16,612 | 3.84% |
| 3'UTR | 40,441 | 4.66% | 26,888 | 6.33% | 27,219 | 6.30% |
| | **Number of indels** | **Relative percent** | **Number of indels** | **Relative percent** | **Number of indels** | **Relative percent** |
| **Indels with gene effects** | **101,475** | | **82,921** | | **83,768** | |
| conservative inframe deletion | 6,045 | 5.96% | 5,052 | 6.09% | 5,076 | 6.06% |
| conservative inframe insertion | 4,852 | 4.78% | 4,011 | 4.84% | 4,087 | 4.88% |
| disruptive inframe deletion | 10,498 | 10.35% | 8,652 | 10.43% | 8,686 | 10.37% |
| disruptive inframe insertion | 4,916 | 4.84% | 4,058 | 4.89% | 4,143 | 4.95% |
| exon loss variant | 78 | 0.08% | 37 | 0.04% | 67 | 0.08% |
| frameshift variant | 75,086 | 73.99% | 61,111 | 73.70% | 61,709 | 73.67% |
| | **Average (in Mb)** | **Range** | **Average (in Mb)** | **Range** | **Average (in Mb)** | **Range** |
| **Total Inserted** | +6.96 | +6.02 to +7.46 | +7.25 | +6.86 to +7.46 | +7.16 | +6.02 to +7.43 |
| **Total Deleted** | -10.97 | -12.80 to -10.55 | -10.72 | -11.15 to -10.62 | -10.74 | -12.34 to -10.55 |
| **Net Total** | -4.01 | -6.32 to -3.21 | -3.46 | -4.28 to -3.21 | -3.58 | -6.32 to -3.28 |

Supplementary Table 6. Nucleotide diversity (pi) by window for the combined species (both), *G. herbaceum* (A1), and *G. arboreum* (A2)

| CHROM | BIN_START | BIN_END | Pi (both) | Pi (A1) | Pi (A2) |
|---|---|---|---|---|---|
| F01 | 1 | 100000 | 0.0019 | 0.0012 | 0.0020 |
| F01 | 20001 | 120000 | 0.0021 | 0.0011 | 0.0022 |
| F01 | 40001 | 140000 | 0.0016 | 0.0005 | 0.0017 |
| F01 | 60001 | 160000 | 0.0013 | 0.0003 | 0.0014 |
| F01 | 80001 | 180000 | 0.0015 | 0.0004 | 0.0016 |
| F01 | 100001 | 200000 | 0.0016 | 0.0004 | 0.0015 |
| F01 | 120001 | 220000 | 0.0016 | 0.0003 | 0.0014 |
| F01 | 140001 | 240000 | 0.0015 | 0.0003 | 0.0013 |
| F01 | 160001 | 260000 | 0.0018 | 0.0003 | 0.0017 |
| F01 | 180001 | 280000 | 0.0017 | 0.0002 | 0.0015 |
| F01 | 200001 | 300000 | 0.0017 | 0.0002 | 0.0015 |
| F01 | 220001 | 320000 | 0.0017 | 0.0003 | 0.0016 |
| F01 | 240001 | 340000 | 0.0017 | 0.0003 | 0.0013 |
| F01 | 260001 | 360000 | 0.0019 | 0.0007 | 0.0013 |
| F01 | 280001 | 380000 | 0.0018 | 0.0006 | 0.0010 |
| F01 | 300001 | 400000 | 0.0019 | 0.0006 | 0.0008 |
| F01 | 320001 | 420000 | 0.0019 | 0.0009 | 0.0008 |
| F01 | 340001 | 440000 | 0.0021 | 0.0014 | 0.0012 |
| F01 | 360001 | 460000 | 0.0017 | 0.0018 | 0.0008 |
| F01 | 380001 | 480000 | 0.0015 | 0.0022 | 0.0008 |
| F01 | 400001 | 500000 | 0.0014 | 0.0028 | 0.0008 |
| F01 | 420001 | 520000 | 0.0012 | 0.0028 | 0.0006 |
| F01 | 440001 | 540000 | 0.0011 | 0.0027 | 0.0004 |
| F01 | 460001 | 560000 | 0.0012 | 0.0024 | 0.0006 |
| F01 | 480001 | 580000 | 0.0018 | 0.0028 | 0.0013 |
| F01 | 500001 | 600000 | 0.0021 | 0.0027 | 0.0017 |
| F01 | 520001 | 620000 | 0.0022 | 0.0026 | 0.0018 |
| F01 | 540001 | 640000 | 0.0026 | 0.0029 | 0.0022 |
| F01 | 560001 | 660000 | 0.0027 | 0.0026 | 0.0024 |
| F01 | 580001 | 680000 | 0.0023 | 0.0019 | 0.0019 |
| F01 | 600001 | 700000 | 0.0021 | 0.0013 | 0.0015 |
| F01 | 620001 | 720000 | 0.0022 | 0.0011 | 0.0016 |
| F01 | 640001 | 740000 | 0.0018 | 0.0013 | 0.0011 |
| F01 | 660001 | 760000 | 0.0015 | 0.0017 | 0.0007 |
| F01 | 680001 | 780000 | 0.0014 | 0.0019 | 0.0005 |
| F01 | 700001 | 800000 | 0.0022 | 0.0028 | 0.0014 |
| F01 | 720001 | 820000 | 0.0022 | 0.0028 | 0.0013 |
| F01 | 740001 | 840000 | 0.0032 | 0.0033 | 0.0024 |
| F01 | 760001 | 860000 | 0.0035 | 0.0029 | 0.0024 |
| F01 | 780001 | 880000 | 0.0049 | 0.0038 | 0.0037 |
| F01 | 800001 | 900000 | 0.0047 | 0.0037 | 0.0036 |
| F01 | 820001 | 920000 | 0.0051 | 0.0041 | 0.0042 |
| F01 | 840001 | 940000 | 0.0045 | 0.0035 | 0.0036 |
| F01 | 860001 | 960000 | 0.0044 | 0.0034 | 0.0036 |
| F01 | 880001 | 980000 | 0.0030 | 0.0022 | 0.0024 |
| F01 | 900001 | 1000000 | 0.0024 | 0.0014 | 0.0016 |
| F01 | 920001 | 1020000 | 0.0019 | 0.0009 | 0.0010 |
| F01 | 940001 | 1040000 | 0.0014 | 0.0002 | 0.0004 |
| F01 | 960001 | 1060000 | 0.0016 | 0.0003 | 0.0005 |
| F01 | 980001 | 1080000 | 0.0019 | 0.0004 | 0.0008 |
| F01 | 1000001 | 1100000 | 0.0020 | 0.0006 | 0.0009 |
| F01 | 1020001 | 1120000 | 0.0023 | 0.0013 | 0.0013 |
| F01 | 1040001 | 1140000 | 0.0027 | 0.0021 | 0.0019 |
| F01 | 1060001 | 1160000 | 0.0025 | 0.0024 | 0.0018 |

| | A1 | A2 | A1+A2 |
|---|---|---|---|
| Minimum | 0.0000 | 0.0000 | 0.0000 |
| Maximum | 0.0147 | 0.0169 | 0.0170 |
| Mean | 0.0022 | 0.0024 | 0.0029 |
| 1st quartile | 0.0014 | 0.0015 | 0.0019 |
| 3rd quartil | 0.0028 | 0.0030 | 0.0036 |

| Chr | Start | End | both | A1 | A2 |
|---|---|---|---|---|---|
| F01 | 1080001 | 1180000 | 0.0023 | 0.0023 | 0.0014 |
| F01 | 1100001 | 1200000 | 0.0022 | 0.0022 | 0.0013 |
| F01 | 1120001 | 1220000 | 0.0018 | 0.0018 | 0.0009 |
| F01 | 1140001 | 1240000 | 0.0012 | 0.0014 | 0.0003 |
| F01 | 1160001 | 1260000 | 0.0012 | 0.0018 | 0.0003 |
| F01 | 1180001 | 1280000 | 0.0013 | 0.0028 | 0.0003 |
| F01 | 1200001 | 1300000 | 0.0025 | 0.0045 | 0.0017 |
| F01 | 1220001 | 1320000 | 0.0039 | 0.0059 | 0.0031 |
| F01 | 1240001 | 1340000 | 0.0048 | 0.0067 | 0.0041 |
| F01 | 1260001 | 1360000 | 0.0048 | 0.0065 | 0.0041 |
| F01 | 1280001 | 1380000 | 0.0050 | 0.0055 | 0.0045 |
| F01 | 1300001 | 1400000 | 0.0039 | 0.0037 | 0.0035 |
| F01 | 1320001 | 1420000 | 0.0029 | 0.0021 | 0.0021 |
| F01 | 1340001 | 1440000 | 0.0024 | 0.0010 | 0.0013 |
| F01 | 1360001 | 1460000 | 0.0027 | 0.0004 | 0.0014 |
| F01 | 1380001 | 1480000 | 0.0024 | 0.0002 | 0.0010 |
| F01 | 1400001 | 1500000 | 0.0026 | 0.0003 | 0.0009 |
| F01 | 1420001 | 1520000 | 0.0024 | 0.0002 | 0.0008 |
| F01 | 1440001 | 1540000 | 0.0022 | 0.0002 | 0.0006 |
| F01 | 1460001 | 1560000 | 0.0022 | 0.0004 | 0.0007 |
| F01 | 1480001 | 1580000 | 0.0023 | 0.0004 | 0.0008 |
| F01 | 1500001 | 1600000 | 0.0023 | 0.0005 | 0.0007 |
| F01 | 1520001 | 1620000 | 0.0021 | 0.0009 | 0.0007 |
| F01 | 1540001 | 1640000 | 0.0019 | 0.0011 | 0.0007 |
| F01 | 1560001 | 1660000 | 0.0014 | 0.0013 | 0.0005 |
| F01 | 1580001 | 1680000 | 0.0015 | 0.0014 | 0.0007 |
| F01 | 1600001 | 1700000 | 0.0013 | 0.0013 | 0.0009 |
| F01 | 1620001 | 1720000 | 0.0011 | 0.0009 | 0.0009 |
| F01 | 1640001 | 1740000 | 0.0011 | 0.0006 | 0.0009 |
| F01 | 1660001 | 1760000 | 0.0010 | 0.0003 | 0.0008 |
| F01 | 1680001 | 1780000 | 0.0006 | 0.0001 | 0.0005 |
| F01 | 1700001 | 1800000 | 0.0007 | 0.0003 | 0.0006 |
| F01 | 1720001 | 1820000 | 0.0010 | 0.0003 | 0.0008 |
| F01 | 1740001 | 1840000 | 0.0015 | 0.0003 | 0.0013 |
| F01 | 1760001 | 1860000 | 0.0018 | 0.0003 | 0.0015 |
| F01 | 1780001 | 1880000 | 0.0020 | 0.0005 | 0.0016 |
| F01 | 1800001 | 1900000 | 0.0019 | 0.0004 | 0.0015 |
| F01 | 1820001 | 1920000 | 0.0017 | 0.0007 | 0.0014 |
| F01 | 1840001 | 1940000 | 0.0015 | 0.0009 | 0.0012 |
| F01 | 1860001 | 1960000 | 0.0012 | 0.0008 | 0.0010 |
| F01 | 1880001 | 1980000 | 0.0009 | 0.0007 | 0.0008 |
| F01 | 1900001 | 2000000 | 0.0005 | 0.0005 | 0.0005 |
| F01 | 1920001 | 2020000 | 0.0005 | 0.0003 | 0.0005 |
| F01 | 1940001 | 2040000 | 0.0004 | 0.0003 | 0.0004 |
| F01 | 1960001 | 2060000 | 0.0004 | 0.0003 | 0.0004 |
| F01 | 1980001 | 2080000 | 0.0007 | 0.0005 | 0.0007 |
| F01 | 2000001 | 2100000 | 0.0014 | 0.0009 | 0.0014 |
| F01 | 2020001 | 2120000 | 0.0017 | 0.0011 | 0.0017 |
| F01 | 2040001 | 2140000 | 0.0019 | 0.0013 | 0.0020 |
| F01 | 2060001 | 2160000 | 0.0021 | 0.0014 | 0.0021 |
| F01 | 2080001 | 2180000 | 0.0023 | 0.0012 | 0.0024 |
| F01 | 2100001 | 2200000 | 0.0021 | 0.0008 | 0.0022 |
| F01 | 2120001 | 2220000 | 0.0023 | 0.0006 | 0.0024 |
| F01 | 2140001 | 2240000 | 0.0021 | 0.0003 | 0.0022 |
| F01 | 2160001 | 2260000 | 0.0023 | 0.0003 | 0.0023 |
| F01 | 2180001 | 2280000 | 0.0021 | 0.0003 | 0.0020 |
| F01 | 2200001 | 2300000 | 0.0019 | 0.0003 | 0.0017 |
| F01 | 2220001 | 2320000 | 0.0016 | 0.0003 | 0.0012 |

| Chr | Start | End | both | A1 | A2 |
|---|---|---|---|---|---|
| F01 | 2240001 | 2340000 | 0.0017 | 0.0005 | 0.0010 |
| F01 | 2260001 | 2360000 | 0.0018 | 0.0009 | 0.0009 |
| F01 | 2280001 | 2380000 | 0.0019 | 0.0011 | 0.0008 |
| F01 | 2300001 | 2400000 | 0.0021 | 0.0014 | 0.0011 |
| F01 | 2320001 | 2420000 | 0.0023 | 0.0015 | 0.0015 |
| F01 | 2340001 | 2440000 | 0.0022 | 0.0013 | 0.0016 |
| F01 | 2360001 | 2460000 | 0.0022 | 0.0010 | 0.0018 |
| F01 | 2380001 | 2480000 | 0.0021 | 0.0008 | 0.0017 |
| F01 | 2400001 | 2500000 | 0.0018 | 0.0008 | 0.0013 |
| F01 | 2420001 | 2520000 | 0.0015 | 0.0010 | 0.0008 |
| F01 | 2440001 | 2540000 | 0.0016 | 0.0014 | 0.0009 |
| F01 | 2460001 | 2560000 | 0.0016 | 0.0019 | 0.0009 |
| F01 | 2480001 | 2580000 | 0.0016 | 0.0024 | 0.0010 |
| F01 | 2500001 | 2600000 | 0.0029 | 0.0036 | 0.0024 |
| F01 | 2520001 | 2620000 | 0.0048 | 0.0053 | 0.0045 |
| F01 | 2540001 | 2640000 | 0.0052 | 0.0057 | 0.0049 |
| F01 | 2560001 | 2660000 | 0.0049 | 0.0053 | 0.0047 |
| F01 | 2580001 | 2680000 | 0.0051 | 0.0052 | 0.0050 |
| F01 | 2600001 | 2700000 | 0.0045 | 0.0046 | 0.0043 |
| F01 | 2620001 | 2720000 | 0.0044 | 0.0044 | 0.0042 |
| F01 | 2640001 | 2740000 | 0.0041 | 0.0042 | 0.0039 |
| F01 | 2660001 | 2760000 | 0.0040 | 0.0042 | 0.0038 |
| F01 | 2680001 | 2780000 | 0.0037 | 0.0042 | 0.0034 |
| F01 | 2700001 | 2800000 | 0.0029 | 0.0035 | 0.0026 |
| F01 | 2720001 | 2820000 | 0.0011 | 0.0020 | 0.0007 |
| F01 | 2740001 | 2840000 | 0.0009 | 0.0019 | 0.0004 |
| F01 | 2760001 | 2860000 | 0.0011 | 0.0021 | 0.0007 |
| F01 | 2780001 | 2880000 | 0.0014 | 0.0022 | 0.0010 |
| F01 | 2800001 | 2900000 | 0.0013 | 0.0020 | 0.0010 |
| F01 | 2820001 | 2920000 | 0.0012 | 0.0017 | 0.0009 |
| F01 | 2840001 | 2940000 | 0.0012 | 0.0016 | 0.0010 |
| F01 | 2860001 | 2960000 | 0.0023 | 0.0025 | 0.0021 |
| F01 | 2880001 | 2980000 | 0.0022 | 0.0021 | 0.0018 |
| F01 | 2900001 | 3000000 | 0.0028 | 0.0024 | 0.0023 |
| F01 | 2920001 | 3020000 | 0.0031 | 0.0025 | 0.0025 |
| F01 | 2940001 | 3040000 | 0.0031 | 0.0022 | 0.0024 |
| F01 | 2960001 | 3060000 | 0.0022 | 0.0011 | 0.0014 |
| F01 | 2980001 | 3080000 | 0.0021 | 0.0011 | 0.0014 |
| F01 | 3000001 | 3100000 | 0.0017 | 0.0010 | 0.0010 |
| F01 | 3020001 | 3120000 | 0.0018 | 0.0015 | 0.0011 |
| F01 | 3040001 | 3140000 | 0.0021 | 0.0021 | 0.0015 |
| F01 | 3060001 | 3160000 | 0.0020 | 0.0023 | 0.0015 |
| F01 | 3080001 | 3180000 | 0.0022 | 0.0025 | 0.0018 |
| F01 | 3100001 | 3200000 | 0.0022 | 0.0025 | 0.0018 |
| F01 | 3120001 | 3220000 | 0.0022 | 0.0022 | 0.0019 |
| F01 | 3140001 | 3240000 | 0.0020 | 0.0018 | 0.0017 |
| F01 | 3160001 | 3260000 | 0.0027 | 0.0025 | 0.0025 |
| F01 | 3180001 | 3280000 | 0.0038 | 0.0034 | 0.0037 |
| F01 | 3200001 | 3300000 | 0.0045 | 0.0039 | 0.0045 |
| F01 | 3220001 | 3320000 | 0.0046 | 0.0041 | 0.0046 |
| F01 | 3240001 | 3340000 | 0.0047 | 0.0042 | 0.0047 |
| F01 | 3260001 | 3360000 | 0.0046 | 0.0041 | 0.0046 |
| F01 | 3280001 | 3380000 | 0.0041 | 0.0036 | 0.0041 |
| F01 | 3300001 | 3400000 | 0.0044 | 0.0037 | 0.0044 |
| F01 | 3320001 | 3420000 | 0.0048 | 0.0040 | 0.0048 |
| F01 | 3340001 | 3440000 | 0.0047 | 0.0038 | 0.0047 |
| F01 | 3360001 | 3460000 | 0.0041 | 0.0032 | 0.0041 |
| F01 | 3380001 | 3480000 | 0.0035 | 0.0029 | 0.0035 |

| Chr | Start | End | both | A1 | A2 |
|---|---|---|---|---|---|
| F01 | 3400001 | 3500000 | 0.0027 | 0.0023 | 0.0026 |
| F01 | 3420001 | 3520000 | 0.0029 | 0.0028 | 0.0028 |
| F01 | 3440001 | 3540000 | 0.0041 | 0.0039 | 0.0039 |
| F01 | 3460001 | 3560000 | 0.0039 | 0.0039 | 0.0037 |
| F01 | 3480001 | 3580000 | 0.0048 | 0.0048 | 0.0046 |
| F01 | 3500001 | 3600000 | 0.0059 | 0.0060 | 0.0056 |
| F01 | 3520001 | 3620000 | 0.0061 | 0.0061 | 0.0058 |
| F01 | 3540001 | 3640000 | 0.0058 | 0.0059 | 0.0056 |
| F01 | 3560001 | 3660000 | 0.0082 | 0.0080 | 0.0079 |
| F01 | 3580001 | 3680000 | 0.0075 | 0.0072 | 0.0073 |
| F01 | 3600001 | 3700000 | 0.0064 | 0.0060 | 0.0063 |
| F01 | 3620001 | 3720000 | 0.0063 | 0.0060 | 0.0061 |
| F01 | 3640001 | 3740000 | 0.0054 | 0.0051 | 0.0053 |
| F01 | 3660001 | 3760000 | 0.0033 | 0.0035 | 0.0032 |
| F01 | 3680001 | 3780000 | 0.0033 | 0.0035 | 0.0031 |
| F01 | 3700001 | 3800000 | 0.0045 | 0.0047 | 0.0043 |
| F01 | 3720001 | 3820000 | 0.0035 | 0.0038 | 0.0033 |
| F01 | 3740001 | 3840000 | 0.0034 | 0.0036 | 0.0031 |
| F01 | 3760001 | 3860000 | 0.0030 | 0.0030 | 0.0028 |
| F01 | 3780001 | 3880000 | 0.0031 | 0.0029 | 0.0029 |
| F01 | 3800001 | 3900000 | 0.0016 | 0.0015 | 0.0015 |
| F01 | 3820001 | 3920000 | 0.0029 | 0.0020 | 0.0029 |
| F01 | 3840001 | 3940000 | 0.0033 | 0.0020 | 0.0033 |
| F01 | 3860001 | 3960000 | 0.0032 | 0.0019 | 0.0033 |
| F01 | 3880001 | 3980000 | 0.0027 | 0.0014 | 0.0028 |
| F01 | 3900001 | 4000000 | 0.0036 | 0.0025 | 0.0037 |
| F01 | 3920001 | 4020000 | 0.0022 | 0.0016 | 0.0022 |
| F01 | 3940001 | 4040000 | 0.0016 | 0.0015 | 0.0016 |
| F01 | 3960001 | 4060000 | 0.0016 | 0.0014 | 0.0015 |
| F01 | 3980001 | 4080000 | 0.0015 | 0.0014 | 0.0015 |
| F01 | 4000001 | 4100000 | 0.0006 | 0.0003 | 0.0007 |
| F01 | 4020001 | 4120000 | 0.0006 | 0.0003 | 0.0007 |
| F01 | 4040001 | 4140000 | 0.0022 | 0.0018 | 0.0022 |
| F01 | 4060001 | 4160000 | 0.0022 | 0.0019 | 0.0022 |
| F01 | 4080001 | 4180000 | 0.0042 | 0.0039 | 0.0042 |
| F01 | 4100001 | 4200000 | 0.0079 | 0.0078 | 0.0078 |
| F01 | 4120001 | 4220000 | 0.0093 | 0.0093 | 0.0092 |
| F01 | 4140001 | 4240000 | 0.0079 | 0.0079 | 0.0079 |
| F01 | 4160001 | 4260000 | 0.0080 | 0.0077 | 0.0080 |
| F01 | 4180001 | 4280000 | 0.0061 | 0.0058 | 0.0061 |
| F01 | 4200001 | 4300000 | 0.0028 | 0.0023 | 0.0029 |
| F01 | 4220001 | 4320000 | 0.0027 | 0.0020 | 0.0028 |
| F01 | 4240001 | 4340000 | 0.0032 | 0.0022 | 0.0032 |
| F01 | 4260001 | 4360000 | 0.0032 | 0.0024 | 0.0033 |
| F01 | 4280001 | 4380000 | 0.0033 | 0.0026 | 0.0033 |
| F01 | 4300001 | 4400000 | 0.0037 | 0.0029 | 0.0036 |
| F01 | 4320001 | 4420000 | 0.0029 | 0.0020 | 0.0027 |
| F01 | 4340001 | 4440000 | 0.0030 | 0.0019 | 0.0027 |
| F01 | 4360001 | 4460000 | 0.0041 | 0.0026 | 0.0038 |
| F01 | 4380001 | 4480000 | 0.0047 | 0.0029 | 0.0044 |
| F01 | 4400001 | 4500000 | 0.0044 | 0.0025 | 0.0041 |
| F01 | 4420001 | 4520000 | 0.0048 | 0.0029 | 0.0046 |
| F01 | 4440001 | 4540000 | 0.0052 | 0.0035 | 0.0049 |
| F01 | 4460001 | 4560000 | 0.0053 | 0.0038 | 0.0051 |
| F01 | 4480001 | 4580000 | 0.0046 | 0.0034 | 0.0044 |
| F01 | 4500001 | 4600000 | 0.0047 | 0.0034 | 0.0043 |
| F01 | 4520001 | 4620000 | 0.0041 | 0.0028 | 0.0036 |
| F01 | 4540001 | 4640000 | 0.0034 | 0.0020 | 0.0027 |

| Chr | Start | End | both | A1 | A2 |
|---|---|---|---|---|---|
| F01 | 4560001 | 4660000 | 0.0021 | 0.0009 | 0.0015 |
| F01 | 4580001 | 4680000 | 0.0027 | 0.0015 | 0.0020 |
| F01 | 4600001 | 4700000 | 0.0022 | 0.0011 | 0.0015 |
| F01 | 4620001 | 4720000 | 0.0022 | 0.0011 | 0.0014 |
| F01 | 4640001 | 4740000 | 0.0032 | 0.0013 | 0.0022 |
| F01 | 4660001 | 4760000 | 0.0042 | 0.0015 | 0.0029 |
| F01 | 4680001 | 4780000 | 0.0038 | 0.0012 | 0.0026 |
| F01 | 4700001 | 4800000 | 0.0054 | 0.0019 | 0.0042 |
| F01 | 4720001 | 4820000 | 0.0059 | 0.0022 | 0.0049 |
| F01 | 4740001 | 4840000 | 0.0053 | 0.0021 | 0.0045 |
| F01 | 4760001 | 4860000 | 0.0043 | 0.0019 | 0.0037 |
| F01 | 4780001 | 4880000 | 0.0048 | 0.0019 | 0.0044 |
| F01 | 4800001 | 4900000 | 0.0035 | 0.0014 | 0.0030 |
| F01 | 4820001 | 4920000 | 0.0029 | 0.0011 | 0.0025 |
| F01 | 4840001 | 4940000 | 0.0023 | 0.0008 | 0.0020 |
| F01 | 4860001 | 4960000 | 0.0025 | 0.0007 | 0.0022 |
| F01 | 4880001 | 4980000 | 0.0021 | 0.0008 | 0.0018 |
| F01 | 4900001 | 5000000 | 0.0025 | 0.0010 | 0.0023 |
| F01 | 4920001 | 5020000 | 0.0027 | 0.0012 | 0.0025 |
| F01 | 4940001 | 5040000 | 0.0044 | 0.0027 | 0.0042 |
| F01 | 4960001 | 5060000 | 0.0045 | 0.0028 | 0.0043 |
| F01 | 4980001 | 5080000 | 0.0051 | 0.0035 | 0.0048 |
| F01 | 5000001 | 5100000 | 0.0048 | 0.0032 | 0.0044 |
| F01 | 5020001 | 5120000 | 0.0046 | 0.0032 | 0.0041 |
| F01 | 5040001 | 5140000 | 0.0031 | 0.0018 | 0.0025 |
| F01 | 5060001 | 5160000 | 0.0032 | 0.0018 | 0.0024 |
| F01 | 5080001 | 5180000 | 0.0021 | 0.0009 | 0.0013 |
| F01 | 5100001 | 5200000 | 0.0020 | 0.0009 | 0.0013 |
| F01 | 5120001 | 5220000 | 0.0032 | 0.0019 | 0.0025 |
| F01 | 5140001 | 5240000 | 0.0044 | 0.0027 | 0.0038 |
| F01 | 5160001 | 5260000 | 0.0052 | 0.0034 | 0.0048 |
| F01 | 5180001 | 5280000 | 0.0062 | 0.0045 | 0.0057 |
| F01 | 5200001 | 5300000 | 0.0061 | 0.0045 | 0.0057 |
| F01 | 5220001 | 5320000 | 0.0054 | 0.0040 | 0.0053 |
| F01 | 5240001 | 5340000 | 0.0050 | 0.0039 | 0.0049 |
| F01 | 5260001 | 5360000 | 0.0064 | 0.0046 | 0.0062 |
| F01 | 5280001 | 5380000 | 0.0070 | 0.0039 | 0.0069 |
| F01 | 5300001 | 5400000 | 0.0072 | 0.0041 | 0.0070 |
| F01 | 5320001 | 5420000 | 0.0069 | 0.0039 | 0.0068 |
| F01 | 5340001 | 5440000 | 0.0067 | 0.0037 | 0.0066 |
| F01 | 5360001 | 5460000 | 0.0048 | 0.0027 | 0.0048 |
| F01 | 5380001 | 5480000 | 0.0044 | 0.0035 | 0.0043 |
| F01 | 5400001 | 5500000 | 0.0043 | 0.0033 | 0.0043 |
| F01 | 5420001 | 5520000 | 0.0049 | 0.0038 | 0.0048 |
| F01 | 5440001 | 5540000 | 0.0048 | 0.0035 | 0.0047 |
| F01 | 5460001 | 5560000 | 0.0044 | 0.0028 | 0.0043 |
| F01 | 5480001 | 5580000 | 0.0034 | 0.0018 | 0.0033 |
| F01 | 5500001 | 5600000 | 0.0032 | 0.0016 | 0.0028 |
| F01 | 5520001 | 5620000 | 0.0023 | 0.0008 | 0.0019 |
| F01 | 5540001 | 5640000 | 0.0025 | 0.0011 | 0.0017 |
| F01 | 5560001 | 5660000 | 0.0022 | 0.0012 | 0.0014 |
| F01 | 5580001 | 5680000 | 0.0027 | 0.0016 | 0.0019 |
| F01 | 5600001 | 5700000 | 0.0035 | 0.0024 | 0.0029 |
| F01 | 5620001 | 5720000 | 0.0039 | 0.0030 | 0.0033 |
| F01 | 5640001 | 5740000 | 0.0032 | 0.0025 | 0.0030 |
| F01 | 5660001 | 5760000 | 0.0031 | 0.0024 | 0.0029 |
| F01 | 5680001 | 5780000 | 0.0032 | 0.0025 | 0.0029 |
| F01 | 5700001 | 5800000 | 0.0032 | 0.0020 | 0.0024 |

| Chr | Start | End | both | A1 | A2 |
|---|---|---|---|---|---|
| F01 | 5720001 | 5820000 | 0.0038 | 0.0023 | 0.0026 |
| F01 | 5740001 | 5840000 | 0.0042 | 0.0022 | 0.0027 |
| F01 | 5760001 | 5860000 | 0.0049 | 0.0026 | 0.0033 |
| F01 | 5780001 | 5880000 | 0.0045 | 0.0023 | 0.0030 |
| F01 | 5800001 | 5900000 | 0.0045 | 0.0027 | 0.0033 |
| F01 | 5820001 | 5920000 | 0.0035 | 0.0021 | 0.0028 |
| F01 | 5840001 | 5940000 | 0.0031 | 0.0021 | 0.0027 |
| F01 | 5860001 | 5960000 | 0.0026 | 0.0018 | 0.0024 |
| F01 | 5880001 | 5980000 | 0.0027 | 0.0015 | 0.0024 |
| F01 | 5900001 | 6000000 | 0.0023 | 0.0011 | 0.0021 |
| F01 | 5920001 | 6020000 | 0.0025 | 0.0011 | 0.0023 |
| F01 | 5940001 | 6040000 | 0.0025 | 0.0010 | 0.0022 |
| F01 | 5960001 | 6060000 | 0.0025 | 0.0010 | 0.0022 |
| F01 | 5980001 | 6080000 | 0.0029 | 0.0017 | 0.0026 |
| F01 | 6000001 | 6100000 | 0.0028 | 0.0018 | 0.0026 |
| F01 | 6020001 | 6120000 | 0.0030 | 0.0021 | 0.0028 |
| F01 | 6040001 | 6140000 | 0.0036 | 0.0030 | 0.0035 |
| F01 | 6060001 | 6160000 | 0.0038 | 0.0033 | 0.0037 |
| F01 | 6080001 | 6180000 | 0.0030 | 0.0026 | 0.0030 |
| F01 | 6100001 | 6200000 | 0.0034 | 0.0029 | 0.0034 |
| F01 | 6120001 | 6220000 | 0.0028 | 0.0024 | 0.0028 |
| F01 | 6140001 | 6240000 | 0.0025 | 0.0020 | 0.0025 |
| F01 | 6160001 | 6260000 | 0.0024 | 0.0019 | 0.0024 |
| F01 | 6180001 | 6280000 | 0.0026 | 0.0019 | 0.0026 |
| F01 | 6200001 | 6300000 | 0.0022 | 0.0012 | 0.0022 |
| F01 | 6220001 | 6320000 | 0.0024 | 0.0012 | 0.0025 |
| F01 | 6240001 | 6340000 | 0.0021 | 0.0008 | 0.0021 |
| F01 | 6260001 | 6360000 | 0.0019 | 0.0007 | 0.0019 |
| F01 | 6280001 | 6380000 | 0.0015 | 0.0006 | 0.0015 |
| F01 | 6300001 | 6400000 | 0.0016 | 0.0010 | 0.0016 |
| F01 | 6320001 | 6420000 | 0.0016 | 0.0011 | 0.0016 |
| F01 | 6340001 | 6440000 | 0.0015 | 0.0010 | 0.0016 |
| F01 | 6360001 | 6460000 | 0.0018 | 0.0009 | 0.0019 |
| F01 | 6380001 | 6480000 | 0.0023 | 0.0011 | 0.0023 |
| F01 | 6400001 | 6500000 | 0.0024 | 0.0008 | 0.0024 |
| F01 | 6420001 | 6520000 | 0.0025 | 0.0009 | 0.0026 |
| F01 | 6440001 | 6540000 | 0.0026 | 0.0010 | 0.0027 |
| F01 | 6460001 | 6560000 | 0.0024 | 0.0010 | 0.0025 |
| F01 | 6480001 | 6580000 | 0.0024 | 0.0008 | 0.0025 |
| F01 | 6500001 | 6600000 | 0.0025 | 0.0012 | 0.0026 |
| F01 | 6520001 | 6620000 | 0.0024 | 0.0009 | 0.0025 |
| F01 | 6540001 | 6640000 | 0.0026 | 0.0011 | 0.0027 |
| F01 | 6560001 | 6660000 | 0.0039 | 0.0026 | 0.0038 |
| F01 | 6580001 | 6680000 | 0.0056 | 0.0045 | 0.0054 |
| F01 | 6600001 | 6700000 | 0.0057 | 0.0045 | 0.0055 |
| F01 | 6620001 | 6720000 | 0.0056 | 0.0046 | 0.0051 |
| F01 | 6640001 | 6740000 | 0.0053 | 0.0044 | 0.0047 |
| F01 | 6660001 | 6760000 | 0.0048 | 0.0039 | 0.0041 |
| F01 | 6680001 | 6780000 | 0.0027 | 0.0019 | 0.0021 |
| F01 | 6700001 | 6800000 | 0.0022 | 0.0015 | 0.0014 |
| F01 | 6720001 | 6820000 | 0.0034 | 0.0026 | 0.0026 |
| F01 | 6740001 | 6840000 | 0.0040 | 0.0031 | 0.0033 |
| F01 | 6760001 | 6860000 | 0.0041 | 0.0030 | 0.0033 |
| F01 | 6780001 | 6880000 | 0.0047 | 0.0036 | 0.0039 |
| F01 | 6800001 | 6900000 | 0.0051 | 0.0040 | 0.0044 |
| F01 | 6820001 | 6920000 | 0.0056 | 0.0038 | 0.0049 |
| F01 | 6840001 | 6940000 | 0.0059 | 0.0038 | 0.0052 |
| F01 | 6860001 | 6960000 | 0.0049 | 0.0030 | 0.0044 |

| Chr | Start | End | both | A1 | A2 |
|---|---|---|---|---|---|
| F01 | 6880001 | 6980000 | 0.0048 | 0.0028 | 0.0042 |
| F01 | 6900001 | 7000000 | 0.0048 | 0.0026 | 0.0040 |
| F01 | 6920001 | 7020000 | 0.0030 | 0.0018 | 0.0023 |
| F01 | 6940001 | 7040000 | 0.0022 | 0.0015 | 0.0016 |
| F01 | 6960001 | 7060000 | 0.0031 | 0.0023 | 0.0025 |
| F01 | 6980001 | 7080000 | 0.0026 | 0.0021 | 0.0021 |
| F01 | 7000001 | 7100000 | 0.0025 | 0.0019 | 0.0021 |
| F01 | 7020001 | 7120000 | 0.0028 | 0.0020 | 0.0022 |
| F01 | 7040001 | 7140000 | 0.0026 | 0.0017 | 0.0019 |
| F01 | 7060001 | 7160000 | 0.0022 | 0.0012 | 0.0015 |
| F01 | 7080001 | 7180000 | 0.0025 | 0.0014 | 0.0016 |
| F01 | 7100001 | 7200000 | 0.0022 | 0.0011 | 0.0012 |
| F01 | 7120001 | 7220000 | 0.0024 | 0.0015 | 0.0016 |
| F01 | 7140001 | 7240000 | 0.0026 | 0.0018 | 0.0018 |
| F01 | 7160001 | 7260000 | 0.0021 | 0.0015 | 0.0014 |
| F01 | 7180001 | 7280000 | 0.0024 | 0.0020 | 0.0019 |
| F01 | 7200001 | 7300000 | 0.0023 | 0.0020 | 0.0020 |
| F01 | 7220001 | 7320000 | 0.0019 | 0.0018 | 0.0015 |
| F01 | 7240001 | 7340000 | 0.0017 | 0.0016 | 0.0013 |
| F01 | 7260001 | 7360000 | 0.0024 | 0.0023 | 0.0019 |
| F01 | 7280001 | 7380000 | 0.0026 | 0.0022 | 0.0019 |
| F01 | 7300001 | 7400000 | 0.0035 | 0.0031 | 0.0027 |
| F01 | 7320001 | 7420000 | 0.0040 | 0.0035 | 0.0033 |
| F01 | 7340001 | 7440000 | 0.0043 | 0.0040 | 0.0034 |
| F01 | 7360001 | 7460000 | 0.0038 | 0.0036 | 0.0028 |
| F01 | 7380001 | 7480000 | 0.0032 | 0.0033 | 0.0023 |
| F01 | 7400001 | 7500000 | 0.0028 | 0.0031 | 0.0019 |
| F01 | 7420001 | 7520000 | 0.0025 | 0.0027 | 0.0013 |
| F01 | 7440001 | 7540000 | 0.0022 | 0.0021 | 0.0012 |
| F01 | 7460001 | 7560000 | 0.0032 | 0.0029 | 0.0023 |
| F01 | 7480001 | 7580000 | 0.0036 | 0.0030 | 0.0028 |
| F01 | 7500001 | 7600000 | 0.0039 | 0.0031 | 0.0031 |
| F01 | 7520001 | 7620000 | 0.0037 | 0.0032 | 0.0032 |
| F01 | 7540001 | 7640000 | 0.0040 | 0.0034 | 0.0035 |
| F01 | 7560001 | 7660000 | 0.0029 | 0.0022 | 0.0022 |
| F01 | 7580001 | 7680000 | 0.0024 | 0.0017 | 0.0017 |
| F01 | 7600001 | 7700000 | 0.0017 | 0.0010 | 0.0009 |
| F01 | 7620001 | 7720000 | 0.0018 | 0.0006 | 0.0006 |
| F01 | 7640001 | 7740000 | 0.0016 | 0.0008 | 0.0002 |
| F01 | 7660001 | 7760000 | 0.0016 | 0.0011 | 0.0004 |
| F01 | 7680001 | 7780000 | 0.0022 | 0.0019 | 0.0010 |
| F01 | 7700001 | 7800000 | 0.0024 | 0.0021 | 0.0013 |
| F01 | 7720001 | 7820000 | 0.0020 | 0.0025 | 0.0013 |
| F01 | 7740001 | 7840000 | 0.0019 | 0.0024 | 0.0013 |
| F01 | 7760001 | 7860000 | 0.0022 | 0.0029 | 0.0015 |
| F01 | 7780001 | 7880000 | 0.0017 | 0.0025 | 0.0009 |
| F01 | 7800001 | 7900000 | 0.0017 | 0.0027 | 0.0009 |
| F01 | 7820001 | 7920000 | 0.0021 | 0.0030 | 0.0014 |
| F01 | 7840001 | 7940000 | 0.0021 | 0.0028 | 0.0014 |
| F01 | 7860001 | 7960000 | 0.0015 | 0.0020 | 0.0010 |
| F01 | 7880001 | 7980000 | 0.0014 | 0.0017 | 0.0011 |
| F01 | 7900001 | 8000000 | 0.0013 | 0.0014 | 0.0011 |
| F01 | 7920001 | 8020000 | 0.0007 | 0.0008 | 0.0006 |
| F01 | 7940001 | 8040000 | 0.0011 | 0.0012 | 0.0010 |
| F01 | 7960001 | 8060000 | 0.0013 | 0.0014 | 0.0012 |
| F01 | 7980001 | 8080000 | 0.0013 | 0.0014 | 0.0011 |
| F01 | 8000001 | 8100000 | 0.0017 | 0.0020 | 0.0014 |
| F01 | 8020001 | 8120000 | 0.0028 | 0.0030 | 0.0025 |

| Chr | Start | End | both | A1 | A2 |
|---|---|---|---|---|---|
| F01 | 8040001 | 8140000 | 0.0028 | 0.0030 | 0.0026 |
| F01 | 8060001 | 8160000 | 0.0029 | 0.0031 | 0.0025 |
| F01 | 8080001 | 8180000 | 0.0039 | 0.0041 | 0.0035 |
| F01 | 8100001 | 8200000 | 0.0036 | 0.0036 | 0.0034 |
| F01 | 8120001 | 8220000 | 0.0040 | 0.0040 | 0.0037 |
| F01 | 8140001 | 8240000 | 0.0042 | 0.0042 | 0.0037 |
| F01 | 8160001 | 8260000 | 0.0042 | 0.0041 | 0.0037 |
| F01 | 8180001 | 8280000 | 0.0033 | 0.0031 | 0.0027 |
| F01 | 8200001 | 8300000 | 0.0039 | 0.0034 | 0.0030 |
| F01 | 8220001 | 8320000 | 0.0028 | 0.0021 | 0.0018 |
| F01 | 8240001 | 8340000 | 0.0026 | 0.0015 | 0.0016 |
| F01 | 8260001 | 8360000 | 0.0028 | 0.0016 | 0.0017 |
| F01 | 8280001 | 8380000 | 0.0029 | 0.0017 | 0.0018 |
| F01 | 8300001 | 8400000 | 0.0026 | 0.0015 | 0.0016 |
| F01 | 8320001 | 8420000 | 0.0029 | 0.0018 | 0.0019 |
| F01 | 8340001 | 8440000 | 0.0029 | 0.0020 | 0.0021 |
| F01 | 8360001 | 8460000 | 0.0033 | 0.0024 | 0.0025 |
| F01 | 8380001 | 8480000 | 0.0034 | 0.0023 | 0.0024 |
| F01 | 8400001 | 8500000 | 0.0037 | 0.0025 | 0.0027 |
| F01 | 8420001 | 8520000 | 0.0043 | 0.0031 | 0.0034 |
| F01 | 8440001 | 8540000 | 0.0043 | 0.0029 | 0.0032 |
| F01 | 8460001 | 8560000 | 0.0040 | 0.0026 | 0.0029 |
| F01 | 8480001 | 8580000 | 0.0035 | 0.0025 | 0.0028 |
| F01 | 8500001 | 8600000 | 0.0028 | 0.0020 | 0.0022 |
| F01 | 8520001 | 8620000 | 0.0022 | 0.0014 | 0.0016 |
| F01 | 8540001 | 8640000 | 0.0031 | 0.0018 | 0.0022 |
| F01 | 8560001 | 8660000 | 0.0031 | 0.0018 | 0.0023 |
| F01 | 8580001 | 8680000 | 0.0044 | 0.0029 | 0.0035 |
| F01 | 8600001 | 8700000 | 0.0048 | 0.0032 | 0.0039 |
| F01 | 8620001 | 8720000 | 0.0053 | 0.0035 | 0.0040 |
| F01 | 8640001 | 8740000 | 0.0044 | 0.0030 | 0.0033 |
| F01 | 8660001 | 8760000 | 0.0044 | 0.0029 | 0.0032 |
| F01 | 8680001 | 8780000 | 0.0034 | 0.0021 | 0.0023 |
| F01 | 8700001 | 8800000 | 0.0035 | 0.0022 | 0.0023 |
| F01 | 8720001 | 8820000 | 0.0030 | 0.0015 | 0.0021 |
| F01 | 8740001 | 8840000 | 0.0033 | 0.0019 | 0.0024 |
| F01 | 8760001 | 8860000 | 0.0032 | 0.0018 | 0.0024 |
| F01 | 8780001 | 8880000 | 0.0031 | 0.0017 | 0.0022 |
| F01 | 8800001 | 8900000 | 0.0027 | 0.0014 | 0.0019 |
| F01 | 8820001 | 8920000 | 0.0022 | 0.0014 | 0.0015 |
| F01 | 8840001 | 8940000 | 0.0023 | 0.0015 | 0.0016 |
| F01 | 8860001 | 8960000 | 0.0021 | 0.0013 | 0.0014 |
| F01 | 8880001 | 8980000 | 0.0029 | 0.0020 | 0.0021 |
| F01 | 8900001 | 9000000 | 0.0027 | 0.0018 | 0.0019 |
| F01 | 8920001 | 9020000 | 0.0026 | 0.0018 | 0.0018 |
| F01 | 8940001 | 9040000 | 0.0019 | 0.0014 | 0.0013 |
| F01 | 8960001 | 9060000 | 0.0021 | 0.0017 | 0.0016 |
| F01 | 8980001 | 9080000 | 0.0012 | 0.0011 | 0.0008 |
| F01 | 9000001 | 9100000 | 0.0013 | 0.0012 | 0.0008 |
| F01 | 9020001 | 9120000 | 0.0013 | 0.0012 | 0.0009 |
| F01 | 9040001 | 9140000 | 0.0019 | 0.0018 | 0.0015 |
| F01 | 9060001 | 9160000 | 0.0021 | 0.0020 | 0.0017 |
| F01 | 9080001 | 9180000 | 0.0032 | 0.0028 | 0.0028 |
| F01 | 9100001 | 9200000 | 0.0042 | 0.0038 | 0.0040 |
| F01 | 9120001 | 9220000 | 0.0045 | 0.0040 | 0.0042 |
| F01 | 9140001 | 9240000 | 0.0042 | 0.0038 | 0.0038 |
| F01 | 9160001 | 9260000 | 0.0042 | 0.0038 | 0.0038 |
| F01 | 9180001 | 9280000 | 0.0033 | 0.0030 | 0.0028 |

| Chr | Start | End | both | A1 | A2 |
|---|---|---|---|---|---|
| F01 | 9200001 | 9300000 | 0.0027 | 0.0024 | 0.0021 |
| F01 | 9220001 | 9320000 | 0.0023 | 0.0020 | 0.0018 |
| F01 | 9240001 | 9340000 | 0.0021 | 0.0017 | 0.0017 |
| F01 | 9260001 | 9360000 | 0.0023 | 0.0019 | 0.0019 |
| F01 | 9280001 | 9380000 | 0.0023 | 0.0019 | 0.0018 |
| F01 | 9300001 | 9400000 | 0.0021 | 0.0019 | 0.0014 |
| F01 | 9320001 | 9420000 | 0.0024 | 0.0022 | 0.0015 |
| F01 | 9340001 | 9440000 | 0.0030 | 0.0027 | 0.0020 |
| F01 | 9360001 | 9460000 | 0.0026 | 0.0024 | 0.0016 |
| F01 | 9380001 | 9480000 | 0.0031 | 0.0029 | 0.0023 |
| F01 | 9400001 | 9500000 | 0.0035 | 0.0033 | 0.0029 |
| F01 | 9420001 | 9520000 | 0.0032 | 0.0032 | 0.0029 |
| F01 | 9440001 | 9540000 | 0.0027 | 0.0026 | 0.0025 |
| F01 | 9460001 | 9560000 | 0.0024 | 0.0023 | 0.0022 |
| F01 | 9480001 | 9580000 | 0.0015 | 0.0015 | 0.0014 |
| F01 | 9500001 | 9600000 | 0.0011 | 0.0009 | 0.0010 |
| F01 | 9520001 | 9620000 | 0.0022 | 0.0018 | 0.0021 |
| F01 | 9540001 | 9640000 | 0.0029 | 0.0024 | 0.0029 |
| F01 | 9560001 | 9660000 | 0.0032 | 0.0027 | 0.0032 |
| F01 | 9580001 | 9680000 | 0.0037 | 0.0030 | 0.0037 |
| F01 | 9600001 | 9700000 | 0.0036 | 0.0029 | 0.0035 |
| F01 | 9620001 | 9720000 | 0.0033 | 0.0027 | 0.0033 |
| F01 | 9640001 | 9740000 | 0.0031 | 0.0025 | 0.0031 |
| F01 | 9660001 | 9760000 | 0.0033 | 0.0028 | 0.0033 |
| F01 | 9680001 | 9780000 | 0.0028 | 0.0025 | 0.0028 |
| F01 | 9700001 | 9800000 | 0.0027 | 0.0023 | 0.0026 |
| F01 | 9720001 | 9820000 | 0.0020 | 0.0019 | 0.0020 |
| F01 | 9740001 | 9840000 | 0.0028 | 0.0026 | 0.0026 |
| F01 | 9760001 | 9860000 | 0.0043 | 0.0039 | 0.0040 |
| F01 | 9780001 | 9880000 | 0.0051 | 0.0047 | 0.0047 |
| F01 | 9800001 | 9900000 | 0.0054 | 0.0049 | 0.0050 |
| F01 | 9820001 | 9920000 | 0.0052 | 0.0049 | 0.0047 |
| F01 | 9840001 | 9940000 | 0.0039 | 0.0036 | 0.0032 |
| F01 | 9860001 | 9960000 | 0.0020 | 0.0019 | 0.0015 |
| F01 | 9880001 | 9980000 | 0.0013 | 0.0012 | 0.0008 |
| F01 | 9900001 | 10000000 | 0.0013 | 0.0011 | 0.0007 |
| F01 | 9920001 | 10020000 | 0.0025 | 0.0017 | 0.0018 |
| F01 | 9940001 | 10040000 | 0.0035 | 0.0027 | 0.0027 |
| F01 | 9960001 | 10060000 | 0.0035 | 0.0027 | 0.0026 |
| F01 | 9980001 | 10080000 | 0.0047 | 0.0037 | 0.0035 |
| F01 | 10000001 | 10100000 | 0.0050 | 0.0038 | 0.0036 |
| F01 | 10020001 | 10120000 | 0.0039 | 0.0027 | 0.0025 |
| F01 | 10040001 | 10140000 | 0.0029 | 0.0018 | 0.0017 |
| F01 | 10060001 | 10160000 | 0.0033 | 0.0021 | 0.0022 |
| F01 | 10080001 | 10180000 | 0.0023 | 0.0010 | 0.0013 |
| F01 | 10100001 | 10200000 | 0.0020 | 0.0009 | 0.0011 |
| F01 | 10120001 | 10220000 | 0.0020 | 0.0008 | 0.0011 |
| F01 | 10140001 | 10240000 | 0.0023 | 0.0011 | 0.0016 |
| F01 | 10160001 | 10260000 | 0.0025 | 0.0014 | 0.0017 |
| F01 | 10180001 | 10280000 | 0.0024 | 0.0014 | 0.0017 |
| F01 | 10200001 | 10300000 | 0.0025 | 0.0013 | 0.0017 |
| F01 | 10220001 | 10320000 | 0.0028 | 0.0015 | 0.0020 |
| F01 | 10240001 | 10340000 | 0.0027 | 0.0016 | 0.0018 |
| F01 | 10260001 | 10360000 | 0.0021 | 0.0013 | 0.0010 |
| F01 | 10280001 | 10380000 | 0.0023 | 0.0019 | 0.0012 |
| F01 | 10300001 | 10400000 | 0.0022 | 0.0024 | 0.0011 |
| F01 | 10320001 | 10420000 | 0.0018 | 0.0025 | 0.0009 |
| F01 | 10340001 | 10440000 | 0.0019 | 0.0027 | 0.0012 |

| Chr | Start | End | both | A1 | A2 |
|---|---|---|---|---|---|
| F01 | 10360001 | 10460000 | 0.0026 | 0.0035 | 0.0020 |
| F01 | 10380001 | 10480000 | 0.0026 | 0.0031 | 0.0021 |
| F01 | 10400001 | 10500000 | 0.0038 | 0.0040 | 0.0035 |
| F01 | 10420001 | 10520000 | 0.0038 | 0.0041 | 0.0034 |
| F01 | 10440001 | 10540000 | 0.0039 | 0.0045 | 0.0035 |
| F01 | 10460001 | 10560000 | 0.0032 | 0.0039 | 0.0027 |
| F01 | 10480001 | 10580000 | 0.0035 | 0.0045 | 0.0030 |
| F01 | 10500001 | 10600000 | 0.0023 | 0.0036 | 0.0017 |
| F01 | 10520001 | 10620000 | 0.0029 | 0.0040 | 0.0024 |
| F01 | 10540001 | 10640000 | 0.0030 | 0.0042 | 0.0025 |
| F01 | 10560001 | 10660000 | 0.0030 | 0.0039 | 0.0025 |
| F01 | 10580001 | 10680000 | 0.0029 | 0.0037 | 0.0025 |
| F01 | 10600001 | 10700000 | 0.0058 | 0.0064 | 0.0054 |
| F01 | 10620001 | 10720000 | 0.0056 | 0.0063 | 0.0051 |
| F01 | 10640001 | 10740000 | 0.0050 | 0.0054 | 0.0046 |
| F01 | 10660001 | 10760000 | 0.0049 | 0.0055 | 0.0045 |
| F01 | 10680001 | 10780000 | 0.0048 | 0.0057 | 0.0043 |
| F01 | 10700001 | 10800000 | 0.0018 | 0.0031 | 0.0013 |
| F01 | 10720001 | 10820000 | 0.0013 | 0.0023 | 0.0009 |
| F01 | 10740001 | 10840000 | 0.0013 | 0.0027 | 0.0008 |
| F01 | 10760001 | 10860000 | 0.0014 | 0.0027 | 0.0008 |
| F01 | 10780001 | 10880000 | 0.0013 | 0.0025 | 0.0009 |
| F01 | 10800001 | 10900000 | 0.0013 | 0.0023 | 0.0010 |
| F01 | 10820001 | 10920000 | 0.0014 | 0.0025 | 0.0010 |
| F01 | 10840001 | 10940000 | 0.0021 | 0.0032 | 0.0016 |
| F01 | 10860001 | 10960000 | 0.0024 | 0.0033 | 0.0019 |
| F01 | 10880001 | 10980000 | 0.0028 | 0.0035 | 0.0025 |
| F01 | 10900001 | 11000000 | 0.0038 | 0.0045 | 0.0034 |
| F01 | 10920001 | 11020000 | 0.0039 | 0.0046 | 0.0035 |
| F01 | 10940001 | 11040000 | 0.0036 | 0.0039 | 0.0032 |
| F01 | 10960001 | 11060000 | 0.0034 | 0.0040 | 0.0030 |
| F01 | 10980001 | 11080000 | 0.0029 | 0.0036 | 0.0024 |
| F01 | 11000001 | 11100000 | 0.0020 | 0.0026 | 0.0017 |
| F01 | 11020001 | 11120000 | 0.0022 | 0.0026 | 0.0018 |
| F01 | 11040001 | 11140000 | 0.0020 | 0.0030 | 0.0015 |
| F01 | 11060001 | 11160000 | 0.0019 | 0.0030 | 0.0014 |
| F01 | 11080001 | 11180000 | 0.0019 | 0.0031 | 0.0013 |
| F01 | 11100001 | 11200000 | 0.0020 | 0.0034 | 0.0012 |
| F01 | 11120001 | 11220000 | 0.0018 | 0.0033 | 0.0010 |
| F01 | 11140001 | 11240000 | 0.0014 | 0.0022 | 0.0008 |
| F01 | 11160001 | 11260000 | 0.0014 | 0.0017 | 0.0008 |
| F01 | 11180001 | 11280000 | 0.0013 | 0.0013 | 0.0007 |
| F01 | 11200001 | 11300000 | 0.0012 | 0.0008 | 0.0007 |
| F01 | 11220001 | 11320000 | 0.0015 | 0.0010 | 0.0010 |
| F01 | 11240001 | 11340000 | 0.0016 | 0.0010 | 0.0010 |
| F01 | 11260001 | 11360000 | 0.0016 | 0.0010 | 0.0010 |
| F01 | 11280001 | 11380000 | 0.0017 | 0.0011 | 0.0012 |
| F01 | 11300001 | 11400000 | 0.0016 | 0.0010 | 0.0010 |
| F01 | 11320001 | 11420000 | 0.0014 | 0.0008 | 0.0010 |
| F01 | 11340001 | 11440000 | 0.0015 | 0.0009 | 0.0011 |
| F01 | 11360001 | 11460000 | 0.0017 | 0.0011 | 0.0013 |
| F01 | 11380001 | 11480000 | 0.0017 | 0.0012 | 0.0013 |
| F01 | 11400001 | 11500000 | 0.0018 | 0.0013 | 0.0015 |
| F01 | 11420001 | 11520000 | 0.0018 | 0.0013 | 0.0014 |
| F01 | 11440001 | 11540000 | 0.0018 | 0.0014 | 0.0015 |
| F01 | 11460001 | 11560000 | 0.0023 | 0.0018 | 0.0020 |
| F01 | 11480001 | 11580000 | 0.0029 | 0.0022 | 0.0025 |
| F01 | 11500001 | 11600000 | 0.0040 | 0.0029 | 0.0034 |

| Chr | Start | End | both | A1 | A2 |
|---|---|---|---|---|---|
| F01 | 11520001 | 11620000 | 0.0042 | 0.0030 | 0.0034 |
| F01 | 11540001 | 11640000 | 0.0050 | 0.0037 | 0.0043 |
| F01 | 11560001 | 11660000 | 0.0047 | 0.0033 | 0.0038 |
| F01 | 11580001 | 11680000 | 0.0043 | 0.0033 | 0.0036 |
| F01 | 11600001 | 11700000 | 0.0032 | 0.0026 | 0.0027 |
| F01 | 11620001 | 11720000 | 0.0029 | 0.0025 | 0.0025 |
| F01 | 11640001 | 11740000 | 0.0024 | 0.0020 | 0.0017 |
| F01 | 11660001 | 11760000 | 0.0027 | 0.0022 | 0.0020 |
| F01 | 11680001 | 11780000 | 0.0030 | 0.0023 | 0.0023 |
| F01 | 11700001 | 11800000 | 0.0032 | 0.0025 | 0.0024 |
| F01 | 11720001 | 11820000 | 0.0035 | 0.0024 | 0.0024 |
| F01 | 11740001 | 11840000 | 0.0030 | 0.0019 | 0.0021 |
| F01 | 11760001 | 11860000 | 0.0026 | 0.0015 | 0.0016 |
| F01 | 11780001 | 11880000 | 0.0027 | 0.0013 | 0.0016 |
| F01 | 11800001 | 11900000 | 0.0032 | 0.0015 | 0.0020 |
| F01 | 11820001 | 11920000 | 0.0032 | 0.0016 | 0.0020 |
| F01 | 11840001 | 11940000 | 0.0035 | 0.0019 | 0.0023 |
| F01 | 11860001 | 11960000 | 0.0037 | 0.0022 | 0.0027 |
| F01 | 11880001 | 11980000 | 0.0030 | 0.0018 | 0.0020 |
| F01 | 11900001 | 12000000 | 0.0027 | 0.0018 | 0.0019 |
| F01 | 11920001 | 12020000 | 0.0034 | 0.0023 | 0.0026 |
| F01 | 11940001 | 12040000 | 0.0031 | 0.0020 | 0.0024 |
| F01 | 11960001 | 12060000 | 0.0034 | 0.0021 | 0.0024 |
| F01 | 11980001 | 12080000 | 0.0031 | 0.0018 | 0.0021 |
| F01 | 12000001 | 12100000 | 0.0027 | 0.0013 | 0.0017 |
| F01 | 12020001 | 12120000 | 0.0017 | 0.0006 | 0.0007 |
| F01 | 12040001 | 12140000 | 0.0023 | 0.0010 | 0.0012 |
| F01 | 12060001 | 12160000 | 0.0022 | 0.0012 | 0.0014 |
| F01 | 12080001 | 12180000 | 0.0024 | 0.0014 | 0.0016 |
| F01 | 12100001 | 12200000 | 0.0023 | 0.0015 | 0.0016 |
| F01 | 12120001 | 12220000 | 0.0025 | 0.0016 | 0.0018 |
| F01 | 12140001 | 12240000 | 0.0021 | 0.0012 | 0.0014 |
| F01 | 12160001 | 12260000 | 0.0017 | 0.0007 | 0.0010 |
| F01 | 12180001 | 12280000 | 0.0019 | 0.0007 | 0.0009 |
| F01 | 12200001 | 12300000 | 0.0021 | 0.0009 | 0.0011 |
| F01 | 12220001 | 12320000 | 0.0027 | 0.0015 | 0.0016 |
| F01 | 12240001 | 12340000 | 0.0029 | 0.0018 | 0.0019 |
| F01 | 12260001 | 12360000 | 0.0027 | 0.0018 | 0.0018 |
| F01 | 12280001 | 12380000 | 0.0024 | 0.0017 | 0.0016 |
| F01 | 12300001 | 12400000 | 0.0022 | 0.0015 | 0.0015 |
| F01 | 12320001 | 12420000 | 0.0020 | 0.0011 | 0.0014 |
| F01 | 12340001 | 12440000 | 0.0019 | 0.0011 | 0.0014 |
| F01 | 12360001 | 12460000 | 0.0020 | 0.0011 | 0.0014 |
| F01 | 12380001 | 12480000 | 0.0026 | 0.0015 | 0.0019 |
| F01 | 12400001 | 12500000 | 0.0027 | 0.0016 | 0.0021 |
| F01 | 12420001 | 12520000 | 0.0023 | 0.0013 | 0.0016 |
| F01 | 12440001 | 12540000 | 0.0021 | 0.0010 | 0.0014 |
| F01 | 12460001 | 12560000 | 0.0023 | 0.0013 | 0.0017 |
| F01 | 12480001 | 12580000 | 0.0020 | 0.0012 | 0.0015 |
| F01 | 12500001 | 12600000 | 0.0021 | 0.0011 | 0.0013 |
| F01 | 12520001 | 12620000 | 0.0021 | 0.0012 | 0.0013 |
| F01 | 12540001 | 12640000 | 0.0027 | 0.0016 | 0.0017 |
| F01 | 12560001 | 12660000 | 0.0024 | 0.0013 | 0.0014 |
| F01 | 12580001 | 12680000 | 0.0024 | 0.0012 | 0.0014 |
| F01 | 12600001 | 12700000 | 0.0025 | 0.0012 | 0.0015 |
| F01 | 12620001 | 12720000 | 0.0044 | 0.0029 | 0.0035 |
| F01 | 12640001 | 12740000 | 0.0039 | 0.0027 | 0.0032 |
| F01 | 12660001 | 12760000 | 0.0041 | 0.0029 | 0.0034 |

| Chr | Start | End | both | A1 | A2 |
|---|---|---|---|---|---|
| F01 | 12680001 | 12780000 | 0.0045 | 0.0034 | 0.0038 |
| F01 | 12700001 | 12800000 | 0.0045 | 0.0036 | 0.0040 |
| F01 | 12720001 | 12820000 | 0.0025 | 0.0021 | 0.0022 |
| F01 | 12740001 | 12840000 | 0.0025 | 0.0020 | 0.0022 |
| F01 | 12760001 | 12860000 | 0.0028 | 0.0022 | 0.0024 |
| F01 | 12780001 | 12880000 | 0.0024 | 0.0018 | 0.0020 |
| F01 | 12800001 | 12900000 | 0.0025 | 0.0019 | 0.0020 |
| F01 | 12820001 | 12920000 | 0.0022 | 0.0015 | 0.0017 |
| F01 | 12840001 | 12940000 | 0.0022 | 0.0015 | 0.0016 |
| F01 | 12860001 | 12960000 | 0.0025 | 0.0017 | 0.0018 |
| F01 | 12880001 | 12980000 | 0.0024 | 0.0016 | 0.0017 |
| F01 | 12900001 | 13000000 | 0.0021 | 0.0014 | 0.0015 |
| F01 | 12920001 | 13020000 | 0.0023 | 0.0017 | 0.0017 |
| F01 | 12940001 | 13040000 | 0.0032 | 0.0028 | 0.0028 |
| F01 | 12960001 | 13060000 | 0.0029 | 0.0023 | 0.0026 |
| F01 | 12980001 | 13080000 | 0.0031 | 0.0025 | 0.0029 |
| F01 | 13000001 | 13100000 | 0.0028 | 0.0022 | 0.0026 |
| F01 | 13020001 | 13120000 | 0.0032 | 0.0024 | 0.0029 |
| F01 | 13040001 | 13140000 | 0.0024 | 0.0015 | 0.0020 |
| F01 | 13060001 | 13160000 | 0.0023 | 0.0016 | 0.0017 |
| F01 | 13080001 | 13180000 | 0.0023 | 0.0016 | 0.0017 |
| F01 | 13100001 | 13200000 | 0.0026 | 0.0019 | 0.0019 |
| F01 | 13120001 | 13220000 | 0.0024 | 0.0018 | 0.0018 |
| F01 | 13140001 | 13240000 | 0.0023 | 0.0018 | 0.0016 |
| F01 | 13160001 | 13260000 | 0.0031 | 0.0022 | 0.0023 |
| F01 | 13180001 | 13280000 | 0.0038 | 0.0026 | 0.0028 |
| F01 | 13200001 | 13300000 | 0.0049 | 0.0029 | 0.0038 |
| F01 | 13220001 | 13320000 | 0.0049 | 0.0027 | 0.0035 |
| F01 | 13240001 | 13340000 | 0.0047 | 0.0023 | 0.0033 |
| F01 | 13260001 | 13360000 | 0.0042 | 0.0021 | 0.0028 |
| F01 | 13280001 | 13380000 | 0.0047 | 0.0027 | 0.0035 |
| F01 | 13300001 | 13400000 | 0.0041 | 0.0028 | 0.0030 |
| F01 | 13320001 | 13420000 | 0.0064 | 0.0051 | 0.0054 |
| F01 | 13340001 | 13440000 | 0.0066 | 0.0055 | 0.0058 |
| F01 | 13360001 | 13460000 | 0.0066 | 0.0055 | 0.0059 |
| F01 | 13380001 | 13480000 | 0.0053 | 0.0042 | 0.0044 |
| F01 | 13400001 | 13500000 | 0.0048 | 0.0037 | 0.0039 |
| F01 | 13420001 | 13520000 | 0.0023 | 0.0013 | 0.0016 |
| F01 | 13440001 | 13540000 | 0.0021 | 0.0011 | 0.0015 |
| F01 | 13460001 | 13560000 | 0.0016 | 0.0008 | 0.0012 |
| F01 | 13480001 | 13580000 | 0.0023 | 0.0016 | 0.0019 |
| F01 | 13500001 | 13600000 | 0.0025 | 0.0018 | 0.0020 |
| F01 | 13520001 | 13620000 | 0.0027 | 0.0020 | 0.0023 |
| F01 | 13540001 | 13640000 | 0.0027 | 0.0021 | 0.0024 |
| F01 | 13560001 | 13660000 | 0.0026 | 0.0019 | 0.0022 |
| F01 | 13580001 | 13680000 | 0.0024 | 0.0016 | 0.0019 |
| F01 | 13600001 | 13700000 | 0.0026 | 0.0016 | 0.0019 |
| F01 | 13620001 | 13720000 | 0.0026 | 0.0016 | 0.0018 |
| F01 | 13640001 | 13740000 | 0.0024 | 0.0014 | 0.0016 |
| F01 | 13660001 | 13760000 | 0.0028 | 0.0017 | 0.0020 |
| F01 | 13680001 | 13780000 | 0.0027 | 0.0019 | 0.0021 |
| F01 | 13700001 | 13800000 | 0.0027 | 0.0022 | 0.0023 |
| F01 | 13720001 | 13820000 | 0.0028 | 0.0022 | 0.0024 |
| F01 | 13740001 | 13840000 | 0.0030 | 0.0024 | 0.0026 |
| F01 | 13760001 | 13860000 | 0.0026 | 0.0021 | 0.0023 |
| F01 | 13780001 | 13880000 | 0.0023 | 0.0017 | 0.0020 |
| F01 | 13800001 | 13900000 | 0.0019 | 0.0013 | 0.0016 |
| F01 | 13820001 | 13920000 | 0.0020 | 0.0014 | 0.0018 |

| Chr | Start | End | both | A1 | A2 |
|---|---|---|---|---|---|
| F01 | 13840001 | 13940000 | 0.0026 | 0.0021 | 0.0025 |
| F01 | 13860001 | 13960000 | 0.0025 | 0.0020 | 0.0023 |
| F01 | 13880001 | 13980000 | 0.0025 | 0.0020 | 0.0023 |
| F01 | 13900001 | 14000000 | 0.0026 | 0.0021 | 0.0024 |
| F01 | 13920001 | 14020000 | 0.0031 | 0.0024 | 0.0028 |
| F01 | 13940001 | 14040000 | 0.0027 | 0.0022 | 0.0024 |
| F01 | 13960001 | 14060000 | 0.0030 | 0.0025 | 0.0027 |
| F01 | 13980001 | 14080000 | 0.0028 | 0.0023 | 0.0025 |
| F01 | 14000001 | 14100000 | 0.0028 | 0.0024 | 0.0026 |
| F01 | 14020001 | 14120000 | 0.0030 | 0.0027 | 0.0027 |
| F01 | 14040001 | 14140000 | 0.0030 | 0.0024 | 0.0027 |
| F01 | 14060001 | 14160000 | 0.0033 | 0.0026 | 0.0030 |
| F01 | 14080001 | 14180000 | 0.0031 | 0.0024 | 0.0028 |
| F01 | 14100001 | 14200000 | 0.0029 | 0.0023 | 0.0026 |
| F01 | 14120001 | 14220000 | 0.0022 | 0.0016 | 0.0020 |
| F01 | 14140001 | 14240000 | 0.0021 | 0.0016 | 0.0019 |
| F01 | 14160001 | 14260000 | 0.0021 | 0.0016 | 0.0018 |
| F01 | 14180001 | 14280000 | 0.0021 | 0.0016 | 0.0019 |
| F01 | 14200001 | 14300000 | 0.0018 | 0.0014 | 0.0016 |
| F01 | 14220001 | 14320000 | 0.0016 | 0.0013 | 0.0015 |
| F01 | 14240001 | 14340000 | 0.0014 | 0.0012 | 0.0014 |
| F01 | 14260001 | 14360000 | 0.0021 | 0.0020 | 0.0021 |
| F01 | 14280001 | 14380000 | 0.0033 | 0.0032 | 0.0033 |
| F01 | 14300001 | 14400000 | 0.0034 | 0.0032 | 0.0033 |
| F01 | 14320001 | 14420000 | 0.0044 | 0.0041 | 0.0041 |
| F01 | 14340001 | 14440000 | 0.0054 | 0.0050 | 0.0050 |
| F01 | 14360001 | 14460000 | 0.0050 | 0.0045 | 0.0046 |
| F01 | 14380001 | 14480000 | 0.0042 | 0.0036 | 0.0039 |
| F01 | 14400001 | 14500000 | 0.0043 | 0.0035 | 0.0038 |
| F01 | 14420001 | 14520000 | 0.0045 | 0.0037 | 0.0040 |
| F01 | 14440001 | 14540000 | 0.0035 | 0.0027 | 0.0032 |
| F01 | 14460001 | 14560000 | 0.0036 | 0.0026 | 0.0033 |
| F01 | 14480001 | 14580000 | 0.0042 | 0.0031 | 0.0039 |
| F01 | 14500001 | 14600000 | 0.0045 | 0.0033 | 0.0042 |
| F01 | 14520001 | 14620000 | 0.0045 | 0.0034 | 0.0042 |
| F01 | 14540001 | 14640000 | 0.0047 | 0.0036 | 0.0044 |
| F01 | 14560001 | 14660000 | 0.0041 | 0.0031 | 0.0037 |
| F01 | 14580001 | 14680000 | 0.0033 | 0.0025 | 0.0029 |
| F01 | 14600001 | 14700000 | 0.0033 | 0.0027 | 0.0030 |
| F01 | 14620001 | 14720000 | 0.0030 | 0.0023 | 0.0026 |
| F01 | 14640001 | 14740000 | 0.0026 | 0.0020 | 0.0023 |
| F01 | 14660001 | 14760000 | 0.0022 | 0.0018 | 0.0020 |
| F01 | 14680001 | 14780000 | 0.0025 | 0.0020 | 0.0022 |
| F01 | 14700001 | 14800000 | 0.0029 | 0.0023 | 0.0026 |
| F01 | 14720001 | 14820000 | 0.0027 | 0.0020 | 0.0024 |
| F01 | 14740001 | 14840000 | 0.0038 | 0.0031 | 0.0035 |
| F01 | 14760001 | 14860000 | 0.0044 | 0.0036 | 0.0040 |
| F01 | 14780001 | 14880000 | 0.0040 | 0.0032 | 0.0037 |
| F01 | 14800001 | 14900000 | 0.0034 | 0.0026 | 0.0028 |
| F01 | 14820001 | 14920000 | 0.0026 | 0.0020 | 0.0021 |
| F01 | 14840001 | 14940000 | 0.0014 | 0.0007 | 0.0008 |
| F01 | 14860001 | 14960000 | 0.0018 | 0.0010 | 0.0012 |
| F01 | 14880001 | 14980000 | 0.0021 | 0.0013 | 0.0016 |
| F01 | 14900001 | 15000000 | 0.0025 | 0.0016 | 0.0021 |
| F01 | 14920001 | 15020000 | 0.0028 | 0.0018 | 0.0023 |
| F01 | 14940001 | 15040000 | 0.0036 | 0.0023 | 0.0030 |
| F01 | 14960001 | 15060000 | 0.0027 | 0.0016 | 0.0022 |
| F01 | 14980001 | 15080000 | 0.0022 | 0.0011 | 0.0017 |

| Chr | Start | End | both | A1 | A2 |
|---|---|---|---|---|---|
| F01 | 15000001 | 15100000 | 0.0022 | 0.0014 | 0.0018 |
| F01 | 15020001 | 15120000 | 0.0027 | 0.0016 | 0.0021 |
| F01 | 15040001 | 15140000 | 0.0022 | 0.0013 | 0.0016 |
| F01 | 15060001 | 15160000 | 0.0024 | 0.0014 | 0.0017 |
| F01 | 15080001 | 15180000 | 0.0032 | 0.0023 | 0.0025 |
| F01 | 15100001 | 15200000 | 0.0027 | 0.0018 | 0.0018 |
| F01 | 15120001 | 15220000 | 0.0027 | 0.0017 | 0.0019 |
| F01 | 15140001 | 15240000 | 0.0031 | 0.0021 | 0.0022 |
| F01 | 15160001 | 15260000 | 0.0030 | 0.0021 | 0.0022 |
| F01 | 15180001 | 15280000 | 0.0033 | 0.0022 | 0.0024 |
| F01 | 15200001 | 15300000 | 0.0038 | 0.0028 | 0.0031 |
| F01 | 15220001 | 15320000 | 0.0040 | 0.0030 | 0.0033 |
| F01 | 15240001 | 15340000 | 0.0038 | 0.0029 | 0.0032 |
| F01 | 15260001 | 15360000 | 0.0041 | 0.0032 | 0.0035 |
| F01 | 15280001 | 15380000 | 0.0034 | 0.0027 | 0.0029 |
| F01 | 15300001 | 15400000 | 0.0030 | 0.0022 | 0.0025 |
| F01 | 15320001 | 15420000 | 0.0026 | 0.0021 | 0.0022 |
| F01 | 15340001 | 15440000 | 0.0029 | 0.0024 | 0.0026 |
| F01 | 15360001 | 15460000 | 0.0026 | 0.0020 | 0.0023 |
| F01 | 15380001 | 15480000 | 0.0023 | 0.0017 | 0.0020 |
| F01 | 15400001 | 15500000 | 0.0024 | 0.0017 | 0.0019 |
| F01 | 15420001 | 15520000 | 0.0022 | 0.0014 | 0.0017 |
| F01 | 15440001 | 15540000 | 0.0017 | 0.0011 | 0.0012 |
| F01 | 15460001 | 15560000 | 0.0018 | 0.0012 | 0.0014 |
| F01 | 15480001 | 15580000 | 0.0019 | 0.0012 | 0.0015 |
| F01 | 15500001 | 15600000 | 0.0029 | 0.0024 | 0.0026 |
| F01 | 15520001 | 15620000 | 0.0040 | 0.0032 | 0.0035 |
| F01 | 15540001 | 15640000 | 0.0045 | 0.0034 | 0.0037 |
| F01 | 15560001 | 15660000 | 0.0046 | 0.0033 | 0.0035 |
| F01 | 15580001 | 15680000 | 0.0058 | 0.0043 | 0.0046 |
| F01 | 15600001 | 15700000 | 0.0052 | 0.0037 | 0.0039 |
| F01 | 15620001 | 15720000 | 0.0049 | 0.0033 | 0.0036 |
| F01 | 15640001 | 15740000 | 0.0045 | 0.0033 | 0.0035 |
| F01 | 15660001 | 15760000 | 0.0045 | 0.0035 | 0.0038 |
| F01 | 15680001 | 15780000 | 0.0036 | 0.0027 | 0.0030 |
| F01 | 15700001 | 15800000 | 0.0032 | 0.0022 | 0.0026 |
| F01 | 15720001 | 15820000 | 0.0023 | 0.0014 | 0.0018 |
| F01 | 15740001 | 15840000 | 0.0023 | 0.0015 | 0.0019 |
| F01 | 15760001 | 15860000 | 0.0021 | 0.0014 | 0.0017 |
| F01 | 15780001 | 15880000 | 0.0024 | 0.0018 | 0.0021 |
| F01 | 15800001 | 15900000 | 0.0025 | 0.0020 | 0.0022 |
| F01 | 15820001 | 15920000 | 0.0029 | 0.0024 | 0.0025 |
| F01 | 15840001 | 15940000 | 0.0028 | 0.0019 | 0.0022 |
| F01 | 15860001 | 15960000 | 0.0035 | 0.0025 | 0.0029 |
| F01 | 15880001 | 15980000 | 0.0035 | 0.0024 | 0.0027 |
| F01 | 15900001 | 16000000 | 0.0039 | 0.0026 | 0.0029 |
| F01 | 15920001 | 16020000 | 0.0038 | 0.0025 | 0.0027 |
| F01 | 15940001 | 16040000 | 0.0043 | 0.0030 | 0.0033 |
| F01 | 15960001 | 16060000 | 0.0042 | 0.0030 | 0.0031 |
| F01 | 15980001 | 16080000 | 0.0041 | 0.0030 | 0.0031 |
| F01 | 16000001 | 16100000 | 0.0045 | 0.0033 | 0.0036 |
| F01 | 16020001 | 16120000 | 0.0043 | 0.0031 | 0.0034 |
| F01 | 16040001 | 16140000 | 0.0035 | 0.0024 | 0.0026 |
| F01 | 16060001 | 16160000 | 0.0031 | 0.0018 | 0.0021 |
| F01 | 16080001 | 16180000 | 0.0031 | 0.0016 | 0.0018 |
| F01 | 16100001 | 16200000 | 0.0025 | 0.0013 | 0.0013 |
| F01 | 16120001 | 16220000 | 0.0031 | 0.0021 | 0.0022 |
| F01 | 16140001 | 16240000 | 0.0038 | 0.0029 | 0.0032 |

| Chr | Start | End | both | A1 | A2 |
|---|---|---|---|---|---|
| F01 | 16160001 | 16260000 | 0.0038 | 0.0031 | 0.0033 |
| F01 | 16180001 | 16280000 | 0.0039 | 0.0034 | 0.0035 |
| F01 | 16200001 | 16300000 | 0.0036 | 0.0030 | 0.0031 |
| F01 | 16220001 | 16320000 | 0.0027 | 0.0020 | 0.0022 |
| F01 | 16240001 | 16340000 | 0.0022 | 0.0014 | 0.0015 |
| F01 | 16260001 | 16360000 | 0.0022 | 0.0013 | 0.0014 |
| F01 | 16280001 | 16380000 | 0.0021 | 0.0011 | 0.0013 |
| F01 | 16300001 | 16400000 | 0.0025 | 0.0014 | 0.0017 |
| F01 | 16320001 | 16420000 | 0.0036 | 0.0023 | 0.0027 |
| F01 | 16340001 | 16440000 | 0.0034 | 0.0023 | 0.0025 |
| F01 | 16360001 | 16460000 | 0.0033 | 0.0021 | 0.0023 |
| F01 | 16380001 | 16480000 | 0.0037 | 0.0025 | 0.0027 |
| F01 | 16400001 | 16500000 | 0.0034 | 0.0021 | 0.0023 |
| F01 | 16420001 | 16520000 | 0.0023 | 0.0012 | 0.0013 |
| F01 | 16440001 | 16540000 | 0.0026 | 0.0015 | 0.0017 |
| F01 | 16460001 | 16560000 | 0.0025 | 0.0014 | 0.0016 |
| F01 | 16480001 | 16580000 | 0.0017 | 0.0007 | 0.0008 |
| F01 | 16500001 | 16600000 | 0.0017 | 0.0009 | 0.0010 |
| F01 | 16520001 | 16620000 | 0.0028 | 0.0018 | 0.0019 |
| F01 | 16540001 | 16640000 | 0.0032 | 0.0022 | 0.0023 |
| F01 | 16560001 | 16660000 | 0.0032 | 0.0023 | 0.0024 |
| F01 | 16580001 | 16680000 | 0.0037 | 0.0027 | 0.0028 |
| F01 | 16600001 | 16700000 | 0.0037 | 0.0026 | 0.0028 |
| F01 | 16620001 | 16720000 | 0.0035 | 0.0025 | 0.0026 |
| F01 | 16640001 | 16740000 | 0.0035 | 0.0026 | 0.0025 |
| F01 | 16660001 | 16760000 | 0.0037 | 0.0026 | 0.0025 |
| F01 | 16680001 | 16780000 | 0.0040 | 0.0027 | 0.0027 |
| F01 | 16700001 | 16800000 | 0.0036 | 0.0024 | 0.0024 |
| F01 | 16720001 | 16820000 | 0.0029 | 0.0017 | 0.0017 |
| F01 | 16740001 | 16840000 | 0.0023 | 0.0011 | 0.0010 |
| F01 | 16760001 | 16860000 | 0.0021 | 0.0010 | 0.0009 |
| F01 | 16780001 | 16880000 | 0.0025 | 0.0011 | 0.0012 |
| F01 | 16800001 | 16900000 | 0.0039 | 0.0022 | 0.0025 |
| F01 | 16820001 | 16920000 | 0.0041 | 0.0023 | 0.0026 |
| F01 | 16840001 | 16940000 | 0.0040 | 0.0021 | 0.0025 |
| F01 | 16860001 | 16960000 | 0.0055 | 0.0038 | 0.0040 |
| F01 | 16880001 | 16980000 | 0.0052 | 0.0038 | 0.0040 |
| F01 | 16900001 | 17000000 | 0.0044 | 0.0033 | 0.0034 |
| F01 | 16920001 | 17020000 | 0.0041 | 0.0032 | 0.0033 |
| F01 | 16940001 | 17040000 | 0.0048 | 0.0041 | 0.0043 |
| F01 | 16960001 | 17060000 | 0.0035 | 0.0027 | 0.0030 |
| F01 | 16980001 | 17080000 | 0.0032 | 0.0023 | 0.0024 |
| F01 | 17000001 | 17100000 | 0.0031 | 0.0020 | 0.0020 |
| F01 | 17020001 | 17120000 | 0.0039 | 0.0027 | 0.0028 |
| F01 | 17040001 | 17140000 | 0.0048 | 0.0034 | 0.0036 |
| F01 | 17060001 | 17160000 | 0.0057 | 0.0043 | 0.0046 |
| F01 | 17080001 | 17180000 | 0.0057 | 0.0047 | 0.0049 |
| F01 | 17100001 | 17200000 | 0.0054 | 0.0047 | 0.0050 |
| F01 | 17120001 | 17220000 | 0.0047 | 0.0041 | 0.0043 |
| F01 | 17140001 | 17240000 | 0.0034 | 0.0027 | 0.0029 |
| F01 | 17160001 | 17260000 | 0.0028 | 0.0021 | 0.0023 |
| F01 | 17180001 | 17280000 | 0.0029 | 0.0020 | 0.0023 |
| F01 | 17200001 | 17300000 | 0.0029 | 0.0019 | 0.0021 |
| F01 | 17220001 | 17320000 | 0.0031 | 0.0021 | 0.0022 |
| F01 | 17240001 | 17340000 | 0.0030 | 0.0021 | 0.0021 |
| F01 | 17260001 | 17360000 | 0.0030 | 0.0020 | 0.0019 |
| F01 | 17280001 | 17380000 | 0.0030 | 0.0020 | 0.0018 |
| F01 | 17300001 | 17400000 | 0.0040 | 0.0029 | 0.0030 |

| Chr | Start | End | both | A1 | A2 |
|---|---|---|---|---|---|
| F01 | 17320001 | 17420000 | 0.0054 | 0.0044 | 0.0045 |
| F01 | 17340001 | 17440000 | 0.0050 | 0.0040 | 0.0042 |
| F01 | 17360001 | 17460000 | 0.0050 | 0.0041 | 0.0043 |
| F01 | 17380001 | 17480000 | 0.0048 | 0.0039 | 0.0040 |
| F01 | 17400001 | 17500000 | 0.0049 | 0.0038 | 0.0039 |
| F01 | 17420001 | 17520000 | 0.0039 | 0.0028 | 0.0029 |
| F01 | 17440001 | 17540000 | 0.0045 | 0.0032 | 0.0032 |
| F01 | 17460001 | 17560000 | 0.0050 | 0.0036 | 0.0037 |
| F01 | 17480001 | 17580000 | 0.0046 | 0.0035 | 0.0036 |
| F01 | 17500001 | 17600000 | 0.0037 | 0.0026 | 0.0026 |
| F01 | 17520001 | 17620000 | 0.0036 | 0.0024 | 0.0023 |
| F01 | 17540001 | 17640000 | 0.0035 | 0.0025 | 0.0024 |
| F01 | 17560001 | 17660000 | 0.0036 | 0.0023 | 0.0024 |
| F01 | 17580001 | 17680000 | 0.0038 | 0.0024 | 0.0025 |
| F01 | 17600001 | 17700000 | 0.0036 | 0.0022 | 0.0024 |
| F01 | 17620001 | 17720000 | 0.0031 | 0.0018 | 0.0020 |
| F01 | 17640001 | 17740000 | 0.0029 | 0.0018 | 0.0021 |
| F01 | 17660001 | 17760000 | 0.0021 | 0.0013 | 0.0015 |
| F01 | 17680001 | 17780000 | 0.0020 | 0.0013 | 0.0015 |
| F01 | 17700001 | 17800000 | 0.0023 | 0.0016 | 0.0018 |
| F01 | 17720001 | 17820000 | 0.0026 | 0.0020 | 0.0022 |
| F01 | 17740001 | 17840000 | 0.0026 | 0.0020 | 0.0022 |
| F01 | 17760001 | 17860000 | 0.0028 | 0.0021 | 0.0022 |
| F01 | 17780001 | 17880000 | 0.0033 | 0.0026 | 0.0027 |
| F01 | 17800001 | 17900000 | 0.0034 | 0.0027 | 0.0028 |
| F01 | 17820001 | 17920000 | 0.0030 | 0.0024 | 0.0025 |
| F01 | 17840001 | 17940000 | 0.0028 | 0.0021 | 0.0022 |
| F01 | 17860001 | 17960000 | 0.0027 | 0.0018 | 0.0019 |
| F01 | 17880001 | 17980000 | 0.0022 | 0.0013 | 0.0013 |
| F01 | 17900001 | 18000000 | 0.0024 | 0.0015 | 0.0016 |
| F01 | 17920001 | 18020000 | 0.0026 | 0.0017 | 0.0018 |
| F01 | 17940001 | 18040000 | 0.0025 | 0.0017 | 0.0019 |
| F01 | 17960001 | 18060000 | 0.0028 | 0.0021 | 0.0025 |
| F01 | 17980001 | 18080000 | 0.0034 | 0.0026 | 0.0031 |
| F01 | 18000001 | 18100000 | 0.0028 | 0.0021 | 0.0025 |
| F01 | 18020001 | 18120000 | 0.0029 | 0.0021 | 0.0026 |
| F01 | 18040001 | 18140000 | 0.0031 | 0.0023 | 0.0027 |
| F01 | 18060001 | 18160000 | 0.0029 | 0.0019 | 0.0022 |
| F01 | 18080001 | 18180000 | 0.0026 | 0.0017 | 0.0018 |
| F01 | 18100001 | 18200000 | 0.0028 | 0.0019 | 0.0020 |
| F01 | 18120001 | 18220000 | 0.0035 | 0.0025 | 0.0028 |
| F01 | 18140001 | 18240000 | 0.0037 | 0.0026 | 0.0027 |
| F01 | 18160001 | 18260000 | 0.0038 | 0.0028 | 0.0029 |
| F01 | 18180001 | 18280000 | 0.0038 | 0.0029 | 0.0030 |
| F01 | 18200001 | 18300000 | 0.0035 | 0.0027 | 0.0028 |
| F01 | 18220001 | 18320000 | 0.0028 | 0.0022 | 0.0020 |
| F01 | 18240001 | 18340000 | 0.0028 | 0.0021 | 0.0021 |
| F01 | 18260001 | 18360000 | 0.0032 | 0.0023 | 0.0024 |
| F01 | 18280001 | 18380000 | 0.0031 | 0.0021 | 0.0022 |
| F01 | 18300001 | 18400000 | 0.0034 | 0.0022 | 0.0023 |
| F01 | 18320001 | 18420000 | 0.0040 | 0.0025 | 0.0027 |
| F01 | 18340001 | 18440000 | 0.0044 | 0.0031 | 0.0031 |
| F01 | 18360001 | 18460000 | 0.0041 | 0.0029 | 0.0029 |
| F01 | 18380001 | 18480000 | 0.0045 | 0.0032 | 0.0032 |
| F01 | 18400001 | 18500000 | 0.0041 | 0.0029 | 0.0030 |
| F01 | 18420001 | 18520000 | 0.0043 | 0.0033 | 0.0034 |
| F01 | 18440001 | 18540000 | 0.0038 | 0.0027 | 0.0029 |
| F01 | 18460001 | 18560000 | 0.0047 | 0.0033 | 0.0037 |

| Chr | Start | End | both | A1 | A2 |
|---|---|---|---|---|---|
| F01 | 18480001 | 18580000 | 0.0052 | 0.0037 | 0.0042 |
| F01 | 18500001 | 18600000 | 0.0055 | 0.0042 | 0.0046 |
| F01 | 18520001 | 18620000 | 0.0055 | 0.0042 | 0.0046 |
| F01 | 18540001 | 18640000 | 0.0064 | 0.0049 | 0.0054 |
| F01 | 18560001 | 18660000 | 0.0053 | 0.0040 | 0.0044 |
| F01 | 18580001 | 18680000 | 0.0050 | 0.0039 | 0.0041 |
| F01 | 18600001 | 18700000 | 0.0050 | 0.0039 | 0.0041 |
| F01 | 18620001 | 18720000 | 0.0044 | 0.0033 | 0.0035 |
| F01 | 18640001 | 18740000 | 0.0031 | 0.0025 | 0.0025 |
| F01 | 18660001 | 18760000 | 0.0029 | 0.0023 | 0.0023 |
| F01 | 18680001 | 18780000 | 0.0022 | 0.0016 | 0.0016 |
| F01 | 18700001 | 18800000 | 0.0029 | 0.0020 | 0.0022 |
| F01 | 18720001 | 18820000 | 0.0034 | 0.0024 | 0.0026 |
| F01 | 18740001 | 18840000 | 0.0046 | 0.0034 | 0.0037 |
| F01 | 18760001 | 18860000 | 0.0051 | 0.0037 | 0.0041 |
| F01 | 18780001 | 18880000 | 0.0057 | 0.0042 | 0.0046 |
| F01 | 18800001 | 18900000 | 0.0052 | 0.0038 | 0.0043 |
| F01 | 18820001 | 18920000 | 0.0045 | 0.0033 | 0.0037 |
| F01 | 18840001 | 18940000 | 0.0043 | 0.0032 | 0.0034 |
| F01 | 18860001 | 18960000 | 0.0043 | 0.0032 | 0.0034 |
| F01 | 18880001 | 18980000 | 0.0039 | 0.0029 | 0.0029 |
| F01 | 18900001 | 19000000 | 0.0037 | 0.0030 | 0.0029 |
| F01 | 18920001 | 19020000 | 0.0050 | 0.0041 | 0.0042 |
| F01 | 18940001 | 19040000 | 0.0046 | 0.0038 | 0.0039 |
| F01 | 18960001 | 19060000 | 0.0050 | 0.0042 | 0.0043 |
| F01 | 18980001 | 19080000 | 0.0051 | 0.0042 | 0.0043 |
| F01 | 19000001 | 19100000 | 0.0056 | 0.0045 | 0.0047 |
| F01 | 19020001 | 19120000 | 0.0045 | 0.0034 | 0.0035 |
| F01 | 19040001 | 19140000 | 0.0045 | 0.0034 | 0.0036 |
| F01 | 19060001 | 19160000 | 0.0035 | 0.0026 | 0.0028 |
| F01 | 19080001 | 19180000 | 0.0032 | 0.0023 | 0.0026 |
| F01 | 19100001 | 19200000 | 0.0025 | 0.0017 | 0.0020 |
| F01 | 19120001 | 19220000 | 0.0021 | 0.0013 | 0.0016 |
| F01 | 19140001 | 19240000 | 0.0022 | 0.0014 | 0.0017 |
| F01 | 19160001 | 19260000 | 0.0024 | 0.0016 | 0.0018 |
| F01 | 19180001 | 19280000 | 0.0024 | 0.0016 | 0.0017 |
| F01 | 19200001 | 19300000 | 0.0029 | 0.0020 | 0.0022 |
| F01 | 19220001 | 19320000 | 0.0034 | 0.0022 | 0.0023 |
| F01 | 19240001 | 19340000 | 0.0046 | 0.0034 | 0.0034 |
| F01 | 19260001 | 19360000 | 0.0054 | 0.0042 | 0.0041 |
| F01 | 19280001 | 19380000 | 0.0057 | 0.0045 | 0.0043 |
| F01 | 19300001 | 19400000 | 0.0059 | 0.0046 | 0.0045 |
| F01 | 19320001 | 19420000 | 0.0054 | 0.0044 | 0.0042 |
| F01 | 19340001 | 19440000 | 0.0043 | 0.0030 | 0.0031 |
| F01 | 19360001 | 19460000 | 0.0031 | 0.0019 | 0.0021 |
| F01 | 19380001 | 19480000 | 0.0030 | 0.0020 | 0.0022 |
| F01 | 19400001 | 19500000 | 0.0028 | 0.0020 | 0.0022 |
| F01 | 19420001 | 19520000 | 0.0029 | 0.0022 | 0.0024 |
| F01 | 19440001 | 19540000 | 0.0027 | 0.0022 | 0.0025 |
| F01 | 19460001 | 19560000 | 0.0035 | 0.0027 | 0.0030 |
| F01 | 19480001 | 19580000 | 0.0037 | 0.0027 | 0.0030 |
| F01 | 19500001 | 19600000 | 0.0032 | 0.0022 | 0.0025 |
| F01 | 19520001 | 19620000 | 0.0033 | 0.0024 | 0.0026 |
| F01 | 19540001 | 19640000 | 0.0032 | 0.0021 | 0.0022 |
| F01 | 19560001 | 19660000 | 0.0033 | 0.0023 | 0.0024 |
| F01 | 19580001 | 19680000 | 0.0042 | 0.0031 | 0.0031 |
| F01 | 19600001 | 19700000 | 0.0042 | 0.0029 | 0.0030 |
| F01 | 19620001 | 19720000 | 0.0043 | 0.0028 | 0.0030 |

| Chr | Start | End | both | A1 | A2 |
|---|---|---|---|---|---|
| F01 | 19640001 | 19740000 | 0.0043 | 0.0029 | 0.0032 |
| F01 | 19660001 | 19760000 | 0.0041 | 0.0028 | 0.0031 |
| F01 | 19680001 | 19780000 | 0.0036 | 0.0024 | 0.0027 |
| F01 | 19700001 | 19800000 | 0.0032 | 0.0024 | 0.0026 |
| F01 | 19720001 | 19820000 | 0.0027 | 0.0021 | 0.0022 |
| F01 | 19740001 | 19840000 | 0.0022 | 0.0019 | 0.0019 |
| F01 | 19760001 | 19860000 | 0.0018 | 0.0014 | 0.0014 |
| F01 | 19780001 | 19880000 | 0.0011 | 0.0006 | 0.0008 |
| F01 | 19800001 | 19900000 | 0.0012 | 0.0005 | 0.0006 |
| F01 | 19820001 | 19920000 | 0.0024 | 0.0015 | 0.0017 |
| F01 | 19840001 | 19940000 | 0.0027 | 0.0017 | 0.0018 |
| F01 | 19860001 | 19960000 | 0.0031 | 0.0021 | 0.0022 |
| F01 | 19880001 | 19980000 | 0.0035 | 0.0025 | 0.0024 |
| F01 | 19900001 | 20000000 | 0.0044 | 0.0032 | 0.0033 |
| F01 | 19920001 | 20020000 | 0.0038 | 0.0029 | 0.0027 |
| F01 | 19940001 | 20040000 | 0.0038 | 0.0030 | 0.0029 |
| F01 | 19960001 | 20060000 | 0.0047 | 0.0038 | 0.0039 |
| F01 | 19980001 | 20080000 | 0.0050 | 0.0041 | 0.0043 |
| F01 | 20000001 | 20100000 | 0.0045 | 0.0037 | 0.0040 |
| F01 | 20020001 | 20120000 | 0.0044 | 0.0035 | 0.0040 |
| F01 | 20040001 | 20140000 | 0.0044 | 0.0034 | 0.0038 |
| F01 | 20060001 | 20160000 | 0.0046 | 0.0032 | 0.0040 |
| F01 | 20080001 | 20180000 | 0.0042 | 0.0029 | 0.0036 |
| F01 | 20100001 | 20200000 | 0.0044 | 0.0030 | 0.0036 |
| F01 | 20120001 | 20220000 | 0.0049 | 0.0035 | 0.0040 |
| F01 | 20140001 | 20240000 | 0.0054 | 0.0040 | 0.0045 |
| F01 | 20160001 | 20260000 | 0.0046 | 0.0036 | 0.0038 |
| F01 | 20180001 | 20280000 | 0.0042 | 0.0034 | 0.0036 |
| F01 | 20200001 | 20300000 | 0.0041 | 0.0034 | 0.0036 |
| F01 | 20220001 | 20320000 | 0.0036 | 0.0031 | 0.0033 |
| F01 | 20240001 | 20340000 | 0.0032 | 0.0026 | 0.0028 |
| F01 | 20260001 | 20360000 | 0.0028 | 0.0024 | 0.0024 |
| F01 | 20280001 | 20380000 | 0.0032 | 0.0026 | 0.0025 |
| F01 | 20300001 | 20400000 | 0.0034 | 0.0024 | 0.0024 |
| F01 | 20320001 | 20420000 | 0.0028 | 0.0019 | 0.0019 |
| F01 | 20340001 | 20440000 | 0.0027 | 0.0016 | 0.0017 |
| F01 | 20360001 | 20460000 | 0.0026 | 0.0012 | 0.0013 |
| F01 | 20380001 | 20480000 | 0.0024 | 0.0011 | 0.0013 |
| F01 | 20400001 | 20500000 | 0.0020 | 0.0011 | 0.0012 |
| F01 | 20420001 | 20520000 | 0.0028 | 0.0018 | 0.0020 |
| F01 | 20440001 | 20540000 | 0.0024 | 0.0016 | 0.0018 |
| F01 | 20460001 | 20560000 | 0.0021 | 0.0016 | 0.0018 |
| F01 | 20480001 | 20580000 | 0.0028 | 0.0023 | 0.0025 |
| F01 | 20500001 | 20600000 | 0.0033 | 0.0028 | 0.0029 |
| F01 | 20520001 | 20620000 | 0.0038 | 0.0030 | 0.0031 |
| F01 | 20540001 | 20640000 | 0.0046 | 0.0036 | 0.0037 |
| F01 | 20560001 | 20660000 | 0.0047 | 0.0037 | 0.0038 |
| F01 | 20580001 | 20680000 | 0.0042 | 0.0031 | 0.0033 |
| F01 | 20600001 | 20700000 | 0.0041 | 0.0029 | 0.0031 |
| F01 | 20620001 | 20720000 | 0.0033 | 0.0023 | 0.0024 |
| F01 | 20640001 | 20740000 | 0.0033 | 0.0020 | 0.0023 |
| F01 | 20660001 | 20760000 | 0.0040 | 0.0026 | 0.0030 |
| F01 | 20680001 | 20780000 | 0.0040 | 0.0026 | 0.0030 |
| F01 | 20700001 | 20800000 | 0.0042 | 0.0028 | 0.0032 |
| F01 | 20720001 | 20820000 | 0.0043 | 0.0029 | 0.0033 |
| F01 | 20740001 | 20840000 | 0.0041 | 0.0028 | 0.0032 |
| F01 | 20760001 | 20860000 | 0.0035 | 0.0023 | 0.0024 |
| F01 | 20780001 | 20880000 | 0.0031 | 0.0019 | 0.0020 |

| Chr | Start | End | both | A1 | A2 |
|---|---|---|---|---|---|
| F01 | 20800001 | 20900000 | 0.0025 | 0.0017 | 0.0014 |
| F01 | 20820001 | 20920000 | 0.0031 | 0.0021 | 0.0019 |
| F01 | 20840001 | 20940000 | 0.0027 | 0.0019 | 0.0017 |
| F01 | 20860001 | 20960000 | 0.0031 | 0.0022 | 0.0022 |
| F01 | 20880001 | 20980000 | 0.0038 | 0.0028 | 0.0029 |
| F01 | 20900001 | 21000000 | 0.0045 | 0.0032 | 0.0036 |
| F01 | 20920001 | 21020000 | 0.0042 | 0.0031 | 0.0033 |
| F01 | 20940001 | 21040000 | 0.0045 | 0.0033 | 0.0035 |
| F01 | 20960001 | 21060000 | 0.0049 | 0.0035 | 0.0037 |
| F01 | 20980001 | 21080000 | 0.0042 | 0.0028 | 0.0031 |
| F01 | 21000001 | 21100000 | 0.0044 | 0.0031 | 0.0033 |
| F01 | 21020001 | 21120000 | 0.0044 | 0.0030 | 0.0032 |
| F01 | 21040001 | 21140000 | 0.0050 | 0.0037 | 0.0039 |
| F01 | 21060001 | 21160000 | 0.0044 | 0.0033 | 0.0034 |
| F01 | 21080001 | 21180000 | 0.0054 | 0.0042 | 0.0045 |
| F01 | 21100001 | 21200000 | 0.0047 | 0.0036 | 0.0039 |
| F01 | 21120001 | 21220000 | 0.0050 | 0.0038 | 0.0042 |
| F01 | 21140001 | 21240000 | 0.0049 | 0.0038 | 0.0043 |
| F01 | 21160001 | 21260000 | 0.0053 | 0.0043 | 0.0048 |
| F01 | 21180001 | 21280000 | 0.0044 | 0.0036 | 0.0040 |
| F01 | 21200001 | 21300000 | 0.0040 | 0.0031 | 0.0034 |
| F01 | 21220001 | 21320000 | 0.0039 | 0.0030 | 0.0033 |
| F01 | 21240001 | 21340000 | 0.0033 | 0.0024 | 0.0026 |
| F01 | 21260001 | 21360000 | 0.0055 | 0.0044 | 0.0048 |
| F01 | 21280001 | 21380000 | 0.0061 | 0.0050 | 0.0052 |
| F01 | 21300001 | 21400000 | 0.0069 | 0.0058 | 0.0061 |
| F01 | 21320001 | 21420000 | 0.0062 | 0.0050 | 0.0053 |
| F01 | 21340001 | 21440000 | 0.0059 | 0.0048 | 0.0051 |
| F01 | 21360001 | 21460000 | 0.0031 | 0.0021 | 0.0023 |
| F01 | 21380001 | 21480000 | 0.0024 | 0.0016 | 0.0019 |
| F01 | 21400001 | 21500000 | 0.0021 | 0.0013 | 0.0016 |
| F01 | 21420001 | 21520000 | 0.0026 | 0.0019 | 0.0023 |
| F01 | 21440001 | 21540000 | 0.0030 | 0.0022 | 0.0025 |
| F01 | 21460001 | 21560000 | 0.0033 | 0.0025 | 0.0029 |
| F01 | 21480001 | 21580000 | 0.0040 | 0.0031 | 0.0036 |
| F01 | 21500001 | 21600000 | 0.0043 | 0.0034 | 0.0037 |
| F01 | 21520001 | 21620000 | 0.0038 | 0.0028 | 0.0029 |
| F01 | 21540001 | 21640000 | 0.0039 | 0.0030 | 0.0032 |
| F01 | 21560001 | 21660000 | 0.0036 | 0.0027 | 0.0029 |
| F01 | 21580001 | 21680000 | 0.0029 | 0.0022 | 0.0022 |
| F01 | 21600001 | 21700000 | 0.0030 | 0.0023 | 0.0024 |
| F01 | 21620001 | 21720000 | 0.0031 | 0.0024 | 0.0025 |
| F01 | 21640001 | 21740000 | 0.0029 | 0.0022 | 0.0021 |
| F01 | 21660001 | 21760000 | 0.0027 | 0.0020 | 0.0019 |
| F01 | 21680001 | 21780000 | 0.0025 | 0.0018 | 0.0017 |
| F01 | 21700001 | 21800000 | 0.0019 | 0.0011 | 0.0010 |
| F01 | 21720001 | 21820000 | 0.0017 | 0.0010 | 0.0010 |
| F01 | 21740001 | 21840000 | 0.0024 | 0.0015 | 0.0017 |
| F01 | 21760001 | 21860000 | 0.0039 | 0.0028 | 0.0030 |
| F01 | 21780001 | 21880000 | 0.0043 | 0.0031 | 0.0032 |
| F01 | 21800001 | 21900000 | 0.0045 | 0.0034 | 0.0035 |
| F01 | 21820001 | 21920000 | 0.0047 | 0.0035 | 0.0036 |
| F01 | 21840001 | 21940000 | 0.0039 | 0.0027 | 0.0028 |
| F01 | 21860001 | 21960000 | 0.0032 | 0.0021 | 0.0021 |
| F01 | 21880001 | 21980000 | 0.0040 | 0.0029 | 0.0030 |
| F01 | 21900001 | 22000000 | 0.0036 | 0.0025 | 0.0026 |
| F01 | 21920001 | 22020000 | 0.0039 | 0.0029 | 0.0029 |
| F01 | 21940001 | 22040000 | 0.0039 | 0.0031 | 0.0032 |

| Chr | Start | End | both | A1 | A2 |
|---|---|---|---|---|---|
| F01 | 21960001 | 22060000 | 0.0049 | 0.0038 | 0.0043 |
| F01 | 21980001 | 22080000 | 0.0041 | 0.0031 | 0.0035 |
| F01 | 22000001 | 22100000 | 0.0047 | 0.0038 | 0.0042 |
| F01 | 22020001 | 22120000 | 0.0046 | 0.0038 | 0.0042 |
| F01 | 22040001 | 22140000 | 0.0052 | 0.0042 | 0.0047 |
| F01 | 22060001 | 22160000 | 0.0044 | 0.0036 | 0.0039 |
| F01 | 22080001 | 22180000 | 0.0051 | 0.0042 | 0.0046 |
| F01 | 22100001 | 22200000 | 0.0047 | 0.0037 | 0.0040 |
| F01 | 22120001 | 22220000 | 0.0048 | 0.0036 | 0.0040 |
| F01 | 22140001 | 22240000 | 0.0043 | 0.0032 | 0.0036 |
| F01 | 22160001 | 22260000 | 0.0035 | 0.0023 | 0.0027 |
| F01 | 22180001 | 22280000 | 0.0027 | 0.0016 | 0.0019 |
| F01 | 22200001 | 22300000 | 0.0023 | 0.0012 | 0.0015 |
| F01 | 22220001 | 22320000 | 0.0038 | 0.0033 | 0.0030 |
| F01 | 22240001 | 22340000 | 0.0041 | 0.0033 | 0.0030 |
| F01 | 22260001 | 22360000 | 0.0055 | 0.0047 | 0.0042 |
| F01 | 22280001 | 22380000 | 0.0053 | 0.0045 | 0.0041 |
| F01 | 22300001 | 22400000 | 0.0063 | 0.0053 | 0.0050 |
| F01 | 22320001 | 22420000 | 0.0044 | 0.0033 | 0.0033 |
| F01 | 22340001 | 22440000 | 0.0044 | 0.0031 | 0.0032 |
| F01 | 22360001 | 22460000 | 0.0029 | 0.0019 | 0.0020 |
| F01 | 22380001 | 22480000 | 0.0037 | 0.0028 | 0.0029 |
| F01 | 22400001 | 22500000 | 0.0032 | 0.0022 | 0.0023 |
| F01 | 22420001 | 22520000 | 0.0039 | 0.0028 | 0.0030 |
| F01 | 22440001 | 22540000 | 0.0041 | 0.0031 | 0.0034 |
| F01 | 22460001 | 22560000 | 0.0043 | 0.0035 | 0.0037 |
| F01 | 22480001 | 22580000 | 0.0042 | 0.0033 | 0.0035 |
| F01 | 22500001 | 22600000 | 0.0039 | 0.0031 | 0.0034 |
| F01 | 22520001 | 22620000 | 0.0035 | 0.0027 | 0.0029 |
| F01 | 22540001 | 22640000 | 0.0036 | 0.0028 | 0.0030 |
| F01 | 22560001 | 22660000 | 0.0037 | 0.0027 | 0.0029 |
| F01 | 22580001 | 22680000 | 0.0032 | 0.0023 | 0.0025 |
| F01 | 22600001 | 22700000 | 0.0032 | 0.0025 | 0.0025 |
| F01 | 22620001 | 22720000 | 0.0035 | 0.0025 | 0.0027 |
| F01 | 22640001 | 22740000 | 0.0031 | 0.0022 | 0.0023 |
| F01 | 22660001 | 22760000 | 0.0034 | 0.0024 | 0.0027 |
| F01 | 22680001 | 22780000 | 0.0034 | 0.0024 | 0.0028 |
| F01 | 22700001 | 22800000 | 0.0062 | 0.0052 | 0.0057 |
| F01 | 22720001 | 22820000 | 0.0064 | 0.0053 | 0.0060 |
| F01 | 22740001 | 22840000 | 0.0061 | 0.0050 | 0.0057 |
| F01 | 22760001 | 22860000 | 0.0055 | 0.0047 | 0.0051 |
| F01 | 22780001 | 22880000 | 0.0060 | 0.0049 | 0.0054 |
| F01 | 22800001 | 22900000 | 0.0033 | 0.0022 | 0.0024 |
| F01 | 22820001 | 22920000 | 0.0031 | 0.0021 | 0.0021 |
| F01 | 22840001 | 22940000 | 0.0030 | 0.0020 | 0.0021 |
| F01 | 22860001 | 22960000 | 0.0031 | 0.0020 | 0.0021 |
| F01 | 22880001 | 22980000 | 0.0023 | 0.0016 | 0.0016 |
| F01 | 22900001 | 23000000 | 0.0021 | 0.0012 | 0.0014 |
| F01 | 22920001 | 23020000 | 0.0030 | 0.0020 | 0.0023 |
| F01 | 22940001 | 23040000 | 0.0035 | 0.0025 | 0.0028 |
| F01 | 22960001 | 23060000 | 0.0043 | 0.0032 | 0.0037 |
| F01 | 22980001 | 23080000 | 0.0043 | 0.0033 | 0.0038 |
| F01 | 23000001 | 23100000 | 0.0049 | 0.0038 | 0.0043 |
| F01 | 23020001 | 23120000 | 0.0033 | 0.0024 | 0.0027 |
| F01 | 23040001 | 23140000 | 0.0039 | 0.0029 | 0.0032 |
| F01 | 23060001 | 23160000 | 0.0038 | 0.0026 | 0.0028 |
| F01 | 23080001 | 23180000 | 0.0045 | 0.0030 | 0.0035 |
| F01 | 23100001 | 23200000 | 0.0054 | 0.0041 | 0.0046 |

| Chr | Start | End | both | A1 | A2 |
|---|---|---|---|---|---|
| F01 | 23120001 | 23220000 | 0.0059 | 0.0046 | 0.0051 |
| F01 | 23140001 | 23240000 | 0.0049 | 0.0037 | 0.0042 |
| F01 | 23160001 | 23260000 | 0.0047 | 0.0038 | 0.0042 |
| F01 | 23180001 | 23280000 | 0.0046 | 0.0038 | 0.0041 |
| F01 | 23200001 | 23300000 | 0.0040 | 0.0033 | 0.0033 |
| F01 | 23220001 | 23320000 | 0.0036 | 0.0030 | 0.0030 |
| F01 | 23240001 | 23340000 | 0.0046 | 0.0039 | 0.0039 |
| F01 | 23260001 | 23360000 | 0.0043 | 0.0034 | 0.0036 |
| F01 | 23280001 | 23380000 | 0.0045 | 0.0037 | 0.0038 |
| F01 | 23300001 | 23400000 | 0.0046 | 0.0035 | 0.0038 |
| F01 | 23320001 | 23420000 | 0.0046 | 0.0034 | 0.0036 |
| F01 | 23340001 | 23440000 | 0.0044 | 0.0029 | 0.0032 |
| F01 | 23360001 | 23460000 | 0.0041 | 0.0028 | 0.0029 |
| F01 | 23380001 | 23480000 | 0.0031 | 0.0019 | 0.0020 |
| F01 | 23400001 | 23500000 | 0.0032 | 0.0019 | 0.0019 |
| F01 | 23420001 | 23520000 | 0.0038 | 0.0024 | 0.0024 |
| F01 | 23440001 | 23540000 | 0.0037 | 0.0023 | 0.0024 |
| F01 | 23460001 | 23560000 | 0.0041 | 0.0026 | 0.0027 |
| F01 | 23480001 | 23580000 | 0.0052 | 0.0036 | 0.0039 |
| F01 | 23500001 | 23600000 | 0.0039 | 0.0027 | 0.0032 |
| F01 | 23520001 | 23620000 | 0.0030 | 0.0022 | 0.0026 |
| F01 | 23540001 | 23640000 | 0.0028 | 0.0022 | 0.0025 |
| F01 | 23560001 | 23660000 | 0.0028 | 0.0023 | 0.0026 |
| F01 | 23580001 | 23680000 | 0.0024 | 0.0019 | 0.0021 |
| F01 | 23600001 | 23700000 | 0.0034 | 0.0028 | 0.0030 |
| F01 | 23620001 | 23720000 | 0.0034 | 0.0028 | 0.0030 |
| F01 | 23640001 | 23740000 | 0.0027 | 0.0023 | 0.0024 |
| F01 | 23660001 | 23760000 | 0.0022 | 0.0019 | 0.0020 |
| F01 | 23680001 | 23780000 | 0.0017 | 0.0016 | 0.0016 |
| F01 | 23700001 | 23800000 | 0.0010 | 0.0009 | 0.0010 |
| F01 | 23720001 | 23820000 | 0.0020 | 0.0017 | 0.0019 |
| F01 | 23740001 | 23840000 | 0.0020 | 0.0017 | 0.0020 |
| F01 | 23760001 | 23860000 | 0.0018 | 0.0016 | 0.0018 |
| F01 | 23780001 | 23880000 | 0.0020 | 0.0017 | 0.0019 |
| F01 | 23800001 | 23900000 | 0.0019 | 0.0017 | 0.0019 |
| F01 | 23820001 | 23920000 | 0.0012 | 0.0010 | 0.0011 |
| F01 | 23840001 | 23940000 | 0.0011 | 0.0009 | 0.0010 |
| F01 | 23860001 | 23960000 | 0.0014 | 0.0012 | 0.0013 |
| F01 | 23880001 | 23980000 | 0.0010 | 0.0008 | 0.0009 |
| F01 | 23900001 | 24000000 | 0.0012 | 0.0009 | 0.0011 |
| F01 | 23920001 | 24020000 | 0.0019 | 0.0017 | 0.0018 |
| F01 | 23940001 | 24040000 | 0.0022 | 0.0018 | 0.0019 |
| F01 | 23960001 | 24060000 | 0.0020 | 0.0017 | 0.0019 |
| F01 | 23980001 | 24080000 | 0.0030 | 0.0025 | 0.0028 |
| F01 | 24000001 | 24100000 | 0.0037 | 0.0033 | 0.0035 |
| F01 | 24020001 | 24120000 | 0.0032 | 0.0028 | 0.0031 |
| F01 | 24040001 | 24140000 | 0.0039 | 0.0033 | 0.0038 |
| F01 | 24060001 | 24160000 | 0.0038 | 0.0033 | 0.0038 |
| F01 | 24080001 | 24180000 | 0.0038 | 0.0032 | 0.0036 |
| F01 | 24100001 | 24200000 | 0.0033 | 0.0027 | 0.0031 |
| F01 | 24120001 | 24220000 | 0.0032 | 0.0025 | 0.0028 |
| F01 | 24140001 | 24240000 | 0.0040 | 0.0034 | 0.0034 |
| F01 | 24160001 | 24260000 | 0.0044 | 0.0036 | 0.0037 |
| F01 | 24180001 | 24280000 | 0.0038 | 0.0031 | 0.0033 |
| F01 | 24200001 | 24300000 | 0.0036 | 0.0030 | 0.0031 |
| F01 | 24220001 | 24320000 | 0.0038 | 0.0033 | 0.0034 |
| F01 | 24240001 | 24340000 | 0.0022 | 0.0018 | 0.0021 |
| F01 | 24260001 | 24360000 | 0.0017 | 0.0017 | 0.0017 |

| Chr | Start | End | both | A1 | A2 |
|---|---|---|---|---|---|
| F01 | 24280001 | 24380000 | 0.0019 | 0.0018 | 0.0018 |
| F01 | 24300001 | 24400000 | 0.0014 | 0.0013 | 0.0014 |
| F01 | 24320001 | 24420000 | 0.0010 | 0.0009 | 0.0009 |
| F01 | 24340001 | 24440000 | 0.0010 | 0.0009 | 0.0009 |
| F01 | 24360001 | 24460000 | 0.0010 | 0.0008 | 0.0010 |
| F01 | 24380001 | 24480000 | 0.0007 | 0.0005 | 0.0007 |
| F01 | 24400001 | 24500000 | 0.0006 | 0.0004 | 0.0006 |
| F01 | 24420001 | 24520000 | 0.0009 | 0.0007 | 0.0009 |
| F01 | 24440001 | 24540000 | 0.0009 | 0.0008 | 0.0009 |
| F01 | 24460001 | 24560000 | 0.0010 | 0.0008 | 0.0010 |
| F01 | 24480001 | 24580000 | 0.0010 | 0.0009 | 0.0010 |
| F01 | 24500001 | 24600000 | 0.0014 | 0.0013 | 0.0014 |
| F01 | 24520001 | 24620000 | 0.0013 | 0.0011 | 0.0013 |
| F01 | 24540001 | 24640000 | 0.0012 | 0.0010 | 0.0012 |
| F01 | 24560001 | 24660000 | 0.0014 | 0.0012 | 0.0013 |
| F01 | 24580001 | 24680000 | 0.0020 | 0.0016 | 0.0016 |
| F01 | 24600001 | 24700000 | 0.0023 | 0.0018 | 0.0018 |
| F01 | 24620001 | 24720000 | 0.0025 | 0.0019 | 0.0020 |
| F01 | 24640001 | 24740000 | 0.0025 | 0.0019 | 0.0019 |
| F01 | 24660001 | 24760000 | 0.0026 | 0.0020 | 0.0021 |
| F01 | 24680001 | 24780000 | 0.0021 | 0.0017 | 0.0019 |
| F01 | 24700001 | 24800000 | 0.0017 | 0.0014 | 0.0016 |
| F01 | 24720001 | 24820000 | 0.0016 | 0.0014 | 0.0016 |
| F01 | 24740001 | 24840000 | 0.0017 | 0.0015 | 0.0017 |
| F01 | 24760001 | 24860000 | 0.0019 | 0.0016 | 0.0018 |
| F01 | 24780001 | 24880000 | 0.0017 | 0.0014 | 0.0017 |
| F01 | 24800001 | 24900000 | 0.0014 | 0.0012 | 0.0014 |
| F01 | 24820001 | 24920000 | 0.0011 | 0.0009 | 0.0011 |
| F01 | 24840001 | 24940000 | 0.0014 | 0.0012 | 0.0014 |
| F01 | 24860001 | 24960000 | 0.0015 | 0.0013 | 0.0015 |
| F01 | 24880001 | 24980000 | 0.0016 | 0.0012 | 0.0013 |
| F01 | 24900001 | 25000000 | 0.0025 | 0.0020 | 0.0021 |
| F01 | 24920001 | 25020000 | 0.0028 | 0.0022 | 0.0024 |
| F01 | 24940001 | 25040000 | 0.0031 | 0.0027 | 0.0027 |
| F01 | 24960001 | 25060000 | 0.0026 | 0.0023 | 0.0023 |
| F01 | 24980001 | 25080000 | 0.0025 | 0.0023 | 0.0024 |
| F01 | 25000001 | 25100000 | 0.0018 | 0.0017 | 0.0017 |
| F01 | 25020001 | 25120000 | 0.0015 | 0.0015 | 0.0014 |
| F01 | 25040001 | 25140000 | 0.0015 | 0.0013 | 0.0014 |
| F01 | 25060001 | 25160000 | 0.0016 | 0.0014 | 0.0015 |
| F01 | 25080001 | 25180000 | 0.0018 | 0.0014 | 0.0015 |
| F01 | 25100001 | 25200000 | 0.0017 | 0.0013 | 0.0015 |
| F01 | 25120001 | 25220000 | 0.0019 | 0.0014 | 0.0017 |
| F01 | 25140001 | 25240000 | 0.0013 | 0.0009 | 0.0011 |
| F01 | 25160001 | 25260000 | 0.0013 | 0.0009 | 0.0011 |
| F01 | 25180001 | 25280000 | 0.0016 | 0.0013 | 0.0016 |
| F01 | 25200001 | 25300000 | 0.0016 | 0.0013 | 0.0015 |
| F01 | 25220001 | 25320000 | 0.0013 | 0.0011 | 0.0012 |
| F01 | 25240001 | 25340000 | 0.0012 | 0.0011 | 0.0012 |
| F01 | 25260001 | 25360000 | 0.0011 | 0.0009 | 0.0010 |
| F01 | 25280001 | 25380000 | 0.0005 | 0.0004 | 0.0005 |
| F01 | 25300001 | 25400000 | 0.0010 | 0.0008 | 0.0010 |
| F01 | 25320001 | 25420000 | 0.0018 | 0.0016 | 0.0018 |
| F01 | 25340001 | 25440000 | 0.0030 | 0.0025 | 0.0029 |
| F01 | 25360001 | 25460000 | 0.0030 | 0.0026 | 0.0029 |
| F01 | 25380001 | 25480000 | 0.0037 | 0.0032 | 0.0036 |
| F01 | 25400001 | 25500000 | 0.0043 | 0.0037 | 0.0042 |
| F01 | 25420001 | 25520000 | 0.0042 | 0.0034 | 0.0040 |

| Chr | Start | End | both | A1 | A2 |
|---|---|---|---|---|---|
| F01 | 25440001 | 25540000 | 0.0033 | 0.0027 | 0.0031 |
| F01 | 25460001 | 25560000 | 0.0033 | 0.0026 | 0.0031 |
| F01 | 25480001 | 25580000 | 0.0034 | 0.0030 | 0.0032 |
| F01 | 25500001 | 25600000 | 0.0038 | 0.0035 | 0.0035 |
| F01 | 25520001 | 25620000 | 0.0037 | 0.0033 | 0.0035 |
| F01 | 25540001 | 25640000 | 0.0034 | 0.0031 | 0.0032 |
| F01 | 25560001 | 25660000 | 0.0048 | 0.0045 | 0.0045 |
| F01 | 25580001 | 25680000 | 0.0050 | 0.0046 | 0.0047 |
| F01 | 25600001 | 25700000 | 0.0042 | 0.0036 | 0.0038 |
| F01 | 25620001 | 25720000 | 0.0039 | 0.0034 | 0.0035 |
| F01 | 25640001 | 25740000 | 0.0043 | 0.0037 | 0.0039 |
| F01 | 25660001 | 25760000 | 0.0032 | 0.0025 | 0.0028 |
| F01 | 25680001 | 25780000 | 0.0026 | 0.0019 | 0.0023 |
| F01 | 25700001 | 25800000 | 0.0031 | 0.0025 | 0.0027 |
| F01 | 25720001 | 25820000 | 0.0035 | 0.0026 | 0.0031 |
| F01 | 25740001 | 25840000 | 0.0043 | 0.0031 | 0.0037 |
| F01 | 25760001 | 25860000 | 0.0042 | 0.0028 | 0.0035 |
| F01 | 25780001 | 25880000 | 0.0045 | 0.0029 | 0.0037 |
| F01 | 25800001 | 25900000 | 0.0042 | 0.0029 | 0.0036 |
| F01 | 25820001 | 25920000 | 0.0041 | 0.0031 | 0.0035 |
| F01 | 25840001 | 25940000 | 0.0031 | 0.0024 | 0.0025 |
| F01 | 25860001 | 25960000 | 0.0032 | 0.0026 | 0.0027 |
| F01 | 25880001 | 25980000 | 0.0031 | 0.0023 | 0.0024 |
| F01 | 25900001 | 26000000 | 0.0029 | 0.0020 | 0.0020 |
| F01 | 25920001 | 26020000 | 0.0024 | 0.0016 | 0.0016 |
| F01 | 25940001 | 26040000 | 0.0026 | 0.0017 | 0.0018 |
| F01 | 25960001 | 26060000 | 0.0036 | 0.0026 | 0.0029 |
| F01 | 25980001 | 26080000 | 0.0042 | 0.0029 | 0.0034 |
| F01 | 26000001 | 26100000 | 0.0038 | 0.0028 | 0.0033 |
| F01 | 26020001 | 26120000 | 0.0038 | 0.0027 | 0.0033 |
| F01 | 26040001 | 26140000 | 0.0036 | 0.0025 | 0.0031 |
| F01 | 26060001 | 26160000 | 0.0030 | 0.0020 | 0.0024 |
| F01 | 26080001 | 26180000 | 0.0024 | 0.0018 | 0.0019 |
| F01 | 26100001 | 26200000 | 0.0024 | 0.0017 | 0.0019 |
| F01 | 26120001 | 26220000 | 0.0027 | 0.0020 | 0.0021 |
| F01 | 26140001 | 26240000 | 0.0026 | 0.0020 | 0.0020 |
| F01 | 26160001 | 26260000 | 0.0022 | 0.0017 | 0.0017 |
| F01 | 26180001 | 26280000 | 0.0018 | 0.0013 | 0.0013 |
| F01 | 26200001 | 26300000 | 0.0013 | 0.0010 | 0.0009 |
| F01 | 26220001 | 26320000 | 0.0009 | 0.0006 | 0.0007 |
| F01 | 26240001 | 26340000 | 0.0010 | 0.0006 | 0.0006 |
| F01 | 26260001 | 26360000 | 0.0017 | 0.0013 | 0.0014 |
| F01 | 26280001 | 26380000 | 0.0023 | 0.0016 | 0.0018 |
| F01 | 26300001 | 26400000 | 0.0029 | 0.0020 | 0.0023 |
| F01 | 26320001 | 26420000 | 0.0030 | 0.0021 | 0.0026 |
| F01 | 26340001 | 26440000 | 0.0033 | 0.0024 | 0.0029 |
| F01 | 26360001 | 26460000 | 0.0021 | 0.0013 | 0.0018 |
| F01 | 26380001 | 26480000 | 0.0017 | 0.0011 | 0.0014 |
| F01 | 26400001 | 26500000 | 0.0020 | 0.0015 | 0.0018 |
| F01 | 26420001 | 26520000 | 0.0022 | 0.0017 | 0.0019 |
| F01 | 26440001 | 26540000 | 0.0021 | 0.0016 | 0.0016 |
| F01 | 26460001 | 26560000 | 0.0026 | 0.0017 | 0.0018 |
| F01 | 26480001 | 26580000 | 0.0028 | 0.0018 | 0.0019 |
| F01 | 26500001 | 26600000 | 0.0020 | 0.0011 | 0.0012 |
| F01 | 26520001 | 26620000 | 0.0023 | 0.0015 | 0.0016 |
| F01 | 26540001 | 26640000 | 0.0019 | 0.0013 | 0.0014 |
| F01 | 26560001 | 26660000 | 0.0016 | 0.0014 | 0.0015 |
| F01 | 26580001 | 26680000 | 0.0020 | 0.0019 | 0.0020 |

| Chr | Start | End | both | A1 | A2 |
|---|---|---|---|---|---|
| F01 | 26600001 | 26700000 | 0.0037 | 0.0034 | 0.0036 |
| F01 | 26620001 | 26720000 | 0.0035 | 0.0031 | 0.0032 |
| F01 | 26640001 | 26740000 | 0.0043 | 0.0036 | 0.0038 |
| F01 | 26660001 | 26760000 | 0.0045 | 0.0037 | 0.0038 |
| F01 | 26680001 | 26780000 | 0.0049 | 0.0039 | 0.0042 |
| F01 | 26700001 | 26800000 | 0.0040 | 0.0031 | 0.0032 |
| F01 | 26720001 | 26820000 | 0.0041 | 0.0032 | 0.0035 |
| F01 | 26740001 | 26840000 | 0.0039 | 0.0030 | 0.0033 |
| F01 | 26760001 | 26860000 | 0.0038 | 0.0028 | 0.0032 |
| F01 | 26780001 | 26880000 | 0.0034 | 0.0025 | 0.0028 |
| F01 | 26800001 | 26900000 | 0.0031 | 0.0022 | 0.0027 |
| F01 | 26820001 | 26920000 | 0.0030 | 0.0020 | 0.0025 |
| F01 | 26840001 | 26940000 | 0.0035 | 0.0026 | 0.0030 |
| F01 | 26860001 | 26960000 | 0.0038 | 0.0031 | 0.0034 |
| F01 | 26880001 | 26980000 | 0.0034 | 0.0026 | 0.0029 |
| F01 | 26900001 | 27000000 | 0.0029 | 0.0022 | 0.0024 |
| F01 | 26920001 | 27020000 | 0.0028 | 0.0022 | 0.0023 |
| F01 | 26940001 | 27040000 | 0.0021 | 0.0014 | 0.0015 |
| F01 | 26960001 | 27060000 | 0.0024 | 0.0016 | 0.0018 |
| F01 | 26980001 | 27080000 | 0.0019 | 0.0013 | 0.0015 |
| F01 | 27000001 | 27100000 | 0.0020 | 0.0015 | 0.0017 |
| F01 | 27020001 | 27120000 | 0.0027 | 0.0020 | 0.0021 |
| F01 | 27040001 | 27140000 | 0.0030 | 0.0022 | 0.0024 |
| F01 | 27060001 | 27160000 | 0.0030 | 0.0021 | 0.0023 |
| F01 | 27080001 | 27180000 | 0.0033 | 0.0023 | 0.0024 |
| F01 | 27100001 | 27200000 | 0.0039 | 0.0028 | 0.0028 |
| F01 | 27120001 | 27220000 | 0.0039 | 0.0030 | 0.0029 |
| F01 | 27140001 | 27240000 | 0.0038 | 0.0029 | 0.0030 |
| F01 | 27160001 | 27260000 | 0.0030 | 0.0022 | 0.0022 |
| F01 | 27180001 | 27280000 | 0.0032 | 0.0024 | 0.0025 |
| F01 | 27200001 | 27300000 | 0.0035 | 0.0026 | 0.0030 |
| F01 | 27220001 | 27320000 | 0.0029 | 0.0021 | 0.0025 |
| F01 | 27240001 | 27340000 | 0.0034 | 0.0025 | 0.0029 |
| F01 | 27260001 | 27360000 | 0.0045 | 0.0034 | 0.0040 |
| F01 | 27280001 | 27380000 | 0.0053 | 0.0041 | 0.0047 |
| F01 | 27300001 | 27400000 | 0.0052 | 0.0041 | 0.0046 |
| F01 | 27320001 | 27420000 | 0.0058 | 0.0047 | 0.0053 |
| F01 | 27340001 | 27440000 | 0.0052 | 0.0043 | 0.0046 |
| F01 | 27360001 | 27460000 | 0.0047 | 0.0038 | 0.0041 |
| F01 | 27380001 | 27480000 | 0.0043 | 0.0034 | 0.0035 |
| F01 | 27400001 | 27500000 | 0.0043 | 0.0034 | 0.0035 |
| F01 | 27420001 | 27520000 | 0.0038 | 0.0028 | 0.0027 |
| F01 | 27440001 | 27540000 | 0.0045 | 0.0034 | 0.0034 |
| F01 | 27460001 | 27560000 | 0.0050 | 0.0039 | 0.0039 |
| F01 | 27480001 | 27580000 | 0.0051 | 0.0039 | 0.0041 |
| F01 | 27500001 | 27600000 | 0.0053 | 0.0040 | 0.0044 |
| F01 | 27520001 | 27620000 | 0.0046 | 0.0036 | 0.0040 |
| F01 | 27540001 | 27640000 | 0.0045 | 0.0032 | 0.0037 |
| F01 | 27560001 | 27660000 | 0.0041 | 0.0027 | 0.0033 |
| F01 | 27580001 | 27680000 | 0.0036 | 0.0023 | 0.0026 |
| F01 | 27600001 | 27700000 | 0.0037 | 0.0026 | 0.0028 |
| F01 | 27620001 | 27720000 | 0.0042 | 0.0030 | 0.0031 |
| F01 | 27640001 | 27740000 | 0.0039 | 0.0028 | 0.0029 |
| F01 | 27660001 | 27760000 | 0.0034 | 0.0025 | 0.0025 |
| F01 | 27680001 | 27780000 | 0.0034 | 0.0025 | 0.0026 |
| F01 | 27700001 | 27800000 | 0.0031 | 0.0023 | 0.0025 |
| F01 | 27720001 | 27820000 | 0.0034 | 0.0026 | 0.0028 |
| F01 | 27740001 | 27840000 | 0.0031 | 0.0025 | 0.0027 |

| Chr | Start | End | both | A1 | A2 |
|---|---|---|---|---|---|
| F01 | 27760001 | 27860000 | 0.0034 | 0.0028 | 0.0030 |
| F01 | 27780001 | 27880000 | 0.0030 | 0.0026 | 0.0028 |
| F01 | 27800001 | 27900000 | 0.0036 | 0.0029 | 0.0031 |
| F01 | 27820001 | 27920000 | 0.0046 | 0.0037 | 0.0041 |
| F01 | 27840001 | 27940000 | 0.0051 | 0.0042 | 0.0046 |
| F01 | 27860001 | 27960000 | 0.0049 | 0.0041 | 0.0044 |
| F01 | 27880001 | 27980000 | 0.0050 | 0.0042 | 0.0046 |
| F01 | 27900001 | 28000000 | 0.0039 | 0.0030 | 0.0034 |
| F01 | 27920001 | 28020000 | 0.0036 | 0.0026 | 0.0030 |
| F01 | 27940001 | 28040000 | 0.0039 | 0.0026 | 0.0031 |
| F01 | 27960001 | 28060000 | 0.0040 | 0.0025 | 0.0031 |
| F01 | 27980001 | 28080000 | 0.0042 | 0.0026 | 0.0030 |
| F01 | 28000001 | 28100000 | 0.0043 | 0.0030 | 0.0032 |
| F01 | 28020001 | 28120000 | 0.0035 | 0.0024 | 0.0026 |
| F01 | 28040001 | 28140000 | 0.0025 | 0.0017 | 0.0016 |
| F01 | 28060001 | 28160000 | 0.0029 | 0.0021 | 0.0021 |
| F01 | 28080001 | 28180000 | 0.0028 | 0.0020 | 0.0021 |
| F01 | 28100001 | 28200000 | 0.0027 | 0.0020 | 0.0021 |
| F01 | 28120001 | 28220000 | 0.0022 | 0.0015 | 0.0016 |
| F01 | 28140001 | 28240000 | 0.0023 | 0.0017 | 0.0018 |
| F01 | 28160001 | 28260000 | 0.0016 | 0.0012 | 0.0011 |
| F01 | 28180001 | 28280000 | 0.0012 | 0.0009 | 0.0009 |
| F01 | 28200001 | 28300000 | 0.0013 | 0.0011 | 0.0012 |
| F01 | 28220001 | 28320000 | 0.0015 | 0.0013 | 0.0015 |
| F01 | 28240001 | 28340000 | 0.0017 | 0.0015 | 0.0016 |
| F01 | 28260001 | 28360000 | 0.0020 | 0.0015 | 0.0017 |
| F01 | 28280001 | 28380000 | 0.0025 | 0.0016 | 0.0017 |
| F01 | 28300001 | 28400000 | 0.0022 | 0.0013 | 0.0014 |
| F01 | 28320001 | 28420000 | 0.0025 | 0.0014 | 0.0015 |
| F01 | 28340001 | 28440000 | 0.0023 | 0.0012 | 0.0013 |
| F01 | 28360001 | 28460000 | 0.0024 | 0.0015 | 0.0017 |
| F01 | 28380001 | 28480000 | 0.0024 | 0.0017 | 0.0019 |
| F01 | 28400001 | 28500000 | 0.0024 | 0.0017 | 0.0018 |
| F01 | 28420001 | 28520000 | 0.0024 | 0.0015 | 0.0017 |
| F01 | 28440001 | 28540000 | 0.0024 | 0.0016 | 0.0017 |
| F01 | 28460001 | 28560000 | 0.0023 | 0.0015 | 0.0015 |
| F01 | 28480001 | 28580000 | 0.0023 | 0.0014 | 0.0014 |
| F01 | 28500001 | 28600000 | 0.0019 | 0.0011 | 0.0012 |
| F01 | 28520001 | 28620000 | 0.0015 | 0.0009 | 0.0010 |
| F01 | 28540001 | 28640000 | 0.0023 | 0.0016 | 0.0014 |
| F01 | 28560001 | 28660000 | 0.0024 | 0.0016 | 0.0014 |
| F01 | 28580001 | 28680000 | 0.0034 | 0.0026 | 0.0024 |
| F01 | 28600001 | 28700000 | 0.0036 | 0.0030 | 0.0028 |
| F01 | 28620001 | 28720000 | 0.0036 | 0.0032 | 0.0029 |
| F01 | 28640001 | 28740000 | 0.0031 | 0.0028 | 0.0027 |
| F01 | 28660001 | 28760000 | 0.0028 | 0.0027 | 0.0025 |
| F01 | 28680001 | 28780000 | 0.0022 | 0.0020 | 0.0018 |
| F01 | 28700001 | 28800000 | 0.0042 | 0.0036 | 0.0035 |
| F01 | 28720001 | 28820000 | 0.0060 | 0.0053 | 0.0051 |
| F01 | 28740001 | 28840000 | 0.0061 | 0.0053 | 0.0050 |
| F01 | 28760001 | 28860000 | 0.0069 | 0.0059 | 0.0057 |
| F01 | 28780001 | 28880000 | 0.0070 | 0.0061 | 0.0061 |
| F01 | 28800001 | 28900000 | 0.0054 | 0.0048 | 0.0048 |
| F01 | 28820001 | 28920000 | 0.0041 | 0.0035 | 0.0036 |
| F01 | 28840001 | 28940000 | 0.0041 | 0.0035 | 0.0038 |
| F01 | 28860001 | 28960000 | 0.0033 | 0.0028 | 0.0030 |
| F01 | 28880001 | 28980000 | 0.0028 | 0.0023 | 0.0025 |
| F01 | 28900001 | 29000000 | 0.0023 | 0.0018 | 0.0020 |

| Chr | Start | End | both | A1 | A2 |
|---|---|---|---|---|---|
| F01 | 28920001 | 29020000 | 0.0023 | 0.0016 | 0.0018 |
| F01 | 28940001 | 29040000 | 0.0025 | 0.0018 | 0.0021 |
| F01 | 28960001 | 29060000 | 0.0025 | 0.0020 | 0.0022 |
| F01 | 28980001 | 29080000 | 0.0025 | 0.0020 | 0.0022 |
| F01 | 29000001 | 29100000 | 0.0033 | 0.0027 | 0.0030 |
| F01 | 29020001 | 29120000 | 0.0031 | 0.0026 | 0.0030 |
| F01 | 29040001 | 29140000 | 0.0028 | 0.0023 | 0.0025 |
| F01 | 29060001 | 29160000 | 0.0027 | 0.0021 | 0.0023 |
| F01 | 29080001 | 29180000 | 0.0034 | 0.0028 | 0.0029 |
| F01 | 29100001 | 29200000 | 0.0031 | 0.0026 | 0.0025 |
| F01 | 29120001 | 29220000 | 0.0027 | 0.0023 | 0.0022 |
| F01 | 29140001 | 29240000 | 0.0023 | 0.0020 | 0.0020 |
| F01 | 29160001 | 29260000 | 0.0026 | 0.0023 | 0.0024 |
| F01 | 29180001 | 29280000 | 0.0019 | 0.0018 | 0.0018 |
| F01 | 29200001 | 29300000 | 0.0018 | 0.0017 | 0.0016 |
| F01 | 29220001 | 29320000 | 0.0019 | 0.0016 | 0.0015 |
| F01 | 29240001 | 29340000 | 0.0021 | 0.0017 | 0.0017 |
| F01 | 29260001 | 29360000 | 0.0021 | 0.0016 | 0.0015 |
| F01 | 29280001 | 29380000 | 0.0022 | 0.0016 | 0.0017 |
| F01 | 29300001 | 29400000 | 0.0020 | 0.0014 | 0.0016 |
| F01 | 29320001 | 29420000 | 0.0024 | 0.0019 | 0.0020 |
| F01 | 29340001 | 29440000 | 0.0025 | 0.0019 | 0.0021 |
| F01 | 29360001 | 29460000 | 0.0025 | 0.0019 | 0.0021 |
| F01 | 29380001 | 29480000 | 0.0029 | 0.0022 | 0.0024 |
| F01 | 29400001 | 29500000 | 0.0029 | 0.0021 | 0.0024 |
| F01 | 29420001 | 29520000 | 0.0038 | 0.0034 | 0.0034 |
| F01 | 29440001 | 29540000 | 0.0041 | 0.0037 | 0.0036 |
| F01 | 29460001 | 29560000 | 0.0045 | 0.0041 | 0.0040 |
| F01 | 29480001 | 29580000 | 0.0041 | 0.0037 | 0.0036 |
| F01 | 29500001 | 29600000 | 0.0044 | 0.0038 | 0.0039 |
| F01 | 29520001 | 29620000 | 0.0035 | 0.0026 | 0.0032 |
| F01 | 29540001 | 29640000 | 0.0036 | 0.0026 | 0.0033 |
| F01 | 29560001 | 29660000 | 0.0027 | 0.0020 | 0.0026 |
| F01 | 29580001 | 29680000 | 0.0023 | 0.0016 | 0.0021 |
| F01 | 29600001 | 29700000 | 0.0016 | 0.0014 | 0.0016 |
| F01 | 29620001 | 29720000 | 0.0009 | 0.0008 | 0.0009 |
| F01 | 29640001 | 29740000 | 0.0005 | 0.0004 | 0.0005 |
| F01 | 29660001 | 29760000 | 0.0007 | 0.0006 | 0.0006 |
| F01 | 29680001 | 29780000 | 0.0010 | 0.0009 | 0.0010 |
| F01 | 29700001 | 29800000 | 0.0013 | 0.0012 | 0.0013 |
| F01 | 29720001 | 29820000 | 0.0013 | 0.0012 | 0.0012 |
| F01 | 29740001 | 29840000 | 0.0021 | 0.0018 | 0.0019 |
| F01 | 29760001 | 29860000 | 0.0023 | 0.0019 | 0.0022 |
| F01 | 29780001 | 29880000 | 0.0031 | 0.0025 | 0.0028 |
| F01 | 29800001 | 29900000 | 0.0030 | 0.0023 | 0.0027 |
| F01 | 29820001 | 29920000 | 0.0031 | 0.0024 | 0.0028 |
| F01 | 29840001 | 29940000 | 0.0025 | 0.0020 | 0.0024 |
| F01 | 29860001 | 29960000 | 0.0031 | 0.0026 | 0.0029 |
| F01 | 29880001 | 29980000 | 0.0025 | 0.0021 | 0.0024 |
| F01 | 29900001 | 30000000 | 0.0024 | 0.0021 | 0.0023 |
| F01 | 29920001 | 30020000 | 0.0037 | 0.0032 | 0.0036 |
| F01 | 29940001 | 30040000 | 0.0040 | 0.0035 | 0.0037 |
| F01 | 29960001 | 30060000 | 0.0037 | 0.0033 | 0.0035 |
| F01 | 29980001 | 30080000 | 0.0031 | 0.0028 | 0.0030 |
| F01 | 30000001 | 30100000 | 0.0030 | 0.0027 | 0.0028 |
| F01 | 30020001 | 30120000 | 0.0023 | 0.0019 | 0.0019 |
| F01 | 30040001 | 30140000 | 0.0023 | 0.0016 | 0.0017 |
| F01 | 30060001 | 30160000 | 0.0023 | 0.0015 | 0.0016 |

| Chr | Start | End | both | A1 | A2 |
|---|---|---|---|---|---|
| F01 | 30080001 | 30180000 | 0.0025 | 0.0016 | 0.0017 |
| F01 | 30100001 | 30200000 | 0.0027 | 0.0018 | 0.0019 |
| F01 | 30120001 | 30220000 | 0.0023 | 0.0016 | 0.0017 |
| F01 | 30140001 | 30240000 | 0.0026 | 0.0022 | 0.0023 |
| F01 | 30160001 | 30260000 | 0.0018 | 0.0016 | 0.0016 |
| F01 | 30180001 | 30280000 | 0.0018 | 0.0016 | 0.0016 |
| F01 | 30200001 | 30300000 | 0.0014 | 0.0013 | 0.0014 |
| F01 | 30220001 | 30320000 | 0.0017 | 0.0015 | 0.0017 |
| F01 | 30240001 | 30340000 | 0.0010 | 0.0008 | 0.0009 |
| F01 | 30260001 | 30360000 | 0.0009 | 0.0007 | 0.0009 |
| F01 | 30280001 | 30380000 | 0.0007 | 0.0006 | 0.0007 |
| F01 | 30300001 | 30400000 | 0.0008 | 0.0007 | 0.0008 |
| F01 | 30320001 | 30420000 | 0.0011 | 0.0010 | 0.0011 |
| F01 | 30340001 | 30440000 | 0.0014 | 0.0012 | 0.0013 |
| F01 | 30360001 | 30460000 | 0.0020 | 0.0018 | 0.0020 |
| F01 | 30380001 | 30480000 | 0.0021 | 0.0019 | 0.0021 |
| F01 | 30400001 | 30500000 | 0.0021 | 0.0018 | 0.0020 |
| F01 | 30420001 | 30520000 | 0.0016 | 0.0014 | 0.0015 |
| F01 | 30440001 | 30540000 | 0.0013 | 0.0012 | 0.0013 |
| F01 | 30460001 | 30560000 | 0.0007 | 0.0006 | 0.0007 |
| F01 | 30480001 | 30580000 | 0.0009 | 0.0007 | 0.0008 |
| F01 | 30500001 | 30600000 | 0.0016 | 0.0012 | 0.0013 |
| F01 | 30520001 | 30620000 | 0.0024 | 0.0020 | 0.0019 |
| F01 | 30540001 | 30640000 | 0.0030 | 0.0025 | 0.0023 |
| F01 | 30560001 | 30660000 | 0.0037 | 0.0030 | 0.0029 |
| F01 | 30580001 | 30680000 | 0.0042 | 0.0034 | 0.0034 |
| F01 | 30600001 | 30700000 | 0.0042 | 0.0034 | 0.0034 |
| F01 | 30620001 | 30720000 | 0.0034 | 0.0024 | 0.0025 |
| F01 | 30640001 | 30740000 | 0.0034 | 0.0025 | 0.0027 |
| F01 | 30660001 | 30760000 | 0.0033 | 0.0025 | 0.0027 |
| F01 | 30680001 | 30780000 | 0.0031 | 0.0024 | 0.0026 |
| F01 | 30700001 | 30800000 | 0.0028 | 0.0022 | 0.0024 |
| F01 | 30720001 | 30820000 | 0.0033 | 0.0027 | 0.0029 |
| F01 | 30740001 | 30840000 | 0.0033 | 0.0027 | 0.0028 |
| F01 | 30760001 | 30860000 | 0.0031 | 0.0027 | 0.0026 |
| F01 | 30780001 | 30880000 | 0.0024 | 0.0021 | 0.0021 |
| F01 | 30800001 | 30900000 | 0.0022 | 0.0020 | 0.0019 |
| F01 | 30820001 | 30920000 | 0.0017 | 0.0015 | 0.0015 |
| F01 | 30840001 | 30940000 | 0.0013 | 0.0011 | 0.0012 |
| F01 | 30860001 | 30960000 | 0.0011 | 0.0009 | 0.0011 |
| F01 | 30880001 | 30980000 | 0.0017 | 0.0015 | 0.0017 |
| F01 | 30900001 | 31000000 | 0.0018 | 0.0016 | 0.0018 |
| F01 | 30920001 | 31020000 | 0.0016 | 0.0014 | 0.0016 |
| F01 | 30940001 | 31040000 | 0.0021 | 0.0018 | 0.0020 |
| F01 | 30960001 | 31060000 | 0.0023 | 0.0019 | 0.0022 |
| F01 | 30980001 | 31080000 | 0.0023 | 0.0019 | 0.0022 |
| F01 | 31000001 | 31100000 | 0.0021 | 0.0018 | 0.0020 |
| F01 | 31020001 | 31120000 | 0.0022 | 0.0019 | 0.0021 |
| F01 | 31040001 | 31140000 | 0.0016 | 0.0013 | 0.0015 |
| F01 | 31060001 | 31160000 | 0.0013 | 0.0011 | 0.0012 |
| F01 | 31080001 | 31180000 | 0.0013 | 0.0011 | 0.0013 |
| F01 | 31100001 | 31200000 | 0.0014 | 0.0012 | 0.0014 |
| F01 | 31120001 | 31220000 | 0.0017 | 0.0013 | 0.0016 |
| F01 | 31140001 | 31240000 | 0.0024 | 0.0020 | 0.0023 |
| F01 | 31160001 | 31260000 | 0.0027 | 0.0022 | 0.0026 |
| F01 | 31180001 | 31280000 | 0.0032 | 0.0026 | 0.0030 |
| F01 | 31200001 | 31300000 | 0.0035 | 0.0029 | 0.0033 |
| F01 | 31220001 | 31320000 | 0.0036 | 0.0030 | 0.0034 |

| Chr | Start | End | both | A1 | A2 |
|---|---|---|---|---|---|
| F01 | 31240001 | 31340000 | 0.0029 | 0.0023 | 0.0027 |
| F01 | 31260001 | 31360000 | 0.0034 | 0.0026 | 0.0031 |
| F01 | 31280001 | 31380000 | 0.0028 | 0.0020 | 0.0024 |
| F01 | 31300001 | 31400000 | 0.0030 | 0.0020 | 0.0025 |
| F01 | 31320001 | 31420000 | 0.0033 | 0.0023 | 0.0026 |
| F01 | 31340001 | 31440000 | 0.0041 | 0.0029 | 0.0032 |
| F01 | 31360001 | 31460000 | 0.0039 | 0.0028 | 0.0029 |
| F01 | 31380001 | 31480000 | 0.0035 | 0.0027 | 0.0027 |
| F01 | 31400001 | 31500000 | 0.0031 | 0.0024 | 0.0023 |
| F01 | 31420001 | 31520000 | 0.0036 | 0.0028 | 0.0028 |
| F01 | 31440001 | 31540000 | 0.0031 | 0.0024 | 0.0025 |
| F01 | 31460001 | 31560000 | 0.0026 | 0.0019 | 0.0019 |
| F01 | 31480001 | 31580000 | 0.0034 | 0.0025 | 0.0026 |
| F01 | 31500001 | 31600000 | 0.0041 | 0.0031 | 0.0033 |
| F01 | 31520001 | 31620000 | 0.0035 | 0.0025 | 0.0029 |
| F01 | 31540001 | 31640000 | 0.0031 | 0.0022 | 0.0026 |
| F01 | 31560001 | 31660000 | 0.0036 | 0.0026 | 0.0030 |
| F01 | 31580001 | 31680000 | 0.0029 | 0.0020 | 0.0023 |
| F01 | 31600001 | 31700000 | 0.0023 | 0.0014 | 0.0017 |
| F01 | 31620001 | 31720000 | 0.0027 | 0.0018 | 0.0020 |
| F01 | 31640001 | 31740000 | 0.0031 | 0.0020 | 0.0021 |
| F01 | 31660001 | 31760000 | 0.0029 | 0.0020 | 0.0020 |
| F01 | 31680001 | 31780000 | 0.0036 | 0.0026 | 0.0027 |
| F01 | 31700001 | 31800000 | 0.0040 | 0.0029 | 0.0031 |
| F01 | 31720001 | 31820000 | 0.0047 | 0.0032 | 0.0036 |
| F01 | 31740001 | 31840000 | 0.0046 | 0.0031 | 0.0036 |
| F01 | 31760001 | 31860000 | 0.0050 | 0.0034 | 0.0038 |
| F01 | 31780001 | 31880000 | 0.0040 | 0.0026 | 0.0030 |
| F01 | 31800001 | 31900000 | 0.0034 | 0.0025 | 0.0027 |
| F01 | 31820001 | 31920000 | 0.0022 | 0.0017 | 0.0018 |
| F01 | 31840001 | 31940000 | 0.0020 | 0.0017 | 0.0018 |
| F01 | 31860001 | 31960000 | 0.0015 | 0.0014 | 0.0013 |
| F01 | 31880001 | 31980000 | 0.0026 | 0.0026 | 0.0023 |
| F01 | 31900001 | 32000000 | 0.0029 | 0.0028 | 0.0026 |
| F01 | 31920001 | 32020000 | 0.0025 | 0.0024 | 0.0022 |
| F01 | 31940001 | 32040000 | 0.0023 | 0.0023 | 0.0020 |
| F01 | 31960001 | 32060000 | 0.0024 | 0.0023 | 0.0022 |
| F01 | 31980001 | 32080000 | 0.0017 | 0.0013 | 0.0016 |
| F01 | 32000001 | 32100000 | 0.0013 | 0.0010 | 0.0012 |
| F01 | 32020001 | 32120000 | 0.0015 | 0.0011 | 0.0014 |
| F01 | 32040001 | 32140000 | 0.0016 | 0.0012 | 0.0014 |
| F01 | 32060001 | 32160000 | 0.0015 | 0.0011 | 0.0013 |
| F01 | 32080001 | 32180000 | 0.0014 | 0.0010 | 0.0011 |
| F01 | 32100001 | 32200000 | 0.0013 | 0.0009 | 0.0009 |
| F01 | 32120001 | 32220000 | 0.0015 | 0.0010 | 0.0011 |
| F01 | 32140001 | 32240000 | 0.0016 | 0.0011 | 0.0012 |
| F01 | 32160001 | 32260000 | 0.0018 | 0.0012 | 0.0014 |
| F01 | 32180001 | 32280000 | 0.0016 | 0.0012 | 0.0014 |
| F01 | 32200001 | 32300000 | 0.0021 | 0.0016 | 0.0019 |
| F01 | 32220001 | 32320000 | 0.0020 | 0.0017 | 0.0019 |
| F01 | 32240001 | 32340000 | 0.0026 | 0.0023 | 0.0025 |
| F01 | 32260001 | 32360000 | 0.0023 | 0.0020 | 0.0022 |
| F01 | 32280001 | 32380000 | 0.0021 | 0.0019 | 0.0021 |
| F01 | 32300001 | 32400000 | 0.0022 | 0.0019 | 0.0022 |
| F01 | 32320001 | 32420000 | 0.0024 | 0.0021 | 0.0023 |
| F01 | 32340001 | 32440000 | 0.0016 | 0.0013 | 0.0016 |
| F01 | 32360001 | 32460000 | 0.0015 | 0.0013 | 0.0015 |
| F01 | 32380001 | 32480000 | 0.0017 | 0.0015 | 0.0017 |

| Chr | Start | End | both | A1 | A2 |
|---|---|---|---|---|---|
| F01 | 32400001 | 32500000 | 0.0011 | 0.0010 | 0.0011 |
| F01 | 32420001 | 32520000 | 0.0008 | 0.0007 | 0.0008 |
| F01 | 32440001 | 32540000 | 0.0012 | 0.0011 | 0.0011 |
| F01 | 32460001 | 32560000 | 0.0011 | 0.0010 | 0.0011 |
| F01 | 32480001 | 32580000 | 0.0008 | 0.0007 | 0.0008 |
| F01 | 32500001 | 32600000 | 0.0008 | 0.0008 | 0.0008 |
| F01 | 32520001 | 32620000 | 0.0010 | 0.0010 | 0.0010 |
| F01 | 32540001 | 32640000 | 0.0009 | 0.0008 | 0.0009 |
| F01 | 32560001 | 32660000 | 0.0009 | 0.0009 | 0.0009 |
| F01 | 32580001 | 32680000 | 0.0011 | 0.0010 | 0.0011 |
| F01 | 32600001 | 32700000 | 0.0010 | 0.0009 | 0.0010 |
| F01 | 32620001 | 32720000 | 0.0007 | 0.0006 | 0.0007 |
| F01 | 32640001 | 32740000 | 0.0003 | 0.0003 | 0.0003 |
| F01 | 32660001 | 32760000 | 0.0004 | 0.0003 | 0.0003 |
| F01 | 32680001 | 32780000 | 0.0002 | 0.0002 | 0.0002 |
| F01 | 32700001 | 32800000 | 0.0003 | 0.0002 | 0.0003 |
| F01 | 32720001 | 32820000 | 0.0005 | 0.0003 | 0.0005 |
| F01 | 32740001 | 32840000 | 0.0006 | 0.0004 | 0.0006 |
| F01 | 32760001 | 32860000 | 0.0005 | 0.0004 | 0.0005 |
| F01 | 32780001 | 32880000 | 0.0007 | 0.0006 | 0.0006 |
| F01 | 32800001 | 32900000 | 0.0009 | 0.0008 | 0.0008 |
| F01 | 32820001 | 32920000 | 0.0013 | 0.0011 | 0.0013 |
| F01 | 32840001 | 32940000 | 0.0012 | 0.0010 | 0.0012 |
| F01 | 32860001 | 32960000 | 0.0014 | 0.0012 | 0.0014 |
| F01 | 32880001 | 32980000 | 0.0013 | 0.0010 | 0.0013 |
| F01 | 32900001 | 33000000 | 0.0012 | 0.0009 | 0.0012 |
| F01 | 32920001 | 33020000 | 0.0008 | 0.0007 | 0.0008 |
| F01 | 32940001 | 33040000 | 0.0012 | 0.0010 | 0.0011 |
| F01 | 32960001 | 33060000 | 0.0010 | 0.0008 | 0.0009 |
| F01 | 32980001 | 33080000 | 0.0009 | 0.0008 | 0.0009 |
| F01 | 33000001 | 33100000 | 0.0011 | 0.0010 | 0.0011 |
| F01 | 33020001 | 33120000 | 0.0012 | 0.0012 | 0.0012 |
| F01 | 33040001 | 33140000 | 0.0011 | 0.0012 | 0.0011 |
| F01 | 33060001 | 33160000 | 0.0018 | 0.0018 | 0.0017 |
| F01 | 33080001 | 33180000 | 0.0023 | 0.0023 | 0.0021 |
| F01 | 33100001 | 33200000 | 0.0023 | 0.0022 | 0.0021 |
| F01 | 33120001 | 33220000 | 0.0020 | 0.0019 | 0.0019 |
| F01 | 33140001 | 33240000 | 0.0018 | 0.0016 | 0.0017 |
| F01 | 33160001 | 33260000 | 0.0018 | 0.0016 | 0.0018 |
| F01 | 33180001 | 33280000 | 0.0015 | 0.0013 | 0.0015 |
| F01 | 33200001 | 33300000 | 0.0015 | 0.0013 | 0.0015 |
| F01 | 33220001 | 33320000 | 0.0018 | 0.0016 | 0.0017 |
| F01 | 33240001 | 33340000 | 0.0030 | 0.0022 | 0.0027 |
| F01 | 33260001 | 33360000 | 0.0031 | 0.0022 | 0.0026 |
| F01 | 33280001 | 33380000 | 0.0033 | 0.0022 | 0.0028 |
| F01 | 33300001 | 33400000 | 0.0037 | 0.0024 | 0.0029 |
| F01 | 33320001 | 33420000 | 0.0039 | 0.0022 | 0.0028 |
| F01 | 33340001 | 33440000 | 0.0028 | 0.0018 | 0.0019 |
| F01 | 33360001 | 33460000 | 0.0029 | 0.0016 | 0.0019 |
| F01 | 33380001 | 33480000 | 0.0032 | 0.0020 | 0.0022 |
| F01 | 33400001 | 33500000 | 0.0035 | 0.0023 | 0.0025 |
| F01 | 33420001 | 33520000 | 0.0031 | 0.0022 | 0.0023 |
| F01 | 33440001 | 33540000 | 0.0035 | 0.0023 | 0.0025 |
| F01 | 33460001 | 33560000 | 0.0042 | 0.0031 | 0.0032 |
| F01 | 33480001 | 33580000 | 0.0041 | 0.0032 | 0.0032 |
| F01 | 33500001 | 33600000 | 0.0041 | 0.0033 | 0.0034 |
| F01 | 33520001 | 33620000 | 0.0043 | 0.0036 | 0.0038 |
| F01 | 33540001 | 33640000 | 0.0041 | 0.0037 | 0.0037 |

| Chr | Start | End | both | A1 | A2 |
|---|---|---|---|---|---|
| F01 | 33560001 | 33660000 | 0.0034 | 0.0030 | 0.0030 |
| F01 | 33580001 | 33680000 | 0.0033 | 0.0028 | 0.0028 |
| F01 | 33600001 | 33700000 | 0.0028 | 0.0023 | 0.0023 |
| F01 | 33620001 | 33720000 | 0.0027 | 0.0021 | 0.0022 |
| F01 | 33640001 | 33740000 | 0.0027 | 0.0020 | 0.0022 |
| F01 | 33660001 | 33760000 | 0.0020 | 0.0015 | 0.0018 |
| F01 | 33680001 | 33780000 | 0.0016 | 0.0012 | 0.0015 |
| F01 | 33700001 | 33800000 | 0.0016 | 0.0011 | 0.0015 |
| F01 | 33720001 | 33820000 | 0.0018 | 0.0013 | 0.0017 |
| F01 | 33740001 | 33840000 | 0.0015 | 0.0011 | 0.0015 |
| F01 | 33760001 | 33860000 | 0.0022 | 0.0017 | 0.0022 |
| F01 | 33780001 | 33880000 | 0.0025 | 0.0020 | 0.0025 |
| F01 | 33800001 | 33900000 | 0.0023 | 0.0018 | 0.0021 |
| F01 | 33820001 | 33920000 | 0.0025 | 0.0018 | 0.0019 |
| F01 | 33840001 | 33940000 | 0.0024 | 0.0018 | 0.0017 |
| F01 | 33860001 | 33960000 | 0.0022 | 0.0015 | 0.0014 |
| F01 | 33880001 | 33980000 | 0.0025 | 0.0016 | 0.0014 |
| F01 | 33900001 | 34000000 | 0.0030 | 0.0021 | 0.0022 |
| F01 | 33920001 | 34020000 | 0.0030 | 0.0022 | 0.0025 |
| F01 | 33940001 | 34040000 | 0.0039 | 0.0032 | 0.0035 |
| F01 | 33960001 | 34060000 | 0.0041 | 0.0032 | 0.0036 |
| F01 | 33980001 | 34080000 | 0.0045 | 0.0035 | 0.0040 |
| F01 | 34000001 | 34100000 | 0.0047 | 0.0036 | 0.0038 |
| F01 | 34020001 | 34120000 | 0.0044 | 0.0033 | 0.0035 |
| F01 | 34040001 | 34140000 | 0.0036 | 0.0024 | 0.0026 |
| F01 | 34060001 | 34160000 | 0.0036 | 0.0024 | 0.0025 |
| F01 | 34080001 | 34180000 | 0.0036 | 0.0024 | 0.0027 |
| F01 | 34100001 | 34200000 | 0.0027 | 0.0018 | 0.0021 |
| F01 | 34120001 | 34220000 | 0.0027 | 0.0019 | 0.0022 |
| F01 | 34140001 | 34240000 | 0.0033 | 0.0020 | 0.0027 |
| F01 | 34160001 | 34260000 | 0.0032 | 0.0021 | 0.0029 |
| F01 | 34180001 | 34280000 | 0.0023 | 0.0014 | 0.0021 |
| F01 | 34200001 | 34300000 | 0.0032 | 0.0023 | 0.0031 |
| F01 | 34220001 | 34320000 | 0.0035 | 0.0025 | 0.0032 |
| F01 | 34240001 | 34340000 | 0.0033 | 0.0029 | 0.0031 |
| F01 | 34260001 | 34360000 | 0.0031 | 0.0027 | 0.0029 |
| F01 | 34280001 | 34380000 | 0.0035 | 0.0032 | 0.0033 |
| F01 | 34300001 | 34400000 | 0.0027 | 0.0025 | 0.0024 |
| F01 | 34320001 | 34420000 | 0.0027 | 0.0026 | 0.0025 |
| F01 | 34340001 | 34440000 | 0.0021 | 0.0021 | 0.0020 |
| F01 | 34360001 | 34460000 | 0.0020 | 0.0019 | 0.0018 |
| F01 | 34380001 | 34480000 | 0.0013 | 0.0012 | 0.0013 |
| F01 | 34400001 | 34500000 | 0.0014 | 0.0012 | 0.0013 |
| F01 | 34420001 | 34520000 | 0.0011 | 0.0009 | 0.0010 |
| F01 | 34440001 | 34540000 | 0.0010 | 0.0009 | 0.0010 |
| F01 | 34460001 | 34560000 | 0.0009 | 0.0007 | 0.0009 |
| F01 | 34480001 | 34580000 | 0.0007 | 0.0006 | 0.0007 |
| F01 | 34500001 | 34600000 | 0.0007 | 0.0005 | 0.0007 |
| F01 | 34520001 | 34620000 | 0.0003 | 0.0003 | 0.0003 |
| F01 | 34540001 | 34640000 | 0.0006 | 0.0005 | 0.0005 |
| F01 | 34560001 | 34660000 | 0.0006 | 0.0005 | 0.0006 |
| F01 | 34580001 | 34680000 | 0.0008 | 0.0007 | 0.0008 |
| F01 | 34600001 | 34700000 | 0.0009 | 0.0008 | 0.0009 |
| F01 | 34620001 | 34720000 | 0.0015 | 0.0013 | 0.0015 |
| F01 | 34640001 | 34740000 | 0.0016 | 0.0013 | 0.0016 |
| F01 | 34660001 | 34760000 | 0.0021 | 0.0018 | 0.0020 |
| F01 | 34680001 | 34780000 | 0.0023 | 0.0020 | 0.0023 |
| F01 | 34700001 | 34800000 | 0.0026 | 0.0022 | 0.0025 |

| Chr | Start | End | both | A1 | A2 |
|---|---|---|---|---|---|
| F01 | 34720001 | 34820000 | 0.0030 | 0.0026 | 0.0029 |
| F01 | 34740001 | 34840000 | 0.0030 | 0.0025 | 0.0028 |
| F01 | 34760001 | 34860000 | 0.0031 | 0.0024 | 0.0028 |
| F01 | 34780001 | 34880000 | 0.0040 | 0.0030 | 0.0035 |
| F01 | 34800001 | 34900000 | 0.0041 | 0.0030 | 0.0035 |
| F01 | 34820001 | 34920000 | 0.0049 | 0.0036 | 0.0041 |
| F01 | 34840001 | 34940000 | 0.0048 | 0.0036 | 0.0041 |
| F01 | 34860001 | 34960000 | 0.0051 | 0.0038 | 0.0043 |
| F01 | 34880001 | 34980000 | 0.0040 | 0.0029 | 0.0032 |
| F01 | 34900001 | 35000000 | 0.0041 | 0.0029 | 0.0031 |
| F01 | 34920001 | 35020000 | 0.0027 | 0.0018 | 0.0018 |
| F01 | 34940001 | 35040000 | 0.0041 | 0.0033 | 0.0031 |
| F01 | 34960001 | 35060000 | 0.0033 | 0.0027 | 0.0026 |
| F01 | 34980001 | 35080000 | 0.0038 | 0.0032 | 0.0030 |
| F01 | 35000001 | 35100000 | 0.0039 | 0.0032 | 0.0031 |
| F01 | 35020001 | 35120000 | 0.0041 | 0.0033 | 0.0032 |
| F01 | 35040001 | 35140000 | 0.0028 | 0.0017 | 0.0017 |
| F01 | 35060001 | 35160000 | 0.0028 | 0.0016 | 0.0016 |
| F01 | 35080001 | 35180000 | 0.0025 | 0.0014 | 0.0015 |
| F01 | 35100001 | 35200000 | 0.0025 | 0.0013 | 0.0015 |
| F01 | 35120001 | 35220000 | 0.0034 | 0.0021 | 0.0021 |
| F01 | 35140001 | 35240000 | 0.0033 | 0.0022 | 0.0022 |
| F01 | 35160001 | 35260000 | 0.0034 | 0.0023 | 0.0024 |
| F01 | 35180001 | 35280000 | 0.0030 | 0.0020 | 0.0020 |
| F01 | 35200001 | 35300000 | 0.0030 | 0.0021 | 0.0021 |
| F01 | 35220001 | 35320000 | 0.0019 | 0.0011 | 0.0012 |
| F01 | 35240001 | 35340000 | 0.0017 | 0.0010 | 0.0011 |
| F01 | 35260001 | 35360000 | 0.0029 | 0.0022 | 0.0023 |
| F01 | 35280001 | 35380000 | 0.0028 | 0.0023 | 0.0024 |
| F01 | 35300001 | 35400000 | 0.0033 | 0.0028 | 0.0029 |
| F01 | 35320001 | 35420000 | 0.0035 | 0.0029 | 0.0029 |
| F01 | 35340001 | 35440000 | 0.0040 | 0.0031 | 0.0033 |
| F01 | 35360001 | 35460000 | 0.0031 | 0.0022 | 0.0023 |
| F01 | 35380001 | 35480000 | 0.0046 | 0.0033 | 0.0035 |
| F01 | 35400001 | 35500000 | 0.0038 | 0.0026 | 0.0028 |
| F01 | 35420001 | 35520000 | 0.0036 | 0.0026 | 0.0029 |
| F01 | 35440001 | 35540000 | 0.0037 | 0.0025 | 0.0026 |
| F01 | 35460001 | 35560000 | 0.0037 | 0.0024 | 0.0025 |
| F01 | 35480001 | 35580000 | 0.0027 | 0.0017 | 0.0017 |
| F01 | 35500001 | 35600000 | 0.0024 | 0.0013 | 0.0013 |
| F01 | 35520001 | 35620000 | 0.0027 | 0.0014 | 0.0013 |
| F01 | 35540001 | 35640000 | 0.0038 | 0.0025 | 0.0025 |
| F01 | 35560001 | 35660000 | 0.0040 | 0.0027 | 0.0029 |
| F01 | 35580001 | 35680000 | 0.0041 | 0.0029 | 0.0032 |
| F01 | 35600001 | 35700000 | 0.0043 | 0.0031 | 0.0035 |
| F01 | 35620001 | 35720000 | 0.0047 | 0.0036 | 0.0042 |
| F01 | 35640001 | 35740000 | 0.0034 | 0.0023 | 0.0029 |
| F01 | 35660001 | 35760000 | 0.0031 | 0.0018 | 0.0023 |
| F01 | 35680001 | 35780000 | 0.0027 | 0.0013 | 0.0017 |
| F01 | 35700001 | 35800000 | 0.0040 | 0.0023 | 0.0027 |
| F01 | 35720001 | 35820000 | 0.0033 | 0.0016 | 0.0018 |
| F01 | 35740001 | 35840000 | 0.0038 | 0.0022 | 0.0025 |
| F01 | 35760001 | 35860000 | 0.0035 | 0.0021 | 0.0024 |
| F01 | 35780001 | 35880000 | 0.0040 | 0.0026 | 0.0030 |
| F01 | 35800001 | 35900000 | 0.0036 | 0.0021 | 0.0025 |
| F01 | 35820001 | 35920000 | 0.0040 | 0.0025 | 0.0030 |
| F01 | 35840001 | 35940000 | 0.0045 | 0.0025 | 0.0034 |
| F01 | 35860001 | 35960000 | 0.0052 | 0.0033 | 0.0042 |

| Chr | Start | End | both | A1 | A2 |
|---|---|---|---|---|---|
| F01 | 35880001 | 35980000 | 0.0047 | 0.0030 | 0.0038 |
| F01 | 35900001 | 36000000 | 0.0037 | 0.0025 | 0.0031 |
| F01 | 35920001 | 36020000 | 0.0037 | 0.0027 | 0.0032 |
| F01 | 35940001 | 36040000 | 0.0036 | 0.0033 | 0.0032 |
| F01 | 35960001 | 36060000 | 0.0040 | 0.0036 | 0.0037 |
| F01 | 35980001 | 36080000 | 0.0048 | 0.0042 | 0.0043 |
| F01 | 36000001 | 36100000 | 0.0054 | 0.0045 | 0.0048 |
| F01 | 36020001 | 36120000 | 0.0053 | 0.0044 | 0.0048 |
| F01 | 36040001 | 36140000 | 0.0041 | 0.0033 | 0.0036 |
| F01 | 36060001 | 36160000 | 0.0030 | 0.0023 | 0.0025 |
| F01 | 36080001 | 36180000 | 0.0023 | 0.0016 | 0.0018 |
| F01 | 36100001 | 36200000 | 0.0018 | 0.0012 | 0.0012 |
| F01 | 36120001 | 36220000 | 0.0021 | 0.0014 | 0.0015 |
| F01 | 36140001 | 36240000 | 0.0019 | 0.0011 | 0.0013 |
| F01 | 36160001 | 36260000 | 0.0022 | 0.0014 | 0.0016 |
| F01 | 36180001 | 36280000 | 0.0020 | 0.0015 | 0.0017 |
| F01 | 36200001 | 36300000 | 0.0020 | 0.0015 | 0.0017 |
| F01 | 36220001 | 36320000 | 0.0017 | 0.0014 | 0.0014 |
| F01 | 36240001 | 36340000 | 0.0020 | 0.0016 | 0.0017 |
| F01 | 36260001 | 36360000 | 0.0020 | 0.0018 | 0.0019 |
| F01 | 36280001 | 36380000 | 0.0019 | 0.0016 | 0.0017 |
| F01 | 36300001 | 36400000 | 0.0020 | 0.0017 | 0.0018 |
| F01 | 36320001 | 36420000 | 0.0016 | 0.0013 | 0.0014 |
| F01 | 36340001 | 36440000 | 0.0018 | 0.0014 | 0.0018 |
| F01 | 36360001 | 36460000 | 0.0014 | 0.0010 | 0.0013 |
| F01 | 36380001 | 36480000 | 0.0013 | 0.0010 | 0.0013 |
| F01 | 36400001 | 36500000 | 0.0009 | 0.0007 | 0.0009 |
| F01 | 36420001 | 36520000 | 0.0008 | 0.0006 | 0.0009 |
| F01 | 36440001 | 36540000 | 0.0004 | 0.0003 | 0.0004 |
| F01 | 36460001 | 36560000 | 0.0008 | 0.0007 | 0.0008 |
| F01 | 36480001 | 36580000 | 0.0015 | 0.0013 | 0.0015 |
| F01 | 36500001 | 36600000 | 0.0020 | 0.0017 | 0.0020 |
| F01 | 36520001 | 36620000 | 0.0022 | 0.0019 | 0.0021 |
| F01 | 36540001 | 36640000 | 0.0022 | 0.0019 | 0.0022 |
| F01 | 36560001 | 36660000 | 0.0019 | 0.0016 | 0.0019 |
| F01 | 36580001 | 36680000 | 0.0011 | 0.0010 | 0.0011 |
| F01 | 36600001 | 36700000 | 0.0008 | 0.0007 | 0.0008 |
| F01 | 36620001 | 36720000 | 0.0005 | 0.0005 | 0.0005 |
| F01 | 36640001 | 36740000 | 0.0005 | 0.0004 | 0.0005 |
| F01 | 36660001 | 36760000 | 0.0006 | 0.0005 | 0.0005 |
| F01 | 36680001 | 36780000 | 0.0009 | 0.0008 | 0.0008 |
| F01 | 36700001 | 36800000 | 0.0011 | 0.0008 | 0.0009 |
| F01 | 36720001 | 36820000 | 0.0011 | 0.0007 | 0.0008 |
| F01 | 36740001 | 36840000 | 0.0016 | 0.0011 | 0.0011 |
| F01 | 36760001 | 36860000 | 0.0018 | 0.0013 | 0.0013 |
| F01 | 36780001 | 36880000 | 0.0016 | 0.0011 | 0.0011 |
| F01 | 36800001 | 36900000 | 0.0018 | 0.0013 | 0.0014 |
| F01 | 36820001 | 36920000 | 0.0025 | 0.0019 | 0.0020 |
| F01 | 36840001 | 36940000 | 0.0024 | 0.0018 | 0.0021 |
| F01 | 36860001 | 36960000 | 0.0020 | 0.0015 | 0.0018 |
| F01 | 36880001 | 36980000 | 0.0023 | 0.0017 | 0.0019 |
| F01 | 36900001 | 37000000 | 0.0018 | 0.0013 | 0.0015 |
| F01 | 36920001 | 37020000 | 0.0013 | 0.0009 | 0.0011 |
| F01 | 36940001 | 37040000 | 0.0012 | 0.0008 | 0.0008 |
| F01 | 36960001 | 37060000 | 0.0012 | 0.0008 | 0.0008 |
| F01 | 36980001 | 37080000 | 0.0013 | 0.0009 | 0.0009 |
| F01 | 37000001 | 37100000 | 0.0015 | 0.0010 | 0.0011 |
| F01 | 37020001 | 37120000 | 0.0015 | 0.0011 | 0.0012 |

| Chr | Start | End | both | A1 | A2 |
|---|---|---|---|---|---|
| F01 | 37040001 | 37140000 | 0.0011 | 0.0008 | 0.0010 |
| F01 | 37060001 | 37160000 | 0.0019 | 0.0015 | 0.0017 |
| F01 | 37080001 | 37180000 | 0.0017 | 0.0012 | 0.0014 |
| F01 | 37100001 | 37200000 | 0.0019 | 0.0014 | 0.0017 |
| F01 | 37120001 | 37220000 | 0.0022 | 0.0016 | 0.0019 |
| F01 | 37140001 | 37240000 | 0.0024 | 0.0018 | 0.0021 |
| F01 | 37160001 | 37260000 | 0.0017 | 0.0012 | 0.0015 |
| F01 | 37180001 | 37280000 | 0.0020 | 0.0014 | 0.0017 |
| F01 | 37200001 | 37300000 | 0.0019 | 0.0014 | 0.0016 |
| F01 | 37220001 | 37320000 | 0.0018 | 0.0011 | 0.0014 |
| F01 | 37240001 | 37340000 | 0.0030 | 0.0020 | 0.0022 |
| F01 | 37260001 | 37360000 | 0.0031 | 0.0020 | 0.0022 |
| F01 | 37280001 | 37380000 | 0.0029 | 0.0018 | 0.0020 |
| F01 | 37300001 | 37400000 | 0.0035 | 0.0023 | 0.0024 |
| F01 | 37320001 | 37420000 | 0.0034 | 0.0023 | 0.0024 |
| F01 | 37340001 | 37440000 | 0.0029 | 0.0021 | 0.0022 |
| F01 | 37360001 | 37460000 | 0.0029 | 0.0021 | 0.0023 |
| F01 | 37380001 | 37480000 | 0.0034 | 0.0028 | 0.0029 |
| F01 | 37400001 | 37500000 | 0.0029 | 0.0024 | 0.0025 |
| F01 | 37420001 | 37520000 | 0.0031 | 0.0025 | 0.0025 |
| F01 | 37440001 | 37540000 | 0.0025 | 0.0019 | 0.0021 |
| F01 | 37460001 | 37560000 | 0.0022 | 0.0018 | 0.0019 |
| F01 | 37480001 | 37580000 | 0.0015 | 0.0012 | 0.0013 |
| F01 | 37500001 | 37600000 | 0.0013 | 0.0010 | 0.0011 |
| F01 | 37520001 | 37620000 | 0.0009 | 0.0008 | 0.0009 |
| F01 | 37540001 | 37640000 | 0.0008 | 0.0007 | 0.0008 |
| F01 | 37560001 | 37660000 | 0.0010 | 0.0008 | 0.0010 |
| F01 | 37580001 | 37680000 | 0.0010 | 0.0008 | 0.0009 |
| F01 | 37600001 | 37700000 | 0.0011 | 0.0008 | 0.0009 |
| F01 | 37620001 | 37720000 | 0.0016 | 0.0012 | 0.0014 |
| F01 | 37640001 | 37740000 | 0.0017 | 0.0013 | 0.0015 |
| F01 | 37660001 | 37760000 | 0.0027 | 0.0020 | 0.0023 |
| F01 | 37680001 | 37780000 | 0.0033 | 0.0024 | 0.0028 |
| F01 | 37700001 | 37800000 | 0.0033 | 0.0025 | 0.0028 |
| F01 | 37720001 | 37820000 | 0.0034 | 0.0026 | 0.0029 |
| F01 | 37740001 | 37840000 | 0.0037 | 0.0028 | 0.0031 |
| F01 | 37760001 | 37860000 | 0.0032 | 0.0025 | 0.0028 |
| F01 | 37780001 | 37880000 | 0.0036 | 0.0030 | 0.0032 |
| F01 | 37800001 | 37900000 | 0.0036 | 0.0030 | 0.0033 |
| F01 | 37820001 | 37920000 | 0.0032 | 0.0027 | 0.0029 |
| F01 | 37840001 | 37940000 | 0.0029 | 0.0024 | 0.0027 |
| F01 | 37860001 | 37960000 | 0.0029 | 0.0025 | 0.0026 |
| F01 | 37880001 | 37980000 | 0.0018 | 0.0015 | 0.0016 |
| F01 | 37900001 | 38000000 | 0.0018 | 0.0015 | 0.0017 |
| F01 | 37920001 | 38020000 | 0.0020 | 0.0017 | 0.0020 |
| F01 | 37940001 | 38040000 | 0.0019 | 0.0017 | 0.0018 |
| F01 | 37960001 | 38060000 | 0.0014 | 0.0011 | 0.0011 |
| F01 | 37980001 | 38080000 | 0.0017 | 0.0012 | 0.0012 |
| F01 | 38000001 | 38100000 | 0.0016 | 0.0011 | 0.0011 |
| F01 | 38020001 | 38120000 | 0.0015 | 0.0011 | 0.0010 |
| F01 | 38040001 | 38140000 | 0.0017 | 0.0012 | 0.0013 |
| F01 | 38060001 | 38160000 | 0.0021 | 0.0017 | 0.0018 |
| F01 | 38080001 | 38180000 | 0.0020 | 0.0017 | 0.0019 |
| F01 | 38100001 | 38200000 | 0.0020 | 0.0017 | 0.0018 |
| F01 | 38120001 | 38220000 | 0.0016 | 0.0014 | 0.0014 |
| F01 | 38140001 | 38240000 | 0.0017 | 0.0013 | 0.0014 |
| F01 | 38160001 | 38260000 | 0.0015 | 0.0012 | 0.0013 |
| F01 | 38180001 | 38280000 | 0.0015 | 0.0012 | 0.0012 |

| Chr | Start | End | both | A1 | A2 |
|---|---|---|---|---|---|
| F01 | 38200001 | 38300000 | 0.0015 | 0.0012 | 0.0013 |
| F01 | 38220001 | 38320000 | 0.0018 | 0.0014 | 0.0015 |
| F01 | 38240001 | 38340000 | 0.0017 | 0.0012 | 0.0012 |
| F01 | 38260001 | 38360000 | 0.0017 | 0.0011 | 0.0011 |
| F01 | 38280001 | 38380000 | 0.0021 | 0.0011 | 0.0013 |
| F01 | 38300001 | 38400000 | 0.0024 | 0.0014 | 0.0016 |
| F01 | 38320001 | 38420000 | 0.0027 | 0.0017 | 0.0018 |
| F01 | 38340001 | 38440000 | 0.0029 | 0.0019 | 0.0020 |
| F01 | 38360001 | 38460000 | 0.0034 | 0.0024 | 0.0026 |
| F01 | 38380001 | 38480000 | 0.0028 | 0.0021 | 0.0022 |
| F01 | 38400001 | 38500000 | 0.0024 | 0.0017 | 0.0018 |
| F01 | 38420001 | 38520000 | 0.0024 | 0.0019 | 0.0019 |
| F01 | 38440001 | 38540000 | 0.0022 | 0.0019 | 0.0020 |
| F01 | 38460001 | 38560000 | 0.0019 | 0.0016 | 0.0017 |
| F01 | 38480001 | 38580000 | 0.0024 | 0.0019 | 0.0021 |
| F01 | 38500001 | 38600000 | 0.0025 | 0.0021 | 0.0021 |
| F01 | 38520001 | 38620000 | 0.0024 | 0.0018 | 0.0020 |
| F01 | 38540001 | 38640000 | 0.0024 | 0.0017 | 0.0019 |
| F01 | 38560001 | 38660000 | 0.0022 | 0.0017 | 0.0017 |
| F01 | 38580001 | 38680000 | 0.0021 | 0.0015 | 0.0017 |
| F01 | 38600001 | 38700000 | 0.0019 | 0.0014 | 0.0015 |
| F01 | 38620001 | 38720000 | 0.0025 | 0.0020 | 0.0022 |
| F01 | 38640001 | 38740000 | 0.0025 | 0.0019 | 0.0019 |
| F01 | 38660001 | 38760000 | 0.0029 | 0.0023 | 0.0024 |
| F01 | 38680001 | 38780000 | 0.0033 | 0.0025 | 0.0026 |
| F01 | 38700001 | 38800000 | 0.0034 | 0.0027 | 0.0027 |
| F01 | 38720001 | 38820000 | 0.0031 | 0.0026 | 0.0024 |
| F01 | 38740001 | 38840000 | 0.0029 | 0.0024 | 0.0023 |
| F01 | 38760001 | 38860000 | 0.0025 | 0.0018 | 0.0019 |
| F01 | 38780001 | 38880000 | 0.0025 | 0.0018 | 0.0019 |
| F01 | 38800001 | 38900000 | 0.0029 | 0.0021 | 0.0021 |
| F01 | 38820001 | 38920000 | 0.0027 | 0.0017 | 0.0019 |
| F01 | 38840001 | 38940000 | 0.0041 | 0.0030 | 0.0032 |
| F01 | 38860001 | 38960000 | 0.0044 | 0.0034 | 0.0035 |
| F01 | 38880001 | 38980000 | 0.0046 | 0.0036 | 0.0037 |
| F01 | 38900001 | 39000000 | 0.0045 | 0.0034 | 0.0036 |
| F01 | 38920001 | 39020000 | 0.0045 | 0.0034 | 0.0034 |
| F01 | 38940001 | 39040000 | 0.0037 | 0.0025 | 0.0027 |
| F01 | 38960001 | 39060000 | 0.0028 | 0.0019 | 0.0021 |
| F01 | 38980001 | 39080000 | 0.0026 | 0.0018 | 0.0020 |
| F01 | 39000001 | 39100000 | 0.0028 | 0.0020 | 0.0024 |
| F01 | 39020001 | 39120000 | 0.0037 | 0.0028 | 0.0033 |
| F01 | 39040001 | 39140000 | 0.0034 | 0.0025 | 0.0029 |
| F01 | 39060001 | 39160000 | 0.0035 | 0.0024 | 0.0028 |
| F01 | 39080001 | 39180000 | 0.0032 | 0.0020 | 0.0023 |
| F01 | 39100001 | 39200000 | 0.0031 | 0.0018 | 0.0019 |
| F01 | 39120001 | 39220000 | 0.0021 | 0.0010 | 0.0011 |
| F01 | 39140001 | 39240000 | 0.0021 | 0.0012 | 0.0013 |
| F01 | 39160001 | 39260000 | 0.0023 | 0.0013 | 0.0014 |
| F01 | 39180001 | 39280000 | 0.0026 | 0.0018 | 0.0019 |
| F01 | 39200001 | 39300000 | 0.0028 | 0.0019 | 0.0021 |
| F01 | 39220001 | 39320000 | 0.0029 | 0.0020 | 0.0021 |
| F01 | 39240001 | 39340000 | 0.0032 | 0.0023 | 0.0023 |
| F01 | 39260001 | 39360000 | 0.0032 | 0.0023 | 0.0023 |
| F01 | 39280001 | 39380000 | 0.0036 | 0.0026 | 0.0027 |
| F01 | 39300001 | 39400000 | 0.0037 | 0.0029 | 0.0029 |
| F01 | 39320001 | 39420000 | 0.0043 | 0.0034 | 0.0035 |
| F01 | 39340001 | 39440000 | 0.0040 | 0.0031 | 0.0032 |

| Chr | Start | End | both | A1 | A2 |
|---|---|---|---|---|---|
| F01 | 39360001 | 39460000 | 0.0044 | 0.0032 | 0.0033 |
| F01 | 39380001 | 39480000 | 0.0038 | 0.0028 | 0.0028 |
| F01 | 39400001 | 39500000 | 0.0033 | 0.0022 | 0.0022 |
| F01 | 39420001 | 39520000 | 0.0027 | 0.0017 | 0.0017 |
| F01 | 39440001 | 39540000 | 0.0026 | 0.0015 | 0.0014 |
| F01 | 39460001 | 39560000 | 0.0025 | 0.0013 | 0.0013 |
| F01 | 39480001 | 39580000 | 0.0026 | 0.0013 | 0.0014 |
| F01 | 39500001 | 39600000 | 0.0025 | 0.0012 | 0.0013 |
| F01 | 39520001 | 39620000 | 0.0023 | 0.0010 | 0.0009 |
| F01 | 39540001 | 39640000 | 0.0020 | 0.0008 | 0.0009 |
| F01 | 39560001 | 39660000 | 0.0017 | 0.0007 | 0.0007 |
| F01 | 39580001 | 39680000 | 0.0015 | 0.0006 | 0.0006 |
| F01 | 39600001 | 39700000 | 0.0014 | 0.0007 | 0.0007 |
| F01 | 39620001 | 39720000 | 0.0021 | 0.0015 | 0.0015 |
| F01 | 39640001 | 39740000 | 0.0027 | 0.0019 | 0.0019 |
| F01 | 39660001 | 39760000 | 0.0029 | 0.0022 | 0.0022 |
| F01 | 39680001 | 39780000 | 0.0030 | 0.0023 | 0.0023 |
| F01 | 39700001 | 39800000 | 0.0039 | 0.0032 | 0.0032 |
| F01 | 39720001 | 39820000 | 0.0031 | 0.0024 | 0.0023 |
| F01 | 39740001 | 39840000 | 0.0032 | 0.0026 | 0.0025 |
| F01 | 39760001 | 39860000 | 0.0033 | 0.0027 | 0.0027 |
| F01 | 39780001 | 39880000 | 0.0035 | 0.0029 | 0.0032 |
| F01 | 39800001 | 39900000 | 0.0024 | 0.0019 | 0.0022 |
| F01 | 39820001 | 39920000 | 0.0027 | 0.0022 | 0.0024 |
| F01 | 39840001 | 39940000 | 0.0030 | 0.0024 | 0.0026 |
| F01 | 39860001 | 39960000 | 0.0026 | 0.0020 | 0.0022 |
| F01 | 39880001 | 39980000 | 0.0024 | 0.0019 | 0.0019 |
| F01 | 39900001 | 40000000 | 0.0026 | 0.0018 | 0.0018 |
| F01 | 39920001 | 40020000 | 0.0022 | 0.0016 | 0.0016 |
| F01 | 39940001 | 40040000 | 0.0021 | 0.0014 | 0.0014 |
| F01 | 39960001 | 40060000 | 0.0032 | 0.0024 | 0.0023 |
| F01 | 39980001 | 40080000 | 0.0033 | 0.0023 | 0.0021 |
| F01 | 40000001 | 40100000 | 0.0032 | 0.0022 | 0.0021 |
| F01 | 40020001 | 40120000 | 0.0038 | 0.0027 | 0.0027 |
| F01 | 40040001 | 40140000 | 0.0033 | 0.0024 | 0.0023 |
| F01 | 40060001 | 40160000 | 0.0022 | 0.0014 | 0.0013 |
| F01 | 40080001 | 40180000 | 0.0026 | 0.0019 | 0.0018 |
| F01 | 40100001 | 40200000 | 0.0035 | 0.0029 | 0.0027 |
| F01 | 40120001 | 40220000 | 0.0029 | 0.0024 | 0.0022 |
| F01 | 40140001 | 40240000 | 0.0034 | 0.0029 | 0.0027 |
| F01 | 40160001 | 40260000 | 0.0036 | 0.0030 | 0.0030 |
| F01 | 40180001 | 40280000 | 0.0026 | 0.0021 | 0.0021 |
| F01 | 40200001 | 40300000 | 0.0017 | 0.0013 | 0.0013 |
| F01 | 40220001 | 40320000 | 0.0020 | 0.0014 | 0.0016 |
| F01 | 40240001 | 40340000 | 0.0016 | 0.0012 | 0.0013 |
| F01 | 40260001 | 40360000 | 0.0014 | 0.0011 | 0.0013 |
| F01 | 40280001 | 40380000 | 0.0018 | 0.0014 | 0.0016 |
| F01 | 40300001 | 40400000 | 0.0020 | 0.0017 | 0.0019 |
| F01 | 40320001 | 40420000 | 0.0019 | 0.0016 | 0.0018 |
| F01 | 40340001 | 40440000 | 0.0020 | 0.0018 | 0.0019 |
| F01 | 40360001 | 40460000 | 0.0020 | 0.0018 | 0.0018 |
| F01 | 40380001 | 40480000 | 0.0018 | 0.0016 | 0.0015 |
| F01 | 40400001 | 40500000 | 0.0014 | 0.0013 | 0.0012 |
| F01 | 40420001 | 40520000 | 0.0015 | 0.0013 | 0.0012 |
| F01 | 40440001 | 40540000 | 0.0011 | 0.0009 | 0.0008 |
| F01 | 40460001 | 40560000 | 0.0010 | 0.0008 | 0.0009 |
| F01 | 40480001 | 40580000 | 0.0016 | 0.0015 | 0.0015 |
| F01 | 40500001 | 40600000 | 0.0021 | 0.0018 | 0.0018 |

| Chr | Start | End | both | A1 | A2 |
|---|---|---|---|---|---|
| F01 | 40520001 | 40620000 | 0.0019 | 0.0017 | 0.0016 |
| F01 | 40540001 | 40640000 | 0.0026 | 0.0022 | 0.0023 |
| F01 | 40560001 | 40660000 | 0.0027 | 0.0021 | 0.0021 |
| F01 | 40580001 | 40680000 | 0.0023 | 0.0017 | 0.0018 |
| F01 | 40600001 | 40700000 | 0.0021 | 0.0015 | 0.0018 |
| F01 | 40620001 | 40720000 | 0.0020 | 0.0014 | 0.0017 |
| F01 | 40640001 | 40740000 | 0.0018 | 0.0015 | 0.0015 |
| F01 | 40660001 | 40760000 | 0.0018 | 0.0017 | 0.0017 |
| F01 | 40680001 | 40780000 | 0.0019 | 0.0018 | 0.0017 |
| F01 | 40700001 | 40800000 | 0.0018 | 0.0018 | 0.0017 |
| F01 | 40720001 | 40820000 | 0.0022 | 0.0022 | 0.0020 |
| F01 | 40740001 | 40840000 | 0.0021 | 0.0020 | 0.0020 |
| F01 | 40760001 | 40860000 | 0.0020 | 0.0018 | 0.0019 |
| F01 | 40780001 | 40880000 | 0.0017 | 0.0015 | 0.0016 |
| F01 | 40800001 | 40900000 | 0.0015 | 0.0013 | 0.0014 |
| F01 | 40820001 | 40920000 | 0.0012 | 0.0009 | 0.0011 |
| F01 | 40840001 | 40940000 | 0.0008 | 0.0005 | 0.0006 |
| F01 | 40860001 | 40960000 | 0.0010 | 0.0006 | 0.0008 |
| F01 | 40880001 | 40980000 | 0.0013 | 0.0010 | 0.0011 |
| F01 | 40900001 | 41000000 | 0.0016 | 0.0013 | 0.0014 |
| F01 | 40920001 | 41020000 | 0.0018 | 0.0015 | 0.0016 |
| F01 | 40940001 | 41040000 | 0.0020 | 0.0017 | 0.0017 |
| F01 | 40960001 | 41060000 | 0.0020 | 0.0017 | 0.0016 |
| F01 | 40980001 | 41080000 | 0.0019 | 0.0016 | 0.0015 |
| F01 | 41000001 | 41100000 | 0.0020 | 0.0018 | 0.0016 |
| F01 | 41020001 | 41120000 | 0.0020 | 0.0018 | 0.0017 |
| F01 | 41040001 | 41140000 | 0.0028 | 0.0026 | 0.0026 |
| F01 | 41060001 | 41160000 | 0.0027 | 0.0025 | 0.0025 |
| F01 | 41080001 | 41180000 | 0.0028 | 0.0024 | 0.0025 |
| F01 | 41100001 | 41200000 | 0.0026 | 0.0021 | 0.0022 |
| F01 | 41120001 | 41220000 | 0.0022 | 0.0018 | 0.0019 |
| F01 | 41140001 | 41240000 | 0.0010 | 0.0007 | 0.0007 |
| F01 | 41160001 | 41260000 | 0.0012 | 0.0009 | 0.0009 |
| F01 | 41180001 | 41280000 | 0.0009 | 0.0008 | 0.0007 |
| F01 | 41200001 | 41300000 | 0.0007 | 0.0006 | 0.0007 |
| F01 | 41220001 | 41320000 | 0.0008 | 0.0006 | 0.0008 |
| F01 | 41240001 | 41340000 | 0.0008 | 0.0006 | 0.0008 |
| F01 | 41260001 | 41360000 | 0.0005 | 0.0004 | 0.0005 |
| F01 | 41280001 | 41380000 | 0.0005 | 0.0004 | 0.0005 |
| F01 | 41300001 | 41400000 | 0.0006 | 0.0006 | 0.0006 |
| F01 | 41320001 | 41420000 | 0.0008 | 0.0007 | 0.0008 |
| F01 | 41340001 | 41440000 | 0.0010 | 0.0009 | 0.0010 |
| F01 | 41360001 | 41460000 | 0.0011 | 0.0010 | 0.0011 |
| F01 | 41380001 | 41480000 | 0.0010 | 0.0008 | 0.0010 |
| F01 | 41400001 | 41500000 | 0.0012 | 0.0010 | 0.0012 |
| F01 | 41420001 | 41520000 | 0.0012 | 0.0009 | 0.0010 |
| F01 | 41440001 | 41540000 | 0.0014 | 0.0010 | 0.0012 |
| F01 | 41460001 | 41560000 | 0.0015 | 0.0011 | 0.0013 |
| F01 | 41480001 | 41580000 | 0.0014 | 0.0010 | 0.0012 |
| F01 | 41500001 | 41600000 | 0.0014 | 0.0011 | 0.0012 |
| F01 | 41520001 | 41620000 | 0.0014 | 0.0012 | 0.0013 |
| F01 | 41540001 | 41640000 | 0.0015 | 0.0013 | 0.0015 |
| F01 | 41560001 | 41660000 | 0.0023 | 0.0020 | 0.0021 |
| F01 | 41580001 | 41680000 | 0.0032 | 0.0028 | 0.0029 |
| F01 | 41600001 | 41700000 | 0.0032 | 0.0027 | 0.0027 |
| F01 | 41620001 | 41720000 | 0.0034 | 0.0027 | 0.0026 |
| F01 | 41640001 | 41740000 | 0.0030 | 0.0023 | 0.0022 |
| F01 | 41660001 | 41760000 | 0.0020 | 0.0015 | 0.0012 |

| Chr | Start | End | both | A1 | A2 |
|---|---|---|---|---|---|
| F01 | 41680001 | 41780000 | 0.0015 | 0.0010 | 0.0009 |
| F01 | 41700001 | 41800000 | 0.0015 | 0.0010 | 0.0010 |
| F01 | 41720001 | 41820000 | 0.0013 | 0.0010 | 0.0010 |
| F01 | 41740001 | 41840000 | 0.0013 | 0.0009 | 0.0009 |
| F01 | 41760001 | 41860000 | 0.0020 | 0.0013 | 0.0015 |
| F01 | 41780001 | 41880000 | 0.0018 | 0.0011 | 0.0013 |
| F01 | 41800001 | 41900000 | 0.0015 | 0.0009 | 0.0011 |
| F01 | 41820001 | 41920000 | 0.0015 | 0.0009 | 0.0012 |
| F01 | 41840001 | 41940000 | 0.0017 | 0.0009 | 0.0012 |
| F01 | 41860001 | 41960000 | 0.0014 | 0.0008 | 0.0009 |
| F01 | 41880001 | 41980000 | 0.0013 | 0.0008 | 0.0008 |
| F01 | 41900001 | 42000000 | 0.0014 | 0.0008 | 0.0009 |
| F01 | 41920001 | 42020000 | 0.0013 | 0.0008 | 0.0008 |
| F01 | 41940001 | 42040000 | 0.0011 | 0.0008 | 0.0009 |
| F01 | 41960001 | 42060000 | 0.0007 | 0.0006 | 0.0007 |
| F01 | 41980001 | 42080000 | 0.0015 | 0.0013 | 0.0015 |
| F01 | 42000001 | 42100000 | 0.0015 | 0.0013 | 0.0015 |
| F01 | 42020001 | 42120000 | 0.0016 | 0.0013 | 0.0016 |
| F01 | 42040001 | 42140000 | 0.0016 | 0.0013 | 0.0015 |
| F01 | 42060001 | 42160000 | 0.0018 | 0.0014 | 0.0018 |
| F01 | 42080001 | 42180000 | 0.0013 | 0.0010 | 0.0014 |
| F01 | 42100001 | 42200000 | 0.0016 | 0.0012 | 0.0016 |
| F01 | 42120001 | 42220000 | 0.0013 | 0.0010 | 0.0013 |
| F01 | 42140001 | 42240000 | 0.0013 | 0.0011 | 0.0014 |
| F01 | 42160001 | 42260000 | 0.0013 | 0.0010 | 0.0013 |
| F01 | 42180001 | 42280000 | 0.0010 | 0.0007 | 0.0010 |
| F01 | 42200001 | 42300000 | 0.0006 | 0.0005 | 0.0006 |
| F01 | 42220001 | 42320000 | 0.0008 | 0.0007 | 0.0008 |
| F01 | 42240001 | 42340000 | 0.0010 | 0.0009 | 0.0010 |
| F01 | 42260001 | 42360000 | 0.0009 | 0.0008 | 0.0009 |
| F01 | 42280001 | 42380000 | 0.0010 | 0.0008 | 0.0010 |
| F01 | 42300001 | 42400000 | 0.0011 | 0.0008 | 0.0011 |
| F01 | 42320001 | 42420000 | 0.0009 | 0.0006 | 0.0010 |
| F01 | 42340001 | 42440000 | 0.0006 | 0.0004 | 0.0007 |
| F01 | 42360001 | 42460000 | 0.0006 | 0.0004 | 0.0006 |
| F01 | 42380001 | 42480000 | 0.0005 | 0.0003 | 0.0005 |
| F01 | 42400001 | 42500000 | 0.0004 | 0.0003 | 0.0004 |
| F01 | 42420001 | 42520000 | 0.0005 | 0.0004 | 0.0005 |
| F01 | 42440001 | 42540000 | 0.0007 | 0.0006 | 0.0007 |
| F01 | 42460001 | 42560000 | 0.0008 | 0.0006 | 0.0008 |
| F01 | 42480001 | 42580000 | 0.0014 | 0.0012 | 0.0014 |
| F01 | 42500001 | 42600000 | 0.0015 | 0.0013 | 0.0015 |
| F01 | 42520001 | 42620000 | 0.0017 | 0.0015 | 0.0017 |
| F01 | 42540001 | 42640000 | 0.0016 | 0.0013 | 0.0015 |
| F01 | 42560001 | 42660000 | 0.0016 | 0.0013 | 0.0014 |
| F01 | 42580001 | 42680000 | 0.0013 | 0.0009 | 0.0010 |
| F01 | 42600001 | 42700000 | 0.0019 | 0.0013 | 0.0012 |
| F01 | 42620001 | 42720000 | 0.0019 | 0.0013 | 0.0010 |
| F01 | 42640001 | 42740000 | 0.0020 | 0.0013 | 0.0011 |
| F01 | 42660001 | 42760000 | 0.0024 | 0.0018 | 0.0015 |
| F01 | 42680001 | 42780000 | 0.0030 | 0.0025 | 0.0021 |
| F01 | 42700001 | 42800000 | 0.0025 | 0.0021 | 0.0020 |
| F01 | 42720001 | 42820000 | 0.0024 | 0.0021 | 0.0021 |
| F01 | 42740001 | 42840000 | 0.0032 | 0.0027 | 0.0027 |
| F01 | 42760001 | 42860000 | 0.0031 | 0.0023 | 0.0023 |
| F01 | 42780001 | 42880000 | 0.0028 | 0.0018 | 0.0018 |
| F01 | 42800001 | 42900000 | 0.0031 | 0.0019 | 0.0018 |
| F01 | 42820001 | 42920000 | 0.0036 | 0.0023 | 0.0022 |

| Chr | Start | End | both | A1 | A2 |
|---|---|---|---|---|---|
| F01 | 42840001 | 42940000 | 0.0027 | 0.0016 | 0.0015 |
| F01 | 42860001 | 42960000 | 0.0029 | 0.0020 | 0.0021 |
| F01 | 42880001 | 42980000 | 0.0026 | 0.0019 | 0.0021 |
| F01 | 42900001 | 43000000 | 0.0021 | 0.0016 | 0.0018 |
| F01 | 42920001 | 43020000 | 0.0023 | 0.0019 | 0.0020 |
| F01 | 42940001 | 43040000 | 0.0034 | 0.0028 | 0.0030 |
| F01 | 42960001 | 43060000 | 0.0036 | 0.0030 | 0.0031 |
| F01 | 42980001 | 43080000 | 0.0038 | 0.0030 | 0.0031 |
| F01 | 43000001 | 43100000 | 0.0037 | 0.0028 | 0.0030 |
| F01 | 43020001 | 43120000 | 0.0033 | 0.0023 | 0.0025 |
| F01 | 43040001 | 43140000 | 0.0028 | 0.0019 | 0.0021 |
| F01 | 43060001 | 43160000 | 0.0021 | 0.0013 | 0.0015 |
| F01 | 43080001 | 43180000 | 0.0023 | 0.0016 | 0.0019 |
| F01 | 43100001 | 43200000 | 0.0032 | 0.0024 | 0.0028 |
| F01 | 43120001 | 43220000 | 0.0030 | 0.0022 | 0.0025 |
| F01 | 43140001 | 43240000 | 0.0029 | 0.0022 | 0.0024 |
| F01 | 43160001 | 43260000 | 0.0032 | 0.0025 | 0.0026 |
| F01 | 43180001 | 43280000 | 0.0036 | 0.0027 | 0.0028 |
| F01 | 43200001 | 43300000 | 0.0034 | 0.0023 | 0.0024 |
| F01 | 43220001 | 43320000 | 0.0043 | 0.0033 | 0.0035 |
| F01 | 43240001 | 43340000 | 0.0043 | 0.0033 | 0.0035 |
| F01 | 43260001 | 43360000 | 0.0045 | 0.0034 | 0.0038 |
| F01 | 43280001 | 43380000 | 0.0037 | 0.0029 | 0.0033 |
| F01 | 43300001 | 43400000 | 0.0034 | 0.0026 | 0.0030 |
| F01 | 43320001 | 43420000 | 0.0025 | 0.0019 | 0.0022 |
| F01 | 43340001 | 43440000 | 0.0022 | 0.0017 | 0.0020 |
| F01 | 43360001 | 43460000 | 0.0021 | 0.0017 | 0.0018 |
| F01 | 43380001 | 43480000 | 0.0021 | 0.0016 | 0.0018 |
| F01 | 43400001 | 43500000 | 0.0023 | 0.0019 | 0.0021 |
| F01 | 43420001 | 43520000 | 0.0028 | 0.0023 | 0.0026 |
| F01 | 43440001 | 43540000 | 0.0026 | 0.0021 | 0.0024 |
| F01 | 43460001 | 43560000 | 0.0025 | 0.0020 | 0.0024 |
| F01 | 43480001 | 43580000 | 0.0021 | 0.0018 | 0.0021 |
| F01 | 43500001 | 43600000 | 0.0022 | 0.0018 | 0.0022 |
| F01 | 43520001 | 43620000 | 0.0018 | 0.0015 | 0.0018 |
| F01 | 43540001 | 43640000 | 0.0021 | 0.0018 | 0.0021 |
| F01 | 43560001 | 43660000 | 0.0018 | 0.0016 | 0.0018 |
| F01 | 43580001 | 43680000 | 0.0021 | 0.0018 | 0.0020 |
| F01 | 43600001 | 43700000 | 0.0018 | 0.0016 | 0.0018 |
| F01 | 43620001 | 43720000 | 0.0015 | 0.0014 | 0.0015 |
| F01 | 43640001 | 43740000 | 0.0014 | 0.0012 | 0.0014 |
| F01 | 43660001 | 43760000 | 0.0013 | 0.0010 | 0.0013 |
| F01 | 43680001 | 43780000 | 0.0019 | 0.0013 | 0.0017 |
| F01 | 43700001 | 43800000 | 0.0018 | 0.0012 | 0.0016 |
| F01 | 43720001 | 43820000 | 0.0019 | 0.0014 | 0.0017 |
| F01 | 43740001 | 43840000 | 0.0025 | 0.0019 | 0.0023 |
| F01 | 43760001 | 43860000 | 0.0029 | 0.0023 | 0.0027 |
| F01 | 43780001 | 43880000 | 0.0021 | 0.0018 | 0.0020 |
| F01 | 43800001 | 43900000 | 0.0017 | 0.0015 | 0.0017 |
| F01 | 43820001 | 43920000 | 0.0017 | 0.0016 | 0.0017 |
| F01 | 43840001 | 43940000 | 0.0015 | 0.0014 | 0.0014 |
| F01 | 43860001 | 43960000 | 0.0011 | 0.0011 | 0.0011 |
| F01 | 43880001 | 43980000 | 0.0013 | 0.0012 | 0.0012 |
| F01 | 43900001 | 44000000 | 0.0017 | 0.0014 | 0.0014 |
| F01 | 43920001 | 44020000 | 0.0015 | 0.0011 | 0.0012 |
| F01 | 43940001 | 44040000 | 0.0011 | 0.0008 | 0.0008 |
| F01 | 43960001 | 44060000 | 0.0010 | 0.0008 | 0.0008 |
| F01 | 43980001 | 44080000 | 0.0009 | 0.0007 | 0.0007 |

| Chr | Start | End | both | A1 | A2 |
|---|---|---|---|---|---|
| F01 | 44000001 | 44100000 | 0.0007 | 0.0006 | 0.0006 |
| F01 | 44020001 | 44120000 | 0.0006 | 0.0005 | 0.0005 |
| F01 | 44040001 | 44140000 | 0.0007 | 0.0005 | 0.0006 |
| F01 | 44060001 | 44160000 | 0.0007 | 0.0006 | 0.0007 |
| F01 | 44080001 | 44180000 | 0.0007 | 0.0005 | 0.0007 |
| F01 | 44100001 | 44200000 | 0.0012 | 0.0008 | 0.0011 |
| F01 | 44120001 | 44220000 | 0.0013 | 0.0009 | 0.0012 |
| F01 | 44140001 | 44240000 | 0.0015 | 0.0011 | 0.0013 |
| F01 | 44160001 | 44260000 | 0.0017 | 0.0011 | 0.0014 |
| F01 | 44180001 | 44280000 | 0.0021 | 0.0014 | 0.0017 |
| F01 | 44200001 | 44300000 | 0.0019 | 0.0012 | 0.0013 |
| F01 | 44220001 | 44320000 | 0.0025 | 0.0017 | 0.0017 |
| F01 | 44240001 | 44340000 | 0.0024 | 0.0014 | 0.0016 |
| F01 | 44260001 | 44360000 | 0.0022 | 0.0013 | 0.0014 |
| F01 | 44280001 | 44380000 | 0.0023 | 0.0014 | 0.0015 |
| F01 | 44300001 | 44400000 | 0.0020 | 0.0013 | 0.0014 |
| F01 | 44320001 | 44420000 | 0.0017 | 0.0010 | 0.0013 |
| F01 | 44340001 | 44440000 | 0.0017 | 0.0013 | 0.0014 |
| F01 | 44360001 | 44460000 | 0.0019 | 0.0016 | 0.0017 |
| F01 | 44380001 | 44480000 | 0.0015 | 0.0013 | 0.0013 |
| F01 | 44400001 | 44500000 | 0.0020 | 0.0016 | 0.0017 |
| F01 | 44420001 | 44520000 | 0.0020 | 0.0016 | 0.0016 |
| F01 | 44440001 | 44540000 | 0.0019 | 0.0014 | 0.0013 |
| F01 | 44460001 | 44560000 | 0.0017 | 0.0011 | 0.0010 |
| F01 | 44480001 | 44580000 | 0.0020 | 0.0013 | 0.0012 |
| F01 | 44500001 | 44600000 | 0.0018 | 0.0010 | 0.0008 |
| F01 | 44520001 | 44620000 | 0.0023 | 0.0014 | 0.0014 |
| F01 | 44540001 | 44640000 | 0.0024 | 0.0015 | 0.0015 |
| F01 | 44560001 | 44660000 | 0.0021 | 0.0014 | 0.0014 |
| F01 | 44580001 | 44680000 | 0.0019 | 0.0012 | 0.0011 |
| F01 | 44600001 | 44700000 | 0.0018 | 0.0012 | 0.0012 |
| F01 | 44620001 | 44720000 | 0.0012 | 0.0006 | 0.0006 |
| F01 | 44640001 | 44740000 | 0.0015 | 0.0009 | 0.0010 |
| F01 | 44660001 | 44760000 | 0.0019 | 0.0010 | 0.0012 |
| F01 | 44680001 | 44780000 | 0.0017 | 0.0010 | 0.0011 |
| F01 | 44700001 | 44800000 | 0.0018 | 0.0011 | 0.0012 |
| F01 | 44720001 | 44820000 | 0.0021 | 0.0013 | 0.0013 |
| F01 | 44740001 | 44840000 | 0.0018 | 0.0010 | 0.0009 |
| F01 | 44760001 | 44860000 | 0.0015 | 0.0009 | 0.0008 |
| F01 | 44780001 | 44880000 | 0.0025 | 0.0018 | 0.0017 |
| F01 | 44800001 | 44900000 | 0.0022 | 0.0016 | 0.0015 |
| F01 | 44820001 | 44920000 | 0.0019 | 0.0015 | 0.0015 |
| F01 | 44840001 | 44940000 | 0.0021 | 0.0017 | 0.0017 |
| F01 | 44860001 | 44960000 | 0.0025 | 0.0020 | 0.0020 |
| F01 | 44880001 | 44980000 | 0.0017 | 0.0012 | 0.0014 |
| F01 | 44900001 | 45000000 | 0.0019 | 0.0013 | 0.0015 |
| F01 | 44920001 | 45020000 | 0.0025 | 0.0016 | 0.0019 |
| F01 | 44940001 | 45040000 | 0.0027 | 0.0019 | 0.0021 |
| F01 | 44960001 | 45060000 | 0.0033 | 0.0026 | 0.0028 |
| F01 | 44980001 | 45080000 | 0.0039 | 0.0031 | 0.0033 |
| F01 | 45000001 | 45100000 | 0.0047 | 0.0038 | 0.0042 |
| F01 | 45020001 | 45120000 | 0.0047 | 0.0038 | 0.0041 |
| F01 | 45040001 | 45140000 | 0.0047 | 0.0037 | 0.0040 |
| F01 | 45060001 | 45160000 | 0.0045 | 0.0036 | 0.0038 |
| F01 | 45080001 | 45180000 | 0.0038 | 0.0030 | 0.0033 |
| F01 | 45100001 | 45200000 | 0.0037 | 0.0030 | 0.0031 |
| F01 | 45120001 | 45220000 | 0.0032 | 0.0025 | 0.0028 |
| F01 | 45140001 | 45240000 | 0.0030 | 0.0023 | 0.0025 |

| Chr | Start | End | both | A1 | A2 |
|---|---|---|---|---|---|
| F01 | 45160001 | 45260000 | 0.0020 | 0.0014 | 0.0016 |
| F01 | 45180001 | 45280000 | 0.0021 | 0.0015 | 0.0017 |
| F01 | 45200001 | 45300000 | 0.0014 | 0.0008 | 0.0010 |
| F01 | 45220001 | 45320000 | 0.0015 | 0.0012 | 0.0013 |
| F01 | 45240001 | 45340000 | 0.0023 | 0.0022 | 0.0023 |
| F01 | 45260001 | 45360000 | 0.0024 | 0.0023 | 0.0023 |
| F01 | 45280001 | 45380000 | 0.0023 | 0.0022 | 0.0022 |
| F01 | 45300001 | 45400000 | 0.0026 | 0.0025 | 0.0025 |
| F01 | 45320001 | 45420000 | 0.0025 | 0.0025 | 0.0022 |
| F01 | 45340001 | 45440000 | 0.0016 | 0.0015 | 0.0012 |
| F01 | 45360001 | 45460000 | 0.0020 | 0.0018 | 0.0016 |
| F01 | 45380001 | 45480000 | 0.0020 | 0.0017 | 0.0015 |
| F01 | 45400001 | 45500000 | 0.0019 | 0.0016 | 0.0013 |
| F01 | 45420001 | 45520000 | 0.0014 | 0.0010 | 0.0010 |
| F01 | 45440001 | 45540000 | 0.0011 | 0.0007 | 0.0008 |
| F01 | 45460001 | 45560000 | 0.0009 | 0.0006 | 0.0007 |
| F01 | 45480001 | 45580000 | 0.0012 | 0.0008 | 0.0008 |
| F01 | 45500001 | 45600000 | 0.0012 | 0.0008 | 0.0009 |
| F01 | 45520001 | 45620000 | 0.0028 | 0.0021 | 0.0025 |
| F01 | 45540001 | 45640000 | 0.0045 | 0.0038 | 0.0040 |
| F01 | 45560001 | 45660000 | 0.0051 | 0.0043 | 0.0045 |
| F01 | 45580001 | 45680000 | 0.0053 | 0.0046 | 0.0048 |
| F01 | 45600001 | 45700000 | 0.0056 | 0.0048 | 0.0050 |
| F01 | 45620001 | 45720000 | 0.0039 | 0.0036 | 0.0035 |
| F01 | 45640001 | 45740000 | 0.0023 | 0.0020 | 0.0020 |
| F01 | 45660001 | 45760000 | 0.0020 | 0.0016 | 0.0018 |
| F01 | 45680001 | 45780000 | 0.0015 | 0.0013 | 0.0014 |
| F01 | 45700001 | 45800000 | 0.0015 | 0.0013 | 0.0013 |
| F01 | 45720001 | 45820000 | 0.0016 | 0.0013 | 0.0013 |
| F01 | 45740001 | 45840000 | 0.0020 | 0.0016 | 0.0015 |
| F01 | 45760001 | 45860000 | 0.0024 | 0.0020 | 0.0018 |
| F01 | 45780001 | 45880000 | 0.0025 | 0.0019 | 0.0017 |
| F01 | 45800001 | 45900000 | 0.0028 | 0.0021 | 0.0019 |
| F01 | 45820001 | 45920000 | 0.0033 | 0.0026 | 0.0024 |
| F01 | 45840001 | 45940000 | 0.0034 | 0.0026 | 0.0026 |
| F01 | 45860001 | 45960000 | 0.0034 | 0.0027 | 0.0028 |
| F01 | 45880001 | 45980000 | 0.0043 | 0.0037 | 0.0039 |
| F01 | 45900001 | 46000000 | 0.0041 | 0.0035 | 0.0038 |
| F01 | 45920001 | 46020000 | 0.0037 | 0.0033 | 0.0035 |
| F01 | 45940001 | 46040000 | 0.0037 | 0.0034 | 0.0034 |
| F01 | 45960001 | 46060000 | 0.0034 | 0.0030 | 0.0031 |
| F01 | 45980001 | 46080000 | 0.0028 | 0.0023 | 0.0024 |
| F01 | 46000001 | 46100000 | 0.0022 | 0.0019 | 0.0020 |
| F01 | 46020001 | 46120000 | 0.0023 | 0.0018 | 0.0020 |
| F01 | 46040001 | 46140000 | 0.0029 | 0.0022 | 0.0025 |
| F01 | 46060001 | 46160000 | 0.0023 | 0.0018 | 0.0020 |
| F01 | 46080001 | 46180000 | 0.0020 | 0.0014 | 0.0016 |
| F01 | 46100001 | 46200000 | 0.0025 | 0.0018 | 0.0020 |
| F01 | 46120001 | 46220000 | 0.0024 | 0.0019 | 0.0021 |
| F01 | 46140001 | 46240000 | 0.0023 | 0.0018 | 0.0019 |
| F01 | 46160001 | 46260000 | 0.0028 | 0.0023 | 0.0023 |
| F01 | 46180001 | 46280000 | 0.0028 | 0.0023 | 0.0024 |
| F01 | 46200001 | 46300000 | 0.0029 | 0.0024 | 0.0024 |
| F01 | 46220001 | 46320000 | 0.0028 | 0.0021 | 0.0021 |
| F01 | 46240001 | 46340000 | 0.0032 | 0.0023 | 0.0023 |
| F01 | 46260001 | 46360000 | 0.0035 | 0.0024 | 0.0026 |
| F01 | 46280001 | 46380000 | 0.0035 | 0.0025 | 0.0027 |
| F01 | 46300001 | 46400000 | 0.0032 | 0.0022 | 0.0025 |

| Chr | Start | End | both | A1 | A2 |
|---|---|---|---|---|---|
| F01 | 46320001 | 46420000 | 0.0035 | 0.0025 | 0.0029 |
| F01 | 46340001 | 46440000 | 0.0028 | 0.0019 | 0.0024 |
| F01 | 46360001 | 46460000 | 0.0026 | 0.0017 | 0.0019 |
| F01 | 46380001 | 46480000 | 0.0025 | 0.0016 | 0.0017 |
| F01 | 46400001 | 46500000 | 0.0022 | 0.0014 | 0.0015 |
| F01 | 46420001 | 46520000 | 0.0019 | 0.0012 | 0.0011 |
| F01 | 46440001 | 46540000 | 0.0017 | 0.0011 | 0.0010 |
| F01 | 46460001 | 46560000 | 0.0013 | 0.0008 | 0.0008 |
| F01 | 46480001 | 46580000 | 0.0015 | 0.0010 | 0.0012 |
| F01 | 46500001 | 46600000 | 0.0012 | 0.0008 | 0.0009 |
| F01 | 46520001 | 46620000 | 0.0014 | 0.0010 | 0.0011 |
| F01 | 46540001 | 46640000 | 0.0013 | 0.0009 | 0.0010 |
| F01 | 46560001 | 46660000 | 0.0014 | 0.0009 | 0.0011 |
| F01 | 46580001 | 46680000 | 0.0015 | 0.0010 | 0.0011 |
| F01 | 46600001 | 46700000 | 0.0022 | 0.0016 | 0.0018 |
| F01 | 46620001 | 46720000 | 0.0035 | 0.0029 | 0.0032 |
| F01 | 46640001 | 46740000 | 0.0037 | 0.0029 | 0.0031 |
| F01 | 46660001 | 46760000 | 0.0040 | 0.0032 | 0.0034 |
| F01 | 46680001 | 46780000 | 0.0038 | 0.0031 | 0.0033 |
| F01 | 46700001 | 46800000 | 0.0043 | 0.0036 | 0.0038 |
| F01 | 46720001 | 46820000 | 0.0030 | 0.0024 | 0.0026 |
| F01 | 46740001 | 46840000 | 0.0028 | 0.0023 | 0.0026 |
| F01 | 46760001 | 46860000 | 0.0030 | 0.0024 | 0.0028 |
| F01 | 46780001 | 46880000 | 0.0032 | 0.0025 | 0.0029 |
| F01 | 46800001 | 46900000 | 0.0028 | 0.0022 | 0.0025 |
| F01 | 46820001 | 46920000 | 0.0030 | 0.0023 | 0.0026 |
| F01 | 46840001 | 46940000 | 0.0034 | 0.0025 | 0.0029 |
| F01 | 46860001 | 46960000 | 0.0035 | 0.0026 | 0.0029 |
| F01 | 46880001 | 46980000 | 0.0033 | 0.0025 | 0.0028 |
| F01 | 46900001 | 47000000 | 0.0028 | 0.0020 | 0.0022 |
| F01 | 46920001 | 47020000 | 0.0027 | 0.0019 | 0.0022 |
| F01 | 46940001 | 47040000 | 0.0021 | 0.0015 | 0.0016 |
| F01 | 46960001 | 47060000 | 0.0016 | 0.0010 | 0.0011 |
| F01 | 46980001 | 47080000 | 0.0016 | 0.0010 | 0.0011 |
| F01 | 47000001 | 47100000 | 0.0018 | 0.0012 | 0.0016 |
| F01 | 47020001 | 47120000 | 0.0013 | 0.0009 | 0.0012 |
| F01 | 47040001 | 47140000 | 0.0013 | 0.0008 | 0.0011 |
| F01 | 47060001 | 47160000 | 0.0012 | 0.0009 | 0.0012 |
| F01 | 47080001 | 47180000 | 0.0017 | 0.0013 | 0.0017 |
| F01 | 47100001 | 47200000 | 0.0018 | 0.0015 | 0.0016 |
| F01 | 47120001 | 47220000 | 0.0020 | 0.0017 | 0.0018 |
| F01 | 47140001 | 47240000 | 0.0024 | 0.0020 | 0.0022 |
| F01 | 47160001 | 47260000 | 0.0025 | 0.0020 | 0.0022 |
| F01 | 47180001 | 47280000 | 0.0019 | 0.0016 | 0.0017 |
| F01 | 47200001 | 47300000 | 0.0019 | 0.0014 | 0.0017 |
| F01 | 47220001 | 47320000 | 0.0020 | 0.0015 | 0.0018 |
| F01 | 47240001 | 47340000 | 0.0018 | 0.0014 | 0.0017 |
| F01 | 47260001 | 47360000 | 0.0019 | 0.0014 | 0.0017 |
| F01 | 47280001 | 47380000 | 0.0016 | 0.0012 | 0.0014 |
| F01 | 47300001 | 47400000 | 0.0016 | 0.0013 | 0.0013 |
| F01 | 47320001 | 47420000 | 0.0018 | 0.0014 | 0.0014 |
| F01 | 47340001 | 47440000 | 0.0020 | 0.0015 | 0.0015 |
| F01 | 47360001 | 47460000 | 0.0020 | 0.0016 | 0.0016 |
| F01 | 47380001 | 47480000 | 0.0021 | 0.0017 | 0.0018 |
| F01 | 47400001 | 47500000 | 0.0023 | 0.0019 | 0.0021 |
| F01 | 47420001 | 47520000 | 0.0029 | 0.0025 | 0.0027 |
| F01 | 47440001 | 47540000 | 0.0030 | 0.0026 | 0.0029 |
| F01 | 47460001 | 47560000 | 0.0028 | 0.0024 | 0.0028 |

| Chr | Start | End | both | A1 | A2 |
|---|---|---|---|---|---|
| F01 | 47480001 | 47580000 | 0.0030 | 0.0026 | 0.0029 |
| F01 | 47500001 | 47600000 | 0.0028 | 0.0023 | 0.0026 |
| F01 | 47520001 | 47620000 | 0.0020 | 0.0017 | 0.0018 |
| F01 | 47540001 | 47640000 | 0.0020 | 0.0015 | 0.0016 |
| F01 | 47560001 | 47660000 | 0.0020 | 0.0016 | 0.0017 |
| F01 | 47580001 | 47680000 | 0.0018 | 0.0013 | 0.0014 |
| F01 | 47600001 | 47700000 | 0.0016 | 0.0011 | 0.0013 |
| F01 | 47620001 | 47720000 | 0.0013 | 0.0009 | 0.0011 |
| F01 | 47640001 | 47740000 | 0.0009 | 0.0007 | 0.0009 |
| F01 | 47660001 | 47760000 | 0.0010 | 0.0008 | 0.0007 |
| F01 | 47680001 | 47780000 | 0.0009 | 0.0007 | 0.0006 |
| F01 | 47700001 | 47800000 | 0.0010 | 0.0008 | 0.0007 |
| F01 | 47720001 | 47820000 | 0.0010 | 0.0008 | 0.0007 |
| F01 | 47740001 | 47840000 | 0.0013 | 0.0009 | 0.0008 |
| F01 | 47760001 | 47860000 | 0.0015 | 0.0011 | 0.0012 |
| F01 | 47780001 | 47880000 | 0.0018 | 0.0012 | 0.0014 |
| F01 | 47800001 | 47900000 | 0.0015 | 0.0010 | 0.0011 |
| F01 | 47820001 | 47920000 | 0.0019 | 0.0013 | 0.0014 |
| F01 | 47840001 | 47940000 | 0.0016 | 0.0012 | 0.0013 |
| F01 | 47860001 | 47960000 | 0.0011 | 0.0009 | 0.0009 |
| F01 | 47880001 | 47980000 | 0.0010 | 0.0008 | 0.0008 |
| F01 | 47900001 | 48000000 | 0.0011 | 0.0010 | 0.0009 |
| F01 | 47920001 | 48020000 | 0.0014 | 0.0012 | 0.0012 |
| F01 | 47940001 | 48040000 | 0.0017 | 0.0013 | 0.0013 |
| F01 | 47960001 | 48060000 | 0.0022 | 0.0016 | 0.0016 |
| F01 | 47980001 | 48080000 | 0.0028 | 0.0022 | 0.0021 |
| F01 | 48000001 | 48100000 | 0.0028 | 0.0021 | 0.0021 |
| F01 | 48020001 | 48120000 | 0.0023 | 0.0017 | 0.0016 |
| F01 | 48040001 | 48140000 | 0.0025 | 0.0019 | 0.0019 |
| F01 | 48060001 | 48160000 | 0.0024 | 0.0019 | 0.0019 |
| F01 | 48080001 | 48180000 | 0.0023 | 0.0018 | 0.0019 |
| F01 | 48100001 | 48200000 | 0.0023 | 0.0017 | 0.0019 |
| F01 | 48120001 | 48220000 | 0.0021 | 0.0016 | 0.0017 |
| F01 | 48140001 | 48240000 | 0.0015 | 0.0013 | 0.0013 |
| F01 | 48160001 | 48260000 | 0.0011 | 0.0009 | 0.0010 |
| F01 | 48180001 | 48280000 | 0.0005 | 0.0004 | 0.0005 |
| F01 | 48200001 | 48300000 | 0.0002 | 0.0002 | 0.0002 |
| F01 | 48220001 | 48320000 | 0.0010 | 0.0008 | 0.0008 |
| F01 | 48240001 | 48340000 | 0.0014 | 0.0013 | 0.0012 |
| F01 | 48260001 | 48360000 | 0.0019 | 0.0016 | 0.0017 |
| F01 | 48280001 | 48380000 | 0.0025 | 0.0020 | 0.0022 |
| F01 | 48300001 | 48400000 | 0.0024 | 0.0020 | 0.0022 |
| F01 | 48320001 | 48420000 | 0.0017 | 0.0014 | 0.0015 |
| F01 | 48340001 | 48440000 | 0.0012 | 0.0010 | 0.0012 |
| F01 | 48360001 | 48460000 | 0.0007 | 0.0005 | 0.0006 |
| F01 | 48380001 | 48480000 | 0.0004 | 0.0004 | 0.0003 |
| F01 | 48400001 | 48500000 | 0.0007 | 0.0007 | 0.0006 |
| F01 | 48420001 | 48520000 | 0.0012 | 0.0011 | 0.0010 |
| F01 | 48440001 | 48540000 | 0.0012 | 0.0011 | 0.0010 |
| F01 | 48460001 | 48560000 | 0.0016 | 0.0014 | 0.0013 |
| F01 | 48480001 | 48580000 | 0.0015 | 0.0012 | 0.0012 |
| F01 | 48500001 | 48600000 | 0.0015 | 0.0011 | 0.0012 |
| F01 | 48520001 | 48620000 | 0.0012 | 0.0009 | 0.0009 |
| F01 | 48540001 | 48640000 | 0.0013 | 0.0009 | 0.0010 |
| F01 | 48560001 | 48660000 | 0.0011 | 0.0009 | 0.0009 |
| F01 | 48580001 | 48680000 | 0.0011 | 0.0009 | 0.0009 |
| F01 | 48600001 | 48700000 | 0.0010 | 0.0008 | 0.0009 |
| F01 | 48620001 | 48720000 | 0.0010 | 0.0008 | 0.0009 |

| Chr | Start | End | both | A1 | A2 |
|---|---|---|---|---|---|
| F01 | 48640001 | 48740000 | 0.0010 | 0.0008 | 0.0008 |
| F01 | 48660001 | 48760000 | 0.0008 | 0.0006 | 0.0007 |
| F01 | 48680001 | 48780000 | 0.0007 | 0.0005 | 0.0006 |
| F01 | 48700001 | 48800000 | 0.0007 | 0.0005 | 0.0005 |
| F01 | 48720001 | 48820000 | 0.0010 | 0.0008 | 0.0009 |
| F01 | 48740001 | 48840000 | 0.0012 | 0.0010 | 0.0011 |
| F01 | 48760001 | 48860000 | 0.0011 | 0.0009 | 0.0010 |
| F01 | 48780001 | 48880000 | 0.0011 | 0.0010 | 0.0010 |
| F01 | 48800001 | 48900000 | 0.0012 | 0.0011 | 0.0012 |
| F01 | 48820001 | 48920000 | 0.0012 | 0.0011 | 0.0011 |
| F01 | 48840001 | 48940000 | 0.0013 | 0.0011 | 0.0010 |
| F01 | 48860001 | 48960000 | 0.0017 | 0.0013 | 0.0013 |
| F01 | 48880001 | 48980000 | 0.0020 | 0.0015 | 0.0015 |
| F01 | 48900001 | 49000000 | 0.0026 | 0.0021 | 0.0020 |
| F01 | 48920001 | 49020000 | 0.0023 | 0.0019 | 0.0018 |
| F01 | 48940001 | 49040000 | 0.0022 | 0.0019 | 0.0019 |
| F01 | 48960001 | 49060000 | 0.0020 | 0.0018 | 0.0019 |
| F01 | 48980001 | 49080000 | 0.0020 | 0.0018 | 0.0019 |
| F01 | 49000001 | 49100000 | 0.0013 | 0.0011 | 0.0013 |
| F01 | 49020001 | 49120000 | 0.0012 | 0.0011 | 0.0012 |
| F01 | 49040001 | 49140000 | 0.0013 | 0.0011 | 0.0013 |
| F01 | 49060001 | 49160000 | 0.0018 | 0.0015 | 0.0016 |
| F01 | 49080001 | 49180000 | 0.0023 | 0.0020 | 0.0021 |
| F01 | 49100001 | 49200000 | 0.0022 | 0.0019 | 0.0019 |
| F01 | 49120001 | 49220000 | 0.0025 | 0.0021 | 0.0021 |
| F01 | 49140001 | 49240000 | 0.0031 | 0.0026 | 0.0026 |
| F01 | 49160001 | 49260000 | 0.0028 | 0.0024 | 0.0025 |
| F01 | 49180001 | 49280000 | 0.0023 | 0.0018 | 0.0020 |
| F01 | 49200001 | 49300000 | 0.0025 | 0.0020 | 0.0023 |
| F01 | 49220001 | 49320000 | 0.0026 | 0.0023 | 0.0025 |
| F01 | 49240001 | 49340000 | 0.0020 | 0.0017 | 0.0020 |
| F01 | 49260001 | 49360000 | 0.0027 | 0.0024 | 0.0025 |
| F01 | 49280001 | 49380000 | 0.0028 | 0.0025 | 0.0025 |
| F01 | 49300001 | 49400000 | 0.0027 | 0.0024 | 0.0025 |
| F01 | 49320001 | 49420000 | 0.0027 | 0.0023 | 0.0024 |
| F01 | 49340001 | 49440000 | 0.0027 | 0.0022 | 0.0021 |
| F01 | 49360001 | 49460000 | 0.0020 | 0.0016 | 0.0016 |
| F01 | 49380001 | 49480000 | 0.0030 | 0.0023 | 0.0023 |
| F01 | 49400001 | 49500000 | 0.0035 | 0.0025 | 0.0025 |
| F01 | 49420001 | 49520000 | 0.0034 | 0.0023 | 0.0024 |
| F01 | 49440001 | 49540000 | 0.0031 | 0.0022 | 0.0024 |
| F01 | 49460001 | 49560000 | 0.0029 | 0.0021 | 0.0023 |
| F01 | 49480001 | 49580000 | 0.0022 | 0.0015 | 0.0017 |
| F01 | 49500001 | 49600000 | 0.0017 | 0.0011 | 0.0013 |
| F01 | 49520001 | 49620000 | 0.0017 | 0.0010 | 0.0012 |
| F01 | 49540001 | 49640000 | 0.0024 | 0.0015 | 0.0018 |
| F01 | 49560001 | 49660000 | 0.0023 | 0.0014 | 0.0017 |
| F01 | 49580001 | 49680000 | 0.0020 | 0.0014 | 0.0016 |
| F01 | 49600001 | 49700000 | 0.0018 | 0.0014 | 0.0016 |
| F01 | 49620001 | 49720000 | 0.0023 | 0.0019 | 0.0021 |
| F01 | 49640001 | 49740000 | 0.0020 | 0.0017 | 0.0019 |
| F01 | 49660001 | 49760000 | 0.0020 | 0.0018 | 0.0019 |
| F01 | 49680001 | 49780000 | 0.0023 | 0.0020 | 0.0022 |
| F01 | 49700001 | 49800000 | 0.0024 | 0.0022 | 0.0023 |
| F01 | 49720001 | 49820000 | 0.0021 | 0.0019 | 0.0019 |
| F01 | 49740001 | 49840000 | 0.0018 | 0.0016 | 0.0016 |
| F01 | 49760001 | 49860000 | 0.0019 | 0.0016 | 0.0017 |
| F01 | 49780001 | 49880000 | 0.0015 | 0.0013 | 0.0013 |

| Chr | Start | End | both | A1 | A2 |
|---|---|---|---|---|---|
| F01 | 49800001 | 49900000 | 0.0015 | 0.0013 | 0.0013 |
| F01 | 49820001 | 49920000 | 0.0023 | 0.0019 | 0.0020 |
| F01 | 49840001 | 49940000 | 0.0027 | 0.0021 | 0.0022 |
| F01 | 49860001 | 49960000 | 0.0027 | 0.0020 | 0.0021 |
| F01 | 49880001 | 49980000 | 0.0032 | 0.0022 | 0.0025 |
| F01 | 49900001 | 50000000 | 0.0037 | 0.0025 | 0.0027 |
| F01 | 49920001 | 50020000 | 0.0035 | 0.0021 | 0.0021 |
| F01 | 49940001 | 50040000 | 0.0033 | 0.0022 | 0.0021 |
| F01 | 49960001 | 50060000 | 0.0030 | 0.0020 | 0.0018 |
| F01 | 49980001 | 50080000 | 0.0031 | 0.0020 | 0.0017 |
| F01 | 50000001 | 50100000 | 0.0034 | 0.0022 | 0.0023 |
| F01 | 50020001 | 50120000 | 0.0029 | 0.0018 | 0.0020 |
| F01 | 50040001 | 50140000 | 0.0034 | 0.0025 | 0.0026 |
| F01 | 50060001 | 50160000 | 0.0032 | 0.0024 | 0.0026 |
| F01 | 50080001 | 50180000 | 0.0036 | 0.0029 | 0.0031 |
| F01 | 50100001 | 50200000 | 0.0035 | 0.0026 | 0.0030 |
| F01 | 50120001 | 50220000 | 0.0034 | 0.0026 | 0.0030 |
| F01 | 50140001 | 50240000 | 0.0029 | 0.0020 | 0.0024 |
| F01 | 50160001 | 50260000 | 0.0037 | 0.0025 | 0.0031 |
| F01 | 50180001 | 50280000 | 0.0036 | 0.0021 | 0.0029 |
| F01 | 50200001 | 50300000 | 0.0030 | 0.0019 | 0.0024 |
| F01 | 50220001 | 50320000 | 0.0033 | 0.0020 | 0.0024 |
| F01 | 50240001 | 50340000 | 0.0031 | 0.0017 | 0.0021 |
| F01 | 50260001 | 50360000 | 0.0038 | 0.0023 | 0.0028 |
| F01 | 50280001 | 50380000 | 0.0032 | 0.0022 | 0.0025 |
| F01 | 50300001 | 50400000 | 0.0035 | 0.0023 | 0.0026 |
| F01 | 50320001 | 50420000 | 0.0035 | 0.0024 | 0.0028 |
| F01 | 50340001 | 50440000 | 0.0042 | 0.0031 | 0.0034 |
| F01 | 50360001 | 50460000 | 0.0030 | 0.0022 | 0.0024 |
| F01 | 50380001 | 50480000 | 0.0029 | 0.0022 | 0.0024 |
| F01 | 50400001 | 50500000 | 0.0022 | 0.0018 | 0.0018 |
| F01 | 50420001 | 50520000 | 0.0029 | 0.0024 | 0.0026 |
| F01 | 50440001 | 50540000 | 0.0021 | 0.0018 | 0.0020 |
| F01 | 50460001 | 50560000 | 0.0019 | 0.0017 | 0.0018 |
| F01 | 50480001 | 50580000 | 0.0018 | 0.0016 | 0.0017 |
| F01 | 50500001 | 50600000 | 0.0022 | 0.0020 | 0.0021 |
| F01 | 50520001 | 50620000 | 0.0017 | 0.0015 | 0.0014 |
| F01 | 50540001 | 50640000 | 0.0016 | 0.0014 | 0.0014 |
| F01 | 50560001 | 50660000 | 0.0016 | 0.0014 | 0.0014 |
| F01 | 50580001 | 50680000 | 0.0013 | 0.0011 | 0.0011 |
| F01 | 50600001 | 50700000 | 0.0011 | 0.0009 | 0.0010 |
| F01 | 50620001 | 50720000 | 0.0010 | 0.0008 | 0.0010 |
| F01 | 50640001 | 50740000 | 0.0019 | 0.0015 | 0.0017 |
| F01 | 50660001 | 50760000 | 0.0018 | 0.0014 | 0.0017 |
| F01 | 50680001 | 50780000 | 0.0024 | 0.0018 | 0.0022 |
| F01 | 50700001 | 50800000 | 0.0030 | 0.0023 | 0.0026 |
| F01 | 50720001 | 50820000 | 0.0030 | 0.0022 | 0.0024 |
| F01 | 50740001 | 50840000 | 0.0023 | 0.0017 | 0.0018 |
| F01 | 50760001 | 50860000 | 0.0028 | 0.0019 | 0.0020 |
| F01 | 50780001 | 50880000 | 0.0027 | 0.0018 | 0.0016 |
| F01 | 50800001 | 50900000 | 0.0029 | 0.0018 | 0.0018 |
| F01 | 50820001 | 50920000 | 0.0032 | 0.0021 | 0.0021 |
| F01 | 50840001 | 50940000 | 0.0034 | 0.0020 | 0.0021 |
| F01 | 50860001 | 50960000 | 0.0040 | 0.0028 | 0.0029 |
| F01 | 50880001 | 50980000 | 0.0044 | 0.0033 | 0.0035 |
| F01 | 50900001 | 51000000 | 0.0039 | 0.0029 | 0.0031 |
| F01 | 50920001 | 51020000 | 0.0035 | 0.0027 | 0.0029 |
| F01 | 50940001 | 51040000 | 0.0031 | 0.0026 | 0.0028 |

| Chr | Start | End | both | A1 | A2 |
|---|---|---|---|---|---|
| F01 | 50960001 | 51060000 | 0.0024 | 0.0020 | 0.0022 |
| F01 | 50980001 | 51080000 | 0.0021 | 0.0017 | 0.0019 |
| F01 | 51000001 | 51100000 | 0.0023 | 0.0021 | 0.0022 |
| F01 | 51020001 | 51120000 | 0.0021 | 0.0019 | 0.0021 |
| F01 | 51040001 | 51140000 | 0.0022 | 0.0020 | 0.0022 |
| F01 | 51060001 | 51160000 | 0.0019 | 0.0017 | 0.0018 |
| F01 | 51080001 | 51180000 | 0.0018 | 0.0016 | 0.0018 |
| F01 | 51100001 | 51200000 | 0.0015 | 0.0014 | 0.0014 |
| F01 | 51120001 | 51220000 | 0.0018 | 0.0016 | 0.0017 |
| F01 | 51140001 | 51240000 | 0.0017 | 0.0016 | 0.0017 |
| F01 | 51160001 | 51260000 | 0.0017 | 0.0015 | 0.0016 |
| F01 | 51180001 | 51280000 | 0.0017 | 0.0015 | 0.0016 |
| F01 | 51200001 | 51300000 | 0.0015 | 0.0012 | 0.0013 |
| F01 | 51220001 | 51320000 | 0.0015 | 0.0010 | 0.0012 |
| F01 | 51240001 | 51340000 | 0.0020 | 0.0014 | 0.0015 |
| F01 | 51260001 | 51360000 | 0.0035 | 0.0027 | 0.0030 |
| F01 | 51280001 | 51380000 | 0.0038 | 0.0029 | 0.0031 |
| F01 | 51300001 | 51400000 | 0.0046 | 0.0036 | 0.0040 |
| F01 | 51320001 | 51420000 | 0.0047 | 0.0037 | 0.0042 |
| F01 | 51340001 | 51440000 | 0.0044 | 0.0034 | 0.0039 |
| F01 | 51360001 | 51460000 | 0.0030 | 0.0022 | 0.0025 |
| F01 | 51380001 | 51480000 | 0.0025 | 0.0018 | 0.0022 |
| F01 | 51400001 | 51500000 | 0.0019 | 0.0014 | 0.0016 |
| F01 | 51420001 | 51520000 | 0.0023 | 0.0018 | 0.0020 |
| F01 | 51440001 | 51540000 | 0.0025 | 0.0021 | 0.0023 |
| F01 | 51460001 | 51560000 | 0.0030 | 0.0025 | 0.0027 |
| F01 | 51480001 | 51580000 | 0.0031 | 0.0026 | 0.0028 |
| F01 | 51500001 | 51600000 | 0.0030 | 0.0024 | 0.0027 |
| F01 | 51520001 | 51620000 | 0.0032 | 0.0027 | 0.0028 |
| F01 | 51540001 | 51640000 | 0.0038 | 0.0033 | 0.0033 |
| F01 | 51560001 | 51660000 | 0.0044 | 0.0039 | 0.0039 |
| F01 | 51580001 | 51680000 | 0.0047 | 0.0042 | 0.0042 |
| F01 | 51600001 | 51700000 | 0.0050 | 0.0045 | 0.0045 |
| F01 | 51620001 | 51720000 | 0.0048 | 0.0041 | 0.0045 |
| F01 | 51640001 | 51740000 | 0.0035 | 0.0030 | 0.0033 |
| F01 | 51660001 | 51760000 | 0.0026 | 0.0022 | 0.0025 |
| F01 | 51680001 | 51780000 | 0.0023 | 0.0019 | 0.0023 |
| F01 | 51700001 | 51800000 | 0.0017 | 0.0014 | 0.0016 |
| F01 | 51720001 | 51820000 | 0.0013 | 0.0011 | 0.0012 |
| F01 | 51740001 | 51840000 | 0.0013 | 0.0011 | 0.0012 |
| F01 | 51760001 | 51860000 | 0.0010 | 0.0008 | 0.0009 |
| F01 | 51780001 | 51880000 | 0.0006 | 0.0005 | 0.0006 |
| F01 | 51800001 | 51900000 | 0.0009 | 0.0007 | 0.0009 |
| F01 | 51820001 | 51920000 | 0.0006 | 0.0005 | 0.0006 |
| F01 | 51840001 | 51940000 | 0.0007 | 0.0006 | 0.0007 |
| F01 | 51860001 | 51960000 | 0.0008 | 0.0007 | 0.0008 |
| F01 | 51880001 | 51980000 | 0.0010 | 0.0009 | 0.0010 |
| F01 | 51900001 | 52000000 | 0.0011 | 0.0010 | 0.0011 |
| F01 | 51920001 | 52020000 | 0.0014 | 0.0013 | 0.0014 |
| F01 | 51940001 | 52040000 | 0.0014 | 0.0012 | 0.0014 |
| F01 | 51960001 | 52060000 | 0.0012 | 0.0010 | 0.0012 |
| F01 | 51980001 | 52080000 | 0.0012 | 0.0010 | 0.0012 |
| F01 | 52000001 | 52100000 | 0.0009 | 0.0006 | 0.0009 |
| F01 | 52020001 | 52120000 | 0.0009 | 0.0007 | 0.0009 |
| F01 | 52040001 | 52140000 | 0.0014 | 0.0010 | 0.0012 |
| F01 | 52060001 | 52160000 | 0.0019 | 0.0013 | 0.0016 |
| F01 | 52080001 | 52180000 | 0.0025 | 0.0020 | 0.0022 |
| F01 | 52100001 | 52200000 | 0.0026 | 0.0021 | 0.0023 |

| Chr | Start | End | both | A1 | A2 |
|---|---|---|---|---|---|
| F01 | 52120001 | 52220000 | 0.0023 | 0.0018 | 0.0020 |
| F01 | 52140001 | 52240000 | 0.0016 | 0.0014 | 0.0015 |
| F01 | 52160001 | 52260000 | 0.0011 | 0.0011 | 0.0011 |
| F01 | 52180001 | 52280000 | 0.0004 | 0.0004 | 0.0004 |
| F01 | 52200001 | 52300000 | 0.0003 | 0.0003 | 0.0003 |
| F01 | 52220001 | 52320000 | 0.0007 | 0.0005 | 0.0007 |
| F01 | 52240001 | 52340000 | 0.0010 | 0.0008 | 0.0010 |
| F01 | 52260001 | 52360000 | 0.0012 | 0.0009 | 0.0012 |
| F01 | 52280001 | 52380000 | 0.0015 | 0.0012 | 0.0015 |
| F01 | 52300001 | 52400000 | 0.0015 | 0.0012 | 0.0015 |
| F01 | 52320001 | 52420000 | 0.0012 | 0.0010 | 0.0012 |
| F01 | 52340001 | 52440000 | 0.0012 | 0.0010 | 0.0013 |
| F01 | 52360001 | 52460000 | 0.0016 | 0.0013 | 0.0015 |
| F01 | 52380001 | 52480000 | 0.0016 | 0.0012 | 0.0014 |
| F01 | 52400001 | 52500000 | 0.0020 | 0.0015 | 0.0017 |
| F01 | 52420001 | 52520000 | 0.0023 | 0.0015 | 0.0018 |
| F01 | 52440001 | 52540000 | 0.0027 | 0.0018 | 0.0020 |
| F01 | 52460001 | 52560000 | 0.0026 | 0.0018 | 0.0020 |
| F01 | 52480001 | 52580000 | 0.0023 | 0.0015 | 0.0018 |
| F01 | 52500001 | 52600000 | 0.0020 | 0.0014 | 0.0016 |
| F01 | 52520001 | 52620000 | 0.0017 | 0.0014 | 0.0016 |
| F01 | 52540001 | 52640000 | 0.0015 | 0.0012 | 0.0014 |
| F01 | 52560001 | 52660000 | 0.0017 | 0.0014 | 0.0015 |
| F01 | 52580001 | 52680000 | 0.0019 | 0.0017 | 0.0017 |
| F01 | 52600001 | 52700000 | 0.0020 | 0.0017 | 0.0018 |
| F01 | 52620001 | 52720000 | 0.0023 | 0.0020 | 0.0022 |
| F01 | 52640001 | 52740000 | 0.0019 | 0.0016 | 0.0018 |
| F01 | 52660001 | 52760000 | 0.0014 | 0.0011 | 0.0013 |
| F01 | 52680001 | 52780000 | 0.0014 | 0.0011 | 0.0014 |
| F01 | 52700001 | 52800000 | 0.0015 | 0.0012 | 0.0014 |
| F01 | 52720001 | 52820000 | 0.0012 | 0.0010 | 0.0011 |
| F01 | 52740001 | 52840000 | 0.0013 | 0.0011 | 0.0012 |
| F01 | 52760001 | 52860000 | 0.0016 | 0.0014 | 0.0015 |
| F01 | 52780001 | 52880000 | 0.0015 | 0.0013 | 0.0014 |
| F01 | 52800001 | 52900000 | 0.0011 | 0.0010 | 0.0010 |
| F01 | 52820001 | 52920000 | 0.0016 | 0.0014 | 0.0016 |
| F01 | 52840001 | 52940000 | 0.0015 | 0.0013 | 0.0014 |
| F01 | 52860001 | 52960000 | 0.0014 | 0.0012 | 0.0013 |
| F01 | 52880001 | 52980000 | 0.0015 | 0.0012 | 0.0014 |
| F01 | 52900001 | 53000000 | 0.0016 | 0.0013 | 0.0015 |
| F01 | 52920001 | 53020000 | 0.0011 | 0.0009 | 0.0010 |
| F01 | 52940001 | 53040000 | 0.0012 | 0.0010 | 0.0011 |
| F01 | 52960001 | 53060000 | 0.0014 | 0.0011 | 0.0014 |
| F01 | 52980001 | 53080000 | 0.0013 | 0.0010 | 0.0012 |
| F01 | 53000001 | 53100000 | 0.0019 | 0.0013 | 0.0015 |
| F01 | 53020001 | 53120000 | 0.0024 | 0.0015 | 0.0016 |
| F01 | 53040001 | 53140000 | 0.0026 | 0.0017 | 0.0018 |
| F01 | 53060001 | 53160000 | 0.0022 | 0.0014 | 0.0013 |
| F01 | 53080001 | 53180000 | 0.0027 | 0.0018 | 0.0018 |
| F01 | 53100001 | 53200000 | 0.0021 | 0.0015 | 0.0015 |
| F01 | 53120001 | 53220000 | 0.0020 | 0.0014 | 0.0014 |
| F01 | 53140001 | 53240000 | 0.0020 | 0.0013 | 0.0013 |
| F01 | 53160001 | 53260000 | 0.0020 | 0.0012 | 0.0012 |
| F01 | 53180001 | 53280000 | 0.0014 | 0.0007 | 0.0008 |
| F01 | 53200001 | 53300000 | 0.0019 | 0.0011 | 0.0011 |
| F01 | 53220001 | 53320000 | 0.0018 | 0.0012 | 0.0012 |
| F01 | 53240001 | 53340000 | 0.0018 | 0.0012 | 0.0012 |
| F01 | 53260001 | 53360000 | 0.0022 | 0.0014 | 0.0014 |

| Chr | Start | End | both | A1 | A2 |
|---|---|---|---|---|---|
| F01 | 53280001 | 53380000 | 0.0026 | 0.0017 | 0.0017 |
| F01 | 53300001 | 53400000 | 0.0024 | 0.0014 | 0.0015 |
| F01 | 53320001 | 53420000 | 0.0025 | 0.0014 | 0.0016 |
| F01 | 53340001 | 53440000 | 0.0026 | 0.0016 | 0.0018 |
| F01 | 53360001 | 53460000 | 0.0025 | 0.0016 | 0.0018 |
| F01 | 53380001 | 53480000 | 0.0033 | 0.0022 | 0.0028 |
| F01 | 53400001 | 53500000 | 0.0036 | 0.0023 | 0.0028 |
| F01 | 53420001 | 53520000 | 0.0039 | 0.0025 | 0.0029 |
| F01 | 53440001 | 53540000 | 0.0041 | 0.0027 | 0.0030 |
| F01 | 53460001 | 53560000 | 0.0046 | 0.0032 | 0.0036 |
| F01 | 53480001 | 53580000 | 0.0036 | 0.0024 | 0.0026 |
| F01 | 53500001 | 53600000 | 0.0030 | 0.0020 | 0.0022 |
| F01 | 53520001 | 53620000 | 0.0026 | 0.0019 | 0.0020 |
| F01 | 53540001 | 53640000 | 0.0022 | 0.0016 | 0.0018 |
| F01 | 53560001 | 53660000 | 0.0016 | 0.0011 | 0.0012 |
| F01 | 53580001 | 53680000 | 0.0021 | 0.0016 | 0.0018 |
| F01 | 53600001 | 53700000 | 0.0025 | 0.0021 | 0.0023 |
| F01 | 53620001 | 53720000 | 0.0022 | 0.0018 | 0.0020 |
| F01 | 53640001 | 53740000 | 0.0022 | 0.0018 | 0.0021 |
| F01 | 53660001 | 53760000 | 0.0030 | 0.0027 | 0.0030 |
| F01 | 53680001 | 53780000 | 0.0024 | 0.0022 | 0.0024 |
| F01 | 53700001 | 53800000 | 0.0021 | 0.0019 | 0.0021 |
| F01 | 53720001 | 53820000 | 0.0021 | 0.0019 | 0.0020 |
| F01 | 53740001 | 53840000 | 0.0021 | 0.0019 | 0.0020 |
| F01 | 53760001 | 53860000 | 0.0009 | 0.0008 | 0.0009 |
| F01 | 53780001 | 53880000 | 0.0008 | 0.0008 | 0.0008 |
| F01 | 53800001 | 53900000 | 0.0010 | 0.0010 | 0.0010 |
| F01 | 53820001 | 53920000 | 0.0017 | 0.0013 | 0.0014 |
| F01 | 53840001 | 53940000 | 0.0024 | 0.0020 | 0.0020 |
| F01 | 53860001 | 53960000 | 0.0026 | 0.0021 | 0.0021 |
| F01 | 53880001 | 53980000 | 0.0025 | 0.0020 | 0.0020 |
| F01 | 53900001 | 54000000 | 0.0027 | 0.0022 | 0.0022 |
| F01 | 53920001 | 54020000 | 0.0022 | 0.0019 | 0.0019 |
| F01 | 53940001 | 54040000 | 0.0014 | 0.0010 | 0.0011 |
| F01 | 53960001 | 54060000 | 0.0017 | 0.0013 | 0.0014 |
| F01 | 53980001 | 54080000 | 0.0020 | 0.0014 | 0.0015 |
| F01 | 54000001 | 54100000 | 0.0019 | 0.0012 | 0.0012 |
| F01 | 54020001 | 54120000 | 0.0018 | 0.0011 | 0.0012 |
| F01 | 54040001 | 54140000 | 0.0024 | 0.0018 | 0.0018 |
| F01 | 54060001 | 54160000 | 0.0027 | 0.0019 | 0.0018 |
| F01 | 54080001 | 54180000 | 0.0030 | 0.0021 | 0.0023 |
| F01 | 54100001 | 54200000 | 0.0031 | 0.0022 | 0.0024 |
| F01 | 54120001 | 54220000 | 0.0033 | 0.0022 | 0.0024 |
| F01 | 54140001 | 54240000 | 0.0029 | 0.0015 | 0.0019 |
| F01 | 54160001 | 54260000 | 0.0027 | 0.0014 | 0.0019 |
| F01 | 54180001 | 54280000 | 0.0029 | 0.0020 | 0.0021 |
| F01 | 54200001 | 54300000 | 0.0032 | 0.0018 | 0.0019 |
| F01 | 54220001 | 54320000 | 0.0038 | 0.0024 | 0.0025 |
| F01 | 54240001 | 54340000 | 0.0040 | 0.0028 | 0.0030 |
| F01 | 54260001 | 54360000 | 0.0042 | 0.0030 | 0.0032 |
| F01 | 54280001 | 54380000 | 0.0038 | 0.0024 | 0.0027 |
| F01 | 54300001 | 54400000 | 0.0036 | 0.0025 | 0.0030 |
| F01 | 54320001 | 54420000 | 0.0029 | 0.0021 | 0.0024 |
| F01 | 54340001 | 54440000 | 0.0022 | 0.0015 | 0.0018 |
| F01 | 54360001 | 54460000 | 0.0018 | 0.0013 | 0.0013 |
| F01 | 54380001 | 54480000 | 0.0019 | 0.0016 | 0.0016 |
| F01 | 54400001 | 54500000 | 0.0013 | 0.0012 | 0.0012 |
| F01 | 54420001 | 54520000 | 0.0015 | 0.0015 | 0.0014 |

| Chr | Start | End | both | A1 | A2 |
|---|---|---|---|---|---|
| F01 | 54440001 | 54540000 | 0.0019 | 0.0017 | 0.0016 |
| F01 | 54460001 | 54560000 | 0.0020 | 0.0017 | 0.0017 |
| F01 | 54480001 | 54580000 | 0.0018 | 0.0014 | 0.0015 |
| F01 | 54500001 | 54600000 | 0.0019 | 0.0016 | 0.0017 |
| F01 | 54520001 | 54620000 | 0.0017 | 0.0013 | 0.0014 |
| F01 | 54540001 | 54640000 | 0.0015 | 0.0011 | 0.0013 |
| F01 | 54560001 | 54660000 | 0.0016 | 0.0012 | 0.0014 |
| F01 | 54580001 | 54680000 | 0.0020 | 0.0013 | 0.0016 |
| F01 | 54600001 | 54700000 | 0.0023 | 0.0015 | 0.0019 |
| F01 | 54620001 | 54720000 | 0.0029 | 0.0022 | 0.0025 |
| F01 | 54640001 | 54740000 | 0.0029 | 0.0022 | 0.0026 |
| F01 | 54660001 | 54760000 | 0.0030 | 0.0024 | 0.0027 |
| F01 | 54680001 | 54780000 | 0.0024 | 0.0020 | 0.0022 |
| F01 | 54700001 | 54800000 | 0.0024 | 0.0020 | 0.0022 |
| F01 | 54720001 | 54820000 | 0.0016 | 0.0012 | 0.0015 |
| F01 | 54740001 | 54840000 | 0.0017 | 0.0013 | 0.0016 |
| F01 | 54760001 | 54860000 | 0.0013 | 0.0010 | 0.0012 |
| F01 | 54780001 | 54880000 | 0.0011 | 0.0009 | 0.0011 |
| F01 | 54800001 | 54900000 | 0.0008 | 0.0007 | 0.0007 |
| F01 | 54820001 | 54920000 | 0.0011 | 0.0009 | 0.0008 |
| F01 | 54840001 | 54940000 | 0.0020 | 0.0018 | 0.0016 |
| F01 | 54860001 | 54960000 | 0.0021 | 0.0020 | 0.0018 |
| F01 | 54880001 | 54980000 | 0.0022 | 0.0020 | 0.0018 |
| F01 | 54900001 | 55000000 | 0.0021 | 0.0018 | 0.0019 |
| F01 | 54920001 | 55020000 | 0.0018 | 0.0016 | 0.0017 |
| F01 | 54940001 | 55040000 | 0.0012 | 0.0009 | 0.0011 |
| F01 | 54960001 | 55060000 | 0.0017 | 0.0012 | 0.0015 |
| F01 | 54980001 | 55080000 | 0.0015 | 0.0011 | 0.0013 |
| F01 | 55000001 | 55100000 | 0.0013 | 0.0010 | 0.0011 |
| F01 | 55020001 | 55120000 | 0.0013 | 0.0010 | 0.0011 |
| F01 | 55040001 | 55140000 | 0.0007 | 0.0006 | 0.0006 |
| F01 | 55060001 | 55160000 | 0.0003 | 0.0002 | 0.0003 |
| F01 | 55080001 | 55180000 | 0.0004 | 0.0003 | 0.0004 |
| F01 | 55100001 | 55200000 | 0.0007 | 0.0006 | 0.0006 |
| F01 | 55120001 | 55220000 | 0.0013 | 0.0011 | 0.0011 |
| F01 | 55140001 | 55240000 | 0.0015 | 0.0012 | 0.0013 |
| F01 | 55160001 | 55260000 | 0.0025 | 0.0021 | 0.0022 |
| F01 | 55180001 | 55280000 | 0.0026 | 0.0021 | 0.0022 |
| F01 | 55200001 | 55300000 | 0.0029 | 0.0023 | 0.0025 |
| F01 | 55220001 | 55320000 | 0.0027 | 0.0022 | 0.0024 |
| F01 | 55240001 | 55340000 | 0.0028 | 0.0024 | 0.0026 |
| F01 | 55260001 | 55360000 | 0.0022 | 0.0019 | 0.0020 |
| F01 | 55280001 | 55380000 | 0.0029 | 0.0025 | 0.0026 |
| F01 | 55300001 | 55400000 | 0.0032 | 0.0026 | 0.0026 |
| F01 | 55320001 | 55420000 | 0.0043 | 0.0037 | 0.0037 |
| F01 | 55340001 | 55440000 | 0.0049 | 0.0042 | 0.0042 |
| F01 | 55360001 | 55460000 | 0.0052 | 0.0044 | 0.0044 |
| F01 | 55380001 | 55480000 | 0.0048 | 0.0040 | 0.0040 |
| F01 | 55400001 | 55500000 | 0.0047 | 0.0040 | 0.0041 |
| F01 | 55420001 | 55520000 | 0.0035 | 0.0028 | 0.0028 |
| F01 | 55440001 | 55540000 | 0.0026 | 0.0019 | 0.0019 |
| F01 | 55460001 | 55560000 | 0.0020 | 0.0014 | 0.0014 |
| F01 | 55480001 | 55580000 | 0.0019 | 0.0013 | 0.0013 |
| F01 | 55500001 | 55600000 | 0.0018 | 0.0012 | 0.0011 |
| F01 | 55520001 | 55620000 | 0.0017 | 0.0012 | 0.0011 |
| F01 | 55540001 | 55640000 | 0.0021 | 0.0014 | 0.0014 |
| F01 | 55560001 | 55660000 | 0.0022 | 0.0014 | 0.0015 |
| F01 | 55580001 | 55680000 | 0.0030 | 0.0023 | 0.0025 |

| Chr | Start | End | both | A1 | A2 |
|---|---|---|---|---|---|
| F01 | 55600001 | 55700000 | 0.0036 | 0.0026 | 0.0030 |
| F01 | 55620001 | 55720000 | 0.0037 | 0.0024 | 0.0030 |
| F01 | 55640001 | 55740000 | 0.0038 | 0.0025 | 0.0030 |
| F01 | 55660001 | 55760000 | 0.0040 | 0.0026 | 0.0031 |
| F01 | 55680001 | 55780000 | 0.0031 | 0.0017 | 0.0022 |
| F01 | 55700001 | 55800000 | 0.0028 | 0.0017 | 0.0020 |
| F01 | 55720001 | 55820000 | 0.0031 | 0.0023 | 0.0026 |
| F01 | 55740001 | 55840000 | 0.0031 | 0.0025 | 0.0028 |
| F01 | 55760001 | 55860000 | 0.0035 | 0.0031 | 0.0033 |
| F01 | 55780001 | 55880000 | 0.0041 | 0.0035 | 0.0037 |
| F01 | 55800001 | 55900000 | 0.0036 | 0.0031 | 0.0032 |
| F01 | 55820001 | 55920000 | 0.0040 | 0.0032 | 0.0032 |
| F01 | 55840001 | 55940000 | 0.0045 | 0.0033 | 0.0035 |
| F01 | 55860001 | 55960000 | 0.0040 | 0.0027 | 0.0030 |
| F01 | 55880001 | 55980000 | 0.0033 | 0.0021 | 0.0022 |
| F01 | 55900001 | 56000000 | 0.0035 | 0.0022 | 0.0024 |
| F01 | 55920001 | 56020000 | 0.0027 | 0.0018 | 0.0019 |
| F01 | 55940001 | 56040000 | 0.0018 | 0.0012 | 0.0011 |
| F01 | 55960001 | 56060000 | 0.0020 | 0.0015 | 0.0016 |
| F01 | 55980001 | 56080000 | 0.0020 | 0.0015 | 0.0018 |
| F01 | 56000001 | 56100000 | 0.0015 | 0.0011 | 0.0014 |
| F01 | 56020001 | 56120000 | 0.0015 | 0.0012 | 0.0015 |
| F01 | 56040001 | 56140000 | 0.0016 | 0.0013 | 0.0015 |
| F01 | 56060001 | 56160000 | 0.0011 | 0.0010 | 0.0011 |
| F01 | 56080001 | 56180000 | 0.0014 | 0.0011 | 0.0011 |
| F01 | 56100001 | 56200000 | 0.0017 | 0.0013 | 0.0014 |
| F01 | 56120001 | 56220000 | 0.0018 | 0.0013 | 0.0014 |
| F01 | 56140001 | 56240000 | 0.0019 | 0.0013 | 0.0015 |
| F01 | 56160001 | 56260000 | 0.0024 | 0.0017 | 0.0021 |
| F01 | 56180001 | 56280000 | 0.0020 | 0.0016 | 0.0019 |
| F01 | 56200001 | 56300000 | 0.0018 | 0.0015 | 0.0017 |
| F01 | 56220001 | 56320000 | 0.0018 | 0.0015 | 0.0017 |
| F01 | 56240001 | 56340000 | 0.0016 | 0.0014 | 0.0015 |
| F01 | 56260001 | 56360000 | 0.0012 | 0.0010 | 0.0010 |
| F01 | 56280001 | 56380000 | 0.0020 | 0.0017 | 0.0018 |
| F01 | 56300001 | 56400000 | 0.0021 | 0.0018 | 0.0018 |
| F01 | 56320001 | 56420000 | 0.0019 | 0.0017 | 0.0018 |
| F01 | 56340001 | 56440000 | 0.0022 | 0.0019 | 0.0021 |
| F01 | 56360001 | 56460000 | 0.0022 | 0.0019 | 0.0021 |
| F01 | 56380001 | 56480000 | 0.0012 | 0.0011 | 0.0012 |
| F01 | 56400001 | 56500000 | 0.0011 | 0.0009 | 0.0010 |
| F01 | 56420001 | 56520000 | 0.0006 | 0.0005 | 0.0006 |
| F01 | 56440001 | 56540000 | 0.0007 | 0.0005 | 0.0007 |
| F01 | 56460001 | 56560000 | 0.0014 | 0.0012 | 0.0014 |
| F01 | 56480001 | 56580000 | 0.0015 | 0.0012 | 0.0015 |
| F01 | 56500001 | 56600000 | 0.0016 | 0.0013 | 0.0016 |
| F01 | 56520001 | 56620000 | 0.0021 | 0.0017 | 0.0021 |
| F01 | 56540001 | 56640000 | 0.0017 | 0.0014 | 0.0017 |
| F01 | 56560001 | 56660000 | 0.0010 | 0.0008 | 0.0009 |
| F01 | 56580001 | 56680000 | 0.0010 | 0.0008 | 0.0010 |
| F01 | 56600001 | 56700000 | 0.0012 | 0.0010 | 0.0012 |
| F01 | 56620001 | 56720000 | 0.0011 | 0.0009 | 0.0010 |
| F01 | 56640001 | 56740000 | 0.0012 | 0.0010 | 0.0012 |
| F01 | 56660001 | 56760000 | 0.0014 | 0.0012 | 0.0014 |
| F01 | 56680001 | 56780000 | 0.0015 | 0.0013 | 0.0015 |
| F01 | 56700001 | 56800000 | 0.0012 | 0.0011 | 0.0012 |
| F01 | 56720001 | 56820000 | 0.0018 | 0.0015 | 0.0018 |
| F01 | 56740001 | 56840000 | 0.0018 | 0.0016 | 0.0018 |

| Chr | Start | End | both | A1 | A2 |
|---|---|---|---|---|---|
| F01 | 56760001 | 56860000 | 0.0021 | 0.0018 | 0.0021 |
| F01 | 56780001 | 56880000 | 0.0020 | 0.0017 | 0.0020 |
| F01 | 56800001 | 56900000 | 0.0019 | 0.0016 | 0.0019 |
| F01 | 56820001 | 56920000 | 0.0014 | 0.0013 | 0.0014 |
| F01 | 56840001 | 56940000 | 0.0015 | 0.0012 | 0.0015 |
| F01 | 56860001 | 56960000 | 0.0012 | 0.0009 | 0.0011 |
| F01 | 56880001 | 56980000 | 0.0018 | 0.0014 | 0.0018 |
| F01 | 56900001 | 57000000 | 0.0025 | 0.0020 | 0.0024 |
| F01 | 56920001 | 57020000 | 0.0022 | 0.0017 | 0.0022 |
| F01 | 56940001 | 57040000 | 0.0019 | 0.0016 | 0.0018 |
| F01 | 56960001 | 57060000 | 0.0021 | 0.0018 | 0.0021 |
| F01 | 56980001 | 57080000 | 0.0015 | 0.0013 | 0.0015 |
| F01 | 57000001 | 57100000 | 0.0011 | 0.0010 | 0.0011 |
| F01 | 57020001 | 57120000 | 0.0010 | 0.0009 | 0.0010 |
| F01 | 57040001 | 57140000 | 0.0010 | 0.0008 | 0.0010 |
| F01 | 57060001 | 57160000 | 0.0007 | 0.0006 | 0.0007 |
| F01 | 57080001 | 57180000 | 0.0006 | 0.0006 | 0.0006 |
| F01 | 57100001 | 57200000 | 0.0008 | 0.0006 | 0.0006 |
| F01 | 57120001 | 57220000 | 0.0016 | 0.0014 | 0.0014 |
| F01 | 57140001 | 57240000 | 0.0019 | 0.0018 | 0.0017 |
| F01 | 57160001 | 57260000 | 0.0021 | 0.0019 | 0.0019 |
| F01 | 57180001 | 57280000 | 0.0023 | 0.0020 | 0.0020 |
| F01 | 57200001 | 57300000 | 0.0024 | 0.0021 | 0.0023 |
| F01 | 57220001 | 57320000 | 0.0030 | 0.0024 | 0.0028 |
| F01 | 57240001 | 57340000 | 0.0032 | 0.0023 | 0.0029 |
| F01 | 57260001 | 57360000 | 0.0030 | 0.0022 | 0.0027 |
| F01 | 57280001 | 57380000 | 0.0035 | 0.0025 | 0.0032 |
| F01 | 57300001 | 57400000 | 0.0039 | 0.0024 | 0.0036 |
| F01 | 57320001 | 57420000 | 0.0033 | 0.0017 | 0.0032 |
| F01 | 57340001 | 57440000 | 0.0033 | 0.0018 | 0.0032 |
| F01 | 57360001 | 57460000 | 0.0040 | 0.0024 | 0.0039 |
| F01 | 57380001 | 57480000 | 0.0039 | 0.0023 | 0.0038 |
| F01 | 57400001 | 57500000 | 0.0033 | 0.0022 | 0.0032 |
| F01 | 57420001 | 57520000 | 0.0026 | 0.0020 | 0.0026 |
| F01 | 57440001 | 57540000 | 0.0028 | 0.0022 | 0.0027 |
| F01 | 57460001 | 57560000 | 0.0024 | 0.0019 | 0.0023 |
| F01 | 57480001 | 57580000 | 0.0019 | 0.0016 | 0.0018 |
| F01 | 57500001 | 57600000 | 0.0016 | 0.0013 | 0.0015 |
| F01 | 57520001 | 57620000 | 0.0017 | 0.0014 | 0.0016 |
| F01 | 57540001 | 57640000 | 0.0011 | 0.0008 | 0.0010 |
| F01 | 57560001 | 57660000 | 0.0008 | 0.0006 | 0.0007 |
| F01 | 57580001 | 57680000 | 0.0013 | 0.0011 | 0.0013 |
| F01 | 57600001 | 57700000 | 0.0014 | 0.0012 | 0.0014 |
| F01 | 57620001 | 57720000 | 0.0014 | 0.0011 | 0.0013 |
| F01 | 57640001 | 57740000 | 0.0017 | 0.0014 | 0.0017 |
| F01 | 57660001 | 57760000 | 0.0020 | 0.0016 | 0.0020 |
| F01 | 57680001 | 57780000 | 0.0015 | 0.0012 | 0.0015 |
| F01 | 57700001 | 57800000 | 0.0016 | 0.0013 | 0.0016 |
| F01 | 57720001 | 57820000 | 0.0014 | 0.0012 | 0.0014 |
| F01 | 57740001 | 57840000 | 0.0010 | 0.0008 | 0.0010 |
| F01 | 57760001 | 57860000 | 0.0007 | 0.0005 | 0.0007 |
| F01 | 57780001 | 57880000 | 0.0008 | 0.0006 | 0.0008 |
| F01 | 57800001 | 57900000 | 0.0005 | 0.0004 | 0.0005 |
| F01 | 57820001 | 57920000 | 0.0006 | 0.0004 | 0.0006 |
| F01 | 57840001 | 57940000 | 0.0006 | 0.0004 | 0.0006 |
| F01 | 57860001 | 57960000 | 0.0005 | 0.0004 | 0.0005 |
| F01 | 57880001 | 57980000 | 0.0002 | 0.0002 | 0.0002 |
| F01 | 57900001 | 58000000 | 0.0005 | 0.0004 | 0.0005 |

| Chr | Start | End | both | A1 | A2 |
|---|---|---|---|---|---|
| F01 | 57920001 | 58020000 | 0.0004 | 0.0003 | 0.0004 |
| F01 | 57940001 | 58040000 | 0.0004 | 0.0004 | 0.0004 |
| F01 | 57960001 | 58060000 | 0.0004 | 0.0004 | 0.0004 |
| F01 | 57980001 | 58080000 | 0.0004 | 0.0004 | 0.0004 |
| F01 | 58000001 | 58100000 | 0.0003 | 0.0002 | 0.0003 |
| F01 | 58020001 | 58120000 | 0.0003 | 0.0002 | 0.0003 |
| F01 | 58040001 | 58140000 | 0.0002 | 0.0001 | 0.0002 |
| F01 | 58060001 | 58160000 | 0.0002 | 0.0001 | 0.0002 |
| F01 | 58080001 | 58180000 | 0.0004 | 0.0004 | 0.0004 |
| F01 | 58100001 | 58200000 | 0.0005 | 0.0004 | 0.0005 |
| F01 | 58120001 | 58220000 | 0.0006 | 0.0005 | 0.0006 |
| F01 | 58140001 | 58240000 | 0.0009 | 0.0008 | 0.0009 |
| F01 | 58160001 | 58260000 | 0.0010 | 0.0009 | 0.0010 |
| F01 | 58180001 | 58280000 | 0.0009 | 0.0008 | 0.0009 |
| F01 | 58200001 | 58300000 | 0.0009 | 0.0008 | 0.0009 |
| F01 | 58220001 | 58320000 | 0.0009 | 0.0009 | 0.0009 |
| F01 | 58240001 | 58340000 | 0.0008 | 0.0007 | 0.0008 |
| F01 | 58260001 | 58360000 | 0.0008 | 0.0007 | 0.0008 |
| F01 | 58280001 | 58380000 | 0.0009 | 0.0007 | 0.0009 |
| F01 | 58300001 | 58400000 | 0.0009 | 0.0007 | 0.0009 |
| F01 | 58320001 | 58420000 | 0.0010 | 0.0008 | 0.0011 |
| F01 | 58340001 | 58440000 | 0.0014 | 0.0009 | 0.0014 |
| F01 | 58360001 | 58460000 | 0.0014 | 0.0009 | 0.0014 |
| F01 | 58380001 | 58480000 | 0.0011 | 0.0007 | 0.0011 |
| F01 | 58400001 | 58500000 | 0.0011 | 0.0008 | 0.0012 |
| F01 | 58420001 | 58520000 | 0.0015 | 0.0012 | 0.0015 |
| F01 | 58440001 | 58540000 | 0.0015 | 0.0013 | 0.0015 |
| F01 | 58460001 | 58560000 | 0.0022 | 0.0019 | 0.0022 |
| F01 | 58480001 | 58580000 | 0.0025 | 0.0022 | 0.0024 |
| F01 | 58500001 | 58600000 | 0.0036 | 0.0032 | 0.0035 |
| F01 | 58520001 | 58620000 | 0.0031 | 0.0028 | 0.0029 |
| F01 | 58540001 | 58640000 | 0.0031 | 0.0029 | 0.0029 |
| F01 | 58560001 | 58660000 | 0.0023 | 0.0022 | 0.0021 |
| F01 | 58580001 | 58680000 | 0.0022 | 0.0020 | 0.0020 |
| F01 | 58600001 | 58700000 | 0.0015 | 0.0014 | 0.0013 |
| F01 | 58620001 | 58720000 | 0.0015 | 0.0014 | 0.0014 |
| F01 | 58640001 | 58740000 | 0.0011 | 0.0010 | 0.0010 |
| F01 | 58660001 | 58760000 | 0.0012 | 0.0011 | 0.0011 |
| F01 | 58680001 | 58780000 | 0.0011 | 0.0010 | 0.0010 |
| F01 | 58700001 | 58800000 | 0.0011 | 0.0010 | 0.0010 |
| F01 | 58720001 | 58820000 | 0.0011 | 0.0008 | 0.0010 |
| F01 | 58740001 | 58840000 | 0.0015 | 0.0012 | 0.0013 |
| F01 | 58760001 | 58860000 | 0.0016 | 0.0013 | 0.0015 |
| F01 | 58780001 | 58880000 | 0.0021 | 0.0018 | 0.0020 |
| F01 | 58800001 | 58900000 | 0.0019 | 0.0015 | 0.0018 |
| F01 | 58820001 | 58920000 | 0.0020 | 0.0018 | 0.0019 |
| F01 | 58840001 | 58940000 | 0.0017 | 0.0016 | 0.0017 |
| F01 | 58860001 | 58960000 | 0.0017 | 0.0015 | 0.0017 |
| F01 | 58880001 | 58980000 | 0.0017 | 0.0015 | 0.0017 |
| F01 | 58900001 | 59000000 | 0.0013 | 0.0012 | 0.0013 |
| F01 | 58920001 | 59020000 | 0.0011 | 0.0009 | 0.0010 |
| F01 | 58940001 | 59040000 | 0.0010 | 0.0008 | 0.0009 |
| F01 | 58960001 | 59060000 | 0.0010 | 0.0007 | 0.0008 |
| F01 | 58980001 | 59080000 | 0.0013 | 0.0009 | 0.0009 |
| F01 | 59000001 | 59100000 | 0.0020 | 0.0016 | 0.0015 |
| F01 | 59020001 | 59120000 | 0.0021 | 0.0017 | 0.0016 |
| F01 | 59040001 | 59140000 | 0.0025 | 0.0019 | 0.0020 |
| F01 | 59060001 | 59160000 | 0.0023 | 0.0019 | 0.0020 |

| Chr | Start | End | both | A1 | A2 |
|---|---|---|---|---|---|
| F01 | 59080001 | 59180000 | 0.0019 | 0.0015 | 0.0017 |
| F01 | 59100001 | 59200000 | 0.0012 | 0.0008 | 0.0010 |
| F01 | 59120001 | 59220000 | 0.0009 | 0.0007 | 0.0009 |
| F01 | 59140001 | 59240000 | 0.0012 | 0.0010 | 0.0012 |
| F01 | 59160001 | 59260000 | 0.0015 | 0.0012 | 0.0015 |
| F01 | 59180001 | 59280000 | 0.0013 | 0.0010 | 0.0013 |
| F01 | 59200001 | 59300000 | 0.0018 | 0.0013 | 0.0018 |
| F01 | 59220001 | 59320000 | 0.0018 | 0.0013 | 0.0017 |
| F01 | 59240001 | 59340000 | 0.0014 | 0.0012 | 0.0014 |
| F01 | 59260001 | 59360000 | 0.0012 | 0.0010 | 0.0012 |
| F01 | 59280001 | 59380000 | 0.0014 | 0.0011 | 0.0013 |
| F01 | 59300001 | 59400000 | 0.0013 | 0.0009 | 0.0012 |
| F01 | 59320001 | 59420000 | 0.0020 | 0.0015 | 0.0020 |
| F01 | 59340001 | 59440000 | 0.0018 | 0.0013 | 0.0018 |
| F01 | 59360001 | 59460000 | 0.0017 | 0.0013 | 0.0017 |
| F01 | 59380001 | 59480000 | 0.0018 | 0.0016 | 0.0018 |
| F01 | 59400001 | 59500000 | 0.0017 | 0.0016 | 0.0017 |
| F01 | 59420001 | 59520000 | 0.0010 | 0.0010 | 0.0010 |
| F01 | 59440001 | 59540000 | 0.0009 | 0.0008 | 0.0009 |
| F01 | 59460001 | 59560000 | 0.0012 | 0.0010 | 0.0011 |
| F01 | 59480001 | 59580000 | 0.0012 | 0.0011 | 0.0012 |
| F01 | 59500001 | 59600000 | 0.0011 | 0.0010 | 0.0011 |
| F01 | 59520001 | 59620000 | 0.0018 | 0.0017 | 0.0018 |
| F01 | 59540001 | 59640000 | 0.0023 | 0.0021 | 0.0023 |
| F01 | 59560001 | 59660000 | 0.0025 | 0.0023 | 0.0024 |
| F01 | 59580001 | 59680000 | 0.0021 | 0.0018 | 0.0020 |
| F01 | 59600001 | 59700000 | 0.0022 | 0.0020 | 0.0022 |
| F01 | 59620001 | 59720000 | 0.0016 | 0.0014 | 0.0016 |
| F01 | 59640001 | 59740000 | 0.0016 | 0.0014 | 0.0014 |
| F01 | 59660001 | 59760000 | 0.0012 | 0.0010 | 0.0009 |
| F01 | 59680001 | 59780000 | 0.0013 | 0.0011 | 0.0010 |
| F01 | 59700001 | 59800000 | 0.0010 | 0.0008 | 0.0007 |
| F01 | 59720001 | 59820000 | 0.0012 | 0.0010 | 0.0010 |
| F01 | 59740001 | 59840000 | 0.0013 | 0.0010 | 0.0011 |
| F01 | 59760001 | 59860000 | 0.0011 | 0.0010 | 0.0011 |
| F01 | 59780001 | 59880000 | 0.0012 | 0.0011 | 0.0012 |
| F01 | 59800001 | 59900000 | 0.0012 | 0.0011 | 0.0012 |
| F01 | 59820001 | 59920000 | 0.0010 | 0.0008 | 0.0009 |
| F01 | 59840001 | 59940000 | 0.0010 | 0.0008 | 0.0010 |
| F01 | 59860001 | 59960000 | 0.0010 | 0.0008 | 0.0010 |
| F01 | 59880001 | 59980000 | 0.0014 | 0.0011 | 0.0014 |
| F01 | 59900001 | 60000000 | 0.0015 | 0.0013 | 0.0015 |
| F01 | 59920001 | 60020000 | 0.0015 | 0.0013 | 0.0015 |
| F01 | 59940001 | 60040000 | 0.0016 | 0.0014 | 0.0015 |
| F01 | 59960001 | 60060000 | 0.0019 | 0.0017 | 0.0018 |
| F01 | 59980001 | 60080000 | 0.0014 | 0.0013 | 0.0013 |
| F01 | 60000001 | 60100000 | 0.0013 | 0.0012 | 0.0013 |
| F01 | 60020001 | 60120000 | 0.0013 | 0.0011 | 0.0012 |
| F01 | 60040001 | 60140000 | 0.0022 | 0.0016 | 0.0022 |
| F01 | 60060001 | 60160000 | 0.0020 | 0.0015 | 0.0021 |
| F01 | 60080001 | 60180000 | 0.0021 | 0.0016 | 0.0022 |
| F01 | 60100001 | 60200000 | 0.0024 | 0.0017 | 0.0024 |
| F01 | 60120001 | 60220000 | 0.0023 | 0.0016 | 0.0023 |
| F01 | 60140001 | 60240000 | 0.0008 | 0.0006 | 0.0008 |
| F01 | 60160001 | 60260000 | 0.0006 | 0.0004 | 0.0006 |
| F01 | 60180001 | 60280000 | 0.0006 | 0.0004 | 0.0006 |
| F01 | 60200001 | 60300000 | 0.0010 | 0.0009 | 0.0009 |
| F01 | 60220001 | 60320000 | 0.0015 | 0.0013 | 0.0013 |

| Chr | Start | End | both | A1 | A2 |
|---|---|---|---|---|---|
| F01 | 60240001 | 60340000 | 0.0020 | 0.0017 | 0.0018 |
| F01 | 60260001 | 60360000 | 0.0024 | 0.0022 | 0.0021 |
| F01 | 60280001 | 60380000 | 0.0023 | 0.0021 | 0.0020 |
| F01 | 60300001 | 60400000 | 0.0029 | 0.0025 | 0.0026 |
| F01 | 60320001 | 60420000 | 0.0035 | 0.0028 | 0.0030 |
| F01 | 60340001 | 60440000 | 0.0032 | 0.0025 | 0.0026 |
| F01 | 60360001 | 60460000 | 0.0031 | 0.0021 | 0.0023 |
| F01 | 60380001 | 60480000 | 0.0033 | 0.0022 | 0.0024 |
| F01 | 60400001 | 60500000 | 0.0028 | 0.0016 | 0.0018 |
| F01 | 60420001 | 60520000 | 0.0022 | 0.0012 | 0.0013 |
| F01 | 60440001 | 60540000 | 0.0023 | 0.0013 | 0.0013 |
| F01 | 60460001 | 60560000 | 0.0024 | 0.0016 | 0.0017 |
| F01 | 60480001 | 60580000 | 0.0023 | 0.0016 | 0.0017 |
| F01 | 60500001 | 60600000 | 0.0018 | 0.0013 | 0.0015 |
| F01 | 60520001 | 60620000 | 0.0020 | 0.0013 | 0.0018 |
| F01 | 60540001 | 60640000 | 0.0018 | 0.0013 | 0.0017 |
| F01 | 60560001 | 60660000 | 0.0016 | 0.0011 | 0.0015 |
| F01 | 60580001 | 60680000 | 0.0015 | 0.0010 | 0.0014 |
| F01 | 60600001 | 60700000 | 0.0016 | 0.0012 | 0.0015 |
| F01 | 60620001 | 60720000 | 0.0009 | 0.0008 | 0.0009 |
| F01 | 60640001 | 60740000 | 0.0012 | 0.0009 | 0.0011 |
| F01 | 60660001 | 60760000 | 0.0015 | 0.0010 | 0.0012 |
| F01 | 60680001 | 60780000 | 0.0023 | 0.0016 | 0.0018 |
| F01 | 60700001 | 60800000 | 0.0025 | 0.0018 | 0.0021 |
| F01 | 60720001 | 60820000 | 0.0032 | 0.0023 | 0.0026 |
| F01 | 60740001 | 60840000 | 0.0033 | 0.0024 | 0.0028 |
| F01 | 60760001 | 60860000 | 0.0030 | 0.0023 | 0.0026 |
| F01 | 60780001 | 60880000 | 0.0020 | 0.0015 | 0.0018 |
| F01 | 60800001 | 60900000 | 0.0014 | 0.0010 | 0.0012 |
| F01 | 60820001 | 60920000 | 0.0008 | 0.0006 | 0.0007 |
| F01 | 60840001 | 60940000 | 0.0007 | 0.0004 | 0.0005 |
| F01 | 60860001 | 60960000 | 0.0006 | 0.0004 | 0.0004 |
| F01 | 60880001 | 60980000 | 0.0008 | 0.0005 | 0.0005 |
| F01 | 60900001 | 61000000 | 0.0012 | 0.0010 | 0.0009 |
| F01 | 60920001 | 61020000 | 0.0014 | 0.0011 | 0.0012 |
| F01 | 60940001 | 61040000 | 0.0014 | 0.0012 | 0.0013 |
| F01 | 60960001 | 61060000 | 0.0020 | 0.0018 | 0.0020 |
| F01 | 60980001 | 61080000 | 0.0025 | 0.0022 | 0.0024 |
| F01 | 61000001 | 61100000 | 0.0021 | 0.0018 | 0.0021 |
| F01 | 61020001 | 61120000 | 0.0018 | 0.0016 | 0.0017 |
| F01 | 61040001 | 61140000 | 0.0019 | 0.0017 | 0.0019 |
| F01 | 61060001 | 61160000 | 0.0012 | 0.0010 | 0.0011 |
| F01 | 61080001 | 61180000 | 0.0007 | 0.0007 | 0.0007 |
| F01 | 61100001 | 61200000 | 0.0007 | 0.0007 | 0.0007 |
| F01 | 61120001 | 61220000 | 0.0008 | 0.0007 | 0.0008 |
| F01 | 61140001 | 61240000 | 0.0005 | 0.0004 | 0.0004 |
| F01 | 61160001 | 61260000 | 0.0004 | 0.0004 | 0.0004 |
| F01 | 61180001 | 61280000 | 0.0005 | 0.0005 | 0.0005 |
| F01 | 61200001 | 61300000 | 0.0007 | 0.0007 | 0.0006 |
| F01 | 61220001 | 61320000 | 0.0011 | 0.0010 | 0.0010 |
| F01 | 61240001 | 61340000 | 0.0012 | 0.0011 | 0.0011 |
| F01 | 61260001 | 61360000 | 0.0013 | 0.0011 | 0.0012 |
| F01 | 61280001 | 61380000 | 0.0014 | 0.0012 | 0.0013 |
| F01 | 61300001 | 61400000 | 0.0015 | 0.0013 | 0.0015 |
| F01 | 61320001 | 61420000 | 0.0018 | 0.0014 | 0.0018 |
| F01 | 61340001 | 61440000 | 0.0016 | 0.0013 | 0.0016 |
| F01 | 61360001 | 61460000 | 0.0015 | 0.0012 | 0.0015 |
| F01 | 61380001 | 61480000 | 0.0014 | 0.0011 | 0.0014 |

| Chr | Start | End | both | A1 | A2 |
|---|---|---|---|---|---|
| F01 | 61400001 | 61500000 | 0.0013 | 0.0010 | 0.0013 |
| F01 | 61420001 | 61520000 | 0.0009 | 0.0007 | 0.0009 |
| F01 | 61440001 | 61540000 | 0.0026 | 0.0021 | 0.0026 |
| F01 | 61460001 | 61560000 | 0.0029 | 0.0023 | 0.0029 |
| F01 | 61480001 | 61580000 | 0.0030 | 0.0024 | 0.0029 |
| F01 | 61500001 | 61600000 | 0.0031 | 0.0025 | 0.0031 |
| F01 | 61520001 | 61620000 | 0.0032 | 0.0025 | 0.0031 |
| F01 | 61540001 | 61640000 | 0.0020 | 0.0013 | 0.0018 |
| F01 | 61560001 | 61660000 | 0.0021 | 0.0014 | 0.0018 |
| F01 | 61580001 | 61680000 | 0.0022 | 0.0014 | 0.0018 |
| F01 | 61600001 | 61700000 | 0.0022 | 0.0012 | 0.0016 |
| F01 | 61620001 | 61720000 | 0.0024 | 0.0010 | 0.0013 |
| F01 | 61640001 | 61740000 | 0.0024 | 0.0010 | 0.0014 |
| F01 | 61660001 | 61760000 | 0.0029 | 0.0016 | 0.0019 |
| F01 | 61680001 | 61780000 | 0.0027 | 0.0015 | 0.0018 |
| F01 | 61700001 | 61800000 | 0.0027 | 0.0018 | 0.0020 |
| F01 | 61720001 | 61820000 | 0.0029 | 0.0024 | 0.0027 |
| F01 | 61740001 | 61840000 | 0.0024 | 0.0021 | 0.0024 |
| F01 | 61760001 | 61860000 | 0.0021 | 0.0019 | 0.0021 |
| F01 | 61780001 | 61880000 | 0.0021 | 0.0019 | 0.0021 |
| F01 | 61800001 | 61900000 | 0.0024 | 0.0020 | 0.0024 |
| F01 | 61820001 | 61920000 | 0.0021 | 0.0017 | 0.0019 |
| F01 | 61840001 | 61940000 | 0.0024 | 0.0019 | 0.0021 |
| F01 | 61860001 | 61960000 | 0.0025 | 0.0019 | 0.0021 |
| F01 | 61880001 | 61980000 | 0.0030 | 0.0022 | 0.0025 |
| F01 | 61900001 | 62000000 | 0.0029 | 0.0022 | 0.0024 |
| F01 | 61920001 | 62020000 | 0.0034 | 0.0029 | 0.0030 |
| F01 | 61940001 | 62040000 | 0.0041 | 0.0033 | 0.0036 |
| F01 | 61960001 | 62060000 | 0.0039 | 0.0030 | 0.0033 |
| F01 | 61980001 | 62080000 | 0.0042 | 0.0032 | 0.0036 |
| F01 | 62000001 | 62100000 | 0.0039 | 0.0030 | 0.0033 |
| F01 | 62020001 | 62120000 | 0.0031 | 0.0021 | 0.0025 |
| F01 | 62040001 | 62140000 | 0.0020 | 0.0015 | 0.0017 |
| F01 | 62060001 | 62160000 | 0.0021 | 0.0017 | 0.0018 |
| F01 | 62080001 | 62180000 | 0.0014 | 0.0012 | 0.0011 |
| F01 | 62100001 | 62200000 | 0.0012 | 0.0009 | 0.0009 |
| F01 | 62120001 | 62220000 | 0.0014 | 0.0010 | 0.0010 |
| F01 | 62140001 | 62240000 | 0.0018 | 0.0011 | 0.0014 |
| F01 | 62160001 | 62260000 | 0.0014 | 0.0007 | 0.0010 |
| F01 | 62180001 | 62280000 | 0.0012 | 0.0006 | 0.0009 |
| F01 | 62200001 | 62300000 | 0.0016 | 0.0010 | 0.0014 |
| F01 | 62220001 | 62320000 | 0.0014 | 0.0009 | 0.0013 |
| F01 | 62240001 | 62340000 | 0.0010 | 0.0007 | 0.0010 |
| F01 | 62260001 | 62360000 | 0.0009 | 0.0007 | 0.0009 |
| F01 | 62280001 | 62380000 | 0.0011 | 0.0009 | 0.0011 |
| F01 | 62300001 | 62400000 | 0.0010 | 0.0007 | 0.0008 |
| F01 | 62320001 | 62420000 | 0.0008 | 0.0005 | 0.0006 |
| F01 | 62340001 | 62440000 | 0.0010 | 0.0007 | 0.0008 |
| F01 | 62360001 | 62460000 | 0.0010 | 0.0007 | 0.0009 |
| F01 | 62380001 | 62480000 | 0.0010 | 0.0008 | 0.0009 |
| F01 | 62400001 | 62500000 | 0.0012 | 0.0009 | 0.0010 |
| F01 | 62420001 | 62520000 | 0.0015 | 0.0011 | 0.0012 |
| F01 | 62440001 | 62540000 | 0.0016 | 0.0012 | 0.0014 |
| F01 | 62460001 | 62560000 | 0.0020 | 0.0014 | 0.0016 |
| F01 | 62480001 | 62580000 | 0.0019 | 0.0014 | 0.0015 |
| F01 | 62500001 | 62600000 | 0.0015 | 0.0012 | 0.0013 |
| F01 | 62520001 | 62620000 | 0.0013 | 0.0010 | 0.0011 |
| F01 | 62540001 | 62640000 | 0.0017 | 0.0013 | 0.0014 |

| Chr | Start | End | both | A1 | A2 |
|---|---|---|---|---|---|
| F01 | 62560001 | 62660000 | 0.0020 | 0.0016 | 0.0019 |
| F01 | 62580001 | 62680000 | 0.0020 | 0.0016 | 0.0019 |
| F01 | 62600001 | 62700000 | 0.0018 | 0.0014 | 0.0017 |
| F01 | 62620001 | 62720000 | 0.0017 | 0.0014 | 0.0017 |
| F01 | 62640001 | 62740000 | 0.0015 | 0.0012 | 0.0012 |
| F01 | 62660001 | 62760000 | 0.0007 | 0.0006 | 0.0005 |
| F01 | 62680001 | 62780000 | 0.0013 | 0.0009 | 0.0008 |
| F01 | 62700001 | 62800000 | 0.0029 | 0.0021 | 0.0021 |
| F01 | 62720001 | 62820000 | 0.0034 | 0.0025 | 0.0024 |
| F01 | 62740001 | 62840000 | 0.0036 | 0.0026 | 0.0026 |
| F01 | 62760001 | 62860000 | 0.0042 | 0.0030 | 0.0030 |
| F01 | 62780001 | 62880000 | 0.0039 | 0.0029 | 0.0030 |
| F01 | 62800001 | 62900000 | 0.0029 | 0.0022 | 0.0022 |
| F01 | 62820001 | 62920000 | 0.0030 | 0.0023 | 0.0024 |
| F01 | 62840001 | 62940000 | 0.0031 | 0.0026 | 0.0027 |
| F01 | 62860001 | 62960000 | 0.0028 | 0.0025 | 0.0025 |
| F01 | 62880001 | 62980000 | 0.0025 | 0.0023 | 0.0023 |
| F01 | 62900001 | 63000000 | 0.0020 | 0.0018 | 0.0018 |
| F01 | 62920001 | 63020000 | 0.0021 | 0.0018 | 0.0018 |
| F01 | 62940001 | 63040000 | 0.0017 | 0.0014 | 0.0014 |
| F01 | 62960001 | 63060000 | 0.0019 | 0.0015 | 0.0016 |
| F01 | 62980001 | 63080000 | 0.0024 | 0.0020 | 0.0022 |
| F01 | 63000001 | 63100000 | 0.0024 | 0.0019 | 0.0021 |
| F01 | 63020001 | 63120000 | 0.0018 | 0.0016 | 0.0016 |
| F01 | 63040001 | 63140000 | 0.0016 | 0.0013 | 0.0014 |
| F01 | 63060001 | 63160000 | 0.0018 | 0.0015 | 0.0017 |
| F01 | 63080001 | 63180000 | 0.0017 | 0.0012 | 0.0014 |
| F01 | 63100001 | 63200000 | 0.0022 | 0.0016 | 0.0018 |
| F01 | 63120001 | 63220000 | 0.0024 | 0.0018 | 0.0021 |
| F01 | 63140001 | 63240000 | 0.0028 | 0.0019 | 0.0022 |
| F01 | 63160001 | 63260000 | 0.0026 | 0.0018 | 0.0019 |
| F01 | 63180001 | 63280000 | 0.0028 | 0.0020 | 0.0022 |
| F01 | 63200001 | 63300000 | 0.0035 | 0.0025 | 0.0029 |
| F01 | 63220001 | 63320000 | 0.0038 | 0.0027 | 0.0032 |
| F01 | 63240001 | 63340000 | 0.0039 | 0.0031 | 0.0035 |
| F01 | 63260001 | 63360000 | 0.0033 | 0.0027 | 0.0030 |
| F01 | 63280001 | 63380000 | 0.0032 | 0.0026 | 0.0030 |
| F01 | 63300001 | 63400000 | 0.0023 | 0.0019 | 0.0021 |
| F01 | 63320001 | 63420000 | 0.0018 | 0.0014 | 0.0015 |
| F01 | 63340001 | 63440000 | 0.0018 | 0.0012 | 0.0014 |
| F01 | 63360001 | 63460000 | 0.0023 | 0.0014 | 0.0017 |
| F01 | 63380001 | 63480000 | 0.0020 | 0.0012 | 0.0013 |
| F01 | 63400001 | 63500000 | 0.0020 | 0.0010 | 0.0012 |
| F01 | 63420001 | 63520000 | 0.0023 | 0.0011 | 0.0013 |
| F01 | 63440001 | 63540000 | 0.0021 | 0.0011 | 0.0012 |
| F01 | 63460001 | 63560000 | 0.0022 | 0.0011 | 0.0013 |
| F01 | 63480001 | 63580000 | 0.0028 | 0.0019 | 0.0020 |
| F01 | 63500001 | 63600000 | 0.0026 | 0.0019 | 0.0020 |
| F01 | 63520001 | 63620000 | 0.0021 | 0.0017 | 0.0018 |
| F01 | 63540001 | 63640000 | 0.0021 | 0.0017 | 0.0019 |
| F01 | 63560001 | 63660000 | 0.0019 | 0.0016 | 0.0018 |
| F01 | 63580001 | 63680000 | 0.0018 | 0.0015 | 0.0017 |
| F01 | 63600001 | 63700000 | 0.0027 | 0.0023 | 0.0026 |
| F01 | 63620001 | 63720000 | 0.0032 | 0.0026 | 0.0030 |
| F01 | 63640001 | 63740000 | 0.0031 | 0.0026 | 0.0030 |
| F01 | 63660001 | 63760000 | 0.0035 | 0.0029 | 0.0033 |
| F01 | 63680001 | 63780000 | 0.0033 | 0.0026 | 0.0030 |
| F01 | 63700001 | 63800000 | 0.0026 | 0.0018 | 0.0022 |

| Chr | Start | End | both | A1 | A2 |
|---|---|---|---|---|---|
| F01 | 63720001 | 63820000 | 0.0028 | 0.0020 | 0.0023 |
| F01 | 63740001 | 63840000 | 0.0026 | 0.0018 | 0.0019 |
| F01 | 63760001 | 63860000 | 0.0021 | 0.0013 | 0.0014 |
| F01 | 63780001 | 63880000 | 0.0024 | 0.0015 | 0.0017 |
| F01 | 63800001 | 63900000 | 0.0029 | 0.0020 | 0.0022 |
| F01 | 63820001 | 63920000 | 0.0024 | 0.0016 | 0.0019 |
| F01 | 63840001 | 63940000 | 0.0037 | 0.0028 | 0.0032 |
| F01 | 63860001 | 63960000 | 0.0037 | 0.0027 | 0.0032 |
| F01 | 63880001 | 63980000 | 0.0030 | 0.0023 | 0.0026 |
| F01 | 63900001 | 64000000 | 0.0025 | 0.0020 | 0.0023 |
| F01 | 63920001 | 64020000 | 0.0028 | 0.0022 | 0.0026 |
| F01 | 63940001 | 64040000 | 0.0018 | 0.0013 | 0.0017 |
| F01 | 63960001 | 64060000 | 0.0016 | 0.0013 | 0.0016 |
| F01 | 63980001 | 64080000 | 0.0021 | 0.0016 | 0.0019 |
| F01 | 64000001 | 64100000 | 0.0023 | 0.0017 | 0.0019 |
| F01 | 64020001 | 64120000 | 0.0023 | 0.0017 | 0.0019 |
| F01 | 64040001 | 64140000 | 0.0021 | 0.0016 | 0.0017 |
| F01 | 64060001 | 64160000 | 0.0022 | 0.0016 | 0.0018 |
| F01 | 64080001 | 64180000 | 0.0015 | 0.0012 | 0.0013 |
| F01 | 64100001 | 64200000 | 0.0009 | 0.0008 | 0.0009 |
| F01 | 64120001 | 64220000 | 0.0009 | 0.0008 | 0.0009 |
| F01 | 64140001 | 64240000 | 0.0009 | 0.0007 | 0.0009 |
| F01 | 64160001 | 64260000 | 0.0010 | 0.0009 | 0.0010 |
| F01 | 64180001 | 64280000 | 0.0011 | 0.0009 | 0.0010 |
| F01 | 64200001 | 64300000 | 0.0010 | 0.0008 | 0.0010 |
| F01 | 64220001 | 64320000 | 0.0006 | 0.0005 | 0.0006 |
| F01 | 64240001 | 64340000 | 0.0004 | 0.0004 | 0.0004 |
| F01 | 64260001 | 64360000 | 0.0002 | 0.0001 | 0.0002 |
| F01 | 64280001 | 64380000 | 0.0003 | 0.0002 | 0.0003 |
| F01 | 64300001 | 64400000 | 0.0004 | 0.0003 | 0.0004 |
| F01 | 64320001 | 64420000 | 0.0004 | 0.0003 | 0.0004 |
| F01 | 64340001 | 64440000 | 0.0005 | 0.0004 | 0.0005 |
| F01 | 64360001 | 64460000 | 0.0004 | 0.0004 | 0.0004 |
| F01 | 64380001 | 64480000 | 0.0004 | 0.0003 | 0.0004 |
| F01 | 64400001 | 64500000 | 0.0004 | 0.0003 | 0.0004 |
| F01 | 64420001 | 64520000 | 0.0005 | 0.0003 | 0.0005 |
| F01 | 64440001 | 64540000 | 0.0008 | 0.0007 | 0.0008 |
| F01 | 64460001 | 64560000 | 0.0009 | 0.0008 | 0.0008 |
| F01 | 64480001 | 64580000 | 0.0010 | 0.0008 | 0.0010 |
| F01 | 64500001 | 64600000 | 0.0013 | 0.0011 | 0.0012 |
| F01 | 64520001 | 64620000 | 0.0014 | 0.0012 | 0.0014 |
| F01 | 64540001 | 64640000 | 0.0017 | 0.0015 | 0.0016 |
| F01 | 64560001 | 64660000 | 0.0018 | 0.0015 | 0.0017 |
| F01 | 64580001 | 64680000 | 0.0015 | 0.0014 | 0.0015 |
| F01 | 64600001 | 64700000 | 0.0014 | 0.0013 | 0.0013 |
| F01 | 64620001 | 64720000 | 0.0012 | 0.0012 | 0.0012 |
| F01 | 64640001 | 64740000 | 0.0007 | 0.0006 | 0.0007 |
| F01 | 64660001 | 64760000 | 0.0010 | 0.0008 | 0.0010 |
| F01 | 64680001 | 64780000 | 0.0011 | 0.0010 | 0.0011 |
| F01 | 64700001 | 64800000 | 0.0014 | 0.0011 | 0.0014 |
| F01 | 64720001 | 64820000 | 0.0019 | 0.0016 | 0.0019 |
| F01 | 64740001 | 64840000 | 0.0018 | 0.0015 | 0.0017 |
| F01 | 64760001 | 64860000 | 0.0020 | 0.0017 | 0.0019 |
| F01 | 64780001 | 64880000 | 0.0030 | 0.0025 | 0.0028 |
| F01 | 64800001 | 64900000 | 0.0030 | 0.0025 | 0.0027 |
| F01 | 64820001 | 64920000 | 0.0031 | 0.0024 | 0.0028 |
| F01 | 64840001 | 64940000 | 0.0035 | 0.0028 | 0.0032 |
| F01 | 64860001 | 64960000 | 0.0031 | 0.0025 | 0.0028 |

| Chr | Start | End | both | A1 | A2 |
|---|---|---|---|---|---|
| F01 | 64880001 | 64980000 | 0.0026 | 0.0019 | 0.0023 |
| F01 | 64900001 | 65000000 | 0.0023 | 0.0017 | 0.0021 |
| F01 | 64920001 | 65020000 | 0.0017 | 0.0014 | 0.0016 |
| F01 | 64940001 | 65040000 | 0.0015 | 0.0012 | 0.0014 |
| F01 | 64960001 | 65060000 | 0.0016 | 0.0012 | 0.0014 |
| F01 | 64980001 | 65080000 | 0.0016 | 0.0014 | 0.0015 |
| F01 | 65000001 | 65100000 | 0.0015 | 0.0013 | 0.0014 |
| F01 | 65020001 | 65120000 | 0.0022 | 0.0019 | 0.0020 |
| F01 | 65040001 | 65140000 | 0.0026 | 0.0024 | 0.0024 |
| F01 | 65060001 | 65160000 | 0.0026 | 0.0024 | 0.0024 |
| F01 | 65080001 | 65180000 | 0.0021 | 0.0019 | 0.0020 |
| F01 | 65100001 | 65200000 | 0.0022 | 0.0020 | 0.0021 |
| F01 | 65120001 | 65220000 | 0.0016 | 0.0014 | 0.0015 |
| F01 | 65140001 | 65240000 | 0.0016 | 0.0014 | 0.0015 |
| F01 | 65160001 | 65260000 | 0.0016 | 0.0015 | 0.0015 |
| F01 | 65180001 | 65280000 | 0.0017 | 0.0016 | 0.0015 |
| F01 | 65200001 | 65300000 | 0.0018 | 0.0017 | 0.0016 |
| F01 | 65220001 | 65320000 | 0.0018 | 0.0017 | 0.0016 |
| F01 | 65240001 | 65340000 | 0.0010 | 0.0010 | 0.0009 |
| F01 | 65260001 | 65360000 | 0.0019 | 0.0016 | 0.0019 |
| F01 | 65280001 | 65380000 | 0.0018 | 0.0014 | 0.0018 |
| F01 | 65300001 | 65400000 | 0.0017 | 0.0013 | 0.0017 |
| F01 | 65320001 | 65420000 | 0.0018 | 0.0013 | 0.0018 |
| F01 | 65340001 | 65440000 | 0.0019 | 0.0014 | 0.0019 |
| F01 | 65360001 | 65460000 | 0.0008 | 0.0007 | 0.0007 |
| F01 | 65380001 | 65480000 | 0.0012 | 0.0010 | 0.0011 |
| F01 | 65400001 | 65500000 | 0.0013 | 0.0012 | 0.0012 |
| F01 | 65420001 | 65520000 | 0.0014 | 0.0013 | 0.0012 |
| F01 | 65440001 | 65540000 | 0.0021 | 0.0018 | 0.0019 |
| F01 | 65460001 | 65560000 | 0.0028 | 0.0024 | 0.0024 |
| F01 | 65480001 | 65580000 | 0.0024 | 0.0021 | 0.0022 |
| F01 | 65500001 | 65600000 | 0.0027 | 0.0024 | 0.0025 |
| F01 | 65520001 | 65620000 | 0.0032 | 0.0024 | 0.0026 |
| F01 | 65540001 | 65640000 | 0.0028 | 0.0021 | 0.0022 |
| F01 | 65560001 | 65660000 | 0.0026 | 0.0019 | 0.0021 |
| F01 | 65580001 | 65680000 | 0.0025 | 0.0018 | 0.0020 |
| F01 | 65600001 | 65700000 | 0.0026 | 0.0019 | 0.0022 |
| F01 | 65620001 | 65720000 | 0.0021 | 0.0018 | 0.0020 |
| F01 | 65640001 | 65740000 | 0.0019 | 0.0014 | 0.0017 |
| F01 | 65660001 | 65760000 | 0.0016 | 0.0012 | 0.0014 |
| F01 | 65680001 | 65780000 | 0.0024 | 0.0018 | 0.0020 |
| F01 | 65700001 | 65800000 | 0.0031 | 0.0023 | 0.0026 |
| F01 | 65720001 | 65820000 | 0.0034 | 0.0025 | 0.0028 |
| F01 | 65740001 | 65840000 | 0.0037 | 0.0029 | 0.0032 |
| F01 | 65760001 | 65860000 | 0.0039 | 0.0029 | 0.0032 |
| F01 | 65780001 | 65880000 | 0.0035 | 0.0025 | 0.0028 |
| F01 | 65800001 | 65900000 | 0.0031 | 0.0024 | 0.0025 |
| F01 | 65820001 | 65920000 | 0.0037 | 0.0031 | 0.0032 |
| F01 | 65840001 | 65940000 | 0.0033 | 0.0027 | 0.0028 |
| F01 | 65860001 | 65960000 | 0.0040 | 0.0034 | 0.0034 |
| F01 | 65880001 | 65980000 | 0.0047 | 0.0041 | 0.0044 |
| F01 | 65900001 | 66000000 | 0.0043 | 0.0037 | 0.0040 |
| F01 | 65920001 | 66020000 | 0.0044 | 0.0038 | 0.0041 |
| F01 | 65940001 | 66040000 | 0.0050 | 0.0041 | 0.0046 |
| F01 | 65960001 | 66060000 | 0.0052 | 0.0042 | 0.0046 |
| F01 | 65980001 | 66080000 | 0.0047 | 0.0037 | 0.0042 |
| F01 | 66000001 | 66100000 | 0.0043 | 0.0033 | 0.0038 |
| F01 | 66020001 | 66120000 | 0.0032 | 0.0024 | 0.0028 |

| Chr | Start | End | both | A1 | A2 |
|---|---|---|---|---|---|
| F01 | 66040001 | 66140000 | 0.0026 | 0.0020 | 0.0023 |
| F01 | 66060001 | 66160000 | 0.0014 | 0.0011 | 0.0014 |
| F01 | 66080001 | 66180000 | 0.0008 | 0.0006 | 0.0008 |
| F01 | 66100001 | 66200000 | 0.0013 | 0.0011 | 0.0012 |
| F01 | 66120001 | 66220000 | 0.0013 | 0.0011 | 0.0012 |
| F01 | 66140001 | 66240000 | 0.0013 | 0.0012 | 0.0013 |
| F01 | 66160001 | 66260000 | 0.0020 | 0.0016 | 0.0016 |
| F01 | 66180001 | 66280000 | 0.0026 | 0.0020 | 0.0022 |
| F01 | 66200001 | 66300000 | 0.0026 | 0.0021 | 0.0022 |
| F01 | 66220001 | 66320000 | 0.0026 | 0.0021 | 0.0022 |
| F01 | 66240001 | 66340000 | 0.0033 | 0.0025 | 0.0027 |
| F01 | 66260001 | 66360000 | 0.0032 | 0.0025 | 0.0028 |
| F01 | 66280001 | 66380000 | 0.0034 | 0.0027 | 0.0029 |
| F01 | 66300001 | 66400000 | 0.0031 | 0.0022 | 0.0025 |
| F01 | 66320001 | 66420000 | 0.0038 | 0.0028 | 0.0032 |
| F01 | 66340001 | 66440000 | 0.0034 | 0.0026 | 0.0029 |
| F01 | 66360001 | 66460000 | 0.0036 | 0.0029 | 0.0031 |
| F01 | 66380001 | 66480000 | 0.0029 | 0.0024 | 0.0026 |
| F01 | 66400001 | 66500000 | 0.0028 | 0.0023 | 0.0026 |
| F01 | 66420001 | 66520000 | 0.0020 | 0.0017 | 0.0019 |
| F01 | 66440001 | 66540000 | 0.0028 | 0.0023 | 0.0026 |
| F01 | 66460001 | 66560000 | 0.0026 | 0.0020 | 0.0023 |
| F01 | 66480001 | 66580000 | 0.0028 | 0.0022 | 0.0025 |
| F01 | 66500001 | 66600000 | 0.0034 | 0.0028 | 0.0031 |
| F01 | 66520001 | 66620000 | 0.0038 | 0.0031 | 0.0034 |
| F01 | 66540001 | 66640000 | 0.0031 | 0.0025 | 0.0027 |
| F01 | 66560001 | 66660000 | 0.0026 | 0.0022 | 0.0024 |
| F01 | 66580001 | 66680000 | 0.0024 | 0.0022 | 0.0022 |
| F01 | 66600001 | 66700000 | 0.0018 | 0.0016 | 0.0017 |
| F01 | 66620001 | 66720000 | 0.0016 | 0.0016 | 0.0014 |
| F01 | 66640001 | 66740000 | 0.0020 | 0.0020 | 0.0018 |
| F01 | 66660001 | 66760000 | 0.0022 | 0.0022 | 0.0020 |
| F01 | 66680001 | 66780000 | 0.0021 | 0.0020 | 0.0018 |
| F01 | 66700001 | 66800000 | 0.0023 | 0.0022 | 0.0020 |
| F01 | 66720001 | 66820000 | 0.0022 | 0.0019 | 0.0019 |
| F01 | 66740001 | 66840000 | 0.0018 | 0.0015 | 0.0015 |
| F01 | 66760001 | 66860000 | 0.0018 | 0.0014 | 0.0015 |
| F01 | 66780001 | 66880000 | 0.0016 | 0.0012 | 0.0013 |
| F01 | 66800001 | 66900000 | 0.0025 | 0.0021 | 0.0023 |
| F01 | 66820001 | 66920000 | 0.0023 | 0.0020 | 0.0022 |
| F01 | 66840001 | 66940000 | 0.0020 | 0.0018 | 0.0020 |
| F01 | 66860001 | 66960000 | 0.0018 | 0.0016 | 0.0018 |
| F01 | 66880001 | 66980000 | 0.0023 | 0.0019 | 0.0020 |
| F01 | 66900001 | 67000000 | 0.0009 | 0.0007 | 0.0006 |
| F01 | 66920001 | 67020000 | 0.0011 | 0.0008 | 0.0009 |
| F01 | 66940001 | 67040000 | 0.0016 | 0.0013 | 0.0013 |
| F01 | 66960001 | 67060000 | 0.0018 | 0.0013 | 0.0014 |
| F01 | 66980001 | 67080000 | 0.0020 | 0.0015 | 0.0016 |
| F01 | 67000001 | 67100000 | 0.0032 | 0.0023 | 0.0026 |
| F01 | 67020001 | 67120000 | 0.0038 | 0.0027 | 0.0031 |
| F01 | 67040001 | 67140000 | 0.0043 | 0.0030 | 0.0035 |
| F01 | 67060001 | 67160000 | 0.0054 | 0.0041 | 0.0046 |
| F01 | 67080001 | 67180000 | 0.0050 | 0.0038 | 0.0043 |
| F01 | 67100001 | 67200000 | 0.0037 | 0.0028 | 0.0033 |
| F01 | 67120001 | 67220000 | 0.0028 | 0.0022 | 0.0025 |
| F01 | 67140001 | 67240000 | 0.0018 | 0.0016 | 0.0016 |
| F01 | 67160001 | 67260000 | 0.0007 | 0.0006 | 0.0007 |
| F01 | 67180001 | 67280000 | 0.0007 | 0.0006 | 0.0006 |

| Chr | Start | End | both | A1 | A2 |
|---|---|---|---|---|---|
| F01 | 67200001 | 67300000 | 0.0017 | 0.0014 | 0.0017 |
| F01 | 67220001 | 67320000 | 0.0018 | 0.0014 | 0.0017 |
| F01 | 67240001 | 67340000 | 0.0027 | 0.0019 | 0.0025 |
| F01 | 67260001 | 67360000 | 0.0027 | 0.0020 | 0.0025 |
| F01 | 67280001 | 67380000 | 0.0027 | 0.0020 | 0.0025 |
| F01 | 67300001 | 67400000 | 0.0023 | 0.0014 | 0.0017 |
| F01 | 67320001 | 67420000 | 0.0024 | 0.0015 | 0.0017 |
| F01 | 67340001 | 67440000 | 0.0016 | 0.0011 | 0.0011 |
| F01 | 67360001 | 67460000 | 0.0021 | 0.0013 | 0.0014 |
| F01 | 67380001 | 67480000 | 0.0027 | 0.0018 | 0.0019 |
| F01 | 67400001 | 67500000 | 0.0025 | 0.0019 | 0.0020 |
| F01 | 67420001 | 67520000 | 0.0035 | 0.0025 | 0.0028 |
| F01 | 67440001 | 67540000 | 0.0039 | 0.0027 | 0.0030 |
| F01 | 67460001 | 67560000 | 0.0033 | 0.0024 | 0.0026 |
| F01 | 67480001 | 67580000 | 0.0029 | 0.0021 | 0.0023 |
| F01 | 67500001 | 67600000 | 0.0033 | 0.0024 | 0.0027 |
| F01 | 67520001 | 67620000 | 0.0031 | 0.0022 | 0.0023 |
| F01 | 67540001 | 67640000 | 0.0032 | 0.0022 | 0.0024 |
| F01 | 67560001 | 67660000 | 0.0036 | 0.0028 | 0.0029 |
| F01 | 67580001 | 67680000 | 0.0035 | 0.0027 | 0.0029 |
| F01 | 67600001 | 67700000 | 0.0031 | 0.0022 | 0.0023 |
| F01 | 67620001 | 67720000 | 0.0022 | 0.0015 | 0.0017 |
| F01 | 67640001 | 67740000 | 0.0026 | 0.0017 | 0.0018 |
| F01 | 67660001 | 67760000 | 0.0025 | 0.0015 | 0.0017 |
| F01 | 67680001 | 67780000 | 0.0030 | 0.0018 | 0.0021 |
| F01 | 67700001 | 67800000 | 0.0029 | 0.0018 | 0.0021 |
| F01 | 67720001 | 67820000 | 0.0038 | 0.0028 | 0.0030 |
| F01 | 67740001 | 67840000 | 0.0030 | 0.0024 | 0.0026 |
| F01 | 67760001 | 67860000 | 0.0027 | 0.0020 | 0.0022 |
| F01 | 67780001 | 67880000 | 0.0022 | 0.0018 | 0.0018 |
| F01 | 67800001 | 67900000 | 0.0023 | 0.0017 | 0.0018 |
| F01 | 67820001 | 67920000 | 0.0015 | 0.0010 | 0.0010 |
| F01 | 67840001 | 67940000 | 0.0023 | 0.0014 | 0.0016 |
| F01 | 67860001 | 67960000 | 0.0023 | 0.0015 | 0.0016 |
| F01 | 67880001 | 67980000 | 0.0023 | 0.0014 | 0.0017 |
| F01 | 67900001 | 68000000 | 0.0031 | 0.0021 | 0.0024 |
| F01 | 67920001 | 68020000 | 0.0033 | 0.0022 | 0.0027 |
| F01 | 67940001 | 68040000 | 0.0032 | 0.0023 | 0.0027 |
| F01 | 67960001 | 68060000 | 0.0041 | 0.0031 | 0.0034 |
| F01 | 67980001 | 68080000 | 0.0038 | 0.0029 | 0.0031 |
| F01 | 68000001 | 68100000 | 0.0036 | 0.0027 | 0.0028 |
| F01 | 68020001 | 68120000 | 0.0034 | 0.0025 | 0.0026 |
| F01 | 68040001 | 68140000 | 0.0028 | 0.0018 | 0.0018 |
| F01 | 68060001 | 68160000 | 0.0027 | 0.0018 | 0.0018 |
| F01 | 68080001 | 68180000 | 0.0027 | 0.0018 | 0.0019 |
| F01 | 68100001 | 68200000 | 0.0025 | 0.0018 | 0.0019 |
| F01 | 68120001 | 68220000 | 0.0025 | 0.0019 | 0.0019 |
| F01 | 68140001 | 68240000 | 0.0026 | 0.0020 | 0.0022 |
| F01 | 68160001 | 68260000 | 0.0019 | 0.0014 | 0.0015 |
| F01 | 68180001 | 68280000 | 0.0020 | 0.0016 | 0.0017 |
| F01 | 68200001 | 68300000 | 0.0025 | 0.0017 | 0.0018 |
| F01 | 68220001 | 68320000 | 0.0027 | 0.0019 | 0.0021 |
| F01 | 68240001 | 68340000 | 0.0034 | 0.0024 | 0.0028 |
| F01 | 68260001 | 68360000 | 0.0032 | 0.0022 | 0.0026 |
| F01 | 68280001 | 68380000 | 0.0031 | 0.0018 | 0.0024 |
| F01 | 68300001 | 68400000 | 0.0030 | 0.0020 | 0.0025 |
| F01 | 68320001 | 68420000 | 0.0025 | 0.0016 | 0.0021 |
| F01 | 68340001 | 68440000 | 0.0015 | 0.0009 | 0.0011 |

| Chr | Start | End | both | A1 | A2 |
|---|---|---|---|---|---|
| F01 | 68360001 | 68460000 | 0.0016 | 0.0010 | 0.0013 |
| F01 | 68380001 | 68480000 | 0.0014 | 0.0011 | 0.0013 |
| F01 | 68400001 | 68500000 | 0.0013 | 0.0008 | 0.0010 |
| F01 | 68420001 | 68520000 | 0.0018 | 0.0012 | 0.0014 |
| F01 | 68440001 | 68540000 | 0.0019 | 0.0013 | 0.0015 |
| F01 | 68460001 | 68560000 | 0.0019 | 0.0014 | 0.0015 |
| F01 | 68480001 | 68580000 | 0.0023 | 0.0017 | 0.0019 |
| F01 | 68500001 | 68600000 | 0.0022 | 0.0016 | 0.0019 |
| F01 | 68520001 | 68620000 | 0.0019 | 0.0014 | 0.0015 |
| F01 | 68540001 | 68640000 | 0.0023 | 0.0016 | 0.0019 |
| F01 | 68560001 | 68660000 | 0.0023 | 0.0016 | 0.0018 |
| F01 | 68580001 | 68680000 | 0.0017 | 0.0012 | 0.0013 |
| F01 | 68600001 | 68700000 | 0.0017 | 0.0013 | 0.0014 |
| F01 | 68620001 | 68720000 | 0.0014 | 0.0012 | 0.0013 |
| F01 | 68640001 | 68740000 | 0.0012 | 0.0011 | 0.0011 |
| F01 | 68660001 | 68760000 | 0.0012 | 0.0011 | 0.0011 |
| F01 | 68680001 | 68780000 | 0.0014 | 0.0012 | 0.0013 |
| F01 | 68700001 | 68800000 | 0.0017 | 0.0015 | 0.0016 |
| F01 | 68720001 | 68820000 | 0.0017 | 0.0015 | 0.0016 |
| F01 | 68740001 | 68840000 | 0.0016 | 0.0013 | 0.0014 |
| F01 | 68760001 | 68860000 | 0.0015 | 0.0012 | 0.0013 |
| F01 | 68780001 | 68880000 | 0.0014 | 0.0012 | 0.0013 |
| F01 | 68800001 | 68900000 | 0.0009 | 0.0007 | 0.0008 |
| F01 | 68820001 | 68920000 | 0.0011 | 0.0008 | 0.0010 |
| F01 | 68840001 | 68940000 | 0.0018 | 0.0013 | 0.0015 |
| F01 | 68860001 | 68960000 | 0.0020 | 0.0014 | 0.0016 |
| F01 | 68880001 | 68980000 | 0.0029 | 0.0020 | 0.0021 |
| F01 | 68900001 | 69000000 | 0.0030 | 0.0020 | 0.0022 |
| F01 | 68920001 | 69020000 | 0.0029 | 0.0020 | 0.0022 |
| F01 | 68940001 | 69040000 | 0.0023 | 0.0015 | 0.0017 |
| F01 | 68960001 | 69060000 | 0.0021 | 0.0013 | 0.0015 |
| F01 | 68980001 | 69080000 | 0.0011 | 0.0007 | 0.0009 |
| F01 | 69000001 | 69100000 | 0.0007 | 0.0005 | 0.0006 |
| F01 | 69020001 | 69120000 | 0.0007 | 0.0005 | 0.0006 |
| F01 | 69040001 | 69140000 | 0.0008 | 0.0007 | 0.0007 |
| F01 | 69060001 | 69160000 | 0.0007 | 0.0006 | 0.0007 |
| F01 | 69080001 | 69180000 | 0.0007 | 0.0006 | 0.0007 |
| F01 | 69100001 | 69200000 | 0.0010 | 0.0009 | 0.0009 |
| F01 | 69120001 | 69220000 | 0.0015 | 0.0014 | 0.0013 |
| F01 | 69140001 | 69240000 | 0.0019 | 0.0016 | 0.0016 |
| F01 | 69160001 | 69260000 | 0.0021 | 0.0016 | 0.0016 |
| F01 | 69180001 | 69280000 | 0.0026 | 0.0018 | 0.0020 |
| F01 | 69200001 | 69300000 | 0.0026 | 0.0018 | 0.0020 |
| F01 | 69220001 | 69320000 | 0.0025 | 0.0015 | 0.0018 |
| F01 | 69240001 | 69340000 | 0.0026 | 0.0014 | 0.0016 |
| F01 | 69260001 | 69360000 | 0.0032 | 0.0019 | 0.0022 |
| F01 | 69280001 | 69380000 | 0.0030 | 0.0018 | 0.0019 |
| F01 | 69300001 | 69400000 | 0.0028 | 0.0017 | 0.0016 |
| F01 | 69320001 | 69420000 | 0.0026 | 0.0017 | 0.0016 |
| F01 | 69340001 | 69440000 | 0.0022 | 0.0016 | 0.0015 |
| F01 | 69360001 | 69460000 | 0.0014 | 0.0011 | 0.0010 |
| F01 | 69380001 | 69480000 | 0.0014 | 0.0014 | 0.0013 |
| F01 | 69400001 | 69500000 | 0.0013 | 0.0012 | 0.0012 |
| F01 | 69420001 | 69520000 | 0.0010 | 0.0010 | 0.0010 |
| F01 | 69440001 | 69540000 | 0.0008 | 0.0007 | 0.0008 |
| F01 | 69460001 | 69560000 | 0.0015 | 0.0013 | 0.0014 |
| F01 | 69480001 | 69580000 | 0.0015 | 0.0010 | 0.0011 |
| F01 | 69500001 | 69600000 | 0.0021 | 0.0014 | 0.0016 |

| Chr | Start | End | both | A1 | A2 |
|---|---|---|---|---|---|
| F01 | 69520001 | 69620000 | 0.0026 | 0.0019 | 0.0021 |
| F01 | 69540001 | 69640000 | 0.0031 | 0.0022 | 0.0026 |
| F01 | 69560001 | 69660000 | 0.0024 | 0.0017 | 0.0021 |
| F01 | 69580001 | 69680000 | 0.0024 | 0.0020 | 0.0023 |
| F01 | 69600001 | 69700000 | 0.0020 | 0.0018 | 0.0019 |
| F01 | 69620001 | 69720000 | 0.0018 | 0.0014 | 0.0016 |
| F01 | 69640001 | 69740000 | 0.0015 | 0.0013 | 0.0013 |
| F01 | 69660001 | 69760000 | 0.0016 | 0.0013 | 0.0014 |
| F01 | 69680001 | 69780000 | 0.0018 | 0.0013 | 0.0015 |
| F01 | 69700001 | 69800000 | 0.0025 | 0.0019 | 0.0022 |
| F01 | 69720001 | 69820000 | 0.0030 | 0.0023 | 0.0026 |
| F01 | 69740001 | 69840000 | 0.0031 | 0.0023 | 0.0026 |
| F01 | 69760001 | 69860000 | 0.0033 | 0.0024 | 0.0026 |
| F01 | 69780001 | 69880000 | 0.0032 | 0.0024 | 0.0026 |
| F01 | 69800001 | 69900000 | 0.0029 | 0.0020 | 0.0022 |
| F01 | 69820001 | 69920000 | 0.0025 | 0.0013 | 0.0017 |
| F01 | 69840001 | 69940000 | 0.0026 | 0.0015 | 0.0019 |
| F01 | 69860001 | 69960000 | 0.0034 | 0.0022 | 0.0027 |
| F01 | 69880001 | 69980000 | 0.0038 | 0.0025 | 0.0031 |
| F01 | 69900001 | 70000000 | 0.0038 | 0.0026 | 0.0031 |
| F01 | 69920001 | 70020000 | 0.0036 | 0.0025 | 0.0029 |
| F01 | 69940001 | 70040000 | 0.0033 | 0.0022 | 0.0026 |
| F01 | 69960001 | 70060000 | 0.0029 | 0.0019 | 0.0024 |
| F01 | 69980001 | 70080000 | 0.0026 | 0.0017 | 0.0020 |
| F01 | 70000001 | 70100000 | 0.0032 | 0.0023 | 0.0026 |
| F01 | 70020001 | 70120000 | 0.0034 | 0.0025 | 0.0029 |
| F01 | 70040001 | 70140000 | 0.0033 | 0.0026 | 0.0030 |
| F01 | 70060001 | 70160000 | 0.0038 | 0.0031 | 0.0034 |
| F01 | 70080001 | 70180000 | 0.0040 | 0.0033 | 0.0036 |
| F01 | 70100001 | 70200000 | 0.0033 | 0.0027 | 0.0030 |
| F01 | 70120001 | 70220000 | 0.0030 | 0.0025 | 0.0029 |
| F01 | 70140001 | 70240000 | 0.0032 | 0.0027 | 0.0030 |
| F01 | 70160001 | 70260000 | 0.0021 | 0.0017 | 0.0021 |
| F01 | 70180001 | 70280000 | 0.0021 | 0.0016 | 0.0019 |
| F01 | 70200001 | 70300000 | 0.0029 | 0.0020 | 0.0024 |
| F01 | 70220001 | 70320000 | 0.0034 | 0.0023 | 0.0027 |
| F01 | 70240001 | 70340000 | 0.0038 | 0.0025 | 0.0029 |
| F01 | 70260001 | 70360000 | 0.0043 | 0.0028 | 0.0033 |
| F01 | 70280001 | 70380000 | 0.0042 | 0.0028 | 0.0033 |
| F01 | 70300001 | 70400000 | 0.0035 | 0.0024 | 0.0028 |
| F01 | 70320001 | 70420000 | 0.0031 | 0.0021 | 0.0026 |
| F01 | 70340001 | 70440000 | 0.0024 | 0.0017 | 0.0021 |
| F01 | 70360001 | 70460000 | 0.0019 | 0.0013 | 0.0017 |
| F01 | 70380001 | 70480000 | 0.0014 | 0.0010 | 0.0013 |
| F01 | 70400001 | 70500000 | 0.0009 | 0.0007 | 0.0009 |
| F01 | 70420001 | 70520000 | 0.0007 | 0.0005 | 0.0007 |
| F01 | 70440001 | 70540000 | 0.0009 | 0.0007 | 0.0009 |
| F01 | 70460001 | 70560000 | 0.0012 | 0.0010 | 0.0012 |
| F01 | 70480001 | 70580000 | 0.0013 | 0.0011 | 0.0013 |
| F01 | 70500001 | 70600000 | 0.0014 | 0.0012 | 0.0014 |
| F01 | 70520001 | 70620000 | 0.0015 | 0.0014 | 0.0014 |
| F01 | 70540001 | 70640000 | 0.0014 | 0.0013 | 0.0013 |
| F01 | 70560001 | 70660000 | 0.0010 | 0.0009 | 0.0009 |
| F01 | 70580001 | 70680000 | 0.0007 | 0.0007 | 0.0007 |
| F01 | 70600001 | 70700000 | 0.0005 | 0.0005 | 0.0005 |
| F01 | 70620001 | 70720000 | 0.0005 | 0.0004 | 0.0004 |
| F01 | 70640001 | 70740000 | 0.0004 | 0.0003 | 0.0004 |
| F01 | 70660001 | 70760000 | 0.0008 | 0.0007 | 0.0007 |

| Chr | Start | End | both | A1 | A2 |
|---|---|---|---|---|---|
| F01 | 70680001 | 70780000 | 0.0009 | 0.0008 | 0.0009 |
| F01 | 70700001 | 70800000 | 0.0015 | 0.0013 | 0.0015 |
| F01 | 70720001 | 70820000 | 0.0016 | 0.0014 | 0.0015 |
| F01 | 70740001 | 70840000 | 0.0021 | 0.0019 | 0.0021 |
| F01 | 70760001 | 70860000 | 0.0021 | 0.0017 | 0.0021 |
| F01 | 70780001 | 70880000 | 0.0024 | 0.0020 | 0.0024 |
| F01 | 70800001 | 70900000 | 0.0020 | 0.0017 | 0.0020 |
| F01 | 70820001 | 70920000 | 0.0025 | 0.0020 | 0.0025 |
| F01 | 70840001 | 70940000 | 0.0029 | 0.0023 | 0.0027 |
| F01 | 70860001 | 70960000 | 0.0033 | 0.0026 | 0.0030 |
| F01 | 70880001 | 70980000 | 0.0041 | 0.0028 | 0.0035 |
| F01 | 70900001 | 71000000 | 0.0043 | 0.0029 | 0.0036 |
| F01 | 70920001 | 71020000 | 0.0035 | 0.0023 | 0.0028 |
| F01 | 70940001 | 71040000 | 0.0028 | 0.0018 | 0.0023 |
| F01 | 70960001 | 71060000 | 0.0019 | 0.0011 | 0.0015 |
| F01 | 70980001 | 71080000 | 0.0007 | 0.0005 | 0.0006 |
| F01 | 71000001 | 71100000 | 0.0004 | 0.0003 | 0.0004 |
| F01 | 71020001 | 71120000 | 0.0005 | 0.0004 | 0.0005 |
| F01 | 71040001 | 71140000 | 0.0009 | 0.0007 | 0.0009 |
| F01 | 71060001 | 71160000 | 0.0011 | 0.0008 | 0.0011 |
| F01 | 71080001 | 71180000 | 0.0014 | 0.0008 | 0.0011 |
| F01 | 71100001 | 71200000 | 0.0020 | 0.0013 | 0.0016 |
| F01 | 71120001 | 71220000 | 0.0021 | 0.0012 | 0.0016 |
| F01 | 71140001 | 71240000 | 0.0026 | 0.0016 | 0.0020 |
| F01 | 71160001 | 71260000 | 0.0033 | 0.0023 | 0.0029 |
| F01 | 71180001 | 71280000 | 0.0039 | 0.0030 | 0.0036 |
| F01 | 71200001 | 71300000 | 0.0049 | 0.0040 | 0.0046 |
| F01 | 71220001 | 71320000 | 0.0059 | 0.0051 | 0.0057 |
| F01 | 71240001 | 71340000 | 0.0051 | 0.0041 | 0.0048 |
| F01 | 71260001 | 71360000 | 0.0053 | 0.0042 | 0.0047 |
| F01 | 71280001 | 71380000 | 0.0048 | 0.0036 | 0.0041 |
| F01 | 71300001 | 71400000 | 0.0045 | 0.0032 | 0.0036 |
| F01 | 71320001 | 71420000 | 0.0039 | 0.0026 | 0.0029 |
| F01 | 71340001 | 71440000 | 0.0049 | 0.0037 | 0.0039 |
| F01 | 71360001 | 71460000 | 0.0043 | 0.0033 | 0.0035 |
| F01 | 71380001 | 71480000 | 0.0053 | 0.0043 | 0.0045 |
| F01 | 71400001 | 71500000 | 0.0043 | 0.0032 | 0.0036 |
| F01 | 71420001 | 71520000 | 0.0041 | 0.0030 | 0.0034 |
| F01 | 71440001 | 71540000 | 0.0031 | 0.0019 | 0.0024 |
| F01 | 71460001 | 71560000 | 0.0029 | 0.0016 | 0.0023 |
| F01 | 71480001 | 71580000 | 0.0026 | 0.0012 | 0.0020 |
| F01 | 71500001 | 71600000 | 0.0028 | 0.0015 | 0.0022 |
| F01 | 71520001 | 71620000 | 0.0029 | 0.0016 | 0.0023 |
| F01 | 71540001 | 71640000 | 0.0031 | 0.0020 | 0.0026 |
| F01 | 71560001 | 71660000 | 0.0030 | 0.0019 | 0.0026 |
| F01 | 71580001 | 71680000 | 0.0026 | 0.0016 | 0.0021 |
| F01 | 71600001 | 71700000 | 0.0025 | 0.0016 | 0.0021 |
| F01 | 71620001 | 71720000 | 0.0026 | 0.0016 | 0.0021 |
| F01 | 71640001 | 71740000 | 0.0033 | 0.0020 | 0.0025 |
| F01 | 71660001 | 71760000 | 0.0042 | 0.0031 | 0.0035 |
| F01 | 71680001 | 71780000 | 0.0049 | 0.0038 | 0.0043 |
| F01 | 71700001 | 71800000 | 0.0049 | 0.0037 | 0.0041 |
| F01 | 71720001 | 71820000 | 0.0048 | 0.0037 | 0.0043 |
| F01 | 71740001 | 71840000 | 0.0044 | 0.0037 | 0.0041 |
| F01 | 71760001 | 71860000 | 0.0031 | 0.0025 | 0.0028 |
| F01 | 71780001 | 71880000 | 0.0020 | 0.0016 | 0.0019 |
| F01 | 71800001 | 71900000 | 0.0020 | 0.0017 | 0.0019 |
| F01 | 71820001 | 71920000 | 0.0024 | 0.0019 | 0.0021 |

| Chr | Start | End | both | A1 | A2 |
|---|---|---|---|---|---|
| F01 | 71840001 | 71940000 | 0.0024 | 0.0017 | 0.0021 |
| F01 | 71860001 | 71960000 | 0.0025 | 0.0018 | 0.0021 |
| F01 | 71880001 | 71980000 | 0.0024 | 0.0017 | 0.0021 |
| F01 | 71900001 | 72000000 | 0.0023 | 0.0017 | 0.0020 |
| F01 | 71920001 | 72020000 | 0.0019 | 0.0016 | 0.0017 |
| F01 | 71940001 | 72040000 | 0.0012 | 0.0010 | 0.0010 |
| F01 | 71960001 | 72060000 | 0.0022 | 0.0018 | 0.0018 |
| F01 | 71980001 | 72080000 | 0.0027 | 0.0022 | 0.0023 |
| F01 | 72000001 | 72100000 | 0.0034 | 0.0028 | 0.0029 |
| F01 | 72020001 | 72120000 | 0.0036 | 0.0027 | 0.0029 |
| F01 | 72040001 | 72140000 | 0.0039 | 0.0029 | 0.0032 |
| F01 | 72060001 | 72160000 | 0.0032 | 0.0024 | 0.0026 |
| F01 | 72080001 | 72180000 | 0.0026 | 0.0019 | 0.0020 |
| F01 | 72100001 | 72200000 | 0.0019 | 0.0014 | 0.0014 |
| F01 | 72120001 | 72220000 | 0.0017 | 0.0013 | 0.0015 |
| F01 | 72140001 | 72240000 | 0.0012 | 0.0009 | 0.0011 |
| F01 | 72160001 | 72260000 | 0.0009 | 0.0007 | 0.0009 |
| F01 | 72180001 | 72280000 | 0.0016 | 0.0013 | 0.0016 |
| F01 | 72200001 | 72300000 | 0.0013 | 0.0011 | 0.0013 |
| F01 | 72220001 | 72320000 | 0.0009 | 0.0008 | 0.0009 |
| F01 | 72240001 | 72340000 | 0.0010 | 0.0009 | 0.0010 |
| F01 | 72260001 | 72360000 | 0.0010 | 0.0009 | 0.0010 |
| F01 | 72280001 | 72380000 | 0.0006 | 0.0005 | 0.0006 |
| F01 | 72300001 | 72400000 | 0.0007 | 0.0006 | 0.0007 |
| F01 | 72320001 | 72420000 | 0.0008 | 0.0007 | 0.0007 |
| F01 | 72340001 | 72440000 | 0.0009 | 0.0008 | 0.0008 |
| F01 | 72360001 | 72460000 | 0.0014 | 0.0011 | 0.0012 |
| F01 | 72380001 | 72480000 | 0.0015 | 0.0009 | 0.0011 |
| F01 | 72400001 | 72500000 | 0.0018 | 0.0010 | 0.0012 |
| F01 | 72420001 | 72520000 | 0.0018 | 0.0010 | 0.0013 |
| F01 | 72440001 | 72540000 | 0.0019 | 0.0010 | 0.0013 |
| F01 | 72460001 | 72560000 | 0.0014 | 0.0007 | 0.0010 |
| F01 | 72480001 | 72580000 | 0.0010 | 0.0006 | 0.0007 |
| F01 | 72500001 | 72600000 | 0.0014 | 0.0011 | 0.0012 |
| F01 | 72520001 | 72620000 | 0.0021 | 0.0016 | 0.0019 |
| F01 | 72540001 | 72640000 | 0.0025 | 0.0019 | 0.0023 |
| F01 | 72560001 | 72660000 | 0.0026 | 0.0020 | 0.0024 |
| F01 | 72580001 | 72680000 | 0.0028 | 0.0021 | 0.0025 |
| F01 | 72600001 | 72700000 | 0.0030 | 0.0023 | 0.0025 |
| F01 | 72620001 | 72720000 | 0.0028 | 0.0020 | 0.0023 |
| F01 | 72640001 | 72740000 | 0.0026 | 0.0018 | 0.0019 |
| F01 | 72660001 | 72760000 | 0.0026 | 0.0017 | 0.0018 |
| F01 | 72680001 | 72780000 | 0.0024 | 0.0016 | 0.0018 |
| F01 | 72700001 | 72800000 | 0.0016 | 0.0009 | 0.0012 |
| F01 | 72720001 | 72820000 | 0.0013 | 0.0007 | 0.0009 |
| F01 | 72740001 | 72840000 | 0.0008 | 0.0004 | 0.0005 |
| F01 | 72760001 | 72860000 | 0.0012 | 0.0009 | 0.0009 |
| F01 | 72780001 | 72880000 | 0.0014 | 0.0012 | 0.0012 |
| F01 | 72800001 | 72900000 | 0.0018 | 0.0016 | 0.0016 |
| F01 | 72820001 | 72920000 | 0.0021 | 0.0017 | 0.0018 |
| F01 | 72840001 | 72940000 | 0.0027 | 0.0021 | 0.0023 |
| F01 | 72860001 | 72960000 | 0.0028 | 0.0020 | 0.0022 |
| F01 | 72880001 | 72980000 | 0.0029 | 0.0019 | 0.0022 |
| F01 | 72900001 | 73000000 | 0.0027 | 0.0017 | 0.0021 |
| F01 | 72920001 | 73020000 | 0.0022 | 0.0015 | 0.0017 |
| F01 | 72940001 | 73040000 | 0.0019 | 0.0013 | 0.0015 |
| F01 | 72960001 | 73060000 | 0.0013 | 0.0010 | 0.0012 |
| F01 | 72980001 | 73080000 | 0.0011 | 0.0009 | 0.0010 |

| Chr | Start | End | both | A1 | A2 |
|---|---|---|---|---|---|
| F01 | 73000001 | 73100000 | 0.0008 | 0.0007 | 0.0008 |
| F01 | 73020001 | 73120000 | 0.0006 | 0.0005 | 0.0006 |
| F01 | 73040001 | 73140000 | 0.0008 | 0.0006 | 0.0009 |
| F01 | 73060001 | 73160000 | 0.0015 | 0.0012 | 0.0015 |
| F01 | 73080001 | 73180000 | 0.0015 | 0.0011 | 0.0014 |
| F01 | 73100001 | 73200000 | 0.0021 | 0.0013 | 0.0018 |
| F01 | 73120001 | 73220000 | 0.0029 | 0.0019 | 0.0024 |
| F01 | 73140001 | 73240000 | 0.0026 | 0.0018 | 0.0022 |
| F01 | 73160001 | 73260000 | 0.0022 | 0.0013 | 0.0016 |
| F01 | 73180001 | 73280000 | 0.0024 | 0.0016 | 0.0020 |
| F01 | 73200001 | 73300000 | 0.0021 | 0.0016 | 0.0017 |
| F01 | 73220001 | 73320000 | 0.0018 | 0.0015 | 0.0017 |
| F01 | 73240001 | 73340000 | 0.0016 | 0.0013 | 0.0015 |
| F01 | 73260001 | 73360000 | 0.0014 | 0.0012 | 0.0014 |
| F01 | 73280001 | 73380000 | 0.0010 | 0.0008 | 0.0009 |
| F01 | 73300001 | 73400000 | 0.0007 | 0.0006 | 0.0007 |
| F01 | 73320001 | 73420000 | 0.0006 | 0.0005 | 0.0006 |
| F01 | 73340001 | 73440000 | 0.0006 | 0.0005 | 0.0006 |
| F01 | 73360001 | 73460000 | 0.0007 | 0.0005 | 0.0007 |
| F01 | 73380001 | 73480000 | 0.0006 | 0.0005 | 0.0006 |
| F01 | 73400001 | 73500000 | 0.0008 | 0.0006 | 0.0008 |
| F01 | 73420001 | 73520000 | 0.0003 | 0.0002 | 0.0003 |
| F01 | 73440001 | 73540000 | 0.0003 | 0.0002 | 0.0003 |
| F01 | 73460001 | 73560000 | 0.0003 | 0.0002 | 0.0003 |
| F01 | 73480001 | 73580000 | 0.0005 | 0.0004 | 0.0005 |
| F01 | 73500001 | 73600000 | 0.0006 | 0.0004 | 0.0006 |
| F01 | 73520001 | 73620000 | 0.0006 | 0.0005 | 0.0006 |
| F01 | 73540001 | 73640000 | 0.0011 | 0.0009 | 0.0011 |
| F01 | 73560001 | 73660000 | 0.0019 | 0.0016 | 0.0019 |
| F01 | 73580001 | 73680000 | 0.0025 | 0.0021 | 0.0025 |
| F01 | 73600001 | 73700000 | 0.0028 | 0.0021 | 0.0025 |
| F01 | 73620001 | 73720000 | 0.0028 | 0.0022 | 0.0026 |
| F01 | 73640001 | 73740000 | 0.0024 | 0.0018 | 0.0022 |
| F01 | 73660001 | 73760000 | 0.0019 | 0.0014 | 0.0017 |
| F01 | 73680001 | 73780000 | 0.0015 | 0.0011 | 0.0013 |
| F01 | 73700001 | 73800000 | 0.0012 | 0.0009 | 0.0011 |
| F01 | 73720001 | 73820000 | 0.0015 | 0.0012 | 0.0015 |
| F01 | 73740001 | 73840000 | 0.0022 | 0.0017 | 0.0021 |
| F01 | 73760001 | 73860000 | 0.0022 | 0.0017 | 0.0019 |
| F01 | 73780001 | 73880000 | 0.0017 | 0.0013 | 0.0015 |
| F01 | 73800001 | 73900000 | 0.0024 | 0.0014 | 0.0021 |
| F01 | 73820001 | 73920000 | 0.0023 | 0.0012 | 0.0017 |
| F01 | 73840001 | 73940000 | 0.0017 | 0.0007 | 0.0012 |
| F01 | 73860001 | 73960000 | 0.0021 | 0.0010 | 0.0016 |
| F01 | 73880001 | 73980000 | 0.0027 | 0.0014 | 0.0020 |
| F01 | 73900001 | 74000000 | 0.0030 | 0.0019 | 0.0024 |
| F01 | 73920001 | 74020000 | 0.0032 | 0.0023 | 0.0028 |
| F01 | 73940001 | 74040000 | 0.0035 | 0.0025 | 0.0030 |
| F01 | 73960001 | 74060000 | 0.0039 | 0.0030 | 0.0035 |
| F01 | 73980001 | 74080000 | 0.0038 | 0.0028 | 0.0034 |
| F01 | 74000001 | 74100000 | 0.0034 | 0.0026 | 0.0030 |
| F01 | 74020001 | 74120000 | 0.0036 | 0.0029 | 0.0032 |
| F01 | 74040001 | 74140000 | 0.0040 | 0.0030 | 0.0035 |
| F01 | 74060001 | 74160000 | 0.0032 | 0.0023 | 0.0026 |
| F01 | 74080001 | 74180000 | 0.0028 | 0.0021 | 0.0024 |
| F01 | 74100001 | 74200000 | 0.0022 | 0.0016 | 0.0019 |
| F01 | 74120001 | 74220000 | 0.0014 | 0.0010 | 0.0011 |
| F01 | 74140001 | 74240000 | 0.0008 | 0.0006 | 0.0007 |

| Chr | Start | End | both | A1 | A2 |
|---|---|---|---|---|---|
| F01 | 74160001 | 74260000 | 0.0007 | 0.0006 | 0.0006 |
| F01 | 74180001 | 74280000 | 0.0007 | 0.0006 | 0.0006 |
| F01 | 74200001 | 74300000 | 0.0010 | 0.0009 | 0.0009 |
| F01 | 74220001 | 74320000 | 0.0010 | 0.0008 | 0.0009 |
| F01 | 74240001 | 74340000 | 0.0010 | 0.0008 | 0.0009 |
| F01 | 74260001 | 74360000 | 0.0009 | 0.0008 | 0.0009 |
| F01 | 74280001 | 74380000 | 0.0010 | 0.0009 | 0.0010 |
| F01 | 74300001 | 74400000 | 0.0010 | 0.0008 | 0.0008 |
| F01 | 74320001 | 74420000 | 0.0015 | 0.0012 | 0.0013 |
| F01 | 74340001 | 74440000 | 0.0017 | 0.0015 | 0.0015 |
| F01 | 74360001 | 74460000 | 0.0025 | 0.0022 | 0.0022 |
| F01 | 74380001 | 74480000 | 0.0031 | 0.0022 | 0.0025 |
| F01 | 74400001 | 74500000 | 0.0031 | 0.0022 | 0.0027 |
| F01 | 74420001 | 74520000 | 0.0031 | 0.0022 | 0.0026 |
| F01 | 74440001 | 74540000 | 0.0033 | 0.0022 | 0.0025 |
| F01 | 74460001 | 74560000 | 0.0026 | 0.0014 | 0.0018 |
| F01 | 74480001 | 74580000 | 0.0023 | 0.0015 | 0.0018 |
| F01 | 74500001 | 74600000 | 0.0021 | 0.0014 | 0.0017 |
| F01 | 74520001 | 74620000 | 0.0021 | 0.0014 | 0.0016 |
| F01 | 74540001 | 74640000 | 0.0022 | 0.0016 | 0.0019 |
| F01 | 74560001 | 74660000 | 0.0023 | 0.0015 | 0.0019 |
| F01 | 74580001 | 74680000 | 0.0025 | 0.0015 | 0.0020 |
| F01 | 74600001 | 74700000 | 0.0029 | 0.0018 | 0.0022 |
| F01 | 74620001 | 74720000 | 0.0038 | 0.0025 | 0.0030 |
| F01 | 74640001 | 74740000 | 0.0033 | 0.0022 | 0.0026 |
| F01 | 74660001 | 74760000 | 0.0033 | 0.0023 | 0.0027 |
| F01 | 74680001 | 74780000 | 0.0027 | 0.0020 | 0.0022 |
| F01 | 74700001 | 74800000 | 0.0027 | 0.0021 | 0.0024 |
| F01 | 74720001 | 74820000 | 0.0016 | 0.0012 | 0.0015 |
| F01 | 74740001 | 74840000 | 0.0019 | 0.0013 | 0.0019 |
| F01 | 74760001 | 74860000 | 0.0024 | 0.0018 | 0.0022 |
| F01 | 74780001 | 74880000 | 0.0029 | 0.0019 | 0.0025 |
| F01 | 74800001 | 74900000 | 0.0028 | 0.0018 | 0.0024 |
| F01 | 74820001 | 74920000 | 0.0026 | 0.0017 | 0.0023 |
| F01 | 74840001 | 74940000 | 0.0020 | 0.0015 | 0.0017 |
| F01 | 74860001 | 74960000 | 0.0018 | 0.0012 | 0.0015 |
| F01 | 74880001 | 74980000 | 0.0016 | 0.0011 | 0.0014 |
| F01 | 74900001 | 75000000 | 0.0013 | 0.0008 | 0.0011 |
| F01 | 74920001 | 75020000 | 0.0014 | 0.0008 | 0.0011 |
| F01 | 74940001 | 75040000 | 0.0029 | 0.0018 | 0.0023 |
| F01 | 74960001 | 75060000 | 0.0030 | 0.0018 | 0.0023 |
| F01 | 74980001 | 75080000 | 0.0033 | 0.0022 | 0.0026 |
| F01 | 75000001 | 75100000 | 0.0039 | 0.0025 | 0.0031 |
| F01 | 75020001 | 75120000 | 0.0046 | 0.0030 | 0.0036 |
| F01 | 75040001 | 75140000 | 0.0039 | 0.0023 | 0.0030 |
| F01 | 75060001 | 75160000 | 0.0034 | 0.0020 | 0.0026 |
| F01 | 75080001 | 75180000 | 0.0029 | 0.0015 | 0.0020 |
| F01 | 75100001 | 75200000 | 0.0027 | 0.0013 | 0.0016 |
| F01 | 75120001 | 75220000 | 0.0020 | 0.0009 | 0.0011 |
| F01 | 75140001 | 75240000 | 0.0016 | 0.0007 | 0.0007 |
| F01 | 75160001 | 75260000 | 0.0020 | 0.0011 | 0.0011 |
| F01 | 75180001 | 75280000 | 0.0026 | 0.0016 | 0.0017 |
| F01 | 75200001 | 75300000 | 0.0023 | 0.0014 | 0.0016 |
| F01 | 75220001 | 75320000 | 0.0028 | 0.0015 | 0.0017 |
| F01 | 75240001 | 75340000 | 0.0025 | 0.0014 | 0.0016 |
| F01 | 75260001 | 75360000 | 0.0026 | 0.0013 | 0.0017 |
| F01 | 75280001 | 75380000 | 0.0024 | 0.0011 | 0.0015 |
| F01 | 75300001 | 75400000 | 0.0026 | 0.0016 | 0.0019 |

| Chr | Start | End | both | A1 | A2 |
|---|---|---|---|---|---|
| F01 | 75320001 | 75420000 | 0.0022 | 0.0014 | 0.0017 |
| F01 | 75340001 | 75440000 | 0.0028 | 0.0019 | 0.0024 |
| F01 | 75360001 | 75460000 | 0.0034 | 0.0027 | 0.0030 |
| F01 | 75380001 | 75480000 | 0.0031 | 0.0025 | 0.0028 |
| F01 | 75400001 | 75500000 | 0.0027 | 0.0022 | 0.0023 |
| F01 | 75420001 | 75520000 | 0.0029 | 0.0023 | 0.0025 |
| F01 | 75440001 | 75540000 | 0.0022 | 0.0018 | 0.0019 |
| F01 | 75460001 | 75560000 | 0.0016 | 0.0013 | 0.0013 |
| F01 | 75480001 | 75580000 | 0.0017 | 0.0011 | 0.0012 |
| F01 | 75500001 | 75600000 | 0.0018 | 0.0011 | 0.0013 |
| F01 | 75520001 | 75620000 | 0.0026 | 0.0019 | 0.0019 |
| F01 | 75540001 | 75640000 | 0.0041 | 0.0033 | 0.0034 |
| F01 | 75560001 | 75660000 | 0.0039 | 0.0029 | 0.0031 |
| F01 | 75580001 | 75680000 | 0.0047 | 0.0035 | 0.0039 |
| F01 | 75600001 | 75700000 | 0.0045 | 0.0034 | 0.0038 |
| F01 | 75620001 | 75720000 | 0.0038 | 0.0026 | 0.0032 |
| F01 | 75640001 | 75740000 | 0.0031 | 0.0017 | 0.0024 |
| F01 | 75660001 | 75760000 | 0.0033 | 0.0020 | 0.0026 |
| F01 | 75680001 | 75780000 | 0.0026 | 0.0017 | 0.0020 |
| F01 | 75700001 | 75800000 | 0.0025 | 0.0017 | 0.0019 |
| F01 | 75720001 | 75820000 | 0.0030 | 0.0022 | 0.0024 |
| F01 | 75740001 | 75840000 | 0.0025 | 0.0019 | 0.0021 |
| F01 | 75760001 | 75860000 | 0.0023 | 0.0016 | 0.0018 |
| F01 | 75780001 | 75880000 | 0.0023 | 0.0014 | 0.0017 |
| F01 | 75800001 | 75900000 | 0.0027 | 0.0017 | 0.0020 |
| F01 | 75820001 | 75920000 | 0.0024 | 0.0014 | 0.0017 |
| F01 | 75840001 | 75940000 | 0.0020 | 0.0012 | 0.0014 |
| F01 | 75860001 | 75960000 | 0.0016 | 0.0010 | 0.0012 |
| F01 | 75880001 | 75980000 | 0.0015 | 0.0011 | 0.0012 |
| F01 | 75900001 | 76000000 | 0.0009 | 0.0007 | 0.0008 |
| F01 | 75920001 | 76020000 | 0.0003 | 0.0003 | 0.0002 |
| F01 | 75940001 | 76040000 | 0.0003 | 0.0003 | 0.0002 |
| F01 | 75960001 | 76060000 | 0.0004 | 0.0003 | 0.0003 |
| F01 | 75980001 | 76080000 | 0.0005 | 0.0003 | 0.0003 |
| F01 | 76000001 | 76100000 | 0.0007 | 0.0003 | 0.0005 |
| F01 | 76020001 | 76120000 | 0.0007 | 0.0003 | 0.0005 |
| F01 | 76040001 | 76140000 | 0.0008 | 0.0004 | 0.0005 |
| F01 | 76060001 | 76160000 | 0.0010 | 0.0006 | 0.0007 |
| F01 | 76080001 | 76180000 | 0.0009 | 0.0006 | 0.0007 |
| F01 | 76100001 | 76200000 | 0.0008 | 0.0007 | 0.0007 |
| F01 | 76120001 | 76220000 | 0.0009 | 0.0007 | 0.0007 |
| F01 | 76140001 | 76240000 | 0.0018 | 0.0014 | 0.0016 |
| F01 | 76160001 | 76260000 | 0.0022 | 0.0017 | 0.0021 |
| F01 | 76180001 | 76280000 | 0.0024 | 0.0019 | 0.0022 |
| F01 | 76200001 | 76300000 | 0.0025 | 0.0020 | 0.0024 |
| F01 | 76220001 | 76320000 | 0.0025 | 0.0021 | 0.0024 |
| F01 | 76240001 | 76340000 | 0.0019 | 0.0017 | 0.0018 |
| F01 | 76260001 | 76360000 | 0.0018 | 0.0016 | 0.0017 |
| F01 | 76280001 | 76380000 | 0.0015 | 0.0014 | 0.0015 |
| F01 | 76300001 | 76400000 | 0.0016 | 0.0015 | 0.0016 |
| F01 | 76320001 | 76420000 | 0.0019 | 0.0017 | 0.0018 |
| F01 | 76340001 | 76440000 | 0.0017 | 0.0015 | 0.0017 |
| F01 | 76360001 | 76460000 | 0.0019 | 0.0016 | 0.0018 |
| F01 | 76380001 | 76480000 | 0.0020 | 0.0018 | 0.0019 |
| F01 | 76400001 | 76500000 | 0.0017 | 0.0015 | 0.0016 |
| F01 | 76420001 | 76520000 | 0.0018 | 0.0015 | 0.0016 |
| F01 | 76440001 | 76540000 | 0.0021 | 0.0017 | 0.0018 |
| F01 | 76460001 | 76560000 | 0.0014 | 0.0011 | 0.0012 |

| Chr | Start | End | both | A1 | A2 |
|---|---|---|---|---|---|
| F01 | 76480001 | 76580000 | 0.0012 | 0.0009 | 0.0011 |
| F01 | 76500001 | 76600000 | 0.0012 | 0.0009 | 0.0011 |
| F01 | 76520001 | 76620000 | 0.0010 | 0.0008 | 0.0009 |
| F01 | 76540001 | 76640000 | 0.0006 | 0.0005 | 0.0006 |
| F01 | 76560001 | 76660000 | 0.0006 | 0.0006 | 0.0006 |
| F01 | 76580001 | 76680000 | 0.0007 | 0.0006 | 0.0007 |
| F01 | 76600001 | 76700000 | 0.0022 | 0.0020 | 0.0021 |
| F01 | 76620001 | 76720000 | 0.0024 | 0.0020 | 0.0022 |
| F01 | 76640001 | 76740000 | 0.0028 | 0.0024 | 0.0026 |
| F01 | 76660001 | 76760000 | 0.0027 | 0.0024 | 0.0026 |
| F01 | 76680001 | 76780000 | 0.0029 | 0.0025 | 0.0027 |
| F01 | 76700001 | 76800000 | 0.0012 | 0.0011 | 0.0011 |
| F01 | 76720001 | 76820000 | 0.0011 | 0.0010 | 0.0010 |
| F01 | 76740001 | 76840000 | 0.0007 | 0.0005 | 0.0005 |
| F01 | 76760001 | 76860000 | 0.0009 | 0.0007 | 0.0007 |
| F01 | 76780001 | 76880000 | 0.0009 | 0.0006 | 0.0007 |
| F01 | 76800001 | 76900000 | 0.0008 | 0.0006 | 0.0007 |
| F01 | 76820001 | 76920000 | 0.0009 | 0.0007 | 0.0008 |
| F01 | 76840001 | 76940000 | 0.0013 | 0.0009 | 0.0012 |
| F01 | 76860001 | 76960000 | 0.0012 | 0.0007 | 0.0010 |
| F01 | 76880001 | 76980000 | 0.0010 | 0.0006 | 0.0008 |
| F01 | 76900001 | 77000000 | 0.0014 | 0.0010 | 0.0011 |
| F01 | 76920001 | 77020000 | 0.0015 | 0.0010 | 0.0012 |
| F01 | 76940001 | 77040000 | 0.0011 | 0.0007 | 0.0008 |
| F01 | 76960001 | 77060000 | 0.0012 | 0.0007 | 0.0009 |
| F01 | 76980001 | 77080000 | 0.0014 | 0.0010 | 0.0011 |
| F01 | 77000001 | 77100000 | 0.0013 | 0.0009 | 0.0011 |
| F01 | 77020001 | 77120000 | 0.0012 | 0.0008 | 0.0009 |
| F01 | 77040001 | 77140000 | 0.0015 | 0.0010 | 0.0011 |
| F01 | 77060001 | 77160000 | 0.0017 | 0.0015 | 0.0015 |
| F01 | 77080001 | 77180000 | 0.0017 | 0.0015 | 0.0016 |
| F01 | 77100001 | 77200000 | 0.0018 | 0.0017 | 0.0017 |
| F01 | 77120001 | 77220000 | 0.0032 | 0.0029 | 0.0030 |
| F01 | 77140001 | 77240000 | 0.0042 | 0.0036 | 0.0038 |
| F01 | 77160001 | 77260000 | 0.0048 | 0.0040 | 0.0043 |
| F01 | 77180001 | 77280000 | 0.0050 | 0.0041 | 0.0043 |
| F01 | 77200001 | 77300000 | 0.0054 | 0.0044 | 0.0048 |
| F01 | 77220001 | 77320000 | 0.0042 | 0.0033 | 0.0037 |
| F01 | 77240001 | 77340000 | 0.0035 | 0.0027 | 0.0029 |
| F01 | 77260001 | 77360000 | 0.0028 | 0.0020 | 0.0022 |
| F01 | 77280001 | 77380000 | 0.0027 | 0.0019 | 0.0022 |
| F01 | 77300001 | 77400000 | 0.0026 | 0.0019 | 0.0022 |
| F01 | 77320001 | 77420000 | 0.0027 | 0.0020 | 0.0022 |
| F01 | 77340001 | 77440000 | 0.0032 | 0.0024 | 0.0028 |
| F01 | 77360001 | 77460000 | 0.0027 | 0.0021 | 0.0024 |
| F01 | 77380001 | 77480000 | 0.0026 | 0.0020 | 0.0023 |
| F01 | 77400001 | 77500000 | 0.0028 | 0.0020 | 0.0024 |
| F01 | 77420001 | 77520000 | 0.0048 | 0.0036 | 0.0042 |
| F01 | 77440001 | 77540000 | 0.0041 | 0.0031 | 0.0037 |
| F01 | 77460001 | 77560000 | 0.0047 | 0.0036 | 0.0043 |
| F01 | 77480001 | 77580000 | 0.0050 | 0.0039 | 0.0046 |
| F01 | 77500001 | 77600000 | 0.0048 | 0.0038 | 0.0046 |
| F01 | 77520001 | 77620000 | 0.0030 | 0.0022 | 0.0027 |
| F01 | 77540001 | 77640000 | 0.0029 | 0.0021 | 0.0027 |
| F01 | 77560001 | 77660000 | 0.0026 | 0.0018 | 0.0024 |
| F01 | 77580001 | 77680000 | 0.0022 | 0.0016 | 0.0020 |
| F01 | 77600001 | 77700000 | 0.0025 | 0.0019 | 0.0022 |
| F01 | 77620001 | 77720000 | 0.0025 | 0.0017 | 0.0019 |

| Chr | Start | End | both | A1 | A2 |
|---|---|---|---|---|---|
| F01 | 77640001 | 77740000 | 0.0033 | 0.0022 | 0.0024 |
| F01 | 77660001 | 77760000 | 0.0036 | 0.0022 | 0.0025 |
| F01 | 77680001 | 77780000 | 0.0038 | 0.0022 | 0.0024 |
| F01 | 77700001 | 77800000 | 0.0029 | 0.0013 | 0.0014 |
| F01 | 77720001 | 77820000 | 0.0026 | 0.0013 | 0.0014 |
| F01 | 77740001 | 77840000 | 0.0020 | 0.0009 | 0.0010 |
| F01 | 77760001 | 77860000 | 0.0019 | 0.0009 | 0.0010 |
| F01 | 77780001 | 77880000 | 0.0019 | 0.0009 | 0.0009 |
| F01 | 77800001 | 77900000 | 0.0022 | 0.0012 | 0.0012 |
| F01 | 77820001 | 77920000 | 0.0021 | 0.0013 | 0.0014 |
| F01 | 77840001 | 77940000 | 0.0019 | 0.0012 | 0.0012 |
| F01 | 77860001 | 77960000 | 0.0021 | 0.0014 | 0.0014 |
| F01 | 77880001 | 77980000 | 0.0021 | 0.0015 | 0.0016 |
| F01 | 77900001 | 78000000 | 0.0022 | 0.0017 | 0.0017 |
| F01 | 77920001 | 78020000 | 0.0034 | 0.0027 | 0.0026 |
| F01 | 77940001 | 78040000 | 0.0049 | 0.0042 | 0.0041 |
| F01 | 77960001 | 78060000 | 0.0046 | 0.0038 | 0.0039 |
| F01 | 77980001 | 78080000 | 0.0052 | 0.0044 | 0.0045 |
| F01 | 78000001 | 78100000 | 0.0054 | 0.0042 | 0.0045 |
| F01 | 78020001 | 78120000 | 0.0043 | 0.0034 | 0.0037 |
| F01 | 78040001 | 78140000 | 0.0031 | 0.0022 | 0.0025 |
| F01 | 78060001 | 78160000 | 0.0034 | 0.0023 | 0.0025 |
| F01 | 78080001 | 78180000 | 0.0031 | 0.0019 | 0.0021 |
| F01 | 78100001 | 78200000 | 0.0032 | 0.0022 | 0.0025 |
| F01 | 78120001 | 78220000 | 0.0035 | 0.0025 | 0.0027 |
| F01 | 78140001 | 78240000 | 0.0030 | 0.0021 | 0.0023 |
| F01 | 78160001 | 78260000 | 0.0025 | 0.0020 | 0.0021 |
| F01 | 78180001 | 78280000 | 0.0022 | 0.0019 | 0.0020 |
| F01 | 78200001 | 78300000 | 0.0020 | 0.0019 | 0.0019 |
| F01 | 78220001 | 78320000 | 0.0014 | 0.0014 | 0.0013 |
| F01 | 78240001 | 78340000 | 0.0018 | 0.0018 | 0.0017 |
| F01 | 78260001 | 78360000 | 0.0024 | 0.0020 | 0.0020 |
| F01 | 78280001 | 78380000 | 0.0030 | 0.0023 | 0.0026 |
| F01 | 78300001 | 78400000 | 0.0025 | 0.0018 | 0.0021 |
| F01 | 78320001 | 78420000 | 0.0023 | 0.0016 | 0.0019 |
| F01 | 78340001 | 78440000 | 0.0020 | 0.0012 | 0.0016 |
| F01 | 78360001 | 78460000 | 0.0014 | 0.0010 | 0.0013 |
| F01 | 78380001 | 78480000 | 0.0003 | 0.0003 | 0.0003 |
| F01 | 78400001 | 78500000 | 0.0004 | 0.0003 | 0.0003 |
| F01 | 78420001 | 78520000 | 0.0004 | 0.0003 | 0.0003 |
| F01 | 78440001 | 78540000 | 0.0008 | 0.0007 | 0.0007 |
| F01 | 78460001 | 78560000 | 0.0009 | 0.0007 | 0.0008 |
| F01 | 78480001 | 78580000 | 0.0011 | 0.0008 | 0.0010 |
| F01 | 78500001 | 78600000 | 0.0011 | 0.0009 | 0.0010 |
| F01 | 78520001 | 78620000 | 0.0020 | 0.0015 | 0.0018 |
| F01 | 78540001 | 78640000 | 0.0029 | 0.0021 | 0.0026 |
| F01 | 78560001 | 78660000 | 0.0031 | 0.0021 | 0.0026 |
| F01 | 78580001 | 78680000 | 0.0036 | 0.0024 | 0.0030 |
| F01 | 78600001 | 78700000 | 0.0040 | 0.0026 | 0.0032 |
| F01 | 78620001 | 78720000 | 0.0037 | 0.0024 | 0.0030 |
| F01 | 78640001 | 78740000 | 0.0032 | 0.0021 | 0.0024 |
| F01 | 78660001 | 78760000 | 0.0032 | 0.0021 | 0.0024 |
| F01 | 78680001 | 78780000 | 0.0029 | 0.0019 | 0.0021 |
| F01 | 78700001 | 78800000 | 0.0027 | 0.0019 | 0.0019 |
| F01 | 78720001 | 78820000 | 0.0025 | 0.0015 | 0.0015 |
| F01 | 78740001 | 78840000 | 0.0020 | 0.0011 | 0.0011 |
| F01 | 78760001 | 78860000 | 0.0029 | 0.0019 | 0.0022 |
| F01 | 78780001 | 78880000 | 0.0028 | 0.0018 | 0.0021 |

| Chr | Start | End | both | A1 | A2 |
|---|---|---|---|---|---|
| F01 | 78800001 | 78900000 | 0.0039 | 0.0029 | 0.0032 |
| F01 | 78820001 | 78920000 | 0.0041 | 0.0033 | 0.0037 |
| F01 | 78840001 | 78940000 | 0.0047 | 0.0039 | 0.0043 |
| F01 | 78860001 | 78960000 | 0.0046 | 0.0038 | 0.0040 |
| F01 | 78880001 | 78980000 | 0.0051 | 0.0045 | 0.0046 |
| F01 | 78900001 | 79000000 | 0.0045 | 0.0037 | 0.0040 |
| F01 | 78920001 | 79020000 | 0.0051 | 0.0042 | 0.0045 |
| F01 | 78940001 | 79040000 | 0.0050 | 0.0038 | 0.0042 |
| F01 | 78960001 | 79060000 | 0.0064 | 0.0053 | 0.0057 |
| F01 | 78980001 | 79080000 | 0.0060 | 0.0047 | 0.0051 |
| F01 | 79000001 | 79100000 | 0.0057 | 0.0046 | 0.0048 |
| F01 | 79020001 | 79120000 | 0.0049 | 0.0039 | 0.0039 |
| F01 | 79040001 | 79140000 | 0.0047 | 0.0036 | 0.0036 |
| F01 | 79060001 | 79160000 | 0.0026 | 0.0015 | 0.0013 |
| F01 | 79080001 | 79180000 | 0.0021 | 0.0014 | 0.0011 |
| F01 | 79100001 | 79200000 | 0.0030 | 0.0022 | 0.0021 |
| F01 | 79120001 | 79220000 | 0.0038 | 0.0027 | 0.0029 |
| F01 | 79140001 | 79240000 | 0.0038 | 0.0029 | 0.0031 |
| F01 | 79160001 | 79260000 | 0.0042 | 0.0032 | 0.0035 |
| F01 | 79180001 | 79280000 | 0.0043 | 0.0032 | 0.0035 |
| F01 | 79200001 | 79300000 | 0.0035 | 0.0024 | 0.0025 |
| F01 | 79220001 | 79320000 | 0.0031 | 0.0021 | 0.0022 |
| F01 | 79240001 | 79340000 | 0.0032 | 0.0020 | 0.0022 |
| F01 | 79260001 | 79360000 | 0.0033 | 0.0024 | 0.0024 |
| F01 | 79280001 | 79380000 | 0.0039 | 0.0027 | 0.0027 |
| F01 | 79300001 | 79400000 | 0.0050 | 0.0039 | 0.0039 |
| F01 | 79320001 | 79420000 | 0.0050 | 0.0038 | 0.0037 |
| F01 | 79340001 | 79440000 | 0.0044 | 0.0032 | 0.0031 |
| F01 | 79360001 | 79460000 | 0.0042 | 0.0030 | 0.0028 |
| F01 | 79380001 | 79480000 | 0.0043 | 0.0035 | 0.0034 |
| F01 | 79400001 | 79500000 | 0.0030 | 0.0022 | 0.0022 |
| F01 | 79420001 | 79520000 | 0.0029 | 0.0021 | 0.0020 |
| F01 | 79440001 | 79540000 | 0.0032 | 0.0022 | 0.0022 |
| F01 | 79460001 | 79560000 | 0.0043 | 0.0032 | 0.0034 |
| F01 | 79480001 | 79580000 | 0.0040 | 0.0030 | 0.0031 |
| F01 | 79500001 | 79600000 | 0.0044 | 0.0033 | 0.0037 |
| F01 | 79520001 | 79620000 | 0.0046 | 0.0037 | 0.0043 |
| F01 | 79540001 | 79640000 | 0.0053 | 0.0047 | 0.0051 |
| F01 | 79560001 | 79660000 | 0.0043 | 0.0037 | 0.0039 |
| F01 | 79580001 | 79680000 | 0.0042 | 0.0034 | 0.0038 |
| F01 | 79600001 | 79700000 | 0.0042 | 0.0035 | 0.0038 |
| F01 | 79620001 | 79720000 | 0.0042 | 0.0035 | 0.0037 |
| F01 | 79640001 | 79740000 | 0.0035 | 0.0027 | 0.0031 |
| F01 | 79660001 | 79760000 | 0.0036 | 0.0030 | 0.0034 |
| F01 | 79680001 | 79780000 | 0.0031 | 0.0027 | 0.0030 |
| F01 | 79700001 | 79800000 | 0.0038 | 0.0034 | 0.0037 |
| F01 | 79720001 | 79820000 | 0.0038 | 0.0033 | 0.0035 |
| F01 | 79740001 | 79840000 | 0.0040 | 0.0031 | 0.0034 |
| F01 | 79760001 | 79860000 | 0.0037 | 0.0027 | 0.0029 |
| F01 | 79780001 | 79880000 | 0.0038 | 0.0026 | 0.0029 |
| F01 | 79800001 | 79900000 | 0.0024 | 0.0013 | 0.0013 |
| F01 | 79820001 | 79920000 | 0.0018 | 0.0007 | 0.0008 |
| F01 | 79840001 | 79940000 | 0.0018 | 0.0010 | 0.0009 |
| F01 | 79860001 | 79960000 | 0.0028 | 0.0021 | 0.0018 |
| F01 | 79880001 | 79980000 | 0.0032 | 0.0026 | 0.0024 |
| F01 | 79900001 | 80000000 | 0.0040 | 0.0034 | 0.0033 |
| F01 | 79920001 | 80020000 | 0.0053 | 0.0045 | 0.0045 |
| F01 | 79940001 | 80040000 | 0.0058 | 0.0051 | 0.0052 |

| Chr | Start | End | both | A1 | A2 |
|---|---|---|---|---|---|
| F01 | 79960001 | 80060000 | 0.0049 | 0.0041 | 0.0044 |
| F01 | 79980001 | 80080000 | 0.0049 | 0.0041 | 0.0044 |
| F01 | 80000001 | 80100000 | 0.0039 | 0.0033 | 0.0035 |
| F01 | 80020001 | 80120000 | 0.0038 | 0.0031 | 0.0033 |
| F01 | 80040001 | 80140000 | 0.0037 | 0.0029 | 0.0031 |
| F01 | 80060001 | 80160000 | 0.0035 | 0.0028 | 0.0029 |
| F01 | 80080001 | 80180000 | 0.0036 | 0.0028 | 0.0028 |
| F01 | 80100001 | 80200000 | 0.0043 | 0.0035 | 0.0034 |
| F01 | 80120001 | 80220000 | 0.0035 | 0.0028 | 0.0028 |
| F01 | 80140001 | 80240000 | 0.0028 | 0.0021 | 0.0021 |
| F01 | 80160001 | 80260000 | 0.0034 | 0.0025 | 0.0024 |
| F01 | 80180001 | 80280000 | 0.0030 | 0.0022 | 0.0023 |
| F01 | 80200001 | 80300000 | 0.0034 | 0.0024 | 0.0027 |
| F01 | 80220001 | 80320000 | 0.0031 | 0.0019 | 0.0023 |
| F01 | 80240001 | 80340000 | 0.0028 | 0.0017 | 0.0021 |
| F01 | 80260001 | 80360000 | 0.0029 | 0.0018 | 0.0023 |
| F01 | 80280001 | 80380000 | 0.0032 | 0.0021 | 0.0025 |
| F01 | 80300001 | 80400000 | 0.0022 | 0.0013 | 0.0015 |
| F01 | 80320001 | 80420000 | 0.0030 | 0.0019 | 0.0020 |
| F01 | 80340001 | 80440000 | 0.0033 | 0.0021 | 0.0022 |
| F01 | 80360001 | 80460000 | 0.0026 | 0.0018 | 0.0017 |
| F01 | 80380001 | 80480000 | 0.0023 | 0.0014 | 0.0014 |
| F01 | 80400001 | 80500000 | 0.0025 | 0.0016 | 0.0016 |
| F01 | 80420001 | 80520000 | 0.0016 | 0.0011 | 0.0010 |
| F01 | 80440001 | 80540000 | 0.0015 | 0.0010 | 0.0010 |
| F01 | 80460001 | 80560000 | 0.0012 | 0.0007 | 0.0007 |
| F01 | 80480001 | 80580000 | 0.0015 | 0.0012 | 0.0012 |
| F01 | 80500001 | 80600000 | 0.0021 | 0.0016 | 0.0017 |
| F01 | 80520001 | 80620000 | 0.0026 | 0.0019 | 0.0020 |
| F01 | 80540001 | 80640000 | 0.0025 | 0.0019 | 0.0020 |
| F01 | 80560001 | 80660000 | 0.0028 | 0.0021 | 0.0023 |
| F01 | 80580001 | 80680000 | 0.0023 | 0.0016 | 0.0017 |
| F01 | 80600001 | 80700000 | 0.0020 | 0.0013 | 0.0014 |
| F01 | 80620001 | 80720000 | 0.0024 | 0.0017 | 0.0016 |
| F01 | 80640001 | 80740000 | 0.0032 | 0.0024 | 0.0025 |
| F01 | 80660001 | 80760000 | 0.0042 | 0.0031 | 0.0032 |
| F01 | 80680001 | 80780000 | 0.0048 | 0.0036 | 0.0037 |
| F01 | 80700001 | 80800000 | 0.0050 | 0.0039 | 0.0041 |
| F01 | 80720001 | 80820000 | 0.0050 | 0.0036 | 0.0040 |
| F01 | 80740001 | 80840000 | 0.0046 | 0.0032 | 0.0035 |
| F01 | 80760001 | 80860000 | 0.0043 | 0.0031 | 0.0034 |
| F01 | 80780001 | 80880000 | 0.0039 | 0.0029 | 0.0031 |
| F01 | 80800001 | 80900000 | 0.0042 | 0.0032 | 0.0035 |
| F01 | 80820001 | 80920000 | 0.0048 | 0.0040 | 0.0042 |
| F01 | 80840001 | 80940000 | 0.0051 | 0.0043 | 0.0045 |
| F01 | 80860001 | 80960000 | 0.0047 | 0.0036 | 0.0040 |
| F01 | 80880001 | 80980000 | 0.0048 | 0.0035 | 0.0039 |
| F01 | 80900001 | 81000000 | 0.0048 | 0.0034 | 0.0038 |
| F01 | 80920001 | 81020000 | 0.0047 | 0.0032 | 0.0036 |
| F01 | 80940001 | 81040000 | 0.0053 | 0.0038 | 0.0041 |
| F01 | 80960001 | 81060000 | 0.0053 | 0.0040 | 0.0042 |
| F01 | 80980001 | 81080000 | 0.0051 | 0.0039 | 0.0042 |
| F01 | 81000001 | 81100000 | 0.0046 | 0.0037 | 0.0039 |
| F01 | 81020001 | 81120000 | 0.0039 | 0.0032 | 0.0034 |
| F01 | 81040001 | 81140000 | 0.0025 | 0.0020 | 0.0021 |
| F01 | 81060001 | 81160000 | 0.0028 | 0.0023 | 0.0025 |
| F01 | 81080001 | 81180000 | 0.0036 | 0.0029 | 0.0032 |
| F01 | 81100001 | 81200000 | 0.0035 | 0.0027 | 0.0029 |

| Chr | Start | End | both | A1 | A2 |
|---|---|---|---|---|---|
| F01 | 81120001 | 81220000 | 0.0039 | 0.0029 | 0.0032 |
| F01 | 81140001 | 81240000 | 0.0044 | 0.0033 | 0.0038 |
| F01 | 81160001 | 81260000 | 0.0039 | 0.0028 | 0.0033 |
| F01 | 81180001 | 81280000 | 0.0036 | 0.0028 | 0.0033 |
| F01 | 81200001 | 81300000 | 0.0037 | 0.0029 | 0.0034 |
| F01 | 81220001 | 81320000 | 0.0040 | 0.0032 | 0.0037 |
| F01 | 81240001 | 81340000 | 0.0037 | 0.0029 | 0.0032 |
| F01 | 81260001 | 81360000 | 0.0038 | 0.0031 | 0.0033 |
| F01 | 81280001 | 81380000 | 0.0036 | 0.0027 | 0.0029 |
| F01 | 81300001 | 81400000 | 0.0031 | 0.0023 | 0.0025 |
| F01 | 81320001 | 81420000 | 0.0024 | 0.0019 | 0.0018 |
| F01 | 81340001 | 81440000 | 0.0024 | 0.0020 | 0.0018 |
| F01 | 81360001 | 81460000 | 0.0034 | 0.0026 | 0.0025 |
| F01 | 81380001 | 81480000 | 0.0035 | 0.0028 | 0.0028 |
| F01 | 81400001 | 81500000 | 0.0042 | 0.0035 | 0.0036 |
| F01 | 81420001 | 81520000 | 0.0039 | 0.0032 | 0.0033 |
| F01 | 81440001 | 81540000 | 0.0037 | 0.0029 | 0.0033 |
| F01 | 81460001 | 81560000 | 0.0027 | 0.0021 | 0.0024 |
| F01 | 81480001 | 81580000 | 0.0023 | 0.0017 | 0.0020 |
| F01 | 81500001 | 81600000 | 0.0024 | 0.0016 | 0.0018 |
| F01 | 81520001 | 81620000 | 0.0023 | 0.0013 | 0.0016 |
| F01 | 81540001 | 81640000 | 0.0030 | 0.0021 | 0.0022 |
| F01 | 81560001 | 81660000 | 0.0037 | 0.0028 | 0.0029 |
| F01 | 81580001 | 81680000 | 0.0040 | 0.0032 | 0.0034 |
| F01 | 81600001 | 81700000 | 0.0036 | 0.0029 | 0.0031 |
| F01 | 81620001 | 81720000 | 0.0043 | 0.0035 | 0.0039 |
| F01 | 81640001 | 81740000 | 0.0045 | 0.0037 | 0.0040 |
| F01 | 81660001 | 81760000 | 0.0044 | 0.0037 | 0.0040 |
| F01 | 81680001 | 81780000 | 0.0048 | 0.0041 | 0.0043 |
| F01 | 81700001 | 81800000 | 0.0054 | 0.0045 | 0.0047 |
| F01 | 81720001 | 81820000 | 0.0046 | 0.0039 | 0.0040 |
| F01 | 81740001 | 81840000 | 0.0037 | 0.0030 | 0.0032 |
| F01 | 81760001 | 81860000 | 0.0036 | 0.0027 | 0.0030 |
| F01 | 81780001 | 81880000 | 0.0030 | 0.0021 | 0.0024 |
| F01 | 81800001 | 81900000 | 0.0024 | 0.0016 | 0.0019 |
| F01 | 81820001 | 81920000 | 0.0033 | 0.0026 | 0.0030 |
| F01 | 81840001 | 81940000 | 0.0034 | 0.0026 | 0.0030 |
| F01 | 81860001 | 81960000 | 0.0031 | 0.0025 | 0.0029 |
| F01 | 81880001 | 81980000 | 0.0036 | 0.0029 | 0.0033 |
| F01 | 81900001 | 82000000 | 0.0040 | 0.0033 | 0.0038 |
| F01 | 81920001 | 82020000 | 0.0032 | 0.0025 | 0.0030 |
| F01 | 81940001 | 82040000 | 0.0030 | 0.0025 | 0.0029 |
| F01 | 81960001 | 82060000 | 0.0029 | 0.0024 | 0.0027 |
| F01 | 81980001 | 82080000 | 0.0027 | 0.0021 | 0.0024 |
| F01 | 82000001 | 82100000 | 0.0028 | 0.0021 | 0.0025 |
| F01 | 82020001 | 82120000 | 0.0033 | 0.0026 | 0.0028 |
| F01 | 82040001 | 82140000 | 0.0031 | 0.0024 | 0.0026 |
| F01 | 82060001 | 82160000 | 0.0037 | 0.0029 | 0.0032 |
| F01 | 82080001 | 82180000 | 0.0038 | 0.0030 | 0.0034 |
| F01 | 82100001 | 82200000 | 0.0040 | 0.0033 | 0.0037 |
| F01 | 82120001 | 82220000 | 0.0037 | 0.0031 | 0.0036 |
| F01 | 82140001 | 82240000 | 0.0039 | 0.0033 | 0.0038 |
| F01 | 82160001 | 82260000 | 0.0034 | 0.0028 | 0.0032 |
| F01 | 82180001 | 82280000 | 0.0043 | 0.0037 | 0.0039 |
| F01 | 82200001 | 82300000 | 0.0034 | 0.0028 | 0.0029 |
| F01 | 82220001 | 82320000 | 0.0039 | 0.0031 | 0.0033 |
| F01 | 82240001 | 82340000 | 0.0041 | 0.0032 | 0.0035 |
| F01 | 82260001 | 82360000 | 0.0046 | 0.0036 | 0.0039 |

| Chr | Start | End | both | A1 | A2 |
|---|---|---|---|---|---|
| F01 | 82280001 | 82380000 | 0.0038 | 0.0029 | 0.0031 |
| F01 | 82300001 | 82400000 | 0.0042 | 0.0032 | 0.0036 |
| F01 | 82320001 | 82420000 | 0.0039 | 0.0031 | 0.0033 |
| F01 | 82340001 | 82440000 | 0.0042 | 0.0034 | 0.0037 |
| F01 | 82360001 | 82460000 | 0.0049 | 0.0039 | 0.0043 |
| F01 | 82380001 | 82480000 | 0.0051 | 0.0041 | 0.0044 |
| F01 | 82400001 | 82500000 | 0.0057 | 0.0045 | 0.0048 |
| F01 | 82420001 | 82520000 | 0.0063 | 0.0049 | 0.0053 |
| F01 | 82440001 | 82540000 | 0.0061 | 0.0045 | 0.0049 |
| F01 | 82460001 | 82560000 | 0.0052 | 0.0038 | 0.0040 |
| F01 | 82480001 | 82580000 | 0.0056 | 0.0040 | 0.0043 |
| F01 | 82500001 | 82600000 | 0.0053 | 0.0038 | 0.0041 |
| F01 | 82520001 | 82620000 | 0.0047 | 0.0032 | 0.0034 |
| F01 | 82540001 | 82640000 | 0.0051 | 0.0038 | 0.0041 |
| F01 | 82560001 | 82660000 | 0.0054 | 0.0041 | 0.0043 |
| F01 | 82580001 | 82680000 | 0.0045 | 0.0033 | 0.0035 |
| F01 | 82600001 | 82700000 | 0.0046 | 0.0033 | 0.0035 |
| F01 | 82620001 | 82720000 | 0.0042 | 0.0030 | 0.0031 |
| F01 | 82640001 | 82740000 | 0.0038 | 0.0026 | 0.0026 |
| F01 | 82660001 | 82760000 | 0.0026 | 0.0018 | 0.0018 |
| F01 | 82680001 | 82780000 | 0.0022 | 0.0014 | 0.0015 |
| F01 | 82700001 | 82800000 | 0.0012 | 0.0009 | 0.0009 |
| F01 | 82720001 | 82820000 | 0.0016 | 0.0013 | 0.0014 |
| F01 | 82740001 | 82840000 | 0.0018 | 0.0015 | 0.0015 |
| F01 | 82760001 | 82860000 | 0.0032 | 0.0024 | 0.0026 |
| F01 | 82780001 | 82880000 | 0.0036 | 0.0027 | 0.0030 |
| F01 | 82800001 | 82900000 | 0.0046 | 0.0033 | 0.0038 |
| F01 | 82820001 | 82920000 | 0.0043 | 0.0030 | 0.0034 |
| F01 | 82840001 | 82940000 | 0.0044 | 0.0031 | 0.0035 |
| F01 | 82860001 | 82960000 | 0.0048 | 0.0037 | 0.0041 |
| F01 | 82880001 | 82980000 | 0.0052 | 0.0038 | 0.0042 |
| F01 | 82900001 | 83000000 | 0.0048 | 0.0036 | 0.0039 |
| F01 | 82920001 | 83020000 | 0.0056 | 0.0044 | 0.0049 |
| F01 | 82940001 | 83040000 | 0.0052 | 0.0039 | 0.0045 |
| F01 | 82960001 | 83060000 | 0.0042 | 0.0030 | 0.0036 |
| F01 | 82980001 | 83080000 | 0.0053 | 0.0043 | 0.0048 |
| F01 | 83000001 | 83100000 | 0.0061 | 0.0048 | 0.0052 |
| F01 | 83020001 | 83120000 | 0.0061 | 0.0048 | 0.0048 |
| F01 | 83040001 | 83140000 | 0.0070 | 0.0059 | 0.0057 |
| F01 | 83060001 | 83160000 | 0.0068 | 0.0058 | 0.0056 |
| F01 | 83080001 | 83180000 | 0.0050 | 0.0040 | 0.0038 |
| F01 | 83100001 | 83200000 | 0.0041 | 0.0032 | 0.0031 |
| F01 | 83120001 | 83220000 | 0.0035 | 0.0025 | 0.0026 |
| F01 | 83140001 | 83240000 | 0.0031 | 0.0019 | 0.0022 |
| F01 | 83160001 | 83260000 | 0.0034 | 0.0020 | 0.0021 |
| F01 | 83180001 | 83280000 | 0.0036 | 0.0022 | 0.0025 |
| F01 | 83200001 | 83300000 | 0.0044 | 0.0030 | 0.0033 |
| F01 | 83220001 | 83320000 | 0.0040 | 0.0028 | 0.0029 |
| F01 | 83240001 | 83340000 | 0.0032 | 0.0023 | 0.0023 |
| F01 | 83260001 | 83360000 | 0.0029 | 0.0022 | 0.0022 |
| F01 | 83280001 | 83380000 | 0.0038 | 0.0030 | 0.0032 |
| F01 | 83300001 | 83400000 | 0.0039 | 0.0033 | 0.0034 |
| F01 | 83320001 | 83420000 | 0.0047 | 0.0039 | 0.0044 |
| F01 | 83340001 | 83440000 | 0.0055 | 0.0044 | 0.0049 |
| F01 | 83360001 | 83460000 | 0.0052 | 0.0041 | 0.0046 |
| F01 | 83380001 | 83480000 | 0.0045 | 0.0036 | 0.0039 |
| F01 | 83400001 | 83500000 | 0.0039 | 0.0029 | 0.0032 |
| F01 | 83420001 | 83520000 | 0.0043 | 0.0034 | 0.0037 |

| Chr | Start | End | both | A1 | A2 |
|---|---|---|---|---|---|
| F01 | 83440001 | 83540000 | 0.0046 | 0.0035 | 0.0040 |
| F01 | 83460001 | 83560000 | 0.0051 | 0.0039 | 0.0045 |
| F01 | 83480001 | 83580000 | 0.0050 | 0.0034 | 0.0043 |
| F01 | 83500001 | 83600000 | 0.0053 | 0.0036 | 0.0044 |
| F01 | 83520001 | 83620000 | 0.0043 | 0.0029 | 0.0034 |
| F01 | 83540001 | 83640000 | 0.0043 | 0.0033 | 0.0035 |
| F01 | 83560001 | 83660000 | 0.0040 | 0.0031 | 0.0031 |
| F01 | 83580001 | 83680000 | 0.0046 | 0.0037 | 0.0037 |
| F01 | 83600001 | 83700000 | 0.0044 | 0.0032 | 0.0033 |
| F01 | 83620001 | 83720000 | 0.0050 | 0.0038 | 0.0038 |
| F01 | 83640001 | 83740000 | 0.0050 | 0.0035 | 0.0036 |
| F01 | 83660001 | 83760000 | 0.0048 | 0.0031 | 0.0034 |
| F01 | 83680001 | 83780000 | 0.0042 | 0.0028 | 0.0031 |
| F01 | 83700001 | 83800000 | 0.0034 | 0.0025 | 0.0026 |
| F01 | 83720001 | 83820000 | 0.0029 | 0.0020 | 0.0022 |
| F01 | 83740001 | 83840000 | 0.0028 | 0.0020 | 0.0021 |
| F01 | 83760001 | 83860000 | 0.0036 | 0.0027 | 0.0027 |
| F01 | 83780001 | 83880000 | 0.0037 | 0.0027 | 0.0028 |
| F01 | 83800001 | 83900000 | 0.0037 | 0.0027 | 0.0028 |
| F01 | 83820001 | 83920000 | 0.0040 | 0.0029 | 0.0032 |
| F01 | 83840001 | 83940000 | 0.0038 | 0.0026 | 0.0031 |
| F01 | 83860001 | 83960000 | 0.0032 | 0.0024 | 0.0029 |
| F01 | 83880001 | 83980000 | 0.0030 | 0.0021 | 0.0026 |
| F01 | 83900001 | 84000000 | 0.0044 | 0.0032 | 0.0038 |
| F01 | 83920001 | 84020000 | 0.0043 | 0.0031 | 0.0035 |
| F01 | 83940001 | 84040000 | 0.0042 | 0.0031 | 0.0034 |
| F01 | 83960001 | 84060000 | 0.0041 | 0.0029 | 0.0031 |
| F01 | 83980001 | 84080000 | 0.0040 | 0.0029 | 0.0031 |
| F01 | 84000001 | 84100000 | 0.0033 | 0.0023 | 0.0025 |
| F01 | 84020001 | 84120000 | 0.0027 | 0.0018 | 0.0020 |
| F01 | 84040001 | 84140000 | 0.0023 | 0.0016 | 0.0018 |
| F01 | 84060001 | 84160000 | 0.0023 | 0.0017 | 0.0020 |
| F01 | 84080001 | 84180000 | 0.0022 | 0.0017 | 0.0020 |
| F01 | 84100001 | 84200000 | 0.0025 | 0.0019 | 0.0021 |
| F01 | 84120001 | 84220000 | 0.0032 | 0.0024 | 0.0025 |
| F01 | 84140001 | 84240000 | 0.0044 | 0.0032 | 0.0034 |
| F01 | 84160001 | 84260000 | 0.0053 | 0.0039 | 0.0041 |
| F01 | 84180001 | 84280000 | 0.0053 | 0.0038 | 0.0039 |
| F01 | 84200001 | 84300000 | 0.0048 | 0.0032 | 0.0034 |
| F01 | 84220001 | 84320000 | 0.0059 | 0.0044 | 0.0048 |
| F01 | 84240001 | 84340000 | 0.0047 | 0.0033 | 0.0036 |
| F01 | 84260001 | 84360000 | 0.0046 | 0.0032 | 0.0035 |
| F01 | 84280001 | 84380000 | 0.0054 | 0.0038 | 0.0040 |
| F01 | 84300001 | 84400000 | 0.0052 | 0.0037 | 0.0039 |
| F01 | 84320001 | 84420000 | 0.0036 | 0.0021 | 0.0023 |
| F01 | 84340001 | 84440000 | 0.0043 | 0.0027 | 0.0029 |
| F01 | 84360001 | 84460000 | 0.0033 | 0.0020 | 0.0021 |
| F01 | 84380001 | 84480000 | 0.0028 | 0.0017 | 0.0018 |
| F01 | 84400001 | 84500000 | 0.0035 | 0.0025 | 0.0024 |
| F01 | 84420001 | 84520000 | 0.0035 | 0.0025 | 0.0024 |
| F01 | 84440001 | 84540000 | 0.0032 | 0.0023 | 0.0022 |
| F01 | 84460001 | 84560000 | 0.0051 | 0.0038 | 0.0041 |
| F01 | 84480001 | 84580000 | 0.0057 | 0.0044 | 0.0045 |
| F01 | 84500001 | 84600000 | 0.0053 | 0.0038 | 0.0041 |
| F01 | 84520001 | 84620000 | 0.0054 | 0.0040 | 0.0044 |
| F01 | 84540001 | 84640000 | 0.0052 | 0.0039 | 0.0042 |
| F01 | 84560001 | 84660000 | 0.0035 | 0.0024 | 0.0024 |
| F01 | 84580001 | 84680000 | 0.0027 | 0.0017 | 0.0018 |

| Chr | Start | End | both | A1 | A2 |
|---|---|---|---|---|---|
| F01 | 84600001 | 84700000 | 0.0027 | 0.0016 | 0.0017 |
| F01 | 84620001 | 84720000 | 0.0026 | 0.0013 | 0.0013 |
| F01 | 84640001 | 84740000 | 0.0027 | 0.0013 | 0.0013 |
| F01 | 84660001 | 84760000 | 0.0035 | 0.0020 | 0.0020 |
| F01 | 84680001 | 84780000 | 0.0036 | 0.0022 | 0.0022 |
| F01 | 84700001 | 84800000 | 0.0043 | 0.0028 | 0.0029 |
| F01 | 84720001 | 84820000 | 0.0049 | 0.0035 | 0.0037 |
| F01 | 84740001 | 84840000 | 0.0050 | 0.0036 | 0.0040 |
| F01 | 84760001 | 84860000 | 0.0040 | 0.0027 | 0.0031 |
| F01 | 84780001 | 84880000 | 0.0041 | 0.0027 | 0.0031 |
| F01 | 84800001 | 84900000 | 0.0032 | 0.0020 | 0.0023 |
| F01 | 84820001 | 84920000 | 0.0025 | 0.0013 | 0.0016 |
| F01 | 84840001 | 84940000 | 0.0028 | 0.0014 | 0.0016 |
| F01 | 84860001 | 84960000 | 0.0029 | 0.0015 | 0.0018 |
| F01 | 84880001 | 84980000 | 0.0025 | 0.0013 | 0.0015 |
| F01 | 84900001 | 85000000 | 0.0023 | 0.0013 | 0.0014 |
| F01 | 84920001 | 85020000 | 0.0032 | 0.0021 | 0.0023 |
| F01 | 84940001 | 85040000 | 0.0035 | 0.0024 | 0.0024 |
| F01 | 84960001 | 85060000 | 0.0041 | 0.0032 | 0.0030 |
| F01 | 84980001 | 85080000 | 0.0041 | 0.0032 | 0.0031 |
| F01 | 85000001 | 85100000 | 0.0044 | 0.0035 | 0.0034 |
| F01 | 85020001 | 85120000 | 0.0036 | 0.0029 | 0.0028 |
| F01 | 85040001 | 85140000 | 0.0026 | 0.0023 | 0.0021 |
| F01 | 85060001 | 85160000 | 0.0019 | 0.0014 | 0.0014 |
| F01 | 85080001 | 85180000 | 0.0019 | 0.0013 | 0.0013 |
| F01 | 85100001 | 85200000 | 0.0016 | 0.0011 | 0.0011 |
| F01 | 85120001 | 85220000 | 0.0018 | 0.0013 | 0.0012 |
| F01 | 85140001 | 85240000 | 0.0020 | 0.0014 | 0.0015 |
| F01 | 85160001 | 85260000 | 0.0019 | 0.0014 | 0.0016 |
| F01 | 85180001 | 85280000 | 0.0031 | 0.0024 | 0.0025 |
| F01 | 85200001 | 85300000 | 0.0035 | 0.0024 | 0.0025 |
| F01 | 85220001 | 85320000 | 0.0036 | 0.0021 | 0.0022 |
| F01 | 85240001 | 85340000 | 0.0034 | 0.0020 | 0.0019 |
| F01 | 85260001 | 85360000 | 0.0041 | 0.0026 | 0.0026 |
| F01 | 85280001 | 85380000 | 0.0030 | 0.0017 | 0.0016 |
| F01 | 85300001 | 85400000 | 0.0028 | 0.0019 | 0.0018 |
| F01 | 85320001 | 85420000 | 0.0029 | 0.0024 | 0.0022 |
| F01 | 85340001 | 85440000 | 0.0035 | 0.0030 | 0.0030 |
| F01 | 85360001 | 85460000 | 0.0027 | 0.0022 | 0.0021 |
| F01 | 85380001 | 85480000 | 0.0026 | 0.0021 | 0.0020 |
| F01 | 85400001 | 85500000 | 0.0025 | 0.0017 | 0.0018 |
| F01 | 85420001 | 85520000 | 0.0031 | 0.0020 | 0.0022 |
| F01 | 85440001 | 85540000 | 0.0022 | 0.0013 | 0.0014 |
| F01 | 85460001 | 85560000 | 0.0034 | 0.0022 | 0.0024 |
| F01 | 85480001 | 85580000 | 0.0039 | 0.0026 | 0.0027 |
| F01 | 85500001 | 85600000 | 0.0044 | 0.0032 | 0.0033 |
| F01 | 85520001 | 85620000 | 0.0045 | 0.0034 | 0.0035 |
| F01 | 85540001 | 85640000 | 0.0047 | 0.0035 | 0.0037 |
| F01 | 85560001 | 85660000 | 0.0041 | 0.0033 | 0.0033 |
| F01 | 85580001 | 85680000 | 0.0041 | 0.0035 | 0.0036 |
| F01 | 85600001 | 85700000 | 0.0040 | 0.0030 | 0.0032 |
| F01 | 85620001 | 85720000 | 0.0031 | 0.0022 | 0.0023 |
| F01 | 85640001 | 85740000 | 0.0031 | 0.0022 | 0.0023 |
| F01 | 85660001 | 85760000 | 0.0034 | 0.0022 | 0.0025 |
| F01 | 85680001 | 85780000 | 0.0031 | 0.0018 | 0.0021 |
| F01 | 85700001 | 85800000 | 0.0029 | 0.0019 | 0.0022 |
| F01 | 85720001 | 85820000 | 0.0026 | 0.0017 | 0.0020 |
| F01 | 85740001 | 85840000 | 0.0026 | 0.0020 | 0.0020 |

| Chr | Start | End | both | A1 | A2 |
|---|---|---|---|---|---|
| F01 | 85760001 | 85860000 | 0.0026 | 0.0020 | 0.0018 |
| F01 | 85780001 | 85880000 | 0.0028 | 0.0021 | 0.0019 |
| F01 | 85800001 | 85900000 | 0.0026 | 0.0019 | 0.0018 |
| F01 | 85820001 | 85920000 | 0.0028 | 0.0020 | 0.0020 |
| F01 | 85840001 | 85940000 | 0.0028 | 0.0018 | 0.0020 |
| F01 | 85860001 | 85960000 | 0.0020 | 0.0011 | 0.0014 |
| F01 | 85880001 | 85980000 | 0.0016 | 0.0011 | 0.0013 |
| F01 | 85900001 | 86000000 | 0.0025 | 0.0017 | 0.0022 |
| F01 | 85920001 | 86020000 | 0.0027 | 0.0019 | 0.0024 |
| F01 | 85940001 | 86040000 | 0.0029 | 0.0022 | 0.0027 |
| F01 | 85960001 | 86060000 | 0.0032 | 0.0025 | 0.0030 |
| F01 | 85980001 | 86080000 | 0.0051 | 0.0038 | 0.0045 |
| F01 | 86000001 | 86100000 | 0.0043 | 0.0032 | 0.0037 |
| F01 | 86020001 | 86120000 | 0.0039 | 0.0030 | 0.0033 |
| F01 | 86040001 | 86140000 | 0.0031 | 0.0023 | 0.0026 |
| F01 | 86060001 | 86160000 | 0.0028 | 0.0020 | 0.0024 |
| F01 | 86080001 | 86180000 | 0.0013 | 0.0011 | 0.0012 |
| F01 | 86100001 | 86200000 | 0.0012 | 0.0010 | 0.0012 |
| F01 | 86120001 | 86220000 | 0.0012 | 0.0010 | 0.0012 |
| F01 | 86140001 | 86240000 | 0.0018 | 0.0015 | 0.0018 |
| F01 | 86160001 | 86260000 | 0.0018 | 0.0014 | 0.0017 |
| F01 | 86180001 | 86280000 | 0.0014 | 0.0012 | 0.0014 |
| F01 | 86200001 | 86300000 | 0.0017 | 0.0014 | 0.0017 |
| F01 | 86220001 | 86320000 | 0.0017 | 0.0014 | 0.0016 |
| F01 | 86240001 | 86340000 | 0.0011 | 0.0010 | 0.0011 |
| F01 | 86260001 | 86360000 | 0.0012 | 0.0011 | 0.0012 |
| F01 | 86280001 | 86380000 | 0.0010 | 0.0009 | 0.0010 |
| F01 | 86300001 | 86400000 | 0.0006 | 0.0005 | 0.0005 |
| F01 | 86320001 | 86420000 | 0.0008 | 0.0006 | 0.0008 |
| F01 | 86340001 | 86440000 | 0.0015 | 0.0012 | 0.0014 |
| F01 | 86360001 | 86460000 | 0.0013 | 0.0010 | 0.0013 |
| F01 | 86380001 | 86480000 | 0.0016 | 0.0013 | 0.0015 |
| F01 | 86400001 | 86500000 | 0.0017 | 0.0013 | 0.0015 |
| F01 | 86420001 | 86520000 | 0.0016 | 0.0012 | 0.0014 |
| F01 | 86440001 | 86540000 | 0.0011 | 0.0009 | 0.0010 |
| F01 | 86460001 | 86560000 | 0.0014 | 0.0011 | 0.0013 |
| F01 | 86480001 | 86580000 | 0.0013 | 0.0011 | 0.0013 |
| F01 | 86500001 | 86600000 | 0.0022 | 0.0019 | 0.0021 |
| F01 | 86520001 | 86620000 | 0.0032 | 0.0024 | 0.0029 |
| F01 | 86540001 | 86640000 | 0.0038 | 0.0029 | 0.0032 |
| F01 | 86560001 | 86660000 | 0.0042 | 0.0031 | 0.0034 |
| F01 | 86580001 | 86680000 | 0.0048 | 0.0036 | 0.0038 |
| F01 | 86600001 | 86700000 | 0.0044 | 0.0030 | 0.0033 |
| F01 | 86620001 | 86720000 | 0.0040 | 0.0029 | 0.0030 |
| F01 | 86640001 | 86740000 | 0.0032 | 0.0023 | 0.0024 |
| F01 | 86660001 | 86760000 | 0.0027 | 0.0020 | 0.0021 |
| F01 | 86680001 | 86780000 | 0.0021 | 0.0014 | 0.0017 |
| F01 | 86700001 | 86800000 | 0.0024 | 0.0020 | 0.0021 |
| F01 | 86720001 | 86820000 | 0.0027 | 0.0022 | 0.0025 |
| F01 | 86740001 | 86840000 | 0.0032 | 0.0026 | 0.0028 |
| F01 | 86760001 | 86860000 | 0.0036 | 0.0027 | 0.0029 |
| F01 | 86780001 | 86880000 | 0.0039 | 0.0029 | 0.0030 |
| F01 | 86800001 | 86900000 | 0.0036 | 0.0024 | 0.0025 |
| F01 | 86820001 | 86920000 | 0.0030 | 0.0019 | 0.0019 |
| F01 | 86840001 | 86940000 | 0.0028 | 0.0015 | 0.0017 |
| F01 | 86860001 | 86960000 | 0.0025 | 0.0016 | 0.0016 |
| F01 | 86880001 | 86980000 | 0.0024 | 0.0016 | 0.0017 |
| F01 | 86900001 | 87000000 | 0.0021 | 0.0014 | 0.0015 |

| Chr | Start | End | both | A1 | A2 |
|---|---|---|---|---|---|
| F01 | 86920001 | 87020000 | 0.0027 | 0.0020 | 0.0021 |
| F01 | 86940001 | 87040000 | 0.0028 | 0.0021 | 0.0022 |
| F01 | 86960001 | 87060000 | 0.0024 | 0.0017 | 0.0019 |
| F01 | 86980001 | 87080000 | 0.0018 | 0.0013 | 0.0015 |
| F01 | 87000001 | 87100000 | 0.0024 | 0.0019 | 0.0021 |
| F01 | 87020001 | 87120000 | 0.0018 | 0.0012 | 0.0013 |
| F01 | 87040001 | 87140000 | 0.0024 | 0.0019 | 0.0019 |
| F01 | 87060001 | 87160000 | 0.0031 | 0.0025 | 0.0023 |
| F01 | 87080001 | 87180000 | 0.0039 | 0.0031 | 0.0029 |
| F01 | 87100001 | 87200000 | 0.0041 | 0.0030 | 0.0030 |
| F01 | 87120001 | 87220000 | 0.0040 | 0.0030 | 0.0029 |
| F01 | 87140001 | 87240000 | 0.0038 | 0.0026 | 0.0027 |
| F01 | 87160001 | 87260000 | 0.0038 | 0.0027 | 0.0028 |
| F01 | 87180001 | 87280000 | 0.0035 | 0.0023 | 0.0024 |
| F01 | 87200001 | 87300000 | 0.0031 | 0.0020 | 0.0019 |
| F01 | 87220001 | 87320000 | 0.0029 | 0.0019 | 0.0019 |
| F01 | 87240001 | 87340000 | 0.0053 | 0.0041 | 0.0043 |
| F01 | 87260001 | 87360000 | 0.0051 | 0.0038 | 0.0043 |
| F01 | 87280001 | 87380000 | 0.0053 | 0.0041 | 0.0045 |
| F01 | 87300001 | 87400000 | 0.0054 | 0.0042 | 0.0048 |
| F01 | 87320001 | 87420000 | 0.0057 | 0.0045 | 0.0053 |
| F01 | 87340001 | 87440000 | 0.0029 | 0.0021 | 0.0024 |
| F01 | 87360001 | 87460000 | 0.0032 | 0.0022 | 0.0026 |
| F01 | 87380001 | 87480000 | 0.0050 | 0.0041 | 0.0046 |
| F01 | 87400001 | 87500000 | 0.0049 | 0.0039 | 0.0044 |
| F01 | 87420001 | 87520000 | 0.0057 | 0.0045 | 0.0050 |
| F01 | 87440001 | 87540000 | 0.0060 | 0.0048 | 0.0052 |
| F01 | 87460001 | 87560000 | 0.0068 | 0.0054 | 0.0058 |
| F01 | 87480001 | 87580000 | 0.0044 | 0.0031 | 0.0034 |
| F01 | 87500001 | 87600000 | 0.0047 | 0.0036 | 0.0036 |
| F01 | 87520001 | 87620000 | 0.0042 | 0.0033 | 0.0033 |
| F01 | 87540001 | 87640000 | 0.0042 | 0.0034 | 0.0035 |
| F01 | 87560001 | 87660000 | 0.0041 | 0.0036 | 0.0034 |
| F01 | 87580001 | 87680000 | 0.0044 | 0.0040 | 0.0039 |
| F01 | 87600001 | 87700000 | 0.0041 | 0.0037 | 0.0037 |
| F01 | 87620001 | 87720000 | 0.0034 | 0.0031 | 0.0031 |
| F01 | 87640001 | 87740000 | 0.0033 | 0.0030 | 0.0028 |
| F01 | 87660001 | 87760000 | 0.0024 | 0.0020 | 0.0021 |
| F01 | 87680001 | 87780000 | 0.0024 | 0.0019 | 0.0020 |
| F01 | 87700001 | 87800000 | 0.0025 | 0.0019 | 0.0020 |
| F01 | 87720001 | 87820000 | 0.0041 | 0.0030 | 0.0033 |
| F01 | 87740001 | 87840000 | 0.0042 | 0.0030 | 0.0034 |
| F01 | 87760001 | 87860000 | 0.0042 | 0.0029 | 0.0033 |
| F01 | 87780001 | 87880000 | 0.0038 | 0.0027 | 0.0031 |
| F01 | 87800001 | 87900000 | 0.0045 | 0.0035 | 0.0038 |
| F01 | 87820001 | 87920000 | 0.0031 | 0.0024 | 0.0025 |
| F01 | 87840001 | 87940000 | 0.0025 | 0.0021 | 0.0021 |
| F01 | 87860001 | 87960000 | 0.0028 | 0.0022 | 0.0023 |
| F01 | 87880001 | 87980000 | 0.0034 | 0.0024 | 0.0026 |
| F01 | 87900001 | 88000000 | 0.0032 | 0.0020 | 0.0023 |
| F01 | 87920001 | 88020000 | 0.0033 | 0.0019 | 0.0022 |
| F01 | 87940001 | 88040000 | 0.0039 | 0.0024 | 0.0027 |
| F01 | 87960001 | 88060000 | 0.0036 | 0.0021 | 0.0023 |
| F01 | 87980001 | 88080000 | 0.0033 | 0.0020 | 0.0021 |
| F01 | 88000001 | 88100000 | 0.0027 | 0.0014 | 0.0016 |
| F01 | 88020001 | 88120000 | 0.0026 | 0.0014 | 0.0016 |
| F01 | 88040001 | 88140000 | 0.0023 | 0.0010 | 0.0012 |
| F01 | 88060001 | 88160000 | 0.0022 | 0.0011 | 0.0011 |

| Chr | Start | End | both | A1 | A2 |
|---|---|---|---|---|---|
| F01 | 88080001 | 88180000 | 0.0021 | 0.0011 | 0.0012 |
| F01 | 88100001 | 88200000 | 0.0027 | 0.0016 | 0.0018 |
| F01 | 88120001 | 88220000 | 0.0029 | 0.0018 | 0.0019 |
| F01 | 88140001 | 88240000 | 0.0029 | 0.0021 | 0.0022 |
| F01 | 88160001 | 88260000 | 0.0031 | 0.0022 | 0.0025 |
| F01 | 88180001 | 88280000 | 0.0032 | 0.0025 | 0.0026 |
| F01 | 88200001 | 88300000 | 0.0027 | 0.0021 | 0.0023 |
| F01 | 88220001 | 88320000 | 0.0023 | 0.0019 | 0.0021 |
| F01 | 88240001 | 88340000 | 0.0024 | 0.0016 | 0.0019 |
| F01 | 88260001 | 88360000 | 0.0037 | 0.0027 | 0.0032 |
| F01 | 88280001 | 88380000 | 0.0037 | 0.0023 | 0.0028 |
| F01 | 88300001 | 88400000 | 0.0034 | 0.0020 | 0.0024 |
| F01 | 88320001 | 88420000 | 0.0041 | 0.0026 | 0.0029 |
| F01 | 88340001 | 88440000 | 0.0044 | 0.0031 | 0.0035 |
| F01 | 88360001 | 88460000 | 0.0034 | 0.0021 | 0.0023 |
| F01 | 88380001 | 88480000 | 0.0036 | 0.0025 | 0.0026 |
| F01 | 88400001 | 88500000 | 0.0040 | 0.0028 | 0.0029 |
| F01 | 88420001 | 88520000 | 0.0036 | 0.0024 | 0.0024 |
| F01 | 88440001 | 88540000 | 0.0033 | 0.0020 | 0.0020 |
| F01 | 88460001 | 88560000 | 0.0036 | 0.0023 | 0.0023 |
| F01 | 88480001 | 88580000 | 0.0032 | 0.0019 | 0.0019 |
| F01 | 88500001 | 88600000 | 0.0035 | 0.0021 | 0.0022 |
| F01 | 88520001 | 88620000 | 0.0038 | 0.0023 | 0.0026 |
| F01 | 88540001 | 88640000 | 0.0039 | 0.0023 | 0.0025 |
| F01 | 88560001 | 88660000 | 0.0037 | 0.0024 | 0.0025 |
| F01 | 88580001 | 88680000 | 0.0041 | 0.0028 | 0.0029 |
| F01 | 88600001 | 88700000 | 0.0037 | 0.0027 | 0.0028 |
| F01 | 88620001 | 88720000 | 0.0037 | 0.0028 | 0.0030 |
| F01 | 88640001 | 88740000 | 0.0037 | 0.0029 | 0.0030 |
| F01 | 88660001 | 88760000 | 0.0032 | 0.0025 | 0.0026 |
| F01 | 88680001 | 88780000 | 0.0028 | 0.0021 | 0.0023 |
| F01 | 88700001 | 88800000 | 0.0033 | 0.0026 | 0.0028 |
| F01 | 88720001 | 88820000 | 0.0032 | 0.0026 | 0.0026 |
| F01 | 88740001 | 88840000 | 0.0033 | 0.0026 | 0.0027 |
| F01 | 88760001 | 88860000 | 0.0034 | 0.0026 | 0.0028 |
| F01 | 88780001 | 88880000 | 0.0032 | 0.0025 | 0.0027 |
| F01 | 88800001 | 88900000 | 0.0024 | 0.0018 | 0.0020 |
| F01 | 88820001 | 88920000 | 0.0017 | 0.0012 | 0.0014 |
| F01 | 88840001 | 88940000 | 0.0024 | 0.0018 | 0.0020 |
| F01 | 88860001 | 88960000 | 0.0025 | 0.0020 | 0.0022 |
| F01 | 88880001 | 88980000 | 0.0029 | 0.0023 | 0.0025 |
| F01 | 88900001 | 89000000 | 0.0027 | 0.0022 | 0.0022 |
| F01 | 88920001 | 89020000 | 0.0031 | 0.0025 | 0.0025 |
| F01 | 88940001 | 89040000 | 0.0030 | 0.0024 | 0.0025 |
| F01 | 88960001 | 89060000 | 0.0030 | 0.0025 | 0.0025 |
| F01 | 88980001 | 89080000 | 0.0028 | 0.0023 | 0.0024 |
| F01 | 89000001 | 89100000 | 0.0031 | 0.0026 | 0.0028 |
| F01 | 89020001 | 89120000 | 0.0030 | 0.0024 | 0.0026 |
| F01 | 89040001 | 89140000 | 0.0019 | 0.0015 | 0.0016 |
| F01 | 89060001 | 89160000 | 0.0019 | 0.0014 | 0.0015 |
| F01 | 89080001 | 89180000 | 0.0019 | 0.0014 | 0.0014 |
| F01 | 89100001 | 89200000 | 0.0020 | 0.0013 | 0.0015 |
| F01 | 89120001 | 89220000 | 0.0022 | 0.0012 | 0.0015 |
| F01 | 89140001 | 89240000 | 0.0021 | 0.0011 | 0.0014 |
| F01 | 89160001 | 89260000 | 0.0018 | 0.0008 | 0.0012 |
| F01 | 89180001 | 89280000 | 0.0017 | 0.0007 | 0.0010 |
| F01 | 89200001 | 89300000 | 0.0013 | 0.0003 | 0.0005 |
| F01 | 89220001 | 89320000 | 0.0015 | 0.0007 | 0.0006 |

| Chr | Start | End | both | A1 | A2 |
|---|---|---|---|---|---|
| F01 | 89240001 | 89340000 | 0.0016 | 0.0008 | 0.0008 |
| F01 | 89260001 | 89360000 | 0.0015 | 0.0008 | 0.0007 |
| F01 | 89280001 | 89380000 | 0.0013 | 0.0009 | 0.0008 |
| F01 | 89300001 | 89400000 | 0.0015 | 0.0012 | 0.0011 |
| F01 | 89320001 | 89420000 | 0.0010 | 0.0008 | 0.0009 |
| F01 | 89340001 | 89440000 | 0.0010 | 0.0008 | 0.0009 |
| F01 | 89360001 | 89460000 | 0.0011 | 0.0009 | 0.0010 |
| F01 | 89380001 | 89480000 | 0.0011 | 0.0009 | 0.0010 |
| F01 | 89400001 | 89500000 | 0.0017 | 0.0014 | 0.0016 |
| F01 | 89420001 | 89520000 | 0.0016 | 0.0012 | 0.0015 |
| F01 | 89440001 | 89540000 | 0.0019 | 0.0015 | 0.0017 |
| F01 | 89460001 | 89560000 | 0.0021 | 0.0017 | 0.0019 |
| F01 | 89480001 | 89580000 | 0.0023 | 0.0020 | 0.0022 |
| F01 | 89500001 | 89600000 | 0.0016 | 0.0015 | 0.0015 |
| F01 | 89520001 | 89620000 | 0.0033 | 0.0032 | 0.0030 |
| F01 | 89540001 | 89640000 | 0.0036 | 0.0035 | 0.0032 |
| F01 | 89560001 | 89660000 | 0.0039 | 0.0038 | 0.0034 |
| F01 | 89580001 | 89680000 | 0.0039 | 0.0037 | 0.0032 |
| F01 | 89600001 | 89700000 | 0.0044 | 0.0039 | 0.0035 |
| F01 | 89620001 | 89720000 | 0.0034 | 0.0027 | 0.0026 |
| F01 | 89640001 | 89740000 | 0.0028 | 0.0022 | 0.0020 |
| F01 | 89660001 | 89760000 | 0.0024 | 0.0014 | 0.0015 |
| F01 | 89680001 | 89780000 | 0.0026 | 0.0016 | 0.0017 |
| F01 | 89700001 | 89800000 | 0.0030 | 0.0019 | 0.0021 |
| F01 | 89720001 | 89820000 | 0.0028 | 0.0017 | 0.0019 |
| F01 | 89740001 | 89840000 | 0.0029 | 0.0018 | 0.0021 |
| F01 | 89760001 | 89860000 | 0.0032 | 0.0022 | 0.0025 |
| F01 | 89780001 | 89880000 | 0.0032 | 0.0022 | 0.0026 |
| F01 | 89800001 | 89900000 | 0.0027 | 0.0021 | 0.0023 |
| F01 | 89820001 | 89920000 | 0.0029 | 0.0022 | 0.0024 |
| F01 | 89840001 | 89940000 | 0.0030 | 0.0021 | 0.0022 |
| F01 | 89860001 | 89960000 | 0.0027 | 0.0018 | 0.0019 |
| F01 | 89880001 | 89980000 | 0.0024 | 0.0015 | 0.0015 |
| F01 | 89900001 | 90000000 | 0.0024 | 0.0013 | 0.0015 |
| F01 | 89920001 | 90020000 | 0.0022 | 0.0012 | 0.0015 |
| F01 | 89940001 | 90040000 | 0.0023 | 0.0014 | 0.0016 |
| F01 | 89960001 | 90060000 | 0.0025 | 0.0016 | 0.0018 |
| F01 | 89980001 | 90080000 | 0.0026 | 0.0017 | 0.0020 |
| F01 | 90000001 | 90100000 | 0.0024 | 0.0014 | 0.0016 |
| F01 | 90020001 | 90120000 | 0.0023 | 0.0012 | 0.0014 |
| F01 | 90040001 | 90140000 | 0.0023 | 0.0014 | 0.0017 |
| F01 | 90060001 | 90160000 | 0.0027 | 0.0019 | 0.0021 |
| F01 | 90080001 | 90180000 | 0.0038 | 0.0029 | 0.0031 |
| F01 | 90100001 | 90200000 | 0.0037 | 0.0029 | 0.0031 |
| F01 | 90120001 | 90220000 | 0.0049 | 0.0040 | 0.0042 |
| F01 | 90140001 | 90240000 | 0.0048 | 0.0038 | 0.0040 |
| F01 | 90160001 | 90260000 | 0.0047 | 0.0035 | 0.0037 |
| F01 | 90180001 | 90280000 | 0.0037 | 0.0024 | 0.0027 |
| F01 | 90200001 | 90300000 | 0.0040 | 0.0027 | 0.0030 |
| F01 | 90220001 | 90320000 | 0.0035 | 0.0024 | 0.0026 |
| F01 | 90240001 | 90340000 | 0.0039 | 0.0025 | 0.0028 |
| F01 | 90260001 | 90360000 | 0.0035 | 0.0020 | 0.0023 |
| F01 | 90280001 | 90380000 | 0.0042 | 0.0025 | 0.0029 |
| F01 | 90300001 | 90400000 | 0.0042 | 0.0024 | 0.0026 |
| F01 | 90320001 | 90420000 | 0.0036 | 0.0018 | 0.0019 |
| F01 | 90340001 | 90440000 | 0.0039 | 0.0023 | 0.0023 |
| F01 | 90360001 | 90460000 | 0.0039 | 0.0025 | 0.0024 |
| F01 | 90380001 | 90480000 | 0.0032 | 0.0023 | 0.0020 |

| Chr | Start | End | both | A1 | A2 |
|---|---|---|---|---|---|
| F01 | 90400001 | 90500000 | 0.0040 | 0.0030 | 0.0026 |
| F01 | 90420001 | 90520000 | 0.0043 | 0.0035 | 0.0030 |
| F01 | 90440001 | 90540000 | 0.0040 | 0.0033 | 0.0029 |
| F01 | 90460001 | 90560000 | 0.0086 | 0.0068 | 0.0076 |
| F01 | 90480001 | 90580000 | 0.0111 | 0.0090 | 0.0103 |
| F01 | 90500001 | 90600000 | 0.0128 | 0.0105 | 0.0123 |
| F01 | 90520001 | 90620000 | 0.0138 | 0.0113 | 0.0134 |
| F01 | 90540001 | 90640000 | 0.0134 | 0.0109 | 0.0131 |
| F01 | 90560001 | 90660000 | 0.0091 | 0.0078 | 0.0088 |
| F01 | 90580001 | 90680000 | 0.0063 | 0.0054 | 0.0059 |
| F01 | 90600001 | 90700000 | 0.0037 | 0.0032 | 0.0033 |
| F01 | 90620001 | 90720000 | 0.0020 | 0.0018 | 0.0017 |
| F01 | 90640001 | 90740000 | 0.0017 | 0.0016 | 0.0012 |
| F01 | 90660001 | 90760000 | 0.0012 | 0.0010 | 0.0007 |
| F01 | 90680001 | 90780000 | 0.0019 | 0.0015 | 0.0012 |
| F01 | 90700001 | 90800000 | 0.0025 | 0.0021 | 0.0017 |
| F01 | 90720001 | 90820000 | 0.0026 | 0.0021 | 0.0018 |
| F01 | 90740001 | 90840000 | 0.0033 | 0.0027 | 0.0025 |
| F01 | 90760001 | 90860000 | 0.0038 | 0.0032 | 0.0031 |
| F01 | 90780001 | 90880000 | 0.0036 | 0.0030 | 0.0030 |
| F01 | 90800001 | 90900000 | 0.0036 | 0.0026 | 0.0031 |
| F01 | 90820001 | 90920000 | 0.0037 | 0.0026 | 0.0032 |
| F01 | 90840001 | 90940000 | 0.0036 | 0.0024 | 0.0032 |
| F01 | 90860001 | 90960000 | 0.0033 | 0.0022 | 0.0030 |
| F01 | 90880001 | 90980000 | 0.0031 | 0.0019 | 0.0028 |
| F01 | 90900001 | 91000000 | 0.0025 | 0.0018 | 0.0022 |
| F01 | 90920001 | 91020000 | 0.0022 | 0.0017 | 0.0020 |
| F01 | 90940001 | 91040000 | 0.0024 | 0.0019 | 0.0022 |
| F01 | 90960001 | 91060000 | 0.0026 | 0.0020 | 0.0023 |
| F01 | 90980001 | 91080000 | 0.0023 | 0.0019 | 0.0021 |
| F01 | 91000001 | 91100000 | 0.0023 | 0.0018 | 0.0022 |
| F01 | 91020001 | 91120000 | 0.0025 | 0.0019 | 0.0022 |
| F01 | 91040001 | 91140000 | 0.0027 | 0.0024 | 0.0023 |
| F01 | 91060001 | 91160000 | 0.0032 | 0.0027 | 0.0026 |
| F01 | 91080001 | 91180000 | 0.0037 | 0.0031 | 0.0029 |
| F01 | 91100001 | 91200000 | 0.0042 | 0.0036 | 0.0034 |
| F01 | 91120001 | 91220000 | 0.0044 | 0.0038 | 0.0036 |
| F01 | 91140001 | 91240000 | 0.0033 | 0.0025 | 0.0027 |
| F01 | 91160001 | 91260000 | 0.0027 | 0.0021 | 0.0023 |
| F01 | 91180001 | 91280000 | 0.0033 | 0.0028 | 0.0031 |
| F01 | 91200001 | 91300000 | 0.0026 | 0.0021 | 0.0023 |
| F01 | 91220001 | 91320000 | 0.0025 | 0.0021 | 0.0021 |
| F01 | 91240001 | 91340000 | 0.0031 | 0.0027 | 0.0026 |
| F01 | 91260001 | 91360000 | 0.0038 | 0.0033 | 0.0031 |
| F01 | 91280001 | 91380000 | 0.0033 | 0.0026 | 0.0024 |
| F01 | 91300001 | 91400000 | 0.0037 | 0.0030 | 0.0029 |
| F01 | 91320001 | 91420000 | 0.0038 | 0.0030 | 0.0031 |
| F01 | 91340001 | 91440000 | 0.0032 | 0.0025 | 0.0025 |
| F01 | 91360001 | 91460000 | 0.0034 | 0.0026 | 0.0026 |
| F01 | 91380001 | 91480000 | 0.0029 | 0.0023 | 0.0023 |
| F01 | 91400001 | 91500000 | 0.0032 | 0.0026 | 0.0024 |
| F01 | 91420001 | 91520000 | 0.0035 | 0.0029 | 0.0029 |
| F01 | 91440001 | 91540000 | 0.0048 | 0.0043 | 0.0041 |
| F01 | 91460001 | 91560000 | 0.0060 | 0.0057 | 0.0052 |
| F01 | 91480001 | 91580000 | 0.0062 | 0.0062 | 0.0053 |
| F01 | 91500001 | 91600000 | 0.0057 | 0.0059 | 0.0049 |
| F01 | 91520001 | 91620000 | 0.0049 | 0.0053 | 0.0041 |
| F01 | 91540001 | 91640000 | 0.0039 | 0.0043 | 0.0034 |

| Chr | Start | End | both | A1 | A2 |
|---|---|---|---|---|---|
| F01 | 91560001 | 91660000 | 0.0022 | 0.0024 | 0.0019 |
| F01 | 91580001 | 91680000 | 0.0024 | 0.0023 | 0.0022 |
| F01 | 91600001 | 91700000 | 0.0026 | 0.0024 | 0.0024 |
| F01 | 91620001 | 91720000 | 0.0032 | 0.0029 | 0.0029 |
| F01 | 91640001 | 91740000 | 0.0029 | 0.0024 | 0.0024 |
| F01 | 91660001 | 91760000 | 0.0028 | 0.0024 | 0.0023 |
| F01 | 91680001 | 91780000 | 0.0027 | 0.0022 | 0.0020 |
| F01 | 91700001 | 91800000 | 0.0029 | 0.0022 | 0.0019 |
| F01 | 91720001 | 91820000 | 0.0029 | 0.0022 | 0.0018 |
| F01 | 91740001 | 91840000 | 0.0029 | 0.0022 | 0.0019 |
| F01 | 91760001 | 91860000 | 0.0032 | 0.0023 | 0.0022 |
| F01 | 91780001 | 91880000 | 0.0028 | 0.0019 | 0.0021 |
| F01 | 91800001 | 91900000 | 0.0030 | 0.0020 | 0.0026 |
| F01 | 91820001 | 91920000 | 0.0029 | 0.0018 | 0.0028 |
| F01 | 91840001 | 91940000 | 0.0030 | 0.0019 | 0.0031 |
| F01 | 91860001 | 91960000 | 0.0036 | 0.0023 | 0.0037 |
| F01 | 91880001 | 91980000 | 0.0041 | 0.0028 | 0.0042 |
| F01 | 91900001 | 92000000 | 0.0040 | 0.0028 | 0.0041 |
| F01 | 91920001 | 92020000 | 0.0044 | 0.0032 | 0.0045 |
| F01 | 91940001 | 92040000 | 0.0046 | 0.0034 | 0.0048 |
| F01 | 91960001 | 92060000 | 0.0039 | 0.0027 | 0.0040 |
| F01 | 91980001 | 92080000 | 0.0036 | 0.0026 | 0.0036 |
| F01 | 92000001 | 92100000 | 0.0040 | 0.0030 | 0.0041 |
| F01 | 92020001 | 92120000 | 0.0058 | 0.0047 | 0.0059 |
| F01 | 92040001 | 92140000 | 0.0068 | 0.0054 | 0.0069 |
| F01 | 92060001 | 92160000 | 0.0065 | 0.0051 | 0.0066 |
| F01 | 92080001 | 92180000 | 0.0075 | 0.0059 | 0.0076 |
| F01 | 92100001 | 92200000 | 0.0072 | 0.0055 | 0.0073 |
| F01 | 92120001 | 92220000 | 0.0047 | 0.0032 | 0.0048 |
| F01 | 92140001 | 92240000 | 0.0037 | 0.0026 | 0.0038 |
| F01 | 92160001 | 92260000 | 0.0045 | 0.0033 | 0.0046 |
| F01 | 92180001 | 92280000 | 0.0039 | 0.0026 | 0.0040 |
| F01 | 92200001 | 92300000 | 0.0039 | 0.0029 | 0.0039 |
| F01 | 92220001 | 92320000 | 0.0042 | 0.0033 | 0.0042 |
| F01 | 92240001 | 92340000 | 0.0042 | 0.0031 | 0.0042 |
| F01 | 92260001 | 92360000 | 0.0033 | 0.0027 | 0.0033 |
| F01 | 92280001 | 92380000 | 0.0035 | 0.0028 | 0.0035 |
| F01 | 92300001 | 92400000 | 0.0035 | 0.0023 | 0.0036 |
| F01 | 92320001 | 92420000 | 0.0030 | 0.0019 | 0.0031 |
| F01 | 92340001 | 92440000 | 0.0030 | 0.0017 | 0.0031 |
| F01 | 92360001 | 92460000 | 0.0028 | 0.0015 | 0.0029 |
| F01 | 92380001 | 92480000 | 0.0024 | 0.0014 | 0.0024 |
| F01 | 92400001 | 92500000 | 0.0027 | 0.0021 | 0.0027 |
| F01 | 92420001 | 92520000 | 0.0032 | 0.0026 | 0.0032 |
| F01 | 92440001 | 92540000 | 0.0034 | 0.0026 | 0.0034 |
| F01 | 92460001 | 92560000 | 0.0040 | 0.0028 | 0.0040 |
| F01 | 92480001 | 92580000 | 0.0038 | 0.0024 | 0.0038 |
| F01 | 92500001 | 92600000 | 0.0030 | 0.0016 | 0.0030 |
| F01 | 92520001 | 92620000 | 0.0034 | 0.0019 | 0.0034 |
| F01 | 92540001 | 92640000 | 0.0034 | 0.0020 | 0.0033 |
| F01 | 92560001 | 92660000 | 0.0030 | 0.0017 | 0.0029 |
| F01 | 92580001 | 92680000 | 0.0032 | 0.0020 | 0.0029 |
| F01 | 92600001 | 92700000 | 0.0032 | 0.0021 | 0.0029 |
| F01 | 92620001 | 92720000 | 0.0025 | 0.0012 | 0.0020 |
| F01 | 92640001 | 92740000 | 0.0021 | 0.0010 | 0.0016 |
| F01 | 92660001 | 92760000 | 0.0026 | 0.0016 | 0.0020 |
| F01 | 92680001 | 92780000 | 0.0026 | 0.0016 | 0.0018 |
| F01 | 92700001 | 92800000 | 0.0032 | 0.0020 | 0.0021 |

| Chr | Start | End | both | A1 | A2 |
|---|---|---|---|---|---|
| F01 | 92720001 | 92820000 | 0.0031 | 0.0021 | 0.0021 |
| F01 | 92740001 | 92840000 | 0.0029 | 0.0021 | 0.0020 |
| F01 | 92760001 | 92860000 | 0.0027 | 0.0020 | 0.0018 |
| F01 | 92780001 | 92880000 | 0.0031 | 0.0025 | 0.0023 |
| F01 | 92800001 | 92900000 | 0.0029 | 0.0025 | 0.0024 |
| F01 | 92820001 | 92920000 | 0.0027 | 0.0023 | 0.0022 |
| F01 | 92840001 | 92940000 | 0.0030 | 0.0024 | 0.0023 |
| F01 | 92860001 | 92960000 | 0.0028 | 0.0021 | 0.0021 |
| F01 | 92880001 | 92980000 | 0.0027 | 0.0018 | 0.0020 |
| F01 | 92900001 | 93000000 | 0.0028 | 0.0018 | 0.0020 |
| F01 | 92920001 | 93020000 | 0.0030 | 0.0019 | 0.0022 |
| F01 | 92940001 | 93040000 | 0.0029 | 0.0018 | 0.0022 |
| F01 | 92960001 | 93060000 | 0.0026 | 0.0016 | 0.0019 |
| F01 | 92980001 | 93080000 | 0.0026 | 0.0016 | 0.0020 |
| F01 | 93000001 | 93100000 | 0.0021 | 0.0012 | 0.0016 |
| F01 | 93020001 | 93120000 | 0.0026 | 0.0019 | 0.0022 |
| F01 | 93040001 | 93140000 | 0.0035 | 0.0026 | 0.0033 |
| F01 | 93060001 | 93160000 | 0.0037 | 0.0027 | 0.0035 |
| F01 | 93080001 | 93180000 | 0.0036 | 0.0024 | 0.0034 |
| F01 | 93100001 | 93200000 | 0.0034 | 0.0022 | 0.0033 |
| F01 | 93120001 | 93220000 | 0.0030 | 0.0018 | 0.0029 |
| F01 | 93140001 | 93240000 | 0.0034 | 0.0022 | 0.0034 |
| F01 | 93160001 | 93260000 | 0.0038 | 0.0027 | 0.0036 |
| F01 | 93180001 | 93280000 | 0.0033 | 0.0026 | 0.0031 |
| F01 | 93200001 | 93300000 | 0.0033 | 0.0027 | 0.0030 |
| F01 | 93220001 | 93320000 | 0.0032 | 0.0026 | 0.0027 |
| F01 | 93240001 | 93340000 | 0.0029 | 0.0024 | 0.0021 |
| F01 | 93260001 | 93360000 | 0.0024 | 0.0019 | 0.0016 |
| F01 | 93280001 | 93380000 | 0.0024 | 0.0019 | 0.0015 |
| F01 | 93300001 | 93400000 | 0.0026 | 0.0019 | 0.0016 |
| F01 | 93320001 | 93420000 | 0.0040 | 0.0033 | 0.0032 |
| F01 | 93340001 | 93440000 | 0.0048 | 0.0042 | 0.0044 |
| F01 | 93360001 | 93460000 | 0.0054 | 0.0048 | 0.0051 |
| F01 | 93380001 | 93480000 | 0.0053 | 0.0047 | 0.0050 |
| F01 | 93400001 | 93500000 | 0.0054 | 0.0047 | 0.0052 |
| F01 | 93420001 | 93520000 | 0.0042 | 0.0033 | 0.0040 |
| F01 | 93440001 | 93540000 | 0.0024 | 0.0015 | 0.0020 |
| F01 | 93460001 | 93560000 | 0.0026 | 0.0016 | 0.0020 |
| F01 | 93480001 | 93580000 | 0.0029 | 0.0017 | 0.0019 |
| F01 | 93500001 | 93600000 | 0.0030 | 0.0019 | 0.0020 |
| F01 | 93520001 | 93620000 | 0.0027 | 0.0017 | 0.0014 |
| F01 | 93540001 | 93640000 | 0.0027 | 0.0017 | 0.0014 |
| F01 | 93560001 | 93660000 | 0.0022 | 0.0013 | 0.0010 |
| F01 | 93580001 | 93680000 | 0.0019 | 0.0012 | 0.0011 |
| F01 | 93600001 | 93700000 | 0.0020 | 0.0014 | 0.0014 |
| F01 | 93620001 | 93720000 | 0.0037 | 0.0030 | 0.0032 |
| F01 | 93640001 | 93740000 | 0.0037 | 0.0030 | 0.0033 |
| F01 | 93660001 | 93760000 | 0.0037 | 0.0030 | 0.0031 |
| F01 | 93680001 | 93780000 | 0.0039 | 0.0031 | 0.0032 |
| F01 | 93700001 | 93800000 | 0.0041 | 0.0035 | 0.0033 |
| F01 | 93720001 | 93820000 | 0.0026 | 0.0020 | 0.0017 |
| F01 | 93740001 | 93840000 | 0.0033 | 0.0028 | 0.0026 |
| F01 | 93760001 | 93860000 | 0.0032 | 0.0026 | 0.0024 |
| F01 | 93780001 | 93880000 | 0.0030 | 0.0024 | 0.0022 |
| F01 | 93800001 | 93900000 | 0.0022 | 0.0015 | 0.0013 |
| F01 | 93820001 | 93920000 | 0.0023 | 0.0016 | 0.0013 |
| F01 | 93840001 | 93940000 | 0.0020 | 0.0014 | 0.0010 |
| F01 | 93860001 | 93960000 | 0.0026 | 0.0018 | 0.0015 |

| Chr | Start | End | both | A1 | A2 |
|---|---|---|---|---|---|
| F01 | 93880001 | 93980000 | 0.0032 | 0.0024 | 0.0020 |
| F01 | 93900001 | 94000000 | 0.0034 | 0.0025 | 0.0021 |
| F01 | 93920001 | 94020000 | 0.0043 | 0.0035 | 0.0032 |
| F01 | 93940001 | 94040000 | 0.0045 | 0.0035 | 0.0035 |
| F01 | 93960001 | 94060000 | 0.0058 | 0.0050 | 0.0050 |
| F01 | 93980001 | 94080000 | 0.0050 | 0.0043 | 0.0045 |
| F01 | 94000001 | 94100000 | 0.0053 | 0.0046 | 0.0048 |
| F01 | 94020001 | 94120000 | 0.0046 | 0.0038 | 0.0039 |
| F01 | 94040001 | 94140000 | 0.0042 | 0.0037 | 0.0036 |
| F01 | 94060001 | 94160000 | 0.0028 | 0.0021 | 0.0020 |
| F01 | 94080001 | 94180000 | 0.0030 | 0.0022 | 0.0021 |
| F01 | 94100001 | 94200000 | 0.0029 | 0.0021 | 0.0020 |
| F01 | 94120001 | 94220000 | 0.0028 | 0.0020 | 0.0020 |
| F01 | 94140001 | 94240000 | 0.0024 | 0.0015 | 0.0016 |
| F01 | 94160001 | 94260000 | 0.0020 | 0.0011 | 0.0013 |
| F01 | 94180001 | 94280000 | 0.0021 | 0.0013 | 0.0016 |
| F01 | 94200001 | 94300000 | 0.0026 | 0.0018 | 0.0021 |
| F01 | 94220001 | 94320000 | 0.0028 | 0.0020 | 0.0022 |
| F01 | 94240001 | 94340000 | 0.0028 | 0.0022 | 0.0023 |
| F01 | 94260001 | 94360000 | 0.0027 | 0.0021 | 0.0023 |
| F01 | 94280001 | 94380000 | 0.0029 | 0.0021 | 0.0024 |
| F01 | 94300001 | 94400000 | 0.0034 | 0.0026 | 0.0030 |
| F01 | 94320001 | 94420000 | 0.0030 | 0.0019 | 0.0027 |
| F01 | 94340001 | 94440000 | 0.0030 | 0.0018 | 0.0028 |
| F01 | 94360001 | 94460000 | 0.0032 | 0.0018 | 0.0030 |
| F01 | 94380001 | 94480000 | 0.0030 | 0.0016 | 0.0030 |
| F01 | 94400001 | 94500000 | 0.0018 | 0.0005 | 0.0019 |
| F01 | 94420001 | 94520000 | 0.0021 | 0.0008 | 0.0022 |
| F01 | 94440001 | 94540000 | 0.0033 | 0.0022 | 0.0034 |
| F01 | 94460001 | 94560000 | 0.0034 | 0.0022 | 0.0035 |
| F01 | 94480001 | 94580000 | 0.0033 | 0.0022 | 0.0034 |
| F01 | 94500001 | 94600000 | 0.0033 | 0.0022 | 0.0034 |
| F01 | 94520001 | 94620000 | 0.0031 | 0.0019 | 0.0032 |
| F01 | 94540001 | 94640000 | 0.0017 | 0.0005 | 0.0018 |
| F01 | 94560001 | 94660000 | 0.0017 | 0.0006 | 0.0018 |
| F01 | 94580001 | 94680000 | 0.0018 | 0.0007 | 0.0017 |
| F01 | 94600001 | 94700000 | 0.0017 | 0.0006 | 0.0014 |
| F01 | 94620001 | 94720000 | 0.0019 | 0.0008 | 0.0013 |
| F01 | 94640001 | 94740000 | 0.0020 | 0.0008 | 0.0012 |
| F01 | 94660001 | 94760000 | 0.0018 | 0.0007 | 0.0009 |
| F01 | 94680001 | 94780000 | 0.0020 | 0.0010 | 0.0011 |
| F01 | 94700001 | 94800000 | 0.0025 | 0.0013 | 0.0016 |
| F01 | 94720001 | 94820000 | 0.0020 | 0.0011 | 0.0013 |
| F01 | 94740001 | 94840000 | 0.0019 | 0.0011 | 0.0012 |
| F01 | 94760001 | 94860000 | 0.0021 | 0.0011 | 0.0014 |
| F01 | 94780001 | 94880000 | 0.0018 | 0.0008 | 0.0012 |
| F01 | 94800001 | 94900000 | 0.0018 | 0.0008 | 0.0011 |
| F01 | 94820001 | 94920000 | 0.0032 | 0.0020 | 0.0023 |
| F01 | 94840001 | 94940000 | 0.0032 | 0.0019 | 0.0022 |
| F01 | 94860001 | 94960000 | 0.0041 | 0.0028 | 0.0032 |
| F01 | 94880001 | 94980000 | 0.0050 | 0.0037 | 0.0040 |
| F01 | 94900001 | 95000000 | 0.0050 | 0.0035 | 0.0039 |
| F01 | 94920001 | 95020000 | 0.0037 | 0.0023 | 0.0029 |
| F01 | 94940001 | 95040000 | 0.0041 | 0.0027 | 0.0031 |
| F01 | 94960001 | 95060000 | 0.0038 | 0.0024 | 0.0026 |
| F01 | 94980001 | 95080000 | 0.0030 | 0.0018 | 0.0019 |
| F01 | 95000001 | 95100000 | 0.0049 | 0.0038 | 0.0039 |
| F01 | 95020001 | 95120000 | 0.0051 | 0.0039 | 0.0040 |

| Chr | Start | End | both | A1 | A2 |
|---|---|---|---|---|---|
| F01 | 95040001 | 95140000 | 0.0048 | 0.0036 | 0.0038 |
| F01 | 95060001 | 95160000 | 0.0041 | 0.0029 | 0.0031 |
| F01 | 95080001 | 95180000 | 0.0041 | 0.0028 | 0.0031 |
| F01 | 95100001 | 95200000 | 0.0018 | 0.0007 | 0.0007 |
| F01 | 95120001 | 95220000 | 0.0018 | 0.0011 | 0.0008 |
| F01 | 95140001 | 95240000 | 0.0020 | 0.0014 | 0.0010 |
| F01 | 95160001 | 95260000 | 0.0021 | 0.0016 | 0.0012 |
| F01 | 95180001 | 95280000 | 0.0024 | 0.0020 | 0.0016 |
| F01 | 95200001 | 95300000 | 0.0027 | 0.0021 | 0.0019 |
| F01 | 95220001 | 95320000 | 0.0026 | 0.0016 | 0.0018 |
| F01 | 95240001 | 95340000 | 0.0025 | 0.0014 | 0.0016 |
| F01 | 95260001 | 95360000 | 0.0027 | 0.0016 | 0.0019 |
| F01 | 95280001 | 95380000 | 0.0026 | 0.0017 | 0.0019 |
| F01 | 95300001 | 95400000 | 0.0032 | 0.0020 | 0.0023 |
| F01 | 95320001 | 95420000 | 0.0032 | 0.0021 | 0.0026 |
| F01 | 95340001 | 95440000 | 0.0031 | 0.0021 | 0.0026 |
| F01 | 95360001 | 95460000 | 0.0035 | 0.0024 | 0.0030 |
| F01 | 95380001 | 95480000 | 0.0037 | 0.0027 | 0.0034 |
| F01 | 95400001 | 95500000 | 0.0031 | 0.0023 | 0.0030 |
| F01 | 95420001 | 95520000 | 0.0030 | 0.0025 | 0.0029 |
| F01 | 95440001 | 95540000 | 0.0032 | 0.0026 | 0.0031 |
| F01 | 95460001 | 95560000 | 0.0026 | 0.0021 | 0.0025 |
| F01 | 95480001 | 95580000 | 0.0027 | 0.0019 | 0.0026 |
| F01 | 95500001 | 95600000 | 0.0030 | 0.0022 | 0.0029 |
| F01 | 95520001 | 95620000 | 0.0039 | 0.0028 | 0.0038 |
| F01 | 95540001 | 95640000 | 0.0045 | 0.0032 | 0.0043 |
| F01 | 95560001 | 95660000 | 0.0047 | 0.0035 | 0.0045 |
| F01 | 95580001 | 95680000 | 0.0043 | 0.0033 | 0.0041 |
| F01 | 95600001 | 95700000 | 0.0041 | 0.0032 | 0.0039 |
| F01 | 95620001 | 95720000 | 0.0036 | 0.0026 | 0.0033 |
| F01 | 95640001 | 95740000 | 0.0033 | 0.0024 | 0.0031 |
| F01 | 95660001 | 95760000 | 0.0039 | 0.0030 | 0.0037 |
| F01 | 95680001 | 95780000 | 0.0043 | 0.0033 | 0.0042 |
| F01 | 95700001 | 95800000 | 0.0051 | 0.0033 | 0.0049 |
| F01 | 95720001 | 95820000 | 0.0050 | 0.0034 | 0.0047 |
| F01 | 95740001 | 95840000 | 0.0045 | 0.0029 | 0.0041 |
| F01 | 95760001 | 95860000 | 0.0042 | 0.0023 | 0.0035 |
| F01 | 95780001 | 95880000 | 0.0038 | 0.0015 | 0.0028 |
| F01 | 95800001 | 95900000 | 0.0038 | 0.0019 | 0.0026 |
| F01 | 95820001 | 95920000 | 0.0035 | 0.0016 | 0.0023 |
| F01 | 95840001 | 95940000 | 0.0037 | 0.0016 | 0.0025 |
| F01 | 95860001 | 95960000 | 0.0033 | 0.0012 | 0.0022 |
| F01 | 95880001 | 95980000 | 0.0028 | 0.0010 | 0.0020 |
| F01 | 95900001 | 96000000 | 0.0019 | 0.0002 | 0.0010 |
| F01 | 95920001 | 96020000 | 0.0019 | 0.0003 | 0.0010 |
| F01 | 95940001 | 96040000 | 0.0017 | 0.0005 | 0.0009 |
| F01 | 95960001 | 96060000 | 0.0018 | 0.0007 | 0.0008 |
| F01 | 95980001 | 96080000 | 0.0018 | 0.0008 | 0.0009 |
| F01 | 96000001 | 96100000 | 0.0018 | 0.0010 | 0.0010 |
| F01 | 96020001 | 96120000 | 0.0021 | 0.0014 | 0.0014 |
| F01 | 96040001 | 96140000 | 0.0028 | 0.0018 | 0.0020 |
| F01 | 96060001 | 96160000 | 0.0023 | 0.0014 | 0.0016 |
| F01 | 96080001 | 96180000 | 0.0027 | 0.0016 | 0.0018 |
| F01 | 96100001 | 96200000 | 0.0026 | 0.0014 | 0.0016 |
| F01 | 96120001 | 96220000 | 0.0028 | 0.0016 | 0.0017 |
| F01 | 96140001 | 96240000 | 0.0026 | 0.0015 | 0.0016 |
| F01 | 96160001 | 96260000 | 0.0029 | 0.0017 | 0.0018 |
| F01 | 96180001 | 96280000 | 0.0025 | 0.0015 | 0.0016 |

| Chr | Start | End | both | A1 | A2 |
|---|---|---|---|---|---|
| F01 | 96200001 | 96300000 | 0.0025 | 0.0018 | 0.0018 |
| F01 | 96220001 | 96320000 | 0.0022 | 0.0015 | 0.0015 |
| F01 | 96240001 | 96340000 | 0.0018 | 0.0016 | 0.0012 |
| F01 | 96260001 | 96360000 | 0.0017 | 0.0019 | 0.0012 |
| F01 | 96280001 | 96380000 | 0.0018 | 0.0023 | 0.0013 |
| F01 | 96300001 | 96400000 | 0.0024 | 0.0028 | 0.0018 |
| F01 | 96320001 | 96420000 | 0.0026 | 0.0033 | 0.0021 |
| F01 | 96340001 | 96440000 | 0.0025 | 0.0031 | 0.0020 |
| F01 | 96360001 | 96460000 | 0.0025 | 0.0025 | 0.0019 |
| F01 | 96380001 | 96480000 | 0.0031 | 0.0028 | 0.0022 |
| F01 | 96400001 | 96500000 | 0.0027 | 0.0022 | 0.0017 |
| F01 | 96420001 | 96520000 | 0.0026 | 0.0017 | 0.0016 |
| F01 | 96440001 | 96540000 | 0.0030 | 0.0015 | 0.0021 |
| F01 | 96460001 | 96560000 | 0.0033 | 0.0015 | 0.0024 |
| F01 | 96480001 | 96580000 | 0.0028 | 0.0008 | 0.0020 |
| F01 | 96500001 | 96600000 | 0.0026 | 0.0007 | 0.0018 |
| F01 | 96520001 | 96620000 | 0.0024 | 0.0005 | 0.0016 |
| F01 | 96540001 | 96640000 | 0.0021 | 0.0005 | 0.0013 |
| F01 | 96560001 | 96660000 | 0.0024 | 0.0011 | 0.0018 |
| F01 | 96580001 | 96680000 | 0.0024 | 0.0010 | 0.0019 |
| F01 | 96600001 | 96700000 | 0.0028 | 0.0011 | 0.0023 |
| F01 | 96620001 | 96720000 | 0.0028 | 0.0011 | 0.0024 |
| F01 | 96640001 | 96740000 | 0.0029 | 0.0012 | 0.0025 |
| F01 | 96660001 | 96760000 | 0.0028 | 0.0013 | 0.0024 |
| F01 | 96680001 | 96780000 | 0.0030 | 0.0015 | 0.0028 |
| F01 | 96700001 | 96800000 | 0.0029 | 0.0015 | 0.0027 |
| F01 | 96720001 | 96820000 | 0.0029 | 0.0014 | 0.0024 |
| F01 | 96740001 | 96840000 | 0.0027 | 0.0012 | 0.0020 |
| F01 | 96760001 | 96860000 | 0.0033 | 0.0016 | 0.0025 |
| F01 | 96780001 | 96880000 | 0.0032 | 0.0015 | 0.0023 |
| F01 | 96800001 | 96900000 | 0.0028 | 0.0015 | 0.0021 |
| F01 | 96820001 | 96920000 | 0.0027 | 0.0014 | 0.0022 |
| F01 | 96840001 | 96940000 | 0.0024 | 0.0014 | 0.0022 |
| F01 | 96860001 | 96960000 | 0.0014 | 0.0005 | 0.0014 |
| F01 | 96880001 | 96980000 | 0.0012 | 0.0003 | 0.0012 |
| F01 | 96900001 | 97000000 | 0.0016 | 0.0005 | 0.0017 |
| F01 | 96920001 | 97020000 | 0.0016 | 0.0006 | 0.0015 |
| F01 | 96940001 | 97040000 | 0.0019 | 0.0006 | 0.0018 |
| F01 | 96960001 | 97060000 | 0.0019 | 0.0004 | 0.0016 |
| F01 | 96980001 | 97080000 | 0.0019 | 0.0008 | 0.0015 |
| F01 | 97000001 | 97100000 | 0.0017 | 0.0012 | 0.0010 |
| F01 | 97020001 | 97120000 | 0.0016 | 0.0014 | 0.0008 |
| F01 | 97040001 | 97140000 | 0.0015 | 0.0017 | 0.0009 |
| F01 | 97060001 | 97160000 | 0.0018 | 0.0021 | 0.0014 |
| F01 | 97080001 | 97180000 | 0.0020 | 0.0018 | 0.0016 |
| F01 | 97100001 | 97200000 | 0.0028 | 0.0023 | 0.0026 |
| F01 | 97120001 | 97220000 | 0.0034 | 0.0025 | 0.0033 |
| F01 | 97140001 | 97240000 | 0.0033 | 0.0022 | 0.0033 |
| F01 | 97160001 | 97260000 | 0.0029 | 0.0018 | 0.0028 |
| F01 | 97180001 | 97280000 | 0.0031 | 0.0022 | 0.0030 |
| F01 | 97200001 | 97300000 | 0.0024 | 0.0018 | 0.0022 |
| F01 | 97220001 | 97320000 | 0.0020 | 0.0019 | 0.0016 |
| F01 | 97240001 | 97340000 | 0.0021 | 0.0023 | 0.0017 |
| F01 | 97260001 | 97360000 | 0.0024 | 0.0024 | 0.0019 |
| F01 | 97280001 | 97380000 | 0.0021 | 0.0019 | 0.0016 |
| F01 | 97300001 | 97400000 | 0.0023 | 0.0017 | 0.0018 |
| F01 | 97320001 | 97420000 | 0.0026 | 0.0012 | 0.0024 |
| F01 | 97340001 | 97440000 | 0.0025 | 0.0008 | 0.0024 |

| Chr | Start | End | both | A1 | A2 |
|---|---|---|---|---|---|
| F01 | 97360001 | 97460000 | 0.0024 | 0.0009 | 0.0022 |
| F01 | 97380001 | 97480000 | 0.0026 | 0.0014 | 0.0023 |
| F01 | 97400001 | 97500000 | 0.0024 | 0.0013 | 0.0020 |
| F01 | 97420001 | 97520000 | 0.0023 | 0.0016 | 0.0018 |
| F01 | 97440001 | 97540000 | 0.0023 | 0.0018 | 0.0018 |
| F01 | 97460001 | 97560000 | 0.0022 | 0.0016 | 0.0017 |
| F01 | 97480001 | 97580000 | 0.0021 | 0.0011 | 0.0016 |
| F01 | 97500001 | 97600000 | 0.0019 | 0.0008 | 0.0014 |
| F01 | 97520001 | 97620000 | 0.0023 | 0.0011 | 0.0018 |
| F01 | 97540001 | 97640000 | 0.0025 | 0.0013 | 0.0020 |
| F01 | 97560001 | 97660000 | 0.0031 | 0.0014 | 0.0027 |
| F01 | 97580001 | 97680000 | 0.0032 | 0.0014 | 0.0028 |
| F01 | 97600001 | 97700000 | 0.0034 | 0.0016 | 0.0031 |
| F01 | 97620001 | 97720000 | 0.0031 | 0.0009 | 0.0027 |
| F01 | 97640001 | 97740000 | 0.0028 | 0.0005 | 0.0022 |
| F01 | 97660001 | 97760000 | 0.0023 | 0.0004 | 0.0014 |
| F01 | 97680001 | 97780000 | 0.0020 | 0.0005 | 0.0011 |
| F01 | 97700001 | 97800000 | 0.0018 | 0.0004 | 0.0010 |
| F01 | 97720001 | 97820000 | 0.0019 | 0.0011 | 0.0011 |
| F01 | 97740001 | 97840000 | 0.0023 | 0.0018 | 0.0017 |
| F01 | 97760001 | 97860000 | 0.0028 | 0.0025 | 0.0025 |
| F01 | 97780001 | 97880000 | 0.0038 | 0.0036 | 0.0037 |
| F01 | 97800001 | 97900000 | 0.0039 | 0.0034 | 0.0039 |
| F01 | 97820001 | 97920000 | 0.0040 | 0.0032 | 0.0040 |
| F01 | 97840001 | 97940000 | 0.0034 | 0.0025 | 0.0035 |
| F01 | 97860001 | 97960000 | 0.0032 | 0.0018 | 0.0033 |
| F01 | 97880001 | 97980000 | 0.0022 | 0.0006 | 0.0023 |
| F01 | 97900001 | 98000000 | 0.0025 | 0.0007 | 0.0025 |
| F01 | 97920001 | 98020000 | 0.0022 | 0.0004 | 0.0021 |
| F01 | 97940001 | 98040000 | 0.0026 | 0.0005 | 0.0023 |
| F01 | 97960001 | 98060000 | 0.0026 | 0.0005 | 0.0022 |
| F01 | 97980001 | 98080000 | 0.0026 | 0.0005 | 0.0022 |
| F01 | 98000001 | 98100000 | 0.0022 | 0.0005 | 0.0018 |
| F01 | 98020001 | 98120000 | 0.0024 | 0.0007 | 0.0020 |
| F01 | 98040001 | 98140000 | 0.0022 | 0.0007 | 0.0018 |
| F01 | 98060001 | 98160000 | 0.0018 | 0.0009 | 0.0014 |
| F01 | 98080001 | 98180000 | 0.0017 | 0.0013 | 0.0012 |
| F01 | 98100001 | 98200000 | 0.0019 | 0.0017 | 0.0013 |
| F01 | 98120001 | 98220000 | 0.0020 | 0.0020 | 0.0016 |
| F01 | 98140001 | 98240000 | 0.0024 | 0.0026 | 0.0021 |
| F01 | 98160001 | 98260000 | 0.0028 | 0.0028 | 0.0025 |
| F01 | 98180001 | 98280000 | 0.0028 | 0.0025 | 0.0024 |
| F01 | 98200001 | 98300000 | 0.0030 | 0.0023 | 0.0027 |
| F01 | 98220001 | 98320000 | 0.0027 | 0.0017 | 0.0023 |
| F01 | 98240001 | 98340000 | 0.0024 | 0.0010 | 0.0019 |
| F01 | 98260001 | 98360000 | 0.0021 | 0.0005 | 0.0017 |
| F01 | 98280001 | 98380000 | 0.0022 | 0.0004 | 0.0016 |
| F01 | 98300001 | 98400000 | 0.0019 | 0.0001 | 0.0013 |
| F01 | 98320001 | 98420000 | 0.0017 | 0.0001 | 0.0011 |
| F01 | 98340001 | 98440000 | 0.0019 | 0.0001 | 0.0013 |
| F01 | 98360001 | 98460000 | 0.0026 | 0.0010 | 0.0022 |
| F01 | 98380001 | 98480000 | 0.0023 | 0.0010 | 0.0020 |
| F01 | 98400001 | 98500000 | 0.0020 | 0.0010 | 0.0018 |
| F01 | 98420001 | 98520000 | 0.0016 | 0.0010 | 0.0015 |
| F01 | 98440001 | 98540000 | 0.0011 | 0.0009 | 0.0011 |
| F01 | 98460001 | 98560000 | 0.0000 | 0.0000 | 0.0000 |
| F02 | 1 | 100000 | 0.0018 | 0.0016 | 0.0015 |
| F02 | 20001 | 120000 | 0.0010 | 0.0008 | 0.0007 |

| Chr | Start | End | both | A1 | A2 |
|---|---|---|---|---|---|
| F02 | 40001 | 140000 | 0.0010 | 0.0007 | 0.0005 |
| F02 | 60001 | 160000 | 0.0011 | 0.0007 | 0.0005 |
| F02 | 80001 | 180000 | 0.0011 | 0.0006 | 0.0005 |
| F02 | 100001 | 200000 | 0.0015 | 0.0007 | 0.0009 |
| F02 | 120001 | 220000 | 0.0017 | 0.0008 | 0.0012 |
| F02 | 140001 | 240000 | 0.0018 | 0.0008 | 0.0013 |
| F02 | 160001 | 260000 | 0.0018 | 0.0007 | 0.0013 |
| F02 | 180001 | 280000 | 0.0021 | 0.0012 | 0.0016 |
| F02 | 200001 | 300000 | 0.0018 | 0.0010 | 0.0012 |
| F02 | 220001 | 320000 | 0.0018 | 0.0010 | 0.0010 |
| F02 | 240001 | 340000 | 0.0019 | 0.0009 | 0.0009 |
| F02 | 260001 | 360000 | 0.0021 | 0.0009 | 0.0012 |
| F02 | 280001 | 380000 | 0.0020 | 0.0008 | 0.0012 |
| F02 | 300001 | 400000 | 0.0022 | 0.0008 | 0.0015 |
| F02 | 320001 | 420000 | 0.0024 | 0.0008 | 0.0018 |
| F02 | 340001 | 440000 | 0.0023 | 0.0009 | 0.0020 |
| F02 | 360001 | 460000 | 0.0025 | 0.0012 | 0.0021 |
| F02 | 380001 | 480000 | 0.0024 | 0.0010 | 0.0019 |
| F02 | 400001 | 500000 | 0.0025 | 0.0011 | 0.0021 |
| F02 | 420001 | 520000 | 0.0024 | 0.0012 | 0.0020 |
| F02 | 440001 | 540000 | 0.0025 | 0.0013 | 0.0021 |
| F02 | 460001 | 560000 | 0.0019 | 0.0011 | 0.0016 |
| F02 | 480001 | 580000 | 0.0016 | 0.0010 | 0.0014 |
| F02 | 500001 | 600000 | 0.0013 | 0.0012 | 0.0011 |
| F02 | 520001 | 620000 | 0.0011 | 0.0013 | 0.0009 |
| F02 | 540001 | 640000 | 0.0008 | 0.0015 | 0.0005 |
| F02 | 560001 | 660000 | 0.0009 | 0.0016 | 0.0005 |
| F02 | 580001 | 680000 | 0.0010 | 0.0017 | 0.0004 |
| F02 | 600001 | 700000 | 0.0011 | 0.0018 | 0.0004 |
| F02 | 620001 | 720000 | 0.0011 | 0.0017 | 0.0003 |
| F02 | 640001 | 740000 | 0.0012 | 0.0013 | 0.0004 |
| F02 | 660001 | 760000 | 0.0014 | 0.0014 | 0.0006 |
| F02 | 680001 | 780000 | 0.0014 | 0.0011 | 0.0009 |
| F02 | 700001 | 800000 | 0.0017 | 0.0008 | 0.0011 |
| F02 | 720001 | 820000 | 0.0018 | 0.0006 | 0.0011 |
| F02 | 740001 | 840000 | 0.0017 | 0.0005 | 0.0010 |
| F02 | 760001 | 860000 | 0.0016 | 0.0004 | 0.0008 |
| F02 | 780001 | 880000 | 0.0015 | 0.0006 | 0.0006 |
| F02 | 800001 | 900000 | 0.0015 | 0.0011 | 0.0005 |
| F02 | 820001 | 920000 | 0.0016 | 0.0011 | 0.0007 |
| F02 | 840001 | 940000 | 0.0018 | 0.0011 | 0.0008 |
| F02 | 860001 | 960000 | 0.0020 | 0.0009 | 0.0011 |
| F02 | 880001 | 980000 | 0.0022 | 0.0006 | 0.0012 |
| F02 | 900001 | 1000000 | 0.0023 | 0.0002 | 0.0016 |
| F02 | 920001 | 1020000 | 0.0023 | 0.0002 | 0.0016 |
| F02 | 940001 | 1040000 | 0.0024 | 0.0003 | 0.0020 |
| F02 | 960001 | 1060000 | 0.0025 | 0.0003 | 0.0019 |
| F02 | 980001 | 1080000 | 0.0025 | 0.0003 | 0.0018 |
| F02 | 1000001 | 1100000 | 0.0023 | 0.0002 | 0.0013 |
| F02 | 1020001 | 1120000 | 0.0022 | 0.0002 | 0.0011 |
| F02 | 1040001 | 1140000 | 0.0027 | 0.0009 | 0.0015 |
| F02 | 1060001 | 1160000 | 0.0034 | 0.0020 | 0.0023 |
| F02 | 1080001 | 1180000 | 0.0035 | 0.0022 | 0.0026 |
| F02 | 1100001 | 1200000 | 0.0036 | 0.0025 | 0.0030 |
| F02 | 1120001 | 1220000 | 0.0039 | 0.0029 | 0.0035 |
| F02 | 1140001 | 1240000 | 0.0034 | 0.0025 | 0.0031 |
| F02 | 1160001 | 1260000 | 0.0023 | 0.0016 | 0.0021 |
| F02 | 1180001 | 1280000 | 0.0022 | 0.0017 | 0.0019 |

| Chr | Start | End | both | A1 | A2 |
|---|---|---|---|---|---|
| F02 | 1200001 | 1300000 | 0.0020 | 0.0019 | 0.0017 |
| F02 | 1220001 | 1320000 | 0.0017 | 0.0019 | 0.0014 |
| F02 | 1240001 | 1340000 | 0.0014 | 0.0020 | 0.0011 |
| F02 | 1260001 | 1360000 | 0.0020 | 0.0026 | 0.0017 |
| F02 | 1280001 | 1380000 | 0.0020 | 0.0026 | 0.0018 |
| F02 | 1300001 | 1400000 | 0.0019 | 0.0026 | 0.0016 |
| F02 | 1320001 | 1420000 | 0.0018 | 0.0026 | 0.0014 |
| F02 | 1340001 | 1440000 | 0.0017 | 0.0025 | 0.0013 |
| F02 | 1360001 | 1460000 | 0.0012 | 0.0022 | 0.0006 |
| F02 | 1380001 | 1480000 | 0.0009 | 0.0020 | 0.0004 |
| F02 | 1400001 | 1500000 | 0.0009 | 0.0019 | 0.0003 |
| F02 | 1420001 | 1520000 | 0.0011 | 0.0021 | 0.0007 |
| F02 | 1440001 | 1540000 | 0.0026 | 0.0035 | 0.0023 |
| F02 | 1460001 | 1560000 | 0.0026 | 0.0037 | 0.0023 |
| F02 | 1480001 | 1580000 | 0.0028 | 0.0037 | 0.0024 |
| F02 | 1500001 | 1600000 | 0.0028 | 0.0036 | 0.0025 |
| F02 | 1520001 | 1620000 | 0.0028 | 0.0035 | 0.0026 |
| F02 | 1540001 | 1640000 | 0.0013 | 0.0019 | 0.0011 |
| F02 | 1560001 | 1660000 | 0.0014 | 0.0017 | 0.0013 |
| F02 | 1580001 | 1680000 | 0.0020 | 0.0021 | 0.0018 |
| F02 | 1600001 | 1700000 | 0.0021 | 0.0023 | 0.0019 |
| F02 | 1620001 | 1720000 | 0.0017 | 0.0021 | 0.0015 |
| F02 | 1640001 | 1740000 | 0.0016 | 0.0020 | 0.0013 |
| F02 | 1660001 | 1760000 | 0.0030 | 0.0034 | 0.0027 |
| F02 | 1680001 | 1780000 | 0.0032 | 0.0038 | 0.0029 |
| F02 | 1700001 | 1800000 | 0.0035 | 0.0039 | 0.0032 |
| F02 | 1720001 | 1820000 | 0.0053 | 0.0054 | 0.0051 |
| F02 | 1740001 | 1840000 | 0.0060 | 0.0058 | 0.0058 |
| F02 | 1760001 | 1860000 | 0.0047 | 0.0045 | 0.0046 |
| F02 | 1780001 | 1880000 | 0.0045 | 0.0041 | 0.0044 |
| F02 | 1800001 | 1900000 | 0.0062 | 0.0058 | 0.0061 |
| F02 | 1820001 | 1920000 | 0.0058 | 0.0056 | 0.0057 |
| F02 | 1840001 | 1940000 | 0.0056 | 0.0056 | 0.0054 |
| F02 | 1860001 | 1960000 | 0.0053 | 0.0057 | 0.0051 |
| F02 | 1880001 | 1980000 | 0.0046 | 0.0050 | 0.0043 |
| F02 | 1900001 | 2000000 | 0.0027 | 0.0034 | 0.0024 |
| F02 | 1920001 | 2020000 | 0.0014 | 0.0025 | 0.0010 |
| F02 | 1940001 | 2040000 | 0.0012 | 0.0022 | 0.0009 |
| F02 | 1960001 | 2060000 | 0.0014 | 0.0021 | 0.0012 |
| F02 | 1980001 | 2080000 | 0.0019 | 0.0025 | 0.0016 |
| F02 | 2000001 | 2100000 | 0.0036 | 0.0040 | 0.0033 |
| F02 | 2020001 | 2120000 | 0.0035 | 0.0038 | 0.0033 |
| F02 | 2040001 | 2140000 | 0.0032 | 0.0039 | 0.0029 |
| F02 | 2060001 | 2160000 | 0.0031 | 0.0040 | 0.0028 |
| F02 | 2080001 | 2180000 | 0.0029 | 0.0039 | 0.0025 |
| F02 | 2100001 | 2200000 | 0.0019 | 0.0030 | 0.0014 |
| F02 | 2120001 | 2220000 | 0.0020 | 0.0031 | 0.0016 |
| F02 | 2140001 | 2240000 | 0.0021 | 0.0027 | 0.0016 |
| F02 | 2160001 | 2260000 | 0.0024 | 0.0023 | 0.0016 |
| F02 | 2180001 | 2280000 | 0.0024 | 0.0019 | 0.0015 |
| F02 | 2200001 | 2300000 | 0.0016 | 0.0008 | 0.0005 |
| F02 | 2220001 | 2320000 | 0.0015 | 0.0006 | 0.0004 |
| F02 | 2240001 | 2340000 | 0.0017 | 0.0011 | 0.0006 |
| F02 | 2260001 | 2360000 | 0.0017 | 0.0016 | 0.0010 |
| F02 | 2280001 | 2380000 | 0.0024 | 0.0025 | 0.0019 |
| F02 | 2300001 | 2400000 | 0.0040 | 0.0042 | 0.0037 |
| F02 | 2320001 | 2420000 | 0.0040 | 0.0042 | 0.0039 |
| F02 | 2340001 | 2440000 | 0.0040 | 0.0041 | 0.0038 |

| Chr | Start | End | both | A1 | A2 |
|---|---|---|---|---|---|
| F02 | 2360001 | 2460000 | 0.0036 | 0.0041 | 0.0033 |
| F02 | 2380001 | 2480000 | 0.0030 | 0.0036 | 0.0026 |
| F02 | 2400001 | 2500000 | 0.0013 | 0.0022 | 0.0009 |
| F02 | 2420001 | 2520000 | 0.0014 | 0.0022 | 0.0008 |
| F02 | 2440001 | 2540000 | 0.0016 | 0.0020 | 0.0008 |
| F02 | 2460001 | 2560000 | 0.0018 | 0.0018 | 0.0011 |
| F02 | 2480001 | 2580000 | 0.0022 | 0.0019 | 0.0016 |
| F02 | 2500001 | 2600000 | 0.0026 | 0.0018 | 0.0019 |
| F02 | 2520001 | 2620000 | 0.0025 | 0.0018 | 0.0021 |
| F02 | 2540001 | 2640000 | 0.0021 | 0.0015 | 0.0020 |
| F02 | 2560001 | 2660000 | 0.0022 | 0.0013 | 0.0022 |
| F02 | 2580001 | 2680000 | 0.0021 | 0.0007 | 0.0021 |
| F02 | 2600001 | 2700000 | 0.0026 | 0.0011 | 0.0025 |
| F02 | 2620001 | 2720000 | 0.0025 | 0.0008 | 0.0023 |
| F02 | 2640001 | 2740000 | 0.0028 | 0.0009 | 0.0023 |
| F02 | 2660001 | 2760000 | 0.0024 | 0.0008 | 0.0017 |
| F02 | 2680001 | 2780000 | 0.0021 | 0.0008 | 0.0012 |
| F02 | 2700001 | 2800000 | 0.0015 | 0.0003 | 0.0005 |
| F02 | 2720001 | 2820000 | 0.0017 | 0.0003 | 0.0005 |
| F02 | 2740001 | 2840000 | 0.0017 | 0.0004 | 0.0005 |
| F02 | 2760001 | 2860000 | 0.0029 | 0.0015 | 0.0018 |
| F02 | 2780001 | 2880000 | 0.0030 | 0.0018 | 0.0021 |
| F02 | 2800001 | 2900000 | 0.0029 | 0.0019 | 0.0021 |
| F02 | 2820001 | 2920000 | 0.0029 | 0.0019 | 0.0022 |
| F02 | 2840001 | 2940000 | 0.0027 | 0.0018 | 0.0021 |
| F02 | 2860001 | 2960000 | 0.0021 | 0.0013 | 0.0013 |
| F02 | 2880001 | 2980000 | 0.0025 | 0.0016 | 0.0017 |
| F02 | 2900001 | 3000000 | 0.0023 | 0.0017 | 0.0017 |
| F02 | 2920001 | 3020000 | 0.0023 | 0.0020 | 0.0019 |
| F02 | 2940001 | 3040000 | 0.0021 | 0.0019 | 0.0019 |
| F02 | 2960001 | 3060000 | 0.0019 | 0.0017 | 0.0017 |
| F02 | 2980001 | 3080000 | 0.0011 | 0.0012 | 0.0010 |
| F02 | 3000001 | 3100000 | 0.0009 | 0.0009 | 0.0009 |
| F02 | 3020001 | 3120000 | 0.0010 | 0.0010 | 0.0009 |
| F02 | 3040001 | 3140000 | 0.0012 | 0.0013 | 0.0011 |
| F02 | 3060001 | 3160000 | 0.0010 | 0.0013 | 0.0009 |
| F02 | 3080001 | 3180000 | 0.0011 | 0.0014 | 0.0009 |
| F02 | 3100001 | 3200000 | 0.0012 | 0.0016 | 0.0010 |
| F02 | 3120001 | 3220000 | 0.0010 | 0.0015 | 0.0008 |
| F02 | 3140001 | 3240000 | 0.0010 | 0.0016 | 0.0008 |
| F02 | 3160001 | 3260000 | 0.0040 | 0.0042 | 0.0037 |
| F02 | 3180001 | 3280000 | 0.0041 | 0.0043 | 0.0038 |
| F02 | 3200001 | 3300000 | 0.0049 | 0.0052 | 0.0046 |
| F02 | 3220001 | 3320000 | 0.0057 | 0.0059 | 0.0054 |
| F02 | 3240001 | 3340000 | 0.0056 | 0.0059 | 0.0054 |
| F02 | 3260001 | 3360000 | 0.0026 | 0.0033 | 0.0023 |
| F02 | 3280001 | 3380000 | 0.0026 | 0.0034 | 0.0023 |
| F02 | 3300001 | 3400000 | 0.0024 | 0.0031 | 0.0021 |
| F02 | 3320001 | 3420000 | 0.0016 | 0.0023 | 0.0014 |
| F02 | 3340001 | 3440000 | 0.0017 | 0.0023 | 0.0014 |
| F02 | 3360001 | 3460000 | 0.0027 | 0.0029 | 0.0025 |
| F02 | 3380001 | 3480000 | 0.0026 | 0.0027 | 0.0025 |
| F02 | 3400001 | 3500000 | 0.0023 | 0.0025 | 0.0022 |
| F02 | 3420001 | 3520000 | 0.0028 | 0.0031 | 0.0026 |
| F02 | 3440001 | 3540000 | 0.0026 | 0.0029 | 0.0024 |
| F02 | 3460001 | 3560000 | 0.0022 | 0.0026 | 0.0020 |
| F02 | 3480001 | 3580000 | 0.0027 | 0.0029 | 0.0025 |
| F02 | 3500001 | 3600000 | 0.0032 | 0.0034 | 0.0029 |

| Chr | Start | End | both | A1 | A2 |
|---|---|---|---|---|---|
| F02 | 3520001 | 3620000 | 0.0026 | 0.0028 | 0.0022 |
| F02 | 3540001 | 3640000 | 0.0028 | 0.0028 | 0.0023 |
| F02 | 3560001 | 3660000 | 0.0024 | 0.0022 | 0.0016 |
| F02 | 3580001 | 3680000 | 0.0028 | 0.0022 | 0.0019 |
| F02 | 3600001 | 3700000 | 0.0029 | 0.0020 | 0.0019 |
| F02 | 3620001 | 3720000 | 0.0035 | 0.0024 | 0.0026 |
| F02 | 3640001 | 3740000 | 0.0036 | 0.0025 | 0.0028 |
| F02 | 3660001 | 3760000 | 0.0039 | 0.0030 | 0.0032 |
| F02 | 3680001 | 3780000 | 0.0035 | 0.0029 | 0.0030 |
| F02 | 3700001 | 3800000 | 0.0035 | 0.0028 | 0.0030 |
| F02 | 3720001 | 3820000 | 0.0033 | 0.0025 | 0.0029 |
| F02 | 3740001 | 3840000 | 0.0041 | 0.0031 | 0.0037 |
| F02 | 3760001 | 3860000 | 0.0039 | 0.0030 | 0.0036 |
| F02 | 3780001 | 3880000 | 0.0037 | 0.0027 | 0.0034 |
| F02 | 3800001 | 3900000 | 0.0030 | 0.0023 | 0.0027 |
| F02 | 3820001 | 3920000 | 0.0026 | 0.0020 | 0.0022 |
| F02 | 3840001 | 3940000 | 0.0016 | 0.0010 | 0.0012 |
| F02 | 3860001 | 3960000 | 0.0015 | 0.0007 | 0.0009 |
| F02 | 3880001 | 3980000 | 0.0014 | 0.0006 | 0.0006 |
| F02 | 3900001 | 4000000 | 0.0014 | 0.0006 | 0.0007 |
| F02 | 3920001 | 4020000 | 0.0014 | 0.0006 | 0.0007 |
| F02 | 3940001 | 4040000 | 0.0015 | 0.0008 | 0.0009 |
| F02 | 3960001 | 4060000 | 0.0015 | 0.0007 | 0.0009 |
| F02 | 3980001 | 4080000 | 0.0016 | 0.0006 | 0.0008 |
| F02 | 4000001 | 4100000 | 0.0026 | 0.0014 | 0.0018 |
| F02 | 4020001 | 4120000 | 0.0031 | 0.0017 | 0.0020 |
| F02 | 4040001 | 4140000 | 0.0030 | 0.0015 | 0.0018 |
| F02 | 4060001 | 4160000 | 0.0039 | 0.0026 | 0.0027 |
| F02 | 4080001 | 4180000 | 0.0042 | 0.0030 | 0.0031 |
| F02 | 4100001 | 4200000 | 0.0032 | 0.0023 | 0.0022 |
| F02 | 4120001 | 4220000 | 0.0029 | 0.0019 | 0.0019 |
| F02 | 4140001 | 4240000 | 0.0028 | 0.0019 | 0.0019 |
| F02 | 4160001 | 4260000 | 0.0017 | 0.0009 | 0.0010 |
| F02 | 4180001 | 4280000 | 0.0015 | 0.0007 | 0.0008 |
| F02 | 4200001 | 4300000 | 0.0015 | 0.0006 | 0.0008 |
| F02 | 4220001 | 4320000 | 0.0014 | 0.0007 | 0.0009 |
| F02 | 4240001 | 4340000 | 0.0014 | 0.0007 | 0.0009 |
| F02 | 4260001 | 4360000 | 0.0020 | 0.0012 | 0.0014 |
| F02 | 4280001 | 4380000 | 0.0019 | 0.0010 | 0.0012 |
| F02 | 4300001 | 4400000 | 0.0018 | 0.0011 | 0.0013 |
| F02 | 4320001 | 4420000 | 0.0020 | 0.0012 | 0.0015 |
| F02 | 4340001 | 4440000 | 0.0020 | 0.0012 | 0.0015 |
| F02 | 4360001 | 4460000 | 0.0014 | 0.0007 | 0.0009 |
| F02 | 4380001 | 4480000 | 0.0020 | 0.0013 | 0.0014 |
| F02 | 4400001 | 4500000 | 0.0020 | 0.0013 | 0.0013 |
| F02 | 4420001 | 4520000 | 0.0025 | 0.0016 | 0.0016 |
| F02 | 4440001 | 4540000 | 0.0027 | 0.0018 | 0.0019 |
| F02 | 4460001 | 4560000 | 0.0029 | 0.0018 | 0.0020 |
| F02 | 4480001 | 4580000 | 0.0029 | 0.0017 | 0.0019 |
| F02 | 4500001 | 4600000 | 0.0032 | 0.0019 | 0.0022 |
| F02 | 4520001 | 4620000 | 0.0025 | 0.0016 | 0.0019 |
| F02 | 4540001 | 4640000 | 0.0029 | 0.0019 | 0.0022 |
| F02 | 4560001 | 4660000 | 0.0034 | 0.0026 | 0.0028 |
| F02 | 4580001 | 4680000 | 0.0031 | 0.0025 | 0.0027 |
| F02 | 4600001 | 4700000 | 0.0037 | 0.0028 | 0.0030 |
| F02 | 4620001 | 4720000 | 0.0042 | 0.0031 | 0.0035 |
| F02 | 4640001 | 4740000 | 0.0041 | 0.0032 | 0.0035 |
| F02 | 4660001 | 4760000 | 0.0037 | 0.0027 | 0.0029 |

| Chr | Start | End | both | A1 | A2 |
|---|---|---|---|---|---|
| F02 | 4680001 | 4780000 | 0.0038 | 0.0028 | 0.0030 |
| F02 | 4700001 | 4800000 | 0.0028 | 0.0022 | 0.0024 |
| F02 | 4720001 | 4820000 | 0.0023 | 0.0017 | 0.0017 |
| F02 | 4740001 | 4840000 | 0.0021 | 0.0013 | 0.0013 |
| F02 | 4760001 | 4860000 | 0.0020 | 0.0012 | 0.0012 |
| F02 | 4780001 | 4880000 | 0.0016 | 0.0008 | 0.0007 |
| F02 | 4800001 | 4900000 | 0.0022 | 0.0013 | 0.0014 |
| F02 | 4820001 | 4920000 | 0.0025 | 0.0015 | 0.0016 |
| F02 | 4840001 | 4940000 | 0.0028 | 0.0017 | 0.0020 |
| F02 | 4860001 | 4960000 | 0.0030 | 0.0019 | 0.0022 |
| F02 | 4880001 | 4980000 | 0.0032 | 0.0019 | 0.0022 |
| F02 | 4900001 | 5000000 | 0.0027 | 0.0014 | 0.0016 |
| F02 | 4920001 | 5020000 | 0.0030 | 0.0018 | 0.0020 |
| F02 | 4940001 | 5040000 | 0.0028 | 0.0018 | 0.0017 |
| F02 | 4960001 | 5060000 | 0.0028 | 0.0019 | 0.0019 |
| F02 | 4980001 | 5080000 | 0.0025 | 0.0019 | 0.0019 |
| F02 | 5000001 | 5100000 | 0.0028 | 0.0020 | 0.0021 |
| F02 | 5020001 | 5120000 | 0.0023 | 0.0014 | 0.0015 |
| F02 | 5040001 | 5140000 | 0.0024 | 0.0012 | 0.0013 |
| F02 | 5060001 | 5160000 | 0.0019 | 0.0009 | 0.0010 |
| F02 | 5080001 | 5180000 | 0.0023 | 0.0014 | 0.0015 |
| F02 | 5100001 | 5200000 | 0.0023 | 0.0015 | 0.0016 |
| F02 | 5120001 | 5220000 | 0.0030 | 0.0026 | 0.0023 |
| F02 | 5140001 | 5240000 | 0.0027 | 0.0025 | 0.0023 |
| F02 | 5160001 | 5260000 | 0.0030 | 0.0028 | 0.0026 |
| F02 | 5180001 | 5280000 | 0.0031 | 0.0029 | 0.0028 |
| F02 | 5200001 | 5300000 | 0.0032 | 0.0031 | 0.0030 |
| F02 | 5220001 | 5320000 | 0.0030 | 0.0026 | 0.0029 |
| F02 | 5240001 | 5340000 | 0.0032 | 0.0026 | 0.0030 |
| F02 | 5260001 | 5360000 | 0.0039 | 0.0028 | 0.0033 |
| F02 | 5280001 | 5380000 | 0.0032 | 0.0021 | 0.0026 |
| F02 | 5300001 | 5400000 | 0.0033 | 0.0020 | 0.0026 |
| F02 | 5320001 | 5420000 | 0.0029 | 0.0015 | 0.0022 |
| F02 | 5340001 | 5440000 | 0.0031 | 0.0017 | 0.0022 |
| F02 | 5360001 | 5460000 | 0.0029 | 0.0019 | 0.0021 |
| F02 | 5380001 | 5480000 | 0.0029 | 0.0019 | 0.0021 |
| F02 | 5400001 | 5500000 | 0.0029 | 0.0020 | 0.0022 |
| F02 | 5420001 | 5520000 | 0.0042 | 0.0033 | 0.0035 |
| F02 | 5440001 | 5540000 | 0.0045 | 0.0035 | 0.0038 |
| F02 | 5460001 | 5560000 | 0.0047 | 0.0034 | 0.0038 |
| F02 | 5480001 | 5580000 | 0.0051 | 0.0037 | 0.0041 |
| F02 | 5500001 | 5600000 | 0.0050 | 0.0036 | 0.0040 |
| F02 | 5520001 | 5620000 | 0.0039 | 0.0026 | 0.0030 |
| F02 | 5540001 | 5640000 | 0.0033 | 0.0022 | 0.0025 |
| F02 | 5560001 | 5660000 | 0.0035 | 0.0028 | 0.0029 |
| F02 | 5580001 | 5680000 | 0.0032 | 0.0026 | 0.0028 |
| F02 | 5600001 | 5700000 | 0.0031 | 0.0025 | 0.0027 |
| F02 | 5620001 | 5720000 | 0.0037 | 0.0030 | 0.0031 |
| F02 | 5640001 | 5740000 | 0.0040 | 0.0034 | 0.0034 |
| F02 | 5660001 | 5760000 | 0.0031 | 0.0023 | 0.0025 |
| F02 | 5680001 | 5780000 | 0.0037 | 0.0028 | 0.0029 |
| F02 | 5700001 | 5800000 | 0.0040 | 0.0031 | 0.0032 |
| F02 | 5720001 | 5820000 | 0.0034 | 0.0025 | 0.0027 |
| F02 | 5740001 | 5840000 | 0.0032 | 0.0021 | 0.0024 |
| F02 | 5760001 | 5860000 | 0.0038 | 0.0028 | 0.0030 |
| F02 | 5780001 | 5880000 | 0.0041 | 0.0032 | 0.0032 |
| F02 | 5800001 | 5900000 | 0.0043 | 0.0034 | 0.0034 |
| F02 | 5820001 | 5920000 | 0.0038 | 0.0030 | 0.0028 |

| Chr | Start | End | both | A1 | A2 |
|---|---|---|---|---|---|
| F02 | 5840001 | 5940000 | 0.0044 | 0.0035 | 0.0034 |
| F02 | 5860001 | 5960000 | 0.0045 | 0.0034 | 0.0034 |
| F02 | 5880001 | 5980000 | 0.0039 | 0.0027 | 0.0028 |
| F02 | 5900001 | 6000000 | 0.0042 | 0.0026 | 0.0028 |
| F02 | 5920001 | 6020000 | 0.0043 | 0.0027 | 0.0029 |
| F02 | 5940001 | 6040000 | 0.0036 | 0.0022 | 0.0023 |
| F02 | 5960001 | 6060000 | 0.0040 | 0.0025 | 0.0027 |
| F02 | 5980001 | 6080000 | 0.0039 | 0.0024 | 0.0026 |
| F02 | 6000001 | 6100000 | 0.0029 | 0.0019 | 0.0020 |
| F02 | 6020001 | 6120000 | 0.0032 | 0.0019 | 0.0020 |
| F02 | 6040001 | 6140000 | 0.0044 | 0.0027 | 0.0030 |
| F02 | 6060001 | 6160000 | 0.0038 | 0.0020 | 0.0024 |
| F02 | 6080001 | 6180000 | 0.0044 | 0.0027 | 0.0029 |
| F02 | 6100001 | 6200000 | 0.0062 | 0.0045 | 0.0047 |
| F02 | 6120001 | 6220000 | 0.0062 | 0.0047 | 0.0050 |
| F02 | 6140001 | 6240000 | 0.0053 | 0.0041 | 0.0043 |
| F02 | 6160001 | 6260000 | 0.0052 | 0.0042 | 0.0044 |
| F02 | 6180001 | 6280000 | 0.0045 | 0.0033 | 0.0037 |
| F02 | 6200001 | 6300000 | 0.0027 | 0.0014 | 0.0019 |
| F02 | 6220001 | 6320000 | 0.0026 | 0.0012 | 0.0018 |
| F02 | 6240001 | 6340000 | 0.0029 | 0.0015 | 0.0022 |
| F02 | 6260001 | 6360000 | 0.0029 | 0.0017 | 0.0023 |
| F02 | 6280001 | 6380000 | 0.0033 | 0.0022 | 0.0027 |
| F02 | 6300001 | 6400000 | 0.0032 | 0.0022 | 0.0026 |
| F02 | 6320001 | 6420000 | 0.0033 | 0.0025 | 0.0028 |
| F02 | 6340001 | 6440000 | 0.0029 | 0.0022 | 0.0024 |
| F02 | 6360001 | 6460000 | 0.0026 | 0.0020 | 0.0022 |
| F02 | 6380001 | 6480000 | 0.0027 | 0.0017 | 0.0022 |
| F02 | 6400001 | 6500000 | 0.0028 | 0.0018 | 0.0023 |
| F02 | 6420001 | 6520000 | 0.0030 | 0.0019 | 0.0023 |
| F02 | 6440001 | 6540000 | 0.0027 | 0.0016 | 0.0020 |
| F02 | 6460001 | 6560000 | 0.0029 | 0.0016 | 0.0019 |
| F02 | 6480001 | 6580000 | 0.0027 | 0.0016 | 0.0017 |
| F02 | 6500001 | 6600000 | 0.0029 | 0.0019 | 0.0019 |
| F02 | 6520001 | 6620000 | 0.0027 | 0.0013 | 0.0015 |
| F02 | 6540001 | 6640000 | 0.0034 | 0.0019 | 0.0022 |
| F02 | 6560001 | 6660000 | 0.0035 | 0.0021 | 0.0025 |
| F02 | 6580001 | 6680000 | 0.0040 | 0.0023 | 0.0028 |
| F02 | 6600001 | 6700000 | 0.0040 | 0.0022 | 0.0029 |
| F02 | 6620001 | 6720000 | 0.0042 | 0.0025 | 0.0033 |
| F02 | 6640001 | 6740000 | 0.0040 | 0.0025 | 0.0032 |
| F02 | 6660001 | 6760000 | 0.0038 | 0.0023 | 0.0031 |
| F02 | 6680001 | 6780000 | 0.0031 | 0.0019 | 0.0026 |
| F02 | 6700001 | 6800000 | 0.0026 | 0.0016 | 0.0021 |
| F02 | 6720001 | 6820000 | 0.0021 | 0.0015 | 0.0018 |
| F02 | 6740001 | 6840000 | 0.0021 | 0.0011 | 0.0015 |
| F02 | 6760001 | 6860000 | 0.0020 | 0.0012 | 0.0015 |
| F02 | 6780001 | 6880000 | 0.0025 | 0.0019 | 0.0022 |
| F02 | 6800001 | 6900000 | 0.0028 | 0.0020 | 0.0023 |
| F02 | 6820001 | 6920000 | 0.0034 | 0.0025 | 0.0028 |
| F02 | 6840001 | 6940000 | 0.0031 | 0.0023 | 0.0024 |
| F02 | 6860001 | 6960000 | 0.0033 | 0.0024 | 0.0026 |
| F02 | 6880001 | 6980000 | 0.0038 | 0.0026 | 0.0029 |
| F02 | 6900001 | 7000000 | 0.0045 | 0.0033 | 0.0037 |
| F02 | 6920001 | 7020000 | 0.0050 | 0.0037 | 0.0042 |
| F02 | 6940001 | 7040000 | 0.0064 | 0.0049 | 0.0056 |
| F02 | 6960001 | 7060000 | 0.0070 | 0.0055 | 0.0061 |
| F02 | 6980001 | 7080000 | 0.0065 | 0.0049 | 0.0055 |

| Chr | Start | End | both | A1 | A2 |
|---|---|---|---|---|---|
| F02 | 7000001 | 7100000 | 0.0057 | 0.0042 | 0.0049 |
| F02 | 7020001 | 7120000 | 0.0048 | 0.0035 | 0.0039 |
| F02 | 7040001 | 7140000 | 0.0035 | 0.0023 | 0.0025 |
| F02 | 7060001 | 7160000 | 0.0031 | 0.0018 | 0.0020 |
| F02 | 7080001 | 7180000 | 0.0029 | 0.0020 | 0.0021 |
| F02 | 7100001 | 7200000 | 0.0030 | 0.0020 | 0.0021 |
| F02 | 7120001 | 7220000 | 0.0031 | 0.0022 | 0.0023 |
| F02 | 7140001 | 7240000 | 0.0031 | 0.0023 | 0.0023 |
| F02 | 7160001 | 7260000 | 0.0027 | 0.0020 | 0.0021 |
| F02 | 7180001 | 7280000 | 0.0020 | 0.0014 | 0.0015 |
| F02 | 7200001 | 7300000 | 0.0017 | 0.0012 | 0.0012 |
| F02 | 7220001 | 7320000 | 0.0015 | 0.0008 | 0.0010 |
| F02 | 7240001 | 7340000 | 0.0015 | 0.0007 | 0.0010 |
| F02 | 7260001 | 7360000 | 0.0012 | 0.0004 | 0.0006 |
| F02 | 7280001 | 7380000 | 0.0017 | 0.0008 | 0.0010 |
| F02 | 7300001 | 7400000 | 0.0021 | 0.0012 | 0.0013 |
| F02 | 7320001 | 7420000 | 0.0025 | 0.0017 | 0.0018 |
| F02 | 7340001 | 7440000 | 0.0027 | 0.0018 | 0.0019 |
| F02 | 7360001 | 7460000 | 0.0040 | 0.0030 | 0.0034 |
| F02 | 7380001 | 7480000 | 0.0043 | 0.0030 | 0.0036 |
| F02 | 7400001 | 7500000 | 0.0041 | 0.0028 | 0.0035 |
| F02 | 7420001 | 7520000 | 0.0040 | 0.0026 | 0.0031 |
| F02 | 7440001 | 7540000 | 0.0044 | 0.0029 | 0.0037 |
| F02 | 7460001 | 7560000 | 0.0038 | 0.0025 | 0.0029 |
| F02 | 7480001 | 7580000 | 0.0042 | 0.0030 | 0.0034 |
| F02 | 7500001 | 7600000 | 0.0048 | 0.0036 | 0.0039 |
| F02 | 7520001 | 7620000 | 0.0044 | 0.0032 | 0.0036 |
| F02 | 7540001 | 7640000 | 0.0047 | 0.0036 | 0.0039 |
| F02 | 7560001 | 7660000 | 0.0043 | 0.0033 | 0.0037 |
| F02 | 7580001 | 7680000 | 0.0032 | 0.0024 | 0.0026 |
| F02 | 7600001 | 7700000 | 0.0024 | 0.0019 | 0.0021 |
| F02 | 7620001 | 7720000 | 0.0027 | 0.0021 | 0.0023 |
| F02 | 7640001 | 7740000 | 0.0023 | 0.0017 | 0.0020 |
| F02 | 7660001 | 7760000 | 0.0021 | 0.0013 | 0.0016 |
| F02 | 7680001 | 7780000 | 0.0022 | 0.0013 | 0.0016 |
| F02 | 7700001 | 7800000 | 0.0021 | 0.0012 | 0.0014 |
| F02 | 7720001 | 7820000 | 0.0018 | 0.0010 | 0.0012 |
| F02 | 7740001 | 7840000 | 0.0009 | 0.0004 | 0.0005 |
| F02 | 7760001 | 7860000 | 0.0011 | 0.0008 | 0.0007 |
| F02 | 7780001 | 7880000 | 0.0011 | 0.0007 | 0.0007 |
| F02 | 7800001 | 7900000 | 0.0010 | 0.0007 | 0.0006 |
| F02 | 7820001 | 7920000 | 0.0010 | 0.0008 | 0.0008 |
| F02 | 7840001 | 7940000 | 0.0013 | 0.0010 | 0.0010 |
| F02 | 7860001 | 7960000 | 0.0011 | 0.0008 | 0.0009 |
| F02 | 7880001 | 7980000 | 0.0009 | 0.0008 | 0.0008 |
| F02 | 7900001 | 8000000 | 0.0014 | 0.0012 | 0.0012 |
| F02 | 7920001 | 8020000 | 0.0020 | 0.0016 | 0.0017 |
| F02 | 7940001 | 8040000 | 0.0028 | 0.0023 | 0.0024 |
| F02 | 7960001 | 8060000 | 0.0036 | 0.0029 | 0.0032 |
| F02 | 7980001 | 8080000 | 0.0042 | 0.0033 | 0.0037 |
| F02 | 8000001 | 8100000 | 0.0050 | 0.0040 | 0.0044 |
| F02 | 8020001 | 8120000 | 0.0048 | 0.0037 | 0.0040 |
| F02 | 8040001 | 8140000 | 0.0047 | 0.0033 | 0.0038 |
| F02 | 8060001 | 8160000 | 0.0039 | 0.0025 | 0.0028 |
| F02 | 8080001 | 8180000 | 0.0034 | 0.0022 | 0.0024 |
| F02 | 8100001 | 8200000 | 0.0021 | 0.0012 | 0.0013 |
| F02 | 8120001 | 8220000 | 0.0022 | 0.0014 | 0.0017 |
| F02 | 8140001 | 8240000 | 0.0024 | 0.0018 | 0.0021 |

| Chr | Start | End | both | A1 | A2 |
|---|---|---|---|---|---|
| F02 | 8160001 | 8260000 | 0.0028 | 0.0021 | 0.0025 |
| F02 | 8180001 | 8280000 | 0.0035 | 0.0025 | 0.0030 |
| F02 | 8200001 | 8300000 | 0.0048 | 0.0035 | 0.0044 |
| F02 | 8220001 | 8320000 | 0.0048 | 0.0036 | 0.0043 |
| F02 | 8240001 | 8340000 | 0.0050 | 0.0038 | 0.0044 |
| F02 | 8260001 | 8360000 | 0.0049 | 0.0038 | 0.0045 |
| F02 | 8280001 | 8380000 | 0.0054 | 0.0044 | 0.0049 |
| F02 | 8300001 | 8400000 | 0.0045 | 0.0036 | 0.0039 |
| F02 | 8320001 | 8420000 | 0.0052 | 0.0042 | 0.0046 |
| F02 | 8340001 | 8440000 | 0.0049 | 0.0040 | 0.0044 |
| F02 | 8360001 | 8460000 | 0.0048 | 0.0038 | 0.0042 |
| F02 | 8380001 | 8480000 | 0.0042 | 0.0033 | 0.0037 |
| F02 | 8400001 | 8500000 | 0.0041 | 0.0035 | 0.0037 |
| F02 | 8420001 | 8520000 | 0.0034 | 0.0029 | 0.0030 |
| F02 | 8440001 | 8540000 | 0.0028 | 0.0023 | 0.0023 |
| F02 | 8460001 | 8560000 | 0.0028 | 0.0023 | 0.0024 |
| F02 | 8480001 | 8580000 | 0.0027 | 0.0022 | 0.0024 |
| F02 | 8500001 | 8600000 | 0.0030 | 0.0022 | 0.0024 |
| F02 | 8520001 | 8620000 | 0.0028 | 0.0020 | 0.0022 |
| F02 | 8540001 | 8640000 | 0.0030 | 0.0021 | 0.0024 |
| F02 | 8560001 | 8660000 | 0.0033 | 0.0024 | 0.0026 |
| F02 | 8580001 | 8680000 | 0.0039 | 0.0028 | 0.0030 |
| F02 | 8600001 | 8700000 | 0.0033 | 0.0026 | 0.0027 |
| F02 | 8620001 | 8720000 | 0.0031 | 0.0023 | 0.0024 |
| F02 | 8640001 | 8740000 | 0.0027 | 0.0019 | 0.0020 |
| F02 | 8660001 | 8760000 | 0.0025 | 0.0015 | 0.0016 |
| F02 | 8680001 | 8780000 | 0.0018 | 0.0011 | 0.0012 |
| F02 | 8700001 | 8800000 | 0.0028 | 0.0018 | 0.0020 |
| F02 | 8720001 | 8820000 | 0.0033 | 0.0024 | 0.0026 |
| F02 | 8740001 | 8840000 | 0.0040 | 0.0029 | 0.0032 |
| F02 | 8760001 | 8860000 | 0.0040 | 0.0030 | 0.0034 |
| F02 | 8780001 | 8880000 | 0.0037 | 0.0027 | 0.0032 |
| F02 | 8800001 | 8900000 | 0.0027 | 0.0019 | 0.0022 |
| F02 | 8820001 | 8920000 | 0.0025 | 0.0015 | 0.0019 |
| F02 | 8840001 | 8940000 | 0.0023 | 0.0016 | 0.0019 |
| F02 | 8860001 | 8960000 | 0.0030 | 0.0023 | 0.0025 |
| F02 | 8880001 | 8980000 | 0.0033 | 0.0025 | 0.0026 |
| F02 | 8900001 | 9000000 | 0.0036 | 0.0028 | 0.0029 |
| F02 | 8920001 | 9020000 | 0.0038 | 0.0032 | 0.0032 |
| F02 | 8940001 | 9040000 | 0.0042 | 0.0035 | 0.0037 |
| F02 | 8960001 | 9060000 | 0.0043 | 0.0033 | 0.0036 |
| F02 | 8980001 | 9080000 | 0.0041 | 0.0033 | 0.0037 |
| F02 | 9000001 | 9100000 | 0.0042 | 0.0033 | 0.0038 |
| F02 | 9020001 | 9120000 | 0.0041 | 0.0032 | 0.0037 |
| F02 | 9040001 | 9140000 | 0.0040 | 0.0030 | 0.0035 |
| F02 | 9060001 | 9160000 | 0.0042 | 0.0033 | 0.0036 |
| F02 | 9080001 | 9180000 | 0.0048 | 0.0036 | 0.0039 |
| F02 | 9100001 | 9200000 | 0.0051 | 0.0038 | 0.0042 |
| F02 | 9120001 | 9220000 | 0.0050 | 0.0035 | 0.0040 |
| F02 | 9140001 | 9240000 | 0.0051 | 0.0038 | 0.0041 |
| F02 | 9160001 | 9260000 | 0.0041 | 0.0029 | 0.0032 |
| F02 | 9180001 | 9280000 | 0.0042 | 0.0033 | 0.0035 |
| F02 | 9200001 | 9300000 | 0.0044 | 0.0036 | 0.0037 |
| F02 | 9220001 | 9320000 | 0.0054 | 0.0045 | 0.0044 |
| F02 | 9240001 | 9340000 | 0.0047 | 0.0036 | 0.0036 |
| F02 | 9260001 | 9360000 | 0.0056 | 0.0044 | 0.0045 |
| F02 | 9280001 | 9380000 | 0.0060 | 0.0047 | 0.0049 |
| F02 | 9300001 | 9400000 | 0.0057 | 0.0045 | 0.0047 |

| Chr | Start | End | both | A1 | A2 |
|---|---|---|---|---|---|
| F02 | 9320001 | 9420000 | 0.0050 | 0.0037 | 0.0041 |
| F02 | 9340001 | 9440000 | 0.0051 | 0.0040 | 0.0044 |
| F02 | 9360001 | 9460000 | 0.0048 | 0.0038 | 0.0041 |
| F02 | 9380001 | 9480000 | 0.0034 | 0.0027 | 0.0029 |
| F02 | 9400001 | 9500000 | 0.0033 | 0.0026 | 0.0028 |
| F02 | 9420001 | 9520000 | 0.0028 | 0.0022 | 0.0023 |
| F02 | 9440001 | 9540000 | 0.0024 | 0.0020 | 0.0021 |
| F02 | 9460001 | 9560000 | 0.0017 | 0.0014 | 0.0016 |
| F02 | 9480001 | 9580000 | 0.0026 | 0.0021 | 0.0024 |
| F02 | 9500001 | 9600000 | 0.0024 | 0.0019 | 0.0023 |
| F02 | 9520001 | 9620000 | 0.0029 | 0.0022 | 0.0025 |
| F02 | 9540001 | 9640000 | 0.0032 | 0.0025 | 0.0028 |
| F02 | 9560001 | 9660000 | 0.0032 | 0.0025 | 0.0027 |
| F02 | 9580001 | 9680000 | 0.0025 | 0.0020 | 0.0022 |
| F02 | 9600001 | 9700000 | 0.0029 | 0.0022 | 0.0024 |
| F02 | 9620001 | 9720000 | 0.0030 | 0.0024 | 0.0028 |
| F02 | 9640001 | 9740000 | 0.0027 | 0.0022 | 0.0025 |
| F02 | 9660001 | 9760000 | 0.0027 | 0.0023 | 0.0025 |
| F02 | 9680001 | 9780000 | 0.0030 | 0.0026 | 0.0028 |
| F02 | 9700001 | 9800000 | 0.0032 | 0.0029 | 0.0031 |
| F02 | 9720001 | 9820000 | 0.0055 | 0.0052 | 0.0053 |
| F02 | 9740001 | 9840000 | 0.0059 | 0.0055 | 0.0056 |
| F02 | 9760001 | 9860000 | 0.0058 | 0.0052 | 0.0055 |
| F02 | 9780001 | 9880000 | 0.0055 | 0.0048 | 0.0052 |
| F02 | 9800001 | 9900000 | 0.0054 | 0.0046 | 0.0051 |
| F02 | 9820001 | 9920000 | 0.0033 | 0.0026 | 0.0031 |
| F02 | 9840001 | 9940000 | 0.0039 | 0.0032 | 0.0037 |
| F02 | 9860001 | 9960000 | 0.0038 | 0.0032 | 0.0036 |
| F02 | 9880001 | 9980000 | 0.0042 | 0.0037 | 0.0040 |
| F02 | 9900001 | 10000000 | 0.0037 | 0.0033 | 0.0035 |
| F02 | 9920001 | 10020000 | 0.0037 | 0.0031 | 0.0035 |
| F02 | 9940001 | 10040000 | 0.0033 | 0.0027 | 0.0031 |
| F02 | 9960001 | 10060000 | 0.0036 | 0.0031 | 0.0035 |
| F02 | 9980001 | 10080000 | 0.0038 | 0.0032 | 0.0036 |
| F02 | 10000001 | 10100000 | 0.0044 | 0.0036 | 0.0042 |
| F02 | 10020001 | 10120000 | 0.0043 | 0.0036 | 0.0041 |
| F02 | 10040001 | 10140000 | 0.0050 | 0.0041 | 0.0048 |
| F02 | 10060001 | 10160000 | 0.0051 | 0.0041 | 0.0048 |
| F02 | 10080001 | 10180000 | 0.0062 | 0.0052 | 0.0061 |
| F02 | 10100001 | 10200000 | 0.0058 | 0.0050 | 0.0057 |
| F02 | 10120001 | 10220000 | 0.0057 | 0.0049 | 0.0055 |
| F02 | 10140001 | 10240000 | 0.0046 | 0.0041 | 0.0045 |
| F02 | 10160001 | 10260000 | 0.0046 | 0.0041 | 0.0044 |
| F02 | 10180001 | 10280000 | 0.0031 | 0.0025 | 0.0028 |
| F02 | 10200001 | 10300000 | 0.0026 | 0.0021 | 0.0023 |
| F02 | 10220001 | 10320000 | 0.0022 | 0.0016 | 0.0019 |
| F02 | 10240001 | 10340000 | 0.0020 | 0.0014 | 0.0018 |
| F02 | 10260001 | 10360000 | 0.0015 | 0.0009 | 0.0013 |
| F02 | 10280001 | 10380000 | 0.0009 | 0.0006 | 0.0009 |
| F02 | 10300001 | 10400000 | 0.0015 | 0.0011 | 0.0014 |
| F02 | 10320001 | 10420000 | 0.0015 | 0.0011 | 0.0014 |
| F02 | 10340001 | 10440000 | 0.0018 | 0.0014 | 0.0015 |
| F02 | 10360001 | 10460000 | 0.0022 | 0.0017 | 0.0019 |
| F02 | 10380001 | 10480000 | 0.0025 | 0.0019 | 0.0022 |
| F02 | 10400001 | 10500000 | 0.0031 | 0.0025 | 0.0026 |
| F02 | 10420001 | 10520000 | 0.0028 | 0.0024 | 0.0025 |
| F02 | 10440001 | 10540000 | 0.0029 | 0.0025 | 0.0027 |
| F02 | 10460001 | 10560000 | 0.0027 | 0.0021 | 0.0023 |

| Chr | Start | End | both | A1 | A2 |
|---|---|---|---|---|---|
| F02 | 10480001 | 10580000 | 0.0029 | 0.0023 | 0.0025 |
| F02 | 10500001 | 10600000 | 0.0023 | 0.0016 | 0.0017 |
| F02 | 10520001 | 10620000 | 0.0030 | 0.0021 | 0.0022 |
| F02 | 10540001 | 10640000 | 0.0034 | 0.0024 | 0.0025 |
| F02 | 10560001 | 10660000 | 0.0039 | 0.0030 | 0.0030 |
| F02 | 10580001 | 10680000 | 0.0038 | 0.0026 | 0.0027 |
| F02 | 10600001 | 10700000 | 0.0042 | 0.0031 | 0.0032 |
| F02 | 10620001 | 10720000 | 0.0042 | 0.0031 | 0.0033 |
| F02 | 10640001 | 10740000 | 0.0034 | 0.0024 | 0.0027 |
| F02 | 10660001 | 10760000 | 0.0031 | 0.0022 | 0.0025 |
| F02 | 10680001 | 10780000 | 0.0034 | 0.0028 | 0.0030 |
| F02 | 10700001 | 10800000 | 0.0027 | 0.0024 | 0.0025 |
| F02 | 10720001 | 10820000 | 0.0029 | 0.0025 | 0.0026 |
| F02 | 10740001 | 10840000 | 0.0036 | 0.0031 | 0.0031 |
| F02 | 10760001 | 10860000 | 0.0036 | 0.0029 | 0.0030 |
| F02 | 10780001 | 10880000 | 0.0032 | 0.0024 | 0.0026 |
| F02 | 10800001 | 10900000 | 0.0032 | 0.0022 | 0.0026 |
| F02 | 10820001 | 10920000 | 0.0026 | 0.0019 | 0.0022 |
| F02 | 10840001 | 10940000 | 0.0023 | 0.0016 | 0.0020 |
| F02 | 10860001 | 10960000 | 0.0021 | 0.0016 | 0.0018 |
| F02 | 10880001 | 10980000 | 0.0021 | 0.0016 | 0.0017 |
| F02 | 10900001 | 11000000 | 0.0016 | 0.0013 | 0.0013 |
| F02 | 10920001 | 11020000 | 0.0029 | 0.0023 | 0.0025 |
| F02 | 10940001 | 11040000 | 0.0028 | 0.0020 | 0.0023 |
| F02 | 10960001 | 11060000 | 0.0033 | 0.0029 | 0.0030 |
| F02 | 10980001 | 11080000 | 0.0042 | 0.0037 | 0.0039 |
| F02 | 11000001 | 11100000 | 0.0046 | 0.0040 | 0.0042 |
| F02 | 11020001 | 11120000 | 0.0035 | 0.0029 | 0.0030 |
| F02 | 11040001 | 11140000 | 0.0036 | 0.0028 | 0.0030 |
| F02 | 11060001 | 11160000 | 0.0029 | 0.0019 | 0.0024 |
| F02 | 11080001 | 11180000 | 0.0025 | 0.0016 | 0.0021 |
| F02 | 11100001 | 11200000 | 0.0028 | 0.0020 | 0.0026 |
| F02 | 11120001 | 11220000 | 0.0023 | 0.0017 | 0.0021 |
| F02 | 11140001 | 11240000 | 0.0019 | 0.0016 | 0.0019 |
| F02 | 11160001 | 11260000 | 0.0023 | 0.0021 | 0.0023 |
| F02 | 11180001 | 11280000 | 0.0016 | 0.0014 | 0.0015 |
| F02 | 11200001 | 11300000 | 0.0010 | 0.0010 | 0.0010 |
| F02 | 11220001 | 11320000 | 0.0011 | 0.0011 | 0.0011 |
| F02 | 11240001 | 11340000 | 0.0019 | 0.0018 | 0.0018 |
| F02 | 11260001 | 11360000 | 0.0020 | 0.0016 | 0.0017 |
| F02 | 11280001 | 11380000 | 0.0025 | 0.0018 | 0.0020 |
| F02 | 11300001 | 11400000 | 0.0037 | 0.0028 | 0.0030 |
| F02 | 11320001 | 11420000 | 0.0050 | 0.0040 | 0.0041 |
| F02 | 11340001 | 11440000 | 0.0056 | 0.0042 | 0.0045 |
| F02 | 11360001 | 11460000 | 0.0055 | 0.0040 | 0.0043 |
| F02 | 11380001 | 11480000 | 0.0052 | 0.0039 | 0.0042 |
| F02 | 11400001 | 11500000 | 0.0044 | 0.0031 | 0.0035 |
| F02 | 11420001 | 11520000 | 0.0038 | 0.0023 | 0.0029 |
| F02 | 11440001 | 11540000 | 0.0027 | 0.0017 | 0.0021 |
| F02 | 11460001 | 11560000 | 0.0021 | 0.0014 | 0.0018 |
| F02 | 11480001 | 11580000 | 0.0025 | 0.0018 | 0.0022 |
| F02 | 11500001 | 11600000 | 0.0025 | 0.0016 | 0.0022 |
| F02 | 11520001 | 11620000 | 0.0026 | 0.0017 | 0.0023 |
| F02 | 11540001 | 11640000 | 0.0028 | 0.0019 | 0.0024 |
| F02 | 11560001 | 11660000 | 0.0033 | 0.0024 | 0.0029 |
| F02 | 11580001 | 11680000 | 0.0030 | 0.0021 | 0.0025 |
| F02 | 11600001 | 11700000 | 0.0024 | 0.0019 | 0.0020 |
| F02 | 11620001 | 11720000 | 0.0026 | 0.0021 | 0.0021 |

| Chr | Start | End | both | A1 | A2 |
|---|---|---|---|---|---|
| F02 | 11640001 | 11740000 | 0.0029 | 0.0022 | 0.0023 |
| F02 | 11660001 | 11760000 | 0.0037 | 0.0029 | 0.0031 |
| F02 | 11680001 | 11780000 | 0.0037 | 0.0028 | 0.0031 |
| F02 | 11700001 | 11800000 | 0.0047 | 0.0036 | 0.0040 |
| F02 | 11720001 | 11820000 | 0.0048 | 0.0037 | 0.0040 |
| F02 | 11740001 | 11840000 | 0.0052 | 0.0040 | 0.0043 |
| F02 | 11760001 | 11860000 | 0.0045 | 0.0034 | 0.0036 |
| F02 | 11780001 | 11880000 | 0.0049 | 0.0038 | 0.0039 |
| F02 | 11800001 | 11900000 | 0.0045 | 0.0035 | 0.0035 |
| F02 | 11820001 | 11920000 | 0.0047 | 0.0037 | 0.0041 |
| F02 | 11840001 | 11940000 | 0.0043 | 0.0033 | 0.0037 |
| F02 | 11860001 | 11960000 | 0.0047 | 0.0035 | 0.0040 |
| F02 | 11880001 | 11980000 | 0.0046 | 0.0033 | 0.0039 |
| F02 | 11900001 | 12000000 | 0.0038 | 0.0027 | 0.0034 |
| F02 | 11920001 | 12020000 | 0.0025 | 0.0017 | 0.0021 |
| F02 | 11940001 | 12040000 | 0.0020 | 0.0012 | 0.0016 |
| F02 | 11960001 | 12060000 | 0.0025 | 0.0017 | 0.0021 |
| F02 | 11980001 | 12080000 | 0.0030 | 0.0023 | 0.0026 |
| F02 | 12000001 | 12100000 | 0.0038 | 0.0029 | 0.0033 |
| F02 | 12020001 | 12120000 | 0.0037 | 0.0029 | 0.0032 |
| F02 | 12040001 | 12140000 | 0.0037 | 0.0029 | 0.0031 |
| F02 | 12060001 | 12160000 | 0.0023 | 0.0018 | 0.0020 |
| F02 | 12080001 | 12180000 | 0.0019 | 0.0012 | 0.0013 |
| F02 | 12100001 | 12200000 | 0.0012 | 0.0007 | 0.0007 |
| F02 | 12120001 | 12220000 | 0.0016 | 0.0009 | 0.0010 |
| F02 | 12140001 | 12240000 | 0.0026 | 0.0017 | 0.0018 |
| F02 | 12160001 | 12260000 | 0.0030 | 0.0018 | 0.0019 |
| F02 | 12180001 | 12280000 | 0.0036 | 0.0022 | 0.0023 |
| F02 | 12200001 | 12300000 | 0.0039 | 0.0025 | 0.0026 |
| F02 | 12220001 | 12320000 | 0.0040 | 0.0024 | 0.0026 |
| F02 | 12240001 | 12340000 | 0.0032 | 0.0019 | 0.0020 |
| F02 | 12260001 | 12360000 | 0.0038 | 0.0028 | 0.0031 |
| F02 | 12280001 | 12380000 | 0.0030 | 0.0024 | 0.0026 |
| F02 | 12300001 | 12400000 | 0.0026 | 0.0021 | 0.0023 |
| F02 | 12320001 | 12420000 | 0.0022 | 0.0019 | 0.0021 |
| F02 | 12340001 | 12440000 | 0.0024 | 0.0019 | 0.0021 |
| F02 | 12360001 | 12460000 | 0.0020 | 0.0014 | 0.0015 |
| F02 | 12380001 | 12480000 | 0.0023 | 0.0016 | 0.0017 |
| F02 | 12400001 | 12500000 | 0.0032 | 0.0021 | 0.0023 |
| F02 | 12420001 | 12520000 | 0.0032 | 0.0021 | 0.0022 |
| F02 | 12440001 | 12540000 | 0.0038 | 0.0029 | 0.0030 |
| F02 | 12460001 | 12560000 | 0.0060 | 0.0047 | 0.0053 |
| F02 | 12480001 | 12580000 | 0.0054 | 0.0043 | 0.0048 |
| F02 | 12500001 | 12600000 | 0.0049 | 0.0040 | 0.0044 |
| F02 | 12520001 | 12620000 | 0.0049 | 0.0040 | 0.0045 |
| F02 | 12540001 | 12640000 | 0.0039 | 0.0032 | 0.0036 |
| F02 | 12560001 | 12660000 | 0.0015 | 0.0012 | 0.0013 |
| F02 | 12580001 | 12680000 | 0.0019 | 0.0015 | 0.0017 |
| F02 | 12600001 | 12700000 | 0.0018 | 0.0014 | 0.0017 |
| F02 | 12620001 | 12720000 | 0.0028 | 0.0022 | 0.0026 |
| F02 | 12640001 | 12740000 | 0.0028 | 0.0022 | 0.0026 |
| F02 | 12660001 | 12760000 | 0.0043 | 0.0034 | 0.0040 |
| F02 | 12680001 | 12780000 | 0.0048 | 0.0039 | 0.0044 |
| F02 | 12700001 | 12800000 | 0.0049 | 0.0040 | 0.0045 |
| F02 | 12720001 | 12820000 | 0.0050 | 0.0040 | 0.0046 |
| F02 | 12740001 | 12840000 | 0.0051 | 0.0038 | 0.0044 |
| F02 | 12760001 | 12860000 | 0.0034 | 0.0023 | 0.0026 |
| F02 | 12780001 | 12880000 | 0.0029 | 0.0018 | 0.0022 |

| Chr | Start | End | both | A1 | A2 |
|---|---|---|---|---|---|
| F02 | 12800001 | 12900000 | 0.0031 | 0.0019 | 0.0023 |
| F02 | 12820001 | 12920000 | 0.0024 | 0.0013 | 0.0018 |
| F02 | 12840001 | 12940000 | 0.0026 | 0.0014 | 0.0020 |
| F02 | 12860001 | 12960000 | 0.0028 | 0.0017 | 0.0021 |
| F02 | 12880001 | 12980000 | 0.0030 | 0.0017 | 0.0022 |
| F02 | 12900001 | 13000000 | 0.0030 | 0.0015 | 0.0019 |
| F02 | 12920001 | 13020000 | 0.0036 | 0.0023 | 0.0025 |
| F02 | 12940001 | 13040000 | 0.0039 | 0.0027 | 0.0029 |
| F02 | 12960001 | 13060000 | 0.0036 | 0.0023 | 0.0026 |
| F02 | 12980001 | 13080000 | 0.0036 | 0.0025 | 0.0027 |
| F02 | 13000001 | 13100000 | 0.0035 | 0.0026 | 0.0027 |
| F02 | 13020001 | 13120000 | 0.0032 | 0.0023 | 0.0024 |
| F02 | 13040001 | 13140000 | 0.0028 | 0.0018 | 0.0020 |
| F02 | 13060001 | 13160000 | 0.0028 | 0.0018 | 0.0020 |
| F02 | 13080001 | 13180000 | 0.0025 | 0.0015 | 0.0017 |
| F02 | 13100001 | 13200000 | 0.0029 | 0.0018 | 0.0021 |
| F02 | 13120001 | 13220000 | 0.0033 | 0.0021 | 0.0023 |
| F02 | 13140001 | 13240000 | 0.0047 | 0.0033 | 0.0036 |
| F02 | 13160001 | 13260000 | 0.0057 | 0.0048 | 0.0045 |
| F02 | 13180001 | 13280000 | 0.0069 | 0.0058 | 0.0056 |
| F02 | 13200001 | 13300000 | 0.0070 | 0.0058 | 0.0056 |
| F02 | 13220001 | 13320000 | 0.0064 | 0.0053 | 0.0051 |
| F02 | 13240001 | 13340000 | 0.0053 | 0.0043 | 0.0040 |
| F02 | 13260001 | 13360000 | 0.0044 | 0.0030 | 0.0033 |
| F02 | 13280001 | 13380000 | 0.0030 | 0.0019 | 0.0021 |
| F02 | 13300001 | 13400000 | 0.0024 | 0.0015 | 0.0016 |
| F02 | 13320001 | 13420000 | 0.0022 | 0.0014 | 0.0016 |
| F02 | 13340001 | 13440000 | 0.0022 | 0.0016 | 0.0016 |
| F02 | 13360001 | 13460000 | 0.0023 | 0.0014 | 0.0014 |
| F02 | 13380001 | 13480000 | 0.0025 | 0.0016 | 0.0016 |
| F02 | 13400001 | 13500000 | 0.0025 | 0.0015 | 0.0016 |
| F02 | 13420001 | 13520000 | 0.0029 | 0.0015 | 0.0016 |
| F02 | 13440001 | 13540000 | 0.0036 | 0.0021 | 0.0024 |
| F02 | 13460001 | 13560000 | 0.0045 | 0.0028 | 0.0034 |
| F02 | 13480001 | 13580000 | 0.0051 | 0.0034 | 0.0039 |
| F02 | 13500001 | 13600000 | 0.0051 | 0.0034 | 0.0039 |
| F02 | 13520001 | 13620000 | 0.0049 | 0.0033 | 0.0037 |
| F02 | 13540001 | 13640000 | 0.0039 | 0.0025 | 0.0029 |
| F02 | 13560001 | 13660000 | 0.0030 | 0.0021 | 0.0021 |
| F02 | 13580001 | 13680000 | 0.0021 | 0.0015 | 0.0016 |
| F02 | 13600001 | 13700000 | 0.0025 | 0.0019 | 0.0020 |
| F02 | 13620001 | 13720000 | 0.0020 | 0.0018 | 0.0018 |
| F02 | 13640001 | 13740000 | 0.0024 | 0.0020 | 0.0021 |
| F02 | 13660001 | 13760000 | 0.0027 | 0.0021 | 0.0023 |
| F02 | 13680001 | 13780000 | 0.0028 | 0.0020 | 0.0023 |
| F02 | 13700001 | 13800000 | 0.0030 | 0.0021 | 0.0025 |
| F02 | 13720001 | 13820000 | 0.0038 | 0.0027 | 0.0031 |
| F02 | 13740001 | 13840000 | 0.0050 | 0.0033 | 0.0043 |
| F02 | 13760001 | 13860000 | 0.0049 | 0.0031 | 0.0041 |
| F02 | 13780001 | 13880000 | 0.0052 | 0.0035 | 0.0044 |
| F02 | 13800001 | 13900000 | 0.0057 | 0.0040 | 0.0047 |
| F02 | 13820001 | 13920000 | 0.0053 | 0.0036 | 0.0042 |
| F02 | 13840001 | 13940000 | 0.0034 | 0.0024 | 0.0023 |
| F02 | 13860001 | 13960000 | 0.0035 | 0.0022 | 0.0021 |
| F02 | 13880001 | 13980000 | 0.0034 | 0.0018 | 0.0017 |
| F02 | 13900001 | 14000000 | 0.0031 | 0.0013 | 0.0015 |
| F02 | 13920001 | 14020000 | 0.0036 | 0.0018 | 0.0021 |
| F02 | 13940001 | 14040000 | 0.0037 | 0.0020 | 0.0023 |

| Chr | Start | End | both | A1 | A2 |
|---|---|---|---|---|---|
| F02 | 13960001 | 14060000 | 0.0036 | 0.0020 | 0.0022 |
| F02 | 13980001 | 14080000 | 0.0038 | 0.0024 | 0.0025 |
| F02 | 14000001 | 14100000 | 0.0031 | 0.0018 | 0.0019 |
| F02 | 14020001 | 14120000 | 0.0026 | 0.0014 | 0.0014 |
| F02 | 14040001 | 14140000 | 0.0034 | 0.0021 | 0.0023 |
| F02 | 14060001 | 14160000 | 0.0041 | 0.0031 | 0.0034 |
| F02 | 14080001 | 14180000 | 0.0042 | 0.0029 | 0.0034 |
| F02 | 14100001 | 14200000 | 0.0041 | 0.0030 | 0.0035 |
| F02 | 14120001 | 14220000 | 0.0038 | 0.0028 | 0.0033 |
| F02 | 14140001 | 14240000 | 0.0029 | 0.0020 | 0.0024 |
| F02 | 14160001 | 14260000 | 0.0017 | 0.0010 | 0.0013 |
| F02 | 14180001 | 14280000 | 0.0017 | 0.0009 | 0.0012 |
| F02 | 14200001 | 14300000 | 0.0016 | 0.0008 | 0.0010 |
| F02 | 14220001 | 14320000 | 0.0013 | 0.0006 | 0.0007 |
| F02 | 14240001 | 14340000 | 0.0020 | 0.0008 | 0.0012 |
| F02 | 14260001 | 14360000 | 0.0027 | 0.0012 | 0.0016 |
| F02 | 14280001 | 14380000 | 0.0025 | 0.0011 | 0.0014 |
| F02 | 14300001 | 14400000 | 0.0032 | 0.0017 | 0.0019 |
| F02 | 14320001 | 14420000 | 0.0037 | 0.0019 | 0.0022 |
| F02 | 14340001 | 14440000 | 0.0032 | 0.0018 | 0.0018 |
| F02 | 14360001 | 14460000 | 0.0034 | 0.0020 | 0.0022 |
| F02 | 14380001 | 14480000 | 0.0034 | 0.0019 | 0.0023 |
| F02 | 14400001 | 14500000 | 0.0035 | 0.0022 | 0.0025 |
| F02 | 14420001 | 14520000 | 0.0040 | 0.0027 | 0.0031 |
| F02 | 14440001 | 14540000 | 0.0041 | 0.0027 | 0.0032 |
| F02 | 14460001 | 14560000 | 0.0037 | 0.0025 | 0.0028 |
| F02 | 14480001 | 14580000 | 0.0035 | 0.0024 | 0.0027 |
| F02 | 14500001 | 14600000 | 0.0026 | 0.0016 | 0.0020 |
| F02 | 14520001 | 14620000 | 0.0018 | 0.0011 | 0.0014 |
| F02 | 14540001 | 14640000 | 0.0016 | 0.0009 | 0.0011 |
| F02 | 14560001 | 14660000 | 0.0017 | 0.0008 | 0.0009 |
| F02 | 14580001 | 14680000 | 0.0023 | 0.0010 | 0.0013 |
| F02 | 14600001 | 14700000 | 0.0036 | 0.0016 | 0.0020 |
| F02 | 14620001 | 14720000 | 0.0043 | 0.0023 | 0.0026 |
| F02 | 14640001 | 14740000 | 0.0049 | 0.0024 | 0.0029 |
| F02 | 14660001 | 14760000 | 0.0048 | 0.0026 | 0.0031 |
| F02 | 14680001 | 14780000 | 0.0045 | 0.0024 | 0.0029 |
| F02 | 14700001 | 14800000 | 0.0043 | 0.0023 | 0.0030 |
| F02 | 14720001 | 14820000 | 0.0036 | 0.0016 | 0.0023 |
| F02 | 14740001 | 14840000 | 0.0040 | 0.0023 | 0.0032 |
| F02 | 14760001 | 14860000 | 0.0048 | 0.0025 | 0.0038 |
| F02 | 14780001 | 14880000 | 0.0047 | 0.0024 | 0.0038 |
| F02 | 14800001 | 14900000 | 0.0035 | 0.0020 | 0.0029 |
| F02 | 14820001 | 14920000 | 0.0037 | 0.0020 | 0.0031 |
| F02 | 14840001 | 14940000 | 0.0028 | 0.0013 | 0.0022 |
| F02 | 14860001 | 14960000 | 0.0019 | 0.0011 | 0.0015 |
| F02 | 14880001 | 14980000 | 0.0018 | 0.0013 | 0.0016 |
| F02 | 14900001 | 15000000 | 0.0021 | 0.0015 | 0.0018 |
| F02 | 14920001 | 15020000 | 0.0027 | 0.0020 | 0.0022 |
| F02 | 14940001 | 15040000 | 0.0032 | 0.0024 | 0.0026 |
| F02 | 14960001 | 15060000 | 0.0040 | 0.0031 | 0.0033 |
| F02 | 14980001 | 15080000 | 0.0043 | 0.0033 | 0.0034 |
| F02 | 15000001 | 15100000 | 0.0038 | 0.0029 | 0.0030 |
| F02 | 15020001 | 15120000 | 0.0035 | 0.0025 | 0.0027 |
| F02 | 15040001 | 15140000 | 0.0036 | 0.0025 | 0.0028 |
| F02 | 15060001 | 15160000 | 0.0025 | 0.0016 | 0.0018 |
| F02 | 15080001 | 15180000 | 0.0023 | 0.0014 | 0.0017 |
| F02 | 15100001 | 15200000 | 0.0036 | 0.0024 | 0.0026 |

| Chr | Start | End | both | A1 | A2 |
|---|---|---|---|---|---|
| F02 | 15120001 | 15220000 | 0.0034 | 0.0022 | 0.0023 |
| F02 | 15140001 | 15240000 | 0.0032 | 0.0019 | 0.0021 |
| F02 | 15160001 | 15260000 | 0.0036 | 0.0022 | 0.0025 |
| F02 | 15180001 | 15280000 | 0.0042 | 0.0026 | 0.0030 |
| F02 | 15200001 | 15300000 | 0.0029 | 0.0018 | 0.0022 |
| F02 | 15220001 | 15320000 | 0.0032 | 0.0022 | 0.0026 |
| F02 | 15240001 | 15340000 | 0.0032 | 0.0027 | 0.0028 |
| F02 | 15260001 | 15360000 | 0.0030 | 0.0025 | 0.0025 |
| F02 | 15280001 | 15380000 | 0.0023 | 0.0019 | 0.0019 |
| F02 | 15300001 | 15400000 | 0.0038 | 0.0031 | 0.0032 |
| F02 | 15320001 | 15420000 | 0.0040 | 0.0031 | 0.0034 |
| F02 | 15340001 | 15440000 | 0.0034 | 0.0025 | 0.0027 |
| F02 | 15360001 | 15460000 | 0.0032 | 0.0024 | 0.0027 |
| F02 | 15380001 | 15480000 | 0.0035 | 0.0026 | 0.0029 |
| F02 | 15400001 | 15500000 | 0.0020 | 0.0014 | 0.0016 |
| F02 | 15420001 | 15520000 | 0.0018 | 0.0011 | 0.0013 |
| F02 | 15440001 | 15540000 | 0.0017 | 0.0010 | 0.0013 |
| F02 | 15460001 | 15560000 | 0.0020 | 0.0012 | 0.0015 |
| F02 | 15480001 | 15580000 | 0.0018 | 0.0011 | 0.0013 |
| F02 | 15500001 | 15600000 | 0.0027 | 0.0017 | 0.0020 |
| F02 | 15520001 | 15620000 | 0.0027 | 0.0016 | 0.0018 |
| F02 | 15540001 | 15640000 | 0.0034 | 0.0024 | 0.0026 |
| F02 | 15560001 | 15660000 | 0.0048 | 0.0037 | 0.0039 |
| F02 | 15580001 | 15680000 | 0.0051 | 0.0039 | 0.0042 |
| F02 | 15600001 | 15700000 | 0.0048 | 0.0036 | 0.0039 |
| F02 | 15620001 | 15720000 | 0.0046 | 0.0036 | 0.0040 |
| F02 | 15640001 | 15740000 | 0.0050 | 0.0036 | 0.0042 |
| F02 | 15660001 | 15760000 | 0.0038 | 0.0024 | 0.0028 |
| F02 | 15680001 | 15780000 | 0.0038 | 0.0022 | 0.0026 |
| F02 | 15700001 | 15800000 | 0.0032 | 0.0020 | 0.0023 |
| F02 | 15720001 | 15820000 | 0.0033 | 0.0021 | 0.0024 |
| F02 | 15740001 | 15840000 | 0.0023 | 0.0013 | 0.0016 |
| F02 | 15760001 | 15860000 | 0.0021 | 0.0014 | 0.0017 |
| F02 | 15780001 | 15880000 | 0.0020 | 0.0016 | 0.0018 |
| F02 | 15800001 | 15900000 | 0.0028 | 0.0021 | 0.0024 |
| F02 | 15820001 | 15920000 | 0.0038 | 0.0032 | 0.0034 |
| F02 | 15840001 | 15940000 | 0.0044 | 0.0037 | 0.0039 |
| F02 | 15860001 | 15960000 | 0.0050 | 0.0043 | 0.0045 |
| F02 | 15880001 | 15980000 | 0.0056 | 0.0045 | 0.0049 |
| F02 | 15900001 | 16000000 | 0.0054 | 0.0044 | 0.0049 |
| F02 | 15920001 | 16020000 | 0.0045 | 0.0035 | 0.0039 |
| F02 | 15940001 | 16040000 | 0.0049 | 0.0038 | 0.0042 |
| F02 | 15960001 | 16060000 | 0.0040 | 0.0030 | 0.0034 |
| F02 | 15980001 | 16080000 | 0.0031 | 0.0025 | 0.0027 |
| F02 | 16000001 | 16100000 | 0.0024 | 0.0019 | 0.0021 |
| F02 | 16020001 | 16120000 | 0.0021 | 0.0016 | 0.0020 |
| F02 | 16040001 | 16140000 | 0.0011 | 0.0009 | 0.0011 |
| F02 | 16060001 | 16160000 | 0.0010 | 0.0008 | 0.0010 |
| F02 | 16080001 | 16180000 | 0.0013 | 0.0009 | 0.0013 |
| F02 | 16100001 | 16200000 | 0.0011 | 0.0009 | 0.0011 |
| F02 | 16120001 | 16220000 | 0.0011 | 0.0008 | 0.0011 |
| F02 | 16140001 | 16240000 | 0.0016 | 0.0012 | 0.0015 |
| F02 | 16160001 | 16260000 | 0.0028 | 0.0022 | 0.0025 |
| F02 | 16180001 | 16280000 | 0.0027 | 0.0022 | 0.0024 |
| F02 | 16200001 | 16300000 | 0.0032 | 0.0025 | 0.0027 |
| F02 | 16220001 | 16320000 | 0.0039 | 0.0029 | 0.0029 |
| F02 | 16240001 | 16340000 | 0.0037 | 0.0027 | 0.0026 |
| F02 | 16260001 | 16360000 | 0.0027 | 0.0019 | 0.0019 |

| Chr | Start | End | both | A1 | A2 |
|---|---|---|---|---|---|
| F02 | 16280001 | 16380000 | 0.0032 | 0.0023 | 0.0023 |
| F02 | 16300001 | 16400000 | 0.0030 | 0.0023 | 0.0023 |
| F02 | 16320001 | 16420000 | 0.0023 | 0.0019 | 0.0019 |
| F02 | 16340001 | 16440000 | 0.0026 | 0.0021 | 0.0024 |
| F02 | 16360001 | 16460000 | 0.0026 | 0.0021 | 0.0024 |
| F02 | 16380001 | 16480000 | 0.0022 | 0.0019 | 0.0022 |
| F02 | 16400001 | 16500000 | 0.0027 | 0.0023 | 0.0027 |
| F02 | 16420001 | 16520000 | 0.0028 | 0.0023 | 0.0027 |
| F02 | 16440001 | 16540000 | 0.0026 | 0.0023 | 0.0023 |
| F02 | 16460001 | 16560000 | 0.0025 | 0.0020 | 0.0021 |
| F02 | 16480001 | 16580000 | 0.0042 | 0.0034 | 0.0037 |
| F02 | 16500001 | 16600000 | 0.0041 | 0.0032 | 0.0035 |
| F02 | 16520001 | 16620000 | 0.0050 | 0.0039 | 0.0042 |
| F02 | 16540001 | 16640000 | 0.0045 | 0.0035 | 0.0038 |
| F02 | 16560001 | 16660000 | 0.0046 | 0.0036 | 0.0039 |
| F02 | 16580001 | 16680000 | 0.0031 | 0.0021 | 0.0023 |
| F02 | 16600001 | 16700000 | 0.0025 | 0.0017 | 0.0019 |
| F02 | 16620001 | 16720000 | 0.0019 | 0.0012 | 0.0013 |
| F02 | 16640001 | 16740000 | 0.0029 | 0.0019 | 0.0021 |
| F02 | 16660001 | 16760000 | 0.0039 | 0.0028 | 0.0030 |
| F02 | 16680001 | 16780000 | 0.0035 | 0.0026 | 0.0027 |
| F02 | 16700001 | 16800000 | 0.0040 | 0.0031 | 0.0030 |
| F02 | 16720001 | 16820000 | 0.0039 | 0.0030 | 0.0029 |
| F02 | 16740001 | 16840000 | 0.0038 | 0.0032 | 0.0029 |
| F02 | 16760001 | 16860000 | 0.0030 | 0.0023 | 0.0020 |
| F02 | 16780001 | 16880000 | 0.0052 | 0.0045 | 0.0041 |
| F02 | 16800001 | 16900000 | 0.0050 | 0.0041 | 0.0039 |
| F02 | 16820001 | 16920000 | 0.0055 | 0.0046 | 0.0045 |
| F02 | 16840001 | 16940000 | 0.0059 | 0.0049 | 0.0049 |
| F02 | 16860001 | 16960000 | 0.0058 | 0.0050 | 0.0050 |
| F02 | 16880001 | 16980000 | 0.0042 | 0.0036 | 0.0035 |
| F02 | 16900001 | 17000000 | 0.0043 | 0.0037 | 0.0036 |
| F02 | 16920001 | 17020000 | 0.0043 | 0.0035 | 0.0034 |
| F02 | 16940001 | 17040000 | 0.0046 | 0.0038 | 0.0038 |
| F02 | 16960001 | 17060000 | 0.0044 | 0.0037 | 0.0036 |
| F02 | 16980001 | 17080000 | 0.0044 | 0.0036 | 0.0038 |
| F02 | 17000001 | 17100000 | 0.0040 | 0.0033 | 0.0036 |
| F02 | 17020001 | 17120000 | 0.0038 | 0.0031 | 0.0034 |
| F02 | 17040001 | 17140000 | 0.0022 | 0.0017 | 0.0019 |
| F02 | 17060001 | 17160000 | 0.0026 | 0.0017 | 0.0020 |
| F02 | 17080001 | 17180000 | 0.0019 | 0.0011 | 0.0013 |
| F02 | 17100001 | 17200000 | 0.0050 | 0.0039 | 0.0044 |
| F02 | 17120001 | 17220000 | 0.0059 | 0.0047 | 0.0052 |
| F02 | 17140001 | 17240000 | 0.0071 | 0.0058 | 0.0064 |
| F02 | 17160001 | 17260000 | 0.0078 | 0.0064 | 0.0071 |
| F02 | 17180001 | 17280000 | 0.0082 | 0.0067 | 0.0072 |
| F02 | 17200001 | 17300000 | 0.0075 | 0.0057 | 0.0066 |
| F02 | 17220001 | 17320000 | 0.0071 | 0.0053 | 0.0063 |
| F02 | 17240001 | 17340000 | 0.0069 | 0.0050 | 0.0062 |
| F02 | 17260001 | 17360000 | 0.0061 | 0.0045 | 0.0056 |
| F02 | 17280001 | 17380000 | 0.0061 | 0.0046 | 0.0057 |
| F02 | 17300001 | 17400000 | 0.0049 | 0.0039 | 0.0046 |
| F02 | 17320001 | 17420000 | 0.0047 | 0.0036 | 0.0042 |
| F02 | 17340001 | 17440000 | 0.0039 | 0.0030 | 0.0033 |
| F02 | 17360001 | 17460000 | 0.0036 | 0.0027 | 0.0030 |
| F02 | 17380001 | 17480000 | 0.0030 | 0.0022 | 0.0026 |
| F02 | 17400001 | 17500000 | 0.0017 | 0.0011 | 0.0014 |
| F02 | 17420001 | 17520000 | 0.0015 | 0.0010 | 0.0013 |

| Chr | Start | End | both | A1 | A2 |
|---|---|---|---|---|---|
| F02 | 17440001 | 17540000 | 0.0017 | 0.0014 | 0.0016 |
| F02 | 17460001 | 17560000 | 0.0020 | 0.0017 | 0.0019 |
| F02 | 17480001 | 17580000 | 0.0026 | 0.0023 | 0.0024 |
| F02 | 17500001 | 17600000 | 0.0037 | 0.0031 | 0.0033 |
| F02 | 17520001 | 17620000 | 0.0041 | 0.0032 | 0.0037 |
| F02 | 17540001 | 17640000 | 0.0043 | 0.0034 | 0.0039 |
| F02 | 17560001 | 17660000 | 0.0037 | 0.0029 | 0.0033 |
| F02 | 17580001 | 17680000 | 0.0034 | 0.0025 | 0.0030 |
| F02 | 17600001 | 17700000 | 0.0027 | 0.0020 | 0.0025 |
| F02 | 17620001 | 17720000 | 0.0023 | 0.0018 | 0.0020 |
| F02 | 17640001 | 17740000 | 0.0024 | 0.0017 | 0.0018 |
| F02 | 17660001 | 17760000 | 0.0032 | 0.0023 | 0.0024 |
| F02 | 17680001 | 17780000 | 0.0031 | 0.0023 | 0.0024 |
| F02 | 17700001 | 17800000 | 0.0029 | 0.0023 | 0.0023 |
| F02 | 17720001 | 17820000 | 0.0024 | 0.0019 | 0.0019 |
| F02 | 17740001 | 17840000 | 0.0018 | 0.0016 | 0.0016 |
| F02 | 17760001 | 17860000 | 0.0016 | 0.0011 | 0.0012 |
| F02 | 17780001 | 17880000 | 0.0017 | 0.0010 | 0.0011 |
| F02 | 17800001 | 17900000 | 0.0020 | 0.0011 | 0.0013 |
| F02 | 17820001 | 17920000 | 0.0029 | 0.0018 | 0.0020 |
| F02 | 17840001 | 17940000 | 0.0037 | 0.0020 | 0.0023 |
| F02 | 17860001 | 17960000 | 0.0036 | 0.0021 | 0.0024 |
| F02 | 17880001 | 17980000 | 0.0041 | 0.0026 | 0.0030 |
| F02 | 17900001 | 18000000 | 0.0041 | 0.0028 | 0.0030 |
| F02 | 17920001 | 18020000 | 0.0032 | 0.0021 | 0.0024 |
| F02 | 17940001 | 18040000 | 0.0023 | 0.0018 | 0.0020 |
| F02 | 17960001 | 18060000 | 0.0027 | 0.0022 | 0.0025 |
| F02 | 17980001 | 18080000 | 0.0023 | 0.0017 | 0.0020 |
| F02 | 18000001 | 18100000 | 0.0025 | 0.0018 | 0.0021 |
| F02 | 18020001 | 18120000 | 0.0041 | 0.0031 | 0.0034 |
| F02 | 18040001 | 18140000 | 0.0047 | 0.0035 | 0.0039 |
| F02 | 18060001 | 18160000 | 0.0039 | 0.0028 | 0.0031 |
| F02 | 18080001 | 18180000 | 0.0037 | 0.0027 | 0.0030 |
| F02 | 18100001 | 18200000 | 0.0037 | 0.0027 | 0.0030 |
| F02 | 18120001 | 18220000 | 0.0022 | 0.0016 | 0.0019 |
| F02 | 18140001 | 18240000 | 0.0020 | 0.0015 | 0.0017 |
| F02 | 18160001 | 18260000 | 0.0028 | 0.0020 | 0.0022 |
| F02 | 18180001 | 18280000 | 0.0036 | 0.0027 | 0.0029 |
| F02 | 18200001 | 18300000 | 0.0035 | 0.0026 | 0.0027 |
| F02 | 18220001 | 18320000 | 0.0045 | 0.0031 | 0.0032 |
| F02 | 18240001 | 18340000 | 0.0043 | 0.0027 | 0.0027 |
| F02 | 18260001 | 18360000 | 0.0041 | 0.0027 | 0.0027 |
| F02 | 18280001 | 18380000 | 0.0041 | 0.0027 | 0.0028 |
| F02 | 18300001 | 18400000 | 0.0036 | 0.0021 | 0.0022 |
| F02 | 18320001 | 18420000 | 0.0033 | 0.0017 | 0.0019 |
| F02 | 18340001 | 18440000 | 0.0035 | 0.0023 | 0.0024 |
| F02 | 18360001 | 18460000 | 0.0046 | 0.0033 | 0.0033 |
| F02 | 18380001 | 18480000 | 0.0041 | 0.0030 | 0.0030 |
| F02 | 18400001 | 18500000 | 0.0045 | 0.0035 | 0.0035 |
| F02 | 18420001 | 18520000 | 0.0041 | 0.0035 | 0.0033 |
| F02 | 18440001 | 18540000 | 0.0047 | 0.0038 | 0.0038 |
| F02 | 18460001 | 18560000 | 0.0037 | 0.0027 | 0.0026 |
| F02 | 18480001 | 18580000 | 0.0048 | 0.0034 | 0.0035 |
| F02 | 18500001 | 18600000 | 0.0049 | 0.0033 | 0.0034 |
| F02 | 18520001 | 18620000 | 0.0050 | 0.0031 | 0.0034 |
| F02 | 18540001 | 18640000 | 0.0056 | 0.0038 | 0.0039 |
| F02 | 18560001 | 18660000 | 0.0054 | 0.0036 | 0.0039 |
| F02 | 18580001 | 18680000 | 0.0061 | 0.0043 | 0.0048 |

| Chr | Start | End | both | A1 | A2 |
|---|---|---|---|---|---|
| F02 | 18600001 | 18700000 | 0.0067 | 0.0049 | 0.0053 |
| F02 | 18620001 | 18720000 | 0.0071 | 0.0055 | 0.0059 |
| F02 | 18640001 | 18740000 | 0.0056 | 0.0040 | 0.0045 |
| F02 | 18660001 | 18760000 | 0.0054 | 0.0038 | 0.0042 |
| F02 | 18680001 | 18780000 | 0.0036 | 0.0025 | 0.0024 |
| F02 | 18700001 | 18800000 | 0.0033 | 0.0021 | 0.0022 |
| F02 | 18720001 | 18820000 | 0.0029 | 0.0017 | 0.0016 |
| F02 | 18740001 | 18840000 | 0.0035 | 0.0023 | 0.0022 |
| F02 | 18760001 | 18860000 | 0.0041 | 0.0030 | 0.0030 |
| F02 | 18780001 | 18880000 | 0.0051 | 0.0037 | 0.0038 |
| F02 | 18800001 | 18900000 | 0.0046 | 0.0033 | 0.0034 |
| F02 | 18820001 | 18920000 | 0.0051 | 0.0037 | 0.0039 |
| F02 | 18840001 | 18940000 | 0.0044 | 0.0032 | 0.0034 |
| F02 | 18860001 | 18960000 | 0.0037 | 0.0026 | 0.0027 |
| F02 | 18880001 | 18980000 | 0.0039 | 0.0028 | 0.0029 |
| F02 | 18900001 | 19000000 | 0.0048 | 0.0035 | 0.0038 |
| F02 | 18920001 | 19020000 | 0.0041 | 0.0031 | 0.0032 |
| F02 | 18940001 | 19040000 | 0.0051 | 0.0036 | 0.0040 |
| F02 | 18960001 | 19060000 | 0.0063 | 0.0048 | 0.0053 |
| F02 | 18980001 | 19080000 | 0.0056 | 0.0041 | 0.0046 |
| F02 | 19000001 | 19100000 | 0.0056 | 0.0039 | 0.0044 |
| F02 | 19020001 | 19120000 | 0.0056 | 0.0038 | 0.0043 |
| F02 | 19040001 | 19140000 | 0.0054 | 0.0039 | 0.0044 |
| F02 | 19060001 | 19160000 | 0.0045 | 0.0030 | 0.0035 |
| F02 | 19080001 | 19180000 | 0.0036 | 0.0024 | 0.0028 |
| F02 | 19100001 | 19200000 | 0.0026 | 0.0019 | 0.0023 |
| F02 | 19120001 | 19220000 | 0.0024 | 0.0019 | 0.0023 |
| F02 | 19140001 | 19240000 | 0.0019 | 0.0015 | 0.0019 |
| F02 | 19160001 | 19260000 | 0.0013 | 0.0010 | 0.0013 |
| F02 | 19180001 | 19280000 | 0.0022 | 0.0017 | 0.0020 |
| F02 | 19200001 | 19300000 | 0.0027 | 0.0021 | 0.0024 |
| F02 | 19220001 | 19320000 | 0.0027 | 0.0021 | 0.0024 |
| F02 | 19240001 | 19340000 | 0.0032 | 0.0026 | 0.0028 |
| F02 | 19260001 | 19360000 | 0.0038 | 0.0032 | 0.0034 |
| F02 | 19280001 | 19380000 | 0.0038 | 0.0032 | 0.0035 |
| F02 | 19300001 | 19400000 | 0.0034 | 0.0029 | 0.0031 |
| F02 | 19320001 | 19420000 | 0.0040 | 0.0035 | 0.0036 |
| F02 | 19340001 | 19440000 | 0.0041 | 0.0034 | 0.0036 |
| F02 | 19360001 | 19460000 | 0.0040 | 0.0031 | 0.0033 |
| F02 | 19380001 | 19480000 | 0.0035 | 0.0026 | 0.0028 |
| F02 | 19400001 | 19500000 | 0.0041 | 0.0031 | 0.0033 |
| F02 | 19420001 | 19520000 | 0.0057 | 0.0044 | 0.0049 |
| F02 | 19440001 | 19540000 | 0.0053 | 0.0039 | 0.0045 |
| F02 | 19460001 | 19560000 | 0.0062 | 0.0048 | 0.0055 |
| F02 | 19480001 | 19580000 | 0.0059 | 0.0047 | 0.0054 |
| F02 | 19500001 | 19600000 | 0.0057 | 0.0045 | 0.0051 |
| F02 | 19520001 | 19620000 | 0.0043 | 0.0035 | 0.0038 |
| F02 | 19540001 | 19640000 | 0.0038 | 0.0031 | 0.0033 |
| F02 | 19560001 | 19660000 | 0.0029 | 0.0020 | 0.0021 |
| F02 | 19580001 | 19680000 | 0.0033 | 0.0022 | 0.0023 |
| F02 | 19600001 | 19700000 | 0.0033 | 0.0023 | 0.0024 |
| F02 | 19620001 | 19720000 | 0.0036 | 0.0024 | 0.0026 |
| F02 | 19640001 | 19740000 | 0.0034 | 0.0023 | 0.0026 |
| F02 | 19660001 | 19760000 | 0.0037 | 0.0026 | 0.0029 |
| F02 | 19680001 | 19780000 | 0.0040 | 0.0030 | 0.0032 |
| F02 | 19700001 | 19800000 | 0.0041 | 0.0030 | 0.0033 |
| F02 | 19720001 | 19820000 | 0.0038 | 0.0027 | 0.0031 |
| F02 | 19740001 | 19840000 | 0.0045 | 0.0033 | 0.0039 |

| Chr | Start | End | both | A1 | A2 |
|---|---|---|---|---|---|
| F02 | 19760001 | 19860000 | 0.0038 | 0.0029 | 0.0034 |
| F02 | 19780001 | 19880000 | 0.0040 | 0.0033 | 0.0038 |
| F02 | 19800001 | 19900000 | 0.0035 | 0.0029 | 0.0034 |
| F02 | 19820001 | 19920000 | 0.0027 | 0.0023 | 0.0027 |
| F02 | 19840001 | 19940000 | 0.0026 | 0.0020 | 0.0026 |
| F02 | 19860001 | 19960000 | 0.0037 | 0.0028 | 0.0038 |
| F02 | 19880001 | 19980000 | 0.0032 | 0.0022 | 0.0032 |
| F02 | 19900001 | 20000000 | 0.0041 | 0.0029 | 0.0041 |
| F02 | 19920001 | 20020000 | 0.0055 | 0.0038 | 0.0056 |
| F02 | 19940001 | 20040000 | 0.0053 | 0.0038 | 0.0053 |
| F02 | 19960001 | 20060000 | 0.0048 | 0.0036 | 0.0048 |
| F02 | 19980001 | 20080000 | 0.0046 | 0.0034 | 0.0046 |
| F02 | 20000001 | 20100000 | 0.0040 | 0.0030 | 0.0040 |
| F02 | 20020001 | 20120000 | 0.0036 | 0.0029 | 0.0035 |
| F02 | 20040001 | 20140000 | 0.0053 | 0.0043 | 0.0051 |
| F02 | 20060001 | 20160000 | 0.0056 | 0.0046 | 0.0055 |
| F02 | 20080001 | 20180000 | 0.0059 | 0.0049 | 0.0057 |
| F02 | 20100001 | 20200000 | 0.0062 | 0.0052 | 0.0060 |
| F02 | 20120001 | 20220000 | 0.0069 | 0.0059 | 0.0067 |
| F02 | 20140001 | 20240000 | 0.0050 | 0.0044 | 0.0049 |
| F02 | 20160001 | 20260000 | 0.0041 | 0.0036 | 0.0040 |
| F02 | 20180001 | 20280000 | 0.0041 | 0.0038 | 0.0041 |
| F02 | 20200001 | 20300000 | 0.0043 | 0.0039 | 0.0042 |
| F02 | 20220001 | 20320000 | 0.0030 | 0.0026 | 0.0027 |
| F02 | 20240001 | 20340000 | 0.0037 | 0.0031 | 0.0032 |
| F02 | 20260001 | 20360000 | 0.0042 | 0.0032 | 0.0035 |
| F02 | 20280001 | 20380000 | 0.0038 | 0.0027 | 0.0030 |
| F02 | 20300001 | 20400000 | 0.0031 | 0.0019 | 0.0022 |
| F02 | 20320001 | 20420000 | 0.0029 | 0.0018 | 0.0022 |
| F02 | 20340001 | 20440000 | 0.0024 | 0.0014 | 0.0019 |
| F02 | 20360001 | 20460000 | 0.0020 | 0.0011 | 0.0015 |
| F02 | 20380001 | 20480000 | 0.0022 | 0.0013 | 0.0017 |
| F02 | 20400001 | 20500000 | 0.0027 | 0.0019 | 0.0021 |
| F02 | 20420001 | 20520000 | 0.0030 | 0.0020 | 0.0021 |
| F02 | 20440001 | 20540000 | 0.0035 | 0.0026 | 0.0025 |
| F02 | 20460001 | 20560000 | 0.0040 | 0.0029 | 0.0028 |
| F02 | 20480001 | 20580000 | 0.0040 | 0.0033 | 0.0027 |
| F02 | 20500001 | 20600000 | 0.0037 | 0.0030 | 0.0024 |
| F02 | 20520001 | 20620000 | 0.0032 | 0.0027 | 0.0021 |
| F02 | 20540001 | 20640000 | 0.0024 | 0.0021 | 0.0016 |
| F02 | 20560001 | 20660000 | 0.0021 | 0.0019 | 0.0015 |
| F02 | 20580001 | 20680000 | 0.0038 | 0.0030 | 0.0034 |
| F02 | 20600001 | 20700000 | 0.0031 | 0.0024 | 0.0029 |
| F02 | 20620001 | 20720000 | 0.0030 | 0.0024 | 0.0029 |
| F02 | 20640001 | 20740000 | 0.0029 | 0.0024 | 0.0028 |
| F02 | 20660001 | 20760000 | 0.0030 | 0.0024 | 0.0029 |
| F02 | 20680001 | 20780000 | 0.0011 | 0.0008 | 0.0011 |
| F02 | 20700001 | 20800000 | 0.0015 | 0.0011 | 0.0015 |
| F02 | 20720001 | 20820000 | 0.0016 | 0.0012 | 0.0016 |
| F02 | 20740001 | 20840000 | 0.0021 | 0.0015 | 0.0021 |
| F02 | 20760001 | 20860000 | 0.0017 | 0.0013 | 0.0017 |
| F02 | 20780001 | 20880000 | 0.0014 | 0.0010 | 0.0014 |
| F02 | 20800001 | 20900000 | 0.0011 | 0.0009 | 0.0011 |
| F02 | 20820001 | 20920000 | 0.0011 | 0.0008 | 0.0011 |
| F02 | 20840001 | 20940000 | 0.0006 | 0.0005 | 0.0005 |
| F02 | 20860001 | 20960000 | 0.0007 | 0.0006 | 0.0007 |
| F02 | 20880001 | 20980000 | 0.0008 | 0.0007 | 0.0008 |
| F02 | 20900001 | 21000000 | 0.0008 | 0.0006 | 0.0009 |

| Chr | Start | End | both | A1 | A2 |
|---|---|---|---|---|---|
| F02 | 20920001 | 21020000 | 0.0011 | 0.0007 | 0.0011 |
| F02 | 20940001 | 21040000 | 0.0012 | 0.0009 | 0.0012 |
| F02 | 20960001 | 21060000 | 0.0016 | 0.0012 | 0.0015 |
| F02 | 20980001 | 21080000 | 0.0015 | 0.0012 | 0.0015 |
| F02 | 21000001 | 21100000 | 0.0014 | 0.0011 | 0.0013 |
| F02 | 21020001 | 21120000 | 0.0013 | 0.0011 | 0.0013 |
| F02 | 21040001 | 21140000 | 0.0013 | 0.0011 | 0.0013 |
| F02 | 21060001 | 21160000 | 0.0009 | 0.0007 | 0.0009 |
| F02 | 21080001 | 21180000 | 0.0010 | 0.0008 | 0.0010 |
| F02 | 21100001 | 21200000 | 0.0010 | 0.0008 | 0.0010 |
| F02 | 21120001 | 21220000 | 0.0009 | 0.0007 | 0.0009 |
| F02 | 21140001 | 21240000 | 0.0010 | 0.0008 | 0.0010 |
| F02 | 21160001 | 21260000 | 0.0018 | 0.0014 | 0.0018 |
| F02 | 21180001 | 21280000 | 0.0015 | 0.0013 | 0.0016 |
| F02 | 21200001 | 21300000 | 0.0015 | 0.0013 | 0.0015 |
| F02 | 21220001 | 21320000 | 0.0016 | 0.0012 | 0.0016 |
| F02 | 21240001 | 21340000 | 0.0015 | 0.0012 | 0.0015 |
| F02 | 21260001 | 21360000 | 0.0014 | 0.0011 | 0.0014 |
| F02 | 21280001 | 21380000 | 0.0026 | 0.0021 | 0.0026 |
| F02 | 21300001 | 21400000 | 0.0041 | 0.0035 | 0.0041 |
| F02 | 21320001 | 21420000 | 0.0044 | 0.0038 | 0.0043 |
| F02 | 21340001 | 21440000 | 0.0050 | 0.0043 | 0.0049 |
| F02 | 21360001 | 21460000 | 0.0045 | 0.0039 | 0.0044 |
| F02 | 21380001 | 21480000 | 0.0050 | 0.0044 | 0.0048 |
| F02 | 21400001 | 21500000 | 0.0041 | 0.0035 | 0.0039 |
| F02 | 21420001 | 21520000 | 0.0036 | 0.0032 | 0.0034 |
| F02 | 21440001 | 21540000 | 0.0030 | 0.0025 | 0.0028 |
| F02 | 21460001 | 21560000 | 0.0029 | 0.0025 | 0.0028 |
| F02 | 21480001 | 21580000 | 0.0015 | 0.0012 | 0.0013 |
| F02 | 21500001 | 21600000 | 0.0011 | 0.0009 | 0.0011 |
| F02 | 21520001 | 21620000 | 0.0027 | 0.0021 | 0.0024 |
| F02 | 21540001 | 21640000 | 0.0034 | 0.0025 | 0.0028 |
| F02 | 21560001 | 21660000 | 0.0032 | 0.0023 | 0.0025 |
| F02 | 21580001 | 21680000 | 0.0038 | 0.0028 | 0.0030 |
| F02 | 21600001 | 21700000 | 0.0045 | 0.0030 | 0.0036 |
| F02 | 21620001 | 21720000 | 0.0039 | 0.0024 | 0.0028 |
| F02 | 21640001 | 21740000 | 0.0042 | 0.0027 | 0.0032 |
| F02 | 21660001 | 21760000 | 0.0058 | 0.0041 | 0.0045 |
| F02 | 21680001 | 21780000 | 0.0052 | 0.0038 | 0.0039 |
| F02 | 21700001 | 21800000 | 0.0044 | 0.0035 | 0.0033 |
| F02 | 21720001 | 21820000 | 0.0038 | 0.0031 | 0.0028 |
| F02 | 21740001 | 21840000 | 0.0028 | 0.0025 | 0.0021 |
| F02 | 21760001 | 21860000 | 0.0013 | 0.0012 | 0.0010 |
| F02 | 21780001 | 21880000 | 0.0014 | 0.0010 | 0.0011 |
| F02 | 21800001 | 21900000 | 0.0015 | 0.0010 | 0.0011 |
| F02 | 21820001 | 21920000 | 0.0019 | 0.0016 | 0.0016 |
| F02 | 21840001 | 21940000 | 0.0028 | 0.0024 | 0.0024 |
| F02 | 21860001 | 21960000 | 0.0037 | 0.0031 | 0.0033 |
| F02 | 21880001 | 21980000 | 0.0034 | 0.0029 | 0.0031 |
| F02 | 21900001 | 22000000 | 0.0033 | 0.0027 | 0.0031 |
| F02 | 21920001 | 22020000 | 0.0024 | 0.0020 | 0.0023 |
| F02 | 21940001 | 22040000 | 0.0022 | 0.0017 | 0.0021 |
| F02 | 21960001 | 22060000 | 0.0012 | 0.0009 | 0.0011 |
| F02 | 21980001 | 22080000 | 0.0011 | 0.0008 | 0.0010 |
| F02 | 22000001 | 22100000 | 0.0010 | 0.0008 | 0.0010 |
| F02 | 22020001 | 22120000 | 0.0011 | 0.0009 | 0.0011 |
| F02 | 22040001 | 22140000 | 0.0004 | 0.0003 | 0.0004 |
| F02 | 22060001 | 22160000 | 0.0005 | 0.0004 | 0.0004 |

| Chr | Start | End | both | A1 | A2 |
|---|---|---|---|---|---|
| F02 | 22080001 | 22180000 | 0.0006 | 0.0005 | 0.0006 |
| F02 | 22100001 | 22200000 | 0.0005 | 0.0004 | 0.0005 |
| F02 | 22120001 | 22220000 | 0.0004 | 0.0003 | 0.0003 |
| F02 | 22140001 | 22240000 | 0.0017 | 0.0014 | 0.0016 |
| F02 | 22160001 | 22260000 | 0.0023 | 0.0020 | 0.0023 |
| F02 | 22180001 | 22280000 | 0.0024 | 0.0020 | 0.0023 |
| F02 | 22200001 | 22300000 | 0.0027 | 0.0022 | 0.0026 |
| F02 | 22220001 | 22320000 | 0.0026 | 0.0023 | 0.0026 |
| F02 | 22240001 | 22340000 | 0.0016 | 0.0013 | 0.0016 |
| F02 | 22260001 | 22360000 | 0.0012 | 0.0011 | 0.0011 |
| F02 | 22280001 | 22380000 | 0.0015 | 0.0013 | 0.0013 |
| F02 | 22300001 | 22400000 | 0.0023 | 0.0021 | 0.0022 |
| F02 | 22320001 | 22420000 | 0.0024 | 0.0022 | 0.0022 |
| F02 | 22340001 | 22440000 | 0.0024 | 0.0021 | 0.0021 |
| F02 | 22360001 | 22460000 | 0.0039 | 0.0031 | 0.0033 |
| F02 | 22380001 | 22480000 | 0.0048 | 0.0037 | 0.0038 |
| F02 | 22400001 | 22500000 | 0.0049 | 0.0035 | 0.0035 |
| F02 | 22420001 | 22520000 | 0.0052 | 0.0035 | 0.0036 |
| F02 | 22440001 | 22540000 | 0.0061 | 0.0042 | 0.0046 |
| F02 | 22460001 | 22560000 | 0.0050 | 0.0033 | 0.0037 |
| F02 | 22480001 | 22580000 | 0.0036 | 0.0023 | 0.0027 |
| F02 | 22500001 | 22600000 | 0.0040 | 0.0028 | 0.0033 |
| F02 | 22520001 | 22620000 | 0.0043 | 0.0033 | 0.0039 |
| F02 | 22540001 | 22640000 | 0.0044 | 0.0036 | 0.0041 |
| F02 | 22560001 | 22660000 | 0.0044 | 0.0037 | 0.0042 |
| F02 | 22580001 | 22680000 | 0.0046 | 0.0038 | 0.0043 |
| F02 | 22600001 | 22700000 | 0.0066 | 0.0056 | 0.0064 |
| F02 | 22620001 | 22720000 | 0.0058 | 0.0049 | 0.0056 |
| F02 | 22640001 | 22740000 | 0.0055 | 0.0048 | 0.0054 |
| F02 | 22660001 | 22760000 | 0.0051 | 0.0045 | 0.0050 |
| F02 | 22680001 | 22780000 | 0.0052 | 0.0046 | 0.0051 |
| F02 | 22700001 | 22800000 | 0.0019 | 0.0018 | 0.0019 |
| F02 | 22720001 | 22820000 | 0.0023 | 0.0020 | 0.0022 |
| F02 | 22740001 | 22840000 | 0.0019 | 0.0017 | 0.0019 |
| F02 | 22760001 | 22860000 | 0.0017 | 0.0015 | 0.0016 |
| F02 | 22780001 | 22880000 | 0.0015 | 0.0013 | 0.0015 |
| F02 | 22800001 | 22900000 | 0.0024 | 0.0021 | 0.0023 |
| F02 | 22820001 | 22920000 | 0.0025 | 0.0021 | 0.0023 |
| F02 | 22840001 | 22940000 | 0.0023 | 0.0019 | 0.0021 |
| F02 | 22860001 | 22960000 | 0.0030 | 0.0025 | 0.0028 |
| F02 | 22880001 | 22980000 | 0.0030 | 0.0026 | 0.0028 |
| F02 | 22900001 | 23000000 | 0.0028 | 0.0023 | 0.0026 |
| F02 | 22920001 | 23020000 | 0.0034 | 0.0027 | 0.0029 |
| F02 | 22940001 | 23040000 | 0.0044 | 0.0035 | 0.0038 |
| F02 | 22960001 | 23060000 | 0.0041 | 0.0030 | 0.0033 |
| F02 | 22980001 | 23080000 | 0.0044 | 0.0031 | 0.0035 |
| F02 | 23000001 | 23100000 | 0.0046 | 0.0031 | 0.0035 |
| F02 | 23020001 | 23120000 | 0.0038 | 0.0027 | 0.0029 |
| F02 | 23040001 | 23140000 | 0.0025 | 0.0017 | 0.0018 |
| F02 | 23060001 | 23160000 | 0.0025 | 0.0018 | 0.0020 |
| F02 | 23080001 | 23180000 | 0.0023 | 0.0018 | 0.0020 |
| F02 | 23100001 | 23200000 | 0.0013 | 0.0012 | 0.0012 |
| F02 | 23120001 | 23220000 | 0.0010 | 0.0009 | 0.0010 |
| F02 | 23140001 | 23240000 | 0.0009 | 0.0008 | 0.0009 |
| F02 | 23160001 | 23260000 | 0.0008 | 0.0007 | 0.0008 |
| F02 | 23180001 | 23280000 | 0.0008 | 0.0007 | 0.0008 |
| F02 | 23200001 | 23300000 | 0.0012 | 0.0009 | 0.0012 |
| F02 | 23220001 | 23320000 | 0.0022 | 0.0019 | 0.0022 |

| Chr | Start | End | both | A1 | A2 |
|---|---|---|---|---|---|
| F02 | 23240001 | 23340000 | 0.0027 | 0.0022 | 0.0027 |
| F02 | 23260001 | 23360000 | 0.0027 | 0.0021 | 0.0027 |
| F02 | 23280001 | 23380000 | 0.0031 | 0.0023 | 0.0026 |
| F02 | 23300001 | 23400000 | 0.0032 | 0.0024 | 0.0025 |
| F02 | 23320001 | 23420000 | 0.0027 | 0.0019 | 0.0019 |
| F02 | 23340001 | 23440000 | 0.0024 | 0.0016 | 0.0016 |
| F02 | 23360001 | 23460000 | 0.0025 | 0.0018 | 0.0017 |
| F02 | 23380001 | 23480000 | 0.0024 | 0.0019 | 0.0021 |
| F02 | 23400001 | 23500000 | 0.0024 | 0.0019 | 0.0022 |
| F02 | 23420001 | 23520000 | 0.0019 | 0.0015 | 0.0018 |
| F02 | 23440001 | 23540000 | 0.0020 | 0.0016 | 0.0019 |
| F02 | 23460001 | 23560000 | 0.0019 | 0.0016 | 0.0019 |
| F02 | 23480001 | 23580000 | 0.0020 | 0.0017 | 0.0019 |
| F02 | 23500001 | 23600000 | 0.0018 | 0.0015 | 0.0018 |
| F02 | 23520001 | 23620000 | 0.0019 | 0.0017 | 0.0019 |
| F02 | 23540001 | 23640000 | 0.0026 | 0.0021 | 0.0025 |
| F02 | 23560001 | 23660000 | 0.0026 | 0.0021 | 0.0025 |
| F02 | 23580001 | 23680000 | 0.0020 | 0.0017 | 0.0020 |
| F02 | 23600001 | 23700000 | 0.0020 | 0.0017 | 0.0020 |
| F02 | 23620001 | 23720000 | 0.0021 | 0.0019 | 0.0021 |
| F02 | 23640001 | 23740000 | 0.0026 | 0.0023 | 0.0026 |
| F02 | 23660001 | 23760000 | 0.0028 | 0.0025 | 0.0028 |
| F02 | 23680001 | 23780000 | 0.0027 | 0.0024 | 0.0027 |
| F02 | 23700001 | 23800000 | 0.0030 | 0.0026 | 0.0029 |
| F02 | 23720001 | 23820000 | 0.0030 | 0.0026 | 0.0029 |
| F02 | 23740001 | 23840000 | 0.0016 | 0.0014 | 0.0015 |
| F02 | 23760001 | 23860000 | 0.0021 | 0.0016 | 0.0018 |
| F02 | 23780001 | 23880000 | 0.0035 | 0.0025 | 0.0029 |
| F02 | 23800001 | 23900000 | 0.0043 | 0.0033 | 0.0037 |
| F02 | 23820001 | 23920000 | 0.0045 | 0.0034 | 0.0038 |
| F02 | 23840001 | 23940000 | 0.0048 | 0.0036 | 0.0042 |
| F02 | 23860001 | 23960000 | 0.0046 | 0.0037 | 0.0041 |
| F02 | 23880001 | 23980000 | 0.0038 | 0.0032 | 0.0035 |
| F02 | 23900001 | 24000000 | 0.0032 | 0.0026 | 0.0030 |
| F02 | 23920001 | 24020000 | 0.0037 | 0.0030 | 0.0032 |
| F02 | 23940001 | 24040000 | 0.0035 | 0.0027 | 0.0029 |
| F02 | 23960001 | 24060000 | 0.0038 | 0.0030 | 0.0033 |
| F02 | 23980001 | 24080000 | 0.0036 | 0.0027 | 0.0029 |
| F02 | 24000001 | 24100000 | 0.0032 | 0.0022 | 0.0023 |
| F02 | 24020001 | 24120000 | 0.0027 | 0.0018 | 0.0020 |
| F02 | 24040001 | 24140000 | 0.0024 | 0.0018 | 0.0019 |
| F02 | 24060001 | 24160000 | 0.0016 | 0.0010 | 0.0011 |
| F02 | 24080001 | 24180000 | 0.0013 | 0.0007 | 0.0008 |
| F02 | 24100001 | 24200000 | 0.0019 | 0.0013 | 0.0015 |
| F02 | 24120001 | 24220000 | 0.0016 | 0.0010 | 0.0012 |
| F02 | 24140001 | 24240000 | 0.0017 | 0.0011 | 0.0013 |
| F02 | 24160001 | 24260000 | 0.0043 | 0.0030 | 0.0036 |
| F02 | 24180001 | 24280000 | 0.0048 | 0.0035 | 0.0043 |
| F02 | 24200001 | 24300000 | 0.0050 | 0.0037 | 0.0044 |
| F02 | 24220001 | 24320000 | 0.0055 | 0.0040 | 0.0048 |
| F02 | 24240001 | 24340000 | 0.0060 | 0.0046 | 0.0053 |
| F02 | 24260001 | 24360000 | 0.0044 | 0.0032 | 0.0035 |
| F02 | 24280001 | 24380000 | 0.0038 | 0.0027 | 0.0029 |
| F02 | 24300001 | 24400000 | 0.0035 | 0.0025 | 0.0029 |
| F02 | 24320001 | 24420000 | 0.0034 | 0.0024 | 0.0028 |
| F02 | 24340001 | 24440000 | 0.0033 | 0.0019 | 0.0022 |
| F02 | 24360001 | 24460000 | 0.0025 | 0.0015 | 0.0018 |
| F02 | 24380001 | 24480000 | 0.0026 | 0.0016 | 0.0019 |

| Chr | Start | End | both | A1 | A2 |
|---|---|---|---|---|---|
| F02 | 24400001 | 24500000 | 0.0016 | 0.0008 | 0.0010 |
| F02 | 24420001 | 24520000 | 0.0013 | 0.0006 | 0.0008 |
| F02 | 24440001 | 24540000 | 0.0016 | 0.0010 | 0.0012 |
| F02 | 24460001 | 24560000 | 0.0015 | 0.0009 | 0.0012 |
| F02 | 24480001 | 24580000 | 0.0019 | 0.0012 | 0.0015 |
| F02 | 24500001 | 24600000 | 0.0035 | 0.0025 | 0.0028 |
| F02 | 24520001 | 24620000 | 0.0038 | 0.0026 | 0.0030 |
| F02 | 24540001 | 24640000 | 0.0041 | 0.0031 | 0.0034 |
| F02 | 24560001 | 24660000 | 0.0052 | 0.0039 | 0.0043 |
| F02 | 24580001 | 24680000 | 0.0053 | 0.0041 | 0.0045 |
| F02 | 24600001 | 24700000 | 0.0039 | 0.0030 | 0.0034 |
| F02 | 24620001 | 24720000 | 0.0037 | 0.0029 | 0.0032 |
| F02 | 24640001 | 24740000 | 0.0033 | 0.0026 | 0.0029 |
| F02 | 24660001 | 24760000 | 0.0025 | 0.0020 | 0.0023 |
| F02 | 24680001 | 24780000 | 0.0032 | 0.0026 | 0.0030 |
| F02 | 24700001 | 24800000 | 0.0041 | 0.0033 | 0.0037 |
| F02 | 24720001 | 24820000 | 0.0050 | 0.0040 | 0.0044 |
| F02 | 24740001 | 24840000 | 0.0052 | 0.0040 | 0.0046 |
| F02 | 24760001 | 24860000 | 0.0052 | 0.0039 | 0.0045 |
| F02 | 24780001 | 24880000 | 0.0042 | 0.0031 | 0.0035 |
| F02 | 24800001 | 24900000 | 0.0036 | 0.0025 | 0.0029 |
| F02 | 24820001 | 24920000 | 0.0038 | 0.0027 | 0.0030 |
| F02 | 24840001 | 24940000 | 0.0033 | 0.0025 | 0.0025 |
| F02 | 24860001 | 24960000 | 0.0039 | 0.0029 | 0.0030 |
| F02 | 24880001 | 24980000 | 0.0035 | 0.0025 | 0.0026 |
| F02 | 24900001 | 25000000 | 0.0032 | 0.0023 | 0.0023 |
| F02 | 24920001 | 25020000 | 0.0022 | 0.0013 | 0.0014 |
| F02 | 24940001 | 25040000 | 0.0021 | 0.0013 | 0.0012 |
| F02 | 24960001 | 25060000 | 0.0019 | 0.0011 | 0.0011 |
| F02 | 24980001 | 25080000 | 0.0023 | 0.0014 | 0.0015 |
| F02 | 25000001 | 25100000 | 0.0026 | 0.0016 | 0.0019 |
| F02 | 25020001 | 25120000 | 0.0026 | 0.0017 | 0.0022 |
| F02 | 25040001 | 25140000 | 0.0026 | 0.0017 | 0.0023 |
| F02 | 25060001 | 25160000 | 0.0020 | 0.0013 | 0.0018 |
| F02 | 25080001 | 25180000 | 0.0017 | 0.0013 | 0.0016 |
| F02 | 25100001 | 25200000 | 0.0016 | 0.0013 | 0.0015 |
| F02 | 25120001 | 25220000 | 0.0017 | 0.0013 | 0.0014 |
| F02 | 25140001 | 25240000 | 0.0018 | 0.0014 | 0.0015 |
| F02 | 25160001 | 25260000 | 0.0020 | 0.0015 | 0.0017 |
| F02 | 25180001 | 25280000 | 0.0022 | 0.0018 | 0.0019 |
| F02 | 25200001 | 25300000 | 0.0025 | 0.0019 | 0.0019 |
| F02 | 25220001 | 25320000 | 0.0027 | 0.0021 | 0.0020 |
| F02 | 25240001 | 25340000 | 0.0028 | 0.0020 | 0.0019 |
| F02 | 25260001 | 25360000 | 0.0029 | 0.0020 | 0.0020 |
| F02 | 25280001 | 25380000 | 0.0027 | 0.0016 | 0.0016 |
| F02 | 25300001 | 25400000 | 0.0024 | 0.0015 | 0.0015 |
| F02 | 25320001 | 25420000 | 0.0036 | 0.0025 | 0.0026 |
| F02 | 25340001 | 25440000 | 0.0039 | 0.0028 | 0.0031 |
| F02 | 25360001 | 25460000 | 0.0040 | 0.0031 | 0.0033 |
| F02 | 25380001 | 25480000 | 0.0045 | 0.0036 | 0.0039 |
| F02 | 25400001 | 25500000 | 0.0044 | 0.0036 | 0.0039 |
| F02 | 25420001 | 25520000 | 0.0036 | 0.0030 | 0.0032 |
| F02 | 25440001 | 25540000 | 0.0035 | 0.0029 | 0.0032 |
| F02 | 25460001 | 25560000 | 0.0033 | 0.0027 | 0.0029 |
| F02 | 25480001 | 25580000 | 0.0041 | 0.0034 | 0.0035 |
| F02 | 25500001 | 25600000 | 0.0044 | 0.0035 | 0.0036 |
| F02 | 25520001 | 25620000 | 0.0038 | 0.0029 | 0.0030 |
| F02 | 25540001 | 25640000 | 0.0033 | 0.0023 | 0.0025 |

| Chr | Start | End | both | A1 | A2 |
|---|---|---|---|---|---|
| F02 | 25560001 | 25660000 | 0.0033 | 0.0021 | 0.0023 |
| F02 | 25580001 | 25680000 | 0.0029 | 0.0016 | 0.0017 |
| F02 | 25600001 | 25700000 | 0.0029 | 0.0016 | 0.0017 |
| F02 | 25620001 | 25720000 | 0.0032 | 0.0017 | 0.0020 |
| F02 | 25640001 | 25740000 | 0.0044 | 0.0026 | 0.0030 |
| F02 | 25660001 | 25760000 | 0.0044 | 0.0027 | 0.0033 |
| F02 | 25680001 | 25780000 | 0.0033 | 0.0020 | 0.0026 |
| F02 | 25700001 | 25800000 | 0.0032 | 0.0020 | 0.0026 |
| F02 | 25720001 | 25820000 | 0.0033 | 0.0023 | 0.0027 |
| F02 | 25740001 | 25840000 | 0.0022 | 0.0015 | 0.0017 |
| F02 | 25760001 | 25860000 | 0.0019 | 0.0014 | 0.0013 |
| F02 | 25780001 | 25880000 | 0.0021 | 0.0015 | 0.0014 |
| F02 | 25800001 | 25900000 | 0.0018 | 0.0013 | 0.0012 |
| F02 | 25820001 | 25920000 | 0.0029 | 0.0020 | 0.0021 |
| F02 | 25840001 | 25940000 | 0.0030 | 0.0021 | 0.0022 |
| F02 | 25860001 | 25960000 | 0.0028 | 0.0020 | 0.0022 |
| F02 | 25880001 | 25980000 | 0.0033 | 0.0023 | 0.0026 |
| F02 | 25900001 | 26000000 | 0.0035 | 0.0023 | 0.0026 |
| F02 | 25920001 | 26020000 | 0.0027 | 0.0016 | 0.0017 |
| F02 | 25940001 | 26040000 | 0.0029 | 0.0014 | 0.0016 |
| F02 | 25960001 | 26060000 | 0.0034 | 0.0017 | 0.0019 |
| F02 | 25980001 | 26080000 | 0.0030 | 0.0014 | 0.0015 |
| F02 | 26000001 | 26100000 | 0.0031 | 0.0018 | 0.0018 |
| F02 | 26020001 | 26120000 | 0.0029 | 0.0017 | 0.0018 |
| F02 | 26040001 | 26140000 | 0.0031 | 0.0021 | 0.0022 |
| F02 | 26060001 | 26160000 | 0.0035 | 0.0025 | 0.0027 |
| F02 | 26080001 | 26180000 | 0.0041 | 0.0030 | 0.0033 |
| F02 | 26100001 | 26200000 | 0.0046 | 0.0034 | 0.0037 |
| F02 | 26120001 | 26220000 | 0.0046 | 0.0036 | 0.0035 |
| F02 | 26140001 | 26240000 | 0.0047 | 0.0039 | 0.0038 |
| F02 | 26160001 | 26260000 | 0.0048 | 0.0039 | 0.0038 |
| F02 | 26180001 | 26280000 | 0.0045 | 0.0037 | 0.0036 |
| F02 | 26200001 | 26300000 | 0.0042 | 0.0034 | 0.0032 |
| F02 | 26220001 | 26320000 | 0.0042 | 0.0031 | 0.0032 |
| F02 | 26240001 | 26340000 | 0.0043 | 0.0031 | 0.0032 |
| F02 | 26260001 | 26360000 | 0.0033 | 0.0024 | 0.0023 |
| F02 | 26280001 | 26380000 | 0.0035 | 0.0026 | 0.0026 |
| F02 | 26300001 | 26400000 | 0.0034 | 0.0026 | 0.0028 |
| F02 | 26320001 | 26420000 | 0.0041 | 0.0033 | 0.0036 |
| F02 | 26340001 | 26440000 | 0.0042 | 0.0033 | 0.0036 |
| F02 | 26360001 | 26460000 | 0.0050 | 0.0036 | 0.0041 |
| F02 | 26380001 | 26480000 | 0.0052 | 0.0038 | 0.0042 |
| F02 | 26400001 | 26500000 | 0.0056 | 0.0039 | 0.0043 |
| F02 | 26420001 | 26520000 | 0.0046 | 0.0030 | 0.0031 |
| F02 | 26440001 | 26540000 | 0.0040 | 0.0027 | 0.0028 |
| F02 | 26460001 | 26560000 | 0.0041 | 0.0029 | 0.0032 |
| F02 | 26480001 | 26580000 | 0.0041 | 0.0029 | 0.0033 |
| F02 | 26500001 | 26600000 | 0.0042 | 0.0034 | 0.0033 |
| F02 | 26520001 | 26620000 | 0.0046 | 0.0037 | 0.0038 |
| F02 | 26540001 | 26640000 | 0.0046 | 0.0036 | 0.0038 |
| F02 | 26560001 | 26660000 | 0.0046 | 0.0040 | 0.0039 |
| F02 | 26580001 | 26680000 | 0.0044 | 0.0037 | 0.0038 |
| F02 | 26600001 | 26700000 | 0.0046 | 0.0036 | 0.0041 |
| F02 | 26620001 | 26720000 | 0.0044 | 0.0034 | 0.0040 |
| F02 | 26640001 | 26740000 | 0.0045 | 0.0035 | 0.0042 |
| F02 | 26660001 | 26760000 | 0.0042 | 0.0031 | 0.0037 |
| F02 | 26680001 | 26780000 | 0.0038 | 0.0028 | 0.0033 |
| F02 | 26700001 | 26800000 | 0.0035 | 0.0024 | 0.0030 |

| Chr | Start | End | both | A1 | A2 |
|---|---|---|---|---|---|
| F02 | 26720001 | 26820000 | 0.0033 | 0.0021 | 0.0027 |
| F02 | 26740001 | 26840000 | 0.0040 | 0.0026 | 0.0031 |
| F02 | 26760001 | 26860000 | 0.0037 | 0.0022 | 0.0027 |
| F02 | 26780001 | 26880000 | 0.0035 | 0.0021 | 0.0026 |
| F02 | 26800001 | 26900000 | 0.0030 | 0.0017 | 0.0019 |
| F02 | 26820001 | 26920000 | 0.0036 | 0.0022 | 0.0025 |
| F02 | 26840001 | 26940000 | 0.0025 | 0.0015 | 0.0018 |
| F02 | 26860001 | 26960000 | 0.0026 | 0.0018 | 0.0021 |
| F02 | 26880001 | 26980000 | 0.0031 | 0.0020 | 0.0022 |
| F02 | 26900001 | 27000000 | 0.0031 | 0.0020 | 0.0022 |
| F02 | 26920001 | 27020000 | 0.0030 | 0.0017 | 0.0020 |
| F02 | 26940001 | 27040000 | 0.0044 | 0.0030 | 0.0034 |
| F02 | 26960001 | 27060000 | 0.0039 | 0.0026 | 0.0030 |
| F02 | 26980001 | 27080000 | 0.0034 | 0.0026 | 0.0030 |
| F02 | 27000001 | 27100000 | 0.0042 | 0.0033 | 0.0038 |
| F02 | 27020001 | 27120000 | 0.0042 | 0.0036 | 0.0039 |
| F02 | 27040001 | 27140000 | 0.0028 | 0.0023 | 0.0025 |
| F02 | 27060001 | 27160000 | 0.0037 | 0.0030 | 0.0033 |
| F02 | 27080001 | 27180000 | 0.0038 | 0.0031 | 0.0034 |
| F02 | 27100001 | 27200000 | 0.0030 | 0.0026 | 0.0027 |
| F02 | 27120001 | 27220000 | 0.0032 | 0.0026 | 0.0029 |
| F02 | 27140001 | 27240000 | 0.0035 | 0.0028 | 0.0031 |
| F02 | 27160001 | 27260000 | 0.0032 | 0.0026 | 0.0028 |
| F02 | 27180001 | 27280000 | 0.0031 | 0.0025 | 0.0027 |
| F02 | 27200001 | 27300000 | 0.0031 | 0.0021 | 0.0023 |
| F02 | 27220001 | 27320000 | 0.0035 | 0.0026 | 0.0028 |
| F02 | 27240001 | 27340000 | 0.0044 | 0.0034 | 0.0036 |
| F02 | 27260001 | 27360000 | 0.0045 | 0.0033 | 0.0037 |
| F02 | 27280001 | 27380000 | 0.0045 | 0.0032 | 0.0037 |
| F02 | 27300001 | 27400000 | 0.0041 | 0.0031 | 0.0036 |
| F02 | 27320001 | 27420000 | 0.0029 | 0.0024 | 0.0026 |
| F02 | 27340001 | 27440000 | 0.0019 | 0.0015 | 0.0016 |
| F02 | 27360001 | 27460000 | 0.0019 | 0.0018 | 0.0018 |
| F02 | 27380001 | 27480000 | 0.0020 | 0.0019 | 0.0019 |
| F02 | 27400001 | 27500000 | 0.0023 | 0.0022 | 0.0022 |
| F02 | 27420001 | 27520000 | 0.0026 | 0.0024 | 0.0025 |
| F02 | 27440001 | 27540000 | 0.0026 | 0.0023 | 0.0025 |
| F02 | 27460001 | 27560000 | 0.0020 | 0.0019 | 0.0020 |
| F02 | 27480001 | 27580000 | 0.0017 | 0.0014 | 0.0015 |
| F02 | 27500001 | 27600000 | 0.0018 | 0.0015 | 0.0016 |
| F02 | 27520001 | 27620000 | 0.0014 | 0.0010 | 0.0012 |
| F02 | 27540001 | 27640000 | 0.0011 | 0.0008 | 0.0009 |
| F02 | 27560001 | 27660000 | 0.0010 | 0.0008 | 0.0009 |
| F02 | 27580001 | 27680000 | 0.0012 | 0.0010 | 0.0011 |
| F02 | 27600001 | 27700000 | 0.0018 | 0.0014 | 0.0016 |
| F02 | 27620001 | 27720000 | 0.0022 | 0.0017 | 0.0019 |
| F02 | 27640001 | 27740000 | 0.0029 | 0.0023 | 0.0025 |
| F02 | 27660001 | 27760000 | 0.0030 | 0.0023 | 0.0025 |
| F02 | 27680001 | 27780000 | 0.0028 | 0.0022 | 0.0025 |
| F02 | 27700001 | 27800000 | 0.0023 | 0.0019 | 0.0021 |
| F02 | 27720001 | 27820000 | 0.0018 | 0.0017 | 0.0017 |
| F02 | 27740001 | 27840000 | 0.0010 | 0.0010 | 0.0010 |
| F02 | 27760001 | 27860000 | 0.0012 | 0.0011 | 0.0011 |
| F02 | 27780001 | 27880000 | 0.0016 | 0.0016 | 0.0015 |
| F02 | 27800001 | 27900000 | 0.0020 | 0.0018 | 0.0017 |
| F02 | 27820001 | 27920000 | 0.0027 | 0.0022 | 0.0022 |
| F02 | 27840001 | 27940000 | 0.0034 | 0.0027 | 0.0029 |
| F02 | 27860001 | 27960000 | 0.0036 | 0.0029 | 0.0031 |

| Chr | Start | End | both | A1 | A2 |
|---|---|---|---|---|---|
| F02 | 27880001 | 27980000 | 0.0042 | 0.0031 | 0.0037 |
| F02 | 27900001 | 28000000 | 0.0041 | 0.0028 | 0.0034 |
| F02 | 27920001 | 28020000 | 0.0038 | 0.0026 | 0.0030 |
| F02 | 27940001 | 28040000 | 0.0047 | 0.0036 | 0.0040 |
| F02 | 27960001 | 28060000 | 0.0053 | 0.0040 | 0.0044 |
| F02 | 27980001 | 28080000 | 0.0048 | 0.0036 | 0.0039 |
| F02 | 28000001 | 28100000 | 0.0042 | 0.0035 | 0.0037 |
| F02 | 28020001 | 28120000 | 0.0044 | 0.0037 | 0.0039 |
| F02 | 28040001 | 28140000 | 0.0036 | 0.0027 | 0.0030 |
| F02 | 28060001 | 28160000 | 0.0034 | 0.0027 | 0.0028 |
| F02 | 28080001 | 28180000 | 0.0032 | 0.0025 | 0.0025 |
| F02 | 28100001 | 28200000 | 0.0032 | 0.0026 | 0.0026 |
| F02 | 28120001 | 28220000 | 0.0030 | 0.0023 | 0.0025 |
| F02 | 28140001 | 28240000 | 0.0025 | 0.0020 | 0.0023 |
| F02 | 28160001 | 28260000 | 0.0017 | 0.0013 | 0.0017 |
| F02 | 28180001 | 28280000 | 0.0016 | 0.0012 | 0.0016 |
| F02 | 28200001 | 28300000 | 0.0014 | 0.0011 | 0.0014 |
| F02 | 28220001 | 28320000 | 0.0011 | 0.0010 | 0.0011 |
| F02 | 28240001 | 28340000 | 0.0013 | 0.0011 | 0.0012 |
| F02 | 28260001 | 28360000 | 0.0025 | 0.0021 | 0.0023 |
| F02 | 28280001 | 28380000 | 0.0025 | 0.0021 | 0.0023 |
| F02 | 28300001 | 28400000 | 0.0028 | 0.0024 | 0.0026 |
| F02 | 28320001 | 28420000 | 0.0031 | 0.0026 | 0.0029 |
| F02 | 28340001 | 28440000 | 0.0029 | 0.0025 | 0.0028 |
| F02 | 28360001 | 28460000 | 0.0024 | 0.0020 | 0.0022 |
| F02 | 28380001 | 28480000 | 0.0025 | 0.0018 | 0.0021 |
| F02 | 28400001 | 28500000 | 0.0029 | 0.0019 | 0.0022 |
| F02 | 28420001 | 28520000 | 0.0030 | 0.0020 | 0.0023 |
| F02 | 28440001 | 28540000 | 0.0028 | 0.0018 | 0.0020 |
| F02 | 28460001 | 28560000 | 0.0023 | 0.0014 | 0.0015 |
| F02 | 28480001 | 28580000 | 0.0020 | 0.0015 | 0.0015 |
| F02 | 28500001 | 28600000 | 0.0013 | 0.0010 | 0.0011 |
| F02 | 28520001 | 28620000 | 0.0008 | 0.0006 | 0.0006 |
| F02 | 28540001 | 28640000 | 0.0008 | 0.0007 | 0.0007 |
| F02 | 28560001 | 28660000 | 0.0010 | 0.0009 | 0.0009 |
| F02 | 28580001 | 28680000 | 0.0009 | 0.0008 | 0.0009 |
| F02 | 28600001 | 28700000 | 0.0011 | 0.0009 | 0.0010 |
| F02 | 28620001 | 28720000 | 0.0020 | 0.0015 | 0.0019 |
| F02 | 28640001 | 28740000 | 0.0027 | 0.0019 | 0.0025 |
| F02 | 28660001 | 28760000 | 0.0027 | 0.0020 | 0.0025 |
| F02 | 28680001 | 28780000 | 0.0033 | 0.0024 | 0.0030 |
| F02 | 28700001 | 28800000 | 0.0032 | 0.0025 | 0.0030 |
| F02 | 28720001 | 28820000 | 0.0024 | 0.0018 | 0.0022 |
| F02 | 28740001 | 28840000 | 0.0019 | 0.0015 | 0.0017 |
| F02 | 28760001 | 28860000 | 0.0020 | 0.0014 | 0.0019 |
| F02 | 28780001 | 28880000 | 0.0019 | 0.0013 | 0.0017 |
| F02 | 28800001 | 28900000 | 0.0024 | 0.0016 | 0.0021 |
| F02 | 28820001 | 28920000 | 0.0026 | 0.0019 | 0.0024 |
| F02 | 28840001 | 28940000 | 0.0023 | 0.0018 | 0.0021 |
| F02 | 28860001 | 28960000 | 0.0023 | 0.0017 | 0.0020 |
| F02 | 28880001 | 28980000 | 0.0017 | 0.0013 | 0.0016 |
| F02 | 28900001 | 29000000 | 0.0014 | 0.0010 | 0.0013 |
| F02 | 28920001 | 29020000 | 0.0011 | 0.0008 | 0.0011 |
| F02 | 28940001 | 29040000 | 0.0015 | 0.0011 | 0.0014 |
| F02 | 28960001 | 29060000 | 0.0017 | 0.0014 | 0.0016 |
| F02 | 28980001 | 29080000 | 0.0023 | 0.0019 | 0.0022 |
| F02 | 29000001 | 29100000 | 0.0023 | 0.0020 | 0.0022 |
| F02 | 29020001 | 29120000 | 0.0023 | 0.0020 | 0.0022 |

| Chr | Start | End | both | A1 | A2 |
|---|---|---|---|---|---|
| F02 | 29040001 | 29140000 | 0.0031 | 0.0028 | 0.0030 |
| F02 | 29060001 | 29160000 | 0.0030 | 0.0027 | 0.0029 |
| F02 | 29080001 | 29180000 | 0.0030 | 0.0026 | 0.0028 |
| F02 | 29100001 | 29200000 | 0.0035 | 0.0028 | 0.0032 |
| F02 | 29120001 | 29220000 | 0.0039 | 0.0031 | 0.0036 |
| F02 | 29140001 | 29240000 | 0.0030 | 0.0020 | 0.0026 |
| F02 | 29160001 | 29260000 | 0.0029 | 0.0019 | 0.0024 |
| F02 | 29180001 | 29280000 | 0.0027 | 0.0019 | 0.0024 |
| F02 | 29200001 | 29300000 | 0.0021 | 0.0015 | 0.0019 |
| F02 | 29220001 | 29320000 | 0.0017 | 0.0012 | 0.0015 |
| F02 | 29240001 | 29340000 | 0.0021 | 0.0016 | 0.0020 |
| F02 | 29260001 | 29360000 | 0.0020 | 0.0015 | 0.0017 |
| F02 | 29280001 | 29380000 | 0.0019 | 0.0014 | 0.0017 |
| F02 | 29300001 | 29400000 | 0.0021 | 0.0016 | 0.0018 |
| F02 | 29320001 | 29420000 | 0.0022 | 0.0016 | 0.0019 |
| F02 | 29340001 | 29440000 | 0.0022 | 0.0018 | 0.0019 |
| F02 | 29360001 | 29460000 | 0.0021 | 0.0017 | 0.0019 |
| F02 | 29380001 | 29480000 | 0.0017 | 0.0015 | 0.0016 |
| F02 | 29400001 | 29500000 | 0.0017 | 0.0014 | 0.0016 |
| F02 | 29420001 | 29520000 | 0.0019 | 0.0017 | 0.0018 |
| F02 | 29440001 | 29540000 | 0.0021 | 0.0019 | 0.0020 |
| F02 | 29460001 | 29560000 | 0.0019 | 0.0017 | 0.0018 |
| F02 | 29480001 | 29580000 | 0.0017 | 0.0016 | 0.0017 |
| F02 | 29500001 | 29600000 | 0.0017 | 0.0016 | 0.0017 |
| F02 | 29520001 | 29620000 | 0.0013 | 0.0012 | 0.0012 |
| F02 | 29540001 | 29640000 | 0.0006 | 0.0004 | 0.0006 |
| F02 | 29560001 | 29660000 | 0.0006 | 0.0004 | 0.0006 |
| F02 | 29580001 | 29680000 | 0.0006 | 0.0005 | 0.0007 |
| F02 | 29600001 | 29700000 | 0.0007 | 0.0005 | 0.0007 |
| F02 | 29620001 | 29720000 | 0.0012 | 0.0009 | 0.0012 |
| F02 | 29640001 | 29740000 | 0.0020 | 0.0016 | 0.0020 |
| F02 | 29660001 | 29760000 | 0.0020 | 0.0016 | 0.0020 |
| F02 | 29680001 | 29780000 | 0.0022 | 0.0017 | 0.0022 |
| F02 | 29700001 | 29800000 | 0.0021 | 0.0016 | 0.0019 |
| F02 | 29720001 | 29820000 | 0.0017 | 0.0013 | 0.0015 |
| F02 | 29740001 | 29840000 | 0.0010 | 0.0006 | 0.0006 |
| F02 | 29760001 | 29860000 | 0.0015 | 0.0010 | 0.0011 |
| F02 | 29780001 | 29880000 | 0.0014 | 0.0010 | 0.0010 |
| F02 | 29800001 | 29900000 | 0.0016 | 0.0011 | 0.0013 |
| F02 | 29820001 | 29920000 | 0.0022 | 0.0017 | 0.0018 |
| F02 | 29840001 | 29940000 | 0.0025 | 0.0021 | 0.0022 |
| F02 | 29860001 | 29960000 | 0.0028 | 0.0021 | 0.0023 |
| F02 | 29880001 | 29980000 | 0.0030 | 0.0021 | 0.0023 |
| F02 | 29900001 | 30000000 | 0.0038 | 0.0029 | 0.0031 |
| F02 | 29920001 | 30020000 | 0.0040 | 0.0030 | 0.0033 |
| F02 | 29940001 | 30040000 | 0.0042 | 0.0031 | 0.0035 |
| F02 | 29960001 | 30060000 | 0.0034 | 0.0027 | 0.0029 |
| F02 | 29980001 | 30080000 | 0.0035 | 0.0028 | 0.0032 |
| F02 | 30000001 | 30100000 | 0.0029 | 0.0022 | 0.0026 |
| F02 | 30020001 | 30120000 | 0.0026 | 0.0020 | 0.0024 |
| F02 | 30040001 | 30140000 | 0.0023 | 0.0017 | 0.0021 |
| F02 | 30060001 | 30160000 | 0.0030 | 0.0023 | 0.0026 |
| F02 | 30080001 | 30180000 | 0.0035 | 0.0027 | 0.0030 |
| F02 | 30100001 | 30200000 | 0.0032 | 0.0025 | 0.0026 |
| F02 | 30120001 | 30220000 | 0.0026 | 0.0020 | 0.0021 |
| F02 | 30140001 | 30240000 | 0.0032 | 0.0024 | 0.0025 |
| F02 | 30160001 | 30260000 | 0.0030 | 0.0021 | 0.0023 |
| F02 | 30180001 | 30280000 | 0.0025 | 0.0018 | 0.0020 |

| Chr | Start | End | both | A1 | A2 |
|---|---|---|---|---|---|
| F02 | 30200001 | 30300000 | 0.0027 | 0.0021 | 0.0023 |
| F02 | 30220001 | 30320000 | 0.0029 | 0.0022 | 0.0025 |
| F02 | 30240001 | 30340000 | 0.0030 | 0.0027 | 0.0028 |
| F02 | 30260001 | 30360000 | 0.0032 | 0.0028 | 0.0029 |
| F02 | 30280001 | 30380000 | 0.0032 | 0.0027 | 0.0028 |
| F02 | 30300001 | 30400000 | 0.0030 | 0.0025 | 0.0026 |
| F02 | 30320001 | 30420000 | 0.0029 | 0.0024 | 0.0026 |
| F02 | 30340001 | 30440000 | 0.0027 | 0.0020 | 0.0022 |
| F02 | 30360001 | 30460000 | 0.0034 | 0.0025 | 0.0028 |
| F02 | 30380001 | 30480000 | 0.0037 | 0.0028 | 0.0030 |
| F02 | 30400001 | 30500000 | 0.0036 | 0.0026 | 0.0027 |
| F02 | 30420001 | 30520000 | 0.0043 | 0.0031 | 0.0033 |
| F02 | 30440001 | 30540000 | 0.0041 | 0.0032 | 0.0032 |
| F02 | 30460001 | 30560000 | 0.0034 | 0.0026 | 0.0026 |
| F02 | 30480001 | 30580000 | 0.0028 | 0.0021 | 0.0022 |
| F02 | 30500001 | 30600000 | 0.0029 | 0.0023 | 0.0025 |
| F02 | 30520001 | 30620000 | 0.0028 | 0.0019 | 0.0025 |
| F02 | 30540001 | 30640000 | 0.0024 | 0.0014 | 0.0023 |
| F02 | 30560001 | 30660000 | 0.0024 | 0.0013 | 0.0022 |
| F02 | 30580001 | 30680000 | 0.0036 | 0.0022 | 0.0033 |
| F02 | 30600001 | 30700000 | 0.0038 | 0.0026 | 0.0034 |
| F02 | 30620001 | 30720000 | 0.0036 | 0.0027 | 0.0031 |
| F02 | 30640001 | 30740000 | 0.0040 | 0.0030 | 0.0033 |
| F02 | 30660001 | 30760000 | 0.0040 | 0.0030 | 0.0032 |
| F02 | 30680001 | 30780000 | 0.0037 | 0.0030 | 0.0030 |
| F02 | 30700001 | 30800000 | 0.0031 | 0.0024 | 0.0024 |
| F02 | 30720001 | 30820000 | 0.0029 | 0.0021 | 0.0022 |
| F02 | 30740001 | 30840000 | 0.0035 | 0.0029 | 0.0030 |
| F02 | 30760001 | 30860000 | 0.0035 | 0.0030 | 0.0032 |
| F02 | 30780001 | 30880000 | 0.0027 | 0.0023 | 0.0025 |
| F02 | 30800001 | 30900000 | 0.0039 | 0.0029 | 0.0034 |
| F02 | 30820001 | 30920000 | 0.0044 | 0.0033 | 0.0039 |
| F02 | 30840001 | 30940000 | 0.0030 | 0.0020 | 0.0026 |
| F02 | 30860001 | 30960000 | 0.0030 | 0.0017 | 0.0023 |
| F02 | 30880001 | 30980000 | 0.0038 | 0.0023 | 0.0028 |
| F02 | 30900001 | 31000000 | 0.0038 | 0.0028 | 0.0030 |
| F02 | 30920001 | 31020000 | 0.0031 | 0.0024 | 0.0024 |
| F02 | 30940001 | 31040000 | 0.0035 | 0.0027 | 0.0027 |
| F02 | 30960001 | 31060000 | 0.0033 | 0.0026 | 0.0027 |
| F02 | 30980001 | 31080000 | 0.0027 | 0.0020 | 0.0022 |
| F02 | 31000001 | 31100000 | 0.0023 | 0.0013 | 0.0016 |
| F02 | 31020001 | 31120000 | 0.0032 | 0.0021 | 0.0026 |
| F02 | 31040001 | 31140000 | 0.0029 | 0.0019 | 0.0023 |
| F02 | 31060001 | 31160000 | 0.0028 | 0.0020 | 0.0024 |
| F02 | 31080001 | 31180000 | 0.0029 | 0.0022 | 0.0027 |
| F02 | 31100001 | 31200000 | 0.0025 | 0.0021 | 0.0024 |
| F02 | 31120001 | 31220000 | 0.0013 | 0.0011 | 0.0013 |
| F02 | 31140001 | 31240000 | 0.0018 | 0.0016 | 0.0018 |
| F02 | 31160001 | 31260000 | 0.0017 | 0.0015 | 0.0017 |
| F02 | 31180001 | 31280000 | 0.0015 | 0.0013 | 0.0015 |
| F02 | 31200001 | 31300000 | 0.0015 | 0.0013 | 0.0015 |
| F02 | 31220001 | 31320000 | 0.0021 | 0.0016 | 0.0019 |
| F02 | 31240001 | 31340000 | 0.0021 | 0.0016 | 0.0020 |
| F02 | 31260001 | 31360000 | 0.0021 | 0.0014 | 0.0018 |
| F02 | 31280001 | 31380000 | 0.0021 | 0.0014 | 0.0018 |
| F02 | 31300001 | 31400000 | 0.0020 | 0.0014 | 0.0017 |
| F02 | 31320001 | 31420000 | 0.0021 | 0.0016 | 0.0019 |
| F02 | 31340001 | 31440000 | 0.0025 | 0.0018 | 0.0020 |

| Chr | Start | End | both | A1 | A2 |
|---|---|---|---|---|---|
| F02 | 31360001 | 31460000 | 0.0027 | 0.0020 | 0.0022 |
| F02 | 31380001 | 31480000 | 0.0031 | 0.0021 | 0.0023 |
| F02 | 31400001 | 31500000 | 0.0036 | 0.0023 | 0.0027 |
| F02 | 31420001 | 31520000 | 0.0037 | 0.0022 | 0.0027 |
| F02 | 31440001 | 31540000 | 0.0030 | 0.0017 | 0.0023 |
| F02 | 31460001 | 31560000 | 0.0029 | 0.0018 | 0.0024 |
| F02 | 31480001 | 31580000 | 0.0025 | 0.0018 | 0.0022 |
| F02 | 31500001 | 31600000 | 0.0019 | 0.0015 | 0.0019 |
| F02 | 31520001 | 31620000 | 0.0015 | 0.0013 | 0.0015 |
| F02 | 31540001 | 31640000 | 0.0014 | 0.0012 | 0.0014 |
| F02 | 31560001 | 31660000 | 0.0013 | 0.0010 | 0.0013 |
| F02 | 31580001 | 31680000 | 0.0018 | 0.0016 | 0.0018 |
| F02 | 31600001 | 31700000 | 0.0019 | 0.0016 | 0.0019 |
| F02 | 31620001 | 31720000 | 0.0024 | 0.0020 | 0.0022 |
| F02 | 31640001 | 31740000 | 0.0026 | 0.0021 | 0.0024 |
| F02 | 31660001 | 31760000 | 0.0036 | 0.0030 | 0.0033 |
| F02 | 31680001 | 31780000 | 0.0037 | 0.0027 | 0.0031 |
| F02 | 31700001 | 31800000 | 0.0047 | 0.0037 | 0.0041 |
| F02 | 31720001 | 31820000 | 0.0041 | 0.0032 | 0.0036 |
| F02 | 31740001 | 31840000 | 0.0036 | 0.0029 | 0.0033 |
| F02 | 31760001 | 31860000 | 0.0030 | 0.0021 | 0.0025 |
| F02 | 31780001 | 31880000 | 0.0034 | 0.0026 | 0.0027 |
| F02 | 31800001 | 31900000 | 0.0036 | 0.0025 | 0.0028 |
| F02 | 31820001 | 31920000 | 0.0036 | 0.0025 | 0.0027 |
| F02 | 31840001 | 31940000 | 0.0038 | 0.0027 | 0.0030 |
| F02 | 31860001 | 31960000 | 0.0043 | 0.0029 | 0.0033 |
| F02 | 31880001 | 31980000 | 0.0036 | 0.0024 | 0.0027 |
| F02 | 31900001 | 32000000 | 0.0034 | 0.0025 | 0.0026 |
| F02 | 31920001 | 32020000 | 0.0038 | 0.0028 | 0.0030 |
| F02 | 31940001 | 32040000 | 0.0039 | 0.0029 | 0.0031 |
| F02 | 31960001 | 32060000 | 0.0033 | 0.0027 | 0.0029 |
| F02 | 31980001 | 32080000 | 0.0036 | 0.0031 | 0.0034 |
| F02 | 32000001 | 32100000 | 0.0030 | 0.0024 | 0.0030 |
| F02 | 32020001 | 32120000 | 0.0028 | 0.0024 | 0.0028 |
| F02 | 32040001 | 32140000 | 0.0036 | 0.0031 | 0.0035 |
| F02 | 32060001 | 32160000 | 0.0036 | 0.0029 | 0.0033 |
| F02 | 32080001 | 32180000 | 0.0043 | 0.0032 | 0.0036 |
| F02 | 32100001 | 32200000 | 0.0047 | 0.0033 | 0.0039 |
| F02 | 32120001 | 32220000 | 0.0050 | 0.0036 | 0.0039 |
| F02 | 32140001 | 32240000 | 0.0057 | 0.0041 | 0.0043 |
| F02 | 32160001 | 32260000 | 0.0068 | 0.0049 | 0.0053 |
| F02 | 32180001 | 32280000 | 0.0059 | 0.0043 | 0.0045 |
| F02 | 32200001 | 32300000 | 0.0052 | 0.0040 | 0.0038 |
| F02 | 32220001 | 32320000 | 0.0061 | 0.0048 | 0.0047 |
| F02 | 32240001 | 32340000 | 0.0046 | 0.0036 | 0.0035 |
| F02 | 32260001 | 32360000 | 0.0037 | 0.0027 | 0.0025 |
| F02 | 32280001 | 32380000 | 0.0042 | 0.0032 | 0.0031 |
| F02 | 32300001 | 32400000 | 0.0039 | 0.0029 | 0.0030 |
| F02 | 32320001 | 32420000 | 0.0036 | 0.0024 | 0.0027 |
| F02 | 32340001 | 32440000 | 0.0049 | 0.0037 | 0.0041 |
| F02 | 32360001 | 32460000 | 0.0054 | 0.0041 | 0.0047 |
| F02 | 32380001 | 32480000 | 0.0050 | 0.0038 | 0.0042 |
| F02 | 32400001 | 32500000 | 0.0051 | 0.0039 | 0.0042 |
| F02 | 32420001 | 32520000 | 0.0043 | 0.0031 | 0.0034 |
| F02 | 32440001 | 32540000 | 0.0026 | 0.0017 | 0.0018 |
| F02 | 32460001 | 32560000 | 0.0019 | 0.0014 | 0.0014 |
| F02 | 32480001 | 32580000 | 0.0012 | 0.0009 | 0.0010 |
| F02 | 32500001 | 32600000 | 0.0012 | 0.0009 | 0.0011 |

| Chr | Start | End | both | A1 | A2 |
|---|---|---|---|---|---|
| F02 | 32520001 | 32620000 | 0.0040 | 0.0034 | 0.0036 |
| F02 | 32540001 | 32640000 | 0.0041 | 0.0034 | 0.0037 |
| F02 | 32560001 | 32660000 | 0.0047 | 0.0037 | 0.0040 |
| F02 | 32580001 | 32680000 | 0.0053 | 0.0042 | 0.0046 |
| F02 | 32600001 | 32700000 | 0.0063 | 0.0049 | 0.0055 |
| F02 | 32620001 | 32720000 | 0.0044 | 0.0034 | 0.0037 |
| F02 | 32640001 | 32740000 | 0.0058 | 0.0044 | 0.0049 |
| F02 | 32660001 | 32760000 | 0.0058 | 0.0045 | 0.0048 |
| F02 | 32680001 | 32780000 | 0.0057 | 0.0043 | 0.0045 |
| F02 | 32700001 | 32800000 | 0.0052 | 0.0037 | 0.0040 |
| F02 | 32720001 | 32820000 | 0.0044 | 0.0030 | 0.0033 |
| F02 | 32740001 | 32840000 | 0.0030 | 0.0020 | 0.0021 |
| F02 | 32760001 | 32860000 | 0.0023 | 0.0017 | 0.0019 |
| F02 | 32780001 | 32880000 | 0.0029 | 0.0024 | 0.0028 |
| F02 | 32800001 | 32900000 | 0.0028 | 0.0026 | 0.0027 |
| F02 | 32820001 | 32920000 | 0.0035 | 0.0030 | 0.0033 |
| F02 | 32840001 | 32940000 | 0.0040 | 0.0033 | 0.0036 |
| F02 | 32860001 | 32960000 | 0.0047 | 0.0037 | 0.0040 |
| F02 | 32880001 | 32980000 | 0.0037 | 0.0027 | 0.0028 |
| F02 | 32900001 | 33000000 | 0.0037 | 0.0026 | 0.0027 |
| F02 | 32920001 | 33020000 | 0.0034 | 0.0021 | 0.0021 |
| F02 | 32940001 | 33040000 | 0.0031 | 0.0019 | 0.0021 |
| F02 | 32960001 | 33060000 | 0.0032 | 0.0021 | 0.0024 |
| F02 | 32980001 | 33080000 | 0.0036 | 0.0023 | 0.0027 |
| F02 | 33000001 | 33100000 | 0.0037 | 0.0023 | 0.0029 |
| F02 | 33020001 | 33120000 | 0.0033 | 0.0023 | 0.0028 |
| F02 | 33040001 | 33140000 | 0.0035 | 0.0021 | 0.0026 |
| F02 | 33060001 | 33160000 | 0.0035 | 0.0022 | 0.0026 |
| F02 | 33080001 | 33180000 | 0.0039 | 0.0028 | 0.0032 |
| F02 | 33100001 | 33200000 | 0.0045 | 0.0034 | 0.0038 |
| F02 | 33120001 | 33220000 | 0.0050 | 0.0038 | 0.0042 |
| F02 | 33140001 | 33240000 | 0.0058 | 0.0049 | 0.0052 |
| F02 | 33160001 | 33260000 | 0.0052 | 0.0041 | 0.0044 |
| F02 | 33180001 | 33280000 | 0.0053 | 0.0038 | 0.0041 |
| F02 | 33200001 | 33300000 | 0.0046 | 0.0031 | 0.0033 |
| F02 | 33220001 | 33320000 | 0.0040 | 0.0025 | 0.0026 |
| F02 | 33240001 | 33340000 | 0.0029 | 0.0015 | 0.0017 |
| F02 | 33260001 | 33360000 | 0.0034 | 0.0021 | 0.0023 |
| F02 | 33280001 | 33380000 | 0.0033 | 0.0024 | 0.0026 |
| F02 | 33300001 | 33400000 | 0.0037 | 0.0031 | 0.0032 |
| F02 | 33320001 | 33420000 | 0.0042 | 0.0038 | 0.0040 |
| F02 | 33340001 | 33440000 | 0.0049 | 0.0045 | 0.0047 |
| F02 | 33360001 | 33460000 | 0.0042 | 0.0039 | 0.0040 |
| F02 | 33380001 | 33480000 | 0.0034 | 0.0032 | 0.0033 |
| F02 | 33400001 | 33500000 | 0.0027 | 0.0024 | 0.0026 |
| F02 | 33420001 | 33520000 | 0.0022 | 0.0020 | 0.0022 |
| F02 | 33440001 | 33540000 | 0.0022 | 0.0019 | 0.0022 |
| F02 | 33460001 | 33560000 | 0.0024 | 0.0020 | 0.0022 |
| F02 | 33480001 | 33580000 | 0.0029 | 0.0024 | 0.0026 |
| F02 | 33500001 | 33600000 | 0.0037 | 0.0030 | 0.0033 |
| F02 | 33520001 | 33620000 | 0.0032 | 0.0025 | 0.0029 |
| F02 | 33540001 | 33640000 | 0.0027 | 0.0021 | 0.0024 |
| F02 | 33560001 | 33660000 | 0.0024 | 0.0020 | 0.0024 |
| F02 | 33580001 | 33680000 | 0.0022 | 0.0018 | 0.0022 |
| F02 | 33600001 | 33700000 | 0.0013 | 0.0010 | 0.0012 |
| F02 | 33620001 | 33720000 | 0.0019 | 0.0014 | 0.0016 |
| F02 | 33640001 | 33740000 | 0.0019 | 0.0015 | 0.0016 |
| F02 | 33660001 | 33760000 | 0.0023 | 0.0017 | 0.0018 |

| Chr | Start | End | both | A1 | A2 |
|---|---|---|---|---|---|
| F02 | 33680001 | 33780000 | 0.0024 | 0.0018 | 0.0019 |
| F02 | 33700001 | 33800000 | 0.0026 | 0.0020 | 0.0020 |
| F02 | 33720001 | 33820000 | 0.0025 | 0.0019 | 0.0019 |
| F02 | 33740001 | 33840000 | 0.0027 | 0.0020 | 0.0022 |
| F02 | 33760001 | 33860000 | 0.0028 | 0.0022 | 0.0024 |
| F02 | 33780001 | 33880000 | 0.0029 | 0.0022 | 0.0025 |
| F02 | 33800001 | 33900000 | 0.0031 | 0.0023 | 0.0027 |
| F02 | 33820001 | 33920000 | 0.0027 | 0.0021 | 0.0025 |
| F02 | 33840001 | 33940000 | 0.0025 | 0.0019 | 0.0023 |
| F02 | 33860001 | 33960000 | 0.0017 | 0.0012 | 0.0016 |
| F02 | 33880001 | 33980000 | 0.0012 | 0.0009 | 0.0012 |
| F02 | 33900001 | 34000000 | 0.0011 | 0.0008 | 0.0011 |
| F02 | 33920001 | 34020000 | 0.0010 | 0.0008 | 0.0010 |
| F02 | 33940001 | 34040000 | 0.0020 | 0.0018 | 0.0019 |
| F02 | 33960001 | 34060000 | 0.0021 | 0.0018 | 0.0020 |
| F02 | 33980001 | 34080000 | 0.0020 | 0.0017 | 0.0018 |
| F02 | 34000001 | 34100000 | 0.0021 | 0.0017 | 0.0019 |
| F02 | 34020001 | 34120000 | 0.0022 | 0.0018 | 0.0020 |
| F02 | 34040001 | 34140000 | 0.0011 | 0.0009 | 0.0010 |
| F02 | 34060001 | 34160000 | 0.0013 | 0.0010 | 0.0011 |
| F02 | 34080001 | 34180000 | 0.0022 | 0.0017 | 0.0019 |
| F02 | 34100001 | 34200000 | 0.0026 | 0.0021 | 0.0023 |
| F02 | 34120001 | 34220000 | 0.0028 | 0.0022 | 0.0026 |
| F02 | 34140001 | 34240000 | 0.0025 | 0.0020 | 0.0023 |
| F02 | 34160001 | 34260000 | 0.0026 | 0.0021 | 0.0025 |
| F02 | 34180001 | 34280000 | 0.0019 | 0.0016 | 0.0019 |
| F02 | 34200001 | 34300000 | 0.0013 | 0.0011 | 0.0013 |
| F02 | 34220001 | 34320000 | 0.0008 | 0.0007 | 0.0008 |
| F02 | 34240001 | 34340000 | 0.0008 | 0.0007 | 0.0008 |
| F02 | 34260001 | 34360000 | 0.0005 | 0.0004 | 0.0005 |
| F02 | 34280001 | 34380000 | 0.0003 | 0.0002 | 0.0003 |
| F02 | 34300001 | 34400000 | 0.0002 | 0.0001 | 0.0002 |
| F02 | 34320001 | 34420000 | 0.0001 | 0.0001 | 0.0001 |
| F02 | 34340001 | 34440000 | 0.0004 | 0.0003 | 0.0004 |
| F02 | 34360001 | 34460000 | 0.0005 | 0.0004 | 0.0005 |
| F02 | 34380001 | 34480000 | 0.0008 | 0.0007 | 0.0007 |
| F02 | 34400001 | 34500000 | 0.0009 | 0.0009 | 0.0009 |
| F02 | 34420001 | 34520000 | 0.0011 | 0.0010 | 0.0011 |
| F02 | 34440001 | 34540000 | 0.0011 | 0.0010 | 0.0011 |
| F02 | 34460001 | 34560000 | 0.0010 | 0.0009 | 0.0010 |
| F02 | 34480001 | 34580000 | 0.0008 | 0.0007 | 0.0008 |
| F02 | 34500001 | 34600000 | 0.0008 | 0.0007 | 0.0008 |
| F02 | 34520001 | 34620000 | 0.0013 | 0.0012 | 0.0013 |
| F02 | 34540001 | 34640000 | 0.0012 | 0.0010 | 0.0011 |
| F02 | 34560001 | 34660000 | 0.0016 | 0.0015 | 0.0015 |
| F02 | 34580001 | 34680000 | 0.0018 | 0.0017 | 0.0018 |
| F02 | 34600001 | 34700000 | 0.0016 | 0.0016 | 0.0016 |
| F02 | 34620001 | 34720000 | 0.0010 | 0.0010 | 0.0010 |
| F02 | 34640001 | 34740000 | 0.0009 | 0.0009 | 0.0009 |
| F02 | 34660001 | 34760000 | 0.0005 | 0.0004 | 0.0005 |
| F02 | 34680001 | 34780000 | 0.0004 | 0.0004 | 0.0004 |
| F02 | 34700001 | 34800000 | 0.0006 | 0.0005 | 0.0006 |
| F02 | 34720001 | 34820000 | 0.0008 | 0.0007 | 0.0008 |
| F02 | 34740001 | 34840000 | 0.0010 | 0.0009 | 0.0010 |
| F02 | 34760001 | 34860000 | 0.0014 | 0.0012 | 0.0014 |
| F02 | 34780001 | 34880000 | 0.0012 | 0.0010 | 0.0012 |
| F02 | 34800001 | 34900000 | 0.0015 | 0.0013 | 0.0015 |
| F02 | 34820001 | 34920000 | 0.0015 | 0.0014 | 0.0015 |

| Chr | Start | End | both | A1 | A2 |
|---|---|---|---|---|---|
| F02 | 34840001 | 34940000 | 0.0013 | 0.0012 | 0.0012 |
| F02 | 34860001 | 34960000 | 0.0009 | 0.0009 | 0.0009 |
| F02 | 34880001 | 34980000 | 0.0009 | 0.0009 | 0.0008 |
| F02 | 34900001 | 35000000 | 0.0005 | 0.0005 | 0.0004 |
| F02 | 34920001 | 35020000 | 0.0004 | 0.0003 | 0.0004 |
| F02 | 34940001 | 35040000 | 0.0005 | 0.0004 | 0.0005 |
| F02 | 34960001 | 35060000 | 0.0007 | 0.0006 | 0.0007 |
| F02 | 34980001 | 35080000 | 0.0008 | 0.0007 | 0.0008 |
| F02 | 35000001 | 35100000 | 0.0011 | 0.0008 | 0.0010 |
| F02 | 35020001 | 35120000 | 0.0012 | 0.0010 | 0.0012 |
| F02 | 35040001 | 35140000 | 0.0016 | 0.0013 | 0.0016 |
| F02 | 35060001 | 35160000 | 0.0020 | 0.0016 | 0.0019 |
| F02 | 35080001 | 35180000 | 0.0018 | 0.0015 | 0.0017 |
| F02 | 35100001 | 35200000 | 0.0020 | 0.0017 | 0.0019 |
| F02 | 35120001 | 35220000 | 0.0025 | 0.0020 | 0.0022 |
| F02 | 35140001 | 35240000 | 0.0025 | 0.0019 | 0.0021 |
| F02 | 35160001 | 35260000 | 0.0029 | 0.0024 | 0.0027 |
| F02 | 35180001 | 35280000 | 0.0034 | 0.0026 | 0.0030 |
| F02 | 35200001 | 35300000 | 0.0034 | 0.0026 | 0.0031 |
| F02 | 35220001 | 35320000 | 0.0032 | 0.0024 | 0.0028 |
| F02 | 35240001 | 35340000 | 0.0029 | 0.0022 | 0.0026 |
| F02 | 35260001 | 35360000 | 0.0027 | 0.0019 | 0.0024 |
| F02 | 35280001 | 35380000 | 0.0024 | 0.0017 | 0.0021 |
| F02 | 35300001 | 35400000 | 0.0024 | 0.0015 | 0.0021 |
| F02 | 35320001 | 35420000 | 0.0019 | 0.0012 | 0.0017 |
| F02 | 35340001 | 35440000 | 0.0020 | 0.0014 | 0.0018 |
| F02 | 35360001 | 35460000 | 0.0021 | 0.0015 | 0.0017 |
| F02 | 35380001 | 35480000 | 0.0024 | 0.0018 | 0.0020 |
| F02 | 35400001 | 35500000 | 0.0019 | 0.0016 | 0.0016 |
| F02 | 35420001 | 35520000 | 0.0019 | 0.0016 | 0.0015 |
| F02 | 35440001 | 35540000 | 0.0018 | 0.0014 | 0.0014 |
| F02 | 35460001 | 35560000 | 0.0010 | 0.0009 | 0.0009 |
| F02 | 35480001 | 35580000 | 0.0010 | 0.0009 | 0.0009 |
| F02 | 35500001 | 35600000 | 0.0017 | 0.0015 | 0.0016 |
| F02 | 35520001 | 35620000 | 0.0019 | 0.0018 | 0.0019 |
| F02 | 35540001 | 35640000 | 0.0020 | 0.0018 | 0.0019 |
| F02 | 35560001 | 35660000 | 0.0025 | 0.0022 | 0.0024 |
| F02 | 35580001 | 35680000 | 0.0025 | 0.0021 | 0.0023 |
| F02 | 35600001 | 35700000 | 0.0020 | 0.0015 | 0.0017 |
| F02 | 35620001 | 35720000 | 0.0023 | 0.0018 | 0.0020 |
| F02 | 35640001 | 35740000 | 0.0035 | 0.0030 | 0.0030 |
| F02 | 35660001 | 35760000 | 0.0035 | 0.0029 | 0.0030 |
| F02 | 35680001 | 35780000 | 0.0042 | 0.0036 | 0.0035 |
| F02 | 35700001 | 35800000 | 0.0050 | 0.0042 | 0.0042 |
| F02 | 35720001 | 35820000 | 0.0058 | 0.0050 | 0.0050 |
| F02 | 35740001 | 35840000 | 0.0043 | 0.0038 | 0.0038 |
| F02 | 35760001 | 35860000 | 0.0039 | 0.0034 | 0.0034 |
| F02 | 35780001 | 35880000 | 0.0036 | 0.0032 | 0.0033 |
| F02 | 35800001 | 35900000 | 0.0043 | 0.0041 | 0.0042 |
| F02 | 35820001 | 35920000 | 0.0037 | 0.0033 | 0.0035 |
| F02 | 35840001 | 35940000 | 0.0047 | 0.0041 | 0.0043 |
| F02 | 35860001 | 35960000 | 0.0053 | 0.0046 | 0.0048 |
| F02 | 35880001 | 35980000 | 0.0064 | 0.0052 | 0.0056 |
| F02 | 35900001 | 36000000 | 0.0054 | 0.0039 | 0.0045 |
| F02 | 35920001 | 36020000 | 0.0053 | 0.0037 | 0.0042 |
| F02 | 35940001 | 36040000 | 0.0049 | 0.0031 | 0.0037 |
| F02 | 35960001 | 36060000 | 0.0046 | 0.0028 | 0.0034 |
| F02 | 35980001 | 36080000 | 0.0031 | 0.0017 | 0.0022 |

| Chr | Start | End | both | A1 | A2 |
|---|---|---|---|---|---|
| F02 | 36000001 | 36100000 | 0.0024 | 0.0015 | 0.0017 |
| F02 | 36020001 | 36120000 | 0.0022 | 0.0015 | 0.0017 |
| F02 | 36040001 | 36140000 | 0.0016 | 0.0012 | 0.0014 |
| F02 | 36060001 | 36160000 | 0.0014 | 0.0011 | 0.0013 |
| F02 | 36080001 | 36180000 | 0.0019 | 0.0013 | 0.0018 |
| F02 | 36100001 | 36200000 | 0.0026 | 0.0017 | 0.0024 |
| F02 | 36120001 | 36220000 | 0.0029 | 0.0020 | 0.0026 |
| F02 | 36140001 | 36240000 | 0.0039 | 0.0028 | 0.0036 |
| F02 | 36160001 | 36260000 | 0.0051 | 0.0038 | 0.0046 |
| F02 | 36180001 | 36280000 | 0.0052 | 0.0040 | 0.0046 |
| F02 | 36200001 | 36300000 | 0.0063 | 0.0049 | 0.0055 |
| F02 | 36220001 | 36320000 | 0.0065 | 0.0048 | 0.0056 |
| F02 | 36240001 | 36340000 | 0.0054 | 0.0039 | 0.0045 |
| F02 | 36260001 | 36360000 | 0.0047 | 0.0030 | 0.0036 |
| F02 | 36280001 | 36380000 | 0.0051 | 0.0035 | 0.0040 |
| F02 | 36300001 | 36400000 | 0.0036 | 0.0023 | 0.0026 |
| F02 | 36320001 | 36420000 | 0.0039 | 0.0023 | 0.0028 |
| F02 | 36340001 | 36440000 | 0.0046 | 0.0028 | 0.0033 |
| F02 | 36360001 | 36460000 | 0.0049 | 0.0034 | 0.0038 |
| F02 | 36380001 | 36480000 | 0.0050 | 0.0032 | 0.0038 |
| F02 | 36400001 | 36500000 | 0.0049 | 0.0033 | 0.0039 |
| F02 | 36420001 | 36520000 | 0.0045 | 0.0033 | 0.0038 |
| F02 | 36440001 | 36540000 | 0.0039 | 0.0029 | 0.0034 |
| F02 | 36460001 | 36560000 | 0.0029 | 0.0023 | 0.0027 |
| F02 | 36480001 | 36580000 | 0.0015 | 0.0012 | 0.0014 |
| F02 | 36500001 | 36600000 | 0.0018 | 0.0016 | 0.0017 |
| F02 | 36520001 | 36620000 | 0.0014 | 0.0013 | 0.0013 |
| F02 | 36540001 | 36640000 | 0.0020 | 0.0018 | 0.0019 |
| F02 | 36560001 | 36660000 | 0.0020 | 0.0017 | 0.0019 |
| F02 | 36580001 | 36680000 | 0.0023 | 0.0019 | 0.0021 |
| F02 | 36600001 | 36700000 | 0.0020 | 0.0016 | 0.0018 |
| F02 | 36620001 | 36720000 | 0.0018 | 0.0014 | 0.0017 |
| F02 | 36640001 | 36740000 | 0.0018 | 0.0014 | 0.0017 |
| F02 | 36660001 | 36760000 | 0.0018 | 0.0014 | 0.0017 |
| F02 | 36680001 | 36780000 | 0.0023 | 0.0017 | 0.0023 |
| F02 | 36700001 | 36800000 | 0.0027 | 0.0022 | 0.0027 |
| F02 | 36720001 | 36820000 | 0.0035 | 0.0028 | 0.0035 |
| F02 | 36740001 | 36840000 | 0.0037 | 0.0029 | 0.0035 |
| F02 | 36760001 | 36860000 | 0.0041 | 0.0032 | 0.0038 |
| F02 | 36780001 | 36880000 | 0.0039 | 0.0030 | 0.0034 |
| F02 | 36800001 | 36900000 | 0.0039 | 0.0029 | 0.0033 |
| F02 | 36820001 | 36920000 | 0.0036 | 0.0027 | 0.0031 |
| F02 | 36840001 | 36940000 | 0.0035 | 0.0026 | 0.0030 |
| F02 | 36860001 | 36960000 | 0.0039 | 0.0028 | 0.0032 |
| F02 | 36880001 | 36980000 | 0.0036 | 0.0026 | 0.0030 |
| F02 | 36900001 | 37000000 | 0.0031 | 0.0023 | 0.0026 |
| F02 | 36920001 | 37020000 | 0.0036 | 0.0027 | 0.0031 |
| F02 | 36940001 | 37040000 | 0.0032 | 0.0024 | 0.0027 |
| F02 | 36960001 | 37060000 | 0.0032 | 0.0024 | 0.0028 |
| F02 | 36980001 | 37080000 | 0.0045 | 0.0039 | 0.0041 |
| F02 | 37000001 | 37100000 | 0.0050 | 0.0043 | 0.0045 |
| F02 | 37020001 | 37120000 | 0.0045 | 0.0039 | 0.0040 |
| F02 | 37040001 | 37140000 | 0.0048 | 0.0042 | 0.0044 |
| F02 | 37060001 | 37160000 | 0.0046 | 0.0043 | 0.0043 |
| F02 | 37080001 | 37180000 | 0.0032 | 0.0028 | 0.0031 |
| F02 | 37100001 | 37200000 | 0.0031 | 0.0027 | 0.0030 |
| F02 | 37120001 | 37220000 | 0.0025 | 0.0021 | 0.0025 |
| F02 | 37140001 | 37240000 | 0.0024 | 0.0020 | 0.0024 |

| Chr | Start | End | both | A1 | A2 |
|---|---|---|---|---|---|
| F02 | 37160001 | 37260000 | 0.0021 | 0.0017 | 0.0021 |
| F02 | 37180001 | 37280000 | 0.0029 | 0.0023 | 0.0028 |
| F02 | 37200001 | 37300000 | 0.0023 | 0.0018 | 0.0022 |
| F02 | 37220001 | 37320000 | 0.0026 | 0.0021 | 0.0025 |
| F02 | 37240001 | 37340000 | 0.0022 | 0.0018 | 0.0021 |
| F02 | 37260001 | 37360000 | 0.0023 | 0.0017 | 0.0020 |
| F02 | 37280001 | 37380000 | 0.0019 | 0.0012 | 0.0014 |
| F02 | 37300001 | 37400000 | 0.0025 | 0.0017 | 0.0019 |
| F02 | 37320001 | 37420000 | 0.0033 | 0.0021 | 0.0023 |
| F02 | 37340001 | 37440000 | 0.0036 | 0.0022 | 0.0023 |
| F02 | 37360001 | 37460000 | 0.0035 | 0.0022 | 0.0023 |
| F02 | 37380001 | 37480000 | 0.0040 | 0.0029 | 0.0032 |
| F02 | 37400001 | 37500000 | 0.0036 | 0.0026 | 0.0029 |
| F02 | 37420001 | 37520000 | 0.0031 | 0.0024 | 0.0027 |
| F02 | 37440001 | 37540000 | 0.0030 | 0.0023 | 0.0028 |
| F02 | 37460001 | 37560000 | 0.0039 | 0.0028 | 0.0034 |
| F02 | 37480001 | 37580000 | 0.0036 | 0.0023 | 0.0028 |
| F02 | 37500001 | 37600000 | 0.0035 | 0.0023 | 0.0028 |
| F02 | 37520001 | 37620000 | 0.0037 | 0.0022 | 0.0028 |
| F02 | 37540001 | 37640000 | 0.0047 | 0.0032 | 0.0038 |
| F02 | 37560001 | 37660000 | 0.0040 | 0.0028 | 0.0034 |
| F02 | 37580001 | 37680000 | 0.0034 | 0.0026 | 0.0031 |
| F02 | 37600001 | 37700000 | 0.0037 | 0.0029 | 0.0034 |
| F02 | 37620001 | 37720000 | 0.0030 | 0.0026 | 0.0029 |
| F02 | 37640001 | 37740000 | 0.0020 | 0.0018 | 0.0020 |
| F02 | 37660001 | 37760000 | 0.0024 | 0.0021 | 0.0023 |
| F02 | 37680001 | 37780000 | 0.0030 | 0.0028 | 0.0029 |
| F02 | 37700001 | 37800000 | 0.0029 | 0.0027 | 0.0028 |
| F02 | 37720001 | 37820000 | 0.0033 | 0.0032 | 0.0032 |
| F02 | 37740001 | 37840000 | 0.0030 | 0.0029 | 0.0029 |
| F02 | 37760001 | 37860000 | 0.0036 | 0.0033 | 0.0033 |
| F02 | 37780001 | 37880000 | 0.0035 | 0.0030 | 0.0030 |
| F02 | 37800001 | 37900000 | 0.0039 | 0.0034 | 0.0033 |
| F02 | 37820001 | 37920000 | 0.0046 | 0.0039 | 0.0040 |
| F02 | 37840001 | 37940000 | 0.0046 | 0.0039 | 0.0040 |
| F02 | 37860001 | 37960000 | 0.0041 | 0.0035 | 0.0036 |
| F02 | 37880001 | 37980000 | 0.0041 | 0.0037 | 0.0038 |
| F02 | 37900001 | 38000000 | 0.0034 | 0.0030 | 0.0033 |
| F02 | 37920001 | 38020000 | 0.0024 | 0.0021 | 0.0024 |
| F02 | 37940001 | 38040000 | 0.0034 | 0.0029 | 0.0034 |
| F02 | 37960001 | 38060000 | 0.0033 | 0.0027 | 0.0033 |
| F02 | 37980001 | 38080000 | 0.0026 | 0.0021 | 0.0026 |
| F02 | 38000001 | 38100000 | 0.0032 | 0.0027 | 0.0031 |
| F02 | 38020001 | 38120000 | 0.0035 | 0.0031 | 0.0035 |
| F02 | 38040001 | 38140000 | 0.0025 | 0.0023 | 0.0025 |
| F02 | 38060001 | 38160000 | 0.0022 | 0.0021 | 0.0021 |
| F02 | 38080001 | 38180000 | 0.0028 | 0.0026 | 0.0027 |
| F02 | 38100001 | 38200000 | 0.0025 | 0.0022 | 0.0024 |
| F02 | 38120001 | 38220000 | 0.0018 | 0.0015 | 0.0017 |
| F02 | 38140001 | 38240000 | 0.0017 | 0.0015 | 0.0017 |
| F02 | 38160001 | 38260000 | 0.0016 | 0.0013 | 0.0017 |
| F02 | 38180001 | 38280000 | 0.0014 | 0.0011 | 0.0014 |
| F02 | 38200001 | 38300000 | 0.0010 | 0.0008 | 0.0010 |
| F02 | 38220001 | 38320000 | 0.0016 | 0.0014 | 0.0015 |
| F02 | 38240001 | 38340000 | 0.0016 | 0.0013 | 0.0016 |
| F02 | 38260001 | 38360000 | 0.0027 | 0.0023 | 0.0027 |
| F02 | 38280001 | 38380000 | 0.0030 | 0.0025 | 0.0030 |
| F02 | 38300001 | 38400000 | 0.0031 | 0.0026 | 0.0030 |

| Chr | Start | End | both | A1 | A2 |
|---|---|---|---|---|---|
| F02 | 38320001 | 38420000 | 0.0031 | 0.0023 | 0.0029 |
| F02 | 38340001 | 38440000 | 0.0038 | 0.0029 | 0.0034 |
| F02 | 38360001 | 38460000 | 0.0032 | 0.0022 | 0.0026 |
| F02 | 38380001 | 38480000 | 0.0027 | 0.0018 | 0.0021 |
| F02 | 38400001 | 38500000 | 0.0027 | 0.0019 | 0.0021 |
| F02 | 38420001 | 38520000 | 0.0024 | 0.0017 | 0.0019 |
| F02 | 38440001 | 38540000 | 0.0017 | 0.0011 | 0.0013 |
| F02 | 38460001 | 38560000 | 0.0010 | 0.0008 | 0.0009 |
| F02 | 38480001 | 38580000 | 0.0011 | 0.0009 | 0.0009 |
| F02 | 38500001 | 38600000 | 0.0011 | 0.0009 | 0.0010 |
| F02 | 38520001 | 38620000 | 0.0015 | 0.0013 | 0.0013 |
| F02 | 38540001 | 38640000 | 0.0016 | 0.0013 | 0.0013 |
| F02 | 38560001 | 38660000 | 0.0017 | 0.0014 | 0.0014 |
| F02 | 38580001 | 38680000 | 0.0015 | 0.0013 | 0.0013 |
| F02 | 38600001 | 38700000 | 0.0013 | 0.0011 | 0.0011 |
| F02 | 38620001 | 38720000 | 0.0008 | 0.0007 | 0.0008 |
| F02 | 38640001 | 38740000 | 0.0013 | 0.0011 | 0.0013 |
| F02 | 38660001 | 38760000 | 0.0011 | 0.0009 | 0.0011 |
| F02 | 38680001 | 38780000 | 0.0015 | 0.0012 | 0.0015 |
| F02 | 38700001 | 38800000 | 0.0016 | 0.0013 | 0.0016 |
| F02 | 38720001 | 38820000 | 0.0017 | 0.0014 | 0.0017 |
| F02 | 38740001 | 38840000 | 0.0017 | 0.0015 | 0.0017 |
| F02 | 38760001 | 38860000 | 0.0022 | 0.0018 | 0.0021 |
| F02 | 38780001 | 38880000 | 0.0028 | 0.0024 | 0.0027 |
| F02 | 38800001 | 38900000 | 0.0029 | 0.0025 | 0.0027 |
| F02 | 38820001 | 38920000 | 0.0028 | 0.0023 | 0.0025 |
| F02 | 38840001 | 38940000 | 0.0026 | 0.0019 | 0.0022 |
| F02 | 38860001 | 38960000 | 0.0029 | 0.0021 | 0.0022 |
| F02 | 38880001 | 38980000 | 0.0027 | 0.0018 | 0.0020 |
| F02 | 38900001 | 39000000 | 0.0028 | 0.0020 | 0.0022 |
| F02 | 38920001 | 39020000 | 0.0028 | 0.0021 | 0.0022 |
| F02 | 38940001 | 39040000 | 0.0024 | 0.0019 | 0.0019 |
| F02 | 38960001 | 39060000 | 0.0017 | 0.0014 | 0.0016 |
| F02 | 38980001 | 39080000 | 0.0010 | 0.0009 | 0.0010 |
| F02 | 39000001 | 39100000 | 0.0007 | 0.0006 | 0.0007 |
| F02 | 39020001 | 39120000 | 0.0005 | 0.0005 | 0.0005 |
| F02 | 39040001 | 39140000 | 0.0007 | 0.0006 | 0.0007 |
| F02 | 39060001 | 39160000 | 0.0011 | 0.0010 | 0.0011 |
| F02 | 39080001 | 39180000 | 0.0016 | 0.0014 | 0.0016 |
| F02 | 39100001 | 39200000 | 0.0017 | 0.0015 | 0.0016 |
| F02 | 39120001 | 39220000 | 0.0017 | 0.0015 | 0.0016 |
| F02 | 39140001 | 39240000 | 0.0018 | 0.0015 | 0.0017 |
| F02 | 39160001 | 39260000 | 0.0017 | 0.0014 | 0.0015 |
| F02 | 39180001 | 39280000 | 0.0017 | 0.0012 | 0.0013 |
| F02 | 39200001 | 39300000 | 0.0016 | 0.0011 | 0.0013 |
| F02 | 39220001 | 39320000 | 0.0020 | 0.0014 | 0.0016 |
| F02 | 39240001 | 39340000 | 0.0020 | 0.0014 | 0.0015 |
| F02 | 39260001 | 39360000 | 0.0018 | 0.0011 | 0.0013 |
| F02 | 39280001 | 39380000 | 0.0024 | 0.0018 | 0.0019 |
| F02 | 39300001 | 39400000 | 0.0027 | 0.0020 | 0.0022 |
| F02 | 39320001 | 39420000 | 0.0031 | 0.0023 | 0.0025 |
| F02 | 39340001 | 39440000 | 0.0043 | 0.0034 | 0.0035 |
| F02 | 39360001 | 39460000 | 0.0048 | 0.0039 | 0.0041 |
| F02 | 39380001 | 39480000 | 0.0044 | 0.0036 | 0.0038 |
| F02 | 39400001 | 39500000 | 0.0047 | 0.0039 | 0.0041 |
| F02 | 39420001 | 39520000 | 0.0045 | 0.0037 | 0.0038 |
| F02 | 39440001 | 39540000 | 0.0032 | 0.0026 | 0.0027 |
| F02 | 39460001 | 39560000 | 0.0025 | 0.0020 | 0.0021 |

| Chr | Start | End | both | A1 | A2 |
|---|---|---|---|---|---|
| F02 | 39480001 | 39580000 | 0.0018 | 0.0014 | 0.0015 |
| F02 | 39500001 | 39600000 | 0.0013 | 0.0009 | 0.0010 |
| F02 | 39520001 | 39620000 | 0.0008 | 0.0006 | 0.0007 |
| F02 | 39540001 | 39640000 | 0.0007 | 0.0005 | 0.0007 |
| F02 | 39560001 | 39660000 | 0.0009 | 0.0008 | 0.0009 |
| F02 | 39580001 | 39680000 | 0.0012 | 0.0010 | 0.0011 |
| F02 | 39600001 | 39700000 | 0.0011 | 0.0009 | 0.0009 |
| F02 | 39620001 | 39720000 | 0.0010 | 0.0008 | 0.0009 |
| F02 | 39640001 | 39740000 | 0.0020 | 0.0017 | 0.0018 |
| F02 | 39660001 | 39760000 | 0.0018 | 0.0014 | 0.0017 |
| F02 | 39680001 | 39780000 | 0.0016 | 0.0013 | 0.0015 |
| F02 | 39700001 | 39800000 | 0.0015 | 0.0012 | 0.0014 |
| F02 | 39720001 | 39820000 | 0.0023 | 0.0021 | 0.0022 |
| F02 | 39740001 | 39840000 | 0.0019 | 0.0016 | 0.0017 |
| F02 | 39760001 | 39860000 | 0.0018 | 0.0016 | 0.0016 |
| F02 | 39780001 | 39880000 | 0.0021 | 0.0018 | 0.0020 |
| F02 | 39800001 | 39900000 | 0.0023 | 0.0018 | 0.0021 |
| F02 | 39820001 | 39920000 | 0.0013 | 0.0009 | 0.0013 |
| F02 | 39840001 | 39940000 | 0.0011 | 0.0008 | 0.0011 |
| F02 | 39860001 | 39960000 | 0.0010 | 0.0008 | 0.0010 |
| F02 | 39880001 | 39980000 | 0.0012 | 0.0009 | 0.0011 |
| F02 | 39900001 | 40000000 | 0.0012 | 0.0010 | 0.0011 |
| F02 | 39920001 | 40020000 | 0.0012 | 0.0011 | 0.0012 |
| F02 | 39940001 | 40040000 | 0.0012 | 0.0010 | 0.0012 |
| F02 | 39960001 | 40060000 | 0.0012 | 0.0010 | 0.0012 |
| F02 | 39980001 | 40080000 | 0.0009 | 0.0007 | 0.0008 |
| F02 | 40000001 | 40100000 | 0.0007 | 0.0005 | 0.0007 |
| F02 | 40020001 | 40120000 | 0.0007 | 0.0005 | 0.0007 |
| F02 | 40040001 | 40140000 | 0.0006 | 0.0005 | 0.0006 |
| F02 | 40060001 | 40160000 | 0.0007 | 0.0005 | 0.0007 |
| F02 | 40080001 | 40180000 | 0.0007 | 0.0006 | 0.0007 |
| F02 | 40100001 | 40200000 | 0.0013 | 0.0010 | 0.0013 |
| F02 | 40120001 | 40220000 | 0.0012 | 0.0009 | 0.0012 |
| F02 | 40140001 | 40240000 | 0.0009 | 0.0006 | 0.0009 |
| F02 | 40160001 | 40260000 | 0.0012 | 0.0010 | 0.0012 |
| F02 | 40180001 | 40280000 | 0.0012 | 0.0010 | 0.0012 |
| F02 | 40200001 | 40300000 | 0.0007 | 0.0007 | 0.0007 |
| F02 | 40220001 | 40320000 | 0.0008 | 0.0007 | 0.0008 |
| F02 | 40240001 | 40340000 | 0.0009 | 0.0009 | 0.0009 |
| F02 | 40260001 | 40360000 | 0.0007 | 0.0006 | 0.0006 |
| F02 | 40280001 | 40380000 | 0.0008 | 0.0006 | 0.0007 |
| F02 | 40300001 | 40400000 | 0.0011 | 0.0009 | 0.0011 |
| F02 | 40320001 | 40420000 | 0.0012 | 0.0010 | 0.0011 |
| F02 | 40340001 | 40440000 | 0.0012 | 0.0010 | 0.0012 |
| F02 | 40360001 | 40460000 | 0.0017 | 0.0015 | 0.0017 |
| F02 | 40380001 | 40480000 | 0.0018 | 0.0016 | 0.0018 |
| F02 | 40400001 | 40500000 | 0.0016 | 0.0014 | 0.0015 |
| F02 | 40420001 | 40520000 | 0.0019 | 0.0017 | 0.0018 |
| F02 | 40440001 | 40540000 | 0.0019 | 0.0017 | 0.0018 |
| F02 | 40460001 | 40560000 | 0.0015 | 0.0013 | 0.0013 |
| F02 | 40480001 | 40580000 | 0.0015 | 0.0011 | 0.0010 |
| F02 | 40500001 | 40600000 | 0.0021 | 0.0016 | 0.0017 |
| F02 | 40520001 | 40620000 | 0.0023 | 0.0017 | 0.0018 |
| F02 | 40540001 | 40640000 | 0.0035 | 0.0025 | 0.0031 |
| F02 | 40560001 | 40660000 | 0.0051 | 0.0038 | 0.0046 |
| F02 | 40580001 | 40680000 | 0.0059 | 0.0046 | 0.0055 |
| F02 | 40600001 | 40700000 | 0.0052 | 0.0040 | 0.0049 |
| F02 | 40620001 | 40720000 | 0.0059 | 0.0044 | 0.0055 |

| Chr | Start | End | both | A1 | A2 |
|---|---|---|---|---|---|
| F02 | 40640001 | 40740000 | 0.0045 | 0.0036 | 0.0042 |
| F02 | 40660001 | 40760000 | 0.0029 | 0.0022 | 0.0026 |
| F02 | 40680001 | 40780000 | 0.0022 | 0.0017 | 0.0021 |
| F02 | 40700001 | 40800000 | 0.0030 | 0.0024 | 0.0029 |
| F02 | 40720001 | 40820000 | 0.0020 | 0.0016 | 0.0019 |
| F02 | 40740001 | 40840000 | 0.0022 | 0.0019 | 0.0022 |
| F02 | 40760001 | 40860000 | 0.0020 | 0.0017 | 0.0020 |
| F02 | 40780001 | 40880000 | 0.0016 | 0.0014 | 0.0016 |
| F02 | 40800001 | 40900000 | 0.0015 | 0.0014 | 0.0015 |
| F02 | 40820001 | 40920000 | 0.0015 | 0.0014 | 0.0015 |
| F02 | 40840001 | 40940000 | 0.0022 | 0.0019 | 0.0020 |
| F02 | 40860001 | 40960000 | 0.0027 | 0.0023 | 0.0023 |
| F02 | 40880001 | 40980000 | 0.0032 | 0.0028 | 0.0028 |
| F02 | 40900001 | 41000000 | 0.0029 | 0.0024 | 0.0024 |
| F02 | 40920001 | 41020000 | 0.0033 | 0.0027 | 0.0028 |
| F02 | 40940001 | 41040000 | 0.0026 | 0.0021 | 0.0023 |
| F02 | 40960001 | 41060000 | 0.0032 | 0.0027 | 0.0029 |
| F02 | 40980001 | 41080000 | 0.0029 | 0.0025 | 0.0027 |
| F02 | 41000001 | 41100000 | 0.0024 | 0.0021 | 0.0023 |
| F02 | 41020001 | 41120000 | 0.0025 | 0.0020 | 0.0022 |
| F02 | 41040001 | 41140000 | 0.0031 | 0.0025 | 0.0028 |
| F02 | 41060001 | 41160000 | 0.0027 | 0.0020 | 0.0024 |
| F02 | 41080001 | 41180000 | 0.0027 | 0.0019 | 0.0024 |
| F02 | 41100001 | 41200000 | 0.0032 | 0.0024 | 0.0029 |
| F02 | 41120001 | 41220000 | 0.0031 | 0.0024 | 0.0030 |
| F02 | 41140001 | 41240000 | 0.0025 | 0.0019 | 0.0024 |
| F02 | 41160001 | 41260000 | 0.0019 | 0.0015 | 0.0019 |
| F02 | 41180001 | 41280000 | 0.0018 | 0.0015 | 0.0018 |
| F02 | 41200001 | 41300000 | 0.0014 | 0.0011 | 0.0014 |
| F02 | 41220001 | 41320000 | 0.0010 | 0.0007 | 0.0010 |
| F02 | 41240001 | 41340000 | 0.0010 | 0.0007 | 0.0009 |
| F02 | 41260001 | 41360000 | 0.0013 | 0.0010 | 0.0011 |
| F02 | 41280001 | 41380000 | 0.0010 | 0.0008 | 0.0009 |
| F02 | 41300001 | 41400000 | 0.0010 | 0.0008 | 0.0009 |
| F02 | 41320001 | 41420000 | 0.0011 | 0.0009 | 0.0010 |
| F02 | 41340001 | 41440000 | 0.0016 | 0.0013 | 0.0016 |
| F02 | 41360001 | 41460000 | 0.0020 | 0.0015 | 0.0020 |
| F02 | 41380001 | 41480000 | 0.0024 | 0.0018 | 0.0024 |
| F02 | 41400001 | 41500000 | 0.0032 | 0.0024 | 0.0032 |
| F02 | 41420001 | 41520000 | 0.0033 | 0.0025 | 0.0033 |
| F02 | 41440001 | 41540000 | 0.0030 | 0.0023 | 0.0030 |
| F02 | 41460001 | 41560000 | 0.0031 | 0.0025 | 0.0030 |
| F02 | 41480001 | 41580000 | 0.0037 | 0.0029 | 0.0036 |
| F02 | 41500001 | 41600000 | 0.0035 | 0.0028 | 0.0032 |
| F02 | 41520001 | 41620000 | 0.0039 | 0.0029 | 0.0036 |
| F02 | 41540001 | 41640000 | 0.0051 | 0.0036 | 0.0047 |
| F02 | 41560001 | 41660000 | 0.0058 | 0.0044 | 0.0053 |
| F02 | 41580001 | 41680000 | 0.0052 | 0.0037 | 0.0045 |
| F02 | 41600001 | 41700000 | 0.0055 | 0.0042 | 0.0049 |
| F02 | 41620001 | 41720000 | 0.0052 | 0.0039 | 0.0044 |
| F02 | 41640001 | 41740000 | 0.0042 | 0.0033 | 0.0034 |
| F02 | 41660001 | 41760000 | 0.0034 | 0.0024 | 0.0025 |
| F02 | 41680001 | 41780000 | 0.0033 | 0.0025 | 0.0027 |
| F02 | 41700001 | 41800000 | 0.0030 | 0.0021 | 0.0024 |
| F02 | 41720001 | 41820000 | 0.0025 | 0.0019 | 0.0022 |
| F02 | 41740001 | 41840000 | 0.0022 | 0.0017 | 0.0020 |
| F02 | 41760001 | 41860000 | 0.0018 | 0.0015 | 0.0017 |
| F02 | 41780001 | 41880000 | 0.0018 | 0.0015 | 0.0018 |

| Chr | Start | End | both | A1 | A2 |
|---|---|---|---|---|---|
| F02 | 41800001 | 41900000 | 0.0014 | 0.0012 | 0.0014 |
| F02 | 41820001 | 41920000 | 0.0014 | 0.0011 | 0.0014 |
| F02 | 41840001 | 41940000 | 0.0015 | 0.0013 | 0.0015 |
| F02 | 41860001 | 41960000 | 0.0026 | 0.0022 | 0.0025 |
| F02 | 41880001 | 41980000 | 0.0027 | 0.0022 | 0.0026 |
| F02 | 41900001 | 42000000 | 0.0028 | 0.0023 | 0.0026 |
| F02 | 41920001 | 42020000 | 0.0030 | 0.0025 | 0.0029 |
| F02 | 41940001 | 42040000 | 0.0027 | 0.0022 | 0.0026 |
| F02 | 41960001 | 42060000 | 0.0019 | 0.0015 | 0.0019 |
| F02 | 41980001 | 42080000 | 0.0014 | 0.0012 | 0.0014 |
| F02 | 42000001 | 42100000 | 0.0011 | 0.0009 | 0.0010 |
| F02 | 42020001 | 42120000 | 0.0007 | 0.0006 | 0.0007 |
| F02 | 42040001 | 42140000 | 0.0009 | 0.0007 | 0.0009 |
| F02 | 42060001 | 42160000 | 0.0004 | 0.0003 | 0.0004 |
| F02 | 42080001 | 42180000 | 0.0005 | 0.0003 | 0.0005 |
| F02 | 42100001 | 42200000 | 0.0007 | 0.0004 | 0.0008 |
| F02 | 42120001 | 42220000 | 0.0008 | 0.0005 | 0.0009 |
| F02 | 42140001 | 42240000 | 0.0011 | 0.0006 | 0.0011 |
| F02 | 42160001 | 42260000 | 0.0011 | 0.0007 | 0.0011 |
| F02 | 42180001 | 42280000 | 0.0011 | 0.0007 | 0.0011 |
| F02 | 42200001 | 42300000 | 0.0009 | 0.0006 | 0.0009 |
| F02 | 42220001 | 42320000 | 0.0009 | 0.0007 | 0.0009 |
| F02 | 42240001 | 42340000 | 0.0012 | 0.0009 | 0.0012 |
| F02 | 42260001 | 42360000 | 0.0014 | 0.0011 | 0.0014 |
| F02 | 42280001 | 42380000 | 0.0018 | 0.0013 | 0.0018 |
| F02 | 42300001 | 42400000 | 0.0020 | 0.0014 | 0.0020 |
| F02 | 42320001 | 42420000 | 0.0020 | 0.0014 | 0.0020 |
| F02 | 42340001 | 42440000 | 0.0014 | 0.0010 | 0.0014 |
| F02 | 42360001 | 42460000 | 0.0015 | 0.0010 | 0.0015 |
| F02 | 42380001 | 42480000 | 0.0010 | 0.0007 | 0.0010 |
| F02 | 42400001 | 42500000 | 0.0014 | 0.0012 | 0.0014 |
| F02 | 42420001 | 42520000 | 0.0014 | 0.0011 | 0.0013 |
| F02 | 42440001 | 42540000 | 0.0015 | 0.0013 | 0.0015 |
| F02 | 42460001 | 42560000 | 0.0017 | 0.0015 | 0.0016 |
| F02 | 42480001 | 42580000 | 0.0018 | 0.0016 | 0.0017 |
| F02 | 42500001 | 42600000 | 0.0013 | 0.0011 | 0.0013 |
| F02 | 42520001 | 42620000 | 0.0016 | 0.0014 | 0.0016 |
| F02 | 42540001 | 42640000 | 0.0023 | 0.0019 | 0.0023 |
| F02 | 42560001 | 42660000 | 0.0023 | 0.0019 | 0.0023 |
| F02 | 42580001 | 42680000 | 0.0023 | 0.0019 | 0.0023 |
| F02 | 42600001 | 42700000 | 0.0022 | 0.0018 | 0.0022 |
| F02 | 42620001 | 42720000 | 0.0017 | 0.0014 | 0.0017 |
| F02 | 42640001 | 42740000 | 0.0009 | 0.0007 | 0.0009 |
| F02 | 42660001 | 42760000 | 0.0004 | 0.0003 | 0.0004 |
| F02 | 42680001 | 42780000 | 0.0003 | 0.0002 | 0.0003 |
| F02 | 42700001 | 42800000 | 0.0010 | 0.0007 | 0.0010 |
| F02 | 42720001 | 42820000 | 0.0011 | 0.0009 | 0.0011 |
| F02 | 42740001 | 42840000 | 0.0010 | 0.0008 | 0.0010 |
| F02 | 42760001 | 42860000 | 0.0009 | 0.0007 | 0.0009 |
| F02 | 42780001 | 42880000 | 0.0010 | 0.0008 | 0.0009 |
| F02 | 42800001 | 42900000 | 0.0009 | 0.0007 | 0.0009 |
| F02 | 42820001 | 42920000 | 0.0011 | 0.0009 | 0.0010 |
| F02 | 42840001 | 42940000 | 0.0024 | 0.0020 | 0.0022 |
| F02 | 42860001 | 42960000 | 0.0037 | 0.0031 | 0.0035 |
| F02 | 42880001 | 42980000 | 0.0043 | 0.0035 | 0.0039 |
| F02 | 42900001 | 43000000 | 0.0046 | 0.0037 | 0.0041 |
| F02 | 42920001 | 43020000 | 0.0048 | 0.0038 | 0.0043 |
| F02 | 42940001 | 43040000 | 0.0042 | 0.0033 | 0.0037 |

| Chr | Start | End | both | A1 | A2 |
|---|---|---|---|---|---|
| F02 | 42960001 | 43060000 | 0.0035 | 0.0026 | 0.0029 |
| F02 | 42980001 | 43080000 | 0.0036 | 0.0028 | 0.0029 |
| F02 | 43000001 | 43100000 | 0.0032 | 0.0025 | 0.0024 |
| F02 | 43020001 | 43120000 | 0.0027 | 0.0021 | 0.0020 |
| F02 | 43040001 | 43140000 | 0.0023 | 0.0018 | 0.0016 |
| F02 | 43060001 | 43160000 | 0.0019 | 0.0014 | 0.0013 |
| F02 | 43080001 | 43180000 | 0.0013 | 0.0011 | 0.0011 |
| F02 | 43100001 | 43200000 | 0.0011 | 0.0009 | 0.0009 |
| F02 | 43120001 | 43220000 | 0.0014 | 0.0012 | 0.0013 |
| F02 | 43140001 | 43240000 | 0.0013 | 0.0012 | 0.0013 |
| F02 | 43160001 | 43260000 | 0.0012 | 0.0011 | 0.0011 |
| F02 | 43180001 | 43280000 | 0.0010 | 0.0009 | 0.0010 |
| F02 | 43200001 | 43300000 | 0.0008 | 0.0006 | 0.0008 |
| F02 | 43220001 | 43320000 | 0.0005 | 0.0004 | 0.0005 |
| F02 | 43240001 | 43340000 | 0.0004 | 0.0003 | 0.0004 |
| F02 | 43260001 | 43360000 | 0.0004 | 0.0003 | 0.0004 |
| F02 | 43280001 | 43380000 | 0.0007 | 0.0005 | 0.0007 |
| F02 | 43300001 | 43400000 | 0.0007 | 0.0005 | 0.0007 |
| F02 | 43320001 | 43420000 | 0.0005 | 0.0004 | 0.0006 |
| F02 | 43340001 | 43440000 | 0.0006 | 0.0003 | 0.0006 |
| F02 | 43360001 | 43460000 | 0.0011 | 0.0007 | 0.0011 |
| F02 | 43380001 | 43480000 | 0.0011 | 0.0007 | 0.0011 |
| F02 | 43400001 | 43500000 | 0.0013 | 0.0009 | 0.0013 |
| F02 | 43420001 | 43520000 | 0.0013 | 0.0009 | 0.0013 |
| F02 | 43440001 | 43540000 | 0.0015 | 0.0010 | 0.0015 |
| F02 | 43460001 | 43560000 | 0.0011 | 0.0008 | 0.0011 |
| F02 | 43480001 | 43580000 | 0.0007 | 0.0005 | 0.0007 |
| F02 | 43500001 | 43600000 | 0.0005 | 0.0004 | 0.0005 |
| F02 | 43520001 | 43620000 | 0.0005 | 0.0003 | 0.0005 |
| F02 | 43540001 | 43640000 | 0.0003 | 0.0002 | 0.0003 |
| F02 | 43560001 | 43660000 | 0.0004 | 0.0002 | 0.0004 |
| F02 | 43580001 | 43680000 | 0.0005 | 0.0003 | 0.0005 |
| F02 | 43600001 | 43700000 | 0.0006 | 0.0004 | 0.0006 |
| F02 | 43620001 | 43720000 | 0.0008 | 0.0006 | 0.0008 |
| F02 | 43640001 | 43740000 | 0.0007 | 0.0006 | 0.0007 |
| F02 | 43660001 | 43760000 | 0.0006 | 0.0005 | 0.0006 |
| F02 | 43680001 | 43780000 | 0.0008 | 0.0006 | 0.0008 |
| F02 | 43700001 | 43800000 | 0.0007 | 0.0006 | 0.0007 |
| F02 | 43720001 | 43820000 | 0.0006 | 0.0005 | 0.0006 |
| F02 | 43740001 | 43840000 | 0.0010 | 0.0008 | 0.0010 |
| F02 | 43760001 | 43860000 | 0.0011 | 0.0008 | 0.0011 |
| F02 | 43780001 | 43880000 | 0.0009 | 0.0007 | 0.0008 |
| F02 | 43800001 | 43900000 | 0.0007 | 0.0005 | 0.0007 |
| F02 | 43820001 | 43920000 | 0.0006 | 0.0004 | 0.0006 |
| F02 | 43840001 | 43940000 | 0.0003 | 0.0002 | 0.0003 |
| F02 | 43860001 | 43960000 | 0.0002 | 0.0002 | 0.0002 |
| F02 | 43880001 | 43980000 | 0.0006 | 0.0005 | 0.0006 |
| F02 | 43900001 | 44000000 | 0.0007 | 0.0005 | 0.0007 |
| F02 | 43920001 | 44020000 | 0.0007 | 0.0005 | 0.0007 |
| F02 | 43940001 | 44040000 | 0.0014 | 0.0011 | 0.0014 |
| F02 | 43960001 | 44060000 | 0.0024 | 0.0021 | 0.0023 |
| F02 | 43980001 | 44080000 | 0.0026 | 0.0022 | 0.0024 |
| F02 | 44000001 | 44100000 | 0.0027 | 0.0024 | 0.0025 |
| F02 | 44020001 | 44120000 | 0.0033 | 0.0028 | 0.0030 |
| F02 | 44040001 | 44140000 | 0.0033 | 0.0028 | 0.0030 |
| F02 | 44060001 | 44160000 | 0.0022 | 0.0018 | 0.0021 |
| F02 | 44080001 | 44180000 | 0.0021 | 0.0018 | 0.0021 |
| F02 | 44100001 | 44200000 | 0.0026 | 0.0021 | 0.0026 |

| Chr | Start | End | both | A1 | A2 |
|---|---|---|---|---|---|
| F02 | 44120001 | 44220000 | 0.0031 | 0.0024 | 0.0027 |
| F02 | 44140001 | 44240000 | 0.0029 | 0.0021 | 0.0024 |
| F02 | 44160001 | 44260000 | 0.0034 | 0.0026 | 0.0029 |
| F02 | 44180001 | 44280000 | 0.0042 | 0.0032 | 0.0036 |
| F02 | 44200001 | 44300000 | 0.0043 | 0.0031 | 0.0034 |
| F02 | 44220001 | 44320000 | 0.0042 | 0.0032 | 0.0036 |
| F02 | 44240001 | 44340000 | 0.0041 | 0.0030 | 0.0033 |
| F02 | 44260001 | 44360000 | 0.0036 | 0.0026 | 0.0029 |
| F02 | 44280001 | 44380000 | 0.0024 | 0.0016 | 0.0017 |
| F02 | 44300001 | 44400000 | 0.0023 | 0.0016 | 0.0017 |
| F02 | 44320001 | 44420000 | 0.0019 | 0.0014 | 0.0014 |
| F02 | 44340001 | 44440000 | 0.0021 | 0.0018 | 0.0018 |
| F02 | 44360001 | 44460000 | 0.0023 | 0.0018 | 0.0019 |
| F02 | 44380001 | 44480000 | 0.0031 | 0.0021 | 0.0023 |
| F02 | 44400001 | 44500000 | 0.0027 | 0.0018 | 0.0021 |
| F02 | 44420001 | 44520000 | 0.0025 | 0.0016 | 0.0019 |
| F02 | 44440001 | 44540000 | 0.0021 | 0.0012 | 0.0015 |
| F02 | 44460001 | 44560000 | 0.0026 | 0.0017 | 0.0022 |
| F02 | 44480001 | 44580000 | 0.0024 | 0.0016 | 0.0021 |
| F02 | 44500001 | 44600000 | 0.0027 | 0.0018 | 0.0024 |
| F02 | 44520001 | 44620000 | 0.0029 | 0.0020 | 0.0026 |
| F02 | 44540001 | 44640000 | 0.0035 | 0.0025 | 0.0032 |
| F02 | 44560001 | 44660000 | 0.0028 | 0.0021 | 0.0025 |
| F02 | 44580001 | 44680000 | 0.0026 | 0.0021 | 0.0024 |
| F02 | 44600001 | 44700000 | 0.0025 | 0.0020 | 0.0023 |
| F02 | 44620001 | 44720000 | 0.0023 | 0.0018 | 0.0021 |
| F02 | 44640001 | 44740000 | 0.0030 | 0.0025 | 0.0028 |
| F02 | 44660001 | 44760000 | 0.0037 | 0.0030 | 0.0034 |
| F02 | 44680001 | 44780000 | 0.0037 | 0.0030 | 0.0035 |
| F02 | 44700001 | 44800000 | 0.0033 | 0.0027 | 0.0031 |
| F02 | 44720001 | 44820000 | 0.0032 | 0.0025 | 0.0028 |
| F02 | 44740001 | 44840000 | 0.0019 | 0.0013 | 0.0016 |
| F02 | 44760001 | 44860000 | 0.0014 | 0.0009 | 0.0011 |
| F02 | 44780001 | 44880000 | 0.0015 | 0.0010 | 0.0013 |
| F02 | 44800001 | 44900000 | 0.0018 | 0.0012 | 0.0016 |
| F02 | 44820001 | 44920000 | 0.0018 | 0.0013 | 0.0017 |
| F02 | 44840001 | 44940000 | 0.0015 | 0.0011 | 0.0015 |
| F02 | 44860001 | 44960000 | 0.0022 | 0.0019 | 0.0022 |
| F02 | 44880001 | 44980000 | 0.0024 | 0.0021 | 0.0023 |
| F02 | 44900001 | 45000000 | 0.0026 | 0.0023 | 0.0025 |
| F02 | 44920001 | 45020000 | 0.0024 | 0.0021 | 0.0024 |
| F02 | 44940001 | 45040000 | 0.0026 | 0.0023 | 0.0025 |
| F02 | 44960001 | 45060000 | 0.0021 | 0.0018 | 0.0021 |
| F02 | 44980001 | 45080000 | 0.0021 | 0.0017 | 0.0021 |
| F02 | 45000001 | 45100000 | 0.0015 | 0.0013 | 0.0015 |
| F02 | 45020001 | 45120000 | 0.0016 | 0.0012 | 0.0016 |
| F02 | 45040001 | 45140000 | 0.0029 | 0.0023 | 0.0028 |
| F02 | 45060001 | 45160000 | 0.0027 | 0.0021 | 0.0026 |
| F02 | 45080001 | 45180000 | 0.0022 | 0.0017 | 0.0022 |
| F02 | 45100001 | 45200000 | 0.0025 | 0.0019 | 0.0025 |
| F02 | 45120001 | 45220000 | 0.0025 | 0.0021 | 0.0025 |
| F02 | 45140001 | 45240000 | 0.0017 | 0.0014 | 0.0017 |
| F02 | 45160001 | 45260000 | 0.0015 | 0.0012 | 0.0015 |
| F02 | 45180001 | 45280000 | 0.0020 | 0.0017 | 0.0020 |
| F02 | 45200001 | 45300000 | 0.0017 | 0.0014 | 0.0017 |
| F02 | 45220001 | 45320000 | 0.0015 | 0.0013 | 0.0015 |
| F02 | 45240001 | 45340000 | 0.0013 | 0.0011 | 0.0013 |
| F02 | 45260001 | 45360000 | 0.0013 | 0.0011 | 0.0013 |

| Chr | Start | End | both | A1 | A2 |
|---|---|---|---|---|---|
| F02 | 45280001 | 45380000 | 0.0009 | 0.0007 | 0.0009 |
| F02 | 45300001 | 45400000 | 0.0015 | 0.0013 | 0.0015 |
| F02 | 45320001 | 45420000 | 0.0021 | 0.0019 | 0.0020 |
| F02 | 45340001 | 45440000 | 0.0033 | 0.0029 | 0.0030 |
| F02 | 45360001 | 45460000 | 0.0041 | 0.0034 | 0.0036 |
| F02 | 45380001 | 45480000 | 0.0054 | 0.0046 | 0.0047 |
| F02 | 45400001 | 45500000 | 0.0054 | 0.0043 | 0.0045 |
| F02 | 45420001 | 45520000 | 0.0056 | 0.0043 | 0.0045 |
| F02 | 45440001 | 45540000 | 0.0050 | 0.0036 | 0.0038 |
| F02 | 45460001 | 45560000 | 0.0047 | 0.0032 | 0.0034 |
| F02 | 45480001 | 45580000 | 0.0038 | 0.0021 | 0.0024 |
| F02 | 45500001 | 45600000 | 0.0040 | 0.0022 | 0.0023 |
| F02 | 45520001 | 45620000 | 0.0042 | 0.0022 | 0.0025 |
| F02 | 45540001 | 45640000 | 0.0041 | 0.0022 | 0.0025 |
| F02 | 45560001 | 45660000 | 0.0039 | 0.0024 | 0.0024 |
| F02 | 45580001 | 45680000 | 0.0036 | 0.0022 | 0.0022 |
| F02 | 45600001 | 45700000 | 0.0041 | 0.0026 | 0.0028 |
| F02 | 45620001 | 45720000 | 0.0033 | 0.0022 | 0.0021 |
| F02 | 45640001 | 45740000 | 0.0030 | 0.0018 | 0.0018 |
| F02 | 45660001 | 45760000 | 0.0029 | 0.0018 | 0.0020 |
| F02 | 45680001 | 45780000 | 0.0029 | 0.0019 | 0.0023 |
| F02 | 45700001 | 45800000 | 0.0025 | 0.0017 | 0.0020 |
| F02 | 45720001 | 45820000 | 0.0030 | 0.0022 | 0.0025 |
| F02 | 45740001 | 45840000 | 0.0033 | 0.0028 | 0.0031 |
| F02 | 45760001 | 45860000 | 0.0033 | 0.0026 | 0.0031 |
| F02 | 45780001 | 45880000 | 0.0038 | 0.0031 | 0.0035 |
| F02 | 45800001 | 45900000 | 0.0038 | 0.0030 | 0.0035 |
| F02 | 45820001 | 45920000 | 0.0033 | 0.0026 | 0.0031 |
| F02 | 45840001 | 45940000 | 0.0025 | 0.0019 | 0.0023 |
| F02 | 45860001 | 45960000 | 0.0021 | 0.0017 | 0.0019 |
| F02 | 45880001 | 45980000 | 0.0017 | 0.0014 | 0.0016 |
| F02 | 45900001 | 46000000 | 0.0020 | 0.0016 | 0.0019 |
| F02 | 45920001 | 46020000 | 0.0018 | 0.0014 | 0.0017 |
| F02 | 45940001 | 46040000 | 0.0019 | 0.0015 | 0.0019 |
| F02 | 45960001 | 46060000 | 0.0022 | 0.0018 | 0.0022 |
| F02 | 45980001 | 46080000 | 0.0021 | 0.0017 | 0.0021 |
| F02 | 46000001 | 46100000 | 0.0012 | 0.0010 | 0.0012 |
| F02 | 46020001 | 46120000 | 0.0016 | 0.0014 | 0.0016 |
| F02 | 46040001 | 46140000 | 0.0016 | 0.0013 | 0.0015 |
| F02 | 46060001 | 46160000 | 0.0019 | 0.0017 | 0.0018 |
| F02 | 46080001 | 46180000 | 0.0019 | 0.0017 | 0.0018 |
| F02 | 46100001 | 46200000 | 0.0017 | 0.0015 | 0.0017 |
| F02 | 46120001 | 46220000 | 0.0014 | 0.0012 | 0.0014 |
| F02 | 46140001 | 46240000 | 0.0013 | 0.0012 | 0.0013 |
| F02 | 46160001 | 46260000 | 0.0008 | 0.0007 | 0.0007 |
| F02 | 46180001 | 46280000 | 0.0008 | 0.0007 | 0.0007 |
| F02 | 46200001 | 46300000 | 0.0010 | 0.0008 | 0.0008 |
| F02 | 46220001 | 46320000 | 0.0011 | 0.0008 | 0.0008 |
| F02 | 46240001 | 46340000 | 0.0010 | 0.0007 | 0.0007 |
| F02 | 46260001 | 46360000 | 0.0017 | 0.0010 | 0.0011 |
| F02 | 46280001 | 46380000 | 0.0018 | 0.0011 | 0.0011 |
| F02 | 46300001 | 46400000 | 0.0019 | 0.0013 | 0.0014 |
| F02 | 46320001 | 46420000 | 0.0026 | 0.0018 | 0.0021 |
| F02 | 46340001 | 46440000 | 0.0029 | 0.0020 | 0.0023 |
| F02 | 46360001 | 46460000 | 0.0029 | 0.0023 | 0.0025 |
| F02 | 46380001 | 46480000 | 0.0033 | 0.0027 | 0.0028 |
| F02 | 46400001 | 46500000 | 0.0036 | 0.0029 | 0.0031 |
| F02 | 46420001 | 46520000 | 0.0033 | 0.0027 | 0.0028 |

| Chr | Start | End | both | A1 | A2 |
|---|---|---|---|---|---|
| F02 | 46440001 | 46540000 | 0.0031 | 0.0025 | 0.0026 |
| F02 | 46460001 | 46560000 | 0.0025 | 0.0021 | 0.0024 |
| F02 | 46480001 | 46580000 | 0.0027 | 0.0024 | 0.0025 |
| F02 | 46500001 | 46600000 | 0.0022 | 0.0020 | 0.0021 |
| F02 | 46520001 | 46620000 | 0.0020 | 0.0018 | 0.0019 |
| F02 | 46540001 | 46640000 | 0.0020 | 0.0017 | 0.0019 |
| F02 | 46560001 | 46660000 | 0.0024 | 0.0019 | 0.0021 |
| F02 | 46580001 | 46680000 | 0.0025 | 0.0019 | 0.0023 |
| F02 | 46600001 | 46700000 | 0.0031 | 0.0023 | 0.0027 |
| F02 | 46620001 | 46720000 | 0.0029 | 0.0022 | 0.0025 |
| F02 | 46640001 | 46740000 | 0.0027 | 0.0021 | 0.0023 |
| F02 | 46660001 | 46760000 | 0.0020 | 0.0017 | 0.0019 |
| F02 | 46680001 | 46780000 | 0.0017 | 0.0014 | 0.0016 |
| F02 | 46700001 | 46800000 | 0.0019 | 0.0018 | 0.0018 |
| F02 | 46720001 | 46820000 | 0.0025 | 0.0024 | 0.0025 |
| F02 | 46740001 | 46840000 | 0.0032 | 0.0030 | 0.0032 |
| F02 | 46760001 | 46860000 | 0.0032 | 0.0029 | 0.0031 |
| F02 | 46780001 | 46880000 | 0.0030 | 0.0028 | 0.0029 |
| F02 | 46800001 | 46900000 | 0.0032 | 0.0028 | 0.0032 |
| F02 | 46820001 | 46920000 | 0.0032 | 0.0028 | 0.0032 |
| F02 | 46840001 | 46940000 | 0.0027 | 0.0024 | 0.0027 |
| F02 | 46860001 | 46960000 | 0.0029 | 0.0026 | 0.0029 |
| F02 | 46880001 | 46980000 | 0.0035 | 0.0031 | 0.0034 |
| F02 | 46900001 | 47000000 | 0.0028 | 0.0024 | 0.0028 |
| F02 | 46920001 | 47020000 | 0.0029 | 0.0026 | 0.0028 |
| F02 | 46940001 | 47040000 | 0.0031 | 0.0029 | 0.0030 |
| F02 | 46960001 | 47060000 | 0.0032 | 0.0030 | 0.0031 |
| F02 | 46980001 | 47080000 | 0.0025 | 0.0024 | 0.0025 |
| F02 | 47000001 | 47100000 | 0.0028 | 0.0026 | 0.0025 |
| F02 | 47020001 | 47120000 | 0.0029 | 0.0025 | 0.0026 |
| F02 | 47040001 | 47140000 | 0.0028 | 0.0026 | 0.0026 |
| F02 | 47060001 | 47160000 | 0.0030 | 0.0027 | 0.0028 |
| F02 | 47080001 | 47180000 | 0.0047 | 0.0044 | 0.0041 |
| F02 | 47100001 | 47200000 | 0.0043 | 0.0040 | 0.0038 |
| F02 | 47120001 | 47220000 | 0.0041 | 0.0039 | 0.0036 |
| F02 | 47140001 | 47240000 | 0.0043 | 0.0040 | 0.0038 |
| F02 | 47160001 | 47260000 | 0.0043 | 0.0039 | 0.0037 |
| F02 | 47180001 | 47280000 | 0.0033 | 0.0024 | 0.0027 |
| F02 | 47200001 | 47300000 | 0.0034 | 0.0026 | 0.0028 |
| F02 | 47220001 | 47320000 | 0.0031 | 0.0023 | 0.0025 |
| F02 | 47240001 | 47340000 | 0.0031 | 0.0023 | 0.0024 |
| F02 | 47260001 | 47360000 | 0.0025 | 0.0019 | 0.0019 |
| F02 | 47280001 | 47380000 | 0.0020 | 0.0016 | 0.0016 |
| F02 | 47300001 | 47400000 | 0.0026 | 0.0020 | 0.0022 |
| F02 | 47320001 | 47420000 | 0.0033 | 0.0025 | 0.0027 |
| F02 | 47340001 | 47440000 | 0.0034 | 0.0025 | 0.0029 |
| F02 | 47360001 | 47460000 | 0.0035 | 0.0025 | 0.0029 |
| F02 | 47380001 | 47480000 | 0.0040 | 0.0030 | 0.0033 |
| F02 | 47400001 | 47500000 | 0.0030 | 0.0022 | 0.0023 |
| F02 | 47420001 | 47520000 | 0.0029 | 0.0020 | 0.0024 |
| F02 | 47440001 | 47540000 | 0.0029 | 0.0020 | 0.0023 |
| F02 | 47460001 | 47560000 | 0.0036 | 0.0025 | 0.0028 |
| F02 | 47480001 | 47580000 | 0.0029 | 0.0020 | 0.0023 |
| F02 | 47500001 | 47600000 | 0.0033 | 0.0021 | 0.0024 |
| F02 | 47520001 | 47620000 | 0.0032 | 0.0021 | 0.0023 |
| F02 | 47540001 | 47640000 | 0.0030 | 0.0017 | 0.0019 |
| F02 | 47560001 | 47660000 | 0.0031 | 0.0016 | 0.0019 |
| F02 | 47580001 | 47680000 | 0.0042 | 0.0026 | 0.0032 |

| Chr | Start | End | both | A1 | A2 |
|---|---|---|---|---|---|
| F02 | 47600001 | 47700000 | 0.0046 | 0.0031 | 0.0038 |
| F02 | 47620001 | 47720000 | 0.0041 | 0.0027 | 0.0033 |
| F02 | 47640001 | 47740000 | 0.0042 | 0.0029 | 0.0035 |
| F02 | 47660001 | 47760000 | 0.0052 | 0.0040 | 0.0046 |
| F02 | 47680001 | 47780000 | 0.0042 | 0.0031 | 0.0035 |
| F02 | 47700001 | 47800000 | 0.0041 | 0.0030 | 0.0032 |
| F02 | 47720001 | 47820000 | 0.0050 | 0.0039 | 0.0043 |
| F02 | 47740001 | 47840000 | 0.0051 | 0.0040 | 0.0044 |
| F02 | 47760001 | 47860000 | 0.0042 | 0.0033 | 0.0035 |
| F02 | 47780001 | 47880000 | 0.0043 | 0.0034 | 0.0037 |
| F02 | 47800001 | 47900000 | 0.0047 | 0.0038 | 0.0042 |
| F02 | 47820001 | 47920000 | 0.0034 | 0.0027 | 0.0030 |
| F02 | 47840001 | 47940000 | 0.0030 | 0.0026 | 0.0029 |
| F02 | 47860001 | 47960000 | 0.0027 | 0.0021 | 0.0023 |
| F02 | 47880001 | 47980000 | 0.0030 | 0.0024 | 0.0024 |
| F02 | 47900001 | 48000000 | 0.0021 | 0.0015 | 0.0015 |
| F02 | 47920001 | 48020000 | 0.0029 | 0.0022 | 0.0022 |
| F02 | 47940001 | 48040000 | 0.0030 | 0.0022 | 0.0022 |
| F02 | 47960001 | 48060000 | 0.0036 | 0.0029 | 0.0030 |
| F02 | 47980001 | 48080000 | 0.0032 | 0.0024 | 0.0028 |
| F02 | 48000001 | 48100000 | 0.0040 | 0.0032 | 0.0035 |
| F02 | 48020001 | 48120000 | 0.0036 | 0.0026 | 0.0029 |
| F02 | 48040001 | 48140000 | 0.0033 | 0.0024 | 0.0027 |
| F02 | 48060001 | 48160000 | 0.0026 | 0.0017 | 0.0019 |
| F02 | 48080001 | 48180000 | 0.0025 | 0.0015 | 0.0017 |
| F02 | 48100001 | 48200000 | 0.0028 | 0.0015 | 0.0018 |
| F02 | 48120001 | 48220000 | 0.0031 | 0.0019 | 0.0022 |
| F02 | 48140001 | 48240000 | 0.0039 | 0.0026 | 0.0029 |
| F02 | 48160001 | 48260000 | 0.0041 | 0.0029 | 0.0032 |
| F02 | 48180001 | 48280000 | 0.0042 | 0.0030 | 0.0032 |
| F02 | 48200001 | 48300000 | 0.0037 | 0.0028 | 0.0030 |
| F02 | 48220001 | 48320000 | 0.0034 | 0.0025 | 0.0027 |
| F02 | 48240001 | 48340000 | 0.0030 | 0.0019 | 0.0022 |
| F02 | 48260001 | 48360000 | 0.0029 | 0.0020 | 0.0022 |
| F02 | 48280001 | 48380000 | 0.0029 | 0.0021 | 0.0023 |
| F02 | 48300001 | 48400000 | 0.0025 | 0.0017 | 0.0020 |
| F02 | 48320001 | 48420000 | 0.0026 | 0.0018 | 0.0020 |
| F02 | 48340001 | 48440000 | 0.0029 | 0.0022 | 0.0025 |
| F02 | 48360001 | 48460000 | 0.0033 | 0.0022 | 0.0027 |
| F02 | 48380001 | 48480000 | 0.0030 | 0.0019 | 0.0024 |
| F02 | 48400001 | 48500000 | 0.0036 | 0.0024 | 0.0027 |
| F02 | 48420001 | 48520000 | 0.0033 | 0.0020 | 0.0024 |
| F02 | 48440001 | 48540000 | 0.0037 | 0.0026 | 0.0028 |
| F02 | 48460001 | 48560000 | 0.0037 | 0.0028 | 0.0029 |
| F02 | 48480001 | 48580000 | 0.0037 | 0.0029 | 0.0030 |
| F02 | 48500001 | 48600000 | 0.0031 | 0.0025 | 0.0027 |
| F02 | 48520001 | 48620000 | 0.0040 | 0.0033 | 0.0037 |
| F02 | 48540001 | 48640000 | 0.0025 | 0.0020 | 0.0023 |
| F02 | 48560001 | 48660000 | 0.0027 | 0.0022 | 0.0026 |
| F02 | 48580001 | 48680000 | 0.0025 | 0.0021 | 0.0025 |
| F02 | 48600001 | 48700000 | 0.0029 | 0.0025 | 0.0029 |
| F02 | 48620001 | 48720000 | 0.0026 | 0.0023 | 0.0025 |
| F02 | 48640001 | 48740000 | 0.0033 | 0.0028 | 0.0030 |
| F02 | 48660001 | 48760000 | 0.0037 | 0.0030 | 0.0033 |
| F02 | 48680001 | 48780000 | 0.0048 | 0.0040 | 0.0043 |
| F02 | 48700001 | 48800000 | 0.0043 | 0.0036 | 0.0038 |
| F02 | 48720001 | 48820000 | 0.0039 | 0.0032 | 0.0035 |
| F02 | 48740001 | 48840000 | 0.0035 | 0.0029 | 0.0032 |

| Chr | Start | End | both | A1 | A2 |
|---|---|---|---|---|---|
| F02 | 48760001 | 48860000 | 0.0028 | 0.0021 | 0.0024 |
| F02 | 48780001 | 48880000 | 0.0020 | 0.0015 | 0.0017 |
| F02 | 48800001 | 48900000 | 0.0022 | 0.0017 | 0.0018 |
| F02 | 48820001 | 48920000 | 0.0023 | 0.0017 | 0.0019 |
| F02 | 48840001 | 48940000 | 0.0031 | 0.0022 | 0.0025 |
| F02 | 48860001 | 48960000 | 0.0032 | 0.0023 | 0.0026 |
| F02 | 48880001 | 48980000 | 0.0035 | 0.0025 | 0.0029 |
| F02 | 48900001 | 49000000 | 0.0035 | 0.0024 | 0.0028 |
| F02 | 48920001 | 49020000 | 0.0040 | 0.0026 | 0.0031 |
| F02 | 48940001 | 49040000 | 0.0036 | 0.0022 | 0.0026 |
| F02 | 48960001 | 49060000 | 0.0032 | 0.0018 | 0.0022 |
| F02 | 48980001 | 49080000 | 0.0039 | 0.0024 | 0.0028 |
| F02 | 49000001 | 49100000 | 0.0044 | 0.0028 | 0.0032 |
| F02 | 49020001 | 49120000 | 0.0044 | 0.0032 | 0.0036 |
| F02 | 49040001 | 49140000 | 0.0041 | 0.0032 | 0.0035 |
| F02 | 49060001 | 49160000 | 0.0041 | 0.0034 | 0.0037 |
| F02 | 49080001 | 49180000 | 0.0033 | 0.0028 | 0.0029 |
| F02 | 49100001 | 49200000 | 0.0038 | 0.0032 | 0.0036 |
| F02 | 49120001 | 49220000 | 0.0034 | 0.0028 | 0.0031 |
| F02 | 49140001 | 49240000 | 0.0036 | 0.0027 | 0.0031 |
| F02 | 49160001 | 49260000 | 0.0035 | 0.0027 | 0.0029 |
| F02 | 49180001 | 49280000 | 0.0035 | 0.0026 | 0.0028 |
| F02 | 49200001 | 49300000 | 0.0023 | 0.0016 | 0.0016 |
| F02 | 49220001 | 49320000 | 0.0020 | 0.0014 | 0.0014 |
| F02 | 49240001 | 49340000 | 0.0016 | 0.0011 | 0.0011 |
| F02 | 49260001 | 49360000 | 0.0013 | 0.0009 | 0.0010 |
| F02 | 49280001 | 49380000 | 0.0010 | 0.0008 | 0.0008 |
| F02 | 49300001 | 49400000 | 0.0014 | 0.0008 | 0.0011 |
| F02 | 49320001 | 49420000 | 0.0015 | 0.0007 | 0.0010 |
| F02 | 49340001 | 49440000 | 0.0023 | 0.0010 | 0.0015 |
| F02 | 49360001 | 49460000 | 0.0028 | 0.0012 | 0.0016 |
| F02 | 49380001 | 49480000 | 0.0028 | 0.0013 | 0.0016 |
| F02 | 49400001 | 49500000 | 0.0024 | 0.0012 | 0.0014 |
| F02 | 49420001 | 49520000 | 0.0022 | 0.0011 | 0.0014 |
| F02 | 49440001 | 49540000 | 0.0013 | 0.0009 | 0.0010 |
| F02 | 49460001 | 49560000 | 0.0011 | 0.0010 | 0.0011 |
| F02 | 49480001 | 49580000 | 0.0011 | 0.0010 | 0.0011 |
| F02 | 49500001 | 49600000 | 0.0014 | 0.0014 | 0.0014 |
| F02 | 49520001 | 49620000 | 0.0015 | 0.0014 | 0.0015 |
| F02 | 49540001 | 49640000 | 0.0023 | 0.0020 | 0.0022 |
| F02 | 49560001 | 49660000 | 0.0022 | 0.0017 | 0.0019 |
| F02 | 49580001 | 49680000 | 0.0022 | 0.0016 | 0.0018 |
| F02 | 49600001 | 49700000 | 0.0021 | 0.0012 | 0.0015 |
| F02 | 49620001 | 49720000 | 0.0025 | 0.0015 | 0.0018 |
| F02 | 49640001 | 49740000 | 0.0024 | 0.0016 | 0.0017 |
| F02 | 49660001 | 49760000 | 0.0031 | 0.0023 | 0.0024 |
| F02 | 49680001 | 49780000 | 0.0032 | 0.0025 | 0.0026 |
| F02 | 49700001 | 49800000 | 0.0031 | 0.0025 | 0.0027 |
| F02 | 49720001 | 49820000 | 0.0029 | 0.0024 | 0.0026 |
| F02 | 49740001 | 49840000 | 0.0028 | 0.0021 | 0.0025 |
| F02 | 49760001 | 49860000 | 0.0021 | 0.0016 | 0.0018 |
| F02 | 49780001 | 49880000 | 0.0037 | 0.0036 | 0.0031 |
| F02 | 49800001 | 49900000 | 0.0050 | 0.0050 | 0.0044 |
| F02 | 49820001 | 49920000 | 0.0055 | 0.0055 | 0.0049 |
| F02 | 49840001 | 49940000 | 0.0056 | 0.0056 | 0.0050 |
| F02 | 49860001 | 49960000 | 0.0059 | 0.0057 | 0.0052 |
| F02 | 49880001 | 49980000 | 0.0040 | 0.0035 | 0.0036 |
| F02 | 49900001 | 50000000 | 0.0032 | 0.0027 | 0.0028 |

| Chr | Start | End | both | A1 | A2 |
|---|---|---|---|---|---|
| F02 | 49920001 | 50020000 | 0.0028 | 0.0023 | 0.0023 |
| F02 | 49940001 | 50040000 | 0.0024 | 0.0020 | 0.0019 |
| F02 | 49960001 | 50060000 | 0.0021 | 0.0020 | 0.0018 |
| F02 | 49980001 | 50080000 | 0.0027 | 0.0024 | 0.0023 |
| F02 | 50000001 | 50100000 | 0.0018 | 0.0015 | 0.0015 |
| F02 | 50020001 | 50120000 | 0.0018 | 0.0015 | 0.0015 |
| F02 | 50040001 | 50140000 | 0.0020 | 0.0016 | 0.0015 |
| F02 | 50060001 | 50160000 | 0.0019 | 0.0014 | 0.0014 |
| F02 | 50080001 | 50180000 | 0.0016 | 0.0013 | 0.0012 |
| F02 | 50100001 | 50200000 | 0.0021 | 0.0018 | 0.0017 |
| F02 | 50120001 | 50220000 | 0.0018 | 0.0015 | 0.0015 |
| F02 | 50140001 | 50240000 | 0.0028 | 0.0025 | 0.0025 |
| F02 | 50160001 | 50260000 | 0.0028 | 0.0025 | 0.0025 |
| F02 | 50180001 | 50280000 | 0.0030 | 0.0023 | 0.0025 |
| F02 | 50200001 | 50300000 | 0.0027 | 0.0019 | 0.0021 |
| F02 | 50220001 | 50320000 | 0.0032 | 0.0023 | 0.0025 |
| F02 | 50240001 | 50340000 | 0.0024 | 0.0014 | 0.0017 |
| F02 | 50260001 | 50360000 | 0.0030 | 0.0019 | 0.0023 |
| F02 | 50280001 | 50380000 | 0.0035 | 0.0025 | 0.0030 |
| F02 | 50300001 | 50400000 | 0.0032 | 0.0023 | 0.0028 |
| F02 | 50320001 | 50420000 | 0.0033 | 0.0027 | 0.0031 |
| F02 | 50340001 | 50440000 | 0.0032 | 0.0025 | 0.0028 |
| F02 | 50360001 | 50460000 | 0.0029 | 0.0021 | 0.0024 |
| F02 | 50380001 | 50480000 | 0.0029 | 0.0019 | 0.0019 |
| F02 | 50400001 | 50500000 | 0.0032 | 0.0021 | 0.0022 |
| F02 | 50420001 | 50520000 | 0.0028 | 0.0016 | 0.0018 |
| F02 | 50440001 | 50540000 | 0.0031 | 0.0021 | 0.0023 |
| F02 | 50460001 | 50560000 | 0.0032 | 0.0024 | 0.0025 |
| F02 | 50480001 | 50580000 | 0.0024 | 0.0018 | 0.0020 |
| F02 | 50500001 | 50600000 | 0.0027 | 0.0022 | 0.0022 |
| F02 | 50520001 | 50620000 | 0.0027 | 0.0022 | 0.0022 |
| F02 | 50540001 | 50640000 | 0.0028 | 0.0020 | 0.0021 |
| F02 | 50560001 | 50660000 | 0.0036 | 0.0028 | 0.0029 |
| F02 | 50580001 | 50680000 | 0.0045 | 0.0036 | 0.0037 |
| F02 | 50600001 | 50700000 | 0.0043 | 0.0034 | 0.0035 |
| F02 | 50620001 | 50720000 | 0.0041 | 0.0032 | 0.0033 |
| F02 | 50640001 | 50740000 | 0.0039 | 0.0030 | 0.0032 |
| F02 | 50660001 | 50760000 | 0.0033 | 0.0024 | 0.0026 |
| F02 | 50680001 | 50780000 | 0.0027 | 0.0019 | 0.0021 |
| F02 | 50700001 | 50800000 | 0.0024 | 0.0017 | 0.0020 |
| F02 | 50720001 | 50820000 | 0.0035 | 0.0027 | 0.0029 |
| F02 | 50740001 | 50840000 | 0.0055 | 0.0047 | 0.0049 |
| F02 | 50760001 | 50860000 | 0.0055 | 0.0044 | 0.0046 |
| F02 | 50780001 | 50880000 | 0.0062 | 0.0046 | 0.0048 |
| F02 | 50800001 | 50900000 | 0.0070 | 0.0052 | 0.0055 |
| F02 | 50820001 | 50920000 | 0.0064 | 0.0046 | 0.0050 |
| F02 | 50840001 | 50940000 | 0.0047 | 0.0031 | 0.0034 |
| F02 | 50860001 | 50960000 | 0.0049 | 0.0034 | 0.0036 |
| F02 | 50880001 | 50980000 | 0.0051 | 0.0038 | 0.0040 |
| F02 | 50900001 | 51000000 | 0.0052 | 0.0040 | 0.0041 |
| F02 | 50920001 | 51020000 | 0.0054 | 0.0039 | 0.0039 |
| F02 | 50940001 | 51040000 | 0.0051 | 0.0035 | 0.0035 |
| F02 | 50960001 | 51060000 | 0.0047 | 0.0033 | 0.0031 |
| F02 | 50980001 | 51080000 | 0.0041 | 0.0029 | 0.0027 |
| F02 | 51000001 | 51100000 | 0.0033 | 0.0019 | 0.0019 |
| F02 | 51020001 | 51120000 | 0.0030 | 0.0019 | 0.0019 |
| F02 | 51040001 | 51140000 | 0.0038 | 0.0025 | 0.0027 |
| F02 | 51060001 | 51160000 | 0.0031 | 0.0020 | 0.0024 |

| Chr | Start | End | both | A1 | A2 |
|---|---|---|---|---|---|
| F02 | 51080001 | 51180000 | 0.0043 | 0.0028 | 0.0034 |
| F02 | 51100001 | 51200000 | 0.0042 | 0.0029 | 0.0033 |
| F02 | 51120001 | 51220000 | 0.0043 | 0.0031 | 0.0035 |
| F02 | 51140001 | 51240000 | 0.0040 | 0.0029 | 0.0031 |
| F02 | 51160001 | 51260000 | 0.0052 | 0.0038 | 0.0042 |
| F02 | 51180001 | 51280000 | 0.0045 | 0.0034 | 0.0036 |
| F02 | 51200001 | 51300000 | 0.0047 | 0.0037 | 0.0040 |
| F02 | 51220001 | 51320000 | 0.0053 | 0.0043 | 0.0046 |
| F02 | 51240001 | 51340000 | 0.0049 | 0.0040 | 0.0043 |
| F02 | 51260001 | 51360000 | 0.0043 | 0.0036 | 0.0039 |
| F02 | 51280001 | 51380000 | 0.0050 | 0.0040 | 0.0046 |
| F02 | 51300001 | 51400000 | 0.0047 | 0.0038 | 0.0043 |
| F02 | 51320001 | 51420000 | 0.0046 | 0.0033 | 0.0040 |
| F02 | 51340001 | 51440000 | 0.0046 | 0.0032 | 0.0040 |
| F02 | 51360001 | 51460000 | 0.0047 | 0.0033 | 0.0040 |
| F02 | 51380001 | 51480000 | 0.0041 | 0.0030 | 0.0033 |
| F02 | 51400001 | 51500000 | 0.0043 | 0.0032 | 0.0035 |
| F02 | 51420001 | 51520000 | 0.0041 | 0.0034 | 0.0032 |
| F02 | 51440001 | 51540000 | 0.0042 | 0.0035 | 0.0033 |
| F02 | 51460001 | 51560000 | 0.0050 | 0.0042 | 0.0042 |
| F02 | 51480001 | 51580000 | 0.0052 | 0.0044 | 0.0045 |
| F02 | 51500001 | 51600000 | 0.0057 | 0.0047 | 0.0048 |
| F02 | 51520001 | 51620000 | 0.0065 | 0.0050 | 0.0054 |
| F02 | 51540001 | 51640000 | 0.0064 | 0.0052 | 0.0054 |
| F02 | 51560001 | 51660000 | 0.0056 | 0.0043 | 0.0046 |
| F02 | 51580001 | 51680000 | 0.0052 | 0.0036 | 0.0038 |
| F02 | 51600001 | 51700000 | 0.0045 | 0.0030 | 0.0033 |
| F02 | 51620001 | 51720000 | 0.0036 | 0.0022 | 0.0025 |
| F02 | 51640001 | 51740000 | 0.0036 | 0.0018 | 0.0022 |
| F02 | 51660001 | 51760000 | 0.0038 | 0.0019 | 0.0022 |
| F02 | 51680001 | 51780000 | 0.0038 | 0.0021 | 0.0024 |
| F02 | 51700001 | 51800000 | 0.0039 | 0.0021 | 0.0024 |
| F02 | 51720001 | 51820000 | 0.0036 | 0.0021 | 0.0024 |
| F02 | 51740001 | 51840000 | 0.0035 | 0.0022 | 0.0026 |
| F02 | 51760001 | 51860000 | 0.0030 | 0.0019 | 0.0022 |
| F02 | 51780001 | 51880000 | 0.0027 | 0.0016 | 0.0019 |
| F02 | 51800001 | 51900000 | 0.0026 | 0.0016 | 0.0018 |
| F02 | 51820001 | 51920000 | 0.0026 | 0.0014 | 0.0017 |
| F02 | 51840001 | 51940000 | 0.0019 | 0.0009 | 0.0011 |
| F02 | 51860001 | 51960000 | 0.0020 | 0.0008 | 0.0009 |
| F02 | 51880001 | 51980000 | 0.0018 | 0.0007 | 0.0008 |
| F02 | 51900001 | 52000000 | 0.0022 | 0.0011 | 0.0011 |
| F02 | 51920001 | 52020000 | 0.0023 | 0.0010 | 0.0010 |
| F02 | 51940001 | 52040000 | 0.0026 | 0.0014 | 0.0015 |
| F02 | 51960001 | 52060000 | 0.0029 | 0.0017 | 0.0017 |
| F02 | 51980001 | 52080000 | 0.0032 | 0.0019 | 0.0020 |
| F02 | 52000001 | 52100000 | 0.0034 | 0.0018 | 0.0020 |
| F02 | 52020001 | 52120000 | 0.0035 | 0.0020 | 0.0021 |
| F02 | 52040001 | 52140000 | 0.0036 | 0.0021 | 0.0022 |
| F02 | 52060001 | 52160000 | 0.0033 | 0.0021 | 0.0022 |
| F02 | 52080001 | 52180000 | 0.0034 | 0.0021 | 0.0023 |
| F02 | 52100001 | 52200000 | 0.0032 | 0.0019 | 0.0023 |
| F02 | 52120001 | 52220000 | 0.0040 | 0.0027 | 0.0031 |
| F02 | 52140001 | 52240000 | 0.0039 | 0.0025 | 0.0030 |
| F02 | 52160001 | 52260000 | 0.0039 | 0.0024 | 0.0030 |
| F02 | 52180001 | 52280000 | 0.0041 | 0.0028 | 0.0034 |
| F02 | 52200001 | 52300000 | 0.0050 | 0.0039 | 0.0043 |
| F02 | 52220001 | 52320000 | 0.0043 | 0.0034 | 0.0038 |

| Chr | Start | End | both | A1 | A2 |
|---|---|---|---|---|---|
| F02 | 52240001 | 52340000 | 0.0049 | 0.0038 | 0.0042 |
| F02 | 52260001 | 52360000 | 0.0056 | 0.0043 | 0.0046 |
| F02 | 52280001 | 52380000 | 0.0046 | 0.0035 | 0.0036 |
| F02 | 52300001 | 52400000 | 0.0040 | 0.0028 | 0.0031 |
| F02 | 52320001 | 52420000 | 0.0050 | 0.0037 | 0.0041 |
| F02 | 52340001 | 52440000 | 0.0044 | 0.0032 | 0.0036 |
| F02 | 52360001 | 52460000 | 0.0036 | 0.0027 | 0.0030 |
| F02 | 52380001 | 52480000 | 0.0038 | 0.0030 | 0.0033 |
| F02 | 52400001 | 52500000 | 0.0031 | 0.0022 | 0.0025 |
| F02 | 52420001 | 52520000 | 0.0020 | 0.0012 | 0.0014 |
| F02 | 52440001 | 52540000 | 0.0018 | 0.0011 | 0.0012 |
| F02 | 52460001 | 52560000 | 0.0023 | 0.0015 | 0.0017 |
| F02 | 52480001 | 52580000 | 0.0023 | 0.0011 | 0.0014 |
| F02 | 52500001 | 52600000 | 0.0024 | 0.0013 | 0.0015 |
| F02 | 52520001 | 52620000 | 0.0031 | 0.0019 | 0.0022 |
| F02 | 52540001 | 52640000 | 0.0041 | 0.0026 | 0.0028 |
| F02 | 52560001 | 52660000 | 0.0038 | 0.0022 | 0.0024 |
| F02 | 52580001 | 52680000 | 0.0037 | 0.0025 | 0.0027 |
| F02 | 52600001 | 52700000 | 0.0039 | 0.0029 | 0.0029 |
| F02 | 52620001 | 52720000 | 0.0032 | 0.0023 | 0.0022 |
| F02 | 52640001 | 52740000 | 0.0024 | 0.0017 | 0.0018 |
| F02 | 52660001 | 52760000 | 0.0032 | 0.0026 | 0.0027 |
| F02 | 52680001 | 52780000 | 0.0038 | 0.0030 | 0.0032 |
| F02 | 52700001 | 52800000 | 0.0038 | 0.0031 | 0.0033 |
| F02 | 52720001 | 52820000 | 0.0042 | 0.0035 | 0.0038 |
| F02 | 52740001 | 52840000 | 0.0042 | 0.0035 | 0.0038 |
| F02 | 52760001 | 52860000 | 0.0037 | 0.0032 | 0.0034 |
| F02 | 52780001 | 52880000 | 0.0032 | 0.0027 | 0.0028 |
| F02 | 52800001 | 52900000 | 0.0030 | 0.0021 | 0.0024 |
| F02 | 52820001 | 52920000 | 0.0033 | 0.0023 | 0.0027 |
| F02 | 52840001 | 52940000 | 0.0042 | 0.0030 | 0.0034 |
| F02 | 52860001 | 52960000 | 0.0043 | 0.0032 | 0.0036 |
| F02 | 52880001 | 52980000 | 0.0057 | 0.0045 | 0.0049 |
| F02 | 52900001 | 53000000 | 0.0060 | 0.0048 | 0.0052 |
| F02 | 52920001 | 53020000 | 0.0054 | 0.0042 | 0.0045 |
| F02 | 52940001 | 53040000 | 0.0043 | 0.0033 | 0.0035 |
| F02 | 52960001 | 53060000 | 0.0046 | 0.0034 | 0.0037 |
| F02 | 52980001 | 53080000 | 0.0033 | 0.0020 | 0.0023 |
| F02 | 53000001 | 53100000 | 0.0035 | 0.0023 | 0.0027 |
| F02 | 53020001 | 53120000 | 0.0033 | 0.0022 | 0.0026 |
| F02 | 53040001 | 53140000 | 0.0039 | 0.0027 | 0.0032 |
| F02 | 53060001 | 53160000 | 0.0036 | 0.0022 | 0.0027 |
| F02 | 53080001 | 53180000 | 0.0041 | 0.0027 | 0.0032 |
| F02 | 53100001 | 53200000 | 0.0037 | 0.0025 | 0.0029 |
| F02 | 53120001 | 53220000 | 0.0036 | 0.0025 | 0.0028 |
| F02 | 53140001 | 53240000 | 0.0032 | 0.0021 | 0.0024 |
| F02 | 53160001 | 53260000 | 0.0032 | 0.0024 | 0.0027 |
| F02 | 53180001 | 53280000 | 0.0034 | 0.0028 | 0.0031 |
| F02 | 53200001 | 53300000 | 0.0042 | 0.0035 | 0.0039 |
| F02 | 53220001 | 53320000 | 0.0044 | 0.0037 | 0.0042 |
| F02 | 53240001 | 53340000 | 0.0042 | 0.0036 | 0.0040 |
| F02 | 53260001 | 53360000 | 0.0043 | 0.0037 | 0.0040 |
| F02 | 53280001 | 53380000 | 0.0036 | 0.0031 | 0.0033 |
| F02 | 53300001 | 53400000 | 0.0025 | 0.0022 | 0.0022 |
| F02 | 53320001 | 53420000 | 0.0022 | 0.0020 | 0.0020 |
| F02 | 53340001 | 53440000 | 0.0019 | 0.0017 | 0.0018 |
| F02 | 53360001 | 53460000 | 0.0014 | 0.0011 | 0.0012 |
| F02 | 53380001 | 53480000 | 0.0014 | 0.0011 | 0.0012 |

| Chr | Start | End | both | A1 | A2 |
|---|---|---|---|---|---|
| F02 | 53400001 | 53500000 | 0.0016 | 0.0011 | 0.0013 |
| F02 | 53420001 | 53520000 | 0.0018 | 0.0012 | 0.0013 |
| F02 | 53440001 | 53540000 | 0.0020 | 0.0013 | 0.0014 |
| F02 | 53460001 | 53560000 | 0.0018 | 0.0012 | 0.0013 |
| F02 | 53480001 | 53580000 | 0.0020 | 0.0012 | 0.0012 |
| F02 | 53500001 | 53600000 | 0.0028 | 0.0017 | 0.0019 |
| F02 | 53520001 | 53620000 | 0.0032 | 0.0021 | 0.0022 |
| F02 | 53540001 | 53640000 | 0.0035 | 0.0023 | 0.0024 |
| F02 | 53560001 | 53660000 | 0.0043 | 0.0030 | 0.0031 |
| F02 | 53580001 | 53680000 | 0.0054 | 0.0042 | 0.0043 |
| F02 | 53600001 | 53700000 | 0.0052 | 0.0042 | 0.0041 |
| F02 | 53620001 | 53720000 | 0.0049 | 0.0039 | 0.0040 |
| F02 | 53640001 | 53740000 | 0.0049 | 0.0041 | 0.0041 |
| F02 | 53660001 | 53760000 | 0.0041 | 0.0034 | 0.0035 |
| F02 | 53680001 | 53780000 | 0.0034 | 0.0025 | 0.0029 |
| F02 | 53700001 | 53800000 | 0.0036 | 0.0028 | 0.0031 |
| F02 | 53720001 | 53820000 | 0.0048 | 0.0041 | 0.0042 |
| F02 | 53740001 | 53840000 | 0.0046 | 0.0037 | 0.0039 |
| F02 | 53760001 | 53860000 | 0.0053 | 0.0041 | 0.0042 |
| F02 | 53780001 | 53880000 | 0.0055 | 0.0043 | 0.0044 |
| F02 | 53800001 | 53900000 | 0.0051 | 0.0035 | 0.0038 |
| F02 | 53820001 | 53920000 | 0.0038 | 0.0021 | 0.0024 |
| F02 | 53840001 | 53940000 | 0.0039 | 0.0022 | 0.0025 |
| F02 | 53860001 | 53960000 | 0.0034 | 0.0019 | 0.0023 |
| F02 | 53880001 | 53980000 | 0.0034 | 0.0020 | 0.0023 |
| F02 | 53900001 | 54000000 | 0.0039 | 0.0032 | 0.0032 |
| F02 | 53920001 | 54020000 | 0.0045 | 0.0038 | 0.0038 |
| F02 | 53940001 | 54040000 | 0.0046 | 0.0040 | 0.0041 |
| F02 | 53960001 | 54060000 | 0.0046 | 0.0039 | 0.0040 |
| F02 | 53980001 | 54080000 | 0.0042 | 0.0034 | 0.0035 |
| F02 | 54000001 | 54100000 | 0.0033 | 0.0023 | 0.0026 |
| F02 | 54020001 | 54120000 | 0.0031 | 0.0024 | 0.0025 |
| F02 | 54040001 | 54140000 | 0.0037 | 0.0027 | 0.0029 |
| F02 | 54060001 | 54160000 | 0.0038 | 0.0028 | 0.0030 |
| F02 | 54080001 | 54180000 | 0.0047 | 0.0035 | 0.0038 |
| F02 | 54100001 | 54200000 | 0.0048 | 0.0038 | 0.0040 |
| F02 | 54120001 | 54220000 | 0.0048 | 0.0036 | 0.0040 |
| F02 | 54140001 | 54240000 | 0.0045 | 0.0035 | 0.0040 |
| F02 | 54160001 | 54260000 | 0.0044 | 0.0032 | 0.0037 |
| F02 | 54180001 | 54280000 | 0.0033 | 0.0023 | 0.0026 |
| F02 | 54200001 | 54300000 | 0.0031 | 0.0020 | 0.0024 |
| F02 | 54220001 | 54320000 | 0.0032 | 0.0019 | 0.0023 |
| F02 | 54240001 | 54340000 | 0.0031 | 0.0020 | 0.0022 |
| F02 | 54260001 | 54360000 | 0.0032 | 0.0022 | 0.0024 |
| F02 | 54280001 | 54380000 | 0.0048 | 0.0039 | 0.0040 |
| F02 | 54300001 | 54400000 | 0.0048 | 0.0036 | 0.0038 |
| F02 | 54320001 | 54420000 | 0.0057 | 0.0045 | 0.0048 |
| F02 | 54340001 | 54440000 | 0.0052 | 0.0041 | 0.0044 |
| F02 | 54360001 | 54460000 | 0.0058 | 0.0044 | 0.0047 |
| F02 | 54380001 | 54480000 | 0.0046 | 0.0033 | 0.0036 |
| F02 | 54400001 | 54500000 | 0.0051 | 0.0039 | 0.0042 |
| F02 | 54420001 | 54520000 | 0.0041 | 0.0031 | 0.0032 |
| F02 | 54440001 | 54540000 | 0.0042 | 0.0032 | 0.0035 |
| F02 | 54460001 | 54560000 | 0.0042 | 0.0036 | 0.0038 |
| F02 | 54480001 | 54580000 | 0.0034 | 0.0029 | 0.0031 |
| F02 | 54500001 | 54600000 | 0.0027 | 0.0023 | 0.0025 |
| F02 | 54520001 | 54620000 | 0.0030 | 0.0028 | 0.0029 |
| F02 | 54540001 | 54640000 | 0.0032 | 0.0029 | 0.0031 |

| Chr | Start | End | both | A1 | A2 |
|---|---|---|---|---|---|
| F02 | 54560001 | 54660000 | 0.0027 | 0.0025 | 0.0026 |
| F02 | 54580001 | 54680000 | 0.0035 | 0.0031 | 0.0033 |
| F02 | 54600001 | 54700000 | 0.0035 | 0.0030 | 0.0032 |
| F02 | 54620001 | 54720000 | 0.0034 | 0.0026 | 0.0030 |
| F02 | 54640001 | 54740000 | 0.0039 | 0.0031 | 0.0035 |
| F02 | 54660001 | 54760000 | 0.0038 | 0.0030 | 0.0033 |
| F02 | 54680001 | 54780000 | 0.0034 | 0.0026 | 0.0030 |
| F02 | 54700001 | 54800000 | 0.0039 | 0.0029 | 0.0035 |
| F02 | 54720001 | 54820000 | 0.0034 | 0.0022 | 0.0027 |
| F02 | 54740001 | 54840000 | 0.0029 | 0.0016 | 0.0021 |
| F02 | 54760001 | 54860000 | 0.0032 | 0.0016 | 0.0023 |
| F02 | 54780001 | 54880000 | 0.0034 | 0.0020 | 0.0025 |
| F02 | 54800001 | 54900000 | 0.0032 | 0.0021 | 0.0024 |
| F02 | 54820001 | 54920000 | 0.0027 | 0.0020 | 0.0023 |
| F02 | 54840001 | 54940000 | 0.0028 | 0.0023 | 0.0025 |
| F02 | 54860001 | 54960000 | 0.0031 | 0.0026 | 0.0029 |
| F02 | 54880001 | 54980000 | 0.0027 | 0.0022 | 0.0026 |
| F02 | 54900001 | 55000000 | 0.0028 | 0.0023 | 0.0027 |
| F02 | 54920001 | 55020000 | 0.0026 | 0.0022 | 0.0025 |
| F02 | 54940001 | 55040000 | 0.0029 | 0.0025 | 0.0028 |
| F02 | 54960001 | 55060000 | 0.0025 | 0.0022 | 0.0023 |
| F02 | 54980001 | 55080000 | 0.0022 | 0.0019 | 0.0020 |
| F02 | 55000001 | 55100000 | 0.0020 | 0.0018 | 0.0019 |
| F02 | 55020001 | 55120000 | 0.0021 | 0.0018 | 0.0019 |
| F02 | 55040001 | 55140000 | 0.0015 | 0.0014 | 0.0015 |
| F02 | 55060001 | 55160000 | 0.0013 | 0.0012 | 0.0013 |
| F02 | 55080001 | 55180000 | 0.0014 | 0.0012 | 0.0013 |
| F02 | 55100001 | 55200000 | 0.0014 | 0.0011 | 0.0013 |
| F02 | 55120001 | 55220000 | 0.0014 | 0.0011 | 0.0013 |
| F02 | 55140001 | 55240000 | 0.0008 | 0.0006 | 0.0007 |
| F02 | 55160001 | 55260000 | 0.0009 | 0.0007 | 0.0009 |
| F02 | 55180001 | 55280000 | 0.0009 | 0.0007 | 0.0008 |
| F02 | 55200001 | 55300000 | 0.0006 | 0.0004 | 0.0006 |
| F02 | 55220001 | 55320000 | 0.0008 | 0.0005 | 0.0007 |
| F02 | 55240001 | 55340000 | 0.0008 | 0.0005 | 0.0006 |
| F02 | 55260001 | 55360000 | 0.0007 | 0.0005 | 0.0005 |
| F02 | 55280001 | 55380000 | 0.0009 | 0.0007 | 0.0007 |
| F02 | 55300001 | 55400000 | 0.0021 | 0.0018 | 0.0019 |
| F02 | 55320001 | 55420000 | 0.0027 | 0.0024 | 0.0025 |
| F02 | 55340001 | 55440000 | 0.0032 | 0.0028 | 0.0029 |
| F02 | 55360001 | 55460000 | 0.0032 | 0.0027 | 0.0028 |
| F02 | 55380001 | 55480000 | 0.0033 | 0.0028 | 0.0030 |
| F02 | 55400001 | 55500000 | 0.0027 | 0.0021 | 0.0024 |
| F02 | 55420001 | 55520000 | 0.0020 | 0.0015 | 0.0018 |
| F02 | 55440001 | 55540000 | 0.0016 | 0.0013 | 0.0015 |
| F02 | 55460001 | 55560000 | 0.0016 | 0.0014 | 0.0016 |
| F02 | 55480001 | 55580000 | 0.0018 | 0.0014 | 0.0017 |
| F02 | 55500001 | 55600000 | 0.0016 | 0.0013 | 0.0015 |
| F02 | 55520001 | 55620000 | 0.0017 | 0.0013 | 0.0016 |
| F02 | 55540001 | 55640000 | 0.0018 | 0.0014 | 0.0017 |
| F02 | 55560001 | 55660000 | 0.0020 | 0.0015 | 0.0019 |
| F02 | 55580001 | 55680000 | 0.0022 | 0.0016 | 0.0019 |
| F02 | 55600001 | 55700000 | 0.0019 | 0.0015 | 0.0016 |
| F02 | 55620001 | 55720000 | 0.0020 | 0.0016 | 0.0018 |
| F02 | 55640001 | 55740000 | 0.0023 | 0.0019 | 0.0021 |
| F02 | 55660001 | 55760000 | 0.0028 | 0.0023 | 0.0026 |
| F02 | 55680001 | 55780000 | 0.0032 | 0.0028 | 0.0031 |
| F02 | 55700001 | 55800000 | 0.0041 | 0.0035 | 0.0038 |

| Chr | Start | End | both | A1 | A2 |
|---|---|---|---|---|---|
| F02 | 55720001 | 55820000 | 0.0042 | 0.0036 | 0.0039 |
| F02 | 55740001 | 55840000 | 0.0040 | 0.0034 | 0.0037 |
| F02 | 55760001 | 55860000 | 0.0035 | 0.0029 | 0.0031 |
| F02 | 55780001 | 55880000 | 0.0035 | 0.0027 | 0.0031 |
| F02 | 55800001 | 55900000 | 0.0029 | 0.0022 | 0.0026 |
| F02 | 55820001 | 55920000 | 0.0032 | 0.0023 | 0.0028 |
| F02 | 55840001 | 55940000 | 0.0034 | 0.0024 | 0.0028 |
| F02 | 55860001 | 55960000 | 0.0031 | 0.0022 | 0.0026 |
| F02 | 55880001 | 55980000 | 0.0031 | 0.0025 | 0.0026 |
| F02 | 55900001 | 56000000 | 0.0034 | 0.0025 | 0.0026 |
| F02 | 55920001 | 56020000 | 0.0032 | 0.0024 | 0.0024 |
| F02 | 55940001 | 56040000 | 0.0039 | 0.0029 | 0.0033 |
| F02 | 55960001 | 56060000 | 0.0044 | 0.0030 | 0.0035 |
| F02 | 55980001 | 56080000 | 0.0039 | 0.0024 | 0.0029 |
| F02 | 56000001 | 56100000 | 0.0038 | 0.0025 | 0.0031 |
| F02 | 56020001 | 56120000 | 0.0035 | 0.0022 | 0.0029 |
| F02 | 56040001 | 56140000 | 0.0030 | 0.0020 | 0.0023 |
| F02 | 56060001 | 56160000 | 0.0030 | 0.0022 | 0.0024 |
| F02 | 56080001 | 56180000 | 0.0035 | 0.0025 | 0.0027 |
| F02 | 56100001 | 56200000 | 0.0036 | 0.0027 | 0.0027 |
| F02 | 56120001 | 56220000 | 0.0033 | 0.0025 | 0.0025 |
| F02 | 56140001 | 56240000 | 0.0028 | 0.0020 | 0.0021 |
| F02 | 56160001 | 56260000 | 0.0021 | 0.0016 | 0.0017 |
| F02 | 56180001 | 56280000 | 0.0016 | 0.0014 | 0.0013 |
| F02 | 56200001 | 56300000 | 0.0021 | 0.0015 | 0.0018 |
| F02 | 56220001 | 56320000 | 0.0024 | 0.0016 | 0.0019 |
| F02 | 56240001 | 56340000 | 0.0029 | 0.0020 | 0.0022 |
| F02 | 56260001 | 56360000 | 0.0038 | 0.0024 | 0.0028 |
| F02 | 56280001 | 56380000 | 0.0037 | 0.0025 | 0.0029 |
| F02 | 56300001 | 56400000 | 0.0033 | 0.0022 | 0.0026 |
| F02 | 56320001 | 56420000 | 0.0032 | 0.0023 | 0.0027 |
| F02 | 56340001 | 56440000 | 0.0027 | 0.0019 | 0.0022 |
| F02 | 56360001 | 56460000 | 0.0025 | 0.0020 | 0.0022 |
| F02 | 56380001 | 56480000 | 0.0024 | 0.0019 | 0.0021 |
| F02 | 56400001 | 56500000 | 0.0021 | 0.0018 | 0.0018 |
| F02 | 56420001 | 56520000 | 0.0028 | 0.0023 | 0.0025 |
| F02 | 56440001 | 56540000 | 0.0029 | 0.0024 | 0.0027 |
| F02 | 56460001 | 56560000 | 0.0023 | 0.0019 | 0.0021 |
| F02 | 56480001 | 56580000 | 0.0024 | 0.0020 | 0.0022 |
| F02 | 56500001 | 56600000 | 0.0021 | 0.0019 | 0.0021 |
| F02 | 56520001 | 56620000 | 0.0020 | 0.0018 | 0.0019 |
| F02 | 56540001 | 56640000 | 0.0028 | 0.0024 | 0.0026 |
| F02 | 56560001 | 56660000 | 0.0032 | 0.0027 | 0.0029 |
| F02 | 56580001 | 56680000 | 0.0030 | 0.0025 | 0.0028 |
| F02 | 56600001 | 56700000 | 0.0034 | 0.0028 | 0.0032 |
| F02 | 56620001 | 56720000 | 0.0033 | 0.0026 | 0.0030 |
| F02 | 56640001 | 56740000 | 0.0031 | 0.0024 | 0.0028 |
| F02 | 56660001 | 56760000 | 0.0035 | 0.0026 | 0.0030 |
| F02 | 56680001 | 56780000 | 0.0037 | 0.0025 | 0.0029 |
| F02 | 56700001 | 56800000 | 0.0035 | 0.0023 | 0.0026 |
| F02 | 56720001 | 56820000 | 0.0038 | 0.0026 | 0.0029 |
| F02 | 56740001 | 56840000 | 0.0033 | 0.0022 | 0.0025 |
| F02 | 56760001 | 56860000 | 0.0030 | 0.0023 | 0.0024 |
| F02 | 56780001 | 56880000 | 0.0028 | 0.0025 | 0.0026 |
| F02 | 56800001 | 56900000 | 0.0030 | 0.0028 | 0.0030 |
| F02 | 56820001 | 56920000 | 0.0027 | 0.0024 | 0.0026 |
| F02 | 56840001 | 56940000 | 0.0023 | 0.0020 | 0.0022 |
| F02 | 56860001 | 56960000 | 0.0019 | 0.0016 | 0.0018 |

| Chr | Start | End | both | A1 | A2 |
|---|---|---|---|---|---|
| F02 | 56880001 | 56980000 | 0.0016 | 0.0014 | 0.0016 |
| F02 | 56900001 | 57000000 | 0.0020 | 0.0016 | 0.0018 |
| F02 | 56920001 | 57020000 | 0.0019 | 0.0014 | 0.0016 |
| F02 | 56940001 | 57040000 | 0.0024 | 0.0015 | 0.0017 |
| F02 | 56960001 | 57060000 | 0.0037 | 0.0022 | 0.0024 |
| F02 | 56980001 | 57080000 | 0.0044 | 0.0024 | 0.0026 |
| F02 | 57000001 | 57100000 | 0.0043 | 0.0022 | 0.0025 |
| F02 | 57020001 | 57120000 | 0.0043 | 0.0024 | 0.0026 |
| F02 | 57040001 | 57140000 | 0.0047 | 0.0031 | 0.0033 |
| F02 | 57060001 | 57160000 | 0.0040 | 0.0025 | 0.0027 |
| F02 | 57080001 | 57180000 | 0.0047 | 0.0036 | 0.0038 |
| F02 | 57100001 | 57200000 | 0.0042 | 0.0031 | 0.0033 |
| F02 | 57120001 | 57220000 | 0.0039 | 0.0027 | 0.0030 |
| F02 | 57140001 | 57240000 | 0.0039 | 0.0023 | 0.0026 |
| F02 | 57160001 | 57260000 | 0.0043 | 0.0028 | 0.0031 |
| F02 | 57180001 | 57280000 | 0.0039 | 0.0024 | 0.0026 |
| F02 | 57200001 | 57300000 | 0.0050 | 0.0036 | 0.0038 |
| F02 | 57220001 | 57320000 | 0.0051 | 0.0036 | 0.0038 |
| F02 | 57240001 | 57340000 | 0.0051 | 0.0036 | 0.0041 |
| F02 | 57260001 | 57360000 | 0.0045 | 0.0032 | 0.0036 |
| F02 | 57280001 | 57380000 | 0.0044 | 0.0033 | 0.0034 |
| F02 | 57300001 | 57400000 | 0.0037 | 0.0024 | 0.0027 |
| F02 | 57320001 | 57420000 | 0.0037 | 0.0026 | 0.0029 |
| F02 | 57340001 | 57440000 | 0.0036 | 0.0029 | 0.0030 |
| F02 | 57360001 | 57460000 | 0.0036 | 0.0028 | 0.0028 |
| F02 | 57380001 | 57480000 | 0.0034 | 0.0023 | 0.0025 |
| F02 | 57400001 | 57500000 | 0.0033 | 0.0023 | 0.0025 |
| F02 | 57420001 | 57520000 | 0.0032 | 0.0022 | 0.0024 |
| F02 | 57440001 | 57540000 | 0.0030 | 0.0017 | 0.0020 |
| F02 | 57460001 | 57560000 | 0.0036 | 0.0025 | 0.0028 |
| F02 | 57480001 | 57580000 | 0.0030 | 0.0021 | 0.0024 |
| F02 | 57500001 | 57600000 | 0.0044 | 0.0033 | 0.0038 |
| F02 | 57520001 | 57620000 | 0.0046 | 0.0034 | 0.0039 |
| F02 | 57540001 | 57640000 | 0.0048 | 0.0039 | 0.0041 |
| F02 | 57560001 | 57660000 | 0.0047 | 0.0036 | 0.0040 |
| F02 | 57580001 | 57680000 | 0.0053 | 0.0041 | 0.0045 |
| F02 | 57600001 | 57700000 | 0.0041 | 0.0030 | 0.0033 |
| F02 | 57620001 | 57720000 | 0.0050 | 0.0040 | 0.0043 |
| F02 | 57640001 | 57740000 | 0.0045 | 0.0036 | 0.0039 |
| F02 | 57660001 | 57760000 | 0.0043 | 0.0032 | 0.0035 |
| F02 | 57680001 | 57780000 | 0.0040 | 0.0029 | 0.0033 |
| F02 | 57700001 | 57800000 | 0.0039 | 0.0029 | 0.0031 |
| F02 | 57720001 | 57820000 | 0.0033 | 0.0022 | 0.0024 |
| F02 | 57740001 | 57840000 | 0.0037 | 0.0025 | 0.0027 |
| F02 | 57760001 | 57860000 | 0.0039 | 0.0027 | 0.0030 |
| F02 | 57780001 | 57880000 | 0.0054 | 0.0042 | 0.0042 |
| F02 | 57800001 | 57900000 | 0.0052 | 0.0041 | 0.0042 |
| F02 | 57820001 | 57920000 | 0.0050 | 0.0037 | 0.0039 |
| F02 | 57840001 | 57940000 | 0.0045 | 0.0033 | 0.0034 |
| F02 | 57860001 | 57960000 | 0.0038 | 0.0028 | 0.0028 |
| F02 | 57880001 | 57980000 | 0.0032 | 0.0020 | 0.0024 |
| F02 | 57900001 | 58000000 | 0.0035 | 0.0021 | 0.0026 |
| F02 | 57920001 | 58020000 | 0.0037 | 0.0022 | 0.0027 |
| F02 | 57940001 | 58040000 | 0.0040 | 0.0025 | 0.0031 |
| F02 | 57960001 | 58060000 | 0.0049 | 0.0032 | 0.0037 |
| F02 | 57980001 | 58080000 | 0.0040 | 0.0026 | 0.0029 |
| F02 | 58000001 | 58100000 | 0.0037 | 0.0022 | 0.0026 |
| F02 | 58020001 | 58120000 | 0.0036 | 0.0023 | 0.0026 |

| Chr | Start | End | both | A1 | A2 |
|---|---|---|---|---|---|
| F02 | 58040001 | 58140000 | 0.0031 | 0.0020 | 0.0023 |
| F02 | 58060001 | 58160000 | 0.0022 | 0.0012 | 0.0016 |
| F02 | 58080001 | 58180000 | 0.0016 | 0.0008 | 0.0011 |
| F02 | 58100001 | 58200000 | 0.0020 | 0.0014 | 0.0016 |
| F02 | 58120001 | 58220000 | 0.0018 | 0.0014 | 0.0015 |
| F02 | 58140001 | 58240000 | 0.0019 | 0.0014 | 0.0016 |
| F02 | 58160001 | 58260000 | 0.0025 | 0.0018 | 0.0020 |
| F02 | 58180001 | 58280000 | 0.0028 | 0.0019 | 0.0021 |
| F02 | 58200001 | 58300000 | 0.0044 | 0.0035 | 0.0038 |
| F02 | 58220001 | 58320000 | 0.0049 | 0.0039 | 0.0041 |
| F02 | 58240001 | 58340000 | 0.0056 | 0.0045 | 0.0047 |
| F02 | 58260001 | 58360000 | 0.0056 | 0.0046 | 0.0049 |
| F02 | 58280001 | 58380000 | 0.0054 | 0.0045 | 0.0049 |
| F02 | 58300001 | 58400000 | 0.0043 | 0.0034 | 0.0036 |
| F02 | 58320001 | 58420000 | 0.0038 | 0.0030 | 0.0032 |
| F02 | 58340001 | 58440000 | 0.0038 | 0.0031 | 0.0033 |
| F02 | 58360001 | 58460000 | 0.0043 | 0.0037 | 0.0040 |
| F02 | 58380001 | 58480000 | 0.0049 | 0.0041 | 0.0044 |
| F02 | 58400001 | 58500000 | 0.0037 | 0.0030 | 0.0034 |
| F02 | 58420001 | 58520000 | 0.0035 | 0.0027 | 0.0033 |
| F02 | 58440001 | 58540000 | 0.0034 | 0.0027 | 0.0031 |
| F02 | 58460001 | 58560000 | 0.0025 | 0.0018 | 0.0020 |
| F02 | 58480001 | 58580000 | 0.0026 | 0.0019 | 0.0021 |
| F02 | 58500001 | 58600000 | 0.0038 | 0.0026 | 0.0027 |
| F02 | 58520001 | 58620000 | 0.0047 | 0.0031 | 0.0033 |
| F02 | 58540001 | 58640000 | 0.0046 | 0.0030 | 0.0032 |
| F02 | 58560001 | 58660000 | 0.0053 | 0.0036 | 0.0037 |
| F02 | 58580001 | 58680000 | 0.0049 | 0.0032 | 0.0034 |
| F02 | 58600001 | 58700000 | 0.0054 | 0.0038 | 0.0041 |
| F02 | 58620001 | 58720000 | 0.0051 | 0.0039 | 0.0041 |
| F02 | 58640001 | 58740000 | 0.0059 | 0.0044 | 0.0047 |
| F02 | 58660001 | 58760000 | 0.0055 | 0.0042 | 0.0045 |
| F02 | 58680001 | 58780000 | 0.0061 | 0.0047 | 0.0051 |
| F02 | 58700001 | 58800000 | 0.0048 | 0.0037 | 0.0039 |
| F02 | 58720001 | 58820000 | 0.0043 | 0.0033 | 0.0036 |
| F02 | 58740001 | 58840000 | 0.0029 | 0.0022 | 0.0024 |
| F02 | 58760001 | 58860000 | 0.0026 | 0.0018 | 0.0020 |
| F02 | 58780001 | 58880000 | 0.0021 | 0.0013 | 0.0014 |
| F02 | 58800001 | 58900000 | 0.0021 | 0.0013 | 0.0014 |
| F02 | 58820001 | 58920000 | 0.0026 | 0.0017 | 0.0016 |
| F02 | 58840001 | 58940000 | 0.0032 | 0.0022 | 0.0021 |
| F02 | 58860001 | 58960000 | 0.0035 | 0.0026 | 0.0025 |
| F02 | 58880001 | 58980000 | 0.0030 | 0.0024 | 0.0024 |
| F02 | 58900001 | 59000000 | 0.0031 | 0.0025 | 0.0026 |
| F02 | 58920001 | 59020000 | 0.0035 | 0.0027 | 0.0032 |
| F02 | 58940001 | 59040000 | 0.0037 | 0.0029 | 0.0035 |
| F02 | 58960001 | 59060000 | 0.0062 | 0.0050 | 0.0058 |
| F02 | 58980001 | 59080000 | 0.0062 | 0.0050 | 0.0058 |
| F02 | 59000001 | 59100000 | 0.0069 | 0.0055 | 0.0062 |
| F02 | 59020001 | 59120000 | 0.0067 | 0.0055 | 0.0060 |
| F02 | 59040001 | 59140000 | 0.0070 | 0.0056 | 0.0060 |
| F02 | 59060001 | 59160000 | 0.0045 | 0.0033 | 0.0035 |
| F02 | 59080001 | 59180000 | 0.0054 | 0.0040 | 0.0042 |
| F02 | 59100001 | 59200000 | 0.0044 | 0.0033 | 0.0036 |
| F02 | 59120001 | 59220000 | 0.0035 | 0.0024 | 0.0027 |
| F02 | 59140001 | 59240000 | 0.0029 | 0.0020 | 0.0023 |
| F02 | 59160001 | 59260000 | 0.0025 | 0.0016 | 0.0019 |
| F02 | 59180001 | 59280000 | 0.0016 | 0.0007 | 0.0011 |

| Chr | Start | End | both | A1 | A2 |
|---|---|---|---|---|---|
| F02 | 59200001 | 59300000 | 0.0025 | 0.0014 | 0.0019 |
| F02 | 59220001 | 59320000 | 0.0028 | 0.0017 | 0.0022 |
| F02 | 59240001 | 59340000 | 0.0029 | 0.0016 | 0.0021 |
| F02 | 59260001 | 59360000 | 0.0031 | 0.0018 | 0.0023 |
| F02 | 59280001 | 59380000 | 0.0041 | 0.0026 | 0.0031 |
| F02 | 59300001 | 59400000 | 0.0030 | 0.0017 | 0.0021 |
| F02 | 59320001 | 59420000 | 0.0026 | 0.0014 | 0.0017 |
| F02 | 59340001 | 59440000 | 0.0028 | 0.0015 | 0.0018 |
| F02 | 59360001 | 59460000 | 0.0030 | 0.0015 | 0.0018 |
| F02 | 59380001 | 59480000 | 0.0026 | 0.0011 | 0.0015 |
| F02 | 59400001 | 59500000 | 0.0035 | 0.0018 | 0.0023 |
| F02 | 59420001 | 59520000 | 0.0046 | 0.0030 | 0.0034 |
| F02 | 59440001 | 59540000 | 0.0044 | 0.0029 | 0.0032 |
| F02 | 59460001 | 59560000 | 0.0041 | 0.0029 | 0.0031 |
| F02 | 59480001 | 59580000 | 0.0033 | 0.0023 | 0.0025 |
| F02 | 59500001 | 59600000 | 0.0027 | 0.0018 | 0.0019 |
| F02 | 59520001 | 59620000 | 0.0022 | 0.0013 | 0.0014 |
| F02 | 59540001 | 59640000 | 0.0022 | 0.0013 | 0.0015 |
| F02 | 59560001 | 59660000 | 0.0029 | 0.0019 | 0.0023 |
| F02 | 59580001 | 59680000 | 0.0041 | 0.0029 | 0.0034 |
| F02 | 59600001 | 59700000 | 0.0048 | 0.0035 | 0.0039 |
| F02 | 59620001 | 59720000 | 0.0043 | 0.0030 | 0.0034 |
| F02 | 59640001 | 59740000 | 0.0040 | 0.0028 | 0.0032 |
| F02 | 59660001 | 59760000 | 0.0039 | 0.0028 | 0.0030 |
| F02 | 59680001 | 59780000 | 0.0036 | 0.0023 | 0.0025 |
| F02 | 59700001 | 59800000 | 0.0029 | 0.0017 | 0.0017 |
| F02 | 59720001 | 59820000 | 0.0034 | 0.0022 | 0.0023 |
| F02 | 59740001 | 59840000 | 0.0034 | 0.0021 | 0.0022 |
| F02 | 59760001 | 59860000 | 0.0034 | 0.0016 | 0.0019 |
| F02 | 59780001 | 59880000 | 0.0033 | 0.0016 | 0.0019 |
| F02 | 59800001 | 59900000 | 0.0042 | 0.0026 | 0.0028 |
| F02 | 59820001 | 59920000 | 0.0042 | 0.0024 | 0.0026 |
| F02 | 59840001 | 59940000 | 0.0042 | 0.0026 | 0.0028 |
| F02 | 59860001 | 59960000 | 0.0034 | 0.0024 | 0.0025 |
| F02 | 59880001 | 59980000 | 0.0033 | 0.0024 | 0.0025 |
| F02 | 59900001 | 60000000 | 0.0027 | 0.0019 | 0.0021 |
| F02 | 59920001 | 60020000 | 0.0019 | 0.0015 | 0.0017 |
| F02 | 59940001 | 60040000 | 0.0021 | 0.0017 | 0.0019 |
| F02 | 59960001 | 60060000 | 0.0021 | 0.0017 | 0.0020 |
| F02 | 59980001 | 60080000 | 0.0016 | 0.0012 | 0.0014 |
| F02 | 60000001 | 60100000 | 0.0016 | 0.0012 | 0.0013 |
| F02 | 60020001 | 60120000 | 0.0028 | 0.0020 | 0.0023 |
| F02 | 60040001 | 60140000 | 0.0032 | 0.0022 | 0.0025 |
| F02 | 60060001 | 60160000 | 0.0045 | 0.0035 | 0.0037 |
| F02 | 60080001 | 60180000 | 0.0056 | 0.0045 | 0.0048 |
| F02 | 60100001 | 60200000 | 0.0065 | 0.0051 | 0.0056 |
| F02 | 60120001 | 60220000 | 0.0057 | 0.0046 | 0.0048 |
| F02 | 60140001 | 60240000 | 0.0055 | 0.0044 | 0.0046 |
| F02 | 60160001 | 60260000 | 0.0048 | 0.0038 | 0.0040 |
| F02 | 60180001 | 60280000 | 0.0040 | 0.0031 | 0.0033 |
| F02 | 60200001 | 60300000 | 0.0033 | 0.0027 | 0.0027 |
| F02 | 60220001 | 60320000 | 0.0030 | 0.0023 | 0.0024 |
| F02 | 60240001 | 60340000 | 0.0029 | 0.0021 | 0.0022 |
| F02 | 60260001 | 60360000 | 0.0025 | 0.0017 | 0.0019 |
| F02 | 60280001 | 60380000 | 0.0022 | 0.0015 | 0.0017 |
| F02 | 60300001 | 60400000 | 0.0020 | 0.0010 | 0.0012 |
| F02 | 60320001 | 60420000 | 0.0024 | 0.0013 | 0.0015 |
| F02 | 60340001 | 60440000 | 0.0026 | 0.0015 | 0.0018 |

| Chr | Start | End | both | A1 | A2 |
|---|---|---|---|---|---|
| F02 | 60360001 | 60460000 | 0.0029 | 0.0017 | 0.0020 |
| F02 | 60380001 | 60480000 | 0.0037 | 0.0025 | 0.0028 |
| F02 | 60400001 | 60500000 | 0.0040 | 0.0029 | 0.0033 |
| F02 | 60420001 | 60520000 | 0.0036 | 0.0028 | 0.0032 |
| F02 | 60440001 | 60540000 | 0.0031 | 0.0024 | 0.0029 |
| F02 | 60460001 | 60560000 | 0.0031 | 0.0022 | 0.0026 |
| F02 | 60480001 | 60580000 | 0.0024 | 0.0013 | 0.0016 |
| F02 | 60500001 | 60600000 | 0.0028 | 0.0017 | 0.0021 |
| F02 | 60520001 | 60620000 | 0.0029 | 0.0017 | 0.0020 |
| F02 | 60540001 | 60640000 | 0.0032 | 0.0021 | 0.0024 |
| F02 | 60560001 | 60660000 | 0.0028 | 0.0019 | 0.0022 |
| F02 | 60580001 | 60680000 | 0.0033 | 0.0025 | 0.0028 |
| F02 | 60600001 | 60700000 | 0.0031 | 0.0024 | 0.0025 |
| F02 | 60620001 | 60720000 | 0.0037 | 0.0028 | 0.0031 |
| F02 | 60640001 | 60740000 | 0.0040 | 0.0028 | 0.0032 |
| F02 | 60660001 | 60760000 | 0.0047 | 0.0036 | 0.0040 |
| F02 | 60680001 | 60780000 | 0.0044 | 0.0033 | 0.0038 |
| F02 | 60700001 | 60800000 | 0.0039 | 0.0028 | 0.0033 |
| F02 | 60720001 | 60820000 | 0.0035 | 0.0026 | 0.0030 |
| F02 | 60740001 | 60840000 | 0.0027 | 0.0021 | 0.0024 |
| F02 | 60760001 | 60860000 | 0.0031 | 0.0021 | 0.0026 |
| F02 | 60780001 | 60880000 | 0.0040 | 0.0029 | 0.0036 |
| F02 | 60800001 | 60900000 | 0.0041 | 0.0029 | 0.0036 |
| F02 | 60820001 | 60920000 | 0.0055 | 0.0042 | 0.0046 |
| F02 | 60840001 | 60940000 | 0.0067 | 0.0050 | 0.0056 |
| F02 | 60860001 | 60960000 | 0.0056 | 0.0042 | 0.0046 |
| F02 | 60880001 | 60980000 | 0.0054 | 0.0041 | 0.0042 |
| F02 | 60900001 | 61000000 | 0.0058 | 0.0048 | 0.0048 |
| F02 | 60920001 | 61020000 | 0.0041 | 0.0032 | 0.0033 |
| F02 | 60940001 | 61040000 | 0.0036 | 0.0029 | 0.0029 |
| F02 | 60960001 | 61060000 | 0.0044 | 0.0035 | 0.0034 |
| F02 | 60980001 | 61080000 | 0.0033 | 0.0024 | 0.0025 |
| F02 | 61000001 | 61100000 | 0.0026 | 0.0018 | 0.0019 |
| F02 | 61020001 | 61120000 | 0.0029 | 0.0018 | 0.0020 |
| F02 | 61040001 | 61140000 | 0.0026 | 0.0017 | 0.0019 |
| F02 | 61060001 | 61160000 | 0.0020 | 0.0013 | 0.0015 |
| F02 | 61080001 | 61180000 | 0.0033 | 0.0025 | 0.0028 |
| F02 | 61100001 | 61200000 | 0.0032 | 0.0024 | 0.0029 |
| F02 | 61120001 | 61220000 | 0.0029 | 0.0024 | 0.0028 |
| F02 | 61140001 | 61240000 | 0.0026 | 0.0021 | 0.0026 |
| F02 | 61160001 | 61260000 | 0.0024 | 0.0019 | 0.0023 |
| F02 | 61180001 | 61280000 | 0.0019 | 0.0013 | 0.0015 |
| F02 | 61200001 | 61300000 | 0.0019 | 0.0014 | 0.0015 |
| F02 | 61220001 | 61320000 | 0.0020 | 0.0014 | 0.0015 |
| F02 | 61240001 | 61340000 | 0.0019 | 0.0013 | 0.0013 |
| F02 | 61260001 | 61360000 | 0.0021 | 0.0015 | 0.0016 |
| F02 | 61280001 | 61380000 | 0.0016 | 0.0009 | 0.0010 |
| F02 | 61300001 | 61400000 | 0.0019 | 0.0007 | 0.0009 |
| F02 | 61320001 | 61420000 | 0.0023 | 0.0010 | 0.0013 |
| F02 | 61340001 | 61440000 | 0.0047 | 0.0031 | 0.0035 |
| F02 | 61360001 | 61460000 | 0.0052 | 0.0036 | 0.0040 |
| F02 | 61380001 | 61480000 | 0.0059 | 0.0044 | 0.0048 |
| F02 | 61400001 | 61500000 | 0.0059 | 0.0047 | 0.0051 |
| F02 | 61420001 | 61520000 | 0.0056 | 0.0044 | 0.0048 |
| F02 | 61440001 | 61540000 | 0.0037 | 0.0027 | 0.0031 |
| F02 | 61460001 | 61560000 | 0.0031 | 0.0021 | 0.0024 |
| F02 | 61480001 | 61580000 | 0.0029 | 0.0019 | 0.0022 |
| F02 | 61500001 | 61600000 | 0.0032 | 0.0021 | 0.0024 |

| Chr | Start | End | both | A1 | A2 |
|---|---|---|---|---|---|
| F02 | 61520001 | 61620000 | 0.0036 | 0.0025 | 0.0028 |
| F02 | 61540001 | 61640000 | 0.0038 | 0.0025 | 0.0027 |
| F02 | 61560001 | 61660000 | 0.0038 | 0.0027 | 0.0029 |
| F02 | 61580001 | 61680000 | 0.0049 | 0.0039 | 0.0042 |
| F02 | 61600001 | 61700000 | 0.0049 | 0.0037 | 0.0042 |
| F02 | 61620001 | 61720000 | 0.0052 | 0.0041 | 0.0045 |
| F02 | 61640001 | 61740000 | 0.0046 | 0.0036 | 0.0041 |
| F02 | 61660001 | 61760000 | 0.0051 | 0.0041 | 0.0046 |
| F02 | 61680001 | 61780000 | 0.0033 | 0.0025 | 0.0029 |
| F02 | 61700001 | 61800000 | 0.0037 | 0.0030 | 0.0035 |
| F02 | 61720001 | 61820000 | 0.0030 | 0.0023 | 0.0027 |
| F02 | 61740001 | 61840000 | 0.0035 | 0.0026 | 0.0031 |
| F02 | 61760001 | 61860000 | 0.0032 | 0.0023 | 0.0027 |
| F02 | 61780001 | 61880000 | 0.0040 | 0.0029 | 0.0033 |
| F02 | 61800001 | 61900000 | 0.0029 | 0.0019 | 0.0020 |
| F02 | 61820001 | 61920000 | 0.0033 | 0.0022 | 0.0024 |
| F02 | 61840001 | 61940000 | 0.0034 | 0.0022 | 0.0025 |
| F02 | 61860001 | 61960000 | 0.0039 | 0.0025 | 0.0029 |
| F02 | 61880001 | 61980000 | 0.0034 | 0.0019 | 0.0023 |
| F02 | 61900001 | 62000000 | 0.0038 | 0.0024 | 0.0028 |
| F02 | 61920001 | 62020000 | 0.0032 | 0.0019 | 0.0022 |
| F02 | 61940001 | 62040000 | 0.0031 | 0.0017 | 0.0020 |
| F02 | 61960001 | 62060000 | 0.0026 | 0.0012 | 0.0015 |
| F02 | 61980001 | 62080000 | 0.0026 | 0.0014 | 0.0016 |
| F02 | 62000001 | 62100000 | 0.0031 | 0.0016 | 0.0019 |
| F02 | 62020001 | 62120000 | 0.0033 | 0.0017 | 0.0021 |
| F02 | 62040001 | 62140000 | 0.0031 | 0.0019 | 0.0022 |
| F02 | 62060001 | 62160000 | 0.0031 | 0.0020 | 0.0022 |
| F02 | 62080001 | 62180000 | 0.0031 | 0.0019 | 0.0022 |
| F02 | 62100001 | 62200000 | 0.0021 | 0.0013 | 0.0014 |
| F02 | 62120001 | 62220000 | 0.0020 | 0.0012 | 0.0014 |
| F02 | 62140001 | 62240000 | 0.0019 | 0.0009 | 0.0011 |
| F02 | 62160001 | 62260000 | 0.0023 | 0.0013 | 0.0014 |
| F02 | 62180001 | 62280000 | 0.0024 | 0.0013 | 0.0015 |
| F02 | 62200001 | 62300000 | 0.0027 | 0.0015 | 0.0016 |
| F02 | 62220001 | 62320000 | 0.0027 | 0.0015 | 0.0016 |
| F02 | 62240001 | 62340000 | 0.0052 | 0.0038 | 0.0042 |
| F02 | 62260001 | 62360000 | 0.0043 | 0.0032 | 0.0035 |
| F02 | 62280001 | 62380000 | 0.0041 | 0.0033 | 0.0036 |
| F02 | 62300001 | 62400000 | 0.0040 | 0.0032 | 0.0038 |
| F02 | 62320001 | 62420000 | 0.0049 | 0.0039 | 0.0045 |
| F02 | 62340001 | 62440000 | 0.0033 | 0.0023 | 0.0026 |
| F02 | 62360001 | 62460000 | 0.0040 | 0.0028 | 0.0031 |
| F02 | 62380001 | 62480000 | 0.0047 | 0.0032 | 0.0034 |
| F02 | 62400001 | 62500000 | 0.0050 | 0.0035 | 0.0037 |
| F02 | 62420001 | 62520000 | 0.0041 | 0.0029 | 0.0029 |
| F02 | 62440001 | 62540000 | 0.0030 | 0.0021 | 0.0021 |
| F02 | 62460001 | 62560000 | 0.0030 | 0.0021 | 0.0022 |
| F02 | 62480001 | 62580000 | 0.0024 | 0.0016 | 0.0016 |
| F02 | 62500001 | 62600000 | 0.0023 | 0.0014 | 0.0013 |
| F02 | 62520001 | 62620000 | 0.0025 | 0.0016 | 0.0015 |
| F02 | 62540001 | 62640000 | 0.0033 | 0.0025 | 0.0023 |
| F02 | 62560001 | 62660000 | 0.0038 | 0.0031 | 0.0028 |
| F02 | 62580001 | 62680000 | 0.0044 | 0.0037 | 0.0036 |
| F02 | 62600001 | 62700000 | 0.0052 | 0.0044 | 0.0043 |
| F02 | 62620001 | 62720000 | 0.0060 | 0.0049 | 0.0050 |
| F02 | 62640001 | 62740000 | 0.0057 | 0.0045 | 0.0047 |
| F02 | 62660001 | 62760000 | 0.0046 | 0.0034 | 0.0035 |

| Chr | Start | End | both | A1 | A2 |
|---|---|---|---|---|---|
| F02 | 62680001 | 62780000 | 0.0040 | 0.0026 | 0.0028 |
| F02 | 62700001 | 62800000 | 0.0032 | 0.0021 | 0.0022 |
| F02 | 62720001 | 62820000 | 0.0029 | 0.0020 | 0.0020 |
| F02 | 62740001 | 62840000 | 0.0032 | 0.0022 | 0.0023 |
| F02 | 62760001 | 62860000 | 0.0040 | 0.0030 | 0.0031 |
| F02 | 62780001 | 62880000 | 0.0044 | 0.0035 | 0.0036 |
| F02 | 62800001 | 62900000 | 0.0048 | 0.0038 | 0.0039 |
| F02 | 62820001 | 62920000 | 0.0054 | 0.0043 | 0.0046 |
| F02 | 62840001 | 62940000 | 0.0050 | 0.0042 | 0.0044 |
| F02 | 62860001 | 62960000 | 0.0045 | 0.0039 | 0.0040 |
| F02 | 62880001 | 62980000 | 0.0043 | 0.0035 | 0.0037 |
| F02 | 62900001 | 63000000 | 0.0044 | 0.0036 | 0.0036 |
| F02 | 62920001 | 63020000 | 0.0035 | 0.0026 | 0.0025 |
| F02 | 62940001 | 63040000 | 0.0037 | 0.0024 | 0.0024 |
| F02 | 62960001 | 63060000 | 0.0042 | 0.0027 | 0.0028 |
| F02 | 62980001 | 63080000 | 0.0050 | 0.0036 | 0.0036 |
| F02 | 63000001 | 63100000 | 0.0044 | 0.0030 | 0.0031 |
| F02 | 63020001 | 63120000 | 0.0044 | 0.0031 | 0.0033 |
| F02 | 63040001 | 63140000 | 0.0048 | 0.0035 | 0.0037 |
| F02 | 63060001 | 63160000 | 0.0052 | 0.0040 | 0.0042 |
| F02 | 63080001 | 63180000 | 0.0049 | 0.0037 | 0.0040 |
| F02 | 63100001 | 63200000 | 0.0053 | 0.0040 | 0.0044 |
| F02 | 63120001 | 63220000 | 0.0053 | 0.0040 | 0.0042 |
| F02 | 63140001 | 63240000 | 0.0052 | 0.0042 | 0.0044 |
| F02 | 63160001 | 63260000 | 0.0043 | 0.0032 | 0.0035 |
| F02 | 63180001 | 63280000 | 0.0047 | 0.0035 | 0.0038 |
| F02 | 63200001 | 63300000 | 0.0052 | 0.0040 | 0.0042 |
| F02 | 63220001 | 63320000 | 0.0064 | 0.0053 | 0.0056 |
| F02 | 63240001 | 63340000 | 0.0070 | 0.0060 | 0.0061 |
| F02 | 63260001 | 63360000 | 0.0074 | 0.0063 | 0.0064 |
| F02 | 63280001 | 63380000 | 0.0064 | 0.0053 | 0.0053 |
| F02 | 63300001 | 63400000 | 0.0062 | 0.0050 | 0.0051 |
| F02 | 63320001 | 63420000 | 0.0052 | 0.0039 | 0.0039 |
| F02 | 63340001 | 63440000 | 0.0042 | 0.0027 | 0.0029 |
| F02 | 63360001 | 63460000 | 0.0038 | 0.0025 | 0.0027 |
| F02 | 63380001 | 63480000 | 0.0037 | 0.0027 | 0.0030 |
| F02 | 63400001 | 63500000 | 0.0042 | 0.0032 | 0.0035 |
| F02 | 63420001 | 63520000 | 0.0035 | 0.0027 | 0.0032 |
| F02 | 63440001 | 63540000 | 0.0034 | 0.0028 | 0.0030 |
| F02 | 63460001 | 63560000 | 0.0035 | 0.0028 | 0.0031 |
| F02 | 63480001 | 63580000 | 0.0038 | 0.0030 | 0.0032 |
| F02 | 63500001 | 63600000 | 0.0037 | 0.0028 | 0.0029 |
| F02 | 63520001 | 63620000 | 0.0040 | 0.0030 | 0.0030 |
| F02 | 63540001 | 63640000 | 0.0041 | 0.0033 | 0.0033 |
| F02 | 63560001 | 63660000 | 0.0041 | 0.0032 | 0.0033 |
| F02 | 63580001 | 63680000 | 0.0042 | 0.0035 | 0.0034 |
| F02 | 63600001 | 63700000 | 0.0041 | 0.0031 | 0.0032 |
| F02 | 63620001 | 63720000 | 0.0040 | 0.0029 | 0.0030 |
| F02 | 63640001 | 63740000 | 0.0035 | 0.0023 | 0.0024 |
| F02 | 63660001 | 63760000 | 0.0034 | 0.0022 | 0.0023 |
| F02 | 63680001 | 63780000 | 0.0031 | 0.0017 | 0.0017 |
| F02 | 63700001 | 63800000 | 0.0025 | 0.0014 | 0.0013 |
| F02 | 63720001 | 63820000 | 0.0025 | 0.0015 | 0.0015 |
| F02 | 63740001 | 63840000 | 0.0027 | 0.0017 | 0.0016 |
| F02 | 63760001 | 63860000 | 0.0030 | 0.0020 | 0.0021 |
| F02 | 63780001 | 63880000 | 0.0030 | 0.0021 | 0.0024 |
| F02 | 63800001 | 63900000 | 0.0027 | 0.0017 | 0.0020 |
| F02 | 63820001 | 63920000 | 0.0030 | 0.0019 | 0.0022 |

| Chr | Start | End | both | A1 | A2 |
|---|---|---|---|---|---|
| F02 | 63840001 | 63940000 | 0.0033 | 0.0023 | 0.0026 |
| F02 | 63860001 | 63960000 | 0.0035 | 0.0026 | 0.0027 |
| F02 | 63880001 | 63980000 | 0.0036 | 0.0027 | 0.0028 |
| F02 | 63900001 | 64000000 | 0.0034 | 0.0028 | 0.0029 |
| F02 | 63920001 | 64020000 | 0.0037 | 0.0032 | 0.0035 |
| F02 | 63940001 | 64040000 | 0.0038 | 0.0032 | 0.0034 |
| F02 | 63960001 | 64060000 | 0.0031 | 0.0023 | 0.0026 |
| F02 | 63980001 | 64080000 | 0.0031 | 0.0023 | 0.0025 |
| F02 | 64000001 | 64100000 | 0.0035 | 0.0023 | 0.0026 |
| F02 | 64020001 | 64120000 | 0.0029 | 0.0018 | 0.0021 |
| F02 | 64040001 | 64140000 | 0.0021 | 0.0012 | 0.0015 |
| F02 | 64060001 | 64160000 | 0.0030 | 0.0022 | 0.0025 |
| F02 | 64080001 | 64180000 | 0.0026 | 0.0019 | 0.0021 |
| F02 | 64100001 | 64200000 | 0.0021 | 0.0019 | 0.0019 |
| F02 | 64120001 | 64220000 | 0.0020 | 0.0017 | 0.0018 |
| F02 | 64140001 | 64240000 | 0.0029 | 0.0024 | 0.0024 |
| F02 | 64160001 | 64260000 | 0.0024 | 0.0018 | 0.0018 |
| F02 | 64180001 | 64280000 | 0.0032 | 0.0023 | 0.0025 |
| F02 | 64200001 | 64300000 | 0.0037 | 0.0024 | 0.0027 |
| F02 | 64220001 | 64320000 | 0.0035 | 0.0022 | 0.0024 |
| F02 | 64240001 | 64340000 | 0.0034 | 0.0021 | 0.0023 |
| F02 | 64260001 | 64360000 | 0.0031 | 0.0016 | 0.0019 |
| F02 | 64280001 | 64380000 | 0.0029 | 0.0017 | 0.0017 |
| F02 | 64300001 | 64400000 | 0.0037 | 0.0027 | 0.0026 |
| F02 | 64320001 | 64420000 | 0.0045 | 0.0035 | 0.0034 |
| F02 | 64340001 | 64440000 | 0.0043 | 0.0033 | 0.0032 |
| F02 | 64360001 | 64460000 | 0.0043 | 0.0034 | 0.0035 |
| F02 | 64380001 | 64480000 | 0.0042 | 0.0032 | 0.0032 |
| F02 | 64400001 | 64500000 | 0.0032 | 0.0022 | 0.0023 |
| F02 | 64420001 | 64520000 | 0.0028 | 0.0018 | 0.0020 |
| F02 | 64440001 | 64540000 | 0.0024 | 0.0016 | 0.0018 |
| F02 | 64460001 | 64560000 | 0.0024 | 0.0014 | 0.0016 |
| F02 | 64480001 | 64580000 | 0.0019 | 0.0011 | 0.0013 |
| F02 | 64500001 | 64600000 | 0.0018 | 0.0010 | 0.0011 |
| F02 | 64520001 | 64620000 | 0.0035 | 0.0024 | 0.0027 |
| F02 | 64540001 | 64640000 | 0.0040 | 0.0026 | 0.0030 |
| F02 | 64560001 | 64660000 | 0.0040 | 0.0028 | 0.0032 |
| F02 | 64580001 | 64680000 | 0.0039 | 0.0028 | 0.0032 |
| F02 | 64600001 | 64700000 | 0.0041 | 0.0030 | 0.0034 |
| F02 | 64620001 | 64720000 | 0.0038 | 0.0027 | 0.0031 |
| F02 | 64640001 | 64740000 | 0.0040 | 0.0028 | 0.0034 |
| F02 | 64660001 | 64760000 | 0.0040 | 0.0026 | 0.0032 |
| F02 | 64680001 | 64780000 | 0.0043 | 0.0028 | 0.0033 |
| F02 | 64700001 | 64800000 | 0.0041 | 0.0027 | 0.0031 |
| F02 | 64720001 | 64820000 | 0.0026 | 0.0013 | 0.0015 |
| F02 | 64740001 | 64840000 | 0.0022 | 0.0011 | 0.0011 |
| F02 | 64760001 | 64860000 | 0.0029 | 0.0017 | 0.0019 |
| F02 | 64780001 | 64880000 | 0.0034 | 0.0021 | 0.0026 |
| F02 | 64800001 | 64900000 | 0.0037 | 0.0023 | 0.0028 |
| F02 | 64820001 | 64920000 | 0.0039 | 0.0027 | 0.0031 |
| F02 | 64840001 | 64940000 | 0.0045 | 0.0033 | 0.0037 |
| F02 | 64860001 | 64960000 | 0.0043 | 0.0031 | 0.0034 |
| F02 | 64880001 | 64980000 | 0.0044 | 0.0033 | 0.0034 |
| F02 | 64900001 | 65000000 | 0.0052 | 0.0038 | 0.0039 |
| F02 | 64920001 | 65020000 | 0.0050 | 0.0036 | 0.0038 |
| F02 | 64940001 | 65040000 | 0.0058 | 0.0044 | 0.0046 |
| F02 | 64960001 | 65060000 | 0.0070 | 0.0055 | 0.0060 |
| F02 | 64980001 | 65080000 | 0.0066 | 0.0053 | 0.0057 |

| Chr | Start | End | both | A1 | A2 |
|---|---|---|---|---|---|
| F02 | 65000001 | 65100000 | 0.0060 | 0.0048 | 0.0054 |
| F02 | 65020001 | 65120000 | 0.0072 | 0.0058 | 0.0066 |
| F02 | 65040001 | 65140000 | 0.0074 | 0.0059 | 0.0066 |
| F02 | 65060001 | 65160000 | 0.0060 | 0.0044 | 0.0049 |
| F02 | 65080001 | 65180000 | 0.0066 | 0.0049 | 0.0054 |
| F02 | 65100001 | 65200000 | 0.0081 | 0.0062 | 0.0066 |
| F02 | 65120001 | 65220000 | 0.0066 | 0.0049 | 0.0052 |
| F02 | 65140001 | 65240000 | 0.0064 | 0.0048 | 0.0052 |
| F02 | 65160001 | 65260000 | 0.0063 | 0.0049 | 0.0054 |
| F02 | 65180001 | 65280000 | 0.0049 | 0.0038 | 0.0042 |
| F02 | 65200001 | 65300000 | 0.0035 | 0.0025 | 0.0029 |
| F02 | 65220001 | 65320000 | 0.0032 | 0.0024 | 0.0028 |
| F02 | 65240001 | 65340000 | 0.0021 | 0.0013 | 0.0016 |
| F02 | 65260001 | 65360000 | 0.0021 | 0.0012 | 0.0014 |
| F02 | 65280001 | 65380000 | 0.0032 | 0.0021 | 0.0022 |
| F02 | 65300001 | 65400000 | 0.0030 | 0.0020 | 0.0021 |
| F02 | 65320001 | 65420000 | 0.0031 | 0.0019 | 0.0020 |
| F02 | 65340001 | 65440000 | 0.0044 | 0.0034 | 0.0036 |
| F02 | 65360001 | 65460000 | 0.0049 | 0.0039 | 0.0042 |
| F02 | 65380001 | 65480000 | 0.0044 | 0.0035 | 0.0039 |
| F02 | 65400001 | 65500000 | 0.0048 | 0.0037 | 0.0041 |
| F02 | 65420001 | 65520000 | 0.0051 | 0.0039 | 0.0044 |
| F02 | 65440001 | 65540000 | 0.0033 | 0.0023 | 0.0025 |
| F02 | 65460001 | 65560000 | 0.0028 | 0.0018 | 0.0019 |
| F02 | 65480001 | 65580000 | 0.0030 | 0.0019 | 0.0019 |
| F02 | 65500001 | 65600000 | 0.0023 | 0.0015 | 0.0014 |
| F02 | 65520001 | 65620000 | 0.0040 | 0.0032 | 0.0030 |
| F02 | 65540001 | 65640000 | 0.0038 | 0.0030 | 0.0028 |
| F02 | 65560001 | 65660000 | 0.0034 | 0.0026 | 0.0025 |
| F02 | 65580001 | 65680000 | 0.0041 | 0.0031 | 0.0034 |
| F02 | 65600001 | 65700000 | 0.0044 | 0.0034 | 0.0037 |
| F02 | 65620001 | 65720000 | 0.0025 | 0.0017 | 0.0020 |
| F02 | 65640001 | 65740000 | 0.0026 | 0.0020 | 0.0023 |
| F02 | 65660001 | 65760000 | 0.0028 | 0.0026 | 0.0025 |
| F02 | 65680001 | 65780000 | 0.0021 | 0.0027 | 0.0017 |
| F02 | 65700001 | 65800000 | 0.0024 | 0.0031 | 0.0019 |
| F02 | 65720001 | 65820000 | 0.0029 | 0.0037 | 0.0024 |
| F02 | 65740001 | 65840000 | 0.0054 | 0.0062 | 0.0048 |
| F02 | 65760001 | 65860000 | 0.0065 | 0.0073 | 0.0059 |
| F02 | 65780001 | 65880000 | 0.0061 | 0.0067 | 0.0055 |
| F02 | 65800001 | 65900000 | 0.0060 | 0.0065 | 0.0055 |
| F02 | 65820001 | 65920000 | 0.0060 | 0.0065 | 0.0055 |
| F02 | 65840001 | 65940000 | 0.0032 | 0.0037 | 0.0029 |
| F02 | 65860001 | 65960000 | 0.0023 | 0.0028 | 0.0021 |
| F02 | 65880001 | 65980000 | 0.0022 | 0.0026 | 0.0019 |
| F02 | 65900001 | 66000000 | 0.0020 | 0.0026 | 0.0017 |
| F02 | 65920001 | 66020000 | 0.0031 | 0.0035 | 0.0029 |
| F02 | 65940001 | 66040000 | 0.0047 | 0.0049 | 0.0044 |
| F02 | 65960001 | 66060000 | 0.0051 | 0.0053 | 0.0047 |
| F02 | 65980001 | 66080000 | 0.0056 | 0.0058 | 0.0053 |
| F02 | 66000001 | 66100000 | 0.0060 | 0.0059 | 0.0057 |
| F02 | 66020001 | 66120000 | 0.0051 | 0.0053 | 0.0048 |
| F02 | 66040001 | 66140000 | 0.0041 | 0.0046 | 0.0038 |
| F02 | 66060001 | 66160000 | 0.0044 | 0.0049 | 0.0041 |
| F02 | 66080001 | 66180000 | 0.0040 | 0.0047 | 0.0037 |
| F02 | 66100001 | 66200000 | 0.0035 | 0.0043 | 0.0031 |
| F02 | 66120001 | 66220000 | 0.0030 | 0.0039 | 0.0025 |
| F02 | 66140001 | 66240000 | 0.0028 | 0.0041 | 0.0022 |

| Chr | Start | End | both | A1 | A2 |
|---|---|---|---|---|---|
| F02 | 66160001 | 66260000 | 0.0035 | 0.0050 | 0.0028 |
| F02 | 66180001 | 66280000 | 0.0034 | 0.0051 | 0.0028 |
| F02 | 66200001 | 66300000 | 0.0038 | 0.0055 | 0.0031 |
| F02 | 66220001 | 66320000 | 0.0038 | 0.0052 | 0.0031 |
| F02 | 66240001 | 66340000 | 0.0041 | 0.0051 | 0.0035 |
| F02 | 66260001 | 66360000 | 0.0027 | 0.0033 | 0.0023 |
| F02 | 66280001 | 66380000 | 0.0028 | 0.0031 | 0.0025 |
| F02 | 66300001 | 66400000 | 0.0027 | 0.0030 | 0.0024 |
| F02 | 66320001 | 66420000 | 0.0028 | 0.0031 | 0.0025 |
| F02 | 66340001 | 66440000 | 0.0028 | 0.0029 | 0.0025 |
| F02 | 66360001 | 66460000 | 0.0023 | 0.0025 | 0.0020 |
| F02 | 66380001 | 66480000 | 0.0021 | 0.0023 | 0.0019 |
| F02 | 66400001 | 66500000 | 0.0020 | 0.0024 | 0.0017 |
| F02 | 66420001 | 66520000 | 0.0019 | 0.0025 | 0.0016 |
| F02 | 66440001 | 66540000 | 0.0026 | 0.0032 | 0.0023 |
| F02 | 66460001 | 66560000 | 0.0033 | 0.0039 | 0.0030 |
| F02 | 66480001 | 66580000 | 0.0032 | 0.0040 | 0.0028 |
| F02 | 66500001 | 66600000 | 0.0034 | 0.0040 | 0.0031 |
| F02 | 66520001 | 66620000 | 0.0031 | 0.0034 | 0.0029 |
| F02 | 66540001 | 66640000 | 0.0019 | 0.0022 | 0.0017 |
| F02 | 66560001 | 66660000 | 0.0018 | 0.0020 | 0.0017 |
| F02 | 66580001 | 66680000 | 0.0029 | 0.0026 | 0.0029 |
| F02 | 66600001 | 66700000 | 0.0034 | 0.0032 | 0.0033 |
| F02 | 66620001 | 66720000 | 0.0033 | 0.0034 | 0.0031 |
| F02 | 66640001 | 66740000 | 0.0039 | 0.0043 | 0.0036 |
| F02 | 66660001 | 66760000 | 0.0034 | 0.0039 | 0.0031 |
| F02 | 66680001 | 66780000 | 0.0025 | 0.0035 | 0.0020 |
| F02 | 66700001 | 66800000 | 0.0019 | 0.0027 | 0.0015 |
| F02 | 66720001 | 66820000 | 0.0029 | 0.0038 | 0.0025 |
| F02 | 66740001 | 66840000 | 0.0029 | 0.0039 | 0.0025 |
| F02 | 66760001 | 66860000 | 0.0033 | 0.0041 | 0.0029 |
| F02 | 66780001 | 66880000 | 0.0034 | 0.0041 | 0.0031 |
| F02 | 66800001 | 66900000 | 0.0033 | 0.0041 | 0.0030 |
| F02 | 66820001 | 66920000 | 0.0029 | 0.0033 | 0.0026 |
| F02 | 66840001 | 66940000 | 0.0025 | 0.0030 | 0.0022 |
| F02 | 66860001 | 66960000 | 0.0025 | 0.0030 | 0.0022 |
| F02 | 66880001 | 66980000 | 0.0024 | 0.0028 | 0.0020 |
| F02 | 66900001 | 67000000 | 0.0028 | 0.0032 | 0.0024 |
| F02 | 66920001 | 67020000 | 0.0033 | 0.0037 | 0.0030 |
| F02 | 66940001 | 67040000 | 0.0031 | 0.0035 | 0.0029 |
| F02 | 66960001 | 67060000 | 0.0032 | 0.0037 | 0.0028 |
| F02 | 66980001 | 67080000 | 0.0032 | 0.0041 | 0.0028 |
| F02 | 67000001 | 67100000 | 0.0030 | 0.0042 | 0.0024 |
| F02 | 67020001 | 67120000 | 0.0020 | 0.0036 | 0.0013 |
| F02 | 67040001 | 67140000 | 0.0020 | 0.0037 | 0.0012 |
| F02 | 67060001 | 67160000 | 0.0027 | 0.0043 | 0.0019 |
| F02 | 67080001 | 67180000 | 0.0026 | 0.0042 | 0.0018 |
| F02 | 67100001 | 67200000 | 0.0030 | 0.0044 | 0.0023 |
| F02 | 67120001 | 67220000 | 0.0033 | 0.0048 | 0.0026 |
| F02 | 67140001 | 67240000 | 0.0036 | 0.0048 | 0.0029 |
| F02 | 67160001 | 67260000 | 0.0029 | 0.0040 | 0.0022 |
| F02 | 67180001 | 67280000 | 0.0031 | 0.0040 | 0.0025 |
| F02 | 67200001 | 67300000 | 0.0028 | 0.0037 | 0.0023 |
| F02 | 67220001 | 67320000 | 0.0026 | 0.0032 | 0.0022 |
| F02 | 67240001 | 67340000 | 0.0027 | 0.0035 | 0.0022 |
| F02 | 67260001 | 67360000 | 0.0030 | 0.0039 | 0.0025 |
| F02 | 67280001 | 67380000 | 0.0031 | 0.0039 | 0.0026 |
| F02 | 67300001 | 67400000 | 0.0030 | 0.0039 | 0.0025 |

| Chr | Start | End | both | A1 | A2 |
|---|---|---|---|---|---|
| F02 | 67320001 | 67420000 | 0.0029 | 0.0038 | 0.0024 |
| F02 | 67340001 | 67440000 | 0.0024 | 0.0030 | 0.0020 |
| F02 | 67360001 | 67460000 | 0.0024 | 0.0028 | 0.0021 |
| F02 | 67380001 | 67480000 | 0.0025 | 0.0028 | 0.0022 |
| F02 | 67400001 | 67500000 | 0.0033 | 0.0035 | 0.0031 |
| F02 | 67420001 | 67520000 | 0.0039 | 0.0043 | 0.0037 |
| F02 | 67440001 | 67540000 | 0.0043 | 0.0052 | 0.0039 |
| F02 | 67460001 | 67560000 | 0.0037 | 0.0048 | 0.0032 |
| F02 | 67480001 | 67580000 | 0.0033 | 0.0047 | 0.0027 |
| F02 | 67500001 | 67600000 | 0.0021 | 0.0034 | 0.0016 |
| F02 | 67520001 | 67620000 | 0.0014 | 0.0024 | 0.0010 |
| F02 | 67540001 | 67640000 | 0.0010 | 0.0018 | 0.0007 |
| F02 | 67560001 | 67660000 | 0.0010 | 0.0016 | 0.0007 |
| F02 | 67580001 | 67680000 | 0.0015 | 0.0019 | 0.0011 |
| F02 | 67600001 | 67700000 | 0.0017 | 0.0020 | 0.0010 |
| F02 | 67620001 | 67720000 | 0.0016 | 0.0019 | 0.0009 |
| F02 | 67640001 | 67740000 | 0.0017 | 0.0018 | 0.0009 |
| F02 | 67660001 | 67760000 | 0.0018 | 0.0017 | 0.0009 |
| F02 | 67680001 | 67780000 | 0.0015 | 0.0013 | 0.0008 |
| F02 | 67700001 | 67800000 | 0.0021 | 0.0018 | 0.0015 |
| F02 | 67720001 | 67820000 | 0.0022 | 0.0018 | 0.0016 |
| F02 | 67740001 | 67840000 | 0.0028 | 0.0024 | 0.0021 |
| F02 | 67760001 | 67860000 | 0.0031 | 0.0026 | 0.0023 |
| F02 | 67780001 | 67880000 | 0.0027 | 0.0023 | 0.0020 |
| F02 | 67800001 | 67900000 | 0.0021 | 0.0017 | 0.0014 |
| F02 | 67820001 | 67920000 | 0.0021 | 0.0017 | 0.0014 |
| F02 | 67840001 | 67940000 | 0.0014 | 0.0011 | 0.0010 |
| F02 | 67860001 | 67960000 | 0.0017 | 0.0014 | 0.0011 |
| F02 | 67880001 | 67980000 | 0.0026 | 0.0022 | 0.0019 |
| F02 | 67900001 | 68000000 | 0.0027 | 0.0023 | 0.0019 |
| F02 | 67920001 | 68020000 | 0.0028 | 0.0023 | 0.0018 |
| F02 | 67940001 | 68040000 | 0.0031 | 0.0026 | 0.0021 |
| F02 | 67960001 | 68060000 | 0.0026 | 0.0021 | 0.0016 |
| F02 | 67980001 | 68080000 | 0.0021 | 0.0017 | 0.0014 |
| F02 | 68000001 | 68100000 | 0.0024 | 0.0020 | 0.0017 |
| F02 | 68020001 | 68120000 | 0.0026 | 0.0022 | 0.0020 |
| F02 | 68040001 | 68140000 | 0.0028 | 0.0025 | 0.0022 |
| F02 | 68060001 | 68160000 | 0.0032 | 0.0029 | 0.0026 |
| F02 | 68080001 | 68180000 | 0.0027 | 0.0024 | 0.0022 |
| F02 | 68100001 | 68200000 | 0.0024 | 0.0022 | 0.0020 |
| F02 | 68120001 | 68220000 | 0.0020 | 0.0019 | 0.0018 |
| F02 | 68140001 | 68240000 | 0.0012 | 0.0011 | 0.0011 |
| F02 | 68160001 | 68260000 | 0.0009 | 0.0008 | 0.0008 |
| F02 | 68180001 | 68280000 | 0.0013 | 0.0011 | 0.0010 |
| F02 | 68200001 | 68300000 | 0.0020 | 0.0018 | 0.0016 |
| F02 | 68220001 | 68320000 | 0.0021 | 0.0019 | 0.0016 |
| F02 | 68240001 | 68340000 | 0.0024 | 0.0021 | 0.0018 |
| F02 | 68260001 | 68360000 | 0.0028 | 0.0025 | 0.0022 |
| F02 | 68280001 | 68380000 | 0.0030 | 0.0028 | 0.0027 |
| F02 | 68300001 | 68400000 | 0.0030 | 0.0026 | 0.0028 |
| F02 | 68320001 | 68420000 | 0.0042 | 0.0038 | 0.0039 |
| F02 | 68340001 | 68440000 | 0.0048 | 0.0042 | 0.0046 |
| F02 | 68360001 | 68460000 | 0.0051 | 0.0044 | 0.0050 |
| F02 | 68380001 | 68480000 | 0.0052 | 0.0046 | 0.0050 |
| F02 | 68400001 | 68500000 | 0.0050 | 0.0045 | 0.0046 |
| F02 | 68420001 | 68520000 | 0.0046 | 0.0041 | 0.0042 |
| F02 | 68440001 | 68540000 | 0.0039 | 0.0038 | 0.0034 |
| F02 | 68460001 | 68560000 | 0.0033 | 0.0034 | 0.0026 |

| Chr | Start | End | both | A1 | A2 |
|---|---|---|---|---|---|
| F02 | 68480001 | 68580000 | 0.0037 | 0.0039 | 0.0030 |
| F02 | 68500001 | 68600000 | 0.0034 | 0.0035 | 0.0028 |
| F02 | 68520001 | 68620000 | 0.0027 | 0.0030 | 0.0020 |
| F02 | 68540001 | 68640000 | 0.0027 | 0.0029 | 0.0020 |
| F02 | 68560001 | 68660000 | 0.0027 | 0.0029 | 0.0020 |
| F02 | 68580001 | 68680000 | 0.0016 | 0.0017 | 0.0008 |
| F02 | 68600001 | 68700000 | 0.0020 | 0.0020 | 0.0012 |
| F02 | 68620001 | 68720000 | 0.0020 | 0.0021 | 0.0013 |
| F02 | 68640001 | 68740000 | 0.0020 | 0.0019 | 0.0013 |
| F02 | 68660001 | 68760000 | 0.0021 | 0.0019 | 0.0014 |
| F02 | 68680001 | 68780000 | 0.0028 | 0.0026 | 0.0022 |
| F02 | 68700001 | 68800000 | 0.0027 | 0.0026 | 0.0021 |
| F02 | 68720001 | 68820000 | 0.0037 | 0.0036 | 0.0030 |
| F02 | 68740001 | 68840000 | 0.0059 | 0.0057 | 0.0052 |
| F02 | 68760001 | 68860000 | 0.0060 | 0.0058 | 0.0052 |
| F02 | 68780001 | 68880000 | 0.0052 | 0.0050 | 0.0044 |
| F02 | 68800001 | 68900000 | 0.0047 | 0.0045 | 0.0039 |
| F02 | 68820001 | 68920000 | 0.0044 | 0.0040 | 0.0037 |
| F02 | 68840001 | 68940000 | 0.0027 | 0.0023 | 0.0022 |
| F02 | 68860001 | 68960000 | 0.0025 | 0.0023 | 0.0021 |
| F02 | 68880001 | 68980000 | 0.0029 | 0.0029 | 0.0024 |
| F02 | 68900001 | 69000000 | 0.0033 | 0.0034 | 0.0026 |
| F02 | 68920001 | 69020000 | 0.0025 | 0.0028 | 0.0018 |
| F02 | 68940001 | 69040000 | 0.0022 | 0.0026 | 0.0013 |
| F02 | 68960001 | 69060000 | 0.0020 | 0.0023 | 0.0012 |
| F02 | 68980001 | 69080000 | 0.0019 | 0.0021 | 0.0010 |
| F02 | 69000001 | 69100000 | 0.0017 | 0.0016 | 0.0008 |
| F02 | 69020001 | 69120000 | 0.0017 | 0.0013 | 0.0008 |
| F02 | 69040001 | 69140000 | 0.0019 | 0.0013 | 0.0010 |
| F02 | 69060001 | 69160000 | 0.0020 | 0.0012 | 0.0010 |
| F02 | 69080001 | 69180000 | 0.0023 | 0.0012 | 0.0012 |
| F02 | 69100001 | 69200000 | 0.0023 | 0.0010 | 0.0011 |
| F02 | 69120001 | 69220000 | 0.0026 | 0.0016 | 0.0015 |
| F02 | 69140001 | 69240000 | 0.0030 | 0.0020 | 0.0019 |
| F02 | 69160001 | 69260000 | 0.0030 | 0.0020 | 0.0020 |
| F02 | 69180001 | 69280000 | 0.0022 | 0.0016 | 0.0015 |
| F02 | 69200001 | 69300000 | 0.0020 | 0.0016 | 0.0015 |
| F02 | 69220001 | 69320000 | 0.0019 | 0.0013 | 0.0013 |
| F02 | 69240001 | 69340000 | 0.0019 | 0.0012 | 0.0012 |
| F02 | 69260001 | 69360000 | 0.0022 | 0.0014 | 0.0014 |
| F02 | 69280001 | 69380000 | 0.0025 | 0.0015 | 0.0014 |
| F02 | 69300001 | 69400000 | 0.0030 | 0.0018 | 0.0017 |
| F02 | 69320001 | 69420000 | 0.0031 | 0.0018 | 0.0018 |
| F02 | 69340001 | 69440000 | 0.0023 | 0.0010 | 0.0009 |
| F02 | 69360001 | 69460000 | 0.0022 | 0.0007 | 0.0007 |
| F02 | 69380001 | 69480000 | 0.0037 | 0.0022 | 0.0024 |
| F02 | 69400001 | 69500000 | 0.0036 | 0.0025 | 0.0027 |
| F02 | 69420001 | 69520000 | 0.0036 | 0.0028 | 0.0031 |
| F02 | 69440001 | 69540000 | 0.0042 | 0.0034 | 0.0039 |
| F02 | 69460001 | 69560000 | 0.0040 | 0.0035 | 0.0039 |
| F02 | 69480001 | 69580000 | 0.0026 | 0.0021 | 0.0024 |
| F02 | 69500001 | 69600000 | 0.0028 | 0.0022 | 0.0025 |
| F02 | 69520001 | 69620000 | 0.0029 | 0.0022 | 0.0024 |
| F02 | 69540001 | 69640000 | 0.0028 | 0.0019 | 0.0020 |
| F02 | 69560001 | 69660000 | 0.0031 | 0.0019 | 0.0020 |
| F02 | 69580001 | 69680000 | 0.0030 | 0.0018 | 0.0018 |
| F02 | 69600001 | 69700000 | 0.0025 | 0.0015 | 0.0014 |
| F02 | 69620001 | 69720000 | 0.0019 | 0.0009 | 0.0009 |

| Chr | Start | End | both | A1 | A2 |
|---|---|---|---|---|---|
| F02 | 69640001 | 69740000 | 0.0016 | 0.0009 | 0.0008 |
| F02 | 69660001 | 69760000 | 0.0016 | 0.0010 | 0.0008 |
| F02 | 69680001 | 69780000 | 0.0014 | 0.0009 | 0.0007 |
| F02 | 69700001 | 69800000 | 0.0015 | 0.0009 | 0.0007 |
| F02 | 69720001 | 69820000 | 0.0017 | 0.0011 | 0.0009 |
| F02 | 69740001 | 69840000 | 0.0017 | 0.0010 | 0.0010 |
| F02 | 69760001 | 69860000 | 0.0018 | 0.0013 | 0.0013 |
| F02 | 69780001 | 69880000 | 0.0026 | 0.0021 | 0.0018 |
| F02 | 69800001 | 69900000 | 0.0026 | 0.0022 | 0.0018 |
| F02 | 69820001 | 69920000 | 0.0026 | 0.0024 | 0.0018 |
| F02 | 69840001 | 69940000 | 0.0026 | 0.0025 | 0.0017 |
| F02 | 69860001 | 69960000 | 0.0030 | 0.0028 | 0.0024 |
| F02 | 69880001 | 69980000 | 0.0041 | 0.0038 | 0.0036 |
| F02 | 69900001 | 70000000 | 0.0043 | 0.0039 | 0.0037 |
| F02 | 69920001 | 70020000 | 0.0044 | 0.0041 | 0.0038 |
| F02 | 69940001 | 70040000 | 0.0048 | 0.0045 | 0.0041 |
| F02 | 69960001 | 70060000 | 0.0047 | 0.0045 | 0.0038 |
| F02 | 69980001 | 70080000 | 0.0041 | 0.0037 | 0.0034 |
| F02 | 70000001 | 70100000 | 0.0037 | 0.0033 | 0.0032 |
| F02 | 70020001 | 70120000 | 0.0041 | 0.0036 | 0.0037 |
| F02 | 70040001 | 70140000 | 0.0034 | 0.0029 | 0.0032 |
| F02 | 70060001 | 70160000 | 0.0040 | 0.0034 | 0.0038 |
| F02 | 70080001 | 70180000 | 0.0037 | 0.0032 | 0.0033 |
| F02 | 70100001 | 70200000 | 0.0042 | 0.0037 | 0.0038 |
| F02 | 70120001 | 70220000 | 0.0041 | 0.0036 | 0.0035 |
| F02 | 70140001 | 70240000 | 0.0046 | 0.0040 | 0.0037 |
| F02 | 70160001 | 70260000 | 0.0037 | 0.0032 | 0.0029 |
| F02 | 70180001 | 70280000 | 0.0030 | 0.0027 | 0.0023 |
| F02 | 70200001 | 70300000 | 0.0032 | 0.0029 | 0.0025 |
| F02 | 70220001 | 70320000 | 0.0033 | 0.0030 | 0.0027 |
| F02 | 70240001 | 70340000 | 0.0030 | 0.0027 | 0.0026 |
| F02 | 70260001 | 70360000 | 0.0032 | 0.0029 | 0.0027 |
| F02 | 70280001 | 70380000 | 0.0033 | 0.0029 | 0.0027 |
| F02 | 70300001 | 70400000 | 0.0036 | 0.0032 | 0.0030 |
| F02 | 70320001 | 70420000 | 0.0032 | 0.0029 | 0.0025 |
| F02 | 70340001 | 70440000 | 0.0037 | 0.0035 | 0.0030 |
| F02 | 70360001 | 70460000 | 0.0037 | 0.0035 | 0.0028 |
| F02 | 70380001 | 70480000 | 0.0046 | 0.0044 | 0.0040 |
| F02 | 70400001 | 70500000 | 0.0038 | 0.0036 | 0.0031 |
| F02 | 70420001 | 70520000 | 0.0036 | 0.0034 | 0.0029 |
| F02 | 70440001 | 70540000 | 0.0031 | 0.0029 | 0.0022 |
| F02 | 70460001 | 70560000 | 0.0033 | 0.0029 | 0.0026 |
| F02 | 70480001 | 70580000 | 0.0027 | 0.0024 | 0.0020 |
| F02 | 70500001 | 70600000 | 0.0026 | 0.0023 | 0.0020 |
| F02 | 70520001 | 70620000 | 0.0039 | 0.0036 | 0.0032 |
| F02 | 70540001 | 70640000 | 0.0040 | 0.0036 | 0.0032 |
| F02 | 70560001 | 70660000 | 0.0033 | 0.0031 | 0.0025 |
| F02 | 70580001 | 70680000 | 0.0028 | 0.0026 | 0.0020 |
| F02 | 70600001 | 70700000 | 0.0028 | 0.0027 | 0.0019 |
| F02 | 70620001 | 70720000 | 0.0016 | 0.0015 | 0.0008 |
| F02 | 70640001 | 70740000 | 0.0016 | 0.0015 | 0.0008 |
| F02 | 70660001 | 70760000 | 0.0018 | 0.0017 | 0.0010 |
| F02 | 70680001 | 70780000 | 0.0018 | 0.0017 | 0.0008 |
| F02 | 70700001 | 70800000 | 0.0021 | 0.0017 | 0.0011 |
| F02 | 70720001 | 70820000 | 0.0023 | 0.0017 | 0.0014 |
| F02 | 70740001 | 70840000 | 0.0023 | 0.0015 | 0.0013 |
| F02 | 70760001 | 70860000 | 0.0025 | 0.0014 | 0.0015 |
| F02 | 70780001 | 70880000 | 0.0030 | 0.0018 | 0.0022 |

| Chr | Start | End | both | A1 | A2 |
|---|---|---|---|---|---|
| F02 | 70800001 | 70900000 | 0.0032 | 0.0022 | 0.0025 |
| F02 | 70820001 | 70920000 | 0.0029 | 0.0020 | 0.0023 |
| F02 | 70840001 | 70940000 | 0.0041 | 0.0033 | 0.0035 |
| F02 | 70860001 | 70960000 | 0.0038 | 0.0031 | 0.0033 |
| F02 | 70880001 | 70980000 | 0.0035 | 0.0030 | 0.0031 |
| F02 | 70900001 | 71000000 | 0.0031 | 0.0025 | 0.0026 |
| F02 | 70920001 | 71020000 | 0.0035 | 0.0029 | 0.0028 |
| F02 | 70940001 | 71040000 | 0.0025 | 0.0020 | 0.0020 |
| F02 | 70960001 | 71060000 | 0.0023 | 0.0018 | 0.0017 |
| F02 | 70980001 | 71080000 | 0.0025 | 0.0017 | 0.0017 |
| F02 | 71000001 | 71100000 | 0.0025 | 0.0016 | 0.0015 |
| F02 | 71020001 | 71120000 | 0.0023 | 0.0013 | 0.0016 |
| F02 | 71040001 | 71140000 | 0.0021 | 0.0011 | 0.0012 |
| F02 | 71060001 | 71160000 | 0.0023 | 0.0014 | 0.0013 |
| F02 | 71080001 | 71180000 | 0.0018 | 0.0014 | 0.0009 |
| F02 | 71100001 | 71200000 | 0.0015 | 0.0017 | 0.0009 |
| F02 | 71120001 | 71220000 | 0.0012 | 0.0018 | 0.0005 |
| F02 | 71140001 | 71240000 | 0.0011 | 0.0023 | 0.0005 |
| F02 | 71160001 | 71260000 | 0.0015 | 0.0028 | 0.0010 |
| F02 | 71180001 | 71280000 | 0.0023 | 0.0033 | 0.0019 |
| F02 | 71200001 | 71300000 | 0.0027 | 0.0038 | 0.0023 |
| F02 | 71220001 | 71320000 | 0.0031 | 0.0039 | 0.0027 |
| F02 | 71240001 | 71340000 | 0.0033 | 0.0038 | 0.0030 |
| F02 | 71260001 | 71360000 | 0.0036 | 0.0038 | 0.0034 |
| F02 | 71280001 | 71380000 | 0.0026 | 0.0031 | 0.0023 |
| F02 | 71300001 | 71400000 | 0.0024 | 0.0029 | 0.0022 |
| F02 | 71320001 | 71420000 | 0.0021 | 0.0026 | 0.0018 |
| F02 | 71340001 | 71440000 | 0.0019 | 0.0029 | 0.0014 |
| F02 | 71360001 | 71460000 | 0.0011 | 0.0025 | 0.0005 |
| F02 | 71380001 | 71480000 | 0.0011 | 0.0024 | 0.0005 |
| F02 | 71400001 | 71500000 | 0.0021 | 0.0029 | 0.0016 |
| F02 | 71420001 | 71520000 | 0.0024 | 0.0032 | 0.0019 |
| F02 | 71440001 | 71540000 | 0.0023 | 0.0027 | 0.0019 |
| F02 | 71460001 | 71560000 | 0.0026 | 0.0029 | 0.0023 |
| F02 | 71480001 | 71580000 | 0.0028 | 0.0031 | 0.0024 |
| F02 | 71500001 | 71600000 | 0.0018 | 0.0023 | 0.0013 |
| F02 | 71520001 | 71620000 | 0.0024 | 0.0026 | 0.0018 |
| F02 | 71540001 | 71640000 | 0.0043 | 0.0040 | 0.0035 |
| F02 | 71560001 | 71660000 | 0.0042 | 0.0036 | 0.0034 |
| F02 | 71580001 | 71680000 | 0.0042 | 0.0034 | 0.0035 |
| F02 | 71600001 | 71700000 | 0.0044 | 0.0034 | 0.0037 |
| F02 | 71620001 | 71720000 | 0.0038 | 0.0028 | 0.0030 |
| F02 | 71640001 | 71740000 | 0.0022 | 0.0014 | 0.0014 |
| F02 | 71660001 | 71760000 | 0.0021 | 0.0013 | 0.0012 |
| F02 | 71680001 | 71780000 | 0.0020 | 0.0011 | 0.0010 |
| F02 | 71700001 | 71800000 | 0.0026 | 0.0016 | 0.0016 |
| F02 | 71720001 | 71820000 | 0.0028 | 0.0016 | 0.0016 |
| F02 | 71740001 | 71840000 | 0.0028 | 0.0015 | 0.0016 |
| F02 | 71760001 | 71860000 | 0.0036 | 0.0025 | 0.0024 |
| F02 | 71780001 | 71880000 | 0.0039 | 0.0028 | 0.0027 |
| F02 | 71800001 | 71900000 | 0.0030 | 0.0018 | 0.0018 |
| F02 | 71820001 | 71920000 | 0.0027 | 0.0018 | 0.0017 |
| F02 | 71840001 | 71940000 | 0.0030 | 0.0022 | 0.0020 |
| F02 | 71860001 | 71960000 | 0.0023 | 0.0016 | 0.0014 |
| F02 | 71880001 | 71980000 | 0.0019 | 0.0014 | 0.0012 |
| F02 | 71900001 | 72000000 | 0.0018 | 0.0016 | 0.0011 |
| F02 | 71920001 | 72020000 | 0.0020 | 0.0019 | 0.0012 |
| F02 | 71940001 | 72040000 | 0.0018 | 0.0018 | 0.0009 |

| Chr | Start | End | both | A1 | A2 |
|---|---|---|---|---|---|
| F02 | 71960001 | 72060000 | 0.0016 | 0.0015 | 0.0006 |
| F02 | 71980001 | 72080000 | 0.0017 | 0.0015 | 0.0007 |
| F02 | 72000001 | 72100000 | 0.0017 | 0.0012 | 0.0006 |
| F02 | 72020001 | 72120000 | 0.0017 | 0.0010 | 0.0006 |
| F02 | 72040001 | 72140000 | 0.0015 | 0.0005 | 0.0004 |
| F02 | 72060001 | 72160000 | 0.0015 | 0.0004 | 0.0004 |
| F02 | 72080001 | 72180000 | 0.0015 | 0.0003 | 0.0004 |
| F02 | 72100001 | 72200000 | 0.0017 | 0.0003 | 0.0005 |
| F02 | 72120001 | 72220000 | 0.0020 | 0.0003 | 0.0006 |
| F02 | 72140001 | 72240000 | 0.0019 | 0.0003 | 0.0007 |
| F02 | 72160001 | 72260000 | 0.0020 | 0.0004 | 0.0007 |
| F02 | 72180001 | 72280000 | 0.0019 | 0.0004 | 0.0007 |
| F02 | 72200001 | 72300000 | 0.0019 | 0.0007 | 0.0007 |
| F02 | 72220001 | 72320000 | 0.0016 | 0.0007 | 0.0005 |
| F02 | 72240001 | 72340000 | 0.0018 | 0.0008 | 0.0007 |
| F02 | 72260001 | 72360000 | 0.0019 | 0.0012 | 0.0009 |
| F02 | 72280001 | 72380000 | 0.0023 | 0.0016 | 0.0014 |
| F02 | 72300001 | 72400000 | 0.0023 | 0.0018 | 0.0014 |
| F02 | 72320001 | 72420000 | 0.0023 | 0.0018 | 0.0013 |
| F02 | 72340001 | 72440000 | 0.0020 | 0.0017 | 0.0011 |
| F02 | 72360001 | 72460000 | 0.0027 | 0.0022 | 0.0017 |
| F02 | 72380001 | 72480000 | 0.0026 | 0.0019 | 0.0015 |
| F02 | 72400001 | 72500000 | 0.0029 | 0.0019 | 0.0019 |
| F02 | 72420001 | 72520000 | 0.0042 | 0.0031 | 0.0033 |
| F02 | 72440001 | 72540000 | 0.0046 | 0.0036 | 0.0037 |
| F02 | 72460001 | 72560000 | 0.0040 | 0.0032 | 0.0030 |
| F02 | 72480001 | 72580000 | 0.0041 | 0.0036 | 0.0031 |
| F02 | 72500001 | 72600000 | 0.0036 | 0.0033 | 0.0026 |
| F02 | 72520001 | 72620000 | 0.0023 | 0.0023 | 0.0013 |
| F02 | 72540001 | 72640000 | 0.0020 | 0.0018 | 0.0009 |
| F02 | 72560001 | 72660000 | 0.0016 | 0.0015 | 0.0007 |
| F02 | 72580001 | 72680000 | 0.0013 | 0.0011 | 0.0003 |
| F02 | 72600001 | 72700000 | 0.0018 | 0.0016 | 0.0007 |
| F02 | 72620001 | 72720000 | 0.0019 | 0.0018 | 0.0009 |
| F02 | 72640001 | 72740000 | 0.0022 | 0.0025 | 0.0012 |
| F02 | 72660001 | 72760000 | 0.0022 | 0.0025 | 0.0013 |
| F02 | 72680001 | 72780000 | 0.0024 | 0.0028 | 0.0014 |
| F02 | 72700001 | 72800000 | 0.0020 | 0.0024 | 0.0011 |
| F02 | 72720001 | 72820000 | 0.0019 | 0.0021 | 0.0010 |
| F02 | 72740001 | 72840000 | 0.0016 | 0.0017 | 0.0007 |
| F02 | 72760001 | 72860000 | 0.0017 | 0.0019 | 0.0010 |
| F02 | 72780001 | 72880000 | 0.0017 | 0.0016 | 0.0011 |
| F02 | 72800001 | 72900000 | 0.0019 | 0.0017 | 0.0014 |
| F02 | 72820001 | 72920000 | 0.0019 | 0.0019 | 0.0015 |
| F02 | 72840001 | 72940000 | 0.0023 | 0.0019 | 0.0020 |
| F02 | 72860001 | 72960000 | 0.0024 | 0.0018 | 0.0020 |
| F02 | 72880001 | 72980000 | 0.0021 | 0.0017 | 0.0019 |
| F02 | 72900001 | 73000000 | 0.0019 | 0.0013 | 0.0016 |
| F02 | 72920001 | 73020000 | 0.0017 | 0.0012 | 0.0013 |
| F02 | 72940001 | 73040000 | 0.0014 | 0.0012 | 0.0010 |
| F02 | 72960001 | 73060000 | 0.0012 | 0.0014 | 0.0007 |
| F02 | 72980001 | 73080000 | 0.0016 | 0.0021 | 0.0011 |
| F02 | 73000001 | 73100000 | 0.0027 | 0.0032 | 0.0022 |
| F02 | 73020001 | 73120000 | 0.0028 | 0.0033 | 0.0026 |
| F02 | 73040001 | 73140000 | 0.0028 | 0.0032 | 0.0026 |
| F02 | 73060001 | 73160000 | 0.0031 | 0.0033 | 0.0029 |
| F02 | 73080001 | 73180000 | 0.0026 | 0.0028 | 0.0024 |
| F02 | 73100001 | 73200000 | 0.0014 | 0.0021 | 0.0011 |

| Chr | Start | End | both | A1 | A2 |
|---|---|---|---|---|---|
| F02 | 73120001 | 73220000 | 0.0012 | 0.0021 | 0.0008 |
| F02 | 73140001 | 73240000 | 0.0011 | 0.0019 | 0.0006 |
| F02 | 73160001 | 73260000 | 0.0013 | 0.0017 | 0.0007 |
| F02 | 73180001 | 73280000 | 0.0014 | 0.0016 | 0.0008 |
| F02 | 73200001 | 73300000 | 0.0021 | 0.0017 | 0.0014 |
| F02 | 73220001 | 73320000 | 0.0026 | 0.0016 | 0.0019 |
| F02 | 73240001 | 73340000 | 0.0027 | 0.0017 | 0.0019 |
| F02 | 73260001 | 73360000 | 0.0024 | 0.0014 | 0.0015 |
| F02 | 73280001 | 73380000 | 0.0026 | 0.0012 | 0.0015 |
| F02 | 73300001 | 73400000 | 0.0022 | 0.0007 | 0.0010 |
| F02 | 73320001 | 73420000 | 0.0017 | 0.0004 | 0.0005 |
| F02 | 73340001 | 73440000 | 0.0019 | 0.0003 | 0.0008 |
| F02 | 73360001 | 73460000 | 0.0020 | 0.0003 | 0.0009 |
| F02 | 73380001 | 73480000 | 0.0023 | 0.0006 | 0.0013 |
| F02 | 73400001 | 73500000 | 0.0027 | 0.0011 | 0.0018 |
| F02 | 73420001 | 73520000 | 0.0026 | 0.0011 | 0.0018 |
| F02 | 73440001 | 73540000 | 0.0024 | 0.0010 | 0.0016 |
| F02 | 73460001 | 73560000 | 0.0024 | 0.0013 | 0.0016 |
| F02 | 73480001 | 73580000 | 0.0019 | 0.0010 | 0.0012 |
| F02 | 73500001 | 73600000 | 0.0012 | 0.0005 | 0.0005 |
| F02 | 73520001 | 73620000 | 0.0014 | 0.0005 | 0.0006 |
| F02 | 73540001 | 73640000 | 0.0015 | 0.0006 | 0.0009 |
| F02 | 73560001 | 73660000 | 0.0016 | 0.0003 | 0.0011 |
| F02 | 73580001 | 73680000 | 0.0024 | 0.0004 | 0.0019 |
| F02 | 73600001 | 73700000 | 0.0033 | 0.0014 | 0.0029 |
| F02 | 73620001 | 73720000 | 0.0042 | 0.0016 | 0.0037 |
| F02 | 73640001 | 73740000 | 0.0051 | 0.0027 | 0.0047 |
| F02 | 73660001 | 73760000 | 0.0052 | 0.0029 | 0.0047 |
| F02 | 73680001 | 73780000 | 0.0064 | 0.0050 | 0.0059 |
| F02 | 73700001 | 73800000 | 0.0060 | 0.0046 | 0.0056 |
| F02 | 73720001 | 73820000 | 0.0052 | 0.0046 | 0.0049 |
| F02 | 73740001 | 73840000 | 0.0047 | 0.0041 | 0.0044 |
| F02 | 73760001 | 73860000 | 0.0045 | 0.0039 | 0.0041 |
| F02 | 73780001 | 73880000 | 0.0030 | 0.0020 | 0.0026 |
| F02 | 73800001 | 73900000 | 0.0026 | 0.0016 | 0.0021 |
| F02 | 73820001 | 73920000 | 0.0027 | 0.0013 | 0.0019 |
| F02 | 73840001 | 73940000 | 0.0027 | 0.0007 | 0.0014 |
| F02 | 73860001 | 73960000 | 0.0028 | 0.0009 | 0.0014 |
| F02 | 73880001 | 73980000 | 0.0031 | 0.0012 | 0.0015 |
| F02 | 73900001 | 74000000 | 0.0041 | 0.0024 | 0.0026 |
| F02 | 73920001 | 74020000 | 0.0048 | 0.0031 | 0.0033 |
| F02 | 73940001 | 74040000 | 0.0047 | 0.0034 | 0.0035 |
| F02 | 73960001 | 74060000 | 0.0044 | 0.0032 | 0.0034 |
| F02 | 73980001 | 74080000 | 0.0037 | 0.0027 | 0.0029 |
| F02 | 74000001 | 74100000 | 0.0027 | 0.0014 | 0.0019 |
| F02 | 74020001 | 74120000 | 0.0021 | 0.0008 | 0.0016 |
| F02 | 74040001 | 74140000 | 0.0021 | 0.0008 | 0.0018 |
| F02 | 74060001 | 74160000 | 0.0025 | 0.0012 | 0.0023 |
| F02 | 74080001 | 74180000 | 0.0029 | 0.0014 | 0.0027 |
| F02 | 74100001 | 74200000 | 0.0032 | 0.0015 | 0.0031 |
| F02 | 74120001 | 74220000 | 0.0034 | 0.0017 | 0.0033 |
| F02 | 74140001 | 74240000 | 0.0029 | 0.0015 | 0.0026 |
| F02 | 74160001 | 74260000 | 0.0027 | 0.0014 | 0.0024 |
| F02 | 74180001 | 74280000 | 0.0030 | 0.0013 | 0.0025 |
| F02 | 74200001 | 74300000 | 0.0026 | 0.0011 | 0.0022 |
| F02 | 74220001 | 74320000 | 0.0026 | 0.0008 | 0.0024 |
| F02 | 74240001 | 74340000 | 0.0036 | 0.0018 | 0.0035 |
| F02 | 74260001 | 74360000 | 0.0034 | 0.0017 | 0.0034 |

| Chr | Start | End | both | A1 | A2 |
|---|---|---|---|---|---|
| F02 | 74280001 | 74380000 | 0.0042 | 0.0030 | 0.0044 |
| F02 | 74300001 | 74400000 | 0.0052 | 0.0039 | 0.0053 |
| F02 | 74320001 | 74420000 | 0.0048 | 0.0041 | 0.0048 |
| F02 | 74340001 | 74440000 | 0.0046 | 0.0037 | 0.0047 |
| F02 | 74360001 | 74460000 | 0.0047 | 0.0035 | 0.0048 |
| F02 | 74380001 | 74480000 | 0.0034 | 0.0022 | 0.0035 |
| F02 | 74400001 | 74500000 | 0.0026 | 0.0015 | 0.0026 |
| F02 | 74420001 | 74520000 | 0.0030 | 0.0016 | 0.0030 |
| F02 | 74440001 | 74540000 | 0.0028 | 0.0017 | 0.0028 |
| F02 | 74460001 | 74560000 | 0.0029 | 0.0016 | 0.0028 |
| F02 | 74480001 | 74580000 | 0.0028 | 0.0016 | 0.0025 |
| F02 | 74500001 | 74600000 | 0.0029 | 0.0014 | 0.0025 |
| F02 | 74520001 | 74620000 | 0.0026 | 0.0011 | 0.0021 |
| F02 | 74540001 | 74640000 | 0.0019 | 0.0004 | 0.0013 |
| F02 | 74560001 | 74660000 | 0.0018 | 0.0003 | 0.0012 |
| F02 | 74580001 | 74680000 | 0.0020 | 0.0003 | 0.0012 |
| F02 | 74600001 | 74700000 | 0.0021 | 0.0003 | 0.0012 |
| F02 | 74620001 | 74720000 | 0.0020 | 0.0003 | 0.0011 |
| F02 | 74640001 | 74740000 | 0.0021 | 0.0003 | 0.0012 |
| F02 | 74660001 | 74760000 | 0.0022 | 0.0003 | 0.0012 |
| F02 | 74680001 | 74780000 | 0.0021 | 0.0004 | 0.0013 |
| F02 | 74700001 | 74800000 | 0.0020 | 0.0005 | 0.0013 |
| F02 | 74720001 | 74820000 | 0.0019 | 0.0004 | 0.0011 |
| F02 | 74740001 | 74840000 | 0.0020 | 0.0006 | 0.0013 |
| F02 | 74760001 | 74860000 | 0.0021 | 0.0006 | 0.0013 |
| F02 | 74780001 | 74880000 | 0.0021 | 0.0005 | 0.0012 |
| F02 | 74800001 | 74900000 | 0.0022 | 0.0004 | 0.0011 |
| F02 | 74820001 | 74920000 | 0.0024 | 0.0005 | 0.0011 |
| F02 | 74840001 | 74940000 | 0.0020 | 0.0004 | 0.0008 |
| F02 | 74860001 | 74960000 | 0.0018 | 0.0005 | 0.0006 |
| F02 | 74880001 | 74980000 | 0.0031 | 0.0014 | 0.0020 |
| F02 | 74900001 | 75000000 | 0.0053 | 0.0030 | 0.0044 |
| F02 | 74920001 | 75020000 | 0.0063 | 0.0037 | 0.0055 |
| F02 | 74940001 | 75040000 | 0.0068 | 0.0040 | 0.0060 |
| F02 | 74960001 | 75060000 | 0.0070 | 0.0042 | 0.0063 |
| F02 | 74980001 | 75080000 | 0.0057 | 0.0037 | 0.0048 |
| F02 | 75000001 | 75100000 | 0.0034 | 0.0027 | 0.0022 |
| F02 | 75020001 | 75120000 | 0.0028 | 0.0027 | 0.0016 |
| F02 | 75040001 | 75140000 | 0.0024 | 0.0027 | 0.0012 |
| F02 | 75060001 | 75160000 | 0.0022 | 0.0027 | 0.0009 |
| F02 | 75080001 | 75180000 | 0.0022 | 0.0029 | 0.0011 |
| F02 | 75100001 | 75200000 | 0.0042 | 0.0044 | 0.0035 |
| F02 | 75120001 | 75220000 | 0.0040 | 0.0045 | 0.0032 |
| F02 | 75140001 | 75240000 | 0.0041 | 0.0046 | 0.0033 |
| F02 | 75160001 | 75260000 | 0.0046 | 0.0052 | 0.0039 |
| F02 | 75180001 | 75280000 | 0.0048 | 0.0053 | 0.0042 |
| F02 | 75200001 | 75300000 | 0.0032 | 0.0037 | 0.0026 |
| F02 | 75220001 | 75320000 | 0.0032 | 0.0035 | 0.0026 |
| F02 | 75240001 | 75340000 | 0.0029 | 0.0030 | 0.0024 |
| F02 | 75260001 | 75360000 | 0.0024 | 0.0023 | 0.0020 |
| F02 | 75280001 | 75380000 | 0.0022 | 0.0019 | 0.0018 |
| F02 | 75300001 | 75400000 | 0.0017 | 0.0016 | 0.0012 |
| F02 | 75320001 | 75420000 | 0.0015 | 0.0012 | 0.0009 |
| F02 | 75340001 | 75440000 | 0.0015 | 0.0011 | 0.0009 |
| F02 | 75360001 | 75460000 | 0.0016 | 0.0010 | 0.0007 |
| F02 | 75380001 | 75480000 | 0.0016 | 0.0009 | 0.0006 |
| F02 | 75400001 | 75500000 | 0.0016 | 0.0007 | 0.0007 |
| F02 | 75420001 | 75520000 | 0.0017 | 0.0010 | 0.0008 |

| Chr | Start | End | both | A1 | A2 |
|---|---|---|---|---|---|
| F02 | 75440001 | 75540000 | 0.0019 | 0.0013 | 0.0011 |
| F02 | 75460001 | 75560000 | 0.0018 | 0.0017 | 0.0011 |
| F02 | 75480001 | 75580000 | 0.0016 | 0.0017 | 0.0009 |
| F02 | 75500001 | 75600000 | 0.0013 | 0.0016 | 0.0008 |
| F02 | 75520001 | 75620000 | 0.0013 | 0.0018 | 0.0007 |
| F02 | 75540001 | 75640000 | 0.0012 | 0.0017 | 0.0006 |
| F02 | 75560001 | 75660000 | 0.0012 | 0.0016 | 0.0006 |
| F02 | 75580001 | 75680000 | 0.0013 | 0.0015 | 0.0006 |
| F02 | 75600001 | 75700000 | 0.0026 | 0.0027 | 0.0018 |
| F02 | 75620001 | 75720000 | 0.0027 | 0.0027 | 0.0019 |
| F02 | 75640001 | 75740000 | 0.0025 | 0.0027 | 0.0018 |
| F02 | 75660001 | 75760000 | 0.0024 | 0.0027 | 0.0018 |
| F02 | 75680001 | 75780000 | 0.0024 | 0.0028 | 0.0018 |
| F02 | 75700001 | 75800000 | 0.0013 | 0.0021 | 0.0006 |
| F02 | 75720001 | 75820000 | 0.0021 | 0.0028 | 0.0016 |
| F02 | 75740001 | 75840000 | 0.0025 | 0.0030 | 0.0019 |
| F02 | 75760001 | 75860000 | 0.0028 | 0.0028 | 0.0023 |
| F02 | 75780001 | 75880000 | 0.0038 | 0.0031 | 0.0033 |
| F02 | 75800001 | 75900000 | 0.0039 | 0.0026 | 0.0034 |
| F02 | 75820001 | 75920000 | 0.0031 | 0.0018 | 0.0025 |
| F02 | 75840001 | 75940000 | 0.0028 | 0.0014 | 0.0021 |
| F02 | 75860001 | 75960000 | 0.0026 | 0.0013 | 0.0018 |
| F02 | 75880001 | 75980000 | 0.0016 | 0.0010 | 0.0009 |
| F02 | 75900001 | 76000000 | 0.0014 | 0.0013 | 0.0007 |
| F02 | 75920001 | 76020000 | 0.0011 | 0.0014 | 0.0005 |
| F02 | 75940001 | 76040000 | 0.0013 | 0.0018 | 0.0008 |
| F02 | 75960001 | 76060000 | 0.0024 | 0.0028 | 0.0018 |
| F02 | 75980001 | 76080000 | 0.0038 | 0.0039 | 0.0031 |
| F02 | 76000001 | 76100000 | 0.0039 | 0.0035 | 0.0031 |
| F02 | 76020001 | 76120000 | 0.0044 | 0.0035 | 0.0034 |
| F02 | 76040001 | 76140000 | 0.0042 | 0.0031 | 0.0032 |
| F02 | 76060001 | 76160000 | 0.0033 | 0.0022 | 0.0023 |
| F02 | 76080001 | 76180000 | 0.0025 | 0.0017 | 0.0015 |
| F02 | 76100001 | 76200000 | 0.0034 | 0.0029 | 0.0028 |
| F02 | 76120001 | 76220000 | 0.0038 | 0.0032 | 0.0034 |
| F02 | 76140001 | 76240000 | 0.0040 | 0.0035 | 0.0038 |
| F02 | 76160001 | 76260000 | 0.0036 | 0.0033 | 0.0036 |
| F02 | 76180001 | 76280000 | 0.0038 | 0.0035 | 0.0038 |
| F02 | 76200001 | 76300000 | 0.0029 | 0.0027 | 0.0029 |
| F02 | 76220001 | 76320000 | 0.0020 | 0.0019 | 0.0019 |
| F02 | 76240001 | 76340000 | 0.0015 | 0.0015 | 0.0014 |
| F02 | 76260001 | 76360000 | 0.0014 | 0.0014 | 0.0014 |
| F02 | 76280001 | 76380000 | 0.0004 | 0.0004 | 0.0004 |
| F03 | 1 | 100000 | 0.0031 | 0.0028 | 0.0030 |
| F03 | 20001 | 120000 | 0.0036 | 0.0034 | 0.0035 |
| F03 | 40001 | 140000 | 0.0033 | 0.0033 | 0.0032 |
| F03 | 60001 | 160000 | 0.0027 | 0.0029 | 0.0026 |
| F03 | 80001 | 180000 | 0.0014 | 0.0017 | 0.0013 |
| F03 | 100001 | 200000 | 0.0010 | 0.0015 | 0.0008 |
| F03 | 120001 | 220000 | 0.0006 | 0.0013 | 0.0004 |
| F03 | 140001 | 240000 | 0.0007 | 0.0014 | 0.0005 |
| F03 | 160001 | 260000 | 0.0007 | 0.0015 | 0.0005 |
| F03 | 180001 | 280000 | 0.0011 | 0.0015 | 0.0007 |
| F03 | 200001 | 300000 | 0.0014 | 0.0014 | 0.0010 |
| F03 | 220001 | 320000 | 0.0016 | 0.0010 | 0.0011 |
| F03 | 240001 | 340000 | 0.0018 | 0.0008 | 0.0013 |
| F03 | 260001 | 360000 | 0.0024 | 0.0007 | 0.0020 |
| F03 | 280001 | 380000 | 0.0024 | 0.0008 | 0.0020 |

| Chr | Start | End | both | A1 | A2 |
|---|---|---|---|---|---|
| F03 | 300001 | 400000 | 0.0023 | 0.0010 | 0.0019 |
| F03 | 320001 | 420000 | 0.0022 | 0.0014 | 0.0019 |
| F03 | 340001 | 440000 | 0.0022 | 0.0020 | 0.0017 |
| F03 | 360001 | 460000 | 0.0019 | 0.0024 | 0.0013 |
| F03 | 380001 | 480000 | 0.0024 | 0.0032 | 0.0019 |
| F03 | 400001 | 500000 | 0.0029 | 0.0039 | 0.0025 |
| F03 | 420001 | 520000 | 0.0040 | 0.0046 | 0.0037 |
| F03 | 440001 | 540000 | 0.0040 | 0.0042 | 0.0038 |
| F03 | 460001 | 560000 | 0.0042 | 0.0043 | 0.0041 |
| F03 | 480001 | 580000 | 0.0043 | 0.0041 | 0.0042 |
| F03 | 500001 | 600000 | 0.0049 | 0.0046 | 0.0049 |
| F03 | 520001 | 620000 | 0.0040 | 0.0040 | 0.0039 |
| F03 | 540001 | 640000 | 0.0041 | 0.0044 | 0.0040 |
| F03 | 560001 | 660000 | 0.0038 | 0.0043 | 0.0035 |
| F03 | 580001 | 680000 | 0.0030 | 0.0041 | 0.0026 |
| F03 | 600001 | 700000 | 0.0017 | 0.0031 | 0.0012 |
| F03 | 620001 | 720000 | 0.0020 | 0.0036 | 0.0014 |
| F03 | 640001 | 740000 | 0.0018 | 0.0034 | 0.0012 |
| F03 | 660001 | 760000 | 0.0032 | 0.0050 | 0.0027 |
| F03 | 680001 | 780000 | 0.0035 | 0.0048 | 0.0030 |
| F03 | 700001 | 800000 | 0.0048 | 0.0052 | 0.0044 |
| F03 | 720001 | 820000 | 0.0058 | 0.0052 | 0.0055 |
| F03 | 740001 | 840000 | 0.0062 | 0.0055 | 0.0060 |
| F03 | 760001 | 860000 | 0.0057 | 0.0046 | 0.0056 |
| F03 | 780001 | 880000 | 0.0079 | 0.0067 | 0.0077 |
| F03 | 800001 | 900000 | 0.0075 | 0.0068 | 0.0073 |
| F03 | 820001 | 920000 | 0.0060 | 0.0061 | 0.0058 |
| F03 | 840001 | 940000 | 0.0056 | 0.0055 | 0.0054 |
| F03 | 860001 | 960000 | 0.0044 | 0.0044 | 0.0043 |
| F03 | 880001 | 980000 | 0.0021 | 0.0021 | 0.0020 |
| F03 | 900001 | 1000000 | 0.0029 | 0.0035 | 0.0026 |
| F03 | 920001 | 1020000 | 0.0052 | 0.0063 | 0.0046 |
| F03 | 940001 | 1040000 | 0.0062 | 0.0074 | 0.0054 |
| F03 | 960001 | 1060000 | 0.0065 | 0.0077 | 0.0056 |
| F03 | 980001 | 1080000 | 0.0063 | 0.0073 | 0.0055 |
| F03 | 1000001 | 1100000 | 0.0057 | 0.0062 | 0.0051 |
| F03 | 1020001 | 1120000 | 0.0035 | 0.0035 | 0.0030 |
| F03 | 1040001 | 1140000 | 0.0025 | 0.0025 | 0.0021 |
| F03 | 1060001 | 1160000 | 0.0026 | 0.0025 | 0.0020 |
| F03 | 1080001 | 1180000 | 0.0027 | 0.0026 | 0.0020 |
| F03 | 1100001 | 1200000 | 0.0019 | 0.0017 | 0.0012 |
| F03 | 1120001 | 1220000 | 0.0034 | 0.0029 | 0.0026 |
| F03 | 1140001 | 1240000 | 0.0034 | 0.0026 | 0.0026 |
| F03 | 1160001 | 1260000 | 0.0036 | 0.0027 | 0.0028 |
| F03 | 1180001 | 1280000 | 0.0042 | 0.0034 | 0.0035 |
| F03 | 1200001 | 1300000 | 0.0039 | 0.0032 | 0.0032 |
| F03 | 1220001 | 1320000 | 0.0029 | 0.0022 | 0.0024 |
| F03 | 1240001 | 1340000 | 0.0035 | 0.0028 | 0.0029 |
| F03 | 1260001 | 1360000 | 0.0034 | 0.0028 | 0.0027 |
| F03 | 1280001 | 1380000 | 0.0031 | 0.0023 | 0.0024 |
| F03 | 1300001 | 1400000 | 0.0040 | 0.0031 | 0.0032 |
| F03 | 1320001 | 1420000 | 0.0037 | 0.0028 | 0.0026 |
| F03 | 1340001 | 1440000 | 0.0030 | 0.0022 | 0.0021 |
| F03 | 1360001 | 1460000 | 0.0034 | 0.0025 | 0.0027 |
| F03 | 1380001 | 1480000 | 0.0048 | 0.0037 | 0.0040 |
| F03 | 1400001 | 1500000 | 0.0052 | 0.0040 | 0.0040 |
| F03 | 1420001 | 1520000 | 0.0063 | 0.0050 | 0.0053 |
| F03 | 1440001 | 1540000 | 0.0085 | 0.0073 | 0.0073 |

| Chr | Start | End | both | A1 | A2 |
|---|---|---|---|---|---|
| F03 | 1460001 | 1560000 | 0.0083 | 0.0073 | 0.0070 |
| F03 | 1480001 | 1580000 | 0.0070 | 0.0064 | 0.0057 |
| F03 | 1500001 | 1600000 | 0.0058 | 0.0055 | 0.0048 |
| F03 | 1520001 | 1620000 | 0.0049 | 0.0048 | 0.0039 |
| F03 | 1540001 | 1640000 | 0.0026 | 0.0023 | 0.0017 |
| F03 | 1560001 | 1660000 | 0.0020 | 0.0017 | 0.0013 |
| F03 | 1580001 | 1680000 | 0.0017 | 0.0012 | 0.0011 |
| F03 | 1600001 | 1700000 | 0.0018 | 0.0011 | 0.0014 |
| F03 | 1620001 | 1720000 | 0.0013 | 0.0006 | 0.0009 |
| F03 | 1640001 | 1740000 | 0.0017 | 0.0008 | 0.0010 |
| F03 | 1660001 | 1760000 | 0.0023 | 0.0012 | 0.0015 |
| F03 | 1680001 | 1780000 | 0.0024 | 0.0014 | 0.0016 |
| F03 | 1700001 | 1800000 | 0.0027 | 0.0017 | 0.0017 |
| F03 | 1720001 | 1820000 | 0.0033 | 0.0020 | 0.0024 |
| F03 | 1740001 | 1840000 | 0.0033 | 0.0020 | 0.0026 |
| F03 | 1760001 | 1860000 | 0.0029 | 0.0016 | 0.0021 |
| F03 | 1780001 | 1880000 | 0.0028 | 0.0015 | 0.0020 |
| F03 | 1800001 | 1900000 | 0.0034 | 0.0020 | 0.0025 |
| F03 | 1820001 | 1920000 | 0.0038 | 0.0026 | 0.0031 |
| F03 | 1840001 | 1940000 | 0.0043 | 0.0032 | 0.0038 |
| F03 | 1860001 | 1960000 | 0.0048 | 0.0037 | 0.0045 |
| F03 | 1880001 | 1980000 | 0.0056 | 0.0045 | 0.0055 |
| F03 | 1900001 | 2000000 | 0.0060 | 0.0050 | 0.0060 |
| F03 | 1920001 | 2020000 | 0.0057 | 0.0049 | 0.0057 |
| F03 | 1940001 | 2040000 | 0.0052 | 0.0046 | 0.0052 |
| F03 | 1960001 | 2060000 | 0.0049 | 0.0044 | 0.0049 |
| F03 | 1980001 | 2080000 | 0.0046 | 0.0039 | 0.0046 |
| F03 | 2000001 | 2100000 | 0.0036 | 0.0026 | 0.0036 |
| F03 | 2020001 | 2120000 | 0.0031 | 0.0021 | 0.0030 |
| F03 | 2040001 | 2140000 | 0.0031 | 0.0021 | 0.0030 |
| F03 | 2060001 | 2160000 | 0.0039 | 0.0028 | 0.0036 |
| F03 | 2080001 | 2180000 | 0.0050 | 0.0043 | 0.0047 |
| F03 | 2100001 | 2200000 | 0.0047 | 0.0044 | 0.0043 |
| F03 | 2120001 | 2220000 | 0.0046 | 0.0045 | 0.0041 |
| F03 | 2140001 | 2240000 | 0.0045 | 0.0045 | 0.0037 |
| F03 | 2160001 | 2260000 | 0.0036 | 0.0038 | 0.0026 |
| F03 | 2180001 | 2280000 | 0.0020 | 0.0023 | 0.0009 |
| F03 | 2200001 | 2300000 | 0.0021 | 0.0022 | 0.0009 |
| F03 | 2220001 | 2320000 | 0.0021 | 0.0020 | 0.0009 |
| F03 | 2240001 | 2340000 | 0.0033 | 0.0031 | 0.0021 |
| F03 | 2260001 | 2360000 | 0.0039 | 0.0036 | 0.0030 |
| F03 | 2280001 | 2380000 | 0.0058 | 0.0051 | 0.0049 |
| F03 | 2300001 | 2400000 | 0.0076 | 0.0067 | 0.0067 |
| F03 | 2320001 | 2420000 | 0.0074 | 0.0065 | 0.0067 |
| F03 | 2340001 | 2440000 | 0.0065 | 0.0055 | 0.0060 |
| F03 | 2360001 | 2460000 | 0.0063 | 0.0052 | 0.0058 |
| F03 | 2380001 | 2480000 | 0.0048 | 0.0037 | 0.0043 |
| F03 | 2400001 | 2500000 | 0.0046 | 0.0031 | 0.0039 |
| F03 | 2420001 | 2520000 | 0.0051 | 0.0035 | 0.0042 |
| F03 | 2440001 | 2540000 | 0.0047 | 0.0034 | 0.0038 |
| F03 | 2460001 | 2560000 | 0.0042 | 0.0037 | 0.0031 |
| F03 | 2480001 | 2580000 | 0.0037 | 0.0038 | 0.0025 |
| F03 | 2500001 | 2600000 | 0.0020 | 0.0039 | 0.0008 |
| F03 | 2520001 | 2620000 | 0.0015 | 0.0038 | 0.0003 |
| F03 | 2540001 | 2640000 | 0.0014 | 0.0035 | 0.0003 |
| F03 | 2560001 | 2660000 | 0.0017 | 0.0031 | 0.0005 |
| F03 | 2580001 | 2680000 | 0.0024 | 0.0029 | 0.0012 |
| F03 | 2600001 | 2700000 | 0.0019 | 0.0015 | 0.0011 |

| Chr | Start | End | both | A1 | A2 |
|---|---|---|---|---|---|
| F03 | 2620001 | 2720000 | 0.0021 | 0.0011 | 0.0014 |
| F03 | 2640001 | 2740000 | 0.0025 | 0.0013 | 0.0017 |
| F03 | 2660001 | 2760000 | 0.0022 | 0.0011 | 0.0017 |
| F03 | 2680001 | 2780000 | 0.0019 | 0.0011 | 0.0014 |
| F03 | 2700001 | 2800000 | 0.0024 | 0.0015 | 0.0017 |
| F03 | 2720001 | 2820000 | 0.0028 | 0.0020 | 0.0021 |
| F03 | 2740001 | 2840000 | 0.0034 | 0.0027 | 0.0026 |
| F03 | 2760001 | 2860000 | 0.0038 | 0.0032 | 0.0032 |
| F03 | 2780001 | 2880000 | 0.0043 | 0.0036 | 0.0036 |
| F03 | 2800001 | 2900000 | 0.0045 | 0.0039 | 0.0041 |
| F03 | 2820001 | 2920000 | 0.0044 | 0.0036 | 0.0039 |
| F03 | 2840001 | 2940000 | 0.0035 | 0.0028 | 0.0032 |
| F03 | 2860001 | 2960000 | 0.0030 | 0.0023 | 0.0028 |
| F03 | 2880001 | 2980000 | 0.0030 | 0.0020 | 0.0026 |
| F03 | 2900001 | 3000000 | 0.0028 | 0.0014 | 0.0024 |
| F03 | 2920001 | 3020000 | 0.0026 | 0.0012 | 0.0023 |
| F03 | 2940001 | 3040000 | 0.0030 | 0.0013 | 0.0027 |
| F03 | 2960001 | 3060000 | 0.0034 | 0.0015 | 0.0033 |
| F03 | 2980001 | 3080000 | 0.0031 | 0.0012 | 0.0031 |
| F03 | 3000001 | 3100000 | 0.0029 | 0.0012 | 0.0029 |
| F03 | 3020001 | 3120000 | 0.0030 | 0.0012 | 0.0027 |
| F03 | 3040001 | 3140000 | 0.0027 | 0.0011 | 0.0021 |
| F03 | 3060001 | 3160000 | 0.0028 | 0.0014 | 0.0019 |
| F03 | 3080001 | 3180000 | 0.0035 | 0.0020 | 0.0023 |
| F03 | 3100001 | 3200000 | 0.0046 | 0.0029 | 0.0032 |
| F03 | 3120001 | 3220000 | 0.0064 | 0.0046 | 0.0050 |
| F03 | 3140001 | 3240000 | 0.0079 | 0.0059 | 0.0067 |
| F03 | 3160001 | 3260000 | 0.0080 | 0.0061 | 0.0068 |
| F03 | 3180001 | 3280000 | 0.0083 | 0.0068 | 0.0072 |
| F03 | 3200001 | 3300000 | 0.0081 | 0.0069 | 0.0071 |
| F03 | 3220001 | 3320000 | 0.0063 | 0.0053 | 0.0052 |
| F03 | 3240001 | 3340000 | 0.0049 | 0.0043 | 0.0038 |
| F03 | 3260001 | 3360000 | 0.0043 | 0.0036 | 0.0031 |
| F03 | 3280001 | 3380000 | 0.0029 | 0.0022 | 0.0019 |
| F03 | 3300001 | 3400000 | 0.0020 | 0.0015 | 0.0011 |
| F03 | 3320001 | 3420000 | 0.0022 | 0.0017 | 0.0015 |
| F03 | 3340001 | 3440000 | 0.0021 | 0.0016 | 0.0015 |
| F03 | 3360001 | 3460000 | 0.0020 | 0.0017 | 0.0016 |
| F03 | 3380001 | 3480000 | 0.0022 | 0.0018 | 0.0016 |
| F03 | 3400001 | 3500000 | 0.0019 | 0.0014 | 0.0012 |
| F03 | 3420001 | 3520000 | 0.0016 | 0.0010 | 0.0008 |
| F03 | 3440001 | 3540000 | 0.0015 | 0.0009 | 0.0005 |
| F03 | 3460001 | 3560000 | 0.0014 | 0.0007 | 0.0003 |
| F03 | 3480001 | 3580000 | 0.0014 | 0.0006 | 0.0003 |
| F03 | 3500001 | 3600000 | 0.0017 | 0.0008 | 0.0004 |
| F03 | 3520001 | 3620000 | 0.0016 | 0.0007 | 0.0003 |
| F03 | 3540001 | 3640000 | 0.0016 | 0.0007 | 0.0003 |
| F03 | 3560001 | 3660000 | 0.0016 | 0.0007 | 0.0003 |
| F03 | 3580001 | 3680000 | 0.0016 | 0.0006 | 0.0003 |
| F03 | 3600001 | 3700000 | 0.0017 | 0.0007 | 0.0005 |
| F03 | 3620001 | 3720000 | 0.0018 | 0.0007 | 0.0006 |
| F03 | 3640001 | 3740000 | 0.0016 | 0.0006 | 0.0006 |
| F03 | 3660001 | 3760000 | 0.0018 | 0.0009 | 0.0011 |
| F03 | 3680001 | 3780000 | 0.0025 | 0.0014 | 0.0019 |
| F03 | 3700001 | 3800000 | 0.0030 | 0.0021 | 0.0026 |
| F03 | 3720001 | 3820000 | 0.0031 | 0.0021 | 0.0027 |
| F03 | 3740001 | 3840000 | 0.0033 | 0.0022 | 0.0028 |
| F03 | 3760001 | 3860000 | 0.0029 | 0.0017 | 0.0024 |

| Chr | Start | End | both | A1 | A2 |
|---|---|---|---|---|---|
| F03 | 3780001 | 3880000 | 0.0021 | 0.0013 | 0.0016 |
| F03 | 3800001 | 3900000 | 0.0017 | 0.0006 | 0.0012 |
| F03 | 3820001 | 3920000 | 0.0016 | 0.0006 | 0.0010 |
| F03 | 3840001 | 3940000 | 0.0017 | 0.0005 | 0.0011 |
| F03 | 3860001 | 3960000 | 0.0018 | 0.0005 | 0.0011 |
| F03 | 3880001 | 3980000 | 0.0022 | 0.0006 | 0.0014 |
| F03 | 3900001 | 4000000 | 0.0026 | 0.0010 | 0.0019 |
| F03 | 3920001 | 4020000 | 0.0031 | 0.0011 | 0.0025 |
| F03 | 3940001 | 4040000 | 0.0033 | 0.0013 | 0.0029 |
| F03 | 3960001 | 4060000 | 0.0036 | 0.0017 | 0.0034 |
| F03 | 3980001 | 4080000 | 0.0036 | 0.0017 | 0.0035 |
| F03 | 4000001 | 4100000 | 0.0031 | 0.0010 | 0.0028 |
| F03 | 4020001 | 4120000 | 0.0027 | 0.0009 | 0.0022 |
| F03 | 4040001 | 4140000 | 0.0026 | 0.0009 | 0.0019 |
| F03 | 4060001 | 4160000 | 0.0027 | 0.0008 | 0.0016 |
| F03 | 4080001 | 4180000 | 0.0024 | 0.0006 | 0.0012 |
| F03 | 4100001 | 4200000 | 0.0022 | 0.0006 | 0.0010 |
| F03 | 4120001 | 4220000 | 0.0023 | 0.0009 | 0.0013 |
| F03 | 4140001 | 4240000 | 0.0021 | 0.0008 | 0.0012 |
| F03 | 4160001 | 4260000 | 0.0021 | 0.0010 | 0.0013 |
| F03 | 4180001 | 4280000 | 0.0025 | 0.0016 | 0.0019 |
| F03 | 4200001 | 4300000 | 0.0027 | 0.0017 | 0.0020 |
| F03 | 4220001 | 4320000 | 0.0031 | 0.0021 | 0.0024 |
| F03 | 4240001 | 4340000 | 0.0036 | 0.0024 | 0.0028 |
| F03 | 4260001 | 4360000 | 0.0030 | 0.0020 | 0.0024 |
| F03 | 4280001 | 4380000 | 0.0029 | 0.0017 | 0.0022 |
| F03 | 4300001 | 4400000 | 0.0029 | 0.0018 | 0.0023 |
| F03 | 4320001 | 4420000 | 0.0021 | 0.0011 | 0.0016 |
| F03 | 4340001 | 4440000 | 0.0018 | 0.0009 | 0.0013 |
| F03 | 4360001 | 4460000 | 0.0021 | 0.0013 | 0.0015 |
| F03 | 4380001 | 4480000 | 0.0023 | 0.0015 | 0.0017 |
| F03 | 4400001 | 4500000 | 0.0020 | 0.0013 | 0.0014 |
| F03 | 4420001 | 4520000 | 0.0024 | 0.0016 | 0.0017 |
| F03 | 4440001 | 4540000 | 0.0022 | 0.0015 | 0.0015 |
| F03 | 4460001 | 4560000 | 0.0017 | 0.0010 | 0.0011 |
| F03 | 4480001 | 4580000 | 0.0013 | 0.0006 | 0.0007 |
| F03 | 4500001 | 4600000 | 0.0019 | 0.0011 | 0.0013 |
| F03 | 4520001 | 4620000 | 0.0014 | 0.0007 | 0.0010 |
| F03 | 4540001 | 4640000 | 0.0017 | 0.0010 | 0.0012 |
| F03 | 4560001 | 4660000 | 0.0023 | 0.0016 | 0.0017 |
| F03 | 4580001 | 4680000 | 0.0028 | 0.0023 | 0.0023 |
| F03 | 4600001 | 4700000 | 0.0026 | 0.0023 | 0.0022 |
| F03 | 4620001 | 4720000 | 0.0031 | 0.0026 | 0.0026 |
| F03 | 4640001 | 4740000 | 0.0031 | 0.0029 | 0.0029 |
| F03 | 4660001 | 4760000 | 0.0042 | 0.0036 | 0.0039 |
| F03 | 4680001 | 4780000 | 0.0038 | 0.0032 | 0.0035 |
| F03 | 4700001 | 4800000 | 0.0039 | 0.0032 | 0.0036 |
| F03 | 4720001 | 4820000 | 0.0038 | 0.0030 | 0.0033 |
| F03 | 4740001 | 4840000 | 0.0036 | 0.0026 | 0.0029 |
| F03 | 4760001 | 4860000 | 0.0022 | 0.0013 | 0.0015 |
| F03 | 4780001 | 4880000 | 0.0020 | 0.0009 | 0.0011 |
| F03 | 4800001 | 4900000 | 0.0019 | 0.0007 | 0.0010 |
| F03 | 4820001 | 4920000 | 0.0016 | 0.0005 | 0.0008 |
| F03 | 4840001 | 4940000 | 0.0017 | 0.0006 | 0.0009 |
| F03 | 4860001 | 4960000 | 0.0017 | 0.0008 | 0.0010 |
| F03 | 4880001 | 4980000 | 0.0028 | 0.0019 | 0.0021 |
| F03 | 4900001 | 5000000 | 0.0023 | 0.0017 | 0.0017 |
| F03 | 4920001 | 5020000 | 0.0023 | 0.0017 | 0.0018 |

| Chr | Start | End | both | A1 | A2 |
| --- | --- | --- | --- | --- | --- |
| F03 | 4940001 | 5040000 | 0.0025 | 0.0021 | 0.0021 |
| F03 | 4960001 | 5060000 | 0.0023 | 0.0019 | 0.0019 |
| F03 | 4980001 | 5080000 | 0.0031 | 0.0027 | 0.0026 |
| F03 | 5000001 | 5100000 | 0.0036 | 0.0032 | 0.0032 |
| F03 | 5020001 | 5120000 | 0.0035 | 0.0032 | 0.0032 |
| F03 | 5040001 | 5140000 | 0.0034 | 0.0029 | 0.0030 |
| F03 | 5060001 | 5160000 | 0.0035 | 0.0028 | 0.0029 |
| F03 | 5080001 | 5180000 | 0.0016 | 0.0009 | 0.0010 |
| F03 | 5100001 | 5200000 | 0.0012 | 0.0005 | 0.0006 |
| F03 | 5120001 | 5220000 | 0.0014 | 0.0006 | 0.0007 |
| F03 | 5140001 | 5240000 | 0.0022 | 0.0014 | 0.0014 |
| F03 | 5160001 | 5260000 | 0.0022 | 0.0014 | 0.0014 |
| F03 | 5180001 | 5280000 | 0.0023 | 0.0014 | 0.0015 |
| F03 | 5200001 | 5300000 | 0.0022 | 0.0012 | 0.0013 |
| F03 | 5220001 | 5320000 | 0.0026 | 0.0014 | 0.0015 |
| F03 | 5240001 | 5340000 | 0.0020 | 0.0010 | 0.0010 |
| F03 | 5260001 | 5360000 | 0.0026 | 0.0015 | 0.0016 |
| F03 | 5280001 | 5380000 | 0.0038 | 0.0030 | 0.0028 |
| F03 | 5300001 | 5400000 | 0.0048 | 0.0041 | 0.0039 |
| F03 | 5320001 | 5420000 | 0.0044 | 0.0039 | 0.0038 |
| F03 | 5340001 | 5440000 | 0.0049 | 0.0044 | 0.0044 |
| F03 | 5360001 | 5460000 | 0.0072 | 0.0074 | 0.0066 |
| F03 | 5380001 | 5480000 | 0.0060 | 0.0061 | 0.0055 |
| F03 | 5400001 | 5500000 | 0.0060 | 0.0062 | 0.0055 |
| F03 | 5420001 | 5520000 | 0.0066 | 0.0068 | 0.0061 |
| F03 | 5440001 | 5540000 | 0.0059 | 0.0061 | 0.0054 |
| F03 | 5460001 | 5560000 | 0.0035 | 0.0030 | 0.0031 |
| F03 | 5480001 | 5580000 | 0.0038 | 0.0032 | 0.0032 |
| F03 | 5500001 | 5600000 | 0.0031 | 0.0021 | 0.0024 |
| F03 | 5520001 | 5620000 | 0.0025 | 0.0013 | 0.0017 |
| F03 | 5540001 | 5640000 | 0.0027 | 0.0015 | 0.0018 |
| F03 | 5560001 | 5660000 | 0.0026 | 0.0013 | 0.0018 |
| F03 | 5580001 | 5680000 | 0.0024 | 0.0010 | 0.0017 |
| F03 | 5600001 | 5700000 | 0.0020 | 0.0009 | 0.0015 |
| F03 | 5620001 | 5720000 | 0.0021 | 0.0010 | 0.0016 |
| F03 | 5640001 | 5740000 | 0.0020 | 0.0008 | 0.0015 |
| F03 | 5660001 | 5760000 | 0.0017 | 0.0005 | 0.0011 |
| F03 | 5680001 | 5780000 | 0.0018 | 0.0005 | 0.0011 |
| F03 | 5700001 | 5800000 | 0.0019 | 0.0005 | 0.0011 |
| F03 | 5720001 | 5820000 | 0.0018 | 0.0004 | 0.0009 |
| F03 | 5740001 | 5840000 | 0.0017 | 0.0004 | 0.0008 |
| F03 | 5760001 | 5860000 | 0.0018 | 0.0006 | 0.0010 |
| F03 | 5780001 | 5880000 | 0.0019 | 0.0010 | 0.0012 |
| F03 | 5800001 | 5900000 | 0.0019 | 0.0012 | 0.0014 |
| F03 | 5820001 | 5920000 | 0.0019 | 0.0012 | 0.0014 |
| F03 | 5840001 | 5940000 | 0.0021 | 0.0013 | 0.0017 |
| F03 | 5860001 | 5960000 | 0.0018 | 0.0012 | 0.0015 |
| F03 | 5880001 | 5980000 | 0.0025 | 0.0017 | 0.0021 |
| F03 | 5900001 | 6000000 | 0.0026 | 0.0016 | 0.0021 |
| F03 | 5920001 | 6020000 | 0.0044 | 0.0030 | 0.0039 |
| F03 | 5940001 | 6040000 | 0.0048 | 0.0034 | 0.0043 |
| F03 | 5960001 | 6060000 | 0.0060 | 0.0044 | 0.0054 |
| F03 | 5980001 | 6080000 | 0.0053 | 0.0035 | 0.0047 |
| F03 | 6000001 | 6100000 | 0.0053 | 0.0034 | 0.0048 |
| F03 | 6020001 | 6120000 | 0.0044 | 0.0025 | 0.0041 |
| F03 | 6040001 | 6140000 | 0.0052 | 0.0030 | 0.0047 |
| F03 | 6060001 | 6160000 | 0.0041 | 0.0021 | 0.0039 |
| F03 | 6080001 | 6180000 | 0.0039 | 0.0022 | 0.0036 |

| Chr | Start | End | both | A1 | A2 |
|---|---|---|---|---|---|
| F03 | 6100001 | 6200000 | 0.0036 | 0.0024 | 0.0034 |
| F03 | 6120001 | 6220000 | 0.0026 | 0.0019 | 0.0023 |
| F03 | 6140001 | 6240000 | 0.0009 | 0.0007 | 0.0009 |
| F03 | 6160001 | 6260000 | 0.0011 | 0.0009 | 0.0010 |
| F03 | 6180001 | 6280000 | 0.0010 | 0.0008 | 0.0010 |
| F03 | 6200001 | 6300000 | 0.0007 | 0.0006 | 0.0007 |
| F03 | 6220001 | 6320000 | 0.0007 | 0.0006 | 0.0007 |
| F03 | 6240001 | 6340000 | 0.0010 | 0.0009 | 0.0010 |
| F03 | 6260001 | 6360000 | 0.0009 | 0.0008 | 0.0008 |
| F03 | 6280001 | 6380000 | 0.0009 | 0.0009 | 0.0009 |
| F03 | 6300001 | 6400000 | 0.0010 | 0.0009 | 0.0009 |
| F03 | 6320001 | 6420000 | 0.0012 | 0.0011 | 0.0012 |
| F03 | 6340001 | 6440000 | 0.0026 | 0.0024 | 0.0026 |
| F03 | 6360001 | 6460000 | 0.0025 | 0.0023 | 0.0025 |
| F03 | 6380001 | 6480000 | 0.0024 | 0.0022 | 0.0023 |
| F03 | 6400001 | 6500000 | 0.0025 | 0.0023 | 0.0024 |
| F03 | 6420001 | 6520000 | 0.0028 | 0.0025 | 0.0027 |
| F03 | 6440001 | 6540000 | 0.0015 | 0.0012 | 0.0013 |
| F03 | 6460001 | 6560000 | 0.0015 | 0.0010 | 0.0012 |
| F03 | 6480001 | 6580000 | 0.0020 | 0.0014 | 0.0016 |
| F03 | 6500001 | 6600000 | 0.0021 | 0.0013 | 0.0015 |
| F03 | 6520001 | 6620000 | 0.0021 | 0.0012 | 0.0016 |
| F03 | 6540001 | 6640000 | 0.0021 | 0.0013 | 0.0017 |
| F03 | 6560001 | 6660000 | 0.0024 | 0.0013 | 0.0018 |
| F03 | 6580001 | 6680000 | 0.0021 | 0.0010 | 0.0015 |
| F03 | 6600001 | 6700000 | 0.0025 | 0.0011 | 0.0016 |
| F03 | 6620001 | 6720000 | 0.0022 | 0.0006 | 0.0011 |
| F03 | 6640001 | 6740000 | 0.0023 | 0.0005 | 0.0010 |
| F03 | 6660001 | 6760000 | 0.0023 | 0.0010 | 0.0013 |
| F03 | 6680001 | 6780000 | 0.0024 | 0.0012 | 0.0015 |
| F03 | 6700001 | 6800000 | 0.0022 | 0.0012 | 0.0015 |
| F03 | 6720001 | 6820000 | 0.0022 | 0.0013 | 0.0016 |
| F03 | 6740001 | 6840000 | 0.0021 | 0.0012 | 0.0014 |
| F03 | 6760001 | 6860000 | 0.0023 | 0.0014 | 0.0015 |
| F03 | 6780001 | 6880000 | 0.0023 | 0.0014 | 0.0016 |
| F03 | 6800001 | 6900000 | 0.0025 | 0.0016 | 0.0019 |
| F03 | 6820001 | 6920000 | 0.0031 | 0.0022 | 0.0025 |
| F03 | 6840001 | 6940000 | 0.0038 | 0.0025 | 0.0035 |
| F03 | 6860001 | 6960000 | 0.0037 | 0.0023 | 0.0035 |
| F03 | 6880001 | 6980000 | 0.0036 | 0.0022 | 0.0033 |
| F03 | 6900001 | 7000000 | 0.0031 | 0.0020 | 0.0030 |
| F03 | 6920001 | 7020000 | 0.0026 | 0.0016 | 0.0025 |
| F03 | 6940001 | 7040000 | 0.0018 | 0.0012 | 0.0014 |
| F03 | 6960001 | 7060000 | 0.0014 | 0.0008 | 0.0010 |
| F03 | 6980001 | 7080000 | 0.0016 | 0.0009 | 0.0010 |
| F03 | 7000001 | 7100000 | 0.0017 | 0.0009 | 0.0010 |
| F03 | 7020001 | 7120000 | 0.0020 | 0.0009 | 0.0010 |
| F03 | 7040001 | 7140000 | 0.0019 | 0.0009 | 0.0010 |
| F03 | 7060001 | 7160000 | 0.0022 | 0.0013 | 0.0013 |
| F03 | 7080001 | 7180000 | 0.0054 | 0.0046 | 0.0047 |
| F03 | 7100001 | 7200000 | 0.0064 | 0.0057 | 0.0058 |
| F03 | 7120001 | 7220000 | 0.0063 | 0.0058 | 0.0059 |
| F03 | 7140001 | 7240000 | 0.0061 | 0.0057 | 0.0057 |
| F03 | 7160001 | 7260000 | 0.0058 | 0.0053 | 0.0054 |
| F03 | 7180001 | 7280000 | 0.0023 | 0.0016 | 0.0021 |
| F03 | 7200001 | 7300000 | 0.0023 | 0.0015 | 0.0019 |
| F03 | 7220001 | 7320000 | 0.0023 | 0.0015 | 0.0020 |
| F03 | 7240001 | 7340000 | 0.0027 | 0.0018 | 0.0026 |

| Chr | Start | End | both | A1 | A2 |
|---|---|---|---|---|---|
| F03 | 7260001 | 7360000 | 0.0031 | 0.0018 | 0.0029 |
| F03 | 7280001 | 7380000 | 0.0034 | 0.0023 | 0.0032 |
| F03 | 7300001 | 7400000 | 0.0031 | 0.0020 | 0.0030 |
| F03 | 7320001 | 7420000 | 0.0034 | 0.0023 | 0.0033 |
| F03 | 7340001 | 7440000 | 0.0041 | 0.0030 | 0.0041 |
| F03 | 7360001 | 7460000 | 0.0042 | 0.0031 | 0.0041 |
| F03 | 7380001 | 7480000 | 0.0042 | 0.0031 | 0.0042 |
| F03 | 7400001 | 7500000 | 0.0038 | 0.0027 | 0.0038 |
| F03 | 7420001 | 7520000 | 0.0042 | 0.0031 | 0.0042 |
| F03 | 7440001 | 7540000 | 0.0030 | 0.0021 | 0.0030 |
| F03 | 7460001 | 7560000 | 0.0030 | 0.0021 | 0.0030 |
| F03 | 7480001 | 7580000 | 0.0025 | 0.0018 | 0.0025 |
| F03 | 7500001 | 7600000 | 0.0024 | 0.0017 | 0.0024 |
| F03 | 7520001 | 7620000 | 0.0018 | 0.0009 | 0.0017 |
| F03 | 7540001 | 7640000 | 0.0021 | 0.0011 | 0.0019 |
| F03 | 7560001 | 7660000 | 0.0025 | 0.0012 | 0.0018 |
| F03 | 7580001 | 7680000 | 0.0031 | 0.0018 | 0.0021 |
| F03 | 7600001 | 7700000 | 0.0028 | 0.0016 | 0.0017 |
| F03 | 7620001 | 7720000 | 0.0025 | 0.0016 | 0.0013 |
| F03 | 7640001 | 7740000 | 0.0026 | 0.0016 | 0.0015 |
| F03 | 7660001 | 7760000 | 0.0026 | 0.0018 | 0.0017 |
| F03 | 7680001 | 7780000 | 0.0030 | 0.0019 | 0.0022 |
| F03 | 7700001 | 7800000 | 0.0036 | 0.0024 | 0.0027 |
| F03 | 7720001 | 7820000 | 0.0036 | 0.0025 | 0.0027 |
| F03 | 7740001 | 7840000 | 0.0033 | 0.0022 | 0.0024 |
| F03 | 7760001 | 7860000 | 0.0028 | 0.0018 | 0.0020 |
| F03 | 7780001 | 7880000 | 0.0020 | 0.0013 | 0.0014 |
| F03 | 7800001 | 7900000 | 0.0017 | 0.0011 | 0.0012 |
| F03 | 7820001 | 7920000 | 0.0015 | 0.0009 | 0.0011 |
| F03 | 7840001 | 7940000 | 0.0014 | 0.0007 | 0.0009 |
| F03 | 7860001 | 7960000 | 0.0016 | 0.0008 | 0.0010 |
| F03 | 7880001 | 7980000 | 0.0015 | 0.0009 | 0.0008 |
| F03 | 7900001 | 8000000 | 0.0014 | 0.0007 | 0.0006 |
| F03 | 7920001 | 8020000 | 0.0031 | 0.0020 | 0.0022 |
| F03 | 7940001 | 8040000 | 0.0038 | 0.0026 | 0.0028 |
| F03 | 7960001 | 8060000 | 0.0038 | 0.0026 | 0.0028 |
| F03 | 7980001 | 8080000 | 0.0038 | 0.0025 | 0.0027 |
| F03 | 8000001 | 8100000 | 0.0036 | 0.0025 | 0.0027 |
| F03 | 8020001 | 8120000 | 0.0022 | 0.0011 | 0.0012 |
| F03 | 8040001 | 8140000 | 0.0015 | 0.0006 | 0.0006 |
| F03 | 8060001 | 8160000 | 0.0014 | 0.0005 | 0.0006 |
| F03 | 8080001 | 8180000 | 0.0013 | 0.0004 | 0.0005 |
| F03 | 8100001 | 8200000 | 0.0015 | 0.0003 | 0.0005 |
| F03 | 8120001 | 8220000 | 0.0017 | 0.0003 | 0.0005 |
| F03 | 8140001 | 8240000 | 0.0017 | 0.0004 | 0.0006 |
| F03 | 8160001 | 8260000 | 0.0020 | 0.0006 | 0.0009 |
| F03 | 8180001 | 8280000 | 0.0021 | 0.0006 | 0.0009 |
| F03 | 8200001 | 8300000 | 0.0018 | 0.0005 | 0.0008 |
| F03 | 8220001 | 8320000 | 0.0016 | 0.0006 | 0.0008 |
| F03 | 8240001 | 8340000 | 0.0020 | 0.0010 | 0.0013 |
| F03 | 8260001 | 8360000 | 0.0019 | 0.0011 | 0.0014 |
| F03 | 8280001 | 8380000 | 0.0019 | 0.0011 | 0.0014 |
| F03 | 8300001 | 8400000 | 0.0022 | 0.0015 | 0.0018 |
| F03 | 8320001 | 8420000 | 0.0022 | 0.0014 | 0.0016 |
| F03 | 8340001 | 8440000 | 0.0023 | 0.0014 | 0.0017 |
| F03 | 8360001 | 8460000 | 0.0018 | 0.0010 | 0.0012 |
| F03 | 8380001 | 8480000 | 0.0017 | 0.0009 | 0.0011 |
| F03 | 8400001 | 8500000 | 0.0014 | 0.0007 | 0.0009 |

| Chr | Start | End | both | A1 | A2 |
|---|---|---|---|---|---|
| F03 | 8420001 | 8520000 | 0.0016 | 0.0010 | 0.0012 |
| F03 | 8440001 | 8540000 | 0.0016 | 0.0011 | 0.0012 |
| F03 | 8460001 | 8560000 | 0.0023 | 0.0016 | 0.0019 |
| F03 | 8480001 | 8580000 | 0.0024 | 0.0017 | 0.0019 |
| F03 | 8500001 | 8600000 | 0.0031 | 0.0022 | 0.0026 |
| F03 | 8520001 | 8620000 | 0.0041 | 0.0029 | 0.0034 |
| F03 | 8540001 | 8640000 | 0.0036 | 0.0026 | 0.0030 |
| F03 | 8560001 | 8660000 | 0.0034 | 0.0021 | 0.0025 |
| F03 | 8580001 | 8680000 | 0.0039 | 0.0026 | 0.0029 |
| F03 | 8600001 | 8700000 | 0.0041 | 0.0026 | 0.0030 |
| F03 | 8620001 | 8720000 | 0.0029 | 0.0016 | 0.0018 |
| F03 | 8640001 | 8740000 | 0.0031 | 0.0015 | 0.0018 |
| F03 | 8660001 | 8760000 | 0.0037 | 0.0022 | 0.0024 |
| F03 | 8680001 | 8780000 | 0.0033 | 0.0019 | 0.0021 |
| F03 | 8700001 | 8800000 | 0.0027 | 0.0015 | 0.0016 |
| F03 | 8720001 | 8820000 | 0.0033 | 0.0022 | 0.0022 |
| F03 | 8740001 | 8840000 | 0.0043 | 0.0035 | 0.0032 |
| F03 | 8760001 | 8860000 | 0.0038 | 0.0031 | 0.0029 |
| F03 | 8780001 | 8880000 | 0.0038 | 0.0032 | 0.0030 |
| F03 | 8800001 | 8900000 | 0.0036 | 0.0029 | 0.0028 |
| F03 | 8820001 | 8920000 | 0.0033 | 0.0023 | 0.0024 |
| F03 | 8840001 | 8940000 | 0.0024 | 0.0011 | 0.0014 |
| F03 | 8860001 | 8960000 | 0.0026 | 0.0012 | 0.0015 |
| F03 | 8880001 | 8980000 | 0.0025 | 0.0012 | 0.0015 |
| F03 | 8900001 | 9000000 | 0.0032 | 0.0019 | 0.0021 |
| F03 | 8920001 | 9020000 | 0.0031 | 0.0022 | 0.0023 |
| F03 | 8940001 | 9040000 | 0.0031 | 0.0024 | 0.0026 |
| F03 | 8960001 | 9060000 | 0.0030 | 0.0024 | 0.0025 |
| F03 | 8980001 | 9080000 | 0.0031 | 0.0024 | 0.0025 |
| F03 | 9000001 | 9100000 | 0.0030 | 0.0024 | 0.0025 |
| F03 | 9020001 | 9120000 | 0.0029 | 0.0023 | 0.0025 |
| F03 | 9040001 | 9140000 | 0.0026 | 0.0020 | 0.0022 |
| F03 | 9060001 | 9160000 | 0.0024 | 0.0018 | 0.0021 |
| F03 | 9080001 | 9180000 | 0.0028 | 0.0020 | 0.0022 |
| F03 | 9100001 | 9200000 | 0.0035 | 0.0024 | 0.0028 |
| F03 | 9120001 | 9220000 | 0.0034 | 0.0023 | 0.0025 |
| F03 | 9140001 | 9240000 | 0.0042 | 0.0025 | 0.0032 |
| F03 | 9160001 | 9260000 | 0.0040 | 0.0022 | 0.0028 |
| F03 | 9180001 | 9280000 | 0.0034 | 0.0018 | 0.0024 |
| F03 | 9200001 | 9300000 | 0.0020 | 0.0008 | 0.0013 |
| F03 | 9220001 | 9320000 | 0.0018 | 0.0008 | 0.0014 |
| F03 | 9240001 | 9340000 | 0.0012 | 0.0006 | 0.0006 |
| F03 | 9260001 | 9360000 | 0.0017 | 0.0011 | 0.0013 |
| F03 | 9280001 | 9380000 | 0.0020 | 0.0014 | 0.0017 |
| F03 | 9300001 | 9400000 | 0.0022 | 0.0015 | 0.0018 |
| F03 | 9320001 | 9420000 | 0.0027 | 0.0018 | 0.0022 |
| F03 | 9340001 | 9440000 | 0.0032 | 0.0024 | 0.0028 |
| F03 | 9360001 | 9460000 | 0.0028 | 0.0021 | 0.0025 |
| F03 | 9380001 | 9480000 | 0.0024 | 0.0017 | 0.0020 |
| F03 | 9400001 | 9500000 | 0.0021 | 0.0015 | 0.0018 |
| F03 | 9420001 | 9520000 | 0.0015 | 0.0010 | 0.0012 |
| F03 | 9440001 | 9540000 | 0.0008 | 0.0004 | 0.0005 |
| F03 | 9460001 | 9560000 | 0.0005 | 0.0003 | 0.0003 |
| F03 | 9480001 | 9580000 | 0.0012 | 0.0009 | 0.0010 |
| F03 | 9500001 | 9600000 | 0.0024 | 0.0022 | 0.0022 |
| F03 | 9520001 | 9620000 | 0.0029 | 0.0024 | 0.0025 |
| F03 | 9540001 | 9640000 | 0.0035 | 0.0027 | 0.0028 |
| F03 | 9560001 | 9660000 | 0.0038 | 0.0027 | 0.0028 |

| Chr | Start | End | both | A1 | A2 |
|---|---|---|---|---|---|
| F03 | 9580001 | 9680000 | 0.0033 | 0.0022 | 0.0022 |
| F03 | 9600001 | 9700000 | 0.0026 | 0.0013 | 0.0015 |
| F03 | 9620001 | 9720000 | 0.0022 | 0.0011 | 0.0012 |
| F03 | 9640001 | 9740000 | 0.0026 | 0.0013 | 0.0014 |
| F03 | 9660001 | 9760000 | 0.0028 | 0.0016 | 0.0015 |
| F03 | 9680001 | 9780000 | 0.0029 | 0.0018 | 0.0018 |
| F03 | 9700001 | 9800000 | 0.0025 | 0.0015 | 0.0016 |
| F03 | 9720001 | 9820000 | 0.0024 | 0.0015 | 0.0016 |
| F03 | 9740001 | 9840000 | 0.0021 | 0.0012 | 0.0014 |
| F03 | 9760001 | 9860000 | 0.0021 | 0.0011 | 0.0015 |
| F03 | 9780001 | 9880000 | 0.0027 | 0.0015 | 0.0018 |
| F03 | 9800001 | 9900000 | 0.0026 | 0.0014 | 0.0017 |
| F03 | 9820001 | 9920000 | 0.0029 | 0.0017 | 0.0021 |
| F03 | 9840001 | 9940000 | 0.0026 | 0.0014 | 0.0018 |
| F03 | 9860001 | 9960000 | 0.0026 | 0.0015 | 0.0019 |
| F03 | 9880001 | 9980000 | 0.0029 | 0.0018 | 0.0024 |
| F03 | 9900001 | 10000000 | 0.0034 | 0.0023 | 0.0029 |
| F03 | 9920001 | 10020000 | 0.0032 | 0.0020 | 0.0026 |
| F03 | 9940001 | 10040000 | 0.0030 | 0.0020 | 0.0025 |
| F03 | 9960001 | 10060000 | 0.0026 | 0.0021 | 0.0024 |
| F03 | 9980001 | 10080000 | 0.0021 | 0.0015 | 0.0017 |
| F03 | 10000001 | 10100000 | 0.0026 | 0.0018 | 0.0020 |
| F03 | 10020001 | 10120000 | 0.0025 | 0.0017 | 0.0019 |
| F03 | 10040001 | 10140000 | 0.0028 | 0.0018 | 0.0020 |
| F03 | 10060001 | 10160000 | 0.0028 | 0.0015 | 0.0018 |
| F03 | 10080001 | 10180000 | 0.0027 | 0.0016 | 0.0019 |
| F03 | 10100001 | 10200000 | 0.0021 | 0.0012 | 0.0013 |
| F03 | 10120001 | 10220000 | 0.0022 | 0.0013 | 0.0014 |
| F03 | 10140001 | 10240000 | 0.0020 | 0.0013 | 0.0014 |
| F03 | 10160001 | 10260000 | 0.0024 | 0.0019 | 0.0019 |
| F03 | 10180001 | 10280000 | 0.0022 | 0.0016 | 0.0017 |
| F03 | 10200001 | 10300000 | 0.0022 | 0.0016 | 0.0017 |
| F03 | 10220001 | 10320000 | 0.0021 | 0.0014 | 0.0015 |
| F03 | 10240001 | 10340000 | 0.0022 | 0.0015 | 0.0017 |
| F03 | 10260001 | 10360000 | 0.0020 | 0.0011 | 0.0013 |
| F03 | 10280001 | 10380000 | 0.0025 | 0.0015 | 0.0019 |
| F03 | 10300001 | 10400000 | 0.0025 | 0.0014 | 0.0018 |
| F03 | 10320001 | 10420000 | 0.0030 | 0.0018 | 0.0023 |
| F03 | 10340001 | 10440000 | 0.0037 | 0.0024 | 0.0028 |
| F03 | 10360001 | 10460000 | 0.0040 | 0.0030 | 0.0032 |
| F03 | 10380001 | 10480000 | 0.0040 | 0.0028 | 0.0029 |
| F03 | 10400001 | 10500000 | 0.0043 | 0.0028 | 0.0031 |
| F03 | 10420001 | 10520000 | 0.0050 | 0.0035 | 0.0038 |
| F03 | 10440001 | 10540000 | 0.0048 | 0.0032 | 0.0035 |
| F03 | 10460001 | 10560000 | 0.0050 | 0.0031 | 0.0036 |
| F03 | 10480001 | 10580000 | 0.0047 | 0.0029 | 0.0032 |
| F03 | 10500001 | 10600000 | 0.0051 | 0.0038 | 0.0037 |
| F03 | 10520001 | 10620000 | 0.0045 | 0.0031 | 0.0031 |
| F03 | 10540001 | 10640000 | 0.0042 | 0.0030 | 0.0030 |
| F03 | 10560001 | 10660000 | 0.0037 | 0.0026 | 0.0025 |
| F03 | 10580001 | 10680000 | 0.0035 | 0.0024 | 0.0024 |
| F03 | 10600001 | 10700000 | 0.0027 | 0.0013 | 0.0015 |
| F03 | 10620001 | 10720000 | 0.0023 | 0.0010 | 0.0012 |
| F03 | 10640001 | 10740000 | 0.0024 | 0.0010 | 0.0013 |
| F03 | 10660001 | 10760000 | 0.0022 | 0.0009 | 0.0012 |
| F03 | 10680001 | 10780000 | 0.0024 | 0.0012 | 0.0014 |
| F03 | 10700001 | 10800000 | 0.0026 | 0.0014 | 0.0015 |
| F03 | 10720001 | 10820000 | 0.0023 | 0.0012 | 0.0013 |

| Chr | Start | End | both | A1 | A2 |
|---|---|---|---|---|---|
| F03 | 10740001 | 10840000 | 0.0019 | 0.0008 | 0.0009 |
| F03 | 10760001 | 10860000 | 0.0020 | 0.0010 | 0.0011 |
| F03 | 10780001 | 10880000 | 0.0022 | 0.0013 | 0.0015 |
| F03 | 10800001 | 10900000 | 0.0024 | 0.0015 | 0.0017 |
| F03 | 10820001 | 10920000 | 0.0035 | 0.0025 | 0.0027 |
| F03 | 10840001 | 10940000 | 0.0042 | 0.0029 | 0.0032 |
| F03 | 10860001 | 10960000 | 0.0040 | 0.0028 | 0.0030 |
| F03 | 10880001 | 10980000 | 0.0038 | 0.0023 | 0.0027 |
| F03 | 10900001 | 11000000 | 0.0034 | 0.0020 | 0.0025 |
| F03 | 10920001 | 11020000 | 0.0029 | 0.0014 | 0.0022 |
| F03 | 10940001 | 11040000 | 0.0035 | 0.0021 | 0.0031 |
| F03 | 10960001 | 11060000 | 0.0036 | 0.0020 | 0.0033 |
| F03 | 10980001 | 11080000 | 0.0031 | 0.0019 | 0.0030 |
| F03 | 11000001 | 11100000 | 0.0035 | 0.0022 | 0.0032 |
| F03 | 11020001 | 11120000 | 0.0031 | 0.0019 | 0.0027 |
| F03 | 11040001 | 11140000 | 0.0021 | 0.0010 | 0.0017 |
| F03 | 11060001 | 11160000 | 0.0022 | 0.0013 | 0.0016 |
| F03 | 11080001 | 11180000 | 0.0026 | 0.0016 | 0.0020 |
| F03 | 11100001 | 11200000 | 0.0031 | 0.0020 | 0.0025 |
| F03 | 11120001 | 11220000 | 0.0038 | 0.0025 | 0.0029 |
| F03 | 11140001 | 11240000 | 0.0036 | 0.0023 | 0.0025 |
| F03 | 11160001 | 11260000 | 0.0039 | 0.0022 | 0.0026 |
| F03 | 11180001 | 11280000 | 0.0041 | 0.0024 | 0.0027 |
| F03 | 11200001 | 11300000 | 0.0036 | 0.0020 | 0.0022 |
| F03 | 11220001 | 11320000 | 0.0034 | 0.0021 | 0.0023 |
| F03 | 11240001 | 11340000 | 0.0036 | 0.0024 | 0.0026 |
| F03 | 11260001 | 11360000 | 0.0034 | 0.0026 | 0.0028 |
| F03 | 11280001 | 11380000 | 0.0032 | 0.0026 | 0.0028 |
| F03 | 11300001 | 11400000 | 0.0035 | 0.0028 | 0.0030 |
| F03 | 11320001 | 11420000 | 0.0035 | 0.0028 | 0.0032 |
| F03 | 11340001 | 11440000 | 0.0032 | 0.0025 | 0.0028 |
| F03 | 11360001 | 11460000 | 0.0040 | 0.0033 | 0.0035 |
| F03 | 11380001 | 11480000 | 0.0040 | 0.0033 | 0.0034 |
| F03 | 11400001 | 11500000 | 0.0041 | 0.0031 | 0.0035 |
| F03 | 11420001 | 11520000 | 0.0036 | 0.0027 | 0.0029 |
| F03 | 11440001 | 11540000 | 0.0037 | 0.0027 | 0.0028 |
| F03 | 11460001 | 11560000 | 0.0025 | 0.0016 | 0.0017 |
| F03 | 11480001 | 11580000 | 0.0022 | 0.0013 | 0.0015 |
| F03 | 11500001 | 11600000 | 0.0016 | 0.0010 | 0.0011 |
| F03 | 11520001 | 11620000 | 0.0014 | 0.0008 | 0.0009 |
| F03 | 11540001 | 11640000 | 0.0023 | 0.0017 | 0.0019 |
| F03 | 11560001 | 11660000 | 0.0020 | 0.0015 | 0.0018 |
| F03 | 11580001 | 11680000 | 0.0022 | 0.0018 | 0.0020 |
| F03 | 11600001 | 11700000 | 0.0024 | 0.0021 | 0.0022 |
| F03 | 11620001 | 11720000 | 0.0024 | 0.0021 | 0.0022 |
| F03 | 11640001 | 11740000 | 0.0019 | 0.0016 | 0.0018 |
| F03 | 11660001 | 11760000 | 0.0018 | 0.0016 | 0.0017 |
| F03 | 11680001 | 11780000 | 0.0020 | 0.0013 | 0.0016 |
| F03 | 11700001 | 11800000 | 0.0018 | 0.0012 | 0.0013 |
| F03 | 11720001 | 11820000 | 0.0019 | 0.0014 | 0.0014 |
| F03 | 11740001 | 11840000 | 0.0025 | 0.0018 | 0.0018 |
| F03 | 11760001 | 11860000 | 0.0027 | 0.0019 | 0.0019 |
| F03 | 11780001 | 11880000 | 0.0027 | 0.0019 | 0.0019 |
| F03 | 11800001 | 11900000 | 0.0028 | 0.0019 | 0.0020 |
| F03 | 11820001 | 11920000 | 0.0040 | 0.0030 | 0.0031 |
| F03 | 11840001 | 11940000 | 0.0031 | 0.0022 | 0.0023 |
| F03 | 11860001 | 11960000 | 0.0035 | 0.0026 | 0.0026 |
| F03 | 11880001 | 11980000 | 0.0034 | 0.0027 | 0.0027 |

| Chr | Start | End | both | A1 | A2 |
|---|---|---|---|---|---|
| F03 | 11900001 | 12000000 | 0.0032 | 0.0025 | 0.0025 |
| F03 | 11920001 | 12020000 | 0.0021 | 0.0012 | 0.0014 |
| F03 | 11940001 | 12040000 | 0.0027 | 0.0015 | 0.0017 |
| F03 | 11960001 | 12060000 | 0.0023 | 0.0012 | 0.0014 |
| F03 | 11980001 | 12080000 | 0.0022 | 0.0009 | 0.0012 |
| F03 | 12000001 | 12100000 | 0.0025 | 0.0012 | 0.0015 |
| F03 | 12020001 | 12120000 | 0.0033 | 0.0020 | 0.0023 |
| F03 | 12040001 | 12140000 | 0.0029 | 0.0020 | 0.0023 |
| F03 | 12060001 | 12160000 | 0.0028 | 0.0019 | 0.0022 |
| F03 | 12080001 | 12180000 | 0.0035 | 0.0024 | 0.0027 |
| F03 | 12100001 | 12200000 | 0.0034 | 0.0023 | 0.0026 |
| F03 | 12120001 | 12220000 | 0.0029 | 0.0014 | 0.0017 |
| F03 | 12140001 | 12240000 | 0.0027 | 0.0012 | 0.0014 |
| F03 | 12160001 | 12260000 | 0.0032 | 0.0015 | 0.0018 |
| F03 | 12180001 | 12280000 | 0.0028 | 0.0013 | 0.0015 |
| F03 | 12200001 | 12300000 | 0.0027 | 0.0012 | 0.0015 |
| F03 | 12220001 | 12320000 | 0.0026 | 0.0015 | 0.0017 |
| F03 | 12240001 | 12340000 | 0.0025 | 0.0016 | 0.0018 |
| F03 | 12260001 | 12360000 | 0.0020 | 0.0014 | 0.0015 |
| F03 | 12280001 | 12380000 | 0.0020 | 0.0015 | 0.0017 |
| F03 | 12300001 | 12400000 | 0.0019 | 0.0015 | 0.0017 |
| F03 | 12320001 | 12420000 | 0.0017 | 0.0011 | 0.0012 |
| F03 | 12340001 | 12440000 | 0.0022 | 0.0014 | 0.0015 |
| F03 | 12360001 | 12460000 | 0.0020 | 0.0012 | 0.0013 |
| F03 | 12380001 | 12480000 | 0.0015 | 0.0008 | 0.0008 |
| F03 | 12400001 | 12500000 | 0.0014 | 0.0008 | 0.0008 |
| F03 | 12420001 | 12520000 | 0.0012 | 0.0008 | 0.0008 |
| F03 | 12440001 | 12540000 | 0.0009 | 0.0007 | 0.0008 |
| F03 | 12460001 | 12560000 | 0.0010 | 0.0007 | 0.0009 |
| F03 | 12480001 | 12580000 | 0.0012 | 0.0010 | 0.0010 |
| F03 | 12500001 | 12600000 | 0.0021 | 0.0019 | 0.0019 |
| F03 | 12520001 | 12620000 | 0.0020 | 0.0019 | 0.0018 |
| F03 | 12540001 | 12640000 | 0.0019 | 0.0017 | 0.0016 |
| F03 | 12560001 | 12660000 | 0.0025 | 0.0022 | 0.0021 |
| F03 | 12580001 | 12680000 | 0.0037 | 0.0029 | 0.0032 |
| F03 | 12600001 | 12700000 | 0.0033 | 0.0021 | 0.0023 |
| F03 | 12620001 | 12720000 | 0.0036 | 0.0022 | 0.0025 |
| F03 | 12640001 | 12740000 | 0.0041 | 0.0024 | 0.0029 |
| F03 | 12660001 | 12760000 | 0.0042 | 0.0024 | 0.0028 |
| F03 | 12680001 | 12780000 | 0.0035 | 0.0015 | 0.0016 |
| F03 | 12700001 | 12800000 | 0.0040 | 0.0020 | 0.0022 |
| F03 | 12720001 | 12820000 | 0.0043 | 0.0025 | 0.0026 |
| F03 | 12740001 | 12840000 | 0.0045 | 0.0027 | 0.0027 |
| F03 | 12760001 | 12860000 | 0.0043 | 0.0026 | 0.0026 |
| F03 | 12780001 | 12880000 | 0.0043 | 0.0027 | 0.0029 |
| F03 | 12800001 | 12900000 | 0.0048 | 0.0034 | 0.0037 |
| F03 | 12820001 | 12920000 | 0.0046 | 0.0033 | 0.0037 |
| F03 | 12840001 | 12940000 | 0.0043 | 0.0033 | 0.0036 |
| F03 | 12860001 | 12960000 | 0.0044 | 0.0035 | 0.0038 |
| F03 | 12880001 | 12980000 | 0.0045 | 0.0039 | 0.0042 |
| F03 | 12900001 | 13000000 | 0.0037 | 0.0032 | 0.0035 |
| F03 | 12920001 | 13020000 | 0.0035 | 0.0030 | 0.0031 |
| F03 | 12940001 | 13040000 | 0.0039 | 0.0030 | 0.0033 |
| F03 | 12960001 | 13060000 | 0.0034 | 0.0026 | 0.0028 |
| F03 | 12980001 | 13080000 | 0.0028 | 0.0021 | 0.0023 |
| F03 | 13000001 | 13100000 | 0.0026 | 0.0018 | 0.0019 |
| F03 | 13020001 | 13120000 | 0.0022 | 0.0015 | 0.0017 |
| F03 | 13040001 | 13140000 | 0.0020 | 0.0015 | 0.0016 |

| Chr | Start | End | both | A1 | A2 |
|---|---|---|---|---|---|
| F03 | 13060001 | 13160000 | 0.0046 | 0.0038 | 0.0040 |
| F03 | 13080001 | 13180000 | 0.0047 | 0.0036 | 0.0039 |
| F03 | 13100001 | 13200000 | 0.0051 | 0.0039 | 0.0040 |
| F03 | 13120001 | 13220000 | 0.0052 | 0.0039 | 0.0042 |
| F03 | 13140001 | 13240000 | 0.0053 | 0.0039 | 0.0042 |
| F03 | 13160001 | 13260000 | 0.0028 | 0.0016 | 0.0018 |
| F03 | 13180001 | 13280000 | 0.0037 | 0.0027 | 0.0028 |
| F03 | 13200001 | 13300000 | 0.0035 | 0.0025 | 0.0028 |
| F03 | 13220001 | 13320000 | 0.0042 | 0.0033 | 0.0035 |
| F03 | 13240001 | 13340000 | 0.0042 | 0.0032 | 0.0035 |
| F03 | 13260001 | 13360000 | 0.0038 | 0.0030 | 0.0032 |
| F03 | 13280001 | 13380000 | 0.0032 | 0.0021 | 0.0024 |
| F03 | 13300001 | 13400000 | 0.0027 | 0.0018 | 0.0020 |
| F03 | 13320001 | 13420000 | 0.0029 | 0.0018 | 0.0021 |
| F03 | 13340001 | 13440000 | 0.0034 | 0.0020 | 0.0025 |
| F03 | 13360001 | 13460000 | 0.0039 | 0.0023 | 0.0028 |
| F03 | 13380001 | 13480000 | 0.0038 | 0.0025 | 0.0030 |
| F03 | 13400001 | 13500000 | 0.0042 | 0.0029 | 0.0034 |
| F03 | 13420001 | 13520000 | 0.0038 | 0.0024 | 0.0028 |
| F03 | 13440001 | 13540000 | 0.0035 | 0.0022 | 0.0024 |
| F03 | 13460001 | 13560000 | 0.0040 | 0.0027 | 0.0027 |
| F03 | 13480001 | 13580000 | 0.0040 | 0.0023 | 0.0023 |
| F03 | 13500001 | 13600000 | 0.0038 | 0.0019 | 0.0019 |
| F03 | 13520001 | 13620000 | 0.0035 | 0.0017 | 0.0018 |
| F03 | 13540001 | 13640000 | 0.0032 | 0.0017 | 0.0016 |
| F03 | 13560001 | 13660000 | 0.0024 | 0.0010 | 0.0011 |
| F03 | 13580001 | 13680000 | 0.0033 | 0.0017 | 0.0021 |
| F03 | 13600001 | 13700000 | 0.0029 | 0.0015 | 0.0019 |
| F03 | 13620001 | 13720000 | 0.0027 | 0.0014 | 0.0019 |
| F03 | 13640001 | 13740000 | 0.0030 | 0.0017 | 0.0021 |
| F03 | 13660001 | 13760000 | 0.0033 | 0.0020 | 0.0024 |
| F03 | 13680001 | 13780000 | 0.0031 | 0.0021 | 0.0021 |
| F03 | 13700001 | 13800000 | 0.0047 | 0.0035 | 0.0037 |
| F03 | 13720001 | 13820000 | 0.0062 | 0.0048 | 0.0052 |
| F03 | 13740001 | 13840000 | 0.0057 | 0.0042 | 0.0047 |
| F03 | 13760001 | 13860000 | 0.0056 | 0.0044 | 0.0049 |
| F03 | 13780001 | 13880000 | 0.0059 | 0.0045 | 0.0051 |
| F03 | 13800001 | 13900000 | 0.0047 | 0.0035 | 0.0039 |
| F03 | 13820001 | 13920000 | 0.0035 | 0.0024 | 0.0026 |
| F03 | 13840001 | 13940000 | 0.0040 | 0.0027 | 0.0030 |
| F03 | 13860001 | 13960000 | 0.0040 | 0.0025 | 0.0029 |
| F03 | 13880001 | 13980000 | 0.0031 | 0.0017 | 0.0021 |
| F03 | 13900001 | 14000000 | 0.0032 | 0.0016 | 0.0020 |
| F03 | 13920001 | 14020000 | 0.0032 | 0.0017 | 0.0020 |
| F03 | 13940001 | 14040000 | 0.0027 | 0.0014 | 0.0017 |
| F03 | 13960001 | 14060000 | 0.0030 | 0.0011 | 0.0021 |
| F03 | 13980001 | 14080000 | 0.0035 | 0.0015 | 0.0024 |
| F03 | 14000001 | 14100000 | 0.0033 | 0.0016 | 0.0024 |
| F03 | 14020001 | 14120000 | 0.0033 | 0.0015 | 0.0022 |
| F03 | 14040001 | 14140000 | 0.0029 | 0.0012 | 0.0019 |
| F03 | 14060001 | 14160000 | 0.0024 | 0.0012 | 0.0012 |
| F03 | 14080001 | 14180000 | 0.0026 | 0.0014 | 0.0016 |
| F03 | 14100001 | 14200000 | 0.0031 | 0.0018 | 0.0021 |
| F03 | 14120001 | 14220000 | 0.0032 | 0.0020 | 0.0024 |
| F03 | 14140001 | 14240000 | 0.0034 | 0.0021 | 0.0026 |
| F03 | 14160001 | 14260000 | 0.0036 | 0.0024 | 0.0028 |
| F03 | 14180001 | 14280000 | 0.0039 | 0.0028 | 0.0031 |
| F03 | 14200001 | 14300000 | 0.0032 | 0.0024 | 0.0025 |

| Chr | Start | End | both | A1 | A2 |
|---|---|---|---|---|---|
| F03 | 14220001 | 14320000 | 0.0032 | 0.0023 | 0.0024 |
| F03 | 14240001 | 14340000 | 0.0038 | 0.0029 | 0.0030 |
| F03 | 14260001 | 14360000 | 0.0036 | 0.0027 | 0.0028 |
| F03 | 14280001 | 14380000 | 0.0028 | 0.0020 | 0.0021 |
| F03 | 14300001 | 14400000 | 0.0030 | 0.0019 | 0.0022 |
| F03 | 14320001 | 14420000 | 0.0036 | 0.0026 | 0.0029 |
| F03 | 14340001 | 14440000 | 0.0034 | 0.0025 | 0.0027 |
| F03 | 14360001 | 14460000 | 0.0033 | 0.0023 | 0.0026 |
| F03 | 14380001 | 14480000 | 0.0029 | 0.0020 | 0.0022 |
| F03 | 14400001 | 14500000 | 0.0035 | 0.0026 | 0.0029 |
| F03 | 14420001 | 14520000 | 0.0023 | 0.0016 | 0.0018 |
| F03 | 14440001 | 14540000 | 0.0016 | 0.0010 | 0.0012 |
| F03 | 14460001 | 14560000 | 0.0016 | 0.0012 | 0.0013 |
| F03 | 14480001 | 14580000 | 0.0016 | 0.0013 | 0.0015 |
| F03 | 14500001 | 14600000 | 0.0007 | 0.0006 | 0.0006 |
| F03 | 14520001 | 14620000 | 0.0013 | 0.0012 | 0.0013 |
| F03 | 14540001 | 14640000 | 0.0014 | 0.0013 | 0.0014 |
| F03 | 14560001 | 14660000 | 0.0014 | 0.0013 | 0.0014 |
| F03 | 14580001 | 14680000 | 0.0012 | 0.0010 | 0.0011 |
| F03 | 14600001 | 14700000 | 0.0019 | 0.0016 | 0.0018 |
| F03 | 14620001 | 14720000 | 0.0013 | 0.0011 | 0.0012 |
| F03 | 14640001 | 14740000 | 0.0023 | 0.0018 | 0.0019 |
| F03 | 14660001 | 14760000 | 0.0026 | 0.0019 | 0.0021 |
| F03 | 14680001 | 14780000 | 0.0031 | 0.0022 | 0.0024 |
| F03 | 14700001 | 14800000 | 0.0028 | 0.0021 | 0.0022 |
| F03 | 14720001 | 14820000 | 0.0029 | 0.0020 | 0.0021 |
| F03 | 14740001 | 14840000 | 0.0025 | 0.0017 | 0.0018 |
| F03 | 14760001 | 14860000 | 0.0035 | 0.0023 | 0.0028 |
| F03 | 14780001 | 14880000 | 0.0042 | 0.0029 | 0.0033 |
| F03 | 14800001 | 14900000 | 0.0047 | 0.0032 | 0.0038 |
| F03 | 14820001 | 14920000 | 0.0053 | 0.0037 | 0.0043 |
| F03 | 14840001 | 14940000 | 0.0054 | 0.0037 | 0.0043 |
| F03 | 14860001 | 14960000 | 0.0055 | 0.0039 | 0.0042 |
| F03 | 14880001 | 14980000 | 0.0045 | 0.0032 | 0.0035 |
| F03 | 14900001 | 15000000 | 0.0035 | 0.0025 | 0.0027 |
| F03 | 14920001 | 15020000 | 0.0031 | 0.0022 | 0.0024 |
| F03 | 14940001 | 15040000 | 0.0026 | 0.0018 | 0.0020 |
| F03 | 14960001 | 15060000 | 0.0033 | 0.0028 | 0.0029 |
| F03 | 14980001 | 15080000 | 0.0034 | 0.0030 | 0.0031 |
| F03 | 15000001 | 15100000 | 0.0035 | 0.0031 | 0.0032 |
| F03 | 15020001 | 15120000 | 0.0037 | 0.0034 | 0.0035 |
| F03 | 15040001 | 15140000 | 0.0034 | 0.0033 | 0.0033 |
| F03 | 15060001 | 15160000 | 0.0027 | 0.0028 | 0.0026 |
| F03 | 15080001 | 15180000 | 0.0030 | 0.0030 | 0.0028 |
| F03 | 15100001 | 15200000 | 0.0029 | 0.0029 | 0.0028 |
| F03 | 15120001 | 15220000 | 0.0028 | 0.0026 | 0.0025 |
| F03 | 15140001 | 15240000 | 0.0032 | 0.0028 | 0.0027 |
| F03 | 15160001 | 15260000 | 0.0020 | 0.0014 | 0.0015 |
| F03 | 15180001 | 15280000 | 0.0023 | 0.0015 | 0.0016 |
| F03 | 15200001 | 15300000 | 0.0030 | 0.0020 | 0.0020 |
| F03 | 15220001 | 15320000 | 0.0033 | 0.0024 | 0.0025 |
| F03 | 15240001 | 15340000 | 0.0031 | 0.0022 | 0.0023 |
| F03 | 15260001 | 15360000 | 0.0027 | 0.0020 | 0.0021 |
| F03 | 15280001 | 15380000 | 0.0019 | 0.0014 | 0.0015 |
| F03 | 15300001 | 15400000 | 0.0017 | 0.0014 | 0.0013 |
| F03 | 15320001 | 15420000 | 0.0016 | 0.0016 | 0.0013 |
| F03 | 15340001 | 15440000 | 0.0030 | 0.0027 | 0.0024 |
| F03 | 15360001 | 15460000 | 0.0034 | 0.0030 | 0.0029 |

| Chr | Start | End | both | A1 | A2 |
|---|---|---|---|---|---|
| F03 | 15380001 | 15480000 | 0.0041 | 0.0036 | 0.0035 |
| F03 | 15400001 | 15500000 | 0.0036 | 0.0030 | 0.0032 |
| F03 | 15420001 | 15520000 | 0.0031 | 0.0025 | 0.0028 |
| F03 | 15440001 | 15540000 | 0.0016 | 0.0013 | 0.0015 |
| F03 | 15460001 | 15560000 | 0.0020 | 0.0013 | 0.0015 |
| F03 | 15480001 | 15580000 | 0.0015 | 0.0008 | 0.0009 |
| F03 | 15500001 | 15600000 | 0.0023 | 0.0012 | 0.0014 |
| F03 | 15520001 | 15620000 | 0.0028 | 0.0015 | 0.0017 |
| F03 | 15540001 | 15640000 | 0.0033 | 0.0019 | 0.0020 |
| F03 | 15560001 | 15660000 | 0.0032 | 0.0020 | 0.0022 |
| F03 | 15580001 | 15680000 | 0.0033 | 0.0021 | 0.0024 |
| F03 | 15600001 | 15700000 | 0.0028 | 0.0018 | 0.0020 |
| F03 | 15620001 | 15720000 | 0.0025 | 0.0016 | 0.0019 |
| F03 | 15640001 | 15740000 | 0.0023 | 0.0015 | 0.0019 |
| F03 | 15660001 | 15760000 | 0.0019 | 0.0011 | 0.0014 |
| F03 | 15680001 | 15780000 | 0.0022 | 0.0013 | 0.0016 |
| F03 | 15700001 | 15800000 | 0.0026 | 0.0017 | 0.0020 |
| F03 | 15720001 | 15820000 | 0.0030 | 0.0019 | 0.0021 |
| F03 | 15740001 | 15840000 | 0.0031 | 0.0020 | 0.0021 |
| F03 | 15760001 | 15860000 | 0.0031 | 0.0020 | 0.0021 |
| F03 | 15780001 | 15880000 | 0.0034 | 0.0023 | 0.0022 |
| F03 | 15800001 | 15900000 | 0.0037 | 0.0026 | 0.0024 |
| F03 | 15820001 | 15920000 | 0.0030 | 0.0022 | 0.0021 |
| F03 | 15840001 | 15940000 | 0.0033 | 0.0025 | 0.0022 |
| F03 | 15860001 | 15960000 | 0.0034 | 0.0024 | 0.0023 |
| F03 | 15880001 | 15980000 | 0.0026 | 0.0018 | 0.0018 |
| F03 | 15900001 | 16000000 | 0.0021 | 0.0012 | 0.0012 |
| F03 | 15920001 | 16020000 | 0.0029 | 0.0015 | 0.0018 |
| F03 | 15940001 | 16040000 | 0.0029 | 0.0013 | 0.0016 |
| F03 | 15960001 | 16060000 | 0.0036 | 0.0021 | 0.0024 |
| F03 | 15980001 | 16080000 | 0.0041 | 0.0025 | 0.0030 |
| F03 | 16000001 | 16100000 | 0.0044 | 0.0026 | 0.0031 |
| F03 | 16020001 | 16120000 | 0.0054 | 0.0038 | 0.0042 |
| F03 | 16040001 | 16140000 | 0.0059 | 0.0045 | 0.0051 |
| F03 | 16060001 | 16160000 | 0.0060 | 0.0044 | 0.0049 |
| F03 | 16080001 | 16180000 | 0.0059 | 0.0042 | 0.0047 |
| F03 | 16100001 | 16200000 | 0.0065 | 0.0051 | 0.0055 |
| F03 | 16120001 | 16220000 | 0.0057 | 0.0043 | 0.0046 |
| F03 | 16140001 | 16240000 | 0.0049 | 0.0033 | 0.0036 |
| F03 | 16160001 | 16260000 | 0.0051 | 0.0038 | 0.0042 |
| F03 | 16180001 | 16280000 | 0.0048 | 0.0037 | 0.0040 |
| F03 | 16200001 | 16300000 | 0.0041 | 0.0029 | 0.0034 |
| F03 | 16220001 | 16320000 | 0.0036 | 0.0026 | 0.0030 |
| F03 | 16240001 | 16340000 | 0.0039 | 0.0032 | 0.0035 |
| F03 | 16260001 | 16360000 | 0.0031 | 0.0024 | 0.0027 |
| F03 | 16280001 | 16380000 | 0.0035 | 0.0027 | 0.0031 |
| F03 | 16300001 | 16400000 | 0.0033 | 0.0026 | 0.0028 |
| F03 | 16320001 | 16420000 | 0.0033 | 0.0026 | 0.0028 |
| F03 | 16340001 | 16440000 | 0.0026 | 0.0021 | 0.0023 |
| F03 | 16360001 | 16460000 | 0.0026 | 0.0022 | 0.0023 |
| F03 | 16380001 | 16480000 | 0.0023 | 0.0019 | 0.0019 |
| F03 | 16400001 | 16500000 | 0.0025 | 0.0021 | 0.0022 |
| F03 | 16420001 | 16520000 | 0.0031 | 0.0025 | 0.0026 |
| F03 | 16440001 | 16540000 | 0.0040 | 0.0033 | 0.0035 |
| F03 | 16460001 | 16560000 | 0.0038 | 0.0031 | 0.0033 |
| F03 | 16480001 | 16580000 | 0.0039 | 0.0034 | 0.0036 |
| F03 | 16500001 | 16600000 | 0.0038 | 0.0032 | 0.0035 |
| F03 | 16520001 | 16620000 | 0.0042 | 0.0036 | 0.0040 |

| Chr | Start | End | both | A1 | A2 |
|---|---|---|---|---|---|
| F03 | 16540001 | 16640000 | 0.0042 | 0.0033 | 0.0039 |
| F03 | 16560001 | 16660000 | 0.0039 | 0.0030 | 0.0036 |
| F03 | 16580001 | 16680000 | 0.0044 | 0.0036 | 0.0041 |
| F03 | 16600001 | 16700000 | 0.0042 | 0.0035 | 0.0040 |
| F03 | 16620001 | 16720000 | 0.0028 | 0.0022 | 0.0026 |
| F03 | 16640001 | 16740000 | 0.0026 | 0.0023 | 0.0024 |
| F03 | 16660001 | 16760000 | 0.0034 | 0.0029 | 0.0029 |
| F03 | 16680001 | 16780000 | 0.0026 | 0.0021 | 0.0021 |
| F03 | 16700001 | 16800000 | 0.0028 | 0.0023 | 0.0024 |
| F03 | 16720001 | 16820000 | 0.0031 | 0.0025 | 0.0026 |
| F03 | 16740001 | 16840000 | 0.0023 | 0.0019 | 0.0020 |
| F03 | 16760001 | 16860000 | 0.0025 | 0.0020 | 0.0023 |
| F03 | 16780001 | 16880000 | 0.0029 | 0.0022 | 0.0025 |
| F03 | 16800001 | 16900000 | 0.0034 | 0.0024 | 0.0028 |
| F03 | 16820001 | 16920000 | 0.0037 | 0.0025 | 0.0029 |
| F03 | 16840001 | 16940000 | 0.0049 | 0.0035 | 0.0040 |
| F03 | 16860001 | 16960000 | 0.0046 | 0.0033 | 0.0036 |
| F03 | 16880001 | 16980000 | 0.0049 | 0.0036 | 0.0040 |
| F03 | 16900001 | 17000000 | 0.0049 | 0.0038 | 0.0041 |
| F03 | 16920001 | 17020000 | 0.0046 | 0.0035 | 0.0038 |
| F03 | 16940001 | 17040000 | 0.0034 | 0.0025 | 0.0027 |
| F03 | 16960001 | 17060000 | 0.0032 | 0.0022 | 0.0024 |
| F03 | 16980001 | 17080000 | 0.0028 | 0.0019 | 0.0019 |
| F03 | 17000001 | 17100000 | 0.0022 | 0.0013 | 0.0014 |
| F03 | 17020001 | 17120000 | 0.0023 | 0.0014 | 0.0015 |
| F03 | 17040001 | 17140000 | 0.0029 | 0.0018 | 0.0020 |
| F03 | 17060001 | 17160000 | 0.0035 | 0.0023 | 0.0026 |
| F03 | 17080001 | 17180000 | 0.0031 | 0.0019 | 0.0024 |
| F03 | 17100001 | 17200000 | 0.0029 | 0.0018 | 0.0022 |
| F03 | 17120001 | 17220000 | 0.0032 | 0.0022 | 0.0025 |
| F03 | 17140001 | 17240000 | 0.0029 | 0.0020 | 0.0023 |
| F03 | 17160001 | 17260000 | 0.0024 | 0.0018 | 0.0020 |
| F03 | 17180001 | 17280000 | 0.0024 | 0.0017 | 0.0019 |
| F03 | 17200001 | 17300000 | 0.0044 | 0.0032 | 0.0036 |
| F03 | 17220001 | 17320000 | 0.0050 | 0.0038 | 0.0042 |
| F03 | 17240001 | 17340000 | 0.0050 | 0.0038 | 0.0042 |
| F03 | 17260001 | 17360000 | 0.0056 | 0.0044 | 0.0048 |
| F03 | 17280001 | 17380000 | 0.0064 | 0.0052 | 0.0057 |
| F03 | 17300001 | 17400000 | 0.0052 | 0.0043 | 0.0046 |
| F03 | 17320001 | 17420000 | 0.0041 | 0.0032 | 0.0035 |
| F03 | 17340001 | 17440000 | 0.0038 | 0.0032 | 0.0034 |
| F03 | 17360001 | 17460000 | 0.0046 | 0.0038 | 0.0040 |
| F03 | 17380001 | 17480000 | 0.0054 | 0.0046 | 0.0048 |
| F03 | 17400001 | 17500000 | 0.0051 | 0.0044 | 0.0047 |
| F03 | 17420001 | 17520000 | 0.0056 | 0.0049 | 0.0052 |
| F03 | 17440001 | 17540000 | 0.0056 | 0.0048 | 0.0051 |
| F03 | 17460001 | 17560000 | 0.0046 | 0.0039 | 0.0040 |
| F03 | 17480001 | 17580000 | 0.0033 | 0.0027 | 0.0029 |
| F03 | 17500001 | 17600000 | 0.0029 | 0.0023 | 0.0025 |
| F03 | 17520001 | 17620000 | 0.0025 | 0.0020 | 0.0022 |
| F03 | 17540001 | 17640000 | 0.0031 | 0.0026 | 0.0028 |
| F03 | 17560001 | 17660000 | 0.0028 | 0.0023 | 0.0026 |
| F03 | 17580001 | 17680000 | 0.0032 | 0.0026 | 0.0029 |
| F03 | 17600001 | 17700000 | 0.0034 | 0.0027 | 0.0031 |
| F03 | 17620001 | 17720000 | 0.0032 | 0.0024 | 0.0028 |
| F03 | 17640001 | 17740000 | 0.0035 | 0.0025 | 0.0031 |
| F03 | 17660001 | 17760000 | 0.0042 | 0.0030 | 0.0037 |
| F03 | 17680001 | 17780000 | 0.0032 | 0.0022 | 0.0027 |

| Chr | Start | End | both | A1 | A2 |
|---|---|---|---|---|---|
| F03 | 17700001 | 17800000 | 0.0033 | 0.0023 | 0.0027 |
| F03 | 17720001 | 17820000 | 0.0036 | 0.0026 | 0.0031 |
| F03 | 17740001 | 17840000 | 0.0029 | 0.0023 | 0.0024 |
| F03 | 17760001 | 17860000 | 0.0024 | 0.0018 | 0.0020 |
| F03 | 17780001 | 17880000 | 0.0029 | 0.0025 | 0.0027 |
| F03 | 17800001 | 17900000 | 0.0030 | 0.0028 | 0.0030 |
| F03 | 17820001 | 17920000 | 0.0029 | 0.0027 | 0.0029 |
| F03 | 17840001 | 17940000 | 0.0026 | 0.0023 | 0.0025 |
| F03 | 17860001 | 17960000 | 0.0019 | 0.0018 | 0.0018 |
| F03 | 17880001 | 17980000 | 0.0015 | 0.0012 | 0.0013 |
| F03 | 17900001 | 18000000 | 0.0009 | 0.0007 | 0.0007 |
| F03 | 17920001 | 18020000 | 0.0007 | 0.0005 | 0.0005 |
| F03 | 17940001 | 18040000 | 0.0018 | 0.0016 | 0.0016 |
| F03 | 17960001 | 18060000 | 0.0029 | 0.0026 | 0.0025 |
| F03 | 17980001 | 18080000 | 0.0031 | 0.0025 | 0.0025 |
| F03 | 18000001 | 18100000 | 0.0038 | 0.0029 | 0.0028 |
| F03 | 18020001 | 18120000 | 0.0050 | 0.0039 | 0.0039 |
| F03 | 18040001 | 18140000 | 0.0048 | 0.0035 | 0.0038 |
| F03 | 18060001 | 18160000 | 0.0043 | 0.0028 | 0.0032 |
| F03 | 18080001 | 18180000 | 0.0045 | 0.0033 | 0.0037 |
| F03 | 18100001 | 18200000 | 0.0049 | 0.0037 | 0.0043 |
| F03 | 18120001 | 18220000 | 0.0046 | 0.0035 | 0.0038 |
| F03 | 18140001 | 18240000 | 0.0040 | 0.0030 | 0.0031 |
| F03 | 18160001 | 18260000 | 0.0036 | 0.0027 | 0.0028 |
| F03 | 18180001 | 18280000 | 0.0031 | 0.0024 | 0.0024 |
| F03 | 18200001 | 18300000 | 0.0026 | 0.0021 | 0.0021 |
| F03 | 18220001 | 18320000 | 0.0020 | 0.0017 | 0.0017 |
| F03 | 18240001 | 18340000 | 0.0018 | 0.0015 | 0.0015 |
| F03 | 18260001 | 18360000 | 0.0019 | 0.0016 | 0.0015 |
| F03 | 18280001 | 18380000 | 0.0019 | 0.0015 | 0.0015 |
| F03 | 18300001 | 18400000 | 0.0015 | 0.0011 | 0.0010 |
| F03 | 18320001 | 18420000 | 0.0019 | 0.0010 | 0.0012 |
| F03 | 18340001 | 18440000 | 0.0019 | 0.0010 | 0.0012 |
| F03 | 18360001 | 18460000 | 0.0025 | 0.0012 | 0.0015 |
| F03 | 18380001 | 18480000 | 0.0038 | 0.0024 | 0.0027 |
| F03 | 18400001 | 18500000 | 0.0047 | 0.0033 | 0.0035 |
| F03 | 18420001 | 18520000 | 0.0047 | 0.0036 | 0.0038 |
| F03 | 18440001 | 18540000 | 0.0054 | 0.0041 | 0.0045 |
| F03 | 18460001 | 18560000 | 0.0052 | 0.0042 | 0.0045 |
| F03 | 18480001 | 18580000 | 0.0038 | 0.0030 | 0.0033 |
| F03 | 18500001 | 18600000 | 0.0031 | 0.0023 | 0.0026 |
| F03 | 18520001 | 18620000 | 0.0029 | 0.0018 | 0.0021 |
| F03 | 18540001 | 18640000 | 0.0028 | 0.0017 | 0.0019 |
| F03 | 18560001 | 18660000 | 0.0035 | 0.0025 | 0.0028 |
| F03 | 18580001 | 18680000 | 0.0035 | 0.0025 | 0.0027 |
| F03 | 18600001 | 18700000 | 0.0033 | 0.0025 | 0.0027 |
| F03 | 18620001 | 18720000 | 0.0035 | 0.0027 | 0.0030 |
| F03 | 18640001 | 18740000 | 0.0051 | 0.0041 | 0.0045 |
| F03 | 18660001 | 18760000 | 0.0046 | 0.0035 | 0.0039 |
| F03 | 18680001 | 18780000 | 0.0053 | 0.0040 | 0.0044 |
| F03 | 18700001 | 18800000 | 0.0057 | 0.0041 | 0.0047 |
| F03 | 18720001 | 18820000 | 0.0050 | 0.0038 | 0.0043 |
| F03 | 18740001 | 18840000 | 0.0028 | 0.0020 | 0.0023 |
| F03 | 18760001 | 18860000 | 0.0021 | 0.0016 | 0.0018 |
| F03 | 18780001 | 18880000 | 0.0020 | 0.0015 | 0.0017 |
| F03 | 18800001 | 18900000 | 0.0014 | 0.0012 | 0.0014 |
| F03 | 18820001 | 18920000 | 0.0014 | 0.0011 | 0.0014 |
| F03 | 18840001 | 18940000 | 0.0019 | 0.0016 | 0.0019 |

| Chr | Start | End | both | A1 | A2 |
|---|---|---|---|---|---|
| F03 | 18860001 | 18960000 | 0.0019 | 0.0015 | 0.0019 |
| F03 | 18880001 | 18980000 | 0.0018 | 0.0013 | 0.0016 |
| F03 | 18900001 | 19000000 | 0.0023 | 0.0014 | 0.0018 |
| F03 | 18920001 | 19020000 | 0.0028 | 0.0018 | 0.0023 |
| F03 | 18940001 | 19040000 | 0.0026 | 0.0014 | 0.0018 |
| F03 | 18960001 | 19060000 | 0.0033 | 0.0018 | 0.0023 |
| F03 | 18980001 | 19080000 | 0.0037 | 0.0021 | 0.0025 |
| F03 | 19000001 | 19100000 | 0.0036 | 0.0022 | 0.0025 |
| F03 | 19020001 | 19120000 | 0.0035 | 0.0022 | 0.0024 |
| F03 | 19040001 | 19140000 | 0.0033 | 0.0022 | 0.0024 |
| F03 | 19060001 | 19160000 | 0.0033 | 0.0024 | 0.0026 |
| F03 | 19080001 | 19180000 | 0.0038 | 0.0030 | 0.0034 |
| F03 | 19100001 | 19200000 | 0.0040 | 0.0031 | 0.0035 |
| F03 | 19120001 | 19220000 | 0.0043 | 0.0033 | 0.0036 |
| F03 | 19140001 | 19240000 | 0.0051 | 0.0040 | 0.0046 |
| F03 | 19160001 | 19260000 | 0.0047 | 0.0035 | 0.0039 |
| F03 | 19180001 | 19280000 | 0.0041 | 0.0027 | 0.0030 |
| F03 | 19200001 | 19300000 | 0.0034 | 0.0023 | 0.0025 |
| F03 | 19220001 | 19320000 | 0.0029 | 0.0021 | 0.0022 |
| F03 | 19240001 | 19340000 | 0.0020 | 0.0013 | 0.0014 |
| F03 | 19260001 | 19360000 | 0.0016 | 0.0013 | 0.0013 |
| F03 | 19280001 | 19380000 | 0.0009 | 0.0009 | 0.0009 |
| F03 | 19300001 | 19400000 | 0.0010 | 0.0009 | 0.0010 |
| F03 | 19320001 | 19420000 | 0.0007 | 0.0006 | 0.0006 |
| F03 | 19340001 | 19440000 | 0.0008 | 0.0008 | 0.0008 |
| F03 | 19360001 | 19460000 | 0.0011 | 0.0009 | 0.0011 |
| F03 | 19380001 | 19480000 | 0.0020 | 0.0017 | 0.0020 |
| F03 | 19400001 | 19500000 | 0.0023 | 0.0020 | 0.0022 |
| F03 | 19420001 | 19520000 | 0.0026 | 0.0021 | 0.0024 |
| F03 | 19440001 | 19540000 | 0.0033 | 0.0026 | 0.0029 |
| F03 | 19460001 | 19560000 | 0.0037 | 0.0031 | 0.0030 |
| F03 | 19480001 | 19580000 | 0.0030 | 0.0024 | 0.0022 |
| F03 | 19500001 | 19600000 | 0.0029 | 0.0022 | 0.0022 |
| F03 | 19520001 | 19620000 | 0.0033 | 0.0025 | 0.0025 |
| F03 | 19540001 | 19640000 | 0.0036 | 0.0027 | 0.0028 |
| F03 | 19560001 | 19660000 | 0.0038 | 0.0028 | 0.0031 |
| F03 | 19580001 | 19680000 | 0.0049 | 0.0039 | 0.0043 |
| F03 | 19600001 | 19700000 | 0.0052 | 0.0042 | 0.0046 |
| F03 | 19620001 | 19720000 | 0.0053 | 0.0042 | 0.0048 |
| F03 | 19640001 | 19740000 | 0.0043 | 0.0034 | 0.0040 |
| F03 | 19660001 | 19760000 | 0.0052 | 0.0042 | 0.0051 |
| F03 | 19680001 | 19780000 | 0.0047 | 0.0037 | 0.0046 |
| F03 | 19700001 | 19800000 | 0.0048 | 0.0037 | 0.0048 |
| F03 | 19720001 | 19820000 | 0.0048 | 0.0038 | 0.0047 |
| F03 | 19740001 | 19840000 | 0.0050 | 0.0040 | 0.0048 |
| F03 | 19760001 | 19860000 | 0.0033 | 0.0026 | 0.0032 |
| F03 | 19780001 | 19880000 | 0.0025 | 0.0020 | 0.0024 |
| F03 | 19800001 | 19900000 | 0.0020 | 0.0017 | 0.0019 |
| F03 | 19820001 | 19920000 | 0.0015 | 0.0013 | 0.0015 |
| F03 | 19840001 | 19940000 | 0.0016 | 0.0013 | 0.0015 |
| F03 | 19860001 | 19960000 | 0.0020 | 0.0016 | 0.0019 |
| F03 | 19880001 | 19980000 | 0.0027 | 0.0021 | 0.0024 |
| F03 | 19900001 | 20000000 | 0.0027 | 0.0021 | 0.0024 |
| F03 | 19920001 | 20020000 | 0.0034 | 0.0027 | 0.0029 |
| F03 | 19940001 | 20040000 | 0.0030 | 0.0023 | 0.0024 |
| F03 | 19960001 | 20060000 | 0.0035 | 0.0028 | 0.0029 |
| F03 | 19980001 | 20080000 | 0.0033 | 0.0027 | 0.0028 |
| F03 | 20000001 | 20100000 | 0.0029 | 0.0025 | 0.0025 |

| Chr | Start | End | both | A1 | A2 |
|---|---|---|---|---|---|
| F03 | 20020001 | 20120000 | 0.0026 | 0.0022 | 0.0023 |
| F03 | 20040001 | 20140000 | 0.0029 | 0.0023 | 0.0027 |
| F03 | 20060001 | 20160000 | 0.0022 | 0.0017 | 0.0021 |
| F03 | 20080001 | 20180000 | 0.0027 | 0.0020 | 0.0026 |
| F03 | 20100001 | 20200000 | 0.0028 | 0.0020 | 0.0027 |
| F03 | 20120001 | 20220000 | 0.0024 | 0.0018 | 0.0024 |
| F03 | 20140001 | 20240000 | 0.0024 | 0.0019 | 0.0024 |
| F03 | 20160001 | 20260000 | 0.0024 | 0.0018 | 0.0023 |
| F03 | 20180001 | 20280000 | 0.0030 | 0.0024 | 0.0026 |
| F03 | 20200001 | 20300000 | 0.0036 | 0.0030 | 0.0032 |
| F03 | 20220001 | 20320000 | 0.0039 | 0.0031 | 0.0034 |
| F03 | 20240001 | 20340000 | 0.0044 | 0.0034 | 0.0037 |
| F03 | 20260001 | 20360000 | 0.0045 | 0.0036 | 0.0039 |
| F03 | 20280001 | 20380000 | 0.0033 | 0.0026 | 0.0029 |
| F03 | 20300001 | 20400000 | 0.0046 | 0.0040 | 0.0042 |
| F03 | 20320001 | 20420000 | 0.0052 | 0.0047 | 0.0049 |
| F03 | 20340001 | 20440000 | 0.0048 | 0.0044 | 0.0045 |
| F03 | 20360001 | 20460000 | 0.0055 | 0.0050 | 0.0052 |
| F03 | 20380001 | 20480000 | 0.0064 | 0.0057 | 0.0059 |
| F03 | 20400001 | 20500000 | 0.0046 | 0.0036 | 0.0040 |
| F03 | 20420001 | 20520000 | 0.0055 | 0.0043 | 0.0049 |
| F03 | 20440001 | 20540000 | 0.0058 | 0.0045 | 0.0053 |
| F03 | 20460001 | 20560000 | 0.0052 | 0.0038 | 0.0045 |
| F03 | 20480001 | 20580000 | 0.0043 | 0.0030 | 0.0037 |
| F03 | 20500001 | 20600000 | 0.0047 | 0.0037 | 0.0042 |
| F03 | 20520001 | 20620000 | 0.0034 | 0.0024 | 0.0026 |
| F03 | 20540001 | 20640000 | 0.0031 | 0.0020 | 0.0021 |
| F03 | 20560001 | 20660000 | 0.0031 | 0.0021 | 0.0022 |
| F03 | 20580001 | 20680000 | 0.0029 | 0.0019 | 0.0020 |
| F03 | 20600001 | 20700000 | 0.0025 | 0.0014 | 0.0017 |
| F03 | 20620001 | 20720000 | 0.0020 | 0.0012 | 0.0015 |
| F03 | 20640001 | 20740000 | 0.0025 | 0.0020 | 0.0022 |
| F03 | 20660001 | 20760000 | 0.0021 | 0.0018 | 0.0019 |
| F03 | 20680001 | 20780000 | 0.0027 | 0.0022 | 0.0023 |
| F03 | 20700001 | 20800000 | 0.0028 | 0.0022 | 0.0023 |
| F03 | 20720001 | 20820000 | 0.0033 | 0.0027 | 0.0028 |
| F03 | 20740001 | 20840000 | 0.0036 | 0.0029 | 0.0031 |
| F03 | 20760001 | 20860000 | 0.0040 | 0.0032 | 0.0036 |
| F03 | 20780001 | 20880000 | 0.0037 | 0.0031 | 0.0034 |
| F03 | 20800001 | 20900000 | 0.0037 | 0.0033 | 0.0036 |
| F03 | 20820001 | 20920000 | 0.0035 | 0.0031 | 0.0034 |
| F03 | 20840001 | 20940000 | 0.0026 | 0.0023 | 0.0026 |
| F03 | 20860001 | 20960000 | 0.0025 | 0.0021 | 0.0025 |
| F03 | 20880001 | 20980000 | 0.0033 | 0.0029 | 0.0033 |
| F03 | 20900001 | 21000000 | 0.0030 | 0.0027 | 0.0030 |
| F03 | 20920001 | 21020000 | 0.0041 | 0.0037 | 0.0040 |
| F03 | 20940001 | 21040000 | 0.0047 | 0.0042 | 0.0045 |
| F03 | 20960001 | 21060000 | 0.0048 | 0.0042 | 0.0044 |
| F03 | 20980001 | 21080000 | 0.0039 | 0.0033 | 0.0035 |
| F03 | 21000001 | 21100000 | 0.0040 | 0.0033 | 0.0035 |
| F03 | 21020001 | 21120000 | 0.0027 | 0.0021 | 0.0023 |
| F03 | 21040001 | 21140000 | 0.0017 | 0.0011 | 0.0013 |
| F03 | 21060001 | 21160000 | 0.0012 | 0.0008 | 0.0009 |
| F03 | 21080001 | 21180000 | 0.0012 | 0.0007 | 0.0007 |
| F03 | 21100001 | 21200000 | 0.0015 | 0.0010 | 0.0011 |
| F03 | 21120001 | 21220000 | 0.0020 | 0.0012 | 0.0014 |
| F03 | 21140001 | 21240000 | 0.0021 | 0.0012 | 0.0015 |
| F03 | 21160001 | 21260000 | 0.0026 | 0.0015 | 0.0019 |

| Chr | Start | End | both | A1 | A2 |
|---|---|---|---|---|---|
| F03 | 21180001 | 21280000 | 0.0029 | 0.0019 | 0.0023 |
| F03 | 21200001 | 21300000 | 0.0034 | 0.0023 | 0.0027 |
| F03 | 21220001 | 21320000 | 0.0035 | 0.0025 | 0.0028 |
| F03 | 21240001 | 21340000 | 0.0035 | 0.0024 | 0.0026 |
| F03 | 21260001 | 21360000 | 0.0031 | 0.0021 | 0.0022 |
| F03 | 21280001 | 21380000 | 0.0029 | 0.0018 | 0.0019 |
| F03 | 21300001 | 21400000 | 0.0029 | 0.0017 | 0.0019 |
| F03 | 21320001 | 21420000 | 0.0030 | 0.0019 | 0.0020 |
| F03 | 21340001 | 21440000 | 0.0041 | 0.0030 | 0.0032 |
| F03 | 21360001 | 21460000 | 0.0051 | 0.0041 | 0.0045 |
| F03 | 21380001 | 21480000 | 0.0054 | 0.0042 | 0.0047 |
| F03 | 21400001 | 21500000 | 0.0047 | 0.0037 | 0.0042 |
| F03 | 21420001 | 21520000 | 0.0043 | 0.0032 | 0.0038 |
| F03 | 21440001 | 21540000 | 0.0036 | 0.0026 | 0.0030 |
| F03 | 21460001 | 21560000 | 0.0029 | 0.0017 | 0.0022 |
| F03 | 21480001 | 21580000 | 0.0030 | 0.0019 | 0.0024 |
| F03 | 21500001 | 21600000 | 0.0027 | 0.0017 | 0.0021 |
| F03 | 21520001 | 21620000 | 0.0025 | 0.0017 | 0.0021 |
| F03 | 21540001 | 21640000 | 0.0027 | 0.0018 | 0.0024 |
| F03 | 21560001 | 21660000 | 0.0023 | 0.0016 | 0.0021 |
| F03 | 21580001 | 21680000 | 0.0015 | 0.0012 | 0.0015 |
| F03 | 21600001 | 21700000 | 0.0017 | 0.0013 | 0.0016 |
| F03 | 21620001 | 21720000 | 0.0019 | 0.0015 | 0.0018 |
| F03 | 21640001 | 21740000 | 0.0011 | 0.0009 | 0.0009 |
| F03 | 21660001 | 21760000 | 0.0013 | 0.0010 | 0.0012 |
| F03 | 21680001 | 21780000 | 0.0023 | 0.0017 | 0.0021 |
| F03 | 21700001 | 21800000 | 0.0034 | 0.0027 | 0.0031 |
| F03 | 21720001 | 21820000 | 0.0039 | 0.0029 | 0.0034 |
| F03 | 21740001 | 21840000 | 0.0056 | 0.0045 | 0.0050 |
| F03 | 21760001 | 21860000 | 0.0055 | 0.0044 | 0.0048 |
| F03 | 21780001 | 21880000 | 0.0048 | 0.0038 | 0.0041 |
| F03 | 21800001 | 21900000 | 0.0038 | 0.0031 | 0.0032 |
| F03 | 21820001 | 21920000 | 0.0033 | 0.0029 | 0.0030 |
| F03 | 21840001 | 21940000 | 0.0017 | 0.0013 | 0.0016 |
| F03 | 21860001 | 21960000 | 0.0014 | 0.0012 | 0.0015 |
| F03 | 21880001 | 21980000 | 0.0014 | 0.0011 | 0.0014 |
| F03 | 21900001 | 22000000 | 0.0014 | 0.0011 | 0.0013 |
| F03 | 21920001 | 22020000 | 0.0012 | 0.0011 | 0.0012 |
| F03 | 21940001 | 22040000 | 0.0014 | 0.0013 | 0.0013 |
| F03 | 21960001 | 22060000 | 0.0013 | 0.0013 | 0.0013 |
| F03 | 21980001 | 22080000 | 0.0014 | 0.0014 | 0.0013 |
| F03 | 22000001 | 22100000 | 0.0011 | 0.0011 | 0.0011 |
| F03 | 22020001 | 22120000 | 0.0016 | 0.0015 | 0.0015 |
| F03 | 22040001 | 22140000 | 0.0022 | 0.0020 | 0.0021 |
| F03 | 22060001 | 22160000 | 0.0024 | 0.0022 | 0.0024 |
| F03 | 22080001 | 22180000 | 0.0025 | 0.0022 | 0.0025 |
| F03 | 22100001 | 22200000 | 0.0035 | 0.0031 | 0.0034 |
| F03 | 22120001 | 22220000 | 0.0034 | 0.0029 | 0.0032 |
| F03 | 22140001 | 22240000 | 0.0027 | 0.0023 | 0.0026 |
| F03 | 22160001 | 22260000 | 0.0024 | 0.0020 | 0.0023 |
| F03 | 22180001 | 22280000 | 0.0025 | 0.0019 | 0.0023 |
| F03 | 22200001 | 22300000 | 0.0027 | 0.0021 | 0.0024 |
| F03 | 22220001 | 22320000 | 0.0031 | 0.0022 | 0.0025 |
| F03 | 22240001 | 22340000 | 0.0035 | 0.0024 | 0.0028 |
| F03 | 22260001 | 22360000 | 0.0038 | 0.0024 | 0.0028 |
| F03 | 22280001 | 22380000 | 0.0040 | 0.0025 | 0.0030 |
| F03 | 22300001 | 22400000 | 0.0033 | 0.0019 | 0.0024 |
| F03 | 22320001 | 22420000 | 0.0031 | 0.0018 | 0.0023 |

| Chr | Start | End | both | A1 | A2 |
|---|---|---|---|---|---|
| F03 | 22340001 | 22440000 | 0.0032 | 0.0021 | 0.0024 |
| F03 | 22360001 | 22460000 | 0.0040 | 0.0029 | 0.0032 |
| F03 | 22380001 | 22480000 | 0.0039 | 0.0028 | 0.0031 |
| F03 | 22400001 | 22500000 | 0.0042 | 0.0032 | 0.0035 |
| F03 | 22420001 | 22520000 | 0.0045 | 0.0037 | 0.0039 |
| F03 | 22440001 | 22540000 | 0.0052 | 0.0044 | 0.0048 |
| F03 | 22460001 | 22560000 | 0.0045 | 0.0039 | 0.0042 |
| F03 | 22480001 | 22580000 | 0.0040 | 0.0035 | 0.0037 |
| F03 | 22500001 | 22600000 | 0.0032 | 0.0028 | 0.0030 |
| F03 | 22520001 | 22620000 | 0.0023 | 0.0020 | 0.0021 |
| F03 | 22540001 | 22640000 | 0.0018 | 0.0013 | 0.0017 |
| F03 | 22560001 | 22660000 | 0.0016 | 0.0011 | 0.0015 |
| F03 | 22580001 | 22680000 | 0.0020 | 0.0015 | 0.0018 |
| F03 | 22600001 | 22700000 | 0.0032 | 0.0023 | 0.0028 |
| F03 | 22620001 | 22720000 | 0.0032 | 0.0023 | 0.0029 |
| F03 | 22640001 | 22740000 | 0.0040 | 0.0032 | 0.0036 |
| F03 | 22660001 | 22760000 | 0.0046 | 0.0038 | 0.0043 |
| F03 | 22680001 | 22780000 | 0.0045 | 0.0037 | 0.0043 |
| F03 | 22700001 | 22800000 | 0.0040 | 0.0035 | 0.0039 |
| F03 | 22720001 | 22820000 | 0.0041 | 0.0034 | 0.0039 |
| F03 | 22740001 | 22840000 | 0.0038 | 0.0031 | 0.0036 |
| F03 | 22760001 | 22860000 | 0.0037 | 0.0028 | 0.0033 |
| F03 | 22780001 | 22880000 | 0.0037 | 0.0028 | 0.0032 |
| F03 | 22800001 | 22900000 | 0.0037 | 0.0027 | 0.0031 |
| F03 | 22820001 | 22920000 | 0.0042 | 0.0030 | 0.0033 |
| F03 | 22840001 | 22940000 | 0.0040 | 0.0028 | 0.0030 |
| F03 | 22860001 | 22960000 | 0.0038 | 0.0027 | 0.0030 |
| F03 | 22880001 | 22980000 | 0.0038 | 0.0028 | 0.0031 |
| F03 | 22900001 | 23000000 | 0.0043 | 0.0033 | 0.0036 |
| F03 | 22920001 | 23020000 | 0.0044 | 0.0037 | 0.0041 |
| F03 | 22940001 | 23040000 | 0.0037 | 0.0030 | 0.0032 |
| F03 | 22960001 | 23060000 | 0.0043 | 0.0034 | 0.0036 |
| F03 | 22980001 | 23080000 | 0.0046 | 0.0035 | 0.0038 |
| F03 | 23000001 | 23100000 | 0.0036 | 0.0025 | 0.0027 |
| F03 | 23020001 | 23120000 | 0.0029 | 0.0019 | 0.0021 |
| F03 | 23040001 | 23140000 | 0.0033 | 0.0025 | 0.0028 |
| F03 | 23060001 | 23160000 | 0.0027 | 0.0019 | 0.0022 |
| F03 | 23080001 | 23180000 | 0.0030 | 0.0022 | 0.0024 |
| F03 | 23100001 | 23200000 | 0.0034 | 0.0027 | 0.0029 |
| F03 | 23120001 | 23220000 | 0.0042 | 0.0033 | 0.0035 |
| F03 | 23140001 | 23240000 | 0.0035 | 0.0026 | 0.0028 |
| F03 | 23160001 | 23260000 | 0.0034 | 0.0028 | 0.0028 |
| F03 | 23180001 | 23280000 | 0.0039 | 0.0032 | 0.0033 |
| F03 | 23200001 | 23300000 | 0.0034 | 0.0028 | 0.0029 |
| F03 | 23220001 | 23320000 | 0.0024 | 0.0021 | 0.0022 |
| F03 | 23240001 | 23340000 | 0.0027 | 0.0025 | 0.0024 |
| F03 | 23260001 | 23360000 | 0.0027 | 0.0024 | 0.0024 |
| F03 | 23280001 | 23380000 | 0.0016 | 0.0014 | 0.0014 |
| F03 | 23300001 | 23400000 | 0.0019 | 0.0017 | 0.0017 |
| F03 | 23320001 | 23420000 | 0.0028 | 0.0025 | 0.0025 |
| F03 | 23340001 | 23440000 | 0.0032 | 0.0026 | 0.0030 |
| F03 | 23360001 | 23460000 | 0.0041 | 0.0033 | 0.0037 |
| F03 | 23380001 | 23480000 | 0.0048 | 0.0038 | 0.0044 |
| F03 | 23400001 | 23500000 | 0.0047 | 0.0036 | 0.0043 |
| F03 | 23420001 | 23520000 | 0.0042 | 0.0030 | 0.0037 |
| F03 | 23440001 | 23540000 | 0.0035 | 0.0026 | 0.0031 |
| F03 | 23460001 | 23560000 | 0.0025 | 0.0017 | 0.0023 |
| F03 | 23480001 | 23580000 | 0.0020 | 0.0013 | 0.0018 |

| Chr | Start | End | both | A1 | A2 |
|---|---|---|---|---|---|
| F03 | 23500001 | 23600000 | 0.0026 | 0.0022 | 0.0024 |
| F03 | 23520001 | 23620000 | 0.0023 | 0.0019 | 0.0022 |
| F03 | 23540001 | 23640000 | 0.0024 | 0.0021 | 0.0023 |
| F03 | 23560001 | 23660000 | 0.0024 | 0.0022 | 0.0024 |
| F03 | 23580001 | 23680000 | 0.0023 | 0.0021 | 0.0022 |
| F03 | 23600001 | 23700000 | 0.0019 | 0.0017 | 0.0019 |
| F03 | 23620001 | 23720000 | 0.0025 | 0.0023 | 0.0024 |
| F03 | 23640001 | 23740000 | 0.0026 | 0.0024 | 0.0025 |
| F03 | 23660001 | 23760000 | 0.0025 | 0.0022 | 0.0023 |
| F03 | 23680001 | 23780000 | 0.0023 | 0.0020 | 0.0022 |
| F03 | 23700001 | 23800000 | 0.0027 | 0.0022 | 0.0024 |
| F03 | 23720001 | 23820000 | 0.0024 | 0.0018 | 0.0020 |
| F03 | 23740001 | 23840000 | 0.0026 | 0.0020 | 0.0022 |
| F03 | 23760001 | 23860000 | 0.0029 | 0.0023 | 0.0026 |
| F03 | 23780001 | 23880000 | 0.0031 | 0.0025 | 0.0028 |
| F03 | 23800001 | 23900000 | 0.0029 | 0.0023 | 0.0027 |
| F03 | 23820001 | 23920000 | 0.0027 | 0.0023 | 0.0027 |
| F03 | 23840001 | 23940000 | 0.0024 | 0.0020 | 0.0024 |
| F03 | 23860001 | 23960000 | 0.0025 | 0.0021 | 0.0025 |
| F03 | 23880001 | 23980000 | 0.0036 | 0.0030 | 0.0035 |
| F03 | 23900001 | 24000000 | 0.0039 | 0.0032 | 0.0037 |
| F03 | 23920001 | 24020000 | 0.0047 | 0.0039 | 0.0046 |
| F03 | 23940001 | 24040000 | 0.0047 | 0.0038 | 0.0045 |
| F03 | 23960001 | 24060000 | 0.0057 | 0.0047 | 0.0055 |
| F03 | 23980001 | 24080000 | 0.0051 | 0.0043 | 0.0049 |
| F03 | 24000001 | 24100000 | 0.0040 | 0.0034 | 0.0040 |
| F03 | 24020001 | 24120000 | 0.0031 | 0.0027 | 0.0031 |
| F03 | 24040001 | 24140000 | 0.0040 | 0.0035 | 0.0040 |
| F03 | 24060001 | 24160000 | 0.0026 | 0.0023 | 0.0026 |
| F03 | 24080001 | 24180000 | 0.0019 | 0.0016 | 0.0019 |
| F03 | 24100001 | 24200000 | 0.0020 | 0.0017 | 0.0019 |
| F03 | 24120001 | 24220000 | 0.0018 | 0.0015 | 0.0018 |
| F03 | 24140001 | 24240000 | 0.0012 | 0.0010 | 0.0012 |
| F03 | 24160001 | 24260000 | 0.0015 | 0.0014 | 0.0015 |
| F03 | 24180001 | 24280000 | 0.0021 | 0.0019 | 0.0021 |
| F03 | 24200001 | 24300000 | 0.0021 | 0.0019 | 0.0020 |
| F03 | 24220001 | 24320000 | 0.0025 | 0.0023 | 0.0024 |
| F03 | 24240001 | 24340000 | 0.0030 | 0.0028 | 0.0029 |
| F03 | 24260001 | 24360000 | 0.0028 | 0.0025 | 0.0028 |
| F03 | 24280001 | 24380000 | 0.0027 | 0.0023 | 0.0026 |
| F03 | 24300001 | 24400000 | 0.0029 | 0.0024 | 0.0027 |
| F03 | 24320001 | 24420000 | 0.0029 | 0.0025 | 0.0027 |
| F03 | 24340001 | 24440000 | 0.0026 | 0.0021 | 0.0024 |
| F03 | 24360001 | 24460000 | 0.0031 | 0.0025 | 0.0028 |
| F03 | 24380001 | 24480000 | 0.0037 | 0.0030 | 0.0034 |
| F03 | 24400001 | 24500000 | 0.0042 | 0.0035 | 0.0039 |
| F03 | 24420001 | 24520000 | 0.0049 | 0.0042 | 0.0046 |
| F03 | 24440001 | 24540000 | 0.0043 | 0.0037 | 0.0040 |
| F03 | 24460001 | 24560000 | 0.0038 | 0.0034 | 0.0036 |
| F03 | 24480001 | 24580000 | 0.0034 | 0.0029 | 0.0030 |
| F03 | 24500001 | 24600000 | 0.0041 | 0.0034 | 0.0038 |
| F03 | 24520001 | 24620000 | 0.0034 | 0.0026 | 0.0030 |
| F03 | 24540001 | 24640000 | 0.0037 | 0.0028 | 0.0033 |
| F03 | 24560001 | 24660000 | 0.0038 | 0.0028 | 0.0034 |
| F03 | 24580001 | 24680000 | 0.0039 | 0.0029 | 0.0035 |
| F03 | 24600001 | 24700000 | 0.0047 | 0.0040 | 0.0043 |
| F03 | 24620001 | 24720000 | 0.0047 | 0.0038 | 0.0042 |
| F03 | 24640001 | 24740000 | 0.0046 | 0.0038 | 0.0041 |

| Chr | Start | End | both | A1 | A2 |
|---|---|---|---|---|---|
| F03 | 24660001 | 24760000 | 0.0045 | 0.0037 | 0.0040 |
| F03 | 24680001 | 24780000 | 0.0042 | 0.0036 | 0.0038 |
| F03 | 24700001 | 24800000 | 0.0027 | 0.0021 | 0.0024 |
| F03 | 24720001 | 24820000 | 0.0022 | 0.0019 | 0.0021 |
| F03 | 24740001 | 24840000 | 0.0018 | 0.0016 | 0.0017 |
| F03 | 24760001 | 24860000 | 0.0023 | 0.0021 | 0.0022 |
| F03 | 24780001 | 24880000 | 0.0021 | 0.0020 | 0.0021 |
| F03 | 24800001 | 24900000 | 0.0030 | 0.0027 | 0.0029 |
| F03 | 24820001 | 24920000 | 0.0035 | 0.0032 | 0.0034 |
| F03 | 24840001 | 24940000 | 0.0036 | 0.0033 | 0.0035 |
| F03 | 24860001 | 24960000 | 0.0039 | 0.0034 | 0.0037 |
| F03 | 24880001 | 24980000 | 0.0051 | 0.0044 | 0.0049 |
| F03 | 24900001 | 25000000 | 0.0040 | 0.0035 | 0.0039 |
| F03 | 24920001 | 25020000 | 0.0038 | 0.0032 | 0.0036 |
| F03 | 24940001 | 25040000 | 0.0047 | 0.0040 | 0.0043 |
| F03 | 24960001 | 25060000 | 0.0044 | 0.0039 | 0.0041 |
| F03 | 24980001 | 25080000 | 0.0039 | 0.0035 | 0.0035 |
| F03 | 25000001 | 25100000 | 0.0042 | 0.0037 | 0.0038 |
| F03 | 25020001 | 25120000 | 0.0058 | 0.0049 | 0.0052 |
| F03 | 25040001 | 25140000 | 0.0062 | 0.0053 | 0.0058 |
| F03 | 25060001 | 25160000 | 0.0058 | 0.0047 | 0.0052 |
| F03 | 25080001 | 25180000 | 0.0073 | 0.0058 | 0.0065 |
| F03 | 25100001 | 25200000 | 0.0064 | 0.0051 | 0.0057 |
| F03 | 25120001 | 25220000 | 0.0052 | 0.0040 | 0.0046 |
| F03 | 25140001 | 25240000 | 0.0042 | 0.0032 | 0.0036 |
| F03 | 25160001 | 25260000 | 0.0040 | 0.0031 | 0.0036 |
| F03 | 25180001 | 25280000 | 0.0027 | 0.0018 | 0.0022 |
| F03 | 25200001 | 25300000 | 0.0028 | 0.0019 | 0.0023 |
| F03 | 25220001 | 25320000 | 0.0027 | 0.0017 | 0.0022 |
| F03 | 25240001 | 25340000 | 0.0026 | 0.0017 | 0.0020 |
| F03 | 25260001 | 25360000 | 0.0040 | 0.0030 | 0.0036 |
| F03 | 25280001 | 25380000 | 0.0044 | 0.0036 | 0.0041 |
| F03 | 25300001 | 25400000 | 0.0048 | 0.0041 | 0.0045 |
| F03 | 25320001 | 25420000 | 0.0049 | 0.0041 | 0.0045 |
| F03 | 25340001 | 25440000 | 0.0055 | 0.0047 | 0.0051 |
| F03 | 25360001 | 25460000 | 0.0046 | 0.0038 | 0.0041 |
| F03 | 25380001 | 25480000 | 0.0042 | 0.0035 | 0.0037 |
| F03 | 25400001 | 25500000 | 0.0042 | 0.0034 | 0.0038 |
| F03 | 25420001 | 25520000 | 0.0048 | 0.0041 | 0.0045 |
| F03 | 25440001 | 25540000 | 0.0039 | 0.0033 | 0.0036 |
| F03 | 25460001 | 25560000 | 0.0040 | 0.0035 | 0.0038 |
| F03 | 25480001 | 25580000 | 0.0035 | 0.0028 | 0.0033 |
| F03 | 25500001 | 25600000 | 0.0034 | 0.0027 | 0.0032 |
| F03 | 25520001 | 25620000 | 0.0025 | 0.0019 | 0.0024 |
| F03 | 25540001 | 25640000 | 0.0033 | 0.0027 | 0.0030 |
| F03 | 25560001 | 25660000 | 0.0036 | 0.0029 | 0.0031 |
| F03 | 25580001 | 25680000 | 0.0035 | 0.0028 | 0.0030 |
| F03 | 25600001 | 25700000 | 0.0037 | 0.0029 | 0.0029 |
| F03 | 25620001 | 25720000 | 0.0039 | 0.0032 | 0.0028 |
| F03 | 25640001 | 25740000 | 0.0038 | 0.0028 | 0.0028 |
| F03 | 25660001 | 25760000 | 0.0035 | 0.0026 | 0.0025 |
| F03 | 25680001 | 25780000 | 0.0052 | 0.0042 | 0.0043 |
| F03 | 25700001 | 25800000 | 0.0052 | 0.0042 | 0.0043 |
| F03 | 25720001 | 25820000 | 0.0052 | 0.0042 | 0.0045 |
| F03 | 25740001 | 25840000 | 0.0054 | 0.0044 | 0.0045 |
| F03 | 25760001 | 25860000 | 0.0049 | 0.0039 | 0.0040 |
| F03 | 25780001 | 25880000 | 0.0035 | 0.0025 | 0.0026 |
| F03 | 25800001 | 25900000 | 0.0034 | 0.0026 | 0.0027 |

| Chr | Start | End | both | A1 | A2 |
|---|---|---|---|---|---|
| F03 | 25820001 | 25920000 | 0.0031 | 0.0024 | 0.0025 |
| F03 | 25840001 | 25940000 | 0.0030 | 0.0024 | 0.0025 |
| F03 | 25860001 | 25960000 | 0.0028 | 0.0022 | 0.0024 |
| F03 | 25880001 | 25980000 | 0.0028 | 0.0022 | 0.0023 |
| F03 | 25900001 | 26000000 | 0.0023 | 0.0017 | 0.0019 |
| F03 | 25920001 | 26020000 | 0.0021 | 0.0015 | 0.0017 |
| F03 | 25940001 | 26040000 | 0.0016 | 0.0011 | 0.0012 |
| F03 | 25960001 | 26060000 | 0.0016 | 0.0011 | 0.0012 |
| F03 | 25980001 | 26080000 | 0.0013 | 0.0010 | 0.0012 |
| F03 | 26000001 | 26100000 | 0.0016 | 0.0014 | 0.0014 |
| F03 | 26020001 | 26120000 | 0.0022 | 0.0018 | 0.0019 |
| F03 | 26040001 | 26140000 | 0.0021 | 0.0016 | 0.0017 |
| F03 | 26060001 | 26160000 | 0.0023 | 0.0017 | 0.0019 |
| F03 | 26080001 | 26180000 | 0.0023 | 0.0019 | 0.0020 |
| F03 | 26100001 | 26200000 | 0.0020 | 0.0015 | 0.0016 |
| F03 | 26120001 | 26220000 | 0.0019 | 0.0017 | 0.0017 |
| F03 | 26140001 | 26240000 | 0.0020 | 0.0017 | 0.0018 |
| F03 | 26160001 | 26260000 | 0.0019 | 0.0015 | 0.0017 |
| F03 | 26180001 | 26280000 | 0.0017 | 0.0012 | 0.0014 |
| F03 | 26200001 | 26300000 | 0.0017 | 0.0011 | 0.0013 |
| F03 | 26220001 | 26320000 | 0.0012 | 0.0006 | 0.0008 |
| F03 | 26240001 | 26340000 | 0.0011 | 0.0006 | 0.0008 |
| F03 | 26260001 | 26360000 | 0.0010 | 0.0007 | 0.0008 |
| F03 | 26280001 | 26380000 | 0.0010 | 0.0007 | 0.0008 |
| F03 | 26300001 | 26400000 | 0.0010 | 0.0007 | 0.0009 |
| F03 | 26320001 | 26420000 | 0.0011 | 0.0009 | 0.0010 |
| F03 | 26340001 | 26440000 | 0.0012 | 0.0010 | 0.0012 |
| F03 | 26360001 | 26460000 | 0.0009 | 0.0008 | 0.0009 |
| F03 | 26380001 | 26480000 | 0.0008 | 0.0007 | 0.0007 |
| F03 | 26400001 | 26500000 | 0.0014 | 0.0012 | 0.0014 |
| F03 | 26420001 | 26520000 | 0.0020 | 0.0015 | 0.0018 |
| F03 | 26440001 | 26540000 | 0.0020 | 0.0015 | 0.0018 |
| F03 | 26460001 | 26560000 | 0.0022 | 0.0017 | 0.0020 |
| F03 | 26480001 | 26580000 | 0.0025 | 0.0018 | 0.0022 |
| F03 | 26500001 | 26600000 | 0.0017 | 0.0013 | 0.0015 |
| F03 | 26520001 | 26620000 | 0.0011 | 0.0009 | 0.0009 |
| F03 | 26540001 | 26640000 | 0.0012 | 0.0010 | 0.0011 |
| F03 | 26560001 | 26660000 | 0.0013 | 0.0011 | 0.0013 |
| F03 | 26580001 | 26680000 | 0.0012 | 0.0010 | 0.0011 |
| F03 | 26600001 | 26700000 | 0.0011 | 0.0009 | 0.0010 |
| F03 | 26620001 | 26720000 | 0.0013 | 0.0011 | 0.0012 |
| F03 | 26640001 | 26740000 | 0.0012 | 0.0010 | 0.0012 |
| F03 | 26660001 | 26760000 | 0.0013 | 0.0011 | 0.0013 |
| F03 | 26680001 | 26780000 | 0.0012 | 0.0010 | 0.0011 |
| F03 | 26700001 | 26800000 | 0.0012 | 0.0010 | 0.0011 |
| F03 | 26720001 | 26820000 | 0.0013 | 0.0011 | 0.0012 |
| F03 | 26740001 | 26840000 | 0.0016 | 0.0013 | 0.0015 |
| F03 | 26760001 | 26860000 | 0.0015 | 0.0013 | 0.0015 |
| F03 | 26780001 | 26880000 | 0.0020 | 0.0018 | 0.0020 |
| F03 | 26800001 | 26900000 | 0.0024 | 0.0021 | 0.0024 |
| F03 | 26820001 | 26920000 | 0.0023 | 0.0019 | 0.0022 |
| F03 | 26840001 | 26940000 | 0.0024 | 0.0020 | 0.0023 |
| F03 | 26860001 | 26960000 | 0.0024 | 0.0019 | 0.0022 |
| F03 | 26880001 | 26980000 | 0.0020 | 0.0015 | 0.0018 |
| F03 | 26900001 | 27000000 | 0.0022 | 0.0017 | 0.0020 |
| F03 | 26920001 | 27020000 | 0.0021 | 0.0016 | 0.0017 |
| F03 | 26940001 | 27040000 | 0.0020 | 0.0015 | 0.0016 |
| F03 | 26960001 | 27060000 | 0.0021 | 0.0018 | 0.0019 |

| Chr | Start | End | both | A1 | A2 |
|---|---|---|---|---|---|
| F03 | 26980001 | 27080000 | 0.0022 | 0.0019 | 0.0020 |
| F03 | 27000001 | 27100000 | 0.0023 | 0.0020 | 0.0021 |
| F03 | 27020001 | 27120000 | 0.0025 | 0.0023 | 0.0023 |
| F03 | 27040001 | 27140000 | 0.0023 | 0.0021 | 0.0022 |
| F03 | 27060001 | 27160000 | 0.0020 | 0.0018 | 0.0019 |
| F03 | 27080001 | 27180000 | 0.0022 | 0.0020 | 0.0020 |
| F03 | 27100001 | 27200000 | 0.0016 | 0.0015 | 0.0016 |
| F03 | 27120001 | 27220000 | 0.0015 | 0.0013 | 0.0014 |
| F03 | 27140001 | 27240000 | 0.0011 | 0.0011 | 0.0011 |
| F03 | 27160001 | 27260000 | 0.0012 | 0.0010 | 0.0011 |
| F03 | 27180001 | 27280000 | 0.0009 | 0.0008 | 0.0009 |
| F03 | 27200001 | 27300000 | 0.0009 | 0.0007 | 0.0008 |
| F03 | 27220001 | 27320000 | 0.0010 | 0.0008 | 0.0010 |
| F03 | 27240001 | 27340000 | 0.0010 | 0.0008 | 0.0010 |
| F03 | 27260001 | 27360000 | 0.0012 | 0.0011 | 0.0011 |
| F03 | 27280001 | 27380000 | 0.0015 | 0.0014 | 0.0015 |
| F03 | 27300001 | 27400000 | 0.0018 | 0.0016 | 0.0018 |
| F03 | 27320001 | 27420000 | 0.0021 | 0.0019 | 0.0020 |
| F03 | 27340001 | 27440000 | 0.0022 | 0.0020 | 0.0022 |
| F03 | 27360001 | 27460000 | 0.0020 | 0.0016 | 0.0020 |
| F03 | 27380001 | 27480000 | 0.0021 | 0.0017 | 0.0020 |
| F03 | 27400001 | 27500000 | 0.0025 | 0.0019 | 0.0022 |
| F03 | 27420001 | 27520000 | 0.0024 | 0.0019 | 0.0021 |
| F03 | 27440001 | 27540000 | 0.0029 | 0.0021 | 0.0025 |
| F03 | 27460001 | 27560000 | 0.0036 | 0.0030 | 0.0032 |
| F03 | 27480001 | 27580000 | 0.0046 | 0.0038 | 0.0042 |
| F03 | 27500001 | 27600000 | 0.0041 | 0.0036 | 0.0039 |
| F03 | 27520001 | 27620000 | 0.0046 | 0.0038 | 0.0042 |
| F03 | 27540001 | 27640000 | 0.0042 | 0.0036 | 0.0039 |
| F03 | 27560001 | 27660000 | 0.0037 | 0.0030 | 0.0033 |
| F03 | 27580001 | 27680000 | 0.0027 | 0.0021 | 0.0023 |
| F03 | 27600001 | 27700000 | 0.0025 | 0.0020 | 0.0021 |
| F03 | 27620001 | 27720000 | 0.0019 | 0.0014 | 0.0016 |
| F03 | 27640001 | 27740000 | 0.0021 | 0.0016 | 0.0018 |
| F03 | 27660001 | 27760000 | 0.0022 | 0.0017 | 0.0020 |
| F03 | 27680001 | 27780000 | 0.0025 | 0.0019 | 0.0019 |
| F03 | 27700001 | 27800000 | 0.0029 | 0.0023 | 0.0022 |
| F03 | 27720001 | 27820000 | 0.0028 | 0.0023 | 0.0022 |
| F03 | 27740001 | 27840000 | 0.0028 | 0.0023 | 0.0022 |
| F03 | 27760001 | 27860000 | 0.0044 | 0.0037 | 0.0037 |
| F03 | 27780001 | 27880000 | 0.0035 | 0.0030 | 0.0033 |
| F03 | 27800001 | 27900000 | 0.0038 | 0.0032 | 0.0035 |
| F03 | 27820001 | 27920000 | 0.0046 | 0.0040 | 0.0043 |
| F03 | 27840001 | 27940000 | 0.0049 | 0.0042 | 0.0046 |
| F03 | 27860001 | 27960000 | 0.0036 | 0.0030 | 0.0033 |
| F03 | 27880001 | 27980000 | 0.0042 | 0.0035 | 0.0040 |
| F03 | 27900001 | 28000000 | 0.0044 | 0.0035 | 0.0041 |
| F03 | 27920001 | 28020000 | 0.0039 | 0.0031 | 0.0037 |
| F03 | 27940001 | 28040000 | 0.0045 | 0.0035 | 0.0042 |
| F03 | 27960001 | 28060000 | 0.0039 | 0.0030 | 0.0036 |
| F03 | 27980001 | 28080000 | 0.0038 | 0.0029 | 0.0034 |
| F03 | 28000001 | 28100000 | 0.0037 | 0.0030 | 0.0034 |
| F03 | 28020001 | 28120000 | 0.0047 | 0.0040 | 0.0043 |
| F03 | 28040001 | 28140000 | 0.0041 | 0.0035 | 0.0037 |
| F03 | 28060001 | 28160000 | 0.0049 | 0.0040 | 0.0044 |
| F03 | 28080001 | 28180000 | 0.0053 | 0.0043 | 0.0048 |
| F03 | 28100001 | 28200000 | 0.0048 | 0.0037 | 0.0043 |
| F03 | 28120001 | 28220000 | 0.0038 | 0.0028 | 0.0034 |

| Chr | Start | End | both | A1 | A2 |
|---|---|---|---|---|---|
| F03 | 28140001 | 28240000 | 0.0031 | 0.0023 | 0.0029 |
| F03 | 28160001 | 28260000 | 0.0027 | 0.0020 | 0.0025 |
| F03 | 28180001 | 28280000 | 0.0026 | 0.0021 | 0.0025 |
| F03 | 28200001 | 28300000 | 0.0023 | 0.0019 | 0.0022 |
| F03 | 28220001 | 28320000 | 0.0022 | 0.0019 | 0.0021 |
| F03 | 28240001 | 28340000 | 0.0031 | 0.0027 | 0.0029 |
| F03 | 28260001 | 28360000 | 0.0035 | 0.0030 | 0.0033 |
| F03 | 28280001 | 28380000 | 0.0034 | 0.0029 | 0.0031 |
| F03 | 28300001 | 28400000 | 0.0039 | 0.0032 | 0.0033 |
| F03 | 28320001 | 28420000 | 0.0042 | 0.0033 | 0.0035 |
| F03 | 28340001 | 28440000 | 0.0037 | 0.0029 | 0.0031 |
| F03 | 28360001 | 28460000 | 0.0032 | 0.0022 | 0.0025 |
| F03 | 28380001 | 28480000 | 0.0031 | 0.0021 | 0.0025 |
| F03 | 28400001 | 28500000 | 0.0036 | 0.0026 | 0.0030 |
| F03 | 28420001 | 28520000 | 0.0029 | 0.0021 | 0.0025 |
| F03 | 28440001 | 28540000 | 0.0035 | 0.0024 | 0.0029 |
| F03 | 28460001 | 28560000 | 0.0034 | 0.0026 | 0.0029 |
| F03 | 28480001 | 28580000 | 0.0036 | 0.0028 | 0.0032 |
| F03 | 28500001 | 28600000 | 0.0042 | 0.0033 | 0.0039 |
| F03 | 28520001 | 28620000 | 0.0043 | 0.0036 | 0.0042 |
| F03 | 28540001 | 28640000 | 0.0036 | 0.0030 | 0.0036 |
| F03 | 28560001 | 28660000 | 0.0040 | 0.0035 | 0.0040 |
| F03 | 28580001 | 28680000 | 0.0030 | 0.0026 | 0.0030 |
| F03 | 28600001 | 28700000 | 0.0015 | 0.0013 | 0.0014 |
| F03 | 28620001 | 28720000 | 0.0013 | 0.0012 | 0.0013 |
| F03 | 28640001 | 28740000 | 0.0021 | 0.0019 | 0.0020 |
| F03 | 28660001 | 28760000 | 0.0018 | 0.0015 | 0.0016 |
| F03 | 28680001 | 28780000 | 0.0028 | 0.0023 | 0.0025 |
| F03 | 28700001 | 28800000 | 0.0031 | 0.0025 | 0.0027 |
| F03 | 28720001 | 28820000 | 0.0041 | 0.0033 | 0.0037 |
| F03 | 28740001 | 28840000 | 0.0040 | 0.0031 | 0.0035 |
| F03 | 28760001 | 28860000 | 0.0038 | 0.0029 | 0.0033 |
| F03 | 28780001 | 28880000 | 0.0038 | 0.0028 | 0.0033 |
| F03 | 28800001 | 28900000 | 0.0041 | 0.0030 | 0.0035 |
| F03 | 28820001 | 28920000 | 0.0037 | 0.0027 | 0.0030 |
| F03 | 28840001 | 28940000 | 0.0031 | 0.0021 | 0.0025 |
| F03 | 28860001 | 28960000 | 0.0034 | 0.0024 | 0.0028 |
| F03 | 28880001 | 28980000 | 0.0024 | 0.0017 | 0.0020 |
| F03 | 28900001 | 29000000 | 0.0022 | 0.0014 | 0.0017 |
| F03 | 28920001 | 29020000 | 0.0021 | 0.0011 | 0.0014 |
| F03 | 28940001 | 29040000 | 0.0024 | 0.0016 | 0.0017 |
| F03 | 28960001 | 29060000 | 0.0034 | 0.0024 | 0.0027 |
| F03 | 28980001 | 29080000 | 0.0045 | 0.0032 | 0.0036 |
| F03 | 29000001 | 29100000 | 0.0061 | 0.0049 | 0.0054 |
| F03 | 29020001 | 29120000 | 0.0061 | 0.0050 | 0.0056 |
| F03 | 29040001 | 29140000 | 0.0057 | 0.0047 | 0.0054 |
| F03 | 29060001 | 29160000 | 0.0048 | 0.0039 | 0.0045 |
| F03 | 29080001 | 29180000 | 0.0041 | 0.0032 | 0.0037 |
| F03 | 29100001 | 29200000 | 0.0028 | 0.0020 | 0.0023 |
| F03 | 29120001 | 29220000 | 0.0040 | 0.0033 | 0.0036 |
| F03 | 29140001 | 29240000 | 0.0042 | 0.0034 | 0.0036 |
| F03 | 29160001 | 29260000 | 0.0041 | 0.0033 | 0.0034 |
| F03 | 29180001 | 29280000 | 0.0043 | 0.0037 | 0.0038 |
| F03 | 29200001 | 29300000 | 0.0038 | 0.0033 | 0.0035 |
| F03 | 29220001 | 29320000 | 0.0022 | 0.0017 | 0.0020 |
| F03 | 29240001 | 29340000 | 0.0023 | 0.0018 | 0.0022 |
| F03 | 29260001 | 29360000 | 0.0023 | 0.0017 | 0.0020 |
| F03 | 29280001 | 29380000 | 0.0020 | 0.0015 | 0.0017 |

| Chr | Start | End | both | A1 | A2 |
|---|---|---|---|---|---|
| F03 | 29300001 | 29400000 | 0.0022 | 0.0017 | 0.0019 |
| F03 | 29320001 | 29420000 | 0.0026 | 0.0018 | 0.0019 |
| F03 | 29340001 | 29440000 | 0.0027 | 0.0019 | 0.0020 |
| F03 | 29360001 | 29460000 | 0.0024 | 0.0018 | 0.0020 |
| F03 | 29380001 | 29480000 | 0.0030 | 0.0023 | 0.0026 |
| F03 | 29400001 | 29500000 | 0.0028 | 0.0022 | 0.0026 |
| F03 | 29420001 | 29520000 | 0.0026 | 0.0022 | 0.0026 |
| F03 | 29440001 | 29540000 | 0.0022 | 0.0019 | 0.0022 |
| F03 | 29460001 | 29560000 | 0.0021 | 0.0018 | 0.0021 |
| F03 | 29480001 | 29580000 | 0.0018 | 0.0016 | 0.0018 |
| F03 | 29500001 | 29600000 | 0.0015 | 0.0013 | 0.0015 |
| F03 | 29520001 | 29620000 | 0.0014 | 0.0012 | 0.0014 |
| F03 | 29540001 | 29640000 | 0.0013 | 0.0011 | 0.0013 |
| F03 | 29560001 | 29660000 | 0.0013 | 0.0010 | 0.0012 |
| F03 | 29580001 | 29680000 | 0.0011 | 0.0010 | 0.0011 |
| F03 | 29600001 | 29700000 | 0.0013 | 0.0012 | 0.0013 |
| F03 | 29620001 | 29720000 | 0.0010 | 0.0011 | 0.0010 |
| F03 | 29640001 | 29740000 | 0.0014 | 0.0014 | 0.0013 |
| F03 | 29660001 | 29760000 | 0.0028 | 0.0027 | 0.0027 |
| F03 | 29680001 | 29780000 | 0.0028 | 0.0024 | 0.0024 |
| F03 | 29700001 | 29800000 | 0.0029 | 0.0024 | 0.0024 |
| F03 | 29720001 | 29820000 | 0.0031 | 0.0025 | 0.0024 |
| F03 | 29740001 | 29840000 | 0.0031 | 0.0024 | 0.0023 |
| F03 | 29760001 | 29860000 | 0.0023 | 0.0017 | 0.0017 |
| F03 | 29780001 | 29880000 | 0.0024 | 0.0019 | 0.0019 |
| F03 | 29800001 | 29900000 | 0.0037 | 0.0031 | 0.0031 |
| F03 | 29820001 | 29920000 | 0.0042 | 0.0035 | 0.0038 |
| F03 | 29840001 | 29940000 | 0.0046 | 0.0040 | 0.0043 |
| F03 | 29860001 | 29960000 | 0.0046 | 0.0036 | 0.0040 |
| F03 | 29880001 | 29980000 | 0.0055 | 0.0046 | 0.0049 |
| F03 | 29900001 | 30000000 | 0.0051 | 0.0041 | 0.0044 |
| F03 | 29920001 | 30020000 | 0.0049 | 0.0040 | 0.0041 |
| F03 | 29940001 | 30040000 | 0.0048 | 0.0039 | 0.0039 |
| F03 | 29960001 | 30060000 | 0.0054 | 0.0047 | 0.0046 |
| F03 | 29980001 | 30080000 | 0.0052 | 0.0045 | 0.0046 |
| F03 | 30000001 | 30100000 | 0.0046 | 0.0041 | 0.0042 |
| F03 | 30020001 | 30120000 | 0.0046 | 0.0041 | 0.0043 |
| F03 | 30040001 | 30140000 | 0.0042 | 0.0036 | 0.0040 |
| F03 | 30060001 | 30160000 | 0.0040 | 0.0032 | 0.0035 |
| F03 | 30080001 | 30180000 | 0.0033 | 0.0023 | 0.0027 |
| F03 | 30100001 | 30200000 | 0.0037 | 0.0027 | 0.0030 |
| F03 | 30120001 | 30220000 | 0.0037 | 0.0025 | 0.0029 |
| F03 | 30140001 | 30240000 | 0.0058 | 0.0044 | 0.0050 |
| F03 | 30160001 | 30260000 | 0.0052 | 0.0040 | 0.0046 |
| F03 | 30180001 | 30280000 | 0.0050 | 0.0039 | 0.0045 |
| F03 | 30200001 | 30300000 | 0.0048 | 0.0037 | 0.0044 |
| F03 | 30220001 | 30320000 | 0.0057 | 0.0048 | 0.0054 |
| F03 | 30240001 | 30340000 | 0.0051 | 0.0044 | 0.0045 |
| F03 | 30260001 | 30360000 | 0.0071 | 0.0064 | 0.0065 |
| F03 | 30280001 | 30380000 | 0.0088 | 0.0078 | 0.0081 |
| F03 | 30300001 | 30400000 | 0.0085 | 0.0076 | 0.0079 |
| F03 | 30320001 | 30420000 | 0.0073 | 0.0065 | 0.0068 |
| F03 | 30340001 | 30440000 | 0.0064 | 0.0058 | 0.0061 |
| F03 | 30360001 | 30460000 | 0.0043 | 0.0037 | 0.0039 |
| F03 | 30380001 | 30480000 | 0.0033 | 0.0029 | 0.0030 |
| F03 | 30400001 | 30500000 | 0.0030 | 0.0026 | 0.0026 |
| F03 | 30420001 | 30520000 | 0.0028 | 0.0024 | 0.0024 |
| F03 | 30440001 | 30540000 | 0.0025 | 0.0017 | 0.0020 |

| Chr | Start | End | both | A1 | A2 |
|---|---|---|---|---|---|
| F03 | 30460001 | 30560000 | 0.0025 | 0.0016 | 0.0020 |
| F03 | 30480001 | 30580000 | 0.0019 | 0.0010 | 0.0012 |
| F03 | 30500001 | 30600000 | 0.0020 | 0.0011 | 0.0013 |
| F03 | 30520001 | 30620000 | 0.0024 | 0.0014 | 0.0017 |
| F03 | 30540001 | 30640000 | 0.0025 | 0.0016 | 0.0019 |
| F03 | 30560001 | 30660000 | 0.0031 | 0.0022 | 0.0025 |
| F03 | 30580001 | 30680000 | 0.0033 | 0.0023 | 0.0026 |
| F03 | 30600001 | 30700000 | 0.0037 | 0.0026 | 0.0031 |
| F03 | 30620001 | 30720000 | 0.0036 | 0.0026 | 0.0030 |
| F03 | 30640001 | 30740000 | 0.0036 | 0.0027 | 0.0030 |
| F03 | 30660001 | 30760000 | 0.0037 | 0.0026 | 0.0031 |
| F03 | 30680001 | 30780000 | 0.0038 | 0.0029 | 0.0033 |
| F03 | 30700001 | 30800000 | 0.0040 | 0.0031 | 0.0033 |
| F03 | 30720001 | 30820000 | 0.0047 | 0.0036 | 0.0039 |
| F03 | 30740001 | 30840000 | 0.0046 | 0.0033 | 0.0038 |
| F03 | 30760001 | 30860000 | 0.0044 | 0.0033 | 0.0036 |
| F03 | 30780001 | 30880000 | 0.0049 | 0.0037 | 0.0040 |
| F03 | 30800001 | 30900000 | 0.0041 | 0.0029 | 0.0032 |
| F03 | 30820001 | 30920000 | 0.0039 | 0.0027 | 0.0031 |
| F03 | 30840001 | 30940000 | 0.0042 | 0.0030 | 0.0034 |
| F03 | 30860001 | 30960000 | 0.0035 | 0.0025 | 0.0028 |
| F03 | 30880001 | 30980000 | 0.0031 | 0.0022 | 0.0025 |
| F03 | 30900001 | 31000000 | 0.0041 | 0.0032 | 0.0038 |
| F03 | 30920001 | 31020000 | 0.0035 | 0.0029 | 0.0033 |
| F03 | 30940001 | 31040000 | 0.0034 | 0.0028 | 0.0031 |
| F03 | 30960001 | 31060000 | 0.0038 | 0.0029 | 0.0033 |
| F03 | 30980001 | 31080000 | 0.0036 | 0.0027 | 0.0031 |
| F03 | 31000001 | 31100000 | 0.0029 | 0.0021 | 0.0024 |
| F03 | 31020001 | 31120000 | 0.0030 | 0.0022 | 0.0026 |
| F03 | 31040001 | 31140000 | 0.0023 | 0.0017 | 0.0021 |
| F03 | 31060001 | 31160000 | 0.0022 | 0.0017 | 0.0021 |
| F03 | 31080001 | 31180000 | 0.0019 | 0.0016 | 0.0018 |
| F03 | 31100001 | 31200000 | 0.0014 | 0.0011 | 0.0014 |
| F03 | 31120001 | 31220000 | 0.0013 | 0.0012 | 0.0013 |
| F03 | 31140001 | 31240000 | 0.0015 | 0.0013 | 0.0015 |
| F03 | 31160001 | 31260000 | 0.0020 | 0.0017 | 0.0020 |
| F03 | 31180001 | 31280000 | 0.0025 | 0.0020 | 0.0025 |
| F03 | 31200001 | 31300000 | 0.0025 | 0.0020 | 0.0025 |
| F03 | 31220001 | 31320000 | 0.0022 | 0.0018 | 0.0022 |
| F03 | 31240001 | 31340000 | 0.0023 | 0.0016 | 0.0021 |
| F03 | 31260001 | 31360000 | 0.0020 | 0.0012 | 0.0017 |
| F03 | 31280001 | 31380000 | 0.0015 | 0.0009 | 0.0012 |
| F03 | 31300001 | 31400000 | 0.0019 | 0.0011 | 0.0015 |
| F03 | 31320001 | 31420000 | 0.0026 | 0.0017 | 0.0021 |
| F03 | 31340001 | 31440000 | 0.0043 | 0.0035 | 0.0039 |
| F03 | 31360001 | 31460000 | 0.0046 | 0.0040 | 0.0042 |
| F03 | 31380001 | 31480000 | 0.0045 | 0.0038 | 0.0040 |
| F03 | 31400001 | 31500000 | 0.0049 | 0.0041 | 0.0045 |
| F03 | 31420001 | 31520000 | 0.0044 | 0.0038 | 0.0040 |
| F03 | 31440001 | 31540000 | 0.0024 | 0.0020 | 0.0023 |
| F03 | 31460001 | 31560000 | 0.0021 | 0.0017 | 0.0021 |
| F03 | 31480001 | 31580000 | 0.0027 | 0.0023 | 0.0027 |
| F03 | 31500001 | 31600000 | 0.0022 | 0.0019 | 0.0022 |
| F03 | 31520001 | 31620000 | 0.0032 | 0.0025 | 0.0029 |
| F03 | 31540001 | 31640000 | 0.0036 | 0.0028 | 0.0031 |
| F03 | 31560001 | 31660000 | 0.0040 | 0.0031 | 0.0033 |
| F03 | 31580001 | 31680000 | 0.0039 | 0.0030 | 0.0032 |
| F03 | 31600001 | 31700000 | 0.0045 | 0.0035 | 0.0038 |

| Chr | Start | End | both | A1 | A2 |
|---|---|---|---|---|---|
| F03 | 31620001 | 31720000 | 0.0036 | 0.0028 | 0.0031 |
| F03 | 31640001 | 31740000 | 0.0037 | 0.0029 | 0.0033 |
| F03 | 31660001 | 31760000 | 0.0035 | 0.0025 | 0.0030 |
| F03 | 31680001 | 31780000 | 0.0031 | 0.0022 | 0.0027 |
| F03 | 31700001 | 31800000 | 0.0030 | 0.0022 | 0.0026 |
| F03 | 31720001 | 31820000 | 0.0034 | 0.0024 | 0.0028 |
| F03 | 31740001 | 31840000 | 0.0033 | 0.0025 | 0.0028 |
| F03 | 31760001 | 31860000 | 0.0027 | 0.0021 | 0.0023 |
| F03 | 31780001 | 31880000 | 0.0024 | 0.0019 | 0.0021 |
| F03 | 31800001 | 31900000 | 0.0016 | 0.0012 | 0.0013 |
| F03 | 31820001 | 31920000 | 0.0008 | 0.0007 | 0.0007 |
| F03 | 31840001 | 31940000 | 0.0004 | 0.0002 | 0.0003 |
| F03 | 31860001 | 31960000 | 0.0008 | 0.0005 | 0.0006 |
| F03 | 31880001 | 31980000 | 0.0011 | 0.0008 | 0.0009 |
| F03 | 31900001 | 32000000 | 0.0013 | 0.0010 | 0.0011 |
| F03 | 31920001 | 32020000 | 0.0018 | 0.0014 | 0.0016 |
| F03 | 31940001 | 32040000 | 0.0027 | 0.0022 | 0.0025 |
| F03 | 31960001 | 32060000 | 0.0027 | 0.0022 | 0.0026 |
| F03 | 31980001 | 32080000 | 0.0028 | 0.0023 | 0.0027 |
| F03 | 32000001 | 32100000 | 0.0030 | 0.0023 | 0.0028 |
| F03 | 32020001 | 32120000 | 0.0030 | 0.0022 | 0.0027 |
| F03 | 32040001 | 32140000 | 0.0024 | 0.0015 | 0.0020 |
| F03 | 32060001 | 32160000 | 0.0024 | 0.0016 | 0.0019 |
| F03 | 32080001 | 32180000 | 0.0022 | 0.0014 | 0.0017 |
| F03 | 32100001 | 32200000 | 0.0021 | 0.0014 | 0.0017 |
| F03 | 32120001 | 32220000 | 0.0023 | 0.0018 | 0.0021 |
| F03 | 32140001 | 32240000 | 0.0022 | 0.0020 | 0.0022 |
| F03 | 32160001 | 32260000 | 0.0019 | 0.0017 | 0.0019 |
| F03 | 32180001 | 32280000 | 0.0021 | 0.0019 | 0.0021 |
| F03 | 32200001 | 32300000 | 0.0025 | 0.0020 | 0.0024 |
| F03 | 32220001 | 32320000 | 0.0028 | 0.0023 | 0.0027 |
| F03 | 32240001 | 32340000 | 0.0035 | 0.0028 | 0.0033 |
| F03 | 32260001 | 32360000 | 0.0038 | 0.0030 | 0.0034 |
| F03 | 32280001 | 32380000 | 0.0039 | 0.0029 | 0.0034 |
| F03 | 32300001 | 32400000 | 0.0037 | 0.0028 | 0.0031 |
| F03 | 32320001 | 32420000 | 0.0027 | 0.0019 | 0.0022 |
| F03 | 32340001 | 32440000 | 0.0018 | 0.0012 | 0.0014 |
| F03 | 32360001 | 32460000 | 0.0026 | 0.0021 | 0.0022 |
| F03 | 32380001 | 32480000 | 0.0026 | 0.0022 | 0.0023 |
| F03 | 32400001 | 32500000 | 0.0028 | 0.0025 | 0.0026 |
| F03 | 32420001 | 32520000 | 0.0034 | 0.0029 | 0.0032 |
| F03 | 32440001 | 32540000 | 0.0043 | 0.0037 | 0.0040 |
| F03 | 32460001 | 32560000 | 0.0038 | 0.0031 | 0.0035 |
| F03 | 32480001 | 32580000 | 0.0035 | 0.0026 | 0.0031 |
| F03 | 32500001 | 32600000 | 0.0033 | 0.0023 | 0.0028 |
| F03 | 32520001 | 32620000 | 0.0040 | 0.0032 | 0.0036 |
| F03 | 32540001 | 32640000 | 0.0030 | 0.0024 | 0.0026 |
| F03 | 32560001 | 32660000 | 0.0029 | 0.0023 | 0.0025 |
| F03 | 32580001 | 32680000 | 0.0026 | 0.0023 | 0.0024 |
| F03 | 32600001 | 32700000 | 0.0022 | 0.0020 | 0.0021 |
| F03 | 32620001 | 32720000 | 0.0014 | 0.0011 | 0.0013 |
| F03 | 32640001 | 32740000 | 0.0016 | 0.0012 | 0.0015 |
| F03 | 32660001 | 32760000 | 0.0013 | 0.0010 | 0.0013 |
| F03 | 32680001 | 32780000 | 0.0014 | 0.0011 | 0.0014 |
| F03 | 32700001 | 32800000 | 0.0015 | 0.0012 | 0.0014 |
| F03 | 32720001 | 32820000 | 0.0011 | 0.0008 | 0.0011 |
| F03 | 32740001 | 32840000 | 0.0008 | 0.0007 | 0.0008 |
| F03 | 32760001 | 32860000 | 0.0008 | 0.0007 | 0.0008 |

| Chr | Start | End | both | A1 | A2 |
|---|---|---|---|---|---|
| F03 | 32780001 | 32880000 | 0.0010 | 0.0009 | 0.0010 |
| F03 | 32800001 | 32900000 | 0.0009 | 0.0008 | 0.0009 |
| F03 | 32820001 | 32920000 | 0.0009 | 0.0008 | 0.0009 |
| F03 | 32840001 | 32940000 | 0.0009 | 0.0008 | 0.0009 |
| F03 | 32860001 | 32960000 | 0.0011 | 0.0009 | 0.0010 |
| F03 | 32880001 | 32980000 | 0.0012 | 0.0009 | 0.0011 |
| F03 | 32900001 | 33000000 | 0.0013 | 0.0011 | 0.0013 |
| F03 | 32920001 | 33020000 | 0.0016 | 0.0014 | 0.0015 |
| F03 | 32940001 | 33040000 | 0.0022 | 0.0019 | 0.0020 |
| F03 | 32960001 | 33060000 | 0.0029 | 0.0024 | 0.0025 |
| F03 | 32980001 | 33080000 | 0.0035 | 0.0028 | 0.0031 |
| F03 | 33000001 | 33100000 | 0.0036 | 0.0030 | 0.0032 |
| F03 | 33020001 | 33120000 | 0.0041 | 0.0032 | 0.0037 |
| F03 | 33040001 | 33140000 | 0.0044 | 0.0032 | 0.0039 |
| F03 | 33060001 | 33160000 | 0.0044 | 0.0032 | 0.0039 |
| F03 | 33080001 | 33180000 | 0.0040 | 0.0029 | 0.0035 |
| F03 | 33100001 | 33200000 | 0.0043 | 0.0033 | 0.0038 |
| F03 | 33120001 | 33220000 | 0.0038 | 0.0029 | 0.0034 |
| F03 | 33140001 | 33240000 | 0.0032 | 0.0025 | 0.0029 |
| F03 | 33160001 | 33260000 | 0.0030 | 0.0024 | 0.0028 |
| F03 | 33180001 | 33280000 | 0.0031 | 0.0025 | 0.0030 |
| F03 | 33200001 | 33300000 | 0.0029 | 0.0022 | 0.0028 |
| F03 | 33220001 | 33320000 | 0.0033 | 0.0027 | 0.0033 |
| F03 | 33240001 | 33340000 | 0.0035 | 0.0029 | 0.0034 |
| F03 | 33260001 | 33360000 | 0.0028 | 0.0022 | 0.0026 |
| F03 | 33280001 | 33380000 | 0.0026 | 0.0019 | 0.0021 |
| F03 | 33300001 | 33400000 | 0.0028 | 0.0020 | 0.0023 |
| F03 | 33320001 | 33420000 | 0.0025 | 0.0015 | 0.0018 |
| F03 | 33340001 | 33440000 | 0.0038 | 0.0026 | 0.0030 |
| F03 | 33360001 | 33460000 | 0.0048 | 0.0034 | 0.0041 |
| F03 | 33380001 | 33480000 | 0.0047 | 0.0033 | 0.0042 |
| F03 | 33400001 | 33500000 | 0.0054 | 0.0037 | 0.0046 |
| F03 | 33420001 | 33520000 | 0.0056 | 0.0039 | 0.0048 |
| F03 | 33440001 | 33540000 | 0.0046 | 0.0029 | 0.0036 |
| F03 | 33460001 | 33560000 | 0.0042 | 0.0028 | 0.0032 |
| F03 | 33480001 | 33580000 | 0.0046 | 0.0034 | 0.0036 |
| F03 | 33500001 | 33600000 | 0.0034 | 0.0026 | 0.0026 |
| F03 | 33520001 | 33620000 | 0.0031 | 0.0024 | 0.0024 |
| F03 | 33540001 | 33640000 | 0.0028 | 0.0023 | 0.0023 |
| F03 | 33560001 | 33660000 | 0.0022 | 0.0017 | 0.0017 |
| F03 | 33580001 | 33680000 | 0.0021 | 0.0015 | 0.0016 |
| F03 | 33600001 | 33700000 | 0.0025 | 0.0021 | 0.0022 |
| F03 | 33620001 | 33720000 | 0.0020 | 0.0017 | 0.0018 |
| F03 | 33640001 | 33740000 | 0.0022 | 0.0019 | 0.0020 |
| F03 | 33660001 | 33760000 | 0.0028 | 0.0022 | 0.0025 |
| F03 | 33680001 | 33780000 | 0.0028 | 0.0020 | 0.0024 |
| F03 | 33700001 | 33800000 | 0.0023 | 0.0016 | 0.0019 |
| F03 | 33720001 | 33820000 | 0.0032 | 0.0022 | 0.0027 |
| F03 | 33740001 | 33840000 | 0.0034 | 0.0023 | 0.0028 |
| F03 | 33760001 | 33860000 | 0.0034 | 0.0023 | 0.0027 |
| F03 | 33780001 | 33880000 | 0.0033 | 0.0022 | 0.0026 |
| F03 | 33800001 | 33900000 | 0.0043 | 0.0030 | 0.0033 |
| F03 | 33820001 | 33920000 | 0.0039 | 0.0025 | 0.0027 |
| F03 | 33840001 | 33940000 | 0.0044 | 0.0031 | 0.0032 |
| F03 | 33860001 | 33960000 | 0.0043 | 0.0031 | 0.0033 |
| F03 | 33880001 | 33980000 | 0.0043 | 0.0032 | 0.0033 |
| F03 | 33900001 | 34000000 | 0.0045 | 0.0034 | 0.0036 |
| F03 | 33920001 | 34020000 | 0.0042 | 0.0033 | 0.0034 |

| Chr | Start | End | both | A1 | A2 |
| --- | --- | --- | --- | --- | --- |
| F03 | 33940001 | 34040000 | 0.0040 | 0.0032 | 0.0035 |
| F03 | 33960001 | 34060000 | 0.0041 | 0.0034 | 0.0036 |
| F03 | 33980001 | 34080000 | 0.0035 | 0.0029 | 0.0031 |
| F03 | 34000001 | 34100000 | 0.0028 | 0.0023 | 0.0025 |
| F03 | 34020001 | 34120000 | 0.0033 | 0.0028 | 0.0030 |
| F03 | 34040001 | 34140000 | 0.0025 | 0.0020 | 0.0022 |
| F03 | 34060001 | 34160000 | 0.0020 | 0.0016 | 0.0018 |
| F03 | 34080001 | 34180000 | 0.0022 | 0.0017 | 0.0020 |
| F03 | 34100001 | 34200000 | 0.0019 | 0.0015 | 0.0017 |
| F03 | 34120001 | 34220000 | 0.0013 | 0.0010 | 0.0011 |
| F03 | 34140001 | 34240000 | 0.0014 | 0.0011 | 0.0013 |
| F03 | 34160001 | 34260000 | 0.0014 | 0.0011 | 0.0013 |
| F03 | 34180001 | 34280000 | 0.0015 | 0.0012 | 0.0013 |
| F03 | 34200001 | 34300000 | 0.0018 | 0.0014 | 0.0017 |
| F03 | 34220001 | 34320000 | 0.0024 | 0.0020 | 0.0022 |
| F03 | 34240001 | 34340000 | 0.0022 | 0.0018 | 0.0020 |
| F03 | 34260001 | 34360000 | 0.0022 | 0.0018 | 0.0020 |
| F03 | 34280001 | 34380000 | 0.0026 | 0.0021 | 0.0023 |
| F03 | 34300001 | 34400000 | 0.0026 | 0.0022 | 0.0022 |
| F03 | 34320001 | 34420000 | 0.0021 | 0.0016 | 0.0017 |
| F03 | 34340001 | 34440000 | 0.0020 | 0.0017 | 0.0017 |
| F03 | 34360001 | 34460000 | 0.0020 | 0.0018 | 0.0017 |
| F03 | 34380001 | 34480000 | 0.0013 | 0.0012 | 0.0011 |
| F03 | 34400001 | 34500000 | 0.0010 | 0.0011 | 0.0009 |
| F03 | 34420001 | 34520000 | 0.0018 | 0.0018 | 0.0016 |
| F03 | 34440001 | 34540000 | 0.0021 | 0.0018 | 0.0018 |
| F03 | 34460001 | 34560000 | 0.0021 | 0.0018 | 0.0018 |
| F03 | 34480001 | 34580000 | 0.0021 | 0.0017 | 0.0018 |
| F03 | 34500001 | 34600000 | 0.0019 | 0.0014 | 0.0015 |
| F03 | 34520001 | 34620000 | 0.0018 | 0.0012 | 0.0013 |
| F03 | 34540001 | 34640000 | 0.0016 | 0.0010 | 0.0012 |
| F03 | 34560001 | 34660000 | 0.0016 | 0.0011 | 0.0013 |
| F03 | 34580001 | 34680000 | 0.0015 | 0.0011 | 0.0012 |
| F03 | 34600001 | 34700000 | 0.0023 | 0.0018 | 0.0020 |
| F03 | 34620001 | 34720000 | 0.0017 | 0.0013 | 0.0015 |
| F03 | 34640001 | 34740000 | 0.0025 | 0.0020 | 0.0021 |
| F03 | 34660001 | 34760000 | 0.0025 | 0.0020 | 0.0021 |
| F03 | 34680001 | 34780000 | 0.0029 | 0.0023 | 0.0023 |
| F03 | 34700001 | 34800000 | 0.0020 | 0.0016 | 0.0016 |
| F03 | 34720001 | 34820000 | 0.0022 | 0.0017 | 0.0017 |
| F03 | 34740001 | 34840000 | 0.0017 | 0.0013 | 0.0014 |
| F03 | 34760001 | 34860000 | 0.0022 | 0.0017 | 0.0019 |
| F03 | 34780001 | 34880000 | 0.0025 | 0.0021 | 0.0022 |
| F03 | 34800001 | 34900000 | 0.0031 | 0.0026 | 0.0027 |
| F03 | 34820001 | 34920000 | 0.0033 | 0.0027 | 0.0029 |
| F03 | 34840001 | 34940000 | 0.0030 | 0.0022 | 0.0024 |
| F03 | 34860001 | 34960000 | 0.0030 | 0.0023 | 0.0024 |
| F03 | 34880001 | 34980000 | 0.0035 | 0.0024 | 0.0027 |
| F03 | 34900001 | 35000000 | 0.0032 | 0.0021 | 0.0023 |
| F03 | 34920001 | 35020000 | 0.0028 | 0.0019 | 0.0022 |
| F03 | 34940001 | 35040000 | 0.0029 | 0.0022 | 0.0025 |
| F03 | 34960001 | 35060000 | 0.0025 | 0.0017 | 0.0021 |
| F03 | 34980001 | 35080000 | 0.0016 | 0.0010 | 0.0013 |
| F03 | 35000001 | 35100000 | 0.0018 | 0.0011 | 0.0015 |
| F03 | 35020001 | 35120000 | 0.0021 | 0.0014 | 0.0019 |
| F03 | 35040001 | 35140000 | 0.0023 | 0.0013 | 0.0018 |
| F03 | 35060001 | 35160000 | 0.0024 | 0.0014 | 0.0018 |
| F03 | 35080001 | 35180000 | 0.0022 | 0.0013 | 0.0017 |

| Chr | Start | End | both | A1 | A2 |
|---|---|---|---|---|---|
| F03 | 35100001 | 35200000 | 0.0023 | 0.0016 | 0.0018 |
| F03 | 35120001 | 35220000 | 0.0024 | 0.0018 | 0.0019 |
| F03 | 35140001 | 35240000 | 0.0023 | 0.0019 | 0.0019 |
| F03 | 35160001 | 35260000 | 0.0021 | 0.0017 | 0.0018 |
| F03 | 35180001 | 35280000 | 0.0023 | 0.0020 | 0.0020 |
| F03 | 35200001 | 35300000 | 0.0021 | 0.0016 | 0.0019 |
| F03 | 35220001 | 35320000 | 0.0021 | 0.0015 | 0.0017 |
| F03 | 35240001 | 35340000 | 0.0020 | 0.0014 | 0.0018 |
| F03 | 35260001 | 35360000 | 0.0020 | 0.0013 | 0.0017 |
| F03 | 35280001 | 35380000 | 0.0020 | 0.0012 | 0.0017 |
| F03 | 35300001 | 35400000 | 0.0020 | 0.0013 | 0.0016 |
| F03 | 35320001 | 35420000 | 0.0018 | 0.0013 | 0.0016 |
| F03 | 35340001 | 35440000 | 0.0018 | 0.0012 | 0.0016 |
| F03 | 35360001 | 35460000 | 0.0014 | 0.0011 | 0.0014 |
| F03 | 35380001 | 35480000 | 0.0014 | 0.0012 | 0.0014 |
| F03 | 35400001 | 35500000 | 0.0014 | 0.0012 | 0.0013 |
| F03 | 35420001 | 35520000 | 0.0018 | 0.0016 | 0.0017 |
| F03 | 35440001 | 35540000 | 0.0020 | 0.0016 | 0.0017 |
| F03 | 35460001 | 35560000 | 0.0022 | 0.0016 | 0.0017 |
| F03 | 35480001 | 35580000 | 0.0020 | 0.0014 | 0.0016 |
| F03 | 35500001 | 35600000 | 0.0019 | 0.0013 | 0.0015 |
| F03 | 35520001 | 35620000 | 0.0011 | 0.0006 | 0.0008 |
| F03 | 35540001 | 35640000 | 0.0014 | 0.0008 | 0.0011 |
| F03 | 35560001 | 35660000 | 0.0014 | 0.0010 | 0.0013 |
| F03 | 35580001 | 35680000 | 0.0018 | 0.0014 | 0.0017 |
| F03 | 35600001 | 35700000 | 0.0019 | 0.0016 | 0.0017 |
| F03 | 35620001 | 35720000 | 0.0021 | 0.0018 | 0.0019 |
| F03 | 35640001 | 35740000 | 0.0016 | 0.0015 | 0.0016 |
| F03 | 35660001 | 35760000 | 0.0016 | 0.0015 | 0.0016 |
| F03 | 35680001 | 35780000 | 0.0013 | 0.0012 | 0.0013 |
| F03 | 35700001 | 35800000 | 0.0013 | 0.0012 | 0.0013 |
| F03 | 35720001 | 35820000 | 0.0011 | 0.0010 | 0.0011 |
| F03 | 35740001 | 35840000 | 0.0015 | 0.0013 | 0.0014 |
| F03 | 35760001 | 35860000 | 0.0020 | 0.0018 | 0.0019 |
| F03 | 35780001 | 35880000 | 0.0022 | 0.0020 | 0.0021 |
| F03 | 35800001 | 35900000 | 0.0025 | 0.0023 | 0.0024 |
| F03 | 35820001 | 35920000 | 0.0038 | 0.0036 | 0.0037 |
| F03 | 35840001 | 35940000 | 0.0038 | 0.0035 | 0.0036 |
| F03 | 35860001 | 35960000 | 0.0034 | 0.0031 | 0.0032 |
| F03 | 35880001 | 35980000 | 0.0034 | 0.0031 | 0.0032 |
| F03 | 35900001 | 36000000 | 0.0038 | 0.0033 | 0.0036 |
| F03 | 35920001 | 36020000 | 0.0030 | 0.0025 | 0.0027 |
| F03 | 35940001 | 36040000 | 0.0035 | 0.0029 | 0.0030 |
| F03 | 35960001 | 36060000 | 0.0040 | 0.0032 | 0.0033 |
| F03 | 35980001 | 36080000 | 0.0045 | 0.0035 | 0.0037 |
| F03 | 36000001 | 36100000 | 0.0040 | 0.0028 | 0.0030 |
| F03 | 36020001 | 36120000 | 0.0045 | 0.0033 | 0.0035 |
| F03 | 36040001 | 36140000 | 0.0042 | 0.0030 | 0.0033 |
| F03 | 36060001 | 36160000 | 0.0038 | 0.0027 | 0.0031 |
| F03 | 36080001 | 36180000 | 0.0035 | 0.0024 | 0.0027 |
| F03 | 36100001 | 36200000 | 0.0032 | 0.0024 | 0.0026 |
| F03 | 36120001 | 36220000 | 0.0033 | 0.0023 | 0.0027 |
| F03 | 36140001 | 36240000 | 0.0027 | 0.0019 | 0.0022 |
| F03 | 36160001 | 36260000 | 0.0027 | 0.0021 | 0.0023 |
| F03 | 36180001 | 36280000 | 0.0032 | 0.0027 | 0.0029 |
| F03 | 36200001 | 36300000 | 0.0033 | 0.0026 | 0.0029 |
| F03 | 36220001 | 36320000 | 0.0025 | 0.0021 | 0.0022 |
| F03 | 36240001 | 36340000 | 0.0030 | 0.0025 | 0.0027 |

| Chr | Start | End | both | A1 | A2 |
|---|---|---|---|---|---|
| F03 | 36260001 | 36360000 | 0.0036 | 0.0031 | 0.0034 |
| F03 | 36280001 | 36380000 | 0.0034 | 0.0030 | 0.0032 |
| F03 | 36300001 | 36400000 | 0.0049 | 0.0043 | 0.0047 |
| F03 | 36320001 | 36420000 | 0.0053 | 0.0048 | 0.0051 |
| F03 | 36340001 | 36440000 | 0.0057 | 0.0049 | 0.0052 |
| F03 | 36360001 | 36460000 | 0.0053 | 0.0046 | 0.0048 |
| F03 | 36380001 | 36480000 | 0.0048 | 0.0041 | 0.0043 |
| F03 | 36400001 | 36500000 | 0.0038 | 0.0031 | 0.0032 |
| F03 | 36420001 | 36520000 | 0.0036 | 0.0025 | 0.0028 |
| F03 | 36440001 | 36540000 | 0.0037 | 0.0026 | 0.0030 |
| F03 | 36460001 | 36560000 | 0.0038 | 0.0025 | 0.0030 |
| F03 | 36480001 | 36580000 | 0.0037 | 0.0023 | 0.0028 |
| F03 | 36500001 | 36600000 | 0.0037 | 0.0023 | 0.0029 |
| F03 | 36520001 | 36620000 | 0.0039 | 0.0026 | 0.0031 |
| F03 | 36540001 | 36640000 | 0.0032 | 0.0019 | 0.0023 |
| F03 | 36560001 | 36660000 | 0.0027 | 0.0017 | 0.0019 |
| F03 | 36580001 | 36680000 | 0.0029 | 0.0020 | 0.0022 |
| F03 | 36600001 | 36700000 | 0.0029 | 0.0020 | 0.0024 |
| F03 | 36620001 | 36720000 | 0.0032 | 0.0024 | 0.0029 |
| F03 | 36640001 | 36740000 | 0.0034 | 0.0028 | 0.0032 |
| F03 | 36660001 | 36760000 | 0.0033 | 0.0028 | 0.0032 |
| F03 | 36680001 | 36780000 | 0.0033 | 0.0027 | 0.0031 |
| F03 | 36700001 | 36800000 | 0.0027 | 0.0023 | 0.0026 |
| F03 | 36720001 | 36820000 | 0.0018 | 0.0016 | 0.0018 |
| F03 | 36740001 | 36840000 | 0.0016 | 0.0013 | 0.0015 |
| F03 | 36760001 | 36860000 | 0.0014 | 0.0012 | 0.0013 |
| F03 | 36780001 | 36880000 | 0.0012 | 0.0011 | 0.0011 |
| F03 | 36800001 | 36900000 | 0.0013 | 0.0011 | 0.0013 |
| F03 | 36820001 | 36920000 | 0.0021 | 0.0017 | 0.0020 |
| F03 | 36840001 | 36940000 | 0.0019 | 0.0016 | 0.0018 |
| F03 | 36860001 | 36960000 | 0.0017 | 0.0013 | 0.0017 |
| F03 | 36880001 | 36980000 | 0.0037 | 0.0030 | 0.0035 |
| F03 | 36900001 | 37000000 | 0.0039 | 0.0031 | 0.0036 |
| F03 | 36920001 | 37020000 | 0.0031 | 0.0024 | 0.0027 |
| F03 | 36940001 | 37040000 | 0.0049 | 0.0039 | 0.0043 |
| F03 | 36960001 | 37060000 | 0.0064 | 0.0051 | 0.0055 |
| F03 | 36980001 | 37080000 | 0.0048 | 0.0036 | 0.0039 |
| F03 | 37000001 | 37100000 | 0.0043 | 0.0032 | 0.0035 |
| F03 | 37020001 | 37120000 | 0.0044 | 0.0033 | 0.0037 |
| F03 | 37040001 | 37140000 | 0.0033 | 0.0021 | 0.0023 |
| F03 | 37060001 | 37160000 | 0.0031 | 0.0020 | 0.0023 |
| F03 | 37080001 | 37180000 | 0.0027 | 0.0019 | 0.0020 |
| F03 | 37100001 | 37200000 | 0.0035 | 0.0023 | 0.0026 |
| F03 | 37120001 | 37220000 | 0.0039 | 0.0025 | 0.0027 |
| F03 | 37140001 | 37240000 | 0.0035 | 0.0024 | 0.0026 |
| F03 | 37160001 | 37260000 | 0.0021 | 0.0013 | 0.0015 |
| F03 | 37180001 | 37280000 | 0.0018 | 0.0012 | 0.0013 |
| F03 | 37200001 | 37300000 | 0.0016 | 0.0012 | 0.0013 |
| F03 | 37220001 | 37320000 | 0.0011 | 0.0009 | 0.0010 |
| F03 | 37240001 | 37340000 | 0.0015 | 0.0011 | 0.0014 |
| F03 | 37260001 | 37360000 | 0.0019 | 0.0014 | 0.0017 |
| F03 | 37280001 | 37380000 | 0.0018 | 0.0013 | 0.0016 |
| F03 | 37300001 | 37400000 | 0.0014 | 0.0010 | 0.0012 |
| F03 | 37320001 | 37420000 | 0.0012 | 0.0009 | 0.0011 |
| F03 | 37340001 | 37440000 | 0.0009 | 0.0007 | 0.0008 |
| F03 | 37360001 | 37460000 | 0.0007 | 0.0006 | 0.0007 |
| F03 | 37380001 | 37480000 | 0.0010 | 0.0007 | 0.0008 |
| F03 | 37400001 | 37500000 | 0.0015 | 0.0010 | 0.0012 |

| Chr | Start | End | both | A1 | A2 |
|---|---|---|---|---|---|
| F03 | 37420001 | 37520000 | 0.0016 | 0.0011 | 0.0013 |
| F03 | 37440001 | 37540000 | 0.0018 | 0.0013 | 0.0015 |
| F03 | 37460001 | 37560000 | 0.0022 | 0.0016 | 0.0019 |
| F03 | 37480001 | 37580000 | 0.0024 | 0.0018 | 0.0022 |
| F03 | 37500001 | 37600000 | 0.0020 | 0.0014 | 0.0017 |
| F03 | 37520001 | 37620000 | 0.0022 | 0.0016 | 0.0019 |
| F03 | 37540001 | 37640000 | 0.0017 | 0.0012 | 0.0014 |
| F03 | 37560001 | 37660000 | 0.0025 | 0.0018 | 0.0020 |
| F03 | 37580001 | 37680000 | 0.0022 | 0.0017 | 0.0018 |
| F03 | 37600001 | 37700000 | 0.0022 | 0.0017 | 0.0019 |
| F03 | 37620001 | 37720000 | 0.0021 | 0.0017 | 0.0019 |
| F03 | 37640001 | 37740000 | 0.0021 | 0.0017 | 0.0019 |
| F03 | 37660001 | 37760000 | 0.0013 | 0.0011 | 0.0013 |
| F03 | 37680001 | 37780000 | 0.0015 | 0.0012 | 0.0014 |
| F03 | 37700001 | 37800000 | 0.0015 | 0.0013 | 0.0014 |
| F03 | 37720001 | 37820000 | 0.0013 | 0.0011 | 0.0012 |
| F03 | 37740001 | 37840000 | 0.0013 | 0.0011 | 0.0012 |
| F03 | 37760001 | 37860000 | 0.0017 | 0.0012 | 0.0014 |
| F03 | 37780001 | 37880000 | 0.0014 | 0.0010 | 0.0012 |
| F03 | 37800001 | 37900000 | 0.0020 | 0.0013 | 0.0015 |
| F03 | 37820001 | 37920000 | 0.0026 | 0.0017 | 0.0020 |
| F03 | 37840001 | 37940000 | 0.0034 | 0.0024 | 0.0028 |
| F03 | 37860001 | 37960000 | 0.0032 | 0.0022 | 0.0025 |
| F03 | 37880001 | 37980000 | 0.0030 | 0.0020 | 0.0023 |
| F03 | 37900001 | 38000000 | 0.0023 | 0.0016 | 0.0019 |
| F03 | 37920001 | 38020000 | 0.0020 | 0.0013 | 0.0016 |
| F03 | 37940001 | 38040000 | 0.0020 | 0.0015 | 0.0017 |
| F03 | 37960001 | 38060000 | 0.0018 | 0.0015 | 0.0017 |
| F03 | 37980001 | 38080000 | 0.0020 | 0.0017 | 0.0019 |
| F03 | 38000001 | 38100000 | 0.0029 | 0.0024 | 0.0028 |
| F03 | 38020001 | 38120000 | 0.0037 | 0.0032 | 0.0035 |
| F03 | 38040001 | 38140000 | 0.0035 | 0.0028 | 0.0031 |
| F03 | 38060001 | 38160000 | 0.0032 | 0.0025 | 0.0028 |
| F03 | 38080001 | 38180000 | 0.0036 | 0.0028 | 0.0031 |
| F03 | 38100001 | 38200000 | 0.0040 | 0.0031 | 0.0035 |
| F03 | 38120001 | 38220000 | 0.0038 | 0.0030 | 0.0032 |
| F03 | 38140001 | 38240000 | 0.0038 | 0.0028 | 0.0030 |
| F03 | 38160001 | 38260000 | 0.0044 | 0.0033 | 0.0035 |
| F03 | 38180001 | 38280000 | 0.0045 | 0.0033 | 0.0035 |
| F03 | 38200001 | 38300000 | 0.0038 | 0.0027 | 0.0028 |
| F03 | 38220001 | 38320000 | 0.0030 | 0.0020 | 0.0021 |
| F03 | 38240001 | 38340000 | 0.0026 | 0.0019 | 0.0020 |
| F03 | 38260001 | 38360000 | 0.0027 | 0.0019 | 0.0021 |
| F03 | 38280001 | 38380000 | 0.0024 | 0.0016 | 0.0020 |
| F03 | 38300001 | 38400000 | 0.0023 | 0.0015 | 0.0019 |
| F03 | 38320001 | 38420000 | 0.0029 | 0.0020 | 0.0024 |
| F03 | 38340001 | 38440000 | 0.0034 | 0.0025 | 0.0029 |
| F03 | 38360001 | 38460000 | 0.0026 | 0.0021 | 0.0023 |
| F03 | 38380001 | 38480000 | 0.0026 | 0.0020 | 0.0022 |
| F03 | 38400001 | 38500000 | 0.0025 | 0.0020 | 0.0020 |
| F03 | 38420001 | 38520000 | 0.0019 | 0.0015 | 0.0016 |
| F03 | 38440001 | 38540000 | 0.0018 | 0.0015 | 0.0015 |
| F03 | 38460001 | 38560000 | 0.0024 | 0.0020 | 0.0020 |
| F03 | 38480001 | 38580000 | 0.0027 | 0.0020 | 0.0022 |
| F03 | 38500001 | 38600000 | 0.0032 | 0.0025 | 0.0027 |
| F03 | 38520001 | 38620000 | 0.0034 | 0.0024 | 0.0027 |
| F03 | 38540001 | 38640000 | 0.0035 | 0.0025 | 0.0027 |
| F03 | 38560001 | 38660000 | 0.0040 | 0.0029 | 0.0030 |

| Chr | Start | End | both | A1 | A2 |
|---|---|---|---|---|---|
| F03 | 38580001 | 38680000 | 0.0044 | 0.0033 | 0.0034 |
| F03 | 38600001 | 38700000 | 0.0034 | 0.0026 | 0.0027 |
| F03 | 38620001 | 38720000 | 0.0031 | 0.0024 | 0.0025 |
| F03 | 38640001 | 38740000 | 0.0031 | 0.0025 | 0.0026 |
| F03 | 38660001 | 38760000 | 0.0023 | 0.0019 | 0.0022 |
| F03 | 38680001 | 38780000 | 0.0016 | 0.0015 | 0.0016 |
| F03 | 38700001 | 38800000 | 0.0024 | 0.0021 | 0.0022 |
| F03 | 38720001 | 38820000 | 0.0042 | 0.0036 | 0.0038 |
| F03 | 38740001 | 38840000 | 0.0037 | 0.0032 | 0.0033 |
| F03 | 38760001 | 38860000 | 0.0041 | 0.0036 | 0.0037 |
| F03 | 38780001 | 38880000 | 0.0054 | 0.0048 | 0.0049 |
| F03 | 38800001 | 38900000 | 0.0052 | 0.0046 | 0.0047 |
| F03 | 38820001 | 38920000 | 0.0042 | 0.0036 | 0.0036 |
| F03 | 38840001 | 38940000 | 0.0053 | 0.0047 | 0.0046 |
| F03 | 38860001 | 38960000 | 0.0049 | 0.0044 | 0.0043 |
| F03 | 38880001 | 38980000 | 0.0050 | 0.0044 | 0.0042 |
| F03 | 38900001 | 39000000 | 0.0052 | 0.0046 | 0.0045 |
| F03 | 38920001 | 39020000 | 0.0047 | 0.0043 | 0.0042 |
| F03 | 38940001 | 39040000 | 0.0053 | 0.0048 | 0.0047 |
| F03 | 38960001 | 39060000 | 0.0056 | 0.0048 | 0.0049 |
| F03 | 38980001 | 39080000 | 0.0048 | 0.0039 | 0.0042 |
| F03 | 39000001 | 39100000 | 0.0049 | 0.0038 | 0.0041 |
| F03 | 39020001 | 39120000 | 0.0055 | 0.0043 | 0.0046 |
| F03 | 39040001 | 39140000 | 0.0042 | 0.0029 | 0.0032 |
| F03 | 39060001 | 39160000 | 0.0035 | 0.0023 | 0.0027 |
| F03 | 39080001 | 39180000 | 0.0031 | 0.0019 | 0.0022 |
| F03 | 39100001 | 39200000 | 0.0023 | 0.0014 | 0.0016 |
| F03 | 39120001 | 39220000 | 0.0018 | 0.0011 | 0.0013 |
| F03 | 39140001 | 39240000 | 0.0013 | 0.0008 | 0.0010 |
| F03 | 39160001 | 39260000 | 0.0025 | 0.0021 | 0.0021 |
| F03 | 39180001 | 39280000 | 0.0027 | 0.0024 | 0.0024 |
| F03 | 39200001 | 39300000 | 0.0031 | 0.0026 | 0.0026 |
| F03 | 39220001 | 39320000 | 0.0034 | 0.0027 | 0.0027 |
| F03 | 39240001 | 39340000 | 0.0033 | 0.0025 | 0.0025 |
| F03 | 39260001 | 39360000 | 0.0024 | 0.0015 | 0.0017 |
| F03 | 39280001 | 39380000 | 0.0025 | 0.0016 | 0.0019 |
| F03 | 39300001 | 39400000 | 0.0026 | 0.0017 | 0.0021 |
| F03 | 39320001 | 39420000 | 0.0021 | 0.0015 | 0.0019 |
| F03 | 39340001 | 39440000 | 0.0022 | 0.0016 | 0.0020 |
| F03 | 39360001 | 39460000 | 0.0021 | 0.0017 | 0.0020 |
| F03 | 39380001 | 39480000 | 0.0020 | 0.0016 | 0.0020 |
| F03 | 39400001 | 39500000 | 0.0019 | 0.0015 | 0.0019 |
| F03 | 39420001 | 39520000 | 0.0020 | 0.0015 | 0.0020 |
| F03 | 39440001 | 39540000 | 0.0026 | 0.0020 | 0.0025 |
| F03 | 39460001 | 39560000 | 0.0021 | 0.0015 | 0.0020 |
| F03 | 39480001 | 39580000 | 0.0020 | 0.0015 | 0.0018 |
| F03 | 39500001 | 39600000 | 0.0015 | 0.0011 | 0.0014 |
| F03 | 39520001 | 39620000 | 0.0014 | 0.0012 | 0.0013 |
| F03 | 39540001 | 39640000 | 0.0012 | 0.0010 | 0.0011 |
| F03 | 39560001 | 39660000 | 0.0013 | 0.0010 | 0.0011 |
| F03 | 39580001 | 39680000 | 0.0010 | 0.0008 | 0.0008 |
| F03 | 39600001 | 39700000 | 0.0010 | 0.0008 | 0.0008 |
| F03 | 39620001 | 39720000 | 0.0013 | 0.0010 | 0.0011 |
| F03 | 39640001 | 39740000 | 0.0008 | 0.0006 | 0.0006 |
| F03 | 39660001 | 39760000 | 0.0009 | 0.0007 | 0.0008 |
| F03 | 39680001 | 39780000 | 0.0010 | 0.0008 | 0.0009 |
| F03 | 39700001 | 39800000 | 0.0016 | 0.0013 | 0.0016 |
| F03 | 39720001 | 39820000 | 0.0014 | 0.0012 | 0.0014 |

| Chr | Start | End | both | A1 | A2 |
|---|---|---|---|---|---|
| F03 | 39740001 | 39840000 | 0.0017 | 0.0015 | 0.0017 |
| F03 | 39760001 | 39860000 | 0.0019 | 0.0016 | 0.0018 |
| F03 | 39780001 | 39880000 | 0.0022 | 0.0018 | 0.0021 |
| F03 | 39800001 | 39900000 | 0.0025 | 0.0020 | 0.0022 |
| F03 | 39820001 | 39920000 | 0.0031 | 0.0024 | 0.0027 |
| F03 | 39840001 | 39940000 | 0.0029 | 0.0021 | 0.0025 |
| F03 | 39860001 | 39960000 | 0.0031 | 0.0023 | 0.0026 |
| F03 | 39880001 | 39980000 | 0.0032 | 0.0025 | 0.0027 |
| F03 | 39900001 | 40000000 | 0.0025 | 0.0019 | 0.0021 |
| F03 | 39920001 | 40020000 | 0.0024 | 0.0019 | 0.0021 |
| F03 | 39940001 | 40040000 | 0.0035 | 0.0030 | 0.0033 |
| F03 | 39960001 | 40060000 | 0.0030 | 0.0025 | 0.0028 |
| F03 | 39980001 | 40080000 | 0.0026 | 0.0023 | 0.0025 |
| F03 | 40000001 | 40100000 | 0.0025 | 0.0023 | 0.0025 |
| F03 | 40020001 | 40120000 | 0.0023 | 0.0020 | 0.0022 |
| F03 | 40040001 | 40140000 | 0.0014 | 0.0010 | 0.0011 |
| F03 | 40060001 | 40160000 | 0.0019 | 0.0014 | 0.0014 |
| F03 | 40080001 | 40180000 | 0.0017 | 0.0011 | 0.0012 |
| F03 | 40100001 | 40200000 | 0.0021 | 0.0015 | 0.0016 |
| F03 | 40120001 | 40220000 | 0.0020 | 0.0015 | 0.0015 |
| F03 | 40140001 | 40240000 | 0.0021 | 0.0016 | 0.0015 |
| F03 | 40160001 | 40260000 | 0.0017 | 0.0012 | 0.0013 |
| F03 | 40180001 | 40280000 | 0.0018 | 0.0013 | 0.0014 |
| F03 | 40200001 | 40300000 | 0.0022 | 0.0016 | 0.0018 |
| F03 | 40220001 | 40320000 | 0.0018 | 0.0013 | 0.0014 |
| F03 | 40240001 | 40340000 | 0.0016 | 0.0012 | 0.0014 |
| F03 | 40260001 | 40360000 | 0.0018 | 0.0015 | 0.0017 |
| F03 | 40280001 | 40380000 | 0.0016 | 0.0014 | 0.0016 |
| F03 | 40300001 | 40400000 | 0.0007 | 0.0007 | 0.0007 |
| F03 | 40320001 | 40420000 | 0.0009 | 0.0008 | 0.0009 |
| F03 | 40340001 | 40440000 | 0.0007 | 0.0006 | 0.0007 |
| F03 | 40360001 | 40460000 | 0.0006 | 0.0005 | 0.0006 |
| F03 | 40380001 | 40480000 | 0.0006 | 0.0005 | 0.0006 |
| F03 | 40400001 | 40500000 | 0.0010 | 0.0008 | 0.0009 |
| F03 | 40420001 | 40520000 | 0.0011 | 0.0010 | 0.0011 |
| F03 | 40440001 | 40540000 | 0.0011 | 0.0009 | 0.0011 |
| F03 | 40460001 | 40560000 | 0.0011 | 0.0009 | 0.0010 |
| F03 | 40480001 | 40580000 | 0.0014 | 0.0011 | 0.0013 |
| F03 | 40500001 | 40600000 | 0.0018 | 0.0014 | 0.0015 |
| F03 | 40520001 | 40620000 | 0.0022 | 0.0017 | 0.0019 |
| F03 | 40540001 | 40640000 | 0.0035 | 0.0027 | 0.0030 |
| F03 | 40560001 | 40660000 | 0.0037 | 0.0030 | 0.0033 |
| F03 | 40580001 | 40680000 | 0.0041 | 0.0032 | 0.0035 |
| F03 | 40600001 | 40700000 | 0.0036 | 0.0028 | 0.0031 |
| F03 | 40620001 | 40720000 | 0.0035 | 0.0027 | 0.0030 |
| F03 | 40640001 | 40740000 | 0.0022 | 0.0018 | 0.0019 |
| F03 | 40660001 | 40760000 | 0.0025 | 0.0018 | 0.0022 |
| F03 | 40680001 | 40780000 | 0.0024 | 0.0016 | 0.0020 |
| F03 | 40700001 | 40800000 | 0.0025 | 0.0017 | 0.0022 |
| F03 | 40720001 | 40820000 | 0.0020 | 0.0013 | 0.0017 |
| F03 | 40740001 | 40840000 | 0.0023 | 0.0015 | 0.0019 |
| F03 | 40760001 | 40860000 | 0.0022 | 0.0016 | 0.0019 |
| F03 | 40780001 | 40880000 | 0.0019 | 0.0015 | 0.0018 |
| F03 | 40800001 | 40900000 | 0.0020 | 0.0016 | 0.0019 |
| F03 | 40820001 | 40920000 | 0.0025 | 0.0019 | 0.0022 |
| F03 | 40840001 | 40940000 | 0.0024 | 0.0016 | 0.0022 |
| F03 | 40860001 | 40960000 | 0.0026 | 0.0016 | 0.0022 |
| F03 | 40880001 | 40980000 | 0.0026 | 0.0015 | 0.0021 |

| Chr | Start | End | both | A1 | A2 |
|---|---|---|---|---|---|
| F03 | 40900001 | 41000000 | 0.0022 | 0.0012 | 0.0017 |
| F03 | 40920001 | 41020000 | 0.0018 | 0.0010 | 0.0015 |
| F03 | 40940001 | 41040000 | 0.0015 | 0.0011 | 0.0013 |
| F03 | 40960001 | 41060000 | 0.0008 | 0.0006 | 0.0007 |
| F03 | 40980001 | 41080000 | 0.0006 | 0.0005 | 0.0006 |
| F03 | 41000001 | 41100000 | 0.0008 | 0.0007 | 0.0008 |
| F03 | 41020001 | 41120000 | 0.0009 | 0.0008 | 0.0009 |
| F03 | 41040001 | 41140000 | 0.0011 | 0.0010 | 0.0011 |
| F03 | 41060001 | 41160000 | 0.0013 | 0.0012 | 0.0012 |
| F03 | 41080001 | 41180000 | 0.0015 | 0.0013 | 0.0014 |
| F03 | 41100001 | 41200000 | 0.0013 | 0.0012 | 0.0012 |
| F03 | 41120001 | 41220000 | 0.0012 | 0.0010 | 0.0010 |
| F03 | 41140001 | 41240000 | 0.0012 | 0.0009 | 0.0010 |
| F03 | 41160001 | 41260000 | 0.0012 | 0.0009 | 0.0010 |
| F03 | 41180001 | 41280000 | 0.0011 | 0.0008 | 0.0009 |
| F03 | 41200001 | 41300000 | 0.0012 | 0.0010 | 0.0011 |
| F03 | 41220001 | 41320000 | 0.0015 | 0.0012 | 0.0014 |
| F03 | 41240001 | 41340000 | 0.0016 | 0.0014 | 0.0015 |
| F03 | 41260001 | 41360000 | 0.0016 | 0.0014 | 0.0016 |
| F03 | 41280001 | 41380000 | 0.0023 | 0.0020 | 0.0023 |
| F03 | 41300001 | 41400000 | 0.0025 | 0.0021 | 0.0024 |
| F03 | 41320001 | 41420000 | 0.0021 | 0.0017 | 0.0020 |
| F03 | 41340001 | 41440000 | 0.0022 | 0.0019 | 0.0021 |
| F03 | 41360001 | 41460000 | 0.0024 | 0.0019 | 0.0022 |
| F03 | 41380001 | 41480000 | 0.0019 | 0.0015 | 0.0017 |
| F03 | 41400001 | 41500000 | 0.0017 | 0.0014 | 0.0015 |
| F03 | 41420001 | 41520000 | 0.0016 | 0.0014 | 0.0015 |
| F03 | 41440001 | 41540000 | 0.0015 | 0.0012 | 0.0014 |
| F03 | 41460001 | 41560000 | 0.0014 | 0.0012 | 0.0014 |
| F03 | 41480001 | 41580000 | 0.0013 | 0.0011 | 0.0013 |
| F03 | 41500001 | 41600000 | 0.0011 | 0.0009 | 0.0011 |
| F03 | 41520001 | 41620000 | 0.0011 | 0.0009 | 0.0011 |
| F03 | 41540001 | 41640000 | 0.0007 | 0.0006 | 0.0008 |
| F03 | 41560001 | 41660000 | 0.0005 | 0.0004 | 0.0005 |
| F03 | 41580001 | 41680000 | 0.0002 | 0.0002 | 0.0002 |
| F03 | 41600001 | 41700000 | 0.0004 | 0.0003 | 0.0004 |
| F03 | 41620001 | 41720000 | 0.0009 | 0.0007 | 0.0009 |
| F03 | 41640001 | 41740000 | 0.0009 | 0.0007 | 0.0009 |
| F03 | 41660001 | 41760000 | 0.0010 | 0.0007 | 0.0009 |
| F03 | 41680001 | 41780000 | 0.0014 | 0.0010 | 0.0013 |
| F03 | 41700001 | 41800000 | 0.0012 | 0.0009 | 0.0012 |
| F03 | 41720001 | 41820000 | 0.0008 | 0.0006 | 0.0008 |
| F03 | 41740001 | 41840000 | 0.0007 | 0.0006 | 0.0007 |
| F03 | 41760001 | 41860000 | 0.0007 | 0.0006 | 0.0007 |
| F03 | 41780001 | 41880000 | 0.0003 | 0.0002 | 0.0003 |
| F03 | 41800001 | 41900000 | 0.0003 | 0.0002 | 0.0003 |
| F03 | 41820001 | 41920000 | 0.0002 | 0.0001 | 0.0002 |
| F03 | 41840001 | 41940000 | 0.0004 | 0.0003 | 0.0003 |
| F03 | 41860001 | 41960000 | 0.0009 | 0.0009 | 0.0006 |
| F03 | 41880001 | 41980000 | 0.0015 | 0.0014 | 0.0011 |
| F03 | 41900001 | 42000000 | 0.0015 | 0.0014 | 0.0012 |
| F03 | 41920001 | 42020000 | 0.0016 | 0.0014 | 0.0012 |
| F03 | 41940001 | 42040000 | 0.0018 | 0.0017 | 0.0015 |
| F03 | 41960001 | 42060000 | 0.0020 | 0.0015 | 0.0018 |
| F03 | 41980001 | 42080000 | 0.0017 | 0.0013 | 0.0014 |
| F03 | 42000001 | 42100000 | 0.0018 | 0.0014 | 0.0016 |
| F03 | 42020001 | 42120000 | 0.0019 | 0.0015 | 0.0017 |
| F03 | 42040001 | 42140000 | 0.0016 | 0.0012 | 0.0014 |

| Chr | Start | End | both | A1 | A2 |
|---|---|---|---|---|---|
| F03 | 42060001 | 42160000 | 0.0007 | 0.0006 | 0.0007 |
| F03 | 42080001 | 42180000 | 0.0008 | 0.0006 | 0.0008 |
| F03 | 42100001 | 42200000 | 0.0007 | 0.0006 | 0.0007 |
| F03 | 42120001 | 42220000 | 0.0011 | 0.0009 | 0.0010 |
| F03 | 42140001 | 42240000 | 0.0010 | 0.0008 | 0.0009 |
| F03 | 42160001 | 42260000 | 0.0018 | 0.0014 | 0.0018 |
| F03 | 42180001 | 42280000 | 0.0016 | 0.0012 | 0.0015 |
| F03 | 42200001 | 42300000 | 0.0017 | 0.0012 | 0.0016 |
| F03 | 42220001 | 42320000 | 0.0013 | 0.0008 | 0.0011 |
| F03 | 42240001 | 42340000 | 0.0018 | 0.0012 | 0.0016 |
| F03 | 42260001 | 42360000 | 0.0013 | 0.0009 | 0.0010 |
| F03 | 42280001 | 42380000 | 0.0014 | 0.0009 | 0.0011 |
| F03 | 42300001 | 42400000 | 0.0015 | 0.0010 | 0.0011 |
| F03 | 42320001 | 42420000 | 0.0016 | 0.0011 | 0.0013 |
| F03 | 42340001 | 42440000 | 0.0014 | 0.0010 | 0.0012 |
| F03 | 42360001 | 42460000 | 0.0013 | 0.0009 | 0.0011 |
| F03 | 42380001 | 42480000 | 0.0013 | 0.0009 | 0.0010 |
| F03 | 42400001 | 42500000 | 0.0018 | 0.0014 | 0.0015 |
| F03 | 42420001 | 42520000 | 0.0022 | 0.0015 | 0.0017 |
| F03 | 42440001 | 42540000 | 0.0024 | 0.0014 | 0.0017 |
| F03 | 42460001 | 42560000 | 0.0033 | 0.0022 | 0.0026 |
| F03 | 42480001 | 42580000 | 0.0038 | 0.0027 | 0.0031 |
| F03 | 42500001 | 42600000 | 0.0039 | 0.0028 | 0.0034 |
| F03 | 42520001 | 42620000 | 0.0042 | 0.0032 | 0.0037 |
| F03 | 42540001 | 42640000 | 0.0039 | 0.0031 | 0.0034 |
| F03 | 42560001 | 42660000 | 0.0034 | 0.0026 | 0.0030 |
| F03 | 42580001 | 42680000 | 0.0047 | 0.0037 | 0.0042 |
| F03 | 42600001 | 42700000 | 0.0046 | 0.0038 | 0.0041 |
| F03 | 42620001 | 42720000 | 0.0042 | 0.0035 | 0.0038 |
| F03 | 42640001 | 42740000 | 0.0045 | 0.0038 | 0.0041 |
| F03 | 42660001 | 42760000 | 0.0051 | 0.0043 | 0.0046 |
| F03 | 42680001 | 42780000 | 0.0037 | 0.0030 | 0.0033 |
| F03 | 42700001 | 42800000 | 0.0032 | 0.0024 | 0.0027 |
| F03 | 42720001 | 42820000 | 0.0032 | 0.0025 | 0.0028 |
| F03 | 42740001 | 42840000 | 0.0032 | 0.0026 | 0.0029 |
| F03 | 42760001 | 42860000 | 0.0024 | 0.0020 | 0.0022 |
| F03 | 42780001 | 42880000 | 0.0021 | 0.0018 | 0.0019 |
| F03 | 42800001 | 42900000 | 0.0019 | 0.0016 | 0.0016 |
| F03 | 42820001 | 42920000 | 0.0017 | 0.0013 | 0.0014 |
| F03 | 42840001 | 42940000 | 0.0012 | 0.0009 | 0.0009 |
| F03 | 42860001 | 42960000 | 0.0014 | 0.0008 | 0.0009 |
| F03 | 42880001 | 42980000 | 0.0016 | 0.0010 | 0.0011 |
| F03 | 42900001 | 43000000 | 0.0020 | 0.0013 | 0.0015 |
| F03 | 42920001 | 43020000 | 0.0020 | 0.0014 | 0.0016 |
| F03 | 42940001 | 43040000 | 0.0022 | 0.0016 | 0.0017 |
| F03 | 42960001 | 43060000 | 0.0021 | 0.0014 | 0.0015 |
| F03 | 42980001 | 43080000 | 0.0024 | 0.0016 | 0.0018 |
| F03 | 43000001 | 43100000 | 0.0019 | 0.0012 | 0.0014 |
| F03 | 43020001 | 43120000 | 0.0020 | 0.0012 | 0.0013 |
| F03 | 43040001 | 43140000 | 0.0020 | 0.0011 | 0.0014 |
| F03 | 43060001 | 43160000 | 0.0019 | 0.0012 | 0.0014 |
| F03 | 43080001 | 43180000 | 0.0017 | 0.0008 | 0.0009 |
| F03 | 43100001 | 43200000 | 0.0027 | 0.0016 | 0.0019 |
| F03 | 43120001 | 43220000 | 0.0032 | 0.0021 | 0.0024 |
| F03 | 43140001 | 43240000 | 0.0038 | 0.0027 | 0.0030 |
| F03 | 43160001 | 43260000 | 0.0039 | 0.0028 | 0.0033 |
| F03 | 43180001 | 43280000 | 0.0040 | 0.0031 | 0.0035 |
| F03 | 43200001 | 43300000 | 0.0045 | 0.0034 | 0.0037 |

| Chr | Start | End | both | A1 | A2 |
|---|---|---|---|---|---|
| F03 | 43220001 | 43320000 | 0.0047 | 0.0034 | 0.0039 |
| F03 | 43240001 | 43340000 | 0.0044 | 0.0032 | 0.0038 |
| F03 | 43260001 | 43360000 | 0.0039 | 0.0029 | 0.0033 |
| F03 | 43280001 | 43380000 | 0.0036 | 0.0027 | 0.0032 |
| F03 | 43300001 | 43400000 | 0.0021 | 0.0016 | 0.0020 |
| F03 | 43320001 | 43420000 | 0.0013 | 0.0011 | 0.0013 |
| F03 | 43340001 | 43440000 | 0.0011 | 0.0009 | 0.0010 |
| F03 | 43360001 | 43460000 | 0.0012 | 0.0010 | 0.0012 |
| F03 | 43380001 | 43480000 | 0.0011 | 0.0008 | 0.0010 |
| F03 | 43400001 | 43500000 | 0.0011 | 0.0008 | 0.0010 |
| F03 | 43420001 | 43520000 | 0.0015 | 0.0011 | 0.0015 |
| F03 | 43440001 | 43540000 | 0.0011 | 0.0008 | 0.0011 |
| F03 | 43460001 | 43560000 | 0.0011 | 0.0007 | 0.0010 |
| F03 | 43480001 | 43580000 | 0.0015 | 0.0012 | 0.0014 |
| F03 | 43500001 | 43600000 | 0.0027 | 0.0022 | 0.0026 |
| F03 | 43520001 | 43620000 | 0.0032 | 0.0026 | 0.0030 |
| F03 | 43540001 | 43640000 | 0.0049 | 0.0040 | 0.0045 |
| F03 | 43560001 | 43660000 | 0.0054 | 0.0044 | 0.0050 |
| F03 | 43580001 | 43680000 | 0.0053 | 0.0041 | 0.0046 |
| F03 | 43600001 | 43700000 | 0.0046 | 0.0035 | 0.0039 |
| F03 | 43620001 | 43720000 | 0.0041 | 0.0031 | 0.0033 |
| F03 | 43640001 | 43740000 | 0.0034 | 0.0023 | 0.0025 |
| F03 | 43660001 | 43760000 | 0.0034 | 0.0021 | 0.0024 |
| F03 | 43680001 | 43780000 | 0.0033 | 0.0022 | 0.0024 |
| F03 | 43700001 | 43800000 | 0.0030 | 0.0018 | 0.0021 |
| F03 | 43720001 | 43820000 | 0.0027 | 0.0015 | 0.0018 |
| F03 | 43740001 | 43840000 | 0.0025 | 0.0017 | 0.0019 |
| F03 | 43760001 | 43860000 | 0.0030 | 0.0022 | 0.0026 |
| F03 | 43780001 | 43880000 | 0.0032 | 0.0022 | 0.0025 |
| F03 | 43800001 | 43900000 | 0.0031 | 0.0021 | 0.0024 |
| F03 | 43820001 | 43920000 | 0.0032 | 0.0024 | 0.0027 |
| F03 | 43840001 | 43940000 | 0.0028 | 0.0018 | 0.0020 |
| F03 | 43860001 | 43960000 | 0.0027 | 0.0018 | 0.0019 |
| F03 | 43880001 | 43980000 | 0.0025 | 0.0019 | 0.0021 |
| F03 | 43900001 | 44000000 | 0.0024 | 0.0020 | 0.0020 |
| F03 | 43920001 | 44020000 | 0.0024 | 0.0018 | 0.0019 |
| F03 | 43940001 | 44040000 | 0.0023 | 0.0017 | 0.0019 |
| F03 | 43960001 | 44060000 | 0.0017 | 0.0012 | 0.0013 |
| F03 | 43980001 | 44080000 | 0.0017 | 0.0011 | 0.0013 |
| F03 | 44000001 | 44100000 | 0.0016 | 0.0011 | 0.0013 |
| F03 | 44020001 | 44120000 | 0.0019 | 0.0011 | 0.0013 |
| F03 | 44040001 | 44140000 | 0.0016 | 0.0009 | 0.0011 |
| F03 | 44060001 | 44160000 | 0.0018 | 0.0010 | 0.0012 |
| F03 | 44080001 | 44180000 | 0.0016 | 0.0008 | 0.0010 |
| F03 | 44100001 | 44200000 | 0.0019 | 0.0010 | 0.0013 |
| F03 | 44120001 | 44220000 | 0.0014 | 0.0009 | 0.0011 |
| F03 | 44140001 | 44240000 | 0.0015 | 0.0010 | 0.0013 |
| F03 | 44160001 | 44260000 | 0.0023 | 0.0017 | 0.0021 |
| F03 | 44180001 | 44280000 | 0.0034 | 0.0027 | 0.0030 |
| F03 | 44200001 | 44300000 | 0.0037 | 0.0027 | 0.0029 |
| F03 | 44220001 | 44320000 | 0.0041 | 0.0032 | 0.0034 |
| F03 | 44240001 | 44340000 | 0.0047 | 0.0035 | 0.0039 |
| F03 | 44260001 | 44360000 | 0.0040 | 0.0030 | 0.0030 |
| F03 | 44280001 | 44380000 | 0.0032 | 0.0023 | 0.0025 |
| F03 | 44300001 | 44400000 | 0.0030 | 0.0021 | 0.0022 |
| F03 | 44320001 | 44420000 | 0.0027 | 0.0019 | 0.0019 |
| F03 | 44340001 | 44440000 | 0.0028 | 0.0018 | 0.0018 |
| F03 | 44360001 | 44460000 | 0.0035 | 0.0023 | 0.0024 |

| Chr | Start | End | both | A1 | A2 |
|---|---|---|---|---|---|
| F03 | 44380001 | 44480000 | 0.0032 | 0.0019 | 0.0020 |
| F03 | 44400001 | 44500000 | 0.0031 | 0.0020 | 0.0021 |
| F03 | 44420001 | 44520000 | 0.0032 | 0.0021 | 0.0023 |
| F03 | 44440001 | 44540000 | 0.0026 | 0.0019 | 0.0020 |
| F03 | 44460001 | 44560000 | 0.0013 | 0.0009 | 0.0010 |
| F03 | 44480001 | 44580000 | 0.0013 | 0.0011 | 0.0012 |
| F03 | 44500001 | 44600000 | 0.0012 | 0.0011 | 0.0012 |
| F03 | 44520001 | 44620000 | 0.0010 | 0.0009 | 0.0010 |
| F03 | 44540001 | 44640000 | 0.0008 | 0.0007 | 0.0008 |
| F03 | 44560001 | 44660000 | 0.0012 | 0.0011 | 0.0012 |
| F03 | 44580001 | 44680000 | 0.0015 | 0.0013 | 0.0015 |
| F03 | 44600001 | 44700000 | 0.0023 | 0.0020 | 0.0023 |
| F03 | 44620001 | 44720000 | 0.0021 | 0.0018 | 0.0020 |
| F03 | 44640001 | 44740000 | 0.0021 | 0.0018 | 0.0021 |
| F03 | 44660001 | 44760000 | 0.0018 | 0.0016 | 0.0018 |
| F03 | 44680001 | 44780000 | 0.0023 | 0.0018 | 0.0023 |
| F03 | 44700001 | 44800000 | 0.0016 | 0.0011 | 0.0016 |
| F03 | 44720001 | 44820000 | 0.0017 | 0.0012 | 0.0017 |
| F03 | 44740001 | 44840000 | 0.0017 | 0.0012 | 0.0017 |
| F03 | 44760001 | 44860000 | 0.0017 | 0.0012 | 0.0017 |
| F03 | 44780001 | 44880000 | 0.0013 | 0.0011 | 0.0013 |
| F03 | 44800001 | 44900000 | 0.0012 | 0.0009 | 0.0012 |
| F03 | 44820001 | 44920000 | 0.0011 | 0.0010 | 0.0011 |
| F03 | 44840001 | 44940000 | 0.0013 | 0.0011 | 0.0013 |
| F03 | 44860001 | 44960000 | 0.0015 | 0.0013 | 0.0015 |
| F03 | 44880001 | 44980000 | 0.0009 | 0.0008 | 0.0009 |
| F03 | 44900001 | 45000000 | 0.0017 | 0.0014 | 0.0016 |
| F03 | 44920001 | 45020000 | 0.0021 | 0.0017 | 0.0020 |
| F03 | 44940001 | 45040000 | 0.0022 | 0.0015 | 0.0020 |
| F03 | 44960001 | 45060000 | 0.0019 | 0.0012 | 0.0017 |
| F03 | 44980001 | 45080000 | 0.0024 | 0.0017 | 0.0022 |
| F03 | 45000001 | 45100000 | 0.0022 | 0.0018 | 0.0020 |
| F03 | 45020001 | 45120000 | 0.0019 | 0.0015 | 0.0017 |
| F03 | 45040001 | 45140000 | 0.0027 | 0.0023 | 0.0025 |
| F03 | 45060001 | 45160000 | 0.0035 | 0.0030 | 0.0032 |
| F03 | 45080001 | 45180000 | 0.0034 | 0.0029 | 0.0031 |
| F03 | 45100001 | 45200000 | 0.0031 | 0.0023 | 0.0027 |
| F03 | 45120001 | 45220000 | 0.0029 | 0.0023 | 0.0026 |
| F03 | 45140001 | 45240000 | 0.0022 | 0.0017 | 0.0019 |
| F03 | 45160001 | 45260000 | 0.0013 | 0.0010 | 0.0012 |
| F03 | 45180001 | 45280000 | 0.0010 | 0.0007 | 0.0009 |
| F03 | 45200001 | 45300000 | 0.0011 | 0.0009 | 0.0011 |
| F03 | 45220001 | 45320000 | 0.0010 | 0.0008 | 0.0010 |
| F03 | 45240001 | 45340000 | 0.0009 | 0.0007 | 0.0009 |
| F03 | 45260001 | 45360000 | 0.0010 | 0.0008 | 0.0010 |
| F03 | 45280001 | 45380000 | 0.0009 | 0.0007 | 0.0009 |
| F03 | 45300001 | 45400000 | 0.0009 | 0.0006 | 0.0008 |
| F03 | 45320001 | 45420000 | 0.0010 | 0.0006 | 0.0009 |
| F03 | 45340001 | 45440000 | 0.0014 | 0.0010 | 0.0012 |
| F03 | 45360001 | 45460000 | 0.0014 | 0.0009 | 0.0011 |
| F03 | 45380001 | 45480000 | 0.0015 | 0.0010 | 0.0012 |
| F03 | 45400001 | 45500000 | 0.0011 | 0.0009 | 0.0009 |
| F03 | 45420001 | 45520000 | 0.0011 | 0.0008 | 0.0009 |
| F03 | 45440001 | 45540000 | 0.0011 | 0.0009 | 0.0010 |
| F03 | 45460001 | 45560000 | 0.0011 | 0.0008 | 0.0010 |
| F03 | 45480001 | 45580000 | 0.0011 | 0.0009 | 0.0011 |
| F03 | 45500001 | 45600000 | 0.0010 | 0.0008 | 0.0011 |
| F03 | 45520001 | 45620000 | 0.0010 | 0.0008 | 0.0010 |

| Chr | Start | End | both | A1 | A2 |
|---|---|---|---|---|---|
| F03 | 45540001 | 45640000 | 0.0004 | 0.0003 | 0.0004 |
| F03 | 45560001 | 45660000 | 0.0007 | 0.0004 | 0.0006 |
| F03 | 45580001 | 45680000 | 0.0011 | 0.0005 | 0.0008 |
| F03 | 45600001 | 45700000 | 0.0012 | 0.0006 | 0.0008 |
| F03 | 45620001 | 45720000 | 0.0017 | 0.0009 | 0.0012 |
| F03 | 45640001 | 45740000 | 0.0023 | 0.0014 | 0.0017 |
| F03 | 45660001 | 45760000 | 0.0021 | 0.0015 | 0.0016 |
| F03 | 45680001 | 45780000 | 0.0016 | 0.0012 | 0.0013 |
| F03 | 45700001 | 45800000 | 0.0019 | 0.0015 | 0.0017 |
| F03 | 45720001 | 45820000 | 0.0015 | 0.0012 | 0.0013 |
| F03 | 45740001 | 45840000 | 0.0013 | 0.0010 | 0.0011 |
| F03 | 45760001 | 45860000 | 0.0016 | 0.0011 | 0.0013 |
| F03 | 45780001 | 45880000 | 0.0021 | 0.0015 | 0.0018 |
| F03 | 45800001 | 45900000 | 0.0020 | 0.0015 | 0.0017 |
| F03 | 45820001 | 45920000 | 0.0025 | 0.0020 | 0.0022 |
| F03 | 45840001 | 45940000 | 0.0027 | 0.0021 | 0.0024 |
| F03 | 45860001 | 45960000 | 0.0040 | 0.0034 | 0.0035 |
| F03 | 45880001 | 45980000 | 0.0041 | 0.0034 | 0.0035 |
| F03 | 45900001 | 46000000 | 0.0045 | 0.0037 | 0.0038 |
| F03 | 45920001 | 46020000 | 0.0060 | 0.0050 | 0.0053 |
| F03 | 45940001 | 46040000 | 0.0067 | 0.0058 | 0.0060 |
| F03 | 45960001 | 46060000 | 0.0055 | 0.0048 | 0.0049 |
| F03 | 45980001 | 46080000 | 0.0057 | 0.0050 | 0.0051 |
| F03 | 46000001 | 46100000 | 0.0056 | 0.0050 | 0.0051 |
| F03 | 46020001 | 46120000 | 0.0054 | 0.0047 | 0.0048 |
| F03 | 46040001 | 46140000 | 0.0043 | 0.0036 | 0.0040 |
| F03 | 46060001 | 46160000 | 0.0046 | 0.0038 | 0.0041 |
| F03 | 46080001 | 46180000 | 0.0043 | 0.0033 | 0.0038 |
| F03 | 46100001 | 46200000 | 0.0045 | 0.0035 | 0.0040 |
| F03 | 46120001 | 46220000 | 0.0033 | 0.0023 | 0.0028 |
| F03 | 46140001 | 46240000 | 0.0036 | 0.0028 | 0.0031 |
| F03 | 46160001 | 46260000 | 0.0032 | 0.0025 | 0.0028 |
| F03 | 46180001 | 46280000 | 0.0031 | 0.0025 | 0.0028 |
| F03 | 46200001 | 46300000 | 0.0035 | 0.0029 | 0.0033 |
| F03 | 46220001 | 46320000 | 0.0041 | 0.0035 | 0.0039 |
| F03 | 46240001 | 46340000 | 0.0041 | 0.0031 | 0.0037 |
| F03 | 46260001 | 46360000 | 0.0043 | 0.0034 | 0.0039 |
| F03 | 46280001 | 46380000 | 0.0038 | 0.0030 | 0.0035 |
| F03 | 46300001 | 46400000 | 0.0030 | 0.0023 | 0.0027 |
| F03 | 46320001 | 46420000 | 0.0020 | 0.0015 | 0.0019 |
| F03 | 46340001 | 46440000 | 0.0017 | 0.0014 | 0.0015 |
| F03 | 46360001 | 46460000 | 0.0017 | 0.0015 | 0.0017 |
| F03 | 46380001 | 46480000 | 0.0017 | 0.0015 | 0.0017 |
| F03 | 46400001 | 46500000 | 0.0016 | 0.0013 | 0.0015 |
| F03 | 46420001 | 46520000 | 0.0018 | 0.0015 | 0.0017 |
| F03 | 46440001 | 46540000 | 0.0017 | 0.0015 | 0.0017 |
| F03 | 46460001 | 46560000 | 0.0019 | 0.0017 | 0.0018 |
| F03 | 46480001 | 46580000 | 0.0021 | 0.0018 | 0.0019 |
| F03 | 46500001 | 46600000 | 0.0020 | 0.0018 | 0.0018 |
| F03 | 46520001 | 46620000 | 0.0020 | 0.0018 | 0.0018 |
| F03 | 46540001 | 46640000 | 0.0020 | 0.0019 | 0.0019 |
| F03 | 46560001 | 46660000 | 0.0015 | 0.0014 | 0.0014 |
| F03 | 46580001 | 46680000 | 0.0016 | 0.0014 | 0.0015 |
| F03 | 46600001 | 46700000 | 0.0024 | 0.0021 | 0.0024 |
| F03 | 46620001 | 46720000 | 0.0025 | 0.0021 | 0.0023 |
| F03 | 46640001 | 46740000 | 0.0035 | 0.0029 | 0.0031 |
| F03 | 46660001 | 46760000 | 0.0032 | 0.0027 | 0.0028 |
| F03 | 46680001 | 46780000 | 0.0033 | 0.0028 | 0.0029 |

| Chr | Start | End | both | A1 | A2 |
|---|---|---|---|---|---|
| F03 | 46700001 | 46800000 | 0.0026 | 0.0021 | 0.0022 |
| F03 | 46720001 | 46820000 | 0.0035 | 0.0030 | 0.0031 |
| F03 | 46740001 | 46840000 | 0.0032 | 0.0025 | 0.0028 |
| F03 | 46760001 | 46860000 | 0.0037 | 0.0027 | 0.0031 |
| F03 | 46780001 | 46880000 | 0.0040 | 0.0029 | 0.0034 |
| F03 | 46800001 | 46900000 | 0.0041 | 0.0028 | 0.0034 |
| F03 | 46820001 | 46920000 | 0.0032 | 0.0020 | 0.0025 |
| F03 | 46840001 | 46940000 | 0.0038 | 0.0026 | 0.0032 |
| F03 | 46860001 | 46960000 | 0.0055 | 0.0041 | 0.0048 |
| F03 | 46880001 | 46980000 | 0.0051 | 0.0038 | 0.0043 |
| F03 | 46900001 | 47000000 | 0.0049 | 0.0038 | 0.0041 |
| F03 | 46920001 | 47020000 | 0.0050 | 0.0037 | 0.0040 |
| F03 | 46940001 | 47040000 | 0.0050 | 0.0038 | 0.0039 |
| F03 | 46960001 | 47060000 | 0.0033 | 0.0023 | 0.0024 |
| F03 | 46980001 | 47080000 | 0.0040 | 0.0031 | 0.0031 |
| F03 | 47000001 | 47100000 | 0.0042 | 0.0033 | 0.0034 |
| F03 | 47020001 | 47120000 | 0.0043 | 0.0036 | 0.0038 |
| F03 | 47040001 | 47140000 | 0.0039 | 0.0032 | 0.0034 |
| F03 | 47060001 | 47160000 | 0.0036 | 0.0030 | 0.0031 |
| F03 | 47080001 | 47180000 | 0.0038 | 0.0031 | 0.0033 |
| F03 | 47100001 | 47200000 | 0.0039 | 0.0029 | 0.0031 |
| F03 | 47120001 | 47220000 | 0.0045 | 0.0035 | 0.0037 |
| F03 | 47140001 | 47240000 | 0.0040 | 0.0029 | 0.0031 |
| F03 | 47160001 | 47260000 | 0.0038 | 0.0029 | 0.0031 |
| F03 | 47180001 | 47280000 | 0.0031 | 0.0026 | 0.0024 |
| F03 | 47200001 | 47300000 | 0.0034 | 0.0031 | 0.0028 |
| F03 | 47220001 | 47320000 | 0.0027 | 0.0024 | 0.0022 |
| F03 | 47240001 | 47340000 | 0.0032 | 0.0030 | 0.0030 |
| F03 | 47260001 | 47360000 | 0.0031 | 0.0029 | 0.0029 |
| F03 | 47280001 | 47380000 | 0.0029 | 0.0024 | 0.0027 |
| F03 | 47300001 | 47400000 | 0.0026 | 0.0021 | 0.0024 |
| F03 | 47320001 | 47420000 | 0.0025 | 0.0020 | 0.0023 |
| F03 | 47340001 | 47440000 | 0.0022 | 0.0018 | 0.0019 |
| F03 | 47360001 | 47460000 | 0.0024 | 0.0019 | 0.0021 |
| F03 | 47380001 | 47480000 | 0.0031 | 0.0024 | 0.0026 |
| F03 | 47400001 | 47500000 | 0.0034 | 0.0027 | 0.0028 |
| F03 | 47420001 | 47520000 | 0.0029 | 0.0022 | 0.0023 |
| F03 | 47440001 | 47540000 | 0.0030 | 0.0021 | 0.0023 |
| F03 | 47460001 | 47560000 | 0.0035 | 0.0025 | 0.0027 |
| F03 | 47480001 | 47580000 | 0.0036 | 0.0025 | 0.0028 |
| F03 | 47500001 | 47600000 | 0.0034 | 0.0024 | 0.0026 |
| F03 | 47520001 | 47620000 | 0.0038 | 0.0025 | 0.0026 |
| F03 | 47540001 | 47640000 | 0.0045 | 0.0032 | 0.0033 |
| F03 | 47560001 | 47660000 | 0.0041 | 0.0027 | 0.0028 |
| F03 | 47580001 | 47680000 | 0.0036 | 0.0023 | 0.0023 |
| F03 | 47600001 | 47700000 | 0.0037 | 0.0023 | 0.0023 |
| F03 | 47620001 | 47720000 | 0.0046 | 0.0034 | 0.0032 |
| F03 | 47640001 | 47740000 | 0.0035 | 0.0024 | 0.0022 |
| F03 | 47660001 | 47760000 | 0.0036 | 0.0026 | 0.0024 |
| F03 | 47680001 | 47780000 | 0.0039 | 0.0029 | 0.0027 |
| F03 | 47700001 | 47800000 | 0.0036 | 0.0027 | 0.0025 |
| F03 | 47720001 | 47820000 | 0.0022 | 0.0014 | 0.0015 |
| F03 | 47740001 | 47840000 | 0.0025 | 0.0016 | 0.0017 |
| F03 | 47760001 | 47860000 | 0.0025 | 0.0015 | 0.0017 |
| F03 | 47780001 | 47880000 | 0.0023 | 0.0012 | 0.0014 |
| F03 | 47800001 | 47900000 | 0.0023 | 0.0013 | 0.0016 |
| F03 | 47820001 | 47920000 | 0.0028 | 0.0017 | 0.0021 |
| F03 | 47840001 | 47940000 | 0.0027 | 0.0016 | 0.0021 |

| Chr | Start | End | both | A1 | A2 |
|---|---|---|---|---|---|
| F03 | 47860001 | 47960000 | 0.0028 | 0.0020 | 0.0024 |
| F03 | 47880001 | 47980000 | 0.0033 | 0.0025 | 0.0029 |
| F03 | 47900001 | 48000000 | 0.0034 | 0.0026 | 0.0029 |
| F03 | 47920001 | 48020000 | 0.0030 | 0.0021 | 0.0024 |
| F03 | 47940001 | 48040000 | 0.0032 | 0.0023 | 0.0024 |
| F03 | 47960001 | 48060000 | 0.0036 | 0.0029 | 0.0029 |
| F03 | 47980001 | 48080000 | 0.0028 | 0.0023 | 0.0023 |
| F03 | 48000001 | 48100000 | 0.0029 | 0.0024 | 0.0025 |
| F03 | 48020001 | 48120000 | 0.0031 | 0.0027 | 0.0027 |
| F03 | 48040001 | 48140000 | 0.0026 | 0.0023 | 0.0024 |
| F03 | 48060001 | 48160000 | 0.0018 | 0.0016 | 0.0016 |
| F03 | 48080001 | 48180000 | 0.0027 | 0.0022 | 0.0025 |
| F03 | 48100001 | 48200000 | 0.0031 | 0.0025 | 0.0030 |
| F03 | 48120001 | 48220000 | 0.0029 | 0.0022 | 0.0027 |
| F03 | 48140001 | 48240000 | 0.0029 | 0.0023 | 0.0028 |
| F03 | 48160001 | 48260000 | 0.0033 | 0.0025 | 0.0032 |
| F03 | 48180001 | 48280000 | 0.0024 | 0.0018 | 0.0022 |
| F03 | 48200001 | 48300000 | 0.0017 | 0.0013 | 0.0015 |
| F03 | 48220001 | 48320000 | 0.0023 | 0.0018 | 0.0020 |
| F03 | 48240001 | 48340000 | 0.0029 | 0.0023 | 0.0025 |
| F03 | 48260001 | 48360000 | 0.0025 | 0.0020 | 0.0022 |
| F03 | 48280001 | 48380000 | 0.0031 | 0.0025 | 0.0027 |
| F03 | 48300001 | 48400000 | 0.0031 | 0.0024 | 0.0026 |
| F03 | 48320001 | 48420000 | 0.0025 | 0.0021 | 0.0022 |
| F03 | 48340001 | 48440000 | 0.0019 | 0.0016 | 0.0016 |
| F03 | 48360001 | 48460000 | 0.0018 | 0.0014 | 0.0014 |
| F03 | 48380001 | 48480000 | 0.0014 | 0.0009 | 0.0010 |
| F03 | 48400001 | 48500000 | 0.0013 | 0.0009 | 0.0009 |
| F03 | 48420001 | 48520000 | 0.0017 | 0.0013 | 0.0012 |
| F03 | 48440001 | 48540000 | 0.0020 | 0.0014 | 0.0015 |
| F03 | 48460001 | 48560000 | 0.0020 | 0.0014 | 0.0015 |
| F03 | 48480001 | 48580000 | 0.0016 | 0.0014 | 0.0015 |
| F03 | 48500001 | 48600000 | 0.0018 | 0.0016 | 0.0017 |
| F03 | 48520001 | 48620000 | 0.0019 | 0.0016 | 0.0017 |
| F03 | 48540001 | 48640000 | 0.0018 | 0.0015 | 0.0016 |
| F03 | 48560001 | 48660000 | 0.0019 | 0.0016 | 0.0017 |
| F03 | 48580001 | 48680000 | 0.0021 | 0.0018 | 0.0019 |
| F03 | 48600001 | 48700000 | 0.0022 | 0.0018 | 0.0020 |
| F03 | 48620001 | 48720000 | 0.0022 | 0.0019 | 0.0021 |
| F03 | 48640001 | 48740000 | 0.0026 | 0.0024 | 0.0024 |
| F03 | 48660001 | 48760000 | 0.0029 | 0.0025 | 0.0027 |
| F03 | 48680001 | 48780000 | 0.0028 | 0.0026 | 0.0026 |
| F03 | 48700001 | 48800000 | 0.0034 | 0.0032 | 0.0031 |
| F03 | 48720001 | 48820000 | 0.0034 | 0.0032 | 0.0032 |
| F03 | 48740001 | 48840000 | 0.0032 | 0.0028 | 0.0029 |
| F03 | 48760001 | 48860000 | 0.0038 | 0.0035 | 0.0035 |
| F03 | 48780001 | 48880000 | 0.0037 | 0.0033 | 0.0034 |
| F03 | 48800001 | 48900000 | 0.0037 | 0.0031 | 0.0035 |
| F03 | 48820001 | 48920000 | 0.0034 | 0.0028 | 0.0031 |
| F03 | 48840001 | 48940000 | 0.0032 | 0.0027 | 0.0029 |
| F03 | 48860001 | 48960000 | 0.0021 | 0.0016 | 0.0019 |
| F03 | 48880001 | 48980000 | 0.0026 | 0.0022 | 0.0024 |
| F03 | 48900001 | 49000000 | 0.0018 | 0.0016 | 0.0017 |
| F03 | 48920001 | 49020000 | 0.0018 | 0.0016 | 0.0017 |
| F03 | 48940001 | 49040000 | 0.0020 | 0.0018 | 0.0019 |
| F03 | 48960001 | 49060000 | 0.0022 | 0.0021 | 0.0021 |
| F03 | 48980001 | 49080000 | 0.0019 | 0.0017 | 0.0019 |
| F03 | 49000001 | 49100000 | 0.0017 | 0.0016 | 0.0016 |

| Chr | Start | End | both | A1 | A2 |
|---|---|---|---|---|---|
| F03 | 49020001 | 49120000 | 0.0013 | 0.0012 | 0.0013 |
| F03 | 49040001 | 49140000 | 0.0017 | 0.0016 | 0.0017 |
| F03 | 49060001 | 49160000 | 0.0018 | 0.0016 | 0.0018 |
| F03 | 49080001 | 49180000 | 0.0023 | 0.0019 | 0.0022 |
| F03 | 49100001 | 49200000 | 0.0025 | 0.0021 | 0.0025 |
| F03 | 49120001 | 49220000 | 0.0026 | 0.0022 | 0.0025 |
| F03 | 49140001 | 49240000 | 0.0020 | 0.0016 | 0.0019 |
| F03 | 49160001 | 49260000 | 0.0021 | 0.0017 | 0.0021 |
| F03 | 49180001 | 49280000 | 0.0013 | 0.0011 | 0.0013 |
| F03 | 49200001 | 49300000 | 0.0011 | 0.0008 | 0.0010 |
| F03 | 49220001 | 49320000 | 0.0015 | 0.0012 | 0.0015 |
| F03 | 49240001 | 49340000 | 0.0021 | 0.0017 | 0.0020 |
| F03 | 49260001 | 49360000 | 0.0020 | 0.0017 | 0.0019 |
| F03 | 49280001 | 49380000 | 0.0025 | 0.0022 | 0.0024 |
| F03 | 49300001 | 49400000 | 0.0029 | 0.0025 | 0.0028 |
| F03 | 49320001 | 49420000 | 0.0026 | 0.0022 | 0.0024 |
| F03 | 49340001 | 49440000 | 0.0041 | 0.0037 | 0.0038 |
| F03 | 49360001 | 49460000 | 0.0038 | 0.0034 | 0.0036 |
| F03 | 49380001 | 49480000 | 0.0036 | 0.0031 | 0.0032 |
| F03 | 49400001 | 49500000 | 0.0034 | 0.0028 | 0.0029 |
| F03 | 49420001 | 49520000 | 0.0031 | 0.0027 | 0.0027 |
| F03 | 49440001 | 49540000 | 0.0008 | 0.0004 | 0.0005 |
| F03 | 49460001 | 49560000 | 0.0008 | 0.0005 | 0.0006 |
| F03 | 49480001 | 49580000 | 0.0005 | 0.0003 | 0.0004 |
| F03 | 49500001 | 49600000 | 0.0008 | 0.0006 | 0.0008 |
| F03 | 49520001 | 49620000 | 0.0014 | 0.0011 | 0.0014 |
| F03 | 49540001 | 49640000 | 0.0022 | 0.0018 | 0.0022 |
| F03 | 49560001 | 49660000 | 0.0020 | 0.0016 | 0.0020 |
| F03 | 49580001 | 49680000 | 0.0024 | 0.0019 | 0.0023 |
| F03 | 49600001 | 49700000 | 0.0026 | 0.0022 | 0.0025 |
| F03 | 49620001 | 49720000 | 0.0026 | 0.0021 | 0.0024 |
| F03 | 49640001 | 49740000 | 0.0020 | 0.0015 | 0.0018 |
| F03 | 49660001 | 49760000 | 0.0022 | 0.0018 | 0.0020 |
| F03 | 49680001 | 49780000 | 0.0026 | 0.0022 | 0.0024 |
| F03 | 49700001 | 49800000 | 0.0023 | 0.0020 | 0.0022 |
| F03 | 49720001 | 49820000 | 0.0019 | 0.0018 | 0.0018 |
| F03 | 49740001 | 49840000 | 0.0021 | 0.0020 | 0.0020 |
| F03 | 49760001 | 49860000 | 0.0027 | 0.0026 | 0.0024 |
| F03 | 49780001 | 49880000 | 0.0025 | 0.0023 | 0.0022 |
| F03 | 49800001 | 49900000 | 0.0023 | 0.0021 | 0.0020 |
| F03 | 49820001 | 49920000 | 0.0028 | 0.0025 | 0.0025 |
| F03 | 49840001 | 49940000 | 0.0026 | 0.0024 | 0.0024 |
| F03 | 49860001 | 49960000 | 0.0020 | 0.0016 | 0.0018 |
| F03 | 49880001 | 49980000 | 0.0015 | 0.0013 | 0.0015 |
| F03 | 49900001 | 50000000 | 0.0013 | 0.0011 | 0.0013 |
| F03 | 49920001 | 50020000 | 0.0010 | 0.0009 | 0.0010 |
| F03 | 49940001 | 50040000 | 0.0011 | 0.0008 | 0.0010 |
| F03 | 49960001 | 50060000 | 0.0014 | 0.0011 | 0.0013 |
| F03 | 49980001 | 50080000 | 0.0014 | 0.0011 | 0.0013 |
| F03 | 50000001 | 50100000 | 0.0014 | 0.0010 | 0.0012 |
| F03 | 50020001 | 50120000 | 0.0011 | 0.0007 | 0.0009 |
| F03 | 50040001 | 50140000 | 0.0011 | 0.0009 | 0.0010 |
| F03 | 50060001 | 50160000 | 0.0008 | 0.0007 | 0.0008 |
| F03 | 50080001 | 50180000 | 0.0009 | 0.0008 | 0.0009 |
| F03 | 50100001 | 50200000 | 0.0010 | 0.0009 | 0.0010 |
| F03 | 50120001 | 50220000 | 0.0013 | 0.0010 | 0.0013 |
| F03 | 50140001 | 50240000 | 0.0009 | 0.0007 | 0.0009 |
| F03 | 50160001 | 50260000 | 0.0017 | 0.0015 | 0.0016 |

| Chr | Start | End | both | A1 | A2 |
|---|---|---|---|---|---|
| F03 | 50180001 | 50280000 | 0.0017 | 0.0016 | 0.0016 |
| F03 | 50200001 | 50300000 | 0.0020 | 0.0020 | 0.0019 |
| F03 | 50220001 | 50320000 | 0.0022 | 0.0022 | 0.0021 |
| F03 | 50240001 | 50340000 | 0.0029 | 0.0029 | 0.0028 |
| F03 | 50260001 | 50360000 | 0.0025 | 0.0024 | 0.0023 |
| F03 | 50280001 | 50380000 | 0.0025 | 0.0024 | 0.0024 |
| F03 | 50300001 | 50400000 | 0.0025 | 0.0021 | 0.0024 |
| F03 | 50320001 | 50420000 | 0.0021 | 0.0018 | 0.0021 |
| F03 | 50340001 | 50440000 | 0.0019 | 0.0015 | 0.0019 |
| F03 | 50360001 | 50460000 | 0.0016 | 0.0012 | 0.0015 |
| F03 | 50380001 | 50480000 | 0.0019 | 0.0015 | 0.0019 |
| F03 | 50400001 | 50500000 | 0.0017 | 0.0014 | 0.0016 |
| F03 | 50420001 | 50520000 | 0.0021 | 0.0018 | 0.0020 |
| F03 | 50440001 | 50540000 | 0.0018 | 0.0014 | 0.0017 |
| F03 | 50460001 | 50560000 | 0.0020 | 0.0017 | 0.0020 |
| F03 | 50480001 | 50580000 | 0.0017 | 0.0015 | 0.0017 |
| F03 | 50500001 | 50600000 | 0.0020 | 0.0017 | 0.0019 |
| F03 | 50520001 | 50620000 | 0.0019 | 0.0017 | 0.0019 |
| F03 | 50540001 | 50640000 | 0.0020 | 0.0019 | 0.0020 |
| F03 | 50560001 | 50660000 | 0.0021 | 0.0020 | 0.0020 |
| F03 | 50580001 | 50680000 | 0.0021 | 0.0019 | 0.0020 |
| F03 | 50600001 | 50700000 | 0.0017 | 0.0016 | 0.0016 |
| F03 | 50620001 | 50720000 | 0.0015 | 0.0012 | 0.0014 |
| F03 | 50640001 | 50740000 | 0.0013 | 0.0010 | 0.0011 |
| F03 | 50660001 | 50760000 | 0.0010 | 0.0006 | 0.0008 |
| F03 | 50680001 | 50780000 | 0.0010 | 0.0007 | 0.0008 |
| F03 | 50700001 | 50800000 | 0.0010 | 0.0007 | 0.0007 |
| F03 | 50720001 | 50820000 | 0.0010 | 0.0008 | 0.0008 |
| F03 | 50740001 | 50840000 | 0.0011 | 0.0009 | 0.0010 |
| F03 | 50760001 | 50860000 | 0.0009 | 0.0008 | 0.0009 |
| F03 | 50780001 | 50880000 | 0.0010 | 0.0009 | 0.0010 |
| F03 | 50800001 | 50900000 | 0.0016 | 0.0014 | 0.0015 |
| F03 | 50820001 | 50920000 | 0.0025 | 0.0021 | 0.0024 |
| F03 | 50840001 | 50940000 | 0.0023 | 0.0019 | 0.0022 |
| F03 | 50860001 | 50960000 | 0.0024 | 0.0020 | 0.0023 |
| F03 | 50880001 | 50980000 | 0.0026 | 0.0021 | 0.0025 |
| F03 | 50900001 | 51000000 | 0.0029 | 0.0024 | 0.0029 |
| F03 | 50920001 | 51020000 | 0.0019 | 0.0016 | 0.0019 |
| F03 | 50940001 | 51040000 | 0.0032 | 0.0029 | 0.0031 |
| F03 | 50960001 | 51060000 | 0.0051 | 0.0049 | 0.0050 |
| F03 | 50980001 | 51080000 | 0.0051 | 0.0049 | 0.0050 |
| F03 | 51000001 | 51100000 | 0.0042 | 0.0040 | 0.0040 |
| F03 | 51020001 | 51120000 | 0.0041 | 0.0039 | 0.0039 |
| F03 | 51040001 | 51140000 | 0.0029 | 0.0028 | 0.0028 |
| F03 | 51060001 | 51160000 | 0.0013 | 0.0011 | 0.0013 |
| F03 | 51080001 | 51180000 | 0.0013 | 0.0010 | 0.0012 |
| F03 | 51100001 | 51200000 | 0.0012 | 0.0009 | 0.0012 |
| F03 | 51120001 | 51220000 | 0.0011 | 0.0009 | 0.0011 |
| F03 | 51140001 | 51240000 | 0.0012 | 0.0010 | 0.0012 |
| F03 | 51160001 | 51260000 | 0.0010 | 0.0009 | 0.0010 |
| F03 | 51180001 | 51280000 | 0.0010 | 0.0008 | 0.0010 |
| F03 | 51200001 | 51300000 | 0.0011 | 0.0009 | 0.0011 |
| F03 | 51220001 | 51320000 | 0.0012 | 0.0010 | 0.0012 |
| F03 | 51240001 | 51340000 | 0.0009 | 0.0007 | 0.0009 |
| F03 | 51260001 | 51360000 | 0.0007 | 0.0005 | 0.0007 |
| F03 | 51280001 | 51380000 | 0.0007 | 0.0006 | 0.0007 |
| F03 | 51300001 | 51400000 | 0.0007 | 0.0006 | 0.0007 |
| F03 | 51320001 | 51420000 | 0.0010 | 0.0009 | 0.0010 |

| Chr | Start | End | both | A1 | A2 |
|---|---|---|---|---|---|
| F03 | 51340001 | 51440000 | 0.0013 | 0.0011 | 0.0013 |
| F03 | 51360001 | 51460000 | 0.0015 | 0.0012 | 0.0015 |
| F03 | 51380001 | 51480000 | 0.0015 | 0.0012 | 0.0015 |
| F03 | 51400001 | 51500000 | 0.0014 | 0.0011 | 0.0014 |
| F03 | 51420001 | 51520000 | 0.0010 | 0.0008 | 0.0010 |
| F03 | 51440001 | 51540000 | 0.0008 | 0.0006 | 0.0008 |
| F03 | 51460001 | 51560000 | 0.0007 | 0.0005 | 0.0007 |
| F03 | 51480001 | 51580000 | 0.0004 | 0.0003 | 0.0004 |
| F03 | 51500001 | 51600000 | 0.0006 | 0.0004 | 0.0006 |
| F03 | 51520001 | 51620000 | 0.0006 | 0.0004 | 0.0006 |
| F03 | 51540001 | 51640000 | 0.0005 | 0.0004 | 0.0005 |
| F03 | 51560001 | 51660000 | 0.0005 | 0.0004 | 0.0005 |
| F03 | 51580001 | 51680000 | 0.0004 | 0.0003 | 0.0004 |
| F03 | 51600001 | 51700000 | 0.0003 | 0.0002 | 0.0003 |
| F03 | 51620001 | 51720000 | 0.0004 | 0.0003 | 0.0004 |
| F03 | 51640001 | 51740000 | 0.0012 | 0.0010 | 0.0012 |
| F03 | 51660001 | 51760000 | 0.0014 | 0.0012 | 0.0014 |
| F03 | 51680001 | 51780000 | 0.0015 | 0.0013 | 0.0015 |
| F03 | 51700001 | 51800000 | 0.0016 | 0.0014 | 0.0016 |
| F03 | 51720001 | 51820000 | 0.0015 | 0.0013 | 0.0015 |
| F03 | 51740001 | 51840000 | 0.0007 | 0.0006 | 0.0007 |
| F03 | 51760001 | 51860000 | 0.0004 | 0.0003 | 0.0004 |
| F03 | 51780001 | 51880000 | 0.0002 | 0.0002 | 0.0003 |
| F03 | 51800001 | 51900000 | 0.0001 | 0.0001 | 0.0001 |
| F03 | 51820001 | 51920000 | 0.0001 | 0.0001 | 0.0001 |
| F03 | 51840001 | 51940000 | 0.0001 | 0.0000 | 0.0001 |
| F03 | 51860001 | 51960000 | 0.0002 | 0.0001 | 0.0002 |
| F03 | 51880001 | 51980000 | 0.0003 | 0.0002 | 0.0003 |
| F03 | 51900001 | 52000000 | 0.0003 | 0.0002 | 0.0003 |
| F03 | 51920001 | 52020000 | 0.0004 | 0.0003 | 0.0004 |
| F03 | 51940001 | 52040000 | 0.0008 | 0.0006 | 0.0008 |
| F03 | 51960001 | 52060000 | 0.0007 | 0.0005 | 0.0007 |
| F03 | 51980001 | 52080000 | 0.0007 | 0.0005 | 0.0007 |
| F03 | 52000001 | 52100000 | 0.0013 | 0.0010 | 0.0013 |
| F03 | 52020001 | 52120000 | 0.0011 | 0.0009 | 0.0011 |
| F03 | 52040001 | 52140000 | 0.0009 | 0.0007 | 0.0009 |
| F03 | 52060001 | 52160000 | 0.0013 | 0.0011 | 0.0013 |
| F03 | 52080001 | 52180000 | 0.0012 | 0.0010 | 0.0012 |
| F03 | 52100001 | 52200000 | 0.0008 | 0.0007 | 0.0008 |
| F03 | 52120001 | 52220000 | 0.0008 | 0.0007 | 0.0008 |
| F03 | 52140001 | 52240000 | 0.0007 | 0.0006 | 0.0007 |
| F03 | 52160001 | 52260000 | 0.0007 | 0.0005 | 0.0007 |
| F03 | 52180001 | 52280000 | 0.0010 | 0.0007 | 0.0010 |
| F03 | 52200001 | 52300000 | 0.0008 | 0.0006 | 0.0009 |
| F03 | 52220001 | 52320000 | 0.0012 | 0.0010 | 0.0012 |
| F03 | 52240001 | 52340000 | 0.0013 | 0.0011 | 0.0013 |
| F03 | 52260001 | 52360000 | 0.0014 | 0.0012 | 0.0014 |
| F03 | 52280001 | 52380000 | 0.0012 | 0.0011 | 0.0012 |
| F03 | 52300001 | 52400000 | 0.0017 | 0.0015 | 0.0016 |
| F03 | 52320001 | 52420000 | 0.0018 | 0.0015 | 0.0017 |
| F03 | 52340001 | 52440000 | 0.0016 | 0.0013 | 0.0015 |
| F03 | 52360001 | 52460000 | 0.0012 | 0.0010 | 0.0012 |
| F03 | 52380001 | 52480000 | 0.0010 | 0.0008 | 0.0010 |
| F03 | 52400001 | 52500000 | 0.0010 | 0.0008 | 0.0009 |
| F03 | 52420001 | 52520000 | 0.0005 | 0.0005 | 0.0005 |
| F03 | 52440001 | 52540000 | 0.0006 | 0.0005 | 0.0005 |
| F03 | 52460001 | 52560000 | 0.0010 | 0.0009 | 0.0010 |
| F03 | 52480001 | 52580000 | 0.0017 | 0.0013 | 0.0017 |

| Chr | Start | End | both | A1 | A2 |
|---|---|---|---|---|---|
| F03 | 52500001 | 52600000 | 0.0021 | 0.0014 | 0.0021 |
| F03 | 52520001 | 52620000 | 0.0028 | 0.0020 | 0.0027 |
| F03 | 52540001 | 52640000 | 0.0032 | 0.0023 | 0.0031 |
| F03 | 52560001 | 52660000 | 0.0030 | 0.0021 | 0.0029 |
| F03 | 52580001 | 52680000 | 0.0033 | 0.0026 | 0.0032 |
| F03 | 52600001 | 52700000 | 0.0032 | 0.0026 | 0.0031 |
| F03 | 52620001 | 52720000 | 0.0030 | 0.0024 | 0.0030 |
| F03 | 52640001 | 52740000 | 0.0032 | 0.0024 | 0.0031 |
| F03 | 52660001 | 52760000 | 0.0029 | 0.0022 | 0.0028 |
| F03 | 52680001 | 52780000 | 0.0021 | 0.0015 | 0.0020 |
| F03 | 52700001 | 52800000 | 0.0016 | 0.0011 | 0.0015 |
| F03 | 52720001 | 52820000 | 0.0028 | 0.0023 | 0.0027 |
| F03 | 52740001 | 52840000 | 0.0030 | 0.0028 | 0.0029 |
| F03 | 52760001 | 52860000 | 0.0030 | 0.0028 | 0.0029 |
| F03 | 52780001 | 52880000 | 0.0028 | 0.0027 | 0.0027 |
| F03 | 52800001 | 52900000 | 0.0029 | 0.0028 | 0.0028 |
| F03 | 52820001 | 52920000 | 0.0021 | 0.0020 | 0.0018 |
| F03 | 52840001 | 52940000 | 0.0014 | 0.0012 | 0.0013 |
| F03 | 52860001 | 52960000 | 0.0016 | 0.0014 | 0.0014 |
| F03 | 52880001 | 52980000 | 0.0023 | 0.0019 | 0.0021 |
| F03 | 52900001 | 53000000 | 0.0021 | 0.0017 | 0.0019 |
| F03 | 52920001 | 53020000 | 0.0015 | 0.0011 | 0.0013 |
| F03 | 52940001 | 53040000 | 0.0016 | 0.0013 | 0.0014 |
| F03 | 52960001 | 53060000 | 0.0016 | 0.0013 | 0.0014 |
| F03 | 52980001 | 53080000 | 0.0016 | 0.0012 | 0.0013 |
| F03 | 53000001 | 53100000 | 0.0016 | 0.0012 | 0.0013 |
| F03 | 53020001 | 53120000 | 0.0014 | 0.0011 | 0.0012 |
| F03 | 53040001 | 53140000 | 0.0014 | 0.0010 | 0.0012 |
| F03 | 53060001 | 53160000 | 0.0015 | 0.0011 | 0.0013 |
| F03 | 53080001 | 53180000 | 0.0014 | 0.0010 | 0.0012 |
| F03 | 53100001 | 53200000 | 0.0014 | 0.0011 | 0.0013 |
| F03 | 53120001 | 53220000 | 0.0020 | 0.0016 | 0.0019 |
| F03 | 53140001 | 53240000 | 0.0019 | 0.0014 | 0.0018 |
| F03 | 53160001 | 53260000 | 0.0017 | 0.0013 | 0.0016 |
| F03 | 53180001 | 53280000 | 0.0012 | 0.0010 | 0.0013 |
| F03 | 53200001 | 53300000 | 0.0014 | 0.0011 | 0.0014 |
| F03 | 53220001 | 53320000 | 0.0009 | 0.0007 | 0.0009 |
| F03 | 53240001 | 53340000 | 0.0012 | 0.0010 | 0.0012 |
| F03 | 53260001 | 53360000 | 0.0011 | 0.0009 | 0.0011 |
| F03 | 53280001 | 53380000 | 0.0010 | 0.0008 | 0.0010 |
| F03 | 53300001 | 53400000 | 0.0010 | 0.0008 | 0.0010 |
| F03 | 53320001 | 53420000 | 0.0011 | 0.0009 | 0.0011 |
| F03 | 53340001 | 53440000 | 0.0010 | 0.0008 | 0.0010 |
| F03 | 53360001 | 53460000 | 0.0013 | 0.0011 | 0.0013 |
| F03 | 53380001 | 53480000 | 0.0013 | 0.0011 | 0.0013 |
| F03 | 53400001 | 53500000 | 0.0014 | 0.0011 | 0.0013 |
| F03 | 53420001 | 53520000 | 0.0018 | 0.0012 | 0.0017 |
| F03 | 53440001 | 53540000 | 0.0018 | 0.0011 | 0.0017 |
| F03 | 53460001 | 53560000 | 0.0015 | 0.0009 | 0.0015 |
| F03 | 53480001 | 53580000 | 0.0020 | 0.0011 | 0.0019 |
| F03 | 53500001 | 53600000 | 0.0017 | 0.0009 | 0.0016 |
| F03 | 53520001 | 53620000 | 0.0015 | 0.0009 | 0.0014 |
| F03 | 53540001 | 53640000 | 0.0016 | 0.0010 | 0.0015 |
| F03 | 53560001 | 53660000 | 0.0022 | 0.0016 | 0.0021 |
| F03 | 53580001 | 53680000 | 0.0019 | 0.0015 | 0.0018 |
| F03 | 53600001 | 53700000 | 0.0025 | 0.0020 | 0.0024 |
| F03 | 53620001 | 53720000 | 0.0021 | 0.0018 | 0.0021 |
| F03 | 53640001 | 53740000 | 0.0019 | 0.0016 | 0.0018 |

| Chr | Start | End | both | A1 | A2 |
|---|---|---|---|---|---|
| F03 | 53660001 | 53760000 | 0.0019 | 0.0016 | 0.0018 |
| F03 | 53680001 | 53780000 | 0.0019 | 0.0015 | 0.0018 |
| F03 | 53700001 | 53800000 | 0.0018 | 0.0015 | 0.0018 |
| F03 | 53720001 | 53820000 | 0.0018 | 0.0015 | 0.0017 |
| F03 | 53740001 | 53840000 | 0.0021 | 0.0017 | 0.0021 |
| F03 | 53760001 | 53860000 | 0.0019 | 0.0016 | 0.0019 |
| F03 | 53780001 | 53880000 | 0.0020 | 0.0017 | 0.0020 |
| F03 | 53800001 | 53900000 | 0.0017 | 0.0013 | 0.0016 |
| F03 | 53820001 | 53920000 | 0.0018 | 0.0014 | 0.0017 |
| F03 | 53840001 | 53940000 | 0.0016 | 0.0013 | 0.0015 |
| F03 | 53860001 | 53960000 | 0.0012 | 0.0009 | 0.0011 |
| F03 | 53880001 | 53980000 | 0.0013 | 0.0010 | 0.0012 |
| F03 | 53900001 | 54000000 | 0.0014 | 0.0011 | 0.0013 |
| F03 | 53920001 | 54020000 | 0.0016 | 0.0013 | 0.0016 |
| F03 | 53940001 | 54040000 | 0.0022 | 0.0018 | 0.0021 |
| F03 | 53960001 | 54060000 | 0.0026 | 0.0022 | 0.0025 |
| F03 | 53980001 | 54080000 | 0.0027 | 0.0023 | 0.0026 |
| F03 | 54000001 | 54100000 | 0.0027 | 0.0024 | 0.0026 |
| F03 | 54020001 | 54120000 | 0.0024 | 0.0022 | 0.0023 |
| F03 | 54040001 | 54140000 | 0.0016 | 0.0014 | 0.0015 |
| F03 | 54060001 | 54160000 | 0.0012 | 0.0010 | 0.0011 |
| F03 | 54080001 | 54180000 | 0.0008 | 0.0007 | 0.0007 |
| F03 | 54100001 | 54200000 | 0.0011 | 0.0008 | 0.0010 |
| F03 | 54120001 | 54220000 | 0.0011 | 0.0007 | 0.0009 |
| F03 | 54140001 | 54240000 | 0.0013 | 0.0008 | 0.0009 |
| F03 | 54160001 | 54260000 | 0.0016 | 0.0011 | 0.0012 |
| F03 | 54180001 | 54280000 | 0.0019 | 0.0013 | 0.0014 |
| F03 | 54200001 | 54300000 | 0.0023 | 0.0016 | 0.0017 |
| F03 | 54220001 | 54320000 | 0.0023 | 0.0018 | 0.0018 |
| F03 | 54240001 | 54340000 | 0.0027 | 0.0022 | 0.0023 |
| F03 | 54260001 | 54360000 | 0.0030 | 0.0023 | 0.0026 |
| F03 | 54280001 | 54380000 | 0.0030 | 0.0024 | 0.0028 |
| F03 | 54300001 | 54400000 | 0.0021 | 0.0016 | 0.0019 |
| F03 | 54320001 | 54420000 | 0.0021 | 0.0017 | 0.0020 |
| F03 | 54340001 | 54440000 | 0.0021 | 0.0015 | 0.0020 |
| F03 | 54360001 | 54460000 | 0.0019 | 0.0014 | 0.0018 |
| F03 | 54380001 | 54480000 | 0.0016 | 0.0010 | 0.0014 |
| F03 | 54400001 | 54500000 | 0.0018 | 0.0012 | 0.0016 |
| F03 | 54420001 | 54520000 | 0.0017 | 0.0011 | 0.0015 |
| F03 | 54440001 | 54540000 | 0.0016 | 0.0012 | 0.0014 |
| F03 | 54460001 | 54560000 | 0.0018 | 0.0015 | 0.0017 |
| F03 | 54480001 | 54580000 | 0.0033 | 0.0028 | 0.0033 |
| F03 | 54500001 | 54600000 | 0.0031 | 0.0026 | 0.0031 |
| F03 | 54520001 | 54620000 | 0.0033 | 0.0028 | 0.0032 |
| F03 | 54540001 | 54640000 | 0.0033 | 0.0028 | 0.0032 |
| F03 | 54560001 | 54660000 | 0.0029 | 0.0025 | 0.0029 |
| F03 | 54580001 | 54680000 | 0.0019 | 0.0018 | 0.0018 |
| F03 | 54600001 | 54700000 | 0.0023 | 0.0021 | 0.0021 |
| F03 | 54620001 | 54720000 | 0.0026 | 0.0020 | 0.0021 |
| F03 | 54640001 | 54740000 | 0.0030 | 0.0024 | 0.0026 |
| F03 | 54660001 | 54760000 | 0.0038 | 0.0031 | 0.0032 |
| F03 | 54680001 | 54780000 | 0.0042 | 0.0032 | 0.0037 |
| F03 | 54700001 | 54800000 | 0.0044 | 0.0034 | 0.0039 |
| F03 | 54720001 | 54820000 | 0.0041 | 0.0034 | 0.0037 |
| F03 | 54740001 | 54840000 | 0.0044 | 0.0035 | 0.0037 |
| F03 | 54760001 | 54860000 | 0.0045 | 0.0035 | 0.0038 |
| F03 | 54780001 | 54880000 | 0.0051 | 0.0044 | 0.0045 |
| F03 | 54800001 | 54900000 | 0.0054 | 0.0043 | 0.0044 |

| Chr | Start | End | both | A1 | A2 |
|---|---|---|---|---|---|
| F03 | 54820001 | 54920000 | 0.0058 | 0.0045 | 0.0047 |
| F03 | 54840001 | 54940000 | 0.0058 | 0.0045 | 0.0048 |
| F03 | 54860001 | 54960000 | 0.0049 | 0.0037 | 0.0039 |
| F03 | 54880001 | 54980000 | 0.0034 | 0.0021 | 0.0024 |
| F03 | 54900001 | 55000000 | 0.0027 | 0.0019 | 0.0021 |
| F03 | 54920001 | 55020000 | 0.0022 | 0.0017 | 0.0018 |
| F03 | 54940001 | 55040000 | 0.0010 | 0.0009 | 0.0008 |
| F03 | 54960001 | 55060000 | 0.0015 | 0.0014 | 0.0014 |
| F03 | 54980001 | 55080000 | 0.0016 | 0.0015 | 0.0014 |
| F03 | 55000001 | 55100000 | 0.0017 | 0.0016 | 0.0016 |
| F03 | 55020001 | 55120000 | 0.0021 | 0.0019 | 0.0020 |
| F03 | 55040001 | 55140000 | 0.0029 | 0.0026 | 0.0029 |
| F03 | 55060001 | 55160000 | 0.0022 | 0.0020 | 0.0022 |
| F03 | 55080001 | 55180000 | 0.0022 | 0.0018 | 0.0020 |
| F03 | 55100001 | 55200000 | 0.0026 | 0.0020 | 0.0023 |
| F03 | 55120001 | 55220000 | 0.0029 | 0.0022 | 0.0025 |
| F03 | 55140001 | 55240000 | 0.0025 | 0.0017 | 0.0018 |
| F03 | 55160001 | 55260000 | 0.0030 | 0.0019 | 0.0020 |
| F03 | 55180001 | 55280000 | 0.0031 | 0.0020 | 0.0021 |
| F03 | 55200001 | 55300000 | 0.0031 | 0.0021 | 0.0021 |
| F03 | 55220001 | 55320000 | 0.0029 | 0.0019 | 0.0020 |
| F03 | 55240001 | 55340000 | 0.0027 | 0.0017 | 0.0019 |
| F03 | 55260001 | 55360000 | 0.0024 | 0.0014 | 0.0016 |
| F03 | 55280001 | 55380000 | 0.0031 | 0.0018 | 0.0021 |
| F03 | 55300001 | 55400000 | 0.0029 | 0.0018 | 0.0020 |
| F03 | 55320001 | 55420000 | 0.0035 | 0.0023 | 0.0026 |
| F03 | 55340001 | 55440000 | 0.0039 | 0.0028 | 0.0031 |
| F03 | 55360001 | 55460000 | 0.0037 | 0.0030 | 0.0032 |
| F03 | 55380001 | 55480000 | 0.0028 | 0.0025 | 0.0027 |
| F03 | 55400001 | 55500000 | 0.0023 | 0.0020 | 0.0022 |
| F03 | 55420001 | 55520000 | 0.0013 | 0.0012 | 0.0013 |
| F03 | 55440001 | 55540000 | 0.0007 | 0.0006 | 0.0007 |
| F03 | 55460001 | 55560000 | 0.0006 | 0.0005 | 0.0006 |
| F03 | 55480001 | 55580000 | 0.0007 | 0.0006 | 0.0006 |
| F03 | 55500001 | 55600000 | 0.0006 | 0.0005 | 0.0005 |
| F03 | 55520001 | 55620000 | 0.0007 | 0.0006 | 0.0007 |
| F03 | 55540001 | 55640000 | 0.0016 | 0.0012 | 0.0014 |
| F03 | 55560001 | 55660000 | 0.0018 | 0.0014 | 0.0017 |
| F03 | 55580001 | 55680000 | 0.0025 | 0.0020 | 0.0023 |
| F03 | 55600001 | 55700000 | 0.0028 | 0.0022 | 0.0026 |
| F03 | 55620001 | 55720000 | 0.0027 | 0.0020 | 0.0024 |
| F03 | 55640001 | 55740000 | 0.0026 | 0.0019 | 0.0022 |
| F03 | 55660001 | 55760000 | 0.0029 | 0.0020 | 0.0023 |
| F03 | 55680001 | 55780000 | 0.0025 | 0.0017 | 0.0020 |
| F03 | 55700001 | 55800000 | 0.0027 | 0.0016 | 0.0020 |
| F03 | 55720001 | 55820000 | 0.0027 | 0.0015 | 0.0019 |
| F03 | 55740001 | 55840000 | 0.0023 | 0.0013 | 0.0015 |
| F03 | 55760001 | 55860000 | 0.0017 | 0.0009 | 0.0011 |
| F03 | 55780001 | 55880000 | 0.0018 | 0.0008 | 0.0011 |
| F03 | 55800001 | 55900000 | 0.0023 | 0.0013 | 0.0017 |
| F03 | 55820001 | 55920000 | 0.0029 | 0.0016 | 0.0021 |
| F03 | 55840001 | 55940000 | 0.0027 | 0.0015 | 0.0021 |
| F03 | 55860001 | 55960000 | 0.0027 | 0.0015 | 0.0021 |
| F03 | 55880001 | 55980000 | 0.0030 | 0.0019 | 0.0023 |
| F03 | 55900001 | 56000000 | 0.0023 | 0.0016 | 0.0018 |
| F03 | 55920001 | 56020000 | 0.0022 | 0.0017 | 0.0018 |
| F03 | 55940001 | 56040000 | 0.0023 | 0.0019 | 0.0020 |
| F03 | 55960001 | 56060000 | 0.0029 | 0.0024 | 0.0024 |

| Chr | Start | End | both | A1 | A2 |
|---|---|---|---|---|---|
| F03 | 55980001 | 56080000 | 0.0022 | 0.0019 | 0.0019 |
| F03 | 56000001 | 56100000 | 0.0031 | 0.0025 | 0.0026 |
| F03 | 56020001 | 56120000 | 0.0033 | 0.0026 | 0.0027 |
| F03 | 56040001 | 56140000 | 0.0037 | 0.0028 | 0.0029 |
| F03 | 56060001 | 56160000 | 0.0043 | 0.0032 | 0.0033 |
| F03 | 56080001 | 56180000 | 0.0053 | 0.0039 | 0.0043 |
| F03 | 56100001 | 56200000 | 0.0043 | 0.0031 | 0.0034 |
| F03 | 56120001 | 56220000 | 0.0037 | 0.0026 | 0.0029 |
| F03 | 56140001 | 56240000 | 0.0037 | 0.0027 | 0.0031 |
| F03 | 56160001 | 56260000 | 0.0031 | 0.0023 | 0.0027 |
| F03 | 56180001 | 56280000 | 0.0020 | 0.0014 | 0.0018 |
| F03 | 56200001 | 56300000 | 0.0022 | 0.0017 | 0.0021 |
| F03 | 56220001 | 56320000 | 0.0022 | 0.0018 | 0.0021 |
| F03 | 56240001 | 56340000 | 0.0016 | 0.0014 | 0.0015 |
| F03 | 56260001 | 56360000 | 0.0019 | 0.0015 | 0.0016 |
| F03 | 56280001 | 56380000 | 0.0020 | 0.0014 | 0.0016 |
| F03 | 56300001 | 56400000 | 0.0024 | 0.0018 | 0.0019 |
| F03 | 56320001 | 56420000 | 0.0031 | 0.0024 | 0.0026 |
| F03 | 56340001 | 56440000 | 0.0035 | 0.0027 | 0.0029 |
| F03 | 56360001 | 56460000 | 0.0034 | 0.0027 | 0.0030 |
| F03 | 56380001 | 56480000 | 0.0037 | 0.0033 | 0.0035 |
| F03 | 56400001 | 56500000 | 0.0030 | 0.0026 | 0.0029 |
| F03 | 56420001 | 56520000 | 0.0024 | 0.0021 | 0.0023 |
| F03 | 56440001 | 56540000 | 0.0019 | 0.0016 | 0.0018 |
| F03 | 56460001 | 56560000 | 0.0014 | 0.0013 | 0.0014 |
| F03 | 56480001 | 56580000 | 0.0012 | 0.0011 | 0.0012 |
| F03 | 56500001 | 56600000 | 0.0011 | 0.0011 | 0.0011 |
| F03 | 56520001 | 56620000 | 0.0009 | 0.0008 | 0.0009 |
| F03 | 56540001 | 56640000 | 0.0010 | 0.0009 | 0.0010 |
| F03 | 56560001 | 56660000 | 0.0009 | 0.0008 | 0.0009 |
| F03 | 56580001 | 56680000 | 0.0006 | 0.0005 | 0.0005 |
| F03 | 56600001 | 56700000 | 0.0007 | 0.0006 | 0.0007 |
| F03 | 56620001 | 56720000 | 0.0017 | 0.0016 | 0.0017 |
| F03 | 56640001 | 56740000 | 0.0020 | 0.0017 | 0.0018 |
| F03 | 56660001 | 56760000 | 0.0023 | 0.0017 | 0.0019 |
| F03 | 56680001 | 56780000 | 0.0025 | 0.0019 | 0.0020 |
| F03 | 56700001 | 56800000 | 0.0028 | 0.0022 | 0.0023 |
| F03 | 56720001 | 56820000 | 0.0022 | 0.0016 | 0.0017 |
| F03 | 56740001 | 56840000 | 0.0028 | 0.0018 | 0.0022 |
| F03 | 56760001 | 56860000 | 0.0025 | 0.0016 | 0.0019 |
| F03 | 56780001 | 56880000 | 0.0032 | 0.0022 | 0.0026 |
| F03 | 56800001 | 56900000 | 0.0035 | 0.0021 | 0.0026 |
| F03 | 56820001 | 56920000 | 0.0037 | 0.0022 | 0.0028 |
| F03 | 56840001 | 56940000 | 0.0035 | 0.0022 | 0.0027 |
| F03 | 56860001 | 56960000 | 0.0049 | 0.0035 | 0.0041 |
| F03 | 56880001 | 56980000 | 0.0044 | 0.0030 | 0.0036 |
| F03 | 56900001 | 57000000 | 0.0038 | 0.0026 | 0.0031 |
| F03 | 56920001 | 57020000 | 0.0032 | 0.0022 | 0.0027 |
| F03 | 56940001 | 57040000 | 0.0025 | 0.0017 | 0.0021 |
| F03 | 56960001 | 57060000 | 0.0011 | 0.0006 | 0.0008 |
| F03 | 56980001 | 57080000 | 0.0006 | 0.0003 | 0.0004 |
| F03 | 57000001 | 57100000 | 0.0003 | 0.0002 | 0.0003 |
| F03 | 57020001 | 57120000 | 0.0003 | 0.0002 | 0.0003 |
| F03 | 57040001 | 57140000 | 0.0004 | 0.0003 | 0.0004 |
| F03 | 57060001 | 57160000 | 0.0005 | 0.0005 | 0.0006 |
| F03 | 57080001 | 57180000 | 0.0007 | 0.0006 | 0.0007 |
| F03 | 57100001 | 57200000 | 0.0007 | 0.0006 | 0.0007 |
| F03 | 57120001 | 57220000 | 0.0010 | 0.0008 | 0.0010 |

| Chr | Start | End | both | A1 | A2 |
|---|---|---|---|---|---|
| F03 | 57140001 | 57240000 | 0.0011 | 0.0009 | 0.0011 |
| F03 | 57160001 | 57260000 | 0.0010 | 0.0008 | 0.0010 |
| F03 | 57180001 | 57280000 | 0.0014 | 0.0011 | 0.0014 |
| F03 | 57200001 | 57300000 | 0.0016 | 0.0013 | 0.0016 |
| F03 | 57220001 | 57320000 | 0.0014 | 0.0012 | 0.0014 |
| F03 | 57240001 | 57340000 | 0.0021 | 0.0018 | 0.0021 |
| F03 | 57260001 | 57360000 | 0.0021 | 0.0019 | 0.0021 |
| F03 | 57280001 | 57380000 | 0.0017 | 0.0015 | 0.0016 |
| F03 | 57300001 | 57400000 | 0.0015 | 0.0013 | 0.0015 |
| F03 | 57320001 | 57420000 | 0.0015 | 0.0013 | 0.0015 |
| F03 | 57340001 | 57440000 | 0.0010 | 0.0008 | 0.0010 |
| F03 | 57360001 | 57460000 | 0.0010 | 0.0009 | 0.0010 |
| F03 | 57380001 | 57480000 | 0.0011 | 0.0009 | 0.0011 |
| F03 | 57400001 | 57500000 | 0.0018 | 0.0015 | 0.0018 |
| F03 | 57420001 | 57520000 | 0.0027 | 0.0021 | 0.0024 |
| F03 | 57440001 | 57540000 | 0.0032 | 0.0024 | 0.0029 |
| F03 | 57460001 | 57560000 | 0.0036 | 0.0028 | 0.0033 |
| F03 | 57480001 | 57580000 | 0.0039 | 0.0029 | 0.0034 |
| F03 | 57500001 | 57600000 | 0.0046 | 0.0031 | 0.0039 |
| F03 | 57520001 | 57620000 | 0.0042 | 0.0027 | 0.0035 |
| F03 | 57540001 | 57640000 | 0.0041 | 0.0025 | 0.0032 |
| F03 | 57560001 | 57660000 | 0.0041 | 0.0024 | 0.0031 |
| F03 | 57580001 | 57680000 | 0.0048 | 0.0031 | 0.0038 |
| F03 | 57600001 | 57700000 | 0.0045 | 0.0031 | 0.0035 |
| F03 | 57620001 | 57720000 | 0.0043 | 0.0032 | 0.0035 |
| F03 | 57640001 | 57740000 | 0.0051 | 0.0042 | 0.0045 |
| F03 | 57660001 | 57760000 | 0.0049 | 0.0041 | 0.0044 |
| F03 | 57680001 | 57780000 | 0.0042 | 0.0034 | 0.0036 |
| F03 | 57700001 | 57800000 | 0.0036 | 0.0030 | 0.0030 |
| F03 | 57720001 | 57820000 | 0.0041 | 0.0033 | 0.0033 |
| F03 | 57740001 | 57840000 | 0.0027 | 0.0020 | 0.0020 |
| F03 | 57760001 | 57860000 | 0.0024 | 0.0017 | 0.0017 |
| F03 | 57780001 | 57880000 | 0.0025 | 0.0018 | 0.0018 |
| F03 | 57800001 | 57900000 | 0.0022 | 0.0016 | 0.0017 |
| F03 | 57820001 | 57920000 | 0.0019 | 0.0013 | 0.0016 |
| F03 | 57840001 | 57940000 | 0.0032 | 0.0025 | 0.0030 |
| F03 | 57860001 | 57960000 | 0.0032 | 0.0026 | 0.0030 |
| F03 | 57880001 | 57980000 | 0.0028 | 0.0023 | 0.0026 |
| F03 | 57900001 | 58000000 | 0.0027 | 0.0022 | 0.0026 |
| F03 | 57920001 | 58020000 | 0.0027 | 0.0022 | 0.0025 |
| F03 | 57940001 | 58040000 | 0.0017 | 0.0012 | 0.0014 |
| F03 | 57960001 | 58060000 | 0.0016 | 0.0012 | 0.0014 |
| F03 | 57980001 | 58080000 | 0.0016 | 0.0012 | 0.0015 |
| F03 | 58000001 | 58100000 | 0.0015 | 0.0011 | 0.0013 |
| F03 | 58020001 | 58120000 | 0.0011 | 0.0009 | 0.0011 |
| F03 | 58040001 | 58140000 | 0.0012 | 0.0010 | 0.0011 |
| F03 | 58060001 | 58160000 | 0.0013 | 0.0010 | 0.0010 |
| F03 | 58080001 | 58180000 | 0.0020 | 0.0013 | 0.0016 |
| F03 | 58100001 | 58200000 | 0.0023 | 0.0013 | 0.0016 |
| F03 | 58120001 | 58220000 | 0.0031 | 0.0018 | 0.0022 |
| F03 | 58140001 | 58240000 | 0.0027 | 0.0013 | 0.0017 |
| F03 | 58160001 | 58260000 | 0.0026 | 0.0013 | 0.0018 |
| F03 | 58180001 | 58280000 | 0.0021 | 0.0010 | 0.0012 |
| F03 | 58200001 | 58300000 | 0.0019 | 0.0011 | 0.0012 |
| F03 | 58220001 | 58320000 | 0.0014 | 0.0005 | 0.0007 |
| F03 | 58240001 | 58340000 | 0.0019 | 0.0007 | 0.0008 |
| F03 | 58260001 | 58360000 | 0.0023 | 0.0010 | 0.0011 |
| F03 | 58280001 | 58380000 | 0.0023 | 0.0010 | 0.0011 |

| Chr | Start | End | both | A1 | A2 |
|---|---|---|---|---|---|
| F03 | 58300001 | 58400000 | 0.0023 | 0.0009 | 0.0012 |
| F03 | 58320001 | 58420000 | 0.0023 | 0.0010 | 0.0011 |
| F03 | 58340001 | 58440000 | 0.0022 | 0.0010 | 0.0013 |
| F03 | 58360001 | 58460000 | 0.0021 | 0.0012 | 0.0014 |
| F03 | 58380001 | 58480000 | 0.0025 | 0.0016 | 0.0018 |
| F03 | 58400001 | 58500000 | 0.0029 | 0.0021 | 0.0022 |
| F03 | 58420001 | 58520000 | 0.0041 | 0.0032 | 0.0035 |
| F03 | 58440001 | 58540000 | 0.0038 | 0.0030 | 0.0032 |
| F03 | 58460001 | 58560000 | 0.0038 | 0.0030 | 0.0030 |
| F03 | 58480001 | 58580000 | 0.0046 | 0.0039 | 0.0040 |
| F03 | 58500001 | 58600000 | 0.0045 | 0.0037 | 0.0038 |
| F03 | 58520001 | 58620000 | 0.0033 | 0.0028 | 0.0027 |
| F03 | 58540001 | 58640000 | 0.0030 | 0.0028 | 0.0025 |
| F03 | 58560001 | 58660000 | 0.0034 | 0.0028 | 0.0030 |
| F03 | 58580001 | 58680000 | 0.0028 | 0.0021 | 0.0022 |
| F03 | 58600001 | 58700000 | 0.0033 | 0.0026 | 0.0027 |
| F03 | 58620001 | 58720000 | 0.0041 | 0.0033 | 0.0035 |
| F03 | 58640001 | 58740000 | 0.0047 | 0.0038 | 0.0041 |
| F03 | 58660001 | 58760000 | 0.0045 | 0.0036 | 0.0037 |
| F03 | 58680001 | 58780000 | 0.0037 | 0.0031 | 0.0032 |
| F03 | 58700001 | 58800000 | 0.0032 | 0.0027 | 0.0027 |
| F03 | 58720001 | 58820000 | 0.0024 | 0.0019 | 0.0019 |
| F03 | 58740001 | 58840000 | 0.0023 | 0.0017 | 0.0016 |
| F03 | 58760001 | 58860000 | 0.0018 | 0.0015 | 0.0014 |
| F03 | 58780001 | 58880000 | 0.0023 | 0.0019 | 0.0019 |
| F03 | 58800001 | 58900000 | 0.0018 | 0.0014 | 0.0015 |
| F03 | 58820001 | 58920000 | 0.0019 | 0.0016 | 0.0017 |
| F03 | 58840001 | 58940000 | 0.0018 | 0.0017 | 0.0018 |
| F03 | 58860001 | 58960000 | 0.0020 | 0.0019 | 0.0019 |
| F03 | 58880001 | 58980000 | 0.0014 | 0.0014 | 0.0014 |
| F03 | 58900001 | 59000000 | 0.0022 | 0.0021 | 0.0021 |
| F03 | 58920001 | 59020000 | 0.0017 | 0.0016 | 0.0017 |
| F03 | 58940001 | 59040000 | 0.0018 | 0.0016 | 0.0018 |
| F03 | 58960001 | 59060000 | 0.0019 | 0.0016 | 0.0018 |
| F03 | 58980001 | 59080000 | 0.0022 | 0.0019 | 0.0022 |
| F03 | 59000001 | 59100000 | 0.0017 | 0.0016 | 0.0017 |
| F03 | 59020001 | 59120000 | 0.0017 | 0.0016 | 0.0017 |
| F03 | 59040001 | 59140000 | 0.0016 | 0.0014 | 0.0016 |
| F03 | 59060001 | 59160000 | 0.0015 | 0.0013 | 0.0014 |
| F03 | 59080001 | 59180000 | 0.0017 | 0.0014 | 0.0016 |
| F03 | 59100001 | 59200000 | 0.0017 | 0.0012 | 0.0014 |
| F03 | 59120001 | 59220000 | 0.0022 | 0.0015 | 0.0016 |
| F03 | 59140001 | 59240000 | 0.0023 | 0.0016 | 0.0016 |
| F03 | 59160001 | 59260000 | 0.0028 | 0.0019 | 0.0019 |
| F03 | 59180001 | 59280000 | 0.0033 | 0.0023 | 0.0023 |
| F03 | 59200001 | 59300000 | 0.0046 | 0.0036 | 0.0038 |
| F03 | 59220001 | 59320000 | 0.0048 | 0.0038 | 0.0042 |
| F03 | 59240001 | 59340000 | 0.0044 | 0.0034 | 0.0038 |
| F03 | 59260001 | 59360000 | 0.0042 | 0.0033 | 0.0037 |
| F03 | 59280001 | 59380000 | 0.0037 | 0.0028 | 0.0032 |
| F03 | 59300001 | 59400000 | 0.0024 | 0.0015 | 0.0017 |
| F03 | 59320001 | 59420000 | 0.0023 | 0.0016 | 0.0017 |
| F03 | 59340001 | 59440000 | 0.0029 | 0.0018 | 0.0021 |
| F03 | 59360001 | 59460000 | 0.0030 | 0.0020 | 0.0023 |
| F03 | 59380001 | 59480000 | 0.0030 | 0.0022 | 0.0024 |
| F03 | 59400001 | 59500000 | 0.0027 | 0.0021 | 0.0023 |
| F03 | 59420001 | 59520000 | 0.0028 | 0.0022 | 0.0024 |
| F03 | 59440001 | 59540000 | 0.0022 | 0.0019 | 0.0020 |

| Chr | Start | End | both | A1 | A2 |
|---|---|---|---|---|---|
| F03 | 59460001 | 59560000 | 0.0017 | 0.0015 | 0.0016 |
| F03 | 59480001 | 59580000 | 0.0012 | 0.0010 | 0.0011 |
| F03 | 59500001 | 59600000 | 0.0016 | 0.0014 | 0.0014 |
| F03 | 59520001 | 59620000 | 0.0008 | 0.0008 | 0.0006 |
| F03 | 59540001 | 59640000 | 0.0014 | 0.0009 | 0.0008 |
| F03 | 59560001 | 59660000 | 0.0012 | 0.0007 | 0.0006 |
| F03 | 59580001 | 59680000 | 0.0014 | 0.0008 | 0.0007 |
| F03 | 59600001 | 59700000 | 0.0012 | 0.0006 | 0.0007 |
| F03 | 59620001 | 59720000 | 0.0013 | 0.0007 | 0.0007 |
| F03 | 59640001 | 59740000 | 0.0008 | 0.0006 | 0.0006 |
| F03 | 59660001 | 59760000 | 0.0010 | 0.0008 | 0.0009 |
| F03 | 59680001 | 59780000 | 0.0007 | 0.0006 | 0.0007 |
| F03 | 59700001 | 59800000 | 0.0007 | 0.0006 | 0.0007 |
| F03 | 59720001 | 59820000 | 0.0010 | 0.0008 | 0.0009 |
| F03 | 59740001 | 59840000 | 0.0015 | 0.0010 | 0.0012 |
| F03 | 59760001 | 59860000 | 0.0015 | 0.0010 | 0.0011 |
| F03 | 59780001 | 59880000 | 0.0015 | 0.0010 | 0.0011 |
| F03 | 59800001 | 59900000 | 0.0017 | 0.0010 | 0.0012 |
| F03 | 59820001 | 59920000 | 0.0018 | 0.0012 | 0.0014 |
| F03 | 59840001 | 59940000 | 0.0017 | 0.0011 | 0.0013 |
| F03 | 59860001 | 59960000 | 0.0023 | 0.0015 | 0.0017 |
| F03 | 59880001 | 59980000 | 0.0036 | 0.0026 | 0.0030 |
| F03 | 59900001 | 60000000 | 0.0033 | 0.0024 | 0.0027 |
| F03 | 59920001 | 60020000 | 0.0035 | 0.0025 | 0.0028 |
| F03 | 59940001 | 60040000 | 0.0039 | 0.0028 | 0.0029 |
| F03 | 59960001 | 60060000 | 0.0034 | 0.0024 | 0.0025 |
| F03 | 59980001 | 60080000 | 0.0039 | 0.0026 | 0.0029 |
| F03 | 60000001 | 60100000 | 0.0050 | 0.0034 | 0.0039 |
| F03 | 60020001 | 60120000 | 0.0047 | 0.0032 | 0.0037 |
| F03 | 60040001 | 60140000 | 0.0044 | 0.0029 | 0.0035 |
| F03 | 60060001 | 60160000 | 0.0046 | 0.0030 | 0.0038 |
| F03 | 60080001 | 60180000 | 0.0036 | 0.0022 | 0.0027 |
| F03 | 60100001 | 60200000 | 0.0031 | 0.0020 | 0.0023 |
| F03 | 60120001 | 60220000 | 0.0029 | 0.0018 | 0.0021 |
| F03 | 60140001 | 60240000 | 0.0028 | 0.0018 | 0.0021 |
| F03 | 60160001 | 60260000 | 0.0022 | 0.0015 | 0.0016 |
| F03 | 60180001 | 60280000 | 0.0021 | 0.0013 | 0.0015 |
| F03 | 60200001 | 60300000 | 0.0016 | 0.0009 | 0.0011 |
| F03 | 60220001 | 60320000 | 0.0022 | 0.0012 | 0.0014 |
| F03 | 60240001 | 60340000 | 0.0028 | 0.0017 | 0.0020 |
| F03 | 60260001 | 60360000 | 0.0036 | 0.0023 | 0.0027 |
| F03 | 60280001 | 60380000 | 0.0036 | 0.0025 | 0.0029 |
| F03 | 60300001 | 60400000 | 0.0034 | 0.0024 | 0.0027 |
| F03 | 60320001 | 60420000 | 0.0039 | 0.0029 | 0.0032 |
| F03 | 60340001 | 60440000 | 0.0036 | 0.0026 | 0.0028 |
| F03 | 60360001 | 60460000 | 0.0030 | 0.0021 | 0.0021 |
| F03 | 60380001 | 60480000 | 0.0029 | 0.0018 | 0.0018 |
| F03 | 60400001 | 60500000 | 0.0042 | 0.0030 | 0.0033 |
| F03 | 60420001 | 60520000 | 0.0039 | 0.0026 | 0.0030 |
| F03 | 60440001 | 60540000 | 0.0035 | 0.0023 | 0.0027 |
| F03 | 60460001 | 60560000 | 0.0034 | 0.0024 | 0.0028 |
| F03 | 60480001 | 60580000 | 0.0029 | 0.0022 | 0.0026 |
| F03 | 60500001 | 60600000 | 0.0014 | 0.0011 | 0.0012 |
| F03 | 60520001 | 60620000 | 0.0006 | 0.0005 | 0.0005 |
| F03 | 60540001 | 60640000 | 0.0005 | 0.0005 | 0.0005 |
| F03 | 60560001 | 60660000 | 0.0004 | 0.0004 | 0.0004 |
| F03 | 60580001 | 60680000 | 0.0009 | 0.0008 | 0.0008 |
| F03 | 60600001 | 60700000 | 0.0020 | 0.0017 | 0.0020 |

| Chr | Start | End | both | A1 | A2 |
|---|---|---|---|---|---|
| F03 | 60620001 | 60720000 | 0.0024 | 0.0020 | 0.0024 |
| F03 | 60640001 | 60740000 | 0.0023 | 0.0020 | 0.0023 |
| F03 | 60660001 | 60760000 | 0.0025 | 0.0021 | 0.0025 |
| F03 | 60680001 | 60780000 | 0.0024 | 0.0020 | 0.0024 |
| F03 | 60700001 | 60800000 | 0.0015 | 0.0013 | 0.0015 |
| F03 | 60720001 | 60820000 | 0.0013 | 0.0012 | 0.0013 |
| F03 | 60740001 | 60840000 | 0.0020 | 0.0017 | 0.0019 |
| F03 | 60760001 | 60860000 | 0.0022 | 0.0019 | 0.0021 |
| F03 | 60780001 | 60880000 | 0.0034 | 0.0029 | 0.0033 |
| F03 | 60800001 | 60900000 | 0.0042 | 0.0036 | 0.0041 |
| F03 | 60820001 | 60920000 | 0.0050 | 0.0044 | 0.0048 |
| F03 | 60840001 | 60940000 | 0.0049 | 0.0043 | 0.0048 |
| F03 | 60860001 | 60960000 | 0.0049 | 0.0042 | 0.0046 |
| F03 | 60880001 | 60980000 | 0.0034 | 0.0030 | 0.0032 |
| F03 | 60900001 | 61000000 | 0.0026 | 0.0022 | 0.0024 |
| F03 | 60920001 | 61020000 | 0.0027 | 0.0021 | 0.0025 |
| F03 | 60940001 | 61040000 | 0.0023 | 0.0016 | 0.0018 |
| F03 | 60960001 | 61060000 | 0.0022 | 0.0015 | 0.0017 |
| F03 | 60980001 | 61080000 | 0.0027 | 0.0018 | 0.0020 |
| F03 | 61000001 | 61100000 | 0.0026 | 0.0017 | 0.0020 |
| F03 | 61020001 | 61120000 | 0.0024 | 0.0017 | 0.0018 |
| F03 | 61040001 | 61140000 | 0.0022 | 0.0017 | 0.0018 |
| F03 | 61060001 | 61160000 | 0.0023 | 0.0019 | 0.0020 |
| F03 | 61080001 | 61180000 | 0.0019 | 0.0016 | 0.0017 |
| F03 | 61100001 | 61200000 | 0.0021 | 0.0019 | 0.0020 |
| F03 | 61120001 | 61220000 | 0.0018 | 0.0013 | 0.0014 |
| F03 | 61140001 | 61240000 | 0.0022 | 0.0017 | 0.0018 |
| F03 | 61160001 | 61260000 | 0.0021 | 0.0016 | 0.0018 |
| F03 | 61180001 | 61280000 | 0.0028 | 0.0020 | 0.0025 |
| F03 | 61200001 | 61300000 | 0.0028 | 0.0020 | 0.0025 |
| F03 | 61220001 | 61320000 | 0.0026 | 0.0021 | 0.0025 |
| F03 | 61240001 | 61340000 | 0.0025 | 0.0020 | 0.0023 |
| F03 | 61260001 | 61360000 | 0.0029 | 0.0023 | 0.0025 |
| F03 | 61280001 | 61380000 | 0.0026 | 0.0023 | 0.0022 |
| F03 | 61300001 | 61400000 | 0.0025 | 0.0018 | 0.0019 |
| F03 | 61320001 | 61420000 | 0.0028 | 0.0020 | 0.0021 |
| F03 | 61340001 | 61440000 | 0.0031 | 0.0023 | 0.0024 |
| F03 | 61360001 | 61460000 | 0.0027 | 0.0021 | 0.0022 |
| F03 | 61380001 | 61480000 | 0.0030 | 0.0023 | 0.0025 |
| F03 | 61400001 | 61500000 | 0.0029 | 0.0024 | 0.0027 |
| F03 | 61420001 | 61520000 | 0.0026 | 0.0022 | 0.0025 |
| F03 | 61440001 | 61540000 | 0.0022 | 0.0019 | 0.0021 |
| F03 | 61460001 | 61560000 | 0.0021 | 0.0018 | 0.0020 |
| F03 | 61480001 | 61580000 | 0.0020 | 0.0017 | 0.0017 |
| F03 | 61500001 | 61600000 | 0.0019 | 0.0016 | 0.0016 |
| F03 | 61520001 | 61620000 | 0.0022 | 0.0019 | 0.0018 |
| F03 | 61540001 | 61640000 | 0.0025 | 0.0022 | 0.0021 |
| F03 | 61560001 | 61660000 | 0.0026 | 0.0021 | 0.0021 |
| F03 | 61580001 | 61680000 | 0.0024 | 0.0019 | 0.0020 |
| F03 | 61600001 | 61700000 | 0.0024 | 0.0019 | 0.0020 |
| F03 | 61620001 | 61720000 | 0.0021 | 0.0017 | 0.0019 |
| F03 | 61640001 | 61740000 | 0.0017 | 0.0013 | 0.0015 |
| F03 | 61660001 | 61760000 | 0.0024 | 0.0020 | 0.0022 |
| F03 | 61680001 | 61780000 | 0.0023 | 0.0020 | 0.0022 |
| F03 | 61700001 | 61800000 | 0.0025 | 0.0021 | 0.0023 |
| F03 | 61720001 | 61820000 | 0.0028 | 0.0020 | 0.0023 |
| F03 | 61740001 | 61840000 | 0.0030 | 0.0021 | 0.0024 |
| F03 | 61760001 | 61860000 | 0.0033 | 0.0021 | 0.0026 |

| Chr | Start | End | both | A1 | A2 |
|---|---|---|---|---|---|
| F03 | 61780001 | 61880000 | 0.0032 | 0.0020 | 0.0025 |
| F03 | 61800001 | 61900000 | 0.0030 | 0.0020 | 0.0024 |
| F03 | 61820001 | 61920000 | 0.0033 | 0.0026 | 0.0030 |
| F03 | 61840001 | 61940000 | 0.0033 | 0.0025 | 0.0028 |
| F03 | 61860001 | 61960000 | 0.0028 | 0.0020 | 0.0023 |
| F03 | 61880001 | 61980000 | 0.0029 | 0.0020 | 0.0022 |
| F03 | 61900001 | 62000000 | 0.0028 | 0.0018 | 0.0021 |
| F03 | 61920001 | 62020000 | 0.0024 | 0.0014 | 0.0016 |
| F03 | 61940001 | 62040000 | 0.0025 | 0.0016 | 0.0018 |
| F03 | 61960001 | 62060000 | 0.0017 | 0.0012 | 0.0012 |
| F03 | 61980001 | 62080000 | 0.0015 | 0.0011 | 0.0011 |
| F03 | 62000001 | 62100000 | 0.0013 | 0.0010 | 0.0011 |
| F03 | 62020001 | 62120000 | 0.0009 | 0.0007 | 0.0008 |
| F03 | 62040001 | 62140000 | 0.0005 | 0.0005 | 0.0005 |
| F03 | 62060001 | 62160000 | 0.0005 | 0.0005 | 0.0005 |
| F03 | 62080001 | 62180000 | 0.0007 | 0.0006 | 0.0007 |
| F03 | 62100001 | 62200000 | 0.0010 | 0.0009 | 0.0010 |
| F03 | 62120001 | 62220000 | 0.0010 | 0.0009 | 0.0010 |
| F03 | 62140001 | 62240000 | 0.0013 | 0.0013 | 0.0013 |
| F03 | 62160001 | 62260000 | 0.0020 | 0.0018 | 0.0019 |
| F03 | 62180001 | 62280000 | 0.0017 | 0.0016 | 0.0016 |
| F03 | 62200001 | 62300000 | 0.0016 | 0.0015 | 0.0016 |
| F03 | 62220001 | 62320000 | 0.0018 | 0.0016 | 0.0017 |
| F03 | 62240001 | 62340000 | 0.0016 | 0.0014 | 0.0015 |
| F03 | 62260001 | 62360000 | 0.0009 | 0.0007 | 0.0008 |
| F03 | 62280001 | 62380000 | 0.0012 | 0.0010 | 0.0010 |
| F03 | 62300001 | 62400000 | 0.0014 | 0.0013 | 0.0012 |
| F03 | 62320001 | 62420000 | 0.0022 | 0.0019 | 0.0017 |
| F03 | 62340001 | 62440000 | 0.0033 | 0.0031 | 0.0027 |
| F03 | 62360001 | 62460000 | 0.0039 | 0.0035 | 0.0032 |
| F03 | 62380001 | 62480000 | 0.0051 | 0.0047 | 0.0045 |
| F03 | 62400001 | 62500000 | 0.0066 | 0.0060 | 0.0060 |
| F03 | 62420001 | 62520000 | 0.0062 | 0.0056 | 0.0057 |
| F03 | 62440001 | 62540000 | 0.0054 | 0.0048 | 0.0050 |
| F03 | 62460001 | 62560000 | 0.0048 | 0.0043 | 0.0045 |
| F03 | 62480001 | 62580000 | 0.0037 | 0.0033 | 0.0034 |
| F03 | 62500001 | 62600000 | 0.0019 | 0.0016 | 0.0017 |
| F03 | 62520001 | 62620000 | 0.0016 | 0.0014 | 0.0015 |
| F03 | 62540001 | 62640000 | 0.0016 | 0.0014 | 0.0015 |
| F03 | 62560001 | 62660000 | 0.0019 | 0.0017 | 0.0018 |
| F03 | 62580001 | 62680000 | 0.0020 | 0.0016 | 0.0019 |
| F03 | 62600001 | 62700000 | 0.0020 | 0.0017 | 0.0019 |
| F03 | 62620001 | 62720000 | 0.0020 | 0.0017 | 0.0019 |
| F03 | 62640001 | 62740000 | 0.0013 | 0.0011 | 0.0012 |
| F03 | 62660001 | 62760000 | 0.0020 | 0.0017 | 0.0020 |
| F03 | 62680001 | 62780000 | 0.0018 | 0.0016 | 0.0018 |
| F03 | 62700001 | 62800000 | 0.0021 | 0.0018 | 0.0020 |
| F03 | 62720001 | 62820000 | 0.0021 | 0.0017 | 0.0020 |
| F03 | 62740001 | 62840000 | 0.0029 | 0.0025 | 0.0028 |
| F03 | 62760001 | 62860000 | 0.0019 | 0.0017 | 0.0018 |
| F03 | 62780001 | 62880000 | 0.0021 | 0.0017 | 0.0018 |
| F03 | 62800001 | 62900000 | 0.0029 | 0.0025 | 0.0026 |
| F03 | 62820001 | 62920000 | 0.0032 | 0.0027 | 0.0029 |
| F03 | 62840001 | 62940000 | 0.0032 | 0.0028 | 0.0029 |
| F03 | 62860001 | 62960000 | 0.0034 | 0.0030 | 0.0031 |
| F03 | 62880001 | 62980000 | 0.0031 | 0.0028 | 0.0029 |
| F03 | 62900001 | 63000000 | 0.0025 | 0.0022 | 0.0023 |
| F03 | 62920001 | 63020000 | 0.0037 | 0.0032 | 0.0035 |

| Chr | Start | End | both | A1 | A2 |
|---|---|---|---|---|---|
| F03 | 62940001 | 63040000 | 0.0033 | 0.0027 | 0.0030 |
| F03 | 62960001 | 63060000 | 0.0035 | 0.0028 | 0.0030 |
| F03 | 62980001 | 63080000 | 0.0043 | 0.0033 | 0.0037 |
| F03 | 63000001 | 63100000 | 0.0045 | 0.0035 | 0.0041 |
| F03 | 63020001 | 63120000 | 0.0033 | 0.0024 | 0.0029 |
| F03 | 63040001 | 63140000 | 0.0041 | 0.0034 | 0.0038 |
| F03 | 63060001 | 63160000 | 0.0038 | 0.0031 | 0.0035 |
| F03 | 63080001 | 63180000 | 0.0029 | 0.0025 | 0.0028 |
| F03 | 63100001 | 63200000 | 0.0019 | 0.0018 | 0.0019 |
| F03 | 63120001 | 63220000 | 0.0015 | 0.0014 | 0.0014 |
| F03 | 63140001 | 63240000 | 0.0002 | 0.0002 | 0.0002 |
| F03 | 63160001 | 63260000 | 0.0004 | 0.0004 | 0.0004 |
| F03 | 63180001 | 63280000 | 0.0004 | 0.0004 | 0.0004 |
| F03 | 63200001 | 63300000 | 0.0007 | 0.0008 | 0.0007 |
| F03 | 63220001 | 63320000 | 0.0020 | 0.0020 | 0.0018 |
| F03 | 63240001 | 63340000 | 0.0027 | 0.0027 | 0.0025 |
| F03 | 63260001 | 63360000 | 0.0034 | 0.0033 | 0.0032 |
| F03 | 63280001 | 63380000 | 0.0036 | 0.0034 | 0.0034 |
| F03 | 63300001 | 63400000 | 0.0036 | 0.0033 | 0.0035 |
| F03 | 63320001 | 63420000 | 0.0027 | 0.0023 | 0.0026 |
| F03 | 63340001 | 63440000 | 0.0020 | 0.0017 | 0.0019 |
| F03 | 63360001 | 63460000 | 0.0012 | 0.0010 | 0.0012 |
| F03 | 63380001 | 63480000 | 0.0015 | 0.0013 | 0.0015 |
| F03 | 63400001 | 63500000 | 0.0023 | 0.0018 | 0.0022 |
| F03 | 63420001 | 63520000 | 0.0020 | 0.0015 | 0.0019 |
| F03 | 63440001 | 63540000 | 0.0025 | 0.0019 | 0.0023 |
| F03 | 63460001 | 63560000 | 0.0032 | 0.0026 | 0.0030 |
| F03 | 63480001 | 63580000 | 0.0036 | 0.0028 | 0.0033 |
| F03 | 63500001 | 63600000 | 0.0024 | 0.0020 | 0.0023 |
| F03 | 63520001 | 63620000 | 0.0024 | 0.0020 | 0.0023 |
| F03 | 63540001 | 63640000 | 0.0019 | 0.0016 | 0.0018 |
| F03 | 63560001 | 63660000 | 0.0015 | 0.0012 | 0.0014 |
| F03 | 63580001 | 63680000 | 0.0010 | 0.0009 | 0.0009 |
| F03 | 63600001 | 63700000 | 0.0019 | 0.0016 | 0.0017 |
| F03 | 63620001 | 63720000 | 0.0020 | 0.0016 | 0.0017 |
| F03 | 63640001 | 63740000 | 0.0023 | 0.0018 | 0.0020 |
| F03 | 63660001 | 63760000 | 0.0022 | 0.0017 | 0.0018 |
| F03 | 63680001 | 63780000 | 0.0026 | 0.0019 | 0.0021 |
| F03 | 63700001 | 63800000 | 0.0020 | 0.0014 | 0.0017 |
| F03 | 63720001 | 63820000 | 0.0026 | 0.0018 | 0.0023 |
| F03 | 63740001 | 63840000 | 0.0024 | 0.0017 | 0.0021 |
| F03 | 63760001 | 63860000 | 0.0024 | 0.0017 | 0.0022 |
| F03 | 63780001 | 63880000 | 0.0027 | 0.0021 | 0.0025 |
| F03 | 63800001 | 63900000 | 0.0024 | 0.0018 | 0.0022 |
| F03 | 63820001 | 63920000 | 0.0017 | 0.0014 | 0.0015 |
| F03 | 63840001 | 63940000 | 0.0025 | 0.0019 | 0.0021 |
| F03 | 63860001 | 63960000 | 0.0028 | 0.0023 | 0.0025 |
| F03 | 63880001 | 63980000 | 0.0023 | 0.0016 | 0.0018 |
| F03 | 63900001 | 64000000 | 0.0025 | 0.0016 | 0.0018 |
| F03 | 63920001 | 64020000 | 0.0027 | 0.0018 | 0.0020 |
| F03 | 63940001 | 64040000 | 0.0023 | 0.0016 | 0.0017 |
| F03 | 63960001 | 64060000 | 0.0019 | 0.0011 | 0.0013 |
| F03 | 63980001 | 64080000 | 0.0019 | 0.0015 | 0.0016 |
| F03 | 64000001 | 64100000 | 0.0023 | 0.0018 | 0.0020 |
| F03 | 64020001 | 64120000 | 0.0020 | 0.0016 | 0.0018 |
| F03 | 64040001 | 64140000 | 0.0016 | 0.0013 | 0.0014 |
| F03 | 64060001 | 64160000 | 0.0014 | 0.0012 | 0.0012 |
| F03 | 64080001 | 64180000 | 0.0014 | 0.0011 | 0.0012 |

| Chr | Start | End | both | A1 | A2 |
|---|---|---|---|---|---|
| F03 | 64100001 | 64200000 | 0.0008 | 0.0007 | 0.0007 |
| F03 | 64120001 | 64220000 | 0.0012 | 0.0010 | 0.0011 |
| F03 | 64140001 | 64240000 | 0.0019 | 0.0015 | 0.0017 |
| F03 | 64160001 | 64260000 | 0.0021 | 0.0016 | 0.0018 |
| F03 | 64180001 | 64280000 | 0.0021 | 0.0015 | 0.0018 |
| F03 | 64200001 | 64300000 | 0.0026 | 0.0019 | 0.0022 |
| F03 | 64220001 | 64320000 | 0.0024 | 0.0015 | 0.0018 |
| F03 | 64240001 | 64340000 | 0.0026 | 0.0018 | 0.0020 |
| F03 | 64260001 | 64360000 | 0.0031 | 0.0023 | 0.0026 |
| F03 | 64280001 | 64380000 | 0.0031 | 0.0024 | 0.0026 |
| F03 | 64300001 | 64400000 | 0.0033 | 0.0026 | 0.0028 |
| F03 | 64320001 | 64420000 | 0.0036 | 0.0031 | 0.0031 |
| F03 | 64340001 | 64440000 | 0.0029 | 0.0023 | 0.0023 |
| F03 | 64360001 | 64460000 | 0.0029 | 0.0021 | 0.0021 |
| F03 | 64380001 | 64480000 | 0.0030 | 0.0021 | 0.0024 |
| F03 | 64400001 | 64500000 | 0.0025 | 0.0016 | 0.0019 |
| F03 | 64420001 | 64520000 | 0.0027 | 0.0016 | 0.0020 |
| F03 | 64440001 | 64540000 | 0.0027 | 0.0017 | 0.0021 |
| F03 | 64460001 | 64560000 | 0.0020 | 0.0013 | 0.0016 |
| F03 | 64480001 | 64580000 | 0.0018 | 0.0011 | 0.0011 |
| F03 | 64500001 | 64600000 | 0.0025 | 0.0016 | 0.0017 |
| F03 | 64520001 | 64620000 | 0.0021 | 0.0013 | 0.0014 |
| F03 | 64540001 | 64640000 | 0.0023 | 0.0014 | 0.0014 |
| F03 | 64560001 | 64660000 | 0.0025 | 0.0013 | 0.0014 |
| F03 | 64580001 | 64680000 | 0.0024 | 0.0014 | 0.0016 |
| F03 | 64600001 | 64700000 | 0.0020 | 0.0011 | 0.0012 |
| F03 | 64620001 | 64720000 | 0.0019 | 0.0011 | 0.0011 |
| F03 | 64640001 | 64740000 | 0.0018 | 0.0011 | 0.0011 |
| F03 | 64660001 | 64760000 | 0.0022 | 0.0017 | 0.0018 |
| F03 | 64680001 | 64780000 | 0.0028 | 0.0021 | 0.0023 |
| F03 | 64700001 | 64800000 | 0.0031 | 0.0023 | 0.0026 |
| F03 | 64720001 | 64820000 | 0.0030 | 0.0022 | 0.0025 |
| F03 | 64740001 | 64840000 | 0.0043 | 0.0032 | 0.0036 |
| F03 | 64760001 | 64860000 | 0.0047 | 0.0035 | 0.0038 |
| F03 | 64780001 | 64880000 | 0.0040 | 0.0030 | 0.0032 |
| F03 | 64800001 | 64900000 | 0.0043 | 0.0034 | 0.0034 |
| F03 | 64820001 | 64920000 | 0.0042 | 0.0033 | 0.0033 |
| F03 | 64840001 | 64940000 | 0.0036 | 0.0030 | 0.0029 |
| F03 | 64860001 | 64960000 | 0.0034 | 0.0029 | 0.0028 |
| F03 | 64880001 | 64980000 | 0.0034 | 0.0029 | 0.0029 |
| F03 | 64900001 | 65000000 | 0.0027 | 0.0025 | 0.0024 |
| F03 | 64920001 | 65020000 | 0.0030 | 0.0029 | 0.0028 |
| F03 | 64940001 | 65040000 | 0.0023 | 0.0022 | 0.0022 |
| F03 | 64960001 | 65060000 | 0.0017 | 0.0017 | 0.0016 |
| F03 | 64980001 | 65080000 | 0.0016 | 0.0016 | 0.0015 |
| F03 | 65000001 | 65100000 | 0.0012 | 0.0011 | 0.0011 |
| F03 | 65020001 | 65120000 | 0.0009 | 0.0008 | 0.0007 |
| F03 | 65040001 | 65140000 | 0.0007 | 0.0005 | 0.0005 |
| F03 | 65060001 | 65160000 | 0.0007 | 0.0006 | 0.0005 |
| F03 | 65080001 | 65180000 | 0.0007 | 0.0006 | 0.0005 |
| F03 | 65100001 | 65200000 | 0.0009 | 0.0008 | 0.0007 |
| F03 | 65120001 | 65220000 | 0.0008 | 0.0008 | 0.0007 |
| F03 | 65140001 | 65240000 | 0.0012 | 0.0012 | 0.0011 |
| F03 | 65160001 | 65260000 | 0.0012 | 0.0011 | 0.0011 |
| F03 | 65180001 | 65280000 | 0.0013 | 0.0012 | 0.0013 |
| F03 | 65200001 | 65300000 | 0.0014 | 0.0012 | 0.0014 |
| F03 | 65220001 | 65320000 | 0.0019 | 0.0016 | 0.0019 |
| F03 | 65240001 | 65340000 | 0.0017 | 0.0014 | 0.0017 |

| Chr | Start | End | both | A1 | A2 |
|---|---|---|---|---|---|
| F03 | 65260001 | 65360000 | 0.0015 | 0.0013 | 0.0015 |
| F03 | 65280001 | 65380000 | 0.0013 | 0.0011 | 0.0013 |
| F03 | 65300001 | 65400000 | 0.0011 | 0.0009 | 0.0011 |
| F03 | 65320001 | 65420000 | 0.0006 | 0.0005 | 0.0006 |
| F03 | 65340001 | 65440000 | 0.0004 | 0.0003 | 0.0004 |
| F03 | 65360001 | 65460000 | 0.0005 | 0.0005 | 0.0005 |
| F03 | 65380001 | 65480000 | 0.0006 | 0.0005 | 0.0006 |
| F03 | 65400001 | 65500000 | 0.0009 | 0.0008 | 0.0008 |
| F03 | 65420001 | 65520000 | 0.0015 | 0.0014 | 0.0015 |
| F03 | 65440001 | 65540000 | 0.0014 | 0.0013 | 0.0014 |
| F03 | 65460001 | 65560000 | 0.0013 | 0.0012 | 0.0012 |
| F03 | 65480001 | 65580000 | 0.0018 | 0.0015 | 0.0016 |
| F03 | 65500001 | 65600000 | 0.0018 | 0.0014 | 0.0015 |
| F03 | 65520001 | 65620000 | 0.0014 | 0.0009 | 0.0011 |
| F03 | 65540001 | 65640000 | 0.0016 | 0.0011 | 0.0012 |
| F03 | 65560001 | 65660000 | 0.0024 | 0.0016 | 0.0018 |
| F03 | 65580001 | 65680000 | 0.0023 | 0.0017 | 0.0017 |
| F03 | 65600001 | 65700000 | 0.0023 | 0.0017 | 0.0017 |
| F03 | 65620001 | 65720000 | 0.0025 | 0.0019 | 0.0019 |
| F03 | 65640001 | 65740000 | 0.0029 | 0.0023 | 0.0023 |
| F03 | 65660001 | 65760000 | 0.0023 | 0.0019 | 0.0018 |
| F03 | 65680001 | 65780000 | 0.0020 | 0.0016 | 0.0017 |
| F03 | 65700001 | 65800000 | 0.0021 | 0.0017 | 0.0018 |
| F03 | 65720001 | 65820000 | 0.0021 | 0.0015 | 0.0016 |
| F03 | 65740001 | 65840000 | 0.0022 | 0.0015 | 0.0018 |
| F03 | 65760001 | 65860000 | 0.0023 | 0.0017 | 0.0020 |
| F03 | 65780001 | 65880000 | 0.0020 | 0.0015 | 0.0018 |
| F03 | 65800001 | 65900000 | 0.0021 | 0.0015 | 0.0018 |
| F03 | 65820001 | 65920000 | 0.0019 | 0.0013 | 0.0016 |
| F03 | 65840001 | 65940000 | 0.0014 | 0.0008 | 0.0011 |
| F03 | 65860001 | 65960000 | 0.0015 | 0.0009 | 0.0012 |
| F03 | 65880001 | 65980000 | 0.0017 | 0.0011 | 0.0015 |
| F03 | 65900001 | 66000000 | 0.0019 | 0.0013 | 0.0017 |
| F03 | 65920001 | 66020000 | 0.0022 | 0.0018 | 0.0021 |
| F03 | 65940001 | 66040000 | 0.0032 | 0.0026 | 0.0029 |
| F03 | 65960001 | 66060000 | 0.0038 | 0.0031 | 0.0035 |
| F03 | 65980001 | 66080000 | 0.0043 | 0.0035 | 0.0037 |
| F03 | 66000001 | 66100000 | 0.0042 | 0.0034 | 0.0035 |
| F03 | 66020001 | 66120000 | 0.0036 | 0.0029 | 0.0030 |
| F03 | 66040001 | 66140000 | 0.0029 | 0.0024 | 0.0024 |
| F03 | 66060001 | 66160000 | 0.0021 | 0.0015 | 0.0016 |
| F03 | 66080001 | 66180000 | 0.0020 | 0.0016 | 0.0018 |
| F03 | 66100001 | 66200000 | 0.0020 | 0.0017 | 0.0019 |
| F03 | 66120001 | 66220000 | 0.0022 | 0.0019 | 0.0022 |
| F03 | 66140001 | 66240000 | 0.0024 | 0.0021 | 0.0024 |
| F03 | 66160001 | 66260000 | 0.0026 | 0.0022 | 0.0026 |
| F03 | 66180001 | 66280000 | 0.0022 | 0.0019 | 0.0021 |
| F03 | 66200001 | 66300000 | 0.0025 | 0.0021 | 0.0024 |
| F03 | 66220001 | 66320000 | 0.0035 | 0.0030 | 0.0031 |
| F03 | 66240001 | 66340000 | 0.0032 | 0.0026 | 0.0026 |
| F03 | 66260001 | 66360000 | 0.0029 | 0.0024 | 0.0024 |
| F03 | 66280001 | 66380000 | 0.0036 | 0.0029 | 0.0031 |
| F03 | 66300001 | 66400000 | 0.0033 | 0.0025 | 0.0027 |
| F03 | 66320001 | 66420000 | 0.0026 | 0.0019 | 0.0022 |
| F03 | 66340001 | 66440000 | 0.0025 | 0.0019 | 0.0022 |
| F03 | 66360001 | 66460000 | 0.0028 | 0.0022 | 0.0026 |
| F03 | 66380001 | 66480000 | 0.0026 | 0.0021 | 0.0024 |
| F03 | 66400001 | 66500000 | 0.0026 | 0.0023 | 0.0025 |

| Chr | Start | End | both | A1 | A2 |
|---|---|---|---|---|---|
| F03 | 66420001 | 66520000 | 0.0029 | 0.0025 | 0.0028 |
| F03 | 66440001 | 66540000 | 0.0031 | 0.0027 | 0.0030 |
| F03 | 66460001 | 66560000 | 0.0027 | 0.0025 | 0.0027 |
| F03 | 66480001 | 66580000 | 0.0020 | 0.0018 | 0.0019 |
| F03 | 66500001 | 66600000 | 0.0014 | 0.0013 | 0.0014 |
| F03 | 66520001 | 66620000 | 0.0005 | 0.0005 | 0.0005 |
| F03 | 66540001 | 66640000 | 0.0004 | 0.0004 | 0.0004 |
| F03 | 66560001 | 66660000 | 0.0011 | 0.0010 | 0.0011 |
| F03 | 66580001 | 66680000 | 0.0013 | 0.0012 | 0.0013 |
| F03 | 66600001 | 66700000 | 0.0017 | 0.0015 | 0.0017 |
| F03 | 66620001 | 66720000 | 0.0021 | 0.0018 | 0.0021 |
| F03 | 66640001 | 66740000 | 0.0024 | 0.0021 | 0.0024 |
| F03 | 66660001 | 66760000 | 0.0015 | 0.0013 | 0.0015 |
| F03 | 66680001 | 66780000 | 0.0031 | 0.0027 | 0.0031 |
| F03 | 66700001 | 66800000 | 0.0036 | 0.0031 | 0.0036 |
| F03 | 66720001 | 66820000 | 0.0039 | 0.0034 | 0.0038 |
| F03 | 66740001 | 66840000 | 0.0037 | 0.0031 | 0.0035 |
| F03 | 66760001 | 66860000 | 0.0041 | 0.0034 | 0.0039 |
| F03 | 66780001 | 66880000 | 0.0027 | 0.0021 | 0.0025 |
| F03 | 66800001 | 66900000 | 0.0025 | 0.0020 | 0.0022 |
| F03 | 66820001 | 66920000 | 0.0022 | 0.0017 | 0.0019 |
| F03 | 66840001 | 66940000 | 0.0019 | 0.0015 | 0.0017 |
| F03 | 66860001 | 66960000 | 0.0013 | 0.0011 | 0.0012 |
| F03 | 66880001 | 66980000 | 0.0018 | 0.0016 | 0.0018 |
| F03 | 66900001 | 67000000 | 0.0015 | 0.0014 | 0.0015 |
| F03 | 66920001 | 67020000 | 0.0014 | 0.0013 | 0.0014 |
| F03 | 66940001 | 67040000 | 0.0018 | 0.0017 | 0.0018 |
| F03 | 66960001 | 67060000 | 0.0022 | 0.0021 | 0.0021 |
| F03 | 66980001 | 67080000 | 0.0018 | 0.0017 | 0.0018 |
| F03 | 67000001 | 67100000 | 0.0019 | 0.0018 | 0.0019 |
| F03 | 67020001 | 67120000 | 0.0022 | 0.0019 | 0.0022 |
| F03 | 67040001 | 67140000 | 0.0022 | 0.0017 | 0.0019 |
| F03 | 67060001 | 67160000 | 0.0022 | 0.0014 | 0.0017 |
| F03 | 67080001 | 67180000 | 0.0023 | 0.0014 | 0.0015 |
| F03 | 67100001 | 67200000 | 0.0023 | 0.0013 | 0.0014 |
| F03 | 67120001 | 67220000 | 0.0023 | 0.0013 | 0.0014 |
| F03 | 67140001 | 67240000 | 0.0027 | 0.0019 | 0.0020 |
| F03 | 67160001 | 67260000 | 0.0031 | 0.0025 | 0.0026 |
| F03 | 67180001 | 67280000 | 0.0025 | 0.0021 | 0.0023 |
| F03 | 67200001 | 67300000 | 0.0026 | 0.0022 | 0.0024 |
| F03 | 67220001 | 67320000 | 0.0026 | 0.0024 | 0.0024 |
| F03 | 67240001 | 67340000 | 0.0019 | 0.0017 | 0.0017 |
| F03 | 67260001 | 67360000 | 0.0017 | 0.0016 | 0.0015 |
| F03 | 67280001 | 67380000 | 0.0026 | 0.0024 | 0.0022 |
| F03 | 67300001 | 67400000 | 0.0021 | 0.0019 | 0.0017 |
| F03 | 67320001 | 67420000 | 0.0018 | 0.0016 | 0.0015 |
| F03 | 67340001 | 67440000 | 0.0025 | 0.0022 | 0.0022 |
| F03 | 67360001 | 67460000 | 0.0032 | 0.0027 | 0.0029 |
| F03 | 67380001 | 67480000 | 0.0029 | 0.0023 | 0.0026 |
| F03 | 67400001 | 67500000 | 0.0036 | 0.0030 | 0.0032 |
| F03 | 67420001 | 67520000 | 0.0042 | 0.0035 | 0.0039 |
| F03 | 67440001 | 67540000 | 0.0042 | 0.0035 | 0.0039 |
| F03 | 67460001 | 67560000 | 0.0035 | 0.0030 | 0.0032 |
| F03 | 67480001 | 67580000 | 0.0036 | 0.0032 | 0.0035 |
| F03 | 67500001 | 67600000 | 0.0032 | 0.0028 | 0.0031 |
| F03 | 67520001 | 67620000 | 0.0034 | 0.0030 | 0.0033 |
| F03 | 67540001 | 67640000 | 0.0035 | 0.0030 | 0.0033 |
| F03 | 67560001 | 67660000 | 0.0032 | 0.0026 | 0.0028 |

| Chr | Start | End | both | A1 | A2 |
|---|---|---|---|---|---|
| F03 | 67580001 | 67680000 | 0.0029 | 0.0023 | 0.0025 |
| F03 | 67600001 | 67700000 | 0.0030 | 0.0023 | 0.0026 |
| F03 | 67620001 | 67720000 | 0.0024 | 0.0019 | 0.0020 |
| F03 | 67640001 | 67740000 | 0.0022 | 0.0017 | 0.0019 |
| F03 | 67660001 | 67760000 | 0.0028 | 0.0024 | 0.0025 |
| F03 | 67680001 | 67780000 | 0.0029 | 0.0024 | 0.0024 |
| F03 | 67700001 | 67800000 | 0.0035 | 0.0029 | 0.0028 |
| F03 | 67720001 | 67820000 | 0.0038 | 0.0031 | 0.0031 |
| F03 | 67740001 | 67840000 | 0.0040 | 0.0032 | 0.0033 |
| F03 | 67760001 | 67860000 | 0.0046 | 0.0036 | 0.0039 |
| F03 | 67780001 | 67880000 | 0.0047 | 0.0036 | 0.0041 |
| F03 | 67800001 | 67900000 | 0.0046 | 0.0037 | 0.0042 |
| F03 | 67820001 | 67920000 | 0.0042 | 0.0035 | 0.0039 |
| F03 | 67840001 | 67940000 | 0.0038 | 0.0030 | 0.0034 |
| F03 | 67860001 | 67960000 | 0.0035 | 0.0028 | 0.0031 |
| F03 | 67880001 | 67980000 | 0.0032 | 0.0027 | 0.0030 |
| F03 | 67900001 | 68000000 | 0.0028 | 0.0023 | 0.0027 |
| F03 | 67920001 | 68020000 | 0.0023 | 0.0017 | 0.0021 |
| F03 | 67940001 | 68040000 | 0.0025 | 0.0021 | 0.0025 |
| F03 | 67960001 | 68060000 | 0.0030 | 0.0023 | 0.0030 |
| F03 | 67980001 | 68080000 | 0.0030 | 0.0022 | 0.0029 |
| F03 | 68000001 | 68100000 | 0.0044 | 0.0034 | 0.0041 |
| F03 | 68020001 | 68120000 | 0.0046 | 0.0036 | 0.0043 |
| F03 | 68040001 | 68140000 | 0.0045 | 0.0034 | 0.0042 |
| F03 | 68060001 | 68160000 | 0.0031 | 0.0024 | 0.0028 |
| F03 | 68080001 | 68180000 | 0.0034 | 0.0027 | 0.0032 |
| F03 | 68100001 | 68200000 | 0.0020 | 0.0015 | 0.0018 |
| F03 | 68120001 | 68220000 | 0.0019 | 0.0014 | 0.0017 |
| F03 | 68140001 | 68240000 | 0.0017 | 0.0012 | 0.0013 |
| F03 | 68160001 | 68260000 | 0.0028 | 0.0022 | 0.0024 |
| F03 | 68180001 | 68280000 | 0.0025 | 0.0019 | 0.0021 |
| F03 | 68200001 | 68300000 | 0.0022 | 0.0018 | 0.0019 |
| F03 | 68220001 | 68320000 | 0.0023 | 0.0018 | 0.0020 |
| F03 | 68240001 | 68340000 | 0.0026 | 0.0022 | 0.0025 |
| F03 | 68260001 | 68360000 | 0.0019 | 0.0016 | 0.0019 |
| F03 | 68280001 | 68380000 | 0.0022 | 0.0019 | 0.0021 |
| F03 | 68300001 | 68400000 | 0.0022 | 0.0018 | 0.0021 |
| F03 | 68320001 | 68420000 | 0.0022 | 0.0018 | 0.0021 |
| F03 | 68340001 | 68440000 | 0.0022 | 0.0018 | 0.0021 |
| F03 | 68360001 | 68460000 | 0.0018 | 0.0015 | 0.0018 |
| F03 | 68380001 | 68480000 | 0.0017 | 0.0013 | 0.0016 |
| F03 | 68400001 | 68500000 | 0.0014 | 0.0012 | 0.0014 |
| F03 | 68420001 | 68520000 | 0.0014 | 0.0012 | 0.0014 |
| F03 | 68440001 | 68540000 | 0.0017 | 0.0014 | 0.0015 |
| F03 | 68460001 | 68560000 | 0.0032 | 0.0027 | 0.0029 |
| F03 | 68480001 | 68580000 | 0.0033 | 0.0029 | 0.0030 |
| F03 | 68500001 | 68600000 | 0.0043 | 0.0036 | 0.0039 |
| F03 | 68520001 | 68620000 | 0.0042 | 0.0034 | 0.0037 |
| F03 | 68540001 | 68640000 | 0.0036 | 0.0030 | 0.0032 |
| F03 | 68560001 | 68660000 | 0.0023 | 0.0016 | 0.0018 |
| F03 | 68580001 | 68680000 | 0.0024 | 0.0015 | 0.0019 |
| F03 | 68600001 | 68700000 | 0.0022 | 0.0013 | 0.0018 |
| F03 | 68620001 | 68720000 | 0.0027 | 0.0018 | 0.0023 |
| F03 | 68640001 | 68740000 | 0.0026 | 0.0017 | 0.0022 |
| F03 | 68660001 | 68760000 | 0.0023 | 0.0017 | 0.0021 |
| F03 | 68680001 | 68780000 | 0.0024 | 0.0020 | 0.0023 |
| F03 | 68700001 | 68800000 | 0.0018 | 0.0017 | 0.0016 |
| F03 | 68720001 | 68820000 | 0.0020 | 0.0018 | 0.0018 |

| Chr | Start | End | both | A1 | A2 |
|---|---|---|---|---|---|
| F03 | 68740001 | 68840000 | 0.0022 | 0.0020 | 0.0020 |
| F03 | 68760001 | 68860000 | 0.0022 | 0.0020 | 0.0020 |
| F03 | 68780001 | 68880000 | 0.0023 | 0.0019 | 0.0019 |
| F03 | 68800001 | 68900000 | 0.0025 | 0.0020 | 0.0020 |
| F03 | 68820001 | 68920000 | 0.0019 | 0.0014 | 0.0015 |
| F03 | 68840001 | 68940000 | 0.0018 | 0.0013 | 0.0014 |
| F03 | 68860001 | 68960000 | 0.0018 | 0.0012 | 0.0013 |
| F03 | 68880001 | 68980000 | 0.0011 | 0.0007 | 0.0008 |
| F03 | 68900001 | 69000000 | 0.0013 | 0.0010 | 0.0011 |
| F03 | 68920001 | 69020000 | 0.0014 | 0.0010 | 0.0012 |
| F03 | 68940001 | 69040000 | 0.0013 | 0.0010 | 0.0011 |
| F03 | 68960001 | 69060000 | 0.0014 | 0.0012 | 0.0013 |
| F03 | 68980001 | 69080000 | 0.0022 | 0.0020 | 0.0021 |
| F03 | 69000001 | 69100000 | 0.0019 | 0.0016 | 0.0017 |
| F03 | 69020001 | 69120000 | 0.0019 | 0.0017 | 0.0016 |
| F03 | 69040001 | 69140000 | 0.0020 | 0.0018 | 0.0017 |
| F03 | 69060001 | 69160000 | 0.0021 | 0.0018 | 0.0019 |
| F03 | 69080001 | 69180000 | 0.0013 | 0.0010 | 0.0011 |
| F03 | 69100001 | 69200000 | 0.0009 | 0.0008 | 0.0009 |
| F03 | 69120001 | 69220000 | 0.0009 | 0.0007 | 0.0009 |
| F03 | 69140001 | 69240000 | 0.0008 | 0.0006 | 0.0007 |
| F03 | 69160001 | 69260000 | 0.0012 | 0.0009 | 0.0011 |
| F03 | 69180001 | 69280000 | 0.0027 | 0.0021 | 0.0025 |
| F03 | 69200001 | 69300000 | 0.0035 | 0.0025 | 0.0030 |
| F03 | 69220001 | 69320000 | 0.0038 | 0.0027 | 0.0033 |
| F03 | 69240001 | 69340000 | 0.0038 | 0.0027 | 0.0033 |
| F03 | 69260001 | 69360000 | 0.0034 | 0.0023 | 0.0028 |
| F03 | 69280001 | 69380000 | 0.0019 | 0.0012 | 0.0014 |
| F03 | 69300001 | 69400000 | 0.0017 | 0.0011 | 0.0014 |
| F03 | 69320001 | 69420000 | 0.0017 | 0.0013 | 0.0014 |
| F03 | 69340001 | 69440000 | 0.0018 | 0.0014 | 0.0014 |
| F03 | 69360001 | 69460000 | 0.0016 | 0.0013 | 0.0013 |
| F03 | 69380001 | 69480000 | 0.0016 | 0.0013 | 0.0014 |
| F03 | 69400001 | 69500000 | 0.0014 | 0.0011 | 0.0011 |
| F03 | 69420001 | 69520000 | 0.0012 | 0.0009 | 0.0010 |
| F03 | 69440001 | 69540000 | 0.0014 | 0.0012 | 0.0013 |
| F03 | 69460001 | 69560000 | 0.0012 | 0.0010 | 0.0011 |
| F03 | 69480001 | 69580000 | 0.0011 | 0.0008 | 0.0010 |
| F03 | 69500001 | 69600000 | 0.0012 | 0.0008 | 0.0012 |
| F03 | 69520001 | 69620000 | 0.0009 | 0.0005 | 0.0009 |
| F03 | 69540001 | 69640000 | 0.0012 | 0.0007 | 0.0012 |
| F03 | 69560001 | 69660000 | 0.0015 | 0.0010 | 0.0015 |
| F03 | 69580001 | 69680000 | 0.0017 | 0.0011 | 0.0017 |
| F03 | 69600001 | 69700000 | 0.0035 | 0.0027 | 0.0035 |
| F03 | 69620001 | 69720000 | 0.0042 | 0.0034 | 0.0041 |
| F03 | 69640001 | 69740000 | 0.0038 | 0.0030 | 0.0037 |
| F03 | 69660001 | 69760000 | 0.0042 | 0.0033 | 0.0041 |
| F03 | 69680001 | 69780000 | 0.0041 | 0.0033 | 0.0040 |
| F03 | 69700001 | 69800000 | 0.0022 | 0.0019 | 0.0022 |
| F03 | 69720001 | 69820000 | 0.0018 | 0.0015 | 0.0018 |
| F03 | 69740001 | 69840000 | 0.0028 | 0.0024 | 0.0027 |
| F03 | 69760001 | 69860000 | 0.0025 | 0.0022 | 0.0024 |
| F03 | 69780001 | 69880000 | 0.0034 | 0.0031 | 0.0030 |
| F03 | 69800001 | 69900000 | 0.0031 | 0.0029 | 0.0028 |
| F03 | 69820001 | 69920000 | 0.0030 | 0.0029 | 0.0027 |
| F03 | 69840001 | 69940000 | 0.0021 | 0.0020 | 0.0018 |
| F03 | 69860001 | 69960000 | 0.0016 | 0.0016 | 0.0014 |
| F03 | 69880001 | 69980000 | 0.0007 | 0.0006 | 0.0006 |

| Chr | Start | End | both | A1 | A2 |
|---|---|---|---|---|---|
| F03 | 69900001 | 70000000 | 0.0008 | 0.0008 | 0.0008 |
| F03 | 69920001 | 70020000 | 0.0009 | 0.0008 | 0.0008 |
| F03 | 69940001 | 70040000 | 0.0007 | 0.0007 | 0.0007 |
| F03 | 69960001 | 70060000 | 0.0008 | 0.0007 | 0.0007 |
| F03 | 69980001 | 70080000 | 0.0007 | 0.0007 | 0.0007 |
| F03 | 70000001 | 70100000 | 0.0005 | 0.0005 | 0.0005 |
| F03 | 70020001 | 70120000 | 0.0002 | 0.0002 | 0.0002 |
| F03 | 70040001 | 70140000 | 0.0004 | 0.0003 | 0.0003 |
| F03 | 70060001 | 70160000 | 0.0007 | 0.0006 | 0.0006 |
| F03 | 70080001 | 70180000 | 0.0013 | 0.0011 | 0.0013 |
| F03 | 70100001 | 70200000 | 0.0017 | 0.0015 | 0.0016 |
| F03 | 70120001 | 70220000 | 0.0021 | 0.0019 | 0.0020 |
| F03 | 70140001 | 70240000 | 0.0025 | 0.0022 | 0.0024 |
| F03 | 70160001 | 70260000 | 0.0021 | 0.0019 | 0.0021 |
| F03 | 70180001 | 70280000 | 0.0018 | 0.0017 | 0.0018 |
| F03 | 70200001 | 70300000 | 0.0020 | 0.0017 | 0.0020 |
| F03 | 70220001 | 70320000 | 0.0021 | 0.0017 | 0.0020 |
| F03 | 70240001 | 70340000 | 0.0017 | 0.0014 | 0.0017 |
| F03 | 70260001 | 70360000 | 0.0019 | 0.0016 | 0.0018 |
| F03 | 70280001 | 70380000 | 0.0018 | 0.0014 | 0.0016 |
| F03 | 70300001 | 70400000 | 0.0013 | 0.0010 | 0.0011 |
| F03 | 70320001 | 70420000 | 0.0017 | 0.0014 | 0.0013 |
| F03 | 70340001 | 70440000 | 0.0017 | 0.0014 | 0.0013 |
| F03 | 70360001 | 70460000 | 0.0022 | 0.0019 | 0.0018 |
| F03 | 70380001 | 70480000 | 0.0024 | 0.0021 | 0.0020 |
| F03 | 70400001 | 70500000 | 0.0036 | 0.0031 | 0.0030 |
| F03 | 70420001 | 70520000 | 0.0034 | 0.0028 | 0.0029 |
| F03 | 70440001 | 70540000 | 0.0041 | 0.0033 | 0.0037 |
| F03 | 70460001 | 70560000 | 0.0038 | 0.0031 | 0.0034 |
| F03 | 70480001 | 70580000 | 0.0042 | 0.0033 | 0.0035 |
| F03 | 70500001 | 70600000 | 0.0036 | 0.0029 | 0.0031 |
| F03 | 70520001 | 70620000 | 0.0033 | 0.0028 | 0.0030 |
| F03 | 70540001 | 70640000 | 0.0025 | 0.0021 | 0.0021 |
| F03 | 70560001 | 70660000 | 0.0026 | 0.0021 | 0.0021 |
| F03 | 70580001 | 70680000 | 0.0032 | 0.0028 | 0.0030 |
| F03 | 70600001 | 70700000 | 0.0033 | 0.0028 | 0.0031 |
| F03 | 70620001 | 70720000 | 0.0032 | 0.0028 | 0.0031 |
| F03 | 70640001 | 70740000 | 0.0036 | 0.0031 | 0.0034 |
| F03 | 70660001 | 70760000 | 0.0031 | 0.0027 | 0.0030 |
| F03 | 70680001 | 70780000 | 0.0020 | 0.0016 | 0.0019 |
| F03 | 70700001 | 70800000 | 0.0022 | 0.0019 | 0.0020 |
| F03 | 70720001 | 70820000 | 0.0042 | 0.0038 | 0.0039 |
| F03 | 70740001 | 70840000 | 0.0049 | 0.0042 | 0.0044 |
| F03 | 70760001 | 70860000 | 0.0056 | 0.0047 | 0.0052 |
| F03 | 70780001 | 70880000 | 0.0064 | 0.0053 | 0.0058 |
| F03 | 70800001 | 70900000 | 0.0060 | 0.0048 | 0.0054 |
| F03 | 70820001 | 70920000 | 0.0049 | 0.0038 | 0.0044 |
| F03 | 70840001 | 70940000 | 0.0039 | 0.0031 | 0.0036 |
| F03 | 70860001 | 70960000 | 0.0034 | 0.0027 | 0.0031 |
| F03 | 70880001 | 70980000 | 0.0026 | 0.0021 | 0.0025 |
| F03 | 70900001 | 71000000 | 0.0021 | 0.0016 | 0.0020 |
| F03 | 70920001 | 71020000 | 0.0011 | 0.0007 | 0.0011 |
| F03 | 70940001 | 71040000 | 0.0022 | 0.0017 | 0.0021 |
| F03 | 70960001 | 71060000 | 0.0020 | 0.0016 | 0.0018 |
| F03 | 70980001 | 71080000 | 0.0025 | 0.0020 | 0.0023 |
| F03 | 71000001 | 71100000 | 0.0027 | 0.0022 | 0.0024 |
| F03 | 71020001 | 71120000 | 0.0029 | 0.0025 | 0.0026 |
| F03 | 71040001 | 71140000 | 0.0025 | 0.0021 | 0.0023 |

| Chr | Start | End | both | A1 | A2 |
|---|---|---|---|---|---|
| F03 | 71060001 | 71160000 | 0.0024 | 0.0021 | 0.0023 |
| F03 | 71080001 | 71180000 | 0.0021 | 0.0015 | 0.0017 |
| F03 | 71100001 | 71200000 | 0.0020 | 0.0013 | 0.0015 |
| F03 | 71120001 | 71220000 | 0.0016 | 0.0009 | 0.0011 |
| F03 | 71140001 | 71240000 | 0.0009 | 0.0003 | 0.0003 |
| F03 | 71160001 | 71260000 | 0.0010 | 0.0004 | 0.0004 |
| F03 | 71180001 | 71280000 | 0.0010 | 0.0006 | 0.0007 |
| F03 | 71200001 | 71300000 | 0.0014 | 0.0008 | 0.0009 |
| F03 | 71220001 | 71320000 | 0.0014 | 0.0009 | 0.0010 |
| F03 | 71240001 | 71340000 | 0.0016 | 0.0011 | 0.0012 |
| F03 | 71260001 | 71360000 | 0.0013 | 0.0009 | 0.0010 |
| F03 | 71280001 | 71380000 | 0.0013 | 0.0010 | 0.0010 |
| F03 | 71300001 | 71400000 | 0.0015 | 0.0014 | 0.0015 |
| F03 | 71320001 | 71420000 | 0.0017 | 0.0015 | 0.0016 |
| F03 | 71340001 | 71440000 | 0.0016 | 0.0015 | 0.0016 |
| F03 | 71360001 | 71460000 | 0.0018 | 0.0016 | 0.0017 |
| F03 | 71380001 | 71480000 | 0.0015 | 0.0014 | 0.0015 |
| F03 | 71400001 | 71500000 | 0.0010 | 0.0009 | 0.0010 |
| F03 | 71420001 | 71520000 | 0.0016 | 0.0015 | 0.0016 |
| F03 | 71440001 | 71540000 | 0.0014 | 0.0013 | 0.0014 |
| F03 | 71460001 | 71560000 | 0.0019 | 0.0017 | 0.0019 |
| F03 | 71480001 | 71580000 | 0.0019 | 0.0017 | 0.0019 |
| F03 | 71500001 | 71600000 | 0.0020 | 0.0016 | 0.0020 |
| F03 | 71520001 | 71620000 | 0.0019 | 0.0014 | 0.0019 |
| F03 | 71540001 | 71640000 | 0.0022 | 0.0017 | 0.0022 |
| F03 | 71560001 | 71660000 | 0.0021 | 0.0016 | 0.0020 |
| F03 | 71580001 | 71680000 | 0.0027 | 0.0021 | 0.0025 |
| F03 | 71600001 | 71700000 | 0.0026 | 0.0021 | 0.0024 |
| F03 | 71620001 | 71720000 | 0.0027 | 0.0022 | 0.0024 |
| F03 | 71640001 | 71740000 | 0.0023 | 0.0018 | 0.0021 |
| F03 | 71660001 | 71760000 | 0.0028 | 0.0025 | 0.0026 |
| F03 | 71680001 | 71780000 | 0.0029 | 0.0026 | 0.0027 |
| F03 | 71700001 | 71800000 | 0.0036 | 0.0032 | 0.0035 |
| F03 | 71720001 | 71820000 | 0.0033 | 0.0027 | 0.0030 |
| F03 | 71740001 | 71840000 | 0.0040 | 0.0033 | 0.0037 |
| F03 | 71760001 | 71860000 | 0.0032 | 0.0024 | 0.0029 |
| F03 | 71780001 | 71880000 | 0.0028 | 0.0020 | 0.0025 |
| F03 | 71800001 | 71900000 | 0.0024 | 0.0017 | 0.0021 |
| F03 | 71820001 | 71920000 | 0.0020 | 0.0015 | 0.0019 |
| F03 | 71840001 | 71940000 | 0.0015 | 0.0012 | 0.0014 |
| F03 | 71860001 | 71960000 | 0.0015 | 0.0012 | 0.0015 |
| F03 | 71880001 | 71980000 | 0.0012 | 0.0010 | 0.0012 |
| F03 | 71900001 | 72000000 | 0.0011 | 0.0010 | 0.0010 |
| F03 | 71920001 | 72020000 | 0.0016 | 0.0013 | 0.0014 |
| F03 | 71940001 | 72040000 | 0.0023 | 0.0019 | 0.0020 |
| F03 | 71960001 | 72060000 | 0.0022 | 0.0019 | 0.0020 |
| F03 | 71980001 | 72080000 | 0.0025 | 0.0021 | 0.0022 |
| F03 | 72000001 | 72100000 | 0.0028 | 0.0020 | 0.0023 |
| F03 | 72020001 | 72120000 | 0.0028 | 0.0019 | 0.0022 |
| F03 | 72040001 | 72140000 | 0.0025 | 0.0016 | 0.0018 |
| F03 | 72060001 | 72160000 | 0.0025 | 0.0016 | 0.0017 |
| F03 | 72080001 | 72180000 | 0.0025 | 0.0017 | 0.0019 |
| F03 | 72100001 | 72200000 | 0.0021 | 0.0017 | 0.0018 |
| F03 | 72120001 | 72220000 | 0.0020 | 0.0017 | 0.0018 |
| F03 | 72140001 | 72240000 | 0.0015 | 0.0014 | 0.0014 |
| F03 | 72160001 | 72260000 | 0.0022 | 0.0018 | 0.0021 |
| F03 | 72180001 | 72280000 | 0.0021 | 0.0018 | 0.0020 |
| F03 | 72200001 | 72300000 | 0.0025 | 0.0021 | 0.0024 |

| Chr | Start | End | both | A1 | A2 |
|---|---|---|---|---|---|
| F03 | 72220001 | 72320000 | 0.0039 | 0.0033 | 0.0036 |
| F03 | 72240001 | 72340000 | 0.0043 | 0.0034 | 0.0037 |
| F03 | 72260001 | 72360000 | 0.0040 | 0.0033 | 0.0034 |
| F03 | 72280001 | 72380000 | 0.0043 | 0.0034 | 0.0034 |
| F03 | 72300001 | 72400000 | 0.0041 | 0.0030 | 0.0029 |
| F03 | 72320001 | 72420000 | 0.0029 | 0.0017 | 0.0015 |
| F03 | 72340001 | 72440000 | 0.0034 | 0.0025 | 0.0023 |
| F03 | 72360001 | 72460000 | 0.0038 | 0.0027 | 0.0026 |
| F03 | 72380001 | 72480000 | 0.0042 | 0.0030 | 0.0031 |
| F03 | 72400001 | 72500000 | 0.0042 | 0.0031 | 0.0033 |
| F03 | 72420001 | 72520000 | 0.0041 | 0.0031 | 0.0035 |
| F03 | 72440001 | 72540000 | 0.0035 | 0.0024 | 0.0029 |
| F03 | 72460001 | 72560000 | 0.0037 | 0.0029 | 0.0032 |
| F03 | 72480001 | 72580000 | 0.0032 | 0.0026 | 0.0029 |
| F03 | 72500001 | 72600000 | 0.0033 | 0.0028 | 0.0029 |
| F03 | 72520001 | 72620000 | 0.0044 | 0.0039 | 0.0039 |
| F03 | 72540001 | 72640000 | 0.0049 | 0.0040 | 0.0042 |
| F03 | 72560001 | 72660000 | 0.0043 | 0.0034 | 0.0037 |
| F03 | 72580001 | 72680000 | 0.0041 | 0.0031 | 0.0035 |
| F03 | 72600001 | 72700000 | 0.0043 | 0.0033 | 0.0038 |
| F03 | 72620001 | 72720000 | 0.0038 | 0.0029 | 0.0033 |
| F03 | 72640001 | 72740000 | 0.0037 | 0.0030 | 0.0033 |
| F03 | 72660001 | 72760000 | 0.0040 | 0.0031 | 0.0035 |
| F03 | 72680001 | 72780000 | 0.0045 | 0.0035 | 0.0039 |
| F03 | 72700001 | 72800000 | 0.0042 | 0.0032 | 0.0034 |
| F03 | 72720001 | 72820000 | 0.0037 | 0.0027 | 0.0028 |
| F03 | 72740001 | 72840000 | 0.0033 | 0.0024 | 0.0024 |
| F03 | 72760001 | 72860000 | 0.0027 | 0.0020 | 0.0020 |
| F03 | 72780001 | 72880000 | 0.0024 | 0.0018 | 0.0018 |
| F03 | 72800001 | 72900000 | 0.0030 | 0.0024 | 0.0024 |
| F03 | 72820001 | 72920000 | 0.0039 | 0.0032 | 0.0035 |
| F03 | 72840001 | 72940000 | 0.0041 | 0.0033 | 0.0035 |
| F03 | 72860001 | 72960000 | 0.0039 | 0.0031 | 0.0033 |
| F03 | 72880001 | 72980000 | 0.0038 | 0.0031 | 0.0033 |
| F03 | 72900001 | 73000000 | 0.0027 | 0.0022 | 0.0024 |
| F03 | 72920001 | 73020000 | 0.0015 | 0.0012 | 0.0013 |
| F03 | 72940001 | 73040000 | 0.0013 | 0.0010 | 0.0013 |
| F03 | 72960001 | 73060000 | 0.0021 | 0.0018 | 0.0021 |
| F03 | 72980001 | 73080000 | 0.0022 | 0.0018 | 0.0021 |
| F03 | 73000001 | 73100000 | 0.0026 | 0.0021 | 0.0023 |
| F03 | 73020001 | 73120000 | 0.0028 | 0.0022 | 0.0023 |
| F03 | 73040001 | 73140000 | 0.0031 | 0.0023 | 0.0024 |
| F03 | 73060001 | 73160000 | 0.0024 | 0.0015 | 0.0015 |
| F03 | 73080001 | 73180000 | 0.0023 | 0.0013 | 0.0013 |
| F03 | 73100001 | 73200000 | 0.0021 | 0.0013 | 0.0013 |
| F03 | 73120001 | 73220000 | 0.0018 | 0.0010 | 0.0011 |
| F03 | 73140001 | 73240000 | 0.0018 | 0.0013 | 0.0013 |
| F03 | 73160001 | 73260000 | 0.0016 | 0.0013 | 0.0013 |
| F03 | 73180001 | 73280000 | 0.0013 | 0.0011 | 0.0012 |
| F03 | 73200001 | 73300000 | 0.0011 | 0.0009 | 0.0009 |
| F03 | 73220001 | 73320000 | 0.0018 | 0.0017 | 0.0018 |
| F03 | 73240001 | 73340000 | 0.0015 | 0.0013 | 0.0015 |
| F03 | 73260001 | 73360000 | 0.0016 | 0.0014 | 0.0016 |
| F03 | 73280001 | 73380000 | 0.0021 | 0.0018 | 0.0021 |
| F03 | 73300001 | 73400000 | 0.0021 | 0.0019 | 0.0021 |
| F03 | 73320001 | 73420000 | 0.0023 | 0.0020 | 0.0021 |
| F03 | 73340001 | 73440000 | 0.0026 | 0.0022 | 0.0024 |
| F03 | 73360001 | 73460000 | 0.0026 | 0.0022 | 0.0023 |

| Chr | Start | End | both | A1 | A2 |
|---|---|---|---|---|---|
| F03 | 73380001 | 73480000 | 0.0025 | 0.0019 | 0.0019 |
| F03 | 73400001 | 73500000 | 0.0033 | 0.0024 | 0.0024 |
| F03 | 73420001 | 73520000 | 0.0030 | 0.0020 | 0.0020 |
| F03 | 73440001 | 73540000 | 0.0026 | 0.0016 | 0.0014 |
| F03 | 73460001 | 73560000 | 0.0031 | 0.0020 | 0.0020 |
| F03 | 73480001 | 73580000 | 0.0028 | 0.0019 | 0.0019 |
| F03 | 73500001 | 73600000 | 0.0028 | 0.0018 | 0.0020 |
| F03 | 73520001 | 73620000 | 0.0035 | 0.0027 | 0.0028 |
| F03 | 73540001 | 73640000 | 0.0048 | 0.0035 | 0.0040 |
| F03 | 73560001 | 73660000 | 0.0048 | 0.0034 | 0.0039 |
| F03 | 73580001 | 73680000 | 0.0050 | 0.0036 | 0.0041 |
| F03 | 73600001 | 73700000 | 0.0050 | 0.0035 | 0.0040 |
| F03 | 73620001 | 73720000 | 0.0038 | 0.0024 | 0.0029 |
| F03 | 73640001 | 73740000 | 0.0024 | 0.0015 | 0.0018 |
| F03 | 73660001 | 73760000 | 0.0027 | 0.0019 | 0.0021 |
| F03 | 73680001 | 73780000 | 0.0035 | 0.0026 | 0.0028 |
| F03 | 73700001 | 73800000 | 0.0031 | 0.0024 | 0.0026 |
| F03 | 73720001 | 73820000 | 0.0036 | 0.0026 | 0.0028 |
| F03 | 73740001 | 73840000 | 0.0038 | 0.0026 | 0.0028 |
| F03 | 73760001 | 73860000 | 0.0031 | 0.0019 | 0.0021 |
| F03 | 73780001 | 73880000 | 0.0024 | 0.0012 | 0.0015 |
| F03 | 73800001 | 73900000 | 0.0037 | 0.0026 | 0.0026 |
| F03 | 73820001 | 73920000 | 0.0033 | 0.0024 | 0.0025 |
| F03 | 73840001 | 73940000 | 0.0041 | 0.0032 | 0.0031 |
| F03 | 73860001 | 73960000 | 0.0046 | 0.0036 | 0.0036 |
| F03 | 73880001 | 73980000 | 0.0049 | 0.0036 | 0.0037 |
| F03 | 73900001 | 74000000 | 0.0043 | 0.0030 | 0.0031 |
| F03 | 73920001 | 74020000 | 0.0043 | 0.0030 | 0.0032 |
| F03 | 73940001 | 74040000 | 0.0041 | 0.0027 | 0.0031 |
| F03 | 73960001 | 74060000 | 0.0038 | 0.0024 | 0.0028 |
| F03 | 73980001 | 74080000 | 0.0037 | 0.0027 | 0.0030 |
| F03 | 74000001 | 74100000 | 0.0027 | 0.0019 | 0.0022 |
| F03 | 74020001 | 74120000 | 0.0030 | 0.0019 | 0.0020 |
| F03 | 74040001 | 74140000 | 0.0024 | 0.0014 | 0.0014 |
| F03 | 74060001 | 74160000 | 0.0030 | 0.0018 | 0.0019 |
| F03 | 74080001 | 74180000 | 0.0038 | 0.0027 | 0.0027 |
| F03 | 74100001 | 74200000 | 0.0044 | 0.0032 | 0.0033 |
| F03 | 74120001 | 74220000 | 0.0047 | 0.0039 | 0.0038 |
| F03 | 74140001 | 74240000 | 0.0048 | 0.0038 | 0.0038 |
| F03 | 74160001 | 74260000 | 0.0046 | 0.0038 | 0.0037 |
| F03 | 74180001 | 74280000 | 0.0035 | 0.0027 | 0.0026 |
| F03 | 74200001 | 74300000 | 0.0036 | 0.0027 | 0.0026 |
| F03 | 74220001 | 74320000 | 0.0032 | 0.0021 | 0.0020 |
| F03 | 74240001 | 74340000 | 0.0038 | 0.0027 | 0.0028 |
| F03 | 74260001 | 74360000 | 0.0045 | 0.0034 | 0.0036 |
| F03 | 74280001 | 74380000 | 0.0042 | 0.0030 | 0.0032 |
| F03 | 74300001 | 74400000 | 0.0035 | 0.0022 | 0.0026 |
| F03 | 74320001 | 74420000 | 0.0033 | 0.0023 | 0.0026 |
| F03 | 74340001 | 74440000 | 0.0031 | 0.0021 | 0.0023 |
| F03 | 74360001 | 74460000 | 0.0025 | 0.0013 | 0.0015 |
| F03 | 74380001 | 74480000 | 0.0037 | 0.0025 | 0.0026 |
| F03 | 74400001 | 74500000 | 0.0036 | 0.0026 | 0.0027 |
| F03 | 74420001 | 74520000 | 0.0041 | 0.0030 | 0.0032 |
| F03 | 74440001 | 74540000 | 0.0039 | 0.0027 | 0.0028 |
| F03 | 74460001 | 74560000 | 0.0034 | 0.0024 | 0.0024 |
| F03 | 74480001 | 74580000 | 0.0029 | 0.0019 | 0.0021 |
| F03 | 74500001 | 74600000 | 0.0030 | 0.0020 | 0.0020 |
| F03 | 74520001 | 74620000 | 0.0021 | 0.0013 | 0.0012 |

| Chr | Start | End | both | A1 | A2 |
|---|---|---|---|---|---|
| F03 | 74540001 | 74640000 | 0.0029 | 0.0020 | 0.0021 |
| F03 | 74560001 | 74660000 | 0.0039 | 0.0028 | 0.0030 |
| F03 | 74580001 | 74680000 | 0.0035 | 0.0022 | 0.0024 |
| F03 | 74600001 | 74700000 | 0.0040 | 0.0026 | 0.0029 |
| F03 | 74620001 | 74720000 | 0.0045 | 0.0027 | 0.0031 |
| F03 | 74640001 | 74740000 | 0.0035 | 0.0019 | 0.0024 |
| F03 | 74660001 | 74760000 | 0.0026 | 0.0014 | 0.0018 |
| F03 | 74680001 | 74780000 | 0.0023 | 0.0015 | 0.0018 |
| F03 | 74700001 | 74800000 | 0.0016 | 0.0010 | 0.0012 |
| F03 | 74720001 | 74820000 | 0.0013 | 0.0011 | 0.0013 |
| F03 | 74740001 | 74840000 | 0.0013 | 0.0012 | 0.0013 |
| F03 | 74760001 | 74860000 | 0.0019 | 0.0017 | 0.0019 |
| F03 | 74780001 | 74880000 | 0.0018 | 0.0015 | 0.0018 |
| F03 | 74800001 | 74900000 | 0.0020 | 0.0018 | 0.0020 |
| F03 | 74820001 | 74920000 | 0.0021 | 0.0018 | 0.0020 |
| F03 | 74840001 | 74940000 | 0.0024 | 0.0019 | 0.0021 |
| F03 | 74860001 | 74960000 | 0.0021 | 0.0015 | 0.0018 |
| F03 | 74880001 | 74980000 | 0.0029 | 0.0023 | 0.0026 |
| F03 | 74900001 | 75000000 | 0.0032 | 0.0024 | 0.0028 |
| F03 | 74920001 | 75020000 | 0.0036 | 0.0027 | 0.0031 |
| F03 | 74940001 | 75040000 | 0.0045 | 0.0035 | 0.0040 |
| F03 | 74960001 | 75060000 | 0.0043 | 0.0032 | 0.0036 |
| F03 | 74980001 | 75080000 | 0.0040 | 0.0026 | 0.0030 |
| F03 | 75000001 | 75100000 | 0.0037 | 0.0024 | 0.0028 |
| F03 | 75020001 | 75120000 | 0.0034 | 0.0022 | 0.0025 |
| F03 | 75040001 | 75140000 | 0.0031 | 0.0018 | 0.0020 |
| F03 | 75060001 | 75160000 | 0.0032 | 0.0020 | 0.0022 |
| F03 | 75080001 | 75180000 | 0.0033 | 0.0021 | 0.0024 |
| F03 | 75100001 | 75200000 | 0.0037 | 0.0025 | 0.0028 |
| F03 | 75120001 | 75220000 | 0.0040 | 0.0029 | 0.0032 |
| F03 | 75140001 | 75240000 | 0.0034 | 0.0026 | 0.0028 |
| F03 | 75160001 | 75260000 | 0.0032 | 0.0025 | 0.0028 |
| F03 | 75180001 | 75280000 | 0.0030 | 0.0026 | 0.0027 |
| F03 | 75200001 | 75300000 | 0.0029 | 0.0025 | 0.0028 |
| F03 | 75220001 | 75320000 | 0.0028 | 0.0023 | 0.0026 |
| F03 | 75240001 | 75340000 | 0.0027 | 0.0022 | 0.0025 |
| F03 | 75260001 | 75360000 | 0.0028 | 0.0022 | 0.0026 |
| F03 | 75280001 | 75380000 | 0.0026 | 0.0022 | 0.0024 |
| F03 | 75300001 | 75400000 | 0.0033 | 0.0026 | 0.0027 |
| F03 | 75320001 | 75420000 | 0.0040 | 0.0033 | 0.0034 |
| F03 | 75340001 | 75440000 | 0.0044 | 0.0035 | 0.0037 |
| F03 | 75360001 | 75460000 | 0.0043 | 0.0034 | 0.0035 |
| F03 | 75380001 | 75480000 | 0.0045 | 0.0035 | 0.0036 |
| F03 | 75400001 | 75500000 | 0.0042 | 0.0034 | 0.0036 |
| F03 | 75420001 | 75520000 | 0.0037 | 0.0028 | 0.0030 |
| F03 | 75440001 | 75540000 | 0.0038 | 0.0031 | 0.0031 |
| F03 | 75460001 | 75560000 | 0.0042 | 0.0034 | 0.0035 |
| F03 | 75480001 | 75580000 | 0.0045 | 0.0037 | 0.0039 |
| F03 | 75500001 | 75600000 | 0.0040 | 0.0031 | 0.0031 |
| F03 | 75520001 | 75620000 | 0.0043 | 0.0033 | 0.0034 |
| F03 | 75540001 | 75640000 | 0.0039 | 0.0029 | 0.0029 |
| F03 | 75560001 | 75660000 | 0.0036 | 0.0027 | 0.0026 |
| F03 | 75580001 | 75680000 | 0.0035 | 0.0027 | 0.0026 |
| F03 | 75600001 | 75700000 | 0.0039 | 0.0031 | 0.0030 |
| F03 | 75620001 | 75720000 | 0.0037 | 0.0027 | 0.0027 |
| F03 | 75640001 | 75740000 | 0.0037 | 0.0025 | 0.0026 |
| F03 | 75660001 | 75760000 | 0.0037 | 0.0024 | 0.0025 |
| F03 | 75680001 | 75780000 | 0.0037 | 0.0022 | 0.0025 |

| Chr | Start | End | both | A1 | A2 |
|---|---|---|---|---|---|
| F03 | 75700001 | 75800000 | 0.0034 | 0.0019 | 0.0021 |
| F03 | 75720001 | 75820000 | 0.0043 | 0.0029 | 0.0030 |
| F03 | 75740001 | 75840000 | 0.0052 | 0.0040 | 0.0040 |
| F03 | 75760001 | 75860000 | 0.0050 | 0.0038 | 0.0038 |
| F03 | 75780001 | 75880000 | 0.0052 | 0.0039 | 0.0039 |
| F03 | 75800001 | 75900000 | 0.0052 | 0.0039 | 0.0040 |
| F03 | 75820001 | 75920000 | 0.0040 | 0.0028 | 0.0029 |
| F03 | 75840001 | 75940000 | 0.0037 | 0.0026 | 0.0029 |
| F03 | 75860001 | 75960000 | 0.0042 | 0.0032 | 0.0036 |
| F03 | 75880001 | 75980000 | 0.0040 | 0.0032 | 0.0036 |
| F03 | 75900001 | 76000000 | 0.0033 | 0.0029 | 0.0033 |
| F03 | 75920001 | 76020000 | 0.0031 | 0.0027 | 0.0030 |
| F03 | 75940001 | 76040000 | 0.0027 | 0.0023 | 0.0027 |
| F03 | 75960001 | 76060000 | 0.0022 | 0.0019 | 0.0022 |
| F03 | 75980001 | 76080000 | 0.0018 | 0.0017 | 0.0018 |
| F03 | 76000001 | 76100000 | 0.0018 | 0.0017 | 0.0018 |
| F03 | 76020001 | 76120000 | 0.0018 | 0.0017 | 0.0018 |
| F03 | 76040001 | 76140000 | 0.0010 | 0.0009 | 0.0010 |
| F03 | 76060001 | 76160000 | 0.0010 | 0.0009 | 0.0010 |
| F03 | 76080001 | 76180000 | 0.0017 | 0.0013 | 0.0016 |
| F03 | 76100001 | 76200000 | 0.0027 | 0.0022 | 0.0026 |
| F03 | 76120001 | 76220000 | 0.0040 | 0.0031 | 0.0036 |
| F03 | 76140001 | 76240000 | 0.0047 | 0.0038 | 0.0043 |
| F03 | 76160001 | 76260000 | 0.0045 | 0.0035 | 0.0039 |
| F03 | 76180001 | 76280000 | 0.0042 | 0.0033 | 0.0036 |
| F03 | 76200001 | 76300000 | 0.0039 | 0.0031 | 0.0033 |
| F03 | 76220001 | 76320000 | 0.0032 | 0.0026 | 0.0028 |
| F03 | 76240001 | 76340000 | 0.0030 | 0.0024 | 0.0025 |
| F03 | 76260001 | 76360000 | 0.0040 | 0.0032 | 0.0034 |
| F03 | 76280001 | 76380000 | 0.0041 | 0.0033 | 0.0034 |
| F03 | 76300001 | 76400000 | 0.0035 | 0.0026 | 0.0028 |
| F03 | 76320001 | 76420000 | 0.0032 | 0.0023 | 0.0024 |
| F03 | 76340001 | 76440000 | 0.0029 | 0.0020 | 0.0021 |
| F03 | 76360001 | 76460000 | 0.0020 | 0.0013 | 0.0014 |
| F03 | 76380001 | 76480000 | 0.0019 | 0.0009 | 0.0010 |
| F03 | 76400001 | 76500000 | 0.0022 | 0.0010 | 0.0011 |
| F03 | 76420001 | 76520000 | 0.0026 | 0.0011 | 0.0012 |
| F03 | 76440001 | 76540000 | 0.0035 | 0.0019 | 0.0020 |
| F03 | 76460001 | 76560000 | 0.0048 | 0.0029 | 0.0029 |
| F03 | 76480001 | 76580000 | 0.0059 | 0.0043 | 0.0042 |
| F03 | 76500001 | 76600000 | 0.0060 | 0.0046 | 0.0045 |
| F03 | 76520001 | 76620000 | 0.0058 | 0.0047 | 0.0047 |
| F03 | 76540001 | 76640000 | 0.0055 | 0.0046 | 0.0045 |
| F03 | 76560001 | 76660000 | 0.0048 | 0.0038 | 0.0038 |
| F03 | 76580001 | 76680000 | 0.0033 | 0.0024 | 0.0024 |
| F03 | 76600001 | 76700000 | 0.0040 | 0.0029 | 0.0030 |
| F03 | 76620001 | 76720000 | 0.0042 | 0.0030 | 0.0031 |
| F03 | 76640001 | 76740000 | 0.0045 | 0.0032 | 0.0032 |
| F03 | 76660001 | 76760000 | 0.0069 | 0.0055 | 0.0056 |
| F03 | 76680001 | 76780000 | 0.0069 | 0.0056 | 0.0057 |
| F03 | 76700001 | 76800000 | 0.0059 | 0.0048 | 0.0049 |
| F03 | 76720001 | 76820000 | 0.0059 | 0.0048 | 0.0049 |
| F03 | 76740001 | 76840000 | 0.0052 | 0.0040 | 0.0042 |
| F03 | 76760001 | 76860000 | 0.0028 | 0.0016 | 0.0017 |
| F03 | 76780001 | 76880000 | 0.0028 | 0.0017 | 0.0017 |
| F03 | 76800001 | 76900000 | 0.0027 | 0.0017 | 0.0017 |
| F03 | 76820001 | 76920000 | 0.0025 | 0.0015 | 0.0016 |
| F03 | 76840001 | 76940000 | 0.0026 | 0.0017 | 0.0019 |

| Chr | Start | End | both | A1 | A2 |
|---|---|---|---|---|---|
| F03 | 76860001 | 76960000 | 0.0021 | 0.0015 | 0.0018 |
| F03 | 76880001 | 76980000 | 0.0023 | 0.0016 | 0.0019 |
| F03 | 76900001 | 77000000 | 0.0021 | 0.0015 | 0.0018 |
| F03 | 76920001 | 77020000 | 0.0024 | 0.0017 | 0.0021 |
| F03 | 76940001 | 77040000 | 0.0031 | 0.0024 | 0.0028 |
| F03 | 76960001 | 77060000 | 0.0035 | 0.0028 | 0.0032 |
| F03 | 76980001 | 77080000 | 0.0035 | 0.0025 | 0.0030 |
| F03 | 77000001 | 77100000 | 0.0040 | 0.0028 | 0.0034 |
| F03 | 77020001 | 77120000 | 0.0038 | 0.0026 | 0.0030 |
| F03 | 77040001 | 77140000 | 0.0038 | 0.0030 | 0.0030 |
| F03 | 77060001 | 77160000 | 0.0043 | 0.0033 | 0.0034 |
| F03 | 77080001 | 77180000 | 0.0052 | 0.0040 | 0.0042 |
| F03 | 77100001 | 77200000 | 0.0054 | 0.0042 | 0.0042 |
| F03 | 77120001 | 77220000 | 0.0064 | 0.0053 | 0.0054 |
| F03 | 77140001 | 77240000 | 0.0061 | 0.0050 | 0.0051 |
| F03 | 77160001 | 77260000 | 0.0057 | 0.0045 | 0.0045 |
| F03 | 77180001 | 77280000 | 0.0047 | 0.0037 | 0.0037 |
| F03 | 77200001 | 77300000 | 0.0038 | 0.0031 | 0.0031 |
| F03 | 77220001 | 77320000 | 0.0032 | 0.0023 | 0.0025 |
| F03 | 77240001 | 77340000 | 0.0027 | 0.0016 | 0.0020 |
| F03 | 77260001 | 77360000 | 0.0027 | 0.0016 | 0.0021 |
| F03 | 77280001 | 77380000 | 0.0029 | 0.0018 | 0.0021 |
| F03 | 77300001 | 77400000 | 0.0031 | 0.0020 | 0.0023 |
| F03 | 77320001 | 77420000 | 0.0029 | 0.0021 | 0.0022 |
| F03 | 77340001 | 77440000 | 0.0039 | 0.0032 | 0.0032 |
| F03 | 77360001 | 77460000 | 0.0040 | 0.0031 | 0.0032 |
| F03 | 77380001 | 77480000 | 0.0045 | 0.0035 | 0.0036 |
| F03 | 77400001 | 77500000 | 0.0044 | 0.0033 | 0.0035 |
| F03 | 77420001 | 77520000 | 0.0043 | 0.0028 | 0.0032 |
| F03 | 77440001 | 77540000 | 0.0031 | 0.0015 | 0.0020 |
| F03 | 77460001 | 77560000 | 0.0035 | 0.0022 | 0.0026 |
| F03 | 77480001 | 77580000 | 0.0035 | 0.0022 | 0.0026 |
| F03 | 77500001 | 77600000 | 0.0041 | 0.0026 | 0.0031 |
| F03 | 77520001 | 77620000 | 0.0042 | 0.0027 | 0.0030 |
| F03 | 77540001 | 77640000 | 0.0050 | 0.0033 | 0.0037 |
| F03 | 77560001 | 77660000 | 0.0045 | 0.0029 | 0.0032 |
| F03 | 77580001 | 77680000 | 0.0040 | 0.0024 | 0.0026 |
| F03 | 77600001 | 77700000 | 0.0045 | 0.0030 | 0.0032 |
| F03 | 77620001 | 77720000 | 0.0042 | 0.0030 | 0.0031 |
| F03 | 77640001 | 77740000 | 0.0042 | 0.0035 | 0.0032 |
| F03 | 77660001 | 77760000 | 0.0040 | 0.0033 | 0.0031 |
| F03 | 77680001 | 77780000 | 0.0038 | 0.0031 | 0.0031 |
| F03 | 77700001 | 77800000 | 0.0032 | 0.0027 | 0.0027 |
| F03 | 77720001 | 77820000 | 0.0033 | 0.0028 | 0.0029 |
| F03 | 77740001 | 77840000 | 0.0024 | 0.0016 | 0.0019 |
| F03 | 77760001 | 77860000 | 0.0042 | 0.0034 | 0.0038 |
| F03 | 77780001 | 77880000 | 0.0051 | 0.0040 | 0.0044 |
| F03 | 77800001 | 77900000 | 0.0052 | 0.0038 | 0.0041 |
| F03 | 77820001 | 77920000 | 0.0046 | 0.0034 | 0.0037 |
| F03 | 77840001 | 77940000 | 0.0055 | 0.0043 | 0.0046 |
| F03 | 77860001 | 77960000 | 0.0045 | 0.0034 | 0.0037 |
| F03 | 77880001 | 77980000 | 0.0038 | 0.0029 | 0.0031 |
| F03 | 77900001 | 78000000 | 0.0038 | 0.0033 | 0.0034 |
| F03 | 77920001 | 78020000 | 0.0050 | 0.0042 | 0.0044 |
| F03 | 77940001 | 78040000 | 0.0044 | 0.0034 | 0.0036 |
| F03 | 77960001 | 78060000 | 0.0044 | 0.0033 | 0.0035 |
| F03 | 77980001 | 78080000 | 0.0043 | 0.0032 | 0.0035 |
| F03 | 78000001 | 78100000 | 0.0039 | 0.0024 | 0.0029 |

| Chr | Start | End | both | A1 | A2 |
|---|---|---|---|---|---|
| F03 | 78020001 | 78120000 | 0.0029 | 0.0016 | 0.0019 |
| F03 | 78040001 | 78140000 | 0.0026 | 0.0015 | 0.0017 |
| F03 | 78060001 | 78160000 | 0.0019 | 0.0007 | 0.0010 |
| F03 | 78080001 | 78180000 | 0.0018 | 0.0007 | 0.0008 |
| F03 | 78100001 | 78200000 | 0.0016 | 0.0007 | 0.0008 |
| F03 | 78120001 | 78220000 | 0.0027 | 0.0017 | 0.0020 |
| F03 | 78140001 | 78240000 | 0.0032 | 0.0022 | 0.0025 |
| F03 | 78160001 | 78260000 | 0.0033 | 0.0024 | 0.0028 |
| F03 | 78180001 | 78280000 | 0.0034 | 0.0026 | 0.0030 |
| F03 | 78200001 | 78300000 | 0.0033 | 0.0026 | 0.0030 |
| F03 | 78220001 | 78320000 | 0.0024 | 0.0020 | 0.0022 |
| F03 | 78240001 | 78340000 | 0.0020 | 0.0017 | 0.0019 |
| F03 | 78260001 | 78360000 | 0.0013 | 0.0011 | 0.0012 |
| F03 | 78280001 | 78380000 | 0.0014 | 0.0009 | 0.0011 |
| F03 | 78300001 | 78400000 | 0.0018 | 0.0014 | 0.0015 |
| F03 | 78320001 | 78420000 | 0.0020 | 0.0014 | 0.0014 |
| F03 | 78340001 | 78440000 | 0.0018 | 0.0012 | 0.0012 |
| F03 | 78360001 | 78460000 | 0.0021 | 0.0015 | 0.0015 |
| F03 | 78380001 | 78480000 | 0.0019 | 0.0015 | 0.0015 |
| F03 | 78400001 | 78500000 | 0.0013 | 0.0009 | 0.0009 |
| F03 | 78420001 | 78520000 | 0.0005 | 0.0005 | 0.0005 |
| F03 | 78440001 | 78540000 | 0.0011 | 0.0010 | 0.0010 |
| F03 | 78460001 | 78560000 | 0.0015 | 0.0015 | 0.0014 |
| F03 | 78480001 | 78580000 | 0.0014 | 0.0014 | 0.0014 |
| F03 | 78500001 | 78600000 | 0.0019 | 0.0017 | 0.0018 |
| F03 | 78520001 | 78620000 | 0.0024 | 0.0021 | 0.0022 |
| F03 | 78540001 | 78640000 | 0.0022 | 0.0019 | 0.0020 |
| F03 | 78560001 | 78660000 | 0.0024 | 0.0019 | 0.0019 |
| F03 | 78580001 | 78680000 | 0.0028 | 0.0022 | 0.0022 |
| F03 | 78600001 | 78700000 | 0.0034 | 0.0027 | 0.0027 |
| F03 | 78620001 | 78720000 | 0.0035 | 0.0028 | 0.0027 |
| F03 | 78640001 | 78740000 | 0.0040 | 0.0034 | 0.0033 |
| F03 | 78660001 | 78760000 | 0.0036 | 0.0031 | 0.0032 |
| F03 | 78680001 | 78780000 | 0.0034 | 0.0029 | 0.0030 |
| F03 | 78700001 | 78800000 | 0.0026 | 0.0023 | 0.0024 |
| F03 | 78720001 | 78820000 | 0.0022 | 0.0020 | 0.0021 |
| F03 | 78740001 | 78840000 | 0.0016 | 0.0013 | 0.0014 |
| F03 | 78760001 | 78860000 | 0.0013 | 0.0009 | 0.0010 |
| F03 | 78780001 | 78880000 | 0.0014 | 0.0009 | 0.0010 |
| F03 | 78800001 | 78900000 | 0.0021 | 0.0016 | 0.0016 |
| F03 | 78820001 | 78920000 | 0.0035 | 0.0027 | 0.0029 |
| F03 | 78840001 | 78940000 | 0.0042 | 0.0031 | 0.0034 |
| F03 | 78860001 | 78960000 | 0.0052 | 0.0041 | 0.0046 |
| F03 | 78880001 | 78980000 | 0.0053 | 0.0041 | 0.0045 |
| F03 | 78900001 | 79000000 | 0.0050 | 0.0036 | 0.0040 |
| F03 | 78920001 | 79020000 | 0.0035 | 0.0024 | 0.0026 |
| F03 | 78940001 | 79040000 | 0.0029 | 0.0022 | 0.0023 |
| F03 | 78960001 | 79060000 | 0.0026 | 0.0019 | 0.0020 |
| F03 | 78980001 | 79080000 | 0.0023 | 0.0019 | 0.0021 |
| F03 | 79000001 | 79100000 | 0.0018 | 0.0017 | 0.0018 |
| F03 | 79020001 | 79120000 | 0.0023 | 0.0020 | 0.0022 |
| F03 | 79040001 | 79140000 | 0.0019 | 0.0017 | 0.0019 |
| F03 | 79060001 | 79160000 | 0.0013 | 0.0012 | 0.0013 |
| F03 | 79080001 | 79180000 | 0.0011 | 0.0010 | 0.0011 |
| F03 | 79100001 | 79200000 | 0.0011 | 0.0009 | 0.0011 |
| F03 | 79120001 | 79220000 | 0.0009 | 0.0008 | 0.0009 |
| F03 | 79140001 | 79240000 | 0.0008 | 0.0007 | 0.0008 |
| F03 | 79160001 | 79260000 | 0.0006 | 0.0006 | 0.0006 |

| Chr | Start | End | both | A1 | A2 |
|---|---|---|---|---|---|
| F03 | 79180001 | 79280000 | 0.0007 | 0.0007 | 0.0007 |
| F03 | 79200001 | 79300000 | 0.0007 | 0.0007 | 0.0007 |
| F03 | 79220001 | 79320000 | 0.0010 | 0.0009 | 0.0009 |
| F03 | 79240001 | 79340000 | 0.0010 | 0.0009 | 0.0010 |
| F03 | 79260001 | 79360000 | 0.0015 | 0.0014 | 0.0015 |
| F03 | 79280001 | 79380000 | 0.0020 | 0.0016 | 0.0018 |
| F03 | 79300001 | 79400000 | 0.0026 | 0.0021 | 0.0022 |
| F03 | 79320001 | 79420000 | 0.0029 | 0.0022 | 0.0024 |
| F03 | 79340001 | 79440000 | 0.0028 | 0.0022 | 0.0023 |
| F03 | 79360001 | 79460000 | 0.0029 | 0.0023 | 0.0024 |
| F03 | 79380001 | 79480000 | 0.0035 | 0.0029 | 0.0030 |
| F03 | 79400001 | 79500000 | 0.0041 | 0.0034 | 0.0037 |
| F03 | 79420001 | 79520000 | 0.0034 | 0.0028 | 0.0030 |
| F03 | 79440001 | 79540000 | 0.0038 | 0.0029 | 0.0032 |
| F03 | 79460001 | 79560000 | 0.0036 | 0.0026 | 0.0028 |
| F03 | 79480001 | 79580000 | 0.0031 | 0.0021 | 0.0024 |
| F03 | 79500001 | 79600000 | 0.0028 | 0.0020 | 0.0022 |
| F03 | 79520001 | 79620000 | 0.0029 | 0.0022 | 0.0024 |
| F03 | 79540001 | 79640000 | 0.0038 | 0.0029 | 0.0033 |
| F03 | 79560001 | 79660000 | 0.0037 | 0.0029 | 0.0033 |
| F03 | 79580001 | 79680000 | 0.0033 | 0.0025 | 0.0029 |
| F03 | 79600001 | 79700000 | 0.0030 | 0.0022 | 0.0026 |
| F03 | 79620001 | 79720000 | 0.0030 | 0.0022 | 0.0025 |
| F03 | 79640001 | 79740000 | 0.0021 | 0.0014 | 0.0017 |
| F03 | 79660001 | 79760000 | 0.0019 | 0.0013 | 0.0014 |
| F03 | 79680001 | 79780000 | 0.0020 | 0.0013 | 0.0014 |
| F03 | 79700001 | 79800000 | 0.0017 | 0.0011 | 0.0011 |
| F03 | 79720001 | 79820000 | 0.0018 | 0.0011 | 0.0012 |
| F03 | 79740001 | 79840000 | 0.0017 | 0.0012 | 0.0013 |
| F03 | 79760001 | 79860000 | 0.0016 | 0.0012 | 0.0013 |
| F03 | 79780001 | 79880000 | 0.0016 | 0.0013 | 0.0014 |
| F03 | 79800001 | 79900000 | 0.0011 | 0.0008 | 0.0011 |
| F03 | 79820001 | 79920000 | 0.0008 | 0.0006 | 0.0008 |
| F03 | 79840001 | 79940000 | 0.0007 | 0.0006 | 0.0007 |
| F03 | 79860001 | 79960000 | 0.0012 | 0.0010 | 0.0012 |
| F03 | 79880001 | 79980000 | 0.0012 | 0.0011 | 0.0012 |
| F03 | 79900001 | 80000000 | 0.0012 | 0.0011 | 0.0012 |
| F03 | 79920001 | 80020000 | 0.0012 | 0.0011 | 0.0012 |
| F03 | 79940001 | 80040000 | 0.0015 | 0.0013 | 0.0015 |
| F03 | 79960001 | 80060000 | 0.0009 | 0.0008 | 0.0009 |
| F03 | 79980001 | 80080000 | 0.0012 | 0.0010 | 0.0011 |
| F03 | 80000001 | 80100000 | 0.0014 | 0.0012 | 0.0014 |
| F03 | 80020001 | 80120000 | 0.0017 | 0.0015 | 0.0017 |
| F03 | 80040001 | 80140000 | 0.0014 | 0.0012 | 0.0014 |
| F03 | 80060001 | 80160000 | 0.0015 | 0.0013 | 0.0015 |
| F03 | 80080001 | 80180000 | 0.0012 | 0.0010 | 0.0012 |
| F03 | 80100001 | 80200000 | 0.0012 | 0.0010 | 0.0012 |
| F03 | 80120001 | 80220000 | 0.0012 | 0.0009 | 0.0011 |
| F03 | 80140001 | 80240000 | 0.0016 | 0.0010 | 0.0013 |
| F03 | 80160001 | 80260000 | 0.0022 | 0.0016 | 0.0018 |
| F03 | 80180001 | 80280000 | 0.0021 | 0.0016 | 0.0018 |
| F03 | 80200001 | 80300000 | 0.0019 | 0.0014 | 0.0016 |
| F03 | 80220001 | 80320000 | 0.0020 | 0.0015 | 0.0016 |
| F03 | 80240001 | 80340000 | 0.0016 | 0.0013 | 0.0013 |
| F03 | 80260001 | 80360000 | 0.0011 | 0.0009 | 0.0010 |
| F03 | 80280001 | 80380000 | 0.0017 | 0.0011 | 0.0014 |
| F03 | 80300001 | 80400000 | 0.0027 | 0.0019 | 0.0020 |
| F03 | 80320001 | 80420000 | 0.0038 | 0.0028 | 0.0033 |

| Chr | Start | End | both | A1 | A2 |
|---|---|---|---|---|---|
| F03 | 80340001 | 80440000 | 0.0043 | 0.0030 | 0.0033 |
| F03 | 80360001 | 80460000 | 0.0045 | 0.0030 | 0.0033 |
| F03 | 80380001 | 80480000 | 0.0048 | 0.0033 | 0.0034 |
| F03 | 80400001 | 80500000 | 0.0045 | 0.0032 | 0.0034 |
| F03 | 80420001 | 80520000 | 0.0033 | 0.0022 | 0.0022 |
| F03 | 80440001 | 80540000 | 0.0032 | 0.0023 | 0.0023 |
| F03 | 80460001 | 80560000 | 0.0034 | 0.0026 | 0.0026 |
| F03 | 80480001 | 80580000 | 0.0024 | 0.0020 | 0.0021 |
| F03 | 80500001 | 80600000 | 0.0030 | 0.0027 | 0.0027 |
| F03 | 80520001 | 80620000 | 0.0038 | 0.0034 | 0.0036 |
| F03 | 80540001 | 80640000 | 0.0044 | 0.0040 | 0.0041 |
| F03 | 80560001 | 80660000 | 0.0044 | 0.0037 | 0.0038 |
| F03 | 80580001 | 80680000 | 0.0048 | 0.0040 | 0.0041 |
| F03 | 80600001 | 80700000 | 0.0038 | 0.0027 | 0.0028 |
| F03 | 80620001 | 80720000 | 0.0035 | 0.0022 | 0.0024 |
| F03 | 80640001 | 80740000 | 0.0036 | 0.0022 | 0.0024 |
| F03 | 80660001 | 80760000 | 0.0040 | 0.0028 | 0.0029 |
| F03 | 80680001 | 80780000 | 0.0042 | 0.0030 | 0.0032 |
| F03 | 80700001 | 80800000 | 0.0042 | 0.0032 | 0.0034 |
| F03 | 80720001 | 80820000 | 0.0046 | 0.0037 | 0.0039 |
| F03 | 80740001 | 80840000 | 0.0044 | 0.0033 | 0.0036 |
| F03 | 80760001 | 80860000 | 0.0041 | 0.0031 | 0.0034 |
| F03 | 80780001 | 80880000 | 0.0034 | 0.0025 | 0.0028 |
| F03 | 80800001 | 80900000 | 0.0034 | 0.0023 | 0.0026 |
| F03 | 80820001 | 80920000 | 0.0026 | 0.0016 | 0.0019 |
| F03 | 80840001 | 80940000 | 0.0036 | 0.0026 | 0.0029 |
| F03 | 80860001 | 80960000 | 0.0031 | 0.0022 | 0.0025 |
| F03 | 80880001 | 80980000 | 0.0034 | 0.0025 | 0.0028 |
| F03 | 80900001 | 81000000 | 0.0036 | 0.0029 | 0.0032 |
| F03 | 80920001 | 81020000 | 0.0042 | 0.0036 | 0.0039 |
| F03 | 80940001 | 81040000 | 0.0030 | 0.0023 | 0.0025 |
| F03 | 80960001 | 81060000 | 0.0036 | 0.0027 | 0.0027 |
| F03 | 80980001 | 81080000 | 0.0034 | 0.0025 | 0.0026 |
| F03 | 81000001 | 81100000 | 0.0041 | 0.0030 | 0.0031 |
| F03 | 81020001 | 81120000 | 0.0034 | 0.0023 | 0.0025 |
| F03 | 81040001 | 81140000 | 0.0030 | 0.0022 | 0.0024 |
| F03 | 81060001 | 81160000 | 0.0029 | 0.0023 | 0.0026 |
| F03 | 81080001 | 81180000 | 0.0032 | 0.0027 | 0.0031 |
| F03 | 81100001 | 81200000 | 0.0034 | 0.0029 | 0.0034 |
| F03 | 81120001 | 81220000 | 0.0038 | 0.0032 | 0.0037 |
| F03 | 81140001 | 81240000 | 0.0043 | 0.0037 | 0.0042 |
| F03 | 81160001 | 81260000 | 0.0037 | 0.0032 | 0.0036 |
| F03 | 81180001 | 81280000 | 0.0035 | 0.0031 | 0.0034 |
| F03 | 81200001 | 81300000 | 0.0028 | 0.0025 | 0.0027 |
| F03 | 81220001 | 81320000 | 0.0020 | 0.0019 | 0.0019 |
| F03 | 81240001 | 81340000 | 0.0021 | 0.0020 | 0.0021 |
| F03 | 81260001 | 81360000 | 0.0023 | 0.0021 | 0.0022 |
| F03 | 81280001 | 81380000 | 0.0023 | 0.0019 | 0.0020 |
| F03 | 81300001 | 81400000 | 0.0022 | 0.0017 | 0.0019 |
| F03 | 81320001 | 81420000 | 0.0025 | 0.0018 | 0.0020 |
| F03 | 81340001 | 81440000 | 0.0023 | 0.0014 | 0.0016 |
| F03 | 81360001 | 81460000 | 0.0026 | 0.0017 | 0.0018 |
| F03 | 81380001 | 81480000 | 0.0027 | 0.0016 | 0.0018 |
| F03 | 81400001 | 81500000 | 0.0023 | 0.0013 | 0.0014 |
| F03 | 81420001 | 81520000 | 0.0025 | 0.0013 | 0.0015 |
| F03 | 81440001 | 81540000 | 0.0026 | 0.0014 | 0.0017 |
| F03 | 81460001 | 81560000 | 0.0032 | 0.0019 | 0.0021 |
| F03 | 81480001 | 81580000 | 0.0048 | 0.0036 | 0.0039 |

| Chr | Start | End | both | A1 | A2 |
|---|---|---|---|---|---|
| F03 | 81500001 | 81600000 | 0.0050 | 0.0039 | 0.0041 |
| F03 | 81520001 | 81620000 | 0.0047 | 0.0039 | 0.0040 |
| F03 | 81540001 | 81640000 | 0.0041 | 0.0036 | 0.0036 |
| F03 | 81560001 | 81660000 | 0.0032 | 0.0030 | 0.0030 |
| F03 | 81580001 | 81680000 | 0.0017 | 0.0016 | 0.0016 |
| F03 | 81600001 | 81700000 | 0.0016 | 0.0015 | 0.0015 |
| F03 | 81620001 | 81720000 | 0.0019 | 0.0016 | 0.0015 |
| F03 | 81640001 | 81740000 | 0.0027 | 0.0023 | 0.0023 |
| F03 | 81660001 | 81760000 | 0.0027 | 0.0021 | 0.0021 |
| F03 | 81680001 | 81780000 | 0.0025 | 0.0019 | 0.0020 |
| F03 | 81700001 | 81800000 | 0.0033 | 0.0026 | 0.0028 |
| F03 | 81720001 | 81820000 | 0.0029 | 0.0024 | 0.0026 |
| F03 | 81740001 | 81840000 | 0.0034 | 0.0029 | 0.0030 |
| F03 | 81760001 | 81860000 | 0.0032 | 0.0028 | 0.0030 |
| F03 | 81780001 | 81880000 | 0.0027 | 0.0024 | 0.0026 |
| F03 | 81800001 | 81900000 | 0.0020 | 0.0017 | 0.0019 |
| F03 | 81820001 | 81920000 | 0.0035 | 0.0029 | 0.0035 |
| F03 | 81840001 | 81940000 | 0.0029 | 0.0022 | 0.0027 |
| F03 | 81860001 | 81960000 | 0.0033 | 0.0025 | 0.0030 |
| F03 | 81880001 | 81980000 | 0.0049 | 0.0040 | 0.0045 |
| F03 | 81900001 | 82000000 | 0.0052 | 0.0044 | 0.0047 |
| F03 | 81920001 | 82020000 | 0.0045 | 0.0038 | 0.0038 |
| F03 | 81940001 | 82040000 | 0.0049 | 0.0041 | 0.0043 |
| F03 | 81960001 | 82060000 | 0.0049 | 0.0042 | 0.0043 |
| F03 | 81980001 | 82080000 | 0.0042 | 0.0032 | 0.0036 |
| F03 | 82000001 | 82100000 | 0.0039 | 0.0028 | 0.0033 |
| F03 | 82020001 | 82120000 | 0.0034 | 0.0024 | 0.0030 |
| F03 | 82040001 | 82140000 | 0.0028 | 0.0019 | 0.0025 |
| F03 | 82060001 | 82160000 | 0.0024 | 0.0015 | 0.0021 |
| F03 | 82080001 | 82180000 | 0.0017 | 0.0012 | 0.0016 |
| F03 | 82100001 | 82200000 | 0.0014 | 0.0011 | 0.0014 |
| F03 | 82120001 | 82220000 | 0.0014 | 0.0013 | 0.0014 |
| F03 | 82140001 | 82240000 | 0.0013 | 0.0012 | 0.0012 |
| F03 | 82160001 | 82260000 | 0.0015 | 0.0013 | 0.0015 |
| F03 | 82180001 | 82280000 | 0.0017 | 0.0015 | 0.0017 |
| F03 | 82200001 | 82300000 | 0.0021 | 0.0018 | 0.0021 |
| F03 | 82220001 | 82320000 | 0.0024 | 0.0020 | 0.0023 |
| F03 | 82240001 | 82340000 | 0.0024 | 0.0019 | 0.0022 |
| F03 | 82260001 | 82360000 | 0.0022 | 0.0019 | 0.0020 |
| F03 | 82280001 | 82380000 | 0.0021 | 0.0017 | 0.0019 |
| F03 | 82300001 | 82400000 | 0.0019 | 0.0016 | 0.0017 |
| F03 | 82320001 | 82420000 | 0.0013 | 0.0012 | 0.0013 |
| F03 | 82340001 | 82440000 | 0.0019 | 0.0018 | 0.0019 |
| F03 | 82360001 | 82460000 | 0.0024 | 0.0021 | 0.0024 |
| F03 | 82380001 | 82480000 | 0.0028 | 0.0025 | 0.0028 |
| F03 | 82400001 | 82500000 | 0.0025 | 0.0023 | 0.0025 |
| F03 | 82420001 | 82520000 | 0.0024 | 0.0021 | 0.0024 |
| F03 | 82440001 | 82540000 | 0.0018 | 0.0015 | 0.0017 |
| F03 | 82460001 | 82560000 | 0.0015 | 0.0013 | 0.0015 |
| F03 | 82480001 | 82580000 | 0.0013 | 0.0011 | 0.0013 |
| F03 | 82500001 | 82600000 | 0.0016 | 0.0013 | 0.0015 |
| F03 | 82520001 | 82620000 | 0.0016 | 0.0014 | 0.0016 |
| F03 | 82540001 | 82640000 | 0.0020 | 0.0018 | 0.0019 |
| F03 | 82560001 | 82660000 | 0.0016 | 0.0014 | 0.0015 |
| F03 | 82580001 | 82680000 | 0.0017 | 0.0016 | 0.0016 |
| F03 | 82600001 | 82700000 | 0.0024 | 0.0021 | 0.0022 |
| F03 | 82620001 | 82720000 | 0.0033 | 0.0029 | 0.0031 |
| F03 | 82640001 | 82740000 | 0.0035 | 0.0030 | 0.0031 |

| Chr | Start | End | both | A1 | A2 |
|---|---|---|---|---|---|
| F03 | 82660001 | 82760000 | 0.0039 | 0.0033 | 0.0034 |
| F03 | 82680001 | 82780000 | 0.0038 | 0.0032 | 0.0034 |
| F03 | 82700001 | 82800000 | 0.0032 | 0.0027 | 0.0029 |
| F03 | 82720001 | 82820000 | 0.0023 | 0.0019 | 0.0020 |
| F03 | 82740001 | 82840000 | 0.0025 | 0.0019 | 0.0022 |
| F03 | 82760001 | 82860000 | 0.0029 | 0.0022 | 0.0025 |
| F03 | 82780001 | 82880000 | 0.0029 | 0.0023 | 0.0025 |
| F03 | 82800001 | 82900000 | 0.0035 | 0.0029 | 0.0031 |
| F03 | 82820001 | 82920000 | 0.0038 | 0.0032 | 0.0034 |
| F03 | 82840001 | 82940000 | 0.0038 | 0.0033 | 0.0035 |
| F03 | 82860001 | 82960000 | 0.0040 | 0.0036 | 0.0038 |
| F03 | 82880001 | 82980000 | 0.0039 | 0.0035 | 0.0036 |
| F03 | 82900001 | 83000000 | 0.0034 | 0.0030 | 0.0032 |
| F03 | 82920001 | 83020000 | 0.0036 | 0.0031 | 0.0032 |
| F03 | 82940001 | 83040000 | 0.0034 | 0.0028 | 0.0030 |
| F03 | 82960001 | 83060000 | 0.0040 | 0.0032 | 0.0037 |
| F03 | 82980001 | 83080000 | 0.0042 | 0.0033 | 0.0039 |
| F03 | 83000001 | 83100000 | 0.0039 | 0.0031 | 0.0036 |
| F03 | 83020001 | 83120000 | 0.0037 | 0.0031 | 0.0035 |
| F03 | 83040001 | 83140000 | 0.0040 | 0.0033 | 0.0037 |
| F03 | 83060001 | 83160000 | 0.0025 | 0.0020 | 0.0021 |
| F03 | 83080001 | 83180000 | 0.0033 | 0.0023 | 0.0026 |
| F03 | 83100001 | 83200000 | 0.0035 | 0.0024 | 0.0026 |
| F03 | 83120001 | 83220000 | 0.0033 | 0.0020 | 0.0025 |
| F03 | 83140001 | 83240000 | 0.0032 | 0.0020 | 0.0024 |
| F03 | 83160001 | 83260000 | 0.0036 | 0.0025 | 0.0030 |
| F03 | 83180001 | 83280000 | 0.0023 | 0.0018 | 0.0020 |
| F03 | 83200001 | 83300000 | 0.0018 | 0.0014 | 0.0017 |
| F03 | 83220001 | 83320000 | 0.0024 | 0.0019 | 0.0023 |
| F03 | 83240001 | 83340000 | 0.0024 | 0.0019 | 0.0023 |
| F03 | 83260001 | 83360000 | 0.0023 | 0.0016 | 0.0020 |
| F03 | 83280001 | 83380000 | 0.0032 | 0.0023 | 0.0028 |
| F03 | 83300001 | 83400000 | 0.0043 | 0.0032 | 0.0037 |
| F03 | 83320001 | 83420000 | 0.0036 | 0.0027 | 0.0031 |
| F03 | 83340001 | 83440000 | 0.0033 | 0.0025 | 0.0028 |
| F03 | 83360001 | 83460000 | 0.0031 | 0.0023 | 0.0027 |
| F03 | 83380001 | 83480000 | 0.0022 | 0.0017 | 0.0019 |
| F03 | 83400001 | 83500000 | 0.0012 | 0.0010 | 0.0011 |
| F03 | 83420001 | 83520000 | 0.0009 | 0.0007 | 0.0008 |
| F03 | 83440001 | 83540000 | 0.0014 | 0.0010 | 0.0011 |
| F03 | 83460001 | 83560000 | 0.0019 | 0.0015 | 0.0016 |
| F03 | 83480001 | 83580000 | 0.0018 | 0.0015 | 0.0016 |
| F03 | 83500001 | 83600000 | 0.0018 | 0.0015 | 0.0016 |
| F03 | 83520001 | 83620000 | 0.0026 | 0.0021 | 0.0023 |
| F03 | 83540001 | 83640000 | 0.0032 | 0.0029 | 0.0031 |
| F03 | 83560001 | 83660000 | 0.0033 | 0.0030 | 0.0033 |
| F03 | 83580001 | 83680000 | 0.0035 | 0.0031 | 0.0034 |
| F03 | 83600001 | 83700000 | 0.0034 | 0.0030 | 0.0034 |
| F03 | 83620001 | 83720000 | 0.0030 | 0.0025 | 0.0030 |
| F03 | 83640001 | 83740000 | 0.0019 | 0.0015 | 0.0019 |
| F03 | 83660001 | 83760000 | 0.0016 | 0.0013 | 0.0016 |
| F03 | 83680001 | 83780000 | 0.0022 | 0.0018 | 0.0020 |
| F03 | 83700001 | 83800000 | 0.0024 | 0.0020 | 0.0022 |
| F03 | 83720001 | 83820000 | 0.0028 | 0.0023 | 0.0024 |
| F03 | 83740001 | 83840000 | 0.0038 | 0.0030 | 0.0034 |
| F03 | 83760001 | 83860000 | 0.0043 | 0.0034 | 0.0038 |
| F03 | 83780001 | 83880000 | 0.0047 | 0.0039 | 0.0041 |
| F03 | 83800001 | 83900000 | 0.0051 | 0.0042 | 0.0045 |

| Chr | Start | End | both | A1 | A2 |
|---|---|---|---|---|---|
| F03 | 83820001 | 83920000 | 0.0055 | 0.0045 | 0.0050 |
| F03 | 83840001 | 83940000 | 0.0045 | 0.0039 | 0.0040 |
| F03 | 83860001 | 83960000 | 0.0036 | 0.0031 | 0.0033 |
| F03 | 83880001 | 83980000 | 0.0032 | 0.0026 | 0.0030 |
| F03 | 83900001 | 84000000 | 0.0034 | 0.0028 | 0.0031 |
| F03 | 83920001 | 84020000 | 0.0026 | 0.0020 | 0.0022 |
| F03 | 83940001 | 84040000 | 0.0026 | 0.0020 | 0.0022 |
| F03 | 83960001 | 84060000 | 0.0033 | 0.0025 | 0.0028 |
| F03 | 83980001 | 84080000 | 0.0035 | 0.0027 | 0.0029 |
| F03 | 84000001 | 84100000 | 0.0034 | 0.0028 | 0.0031 |
| F03 | 84020001 | 84120000 | 0.0041 | 0.0034 | 0.0037 |
| F03 | 84040001 | 84140000 | 0.0052 | 0.0044 | 0.0049 |
| F03 | 84060001 | 84160000 | 0.0048 | 0.0041 | 0.0047 |
| F03 | 84080001 | 84180000 | 0.0043 | 0.0037 | 0.0042 |
| F03 | 84100001 | 84200000 | 0.0036 | 0.0031 | 0.0034 |
| F03 | 84120001 | 84220000 | 0.0031 | 0.0027 | 0.0030 |
| F03 | 84140001 | 84240000 | 0.0023 | 0.0021 | 0.0022 |
| F03 | 84160001 | 84260000 | 0.0017 | 0.0016 | 0.0016 |
| F03 | 84180001 | 84280000 | 0.0020 | 0.0019 | 0.0020 |
| F03 | 84200001 | 84300000 | 0.0025 | 0.0023 | 0.0025 |
| F03 | 84220001 | 84320000 | 0.0040 | 0.0034 | 0.0039 |
| F03 | 84240001 | 84340000 | 0.0040 | 0.0033 | 0.0037 |
| F03 | 84260001 | 84360000 | 0.0043 | 0.0035 | 0.0039 |
| F03 | 84280001 | 84380000 | 0.0057 | 0.0046 | 0.0050 |
| F03 | 84300001 | 84400000 | 0.0064 | 0.0051 | 0.0055 |
| F03 | 84320001 | 84420000 | 0.0048 | 0.0037 | 0.0038 |
| F03 | 84340001 | 84440000 | 0.0044 | 0.0034 | 0.0036 |
| F03 | 84360001 | 84460000 | 0.0054 | 0.0043 | 0.0043 |
| F03 | 84380001 | 84480000 | 0.0048 | 0.0036 | 0.0037 |
| F03 | 84400001 | 84500000 | 0.0048 | 0.0035 | 0.0037 |
| F03 | 84420001 | 84520000 | 0.0049 | 0.0037 | 0.0039 |
| F03 | 84440001 | 84540000 | 0.0052 | 0.0039 | 0.0041 |
| F03 | 84460001 | 84560000 | 0.0047 | 0.0034 | 0.0039 |
| F03 | 84480001 | 84580000 | 0.0034 | 0.0026 | 0.0030 |
| F03 | 84500001 | 84600000 | 0.0024 | 0.0016 | 0.0017 |
| F03 | 84520001 | 84620000 | 0.0028 | 0.0016 | 0.0019 |
| F03 | 84540001 | 84640000 | 0.0034 | 0.0020 | 0.0022 |
| F03 | 84560001 | 84660000 | 0.0040 | 0.0027 | 0.0027 |
| F03 | 84580001 | 84680000 | 0.0039 | 0.0026 | 0.0027 |
| F03 | 84600001 | 84700000 | 0.0038 | 0.0029 | 0.0030 |
| F03 | 84620001 | 84720000 | 0.0031 | 0.0025 | 0.0026 |
| F03 | 84640001 | 84740000 | 0.0022 | 0.0020 | 0.0020 |
| F03 | 84660001 | 84760000 | 0.0017 | 0.0017 | 0.0015 |
| F03 | 84680001 | 84780000 | 0.0022 | 0.0020 | 0.0019 |
| F03 | 84700001 | 84800000 | 0.0026 | 0.0023 | 0.0021 |
| F03 | 84720001 | 84820000 | 0.0030 | 0.0024 | 0.0022 |
| F03 | 84740001 | 84840000 | 0.0034 | 0.0025 | 0.0023 |
| F03 | 84760001 | 84860000 | 0.0038 | 0.0025 | 0.0028 |
| F03 | 84780001 | 84880000 | 0.0038 | 0.0025 | 0.0026 |
| F03 | 84800001 | 84900000 | 0.0036 | 0.0021 | 0.0024 |
| F03 | 84820001 | 84920000 | 0.0038 | 0.0023 | 0.0025 |
| F03 | 84840001 | 84940000 | 0.0046 | 0.0032 | 0.0034 |
| F03 | 84860001 | 84960000 | 0.0038 | 0.0026 | 0.0027 |
| F03 | 84880001 | 84980000 | 0.0036 | 0.0025 | 0.0027 |
| F03 | 84900001 | 85000000 | 0.0034 | 0.0025 | 0.0026 |
| F03 | 84920001 | 85020000 | 0.0038 | 0.0027 | 0.0031 |
| F03 | 84940001 | 85040000 | 0.0027 | 0.0018 | 0.0023 |
| F03 | 84960001 | 85060000 | 0.0032 | 0.0020 | 0.0025 |

| Chr | Start | End | both | A1 | A2 |
|---|---|---|---|---|---|
| F03 | 84980001 | 85080000 | 0.0037 | 0.0023 | 0.0027 |
| F03 | 85000001 | 85100000 | 0.0039 | 0.0022 | 0.0029 |
| F03 | 85020001 | 85120000 | 0.0045 | 0.0029 | 0.0035 |
| F03 | 85040001 | 85140000 | 0.0052 | 0.0035 | 0.0041 |
| F03 | 85060001 | 85160000 | 0.0048 | 0.0033 | 0.0039 |
| F03 | 85080001 | 85180000 | 0.0044 | 0.0032 | 0.0035 |
| F03 | 85100001 | 85200000 | 0.0041 | 0.0032 | 0.0033 |
| F03 | 85120001 | 85220000 | 0.0036 | 0.0026 | 0.0028 |
| F03 | 85140001 | 85240000 | 0.0035 | 0.0027 | 0.0028 |
| F03 | 85160001 | 85260000 | 0.0037 | 0.0028 | 0.0028 |
| F03 | 85180001 | 85280000 | 0.0039 | 0.0028 | 0.0030 |
| F03 | 85200001 | 85300000 | 0.0044 | 0.0032 | 0.0034 |
| F03 | 85220001 | 85320000 | 0.0037 | 0.0025 | 0.0027 |
| F03 | 85240001 | 85340000 | 0.0035 | 0.0025 | 0.0026 |
| F03 | 85260001 | 85360000 | 0.0035 | 0.0025 | 0.0027 |
| F03 | 85280001 | 85380000 | 0.0039 | 0.0029 | 0.0031 |
| F03 | 85300001 | 85400000 | 0.0036 | 0.0025 | 0.0026 |
| F03 | 85320001 | 85420000 | 0.0040 | 0.0028 | 0.0029 |
| F03 | 85340001 | 85440000 | 0.0041 | 0.0025 | 0.0025 |
| F03 | 85360001 | 85460000 | 0.0038 | 0.0022 | 0.0024 |
| F03 | 85380001 | 85480000 | 0.0033 | 0.0018 | 0.0019 |
| F03 | 85400001 | 85500000 | 0.0037 | 0.0021 | 0.0025 |
| F03 | 85420001 | 85520000 | 0.0042 | 0.0028 | 0.0031 |
| F03 | 85440001 | 85540000 | 0.0045 | 0.0035 | 0.0038 |
| F03 | 85460001 | 85560000 | 0.0061 | 0.0051 | 0.0054 |
| F03 | 85480001 | 85580000 | 0.0061 | 0.0051 | 0.0053 |
| F03 | 85500001 | 85600000 | 0.0059 | 0.0049 | 0.0051 |
| F03 | 85520001 | 85620000 | 0.0066 | 0.0052 | 0.0056 |
| F03 | 85540001 | 85640000 | 0.0058 | 0.0043 | 0.0047 |
| F03 | 85560001 | 85660000 | 0.0041 | 0.0024 | 0.0028 |
| F03 | 85580001 | 85680000 | 0.0045 | 0.0026 | 0.0031 |
| F03 | 85600001 | 85700000 | 0.0045 | 0.0027 | 0.0033 |
| F03 | 85620001 | 85720000 | 0.0036 | 0.0018 | 0.0022 |
| F03 | 85640001 | 85740000 | 0.0035 | 0.0020 | 0.0024 |
| F03 | 85660001 | 85760000 | 0.0041 | 0.0026 | 0.0029 |
| F03 | 85680001 | 85780000 | 0.0041 | 0.0026 | 0.0028 |
| F03 | 85700001 | 85800000 | 0.0050 | 0.0036 | 0.0037 |
| F03 | 85720001 | 85820000 | 0.0047 | 0.0036 | 0.0037 |
| F03 | 85740001 | 85840000 | 0.0058 | 0.0046 | 0.0047 |
| F03 | 85760001 | 85860000 | 0.0053 | 0.0043 | 0.0046 |
| F03 | 85780001 | 85880000 | 0.0050 | 0.0040 | 0.0042 |
| F03 | 85800001 | 85900000 | 0.0036 | 0.0027 | 0.0028 |
| F03 | 85820001 | 85920000 | 0.0034 | 0.0025 | 0.0026 |
| F03 | 85840001 | 85940000 | 0.0025 | 0.0014 | 0.0015 |
| F03 | 85860001 | 85960000 | 0.0021 | 0.0012 | 0.0014 |
| F03 | 85880001 | 85980000 | 0.0022 | 0.0017 | 0.0018 |
| F03 | 85900001 | 86000000 | 0.0025 | 0.0018 | 0.0021 |
| F03 | 85920001 | 86020000 | 0.0028 | 0.0020 | 0.0023 |
| F03 | 85940001 | 86040000 | 0.0026 | 0.0019 | 0.0022 |
| F03 | 85960001 | 86060000 | 0.0025 | 0.0019 | 0.0020 |
| F03 | 85980001 | 86080000 | 0.0023 | 0.0016 | 0.0018 |
| F03 | 86000001 | 86100000 | 0.0022 | 0.0013 | 0.0015 |
| F03 | 86020001 | 86120000 | 0.0024 | 0.0015 | 0.0017 |
| F03 | 86040001 | 86140000 | 0.0024 | 0.0015 | 0.0016 |
| F03 | 86060001 | 86160000 | 0.0031 | 0.0020 | 0.0023 |
| F03 | 86080001 | 86180000 | 0.0036 | 0.0024 | 0.0027 |
| F03 | 86100001 | 86200000 | 0.0039 | 0.0030 | 0.0032 |
| F03 | 86120001 | 86220000 | 0.0048 | 0.0037 | 0.0040 |

| Chr | Start | End | both | A1 | A2 |
|---|---|---|---|---|---|
| F03 | 86140001 | 86240000 | 0.0052 | 0.0043 | 0.0046 |
| F03 | 86160001 | 86260000 | 0.0046 | 0.0038 | 0.0040 |
| F03 | 86180001 | 86280000 | 0.0042 | 0.0035 | 0.0037 |
| F03 | 86200001 | 86300000 | 0.0035 | 0.0029 | 0.0031 |
| F03 | 86220001 | 86320000 | 0.0022 | 0.0018 | 0.0019 |
| F03 | 86240001 | 86340000 | 0.0020 | 0.0014 | 0.0016 |
| F03 | 86260001 | 86360000 | 0.0019 | 0.0014 | 0.0016 |
| F03 | 86280001 | 86380000 | 0.0017 | 0.0013 | 0.0013 |
| F03 | 86300001 | 86400000 | 0.0017 | 0.0014 | 0.0014 |
| F03 | 86320001 | 86420000 | 0.0015 | 0.0012 | 0.0012 |
| F03 | 86340001 | 86440000 | 0.0018 | 0.0013 | 0.0014 |
| F03 | 86360001 | 86460000 | 0.0021 | 0.0015 | 0.0016 |
| F03 | 86380001 | 86480000 | 0.0030 | 0.0018 | 0.0021 |
| F03 | 86400001 | 86500000 | 0.0036 | 0.0021 | 0.0026 |
| F03 | 86420001 | 86520000 | 0.0052 | 0.0033 | 0.0039 |
| F03 | 86440001 | 86540000 | 0.0045 | 0.0029 | 0.0034 |
| F03 | 86460001 | 86560000 | 0.0041 | 0.0025 | 0.0030 |
| F03 | 86480001 | 86580000 | 0.0038 | 0.0027 | 0.0031 |
| F03 | 86500001 | 86600000 | 0.0040 | 0.0028 | 0.0032 |
| F03 | 86520001 | 86620000 | 0.0030 | 0.0020 | 0.0023 |
| F03 | 86540001 | 86640000 | 0.0047 | 0.0034 | 0.0037 |
| F03 | 86560001 | 86660000 | 0.0046 | 0.0035 | 0.0037 |
| F03 | 86580001 | 86680000 | 0.0038 | 0.0026 | 0.0029 |
| F03 | 86600001 | 86700000 | 0.0032 | 0.0022 | 0.0025 |
| F03 | 86620001 | 86720000 | 0.0028 | 0.0021 | 0.0024 |
| F03 | 86640001 | 86740000 | 0.0018 | 0.0010 | 0.0014 |
| F03 | 86660001 | 86760000 | 0.0027 | 0.0017 | 0.0020 |
| F03 | 86680001 | 86780000 | 0.0036 | 0.0025 | 0.0029 |
| F03 | 86700001 | 86800000 | 0.0039 | 0.0025 | 0.0028 |
| F03 | 86720001 | 86820000 | 0.0042 | 0.0027 | 0.0030 |
| F03 | 86740001 | 86840000 | 0.0038 | 0.0025 | 0.0027 |
| F03 | 86760001 | 86860000 | 0.0035 | 0.0023 | 0.0024 |
| F03 | 86780001 | 86880000 | 0.0031 | 0.0020 | 0.0021 |
| F03 | 86800001 | 86900000 | 0.0033 | 0.0026 | 0.0028 |
| F03 | 86820001 | 86920000 | 0.0028 | 0.0022 | 0.0024 |
| F03 | 86840001 | 86940000 | 0.0024 | 0.0020 | 0.0022 |
| F03 | 86860001 | 86960000 | 0.0042 | 0.0038 | 0.0041 |
| F03 | 86880001 | 86980000 | 0.0049 | 0.0042 | 0.0046 |
| F03 | 86900001 | 87000000 | 0.0045 | 0.0038 | 0.0042 |
| F03 | 86920001 | 87020000 | 0.0050 | 0.0040 | 0.0044 |
| F03 | 86940001 | 87040000 | 0.0063 | 0.0052 | 0.0056 |
| F03 | 86960001 | 87060000 | 0.0048 | 0.0035 | 0.0041 |
| F03 | 86980001 | 87080000 | 0.0040 | 0.0031 | 0.0034 |
| F03 | 87000001 | 87100000 | 0.0054 | 0.0045 | 0.0049 |
| F03 | 87020001 | 87120000 | 0.0056 | 0.0046 | 0.0051 |
| F03 | 87040001 | 87140000 | 0.0055 | 0.0044 | 0.0048 |
| F03 | 87060001 | 87160000 | 0.0049 | 0.0040 | 0.0042 |
| F03 | 87080001 | 87180000 | 0.0061 | 0.0049 | 0.0050 |
| F03 | 87100001 | 87200000 | 0.0062 | 0.0047 | 0.0048 |
| F03 | 87120001 | 87220000 | 0.0073 | 0.0058 | 0.0057 |
| F03 | 87140001 | 87240000 | 0.0078 | 0.0063 | 0.0064 |
| F03 | 87160001 | 87260000 | 0.0081 | 0.0065 | 0.0066 |
| F03 | 87180001 | 87280000 | 0.0067 | 0.0051 | 0.0054 |
| F03 | 87200001 | 87300000 | 0.0059 | 0.0045 | 0.0048 |
| F03 | 87220001 | 87320000 | 0.0050 | 0.0033 | 0.0038 |
| F03 | 87240001 | 87340000 | 0.0039 | 0.0024 | 0.0027 |
| F03 | 87260001 | 87360000 | 0.0038 | 0.0020 | 0.0023 |
| F03 | 87280001 | 87380000 | 0.0039 | 0.0020 | 0.0022 |

| Chr | Start | End | both | A1 | A2 |
|---|---|---|---|---|---|
| F03 | 87300001 | 87400000 | 0.0032 | 0.0011 | 0.0013 |
| F03 | 87320001 | 87420000 | 0.0028 | 0.0009 | 0.0010 |
| F03 | 87340001 | 87440000 | 0.0030 | 0.0010 | 0.0013 |
| F03 | 87360001 | 87460000 | 0.0025 | 0.0010 | 0.0013 |
| F03 | 87380001 | 87480000 | 0.0026 | 0.0013 | 0.0016 |
| F03 | 87400001 | 87500000 | 0.0027 | 0.0015 | 0.0018 |
| F03 | 87420001 | 87520000 | 0.0025 | 0.0015 | 0.0018 |
| F03 | 87440001 | 87540000 | 0.0022 | 0.0013 | 0.0016 |
| F03 | 87460001 | 87560000 | 0.0025 | 0.0013 | 0.0016 |
| F03 | 87480001 | 87580000 | 0.0024 | 0.0012 | 0.0014 |
| F03 | 87500001 | 87600000 | 0.0024 | 0.0015 | 0.0017 |
| F03 | 87520001 | 87620000 | 0.0024 | 0.0015 | 0.0017 |
| F03 | 87540001 | 87640000 | 0.0029 | 0.0019 | 0.0020 |
| F03 | 87560001 | 87660000 | 0.0033 | 0.0023 | 0.0025 |
| F03 | 87580001 | 87680000 | 0.0030 | 0.0021 | 0.0024 |
| F03 | 87600001 | 87700000 | 0.0026 | 0.0017 | 0.0019 |
| F03 | 87620001 | 87720000 | 0.0025 | 0.0016 | 0.0018 |
| F03 | 87640001 | 87740000 | 0.0025 | 0.0015 | 0.0018 |
| F03 | 87660001 | 87760000 | 0.0025 | 0.0018 | 0.0020 |
| F03 | 87680001 | 87780000 | 0.0031 | 0.0020 | 0.0023 |
| F03 | 87700001 | 87800000 | 0.0039 | 0.0027 | 0.0030 |
| F03 | 87720001 | 87820000 | 0.0038 | 0.0026 | 0.0029 |
| F03 | 87740001 | 87840000 | 0.0037 | 0.0023 | 0.0026 |
| F03 | 87760001 | 87860000 | 0.0037 | 0.0018 | 0.0023 |
| F03 | 87780001 | 87880000 | 0.0037 | 0.0021 | 0.0026 |
| F03 | 87800001 | 87900000 | 0.0036 | 0.0022 | 0.0026 |
| F03 | 87820001 | 87920000 | 0.0038 | 0.0024 | 0.0029 |
| F03 | 87840001 | 87940000 | 0.0038 | 0.0024 | 0.0029 |
| F03 | 87860001 | 87960000 | 0.0036 | 0.0025 | 0.0029 |
| F03 | 87880001 | 87980000 | 0.0030 | 0.0020 | 0.0023 |
| F03 | 87900001 | 88000000 | 0.0026 | 0.0018 | 0.0021 |
| F03 | 87920001 | 88020000 | 0.0029 | 0.0019 | 0.0022 |
| F03 | 87940001 | 88040000 | 0.0029 | 0.0018 | 0.0021 |
| F03 | 87960001 | 88060000 | 0.0036 | 0.0023 | 0.0026 |
| F03 | 87980001 | 88080000 | 0.0042 | 0.0026 | 0.0030 |
| F03 | 88000001 | 88100000 | 0.0040 | 0.0023 | 0.0027 |
| F03 | 88020001 | 88120000 | 0.0037 | 0.0022 | 0.0025 |
| F03 | 88040001 | 88140000 | 0.0035 | 0.0023 | 0.0026 |
| F03 | 88060001 | 88160000 | 0.0025 | 0.0016 | 0.0018 |
| F03 | 88080001 | 88180000 | 0.0021 | 0.0015 | 0.0016 |
| F03 | 88100001 | 88200000 | 0.0020 | 0.0015 | 0.0016 |
| F03 | 88120001 | 88220000 | 0.0023 | 0.0017 | 0.0020 |
| F03 | 88140001 | 88240000 | 0.0039 | 0.0029 | 0.0034 |
| F03 | 88160001 | 88260000 | 0.0042 | 0.0032 | 0.0037 |
| F03 | 88180001 | 88280000 | 0.0054 | 0.0044 | 0.0049 |
| F03 | 88200001 | 88300000 | 0.0060 | 0.0050 | 0.0055 |
| F03 | 88220001 | 88320000 | 0.0059 | 0.0050 | 0.0054 |
| F03 | 88240001 | 88340000 | 0.0043 | 0.0035 | 0.0037 |
| F03 | 88260001 | 88360000 | 0.0049 | 0.0042 | 0.0042 |
| F03 | 88280001 | 88380000 | 0.0045 | 0.0036 | 0.0036 |
| F03 | 88300001 | 88400000 | 0.0044 | 0.0035 | 0.0035 |
| F03 | 88320001 | 88420000 | 0.0048 | 0.0038 | 0.0037 |
| F03 | 88340001 | 88440000 | 0.0050 | 0.0039 | 0.0038 |
| F03 | 88360001 | 88460000 | 0.0068 | 0.0059 | 0.0055 |
| F03 | 88380001 | 88480000 | 0.0068 | 0.0058 | 0.0055 |
| F03 | 88400001 | 88500000 | 0.0071 | 0.0059 | 0.0057 |
| F03 | 88420001 | 88520000 | 0.0061 | 0.0051 | 0.0050 |
| F03 | 88440001 | 88540000 | 0.0060 | 0.0053 | 0.0051 |

| Chr | Start | End | both | A1 | A2 |
|---|---|---|---|---|---|
| F03 | 88460001 | 88560000 | 0.0033 | 0.0026 | 0.0027 |
| F03 | 88480001 | 88580000 | 0.0033 | 0.0026 | 0.0030 |
| F03 | 88500001 | 88600000 | 0.0024 | 0.0020 | 0.0023 |
| F03 | 88520001 | 88620000 | 0.0033 | 0.0027 | 0.0031 |
| F03 | 88540001 | 88640000 | 0.0037 | 0.0028 | 0.0034 |
| F03 | 88560001 | 88660000 | 0.0039 | 0.0027 | 0.0034 |
| F03 | 88580001 | 88680000 | 0.0038 | 0.0025 | 0.0028 |
| F03 | 88600001 | 88700000 | 0.0052 | 0.0038 | 0.0041 |
| F03 | 88620001 | 88720000 | 0.0055 | 0.0038 | 0.0042 |
| F03 | 88640001 | 88740000 | 0.0055 | 0.0038 | 0.0040 |
| F03 | 88660001 | 88760000 | 0.0055 | 0.0038 | 0.0040 |
| F03 | 88680001 | 88780000 | 0.0053 | 0.0038 | 0.0041 |
| F03 | 88700001 | 88800000 | 0.0044 | 0.0028 | 0.0033 |
| F03 | 88720001 | 88820000 | 0.0043 | 0.0025 | 0.0031 |
| F03 | 88740001 | 88840000 | 0.0041 | 0.0025 | 0.0032 |
| F03 | 88760001 | 88860000 | 0.0050 | 0.0031 | 0.0038 |
| F03 | 88780001 | 88880000 | 0.0052 | 0.0031 | 0.0039 |
| F03 | 88800001 | 88900000 | 0.0060 | 0.0037 | 0.0045 |
| F03 | 88820001 | 88920000 | 0.0060 | 0.0042 | 0.0047 |
| F03 | 88840001 | 88940000 | 0.0063 | 0.0044 | 0.0046 |
| F03 | 88860001 | 88960000 | 0.0058 | 0.0044 | 0.0045 |
| F03 | 88880001 | 88980000 | 0.0056 | 0.0044 | 0.0044 |
| F03 | 88900001 | 89000000 | 0.0051 | 0.0040 | 0.0041 |
| F03 | 88920001 | 89020000 | 0.0050 | 0.0040 | 0.0043 |
| F03 | 88940001 | 89040000 | 0.0042 | 0.0035 | 0.0040 |
| F03 | 88960001 | 89060000 | 0.0033 | 0.0028 | 0.0031 |
| F03 | 88980001 | 89080000 | 0.0029 | 0.0024 | 0.0027 |
| F03 | 89000001 | 89100000 | 0.0023 | 0.0020 | 0.0020 |
| F03 | 89020001 | 89120000 | 0.0019 | 0.0015 | 0.0016 |
| F03 | 89040001 | 89140000 | 0.0025 | 0.0021 | 0.0022 |
| F03 | 89060001 | 89160000 | 0.0026 | 0.0022 | 0.0024 |
| F03 | 89080001 | 89180000 | 0.0033 | 0.0029 | 0.0031 |
| F03 | 89100001 | 89200000 | 0.0031 | 0.0026 | 0.0030 |
| F03 | 89120001 | 89220000 | 0.0025 | 0.0024 | 0.0024 |
| F03 | 89140001 | 89240000 | 0.0018 | 0.0016 | 0.0017 |
| F03 | 89160001 | 89260000 | 0.0020 | 0.0018 | 0.0018 |
| F03 | 89180001 | 89280000 | 0.0026 | 0.0023 | 0.0025 |
| F03 | 89200001 | 89300000 | 0.0039 | 0.0033 | 0.0035 |
| F03 | 89220001 | 89320000 | 0.0047 | 0.0040 | 0.0042 |
| F03 | 89240001 | 89340000 | 0.0052 | 0.0044 | 0.0047 |
| F03 | 89260001 | 89360000 | 0.0052 | 0.0045 | 0.0048 |
| F03 | 89280001 | 89380000 | 0.0041 | 0.0032 | 0.0034 |
| F03 | 89300001 | 89400000 | 0.0038 | 0.0029 | 0.0032 |
| F03 | 89320001 | 89420000 | 0.0033 | 0.0024 | 0.0028 |
| F03 | 89340001 | 89440000 | 0.0028 | 0.0021 | 0.0024 |
| F03 | 89360001 | 89460000 | 0.0027 | 0.0020 | 0.0023 |
| F03 | 89380001 | 89480000 | 0.0024 | 0.0020 | 0.0023 |
| F03 | 89400001 | 89500000 | 0.0023 | 0.0020 | 0.0023 |
| F03 | 89420001 | 89520000 | 0.0027 | 0.0023 | 0.0026 |
| F03 | 89440001 | 89540000 | 0.0035 | 0.0030 | 0.0034 |
| F03 | 89460001 | 89560000 | 0.0032 | 0.0029 | 0.0032 |
| F03 | 89480001 | 89580000 | 0.0036 | 0.0033 | 0.0035 |
| F03 | 89500001 | 89600000 | 0.0030 | 0.0028 | 0.0030 |
| F03 | 89520001 | 89620000 | 0.0022 | 0.0020 | 0.0022 |
| F03 | 89540001 | 89640000 | 0.0015 | 0.0014 | 0.0015 |
| F03 | 89560001 | 89660000 | 0.0017 | 0.0017 | 0.0017 |
| F03 | 89580001 | 89680000 | 0.0012 | 0.0011 | 0.0012 |
| F03 | 89600001 | 89700000 | 0.0013 | 0.0013 | 0.0012 |

| Chr | Start | End | both | A1 | A2 |
|---|---|---|---|---|---|
| F03 | 89620001 | 89720000 | 0.0015 | 0.0014 | 0.0014 |
| F03 | 89640001 | 89740000 | 0.0031 | 0.0027 | 0.0029 |
| F03 | 89660001 | 89760000 | 0.0032 | 0.0026 | 0.0027 |
| F03 | 89680001 | 89780000 | 0.0042 | 0.0031 | 0.0034 |
| F03 | 89700001 | 89800000 | 0.0048 | 0.0033 | 0.0040 |
| F03 | 89720001 | 89820000 | 0.0063 | 0.0045 | 0.0053 |
| F03 | 89740001 | 89840000 | 0.0048 | 0.0030 | 0.0037 |
| F03 | 89760001 | 89860000 | 0.0054 | 0.0038 | 0.0045 |
| F03 | 89780001 | 89880000 | 0.0049 | 0.0034 | 0.0039 |
| F03 | 89800001 | 89900000 | 0.0040 | 0.0029 | 0.0031 |
| F03 | 89820001 | 89920000 | 0.0034 | 0.0023 | 0.0025 |
| F03 | 89840001 | 89940000 | 0.0034 | 0.0024 | 0.0026 |
| F03 | 89860001 | 89960000 | 0.0027 | 0.0016 | 0.0019 |
| F03 | 89880001 | 89980000 | 0.0032 | 0.0025 | 0.0027 |
| F03 | 89900001 | 90000000 | 0.0046 | 0.0037 | 0.0042 |
| F03 | 89920001 | 90020000 | 0.0045 | 0.0037 | 0.0042 |
| F03 | 89940001 | 90040000 | 0.0051 | 0.0041 | 0.0045 |
| F03 | 89960001 | 90060000 | 0.0051 | 0.0041 | 0.0045 |
| F03 | 89980001 | 90080000 | 0.0043 | 0.0033 | 0.0037 |
| F03 | 90000001 | 90100000 | 0.0031 | 0.0023 | 0.0025 |
| F03 | 90020001 | 90120000 | 0.0024 | 0.0017 | 0.0019 |
| F03 | 90040001 | 90140000 | 0.0020 | 0.0015 | 0.0018 |
| F03 | 90060001 | 90160000 | 0.0024 | 0.0018 | 0.0021 |
| F03 | 90080001 | 90180000 | 0.0034 | 0.0026 | 0.0029 |
| F03 | 90100001 | 90200000 | 0.0037 | 0.0025 | 0.0030 |
| F03 | 90120001 | 90220000 | 0.0043 | 0.0030 | 0.0035 |
| F03 | 90140001 | 90240000 | 0.0044 | 0.0029 | 0.0033 |
| F03 | 90160001 | 90260000 | 0.0043 | 0.0028 | 0.0032 |
| F03 | 90180001 | 90280000 | 0.0035 | 0.0021 | 0.0026 |
| F03 | 90200001 | 90300000 | 0.0037 | 0.0025 | 0.0029 |
| F03 | 90220001 | 90320000 | 0.0030 | 0.0021 | 0.0024 |
| F03 | 90240001 | 90340000 | 0.0037 | 0.0029 | 0.0033 |
| F03 | 90260001 | 90360000 | 0.0034 | 0.0026 | 0.0031 |
| F03 | 90280001 | 90380000 | 0.0037 | 0.0030 | 0.0033 |
| F03 | 90300001 | 90400000 | 0.0032 | 0.0025 | 0.0027 |
| F03 | 90320001 | 90420000 | 0.0040 | 0.0029 | 0.0033 |
| F03 | 90340001 | 90440000 | 0.0034 | 0.0023 | 0.0028 |
| F03 | 90360001 | 90460000 | 0.0040 | 0.0031 | 0.0035 |
| F03 | 90380001 | 90480000 | 0.0041 | 0.0029 | 0.0036 |
| F03 | 90400001 | 90500000 | 0.0047 | 0.0035 | 0.0043 |
| F03 | 90420001 | 90520000 | 0.0038 | 0.0029 | 0.0036 |
| F03 | 90440001 | 90540000 | 0.0043 | 0.0031 | 0.0038 |
| F03 | 90460001 | 90560000 | 0.0036 | 0.0025 | 0.0031 |
| F03 | 90480001 | 90580000 | 0.0028 | 0.0021 | 0.0025 |
| F03 | 90500001 | 90600000 | 0.0025 | 0.0018 | 0.0022 |
| F03 | 90520001 | 90620000 | 0.0030 | 0.0021 | 0.0026 |
| F03 | 90540001 | 90640000 | 0.0029 | 0.0025 | 0.0028 |
| F03 | 90560001 | 90660000 | 0.0038 | 0.0031 | 0.0035 |
| F03 | 90580001 | 90680000 | 0.0044 | 0.0036 | 0.0040 |
| F03 | 90600001 | 90700000 | 0.0061 | 0.0049 | 0.0056 |
| F03 | 90620001 | 90720000 | 0.0062 | 0.0046 | 0.0053 |
| F03 | 90640001 | 90740000 | 0.0065 | 0.0046 | 0.0054 |
| F03 | 90660001 | 90760000 | 0.0052 | 0.0037 | 0.0044 |
| F03 | 90680001 | 90780000 | 0.0058 | 0.0040 | 0.0048 |
| F03 | 90700001 | 90800000 | 0.0039 | 0.0023 | 0.0028 |
| F03 | 90720001 | 90820000 | 0.0041 | 0.0027 | 0.0032 |
| F03 | 90740001 | 90840000 | 0.0037 | 0.0023 | 0.0028 |
| F03 | 90760001 | 90860000 | 0.0038 | 0.0022 | 0.0028 |

| Chr | Start | End | both | A1 | A2 |
|---|---|---|---|---|---|
| F03 | 90780001 | 90880000 | 0.0031 | 0.0018 | 0.0023 |
| F03 | 90800001 | 90900000 | 0.0030 | 0.0019 | 0.0023 |
| F03 | 90820001 | 90920000 | 0.0024 | 0.0016 | 0.0019 |
| F03 | 90840001 | 90940000 | 0.0020 | 0.0014 | 0.0017 |
| F03 | 90860001 | 90960000 | 0.0033 | 0.0025 | 0.0027 |
| F03 | 90880001 | 90980000 | 0.0039 | 0.0030 | 0.0033 |
| F03 | 90900001 | 91000000 | 0.0046 | 0.0038 | 0.0041 |
| F03 | 90920001 | 91020000 | 0.0049 | 0.0037 | 0.0042 |
| F03 | 90940001 | 91040000 | 0.0053 | 0.0039 | 0.0044 |
| F03 | 90960001 | 91060000 | 0.0054 | 0.0039 | 0.0044 |
| F03 | 90980001 | 91080000 | 0.0055 | 0.0040 | 0.0046 |
| F03 | 91000001 | 91100000 | 0.0054 | 0.0037 | 0.0045 |
| F03 | 91020001 | 91120000 | 0.0053 | 0.0039 | 0.0046 |
| F03 | 91040001 | 91140000 | 0.0050 | 0.0036 | 0.0043 |
| F03 | 91060001 | 91160000 | 0.0050 | 0.0037 | 0.0045 |
| F03 | 91080001 | 91180000 | 0.0048 | 0.0037 | 0.0044 |
| F03 | 91100001 | 91200000 | 0.0064 | 0.0053 | 0.0060 |
| F03 | 91120001 | 91220000 | 0.0065 | 0.0053 | 0.0059 |
| F03 | 91140001 | 91240000 | 0.0062 | 0.0051 | 0.0057 |
| F03 | 91160001 | 91260000 | 0.0049 | 0.0040 | 0.0045 |
| F03 | 91180001 | 91280000 | 0.0045 | 0.0034 | 0.0039 |
| F03 | 91200001 | 91300000 | 0.0027 | 0.0015 | 0.0020 |
| F03 | 91220001 | 91320000 | 0.0030 | 0.0017 | 0.0022 |
| F03 | 91240001 | 91340000 | 0.0046 | 0.0030 | 0.0036 |
| F03 | 91260001 | 91360000 | 0.0051 | 0.0033 | 0.0039 |
| F03 | 91280001 | 91380000 | 0.0057 | 0.0041 | 0.0047 |
| F03 | 91300001 | 91400000 | 0.0065 | 0.0047 | 0.0055 |
| F03 | 91320001 | 91420000 | 0.0059 | 0.0045 | 0.0053 |
| F03 | 91340001 | 91440000 | 0.0046 | 0.0033 | 0.0039 |
| F03 | 91360001 | 91460000 | 0.0049 | 0.0037 | 0.0044 |
| F03 | 91380001 | 91480000 | 0.0040 | 0.0028 | 0.0035 |
| F03 | 91400001 | 91500000 | 0.0027 | 0.0018 | 0.0022 |
| F03 | 91420001 | 91520000 | 0.0026 | 0.0016 | 0.0020 |
| F03 | 91440001 | 91540000 | 0.0030 | 0.0020 | 0.0023 |
| F03 | 91460001 | 91560000 | 0.0023 | 0.0013 | 0.0016 |
| F03 | 91480001 | 91580000 | 0.0029 | 0.0020 | 0.0024 |
| F03 | 91500001 | 91600000 | 0.0038 | 0.0029 | 0.0033 |
| F03 | 91520001 | 91620000 | 0.0043 | 0.0033 | 0.0038 |
| F03 | 91540001 | 91640000 | 0.0042 | 0.0032 | 0.0037 |
| F03 | 91560001 | 91660000 | 0.0048 | 0.0037 | 0.0042 |
| F03 | 91580001 | 91680000 | 0.0047 | 0.0037 | 0.0041 |
| F03 | 91600001 | 91700000 | 0.0051 | 0.0040 | 0.0044 |
| F03 | 91620001 | 91720000 | 0.0045 | 0.0035 | 0.0038 |
| F03 | 91640001 | 91740000 | 0.0046 | 0.0035 | 0.0038 |
| F03 | 91660001 | 91760000 | 0.0040 | 0.0029 | 0.0033 |
| F03 | 91680001 | 91780000 | 0.0038 | 0.0026 | 0.0029 |
| F03 | 91700001 | 91800000 | 0.0028 | 0.0016 | 0.0018 |
| F03 | 91720001 | 91820000 | 0.0032 | 0.0019 | 0.0020 |
| F03 | 91740001 | 91840000 | 0.0033 | 0.0022 | 0.0024 |
| F03 | 91760001 | 91860000 | 0.0042 | 0.0030 | 0.0031 |
| F03 | 91780001 | 91880000 | 0.0048 | 0.0036 | 0.0037 |
| F03 | 91800001 | 91900000 | 0.0059 | 0.0047 | 0.0049 |
| F03 | 91820001 | 91920000 | 0.0060 | 0.0049 | 0.0051 |
| F03 | 91840001 | 91940000 | 0.0057 | 0.0044 | 0.0047 |
| F03 | 91860001 | 91960000 | 0.0050 | 0.0039 | 0.0043 |
| F03 | 91880001 | 91980000 | 0.0043 | 0.0032 | 0.0035 |
| F03 | 91900001 | 92000000 | 0.0034 | 0.0025 | 0.0027 |
| F03 | 91920001 | 92020000 | 0.0038 | 0.0029 | 0.0033 |

| Chr | Start | End | both | A1 | A2 |
|---|---|---|---|---|---|
| F03 | 91940001 | 92040000 | 0.0040 | 0.0033 | 0.0036 |
| F03 | 91960001 | 92060000 | 0.0042 | 0.0032 | 0.0036 |
| F03 | 91980001 | 92080000 | 0.0043 | 0.0033 | 0.0037 |
| F03 | 92000001 | 92100000 | 0.0041 | 0.0030 | 0.0035 |
| F03 | 92020001 | 92120000 | 0.0031 | 0.0021 | 0.0024 |
| F03 | 92040001 | 92140000 | 0.0029 | 0.0019 | 0.0021 |
| F03 | 92060001 | 92160000 | 0.0035 | 0.0027 | 0.0028 |
| F03 | 92080001 | 92180000 | 0.0039 | 0.0029 | 0.0032 |
| F03 | 92100001 | 92200000 | 0.0045 | 0.0036 | 0.0039 |
| F03 | 92120001 | 92220000 | 0.0047 | 0.0037 | 0.0039 |
| F03 | 92140001 | 92240000 | 0.0046 | 0.0032 | 0.0037 |
| F03 | 92160001 | 92260000 | 0.0041 | 0.0027 | 0.0031 |
| F03 | 92180001 | 92280000 | 0.0039 | 0.0026 | 0.0028 |
| F03 | 92200001 | 92300000 | 0.0033 | 0.0020 | 0.0022 |
| F03 | 92220001 | 92320000 | 0.0037 | 0.0024 | 0.0027 |
| F03 | 92240001 | 92340000 | 0.0035 | 0.0023 | 0.0026 |
| F03 | 92260001 | 92360000 | 0.0027 | 0.0016 | 0.0019 |
| F03 | 92280001 | 92380000 | 0.0030 | 0.0017 | 0.0021 |
| F03 | 92300001 | 92400000 | 0.0032 | 0.0017 | 0.0022 |
| F03 | 92320001 | 92420000 | 0.0039 | 0.0023 | 0.0029 |
| F03 | 92340001 | 92440000 | 0.0049 | 0.0033 | 0.0039 |
| F03 | 92360001 | 92460000 | 0.0051 | 0.0037 | 0.0043 |
| F03 | 92380001 | 92480000 | 0.0042 | 0.0033 | 0.0036 |
| F03 | 92400001 | 92500000 | 0.0037 | 0.0030 | 0.0033 |
| F03 | 92420001 | 92520000 | 0.0028 | 0.0022 | 0.0025 |
| F03 | 92440001 | 92540000 | 0.0019 | 0.0013 | 0.0015 |
| F03 | 92460001 | 92560000 | 0.0022 | 0.0015 | 0.0017 |
| F03 | 92480001 | 92580000 | 0.0037 | 0.0029 | 0.0031 |
| F03 | 92500001 | 92600000 | 0.0038 | 0.0028 | 0.0030 |
| F03 | 92520001 | 92620000 | 0.0042 | 0.0032 | 0.0033 |
| F03 | 92540001 | 92640000 | 0.0042 | 0.0034 | 0.0034 |
| F03 | 92560001 | 92660000 | 0.0041 | 0.0029 | 0.0031 |
| F03 | 92580001 | 92680000 | 0.0029 | 0.0015 | 0.0018 |
| F03 | 92600001 | 92700000 | 0.0034 | 0.0022 | 0.0024 |
| F03 | 92620001 | 92720000 | 0.0047 | 0.0033 | 0.0037 |
| F03 | 92640001 | 92740000 | 0.0050 | 0.0036 | 0.0041 |
| F03 | 92660001 | 92760000 | 0.0050 | 0.0038 | 0.0042 |
| F03 | 92680001 | 92780000 | 0.0056 | 0.0043 | 0.0049 |
| F03 | 92700001 | 92800000 | 0.0048 | 0.0034 | 0.0041 |
| F03 | 92720001 | 92820000 | 0.0034 | 0.0021 | 0.0027 |
| F03 | 92740001 | 92840000 | 0.0030 | 0.0017 | 0.0023 |
| F03 | 92760001 | 92860000 | 0.0027 | 0.0014 | 0.0022 |
| F03 | 92780001 | 92880000 | 0.0021 | 0.0008 | 0.0013 |
| F03 | 92800001 | 92900000 | 0.0028 | 0.0015 | 0.0020 |
| F03 | 92820001 | 92920000 | 0.0023 | 0.0012 | 0.0017 |
| F03 | 92840001 | 92940000 | 0.0023 | 0.0014 | 0.0018 |
| F03 | 92860001 | 92960000 | 0.0030 | 0.0022 | 0.0026 |
| F03 | 92880001 | 92980000 | 0.0037 | 0.0031 | 0.0036 |
| F03 | 92900001 | 93000000 | 0.0032 | 0.0027 | 0.0032 |
| F03 | 92920001 | 93020000 | 0.0042 | 0.0035 | 0.0040 |
| F03 | 92940001 | 93040000 | 0.0049 | 0.0038 | 0.0043 |
| F03 | 92960001 | 93060000 | 0.0047 | 0.0033 | 0.0038 |
| F03 | 92980001 | 93080000 | 0.0044 | 0.0026 | 0.0031 |
| F03 | 93000001 | 93100000 | 0.0056 | 0.0034 | 0.0039 |
| F03 | 93020001 | 93120000 | 0.0051 | 0.0029 | 0.0034 |
| F03 | 93040001 | 93140000 | 0.0043 | 0.0024 | 0.0028 |
| F03 | 93060001 | 93160000 | 0.0039 | 0.0022 | 0.0025 |
| F03 | 93080001 | 93180000 | 0.0038 | 0.0024 | 0.0026 |

| Chr | Start | End | both | A1 | A2 |
|---|---|---|---|---|---|
| F03 | 93100001 | 93200000 | 0.0038 | 0.0026 | 0.0028 |
| F03 | 93120001 | 93220000 | 0.0042 | 0.0029 | 0.0031 |
| F03 | 93140001 | 93240000 | 0.0045 | 0.0031 | 0.0034 |
| F03 | 93160001 | 93260000 | 0.0046 | 0.0034 | 0.0037 |
| F03 | 93180001 | 93280000 | 0.0044 | 0.0032 | 0.0036 |
| F03 | 93200001 | 93300000 | 0.0033 | 0.0020 | 0.0025 |
| F03 | 93220001 | 93320000 | 0.0030 | 0.0019 | 0.0022 |
| F03 | 93240001 | 93340000 | 0.0034 | 0.0022 | 0.0025 |
| F03 | 93260001 | 93360000 | 0.0036 | 0.0025 | 0.0026 |
| F03 | 93280001 | 93380000 | 0.0037 | 0.0028 | 0.0027 |
| F03 | 93300001 | 93400000 | 0.0035 | 0.0027 | 0.0025 |
| F03 | 93320001 | 93420000 | 0.0037 | 0.0029 | 0.0028 |
| F03 | 93340001 | 93440000 | 0.0035 | 0.0027 | 0.0029 |
| F03 | 93360001 | 93460000 | 0.0034 | 0.0026 | 0.0029 |
| F03 | 93380001 | 93480000 | 0.0032 | 0.0024 | 0.0028 |
| F03 | 93400001 | 93500000 | 0.0033 | 0.0024 | 0.0028 |
| F03 | 93420001 | 93520000 | 0.0025 | 0.0018 | 0.0021 |
| F03 | 93440001 | 93540000 | 0.0026 | 0.0018 | 0.0019 |
| F03 | 93460001 | 93560000 | 0.0024 | 0.0015 | 0.0015 |
| F03 | 93480001 | 93580000 | 0.0025 | 0.0014 | 0.0014 |
| F03 | 93500001 | 93600000 | 0.0026 | 0.0014 | 0.0015 |
| F03 | 93520001 | 93620000 | 0.0032 | 0.0017 | 0.0019 |
| F03 | 93540001 | 93640000 | 0.0031 | 0.0016 | 0.0017 |
| F03 | 93560001 | 93660000 | 0.0029 | 0.0015 | 0.0017 |
| F03 | 93580001 | 93680000 | 0.0033 | 0.0020 | 0.0022 |
| F03 | 93600001 | 93700000 | 0.0035 | 0.0022 | 0.0024 |
| F03 | 93620001 | 93720000 | 0.0033 | 0.0021 | 0.0023 |
| F03 | 93640001 | 93740000 | 0.0033 | 0.0020 | 0.0023 |
| F03 | 93660001 | 93760000 | 0.0035 | 0.0021 | 0.0025 |
| F03 | 93680001 | 93780000 | 0.0030 | 0.0016 | 0.0019 |
| F03 | 93700001 | 93800000 | 0.0029 | 0.0015 | 0.0019 |
| F03 | 93720001 | 93820000 | 0.0024 | 0.0013 | 0.0015 |
| F03 | 93740001 | 93840000 | 0.0019 | 0.0012 | 0.0013 |
| F03 | 93760001 | 93860000 | 0.0023 | 0.0015 | 0.0016 |
| F03 | 93780001 | 93880000 | 0.0032 | 0.0024 | 0.0025 |
| F03 | 93800001 | 93900000 | 0.0033 | 0.0025 | 0.0027 |
| F03 | 93820001 | 93920000 | 0.0034 | 0.0027 | 0.0030 |
| F03 | 93840001 | 93940000 | 0.0036 | 0.0027 | 0.0030 |
| F03 | 93860001 | 93960000 | 0.0040 | 0.0031 | 0.0036 |
| F03 | 93880001 | 93980000 | 0.0032 | 0.0023 | 0.0027 |
| F03 | 93900001 | 94000000 | 0.0026 | 0.0019 | 0.0022 |
| F03 | 93920001 | 94020000 | 0.0035 | 0.0024 | 0.0027 |
| F03 | 93940001 | 94040000 | 0.0042 | 0.0029 | 0.0033 |
| F03 | 93960001 | 94060000 | 0.0041 | 0.0026 | 0.0029 |
| F03 | 93980001 | 94080000 | 0.0044 | 0.0029 | 0.0033 |
| F03 | 94000001 | 94100000 | 0.0045 | 0.0029 | 0.0032 |
| F03 | 94020001 | 94120000 | 0.0039 | 0.0026 | 0.0028 |
| F03 | 94040001 | 94140000 | 0.0036 | 0.0027 | 0.0028 |
| F03 | 94060001 | 94160000 | 0.0027 | 0.0020 | 0.0022 |
| F03 | 94080001 | 94180000 | 0.0024 | 0.0018 | 0.0021 |
| F03 | 94100001 | 94200000 | 0.0029 | 0.0023 | 0.0027 |
| F03 | 94120001 | 94220000 | 0.0027 | 0.0022 | 0.0027 |
| F03 | 94140001 | 94240000 | 0.0032 | 0.0026 | 0.0031 |
| F03 | 94160001 | 94260000 | 0.0029 | 0.0024 | 0.0028 |
| F03 | 94180001 | 94280000 | 0.0025 | 0.0020 | 0.0023 |
| F03 | 94200001 | 94300000 | 0.0023 | 0.0017 | 0.0021 |
| F03 | 94220001 | 94320000 | 0.0027 | 0.0020 | 0.0023 |
| F03 | 94240001 | 94340000 | 0.0019 | 0.0014 | 0.0016 |

| Chr | Start | End | both | A1 | A2 |
|---|---|---|---|---|---|
| F03 | 94260001 | 94360000 | 0.0024 | 0.0015 | 0.0020 |
| F03 | 94280001 | 94380000 | 0.0027 | 0.0019 | 0.0025 |
| F03 | 94300001 | 94400000 | 0.0032 | 0.0023 | 0.0029 |
| F03 | 94320001 | 94420000 | 0.0028 | 0.0020 | 0.0026 |
| F03 | 94340001 | 94440000 | 0.0028 | 0.0018 | 0.0026 |
| F03 | 94360001 | 94460000 | 0.0043 | 0.0032 | 0.0039 |
| F03 | 94380001 | 94480000 | 0.0048 | 0.0036 | 0.0044 |
| F03 | 94400001 | 94500000 | 0.0053 | 0.0038 | 0.0047 |
| F03 | 94420001 | 94520000 | 0.0058 | 0.0040 | 0.0049 |
| F03 | 94440001 | 94540000 | 0.0056 | 0.0040 | 0.0047 |
| F03 | 94460001 | 94560000 | 0.0039 | 0.0027 | 0.0030 |
| F03 | 94480001 | 94580000 | 0.0041 | 0.0030 | 0.0032 |
| F03 | 94500001 | 94600000 | 0.0036 | 0.0027 | 0.0028 |
| F03 | 94520001 | 94620000 | 0.0041 | 0.0033 | 0.0035 |
| F03 | 94540001 | 94640000 | 0.0041 | 0.0033 | 0.0033 |
| F03 | 94560001 | 94660000 | 0.0041 | 0.0033 | 0.0034 |
| F03 | 94580001 | 94680000 | 0.0034 | 0.0026 | 0.0027 |
| F03 | 94600001 | 94700000 | 0.0028 | 0.0022 | 0.0022 |
| F03 | 94620001 | 94720000 | 0.0020 | 0.0013 | 0.0014 |
| F03 | 94640001 | 94740000 | 0.0019 | 0.0012 | 0.0014 |
| F03 | 94660001 | 94760000 | 0.0024 | 0.0017 | 0.0018 |
| F03 | 94680001 | 94780000 | 0.0025 | 0.0019 | 0.0019 |
| F03 | 94700001 | 94800000 | 0.0025 | 0.0017 | 0.0017 |
| F03 | 94720001 | 94820000 | 0.0029 | 0.0022 | 0.0023 |
| F03 | 94740001 | 94840000 | 0.0032 | 0.0024 | 0.0025 |
| F03 | 94760001 | 94860000 | 0.0027 | 0.0019 | 0.0020 |
| F03 | 94780001 | 94880000 | 0.0024 | 0.0016 | 0.0017 |
| F03 | 94800001 | 94900000 | 0.0025 | 0.0016 | 0.0018 |
| F03 | 94820001 | 94920000 | 0.0020 | 0.0011 | 0.0012 |
| F03 | 94840001 | 94940000 | 0.0015 | 0.0007 | 0.0008 |
| F03 | 94860001 | 94960000 | 0.0015 | 0.0008 | 0.0009 |
| F03 | 94880001 | 94980000 | 0.0012 | 0.0007 | 0.0008 |
| F03 | 94900001 | 95000000 | 0.0010 | 0.0007 | 0.0008 |
| F03 | 94920001 | 95020000 | 0.0012 | 0.0009 | 0.0011 |
| F03 | 94940001 | 95040000 | 0.0015 | 0.0013 | 0.0015 |
| F03 | 94960001 | 95060000 | 0.0020 | 0.0017 | 0.0019 |
| F03 | 94980001 | 95080000 | 0.0036 | 0.0031 | 0.0034 |
| F03 | 95000001 | 95100000 | 0.0038 | 0.0030 | 0.0034 |
| F03 | 95020001 | 95120000 | 0.0037 | 0.0027 | 0.0030 |
| F03 | 95040001 | 95140000 | 0.0031 | 0.0022 | 0.0024 |
| F03 | 95060001 | 95160000 | 0.0025 | 0.0016 | 0.0019 |
| F03 | 95080001 | 95180000 | 0.0013 | 0.0007 | 0.0008 |
| F03 | 95100001 | 95200000 | 0.0013 | 0.0006 | 0.0007 |
| F03 | 95120001 | 95220000 | 0.0012 | 0.0009 | 0.0009 |
| F03 | 95140001 | 95240000 | 0.0026 | 0.0021 | 0.0022 |
| F03 | 95160001 | 95260000 | 0.0037 | 0.0030 | 0.0033 |
| F03 | 95180001 | 95280000 | 0.0039 | 0.0031 | 0.0034 |
| F03 | 95200001 | 95300000 | 0.0038 | 0.0032 | 0.0034 |
| F03 | 95220001 | 95320000 | 0.0041 | 0.0033 | 0.0036 |
| F03 | 95240001 | 95340000 | 0.0037 | 0.0029 | 0.0033 |
| F03 | 95260001 | 95360000 | 0.0030 | 0.0022 | 0.0026 |
| F03 | 95280001 | 95380000 | 0.0030 | 0.0021 | 0.0024 |
| F03 | 95300001 | 95400000 | 0.0032 | 0.0022 | 0.0025 |
| F03 | 95320001 | 95420000 | 0.0029 | 0.0019 | 0.0023 |
| F03 | 95340001 | 95440000 | 0.0022 | 0.0012 | 0.0014 |
| F03 | 95360001 | 95460000 | 0.0024 | 0.0013 | 0.0016 |
| F03 | 95380001 | 95480000 | 0.0021 | 0.0011 | 0.0013 |
| F03 | 95400001 | 95500000 | 0.0029 | 0.0019 | 0.0021 |

| Chr | Start | End | both | A1 | A2 |
|---|---|---|---|---|---|
| F03 | 95420001 | 95520000 | 0.0028 | 0.0018 | 0.0020 |
| F03 | 95440001 | 95540000 | 0.0028 | 0.0018 | 0.0019 |
| F03 | 95460001 | 95560000 | 0.0026 | 0.0017 | 0.0018 |
| F03 | 95480001 | 95580000 | 0.0036 | 0.0027 | 0.0029 |
| F03 | 95500001 | 95600000 | 0.0033 | 0.0026 | 0.0026 |
| F03 | 95520001 | 95620000 | 0.0041 | 0.0034 | 0.0036 |
| F03 | 95540001 | 95640000 | 0.0040 | 0.0033 | 0.0035 |
| F03 | 95560001 | 95660000 | 0.0040 | 0.0033 | 0.0034 |
| F03 | 95580001 | 95680000 | 0.0036 | 0.0026 | 0.0029 |
| F03 | 95600001 | 95700000 | 0.0034 | 0.0024 | 0.0028 |
| F03 | 95620001 | 95720000 | 0.0027 | 0.0018 | 0.0021 |
| F03 | 95640001 | 95740000 | 0.0034 | 0.0023 | 0.0026 |
| F03 | 95660001 | 95760000 | 0.0034 | 0.0023 | 0.0025 |
| F03 | 95680001 | 95780000 | 0.0030 | 0.0018 | 0.0020 |
| F03 | 95700001 | 95800000 | 0.0027 | 0.0014 | 0.0016 |
| F03 | 95720001 | 95820000 | 0.0032 | 0.0019 | 0.0022 |
| F03 | 95740001 | 95840000 | 0.0026 | 0.0014 | 0.0017 |
| F03 | 95760001 | 95860000 | 0.0024 | 0.0013 | 0.0016 |
| F03 | 95780001 | 95880000 | 0.0026 | 0.0019 | 0.0020 |
| F03 | 95800001 | 95900000 | 0.0025 | 0.0018 | 0.0020 |
| F03 | 95820001 | 95920000 | 0.0021 | 0.0015 | 0.0018 |
| F03 | 95840001 | 95940000 | 0.0031 | 0.0026 | 0.0029 |
| F03 | 95860001 | 95960000 | 0.0039 | 0.0034 | 0.0038 |
| F03 | 95880001 | 95980000 | 0.0035 | 0.0029 | 0.0032 |
| F03 | 95900001 | 96000000 | 0.0042 | 0.0034 | 0.0039 |
| F03 | 95920001 | 96020000 | 0.0042 | 0.0031 | 0.0035 |
| F03 | 95940001 | 96040000 | 0.0034 | 0.0024 | 0.0028 |
| F03 | 95960001 | 96060000 | 0.0025 | 0.0016 | 0.0019 |
| F03 | 95980001 | 96080000 | 0.0021 | 0.0014 | 0.0017 |
| F03 | 96000001 | 96100000 | 0.0023 | 0.0013 | 0.0017 |
| F03 | 96020001 | 96120000 | 0.0022 | 0.0016 | 0.0019 |
| F03 | 96040001 | 96140000 | 0.0029 | 0.0021 | 0.0026 |
| F03 | 96060001 | 96160000 | 0.0029 | 0.0020 | 0.0024 |
| F03 | 96080001 | 96180000 | 0.0030 | 0.0021 | 0.0025 |
| F03 | 96100001 | 96200000 | 0.0022 | 0.0017 | 0.0018 |
| F03 | 96120001 | 96220000 | 0.0027 | 0.0018 | 0.0021 |
| F03 | 96140001 | 96240000 | 0.0017 | 0.0010 | 0.0012 |
| F03 | 96160001 | 96260000 | 0.0025 | 0.0017 | 0.0020 |
| F03 | 96180001 | 96280000 | 0.0025 | 0.0016 | 0.0019 |
| F03 | 96200001 | 96300000 | 0.0030 | 0.0017 | 0.0022 |
| F03 | 96220001 | 96320000 | 0.0027 | 0.0016 | 0.0019 |
| F03 | 96240001 | 96340000 | 0.0031 | 0.0019 | 0.0022 |
| F03 | 96260001 | 96360000 | 0.0030 | 0.0018 | 0.0022 |
| F03 | 96280001 | 96380000 | 0.0046 | 0.0030 | 0.0038 |
| F03 | 96300001 | 96400000 | 0.0043 | 0.0031 | 0.0037 |
| F03 | 96320001 | 96420000 | 0.0047 | 0.0036 | 0.0041 |
| F03 | 96340001 | 96440000 | 0.0048 | 0.0037 | 0.0043 |
| F03 | 96360001 | 96460000 | 0.0042 | 0.0031 | 0.0036 |
| F03 | 96380001 | 96480000 | 0.0028 | 0.0022 | 0.0022 |
| F03 | 96400001 | 96500000 | 0.0031 | 0.0024 | 0.0024 |
| F03 | 96420001 | 96520000 | 0.0031 | 0.0022 | 0.0022 |
| F03 | 96440001 | 96540000 | 0.0026 | 0.0016 | 0.0016 |
| F03 | 96460001 | 96560000 | 0.0033 | 0.0022 | 0.0024 |
| F03 | 96480001 | 96580000 | 0.0034 | 0.0024 | 0.0027 |
| F03 | 96500001 | 96600000 | 0.0030 | 0.0022 | 0.0025 |
| F03 | 96520001 | 96620000 | 0.0022 | 0.0017 | 0.0019 |
| F03 | 96540001 | 96640000 | 0.0027 | 0.0021 | 0.0024 |
| F03 | 96560001 | 96660000 | 0.0024 | 0.0018 | 0.0021 |

| Chr | Start | End | both | A1 | A2 |
|---|---|---|---|---|---|
| F03 | 96580001 | 96680000 | 0.0024 | 0.0018 | 0.0019 |
| F03 | 96600001 | 96700000 | 0.0027 | 0.0020 | 0.0020 |
| F03 | 96620001 | 96720000 | 0.0030 | 0.0022 | 0.0022 |
| F03 | 96640001 | 96740000 | 0.0037 | 0.0029 | 0.0029 |
| F03 | 96660001 | 96760000 | 0.0034 | 0.0026 | 0.0027 |
| F03 | 96680001 | 96780000 | 0.0039 | 0.0030 | 0.0031 |
| F03 | 96700001 | 96800000 | 0.0037 | 0.0029 | 0.0032 |
| F03 | 96720001 | 96820000 | 0.0034 | 0.0027 | 0.0029 |
| F03 | 96740001 | 96840000 | 0.0023 | 0.0017 | 0.0018 |
| F03 | 96760001 | 96860000 | 0.0023 | 0.0016 | 0.0017 |
| F03 | 96780001 | 96880000 | 0.0020 | 0.0013 | 0.0015 |
| F03 | 96800001 | 96900000 | 0.0023 | 0.0013 | 0.0014 |
| F03 | 96820001 | 96920000 | 0.0025 | 0.0013 | 0.0015 |
| F03 | 96840001 | 96940000 | 0.0030 | 0.0017 | 0.0019 |
| F03 | 96860001 | 96960000 | 0.0041 | 0.0027 | 0.0031 |
| F03 | 96880001 | 96980000 | 0.0038 | 0.0024 | 0.0027 |
| F03 | 96900001 | 97000000 | 0.0036 | 0.0024 | 0.0028 |
| F03 | 96920001 | 97020000 | 0.0040 | 0.0028 | 0.0030 |
| F03 | 96940001 | 97040000 | 0.0039 | 0.0026 | 0.0028 |
| F03 | 96960001 | 97060000 | 0.0032 | 0.0021 | 0.0020 |
| F03 | 96980001 | 97080000 | 0.0030 | 0.0019 | 0.0018 |
| F03 | 97000001 | 97100000 | 0.0025 | 0.0016 | 0.0015 |
| F03 | 97020001 | 97120000 | 0.0020 | 0.0011 | 0.0012 |
| F03 | 97040001 | 97140000 | 0.0017 | 0.0009 | 0.0010 |
| F03 | 97060001 | 97160000 | 0.0012 | 0.0005 | 0.0006 |
| F03 | 97080001 | 97180000 | 0.0013 | 0.0007 | 0.0007 |
| F03 | 97100001 | 97200000 | 0.0016 | 0.0008 | 0.0009 |
| F03 | 97120001 | 97220000 | 0.0020 | 0.0010 | 0.0012 |
| F03 | 97140001 | 97240000 | 0.0021 | 0.0012 | 0.0013 |
| F03 | 97160001 | 97260000 | 0.0024 | 0.0015 | 0.0015 |
| F03 | 97180001 | 97280000 | 0.0037 | 0.0028 | 0.0027 |
| F03 | 97200001 | 97300000 | 0.0037 | 0.0029 | 0.0027 |
| F03 | 97220001 | 97320000 | 0.0050 | 0.0042 | 0.0041 |
| F03 | 97240001 | 97340000 | 0.0052 | 0.0045 | 0.0044 |
| F03 | 97260001 | 97360000 | 0.0051 | 0.0043 | 0.0043 |
| F03 | 97280001 | 97380000 | 0.0045 | 0.0035 | 0.0038 |
| F03 | 97300001 | 97400000 | 0.0068 | 0.0056 | 0.0062 |
| F03 | 97320001 | 97420000 | 0.0058 | 0.0046 | 0.0051 |
| F03 | 97340001 | 97440000 | 0.0055 | 0.0044 | 0.0049 |
| F03 | 97360001 | 97460000 | 0.0055 | 0.0045 | 0.0050 |
| F03 | 97380001 | 97480000 | 0.0050 | 0.0042 | 0.0046 |
| F03 | 97400001 | 97500000 | 0.0028 | 0.0024 | 0.0026 |
| F03 | 97420001 | 97520000 | 0.0029 | 0.0025 | 0.0028 |
| F03 | 97440001 | 97540000 | 0.0027 | 0.0024 | 0.0027 |
| F03 | 97460001 | 97560000 | 0.0030 | 0.0027 | 0.0030 |
| F03 | 97480001 | 97580000 | 0.0026 | 0.0023 | 0.0026 |
| F03 | 97500001 | 97600000 | 0.0022 | 0.0019 | 0.0021 |
| F03 | 97520001 | 97620000 | 0.0016 | 0.0014 | 0.0016 |
| F03 | 97540001 | 97640000 | 0.0027 | 0.0024 | 0.0026 |
| F03 | 97560001 | 97660000 | 0.0030 | 0.0026 | 0.0028 |
| F03 | 97580001 | 97680000 | 0.0035 | 0.0029 | 0.0031 |
| F03 | 97600001 | 97700000 | 0.0040 | 0.0030 | 0.0033 |
| F03 | 97620001 | 97720000 | 0.0039 | 0.0028 | 0.0031 |
| F03 | 97640001 | 97740000 | 0.0039 | 0.0027 | 0.0031 |
| F03 | 97660001 | 97760000 | 0.0036 | 0.0023 | 0.0028 |
| F03 | 97680001 | 97780000 | 0.0032 | 0.0021 | 0.0026 |
| F03 | 97700001 | 97800000 | 0.0036 | 0.0025 | 0.0029 |
| F03 | 97720001 | 97820000 | 0.0038 | 0.0027 | 0.0032 |

| Chr | Start | End | both | A1 | A2 |
|---|---|---|---|---|---|
| F03 | 97740001 | 97840000 | 0.0032 | 0.0024 | 0.0027 |
| F03 | 97760001 | 97860000 | 0.0032 | 0.0024 | 0.0028 |
| F03 | 97780001 | 97880000 | 0.0035 | 0.0025 | 0.0029 |
| F03 | 97800001 | 97900000 | 0.0030 | 0.0022 | 0.0026 |
| F03 | 97820001 | 97920000 | 0.0032 | 0.0024 | 0.0028 |
| F03 | 97840001 | 97940000 | 0.0033 | 0.0022 | 0.0026 |
| F03 | 97860001 | 97960000 | 0.0036 | 0.0023 | 0.0028 |
| F03 | 97880001 | 97980000 | 0.0044 | 0.0032 | 0.0038 |
| F03 | 97900001 | 98000000 | 0.0045 | 0.0034 | 0.0039 |
| F03 | 97920001 | 98020000 | 0.0041 | 0.0032 | 0.0036 |
| F03 | 97940001 | 98040000 | 0.0039 | 0.0033 | 0.0037 |
| F03 | 97960001 | 98060000 | 0.0035 | 0.0032 | 0.0034 |
| F03 | 97980001 | 98080000 | 0.0022 | 0.0020 | 0.0021 |
| F03 | 98000001 | 98100000 | 0.0025 | 0.0022 | 0.0024 |
| F03 | 98020001 | 98120000 | 0.0026 | 0.0023 | 0.0025 |
| F03 | 98040001 | 98140000 | 0.0021 | 0.0018 | 0.0020 |
| F03 | 98060001 | 98160000 | 0.0025 | 0.0020 | 0.0022 |
| F03 | 98080001 | 98180000 | 0.0027 | 0.0020 | 0.0022 |
| F03 | 98100001 | 98200000 | 0.0018 | 0.0011 | 0.0012 |
| F03 | 98120001 | 98220000 | 0.0017 | 0.0010 | 0.0010 |
| F03 | 98140001 | 98240000 | 0.0026 | 0.0015 | 0.0018 |
| F03 | 98160001 | 98260000 | 0.0033 | 0.0023 | 0.0027 |
| F03 | 98180001 | 98280000 | 0.0043 | 0.0034 | 0.0039 |
| F03 | 98200001 | 98300000 | 0.0054 | 0.0043 | 0.0049 |
| F03 | 98220001 | 98320000 | 0.0060 | 0.0047 | 0.0054 |
| F03 | 98240001 | 98340000 | 0.0054 | 0.0042 | 0.0047 |
| F03 | 98260001 | 98360000 | 0.0049 | 0.0039 | 0.0043 |
| F03 | 98280001 | 98380000 | 0.0043 | 0.0031 | 0.0035 |
| F03 | 98300001 | 98400000 | 0.0040 | 0.0029 | 0.0032 |
| F03 | 98320001 | 98420000 | 0.0040 | 0.0029 | 0.0034 |
| F03 | 98340001 | 98440000 | 0.0045 | 0.0036 | 0.0041 |
| F03 | 98360001 | 98460000 | 0.0039 | 0.0029 | 0.0034 |
| F03 | 98380001 | 98480000 | 0.0043 | 0.0034 | 0.0040 |
| F03 | 98400001 | 98500000 | 0.0039 | 0.0032 | 0.0037 |
| F03 | 98420001 | 98520000 | 0.0044 | 0.0036 | 0.0042 |
| F03 | 98440001 | 98540000 | 0.0036 | 0.0028 | 0.0034 |
| F03 | 98460001 | 98560000 | 0.0039 | 0.0031 | 0.0037 |
| F03 | 98480001 | 98580000 | 0.0034 | 0.0026 | 0.0031 |
| F03 | 98500001 | 98600000 | 0.0036 | 0.0025 | 0.0032 |
| F03 | 98520001 | 98620000 | 0.0024 | 0.0017 | 0.0020 |
| F03 | 98540001 | 98640000 | 0.0032 | 0.0024 | 0.0027 |
| F03 | 98560001 | 98660000 | 0.0029 | 0.0022 | 0.0025 |
| F03 | 98580001 | 98680000 | 0.0028 | 0.0020 | 0.0022 |
| F03 | 98600001 | 98700000 | 0.0023 | 0.0016 | 0.0017 |
| F03 | 98620001 | 98720000 | 0.0025 | 0.0016 | 0.0019 |
| F03 | 98640001 | 98740000 | 0.0019 | 0.0009 | 0.0011 |
| F03 | 98660001 | 98760000 | 0.0020 | 0.0009 | 0.0010 |
| F03 | 98680001 | 98780000 | 0.0020 | 0.0008 | 0.0010 |
| F03 | 98700001 | 98800000 | 0.0021 | 0.0009 | 0.0010 |
| F03 | 98720001 | 98820000 | 0.0028 | 0.0016 | 0.0018 |
| F03 | 98740001 | 98840000 | 0.0033 | 0.0023 | 0.0024 |
| F03 | 98760001 | 98860000 | 0.0030 | 0.0020 | 0.0022 |
| F03 | 98780001 | 98880000 | 0.0039 | 0.0029 | 0.0033 |
| F03 | 98800001 | 98900000 | 0.0044 | 0.0034 | 0.0039 |
| F03 | 98820001 | 98920000 | 0.0035 | 0.0025 | 0.0028 |
| F03 | 98840001 | 98940000 | 0.0036 | 0.0026 | 0.0030 |
| F03 | 98860001 | 98960000 | 0.0039 | 0.0030 | 0.0034 |
| F03 | 98880001 | 98980000 | 0.0033 | 0.0025 | 0.0025 |

| Chr | Start | End | both | A1 | A2 |
|---|---|---|---|---|---|
| F03 | 98900001 | 99000000 | 0.0034 | 0.0026 | 0.0027 |
| F03 | 98920001 | 99020000 | 0.0038 | 0.0029 | 0.0032 |
| F03 | 98940001 | 99040000 | 0.0041 | 0.0032 | 0.0035 |
| F03 | 98960001 | 99060000 | 0.0038 | 0.0028 | 0.0031 |
| F03 | 98980001 | 99080000 | 0.0034 | 0.0025 | 0.0028 |
| F03 | 99000001 | 99100000 | 0.0030 | 0.0021 | 0.0023 |
| F03 | 99020001 | 99120000 | 0.0029 | 0.0022 | 0.0023 |
| F03 | 99040001 | 99140000 | 0.0019 | 0.0012 | 0.0013 |
| F03 | 99060001 | 99160000 | 0.0022 | 0.0016 | 0.0016 |
| F03 | 99080001 | 99180000 | 0.0029 | 0.0022 | 0.0023 |
| F03 | 99100001 | 99200000 | 0.0031 | 0.0023 | 0.0024 |
| F03 | 99120001 | 99220000 | 0.0028 | 0.0019 | 0.0020 |
| F03 | 99140001 | 99240000 | 0.0031 | 0.0023 | 0.0024 |
| F03 | 99160001 | 99260000 | 0.0033 | 0.0024 | 0.0026 |
| F03 | 99180001 | 99280000 | 0.0028 | 0.0018 | 0.0020 |
| F03 | 99200001 | 99300000 | 0.0026 | 0.0016 | 0.0018 |
| F03 | 99220001 | 99320000 | 0.0027 | 0.0017 | 0.0019 |
| F03 | 99240001 | 99340000 | 0.0027 | 0.0017 | 0.0020 |
| F03 | 99260001 | 99360000 | 0.0023 | 0.0014 | 0.0016 |
| F03 | 99280001 | 99380000 | 0.0029 | 0.0021 | 0.0023 |
| F03 | 99300001 | 99400000 | 0.0034 | 0.0025 | 0.0027 |
| F03 | 99320001 | 99420000 | 0.0031 | 0.0023 | 0.0025 |
| F03 | 99340001 | 99440000 | 0.0031 | 0.0023 | 0.0025 |
| F03 | 99360001 | 99460000 | 0.0034 | 0.0025 | 0.0028 |
| F03 | 99380001 | 99480000 | 0.0029 | 0.0019 | 0.0023 |
| F03 | 99400001 | 99500000 | 0.0029 | 0.0019 | 0.0025 |
| F03 | 99420001 | 99520000 | 0.0030 | 0.0021 | 0.0027 |
| F03 | 99440001 | 99540000 | 0.0030 | 0.0018 | 0.0025 |
| F03 | 99460001 | 99560000 | 0.0029 | 0.0016 | 0.0022 |
| F03 | 99480001 | 99580000 | 0.0024 | 0.0013 | 0.0019 |
| F03 | 99500001 | 99600000 | 0.0024 | 0.0012 | 0.0017 |
| F03 | 99520001 | 99620000 | 0.0028 | 0.0014 | 0.0019 |
| F03 | 99540001 | 99640000 | 0.0027 | 0.0012 | 0.0017 |
| F03 | 99560001 | 99660000 | 0.0026 | 0.0013 | 0.0017 |
| F03 | 99580001 | 99680000 | 0.0030 | 0.0013 | 0.0018 |
| F03 | 99600001 | 99700000 | 0.0025 | 0.0009 | 0.0013 |
| F03 | 99620001 | 99720000 | 0.0020 | 0.0006 | 0.0011 |
| F03 | 99640001 | 99740000 | 0.0024 | 0.0012 | 0.0017 |
| F03 | 99660001 | 99760000 | 0.0027 | 0.0016 | 0.0021 |
| F03 | 99680001 | 99780000 | 0.0030 | 0.0021 | 0.0025 |
| F03 | 99700001 | 99800000 | 0.0031 | 0.0022 | 0.0026 |
| F03 | 99720001 | 99820000 | 0.0032 | 0.0021 | 0.0025 |
| F03 | 99740001 | 99840000 | 0.0026 | 0.0016 | 0.0019 |
| F03 | 99760001 | 99860000 | 0.0023 | 0.0014 | 0.0016 |
| F03 | 99780001 | 99880000 | 0.0017 | 0.0010 | 0.0011 |
| F03 | 99800001 | 99900000 | 0.0017 | 0.0009 | 0.0011 |
| F03 | 99820001 | 99920000 | 0.0014 | 0.0009 | 0.0010 |
| F03 | 99840001 | 99940000 | 0.0017 | 0.0013 | 0.0014 |
| F03 | 99860001 | 99960000 | 0.0016 | 0.0011 | 0.0013 |
| F03 | 99880001 | 99980000 | 0.0023 | 0.0016 | 0.0018 |
| F03 | 99900001 | 1E+08 | 0.0027 | 0.0018 | 0.0019 |
| F03 | 99920001 | 1E+08 | 0.0028 | 0.0018 | 0.0019 |
| F03 | 99940001 | 1E+08 | 0.0024 | 0.0013 | 0.0015 |
| F03 | 99960001 | 1E+08 | 0.0026 | 0.0015 | 0.0017 |
| F03 | 99980001 | 1E+08 | 0.0028 | 0.0016 | 0.0017 |
| F03 | 100000001 | 1E+08 | 0.0029 | 0.0019 | 0.0020 |
| F03 | 100020001 | 1E+08 | 0.0031 | 0.0020 | 0.0022 |
| F03 | 100040001 | 1E+08 | 0.0040 | 0.0027 | 0.0030 |

| Chr | Start | End | both | A1 | A2 |
|---|---|---|---|---|---|
| F03 | 100060001 | 1E+08 | 0.0044 | 0.0031 | 0.0035 |
| F03 | 100080001 | 1E+08 | 0.0046 | 0.0034 | 0.0039 |
| F03 | 100100001 | 1E+08 | 0.0046 | 0.0033 | 0.0036 |
| F03 | 100120001 | 1E+08 | 0.0048 | 0.0035 | 0.0038 |
| F03 | 100140001 | 1E+08 | 0.0043 | 0.0031 | 0.0032 |
| F03 | 100160001 | 1E+08 | 0.0045 | 0.0031 | 0.0032 |
| F03 | 100180001 | 1E+08 | 0.0042 | 0.0029 | 0.0031 |
| F03 | 100200001 | 1E+08 | 0.0038 | 0.0027 | 0.0031 |
| F03 | 100220001 | 1E+08 | 0.0033 | 0.0024 | 0.0029 |
| F03 | 100240001 | 1E+08 | 0.0039 | 0.0029 | 0.0035 |
| F03 | 100260001 | 1E+08 | 0.0029 | 0.0023 | 0.0028 |
| F03 | 100280001 | 1E+08 | 0.0020 | 0.0016 | 0.0020 |
| F03 | 100300001 | 1E+08 | 0.0019 | 0.0016 | 0.0019 |
| F03 | 100320001 | 1E+08 | 0.0024 | 0.0020 | 0.0023 |
| F03 | 100340001 | 1E+08 | 0.0015 | 0.0012 | 0.0014 |
| F03 | 100360001 | 1E+08 | 0.0030 | 0.0025 | 0.0029 |
| F03 | 100380001 | 1E+08 | 0.0040 | 0.0033 | 0.0037 |
| F03 | 100400001 | 1.01E+08 | 0.0038 | 0.0031 | 0.0036 |
| F03 | 100420001 | 1.01E+08 | 0.0039 | 0.0030 | 0.0034 |
| F03 | 100440001 | 1.01E+08 | 0.0042 | 0.0033 | 0.0037 |
| F03 | 100460001 | 1.01E+08 | 0.0033 | 0.0024 | 0.0026 |
| F03 | 100480001 | 1.01E+08 | 0.0029 | 0.0018 | 0.0021 |
| F03 | 100500001 | 1.01E+08 | 0.0035 | 0.0023 | 0.0026 |
| F03 | 100520001 | 1.01E+08 | 0.0030 | 0.0020 | 0.0023 |
| F03 | 100540001 | 1.01E+08 | 0.0038 | 0.0026 | 0.0031 |
| F03 | 100560001 | 1.01E+08 | 0.0057 | 0.0045 | 0.0049 |
| F03 | 100580001 | 1.01E+08 | 0.0058 | 0.0050 | 0.0051 |
| F03 | 100600001 | 1.01E+08 | 0.0055 | 0.0046 | 0.0047 |
| F03 | 100620001 | 1.01E+08 | 0.0069 | 0.0054 | 0.0059 |
| F03 | 100640001 | 1.01E+08 | 0.0066 | 0.0050 | 0.0054 |
| F03 | 100660001 | 1.01E+08 | 0.0046 | 0.0033 | 0.0037 |
| F03 | 100680001 | 1.01E+08 | 0.0045 | 0.0030 | 0.0036 |
| F03 | 100700001 | 1.01E+08 | 0.0043 | 0.0031 | 0.0037 |
| F03 | 100720001 | 1.01E+08 | 0.0041 | 0.0033 | 0.0036 |
| F03 | 100740001 | 1.01E+08 | 0.0035 | 0.0029 | 0.0030 |
| F03 | 100760001 | 1.01E+08 | 0.0038 | 0.0032 | 0.0034 |
| F03 | 100780001 | 1.01E+08 | 0.0037 | 0.0030 | 0.0032 |
| F03 | 100800001 | 1.01E+08 | 0.0037 | 0.0030 | 0.0033 |
| F03 | 100820001 | 1.01E+08 | 0.0027 | 0.0020 | 0.0023 |
| F03 | 100840001 | 1.01E+08 | 0.0028 | 0.0021 | 0.0024 |
| F03 | 100860001 | 1.01E+08 | 0.0021 | 0.0014 | 0.0016 |
| F03 | 100880001 | 1.01E+08 | 0.0030 | 0.0020 | 0.0021 |
| F03 | 100900001 | 1.01E+08 | 0.0028 | 0.0016 | 0.0017 |
| F03 | 100920001 | 1.01E+08 | 0.0030 | 0.0019 | 0.0019 |
| F03 | 100940001 | 1.01E+08 | 0.0038 | 0.0026 | 0.0025 |
| F03 | 100960001 | 1.01E+08 | 0.0039 | 0.0026 | 0.0026 |
| F03 | 100980001 | 1.01E+08 | 0.0030 | 0.0019 | 0.0019 |
| F03 | 101000001 | 1.01E+08 | 0.0031 | 0.0020 | 0.0021 |
| F03 | 101020001 | 1.01E+08 | 0.0029 | 0.0016 | 0.0017 |
| F03 | 101040001 | 1.01E+08 | 0.0021 | 0.0010 | 0.0012 |
| F03 | 101060001 | 1.01E+08 | 0.0018 | 0.0009 | 0.0011 |
| F03 | 101080001 | 1.01E+08 | 0.0022 | 0.0013 | 0.0015 |
| F03 | 101100001 | 1.01E+08 | 0.0021 | 0.0011 | 0.0014 |
| F03 | 101120001 | 1.01E+08 | 0.0019 | 0.0010 | 0.0013 |
| F03 | 101140001 | 1.01E+08 | 0.0025 | 0.0015 | 0.0019 |
| F03 | 101160001 | 1.01E+08 | 0.0031 | 0.0021 | 0.0026 |
| F03 | 101180001 | 1.01E+08 | 0.0032 | 0.0022 | 0.0026 |
| F03 | 101200001 | 1.01E+08 | 0.0037 | 0.0027 | 0.0031 |

| Chr | Start | End | both | A1 | A2 |
|---|---|---|---|---|---|
| F03 | 101220001 | 1.01E+08 | 0.0038 | 0.0029 | 0.0033 |
| F03 | 101240001 | 1.01E+08 | 0.0029 | 0.0021 | 0.0024 |
| F03 | 101260001 | 1.01E+08 | 0.0024 | 0.0016 | 0.0018 |
| F03 | 101280001 | 1.01E+08 | 0.0019 | 0.0011 | 0.0012 |
| F03 | 101300001 | 1.01E+08 | 0.0016 | 0.0009 | 0.0009 |
| F03 | 101320001 | 1.01E+08 | 0.0021 | 0.0013 | 0.0014 |
| F03 | 101340001 | 1.01E+08 | 0.0028 | 0.0019 | 0.0020 |
| F03 | 101360001 | 1.01E+08 | 0.0029 | 0.0020 | 0.0021 |
| F03 | 101380001 | 1.01E+08 | 0.0028 | 0.0018 | 0.0020 |
| F03 | 101400001 | 1.02E+08 | 0.0032 | 0.0020 | 0.0022 |
| F03 | 101420001 | 1.02E+08 | 0.0029 | 0.0018 | 0.0019 |
| F03 | 101440001 | 1.02E+08 | 0.0021 | 0.0012 | 0.0013 |
| F03 | 101460001 | 1.02E+08 | 0.0019 | 0.0011 | 0.0012 |
| F03 | 101480001 | 1.02E+08 | 0.0019 | 0.0011 | 0.0012 |
| F03 | 101500001 | 1.02E+08 | 0.0014 | 0.0008 | 0.0008 |
| F03 | 101520001 | 1.02E+08 | 0.0011 | 0.0004 | 0.0004 |
| F03 | 101540001 | 1.02E+08 | 0.0022 | 0.0014 | 0.0014 |
| F03 | 101560001 | 1.02E+08 | 0.0030 | 0.0021 | 0.0021 |
| F03 | 101580001 | 1.02E+08 | 0.0038 | 0.0030 | 0.0030 |
| F03 | 101600001 | 1.02E+08 | 0.0041 | 0.0035 | 0.0035 |
| F03 | 101620001 | 1.02E+08 | 0.0053 | 0.0048 | 0.0048 |
| F03 | 101640001 | 1.02E+08 | 0.0044 | 0.0039 | 0.0039 |
| F03 | 101660001 | 1.02E+08 | 0.0041 | 0.0036 | 0.0037 |
| F03 | 101680001 | 1.02E+08 | 0.0042 | 0.0035 | 0.0038 |
| F03 | 101700001 | 1.02E+08 | 0.0037 | 0.0029 | 0.0032 |
| F03 | 101720001 | 1.02E+08 | 0.0031 | 0.0023 | 0.0026 |
| F03 | 101740001 | 1.02E+08 | 0.0032 | 0.0024 | 0.0028 |
| F03 | 101760001 | 1.02E+08 | 0.0028 | 0.0022 | 0.0024 |
| F03 | 101780001 | 1.02E+08 | 0.0021 | 0.0014 | 0.0015 |
| F03 | 101800001 | 1.02E+08 | 0.0032 | 0.0025 | 0.0025 |
| F03 | 101820001 | 1.02E+08 | 0.0027 | 0.0021 | 0.0019 |
| F03 | 101840001 | 1.02E+08 | 0.0025 | 0.0021 | 0.0018 |
| F03 | 101860001 | 1.02E+08 | 0.0025 | 0.0020 | 0.0018 |
| F03 | 101880001 | 1.02E+08 | 0.0027 | 0.0021 | 0.0018 |
| F03 | 101900001 | 1.02E+08 | 0.0021 | 0.0014 | 0.0014 |
| F03 | 101920001 | 1.02E+08 | 0.0022 | 0.0014 | 0.0014 |
| F03 | 101940001 | 1.02E+08 | 0.0035 | 0.0025 | 0.0026 |
| F03 | 101960001 | 1.02E+08 | 0.0049 | 0.0038 | 0.0039 |
| F03 | 101980001 | 1.02E+08 | 0.0053 | 0.0040 | 0.0044 |
| F03 | 102000001 | 1.02E+08 | 0.0053 | 0.0041 | 0.0045 |
| F03 | 102020001 | 1.02E+08 | 0.0053 | 0.0041 | 0.0045 |
| F03 | 102040001 | 1.02E+08 | 0.0042 | 0.0032 | 0.0036 |
| F03 | 102060001 | 1.02E+08 | 0.0031 | 0.0022 | 0.0026 |
| F03 | 102080001 | 1.02E+08 | 0.0029 | 0.0023 | 0.0025 |
| F03 | 102100001 | 1.02E+08 | 0.0025 | 0.0020 | 0.0021 |
| F03 | 102120001 | 1.02E+08 | 0.0030 | 0.0026 | 0.0026 |
| F03 | 102140001 | 1.02E+08 | 0.0029 | 0.0023 | 0.0023 |
| F03 | 102160001 | 1.02E+08 | 0.0027 | 0.0018 | 0.0020 |
| F03 | 102180001 | 1.02E+08 | 0.0042 | 0.0032 | 0.0033 |
| F03 | 102200001 | 1.02E+08 | 0.0039 | 0.0029 | 0.0030 |
| F03 | 102220001 | 1.02E+08 | 0.0036 | 0.0024 | 0.0026 |
| F03 | 102240001 | 1.02E+08 | 0.0039 | 0.0026 | 0.0028 |
| F03 | 102260001 | 1.02E+08 | 0.0043 | 0.0031 | 0.0032 |
| F03 | 102280001 | 1.02E+08 | 0.0029 | 0.0018 | 0.0019 |
| F03 | 102300001 | 1.02E+08 | 0.0031 | 0.0019 | 0.0021 |
| F03 | 102320001 | 1.02E+08 | 0.0048 | 0.0037 | 0.0038 |
| F03 | 102340001 | 1.02E+08 | 0.0044 | 0.0035 | 0.0035 |
| F03 | 102360001 | 1.02E+08 | 0.0048 | 0.0039 | 0.0039 |

| Chr | Start | End | both | A1 | A2 |
|---|---|---|---|---|---|
| F03 | 102380001 | 1.02E+08 | 0.0049 | 0.0040 | 0.0040 |
| F03 | 102400001 | 1.03E+08 | 0.0053 | 0.0043 | 0.0045 |
| F03 | 102420001 | 1.03E+08 | 0.0032 | 0.0024 | 0.0025 |
| F03 | 102440001 | 1.03E+08 | 0.0034 | 0.0025 | 0.0026 |
| F03 | 102460001 | 1.03E+08 | 0.0039 | 0.0030 | 0.0030 |
| F03 | 102480001 | 1.03E+08 | 0.0044 | 0.0034 | 0.0037 |
| F03 | 102500001 | 1.03E+08 | 0.0039 | 0.0031 | 0.0033 |
| F03 | 102520001 | 1.03E+08 | 0.0047 | 0.0036 | 0.0040 |
| F03 | 102540001 | 1.03E+08 | 0.0052 | 0.0040 | 0.0045 |
| F03 | 102560001 | 1.03E+08 | 0.0038 | 0.0028 | 0.0032 |
| F03 | 102580001 | 1.03E+08 | 0.0030 | 0.0022 | 0.0024 |
| F03 | 102600001 | 1.03E+08 | 0.0034 | 0.0025 | 0.0028 |
| F03 | 102620001 | 1.03E+08 | 0.0027 | 0.0019 | 0.0020 |
| F03 | 102640001 | 1.03E+08 | 0.0026 | 0.0018 | 0.0020 |
| F03 | 102660001 | 1.03E+08 | 0.0032 | 0.0023 | 0.0026 |
| F03 | 102680001 | 1.03E+08 | 0.0033 | 0.0024 | 0.0027 |
| F03 | 102700001 | 1.03E+08 | 0.0049 | 0.0038 | 0.0042 |
| F03 | 102720001 | 1.03E+08 | 0.0047 | 0.0037 | 0.0041 |
| F03 | 102740001 | 1.03E+08 | 0.0044 | 0.0035 | 0.0039 |
| F03 | 102760001 | 1.03E+08 | 0.0042 | 0.0034 | 0.0036 |
| F03 | 102780001 | 1.03E+08 | 0.0036 | 0.0028 | 0.0030 |
| F03 | 102800001 | 1.03E+08 | 0.0019 | 0.0012 | 0.0012 |
| F03 | 102820001 | 1.03E+08 | 0.0019 | 0.0011 | 0.0012 |
| F03 | 102840001 | 1.03E+08 | 0.0018 | 0.0010 | 0.0011 |
| F03 | 102860001 | 1.03E+08 | 0.0016 | 0.0008 | 0.0009 |
| F03 | 102880001 | 1.03E+08 | 0.0015 | 0.0007 | 0.0008 |
| F03 | 102900001 | 1.03E+08 | 0.0017 | 0.0009 | 0.0010 |
| F03 | 102920001 | 1.03E+08 | 0.0017 | 0.0010 | 0.0011 |
| F03 | 102940001 | 1.03E+08 | 0.0016 | 0.0008 | 0.0009 |
| F03 | 102960001 | 1.03E+08 | 0.0017 | 0.0008 | 0.0009 |
| F03 | 102980001 | 1.03E+08 | 0.0019 | 0.0008 | 0.0009 |
| F03 | 103000001 | 1.03E+08 | 0.0018 | 0.0006 | 0.0008 |
| F03 | 103020001 | 1.03E+08 | 0.0017 | 0.0006 | 0.0008 |
| F03 | 103040001 | 1.03E+08 | 0.0018 | 0.0006 | 0.0008 |
| F03 | 103060001 | 1.03E+08 | 0.0016 | 0.0004 | 0.0007 |
| F03 | 103080001 | 1.03E+08 | 0.0022 | 0.0010 | 0.0015 |
| F03 | 103100001 | 1.03E+08 | 0.0023 | 0.0010 | 0.0016 |
| F03 | 103120001 | 1.03E+08 | 0.0029 | 0.0015 | 0.0022 |
| F03 | 103140001 | 1.03E+08 | 0.0035 | 0.0020 | 0.0029 |
| F03 | 103160001 | 1.03E+08 | 0.0041 | 0.0026 | 0.0036 |
| F03 | 103180001 | 1.03E+08 | 0.0037 | 0.0022 | 0.0033 |
| F03 | 103200001 | 1.03E+08 | 0.0037 | 0.0022 | 0.0033 |
| F03 | 103220001 | 1.03E+08 | 0.0034 | 0.0020 | 0.0031 |
| F03 | 103240001 | 1.03E+08 | 0.0028 | 0.0014 | 0.0024 |
| F03 | 103260001 | 1.03E+08 | 0.0026 | 0.0014 | 0.0023 |
| F03 | 103280001 | 1.03E+08 | 0.0024 | 0.0012 | 0.0021 |
| F03 | 103300001 | 1.03E+08 | 0.0026 | 0.0014 | 0.0023 |
| F03 | 103320001 | 1.03E+08 | 0.0027 | 0.0016 | 0.0024 |
| F03 | 103340001 | 1.03E+08 | 0.0029 | 0.0019 | 0.0027 |
| F03 | 103360001 | 1.03E+08 | 0.0031 | 0.0021 | 0.0029 |
| F03 | 103380001 | 1.03E+08 | 0.0040 | 0.0030 | 0.0037 |
| F03 | 103400001 | 1.04E+08 | 0.0035 | 0.0026 | 0.0032 |
| F03 | 103420001 | 1.04E+08 | 0.0038 | 0.0026 | 0.0033 |
| F03 | 103440001 | 1.04E+08 | 0.0040 | 0.0028 | 0.0035 |
| F03 | 103460001 | 1.04E+08 | 0.0036 | 0.0023 | 0.0030 |
| F03 | 103480001 | 1.04E+08 | 0.0027 | 0.0015 | 0.0021 |
| F03 | 103500001 | 1.04E+08 | 0.0025 | 0.0013 | 0.0019 |
| F03 | 103520001 | 1.04E+08 | 0.0022 | 0.0014 | 0.0017 |

| Chr | Start | End | both | A1 | A2 |
|---|---|---|---|---|---|
| F03 | 103540001 | 1.04E+08 | 0.0020 | 0.0012 | 0.0016 |
| F03 | 103560001 | 1.04E+08 | 0.0019 | 0.0013 | 0.0016 |
| F03 | 103580001 | 1.04E+08 | 0.0018 | 0.0013 | 0.0016 |
| F03 | 103600001 | 1.04E+08 | 0.0020 | 0.0016 | 0.0018 |
| F03 | 103620001 | 1.04E+08 | 0.0021 | 0.0018 | 0.0020 |
| F03 | 103640001 | 1.04E+08 | 0.0021 | 0.0017 | 0.0018 |
| F03 | 103660001 | 1.04E+08 | 0.0023 | 0.0019 | 0.0020 |
| F03 | 103680001 | 1.04E+08 | 0.0028 | 0.0023 | 0.0025 |
| F03 | 103700001 | 1.04E+08 | 0.0034 | 0.0028 | 0.0030 |
| F03 | 103720001 | 1.04E+08 | 0.0029 | 0.0022 | 0.0025 |
| F03 | 103740001 | 1.04E+08 | 0.0027 | 0.0021 | 0.0024 |
| F03 | 103760001 | 1.04E+08 | 0.0023 | 0.0017 | 0.0018 |
| F03 | 103780001 | 1.04E+08 | 0.0020 | 0.0012 | 0.0013 |
| F03 | 103800001 | 1.04E+08 | 0.0018 | 0.0010 | 0.0011 |
| F03 | 103820001 | 1.04E+08 | 0.0022 | 0.0014 | 0.0014 |
| F03 | 103840001 | 1.04E+08 | 0.0022 | 0.0013 | 0.0012 |
| F03 | 103860001 | 1.04E+08 | 0.0023 | 0.0011 | 0.0011 |
| F03 | 103880001 | 1.04E+08 | 0.0025 | 0.0014 | 0.0015 |
| F03 | 103900001 | 1.04E+08 | 0.0028 | 0.0016 | 0.0017 |
| F03 | 103920001 | 1.04E+08 | 0.0025 | 0.0014 | 0.0015 |
| F03 | 103940001 | 1.04E+08 | 0.0028 | 0.0019 | 0.0020 |
| F03 | 103960001 | 1.04E+08 | 0.0030 | 0.0023 | 0.0023 |
| F03 | 103980001 | 1.04E+08 | 0.0029 | 0.0023 | 0.0023 |
| F03 | 104000001 | 1.04E+08 | 0.0030 | 0.0023 | 0.0023 |
| F03 | 104020001 | 1.04E+08 | 0.0032 | 0.0024 | 0.0025 |
| F03 | 104040001 | 1.04E+08 | 0.0031 | 0.0021 | 0.0023 |
| F03 | 104060001 | 1.04E+08 | 0.0032 | 0.0023 | 0.0025 |
| F03 | 104080001 | 1.04E+08 | 0.0028 | 0.0021 | 0.0023 |
| F03 | 104100001 | 1.04E+08 | 0.0020 | 0.0015 | 0.0016 |
| F03 | 104120001 | 1.04E+08 | 0.0018 | 0.0014 | 0.0014 |
| F03 | 104140001 | 1.04E+08 | 0.0031 | 0.0025 | 0.0029 |
| F03 | 104160001 | 1.04E+08 | 0.0036 | 0.0030 | 0.0033 |
| F03 | 104180001 | 1.04E+08 | 0.0038 | 0.0031 | 0.0035 |
| F03 | 104200001 | 1.04E+08 | 0.0054 | 0.0046 | 0.0051 |
| F03 | 104220001 | 1.04E+08 | 0.0058 | 0.0051 | 0.0055 |
| F03 | 104240001 | 1.04E+08 | 0.0051 | 0.0047 | 0.0048 |
| F03 | 104260001 | 1.04E+08 | 0.0045 | 0.0042 | 0.0044 |
| F03 | 104280001 | 1.04E+08 | 0.0045 | 0.0043 | 0.0044 |
| F03 | 104300001 | 1.04E+08 | 0.0032 | 0.0030 | 0.0031 |
| F03 | 104320001 | 1.04E+08 | 0.0032 | 0.0030 | 0.0031 |
| F03 | 104340001 | 1.04E+08 | 0.0024 | 0.0023 | 0.0022 |
| F03 | 104360001 | 1.04E+08 | 0.0021 | 0.0017 | 0.0018 |
| F03 | 104380001 | 1.04E+08 | 0.0027 | 0.0023 | 0.0023 |
| F03 | 104400001 | 1.05E+08 | 0.0027 | 0.0022 | 0.0023 |
| F03 | 104420001 | 1.05E+08 | 0.0025 | 0.0020 | 0.0021 |
| F03 | 104440001 | 1.05E+08 | 0.0026 | 0.0021 | 0.0022 |
| F03 | 104460001 | 1.05E+08 | 0.0026 | 0.0021 | 0.0023 |
| F03 | 104480001 | 1.05E+08 | 0.0021 | 0.0016 | 0.0019 |
| F03 | 104500001 | 1.05E+08 | 0.0021 | 0.0016 | 0.0019 |
| F03 | 104520001 | 1.05E+08 | 0.0022 | 0.0015 | 0.0020 |
| F03 | 104540001 | 1.05E+08 | 0.0020 | 0.0012 | 0.0017 |
| F03 | 104560001 | 1.05E+08 | 0.0020 | 0.0013 | 0.0018 |
| F03 | 104580001 | 1.05E+08 | 0.0016 | 0.0008 | 0.0013 |
| F03 | 104600001 | 1.05E+08 | 0.0016 | 0.0006 | 0.0010 |
| F03 | 104620001 | 1.05E+08 | 0.0013 | 0.0002 | 0.0006 |
| F03 | 104640001 | 1.05E+08 | 0.0014 | 0.0002 | 0.0006 |
| F03 | 104660001 | 1.05E+08 | 0.0014 | 0.0003 | 0.0006 |
| F03 | 104680001 | 1.05E+08 | 0.0017 | 0.0005 | 0.0008 |

| Chr | Start | End | both | A1 | A2 |
|---|---|---|---|---|---|
| F03 | 104700001 | 1.05E+08 | 0.0015 | 0.0005 | 0.0007 |
| F03 | 104720001 | 1.05E+08 | 0.0015 | 0.0006 | 0.0007 |
| F03 | 104740001 | 1.05E+08 | 0.0017 | 0.0007 | 0.0008 |
| F03 | 104760001 | 1.05E+08 | 0.0030 | 0.0018 | 0.0020 |
| F03 | 104780001 | 1.05E+08 | 0.0028 | 0.0016 | 0.0017 |
| F03 | 104800001 | 1.05E+08 | 0.0038 | 0.0027 | 0.0027 |
| F03 | 104820001 | 1.05E+08 | 0.0041 | 0.0029 | 0.0029 |
| F03 | 104840001 | 1.05E+08 | 0.0045 | 0.0036 | 0.0035 |
| F03 | 104860001 | 1.05E+08 | 0.0052 | 0.0042 | 0.0042 |
| F03 | 104880001 | 1.05E+08 | 0.0054 | 0.0046 | 0.0045 |
| F03 | 104900001 | 1.05E+08 | 0.0050 | 0.0041 | 0.0040 |
| F03 | 104920001 | 1.05E+08 | 0.0049 | 0.0040 | 0.0039 |
| F03 | 104940001 | 1.05E+08 | 0.0044 | 0.0033 | 0.0033 |
| F03 | 104960001 | 1.05E+08 | 0.0024 | 0.0014 | 0.0014 |
| F03 | 104980001 | 1.05E+08 | 0.0022 | 0.0010 | 0.0010 |
| F03 | 105000001 | 1.05E+08 | 0.0016 | 0.0006 | 0.0007 |
| F03 | 105020001 | 1.05E+08 | 0.0018 | 0.0007 | 0.0008 |
| F03 | 105040001 | 1.05E+08 | 0.0018 | 0.0008 | 0.0009 |
| F03 | 105060001 | 1.05E+08 | 0.0020 | 0.0010 | 0.0012 |
| F03 | 105080001 | 1.05E+08 | 0.0021 | 0.0011 | 0.0013 |
| F03 | 105100001 | 1.05E+08 | 0.0023 | 0.0010 | 0.0012 |
| F03 | 105120001 | 1.05E+08 | 0.0023 | 0.0011 | 0.0014 |
| F03 | 105140001 | 1.05E+08 | 0.0025 | 0.0013 | 0.0015 |
| F03 | 105160001 | 1.05E+08 | 0.0024 | 0.0011 | 0.0014 |
| F03 | 105180001 | 1.05E+08 | 0.0021 | 0.0010 | 0.0013 |
| F03 | 105200001 | 1.05E+08 | 0.0020 | 0.0009 | 0.0011 |
| F03 | 105220001 | 1.05E+08 | 0.0023 | 0.0011 | 0.0014 |
| F03 | 105240001 | 1.05E+08 | 0.0023 | 0.0011 | 0.0014 |
| F03 | 105260001 | 1.05E+08 | 0.0028 | 0.0018 | 0.0020 |
| F03 | 105280001 | 1.05E+08 | 0.0029 | 0.0020 | 0.0022 |
| F03 | 105300001 | 1.05E+08 | 0.0032 | 0.0023 | 0.0025 |
| F03 | 105320001 | 1.05E+08 | 0.0027 | 0.0020 | 0.0021 |
| F03 | 105340001 | 1.05E+08 | 0.0028 | 0.0022 | 0.0023 |
| F03 | 105360001 | 1.05E+08 | 0.0024 | 0.0017 | 0.0018 |
| F03 | 105380001 | 1.05E+08 | 0.0024 | 0.0016 | 0.0016 |
| F03 | 105400001 | 1.06E+08 | 0.0026 | 0.0018 | 0.0021 |
| F03 | 105420001 | 1.06E+08 | 0.0025 | 0.0015 | 0.0018 |
| F03 | 105440001 | 1.06E+08 | 0.0020 | 0.0008 | 0.0010 |
| F03 | 105460001 | 1.06E+08 | 0.0022 | 0.0008 | 0.0011 |
| F03 | 105480001 | 1.06E+08 | 0.0023 | 0.0008 | 0.0011 |
| F03 | 105500001 | 1.06E+08 | 0.0019 | 0.0004 | 0.0005 |
| F03 | 105520001 | 1.06E+08 | 0.0019 | 0.0006 | 0.0007 |
| F03 | 105540001 | 1.06E+08 | 0.0021 | 0.0009 | 0.0011 |
| F03 | 105560001 | 1.06E+08 | 0.0026 | 0.0015 | 0.0015 |
| F03 | 105580001 | 1.06E+08 | 0.0027 | 0.0015 | 0.0016 |
| F03 | 105600001 | 1.06E+08 | 0.0029 | 0.0019 | 0.0021 |
| F03 | 105620001 | 1.06E+08 | 0.0033 | 0.0023 | 0.0026 |
| F03 | 105640001 | 1.06E+08 | 0.0031 | 0.0022 | 0.0023 |
| F03 | 105660001 | 1.06E+08 | 0.0038 | 0.0029 | 0.0033 |
| F03 | 105680001 | 1.06E+08 | 0.0039 | 0.0032 | 0.0035 |
| F03 | 105700001 | 1.06E+08 | 0.0036 | 0.0028 | 0.0031 |
| F03 | 105720001 | 1.06E+08 | 0.0031 | 0.0023 | 0.0025 |
| F03 | 105740001 | 1.06E+08 | 0.0033 | 0.0022 | 0.0026 |
| F03 | 105760001 | 1.06E+08 | 0.0020 | 0.0009 | 0.0010 |
| F03 | 105780001 | 1.06E+08 | 0.0018 | 0.0006 | 0.0008 |
| F03 | 105800001 | 1.06E+08 | 0.0018 | 0.0006 | 0.0006 |
| F03 | 105820001 | 1.06E+08 | 0.0019 | 0.0008 | 0.0008 |
| F03 | 105840001 | 1.06E+08 | 0.0018 | 0.0008 | 0.0006 |

| Chr | Start | End | both | A1 | A2 |
|---|---|---|---|---|---|
| F03 | 105860001 | 1.06E+08 | 0.0026 | 0.0014 | 0.0014 |
| F03 | 105880001 | 1.06E+08 | 0.0028 | 0.0015 | 0.0016 |
| F03 | 105900001 | 1.06E+08 | 0.0028 | 0.0014 | 0.0015 |
| F03 | 105920001 | 1.06E+08 | 0.0025 | 0.0011 | 0.0013 |
| F03 | 105940001 | 1.06E+08 | 0.0026 | 0.0011 | 0.0013 |
| F03 | 105960001 | 1.06E+08 | 0.0019 | 0.0005 | 0.0007 |
| F03 | 105980001 | 1.06E+08 | 0.0023 | 0.0011 | 0.0014 |
| F03 | 106000001 | 1.06E+08 | 0.0024 | 0.0012 | 0.0015 |
| F03 | 106020001 | 1.06E+08 | 0.0025 | 0.0014 | 0.0017 |
| F03 | 106040001 | 1.06E+08 | 0.0024 | 0.0015 | 0.0017 |
| F03 | 106060001 | 1.06E+08 | 0.0020 | 0.0013 | 0.0014 |
| F03 | 106080001 | 1.06E+08 | 0.0013 | 0.0005 | 0.0006 |
| F03 | 106100001 | 1.06E+08 | 0.0011 | 0.0005 | 0.0005 |
| F03 | 106120001 | 1.06E+08 | 0.0019 | 0.0010 | 0.0012 |
| F03 | 106140001 | 1.06E+08 | 0.0021 | 0.0010 | 0.0012 |
| F03 | 106160001 | 1.06E+08 | 0.0022 | 0.0011 | 0.0012 |
| F03 | 106180001 | 1.06E+08 | 0.0023 | 0.0011 | 0.0012 |
| F03 | 106200001 | 1.06E+08 | 0.0025 | 0.0010 | 0.0013 |
| F03 | 106220001 | 1.06E+08 | 0.0015 | 0.0002 | 0.0005 |
| F03 | 106240001 | 1.06E+08 | 0.0017 | 0.0003 | 0.0009 |
| F03 | 106260001 | 1.06E+08 | 0.0021 | 0.0005 | 0.0013 |
| F03 | 106280001 | 1.06E+08 | 0.0025 | 0.0012 | 0.0020 |
| F03 | 106300001 | 1.06E+08 | 0.0029 | 0.0018 | 0.0027 |
| F03 | 106320001 | 1.06E+08 | 0.0031 | 0.0019 | 0.0028 |
| F03 | 106340001 | 1.06E+08 | 0.0027 | 0.0018 | 0.0024 |
| F03 | 106360001 | 1.06E+08 | 0.0027 | 0.0017 | 0.0022 |
| F03 | 106380001 | 1.06E+08 | 0.0023 | 0.0012 | 0.0017 |
| F03 | 106400001 | 1.07E+08 | 0.0023 | 0.0011 | 0.0015 |
| F03 | 106420001 | 1.07E+08 | 0.0033 | 0.0022 | 0.0027 |
| F03 | 106440001 | 1.07E+08 | 0.0034 | 0.0022 | 0.0026 |
| F03 | 106460001 | 1.07E+08 | 0.0035 | 0.0024 | 0.0028 |
| F03 | 106480001 | 1.07E+08 | 0.0034 | 0.0023 | 0.0028 |
| F03 | 106500001 | 1.07E+08 | 0.0029 | 0.0018 | 0.0024 |
| F03 | 106520001 | 1.07E+08 | 0.0018 | 0.0006 | 0.0013 |
| F03 | 106540001 | 1.07E+08 | 0.0018 | 0.0009 | 0.0015 |
| F03 | 106560001 | 1.07E+08 | 0.0013 | 0.0005 | 0.0011 |
| F03 | 106580001 | 1.07E+08 | 0.0014 | 0.0008 | 0.0013 |
| F03 | 106600001 | 1.07E+08 | 0.0015 | 0.0011 | 0.0015 |
| F03 | 106620001 | 1.07E+08 | 0.0014 | 0.0012 | 0.0014 |
| F03 | 106640001 | 1.07E+08 | 0.0012 | 0.0010 | 0.0012 |
| F03 | 106660001 | 1.07E+08 | 0.0018 | 0.0016 | 0.0018 |
| F03 | 106680001 | 1.07E+08 | 0.0018 | 0.0015 | 0.0018 |
| F03 | 106700001 | 1.07E+08 | 0.0020 | 0.0017 | 0.0020 |
| F03 | 106720001 | 1.07E+08 | 0.0026 | 0.0022 | 0.0027 |
| F03 | 106740001 | 1.07E+08 | 0.0027 | 0.0023 | 0.0027 |
| F03 | 106760001 | 1.07E+08 | 0.0021 | 0.0018 | 0.0021 |
| F03 | 106780001 | 1.07E+08 | 0.0023 | 0.0020 | 0.0023 |
| F03 | 106800001 | 1.07E+08 | 0.0018 | 0.0016 | 0.0017 |
| F03 | 106820001 | 1.07E+08 | 0.0016 | 0.0014 | 0.0016 |
| F03 | 106840001 | 1.07E+08 | 0.0021 | 0.0017 | 0.0021 |
| F03 | 106860001 | 1.07E+08 | 0.0024 | 0.0018 | 0.0023 |
| F03 | 106880001 | 1.07E+08 | 0.0021 | 0.0014 | 0.0020 |
| F03 | 106900001 | 1.07E+08 | 0.0022 | 0.0014 | 0.0021 |
| F03 | 106920001 | 1.07E+08 | 0.0025 | 0.0017 | 0.0022 |
| F03 | 106940001 | 1.07E+08 | 0.0021 | 0.0013 | 0.0017 |
| F03 | 106960001 | 1.07E+08 | 0.0021 | 0.0013 | 0.0015 |
| F03 | 106980001 | 1.07E+08 | 0.0020 | 0.0014 | 0.0013 |
| F03 | 107000001 | 1.07E+08 | 0.0022 | 0.0016 | 0.0015 |

| Chr | Start | End | both | A1 | A2 |
|---|---|---|---|---|---|
| F03 | 107020001 | 1.07E+08 | 0.0016 | 0.0011 | 0.0008 |
| F03 | 107040001 | 1.07E+08 | 0.0017 | 0.0012 | 0.0007 |
| F03 | 107060001 | 1.07E+08 | 0.0016 | 0.0011 | 0.0007 |
| F03 | 107080001 | 1.07E+08 | 0.0015 | 0.0009 | 0.0007 |
| F03 | 107100001 | 1.07E+08 | 0.0013 | 0.0006 | 0.0004 |
| F03 | 107120001 | 1.07E+08 | 0.0013 | 0.0006 | 0.0004 |
| F03 | 107140001 | 1.07E+08 | 0.0013 | 0.0005 | 0.0003 |
| F03 | 107160001 | 1.07E+08 | 0.0016 | 0.0007 | 0.0005 |
| F03 | 107180001 | 1.07E+08 | 0.0017 | 0.0006 | 0.0004 |
| F03 | 107200001 | 1.07E+08 | 0.0016 | 0.0005 | 0.0003 |
| F03 | 107220001 | 1.07E+08 | 0.0016 | 0.0004 | 0.0005 |
| F03 | 107240001 | 1.07E+08 | 0.0015 | 0.0003 | 0.0007 |
| F03 | 107260001 | 1.07E+08 | 0.0013 | 0.0001 | 0.0006 |
| F03 | 107280001 | 1.07E+08 | 0.0020 | 0.0009 | 0.0014 |
| F03 | 107300001 | 1.07E+08 | 0.0036 | 0.0023 | 0.0029 |
| F03 | 107320001 | 1.07E+08 | 0.0038 | 0.0024 | 0.0031 |
| F03 | 107340001 | 1.07E+08 | 0.0039 | 0.0027 | 0.0032 |
| F03 | 107360001 | 1.07E+08 | 0.0042 | 0.0029 | 0.0036 |
| F03 | 107380001 | 1.07E+08 | 0.0038 | 0.0024 | 0.0033 |
| F03 | 107400001 | 1.08E+08 | 0.0024 | 0.0010 | 0.0023 |
| F03 | 107420001 | 1.08E+08 | 0.0024 | 0.0009 | 0.0024 |
| F03 | 107440001 | 1.08E+08 | 0.0025 | 0.0008 | 0.0025 |
| F03 | 107460001 | 1.08E+08 | 0.0025 | 0.0005 | 0.0025 |
| F03 | 107480001 | 1.08E+08 | 0.0021 | 0.0003 | 0.0021 |
| F03 | 107500001 | 1.08E+08 | 0.0022 | 0.0004 | 0.0022 |
| F03 | 107520001 | 1.08E+08 | 0.0023 | 0.0004 | 0.0022 |
| F03 | 107540001 | 1.08E+08 | 0.0025 | 0.0006 | 0.0022 |
| F03 | 107560001 | 1.08E+08 | 0.0024 | 0.0006 | 0.0019 |
| F03 | 107580001 | 1.08E+08 | 0.0026 | 0.0006 | 0.0018 |
| F03 | 107600001 | 1.08E+08 | 0.0024 | 0.0006 | 0.0015 |
| F03 | 107620001 | 1.08E+08 | 0.0022 | 0.0006 | 0.0012 |
| F03 | 107640001 | 1.08E+08 | 0.0020 | 0.0003 | 0.0009 |
| F03 | 107660001 | 1.08E+08 | 0.0018 | 0.0003 | 0.0008 |
| F03 | 107680001 | 1.08E+08 | 0.0020 | 0.0005 | 0.0010 |
| F03 | 107700001 | 1.08E+08 | 0.0020 | 0.0006 | 0.0009 |
| F03 | 107720001 | 1.08E+08 | 0.0022 | 0.0008 | 0.0010 |
| F03 | 107740001 | 1.08E+08 | 0.0029 | 0.0016 | 0.0019 |
| F03 | 107760001 | 1.08E+08 | 0.0029 | 0.0016 | 0.0021 |
| F03 | 107780001 | 1.08E+08 | 0.0028 | 0.0014 | 0.0022 |
| F03 | 107800001 | 1.08E+08 | 0.0030 | 0.0013 | 0.0024 |
| F03 | 107820001 | 1.08E+08 | 0.0027 | 0.0009 | 0.0022 |
| F03 | 107840001 | 1.08E+08 | 0.0019 | 0.0001 | 0.0014 |
| F03 | 107860001 | 1.08E+08 | 0.0019 | 0.0001 | 0.0014 |
| F03 | 107880001 | 1.08E+08 | 0.0018 | 0.0002 | 0.0013 |
| F03 | 107900001 | 1.08E+08 | 0.0019 | 0.0002 | 0.0013 |
| F03 | 107920001 | 1.08E+08 | 0.0018 | 0.0003 | 0.0013 |
| F03 | 107940001 | 1.08E+08 | 0.0020 | 0.0004 | 0.0016 |
| F03 | 107960001 | 1.08E+08 | 0.0023 | 0.0005 | 0.0018 |
| F03 | 107980001 | 1.08E+08 | 0.0025 | 0.0005 | 0.0020 |
| F03 | 108000001 | 1.08E+08 | 0.0025 | 0.0005 | 0.0019 |
| F03 | 108020001 | 1.08E+08 | 0.0026 | 0.0005 | 0.0020 |
| F03 | 108040001 | 1.08E+08 | 0.0025 | 0.0004 | 0.0020 |
| F03 | 108060001 | 1.08E+08 | 0.0024 | 0.0003 | 0.0019 |
| F03 | 108080001 | 1.08E+08 | 0.0023 | 0.0003 | 0.0017 |
| F03 | 108100001 | 1.08E+08 | 0.0024 | 0.0003 | 0.0018 |
| F03 | 108120001 | 1.08E+08 | 0.0025 | 0.0003 | 0.0019 |
| F03 | 108140001 | 1.08E+08 | 0.0023 | 0.0002 | 0.0017 |
| F03 | 108160001 | 1.08E+08 | 0.0023 | 0.0003 | 0.0016 |

| Chr | Start | End | both | A1 | A2 |
|---|---|---|---|---|---|
| F03 | 108180001 | 1.08E+08 | 0.0021 | 0.0001 | 0.0014 |
| F03 | 108200001 | 1.08E+08 | 0.0024 | 0.0007 | 0.0018 |
| F03 | 108220001 | 1.08E+08 | 0.0026 | 0.0010 | 0.0019 |
| F03 | 108240001 | 1.08E+08 | 0.0026 | 0.0011 | 0.0018 |
| F03 | 108260001 | 1.08E+08 | 0.0030 | 0.0015 | 0.0023 |
| F03 | 108280001 | 1.08E+08 | 0.0032 | 0.0018 | 0.0025 |
| F03 | 108300001 | 1.08E+08 | 0.0028 | 0.0014 | 0.0018 |
| F03 | 108320001 | 1.08E+08 | 0.0029 | 0.0015 | 0.0019 |
| F03 | 108340001 | 1.08E+08 | 0.0028 | 0.0016 | 0.0018 |
| F03 | 108360001 | 1.08E+08 | 0.0024 | 0.0013 | 0.0013 |
| F03 | 108380001 | 1.08E+08 | 0.0023 | 0.0012 | 0.0012 |
| F03 | 108400001 | 1.09E+08 | 0.0022 | 0.0011 | 0.0011 |
| F03 | 108420001 | 1.09E+08 | 0.0020 | 0.0010 | 0.0009 |
| F03 | 108440001 | 1.09E+08 | 0.0020 | 0.0015 | 0.0009 |
| F03 | 108460001 | 1.09E+08 | 0.0018 | 0.0020 | 0.0008 |
| F03 | 108480001 | 1.09E+08 | 0.0017 | 0.0020 | 0.0008 |
| F03 | 108500001 | 1.09E+08 | 0.0018 | 0.0022 | 0.0010 |
| F03 | 108520001 | 1.09E+08 | 0.0018 | 0.0021 | 0.0009 |
| F03 | 108540001 | 1.09E+08 | 0.0019 | 0.0015 | 0.0011 |
| F03 | 108560001 | 1.09E+08 | 0.0017 | 0.0010 | 0.0011 |
| F03 | 108580001 | 1.09E+08 | 0.0017 | 0.0009 | 0.0010 |
| F03 | 108600001 | 1.09E+08 | 0.0018 | 0.0011 | 0.0009 |
| F03 | 108620001 | 1.09E+08 | 0.0016 | 0.0013 | 0.0007 |
| F03 | 108640001 | 1.09E+08 | 0.0014 | 0.0017 | 0.0005 |
| F03 | 108660001 | 1.09E+08 | 0.0018 | 0.0021 | 0.0009 |
| F03 | 108680001 | 1.09E+08 | 0.0018 | 0.0022 | 0.0011 |
| F03 | 108700001 | 1.09E+08 | 0.0018 | 0.0021 | 0.0012 |
| F03 | 108720001 | 1.09E+08 | 0.0021 | 0.0020 | 0.0016 |
| F03 | 108740001 | 1.09E+08 | 0.0023 | 0.0016 | 0.0017 |
| F03 | 108760001 | 1.09E+08 | 0.0022 | 0.0011 | 0.0016 |
| F03 | 108780001 | 1.09E+08 | 0.0023 | 0.0011 | 0.0016 |
| F03 | 108800001 | 1.09E+08 | 0.0021 | 0.0008 | 0.0012 |
| F03 | 108820001 | 1.09E+08 | 0.0017 | 0.0005 | 0.0007 |
| F03 | 108840001 | 1.09E+08 | 0.0017 | 0.0006 | 0.0007 |
| F03 | 108860001 | 1.09E+08 | 0.0015 | 0.0005 | 0.0005 |
| F03 | 108880001 | 1.09E+08 | 0.0015 | 0.0007 | 0.0005 |
| F03 | 108900001 | 1.09E+08 | 0.0020 | 0.0013 | 0.0012 |
| F03 | 108920001 | 1.09E+08 | 0.0024 | 0.0019 | 0.0018 |
| F03 | 108940001 | 1.09E+08 | 0.0030 | 0.0026 | 0.0026 |
| F03 | 108960001 | 1.09E+08 | 0.0048 | 0.0045 | 0.0045 |
| F03 | 108980001 | 1.09E+08 | 0.0049 | 0.0043 | 0.0047 |
| F03 | 109000001 | 1.09E+08 | 0.0047 | 0.0041 | 0.0046 |
| F03 | 109020001 | 1.09E+08 | 0.0045 | 0.0041 | 0.0044 |
| F03 | 109040001 | 1.09E+08 | 0.0041 | 0.0039 | 0.0039 |
| F03 | 109060001 | 1.09E+08 | 0.0024 | 0.0019 | 0.0022 |
| F03 | 109080001 | 1.09E+08 | 0.0023 | 0.0017 | 0.0021 |
| F03 | 109100001 | 1.09E+08 | 0.0023 | 0.0012 | 0.0020 |
| F03 | 109120001 | 1.09E+08 | 0.0022 | 0.0007 | 0.0020 |
| F03 | 109140001 | 1.09E+08 | 0.0021 | 0.0001 | 0.0020 |
| F03 | 109160001 | 1.09E+08 | 0.0026 | 0.0005 | 0.0026 |
| F03 | 109180001 | 1.09E+08 | 0.0028 | 0.0006 | 0.0028 |
| F03 | 109200001 | 1.09E+08 | 0.0027 | 0.0006 | 0.0026 |
| F03 | 109220001 | 1.09E+08 | 0.0027 | 0.0007 | 0.0024 |
| F03 | 109240001 | 1.09E+08 | 0.0029 | 0.0010 | 0.0024 |
| F03 | 109260001 | 1.09E+08 | 0.0022 | 0.0009 | 0.0016 |
| F03 | 109280001 | 1.09E+08 | 0.0018 | 0.0010 | 0.0013 |
| F03 | 109300001 | 1.09E+08 | 0.0017 | 0.0015 | 0.0012 |
| F03 | 109320001 | 1.09E+08 | 0.0014 | 0.0017 | 0.0010 |

| Chr | Start | End | both | A1 | A2 |
|---|---|---|---|---|---|
| F03 | 109340001 | 1.09E+08 | 0.0010 | 0.0019 | 0.0006 |
| F03 | 109360001 | 1.09E+08 | 0.0010 | 0.0020 | 0.0006 |
| F03 | 109380001 | 1.09E+08 | 0.0016 | 0.0028 | 0.0011 |
| F03 | 109400001 | 1.1E+08 | 0.0014 | 0.0025 | 0.0010 |
| F03 | 109420001 | 1.1E+08 | 0.0026 | 0.0037 | 0.0022 |
| F03 | 109440001 | 1.1E+08 | 0.0027 | 0.0035 | 0.0024 |
| F03 | 109460001 | 1.1E+08 | 0.0050 | 0.0056 | 0.0046 |
| F03 | 109480001 | 1.1E+08 | 0.0053 | 0.0061 | 0.0050 |
| F03 | 109500001 | 1.1E+08 | 0.0053 | 0.0063 | 0.0049 |
| F03 | 109520001 | 1.1E+08 | 0.0041 | 0.0052 | 0.0037 |
| F03 | 109540001 | 1.1E+08 | 0.0040 | 0.0053 | 0.0035 |
| F03 | 109560001 | 1.1E+08 | 0.0018 | 0.0033 | 0.0011 |
| F03 | 109580001 | 1.1E+08 | 0.0008 | 0.0022 | 0.0001 |
| F03 | 109600001 | 1.1E+08 | 0.0008 | 0.0021 | 0.0002 |
| F03 | 109620001 | 1.1E+08 | 0.0009 | 0.0021 | 0.0002 |
| F03 | 109640001 | 1.1E+08 | 0.0008 | 0.0017 | 0.0003 |
| F03 | 109660001 | 1.1E+08 | 0.0008 | 0.0016 | 0.0004 |
| F03 | 109680001 | 1.1E+08 | 0.0007 | 0.0013 | 0.0003 |
| F03 | 109700001 | 1.1E+08 | 0.0007 | 0.0012 | 0.0004 |
| F03 | 109720001 | 1.1E+08 | 0.0007 | 0.0012 | 0.0005 |
| F03 | 109740001 | 1.1E+08 | 0.0010 | 0.0013 | 0.0006 |
| F03 | 109760001 | 1.1E+08 | 0.0010 | 0.0012 | 0.0005 |
| F03 | 109780001 | 1.1E+08 | 0.0012 | 0.0013 | 0.0005 |
| F03 | 109800001 | 1.1E+08 | 0.0020 | 0.0021 | 0.0011 |
| F03 | 109820001 | 1.1E+08 | 0.0025 | 0.0025 | 0.0015 |
| F03 | 109840001 | 1.1E+08 | 0.0026 | 0.0028 | 0.0019 |
| F03 | 109860001 | 1.1E+08 | 0.0029 | 0.0029 | 0.0022 |
| F03 | 109880001 | 1.1E+08 | 0.0028 | 0.0027 | 0.0022 |
| F03 | 109900001 | 1.1E+08 | 0.0021 | 0.0019 | 0.0016 |
| F03 | 109920001 | 1.1E+08 | 0.0019 | 0.0014 | 0.0013 |
| F03 | 109940001 | 1.1E+08 | 0.0017 | 0.0010 | 0.0009 |
| F03 | 109960001 | 1.1E+08 | 0.0015 | 0.0006 | 0.0007 |
| F03 | 109980001 | 1.1E+08 | 0.0017 | 0.0006 | 0.0008 |
| F03 | 110000001 | 1.1E+08 | 0.0017 | 0.0005 | 0.0010 |
| F03 | 110020001 | 1.1E+08 | 0.0020 | 0.0007 | 0.0014 |
| F03 | 110040001 | 1.1E+08 | 0.0021 | 0.0008 | 0.0018 |
| F03 | 110060001 | 1.1E+08 | 0.0023 | 0.0010 | 0.0021 |
| F03 | 110080001 | 1.1E+08 | 0.0023 | 0.0010 | 0.0023 |
| F03 | 110100001 | 1.1E+08 | 0.0024 | 0.0011 | 0.0024 |
| F03 | 110120001 | 1.1E+08 | 0.0027 | 0.0017 | 0.0028 |
| F03 | 110140001 | 1.1E+08 | 0.0027 | 0.0019 | 0.0027 |
| F03 | 110160001 | 1.1E+08 | 0.0023 | 0.0018 | 0.0023 |
| F03 | 110180001 | 1.1E+08 | 0.0019 | 0.0016 | 0.0019 |
| F03 | 110200001 | 1.1E+08 | 0.0015 | 0.0015 | 0.0014 |
| F03 | 110220001 | 1.1E+08 | 0.0004 | 0.0004 | 0.0003 |
| F04 | 1 | 100000 | 0.0051 | 0.0050 | 0.0045 |
| F04 | 20001 | 120000 | 0.0059 | 0.0037 | 0.0052 |
| F04 | 40001 | 140000 | 0.0048 | 0.0028 | 0.0039 |
| F04 | 60001 | 160000 | 0.0043 | 0.0024 | 0.0033 |
| F04 | 80001 | 180000 | 0.0038 | 0.0021 | 0.0029 |
| F04 | 100001 | 200000 | 0.0038 | 0.0016 | 0.0026 |
| F04 | 120001 | 220000 | 0.0032 | 0.0016 | 0.0020 |
| F04 | 140001 | 240000 | 0.0029 | 0.0017 | 0.0015 |
| F04 | 160001 | 260000 | 0.0026 | 0.0016 | 0.0013 |
| F04 | 180001 | 280000 | 0.0024 | 0.0021 | 0.0011 |
| F04 | 200001 | 300000 | 0.0024 | 0.0022 | 0.0012 |
| F04 | 220001 | 320000 | 0.0023 | 0.0022 | 0.0010 |
| F04 | 240001 | 340000 | 0.0021 | 0.0021 | 0.0009 |

| Chr | Start | End | both | A1 | A2 |
|---|---|---|---|---|---|
| F04 | 260001 | 360000 | 0.0020 | 0.0020 | 0.0009 |
| F04 | 280001 | 380000 | 0.0020 | 0.0014 | 0.0012 |
| F04 | 300001 | 400000 | 0.0019 | 0.0012 | 0.0013 |
| F04 | 320001 | 420000 | 0.0022 | 0.0007 | 0.0016 |
| F04 | 340001 | 440000 | 0.0021 | 0.0011 | 0.0016 |
| F04 | 360001 | 460000 | 0.0032 | 0.0023 | 0.0025 |
| F04 | 380001 | 480000 | 0.0043 | 0.0031 | 0.0035 |
| F04 | 400001 | 500000 | 0.0042 | 0.0036 | 0.0033 |
| F04 | 420001 | 520000 | 0.0040 | 0.0038 | 0.0033 |
| F04 | 440001 | 540000 | 0.0038 | 0.0037 | 0.0032 |
| F04 | 460001 | 560000 | 0.0028 | 0.0023 | 0.0022 |
| F04 | 480001 | 580000 | 0.0018 | 0.0017 | 0.0010 |
| F04 | 500001 | 600000 | 0.0021 | 0.0015 | 0.0013 |
| F04 | 520001 | 620000 | 0.0023 | 0.0019 | 0.0013 |
| F04 | 540001 | 640000 | 0.0030 | 0.0019 | 0.0019 |
| F04 | 560001 | 660000 | 0.0031 | 0.0021 | 0.0020 |
| F04 | 580001 | 680000 | 0.0031 | 0.0019 | 0.0023 |
| F04 | 600001 | 700000 | 0.0028 | 0.0015 | 0.0020 |
| F04 | 620001 | 720000 | 0.0025 | 0.0009 | 0.0019 |
| F04 | 640001 | 740000 | 0.0020 | 0.0010 | 0.0014 |
| F04 | 660001 | 760000 | 0.0027 | 0.0007 | 0.0020 |
| F04 | 680001 | 780000 | 0.0026 | 0.0010 | 0.0019 |
| F04 | 700001 | 800000 | 0.0024 | 0.0012 | 0.0016 |
| F04 | 720001 | 820000 | 0.0023 | 0.0012 | 0.0015 |
| F04 | 740001 | 840000 | 0.0021 | 0.0008 | 0.0014 |
| F04 | 760001 | 860000 | 0.0014 | 0.0014 | 0.0007 |
| F04 | 780001 | 880000 | 0.0018 | 0.0012 | 0.0011 |
| F04 | 800001 | 900000 | 0.0018 | 0.0011 | 0.0011 |
| F04 | 820001 | 920000 | 0.0019 | 0.0012 | 0.0010 |
| F04 | 840001 | 940000 | 0.0020 | 0.0013 | 0.0011 |
| F04 | 860001 | 960000 | 0.0024 | 0.0017 | 0.0015 |
| F04 | 880001 | 980000 | 0.0027 | 0.0016 | 0.0020 |
| F04 | 900001 | 1000000 | 0.0026 | 0.0014 | 0.0020 |
| F04 | 920001 | 1020000 | 0.0026 | 0.0018 | 0.0019 |
| F04 | 940001 | 1040000 | 0.0034 | 0.0018 | 0.0023 |
| F04 | 960001 | 1060000 | 0.0033 | 0.0013 | 0.0021 |
| F04 | 980001 | 1080000 | 0.0028 | 0.0020 | 0.0016 |
| F04 | 1000001 | 1100000 | 0.0036 | 0.0020 | 0.0023 |
| F04 | 1020001 | 1120000 | 0.0036 | 0.0016 | 0.0023 |
| F04 | 1040001 | 1140000 | 0.0029 | 0.0016 | 0.0018 |
| F04 | 1060001 | 1160000 | 0.0028 | 0.0014 | 0.0019 |
| F04 | 1080001 | 1180000 | 0.0027 | 0.0008 | 0.0017 |
| F04 | 1100001 | 1200000 | 0.0021 | 0.0010 | 0.0012 |
| F04 | 1120001 | 1220000 | 0.0024 | 0.0010 | 0.0013 |
| F04 | 1140001 | 1240000 | 0.0024 | 0.0010 | 0.0013 |
| F04 | 1160001 | 1260000 | 0.0026 | 0.0009 | 0.0012 |
| F04 | 1180001 | 1280000 | 0.0024 | 0.0008 | 0.0010 |
| F04 | 1200001 | 1300000 | 0.0023 | 0.0005 | 0.0009 |
| F04 | 1220001 | 1320000 | 0.0020 | 0.0005 | 0.0007 |
| F04 | 1240001 | 1340000 | 0.0021 | 0.0004 | 0.0009 |
| F04 | 1260001 | 1360000 | 0.0018 | 0.0005 | 0.0008 |
| F04 | 1280001 | 1380000 | 0.0019 | 0.0007 | 0.0011 |
| F04 | 1300001 | 1400000 | 0.0023 | 0.0009 | 0.0014 |
| F04 | 1320001 | 1420000 | 0.0025 | 0.0012 | 0.0016 |
| F04 | 1340001 | 1440000 | 0.0026 | 0.0015 | 0.0019 |
| F04 | 1360001 | 1460000 | 0.0027 | 0.0013 | 0.0020 |
| F04 | 1380001 | 1480000 | 0.0027 | 0.0011 | 0.0018 |
| F04 | 1400001 | 1500000 | 0.0024 | 0.0009 | 0.0016 |

| Chr | Start | End | both | A1 | A2 |
|---|---|---|---|---|---|
| F04 | 1420001 | 1520000 | 0.0021 | 0.0005 | 0.0015 |
| F04 | 1440001 | 1540000 | 0.0017 | 0.0003 | 0.0011 |
| F04 | 1460001 | 1560000 | 0.0015 | 0.0001 | 0.0011 |
| F04 | 1480001 | 1580000 | 0.0014 | 0.0001 | 0.0013 |
| F04 | 1500001 | 1600000 | 0.0012 | 0.0003 | 0.0013 |
| F04 | 1520001 | 1620000 | 0.0014 | 0.0003 | 0.0013 |
| F04 | 1540001 | 1640000 | 0.0016 | 0.0003 | 0.0015 |
| F04 | 1560001 | 1660000 | 0.0019 | 0.0006 | 0.0018 |
| F04 | 1580001 | 1680000 | 0.0023 | 0.0008 | 0.0022 |
| F04 | 1600001 | 1700000 | 0.0025 | 0.0007 | 0.0025 |
| F04 | 1620001 | 1720000 | 0.0024 | 0.0010 | 0.0025 |
| F04 | 1640001 | 1740000 | 0.0027 | 0.0020 | 0.0028 |
| F04 | 1660001 | 1760000 | 0.0033 | 0.0020 | 0.0034 |
| F04 | 1680001 | 1780000 | 0.0033 | 0.0022 | 0.0034 |
| F04 | 1700001 | 1800000 | 0.0035 | 0.0025 | 0.0035 |
| F04 | 1720001 | 1820000 | 0.0038 | 0.0029 | 0.0039 |
| F04 | 1740001 | 1840000 | 0.0042 | 0.0024 | 0.0042 |
| F04 | 1760001 | 1860000 | 0.0036 | 0.0023 | 0.0036 |
| F04 | 1780001 | 1880000 | 0.0035 | 0.0022 | 0.0035 |
| F04 | 1800001 | 1900000 | 0.0033 | 0.0023 | 0.0034 |
| F04 | 1820001 | 1920000 | 0.0030 | 0.0021 | 0.0031 |
| F04 | 1840001 | 1940000 | 0.0026 | 0.0019 | 0.0026 |
| F04 | 1860001 | 1960000 | 0.0024 | 0.0022 | 0.0025 |
| F04 | 1880001 | 1980000 | 0.0027 | 0.0021 | 0.0028 |
| F04 | 1900001 | 2000000 | 0.0028 | 0.0021 | 0.0028 |
| F04 | 1920001 | 2020000 | 0.0028 | 0.0016 | 0.0029 |
| F04 | 1940001 | 2040000 | 0.0026 | 0.0014 | 0.0027 |
| F04 | 1960001 | 2060000 | 0.0024 | 0.0009 | 0.0025 |
| F04 | 1980001 | 2080000 | 0.0018 | 0.0012 | 0.0019 |
| F04 | 2000001 | 2100000 | 0.0020 | 0.0010 | 0.0021 |
| F04 | 2020001 | 2120000 | 0.0018 | 0.0017 | 0.0020 |
| F04 | 2040001 | 2140000 | 0.0027 | 0.0018 | 0.0028 |
| F04 | 2060001 | 2160000 | 0.0028 | 0.0017 | 0.0029 |
| F04 | 2080001 | 2180000 | 0.0028 | 0.0018 | 0.0028 |
| F04 | 2100001 | 2200000 | 0.0030 | 0.0017 | 0.0028 |
| F04 | 2120001 | 2220000 | 0.0031 | 0.0009 | 0.0029 |
| F04 | 2140001 | 2240000 | 0.0021 | 0.0008 | 0.0019 |
| F04 | 2160001 | 2260000 | 0.0020 | 0.0012 | 0.0018 |
| F04 | 2180001 | 2280000 | 0.0026 | 0.0009 | 0.0022 |
| F04 | 2200001 | 2300000 | 0.0021 | 0.0012 | 0.0019 |
| F04 | 2220001 | 2320000 | 0.0024 | 0.0013 | 0.0020 |
| F04 | 2240001 | 2340000 | 0.0027 | 0.0012 | 0.0022 |
| F04 | 2260001 | 2360000 | 0.0027 | 0.0010 | 0.0021 |
| F04 | 2280001 | 2380000 | 0.0022 | 0.0009 | 0.0017 |
| F04 | 2300001 | 2400000 | 0.0026 | 0.0011 | 0.0020 |
| F04 | 2320001 | 2420000 | 0.0029 | 0.0016 | 0.0024 |
| F04 | 2340001 | 2440000 | 0.0034 | 0.0027 | 0.0031 |
| F04 | 2360001 | 2460000 | 0.0047 | 0.0026 | 0.0044 |
| F04 | 2380001 | 2480000 | 0.0049 | 0.0023 | 0.0047 |
| F04 | 2400001 | 2500000 | 0.0044 | 0.0022 | 0.0042 |
| F04 | 2420001 | 2520000 | 0.0044 | 0.0023 | 0.0043 |
| F04 | 2440001 | 2540000 | 0.0041 | 0.0012 | 0.0041 |
| F04 | 2460001 | 2560000 | 0.0030 | 0.0015 | 0.0030 |
| F04 | 2480001 | 2580000 | 0.0030 | 0.0017 | 0.0030 |
| F04 | 2500001 | 2600000 | 0.0032 | 0.0014 | 0.0032 |
| F04 | 2520001 | 2620000 | 0.0026 | 0.0013 | 0.0026 |
| F04 | 2540001 | 2640000 | 0.0025 | 0.0015 | 0.0025 |
| F04 | 2560001 | 2660000 | 0.0025 | 0.0016 | 0.0025 |

| Chr | Start | End | both | A1 | A2 |
|---|---|---|---|---|---|
| F04 | 2580001 | 2680000 | 0.0027 | 0.0015 | 0.0027 |
| F04 | 2600001 | 2700000 | 0.0024 | 0.0016 | 0.0024 |
| F04 | 2620001 | 2720000 | 0.0026 | 0.0020 | 0.0024 |
| F04 | 2640001 | 2740000 | 0.0030 | 0.0034 | 0.0026 |
| F04 | 2660001 | 2760000 | 0.0038 | 0.0032 | 0.0031 |
| F04 | 2680001 | 2780000 | 0.0035 | 0.0038 | 0.0026 |
| F04 | 2700001 | 2800000 | 0.0043 | 0.0038 | 0.0033 |
| F04 | 2720001 | 2820000 | 0.0043 | 0.0031 | 0.0032 |
| F04 | 2740001 | 2840000 | 0.0037 | 0.0018 | 0.0026 |
| F04 | 2760001 | 2860000 | 0.0029 | 0.0026 | 0.0020 |
| F04 | 2780001 | 2880000 | 0.0039 | 0.0020 | 0.0030 |
| F04 | 2800001 | 2900000 | 0.0035 | 0.0021 | 0.0023 |
| F04 | 2820001 | 2920000 | 0.0036 | 0.0018 | 0.0024 |
| F04 | 2840001 | 2940000 | 0.0032 | 0.0016 | 0.0021 |
| F04 | 2860001 | 2960000 | 0.0031 | 0.0011 | 0.0020 |
| F04 | 2880001 | 2980000 | 0.0025 | 0.0011 | 0.0014 |
| F04 | 2900001 | 3000000 | 0.0022 | 0.0008 | 0.0014 |
| F04 | 2920001 | 3020000 | 0.0019 | 0.0008 | 0.0011 |
| F04 | 2940001 | 3040000 | 0.0020 | 0.0010 | 0.0013 |
| F04 | 2960001 | 3060000 | 0.0025 | 0.0004 | 0.0019 |
| F04 | 2980001 | 3080000 | 0.0019 | 0.0004 | 0.0015 |
| F04 | 3000001 | 3100000 | 0.0018 | 0.0015 | 0.0015 |
| F04 | 3020001 | 3120000 | 0.0029 | 0.0022 | 0.0027 |
| F04 | 3040001 | 3140000 | 0.0035 | 0.0021 | 0.0034 |
| F04 | 3060001 | 3160000 | 0.0031 | 0.0022 | 0.0030 |
| F04 | 3080001 | 3180000 | 0.0032 | 0.0022 | 0.0031 |
| F04 | 3100001 | 3200000 | 0.0034 | 0.0012 | 0.0033 |
| F04 | 3120001 | 3220000 | 0.0025 | 0.0009 | 0.0022 |
| F04 | 3140001 | 3240000 | 0.0023 | 0.0012 | 0.0019 |
| F04 | 3160001 | 3260000 | 0.0024 | 0.0011 | 0.0019 |
| F04 | 3180001 | 3280000 | 0.0022 | 0.0012 | 0.0017 |
| F04 | 3200001 | 3300000 | 0.0020 | 0.0014 | 0.0015 |
| F04 | 3220001 | 3320000 | 0.0019 | 0.0012 | 0.0016 |
| F04 | 3240001 | 3340000 | 0.0020 | 0.0012 | 0.0017 |
| F04 | 3260001 | 3360000 | 0.0019 | 0.0019 | 0.0017 |
| F04 | 3280001 | 3380000 | 0.0026 | 0.0024 | 0.0024 |
| F04 | 3300001 | 3400000 | 0.0031 | 0.0023 | 0.0029 |
| F04 | 3320001 | 3420000 | 0.0033 | 0.0028 | 0.0030 |
| F04 | 3340001 | 3440000 | 0.0035 | 0.0028 | 0.0033 |
| F04 | 3360001 | 3460000 | 0.0037 | 0.0024 | 0.0034 |
| F04 | 3380001 | 3480000 | 0.0036 | 0.0021 | 0.0032 |
| F04 | 3400001 | 3500000 | 0.0036 | 0.0022 | 0.0032 |
| F04 | 3420001 | 3520000 | 0.0041 | 0.0014 | 0.0038 |
| F04 | 3440001 | 3540000 | 0.0032 | 0.0011 | 0.0030 |
| F04 | 3460001 | 3560000 | 0.0027 | 0.0007 | 0.0027 |
| F04 | 3480001 | 3580000 | 0.0024 | 0.0009 | 0.0025 |
| F04 | 3500001 | 3600000 | 0.0023 | 0.0014 | 0.0024 |
| F04 | 3520001 | 3620000 | 0.0023 | 0.0016 | 0.0025 |
| F04 | 3540001 | 3640000 | 0.0025 | 0.0017 | 0.0027 |
| F04 | 3560001 | 3660000 | 0.0026 | 0.0017 | 0.0027 |
| F04 | 3580001 | 3680000 | 0.0024 | 0.0015 | 0.0025 |
| F04 | 3600001 | 3700000 | 0.0026 | 0.0015 | 0.0027 |
| F04 | 3620001 | 3720000 | 0.0027 | 0.0021 | 0.0028 |
| F04 | 3640001 | 3740000 | 0.0036 | 0.0027 | 0.0036 |
| F04 | 3660001 | 3760000 | 0.0041 | 0.0047 | 0.0042 |
| F04 | 3680001 | 3780000 | 0.0062 | 0.0058 | 0.0061 |
| F04 | 3700001 | 3800000 | 0.0068 | 0.0069 | 0.0067 |
| F04 | 3720001 | 3820000 | 0.0077 | 0.0065 | 0.0075 |

| Chr | Start | End | both | A1 | A2 |
|---|---|---|---|---|---|
| F04 | 3740001 | 3840000 | 0.0072 | 0.0058 | 0.0070 |
| F04 | 3760001 | 3860000 | 0.0067 | 0.0037 | 0.0064 |
| F04 | 3780001 | 3880000 | 0.0050 | 0.0035 | 0.0047 |
| F04 | 3800001 | 3900000 | 0.0049 | 0.0018 | 0.0045 |
| F04 | 3820001 | 3920000 | 0.0035 | 0.0017 | 0.0032 |
| F04 | 3840001 | 3940000 | 0.0032 | 0.0031 | 0.0030 |
| F04 | 3860001 | 3960000 | 0.0044 | 0.0039 | 0.0042 |
| F04 | 3880001 | 3980000 | 0.0048 | 0.0052 | 0.0047 |
| F04 | 3900001 | 4000000 | 0.0064 | 0.0065 | 0.0064 |
| F04 | 3920001 | 4020000 | 0.0079 | 0.0080 | 0.0079 |
| F04 | 3940001 | 4040000 | 0.0092 | 0.0077 | 0.0092 |
| F04 | 3960001 | 4060000 | 0.0087 | 0.0083 | 0.0087 |
| F04 | 3980001 | 4080000 | 0.0094 | 0.0068 | 0.0094 |
| F04 | 4000001 | 4100000 | 0.0076 | 0.0055 | 0.0076 |
| F04 | 4020001 | 4120000 | 0.0057 | 0.0038 | 0.0056 |
| F04 | 4040001 | 4140000 | 0.0045 | 0.0029 | 0.0045 |
| F04 | 4060001 | 4160000 | 0.0040 | 0.0022 | 0.0041 |
| F04 | 4080001 | 4180000 | 0.0033 | 0.0015 | 0.0035 |
| F04 | 4100001 | 4200000 | 0.0030 | 0.0015 | 0.0032 |
| F04 | 4120001 | 4220000 | 0.0032 | 0.0015 | 0.0033 |
| F04 | 4140001 | 4240000 | 0.0029 | 0.0013 | 0.0031 |
| F04 | 4160001 | 4260000 | 0.0025 | 0.0007 | 0.0026 |
| F04 | 4180001 | 4280000 | 0.0022 | 0.0013 | 0.0022 |
| F04 | 4200001 | 4300000 | 0.0026 | 0.0019 | 0.0026 |
| F04 | 4220001 | 4320000 | 0.0031 | 0.0036 | 0.0030 |
| F04 | 4240001 | 4340000 | 0.0047 | 0.0053 | 0.0046 |
| F04 | 4260001 | 4360000 | 0.0064 | 0.0051 | 0.0064 |
| F04 | 4280001 | 4380000 | 0.0063 | 0.0044 | 0.0063 |
| F04 | 4300001 | 4400000 | 0.0056 | 0.0039 | 0.0056 |
| F04 | 4320001 | 4420000 | 0.0056 | 0.0023 | 0.0055 |
| F04 | 4340001 | 4440000 | 0.0040 | 0.0008 | 0.0040 |
| F04 | 4360001 | 4460000 | 0.0027 | 0.0009 | 0.0027 |
| F04 | 4380001 | 4480000 | 0.0027 | 0.0010 | 0.0025 |
| F04 | 4400001 | 4500000 | 0.0027 | 0.0011 | 0.0025 |
| F04 | 4420001 | 4520000 | 0.0029 | 0.0013 | 0.0028 |
| F04 | 4440001 | 4540000 | 0.0032 | 0.0020 | 0.0029 |
| F04 | 4460001 | 4560000 | 0.0036 | 0.0021 | 0.0033 |
| F04 | 4480001 | 4580000 | 0.0038 | 0.0030 | 0.0033 |
| F04 | 4500001 | 4600000 | 0.0046 | 0.0029 | 0.0040 |
| F04 | 4520001 | 4620000 | 0.0039 | 0.0026 | 0.0034 |
| F04 | 4540001 | 4640000 | 0.0037 | 0.0020 | 0.0032 |
| F04 | 4560001 | 4660000 | 0.0028 | 0.0017 | 0.0024 |
| F04 | 4580001 | 4680000 | 0.0027 | 0.0008 | 0.0024 |
| F04 | 4600001 | 4700000 | 0.0019 | 0.0006 | 0.0017 |
| F04 | 4620001 | 4720000 | 0.0019 | 0.0005 | 0.0017 |
| F04 | 4640001 | 4740000 | 0.0017 | 0.0004 | 0.0017 |
| F04 | 4660001 | 4760000 | 0.0019 | 0.0007 | 0.0019 |
| F04 | 4680001 | 4780000 | 0.0020 | 0.0012 | 0.0020 |
| F04 | 4700001 | 4800000 | 0.0029 | 0.0013 | 0.0029 |
| F04 | 4720001 | 4820000 | 0.0030 | 0.0013 | 0.0030 |
| F04 | 4740001 | 4840000 | 0.0030 | 0.0013 | 0.0028 |
| F04 | 4760001 | 4860000 | 0.0030 | 0.0015 | 0.0027 |
| F04 | 4780001 | 4880000 | 0.0030 | 0.0012 | 0.0027 |
| F04 | 4800001 | 4900000 | 0.0021 | 0.0010 | 0.0017 |
| F04 | 4820001 | 4920000 | 0.0022 | 0.0010 | 0.0016 |
| F04 | 4840001 | 4940000 | 0.0022 | 0.0013 | 0.0016 |
| F04 | 4860001 | 4960000 | 0.0024 | 0.0012 | 0.0020 |
| F04 | 4880001 | 4980000 | 0.0024 | 0.0010 | 0.0020 |

| Chr | Start | End | both | A1 | A2 |
|---|---|---|---|---|---|
| F04 | 4900001 | 5000000 | 0.0020 | 0.0015 | 0.0018 |
| F04 | 4920001 | 5020000 | 0.0026 | 0.0018 | 0.0024 |
| F04 | 4940001 | 5040000 | 0.0028 | 0.0022 | 0.0027 |
| F04 | 4960001 | 5060000 | 0.0034 | 0.0021 | 0.0033 |
| F04 | 4980001 | 5080000 | 0.0037 | 0.0022 | 0.0034 |
| F04 | 5000001 | 5100000 | 0.0039 | 0.0026 | 0.0035 |
| F04 | 5020001 | 5120000 | 0.0043 | 0.0024 | 0.0037 |
| F04 | 5040001 | 5140000 | 0.0041 | 0.0016 | 0.0033 |
| F04 | 5060001 | 5160000 | 0.0030 | 0.0018 | 0.0021 |
| F04 | 5080001 | 5180000 | 0.0029 | 0.0020 | 0.0019 |
| F04 | 5100001 | 5200000 | 0.0029 | 0.0012 | 0.0021 |
| F04 | 5120001 | 5220000 | 0.0018 | 0.0011 | 0.0012 |
| F04 | 5140001 | 5240000 | 0.0018 | 0.0013 | 0.0010 |
| F04 | 5160001 | 5260000 | 0.0019 | 0.0012 | 0.0012 |
| F04 | 5180001 | 5280000 | 0.0018 | 0.0013 | 0.0012 |
| F04 | 5200001 | 5300000 | 0.0024 | 0.0011 | 0.0016 |
| F04 | 5220001 | 5320000 | 0.0025 | 0.0016 | 0.0016 |
| F04 | 5240001 | 5340000 | 0.0030 | 0.0023 | 0.0023 |
| F04 | 5260001 | 5360000 | 0.0038 | 0.0020 | 0.0031 |
| F04 | 5280001 | 5380000 | 0.0035 | 0.0020 | 0.0028 |
| F04 | 5300001 | 5400000 | 0.0034 | 0.0022 | 0.0028 |
| F04 | 5320001 | 5420000 | 0.0033 | 0.0023 | 0.0029 |
| F04 | 5340001 | 5440000 | 0.0036 | 0.0015 | 0.0031 |
| F04 | 5360001 | 5460000 | 0.0027 | 0.0019 | 0.0023 |
| F04 | 5380001 | 5480000 | 0.0031 | 0.0018 | 0.0027 |
| F04 | 5400001 | 5500000 | 0.0032 | 0.0018 | 0.0028 |
| F04 | 5420001 | 5520000 | 0.0035 | 0.0014 | 0.0030 |
| F04 | 5440001 | 5540000 | 0.0028 | 0.0024 | 0.0023 |
| F04 | 5460001 | 5560000 | 0.0039 | 0.0023 | 0.0034 |
| F04 | 5480001 | 5580000 | 0.0036 | 0.0023 | 0.0031 |
| F04 | 5500001 | 5600000 | 0.0030 | 0.0021 | 0.0027 |
| F04 | 5520001 | 5620000 | 0.0026 | 0.0026 | 0.0023 |
| F04 | 5540001 | 5640000 | 0.0032 | 0.0018 | 0.0027 |
| F04 | 5560001 | 5660000 | 0.0023 | 0.0021 | 0.0017 |
| F04 | 5580001 | 5680000 | 0.0027 | 0.0030 | 0.0018 |
| F04 | 5600001 | 5700000 | 0.0036 | 0.0029 | 0.0024 |
| F04 | 5620001 | 5720000 | 0.0036 | 0.0024 | 0.0024 |
| F04 | 5640001 | 5740000 | 0.0032 | 0.0022 | 0.0019 |
| F04 | 5660001 | 5760000 | 0.0030 | 0.0030 | 0.0018 |
| F04 | 5680001 | 5780000 | 0.0040 | 0.0023 | 0.0029 |
| F04 | 5700001 | 5800000 | 0.0032 | 0.0029 | 0.0024 |
| F04 | 5720001 | 5820000 | 0.0037 | 0.0037 | 0.0030 |
| F04 | 5740001 | 5840000 | 0.0045 | 0.0040 | 0.0040 |
| F04 | 5760001 | 5860000 | 0.0047 | 0.0031 | 0.0042 |
| F04 | 5780001 | 5880000 | 0.0036 | 0.0034 | 0.0033 |
| F04 | 5800001 | 5900000 | 0.0040 | 0.0035 | 0.0037 |
| F04 | 5820001 | 5920000 | 0.0041 | 0.0026 | 0.0037 |
| F04 | 5840001 | 5940000 | 0.0031 | 0.0024 | 0.0028 |
| F04 | 5860001 | 5960000 | 0.0030 | 0.0030 | 0.0027 |
| F04 | 5880001 | 5980000 | 0.0037 | 0.0023 | 0.0033 |
| F04 | 5900001 | 6000000 | 0.0028 | 0.0025 | 0.0025 |
| F04 | 5920001 | 6020000 | 0.0030 | 0.0039 | 0.0027 |
| F04 | 5940001 | 6040000 | 0.0043 | 0.0041 | 0.0039 |
| F04 | 5960001 | 6060000 | 0.0046 | 0.0034 | 0.0040 |
| F04 | 5980001 | 6080000 | 0.0036 | 0.0039 | 0.0030 |
| F04 | 6000001 | 6100000 | 0.0041 | 0.0037 | 0.0032 |
| F04 | 6020001 | 6120000 | 0.0037 | 0.0024 | 0.0028 |
| F04 | 6040001 | 6140000 | 0.0026 | 0.0018 | 0.0016 |

| Chr | Start | End | both | A1 | A2 |
|---|---|---|---|---|---|
| F04 | 6060001 | 6160000 | 0.0021 | 0.0018 | 0.0011 |
| F04 | 6080001 | 6180000 | 0.0022 | 0.0013 | 0.0013 |
| F04 | 6100001 | 6200000 | 0.0019 | 0.0008 | 0.0011 |
| F04 | 6120001 | 6220000 | 0.0018 | 0.0032 | 0.0010 |
| F04 | 6140001 | 6240000 | 0.0042 | 0.0033 | 0.0035 |
| F04 | 6160001 | 6260000 | 0.0043 | 0.0036 | 0.0035 |
| F04 | 6180001 | 6280000 | 0.0045 | 0.0039 | 0.0037 |
| F04 | 6200001 | 6300000 | 0.0047 | 0.0036 | 0.0039 |
| F04 | 6220001 | 6320000 | 0.0043 | 0.0016 | 0.0036 |
| F04 | 6240001 | 6340000 | 0.0022 | 0.0024 | 0.0015 |
| F04 | 6260001 | 6360000 | 0.0031 | 0.0023 | 0.0023 |
| F04 | 6280001 | 6380000 | 0.0030 | 0.0025 | 0.0020 |
| F04 | 6300001 | 6400000 | 0.0032 | 0.0030 | 0.0020 |
| F04 | 6320001 | 6420000 | 0.0038 | 0.0025 | 0.0026 |
| F04 | 6340001 | 6440000 | 0.0034 | 0.0020 | 0.0023 |
| F04 | 6360001 | 6460000 | 0.0029 | 0.0016 | 0.0018 |
| F04 | 6380001 | 6480000 | 0.0025 | 0.0014 | 0.0018 |
| F04 | 6400001 | 6500000 | 0.0024 | 0.0010 | 0.0017 |
| F04 | 6420001 | 6520000 | 0.0020 | 0.0025 | 0.0013 |
| F04 | 6440001 | 6540000 | 0.0036 | 0.0032 | 0.0026 |
| F04 | 6460001 | 6560000 | 0.0046 | 0.0036 | 0.0035 |
| F04 | 6480001 | 6580000 | 0.0051 | 0.0034 | 0.0038 |
| F04 | 6500001 | 6600000 | 0.0049 | 0.0035 | 0.0037 |
| F04 | 6520001 | 6620000 | 0.0052 | 0.0021 | 0.0038 |
| F04 | 6540001 | 6640000 | 0.0036 | 0.0010 | 0.0025 |
| F04 | 6560001 | 6660000 | 0.0023 | 0.0008 | 0.0012 |
| F04 | 6580001 | 6680000 | 0.0021 | 0.0011 | 0.0011 |
| F04 | 6600001 | 6700000 | 0.0024 | 0.0013 | 0.0014 |
| F04 | 6620001 | 6720000 | 0.0025 | 0.0021 | 0.0016 |
| F04 | 6640001 | 6740000 | 0.0036 | 0.0024 | 0.0026 |
| F04 | 6660001 | 6760000 | 0.0038 | 0.0028 | 0.0030 |
| F04 | 6680001 | 6780000 | 0.0042 | 0.0029 | 0.0032 |
| F04 | 6700001 | 6800000 | 0.0042 | 0.0027 | 0.0033 |
| F04 | 6720001 | 6820000 | 0.0037 | 0.0018 | 0.0029 |
| F04 | 6740001 | 6840000 | 0.0028 | 0.0016 | 0.0019 |
| F04 | 6760001 | 6860000 | 0.0026 | 0.0014 | 0.0016 |
| F04 | 6780001 | 6880000 | 0.0024 | 0.0009 | 0.0014 |
| F04 | 6800001 | 6900000 | 0.0018 | 0.0016 | 0.0009 |
| F04 | 6820001 | 6920000 | 0.0026 | 0.0024 | 0.0015 |
| F04 | 6840001 | 6940000 | 0.0034 | 0.0023 | 0.0023 |
| F04 | 6860001 | 6960000 | 0.0034 | 0.0029 | 0.0021 |
| F04 | 6880001 | 6980000 | 0.0039 | 0.0032 | 0.0026 |
| F04 | 6900001 | 7000000 | 0.0044 | 0.0028 | 0.0029 |
| F04 | 6920001 | 7020000 | 0.0038 | 0.0023 | 0.0024 |
| F04 | 6940001 | 7040000 | 0.0035 | 0.0034 | 0.0021 |
| F04 | 6960001 | 7060000 | 0.0044 | 0.0031 | 0.0031 |
| F04 | 6980001 | 7080000 | 0.0039 | 0.0034 | 0.0028 |
| F04 | 7000001 | 7100000 | 0.0041 | 0.0038 | 0.0032 |
| F04 | 7020001 | 7120000 | 0.0045 | 0.0036 | 0.0037 |
| F04 | 7040001 | 7140000 | 0.0041 | 0.0029 | 0.0036 |
| F04 | 7060001 | 7160000 | 0.0034 | 0.0025 | 0.0030 |
| F04 | 7080001 | 7180000 | 0.0031 | 0.0045 | 0.0027 |
| F04 | 7100001 | 7200000 | 0.0054 | 0.0037 | 0.0049 |
| F04 | 7120001 | 7220000 | 0.0048 | 0.0035 | 0.0042 |
| F04 | 7140001 | 7240000 | 0.0045 | 0.0032 | 0.0039 |
| F04 | 7160001 | 7260000 | 0.0043 | 0.0032 | 0.0036 |
| F04 | 7180001 | 7280000 | 0.0043 | 0.0006 | 0.0035 |
| F04 | 7200001 | 7300000 | 0.0015 | 0.0006 | 0.0007 |

| Chr | Start | End | both | A1 | A2 |
|---|---|---|---|---|---|
| F04 | 7220001 | 7320000 | 0.0014 | 0.0007 | 0.0006 |
| F04 | 7240001 | 7340000 | 0.0015 | 0.0010 | 0.0006 |
| F04 | 7260001 | 7360000 | 0.0017 | 0.0010 | 0.0009 |
| F04 | 7280001 | 7380000 | 0.0019 | 0.0010 | 0.0010 |
| F04 | 7300001 | 7400000 | 0.0019 | 0.0018 | 0.0010 |
| F04 | 7320001 | 7420000 | 0.0028 | 0.0018 | 0.0019 |
| F04 | 7340001 | 7440000 | 0.0028 | 0.0020 | 0.0019 |
| F04 | 7360001 | 7460000 | 0.0030 | 0.0020 | 0.0021 |
| F04 | 7380001 | 7480000 | 0.0028 | 0.0029 | 0.0019 |
| F04 | 7400001 | 7500000 | 0.0038 | 0.0032 | 0.0030 |
| F04 | 7420001 | 7520000 | 0.0038 | 0.0036 | 0.0031 |
| F04 | 7440001 | 7540000 | 0.0043 | 0.0035 | 0.0036 |
| F04 | 7460001 | 7560000 | 0.0041 | 0.0034 | 0.0035 |
| F04 | 7480001 | 7580000 | 0.0039 | 0.0027 | 0.0035 |
| F04 | 7500001 | 7600000 | 0.0029 | 0.0018 | 0.0025 |
| F04 | 7520001 | 7620000 | 0.0020 | 0.0018 | 0.0016 |
| F04 | 7540001 | 7640000 | 0.0018 | 0.0015 | 0.0016 |
| F04 | 7560001 | 7660000 | 0.0014 | 0.0020 | 0.0012 |
| F04 | 7580001 | 7680000 | 0.0018 | 0.0024 | 0.0014 |
| F04 | 7600001 | 7700000 | 0.0022 | 0.0030 | 0.0018 |
| F04 | 7620001 | 7720000 | 0.0029 | 0.0035 | 0.0025 |
| F04 | 7640001 | 7740000 | 0.0034 | 0.0039 | 0.0029 |
| F04 | 7660001 | 7760000 | 0.0036 | 0.0041 | 0.0028 |
| F04 | 7680001 | 7780000 | 0.0037 | 0.0039 | 0.0028 |
| F04 | 7700001 | 7800000 | 0.0038 | 0.0039 | 0.0029 |
| F04 | 7720001 | 7820000 | 0.0037 | 0.0030 | 0.0027 |
| F04 | 7740001 | 7840000 | 0.0027 | 0.0028 | 0.0019 |
| F04 | 7760001 | 7860000 | 0.0028 | 0.0026 | 0.0021 |
| F04 | 7780001 | 7880000 | 0.0027 | 0.0027 | 0.0022 |
| F04 | 7800001 | 7900000 | 0.0022 | 0.0022 | 0.0016 |
| F04 | 7820001 | 7920000 | 0.0016 | 0.0024 | 0.0011 |
| F04 | 7840001 | 7940000 | 0.0019 | 0.0019 | 0.0014 |
| F04 | 7860001 | 7960000 | 0.0014 | 0.0017 | 0.0008 |
| F04 | 7880001 | 7980000 | 0.0015 | 0.0021 | 0.0009 |
| F04 | 7900001 | 8000000 | 0.0023 | 0.0025 | 0.0017 |
| F04 | 7920001 | 8020000 | 0.0028 | 0.0028 | 0.0021 |
| F04 | 7940001 | 8040000 | 0.0028 | 0.0035 | 0.0020 |
| F04 | 7960001 | 8060000 | 0.0036 | 0.0044 | 0.0028 |
| F04 | 7980001 | 8080000 | 0.0041 | 0.0039 | 0.0031 |
| F04 | 8000001 | 8100000 | 0.0034 | 0.0037 | 0.0022 |
| F04 | 8020001 | 8120000 | 0.0032 | 0.0030 | 0.0020 |
| F04 | 8040001 | 8140000 | 0.0027 | 0.0026 | 0.0017 |
| F04 | 8060001 | 8160000 | 0.0024 | 0.0016 | 0.0013 |
| F04 | 8080001 | 8180000 | 0.0016 | 0.0011 | 0.0008 |
| F04 | 8100001 | 8200000 | 0.0013 | 0.0007 | 0.0007 |
| F04 | 8120001 | 8220000 | 0.0011 | 0.0014 | 0.0007 |
| F04 | 8140001 | 8240000 | 0.0020 | 0.0011 | 0.0013 |
| F04 | 8160001 | 8260000 | 0.0018 | 0.0016 | 0.0011 |
| F04 | 8180001 | 8280000 | 0.0024 | 0.0025 | 0.0016 |
| F04 | 8200001 | 8300000 | 0.0033 | 0.0026 | 0.0027 |
| F04 | 8220001 | 8320000 | 0.0035 | 0.0023 | 0.0028 |
| F04 | 8240001 | 8340000 | 0.0031 | 0.0037 | 0.0025 |
| F04 | 8260001 | 8360000 | 0.0046 | 0.0032 | 0.0041 |
| F04 | 8280001 | 8380000 | 0.0043 | 0.0025 | 0.0036 |
| F04 | 8300001 | 8400000 | 0.0034 | 0.0030 | 0.0026 |
| F04 | 8320001 | 8420000 | 0.0038 | 0.0030 | 0.0032 |
| F04 | 8340001 | 8440000 | 0.0037 | 0.0018 | 0.0032 |
| F04 | 8360001 | 8460000 | 0.0025 | 0.0021 | 0.0020 |

| Chr | Start | End | both | A1 | A2 |
|---|---|---|---|---|---|
| F04 | 8380001 | 8480000 | 0.0027 | 0.0028 | 0.0024 |
| F04 | 8400001 | 8500000 | 0.0033 | 0.0024 | 0.0030 |
| F04 | 8420001 | 8520000 | 0.0029 | 0.0021 | 0.0026 |
| F04 | 8440001 | 8540000 | 0.0026 | 0.0020 | 0.0023 |
| F04 | 8460001 | 8560000 | 0.0025 | 0.0018 | 0.0021 |
| F04 | 8480001 | 8580000 | 0.0023 | 0.0013 | 0.0017 |
| F04 | 8500001 | 8600000 | 0.0019 | 0.0022 | 0.0014 |
| F04 | 8520001 | 8620000 | 0.0029 | 0.0031 | 0.0022 |
| F04 | 8540001 | 8640000 | 0.0040 | 0.0030 | 0.0032 |
| F04 | 8560001 | 8660000 | 0.0040 | 0.0031 | 0.0032 |
| F04 | 8580001 | 8680000 | 0.0040 | 0.0029 | 0.0033 |
| F04 | 8600001 | 8700000 | 0.0040 | 0.0018 | 0.0032 |
| F04 | 8620001 | 8720000 | 0.0027 | 0.0018 | 0.0021 |
| F04 | 8640001 | 8740000 | 0.0027 | 0.0024 | 0.0021 |
| F04 | 8660001 | 8760000 | 0.0032 | 0.0025 | 0.0027 |
| F04 | 8680001 | 8780000 | 0.0031 | 0.0029 | 0.0026 |
| F04 | 8700001 | 8800000 | 0.0035 | 0.0029 | 0.0030 |
| F04 | 8720001 | 8820000 | 0.0036 | 0.0021 | 0.0030 |
| F04 | 8740001 | 8840000 | 0.0028 | 0.0013 | 0.0023 |
| F04 | 8760001 | 8860000 | 0.0019 | 0.0012 | 0.0015 |
| F04 | 8780001 | 8880000 | 0.0019 | 0.0011 | 0.0015 |
| F04 | 8800001 | 8900000 | 0.0015 | 0.0012 | 0.0013 |
| F04 | 8820001 | 8920000 | 0.0019 | 0.0013 | 0.0015 |
| F04 | 8840001 | 8940000 | 0.0020 | 0.0019 | 0.0015 |
| F04 | 8860001 | 8960000 | 0.0030 | 0.0019 | 0.0022 |
| F04 | 8880001 | 8980000 | 0.0033 | 0.0016 | 0.0023 |
| F04 | 8900001 | 9000000 | 0.0027 | 0.0021 | 0.0017 |
| F04 | 8920001 | 9020000 | 0.0027 | 0.0027 | 0.0017 |
| F04 | 8940001 | 9040000 | 0.0030 | 0.0028 | 0.0022 |
| F04 | 8960001 | 9060000 | 0.0025 | 0.0036 | 0.0020 |
| F04 | 8980001 | 9080000 | 0.0029 | 0.0040 | 0.0026 |
| F04 | 9000001 | 9100000 | 0.0035 | 0.0041 | 0.0032 |
| F04 | 9020001 | 9120000 | 0.0040 | 0.0038 | 0.0038 |
| F04 | 9040001 | 9140000 | 0.0037 | 0.0034 | 0.0036 |
| F04 | 9060001 | 9160000 | 0.0035 | 0.0029 | 0.0034 |
| F04 | 9080001 | 9180000 | 0.0031 | 0.0033 | 0.0031 |
| F04 | 9100001 | 9200000 | 0.0033 | 0.0029 | 0.0031 |
| F04 | 9120001 | 9220000 | 0.0028 | 0.0035 | 0.0027 |
| F04 | 9140001 | 9240000 | 0.0034 | 0.0039 | 0.0033 |
| F04 | 9160001 | 9260000 | 0.0038 | 0.0037 | 0.0036 |
| F04 | 9180001 | 9280000 | 0.0035 | 0.0031 | 0.0033 |
| F04 | 9200001 | 9300000 | 0.0027 | 0.0032 | 0.0025 |
| F04 | 9220001 | 9320000 | 0.0026 | 0.0023 | 0.0023 |
| F04 | 9240001 | 9340000 | 0.0014 | 0.0017 | 0.0011 |
| F04 | 9260001 | 9360000 | 0.0008 | 0.0017 | 0.0004 |
| F04 | 9280001 | 9380000 | 0.0008 | 0.0018 | 0.0005 |
| F04 | 9300001 | 9400000 | 0.0010 | 0.0017 | 0.0007 |
| F04 | 9320001 | 9420000 | 0.0011 | 0.0020 | 0.0009 |
| F04 | 9340001 | 9440000 | 0.0015 | 0.0027 | 0.0012 |
| F04 | 9360001 | 9460000 | 0.0020 | 0.0041 | 0.0017 |
| F04 | 9380001 | 9480000 | 0.0038 | 0.0039 | 0.0036 |
| F04 | 9400001 | 9500000 | 0.0038 | 0.0043 | 0.0036 |
| F04 | 9420001 | 9520000 | 0.0043 | 0.0046 | 0.0040 |
| F04 | 9440001 | 9540000 | 0.0046 | 0.0043 | 0.0044 |
| F04 | 9460001 | 9560000 | 0.0043 | 0.0032 | 0.0041 |
| F04 | 9480001 | 9580000 | 0.0026 | 0.0042 | 0.0023 |
| F04 | 9500001 | 9600000 | 0.0035 | 0.0042 | 0.0031 |
| F04 | 9520001 | 9620000 | 0.0033 | 0.0037 | 0.0029 |

| Chr | Start | End | both | A1 | A2 |
|---|---|---|---|---|---|
| F04 | 9540001 | 9640000 | 0.0029 | 0.0037 | 0.0026 |
| F04 | 9560001 | 9660000 | 0.0035 | 0.0030 | 0.0031 |
| F04 | 9580001 | 9680000 | 0.0034 | 0.0021 | 0.0031 |
| F04 | 9600001 | 9700000 | 0.0027 | 0.0015 | 0.0025 |
| F04 | 9620001 | 9720000 | 0.0028 | 0.0016 | 0.0027 |
| F04 | 9640001 | 9740000 | 0.0031 | 0.0014 | 0.0030 |
| F04 | 9660001 | 9760000 | 0.0030 | 0.0026 | 0.0030 |
| F04 | 9680001 | 9780000 | 0.0040 | 0.0023 | 0.0039 |
| F04 | 9700001 | 9800000 | 0.0038 | 0.0025 | 0.0036 |
| F04 | 9720001 | 9820000 | 0.0037 | 0.0021 | 0.0035 |
| F04 | 9740001 | 9840000 | 0.0033 | 0.0021 | 0.0029 |
| F04 | 9760001 | 9860000 | 0.0030 | 0.0013 | 0.0024 |
| F04 | 9780001 | 9880000 | 0.0021 | 0.0014 | 0.0015 |
| F04 | 9800001 | 9900000 | 0.0021 | 0.0021 | 0.0016 |
| F04 | 9820001 | 9920000 | 0.0027 | 0.0024 | 0.0022 |
| F04 | 9840001 | 9940000 | 0.0029 | 0.0030 | 0.0024 |
| F04 | 9860001 | 9960000 | 0.0034 | 0.0029 | 0.0030 |
| F04 | 9880001 | 9980000 | 0.0036 | 0.0029 | 0.0029 |
| F04 | 9900001 | 10000000 | 0.0036 | 0.0021 | 0.0028 |
| F04 | 9920001 | 10020000 | 0.0028 | 0.0019 | 0.0020 |
| F04 | 9940001 | 10040000 | 0.0026 | 0.0011 | 0.0018 |
| F04 | 9960001 | 10060000 | 0.0018 | 0.0010 | 0.0012 |
| F04 | 9980001 | 10080000 | 0.0016 | 0.0015 | 0.0010 |
| F04 | 10000001 | 10100000 | 0.0021 | 0.0018 | 0.0014 |
| F04 | 10020001 | 10120000 | 0.0026 | 0.0020 | 0.0018 |
| F04 | 10040001 | 10140000 | 0.0028 | 0.0022 | 0.0020 |
| F04 | 10060001 | 10160000 | 0.0032 | 0.0022 | 0.0022 |
| F04 | 10080001 | 10180000 | 0.0032 | 0.0017 | 0.0022 |
| F04 | 10100001 | 10200000 | 0.0029 | 0.0015 | 0.0019 |
| F04 | 10120001 | 10220000 | 0.0027 | 0.0018 | 0.0015 |
| F04 | 10140001 | 10240000 | 0.0033 | 0.0017 | 0.0019 |
| F04 | 10160001 | 10260000 | 0.0031 | 0.0018 | 0.0017 |
| F04 | 10180001 | 10280000 | 0.0030 | 0.0021 | 0.0018 |
| F04 | 10200001 | 10300000 | 0.0032 | 0.0027 | 0.0021 |
| F04 | 10220001 | 10320000 | 0.0036 | 0.0023 | 0.0028 |
| F04 | 10240001 | 10340000 | 0.0028 | 0.0021 | 0.0024 |
| F04 | 10260001 | 10360000 | 0.0026 | 0.0028 | 0.0023 |
| F04 | 10280001 | 10380000 | 0.0034 | 0.0028 | 0.0028 |
| F04 | 10300001 | 10400000 | 0.0034 | 0.0024 | 0.0027 |
| F04 | 10320001 | 10420000 | 0.0030 | 0.0023 | 0.0023 |
| F04 | 10340001 | 10440000 | 0.0029 | 0.0022 | 0.0021 |
| F04 | 10360001 | 10460000 | 0.0028 | 0.0019 | 0.0021 |
| F04 | 10380001 | 10480000 | 0.0025 | 0.0015 | 0.0019 |
| F04 | 10400001 | 10500000 | 0.0022 | 0.0016 | 0.0015 |
| F04 | 10420001 | 10520000 | 0.0024 | 0.0029 | 0.0016 |
| F04 | 10440001 | 10540000 | 0.0043 | 0.0038 | 0.0033 |
| F04 | 10460001 | 10560000 | 0.0051 | 0.0040 | 0.0042 |
| F04 | 10480001 | 10580000 | 0.0053 | 0.0044 | 0.0043 |
| F04 | 10500001 | 10600000 | 0.0057 | 0.0043 | 0.0047 |
| F04 | 10520001 | 10620000 | 0.0056 | 0.0036 | 0.0049 |
| F04 | 10540001 | 10640000 | 0.0045 | 0.0037 | 0.0037 |
| F04 | 10560001 | 10660000 | 0.0049 | 0.0033 | 0.0040 |
| F04 | 10580001 | 10680000 | 0.0043 | 0.0030 | 0.0036 |
| F04 | 10600001 | 10700000 | 0.0040 | 0.0026 | 0.0033 |
| F04 | 10620001 | 10720000 | 0.0036 | 0.0025 | 0.0027 |
| F04 | 10640001 | 10740000 | 0.0034 | 0.0017 | 0.0027 |
| F04 | 10660001 | 10760000 | 0.0026 | 0.0018 | 0.0017 |
| F04 | 10680001 | 10780000 | 0.0028 | 0.0024 | 0.0017 |

| Chr | Start | End | both | A1 | A2 |
|---|---|---|---|---|---|
| F04 | 10700001 | 10800000 | 0.0034 | 0.0028 | 0.0025 |
| F04 | 10720001 | 10820000 | 0.0038 | 0.0025 | 0.0030 |
| F04 | 10740001 | 10840000 | 0.0037 | 0.0026 | 0.0027 |
| F04 | 10760001 | 10860000 | 0.0036 | 0.0023 | 0.0028 |
| F04 | 10780001 | 10880000 | 0.0034 | 0.0017 | 0.0025 |
| F04 | 10800001 | 10900000 | 0.0028 | 0.0011 | 0.0020 |
| F04 | 10820001 | 10920000 | 0.0021 | 0.0012 | 0.0013 |
| F04 | 10840001 | 10940000 | 0.0022 | 0.0014 | 0.0015 |
| F04 | 10860001 | 10960000 | 0.0025 | 0.0016 | 0.0016 |
| F04 | 10880001 | 10980000 | 0.0027 | 0.0019 | 0.0018 |
| F04 | 10900001 | 11000000 | 0.0028 | 0.0025 | 0.0019 |
| F04 | 10920001 | 11020000 | 0.0035 | 0.0025 | 0.0026 |
| F04 | 10940001 | 11040000 | 0.0035 | 0.0027 | 0.0025 |
| F04 | 10960001 | 11060000 | 0.0038 | 0.0027 | 0.0028 |
| F04 | 10980001 | 11080000 | 0.0039 | 0.0026 | 0.0029 |
| F04 | 11000001 | 11100000 | 0.0039 | 0.0024 | 0.0027 |
| F04 | 11020001 | 11120000 | 0.0038 | 0.0025 | 0.0025 |
| F04 | 11040001 | 11140000 | 0.0038 | 0.0030 | 0.0026 |
| F04 | 11060001 | 11160000 | 0.0042 | 0.0030 | 0.0031 |
| F04 | 11080001 | 11180000 | 0.0041 | 0.0031 | 0.0030 |
| F04 | 11100001 | 11200000 | 0.0040 | 0.0032 | 0.0032 |
| F04 | 11120001 | 11220000 | 0.0040 | 0.0029 | 0.0033 |
| F04 | 11140001 | 11240000 | 0.0037 | 0.0023 | 0.0030 |
| F04 | 11160001 | 11260000 | 0.0030 | 0.0033 | 0.0024 |
| F04 | 11180001 | 11280000 | 0.0041 | 0.0030 | 0.0035 |
| F04 | 11200001 | 11300000 | 0.0037 | 0.0025 | 0.0031 |
| F04 | 11220001 | 11320000 | 0.0031 | 0.0040 | 0.0026 |
| F04 | 11240001 | 11340000 | 0.0048 | 0.0042 | 0.0042 |
| F04 | 11260001 | 11360000 | 0.0052 | 0.0036 | 0.0043 |
| F04 | 11280001 | 11380000 | 0.0045 | 0.0042 | 0.0036 |
| F04 | 11300001 | 11400000 | 0.0054 | 0.0042 | 0.0043 |
| F04 | 11320001 | 11420000 | 0.0052 | 0.0039 | 0.0044 |
| F04 | 11340001 | 11440000 | 0.0049 | 0.0034 | 0.0040 |
| F04 | 11360001 | 11460000 | 0.0044 | 0.0030 | 0.0034 |
| F04 | 11380001 | 11480000 | 0.0039 | 0.0024 | 0.0030 |
| F04 | 11400001 | 11500000 | 0.0034 | 0.0024 | 0.0023 |
| F04 | 11420001 | 11520000 | 0.0035 | 0.0011 | 0.0022 |
| F04 | 11440001 | 11540000 | 0.0021 | 0.0012 | 0.0009 |
| F04 | 11460001 | 11560000 | 0.0023 | 0.0014 | 0.0010 |
| F04 | 11480001 | 11580000 | 0.0027 | 0.0018 | 0.0012 |
| F04 | 11500001 | 11600000 | 0.0030 | 0.0022 | 0.0016 |
| F04 | 11520001 | 11620000 | 0.0032 | 0.0023 | 0.0020 |
| F04 | 11540001 | 11640000 | 0.0033 | 0.0021 | 0.0022 |
| F04 | 11560001 | 11660000 | 0.0030 | 0.0020 | 0.0019 |
| F04 | 11580001 | 11680000 | 0.0029 | 0.0018 | 0.0019 |
| F04 | 11600001 | 11700000 | 0.0028 | 0.0018 | 0.0017 |
| F04 | 11620001 | 11720000 | 0.0031 | 0.0024 | 0.0016 |
| F04 | 11640001 | 11740000 | 0.0038 | 0.0026 | 0.0020 |
| F04 | 11660001 | 11760000 | 0.0041 | 0.0042 | 0.0023 |
| F04 | 11680001 | 11780000 | 0.0058 | 0.0040 | 0.0040 |
| F04 | 11700001 | 11800000 | 0.0057 | 0.0048 | 0.0039 |
| F04 | 11720001 | 11820000 | 0.0065 | 0.0053 | 0.0049 |
| F04 | 11740001 | 11840000 | 0.0070 | 0.0049 | 0.0056 |
| F04 | 11760001 | 11860000 | 0.0064 | 0.0034 | 0.0051 |
| F04 | 11780001 | 11880000 | 0.0049 | 0.0034 | 0.0035 |
| F04 | 11800001 | 11900000 | 0.0048 | 0.0025 | 0.0034 |
| F04 | 11820001 | 11920000 | 0.0037 | 0.0019 | 0.0023 |
| F04 | 11840001 | 11940000 | 0.0031 | 0.0018 | 0.0015 |

| Chr | Start | End | both | A1 | A2 |
|---|---|---|---|---|---|
| F04 | 11860001 | 11960000 | 0.0032 | 0.0014 | 0.0014 |
| F04 | 11880001 | 11980000 | 0.0025 | 0.0015 | 0.0011 |
| F04 | 11900001 | 12000000 | 0.0026 | 0.0016 | 0.0013 |
| F04 | 11920001 | 12020000 | 0.0025 | 0.0013 | 0.0014 |
| F04 | 11940001 | 12040000 | 0.0022 | 0.0013 | 0.0011 |
| F04 | 11960001 | 12060000 | 0.0023 | 0.0019 | 0.0012 |
| F04 | 11980001 | 12080000 | 0.0032 | 0.0020 | 0.0018 |
| F04 | 12000001 | 12100000 | 0.0033 | 0.0019 | 0.0020 |
| F04 | 12020001 | 12120000 | 0.0032 | 0.0020 | 0.0019 |
| F04 | 12040001 | 12140000 | 0.0033 | 0.0023 | 0.0020 |
| F04 | 12060001 | 12160000 | 0.0036 | 0.0019 | 0.0023 |
| F04 | 12080001 | 12180000 | 0.0030 | 0.0016 | 0.0020 |
| F04 | 12100001 | 12200000 | 0.0027 | 0.0019 | 0.0016 |
| F04 | 12120001 | 12220000 | 0.0030 | 0.0018 | 0.0020 |
| F04 | 12140001 | 12240000 | 0.0028 | 0.0018 | 0.0018 |
| F04 | 12160001 | 12260000 | 0.0028 | 0.0016 | 0.0018 |
| F04 | 12180001 | 12280000 | 0.0025 | 0.0017 | 0.0016 |
| F04 | 12200001 | 12300000 | 0.0025 | 0.0019 | 0.0017 |
| F04 | 12220001 | 12320000 | 0.0028 | 0.0018 | 0.0019 |
| F04 | 12240001 | 12340000 | 0.0026 | 0.0014 | 0.0018 |
| F04 | 12260001 | 12360000 | 0.0022 | 0.0020 | 0.0015 |
| F04 | 12280001 | 12380000 | 0.0027 | 0.0022 | 0.0020 |
| F04 | 12300001 | 12400000 | 0.0032 | 0.0017 | 0.0023 |
| F04 | 12320001 | 12420000 | 0.0025 | 0.0021 | 0.0018 |
| F04 | 12340001 | 12440000 | 0.0029 | 0.0023 | 0.0022 |
| F04 | 12360001 | 12460000 | 0.0030 | 0.0021 | 0.0023 |
| F04 | 12380001 | 12480000 | 0.0030 | 0.0018 | 0.0024 |
| F04 | 12400001 | 12500000 | 0.0026 | 0.0018 | 0.0020 |
| F04 | 12420001 | 12520000 | 0.0026 | 0.0014 | 0.0020 |
| F04 | 12440001 | 12540000 | 0.0024 | 0.0016 | 0.0017 |
| F04 | 12460001 | 12560000 | 0.0027 | 0.0016 | 0.0020 |
| F04 | 12480001 | 12580000 | 0.0025 | 0.0016 | 0.0018 |
| F04 | 12500001 | 12600000 | 0.0025 | 0.0015 | 0.0019 |
| F04 | 12520001 | 12620000 | 0.0022 | 0.0018 | 0.0018 |
| F04 | 12540001 | 12640000 | 0.0023 | 0.0015 | 0.0020 |
| F04 | 12560001 | 12660000 | 0.0023 | 0.0012 | 0.0017 |
| F04 | 12580001 | 12680000 | 0.0022 | 0.0012 | 0.0014 |
| F04 | 12600001 | 12700000 | 0.0024 | 0.0020 | 0.0013 |
| F04 | 12620001 | 12720000 | 0.0035 | 0.0023 | 0.0021 |
| F04 | 12640001 | 12740000 | 0.0039 | 0.0024 | 0.0024 |
| F04 | 12660001 | 12760000 | 0.0038 | 0.0027 | 0.0025 |
| F04 | 12680001 | 12780000 | 0.0041 | 0.0027 | 0.0029 |
| F04 | 12700001 | 12800000 | 0.0039 | 0.0021 | 0.0029 |
| F04 | 12720001 | 12820000 | 0.0031 | 0.0013 | 0.0024 |
| F04 | 12740001 | 12840000 | 0.0022 | 0.0017 | 0.0015 |
| F04 | 12760001 | 12860000 | 0.0026 | 0.0015 | 0.0020 |
| F04 | 12780001 | 12880000 | 0.0025 | 0.0017 | 0.0019 |
| F04 | 12800001 | 12900000 | 0.0028 | 0.0015 | 0.0020 |
| F04 | 12820001 | 12920000 | 0.0027 | 0.0022 | 0.0018 |
| F04 | 12840001 | 12940000 | 0.0036 | 0.0024 | 0.0027 |
| F04 | 12860001 | 12960000 | 0.0038 | 0.0023 | 0.0029 |
| F04 | 12880001 | 12980000 | 0.0035 | 0.0023 | 0.0025 |
| F04 | 12900001 | 13000000 | 0.0034 | 0.0022 | 0.0025 |
| F04 | 12920001 | 13020000 | 0.0030 | 0.0016 | 0.0024 |
| F04 | 12940001 | 13040000 | 0.0023 | 0.0018 | 0.0017 |
| F04 | 12960001 | 13060000 | 0.0024 | 0.0023 | 0.0019 |
| F04 | 12980001 | 13080000 | 0.0030 | 0.0020 | 0.0024 |
| F04 | 13000001 | 13100000 | 0.0028 | 0.0027 | 0.0021 |

| Chr | Start | End | both | A1 | A2 |
|---|---|---|---|---|---|
| F04 | 13020001 | 13120000 | 0.0037 | 0.0029 | 0.0030 |
| F04 | 13040001 | 13140000 | 0.0039 | 0.0031 | 0.0033 |
| F04 | 13060001 | 13160000 | 0.0039 | 0.0049 | 0.0033 |
| F04 | 13080001 | 13180000 | 0.0057 | 0.0049 | 0.0052 |
| F04 | 13100001 | 13200000 | 0.0058 | 0.0047 | 0.0052 |
| F04 | 13120001 | 13220000 | 0.0055 | 0.0044 | 0.0047 |
| F04 | 13140001 | 13240000 | 0.0053 | 0.0035 | 0.0043 |
| F04 | 13160001 | 13260000 | 0.0046 | 0.0017 | 0.0035 |
| F04 | 13180001 | 13280000 | 0.0026 | 0.0020 | 0.0016 |
| F04 | 13200001 | 13300000 | 0.0029 | 0.0022 | 0.0019 |
| F04 | 13220001 | 13320000 | 0.0032 | 0.0022 | 0.0023 |
| F04 | 13240001 | 13340000 | 0.0032 | 0.0024 | 0.0024 |
| F04 | 13260001 | 13360000 | 0.0031 | 0.0022 | 0.0025 |
| F04 | 13280001 | 13380000 | 0.0030 | 0.0022 | 0.0024 |
| F04 | 13300001 | 13400000 | 0.0029 | 0.0017 | 0.0025 |
| F04 | 13320001 | 13420000 | 0.0022 | 0.0020 | 0.0018 |
| F04 | 13340001 | 13440000 | 0.0025 | 0.0019 | 0.0021 |
| F04 | 13360001 | 13460000 | 0.0024 | 0.0015 | 0.0020 |
| F04 | 13380001 | 13480000 | 0.0018 | 0.0018 | 0.0015 |
| F04 | 13400001 | 13500000 | 0.0019 | 0.0016 | 0.0018 |
| F04 | 13420001 | 13520000 | 0.0018 | 0.0010 | 0.0016 |
| F04 | 13440001 | 13540000 | 0.0011 | 0.0008 | 0.0011 |
| F04 | 13460001 | 13560000 | 0.0008 | 0.0012 | 0.0008 |
| F04 | 13480001 | 13580000 | 0.0013 | 0.0007 | 0.0013 |
| F04 | 13500001 | 13600000 | 0.0008 | 0.0009 | 0.0007 |
| F04 | 13520001 | 13620000 | 0.0012 | 0.0010 | 0.0011 |
| F04 | 13540001 | 13640000 | 0.0016 | 0.0013 | 0.0013 |
| F04 | 13560001 | 13660000 | 0.0023 | 0.0015 | 0.0018 |
| F04 | 13580001 | 13680000 | 0.0026 | 0.0014 | 0.0019 |
| F04 | 13600001 | 13700000 | 0.0027 | 0.0021 | 0.0018 |
| F04 | 13620001 | 13720000 | 0.0034 | 0.0026 | 0.0024 |
| F04 | 13640001 | 13740000 | 0.0038 | 0.0033 | 0.0029 |
| F04 | 13660001 | 13760000 | 0.0045 | 0.0028 | 0.0035 |
| F04 | 13680001 | 13780000 | 0.0043 | 0.0028 | 0.0032 |
| F04 | 13700001 | 13800000 | 0.0041 | 0.0021 | 0.0032 |
| F04 | 13720001 | 13820000 | 0.0033 | 0.0018 | 0.0024 |
| F04 | 13740001 | 13840000 | 0.0030 | 0.0014 | 0.0021 |
| F04 | 13760001 | 13860000 | 0.0026 | 0.0016 | 0.0016 |
| F04 | 13780001 | 13880000 | 0.0025 | 0.0016 | 0.0017 |
| F04 | 13800001 | 13900000 | 0.0025 | 0.0017 | 0.0016 |
| F04 | 13820001 | 13920000 | 0.0026 | 0.0020 | 0.0019 |
| F04 | 13840001 | 13940000 | 0.0029 | 0.0024 | 0.0021 |
| F04 | 13860001 | 13960000 | 0.0032 | 0.0031 | 0.0026 |
| F04 | 13880001 | 13980000 | 0.0041 | 0.0033 | 0.0035 |
| F04 | 13900001 | 14000000 | 0.0044 | 0.0031 | 0.0039 |
| F04 | 13920001 | 14020000 | 0.0043 | 0.0029 | 0.0035 |
| F04 | 13940001 | 14040000 | 0.0039 | 0.0028 | 0.0034 |
| F04 | 13960001 | 14060000 | 0.0037 | 0.0023 | 0.0032 |
| F04 | 13980001 | 14080000 | 0.0032 | 0.0024 | 0.0026 |
| F04 | 14000001 | 14100000 | 0.0032 | 0.0033 | 0.0027 |
| F04 | 14020001 | 14120000 | 0.0043 | 0.0041 | 0.0037 |
| F04 | 14040001 | 14140000 | 0.0053 | 0.0034 | 0.0047 |
| F04 | 14060001 | 14160000 | 0.0044 | 0.0036 | 0.0038 |
| F04 | 14080001 | 14180000 | 0.0047 | 0.0035 | 0.0040 |
| F04 | 14100001 | 14200000 | 0.0045 | 0.0025 | 0.0038 |
| F04 | 14120001 | 14220000 | 0.0035 | 0.0018 | 0.0028 |
| F04 | 14140001 | 14240000 | 0.0027 | 0.0020 | 0.0020 |
| F04 | 14160001 | 14260000 | 0.0030 | 0.0015 | 0.0023 |

| Chr | Start | End | both | A1 | A2 |
|---|---|---|---|---|---|
| F04 | 14180001 | 14280000 | 0.0024 | 0.0015 | 0.0019 |
| F04 | 14200001 | 14300000 | 0.0024 | 0.0017 | 0.0018 |
| F04 | 14220001 | 14320000 | 0.0024 | 0.0022 | 0.0020 |
| F04 | 14240001 | 14340000 | 0.0028 | 0.0020 | 0.0025 |
| F04 | 14260001 | 14360000 | 0.0024 | 0.0020 | 0.0022 |
| F04 | 14280001 | 14380000 | 0.0024 | 0.0019 | 0.0021 |
| F04 | 14300001 | 14400000 | 0.0022 | 0.0018 | 0.0020 |
| F04 | 14320001 | 14420000 | 0.0021 | 0.0013 | 0.0019 |
| F04 | 14340001 | 14440000 | 0.0016 | 0.0017 | 0.0013 |
| F04 | 14360001 | 14460000 | 0.0020 | 0.0014 | 0.0016 |
| F04 | 14380001 | 14480000 | 0.0020 | 0.0019 | 0.0015 |
| F04 | 14400001 | 14500000 | 0.0027 | 0.0020 | 0.0022 |
| F04 | 14420001 | 14520000 | 0.0031 | 0.0024 | 0.0026 |
| F04 | 14440001 | 14540000 | 0.0038 | 0.0022 | 0.0034 |
| F04 | 14460001 | 14560000 | 0.0038 | 0.0024 | 0.0035 |
| F04 | 14480001 | 14580000 | 0.0041 | 0.0021 | 0.0040 |
| F04 | 14500001 | 14600000 | 0.0038 | 0.0019 | 0.0037 |
| F04 | 14520001 | 14620000 | 0.0034 | 0.0016 | 0.0034 |
| F04 | 14540001 | 14640000 | 0.0034 | 0.0015 | 0.0033 |
| F04 | 14560001 | 14660000 | 0.0033 | 0.0012 | 0.0033 |
| F04 | 14580001 | 14680000 | 0.0029 | 0.0010 | 0.0028 |
| F04 | 14600001 | 14700000 | 0.0025 | 0.0012 | 0.0025 |
| F04 | 14620001 | 14720000 | 0.0025 | 0.0008 | 0.0024 |
| F04 | 14640001 | 14740000 | 0.0015 | 0.0012 | 0.0015 |
| F04 | 14660001 | 14760000 | 0.0015 | 0.0014 | 0.0015 |
| F04 | 14680001 | 14780000 | 0.0017 | 0.0016 | 0.0017 |
| F04 | 14700001 | 14800000 | 0.0019 | 0.0025 | 0.0018 |
| F04 | 14720001 | 14820000 | 0.0030 | 0.0026 | 0.0029 |
| F04 | 14740001 | 14840000 | 0.0033 | 0.0028 | 0.0033 |
| F04 | 14760001 | 14860000 | 0.0039 | 0.0028 | 0.0038 |
| F04 | 14780001 | 14880000 | 0.0042 | 0.0028 | 0.0041 |
| F04 | 14800001 | 14900000 | 0.0046 | 0.0031 | 0.0045 |
| F04 | 14820001 | 14920000 | 0.0050 | 0.0032 | 0.0049 |
| F04 | 14840001 | 14940000 | 0.0051 | 0.0027 | 0.0050 |
| F04 | 14860001 | 14960000 | 0.0045 | 0.0024 | 0.0045 |
| F04 | 14880001 | 14980000 | 0.0044 | 0.0023 | 0.0043 |
| F04 | 14900001 | 15000000 | 0.0042 | 0.0010 | 0.0039 |
| F04 | 14920001 | 15020000 | 0.0029 | 0.0008 | 0.0025 |
| F04 | 14940001 | 15040000 | 0.0027 | 0.0012 | 0.0021 |
| F04 | 14960001 | 15060000 | 0.0029 | 0.0012 | 0.0022 |
| F04 | 14980001 | 15080000 | 0.0026 | 0.0011 | 0.0018 |
| F04 | 15000001 | 15100000 | 0.0025 | 0.0015 | 0.0017 |
| F04 | 15020001 | 15120000 | 0.0028 | 0.0018 | 0.0020 |
| F04 | 15040001 | 15140000 | 0.0029 | 0.0016 | 0.0021 |
| F04 | 15060001 | 15160000 | 0.0028 | 0.0018 | 0.0020 |
| F04 | 15080001 | 15180000 | 0.0029 | 0.0023 | 0.0021 |
| F04 | 15100001 | 15200000 | 0.0033 | 0.0028 | 0.0026 |
| F04 | 15120001 | 15220000 | 0.0039 | 0.0029 | 0.0030 |
| F04 | 15140001 | 15240000 | 0.0039 | 0.0027 | 0.0030 |
| F04 | 15160001 | 15260000 | 0.0034 | 0.0034 | 0.0027 |
| F04 | 15180001 | 15280000 | 0.0041 | 0.0028 | 0.0034 |
| F04 | 15200001 | 15300000 | 0.0034 | 0.0022 | 0.0027 |
| F04 | 15220001 | 15320000 | 0.0028 | 0.0026 | 0.0023 |
| F04 | 15240001 | 15340000 | 0.0032 | 0.0034 | 0.0028 |
| F04 | 15260001 | 15360000 | 0.0042 | 0.0027 | 0.0039 |
| F04 | 15280001 | 15380000 | 0.0037 | 0.0033 | 0.0034 |
| F04 | 15300001 | 15400000 | 0.0043 | 0.0032 | 0.0040 |
| F04 | 15320001 | 15420000 | 0.0040 | 0.0026 | 0.0037 |

| Chr | Start | End | both | A1 | A2 |
|---|---|---|---|---|---|
| F04 | 15340001 | 15440000 | 0.0035 | 0.0022 | 0.0030 |
| F04 | 15360001 | 15460000 | 0.0030 | 0.0025 | 0.0023 |
| F04 | 15380001 | 15480000 | 0.0034 | 0.0020 | 0.0027 |
| F04 | 15400001 | 15500000 | 0.0030 | 0.0023 | 0.0022 |
| F04 | 15420001 | 15520000 | 0.0033 | 0.0029 | 0.0026 |
| F04 | 15440001 | 15540000 | 0.0038 | 0.0027 | 0.0032 |
| F04 | 15460001 | 15560000 | 0.0035 | 0.0026 | 0.0031 |
| F04 | 15480001 | 15580000 | 0.0035 | 0.0026 | 0.0031 |
| F04 | 15500001 | 15600000 | 0.0034 | 0.0024 | 0.0031 |
| F04 | 15520001 | 15620000 | 0.0035 | 0.0021 | 0.0032 |
| F04 | 15540001 | 15640000 | 0.0036 | 0.0019 | 0.0033 |
| F04 | 15560001 | 15660000 | 0.0040 | 0.0020 | 0.0037 |
| F04 | 15580001 | 15680000 | 0.0042 | 0.0019 | 0.0039 |
| F04 | 15600001 | 15700000 | 0.0043 | 0.0024 | 0.0039 |
| F04 | 15620001 | 15720000 | 0.0048 | 0.0022 | 0.0044 |
| F04 | 15640001 | 15740000 | 0.0044 | 0.0024 | 0.0040 |
| F04 | 15660001 | 15760000 | 0.0039 | 0.0020 | 0.0037 |
| F04 | 15680001 | 15780000 | 0.0032 | 0.0022 | 0.0030 |
| F04 | 15700001 | 15800000 | 0.0031 | 0.0016 | 0.0029 |
| F04 | 15720001 | 15820000 | 0.0027 | 0.0015 | 0.0025 |
| F04 | 15740001 | 15840000 | 0.0024 | 0.0019 | 0.0023 |
| F04 | 15760001 | 15860000 | 0.0027 | 0.0017 | 0.0026 |
| F04 | 15780001 | 15880000 | 0.0025 | 0.0021 | 0.0024 |
| F04 | 15800001 | 15900000 | 0.0032 | 0.0021 | 0.0030 |
| F04 | 15820001 | 15920000 | 0.0030 | 0.0022 | 0.0029 |
| F04 | 15840001 | 15940000 | 0.0031 | 0.0022 | 0.0030 |
| F04 | 15860001 | 15960000 | 0.0036 | 0.0024 | 0.0034 |
| F04 | 15880001 | 15980000 | 0.0038 | 0.0022 | 0.0036 |
| F04 | 15900001 | 16000000 | 0.0035 | 0.0020 | 0.0033 |
| F04 | 15920001 | 16020000 | 0.0033 | 0.0029 | 0.0031 |
| F04 | 15940001 | 16040000 | 0.0045 | 0.0032 | 0.0043 |
| F04 | 15960001 | 16060000 | 0.0044 | 0.0030 | 0.0043 |
| F04 | 15980001 | 16080000 | 0.0046 | 0.0032 | 0.0044 |
| F04 | 16000001 | 16100000 | 0.0048 | 0.0032 | 0.0047 |
| F04 | 16020001 | 16120000 | 0.0044 | 0.0025 | 0.0043 |
| F04 | 16040001 | 16140000 | 0.0035 | 0.0020 | 0.0035 |
| F04 | 16060001 | 16160000 | 0.0033 | 0.0019 | 0.0033 |
| F04 | 16080001 | 16180000 | 0.0032 | 0.0014 | 0.0032 |
| F04 | 16100001 | 16200000 | 0.0027 | 0.0027 | 0.0027 |
| F04 | 16120001 | 16220000 | 0.0045 | 0.0034 | 0.0045 |
| F04 | 16140001 | 16240000 | 0.0052 | 0.0033 | 0.0052 |
| F04 | 16160001 | 16260000 | 0.0046 | 0.0034 | 0.0047 |
| F04 | 16180001 | 16280000 | 0.0049 | 0.0055 | 0.0049 |
| F04 | 16200001 | 16300000 | 0.0070 | 0.0045 | 0.0070 |
| F04 | 16220001 | 16320000 | 0.0057 | 0.0043 | 0.0057 |
| F04 | 16240001 | 16340000 | 0.0055 | 0.0045 | 0.0055 |
| F04 | 16260001 | 16360000 | 0.0056 | 0.0047 | 0.0056 |
| F04 | 16280001 | 16380000 | 0.0055 | 0.0028 | 0.0055 |
| F04 | 16300001 | 16400000 | 0.0033 | 0.0031 | 0.0033 |
| F04 | 16320001 | 16420000 | 0.0038 | 0.0024 | 0.0038 |
| F04 | 16340001 | 16440000 | 0.0031 | 0.0022 | 0.0030 |
| F04 | 16360001 | 16460000 | 0.0029 | 0.0019 | 0.0028 |
| F04 | 16380001 | 16480000 | 0.0030 | 0.0023 | 0.0030 |
| F04 | 16400001 | 16500000 | 0.0037 | 0.0032 | 0.0037 |
| F04 | 16420001 | 16520000 | 0.0044 | 0.0039 | 0.0044 |
| F04 | 16440001 | 16540000 | 0.0054 | 0.0042 | 0.0054 |
| F04 | 16460001 | 16560000 | 0.0060 | 0.0043 | 0.0060 |
| F04 | 16480001 | 16580000 | 0.0060 | 0.0049 | 0.0061 |

| Chr | Start | End | both | A1 | A2 |
|---|---|---|---|---|---|
| F04 | 16500001 | 16600000 | 0.0067 | 0.0034 | 0.0067 |
| F04 | 16520001 | 16620000 | 0.0051 | 0.0027 | 0.0051 |
| F04 | 16540001 | 16640000 | 0.0042 | 0.0023 | 0.0042 |
| F04 | 16560001 | 16660000 | 0.0035 | 0.0020 | 0.0034 |
| F04 | 16580001 | 16680000 | 0.0034 | 0.0014 | 0.0033 |
| F04 | 16600001 | 16700000 | 0.0029 | 0.0016 | 0.0028 |
| F04 | 16620001 | 16720000 | 0.0032 | 0.0014 | 0.0032 |
| F04 | 16640001 | 16740000 | 0.0029 | 0.0020 | 0.0029 |
| F04 | 16660001 | 16760000 | 0.0037 | 0.0021 | 0.0037 |
| F04 | 16680001 | 16780000 | 0.0032 | 0.0023 | 0.0032 |
| F04 | 16700001 | 16800000 | 0.0032 | 0.0025 | 0.0032 |
| F04 | 16720001 | 16820000 | 0.0038 | 0.0028 | 0.0038 |
| F04 | 16740001 | 16840000 | 0.0041 | 0.0031 | 0.0041 |
| F04 | 16760001 | 16860000 | 0.0043 | 0.0030 | 0.0043 |
| F04 | 16780001 | 16880000 | 0.0042 | 0.0030 | 0.0042 |
| F04 | 16800001 | 16900000 | 0.0042 | 0.0036 | 0.0042 |
| F04 | 16820001 | 16920000 | 0.0041 | 0.0036 | 0.0040 |
| F04 | 16840001 | 16940000 | 0.0039 | 0.0029 | 0.0037 |
| F04 | 16860001 | 16960000 | 0.0030 | 0.0033 | 0.0028 |
| F04 | 16880001 | 16980000 | 0.0035 | 0.0025 | 0.0033 |
| F04 | 16900001 | 17000000 | 0.0026 | 0.0016 | 0.0024 |
| F04 | 16920001 | 17020000 | 0.0018 | 0.0016 | 0.0016 |
| F04 | 16940001 | 17040000 | 0.0021 | 0.0016 | 0.0021 |
| F04 | 16960001 | 17060000 | 0.0021 | 0.0016 | 0.0021 |
| F04 | 16980001 | 17080000 | 0.0020 | 0.0020 | 0.0020 |
| F04 | 17000001 | 17100000 | 0.0027 | 0.0021 | 0.0027 |
| F04 | 17020001 | 17120000 | 0.0032 | 0.0021 | 0.0032 |
| F04 | 17040001 | 17140000 | 0.0031 | 0.0031 | 0.0031 |
| F04 | 17060001 | 17160000 | 0.0045 | 0.0030 | 0.0045 |
| F04 | 17080001 | 17180000 | 0.0044 | 0.0028 | 0.0045 |
| F04 | 17100001 | 17200000 | 0.0041 | 0.0038 | 0.0042 |
| F04 | 17120001 | 17220000 | 0.0053 | 0.0035 | 0.0053 |
| F04 | 17140001 | 17240000 | 0.0052 | 0.0026 | 0.0052 |
| F04 | 17160001 | 17260000 | 0.0039 | 0.0027 | 0.0039 |
| F04 | 17180001 | 17280000 | 0.0040 | 0.0025 | 0.0039 |
| F04 | 17200001 | 17300000 | 0.0037 | 0.0016 | 0.0037 |
| F04 | 17220001 | 17320000 | 0.0024 | 0.0021 | 0.0024 |
| F04 | 17240001 | 17340000 | 0.0028 | 0.0020 | 0.0025 |
| F04 | 17260001 | 17360000 | 0.0027 | 0.0020 | 0.0023 |
| F04 | 17280001 | 17380000 | 0.0030 | 0.0023 | 0.0025 |
| F04 | 17300001 | 17400000 | 0.0036 | 0.0027 | 0.0027 |
| F04 | 17320001 | 17420000 | 0.0041 | 0.0024 | 0.0031 |
| F04 | 17340001 | 17440000 | 0.0038 | 0.0026 | 0.0029 |
| F04 | 17360001 | 17460000 | 0.0040 | 0.0022 | 0.0030 |
| F04 | 17380001 | 17480000 | 0.0035 | 0.0018 | 0.0024 |
| F04 | 17400001 | 17500000 | 0.0029 | 0.0014 | 0.0020 |
| F04 | 17420001 | 17520000 | 0.0024 | 0.0020 | 0.0016 |
| F04 | 17440001 | 17540000 | 0.0034 | 0.0026 | 0.0024 |
| F04 | 17460001 | 17560000 | 0.0040 | 0.0029 | 0.0031 |
| F04 | 17480001 | 17580000 | 0.0044 | 0.0031 | 0.0034 |
| F04 | 17500001 | 17600000 | 0.0047 | 0.0030 | 0.0037 |
| F04 | 17520001 | 17620000 | 0.0050 | 0.0024 | 0.0039 |
| F04 | 17540001 | 17640000 | 0.0041 | 0.0018 | 0.0030 |
| F04 | 17560001 | 17660000 | 0.0037 | 0.0016 | 0.0024 |
| F04 | 17580001 | 17680000 | 0.0034 | 0.0013 | 0.0021 |
| F04 | 17600001 | 17700000 | 0.0031 | 0.0011 | 0.0019 |
| F04 | 17620001 | 17720000 | 0.0029 | 0.0019 | 0.0017 |
| F04 | 17640001 | 17740000 | 0.0037 | 0.0019 | 0.0028 |

| Chr | Start | End | both | A1 | A2 |
|---|---|---|---|---|---|
| F04 | 17660001 | 17760000 | 0.0033 | 0.0021 | 0.0027 |
| F04 | 17680001 | 17780000 | 0.0034 | 0.0023 | 0.0031 |
| F04 | 17700001 | 17800000 | 0.0036 | 0.0022 | 0.0033 |
| F04 | 17720001 | 17820000 | 0.0030 | 0.0014 | 0.0030 |
| F04 | 17740001 | 17840000 | 0.0021 | 0.0015 | 0.0021 |
| F04 | 17760001 | 17860000 | 0.0024 | 0.0018 | 0.0023 |
| F04 | 17780001 | 17880000 | 0.0029 | 0.0016 | 0.0029 |
| F04 | 17800001 | 17900000 | 0.0030 | 0.0021 | 0.0030 |
| F04 | 17820001 | 17920000 | 0.0042 | 0.0024 | 0.0042 |
| F04 | 17840001 | 17940000 | 0.0046 | 0.0020 | 0.0046 |
| F04 | 17860001 | 17960000 | 0.0040 | 0.0015 | 0.0040 |
| F04 | 17880001 | 17980000 | 0.0031 | 0.0017 | 0.0031 |
| F04 | 17900001 | 18000000 | 0.0028 | 0.0013 | 0.0027 |
| F04 | 17920001 | 18020000 | 0.0018 | 0.0012 | 0.0017 |
| F04 | 17940001 | 18040000 | 0.0016 | 0.0022 | 0.0014 |
| F04 | 17960001 | 18060000 | 0.0027 | 0.0023 | 0.0021 |
| F04 | 17980001 | 18080000 | 0.0029 | 0.0025 | 0.0021 |
| F04 | 18000001 | 18100000 | 0.0034 | 0.0025 | 0.0024 |
| F04 | 18020001 | 18120000 | 0.0034 | 0.0030 | 0.0023 |
| F04 | 18040001 | 18140000 | 0.0038 | 0.0024 | 0.0027 |
| F04 | 18060001 | 18160000 | 0.0030 | 0.0030 | 0.0023 |
| F04 | 18080001 | 18180000 | 0.0038 | 0.0031 | 0.0031 |
| F04 | 18100001 | 18200000 | 0.0039 | 0.0029 | 0.0034 |
| F04 | 18120001 | 18220000 | 0.0036 | 0.0027 | 0.0032 |
| F04 | 18140001 | 18240000 | 0.0035 | 0.0025 | 0.0030 |
| F04 | 18160001 | 18260000 | 0.0032 | 0.0020 | 0.0027 |
| F04 | 18180001 | 18280000 | 0.0029 | 0.0017 | 0.0024 |
| F04 | 18200001 | 18300000 | 0.0025 | 0.0019 | 0.0019 |
| F04 | 18220001 | 18320000 | 0.0029 | 0.0015 | 0.0021 |
| F04 | 18240001 | 18340000 | 0.0024 | 0.0014 | 0.0018 |
| F04 | 18260001 | 18360000 | 0.0024 | 0.0010 | 0.0017 |
| F04 | 18280001 | 18380000 | 0.0018 | 0.0008 | 0.0011 |
| F04 | 18300001 | 18400000 | 0.0018 | 0.0010 | 0.0010 |
| F04 | 18320001 | 18420000 | 0.0020 | 0.0013 | 0.0012 |
| F04 | 18340001 | 18440000 | 0.0025 | 0.0019 | 0.0015 |
| F04 | 18360001 | 18460000 | 0.0031 | 0.0021 | 0.0022 |
| F04 | 18380001 | 18480000 | 0.0033 | 0.0023 | 0.0026 |
| F04 | 18400001 | 18500000 | 0.0034 | 0.0020 | 0.0028 |
| F04 | 18420001 | 18520000 | 0.0030 | 0.0021 | 0.0025 |
| F04 | 18440001 | 18540000 | 0.0031 | 0.0017 | 0.0025 |
| F04 | 18460001 | 18560000 | 0.0025 | 0.0016 | 0.0020 |
| F04 | 18480001 | 18580000 | 0.0023 | 0.0017 | 0.0019 |
| F04 | 18500001 | 18600000 | 0.0024 | 0.0017 | 0.0018 |
| F04 | 18520001 | 18620000 | 0.0025 | 0.0013 | 0.0019 |
| F04 | 18540001 | 18640000 | 0.0021 | 0.0024 | 0.0015 |
| F04 | 18560001 | 18660000 | 0.0035 | 0.0022 | 0.0028 |
| F04 | 18580001 | 18680000 | 0.0035 | 0.0023 | 0.0026 |
| F04 | 18600001 | 18700000 | 0.0034 | 0.0025 | 0.0026 |
| F04 | 18620001 | 18720000 | 0.0035 | 0.0031 | 0.0027 |
| F04 | 18640001 | 18740000 | 0.0043 | 0.0025 | 0.0034 |
| F04 | 18660001 | 18760000 | 0.0036 | 0.0027 | 0.0026 |
| F04 | 18680001 | 18780000 | 0.0037 | 0.0025 | 0.0028 |
| F04 | 18700001 | 18800000 | 0.0037 | 0.0026 | 0.0027 |
| F04 | 18720001 | 18820000 | 0.0037 | 0.0023 | 0.0027 |
| F04 | 18740001 | 18840000 | 0.0031 | 0.0021 | 0.0024 |
| F04 | 18760001 | 18860000 | 0.0029 | 0.0024 | 0.0023 |
| F04 | 18780001 | 18880000 | 0.0034 | 0.0025 | 0.0028 |
| F04 | 18800001 | 18900000 | 0.0034 | 0.0027 | 0.0027 |

| Chr | Start | End | both | A1 | A2 |
|---|---|---|---|---|---|
| F04 | 18820001 | 18920000 | 0.0038 | 0.0024 | 0.0030 |
| F04 | 18840001 | 18940000 | 0.0034 | 0.0019 | 0.0025 |
| F04 | 18860001 | 18960000 | 0.0030 | 0.0019 | 0.0022 |
| F04 | 18880001 | 18980000 | 0.0027 | 0.0026 | 0.0020 |
| F04 | 18900001 | 19000000 | 0.0035 | 0.0020 | 0.0029 |
| F04 | 18920001 | 19020000 | 0.0027 | 0.0032 | 0.0023 |
| F04 | 18940001 | 19040000 | 0.0043 | 0.0032 | 0.0036 |
| F04 | 18960001 | 19060000 | 0.0045 | 0.0032 | 0.0036 |
| F04 | 18980001 | 19080000 | 0.0046 | 0.0030 | 0.0036 |
| F04 | 19000001 | 19100000 | 0.0044 | 0.0041 | 0.0034 |
| F04 | 19020001 | 19120000 | 0.0055 | 0.0029 | 0.0044 |
| F04 | 19040001 | 19140000 | 0.0039 | 0.0032 | 0.0031 |
| F04 | 19060001 | 19160000 | 0.0040 | 0.0027 | 0.0033 |
| F04 | 19080001 | 19180000 | 0.0033 | 0.0020 | 0.0027 |
| F04 | 19100001 | 19200000 | 0.0027 | 0.0014 | 0.0020 |
| F04 | 19120001 | 19220000 | 0.0019 | 0.0023 | 0.0015 |
| F04 | 19140001 | 19240000 | 0.0030 | 0.0027 | 0.0026 |
| F04 | 19160001 | 19260000 | 0.0034 | 0.0032 | 0.0029 |
| F04 | 19180001 | 19280000 | 0.0040 | 0.0031 | 0.0035 |
| F04 | 19200001 | 19300000 | 0.0040 | 0.0031 | 0.0034 |
| F04 | 19220001 | 19320000 | 0.0040 | 0.0028 | 0.0034 |
| F04 | 19240001 | 19340000 | 0.0036 | 0.0023 | 0.0029 |
| F04 | 19260001 | 19360000 | 0.0031 | 0.0021 | 0.0025 |
| F04 | 19280001 | 19380000 | 0.0032 | 0.0027 | 0.0023 |
| F04 | 19300001 | 19400000 | 0.0038 | 0.0026 | 0.0031 |
| F04 | 19320001 | 19420000 | 0.0035 | 0.0019 | 0.0028 |
| F04 | 19340001 | 19440000 | 0.0029 | 0.0020 | 0.0022 |
| F04 | 19360001 | 19460000 | 0.0030 | 0.0022 | 0.0023 |
| F04 | 19380001 | 19480000 | 0.0031 | 0.0016 | 0.0024 |
| F04 | 19400001 | 19500000 | 0.0023 | 0.0017 | 0.0015 |
| F04 | 19420001 | 19520000 | 0.0026 | 0.0023 | 0.0017 |
| F04 | 19440001 | 19540000 | 0.0033 | 0.0026 | 0.0024 |
| F04 | 19460001 | 19560000 | 0.0036 | 0.0026 | 0.0027 |
| F04 | 19480001 | 19580000 | 0.0036 | 0.0028 | 0.0028 |
| F04 | 19500001 | 19600000 | 0.0037 | 0.0026 | 0.0032 |
| F04 | 19520001 | 19620000 | 0.0037 | 0.0031 | 0.0030 |
| F04 | 19540001 | 19640000 | 0.0042 | 0.0031 | 0.0035 |
| F04 | 19560001 | 19660000 | 0.0042 | 0.0026 | 0.0034 |
| F04 | 19580001 | 19680000 | 0.0041 | 0.0031 | 0.0030 |
| F04 | 19600001 | 19700000 | 0.0046 | 0.0035 | 0.0035 |
| F04 | 19620001 | 19720000 | 0.0048 | 0.0031 | 0.0039 |
| F04 | 19640001 | 19740000 | 0.0044 | 0.0026 | 0.0036 |
| F04 | 19660001 | 19760000 | 0.0038 | 0.0025 | 0.0031 |
| F04 | 19680001 | 19780000 | 0.0034 | 0.0018 | 0.0029 |
| F04 | 19700001 | 19800000 | 0.0028 | 0.0014 | 0.0022 |
| F04 | 19720001 | 19820000 | 0.0027 | 0.0014 | 0.0018 |
| F04 | 19740001 | 19840000 | 0.0025 | 0.0014 | 0.0017 |
| F04 | 19760001 | 19860000 | 0.0025 | 0.0017 | 0.0017 |
| F04 | 19780001 | 19880000 | 0.0030 | 0.0022 | 0.0020 |
| F04 | 19800001 | 19900000 | 0.0033 | 0.0022 | 0.0025 |
| F04 | 19820001 | 19920000 | 0.0032 | 0.0031 | 0.0025 |
| F04 | 19840001 | 19940000 | 0.0043 | 0.0041 | 0.0036 |
| F04 | 19860001 | 19960000 | 0.0054 | 0.0045 | 0.0048 |
| F04 | 19880001 | 19980000 | 0.0062 | 0.0048 | 0.0054 |
| F04 | 19900001 | 20000000 | 0.0066 | 0.0050 | 0.0056 |
| F04 | 19920001 | 20020000 | 0.0067 | 0.0037 | 0.0058 |
| F04 | 19940001 | 20040000 | 0.0050 | 0.0032 | 0.0043 |
| F04 | 19960001 | 20060000 | 0.0044 | 0.0029 | 0.0037 |

| Chr | Start | End | both | A1 | A2 |
|---|---|---|---|---|---|
| F04 | 19980001 | 20080000 | 0.0037 | 0.0026 | 0.0033 |
| F04 | 20000001 | 20100000 | 0.0033 | 0.0032 | 0.0030 |
| F04 | 20020001 | 20120000 | 0.0039 | 0.0030 | 0.0035 |
| F04 | 20040001 | 20140000 | 0.0039 | 0.0026 | 0.0032 |
| F04 | 20060001 | 20160000 | 0.0037 | 0.0021 | 0.0027 |
| F04 | 20080001 | 20180000 | 0.0034 | 0.0019 | 0.0022 |
| F04 | 20100001 | 20200000 | 0.0029 | 0.0012 | 0.0019 |
| F04 | 20120001 | 20220000 | 0.0023 | 0.0010 | 0.0012 |
| F04 | 20140001 | 20240000 | 0.0021 | 0.0007 | 0.0010 |
| F04 | 20160001 | 20260000 | 0.0020 | 0.0010 | 0.0010 |
| F04 | 20180001 | 20280000 | 0.0023 | 0.0010 | 0.0012 |
| F04 | 20200001 | 20300000 | 0.0022 | 0.0011 | 0.0011 |
| F04 | 20220001 | 20320000 | 0.0023 | 0.0015 | 0.0013 |
| F04 | 20240001 | 20340000 | 0.0026 | 0.0016 | 0.0017 |
| F04 | 20260001 | 20360000 | 0.0028 | 0.0014 | 0.0018 |
| F04 | 20280001 | 20380000 | 0.0026 | 0.0020 | 0.0016 |
| F04 | 20300001 | 20400000 | 0.0036 | 0.0026 | 0.0025 |
| F04 | 20320001 | 20420000 | 0.0040 | 0.0031 | 0.0030 |
| F04 | 20340001 | 20440000 | 0.0046 | 0.0048 | 0.0035 |
| F04 | 20360001 | 20460000 | 0.0059 | 0.0048 | 0.0051 |
| F04 | 20380001 | 20480000 | 0.0056 | 0.0043 | 0.0050 |
| F04 | 20400001 | 20500000 | 0.0050 | 0.0046 | 0.0045 |
| F04 | 20420001 | 20520000 | 0.0052 | 0.0046 | 0.0046 |
| F04 | 20440001 | 20540000 | 0.0053 | 0.0037 | 0.0046 |
| F04 | 20460001 | 20560000 | 0.0046 | 0.0041 | 0.0038 |
| F04 | 20480001 | 20580000 | 0.0049 | 0.0039 | 0.0041 |
| F04 | 20500001 | 20600000 | 0.0046 | 0.0031 | 0.0039 |
| F04 | 20520001 | 20620000 | 0.0040 | 0.0024 | 0.0034 |
| F04 | 20540001 | 20640000 | 0.0029 | 0.0015 | 0.0024 |
| F04 | 20560001 | 20660000 | 0.0019 | 0.0021 | 0.0015 |
| F04 | 20580001 | 20680000 | 0.0027 | 0.0019 | 0.0023 |
| F04 | 20600001 | 20700000 | 0.0028 | 0.0016 | 0.0021 |
| F04 | 20620001 | 20720000 | 0.0030 | 0.0017 | 0.0019 |
| F04 | 20640001 | 20740000 | 0.0031 | 0.0019 | 0.0021 |
| F04 | 20660001 | 20760000 | 0.0033 | 0.0020 | 0.0022 |
| F04 | 20680001 | 20780000 | 0.0036 | 0.0031 | 0.0025 |
| F04 | 20700001 | 20800000 | 0.0045 | 0.0037 | 0.0036 |
| F04 | 20720001 | 20820000 | 0.0046 | 0.0040 | 0.0040 |
| F04 | 20740001 | 20840000 | 0.0052 | 0.0052 | 0.0045 |
| F04 | 20760001 | 20860000 | 0.0061 | 0.0045 | 0.0054 |
| F04 | 20780001 | 20880000 | 0.0054 | 0.0041 | 0.0046 |
| F04 | 20800001 | 20900000 | 0.0052 | 0.0038 | 0.0042 |
| F04 | 20820001 | 20920000 | 0.0049 | 0.0041 | 0.0039 |
| F04 | 20840001 | 20940000 | 0.0052 | 0.0036 | 0.0042 |
| F04 | 20860001 | 20960000 | 0.0048 | 0.0037 | 0.0038 |
| F04 | 20880001 | 20980000 | 0.0049 | 0.0033 | 0.0041 |
| F04 | 20900001 | 21000000 | 0.0044 | 0.0029 | 0.0036 |
| F04 | 20920001 | 21020000 | 0.0039 | 0.0024 | 0.0031 |
| F04 | 20940001 | 21040000 | 0.0032 | 0.0026 | 0.0025 |
| F04 | 20960001 | 21060000 | 0.0035 | 0.0024 | 0.0028 |
| F04 | 20980001 | 21080000 | 0.0033 | 0.0028 | 0.0025 |
| F04 | 21000001 | 21100000 | 0.0037 | 0.0035 | 0.0028 |
| F04 | 21020001 | 21120000 | 0.0046 | 0.0032 | 0.0035 |
| F04 | 21040001 | 21140000 | 0.0045 | 0.0028 | 0.0034 |
| F04 | 21060001 | 21160000 | 0.0041 | 0.0026 | 0.0028 |
| F04 | 21080001 | 21180000 | 0.0038 | 0.0025 | 0.0026 |
| F04 | 21100001 | 21200000 | 0.0036 | 0.0019 | 0.0026 |
| F04 | 21120001 | 21220000 | 0.0029 | 0.0025 | 0.0021 |

| Chr | Start | End | both | A1 | A2 |
|---|---|---|---|---|---|
| F04 | 21140001 | 21240000 | 0.0034 | 0.0026 | 0.0026 |
| F04 | 21160001 | 21260000 | 0.0035 | 0.0032 | 0.0031 |
| F04 | 21180001 | 21280000 | 0.0042 | 0.0034 | 0.0038 |
| F04 | 21200001 | 21300000 | 0.0045 | 0.0036 | 0.0039 |
| F04 | 21220001 | 21320000 | 0.0046 | 0.0037 | 0.0041 |
| F04 | 21240001 | 21340000 | 0.0046 | 0.0048 | 0.0041 |
| F04 | 21260001 | 21360000 | 0.0056 | 0.0052 | 0.0051 |
| F04 | 21280001 | 21380000 | 0.0063 | 0.0054 | 0.0055 |
| F04 | 21300001 | 21400000 | 0.0064 | 0.0062 | 0.0059 |
| F04 | 21320001 | 21420000 | 0.0073 | 0.0058 | 0.0067 |
| F04 | 21340001 | 21440000 | 0.0070 | 0.0045 | 0.0062 |
| F04 | 21360001 | 21460000 | 0.0057 | 0.0041 | 0.0048 |
| F04 | 21380001 | 21480000 | 0.0052 | 0.0034 | 0.0046 |
| F04 | 21400001 | 21500000 | 0.0044 | 0.0024 | 0.0037 |
| F04 | 21420001 | 21520000 | 0.0033 | 0.0026 | 0.0028 |
| F04 | 21440001 | 21540000 | 0.0034 | 0.0023 | 0.0031 |
| F04 | 21460001 | 21560000 | 0.0029 | 0.0018 | 0.0028 |
| F04 | 21480001 | 21580000 | 0.0022 | 0.0019 | 0.0021 |
| F04 | 21500001 | 21600000 | 0.0022 | 0.0027 | 0.0022 |
| F04 | 21520001 | 21620000 | 0.0031 | 0.0022 | 0.0030 |
| F04 | 21540001 | 21640000 | 0.0026 | 0.0020 | 0.0026 |
| F04 | 21560001 | 21660000 | 0.0024 | 0.0017 | 0.0024 |
| F04 | 21580001 | 21680000 | 0.0020 | 0.0019 | 0.0019 |
| F04 | 21600001 | 21700000 | 0.0021 | 0.0024 | 0.0021 |
| F04 | 21620001 | 21720000 | 0.0026 | 0.0026 | 0.0026 |
| F04 | 21640001 | 21740000 | 0.0030 | 0.0025 | 0.0028 |
| F04 | 21660001 | 21760000 | 0.0030 | 0.0028 | 0.0027 |
| F04 | 21680001 | 21780000 | 0.0035 | 0.0033 | 0.0030 |
| F04 | 21700001 | 21800000 | 0.0041 | 0.0026 | 0.0035 |
| F04 | 21720001 | 21820000 | 0.0035 | 0.0026 | 0.0027 |
| F04 | 21740001 | 21840000 | 0.0034 | 0.0029 | 0.0028 |
| F04 | 21760001 | 21860000 | 0.0037 | 0.0031 | 0.0031 |
| F04 | 21780001 | 21880000 | 0.0040 | 0.0026 | 0.0034 |
| F04 | 21800001 | 21900000 | 0.0035 | 0.0026 | 0.0029 |
| F04 | 21820001 | 21920000 | 0.0035 | 0.0030 | 0.0028 |
| F04 | 21840001 | 21940000 | 0.0040 | 0.0032 | 0.0033 |
| F04 | 21860001 | 21960000 | 0.0045 | 0.0033 | 0.0039 |
| F04 | 21880001 | 21980000 | 0.0045 | 0.0031 | 0.0039 |
| F04 | 21900001 | 22000000 | 0.0044 | 0.0026 | 0.0037 |
| F04 | 21920001 | 22020000 | 0.0040 | 0.0034 | 0.0033 |
| F04 | 21940001 | 22040000 | 0.0050 | 0.0035 | 0.0043 |
| F04 | 21960001 | 22060000 | 0.0050 | 0.0033 | 0.0040 |
| F04 | 21980001 | 22080000 | 0.0048 | 0.0034 | 0.0036 |
| F04 | 22000001 | 22100000 | 0.0048 | 0.0036 | 0.0038 |
| F04 | 22020001 | 22120000 | 0.0052 | 0.0028 | 0.0041 |
| F04 | 22040001 | 22140000 | 0.0039 | 0.0025 | 0.0029 |
| F04 | 22060001 | 22160000 | 0.0036 | 0.0024 | 0.0027 |
| F04 | 22080001 | 22180000 | 0.0035 | 0.0023 | 0.0028 |
| F04 | 22100001 | 22200000 | 0.0032 | 0.0030 | 0.0026 |
| F04 | 22120001 | 22220000 | 0.0037 | 0.0023 | 0.0033 |
| F04 | 22140001 | 22240000 | 0.0031 | 0.0021 | 0.0027 |
| F04 | 22160001 | 22260000 | 0.0026 | 0.0025 | 0.0024 |
| F04 | 22180001 | 22280000 | 0.0031 | 0.0029 | 0.0028 |
| F04 | 22200001 | 22300000 | 0.0036 | 0.0022 | 0.0032 |
| F04 | 22220001 | 22320000 | 0.0030 | 0.0022 | 0.0024 |
| F04 | 22240001 | 22340000 | 0.0032 | 0.0024 | 0.0024 |
| F04 | 22260001 | 22360000 | 0.0036 | 0.0017 | 0.0027 |
| F04 | 22280001 | 22380000 | 0.0028 | 0.0017 | 0.0019 |

| Chr | Start | End | both | A1 | A2 |
|---|---|---|---|---|---|
| F04 | 22300001 | 22400000 | 0.0028 | 0.0018 | 0.0019 |
| F04 | 22320001 | 22420000 | 0.0027 | 0.0023 | 0.0018 |
| F04 | 22340001 | 22440000 | 0.0032 | 0.0022 | 0.0023 |
| F04 | 22360001 | 22460000 | 0.0030 | 0.0021 | 0.0022 |
| F04 | 22380001 | 22480000 | 0.0030 | 0.0019 | 0.0020 |
| F04 | 22400001 | 22500000 | 0.0027 | 0.0018 | 0.0018 |
| F04 | 22420001 | 22520000 | 0.0026 | 0.0018 | 0.0019 |
| F04 | 22440001 | 22540000 | 0.0024 | 0.0021 | 0.0020 |
| F04 | 22460001 | 22560000 | 0.0026 | 0.0024 | 0.0023 |
| F04 | 22480001 | 22580000 | 0.0029 | 0.0024 | 0.0027 |
| F04 | 22500001 | 22600000 | 0.0029 | 0.0020 | 0.0026 |
| F04 | 22520001 | 22620000 | 0.0028 | 0.0016 | 0.0023 |
| F04 | 22540001 | 22640000 | 0.0024 | 0.0015 | 0.0018 |
| F04 | 22560001 | 22660000 | 0.0023 | 0.0013 | 0.0018 |
| F04 | 22580001 | 22680000 | 0.0022 | 0.0015 | 0.0015 |
| F04 | 22600001 | 22700000 | 0.0023 | 0.0013 | 0.0018 |
| F04 | 22620001 | 22720000 | 0.0019 | 0.0016 | 0.0015 |
| F04 | 22640001 | 22740000 | 0.0023 | 0.0015 | 0.0018 |
| F04 | 22660001 | 22760000 | 0.0022 | 0.0015 | 0.0017 |
| F04 | 22680001 | 22780000 | 0.0021 | 0.0016 | 0.0017 |
| F04 | 22700001 | 22800000 | 0.0023 | 0.0016 | 0.0018 |
| F04 | 22720001 | 22820000 | 0.0023 | 0.0017 | 0.0019 |
| F04 | 22740001 | 22840000 | 0.0022 | 0.0021 | 0.0019 |
| F04 | 22760001 | 22860000 | 0.0026 | 0.0027 | 0.0023 |
| F04 | 22780001 | 22880000 | 0.0034 | 0.0023 | 0.0031 |
| F04 | 22800001 | 22900000 | 0.0027 | 0.0023 | 0.0027 |
| F04 | 22820001 | 22920000 | 0.0028 | 0.0020 | 0.0027 |
| F04 | 22840001 | 22940000 | 0.0025 | 0.0013 | 0.0024 |
| F04 | 22860001 | 22960000 | 0.0018 | 0.0009 | 0.0016 |
| F04 | 22880001 | 22980000 | 0.0013 | 0.0011 | 0.0010 |
| F04 | 22900001 | 23000000 | 0.0018 | 0.0014 | 0.0013 |
| F04 | 22920001 | 23020000 | 0.0021 | 0.0023 | 0.0015 |
| F04 | 22940001 | 23040000 | 0.0031 | 0.0023 | 0.0025 |
| F04 | 22960001 | 23060000 | 0.0030 | 0.0025 | 0.0024 |
| F04 | 22980001 | 23080000 | 0.0031 | 0.0028 | 0.0027 |
| F04 | 23000001 | 23100000 | 0.0032 | 0.0025 | 0.0029 |
| F04 | 23020001 | 23120000 | 0.0028 | 0.0015 | 0.0026 |
| F04 | 23040001 | 23140000 | 0.0017 | 0.0016 | 0.0016 |
| F04 | 23060001 | 23160000 | 0.0018 | 0.0013 | 0.0016 |
| F04 | 23080001 | 23180000 | 0.0018 | 0.0014 | 0.0015 |
| F04 | 23100001 | 23200000 | 0.0020 | 0.0020 | 0.0015 |
| F04 | 23120001 | 23220000 | 0.0027 | 0.0029 | 0.0022 |
| F04 | 23140001 | 23240000 | 0.0036 | 0.0033 | 0.0031 |
| F04 | 23160001 | 23260000 | 0.0043 | 0.0033 | 0.0036 |
| F04 | 23180001 | 23280000 | 0.0043 | 0.0028 | 0.0037 |
| F04 | 23200001 | 23300000 | 0.0037 | 0.0021 | 0.0031 |
| F04 | 23220001 | 23320000 | 0.0031 | 0.0011 | 0.0023 |
| F04 | 23240001 | 23340000 | 0.0022 | 0.0009 | 0.0014 |
| F04 | 23260001 | 23360000 | 0.0018 | 0.0006 | 0.0011 |
| F04 | 23280001 | 23380000 | 0.0011 | 0.0005 | 0.0007 |
| F04 | 23300001 | 23400000 | 0.0011 | 0.0010 | 0.0006 |
| F04 | 23320001 | 23420000 | 0.0016 | 0.0013 | 0.0011 |
| F04 | 23340001 | 23440000 | 0.0019 | 0.0013 | 0.0014 |
| F04 | 23360001 | 23460000 | 0.0020 | 0.0018 | 0.0015 |
| F04 | 23380001 | 23480000 | 0.0025 | 0.0018 | 0.0019 |
| F04 | 23400001 | 23500000 | 0.0025 | 0.0021 | 0.0020 |
| F04 | 23420001 | 23520000 | 0.0026 | 0.0023 | 0.0023 |
| F04 | 23440001 | 23540000 | 0.0027 | 0.0022 | 0.0025 |

| Chr | Start | End | both | A1 | A2 |
|---|---|---|---|---|---|
| F04 | 23460001 | 23560000 | 0.0025 | 0.0017 | 0.0024 |
| F04 | 23480001 | 23580000 | 0.0020 | 0.0024 | 0.0020 |
| F04 | 23500001 | 23600000 | 0.0027 | 0.0017 | 0.0027 |
| F04 | 23520001 | 23620000 | 0.0020 | 0.0013 | 0.0019 |
| F04 | 23540001 | 23640000 | 0.0016 | 0.0014 | 0.0015 |
| F04 | 23560001 | 23660000 | 0.0017 | 0.0016 | 0.0016 |
| F04 | 23580001 | 23680000 | 0.0019 | 0.0011 | 0.0018 |
| F04 | 23600001 | 23700000 | 0.0014 | 0.0012 | 0.0012 |
| F04 | 23620001 | 23720000 | 0.0017 | 0.0011 | 0.0014 |
| F04 | 23640001 | 23740000 | 0.0016 | 0.0013 | 0.0013 |
| F04 | 23660001 | 23760000 | 0.0018 | 0.0011 | 0.0016 |
| F04 | 23680001 | 23780000 | 0.0017 | 0.0013 | 0.0014 |
| F04 | 23700001 | 23800000 | 0.0018 | 0.0015 | 0.0017 |
| F04 | 23720001 | 23820000 | 0.0018 | 0.0016 | 0.0017 |
| F04 | 23740001 | 23840000 | 0.0022 | 0.0012 | 0.0020 |
| F04 | 23760001 | 23860000 | 0.0019 | 0.0012 | 0.0014 |
| F04 | 23780001 | 23880000 | 0.0021 | 0.0015 | 0.0016 |
| F04 | 23800001 | 23900000 | 0.0024 | 0.0011 | 0.0016 |
| F04 | 23820001 | 23920000 | 0.0022 | 0.0011 | 0.0013 |
| F04 | 23840001 | 23940000 | 0.0019 | 0.0016 | 0.0011 |
| F04 | 23860001 | 23960000 | 0.0022 | 0.0019 | 0.0017 |
| F04 | 23880001 | 23980000 | 0.0024 | 0.0022 | 0.0020 |
| F04 | 23900001 | 24000000 | 0.0028 | 0.0023 | 0.0026 |
| F04 | 23920001 | 24020000 | 0.0028 | 0.0024 | 0.0027 |
| F04 | 23940001 | 24040000 | 0.0029 | 0.0018 | 0.0029 |
| F04 | 23960001 | 24060000 | 0.0024 | 0.0016 | 0.0023 |
| F04 | 23980001 | 24080000 | 0.0022 | 0.0013 | 0.0020 |
| F04 | 24000001 | 24100000 | 0.0019 | 0.0020 | 0.0016 |
| F04 | 24020001 | 24120000 | 0.0029 | 0.0022 | 0.0025 |
| F04 | 24040001 | 24140000 | 0.0031 | 0.0026 | 0.0027 |
| F04 | 24060001 | 24160000 | 0.0034 | 0.0025 | 0.0031 |
| F04 | 24080001 | 24180000 | 0.0033 | 0.0021 | 0.0031 |
| F04 | 24100001 | 24200000 | 0.0026 | 0.0014 | 0.0026 |
| F04 | 24120001 | 24220000 | 0.0017 | 0.0012 | 0.0016 |
| F04 | 24140001 | 24240000 | 0.0015 | 0.0012 | 0.0014 |
| F04 | 24160001 | 24260000 | 0.0015 | 0.0015 | 0.0014 |
| F04 | 24180001 | 24280000 | 0.0018 | 0.0017 | 0.0017 |
| F04 | 24200001 | 24300000 | 0.0020 | 0.0015 | 0.0019 |
| F04 | 24220001 | 24320000 | 0.0018 | 0.0017 | 0.0017 |
| F04 | 24240001 | 24340000 | 0.0022 | 0.0015 | 0.0020 |
| F04 | 24260001 | 24360000 | 0.0019 | 0.0011 | 0.0017 |
| F04 | 24280001 | 24380000 | 0.0014 | 0.0014 | 0.0012 |
| F04 | 24300001 | 24400000 | 0.0018 | 0.0017 | 0.0016 |
| F04 | 24320001 | 24420000 | 0.0020 | 0.0012 | 0.0018 |
| F04 | 24340001 | 24440000 | 0.0013 | 0.0016 | 0.0013 |
| F04 | 24360001 | 24460000 | 0.0018 | 0.0017 | 0.0017 |
| F04 | 24380001 | 24480000 | 0.0021 | 0.0015 | 0.0018 |
| F04 | 24400001 | 24500000 | 0.0018 | 0.0013 | 0.0015 |
| F04 | 24420001 | 24520000 | 0.0016 | 0.0014 | 0.0013 |
| F04 | 24440001 | 24540000 | 0.0018 | 0.0011 | 0.0016 |
| F04 | 24460001 | 24560000 | 0.0014 | 0.0008 | 0.0012 |
| F04 | 24480001 | 24580000 | 0.0010 | 0.0005 | 0.0010 |
| F04 | 24500001 | 24600000 | 0.0007 | 0.0008 | 0.0007 |
| F04 | 24520001 | 24620000 | 0.0011 | 0.0011 | 0.0011 |
| F04 | 24540001 | 24640000 | 0.0012 | 0.0009 | 0.0012 |
| F04 | 24560001 | 24660000 | 0.0011 | 0.0010 | 0.0010 |
| F04 | 24580001 | 24680000 | 0.0012 | 0.0010 | 0.0012 |
| F04 | 24600001 | 24700000 | 0.0012 | 0.0006 | 0.0012 |

| Chr | Start | End | both | A1 | A2 |
|---|---|---|---|---|---|
| F04 | 24620001 | 24720000 | 0.0008 | 0.0003 | 0.0008 |
| F04 | 24640001 | 24740000 | 0.0004 | 0.0003 | 0.0004 |
| F04 | 24660001 | 24760000 | 0.0004 | 0.0003 | 0.0004 |
| F04 | 24680001 | 24780000 | 0.0004 | 0.0004 | 0.0004 |
| F04 | 24700001 | 24800000 | 0.0006 | 0.0006 | 0.0006 |
| F04 | 24720001 | 24820000 | 0.0008 | 0.0008 | 0.0008 |
| F04 | 24740001 | 24840000 | 0.0011 | 0.0011 | 0.0011 |
| F04 | 24760001 | 24860000 | 0.0013 | 0.0010 | 0.0013 |
| F04 | 24780001 | 24880000 | 0.0011 | 0.0008 | 0.0011 |
| F04 | 24800001 | 24900000 | 0.0009 | 0.0009 | 0.0009 |
| F04 | 24820001 | 24920000 | 0.0010 | 0.0009 | 0.0010 |
| F04 | 24840001 | 24940000 | 0.0009 | 0.0012 | 0.0009 |
| F04 | 24860001 | 24960000 | 0.0012 | 0.0014 | 0.0012 |
| F04 | 24880001 | 24980000 | 0.0015 | 0.0014 | 0.0015 |
| F04 | 24900001 | 25000000 | 0.0015 | 0.0011 | 0.0015 |
| F04 | 24920001 | 25020000 | 0.0013 | 0.0016 | 0.0012 |
| F04 | 24940001 | 25040000 | 0.0018 | 0.0012 | 0.0018 |
| F04 | 24960001 | 25060000 | 0.0014 | 0.0010 | 0.0013 |
| F04 | 24980001 | 25080000 | 0.0011 | 0.0011 | 0.0010 |
| F04 | 25000001 | 25100000 | 0.0012 | 0.0014 | 0.0011 |
| F04 | 25020001 | 25120000 | 0.0015 | 0.0014 | 0.0015 |
| F04 | 25040001 | 25140000 | 0.0018 | 0.0015 | 0.0017 |
| F04 | 25060001 | 25160000 | 0.0020 | 0.0017 | 0.0018 |
| F04 | 25080001 | 25180000 | 0.0022 | 0.0023 | 0.0021 |
| F04 | 25100001 | 25200000 | 0.0031 | 0.0027 | 0.0028 |
| F04 | 25120001 | 25220000 | 0.0035 | 0.0020 | 0.0031 |
| F04 | 25140001 | 25240000 | 0.0025 | 0.0022 | 0.0022 |
| F04 | 25160001 | 25260000 | 0.0027 | 0.0022 | 0.0023 |
| F04 | 25180001 | 25280000 | 0.0027 | 0.0022 | 0.0023 |
| F04 | 25200001 | 25300000 | 0.0027 | 0.0017 | 0.0023 |
| F04 | 25220001 | 25320000 | 0.0022 | 0.0019 | 0.0019 |
| F04 | 25240001 | 25340000 | 0.0024 | 0.0016 | 0.0021 |
| F04 | 25260001 | 25360000 | 0.0022 | 0.0015 | 0.0019 |
| F04 | 25280001 | 25380000 | 0.0021 | 0.0011 | 0.0018 |
| F04 | 25300001 | 25400000 | 0.0016 | 0.0014 | 0.0013 |
| F04 | 25320001 | 25420000 | 0.0019 | 0.0017 | 0.0015 |
| F04 | 25340001 | 25440000 | 0.0023 | 0.0019 | 0.0019 |
| F04 | 25360001 | 25460000 | 0.0025 | 0.0020 | 0.0022 |
| F04 | 25380001 | 25480000 | 0.0023 | 0.0018 | 0.0022 |
| F04 | 25400001 | 25500000 | 0.0021 | 0.0013 | 0.0020 |
| F04 | 25420001 | 25520000 | 0.0015 | 0.0007 | 0.0015 |
| F04 | 25440001 | 25540000 | 0.0009 | 0.0009 | 0.0009 |
| F04 | 25460001 | 25560000 | 0.0009 | 0.0008 | 0.0009 |
| F04 | 25480001 | 25580000 | 0.0008 | 0.0007 | 0.0008 |
| F04 | 25500001 | 25600000 | 0.0008 | 0.0007 | 0.0008 |
| F04 | 25520001 | 25620000 | 0.0007 | 0.0009 | 0.0007 |
| F04 | 25540001 | 25640000 | 0.0009 | 0.0003 | 0.0009 |
| F04 | 25560001 | 25660000 | 0.0004 | 0.0008 | 0.0004 |
| F04 | 25580001 | 25680000 | 0.0009 | 0.0018 | 0.0009 |
| F04 | 25600001 | 25700000 | 0.0019 | 0.0022 | 0.0019 |
| F04 | 25620001 | 25720000 | 0.0024 | 0.0020 | 0.0024 |
| F04 | 25640001 | 25740000 | 0.0024 | 0.0023 | 0.0022 |
| F04 | 25660001 | 25760000 | 0.0028 | 0.0021 | 0.0024 |
| F04 | 25680001 | 25780000 | 0.0027 | 0.0018 | 0.0021 |
| F04 | 25700001 | 25800000 | 0.0027 | 0.0018 | 0.0021 |
| F04 | 25720001 | 25820000 | 0.0026 | 0.0021 | 0.0020 |
| F04 | 25740001 | 25840000 | 0.0029 | 0.0023 | 0.0024 |
| F04 | 25760001 | 25860000 | 0.0030 | 0.0037 | 0.0027 |

| Chr | Start | End | both | A1 | A2 |
|---|---|---|---|---|---|
| F04 | 25780001 | 25880000 | 0.0044 | 0.0030 | 0.0042 |
| F04 | 25800001 | 25900000 | 0.0036 | 0.0031 | 0.0033 |
| F04 | 25820001 | 25920000 | 0.0037 | 0.0027 | 0.0033 |
| F04 | 25840001 | 25940000 | 0.0033 | 0.0022 | 0.0029 |
| F04 | 25860001 | 25960000 | 0.0027 | 0.0008 | 0.0024 |
| F04 | 25880001 | 25980000 | 0.0011 | 0.0007 | 0.0008 |
| F04 | 25900001 | 26000000 | 0.0008 | 0.0004 | 0.0007 |
| F04 | 25920001 | 26020000 | 0.0005 | 0.0008 | 0.0004 |
| F04 | 25940001 | 26040000 | 0.0008 | 0.0009 | 0.0008 |
| F04 | 25960001 | 26060000 | 0.0009 | 0.0010 | 0.0008 |
| F04 | 25980001 | 26080000 | 0.0012 | 0.0012 | 0.0010 |
| F04 | 26000001 | 26100000 | 0.0014 | 0.0012 | 0.0012 |
| F04 | 26020001 | 26120000 | 0.0013 | 0.0007 | 0.0012 |
| F04 | 26040001 | 26140000 | 0.0009 | 0.0008 | 0.0007 |
| F04 | 26060001 | 26160000 | 0.0009 | 0.0005 | 0.0008 |
| F04 | 26080001 | 26180000 | 0.0006 | 0.0004 | 0.0006 |
| F04 | 26100001 | 26200000 | 0.0004 | 0.0003 | 0.0004 |
| F04 | 26120001 | 26220000 | 0.0003 | 0.0003 | 0.0003 |
| F04 | 26140001 | 26240000 | 0.0004 | 0.0005 | 0.0003 |
| F04 | 26160001 | 26260000 | 0.0006 | 0.0005 | 0.0006 |
| F04 | 26180001 | 26280000 | 0.0006 | 0.0007 | 0.0006 |
| F04 | 26200001 | 26300000 | 0.0008 | 0.0007 | 0.0008 |
| F04 | 26220001 | 26320000 | 0.0008 | 0.0006 | 0.0008 |
| F04 | 26240001 | 26340000 | 0.0008 | 0.0004 | 0.0008 |
| F04 | 26260001 | 26360000 | 0.0004 | 0.0002 | 0.0004 |
| F04 | 26280001 | 26380000 | 0.0002 | 0.0005 | 0.0002 |
| F04 | 26300001 | 26400000 | 0.0008 | 0.0011 | 0.0007 |
| F04 | 26320001 | 26420000 | 0.0015 | 0.0014 | 0.0014 |
| F04 | 26340001 | 26440000 | 0.0018 | 0.0016 | 0.0017 |
| F04 | 26360001 | 26460000 | 0.0021 | 0.0016 | 0.0020 |
| F04 | 26380001 | 26480000 | 0.0021 | 0.0015 | 0.0019 |
| F04 | 26400001 | 26500000 | 0.0018 | 0.0012 | 0.0017 |
| F04 | 26420001 | 26520000 | 0.0014 | 0.0010 | 0.0013 |
| F04 | 26440001 | 26540000 | 0.0011 | 0.0010 | 0.0010 |
| F04 | 26460001 | 26560000 | 0.0012 | 0.0013 | 0.0011 |
| F04 | 26480001 | 26580000 | 0.0016 | 0.0011 | 0.0014 |
| F04 | 26500001 | 26600000 | 0.0014 | 0.0008 | 0.0013 |
| F04 | 26520001 | 26620000 | 0.0011 | 0.0009 | 0.0010 |
| F04 | 26540001 | 26640000 | 0.0012 | 0.0008 | 0.0011 |
| F04 | 26560001 | 26660000 | 0.0012 | 0.0009 | 0.0011 |
| F04 | 26580001 | 26680000 | 0.0011 | 0.0007 | 0.0011 |
| F04 | 26600001 | 26700000 | 0.0009 | 0.0008 | 0.0009 |
| F04 | 26620001 | 26720000 | 0.0010 | 0.0008 | 0.0009 |
| F04 | 26640001 | 26740000 | 0.0009 | 0.0006 | 0.0008 |
| F04 | 26660001 | 26760000 | 0.0006 | 0.0004 | 0.0005 |
| F04 | 26680001 | 26780000 | 0.0004 | 0.0003 | 0.0004 |
| F04 | 26700001 | 26800000 | 0.0003 | 0.0002 | 0.0003 |
| F04 | 26720001 | 26820000 | 0.0002 | 0.0003 | 0.0002 |
| F04 | 26740001 | 26840000 | 0.0003 | 0.0004 | 0.0003 |
| F04 | 26760001 | 26860000 | 0.0006 | 0.0004 | 0.0005 |
| F04 | 26780001 | 26880000 | 0.0005 | 0.0011 | 0.0004 |
| F04 | 26800001 | 26900000 | 0.0012 | 0.0011 | 0.0011 |
| F04 | 26820001 | 26920000 | 0.0013 | 0.0013 | 0.0012 |
| F04 | 26840001 | 26940000 | 0.0015 | 0.0013 | 0.0013 |
| F04 | 26860001 | 26960000 | 0.0014 | 0.0016 | 0.0014 |
| F04 | 26880001 | 26980000 | 0.0018 | 0.0012 | 0.0017 |
| F04 | 26900001 | 27000000 | 0.0015 | 0.0016 | 0.0013 |
| F04 | 26920001 | 27020000 | 0.0019 | 0.0017 | 0.0017 |

| Chr | Start | End | both | A1 | A2 |
|---|---|---|---|---|---|
| F04 | 26940001 | 27040000 | 0.0020 | 0.0015 | 0.0019 |
| F04 | 26960001 | 27060000 | 0.0018 | 0.0014 | 0.0017 |
| F04 | 26980001 | 27080000 | 0.0017 | 0.0013 | 0.0014 |
| F04 | 27000001 | 27100000 | 0.0015 | 0.0009 | 0.0013 |
| F04 | 27020001 | 27120000 | 0.0011 | 0.0006 | 0.0010 |
| F04 | 27040001 | 27140000 | 0.0008 | 0.0007 | 0.0007 |
| F04 | 27060001 | 27160000 | 0.0009 | 0.0006 | 0.0008 |
| F04 | 27080001 | 27180000 | 0.0008 | 0.0010 | 0.0008 |
| F04 | 27100001 | 27200000 | 0.0013 | 0.0014 | 0.0012 |
| F04 | 27120001 | 27220000 | 0.0016 | 0.0014 | 0.0016 |
| F04 | 27140001 | 27240000 | 0.0016 | 0.0013 | 0.0016 |
| F04 | 27160001 | 27260000 | 0.0016 | 0.0013 | 0.0015 |
| F04 | 27180001 | 27280000 | 0.0015 | 0.0009 | 0.0014 |
| F04 | 27200001 | 27300000 | 0.0011 | 0.0004 | 0.0010 |
| F04 | 27220001 | 27320000 | 0.0005 | 0.0006 | 0.0004 |
| F04 | 27240001 | 27340000 | 0.0007 | 0.0006 | 0.0006 |
| F04 | 27260001 | 27360000 | 0.0008 | 0.0007 | 0.0007 |
| F04 | 27280001 | 27380000 | 0.0008 | 0.0005 | 0.0007 |
| F04 | 27300001 | 27400000 | 0.0006 | 0.0007 | 0.0006 |
| F04 | 27320001 | 27420000 | 0.0009 | 0.0005 | 0.0009 |
| F04 | 27340001 | 27440000 | 0.0006 | 0.0005 | 0.0006 |
| F04 | 27360001 | 27460000 | 0.0006 | 0.0007 | 0.0006 |
| F04 | 27380001 | 27480000 | 0.0008 | 0.0009 | 0.0008 |
| F04 | 27400001 | 27500000 | 0.0010 | 0.0007 | 0.0010 |
| F04 | 27420001 | 27520000 | 0.0008 | 0.0007 | 0.0007 |
| F04 | 27440001 | 27540000 | 0.0008 | 0.0006 | 0.0008 |
| F04 | 27460001 | 27560000 | 0.0007 | 0.0006 | 0.0007 |
| F04 | 27480001 | 27580000 | 0.0008 | 0.0005 | 0.0008 |
| F04 | 27500001 | 27600000 | 0.0007 | 0.0013 | 0.0007 |
| F04 | 27520001 | 27620000 | 0.0015 | 0.0015 | 0.0014 |
| F04 | 27540001 | 27640000 | 0.0016 | 0.0018 | 0.0015 |
| F04 | 27560001 | 27660000 | 0.0020 | 0.0016 | 0.0019 |
| F04 | 27580001 | 27680000 | 0.0016 | 0.0014 | 0.0015 |
| F04 | 27600001 | 27700000 | 0.0015 | 0.0006 | 0.0014 |
| F04 | 27620001 | 27720000 | 0.0007 | 0.0005 | 0.0007 |
| F04 | 27640001 | 27740000 | 0.0005 | 0.0001 | 0.0005 |
| F04 | 27660001 | 27760000 | 0.0001 | 0.0001 | 0.0001 |
| F04 | 27680001 | 27780000 | 0.0001 | 0.0001 | 0.0001 |
| F04 | 27700001 | 27800000 | 0.0002 | 0.0002 | 0.0002 |
| F04 | 27720001 | 27820000 | 0.0004 | 0.0005 | 0.0002 |
| F04 | 27740001 | 27840000 | 0.0007 | 0.0010 | 0.0005 |
| F04 | 27760001 | 27860000 | 0.0013 | 0.0014 | 0.0011 |
| F04 | 27780001 | 27880000 | 0.0017 | 0.0016 | 0.0015 |
| F04 | 27800001 | 27900000 | 0.0019 | 0.0017 | 0.0017 |
| F04 | 27820001 | 27920000 | 0.0019 | 0.0014 | 0.0018 |
| F04 | 27840001 | 27940000 | 0.0016 | 0.0010 | 0.0015 |
| F04 | 27860001 | 27960000 | 0.0011 | 0.0007 | 0.0010 |
| F04 | 27880001 | 27980000 | 0.0008 | 0.0008 | 0.0008 |
| F04 | 27900001 | 28000000 | 0.0010 | 0.0007 | 0.0009 |
| F04 | 27920001 | 28020000 | 0.0009 | 0.0007 | 0.0008 |
| F04 | 27940001 | 28040000 | 0.0008 | 0.0006 | 0.0008 |
| F04 | 27960001 | 28060000 | 0.0008 | 0.0007 | 0.0007 |
| F04 | 27980001 | 28080000 | 0.0008 | 0.0005 | 0.0007 |
| F04 | 28000001 | 28100000 | 0.0006 | 0.0005 | 0.0006 |
| F04 | 28020001 | 28120000 | 0.0005 | 0.0006 | 0.0005 |
| F04 | 28040001 | 28140000 | 0.0008 | 0.0007 | 0.0008 |
| F04 | 28060001 | 28160000 | 0.0008 | 0.0006 | 0.0008 |
| F04 | 28080001 | 28180000 | 0.0007 | 0.0006 | 0.0007 |

| Chr | Start | End | both | A1 | A2 |
|---|---|---|---|---|---|
| F04 | 28100001 | 28200000 | 0.0007 | 0.0006 | 0.0006 |
| F04 | 28120001 | 28220000 | 0.0007 | 0.0005 | 0.0007 |
| F04 | 28140001 | 28240000 | 0.0005 | 0.0005 | 0.0005 |
| F04 | 28160001 | 28260000 | 0.0006 | 0.0010 | 0.0005 |
| F04 | 28180001 | 28280000 | 0.0010 | 0.0015 | 0.0009 |
| F04 | 28200001 | 28300000 | 0.0016 | 0.0016 | 0.0015 |
| F04 | 28220001 | 28320000 | 0.0018 | 0.0015 | 0.0017 |
| F04 | 28240001 | 28340000 | 0.0017 | 0.0016 | 0.0016 |
| F04 | 28260001 | 28360000 | 0.0017 | 0.0012 | 0.0017 |
| F04 | 28280001 | 28380000 | 0.0013 | 0.0006 | 0.0013 |
| F04 | 28300001 | 28400000 | 0.0006 | 0.0005 | 0.0006 |
| F04 | 28320001 | 28420000 | 0.0006 | 0.0007 | 0.0006 |
| F04 | 28340001 | 28440000 | 0.0010 | 0.0009 | 0.0009 |
| F04 | 28360001 | 28460000 | 0.0012 | 0.0010 | 0.0010 |
| F04 | 28380001 | 28480000 | 0.0015 | 0.0010 | 0.0012 |
| F04 | 28400001 | 28500000 | 0.0015 | 0.0009 | 0.0012 |
| F04 | 28420001 | 28520000 | 0.0013 | 0.0009 | 0.0010 |
| F04 | 28440001 | 28540000 | 0.0011 | 0.0007 | 0.0010 |
| F04 | 28460001 | 28560000 | 0.0009 | 0.0006 | 0.0008 |
| F04 | 28480001 | 28580000 | 0.0008 | 0.0005 | 0.0007 |
| F04 | 28500001 | 28600000 | 0.0007 | 0.0006 | 0.0006 |
| F04 | 28520001 | 28620000 | 0.0008 | 0.0005 | 0.0007 |
| F04 | 28540001 | 28640000 | 0.0007 | 0.0007 | 0.0006 |
| F04 | 28560001 | 28660000 | 0.0008 | 0.0007 | 0.0007 |
| F04 | 28580001 | 28680000 | 0.0009 | 0.0009 | 0.0009 |
| F04 | 28600001 | 28700000 | 0.0010 | 0.0009 | 0.0010 |
| F04 | 28620001 | 28720000 | 0.0010 | 0.0009 | 0.0011 |
| F04 | 28640001 | 28740000 | 0.0011 | 0.0006 | 0.0011 |
| F04 | 28660001 | 28760000 | 0.0007 | 0.0004 | 0.0008 |
| F04 | 28680001 | 28780000 | 0.0005 | 0.0003 | 0.0005 |
| F04 | 28700001 | 28800000 | 0.0004 | 0.0007 | 0.0004 |
| F04 | 28720001 | 28820000 | 0.0008 | 0.0011 | 0.0008 |
| F04 | 28740001 | 28840000 | 0.0013 | 0.0014 | 0.0012 |
| F04 | 28760001 | 28860000 | 0.0018 | 0.0023 | 0.0017 |
| F04 | 28780001 | 28880000 | 0.0029 | 0.0025 | 0.0027 |
| F04 | 28800001 | 28900000 | 0.0032 | 0.0022 | 0.0030 |
| F04 | 28820001 | 28920000 | 0.0028 | 0.0021 | 0.0027 |
| F04 | 28840001 | 28940000 | 0.0028 | 0.0020 | 0.0026 |
| F04 | 28860001 | 28960000 | 0.0028 | 0.0014 | 0.0026 |
| F04 | 28880001 | 28980000 | 0.0021 | 0.0012 | 0.0020 |
| F04 | 28900001 | 29000000 | 0.0019 | 0.0012 | 0.0018 |
| F04 | 28920001 | 29020000 | 0.0018 | 0.0010 | 0.0017 |
| F04 | 28940001 | 29040000 | 0.0016 | 0.0008 | 0.0014 |
| F04 | 28960001 | 29060000 | 0.0011 | 0.0007 | 0.0010 |
| F04 | 28980001 | 29080000 | 0.0009 | 0.0008 | 0.0009 |
| F04 | 29000001 | 29100000 | 0.0010 | 0.0012 | 0.0010 |
| F04 | 29020001 | 29120000 | 0.0014 | 0.0010 | 0.0014 |
| F04 | 29040001 | 29140000 | 0.0011 | 0.0010 | 0.0011 |
| F04 | 29060001 | 29160000 | 0.0012 | 0.0008 | 0.0012 |
| F04 | 29080001 | 29180000 | 0.0009 | 0.0008 | 0.0009 |
| F04 | 29100001 | 29200000 | 0.0010 | 0.0005 | 0.0010 |
| F04 | 29120001 | 29220000 | 0.0005 | 0.0006 | 0.0005 |
| F04 | 29140001 | 29240000 | 0.0006 | 0.0007 | 0.0006 |
| F04 | 29160001 | 29260000 | 0.0007 | 0.0007 | 0.0007 |
| F04 | 29180001 | 29280000 | 0.0008 | 0.0007 | 0.0008 |
| F04 | 29200001 | 29300000 | 0.0007 | 0.0006 | 0.0007 |
| F04 | 29220001 | 29320000 | 0.0007 | 0.0007 | 0.0006 |
| F04 | 29240001 | 29340000 | 0.0008 | 0.0011 | 0.0007 |

| Chr | Start | End | both | A1 | A2 |
|---|---|---|---|---|---|
| F04 | 29260001 | 29360000 | 0.0012 | 0.0012 | 0.0012 |
| F04 | 29280001 | 29380000 | 0.0013 | 0.0009 | 0.0013 |
| F04 | 29300001 | 29400000 | 0.0011 | 0.0010 | 0.0011 |
| F04 | 29320001 | 29420000 | 0.0012 | 0.0009 | 0.0012 |
| F04 | 29340001 | 29440000 | 0.0011 | 0.0006 | 0.0011 |
| F04 | 29360001 | 29460000 | 0.0008 | 0.0012 | 0.0008 |
| F04 | 29380001 | 29480000 | 0.0013 | 0.0013 | 0.0013 |
| F04 | 29400001 | 29500000 | 0.0015 | 0.0012 | 0.0014 |
| F04 | 29420001 | 29520000 | 0.0013 | 0.0012 | 0.0013 |
| F04 | 29440001 | 29540000 | 0.0013 | 0.0010 | 0.0012 |
| F04 | 29460001 | 29560000 | 0.0010 | 0.0006 | 0.0010 |
| F04 | 29480001 | 29580000 | 0.0006 | 0.0009 | 0.0006 |
| F04 | 29500001 | 29600000 | 0.0011 | 0.0010 | 0.0011 |
| F04 | 29520001 | 29620000 | 0.0012 | 0.0010 | 0.0012 |
| F04 | 29540001 | 29640000 | 0.0012 | 0.0010 | 0.0012 |
| F04 | 29560001 | 29660000 | 0.0012 | 0.0008 | 0.0012 |
| F04 | 29580001 | 29680000 | 0.0011 | 0.0005 | 0.0011 |
| F04 | 29600001 | 29700000 | 0.0006 | 0.0003 | 0.0006 |
| F04 | 29620001 | 29720000 | 0.0004 | 0.0003 | 0.0004 |
| F04 | 29640001 | 29740000 | 0.0004 | 0.0003 | 0.0004 |
| F04 | 29660001 | 29760000 | 0.0004 | 0.0003 | 0.0004 |
| F04 | 29680001 | 29780000 | 0.0003 | 0.0002 | 0.0003 |
| F04 | 29700001 | 29800000 | 0.0002 | 0.0002 | 0.0002 |
| F04 | 29720001 | 29820000 | 0.0002 | 0.0002 | 0.0002 |
| F04 | 29740001 | 29840000 | 0.0003 | 0.0003 | 0.0003 |
| F04 | 29760001 | 29860000 | 0.0003 | 0.0003 | 0.0003 |
| F04 | 29780001 | 29880000 | 0.0003 | 0.0006 | 0.0003 |
| F04 | 29800001 | 29900000 | 0.0007 | 0.0008 | 0.0006 |
| F04 | 29820001 | 29920000 | 0.0008 | 0.0009 | 0.0008 |
| F04 | 29840001 | 29940000 | 0.0009 | 0.0012 | 0.0008 |
| F04 | 29860001 | 29960000 | 0.0012 | 0.0011 | 0.0011 |
| F04 | 29880001 | 29980000 | 0.0012 | 0.0015 | 0.0011 |
| F04 | 29900001 | 30000000 | 0.0016 | 0.0014 | 0.0016 |
| F04 | 29920001 | 30020000 | 0.0015 | 0.0013 | 0.0014 |
| F04 | 29940001 | 30040000 | 0.0013 | 0.0011 | 0.0013 |
| F04 | 29960001 | 30060000 | 0.0012 | 0.0013 | 0.0012 |
| F04 | 29980001 | 30080000 | 0.0014 | 0.0007 | 0.0013 |
| F04 | 30000001 | 30100000 | 0.0007 | 0.0008 | 0.0007 |
| F04 | 30020001 | 30120000 | 0.0008 | 0.0008 | 0.0008 |
| F04 | 30040001 | 30140000 | 0.0008 | 0.0009 | 0.0008 |
| F04 | 30060001 | 30160000 | 0.0009 | 0.0007 | 0.0009 |
| F04 | 30080001 | 30180000 | 0.0008 | 0.0006 | 0.0008 |
| F04 | 30100001 | 30200000 | 0.0008 | 0.0007 | 0.0008 |
| F04 | 30120001 | 30220000 | 0.0009 | 0.0007 | 0.0009 |
| F04 | 30140001 | 30240000 | 0.0009 | 0.0007 | 0.0009 |
| F04 | 30160001 | 30260000 | 0.0009 | 0.0013 | 0.0008 |
| F04 | 30180001 | 30280000 | 0.0014 | 0.0016 | 0.0013 |
| F04 | 30200001 | 30300000 | 0.0018 | 0.0025 | 0.0016 |
| F04 | 30220001 | 30320000 | 0.0030 | 0.0046 | 0.0026 |
| F04 | 30240001 | 30340000 | 0.0050 | 0.0047 | 0.0045 |
| F04 | 30260001 | 30360000 | 0.0052 | 0.0049 | 0.0047 |
| F04 | 30280001 | 30380000 | 0.0057 | 0.0054 | 0.0050 |
| F04 | 30300001 | 30400000 | 0.0061 | 0.0048 | 0.0055 |
| F04 | 30320001 | 30420000 | 0.0053 | 0.0028 | 0.0047 |
| F04 | 30340001 | 30440000 | 0.0034 | 0.0025 | 0.0029 |
| F04 | 30360001 | 30460000 | 0.0031 | 0.0017 | 0.0026 |
| F04 | 30380001 | 30480000 | 0.0021 | 0.0011 | 0.0018 |
| F04 | 30400001 | 30500000 | 0.0015 | 0.0006 | 0.0012 |

| Chr | Start | End | both | A1 | A2 |
|---|---|---|---|---|---|
| F04 | 30420001 | 30520000 | 0.0008 | 0.0009 | 0.0007 |
| F04 | 30440001 | 30540000 | 0.0012 | 0.0021 | 0.0010 |
| F04 | 30460001 | 30560000 | 0.0026 | 0.0028 | 0.0023 |
| F04 | 30480001 | 30580000 | 0.0035 | 0.0026 | 0.0030 |
| F04 | 30500001 | 30600000 | 0.0036 | 0.0028 | 0.0029 |
| F04 | 30520001 | 30620000 | 0.0038 | 0.0030 | 0.0031 |
| F04 | 30540001 | 30640000 | 0.0039 | 0.0017 | 0.0033 |
| F04 | 30560001 | 30660000 | 0.0024 | 0.0010 | 0.0019 |
| F04 | 30580001 | 30680000 | 0.0015 | 0.0009 | 0.0012 |
| F04 | 30600001 | 30700000 | 0.0010 | 0.0011 | 0.0010 |
| F04 | 30620001 | 30720000 | 0.0011 | 0.0006 | 0.0011 |
| F04 | 30640001 | 30740000 | 0.0006 | 0.0008 | 0.0006 |
| F04 | 30660001 | 30760000 | 0.0010 | 0.0012 | 0.0009 |
| F04 | 30680001 | 30780000 | 0.0014 | 0.0013 | 0.0014 |
| F04 | 30700001 | 30800000 | 0.0015 | 0.0012 | 0.0015 |
| F04 | 30720001 | 30820000 | 0.0014 | 0.0011 | 0.0014 |
| F04 | 30740001 | 30840000 | 0.0014 | 0.0014 | 0.0014 |
| F04 | 30760001 | 30860000 | 0.0016 | 0.0012 | 0.0016 |
| F04 | 30780001 | 30880000 | 0.0013 | 0.0014 | 0.0012 |
| F04 | 30800001 | 30900000 | 0.0014 | 0.0013 | 0.0014 |
| F04 | 30820001 | 30920000 | 0.0013 | 0.0013 | 0.0012 |
| F04 | 30840001 | 30940000 | 0.0013 | 0.0009 | 0.0012 |
| F04 | 30860001 | 30960000 | 0.0009 | 0.0007 | 0.0008 |
| F04 | 30880001 | 30980000 | 0.0008 | 0.0011 | 0.0007 |
| F04 | 30900001 | 31000000 | 0.0011 | 0.0009 | 0.0010 |
| F04 | 30920001 | 31020000 | 0.0010 | 0.0016 | 0.0009 |
| F04 | 30940001 | 31040000 | 0.0018 | 0.0016 | 0.0018 |
| F04 | 30960001 | 31060000 | 0.0018 | 0.0017 | 0.0017 |
| F04 | 30980001 | 31080000 | 0.0018 | 0.0015 | 0.0017 |
| F04 | 31000001 | 31100000 | 0.0018 | 0.0015 | 0.0017 |
| F04 | 31020001 | 31120000 | 0.0018 | 0.0019 | 0.0017 |
| F04 | 31040001 | 31140000 | 0.0021 | 0.0019 | 0.0020 |
| F04 | 31060001 | 31160000 | 0.0023 | 0.0021 | 0.0021 |
| F04 | 31080001 | 31180000 | 0.0032 | 0.0024 | 0.0028 |
| F04 | 31100001 | 31200000 | 0.0035 | 0.0029 | 0.0029 |
| F04 | 31120001 | 31220000 | 0.0040 | 0.0019 | 0.0034 |
| F04 | 31140001 | 31240000 | 0.0030 | 0.0020 | 0.0024 |
| F04 | 31160001 | 31260000 | 0.0029 | 0.0017 | 0.0025 |
| F04 | 31180001 | 31280000 | 0.0020 | 0.0010 | 0.0018 |
| F04 | 31200001 | 31300000 | 0.0013 | 0.0009 | 0.0011 |
| F04 | 31220001 | 31320000 | 0.0012 | 0.0015 | 0.0011 |
| F04 | 31240001 | 31340000 | 0.0020 | 0.0015 | 0.0017 |
| F04 | 31260001 | 31360000 | 0.0023 | 0.0025 | 0.0018 |
| F04 | 31280001 | 31380000 | 0.0036 | 0.0028 | 0.0031 |
| F04 | 31300001 | 31400000 | 0.0039 | 0.0029 | 0.0034 |
| F04 | 31320001 | 31420000 | 0.0041 | 0.0036 | 0.0035 |
| F04 | 31340001 | 31440000 | 0.0048 | 0.0037 | 0.0042 |
| F04 | 31360001 | 31460000 | 0.0045 | 0.0031 | 0.0041 |
| F04 | 31380001 | 31480000 | 0.0038 | 0.0029 | 0.0034 |
| F04 | 31400001 | 31500000 | 0.0037 | 0.0034 | 0.0031 |
| F04 | 31420001 | 31520000 | 0.0040 | 0.0024 | 0.0034 |
| F04 | 31440001 | 31540000 | 0.0030 | 0.0030 | 0.0024 |
| F04 | 31460001 | 31560000 | 0.0035 | 0.0024 | 0.0029 |
| F04 | 31480001 | 31580000 | 0.0027 | 0.0027 | 0.0023 |
| F04 | 31500001 | 31600000 | 0.0029 | 0.0020 | 0.0026 |
| F04 | 31520001 | 31620000 | 0.0024 | 0.0021 | 0.0021 |
| F04 | 31540001 | 31640000 | 0.0026 | 0.0020 | 0.0024 |
| F04 | 31560001 | 31660000 | 0.0029 | 0.0022 | 0.0025 |

| Chr | Start | End | both | A1 | A2 |
|---|---|---|---|---|---|
| F04 | 31580001 | 31680000 | 0.0031 | 0.0025 | 0.0027 |
| F04 | 31600001 | 31700000 | 0.0034 | 0.0023 | 0.0029 |
| F04 | 31620001 | 31720000 | 0.0031 | 0.0017 | 0.0027 |
| F04 | 31640001 | 31740000 | 0.0024 | 0.0013 | 0.0020 |
| F04 | 31660001 | 31760000 | 0.0017 | 0.0018 | 0.0015 |
| F04 | 31680001 | 31780000 | 0.0021 | 0.0016 | 0.0019 |
| F04 | 31700001 | 31800000 | 0.0019 | 0.0021 | 0.0017 |
| F04 | 31720001 | 31820000 | 0.0025 | 0.0020 | 0.0023 |
| F04 | 31740001 | 31840000 | 0.0025 | 0.0017 | 0.0021 |
| F04 | 31760001 | 31860000 | 0.0021 | 0.0016 | 0.0017 |
| F04 | 31780001 | 31880000 | 0.0021 | 0.0018 | 0.0018 |
| F04 | 31800001 | 31900000 | 0.0025 | 0.0014 | 0.0021 |
| F04 | 31820001 | 31920000 | 0.0021 | 0.0018 | 0.0017 |
| F04 | 31840001 | 31940000 | 0.0024 | 0.0018 | 0.0022 |
| F04 | 31860001 | 31960000 | 0.0024 | 0.0016 | 0.0021 |
| F04 | 31880001 | 31980000 | 0.0021 | 0.0009 | 0.0018 |
| F04 | 31900001 | 32000000 | 0.0013 | 0.0008 | 0.0011 |
| F04 | 31920001 | 32020000 | 0.0011 | 0.0004 | 0.0010 |
| F04 | 31940001 | 32040000 | 0.0006 | 0.0005 | 0.0005 |
| F04 | 31960001 | 32060000 | 0.0007 | 0.0016 | 0.0006 |
| F04 | 31980001 | 32080000 | 0.0019 | 0.0025 | 0.0019 |
| F04 | 32000001 | 32100000 | 0.0029 | 0.0028 | 0.0028 |
| F04 | 32020001 | 32120000 | 0.0032 | 0.0032 | 0.0031 |
| F04 | 32040001 | 32140000 | 0.0036 | 0.0043 | 0.0035 |
| F04 | 32060001 | 32160000 | 0.0050 | 0.0031 | 0.0048 |
| F04 | 32080001 | 32180000 | 0.0037 | 0.0021 | 0.0035 |
| F04 | 32100001 | 32200000 | 0.0027 | 0.0020 | 0.0025 |
| F04 | 32120001 | 32220000 | 0.0026 | 0.0022 | 0.0024 |
| F04 | 32140001 | 32240000 | 0.0027 | 0.0013 | 0.0026 |
| F04 | 32160001 | 32260000 | 0.0016 | 0.0016 | 0.0015 |
| F04 | 32180001 | 32280000 | 0.0019 | 0.0016 | 0.0018 |
| F04 | 32200001 | 32300000 | 0.0019 | 0.0013 | 0.0018 |
| F04 | 32220001 | 32320000 | 0.0016 | 0.0009 | 0.0015 |
| F04 | 32240001 | 32340000 | 0.0011 | 0.0007 | 0.0011 |
| F04 | 32260001 | 32360000 | 0.0009 | 0.0006 | 0.0009 |
| F04 | 32280001 | 32380000 | 0.0008 | 0.0010 | 0.0008 |
| F04 | 32300001 | 32400000 | 0.0011 | 0.0014 | 0.0011 |
| F04 | 32320001 | 32420000 | 0.0015 | 0.0020 | 0.0015 |
| F04 | 32340001 | 32440000 | 0.0025 | 0.0028 | 0.0020 |
| F04 | 32360001 | 32460000 | 0.0037 | 0.0033 | 0.0029 |
| F04 | 32380001 | 32480000 | 0.0044 | 0.0029 | 0.0035 |
| F04 | 32400001 | 32500000 | 0.0041 | 0.0028 | 0.0033 |
| F04 | 32420001 | 32520000 | 0.0042 | 0.0027 | 0.0032 |
| F04 | 32440001 | 32540000 | 0.0040 | 0.0021 | 0.0032 |
| F04 | 32460001 | 32560000 | 0.0032 | 0.0014 | 0.0027 |
| F04 | 32480001 | 32580000 | 0.0022 | 0.0014 | 0.0017 |
| F04 | 32500001 | 32600000 | 0.0021 | 0.0013 | 0.0017 |
| F04 | 32520001 | 32620000 | 0.0020 | 0.0015 | 0.0016 |
| F04 | 32540001 | 32640000 | 0.0019 | 0.0013 | 0.0017 |
| F04 | 32560001 | 32660000 | 0.0016 | 0.0016 | 0.0014 |
| F04 | 32580001 | 32680000 | 0.0019 | 0.0020 | 0.0016 |
| F04 | 32600001 | 32700000 | 0.0025 | 0.0018 | 0.0021 |
| F04 | 32620001 | 32720000 | 0.0022 | 0.0011 | 0.0019 |
| F04 | 32640001 | 32740000 | 0.0016 | 0.0009 | 0.0012 |
| F04 | 32660001 | 32760000 | 0.0018 | 0.0007 | 0.0011 |
| F04 | 32680001 | 32780000 | 0.0015 | 0.0013 | 0.0009 |
| F04 | 32700001 | 32800000 | 0.0020 | 0.0016 | 0.0013 |
| F04 | 32720001 | 32820000 | 0.0025 | 0.0022 | 0.0016 |

| Chr | Start | End | both | A1 | A2 |
|---|---|---|---|---|---|
| F04 | 32740001 | 32840000 | 0.0032 | 0.0026 | 0.0024 |
| F04 | 32760001 | 32860000 | 0.0033 | 0.0026 | 0.0027 |
| F04 | 32780001 | 32880000 | 0.0031 | 0.0021 | 0.0027 |
| F04 | 32800001 | 32900000 | 0.0027 | 0.0018 | 0.0023 |
| F04 | 32820001 | 32920000 | 0.0022 | 0.0011 | 0.0021 |
| F04 | 32840001 | 32940000 | 0.0014 | 0.0008 | 0.0013 |
| F04 | 32860001 | 32960000 | 0.0009 | 0.0010 | 0.0009 |
| F04 | 32880001 | 32980000 | 0.0012 | 0.0006 | 0.0012 |
| F04 | 32900001 | 33000000 | 0.0007 | 0.0011 | 0.0007 |
| F04 | 32920001 | 33020000 | 0.0012 | 0.0016 | 0.0012 |
| F04 | 32940001 | 33040000 | 0.0020 | 0.0016 | 0.0020 |
| F04 | 32960001 | 33060000 | 0.0020 | 0.0014 | 0.0020 |
| F04 | 32980001 | 33080000 | 0.0017 | 0.0015 | 0.0017 |
| F04 | 33000001 | 33100000 | 0.0018 | 0.0012 | 0.0018 |
| F04 | 33020001 | 33120000 | 0.0016 | 0.0009 | 0.0016 |
| F04 | 33040001 | 33140000 | 0.0011 | 0.0012 | 0.0011 |
| F04 | 33060001 | 33160000 | 0.0014 | 0.0017 | 0.0014 |
| F04 | 33080001 | 33180000 | 0.0019 | 0.0021 | 0.0019 |
| F04 | 33100001 | 33200000 | 0.0023 | 0.0018 | 0.0023 |
| F04 | 33120001 | 33220000 | 0.0020 | 0.0016 | 0.0020 |
| F04 | 33140001 | 33240000 | 0.0018 | 0.0015 | 0.0018 |
| F04 | 33160001 | 33260000 | 0.0017 | 0.0011 | 0.0016 |
| F04 | 33180001 | 33280000 | 0.0012 | 0.0005 | 0.0012 |
| F04 | 33200001 | 33300000 | 0.0006 | 0.0004 | 0.0005 |
| F04 | 33220001 | 33320000 | 0.0004 | 0.0007 | 0.0004 |
| F04 | 33240001 | 33340000 | 0.0008 | 0.0009 | 0.0008 |
| F04 | 33260001 | 33360000 | 0.0011 | 0.0009 | 0.0011 |
| F04 | 33280001 | 33380000 | 0.0012 | 0.0012 | 0.0012 |
| F04 | 33300001 | 33400000 | 0.0015 | 0.0015 | 0.0015 |
| F04 | 33320001 | 33420000 | 0.0018 | 0.0012 | 0.0018 |
| F04 | 33340001 | 33440000 | 0.0014 | 0.0009 | 0.0014 |
| F04 | 33360001 | 33460000 | 0.0010 | 0.0007 | 0.0009 |
| F04 | 33380001 | 33480000 | 0.0008 | 0.0005 | 0.0008 |
| F04 | 33400001 | 33500000 | 0.0005 | 0.0002 | 0.0005 |
| F04 | 33420001 | 33520000 | 0.0003 | 0.0002 | 0.0003 |
| F04 | 33440001 | 33540000 | 0.0002 | 0.0003 | 0.0002 |
| F04 | 33460001 | 33560000 | 0.0003 | 0.0003 | 0.0003 |
| F04 | 33480001 | 33580000 | 0.0004 | 0.0004 | 0.0004 |
| F04 | 33500001 | 33600000 | 0.0004 | 0.0007 | 0.0004 |
| F04 | 33520001 | 33620000 | 0.0009 | 0.0009 | 0.0009 |
| F04 | 33540001 | 33640000 | 0.0010 | 0.0007 | 0.0011 |
| F04 | 33560001 | 33660000 | 0.0009 | 0.0011 | 0.0009 |
| F04 | 33580001 | 33680000 | 0.0013 | 0.0011 | 0.0013 |
| F04 | 33600001 | 33700000 | 0.0014 | 0.0007 | 0.0014 |
| F04 | 33620001 | 33720000 | 0.0009 | 0.0008 | 0.0009 |
| F04 | 33640001 | 33740000 | 0.0009 | 0.0012 | 0.0009 |
| F04 | 33660001 | 33760000 | 0.0015 | 0.0011 | 0.0015 |
| F04 | 33680001 | 33780000 | 0.0013 | 0.0014 | 0.0013 |
| F04 | 33700001 | 33800000 | 0.0017 | 0.0019 | 0.0017 |
| F04 | 33720001 | 33820000 | 0.0023 | 0.0023 | 0.0023 |
| F04 | 33740001 | 33840000 | 0.0027 | 0.0029 | 0.0026 |
| F04 | 33760001 | 33860000 | 0.0034 | 0.0029 | 0.0032 |
| F04 | 33780001 | 33880000 | 0.0035 | 0.0028 | 0.0032 |
| F04 | 33800001 | 33900000 | 0.0035 | 0.0037 | 0.0032 |
| F04 | 33820001 | 33920000 | 0.0045 | 0.0035 | 0.0042 |
| F04 | 33840001 | 33940000 | 0.0041 | 0.0029 | 0.0037 |
| F04 | 33860001 | 33960000 | 0.0034 | 0.0027 | 0.0031 |
| F04 | 33880001 | 33980000 | 0.0033 | 0.0027 | 0.0029 |

| Chr | Start | End | both | A1 | A2 |
|---|---|---|---|---|---|
| F04 | 33900001 | 34000000 | 0.0032 | 0.0019 | 0.0027 |
| F04 | 33920001 | 34020000 | 0.0025 | 0.0017 | 0.0017 |
| F04 | 33940001 | 34040000 | 0.0025 | 0.0019 | 0.0018 |
| F04 | 33960001 | 34060000 | 0.0027 | 0.0021 | 0.0021 |
| F04 | 33980001 | 34080000 | 0.0028 | 0.0023 | 0.0022 |
| F04 | 34000001 | 34100000 | 0.0032 | 0.0020 | 0.0027 |
| F04 | 34020001 | 34120000 | 0.0027 | 0.0023 | 0.0025 |
| F04 | 34040001 | 34140000 | 0.0030 | 0.0019 | 0.0028 |
| F04 | 34060001 | 34160000 | 0.0024 | 0.0017 | 0.0023 |
| F04 | 34080001 | 34180000 | 0.0023 | 0.0016 | 0.0022 |
| F04 | 34100001 | 34200000 | 0.0020 | 0.0020 | 0.0019 |
| F04 | 34120001 | 34220000 | 0.0024 | 0.0016 | 0.0023 |
| F04 | 34140001 | 34240000 | 0.0020 | 0.0020 | 0.0018 |
| F04 | 34160001 | 34260000 | 0.0025 | 0.0020 | 0.0021 |
| F04 | 34180001 | 34280000 | 0.0024 | 0.0016 | 0.0021 |
| F04 | 34200001 | 34300000 | 0.0020 | 0.0011 | 0.0018 |
| F04 | 34220001 | 34320000 | 0.0015 | 0.0011 | 0.0013 |
| F04 | 34240001 | 34340000 | 0.0015 | 0.0006 | 0.0014 |
| F04 | 34260001 | 34360000 | 0.0009 | 0.0011 | 0.0009 |
| F04 | 34280001 | 34380000 | 0.0014 | 0.0013 | 0.0014 |
| F04 | 34300001 | 34400000 | 0.0015 | 0.0016 | 0.0014 |
| F04 | 34320001 | 34420000 | 0.0018 | 0.0020 | 0.0018 |
| F04 | 34340001 | 34440000 | 0.0023 | 0.0021 | 0.0023 |
| F04 | 34360001 | 34460000 | 0.0025 | 0.0017 | 0.0025 |
| F04 | 34380001 | 34480000 | 0.0022 | 0.0016 | 0.0022 |
| F04 | 34400001 | 34500000 | 0.0022 | 0.0011 | 0.0022 |
| F04 | 34420001 | 34520000 | 0.0016 | 0.0007 | 0.0015 |
| F04 | 34440001 | 34540000 | 0.0011 | 0.0006 | 0.0011 |
| F04 | 34460001 | 34560000 | 0.0010 | 0.0005 | 0.0009 |
| F04 | 34480001 | 34580000 | 0.0006 | 0.0005 | 0.0006 |
| F04 | 34500001 | 34600000 | 0.0006 | 0.0007 | 0.0006 |
| F04 | 34520001 | 34620000 | 0.0008 | 0.0008 | 0.0008 |
| F04 | 34540001 | 34640000 | 0.0009 | 0.0009 | 0.0008 |
| F04 | 34560001 | 34660000 | 0.0010 | 0.0015 | 0.0010 |
| F04 | 34580001 | 34680000 | 0.0016 | 0.0014 | 0.0015 |
| F04 | 34600001 | 34700000 | 0.0015 | 0.0012 | 0.0014 |
| F04 | 34620001 | 34720000 | 0.0013 | 0.0010 | 0.0012 |
| F04 | 34640001 | 34740000 | 0.0012 | 0.0009 | 0.0011 |
| F04 | 34660001 | 34760000 | 0.0010 | 0.0003 | 0.0010 |
| F04 | 34680001 | 34780000 | 0.0004 | 0.0003 | 0.0004 |
| F04 | 34700001 | 34800000 | 0.0004 | 0.0007 | 0.0004 |
| F04 | 34720001 | 34820000 | 0.0009 | 0.0007 | 0.0009 |
| F04 | 34740001 | 34840000 | 0.0009 | 0.0007 | 0.0009 |
| F04 | 34760001 | 34860000 | 0.0008 | 0.0007 | 0.0008 |
| F04 | 34780001 | 34880000 | 0.0009 | 0.0007 | 0.0008 |
| F04 | 34800001 | 34900000 | 0.0008 | 0.0003 | 0.0007 |
| F04 | 34820001 | 34920000 | 0.0003 | 0.0002 | 0.0003 |
| F04 | 34840001 | 34940000 | 0.0002 | 0.0003 | 0.0002 |
| F04 | 34860001 | 34960000 | 0.0004 | 0.0003 | 0.0004 |
| F04 | 34880001 | 34980000 | 0.0004 | 0.0003 | 0.0004 |
| F04 | 34900001 | 35000000 | 0.0003 | 0.0002 | 0.0003 |
| F04 | 34920001 | 35020000 | 0.0003 | 0.0002 | 0.0003 |
| F04 | 34940001 | 35040000 | 0.0003 | 0.0000 | 0.0003 |
| F04 | 34960001 | 35060000 | 0.0001 | 0.0000 | 0.0001 |
| F04 | 34980001 | 35080000 | 0.0001 | 0.0001 | 0.0001 |
| F04 | 35000001 | 35100000 | 0.0002 | 0.0002 | 0.0002 |
| F04 | 35020001 | 35120000 | 0.0003 | 0.0002 | 0.0003 |
| F04 | 35040001 | 35140000 | 0.0003 | 0.0003 | 0.0003 |

| Chr | Start | End | both | A1 | A2 |
|---|---|---|---|---|---|
| F04 | 35060001 | 35160000 | 0.0004 | 0.0003 | 0.0004 |
| F04 | 35080001 | 35180000 | 0.0004 | 0.0002 | 0.0004 |
| F04 | 35100001 | 35200000 | 0.0003 | 0.0002 | 0.0003 |
| F04 | 35120001 | 35220000 | 0.0002 | 0.0004 | 0.0002 |
| F04 | 35140001 | 35240000 | 0.0004 | 0.0005 | 0.0004 |
| F04 | 35160001 | 35260000 | 0.0006 | 0.0006 | 0.0006 |
| F04 | 35180001 | 35280000 | 0.0007 | 0.0007 | 0.0007 |
| F04 | 35200001 | 35300000 | 0.0008 | 0.0007 | 0.0008 |
| F04 | 35220001 | 35320000 | 0.0009 | 0.0006 | 0.0009 |
| F04 | 35240001 | 35340000 | 0.0008 | 0.0021 | 0.0008 |
| F04 | 35260001 | 35360000 | 0.0024 | 0.0022 | 0.0024 |
| F04 | 35280001 | 35380000 | 0.0025 | 0.0027 | 0.0025 |
| F04 | 35300001 | 35400000 | 0.0032 | 0.0026 | 0.0031 |
| F04 | 35320001 | 35420000 | 0.0030 | 0.0029 | 0.0030 |
| F04 | 35340001 | 35440000 | 0.0033 | 0.0012 | 0.0033 |
| F04 | 35360001 | 35460000 | 0.0014 | 0.0010 | 0.0014 |
| F04 | 35380001 | 35480000 | 0.0012 | 0.0005 | 0.0011 |
| F04 | 35400001 | 35500000 | 0.0005 | 0.0005 | 0.0005 |
| F04 | 35420001 | 35520000 | 0.0005 | 0.0002 | 0.0005 |
| F04 | 35440001 | 35540000 | 0.0002 | 0.0002 | 0.0002 |
| F04 | 35460001 | 35560000 | 0.0003 | 0.0002 | 0.0003 |
| F04 | 35480001 | 35580000 | 0.0003 | 0.0004 | 0.0003 |
| F04 | 35500001 | 35600000 | 0.0005 | 0.0004 | 0.0005 |
| F04 | 35520001 | 35620000 | 0.0005 | 0.0005 | 0.0005 |
| F04 | 35540001 | 35640000 | 0.0007 | 0.0007 | 0.0007 |
| F04 | 35560001 | 35660000 | 0.0009 | 0.0007 | 0.0009 |
| F04 | 35580001 | 35680000 | 0.0010 | 0.0005 | 0.0010 |
| F04 | 35600001 | 35700000 | 0.0007 | 0.0011 | 0.0007 |
| F04 | 35620001 | 35720000 | 0.0018 | 0.0019 | 0.0018 |
| F04 | 35640001 | 35740000 | 0.0028 | 0.0020 | 0.0028 |
| F04 | 35660001 | 35760000 | 0.0029 | 0.0021 | 0.0028 |
| F04 | 35680001 | 35780000 | 0.0031 | 0.0021 | 0.0030 |
| F04 | 35700001 | 35800000 | 0.0031 | 0.0016 | 0.0031 |
| F04 | 35720001 | 35820000 | 0.0022 | 0.0013 | 0.0021 |
| F04 | 35740001 | 35840000 | 0.0018 | 0.0013 | 0.0018 |
| F04 | 35760001 | 35860000 | 0.0018 | 0.0014 | 0.0018 |
| F04 | 35780001 | 35880000 | 0.0018 | 0.0015 | 0.0018 |
| F04 | 35800001 | 35900000 | 0.0018 | 0.0017 | 0.0018 |
| F04 | 35820001 | 35920000 | 0.0021 | 0.0015 | 0.0021 |
| F04 | 35840001 | 35940000 | 0.0018 | 0.0014 | 0.0018 |
| F04 | 35860001 | 35960000 | 0.0016 | 0.0013 | 0.0016 |
| F04 | 35880001 | 35980000 | 0.0015 | 0.0016 | 0.0015 |
| F04 | 35900001 | 36000000 | 0.0019 | 0.0022 | 0.0019 |
| F04 | 35920001 | 36020000 | 0.0024 | 0.0020 | 0.0022 |
| F04 | 35940001 | 36040000 | 0.0020 | 0.0018 | 0.0019 |
| F04 | 35960001 | 36060000 | 0.0018 | 0.0019 | 0.0017 |
| F04 | 35980001 | 36080000 | 0.0018 | 0.0020 | 0.0017 |
| F04 | 36000001 | 36100000 | 0.0020 | 0.0013 | 0.0018 |
| F04 | 36020001 | 36120000 | 0.0016 | 0.0013 | 0.0015 |
| F04 | 36040001 | 36140000 | 0.0016 | 0.0015 | 0.0015 |
| F04 | 36060001 | 36160000 | 0.0018 | 0.0015 | 0.0018 |
| F04 | 36080001 | 36180000 | 0.0018 | 0.0016 | 0.0017 |
| F04 | 36100001 | 36200000 | 0.0019 | 0.0019 | 0.0017 |
| F04 | 36120001 | 36220000 | 0.0022 | 0.0020 | 0.0019 |
| F04 | 36140001 | 36240000 | 0.0025 | 0.0026 | 0.0023 |
| F04 | 36160001 | 36260000 | 0.0033 | 0.0030 | 0.0030 |
| F04 | 36180001 | 36280000 | 0.0039 | 0.0029 | 0.0034 |
| F04 | 36200001 | 36300000 | 0.0041 | 0.0028 | 0.0036 |

| Chr | Start | End | both | A1 | A2 |
|---|---|---|---|---|---|
| F04 | 36220001 | 36320000 | 0.0038 | 0.0032 | 0.0034 |
| F04 | 36240001 | 36340000 | 0.0041 | 0.0028 | 0.0037 |
| F04 | 36260001 | 36360000 | 0.0037 | 0.0023 | 0.0032 |
| F04 | 36280001 | 36380000 | 0.0030 | 0.0020 | 0.0029 |
| F04 | 36300001 | 36400000 | 0.0025 | 0.0017 | 0.0025 |
| F04 | 36320001 | 36420000 | 0.0021 | 0.0013 | 0.0021 |
| F04 | 36340001 | 36440000 | 0.0016 | 0.0023 | 0.0015 |
| F04 | 36360001 | 36460000 | 0.0027 | 0.0025 | 0.0027 |
| F04 | 36380001 | 36480000 | 0.0030 | 0.0025 | 0.0029 |
| F04 | 36400001 | 36500000 | 0.0030 | 0.0026 | 0.0029 |
| F04 | 36420001 | 36520000 | 0.0031 | 0.0024 | 0.0030 |
| F04 | 36440001 | 36540000 | 0.0029 | 0.0011 | 0.0029 |
| F04 | 36460001 | 36560000 | 0.0013 | 0.0009 | 0.0012 |
| F04 | 36480001 | 36580000 | 0.0010 | 0.0007 | 0.0010 |
| F04 | 36500001 | 36600000 | 0.0008 | 0.0006 | 0.0008 |
| F04 | 36520001 | 36620000 | 0.0006 | 0.0004 | 0.0006 |
| F04 | 36540001 | 36640000 | 0.0004 | 0.0010 | 0.0004 |
| F04 | 36560001 | 36660000 | 0.0011 | 0.0010 | 0.0010 |
| F04 | 36580001 | 36680000 | 0.0012 | 0.0013 | 0.0011 |
| F04 | 36600001 | 36700000 | 0.0015 | 0.0014 | 0.0014 |
| F04 | 36620001 | 36720000 | 0.0016 | 0.0016 | 0.0016 |
| F04 | 36640001 | 36740000 | 0.0019 | 0.0012 | 0.0018 |
| F04 | 36660001 | 36760000 | 0.0014 | 0.0012 | 0.0013 |
| F04 | 36680001 | 36780000 | 0.0014 | 0.0010 | 0.0013 |
| F04 | 36700001 | 36800000 | 0.0011 | 0.0010 | 0.0010 |
| F04 | 36720001 | 36820000 | 0.0011 | 0.0009 | 0.0010 |
| F04 | 36740001 | 36840000 | 0.0010 | 0.0008 | 0.0010 |
| F04 | 36760001 | 36860000 | 0.0009 | 0.0008 | 0.0009 |
| F04 | 36780001 | 36880000 | 0.0009 | 0.0007 | 0.0009 |
| F04 | 36800001 | 36900000 | 0.0010 | 0.0010 | 0.0010 |
| F04 | 36820001 | 36920000 | 0.0013 | 0.0010 | 0.0013 |
| F04 | 36840001 | 36940000 | 0.0013 | 0.0010 | 0.0013 |
| F04 | 36860001 | 36960000 | 0.0013 | 0.0012 | 0.0013 |
| F04 | 36880001 | 36980000 | 0.0015 | 0.0011 | 0.0014 |
| F04 | 36900001 | 37000000 | 0.0013 | 0.0008 | 0.0012 |
| F04 | 36920001 | 37020000 | 0.0009 | 0.0010 | 0.0009 |
| F04 | 36940001 | 37040000 | 0.0011 | 0.0008 | 0.0010 |
| F04 | 36960001 | 37060000 | 0.0009 | 0.0007 | 0.0008 |
| F04 | 36980001 | 37080000 | 0.0008 | 0.0008 | 0.0007 |
| F04 | 37000001 | 37100000 | 0.0009 | 0.0012 | 0.0008 |
| F04 | 37020001 | 37120000 | 0.0013 | 0.0009 | 0.0013 |
| F04 | 37040001 | 37140000 | 0.0011 | 0.0010 | 0.0010 |
| F04 | 37060001 | 37160000 | 0.0013 | 0.0017 | 0.0012 |
| F04 | 37080001 | 37180000 | 0.0020 | 0.0025 | 0.0018 |
| F04 | 37100001 | 37200000 | 0.0028 | 0.0023 | 0.0027 |
| F04 | 37120001 | 37220000 | 0.0026 | 0.0026 | 0.0024 |
| F04 | 37140001 | 37240000 | 0.0031 | 0.0027 | 0.0029 |
| F04 | 37160001 | 37260000 | 0.0031 | 0.0025 | 0.0030 |
| F04 | 37180001 | 37280000 | 0.0030 | 0.0019 | 0.0029 |
| F04 | 37200001 | 37300000 | 0.0024 | 0.0018 | 0.0023 |
| F04 | 37220001 | 37320000 | 0.0023 | 0.0017 | 0.0022 |
| F04 | 37240001 | 37340000 | 0.0022 | 0.0019 | 0.0021 |
| F04 | 37260001 | 37360000 | 0.0024 | 0.0016 | 0.0023 |
| F04 | 37280001 | 37380000 | 0.0022 | 0.0017 | 0.0021 |
| F04 | 37300001 | 37400000 | 0.0023 | 0.0018 | 0.0022 |
| F04 | 37320001 | 37420000 | 0.0025 | 0.0020 | 0.0024 |
| F04 | 37340001 | 37440000 | 0.0029 | 0.0019 | 0.0025 |
| F04 | 37360001 | 37460000 | 0.0030 | 0.0017 | 0.0024 |

| Chr | Start | End | both | A1 | A2 |
|---|---|---|---|---|---|
| F04 | 37380001 | 37480000 | 0.0026 | 0.0014 | 0.0021 |
| F04 | 37400001 | 37500000 | 0.0023 | 0.0017 | 0.0017 |
| F04 | 37420001 | 37520000 | 0.0026 | 0.0011 | 0.0021 |
| F04 | 37440001 | 37540000 | 0.0017 | 0.0011 | 0.0014 |
| F04 | 37460001 | 37560000 | 0.0013 | 0.0011 | 0.0013 |
| F04 | 37480001 | 37580000 | 0.0016 | 0.0013 | 0.0014 |
| F04 | 37500001 | 37600000 | 0.0019 | 0.0012 | 0.0016 |
| F04 | 37520001 | 37620000 | 0.0019 | 0.0015 | 0.0014 |
| F04 | 37540001 | 37640000 | 0.0024 | 0.0019 | 0.0018 |
| F04 | 37560001 | 37660000 | 0.0030 | 0.0027 | 0.0023 |
| F04 | 37580001 | 37680000 | 0.0036 | 0.0035 | 0.0030 |
| F04 | 37600001 | 37700000 | 0.0045 | 0.0035 | 0.0039 |
| F04 | 37620001 | 37720000 | 0.0043 | 0.0035 | 0.0039 |
| F04 | 37640001 | 37740000 | 0.0045 | 0.0037 | 0.0041 |
| F04 | 37660001 | 37760000 | 0.0047 | 0.0030 | 0.0041 |
| F04 | 37680001 | 37780000 | 0.0040 | 0.0023 | 0.0034 |
| F04 | 37700001 | 37800000 | 0.0034 | 0.0029 | 0.0028 |
| F04 | 37720001 | 37820000 | 0.0039 | 0.0028 | 0.0033 |
| F04 | 37740001 | 37840000 | 0.0037 | 0.0024 | 0.0031 |
| F04 | 37760001 | 37860000 | 0.0030 | 0.0022 | 0.0027 |
| F04 | 37780001 | 37880000 | 0.0026 | 0.0017 | 0.0024 |
| F04 | 37800001 | 37900000 | 0.0020 | 0.0008 | 0.0017 |
| F04 | 37820001 | 37920000 | 0.0010 | 0.0007 | 0.0008 |
| F04 | 37840001 | 37940000 | 0.0008 | 0.0018 | 0.0007 |
| F04 | 37860001 | 37960000 | 0.0020 | 0.0017 | 0.0018 |
| F04 | 37880001 | 37980000 | 0.0020 | 0.0020 | 0.0018 |
| F04 | 37900001 | 38000000 | 0.0023 | 0.0024 | 0.0021 |
| F04 | 37920001 | 38020000 | 0.0028 | 0.0029 | 0.0026 |
| F04 | 37940001 | 38040000 | 0.0033 | 0.0018 | 0.0031 |
| F04 | 37960001 | 38060000 | 0.0023 | 0.0023 | 0.0021 |
| F04 | 37980001 | 38080000 | 0.0028 | 0.0023 | 0.0025 |
| F04 | 38000001 | 38100000 | 0.0029 | 0.0025 | 0.0026 |
| F04 | 38020001 | 38120000 | 0.0030 | 0.0018 | 0.0026 |
| F04 | 38040001 | 38140000 | 0.0023 | 0.0015 | 0.0019 |
| F04 | 38060001 | 38160000 | 0.0017 | 0.0011 | 0.0016 |
| F04 | 38080001 | 38180000 | 0.0014 | 0.0007 | 0.0012 |
| F04 | 38100001 | 38200000 | 0.0008 | 0.0009 | 0.0008 |
| F04 | 38120001 | 38220000 | 0.0011 | 0.0013 | 0.0010 |
| F04 | 38140001 | 38240000 | 0.0016 | 0.0018 | 0.0015 |
| F04 | 38160001 | 38260000 | 0.0022 | 0.0022 | 0.0021 |
| F04 | 38180001 | 38280000 | 0.0027 | 0.0022 | 0.0024 |
| F04 | 38200001 | 38300000 | 0.0029 | 0.0016 | 0.0025 |
| F04 | 38220001 | 38320000 | 0.0022 | 0.0018 | 0.0018 |
| F04 | 38240001 | 38340000 | 0.0025 | 0.0014 | 0.0021 |
| F04 | 38260001 | 38360000 | 0.0021 | 0.0015 | 0.0017 |
| F04 | 38280001 | 38380000 | 0.0024 | 0.0023 | 0.0021 |
| F04 | 38300001 | 38400000 | 0.0033 | 0.0028 | 0.0032 |
| F04 | 38320001 | 38420000 | 0.0037 | 0.0023 | 0.0036 |
| F04 | 38340001 | 38440000 | 0.0031 | 0.0023 | 0.0030 |
| F04 | 38360001 | 38460000 | 0.0029 | 0.0031 | 0.0027 |
| F04 | 38380001 | 38480000 | 0.0033 | 0.0023 | 0.0031 |
| F04 | 38400001 | 38500000 | 0.0023 | 0.0021 | 0.0021 |
| F04 | 38420001 | 38520000 | 0.0021 | 0.0021 | 0.0019 |
| F04 | 38440001 | 38540000 | 0.0024 | 0.0024 | 0.0020 |
| F04 | 38460001 | 38560000 | 0.0029 | 0.0017 | 0.0025 |
| F04 | 38480001 | 38580000 | 0.0023 | 0.0023 | 0.0019 |
| F04 | 38500001 | 38600000 | 0.0029 | 0.0024 | 0.0025 |
| F04 | 38520001 | 38620000 | 0.0032 | 0.0030 | 0.0026 |

| Chr | Start | End | both | A1 | A2 |
|---|---|---|---|---|---|
| F04 | 38540001 | 38640000 | 0.0036 | 0.0026 | 0.0030 |
| F04 | 38560001 | 38660000 | 0.0031 | 0.0023 | 0.0026 |
| F04 | 38580001 | 38680000 | 0.0027 | 0.0021 | 0.0022 |
| F04 | 38600001 | 38700000 | 0.0025 | 0.0020 | 0.0020 |
| F04 | 38620001 | 38720000 | 0.0024 | 0.0017 | 0.0022 |
| F04 | 38640001 | 38740000 | 0.0022 | 0.0020 | 0.0020 |
| F04 | 38660001 | 38760000 | 0.0026 | 0.0020 | 0.0022 |
| F04 | 38680001 | 38780000 | 0.0026 | 0.0025 | 0.0022 |
| F04 | 38700001 | 38800000 | 0.0032 | 0.0022 | 0.0028 |
| F04 | 38720001 | 38820000 | 0.0028 | 0.0022 | 0.0024 |
| F04 | 38740001 | 38840000 | 0.0026 | 0.0022 | 0.0023 |
| F04 | 38760001 | 38860000 | 0.0026 | 0.0028 | 0.0024 |
| F04 | 38780001 | 38880000 | 0.0032 | 0.0018 | 0.0030 |
| F04 | 38800001 | 38900000 | 0.0020 | 0.0022 | 0.0019 |
| F04 | 38820001 | 38920000 | 0.0023 | 0.0018 | 0.0022 |
| F04 | 38840001 | 38940000 | 0.0020 | 0.0014 | 0.0019 |
| F04 | 38860001 | 38960000 | 0.0016 | 0.0015 | 0.0014 |
| F04 | 38880001 | 38980000 | 0.0020 | 0.0024 | 0.0017 |
| F04 | 38900001 | 39000000 | 0.0030 | 0.0032 | 0.0025 |
| F04 | 38920001 | 39020000 | 0.0039 | 0.0034 | 0.0034 |
| F04 | 38940001 | 39040000 | 0.0040 | 0.0033 | 0.0035 |
| F04 | 38960001 | 39060000 | 0.0038 | 0.0026 | 0.0035 |
| F04 | 38980001 | 39080000 | 0.0030 | 0.0021 | 0.0026 |
| F04 | 39000001 | 39100000 | 0.0025 | 0.0007 | 0.0022 |
| F04 | 39020001 | 39120000 | 0.0011 | 0.0007 | 0.0009 |
| F04 | 39040001 | 39140000 | 0.0013 | 0.0010 | 0.0008 |
| F04 | 39060001 | 39160000 | 0.0016 | 0.0013 | 0.0011 |
| F04 | 39080001 | 39180000 | 0.0018 | 0.0015 | 0.0013 |
| F04 | 39100001 | 39200000 | 0.0023 | 0.0016 | 0.0017 |
| F04 | 39120001 | 39220000 | 0.0024 | 0.0019 | 0.0017 |
| F04 | 39140001 | 39240000 | 0.0026 | 0.0020 | 0.0021 |
| F04 | 39160001 | 39260000 | 0.0027 | 0.0017 | 0.0022 |
| F04 | 39180001 | 39280000 | 0.0025 | 0.0012 | 0.0019 |
| F04 | 39200001 | 39300000 | 0.0016 | 0.0018 | 0.0013 |
| F04 | 39220001 | 39320000 | 0.0023 | 0.0016 | 0.0020 |
| F04 | 39240001 | 39340000 | 0.0020 | 0.0012 | 0.0018 |
| F04 | 39260001 | 39360000 | 0.0016 | 0.0013 | 0.0014 |
| F04 | 39280001 | 39380000 | 0.0015 | 0.0013 | 0.0014 |
| F04 | 39300001 | 39400000 | 0.0015 | 0.0008 | 0.0015 |
| F04 | 39320001 | 39420000 | 0.0009 | 0.0005 | 0.0008 |
| F04 | 39340001 | 39440000 | 0.0006 | 0.0006 | 0.0006 |
| F04 | 39360001 | 39460000 | 0.0007 | 0.0006 | 0.0006 |
| F04 | 39380001 | 39480000 | 0.0006 | 0.0004 | 0.0006 |
| F04 | 39400001 | 39500000 | 0.0004 | 0.0004 | 0.0004 |
| F04 | 39420001 | 39520000 | 0.0004 | 0.0006 | 0.0004 |
| F04 | 39440001 | 39540000 | 0.0007 | 0.0006 | 0.0007 |
| F04 | 39460001 | 39560000 | 0.0007 | 0.0005 | 0.0007 |
| F04 | 39480001 | 39580000 | 0.0006 | 0.0006 | 0.0006 |
| F04 | 39500001 | 39600000 | 0.0007 | 0.0006 | 0.0007 |
| F04 | 39520001 | 39620000 | 0.0008 | 0.0005 | 0.0007 |
| F04 | 39540001 | 39640000 | 0.0007 | 0.0005 | 0.0006 |
| F04 | 39560001 | 39660000 | 0.0007 | 0.0008 | 0.0006 |
| F04 | 39580001 | 39680000 | 0.0010 | 0.0007 | 0.0009 |
| F04 | 39600001 | 39700000 | 0.0010 | 0.0008 | 0.0009 |
| F04 | 39620001 | 39720000 | 0.0010 | 0.0007 | 0.0009 |
| F04 | 39640001 | 39740000 | 0.0009 | 0.0006 | 0.0008 |
| F04 | 39660001 | 39760000 | 0.0008 | 0.0006 | 0.0008 |
| F04 | 39680001 | 39780000 | 0.0008 | 0.0013 | 0.0008 |

| Chr | Start | End | both | A1 | A2 |
|---|---|---|---|---|---|
| F04 | 39700001 | 39800000 | 0.0016 | 0.0016 | 0.0016 |
| F04 | 39720001 | 39820000 | 0.0020 | 0.0016 | 0.0020 |
| F04 | 39740001 | 39840000 | 0.0019 | 0.0016 | 0.0019 |
| F04 | 39760001 | 39860000 | 0.0020 | 0.0015 | 0.0020 |
| F04 | 39780001 | 39880000 | 0.0020 | 0.0010 | 0.0018 |
| F04 | 39800001 | 39900000 | 0.0014 | 0.0006 | 0.0012 |
| F04 | 39820001 | 39920000 | 0.0010 | 0.0009 | 0.0007 |
| F04 | 39840001 | 39940000 | 0.0013 | 0.0010 | 0.0010 |
| F04 | 39860001 | 39960000 | 0.0014 | 0.0012 | 0.0011 |
| F04 | 39880001 | 39980000 | 0.0015 | 0.0010 | 0.0012 |
| F04 | 39900001 | 40000000 | 0.0012 | 0.0010 | 0.0010 |
| F04 | 39920001 | 40020000 | 0.0011 | 0.0011 | 0.0011 |
| F04 | 39940001 | 40040000 | 0.0012 | 0.0013 | 0.0012 |
| F04 | 39960001 | 40060000 | 0.0014 | 0.0013 | 0.0013 |
| F04 | 39980001 | 40080000 | 0.0013 | 0.0018 | 0.0013 |
| F04 | 40000001 | 40100000 | 0.0019 | 0.0022 | 0.0018 |
| F04 | 40020001 | 40120000 | 0.0024 | 0.0020 | 0.0022 |
| F04 | 40040001 | 40140000 | 0.0026 | 0.0019 | 0.0020 |
| F04 | 40060001 | 40160000 | 0.0025 | 0.0023 | 0.0020 |
| F04 | 40080001 | 40180000 | 0.0029 | 0.0021 | 0.0024 |
| F04 | 40100001 | 40200000 | 0.0028 | 0.0017 | 0.0022 |
| F04 | 40120001 | 40220000 | 0.0022 | 0.0014 | 0.0017 |
| F04 | 40140001 | 40240000 | 0.0016 | 0.0013 | 0.0015 |
| F04 | 40160001 | 40260000 | 0.0015 | 0.0008 | 0.0013 |
| F04 | 40180001 | 40280000 | 0.0010 | 0.0005 | 0.0009 |
| F04 | 40200001 | 40300000 | 0.0007 | 0.0010 | 0.0006 |
| F04 | 40220001 | 40320000 | 0.0015 | 0.0013 | 0.0012 |
| F04 | 40240001 | 40340000 | 0.0020 | 0.0017 | 0.0018 |
| F04 | 40260001 | 40360000 | 0.0024 | 0.0017 | 0.0022 |
| F04 | 40280001 | 40380000 | 0.0024 | 0.0025 | 0.0022 |
| F04 | 40300001 | 40400000 | 0.0033 | 0.0021 | 0.0031 |
| F04 | 40320001 | 40420000 | 0.0028 | 0.0019 | 0.0026 |
| F04 | 40340001 | 40440000 | 0.0024 | 0.0015 | 0.0021 |
| F04 | 40360001 | 40460000 | 0.0020 | 0.0012 | 0.0017 |
| F04 | 40380001 | 40480000 | 0.0017 | 0.0004 | 0.0014 |
| F04 | 40400001 | 40500000 | 0.0008 | 0.0003 | 0.0005 |
| F04 | 40420001 | 40520000 | 0.0006 | 0.0003 | 0.0003 |
| F04 | 40440001 | 40540000 | 0.0006 | 0.0004 | 0.0003 |
| F04 | 40460001 | 40560000 | 0.0008 | 0.0006 | 0.0006 |
| F04 | 40480001 | 40580000 | 0.0012 | 0.0008 | 0.0008 |
| F04 | 40500001 | 40600000 | 0.0014 | 0.0008 | 0.0010 |
| F04 | 40520001 | 40620000 | 0.0013 | 0.0007 | 0.0011 |
| F04 | 40540001 | 40640000 | 0.0012 | 0.0006 | 0.0010 |
| F04 | 40560001 | 40660000 | 0.0009 | 0.0004 | 0.0007 |
| F04 | 40580001 | 40680000 | 0.0007 | 0.0005 | 0.0006 |
| F04 | 40600001 | 40700000 | 0.0008 | 0.0010 | 0.0006 |
| F04 | 40620001 | 40720000 | 0.0014 | 0.0012 | 0.0011 |
| F04 | 40640001 | 40740000 | 0.0016 | 0.0012 | 0.0013 |
| F04 | 40660001 | 40760000 | 0.0017 | 0.0013 | 0.0014 |
| F04 | 40680001 | 40780000 | 0.0018 | 0.0013 | 0.0016 |
| F04 | 40700001 | 40800000 | 0.0016 | 0.0014 | 0.0014 |
| F04 | 40720001 | 40820000 | 0.0016 | 0.0015 | 0.0015 |
| F04 | 40740001 | 40840000 | 0.0018 | 0.0018 | 0.0017 |
| F04 | 40760001 | 40860000 | 0.0020 | 0.0022 | 0.0020 |
| F04 | 40780001 | 40880000 | 0.0024 | 0.0022 | 0.0023 |
| F04 | 40800001 | 40900000 | 0.0025 | 0.0021 | 0.0024 |
| F04 | 40820001 | 40920000 | 0.0024 | 0.0031 | 0.0024 |
| F04 | 40840001 | 40940000 | 0.0036 | 0.0027 | 0.0035 |

| Chr | Start | End | both | A1 | A2 |
|---|---|---|---|---|---|
| F04 | 40860001 | 40960000 | 0.0034 | 0.0023 | 0.0031 |
| F04 | 40880001 | 40980000 | 0.0032 | 0.0025 | 0.0027 |
| F04 | 40900001 | 41000000 | 0.0034 | 0.0023 | 0.0029 |
| F04 | 40920001 | 41020000 | 0.0030 | 0.0010 | 0.0026 |
| F04 | 40940001 | 41040000 | 0.0015 | 0.0013 | 0.0011 |
| F04 | 40960001 | 41060000 | 0.0016 | 0.0012 | 0.0014 |
| F04 | 40980001 | 41080000 | 0.0015 | 0.0010 | 0.0014 |
| F04 | 41000001 | 41100000 | 0.0011 | 0.0009 | 0.0011 |
| F04 | 41020001 | 41120000 | 0.0010 | 0.0009 | 0.0010 |
| F04 | 41040001 | 41140000 | 0.0011 | 0.0008 | 0.0010 |
| F04 | 41060001 | 41160000 | 0.0010 | 0.0010 | 0.0009 |
| F04 | 41080001 | 41180000 | 0.0013 | 0.0012 | 0.0011 |
| F04 | 41100001 | 41200000 | 0.0015 | 0.0010 | 0.0013 |
| F04 | 41120001 | 41220000 | 0.0013 | 0.0010 | 0.0011 |
| F04 | 41140001 | 41240000 | 0.0013 | 0.0009 | 0.0011 |
| F04 | 41160001 | 41260000 | 0.0011 | 0.0006 | 0.0011 |
| F04 | 41180001 | 41280000 | 0.0007 | 0.0011 | 0.0007 |
| F04 | 41200001 | 41300000 | 0.0012 | 0.0013 | 0.0012 |
| F04 | 41220001 | 41320000 | 0.0014 | 0.0012 | 0.0014 |
| F04 | 41240001 | 41340000 | 0.0014 | 0.0012 | 0.0013 |
| F04 | 41260001 | 41360000 | 0.0014 | 0.0013 | 0.0014 |
| F04 | 41280001 | 41380000 | 0.0014 | 0.0008 | 0.0014 |
| F04 | 41300001 | 41400000 | 0.0010 | 0.0010 | 0.0010 |
| F04 | 41320001 | 41420000 | 0.0011 | 0.0018 | 0.0011 |
| F04 | 41340001 | 41440000 | 0.0021 | 0.0018 | 0.0021 |
| F04 | 41360001 | 41460000 | 0.0020 | 0.0027 | 0.0019 |
| F04 | 41380001 | 41480000 | 0.0031 | 0.0031 | 0.0029 |
| F04 | 41400001 | 41500000 | 0.0036 | 0.0029 | 0.0034 |
| F04 | 41420001 | 41520000 | 0.0035 | 0.0021 | 0.0032 |
| F04 | 41440001 | 41540000 | 0.0026 | 0.0026 | 0.0023 |
| F04 | 41460001 | 41560000 | 0.0032 | 0.0019 | 0.0029 |
| F04 | 41480001 | 41580000 | 0.0022 | 0.0017 | 0.0021 |
| F04 | 41500001 | 41600000 | 0.0021 | 0.0017 | 0.0018 |
| F04 | 41520001 | 41620000 | 0.0020 | 0.0019 | 0.0018 |
| F04 | 41540001 | 41640000 | 0.0026 | 0.0018 | 0.0023 |
| F04 | 41560001 | 41660000 | 0.0024 | 0.0017 | 0.0022 |
| F04 | 41580001 | 41680000 | 0.0023 | 0.0012 | 0.0021 |
| F04 | 41600001 | 41700000 | 0.0016 | 0.0013 | 0.0015 |
| F04 | 41620001 | 41720000 | 0.0019 | 0.0024 | 0.0018 |
| F04 | 41640001 | 41740000 | 0.0028 | 0.0020 | 0.0027 |
| F04 | 41660001 | 41760000 | 0.0023 | 0.0020 | 0.0022 |
| F04 | 41680001 | 41780000 | 0.0024 | 0.0021 | 0.0023 |
| F04 | 41700001 | 41800000 | 0.0025 | 0.0020 | 0.0024 |
| F04 | 41720001 | 41820000 | 0.0022 | 0.0022 | 0.0022 |
| F04 | 41740001 | 41840000 | 0.0023 | 0.0029 | 0.0022 |
| F04 | 41760001 | 41860000 | 0.0030 | 0.0031 | 0.0028 |
| F04 | 41780001 | 41880000 | 0.0032 | 0.0033 | 0.0030 |
| F04 | 41800001 | 41900000 | 0.0035 | 0.0034 | 0.0032 |
| F04 | 41820001 | 41920000 | 0.0037 | 0.0022 | 0.0034 |
| F04 | 41840001 | 41940000 | 0.0027 | 0.0016 | 0.0026 |
| F04 | 41860001 | 41960000 | 0.0021 | 0.0017 | 0.0020 |
| F04 | 41880001 | 41980000 | 0.0022 | 0.0016 | 0.0021 |
| F04 | 41900001 | 42000000 | 0.0020 | 0.0014 | 0.0020 |
| F04 | 41920001 | 42020000 | 0.0017 | 0.0016 | 0.0017 |
| F04 | 41940001 | 42040000 | 0.0019 | 0.0029 | 0.0017 |
| F04 | 41960001 | 42060000 | 0.0031 | 0.0026 | 0.0029 |
| F04 | 41980001 | 42080000 | 0.0029 | 0.0035 | 0.0026 |
| F04 | 42000001 | 42100000 | 0.0038 | 0.0038 | 0.0034 |

| Chr | Start | End | both | A1 | A2 |
|---|---|---|---|---|---|
| F04 | 42020001 | 42120000 | 0.0041 | 0.0038 | 0.0037 |
| F04 | 42040001 | 42140000 | 0.0041 | 0.0028 | 0.0037 |
| F04 | 42060001 | 42160000 | 0.0031 | 0.0027 | 0.0028 |
| F04 | 42080001 | 42180000 | 0.0030 | 0.0022 | 0.0026 |
| F04 | 42100001 | 42200000 | 0.0027 | 0.0020 | 0.0024 |
| F04 | 42120001 | 42220000 | 0.0025 | 0.0014 | 0.0022 |
| F04 | 42140001 | 42240000 | 0.0018 | 0.0021 | 0.0016 |
| F04 | 42160001 | 42260000 | 0.0025 | 0.0022 | 0.0024 |
| F04 | 42180001 | 42280000 | 0.0026 | 0.0016 | 0.0024 |
| F04 | 42200001 | 42300000 | 0.0021 | 0.0021 | 0.0018 |
| F04 | 42220001 | 42320000 | 0.0029 | 0.0022 | 0.0025 |
| F04 | 42240001 | 42340000 | 0.0031 | 0.0013 | 0.0026 |
| F04 | 42260001 | 42360000 | 0.0021 | 0.0011 | 0.0016 |
| F04 | 42280001 | 42380000 | 0.0018 | 0.0014 | 0.0014 |
| F04 | 42300001 | 42400000 | 0.0019 | 0.0007 | 0.0017 |
| F04 | 42320001 | 42420000 | 0.0009 | 0.0009 | 0.0007 |
| F04 | 42340001 | 42440000 | 0.0012 | 0.0009 | 0.0011 |
| F04 | 42360001 | 42460000 | 0.0011 | 0.0009 | 0.0011 |
| F04 | 42380001 | 42480000 | 0.0011 | 0.0008 | 0.0011 |
| F04 | 42400001 | 42500000 | 0.0010 | 0.0015 | 0.0010 |
| F04 | 42420001 | 42520000 | 0.0018 | 0.0013 | 0.0018 |
| F04 | 42440001 | 42540000 | 0.0017 | 0.0014 | 0.0017 |
| F04 | 42460001 | 42560000 | 0.0017 | 0.0016 | 0.0017 |
| F04 | 42480001 | 42580000 | 0.0020 | 0.0018 | 0.0020 |
| F04 | 42500001 | 42600000 | 0.0022 | 0.0014 | 0.0022 |
| F04 | 42520001 | 42620000 | 0.0017 | 0.0013 | 0.0016 |
| F04 | 42540001 | 42640000 | 0.0015 | 0.0014 | 0.0015 |
| F04 | 42560001 | 42660000 | 0.0016 | 0.0012 | 0.0015 |
| F04 | 42580001 | 42680000 | 0.0014 | 0.0013 | 0.0014 |
| F04 | 42600001 | 42700000 | 0.0016 | 0.0012 | 0.0016 |
| F04 | 42620001 | 42720000 | 0.0014 | 0.0014 | 0.0014 |
| F04 | 42640001 | 42740000 | 0.0017 | 0.0017 | 0.0017 |
| F04 | 42660001 | 42760000 | 0.0021 | 0.0015 | 0.0019 |
| F04 | 42680001 | 42780000 | 0.0021 | 0.0012 | 0.0018 |
| F04 | 42700001 | 42800000 | 0.0017 | 0.0016 | 0.0014 |
| F04 | 42720001 | 42820000 | 0.0021 | 0.0015 | 0.0018 |
| F04 | 42740001 | 42840000 | 0.0020 | 0.0011 | 0.0016 |
| F04 | 42760001 | 42860000 | 0.0018 | 0.0019 | 0.0014 |
| F04 | 42780001 | 42880000 | 0.0025 | 0.0018 | 0.0022 |
| F04 | 42800001 | 42900000 | 0.0024 | 0.0014 | 0.0020 |
| F04 | 42820001 | 42920000 | 0.0019 | 0.0018 | 0.0016 |
| F04 | 42840001 | 42940000 | 0.0023 | 0.0026 | 0.0019 |
| F04 | 42860001 | 42960000 | 0.0031 | 0.0020 | 0.0026 |
| F04 | 42880001 | 42980000 | 0.0024 | 0.0021 | 0.0020 |
| F04 | 42900001 | 43000000 | 0.0024 | 0.0020 | 0.0022 |
| F04 | 42920001 | 43020000 | 0.0024 | 0.0015 | 0.0021 |
| F04 | 42940001 | 43040000 | 0.0018 | 0.0008 | 0.0016 |
| F04 | 42960001 | 43060000 | 0.0010 | 0.0006 | 0.0009 |
| F04 | 42980001 | 43080000 | 0.0008 | 0.0006 | 0.0007 |
| F04 | 43000001 | 43100000 | 0.0008 | 0.0012 | 0.0007 |
| F04 | 43020001 | 43120000 | 0.0013 | 0.0014 | 0.0013 |
| F04 | 43040001 | 43140000 | 0.0016 | 0.0013 | 0.0016 |
| F04 | 43060001 | 43160000 | 0.0016 | 0.0013 | 0.0016 |
| F04 | 43080001 | 43180000 | 0.0017 | 0.0012 | 0.0016 |
| F04 | 43100001 | 43200000 | 0.0015 | 0.0008 | 0.0014 |
| F04 | 43120001 | 43220000 | 0.0011 | 0.0007 | 0.0011 |
| F04 | 43140001 | 43240000 | 0.0009 | 0.0009 | 0.0009 |
| F04 | 43160001 | 43260000 | 0.0012 | 0.0013 | 0.0011 |

| Chr | Start | End | both | A1 | A2 |
|---|---|---|---|---|---|
| F04 | 43180001 | 43280000 | 0.0018 | 0.0016 | 0.0015 |
| F04 | 43200001 | 43300000 | 0.0021 | 0.0017 | 0.0018 |
| F04 | 43220001 | 43320000 | 0.0022 | 0.0019 | 0.0017 |
| F04 | 43240001 | 43340000 | 0.0025 | 0.0021 | 0.0021 |
| F04 | 43260001 | 43360000 | 0.0027 | 0.0016 | 0.0023 |
| F04 | 43280001 | 43380000 | 0.0021 | 0.0016 | 0.0018 |
| F04 | 43300001 | 43400000 | 0.0023 | 0.0014 | 0.0019 |
| F04 | 43320001 | 43420000 | 0.0021 | 0.0011 | 0.0017 |
| F04 | 43340001 | 43440000 | 0.0018 | 0.0013 | 0.0013 |
| F04 | 43360001 | 43460000 | 0.0019 | 0.0017 | 0.0016 |
| F04 | 43380001 | 43480000 | 0.0023 | 0.0017 | 0.0019 |
| F04 | 43400001 | 43500000 | 0.0023 | 0.0027 | 0.0018 |
| F04 | 43420001 | 43520000 | 0.0035 | 0.0035 | 0.0029 |
| F04 | 43440001 | 43540000 | 0.0043 | 0.0030 | 0.0036 |
| F04 | 43460001 | 43560000 | 0.0037 | 0.0026 | 0.0030 |
| F04 | 43480001 | 43580000 | 0.0033 | 0.0024 | 0.0027 |
| F04 | 43500001 | 43600000 | 0.0030 | 0.0015 | 0.0025 |
| F04 | 43520001 | 43620000 | 0.0017 | 0.0008 | 0.0015 |
| F04 | 43540001 | 43640000 | 0.0009 | 0.0006 | 0.0008 |
| F04 | 43560001 | 43660000 | 0.0006 | 0.0005 | 0.0006 |
| F04 | 43580001 | 43680000 | 0.0005 | 0.0004 | 0.0005 |
| F04 | 43600001 | 43700000 | 0.0004 | 0.0003 | 0.0004 |
| F04 | 43620001 | 43720000 | 0.0003 | 0.0002 | 0.0003 |
| F04 | 43640001 | 43740000 | 0.0002 | 0.0002 | 0.0002 |
| F04 | 43660001 | 43760000 | 0.0002 | 0.0002 | 0.0002 |
| F04 | 43680001 | 43780000 | 0.0002 | 0.0003 | 0.0002 |
| F04 | 43700001 | 43800000 | 0.0004 | 0.0004 | 0.0003 |
| F04 | 43720001 | 43820000 | 0.0005 | 0.0003 | 0.0004 |
| F04 | 43740001 | 43840000 | 0.0004 | 0.0004 | 0.0003 |
| F04 | 43760001 | 43860000 | 0.0006 | 0.0004 | 0.0005 |
| F04 | 43780001 | 43880000 | 0.0007 | 0.0005 | 0.0005 |
| F04 | 43800001 | 43900000 | 0.0008 | 0.0007 | 0.0006 |
| F04 | 43820001 | 43920000 | 0.0008 | 0.0009 | 0.0007 |
| F04 | 43840001 | 43940000 | 0.0011 | 0.0011 | 0.0010 |
| F04 | 43860001 | 43960000 | 0.0014 | 0.0011 | 0.0012 |
| F04 | 43880001 | 43980000 | 0.0014 | 0.0009 | 0.0012 |
| F04 | 43900001 | 44000000 | 0.0011 | 0.0010 | 0.0010 |
| F04 | 43920001 | 44020000 | 0.0012 | 0.0008 | 0.0011 |
| F04 | 43940001 | 44040000 | 0.0009 | 0.0008 | 0.0009 |
| F04 | 43960001 | 44060000 | 0.0009 | 0.0009 | 0.0009 |
| F04 | 43980001 | 44080000 | 0.0010 | 0.0010 | 0.0010 |
| F04 | 44000001 | 44100000 | 0.0012 | 0.0009 | 0.0012 |
| F04 | 44020001 | 44120000 | 0.0011 | 0.0008 | 0.0011 |
| F04 | 44040001 | 44140000 | 0.0010 | 0.0005 | 0.0010 |
| F04 | 44060001 | 44160000 | 0.0006 | 0.0004 | 0.0007 |
| F04 | 44080001 | 44180000 | 0.0005 | 0.0003 | 0.0005 |
| F04 | 44100001 | 44200000 | 0.0005 | 0.0004 | 0.0005 |
| F04 | 44120001 | 44220000 | 0.0004 | 0.0005 | 0.0004 |
| F04 | 44140001 | 44240000 | 0.0006 | 0.0005 | 0.0006 |
| F04 | 44160001 | 44260000 | 0.0006 | 0.0006 | 0.0006 |
| F04 | 44180001 | 44280000 | 0.0008 | 0.0006 | 0.0008 |
| F04 | 44200001 | 44300000 | 0.0007 | 0.0005 | 0.0007 |
| F04 | 44220001 | 44320000 | 0.0006 | 0.0003 | 0.0006 |
| F04 | 44240001 | 44340000 | 0.0004 | 0.0004 | 0.0004 |
| F04 | 44260001 | 44360000 | 0.0005 | 0.0003 | 0.0005 |
| F04 | 44280001 | 44380000 | 0.0004 | 0.0003 | 0.0004 |
| F04 | 44300001 | 44400000 | 0.0004 | 0.0007 | 0.0004 |
| F04 | 44320001 | 44420000 | 0.0007 | 0.0007 | 0.0007 |

| Chr | Start | End | both | A1 | A2 |
|---|---|---|---|---|---|
| F04 | 44340001 | 44440000 | 0.0007 | 0.0006 | 0.0007 |
| F04 | 44360001 | 44460000 | 0.0006 | 0.0009 | 0.0006 |
| F04 | 44380001 | 44480000 | 0.0010 | 0.0008 | 0.0010 |
| F04 | 44400001 | 44500000 | 0.0009 | 0.0004 | 0.0009 |
| F04 | 44420001 | 44520000 | 0.0005 | 0.0004 | 0.0005 |
| F04 | 44440001 | 44540000 | 0.0005 | 0.0005 | 0.0005 |
| F04 | 44460001 | 44560000 | 0.0006 | 0.0003 | 0.0006 |
| F04 | 44480001 | 44580000 | 0.0003 | 0.0003 | 0.0003 |
| F04 | 44500001 | 44600000 | 0.0003 | 0.0003 | 0.0003 |
| F04 | 44520001 | 44620000 | 0.0004 | 0.0005 | 0.0004 |
| F04 | 44540001 | 44640000 | 0.0006 | 0.0010 | 0.0007 |
| F04 | 44560001 | 44660000 | 0.0012 | 0.0011 | 0.0012 |
| F04 | 44580001 | 44680000 | 0.0013 | 0.0014 | 0.0012 |
| F04 | 44600001 | 44700000 | 0.0016 | 0.0015 | 0.0015 |
| F04 | 44620001 | 44720000 | 0.0017 | 0.0013 | 0.0016 |
| F04 | 44640001 | 44740000 | 0.0014 | 0.0008 | 0.0014 |
| F04 | 44660001 | 44760000 | 0.0008 | 0.0008 | 0.0008 |
| F04 | 44680001 | 44780000 | 0.0010 | 0.0005 | 0.0010 |
| F04 | 44700001 | 44800000 | 0.0007 | 0.0005 | 0.0007 |
| F04 | 44720001 | 44820000 | 0.0007 | 0.0007 | 0.0007 |
| F04 | 44740001 | 44840000 | 0.0010 | 0.0010 | 0.0010 |
| F04 | 44760001 | 44860000 | 0.0013 | 0.0008 | 0.0012 |
| F04 | 44780001 | 44880000 | 0.0010 | 0.0008 | 0.0009 |
| F04 | 44800001 | 44900000 | 0.0010 | 0.0008 | 0.0009 |
| F04 | 44820001 | 44920000 | 0.0010 | 0.0007 | 0.0009 |
| F04 | 44840001 | 44940000 | 0.0009 | 0.0009 | 0.0009 |
| F04 | 44860001 | 44960000 | 0.0013 | 0.0009 | 0.0012 |
| F04 | 44880001 | 44980000 | 0.0012 | 0.0009 | 0.0012 |
| F04 | 44900001 | 45000000 | 0.0013 | 0.0009 | 0.0013 |
| F04 | 44920001 | 45020000 | 0.0013 | 0.0007 | 0.0013 |
| F04 | 44940001 | 45040000 | 0.0011 | 0.0003 | 0.0011 |
| F04 | 44960001 | 45060000 | 0.0004 | 0.0004 | 0.0004 |
| F04 | 44980001 | 45080000 | 0.0005 | 0.0009 | 0.0005 |
| F04 | 45000001 | 45100000 | 0.0011 | 0.0015 | 0.0011 |
| F04 | 45020001 | 45120000 | 0.0019 | 0.0017 | 0.0017 |
| F04 | 45040001 | 45140000 | 0.0021 | 0.0018 | 0.0018 |
| F04 | 45060001 | 45160000 | 0.0024 | 0.0018 | 0.0020 |
| F04 | 45080001 | 45180000 | 0.0024 | 0.0013 | 0.0019 |
| F04 | 45100001 | 45200000 | 0.0019 | 0.0009 | 0.0013 |
| F04 | 45120001 | 45220000 | 0.0015 | 0.0014 | 0.0010 |
| F04 | 45140001 | 45240000 | 0.0021 | 0.0015 | 0.0016 |
| F04 | 45160001 | 45260000 | 0.0023 | 0.0018 | 0.0017 |
| F04 | 45180001 | 45280000 | 0.0026 | 0.0018 | 0.0021 |
| F04 | 45200001 | 45300000 | 0.0027 | 0.0017 | 0.0021 |
| F04 | 45220001 | 45320000 | 0.0026 | 0.0010 | 0.0020 |
| F04 | 45240001 | 45340000 | 0.0017 | 0.0008 | 0.0011 |
| F04 | 45260001 | 45360000 | 0.0014 | 0.0008 | 0.0009 |
| F04 | 45280001 | 45380000 | 0.0015 | 0.0011 | 0.0010 |
| F04 | 45300001 | 45400000 | 0.0015 | 0.0012 | 0.0012 |
| F04 | 45320001 | 45420000 | 0.0015 | 0.0015 | 0.0012 |
| F04 | 45340001 | 45440000 | 0.0018 | 0.0020 | 0.0015 |
| F04 | 45360001 | 45460000 | 0.0024 | 0.0024 | 0.0020 |
| F04 | 45380001 | 45480000 | 0.0028 | 0.0031 | 0.0025 |
| F04 | 45400001 | 45500000 | 0.0036 | 0.0031 | 0.0033 |
| F04 | 45420001 | 45520000 | 0.0036 | 0.0033 | 0.0033 |
| F04 | 45440001 | 45540000 | 0.0040 | 0.0040 | 0.0037 |
| F04 | 45460001 | 45560000 | 0.0045 | 0.0040 | 0.0044 |
| F04 | 45480001 | 45580000 | 0.0048 | 0.0037 | 0.0044 |

| Chr | Start | End | both | A1 | A2 |
|---|---|---|---|---|---|
| F04 | 45500001 | 45600000 | 0.0047 | 0.0049 | 0.0041 |
| F04 | 45520001 | 45620000 | 0.0060 | 0.0043 | 0.0052 |
| F04 | 45540001 | 45640000 | 0.0055 | 0.0033 | 0.0046 |
| F04 | 45560001 | 45660000 | 0.0043 | 0.0025 | 0.0034 |
| F04 | 45580001 | 45680000 | 0.0032 | 0.0020 | 0.0027 |
| F04 | 45600001 | 45700000 | 0.0029 | 0.0010 | 0.0022 |
| F04 | 45620001 | 45720000 | 0.0020 | 0.0018 | 0.0014 |
| F04 | 45640001 | 45740000 | 0.0030 | 0.0023 | 0.0024 |
| F04 | 45660001 | 45760000 | 0.0034 | 0.0023 | 0.0029 |
| F04 | 45680001 | 45780000 | 0.0036 | 0.0025 | 0.0029 |
| F04 | 45700001 | 45800000 | 0.0035 | 0.0030 | 0.0029 |
| F04 | 45720001 | 45820000 | 0.0037 | 0.0027 | 0.0029 |
| F04 | 45740001 | 45840000 | 0.0034 | 0.0029 | 0.0025 |
| F04 | 45760001 | 45860000 | 0.0037 | 0.0028 | 0.0027 |
| F04 | 45780001 | 45880000 | 0.0033 | 0.0027 | 0.0025 |
| F04 | 45800001 | 45900000 | 0.0031 | 0.0027 | 0.0024 |
| F04 | 45820001 | 45920000 | 0.0035 | 0.0025 | 0.0030 |
| F04 | 45840001 | 45940000 | 0.0031 | 0.0024 | 0.0027 |
| F04 | 45860001 | 45960000 | 0.0029 | 0.0024 | 0.0026 |
| F04 | 45880001 | 45980000 | 0.0030 | 0.0026 | 0.0025 |
| F04 | 45900001 | 46000000 | 0.0033 | 0.0017 | 0.0028 |
| F04 | 45920001 | 46020000 | 0.0021 | 0.0014 | 0.0019 |
| F04 | 45940001 | 46040000 | 0.0017 | 0.0010 | 0.0015 |
| F04 | 45960001 | 46060000 | 0.0013 | 0.0013 | 0.0011 |
| F04 | 45980001 | 46080000 | 0.0016 | 0.0009 | 0.0015 |
| F04 | 46000001 | 46100000 | 0.0012 | 0.0011 | 0.0011 |
| F04 | 46020001 | 46120000 | 0.0015 | 0.0012 | 0.0013 |
| F04 | 46040001 | 46140000 | 0.0016 | 0.0015 | 0.0014 |
| F04 | 46060001 | 46160000 | 0.0020 | 0.0011 | 0.0016 |
| F04 | 46080001 | 46180000 | 0.0016 | 0.0009 | 0.0012 |
| F04 | 46100001 | 46200000 | 0.0013 | 0.0008 | 0.0010 |
| F04 | 46120001 | 46220000 | 0.0011 | 0.0009 | 0.0007 |
| F04 | 46140001 | 46240000 | 0.0013 | 0.0009 | 0.0009 |
| F04 | 46160001 | 46260000 | 0.0013 | 0.0012 | 0.0011 |
| F04 | 46180001 | 46280000 | 0.0016 | 0.0015 | 0.0015 |
| F04 | 46200001 | 46300000 | 0.0019 | 0.0015 | 0.0019 |
| F04 | 46220001 | 46320000 | 0.0020 | 0.0018 | 0.0020 |
| F04 | 46240001 | 46340000 | 0.0024 | 0.0016 | 0.0024 |
| F04 | 46260001 | 46360000 | 0.0020 | 0.0017 | 0.0020 |
| F04 | 46280001 | 46380000 | 0.0021 | 0.0018 | 0.0021 |
| F04 | 46300001 | 46400000 | 0.0021 | 0.0019 | 0.0021 |
| F04 | 46320001 | 46420000 | 0.0023 | 0.0016 | 0.0023 |
| F04 | 46340001 | 46440000 | 0.0019 | 0.0015 | 0.0018 |
| F04 | 46360001 | 46460000 | 0.0017 | 0.0014 | 0.0017 |
| F04 | 46380001 | 46480000 | 0.0016 | 0.0011 | 0.0014 |
| F04 | 46400001 | 46500000 | 0.0014 | 0.0010 | 0.0012 |
| F04 | 46420001 | 46520000 | 0.0012 | 0.0009 | 0.0010 |
| F04 | 46440001 | 46540000 | 0.0011 | 0.0019 | 0.0009 |
| F04 | 46460001 | 46560000 | 0.0023 | 0.0015 | 0.0021 |
| F04 | 46480001 | 46580000 | 0.0018 | 0.0015 | 0.0017 |
| F04 | 46500001 | 46600000 | 0.0018 | 0.0017 | 0.0018 |
| F04 | 46520001 | 46620000 | 0.0021 | 0.0016 | 0.0020 |
| F04 | 46540001 | 46640000 | 0.0020 | 0.0013 | 0.0019 |
| F04 | 46560001 | 46660000 | 0.0015 | 0.0014 | 0.0014 |
| F04 | 46580001 | 46680000 | 0.0018 | 0.0015 | 0.0016 |
| F04 | 46600001 | 46700000 | 0.0018 | 0.0016 | 0.0016 |
| F04 | 46620001 | 46720000 | 0.0018 | 0.0020 | 0.0016 |
| F04 | 46640001 | 46740000 | 0.0023 | 0.0014 | 0.0019 |

| Chr | Start | End | both | A1 | A2 |
|---|---|---|---|---|---|
| F04 | 46660001 | 46760000 | 0.0017 | 0.0018 | 0.0013 |
| F04 | 46680001 | 46780000 | 0.0020 | 0.0019 | 0.0017 |
| F04 | 46700001 | 46800000 | 0.0021 | 0.0017 | 0.0019 |
| F04 | 46720001 | 46820000 | 0.0020 | 0.0016 | 0.0017 |
| F04 | 46740001 | 46840000 | 0.0019 | 0.0018 | 0.0017 |
| F04 | 46760001 | 46860000 | 0.0019 | 0.0015 | 0.0018 |
| F04 | 46780001 | 46880000 | 0.0017 | 0.0011 | 0.0015 |
| F04 | 46800001 | 46900000 | 0.0013 | 0.0014 | 0.0012 |
| F04 | 46820001 | 46920000 | 0.0017 | 0.0013 | 0.0016 |
| F04 | 46840001 | 46940000 | 0.0019 | 0.0017 | 0.0016 |
| F04 | 46860001 | 46960000 | 0.0026 | 0.0017 | 0.0021 |
| F04 | 46880001 | 46980000 | 0.0026 | 0.0017 | 0.0021 |
| F04 | 46900001 | 47000000 | 0.0027 | 0.0013 | 0.0021 |
| F04 | 46920001 | 47020000 | 0.0022 | 0.0013 | 0.0016 |
| F04 | 46940001 | 47040000 | 0.0019 | 0.0017 | 0.0015 |
| F04 | 46960001 | 47060000 | 0.0022 | 0.0022 | 0.0019 |
| F04 | 46980001 | 47080000 | 0.0026 | 0.0024 | 0.0021 |
| F04 | 47000001 | 47100000 | 0.0031 | 0.0026 | 0.0023 |
| F04 | 47020001 | 47120000 | 0.0033 | 0.0027 | 0.0025 |
| F04 | 47040001 | 47140000 | 0.0034 | 0.0023 | 0.0027 |
| F04 | 47060001 | 47160000 | 0.0031 | 0.0024 | 0.0021 |
| F04 | 47080001 | 47180000 | 0.0032 | 0.0024 | 0.0025 |
| F04 | 47100001 | 47200000 | 0.0032 | 0.0029 | 0.0025 |
| F04 | 47120001 | 47220000 | 0.0037 | 0.0025 | 0.0029 |
| F04 | 47140001 | 47240000 | 0.0033 | 0.0024 | 0.0025 |
| F04 | 47160001 | 47260000 | 0.0031 | 0.0017 | 0.0025 |
| F04 | 47180001 | 47280000 | 0.0024 | 0.0017 | 0.0017 |
| F04 | 47200001 | 47300000 | 0.0022 | 0.0013 | 0.0017 |
| F04 | 47220001 | 47320000 | 0.0019 | 0.0016 | 0.0014 |
| F04 | 47240001 | 47340000 | 0.0023 | 0.0011 | 0.0015 |
| F04 | 47260001 | 47360000 | 0.0018 | 0.0011 | 0.0011 |
| F04 | 47280001 | 47380000 | 0.0018 | 0.0013 | 0.0011 |
| F04 | 47300001 | 47400000 | 0.0021 | 0.0015 | 0.0013 |
| F04 | 47320001 | 47420000 | 0.0021 | 0.0017 | 0.0013 |
| F04 | 47340001 | 47440000 | 0.0022 | 0.0024 | 0.0016 |
| F04 | 47360001 | 47460000 | 0.0033 | 0.0034 | 0.0024 |
| F04 | 47380001 | 47480000 | 0.0044 | 0.0036 | 0.0036 |
| F04 | 47400001 | 47500000 | 0.0045 | 0.0045 | 0.0037 |
| F04 | 47420001 | 47520000 | 0.0054 | 0.0046 | 0.0048 |
| F04 | 47440001 | 47540000 | 0.0056 | 0.0040 | 0.0050 |
| F04 | 47460001 | 47560000 | 0.0045 | 0.0032 | 0.0042 |
| F04 | 47480001 | 47580000 | 0.0035 | 0.0026 | 0.0033 |
| F04 | 47500001 | 47600000 | 0.0030 | 0.0014 | 0.0028 |
| F04 | 47520001 | 47620000 | 0.0018 | 0.0012 | 0.0016 |
| F04 | 47540001 | 47640000 | 0.0017 | 0.0015 | 0.0014 |
| F04 | 47560001 | 47660000 | 0.0024 | 0.0014 | 0.0016 |
| F04 | 47580001 | 47680000 | 0.0025 | 0.0017 | 0.0016 |
| F04 | 47600001 | 47700000 | 0.0030 | 0.0020 | 0.0018 |
| F04 | 47620001 | 47720000 | 0.0032 | 0.0016 | 0.0021 |
| F04 | 47640001 | 47740000 | 0.0026 | 0.0014 | 0.0017 |
| F04 | 47660001 | 47760000 | 0.0022 | 0.0016 | 0.0015 |
| F04 | 47680001 | 47780000 | 0.0022 | 0.0019 | 0.0015 |
| F04 | 47700001 | 47800000 | 0.0025 | 0.0016 | 0.0019 |
| F04 | 47720001 | 47820000 | 0.0022 | 0.0016 | 0.0015 |
| F04 | 47740001 | 47840000 | 0.0023 | 0.0023 | 0.0016 |
| F04 | 47760001 | 47860000 | 0.0030 | 0.0021 | 0.0024 |
| F04 | 47780001 | 47880000 | 0.0027 | 0.0019 | 0.0022 |
| F04 | 47800001 | 47900000 | 0.0025 | 0.0021 | 0.0019 |

| Chr | Start | End | both | A1 | A2 |
|---|---|---|---|---|---|
| F04 | 47820001 | 47920000 | 0.0026 | 0.0024 | 0.0021 |
| F04 | 47840001 | 47940000 | 0.0030 | 0.0027 | 0.0024 |
| F04 | 47860001 | 47960000 | 0.0033 | 0.0034 | 0.0027 |
| F04 | 47880001 | 47980000 | 0.0042 | 0.0038 | 0.0034 |
| F04 | 47900001 | 48000000 | 0.0049 | 0.0048 | 0.0039 |
| F04 | 47920001 | 48020000 | 0.0059 | 0.0045 | 0.0050 |
| F04 | 47940001 | 48040000 | 0.0054 | 0.0037 | 0.0046 |
| F04 | 47960001 | 48060000 | 0.0046 | 0.0028 | 0.0038 |
| F04 | 47980001 | 48080000 | 0.0035 | 0.0020 | 0.0029 |
| F04 | 48000001 | 48100000 | 0.0026 | 0.0010 | 0.0020 |
| F04 | 48020001 | 48120000 | 0.0016 | 0.0013 | 0.0010 |
| F04 | 48040001 | 48140000 | 0.0023 | 0.0014 | 0.0016 |
| F04 | 48060001 | 48160000 | 0.0023 | 0.0016 | 0.0017 |
| F04 | 48080001 | 48180000 | 0.0023 | 0.0023 | 0.0019 |
| F04 | 48100001 | 48200000 | 0.0028 | 0.0024 | 0.0026 |
| F04 | 48120001 | 48220000 | 0.0028 | 0.0020 | 0.0025 |
| F04 | 48140001 | 48240000 | 0.0022 | 0.0016 | 0.0021 |
| F04 | 48160001 | 48260000 | 0.0019 | 0.0023 | 0.0016 |
| F04 | 48180001 | 48280000 | 0.0028 | 0.0017 | 0.0025 |
| F04 | 48200001 | 48300000 | 0.0025 | 0.0015 | 0.0020 |
| F04 | 48220001 | 48320000 | 0.0024 | 0.0015 | 0.0019 |
| F04 | 48240001 | 48340000 | 0.0029 | 0.0023 | 0.0020 |
| F04 | 48260001 | 48360000 | 0.0038 | 0.0017 | 0.0027 |
| F04 | 48280001 | 48380000 | 0.0029 | 0.0018 | 0.0019 |
| F04 | 48300001 | 48400000 | 0.0029 | 0.0021 | 0.0020 |
| F04 | 48320001 | 48420000 | 0.0031 | 0.0021 | 0.0024 |
| F04 | 48340001 | 48440000 | 0.0026 | 0.0014 | 0.0021 |
| F04 | 48360001 | 48460000 | 0.0016 | 0.0013 | 0.0016 |
| F04 | 48380001 | 48480000 | 0.0016 | 0.0012 | 0.0015 |
| F04 | 48400001 | 48500000 | 0.0018 | 0.0014 | 0.0014 |
| F04 | 48420001 | 48520000 | 0.0020 | 0.0017 | 0.0016 |
| F04 | 48440001 | 48540000 | 0.0023 | 0.0020 | 0.0018 |
| F04 | 48460001 | 48560000 | 0.0027 | 0.0024 | 0.0022 |
| F04 | 48480001 | 48580000 | 0.0030 | 0.0023 | 0.0026 |
| F04 | 48500001 | 48600000 | 0.0026 | 0.0016 | 0.0024 |
| F04 | 48520001 | 48620000 | 0.0019 | 0.0018 | 0.0018 |
| F04 | 48540001 | 48640000 | 0.0022 | 0.0020 | 0.0021 |
| F04 | 48560001 | 48660000 | 0.0024 | 0.0018 | 0.0021 |
| F04 | 48580001 | 48680000 | 0.0022 | 0.0022 | 0.0019 |
| F04 | 48600001 | 48700000 | 0.0027 | 0.0025 | 0.0022 |
| F04 | 48620001 | 48720000 | 0.0031 | 0.0025 | 0.0026 |
| F04 | 48640001 | 48740000 | 0.0032 | 0.0019 | 0.0025 |
| F04 | 48660001 | 48760000 | 0.0027 | 0.0020 | 0.0019 |
| F04 | 48680001 | 48780000 | 0.0028 | 0.0015 | 0.0020 |
| F04 | 48700001 | 48800000 | 0.0023 | 0.0012 | 0.0015 |
| F04 | 48720001 | 48820000 | 0.0019 | 0.0008 | 0.0011 |
| F04 | 48740001 | 48840000 | 0.0013 | 0.0011 | 0.0008 |
| F04 | 48760001 | 48860000 | 0.0015 | 0.0009 | 0.0012 |
| F04 | 48780001 | 48880000 | 0.0012 | 0.0015 | 0.0011 |
| F04 | 48800001 | 48900000 | 0.0019 | 0.0016 | 0.0016 |
| F04 | 48820001 | 48920000 | 0.0020 | 0.0021 | 0.0017 |
| F04 | 48840001 | 48940000 | 0.0026 | 0.0022 | 0.0023 |
| F04 | 48860001 | 48960000 | 0.0030 | 0.0022 | 0.0025 |
| F04 | 48880001 | 48980000 | 0.0031 | 0.0017 | 0.0024 |
| F04 | 48900001 | 49000000 | 0.0026 | 0.0020 | 0.0020 |
| F04 | 48920001 | 49020000 | 0.0029 | 0.0029 | 0.0022 |
| F04 | 48940001 | 49040000 | 0.0038 | 0.0027 | 0.0029 |
| F04 | 48960001 | 49060000 | 0.0035 | 0.0037 | 0.0028 |

| Chr | Start | End | both | A1 | A2 |
|---|---|---|---|---|---|
| F04 | 48980001 | 49080000 | 0.0046 | 0.0041 | 0.0040 |
| F04 | 49000001 | 49100000 | 0.0049 | 0.0044 | 0.0045 |
| F04 | 49020001 | 49120000 | 0.0052 | 0.0028 | 0.0048 |
| F04 | 49040001 | 49140000 | 0.0036 | 0.0026 | 0.0033 |
| F04 | 49060001 | 49160000 | 0.0034 | 0.0016 | 0.0030 |
| F04 | 49080001 | 49180000 | 0.0022 | 0.0012 | 0.0019 |
| F04 | 49100001 | 49200000 | 0.0017 | 0.0007 | 0.0014 |
| F04 | 49120001 | 49220000 | 0.0014 | 0.0014 | 0.0010 |
| F04 | 49140001 | 49240000 | 0.0019 | 0.0017 | 0.0016 |
| F04 | 49160001 | 49260000 | 0.0021 | 0.0016 | 0.0019 |
| F04 | 49180001 | 49280000 | 0.0019 | 0.0018 | 0.0017 |
| F04 | 49200001 | 49300000 | 0.0020 | 0.0019 | 0.0019 |
| F04 | 49220001 | 49320000 | 0.0021 | 0.0014 | 0.0020 |
| F04 | 49240001 | 49340000 | 0.0018 | 0.0015 | 0.0016 |
| F04 | 49260001 | 49360000 | 0.0020 | 0.0022 | 0.0016 |
| F04 | 49280001 | 49380000 | 0.0029 | 0.0020 | 0.0024 |
| F04 | 49300001 | 49400000 | 0.0027 | 0.0018 | 0.0021 |
| F04 | 49320001 | 49420000 | 0.0024 | 0.0016 | 0.0018 |
| F04 | 49340001 | 49440000 | 0.0021 | 0.0016 | 0.0017 |
| F04 | 49360001 | 49460000 | 0.0022 | 0.0012 | 0.0019 |
| F04 | 49380001 | 49480000 | 0.0017 | 0.0014 | 0.0014 |
| F04 | 49400001 | 49500000 | 0.0019 | 0.0024 | 0.0017 |
| F04 | 49420001 | 49520000 | 0.0030 | 0.0028 | 0.0028 |
| F04 | 49440001 | 49540000 | 0.0035 | 0.0024 | 0.0032 |
| F04 | 49460001 | 49560000 | 0.0029 | 0.0031 | 0.0027 |
| F04 | 49480001 | 49580000 | 0.0036 | 0.0029 | 0.0034 |
| F04 | 49500001 | 49600000 | 0.0034 | 0.0020 | 0.0032 |
| F04 | 49520001 | 49620000 | 0.0025 | 0.0017 | 0.0022 |
| F04 | 49540001 | 49640000 | 0.0022 | 0.0017 | 0.0019 |
| F04 | 49560001 | 49660000 | 0.0021 | 0.0009 | 0.0018 |
| F04 | 49580001 | 49680000 | 0.0012 | 0.0008 | 0.0010 |
| F04 | 49600001 | 49700000 | 0.0011 | 0.0006 | 0.0009 |
| F04 | 49620001 | 49720000 | 0.0007 | 0.0007 | 0.0007 |
| F04 | 49640001 | 49740000 | 0.0008 | 0.0008 | 0.0008 |
| F04 | 49660001 | 49760000 | 0.0009 | 0.0006 | 0.0009 |
| F04 | 49680001 | 49780000 | 0.0007 | 0.0007 | 0.0007 |
| F04 | 49700001 | 49800000 | 0.0007 | 0.0007 | 0.0007 |
| F04 | 49720001 | 49820000 | 0.0008 | 0.0005 | 0.0008 |
| F04 | 49740001 | 49840000 | 0.0006 | 0.0004 | 0.0006 |
| F04 | 49760001 | 49860000 | 0.0005 | 0.0003 | 0.0005 |
| F04 | 49780001 | 49880000 | 0.0004 | 0.0002 | 0.0005 |
| F04 | 49800001 | 49900000 | 0.0003 | 0.0004 | 0.0003 |
| F04 | 49820001 | 49920000 | 0.0005 | 0.0007 | 0.0005 |
| F04 | 49840001 | 49940000 | 0.0009 | 0.0009 | 0.0009 |
| F04 | 49860001 | 49960000 | 0.0012 | 0.0009 | 0.0011 |
| F04 | 49880001 | 49980000 | 0.0011 | 0.0010 | 0.0011 |
| F04 | 49900001 | 50000000 | 0.0013 | 0.0007 | 0.0013 |
| F04 | 49920001 | 50020000 | 0.0010 | 0.0006 | 0.0010 |
| F04 | 49940001 | 50040000 | 0.0008 | 0.0004 | 0.0008 |
| F04 | 49960001 | 50060000 | 0.0007 | 0.0007 | 0.0007 |
| F04 | 49980001 | 50080000 | 0.0009 | 0.0007 | 0.0009 |
| F04 | 50000001 | 50100000 | 0.0010 | 0.0009 | 0.0010 |
| F04 | 50020001 | 50120000 | 0.0012 | 0.0012 | 0.0012 |
| F04 | 50040001 | 50140000 | 0.0014 | 0.0014 | 0.0013 |
| F04 | 50060001 | 50160000 | 0.0014 | 0.0015 | 0.0013 |
| F04 | 50080001 | 50180000 | 0.0016 | 0.0021 | 0.0015 |
| F04 | 50100001 | 50200000 | 0.0022 | 0.0019 | 0.0020 |
| F04 | 50120001 | 50220000 | 0.0019 | 0.0016 | 0.0018 |

| Chr | Start | End | both | A1 | A2 |
|---|---|---|---|---|---|
| F04 | 50140001 | 50240000 | 0.0017 | 0.0014 | 0.0015 |
| F04 | 50160001 | 50260000 | 0.0016 | 0.0011 | 0.0014 |
| F04 | 50180001 | 50280000 | 0.0013 | 0.0004 | 0.0011 |
| F04 | 50200001 | 50300000 | 0.0006 | 0.0005 | 0.0004 |
| F04 | 50220001 | 50320000 | 0.0006 | 0.0004 | 0.0004 |
| F04 | 50240001 | 50340000 | 0.0005 | 0.0004 | 0.0004 |
| F04 | 50260001 | 50360000 | 0.0004 | 0.0004 | 0.0004 |
| F04 | 50280001 | 50380000 | 0.0005 | 0.0004 | 0.0004 |
| F04 | 50300001 | 50400000 | 0.0004 | 0.0007 | 0.0004 |
| F04 | 50320001 | 50420000 | 0.0009 | 0.0007 | 0.0009 |
| F04 | 50340001 | 50440000 | 0.0012 | 0.0010 | 0.0009 |
| F04 | 50360001 | 50460000 | 0.0017 | 0.0013 | 0.0013 |
| F04 | 50380001 | 50480000 | 0.0022 | 0.0016 | 0.0016 |
| F04 | 50400001 | 50500000 | 0.0025 | 0.0018 | 0.0019 |
| F04 | 50420001 | 50520000 | 0.0026 | 0.0022 | 0.0020 |
| F04 | 50440001 | 50540000 | 0.0030 | 0.0022 | 0.0025 |
| F04 | 50460001 | 50560000 | 0.0027 | 0.0019 | 0.0024 |
| F04 | 50480001 | 50580000 | 0.0023 | 0.0018 | 0.0022 |
| F04 | 50500001 | 50600000 | 0.0023 | 0.0018 | 0.0021 |
| F04 | 50520001 | 50620000 | 0.0023 | 0.0015 | 0.0022 |
| F04 | 50540001 | 50640000 | 0.0019 | 0.0016 | 0.0018 |
| F04 | 50560001 | 50660000 | 0.0019 | 0.0030 | 0.0019 |
| F04 | 50580001 | 50680000 | 0.0035 | 0.0033 | 0.0035 |
| F04 | 50600001 | 50700000 | 0.0038 | 0.0035 | 0.0036 |
| F04 | 50620001 | 50720000 | 0.0040 | 0.0035 | 0.0038 |
| F04 | 50640001 | 50740000 | 0.0041 | 0.0036 | 0.0039 |
| F04 | 50660001 | 50760000 | 0.0041 | 0.0020 | 0.0039 |
| F04 | 50680001 | 50780000 | 0.0025 | 0.0016 | 0.0023 |
| F04 | 50700001 | 50800000 | 0.0021 | 0.0012 | 0.0020 |
| F04 | 50720001 | 50820000 | 0.0015 | 0.0015 | 0.0014 |
| F04 | 50740001 | 50840000 | 0.0020 | 0.0013 | 0.0018 |
| F04 | 50760001 | 50860000 | 0.0020 | 0.0015 | 0.0018 |
| F04 | 50780001 | 50880000 | 0.0024 | 0.0013 | 0.0019 |
| F04 | 50800001 | 50900000 | 0.0022 | 0.0015 | 0.0016 |
| F04 | 50820001 | 50920000 | 0.0024 | 0.0014 | 0.0020 |
| F04 | 50840001 | 50940000 | 0.0024 | 0.0015 | 0.0017 |
| F04 | 50860001 | 50960000 | 0.0024 | 0.0014 | 0.0018 |
| F04 | 50880001 | 50980000 | 0.0021 | 0.0019 | 0.0016 |
| F04 | 50900001 | 51000000 | 0.0027 | 0.0022 | 0.0022 |
| F04 | 50920001 | 51020000 | 0.0031 | 0.0021 | 0.0025 |
| F04 | 50940001 | 51040000 | 0.0027 | 0.0023 | 0.0024 |
| F04 | 50960001 | 51060000 | 0.0029 | 0.0027 | 0.0025 |
| F04 | 50980001 | 51080000 | 0.0034 | 0.0023 | 0.0029 |
| F04 | 51000001 | 51100000 | 0.0029 | 0.0016 | 0.0024 |
| F04 | 51020001 | 51120000 | 0.0021 | 0.0013 | 0.0016 |
| F04 | 51040001 | 51140000 | 0.0018 | 0.0013 | 0.0013 |
| F04 | 51060001 | 51160000 | 0.0017 | 0.0013 | 0.0014 |
| F04 | 51080001 | 51180000 | 0.0015 | 0.0017 | 0.0013 |
| F04 | 51100001 | 51200000 | 0.0018 | 0.0023 | 0.0018 |
| F04 | 51120001 | 51220000 | 0.0026 | 0.0030 | 0.0024 |
| F04 | 51140001 | 51240000 | 0.0033 | 0.0029 | 0.0029 |
| F04 | 51160001 | 51260000 | 0.0032 | 0.0023 | 0.0028 |
| F04 | 51180001 | 51280000 | 0.0026 | 0.0033 | 0.0022 |
| F04 | 51200001 | 51300000 | 0.0036 | 0.0031 | 0.0032 |
| F04 | 51220001 | 51320000 | 0.0035 | 0.0025 | 0.0031 |
| F04 | 51240001 | 51340000 | 0.0029 | 0.0024 | 0.0027 |
| F04 | 51260001 | 51360000 | 0.0035 | 0.0040 | 0.0033 |
| F04 | 51280001 | 51380000 | 0.0053 | 0.0031 | 0.0050 |

| Chr | Start | End | both | A1 | A2 |
|---|---|---|---|---|---|
| F04 | 51300001 | 51400000 | 0.0044 | 0.0026 | 0.0041 |
| F04 | 51320001 | 51420000 | 0.0039 | 0.0031 | 0.0036 |
| F04 | 51340001 | 51440000 | 0.0044 | 0.0029 | 0.0040 |
| F04 | 51360001 | 51460000 | 0.0037 | 0.0018 | 0.0031 |
| F04 | 51380001 | 51480000 | 0.0027 | 0.0017 | 0.0021 |
| F04 | 51400001 | 51500000 | 0.0026 | 0.0020 | 0.0021 |
| F04 | 51420001 | 51520000 | 0.0029 | 0.0018 | 0.0025 |
| F04 | 51440001 | 51540000 | 0.0027 | 0.0017 | 0.0024 |
| F04 | 51460001 | 51560000 | 0.0024 | 0.0014 | 0.0023 |
| F04 | 51480001 | 51580000 | 0.0018 | 0.0015 | 0.0017 |
| F04 | 51500001 | 51600000 | 0.0019 | 0.0013 | 0.0018 |
| F04 | 51520001 | 51620000 | 0.0017 | 0.0010 | 0.0016 |
| F04 | 51540001 | 51640000 | 0.0014 | 0.0013 | 0.0013 |
| F04 | 51560001 | 51660000 | 0.0017 | 0.0017 | 0.0015 |
| F04 | 51580001 | 51680000 | 0.0023 | 0.0013 | 0.0021 |
| F04 | 51600001 | 51700000 | 0.0017 | 0.0011 | 0.0015 |
| F04 | 51620001 | 51720000 | 0.0014 | 0.0013 | 0.0013 |
| F04 | 51640001 | 51740000 | 0.0017 | 0.0012 | 0.0016 |
| F04 | 51660001 | 51760000 | 0.0016 | 0.0006 | 0.0015 |
| F04 | 51680001 | 51780000 | 0.0012 | 0.0011 | 0.0009 |
| F04 | 51700001 | 51800000 | 0.0017 | 0.0019 | 0.0013 |
| F04 | 51720001 | 51820000 | 0.0027 | 0.0017 | 0.0021 |
| F04 | 51740001 | 51840000 | 0.0025 | 0.0018 | 0.0018 |
| F04 | 51760001 | 51860000 | 0.0026 | 0.0018 | 0.0019 |
| F04 | 51780001 | 51880000 | 0.0023 | 0.0017 | 0.0019 |
| F04 | 51800001 | 51900000 | 0.0022 | 0.0013 | 0.0017 |
| F04 | 51820001 | 51920000 | 0.0016 | 0.0015 | 0.0013 |
| F04 | 51840001 | 51940000 | 0.0017 | 0.0015 | 0.0015 |
| F04 | 51860001 | 51960000 | 0.0018 | 0.0016 | 0.0016 |
| F04 | 51880001 | 51980000 | 0.0020 | 0.0016 | 0.0017 |
| F04 | 51900001 | 52000000 | 0.0018 | 0.0017 | 0.0017 |
| F04 | 51920001 | 52020000 | 0.0020 | 0.0017 | 0.0019 |
| F04 | 51940001 | 52040000 | 0.0020 | 0.0017 | 0.0018 |
| F04 | 51960001 | 52060000 | 0.0018 | 0.0019 | 0.0018 |
| F04 | 51980001 | 52080000 | 0.0021 | 0.0018 | 0.0021 |
| F04 | 52000001 | 52100000 | 0.0021 | 0.0014 | 0.0020 |
| F04 | 52020001 | 52120000 | 0.0016 | 0.0014 | 0.0016 |
| F04 | 52040001 | 52140000 | 0.0016 | 0.0015 | 0.0016 |
| F04 | 52060001 | 52160000 | 0.0017 | 0.0012 | 0.0016 |
| F04 | 52080001 | 52180000 | 0.0012 | 0.0010 | 0.0012 |
| F04 | 52100001 | 52200000 | 0.0011 | 0.0011 | 0.0010 |
| F04 | 52120001 | 52220000 | 0.0013 | 0.0012 | 0.0011 |
| F04 | 52140001 | 52240000 | 0.0017 | 0.0014 | 0.0011 |
| F04 | 52160001 | 52260000 | 0.0021 | 0.0017 | 0.0014 |
| F04 | 52180001 | 52280000 | 0.0026 | 0.0020 | 0.0017 |
| F04 | 52200001 | 52300000 | 0.0029 | 0.0026 | 0.0021 |
| F04 | 52220001 | 52320000 | 0.0034 | 0.0024 | 0.0025 |
| F04 | 52240001 | 52340000 | 0.0032 | 0.0019 | 0.0025 |
| F04 | 52260001 | 52360000 | 0.0024 | 0.0019 | 0.0019 |
| F04 | 52280001 | 52380000 | 0.0024 | 0.0017 | 0.0019 |
| F04 | 52300001 | 52400000 | 0.0024 | 0.0011 | 0.0018 |
| F04 | 52320001 | 52420000 | 0.0019 | 0.0018 | 0.0011 |
| F04 | 52340001 | 52440000 | 0.0026 | 0.0019 | 0.0020 |
| F04 | 52360001 | 52460000 | 0.0027 | 0.0017 | 0.0020 |
| F04 | 52380001 | 52480000 | 0.0025 | 0.0019 | 0.0019 |
| F04 | 52400001 | 52500000 | 0.0025 | 0.0019 | 0.0021 |
| F04 | 52420001 | 52520000 | 0.0024 | 0.0010 | 0.0021 |
| F04 | 52440001 | 52540000 | 0.0016 | 0.0016 | 0.0012 |

| Chr | Start | End | both | A1 | A2 |
|---|---|---|---|---|---|
| F04 | 52460001 | 52560000 | 0.0024 | 0.0016 | 0.0018 |
| F04 | 52480001 | 52580000 | 0.0025 | 0.0020 | 0.0017 |
| F04 | 52500001 | 52600000 | 0.0031 | 0.0025 | 0.0024 |
| F04 | 52520001 | 52620000 | 0.0036 | 0.0038 | 0.0029 |
| F04 | 52540001 | 52640000 | 0.0048 | 0.0034 | 0.0040 |
| F04 | 52560001 | 52660000 | 0.0041 | 0.0037 | 0.0035 |
| F04 | 52580001 | 52680000 | 0.0044 | 0.0029 | 0.0038 |
| F04 | 52600001 | 52700000 | 0.0035 | 0.0023 | 0.0028 |
| F04 | 52620001 | 52720000 | 0.0032 | 0.0013 | 0.0023 |
| F04 | 52640001 | 52740000 | 0.0023 | 0.0013 | 0.0016 |
| F04 | 52660001 | 52760000 | 0.0022 | 0.0008 | 0.0016 |
| F04 | 52680001 | 52780000 | 0.0016 | 0.0011 | 0.0011 |
| F04 | 52700001 | 52800000 | 0.0016 | 0.0015 | 0.0013 |
| F04 | 52720001 | 52820000 | 0.0018 | 0.0012 | 0.0018 |
| F04 | 52740001 | 52840000 | 0.0014 | 0.0014 | 0.0013 |
| F04 | 52760001 | 52860000 | 0.0017 | 0.0016 | 0.0016 |
| F04 | 52780001 | 52880000 | 0.0019 | 0.0017 | 0.0018 |
| F04 | 52800001 | 52900000 | 0.0019 | 0.0014 | 0.0019 |
| F04 | 52820001 | 52920000 | 0.0016 | 0.0017 | 0.0015 |
| F04 | 52840001 | 52940000 | 0.0019 | 0.0013 | 0.0018 |
| F04 | 52860001 | 52960000 | 0.0015 | 0.0013 | 0.0015 |
| F04 | 52880001 | 52980000 | 0.0014 | 0.0013 | 0.0014 |
| F04 | 52900001 | 53000000 | 0.0014 | 0.0015 | 0.0013 |
| F04 | 52920001 | 53020000 | 0.0014 | 0.0013 | 0.0013 |
| F04 | 52940001 | 53040000 | 0.0013 | 0.0015 | 0.0012 |
| F04 | 52960001 | 53060000 | 0.0014 | 0.0013 | 0.0013 |
| F04 | 52980001 | 53080000 | 0.0013 | 0.0014 | 0.0012 |
| F04 | 53000001 | 53100000 | 0.0014 | 0.0010 | 0.0013 |
| F04 | 53020001 | 53120000 | 0.0013 | 0.0010 | 0.0012 |
| F04 | 53040001 | 53140000 | 0.0012 | 0.0009 | 0.0011 |
| F04 | 53060001 | 53160000 | 0.0011 | 0.0011 | 0.0011 |
| F04 | 53080001 | 53180000 | 0.0017 | 0.0009 | 0.0013 |
| F04 | 53100001 | 53200000 | 0.0017 | 0.0011 | 0.0012 |
| F04 | 53120001 | 53220000 | 0.0019 | 0.0013 | 0.0013 |
| F04 | 53140001 | 53240000 | 0.0022 | 0.0015 | 0.0015 |
| F04 | 53160001 | 53260000 | 0.0024 | 0.0016 | 0.0017 |
| F04 | 53180001 | 53280000 | 0.0022 | 0.0015 | 0.0017 |
| F04 | 53200001 | 53300000 | 0.0020 | 0.0013 | 0.0016 |
| F04 | 53220001 | 53320000 | 0.0017 | 0.0016 | 0.0013 |
| F04 | 53240001 | 53340000 | 0.0021 | 0.0015 | 0.0016 |
| F04 | 53260001 | 53360000 | 0.0023 | 0.0012 | 0.0016 |
| F04 | 53280001 | 53380000 | 0.0021 | 0.0011 | 0.0013 |
| F04 | 53300001 | 53400000 | 0.0019 | 0.0015 | 0.0012 |
| F04 | 53320001 | 53420000 | 0.0024 | 0.0013 | 0.0017 |
| F04 | 53340001 | 53440000 | 0.0020 | 0.0014 | 0.0016 |
| F04 | 53360001 | 53460000 | 0.0017 | 0.0018 | 0.0016 |
| F04 | 53380001 | 53480000 | 0.0021 | 0.0018 | 0.0020 |
| F04 | 53400001 | 53500000 | 0.0022 | 0.0014 | 0.0021 |
| F04 | 53420001 | 53520000 | 0.0017 | 0.0013 | 0.0016 |
| F04 | 53440001 | 53540000 | 0.0017 | 0.0011 | 0.0016 |
| F04 | 53460001 | 53560000 | 0.0015 | 0.0009 | 0.0014 |
| F04 | 53480001 | 53580000 | 0.0013 | 0.0009 | 0.0012 |
| F04 | 53500001 | 53600000 | 0.0011 | 0.0015 | 0.0011 |
| F04 | 53520001 | 53620000 | 0.0018 | 0.0015 | 0.0017 |
| F04 | 53540001 | 53640000 | 0.0018 | 0.0017 | 0.0016 |
| F04 | 53560001 | 53660000 | 0.0022 | 0.0017 | 0.0019 |
| F04 | 53580001 | 53680000 | 0.0022 | 0.0021 | 0.0018 |
| F04 | 53600001 | 53700000 | 0.0027 | 0.0017 | 0.0023 |

| Chr | Start | End | both | A1 | A2 |
|---|---|---|---|---|---|
| F04 | 53620001 | 53720000 | 0.0023 | 0.0015 | 0.0018 |
| F04 | 53640001 | 53740000 | 0.0019 | 0.0013 | 0.0017 |
| F04 | 53660001 | 53760000 | 0.0017 | 0.0010 | 0.0015 |
| F04 | 53680001 | 53780000 | 0.0013 | 0.0007 | 0.0012 |
| F04 | 53700001 | 53800000 | 0.0009 | 0.0005 | 0.0008 |
| F04 | 53720001 | 53820000 | 0.0007 | 0.0006 | 0.0007 |
| F04 | 53740001 | 53840000 | 0.0006 | 0.0005 | 0.0006 |
| F04 | 53760001 | 53860000 | 0.0005 | 0.0008 | 0.0005 |
| F04 | 53780001 | 53880000 | 0.0008 | 0.0011 | 0.0008 |
| F04 | 53800001 | 53900000 | 0.0013 | 0.0013 | 0.0011 |
| F04 | 53820001 | 53920000 | 0.0016 | 0.0016 | 0.0014 |
| F04 | 53840001 | 53940000 | 0.0021 | 0.0017 | 0.0017 |
| F04 | 53860001 | 53960000 | 0.0023 | 0.0021 | 0.0019 |
| F04 | 53880001 | 53980000 | 0.0028 | 0.0021 | 0.0024 |
| F04 | 53900001 | 54000000 | 0.0027 | 0.0019 | 0.0024 |
| F04 | 53920001 | 54020000 | 0.0026 | 0.0018 | 0.0021 |
| F04 | 53940001 | 54040000 | 0.0026 | 0.0020 | 0.0021 |
| F04 | 53960001 | 54060000 | 0.0029 | 0.0015 | 0.0023 |
| F04 | 53980001 | 54080000 | 0.0023 | 0.0013 | 0.0017 |
| F04 | 54000001 | 54100000 | 0.0021 | 0.0014 | 0.0014 |
| F04 | 54020001 | 54120000 | 0.0020 | 0.0012 | 0.0015 |
| F04 | 54040001 | 54140000 | 0.0015 | 0.0007 | 0.0013 |
| F04 | 54060001 | 54160000 | 0.0010 | 0.0006 | 0.0008 |
| F04 | 54080001 | 54180000 | 0.0008 | 0.0011 | 0.0007 |
| F04 | 54100001 | 54200000 | 0.0013 | 0.0010 | 0.0013 |
| F04 | 54120001 | 54220000 | 0.0012 | 0.0009 | 0.0012 |
| F04 | 54140001 | 54240000 | 0.0012 | 0.0010 | 0.0012 |
| F04 | 54160001 | 54260000 | 0.0012 | 0.0013 | 0.0011 |
| F04 | 54180001 | 54280000 | 0.0016 | 0.0008 | 0.0016 |
| F04 | 54200001 | 54300000 | 0.0010 | 0.0010 | 0.0010 |
| F04 | 54220001 | 54320000 | 0.0013 | 0.0017 | 0.0013 |
| F04 | 54240001 | 54340000 | 0.0022 | 0.0022 | 0.0021 |
| F04 | 54260001 | 54360000 | 0.0027 | 0.0023 | 0.0026 |
| F04 | 54280001 | 54380000 | 0.0029 | 0.0024 | 0.0027 |
| F04 | 54300001 | 54400000 | 0.0031 | 0.0023 | 0.0029 |
| F04 | 54320001 | 54420000 | 0.0030 | 0.0019 | 0.0027 |
| F04 | 54340001 | 54440000 | 0.0023 | 0.0016 | 0.0021 |
| F04 | 54360001 | 54460000 | 0.0020 | 0.0019 | 0.0018 |
| F04 | 54380001 | 54480000 | 0.0021 | 0.0019 | 0.0020 |
| F04 | 54400001 | 54500000 | 0.0022 | 0.0021 | 0.0020 |
| F04 | 54420001 | 54520000 | 0.0025 | 0.0020 | 0.0023 |
| F04 | 54440001 | 54540000 | 0.0023 | 0.0023 | 0.0021 |
| F04 | 54460001 | 54560000 | 0.0027 | 0.0016 | 0.0025 |
| F04 | 54480001 | 54580000 | 0.0020 | 0.0015 | 0.0018 |
| F04 | 54500001 | 54600000 | 0.0018 | 0.0014 | 0.0016 |
| F04 | 54520001 | 54620000 | 0.0017 | 0.0015 | 0.0015 |
| F04 | 54540001 | 54640000 | 0.0021 | 0.0010 | 0.0018 |
| F04 | 54560001 | 54660000 | 0.0015 | 0.0012 | 0.0012 |
| F04 | 54580001 | 54680000 | 0.0018 | 0.0010 | 0.0015 |
| F04 | 54600001 | 54700000 | 0.0016 | 0.0014 | 0.0013 |
| F04 | 54620001 | 54720000 | 0.0019 | 0.0011 | 0.0016 |
| F04 | 54640001 | 54740000 | 0.0014 | 0.0016 | 0.0012 |
| F04 | 54660001 | 54760000 | 0.0020 | 0.0015 | 0.0017 |
| F04 | 54680001 | 54780000 | 0.0019 | 0.0022 | 0.0017 |
| F04 | 54700001 | 54800000 | 0.0027 | 0.0018 | 0.0025 |
| F04 | 54720001 | 54820000 | 0.0023 | 0.0018 | 0.0021 |
| F04 | 54740001 | 54840000 | 0.0024 | 0.0017 | 0.0023 |
| F04 | 54760001 | 54860000 | 0.0022 | 0.0017 | 0.0022 |

| Chr | Start | End | both | A1 | A2 |
|---|---|---|---|---|---|
| F04 | 54780001 | 54880000 | 0.0021 | 0.0012 | 0.0020 |
| F04 | 54800001 | 54900000 | 0.0014 | 0.0011 | 0.0013 |
| F04 | 54820001 | 54920000 | 0.0013 | 0.0014 | 0.0012 |
| F04 | 54840001 | 54940000 | 0.0017 | 0.0014 | 0.0016 |
| F04 | 54860001 | 54960000 | 0.0017 | 0.0015 | 0.0017 |
| F04 | 54880001 | 54980000 | 0.0018 | 0.0016 | 0.0018 |
| F04 | 54900001 | 55000000 | 0.0020 | 0.0019 | 0.0020 |
| F04 | 54920001 | 55020000 | 0.0025 | 0.0015 | 0.0023 |
| F04 | 54940001 | 55040000 | 0.0019 | 0.0011 | 0.0016 |
| F04 | 54960001 | 55060000 | 0.0015 | 0.0010 | 0.0012 |
| F04 | 54980001 | 55080000 | 0.0013 | 0.0017 | 0.0011 |
| F04 | 55000001 | 55100000 | 0.0021 | 0.0019 | 0.0017 |
| F04 | 55020001 | 55120000 | 0.0022 | 0.0021 | 0.0019 |
| F04 | 55040001 | 55140000 | 0.0024 | 0.0019 | 0.0022 |
| F04 | 55060001 | 55160000 | 0.0022 | 0.0021 | 0.0020 |
| F04 | 55080001 | 55180000 | 0.0023 | 0.0012 | 0.0021 |
| F04 | 55100001 | 55200000 | 0.0013 | 0.0009 | 0.0013 |
| F04 | 55120001 | 55220000 | 0.0011 | 0.0012 | 0.0009 |
| F04 | 55140001 | 55240000 | 0.0017 | 0.0012 | 0.0015 |
| F04 | 55160001 | 55260000 | 0.0016 | 0.0011 | 0.0014 |
| F04 | 55180001 | 55280000 | 0.0016 | 0.0011 | 0.0014 |
| F04 | 55200001 | 55300000 | 0.0018 | 0.0009 | 0.0015 |
| F04 | 55220001 | 55320000 | 0.0016 | 0.0006 | 0.0013 |
| F04 | 55240001 | 55340000 | 0.0010 | 0.0015 | 0.0008 |
| F04 | 55260001 | 55360000 | 0.0020 | 0.0018 | 0.0017 |
| F04 | 55280001 | 55380000 | 0.0024 | 0.0020 | 0.0020 |
| F04 | 55300001 | 55400000 | 0.0025 | 0.0020 | 0.0022 |
| F04 | 55320001 | 55420000 | 0.0026 | 0.0020 | 0.0023 |
| F04 | 55340001 | 55440000 | 0.0025 | 0.0015 | 0.0022 |
| F04 | 55360001 | 55460000 | 0.0020 | 0.0016 | 0.0016 |
| F04 | 55380001 | 55480000 | 0.0020 | 0.0013 | 0.0017 |
| F04 | 55400001 | 55500000 | 0.0016 | 0.0012 | 0.0013 |
| F04 | 55420001 | 55520000 | 0.0017 | 0.0014 | 0.0012 |
| F04 | 55440001 | 55540000 | 0.0019 | 0.0014 | 0.0014 |
| F04 | 55460001 | 55560000 | 0.0021 | 0.0012 | 0.0017 |
| F04 | 55480001 | 55580000 | 0.0019 | 0.0016 | 0.0015 |
| F04 | 55500001 | 55600000 | 0.0023 | 0.0023 | 0.0019 |
| F04 | 55520001 | 55620000 | 0.0030 | 0.0026 | 0.0027 |
| F04 | 55540001 | 55640000 | 0.0031 | 0.0025 | 0.0029 |
| F04 | 55560001 | 55660000 | 0.0028 | 0.0026 | 0.0027 |
| F04 | 55580001 | 55680000 | 0.0030 | 0.0026 | 0.0028 |
| F04 | 55600001 | 55700000 | 0.0032 | 0.0020 | 0.0028 |
| F04 | 55620001 | 55720000 | 0.0026 | 0.0019 | 0.0022 |
| F04 | 55640001 | 55740000 | 0.0026 | 0.0024 | 0.0021 |
| F04 | 55660001 | 55760000 | 0.0031 | 0.0023 | 0.0025 |
| F04 | 55680001 | 55780000 | 0.0028 | 0.0018 | 0.0024 |
| F04 | 55700001 | 55800000 | 0.0023 | 0.0016 | 0.0021 |
| F04 | 55720001 | 55820000 | 0.0020 | 0.0014 | 0.0018 |
| F04 | 55740001 | 55840000 | 0.0017 | 0.0006 | 0.0016 |
| F04 | 55760001 | 55860000 | 0.0008 | 0.0006 | 0.0008 |
| F04 | 55780001 | 55880000 | 0.0008 | 0.0008 | 0.0008 |
| F04 | 55800001 | 55900000 | 0.0009 | 0.0014 | 0.0008 |
| F04 | 55820001 | 55920000 | 0.0016 | 0.0013 | 0.0015 |
| F04 | 55840001 | 55940000 | 0.0015 | 0.0014 | 0.0014 |
| F04 | 55860001 | 55960000 | 0.0016 | 0.0013 | 0.0015 |
| F04 | 55880001 | 55980000 | 0.0016 | 0.0015 | 0.0015 |
| F04 | 55900001 | 56000000 | 0.0018 | 0.0017 | 0.0018 |
| F04 | 55920001 | 56020000 | 0.0019 | 0.0016 | 0.0018 |

| Chr | Start | End | both | A1 | A2 |
|---|---|---|---|---|---|
| F04 | 55940001 | 56040000 | 0.0017 | 0.0017 | 0.0017 |
| F04 | 55960001 | 56060000 | 0.0018 | 0.0018 | 0.0017 |
| F04 | 55980001 | 56080000 | 0.0019 | 0.0015 | 0.0018 |
| F04 | 56000001 | 56100000 | 0.0015 | 0.0006 | 0.0014 |
| F04 | 56020001 | 56120000 | 0.0007 | 0.0014 | 0.0006 |
| F04 | 56040001 | 56140000 | 0.0016 | 0.0015 | 0.0014 |
| F04 | 56060001 | 56160000 | 0.0016 | 0.0014 | 0.0014 |
| F04 | 56080001 | 56180000 | 0.0015 | 0.0015 | 0.0013 |
| F04 | 56100001 | 56200000 | 0.0017 | 0.0020 | 0.0015 |
| F04 | 56120001 | 56220000 | 0.0021 | 0.0012 | 0.0019 |
| F04 | 56140001 | 56240000 | 0.0012 | 0.0013 | 0.0011 |
| F04 | 56160001 | 56260000 | 0.0014 | 0.0014 | 0.0014 |
| F04 | 56180001 | 56280000 | 0.0014 | 0.0013 | 0.0014 |
| F04 | 56200001 | 56300000 | 0.0013 | 0.0008 | 0.0013 |
| F04 | 56220001 | 56320000 | 0.0009 | 0.0009 | 0.0009 |
| F04 | 56240001 | 56340000 | 0.0010 | 0.0006 | 0.0010 |
| F04 | 56260001 | 56360000 | 0.0007 | 0.0006 | 0.0006 |
| F04 | 56280001 | 56380000 | 0.0007 | 0.0010 | 0.0007 |
| F04 | 56300001 | 56400000 | 0.0011 | 0.0013 | 0.0011 |
| F04 | 56320001 | 56420000 | 0.0015 | 0.0026 | 0.0015 |
| F04 | 56340001 | 56440000 | 0.0029 | 0.0026 | 0.0028 |
| F04 | 56360001 | 56460000 | 0.0029 | 0.0024 | 0.0028 |
| F04 | 56380001 | 56480000 | 0.0028 | 0.0021 | 0.0027 |
| F04 | 56400001 | 56500000 | 0.0024 | 0.0020 | 0.0023 |
| F04 | 56420001 | 56520000 | 0.0024 | 0.0009 | 0.0023 |
| F04 | 56440001 | 56540000 | 0.0013 | 0.0011 | 0.0012 |
| F04 | 56460001 | 56560000 | 0.0016 | 0.0014 | 0.0014 |
| F04 | 56480001 | 56580000 | 0.0018 | 0.0018 | 0.0017 |
| F04 | 56500001 | 56600000 | 0.0023 | 0.0019 | 0.0022 |
| F04 | 56520001 | 56620000 | 0.0025 | 0.0017 | 0.0024 |
| F04 | 56540001 | 56640000 | 0.0024 | 0.0015 | 0.0024 |
| F04 | 56560001 | 56660000 | 0.0020 | 0.0014 | 0.0019 |
| F04 | 56580001 | 56680000 | 0.0017 | 0.0009 | 0.0017 |
| F04 | 56600001 | 56700000 | 0.0012 | 0.0006 | 0.0011 |
| F04 | 56620001 | 56720000 | 0.0007 | 0.0005 | 0.0007 |
| F04 | 56640001 | 56740000 | 0.0006 | 0.0007 | 0.0005 |
| F04 | 56660001 | 56760000 | 0.0008 | 0.0008 | 0.0007 |
| F04 | 56680001 | 56780000 | 0.0009 | 0.0010 | 0.0009 |
| F04 | 56700001 | 56800000 | 0.0011 | 0.0011 | 0.0011 |
| F04 | 56720001 | 56820000 | 0.0013 | 0.0014 | 0.0012 |
| F04 | 56740001 | 56840000 | 0.0018 | 0.0013 | 0.0016 |
| F04 | 56760001 | 56860000 | 0.0017 | 0.0010 | 0.0014 |
| F04 | 56780001 | 56880000 | 0.0014 | 0.0009 | 0.0011 |
| F04 | 56800001 | 56900000 | 0.0013 | 0.0010 | 0.0010 |
| F04 | 56820001 | 56920000 | 0.0012 | 0.0006 | 0.0011 |
| F04 | 56840001 | 56940000 | 0.0008 | 0.0008 | 0.0006 |
| F04 | 56860001 | 56960000 | 0.0010 | 0.0007 | 0.0007 |
| F04 | 56880001 | 56980000 | 0.0011 | 0.0010 | 0.0007 |
| F04 | 56900001 | 57000000 | 0.0013 | 0.0008 | 0.0010 |
| F04 | 56920001 | 57020000 | 0.0011 | 0.0013 | 0.0008 |
| F04 | 56940001 | 57040000 | 0.0014 | 0.0014 | 0.0013 |
| F04 | 56960001 | 57060000 | 0.0017 | 0.0013 | 0.0015 |
| F04 | 56980001 | 57080000 | 0.0017 | 0.0011 | 0.0015 |
| F04 | 57000001 | 57100000 | 0.0014 | 0.0011 | 0.0012 |
| F04 | 57020001 | 57120000 | 0.0015 | 0.0009 | 0.0012 |
| F04 | 57040001 | 57140000 | 0.0013 | 0.0007 | 0.0010 |
| F04 | 57060001 | 57160000 | 0.0011 | 0.0008 | 0.0008 |
| F04 | 57080001 | 57180000 | 0.0010 | 0.0012 | 0.0009 |

| Chr | Start | End | both | A1 | A2 |
|---|---|---|---|---|---|
| F04 | 57100001 | 57200000 | 0.0014 | 0.0013 | 0.0013 |
| F04 | 57120001 | 57220000 | 0.0015 | 0.0011 | 0.0014 |
| F04 | 57140001 | 57240000 | 0.0012 | 0.0009 | 0.0012 |
| F04 | 57160001 | 57260000 | 0.0011 | 0.0009 | 0.0010 |
| F04 | 57180001 | 57280000 | 0.0011 | 0.0005 | 0.0010 |
| F04 | 57200001 | 57300000 | 0.0008 | 0.0009 | 0.0006 |
| F04 | 57220001 | 57320000 | 0.0013 | 0.0010 | 0.0010 |
| F04 | 57240001 | 57340000 | 0.0014 | 0.0015 | 0.0010 |
| F04 | 57260001 | 57360000 | 0.0019 | 0.0016 | 0.0016 |
| F04 | 57280001 | 57380000 | 0.0023 | 0.0021 | 0.0019 |
| F04 | 57300001 | 57400000 | 0.0029 | 0.0018 | 0.0023 |
| F04 | 57320001 | 57420000 | 0.0027 | 0.0020 | 0.0019 |
| F04 | 57340001 | 57440000 | 0.0031 | 0.0019 | 0.0021 |
| F04 | 57360001 | 57460000 | 0.0033 | 0.0028 | 0.0021 |
| F04 | 57380001 | 57480000 | 0.0041 | 0.0024 | 0.0030 |
| F04 | 57400001 | 57500000 | 0.0036 | 0.0021 | 0.0028 |
| F04 | 57420001 | 57520000 | 0.0032 | 0.0020 | 0.0025 |
| F04 | 57440001 | 57540000 | 0.0028 | 0.0017 | 0.0025 |
| F04 | 57460001 | 57560000 | 0.0022 | 0.0007 | 0.0021 |
| F04 | 57480001 | 57580000 | 0.0010 | 0.0007 | 0.0009 |
| F04 | 57500001 | 57600000 | 0.0008 | 0.0009 | 0.0008 |
| F04 | 57520001 | 57620000 | 0.0010 | 0.0008 | 0.0010 |
| F04 | 57540001 | 57640000 | 0.0010 | 0.0009 | 0.0010 |
| F04 | 57560001 | 57660000 | 0.0011 | 0.0013 | 0.0010 |
| F04 | 57580001 | 57680000 | 0.0015 | 0.0012 | 0.0015 |
| F04 | 57600001 | 57700000 | 0.0014 | 0.0011 | 0.0014 |
| F04 | 57620001 | 57720000 | 0.0014 | 0.0012 | 0.0013 |
| F04 | 57640001 | 57740000 | 0.0014 | 0.0015 | 0.0014 |
| F04 | 57660001 | 57760000 | 0.0017 | 0.0011 | 0.0016 |
| F04 | 57680001 | 57780000 | 0.0013 | 0.0011 | 0.0012 |
| F04 | 57700001 | 57800000 | 0.0013 | 0.0013 | 0.0012 |
| F04 | 57720001 | 57820000 | 0.0016 | 0.0018 | 0.0014 |
| F04 | 57740001 | 57840000 | 0.0023 | 0.0016 | 0.0020 |
| F04 | 57760001 | 57860000 | 0.0021 | 0.0020 | 0.0017 |
| F04 | 57780001 | 57880000 | 0.0026 | 0.0024 | 0.0021 |
| F04 | 57800001 | 57900000 | 0.0030 | 0.0021 | 0.0025 |
| F04 | 57820001 | 57920000 | 0.0027 | 0.0017 | 0.0022 |
| F04 | 57840001 | 57940000 | 0.0023 | 0.0016 | 0.0019 |
| F04 | 57860001 | 57960000 | 0.0021 | 0.0015 | 0.0018 |
| F04 | 57880001 | 57980000 | 0.0021 | 0.0014 | 0.0017 |
| F04 | 57900001 | 58000000 | 0.0020 | 0.0015 | 0.0016 |
| F04 | 57920001 | 58020000 | 0.0020 | 0.0012 | 0.0017 |
| F04 | 57940001 | 58040000 | 0.0016 | 0.0012 | 0.0013 |
| F04 | 57960001 | 58060000 | 0.0015 | 0.0010 | 0.0013 |
| F04 | 57980001 | 58080000 | 0.0012 | 0.0010 | 0.0011 |
| F04 | 58000001 | 58100000 | 0.0012 | 0.0010 | 0.0011 |
| F04 | 58020001 | 58120000 | 0.0012 | 0.0013 | 0.0012 |
| F04 | 58040001 | 58140000 | 0.0015 | 0.0013 | 0.0014 |
| F04 | 58060001 | 58160000 | 0.0016 | 0.0013 | 0.0014 |
| F04 | 58080001 | 58180000 | 0.0016 | 0.0016 | 0.0014 |
| F04 | 58100001 | 58200000 | 0.0019 | 0.0017 | 0.0018 |
| F04 | 58120001 | 58220000 | 0.0022 | 0.0019 | 0.0020 |
| F04 | 58140001 | 58240000 | 0.0023 | 0.0018 | 0.0022 |
| F04 | 58160001 | 58260000 | 0.0021 | 0.0016 | 0.0020 |
| F04 | 58180001 | 58280000 | 0.0018 | 0.0010 | 0.0018 |
| F04 | 58200001 | 58300000 | 0.0011 | 0.0007 | 0.0011 |
| F04 | 58220001 | 58320000 | 0.0007 | 0.0001 | 0.0007 |
| F04 | 58240001 | 58340000 | 0.0002 | 0.0003 | 0.0002 |

| Chr | Start | End | both | A1 | A2 |
|---|---|---|---|---|---|
| F04 | 58260001 | 58360000 | 0.0004 | 0.0007 | 0.0004 |
| F04 | 58280001 | 58380000 | 0.0008 | 0.0009 | 0.0008 |
| F04 | 58300001 | 58400000 | 0.0011 | 0.0014 | 0.0011 |
| F04 | 58320001 | 58420000 | 0.0014 | 0.0014 | 0.0014 |
| F04 | 58340001 | 58440000 | 0.0015 | 0.0016 | 0.0014 |
| F04 | 58360001 | 58460000 | 0.0016 | 0.0012 | 0.0015 |
| F04 | 58380001 | 58480000 | 0.0012 | 0.0010 | 0.0012 |
| F04 | 58400001 | 58500000 | 0.0010 | 0.0006 | 0.0010 |
| F04 | 58420001 | 58520000 | 0.0007 | 0.0005 | 0.0007 |
| F04 | 58440001 | 58540000 | 0.0006 | 0.0004 | 0.0006 |
| F04 | 58460001 | 58560000 | 0.0005 | 0.0006 | 0.0005 |
| F04 | 58480001 | 58580000 | 0.0008 | 0.0010 | 0.0007 |
| F04 | 58500001 | 58600000 | 0.0012 | 0.0016 | 0.0011 |
| F04 | 58520001 | 58620000 | 0.0018 | 0.0017 | 0.0017 |
| F04 | 58540001 | 58640000 | 0.0020 | 0.0016 | 0.0019 |
| F04 | 58560001 | 58660000 | 0.0022 | 0.0023 | 0.0021 |
| F04 | 58580001 | 58680000 | 0.0031 | 0.0020 | 0.0028 |
| F04 | 58600001 | 58700000 | 0.0027 | 0.0015 | 0.0025 |
| F04 | 58620001 | 58720000 | 0.0021 | 0.0014 | 0.0019 |
| F04 | 58640001 | 58740000 | 0.0020 | 0.0014 | 0.0018 |
| F04 | 58660001 | 58760000 | 0.0017 | 0.0016 | 0.0015 |
| F04 | 58680001 | 58780000 | 0.0017 | 0.0018 | 0.0014 |
| F04 | 58700001 | 58800000 | 0.0021 | 0.0017 | 0.0017 |
| F04 | 58720001 | 58820000 | 0.0024 | 0.0019 | 0.0016 |
| F04 | 58740001 | 58840000 | 0.0027 | 0.0019 | 0.0018 |
| F04 | 58760001 | 58860000 | 0.0028 | 0.0009 | 0.0018 |
| F04 | 58780001 | 58880000 | 0.0019 | 0.0010 | 0.0009 |
| F04 | 58800001 | 58900000 | 0.0019 | 0.0010 | 0.0011 |
| F04 | 58820001 | 58920000 | 0.0015 | 0.0013 | 0.0011 |
| F04 | 58840001 | 58940000 | 0.0019 | 0.0021 | 0.0014 |
| F04 | 58860001 | 58960000 | 0.0028 | 0.0023 | 0.0024 |
| F04 | 58880001 | 58980000 | 0.0029 | 0.0023 | 0.0027 |
| F04 | 58900001 | 59000000 | 0.0028 | 0.0025 | 0.0026 |
| F04 | 58920001 | 59020000 | 0.0030 | 0.0022 | 0.0028 |
| F04 | 58940001 | 59040000 | 0.0025 | 0.0018 | 0.0024 |
| F04 | 58960001 | 59060000 | 0.0021 | 0.0015 | 0.0019 |
| F04 | 58980001 | 59080000 | 0.0021 | 0.0013 | 0.0016 |
| F04 | 59000001 | 59100000 | 0.0019 | 0.0015 | 0.0014 |
| F04 | 59020001 | 59120000 | 0.0022 | 0.0014 | 0.0015 |
| F04 | 59040001 | 59140000 | 0.0021 | 0.0014 | 0.0014 |
| F04 | 59060001 | 59160000 | 0.0020 | 0.0014 | 0.0013 |
| F04 | 59080001 | 59180000 | 0.0018 | 0.0019 | 0.0013 |
| F04 | 59100001 | 59200000 | 0.0024 | 0.0022 | 0.0019 |
| F04 | 59120001 | 59220000 | 0.0027 | 0.0028 | 0.0022 |
| F04 | 59140001 | 59240000 | 0.0037 | 0.0030 | 0.0031 |
| F04 | 59160001 | 59260000 | 0.0042 | 0.0029 | 0.0034 |
| F04 | 59180001 | 59280000 | 0.0040 | 0.0025 | 0.0033 |
| F04 | 59200001 | 59300000 | 0.0036 | 0.0024 | 0.0029 |
| F04 | 59220001 | 59320000 | 0.0033 | 0.0028 | 0.0029 |
| F04 | 59240001 | 59340000 | 0.0033 | 0.0024 | 0.0029 |
| F04 | 59260001 | 59360000 | 0.0027 | 0.0026 | 0.0025 |
| F04 | 59280001 | 59380000 | 0.0029 | 0.0027 | 0.0027 |
| F04 | 59300001 | 59400000 | 0.0030 | 0.0021 | 0.0028 |
| F04 | 59320001 | 59420000 | 0.0024 | 0.0012 | 0.0023 |
| F04 | 59340001 | 59440000 | 0.0015 | 0.0012 | 0.0014 |
| F04 | 59360001 | 59460000 | 0.0015 | 0.0012 | 0.0014 |
| F04 | 59380001 | 59480000 | 0.0015 | 0.0012 | 0.0014 |
| F04 | 59400001 | 59500000 | 0.0016 | 0.0017 | 0.0014 |

| Chr | Start | End | both | A1 | A2 |
|---|---|---|---|---|---|
| F04 | 59420001 | 59520000 | 0.0021 | 0.0016 | 0.0019 |
| F04 | 59440001 | 59540000 | 0.0022 | 0.0017 | 0.0018 |
| F04 | 59460001 | 59560000 | 0.0025 | 0.0018 | 0.0019 |
| F04 | 59480001 | 59580000 | 0.0027 | 0.0016 | 0.0020 |
| F04 | 59500001 | 59600000 | 0.0025 | 0.0012 | 0.0019 |
| F04 | 59520001 | 59620000 | 0.0022 | 0.0012 | 0.0015 |
| F04 | 59540001 | 59640000 | 0.0019 | 0.0010 | 0.0014 |
| F04 | 59560001 | 59660000 | 0.0016 | 0.0007 | 0.0012 |
| F04 | 59580001 | 59680000 | 0.0013 | 0.0008 | 0.0009 |
| F04 | 59600001 | 59700000 | 0.0015 | 0.0011 | 0.0010 |
| F04 | 59620001 | 59720000 | 0.0017 | 0.0010 | 0.0013 |
| F04 | 59640001 | 59740000 | 0.0018 | 0.0015 | 0.0012 |
| F04 | 59660001 | 59760000 | 0.0023 | 0.0020 | 0.0018 |
| F04 | 59680001 | 59780000 | 0.0027 | 0.0021 | 0.0023 |
| F04 | 59700001 | 59800000 | 0.0027 | 0.0024 | 0.0024 |
| F04 | 59720001 | 59820000 | 0.0031 | 0.0026 | 0.0028 |
| F04 | 59740001 | 59840000 | 0.0031 | 0.0021 | 0.0030 |
| F04 | 59760001 | 59860000 | 0.0025 | 0.0018 | 0.0024 |
| F04 | 59780001 | 59880000 | 0.0023 | 0.0013 | 0.0019 |
| F04 | 59800001 | 59900000 | 0.0019 | 0.0009 | 0.0014 |
| F04 | 59820001 | 59920000 | 0.0013 | 0.0022 | 0.0009 |
| F04 | 59840001 | 59940000 | 0.0029 | 0.0026 | 0.0023 |
| F04 | 59860001 | 59960000 | 0.0035 | 0.0027 | 0.0026 |
| F04 | 59880001 | 59980000 | 0.0036 | 0.0028 | 0.0028 |
| F04 | 59900001 | 60000000 | 0.0037 | 0.0028 | 0.0030 |
| F04 | 59920001 | 60020000 | 0.0041 | 0.0017 | 0.0031 |
| F04 | 59940001 | 60040000 | 0.0030 | 0.0021 | 0.0021 |
| F04 | 59960001 | 60060000 | 0.0034 | 0.0021 | 0.0025 |
| F04 | 59980001 | 60080000 | 0.0033 | 0.0031 | 0.0025 |
| F04 | 60000001 | 60100000 | 0.0044 | 0.0038 | 0.0036 |
| F04 | 60020001 | 60120000 | 0.0047 | 0.0037 | 0.0041 |
| F04 | 60040001 | 60140000 | 0.0044 | 0.0038 | 0.0040 |
| F04 | 60060001 | 60160000 | 0.0047 | 0.0039 | 0.0042 |
| F04 | 60080001 | 60180000 | 0.0048 | 0.0031 | 0.0042 |
| F04 | 60100001 | 60200000 | 0.0039 | 0.0023 | 0.0033 |
| F04 | 60120001 | 60220000 | 0.0031 | 0.0022 | 0.0026 |
| F04 | 60140001 | 60240000 | 0.0028 | 0.0013 | 0.0024 |
| F04 | 60160001 | 60260000 | 0.0015 | 0.0010 | 0.0014 |
| F04 | 60180001 | 60280000 | 0.0013 | 0.0018 | 0.0012 |
| F04 | 60200001 | 60300000 | 0.0023 | 0.0024 | 0.0021 |
| F04 | 60220001 | 60320000 | 0.0029 | 0.0025 | 0.0025 |
| F04 | 60240001 | 60340000 | 0.0033 | 0.0027 | 0.0027 |
| F04 | 60260001 | 60360000 | 0.0035 | 0.0031 | 0.0030 |
| F04 | 60280001 | 60380000 | 0.0040 | 0.0022 | 0.0033 |
| F04 | 60300001 | 60400000 | 0.0030 | 0.0020 | 0.0023 |
| F04 | 60320001 | 60420000 | 0.0027 | 0.0023 | 0.0021 |
| F04 | 60340001 | 60440000 | 0.0032 | 0.0022 | 0.0025 |
| F04 | 60360001 | 60460000 | 0.0033 | 0.0018 | 0.0023 |
| F04 | 60380001 | 60480000 | 0.0028 | 0.0017 | 0.0019 |
| F04 | 60400001 | 60500000 | 0.0025 | 0.0012 | 0.0018 |
| F04 | 60420001 | 60520000 | 0.0020 | 0.0010 | 0.0013 |
| F04 | 60440001 | 60540000 | 0.0014 | 0.0014 | 0.0010 |
| F04 | 60460001 | 60560000 | 0.0016 | 0.0016 | 0.0015 |
| F04 | 60480001 | 60580000 | 0.0019 | 0.0017 | 0.0018 |
| F04 | 60500001 | 60600000 | 0.0022 | 0.0017 | 0.0021 |
| F04 | 60520001 | 60620000 | 0.0021 | 0.0015 | 0.0021 |
| F04 | 60540001 | 60640000 | 0.0019 | 0.0016 | 0.0019 |
| F04 | 60560001 | 60660000 | 0.0020 | 0.0017 | 0.0018 |

| Chr | Start | End | both | A1 | A2 |
|---|---|---|---|---|---|
| F04 | 60580001 | 60680000 | 0.0021 | 0.0024 | 0.0016 |
| F04 | 60600001 | 60700000 | 0.0029 | 0.0032 | 0.0022 |
| F04 | 60620001 | 60720000 | 0.0037 | 0.0035 | 0.0030 |
| F04 | 60640001 | 60740000 | 0.0040 | 0.0031 | 0.0033 |
| F04 | 60660001 | 60760000 | 0.0035 | 0.0029 | 0.0030 |
| F04 | 60680001 | 60780000 | 0.0033 | 0.0018 | 0.0029 |
| F04 | 60700001 | 60800000 | 0.0022 | 0.0011 | 0.0019 |
| F04 | 60720001 | 60820000 | 0.0016 | 0.0008 | 0.0012 |
| F04 | 60740001 | 60840000 | 0.0015 | 0.0011 | 0.0009 |
| F04 | 60760001 | 60860000 | 0.0019 | 0.0015 | 0.0013 |
| F04 | 60780001 | 60880000 | 0.0024 | 0.0029 | 0.0017 |
| F04 | 60800001 | 60900000 | 0.0039 | 0.0029 | 0.0032 |
| F04 | 60820001 | 60920000 | 0.0039 | 0.0030 | 0.0033 |
| F04 | 60840001 | 60940000 | 0.0037 | 0.0024 | 0.0034 |
| F04 | 60860001 | 60960000 | 0.0030 | 0.0018 | 0.0028 |
| F04 | 60880001 | 60980000 | 0.0021 | 0.0004 | 0.0021 |
| F04 | 60900001 | 61000000 | 0.0006 | 0.0005 | 0.0006 |
| F04 | 60920001 | 61020000 | 0.0007 | 0.0007 | 0.0007 |
| F04 | 60940001 | 61040000 | 0.0008 | 0.0009 | 0.0008 |
| F04 | 60960001 | 61060000 | 0.0012 | 0.0018 | 0.0012 |
| F04 | 60980001 | 61080000 | 0.0022 | 0.0024 | 0.0022 |
| F04 | 61000001 | 61100000 | 0.0029 | 0.0029 | 0.0027 |
| F04 | 61020001 | 61120000 | 0.0034 | 0.0028 | 0.0032 |
| F04 | 61040001 | 61140000 | 0.0036 | 0.0031 | 0.0033 |
| F04 | 61060001 | 61160000 | 0.0040 | 0.0023 | 0.0036 |
| F04 | 61080001 | 61180000 | 0.0035 | 0.0020 | 0.0029 |
| F04 | 61100001 | 61200000 | 0.0032 | 0.0022 | 0.0026 |
| F04 | 61120001 | 61220000 | 0.0036 | 0.0024 | 0.0028 |
| F04 | 61140001 | 61240000 | 0.0038 | 0.0021 | 0.0028 |
| F04 | 61160001 | 61260000 | 0.0033 | 0.0026 | 0.0024 |
| F04 | 61180001 | 61280000 | 0.0036 | 0.0030 | 0.0027 |
| F04 | 61200001 | 61300000 | 0.0039 | 0.0021 | 0.0030 |
| F04 | 61220001 | 61320000 | 0.0029 | 0.0023 | 0.0021 |
| F04 | 61240001 | 61340000 | 0.0029 | 0.0024 | 0.0022 |
| F04 | 61260001 | 61360000 | 0.0030 | 0.0021 | 0.0022 |
| F04 | 61280001 | 61380000 | 0.0027 | 0.0017 | 0.0020 |
| F04 | 61300001 | 61400000 | 0.0021 | 0.0021 | 0.0017 |
| F04 | 61320001 | 61420000 | 0.0026 | 0.0015 | 0.0023 |
| F04 | 61340001 | 61440000 | 0.0021 | 0.0016 | 0.0019 |
| F04 | 61360001 | 61460000 | 0.0023 | 0.0015 | 0.0021 |
| F04 | 61380001 | 61480000 | 0.0022 | 0.0013 | 0.0021 |
| F04 | 61400001 | 61500000 | 0.0022 | 0.0011 | 0.0019 |
| F04 | 61420001 | 61520000 | 0.0019 | 0.0015 | 0.0014 |
| F04 | 61440001 | 61540000 | 0.0025 | 0.0016 | 0.0019 |
| F04 | 61460001 | 61560000 | 0.0026 | 0.0018 | 0.0018 |
| F04 | 61480001 | 61580000 | 0.0029 | 0.0019 | 0.0019 |
| F04 | 61500001 | 61600000 | 0.0031 | 0.0022 | 0.0021 |
| F04 | 61520001 | 61620000 | 0.0037 | 0.0017 | 0.0026 |
| F04 | 61540001 | 61640000 | 0.0030 | 0.0017 | 0.0020 |
| F04 | 61560001 | 61660000 | 0.0030 | 0.0018 | 0.0021 |
| F04 | 61580001 | 61680000 | 0.0030 | 0.0023 | 0.0022 |
| F04 | 61600001 | 61700000 | 0.0035 | 0.0023 | 0.0027 |
| F04 | 61620001 | 61720000 | 0.0031 | 0.0027 | 0.0024 |
| F04 | 61640001 | 61740000 | 0.0036 | 0.0029 | 0.0029 |
| F04 | 61660001 | 61760000 | 0.0036 | 0.0032 | 0.0030 |
| F04 | 61680001 | 61780000 | 0.0036 | 0.0037 | 0.0031 |
| F04 | 61700001 | 61800000 | 0.0040 | 0.0039 | 0.0036 |
| F04 | 61720001 | 61820000 | 0.0042 | 0.0041 | 0.0040 |

| Chr | Start | End | both | A1 | A2 |
|---|---|---|---|---|---|
| F04 | 61740001 | 61840000 | 0.0042 | 0.0041 | 0.0040 |
| F04 | 61760001 | 61860000 | 0.0044 | 0.0037 | 0.0041 |
| F04 | 61780001 | 61880000 | 0.0043 | 0.0026 | 0.0038 |
| F04 | 61800001 | 61900000 | 0.0033 | 0.0024 | 0.0027 |
| F04 | 61820001 | 61920000 | 0.0032 | 0.0019 | 0.0025 |
| F04 | 61840001 | 61940000 | 0.0028 | 0.0018 | 0.0020 |
| F04 | 61860001 | 61960000 | 0.0027 | 0.0019 | 0.0019 |
| F04 | 61880001 | 61980000 | 0.0025 | 0.0020 | 0.0020 |
| F04 | 61900001 | 62000000 | 0.0025 | 0.0019 | 0.0021 |
| F04 | 61920001 | 62020000 | 0.0023 | 0.0020 | 0.0019 |
| F04 | 61940001 | 62040000 | 0.0023 | 0.0018 | 0.0021 |
| F04 | 61960001 | 62060000 | 0.0021 | 0.0016 | 0.0019 |
| F04 | 61980001 | 62080000 | 0.0020 | 0.0016 | 0.0018 |
| F04 | 62000001 | 62100000 | 0.0020 | 0.0017 | 0.0018 |
| F04 | 62020001 | 62120000 | 0.0021 | 0.0019 | 0.0020 |
| F04 | 62040001 | 62140000 | 0.0024 | 0.0020 | 0.0021 |
| F04 | 62060001 | 62160000 | 0.0027 | 0.0024 | 0.0022 |
| F04 | 62080001 | 62180000 | 0.0032 | 0.0026 | 0.0026 |
| F04 | 62100001 | 62200000 | 0.0036 | 0.0028 | 0.0028 |
| F04 | 62120001 | 62220000 | 0.0038 | 0.0036 | 0.0029 |
| F04 | 62140001 | 62240000 | 0.0047 | 0.0036 | 0.0038 |
| F04 | 62160001 | 62260000 | 0.0047 | 0.0032 | 0.0040 |
| F04 | 62180001 | 62280000 | 0.0040 | 0.0031 | 0.0034 |
| F04 | 62200001 | 62300000 | 0.0037 | 0.0028 | 0.0033 |
| F04 | 62220001 | 62320000 | 0.0034 | 0.0019 | 0.0032 |
| F04 | 62240001 | 62340000 | 0.0024 | 0.0018 | 0.0022 |
| F04 | 62260001 | 62360000 | 0.0022 | 0.0019 | 0.0020 |
| F04 | 62280001 | 62380000 | 0.0025 | 0.0016 | 0.0020 |
| F04 | 62300001 | 62400000 | 0.0023 | 0.0020 | 0.0017 |
| F04 | 62320001 | 62420000 | 0.0028 | 0.0022 | 0.0021 |
| F04 | 62340001 | 62440000 | 0.0029 | 0.0019 | 0.0023 |
| F04 | 62360001 | 62460000 | 0.0026 | 0.0022 | 0.0021 |
| F04 | 62380001 | 62480000 | 0.0026 | 0.0021 | 0.0023 |
| F04 | 62400001 | 62500000 | 0.0024 | 0.0016 | 0.0022 |
| F04 | 62420001 | 62520000 | 0.0021 | 0.0011 | 0.0017 |
| F04 | 62440001 | 62540000 | 0.0016 | 0.0014 | 0.0012 |
| F04 | 62460001 | 62560000 | 0.0019 | 0.0016 | 0.0014 |
| F04 | 62480001 | 62580000 | 0.0023 | 0.0026 | 0.0019 |
| F04 | 62500001 | 62600000 | 0.0035 | 0.0022 | 0.0029 |
| F04 | 62520001 | 62620000 | 0.0029 | 0.0034 | 0.0025 |
| F04 | 62540001 | 62640000 | 0.0041 | 0.0039 | 0.0036 |
| F04 | 62560001 | 62660000 | 0.0046 | 0.0033 | 0.0040 |
| F04 | 62580001 | 62680000 | 0.0039 | 0.0026 | 0.0033 |
| F04 | 62600001 | 62700000 | 0.0028 | 0.0026 | 0.0025 |
| F04 | 62620001 | 62720000 | 0.0028 | 0.0017 | 0.0025 |
| F04 | 62640001 | 62740000 | 0.0017 | 0.0015 | 0.0015 |
| F04 | 62660001 | 62760000 | 0.0017 | 0.0020 | 0.0016 |
| F04 | 62680001 | 62780000 | 0.0022 | 0.0021 | 0.0022 |
| F04 | 62700001 | 62800000 | 0.0023 | 0.0022 | 0.0023 |
| F04 | 62720001 | 62820000 | 0.0026 | 0.0028 | 0.0024 |
| F04 | 62740001 | 62840000 | 0.0033 | 0.0024 | 0.0030 |
| F04 | 62760001 | 62860000 | 0.0028 | 0.0022 | 0.0023 |
| F04 | 62780001 | 62880000 | 0.0026 | 0.0023 | 0.0020 |
| F04 | 62800001 | 62900000 | 0.0028 | 0.0023 | 0.0023 |
| F04 | 62820001 | 62920000 | 0.0028 | 0.0019 | 0.0022 |
| F04 | 62840001 | 62940000 | 0.0024 | 0.0015 | 0.0017 |
| F04 | 62860001 | 62960000 | 0.0020 | 0.0015 | 0.0015 |
| F04 | 62880001 | 62980000 | 0.0020 | 0.0015 | 0.0015 |

| Chr | Start | End | both | A1 | A2 |
|---|---|---|---|---|---|
| F04 | 62900001 | 63000000 | 0.0019 | 0.0014 | 0.0014 |
| F04 | 62920001 | 63020000 | 0.0020 | 0.0015 | 0.0014 |
| F04 | 62940001 | 63040000 | 0.0022 | 0.0016 | 0.0017 |
| F04 | 62960001 | 63060000 | 0.0023 | 0.0016 | 0.0018 |
| F04 | 62980001 | 63080000 | 0.0025 | 0.0018 | 0.0019 |
| F04 | 63000001 | 63100000 | 0.0025 | 0.0021 | 0.0021 |
| F04 | 63020001 | 63120000 | 0.0027 | 0.0017 | 0.0024 |
| F04 | 63040001 | 63140000 | 0.0020 | 0.0016 | 0.0019 |
| F04 | 63060001 | 63160000 | 0.0019 | 0.0011 | 0.0018 |
| F04 | 63080001 | 63180000 | 0.0012 | 0.0011 | 0.0012 |
| F04 | 63100001 | 63200000 | 0.0012 | 0.0009 | 0.0012 |
| F04 | 63120001 | 63220000 | 0.0010 | 0.0011 | 0.0010 |
| F04 | 63140001 | 63240000 | 0.0012 | 0.0012 | 0.0011 |
| F04 | 63160001 | 63260000 | 0.0013 | 0.0015 | 0.0013 |
| F04 | 63180001 | 63280000 | 0.0016 | 0.0013 | 0.0016 |
| F04 | 63200001 | 63300000 | 0.0015 | 0.0015 | 0.0014 |
| F04 | 63220001 | 63320000 | 0.0017 | 0.0017 | 0.0017 |
| F04 | 63240001 | 63340000 | 0.0019 | 0.0015 | 0.0018 |
| F04 | 63260001 | 63360000 | 0.0018 | 0.0013 | 0.0017 |
| F04 | 63280001 | 63380000 | 0.0016 | 0.0012 | 0.0016 |
| F04 | 63300001 | 63400000 | 0.0014 | 0.0008 | 0.0014 |
| F04 | 63320001 | 63420000 | 0.0009 | 0.0005 | 0.0009 |
| F04 | 63340001 | 63440000 | 0.0007 | 0.0005 | 0.0007 |
| F04 | 63360001 | 63460000 | 0.0008 | 0.0004 | 0.0007 |
| F04 | 63380001 | 63480000 | 0.0007 | 0.0003 | 0.0005 |
| F04 | 63400001 | 63500000 | 0.0006 | 0.0009 | 0.0004 |
| F04 | 63420001 | 63520000 | 0.0013 | 0.0016 | 0.0011 |
| F04 | 63440001 | 63540000 | 0.0020 | 0.0026 | 0.0019 |
| F04 | 63460001 | 63560000 | 0.0031 | 0.0036 | 0.0030 |
| F04 | 63480001 | 63580000 | 0.0043 | 0.0036 | 0.0041 |
| F04 | 63500001 | 63600000 | 0.0045 | 0.0034 | 0.0041 |
| F04 | 63520001 | 63620000 | 0.0043 | 0.0028 | 0.0038 |
| F04 | 63540001 | 63640000 | 0.0038 | 0.0021 | 0.0032 |
| F04 | 63560001 | 63660000 | 0.0029 | 0.0011 | 0.0022 |
| F04 | 63580001 | 63680000 | 0.0017 | 0.0012 | 0.0012 |
| F04 | 63600001 | 63700000 | 0.0017 | 0.0008 | 0.0013 |
| F04 | 63620001 | 63720000 | 0.0012 | 0.0006 | 0.0009 |
| F04 | 63640001 | 63740000 | 0.0007 | 0.0005 | 0.0006 |
| F04 | 63660001 | 63760000 | 0.0006 | 0.0005 | 0.0006 |
| F04 | 63680001 | 63780000 | 0.0006 | 0.0005 | 0.0006 |
| F04 | 63700001 | 63800000 | 0.0006 | 0.0008 | 0.0005 |
| F04 | 63720001 | 63820000 | 0.0010 | 0.0010 | 0.0010 |
| F04 | 63740001 | 63840000 | 0.0013 | 0.0011 | 0.0012 |
| F04 | 63760001 | 63860000 | 0.0014 | 0.0014 | 0.0013 |
| F04 | 63780001 | 63880000 | 0.0018 | 0.0019 | 0.0016 |
| F04 | 63800001 | 63900000 | 0.0024 | 0.0016 | 0.0022 |
| F04 | 63820001 | 63920000 | 0.0020 | 0.0019 | 0.0018 |
| F04 | 63840001 | 63940000 | 0.0023 | 0.0025 | 0.0021 |
| F04 | 63860001 | 63960000 | 0.0029 | 0.0027 | 0.0028 |
| F04 | 63880001 | 63980000 | 0.0031 | 0.0022 | 0.0029 |
| F04 | 63900001 | 64000000 | 0.0025 | 0.0024 | 0.0023 |
| F04 | 63920001 | 64020000 | 0.0026 | 0.0021 | 0.0025 |
| F04 | 63940001 | 64040000 | 0.0023 | 0.0014 | 0.0022 |
| F04 | 63960001 | 64060000 | 0.0016 | 0.0012 | 0.0015 |
| F04 | 63980001 | 64080000 | 0.0014 | 0.0012 | 0.0013 |
| F04 | 64000001 | 64100000 | 0.0015 | 0.0010 | 0.0014 |
| F04 | 64020001 | 64120000 | 0.0012 | 0.0015 | 0.0011 |
| F04 | 64040001 | 64140000 | 0.0019 | 0.0017 | 0.0017 |

| Chr | Start | End | both | A1 | A2 |
|---|---|---|---|---|---|
| F04 | 64060001 | 64160000 | 0.0022 | 0.0017 | 0.0019 |
| F04 | 64080001 | 64180000 | 0.0023 | 0.0016 | 0.0020 |
| F04 | 64100001 | 64200000 | 0.0023 | 0.0018 | 0.0019 |
| F04 | 64120001 | 64220000 | 0.0026 | 0.0017 | 0.0021 |
| F04 | 64140001 | 64240000 | 0.0024 | 0.0017 | 0.0019 |
| F04 | 64160001 | 64260000 | 0.0024 | 0.0022 | 0.0019 |
| F04 | 64180001 | 64280000 | 0.0030 | 0.0031 | 0.0024 |
| F04 | 64200001 | 64300000 | 0.0040 | 0.0036 | 0.0034 |
| F04 | 64220001 | 64320000 | 0.0045 | 0.0041 | 0.0039 |
| F04 | 64240001 | 64340000 | 0.0050 | 0.0042 | 0.0044 |
| F04 | 64260001 | 64360000 | 0.0050 | 0.0040 | 0.0045 |
| F04 | 64280001 | 64380000 | 0.0045 | 0.0038 | 0.0042 |
| F04 | 64300001 | 64400000 | 0.0043 | 0.0042 | 0.0038 |
| F04 | 64320001 | 64420000 | 0.0048 | 0.0041 | 0.0044 |
| F04 | 64340001 | 64440000 | 0.0048 | 0.0042 | 0.0045 |
| F04 | 64360001 | 64460000 | 0.0051 | 0.0046 | 0.0045 |
| F04 | 64380001 | 64480000 | 0.0056 | 0.0038 | 0.0049 |
| F04 | 64400001 | 64500000 | 0.0047 | 0.0033 | 0.0041 |
| F04 | 64420001 | 64520000 | 0.0045 | 0.0034 | 0.0036 |
| F04 | 64440001 | 64540000 | 0.0046 | 0.0031 | 0.0037 |
| F04 | 64460001 | 64560000 | 0.0043 | 0.0024 | 0.0034 |
| F04 | 64480001 | 64580000 | 0.0036 | 0.0034 | 0.0028 |
| F04 | 64500001 | 64600000 | 0.0054 | 0.0028 | 0.0047 |
| F04 | 64520001 | 64620000 | 0.0045 | 0.0020 | 0.0041 |
| F04 | 64540001 | 64640000 | 0.0035 | 0.0020 | 0.0031 |
| F04 | 64560001 | 64660000 | 0.0034 | 0.0019 | 0.0031 |
| F04 | 64580001 | 64680000 | 0.0030 | 0.0017 | 0.0028 |
| F04 | 64600001 | 64700000 | 0.0022 | 0.0017 | 0.0020 |
| F04 | 64620001 | 64720000 | 0.0025 | 0.0016 | 0.0022 |
| F04 | 64640001 | 64740000 | 0.0026 | 0.0019 | 0.0022 |
| F04 | 64660001 | 64760000 | 0.0030 | 0.0031 | 0.0026 |
| F04 | 64680001 | 64780000 | 0.0045 | 0.0024 | 0.0041 |
| F04 | 64700001 | 64800000 | 0.0036 | 0.0029 | 0.0032 |
| F04 | 64720001 | 64820000 | 0.0041 | 0.0041 | 0.0038 |
| F04 | 64740001 | 64840000 | 0.0054 | 0.0035 | 0.0050 |
| F04 | 64760001 | 64860000 | 0.0047 | 0.0029 | 0.0042 |
| F04 | 64780001 | 64880000 | 0.0039 | 0.0029 | 0.0033 |
| F04 | 64800001 | 64900000 | 0.0041 | 0.0033 | 0.0033 |
| F04 | 64820001 | 64920000 | 0.0046 | 0.0026 | 0.0036 |
| F04 | 64840001 | 64940000 | 0.0036 | 0.0035 | 0.0027 |
| F04 | 64860001 | 64960000 | 0.0047 | 0.0044 | 0.0038 |
| F04 | 64880001 | 64980000 | 0.0055 | 0.0051 | 0.0046 |
| F04 | 64900001 | 65000000 | 0.0061 | 0.0045 | 0.0054 |
| F04 | 64920001 | 65020000 | 0.0054 | 0.0040 | 0.0047 |
| F04 | 64940001 | 65040000 | 0.0050 | 0.0030 | 0.0041 |
| F04 | 64960001 | 65060000 | 0.0042 | 0.0021 | 0.0031 |
| F04 | 64980001 | 65080000 | 0.0032 | 0.0021 | 0.0021 |
| F04 | 65000001 | 65100000 | 0.0034 | 0.0025 | 0.0022 |
| F04 | 65020001 | 65120000 | 0.0038 | 0.0030 | 0.0029 |
| F04 | 65040001 | 65140000 | 0.0041 | 0.0031 | 0.0034 |
| F04 | 65060001 | 65160000 | 0.0040 | 0.0034 | 0.0035 |
| F04 | 65080001 | 65180000 | 0.0045 | 0.0033 | 0.0041 |
| F04 | 65100001 | 65200000 | 0.0042 | 0.0031 | 0.0041 |
| F04 | 65120001 | 65220000 | 0.0040 | 0.0030 | 0.0036 |
| F04 | 65140001 | 65240000 | 0.0040 | 0.0038 | 0.0035 |
| F04 | 65160001 | 65260000 | 0.0050 | 0.0036 | 0.0044 |
| F04 | 65180001 | 65280000 | 0.0044 | 0.0036 | 0.0039 |
| F04 | 65200001 | 65300000 | 0.0043 | 0.0033 | 0.0038 |

| Chr | Start | End | both | A1 | A2 |
|---|---|---|---|---|---|
| F04 | 65220001 | 65320000 | 0.0039 | 0.0034 | 0.0035 |
| F04 | 65240001 | 65340000 | 0.0040 | 0.0026 | 0.0037 |
| F04 | 65260001 | 65360000 | 0.0029 | 0.0023 | 0.0028 |
| F04 | 65280001 | 65380000 | 0.0027 | 0.0019 | 0.0025 |
| F04 | 65300001 | 65400000 | 0.0023 | 0.0018 | 0.0019 |
| F04 | 65320001 | 65420000 | 0.0024 | 0.0013 | 0.0019 |
| F04 | 65340001 | 65440000 | 0.0020 | 0.0015 | 0.0013 |
| F04 | 65360001 | 65460000 | 0.0023 | 0.0012 | 0.0014 |
| F04 | 65380001 | 65480000 | 0.0020 | 0.0009 | 0.0012 |
| F04 | 65400001 | 65500000 | 0.0019 | 0.0007 | 0.0010 |
| F04 | 65420001 | 65520000 | 0.0017 | 0.0012 | 0.0008 |
| F04 | 65440001 | 65540000 | 0.0022 | 0.0012 | 0.0014 |
| F04 | 65460001 | 65560000 | 0.0022 | 0.0013 | 0.0014 |
| F04 | 65480001 | 65580000 | 0.0024 | 0.0016 | 0.0016 |
| F04 | 65500001 | 65600000 | 0.0023 | 0.0019 | 0.0018 |
| F04 | 65520001 | 65620000 | 0.0026 | 0.0021 | 0.0023 |
| F04 | 65540001 | 65640000 | 0.0029 | 0.0022 | 0.0025 |
| F04 | 65560001 | 65660000 | 0.0028 | 0.0022 | 0.0025 |
| F04 | 65580001 | 65680000 | 0.0027 | 0.0024 | 0.0025 |
| F04 | 65600001 | 65700000 | 0.0030 | 0.0027 | 0.0027 |
| F04 | 65620001 | 65720000 | 0.0035 | 0.0021 | 0.0031 |
| F04 | 65640001 | 65740000 | 0.0027 | 0.0021 | 0.0023 |
| F04 | 65660001 | 65760000 | 0.0028 | 0.0020 | 0.0022 |
| F04 | 65680001 | 65780000 | 0.0028 | 0.0019 | 0.0021 |
| F04 | 65700001 | 65800000 | 0.0028 | 0.0016 | 0.0020 |
| F04 | 65720001 | 65820000 | 0.0025 | 0.0016 | 0.0016 |
| F04 | 65740001 | 65840000 | 0.0024 | 0.0013 | 0.0017 |
| F04 | 65760001 | 65860000 | 0.0020 | 0.0013 | 0.0014 |
| F04 | 65780001 | 65880000 | 0.0018 | 0.0013 | 0.0013 |
| F04 | 65800001 | 65900000 | 0.0016 | 0.0014 | 0.0012 |
| F04 | 65820001 | 65920000 | 0.0016 | 0.0014 | 0.0015 |
| F04 | 65840001 | 65940000 | 0.0016 | 0.0016 | 0.0015 |
| F04 | 65860001 | 65960000 | 0.0021 | 0.0025 | 0.0019 |
| F04 | 65880001 | 65980000 | 0.0031 | 0.0025 | 0.0027 |
| F04 | 65900001 | 66000000 | 0.0033 | 0.0027 | 0.0028 |
| F04 | 65920001 | 66020000 | 0.0035 | 0.0026 | 0.0030 |
| F04 | 65940001 | 66040000 | 0.0034 | 0.0028 | 0.0028 |
| F04 | 65960001 | 66060000 | 0.0036 | 0.0025 | 0.0029 |
| F04 | 65980001 | 66080000 | 0.0033 | 0.0035 | 0.0026 |
| F04 | 66000001 | 66100000 | 0.0045 | 0.0029 | 0.0038 |
| F04 | 66020001 | 66120000 | 0.0038 | 0.0036 | 0.0030 |
| F04 | 66040001 | 66140000 | 0.0047 | 0.0037 | 0.0039 |
| F04 | 66060001 | 66160000 | 0.0047 | 0.0041 | 0.0041 |
| F04 | 66080001 | 66180000 | 0.0051 | 0.0042 | 0.0044 |
| F04 | 66100001 | 66200000 | 0.0050 | 0.0045 | 0.0042 |
| F04 | 66120001 | 66220000 | 0.0054 | 0.0040 | 0.0046 |
| F04 | 66140001 | 66240000 | 0.0047 | 0.0045 | 0.0040 |
| F04 | 66160001 | 66260000 | 0.0054 | 0.0037 | 0.0047 |
| F04 | 66180001 | 66280000 | 0.0046 | 0.0025 | 0.0039 |
| F04 | 66200001 | 66300000 | 0.0033 | 0.0031 | 0.0027 |
| F04 | 66220001 | 66320000 | 0.0042 | 0.0032 | 0.0035 |
| F04 | 66240001 | 66340000 | 0.0044 | 0.0027 | 0.0036 |
| F04 | 66260001 | 66360000 | 0.0040 | 0.0031 | 0.0029 |
| F04 | 66280001 | 66380000 | 0.0043 | 0.0034 | 0.0032 |
| F04 | 66300001 | 66400000 | 0.0050 | 0.0031 | 0.0037 |
| F04 | 66320001 | 66420000 | 0.0043 | 0.0039 | 0.0032 |
| F04 | 66340001 | 66440000 | 0.0049 | 0.0036 | 0.0041 |
| F04 | 66360001 | 66460000 | 0.0044 | 0.0031 | 0.0038 |

| Chr | Start | End | both | A1 | A2 |
|---|---|---|---|---|---|
| F04 | 66380001 | 66480000 | 0.0040 | 0.0025 | 0.0035 |
| F04 | 66400001 | 66500000 | 0.0033 | 0.0024 | 0.0027 |
| F04 | 66420001 | 66520000 | 0.0035 | 0.0020 | 0.0027 |
| F04 | 66440001 | 66540000 | 0.0033 | 0.0021 | 0.0021 |
| F04 | 66460001 | 66560000 | 0.0036 | 0.0027 | 0.0022 |
| F04 | 66480001 | 66580000 | 0.0040 | 0.0028 | 0.0028 |
| F04 | 66500001 | 66600000 | 0.0039 | 0.0026 | 0.0030 |
| F04 | 66520001 | 66620000 | 0.0035 | 0.0019 | 0.0027 |
| F04 | 66540001 | 66640000 | 0.0026 | 0.0015 | 0.0020 |
| F04 | 66560001 | 66660000 | 0.0022 | 0.0017 | 0.0018 |
| F04 | 66580001 | 66680000 | 0.0024 | 0.0015 | 0.0019 |
| F04 | 66600001 | 66700000 | 0.0022 | 0.0011 | 0.0017 |
| F04 | 66620001 | 66720000 | 0.0017 | 0.0013 | 0.0012 |
| F04 | 66640001 | 66740000 | 0.0020 | 0.0013 | 0.0014 |
| F04 | 66660001 | 66760000 | 0.0019 | 0.0008 | 0.0014 |
| F04 | 66680001 | 66780000 | 0.0012 | 0.0011 | 0.0009 |
| F04 | 66700001 | 66800000 | 0.0017 | 0.0015 | 0.0013 |
| F04 | 66720001 | 66820000 | 0.0023 | 0.0019 | 0.0018 |
| F04 | 66740001 | 66840000 | 0.0027 | 0.0020 | 0.0021 |
| F04 | 66760001 | 66860000 | 0.0030 | 0.0020 | 0.0022 |
| F04 | 66780001 | 66880000 | 0.0033 | 0.0024 | 0.0022 |
| F04 | 66800001 | 66900000 | 0.0039 | 0.0023 | 0.0027 |
| F04 | 66820001 | 66920000 | 0.0037 | 0.0024 | 0.0026 |
| F04 | 66840001 | 66940000 | 0.0037 | 0.0026 | 0.0026 |
| F04 | 66860001 | 66960000 | 0.0040 | 0.0030 | 0.0030 |
| F04 | 66880001 | 66980000 | 0.0042 | 0.0026 | 0.0033 |
| F04 | 66900001 | 67000000 | 0.0036 | 0.0023 | 0.0030 |
| F04 | 66920001 | 67020000 | 0.0031 | 0.0021 | 0.0027 |
| F04 | 66940001 | 67040000 | 0.0027 | 0.0021 | 0.0025 |
| F04 | 66960001 | 67060000 | 0.0025 | 0.0021 | 0.0025 |
| F04 | 66980001 | 67080000 | 0.0027 | 0.0020 | 0.0026 |
| F04 | 67000001 | 67100000 | 0.0026 | 0.0024 | 0.0024 |
| F04 | 67020001 | 67120000 | 0.0030 | 0.0026 | 0.0028 |
| F04 | 67040001 | 67140000 | 0.0034 | 0.0021 | 0.0032 |
| F04 | 67060001 | 67160000 | 0.0030 | 0.0022 | 0.0028 |
| F04 | 67080001 | 67180000 | 0.0029 | 0.0024 | 0.0026 |
| F04 | 67100001 | 67200000 | 0.0031 | 0.0030 | 0.0029 |
| F04 | 67120001 | 67220000 | 0.0037 | 0.0029 | 0.0034 |
| F04 | 67140001 | 67240000 | 0.0034 | 0.0028 | 0.0031 |
| F04 | 67160001 | 67260000 | 0.0034 | 0.0027 | 0.0029 |
| F04 | 67180001 | 67280000 | 0.0032 | 0.0023 | 0.0027 |
| F04 | 67200001 | 67300000 | 0.0027 | 0.0014 | 0.0022 |
| F04 | 67220001 | 67320000 | 0.0018 | 0.0013 | 0.0013 |
| F04 | 67240001 | 67340000 | 0.0018 | 0.0019 | 0.0012 |
| F04 | 67260001 | 67360000 | 0.0026 | 0.0024 | 0.0021 |
| F04 | 67280001 | 67380000 | 0.0032 | 0.0024 | 0.0027 |
| F04 | 67300001 | 67400000 | 0.0032 | 0.0025 | 0.0027 |
| F04 | 67320001 | 67420000 | 0.0034 | 0.0028 | 0.0030 |
| F04 | 67340001 | 67440000 | 0.0036 | 0.0029 | 0.0033 |
| F04 | 67360001 | 67460000 | 0.0035 | 0.0024 | 0.0030 |
| F04 | 67380001 | 67480000 | 0.0030 | 0.0032 | 0.0025 |
| F04 | 67400001 | 67500000 | 0.0040 | 0.0044 | 0.0035 |
| F04 | 67420001 | 67520000 | 0.0054 | 0.0042 | 0.0048 |
| F04 | 67440001 | 67540000 | 0.0053 | 0.0036 | 0.0046 |
| F04 | 67460001 | 67560000 | 0.0047 | 0.0032 | 0.0041 |
| F04 | 67480001 | 67580000 | 0.0041 | 0.0026 | 0.0036 |
| F04 | 67500001 | 67600000 | 0.0036 | 0.0013 | 0.0029 |
| F04 | 67520001 | 67620000 | 0.0020 | 0.0010 | 0.0014 |

| Chr | Start | End | both | A1 | A2 |
|---|---|---|---|---|---|
| F04 | 67540001 | 67640000 | 0.0015 | 0.0015 | 0.0012 |
| F04 | 67560001 | 67660000 | 0.0019 | 0.0015 | 0.0017 |
| F04 | 67580001 | 67680000 | 0.0019 | 0.0011 | 0.0018 |
| F04 | 67600001 | 67700000 | 0.0014 | 0.0015 | 0.0014 |
| F04 | 67620001 | 67720000 | 0.0020 | 0.0021 | 0.0019 |
| F04 | 67640001 | 67740000 | 0.0028 | 0.0016 | 0.0025 |
| F04 | 67660001 | 67760000 | 0.0023 | 0.0018 | 0.0019 |
| F04 | 67680001 | 67780000 | 0.0025 | 0.0028 | 0.0021 |
| F04 | 67700001 | 67800000 | 0.0037 | 0.0043 | 0.0032 |
| F04 | 67720001 | 67820000 | 0.0054 | 0.0045 | 0.0049 |
| F04 | 67740001 | 67840000 | 0.0057 | 0.0046 | 0.0051 |
| F04 | 67760001 | 67860000 | 0.0059 | 0.0047 | 0.0053 |
| F04 | 67780001 | 67880000 | 0.0059 | 0.0037 | 0.0053 |
| F04 | 67800001 | 67900000 | 0.0047 | 0.0027 | 0.0041 |
| F04 | 67820001 | 67920000 | 0.0035 | 0.0020 | 0.0029 |
| F04 | 67840001 | 67940000 | 0.0027 | 0.0020 | 0.0021 |
| F04 | 67860001 | 67960000 | 0.0025 | 0.0019 | 0.0021 |
| F04 | 67880001 | 67980000 | 0.0024 | 0.0020 | 0.0020 |
| F04 | 67900001 | 68000000 | 0.0028 | 0.0009 | 0.0021 |
| F04 | 67920001 | 68020000 | 0.0018 | 0.0009 | 0.0010 |
| F04 | 67940001 | 68040000 | 0.0018 | 0.0009 | 0.0010 |
| F04 | 67960001 | 68060000 | 0.0020 | 0.0008 | 0.0009 |
| F04 | 67980001 | 68080000 | 0.0021 | 0.0011 | 0.0009 |
| F04 | 68000001 | 68100000 | 0.0022 | 0.0018 | 0.0012 |
| F04 | 68020001 | 68120000 | 0.0028 | 0.0016 | 0.0021 |
| F04 | 68040001 | 68140000 | 0.0024 | 0.0015 | 0.0018 |
| F04 | 68060001 | 68160000 | 0.0021 | 0.0017 | 0.0017 |
| F04 | 68080001 | 68180000 | 0.0027 | 0.0015 | 0.0020 |
| F04 | 68100001 | 68200000 | 0.0026 | 0.0011 | 0.0017 |
| F04 | 68120001 | 68220000 | 0.0021 | 0.0014 | 0.0013 |
| F04 | 68140001 | 68240000 | 0.0023 | 0.0021 | 0.0016 |
| F04 | 68160001 | 68260000 | 0.0033 | 0.0025 | 0.0024 |
| F04 | 68180001 | 68280000 | 0.0035 | 0.0024 | 0.0030 |
| F04 | 68200001 | 68300000 | 0.0032 | 0.0028 | 0.0030 |
| F04 | 68220001 | 68320000 | 0.0037 | 0.0029 | 0.0034 |
| F04 | 68240001 | 68340000 | 0.0037 | 0.0029 | 0.0035 |
| F04 | 68260001 | 68360000 | 0.0036 | 0.0025 | 0.0033 |
| F04 | 68280001 | 68380000 | 0.0031 | 0.0026 | 0.0027 |
| F04 | 68300001 | 68400000 | 0.0035 | 0.0033 | 0.0029 |
| F04 | 68320001 | 68420000 | 0.0042 | 0.0029 | 0.0035 |
| F04 | 68340001 | 68440000 | 0.0037 | 0.0026 | 0.0031 |
| F04 | 68360001 | 68460000 | 0.0035 | 0.0031 | 0.0029 |
| F04 | 68380001 | 68480000 | 0.0041 | 0.0034 | 0.0034 |
| F04 | 68400001 | 68500000 | 0.0041 | 0.0021 | 0.0036 |
| F04 | 68420001 | 68520000 | 0.0027 | 0.0022 | 0.0022 |
| F04 | 68440001 | 68540000 | 0.0030 | 0.0022 | 0.0024 |
| F04 | 68460001 | 68560000 | 0.0028 | 0.0019 | 0.0024 |
| F04 | 68480001 | 68580000 | 0.0025 | 0.0017 | 0.0021 |
| F04 | 68500001 | 68600000 | 0.0025 | 0.0019 | 0.0019 |
| F04 | 68520001 | 68620000 | 0.0029 | 0.0022 | 0.0021 |
| F04 | 68540001 | 68640000 | 0.0031 | 0.0021 | 0.0023 |
| F04 | 68560001 | 68660000 | 0.0030 | 0.0017 | 0.0022 |
| F04 | 68580001 | 68680000 | 0.0025 | 0.0016 | 0.0017 |
| F04 | 68600001 | 68700000 | 0.0024 | 0.0017 | 0.0018 |
| F04 | 68620001 | 68720000 | 0.0022 | 0.0015 | 0.0018 |
| F04 | 68640001 | 68740000 | 0.0021 | 0.0012 | 0.0018 |
| F04 | 68660001 | 68760000 | 0.0018 | 0.0015 | 0.0015 |
| F04 | 68680001 | 68780000 | 0.0020 | 0.0014 | 0.0019 |

| Chr | Start | End | both | A1 | A2 |
|---|---|---|---|---|---|
| F04 | 68700001 | 68800000 | 0.0019 | 0.0011 | 0.0017 |
| F04 | 68720001 | 68820000 | 0.0015 | 0.0018 | 0.0013 |
| F04 | 68740001 | 68840000 | 0.0023 | 0.0024 | 0.0020 |
| F04 | 68760001 | 68860000 | 0.0030 | 0.0021 | 0.0028 |
| F04 | 68780001 | 68880000 | 0.0030 | 0.0027 | 0.0025 |
| F04 | 68800001 | 68900000 | 0.0037 | 0.0032 | 0.0032 |
| F04 | 68820001 | 68920000 | 0.0043 | 0.0036 | 0.0037 |
| F04 | 68840001 | 68940000 | 0.0050 | 0.0040 | 0.0044 |
| F04 | 68860001 | 68960000 | 0.0054 | 0.0047 | 0.0046 |
| F04 | 68880001 | 68980000 | 0.0060 | 0.0045 | 0.0054 |
| F04 | 68900001 | 69000000 | 0.0058 | 0.0041 | 0.0050 |
| F04 | 68920001 | 69020000 | 0.0054 | 0.0034 | 0.0045 |
| F04 | 68940001 | 69040000 | 0.0043 | 0.0032 | 0.0035 |
| F04 | 68960001 | 69060000 | 0.0042 | 0.0036 | 0.0032 |
| F04 | 68980001 | 69080000 | 0.0047 | 0.0030 | 0.0037 |
| F04 | 69000001 | 69100000 | 0.0039 | 0.0029 | 0.0031 |
| F04 | 69020001 | 69120000 | 0.0037 | 0.0025 | 0.0030 |
| F04 | 69040001 | 69140000 | 0.0032 | 0.0022 | 0.0025 |
| F04 | 69060001 | 69160000 | 0.0030 | 0.0015 | 0.0024 |
| F04 | 69080001 | 69180000 | 0.0024 | 0.0017 | 0.0015 |
| F04 | 69100001 | 69200000 | 0.0026 | 0.0018 | 0.0017 |
| F04 | 69120001 | 69220000 | 0.0027 | 0.0016 | 0.0018 |
| F04 | 69140001 | 69240000 | 0.0026 | 0.0022 | 0.0017 |
| F04 | 69160001 | 69260000 | 0.0030 | 0.0018 | 0.0021 |
| F04 | 69180001 | 69280000 | 0.0026 | 0.0020 | 0.0019 |
| F04 | 69200001 | 69300000 | 0.0030 | 0.0022 | 0.0021 |
| F04 | 69220001 | 69320000 | 0.0036 | 0.0028 | 0.0023 |
| F04 | 69240001 | 69340000 | 0.0043 | 0.0021 | 0.0029 |
| F04 | 69260001 | 69360000 | 0.0036 | 0.0025 | 0.0022 |
| F04 | 69280001 | 69380000 | 0.0040 | 0.0029 | 0.0028 |
| F04 | 69300001 | 69400000 | 0.0044 | 0.0032 | 0.0031 |
| F04 | 69320001 | 69420000 | 0.0043 | 0.0030 | 0.0034 |
| F04 | 69340001 | 69440000 | 0.0041 | 0.0032 | 0.0032 |
| F04 | 69360001 | 69460000 | 0.0042 | 0.0041 | 0.0034 |
| F04 | 69380001 | 69480000 | 0.0053 | 0.0036 | 0.0044 |
| F04 | 69400001 | 69500000 | 0.0049 | 0.0031 | 0.0040 |
| F04 | 69420001 | 69520000 | 0.0043 | 0.0026 | 0.0035 |
| F04 | 69440001 | 69540000 | 0.0037 | 0.0027 | 0.0030 |
| F04 | 69460001 | 69560000 | 0.0036 | 0.0020 | 0.0030 |
| F04 | 69480001 | 69580000 | 0.0027 | 0.0029 | 0.0019 |
| F04 | 69500001 | 69600000 | 0.0036 | 0.0032 | 0.0028 |
| F04 | 69520001 | 69620000 | 0.0040 | 0.0035 | 0.0031 |
| F04 | 69540001 | 69640000 | 0.0045 | 0.0031 | 0.0034 |
| F04 | 69560001 | 69660000 | 0.0042 | 0.0023 | 0.0031 |
| F04 | 69580001 | 69680000 | 0.0034 | 0.0020 | 0.0024 |
| F04 | 69600001 | 69700000 | 0.0031 | 0.0019 | 0.0024 |
| F04 | 69620001 | 69720000 | 0.0031 | 0.0023 | 0.0022 |
| F04 | 69640001 | 69740000 | 0.0036 | 0.0026 | 0.0026 |
| F04 | 69660001 | 69760000 | 0.0037 | 0.0025 | 0.0028 |
| F04 | 69680001 | 69780000 | 0.0035 | 0.0019 | 0.0027 |
| F04 | 69700001 | 69800000 | 0.0028 | 0.0036 | 0.0019 |
| F04 | 69720001 | 69820000 | 0.0044 | 0.0034 | 0.0037 |
| F04 | 69740001 | 69840000 | 0.0042 | 0.0043 | 0.0036 |
| F04 | 69760001 | 69860000 | 0.0052 | 0.0049 | 0.0047 |
| F04 | 69780001 | 69880000 | 0.0057 | 0.0049 | 0.0053 |
| F04 | 69800001 | 69900000 | 0.0056 | 0.0036 | 0.0053 |
| F04 | 69820001 | 69920000 | 0.0042 | 0.0032 | 0.0039 |
| F04 | 69840001 | 69940000 | 0.0038 | 0.0023 | 0.0034 |

| Chr | Start | End | both | A1 | A2 |
|---|---|---|---|---|---|
| F04 | 69860001 | 69960000 | 0.0031 | 0.0019 | 0.0025 |
| F04 | 69880001 | 69980000 | 0.0027 | 0.0021 | 0.0020 |
| F04 | 69900001 | 70000000 | 0.0030 | 0.0019 | 0.0022 |
| F04 | 69920001 | 70020000 | 0.0029 | 0.0017 | 0.0020 |
| F04 | 69940001 | 70040000 | 0.0025 | 0.0013 | 0.0018 |
| F04 | 69960001 | 70060000 | 0.0018 | 0.0014 | 0.0012 |
| F04 | 69980001 | 70080000 | 0.0022 | 0.0016 | 0.0014 |
| F04 | 70000001 | 70100000 | 0.0025 | 0.0014 | 0.0019 |
| F04 | 70020001 | 70120000 | 0.0022 | 0.0018 | 0.0017 |
| F04 | 70040001 | 70140000 | 0.0026 | 0.0021 | 0.0022 |
| F04 | 70060001 | 70160000 | 0.0030 | 0.0024 | 0.0025 |
| F04 | 70080001 | 70180000 | 0.0030 | 0.0026 | 0.0024 |
| F04 | 70100001 | 70200000 | 0.0031 | 0.0037 | 0.0025 |
| F04 | 70120001 | 70220000 | 0.0044 | 0.0042 | 0.0037 |
| F04 | 70140001 | 70240000 | 0.0051 | 0.0042 | 0.0042 |
| F04 | 70160001 | 70260000 | 0.0050 | 0.0036 | 0.0043 |
| F04 | 70180001 | 70280000 | 0.0046 | 0.0030 | 0.0040 |
| F04 | 70200001 | 70300000 | 0.0039 | 0.0026 | 0.0032 |
| F04 | 70220001 | 70320000 | 0.0035 | 0.0021 | 0.0027 |
| F04 | 70240001 | 70340000 | 0.0031 | 0.0022 | 0.0023 |
| F04 | 70260001 | 70360000 | 0.0034 | 0.0025 | 0.0024 |
| F04 | 70280001 | 70380000 | 0.0035 | 0.0034 | 0.0027 |
| F04 | 70300001 | 70400000 | 0.0046 | 0.0032 | 0.0036 |
| F04 | 70320001 | 70420000 | 0.0043 | 0.0030 | 0.0033 |
| F04 | 70340001 | 70440000 | 0.0037 | 0.0024 | 0.0030 |
| F04 | 70360001 | 70460000 | 0.0030 | 0.0023 | 0.0025 |
| F04 | 70380001 | 70480000 | 0.0028 | 0.0013 | 0.0024 |
| F04 | 70400001 | 70500000 | 0.0016 | 0.0014 | 0.0014 |
| F04 | 70420001 | 70520000 | 0.0020 | 0.0014 | 0.0016 |
| F04 | 70440001 | 70540000 | 0.0020 | 0.0017 | 0.0016 |
| F04 | 70460001 | 70560000 | 0.0025 | 0.0018 | 0.0020 |
| F04 | 70480001 | 70580000 | 0.0027 | 0.0020 | 0.0022 |
| F04 | 70500001 | 70600000 | 0.0028 | 0.0013 | 0.0024 |
| F04 | 70520001 | 70620000 | 0.0019 | 0.0012 | 0.0015 |
| F04 | 70540001 | 70640000 | 0.0020 | 0.0010 | 0.0015 |
| F04 | 70560001 | 70660000 | 0.0018 | 0.0012 | 0.0012 |
| F04 | 70580001 | 70680000 | 0.0021 | 0.0028 | 0.0015 |
| F04 | 70600001 | 70700000 | 0.0039 | 0.0036 | 0.0030 |
| F04 | 70620001 | 70720000 | 0.0047 | 0.0039 | 0.0037 |
| F04 | 70640001 | 70740000 | 0.0050 | 0.0042 | 0.0040 |
| F04 | 70660001 | 70760000 | 0.0052 | 0.0054 | 0.0044 |
| F04 | 70680001 | 70780000 | 0.0065 | 0.0043 | 0.0056 |
| F04 | 70700001 | 70800000 | 0.0054 | 0.0040 | 0.0045 |
| F04 | 70720001 | 70820000 | 0.0050 | 0.0043 | 0.0044 |
| F04 | 70740001 | 70840000 | 0.0052 | 0.0051 | 0.0047 |
| F04 | 70760001 | 70860000 | 0.0061 | 0.0036 | 0.0054 |
| F04 | 70780001 | 70880000 | 0.0046 | 0.0031 | 0.0038 |
| F04 | 70800001 | 70900000 | 0.0038 | 0.0026 | 0.0033 |
| F04 | 70820001 | 70920000 | 0.0036 | 0.0023 | 0.0028 |
| F04 | 70840001 | 70940000 | 0.0034 | 0.0016 | 0.0026 |
| F04 | 70860001 | 70960000 | 0.0024 | 0.0016 | 0.0019 |
| F04 | 70880001 | 70980000 | 0.0025 | 0.0024 | 0.0018 |
| F04 | 70900001 | 71000000 | 0.0034 | 0.0031 | 0.0027 |
| F04 | 70920001 | 71020000 | 0.0040 | 0.0032 | 0.0033 |
| F04 | 70940001 | 71040000 | 0.0040 | 0.0033 | 0.0032 |
| F04 | 70960001 | 71060000 | 0.0041 | 0.0040 | 0.0033 |
| F04 | 70980001 | 71080000 | 0.0046 | 0.0040 | 0.0039 |
| F04 | 71000001 | 71100000 | 0.0047 | 0.0040 | 0.0039 |

| Chr | Start | End | both | A1 | A2 |
|---|---|---|---|---|---|
| F04 | 71020001 | 71120000 | 0.0048 | 0.0036 | 0.0040 |
| F04 | 71040001 | 71140000 | 0.0044 | 0.0031 | 0.0036 |
| F04 | 71060001 | 71160000 | 0.0040 | 0.0025 | 0.0032 |
| F04 | 71080001 | 71180000 | 0.0035 | 0.0018 | 0.0027 |
| F04 | 71100001 | 71200000 | 0.0026 | 0.0015 | 0.0019 |
| F04 | 71120001 | 71220000 | 0.0022 | 0.0018 | 0.0015 |
| F04 | 71140001 | 71240000 | 0.0023 | 0.0020 | 0.0018 |
| F04 | 71160001 | 71260000 | 0.0028 | 0.0017 | 0.0022 |
| F04 | 71180001 | 71280000 | 0.0023 | 0.0017 | 0.0018 |
| F04 | 71200001 | 71300000 | 0.0024 | 0.0012 | 0.0019 |
| F04 | 71220001 | 71320000 | 0.0018 | 0.0008 | 0.0014 |
| F04 | 71240001 | 71340000 | 0.0013 | 0.0006 | 0.0010 |
| F04 | 71260001 | 71360000 | 0.0008 | 0.0007 | 0.0007 |
| F04 | 71280001 | 71380000 | 0.0009 | 0.0004 | 0.0009 |
| F04 | 71300001 | 71400000 | 0.0005 | 0.0004 | 0.0005 |
| F04 | 71320001 | 71420000 | 0.0004 | 0.0007 | 0.0004 |
| F04 | 71340001 | 71440000 | 0.0008 | 0.0006 | 0.0008 |
| F04 | 71360001 | 71460000 | 0.0007 | 0.0007 | 0.0007 |
| F04 | 71380001 | 71480000 | 0.0008 | 0.0008 | 0.0007 |
| F04 | 71400001 | 71500000 | 0.0009 | 0.0008 | 0.0008 |
| F04 | 71420001 | 71520000 | 0.0008 | 0.0004 | 0.0008 |
| F04 | 71440001 | 71540000 | 0.0005 | 0.0005 | 0.0005 |
| F04 | 71460001 | 71560000 | 0.0005 | 0.0006 | 0.0005 |
| F04 | 71480001 | 71580000 | 0.0006 | 0.0008 | 0.0006 |
| F04 | 71500001 | 71600000 | 0.0008 | 0.0008 | 0.0007 |
| F04 | 71520001 | 71620000 | 0.0008 | 0.0009 | 0.0007 |
| F04 | 71540001 | 71640000 | 0.0009 | 0.0011 | 0.0009 |
| F04 | 71560001 | 71660000 | 0.0011 | 0.0011 | 0.0011 |
| F04 | 71580001 | 71680000 | 0.0012 | 0.0012 | 0.0012 |
| F04 | 71600001 | 71700000 | 0.0013 | 0.0015 | 0.0013 |
| F04 | 71620001 | 71720000 | 0.0018 | 0.0018 | 0.0017 |
| F04 | 71640001 | 71740000 | 0.0022 | 0.0016 | 0.0020 |
| F04 | 71660001 | 71760000 | 0.0021 | 0.0015 | 0.0019 |
| F04 | 71680001 | 71780000 | 0.0021 | 0.0016 | 0.0017 |
| F04 | 71700001 | 71800000 | 0.0024 | 0.0015 | 0.0019 |
| F04 | 71720001 | 71820000 | 0.0022 | 0.0010 | 0.0017 |
| F04 | 71740001 | 71840000 | 0.0018 | 0.0016 | 0.0013 |
| F04 | 71760001 | 71860000 | 0.0025 | 0.0021 | 0.0019 |
| F04 | 71780001 | 71880000 | 0.0029 | 0.0020 | 0.0025 |
| F04 | 71800001 | 71900000 | 0.0026 | 0.0021 | 0.0022 |
| F04 | 71820001 | 71920000 | 0.0026 | 0.0024 | 0.0023 |
| F04 | 71840001 | 71940000 | 0.0029 | 0.0025 | 0.0027 |
| F04 | 71860001 | 71960000 | 0.0031 | 0.0018 | 0.0029 |
| F04 | 71880001 | 71980000 | 0.0022 | 0.0022 | 0.0021 |
| F04 | 71900001 | 72000000 | 0.0029 | 0.0021 | 0.0025 |
| F04 | 71920001 | 72020000 | 0.0029 | 0.0019 | 0.0024 |
| F04 | 71940001 | 72040000 | 0.0029 | 0.0012 | 0.0022 |
| F04 | 71960001 | 72060000 | 0.0020 | 0.0025 | 0.0013 |
| F04 | 71980001 | 72080000 | 0.0034 | 0.0023 | 0.0024 |
| F04 | 72000001 | 72100000 | 0.0034 | 0.0025 | 0.0023 |
| F04 | 72020001 | 72120000 | 0.0035 | 0.0029 | 0.0024 |
| F04 | 72040001 | 72140000 | 0.0038 | 0.0030 | 0.0029 |
| F04 | 72060001 | 72160000 | 0.0041 | 0.0023 | 0.0030 |
| F04 | 72080001 | 72180000 | 0.0036 | 0.0020 | 0.0023 |
| F04 | 72100001 | 72200000 | 0.0033 | 0.0017 | 0.0019 |
| F04 | 72120001 | 72220000 | 0.0032 | 0.0020 | 0.0017 |
| F04 | 72140001 | 72240000 | 0.0037 | 0.0022 | 0.0020 |
| F04 | 72160001 | 72260000 | 0.0035 | 0.0019 | 0.0021 |

| Chr | Start | End | both | A1 | A2 |
|---|---|---|---|---|---|
| F04 | 72180001 | 72280000 | 0.0032 | 0.0018 | 0.0018 |
| F04 | 72200001 | 72300000 | 0.0029 | 0.0029 | 0.0017 |
| F04 | 72220001 | 72320000 | 0.0040 | 0.0022 | 0.0029 |
| F04 | 72240001 | 72340000 | 0.0033 | 0.0020 | 0.0023 |
| F04 | 72260001 | 72360000 | 0.0033 | 0.0018 | 0.0021 |
| F04 | 72280001 | 72380000 | 0.0031 | 0.0019 | 0.0020 |
| F04 | 72300001 | 72400000 | 0.0030 | 0.0010 | 0.0021 |
| F04 | 72320001 | 72420000 | 0.0021 | 0.0012 | 0.0012 |
| F04 | 72340001 | 72440000 | 0.0021 | 0.0024 | 0.0013 |
| F04 | 72360001 | 72460000 | 0.0033 | 0.0032 | 0.0027 |
| F04 | 72380001 | 72480000 | 0.0039 | 0.0033 | 0.0035 |
| F04 | 72400001 | 72500000 | 0.0042 | 0.0032 | 0.0036 |
| F04 | 72420001 | 72520000 | 0.0040 | 0.0042 | 0.0035 |
| F04 | 72440001 | 72540000 | 0.0051 | 0.0031 | 0.0045 |
| F04 | 72460001 | 72560000 | 0.0038 | 0.0021 | 0.0031 |
| F04 | 72480001 | 72580000 | 0.0027 | 0.0026 | 0.0022 |
| F04 | 72500001 | 72600000 | 0.0033 | 0.0042 | 0.0029 |
| F04 | 72520001 | 72620000 | 0.0052 | 0.0031 | 0.0045 |
| F04 | 72540001 | 72640000 | 0.0042 | 0.0035 | 0.0035 |
| F04 | 72560001 | 72660000 | 0.0045 | 0.0038 | 0.0038 |
| F04 | 72580001 | 72680000 | 0.0051 | 0.0035 | 0.0041 |
| F04 | 72600001 | 72700000 | 0.0047 | 0.0024 | 0.0037 |
| F04 | 72620001 | 72720000 | 0.0034 | 0.0021 | 0.0026 |
| F04 | 72640001 | 72740000 | 0.0029 | 0.0022 | 0.0021 |
| F04 | 72660001 | 72760000 | 0.0031 | 0.0025 | 0.0024 |
| F04 | 72680001 | 72780000 | 0.0033 | 0.0024 | 0.0027 |
| F04 | 72700001 | 72800000 | 0.0030 | 0.0033 | 0.0024 |
| F04 | 72720001 | 72820000 | 0.0039 | 0.0045 | 0.0032 |
| F04 | 72740001 | 72840000 | 0.0052 | 0.0044 | 0.0045 |
| F04 | 72760001 | 72860000 | 0.0051 | 0.0058 | 0.0046 |
| F04 | 72780001 | 72880000 | 0.0064 | 0.0065 | 0.0057 |
| F04 | 72800001 | 72900000 | 0.0072 | 0.0053 | 0.0064 |
| F04 | 72820001 | 72920000 | 0.0062 | 0.0049 | 0.0053 |
| F04 | 72840001 | 72940000 | 0.0059 | 0.0051 | 0.0049 |
| F04 | 72860001 | 72960000 | 0.0061 | 0.0039 | 0.0050 |
| F04 | 72880001 | 72980000 | 0.0049 | 0.0036 | 0.0039 |
| F04 | 72900001 | 73000000 | 0.0045 | 0.0034 | 0.0037 |
| F04 | 72920001 | 73020000 | 0.0042 | 0.0025 | 0.0036 |
| F04 | 72940001 | 73040000 | 0.0033 | 0.0022 | 0.0026 |
| F04 | 72960001 | 73060000 | 0.0030 | 0.0021 | 0.0022 |
| F04 | 72980001 | 73080000 | 0.0030 | 0.0016 | 0.0022 |
| F04 | 73000001 | 73100000 | 0.0026 | 0.0015 | 0.0017 |
| F04 | 73020001 | 73120000 | 0.0025 | 0.0024 | 0.0016 |
| F04 | 73040001 | 73140000 | 0.0034 | 0.0031 | 0.0025 |
| F04 | 73060001 | 73160000 | 0.0040 | 0.0029 | 0.0033 |
| F04 | 73080001 | 73180000 | 0.0037 | 0.0030 | 0.0033 |
| F04 | 73100001 | 73200000 | 0.0039 | 0.0033 | 0.0034 |
| F04 | 73120001 | 73220000 | 0.0041 | 0.0030 | 0.0037 |
| F04 | 73140001 | 73240000 | 0.0036 | 0.0023 | 0.0032 |
| F04 | 73160001 | 73260000 | 0.0029 | 0.0025 | 0.0024 |
| F04 | 73180001 | 73280000 | 0.0033 | 0.0023 | 0.0028 |
| F04 | 73200001 | 73300000 | 0.0034 | 0.0019 | 0.0027 |
| F04 | 73220001 | 73320000 | 0.0032 | 0.0017 | 0.0023 |
| F04 | 73240001 | 73340000 | 0.0033 | 0.0015 | 0.0023 |
| F04 | 73260001 | 73360000 | 0.0031 | 0.0015 | 0.0021 |
| F04 | 73280001 | 73380000 | 0.0030 | 0.0022 | 0.0019 |
| F04 | 73300001 | 73400000 | 0.0033 | 0.0026 | 0.0024 |
| F04 | 73320001 | 73420000 | 0.0038 | 0.0023 | 0.0030 |

| Chr | Start | End | both | A1 | A2 |
|---|---|---|---|---|---|
| F04 | 73340001 | 73440000 | 0.0033 | 0.0024 | 0.0027 |
| F04 | 73360001 | 73460000 | 0.0033 | 0.0022 | 0.0028 |
| F04 | 73380001 | 73480000 | 0.0028 | 0.0015 | 0.0024 |
| F04 | 73400001 | 73500000 | 0.0021 | 0.0022 | 0.0019 |
| F04 | 73420001 | 73520000 | 0.0029 | 0.0028 | 0.0028 |
| F04 | 73440001 | 73540000 | 0.0036 | 0.0030 | 0.0034 |
| F04 | 73460001 | 73560000 | 0.0040 | 0.0040 | 0.0037 |
| F04 | 73480001 | 73580000 | 0.0054 | 0.0041 | 0.0048 |
| F04 | 73500001 | 73600000 | 0.0057 | 0.0028 | 0.0048 |
| F04 | 73520001 | 73620000 | 0.0040 | 0.0025 | 0.0030 |
| F04 | 73540001 | 73640000 | 0.0038 | 0.0023 | 0.0026 |
| F04 | 73560001 | 73660000 | 0.0037 | 0.0015 | 0.0024 |
| F04 | 73580001 | 73680000 | 0.0027 | 0.0017 | 0.0015 |
| F04 | 73600001 | 73700000 | 0.0027 | 0.0017 | 0.0017 |
| F04 | 73620001 | 73720000 | 0.0029 | 0.0020 | 0.0017 |
| F04 | 73640001 | 73740000 | 0.0034 | 0.0017 | 0.0021 |
| F04 | 73660001 | 73760000 | 0.0028 | 0.0013 | 0.0017 |
| F04 | 73680001 | 73780000 | 0.0022 | 0.0008 | 0.0012 |
| F04 | 73700001 | 73800000 | 0.0017 | 0.0010 | 0.0008 |
| F04 | 73720001 | 73820000 | 0.0019 | 0.0014 | 0.0011 |
| F04 | 73740001 | 73840000 | 0.0023 | 0.0018 | 0.0016 |
| F04 | 73760001 | 73860000 | 0.0027 | 0.0023 | 0.0021 |
| F04 | 73780001 | 73880000 | 0.0032 | 0.0024 | 0.0026 |
| F04 | 73800001 | 73900000 | 0.0034 | 0.0029 | 0.0028 |
| F04 | 73820001 | 73920000 | 0.0038 | 0.0021 | 0.0032 |
| F04 | 73840001 | 73940000 | 0.0028 | 0.0019 | 0.0023 |
| F04 | 73860001 | 73960000 | 0.0024 | 0.0015 | 0.0020 |
| F04 | 73880001 | 73980000 | 0.0020 | 0.0030 | 0.0017 |
| F04 | 73900001 | 74000000 | 0.0037 | 0.0026 | 0.0032 |
| F04 | 73920001 | 74020000 | 0.0035 | 0.0031 | 0.0030 |
| F04 | 73940001 | 74040000 | 0.0041 | 0.0033 | 0.0034 |
| F04 | 73960001 | 74060000 | 0.0043 | 0.0033 | 0.0036 |
| F04 | 73980001 | 74080000 | 0.0044 | 0.0019 | 0.0036 |
| F04 | 74000001 | 74100000 | 0.0029 | 0.0026 | 0.0021 |
| F04 | 74020001 | 74120000 | 0.0036 | 0.0025 | 0.0027 |
| F04 | 74040001 | 74140000 | 0.0033 | 0.0027 | 0.0026 |
| F04 | 74060001 | 74160000 | 0.0036 | 0.0034 | 0.0027 |
| F04 | 74080001 | 74180000 | 0.0042 | 0.0039 | 0.0034 |
| F04 | 74100001 | 74200000 | 0.0047 | 0.0030 | 0.0039 |
| F04 | 74120001 | 74220000 | 0.0039 | 0.0025 | 0.0032 |
| F04 | 74140001 | 74240000 | 0.0035 | 0.0022 | 0.0027 |
| F04 | 74160001 | 74260000 | 0.0031 | 0.0020 | 0.0025 |
| F04 | 74180001 | 74280000 | 0.0027 | 0.0013 | 0.0021 |
| F04 | 74200001 | 74300000 | 0.0021 | 0.0014 | 0.0017 |
| F04 | 74220001 | 74320000 | 0.0020 | 0.0018 | 0.0017 |
| F04 | 74240001 | 74340000 | 0.0026 | 0.0016 | 0.0023 |
| F04 | 74260001 | 74360000 | 0.0023 | 0.0014 | 0.0020 |
| F04 | 74280001 | 74380000 | 0.0023 | 0.0020 | 0.0018 |
| F04 | 74300001 | 74400000 | 0.0029 | 0.0020 | 0.0023 |
| F04 | 74320001 | 74420000 | 0.0029 | 0.0018 | 0.0022 |
| F04 | 74340001 | 74440000 | 0.0026 | 0.0020 | 0.0020 |
| F04 | 74360001 | 74460000 | 0.0028 | 0.0018 | 0.0022 |
| F04 | 74380001 | 74480000 | 0.0024 | 0.0014 | 0.0019 |
| F04 | 74400001 | 74500000 | 0.0019 | 0.0012 | 0.0016 |
| F04 | 74420001 | 74520000 | 0.0017 | 0.0009 | 0.0014 |
| F04 | 74440001 | 74540000 | 0.0012 | 0.0008 | 0.0010 |
| F04 | 74460001 | 74560000 | 0.0010 | 0.0014 | 0.0009 |
| F04 | 74480001 | 74580000 | 0.0017 | 0.0015 | 0.0016 |

| Chr | Start | End | both | A1 | A2 |
|---|---|---|---|---|---|
| F04 | 74500001 | 74600000 | 0.0016 | 0.0015 | 0.0015 |
| F04 | 74520001 | 74620000 | 0.0018 | 0.0015 | 0.0015 |
| F04 | 74540001 | 74640000 | 0.0019 | 0.0023 | 0.0015 |
| F04 | 74560001 | 74660000 | 0.0029 | 0.0019 | 0.0024 |
| F04 | 74580001 | 74680000 | 0.0024 | 0.0021 | 0.0019 |
| F04 | 74600001 | 74700000 | 0.0028 | 0.0024 | 0.0020 |
| F04 | 74620001 | 74720000 | 0.0031 | 0.0028 | 0.0023 |
| F04 | 74640001 | 74740000 | 0.0034 | 0.0021 | 0.0026 |
| F04 | 74660001 | 74760000 | 0.0028 | 0.0025 | 0.0018 |
| F04 | 74680001 | 74780000 | 0.0034 | 0.0020 | 0.0022 |
| F04 | 74700001 | 74800000 | 0.0029 | 0.0022 | 0.0019 |
| F04 | 74720001 | 74820000 | 0.0031 | 0.0018 | 0.0023 |
| F04 | 74740001 | 74840000 | 0.0029 | 0.0019 | 0.0020 |
| F04 | 74760001 | 74860000 | 0.0029 | 0.0019 | 0.0021 |
| F04 | 74780001 | 74880000 | 0.0027 | 0.0018 | 0.0021 |
| F04 | 74800001 | 74900000 | 0.0027 | 0.0014 | 0.0020 |
| F04 | 74820001 | 74920000 | 0.0026 | 0.0019 | 0.0014 |
| F04 | 74840001 | 74940000 | 0.0029 | 0.0024 | 0.0020 |
| F04 | 74860001 | 74960000 | 0.0034 | 0.0024 | 0.0025 |
| F04 | 74880001 | 74980000 | 0.0035 | 0.0026 | 0.0024 |
| F04 | 74900001 | 75000000 | 0.0038 | 0.0034 | 0.0026 |
| F04 | 74920001 | 75020000 | 0.0042 | 0.0033 | 0.0034 |
| F04 | 74940001 | 75040000 | 0.0041 | 0.0030 | 0.0032 |
| F04 | 74960001 | 75060000 | 0.0041 | 0.0026 | 0.0030 |
| F04 | 74980001 | 75080000 | 0.0039 | 0.0029 | 0.0027 |
| F04 | 75000001 | 75100000 | 0.0039 | 0.0023 | 0.0032 |
| F04 | 75020001 | 75120000 | 0.0032 | 0.0020 | 0.0026 |
| F04 | 75040001 | 75140000 | 0.0029 | 0.0017 | 0.0023 |
| F04 | 75060001 | 75160000 | 0.0027 | 0.0018 | 0.0019 |
| F04 | 75080001 | 75180000 | 0.0025 | 0.0012 | 0.0018 |
| F04 | 75100001 | 75200000 | 0.0019 | 0.0009 | 0.0012 |
| F04 | 75120001 | 75220000 | 0.0017 | 0.0010 | 0.0009 |
| F04 | 75140001 | 75240000 | 0.0018 | 0.0009 | 0.0012 |
| F04 | 75160001 | 75260000 | 0.0014 | 0.0007 | 0.0012 |
| F04 | 75180001 | 75280000 | 0.0011 | 0.0014 | 0.0010 |
| F04 | 75200001 | 75300000 | 0.0019 | 0.0022 | 0.0018 |
| F04 | 75220001 | 75320000 | 0.0027 | 0.0022 | 0.0026 |
| F04 | 75240001 | 75340000 | 0.0027 | 0.0026 | 0.0025 |
| F04 | 75260001 | 75360000 | 0.0032 | 0.0025 | 0.0029 |
| F04 | 75280001 | 75380000 | 0.0032 | 0.0020 | 0.0028 |
| F04 | 75300001 | 75400000 | 0.0025 | 0.0023 | 0.0021 |
| F04 | 75320001 | 75420000 | 0.0028 | 0.0022 | 0.0024 |
| F04 | 75340001 | 75440000 | 0.0026 | 0.0018 | 0.0022 |
| F04 | 75360001 | 75460000 | 0.0021 | 0.0027 | 0.0019 |
| F04 | 75380001 | 75480000 | 0.0032 | 0.0027 | 0.0028 |
| F04 | 75400001 | 75500000 | 0.0033 | 0.0015 | 0.0029 |
| F04 | 75420001 | 75520000 | 0.0020 | 0.0018 | 0.0017 |
| F04 | 75440001 | 75540000 | 0.0024 | 0.0017 | 0.0020 |
| F04 | 75460001 | 75560000 | 0.0022 | 0.0008 | 0.0019 |
| F04 | 75480001 | 75580000 | 0.0011 | 0.0009 | 0.0009 |
| F04 | 75500001 | 75600000 | 0.0011 | 0.0012 | 0.0010 |
| F04 | 75520001 | 75620000 | 0.0015 | 0.0010 | 0.0013 |
| F04 | 75540001 | 75640000 | 0.0013 | 0.0013 | 0.0012 |
| F04 | 75560001 | 75660000 | 0.0017 | 0.0020 | 0.0014 |
| F04 | 75580001 | 75680000 | 0.0025 | 0.0018 | 0.0022 |
| F04 | 75600001 | 75700000 | 0.0024 | 0.0030 | 0.0020 |
| F04 | 75620001 | 75720000 | 0.0036 | 0.0030 | 0.0032 |
| F04 | 75640001 | 75740000 | 0.0037 | 0.0033 | 0.0033 |

| Chr | Start | End | both | A1 | A2 |
|---|---|---|---|---|---|
| F04 | 75660001 | 75760000 | 0.0039 | 0.0025 | 0.0037 |
| F04 | 75680001 | 75780000 | 0.0031 | 0.0025 | 0.0029 |
| F04 | 75700001 | 75800000 | 0.0030 | 0.0012 | 0.0029 |
| F04 | 75720001 | 75820000 | 0.0014 | 0.0015 | 0.0014 |
| F04 | 75740001 | 75840000 | 0.0017 | 0.0011 | 0.0017 |
| F04 | 75760001 | 75860000 | 0.0012 | 0.0011 | 0.0012 |
| F04 | 75780001 | 75880000 | 0.0012 | 0.0011 | 0.0012 |
| F04 | 75800001 | 75900000 | 0.0013 | 0.0012 | 0.0013 |
| F04 | 75820001 | 75920000 | 0.0015 | 0.0012 | 0.0015 |
| F04 | 75840001 | 75940000 | 0.0015 | 0.0011 | 0.0015 |
| F04 | 75860001 | 75960000 | 0.0014 | 0.0012 | 0.0014 |
| F04 | 75880001 | 75980000 | 0.0015 | 0.0011 | 0.0015 |
| F04 | 75900001 | 76000000 | 0.0014 | 0.0016 | 0.0014 |
| F04 | 75920001 | 76020000 | 0.0020 | 0.0012 | 0.0020 |
| F04 | 75940001 | 76040000 | 0.0015 | 0.0019 | 0.0015 |
| F04 | 75960001 | 76060000 | 0.0021 | 0.0019 | 0.0021 |
| F04 | 75980001 | 76080000 | 0.0021 | 0.0021 | 0.0021 |
| F04 | 76000001 | 76100000 | 0.0024 | 0.0016 | 0.0024 |
| F04 | 76020001 | 76120000 | 0.0018 | 0.0016 | 0.0018 |
| F04 | 76040001 | 76140000 | 0.0018 | 0.0021 | 0.0018 |
| F04 | 76060001 | 76160000 | 0.0026 | 0.0020 | 0.0025 |
| F04 | 76080001 | 76180000 | 0.0025 | 0.0017 | 0.0024 |
| F04 | 76100001 | 76200000 | 0.0020 | 0.0016 | 0.0020 |
| F04 | 76120001 | 76220000 | 0.0020 | 0.0015 | 0.0019 |
| F04 | 76140001 | 76240000 | 0.0022 | 0.0008 | 0.0018 |
| F04 | 76160001 | 76260000 | 0.0014 | 0.0008 | 0.0008 |
| F04 | 76180001 | 76280000 | 0.0018 | 0.0010 | 0.0009 |
| F04 | 76200001 | 76300000 | 0.0021 | 0.0010 | 0.0011 |
| F04 | 76220001 | 76320000 | 0.0020 | 0.0010 | 0.0011 |
| F04 | 76240001 | 76340000 | 0.0022 | 0.0006 | 0.0012 |
| F04 | 76260001 | 76360000 | 0.0015 | 0.0012 | 0.0007 |
| F04 | 76280001 | 76380000 | 0.0018 | 0.0014 | 0.0013 |
| F04 | 76300001 | 76400000 | 0.0018 | 0.0020 | 0.0015 |
| F04 | 76320001 | 76420000 | 0.0025 | 0.0022 | 0.0021 |
| F04 | 76340001 | 76440000 | 0.0027 | 0.0029 | 0.0023 |
| F04 | 76360001 | 76460000 | 0.0036 | 0.0025 | 0.0030 |
| F04 | 76380001 | 76480000 | 0.0033 | 0.0027 | 0.0027 |
| F04 | 76400001 | 76500000 | 0.0037 | 0.0028 | 0.0029 |
| F04 | 76420001 | 76520000 | 0.0040 | 0.0026 | 0.0031 |
| F04 | 76440001 | 76540000 | 0.0035 | 0.0027 | 0.0029 |
| F04 | 76460001 | 76560000 | 0.0037 | 0.0024 | 0.0030 |
| F04 | 76480001 | 76580000 | 0.0035 | 0.0023 | 0.0027 |
| F04 | 76500001 | 76600000 | 0.0032 | 0.0025 | 0.0026 |
| F04 | 76520001 | 76620000 | 0.0033 | 0.0026 | 0.0028 |
| F04 | 76540001 | 76640000 | 0.0036 | 0.0032 | 0.0027 |
| F04 | 76560001 | 76660000 | 0.0043 | 0.0033 | 0.0037 |
| F04 | 76580001 | 76680000 | 0.0045 | 0.0033 | 0.0038 |
| F04 | 76600001 | 76700000 | 0.0046 | 0.0034 | 0.0037 |
| F04 | 76620001 | 76720000 | 0.0050 | 0.0038 | 0.0040 |
| F04 | 76640001 | 76740000 | 0.0053 | 0.0025 | 0.0043 |
| F04 | 76660001 | 76760000 | 0.0039 | 0.0026 | 0.0027 |
| F04 | 76680001 | 76780000 | 0.0041 | 0.0024 | 0.0028 |
| F04 | 76700001 | 76800000 | 0.0041 | 0.0019 | 0.0027 |
| F04 | 76720001 | 76820000 | 0.0032 | 0.0015 | 0.0020 |
| F04 | 76740001 | 76840000 | 0.0030 | 0.0015 | 0.0015 |
| F04 | 76760001 | 76860000 | 0.0027 | 0.0019 | 0.0014 |
| F04 | 76780001 | 76880000 | 0.0031 | 0.0018 | 0.0020 |
| F04 | 76800001 | 76900000 | 0.0031 | 0.0016 | 0.0019 |

| Chr | Start | End | both | A1 | A2 |
|---|---|---|---|---|---|
| F04 | 76820001 | 76920000 | 0.0031 | 0.0025 | 0.0019 |
| F04 | 76840001 | 76940000 | 0.0037 | 0.0033 | 0.0027 |
| F04 | 76860001 | 76960000 | 0.0044 | 0.0030 | 0.0036 |
| F04 | 76880001 | 76980000 | 0.0041 | 0.0037 | 0.0031 |
| F04 | 76900001 | 77000000 | 0.0048 | 0.0042 | 0.0040 |
| F04 | 76920001 | 77020000 | 0.0051 | 0.0041 | 0.0040 |
| F04 | 76940001 | 77040000 | 0.0051 | 0.0038 | 0.0042 |
| F04 | 76960001 | 77060000 | 0.0053 | 0.0038 | 0.0044 |
| F04 | 76980001 | 77080000 | 0.0051 | 0.0048 | 0.0043 |
| F04 | 77000001 | 77100000 | 0.0060 | 0.0051 | 0.0053 |
| F04 | 77020001 | 77120000 | 0.0063 | 0.0048 | 0.0058 |
| F04 | 77040001 | 77140000 | 0.0061 | 0.0048 | 0.0054 |
| F04 | 77060001 | 77160000 | 0.0058 | 0.0057 | 0.0049 |
| F04 | 77080001 | 77180000 | 0.0068 | 0.0041 | 0.0059 |
| F04 | 77100001 | 77200000 | 0.0053 | 0.0031 | 0.0042 |
| F04 | 77120001 | 77220000 | 0.0045 | 0.0028 | 0.0033 |
| F04 | 77140001 | 77240000 | 0.0040 | 0.0027 | 0.0029 |
| F04 | 77160001 | 77260000 | 0.0038 | 0.0016 | 0.0029 |
| F04 | 77180001 | 77280000 | 0.0024 | 0.0017 | 0.0017 |
| F04 | 77200001 | 77300000 | 0.0022 | 0.0017 | 0.0018 |
| F04 | 77220001 | 77320000 | 0.0023 | 0.0023 | 0.0019 |
| F04 | 77240001 | 77340000 | 0.0029 | 0.0023 | 0.0023 |
| F04 | 77260001 | 77360000 | 0.0030 | 0.0024 | 0.0024 |
| F04 | 77280001 | 77380000 | 0.0033 | 0.0030 | 0.0025 |
| F04 | 77300001 | 77400000 | 0.0042 | 0.0034 | 0.0033 |
| F04 | 77320001 | 77420000 | 0.0047 | 0.0027 | 0.0036 |
| F04 | 77340001 | 77440000 | 0.0040 | 0.0046 | 0.0029 |
| F04 | 77360001 | 77460000 | 0.0060 | 0.0047 | 0.0048 |
| F04 | 77380001 | 77480000 | 0.0062 | 0.0044 | 0.0050 |
| F04 | 77400001 | 77500000 | 0.0058 | 0.0041 | 0.0047 |
| F04 | 77420001 | 77520000 | 0.0053 | 0.0057 | 0.0046 |
| F04 | 77440001 | 77540000 | 0.0072 | 0.0039 | 0.0064 |
| F04 | 77460001 | 77560000 | 0.0050 | 0.0039 | 0.0043 |
| F04 | 77480001 | 77580000 | 0.0049 | 0.0033 | 0.0043 |
| F04 | 77500001 | 77600000 | 0.0043 | 0.0037 | 0.0036 |
| F04 | 77520001 | 77620000 | 0.0049 | 0.0022 | 0.0041 |
| F04 | 77540001 | 77640000 | 0.0030 | 0.0018 | 0.0024 |
| F04 | 77560001 | 77660000 | 0.0027 | 0.0019 | 0.0021 |
| F04 | 77580001 | 77680000 | 0.0028 | 0.0021 | 0.0021 |
| F04 | 77600001 | 77700000 | 0.0029 | 0.0020 | 0.0024 |
| F04 | 77620001 | 77720000 | 0.0027 | 0.0026 | 0.0021 |
| F04 | 77640001 | 77740000 | 0.0034 | 0.0029 | 0.0029 |
| F04 | 77660001 | 77760000 | 0.0037 | 0.0029 | 0.0031 |
| F04 | 77680001 | 77780000 | 0.0038 | 0.0033 | 0.0033 |
| F04 | 77700001 | 77800000 | 0.0041 | 0.0028 | 0.0035 |
| F04 | 77720001 | 77820000 | 0.0039 | 0.0031 | 0.0032 |
| F04 | 77740001 | 77840000 | 0.0039 | 0.0033 | 0.0030 |
| F04 | 77760001 | 77860000 | 0.0040 | 0.0028 | 0.0031 |
| F04 | 77780001 | 77880000 | 0.0034 | 0.0025 | 0.0026 |
| F04 | 77800001 | 77900000 | 0.0032 | 0.0034 | 0.0024 |
| F04 | 77820001 | 77920000 | 0.0039 | 0.0029 | 0.0032 |
| F04 | 77840001 | 77940000 | 0.0036 | 0.0034 | 0.0031 |
| F04 | 77860001 | 77960000 | 0.0041 | 0.0037 | 0.0037 |
| F04 | 77880001 | 77980000 | 0.0044 | 0.0041 | 0.0041 |
| F04 | 77900001 | 78000000 | 0.0046 | 0.0033 | 0.0044 |
| F04 | 77920001 | 78020000 | 0.0040 | 0.0031 | 0.0037 |
| F04 | 77940001 | 78040000 | 0.0038 | 0.0025 | 0.0035 |
| F04 | 77960001 | 78060000 | 0.0030 | 0.0032 | 0.0025 |

| Chr | Start | End | both | A1 | A2 |
|---|---|---|---|---|---|
| F04 | 77980001 | 78080000 | 0.0038 | 0.0024 | 0.0031 |
| F04 | 78000001 | 78100000 | 0.0032 | 0.0025 | 0.0024 |
| F04 | 78020001 | 78120000 | 0.0033 | 0.0030 | 0.0026 |
| F04 | 78040001 | 78140000 | 0.0036 | 0.0030 | 0.0029 |
| F04 | 78060001 | 78160000 | 0.0038 | 0.0023 | 0.0033 |
| F04 | 78080001 | 78180000 | 0.0031 | 0.0027 | 0.0026 |
| F04 | 78100001 | 78200000 | 0.0034 | 0.0025 | 0.0030 |
| F04 | 78120001 | 78220000 | 0.0032 | 0.0021 | 0.0026 |
| F04 | 78140001 | 78240000 | 0.0028 | 0.0025 | 0.0022 |
| F04 | 78160001 | 78260000 | 0.0032 | 0.0041 | 0.0025 |
| F04 | 78180001 | 78280000 | 0.0049 | 0.0042 | 0.0041 |
| F04 | 78200001 | 78300000 | 0.0051 | 0.0039 | 0.0042 |
| F04 | 78220001 | 78320000 | 0.0049 | 0.0042 | 0.0041 |
| F04 | 78240001 | 78340000 | 0.0051 | 0.0033 | 0.0042 |
| F04 | 78260001 | 78360000 | 0.0041 | 0.0014 | 0.0035 |
| F04 | 78280001 | 78380000 | 0.0019 | 0.0012 | 0.0015 |
| F04 | 78300001 | 78400000 | 0.0016 | 0.0015 | 0.0013 |
| F04 | 78320001 | 78420000 | 0.0019 | 0.0010 | 0.0015 |
| F04 | 78340001 | 78440000 | 0.0015 | 0.0015 | 0.0012 |
| F04 | 78360001 | 78460000 | 0.0022 | 0.0016 | 0.0017 |
| F04 | 78380001 | 78480000 | 0.0024 | 0.0014 | 0.0019 |
| F04 | 78400001 | 78500000 | 0.0022 | 0.0014 | 0.0017 |
| F04 | 78420001 | 78520000 | 0.0025 | 0.0018 | 0.0017 |
| F04 | 78440001 | 78540000 | 0.0028 | 0.0014 | 0.0021 |
| F04 | 78460001 | 78560000 | 0.0027 | 0.0015 | 0.0018 |
| F04 | 78480001 | 78580000 | 0.0031 | 0.0015 | 0.0020 |
| F04 | 78500001 | 78600000 | 0.0030 | 0.0016 | 0.0020 |
| F04 | 78520001 | 78620000 | 0.0029 | 0.0014 | 0.0021 |
| F04 | 78540001 | 78640000 | 0.0025 | 0.0017 | 0.0018 |
| F04 | 78560001 | 78660000 | 0.0024 | 0.0015 | 0.0019 |
| F04 | 78580001 | 78680000 | 0.0021 | 0.0017 | 0.0017 |
| F04 | 78600001 | 78700000 | 0.0023 | 0.0018 | 0.0018 |
| F04 | 78620001 | 78720000 | 0.0025 | 0.0015 | 0.0020 |
| F04 | 78640001 | 78740000 | 0.0021 | 0.0014 | 0.0017 |
| F04 | 78660001 | 78760000 | 0.0020 | 0.0020 | 0.0016 |
| F04 | 78680001 | 78780000 | 0.0025 | 0.0019 | 0.0022 |
| F04 | 78700001 | 78800000 | 0.0023 | 0.0014 | 0.0021 |
| F04 | 78720001 | 78820000 | 0.0016 | 0.0015 | 0.0015 |
| F04 | 78740001 | 78840000 | 0.0019 | 0.0013 | 0.0016 |
| F04 | 78760001 | 78860000 | 0.0018 | 0.0013 | 0.0014 |
| F04 | 78780001 | 78880000 | 0.0020 | 0.0018 | 0.0014 |
| F04 | 78800001 | 78900000 | 0.0028 | 0.0023 | 0.0019 |
| F04 | 78820001 | 78920000 | 0.0032 | 0.0020 | 0.0023 |
| F04 | 78840001 | 78940000 | 0.0029 | 0.0021 | 0.0022 |
| F04 | 78860001 | 78960000 | 0.0029 | 0.0014 | 0.0022 |
| F04 | 78880001 | 78980000 | 0.0022 | 0.0012 | 0.0015 |
| F04 | 78900001 | 79000000 | 0.0019 | 0.0011 | 0.0012 |
| F04 | 78920001 | 79020000 | 0.0021 | 0.0014 | 0.0013 |
| F04 | 78940001 | 79040000 | 0.0026 | 0.0014 | 0.0016 |
| F04 | 78960001 | 79060000 | 0.0028 | 0.0015 | 0.0017 |
| F04 | 78980001 | 79080000 | 0.0027 | 0.0014 | 0.0018 |
| F04 | 79000001 | 79100000 | 0.0025 | 0.0009 | 0.0016 |
| F04 | 79020001 | 79120000 | 0.0019 | 0.0012 | 0.0010 |
| F04 | 79040001 | 79140000 | 0.0021 | 0.0013 | 0.0014 |
| F04 | 79060001 | 79160000 | 0.0022 | 0.0013 | 0.0015 |
| F04 | 79080001 | 79180000 | 0.0025 | 0.0015 | 0.0015 |
| F04 | 79100001 | 79200000 | 0.0027 | 0.0021 | 0.0017 |
| F04 | 79120001 | 79220000 | 0.0034 | 0.0016 | 0.0024 |

| Chr | Start | End | both | A1 | A2 |
|---|---|---|---|---|---|
| F04 | 79140001 | 79240000 | 0.0028 | 0.0018 | 0.0018 |
| F04 | 79160001 | 79260000 | 0.0031 | 0.0020 | 0.0020 |
| F04 | 79180001 | 79280000 | 0.0032 | 0.0019 | 0.0023 |
| F04 | 79200001 | 79300000 | 0.0032 | 0.0017 | 0.0022 |
| F04 | 79220001 | 79320000 | 0.0029 | 0.0020 | 0.0021 |
| F04 | 79240001 | 79340000 | 0.0031 | 0.0020 | 0.0023 |
| F04 | 79260001 | 79360000 | 0.0031 | 0.0015 | 0.0023 |
| F04 | 79280001 | 79380000 | 0.0028 | 0.0018 | 0.0019 |
| F04 | 79300001 | 79400000 | 0.0030 | 0.0026 | 0.0021 |
| F04 | 79320001 | 79420000 | 0.0036 | 0.0031 | 0.0029 |
| F04 | 79340001 | 79440000 | 0.0041 | 0.0027 | 0.0035 |
| F04 | 79360001 | 79460000 | 0.0034 | 0.0035 | 0.0030 |
| F04 | 79380001 | 79480000 | 0.0039 | 0.0036 | 0.0036 |
| F04 | 79400001 | 79500000 | 0.0040 | 0.0030 | 0.0039 |
| F04 | 79420001 | 79520000 | 0.0035 | 0.0026 | 0.0034 |
| F04 | 79440001 | 79540000 | 0.0029 | 0.0029 | 0.0028 |
| F04 | 79460001 | 79560000 | 0.0033 | 0.0025 | 0.0032 |
| F04 | 79480001 | 79580000 | 0.0029 | 0.0021 | 0.0028 |
| F04 | 79500001 | 79600000 | 0.0025 | 0.0024 | 0.0024 |
| F04 | 79520001 | 79620000 | 0.0027 | 0.0024 | 0.0026 |
| F04 | 79540001 | 79640000 | 0.0030 | 0.0022 | 0.0027 |
| F04 | 79560001 | 79660000 | 0.0027 | 0.0018 | 0.0024 |
| F04 | 79580001 | 79680000 | 0.0024 | 0.0018 | 0.0021 |
| F04 | 79600001 | 79700000 | 0.0024 | 0.0010 | 0.0021 |
| F04 | 79620001 | 79720000 | 0.0018 | 0.0008 | 0.0014 |
| F04 | 79640001 | 79740000 | 0.0012 | 0.0014 | 0.0009 |
| F04 | 79660001 | 79760000 | 0.0018 | 0.0019 | 0.0015 |
| F04 | 79680001 | 79780000 | 0.0024 | 0.0025 | 0.0021 |
| F04 | 79700001 | 79800000 | 0.0030 | 0.0029 | 0.0027 |
| F04 | 79720001 | 79820000 | 0.0033 | 0.0042 | 0.0030 |
| F04 | 79740001 | 79840000 | 0.0047 | 0.0040 | 0.0043 |
| F04 | 79760001 | 79860000 | 0.0046 | 0.0040 | 0.0041 |
| F04 | 79780001 | 79880000 | 0.0044 | 0.0032 | 0.0039 |
| F04 | 79800001 | 79900000 | 0.0036 | 0.0035 | 0.0032 |
| F04 | 79820001 | 79920000 | 0.0040 | 0.0024 | 0.0036 |
| F04 | 79840001 | 79940000 | 0.0029 | 0.0021 | 0.0026 |
| F04 | 79860001 | 79960000 | 0.0027 | 0.0019 | 0.0023 |
| F04 | 79880001 | 79980000 | 0.0028 | 0.0029 | 0.0021 |
| F04 | 79900001 | 80000000 | 0.0042 | 0.0021 | 0.0033 |
| F04 | 79920001 | 80020000 | 0.0033 | 0.0020 | 0.0024 |
| F04 | 79940001 | 80040000 | 0.0033 | 0.0026 | 0.0024 |
| F04 | 79960001 | 80060000 | 0.0038 | 0.0025 | 0.0031 |
| F04 | 79980001 | 80080000 | 0.0034 | 0.0021 | 0.0028 |
| F04 | 80000001 | 80100000 | 0.0028 | 0.0027 | 0.0023 |
| F04 | 80020001 | 80120000 | 0.0034 | 0.0026 | 0.0026 |
| F04 | 80040001 | 80140000 | 0.0034 | 0.0020 | 0.0025 |
| F04 | 80060001 | 80160000 | 0.0030 | 0.0023 | 0.0020 |
| F04 | 80080001 | 80180000 | 0.0033 | 0.0021 | 0.0023 |
| F04 | 80100001 | 80200000 | 0.0031 | 0.0017 | 0.0023 |
| F04 | 80120001 | 80220000 | 0.0025 | 0.0019 | 0.0021 |
| F04 | 80140001 | 80240000 | 0.0025 | 0.0021 | 0.0022 |
| F04 | 80160001 | 80260000 | 0.0024 | 0.0018 | 0.0023 |
| F04 | 80180001 | 80280000 | 0.0020 | 0.0012 | 0.0020 |
| F04 | 80200001 | 80300000 | 0.0014 | 0.0013 | 0.0014 |
| F04 | 80220001 | 80320000 | 0.0015 | 0.0011 | 0.0015 |
| F04 | 80240001 | 80340000 | 0.0013 | 0.0015 | 0.0013 |
| F04 | 80260001 | 80360000 | 0.0018 | 0.0016 | 0.0017 |
| F04 | 80280001 | 80380000 | 0.0022 | 0.0026 | 0.0020 |

| Chr | Start | End | both | A1 | A2 |
|---|---|---|---|---|---|
| F04 | 80300001 | 80400000 | 0.0033 | 0.0025 | 0.0031 |
| F04 | 80320001 | 80420000 | 0.0035 | 0.0022 | 0.0030 |
| F04 | 80340001 | 80440000 | 0.0032 | 0.0019 | 0.0027 |
| F04 | 80360001 | 80460000 | 0.0027 | 0.0022 | 0.0022 |
| F04 | 80380001 | 80480000 | 0.0032 | 0.0016 | 0.0025 |
| F04 | 80400001 | 80500000 | 0.0027 | 0.0020 | 0.0019 |
| F04 | 80420001 | 80520000 | 0.0031 | 0.0026 | 0.0023 |
| F04 | 80440001 | 80540000 | 0.0040 | 0.0022 | 0.0029 |
| F04 | 80460001 | 80560000 | 0.0035 | 0.0021 | 0.0025 |
| F04 | 80480001 | 80580000 | 0.0031 | 0.0019 | 0.0023 |
| F04 | 80500001 | 80600000 | 0.0026 | 0.0014 | 0.0020 |
| F04 | 80520001 | 80620000 | 0.0019 | 0.0010 | 0.0014 |
| F04 | 80540001 | 80640000 | 0.0015 | 0.0015 | 0.0010 |
| F04 | 80560001 | 80660000 | 0.0021 | 0.0013 | 0.0016 |
| F04 | 80580001 | 80680000 | 0.0022 | 0.0021 | 0.0016 |
| F04 | 80600001 | 80700000 | 0.0030 | 0.0022 | 0.0023 |
| F04 | 80620001 | 80720000 | 0.0034 | 0.0026 | 0.0024 |
| F04 | 80640001 | 80740000 | 0.0038 | 0.0024 | 0.0029 |
| F04 | 80660001 | 80760000 | 0.0038 | 0.0021 | 0.0026 |
| F04 | 80680001 | 80780000 | 0.0036 | 0.0013 | 0.0022 |
| F04 | 80700001 | 80800000 | 0.0030 | 0.0015 | 0.0016 |
| F04 | 80720001 | 80820000 | 0.0032 | 0.0016 | 0.0018 |
| F04 | 80740001 | 80840000 | 0.0034 | 0.0016 | 0.0018 |
| F04 | 80760001 | 80860000 | 0.0032 | 0.0020 | 0.0019 |
| F04 | 80780001 | 80880000 | 0.0033 | 0.0023 | 0.0023 |
| F04 | 80800001 | 80900000 | 0.0036 | 0.0024 | 0.0026 |
| F04 | 80820001 | 80920000 | 0.0035 | 0.0022 | 0.0027 |
| F04 | 80840001 | 80940000 | 0.0031 | 0.0025 | 0.0027 |
| F04 | 80860001 | 80960000 | 0.0033 | 0.0022 | 0.0029 |
| F04 | 80880001 | 80980000 | 0.0029 | 0.0020 | 0.0026 |
| F04 | 80900001 | 81000000 | 0.0028 | 0.0021 | 0.0025 |
| F04 | 80920001 | 81020000 | 0.0031 | 0.0018 | 0.0027 |
| F04 | 80940001 | 81040000 | 0.0030 | 0.0018 | 0.0024 |
| F04 | 80960001 | 81060000 | 0.0032 | 0.0020 | 0.0025 |
| F04 | 80980001 | 81080000 | 0.0038 | 0.0022 | 0.0028 |
| F04 | 81000001 | 81100000 | 0.0038 | 0.0020 | 0.0027 |
| F04 | 81020001 | 81120000 | 0.0035 | 0.0019 | 0.0024 |
| F04 | 81040001 | 81140000 | 0.0032 | 0.0017 | 0.0022 |
| F04 | 81060001 | 81160000 | 0.0031 | 0.0016 | 0.0019 |
| F04 | 81080001 | 81180000 | 0.0027 | 0.0012 | 0.0017 |
| F04 | 81100001 | 81200000 | 0.0023 | 0.0021 | 0.0014 |
| F04 | 81120001 | 81220000 | 0.0031 | 0.0022 | 0.0023 |
| F04 | 81140001 | 81240000 | 0.0035 | 0.0020 | 0.0023 |
| F04 | 81160001 | 81260000 | 0.0029 | 0.0022 | 0.0021 |
| F04 | 81180001 | 81280000 | 0.0035 | 0.0023 | 0.0025 |
| F04 | 81200001 | 81300000 | 0.0035 | 0.0014 | 0.0026 |
| F04 | 81220001 | 81320000 | 0.0025 | 0.0014 | 0.0016 |
| F04 | 81240001 | 81340000 | 0.0023 | 0.0018 | 0.0016 |
| F04 | 81260001 | 81360000 | 0.0026 | 0.0015 | 0.0020 |
| F04 | 81280001 | 81380000 | 0.0020 | 0.0016 | 0.0015 |
| F04 | 81300001 | 81400000 | 0.0024 | 0.0018 | 0.0017 |
| F04 | 81320001 | 81420000 | 0.0025 | 0.0020 | 0.0019 |
| F04 | 81340001 | 81440000 | 0.0031 | 0.0021 | 0.0020 |
| F04 | 81360001 | 81460000 | 0.0036 | 0.0030 | 0.0023 |
| F04 | 81380001 | 81480000 | 0.0046 | 0.0034 | 0.0032 |
| F04 | 81400001 | 81500000 | 0.0051 | 0.0035 | 0.0037 |
| F04 | 81420001 | 81520000 | 0.0054 | 0.0034 | 0.0039 |
| F04 | 81440001 | 81540000 | 0.0051 | 0.0030 | 0.0039 |

| Chr | Start | End | both | A1 | A2 |
|---|---|---|---|---|---|
| F04 | 81460001 | 81560000 | 0.0044 | 0.0022 | 0.0033 |
| F04 | 81480001 | 81580000 | 0.0033 | 0.0016 | 0.0026 |
| F04 | 81500001 | 81600000 | 0.0025 | 0.0011 | 0.0018 |
| F04 | 81520001 | 81620000 | 0.0017 | 0.0010 | 0.0013 |
| F04 | 81540001 | 81640000 | 0.0015 | 0.0010 | 0.0010 |
| F04 | 81560001 | 81660000 | 0.0016 | 0.0012 | 0.0010 |
| F04 | 81580001 | 81680000 | 0.0019 | 0.0012 | 0.0012 |
| F04 | 81600001 | 81700000 | 0.0020 | 0.0015 | 0.0012 |
| F04 | 81620001 | 81720000 | 0.0022 | 0.0015 | 0.0015 |
| F04 | 81640001 | 81740000 | 0.0022 | 0.0019 | 0.0015 |
| F04 | 81660001 | 81760000 | 0.0025 | 0.0017 | 0.0019 |
| F04 | 81680001 | 81780000 | 0.0021 | 0.0017 | 0.0016 |
| F04 | 81700001 | 81800000 | 0.0021 | 0.0016 | 0.0017 |
| F04 | 81720001 | 81820000 | 0.0022 | 0.0017 | 0.0016 |
| F04 | 81740001 | 81840000 | 0.0025 | 0.0014 | 0.0018 |
| F04 | 81760001 | 81860000 | 0.0023 | 0.0014 | 0.0017 |
| F04 | 81780001 | 81880000 | 0.0025 | 0.0014 | 0.0018 |
| F04 | 81800001 | 81900000 | 0.0025 | 0.0017 | 0.0017 |
| F04 | 81820001 | 81920000 | 0.0028 | 0.0021 | 0.0021 |
| F04 | 81840001 | 81940000 | 0.0030 | 0.0027 | 0.0023 |
| F04 | 81860001 | 81960000 | 0.0037 | 0.0026 | 0.0028 |
| F04 | 81880001 | 81980000 | 0.0037 | 0.0038 | 0.0027 |
| F04 | 81900001 | 82000000 | 0.0049 | 0.0034 | 0.0039 |
| F04 | 81920001 | 82020000 | 0.0044 | 0.0027 | 0.0035 |
| F04 | 81940001 | 82040000 | 0.0037 | 0.0020 | 0.0028 |
| F04 | 81960001 | 82060000 | 0.0029 | 0.0025 | 0.0021 |
| F04 | 81980001 | 82080000 | 0.0039 | 0.0013 | 0.0031 |
| F04 | 82000001 | 82100000 | 0.0029 | 0.0014 | 0.0021 |
| F04 | 82020001 | 82120000 | 0.0032 | 0.0020 | 0.0025 |
| F04 | 82040001 | 82140000 | 0.0040 | 0.0021 | 0.0034 |
| F04 | 82060001 | 82160000 | 0.0041 | 0.0015 | 0.0037 |
| F04 | 82080001 | 82180000 | 0.0032 | 0.0017 | 0.0028 |
| F04 | 82100001 | 82200000 | 0.0035 | 0.0015 | 0.0030 |
| F04 | 82120001 | 82220000 | 0.0031 | 0.0012 | 0.0027 |
| F04 | 82140001 | 82240000 | 0.0027 | 0.0012 | 0.0023 |
| F04 | 82160001 | 82260000 | 0.0027 | 0.0012 | 0.0023 |
| F04 | 82180001 | 82280000 | 0.0027 | 0.0010 | 0.0023 |
| F04 | 82200001 | 82300000 | 0.0028 | 0.0015 | 0.0024 |
| F04 | 82220001 | 82320000 | 0.0034 | 0.0016 | 0.0031 |
| F04 | 82240001 | 82340000 | 0.0038 | 0.0017 | 0.0035 |
| F04 | 82260001 | 82360000 | 0.0039 | 0.0017 | 0.0034 |
| F04 | 82280001 | 82380000 | 0.0038 | 0.0015 | 0.0031 |
| F04 | 82300001 | 82400000 | 0.0031 | 0.0015 | 0.0024 |
| F04 | 82320001 | 82420000 | 0.0030 | 0.0012 | 0.0021 |
| F04 | 82340001 | 82440000 | 0.0023 | 0.0011 | 0.0014 |
| F04 | 82360001 | 82460000 | 0.0020 | 0.0010 | 0.0011 |
| F04 | 82380001 | 82480000 | 0.0021 | 0.0011 | 0.0012 |
| F04 | 82400001 | 82500000 | 0.0021 | 0.0009 | 0.0013 |
| F04 | 82420001 | 82520000 | 0.0018 | 0.0012 | 0.0011 |
| F04 | 82440001 | 82540000 | 0.0021 | 0.0019 | 0.0014 |
| F04 | 82460001 | 82560000 | 0.0031 | 0.0019 | 0.0022 |
| F04 | 82480001 | 82580000 | 0.0029 | 0.0029 | 0.0021 |
| F04 | 82500001 | 82600000 | 0.0038 | 0.0026 | 0.0032 |
| F04 | 82520001 | 82620000 | 0.0035 | 0.0025 | 0.0029 |
| F04 | 82540001 | 82640000 | 0.0034 | 0.0018 | 0.0029 |
| F04 | 82560001 | 82660000 | 0.0025 | 0.0020 | 0.0020 |
| F04 | 82580001 | 82680000 | 0.0027 | 0.0015 | 0.0023 |
| F04 | 82600001 | 82700000 | 0.0022 | 0.0019 | 0.0016 |

| Chr | Start | End | both | A1 | A2 |
|---|---|---|---|---|---|
| F04 | 82620001 | 82720000 | 0.0026 | 0.0019 | 0.0020 |
| F04 | 82640001 | 82740000 | 0.0024 | 0.0029 | 0.0018 |
| F04 | 82660001 | 82760000 | 0.0036 | 0.0029 | 0.0031 |
| F04 | 82680001 | 82780000 | 0.0037 | 0.0026 | 0.0030 |
| F04 | 82700001 | 82800000 | 0.0033 | 0.0025 | 0.0028 |
| F04 | 82720001 | 82820000 | 0.0033 | 0.0037 | 0.0027 |
| F04 | 82740001 | 82840000 | 0.0045 | 0.0026 | 0.0040 |
| F04 | 82760001 | 82860000 | 0.0034 | 0.0026 | 0.0028 |
| F04 | 82780001 | 82880000 | 0.0034 | 0.0026 | 0.0028 |
| F04 | 82800001 | 82900000 | 0.0035 | 0.0023 | 0.0028 |
| F04 | 82820001 | 82920000 | 0.0031 | 0.0015 | 0.0025 |
| F04 | 82840001 | 82940000 | 0.0025 | 0.0016 | 0.0016 |
| F04 | 82860001 | 82960000 | 0.0028 | 0.0018 | 0.0017 |
| F04 | 82880001 | 82980000 | 0.0025 | 0.0022 | 0.0018 |
| F04 | 82900001 | 83000000 | 0.0030 | 0.0025 | 0.0024 |
| F04 | 82920001 | 83020000 | 0.0033 | 0.0022 | 0.0027 |
| F04 | 82940001 | 83040000 | 0.0029 | 0.0029 | 0.0025 |
| F04 | 82960001 | 83060000 | 0.0034 | 0.0038 | 0.0030 |
| F04 | 82980001 | 83080000 | 0.0045 | 0.0033 | 0.0041 |
| F04 | 83000001 | 83100000 | 0.0042 | 0.0033 | 0.0035 |
| F04 | 83020001 | 83120000 | 0.0042 | 0.0033 | 0.0035 |
| F04 | 83040001 | 83140000 | 0.0041 | 0.0034 | 0.0034 |
| F04 | 83060001 | 83160000 | 0.0044 | 0.0025 | 0.0037 |
| F04 | 83080001 | 83180000 | 0.0035 | 0.0024 | 0.0027 |
| F04 | 83100001 | 83200000 | 0.0033 | 0.0027 | 0.0024 |
| F04 | 83120001 | 83220000 | 0.0037 | 0.0031 | 0.0027 |
| F04 | 83140001 | 83240000 | 0.0041 | 0.0026 | 0.0031 |
| F04 | 83160001 | 83260000 | 0.0034 | 0.0026 | 0.0024 |
| F04 | 83180001 | 83280000 | 0.0033 | 0.0033 | 0.0024 |
| F04 | 83200001 | 83300000 | 0.0039 | 0.0030 | 0.0030 |
| F04 | 83220001 | 83320000 | 0.0037 | 0.0023 | 0.0029 |
| F04 | 83240001 | 83340000 | 0.0029 | 0.0023 | 0.0022 |
| F04 | 83260001 | 83360000 | 0.0029 | 0.0026 | 0.0021 |
| F04 | 83280001 | 83380000 | 0.0033 | 0.0020 | 0.0025 |
| F04 | 83300001 | 83400000 | 0.0028 | 0.0020 | 0.0019 |
| F04 | 83320001 | 83420000 | 0.0027 | 0.0019 | 0.0017 |
| F04 | 83340001 | 83440000 | 0.0028 | 0.0017 | 0.0018 |
| F04 | 83360001 | 83460000 | 0.0025 | 0.0012 | 0.0016 |
| F04 | 83380001 | 83480000 | 0.0020 | 0.0014 | 0.0011 |
| F04 | 83400001 | 83500000 | 0.0022 | 0.0018 | 0.0013 |
| F04 | 83420001 | 83520000 | 0.0027 | 0.0025 | 0.0018 |
| F04 | 83440001 | 83540000 | 0.0035 | 0.0026 | 0.0028 |
| F04 | 83460001 | 83560000 | 0.0037 | 0.0032 | 0.0028 |
| F04 | 83480001 | 83580000 | 0.0044 | 0.0033 | 0.0033 |
| F04 | 83500001 | 83600000 | 0.0045 | 0.0034 | 0.0034 |
| F04 | 83520001 | 83620000 | 0.0044 | 0.0028 | 0.0033 |
| F04 | 83540001 | 83640000 | 0.0036 | 0.0029 | 0.0025 |
| F04 | 83560001 | 83660000 | 0.0036 | 0.0023 | 0.0025 |
| F04 | 83580001 | 83680000 | 0.0031 | 0.0022 | 0.0022 |
| F04 | 83600001 | 83700000 | 0.0028 | 0.0028 | 0.0020 |
| F04 | 83620001 | 83720000 | 0.0036 | 0.0031 | 0.0027 |
| F04 | 83640001 | 83740000 | 0.0040 | 0.0035 | 0.0030 |
| F04 | 83660001 | 83760000 | 0.0043 | 0.0037 | 0.0034 |
| F04 | 83680001 | 83780000 | 0.0044 | 0.0047 | 0.0036 |
| F04 | 83700001 | 83800000 | 0.0057 | 0.0037 | 0.0050 |
| F04 | 83720001 | 83820000 | 0.0046 | 0.0034 | 0.0041 |
| F04 | 83740001 | 83840000 | 0.0043 | 0.0033 | 0.0038 |
| F04 | 83760001 | 83860000 | 0.0042 | 0.0056 | 0.0036 |

| Chr | Start | End | both | A1 | A2 |
|---|---|---|---|---|---|
| F04 | 83780001 | 83880000 | 0.0068 | 0.0046 | 0.0062 |
| F04 | 83800001 | 83900000 | 0.0056 | 0.0048 | 0.0049 |
| F04 | 83820001 | 83920000 | 0.0058 | 0.0047 | 0.0052 |
| F04 | 83840001 | 83940000 | 0.0056 | 0.0050 | 0.0052 |
| F04 | 83860001 | 83960000 | 0.0060 | 0.0039 | 0.0056 |
| F04 | 83880001 | 83980000 | 0.0047 | 0.0036 | 0.0042 |
| F04 | 83900001 | 84000000 | 0.0046 | 0.0033 | 0.0042 |
| F04 | 83920001 | 84020000 | 0.0042 | 0.0032 | 0.0038 |
| F04 | 83940001 | 84040000 | 0.0040 | 0.0025 | 0.0036 |
| F04 | 83960001 | 84060000 | 0.0033 | 0.0009 | 0.0031 |
| F04 | 83980001 | 84080000 | 0.0016 | 0.0018 | 0.0014 |
| F04 | 84000001 | 84100000 | 0.0025 | 0.0031 | 0.0021 |
| F04 | 84020001 | 84120000 | 0.0039 | 0.0032 | 0.0035 |
| F04 | 84040001 | 84140000 | 0.0042 | 0.0032 | 0.0035 |
| F04 | 84060001 | 84160000 | 0.0042 | 0.0042 | 0.0033 |
| F04 | 84080001 | 84180000 | 0.0053 | 0.0032 | 0.0042 |
| F04 | 84100001 | 84200000 | 0.0042 | 0.0018 | 0.0032 |
| F04 | 84120001 | 84220000 | 0.0027 | 0.0018 | 0.0017 |
| F04 | 84140001 | 84240000 | 0.0026 | 0.0019 | 0.0017 |
| F04 | 84160001 | 84260000 | 0.0026 | 0.0013 | 0.0019 |
| F04 | 84180001 | 84280000 | 0.0019 | 0.0017 | 0.0012 |
| F04 | 84200001 | 84300000 | 0.0022 | 0.0018 | 0.0016 |
| F04 | 84220001 | 84320000 | 0.0022 | 0.0018 | 0.0016 |
| F04 | 84240001 | 84340000 | 0.0022 | 0.0015 | 0.0016 |
| F04 | 84260001 | 84360000 | 0.0020 | 0.0014 | 0.0013 |
| F04 | 84280001 | 84380000 | 0.0019 | 0.0016 | 0.0012 |
| F04 | 84300001 | 84400000 | 0.0022 | 0.0023 | 0.0015 |
| F04 | 84320001 | 84420000 | 0.0030 | 0.0032 | 0.0023 |
| F04 | 84340001 | 84440000 | 0.0039 | 0.0031 | 0.0031 |
| F04 | 84360001 | 84460000 | 0.0039 | 0.0041 | 0.0030 |
| F04 | 84380001 | 84480000 | 0.0049 | 0.0035 | 0.0042 |
| F04 | 84400001 | 84500000 | 0.0043 | 0.0028 | 0.0036 |
| F04 | 84420001 | 84520000 | 0.0035 | 0.0028 | 0.0029 |
| F04 | 84440001 | 84540000 | 0.0036 | 0.0033 | 0.0030 |
| F04 | 84460001 | 84560000 | 0.0038 | 0.0024 | 0.0031 |
| F04 | 84480001 | 84580000 | 0.0026 | 0.0022 | 0.0020 |
| F04 | 84500001 | 84600000 | 0.0025 | 0.0025 | 0.0018 |
| F04 | 84520001 | 84620000 | 0.0031 | 0.0022 | 0.0019 |
| F04 | 84540001 | 84640000 | 0.0030 | 0.0021 | 0.0017 |
| F04 | 84560001 | 84660000 | 0.0033 | 0.0021 | 0.0023 |
| F04 | 84580001 | 84680000 | 0.0032 | 0.0022 | 0.0022 |
| F04 | 84600001 | 84700000 | 0.0034 | 0.0025 | 0.0022 |
| F04 | 84620001 | 84720000 | 0.0035 | 0.0022 | 0.0028 |
| F04 | 84640001 | 84740000 | 0.0028 | 0.0026 | 0.0024 |
| F04 | 84660001 | 84760000 | 0.0033 | 0.0029 | 0.0025 |
| F04 | 84680001 | 84780000 | 0.0036 | 0.0031 | 0.0028 |
| F04 | 84700001 | 84800000 | 0.0038 | 0.0024 | 0.0031 |
| F04 | 84720001 | 84820000 | 0.0031 | 0.0030 | 0.0022 |
| F04 | 84740001 | 84840000 | 0.0038 | 0.0027 | 0.0029 |
| F04 | 84760001 | 84860000 | 0.0033 | 0.0024 | 0.0026 |
| F04 | 84780001 | 84880000 | 0.0030 | 0.0022 | 0.0024 |
| F04 | 84800001 | 84900000 | 0.0029 | 0.0025 | 0.0023 |
| F04 | 84820001 | 84920000 | 0.0032 | 0.0014 | 0.0027 |
| F04 | 84840001 | 84940000 | 0.0021 | 0.0012 | 0.0017 |
| F04 | 84860001 | 84960000 | 0.0020 | 0.0011 | 0.0017 |
| F04 | 84880001 | 84980000 | 0.0021 | 0.0014 | 0.0017 |
| F04 | 84900001 | 85000000 | 0.0023 | 0.0019 | 0.0019 |
| F04 | 84920001 | 85020000 | 0.0029 | 0.0039 | 0.0026 |

| Chr | Start | End | both | A1 | A2 |
|---|---|---|---|---|---|
| F04 | 84940001 | 85040000 | 0.0049 | 0.0037 | 0.0045 |
| F04 | 84960001 | 85060000 | 0.0044 | 0.0044 | 0.0041 |
| F04 | 84980001 | 85080000 | 0.0050 | 0.0044 | 0.0048 |
| F04 | 85000001 | 85100000 | 0.0049 | 0.0038 | 0.0047 |
| F04 | 85020001 | 85120000 | 0.0043 | 0.0022 | 0.0041 |
| F04 | 85040001 | 85140000 | 0.0028 | 0.0026 | 0.0024 |
| F04 | 85060001 | 85160000 | 0.0032 | 0.0020 | 0.0027 |
| F04 | 85080001 | 85180000 | 0.0027 | 0.0020 | 0.0019 |
| F04 | 85100001 | 85200000 | 0.0030 | 0.0020 | 0.0021 |
| F04 | 85120001 | 85220000 | 0.0030 | 0.0023 | 0.0020 |
| F04 | 85140001 | 85240000 | 0.0032 | 0.0019 | 0.0023 |
| F04 | 85160001 | 85260000 | 0.0027 | 0.0019 | 0.0019 |
| F04 | 85180001 | 85280000 | 0.0025 | 0.0016 | 0.0019 |
| F04 | 85200001 | 85300000 | 0.0021 | 0.0012 | 0.0017 |
| F04 | 85220001 | 85320000 | 0.0017 | 0.0007 | 0.0015 |
| F04 | 85240001 | 85340000 | 0.0012 | 0.0006 | 0.0012 |
| F04 | 85260001 | 85360000 | 0.0013 | 0.0005 | 0.0013 |
| F04 | 85280001 | 85380000 | 0.0015 | 0.0004 | 0.0015 |
| F04 | 85300001 | 85400000 | 0.0017 | 0.0007 | 0.0017 |
| F04 | 85320001 | 85420000 | 0.0020 | 0.0010 | 0.0021 |
| F04 | 85340001 | 85440000 | 0.0027 | 0.0015 | 0.0027 |
| F04 | 85360001 | 85460000 | 0.0035 | 0.0020 | 0.0034 |
| F04 | 85380001 | 85480000 | 0.0040 | 0.0021 | 0.0039 |
| F04 | 85400001 | 85500000 | 0.0037 | 0.0022 | 0.0037 |
| F04 | 85420001 | 85520000 | 0.0037 | 0.0019 | 0.0037 |
| F04 | 85440001 | 85540000 | 0.0030 | 0.0015 | 0.0030 |
| F04 | 85460001 | 85560000 | 0.0024 | 0.0011 | 0.0024 |
| F04 | 85480001 | 85580000 | 0.0018 | 0.0011 | 0.0018 |
| F04 | 85500001 | 85600000 | 0.0019 | 0.0007 | 0.0019 |
| F04 | 85520001 | 85620000 | 0.0015 | 0.0008 | 0.0016 |
| F04 | 85540001 | 85640000 | 0.0018 | 0.0016 | 0.0019 |
| F04 | 85560001 | 85660000 | 0.0024 | 0.0017 | 0.0025 |
| F04 | 85580001 | 85680000 | 0.0025 | 0.0023 | 0.0025 |
| F04 | 85600001 | 85700000 | 0.0030 | 0.0025 | 0.0030 |
| F04 | 85620001 | 85720000 | 0.0036 | 0.0029 | 0.0036 |
| F04 | 85640001 | 85740000 | 0.0041 | 0.0026 | 0.0041 |
| F04 | 85660001 | 85760000 | 0.0038 | 0.0036 | 0.0039 |
| F04 | 85680001 | 85780000 | 0.0048 | 0.0038 | 0.0049 |
| F04 | 85700001 | 85800000 | 0.0050 | 0.0043 | 0.0051 |
| F04 | 85720001 | 85820000 | 0.0055 | 0.0044 | 0.0056 |
| F04 | 85740001 | 85840000 | 0.0055 | 0.0043 | 0.0056 |
| F04 | 85760001 | 85860000 | 0.0054 | 0.0035 | 0.0054 |
| F04 | 85780001 | 85880000 | 0.0046 | 0.0026 | 0.0047 |
| F04 | 85800001 | 85900000 | 0.0039 | 0.0025 | 0.0040 |
| F04 | 85820001 | 85920000 | 0.0034 | 0.0022 | 0.0035 |
| F04 | 85840001 | 85940000 | 0.0032 | 0.0018 | 0.0032 |
| F04 | 85860001 | 85960000 | 0.0028 | 0.0016 | 0.0029 |
| F04 | 85880001 | 85980000 | 0.0026 | 0.0015 | 0.0026 |
| F04 | 85900001 | 86000000 | 0.0026 | 0.0011 | 0.0026 |
| F04 | 85920001 | 86020000 | 0.0024 | 0.0006 | 0.0022 |
| F04 | 85940001 | 86040000 | 0.0019 | 0.0009 | 0.0016 |
| F04 | 85960001 | 86060000 | 0.0021 | 0.0011 | 0.0017 |
| F04 | 85980001 | 86080000 | 0.0022 | 0.0019 | 0.0018 |
| F04 | 86000001 | 86100000 | 0.0029 | 0.0017 | 0.0025 |
| F04 | 86020001 | 86120000 | 0.0027 | 0.0022 | 0.0022 |
| F04 | 86040001 | 86140000 | 0.0033 | 0.0034 | 0.0027 |
| F04 | 86060001 | 86160000 | 0.0039 | 0.0034 | 0.0031 |
| F04 | 86080001 | 86180000 | 0.0038 | 0.0035 | 0.0030 |

| Chr | Start | End | both | A1 | A2 |
|---|---|---|---|---|---|
| F04 | 86100001 | 86200000 | 0.0037 | 0.0037 | 0.0028 |
| F04 | 86120001 | 86220000 | 0.0036 | 0.0032 | 0.0030 |
| F04 | 86140001 | 86240000 | 0.0028 | 0.0016 | 0.0024 |
| F04 | 86160001 | 86260000 | 0.0017 | 0.0015 | 0.0016 |
| F04 | 86180001 | 86280000 | 0.0015 | 0.0008 | 0.0014 |
| F04 | 86200001 | 86300000 | 0.0007 | 0.0006 | 0.0007 |
| F04 | 86220001 | 86320000 | 0.0006 | 0.0009 | 0.0005 |
| F04 | 86240001 | 86340000 | 0.0009 | 0.0019 | 0.0008 |
| F04 | 86260001 | 86360000 | 0.0022 | 0.0021 | 0.0022 |
| F04 | 86280001 | 86380000 | 0.0026 | 0.0025 | 0.0025 |
| F04 | 86300001 | 86400000 | 0.0031 | 0.0027 | 0.0029 |
| F04 | 86320001 | 86420000 | 0.0034 | 0.0024 | 0.0031 |
| F04 | 86340001 | 86440000 | 0.0034 | 0.0015 | 0.0030 |
| F04 | 86360001 | 86460000 | 0.0024 | 0.0015 | 0.0018 |
| F04 | 86380001 | 86480000 | 0.0025 | 0.0011 | 0.0018 |
| F04 | 86400001 | 86500000 | 0.0023 | 0.0009 | 0.0016 |
| F04 | 86420001 | 86520000 | 0.0021 | 0.0009 | 0.0013 |
| F04 | 86440001 | 86540000 | 0.0019 | 0.0009 | 0.0012 |
| F04 | 86460001 | 86560000 | 0.0021 | 0.0007 | 0.0012 |
| F04 | 86480001 | 86580000 | 0.0016 | 0.0008 | 0.0009 |
| F04 | 86500001 | 86600000 | 0.0015 | 0.0009 | 0.0009 |
| F04 | 86520001 | 86620000 | 0.0015 | 0.0008 | 0.0009 |
| F04 | 86540001 | 86640000 | 0.0015 | 0.0008 | 0.0009 |
| F04 | 86560001 | 86660000 | 0.0014 | 0.0008 | 0.0008 |
| F04 | 86580001 | 86680000 | 0.0016 | 0.0007 | 0.0009 |
| F04 | 86600001 | 86700000 | 0.0016 | 0.0007 | 0.0006 |
| F04 | 86620001 | 86720000 | 0.0016 | 0.0008 | 0.0006 |
| F04 | 86640001 | 86740000 | 0.0017 | 0.0009 | 0.0006 |
| F04 | 86660001 | 86760000 | 0.0017 | 0.0010 | 0.0008 |
| F04 | 86680001 | 86780000 | 0.0018 | 0.0009 | 0.0009 |
| F04 | 86700001 | 86800000 | 0.0017 | 0.0008 | 0.0009 |
| F04 | 86720001 | 86820000 | 0.0016 | 0.0007 | 0.0009 |
| F04 | 86740001 | 86840000 | 0.0018 | 0.0006 | 0.0011 |
| F04 | 86760001 | 86860000 | 0.0018 | 0.0011 | 0.0010 |
| F04 | 86780001 | 86880000 | 0.0021 | 0.0011 | 0.0014 |
| F04 | 86800001 | 86900000 | 0.0021 | 0.0012 | 0.0014 |
| F04 | 86820001 | 86920000 | 0.0023 | 0.0012 | 0.0015 |
| F04 | 86840001 | 86940000 | 0.0021 | 0.0016 | 0.0014 |
| F04 | 86860001 | 86960000 | 0.0024 | 0.0010 | 0.0018 |
| F04 | 86880001 | 86980000 | 0.0018 | 0.0009 | 0.0012 |
| F04 | 86900001 | 87000000 | 0.0018 | 0.0008 | 0.0012 |
| F04 | 86920001 | 87020000 | 0.0016 | 0.0015 | 0.0011 |
| F04 | 86940001 | 87040000 | 0.0025 | 0.0011 | 0.0019 |
| F04 | 86960001 | 87060000 | 0.0024 | 0.0011 | 0.0014 |
| F04 | 86980001 | 87080000 | 0.0023 | 0.0012 | 0.0015 |
| F04 | 87000001 | 87100000 | 0.0023 | 0.0014 | 0.0016 |
| F04 | 87020001 | 87120000 | 0.0025 | 0.0006 | 0.0018 |
| F04 | 87040001 | 87140000 | 0.0017 | 0.0006 | 0.0009 |
| F04 | 87060001 | 87160000 | 0.0016 | 0.0005 | 0.0010 |
| F04 | 87080001 | 87180000 | 0.0018 | 0.0008 | 0.0010 |
| F04 | 87100001 | 87200000 | 0.0022 | 0.0007 | 0.0013 |
| F04 | 87120001 | 87220000 | 0.0020 | 0.0013 | 0.0012 |
| F04 | 87140001 | 87240000 | 0.0025 | 0.0013 | 0.0018 |
| F04 | 87160001 | 87260000 | 0.0022 | 0.0013 | 0.0017 |
| F04 | 87180001 | 87280000 | 0.0019 | 0.0010 | 0.0016 |
| F04 | 87200001 | 87300000 | 0.0015 | 0.0010 | 0.0012 |
| F04 | 87220001 | 87320000 | 0.0017 | 0.0011 | 0.0013 |
| F04 | 87240001 | 87340000 | 0.0017 | 0.0022 | 0.0013 |

| Chr | Start | End | both | A1 | A2 |
|---|---|---|---|---|---|
| F04 | 87260001 | 87360000 | 0.0031 | 0.0022 | 0.0027 |
| F04 | 87280001 | 87380000 | 0.0033 | 0.0026 | 0.0028 |
| F04 | 87300001 | 87400000 | 0.0038 | 0.0029 | 0.0033 |
| F04 | 87320001 | 87420000 | 0.0042 | 0.0029 | 0.0036 |
| F04 | 87340001 | 87440000 | 0.0041 | 0.0018 | 0.0036 |
| F04 | 87360001 | 87460000 | 0.0028 | 0.0019 | 0.0022 |
| F04 | 87380001 | 87480000 | 0.0030 | 0.0015 | 0.0022 |
| F04 | 87400001 | 87500000 | 0.0027 | 0.0020 | 0.0018 |
| F04 | 87420001 | 87520000 | 0.0029 | 0.0030 | 0.0022 |
| F04 | 87440001 | 87540000 | 0.0040 | 0.0041 | 0.0034 |
| F04 | 87460001 | 87560000 | 0.0051 | 0.0050 | 0.0044 |
| F04 | 87480001 | 87580000 | 0.0062 | 0.0050 | 0.0054 |
| F04 | 87500001 | 87600000 | 0.0060 | 0.0054 | 0.0054 |
| F04 | 87520001 | 87620000 | 0.0065 | 0.0045 | 0.0058 |
| F04 | 87540001 | 87640000 | 0.0053 | 0.0037 | 0.0046 |
| F04 | 87560001 | 87660000 | 0.0046 | 0.0026 | 0.0040 |
| F04 | 87580001 | 87680000 | 0.0030 | 0.0031 | 0.0028 |
| F04 | 87600001 | 87700000 | 0.0036 | 0.0020 | 0.0034 |
| F04 | 87620001 | 87720000 | 0.0026 | 0.0016 | 0.0022 |
| F04 | 87640001 | 87740000 | 0.0024 | 0.0021 | 0.0019 |
| F04 | 87660001 | 87760000 | 0.0031 | 0.0026 | 0.0025 |
| F04 | 87680001 | 87780000 | 0.0038 | 0.0023 | 0.0031 |
| F04 | 87700001 | 87800000 | 0.0038 | 0.0022 | 0.0029 |
| F04 | 87720001 | 87820000 | 0.0037 | 0.0020 | 0.0029 |
| F04 | 87740001 | 87840000 | 0.0035 | 0.0011 | 0.0026 |
| F04 | 87760001 | 87860000 | 0.0024 | 0.0007 | 0.0017 |
| F04 | 87780001 | 87880000 | 0.0018 | 0.0004 | 0.0010 |
| F04 | 87800001 | 87900000 | 0.0012 | 0.0008 | 0.0006 |
| F04 | 87820001 | 87920000 | 0.0015 | 0.0010 | 0.0010 |
| F04 | 87840001 | 87940000 | 0.0016 | 0.0010 | 0.0012 |
| F04 | 87860001 | 87960000 | 0.0018 | 0.0028 | 0.0013 |
| F04 | 87880001 | 87980000 | 0.0037 | 0.0030 | 0.0031 |
| F04 | 87900001 | 88000000 | 0.0041 | 0.0031 | 0.0033 |
| F04 | 87920001 | 88020000 | 0.0041 | 0.0031 | 0.0033 |
| F04 | 87940001 | 88040000 | 0.0040 | 0.0032 | 0.0033 |
| F04 | 87960001 | 88060000 | 0.0040 | 0.0015 | 0.0034 |
| F04 | 87980001 | 88080000 | 0.0024 | 0.0015 | 0.0017 |
| F04 | 88000001 | 88100000 | 0.0023 | 0.0011 | 0.0017 |
| F04 | 88020001 | 88120000 | 0.0020 | 0.0010 | 0.0012 |
| F04 | 88040001 | 88140000 | 0.0020 | 0.0009 | 0.0009 |
| F04 | 88060001 | 88160000 | 0.0020 | 0.0010 | 0.0008 |
| F04 | 88080001 | 88180000 | 0.0021 | 0.0010 | 0.0009 |
| F04 | 88100001 | 88200000 | 0.0021 | 0.0009 | 0.0006 |
| F04 | 88120001 | 88220000 | 0.0020 | 0.0010 | 0.0006 |
| F04 | 88140001 | 88240000 | 0.0021 | 0.0010 | 0.0009 |
| F04 | 88160001 | 88260000 | 0.0020 | 0.0009 | 0.0009 |
| F04 | 88180001 | 88280000 | 0.0020 | 0.0008 | 0.0009 |
| F04 | 88200001 | 88300000 | 0.0019 | 0.0008 | 0.0009 |
| F04 | 88220001 | 88320000 | 0.0018 | 0.0006 | 0.0009 |
| F04 | 88240001 | 88340000 | 0.0017 | 0.0006 | 0.0007 |
| F04 | 88260001 | 88360000 | 0.0016 | 0.0005 | 0.0005 |
| F04 | 88280001 | 88380000 | 0.0014 | 0.0006 | 0.0004 |
| F04 | 88300001 | 88400000 | 0.0013 | 0.0010 | 0.0006 |
| F04 | 88320001 | 88420000 | 0.0016 | 0.0011 | 0.0009 |
| F04 | 88340001 | 88440000 | 0.0016 | 0.0012 | 0.0011 |
| F04 | 88360001 | 88460000 | 0.0016 | 0.0024 | 0.0011 |
| F04 | 88380001 | 88480000 | 0.0028 | 0.0024 | 0.0023 |
| F04 | 88400001 | 88500000 | 0.0028 | 0.0020 | 0.0022 |

| Chr | Start | End | both | A1 | A2 |
|---|---|---|---|---|---|
| F04 | 88420001 | 88520000 | 0.0026 | 0.0019 | 0.0018 |
| F04 | 88440001 | 88540000 | 0.0027 | 0.0022 | 0.0017 |
| F04 | 88460001 | 88560000 | 0.0029 | 0.0011 | 0.0019 |
| F04 | 88480001 | 88580000 | 0.0019 | 0.0009 | 0.0008 |
| F04 | 88500001 | 88600000 | 0.0019 | 0.0011 | 0.0007 |
| F04 | 88520001 | 88620000 | 0.0019 | 0.0013 | 0.0008 |
| F04 | 88540001 | 88640000 | 0.0018 | 0.0013 | 0.0008 |
| F04 | 88560001 | 88660000 | 0.0016 | 0.0016 | 0.0006 |
| F04 | 88580001 | 88680000 | 0.0014 | 0.0018 | 0.0007 |
| F04 | 88600001 | 88700000 | 0.0013 | 0.0024 | 0.0007 |
| F04 | 88620001 | 88720000 | 0.0018 | 0.0025 | 0.0013 |
| F04 | 88640001 | 88740000 | 0.0016 | 0.0028 | 0.0012 |
| F04 | 88660001 | 88760000 | 0.0017 | 0.0027 | 0.0013 |
| F04 | 88680001 | 88780000 | 0.0016 | 0.0029 | 0.0011 |
| F04 | 88700001 | 88800000 | 0.0016 | 0.0024 | 0.0011 |
| F04 | 88720001 | 88820000 | 0.0012 | 0.0030 | 0.0008 |
| F04 | 88740001 | 88840000 | 0.0021 | 0.0035 | 0.0018 |
| F04 | 88760001 | 88860000 | 0.0030 | 0.0037 | 0.0027 |
| F04 | 88780001 | 88880000 | 0.0033 | 0.0038 | 0.0030 |
| F04 | 88800001 | 88900000 | 0.0034 | 0.0037 | 0.0031 |
| F04 | 88820001 | 88920000 | 0.0033 | 0.0030 | 0.0030 |
| F04 | 88840001 | 88940000 | 0.0024 | 0.0021 | 0.0020 |
| F04 | 88860001 | 88960000 | 0.0013 | 0.0022 | 0.0009 |
| F04 | 88880001 | 88980000 | 0.0012 | 0.0022 | 0.0008 |
| F04 | 88900001 | 89000000 | 0.0014 | 0.0030 | 0.0011 |
| F04 | 88920001 | 89020000 | 0.0023 | 0.0039 | 0.0020 |
| F04 | 88940001 | 89040000 | 0.0032 | 0.0048 | 0.0029 |
| F04 | 88960001 | 89060000 | 0.0045 | 0.0056 | 0.0043 |
| F04 | 88980001 | 89080000 | 0.0053 | 0.0054 | 0.0051 |
| F04 | 89000001 | 89100000 | 0.0051 | 0.0046 | 0.0049 |
| F04 | 89020001 | 89120000 | 0.0042 | 0.0046 | 0.0040 |
| F04 | 89040001 | 89140000 | 0.0042 | 0.0035 | 0.0041 |
| F04 | 89060001 | 89160000 | 0.0034 | 0.0025 | 0.0033 |
| F04 | 89080001 | 89180000 | 0.0026 | 0.0022 | 0.0026 |
| F04 | 89100001 | 89200000 | 0.0026 | 0.0022 | 0.0027 |
| F04 | 89120001 | 89220000 | 0.0031 | 0.0012 | 0.0032 |
| F04 | 89140001 | 89240000 | 0.0023 | 0.0019 | 0.0025 |
| F04 | 89160001 | 89260000 | 0.0034 | 0.0030 | 0.0035 |
| F04 | 89180001 | 89280000 | 0.0045 | 0.0030 | 0.0045 |
| F04 | 89200001 | 89300000 | 0.0048 | 0.0028 | 0.0048 |
| F04 | 89220001 | 89320000 | 0.0045 | 0.0028 | 0.0044 |
| F04 | 89240001 | 89340000 | 0.0046 | 0.0023 | 0.0042 |
| F04 | 89260001 | 89360000 | 0.0037 | 0.0010 | 0.0029 |
| F04 | 89280001 | 89380000 | 0.0024 | 0.0011 | 0.0016 |
| F04 | 89300001 | 89400000 | 0.0023 | 0.0011 | 0.0013 |
| F04 | 89320001 | 89420000 | 0.0022 | 0.0016 | 0.0010 |
| F04 | 89340001 | 89440000 | 0.0026 | 0.0011 | 0.0015 |
| F04 | 89360001 | 89460000 | 0.0022 | 0.0014 | 0.0013 |
| F04 | 89380001 | 89480000 | 0.0023 | 0.0017 | 0.0013 |
| F04 | 89400001 | 89500000 | 0.0024 | 0.0019 | 0.0016 |
| F04 | 89420001 | 89520000 | 0.0022 | 0.0019 | 0.0016 |
| F04 | 89440001 | 89540000 | 0.0019 | 0.0020 | 0.0014 |
| F04 | 89460001 | 89560000 | 0.0017 | 0.0023 | 0.0014 |
| F04 | 89480001 | 89580000 | 0.0019 | 0.0028 | 0.0018 |
| F04 | 89500001 | 89600000 | 0.0024 | 0.0030 | 0.0021 |
| F04 | 89520001 | 89620000 | 0.0027 | 0.0033 | 0.0025 |
| F04 | 89540001 | 89640000 | 0.0032 | 0.0039 | 0.0029 |
| F04 | 89560001 | 89660000 | 0.0042 | 0.0047 | 0.0039 |

| Chr | Start | End | both | A1 | A2 |
|---|---|---|---|---|---|
| F04 | 89580001 | 89680000 | 0.0054 | 0.0050 | 0.0052 |
| F04 | 89600001 | 89700000 | 0.0059 | 0.0046 | 0.0057 |
| F04 | 89620001 | 89720000 | 0.0059 | 0.0047 | 0.0056 |
| F04 | 89640001 | 89740000 | 0.0061 | 0.0041 | 0.0058 |
| F04 | 89660001 | 89760000 | 0.0053 | 0.0029 | 0.0049 |
| F04 | 89680001 | 89780000 | 0.0039 | 0.0015 | 0.0032 |
| F04 | 89700001 | 89800000 | 0.0028 | 0.0016 | 0.0020 |
| F04 | 89720001 | 89820000 | 0.0028 | 0.0008 | 0.0019 |
| F04 | 89740001 | 89840000 | 0.0026 | 0.0007 | 0.0015 |
| F04 | 89760001 | 89860000 | 0.0024 | 0.0006 | 0.0014 |
| F04 | 89780001 | 89880000 | 0.0023 | 0.0006 | 0.0014 |
| F04 | 89800001 | 89900000 | 0.0022 | 0.0008 | 0.0013 |
| F04 | 89820001 | 89920000 | 0.0019 | 0.0014 | 0.0010 |
| F04 | 89840001 | 89940000 | 0.0017 | 0.0017 | 0.0009 |
| F04 | 89860001 | 89960000 | 0.0018 | 0.0018 | 0.0009 |
| F04 | 89880001 | 89980000 | 0.0016 | 0.0021 | 0.0008 |
| F04 | 89900001 | 90000000 | 0.0017 | 0.0020 | 0.0008 |
| F04 | 89920001 | 90020000 | 0.0017 | 0.0015 | 0.0008 |
| F04 | 89940001 | 90040000 | 0.0012 | 0.0013 | 0.0003 |
| F04 | 89960001 | 90060000 | 0.0013 | 0.0013 | 0.0003 |
| F04 | 89980001 | 90080000 | 0.0014 | 0.0010 | 0.0005 |
| F04 | 90000001 | 90100000 | 0.0015 | 0.0008 | 0.0006 |
| F04 | 90020001 | 90120000 | 0.0018 | 0.0007 | 0.0009 |
| F04 | 90040001 | 90140000 | 0.0019 | 0.0004 | 0.0009 |
| F04 | 90060001 | 90160000 | 0.0019 | 0.0003 | 0.0009 |
| F04 | 90080001 | 90180000 | 0.0023 | 0.0005 | 0.0011 |
| F04 | 90100001 | 90200000 | 0.0026 | 0.0008 | 0.0011 |
| F04 | 90120001 | 90220000 | 0.0025 | 0.0010 | 0.0008 |
| F04 | 90140001 | 90240000 | 0.0024 | 0.0011 | 0.0009 |
| F04 | 90160001 | 90260000 | 0.0024 | 0.0038 | 0.0009 |
| F04 | 90180001 | 90280000 | 0.0046 | 0.0050 | 0.0033 |
| F04 | 90200001 | 90300000 | 0.0056 | 0.0076 | 0.0045 |
| F04 | 90220001 | 90320000 | 0.0082 | 0.0089 | 0.0072 |
| F04 | 90240001 | 90340000 | 0.0096 | 0.0114 | 0.0082 |
| F04 | 90260001 | 90360000 | 0.0119 | 0.0102 | 0.0100 |
| F04 | 90280001 | 90380000 | 0.0109 | 0.0102 | 0.0086 |
| F04 | 90300001 | 90400000 | 0.0118 | 0.0077 | 0.0095 |
| F04 | 90320001 | 90420000 | 0.0095 | 0.0066 | 0.0074 |
| F04 | 90340001 | 90440000 | 0.0092 | 0.0041 | 0.0070 |
| F04 | 90360001 | 90460000 | 0.0072 | 0.0026 | 0.0053 |
| F04 | 90380001 | 90480000 | 0.0057 | 0.0010 | 0.0040 |
| F04 | 90400001 | 90500000 | 0.0036 | 0.0008 | 0.0020 |
| F04 | 90420001 | 90520000 | 0.0032 | 0.0006 | 0.0014 |
| F04 | 90440001 | 90540000 | 0.0019 | 0.0006 | 0.0007 |
| F04 | 90460001 | 90560000 | 0.0019 | 0.0005 | 0.0007 |
| F04 | 90480001 | 90580000 | 0.0018 | 0.0005 | 0.0007 |
| F04 | 90500001 | 90600000 | 0.0019 | 0.0003 | 0.0007 |
| F04 | 90520001 | 90620000 | 0.0019 | 0.0002 | 0.0007 |
| F04 | 90540001 | 90640000 | 0.0020 | 0.0003 | 0.0008 |
| F04 | 90560001 | 90660000 | 0.0024 | 0.0003 | 0.0013 |
| F04 | 90580001 | 90680000 | 0.0026 | 0.0010 | 0.0018 |
| F04 | 90600001 | 90700000 | 0.0036 | 0.0014 | 0.0029 |
| F04 | 90620001 | 90720000 | 0.0041 | 0.0017 | 0.0036 |
| F04 | 90640001 | 90740000 | 0.0049 | 0.0016 | 0.0046 |
| F04 | 90660001 | 90760000 | 0.0045 | 0.0015 | 0.0043 |
| F04 | 90680001 | 90780000 | 0.0042 | 0.0015 | 0.0040 |
| F04 | 90700001 | 90800000 | 0.0031 | 0.0019 | 0.0030 |
| F04 | 90720001 | 90820000 | 0.0030 | 0.0023 | 0.0028 |

| Chr | Start | End | both | A1 | A2 |
|---|---|---|---|---|---|
| F04 | 90740001 | 90840000 | 0.0025 | 0.0031 | 0.0023 |
| F04 | 90760001 | 90860000 | 0.0029 | 0.0033 | 0.0025 |
| F04 | 90780001 | 90880000 | 0.0028 | 0.0027 | 0.0022 |
| F04 | 90800001 | 90900000 | 0.0026 | 0.0019 | 0.0020 |
| F04 | 90820001 | 90920000 | 0.0021 | 0.0013 | 0.0013 |
| F04 | 90840001 | 90940000 | 0.0017 | 0.0007 | 0.0007 |
| F04 | 90860001 | 90960000 | 0.0012 | 0.0008 | 0.0003 |
| F04 | 90880001 | 90980000 | 0.0011 | 0.0012 | 0.0003 |
| F04 | 90900001 | 91000000 | 0.0013 | 0.0018 | 0.0005 |
| F04 | 90920001 | 91020000 | 0.0018 | 0.0018 | 0.0013 |
| F04 | 90940001 | 91040000 | 0.0016 | 0.0015 | 0.0013 |
| F04 | 90960001 | 91060000 | 0.0013 | 0.0012 | 0.0011 |
| F04 | 90980001 | 91080000 | 0.0012 | 0.0008 | 0.0011 |
| F04 | 91000001 | 91100000 | 0.0008 | 0.0000 | 0.0008 |
| F04 | 91020001 | 91120000 | 0.0000 | 0.0036 | 0.0000 |
| F05 | 1 | 100000 | 0.0017 | 0.0010 | 0.0014 |
| F05 | 20001 | 120000 | 0.0014 | 0.0005 | 0.0010 |
| F05 | 40001 | 140000 | 0.0011 | 0.0002 | 0.0005 |
| F05 | 60001 | 160000 | 0.0009 | 0.0004 | 0.0003 |
| F05 | 80001 | 180000 | 0.0010 | 0.0005 | 0.0004 |
| F05 | 100001 | 200000 | 0.0013 | 0.0008 | 0.0006 |
| F05 | 120001 | 220000 | 0.0016 | 0.0008 | 0.0009 |
| F05 | 140001 | 240000 | 0.0016 | 0.0012 | 0.0009 |
| F05 | 160001 | 260000 | 0.0021 | 0.0011 | 0.0014 |
| F05 | 180001 | 280000 | 0.0019 | 0.0009 | 0.0012 |
| F05 | 200001 | 300000 | 0.0018 | 0.0006 | 0.0010 |
| F05 | 220001 | 320000 | 0.0015 | 0.0009 | 0.0008 |
| F05 | 240001 | 340000 | 0.0016 | 0.0004 | 0.0008 |
| F05 | 260001 | 360000 | 0.0012 | 0.0003 | 0.0003 |
| F05 | 280001 | 380000 | 0.0012 | 0.0003 | 0.0002 |
| F05 | 300001 | 400000 | 0.0013 | 0.0004 | 0.0002 |
| F05 | 320001 | 420000 | 0.0013 | 0.0002 | 0.0003 |
| F05 | 340001 | 440000 | 0.0012 | 0.0003 | 0.0002 |
| F05 | 360001 | 460000 | 0.0012 | 0.0005 | 0.0003 |
| F05 | 380001 | 480000 | 0.0014 | 0.0005 | 0.0006 |
| F05 | 400001 | 500000 | 0.0013 | 0.0004 | 0.0006 |
| F05 | 420001 | 520000 | 0.0013 | 0.0004 | 0.0005 |
| F05 | 440001 | 540000 | 0.0013 | 0.0003 | 0.0006 |
| F05 | 460001 | 560000 | 0.0013 | 0.0002 | 0.0005 |
| F05 | 480001 | 580000 | 0.0011 | 0.0002 | 0.0003 |
| F05 | 500001 | 600000 | 0.0010 | 0.0002 | 0.0003 |
| F05 | 520001 | 620000 | 0.0010 | 0.0002 | 0.0003 |
| F05 | 540001 | 640000 | 0.0009 | 0.0002 | 0.0002 |
| F05 | 560001 | 660000 | 0.0010 | 0.0002 | 0.0002 |
| F05 | 580001 | 680000 | 0.0011 | 0.0003 | 0.0002 |
| F05 | 600001 | 700000 | 0.0013 | 0.0003 | 0.0002 |
| F05 | 620001 | 720000 | 0.0013 | 0.0003 | 0.0002 |
| F05 | 640001 | 740000 | 0.0014 | 0.0003 | 0.0002 |
| F05 | 660001 | 760000 | 0.0012 | 0.0003 | 0.0002 |
| F05 | 680001 | 780000 | 0.0011 | 0.0002 | 0.0002 |
| F05 | 700001 | 800000 | 0.0011 | 0.0001 | 0.0002 |
| F05 | 720001 | 820000 | 0.0013 | 0.0001 | 0.0002 |
| F05 | 740001 | 840000 | 0.0014 | 0.0001 | 0.0002 |
| F05 | 760001 | 860000 | 0.0016 | 0.0002 | 0.0004 |
| F05 | 780001 | 880000 | 0.0015 | 0.0002 | 0.0004 |
| F05 | 800001 | 900000 | 0.0015 | 0.0003 | 0.0005 |
| F05 | 820001 | 920000 | 0.0015 | 0.0003 | 0.0007 |
| F05 | 840001 | 940000 | 0.0013 | 0.0005 | 0.0007 |

| Chr | Start | End | both | A1 | A2 |
|---|---|---|---|---|---|
| F05 | 860001 | 960000 | 0.0017 | 0.0004 | 0.0011 |
| F05 | 880001 | 980000 | 0.0020 | 0.0003 | 0.0014 |
| F05 | 900001 | 1000000 | 0.0020 | 0.0003 | 0.0013 |
| F05 | 920001 | 1020000 | 0.0019 | 0.0003 | 0.0012 |
| F05 | 940001 | 1040000 | 0.0020 | 0.0003 | 0.0012 |
| F05 | 960001 | 1060000 | 0.0017 | 0.0003 | 0.0008 |
| F05 | 980001 | 1080000 | 0.0014 | 0.0003 | 0.0006 |
| F05 | 1000001 | 1100000 | 0.0014 | 0.0004 | 0.0006 |
| F05 | 1020001 | 1120000 | 0.0014 | 0.0004 | 0.0006 |
| F05 | 1040001 | 1140000 | 0.0014 | 0.0003 | 0.0005 |
| F05 | 1060001 | 1160000 | 0.0012 | 0.0004 | 0.0003 |
| F05 | 1080001 | 1180000 | 0.0012 | 0.0004 | 0.0003 |
| F05 | 1100001 | 1200000 | 0.0012 | 0.0003 | 0.0003 |
| F05 | 1120001 | 1220000 | 0.0011 | 0.0003 | 0.0002 |
| F05 | 1140001 | 1240000 | 0.0011 | 0.0004 | 0.0003 |
| F05 | 1160001 | 1260000 | 0.0012 | 0.0004 | 0.0003 |
| F05 | 1180001 | 1280000 | 0.0013 | 0.0004 | 0.0003 |
| F05 | 1200001 | 1300000 | 0.0014 | 0.0003 | 0.0004 |
| F05 | 1220001 | 1320000 | 0.0014 | 0.0006 | 0.0004 |
| F05 | 1240001 | 1340000 | 0.0017 | 0.0005 | 0.0007 |
| F05 | 1260001 | 1360000 | 0.0015 | 0.0005 | 0.0006 |
| F05 | 1280001 | 1380000 | 0.0014 | 0.0008 | 0.0006 |
| F05 | 1300001 | 1400000 | 0.0017 | 0.0008 | 0.0008 |
| F05 | 1320001 | 1420000 | 0.0016 | 0.0005 | 0.0009 |
| F05 | 1340001 | 1440000 | 0.0013 | 0.0005 | 0.0006 |
| F05 | 1360001 | 1460000 | 0.0014 | 0.0007 | 0.0007 |
| F05 | 1380001 | 1480000 | 0.0016 | 0.0004 | 0.0009 |
| F05 | 1400001 | 1500000 | 0.0012 | 0.0003 | 0.0006 |
| F05 | 1420001 | 1520000 | 0.0012 | 0.0004 | 0.0005 |
| F05 | 1440001 | 1540000 | 0.0013 | 0.0003 | 0.0006 |
| F05 | 1460001 | 1560000 | 0.0013 | 0.0005 | 0.0006 |
| F05 | 1480001 | 1580000 | 0.0014 | 0.0005 | 0.0007 |
| F05 | 1500001 | 1600000 | 0.0015 | 0.0006 | 0.0008 |
| F05 | 1520001 | 1620000 | 0.0016 | 0.0013 | 0.0008 |
| F05 | 1540001 | 1640000 | 0.0022 | 0.0016 | 0.0014 |
| F05 | 1560001 | 1660000 | 0.0023 | 0.0013 | 0.0015 |
| F05 | 1580001 | 1680000 | 0.0021 | 0.0013 | 0.0013 |
| F05 | 1600001 | 1700000 | 0.0020 | 0.0012 | 0.0013 |
| F05 | 1620001 | 1720000 | 0.0019 | 0.0005 | 0.0012 |
| F05 | 1640001 | 1740000 | 0.0012 | 0.0003 | 0.0006 |
| F05 | 1660001 | 1760000 | 0.0011 | 0.0003 | 0.0004 |
| F05 | 1680001 | 1780000 | 0.0011 | 0.0002 | 0.0003 |
| F05 | 1700001 | 1800000 | 0.0011 | 0.0002 | 0.0003 |
| F05 | 1720001 | 1820000 | 0.0012 | 0.0002 | 0.0003 |
| F05 | 1740001 | 1840000 | 0.0012 | 0.0001 | 0.0004 |
| F05 | 1760001 | 1860000 | 0.0011 | 0.0003 | 0.0003 |
| F05 | 1780001 | 1880000 | 0.0013 | 0.0005 | 0.0005 |
| F05 | 1800001 | 1900000 | 0.0014 | 0.0005 | 0.0006 |
| F05 | 1820001 | 1920000 | 0.0014 | 0.0008 | 0.0007 |
| F05 | 1840001 | 1940000 | 0.0017 | 0.0008 | 0.0008 |
| F05 | 1860001 | 1960000 | 0.0017 | 0.0005 | 0.0008 |
| F05 | 1880001 | 1980000 | 0.0016 | 0.0005 | 0.0006 |
| F05 | 1900001 | 2000000 | 0.0018 | 0.0007 | 0.0007 |
| F05 | 1920001 | 2020000 | 0.0019 | 0.0005 | 0.0008 |
| F05 | 1940001 | 2040000 | 0.0017 | 0.0005 | 0.0006 |
| F05 | 1960001 | 2060000 | 0.0017 | 0.0005 | 0.0006 |
| F05 | 1980001 | 2080000 | 0.0015 | 0.0003 | 0.0005 |
| F05 | 2000001 | 2100000 | 0.0015 | 0.0005 | 0.0004 |

| Chr | Start | End | both | A1 | A2 |
|---|---|---|---|---|---|
| F05 | 2020001 | 2120000 | 0.0016 | 0.0005 | 0.0006 |
| F05 | 2040001 | 2140000 | 0.0016 | 0.0006 | 0.0006 |
| F05 | 2060001 | 2160000 | 0.0017 | 0.0006 | 0.0006 |
| F05 | 2080001 | 2180000 | 0.0018 | 0.0010 | 0.0007 |
| F05 | 2100001 | 2200000 | 0.0022 | 0.0006 | 0.0011 |
| F05 | 2120001 | 2220000 | 0.0018 | 0.0006 | 0.0007 |
| F05 | 2140001 | 2240000 | 0.0018 | 0.0006 | 0.0007 |
| F05 | 2160001 | 2260000 | 0.0016 | 0.0005 | 0.0006 |
| F05 | 2180001 | 2280000 | 0.0016 | 0.0002 | 0.0005 |
| F05 | 2200001 | 2300000 | 0.0012 | 0.0005 | 0.0002 |
| F05 | 2220001 | 2320000 | 0.0014 | 0.0005 | 0.0005 |
| F05 | 2240001 | 2340000 | 0.0014 | 0.0005 | 0.0005 |
| F05 | 2260001 | 2360000 | 0.0013 | 0.0005 | 0.0005 |
| F05 | 2280001 | 2380000 | 0.0014 | 0.0007 | 0.0005 |
| F05 | 2300001 | 2400000 | 0.0017 | 0.0005 | 0.0008 |
| F05 | 2320001 | 2420000 | 0.0016 | 0.0005 | 0.0006 |
| F05 | 2340001 | 2440000 | 0.0017 | 0.0007 | 0.0006 |
| F05 | 2360001 | 2460000 | 0.0019 | 0.0007 | 0.0008 |
| F05 | 2380001 | 2480000 | 0.0019 | 0.0005 | 0.0008 |
| F05 | 2400001 | 2500000 | 0.0017 | 0.0004 | 0.0006 |
| F05 | 2420001 | 2520000 | 0.0016 | 0.0005 | 0.0006 |
| F05 | 2440001 | 2540000 | 0.0016 | 0.0003 | 0.0006 |
| F05 | 2460001 | 2560000 | 0.0014 | 0.0003 | 0.0004 |
| F05 | 2480001 | 2580000 | 0.0012 | 0.0002 | 0.0004 |
| F05 | 2500001 | 2600000 | 0.0010 | 0.0003 | 0.0003 |
| F05 | 2520001 | 2620000 | 0.0010 | 0.0006 | 0.0004 |
| F05 | 2540001 | 2640000 | 0.0013 | 0.0007 | 0.0007 |
| F05 | 2560001 | 2660000 | 0.0015 | 0.0010 | 0.0008 |
| F05 | 2580001 | 2680000 | 0.0018 | 0.0009 | 0.0011 |
| F05 | 2600001 | 2700000 | 0.0018 | 0.0015 | 0.0010 |
| F05 | 2620001 | 2720000 | 0.0025 | 0.0016 | 0.0017 |
| F05 | 2640001 | 2740000 | 0.0026 | 0.0015 | 0.0018 |
| F05 | 2660001 | 2760000 | 0.0026 | 0.0013 | 0.0019 |
| F05 | 2680001 | 2780000 | 0.0025 | 0.0013 | 0.0016 |
| F05 | 2700001 | 2800000 | 0.0026 | 0.0008 | 0.0016 |
| F05 | 2720001 | 2820000 | 0.0020 | 0.0004 | 0.0010 |
| F05 | 2740001 | 2840000 | 0.0016 | 0.0004 | 0.0007 |
| F05 | 2760001 | 2860000 | 0.0014 | 0.0004 | 0.0005 |
| F05 | 2780001 | 2880000 | 0.0012 | 0.0005 | 0.0004 |
| F05 | 2800001 | 2900000 | 0.0012 | 0.0004 | 0.0006 |
| F05 | 2820001 | 2920000 | 0.0011 | 0.0004 | 0.0004 |
| F05 | 2840001 | 2940000 | 0.0012 | 0.0008 | 0.0003 |
| F05 | 2860001 | 2960000 | 0.0015 | 0.0010 | 0.0008 |
| F05 | 2880001 | 2980000 | 0.0019 | 0.0011 | 0.0011 |
| F05 | 2900001 | 3000000 | 0.0021 | 0.0011 | 0.0012 |
| F05 | 2920001 | 3020000 | 0.0020 | 0.0009 | 0.0011 |
| F05 | 2940001 | 3040000 | 0.0020 | 0.0005 | 0.0011 |
| F05 | 2960001 | 3060000 | 0.0017 | 0.0002 | 0.0006 |
| F05 | 2980001 | 3080000 | 0.0014 | 0.0008 | 0.0003 |
| F05 | 3000001 | 3100000 | 0.0019 | 0.0008 | 0.0009 |
| F05 | 3020001 | 3120000 | 0.0019 | 0.0008 | 0.0009 |
| F05 | 3040001 | 3140000 | 0.0019 | 0.0010 | 0.0009 |
| F05 | 3060001 | 3160000 | 0.0018 | 0.0013 | 0.0009 |
| F05 | 3080001 | 3180000 | 0.0020 | 0.0005 | 0.0012 |
| F05 | 3100001 | 3200000 | 0.0012 | 0.0005 | 0.0004 |
| F05 | 3120001 | 3220000 | 0.0012 | 0.0007 | 0.0004 |
| F05 | 3140001 | 3240000 | 0.0013 | 0.0008 | 0.0006 |
| F05 | 3160001 | 3260000 | 0.0015 | 0.0009 | 0.0008 |

| Chr | Start | End | both | A1 | A2 |
|---|---|---|---|---|---|
| F05 | 3180001 | 3280000 | 0.0016 | 0.0009 | 0.0009 |
| F05 | 3200001 | 3300000 | 0.0017 | 0.0010 | 0.0009 |
| F05 | 3220001 | 3320000 | 0.0018 | 0.0008 | 0.0010 |
| F05 | 3240001 | 3340000 | 0.0017 | 0.0006 | 0.0008 |
| F05 | 3260001 | 3360000 | 0.0015 | 0.0002 | 0.0006 |
| F05 | 3280001 | 3380000 | 0.0012 | 0.0004 | 0.0002 |
| F05 | 3300001 | 3400000 | 0.0013 | 0.0002 | 0.0004 |
| F05 | 3320001 | 3420000 | 0.0011 | 0.0002 | 0.0002 |
| F05 | 3340001 | 3440000 | 0.0011 | 0.0005 | 0.0002 |
| F05 | 3360001 | 3460000 | 0.0013 | 0.0005 | 0.0005 |
| F05 | 3380001 | 3480000 | 0.0014 | 0.0005 | 0.0005 |
| F05 | 3400001 | 3500000 | 0.0014 | 0.0009 | 0.0006 |
| F05 | 3420001 | 3520000 | 0.0019 | 0.0011 | 0.0010 |
| F05 | 3440001 | 3540000 | 0.0020 | 0.0011 | 0.0012 |
| F05 | 3460001 | 3560000 | 0.0020 | 0.0011 | 0.0012 |
| F05 | 3480001 | 3580000 | 0.0020 | 0.0009 | 0.0012 |
| F05 | 3500001 | 3600000 | 0.0021 | 0.0005 | 0.0010 |
| F05 | 3520001 | 3620000 | 0.0017 | 0.0003 | 0.0006 |
| F05 | 3540001 | 3640000 | 0.0016 | 0.0001 | 0.0004 |
| F05 | 3560001 | 3660000 | 0.0014 | 0.0003 | 0.0002 |
| F05 | 3580001 | 3680000 | 0.0014 | 0.0008 | 0.0004 |
| F05 | 3600001 | 3700000 | 0.0016 | 0.0015 | 0.0009 |
| F05 | 3620001 | 3720000 | 0.0023 | 0.0015 | 0.0016 |
| F05 | 3640001 | 3740000 | 0.0023 | 0.0018 | 0.0016 |
| F05 | 3660001 | 3760000 | 0.0028 | 0.0023 | 0.0020 |
| F05 | 3680001 | 3780000 | 0.0034 | 0.0018 | 0.0025 |
| F05 | 3700001 | 3800000 | 0.0031 | 0.0011 | 0.0020 |
| F05 | 3720001 | 3820000 | 0.0024 | 0.0012 | 0.0014 |
| F05 | 3740001 | 3840000 | 0.0025 | 0.0011 | 0.0015 |
| F05 | 3760001 | 3860000 | 0.0023 | 0.0005 | 0.0014 |
| F05 | 3780001 | 3880000 | 0.0016 | 0.0005 | 0.0008 |
| F05 | 3800001 | 3900000 | 0.0016 | 0.0005 | 0.0008 |
| F05 | 3820001 | 3920000 | 0.0015 | 0.0005 | 0.0007 |
| F05 | 3840001 | 3940000 | 0.0015 | 0.0003 | 0.0007 |
| F05 | 3860001 | 3960000 | 0.0013 | 0.0003 | 0.0005 |
| F05 | 3880001 | 3980000 | 0.0013 | 0.0005 | 0.0006 |
| F05 | 3900001 | 4000000 | 0.0013 | 0.0004 | 0.0007 |
| F05 | 3920001 | 4020000 | 0.0014 | 0.0004 | 0.0007 |
| F05 | 3940001 | 4040000 | 0.0013 | 0.0004 | 0.0007 |
| F05 | 3960001 | 4060000 | 0.0012 | 0.0003 | 0.0006 |
| F05 | 3980001 | 4080000 | 0.0011 | 0.0002 | 0.0004 |
| F05 | 4000001 | 4100000 | 0.0010 | 0.0002 | 0.0002 |
| F05 | 4020001 | 4120000 | 0.0010 | 0.0005 | 0.0002 |
| F05 | 4040001 | 4140000 | 0.0012 | 0.0005 | 0.0004 |
| F05 | 4060001 | 4160000 | 0.0014 | 0.0008 | 0.0005 |
| F05 | 4080001 | 4180000 | 0.0017 | 0.0016 | 0.0008 |
| F05 | 4100001 | 4200000 | 0.0025 | 0.0019 | 0.0015 |
| F05 | 4120001 | 4220000 | 0.0029 | 0.0021 | 0.0018 |
| F05 | 4140001 | 4240000 | 0.0031 | 0.0020 | 0.0021 |
| F05 | 4160001 | 4260000 | 0.0032 | 0.0019 | 0.0020 |
| F05 | 4180001 | 4280000 | 0.0031 | 0.0014 | 0.0020 |
| F05 | 4200001 | 4300000 | 0.0027 | 0.0012 | 0.0016 |
| F05 | 4220001 | 4320000 | 0.0024 | 0.0012 | 0.0014 |
| F05 | 4240001 | 4340000 | 0.0023 | 0.0019 | 0.0013 |
| F05 | 4260001 | 4360000 | 0.0028 | 0.0022 | 0.0020 |
| F05 | 4280001 | 4380000 | 0.0030 | 0.0021 | 0.0024 |
| F05 | 4300001 | 4400000 | 0.0028 | 0.0020 | 0.0021 |
| F05 | 4320001 | 4420000 | 0.0027 | 0.0016 | 0.0020 |

| Chr | Start | End | both | A1 | A2 |
|---|---|---|---|---|---|
| F05 | 4340001 | 4440000 | 0.0023 | 0.0011 | 0.0016 |
| F05 | 4360001 | 4460000 | 0.0018 | 0.0009 | 0.0009 |
| F05 | 4380001 | 4480000 | 0.0016 | 0.0010 | 0.0005 |
| F05 | 4400001 | 4500000 | 0.0017 | 0.0010 | 0.0006 |
| F05 | 4420001 | 4520000 | 0.0016 | 0.0010 | 0.0006 |
| F05 | 4440001 | 4540000 | 0.0015 | 0.0008 | 0.0007 |
| F05 | 4460001 | 4560000 | 0.0014 | 0.0005 | 0.0006 |
| F05 | 4480001 | 4580000 | 0.0011 | 0.0004 | 0.0002 |
| F05 | 4500001 | 4600000 | 0.0009 | 0.0007 | 0.0002 |
| F05 | 4520001 | 4620000 | 0.0012 | 0.0008 | 0.0005 |
| F05 | 4540001 | 4640000 | 0.0014 | 0.0008 | 0.0006 |
| F05 | 4560001 | 4660000 | 0.0015 | 0.0009 | 0.0007 |
| F05 | 4580001 | 4680000 | 0.0017 | 0.0011 | 0.0008 |
| F05 | 4600001 | 4700000 | 0.0019 | 0.0012 | 0.0010 |
| F05 | 4620001 | 4720000 | 0.0020 | 0.0013 | 0.0011 |
| F05 | 4640001 | 4740000 | 0.0022 | 0.0013 | 0.0012 |
| F05 | 4660001 | 4760000 | 0.0020 | 0.0014 | 0.0011 |
| F05 | 4680001 | 4780000 | 0.0020 | 0.0012 | 0.0012 |
| F05 | 4700001 | 4800000 | 0.0021 | 0.0008 | 0.0011 |
| F05 | 4720001 | 4820000 | 0.0016 | 0.0009 | 0.0007 |
| F05 | 4740001 | 4840000 | 0.0016 | 0.0012 | 0.0006 |
| F05 | 4760001 | 4860000 | 0.0020 | 0.0010 | 0.0009 |
| F05 | 4780001 | 4880000 | 0.0018 | 0.0010 | 0.0007 |
| F05 | 4800001 | 4900000 | 0.0017 | 0.0010 | 0.0006 |
| F05 | 4820001 | 4920000 | 0.0018 | 0.0008 | 0.0007 |
| F05 | 4840001 | 4940000 | 0.0015 | 0.0009 | 0.0005 |
| F05 | 4860001 | 4960000 | 0.0016 | 0.0010 | 0.0007 |
| F05 | 4880001 | 4980000 | 0.0017 | 0.0009 | 0.0007 |
| F05 | 4900001 | 5000000 | 0.0017 | 0.0009 | 0.0007 |
| F05 | 4920001 | 5020000 | 0.0017 | 0.0009 | 0.0006 |
| F05 | 4940001 | 5040000 | 0.0016 | 0.0005 | 0.0006 |
| F05 | 4960001 | 5060000 | 0.0012 | 0.0006 | 0.0002 |
| F05 | 4980001 | 5080000 | 0.0012 | 0.0006 | 0.0003 |
| F05 | 5000001 | 5100000 | 0.0012 | 0.0005 | 0.0003 |
| F05 | 5020001 | 5120000 | 0.0012 | 0.0006 | 0.0003 |
| F05 | 5040001 | 5140000 | 0.0013 | 0.0006 | 0.0003 |
| F05 | 5060001 | 5160000 | 0.0013 | 0.0004 | 0.0004 |
| F05 | 5080001 | 5180000 | 0.0013 | 0.0007 | 0.0003 |
| F05 | 5100001 | 5200000 | 0.0017 | 0.0010 | 0.0008 |
| F05 | 5120001 | 5220000 | 0.0020 | 0.0010 | 0.0011 |
| F05 | 5140001 | 5240000 | 0.0019 | 0.0011 | 0.0011 |
| F05 | 5160001 | 5260000 | 0.0020 | 0.0013 | 0.0012 |
| F05 | 5180001 | 5280000 | 0.0021 | 0.0009 | 0.0014 |
| F05 | 5200001 | 5300000 | 0.0016 | 0.0007 | 0.0010 |
| F05 | 5220001 | 5320000 | 0.0014 | 0.0007 | 0.0007 |
| F05 | 5240001 | 5340000 | 0.0013 | 0.0006 | 0.0007 |
| F05 | 5260001 | 5360000 | 0.0012 | 0.0004 | 0.0005 |
| F05 | 5280001 | 5380000 | 0.0009 | 0.0005 | 0.0002 |
| F05 | 5300001 | 5400000 | 0.0008 | 0.0012 | 0.0002 |
| F05 | 5320001 | 5420000 | 0.0014 | 0.0012 | 0.0009 |
| F05 | 5340001 | 5440000 | 0.0013 | 0.0014 | 0.0009 |
| F05 | 5360001 | 5460000 | 0.0013 | 0.0022 | 0.0011 |
| F05 | 5380001 | 5480000 | 0.0020 | 0.0021 | 0.0018 |
| F05 | 5400001 | 5500000 | 0.0020 | 0.0015 | 0.0019 |
| F05 | 5420001 | 5520000 | 0.0012 | 0.0027 | 0.0011 |
| F05 | 5440001 | 5540000 | 0.0024 | 0.0026 | 0.0023 |
| F05 | 5460001 | 5560000 | 0.0023 | 0.0021 | 0.0021 |
| F05 | 5480001 | 5580000 | 0.0016 | 0.0023 | 0.0014 |

| Chr | Start | End | both | A1 | A2 |
|---|---|---|---|---|---|
| F05 | 5500001 | 5600000 | 0.0017 | 0.0024 | 0.0014 |
| F05 | 5520001 | 5620000 | 0.0017 | 0.0012 | 0.0013 |
| F05 | 5540001 | 5640000 | 0.0005 | 0.0014 | 0.0002 |
| F05 | 5560001 | 5660000 | 0.0006 | 0.0019 | 0.0004 |
| F05 | 5580001 | 5680000 | 0.0010 | 0.0020 | 0.0007 |
| F05 | 5600001 | 5700000 | 0.0011 | 0.0021 | 0.0007 |
| F05 | 5620001 | 5720000 | 0.0014 | 0.0022 | 0.0009 |
| F05 | 5640001 | 5740000 | 0.0015 | 0.0021 | 0.0009 |
| F05 | 5660001 | 5760000 | 0.0017 | 0.0017 | 0.0010 |
| F05 | 5680001 | 5780000 | 0.0017 | 0.0015 | 0.0010 |
| F05 | 5700001 | 5800000 | 0.0018 | 0.0012 | 0.0012 |
| F05 | 5720001 | 5820000 | 0.0018 | 0.0009 | 0.0012 |
| F05 | 5740001 | 5840000 | 0.0019 | 0.0008 | 0.0013 |
| F05 | 5760001 | 5860000 | 0.0018 | 0.0005 | 0.0014 |
| F05 | 5780001 | 5880000 | 0.0014 | 0.0005 | 0.0010 |
| F05 | 5800001 | 5900000 | 0.0011 | 0.0004 | 0.0008 |
| F05 | 5820001 | 5920000 | 0.0010 | 0.0008 | 0.0007 |
| F05 | 5840001 | 5940000 | 0.0013 | 0.0017 | 0.0009 |
| F05 | 5860001 | 5960000 | 0.0022 | 0.0019 | 0.0017 |
| F05 | 5880001 | 5980000 | 0.0023 | 0.0019 | 0.0018 |
| F05 | 5900001 | 6000000 | 0.0024 | 0.0019 | 0.0018 |
| F05 | 5920001 | 6020000 | 0.0023 | 0.0015 | 0.0017 |
| F05 | 5940001 | 6040000 | 0.0020 | 0.0008 | 0.0013 |
| F05 | 5960001 | 6060000 | 0.0012 | 0.0006 | 0.0006 |
| F05 | 5980001 | 6080000 | 0.0010 | 0.0005 | 0.0004 |
| F05 | 6000001 | 6100000 | 0.0008 | 0.0004 | 0.0004 |
| F05 | 6020001 | 6120000 | 0.0007 | 0.0004 | 0.0004 |
| F05 | 6040001 | 6140000 | 0.0007 | 0.0003 | 0.0004 |
| F05 | 6060001 | 6160000 | 0.0005 | 0.0003 | 0.0002 |
| F05 | 6080001 | 6180000 | 0.0007 | 0.0005 | 0.0002 |
| F05 | 6100001 | 6200000 | 0.0010 | 0.0006 | 0.0004 |
| F05 | 6120001 | 6220000 | 0.0013 | 0.0006 | 0.0004 |
| F05 | 6140001 | 6240000 | 0.0012 | 0.0007 | 0.0004 |
| F05 | 6160001 | 6260000 | 0.0015 | 0.0007 | 0.0006 |
| F05 | 6180001 | 6280000 | 0.0014 | 0.0006 | 0.0006 |
| F05 | 6200001 | 6300000 | 0.0013 | 0.0006 | 0.0005 |
| F05 | 6220001 | 6320000 | 0.0013 | 0.0007 | 0.0004 |
| F05 | 6240001 | 6340000 | 0.0013 | 0.0008 | 0.0005 |
| F05 | 6260001 | 6360000 | 0.0015 | 0.0011 | 0.0007 |
| F05 | 6280001 | 6380000 | 0.0017 | 0.0012 | 0.0010 |
| F05 | 6300001 | 6400000 | 0.0017 | 0.0022 | 0.0012 |
| F05 | 6320001 | 6420000 | 0.0026 | 0.0023 | 0.0023 |
| F05 | 6340001 | 6440000 | 0.0026 | 0.0021 | 0.0023 |
| F05 | 6360001 | 6460000 | 0.0023 | 0.0017 | 0.0020 |
| F05 | 6380001 | 6480000 | 0.0020 | 0.0017 | 0.0016 |
| F05 | 6400001 | 6500000 | 0.0021 | 0.0010 | 0.0016 |
| F05 | 6420001 | 6520000 | 0.0015 | 0.0008 | 0.0009 |
| F05 | 6440001 | 6540000 | 0.0014 | 0.0007 | 0.0008 |
| F05 | 6460001 | 6560000 | 0.0013 | 0.0009 | 0.0006 |
| F05 | 6480001 | 6580000 | 0.0015 | 0.0014 | 0.0007 |
| F05 | 6500001 | 6600000 | 0.0021 | 0.0015 | 0.0011 |
| F05 | 6520001 | 6620000 | 0.0021 | 0.0043 | 0.0010 |
| F05 | 6540001 | 6640000 | 0.0049 | 0.0043 | 0.0038 |
| F05 | 6560001 | 6660000 | 0.0049 | 0.0063 | 0.0038 |
| F05 | 6580001 | 6680000 | 0.0071 | 0.0057 | 0.0060 |
| F05 | 6600001 | 6700000 | 0.0066 | 0.0057 | 0.0055 |
| F05 | 6620001 | 6720000 | 0.0065 | 0.0028 | 0.0056 |
| F05 | 6640001 | 6740000 | 0.0036 | 0.0029 | 0.0029 |

| Chr | Start | End | both | A1 | A2 |
|---|---|---|---|---|---|
| F05 | 6660001 | 6760000 | 0.0038 | 0.0009 | 0.0029 |
| F05 | 6680001 | 6780000 | 0.0014 | 0.0009 | 0.0006 |
| F05 | 6700001 | 6800000 | 0.0014 | 0.0010 | 0.0007 |
| F05 | 6720001 | 6820000 | 0.0015 | 0.0010 | 0.0008 |
| F05 | 6740001 | 6840000 | 0.0016 | 0.0009 | 0.0008 |
| F05 | 6760001 | 6860000 | 0.0014 | 0.0009 | 0.0008 |
| F05 | 6780001 | 6880000 | 0.0014 | 0.0010 | 0.0007 |
| F05 | 6800001 | 6900000 | 0.0016 | 0.0007 | 0.0008 |
| F05 | 6820001 | 6920000 | 0.0013 | 0.0007 | 0.0004 |
| F05 | 6840001 | 6940000 | 0.0013 | 0.0009 | 0.0005 |
| F05 | 6860001 | 6960000 | 0.0016 | 0.0009 | 0.0006 |
| F05 | 6880001 | 6980000 | 0.0014 | 0.0006 | 0.0006 |
| F05 | 6900001 | 7000000 | 0.0011 | 0.0006 | 0.0004 |
| F05 | 6920001 | 7020000 | 0.0012 | 0.0009 | 0.0004 |
| F05 | 6940001 | 7040000 | 0.0014 | 0.0008 | 0.0006 |
| F05 | 6960001 | 7060000 | 0.0012 | 0.0009 | 0.0005 |
| F05 | 6980001 | 7080000 | 0.0014 | 0.0009 | 0.0005 |
| F05 | 7000001 | 7100000 | 0.0014 | 0.0013 | 0.0005 |
| F05 | 7020001 | 7120000 | 0.0016 | 0.0012 | 0.0009 |
| F05 | 7040001 | 7140000 | 0.0016 | 0.0011 | 0.0009 |
| F05 | 7060001 | 7160000 | 0.0016 | 0.0010 | 0.0008 |
| F05 | 7080001 | 7180000 | 0.0016 | 0.0010 | 0.0008 |
| F05 | 7100001 | 7200000 | 0.0016 | 0.0006 | 0.0007 |
| F05 | 7120001 | 7220000 | 0.0012 | 0.0004 | 0.0003 |
| F05 | 7140001 | 7240000 | 0.0010 | 0.0004 | 0.0001 |
| F05 | 7160001 | 7260000 | 0.0009 | 0.0004 | 0.0001 |
| F05 | 7180001 | 7280000 | 0.0009 | 0.0003 | 0.0001 |
| F05 | 7200001 | 7300000 | 0.0009 | 0.0004 | 0.0001 |
| F05 | 7220001 | 7320000 | 0.0010 | 0.0004 | 0.0001 |
| F05 | 7240001 | 7340000 | 0.0010 | 0.0006 | 0.0001 |
| F05 | 7260001 | 7360000 | 0.0013 | 0.0007 | 0.0003 |
| F05 | 7280001 | 7380000 | 0.0014 | 0.0007 | 0.0004 |
| F05 | 7300001 | 7400000 | 0.0013 | 0.0007 | 0.0004 |
| F05 | 7320001 | 7420000 | 0.0012 | 0.0008 | 0.0005 |
| F05 | 7340001 | 7440000 | 0.0013 | 0.0010 | 0.0006 |
| F05 | 7360001 | 7460000 | 0.0015 | 0.0009 | 0.0008 |
| F05 | 7380001 | 7480000 | 0.0015 | 0.0009 | 0.0007 |
| F05 | 7400001 | 7500000 | 0.0015 | 0.0009 | 0.0007 |
| F05 | 7420001 | 7520000 | 0.0016 | 0.0010 | 0.0007 |
| F05 | 7440001 | 7540000 | 0.0017 | 0.0006 | 0.0008 |
| F05 | 7460001 | 7560000 | 0.0013 | 0.0006 | 0.0004 |
| F05 | 7480001 | 7580000 | 0.0014 | 0.0009 | 0.0004 |
| F05 | 7500001 | 7600000 | 0.0016 | 0.0010 | 0.0006 |
| F05 | 7520001 | 7620000 | 0.0016 | 0.0011 | 0.0007 |
| F05 | 7540001 | 7640000 | 0.0016 | 0.0011 | 0.0007 |
| F05 | 7560001 | 7660000 | 0.0017 | 0.0012 | 0.0008 |
| F05 | 7580001 | 7680000 | 0.0018 | 0.0010 | 0.0009 |
| F05 | 7600001 | 7700000 | 0.0016 | 0.0009 | 0.0006 |
| F05 | 7620001 | 7720000 | 0.0017 | 0.0011 | 0.0007 |
| F05 | 7640001 | 7740000 | 0.0018 | 0.0010 | 0.0008 |
| F05 | 7660001 | 7760000 | 0.0017 | 0.0009 | 0.0009 |
| F05 | 7680001 | 7780000 | 0.0016 | 0.0014 | 0.0008 |
| F05 | 7700001 | 7800000 | 0.0020 | 0.0013 | 0.0013 |
| F05 | 7720001 | 7820000 | 0.0019 | 0.0010 | 0.0011 |
| F05 | 7740001 | 7840000 | 0.0016 | 0.0010 | 0.0007 |
| F05 | 7760001 | 7860000 | 0.0016 | 0.0021 | 0.0008 |
| F05 | 7780001 | 7880000 | 0.0026 | 0.0016 | 0.0017 |
| F05 | 7800001 | 7900000 | 0.0021 | 0.0015 | 0.0013 |

| Chr | Start | End | both | A1 | A2 |
|---|---|---|---|---|---|
| F05 | 7820001 | 7920000 | 0.0021 | 0.0018 | 0.0013 |
| F05 | 7840001 | 7940000 | 0.0023 | 0.0017 | 0.0016 |
| F05 | 7860001 | 7960000 | 0.0022 | 0.0007 | 0.0015 |
| F05 | 7880001 | 7980000 | 0.0012 | 0.0013 | 0.0006 |
| F05 | 7900001 | 8000000 | 0.0017 | 0.0014 | 0.0010 |
| F05 | 7920001 | 8020000 | 0.0018 | 0.0017 | 0.0010 |
| F05 | 7940001 | 8040000 | 0.0021 | 0.0018 | 0.0012 |
| F05 | 7960001 | 8060000 | 0.0021 | 0.0020 | 0.0012 |
| F05 | 7980001 | 8080000 | 0.0021 | 0.0014 | 0.0013 |
| F05 | 8000001 | 8100000 | 0.0015 | 0.0016 | 0.0009 |
| F05 | 8020001 | 8120000 | 0.0014 | 0.0010 | 0.0010 |
| F05 | 8040001 | 8140000 | 0.0007 | 0.0010 | 0.0005 |
| F05 | 8060001 | 8160000 | 0.0007 | 0.0009 | 0.0005 |
| F05 | 8080001 | 8180000 | 0.0007 | 0.0010 | 0.0005 |
| F05 | 8100001 | 8200000 | 0.0008 | 0.0008 | 0.0006 |
| F05 | 8120001 | 8220000 | 0.0007 | 0.0008 | 0.0004 |
| F05 | 8140001 | 8240000 | 0.0008 | 0.0008 | 0.0004 |
| F05 | 8160001 | 8260000 | 0.0008 | 0.0008 | 0.0004 |
| F05 | 8180001 | 8280000 | 0.0008 | 0.0008 | 0.0003 |
| F05 | 8200001 | 8300000 | 0.0006 | 0.0011 | 0.0001 |
| F05 | 8220001 | 8320000 | 0.0007 | 0.0015 | 0.0002 |
| F05 | 8240001 | 8340000 | 0.0008 | 0.0016 | 0.0002 |
| F05 | 8260001 | 8360000 | 0.0009 | 0.0016 | 0.0003 |
| F05 | 8280001 | 8380000 | 0.0011 | 0.0019 | 0.0004 |
| F05 | 8300001 | 8400000 | 0.0013 | 0.0019 | 0.0006 |
| F05 | 8320001 | 8420000 | 0.0014 | 0.0021 | 0.0007 |
| F05 | 8340001 | 8440000 | 0.0016 | 0.0022 | 0.0009 |
| F05 | 8360001 | 8460000 | 0.0016 | 0.0022 | 0.0010 |
| F05 | 8380001 | 8480000 | 0.0015 | 0.0019 | 0.0009 |
| F05 | 8400001 | 8500000 | 0.0012 | 0.0019 | 0.0007 |
| F05 | 8420001 | 8520000 | 0.0013 | 0.0020 | 0.0009 |
| F05 | 8440001 | 8540000 | 0.0014 | 0.0019 | 0.0012 |
| F05 | 8460001 | 8560000 | 0.0014 | 0.0019 | 0.0012 |
| F05 | 8480001 | 8580000 | 0.0014 | 0.0020 | 0.0011 |
| F05 | 8500001 | 8600000 | 0.0015 | 0.0020 | 0.0011 |
| F05 | 8520001 | 8620000 | 0.0013 | 0.0017 | 0.0009 |
| F05 | 8540001 | 8640000 | 0.0012 | 0.0020 | 0.0008 |
| F05 | 8560001 | 8660000 | 0.0013 | 0.0023 | 0.0009 |
| F05 | 8580001 | 8680000 | 0.0017 | 0.0025 | 0.0014 |
| F05 | 8600001 | 8700000 | 0.0019 | 0.0025 | 0.0015 |
| F05 | 8620001 | 8720000 | 0.0019 | 0.0025 | 0.0015 |
| F05 | 8640001 | 8740000 | 0.0017 | 0.0024 | 0.0013 |
| F05 | 8660001 | 8760000 | 0.0016 | 0.0023 | 0.0012 |
| F05 | 8680001 | 8780000 | 0.0015 | 0.0021 | 0.0011 |
| F05 | 8700001 | 8800000 | 0.0013 | 0.0020 | 0.0009 |
| F05 | 8720001 | 8820000 | 0.0013 | 0.0021 | 0.0009 |
| F05 | 8740001 | 8840000 | 0.0014 | 0.0020 | 0.0009 |
| F05 | 8760001 | 8860000 | 0.0012 | 0.0020 | 0.0007 |
| F05 | 8780001 | 8880000 | 0.0010 | 0.0021 | 0.0005 |
| F05 | 8800001 | 8900000 | 0.0010 | 0.0022 | 0.0005 |
| F05 | 8820001 | 8920000 | 0.0011 | 0.0021 | 0.0007 |
| F05 | 8840001 | 8940000 | 0.0010 | 0.0021 | 0.0005 |
| F05 | 8860001 | 8960000 | 0.0009 | 0.0019 | 0.0005 |
| F05 | 8880001 | 8980000 | 0.0007 | 0.0019 | 0.0003 |
| F05 | 8900001 | 9000000 | 0.0007 | 0.0017 | 0.0003 |
| F05 | 8920001 | 9020000 | 0.0005 | 0.0019 | 0.0001 |
| F05 | 8940001 | 9040000 | 0.0008 | 0.0019 | 0.0004 |
| F05 | 8960001 | 9060000 | 0.0011 | 0.0019 | 0.0008 |

| Chr | Start | End | both | A1 | A2 |
|---|---|---|---|---|---|
| F05 | 8980001 | 9080000 | 0.0011 | 0.0020 | 0.0008 |
| F05 | 9000001 | 9100000 | 0.0013 | 0.0018 | 0.0011 |
| F05 | 9020001 | 9120000 | 0.0015 | 0.0015 | 0.0011 |
| F05 | 9040001 | 9140000 | 0.0013 | 0.0011 | 0.0007 |
| F05 | 9060001 | 9160000 | 0.0012 | 0.0010 | 0.0004 |
| F05 | 9080001 | 9180000 | 0.0013 | 0.0006 | 0.0004 |
| F05 | 9100001 | 9200000 | 0.0013 | 0.0008 | 0.0001 |
| F05 | 9120001 | 9220000 | 0.0014 | 0.0011 | 0.0003 |
| F05 | 9140001 | 9240000 | 0.0018 | 0.0011 | 0.0007 |
| F05 | 9160001 | 9260000 | 0.0017 | 0.0011 | 0.0007 |
| F05 | 9180001 | 9280000 | 0.0017 | 0.0010 | 0.0007 |
| F05 | 9200001 | 9300000 | 0.0017 | 0.0009 | 0.0007 |
| F05 | 9220001 | 9320000 | 0.0015 | 0.0005 | 0.0006 |
| F05 | 9240001 | 9340000 | 0.0011 | 0.0005 | 0.0002 |
| F05 | 9260001 | 9360000 | 0.0011 | 0.0004 | 0.0002 |
| F05 | 9280001 | 9380000 | 0.0011 | 0.0005 | 0.0002 |
| F05 | 9300001 | 9400000 | 0.0013 | 0.0004 | 0.0002 |
| F05 | 9320001 | 9420000 | 0.0013 | 0.0004 | 0.0003 |
| F05 | 9340001 | 9440000 | 0.0012 | 0.0004 | 0.0004 |
| F05 | 9360001 | 9460000 | 0.0012 | 0.0004 | 0.0004 |
| F05 | 9380001 | 9480000 | 0.0011 | 0.0003 | 0.0004 |
| F05 | 9400001 | 9500000 | 0.0011 | 0.0004 | 0.0003 |
| F05 | 9420001 | 9520000 | 0.0012 | 0.0006 | 0.0004 |
| F05 | 9440001 | 9540000 | 0.0012 | 0.0009 | 0.0003 |
| F05 | 9460001 | 9560000 | 0.0012 | 0.0011 | 0.0004 |
| F05 | 9480001 | 9580000 | 0.0013 | 0.0014 | 0.0006 |
| F05 | 9500001 | 9600000 | 0.0013 | 0.0016 | 0.0008 |
| F05 | 9520001 | 9620000 | 0.0012 | 0.0016 | 0.0010 |
| F05 | 9540001 | 9640000 | 0.0013 | 0.0017 | 0.0012 |
| F05 | 9560001 | 9660000 | 0.0015 | 0.0022 | 0.0014 |
| F05 | 9580001 | 9680000 | 0.0019 | 0.0023 | 0.0018 |
| F05 | 9600001 | 9700000 | 0.0020 | 0.0021 | 0.0018 |
| F05 | 9620001 | 9720000 | 0.0021 | 0.0018 | 0.0020 |
| F05 | 9640001 | 9740000 | 0.0019 | 0.0015 | 0.0018 |
| F05 | 9660001 | 9760000 | 0.0019 | 0.0008 | 0.0019 |
| F05 | 9680001 | 9780000 | 0.0016 | 0.0005 | 0.0015 |
| F05 | 9700001 | 9800000 | 0.0018 | 0.0006 | 0.0014 |
| F05 | 9720001 | 9820000 | 0.0019 | 0.0007 | 0.0013 |
| F05 | 9740001 | 9840000 | 0.0019 | 0.0006 | 0.0010 |
| F05 | 9760001 | 9860000 | 0.0017 | 0.0006 | 0.0007 |
| F05 | 9780001 | 9880000 | 0.0015 | 0.0012 | 0.0005 |
| F05 | 9800001 | 9900000 | 0.0019 | 0.0010 | 0.0011 |
| F05 | 9820001 | 9920000 | 0.0017 | 0.0009 | 0.0011 |
| F05 | 9840001 | 9940000 | 0.0016 | 0.0009 | 0.0012 |
| F05 | 9860001 | 9960000 | 0.0014 | 0.0008 | 0.0012 |
| F05 | 9880001 | 9980000 | 0.0013 | 0.0001 | 0.0011 |
| F05 | 9900001 | 10000000 | 0.0006 | 0.0000 | 0.0006 |
| F05 | 9920001 | 10020000 | 0.0004 | 0.0005 | 0.0004 |
| F05 | 9940001 | 10040000 | 0.0010 | 0.0005 | 0.0011 |
| F05 | 9960001 | 10060000 | 0.0011 | 0.0006 | 0.0011 |
| F05 | 9980001 | 10080000 | 0.0012 | 0.0007 | 0.0013 |
| F05 | 10000001 | 10100000 | 0.0012 | 0.0007 | 0.0013 |
| F05 | 10020001 | 10120000 | 0.0013 | 0.0002 | 0.0014 |
| F05 | 10040001 | 10140000 | 0.0009 | 0.0005 | 0.0009 |
| F05 | 10060001 | 10160000 | 0.0011 | 0.0004 | 0.0011 |
| F05 | 10080001 | 10180000 | 0.0013 | 0.0004 | 0.0012 |
| F05 | 10100001 | 10200000 | 0.0014 | 0.0004 | 0.0010 |
| F05 | 10120001 | 10220000 | 0.0014 | 0.0004 | 0.0008 |

| Chr | Start | End | both | A1 | A2 |
|---|---|---|---|---|---|
| F05 | 10140001 | 10240000 | 0.0015 | 0.0002 | 0.0006 |
| F05 | 10160001 | 10260000 | 0.0015 | 0.0004 | 0.0004 |
| F05 | 10180001 | 10280000 | 0.0016 | 0.0004 | 0.0005 |
| F05 | 10200001 | 10300000 | 0.0015 | 0.0004 | 0.0005 |
| F05 | 10220001 | 10320000 | 0.0016 | 0.0004 | 0.0006 |
| F05 | 10240001 | 10340000 | 0.0016 | 0.0003 | 0.0006 |
| F05 | 10260001 | 10360000 | 0.0017 | 0.0003 | 0.0006 |
| F05 | 10280001 | 10380000 | 0.0016 | 0.0004 | 0.0005 |
| F05 | 10300001 | 10400000 | 0.0017 | 0.0003 | 0.0006 |
| F05 | 10320001 | 10420000 | 0.0016 | 0.0003 | 0.0005 |
| F05 | 10340001 | 10440000 | 0.0015 | 0.0003 | 0.0005 |
| F05 | 10360001 | 10460000 | 0.0014 | 0.0014 | 0.0006 |
| F05 | 10380001 | 10480000 | 0.0027 | 0.0015 | 0.0018 |
| F05 | 10400001 | 10500000 | 0.0029 | 0.0015 | 0.0019 |
| F05 | 10420001 | 10520000 | 0.0029 | 0.0016 | 0.0019 |
| F05 | 10440001 | 10540000 | 0.0029 | 0.0019 | 0.0019 |
| F05 | 10460001 | 10560000 | 0.0031 | 0.0006 | 0.0021 |
| F05 | 10480001 | 10580000 | 0.0018 | 0.0005 | 0.0008 |
| F05 | 10500001 | 10600000 | 0.0017 | 0.0004 | 0.0006 |
| F05 | 10520001 | 10620000 | 0.0016 | 0.0004 | 0.0006 |
| F05 | 10540001 | 10640000 | 0.0016 | 0.0001 | 0.0006 |
| F05 | 10560001 | 10660000 | 0.0014 | 0.0001 | 0.0004 |
| F05 | 10580001 | 10680000 | 0.0013 | 0.0001 | 0.0003 |
| F05 | 10600001 | 10700000 | 0.0013 | 0.0001 | 0.0003 |
| F05 | 10620001 | 10720000 | 0.0012 | 0.0002 | 0.0003 |
| F05 | 10640001 | 10740000 | 0.0013 | 0.0002 | 0.0003 |
| F05 | 10660001 | 10760000 | 0.0011 | 0.0003 | 0.0002 |
| F05 | 10680001 | 10780000 | 0.0013 | 0.0003 | 0.0004 |
| F05 | 10700001 | 10800000 | 0.0012 | 0.0003 | 0.0004 |
| F05 | 10720001 | 10820000 | 0.0013 | 0.0003 | 0.0005 |
| F05 | 10740001 | 10840000 | 0.0014 | 0.0003 | 0.0005 |
| F05 | 10760001 | 10860000 | 0.0014 | 0.0002 | 0.0005 |
| F05 | 10780001 | 10880000 | 0.0013 | 0.0002 | 0.0003 |
| F05 | 10800001 | 10900000 | 0.0013 | 0.0002 | 0.0004 |
| F05 | 10820001 | 10920000 | 0.0012 | 0.0002 | 0.0003 |
| F05 | 10840001 | 10940000 | 0.0012 | 0.0002 | 0.0003 |
| F05 | 10860001 | 10960000 | 0.0012 | 0.0002 | 0.0004 |
| F05 | 10880001 | 10980000 | 0.0012 | 0.0002 | 0.0004 |
| F05 | 10900001 | 11000000 | 0.0013 | 0.0001 | 0.0004 |
| F05 | 10920001 | 11020000 | 0.0012 | 0.0002 | 0.0003 |
| F05 | 10940001 | 11040000 | 0.0013 | 0.0002 | 0.0004 |
| F05 | 10960001 | 11060000 | 0.0012 | 0.0001 | 0.0003 |
| F05 | 10980001 | 11080000 | 0.0010 | 0.0001 | 0.0003 |
| F05 | 11000001 | 11100000 | 0.0008 | 0.0002 | 0.0002 |
| F05 | 11020001 | 11120000 | 0.0009 | 0.0001 | 0.0002 |
| F05 | 11040001 | 11140000 | 0.0007 | 0.0002 | 0.0002 |
| F05 | 11060001 | 11160000 | 0.0007 | 0.0003 | 0.0002 |
| F05 | 11080001 | 11180000 | 0.0010 | 0.0005 | 0.0003 |
| F05 | 11100001 | 11200000 | 0.0013 | 0.0004 | 0.0004 |
| F05 | 11120001 | 11220000 | 0.0013 | 0.0005 | 0.0003 |
| F05 | 11140001 | 11240000 | 0.0014 | 0.0005 | 0.0004 |
| F05 | 11160001 | 11260000 | 0.0016 | 0.0004 | 0.0005 |
| F05 | 11180001 | 11280000 | 0.0014 | 0.0002 | 0.0005 |
| F05 | 11200001 | 11300000 | 0.0012 | 0.0003 | 0.0004 |
| F05 | 11220001 | 11320000 | 0.0011 | 0.0003 | 0.0005 |
| F05 | 11240001 | 11340000 | 0.0012 | 0.0003 | 0.0006 |
| F05 | 11260001 | 11360000 | 0.0012 | 0.0003 | 0.0008 |
| F05 | 11280001 | 11380000 | 0.0013 | 0.0003 | 0.0009 |

| Chr | Start | End | both | A1 | A2 |
|---|---|---|---|---|---|
| F05 | 11300001 | 11400000 | 0.0014 | 0.0003 | 0.0009 |
| F05 | 11320001 | 11420000 | 0.0015 | 0.0002 | 0.0008 |
| F05 | 11340001 | 11440000 | 0.0014 | 0.0003 | 0.0007 |
| F05 | 11360001 | 11460000 | 0.0014 | 0.0003 | 0.0006 |
| F05 | 11380001 | 11480000 | 0.0013 | 0.0003 | 0.0006 |
| F05 | 11400001 | 11500000 | 0.0014 | 0.0005 | 0.0006 |
| F05 | 11420001 | 11520000 | 0.0016 | 0.0005 | 0.0009 |
| F05 | 11440001 | 11540000 | 0.0017 | 0.0004 | 0.0010 |
| F05 | 11460001 | 11560000 | 0.0016 | 0.0004 | 0.0009 |
| F05 | 11480001 | 11580000 | 0.0016 | 0.0005 | 0.0010 |
| F05 | 11500001 | 11600000 | 0.0017 | 0.0005 | 0.0012 |
| F05 | 11520001 | 11620000 | 0.0017 | 0.0005 | 0.0012 |
| F05 | 11540001 | 11640000 | 0.0017 | 0.0004 | 0.0011 |
| F05 | 11560001 | 11660000 | 0.0017 | 0.0005 | 0.0011 |
| F05 | 11580001 | 11680000 | 0.0018 | 0.0005 | 0.0011 |
| F05 | 11600001 | 11700000 | 0.0017 | 0.0003 | 0.0011 |
| F05 | 11620001 | 11720000 | 0.0015 | 0.0005 | 0.0009 |
| F05 | 11640001 | 11740000 | 0.0018 | 0.0005 | 0.0012 |
| F05 | 11660001 | 11760000 | 0.0018 | 0.0006 | 0.0010 |
| F05 | 11680001 | 11780000 | 0.0016 | 0.0007 | 0.0008 |
| F05 | 11700001 | 11800000 | 0.0017 | 0.0007 | 0.0006 |
| F05 | 11720001 | 11820000 | 0.0016 | 0.0006 | 0.0005 |
| F05 | 11740001 | 11840000 | 0.0013 | 0.0006 | 0.0002 |
| F05 | 11760001 | 11860000 | 0.0013 | 0.0006 | 0.0002 |
| F05 | 11780001 | 11880000 | 0.0013 | 0.0007 | 0.0003 |
| F05 | 11800001 | 11900000 | 0.0014 | 0.0008 | 0.0005 |
| F05 | 11820001 | 11920000 | 0.0015 | 0.0008 | 0.0007 |
| F05 | 11840001 | 11940000 | 0.0015 | 0.0011 | 0.0007 |
| F05 | 11860001 | 11960000 | 0.0018 | 0.0011 | 0.0010 |
| F05 | 11880001 | 11980000 | 0.0018 | 0.0010 | 0.0009 |
| F05 | 11900001 | 12000000 | 0.0018 | 0.0014 | 0.0009 |
| F05 | 11920001 | 12020000 | 0.0022 | 0.0013 | 0.0013 |
| F05 | 11940001 | 12040000 | 0.0021 | 0.0010 | 0.0013 |
| F05 | 11960001 | 12060000 | 0.0017 | 0.0011 | 0.0010 |
| F05 | 11980001 | 12080000 | 0.0018 | 0.0009 | 0.0011 |
| F05 | 12000001 | 12100000 | 0.0015 | 0.0005 | 0.0008 |
| F05 | 12020001 | 12120000 | 0.0012 | 0.0007 | 0.0004 |
| F05 | 12040001 | 12140000 | 0.0014 | 0.0008 | 0.0004 |
| F05 | 12060001 | 12160000 | 0.0015 | 0.0009 | 0.0005 |
| F05 | 12080001 | 12180000 | 0.0016 | 0.0012 | 0.0006 |
| F05 | 12100001 | 12200000 | 0.0020 | 0.0013 | 0.0009 |
| F05 | 12120001 | 12220000 | 0.0021 | 0.0014 | 0.0012 |
| F05 | 12140001 | 12240000 | 0.0020 | 0.0015 | 0.0011 |
| F05 | 12160001 | 12260000 | 0.0019 | 0.0015 | 0.0010 |
| F05 | 12180001 | 12280000 | 0.0017 | 0.0015 | 0.0008 |
| F05 | 12200001 | 12300000 | 0.0012 | 0.0017 | 0.0005 |
| F05 | 12220001 | 12320000 | 0.0011 | 0.0017 | 0.0005 |
| F05 | 12240001 | 12340000 | 0.0010 | 0.0016 | 0.0006 |
| F05 | 12260001 | 12360000 | 0.0009 | 0.0015 | 0.0007 |
| F05 | 12280001 | 12380000 | 0.0009 | 0.0014 | 0.0007 |
| F05 | 12300001 | 12400000 | 0.0009 | 0.0011 | 0.0007 |
| F05 | 12320001 | 12420000 | 0.0007 | 0.0014 | 0.0005 |
| F05 | 12340001 | 12440000 | 0.0009 | 0.0016 | 0.0007 |
| F05 | 12360001 | 12460000 | 0.0010 | 0.0017 | 0.0007 |
| F05 | 12380001 | 12480000 | 0.0010 | 0.0019 | 0.0006 |
| F05 | 12400001 | 12500000 | 0.0010 | 0.0024 | 0.0005 |
| F05 | 12420001 | 12520000 | 0.0013 | 0.0022 | 0.0007 |
| F05 | 12440001 | 12540000 | 0.0011 | 0.0021 | 0.0006 |

| Chr | Start | End | both | A1 | A2 |
|---|---|---|---|---|---|
| F05 | 12460001 | 12560000 | 0.0011 | 0.0032 | 0.0006 |
| F05 | 12480001 | 12580000 | 0.0024 | 0.0031 | 0.0020 |
| F05 | 12500001 | 12600000 | 0.0024 | 0.0027 | 0.0020 |
| F05 | 12520001 | 12620000 | 0.0021 | 0.0025 | 0.0019 |
| F05 | 12540001 | 12640000 | 0.0019 | 0.0028 | 0.0017 |
| F05 | 12560001 | 12660000 | 0.0023 | 0.0015 | 0.0020 |
| F05 | 12580001 | 12680000 | 0.0009 | 0.0017 | 0.0006 |
| F05 | 12600001 | 12700000 | 0.0012 | 0.0020 | 0.0010 |
| F05 | 12620001 | 12720000 | 0.0016 | 0.0020 | 0.0014 |
| F05 | 12640001 | 12740000 | 0.0016 | 0.0017 | 0.0014 |
| F05 | 12660001 | 12760000 | 0.0013 | 0.0017 | 0.0011 |
| F05 | 12680001 | 12780000 | 0.0013 | 0.0016 | 0.0011 |
| F05 | 12700001 | 12800000 | 0.0011 | 0.0014 | 0.0008 |
| F05 | 12720001 | 12820000 | 0.0007 | 0.0022 | 0.0004 |
| F05 | 12740001 | 12840000 | 0.0016 | 0.0024 | 0.0013 |
| F05 | 12760001 | 12860000 | 0.0017 | 0.0026 | 0.0014 |
| F05 | 12780001 | 12880000 | 0.0021 | 0.0024 | 0.0017 |
| F05 | 12800001 | 12900000 | 0.0021 | 0.0022 | 0.0015 |
| F05 | 12820001 | 12920000 | 0.0021 | 0.0014 | 0.0015 |
| F05 | 12840001 | 12940000 | 0.0014 | 0.0013 | 0.0008 |
| F05 | 12860001 | 12960000 | 0.0014 | 0.0010 | 0.0007 |
| F05 | 12880001 | 12980000 | 0.0012 | 0.0011 | 0.0004 |
| F05 | 12900001 | 13000000 | 0.0013 | 0.0011 | 0.0004 |
| F05 | 12920001 | 13020000 | 0.0015 | 0.0010 | 0.0004 |
| F05 | 12940001 | 13040000 | 0.0014 | 0.0010 | 0.0003 |
| F05 | 12960001 | 13060000 | 0.0014 | 0.0010 | 0.0004 |
| F05 | 12980001 | 13080000 | 0.0015 | 0.0010 | 0.0006 |
| F05 | 13000001 | 13100000 | 0.0014 | 0.0019 | 0.0006 |
| F05 | 13020001 | 13120000 | 0.0025 | 0.0025 | 0.0018 |
| F05 | 13040001 | 13140000 | 0.0032 | 0.0023 | 0.0025 |
| F05 | 13060001 | 13160000 | 0.0030 | 0.0023 | 0.0023 |
| F05 | 13080001 | 13180000 | 0.0029 | 0.0023 | 0.0022 |
| F05 | 13100001 | 13200000 | 0.0028 | 0.0015 | 0.0023 |
| F05 | 13120001 | 13220000 | 0.0017 | 0.0011 | 0.0012 |
| F05 | 13140001 | 13240000 | 0.0011 | 0.0013 | 0.0006 |
| F05 | 13160001 | 13260000 | 0.0011 | 0.0014 | 0.0006 |
| F05 | 13180001 | 13280000 | 0.0010 | 0.0017 | 0.0006 |
| F05 | 13200001 | 13300000 | 0.0012 | 0.0016 | 0.0007 |
| F05 | 13220001 | 13320000 | 0.0011 | 0.0016 | 0.0006 |
| F05 | 13240001 | 13340000 | 0.0011 | 0.0018 | 0.0005 |
| F05 | 13260001 | 13360000 | 0.0013 | 0.0019 | 0.0007 |
| F05 | 13280001 | 13380000 | 0.0015 | 0.0018 | 0.0009 |
| F05 | 13300001 | 13400000 | 0.0013 | 0.0020 | 0.0007 |
| F05 | 13320001 | 13420000 | 0.0015 | 0.0018 | 0.0008 |
| F05 | 13340001 | 13440000 | 0.0014 | 0.0014 | 0.0007 |
| F05 | 13360001 | 13460000 | 0.0012 | 0.0014 | 0.0005 |
| F05 | 13380001 | 13480000 | 0.0011 | 0.0013 | 0.0004 |
| F05 | 13400001 | 13500000 | 0.0012 | 0.0014 | 0.0006 |
| F05 | 13420001 | 13520000 | 0.0014 | 0.0015 | 0.0009 |
| F05 | 13440001 | 13540000 | 0.0016 | 0.0024 | 0.0011 |
| F05 | 13460001 | 13560000 | 0.0025 | 0.0021 | 0.0021 |
| F05 | 13480001 | 13580000 | 0.0024 | 0.0021 | 0.0019 |
| F05 | 13500001 | 13600000 | 0.0025 | 0.0022 | 0.0020 |
| F05 | 13520001 | 13620000 | 0.0026 | 0.0023 | 0.0020 |
| F05 | 13540001 | 13640000 | 0.0028 | 0.0014 | 0.0022 |
| F05 | 13560001 | 13660000 | 0.0018 | 0.0014 | 0.0012 |
| F05 | 13580001 | 13680000 | 0.0018 | 0.0013 | 0.0012 |
| F05 | 13600001 | 13700000 | 0.0016 | 0.0010 | 0.0009 |

| Chr | Start | End | both | A1 | A2 |
|---|---|---|---|---|---|
| F05 | 13620001 | 13720000 | 0.0013 | 0.0007 | 0.0006 |
| F05 | 13640001 | 13740000 | 0.0009 | 0.0010 | 0.0002 |
| F05 | 13660001 | 13760000 | 0.0013 | 0.0012 | 0.0006 |
| F05 | 13680001 | 13780000 | 0.0015 | 0.0012 | 0.0008 |
| F05 | 13700001 | 13800000 | 0.0016 | 0.0010 | 0.0008 |
| F05 | 13720001 | 13820000 | 0.0015 | 0.0011 | 0.0007 |
| F05 | 13740001 | 13840000 | 0.0016 | 0.0011 | 0.0008 |
| F05 | 13760001 | 13860000 | 0.0017 | 0.0009 | 0.0009 |
| F05 | 13780001 | 13880000 | 0.0015 | 0.0011 | 0.0007 |
| F05 | 13800001 | 13900000 | 0.0017 | 0.0011 | 0.0010 |
| F05 | 13820001 | 13920000 | 0.0017 | 0.0012 | 0.0010 |
| F05 | 13840001 | 13940000 | 0.0017 | 0.0009 | 0.0011 |
| F05 | 13860001 | 13960000 | 0.0014 | 0.0009 | 0.0007 |
| F05 | 13880001 | 13980000 | 0.0013 | 0.0006 | 0.0007 |
| F05 | 13900001 | 14000000 | 0.0010 | 0.0007 | 0.0004 |
| F05 | 13920001 | 14020000 | 0.0013 | 0.0008 | 0.0005 |
| F05 | 13940001 | 14040000 | 0.0013 | 0.0007 | 0.0006 |
| F05 | 13960001 | 14060000 | 0.0013 | 0.0007 | 0.0006 |
| F05 | 13980001 | 14080000 | 0.0013 | 0.0008 | 0.0006 |
| F05 | 14000001 | 14100000 | 0.0013 | 0.0007 | 0.0006 |
| F05 | 14020001 | 14120000 | 0.0012 | 0.0006 | 0.0005 |
| F05 | 14040001 | 14140000 | 0.0013 | 0.0006 | 0.0004 |
| F05 | 14060001 | 14160000 | 0.0014 | 0.0005 | 0.0005 |
| F05 | 14080001 | 14180000 | 0.0015 | 0.0004 | 0.0005 |
| F05 | 14100001 | 14200000 | 0.0015 | 0.0004 | 0.0005 |
| F05 | 14120001 | 14220000 | 0.0015 | 0.0005 | 0.0005 |
| F05 | 14140001 | 14240000 | 0.0016 | 0.0005 | 0.0007 |
| F05 | 14160001 | 14260000 | 0.0016 | 0.0005 | 0.0008 |
| F05 | 14180001 | 14280000 | 0.0017 | 0.0005 | 0.0010 |
| F05 | 14200001 | 14300000 | 0.0017 | 0.0005 | 0.0013 |
| F05 | 14220001 | 14320000 | 0.0016 | 0.0003 | 0.0013 |
| F05 | 14240001 | 14340000 | 0.0016 | 0.0003 | 0.0013 |
| F05 | 14260001 | 14360000 | 0.0016 | 0.0003 | 0.0013 |
| F05 | 14280001 | 14380000 | 0.0016 | 0.0003 | 0.0012 |
| F05 | 14300001 | 14400000 | 0.0016 | 0.0004 | 0.0011 |
| F05 | 14320001 | 14420000 | 0.0017 | 0.0004 | 0.0010 |
| F05 | 14340001 | 14440000 | 0.0015 | 0.0004 | 0.0007 |
| F05 | 14360001 | 14460000 | 0.0015 | 0.0005 | 0.0005 |
| F05 | 14380001 | 14480000 | 0.0016 | 0.0006 | 0.0006 |
| F05 | 14400001 | 14500000 | 0.0015 | 0.0007 | 0.0005 |
| F05 | 14420001 | 14520000 | 0.0014 | 0.0011 | 0.0004 |
| F05 | 14440001 | 14540000 | 0.0017 | 0.0012 | 0.0008 |
| F05 | 14460001 | 14560000 | 0.0017 | 0.0012 | 0.0008 |
| F05 | 14480001 | 14580000 | 0.0016 | 0.0010 | 0.0007 |
| F05 | 14500001 | 14600000 | 0.0017 | 0.0015 | 0.0008 |
| F05 | 14520001 | 14620000 | 0.0024 | 0.0011 | 0.0016 |
| F05 | 14540001 | 14640000 | 0.0022 | 0.0009 | 0.0013 |
| F05 | 14560001 | 14660000 | 0.0021 | 0.0008 | 0.0013 |
| F05 | 14580001 | 14680000 | 0.0020 | 0.0009 | 0.0012 |
| F05 | 14600001 | 14700000 | 0.0018 | 0.0001 | 0.0011 |
| F05 | 14620001 | 14720000 | 0.0010 | 0.0005 | 0.0003 |
| F05 | 14640001 | 14740000 | 0.0014 | 0.0007 | 0.0006 |
| F05 | 14660001 | 14760000 | 0.0014 | 0.0013 | 0.0006 |
| F05 | 14680001 | 14780000 | 0.0016 | 0.0021 | 0.0008 |
| F05 | 14700001 | 14800000 | 0.0021 | 0.0026 | 0.0014 |
| F05 | 14720001 | 14820000 | 0.0022 | 0.0025 | 0.0014 |
| F05 | 14740001 | 14840000 | 0.0018 | 0.0026 | 0.0011 |
| F05 | 14760001 | 14860000 | 0.0017 | 0.0024 | 0.0011 |

| Chr | Start | End | both | A1 | A2 |
|---|---|---|---|---|---|
| F05 | 14780001 | 14880000 | 0.0017 | 0.0021 | 0.0011 |
| F05 | 14800001 | 14900000 | 0.0013 | 0.0019 | 0.0006 |
| F05 | 14820001 | 14920000 | 0.0012 | 0.0019 | 0.0005 |
| F05 | 14840001 | 14940000 | 0.0013 | 0.0024 | 0.0006 |
| F05 | 14860001 | 14960000 | 0.0018 | 0.0023 | 0.0010 |
| F05 | 14880001 | 14980000 | 0.0016 | 0.0024 | 0.0008 |
| F05 | 14900001 | 15000000 | 0.0018 | 0.0027 | 0.0011 |
| F05 | 14920001 | 15020000 | 0.0022 | 0.0027 | 0.0015 |
| F05 | 14940001 | 15040000 | 0.0021 | 0.0024 | 0.0015 |
| F05 | 14960001 | 15060000 | 0.0016 | 0.0026 | 0.0011 |
| F05 | 14980001 | 15080000 | 0.0020 | 0.0024 | 0.0016 |
| F05 | 15000001 | 15100000 | 0.0018 | 0.0024 | 0.0014 |
| F05 | 15020001 | 15120000 | 0.0019 | 0.0032 | 0.0014 |
| F05 | 15040001 | 15140000 | 0.0028 | 0.0037 | 0.0024 |
| F05 | 15060001 | 15160000 | 0.0036 | 0.0035 | 0.0033 |
| F05 | 15080001 | 15180000 | 0.0034 | 0.0043 | 0.0031 |
| F05 | 15100001 | 15200000 | 0.0040 | 0.0040 | 0.0036 |
| F05 | 15120001 | 15220000 | 0.0035 | 0.0034 | 0.0031 |
| F05 | 15140001 | 15240000 | 0.0029 | 0.0029 | 0.0024 |
| F05 | 15160001 | 15260000 | 0.0020 | 0.0037 | 0.0015 |
| F05 | 15180001 | 15280000 | 0.0026 | 0.0032 | 0.0021 |
| F05 | 15200001 | 15300000 | 0.0023 | 0.0032 | 0.0019 |
| F05 | 15220001 | 15320000 | 0.0024 | 0.0030 | 0.0020 |
| F05 | 15240001 | 15340000 | 0.0023 | 0.0031 | 0.0019 |
| F05 | 15260001 | 15360000 | 0.0025 | 0.0020 | 0.0021 |
| F05 | 15280001 | 15380000 | 0.0017 | 0.0020 | 0.0012 |
| F05 | 15300001 | 15400000 | 0.0016 | 0.0023 | 0.0010 |
| F05 | 15320001 | 15420000 | 0.0020 | 0.0023 | 0.0013 |
| F05 | 15340001 | 15440000 | 0.0020 | 0.0020 | 0.0014 |
| F05 | 15360001 | 15460000 | 0.0019 | 0.0023 | 0.0012 |
| F05 | 15380001 | 15480000 | 0.0022 | 0.0023 | 0.0014 |
| F05 | 15400001 | 15500000 | 0.0020 | 0.0028 | 0.0013 |
| F05 | 15420001 | 15520000 | 0.0025 | 0.0033 | 0.0017 |
| F05 | 15440001 | 15540000 | 0.0030 | 0.0035 | 0.0023 |
| F05 | 15460001 | 15560000 | 0.0031 | 0.0033 | 0.0025 |
| F05 | 15480001 | 15580000 | 0.0027 | 0.0033 | 0.0022 |
| F05 | 15500001 | 15600000 | 0.0027 | 0.0024 | 0.0023 |
| F05 | 15520001 | 15620000 | 0.0018 | 0.0019 | 0.0015 |
| F05 | 15540001 | 15640000 | 0.0011 | 0.0018 | 0.0007 |
| F05 | 15560001 | 15660000 | 0.0009 | 0.0017 | 0.0004 |
| F05 | 15580001 | 15680000 | 0.0009 | 0.0015 | 0.0004 |
| F05 | 15600001 | 15700000 | 0.0007 | 0.0014 | 0.0002 |
| F05 | 15620001 | 15720000 | 0.0007 | 0.0013 | 0.0001 |
| F05 | 15640001 | 15740000 | 0.0008 | 0.0014 | 0.0001 |
| F05 | 15660001 | 15760000 | 0.0014 | 0.0011 | 0.0006 |
| F05 | 15680001 | 15780000 | 0.0015 | 0.0010 | 0.0006 |
| F05 | 15700001 | 15800000 | 0.0016 | 0.0011 | 0.0007 |
| F05 | 15720001 | 15820000 | 0.0020 | 0.0013 | 0.0009 |
| F05 | 15740001 | 15840000 | 0.0023 | 0.0011 | 0.0012 |
| F05 | 15760001 | 15860000 | 0.0019 | 0.0011 | 0.0010 |
| F05 | 15780001 | 15880000 | 0.0019 | 0.0010 | 0.0010 |
| F05 | 15800001 | 15900000 | 0.0017 | 0.0010 | 0.0009 |
| F05 | 15820001 | 15920000 | 0.0018 | 0.0007 | 0.0010 |
| F05 | 15840001 | 15940000 | 0.0017 | 0.0009 | 0.0009 |
| F05 | 15860001 | 15960000 | 0.0018 | 0.0010 | 0.0011 |
| F05 | 15880001 | 15980000 | 0.0018 | 0.0010 | 0.0012 |
| F05 | 15900001 | 16000000 | 0.0019 | 0.0008 | 0.0012 |
| F05 | 15920001 | 16020000 | 0.0016 | 0.0009 | 0.0009 |

| Chr | Start | End | both | A1 | A2 |
|---|---|---|---|---|---|
| F05 | 15940001 | 16040000 | 0.0018 | 0.0008 | 0.0010 |
| F05 | 15960001 | 16060000 | 0.0016 | 0.0007 | 0.0009 |
| F05 | 15980001 | 16080000 | 0.0017 | 0.0008 | 0.0009 |
| F05 | 16000001 | 16100000 | 0.0017 | 0.0008 | 0.0010 |
| F05 | 16020001 | 16120000 | 0.0019 | 0.0014 | 0.0011 |
| F05 | 16040001 | 16140000 | 0.0024 | 0.0013 | 0.0017 |
| F05 | 16060001 | 16160000 | 0.0023 | 0.0014 | 0.0015 |
| F05 | 16080001 | 16180000 | 0.0024 | 0.0014 | 0.0016 |
| F05 | 16100001 | 16200000 | 0.0024 | 0.0017 | 0.0015 |
| F05 | 16120001 | 16220000 | 0.0027 | 0.0008 | 0.0018 |
| F05 | 16140001 | 16240000 | 0.0019 | 0.0009 | 0.0008 |
| F05 | 16160001 | 16260000 | 0.0020 | 0.0008 | 0.0010 |
| F05 | 16180001 | 16280000 | 0.0018 | 0.0008 | 0.0008 |
| F05 | 16200001 | 16300000 | 0.0017 | 0.0003 | 0.0009 |
| F05 | 16220001 | 16320000 | 0.0011 | 0.0006 | 0.0004 |
| F05 | 16240001 | 16340000 | 0.0014 | 0.0006 | 0.0007 |
| F05 | 16260001 | 16360000 | 0.0014 | 0.0008 | 0.0007 |
| F05 | 16280001 | 16380000 | 0.0014 | 0.0011 | 0.0008 |
| F05 | 16300001 | 16400000 | 0.0018 | 0.0016 | 0.0012 |
| F05 | 16320001 | 16420000 | 0.0023 | 0.0025 | 0.0018 |
| F05 | 16340001 | 16440000 | 0.0031 | 0.0030 | 0.0029 |
| F05 | 16360001 | 16460000 | 0.0035 | 0.0034 | 0.0034 |
| F05 | 16380001 | 16480000 | 0.0039 | 0.0035 | 0.0037 |
| F05 | 16400001 | 16500000 | 0.0040 | 0.0038 | 0.0039 |
| F05 | 16420001 | 16520000 | 0.0043 | 0.0040 | 0.0041 |
| F05 | 16440001 | 16540000 | 0.0045 | 0.0033 | 0.0043 |
| F05 | 16460001 | 16560000 | 0.0039 | 0.0031 | 0.0036 |
| F05 | 16480001 | 16580000 | 0.0037 | 0.0025 | 0.0034 |
| F05 | 16500001 | 16600000 | 0.0032 | 0.0022 | 0.0027 |
| F05 | 16520001 | 16620000 | 0.0031 | 0.0009 | 0.0025 |
| F05 | 16540001 | 16640000 | 0.0017 | 0.0009 | 0.0010 |
| F05 | 16560001 | 16660000 | 0.0017 | 0.0009 | 0.0010 |
| F05 | 16580001 | 16680000 | 0.0017 | 0.0010 | 0.0010 |
| F05 | 16600001 | 16700000 | 0.0018 | 0.0008 | 0.0011 |
| F05 | 16620001 | 16720000 | 0.0015 | 0.0008 | 0.0009 |
| F05 | 16640001 | 16740000 | 0.0015 | 0.0008 | 0.0009 |
| F05 | 16660001 | 16760000 | 0.0014 | 0.0016 | 0.0009 |
| F05 | 16680001 | 16780000 | 0.0024 | 0.0020 | 0.0018 |
| F05 | 16700001 | 16800000 | 0.0029 | 0.0022 | 0.0023 |
| F05 | 16720001 | 16820000 | 0.0032 | 0.0024 | 0.0025 |
| F05 | 16740001 | 16840000 | 0.0032 | 0.0029 | 0.0027 |
| F05 | 16760001 | 16860000 | 0.0039 | 0.0021 | 0.0035 |
| F05 | 16780001 | 16880000 | 0.0028 | 0.0019 | 0.0024 |
| F05 | 16800001 | 16900000 | 0.0026 | 0.0014 | 0.0020 |
| F05 | 16820001 | 16920000 | 0.0020 | 0.0013 | 0.0015 |
| F05 | 16840001 | 16940000 | 0.0021 | 0.0007 | 0.0014 |
| F05 | 16860001 | 16960000 | 0.0015 | 0.0010 | 0.0006 |
| F05 | 16880001 | 16980000 | 0.0019 | 0.0017 | 0.0010 |
| F05 | 16900001 | 17000000 | 0.0025 | 0.0018 | 0.0017 |
| F05 | 16920001 | 17020000 | 0.0026 | 0.0017 | 0.0018 |
| F05 | 16940001 | 17040000 | 0.0025 | 0.0019 | 0.0017 |
| F05 | 16960001 | 17060000 | 0.0026 | 0.0014 | 0.0019 |
| F05 | 16980001 | 17080000 | 0.0022 | 0.0005 | 0.0013 |
| F05 | 17000001 | 17100000 | 0.0014 | 0.0009 | 0.0004 |
| F05 | 17020001 | 17120000 | 0.0016 | 0.0010 | 0.0007 |
| F05 | 17040001 | 17140000 | 0.0017 | 0.0012 | 0.0008 |
| F05 | 17060001 | 17160000 | 0.0018 | 0.0013 | 0.0010 |
| F05 | 17080001 | 17180000 | 0.0019 | 0.0013 | 0.0011 |

| Chr | Start | End | both | A1 | A2 |
|---|---|---|---|---|---|
| F05 | 17100001 | 17200000 | 0.0020 | 0.0011 | 0.0012 |
| F05 | 17120001 | 17220000 | 0.0019 | 0.0012 | 0.0010 |
| F05 | 17140001 | 17240000 | 0.0021 | 0.0008 | 0.0012 |
| F05 | 17160001 | 17260000 | 0.0019 | 0.0008 | 0.0009 |
| F05 | 17180001 | 17280000 | 0.0019 | 0.0007 | 0.0009 |
| F05 | 17200001 | 17300000 | 0.0018 | 0.0007 | 0.0007 |
| F05 | 17220001 | 17320000 | 0.0018 | 0.0006 | 0.0008 |
| F05 | 17240001 | 17340000 | 0.0016 | 0.0007 | 0.0008 |
| F05 | 17260001 | 17360000 | 0.0017 | 0.0009 | 0.0009 |
| F05 | 17280001 | 17380000 | 0.0018 | 0.0009 | 0.0010 |
| F05 | 17300001 | 17400000 | 0.0019 | 0.0008 | 0.0010 |
| F05 | 17320001 | 17420000 | 0.0017 | 0.0007 | 0.0009 |
| F05 | 17340001 | 17440000 | 0.0016 | 0.0009 | 0.0008 |
| F05 | 17360001 | 17460000 | 0.0018 | 0.0009 | 0.0012 |
| F05 | 17380001 | 17480000 | 0.0017 | 0.0012 | 0.0011 |
| F05 | 17400001 | 17500000 | 0.0021 | 0.0015 | 0.0014 |
| F05 | 17420001 | 17520000 | 0.0024 | 0.0014 | 0.0016 |
| F05 | 17440001 | 17540000 | 0.0022 | 0.0012 | 0.0015 |
| F05 | 17460001 | 17560000 | 0.0020 | 0.0011 | 0.0011 |
| F05 | 17480001 | 17580000 | 0.0020 | 0.0008 | 0.0010 |
| F05 | 17500001 | 17600000 | 0.0016 | 0.0004 | 0.0008 |
| F05 | 17520001 | 17620000 | 0.0014 | 0.0003 | 0.0005 |
| F05 | 17540001 | 17640000 | 0.0013 | 0.0002 | 0.0004 |
| F05 | 17560001 | 17660000 | 0.0013 | 0.0002 | 0.0003 |
| F05 | 17580001 | 17680000 | 0.0013 | 0.0001 | 0.0002 |
| F05 | 17600001 | 17700000 | 0.0012 | 0.0005 | 0.0001 |
| F05 | 17620001 | 17720000 | 0.0016 | 0.0005 | 0.0005 |
| F05 | 17640001 | 17740000 | 0.0017 | 0.0007 | 0.0005 |
| F05 | 17660001 | 17760000 | 0.0018 | 0.0008 | 0.0007 |
| F05 | 17680001 | 17780000 | 0.0019 | 0.0011 | 0.0008 |
| F05 | 17700001 | 17800000 | 0.0022 | 0.0007 | 0.0012 |
| F05 | 17720001 | 17820000 | 0.0018 | 0.0009 | 0.0008 |
| F05 | 17740001 | 17840000 | 0.0019 | 0.0007 | 0.0009 |
| F05 | 17760001 | 17860000 | 0.0018 | 0.0006 | 0.0007 |
| F05 | 17780001 | 17880000 | 0.0018 | 0.0003 | 0.0006 |
| F05 | 17800001 | 17900000 | 0.0014 | 0.0003 | 0.0003 |
| F05 | 17820001 | 17920000 | 0.0015 | 0.0005 | 0.0003 |
| F05 | 17840001 | 17940000 | 0.0017 | 0.0010 | 0.0006 |
| F05 | 17860001 | 17960000 | 0.0020 | 0.0010 | 0.0011 |
| F05 | 17880001 | 17980000 | 0.0021 | 0.0010 | 0.0011 |
| F05 | 17900001 | 18000000 | 0.0021 | 0.0010 | 0.0011 |
| F05 | 17920001 | 18020000 | 0.0021 | 0.0006 | 0.0011 |
| F05 | 17940001 | 18040000 | 0.0018 | 0.0002 | 0.0007 |
| F05 | 17960001 | 18060000 | 0.0014 | 0.0002 | 0.0003 |
| F05 | 17980001 | 18080000 | 0.0013 | 0.0002 | 0.0003 |
| F05 | 18000001 | 18100000 | 0.0013 | 0.0002 | 0.0002 |
| F05 | 18020001 | 18120000 | 0.0013 | 0.0003 | 0.0003 |
| F05 | 18040001 | 18140000 | 0.0013 | 0.0006 | 0.0003 |
| F05 | 18060001 | 18160000 | 0.0014 | 0.0007 | 0.0005 |
| F05 | 18080001 | 18180000 | 0.0013 | 0.0008 | 0.0005 |
| F05 | 18100001 | 18200000 | 0.0014 | 0.0012 | 0.0004 |
| F05 | 18120001 | 18220000 | 0.0017 | 0.0011 | 0.0005 |
| F05 | 18140001 | 18240000 | 0.0017 | 0.0010 | 0.0004 |
| F05 | 18160001 | 18260000 | 0.0017 | 0.0008 | 0.0004 |
| F05 | 18180001 | 18280000 | 0.0018 | 0.0008 | 0.0004 |
| F05 | 18200001 | 18300000 | 0.0018 | 0.0008 | 0.0005 |
| F05 | 18220001 | 18320000 | 0.0017 | 0.0006 | 0.0007 |
| F05 | 18240001 | 18340000 | 0.0015 | 0.0006 | 0.0007 |

| Chr | Start | End | both | A1 | A2 |
|---|---|---|---|---|---|
| F05 | 18260001 | 18360000 | 0.0013 | 0.0006 | 0.0006 |
| F05 | 18280001 | 18380000 | 0.0013 | 0.0006 | 0.0005 |
| F05 | 18300001 | 18400000 | 0.0014 | 0.0002 | 0.0006 |
| F05 | 18320001 | 18420000 | 0.0012 | 0.0005 | 0.0003 |
| F05 | 18340001 | 18440000 | 0.0017 | 0.0008 | 0.0007 |
| F05 | 18360001 | 18460000 | 0.0019 | 0.0008 | 0.0009 |
| F05 | 18380001 | 18480000 | 0.0018 | 0.0009 | 0.0010 |
| F05 | 18400001 | 18500000 | 0.0017 | 0.0009 | 0.0009 |
| F05 | 18420001 | 18520000 | 0.0016 | 0.0006 | 0.0009 |
| F05 | 18440001 | 18540000 | 0.0012 | 0.0003 | 0.0005 |
| F05 | 18460001 | 18560000 | 0.0011 | 0.0002 | 0.0003 |
| F05 | 18480001 | 18580000 | 0.0011 | 0.0002 | 0.0003 |
| F05 | 18500001 | 18600000 | 0.0013 | 0.0002 | 0.0003 |
| F05 | 18520001 | 18620000 | 0.0015 | 0.0005 | 0.0004 |
| F05 | 18540001 | 18640000 | 0.0016 | 0.0007 | 0.0005 |
| F05 | 18560001 | 18660000 | 0.0017 | 0.0008 | 0.0008 |
| F05 | 18580001 | 18680000 | 0.0017 | 0.0009 | 0.0008 |
| F05 | 18600001 | 18700000 | 0.0017 | 0.0010 | 0.0010 |
| F05 | 18620001 | 18720000 | 0.0018 | 0.0011 | 0.0010 |
| F05 | 18640001 | 18740000 | 0.0019 | 0.0012 | 0.0013 |
| F05 | 18660001 | 18760000 | 0.0021 | 0.0013 | 0.0013 |
| F05 | 18680001 | 18780000 | 0.0022 | 0.0012 | 0.0013 |
| F05 | 18700001 | 18800000 | 0.0020 | 0.0013 | 0.0011 |
| F05 | 18720001 | 18820000 | 0.0020 | 0.0012 | 0.0012 |
| F05 | 18740001 | 18840000 | 0.0018 | 0.0010 | 0.0009 |
| F05 | 18760001 | 18860000 | 0.0016 | 0.0009 | 0.0006 |
| F05 | 18780001 | 18880000 | 0.0015 | 0.0008 | 0.0006 |
| F05 | 18800001 | 18900000 | 0.0015 | 0.0007 | 0.0005 |
| F05 | 18820001 | 18920000 | 0.0015 | 0.0007 | 0.0006 |
| F05 | 18840001 | 18940000 | 0.0016 | 0.0011 | 0.0007 |
| F05 | 18860001 | 18960000 | 0.0020 | 0.0010 | 0.0011 |
| F05 | 18880001 | 18980000 | 0.0020 | 0.0015 | 0.0012 |
| F05 | 18900001 | 19000000 | 0.0024 | 0.0016 | 0.0018 |
| F05 | 18920001 | 19020000 | 0.0026 | 0.0014 | 0.0019 |
| F05 | 18940001 | 19040000 | 0.0023 | 0.0010 | 0.0016 |
| F05 | 18960001 | 19060000 | 0.0020 | 0.0014 | 0.0012 |
| F05 | 18980001 | 19080000 | 0.0022 | 0.0009 | 0.0014 |
| F05 | 19000001 | 19100000 | 0.0017 | 0.0007 | 0.0007 |
| F05 | 19020001 | 19120000 | 0.0015 | 0.0008 | 0.0005 |
| F05 | 19040001 | 19140000 | 0.0016 | 0.0007 | 0.0006 |
| F05 | 19060001 | 19160000 | 0.0016 | 0.0008 | 0.0006 |
| F05 | 19080001 | 19180000 | 0.0016 | 0.0010 | 0.0006 |
| F05 | 19100001 | 19200000 | 0.0018 | 0.0012 | 0.0009 |
| F05 | 19120001 | 19220000 | 0.0022 | 0.0013 | 0.0015 |
| F05 | 19140001 | 19240000 | 0.0022 | 0.0016 | 0.0017 |
| F05 | 19160001 | 19260000 | 0.0025 | 0.0015 | 0.0021 |
| F05 | 19180001 | 19280000 | 0.0023 | 0.0019 | 0.0019 |
| F05 | 19200001 | 19300000 | 0.0025 | 0.0021 | 0.0020 |
| F05 | 19220001 | 19320000 | 0.0022 | 0.0021 | 0.0015 |
| F05 | 19240001 | 19340000 | 0.0019 | 0.0020 | 0.0013 |
| F05 | 19260001 | 19360000 | 0.0016 | 0.0020 | 0.0009 |
| F05 | 19280001 | 19380000 | 0.0015 | 0.0016 | 0.0008 |
| F05 | 19300001 | 19400000 | 0.0011 | 0.0015 | 0.0004 |
| F05 | 19320001 | 19420000 | 0.0010 | 0.0018 | 0.0003 |
| F05 | 19340001 | 19440000 | 0.0012 | 0.0018 | 0.0003 |
| F05 | 19360001 | 19460000 | 0.0012 | 0.0028 | 0.0003 |
| F05 | 19380001 | 19480000 | 0.0024 | 0.0029 | 0.0017 |
| F05 | 19400001 | 19500000 | 0.0025 | 0.0025 | 0.0018 |

| Chr | Start | End | both | A1 | A2 |
|---|---|---|---|---|---|
| F05 | 19420001 | 19520000 | 0.0024 | 0.0022 | 0.0016 |
| F05 | 19440001 | 19540000 | 0.0024 | 0.0022 | 0.0017 |
| F05 | 19460001 | 19560000 | 0.0025 | 0.0014 | 0.0019 |
| F05 | 19480001 | 19580000 | 0.0018 | 0.0018 | 0.0013 |
| F05 | 19500001 | 19600000 | 0.0023 | 0.0021 | 0.0018 |
| F05 | 19520001 | 19620000 | 0.0025 | 0.0022 | 0.0022 |
| F05 | 19540001 | 19640000 | 0.0027 | 0.0027 | 0.0022 |
| F05 | 19560001 | 19660000 | 0.0034 | 0.0032 | 0.0027 |
| F05 | 19580001 | 19680000 | 0.0039 | 0.0028 | 0.0032 |
| F05 | 19600001 | 19700000 | 0.0036 | 0.0026 | 0.0028 |
| F05 | 19620001 | 19720000 | 0.0037 | 0.0030 | 0.0027 |
| F05 | 19640001 | 19740000 | 0.0040 | 0.0024 | 0.0032 |
| F05 | 19660001 | 19760000 | 0.0033 | 0.0012 | 0.0027 |
| F05 | 19680001 | 19780000 | 0.0022 | 0.0013 | 0.0014 |
| F05 | 19700001 | 19800000 | 0.0023 | 0.0011 | 0.0016 |
| F05 | 19720001 | 19820000 | 0.0021 | 0.0009 | 0.0014 |
| F05 | 19740001 | 19840000 | 0.0019 | 0.0017 | 0.0012 |
| F05 | 19760001 | 19860000 | 0.0027 | 0.0017 | 0.0020 |
| F05 | 19780001 | 19880000 | 0.0027 | 0.0016 | 0.0021 |
| F05 | 19800001 | 19900000 | 0.0025 | 0.0020 | 0.0019 |
| F05 | 19820001 | 19920000 | 0.0026 | 0.0025 | 0.0021 |
| F05 | 19840001 | 19940000 | 0.0030 | 0.0020 | 0.0025 |
| F05 | 19860001 | 19960000 | 0.0024 | 0.0021 | 0.0019 |
| F05 | 19880001 | 19980000 | 0.0023 | 0.0020 | 0.0020 |
| F05 | 19900001 | 20000000 | 0.0023 | 0.0019 | 0.0020 |
| F05 | 19920001 | 20020000 | 0.0022 | 0.0010 | 0.0019 |
| F05 | 19940001 | 20040000 | 0.0016 | 0.0006 | 0.0011 |
| F05 | 19960001 | 20060000 | 0.0014 | 0.0004 | 0.0008 |
| F05 | 19980001 | 20080000 | 0.0015 | 0.0006 | 0.0008 |
| F05 | 20000001 | 20100000 | 0.0017 | 0.0005 | 0.0011 |
| F05 | 20020001 | 20120000 | 0.0017 | 0.0008 | 0.0011 |
| F05 | 20040001 | 20140000 | 0.0021 | 0.0010 | 0.0014 |
| F05 | 20060001 | 20160000 | 0.0022 | 0.0010 | 0.0015 |
| F05 | 20080001 | 20180000 | 0.0021 | 0.0008 | 0.0014 |
| F05 | 20100001 | 20200000 | 0.0018 | 0.0007 | 0.0010 |
| F05 | 20120001 | 20220000 | 0.0018 | 0.0004 | 0.0009 |
| F05 | 20140001 | 20240000 | 0.0014 | 0.0006 | 0.0005 |
| F05 | 20160001 | 20260000 | 0.0016 | 0.0007 | 0.0007 |
| F05 | 20180001 | 20280000 | 0.0018 | 0.0007 | 0.0008 |
| F05 | 20200001 | 20300000 | 0.0020 | 0.0007 | 0.0010 |
| F05 | 20220001 | 20320000 | 0.0020 | 0.0008 | 0.0011 |
| F05 | 20240001 | 20340000 | 0.0023 | 0.0005 | 0.0015 |
| F05 | 20260001 | 20360000 | 0.0021 | 0.0005 | 0.0014 |
| F05 | 20280001 | 20380000 | 0.0019 | 0.0007 | 0.0012 |
| F05 | 20300001 | 20400000 | 0.0018 | 0.0011 | 0.0012 |
| F05 | 20320001 | 20420000 | 0.0020 | 0.0011 | 0.0014 |
| F05 | 20340001 | 20440000 | 0.0017 | 0.0013 | 0.0011 |
| F05 | 20360001 | 20460000 | 0.0015 | 0.0015 | 0.0009 |
| F05 | 20380001 | 20480000 | 0.0017 | 0.0015 | 0.0010 |
| F05 | 20400001 | 20500000 | 0.0015 | 0.0013 | 0.0009 |
| F05 | 20420001 | 20520000 | 0.0013 | 0.0013 | 0.0006 |
| F05 | 20440001 | 20540000 | 0.0013 | 0.0015 | 0.0005 |
| F05 | 20460001 | 20560000 | 0.0014 | 0.0015 | 0.0006 |
| F05 | 20480001 | 20580000 | 0.0014 | 0.0015 | 0.0007 |
| F05 | 20500001 | 20600000 | 0.0014 | 0.0016 | 0.0008 |
| F05 | 20520001 | 20620000 | 0.0015 | 0.0020 | 0.0008 |
| F05 | 20540001 | 20640000 | 0.0018 | 0.0020 | 0.0011 |
| F05 | 20560001 | 20660000 | 0.0018 | 0.0018 | 0.0010 |

| Chr | Start | End | both | A1 | A2 |
|---|---|---|---|---|---|
| F05 | 20580001 | 20680000 | 0.0016 | 0.0015 | 0.0008 |
| F05 | 20600001 | 20700000 | 0.0015 | 0.0014 | 0.0007 |
| F05 | 20620001 | 20720000 | 0.0018 | 0.0008 | 0.0009 |
| F05 | 20640001 | 20740000 | 0.0015 | 0.0007 | 0.0006 |
| F05 | 20660001 | 20760000 | 0.0018 | 0.0006 | 0.0008 |
| F05 | 20680001 | 20780000 | 0.0020 | 0.0014 | 0.0009 |
| F05 | 20700001 | 20800000 | 0.0027 | 0.0012 | 0.0016 |
| F05 | 20720001 | 20820000 | 0.0024 | 0.0015 | 0.0015 |
| F05 | 20740001 | 20840000 | 0.0027 | 0.0014 | 0.0017 |
| F05 | 20760001 | 20860000 | 0.0027 | 0.0018 | 0.0016 |
| F05 | 20780001 | 20880000 | 0.0030 | 0.0012 | 0.0020 |
| F05 | 20800001 | 20900000 | 0.0024 | 0.0013 | 0.0014 |
| F05 | 20820001 | 20920000 | 0.0024 | 0.0010 | 0.0015 |
| F05 | 20840001 | 20940000 | 0.0019 | 0.0012 | 0.0012 |
| F05 | 20860001 | 20960000 | 0.0017 | 0.0011 | 0.0012 |
| F05 | 20880001 | 20980000 | 0.0016 | 0.0011 | 0.0012 |
| F05 | 20900001 | 21000000 | 0.0015 | 0.0010 | 0.0011 |
| F05 | 20920001 | 21020000 | 0.0014 | 0.0013 | 0.0011 |
| F05 | 20940001 | 21040000 | 0.0018 | 0.0011 | 0.0014 |
| F05 | 20960001 | 21060000 | 0.0016 | 0.0007 | 0.0012 |
| F05 | 20980001 | 21080000 | 0.0011 | 0.0008 | 0.0006 |
| F05 | 21000001 | 21100000 | 0.0012 | 0.0020 | 0.0009 |
| F05 | 21020001 | 21120000 | 0.0025 | 0.0025 | 0.0021 |
| F05 | 21040001 | 21140000 | 0.0032 | 0.0027 | 0.0028 |
| F05 | 21060001 | 21160000 | 0.0034 | 0.0029 | 0.0030 |
| F05 | 21080001 | 21180000 | 0.0037 | 0.0027 | 0.0033 |
| F05 | 21100001 | 21200000 | 0.0036 | 0.0017 | 0.0031 |
| F05 | 21120001 | 21220000 | 0.0025 | 0.0012 | 0.0020 |
| F05 | 21140001 | 21240000 | 0.0019 | 0.0012 | 0.0014 |
| F05 | 21160001 | 21260000 | 0.0020 | 0.0011 | 0.0014 |
| F05 | 21180001 | 21280000 | 0.0019 | 0.0013 | 0.0013 |
| F05 | 21200001 | 21300000 | 0.0020 | 0.0017 | 0.0013 |
| F05 | 21220001 | 21320000 | 0.0024 | 0.0012 | 0.0017 |
| F05 | 21240001 | 21340000 | 0.0019 | 0.0012 | 0.0013 |
| F05 | 21260001 | 21360000 | 0.0019 | 0.0009 | 0.0012 |
| F05 | 21280001 | 21380000 | 0.0015 | 0.0011 | 0.0009 |
| F05 | 21300001 | 21400000 | 0.0017 | 0.0008 | 0.0012 |
| F05 | 21320001 | 21420000 | 0.0015 | 0.0020 | 0.0010 |
| F05 | 21340001 | 21440000 | 0.0026 | 0.0026 | 0.0021 |
| F05 | 21360001 | 21460000 | 0.0029 | 0.0029 | 0.0027 |
| F05 | 21380001 | 21480000 | 0.0032 | 0.0031 | 0.0031 |
| F05 | 21400001 | 21500000 | 0.0034 | 0.0041 | 0.0033 |
| F05 | 21420001 | 21520000 | 0.0045 | 0.0030 | 0.0044 |
| F05 | 21440001 | 21540000 | 0.0034 | 0.0029 | 0.0033 |
| F05 | 21460001 | 21560000 | 0.0032 | 0.0029 | 0.0032 |
| F05 | 21480001 | 21580000 | 0.0033 | 0.0027 | 0.0031 |
| F05 | 21500001 | 21600000 | 0.0031 | 0.0014 | 0.0027 |
| F05 | 21520001 | 21620000 | 0.0018 | 0.0017 | 0.0014 |
| F05 | 21540001 | 21640000 | 0.0023 | 0.0012 | 0.0017 |
| F05 | 21560001 | 21660000 | 0.0020 | 0.0013 | 0.0013 |
| F05 | 21580001 | 21680000 | 0.0023 | 0.0012 | 0.0014 |
| F05 | 21600001 | 21700000 | 0.0020 | 0.0014 | 0.0013 |
| F05 | 21620001 | 21720000 | 0.0021 | 0.0012 | 0.0013 |
| F05 | 21640001 | 21740000 | 0.0017 | 0.0011 | 0.0010 |
| F05 | 21660001 | 21760000 | 0.0017 | 0.0008 | 0.0010 |
| F05 | 21680001 | 21780000 | 0.0012 | 0.0014 | 0.0006 |
| F05 | 21700001 | 21800000 | 0.0020 | 0.0012 | 0.0012 |
| F05 | 21720001 | 21820000 | 0.0020 | 0.0011 | 0.0012 |

| Chr | Start | End | both | A1 | A2 |
|---|---|---|---|---|---|
| F05 | 21740001 | 21840000 | 0.0019 | 0.0013 | 0.0012 |
| F05 | 21760001 | 21860000 | 0.0020 | 0.0013 | 0.0015 |
| F05 | 21780001 | 21880000 | 0.0022 | 0.0005 | 0.0016 |
| F05 | 21800001 | 21900000 | 0.0013 | 0.0003 | 0.0008 |
| F05 | 21820001 | 21920000 | 0.0013 | 0.0006 | 0.0009 |
| F05 | 21840001 | 21940000 | 0.0018 | 0.0004 | 0.0016 |
| F05 | 21860001 | 21960000 | 0.0017 | 0.0011 | 0.0013 |
| F05 | 21880001 | 21980000 | 0.0024 | 0.0012 | 0.0020 |
| F05 | 21900001 | 22000000 | 0.0028 | 0.0016 | 0.0021 |
| F05 | 21920001 | 22020000 | 0.0030 | 0.0014 | 0.0022 |
| F05 | 21940001 | 22040000 | 0.0026 | 0.0016 | 0.0017 |
| F05 | 21960001 | 22060000 | 0.0028 | 0.0011 | 0.0019 |
| F05 | 21980001 | 22080000 | 0.0021 | 0.0016 | 0.0013 |
| F05 | 22000001 | 22100000 | 0.0024 | 0.0028 | 0.0017 |
| F05 | 22020001 | 22120000 | 0.0035 | 0.0034 | 0.0027 |
| F05 | 22040001 | 22140000 | 0.0041 | 0.0038 | 0.0033 |
| F05 | 22060001 | 22160000 | 0.0045 | 0.0041 | 0.0037 |
| F05 | 22080001 | 22180000 | 0.0048 | 0.0042 | 0.0041 |
| F05 | 22100001 | 22200000 | 0.0050 | 0.0032 | 0.0044 |
| F05 | 22120001 | 22220000 | 0.0042 | 0.0030 | 0.0036 |
| F05 | 22140001 | 22240000 | 0.0040 | 0.0030 | 0.0035 |
| F05 | 22160001 | 22260000 | 0.0040 | 0.0027 | 0.0035 |
| F05 | 22180001 | 22280000 | 0.0035 | 0.0021 | 0.0031 |
| F05 | 22200001 | 22300000 | 0.0027 | 0.0020 | 0.0024 |
| F05 | 22220001 | 22320000 | 0.0023 | 0.0019 | 0.0022 |
| F05 | 22240001 | 22340000 | 0.0022 | 0.0016 | 0.0022 |
| F05 | 22260001 | 22360000 | 0.0022 | 0.0021 | 0.0022 |
| F05 | 22280001 | 22380000 | 0.0029 | 0.0027 | 0.0029 |
| F05 | 22300001 | 22400000 | 0.0036 | 0.0031 | 0.0036 |
| F05 | 22320001 | 22420000 | 0.0040 | 0.0029 | 0.0040 |
| F05 | 22340001 | 22440000 | 0.0037 | 0.0031 | 0.0037 |
| F05 | 22360001 | 22460000 | 0.0037 | 0.0027 | 0.0037 |
| F05 | 22380001 | 22480000 | 0.0032 | 0.0025 | 0.0032 |
| F05 | 22400001 | 22500000 | 0.0029 | 0.0024 | 0.0028 |
| F05 | 22420001 | 22520000 | 0.0027 | 0.0025 | 0.0025 |
| F05 | 22440001 | 22540000 | 0.0029 | 0.0026 | 0.0026 |
| F05 | 22460001 | 22560000 | 0.0031 | 0.0023 | 0.0026 |
| F05 | 22480001 | 22580000 | 0.0028 | 0.0018 | 0.0023 |
| F05 | 22500001 | 22600000 | 0.0022 | 0.0015 | 0.0017 |
| F05 | 22520001 | 22620000 | 0.0023 | 0.0011 | 0.0016 |
| F05 | 22540001 | 22640000 | 0.0019 | 0.0005 | 0.0013 |
| F05 | 22560001 | 22660000 | 0.0014 | 0.0005 | 0.0006 |
| F05 | 22580001 | 22680000 | 0.0015 | 0.0005 | 0.0007 |
| F05 | 22600001 | 22700000 | 0.0016 | 0.0003 | 0.0007 |
| F05 | 22620001 | 22720000 | 0.0013 | 0.0003 | 0.0003 |
| F05 | 22640001 | 22740000 | 0.0012 | 0.0003 | 0.0004 |
| F05 | 22660001 | 22760000 | 0.0012 | 0.0005 | 0.0004 |
| F05 | 22680001 | 22780000 | 0.0016 | 0.0009 | 0.0006 |
| F05 | 22700001 | 22800000 | 0.0020 | 0.0008 | 0.0012 |
| F05 | 22720001 | 22820000 | 0.0019 | 0.0010 | 0.0011 |
| F05 | 22740001 | 22840000 | 0.0021 | 0.0023 | 0.0013 |
| F05 | 22760001 | 22860000 | 0.0034 | 0.0023 | 0.0026 |
| F05 | 22780001 | 22880000 | 0.0033 | 0.0020 | 0.0025 |
| F05 | 22800001 | 22900000 | 0.0028 | 0.0029 | 0.0021 |
| F05 | 22820001 | 22920000 | 0.0037 | 0.0036 | 0.0031 |
| F05 | 22840001 | 22940000 | 0.0043 | 0.0029 | 0.0039 |
| F05 | 22860001 | 22960000 | 0.0035 | 0.0029 | 0.0032 |
| F05 | 22880001 | 22980000 | 0.0035 | 0.0028 | 0.0032 |

| Chr | Start | End | both | A1 | A2 |
|---|---|---|---|---|---|
| F05 | 22900001 | 23000000 | 0.0035 | 0.0020 | 0.0032 |
| F05 | 22920001 | 23020000 | 0.0028 | 0.0010 | 0.0024 |
| F05 | 22940001 | 23040000 | 0.0021 | 0.0005 | 0.0017 |
| F05 | 22960001 | 23060000 | 0.0017 | 0.0003 | 0.0014 |
| F05 | 22980001 | 23080000 | 0.0017 | 0.0009 | 0.0016 |
| F05 | 23000001 | 23100000 | 0.0026 | 0.0009 | 0.0026 |
| F05 | 23020001 | 23120000 | 0.0027 | 0.0010 | 0.0027 |
| F05 | 23040001 | 23140000 | 0.0027 | 0.0011 | 0.0024 |
| F05 | 23060001 | 23160000 | 0.0028 | 0.0011 | 0.0025 |
| F05 | 23080001 | 23180000 | 0.0026 | 0.0005 | 0.0021 |
| F05 | 23100001 | 23200000 | 0.0017 | 0.0005 | 0.0010 |
| F05 | 23120001 | 23220000 | 0.0016 | 0.0004 | 0.0008 |
| F05 | 23140001 | 23240000 | 0.0017 | 0.0002 | 0.0007 |
| F05 | 23160001 | 23260000 | 0.0017 | 0.0002 | 0.0004 |
| F05 | 23180001 | 23280000 | 0.0016 | 0.0002 | 0.0004 |
| F05 | 23200001 | 23300000 | 0.0017 | 0.0006 | 0.0004 |
| F05 | 23220001 | 23320000 | 0.0021 | 0.0007 | 0.0009 |
| F05 | 23240001 | 23340000 | 0.0019 | 0.0006 | 0.0010 |
| F05 | 23260001 | 23360000 | 0.0018 | 0.0012 | 0.0012 |
| F05 | 23280001 | 23380000 | 0.0025 | 0.0013 | 0.0020 |
| F05 | 23300001 | 23400000 | 0.0026 | 0.0010 | 0.0024 |
| F05 | 23320001 | 23420000 | 0.0023 | 0.0018 | 0.0023 |
| F05 | 23340001 | 23440000 | 0.0031 | 0.0020 | 0.0032 |
| F05 | 23360001 | 23460000 | 0.0034 | 0.0015 | 0.0035 |
| F05 | 23380001 | 23480000 | 0.0030 | 0.0015 | 0.0030 |
| F05 | 23400001 | 23500000 | 0.0029 | 0.0015 | 0.0029 |
| F05 | 23420001 | 23520000 | 0.0029 | 0.0006 | 0.0029 |
| F05 | 23440001 | 23540000 | 0.0021 | 0.0004 | 0.0021 |
| F05 | 23460001 | 23560000 | 0.0018 | 0.0010 | 0.0015 |
| F05 | 23480001 | 23580000 | 0.0022 | 0.0009 | 0.0018 |
| F05 | 23500001 | 23600000 | 0.0019 | 0.0009 | 0.0014 |
| F05 | 23520001 | 23620000 | 0.0019 | 0.0009 | 0.0012 |
| F05 | 23540001 | 23640000 | 0.0017 | 0.0011 | 0.0010 |
| F05 | 23560001 | 23660000 | 0.0019 | 0.0005 | 0.0012 |
| F05 | 23580001 | 23680000 | 0.0013 | 0.0008 | 0.0005 |
| F05 | 23600001 | 23700000 | 0.0019 | 0.0007 | 0.0010 |
| F05 | 23620001 | 23720000 | 0.0018 | 0.0008 | 0.0009 |
| F05 | 23640001 | 23740000 | 0.0020 | 0.0009 | 0.0009 |
| F05 | 23660001 | 23760000 | 0.0018 | 0.0009 | 0.0010 |
| F05 | 23680001 | 23780000 | 0.0018 | 0.0007 | 0.0011 |
| F05 | 23700001 | 23800000 | 0.0015 | 0.0012 | 0.0008 |
| F05 | 23720001 | 23820000 | 0.0021 | 0.0016 | 0.0015 |
| F05 | 23740001 | 23840000 | 0.0023 | 0.0014 | 0.0019 |
| F05 | 23760001 | 23860000 | 0.0023 | 0.0018 | 0.0018 |
| F05 | 23780001 | 23880000 | 0.0026 | 0.0018 | 0.0021 |
| F05 | 23800001 | 23900000 | 0.0025 | 0.0012 | 0.0019 |
| F05 | 23820001 | 23920000 | 0.0019 | 0.0010 | 0.0012 |
| F05 | 23840001 | 23940000 | 0.0018 | 0.0011 | 0.0010 |
| F05 | 23860001 | 23960000 | 0.0019 | 0.0010 | 0.0010 |
| F05 | 23880001 | 23980000 | 0.0020 | 0.0022 | 0.0011 |
| F05 | 23900001 | 24000000 | 0.0031 | 0.0024 | 0.0024 |
| F05 | 23920001 | 24020000 | 0.0032 | 0.0023 | 0.0026 |
| F05 | 23940001 | 24040000 | 0.0029 | 0.0024 | 0.0025 |
| F05 | 23960001 | 24060000 | 0.0030 | 0.0030 | 0.0027 |
| F05 | 23980001 | 24080000 | 0.0036 | 0.0019 | 0.0033 |
| F05 | 24000001 | 24100000 | 0.0026 | 0.0026 | 0.0021 |
| F05 | 24020001 | 24120000 | 0.0034 | 0.0024 | 0.0029 |
| F05 | 24040001 | 24140000 | 0.0032 | 0.0023 | 0.0027 |

| Chr | Start | End | both | A1 | A2 |
|---|---|---|---|---|---|
| F05 | 24060001 | 24160000 | 0.0032 | 0.0016 | 0.0024 |
| F05 | 24080001 | 24180000 | 0.0024 | 0.0017 | 0.0017 |
| F05 | 24100001 | 24200000 | 0.0024 | 0.0013 | 0.0018 |
| F05 | 24120001 | 24220000 | 0.0021 | 0.0013 | 0.0014 |
| F05 | 24140001 | 24240000 | 0.0024 | 0.0018 | 0.0015 |
| F05 | 24160001 | 24260000 | 0.0028 | 0.0016 | 0.0021 |
| F05 | 24180001 | 24280000 | 0.0026 | 0.0016 | 0.0019 |
| F05 | 24200001 | 24300000 | 0.0024 | 0.0010 | 0.0018 |
| F05 | 24220001 | 24320000 | 0.0018 | 0.0012 | 0.0012 |
| F05 | 24240001 | 24340000 | 0.0020 | 0.0022 | 0.0013 |
| F05 | 24260001 | 24360000 | 0.0031 | 0.0026 | 0.0023 |
| F05 | 24280001 | 24380000 | 0.0037 | 0.0027 | 0.0028 |
| F05 | 24300001 | 24400000 | 0.0038 | 0.0029 | 0.0031 |
| F05 | 24320001 | 24420000 | 0.0040 | 0.0028 | 0.0032 |
| F05 | 24340001 | 24440000 | 0.0039 | 0.0012 | 0.0032 |
| F05 | 24360001 | 24460000 | 0.0021 | 0.0006 | 0.0016 |
| F05 | 24380001 | 24480000 | 0.0013 | 0.0009 | 0.0008 |
| F05 | 24400001 | 24500000 | 0.0017 | 0.0009 | 0.0012 |
| F05 | 24420001 | 24520000 | 0.0017 | 0.0013 | 0.0012 |
| F05 | 24440001 | 24540000 | 0.0021 | 0.0013 | 0.0015 |
| F05 | 24460001 | 24560000 | 0.0021 | 0.0018 | 0.0015 |
| F05 | 24480001 | 24580000 | 0.0028 | 0.0014 | 0.0021 |
| F05 | 24500001 | 24600000 | 0.0023 | 0.0021 | 0.0016 |
| F05 | 24520001 | 24620000 | 0.0033 | 0.0018 | 0.0024 |
| F05 | 24540001 | 24640000 | 0.0028 | 0.0019 | 0.0020 |
| F05 | 24560001 | 24660000 | 0.0030 | 0.0014 | 0.0022 |
| F05 | 24580001 | 24680000 | 0.0024 | 0.0021 | 0.0016 |
| F05 | 24600001 | 24700000 | 0.0031 | 0.0012 | 0.0023 |
| F05 | 24620001 | 24720000 | 0.0020 | 0.0010 | 0.0013 |
| F05 | 24640001 | 24740000 | 0.0018 | 0.0008 | 0.0011 |
| F05 | 24660001 | 24760000 | 0.0015 | 0.0008 | 0.0009 |
| F05 | 24680001 | 24780000 | 0.0015 | 0.0002 | 0.0009 |
| F05 | 24700001 | 24800000 | 0.0010 | 0.0005 | 0.0003 |
| F05 | 24720001 | 24820000 | 0.0013 | 0.0007 | 0.0006 |
| F05 | 24740001 | 24840000 | 0.0015 | 0.0012 | 0.0008 |
| F05 | 24760001 | 24860000 | 0.0019 | 0.0016 | 0.0012 |
| F05 | 24780001 | 24880000 | 0.0022 | 0.0016 | 0.0015 |
| F05 | 24800001 | 24900000 | 0.0021 | 0.0014 | 0.0015 |
| F05 | 24820001 | 24920000 | 0.0017 | 0.0012 | 0.0013 |
| F05 | 24840001 | 24940000 | 0.0016 | 0.0012 | 0.0011 |
| F05 | 24860001 | 24960000 | 0.0015 | 0.0007 | 0.0011 |
| F05 | 24880001 | 24980000 | 0.0014 | 0.0009 | 0.0008 |
| F05 | 24900001 | 25000000 | 0.0015 | 0.0009 | 0.0010 |
| F05 | 24920001 | 25020000 | 0.0017 | 0.0010 | 0.0010 |
| F05 | 24940001 | 25040000 | 0.0019 | 0.0010 | 0.0012 |
| F05 | 24960001 | 25060000 | 0.0021 | 0.0010 | 0.0014 |
| F05 | 24980001 | 25080000 | 0.0021 | 0.0008 | 0.0014 |
| F05 | 25000001 | 25100000 | 0.0020 | 0.0009 | 0.0013 |
| F05 | 25020001 | 25120000 | 0.0020 | 0.0008 | 0.0014 |
| F05 | 25040001 | 25140000 | 0.0019 | 0.0005 | 0.0013 |
| F05 | 25060001 | 25160000 | 0.0014 | 0.0006 | 0.0008 |
| F05 | 25080001 | 25180000 | 0.0015 | 0.0007 | 0.0009 |
| F05 | 25100001 | 25200000 | 0.0015 | 0.0030 | 0.0009 |
| F05 | 25120001 | 25220000 | 0.0041 | 0.0033 | 0.0034 |
| F05 | 25140001 | 25240000 | 0.0043 | 0.0032 | 0.0037 |
| F05 | 25160001 | 25260000 | 0.0043 | 0.0031 | 0.0037 |
| F05 | 25180001 | 25280000 | 0.0042 | 0.0032 | 0.0037 |
| F05 | 25200001 | 25300000 | 0.0044 | 0.0008 | 0.0039 |

| Chr | Start | End | both | A1 | A2 |
|---|---|---|---|---|---|
| F05 | 25220001 | 25320000 | 0.0018 | 0.0005 | 0.0013 |
| F05 | 25240001 | 25340000 | 0.0016 | 0.0019 | 0.0010 |
| F05 | 25260001 | 25360000 | 0.0029 | 0.0049 | 0.0023 |
| F05 | 25280001 | 25380000 | 0.0060 | 0.0047 | 0.0052 |
| F05 | 25300001 | 25400000 | 0.0058 | 0.0047 | 0.0049 |
| F05 | 25320001 | 25420000 | 0.0059 | 0.0064 | 0.0049 |
| F05 | 25340001 | 25440000 | 0.0076 | 0.0057 | 0.0068 |
| F05 | 25360001 | 25460000 | 0.0070 | 0.0027 | 0.0062 |
| F05 | 25380001 | 25480000 | 0.0040 | 0.0034 | 0.0032 |
| F05 | 25400001 | 25500000 | 0.0048 | 0.0036 | 0.0039 |
| F05 | 25420001 | 25520000 | 0.0050 | 0.0020 | 0.0041 |
| F05 | 25440001 | 25540000 | 0.0032 | 0.0014 | 0.0023 |
| F05 | 25460001 | 25560000 | 0.0023 | 0.0014 | 0.0015 |
| F05 | 25480001 | 25580000 | 0.0023 | 0.0008 | 0.0016 |
| F05 | 25500001 | 25600000 | 0.0015 | 0.0005 | 0.0011 |
| F05 | 25520001 | 25620000 | 0.0012 | 0.0005 | 0.0008 |
| F05 | 25540001 | 25640000 | 0.0013 | 0.0004 | 0.0009 |
| F05 | 25560001 | 25660000 | 0.0014 | 0.0009 | 0.0009 |
| F05 | 25580001 | 25680000 | 0.0020 | 0.0014 | 0.0014 |
| F05 | 25600001 | 25700000 | 0.0023 | 0.0014 | 0.0018 |
| F05 | 25620001 | 25720000 | 0.0024 | 0.0016 | 0.0018 |
| F05 | 25640001 | 25740000 | 0.0023 | 0.0018 | 0.0019 |
| F05 | 25660001 | 25760000 | 0.0023 | 0.0014 | 0.0020 |
| F05 | 25680001 | 25780000 | 0.0018 | 0.0019 | 0.0015 |
| F05 | 25700001 | 25800000 | 0.0021 | 0.0020 | 0.0016 |
| F05 | 25720001 | 25820000 | 0.0020 | 0.0022 | 0.0016 |
| F05 | 25740001 | 25840000 | 0.0025 | 0.0020 | 0.0019 |
| F05 | 25760001 | 25860000 | 0.0024 | 0.0024 | 0.0016 |
| F05 | 25780001 | 25880000 | 0.0029 | 0.0033 | 0.0020 |
| F05 | 25800001 | 25900000 | 0.0042 | 0.0040 | 0.0033 |
| F05 | 25820001 | 25920000 | 0.0050 | 0.0037 | 0.0040 |
| F05 | 25840001 | 25940000 | 0.0046 | 0.0039 | 0.0036 |
| F05 | 25860001 | 25960000 | 0.0045 | 0.0035 | 0.0038 |
| F05 | 25880001 | 25980000 | 0.0040 | 0.0016 | 0.0035 |
| F05 | 25900001 | 26000000 | 0.0018 | 0.0011 | 0.0015 |
| F05 | 25920001 | 26020000 | 0.0012 | 0.0013 | 0.0009 |
| F05 | 25940001 | 26040000 | 0.0016 | 0.0012 | 0.0013 |
| F05 | 25960001 | 26060000 | 0.0015 | 0.0020 | 0.0011 |
| F05 | 25980001 | 26080000 | 0.0026 | 0.0033 | 0.0020 |
| F05 | 26000001 | 26100000 | 0.0039 | 0.0032 | 0.0032 |
| F05 | 26020001 | 26120000 | 0.0036 | 0.0029 | 0.0029 |
| F05 | 26040001 | 26140000 | 0.0032 | 0.0032 | 0.0025 |
| F05 | 26060001 | 26160000 | 0.0035 | 0.0024 | 0.0026 |
| F05 | 26080001 | 26180000 | 0.0025 | 0.0015 | 0.0017 |
| F05 | 26100001 | 26200000 | 0.0016 | 0.0016 | 0.0009 |
| F05 | 26120001 | 26220000 | 0.0019 | 0.0014 | 0.0012 |
| F05 | 26140001 | 26240000 | 0.0017 | 0.0014 | 0.0012 |
| F05 | 26160001 | 26260000 | 0.0017 | 0.0015 | 0.0011 |
| F05 | 26180001 | 26280000 | 0.0017 | 0.0012 | 0.0012 |
| F05 | 26200001 | 26300000 | 0.0014 | 0.0010 | 0.0008 |
| F05 | 26220001 | 26320000 | 0.0011 | 0.0015 | 0.0005 |
| F05 | 26240001 | 26340000 | 0.0017 | 0.0017 | 0.0010 |
| F05 | 26260001 | 26360000 | 0.0019 | 0.0016 | 0.0013 |
| F05 | 26280001 | 26380000 | 0.0019 | 0.0024 | 0.0013 |
| F05 | 26300001 | 26400000 | 0.0027 | 0.0025 | 0.0019 |
| F05 | 26320001 | 26420000 | 0.0028 | 0.0021 | 0.0020 |
| F05 | 26340001 | 26440000 | 0.0023 | 0.0018 | 0.0015 |
| F05 | 26360001 | 26460000 | 0.0018 | 0.0021 | 0.0011 |

| Chr | Start | End | both | A1 | A2 |
|---|---|---|---|---|---|
| F05 | 26380001 | 26480000 | 0.0020 | 0.0022 | 0.0014 |
| F05 | 26400001 | 26500000 | 0.0022 | 0.0025 | 0.0016 |
| F05 | 26420001 | 26520000 | 0.0024 | 0.0029 | 0.0019 |
| F05 | 26440001 | 26540000 | 0.0026 | 0.0030 | 0.0020 |
| F05 | 26460001 | 26560000 | 0.0028 | 0.0030 | 0.0021 |
| F05 | 26480001 | 26580000 | 0.0026 | 0.0023 | 0.0020 |
| F05 | 26500001 | 26600000 | 0.0019 | 0.0027 | 0.0014 |
| F05 | 26520001 | 26620000 | 0.0020 | 0.0026 | 0.0015 |
| F05 | 26540001 | 26640000 | 0.0020 | 0.0028 | 0.0014 |
| F05 | 26560001 | 26660000 | 0.0020 | 0.0028 | 0.0014 |
| F05 | 26580001 | 26680000 | 0.0022 | 0.0026 | 0.0017 |
| F05 | 26600001 | 26700000 | 0.0021 | 0.0017 | 0.0016 |
| F05 | 26620001 | 26720000 | 0.0014 | 0.0014 | 0.0011 |
| F05 | 26640001 | 26740000 | 0.0013 | 0.0015 | 0.0011 |
| F05 | 26660001 | 26760000 | 0.0014 | 0.0011 | 0.0013 |
| F05 | 26680001 | 26780000 | 0.0011 | 0.0018 | 0.0010 |
| F05 | 26700001 | 26800000 | 0.0015 | 0.0021 | 0.0011 |
| F05 | 26720001 | 26820000 | 0.0016 | 0.0027 | 0.0011 |
| F05 | 26740001 | 26840000 | 0.0023 | 0.0024 | 0.0018 |
| F05 | 26760001 | 26860000 | 0.0020 | 0.0025 | 0.0016 |
| F05 | 26780001 | 26880000 | 0.0020 | 0.0026 | 0.0015 |
| F05 | 26800001 | 26900000 | 0.0022 | 0.0025 | 0.0019 |
| F05 | 26820001 | 26920000 | 0.0022 | 0.0017 | 0.0020 |
| F05 | 26840001 | 26940000 | 0.0015 | 0.0020 | 0.0012 |
| F05 | 26860001 | 26960000 | 0.0017 | 0.0017 | 0.0014 |
| F05 | 26880001 | 26980000 | 0.0015 | 0.0012 | 0.0013 |
| F05 | 26900001 | 27000000 | 0.0009 | 0.0015 | 0.0006 |
| F05 | 26920001 | 27020000 | 0.0010 | 0.0023 | 0.0006 |
| F05 | 26940001 | 27040000 | 0.0019 | 0.0019 | 0.0014 |
| F05 | 26960001 | 27060000 | 0.0017 | 0.0028 | 0.0013 |
| F05 | 26980001 | 27080000 | 0.0026 | 0.0027 | 0.0022 |
| F05 | 27000001 | 27100000 | 0.0026 | 0.0025 | 0.0023 |
| F05 | 27020001 | 27120000 | 0.0027 | 0.0018 | 0.0026 |
| F05 | 27040001 | 27140000 | 0.0019 | 0.0022 | 0.0018 |
| F05 | 27060001 | 27160000 | 0.0024 | 0.0014 | 0.0023 |
| F05 | 27080001 | 27180000 | 0.0016 | 0.0016 | 0.0015 |
| F05 | 27100001 | 27200000 | 0.0017 | 0.0020 | 0.0016 |
| F05 | 27120001 | 27220000 | 0.0022 | 0.0026 | 0.0021 |
| F05 | 27140001 | 27240000 | 0.0024 | 0.0030 | 0.0020 |
| F05 | 27160001 | 27260000 | 0.0025 | 0.0035 | 0.0020 |
| F05 | 27180001 | 27280000 | 0.0030 | 0.0039 | 0.0023 |
| F05 | 27200001 | 27300000 | 0.0035 | 0.0040 | 0.0027 |
| F05 | 27220001 | 27320000 | 0.0037 | 0.0045 | 0.0027 |
| F05 | 27240001 | 27340000 | 0.0049 | 0.0039 | 0.0040 |
| F05 | 27260001 | 27360000 | 0.0046 | 0.0033 | 0.0037 |
| F05 | 27280001 | 27380000 | 0.0043 | 0.0026 | 0.0034 |
| F05 | 27300001 | 27400000 | 0.0038 | 0.0023 | 0.0028 |
| F05 | 27320001 | 27420000 | 0.0032 | 0.0025 | 0.0025 |
| F05 | 27340001 | 27440000 | 0.0035 | 0.0022 | 0.0028 |
| F05 | 27360001 | 27460000 | 0.0031 | 0.0021 | 0.0024 |
| F05 | 27380001 | 27480000 | 0.0029 | 0.0030 | 0.0024 |
| F05 | 27400001 | 27500000 | 0.0037 | 0.0030 | 0.0033 |
| F05 | 27420001 | 27520000 | 0.0038 | 0.0017 | 0.0033 |
| F05 | 27440001 | 27540000 | 0.0023 | 0.0018 | 0.0017 |
| F05 | 27460001 | 27560000 | 0.0028 | 0.0019 | 0.0021 |
| F05 | 27480001 | 27580000 | 0.0030 | 0.0010 | 0.0022 |
| F05 | 27500001 | 27600000 | 0.0024 | 0.0008 | 0.0015 |
| F05 | 27520001 | 27620000 | 0.0021 | 0.0008 | 0.0013 |

| Chr | Start | End | both | A1 | A2 |
|---|---|---|---|---|---|
| F05 | 27540001 | 27640000 | 0.0020 | 0.0014 | 0.0013 |
| F05 | 27560001 | 27660000 | 0.0023 | 0.0014 | 0.0017 |
| F05 | 27580001 | 27680000 | 0.0020 | 0.0013 | 0.0016 |
| F05 | 27600001 | 27700000 | 0.0016 | 0.0014 | 0.0015 |
| F05 | 27620001 | 27720000 | 0.0017 | 0.0019 | 0.0015 |
| F05 | 27640001 | 27740000 | 0.0024 | 0.0017 | 0.0021 |
| F05 | 27660001 | 27760000 | 0.0023 | 0.0018 | 0.0021 |
| F05 | 27680001 | 27780000 | 0.0025 | 0.0019 | 0.0022 |
| F05 | 27700001 | 27800000 | 0.0026 | 0.0017 | 0.0022 |
| F05 | 27720001 | 27820000 | 0.0026 | 0.0023 | 0.0020 |
| F05 | 27740001 | 27840000 | 0.0034 | 0.0018 | 0.0029 |
| F05 | 27760001 | 27860000 | 0.0029 | 0.0017 | 0.0023 |
| F05 | 27780001 | 27880000 | 0.0028 | 0.0019 | 0.0023 |
| F05 | 27800001 | 27900000 | 0.0030 | 0.0022 | 0.0025 |
| F05 | 27820001 | 27920000 | 0.0031 | 0.0009 | 0.0026 |
| F05 | 27840001 | 27940000 | 0.0015 | 0.0008 | 0.0011 |
| F05 | 27860001 | 27960000 | 0.0014 | 0.0007 | 0.0009 |
| F05 | 27880001 | 27980000 | 0.0013 | 0.0005 | 0.0009 |
| F05 | 27900001 | 28000000 | 0.0011 | 0.0005 | 0.0005 |
| F05 | 27920001 | 28020000 | 0.0012 | 0.0005 | 0.0005 |
| F05 | 27940001 | 28040000 | 0.0012 | 0.0009 | 0.0005 |
| F05 | 27960001 | 28060000 | 0.0015 | 0.0011 | 0.0008 |
| F05 | 27980001 | 28080000 | 0.0017 | 0.0011 | 0.0011 |
| F05 | 28000001 | 28100000 | 0.0017 | 0.0016 | 0.0011 |
| F05 | 28020001 | 28120000 | 0.0024 | 0.0016 | 0.0019 |
| F05 | 28040001 | 28140000 | 0.0025 | 0.0015 | 0.0020 |
| F05 | 28060001 | 28160000 | 0.0026 | 0.0014 | 0.0019 |
| F05 | 28080001 | 28180000 | 0.0023 | 0.0013 | 0.0018 |
| F05 | 28100001 | 28200000 | 0.0025 | 0.0006 | 0.0017 |
| F05 | 28120001 | 28220000 | 0.0017 | 0.0008 | 0.0008 |
| F05 | 28140001 | 28240000 | 0.0020 | 0.0008 | 0.0011 |
| F05 | 28160001 | 28260000 | 0.0022 | 0.0010 | 0.0012 |
| F05 | 28180001 | 28280000 | 0.0024 | 0.0015 | 0.0014 |
| F05 | 28200001 | 28300000 | 0.0026 | 0.0017 | 0.0018 |
| F05 | 28220001 | 28320000 | 0.0027 | 0.0016 | 0.0020 |
| F05 | 28240001 | 28340000 | 0.0025 | 0.0016 | 0.0018 |
| F05 | 28260001 | 28360000 | 0.0024 | 0.0015 | 0.0017 |
| F05 | 28280001 | 28380000 | 0.0024 | 0.0023 | 0.0017 |
| F05 | 28300001 | 28400000 | 0.0033 | 0.0022 | 0.0026 |
| F05 | 28320001 | 28420000 | 0.0030 | 0.0022 | 0.0025 |
| F05 | 28340001 | 28440000 | 0.0031 | 0.0028 | 0.0026 |
| F05 | 28360001 | 28460000 | 0.0036 | 0.0028 | 0.0033 |
| F05 | 28380001 | 28480000 | 0.0037 | 0.0019 | 0.0032 |
| F05 | 28400001 | 28500000 | 0.0027 | 0.0030 | 0.0023 |
| F05 | 28420001 | 28520000 | 0.0037 | 0.0031 | 0.0031 |
| F05 | 28440001 | 28540000 | 0.0037 | 0.0031 | 0.0031 |
| F05 | 28460001 | 28560000 | 0.0037 | 0.0038 | 0.0031 |
| F05 | 28480001 | 28580000 | 0.0042 | 0.0047 | 0.0037 |
| F05 | 28500001 | 28600000 | 0.0051 | 0.0038 | 0.0047 |
| F05 | 28520001 | 28620000 | 0.0044 | 0.0043 | 0.0040 |
| F05 | 28540001 | 28640000 | 0.0050 | 0.0037 | 0.0046 |
| F05 | 28560001 | 28660000 | 0.0045 | 0.0029 | 0.0041 |
| F05 | 28580001 | 28680000 | 0.0040 | 0.0019 | 0.0035 |
| F05 | 28600001 | 28700000 | 0.0028 | 0.0017 | 0.0021 |
| F05 | 28620001 | 28720000 | 0.0026 | 0.0010 | 0.0019 |
| F05 | 28640001 | 28740000 | 0.0019 | 0.0007 | 0.0011 |
| F05 | 28660001 | 28760000 | 0.0015 | 0.0010 | 0.0007 |
| F05 | 28680001 | 28780000 | 0.0016 | 0.0017 | 0.0009 |

| Chr | Start | End | both | A1 | A2 |
|---|---|---|---|---|---|
| F05 | 28700001 | 28800000 | 0.0022 | 0.0020 | 0.0016 |
| F05 | 28720001 | 28820000 | 0.0025 | 0.0031 | 0.0019 |
| F05 | 28740001 | 28840000 | 0.0036 | 0.0032 | 0.0030 |
| F05 | 28760001 | 28860000 | 0.0037 | 0.0032 | 0.0031 |
| F05 | 28780001 | 28880000 | 0.0038 | 0.0026 | 0.0032 |
| F05 | 28800001 | 28900000 | 0.0032 | 0.0031 | 0.0025 |
| F05 | 28820001 | 28920000 | 0.0040 | 0.0020 | 0.0033 |
| F05 | 28840001 | 28940000 | 0.0028 | 0.0023 | 0.0022 |
| F05 | 28860001 | 28960000 | 0.0030 | 0.0023 | 0.0025 |
| F05 | 28880001 | 28980000 | 0.0034 | 0.0024 | 0.0026 |
| F05 | 28900001 | 29000000 | 0.0036 | 0.0017 | 0.0030 |
| F05 | 28920001 | 29020000 | 0.0027 | 0.0020 | 0.0021 |
| F05 | 28940001 | 29040000 | 0.0029 | 0.0022 | 0.0023 |
| F05 | 28960001 | 29060000 | 0.0031 | 0.0024 | 0.0026 |
| F05 | 28980001 | 29080000 | 0.0029 | 0.0021 | 0.0025 |
| F05 | 29000001 | 29100000 | 0.0024 | 0.0022 | 0.0021 |
| F05 | 29020001 | 29120000 | 0.0023 | 0.0019 | 0.0022 |
| F05 | 29040001 | 29140000 | 0.0020 | 0.0019 | 0.0019 |
| F05 | 29060001 | 29160000 | 0.0019 | 0.0015 | 0.0019 |
| F05 | 29080001 | 29180000 | 0.0016 | 0.0015 | 0.0016 |
| F05 | 29100001 | 29200000 | 0.0016 | 0.0017 | 0.0015 |
| F05 | 29120001 | 29220000 | 0.0017 | 0.0044 | 0.0016 |
| F05 | 29140001 | 29240000 | 0.0046 | 0.0037 | 0.0044 |
| F05 | 29160001 | 29260000 | 0.0039 | 0.0037 | 0.0036 |
| F05 | 29180001 | 29280000 | 0.0039 | 0.0040 | 0.0037 |
| F05 | 29200001 | 29300000 | 0.0045 | 0.0039 | 0.0043 |
| F05 | 29220001 | 29320000 | 0.0044 | 0.0024 | 0.0042 |
| F05 | 29240001 | 29340000 | 0.0028 | 0.0029 | 0.0027 |
| F05 | 29260001 | 29360000 | 0.0033 | 0.0031 | 0.0033 |
| F05 | 29280001 | 29380000 | 0.0036 | 0.0027 | 0.0035 |
| F05 | 29300001 | 29400000 | 0.0031 | 0.0025 | 0.0029 |
| F05 | 29320001 | 29420000 | 0.0031 | 0.0013 | 0.0028 |
| F05 | 29340001 | 29440000 | 0.0020 | 0.0017 | 0.0016 |
| F05 | 29360001 | 29460000 | 0.0025 | 0.0022 | 0.0018 |
| F05 | 29380001 | 29480000 | 0.0033 | 0.0027 | 0.0024 |
| F05 | 29400001 | 29500000 | 0.0039 | 0.0028 | 0.0031 |
| F05 | 29420001 | 29520000 | 0.0037 | 0.0036 | 0.0031 |
| F05 | 29440001 | 29540000 | 0.0043 | 0.0036 | 0.0038 |
| F05 | 29460001 | 29560000 | 0.0043 | 0.0029 | 0.0040 |
| F05 | 29480001 | 29580000 | 0.0034 | 0.0027 | 0.0032 |
| F05 | 29500001 | 29600000 | 0.0031 | 0.0028 | 0.0029 |
| F05 | 29520001 | 29620000 | 0.0032 | 0.0023 | 0.0030 |
| F05 | 29540001 | 29640000 | 0.0027 | 0.0018 | 0.0024 |
| F05 | 29560001 | 29660000 | 0.0023 | 0.0024 | 0.0021 |
| F05 | 29580001 | 29680000 | 0.0030 | 0.0019 | 0.0027 |
| F05 | 29600001 | 29700000 | 0.0024 | 0.0021 | 0.0022 |
| F05 | 29620001 | 29720000 | 0.0026 | 0.0021 | 0.0024 |
| F05 | 29640001 | 29740000 | 0.0025 | 0.0017 | 0.0024 |
| F05 | 29660001 | 29760000 | 0.0020 | 0.0008 | 0.0019 |
| F05 | 29680001 | 29780000 | 0.0012 | 0.0012 | 0.0010 |
| F05 | 29700001 | 29800000 | 0.0017 | 0.0015 | 0.0015 |
| F05 | 29720001 | 29820000 | 0.0022 | 0.0016 | 0.0018 |
| F05 | 29740001 | 29840000 | 0.0022 | 0.0015 | 0.0018 |
| F05 | 29760001 | 29860000 | 0.0021 | 0.0017 | 0.0017 |
| F05 | 29780001 | 29880000 | 0.0022 | 0.0016 | 0.0018 |
| F05 | 29800001 | 29900000 | 0.0019 | 0.0010 | 0.0016 |
| F05 | 29820001 | 29920000 | 0.0011 | 0.0008 | 0.0009 |
| F05 | 29840001 | 29940000 | 0.0009 | 0.0012 | 0.0008 |

| Chr | Start | End | both | A1 | A2 |
|---|---|---|---|---|---|
| F05 | 29860001 | 29960000 | 0.0013 | 0.0009 | 0.0012 |
| F05 | 29880001 | 29980000 | 0.0012 | 0.0012 | 0.0011 |
| F05 | 29900001 | 30000000 | 0.0018 | 0.0014 | 0.0017 |
| F05 | 29920001 | 30020000 | 0.0022 | 0.0021 | 0.0021 |
| F05 | 29940001 | 30040000 | 0.0030 | 0.0018 | 0.0028 |
| F05 | 29960001 | 30060000 | 0.0030 | 0.0036 | 0.0027 |
| F05 | 29980001 | 30080000 | 0.0050 | 0.0044 | 0.0047 |
| F05 | 30000001 | 30100000 | 0.0057 | 0.0055 | 0.0055 |
| F05 | 30020001 | 30120000 | 0.0071 | 0.0062 | 0.0068 |
| F05 | 30040001 | 30140000 | 0.0080 | 0.0073 | 0.0076 |
| F05 | 30060001 | 30160000 | 0.0092 | 0.0068 | 0.0089 |
| F05 | 30080001 | 30180000 | 0.0082 | 0.0059 | 0.0079 |
| F05 | 30100001 | 30200000 | 0.0070 | 0.0047 | 0.0067 |
| F05 | 30120001 | 30220000 | 0.0054 | 0.0043 | 0.0052 |
| F05 | 30140001 | 30240000 | 0.0050 | 0.0031 | 0.0049 |
| F05 | 30160001 | 30260000 | 0.0038 | 0.0033 | 0.0036 |
| F05 | 30180001 | 30280000 | 0.0042 | 0.0035 | 0.0040 |
| F05 | 30200001 | 30300000 | 0.0045 | 0.0037 | 0.0042 |
| F05 | 30220001 | 30320000 | 0.0048 | 0.0031 | 0.0044 |
| F05 | 30240001 | 30340000 | 0.0040 | 0.0042 | 0.0035 |
| F05 | 30260001 | 30360000 | 0.0052 | 0.0035 | 0.0049 |
| F05 | 30280001 | 30380000 | 0.0044 | 0.0033 | 0.0041 |
| F05 | 30300001 | 30400000 | 0.0041 | 0.0036 | 0.0039 |
| F05 | 30320001 | 30420000 | 0.0043 | 0.0044 | 0.0040 |
| F05 | 30340001 | 30440000 | 0.0048 | 0.0034 | 0.0045 |
| F05 | 30360001 | 30460000 | 0.0037 | 0.0031 | 0.0033 |
| F05 | 30380001 | 30480000 | 0.0035 | 0.0030 | 0.0030 |
| F05 | 30400001 | 30500000 | 0.0034 | 0.0028 | 0.0029 |
| F05 | 30420001 | 30520000 | 0.0034 | 0.0023 | 0.0027 |
| F05 | 30440001 | 30540000 | 0.0031 | 0.0022 | 0.0023 |
| F05 | 30460001 | 30560000 | 0.0029 | 0.0019 | 0.0022 |
| F05 | 30480001 | 30580000 | 0.0025 | 0.0023 | 0.0018 |
| F05 | 30500001 | 30600000 | 0.0030 | 0.0016 | 0.0022 |
| F05 | 30520001 | 30620000 | 0.0022 | 0.0011 | 0.0017 |
| F05 | 30540001 | 30640000 | 0.0016 | 0.0016 | 0.0013 |
| F05 | 30560001 | 30660000 | 0.0021 | 0.0022 | 0.0018 |
| F05 | 30580001 | 30680000 | 0.0030 | 0.0019 | 0.0025 |
| F05 | 30600001 | 30700000 | 0.0027 | 0.0020 | 0.0022 |
| F05 | 30620001 | 30720000 | 0.0029 | 0.0021 | 0.0024 |
| F05 | 30640001 | 30740000 | 0.0030 | 0.0021 | 0.0024 |
| F05 | 30660001 | 30760000 | 0.0030 | 0.0016 | 0.0023 |
| F05 | 30680001 | 30780000 | 0.0025 | 0.0016 | 0.0018 |
| F05 | 30700001 | 30800000 | 0.0021 | 0.0016 | 0.0016 |
| F05 | 30720001 | 30820000 | 0.0021 | 0.0017 | 0.0017 |
| F05 | 30740001 | 30840000 | 0.0026 | 0.0012 | 0.0019 |
| F05 | 30760001 | 30860000 | 0.0023 | 0.0010 | 0.0016 |
| F05 | 30780001 | 30880000 | 0.0020 | 0.0009 | 0.0015 |
| F05 | 30800001 | 30900000 | 0.0022 | 0.0007 | 0.0015 |
| F05 | 30820001 | 30920000 | 0.0020 | 0.0005 | 0.0014 |
| F05 | 30840001 | 30940000 | 0.0018 | 0.0004 | 0.0013 |
| F05 | 30860001 | 30960000 | 0.0017 | 0.0009 | 0.0013 |
| F05 | 30880001 | 30980000 | 0.0022 | 0.0016 | 0.0018 |
| F05 | 30900001 | 31000000 | 0.0030 | 0.0021 | 0.0027 |
| F05 | 30920001 | 31020000 | 0.0036 | 0.0021 | 0.0034 |
| F05 | 30940001 | 31040000 | 0.0037 | 0.0032 | 0.0035 |
| F05 | 30960001 | 31060000 | 0.0047 | 0.0029 | 0.0045 |
| F05 | 30980001 | 31080000 | 0.0042 | 0.0023 | 0.0041 |
| F05 | 31000001 | 31100000 | 0.0037 | 0.0018 | 0.0037 |

| Chr | Start | End | both | A1 | A2 |
|---|---|---|---|---|---|
| F05 | 31020001 | 31120000 | 0.0033 | 0.0019 | 0.0033 |
| F05 | 31040001 | 31140000 | 0.0032 | 0.0009 | 0.0032 |
| F05 | 31060001 | 31160000 | 0.0025 | 0.0010 | 0.0024 |
| F05 | 31080001 | 31180000 | 0.0027 | 0.0011 | 0.0027 |
| F05 | 31100001 | 31200000 | 0.0029 | 0.0015 | 0.0028 |
| F05 | 31120001 | 31220000 | 0.0031 | 0.0013 | 0.0030 |
| F05 | 31140001 | 31240000 | 0.0030 | 0.0015 | 0.0028 |
| F05 | 31160001 | 31260000 | 0.0032 | 0.0012 | 0.0031 |
| F05 | 31180001 | 31280000 | 0.0030 | 0.0013 | 0.0028 |
| F05 | 31200001 | 31300000 | 0.0028 | 0.0012 | 0.0027 |
| F05 | 31220001 | 31320000 | 0.0028 | 0.0018 | 0.0027 |
| F05 | 31240001 | 31340000 | 0.0031 | 0.0014 | 0.0031 |
| F05 | 31260001 | 31360000 | 0.0024 | 0.0018 | 0.0024 |
| F05 | 31280001 | 31380000 | 0.0029 | 0.0017 | 0.0028 |
| F05 | 31300001 | 31400000 | 0.0026 | 0.0016 | 0.0026 |
| F05 | 31320001 | 31420000 | 0.0023 | 0.0016 | 0.0023 |
| F05 | 31340001 | 31440000 | 0.0023 | 0.0019 | 0.0022 |
| F05 | 31360001 | 31460000 | 0.0025 | 0.0017 | 0.0025 |
| F05 | 31380001 | 31480000 | 0.0023 | 0.0016 | 0.0023 |
| F05 | 31400001 | 31500000 | 0.0021 | 0.0015 | 0.0021 |
| F05 | 31420001 | 31520000 | 0.0020 | 0.0010 | 0.0020 |
| F05 | 31440001 | 31540000 | 0.0015 | 0.0010 | 0.0016 |
| F05 | 31460001 | 31560000 | 0.0014 | 0.0009 | 0.0015 |
| F05 | 31480001 | 31580000 | 0.0013 | 0.0013 | 0.0014 |
| F05 | 31500001 | 31600000 | 0.0018 | 0.0014 | 0.0018 |
| F05 | 31520001 | 31620000 | 0.0019 | 0.0014 | 0.0019 |
| F05 | 31540001 | 31640000 | 0.0019 | 0.0013 | 0.0019 |
| F05 | 31560001 | 31660000 | 0.0019 | 0.0011 | 0.0019 |
| F05 | 31580001 | 31680000 | 0.0016 | 0.0006 | 0.0017 |
| F05 | 31600001 | 31700000 | 0.0013 | 0.0004 | 0.0013 |
| F05 | 31620001 | 31720000 | 0.0011 | 0.0008 | 0.0010 |
| F05 | 31640001 | 31740000 | 0.0015 | 0.0008 | 0.0013 |
| F05 | 31660001 | 31760000 | 0.0014 | 0.0009 | 0.0011 |
| F05 | 31680001 | 31780000 | 0.0015 | 0.0012 | 0.0011 |
| F05 | 31700001 | 31800000 | 0.0017 | 0.0023 | 0.0013 |
| F05 | 31720001 | 31820000 | 0.0029 | 0.0025 | 0.0025 |
| F05 | 31740001 | 31840000 | 0.0033 | 0.0025 | 0.0029 |
| F05 | 31760001 | 31860000 | 0.0034 | 0.0027 | 0.0030 |
| F05 | 31780001 | 31880000 | 0.0035 | 0.0026 | 0.0031 |
| F05 | 31800001 | 31900000 | 0.0033 | 0.0016 | 0.0030 |
| F05 | 31820001 | 31920000 | 0.0022 | 0.0016 | 0.0019 |
| F05 | 31840001 | 31940000 | 0.0021 | 0.0023 | 0.0017 |
| F05 | 31860001 | 31960000 | 0.0029 | 0.0023 | 0.0024 |
| F05 | 31880001 | 31980000 | 0.0029 | 0.0024 | 0.0025 |
| F05 | 31900001 | 32000000 | 0.0032 | 0.0025 | 0.0026 |
| F05 | 31920001 | 32020000 | 0.0033 | 0.0041 | 0.0028 |
| F05 | 31940001 | 32040000 | 0.0050 | 0.0035 | 0.0046 |
| F05 | 31960001 | 32060000 | 0.0044 | 0.0039 | 0.0040 |
| F05 | 31980001 | 32080000 | 0.0050 | 0.0038 | 0.0043 |
| F05 | 32000001 | 32100000 | 0.0047 | 0.0040 | 0.0041 |
| F05 | 32020001 | 32120000 | 0.0051 | 0.0024 | 0.0044 |
| F05 | 32040001 | 32140000 | 0.0032 | 0.0027 | 0.0023 |
| F05 | 32060001 | 32160000 | 0.0034 | 0.0020 | 0.0028 |
| F05 | 32080001 | 32180000 | 0.0025 | 0.0020 | 0.0021 |
| F05 | 32100001 | 32200000 | 0.0027 | 0.0020 | 0.0022 |
| F05 | 32120001 | 32220000 | 0.0025 | 0.0017 | 0.0022 |
| F05 | 32140001 | 32240000 | 0.0023 | 0.0019 | 0.0019 |
| F05 | 32160001 | 32260000 | 0.0025 | 0.0027 | 0.0020 |

| Chr | Start | End | both | A1 | A2 |
|---|---|---|---|---|---|
| F05 | 32180001 | 32280000 | 0.0034 | 0.0024 | 0.0029 |
| F05 | 32200001 | 32300000 | 0.0030 | 0.0024 | 0.0025 |
| F05 | 32220001 | 32320000 | 0.0032 | 0.0029 | 0.0027 |
| F05 | 32240001 | 32340000 | 0.0039 | 0.0028 | 0.0036 |
| F05 | 32260001 | 32360000 | 0.0039 | 0.0023 | 0.0035 |
| F05 | 32280001 | 32380000 | 0.0035 | 0.0023 | 0.0029 |
| F05 | 32300001 | 32400000 | 0.0036 | 0.0019 | 0.0029 |
| F05 | 32320001 | 32420000 | 0.0031 | 0.0014 | 0.0025 |
| F05 | 32340001 | 32440000 | 0.0021 | 0.0012 | 0.0015 |
| F05 | 32360001 | 32460000 | 0.0018 | 0.0015 | 0.0013 |
| F05 | 32380001 | 32480000 | 0.0021 | 0.0021 | 0.0016 |
| F05 | 32400001 | 32500000 | 0.0027 | 0.0026 | 0.0022 |
| F05 | 32420001 | 32520000 | 0.0032 | 0.0027 | 0.0027 |
| F05 | 32440001 | 32540000 | 0.0036 | 0.0027 | 0.0027 |
| F05 | 32460001 | 32560000 | 0.0038 | 0.0022 | 0.0028 |
| F05 | 32480001 | 32580000 | 0.0034 | 0.0017 | 0.0023 |
| F05 | 32500001 | 32600000 | 0.0028 | 0.0015 | 0.0018 |
| F05 | 32520001 | 32620000 | 0.0027 | 0.0018 | 0.0015 |
| F05 | 32540001 | 32640000 | 0.0029 | 0.0015 | 0.0019 |
| F05 | 32560001 | 32660000 | 0.0023 | 0.0023 | 0.0015 |
| F05 | 32580001 | 32680000 | 0.0030 | 0.0021 | 0.0024 |
| F05 | 32600001 | 32700000 | 0.0030 | 0.0019 | 0.0024 |
| F05 | 32620001 | 32720000 | 0.0028 | 0.0014 | 0.0021 |
| F05 | 32640001 | 32740000 | 0.0024 | 0.0018 | 0.0015 |
| F05 | 32660001 | 32760000 | 0.0032 | 0.0011 | 0.0020 |
| F05 | 32680001 | 32780000 | 0.0024 | 0.0013 | 0.0012 |
| F05 | 32700001 | 32800000 | 0.0024 | 0.0013 | 0.0014 |
| F05 | 32720001 | 32820000 | 0.0022 | 0.0018 | 0.0015 |
| F05 | 32740001 | 32840000 | 0.0026 | 0.0020 | 0.0020 |
| F05 | 32760001 | 32860000 | 0.0025 | 0.0028 | 0.0021 |
| F05 | 32780001 | 32880000 | 0.0034 | 0.0027 | 0.0030 |
| F05 | 32800001 | 32900000 | 0.0034 | 0.0026 | 0.0029 |
| F05 | 32820001 | 32920000 | 0.0035 | 0.0026 | 0.0030 |
| F05 | 32840001 | 32940000 | 0.0033 | 0.0019 | 0.0029 |
| F05 | 32860001 | 32960000 | 0.0027 | 0.0010 | 0.0022 |
| F05 | 32880001 | 32980000 | 0.0018 | 0.0014 | 0.0012 |
| F05 | 32900001 | 33000000 | 0.0021 | 0.0014 | 0.0016 |
| F05 | 32920001 | 33020000 | 0.0026 | 0.0010 | 0.0018 |
| F05 | 32940001 | 33040000 | 0.0025 | 0.0010 | 0.0015 |
| F05 | 32960001 | 33060000 | 0.0026 | 0.0011 | 0.0015 |
| F05 | 32980001 | 33080000 | 0.0026 | 0.0010 | 0.0016 |
| F05 | 33000001 | 33100000 | 0.0024 | 0.0009 | 0.0014 |
| F05 | 33020001 | 33120000 | 0.0017 | 0.0009 | 0.0011 |
| F05 | 33040001 | 33140000 | 0.0015 | 0.0009 | 0.0010 |
| F05 | 33060001 | 33160000 | 0.0013 | 0.0008 | 0.0009 |
| F05 | 33080001 | 33180000 | 0.0012 | 0.0009 | 0.0009 |
| F05 | 33100001 | 33200000 | 0.0013 | 0.0009 | 0.0010 |
| F05 | 33120001 | 33220000 | 0.0013 | 0.0009 | 0.0010 |
| F05 | 33140001 | 33240000 | 0.0012 | 0.0010 | 0.0011 |
| F05 | 33160001 | 33260000 | 0.0013 | 0.0016 | 0.0011 |
| F05 | 33180001 | 33280000 | 0.0021 | 0.0017 | 0.0020 |
| F05 | 33200001 | 33300000 | 0.0021 | 0.0018 | 0.0020 |
| F05 | 33220001 | 33320000 | 0.0022 | 0.0018 | 0.0021 |
| F05 | 33240001 | 33340000 | 0.0022 | 0.0018 | 0.0021 |
| F05 | 33260001 | 33360000 | 0.0023 | 0.0013 | 0.0022 |
| F05 | 33280001 | 33380000 | 0.0017 | 0.0011 | 0.0015 |
| F05 | 33300001 | 33400000 | 0.0015 | 0.0009 | 0.0012 |
| F05 | 33320001 | 33420000 | 0.0013 | 0.0011 | 0.0011 |

| Chr | Start | End | both | A1 | A2 |
|---|---|---|---|---|---|
| F05 | 33340001 | 33440000 | 0.0015 | 0.0013 | 0.0013 |
| F05 | 33360001 | 33460000 | 0.0017 | 0.0017 | 0.0013 |
| F05 | 33380001 | 33480000 | 0.0022 | 0.0014 | 0.0017 |
| F05 | 33400001 | 33500000 | 0.0018 | 0.0015 | 0.0015 |
| F05 | 33420001 | 33520000 | 0.0020 | 0.0012 | 0.0016 |
| F05 | 33440001 | 33540000 | 0.0017 | 0.0009 | 0.0012 |
| F05 | 33460001 | 33560000 | 0.0013 | 0.0005 | 0.0009 |
| F05 | 33480001 | 33580000 | 0.0012 | 0.0011 | 0.0007 |
| F05 | 33500001 | 33600000 | 0.0019 | 0.0027 | 0.0014 |
| F05 | 33520001 | 33620000 | 0.0035 | 0.0027 | 0.0029 |
| F05 | 33540001 | 33640000 | 0.0038 | 0.0029 | 0.0031 |
| F05 | 33560001 | 33660000 | 0.0041 | 0.0031 | 0.0033 |
| F05 | 33580001 | 33680000 | 0.0041 | 0.0033 | 0.0033 |
| F05 | 33600001 | 33700000 | 0.0043 | 0.0022 | 0.0036 |
| F05 | 33620001 | 33720000 | 0.0032 | 0.0024 | 0.0024 |
| F05 | 33640001 | 33740000 | 0.0031 | 0.0029 | 0.0023 |
| F05 | 33660001 | 33760000 | 0.0034 | 0.0032 | 0.0025 |
| F05 | 33680001 | 33780000 | 0.0035 | 0.0027 | 0.0028 |
| F05 | 33700001 | 33800000 | 0.0028 | 0.0033 | 0.0023 |
| F05 | 33720001 | 33820000 | 0.0033 | 0.0030 | 0.0030 |
| F05 | 33740001 | 33840000 | 0.0031 | 0.0027 | 0.0029 |
| F05 | 33760001 | 33860000 | 0.0029 | 0.0022 | 0.0028 |
| F05 | 33780001 | 33880000 | 0.0024 | 0.0021 | 0.0022 |
| F05 | 33800001 | 33900000 | 0.0023 | 0.0012 | 0.0020 |
| F05 | 33820001 | 33920000 | 0.0012 | 0.0015 | 0.0009 |
| F05 | 33840001 | 33940000 | 0.0015 | 0.0038 | 0.0011 |
| F05 | 33860001 | 33960000 | 0.0038 | 0.0039 | 0.0032 |
| F05 | 33880001 | 33980000 | 0.0038 | 0.0039 | 0.0032 |
| F05 | 33900001 | 34000000 | 0.0036 | 0.0047 | 0.0030 |
| F05 | 33920001 | 34020000 | 0.0045 | 0.0051 | 0.0039 |
| F05 | 33940001 | 34040000 | 0.0050 | 0.0026 | 0.0045 |
| F05 | 33960001 | 34060000 | 0.0025 | 0.0034 | 0.0020 |
| F05 | 33980001 | 34080000 | 0.0035 | 0.0039 | 0.0031 |
| F05 | 34000001 | 34100000 | 0.0042 | 0.0031 | 0.0039 |
| F05 | 34020001 | 34120000 | 0.0033 | 0.0025 | 0.0029 |
| F05 | 34040001 | 34140000 | 0.0025 | 0.0026 | 0.0022 |
| F05 | 34060001 | 34160000 | 0.0027 | 0.0020 | 0.0024 |
| F05 | 34080001 | 34180000 | 0.0020 | 0.0017 | 0.0015 |
| F05 | 34100001 | 34200000 | 0.0013 | 0.0021 | 0.0007 |
| F05 | 34120001 | 34220000 | 0.0018 | 0.0038 | 0.0012 |
| F05 | 34140001 | 34240000 | 0.0034 | 0.0037 | 0.0028 |
| F05 | 34160001 | 34260000 | 0.0033 | 0.0037 | 0.0025 |
| F05 | 34180001 | 34280000 | 0.0034 | 0.0036 | 0.0026 |
| F05 | 34200001 | 34300000 | 0.0035 | 0.0040 | 0.0029 |
| F05 | 34220001 | 34320000 | 0.0039 | 0.0046 | 0.0031 |
| F05 | 34240001 | 34340000 | 0.0046 | 0.0050 | 0.0034 |
| F05 | 34260001 | 34360000 | 0.0049 | 0.0048 | 0.0038 |
| F05 | 34280001 | 34380000 | 0.0049 | 0.0046 | 0.0038 |
| F05 | 34300001 | 34400000 | 0.0045 | 0.0042 | 0.0036 |
| F05 | 34320001 | 34420000 | 0.0041 | 0.0024 | 0.0033 |
| F05 | 34340001 | 34440000 | 0.0021 | 0.0020 | 0.0016 |
| F05 | 34360001 | 34460000 | 0.0017 | 0.0026 | 0.0011 |
| F05 | 34380001 | 34480000 | 0.0021 | 0.0032 | 0.0013 |
| F05 | 34400001 | 34500000 | 0.0028 | 0.0036 | 0.0016 |
| F05 | 34420001 | 34520000 | 0.0031 | 0.0034 | 0.0017 |
| F05 | 34440001 | 34540000 | 0.0031 | 0.0032 | 0.0017 |
| F05 | 34460001 | 34560000 | 0.0030 | 0.0028 | 0.0017 |
| F05 | 34480001 | 34580000 | 0.0027 | 0.0031 | 0.0015 |

| Chr | Start | End | both | A1 | A2 |
|---|---|---|---|---|---|
| F05 | 34500001 | 34600000 | 0.0028 | 0.0029 | 0.0018 |
| F05 | 34520001 | 34620000 | 0.0026 | 0.0032 | 0.0015 |
| F05 | 34540001 | 34640000 | 0.0029 | 0.0035 | 0.0019 |
| F05 | 34560001 | 34660000 | 0.0035 | 0.0046 | 0.0027 |
| F05 | 34580001 | 34680000 | 0.0042 | 0.0041 | 0.0032 |
| F05 | 34600001 | 34700000 | 0.0035 | 0.0050 | 0.0027 |
| F05 | 34620001 | 34720000 | 0.0038 | 0.0047 | 0.0029 |
| F05 | 34640001 | 34740000 | 0.0035 | 0.0043 | 0.0025 |
| F05 | 34660001 | 34760000 | 0.0029 | 0.0036 | 0.0016 |
| F05 | 34680001 | 34780000 | 0.0026 | 0.0033 | 0.0015 |
| F05 | 34700001 | 34800000 | 0.0025 | 0.0019 | 0.0014 |
| F05 | 34720001 | 34820000 | 0.0018 | 0.0017 | 0.0012 |
| F05 | 34740001 | 34840000 | 0.0017 | 0.0014 | 0.0012 |
| F05 | 34760001 | 34860000 | 0.0014 | 0.0006 | 0.0012 |
| F05 | 34780001 | 34880000 | 0.0006 | 0.0005 | 0.0004 |
| F05 | 34800001 | 34900000 | 0.0006 | 0.0007 | 0.0004 |
| F05 | 34820001 | 34920000 | 0.0006 | 0.0011 | 0.0002 |
| F05 | 34840001 | 34940000 | 0.0010 | 0.0015 | 0.0004 |
| F05 | 34860001 | 34960000 | 0.0013 | 0.0017 | 0.0006 |
| F05 | 34880001 | 34980000 | 0.0014 | 0.0020 | 0.0006 |
| F05 | 34900001 | 35000000 | 0.0018 | 0.0026 | 0.0010 |
| F05 | 34920001 | 35020000 | 0.0025 | 0.0028 | 0.0018 |
| F05 | 34940001 | 35040000 | 0.0027 | 0.0029 | 0.0021 |
| F05 | 34960001 | 35060000 | 0.0028 | 0.0039 | 0.0024 |
| F05 | 34980001 | 35080000 | 0.0038 | 0.0035 | 0.0034 |
| F05 | 35000001 | 35100000 | 0.0034 | 0.0026 | 0.0031 |
| F05 | 35020001 | 35120000 | 0.0025 | 0.0020 | 0.0021 |
| F05 | 35040001 | 35140000 | 0.0019 | 0.0015 | 0.0016 |
| F05 | 35060001 | 35160000 | 0.0015 | 0.0009 | 0.0012 |
| F05 | 35080001 | 35180000 | 0.0007 | 0.0015 | 0.0003 |
| F05 | 35100001 | 35200000 | 0.0013 | 0.0015 | 0.0004 |
| F05 | 35120001 | 35220000 | 0.0013 | 0.0016 | 0.0005 |
| F05 | 35140001 | 35240000 | 0.0015 | 0.0026 | 0.0006 |
| F05 | 35160001 | 35260000 | 0.0023 | 0.0030 | 0.0011 |
| F05 | 35180001 | 35280000 | 0.0028 | 0.0030 | 0.0016 |
| F05 | 35200001 | 35300000 | 0.0027 | 0.0039 | 0.0015 |
| F05 | 35220001 | 35320000 | 0.0035 | 0.0049 | 0.0018 |
| F05 | 35240001 | 35340000 | 0.0043 | 0.0045 | 0.0020 |
| F05 | 35260001 | 35360000 | 0.0039 | 0.0047 | 0.0016 |
| F05 | 35280001 | 35380000 | 0.0039 | 0.0048 | 0.0017 |
| F05 | 35300001 | 35400000 | 0.0042 | 0.0043 | 0.0023 |
| F05 | 35320001 | 35420000 | 0.0037 | 0.0040 | 0.0020 |
| F05 | 35340001 | 35440000 | 0.0035 | 0.0041 | 0.0020 |
| F05 | 35360001 | 35460000 | 0.0036 | 0.0039 | 0.0022 |
| F05 | 35380001 | 35480000 | 0.0035 | 0.0035 | 0.0023 |
| F05 | 35400001 | 35500000 | 0.0032 | 0.0034 | 0.0019 |
| F05 | 35420001 | 35520000 | 0.0031 | 0.0027 | 0.0019 |
| F05 | 35440001 | 35540000 | 0.0025 | 0.0021 | 0.0016 |
| F05 | 35460001 | 35560000 | 0.0021 | 0.0016 | 0.0013 |
| F05 | 35480001 | 35580000 | 0.0017 | 0.0014 | 0.0011 |
| F05 | 35500001 | 35600000 | 0.0018 | 0.0024 | 0.0010 |
| F05 | 35520001 | 35620000 | 0.0030 | 0.0025 | 0.0022 |
| F05 | 35540001 | 35640000 | 0.0034 | 0.0025 | 0.0026 |
| F05 | 35560001 | 35660000 | 0.0036 | 0.0020 | 0.0028 |
| F05 | 35580001 | 35680000 | 0.0035 | 0.0020 | 0.0024 |
| F05 | 35600001 | 35700000 | 0.0035 | 0.0009 | 0.0024 |
| F05 | 35620001 | 35720000 | 0.0025 | 0.0008 | 0.0011 |
| F05 | 35640001 | 35740000 | 0.0023 | 0.0012 | 0.0009 |

| Chr | Start | End | both | A1 | A2 |
|---|---|---|---|---|---|
| F05 | 35660001 | 35760000 | 0.0028 | 0.0015 | 0.0013 |
| F05 | 35680001 | 35780000 | 0.0029 | 0.0016 | 0.0016 |
| F05 | 35700001 | 35800000 | 0.0032 | 0.0016 | 0.0016 |
| F05 | 35720001 | 35820000 | 0.0035 | 0.0018 | 0.0018 |
| F05 | 35740001 | 35840000 | 0.0038 | 0.0014 | 0.0019 |
| F05 | 35760001 | 35860000 | 0.0032 | 0.0013 | 0.0015 |
| F05 | 35780001 | 35880000 | 0.0029 | 0.0012 | 0.0013 |
| F05 | 35800001 | 35900000 | 0.0027 | 0.0016 | 0.0013 |
| F05 | 35820001 | 35920000 | 0.0026 | 0.0017 | 0.0017 |
| F05 | 35840001 | 35940000 | 0.0026 | 0.0021 | 0.0020 |
| F05 | 35860001 | 35960000 | 0.0029 | 0.0019 | 0.0024 |
| F05 | 35880001 | 35980000 | 0.0028 | 0.0020 | 0.0023 |
| F05 | 35900001 | 36000000 | 0.0030 | 0.0019 | 0.0023 |
| F05 | 35920001 | 36020000 | 0.0030 | 0.0017 | 0.0023 |
| F05 | 35940001 | 36040000 | 0.0029 | 0.0014 | 0.0019 |
| F05 | 35960001 | 36060000 | 0.0026 | 0.0019 | 0.0016 |
| F05 | 35980001 | 36080000 | 0.0032 | 0.0022 | 0.0021 |
| F05 | 36000001 | 36100000 | 0.0033 | 0.0024 | 0.0024 |
| F05 | 36020001 | 36120000 | 0.0034 | 0.0023 | 0.0024 |
| F05 | 36040001 | 36140000 | 0.0033 | 0.0031 | 0.0024 |
| F05 | 36060001 | 36160000 | 0.0043 | 0.0033 | 0.0033 |
| F05 | 36080001 | 36180000 | 0.0045 | 0.0032 | 0.0035 |
| F05 | 36100001 | 36200000 | 0.0043 | 0.0028 | 0.0035 |
| F05 | 36120001 | 36220000 | 0.0038 | 0.0031 | 0.0031 |
| F05 | 36140001 | 36240000 | 0.0042 | 0.0036 | 0.0035 |
| F05 | 36160001 | 36260000 | 0.0046 | 0.0038 | 0.0038 |
| F05 | 36180001 | 36280000 | 0.0046 | 0.0043 | 0.0039 |
| F05 | 36200001 | 36300000 | 0.0052 | 0.0049 | 0.0043 |
| F05 | 36220001 | 36320000 | 0.0059 | 0.0045 | 0.0052 |
| F05 | 36240001 | 36340000 | 0.0054 | 0.0032 | 0.0048 |
| F05 | 36260001 | 36360000 | 0.0039 | 0.0023 | 0.0036 |
| F05 | 36280001 | 36380000 | 0.0034 | 0.0025 | 0.0031 |
| F05 | 36300001 | 36400000 | 0.0042 | 0.0026 | 0.0042 |
| F05 | 36320001 | 36420000 | 0.0043 | 0.0030 | 0.0043 |
| F05 | 36340001 | 36440000 | 0.0053 | 0.0045 | 0.0052 |
| F05 | 36360001 | 36460000 | 0.0071 | 0.0046 | 0.0070 |
| F05 | 36380001 | 36480000 | 0.0073 | 0.0035 | 0.0071 |
| F05 | 36400001 | 36500000 | 0.0054 | 0.0027 | 0.0053 |
| F05 | 36420001 | 36520000 | 0.0043 | 0.0023 | 0.0041 |
| F05 | 36440001 | 36540000 | 0.0032 | 0.0015 | 0.0031 |
| F05 | 36460001 | 36560000 | 0.0022 | 0.0016 | 0.0021 |
| F05 | 36480001 | 36580000 | 0.0020 | 0.0022 | 0.0019 |
| F05 | 36500001 | 36600000 | 0.0028 | 0.0022 | 0.0028 |
| F05 | 36520001 | 36620000 | 0.0029 | 0.0026 | 0.0028 |
| F05 | 36540001 | 36640000 | 0.0033 | 0.0022 | 0.0032 |
| F05 | 36560001 | 36660000 | 0.0030 | 0.0026 | 0.0029 |
| F05 | 36580001 | 36680000 | 0.0034 | 0.0028 | 0.0033 |
| F05 | 36600001 | 36700000 | 0.0034 | 0.0034 | 0.0030 |
| F05 | 36620001 | 36720000 | 0.0042 | 0.0033 | 0.0035 |
| F05 | 36640001 | 36740000 | 0.0040 | 0.0040 | 0.0034 |
| F05 | 36660001 | 36760000 | 0.0048 | 0.0036 | 0.0041 |
| F05 | 36680001 | 36780000 | 0.0044 | 0.0031 | 0.0036 |
| F05 | 36700001 | 36800000 | 0.0040 | 0.0024 | 0.0033 |
| F05 | 36720001 | 36820000 | 0.0035 | 0.0023 | 0.0027 |
| F05 | 36740001 | 36840000 | 0.0035 | 0.0013 | 0.0027 |
| F05 | 36760001 | 36860000 | 0.0027 | 0.0017 | 0.0015 |
| F05 | 36780001 | 36880000 | 0.0031 | 0.0014 | 0.0020 |
| F05 | 36800001 | 36900000 | 0.0026 | 0.0015 | 0.0016 |

| Chr | Start | End | both | A1 | A2 |
|---|---|---|---|---|---|
| F05 | 36820001 | 36920000 | 0.0027 | 0.0014 | 0.0018 |
| F05 | 36840001 | 36940000 | 0.0025 | 0.0013 | 0.0016 |
| F05 | 36860001 | 36960000 | 0.0021 | 0.0011 | 0.0015 |
| F05 | 36880001 | 36980000 | 0.0017 | 0.0011 | 0.0013 |
| F05 | 36900001 | 37000000 | 0.0019 | 0.0011 | 0.0013 |
| F05 | 36920001 | 37020000 | 0.0019 | 0.0012 | 0.0012 |
| F05 | 36940001 | 37040000 | 0.0021 | 0.0016 | 0.0013 |
| F05 | 36960001 | 37060000 | 0.0024 | 0.0014 | 0.0017 |
| F05 | 36980001 | 37080000 | 0.0024 | 0.0018 | 0.0015 |
| F05 | 37000001 | 37100000 | 0.0026 | 0.0017 | 0.0019 |
| F05 | 37020001 | 37120000 | 0.0023 | 0.0018 | 0.0018 |
| F05 | 37040001 | 37140000 | 0.0024 | 0.0022 | 0.0018 |
| F05 | 37060001 | 37160000 | 0.0032 | 0.0022 | 0.0023 |
| F05 | 37080001 | 37180000 | 0.0031 | 0.0029 | 0.0022 |
| F05 | 37100001 | 37200000 | 0.0040 | 0.0031 | 0.0030 |
| F05 | 37120001 | 37220000 | 0.0043 | 0.0032 | 0.0033 |
| F05 | 37140001 | 37240000 | 0.0042 | 0.0024 | 0.0033 |
| F05 | 37160001 | 37260000 | 0.0032 | 0.0028 | 0.0023 |
| F05 | 37180001 | 37280000 | 0.0038 | 0.0018 | 0.0028 |
| F05 | 37200001 | 37300000 | 0.0029 | 0.0020 | 0.0018 |
| F05 | 37220001 | 37320000 | 0.0031 | 0.0032 | 0.0020 |
| F05 | 37240001 | 37340000 | 0.0044 | 0.0040 | 0.0033 |
| F05 | 37260001 | 37360000 | 0.0050 | 0.0038 | 0.0040 |
| F05 | 37280001 | 37380000 | 0.0047 | 0.0040 | 0.0037 |
| F05 | 37300001 | 37400000 | 0.0046 | 0.0036 | 0.0039 |
| F05 | 37320001 | 37420000 | 0.0044 | 0.0023 | 0.0035 |
| F05 | 37340001 | 37440000 | 0.0032 | 0.0017 | 0.0021 |
| F05 | 37360001 | 37460000 | 0.0031 | 0.0020 | 0.0018 |
| F05 | 37380001 | 37480000 | 0.0033 | 0.0018 | 0.0021 |
| F05 | 37400001 | 37500000 | 0.0030 | 0.0022 | 0.0018 |
| F05 | 37420001 | 37520000 | 0.0034 | 0.0026 | 0.0023 |
| F05 | 37440001 | 37540000 | 0.0039 | 0.0031 | 0.0029 |
| F05 | 37460001 | 37560000 | 0.0041 | 0.0034 | 0.0032 |
| F05 | 37480001 | 37580000 | 0.0043 | 0.0038 | 0.0034 |
| F05 | 37500001 | 37600000 | 0.0049 | 0.0036 | 0.0038 |
| F05 | 37520001 | 37620000 | 0.0045 | 0.0030 | 0.0036 |
| F05 | 37540001 | 37640000 | 0.0036 | 0.0032 | 0.0029 |
| F05 | 37560001 | 37660000 | 0.0038 | 0.0023 | 0.0031 |
| F05 | 37580001 | 37680000 | 0.0029 | 0.0024 | 0.0023 |
| F05 | 37600001 | 37700000 | 0.0032 | 0.0021 | 0.0026 |
| F05 | 37620001 | 37720000 | 0.0028 | 0.0021 | 0.0022 |
| F05 | 37640001 | 37740000 | 0.0029 | 0.0019 | 0.0022 |
| F05 | 37660001 | 37760000 | 0.0027 | 0.0019 | 0.0020 |
| F05 | 37680001 | 37780000 | 0.0027 | 0.0013 | 0.0020 |
| F05 | 37700001 | 37800000 | 0.0018 | 0.0014 | 0.0013 |
| F05 | 37720001 | 37820000 | 0.0022 | 0.0017 | 0.0015 |
| F05 | 37740001 | 37840000 | 0.0025 | 0.0013 | 0.0018 |
| F05 | 37760001 | 37860000 | 0.0022 | 0.0016 | 0.0018 |
| F05 | 37780001 | 37880000 | 0.0025 | 0.0019 | 0.0022 |
| F05 | 37800001 | 37900000 | 0.0029 | 0.0020 | 0.0025 |
| F05 | 37820001 | 37920000 | 0.0029 | 0.0020 | 0.0027 |
| F05 | 37840001 | 37940000 | 0.0029 | 0.0017 | 0.0026 |
| F05 | 37860001 | 37960000 | 0.0025 | 0.0017 | 0.0019 |
| F05 | 37880001 | 37980000 | 0.0025 | 0.0023 | 0.0018 |
| F05 | 37900001 | 38000000 | 0.0033 | 0.0021 | 0.0026 |
| F05 | 37920001 | 38020000 | 0.0031 | 0.0020 | 0.0023 |
| F05 | 37940001 | 38040000 | 0.0031 | 0.0025 | 0.0022 |
| F05 | 37960001 | 38060000 | 0.0036 | 0.0027 | 0.0027 |

| Chr | Start | End | both | A1 | A2 |
|---|---|---|---|---|---|
| F05 | 37980001 | 38080000 | 0.0038 | 0.0019 | 0.0029 |
| F05 | 38000001 | 38100000 | 0.0030 | 0.0029 | 0.0019 |
| F05 | 38020001 | 38120000 | 0.0039 | 0.0033 | 0.0029 |
| F05 | 38040001 | 38140000 | 0.0042 | 0.0036 | 0.0033 |
| F05 | 38060001 | 38160000 | 0.0047 | 0.0037 | 0.0037 |
| F05 | 38080001 | 38180000 | 0.0050 | 0.0035 | 0.0039 |
| F05 | 38100001 | 38200000 | 0.0047 | 0.0027 | 0.0037 |
| F05 | 38120001 | 38220000 | 0.0039 | 0.0021 | 0.0029 |
| F05 | 38140001 | 38240000 | 0.0036 | 0.0012 | 0.0027 |
| F05 | 38160001 | 38260000 | 0.0025 | 0.0007 | 0.0016 |
| F05 | 38180001 | 38280000 | 0.0018 | 0.0007 | 0.0011 |
| F05 | 38200001 | 38300000 | 0.0019 | 0.0004 | 0.0011 |
| F05 | 38220001 | 38320000 | 0.0017 | 0.0004 | 0.0008 |
| F05 | 38240001 | 38340000 | 0.0013 | 0.0005 | 0.0005 |
| F05 | 38260001 | 38360000 | 0.0014 | 0.0008 | 0.0005 |
| F05 | 38280001 | 38380000 | 0.0016 | 0.0009 | 0.0009 |
| F05 | 38300001 | 38400000 | 0.0016 | 0.0009 | 0.0011 |
| F05 | 38320001 | 38420000 | 0.0018 | 0.0010 | 0.0011 |
| F05 | 38340001 | 38440000 | 0.0021 | 0.0010 | 0.0013 |
| F05 | 38360001 | 38460000 | 0.0021 | 0.0007 | 0.0014 |
| F05 | 38380001 | 38480000 | 0.0018 | 0.0011 | 0.0010 |
| F05 | 38400001 | 38500000 | 0.0021 | 0.0019 | 0.0013 |
| F05 | 38420001 | 38520000 | 0.0028 | 0.0017 | 0.0022 |
| F05 | 38440001 | 38540000 | 0.0026 | 0.0019 | 0.0020 |
| F05 | 38460001 | 38560000 | 0.0028 | 0.0018 | 0.0023 |
| F05 | 38480001 | 38580000 | 0.0027 | 0.0016 | 0.0023 |
| F05 | 38500001 | 38600000 | 0.0028 | 0.0017 | 0.0021 |
| F05 | 38520001 | 38620000 | 0.0031 | 0.0017 | 0.0022 |
| F05 | 38540001 | 38640000 | 0.0031 | 0.0023 | 0.0022 |
| F05 | 38560001 | 38660000 | 0.0036 | 0.0031 | 0.0025 |
| F05 | 38580001 | 38680000 | 0.0044 | 0.0031 | 0.0032 |
| F05 | 38600001 | 38700000 | 0.0042 | 0.0025 | 0.0031 |
| F05 | 38620001 | 38720000 | 0.0038 | 0.0027 | 0.0024 |
| F05 | 38640001 | 38740000 | 0.0042 | 0.0020 | 0.0028 |
| F05 | 38660001 | 38760000 | 0.0033 | 0.0017 | 0.0021 |
| F05 | 38680001 | 38780000 | 0.0030 | 0.0018 | 0.0019 |
| F05 | 38700001 | 38800000 | 0.0030 | 0.0020 | 0.0020 |
| F05 | 38720001 | 38820000 | 0.0028 | 0.0025 | 0.0022 |
| F05 | 38740001 | 38840000 | 0.0033 | 0.0036 | 0.0026 |
| F05 | 38760001 | 38860000 | 0.0046 | 0.0035 | 0.0038 |
| F05 | 38780001 | 38880000 | 0.0043 | 0.0033 | 0.0037 |
| F05 | 38800001 | 38900000 | 0.0040 | 0.0031 | 0.0034 |
| F05 | 38820001 | 38920000 | 0.0038 | 0.0028 | 0.0032 |
| F05 | 38840001 | 38940000 | 0.0034 | 0.0015 | 0.0030 |
| F05 | 38860001 | 38960000 | 0.0022 | 0.0021 | 0.0016 |
| F05 | 38880001 | 38980000 | 0.0028 | 0.0021 | 0.0021 |
| F05 | 38900001 | 39000000 | 0.0029 | 0.0025 | 0.0022 |
| F05 | 38920001 | 39020000 | 0.0041 | 0.0025 | 0.0032 |
| F05 | 38940001 | 39040000 | 0.0040 | 0.0033 | 0.0031 |
| F05 | 38960001 | 39060000 | 0.0048 | 0.0030 | 0.0040 |
| F05 | 38980001 | 39080000 | 0.0045 | 0.0036 | 0.0038 |
| F05 | 39000001 | 39100000 | 0.0051 | 0.0033 | 0.0044 |
| F05 | 39020001 | 39120000 | 0.0044 | 0.0029 | 0.0036 |
| F05 | 39040001 | 39140000 | 0.0040 | 0.0023 | 0.0031 |
| F05 | 39060001 | 39160000 | 0.0034 | 0.0018 | 0.0025 |
| F05 | 39080001 | 39180000 | 0.0030 | 0.0019 | 0.0021 |
| F05 | 39100001 | 39200000 | 0.0031 | 0.0015 | 0.0023 |
| F05 | 39120001 | 39220000 | 0.0023 | 0.0020 | 0.0018 |

| Chr | Start | End | both | A1 | A2 |
|---|---|---|---|---|---|
| F05 | 39140001 | 39240000 | 0.0028 | 0.0019 | 0.0024 |
| F05 | 39160001 | 39260000 | 0.0025 | 0.0026 | 0.0023 |
| F05 | 39180001 | 39280000 | 0.0032 | 0.0027 | 0.0029 |
| F05 | 39200001 | 39300000 | 0.0035 | 0.0039 | 0.0030 |
| F05 | 39220001 | 39320000 | 0.0048 | 0.0034 | 0.0042 |
| F05 | 39240001 | 39340000 | 0.0045 | 0.0033 | 0.0038 |
| F05 | 39260001 | 39360000 | 0.0043 | 0.0027 | 0.0037 |
| F05 | 39280001 | 39380000 | 0.0037 | 0.0020 | 0.0031 |
| F05 | 39300001 | 39400000 | 0.0026 | 0.0013 | 0.0022 |
| F05 | 39320001 | 39420000 | 0.0020 | 0.0014 | 0.0014 |
| F05 | 39340001 | 39440000 | 0.0020 | 0.0019 | 0.0014 |
| F05 | 39360001 | 39460000 | 0.0027 | 0.0021 | 0.0019 |
| F05 | 39380001 | 39480000 | 0.0031 | 0.0022 | 0.0022 |
| F05 | 39400001 | 39500000 | 0.0032 | 0.0030 | 0.0023 |
| F05 | 39420001 | 39520000 | 0.0039 | 0.0032 | 0.0032 |
| F05 | 39440001 | 39540000 | 0.0043 | 0.0032 | 0.0035 |
| F05 | 39460001 | 39560000 | 0.0042 | 0.0032 | 0.0034 |
| F05 | 39480001 | 39580000 | 0.0041 | 0.0033 | 0.0033 |
| F05 | 39500001 | 39600000 | 0.0043 | 0.0022 | 0.0034 |
| F05 | 39520001 | 39620000 | 0.0028 | 0.0023 | 0.0022 |
| F05 | 39540001 | 39640000 | 0.0029 | 0.0020 | 0.0022 |
| F05 | 39560001 | 39660000 | 0.0027 | 0.0016 | 0.0020 |
| F05 | 39580001 | 39680000 | 0.0024 | 0.0014 | 0.0017 |
| F05 | 39600001 | 39700000 | 0.0022 | 0.0020 | 0.0014 |
| F05 | 39620001 | 39720000 | 0.0030 | 0.0017 | 0.0022 |
| F05 | 39640001 | 39740000 | 0.0027 | 0.0023 | 0.0020 |
| F05 | 39660001 | 39760000 | 0.0032 | 0.0023 | 0.0026 |
| F05 | 39680001 | 39780000 | 0.0032 | 0.0023 | 0.0026 |
| F05 | 39700001 | 39800000 | 0.0033 | 0.0017 | 0.0029 |
| F05 | 39720001 | 39820000 | 0.0026 | 0.0022 | 0.0020 |
| F05 | 39740001 | 39840000 | 0.0031 | 0.0016 | 0.0026 |
| F05 | 39760001 | 39860000 | 0.0026 | 0.0016 | 0.0021 |
| F05 | 39780001 | 39880000 | 0.0027 | 0.0019 | 0.0021 |
| F05 | 39800001 | 39900000 | 0.0032 | 0.0018 | 0.0025 |
| F05 | 39820001 | 39920000 | 0.0031 | 0.0015 | 0.0023 |
| F05 | 39840001 | 39940000 | 0.0027 | 0.0017 | 0.0020 |
| F05 | 39860001 | 39960000 | 0.0030 | 0.0029 | 0.0024 |
| F05 | 39880001 | 39980000 | 0.0042 | 0.0027 | 0.0036 |
| F05 | 39900001 | 40000000 | 0.0038 | 0.0030 | 0.0032 |
| F05 | 39920001 | 40020000 | 0.0040 | 0.0026 | 0.0035 |
| F05 | 39940001 | 40040000 | 0.0037 | 0.0026 | 0.0031 |
| F05 | 39960001 | 40060000 | 0.0036 | 0.0013 | 0.0029 |
| F05 | 39980001 | 40080000 | 0.0022 | 0.0021 | 0.0015 |
| F05 | 40000001 | 40100000 | 0.0031 | 0.0020 | 0.0024 |
| F05 | 40020001 | 40120000 | 0.0029 | 0.0020 | 0.0023 |
| F05 | 40040001 | 40140000 | 0.0028 | 0.0016 | 0.0023 |
| F05 | 40060001 | 40160000 | 0.0022 | 0.0017 | 0.0018 |
| F05 | 40080001 | 40180000 | 0.0022 | 0.0009 | 0.0020 |
| F05 | 40100001 | 40200000 | 0.0011 | 0.0008 | 0.0011 |
| F05 | 40120001 | 40220000 | 0.0010 | 0.0010 | 0.0010 |
| F05 | 40140001 | 40240000 | 0.0014 | 0.0010 | 0.0014 |
| F05 | 40160001 | 40260000 | 0.0013 | 0.0007 | 0.0013 |
| F05 | 40180001 | 40280000 | 0.0010 | 0.0007 | 0.0010 |
| F05 | 40200001 | 40300000 | 0.0009 | 0.0008 | 0.0009 |
| F05 | 40220001 | 40320000 | 0.0010 | 0.0010 | 0.0010 |
| F05 | 40240001 | 40340000 | 0.0011 | 0.0014 | 0.0010 |
| F05 | 40260001 | 40360000 | 0.0016 | 0.0019 | 0.0014 |
| F05 | 40280001 | 40380000 | 0.0021 | 0.0024 | 0.0019 |

| Chr | Start | End | both | A1 | A2 |
|---|---|---|---|---|---|
| F05 | 40300001 | 40400000 | 0.0027 | 0.0026 | 0.0024 |
| F05 | 40320001 | 40420000 | 0.0030 | 0.0026 | 0.0027 |
| F05 | 40340001 | 40440000 | 0.0031 | 0.0031 | 0.0028 |
| F05 | 40360001 | 40460000 | 0.0037 | 0.0029 | 0.0034 |
| F05 | 40380001 | 40480000 | 0.0036 | 0.0026 | 0.0031 |
| F05 | 40400001 | 40500000 | 0.0033 | 0.0024 | 0.0029 |
| F05 | 40420001 | 40520000 | 0.0030 | 0.0023 | 0.0027 |
| F05 | 40440001 | 40540000 | 0.0028 | 0.0020 | 0.0025 |
| F05 | 40460001 | 40560000 | 0.0025 | 0.0030 | 0.0021 |
| F05 | 40480001 | 40580000 | 0.0037 | 0.0035 | 0.0035 |
| F05 | 40500001 | 40600000 | 0.0042 | 0.0033 | 0.0040 |
| F05 | 40520001 | 40620000 | 0.0040 | 0.0031 | 0.0038 |
| F05 | 40540001 | 40640000 | 0.0039 | 0.0025 | 0.0036 |
| F05 | 40560001 | 40660000 | 0.0034 | 0.0015 | 0.0031 |
| F05 | 40580001 | 40680000 | 0.0023 | 0.0013 | 0.0019 |
| F05 | 40600001 | 40700000 | 0.0021 | 0.0013 | 0.0017 |
| F05 | 40620001 | 40720000 | 0.0022 | 0.0020 | 0.0018 |
| F05 | 40640001 | 40740000 | 0.0032 | 0.0019 | 0.0026 |
| F05 | 40660001 | 40760000 | 0.0029 | 0.0026 | 0.0025 |
| F05 | 40680001 | 40780000 | 0.0035 | 0.0030 | 0.0030 |
| F05 | 40700001 | 40800000 | 0.0039 | 0.0032 | 0.0034 |
| F05 | 40720001 | 40820000 | 0.0040 | 0.0029 | 0.0035 |
| F05 | 40740001 | 40840000 | 0.0034 | 0.0034 | 0.0031 |
| F05 | 40760001 | 40860000 | 0.0040 | 0.0039 | 0.0036 |
| F05 | 40780001 | 40880000 | 0.0047 | 0.0032 | 0.0042 |
| F05 | 40800001 | 40900000 | 0.0042 | 0.0033 | 0.0034 |
| F05 | 40820001 | 40920000 | 0.0044 | 0.0040 | 0.0035 |
| F05 | 40840001 | 40940000 | 0.0051 | 0.0038 | 0.0043 |
| F05 | 40860001 | 40960000 | 0.0051 | 0.0025 | 0.0042 |
| F05 | 40880001 | 40980000 | 0.0039 | 0.0025 | 0.0030 |
| F05 | 40900001 | 41000000 | 0.0038 | 0.0025 | 0.0031 |
| F05 | 40920001 | 41020000 | 0.0038 | 0.0023 | 0.0030 |
| F05 | 40940001 | 41040000 | 0.0037 | 0.0022 | 0.0028 |
| F05 | 40960001 | 41060000 | 0.0033 | 0.0022 | 0.0025 |
| F05 | 40980001 | 41080000 | 0.0031 | 0.0020 | 0.0025 |
| F05 | 41000001 | 41100000 | 0.0027 | 0.0018 | 0.0022 |
| F05 | 41020001 | 41120000 | 0.0024 | 0.0007 | 0.0020 |
| F05 | 41040001 | 41140000 | 0.0011 | 0.0011 | 0.0009 |
| F05 | 41060001 | 41160000 | 0.0015 | 0.0011 | 0.0013 |
| F05 | 41080001 | 41180000 | 0.0014 | 0.0011 | 0.0012 |
| F05 | 41100001 | 41200000 | 0.0017 | 0.0009 | 0.0013 |
| F05 | 41120001 | 41220000 | 0.0014 | 0.0016 | 0.0011 |
| F05 | 41140001 | 41240000 | 0.0022 | 0.0013 | 0.0018 |
| F05 | 41160001 | 41260000 | 0.0018 | 0.0018 | 0.0014 |
| F05 | 41180001 | 41280000 | 0.0024 | 0.0021 | 0.0020 |
| F05 | 41200001 | 41300000 | 0.0026 | 0.0021 | 0.0023 |
| F05 | 41220001 | 41320000 | 0.0027 | 0.0017 | 0.0023 |
| F05 | 41240001 | 41340000 | 0.0022 | 0.0022 | 0.0019 |
| F05 | 41260001 | 41360000 | 0.0027 | 0.0015 | 0.0024 |
| F05 | 41280001 | 41380000 | 0.0019 | 0.0014 | 0.0016 |
| F05 | 41300001 | 41400000 | 0.0020 | 0.0019 | 0.0017 |
| F05 | 41320001 | 41420000 | 0.0026 | 0.0017 | 0.0022 |
| F05 | 41340001 | 41440000 | 0.0024 | 0.0016 | 0.0020 |
| F05 | 41360001 | 41460000 | 0.0023 | 0.0022 | 0.0020 |
| F05 | 41380001 | 41480000 | 0.0031 | 0.0023 | 0.0027 |
| F05 | 41400001 | 41500000 | 0.0029 | 0.0018 | 0.0025 |
| F05 | 41420001 | 41520000 | 0.0023 | 0.0021 | 0.0021 |
| F05 | 41440001 | 41540000 | 0.0026 | 0.0020 | 0.0024 |

| Chr | Start | End | both | A1 | A2 |
|---|---|---|---|---|---|
| F05 | 41460001 | 41560000 | 0.0027 | 0.0016 | 0.0022 |
| F05 | 41480001 | 41580000 | 0.0021 | 0.0013 | 0.0017 |
| F05 | 41500001 | 41600000 | 0.0020 | 0.0014 | 0.0014 |
| F05 | 41520001 | 41620000 | 0.0022 | 0.0019 | 0.0015 |
| F05 | 41540001 | 41640000 | 0.0027 | 0.0018 | 0.0020 |
| F05 | 41560001 | 41660000 | 0.0027 | 0.0016 | 0.0020 |
| F05 | 41580001 | 41680000 | 0.0026 | 0.0016 | 0.0018 |
| F05 | 41600001 | 41700000 | 0.0024 | 0.0015 | 0.0018 |
| F05 | 41620001 | 41720000 | 0.0022 | 0.0010 | 0.0017 |
| F05 | 41640001 | 41740000 | 0.0019 | 0.0009 | 0.0013 |
| F05 | 41660001 | 41760000 | 0.0020 | 0.0009 | 0.0013 |
| F05 | 41680001 | 41780000 | 0.0019 | 0.0019 | 0.0012 |
| F05 | 41700001 | 41800000 | 0.0032 | 0.0020 | 0.0023 |
| F05 | 41720001 | 41820000 | 0.0035 | 0.0019 | 0.0025 |
| F05 | 41740001 | 41840000 | 0.0032 | 0.0023 | 0.0024 |
| F05 | 41760001 | 41860000 | 0.0034 | 0.0027 | 0.0026 |
| F05 | 41780001 | 41880000 | 0.0039 | 0.0017 | 0.0031 |
| F05 | 41800001 | 41900000 | 0.0029 | 0.0015 | 0.0020 |
| F05 | 41820001 | 41920000 | 0.0028 | 0.0019 | 0.0018 |
| F05 | 41840001 | 41940000 | 0.0034 | 0.0017 | 0.0022 |
| F05 | 41860001 | 41960000 | 0.0033 | 0.0014 | 0.0021 |
| F05 | 41880001 | 41980000 | 0.0028 | 0.0014 | 0.0017 |
| F05 | 41900001 | 42000000 | 0.0027 | 0.0014 | 0.0017 |
| F05 | 41920001 | 42020000 | 0.0023 | 0.0010 | 0.0016 |
| F05 | 41940001 | 42040000 | 0.0017 | 0.0008 | 0.0011 |
| F05 | 41960001 | 42060000 | 0.0013 | 0.0011 | 0.0009 |
| F05 | 41980001 | 42080000 | 0.0018 | 0.0020 | 0.0012 |
| F05 | 42000001 | 42100000 | 0.0027 | 0.0022 | 0.0021 |
| F05 | 42020001 | 42120000 | 0.0031 | 0.0024 | 0.0022 |
| F05 | 42040001 | 42140000 | 0.0033 | 0.0021 | 0.0025 |
| F05 | 42060001 | 42160000 | 0.0030 | 0.0019 | 0.0021 |
| F05 | 42080001 | 42180000 | 0.0027 | 0.0012 | 0.0020 |
| F05 | 42100001 | 42200000 | 0.0020 | 0.0012 | 0.0014 |
| F05 | 42120001 | 42220000 | 0.0019 | 0.0013 | 0.0014 |
| F05 | 42140001 | 42240000 | 0.0019 | 0.0016 | 0.0015 |
| F05 | 42160001 | 42260000 | 0.0022 | 0.0015 | 0.0018 |
| F05 | 42180001 | 42280000 | 0.0020 | 0.0013 | 0.0016 |
| F05 | 42200001 | 42300000 | 0.0018 | 0.0012 | 0.0014 |
| F05 | 42220001 | 42320000 | 0.0018 | 0.0011 | 0.0013 |
| F05 | 42240001 | 42340000 | 0.0016 | 0.0008 | 0.0011 |
| F05 | 42260001 | 42360000 | 0.0012 | 0.0009 | 0.0008 |
| F05 | 42280001 | 42380000 | 0.0012 | 0.0011 | 0.0008 |
| F05 | 42300001 | 42400000 | 0.0013 | 0.0018 | 0.0011 |
| F05 | 42320001 | 42420000 | 0.0020 | 0.0015 | 0.0019 |
| F05 | 42340001 | 42440000 | 0.0019 | 0.0018 | 0.0017 |
| F05 | 42360001 | 42460000 | 0.0021 | 0.0019 | 0.0020 |
| F05 | 42380001 | 42480000 | 0.0024 | 0.0021 | 0.0023 |
| F05 | 42400001 | 42500000 | 0.0026 | 0.0014 | 0.0024 |
| F05 | 42420001 | 42520000 | 0.0019 | 0.0015 | 0.0016 |
| F05 | 42440001 | 42540000 | 0.0021 | 0.0013 | 0.0016 |
| F05 | 42460001 | 42560000 | 0.0020 | 0.0012 | 0.0015 |
| F05 | 42480001 | 42580000 | 0.0018 | 0.0012 | 0.0012 |
| F05 | 42500001 | 42600000 | 0.0018 | 0.0019 | 0.0013 |
| F05 | 42520001 | 42620000 | 0.0026 | 0.0023 | 0.0019 |
| F05 | 42540001 | 42640000 | 0.0030 | 0.0028 | 0.0024 |
| F05 | 42560001 | 42660000 | 0.0034 | 0.0027 | 0.0030 |
| F05 | 42580001 | 42680000 | 0.0032 | 0.0023 | 0.0029 |
| F05 | 42600001 | 42700000 | 0.0029 | 0.0028 | 0.0025 |

| Chr | Start | End | both | A1 | A2 |
|---|---|---|---|---|---|
| F05 | 42620001 | 42720000 | 0.0032 | 0.0027 | 0.0027 |
| F05 | 42640001 | 42740000 | 0.0031 | 0.0027 | 0.0025 |
| F05 | 42660001 | 42760000 | 0.0032 | 0.0028 | 0.0025 |
| F05 | 42680001 | 42780000 | 0.0036 | 0.0028 | 0.0026 |
| F05 | 42700001 | 42800000 | 0.0036 | 0.0019 | 0.0026 |
| F05 | 42720001 | 42820000 | 0.0027 | 0.0013 | 0.0020 |
| F05 | 42740001 | 42840000 | 0.0020 | 0.0009 | 0.0015 |
| F05 | 42760001 | 42860000 | 0.0014 | 0.0010 | 0.0010 |
| F05 | 42780001 | 42880000 | 0.0013 | 0.0009 | 0.0011 |
| F05 | 42800001 | 42900000 | 0.0011 | 0.0011 | 0.0010 |
| F05 | 42820001 | 42920000 | 0.0012 | 0.0013 | 0.0012 |
| F05 | 42840001 | 42940000 | 0.0014 | 0.0012 | 0.0014 |
| F05 | 42860001 | 42960000 | 0.0014 | 0.0013 | 0.0014 |
| F05 | 42880001 | 42980000 | 0.0015 | 0.0013 | 0.0014 |
| F05 | 42900001 | 43000000 | 0.0016 | 0.0011 | 0.0015 |
| F05 | 42920001 | 43020000 | 0.0013 | 0.0011 | 0.0011 |
| F05 | 42940001 | 43040000 | 0.0012 | 0.0016 | 0.0010 |
| F05 | 42960001 | 43060000 | 0.0019 | 0.0012 | 0.0017 |
| F05 | 42980001 | 43080000 | 0.0013 | 0.0011 | 0.0012 |
| F05 | 43000001 | 43100000 | 0.0012 | 0.0008 | 0.0011 |
| F05 | 43020001 | 43120000 | 0.0010 | 0.0007 | 0.0010 |
| F05 | 43040001 | 43140000 | 0.0009 | 0.0004 | 0.0009 |
| F05 | 43060001 | 43160000 | 0.0004 | 0.0004 | 0.0004 |
| F05 | 43080001 | 43180000 | 0.0005 | 0.0005 | 0.0005 |
| F05 | 43100001 | 43200000 | 0.0006 | 0.0009 | 0.0006 |
| F05 | 43120001 | 43220000 | 0.0012 | 0.0009 | 0.0012 |
| F05 | 43140001 | 43240000 | 0.0012 | 0.0008 | 0.0012 |
| F05 | 43160001 | 43260000 | 0.0010 | 0.0008 | 0.0010 |
| F05 | 43180001 | 43280000 | 0.0012 | 0.0009 | 0.0012 |
| F05 | 43200001 | 43300000 | 0.0013 | 0.0008 | 0.0013 |
| F05 | 43220001 | 43320000 | 0.0011 | 0.0010 | 0.0010 |
| F05 | 43240001 | 43340000 | 0.0013 | 0.0015 | 0.0013 |
| F05 | 43260001 | 43360000 | 0.0019 | 0.0015 | 0.0019 |
| F05 | 43280001 | 43380000 | 0.0019 | 0.0015 | 0.0018 |
| F05 | 43300001 | 43400000 | 0.0019 | 0.0012 | 0.0017 |
| F05 | 43320001 | 43420000 | 0.0017 | 0.0012 | 0.0014 |
| F05 | 43340001 | 43440000 | 0.0018 | 0.0006 | 0.0014 |
| F05 | 43360001 | 43460000 | 0.0012 | 0.0009 | 0.0007 |
| F05 | 43380001 | 43480000 | 0.0015 | 0.0009 | 0.0011 |
| F05 | 43400001 | 43500000 | 0.0013 | 0.0011 | 0.0011 |
| F05 | 43420001 | 43520000 | 0.0016 | 0.0018 | 0.0014 |
| F05 | 43440001 | 43540000 | 0.0022 | 0.0022 | 0.0021 |
| F05 | 43460001 | 43560000 | 0.0026 | 0.0019 | 0.0025 |
| F05 | 43480001 | 43580000 | 0.0023 | 0.0018 | 0.0021 |
| F05 | 43500001 | 43600000 | 0.0022 | 0.0021 | 0.0020 |
| F05 | 43520001 | 43620000 | 0.0024 | 0.0015 | 0.0022 |
| F05 | 43540001 | 43640000 | 0.0018 | 0.0016 | 0.0016 |
| F05 | 43560001 | 43660000 | 0.0021 | 0.0018 | 0.0019 |
| F05 | 43580001 | 43680000 | 0.0022 | 0.0019 | 0.0021 |
| F05 | 43600001 | 43700000 | 0.0023 | 0.0014 | 0.0022 |
| F05 | 43620001 | 43720000 | 0.0018 | 0.0011 | 0.0017 |
| F05 | 43640001 | 43740000 | 0.0015 | 0.0008 | 0.0014 |
| F05 | 43660001 | 43760000 | 0.0009 | 0.0006 | 0.0009 |
| F05 | 43680001 | 43780000 | 0.0007 | 0.0005 | 0.0006 |
| F05 | 43700001 | 43800000 | 0.0006 | 0.0006 | 0.0006 |
| F05 | 43720001 | 43820000 | 0.0008 | 0.0006 | 0.0008 |
| F05 | 43740001 | 43840000 | 0.0007 | 0.0004 | 0.0007 |
| F05 | 43760001 | 43860000 | 0.0005 | 0.0004 | 0.0005 |

| Chr | Start | End | both | A1 | A2 |
|---|---|---|---|---|---|
| F05 | 43780001 | 43880000 | 0.0005 | 0.0007 | 0.0005 |
| F05 | 43800001 | 43900000 | 0.0009 | 0.0007 | 0.0008 |
| F05 | 43820001 | 43920000 | 0.0009 | 0.0007 | 0.0008 |
| F05 | 43840001 | 43940000 | 0.0009 | 0.0008 | 0.0008 |
| F05 | 43860001 | 43960000 | 0.0010 | 0.0008 | 0.0008 |
| F05 | 43880001 | 43980000 | 0.0010 | 0.0005 | 0.0009 |
| F05 | 43900001 | 44000000 | 0.0007 | 0.0006 | 0.0007 |
| F05 | 43920001 | 44020000 | 0.0008 | 0.0008 | 0.0008 |
| F05 | 43940001 | 44040000 | 0.0009 | 0.0008 | 0.0009 |
| F05 | 43960001 | 44060000 | 0.0010 | 0.0007 | 0.0010 |
| F05 | 43980001 | 44080000 | 0.0009 | 0.0007 | 0.0009 |
| F05 | 44000001 | 44100000 | 0.0008 | 0.0005 | 0.0008 |
| F05 | 44020001 | 44120000 | 0.0006 | 0.0003 | 0.0006 |
| F05 | 44040001 | 44140000 | 0.0004 | 0.0003 | 0.0004 |
| F05 | 44060001 | 44160000 | 0.0004 | 0.0004 | 0.0004 |
| F05 | 44080001 | 44180000 | 0.0007 | 0.0004 | 0.0004 |
| F05 | 44100001 | 44200000 | 0.0007 | 0.0004 | 0.0004 |
| F05 | 44120001 | 44220000 | 0.0007 | 0.0011 | 0.0005 |
| F05 | 44140001 | 44240000 | 0.0016 | 0.0014 | 0.0013 |
| F05 | 44160001 | 44260000 | 0.0018 | 0.0013 | 0.0016 |
| F05 | 44180001 | 44280000 | 0.0015 | 0.0011 | 0.0015 |
| F05 | 44200001 | 44300000 | 0.0014 | 0.0012 | 0.0013 |
| F05 | 44220001 | 44320000 | 0.0014 | 0.0008 | 0.0013 |
| F05 | 44240001 | 44340000 | 0.0009 | 0.0011 | 0.0009 |
| F05 | 44260001 | 44360000 | 0.0013 | 0.0015 | 0.0013 |
| F05 | 44280001 | 44380000 | 0.0017 | 0.0019 | 0.0017 |
| F05 | 44300001 | 44400000 | 0.0022 | 0.0019 | 0.0021 |
| F05 | 44320001 | 44420000 | 0.0023 | 0.0017 | 0.0023 |
| F05 | 44340001 | 44440000 | 0.0021 | 0.0016 | 0.0020 |
| F05 | 44360001 | 44460000 | 0.0020 | 0.0013 | 0.0019 |
| F05 | 44380001 | 44480000 | 0.0018 | 0.0010 | 0.0016 |
| F05 | 44400001 | 44500000 | 0.0015 | 0.0010 | 0.0013 |
| F05 | 44420001 | 44520000 | 0.0014 | 0.0010 | 0.0012 |
| F05 | 44440001 | 44540000 | 0.0012 | 0.0008 | 0.0011 |
| F05 | 44460001 | 44560000 | 0.0011 | 0.0009 | 0.0009 |
| F05 | 44480001 | 44580000 | 0.0011 | 0.0012 | 0.0009 |
| F05 | 44500001 | 44600000 | 0.0014 | 0.0012 | 0.0012 |
| F05 | 44520001 | 44620000 | 0.0015 | 0.0015 | 0.0013 |
| F05 | 44540001 | 44640000 | 0.0018 | 0.0012 | 0.0017 |
| F05 | 44560001 | 44660000 | 0.0015 | 0.0011 | 0.0014 |
| F05 | 44580001 | 44680000 | 0.0013 | 0.0008 | 0.0013 |
| F05 | 44600001 | 44700000 | 0.0010 | 0.0006 | 0.0009 |
| F05 | 44620001 | 44720000 | 0.0008 | 0.0007 | 0.0007 |
| F05 | 44640001 | 44740000 | 0.0009 | 0.0016 | 0.0008 |
| F05 | 44660001 | 44760000 | 0.0019 | 0.0021 | 0.0017 |
| F05 | 44680001 | 44780000 | 0.0025 | 0.0021 | 0.0023 |
| F05 | 44700001 | 44800000 | 0.0024 | 0.0021 | 0.0023 |
| F05 | 44720001 | 44820000 | 0.0025 | 0.0022 | 0.0024 |
| F05 | 44740001 | 44840000 | 0.0025 | 0.0015 | 0.0024 |
| F05 | 44760001 | 44860000 | 0.0018 | 0.0012 | 0.0018 |
| F05 | 44780001 | 44880000 | 0.0015 | 0.0012 | 0.0014 |
| F05 | 44800001 | 44900000 | 0.0016 | 0.0012 | 0.0014 |
| F05 | 44820001 | 44920000 | 0.0015 | 0.0006 | 0.0013 |
| F05 | 44840001 | 44940000 | 0.0010 | 0.0004 | 0.0008 |
| F05 | 44860001 | 44960000 | 0.0007 | 0.0003 | 0.0004 |
| F05 | 44880001 | 44980000 | 0.0006 | 0.0003 | 0.0005 |
| F05 | 44900001 | 45000000 | 0.0006 | 0.0004 | 0.0005 |
| F05 | 44920001 | 45020000 | 0.0007 | 0.0004 | 0.0006 |

| Chr | Start | End | both | A1 | A2 |
|---|---|---|---|---|---|
| F05 | 44940001 | 45040000 | 0.0007 | 0.0004 | 0.0006 |
| F05 | 44960001 | 45060000 | 0.0006 | 0.0005 | 0.0006 |
| F05 | 44980001 | 45080000 | 0.0007 | 0.0006 | 0.0006 |
| F05 | 45000001 | 45100000 | 0.0007 | 0.0006 | 0.0007 |
| F05 | 45020001 | 45120000 | 0.0008 | 0.0010 | 0.0008 |
| F05 | 45040001 | 45140000 | 0.0014 | 0.0011 | 0.0014 |
| F05 | 45060001 | 45160000 | 0.0014 | 0.0013 | 0.0014 |
| F05 | 45080001 | 45180000 | 0.0017 | 0.0013 | 0.0016 |
| F05 | 45100001 | 45200000 | 0.0016 | 0.0017 | 0.0015 |
| F05 | 45120001 | 45220000 | 0.0020 | 0.0016 | 0.0019 |
| F05 | 45140001 | 45240000 | 0.0018 | 0.0019 | 0.0017 |
| F05 | 45160001 | 45260000 | 0.0024 | 0.0020 | 0.0023 |
| F05 | 45180001 | 45280000 | 0.0025 | 0.0021 | 0.0024 |
| F05 | 45200001 | 45300000 | 0.0026 | 0.0025 | 0.0025 |
| F05 | 45220001 | 45320000 | 0.0028 | 0.0025 | 0.0027 |
| F05 | 45240001 | 45340000 | 0.0029 | 0.0023 | 0.0027 |
| F05 | 45260001 | 45360000 | 0.0028 | 0.0021 | 0.0024 |
| F05 | 45280001 | 45380000 | 0.0027 | 0.0022 | 0.0022 |
| F05 | 45300001 | 45400000 | 0.0030 | 0.0014 | 0.0023 |
| F05 | 45320001 | 45420000 | 0.0023 | 0.0023 | 0.0016 |
| F05 | 45340001 | 45440000 | 0.0032 | 0.0025 | 0.0027 |
| F05 | 45360001 | 45460000 | 0.0032 | 0.0023 | 0.0029 |
| F05 | 45380001 | 45480000 | 0.0029 | 0.0023 | 0.0027 |
| F05 | 45400001 | 45500000 | 0.0028 | 0.0024 | 0.0028 |
| F05 | 45420001 | 45520000 | 0.0028 | 0.0018 | 0.0028 |
| F05 | 45440001 | 45540000 | 0.0021 | 0.0019 | 0.0020 |
| F05 | 45460001 | 45560000 | 0.0023 | 0.0028 | 0.0021 |
| F05 | 45480001 | 45580000 | 0.0032 | 0.0025 | 0.0030 |
| F05 | 45500001 | 45600000 | 0.0029 | 0.0024 | 0.0027 |
| F05 | 45520001 | 45620000 | 0.0028 | 0.0022 | 0.0026 |
| F05 | 45540001 | 45640000 | 0.0028 | 0.0021 | 0.0025 |
| F05 | 45560001 | 45660000 | 0.0024 | 0.0017 | 0.0023 |
| F05 | 45580001 | 45680000 | 0.0022 | 0.0020 | 0.0020 |
| F05 | 45600001 | 45700000 | 0.0025 | 0.0021 | 0.0023 |
| F05 | 45620001 | 45720000 | 0.0027 | 0.0023 | 0.0023 |
| F05 | 45640001 | 45740000 | 0.0029 | 0.0027 | 0.0024 |
| F05 | 45660001 | 45760000 | 0.0037 | 0.0021 | 0.0029 |
| F05 | 45680001 | 45780000 | 0.0030 | 0.0024 | 0.0023 |
| F05 | 45700001 | 45800000 | 0.0035 | 0.0028 | 0.0027 |
| F05 | 45720001 | 45820000 | 0.0037 | 0.0025 | 0.0031 |
| F05 | 45740001 | 45840000 | 0.0033 | 0.0026 | 0.0028 |
| F05 | 45760001 | 45860000 | 0.0032 | 0.0025 | 0.0027 |
| F05 | 45780001 | 45880000 | 0.0033 | 0.0020 | 0.0027 |
| F05 | 45800001 | 45900000 | 0.0028 | 0.0019 | 0.0022 |
| F05 | 45820001 | 45920000 | 0.0029 | 0.0023 | 0.0022 |
| F05 | 45840001 | 45940000 | 0.0032 | 0.0021 | 0.0027 |
| F05 | 45860001 | 45960000 | 0.0030 | 0.0022 | 0.0025 |
| F05 | 45880001 | 45980000 | 0.0030 | 0.0025 | 0.0025 |
| F05 | 45900001 | 46000000 | 0.0035 | 0.0021 | 0.0030 |
| F05 | 45920001 | 46020000 | 0.0030 | 0.0015 | 0.0025 |
| F05 | 45940001 | 46040000 | 0.0026 | 0.0012 | 0.0017 |
| F05 | 45960001 | 46060000 | 0.0024 | 0.0015 | 0.0016 |
| F05 | 45980001 | 46080000 | 0.0027 | 0.0011 | 0.0019 |
| F05 | 46000001 | 46100000 | 0.0021 | 0.0013 | 0.0014 |
| F05 | 46020001 | 46120000 | 0.0022 | 0.0017 | 0.0016 |
| F05 | 46040001 | 46140000 | 0.0023 | 0.0017 | 0.0020 |
| F05 | 46060001 | 46160000 | 0.0022 | 0.0018 | 0.0019 |
| F05 | 46080001 | 46180000 | 0.0021 | 0.0028 | 0.0019 |

| Chr | Start | End | both | A1 | A2 |
|---|---|---|---|---|---|
| F05 | 46100001 | 46200000 | 0.0032 | 0.0035 | 0.0030 |
| F05 | 46120001 | 46220000 | 0.0041 | 0.0038 | 0.0039 |
| F05 | 46140001 | 46240000 | 0.0045 | 0.0040 | 0.0043 |
| F05 | 46160001 | 46260000 | 0.0051 | 0.0041 | 0.0045 |
| F05 | 46180001 | 46280000 | 0.0057 | 0.0038 | 0.0051 |
| F05 | 46200001 | 46300000 | 0.0057 | 0.0037 | 0.0048 |
| F05 | 46220001 | 46320000 | 0.0056 | 0.0037 | 0.0047 |
| F05 | 46240001 | 46340000 | 0.0053 | 0.0035 | 0.0044 |
| F05 | 46260001 | 46360000 | 0.0048 | 0.0031 | 0.0043 |
| F05 | 46280001 | 46380000 | 0.0039 | 0.0027 | 0.0033 |
| F05 | 46300001 | 46400000 | 0.0034 | 0.0024 | 0.0029 |
| F05 | 46320001 | 46420000 | 0.0029 | 0.0019 | 0.0023 |
| F05 | 46340001 | 46440000 | 0.0024 | 0.0034 | 0.0018 |
| F05 | 46360001 | 46460000 | 0.0040 | 0.0046 | 0.0034 |
| F05 | 46380001 | 46480000 | 0.0056 | 0.0043 | 0.0050 |
| F05 | 46400001 | 46500000 | 0.0050 | 0.0044 | 0.0046 |
| F05 | 46420001 | 46520000 | 0.0054 | 0.0053 | 0.0050 |
| F05 | 46440001 | 46540000 | 0.0064 | 0.0039 | 0.0060 |
| F05 | 46460001 | 46560000 | 0.0047 | 0.0022 | 0.0044 |
| F05 | 46480001 | 46580000 | 0.0026 | 0.0024 | 0.0025 |
| F05 | 46500001 | 46600000 | 0.0029 | 0.0017 | 0.0028 |
| F05 | 46520001 | 46620000 | 0.0020 | 0.0006 | 0.0020 |
| F05 | 46540001 | 46640000 | 0.0008 | 0.0010 | 0.0007 |
| F05 | 46560001 | 46660000 | 0.0012 | 0.0012 | 0.0011 |
| F05 | 46580001 | 46680000 | 0.0016 | 0.0013 | 0.0014 |
| F05 | 46600001 | 46700000 | 0.0017 | 0.0014 | 0.0015 |
| F05 | 46620001 | 46720000 | 0.0020 | 0.0017 | 0.0015 |
| F05 | 46640001 | 46740000 | 0.0025 | 0.0021 | 0.0019 |
| F05 | 46660001 | 46760000 | 0.0032 | 0.0021 | 0.0024 |
| F05 | 46680001 | 46780000 | 0.0032 | 0.0025 | 0.0024 |
| F05 | 46700001 | 46800000 | 0.0035 | 0.0031 | 0.0028 |
| F05 | 46720001 | 46820000 | 0.0042 | 0.0033 | 0.0035 |
| F05 | 46740001 | 46840000 | 0.0045 | 0.0026 | 0.0037 |
| F05 | 46760001 | 46860000 | 0.0039 | 0.0027 | 0.0031 |
| F05 | 46780001 | 46880000 | 0.0039 | 0.0029 | 0.0032 |
| F05 | 46800001 | 46900000 | 0.0041 | 0.0030 | 0.0033 |
| F05 | 46820001 | 46920000 | 0.0040 | 0.0031 | 0.0032 |
| F05 | 46840001 | 46940000 | 0.0041 | 0.0033 | 0.0035 |
| F05 | 46860001 | 46960000 | 0.0041 | 0.0031 | 0.0036 |
| F05 | 46880001 | 46980000 | 0.0039 | 0.0027 | 0.0034 |
| F05 | 46900001 | 47000000 | 0.0034 | 0.0025 | 0.0029 |
| F05 | 46920001 | 47020000 | 0.0031 | 0.0027 | 0.0028 |
| F05 | 46940001 | 47040000 | 0.0034 | 0.0025 | 0.0029 |
| F05 | 46960001 | 47060000 | 0.0031 | 0.0029 | 0.0027 |
| F05 | 46980001 | 47080000 | 0.0036 | 0.0030 | 0.0033 |
| F05 | 47000001 | 47100000 | 0.0037 | 0.0028 | 0.0034 |
| F05 | 47020001 | 47120000 | 0.0035 | 0.0022 | 0.0032 |
| F05 | 47040001 | 47140000 | 0.0029 | 0.0024 | 0.0027 |
| F05 | 47060001 | 47160000 | 0.0029 | 0.0022 | 0.0027 |
| F05 | 47080001 | 47180000 | 0.0028 | 0.0019 | 0.0025 |
| F05 | 47100001 | 47200000 | 0.0024 | 0.0019 | 0.0021 |
| F05 | 47120001 | 47220000 | 0.0023 | 0.0017 | 0.0022 |
| F05 | 47140001 | 47240000 | 0.0019 | 0.0016 | 0.0018 |
| F05 | 47160001 | 47260000 | 0.0019 | 0.0020 | 0.0019 |
| F05 | 47180001 | 47280000 | 0.0023 | 0.0027 | 0.0023 |
| F05 | 47200001 | 47300000 | 0.0031 | 0.0025 | 0.0030 |
| F05 | 47220001 | 47320000 | 0.0030 | 0.0027 | 0.0028 |
| F05 | 47240001 | 47340000 | 0.0033 | 0.0026 | 0.0030 |

| Chr | Start | End | both | A1 | A2 |
|---|---|---|---|---|---|
| F05 | 47260001 | 47360000 | 0.0032 | 0.0017 | 0.0028 |
| F05 | 47280001 | 47380000 | 0.0022 | 0.0009 | 0.0018 |
| F05 | 47300001 | 47400000 | 0.0016 | 0.0011 | 0.0011 |
| F05 | 47320001 | 47420000 | 0.0017 | 0.0013 | 0.0012 |
| F05 | 47340001 | 47440000 | 0.0018 | 0.0010 | 0.0015 |
| F05 | 47360001 | 47460000 | 0.0015 | 0.0011 | 0.0012 |
| F05 | 47380001 | 47480000 | 0.0015 | 0.0015 | 0.0013 |
| F05 | 47400001 | 47500000 | 0.0020 | 0.0015 | 0.0018 |
| F05 | 47420001 | 47520000 | 0.0020 | 0.0014 | 0.0019 |
| F05 | 47440001 | 47540000 | 0.0019 | 0.0020 | 0.0017 |
| F05 | 47460001 | 47560000 | 0.0026 | 0.0020 | 0.0025 |
| F05 | 47480001 | 47580000 | 0.0026 | 0.0021 | 0.0024 |
| F05 | 47500001 | 47600000 | 0.0025 | 0.0020 | 0.0023 |
| F05 | 47520001 | 47620000 | 0.0024 | 0.0019 | 0.0022 |
| F05 | 47540001 | 47640000 | 0.0024 | 0.0015 | 0.0022 |
| F05 | 47560001 | 47660000 | 0.0018 | 0.0019 | 0.0017 |
| F05 | 47580001 | 47680000 | 0.0024 | 0.0015 | 0.0023 |
| F05 | 47600001 | 47700000 | 0.0021 | 0.0017 | 0.0020 |
| F05 | 47620001 | 47720000 | 0.0024 | 0.0017 | 0.0023 |
| F05 | 47640001 | 47740000 | 0.0024 | 0.0016 | 0.0022 |
| F05 | 47660001 | 47760000 | 0.0022 | 0.0015 | 0.0020 |
| F05 | 47680001 | 47780000 | 0.0021 | 0.0015 | 0.0019 |
| F05 | 47700001 | 47800000 | 0.0020 | 0.0017 | 0.0019 |
| F05 | 47720001 | 47820000 | 0.0021 | 0.0019 | 0.0020 |
| F05 | 47740001 | 47840000 | 0.0023 | 0.0031 | 0.0022 |
| F05 | 47760001 | 47860000 | 0.0036 | 0.0030 | 0.0035 |
| F05 | 47780001 | 47880000 | 0.0037 | 0.0027 | 0.0035 |
| F05 | 47800001 | 47900000 | 0.0034 | 0.0027 | 0.0033 |
| F05 | 47820001 | 47920000 | 0.0033 | 0.0023 | 0.0032 |
| F05 | 47840001 | 47940000 | 0.0030 | 0.0017 | 0.0029 |
| F05 | 47860001 | 47960000 | 0.0023 | 0.0014 | 0.0023 |
| F05 | 47880001 | 47980000 | 0.0018 | 0.0015 | 0.0018 |
| F05 | 47900001 | 48000000 | 0.0018 | 0.0010 | 0.0017 |
| F05 | 47920001 | 48020000 | 0.0013 | 0.0013 | 0.0012 |
| F05 | 47940001 | 48040000 | 0.0015 | 0.0011 | 0.0014 |
| F05 | 47960001 | 48060000 | 0.0013 | 0.0012 | 0.0012 |
| F05 | 47980001 | 48080000 | 0.0014 | 0.0015 | 0.0013 |
| F05 | 48000001 | 48100000 | 0.0017 | 0.0016 | 0.0017 |
| F05 | 48020001 | 48120000 | 0.0019 | 0.0013 | 0.0019 |
| F05 | 48040001 | 48140000 | 0.0016 | 0.0008 | 0.0016 |
| F05 | 48060001 | 48160000 | 0.0011 | 0.0008 | 0.0011 |
| F05 | 48080001 | 48180000 | 0.0011 | 0.0005 | 0.0011 |
| F05 | 48100001 | 48200000 | 0.0008 | 0.0006 | 0.0007 |
| F05 | 48120001 | 48220000 | 0.0008 | 0.0007 | 0.0008 |
| F05 | 48140001 | 48240000 | 0.0009 | 0.0010 | 0.0009 |
| F05 | 48160001 | 48260000 | 0.0012 | 0.0010 | 0.0011 |
| F05 | 48180001 | 48280000 | 0.0011 | 0.0009 | 0.0010 |
| F05 | 48200001 | 48300000 | 0.0009 | 0.0006 | 0.0009 |
| F05 | 48220001 | 48320000 | 0.0007 | 0.0006 | 0.0007 |
| F05 | 48240001 | 48340000 | 0.0007 | 0.0002 | 0.0006 |
| F05 | 48260001 | 48360000 | 0.0003 | 0.0002 | 0.0003 |
| F05 | 48280001 | 48380000 | 0.0002 | 0.0001 | 0.0002 |
| F05 | 48300001 | 48400000 | 0.0002 | 0.0003 | 0.0002 |
| F05 | 48320001 | 48420000 | 0.0003 | 0.0003 | 0.0003 |
| F05 | 48340001 | 48440000 | 0.0003 | 0.0005 | 0.0003 |
| F05 | 48360001 | 48460000 | 0.0005 | 0.0009 | 0.0005 |
| F05 | 48380001 | 48480000 | 0.0010 | 0.0009 | 0.0010 |
| F05 | 48400001 | 48500000 | 0.0010 | 0.0009 | 0.0010 |

| Chr | Start | End | both | A1 | A2 |
|---|---|---|---|---|---|
| F05 | 48420001 | 48520000 | 0.0011 | 0.0010 | 0.0010 |
| F05 | 48440001 | 48540000 | 0.0012 | 0.0009 | 0.0012 |
| F05 | 48460001 | 48560000 | 0.0012 | 0.0009 | 0.0012 |
| F05 | 48480001 | 48580000 | 0.0012 | 0.0011 | 0.0011 |
| F05 | 48500001 | 48600000 | 0.0015 | 0.0009 | 0.0015 |
| F05 | 48520001 | 48620000 | 0.0013 | 0.0007 | 0.0013 |
| F05 | 48540001 | 48640000 | 0.0011 | 0.0008 | 0.0011 |
| F05 | 48560001 | 48660000 | 0.0012 | 0.0010 | 0.0012 |
| F05 | 48580001 | 48680000 | 0.0013 | 0.0010 | 0.0013 |
| F05 | 48600001 | 48700000 | 0.0013 | 0.0010 | 0.0013 |
| F05 | 48620001 | 48720000 | 0.0013 | 0.0010 | 0.0013 |
| F05 | 48640001 | 48740000 | 0.0013 | 0.0008 | 0.0013 |
| F05 | 48660001 | 48760000 | 0.0011 | 0.0005 | 0.0011 |
| F05 | 48680001 | 48780000 | 0.0007 | 0.0006 | 0.0007 |
| F05 | 48700001 | 48800000 | 0.0007 | 0.0009 | 0.0007 |
| F05 | 48720001 | 48820000 | 0.0010 | 0.0010 | 0.0010 |
| F05 | 48740001 | 48840000 | 0.0011 | 0.0011 | 0.0011 |
| F05 | 48760001 | 48860000 | 0.0012 | 0.0011 | 0.0012 |
| F05 | 48780001 | 48880000 | 0.0012 | 0.0010 | 0.0012 |
| F05 | 48800001 | 48900000 | 0.0012 | 0.0009 | 0.0012 |
| F05 | 48820001 | 48920000 | 0.0012 | 0.0022 | 0.0011 |
| F05 | 48840001 | 48940000 | 0.0028 | 0.0021 | 0.0025 |
| F05 | 48860001 | 48960000 | 0.0028 | 0.0022 | 0.0024 |
| F05 | 48880001 | 48980000 | 0.0030 | 0.0023 | 0.0026 |
| F05 | 48900001 | 49000000 | 0.0030 | 0.0027 | 0.0026 |
| F05 | 48920001 | 49020000 | 0.0034 | 0.0016 | 0.0030 |
| F05 | 48940001 | 49040000 | 0.0021 | 0.0017 | 0.0018 |
| F05 | 48960001 | 49060000 | 0.0021 | 0.0017 | 0.0019 |
| F05 | 48980001 | 49080000 | 0.0019 | 0.0013 | 0.0017 |
| F05 | 49000001 | 49100000 | 0.0016 | 0.0014 | 0.0015 |
| F05 | 49020001 | 49120000 | 0.0018 | 0.0017 | 0.0016 |
| F05 | 49040001 | 49140000 | 0.0023 | 0.0022 | 0.0019 |
| F05 | 49060001 | 49160000 | 0.0030 | 0.0021 | 0.0025 |
| F05 | 49080001 | 49180000 | 0.0032 | 0.0023 | 0.0025 |
| F05 | 49100001 | 49200000 | 0.0033 | 0.0024 | 0.0026 |
| F05 | 49120001 | 49220000 | 0.0034 | 0.0028 | 0.0027 |
| F05 | 49140001 | 49240000 | 0.0037 | 0.0024 | 0.0033 |
| F05 | 49160001 | 49260000 | 0.0035 | 0.0025 | 0.0030 |
| F05 | 49180001 | 49280000 | 0.0038 | 0.0033 | 0.0031 |
| F05 | 49200001 | 49300000 | 0.0045 | 0.0028 | 0.0039 |
| F05 | 49220001 | 49320000 | 0.0039 | 0.0023 | 0.0033 |
| F05 | 49240001 | 49340000 | 0.0033 | 0.0024 | 0.0028 |
| F05 | 49260001 | 49360000 | 0.0032 | 0.0028 | 0.0028 |
| F05 | 49280001 | 49380000 | 0.0034 | 0.0019 | 0.0032 |
| F05 | 49300001 | 49400000 | 0.0025 | 0.0026 | 0.0023 |
| F05 | 49320001 | 49420000 | 0.0031 | 0.0023 | 0.0029 |
| F05 | 49340001 | 49440000 | 0.0028 | 0.0023 | 0.0026 |
| F05 | 49360001 | 49460000 | 0.0027 | 0.0020 | 0.0025 |
| F05 | 49380001 | 49480000 | 0.0026 | 0.0026 | 0.0023 |
| F05 | 49400001 | 49500000 | 0.0033 | 0.0017 | 0.0029 |
| F05 | 49420001 | 49520000 | 0.0024 | 0.0016 | 0.0020 |
| F05 | 49440001 | 49540000 | 0.0024 | 0.0015 | 0.0019 |
| F05 | 49460001 | 49560000 | 0.0021 | 0.0016 | 0.0016 |
| F05 | 49480001 | 49580000 | 0.0022 | 0.0012 | 0.0018 |
| F05 | 49500001 | 49600000 | 0.0017 | 0.0014 | 0.0014 |
| F05 | 49520001 | 49620000 | 0.0018 | 0.0014 | 0.0016 |
| F05 | 49540001 | 49640000 | 0.0020 | 0.0014 | 0.0018 |
| F05 | 49560001 | 49660000 | 0.0021 | 0.0015 | 0.0019 |

| Chr | Start | End | both | A1 | A2 |
|---|---|---|---|---|---|
| F05 | 49580001 | 49680000 | 0.0021 | 0.0018 | 0.0019 |
| F05 | 49600001 | 49700000 | 0.0025 | 0.0027 | 0.0023 |
| F05 | 49620001 | 49720000 | 0.0034 | 0.0027 | 0.0033 |
| F05 | 49640001 | 49740000 | 0.0034 | 0.0026 | 0.0033 |
| F05 | 49660001 | 49760000 | 0.0034 | 0.0022 | 0.0031 |
| F05 | 49680001 | 49780000 | 0.0030 | 0.0017 | 0.0027 |
| F05 | 49700001 | 49800000 | 0.0024 | 0.0008 | 0.0021 |
| F05 | 49720001 | 49820000 | 0.0014 | 0.0013 | 0.0011 |
| F05 | 49740001 | 49840000 | 0.0019 | 0.0013 | 0.0016 |
| F05 | 49760001 | 49860000 | 0.0017 | 0.0014 | 0.0015 |
| F05 | 49780001 | 49880000 | 0.0017 | 0.0014 | 0.0015 |
| F05 | 49800001 | 49900000 | 0.0018 | 0.0016 | 0.0016 |
| F05 | 49820001 | 49920000 | 0.0021 | 0.0017 | 0.0018 |
| F05 | 49840001 | 49940000 | 0.0023 | 0.0021 | 0.0019 |
| F05 | 49860001 | 49960000 | 0.0030 | 0.0025 | 0.0026 |
| F05 | 49880001 | 49980000 | 0.0035 | 0.0029 | 0.0031 |
| F05 | 49900001 | 50000000 | 0.0038 | 0.0033 | 0.0035 |
| F05 | 49920001 | 50020000 | 0.0042 | 0.0038 | 0.0039 |
| F05 | 49940001 | 50040000 | 0.0044 | 0.0035 | 0.0043 |
| F05 | 49960001 | 50060000 | 0.0039 | 0.0036 | 0.0038 |
| F05 | 49980001 | 50080000 | 0.0041 | 0.0037 | 0.0040 |
| F05 | 50000001 | 50100000 | 0.0043 | 0.0030 | 0.0041 |
| F05 | 50020001 | 50120000 | 0.0036 | 0.0019 | 0.0034 |
| F05 | 50040001 | 50140000 | 0.0025 | 0.0019 | 0.0022 |
| F05 | 50060001 | 50160000 | 0.0025 | 0.0016 | 0.0022 |
| F05 | 50080001 | 50180000 | 0.0020 | 0.0018 | 0.0017 |
| F05 | 50100001 | 50200000 | 0.0021 | 0.0025 | 0.0019 |
| F05 | 50120001 | 50220000 | 0.0027 | 0.0024 | 0.0025 |
| F05 | 50140001 | 50240000 | 0.0026 | 0.0029 | 0.0025 |
| F05 | 50160001 | 50260000 | 0.0031 | 0.0029 | 0.0030 |
| F05 | 50180001 | 50280000 | 0.0032 | 0.0022 | 0.0030 |
| F05 | 50200001 | 50300000 | 0.0025 | 0.0019 | 0.0023 |
| F05 | 50220001 | 50320000 | 0.0024 | 0.0019 | 0.0023 |
| F05 | 50240001 | 50340000 | 0.0025 | 0.0013 | 0.0023 |
| F05 | 50260001 | 50360000 | 0.0018 | 0.0012 | 0.0016 |
| F05 | 50280001 | 50380000 | 0.0016 | 0.0012 | 0.0015 |
| F05 | 50300001 | 50400000 | 0.0017 | 0.0008 | 0.0016 |
| F05 | 50320001 | 50420000 | 0.0010 | 0.0008 | 0.0010 |
| F05 | 50340001 | 50440000 | 0.0010 | 0.0008 | 0.0010 |
| F05 | 50360001 | 50460000 | 0.0009 | 0.0004 | 0.0009 |
| F05 | 50380001 | 50480000 | 0.0005 | 0.0005 | 0.0005 |
| F05 | 50400001 | 50500000 | 0.0006 | 0.0006 | 0.0006 |
| F05 | 50420001 | 50520000 | 0.0007 | 0.0005 | 0.0007 |
| F05 | 50440001 | 50540000 | 0.0007 | 0.0006 | 0.0007 |
| F05 | 50460001 | 50560000 | 0.0008 | 0.0009 | 0.0008 |
| F05 | 50480001 | 50580000 | 0.0011 | 0.0011 | 0.0012 |
| F05 | 50500001 | 50600000 | 0.0013 | 0.0012 | 0.0014 |
| F05 | 50520001 | 50620000 | 0.0015 | 0.0012 | 0.0015 |
| F05 | 50540001 | 50640000 | 0.0014 | 0.0012 | 0.0014 |
| F05 | 50560001 | 50660000 | 0.0016 | 0.0013 | 0.0015 |
| F05 | 50580001 | 50680000 | 0.0018 | 0.0017 | 0.0016 |
| F05 | 50600001 | 50700000 | 0.0024 | 0.0020 | 0.0021 |
| F05 | 50620001 | 50720000 | 0.0028 | 0.0022 | 0.0024 |
| F05 | 50640001 | 50740000 | 0.0030 | 0.0024 | 0.0027 |
| F05 | 50660001 | 50760000 | 0.0030 | 0.0023 | 0.0028 |
| F05 | 50680001 | 50780000 | 0.0029 | 0.0018 | 0.0027 |
| F05 | 50700001 | 50800000 | 0.0023 | 0.0018 | 0.0021 |
| F05 | 50720001 | 50820000 | 0.0023 | 0.0020 | 0.0021 |

| Chr | Start | End | both | A1 | A2 |
|---|---|---|---|---|---|
| F05 | 50740001 | 50840000 | 0.0025 | 0.0016 | 0.0022 |
| F05 | 50760001 | 50860000 | 0.0021 | 0.0022 | 0.0019 |
| F05 | 50780001 | 50880000 | 0.0027 | 0.0020 | 0.0024 |
| F05 | 50800001 | 50900000 | 0.0022 | 0.0025 | 0.0021 |
| F05 | 50820001 | 50920000 | 0.0027 | 0.0030 | 0.0025 |
| F05 | 50840001 | 50940000 | 0.0033 | 0.0038 | 0.0031 |
| F05 | 50860001 | 50960000 | 0.0043 | 0.0042 | 0.0039 |
| F05 | 50880001 | 50980000 | 0.0050 | 0.0048 | 0.0046 |
| F05 | 50900001 | 51000000 | 0.0060 | 0.0046 | 0.0053 |
| F05 | 50920001 | 51020000 | 0.0060 | 0.0038 | 0.0052 |
| F05 | 50940001 | 51040000 | 0.0052 | 0.0031 | 0.0043 |
| F05 | 50960001 | 51060000 | 0.0047 | 0.0027 | 0.0037 |
| F05 | 50980001 | 51080000 | 0.0042 | 0.0029 | 0.0030 |
| F05 | 51000001 | 51100000 | 0.0041 | 0.0040 | 0.0031 |
| F05 | 51020001 | 51120000 | 0.0052 | 0.0046 | 0.0041 |
| F05 | 51040001 | 51140000 | 0.0057 | 0.0050 | 0.0047 |
| F05 | 51060001 | 51160000 | 0.0056 | 0.0046 | 0.0050 |
| F05 | 51080001 | 51180000 | 0.0052 | 0.0040 | 0.0048 |
| F05 | 51100001 | 51200000 | 0.0047 | 0.0021 | 0.0044 |
| F05 | 51120001 | 51220000 | 0.0029 | 0.0018 | 0.0027 |
| F05 | 51140001 | 51240000 | 0.0031 | 0.0014 | 0.0027 |
| F05 | 51160001 | 51260000 | 0.0029 | 0.0012 | 0.0024 |
| F05 | 51180001 | 51280000 | 0.0027 | 0.0015 | 0.0022 |
| F05 | 51200001 | 51300000 | 0.0028 | 0.0015 | 0.0023 |
| F05 | 51220001 | 51320000 | 0.0026 | 0.0017 | 0.0022 |
| F05 | 51240001 | 51340000 | 0.0024 | 0.0018 | 0.0022 |
| F05 | 51260001 | 51360000 | 0.0022 | 0.0024 | 0.0021 |
| F05 | 51280001 | 51380000 | 0.0028 | 0.0024 | 0.0028 |
| F05 | 51300001 | 51400000 | 0.0027 | 0.0024 | 0.0026 |
| F05 | 51320001 | 51420000 | 0.0028 | 0.0019 | 0.0027 |
| F05 | 51340001 | 51440000 | 0.0022 | 0.0020 | 0.0022 |
| F05 | 51360001 | 51460000 | 0.0024 | 0.0012 | 0.0023 |
| F05 | 51380001 | 51480000 | 0.0016 | 0.0008 | 0.0016 |
| F05 | 51400001 | 51500000 | 0.0010 | 0.0007 | 0.0011 |
| F05 | 51420001 | 51520000 | 0.0009 | 0.0008 | 0.0009 |
| F05 | 51440001 | 51540000 | 0.0011 | 0.0008 | 0.0011 |
| F05 | 51460001 | 51560000 | 0.0010 | 0.0012 | 0.0010 |
| F05 | 51480001 | 51580000 | 0.0016 | 0.0018 | 0.0016 |
| F05 | 51500001 | 51600000 | 0.0023 | 0.0020 | 0.0023 |
| F05 | 51520001 | 51620000 | 0.0025 | 0.0021 | 0.0025 |
| F05 | 51540001 | 51640000 | 0.0027 | 0.0018 | 0.0027 |
| F05 | 51560001 | 51660000 | 0.0024 | 0.0017 | 0.0024 |
| F05 | 51580001 | 51680000 | 0.0020 | 0.0017 | 0.0020 |
| F05 | 51600001 | 51700000 | 0.0021 | 0.0016 | 0.0020 |
| F05 | 51620001 | 51720000 | 0.0020 | 0.0018 | 0.0019 |
| F05 | 51640001 | 51740000 | 0.0023 | 0.0023 | 0.0022 |
| F05 | 51660001 | 51760000 | 0.0029 | 0.0022 | 0.0028 |
| F05 | 51680001 | 51780000 | 0.0029 | 0.0017 | 0.0028 |
| F05 | 51700001 | 51800000 | 0.0022 | 0.0022 | 0.0022 |
| F05 | 51720001 | 51820000 | 0.0027 | 0.0016 | 0.0026 |
| F05 | 51740001 | 51840000 | 0.0020 | 0.0013 | 0.0020 |
| F05 | 51760001 | 51860000 | 0.0015 | 0.0012 | 0.0015 |
| F05 | 51780001 | 51880000 | 0.0015 | 0.0014 | 0.0014 |
| F05 | 51800001 | 51900000 | 0.0016 | 0.0009 | 0.0016 |
| F05 | 51820001 | 51920000 | 0.0011 | 0.0008 | 0.0011 |
| F05 | 51840001 | 51940000 | 0.0009 | 0.0007 | 0.0009 |
| F05 | 51860001 | 51960000 | 0.0009 | 0.0004 | 0.0009 |
| F05 | 51880001 | 51980000 | 0.0005 | 0.0004 | 0.0005 |

| Chr | Start | End | both | A1 | A2 |
|---|---|---|---|---|---|
| F05 | 51900001 | 52000000 | 0.0005 | 0.0004 | 0.0004 |
| F05 | 51920001 | 52020000 | 0.0005 | 0.0003 | 0.0005 |
| F05 | 51940001 | 52040000 | 0.0004 | 0.0004 | 0.0004 |
| F05 | 51960001 | 52060000 | 0.0006 | 0.0005 | 0.0006 |
| F05 | 51980001 | 52080000 | 0.0006 | 0.0003 | 0.0006 |
| F05 | 52000001 | 52100000 | 0.0004 | 0.0003 | 0.0004 |
| F05 | 52020001 | 52120000 | 0.0004 | 0.0003 | 0.0004 |
| F05 | 52040001 | 52140000 | 0.0004 | 0.0004 | 0.0004 |
| F05 | 52060001 | 52160000 | 0.0005 | 0.0003 | 0.0005 |
| F05 | 52080001 | 52180000 | 0.0004 | 0.0003 | 0.0004 |
| F05 | 52100001 | 52200000 | 0.0005 | 0.0003 | 0.0005 |
| F05 | 52120001 | 52220000 | 0.0004 | 0.0004 | 0.0004 |
| F05 | 52140001 | 52240000 | 0.0006 | 0.0003 | 0.0006 |
| F05 | 52160001 | 52260000 | 0.0005 | 0.0004 | 0.0005 |
| F05 | 52180001 | 52280000 | 0.0005 | 0.0004 | 0.0006 |
| F05 | 52200001 | 52300000 | 0.0005 | 0.0004 | 0.0005 |
| F05 | 52220001 | 52320000 | 0.0006 | 0.0003 | 0.0006 |
| F05 | 52240001 | 52340000 | 0.0005 | 0.0005 | 0.0005 |
| F05 | 52260001 | 52360000 | 0.0007 | 0.0006 | 0.0007 |
| F05 | 52280001 | 52380000 | 0.0008 | 0.0006 | 0.0008 |
| F05 | 52300001 | 52400000 | 0.0007 | 0.0007 | 0.0007 |
| F05 | 52320001 | 52420000 | 0.0008 | 0.0009 | 0.0008 |
| F05 | 52340001 | 52440000 | 0.0010 | 0.0010 | 0.0010 |
| F05 | 52360001 | 52460000 | 0.0011 | 0.0011 | 0.0011 |
| F05 | 52380001 | 52480000 | 0.0013 | 0.0017 | 0.0013 |
| F05 | 52400001 | 52500000 | 0.0019 | 0.0021 | 0.0018 |
| F05 | 52420001 | 52520000 | 0.0026 | 0.0019 | 0.0025 |
| F05 | 52440001 | 52540000 | 0.0023 | 0.0016 | 0.0023 |
| F05 | 52460001 | 52560000 | 0.0020 | 0.0015 | 0.0020 |
| F05 | 52480001 | 52580000 | 0.0018 | 0.0010 | 0.0018 |
| F05 | 52500001 | 52600000 | 0.0014 | 0.0008 | 0.0014 |
| F05 | 52520001 | 52620000 | 0.0010 | 0.0008 | 0.0010 |
| F05 | 52540001 | 52640000 | 0.0010 | 0.0017 | 0.0010 |
| F05 | 52560001 | 52660000 | 0.0019 | 0.0020 | 0.0019 |
| F05 | 52580001 | 52680000 | 0.0023 | 0.0019 | 0.0022 |
| F05 | 52600001 | 52700000 | 0.0021 | 0.0019 | 0.0021 |
| F05 | 52620001 | 52720000 | 0.0022 | 0.0028 | 0.0021 |
| F05 | 52640001 | 52740000 | 0.0032 | 0.0024 | 0.0032 |
| F05 | 52660001 | 52760000 | 0.0029 | 0.0025 | 0.0029 |
| F05 | 52680001 | 52780000 | 0.0031 | 0.0034 | 0.0031 |
| F05 | 52700001 | 52800000 | 0.0040 | 0.0033 | 0.0040 |
| F05 | 52720001 | 52820000 | 0.0039 | 0.0044 | 0.0039 |
| F05 | 52740001 | 52840000 | 0.0051 | 0.0045 | 0.0050 |
| F05 | 52760001 | 52860000 | 0.0051 | 0.0039 | 0.0050 |
| F05 | 52780001 | 52880000 | 0.0044 | 0.0031 | 0.0044 |
| F05 | 52800001 | 52900000 | 0.0035 | 0.0031 | 0.0035 |
| F05 | 52820001 | 52920000 | 0.0035 | 0.0013 | 0.0034 |
| F05 | 52840001 | 52940000 | 0.0014 | 0.0007 | 0.0013 |
| F05 | 52860001 | 52960000 | 0.0008 | 0.0008 | 0.0008 |
| F05 | 52880001 | 52980000 | 0.0009 | 0.0011 | 0.0008 |
| F05 | 52900001 | 53000000 | 0.0012 | 0.0008 | 0.0012 |
| F05 | 52920001 | 53020000 | 0.0010 | 0.0008 | 0.0010 |
| F05 | 52940001 | 53040000 | 0.0010 | 0.0012 | 0.0010 |
| F05 | 52960001 | 53060000 | 0.0015 | 0.0013 | 0.0015 |
| F05 | 52980001 | 53080000 | 0.0016 | 0.0013 | 0.0016 |
| F05 | 53000001 | 53100000 | 0.0016 | 0.0013 | 0.0017 |
| F05 | 53020001 | 53120000 | 0.0015 | 0.0015 | 0.0015 |
| F05 | 53040001 | 53140000 | 0.0019 | 0.0018 | 0.0019 |

| Chr | Start | End | both | A1 | A2 |
|---|---|---|---|---|---|
| F05 | 53060001 | 53160000 | 0.0020 | 0.0018 | 0.0020 |
| F05 | 53080001 | 53180000 | 0.0020 | 0.0019 | 0.0020 |
| F05 | 53100001 | 53200000 | 0.0020 | 0.0032 | 0.0019 |
| F05 | 53120001 | 53220000 | 0.0033 | 0.0031 | 0.0033 |
| F05 | 53140001 | 53240000 | 0.0032 | 0.0032 | 0.0031 |
| F05 | 53160001 | 53260000 | 0.0035 | 0.0033 | 0.0034 |
| F05 | 53180001 | 53280000 | 0.0036 | 0.0035 | 0.0035 |
| F05 | 53200001 | 53300000 | 0.0040 | 0.0025 | 0.0039 |
| F05 | 53220001 | 53320000 | 0.0030 | 0.0037 | 0.0029 |
| F05 | 53240001 | 53340000 | 0.0044 | 0.0031 | 0.0042 |
| F05 | 53260001 | 53360000 | 0.0039 | 0.0031 | 0.0035 |
| F05 | 53280001 | 53380000 | 0.0040 | 0.0031 | 0.0036 |
| F05 | 53300001 | 53400000 | 0.0038 | 0.0031 | 0.0034 |
| F05 | 53320001 | 53420000 | 0.0039 | 0.0022 | 0.0034 |
| F05 | 53340001 | 53440000 | 0.0031 | 0.0030 | 0.0024 |
| F05 | 53360001 | 53460000 | 0.0040 | 0.0030 | 0.0034 |
| F05 | 53380001 | 53480000 | 0.0041 | 0.0037 | 0.0033 |
| F05 | 53400001 | 53500000 | 0.0048 | 0.0039 | 0.0040 |
| F05 | 53420001 | 53520000 | 0.0051 | 0.0037 | 0.0043 |
| F05 | 53440001 | 53540000 | 0.0047 | 0.0032 | 0.0041 |
| F05 | 53460001 | 53560000 | 0.0040 | 0.0031 | 0.0036 |
| F05 | 53480001 | 53580000 | 0.0038 | 0.0017 | 0.0034 |
| F05 | 53500001 | 53600000 | 0.0025 | 0.0022 | 0.0020 |
| F05 | 53520001 | 53620000 | 0.0029 | 0.0021 | 0.0025 |
| F05 | 53540001 | 53640000 | 0.0033 | 0.0027 | 0.0028 |
| F05 | 53560001 | 53660000 | 0.0046 | 0.0034 | 0.0039 |
| F05 | 53580001 | 53680000 | 0.0054 | 0.0047 | 0.0049 |
| F05 | 53600001 | 53700000 | 0.0067 | 0.0041 | 0.0063 |
| F05 | 53620001 | 53720000 | 0.0060 | 0.0041 | 0.0057 |
| F05 | 53640001 | 53740000 | 0.0056 | 0.0033 | 0.0054 |
| F05 | 53660001 | 53760000 | 0.0040 | 0.0032 | 0.0039 |
| F05 | 53680001 | 53780000 | 0.0039 | 0.0035 | 0.0038 |
| F05 | 53700001 | 53800000 | 0.0041 | 0.0042 | 0.0040 |
| F05 | 53720001 | 53820000 | 0.0049 | 0.0042 | 0.0048 |
| F05 | 53740001 | 53840000 | 0.0050 | 0.0039 | 0.0049 |
| F05 | 53760001 | 53860000 | 0.0046 | 0.0041 | 0.0045 |
| F05 | 53780001 | 53880000 | 0.0049 | 0.0032 | 0.0048 |
| F05 | 53800001 | 53900000 | 0.0041 | 0.0024 | 0.0041 |
| F05 | 53820001 | 53920000 | 0.0032 | 0.0029 | 0.0032 |
| F05 | 53840001 | 53940000 | 0.0035 | 0.0034 | 0.0034 |
| F05 | 53860001 | 53960000 | 0.0041 | 0.0029 | 0.0041 |
| F05 | 53880001 | 53980000 | 0.0034 | 0.0039 | 0.0033 |
| F05 | 53900001 | 54000000 | 0.0045 | 0.0058 | 0.0044 |
| F05 | 53920001 | 54020000 | 0.0069 | 0.0056 | 0.0067 |
| F05 | 53940001 | 54040000 | 0.0067 | 0.0060 | 0.0066 |
| F05 | 53960001 | 54060000 | 0.0074 | 0.0057 | 0.0070 |
| F05 | 53980001 | 54080000 | 0.0072 | 0.0047 | 0.0068 |
| F05 | 54000001 | 54100000 | 0.0061 | 0.0032 | 0.0056 |
| F05 | 54020001 | 54120000 | 0.0044 | 0.0043 | 0.0038 |
| F05 | 54040001 | 54140000 | 0.0059 | 0.0040 | 0.0052 |
| F05 | 54060001 | 54160000 | 0.0053 | 0.0047 | 0.0048 |
| F05 | 54080001 | 54180000 | 0.0060 | 0.0043 | 0.0055 |
| F05 | 54100001 | 54200000 | 0.0060 | 0.0037 | 0.0050 |
| F05 | 54120001 | 54220000 | 0.0054 | 0.0028 | 0.0044 |
| F05 | 54140001 | 54240000 | 0.0042 | 0.0034 | 0.0031 |
| F05 | 54160001 | 54260000 | 0.0046 | 0.0032 | 0.0035 |
| F05 | 54180001 | 54280000 | 0.0044 | 0.0034 | 0.0033 |
| F05 | 54200001 | 54300000 | 0.0041 | 0.0040 | 0.0035 |

| Chr | Start | End | both | A1 | A2 |
|---|---|---|---|---|---|
| F05 | 54220001 | 54320000 | 0.0047 | 0.0038 | 0.0042 |
| F05 | 54240001 | 54340000 | 0.0045 | 0.0026 | 0.0041 |
| F05 | 54260001 | 54360000 | 0.0034 | 0.0019 | 0.0031 |
| F05 | 54280001 | 54380000 | 0.0026 | 0.0018 | 0.0024 |
| F05 | 54300001 | 54400000 | 0.0024 | 0.0019 | 0.0024 |
| F05 | 54320001 | 54420000 | 0.0025 | 0.0025 | 0.0025 |
| F05 | 54340001 | 54440000 | 0.0029 | 0.0029 | 0.0029 |
| F05 | 54360001 | 54460000 | 0.0034 | 0.0030 | 0.0034 |
| F05 | 54380001 | 54480000 | 0.0036 | 0.0032 | 0.0036 |
| F05 | 54400001 | 54500000 | 0.0037 | 0.0029 | 0.0037 |
| F05 | 54420001 | 54520000 | 0.0032 | 0.0019 | 0.0031 |
| F05 | 54440001 | 54540000 | 0.0021 | 0.0016 | 0.0021 |
| F05 | 54460001 | 54560000 | 0.0019 | 0.0018 | 0.0018 |
| F05 | 54480001 | 54580000 | 0.0019 | 0.0026 | 0.0019 |
| F05 | 54500001 | 54600000 | 0.0027 | 0.0028 | 0.0026 |
| F05 | 54520001 | 54620000 | 0.0031 | 0.0038 | 0.0030 |
| F05 | 54540001 | 54640000 | 0.0041 | 0.0043 | 0.0040 |
| F05 | 54560001 | 54660000 | 0.0047 | 0.0045 | 0.0045 |
| F05 | 54580001 | 54680000 | 0.0049 | 0.0035 | 0.0047 |
| F05 | 54600001 | 54700000 | 0.0040 | 0.0033 | 0.0039 |
| F05 | 54620001 | 54720000 | 0.0038 | 0.0033 | 0.0036 |
| F05 | 54640001 | 54740000 | 0.0037 | 0.0058 | 0.0035 |
| F05 | 54660001 | 54760000 | 0.0065 | 0.0061 | 0.0063 |
| F05 | 54680001 | 54780000 | 0.0069 | 0.0062 | 0.0067 |
| F05 | 54700001 | 54800000 | 0.0071 | 0.0058 | 0.0069 |
| F05 | 54720001 | 54820000 | 0.0065 | 0.0053 | 0.0064 |
| F05 | 54740001 | 54840000 | 0.0062 | 0.0026 | 0.0061 |
| F05 | 54760001 | 54860000 | 0.0032 | 0.0025 | 0.0032 |
| F05 | 54780001 | 54880000 | 0.0032 | 0.0025 | 0.0032 |
| F05 | 54800001 | 54900000 | 0.0032 | 0.0036 | 0.0031 |
| F05 | 54820001 | 54920000 | 0.0045 | 0.0031 | 0.0044 |
| F05 | 54840001 | 54940000 | 0.0039 | 0.0028 | 0.0037 |
| F05 | 54860001 | 54960000 | 0.0034 | 0.0023 | 0.0033 |
| F05 | 54880001 | 54980000 | 0.0029 | 0.0019 | 0.0028 |
| F05 | 54900001 | 55000000 | 0.0023 | 0.0007 | 0.0022 |
| F05 | 54920001 | 55020000 | 0.0009 | 0.0012 | 0.0009 |
| F05 | 54940001 | 55040000 | 0.0015 | 0.0014 | 0.0014 |
| F05 | 54960001 | 55060000 | 0.0017 | 0.0015 | 0.0016 |
| F05 | 54980001 | 55080000 | 0.0018 | 0.0032 | 0.0017 |
| F05 | 55000001 | 55100000 | 0.0036 | 0.0034 | 0.0034 |
| F05 | 55020001 | 55120000 | 0.0041 | 0.0035 | 0.0037 |
| F05 | 55040001 | 55140000 | 0.0042 | 0.0040 | 0.0038 |
| F05 | 55060001 | 55160000 | 0.0047 | 0.0036 | 0.0043 |
| F05 | 55080001 | 55180000 | 0.0043 | 0.0022 | 0.0040 |
| F05 | 55100001 | 55200000 | 0.0028 | 0.0028 | 0.0026 |
| F05 | 55120001 | 55220000 | 0.0033 | 0.0026 | 0.0032 |
| F05 | 55140001 | 55240000 | 0.0030 | 0.0020 | 0.0030 |
| F05 | 55160001 | 55260000 | 0.0022 | 0.0030 | 0.0021 |
| F05 | 55180001 | 55280000 | 0.0033 | 0.0038 | 0.0032 |
| F05 | 55200001 | 55300000 | 0.0043 | 0.0035 | 0.0042 |
| F05 | 55220001 | 55320000 | 0.0039 | 0.0037 | 0.0037 |
| F05 | 55240001 | 55340000 | 0.0043 | 0.0038 | 0.0040 |
| F05 | 55260001 | 55360000 | 0.0045 | 0.0032 | 0.0042 |
| F05 | 55280001 | 55380000 | 0.0038 | 0.0029 | 0.0035 |
| F05 | 55300001 | 55400000 | 0.0034 | 0.0031 | 0.0032 |
| F05 | 55320001 | 55420000 | 0.0037 | 0.0030 | 0.0035 |
| F05 | 55340001 | 55440000 | 0.0038 | 0.0032 | 0.0035 |
| F05 | 55360001 | 55460000 | 0.0041 | 0.0031 | 0.0038 |

| Chr | Start | End | both | A1 | A2 |
|---|---|---|---|---|---|
| F05 | 55380001 | 55480000 | 0.0038 | 0.0027 | 0.0035 |
| F05 | 55400001 | 55500000 | 0.0032 | 0.0020 | 0.0029 |
| F05 | 55420001 | 55520000 | 0.0024 | 0.0030 | 0.0021 |
| F05 | 55440001 | 55540000 | 0.0034 | 0.0039 | 0.0032 |
| F05 | 55460001 | 55560000 | 0.0043 | 0.0043 | 0.0041 |
| F05 | 55480001 | 55580000 | 0.0048 | 0.0045 | 0.0047 |
| F05 | 55500001 | 55600000 | 0.0051 | 0.0051 | 0.0050 |
| F05 | 55520001 | 55620000 | 0.0058 | 0.0037 | 0.0057 |
| F05 | 55540001 | 55640000 | 0.0043 | 0.0029 | 0.0043 |
| F05 | 55560001 | 55660000 | 0.0034 | 0.0032 | 0.0034 |
| F05 | 55580001 | 55680000 | 0.0039 | 0.0034 | 0.0039 |
| F05 | 55600001 | 55700000 | 0.0041 | 0.0028 | 0.0041 |
| F05 | 55620001 | 55720000 | 0.0033 | 0.0029 | 0.0033 |
| F05 | 55640001 | 55740000 | 0.0035 | 0.0025 | 0.0035 |
| F05 | 55660001 | 55760000 | 0.0031 | 0.0029 | 0.0031 |
| F05 | 55680001 | 55780000 | 0.0034 | 0.0025 | 0.0034 |
| F05 | 55700001 | 55800000 | 0.0030 | 0.0025 | 0.0029 |
| F05 | 55720001 | 55820000 | 0.0031 | 0.0027 | 0.0030 |
| F05 | 55740001 | 55840000 | 0.0033 | 0.0031 | 0.0032 |
| F05 | 55760001 | 55860000 | 0.0038 | 0.0043 | 0.0037 |
| F05 | 55780001 | 55880000 | 0.0051 | 0.0050 | 0.0050 |
| F05 | 55800001 | 55900000 | 0.0060 | 0.0053 | 0.0059 |
| F05 | 55820001 | 55920000 | 0.0064 | 0.0053 | 0.0062 |
| F05 | 55840001 | 55940000 | 0.0065 | 0.0058 | 0.0062 |
| F05 | 55860001 | 55960000 | 0.0068 | 0.0065 | 0.0065 |
| F05 | 55880001 | 55980000 | 0.0075 | 0.0060 | 0.0070 |
| F05 | 55900001 | 56000000 | 0.0070 | 0.0059 | 0.0064 |
| F05 | 55920001 | 56020000 | 0.0068 | 0.0058 | 0.0064 |
| F05 | 55940001 | 56040000 | 0.0067 | 0.0062 | 0.0062 |
| F05 | 55960001 | 56060000 | 0.0073 | 0.0033 | 0.0067 |
| F05 | 55980001 | 56080000 | 0.0043 | 0.0037 | 0.0038 |
| F05 | 56000001 | 56100000 | 0.0046 | 0.0044 | 0.0042 |
| F05 | 56020001 | 56120000 | 0.0054 | 0.0052 | 0.0051 |
| F05 | 56040001 | 56140000 | 0.0063 | 0.0052 | 0.0061 |
| F05 | 56060001 | 56160000 | 0.0063 | 0.0057 | 0.0062 |
| F05 | 56080001 | 56180000 | 0.0068 | 0.0074 | 0.0066 |
| F05 | 56100001 | 56200000 | 0.0086 | 0.0080 | 0.0084 |
| F05 | 56120001 | 56220000 | 0.0093 | 0.0075 | 0.0090 |
| F05 | 56140001 | 56240000 | 0.0086 | 0.0073 | 0.0082 |
| F05 | 56160001 | 56260000 | 0.0082 | 0.0077 | 0.0078 |
| F05 | 56180001 | 56280000 | 0.0086 | 0.0049 | 0.0081 |
| F05 | 56200001 | 56300000 | 0.0056 | 0.0037 | 0.0053 |
| F05 | 56220001 | 56320000 | 0.0041 | 0.0049 | 0.0039 |
| F05 | 56240001 | 56340000 | 0.0055 | 0.0046 | 0.0053 |
| F05 | 56260001 | 56360000 | 0.0054 | 0.0039 | 0.0052 |
| F05 | 56280001 | 56380000 | 0.0046 | 0.0043 | 0.0045 |
| F05 | 56300001 | 56400000 | 0.0050 | 0.0054 | 0.0048 |
| F05 | 56320001 | 56420000 | 0.0061 | 0.0054 | 0.0058 |
| F05 | 56340001 | 56440000 | 0.0062 | 0.0052 | 0.0058 |
| F05 | 56360001 | 56460000 | 0.0059 | 0.0051 | 0.0054 |
| F05 | 56380001 | 56480000 | 0.0057 | 0.0064 | 0.0052 |
| F05 | 56400001 | 56500000 | 0.0072 | 0.0058 | 0.0068 |
| F05 | 56420001 | 56520000 | 0.0067 | 0.0046 | 0.0065 |
| F05 | 56440001 | 56540000 | 0.0051 | 0.0045 | 0.0049 |
| F05 | 56460001 | 56560000 | 0.0052 | 0.0047 | 0.0050 |
| F05 | 56480001 | 56580000 | 0.0054 | 0.0037 | 0.0052 |
| F05 | 56500001 | 56600000 | 0.0041 | 0.0031 | 0.0039 |
| F05 | 56520001 | 56620000 | 0.0032 | 0.0039 | 0.0030 |

| Chr | Start | End | both | A1 | A2 |
|---|---|---|---|---|---|
| F05 | 56540001 | 56640000 | 0.0041 | 0.0047 | 0.0038 |
| F05 | 56560001 | 56660000 | 0.0051 | 0.0043 | 0.0049 |
| F05 | 56580001 | 56680000 | 0.0048 | 0.0042 | 0.0046 |
| F05 | 56600001 | 56700000 | 0.0047 | 0.0043 | 0.0046 |
| F05 | 56620001 | 56720000 | 0.0050 | 0.0033 | 0.0049 |
| F05 | 56640001 | 56740000 | 0.0040 | 0.0026 | 0.0040 |
| F05 | 56660001 | 56760000 | 0.0030 | 0.0028 | 0.0030 |
| F05 | 56680001 | 56780000 | 0.0032 | 0.0026 | 0.0032 |
| F05 | 56700001 | 56800000 | 0.0031 | 0.0026 | 0.0031 |
| F05 | 56720001 | 56820000 | 0.0030 | 0.0022 | 0.0030 |
| F05 | 56740001 | 56840000 | 0.0025 | 0.0014 | 0.0025 |
| F05 | 56760001 | 56860000 | 0.0016 | 0.0019 | 0.0016 |
| F05 | 56780001 | 56880000 | 0.0018 | 0.0056 | 0.0016 |
| F05 | 56800001 | 56900000 | 0.0053 | 0.0062 | 0.0047 |
| F05 | 56820001 | 56920000 | 0.0060 | 0.0064 | 0.0054 |
| F05 | 56840001 | 56940000 | 0.0063 | 0.0064 | 0.0057 |
| F05 | 56860001 | 56960000 | 0.0063 | 0.0057 | 0.0058 |
| F05 | 56880001 | 56980000 | 0.0060 | 0.0015 | 0.0056 |
| F05 | 56900001 | 57000000 | 0.0019 | 0.0016 | 0.0018 |
| F05 | 56920001 | 57020000 | 0.0021 | 0.0014 | 0.0020 |
| F05 | 56940001 | 57040000 | 0.0020 | 0.0020 | 0.0016 |
| F05 | 56960001 | 57060000 | 0.0027 | 0.0021 | 0.0023 |
| F05 | 56980001 | 57080000 | 0.0028 | 0.0026 | 0.0023 |
| F05 | 57000001 | 57100000 | 0.0035 | 0.0023 | 0.0030 |
| F05 | 57020001 | 57120000 | 0.0031 | 0.0031 | 0.0028 |
| F05 | 57040001 | 57140000 | 0.0038 | 0.0028 | 0.0036 |
| F05 | 57060001 | 57160000 | 0.0034 | 0.0027 | 0.0032 |
| F05 | 57080001 | 57180000 | 0.0033 | 0.0022 | 0.0032 |
| F05 | 57100001 | 57200000 | 0.0027 | 0.0017 | 0.0025 |
| F05 | 57120001 | 57220000 | 0.0020 | 0.0014 | 0.0017 |
| F05 | 57140001 | 57240000 | 0.0018 | 0.0015 | 0.0015 |
| F05 | 57160001 | 57260000 | 0.0020 | 0.0015 | 0.0015 |
| F05 | 57180001 | 57280000 | 0.0021 | 0.0016 | 0.0016 |
| F05 | 57200001 | 57300000 | 0.0023 | 0.0016 | 0.0017 |
| F05 | 57220001 | 57320000 | 0.0023 | 0.0011 | 0.0017 |
| F05 | 57240001 | 57340000 | 0.0019 | 0.0008 | 0.0012 |
| F05 | 57260001 | 57360000 | 0.0014 | 0.0012 | 0.0008 |
| F05 | 57280001 | 57380000 | 0.0018 | 0.0014 | 0.0013 |
| F05 | 57300001 | 57400000 | 0.0018 | 0.0012 | 0.0015 |
| F05 | 57320001 | 57420000 | 0.0016 | 0.0019 | 0.0013 |
| F05 | 57340001 | 57440000 | 0.0023 | 0.0019 | 0.0021 |
| F05 | 57360001 | 57460000 | 0.0023 | 0.0014 | 0.0021 |
| F05 | 57380001 | 57480000 | 0.0018 | 0.0019 | 0.0015 |
| F05 | 57400001 | 57500000 | 0.0024 | 0.0031 | 0.0022 |
| F05 | 57420001 | 57520000 | 0.0038 | 0.0025 | 0.0035 |
| F05 | 57440001 | 57540000 | 0.0030 | 0.0030 | 0.0027 |
| F05 | 57460001 | 57560000 | 0.0035 | 0.0031 | 0.0033 |
| F05 | 57480001 | 57580000 | 0.0036 | 0.0024 | 0.0034 |
| F05 | 57500001 | 57600000 | 0.0030 | 0.0021 | 0.0027 |
| F05 | 57520001 | 57620000 | 0.0026 | 0.0025 | 0.0023 |
| F05 | 57540001 | 57640000 | 0.0032 | 0.0025 | 0.0029 |
| F05 | 57560001 | 57660000 | 0.0034 | 0.0036 | 0.0031 |
| F05 | 57580001 | 57680000 | 0.0047 | 0.0037 | 0.0043 |
| F05 | 57600001 | 57700000 | 0.0046 | 0.0028 | 0.0044 |
| F05 | 57620001 | 57720000 | 0.0036 | 0.0032 | 0.0035 |
| F05 | 57640001 | 57740000 | 0.0040 | 0.0027 | 0.0038 |
| F05 | 57660001 | 57760000 | 0.0033 | 0.0018 | 0.0032 |
| F05 | 57680001 | 57780000 | 0.0021 | 0.0027 | 0.0020 |

| Chr | Start | End | both | A1 | A2 |
|---|---|---|---|---|---|
| F05 | 57700001 | 57800000 | 0.0032 | 0.0039 | 0.0031 |
| F05 | 57720001 | 57820000 | 0.0046 | 0.0039 | 0.0045 |
| F05 | 57740001 | 57840000 | 0.0047 | 0.0044 | 0.0046 |
| F05 | 57760001 | 57860000 | 0.0053 | 0.0043 | 0.0051 |
| F05 | 57780001 | 57880000 | 0.0051 | 0.0037 | 0.0050 |
| F05 | 57800001 | 57900000 | 0.0044 | 0.0027 | 0.0043 |
| F05 | 57820001 | 57920000 | 0.0030 | 0.0019 | 0.0028 |
| F05 | 57840001 | 57940000 | 0.0022 | 0.0021 | 0.0021 |
| F05 | 57860001 | 57960000 | 0.0022 | 0.0033 | 0.0022 |
| F05 | 57880001 | 57980000 | 0.0035 | 0.0038 | 0.0034 |
| F05 | 57900001 | 58000000 | 0.0040 | 0.0039 | 0.0039 |
| F05 | 57920001 | 58020000 | 0.0042 | 0.0036 | 0.0041 |
| F05 | 57940001 | 58040000 | 0.0039 | 0.0028 | 0.0038 |
| F05 | 57960001 | 58060000 | 0.0032 | 0.0017 | 0.0031 |
| F05 | 57980001 | 58080000 | 0.0020 | 0.0008 | 0.0020 |
| F05 | 58000001 | 58100000 | 0.0010 | 0.0008 | 0.0010 |
| F05 | 58020001 | 58120000 | 0.0011 | 0.0009 | 0.0011 |
| F05 | 58040001 | 58140000 | 0.0012 | 0.0011 | 0.0012 |
| F05 | 58060001 | 58160000 | 0.0014 | 0.0011 | 0.0014 |
| F05 | 58080001 | 58180000 | 0.0013 | 0.0019 | 0.0013 |
| F05 | 58100001 | 58200000 | 0.0023 | 0.0026 | 0.0023 |
| F05 | 58120001 | 58220000 | 0.0029 | 0.0033 | 0.0029 |
| F05 | 58140001 | 58240000 | 0.0038 | 0.0035 | 0.0037 |
| F05 | 58160001 | 58260000 | 0.0042 | 0.0048 | 0.0038 |
| F05 | 58180001 | 58280000 | 0.0056 | 0.0042 | 0.0050 |
| F05 | 58200001 | 58300000 | 0.0051 | 0.0042 | 0.0042 |
| F05 | 58220001 | 58320000 | 0.0052 | 0.0034 | 0.0042 |
| F05 | 58240001 | 58340000 | 0.0045 | 0.0039 | 0.0034 |
| F05 | 58260001 | 58360000 | 0.0049 | 0.0028 | 0.0040 |
| F05 | 58280001 | 58380000 | 0.0038 | 0.0028 | 0.0030 |
| F05 | 58300001 | 58400000 | 0.0036 | 0.0028 | 0.0031 |
| F05 | 58320001 | 58420000 | 0.0036 | 0.0028 | 0.0030 |
| F05 | 58340001 | 58440000 | 0.0035 | 0.0022 | 0.0031 |
| F05 | 58360001 | 58460000 | 0.0028 | 0.0019 | 0.0025 |
| F05 | 58380001 | 58480000 | 0.0023 | 0.0020 | 0.0021 |
| F05 | 58400001 | 58500000 | 0.0027 | 0.0017 | 0.0024 |
| F05 | 58420001 | 58520000 | 0.0023 | 0.0016 | 0.0021 |
| F05 | 58440001 | 58540000 | 0.0022 | 0.0016 | 0.0020 |
| F05 | 58460001 | 58560000 | 0.0021 | 0.0021 | 0.0020 |
| F05 | 58480001 | 58580000 | 0.0026 | 0.0028 | 0.0026 |
| F05 | 58500001 | 58600000 | 0.0031 | 0.0026 | 0.0030 |
| F05 | 58520001 | 58620000 | 0.0029 | 0.0026 | 0.0027 |
| F05 | 58540001 | 58640000 | 0.0028 | 0.0027 | 0.0025 |
| F05 | 58560001 | 58660000 | 0.0029 | 0.0025 | 0.0027 |
| F05 | 58580001 | 58680000 | 0.0028 | 0.0013 | 0.0025 |
| F05 | 58600001 | 58700000 | 0.0018 | 0.0011 | 0.0014 |
| F05 | 58620001 | 58720000 | 0.0017 | 0.0014 | 0.0012 |
| F05 | 58640001 | 58740000 | 0.0020 | 0.0020 | 0.0017 |
| F05 | 58660001 | 58760000 | 0.0026 | 0.0021 | 0.0022 |
| F05 | 58680001 | 58780000 | 0.0026 | 0.0020 | 0.0022 |
| F05 | 58700001 | 58800000 | 0.0023 | 0.0023 | 0.0021 |
| F05 | 58720001 | 58820000 | 0.0026 | 0.0025 | 0.0025 |
| F05 | 58740001 | 58840000 | 0.0027 | 0.0017 | 0.0025 |
| F05 | 58760001 | 58860000 | 0.0018 | 0.0014 | 0.0018 |
| F05 | 58780001 | 58880000 | 0.0015 | 0.0022 | 0.0015 |
| F05 | 58800001 | 58900000 | 0.0024 | 0.0023 | 0.0024 |
| F05 | 58820001 | 58920000 | 0.0025 | 0.0027 | 0.0024 |
| F05 | 58840001 | 58940000 | 0.0030 | 0.0032 | 0.0028 |

| Chr | Start | End | both | A1 | A2 |
|---|---|---|---|---|---|
| F05 | 58860001 | 58960000 | 0.0036 | 0.0032 | 0.0034 |
| F05 | 58880001 | 58980000 | 0.0036 | 0.0030 | 0.0034 |
| F05 | 58900001 | 59000000 | 0.0033 | 0.0026 | 0.0031 |
| F05 | 58920001 | 59020000 | 0.0029 | 0.0019 | 0.0027 |
| F05 | 58940001 | 59040000 | 0.0023 | 0.0019 | 0.0020 |
| F05 | 58960001 | 59060000 | 0.0024 | 0.0020 | 0.0019 |
| F05 | 58980001 | 59080000 | 0.0025 | 0.0016 | 0.0019 |
| F05 | 59000001 | 59100000 | 0.0023 | 0.0017 | 0.0017 |
| F05 | 59020001 | 59120000 | 0.0024 | 0.0020 | 0.0018 |
| F05 | 59040001 | 59140000 | 0.0025 | 0.0014 | 0.0021 |
| F05 | 59060001 | 59160000 | 0.0019 | 0.0022 | 0.0016 |
| F05 | 59080001 | 59180000 | 0.0027 | 0.0020 | 0.0024 |
| F05 | 59100001 | 59200000 | 0.0025 | 0.0021 | 0.0022 |
| F05 | 59120001 | 59220000 | 0.0028 | 0.0017 | 0.0023 |
| F05 | 59140001 | 59240000 | 0.0023 | 0.0021 | 0.0019 |
| F05 | 59160001 | 59260000 | 0.0027 | 0.0013 | 0.0023 |
| F05 | 59180001 | 59280000 | 0.0017 | 0.0014 | 0.0015 |
| F05 | 59200001 | 59300000 | 0.0018 | 0.0015 | 0.0016 |
| F05 | 59220001 | 59320000 | 0.0022 | 0.0017 | 0.0018 |
| F05 | 59240001 | 59340000 | 0.0028 | 0.0016 | 0.0020 |
| F05 | 59260001 | 59360000 | 0.0028 | 0.0048 | 0.0018 |
| F05 | 59280001 | 59380000 | 0.0063 | 0.0065 | 0.0050 |
| F05 | 59300001 | 59400000 | 0.0083 | 0.0062 | 0.0070 |
| F05 | 59320001 | 59420000 | 0.0079 | 0.0066 | 0.0067 |
| F05 | 59340001 | 59440000 | 0.0081 | 0.0077 | 0.0072 |
| F05 | 59360001 | 59460000 | 0.0095 | 0.0049 | 0.0085 |
| F05 | 59380001 | 59480000 | 0.0066 | 0.0034 | 0.0056 |
| F05 | 59400001 | 59500000 | 0.0049 | 0.0039 | 0.0038 |
| F05 | 59420001 | 59520000 | 0.0054 | 0.0039 | 0.0043 |
| F05 | 59440001 | 59540000 | 0.0055 | 0.0027 | 0.0042 |
| F05 | 59460001 | 59560000 | 0.0040 | 0.0027 | 0.0028 |
| F05 | 59480001 | 59580000 | 0.0039 | 0.0027 | 0.0028 |
| F05 | 59500001 | 59600000 | 0.0038 | 0.0025 | 0.0028 |
| F05 | 59520001 | 59620000 | 0.0033 | 0.0020 | 0.0025 |
| F05 | 59540001 | 59640000 | 0.0027 | 0.0023 | 0.0021 |
| F05 | 59560001 | 59660000 | 0.0032 | 0.0022 | 0.0027 |
| F05 | 59580001 | 59680000 | 0.0032 | 0.0023 | 0.0028 |
| F05 | 59600001 | 59700000 | 0.0034 | 0.0035 | 0.0029 |
| F05 | 59620001 | 59720000 | 0.0047 | 0.0037 | 0.0040 |
| F05 | 59640001 | 59740000 | 0.0049 | 0.0037 | 0.0042 |
| F05 | 59660001 | 59760000 | 0.0047 | 0.0041 | 0.0039 |
| F05 | 59680001 | 59780000 | 0.0051 | 0.0044 | 0.0042 |
| F05 | 59700001 | 59800000 | 0.0055 | 0.0031 | 0.0045 |
| F05 | 59720001 | 59820000 | 0.0043 | 0.0027 | 0.0033 |
| F05 | 59740001 | 59840000 | 0.0038 | 0.0028 | 0.0029 |
| F05 | 59760001 | 59860000 | 0.0040 | 0.0025 | 0.0031 |
| F05 | 59780001 | 59880000 | 0.0037 | 0.0024 | 0.0029 |
| F05 | 59800001 | 59900000 | 0.0036 | 0.0028 | 0.0027 |
| F05 | 59820001 | 59920000 | 0.0040 | 0.0034 | 0.0030 |
| F05 | 59840001 | 59940000 | 0.0046 | 0.0048 | 0.0036 |
| F05 | 59860001 | 59960000 | 0.0063 | 0.0053 | 0.0054 |
| F05 | 59880001 | 59980000 | 0.0068 | 0.0047 | 0.0059 |
| F05 | 59900001 | 60000000 | 0.0060 | 0.0047 | 0.0053 |
| F05 | 59920001 | 60020000 | 0.0058 | 0.0041 | 0.0053 |
| F05 | 59940001 | 60040000 | 0.0055 | 0.0032 | 0.0050 |
| F05 | 59960001 | 60060000 | 0.0043 | 0.0032 | 0.0038 |
| F05 | 59980001 | 60080000 | 0.0044 | 0.0037 | 0.0039 |
| F05 | 60000001 | 60100000 | 0.0048 | 0.0032 | 0.0043 |

| Chr | Start | End | both | A1 | A2 |
|---|---|---|---|---|---|
| F05 | 60020001 | 60120000 | 0.0043 | 0.0035 | 0.0038 |
| F05 | 60040001 | 60140000 | 0.0042 | 0.0026 | 0.0038 |
| F05 | 60060001 | 60160000 | 0.0032 | 0.0023 | 0.0030 |
| F05 | 60080001 | 60180000 | 0.0029 | 0.0021 | 0.0024 |
| F05 | 60100001 | 60200000 | 0.0030 | 0.0020 | 0.0024 |
| F05 | 60120001 | 60220000 | 0.0030 | 0.0019 | 0.0022 |
| F05 | 60140001 | 60240000 | 0.0030 | 0.0016 | 0.0021 |
| F05 | 60160001 | 60260000 | 0.0025 | 0.0010 | 0.0017 |
| F05 | 60180001 | 60280000 | 0.0016 | 0.0012 | 0.0011 |
| F05 | 60200001 | 60300000 | 0.0016 | 0.0014 | 0.0014 |
| F05 | 60220001 | 60320000 | 0.0017 | 0.0012 | 0.0016 |
| F05 | 60240001 | 60340000 | 0.0014 | 0.0015 | 0.0013 |
| F05 | 60260001 | 60360000 | 0.0018 | 0.0015 | 0.0017 |
| F05 | 60280001 | 60380000 | 0.0018 | 0.0017 | 0.0017 |
| F05 | 60300001 | 60400000 | 0.0019 | 0.0021 | 0.0018 |
| F05 | 60320001 | 60420000 | 0.0025 | 0.0023 | 0.0022 |
| F05 | 60340001 | 60440000 | 0.0028 | 0.0027 | 0.0025 |
| F05 | 60360001 | 60460000 | 0.0031 | 0.0030 | 0.0028 |
| F05 | 60380001 | 60480000 | 0.0033 | 0.0028 | 0.0029 |
| F05 | 60400001 | 60500000 | 0.0030 | 0.0028 | 0.0027 |
| F05 | 60420001 | 60520000 | 0.0029 | 0.0037 | 0.0026 |
| F05 | 60440001 | 60540000 | 0.0039 | 0.0041 | 0.0036 |
| F05 | 60460001 | 60560000 | 0.0043 | 0.0055 | 0.0040 |
| F05 | 60480001 | 60580000 | 0.0060 | 0.0054 | 0.0055 |
| F05 | 60500001 | 60600000 | 0.0061 | 0.0055 | 0.0057 |
| F05 | 60520001 | 60620000 | 0.0065 | 0.0059 | 0.0061 |
| F05 | 60540001 | 60640000 | 0.0070 | 0.0052 | 0.0064 |
| F05 | 60560001 | 60660000 | 0.0062 | 0.0036 | 0.0058 |
| F05 | 60580001 | 60680000 | 0.0043 | 0.0034 | 0.0041 |
| F05 | 60600001 | 60700000 | 0.0040 | 0.0036 | 0.0038 |
| F05 | 60620001 | 60720000 | 0.0041 | 0.0021 | 0.0039 |
| F05 | 60640001 | 60740000 | 0.0026 | 0.0022 | 0.0024 |
| F05 | 60660001 | 60760000 | 0.0027 | 0.0025 | 0.0024 |
| F05 | 60680001 | 60780000 | 0.0032 | 0.0026 | 0.0028 |
| F05 | 60700001 | 60800000 | 0.0033 | 0.0021 | 0.0028 |
| F05 | 60720001 | 60820000 | 0.0027 | 0.0021 | 0.0022 |
| F05 | 60740001 | 60840000 | 0.0025 | 0.0023 | 0.0021 |
| F05 | 60760001 | 60860000 | 0.0028 | 0.0020 | 0.0025 |
| F05 | 60780001 | 60880000 | 0.0026 | 0.0021 | 0.0021 |
| F05 | 60800001 | 60900000 | 0.0030 | 0.0022 | 0.0026 |
| F05 | 60820001 | 60920000 | 0.0033 | 0.0029 | 0.0026 |
| F05 | 60840001 | 60940000 | 0.0040 | 0.0023 | 0.0034 |
| F05 | 60860001 | 60960000 | 0.0034 | 0.0024 | 0.0027 |
| F05 | 60880001 | 60980000 | 0.0035 | 0.0019 | 0.0029 |
| F05 | 60900001 | 61000000 | 0.0028 | 0.0016 | 0.0023 |
| F05 | 60920001 | 61020000 | 0.0024 | 0.0018 | 0.0019 |
| F05 | 60940001 | 61040000 | 0.0027 | 0.0023 | 0.0021 |
| F05 | 60960001 | 61060000 | 0.0032 | 0.0023 | 0.0026 |
| F05 | 60980001 | 61080000 | 0.0030 | 0.0023 | 0.0025 |
| F05 | 61000001 | 61100000 | 0.0030 | 0.0028 | 0.0025 |
| F05 | 61020001 | 61120000 | 0.0035 | 0.0022 | 0.0030 |
| F05 | 61040001 | 61140000 | 0.0028 | 0.0025 | 0.0025 |
| F05 | 61060001 | 61160000 | 0.0032 | 0.0022 | 0.0030 |
| F05 | 61080001 | 61180000 | 0.0029 | 0.0026 | 0.0027 |
| F05 | 61100001 | 61200000 | 0.0033 | 0.0021 | 0.0029 |
| F05 | 61120001 | 61220000 | 0.0027 | 0.0027 | 0.0023 |
| F05 | 61140001 | 61240000 | 0.0031 | 0.0027 | 0.0027 |
| F05 | 61160001 | 61260000 | 0.0030 | 0.0029 | 0.0025 |

| Chr | Start | End | both | A1 | A2 |
| --- | --- | --- | --- | --- | --- |
| F05 | 61180001 | 61280000 | 0.0033 | 0.0033 | 0.0027 |
| F05 | 61200001 | 61300000 | 0.0039 | 0.0032 | 0.0034 |
| F05 | 61220001 | 61320000 | 0.0038 | 0.0022 | 0.0034 |
| F05 | 61240001 | 61340000 | 0.0027 | 0.0015 | 0.0023 |
| F05 | 61260001 | 61360000 | 0.0018 | 0.0020 | 0.0016 |
| F05 | 61280001 | 61380000 | 0.0023 | 0.0013 | 0.0021 |
| F05 | 61300001 | 61400000 | 0.0015 | 0.0020 | 0.0013 |
| F05 | 61320001 | 61420000 | 0.0021 | 0.0025 | 0.0019 |
| F05 | 61340001 | 61440000 | 0.0028 | 0.0027 | 0.0024 |
| F05 | 61360001 | 61460000 | 0.0032 | 0.0029 | 0.0027 |
| F05 | 61380001 | 61480000 | 0.0035 | 0.0028 | 0.0030 |
| F05 | 61400001 | 61500000 | 0.0036 | 0.0025 | 0.0030 |
| F05 | 61420001 | 61520000 | 0.0035 | 0.0028 | 0.0027 |
| F05 | 61440001 | 61540000 | 0.0036 | 0.0026 | 0.0032 |
| F05 | 61460001 | 61560000 | 0.0032 | 0.0017 | 0.0029 |
| F05 | 61480001 | 61580000 | 0.0022 | 0.0020 | 0.0020 |
| F05 | 61500001 | 61600000 | 0.0024 | 0.0019 | 0.0020 |
| F05 | 61520001 | 61620000 | 0.0022 | 0.0019 | 0.0019 |
| F05 | 61540001 | 61640000 | 0.0023 | 0.0021 | 0.0021 |
| F05 | 61560001 | 61660000 | 0.0026 | 0.0024 | 0.0023 |
| F05 | 61580001 | 61680000 | 0.0032 | 0.0039 | 0.0026 |
| F05 | 61600001 | 61700000 | 0.0048 | 0.0048 | 0.0041 |
| F05 | 61620001 | 61720000 | 0.0057 | 0.0042 | 0.0050 |
| F05 | 61640001 | 61740000 | 0.0051 | 0.0043 | 0.0043 |
| F05 | 61660001 | 61760000 | 0.0053 | 0.0046 | 0.0044 |
| F05 | 61680001 | 61780000 | 0.0058 | 0.0041 | 0.0048 |
| F05 | 61700001 | 61800000 | 0.0051 | 0.0037 | 0.0042 |
| F05 | 61720001 | 61820000 | 0.0046 | 0.0042 | 0.0035 |
| F05 | 61740001 | 61840000 | 0.0052 | 0.0052 | 0.0040 |
| F05 | 61760001 | 61860000 | 0.0062 | 0.0053 | 0.0051 |
| F05 | 61780001 | 61880000 | 0.0060 | 0.0044 | 0.0052 |
| F05 | 61800001 | 61900000 | 0.0051 | 0.0037 | 0.0045 |
| F05 | 61820001 | 61920000 | 0.0045 | 0.0035 | 0.0042 |
| F05 | 61840001 | 61940000 | 0.0040 | 0.0021 | 0.0039 |
| F05 | 61860001 | 61960000 | 0.0024 | 0.0026 | 0.0024 |
| F05 | 61880001 | 61980000 | 0.0031 | 0.0023 | 0.0031 |
| F05 | 61900001 | 62000000 | 0.0027 | 0.0031 | 0.0027 |
| F05 | 61920001 | 62020000 | 0.0036 | 0.0033 | 0.0036 |
| F05 | 61940001 | 62040000 | 0.0038 | 0.0034 | 0.0037 |
| F05 | 61960001 | 62060000 | 0.0039 | 0.0024 | 0.0037 |
| F05 | 61980001 | 62080000 | 0.0030 | 0.0033 | 0.0026 |
| F05 | 62000001 | 62100000 | 0.0038 | 0.0028 | 0.0033 |
| F05 | 62020001 | 62120000 | 0.0038 | 0.0035 | 0.0029 |
| F05 | 62040001 | 62140000 | 0.0047 | 0.0049 | 0.0038 |
| F05 | 62060001 | 62160000 | 0.0063 | 0.0050 | 0.0055 |
| F05 | 62080001 | 62180000 | 0.0064 | 0.0041 | 0.0056 |
| F05 | 62100001 | 62200000 | 0.0054 | 0.0038 | 0.0048 |
| F05 | 62120001 | 62220000 | 0.0047 | 0.0029 | 0.0042 |
| F05 | 62140001 | 62240000 | 0.0039 | 0.0024 | 0.0032 |
| F05 | 62160001 | 62260000 | 0.0033 | 0.0023 | 0.0026 |
| F05 | 62180001 | 62280000 | 0.0031 | 0.0032 | 0.0026 |
| F05 | 62200001 | 62300000 | 0.0041 | 0.0031 | 0.0036 |
| F05 | 62220001 | 62320000 | 0.0040 | 0.0031 | 0.0035 |
| F05 | 62240001 | 62340000 | 0.0039 | 0.0028 | 0.0035 |
| F05 | 62260001 | 62360000 | 0.0035 | 0.0033 | 0.0032 |
| F05 | 62280001 | 62380000 | 0.0039 | 0.0029 | 0.0035 |
| F05 | 62300001 | 62400000 | 0.0035 | 0.0029 | 0.0032 |
| F05 | 62320001 | 62420000 | 0.0034 | 0.0025 | 0.0032 |

| Chr | Start | End | both | A1 | A2 |
|---|---|---|---|---|---|
| F05 | 62340001 | 62440000 | 0.0030 | 0.0023 | 0.0029 |
| F05 | 62360001 | 62460000 | 0.0029 | 0.0029 | 0.0027 |
| F05 | 62380001 | 62480000 | 0.0036 | 0.0023 | 0.0035 |
| F05 | 62400001 | 62500000 | 0.0029 | 0.0021 | 0.0028 |
| F05 | 62420001 | 62520000 | 0.0026 | 0.0022 | 0.0025 |
| F05 | 62440001 | 62540000 | 0.0028 | 0.0022 | 0.0026 |
| F05 | 62460001 | 62560000 | 0.0028 | 0.0010 | 0.0024 |
| F05 | 62480001 | 62580000 | 0.0016 | 0.0012 | 0.0010 |
| F05 | 62500001 | 62600000 | 0.0022 | 0.0013 | 0.0013 |
| F05 | 62520001 | 62620000 | 0.0024 | 0.0011 | 0.0014 |
| F05 | 62540001 | 62640000 | 0.0024 | 0.0016 | 0.0013 |
| F05 | 62560001 | 62660000 | 0.0027 | 0.0024 | 0.0017 |
| F05 | 62580001 | 62680000 | 0.0034 | 0.0024 | 0.0024 |
| F05 | 62600001 | 62700000 | 0.0031 | 0.0026 | 0.0023 |
| F05 | 62620001 | 62720000 | 0.0036 | 0.0043 | 0.0025 |
| F05 | 62640001 | 62740000 | 0.0049 | 0.0039 | 0.0041 |
| F05 | 62660001 | 62760000 | 0.0046 | 0.0031 | 0.0038 |
| F05 | 62680001 | 62780000 | 0.0037 | 0.0037 | 0.0031 |
| F05 | 62700001 | 62800000 | 0.0045 | 0.0036 | 0.0039 |
| F05 | 62720001 | 62820000 | 0.0042 | 0.0025 | 0.0038 |
| F05 | 62740001 | 62840000 | 0.0032 | 0.0022 | 0.0029 |
| F05 | 62760001 | 62860000 | 0.0027 | 0.0025 | 0.0025 |
| F05 | 62780001 | 62880000 | 0.0031 | 0.0020 | 0.0029 |
| F05 | 62800001 | 62900000 | 0.0025 | 0.0021 | 0.0024 |
| F05 | 62820001 | 62920000 | 0.0024 | 0.0022 | 0.0024 |
| F05 | 62840001 | 62940000 | 0.0027 | 0.0025 | 0.0027 |
| F05 | 62860001 | 62960000 | 0.0031 | 0.0028 | 0.0031 |
| F05 | 62880001 | 62980000 | 0.0034 | 0.0028 | 0.0034 |
| F05 | 62900001 | 63000000 | 0.0035 | 0.0032 | 0.0033 |
| F05 | 62920001 | 63020000 | 0.0040 | 0.0039 | 0.0036 |
| F05 | 62940001 | 63040000 | 0.0047 | 0.0038 | 0.0041 |
| F05 | 62960001 | 63060000 | 0.0045 | 0.0035 | 0.0039 |
| F05 | 62980001 | 63080000 | 0.0042 | 0.0031 | 0.0036 |
| F05 | 63000001 | 63100000 | 0.0037 | 0.0025 | 0.0032 |
| F05 | 63020001 | 63120000 | 0.0028 | 0.0013 | 0.0026 |
| F05 | 63040001 | 63140000 | 0.0016 | 0.0014 | 0.0016 |
| F05 | 63060001 | 63160000 | 0.0016 | 0.0015 | 0.0015 |
| F05 | 63080001 | 63180000 | 0.0020 | 0.0028 | 0.0017 |
| F05 | 63100001 | 63200000 | 0.0035 | 0.0036 | 0.0030 |
| F05 | 63120001 | 63220000 | 0.0047 | 0.0041 | 0.0039 |
| F05 | 63140001 | 63240000 | 0.0054 | 0.0041 | 0.0045 |
| F05 | 63160001 | 63260000 | 0.0055 | 0.0038 | 0.0043 |
| F05 | 63180001 | 63280000 | 0.0050 | 0.0031 | 0.0039 |
| F05 | 63200001 | 63300000 | 0.0044 | 0.0027 | 0.0034 |
| F05 | 63220001 | 63320000 | 0.0039 | 0.0025 | 0.0029 |
| F05 | 63240001 | 63340000 | 0.0039 | 0.0030 | 0.0030 |
| F05 | 63260001 | 63360000 | 0.0042 | 0.0028 | 0.0035 |
| F05 | 63280001 | 63380000 | 0.0040 | 0.0025 | 0.0033 |
| F05 | 63300001 | 63400000 | 0.0035 | 0.0023 | 0.0029 |
| F05 | 63320001 | 63420000 | 0.0032 | 0.0021 | 0.0025 |
| F05 | 63340001 | 63440000 | 0.0026 | 0.0020 | 0.0020 |
| F05 | 63360001 | 63460000 | 0.0026 | 0.0022 | 0.0021 |
| F05 | 63380001 | 63480000 | 0.0028 | 0.0028 | 0.0023 |
| F05 | 63400001 | 63500000 | 0.0036 | 0.0027 | 0.0032 |
| F05 | 63420001 | 63520000 | 0.0033 | 0.0023 | 0.0031 |
| F05 | 63440001 | 63540000 | 0.0029 | 0.0017 | 0.0027 |
| F05 | 63460001 | 63560000 | 0.0023 | 0.0015 | 0.0021 |
| F05 | 63480001 | 63580000 | 0.0022 | 0.0006 | 0.0019 |

| Chr | Start | End | both | A1 | A2 |
|---|---|---|---|---|---|
| F05 | 63500001 | 63600000 | 0.0011 | 0.0008 | 0.0007 |
| F05 | 63520001 | 63620000 | 0.0013 | 0.0010 | 0.0009 |
| F05 | 63540001 | 63640000 | 0.0016 | 0.0013 | 0.0012 |
| F05 | 63560001 | 63660000 | 0.0022 | 0.0013 | 0.0015 |
| F05 | 63580001 | 63680000 | 0.0021 | 0.0026 | 0.0014 |
| F05 | 63600001 | 63700000 | 0.0037 | 0.0031 | 0.0030 |
| F05 | 63620001 | 63720000 | 0.0043 | 0.0031 | 0.0034 |
| F05 | 63640001 | 63740000 | 0.0041 | 0.0028 | 0.0034 |
| F05 | 63660001 | 63760000 | 0.0037 | 0.0030 | 0.0031 |
| F05 | 63680001 | 63780000 | 0.0039 | 0.0018 | 0.0032 |
| F05 | 63700001 | 63800000 | 0.0026 | 0.0012 | 0.0019 |
| F05 | 63720001 | 63820000 | 0.0020 | 0.0009 | 0.0012 |
| F05 | 63740001 | 63840000 | 0.0018 | 0.0009 | 0.0009 |
| F05 | 63760001 | 63860000 | 0.0019 | 0.0007 | 0.0010 |
| F05 | 63780001 | 63880000 | 0.0017 | 0.0005 | 0.0009 |
| F05 | 63800001 | 63900000 | 0.0013 | 0.0010 | 0.0006 |
| F05 | 63820001 | 63920000 | 0.0019 | 0.0011 | 0.0012 |
| F05 | 63840001 | 63940000 | 0.0020 | 0.0010 | 0.0013 |
| F05 | 63860001 | 63960000 | 0.0018 | 0.0013 | 0.0012 |
| F05 | 63880001 | 63980000 | 0.0019 | 0.0016 | 0.0013 |
| F05 | 63900001 | 64000000 | 0.0022 | 0.0012 | 0.0016 |
| F05 | 63920001 | 64020000 | 0.0017 | 0.0017 | 0.0012 |
| F05 | 63940001 | 64040000 | 0.0023 | 0.0022 | 0.0018 |
| F05 | 63960001 | 64060000 | 0.0033 | 0.0021 | 0.0026 |
| F05 | 63980001 | 64080000 | 0.0034 | 0.0023 | 0.0025 |
| F05 | 64000001 | 64100000 | 0.0037 | 0.0025 | 0.0029 |
| F05 | 64020001 | 64120000 | 0.0042 | 0.0020 | 0.0033 |
| F05 | 64040001 | 64140000 | 0.0035 | 0.0014 | 0.0027 |
| F05 | 64060001 | 64160000 | 0.0022 | 0.0014 | 0.0017 |
| F05 | 64080001 | 64180000 | 0.0022 | 0.0011 | 0.0019 |
| F05 | 64100001 | 64200000 | 0.0022 | 0.0009 | 0.0016 |
| F05 | 64120001 | 64220000 | 0.0018 | 0.0008 | 0.0011 |
| F05 | 64140001 | 64240000 | 0.0019 | 0.0009 | 0.0011 |
| F05 | 64160001 | 64260000 | 0.0022 | 0.0009 | 0.0014 |
| F05 | 64180001 | 64280000 | 0.0022 | 0.0011 | 0.0013 |
| F05 | 64200001 | 64300000 | 0.0021 | 0.0016 | 0.0014 |
| F05 | 64220001 | 64320000 | 0.0023 | 0.0017 | 0.0018 |
| F05 | 64240001 | 64340000 | 0.0023 | 0.0019 | 0.0019 |
| F05 | 64260001 | 64360000 | 0.0026 | 0.0020 | 0.0020 |
| F05 | 64280001 | 64380000 | 0.0030 | 0.0020 | 0.0022 |
| F05 | 64300001 | 64400000 | 0.0033 | 0.0015 | 0.0022 |
| F05 | 64320001 | 64420000 | 0.0032 | 0.0017 | 0.0018 |
| F05 | 64340001 | 64440000 | 0.0036 | 0.0021 | 0.0020 |
| F05 | 64360001 | 64460000 | 0.0038 | 0.0025 | 0.0022 |
| F05 | 64380001 | 64480000 | 0.0039 | 0.0025 | 0.0026 |
| F05 | 64400001 | 64500000 | 0.0037 | 0.0027 | 0.0027 |
| F05 | 64420001 | 64520000 | 0.0037 | 0.0024 | 0.0030 |
| F05 | 64440001 | 64540000 | 0.0031 | 0.0024 | 0.0027 |
| F05 | 64460001 | 64560000 | 0.0034 | 0.0025 | 0.0028 |
| F05 | 64480001 | 64580000 | 0.0035 | 0.0024 | 0.0028 |
| F05 | 64500001 | 64600000 | 0.0036 | 0.0022 | 0.0027 |
| F05 | 64520001 | 64620000 | 0.0035 | 0.0026 | 0.0025 |
| F05 | 64540001 | 64640000 | 0.0042 | 0.0033 | 0.0029 |
| F05 | 64560001 | 64660000 | 0.0048 | 0.0030 | 0.0035 |
| F05 | 64580001 | 64680000 | 0.0045 | 0.0033 | 0.0032 |
| F05 | 64600001 | 64700000 | 0.0049 | 0.0044 | 0.0037 |
| F05 | 64620001 | 64720000 | 0.0062 | 0.0046 | 0.0048 |
| F05 | 64640001 | 64740000 | 0.0064 | 0.0042 | 0.0050 |

| Chr | Start | End | both | A1 | A2 |
|---|---|---|---|---|---|
| F05 | 64660001 | 64760000 | 0.0061 | 0.0038 | 0.0048 |
| F05 | 64680001 | 64780000 | 0.0056 | 0.0044 | 0.0043 |
| F05 | 64700001 | 64800000 | 0.0059 | 0.0035 | 0.0046 |
| F05 | 64720001 | 64820000 | 0.0047 | 0.0035 | 0.0037 |
| F05 | 64740001 | 64840000 | 0.0044 | 0.0031 | 0.0036 |
| F05 | 64760001 | 64860000 | 0.0035 | 0.0030 | 0.0029 |
| F05 | 64780001 | 64880000 | 0.0036 | 0.0022 | 0.0031 |
| F05 | 64800001 | 64900000 | 0.0027 | 0.0020 | 0.0023 |
| F05 | 64820001 | 64920000 | 0.0030 | 0.0019 | 0.0024 |
| F05 | 64840001 | 64940000 | 0.0029 | 0.0027 | 0.0024 |
| F05 | 64860001 | 64960000 | 0.0037 | 0.0040 | 0.0033 |
| F05 | 64880001 | 64980000 | 0.0050 | 0.0039 | 0.0043 |
| F05 | 64900001 | 65000000 | 0.0051 | 0.0037 | 0.0042 |
| F05 | 64920001 | 65020000 | 0.0046 | 0.0041 | 0.0039 |
| F05 | 64940001 | 65040000 | 0.0048 | 0.0035 | 0.0041 |
| F05 | 64960001 | 65060000 | 0.0039 | 0.0024 | 0.0031 |
| F05 | 64980001 | 65080000 | 0.0027 | 0.0027 | 0.0022 |
| F05 | 65000001 | 65100000 | 0.0028 | 0.0030 | 0.0024 |
| F05 | 65020001 | 65120000 | 0.0030 | 0.0036 | 0.0026 |
| F05 | 65040001 | 65140000 | 0.0039 | 0.0035 | 0.0037 |
| F05 | 65060001 | 65160000 | 0.0039 | 0.0035 | 0.0037 |
| F05 | 65080001 | 65180000 | 0.0039 | 0.0033 | 0.0037 |
| F05 | 65100001 | 65200000 | 0.0038 | 0.0029 | 0.0034 |
| F05 | 65120001 | 65220000 | 0.0034 | 0.0017 | 0.0031 |
| F05 | 65140001 | 65240000 | 0.0020 | 0.0022 | 0.0017 |
| F05 | 65160001 | 65260000 | 0.0026 | 0.0032 | 0.0022 |
| F05 | 65180001 | 65280000 | 0.0037 | 0.0032 | 0.0032 |
| F05 | 65200001 | 65300000 | 0.0035 | 0.0035 | 0.0032 |
| F05 | 65220001 | 65320000 | 0.0039 | 0.0035 | 0.0036 |
| F05 | 65240001 | 65340000 | 0.0039 | 0.0030 | 0.0036 |
| F05 | 65260001 | 65360000 | 0.0033 | 0.0018 | 0.0031 |
| F05 | 65280001 | 65380000 | 0.0020 | 0.0016 | 0.0018 |
| F05 | 65300001 | 65400000 | 0.0019 | 0.0013 | 0.0016 |
| F05 | 65320001 | 65420000 | 0.0015 | 0.0016 | 0.0013 |
| F05 | 65340001 | 65440000 | 0.0019 | 0.0015 | 0.0016 |
| F05 | 65360001 | 65460000 | 0.0017 | 0.0020 | 0.0013 |
| F05 | 65380001 | 65480000 | 0.0022 | 0.0030 | 0.0018 |
| F05 | 65400001 | 65500000 | 0.0034 | 0.0032 | 0.0028 |
| F05 | 65420001 | 65520000 | 0.0037 | 0.0033 | 0.0030 |
| F05 | 65440001 | 65540000 | 0.0038 | 0.0039 | 0.0032 |
| F05 | 65460001 | 65560000 | 0.0049 | 0.0036 | 0.0042 |
| F05 | 65480001 | 65580000 | 0.0045 | 0.0032 | 0.0039 |
| F05 | 65500001 | 65600000 | 0.0041 | 0.0036 | 0.0035 |
| F05 | 65520001 | 65620000 | 0.0046 | 0.0049 | 0.0041 |
| F05 | 65540001 | 65640000 | 0.0057 | 0.0040 | 0.0051 |
| F05 | 65560001 | 65660000 | 0.0046 | 0.0038 | 0.0042 |
| F05 | 65580001 | 65680000 | 0.0043 | 0.0031 | 0.0040 |
| F05 | 65600001 | 65700000 | 0.0034 | 0.0024 | 0.0033 |
| F05 | 65620001 | 65720000 | 0.0028 | 0.0012 | 0.0024 |
| F05 | 65640001 | 65740000 | 0.0018 | 0.0012 | 0.0010 |
| F05 | 65660001 | 65760000 | 0.0021 | 0.0022 | 0.0011 |
| F05 | 65680001 | 65780000 | 0.0032 | 0.0031 | 0.0022 |
| F05 | 65700001 | 65800000 | 0.0042 | 0.0036 | 0.0029 |
| F05 | 65720001 | 65820000 | 0.0045 | 0.0033 | 0.0032 |
| F05 | 65740001 | 65840000 | 0.0040 | 0.0032 | 0.0032 |
| F05 | 65760001 | 65860000 | 0.0037 | 0.0023 | 0.0031 |
| F05 | 65780001 | 65880000 | 0.0028 | 0.0012 | 0.0020 |
| F05 | 65800001 | 65900000 | 0.0019 | 0.0008 | 0.0012 |

| Chr | Start | End | both | A1 | A2 |
|---|---|---|---|---|---|
| F05 | 65820001 | 65920000 | 0.0014 | 0.0004 | 0.0008 |
| F05 | 65840001 | 65940000 | 0.0010 | 0.0006 | 0.0004 |
| F05 | 65860001 | 65960000 | 0.0012 | 0.0010 | 0.0006 |
| F05 | 65880001 | 65980000 | 0.0016 | 0.0016 | 0.0011 |
| F05 | 65900001 | 66000000 | 0.0021 | 0.0019 | 0.0017 |
| F05 | 65920001 | 66020000 | 0.0028 | 0.0031 | 0.0022 |
| F05 | 65940001 | 66040000 | 0.0039 | 0.0035 | 0.0033 |
| F05 | 65960001 | 66060000 | 0.0043 | 0.0035 | 0.0036 |
| F05 | 65980001 | 66080000 | 0.0045 | 0.0038 | 0.0035 |
| F05 | 66000001 | 66100000 | 0.0048 | 0.0034 | 0.0036 |
| F05 | 66020001 | 66120000 | 0.0041 | 0.0031 | 0.0032 |
| F05 | 66040001 | 66140000 | 0.0038 | 0.0029 | 0.0030 |
| F05 | 66060001 | 66160000 | 0.0036 | 0.0026 | 0.0029 |
| F05 | 66080001 | 66180000 | 0.0031 | 0.0019 | 0.0027 |
| F05 | 66100001 | 66200000 | 0.0022 | 0.0022 | 0.0019 |
| F05 | 66120001 | 66220000 | 0.0025 | 0.0026 | 0.0021 |
| F05 | 66140001 | 66240000 | 0.0028 | 0.0033 | 0.0023 |
| F05 | 66160001 | 66260000 | 0.0035 | 0.0035 | 0.0030 |
| F05 | 66180001 | 66280000 | 0.0038 | 0.0044 | 0.0033 |
| F05 | 66200001 | 66300000 | 0.0047 | 0.0048 | 0.0042 |
| F05 | 66220001 | 66320000 | 0.0051 | 0.0043 | 0.0045 |
| F05 | 66240001 | 66340000 | 0.0048 | 0.0053 | 0.0042 |
| F05 | 66260001 | 66360000 | 0.0056 | 0.0062 | 0.0050 |
| F05 | 66280001 | 66380000 | 0.0066 | 0.0057 | 0.0061 |
| F05 | 66300001 | 66400000 | 0.0062 | 0.0051 | 0.0058 |
| F05 | 66320001 | 66420000 | 0.0059 | 0.0045 | 0.0055 |
| F05 | 66340001 | 66440000 | 0.0054 | 0.0031 | 0.0048 |
| F05 | 66360001 | 66460000 | 0.0041 | 0.0017 | 0.0035 |
| F05 | 66380001 | 66480000 | 0.0025 | 0.0014 | 0.0020 |
| F05 | 66400001 | 66500000 | 0.0021 | 0.0011 | 0.0017 |
| F05 | 66420001 | 66520000 | 0.0016 | 0.0008 | 0.0014 |
| F05 | 66440001 | 66540000 | 0.0012 | 0.0008 | 0.0011 |
| F05 | 66460001 | 66560000 | 0.0010 | 0.0011 | 0.0010 |
| F05 | 66480001 | 66580000 | 0.0013 | 0.0009 | 0.0012 |
| F05 | 66500001 | 66600000 | 0.0012 | 0.0014 | 0.0010 |
| F05 | 66520001 | 66620000 | 0.0021 | 0.0014 | 0.0014 |
| F05 | 66540001 | 66640000 | 0.0023 | 0.0012 | 0.0015 |
| F05 | 66560001 | 66660000 | 0.0022 | 0.0017 | 0.0013 |
| F05 | 66580001 | 66680000 | 0.0029 | 0.0020 | 0.0018 |
| F05 | 66600001 | 66700000 | 0.0032 | 0.0018 | 0.0021 |
| F05 | 66620001 | 66720000 | 0.0026 | 0.0023 | 0.0020 |
| F05 | 66640001 | 66740000 | 0.0031 | 0.0020 | 0.0026 |
| F05 | 66660001 | 66760000 | 0.0027 | 0.0031 | 0.0022 |
| F05 | 66680001 | 66780000 | 0.0039 | 0.0034 | 0.0035 |
| F05 | 66700001 | 66800000 | 0.0042 | 0.0037 | 0.0039 |
| F05 | 66720001 | 66820000 | 0.0044 | 0.0032 | 0.0040 |
| F05 | 66740001 | 66840000 | 0.0041 | 0.0035 | 0.0035 |
| F05 | 66760001 | 66860000 | 0.0043 | 0.0029 | 0.0038 |
| F05 | 66780001 | 66880000 | 0.0037 | 0.0034 | 0.0030 |
| F05 | 66800001 | 66900000 | 0.0043 | 0.0034 | 0.0035 |
| F05 | 66820001 | 66920000 | 0.0044 | 0.0038 | 0.0035 |
| F05 | 66840001 | 66940000 | 0.0046 | 0.0037 | 0.0039 |
| F05 | 66860001 | 66960000 | 0.0047 | 0.0031 | 0.0039 |
| F05 | 66880001 | 66980000 | 0.0041 | 0.0024 | 0.0033 |
| F05 | 66900001 | 67000000 | 0.0031 | 0.0019 | 0.0024 |
| F05 | 66920001 | 67020000 | 0.0029 | 0.0020 | 0.0020 |
| F05 | 66940001 | 67040000 | 0.0032 | 0.0026 | 0.0024 |
| F05 | 66960001 | 67060000 | 0.0035 | 0.0028 | 0.0027 |

| Chr | Start | End | both | A1 | A2 |
|---|---|---|---|---|---|
| F05 | 66980001 | 67080000 | 0.0038 | 0.0028 | 0.0030 |
| F05 | 67000001 | 67100000 | 0.0039 | 0.0030 | 0.0029 |
| F05 | 67020001 | 67120000 | 0.0040 | 0.0028 | 0.0031 |
| F05 | 67040001 | 67140000 | 0.0038 | 0.0024 | 0.0028 |
| F05 | 67060001 | 67160000 | 0.0035 | 0.0022 | 0.0024 |
| F05 | 67080001 | 67180000 | 0.0032 | 0.0032 | 0.0020 |
| F05 | 67100001 | 67200000 | 0.0034 | 0.0037 | 0.0021 |
| F05 | 67120001 | 67220000 | 0.0036 | 0.0034 | 0.0023 |
| F05 | 67140001 | 67240000 | 0.0034 | 0.0034 | 0.0019 |
| F05 | 67160001 | 67260000 | 0.0036 | 0.0033 | 0.0021 |
| F05 | 67180001 | 67280000 | 0.0037 | 0.0026 | 0.0022 |
| F05 | 67200001 | 67300000 | 0.0036 | 0.0030 | 0.0023 |
| F05 | 67220001 | 67320000 | 0.0043 | 0.0035 | 0.0033 |
| F05 | 67240001 | 67340000 | 0.0049 | 0.0037 | 0.0038 |
| F05 | 67260001 | 67360000 | 0.0050 | 0.0045 | 0.0041 |
| F05 | 67280001 | 67380000 | 0.0058 | 0.0047 | 0.0050 |
| F05 | 67300001 | 67400000 | 0.0060 | 0.0047 | 0.0054 |
| F05 | 67320001 | 67420000 | 0.0060 | 0.0054 | 0.0054 |
| F05 | 67340001 | 67440000 | 0.0065 | 0.0055 | 0.0062 |
| F05 | 67360001 | 67460000 | 0.0064 | 0.0042 | 0.0062 |
| F05 | 67380001 | 67480000 | 0.0049 | 0.0038 | 0.0048 |
| F05 | 67400001 | 67500000 | 0.0044 | 0.0032 | 0.0042 |
| F05 | 67420001 | 67520000 | 0.0037 | 0.0042 | 0.0034 |
| F05 | 67440001 | 67540000 | 0.0045 | 0.0035 | 0.0043 |
| F05 | 67460001 | 67560000 | 0.0038 | 0.0036 | 0.0036 |
| F05 | 67480001 | 67580000 | 0.0038 | 0.0047 | 0.0036 |
| F05 | 67500001 | 67600000 | 0.0053 | 0.0046 | 0.0049 |
| F05 | 67520001 | 67620000 | 0.0054 | 0.0031 | 0.0050 |
| F05 | 67540001 | 67640000 | 0.0043 | 0.0031 | 0.0036 |
| F05 | 67560001 | 67660000 | 0.0046 | 0.0032 | 0.0038 |
| F05 | 67580001 | 67680000 | 0.0051 | 0.0019 | 0.0040 |
| F05 | 67600001 | 67700000 | 0.0040 | 0.0013 | 0.0029 |
| F05 | 67620001 | 67720000 | 0.0032 | 0.0008 | 0.0021 |
| F05 | 67640001 | 67740000 | 0.0024 | 0.0007 | 0.0014 |
| F05 | 67660001 | 67760000 | 0.0021 | 0.0007 | 0.0012 |
| F05 | 67680001 | 67780000 | 0.0018 | 0.0006 | 0.0009 |
| F05 | 67700001 | 67800000 | 0.0012 | 0.0009 | 0.0006 |
| F05 | 67720001 | 67820000 | 0.0017 | 0.0020 | 0.0010 |
| F05 | 67740001 | 67840000 | 0.0029 | 0.0023 | 0.0021 |
| F05 | 67760001 | 67860000 | 0.0037 | 0.0025 | 0.0026 |
| F05 | 67780001 | 67880000 | 0.0039 | 0.0029 | 0.0028 |
| F05 | 67800001 | 67900000 | 0.0043 | 0.0030 | 0.0031 |
| F05 | 67820001 | 67920000 | 0.0042 | 0.0024 | 0.0028 |
| F05 | 67840001 | 67940000 | 0.0038 | 0.0028 | 0.0025 |
| F05 | 67860001 | 67960000 | 0.0043 | 0.0044 | 0.0032 |
| F05 | 67880001 | 67980000 | 0.0063 | 0.0048 | 0.0050 |
| F05 | 67900001 | 68000000 | 0.0068 | 0.0054 | 0.0052 |
| F05 | 67920001 | 68020000 | 0.0073 | 0.0050 | 0.0059 |
| F05 | 67940001 | 68040000 | 0.0066 | 0.0043 | 0.0052 |
| F05 | 67960001 | 68060000 | 0.0051 | 0.0023 | 0.0040 |
| F05 | 67980001 | 68080000 | 0.0025 | 0.0021 | 0.0018 |
| F05 | 68000001 | 68100000 | 0.0021 | 0.0015 | 0.0017 |
| F05 | 68020001 | 68120000 | 0.0014 | 0.0020 | 0.0013 |
| F05 | 68040001 | 68140000 | 0.0018 | 0.0023 | 0.0016 |
| F05 | 68060001 | 68160000 | 0.0021 | 0.0028 | 0.0019 |
| F05 | 68080001 | 68180000 | 0.0028 | 0.0035 | 0.0026 |
| F05 | 68100001 | 68200000 | 0.0039 | 0.0082 | 0.0038 |
| F05 | 68120001 | 68220000 | 0.0086 | 0.0106 | 0.0082 |

| Chr | Start | End | both | A1 | A2 |
|---|---|---|---|---|---|
| F05 | 68140001 | 68240000 | 0.0112 | 0.0130 | 0.0106 |
| F05 | 68160001 | 68260000 | 0.0135 | 0.0144 | 0.0126 |
| F05 | 68180001 | 68280000 | 0.0147 | 0.0139 | 0.0136 |
| F05 | 68200001 | 68300000 | 0.0140 | 0.0118 | 0.0128 |
| F05 | 68220001 | 68320000 | 0.0120 | 0.0101 | 0.0108 |
| F05 | 68240001 | 68340000 | 0.0102 | 0.0085 | 0.0092 |
| F05 | 68260001 | 68360000 | 0.0088 | 0.0082 | 0.0079 |
| F05 | 68280001 | 68380000 | 0.0087 | 0.0082 | 0.0076 |
| F05 | 68300001 | 68400000 | 0.0088 | 0.0054 | 0.0074 |
| F05 | 68320001 | 68420000 | 0.0059 | 0.0042 | 0.0046 |
| F05 | 68340001 | 68440000 | 0.0046 | 0.0033 | 0.0032 |
| F05 | 68360001 | 68460000 | 0.0036 | 0.0021 | 0.0023 |
| F05 | 68380001 | 68480000 | 0.0022 | 0.0017 | 0.0012 |
| F05 | 68400001 | 68500000 | 0.0017 | 0.0019 | 0.0009 |
| F05 | 68420001 | 68520000 | 0.0020 | 0.0020 | 0.0011 |
| F05 | 68440001 | 68540000 | 0.0021 | 0.0021 | 0.0012 |
| F05 | 68460001 | 68560000 | 0.0021 | 0.0019 | 0.0012 |
| F05 | 68480001 | 68580000 | 0.0018 | 0.0018 | 0.0011 |
| F05 | 68500001 | 68600000 | 0.0019 | 0.0021 | 0.0013 |
| F05 | 68520001 | 68620000 | 0.0021 | 0.0030 | 0.0017 |
| F05 | 68540001 | 68640000 | 0.0031 | 0.0031 | 0.0029 |
| F05 | 68560001 | 68660000 | 0.0034 | 0.0033 | 0.0031 |
| F05 | 68580001 | 68680000 | 0.0039 | 0.0049 | 0.0036 |
| F05 | 68600001 | 68700000 | 0.0061 | 0.0054 | 0.0054 |
| F05 | 68620001 | 68720000 | 0.0070 | 0.0062 | 0.0058 |
| F05 | 68640001 | 68740000 | 0.0080 | 0.0073 | 0.0059 |
| F05 | 68660001 | 68760000 | 0.0090 | 0.0085 | 0.0067 |
| F05 | 68680001 | 68780000 | 0.0102 | 0.0079 | 0.0080 |
| F05 | 68700001 | 68800000 | 0.0089 | 0.0080 | 0.0069 |
| F05 | 68720001 | 68820000 | 0.0087 | 0.0084 | 0.0071 |
| F05 | 68740001 | 68840000 | 0.0091 | 0.0082 | 0.0081 |
| F05 | 68760001 | 68860000 | 0.0090 | 0.0086 | 0.0079 |
| F05 | 68780001 | 68880000 | 0.0090 | 0.0084 | 0.0078 |
| F05 | 68800001 | 68900000 | 0.0091 | 0.0083 | 0.0080 |
| F05 | 68820001 | 68920000 | 0.0087 | 0.0060 | 0.0073 |
| F05 | 68840001 | 68940000 | 0.0061 | 0.0049 | 0.0050 |
| F05 | 68860001 | 68960000 | 0.0055 | 0.0033 | 0.0042 |
| F05 | 68880001 | 68980000 | 0.0043 | 0.0031 | 0.0025 |
| F05 | 68900001 | 69000000 | 0.0043 | 0.0032 | 0.0019 |
| F05 | 68920001 | 69020000 | 0.0045 | 0.0040 | 0.0024 |
| F05 | 68940001 | 69040000 | 0.0053 | 0.0044 | 0.0032 |
| F05 | 68960001 | 69060000 | 0.0056 | 0.0046 | 0.0038 |
| F05 | 68980001 | 69080000 | 0.0055 | 0.0045 | 0.0043 |
| F05 | 69000001 | 69100000 | 0.0049 | 0.0047 | 0.0041 |
| F05 | 69020001 | 69120000 | 0.0050 | 0.0043 | 0.0041 |
| F05 | 69040001 | 69140000 | 0.0046 | 0.0039 | 0.0035 |
| F05 | 69060001 | 69160000 | 0.0042 | 0.0035 | 0.0031 |
| F05 | 69080001 | 69180000 | 0.0038 | 0.0028 | 0.0028 |
| F05 | 69100001 | 69200000 | 0.0034 | 0.0015 | 0.0024 |
| F05 | 69120001 | 69220000 | 0.0022 | 0.0031 | 0.0014 |
| F05 | 69140001 | 69240000 | 0.0041 | 0.0033 | 0.0032 |
| F05 | 69160001 | 69260000 | 0.0047 | 0.0038 | 0.0037 |
| F05 | 69180001 | 69280000 | 0.0051 | 0.0038 | 0.0042 |
| F05 | 69200001 | 69300000 | 0.0048 | 0.0036 | 0.0041 |
| F05 | 69220001 | 69320000 | 0.0044 | 0.0019 | 0.0040 |
| F05 | 69240001 | 69340000 | 0.0023 | 0.0027 | 0.0020 |
| F05 | 69260001 | 69360000 | 0.0032 | 0.0022 | 0.0027 |
| F05 | 69280001 | 69380000 | 0.0027 | 0.0023 | 0.0022 |

| Chr | Start | End | both | A1 | A2 |
|---|---|---|---|---|---|
| F05 | 69300001 | 69400000 | 0.0028 | 0.0029 | 0.0023 |
| F05 | 69320001 | 69420000 | 0.0035 | 0.0028 | 0.0030 |
| F05 | 69340001 | 69440000 | 0.0039 | 0.0025 | 0.0030 |
| F05 | 69360001 | 69460000 | 0.0033 | 0.0049 | 0.0024 |
| F05 | 69380001 | 69480000 | 0.0059 | 0.0066 | 0.0048 |
| F05 | 69400001 | 69500000 | 0.0074 | 0.0061 | 0.0061 |
| F05 | 69420001 | 69520000 | 0.0069 | 0.0070 | 0.0058 |
| F05 | 69440001 | 69540000 | 0.0077 | 0.0077 | 0.0070 |
| F05 | 69460001 | 69560000 | 0.0083 | 0.0069 | 0.0078 |
| F05 | 69480001 | 69580000 | 0.0073 | 0.0055 | 0.0068 |
| F05 | 69500001 | 69600000 | 0.0061 | 0.0066 | 0.0058 |
| F05 | 69520001 | 69620000 | 0.0070 | 0.0069 | 0.0066 |
| F05 | 69540001 | 69640000 | 0.0072 | 0.0057 | 0.0068 |
| F05 | 69560001 | 69660000 | 0.0059 | 0.0053 | 0.0055 |
| F05 | 69580001 | 69680000 | 0.0057 | 0.0064 | 0.0053 |
| F05 | 69600001 | 69700000 | 0.0071 | 0.0059 | 0.0067 |
| F05 | 69620001 | 69720000 | 0.0068 | 0.0062 | 0.0065 |
| F05 | 69640001 | 69740000 | 0.0071 | 0.0058 | 0.0067 |
| F05 | 69660001 | 69760000 | 0.0069 | 0.0046 | 0.0066 |
| F05 | 69680001 | 69780000 | 0.0054 | 0.0034 | 0.0051 |
| F05 | 69700001 | 69800000 | 0.0040 | 0.0040 | 0.0036 |
| F05 | 69720001 | 69820000 | 0.0042 | 0.0046 | 0.0037 |
| F05 | 69740001 | 69840000 | 0.0045 | 0.0060 | 0.0039 |
| F05 | 69760001 | 69860000 | 0.0060 | 0.0063 | 0.0051 |
| F05 | 69780001 | 69880000 | 0.0063 | 0.0061 | 0.0053 |
| F05 | 69800001 | 69900000 | 0.0060 | 0.0051 | 0.0052 |
| F05 | 69820001 | 69920000 | 0.0052 | 0.0037 | 0.0046 |
| F05 | 69840001 | 69940000 | 0.0039 | 0.0040 | 0.0034 |
| F05 | 69860001 | 69960000 | 0.0037 | 0.0038 | 0.0031 |
| F05 | 69880001 | 69980000 | 0.0037 | 0.0042 | 0.0028 |
| F05 | 69900001 | 70000000 | 0.0045 | 0.0048 | 0.0035 |
| F05 | 69920001 | 70020000 | 0.0056 | 0.0042 | 0.0042 |
| F05 | 69940001 | 70040000 | 0.0053 | 0.0023 | 0.0037 |
| F05 | 69960001 | 70060000 | 0.0040 | 0.0025 | 0.0025 |
| F05 | 69980001 | 70080000 | 0.0044 | 0.0031 | 0.0027 |
| F05 | 70000001 | 70100000 | 0.0048 | 0.0028 | 0.0030 |
| F05 | 70020001 | 70120000 | 0.0040 | 0.0035 | 0.0026 |
| F05 | 70040001 | 70140000 | 0.0052 | 0.0038 | 0.0039 |
| F05 | 70060001 | 70160000 | 0.0055 | 0.0039 | 0.0045 |
| F05 | 70080001 | 70180000 | 0.0055 | 0.0030 | 0.0047 |
| F05 | 70100001 | 70200000 | 0.0051 | 0.0030 | 0.0037 |
| F05 | 70120001 | 70220000 | 0.0051 | 0.0021 | 0.0038 |
| F05 | 70140001 | 70240000 | 0.0037 | 0.0021 | 0.0022 |
| F05 | 70160001 | 70260000 | 0.0039 | 0.0022 | 0.0017 |
| F05 | 70180001 | 70280000 | 0.0039 | 0.0021 | 0.0017 |
| F05 | 70200001 | 70300000 | 0.0034 | 0.0019 | 0.0016 |
| F05 | 70220001 | 70320000 | 0.0033 | 0.0028 | 0.0011 |
| F05 | 70240001 | 70340000 | 0.0043 | 0.0030 | 0.0022 |
| F05 | 70260001 | 70360000 | 0.0041 | 0.0026 | 0.0026 |
| F05 | 70280001 | 70380000 | 0.0035 | 0.0027 | 0.0023 |
| F05 | 70300001 | 70400000 | 0.0033 | 0.0029 | 0.0024 |
| F05 | 70320001 | 70420000 | 0.0034 | 0.0021 | 0.0028 |
| F05 | 70340001 | 70440000 | 0.0023 | 0.0019 | 0.0017 |
| F05 | 70360001 | 70460000 | 0.0021 | 0.0047 | 0.0016 |
| F05 | 70380001 | 70480000 | 0.0050 | 0.0056 | 0.0043 |
| F05 | 70400001 | 70500000 | 0.0065 | 0.0050 | 0.0059 |
| F05 | 70420001 | 70520000 | 0.0059 | 0.0082 | 0.0054 |
| F05 | 70440001 | 70540000 | 0.0094 | 0.0103 | 0.0089 |

| Chr | Start | End | both | A1 | A2 |
|---|---|---|---|---|---|
| F05 | 70460001 | 70560000 | 0.0111 | 0.0095 | 0.0103 |
| F05 | 70480001 | 70580000 | 0.0102 | 0.0083 | 0.0095 |
| F05 | 70500001 | 70600000 | 0.0085 | 0.0084 | 0.0078 |
| F05 | 70520001 | 70620000 | 0.0087 | 0.0058 | 0.0080 |
| F05 | 70540001 | 70640000 | 0.0057 | 0.0044 | 0.0052 |
| F05 | 70560001 | 70660000 | 0.0044 | 0.0027 | 0.0041 |
| F05 | 70580001 | 70680000 | 0.0027 | 0.0031 | 0.0025 |
| F05 | 70600001 | 70700000 | 0.0030 | 0.0042 | 0.0026 |
| F05 | 70620001 | 70720000 | 0.0040 | 0.0047 | 0.0034 |
| F05 | 70640001 | 70740000 | 0.0044 | 0.0058 | 0.0037 |
| F05 | 70660001 | 70760000 | 0.0056 | 0.0059 | 0.0049 |
| F05 | 70680001 | 70780000 | 0.0058 | 0.0060 | 0.0051 |
| F05 | 70700001 | 70800000 | 0.0062 | 0.0056 | 0.0057 |
| F05 | 70720001 | 70820000 | 0.0060 | 0.0045 | 0.0055 |
| F05 | 70740001 | 70840000 | 0.0050 | 0.0026 | 0.0047 |
| F05 | 70760001 | 70860000 | 0.0031 | 0.0045 | 0.0029 |
| F05 | 70780001 | 70880000 | 0.0049 | 0.0046 | 0.0047 |
| F05 | 70800001 | 70900000 | 0.0050 | 0.0041 | 0.0048 |
| F05 | 70820001 | 70920000 | 0.0042 | 0.0042 | 0.0041 |
| F05 | 70840001 | 70940000 | 0.0042 | 0.0045 | 0.0041 |
| F05 | 70860001 | 70960000 | 0.0043 | 0.0022 | 0.0041 |
| F05 | 70880001 | 70980000 | 0.0019 | 0.0021 | 0.0018 |
| F05 | 70900001 | 71000000 | 0.0016 | 0.0024 | 0.0015 |
| F05 | 70920001 | 71020000 | 0.0019 | 0.0031 | 0.0018 |
| F05 | 70940001 | 71040000 | 0.0028 | 0.0031 | 0.0027 |
| F05 | 70960001 | 71060000 | 0.0030 | 0.0032 | 0.0029 |
| F05 | 70980001 | 71080000 | 0.0032 | 0.0034 | 0.0031 |
| F05 | 71000001 | 71100000 | 0.0035 | 0.0033 | 0.0034 |
| F05 | 71020001 | 71120000 | 0.0036 | 0.0030 | 0.0035 |
| F05 | 71040001 | 71140000 | 0.0033 | 0.0046 | 0.0032 |
| F05 | 71060001 | 71160000 | 0.0051 | 0.0046 | 0.0050 |
| F05 | 71080001 | 71180000 | 0.0052 | 0.0038 | 0.0051 |
| F05 | 71100001 | 71200000 | 0.0044 | 0.0032 | 0.0044 |
| F05 | 71120001 | 71220000 | 0.0038 | 0.0032 | 0.0038 |
| F05 | 71140001 | 71240000 | 0.0038 | 0.0017 | 0.0038 |
| F05 | 71160001 | 71260000 | 0.0020 | 0.0021 | 0.0019 |
| F05 | 71180001 | 71280000 | 0.0019 | 0.0024 | 0.0017 |
| F05 | 71200001 | 71300000 | 0.0021 | 0.0025 | 0.0019 |
| F05 | 71220001 | 71320000 | 0.0022 | 0.0022 | 0.0020 |
| F05 | 71240001 | 71340000 | 0.0019 | 0.0023 | 0.0017 |
| F05 | 71260001 | 71360000 | 0.0021 | 0.0023 | 0.0019 |
| F05 | 71280001 | 71380000 | 0.0024 | 0.0024 | 0.0023 |
| F05 | 71300001 | 71400000 | 0.0027 | 0.0028 | 0.0022 |
| F05 | 71320001 | 71420000 | 0.0033 | 0.0026 | 0.0025 |
| F05 | 71340001 | 71440000 | 0.0030 | 0.0023 | 0.0020 |
| F05 | 71360001 | 71460000 | 0.0029 | 0.0019 | 0.0017 |
| F05 | 71380001 | 71480000 | 0.0027 | 0.0018 | 0.0013 |
| F05 | 71400001 | 71500000 | 0.0024 | 0.0024 | 0.0014 |
| F05 | 71420001 | 71520000 | 0.0029 | 0.0033 | 0.0020 |
| F05 | 71440001 | 71540000 | 0.0045 | 0.0043 | 0.0031 |
| F05 | 71460001 | 71560000 | 0.0063 | 0.0040 | 0.0048 |
| F05 | 71480001 | 71580000 | 0.0058 | 0.0039 | 0.0046 |
| F05 | 71500001 | 71600000 | 0.0057 | 0.0034 | 0.0045 |
| F05 | 71520001 | 71620000 | 0.0053 | 0.0047 | 0.0043 |
| F05 | 71540001 | 71640000 | 0.0061 | 0.0037 | 0.0055 |
| F05 | 71560001 | 71660000 | 0.0043 | 0.0043 | 0.0041 |
| F05 | 71580001 | 71680000 | 0.0053 | 0.0047 | 0.0048 |
| F05 | 71600001 | 71700000 | 0.0058 | 0.0044 | 0.0051 |

| Chr | Start | End | both | A1 | A2 |
|---|---|---|---|---|---|
| F05 | 71620001 | 71720000 | 0.0054 | 0.0029 | 0.0046 |
| F05 | 71640001 | 71740000 | 0.0039 | 0.0030 | 0.0031 |
| F05 | 71660001 | 71760000 | 0.0040 | 0.0032 | 0.0031 |
| F05 | 71680001 | 71780000 | 0.0039 | 0.0029 | 0.0031 |
| F05 | 71700001 | 71800000 | 0.0036 | 0.0047 | 0.0027 |
| F05 | 71720001 | 71820000 | 0.0055 | 0.0041 | 0.0047 |
| F05 | 71740001 | 71840000 | 0.0048 | 0.0053 | 0.0040 |
| F05 | 71760001 | 71860000 | 0.0059 | 0.0059 | 0.0053 |
| F05 | 71780001 | 71880000 | 0.0064 | 0.0063 | 0.0059 |
| F05 | 71800001 | 71900000 | 0.0068 | 0.0049 | 0.0065 |
| F05 | 71820001 | 71920000 | 0.0052 | 0.0047 | 0.0051 |
| F05 | 71840001 | 71940000 | 0.0051 | 0.0050 | 0.0049 |
| F05 | 71860001 | 71960000 | 0.0054 | 0.0054 | 0.0053 |
| F05 | 71880001 | 71980000 | 0.0060 | 0.0064 | 0.0058 |
| F05 | 71900001 | 72000000 | 0.0071 | 0.0067 | 0.0070 |
| F05 | 71920001 | 72020000 | 0.0075 | 0.0067 | 0.0073 |
| F05 | 71940001 | 72040000 | 0.0076 | 0.0050 | 0.0074 |
| F05 | 71960001 | 72060000 | 0.0058 | 0.0033 | 0.0056 |
| F05 | 71980001 | 72080000 | 0.0044 | 0.0027 | 0.0042 |
| F05 | 72000001 | 72100000 | 0.0039 | 0.0021 | 0.0037 |
| F05 | 72020001 | 72120000 | 0.0035 | 0.0022 | 0.0033 |
| F05 | 72040001 | 72140000 | 0.0038 | 0.0022 | 0.0037 |
| F05 | 72060001 | 72160000 | 0.0040 | 0.0021 | 0.0039 |
| F05 | 72080001 | 72180000 | 0.0038 | 0.0013 | 0.0038 |
| F05 | 72100001 | 72200000 | 0.0029 | 0.0011 | 0.0027 |
| F05 | 72120001 | 72220000 | 0.0024 | 0.0010 | 0.0020 |
| F05 | 72140001 | 72240000 | 0.0020 | 0.0009 | 0.0015 |
| F05 | 72160001 | 72260000 | 0.0024 | 0.0010 | 0.0014 |
| F05 | 72180001 | 72280000 | 0.0023 | 0.0018 | 0.0012 |
| F05 | 72200001 | 72300000 | 0.0033 | 0.0020 | 0.0022 |
| F05 | 72220001 | 72320000 | 0.0036 | 0.0022 | 0.0027 |
| F05 | 72240001 | 72340000 | 0.0039 | 0.0028 | 0.0030 |
| F05 | 72260001 | 72360000 | 0.0038 | 0.0032 | 0.0031 |
| F05 | 72280001 | 72380000 | 0.0042 | 0.0023 | 0.0034 |
| F05 | 72300001 | 72400000 | 0.0033 | 0.0027 | 0.0024 |
| F05 | 72320001 | 72420000 | 0.0037 | 0.0036 | 0.0027 |
| F05 | 72340001 | 72440000 | 0.0045 | 0.0040 | 0.0034 |
| F05 | 72360001 | 72460000 | 0.0049 | 0.0042 | 0.0038 |
| F05 | 72380001 | 72480000 | 0.0051 | 0.0041 | 0.0040 |
| F05 | 72400001 | 72500000 | 0.0047 | 0.0038 | 0.0038 |
| F05 | 72420001 | 72520000 | 0.0044 | 0.0035 | 0.0036 |
| F05 | 72440001 | 72540000 | 0.0042 | 0.0027 | 0.0034 |
| F05 | 72460001 | 72560000 | 0.0032 | 0.0023 | 0.0026 |
| F05 | 72480001 | 72580000 | 0.0027 | 0.0026 | 0.0023 |
| F05 | 72500001 | 72600000 | 0.0031 | 0.0040 | 0.0027 |
| F05 | 72520001 | 72620000 | 0.0046 | 0.0035 | 0.0042 |
| F05 | 72540001 | 72640000 | 0.0040 | 0.0043 | 0.0037 |
| F05 | 72560001 | 72660000 | 0.0052 | 0.0043 | 0.0048 |
| F05 | 72580001 | 72680000 | 0.0054 | 0.0038 | 0.0049 |
| F05 | 72600001 | 72700000 | 0.0052 | 0.0018 | 0.0044 |
| F05 | 72620001 | 72720000 | 0.0033 | 0.0019 | 0.0023 |
| F05 | 72640001 | 72740000 | 0.0034 | 0.0013 | 0.0024 |
| F05 | 72660001 | 72760000 | 0.0028 | 0.0026 | 0.0016 |
| F05 | 72680001 | 72780000 | 0.0041 | 0.0038 | 0.0028 |
| F05 | 72700001 | 72800000 | 0.0056 | 0.0066 | 0.0043 |
| F05 | 72720001 | 72820000 | 0.0085 | 0.0072 | 0.0073 |
| F05 | 72740001 | 72840000 | 0.0094 | 0.0081 | 0.0080 |
| F05 | 72760001 | 72860000 | 0.0101 | 0.0071 | 0.0091 |

| Chr | Start | End | both | A1 | A2 |
|---|---|---|---|---|---|
| F05 | 72780001 | 72880000 | 0.0088 | 0.0078 | 0.0079 |
| F05 | 72800001 | 72900000 | 0.0091 | 0.0055 | 0.0081 |
| F05 | 72820001 | 72920000 | 0.0065 | 0.0045 | 0.0056 |
| F05 | 72840001 | 72940000 | 0.0054 | 0.0032 | 0.0044 |
| F05 | 72860001 | 72960000 | 0.0042 | 0.0030 | 0.0031 |
| F05 | 72880001 | 72980000 | 0.0043 | 0.0012 | 0.0034 |
| F05 | 72900001 | 73000000 | 0.0023 | 0.0009 | 0.0016 |
| F05 | 72920001 | 73020000 | 0.0020 | 0.0010 | 0.0016 |
| F05 | 72940001 | 73040000 | 0.0018 | 0.0012 | 0.0017 |
| F05 | 72960001 | 73060000 | 0.0018 | 0.0011 | 0.0018 |
| F05 | 72980001 | 73080000 | 0.0016 | 0.0021 | 0.0016 |
| F05 | 73000001 | 73100000 | 0.0027 | 0.0021 | 0.0027 |
| F05 | 73020001 | 73120000 | 0.0029 | 0.0020 | 0.0027 |
| F05 | 73040001 | 73140000 | 0.0031 | 0.0023 | 0.0027 |
| F05 | 73060001 | 73160000 | 0.0034 | 0.0028 | 0.0029 |
| F05 | 73080001 | 73180000 | 0.0037 | 0.0020 | 0.0032 |
| F05 | 73100001 | 73200000 | 0.0028 | 0.0026 | 0.0024 |
| F05 | 73120001 | 73220000 | 0.0030 | 0.0036 | 0.0026 |
| F05 | 73140001 | 73240000 | 0.0036 | 0.0032 | 0.0034 |
| F05 | 73160001 | 73260000 | 0.0033 | 0.0026 | 0.0032 |
| F05 | 73180001 | 73280000 | 0.0027 | 0.0036 | 0.0026 |
| F05 | 73200001 | 73300000 | 0.0037 | 0.0034 | 0.0035 |
| F05 | 73220001 | 73320000 | 0.0037 | 0.0029 | 0.0036 |
| F05 | 73240001 | 73340000 | 0.0033 | 0.0036 | 0.0032 |
| F05 | 73260001 | 73360000 | 0.0044 | 0.0043 | 0.0042 |
| F05 | 73280001 | 73380000 | 0.0053 | 0.0034 | 0.0051 |
| F05 | 73300001 | 73400000 | 0.0051 | 0.0031 | 0.0047 |
| F05 | 73320001 | 73420000 | 0.0052 | 0.0031 | 0.0047 |
| F05 | 73340001 | 73440000 | 0.0057 | 0.0030 | 0.0051 |
| F05 | 73360001 | 73460000 | 0.0062 | 0.0037 | 0.0055 |
| F05 | 73380001 | 73480000 | 0.0078 | 0.0034 | 0.0068 |
| F05 | 73400001 | 73500000 | 0.0071 | 0.0034 | 0.0062 |
| F05 | 73420001 | 73520000 | 0.0067 | 0.0029 | 0.0059 |
| F05 | 73440001 | 73540000 | 0.0058 | 0.0034 | 0.0052 |
| F05 | 73460001 | 73560000 | 0.0055 | 0.0019 | 0.0050 |
| F05 | 73480001 | 73580000 | 0.0033 | 0.0030 | 0.0031 |
| F05 | 73500001 | 73600000 | 0.0042 | 0.0028 | 0.0039 |
| F05 | 73520001 | 73620000 | 0.0037 | 0.0036 | 0.0035 |
| F05 | 73540001 | 73640000 | 0.0042 | 0.0030 | 0.0039 |
| F05 | 73560001 | 73660000 | 0.0032 | 0.0030 | 0.0029 |
| F05 | 73580001 | 73680000 | 0.0028 | 0.0025 | 0.0025 |
| F05 | 73600001 | 73700000 | 0.0022 | 0.0024 | 0.0019 |
| F05 | 73620001 | 73720000 | 0.0022 | 0.0028 | 0.0020 |
| F05 | 73640001 | 73740000 | 0.0027 | 0.0025 | 0.0025 |
| F05 | 73660001 | 73760000 | 0.0025 | 0.0038 | 0.0024 |
| F05 | 73680001 | 73780000 | 0.0041 | 0.0043 | 0.0039 |
| F05 | 73700001 | 73800000 | 0.0047 | 0.0045 | 0.0046 |
| F05 | 73720001 | 73820000 | 0.0049 | 0.0041 | 0.0048 |
| F05 | 73740001 | 73840000 | 0.0043 | 0.0039 | 0.0041 |
| F05 | 73760001 | 73860000 | 0.0042 | 0.0029 | 0.0039 |
| F05 | 73780001 | 73880000 | 0.0029 | 0.0022 | 0.0027 |
| F05 | 73800001 | 73900000 | 0.0022 | 0.0021 | 0.0019 |
| F05 | 73820001 | 73920000 | 0.0022 | 0.0015 | 0.0019 |
| F05 | 73840001 | 73940000 | 0.0015 | 0.0023 | 0.0014 |
| F05 | 73860001 | 73960000 | 0.0023 | 0.0025 | 0.0022 |
| F05 | 73880001 | 73980000 | 0.0029 | 0.0031 | 0.0028 |
| F05 | 73900001 | 74000000 | 0.0036 | 0.0032 | 0.0036 |
| F05 | 73920001 | 74020000 | 0.0038 | 0.0033 | 0.0038 |

| Chr | Start | End | both | A1 | A2 |
|---|---|---|---|---|---|
| F05 | 73940001 | 74040000 | 0.0040 | 0.0030 | 0.0039 |
| F05 | 73960001 | 74060000 | 0.0037 | 0.0033 | 0.0037 |
| F05 | 73980001 | 74080000 | 0.0039 | 0.0026 | 0.0038 |
| F05 | 74000001 | 74100000 | 0.0030 | 0.0026 | 0.0030 |
| F05 | 74020001 | 74120000 | 0.0030 | 0.0027 | 0.0029 |
| F05 | 74040001 | 74140000 | 0.0031 | 0.0022 | 0.0029 |
| F05 | 74060001 | 74160000 | 0.0024 | 0.0032 | 0.0023 |
| F05 | 74080001 | 74180000 | 0.0035 | 0.0066 | 0.0031 |
| F05 | 74100001 | 74200000 | 0.0064 | 0.0082 | 0.0053 |
| F05 | 74120001 | 74220000 | 0.0077 | 0.0083 | 0.0060 |
| F05 | 74140001 | 74240000 | 0.0078 | 0.0084 | 0.0059 |
| F05 | 74160001 | 74260000 | 0.0085 | 0.0067 | 0.0067 |
| F05 | 74180001 | 74280000 | 0.0070 | 0.0035 | 0.0053 |
| F05 | 74200001 | 74300000 | 0.0043 | 0.0033 | 0.0032 |
| F05 | 74220001 | 74320000 | 0.0044 | 0.0034 | 0.0039 |
| F05 | 74240001 | 74340000 | 0.0047 | 0.0033 | 0.0042 |
| F05 | 74260001 | 74360000 | 0.0044 | 0.0030 | 0.0034 |
| F05 | 74280001 | 74380000 | 0.0041 | 0.0037 | 0.0030 |
| F05 | 74300001 | 74400000 | 0.0053 | 0.0034 | 0.0039 |
| F05 | 74320001 | 74420000 | 0.0052 | 0.0050 | 0.0037 |
| F05 | 74340001 | 74440000 | 0.0066 | 0.0060 | 0.0051 |
| F05 | 74360001 | 74460000 | 0.0083 | 0.0075 | 0.0070 |
| F05 | 74380001 | 74480000 | 0.0110 | 0.0067 | 0.0099 |
| F05 | 74400001 | 74500000 | 0.0100 | 0.0063 | 0.0090 |
| F05 | 74420001 | 74520000 | 0.0097 | 0.0055 | 0.0087 |
| F05 | 74440001 | 74540000 | 0.0090 | 0.0059 | 0.0084 |
| F05 | 74460001 | 74560000 | 0.0085 | 0.0050 | 0.0080 |
| F05 | 74480001 | 74580000 | 0.0062 | 0.0048 | 0.0058 |
| F05 | 74500001 | 74600000 | 0.0058 | 0.0040 | 0.0055 |
| F05 | 74520001 | 74620000 | 0.0047 | 0.0029 | 0.0046 |
| F05 | 74540001 | 74640000 | 0.0034 | 0.0015 | 0.0033 |
| F05 | 74560001 | 74660000 | 0.0018 | 0.0013 | 0.0017 |
| F05 | 74580001 | 74680000 | 0.0016 | 0.0014 | 0.0015 |
| F05 | 74600001 | 74700000 | 0.0016 | 0.0014 | 0.0015 |
| F05 | 74620001 | 74720000 | 0.0017 | 0.0015 | 0.0015 |
| F05 | 74640001 | 74740000 | 0.0017 | 0.0025 | 0.0015 |
| F05 | 74660001 | 74760000 | 0.0030 | 0.0023 | 0.0028 |
| F05 | 74680001 | 74780000 | 0.0027 | 0.0032 | 0.0025 |
| F05 | 74700001 | 74800000 | 0.0038 | 0.0036 | 0.0036 |
| F05 | 74720001 | 74820000 | 0.0044 | 0.0047 | 0.0041 |
| F05 | 74740001 | 74840000 | 0.0054 | 0.0036 | 0.0049 |
| F05 | 74760001 | 74860000 | 0.0050 | 0.0039 | 0.0038 |
| F05 | 74780001 | 74880000 | 0.0057 | 0.0026 | 0.0041 |
| F05 | 74800001 | 74900000 | 0.0045 | 0.0018 | 0.0027 |
| F05 | 74820001 | 74920000 | 0.0038 | 0.0006 | 0.0018 |
| F05 | 74840001 | 74940000 | 0.0030 | 0.0006 | 0.0012 |
| F05 | 74860001 | 74960000 | 0.0025 | 0.0004 | 0.0014 |
| F05 | 74880001 | 74980000 | 0.0020 | 0.0008 | 0.0011 |
| F05 | 74900001 | 75000000 | 0.0022 | 0.0015 | 0.0016 |
| F05 | 74920001 | 75020000 | 0.0031 | 0.0017 | 0.0025 |
| F05 | 74940001 | 75040000 | 0.0030 | 0.0017 | 0.0024 |
| F05 | 74960001 | 75060000 | 0.0029 | 0.0015 | 0.0021 |
| F05 | 74980001 | 75080000 | 0.0030 | 0.0017 | 0.0020 |
| F05 | 75000001 | 75100000 | 0.0034 | 0.0020 | 0.0022 |
| F05 | 75020001 | 75120000 | 0.0035 | 0.0030 | 0.0025 |
| F05 | 75040001 | 75140000 | 0.0046 | 0.0032 | 0.0037 |
| F05 | 75060001 | 75160000 | 0.0053 | 0.0034 | 0.0042 |
| F05 | 75080001 | 75180000 | 0.0065 | 0.0030 | 0.0048 |

| Chr | Start | End | both | A1 | A2 |
|---|---|---|---|---|---|
| F05 | 75100001 | 75200000 | 0.0061 | 0.0024 | 0.0043 |
| F05 | 75120001 | 75220000 | 0.0057 | 0.0013 | 0.0037 |
| F05 | 75140001 | 75240000 | 0.0046 | 0.0014 | 0.0026 |
| F05 | 75160001 | 75260000 | 0.0044 | 0.0014 | 0.0025 |
| F05 | 75180001 | 75280000 | 0.0035 | 0.0014 | 0.0023 |
| F05 | 75200001 | 75300000 | 0.0034 | 0.0008 | 0.0025 |
| F05 | 75220001 | 75320000 | 0.0032 | 0.0007 | 0.0022 |
| F05 | 75240001 | 75340000 | 0.0032 | 0.0006 | 0.0021 |
| F05 | 75260001 | 75360000 | 0.0032 | 0.0004 | 0.0019 |
| F05 | 75280001 | 75380000 | 0.0026 | 0.0009 | 0.0013 |
| F05 | 75300001 | 75400000 | 0.0030 | 0.0011 | 0.0016 |
| F05 | 75320001 | 75420000 | 0.0026 | 0.0019 | 0.0014 |
| F05 | 75340001 | 75440000 | 0.0031 | 0.0031 | 0.0022 |
| F05 | 75360001 | 75460000 | 0.0043 | 0.0036 | 0.0038 |
| F05 | 75380001 | 75480000 | 0.0047 | 0.0039 | 0.0044 |
| F05 | 75400001 | 75500000 | 0.0051 | 0.0060 | 0.0050 |
| F05 | 75420001 | 75520000 | 0.0075 | 0.0059 | 0.0074 |
| F05 | 75440001 | 75540000 | 0.0077 | 0.0047 | 0.0075 |
| F05 | 75460001 | 75560000 | 0.0063 | 0.0042 | 0.0060 |
| F05 | 75480001 | 75580000 | 0.0057 | 0.0043 | 0.0055 |
| F05 | 75500001 | 75600000 | 0.0057 | 0.0035 | 0.0053 |
| F05 | 75520001 | 75620000 | 0.0046 | 0.0039 | 0.0041 |
| F05 | 75540001 | 75640000 | 0.0042 | 0.0050 | 0.0035 |
| F05 | 75560001 | 75660000 | 0.0049 | 0.0064 | 0.0041 |
| F05 | 75580001 | 75680000 | 0.0061 | 0.0058 | 0.0051 |
| F05 | 75600001 | 75700000 | 0.0054 | 0.0057 | 0.0045 |
| F05 | 75620001 | 75720000 | 0.0054 | 0.0060 | 0.0045 |
| F05 | 75640001 | 75740000 | 0.0059 | 0.0055 | 0.0049 |
| F05 | 75660001 | 75760000 | 0.0058 | 0.0040 | 0.0048 |
| F05 | 75680001 | 75780000 | 0.0052 | 0.0037 | 0.0045 |
| F05 | 75700001 | 75800000 | 0.0055 | 0.0026 | 0.0047 |
| F05 | 75720001 | 75820000 | 0.0045 | 0.0021 | 0.0034 |
| F05 | 75740001 | 75840000 | 0.0044 | 0.0024 | 0.0033 |
| F05 | 75760001 | 75860000 | 0.0047 | 0.0025 | 0.0036 |
| F05 | 75780001 | 75880000 | 0.0048 | 0.0025 | 0.0031 |
| F05 | 75800001 | 75900000 | 0.0047 | 0.0024 | 0.0029 |
| F05 | 75820001 | 75920000 | 0.0056 | 0.0016 | 0.0040 |
| F05 | 75840001 | 75940000 | 0.0050 | 0.0006 | 0.0035 |
| F05 | 75860001 | 75960000 | 0.0040 | 0.0004 | 0.0023 |
| F05 | 75880001 | 75980000 | 0.0036 | 0.0011 | 0.0023 |
| F05 | 75900001 | 76000000 | 0.0036 | 0.0012 | 0.0022 |
| F05 | 75920001 | 76020000 | 0.0030 | 0.0013 | 0.0014 |
| F05 | 75940001 | 76040000 | 0.0031 | 0.0027 | 0.0015 |
| F05 | 75960001 | 76060000 | 0.0046 | 0.0038 | 0.0031 |
| F05 | 75980001 | 76080000 | 0.0052 | 0.0031 | 0.0039 |
| F05 | 76000001 | 76100000 | 0.0053 | 0.0034 | 0.0036 |
| F05 | 76020001 | 76120000 | 0.0053 | 0.0042 | 0.0037 |
| F05 | 76040001 | 76140000 | 0.0060 | 0.0035 | 0.0042 |
| F05 | 76060001 | 76160000 | 0.0050 | 0.0033 | 0.0032 |
| F05 | 76080001 | 76180000 | 0.0048 | 0.0040 | 0.0032 |
| F05 | 76100001 | 76200000 | 0.0051 | 0.0042 | 0.0041 |
| F05 | 76120001 | 76220000 | 0.0049 | 0.0034 | 0.0038 |
| F05 | 76140001 | 76240000 | 0.0039 | 0.0030 | 0.0031 |
| F05 | 76160001 | 76260000 | 0.0032 | 0.0025 | 0.0026 |
| F05 | 76180001 | 76280000 | 0.0026 | 0.0022 | 0.0017 |
| F05 | 76200001 | 76300000 | 0.0023 | 0.0018 | 0.0012 |
| F05 | 76220001 | 76320000 | 0.0020 | 0.0018 | 0.0010 |
| F05 | 76240001 | 76340000 | 0.0019 | 0.0021 | 0.0008 |

| Chr | Start | End | both | A1 | A2 |
|---|---|---|---|---|---|
| F05 | 76260001 | 76360000 | 0.0024 | 0.0019 | 0.0012 |
| F05 | 76280001 | 76380000 | 0.0020 | 0.0018 | 0.0012 |
| F05 | 76300001 | 76400000 | 0.0018 | 0.0015 | 0.0011 |
| F05 | 76320001 | 76420000 | 0.0015 | 0.0013 | 0.0011 |
| F05 | 76340001 | 76440000 | 0.0013 | 0.0014 | 0.0011 |
| F05 | 76360001 | 76460000 | 0.0016 | 0.0018 | 0.0014 |
| F05 | 76380001 | 76480000 | 0.0020 | 0.0014 | 0.0019 |
| F05 | 76400001 | 76500000 | 0.0015 | 0.0012 | 0.0015 |
| F05 | 76420001 | 76520000 | 0.0014 | 0.0011 | 0.0014 |
| F05 | 76440001 | 76540000 | 0.0013 | 0.0004 | 0.0013 |
| F05 | 76460001 | 76560000 | 0.0005 | 0.0000 | 0.0005 |
| F05 | 76480001 | 76580000 | 0.0000 | 0.0007 | 0.0000 |
| F06 | 1 | 100000 | 0.0017 | 0.0006 | 0.0010 |
| F06 | 20001 | 120000 | 0.0015 | 0.0005 | 0.0008 |
| F06 | 40001 | 140000 | 0.0016 | 0.0004 | 0.0007 |
| F06 | 60001 | 160000 | 0.0017 | 0.0006 | 0.0007 |
| F06 | 80001 | 180000 | 0.0015 | 0.0007 | 0.0006 |
| F06 | 100001 | 200000 | 0.0012 | 0.0011 | 0.0004 |
| F06 | 120001 | 220000 | 0.0013 | 0.0014 | 0.0007 |
| F06 | 140001 | 240000 | 0.0012 | 0.0016 | 0.0007 |
| F06 | 160001 | 260000 | 0.0010 | 0.0016 | 0.0007 |
| F06 | 180001 | 280000 | 0.0010 | 0.0017 | 0.0007 |
| F06 | 200001 | 300000 | 0.0010 | 0.0017 | 0.0007 |
| F06 | 220001 | 320000 | 0.0010 | 0.0016 | 0.0007 |
| F06 | 240001 | 340000 | 0.0010 | 0.0017 | 0.0008 |
| F06 | 260001 | 360000 | 0.0010 | 0.0026 | 0.0008 |
| F06 | 280001 | 380000 | 0.0021 | 0.0028 | 0.0018 |
| F06 | 300001 | 400000 | 0.0021 | 0.0026 | 0.0018 |
| F06 | 320001 | 420000 | 0.0018 | 0.0028 | 0.0015 |
| F06 | 340001 | 440000 | 0.0019 | 0.0026 | 0.0015 |
| F06 | 360001 | 460000 | 0.0019 | 0.0013 | 0.0015 |
| F06 | 380001 | 480000 | 0.0010 | 0.0009 | 0.0005 |
| F06 | 400001 | 500000 | 0.0012 | 0.0007 | 0.0006 |
| F06 | 420001 | 520000 | 0.0015 | 0.0005 | 0.0010 |
| F06 | 440001 | 540000 | 0.0022 | 0.0006 | 0.0015 |
| F06 | 460001 | 560000 | 0.0025 | 0.0016 | 0.0017 |
| F06 | 480001 | 580000 | 0.0034 | 0.0017 | 0.0025 |
| F06 | 500001 | 600000 | 0.0034 | 0.0019 | 0.0025 |
| F06 | 520001 | 620000 | 0.0035 | 0.0016 | 0.0021 |
| F06 | 540001 | 640000 | 0.0028 | 0.0018 | 0.0015 |
| F06 | 560001 | 660000 | 0.0031 | 0.0020 | 0.0016 |
| F06 | 580001 | 680000 | 0.0041 | 0.0018 | 0.0025 |
| F06 | 600001 | 700000 | 0.0041 | 0.0017 | 0.0027 |
| F06 | 620001 | 720000 | 0.0043 | 0.0019 | 0.0029 |
| F06 | 640001 | 740000 | 0.0044 | 0.0022 | 0.0031 |
| F06 | 660001 | 760000 | 0.0043 | 0.0011 | 0.0034 |
| F06 | 680001 | 780000 | 0.0023 | 0.0013 | 0.0017 |
| F06 | 700001 | 800000 | 0.0022 | 0.0013 | 0.0016 |
| F06 | 720001 | 820000 | 0.0019 | 0.0012 | 0.0015 |
| F06 | 740001 | 840000 | 0.0017 | 0.0016 | 0.0014 |
| F06 | 760001 | 860000 | 0.0021 | 0.0022 | 0.0017 |
| F06 | 780001 | 880000 | 0.0025 | 0.0023 | 0.0020 |
| F06 | 800001 | 900000 | 0.0025 | 0.0027 | 0.0020 |
| F06 | 820001 | 920000 | 0.0027 | 0.0030 | 0.0024 |
| F06 | 840001 | 940000 | 0.0031 | 0.0021 | 0.0027 |
| F06 | 860001 | 960000 | 0.0023 | 0.0017 | 0.0019 |
| F06 | 880001 | 980000 | 0.0020 | 0.0014 | 0.0018 |
| F06 | 900001 | 1000000 | 0.0018 | 0.0012 | 0.0015 |

| Chr | Start | End | both | A1 | A2 |
|---|---|---|---|---|---|
| F06 | 920001 | 1020000 | 0.0016 | 0.0008 | 0.0013 |
| F06 | 940001 | 1040000 | 0.0014 | 0.0007 | 0.0008 |
| F06 | 960001 | 1060000 | 0.0014 | 0.0005 | 0.0007 |
| F06 | 980001 | 1080000 | 0.0016 | 0.0007 | 0.0005 |
| F06 | 1000001 | 1100000 | 0.0019 | 0.0006 | 0.0006 |
| F06 | 1020001 | 1120000 | 0.0020 | 0.0005 | 0.0005 |
| F06 | 1040001 | 1140000 | 0.0018 | 0.0005 | 0.0004 |
| F06 | 1060001 | 1160000 | 0.0020 | 0.0006 | 0.0005 |
| F06 | 1080001 | 1180000 | 0.0018 | 0.0004 | 0.0007 |
| F06 | 1100001 | 1200000 | 0.0017 | 0.0003 | 0.0006 |
| F06 | 1120001 | 1220000 | 0.0017 | 0.0004 | 0.0007 |
| F06 | 1140001 | 1240000 | 0.0020 | 0.0010 | 0.0009 |
| F06 | 1160001 | 1260000 | 0.0025 | 0.0011 | 0.0015 |
| F06 | 1180001 | 1280000 | 0.0027 | 0.0011 | 0.0016 |
| F06 | 1200001 | 1300000 | 0.0026 | 0.0012 | 0.0015 |
| F06 | 1220001 | 1320000 | 0.0024 | 0.0010 | 0.0014 |
| F06 | 1240001 | 1340000 | 0.0022 | 0.0003 | 0.0012 |
| F06 | 1260001 | 1360000 | 0.0016 | 0.0001 | 0.0005 |
| F06 | 1280001 | 1380000 | 0.0015 | 0.0002 | 0.0002 |
| F06 | 1300001 | 1400000 | 0.0016 | 0.0002 | 0.0002 |
| F06 | 1320001 | 1420000 | 0.0017 | 0.0003 | 0.0003 |
| F06 | 1340001 | 1440000 | 0.0016 | 0.0004 | 0.0002 |
| F06 | 1360001 | 1460000 | 0.0016 | 0.0006 | 0.0002 |
| F06 | 1380001 | 1480000 | 0.0016 | 0.0008 | 0.0003 |
| F06 | 1400001 | 1500000 | 0.0016 | 0.0009 | 0.0005 |
| F06 | 1420001 | 1520000 | 0.0017 | 0.0008 | 0.0006 |
| F06 | 1440001 | 1540000 | 0.0019 | 0.0009 | 0.0006 |
| F06 | 1460001 | 1560000 | 0.0020 | 0.0009 | 0.0009 |
| F06 | 1480001 | 1580000 | 0.0019 | 0.0009 | 0.0009 |
| F06 | 1500001 | 1600000 | 0.0016 | 0.0008 | 0.0007 |
| F06 | 1520001 | 1620000 | 0.0014 | 0.0008 | 0.0005 |
| F06 | 1540001 | 1640000 | 0.0012 | 0.0007 | 0.0005 |
| F06 | 1560001 | 1660000 | 0.0011 | 0.0006 | 0.0004 |
| F06 | 1580001 | 1680000 | 0.0014 | 0.0005 | 0.0004 |
| F06 | 1600001 | 1700000 | 0.0015 | 0.0004 | 0.0004 |
| F06 | 1620001 | 1720000 | 0.0016 | 0.0011 | 0.0004 |
| F06 | 1640001 | 1740000 | 0.0021 | 0.0016 | 0.0011 |
| F06 | 1660001 | 1760000 | 0.0024 | 0.0016 | 0.0015 |
| F06 | 1680001 | 1780000 | 0.0020 | 0.0017 | 0.0014 |
| F06 | 1700001 | 1800000 | 0.0018 | 0.0022 | 0.0015 |
| F06 | 1720001 | 1820000 | 0.0021 | 0.0016 | 0.0020 |
| F06 | 1740001 | 1840000 | 0.0015 | 0.0013 | 0.0013 |
| F06 | 1760001 | 1860000 | 0.0011 | 0.0014 | 0.0010 |
| F06 | 1780001 | 1880000 | 0.0011 | 0.0015 | 0.0009 |
| F06 | 1800001 | 1900000 | 0.0011 | 0.0011 | 0.0009 |
| F06 | 1820001 | 1920000 | 0.0009 | 0.0013 | 0.0005 |
| F06 | 1840001 | 1940000 | 0.0013 | 0.0012 | 0.0008 |
| F06 | 1860001 | 1960000 | 0.0014 | 0.0011 | 0.0007 |
| F06 | 1880001 | 1980000 | 0.0016 | 0.0011 | 0.0007 |
| F06 | 1900001 | 2000000 | 0.0018 | 0.0012 | 0.0008 |
| F06 | 1920001 | 2020000 | 0.0020 | 0.0009 | 0.0010 |
| F06 | 1940001 | 2040000 | 0.0017 | 0.0009 | 0.0007 |
| F06 | 1960001 | 2060000 | 0.0017 | 0.0007 | 0.0007 |
| F06 | 1980001 | 2080000 | 0.0016 | 0.0005 | 0.0007 |
| F06 | 2000001 | 2100000 | 0.0016 | 0.0002 | 0.0008 |
| F06 | 2020001 | 2120000 | 0.0014 | 0.0002 | 0.0005 |
| F06 | 2040001 | 2140000 | 0.0013 | 0.0008 | 0.0006 |
| F06 | 2060001 | 2160000 | 0.0021 | 0.0009 | 0.0012 |

| Chr | Start | End | both | A1 | A2 |
|---|---|---|---|---|---|
| F06 | 2080001 | 2180000 | 0.0022 | 0.0010 | 0.0012 |
| F06 | 2100001 | 2200000 | 0.0023 | 0.0010 | 0.0013 |
| F06 | 2120001 | 2220000 | 0.0023 | 0.0009 | 0.0012 |
| F06 | 2140001 | 2240000 | 0.0022 | 0.0003 | 0.0011 |
| F06 | 2160001 | 2260000 | 0.0016 | 0.0009 | 0.0005 |
| F06 | 2180001 | 2280000 | 0.0022 | 0.0008 | 0.0009 |
| F06 | 2200001 | 2300000 | 0.0021 | 0.0008 | 0.0008 |
| F06 | 2220001 | 2320000 | 0.0020 | 0.0008 | 0.0008 |
| F06 | 2240001 | 2340000 | 0.0020 | 0.0008 | 0.0008 |
| F06 | 2260001 | 2360000 | 0.0019 | 0.0004 | 0.0008 |
| F06 | 2280001 | 2380000 | 0.0012 | 0.0004 | 0.0004 |
| F06 | 2300001 | 2400000 | 0.0011 | 0.0007 | 0.0004 |
| F06 | 2320001 | 2420000 | 0.0013 | 0.0015 | 0.0008 |
| F06 | 2340001 | 2440000 | 0.0016 | 0.0018 | 0.0010 |
| F06 | 2360001 | 2460000 | 0.0017 | 0.0027 | 0.0011 |
| F06 | 2380001 | 2480000 | 0.0024 | 0.0029 | 0.0017 |
| F06 | 2400001 | 2500000 | 0.0027 | 0.0032 | 0.0020 |
| F06 | 2420001 | 2520000 | 0.0029 | 0.0033 | 0.0019 |
| F06 | 2440001 | 2540000 | 0.0032 | 0.0032 | 0.0024 |
| F06 | 2460001 | 2560000 | 0.0032 | 0.0027 | 0.0025 |
| F06 | 2480001 | 2580000 | 0.0027 | 0.0026 | 0.0018 |
| F06 | 2500001 | 2600000 | 0.0024 | 0.0021 | 0.0015 |
| F06 | 2520001 | 2620000 | 0.0019 | 0.0015 | 0.0012 |
| F06 | 2540001 | 2640000 | 0.0013 | 0.0012 | 0.0005 |
| F06 | 2560001 | 2660000 | 0.0012 | 0.0008 | 0.0003 |
| F06 | 2580001 | 2680000 | 0.0009 | 0.0007 | 0.0003 |
| F06 | 2600001 | 2700000 | 0.0010 | 0.0009 | 0.0004 |
| F06 | 2620001 | 2720000 | 0.0013 | 0.0006 | 0.0008 |
| F06 | 2640001 | 2740000 | 0.0013 | 0.0005 | 0.0008 |
| F06 | 2660001 | 2760000 | 0.0012 | 0.0006 | 0.0007 |
| F06 | 2680001 | 2780000 | 0.0014 | 0.0004 | 0.0008 |
| F06 | 2700001 | 2800000 | 0.0015 | 0.0001 | 0.0007 |
| F06 | 2720001 | 2820000 | 0.0011 | 0.0001 | 0.0004 |
| F06 | 2740001 | 2840000 | 0.0012 | 0.0001 | 0.0004 |
| F06 | 2760001 | 2860000 | 0.0012 | 0.0007 | 0.0004 |
| F06 | 2780001 | 2880000 | 0.0018 | 0.0007 | 0.0010 |
| F06 | 2800001 | 2900000 | 0.0017 | 0.0007 | 0.0010 |
| F06 | 2820001 | 2920000 | 0.0017 | 0.0007 | 0.0011 |
| F06 | 2840001 | 2940000 | 0.0016 | 0.0008 | 0.0009 |
| F06 | 2860001 | 2960000 | 0.0019 | 0.0005 | 0.0010 |
| F06 | 2880001 | 2980000 | 0.0012 | 0.0010 | 0.0003 |
| F06 | 2900001 | 3000000 | 0.0011 | 0.0010 | 0.0003 |
| F06 | 2920001 | 3020000 | 0.0010 | 0.0022 | 0.0002 |
| F06 | 2940001 | 3040000 | 0.0016 | 0.0026 | 0.0008 |
| F06 | 2960001 | 3060000 | 0.0013 | 0.0027 | 0.0008 |
| F06 | 2980001 | 3080000 | 0.0013 | 0.0025 | 0.0008 |
| F06 | 3000001 | 3100000 | 0.0012 | 0.0028 | 0.0006 |
| F06 | 3020001 | 3120000 | 0.0013 | 0.0020 | 0.0007 |
| F06 | 3040001 | 3140000 | 0.0008 | 0.0019 | 0.0002 |
| F06 | 3060001 | 3160000 | 0.0010 | 0.0015 | 0.0003 |
| F06 | 3080001 | 3180000 | 0.0011 | 0.0011 | 0.0003 |
| F06 | 3100001 | 3200000 | 0.0012 | 0.0009 | 0.0004 |
| F06 | 3120001 | 3220000 | 0.0015 | 0.0006 | 0.0006 |
| F06 | 3140001 | 3240000 | 0.0014 | 0.0005 | 0.0005 |
| F06 | 3160001 | 3260000 | 0.0016 | 0.0006 | 0.0007 |
| F06 | 3180001 | 3280000 | 0.0016 | 0.0006 | 0.0008 |
| F06 | 3200001 | 3300000 | 0.0017 | 0.0021 | 0.0007 |
| F06 | 3220001 | 3320000 | 0.0034 | 0.0022 | 0.0024 |

| Chr | Start | End | both | A1 | A2 |
|---|---|---|---|---|---|
| F06 | 3240001 | 3340000 | 0.0035 | 0.0019 | 0.0025 |
| F06 | 3260001 | 3360000 | 0.0031 | 0.0022 | 0.0023 |
| F06 | 3280001 | 3380000 | 0.0031 | 0.0023 | 0.0025 |
| F06 | 3300001 | 3400000 | 0.0030 | 0.0009 | 0.0026 |
| F06 | 3320001 | 3420000 | 0.0013 | 0.0008 | 0.0010 |
| F06 | 3340001 | 3440000 | 0.0014 | 0.0008 | 0.0009 |
| F06 | 3360001 | 3460000 | 0.0014 | 0.0010 | 0.0009 |
| F06 | 3380001 | 3480000 | 0.0017 | 0.0009 | 0.0011 |
| F06 | 3400001 | 3500000 | 0.0018 | 0.0009 | 0.0010 |
| F06 | 3420001 | 3520000 | 0.0017 | 0.0011 | 0.0009 |
| F06 | 3440001 | 3540000 | 0.0017 | 0.0014 | 0.0010 |
| F06 | 3460001 | 3560000 | 0.0022 | 0.0013 | 0.0013 |
| F06 | 3480001 | 3580000 | 0.0023 | 0.0017 | 0.0012 |
| F06 | 3500001 | 3600000 | 0.0026 | 0.0022 | 0.0017 |
| F06 | 3520001 | 3620000 | 0.0029 | 0.0022 | 0.0019 |
| F06 | 3540001 | 3640000 | 0.0029 | 0.0019 | 0.0020 |
| F06 | 3560001 | 3660000 | 0.0025 | 0.0016 | 0.0018 |
| F06 | 3580001 | 3680000 | 0.0019 | 0.0013 | 0.0014 |
| F06 | 3600001 | 3700000 | 0.0015 | 0.0007 | 0.0011 |
| F06 | 3620001 | 3720000 | 0.0011 | 0.0007 | 0.0008 |
| F06 | 3640001 | 3740000 | 0.0012 | 0.0006 | 0.0008 |
| F06 | 3660001 | 3760000 | 0.0011 | 0.0010 | 0.0007 |
| F06 | 3680001 | 3780000 | 0.0016 | 0.0010 | 0.0011 |
| F06 | 3700001 | 3800000 | 0.0018 | 0.0009 | 0.0011 |
| F06 | 3720001 | 3820000 | 0.0018 | 0.0013 | 0.0009 |
| F06 | 3740001 | 3840000 | 0.0021 | 0.0016 | 0.0012 |
| F06 | 3760001 | 3860000 | 0.0025 | 0.0013 | 0.0015 |
| F06 | 3780001 | 3880000 | 0.0022 | 0.0013 | 0.0012 |
| F06 | 3800001 | 3900000 | 0.0021 | 0.0014 | 0.0013 |
| F06 | 3820001 | 3920000 | 0.0019 | 0.0014 | 0.0014 |
| F06 | 3840001 | 3940000 | 0.0019 | 0.0020 | 0.0016 |
| F06 | 3860001 | 3960000 | 0.0024 | 0.0022 | 0.0022 |
| F06 | 3880001 | 3980000 | 0.0025 | 0.0020 | 0.0025 |
| F06 | 3900001 | 4000000 | 0.0024 | 0.0019 | 0.0023 |
| F06 | 3920001 | 4020000 | 0.0022 | 0.0014 | 0.0022 |
| F06 | 3940001 | 4040000 | 0.0016 | 0.0010 | 0.0016 |
| F06 | 3960001 | 4060000 | 0.0013 | 0.0007 | 0.0013 |
| F06 | 3980001 | 4080000 | 0.0010 | 0.0013 | 0.0010 |
| F06 | 4000001 | 4100000 | 0.0016 | 0.0013 | 0.0016 |
| F06 | 4020001 | 4120000 | 0.0015 | 0.0015 | 0.0015 |
| F06 | 4040001 | 4140000 | 0.0019 | 0.0018 | 0.0018 |
| F06 | 4060001 | 4160000 | 0.0022 | 0.0027 | 0.0018 |
| F06 | 4080001 | 4180000 | 0.0032 | 0.0023 | 0.0025 |
| F06 | 4100001 | 4200000 | 0.0028 | 0.0023 | 0.0020 |
| F06 | 4120001 | 4220000 | 0.0030 | 0.0020 | 0.0021 |
| F06 | 4140001 | 4240000 | 0.0027 | 0.0017 | 0.0017 |
| F06 | 4160001 | 4260000 | 0.0025 | 0.0008 | 0.0016 |
| F06 | 4180001 | 4280000 | 0.0017 | 0.0007 | 0.0010 |
| F06 | 4200001 | 4300000 | 0.0017 | 0.0011 | 0.0009 |
| F06 | 4220001 | 4320000 | 0.0020 | 0.0019 | 0.0013 |
| F06 | 4240001 | 4340000 | 0.0029 | 0.0017 | 0.0021 |
| F06 | 4260001 | 4360000 | 0.0027 | 0.0021 | 0.0019 |
| F06 | 4280001 | 4380000 | 0.0029 | 0.0027 | 0.0022 |
| F06 | 4300001 | 4400000 | 0.0036 | 0.0048 | 0.0028 |
| F06 | 4320001 | 4420000 | 0.0060 | 0.0045 | 0.0051 |
| F06 | 4340001 | 4440000 | 0.0058 | 0.0045 | 0.0049 |
| F06 | 4360001 | 4460000 | 0.0056 | 0.0043 | 0.0048 |
| F06 | 4380001 | 4480000 | 0.0054 | 0.0040 | 0.0046 |

| Chr | Start | End | both | A1 | A2 |
|---|---|---|---|---|---|
| F06 | 4400001 | 4500000 | 0.0048 | 0.0018 | 0.0044 |
| F06 | 4420001 | 4520000 | 0.0021 | 0.0014 | 0.0018 |
| F06 | 4440001 | 4540000 | 0.0018 | 0.0014 | 0.0015 |
| F06 | 4460001 | 4560000 | 0.0021 | 0.0010 | 0.0014 |
| F06 | 4480001 | 4580000 | 0.0019 | 0.0008 | 0.0010 |
| F06 | 4500001 | 4600000 | 0.0018 | 0.0009 | 0.0008 |
| F06 | 4520001 | 4620000 | 0.0022 | 0.0008 | 0.0010 |
| F06 | 4540001 | 4640000 | 0.0018 | 0.0011 | 0.0009 |
| F06 | 4560001 | 4660000 | 0.0019 | 0.0013 | 0.0013 |
| F06 | 4580001 | 4680000 | 0.0021 | 0.0012 | 0.0016 |
| F06 | 4600001 | 4700000 | 0.0020 | 0.0016 | 0.0015 |
| F06 | 4620001 | 4720000 | 0.0020 | 0.0014 | 0.0018 |
| F06 | 4640001 | 4740000 | 0.0020 | 0.0011 | 0.0016 |
| F06 | 4660001 | 4760000 | 0.0017 | 0.0014 | 0.0012 |
| F06 | 4680001 | 4780000 | 0.0018 | 0.0018 | 0.0014 |
| F06 | 4700001 | 4800000 | 0.0021 | 0.0013 | 0.0017 |
| F06 | 4720001 | 4820000 | 0.0018 | 0.0014 | 0.0013 |
| F06 | 4740001 | 4840000 | 0.0019 | 0.0016 | 0.0014 |
| F06 | 4760001 | 4860000 | 0.0021 | 0.0015 | 0.0017 |
| F06 | 4780001 | 4880000 | 0.0021 | 0.0013 | 0.0015 |
| F06 | 4800001 | 4900000 | 0.0020 | 0.0015 | 0.0014 |
| F06 | 4820001 | 4920000 | 0.0022 | 0.0018 | 0.0015 |
| F06 | 4840001 | 4940000 | 0.0024 | 0.0022 | 0.0018 |
| F06 | 4860001 | 4960000 | 0.0028 | 0.0026 | 0.0024 |
| F06 | 4880001 | 4980000 | 0.0031 | 0.0030 | 0.0028 |
| F06 | 4900001 | 5000000 | 0.0035 | 0.0036 | 0.0032 |
| F06 | 4920001 | 5020000 | 0.0040 | 0.0037 | 0.0038 |
| F06 | 4940001 | 5040000 | 0.0042 | 0.0032 | 0.0041 |
| F06 | 4960001 | 5060000 | 0.0036 | 0.0024 | 0.0034 |
| F06 | 4980001 | 5080000 | 0.0028 | 0.0018 | 0.0026 |
| F06 | 5000001 | 5100000 | 0.0023 | 0.0012 | 0.0019 |
| F06 | 5020001 | 5120000 | 0.0016 | 0.0015 | 0.0012 |
| F06 | 5040001 | 5140000 | 0.0017 | 0.0015 | 0.0014 |
| F06 | 5060001 | 5160000 | 0.0019 | 0.0021 | 0.0015 |
| F06 | 5080001 | 5180000 | 0.0024 | 0.0022 | 0.0018 |
| F06 | 5100001 | 5200000 | 0.0025 | 0.0040 | 0.0020 |
| F06 | 5120001 | 5220000 | 0.0044 | 0.0044 | 0.0039 |
| F06 | 5140001 | 5240000 | 0.0047 | 0.0047 | 0.0041 |
| F06 | 5160001 | 5260000 | 0.0051 | 0.0044 | 0.0044 |
| F06 | 5180001 | 5280000 | 0.0050 | 0.0043 | 0.0043 |
| F06 | 5200001 | 5300000 | 0.0049 | 0.0023 | 0.0042 |
| F06 | 5220001 | 5320000 | 0.0031 | 0.0026 | 0.0023 |
| F06 | 5240001 | 5340000 | 0.0035 | 0.0026 | 0.0027 |
| F06 | 5260001 | 5360000 | 0.0033 | 0.0024 | 0.0027 |
| F06 | 5280001 | 5380000 | 0.0032 | 0.0025 | 0.0025 |
| F06 | 5300001 | 5400000 | 0.0034 | 0.0026 | 0.0025 |
| F06 | 5320001 | 5420000 | 0.0035 | 0.0013 | 0.0025 |
| F06 | 5340001 | 5440000 | 0.0022 | 0.0011 | 0.0013 |
| F06 | 5360001 | 5460000 | 0.0020 | 0.0020 | 0.0011 |
| F06 | 5380001 | 5480000 | 0.0027 | 0.0019 | 0.0021 |
| F06 | 5400001 | 5500000 | 0.0026 | 0.0017 | 0.0021 |
| F06 | 5420001 | 5520000 | 0.0024 | 0.0013 | 0.0019 |
| F06 | 5440001 | 5540000 | 0.0022 | 0.0012 | 0.0016 |
| F06 | 5460001 | 5560000 | 0.0022 | 0.0003 | 0.0015 |
| F06 | 5480001 | 5580000 | 0.0013 | 0.0007 | 0.0005 |
| F06 | 5500001 | 5600000 | 0.0019 | 0.0011 | 0.0009 |
| F06 | 5520001 | 5620000 | 0.0022 | 0.0011 | 0.0012 |
| F06 | 5540001 | 5640000 | 0.0023 | 0.0011 | 0.0012 |

| Chr | Start | End | both | A1 | A2 |
|---|---|---|---|---|---|
| F06 | 5560001 | 5660000 | 0.0022 | 0.0014 | 0.0011 |
| F06 | 5580001 | 5680000 | 0.0027 | 0.0010 | 0.0016 |
| F06 | 5600001 | 5700000 | 0.0023 | 0.0007 | 0.0014 |
| F06 | 5620001 | 5720000 | 0.0021 | 0.0007 | 0.0014 |
| F06 | 5640001 | 5740000 | 0.0021 | 0.0007 | 0.0015 |
| F06 | 5660001 | 5760000 | 0.0021 | 0.0005 | 0.0017 |
| F06 | 5680001 | 5780000 | 0.0018 | 0.0006 | 0.0015 |
| F06 | 5700001 | 5800000 | 0.0020 | 0.0006 | 0.0018 |
| F06 | 5720001 | 5820000 | 0.0021 | 0.0011 | 0.0020 |
| F06 | 5740001 | 5840000 | 0.0026 | 0.0010 | 0.0025 |
| F06 | 5760001 | 5860000 | 0.0025 | 0.0011 | 0.0024 |
| F06 | 5780001 | 5880000 | 0.0026 | 0.0010 | 0.0026 |
| F06 | 5800001 | 5900000 | 0.0024 | 0.0013 | 0.0024 |
| F06 | 5820001 | 5920000 | 0.0023 | 0.0013 | 0.0023 |
| F06 | 5840001 | 5940000 | 0.0025 | 0.0022 | 0.0025 |
| F06 | 5860001 | 5960000 | 0.0033 | 0.0023 | 0.0033 |
| F06 | 5880001 | 5980000 | 0.0036 | 0.0035 | 0.0036 |
| F06 | 5900001 | 6000000 | 0.0049 | 0.0034 | 0.0050 |
| F06 | 5920001 | 6020000 | 0.0051 | 0.0031 | 0.0052 |
| F06 | 5940001 | 6040000 | 0.0046 | 0.0027 | 0.0046 |
| F06 | 5960001 | 6060000 | 0.0042 | 0.0027 | 0.0042 |
| F06 | 5980001 | 6080000 | 0.0041 | 0.0015 | 0.0041 |
| F06 | 6000001 | 6100000 | 0.0028 | 0.0013 | 0.0027 |
| F06 | 6020001 | 6120000 | 0.0025 | 0.0025 | 0.0022 |
| F06 | 6040001 | 6140000 | 0.0035 | 0.0035 | 0.0029 |
| F06 | 6060001 | 6160000 | 0.0051 | 0.0036 | 0.0038 |
| F06 | 6080001 | 6180000 | 0.0056 | 0.0048 | 0.0037 |
| F06 | 6100001 | 6200000 | 0.0075 | 0.0052 | 0.0047 |
| F06 | 6120001 | 6220000 | 0.0084 | 0.0052 | 0.0052 |
| F06 | 6140001 | 6240000 | 0.0088 | 0.0041 | 0.0057 |
| F06 | 6160001 | 6260000 | 0.0073 | 0.0045 | 0.0046 |
| F06 | 6180001 | 6280000 | 0.0071 | 0.0033 | 0.0050 |
| F06 | 6200001 | 6300000 | 0.0048 | 0.0028 | 0.0037 |
| F06 | 6220001 | 6320000 | 0.0039 | 0.0014 | 0.0033 |
| F06 | 6240001 | 6340000 | 0.0023 | 0.0016 | 0.0018 |
| F06 | 6260001 | 6360000 | 0.0023 | 0.0015 | 0.0020 |
| F06 | 6280001 | 6380000 | 0.0023 | 0.0015 | 0.0019 |
| F06 | 6300001 | 6400000 | 0.0025 | 0.0018 | 0.0020 |
| F06 | 6320001 | 6420000 | 0.0027 | 0.0023 | 0.0024 |
| F06 | 6340001 | 6440000 | 0.0034 | 0.0021 | 0.0031 |
| F06 | 6360001 | 6460000 | 0.0031 | 0.0014 | 0.0029 |
| F06 | 6380001 | 6480000 | 0.0025 | 0.0015 | 0.0025 |
| F06 | 6400001 | 6500000 | 0.0027 | 0.0014 | 0.0027 |
| F06 | 6420001 | 6520000 | 0.0029 | 0.0011 | 0.0029 |
| F06 | 6440001 | 6540000 | 0.0028 | 0.0007 | 0.0027 |
| F06 | 6460001 | 6560000 | 0.0024 | 0.0006 | 0.0023 |
| F06 | 6480001 | 6580000 | 0.0023 | 0.0027 | 0.0019 |
| F06 | 6500001 | 6600000 | 0.0043 | 0.0024 | 0.0037 |
| F06 | 6520001 | 6620000 | 0.0038 | 0.0023 | 0.0030 |
| F06 | 6540001 | 6640000 | 0.0034 | 0.0024 | 0.0025 |
| F06 | 6560001 | 6660000 | 0.0034 | 0.0063 | 0.0025 |
| F06 | 6580001 | 6680000 | 0.0078 | 0.0042 | 0.0070 |
| F06 | 6600001 | 6700000 | 0.0057 | 0.0047 | 0.0049 |
| F06 | 6620001 | 6720000 | 0.0062 | 0.0063 | 0.0054 |
| F06 | 6640001 | 6740000 | 0.0077 | 0.0064 | 0.0070 |
| F06 | 6660001 | 6760000 | 0.0077 | 0.0025 | 0.0071 |
| F06 | 6680001 | 6780000 | 0.0032 | 0.0026 | 0.0026 |
| F06 | 6700001 | 6800000 | 0.0032 | 0.0025 | 0.0027 |

| Chr | Start | End | both | A1 | A2 |
|---|---|---|---|---|---|
| F06 | 6720001 | 6820000 | 0.0032 | 0.0014 | 0.0026 |
| F06 | 6740001 | 6840000 | 0.0023 | 0.0017 | 0.0014 |
| F06 | 6760001 | 6860000 | 0.0029 | 0.0018 | 0.0019 |
| F06 | 6780001 | 6880000 | 0.0030 | 0.0023 | 0.0020 |
| F06 | 6800001 | 6900000 | 0.0035 | 0.0020 | 0.0024 |
| F06 | 6820001 | 6920000 | 0.0032 | 0.0017 | 0.0022 |
| F06 | 6840001 | 6940000 | 0.0026 | 0.0013 | 0.0018 |
| F06 | 6860001 | 6960000 | 0.0020 | 0.0014 | 0.0015 |
| F06 | 6880001 | 6980000 | 0.0020 | 0.0014 | 0.0015 |
| F06 | 6900001 | 7000000 | 0.0019 | 0.0013 | 0.0016 |
| F06 | 6920001 | 7020000 | 0.0019 | 0.0012 | 0.0015 |
| F06 | 6940001 | 7040000 | 0.0018 | 0.0013 | 0.0014 |
| F06 | 6960001 | 7060000 | 0.0020 | 0.0012 | 0.0014 |
| F06 | 6980001 | 7080000 | 0.0020 | 0.0007 | 0.0014 |
| F06 | 7000001 | 7100000 | 0.0015 | 0.0007 | 0.0007 |
| F06 | 7020001 | 7120000 | 0.0014 | 0.0008 | 0.0007 |
| F06 | 7040001 | 7140000 | 0.0014 | 0.0007 | 0.0008 |
| F06 | 7060001 | 7160000 | 0.0012 | 0.0017 | 0.0006 |
| F06 | 7080001 | 7180000 | 0.0021 | 0.0018 | 0.0016 |
| F06 | 7100001 | 7200000 | 0.0020 | 0.0022 | 0.0017 |
| F06 | 7120001 | 7220000 | 0.0025 | 0.0024 | 0.0020 |
| F06 | 7140001 | 7240000 | 0.0025 | 0.0028 | 0.0020 |
| F06 | 7160001 | 7260000 | 0.0029 | 0.0018 | 0.0023 |
| F06 | 7180001 | 7280000 | 0.0018 | 0.0021 | 0.0012 |
| F06 | 7200001 | 7300000 | 0.0022 | 0.0020 | 0.0014 |
| F06 | 7220001 | 7320000 | 0.0020 | 0.0023 | 0.0012 |
| F06 | 7240001 | 7340000 | 0.0023 | 0.0023 | 0.0014 |
| F06 | 7260001 | 7360000 | 0.0023 | 0.0025 | 0.0015 |
| F06 | 7280001 | 7380000 | 0.0024 | 0.0021 | 0.0016 |
| F06 | 7300001 | 7400000 | 0.0021 | 0.0022 | 0.0015 |
| F06 | 7320001 | 7420000 | 0.0022 | 0.0021 | 0.0016 |
| F06 | 7340001 | 7440000 | 0.0021 | 0.0028 | 0.0014 |
| F06 | 7360001 | 7460000 | 0.0029 | 0.0028 | 0.0022 |
| F06 | 7380001 | 7480000 | 0.0028 | 0.0031 | 0.0021 |
| F06 | 7400001 | 7500000 | 0.0031 | 0.0027 | 0.0025 |
| F06 | 7420001 | 7520000 | 0.0028 | 0.0024 | 0.0022 |
| F06 | 7440001 | 7540000 | 0.0025 | 0.0014 | 0.0022 |
| F06 | 7460001 | 7560000 | 0.0015 | 0.0016 | 0.0012 |
| F06 | 7480001 | 7580000 | 0.0017 | 0.0021 | 0.0014 |
| F06 | 7500001 | 7600000 | 0.0024 | 0.0025 | 0.0019 |
| F06 | 7520001 | 7620000 | 0.0027 | 0.0028 | 0.0020 |
| F06 | 7540001 | 7640000 | 0.0030 | 0.0028 | 0.0023 |
| F06 | 7560001 | 7660000 | 0.0029 | 0.0030 | 0.0021 |
| F06 | 7580001 | 7680000 | 0.0030 | 0.0025 | 0.0023 |
| F06 | 7600001 | 7700000 | 0.0024 | 0.0023 | 0.0018 |
| F06 | 7620001 | 7720000 | 0.0022 | 0.0019 | 0.0017 |
| F06 | 7640001 | 7740000 | 0.0018 | 0.0020 | 0.0014 |
| F06 | 7660001 | 7760000 | 0.0020 | 0.0024 | 0.0016 |
| F06 | 7680001 | 7780000 | 0.0030 | 0.0024 | 0.0026 |
| F06 | 7700001 | 7800000 | 0.0032 | 0.0026 | 0.0028 |
| F06 | 7720001 | 7820000 | 0.0034 | 0.0032 | 0.0030 |
| F06 | 7740001 | 7840000 | 0.0039 | 0.0034 | 0.0033 |
| F06 | 7760001 | 7860000 | 0.0040 | 0.0026 | 0.0034 |
| F06 | 7780001 | 7880000 | 0.0028 | 0.0032 | 0.0021 |
| F06 | 7800001 | 7900000 | 0.0033 | 0.0028 | 0.0025 |
| F06 | 7820001 | 7920000 | 0.0029 | 0.0027 | 0.0023 |
| F06 | 7840001 | 7940000 | 0.0028 | 0.0027 | 0.0023 |
| F06 | 7860001 | 7960000 | 0.0028 | 0.0031 | 0.0022 |

| Chr | Start | End | both | A1 | A2 |
|---|---|---|---|---|---|
| F06 | 7880001 | 7980000 | 0.0033 | 0.0024 | 0.0025 |
| F06 | 7900001 | 8000000 | 0.0025 | 0.0028 | 0.0018 |
| F06 | 7920001 | 8020000 | 0.0030 | 0.0025 | 0.0023 |
| F06 | 7940001 | 8040000 | 0.0027 | 0.0023 | 0.0019 |
| F06 | 7960001 | 8060000 | 0.0027 | 0.0019 | 0.0020 |
| F06 | 7980001 | 8080000 | 0.0024 | 0.0017 | 0.0019 |
| F06 | 8000001 | 8100000 | 0.0023 | 0.0018 | 0.0017 |
| F06 | 8020001 | 8120000 | 0.0027 | 0.0019 | 0.0018 |
| F06 | 8040001 | 8140000 | 0.0030 | 0.0018 | 0.0021 |
| F06 | 8060001 | 8160000 | 0.0029 | 0.0015 | 0.0019 |
| F06 | 8080001 | 8180000 | 0.0025 | 0.0016 | 0.0017 |
| F06 | 8100001 | 8200000 | 0.0027 | 0.0008 | 0.0017 |
| F06 | 8120001 | 8220000 | 0.0017 | 0.0009 | 0.0009 |
| F06 | 8140001 | 8240000 | 0.0019 | 0.0009 | 0.0011 |
| F06 | 8160001 | 8260000 | 0.0018 | 0.0020 | 0.0011 |
| F06 | 8180001 | 8280000 | 0.0030 | 0.0023 | 0.0022 |
| F06 | 8200001 | 8300000 | 0.0032 | 0.0029 | 0.0026 |
| F06 | 8220001 | 8320000 | 0.0039 | 0.0029 | 0.0032 |
| F06 | 8240001 | 8340000 | 0.0038 | 0.0029 | 0.0031 |
| F06 | 8260001 | 8360000 | 0.0038 | 0.0025 | 0.0030 |
| F06 | 8280001 | 8380000 | 0.0035 | 0.0027 | 0.0027 |
| F06 | 8300001 | 8400000 | 0.0038 | 0.0027 | 0.0028 |
| F06 | 8320001 | 8420000 | 0.0037 | 0.0023 | 0.0027 |
| F06 | 8340001 | 8440000 | 0.0035 | 0.0028 | 0.0025 |
| F06 | 8360001 | 8460000 | 0.0040 | 0.0023 | 0.0030 |
| F06 | 8380001 | 8480000 | 0.0033 | 0.0018 | 0.0024 |
| F06 | 8400001 | 8500000 | 0.0027 | 0.0014 | 0.0020 |
| F06 | 8420001 | 8520000 | 0.0023 | 0.0019 | 0.0015 |
| F06 | 8440001 | 8540000 | 0.0030 | 0.0015 | 0.0022 |
| F06 | 8460001 | 8560000 | 0.0025 | 0.0012 | 0.0017 |
| F06 | 8480001 | 8580000 | 0.0022 | 0.0019 | 0.0014 |
| F06 | 8500001 | 8600000 | 0.0030 | 0.0018 | 0.0024 |
| F06 | 8520001 | 8620000 | 0.0029 | 0.0018 | 0.0023 |
| F06 | 8540001 | 8640000 | 0.0028 | 0.0018 | 0.0022 |
| F06 | 8560001 | 8660000 | 0.0030 | 0.0021 | 0.0023 |
| F06 | 8580001 | 8680000 | 0.0034 | 0.0017 | 0.0025 |
| F06 | 8600001 | 8700000 | 0.0032 | 0.0018 | 0.0022 |
| F06 | 8620001 | 8720000 | 0.0034 | 0.0014 | 0.0025 |
| F06 | 8640001 | 8740000 | 0.0031 | 0.0014 | 0.0021 |
| F06 | 8660001 | 8760000 | 0.0027 | 0.0013 | 0.0019 |
| F06 | 8680001 | 8780000 | 0.0029 | 0.0008 | 0.0019 |
| F06 | 8700001 | 8800000 | 0.0022 | 0.0008 | 0.0012 |
| F06 | 8720001 | 8820000 | 0.0023 | 0.0007 | 0.0012 |
| F06 | 8740001 | 8840000 | 0.0021 | 0.0009 | 0.0010 |
| F06 | 8760001 | 8860000 | 0.0026 | 0.0013 | 0.0013 |
| F06 | 8780001 | 8880000 | 0.0031 | 0.0015 | 0.0016 |
| F06 | 8800001 | 8900000 | 0.0035 | 0.0024 | 0.0018 |
| F06 | 8820001 | 8920000 | 0.0043 | 0.0026 | 0.0028 |
| F06 | 8840001 | 8940000 | 0.0042 | 0.0022 | 0.0028 |
| F06 | 8860001 | 8960000 | 0.0035 | 0.0019 | 0.0023 |
| F06 | 8880001 | 8980000 | 0.0029 | 0.0021 | 0.0021 |
| F06 | 8900001 | 9000000 | 0.0028 | 0.0021 | 0.0022 |
| F06 | 8920001 | 9020000 | 0.0025 | 0.0022 | 0.0021 |
| F06 | 8940001 | 9040000 | 0.0030 | 0.0036 | 0.0025 |
| F06 | 8960001 | 9060000 | 0.0046 | 0.0035 | 0.0042 |
| F06 | 8980001 | 9080000 | 0.0045 | 0.0031 | 0.0040 |
| F06 | 9000001 | 9100000 | 0.0042 | 0.0022 | 0.0037 |
| F06 | 9020001 | 9120000 | 0.0035 | 0.0020 | 0.0027 |

| Chr | Start | End | both | A1 | A2 |
|---|---|---|---|---|---|
| F06 | 9040001 | 9140000 | 0.0031 | 0.0010 | 0.0024 |
| F06 | 9060001 | 9160000 | 0.0021 | 0.0027 | 0.0011 |
| F06 | 9080001 | 9180000 | 0.0038 | 0.0028 | 0.0030 |
| F06 | 9100001 | 9200000 | 0.0040 | 0.0031 | 0.0031 |
| F06 | 9120001 | 9220000 | 0.0042 | 0.0032 | 0.0034 |
| F06 | 9140001 | 9240000 | 0.0044 | 0.0027 | 0.0034 |
| F06 | 9160001 | 9260000 | 0.0041 | 0.0009 | 0.0030 |
| F06 | 9180001 | 9280000 | 0.0022 | 0.0009 | 0.0010 |
| F06 | 9200001 | 9300000 | 0.0023 | 0.0007 | 0.0011 |
| F06 | 9220001 | 9320000 | 0.0022 | 0.0006 | 0.0009 |
| F06 | 9240001 | 9340000 | 0.0018 | 0.0006 | 0.0007 |
| F06 | 9260001 | 9360000 | 0.0017 | 0.0008 | 0.0008 |
| F06 | 9280001 | 9380000 | 0.0017 | 0.0010 | 0.0010 |
| F06 | 9300001 | 9400000 | 0.0018 | 0.0008 | 0.0012 |
| F06 | 9320001 | 9420000 | 0.0017 | 0.0008 | 0.0011 |
| F06 | 9340001 | 9440000 | 0.0022 | 0.0012 | 0.0015 |
| F06 | 9360001 | 9460000 | 0.0025 | 0.0009 | 0.0018 |
| F06 | 9380001 | 9480000 | 0.0024 | 0.0006 | 0.0016 |
| F06 | 9400001 | 9500000 | 0.0019 | 0.0008 | 0.0012 |
| F06 | 9420001 | 9520000 | 0.0024 | 0.0009 | 0.0015 |
| F06 | 9440001 | 9540000 | 0.0022 | 0.0006 | 0.0013 |
| F06 | 9460001 | 9560000 | 0.0018 | 0.0011 | 0.0009 |
| F06 | 9480001 | 9580000 | 0.0021 | 0.0015 | 0.0013 |
| F06 | 9500001 | 9600000 | 0.0028 | 0.0015 | 0.0018 |
| F06 | 9520001 | 9620000 | 0.0022 | 0.0017 | 0.0016 |
| F06 | 9540001 | 9640000 | 0.0027 | 0.0021 | 0.0021 |
| F06 | 9560001 | 9660000 | 0.0032 | 0.0023 | 0.0025 |
| F06 | 9580001 | 9680000 | 0.0035 | 0.0018 | 0.0028 |
| F06 | 9600001 | 9700000 | 0.0031 | 0.0023 | 0.0024 |
| F06 | 9620001 | 9720000 | 0.0036 | 0.0019 | 0.0028 |
| F06 | 9640001 | 9740000 | 0.0036 | 0.0019 | 0.0024 |
| F06 | 9660001 | 9760000 | 0.0037 | 0.0016 | 0.0024 |
| F06 | 9680001 | 9780000 | 0.0034 | 0.0018 | 0.0021 |
| F06 | 9700001 | 9800000 | 0.0036 | 0.0018 | 0.0022 |
| F06 | 9720001 | 9820000 | 0.0034 | 0.0017 | 0.0023 |
| F06 | 9740001 | 9840000 | 0.0027 | 0.0013 | 0.0019 |
| F06 | 9760001 | 9860000 | 0.0023 | 0.0017 | 0.0015 |
| F06 | 9780001 | 9880000 | 0.0024 | 0.0018 | 0.0018 |
| F06 | 9800001 | 9900000 | 0.0024 | 0.0014 | 0.0019 |
| F06 | 9820001 | 9920000 | 0.0020 | 0.0020 | 0.0015 |
| F06 | 9840001 | 9940000 | 0.0027 | 0.0023 | 0.0021 |
| F06 | 9860001 | 9960000 | 0.0030 | 0.0022 | 0.0025 |
| F06 | 9880001 | 9980000 | 0.0028 | 0.0024 | 0.0023 |
| F06 | 9900001 | 10000000 | 0.0030 | 0.0024 | 0.0025 |
| F06 | 9920001 | 10020000 | 0.0030 | 0.0018 | 0.0025 |
| F06 | 9940001 | 10040000 | 0.0023 | 0.0015 | 0.0019 |
| F06 | 9960001 | 10060000 | 0.0023 | 0.0019 | 0.0017 |
| F06 | 9980001 | 10080000 | 0.0028 | 0.0014 | 0.0020 |
| F06 | 10000001 | 10100000 | 0.0023 | 0.0018 | 0.0016 |
| F06 | 10020001 | 10120000 | 0.0027 | 0.0022 | 0.0020 |
| F06 | 10040001 | 10140000 | 0.0034 | 0.0021 | 0.0026 |
| F06 | 10060001 | 10160000 | 0.0032 | 0.0013 | 0.0025 |
| F06 | 10080001 | 10180000 | 0.0024 | 0.0017 | 0.0017 |
| F06 | 10100001 | 10200000 | 0.0027 | 0.0014 | 0.0022 |
| F06 | 10120001 | 10220000 | 0.0025 | 0.0011 | 0.0019 |
| F06 | 10140001 | 10240000 | 0.0021 | 0.0015 | 0.0014 |
| F06 | 10160001 | 10260000 | 0.0026 | 0.0017 | 0.0018 |
| F06 | 10180001 | 10280000 | 0.0029 | 0.0015 | 0.0022 |

| Chr | Start | End | both | A1 | A2 |
|---|---|---|---|---|---|
| F06 | 10200001 | 10300000 | 0.0026 | 0.0014 | 0.0019 |
| F06 | 10220001 | 10320000 | 0.0024 | 0.0014 | 0.0017 |
| F06 | 10240001 | 10340000 | 0.0024 | 0.0014 | 0.0017 |
| F06 | 10260001 | 10360000 | 0.0024 | 0.0013 | 0.0016 |
| F06 | 10280001 | 10380000 | 0.0023 | 0.0012 | 0.0014 |
| F06 | 10300001 | 10400000 | 0.0022 | 0.0014 | 0.0014 |
| F06 | 10320001 | 10420000 | 0.0026 | 0.0018 | 0.0015 |
| F06 | 10340001 | 10440000 | 0.0029 | 0.0017 | 0.0019 |
| F06 | 10360001 | 10460000 | 0.0026 | 0.0016 | 0.0018 |
| F06 | 10380001 | 10480000 | 0.0025 | 0.0016 | 0.0018 |
| F06 | 10400001 | 10500000 | 0.0023 | 0.0026 | 0.0017 |
| F06 | 10420001 | 10520000 | 0.0033 | 0.0025 | 0.0027 |
| F06 | 10440001 | 10540000 | 0.0032 | 0.0034 | 0.0027 |
| F06 | 10460001 | 10560000 | 0.0043 | 0.0037 | 0.0037 |
| F06 | 10480001 | 10580000 | 0.0045 | 0.0041 | 0.0040 |
| F06 | 10500001 | 10600000 | 0.0049 | 0.0027 | 0.0045 |
| F06 | 10520001 | 10620000 | 0.0033 | 0.0026 | 0.0031 |
| F06 | 10540001 | 10640000 | 0.0032 | 0.0017 | 0.0031 |
| F06 | 10560001 | 10660000 | 0.0021 | 0.0020 | 0.0021 |
| F06 | 10580001 | 10680000 | 0.0023 | 0.0016 | 0.0023 |
| F06 | 10600001 | 10700000 | 0.0018 | 0.0017 | 0.0018 |
| F06 | 10620001 | 10720000 | 0.0019 | 0.0014 | 0.0019 |
| F06 | 10640001 | 10740000 | 0.0017 | 0.0021 | 0.0016 |
| F06 | 10660001 | 10760000 | 0.0025 | 0.0022 | 0.0022 |
| F06 | 10680001 | 10780000 | 0.0028 | 0.0027 | 0.0024 |
| F06 | 10700001 | 10800000 | 0.0034 | 0.0029 | 0.0028 |
| F06 | 10720001 | 10820000 | 0.0037 | 0.0034 | 0.0031 |
| F06 | 10740001 | 10840000 | 0.0043 | 0.0024 | 0.0036 |
| F06 | 10760001 | 10860000 | 0.0031 | 0.0023 | 0.0026 |
| F06 | 10780001 | 10880000 | 0.0029 | 0.0020 | 0.0024 |
| F06 | 10800001 | 10900000 | 0.0024 | 0.0016 | 0.0022 |
| F06 | 10820001 | 10920000 | 0.0020 | 0.0013 | 0.0018 |
| F06 | 10840001 | 10940000 | 0.0015 | 0.0022 | 0.0013 |
| F06 | 10860001 | 10960000 | 0.0027 | 0.0014 | 0.0024 |
| F06 | 10880001 | 10980000 | 0.0018 | 0.0014 | 0.0016 |
| F06 | 10900001 | 11000000 | 0.0018 | 0.0017 | 0.0015 |
| F06 | 10920001 | 11020000 | 0.0022 | 0.0019 | 0.0020 |
| F06 | 10940001 | 11040000 | 0.0025 | 0.0013 | 0.0022 |
| F06 | 10960001 | 11060000 | 0.0018 | 0.0019 | 0.0016 |
| F06 | 10980001 | 11080000 | 0.0023 | 0.0023 | 0.0022 |
| F06 | 11000001 | 11100000 | 0.0029 | 0.0027 | 0.0026 |
| F06 | 11020001 | 11120000 | 0.0033 | 0.0030 | 0.0030 |
| F06 | 11040001 | 11140000 | 0.0037 | 0.0034 | 0.0033 |
| F06 | 11060001 | 11160000 | 0.0043 | 0.0031 | 0.0038 |
| F06 | 11080001 | 11180000 | 0.0041 | 0.0029 | 0.0036 |
| F06 | 11100001 | 11200000 | 0.0037 | 0.0022 | 0.0034 |
| F06 | 11120001 | 11220000 | 0.0029 | 0.0023 | 0.0027 |
| F06 | 11140001 | 11240000 | 0.0029 | 0.0017 | 0.0027 |
| F06 | 11160001 | 11260000 | 0.0020 | 0.0015 | 0.0019 |
| F06 | 11180001 | 11280000 | 0.0018 | 0.0016 | 0.0018 |
| F06 | 11200001 | 11300000 | 0.0019 | 0.0016 | 0.0018 |
| F06 | 11220001 | 11320000 | 0.0021 | 0.0009 | 0.0018 |
| F06 | 11240001 | 11340000 | 0.0013 | 0.0007 | 0.0010 |
| F06 | 11260001 | 11360000 | 0.0011 | 0.0006 | 0.0008 |
| F06 | 11280001 | 11380000 | 0.0010 | 0.0003 | 0.0007 |
| F06 | 11300001 | 11400000 | 0.0005 | 0.0003 | 0.0004 |
| F06 | 11320001 | 11420000 | 0.0005 | 0.0003 | 0.0005 |
| F06 | 11340001 | 11440000 | 0.0004 | 0.0005 | 0.0004 |

| Chr | Start | End | both | A1 | A2 |
|---|---|---|---|---|---|
| F06 | 11360001 | 11460000 | 0.0006 | 0.0009 | 0.0006 |
| F06 | 11380001 | 11480000 | 0.0011 | 0.0011 | 0.0010 |
| F06 | 11400001 | 11500000 | 0.0012 | 0.0011 | 0.0012 |
| F06 | 11420001 | 11520000 | 0.0014 | 0.0011 | 0.0013 |
| F06 | 11440001 | 11540000 | 0.0015 | 0.0013 | 0.0013 |
| F06 | 11460001 | 11560000 | 0.0019 | 0.0008 | 0.0015 |
| F06 | 11480001 | 11580000 | 0.0016 | 0.0009 | 0.0011 |
| F06 | 11500001 | 11600000 | 0.0019 | 0.0009 | 0.0014 |
| F06 | 11520001 | 11620000 | 0.0017 | 0.0014 | 0.0013 |
| F06 | 11540001 | 11640000 | 0.0023 | 0.0010 | 0.0018 |
| F06 | 11560001 | 11660000 | 0.0019 | 0.0012 | 0.0013 |
| F06 | 11580001 | 11680000 | 0.0020 | 0.0018 | 0.0014 |
| F06 | 11600001 | 11700000 | 0.0027 | 0.0022 | 0.0020 |
| F06 | 11620001 | 11720000 | 0.0034 | 0.0022 | 0.0026 |
| F06 | 11640001 | 11740000 | 0.0033 | 0.0032 | 0.0026 |
| F06 | 11660001 | 11760000 | 0.0046 | 0.0038 | 0.0036 |
| F06 | 11680001 | 11780000 | 0.0052 | 0.0032 | 0.0042 |
| F06 | 11700001 | 11800000 | 0.0045 | 0.0027 | 0.0035 |
| F06 | 11720001 | 11820000 | 0.0037 | 0.0023 | 0.0029 |
| F06 | 11740001 | 11840000 | 0.0032 | 0.0014 | 0.0025 |
| F06 | 11760001 | 11860000 | 0.0019 | 0.0009 | 0.0015 |
| F06 | 11780001 | 11880000 | 0.0012 | 0.0011 | 0.0011 |
| F06 | 11800001 | 11900000 | 0.0013 | 0.0011 | 0.0013 |
| F06 | 11820001 | 11920000 | 0.0013 | 0.0015 | 0.0013 |
| F06 | 11840001 | 11940000 | 0.0016 | 0.0017 | 0.0016 |
| F06 | 11860001 | 11960000 | 0.0018 | 0.0015 | 0.0018 |
| F06 | 11880001 | 11980000 | 0.0016 | 0.0012 | 0.0016 |
| F06 | 11900001 | 12000000 | 0.0013 | 0.0013 | 0.0013 |
| F06 | 11920001 | 12020000 | 0.0015 | 0.0009 | 0.0015 |
| F06 | 11940001 | 12040000 | 0.0011 | 0.0009 | 0.0011 |
| F06 | 11960001 | 12060000 | 0.0011 | 0.0011 | 0.0011 |
| F06 | 11980001 | 12080000 | 0.0014 | 0.0013 | 0.0014 |
| F06 | 12000001 | 12100000 | 0.0016 | 0.0015 | 0.0015 |
| F06 | 12020001 | 12120000 | 0.0019 | 0.0021 | 0.0017 |
| F06 | 12040001 | 12140000 | 0.0027 | 0.0019 | 0.0024 |
| F06 | 12060001 | 12160000 | 0.0026 | 0.0015 | 0.0023 |
| F06 | 12080001 | 12180000 | 0.0021 | 0.0015 | 0.0018 |
| F06 | 12100001 | 12200000 | 0.0021 | 0.0015 | 0.0018 |
| F06 | 12120001 | 12220000 | 0.0019 | 0.0011 | 0.0017 |
| F06 | 12140001 | 12240000 | 0.0013 | 0.0015 | 0.0012 |
| F06 | 12160001 | 12260000 | 0.0017 | 0.0022 | 0.0016 |
| F06 | 12180001 | 12280000 | 0.0024 | 0.0021 | 0.0023 |
| F06 | 12200001 | 12300000 | 0.0022 | 0.0020 | 0.0022 |
| F06 | 12220001 | 12320000 | 0.0023 | 0.0019 | 0.0023 |
| F06 | 12240001 | 12340000 | 0.0022 | 0.0020 | 0.0021 |
| F06 | 12260001 | 12360000 | 0.0024 | 0.0016 | 0.0023 |
| F06 | 12280001 | 12380000 | 0.0020 | 0.0015 | 0.0019 |
| F06 | 12300001 | 12400000 | 0.0019 | 0.0012 | 0.0018 |
| F06 | 12320001 | 12420000 | 0.0016 | 0.0013 | 0.0015 |
| F06 | 12340001 | 12440000 | 0.0018 | 0.0009 | 0.0017 |
| F06 | 12360001 | 12460000 | 0.0014 | 0.0010 | 0.0012 |
| F06 | 12380001 | 12480000 | 0.0016 | 0.0009 | 0.0012 |
| F06 | 12400001 | 12500000 | 0.0015 | 0.0011 | 0.0012 |
| F06 | 12420001 | 12520000 | 0.0017 | 0.0011 | 0.0013 |
| F06 | 12440001 | 12540000 | 0.0016 | 0.0011 | 0.0012 |
| F06 | 12460001 | 12560000 | 0.0014 | 0.0012 | 0.0012 |
| F06 | 12480001 | 12580000 | 0.0014 | 0.0013 | 0.0013 |
| F06 | 12500001 | 12600000 | 0.0015 | 0.0011 | 0.0014 |

| Chr | Start | End | both | A1 | A2 |
|---|---|---|---|---|---|
| F06 | 12520001 | 12620000 | 0.0013 | 0.0025 | 0.0013 |
| F06 | 12540001 | 12640000 | 0.0028 | 0.0024 | 0.0026 |
| F06 | 12560001 | 12660000 | 0.0029 | 0.0031 | 0.0026 |
| F06 | 12580001 | 12680000 | 0.0037 | 0.0043 | 0.0033 |
| F06 | 12600001 | 12700000 | 0.0050 | 0.0044 | 0.0045 |
| F06 | 12620001 | 12720000 | 0.0053 | 0.0056 | 0.0048 |
| F06 | 12640001 | 12740000 | 0.0065 | 0.0058 | 0.0059 |
| F06 | 12660001 | 12760000 | 0.0068 | 0.0050 | 0.0062 |
| F06 | 12680001 | 12780000 | 0.0059 | 0.0047 | 0.0054 |
| F06 | 12700001 | 12800000 | 0.0055 | 0.0049 | 0.0051 |
| F06 | 12720001 | 12820000 | 0.0057 | 0.0027 | 0.0052 |
| F06 | 12740001 | 12840000 | 0.0033 | 0.0028 | 0.0029 |
| F06 | 12760001 | 12860000 | 0.0033 | 0.0024 | 0.0029 |
| F06 | 12780001 | 12880000 | 0.0029 | 0.0014 | 0.0025 |
| F06 | 12800001 | 12900000 | 0.0019 | 0.0018 | 0.0016 |
| F06 | 12820001 | 12920000 | 0.0021 | 0.0016 | 0.0019 |
| F06 | 12840001 | 12940000 | 0.0023 | 0.0019 | 0.0020 |
| F06 | 12860001 | 12960000 | 0.0025 | 0.0022 | 0.0023 |
| F06 | 12880001 | 12980000 | 0.0029 | 0.0024 | 0.0027 |
| F06 | 12900001 | 13000000 | 0.0031 | 0.0018 | 0.0030 |
| F06 | 12920001 | 13020000 | 0.0025 | 0.0019 | 0.0023 |
| F06 | 12940001 | 13040000 | 0.0025 | 0.0025 | 0.0024 |
| F06 | 12960001 | 13060000 | 0.0032 | 0.0023 | 0.0030 |
| F06 | 12980001 | 13080000 | 0.0032 | 0.0027 | 0.0028 |
| F06 | 13000001 | 13100000 | 0.0035 | 0.0026 | 0.0031 |
| F06 | 13020001 | 13120000 | 0.0034 | 0.0023 | 0.0029 |
| F06 | 13040001 | 13140000 | 0.0033 | 0.0015 | 0.0026 |
| F06 | 13060001 | 13160000 | 0.0023 | 0.0020 | 0.0017 |
| F06 | 13080001 | 13180000 | 0.0027 | 0.0022 | 0.0021 |
| F06 | 13100001 | 13200000 | 0.0031 | 0.0030 | 0.0024 |
| F06 | 13120001 | 13220000 | 0.0042 | 0.0033 | 0.0034 |
| F06 | 13140001 | 13240000 | 0.0045 | 0.0068 | 0.0037 |
| F06 | 13160001 | 13260000 | 0.0082 | 0.0062 | 0.0072 |
| F06 | 13180001 | 13280000 | 0.0075 | 0.0055 | 0.0066 |
| F06 | 13200001 | 13300000 | 0.0067 | 0.0050 | 0.0059 |
| F06 | 13220001 | 13320000 | 0.0061 | 0.0048 | 0.0052 |
| F06 | 13240001 | 13340000 | 0.0060 | 0.0018 | 0.0052 |
| F06 | 13260001 | 13360000 | 0.0028 | 0.0018 | 0.0020 |
| F06 | 13280001 | 13380000 | 0.0031 | 0.0016 | 0.0021 |
| F06 | 13300001 | 13400000 | 0.0029 | 0.0015 | 0.0020 |
| F06 | 13320001 | 13420000 | 0.0028 | 0.0013 | 0.0019 |
| F06 | 13340001 | 13440000 | 0.0024 | 0.0006 | 0.0016 |
| F06 | 13360001 | 13460000 | 0.0018 | 0.0006 | 0.0009 |
| F06 | 13380001 | 13480000 | 0.0015 | 0.0012 | 0.0007 |
| F06 | 13400001 | 13500000 | 0.0022 | 0.0012 | 0.0013 |
| F06 | 13420001 | 13520000 | 0.0023 | 0.0016 | 0.0013 |
| F06 | 13440001 | 13540000 | 0.0027 | 0.0014 | 0.0017 |
| F06 | 13460001 | 13560000 | 0.0025 | 0.0021 | 0.0016 |
| F06 | 13480001 | 13580000 | 0.0034 | 0.0017 | 0.0024 |
| F06 | 13500001 | 13600000 | 0.0028 | 0.0015 | 0.0020 |
| F06 | 13520001 | 13620000 | 0.0026 | 0.0013 | 0.0019 |
| F06 | 13540001 | 13640000 | 0.0024 | 0.0019 | 0.0017 |
| F06 | 13560001 | 13660000 | 0.0031 | 0.0021 | 0.0022 |
| F06 | 13580001 | 13680000 | 0.0033 | 0.0023 | 0.0024 |
| F06 | 13600001 | 13700000 | 0.0036 | 0.0027 | 0.0026 |
| F06 | 13620001 | 13720000 | 0.0040 | 0.0030 | 0.0031 |
| F06 | 13640001 | 13740000 | 0.0041 | 0.0027 | 0.0032 |
| F06 | 13660001 | 13760000 | 0.0037 | 0.0025 | 0.0030 |

| Chr | Start | End | both | A1 | A2 |
|---|---|---|---|---|---|
| F06 | 13680001 | 13780000 | 0.0033 | 0.0024 | 0.0027 |
| F06 | 13700001 | 13800000 | 0.0034 | 0.0035 | 0.0027 |
| F06 | 13720001 | 13820000 | 0.0048 | 0.0045 | 0.0039 |
| F06 | 13740001 | 13840000 | 0.0060 | 0.0044 | 0.0050 |
| F06 | 13760001 | 13860000 | 0.0059 | 0.0049 | 0.0049 |
| F06 | 13780001 | 13880000 | 0.0064 | 0.0051 | 0.0054 |
| F06 | 13800001 | 13900000 | 0.0064 | 0.0039 | 0.0055 |
| F06 | 13820001 | 13920000 | 0.0051 | 0.0028 | 0.0044 |
| F06 | 13840001 | 13940000 | 0.0039 | 0.0027 | 0.0033 |
| F06 | 13860001 | 13960000 | 0.0039 | 0.0015 | 0.0033 |
| F06 | 13880001 | 13980000 | 0.0028 | 0.0014 | 0.0021 |
| F06 | 13900001 | 14000000 | 0.0026 | 0.0009 | 0.0019 |
| F06 | 13920001 | 14020000 | 0.0026 | 0.0006 | 0.0018 |
| F06 | 13940001 | 14040000 | 0.0022 | 0.0006 | 0.0016 |
| F06 | 13960001 | 14060000 | 0.0025 | 0.0008 | 0.0020 |
| F06 | 13980001 | 14080000 | 0.0028 | 0.0008 | 0.0023 |
| F06 | 14000001 | 14100000 | 0.0029 | 0.0009 | 0.0025 |
| F06 | 14020001 | 14120000 | 0.0028 | 0.0012 | 0.0024 |
| F06 | 14040001 | 14140000 | 0.0029 | 0.0014 | 0.0025 |
| F06 | 14060001 | 14160000 | 0.0026 | 0.0017 | 0.0023 |
| F06 | 14080001 | 14180000 | 0.0027 | 0.0020 | 0.0024 |
| F06 | 14100001 | 14200000 | 0.0026 | 0.0022 | 0.0024 |
| F06 | 14120001 | 14220000 | 0.0025 | 0.0023 | 0.0023 |
| F06 | 14140001 | 14240000 | 0.0027 | 0.0031 | 0.0024 |
| F06 | 14160001 | 14260000 | 0.0036 | 0.0025 | 0.0033 |
| F06 | 14180001 | 14280000 | 0.0031 | 0.0022 | 0.0028 |
| F06 | 14200001 | 14300000 | 0.0030 | 0.0021 | 0.0028 |
| F06 | 14220001 | 14320000 | 0.0029 | 0.0016 | 0.0027 |
| F06 | 14240001 | 14340000 | 0.0023 | 0.0009 | 0.0022 |
| F06 | 14260001 | 14360000 | 0.0015 | 0.0011 | 0.0014 |
| F06 | 14280001 | 14380000 | 0.0017 | 0.0022 | 0.0017 |
| F06 | 14300001 | 14400000 | 0.0026 | 0.0019 | 0.0025 |
| F06 | 14320001 | 14420000 | 0.0026 | 0.0026 | 0.0025 |
| F06 | 14340001 | 14440000 | 0.0035 | 0.0033 | 0.0033 |
| F06 | 14360001 | 14460000 | 0.0042 | 0.0034 | 0.0040 |
| F06 | 14380001 | 14480000 | 0.0044 | 0.0026 | 0.0041 |
| F06 | 14400001 | 14500000 | 0.0036 | 0.0030 | 0.0034 |
| F06 | 14420001 | 14520000 | 0.0040 | 0.0024 | 0.0037 |
| F06 | 14440001 | 14540000 | 0.0034 | 0.0014 | 0.0031 |
| F06 | 14460001 | 14560000 | 0.0028 | 0.0011 | 0.0024 |
| F06 | 14480001 | 14580000 | 0.0026 | 0.0008 | 0.0022 |
| F06 | 14500001 | 14600000 | 0.0025 | 0.0004 | 0.0021 |
| F06 | 14520001 | 14620000 | 0.0021 | 0.0003 | 0.0019 |
| F06 | 14540001 | 14640000 | 0.0023 | 0.0008 | 0.0021 |
| F06 | 14560001 | 14660000 | 0.0028 | 0.0011 | 0.0028 |
| F06 | 14580001 | 14680000 | 0.0030 | 0.0013 | 0.0030 |
| F06 | 14600001 | 14700000 | 0.0030 | 0.0015 | 0.0030 |
| F06 | 14620001 | 14720000 | 0.0028 | 0.0015 | 0.0028 |
| F06 | 14640001 | 14740000 | 0.0032 | 0.0014 | 0.0033 |
| F06 | 14660001 | 14760000 | 0.0035 | 0.0015 | 0.0035 |
| F06 | 14680001 | 14780000 | 0.0042 | 0.0024 | 0.0043 |
| F06 | 14700001 | 14800000 | 0.0054 | 0.0024 | 0.0055 |
| F06 | 14720001 | 14820000 | 0.0055 | 0.0024 | 0.0056 |
| F06 | 14740001 | 14840000 | 0.0050 | 0.0020 | 0.0051 |
| F06 | 14760001 | 14860000 | 0.0040 | 0.0015 | 0.0040 |
| F06 | 14780001 | 14880000 | 0.0032 | 0.0004 | 0.0033 |
| F06 | 14800001 | 14900000 | 0.0018 | 0.0011 | 0.0019 |
| F06 | 14820001 | 14920000 | 0.0025 | 0.0014 | 0.0026 |

| Chr | Start | End | both | A1 | A2 |
|---|---|---|---|---|---|
| F06 | 14840001 | 14940000 | 0.0026 | 0.0019 | 0.0026 |
| F06 | 14860001 | 14960000 | 0.0031 | 0.0024 | 0.0031 |
| F06 | 14880001 | 14980000 | 0.0036 | 0.0028 | 0.0036 |
| F06 | 14900001 | 15000000 | 0.0043 | 0.0022 | 0.0044 |
| F06 | 14920001 | 15020000 | 0.0039 | 0.0022 | 0.0039 |
| F06 | 14940001 | 15040000 | 0.0041 | 0.0017 | 0.0041 |
| F06 | 14960001 | 15060000 | 0.0037 | 0.0013 | 0.0037 |
| F06 | 14980001 | 15080000 | 0.0028 | 0.0009 | 0.0028 |
| F06 | 15000001 | 15100000 | 0.0025 | 0.0013 | 0.0024 |
| F06 | 15020001 | 15120000 | 0.0029 | 0.0012 | 0.0028 |
| F06 | 15040001 | 15140000 | 0.0032 | 0.0014 | 0.0031 |
| F06 | 15060001 | 15160000 | 0.0035 | 0.0019 | 0.0033 |
| F06 | 15080001 | 15180000 | 0.0044 | 0.0026 | 0.0041 |
| F06 | 15100001 | 15200000 | 0.0046 | 0.0024 | 0.0044 |
| F06 | 15120001 | 15220000 | 0.0044 | 0.0023 | 0.0040 |
| F06 | 15140001 | 15240000 | 0.0038 | 0.0023 | 0.0032 |
| F06 | 15160001 | 15260000 | 0.0038 | 0.0026 | 0.0031 |
| F06 | 15180001 | 15280000 | 0.0036 | 0.0022 | 0.0028 |
| F06 | 15200001 | 15300000 | 0.0033 | 0.0018 | 0.0024 |
| F06 | 15220001 | 15320000 | 0.0028 | 0.0021 | 0.0019 |
| F06 | 15240001 | 15340000 | 0.0031 | 0.0024 | 0.0022 |
| F06 | 15260001 | 15360000 | 0.0033 | 0.0020 | 0.0025 |
| F06 | 15280001 | 15380000 | 0.0031 | 0.0016 | 0.0023 |
| F06 | 15300001 | 15400000 | 0.0026 | 0.0019 | 0.0020 |
| F06 | 15320001 | 15420000 | 0.0029 | 0.0023 | 0.0023 |
| F06 | 15340001 | 15440000 | 0.0032 | 0.0020 | 0.0026 |
| F06 | 15360001 | 15460000 | 0.0030 | 0.0025 | 0.0023 |
| F06 | 15380001 | 15480000 | 0.0033 | 0.0026 | 0.0026 |
| F06 | 15400001 | 15500000 | 0.0036 | 0.0040 | 0.0027 |
| F06 | 15420001 | 15520000 | 0.0050 | 0.0035 | 0.0042 |
| F06 | 15440001 | 15540000 | 0.0046 | 0.0039 | 0.0037 |
| F06 | 15460001 | 15560000 | 0.0050 | 0.0036 | 0.0042 |
| F06 | 15480001 | 15580000 | 0.0050 | 0.0040 | 0.0040 |
| F06 | 15500001 | 15600000 | 0.0053 | 0.0032 | 0.0046 |
| F06 | 15520001 | 15620000 | 0.0045 | 0.0038 | 0.0036 |
| F06 | 15540001 | 15640000 | 0.0049 | 0.0034 | 0.0042 |
| F06 | 15560001 | 15660000 | 0.0045 | 0.0027 | 0.0037 |
| F06 | 15580001 | 15680000 | 0.0037 | 0.0025 | 0.0030 |
| F06 | 15600001 | 15700000 | 0.0036 | 0.0025 | 0.0028 |
| F06 | 15620001 | 15720000 | 0.0033 | 0.0020 | 0.0026 |
| F06 | 15640001 | 15740000 | 0.0029 | 0.0024 | 0.0019 |
| F06 | 15660001 | 15760000 | 0.0035 | 0.0024 | 0.0023 |
| F06 | 15680001 | 15780000 | 0.0034 | 0.0027 | 0.0023 |
| F06 | 15700001 | 15800000 | 0.0038 | 0.0022 | 0.0026 |
| F06 | 15720001 | 15820000 | 0.0036 | 0.0021 | 0.0023 |
| F06 | 15740001 | 15840000 | 0.0038 | 0.0018 | 0.0022 |
| F06 | 15760001 | 15860000 | 0.0033 | 0.0026 | 0.0019 |
| F06 | 15780001 | 15880000 | 0.0043 | 0.0021 | 0.0027 |
| F06 | 15800001 | 15900000 | 0.0037 | 0.0020 | 0.0022 |
| F06 | 15820001 | 15920000 | 0.0035 | 0.0017 | 0.0021 |
| F06 | 15840001 | 15940000 | 0.0029 | 0.0021 | 0.0019 |
| F06 | 15860001 | 15960000 | 0.0031 | 0.0018 | 0.0022 |
| F06 | 15880001 | 15980000 | 0.0027 | 0.0021 | 0.0020 |
| F06 | 15900001 | 16000000 | 0.0031 | 0.0026 | 0.0025 |
| F06 | 15920001 | 16020000 | 0.0035 | 0.0034 | 0.0029 |
| F06 | 15940001 | 16040000 | 0.0044 | 0.0036 | 0.0036 |
| F06 | 15960001 | 16060000 | 0.0047 | 0.0042 | 0.0038 |
| F06 | 15980001 | 16080000 | 0.0054 | 0.0050 | 0.0045 |

| Chr | Start | End | both | A1 | A2 |
|---|---|---|---|---|---|
| F06 | 16000001 | 16100000 | 0.0062 | 0.0048 | 0.0053 |
| F06 | 16020001 | 16120000 | 0.0063 | 0.0046 | 0.0052 |
| F06 | 16040001 | 16140000 | 0.0062 | 0.0043 | 0.0051 |
| F06 | 16060001 | 16160000 | 0.0057 | 0.0036 | 0.0048 |
| F06 | 16080001 | 16180000 | 0.0054 | 0.0024 | 0.0042 |
| F06 | 16100001 | 16200000 | 0.0043 | 0.0028 | 0.0030 |
| F06 | 16120001 | 16220000 | 0.0044 | 0.0024 | 0.0033 |
| F06 | 16140001 | 16240000 | 0.0044 | 0.0024 | 0.0030 |
| F06 | 16160001 | 16260000 | 0.0042 | 0.0028 | 0.0029 |
| F06 | 16180001 | 16280000 | 0.0042 | 0.0030 | 0.0030 |
| F06 | 16200001 | 16300000 | 0.0047 | 0.0030 | 0.0035 |
| F06 | 16220001 | 16320000 | 0.0045 | 0.0036 | 0.0034 |
| F06 | 16240001 | 16340000 | 0.0049 | 0.0036 | 0.0040 |
| F06 | 16260001 | 16360000 | 0.0050 | 0.0028 | 0.0042 |
| F06 | 16280001 | 16380000 | 0.0043 | 0.0028 | 0.0034 |
| F06 | 16300001 | 16400000 | 0.0038 | 0.0025 | 0.0031 |
| F06 | 16320001 | 16420000 | 0.0037 | 0.0019 | 0.0029 |
| F06 | 16340001 | 16440000 | 0.0034 | 0.0016 | 0.0023 |
| F06 | 16360001 | 16460000 | 0.0034 | 0.0024 | 0.0022 |
| F06 | 16380001 | 16480000 | 0.0041 | 0.0021 | 0.0029 |
| F06 | 16400001 | 16500000 | 0.0043 | 0.0024 | 0.0027 |
| F06 | 16420001 | 16520000 | 0.0045 | 0.0033 | 0.0028 |
| F06 | 16440001 | 16540000 | 0.0050 | 0.0033 | 0.0038 |
| F06 | 16460001 | 16560000 | 0.0048 | 0.0031 | 0.0037 |
| F06 | 16480001 | 16580000 | 0.0045 | 0.0044 | 0.0035 |
| F06 | 16500001 | 16600000 | 0.0056 | 0.0040 | 0.0051 |
| F06 | 16520001 | 16620000 | 0.0052 | 0.0032 | 0.0047 |
| F06 | 16540001 | 16640000 | 0.0045 | 0.0030 | 0.0037 |
| F06 | 16560001 | 16660000 | 0.0042 | 0.0031 | 0.0034 |
| F06 | 16580001 | 16680000 | 0.0044 | 0.0023 | 0.0035 |
| F06 | 16600001 | 16700000 | 0.0037 | 0.0026 | 0.0026 |
| F06 | 16620001 | 16720000 | 0.0038 | 0.0024 | 0.0030 |
| F06 | 16640001 | 16740000 | 0.0033 | 0.0027 | 0.0027 |
| F06 | 16660001 | 16760000 | 0.0034 | 0.0025 | 0.0030 |
| F06 | 16680001 | 16780000 | 0.0033 | 0.0030 | 0.0027 |
| F06 | 16700001 | 16800000 | 0.0034 | 0.0035 | 0.0029 |
| F06 | 16720001 | 16820000 | 0.0042 | 0.0045 | 0.0034 |
| F06 | 16740001 | 16840000 | 0.0054 | 0.0045 | 0.0043 |
| F06 | 16760001 | 16860000 | 0.0055 | 0.0045 | 0.0044 |
| F06 | 16780001 | 16880000 | 0.0056 | 0.0037 | 0.0043 |
| F06 | 16800001 | 16900000 | 0.0051 | 0.0030 | 0.0038 |
| F06 | 16820001 | 16920000 | 0.0042 | 0.0021 | 0.0031 |
| F06 | 16840001 | 16940000 | 0.0032 | 0.0022 | 0.0023 |
| F06 | 16860001 | 16960000 | 0.0034 | 0.0020 | 0.0024 |
| F06 | 16880001 | 16980000 | 0.0028 | 0.0025 | 0.0022 |
| F06 | 16900001 | 17000000 | 0.0031 | 0.0019 | 0.0026 |
| F06 | 16920001 | 17020000 | 0.0029 | 0.0017 | 0.0022 |
| F06 | 16940001 | 17040000 | 0.0028 | 0.0023 | 0.0019 |
| F06 | 16960001 | 17060000 | 0.0032 | 0.0022 | 0.0025 |
| F06 | 16980001 | 17080000 | 0.0032 | 0.0019 | 0.0025 |
| F06 | 17000001 | 17100000 | 0.0030 | 0.0023 | 0.0022 |
| F06 | 17020001 | 17120000 | 0.0033 | 0.0042 | 0.0026 |
| F06 | 17040001 | 17140000 | 0.0048 | 0.0042 | 0.0040 |
| F06 | 17060001 | 17160000 | 0.0046 | 0.0046 | 0.0038 |
| F06 | 17080001 | 17180000 | 0.0050 | 0.0042 | 0.0043 |
| F06 | 17100001 | 17200000 | 0.0044 | 0.0044 | 0.0038 |
| F06 | 17120001 | 17220000 | 0.0042 | 0.0036 | 0.0038 |
| F06 | 17140001 | 17240000 | 0.0038 | 0.0038 | 0.0036 |

| Chr | Start | End | both | A1 | A2 |
|---|---|---|---|---|---|
| F06 | 17160001 | 17260000 | 0.0039 | 0.0038 | 0.0037 |
| F06 | 17180001 | 17280000 | 0.0037 | 0.0045 | 0.0033 |
| F06 | 17200001 | 17300000 | 0.0042 | 0.0046 | 0.0036 |
| F06 | 17220001 | 17320000 | 0.0042 | 0.0039 | 0.0036 |
| F06 | 17240001 | 17340000 | 0.0034 | 0.0031 | 0.0028 |
| F06 | 17260001 | 17360000 | 0.0027 | 0.0030 | 0.0023 |
| F06 | 17280001 | 17380000 | 0.0028 | 0.0025 | 0.0025 |
| F06 | 17300001 | 17400000 | 0.0024 | 0.0024 | 0.0023 |
| F06 | 17320001 | 17420000 | 0.0024 | 0.0030 | 0.0023 |
| F06 | 17340001 | 17440000 | 0.0029 | 0.0036 | 0.0026 |
| F06 | 17360001 | 17460000 | 0.0031 | 0.0035 | 0.0027 |
| F06 | 17380001 | 17480000 | 0.0032 | 0.0038 | 0.0027 |
| F06 | 17400001 | 17500000 | 0.0034 | 0.0037 | 0.0027 |
| F06 | 17420001 | 17520000 | 0.0034 | 0.0033 | 0.0026 |
| F06 | 17440001 | 17540000 | 0.0030 | 0.0025 | 0.0022 |
| F06 | 17460001 | 17560000 | 0.0025 | 0.0025 | 0.0019 |
| F06 | 17480001 | 17580000 | 0.0023 | 0.0025 | 0.0017 |
| F06 | 17500001 | 17600000 | 0.0026 | 0.0034 | 0.0020 |
| F06 | 17520001 | 17620000 | 0.0035 | 0.0047 | 0.0028 |
| F06 | 17540001 | 17640000 | 0.0046 | 0.0051 | 0.0039 |
| F06 | 17560001 | 17660000 | 0.0050 | 0.0051 | 0.0042 |
| F06 | 17580001 | 17680000 | 0.0049 | 0.0048 | 0.0042 |
| F06 | 17600001 | 17700000 | 0.0045 | 0.0041 | 0.0038 |
| F06 | 17620001 | 17720000 | 0.0038 | 0.0029 | 0.0032 |
| F06 | 17640001 | 17740000 | 0.0027 | 0.0033 | 0.0022 |
| F06 | 17660001 | 17760000 | 0.0034 | 0.0039 | 0.0030 |
| F06 | 17680001 | 17780000 | 0.0041 | 0.0046 | 0.0037 |
| F06 | 17700001 | 17800000 | 0.0051 | 0.0043 | 0.0048 |
| F06 | 17720001 | 17820000 | 0.0046 | 0.0039 | 0.0042 |
| F06 | 17740001 | 17840000 | 0.0042 | 0.0037 | 0.0039 |
| F06 | 17760001 | 17860000 | 0.0036 | 0.0033 | 0.0031 |
| F06 | 17780001 | 17880000 | 0.0033 | 0.0032 | 0.0029 |
| F06 | 17800001 | 17900000 | 0.0030 | 0.0037 | 0.0025 |
| F06 | 17820001 | 17920000 | 0.0033 | 0.0040 | 0.0026 |
| F06 | 17840001 | 17940000 | 0.0035 | 0.0041 | 0.0027 |
| F06 | 17860001 | 17960000 | 0.0037 | 0.0039 | 0.0030 |
| F06 | 17880001 | 17980000 | 0.0033 | 0.0031 | 0.0025 |
| F06 | 17900001 | 18000000 | 0.0027 | 0.0026 | 0.0021 |
| F06 | 17920001 | 18020000 | 0.0024 | 0.0035 | 0.0019 |
| F06 | 17940001 | 18040000 | 0.0034 | 0.0037 | 0.0029 |
| F06 | 17960001 | 18060000 | 0.0033 | 0.0036 | 0.0026 |
| F06 | 17980001 | 18080000 | 0.0032 | 0.0037 | 0.0025 |
| F06 | 18000001 | 18100000 | 0.0033 | 0.0043 | 0.0025 |
| F06 | 18020001 | 18120000 | 0.0037 | 0.0044 | 0.0028 |
| F06 | 18040001 | 18140000 | 0.0040 | 0.0039 | 0.0032 |
| F06 | 18060001 | 18160000 | 0.0039 | 0.0048 | 0.0034 |
| F06 | 18080001 | 18180000 | 0.0050 | 0.0047 | 0.0044 |
| F06 | 18100001 | 18200000 | 0.0049 | 0.0042 | 0.0043 |
| F06 | 18120001 | 18220000 | 0.0048 | 0.0027 | 0.0044 |
| F06 | 18140001 | 18240000 | 0.0032 | 0.0021 | 0.0028 |
| F06 | 18160001 | 18260000 | 0.0025 | 0.0012 | 0.0022 |
| F06 | 18180001 | 18280000 | 0.0015 | 0.0009 | 0.0013 |
| F06 | 18200001 | 18300000 | 0.0012 | 0.0006 | 0.0011 |
| F06 | 18220001 | 18320000 | 0.0008 | 0.0009 | 0.0008 |
| F06 | 18240001 | 18340000 | 0.0010 | 0.0011 | 0.0010 |
| F06 | 18260001 | 18360000 | 0.0014 | 0.0009 | 0.0013 |
| F06 | 18280001 | 18380000 | 0.0013 | 0.0013 | 0.0011 |
| F06 | 18300001 | 18400000 | 0.0020 | 0.0019 | 0.0015 |

| Chr | Start | End | both | A1 | A2 |
|---|---|---|---|---|---|
| F06 | 18320001 | 18420000 | 0.0029 | 0.0017 | 0.0023 |
| F06 | 18340001 | 18440000 | 0.0029 | 0.0023 | 0.0022 |
| F06 | 18360001 | 18460000 | 0.0037 | 0.0023 | 0.0029 |
| F06 | 18380001 | 18480000 | 0.0039 | 0.0034 | 0.0029 |
| F06 | 18400001 | 18500000 | 0.0050 | 0.0032 | 0.0041 |
| F06 | 18420001 | 18520000 | 0.0049 | 0.0038 | 0.0037 |
| F06 | 18440001 | 18540000 | 0.0053 | 0.0030 | 0.0042 |
| F06 | 18460001 | 18560000 | 0.0043 | 0.0029 | 0.0033 |
| F06 | 18480001 | 18580000 | 0.0039 | 0.0015 | 0.0031 |
| F06 | 18500001 | 18600000 | 0.0023 | 0.0020 | 0.0015 |
| F06 | 18520001 | 18620000 | 0.0027 | 0.0014 | 0.0022 |
| F06 | 18540001 | 18640000 | 0.0022 | 0.0019 | 0.0017 |
| F06 | 18560001 | 18660000 | 0.0025 | 0.0020 | 0.0022 |
| F06 | 18580001 | 18680000 | 0.0026 | 0.0025 | 0.0023 |
| F06 | 18600001 | 18700000 | 0.0032 | 0.0019 | 0.0029 |
| F06 | 18620001 | 18720000 | 0.0025 | 0.0021 | 0.0022 |
| F06 | 18640001 | 18740000 | 0.0028 | 0.0022 | 0.0025 |
| F06 | 18660001 | 18760000 | 0.0029 | 0.0039 | 0.0026 |
| F06 | 18680001 | 18780000 | 0.0049 | 0.0032 | 0.0043 |
| F06 | 18700001 | 18800000 | 0.0041 | 0.0032 | 0.0036 |
| F06 | 18720001 | 18820000 | 0.0041 | 0.0036 | 0.0035 |
| F06 | 18740001 | 18840000 | 0.0046 | 0.0033 | 0.0039 |
| F06 | 18760001 | 18860000 | 0.0042 | 0.0030 | 0.0034 |
| F06 | 18780001 | 18880000 | 0.0038 | 0.0041 | 0.0032 |
| F06 | 18800001 | 18900000 | 0.0051 | 0.0043 | 0.0044 |
| F06 | 18820001 | 18920000 | 0.0052 | 0.0038 | 0.0045 |
| F06 | 18840001 | 18940000 | 0.0044 | 0.0041 | 0.0040 |
| F06 | 18860001 | 18960000 | 0.0047 | 0.0029 | 0.0044 |
| F06 | 18880001 | 18980000 | 0.0034 | 0.0020 | 0.0031 |
| F06 | 18900001 | 19000000 | 0.0023 | 0.0019 | 0.0022 |
| F06 | 18920001 | 19020000 | 0.0023 | 0.0025 | 0.0023 |
| F06 | 18940001 | 19040000 | 0.0030 | 0.0027 | 0.0030 |
| F06 | 18960001 | 19060000 | 0.0034 | 0.0027 | 0.0034 |
| F06 | 18980001 | 19080000 | 0.0033 | 0.0036 | 0.0033 |
| F06 | 19000001 | 19100000 | 0.0044 | 0.0033 | 0.0041 |
| F06 | 19020001 | 19120000 | 0.0040 | 0.0028 | 0.0037 |
| F06 | 19040001 | 19140000 | 0.0036 | 0.0024 | 0.0032 |
| F06 | 19060001 | 19160000 | 0.0033 | 0.0022 | 0.0026 |
| F06 | 19080001 | 19180000 | 0.0035 | 0.0020 | 0.0024 |
| F06 | 19100001 | 19200000 | 0.0033 | 0.0023 | 0.0022 |
| F06 | 19120001 | 19220000 | 0.0037 | 0.0025 | 0.0024 |
| F06 | 19140001 | 19240000 | 0.0038 | 0.0023 | 0.0026 |
| F06 | 19160001 | 19260000 | 0.0034 | 0.0024 | 0.0025 |
| F06 | 19180001 | 19280000 | 0.0032 | 0.0016 | 0.0026 |
| F06 | 19200001 | 19300000 | 0.0021 | 0.0017 | 0.0018 |
| F06 | 19220001 | 19320000 | 0.0023 | 0.0014 | 0.0021 |
| F06 | 19240001 | 19340000 | 0.0021 | 0.0023 | 0.0018 |
| F06 | 19260001 | 19360000 | 0.0030 | 0.0024 | 0.0027 |
| F06 | 19280001 | 19380000 | 0.0031 | 0.0022 | 0.0027 |
| F06 | 19300001 | 19400000 | 0.0029 | 0.0027 | 0.0025 |
| F06 | 19320001 | 19420000 | 0.0031 | 0.0029 | 0.0027 |
| F06 | 19340001 | 19440000 | 0.0032 | 0.0022 | 0.0030 |
| F06 | 19360001 | 19460000 | 0.0027 | 0.0026 | 0.0025 |
| F06 | 19380001 | 19480000 | 0.0032 | 0.0029 | 0.0029 |
| F06 | 19400001 | 19500000 | 0.0035 | 0.0019 | 0.0033 |
| F06 | 19420001 | 19520000 | 0.0027 | 0.0026 | 0.0022 |
| F06 | 19440001 | 19540000 | 0.0037 | 0.0023 | 0.0031 |
| F06 | 19460001 | 19560000 | 0.0033 | 0.0024 | 0.0027 |

| Chr | Start | End | both | A1 | A2 |
|---|---|---|---|---|---|
| F06 | 19480001 | 19580000 | 0.0032 | 0.0030 | 0.0026 |
| F06 | 19500001 | 19600000 | 0.0039 | 0.0035 | 0.0033 |
| F06 | 19520001 | 19620000 | 0.0042 | 0.0026 | 0.0040 |
| F06 | 19540001 | 19640000 | 0.0029 | 0.0024 | 0.0028 |
| F06 | 19560001 | 19660000 | 0.0027 | 0.0017 | 0.0026 |
| F06 | 19580001 | 19680000 | 0.0021 | 0.0013 | 0.0021 |
| F06 | 19600001 | 19700000 | 0.0016 | 0.0016 | 0.0016 |
| F06 | 19620001 | 19720000 | 0.0020 | 0.0017 | 0.0019 |
| F06 | 19640001 | 19740000 | 0.0023 | 0.0024 | 0.0020 |
| F06 | 19660001 | 19760000 | 0.0035 | 0.0035 | 0.0029 |
| F06 | 19680001 | 19780000 | 0.0048 | 0.0034 | 0.0042 |
| F06 | 19700001 | 19800000 | 0.0046 | 0.0027 | 0.0040 |
| F06 | 19720001 | 19820000 | 0.0038 | 0.0033 | 0.0032 |
| F06 | 19740001 | 19840000 | 0.0043 | 0.0029 | 0.0040 |
| F06 | 19760001 | 19860000 | 0.0036 | 0.0021 | 0.0034 |
| F06 | 19780001 | 19880000 | 0.0028 | 0.0028 | 0.0024 |
| F06 | 19800001 | 19900000 | 0.0037 | 0.0034 | 0.0032 |
| F06 | 19820001 | 19920000 | 0.0042 | 0.0038 | 0.0037 |
| F06 | 19840001 | 19940000 | 0.0047 | 0.0042 | 0.0040 |
| F06 | 19860001 | 19960000 | 0.0050 | 0.0044 | 0.0042 |
| F06 | 19880001 | 19980000 | 0.0054 | 0.0039 | 0.0045 |
| F06 | 19900001 | 20000000 | 0.0052 | 0.0036 | 0.0042 |
| F06 | 19920001 | 20020000 | 0.0051 | 0.0028 | 0.0041 |
| F06 | 19940001 | 20040000 | 0.0040 | 0.0024 | 0.0031 |
| F06 | 19960001 | 20060000 | 0.0034 | 0.0017 | 0.0027 |
| F06 | 19980001 | 20080000 | 0.0025 | 0.0014 | 0.0022 |
| F06 | 20000001 | 20100000 | 0.0017 | 0.0012 | 0.0015 |
| F06 | 20020001 | 20120000 | 0.0014 | 0.0014 | 0.0012 |
| F06 | 20040001 | 20140000 | 0.0016 | 0.0012 | 0.0015 |
| F06 | 20060001 | 20160000 | 0.0015 | 0.0017 | 0.0014 |
| F06 | 20080001 | 20180000 | 0.0020 | 0.0024 | 0.0019 |
| F06 | 20100001 | 20200000 | 0.0029 | 0.0024 | 0.0027 |
| F06 | 20120001 | 20220000 | 0.0030 | 0.0025 | 0.0028 |
| F06 | 20140001 | 20240000 | 0.0031 | 0.0034 | 0.0028 |
| F06 | 20160001 | 20260000 | 0.0040 | 0.0046 | 0.0037 |
| F06 | 20180001 | 20280000 | 0.0052 | 0.0055 | 0.0049 |
| F06 | 20200001 | 20300000 | 0.0061 | 0.0055 | 0.0057 |
| F06 | 20220001 | 20320000 | 0.0062 | 0.0056 | 0.0057 |
| F06 | 20240001 | 20340000 | 0.0064 | 0.0052 | 0.0060 |
| F06 | 20260001 | 20360000 | 0.0061 | 0.0044 | 0.0056 |
| F06 | 20280001 | 20380000 | 0.0052 | 0.0030 | 0.0047 |
| F06 | 20300001 | 20400000 | 0.0038 | 0.0030 | 0.0032 |
| F06 | 20320001 | 20420000 | 0.0037 | 0.0033 | 0.0032 |
| F06 | 20340001 | 20440000 | 0.0041 | 0.0030 | 0.0034 |
| F06 | 20360001 | 20460000 | 0.0038 | 0.0027 | 0.0029 |
| F06 | 20380001 | 20480000 | 0.0037 | 0.0026 | 0.0028 |
| F06 | 20400001 | 20500000 | 0.0036 | 0.0023 | 0.0026 |
| F06 | 20420001 | 20520000 | 0.0033 | 0.0014 | 0.0024 |
| F06 | 20440001 | 20540000 | 0.0022 | 0.0013 | 0.0015 |
| F06 | 20460001 | 20560000 | 0.0020 | 0.0012 | 0.0015 |
| F06 | 20480001 | 20580000 | 0.0020 | 0.0012 | 0.0014 |
| F06 | 20500001 | 20600000 | 0.0023 | 0.0018 | 0.0014 |
| F06 | 20520001 | 20620000 | 0.0028 | 0.0020 | 0.0020 |
| F06 | 20540001 | 20640000 | 0.0032 | 0.0025 | 0.0022 |
| F06 | 20560001 | 20660000 | 0.0035 | 0.0025 | 0.0026 |
| F06 | 20580001 | 20680000 | 0.0034 | 0.0028 | 0.0026 |
| F06 | 20600001 | 20700000 | 0.0033 | 0.0032 | 0.0027 |
| F06 | 20620001 | 20720000 | 0.0038 | 0.0030 | 0.0032 |

| Chr | Start | End | both | A1 | A2 |
|---|---|---|---|---|---|
| F06 | 20640001 | 20740000 | 0.0038 | 0.0023 | 0.0030 |
| F06 | 20660001 | 20760000 | 0.0033 | 0.0019 | 0.0024 |
| F06 | 20680001 | 20780000 | 0.0030 | 0.0023 | 0.0019 |
| F06 | 20700001 | 20800000 | 0.0035 | 0.0018 | 0.0023 |
| F06 | 20720001 | 20820000 | 0.0029 | 0.0022 | 0.0018 |
| F06 | 20740001 | 20840000 | 0.0033 | 0.0023 | 0.0022 |
| F06 | 20760001 | 20860000 | 0.0033 | 0.0023 | 0.0023 |
| F06 | 20780001 | 20880000 | 0.0029 | 0.0019 | 0.0024 |
| F06 | 20800001 | 20900000 | 0.0024 | 0.0013 | 0.0021 |
| F06 | 20820001 | 20920000 | 0.0017 | 0.0020 | 0.0014 |
| F06 | 20840001 | 20940000 | 0.0021 | 0.0029 | 0.0020 |
| F06 | 20860001 | 20960000 | 0.0032 | 0.0034 | 0.0031 |
| F06 | 20880001 | 20980000 | 0.0039 | 0.0033 | 0.0036 |
| F06 | 20900001 | 21000000 | 0.0040 | 0.0035 | 0.0035 |
| F06 | 20920001 | 21020000 | 0.0043 | 0.0028 | 0.0037 |
| F06 | 20940001 | 21040000 | 0.0038 | 0.0031 | 0.0031 |
| F06 | 20960001 | 21060000 | 0.0040 | 0.0033 | 0.0032 |
| F06 | 20980001 | 21080000 | 0.0041 | 0.0039 | 0.0035 |
| F06 | 21000001 | 21100000 | 0.0048 | 0.0038 | 0.0042 |
| F06 | 21020001 | 21120000 | 0.0046 | 0.0035 | 0.0041 |
| F06 | 21040001 | 21140000 | 0.0042 | 0.0024 | 0.0037 |
| F06 | 21060001 | 21160000 | 0.0030 | 0.0019 | 0.0026 |
| F06 | 21080001 | 21180000 | 0.0024 | 0.0018 | 0.0021 |
| F06 | 21100001 | 21200000 | 0.0019 | 0.0019 | 0.0018 |
| F06 | 21120001 | 21220000 | 0.0023 | 0.0024 | 0.0022 |
| F06 | 21140001 | 21240000 | 0.0027 | 0.0022 | 0.0027 |
| F06 | 21160001 | 21260000 | 0.0026 | 0.0023 | 0.0026 |
| F06 | 21180001 | 21280000 | 0.0027 | 0.0025 | 0.0027 |
| F06 | 21200001 | 21300000 | 0.0028 | 0.0023 | 0.0028 |
| F06 | 21220001 | 21320000 | 0.0026 | 0.0019 | 0.0025 |
| F06 | 21240001 | 21340000 | 0.0020 | 0.0024 | 0.0020 |
| F06 | 21260001 | 21360000 | 0.0026 | 0.0021 | 0.0026 |
| F06 | 21280001 | 21380000 | 0.0024 | 0.0014 | 0.0023 |
| F06 | 21300001 | 21400000 | 0.0017 | 0.0015 | 0.0017 |
| F06 | 21320001 | 21420000 | 0.0017 | 0.0016 | 0.0017 |
| F06 | 21340001 | 21440000 | 0.0018 | 0.0015 | 0.0018 |
| F06 | 21360001 | 21460000 | 0.0018 | 0.0014 | 0.0017 |
| F06 | 21380001 | 21480000 | 0.0019 | 0.0020 | 0.0016 |
| F06 | 21400001 | 21500000 | 0.0025 | 0.0025 | 0.0022 |
| F06 | 21420001 | 21520000 | 0.0032 | 0.0031 | 0.0029 |
| F06 | 21440001 | 21540000 | 0.0040 | 0.0028 | 0.0034 |
| F06 | 21460001 | 21560000 | 0.0038 | 0.0041 | 0.0031 |
| F06 | 21480001 | 21580000 | 0.0052 | 0.0047 | 0.0043 |
| F06 | 21500001 | 21600000 | 0.0057 | 0.0041 | 0.0046 |
| F06 | 21520001 | 21620000 | 0.0051 | 0.0042 | 0.0040 |
| F06 | 21540001 | 21640000 | 0.0050 | 0.0038 | 0.0041 |
| F06 | 21560001 | 21660000 | 0.0044 | 0.0027 | 0.0037 |
| F06 | 21580001 | 21680000 | 0.0031 | 0.0019 | 0.0028 |
| F06 | 21600001 | 21700000 | 0.0021 | 0.0016 | 0.0020 |
| F06 | 21620001 | 21720000 | 0.0017 | 0.0008 | 0.0016 |
| F06 | 21640001 | 21740000 | 0.0008 | 0.0009 | 0.0008 |
| F06 | 21660001 | 21760000 | 0.0009 | 0.0012 | 0.0009 |
| F06 | 21680001 | 21780000 | 0.0013 | 0.0011 | 0.0013 |
| F06 | 21700001 | 21800000 | 0.0012 | 0.0011 | 0.0012 |
| F06 | 21720001 | 21820000 | 0.0012 | 0.0012 | 0.0012 |
| F06 | 21740001 | 21840000 | 0.0014 | 0.0013 | 0.0013 |
| F06 | 21760001 | 21860000 | 0.0015 | 0.0020 | 0.0014 |
| F06 | 21780001 | 21880000 | 0.0022 | 0.0022 | 0.0019 |

| Chr | Start | End | both | A1 | A2 |
|---|---|---|---|---|---|
| F06 | 21800001 | 21900000 | 0.0027 | 0.0024 | 0.0023 |
| F06 | 21820001 | 21920000 | 0.0032 | 0.0029 | 0.0025 |
| F06 | 21840001 | 21940000 | 0.0036 | 0.0031 | 0.0029 |
| F06 | 21860001 | 21960000 | 0.0039 | 0.0027 | 0.0031 |
| F06 | 21880001 | 21980000 | 0.0034 | 0.0027 | 0.0027 |
| F06 | 21900001 | 22000000 | 0.0033 | 0.0030 | 0.0027 |
| F06 | 21920001 | 22020000 | 0.0038 | 0.0034 | 0.0033 |
| F06 | 21940001 | 22040000 | 0.0042 | 0.0036 | 0.0038 |
| F06 | 21960001 | 22060000 | 0.0044 | 0.0028 | 0.0039 |
| F06 | 21980001 | 22080000 | 0.0037 | 0.0030 | 0.0032 |
| F06 | 22000001 | 22100000 | 0.0038 | 0.0026 | 0.0033 |
| F06 | 22020001 | 22120000 | 0.0031 | 0.0021 | 0.0028 |
| F06 | 22040001 | 22140000 | 0.0026 | 0.0018 | 0.0022 |
| F06 | 22060001 | 22160000 | 0.0022 | 0.0024 | 0.0020 |
| F06 | 22080001 | 22180000 | 0.0029 | 0.0021 | 0.0028 |
| F06 | 22100001 | 22200000 | 0.0024 | 0.0023 | 0.0023 |
| F06 | 22120001 | 22220000 | 0.0026 | 0.0022 | 0.0025 |
| F06 | 22140001 | 22240000 | 0.0024 | 0.0019 | 0.0024 |
| F06 | 22160001 | 22260000 | 0.0021 | 0.0016 | 0.0021 |
| F06 | 22180001 | 22280000 | 0.0019 | 0.0016 | 0.0018 |
| F06 | 22200001 | 22300000 | 0.0020 | 0.0020 | 0.0018 |
| F06 | 22220001 | 22320000 | 0.0026 | 0.0022 | 0.0023 |
| F06 | 22240001 | 22340000 | 0.0029 | 0.0025 | 0.0025 |
| F06 | 22260001 | 22360000 | 0.0035 | 0.0024 | 0.0029 |
| F06 | 22280001 | 22380000 | 0.0033 | 0.0023 | 0.0028 |
| F06 | 22300001 | 22400000 | 0.0032 | 0.0019 | 0.0026 |
| F06 | 22320001 | 22420000 | 0.0027 | 0.0017 | 0.0023 |
| F06 | 22340001 | 22440000 | 0.0024 | 0.0014 | 0.0021 |
| F06 | 22360001 | 22460000 | 0.0019 | 0.0014 | 0.0017 |
| F06 | 22380001 | 22480000 | 0.0017 | 0.0013 | 0.0016 |
| F06 | 22400001 | 22500000 | 0.0015 | 0.0010 | 0.0015 |
| F06 | 22420001 | 22520000 | 0.0012 | 0.0016 | 0.0011 |
| F06 | 22440001 | 22540000 | 0.0017 | 0.0018 | 0.0017 |
| F06 | 22460001 | 22560000 | 0.0020 | 0.0017 | 0.0019 |
| F06 | 22480001 | 22580000 | 0.0019 | 0.0021 | 0.0018 |
| F06 | 22500001 | 22600000 | 0.0024 | 0.0023 | 0.0023 |
| F06 | 22520001 | 22620000 | 0.0026 | 0.0025 | 0.0025 |
| F06 | 22540001 | 22640000 | 0.0028 | 0.0035 | 0.0026 |
| F06 | 22560001 | 22660000 | 0.0040 | 0.0043 | 0.0038 |
| F06 | 22580001 | 22680000 | 0.0049 | 0.0046 | 0.0047 |
| F06 | 22600001 | 22700000 | 0.0053 | 0.0047 | 0.0051 |
| F06 | 22620001 | 22720000 | 0.0054 | 0.0041 | 0.0053 |
| F06 | 22640001 | 22740000 | 0.0048 | 0.0036 | 0.0047 |
| F06 | 22660001 | 22760000 | 0.0043 | 0.0033 | 0.0041 |
| F06 | 22680001 | 22780000 | 0.0040 | 0.0029 | 0.0037 |
| F06 | 22700001 | 22800000 | 0.0034 | 0.0028 | 0.0031 |
| F06 | 22720001 | 22820000 | 0.0033 | 0.0029 | 0.0030 |
| F06 | 22740001 | 22840000 | 0.0034 | 0.0020 | 0.0031 |
| F06 | 22760001 | 22860000 | 0.0023 | 0.0018 | 0.0022 |
| F06 | 22780001 | 22880000 | 0.0020 | 0.0018 | 0.0018 |
| F06 | 22800001 | 22900000 | 0.0020 | 0.0018 | 0.0017 |
| F06 | 22820001 | 22920000 | 0.0021 | 0.0023 | 0.0017 |
| F06 | 22840001 | 22940000 | 0.0028 | 0.0025 | 0.0023 |
| F06 | 22860001 | 22960000 | 0.0032 | 0.0030 | 0.0026 |
| F06 | 22880001 | 22980000 | 0.0040 | 0.0034 | 0.0033 |
| F06 | 22900001 | 23000000 | 0.0044 | 0.0037 | 0.0038 |
| F06 | 22920001 | 23020000 | 0.0048 | 0.0035 | 0.0040 |
| F06 | 22940001 | 23040000 | 0.0050 | 0.0039 | 0.0039 |

| Chr | Start | End | both | A1 | A2 |
|---|---|---|---|---|---|
| F06 | 22960001 | 23060000 | 0.0054 | 0.0030 | 0.0040 |
| F06 | 22980001 | 23080000 | 0.0043 | 0.0026 | 0.0032 |
| F06 | 23000001 | 23100000 | 0.0040 | 0.0025 | 0.0027 |
| F06 | 23020001 | 23120000 | 0.0039 | 0.0019 | 0.0026 |
| F06 | 23040001 | 23140000 | 0.0028 | 0.0021 | 0.0019 |
| F06 | 23060001 | 23160000 | 0.0029 | 0.0021 | 0.0024 |
| F06 | 23080001 | 23180000 | 0.0028 | 0.0020 | 0.0024 |
| F06 | 23100001 | 23200000 | 0.0025 | 0.0016 | 0.0023 |
| F06 | 23120001 | 23220000 | 0.0019 | 0.0014 | 0.0019 |
| F06 | 23140001 | 23240000 | 0.0016 | 0.0008 | 0.0016 |
| F06 | 23160001 | 23260000 | 0.0010 | 0.0008 | 0.0010 |
| F06 | 23180001 | 23280000 | 0.0009 | 0.0009 | 0.0008 |
| F06 | 23200001 | 23300000 | 0.0010 | 0.0008 | 0.0009 |
| F06 | 23220001 | 23320000 | 0.0010 | 0.0008 | 0.0009 |
| F06 | 23240001 | 23340000 | 0.0009 | 0.0008 | 0.0009 |
| F06 | 23260001 | 23360000 | 0.0009 | 0.0009 | 0.0008 |
| F06 | 23280001 | 23380000 | 0.0010 | 0.0008 | 0.0009 |
| F06 | 23300001 | 23400000 | 0.0008 | 0.0008 | 0.0008 |
| F06 | 23320001 | 23420000 | 0.0009 | 0.0008 | 0.0009 |
| F06 | 23340001 | 23440000 | 0.0009 | 0.0011 | 0.0008 |
| F06 | 23360001 | 23460000 | 0.0012 | 0.0009 | 0.0012 |
| F06 | 23380001 | 23480000 | 0.0010 | 0.0021 | 0.0010 |
| F06 | 23400001 | 23500000 | 0.0023 | 0.0022 | 0.0022 |
| F06 | 23420001 | 23520000 | 0.0024 | 0.0023 | 0.0024 |
| F06 | 23440001 | 23540000 | 0.0025 | 0.0021 | 0.0025 |
| F06 | 23460001 | 23560000 | 0.0023 | 0.0021 | 0.0023 |
| F06 | 23480001 | 23580000 | 0.0024 | 0.0012 | 0.0023 |
| F06 | 23500001 | 23600000 | 0.0014 | 0.0010 | 0.0014 |
| F06 | 23520001 | 23620000 | 0.0011 | 0.0010 | 0.0011 |
| F06 | 23540001 | 23640000 | 0.0011 | 0.0009 | 0.0010 |
| F06 | 23560001 | 23660000 | 0.0010 | 0.0009 | 0.0010 |
| F06 | 23580001 | 23680000 | 0.0010 | 0.0006 | 0.0010 |
| F06 | 23600001 | 23700000 | 0.0007 | 0.0013 | 0.0006 |
| F06 | 23620001 | 23720000 | 0.0015 | 0.0016 | 0.0015 |
| F06 | 23640001 | 23740000 | 0.0018 | 0.0016 | 0.0018 |
| F06 | 23660001 | 23760000 | 0.0017 | 0.0019 | 0.0017 |
| F06 | 23680001 | 23780000 | 0.0021 | 0.0018 | 0.0020 |
| F06 | 23700001 | 23800000 | 0.0020 | 0.0013 | 0.0020 |
| F06 | 23720001 | 23820000 | 0.0014 | 0.0010 | 0.0014 |
| F06 | 23740001 | 23840000 | 0.0012 | 0.0011 | 0.0011 |
| F06 | 23760001 | 23860000 | 0.0013 | 0.0010 | 0.0013 |
| F06 | 23780001 | 23880000 | 0.0012 | 0.0010 | 0.0012 |
| F06 | 23800001 | 23900000 | 0.0014 | 0.0013 | 0.0012 |
| F06 | 23820001 | 23920000 | 0.0017 | 0.0011 | 0.0015 |
| F06 | 23840001 | 23940000 | 0.0016 | 0.0014 | 0.0014 |
| F06 | 23860001 | 23960000 | 0.0020 | 0.0021 | 0.0017 |
| F06 | 23880001 | 23980000 | 0.0028 | 0.0026 | 0.0025 |
| F06 | 23900001 | 24000000 | 0.0034 | 0.0023 | 0.0032 |
| F06 | 23920001 | 24020000 | 0.0032 | 0.0030 | 0.0028 |
| F06 | 23940001 | 24040000 | 0.0042 | 0.0029 | 0.0037 |
| F06 | 23960001 | 24060000 | 0.0040 | 0.0025 | 0.0034 |
| F06 | 23980001 | 24080000 | 0.0038 | 0.0024 | 0.0028 |
| F06 | 24000001 | 24100000 | 0.0039 | 0.0028 | 0.0028 |
| F06 | 24020001 | 24120000 | 0.0042 | 0.0023 | 0.0030 |
| F06 | 24040001 | 24140000 | 0.0036 | 0.0020 | 0.0023 |
| F06 | 24060001 | 24160000 | 0.0033 | 0.0021 | 0.0019 |
| F06 | 24080001 | 24180000 | 0.0034 | 0.0023 | 0.0021 |
| F06 | 24100001 | 24200000 | 0.0036 | 0.0027 | 0.0020 |

| Chr | Start | End | both | A1 | A2 |
|---|---|---|---|---|---|
| F06 | 24120001 | 24220000 | 0.0041 | 0.0038 | 0.0024 |
| F06 | 24140001 | 24240000 | 0.0052 | 0.0042 | 0.0037 |
| F06 | 24160001 | 24260000 | 0.0059 | 0.0037 | 0.0040 |
| F06 | 24180001 | 24280000 | 0.0054 | 0.0031 | 0.0035 |
| F06 | 24200001 | 24300000 | 0.0043 | 0.0034 | 0.0029 |
| F06 | 24220001 | 24320000 | 0.0047 | 0.0026 | 0.0034 |
| F06 | 24240001 | 24340000 | 0.0038 | 0.0030 | 0.0027 |
| F06 | 24260001 | 24360000 | 0.0040 | 0.0033 | 0.0030 |
| F06 | 24280001 | 24380000 | 0.0043 | 0.0034 | 0.0034 |
| F06 | 24300001 | 24400000 | 0.0044 | 0.0026 | 0.0035 |
| F06 | 24320001 | 24420000 | 0.0033 | 0.0024 | 0.0026 |
| F06 | 24340001 | 24440000 | 0.0031 | 0.0022 | 0.0023 |
| F06 | 24360001 | 24460000 | 0.0027 | 0.0021 | 0.0023 |
| F06 | 24380001 | 24480000 | 0.0026 | 0.0025 | 0.0021 |
| F06 | 24400001 | 24500000 | 0.0029 | 0.0023 | 0.0025 |
| F06 | 24420001 | 24520000 | 0.0029 | 0.0022 | 0.0024 |
| F06 | 24440001 | 24540000 | 0.0031 | 0.0021 | 0.0025 |
| F06 | 24460001 | 24560000 | 0.0032 | 0.0021 | 0.0024 |
| F06 | 24480001 | 24580000 | 0.0033 | 0.0032 | 0.0025 |
| F06 | 24500001 | 24600000 | 0.0047 | 0.0036 | 0.0039 |
| F06 | 24520001 | 24620000 | 0.0052 | 0.0034 | 0.0042 |
| F06 | 24540001 | 24640000 | 0.0047 | 0.0029 | 0.0038 |
| F06 | 24560001 | 24660000 | 0.0043 | 0.0031 | 0.0033 |
| F06 | 24580001 | 24680000 | 0.0046 | 0.0016 | 0.0035 |
| F06 | 24600001 | 24700000 | 0.0029 | 0.0014 | 0.0017 |
| F06 | 24620001 | 24720000 | 0.0026 | 0.0016 | 0.0016 |
| F06 | 24640001 | 24740000 | 0.0028 | 0.0017 | 0.0019 |
| F06 | 24660001 | 24760000 | 0.0029 | 0.0012 | 0.0019 |
| F06 | 24680001 | 24780000 | 0.0022 | 0.0011 | 0.0013 |
| F06 | 24700001 | 24800000 | 0.0020 | 0.0011 | 0.0013 |
| F06 | 24720001 | 24820000 | 0.0020 | 0.0011 | 0.0011 |
| F06 | 24740001 | 24840000 | 0.0021 | 0.0010 | 0.0012 |
| F06 | 24760001 | 24860000 | 0.0022 | 0.0013 | 0.0012 |
| F06 | 24780001 | 24880000 | 0.0024 | 0.0019 | 0.0014 |
| F06 | 24800001 | 24900000 | 0.0033 | 0.0019 | 0.0021 |
| F06 | 24820001 | 24920000 | 0.0032 | 0.0017 | 0.0021 |
| F06 | 24840001 | 24940000 | 0.0033 | 0.0018 | 0.0019 |
| F06 | 24860001 | 24960000 | 0.0033 | 0.0018 | 0.0020 |
| F06 | 24880001 | 24980000 | 0.0032 | 0.0012 | 0.0021 |
| F06 | 24900001 | 25000000 | 0.0022 | 0.0011 | 0.0014 |
| F06 | 24920001 | 25020000 | 0.0020 | 0.0014 | 0.0013 |
| F06 | 24940001 | 25040000 | 0.0021 | 0.0017 | 0.0014 |
| F06 | 24960001 | 25060000 | 0.0026 | 0.0016 | 0.0019 |
| F06 | 24980001 | 25080000 | 0.0028 | 0.0020 | 0.0018 |
| F06 | 25000001 | 25100000 | 0.0037 | 0.0020 | 0.0023 |
| F06 | 25020001 | 25120000 | 0.0038 | 0.0020 | 0.0023 |
| F06 | 25040001 | 25140000 | 0.0040 | 0.0018 | 0.0025 |
| F06 | 25060001 | 25160000 | 0.0035 | 0.0017 | 0.0020 |
| F06 | 25080001 | 25180000 | 0.0032 | 0.0016 | 0.0018 |
| F06 | 25100001 | 25200000 | 0.0032 | 0.0021 | 0.0018 |
| F06 | 25120001 | 25220000 | 0.0037 | 0.0023 | 0.0023 |
| F06 | 25140001 | 25240000 | 0.0035 | 0.0027 | 0.0024 |
| F06 | 25160001 | 25260000 | 0.0041 | 0.0033 | 0.0030 |
| F06 | 25180001 | 25280000 | 0.0046 | 0.0036 | 0.0037 |
| F06 | 25200001 | 25300000 | 0.0047 | 0.0034 | 0.0039 |
| F06 | 25220001 | 25320000 | 0.0046 | 0.0030 | 0.0036 |
| F06 | 25240001 | 25340000 | 0.0041 | 0.0027 | 0.0031 |
| F06 | 25260001 | 25360000 | 0.0038 | 0.0029 | 0.0026 |

| Chr | Start | End | both | A1 | A2 |
|---|---|---|---|---|---|
| F06 | 25280001 | 25380000 | 0.0043 | 0.0028 | 0.0028 |
| F06 | 25300001 | 25400000 | 0.0042 | 0.0026 | 0.0028 |
| F06 | 25320001 | 25420000 | 0.0038 | 0.0029 | 0.0024 |
| F06 | 25340001 | 25440000 | 0.0044 | 0.0026 | 0.0028 |
| F06 | 25360001 | 25460000 | 0.0039 | 0.0022 | 0.0026 |
| F06 | 25380001 | 25480000 | 0.0033 | 0.0019 | 0.0022 |
| F06 | 25400001 | 25500000 | 0.0029 | 0.0031 | 0.0019 |
| F06 | 25420001 | 25520000 | 0.0042 | 0.0029 | 0.0033 |
| F06 | 25440001 | 25540000 | 0.0036 | 0.0027 | 0.0032 |
| F06 | 25460001 | 25560000 | 0.0035 | 0.0043 | 0.0030 |
| F06 | 25480001 | 25580000 | 0.0051 | 0.0045 | 0.0045 |
| F06 | 25500001 | 25600000 | 0.0054 | 0.0031 | 0.0048 |
| F06 | 25520001 | 25620000 | 0.0039 | 0.0030 | 0.0032 |
| F06 | 25540001 | 25640000 | 0.0039 | 0.0034 | 0.0030 |
| F06 | 25560001 | 25660000 | 0.0043 | 0.0015 | 0.0035 |
| F06 | 25580001 | 25680000 | 0.0022 | 0.0017 | 0.0016 |
| F06 | 25600001 | 25700000 | 0.0026 | 0.0017 | 0.0019 |
| F06 | 25620001 | 25720000 | 0.0027 | 0.0024 | 0.0018 |
| F06 | 25640001 | 25740000 | 0.0034 | 0.0026 | 0.0025 |
| F06 | 25660001 | 25760000 | 0.0036 | 0.0033 | 0.0027 |
| F06 | 25680001 | 25780000 | 0.0045 | 0.0032 | 0.0034 |
| F06 | 25700001 | 25800000 | 0.0042 | 0.0043 | 0.0034 |
| F06 | 25720001 | 25820000 | 0.0051 | 0.0039 | 0.0044 |
| F06 | 25740001 | 25840000 | 0.0049 | 0.0041 | 0.0042 |
| F06 | 25760001 | 25860000 | 0.0055 | 0.0046 | 0.0044 |
| F06 | 25780001 | 25880000 | 0.0061 | 0.0052 | 0.0049 |
| F06 | 25800001 | 25900000 | 0.0068 | 0.0042 | 0.0055 |
| F06 | 25820001 | 25920000 | 0.0056 | 0.0036 | 0.0045 |
| F06 | 25840001 | 25940000 | 0.0048 | 0.0030 | 0.0039 |
| F06 | 25860001 | 25960000 | 0.0037 | 0.0023 | 0.0032 |
| F06 | 25880001 | 25980000 | 0.0027 | 0.0013 | 0.0025 |
| F06 | 25900001 | 26000000 | 0.0015 | 0.0016 | 0.0014 |
| F06 | 25920001 | 26020000 | 0.0018 | 0.0021 | 0.0018 |
| F06 | 25940001 | 26040000 | 0.0024 | 0.0023 | 0.0023 |
| F06 | 25960001 | 26060000 | 0.0028 | 0.0020 | 0.0025 |
| F06 | 25980001 | 26080000 | 0.0026 | 0.0021 | 0.0022 |
| F06 | 26000001 | 26100000 | 0.0027 | 0.0016 | 0.0023 |
| F06 | 26020001 | 26120000 | 0.0024 | 0.0016 | 0.0018 |
| F06 | 26040001 | 26140000 | 0.0023 | 0.0016 | 0.0018 |
| F06 | 26060001 | 26160000 | 0.0024 | 0.0019 | 0.0018 |
| F06 | 26080001 | 26180000 | 0.0028 | 0.0017 | 0.0020 |
| F06 | 26100001 | 26200000 | 0.0026 | 0.0020 | 0.0018 |
| F06 | 26120001 | 26220000 | 0.0028 | 0.0019 | 0.0020 |
| F06 | 26140001 | 26240000 | 0.0029 | 0.0019 | 0.0019 |
| F06 | 26160001 | 26260000 | 0.0028 | 0.0014 | 0.0019 |
| F06 | 26180001 | 26280000 | 0.0023 | 0.0013 | 0.0014 |
| F06 | 26200001 | 26300000 | 0.0023 | 0.0012 | 0.0013 |
| F06 | 26220001 | 26320000 | 0.0020 | 0.0010 | 0.0012 |
| F06 | 26240001 | 26340000 | 0.0016 | 0.0007 | 0.0010 |
| F06 | 26260001 | 26360000 | 0.0012 | 0.0008 | 0.0007 |
| F06 | 26280001 | 26380000 | 0.0013 | 0.0011 | 0.0009 |
| F06 | 26300001 | 26400000 | 0.0019 | 0.0011 | 0.0014 |
| F06 | 26320001 | 26420000 | 0.0022 | 0.0010 | 0.0015 |
| F06 | 26340001 | 26440000 | 0.0021 | 0.0013 | 0.0015 |
| F06 | 26360001 | 26460000 | 0.0023 | 0.0019 | 0.0017 |
| F06 | 26380001 | 26480000 | 0.0029 | 0.0017 | 0.0024 |
| F06 | 26400001 | 26500000 | 0.0025 | 0.0019 | 0.0019 |
| F06 | 26420001 | 26520000 | 0.0025 | 0.0027 | 0.0020 |

| Chr | Start | End | both | A1 | A2 |
|---|---|---|---|---|---|
| F06 | 26440001 | 26540000 | 0.0034 | 0.0026 | 0.0027 |
| F06 | 26460001 | 26560000 | 0.0034 | 0.0022 | 0.0026 |
| F06 | 26480001 | 26580000 | 0.0028 | 0.0023 | 0.0020 |
| F06 | 26500001 | 26600000 | 0.0029 | 0.0026 | 0.0024 |
| F06 | 26520001 | 26620000 | 0.0035 | 0.0024 | 0.0030 |
| F06 | 26540001 | 26640000 | 0.0032 | 0.0022 | 0.0027 |
| F06 | 26560001 | 26660000 | 0.0030 | 0.0023 | 0.0025 |
| F06 | 26580001 | 26680000 | 0.0031 | 0.0021 | 0.0026 |
| F06 | 26600001 | 26700000 | 0.0027 | 0.0023 | 0.0023 |
| F06 | 26620001 | 26720000 | 0.0029 | 0.0025 | 0.0023 |
| F06 | 26640001 | 26740000 | 0.0032 | 0.0026 | 0.0027 |
| F06 | 26660001 | 26760000 | 0.0032 | 0.0025 | 0.0028 |
| F06 | 26680001 | 26780000 | 0.0031 | 0.0028 | 0.0027 |
| F06 | 26700001 | 26800000 | 0.0035 | 0.0019 | 0.0032 |
| F06 | 26720001 | 26820000 | 0.0024 | 0.0015 | 0.0022 |
| F06 | 26740001 | 26840000 | 0.0017 | 0.0015 | 0.0017 |
| F06 | 26760001 | 26860000 | 0.0017 | 0.0012 | 0.0017 |
| F06 | 26780001 | 26880000 | 0.0015 | 0.0008 | 0.0015 |
| F06 | 26800001 | 26900000 | 0.0009 | 0.0014 | 0.0009 |
| F06 | 26820001 | 26920000 | 0.0015 | 0.0014 | 0.0014 |
| F06 | 26840001 | 26940000 | 0.0017 | 0.0022 | 0.0014 |
| F06 | 26860001 | 26960000 | 0.0028 | 0.0028 | 0.0023 |
| F06 | 26880001 | 26980000 | 0.0034 | 0.0030 | 0.0030 |
| F06 | 26900001 | 27000000 | 0.0041 | 0.0028 | 0.0034 |
| F06 | 26920001 | 27020000 | 0.0042 | 0.0027 | 0.0033 |
| F06 | 26940001 | 27040000 | 0.0041 | 0.0030 | 0.0032 |
| F06 | 26960001 | 27060000 | 0.0042 | 0.0037 | 0.0034 |
| F06 | 26980001 | 27080000 | 0.0050 | 0.0043 | 0.0041 |
| F06 | 27000001 | 27100000 | 0.0053 | 0.0041 | 0.0044 |
| F06 | 27020001 | 27120000 | 0.0049 | 0.0042 | 0.0042 |
| F06 | 27040001 | 27140000 | 0.0048 | 0.0032 | 0.0043 |
| F06 | 27060001 | 27160000 | 0.0038 | 0.0028 | 0.0032 |
| F06 | 27080001 | 27180000 | 0.0035 | 0.0023 | 0.0029 |
| F06 | 27100001 | 27200000 | 0.0029 | 0.0022 | 0.0023 |
| F06 | 27120001 | 27220000 | 0.0029 | 0.0019 | 0.0022 |
| F06 | 27140001 | 27240000 | 0.0027 | 0.0021 | 0.0020 |
| F06 | 27160001 | 27260000 | 0.0028 | 0.0024 | 0.0022 |
| F06 | 27180001 | 27280000 | 0.0033 | 0.0029 | 0.0028 |
| F06 | 27200001 | 27300000 | 0.0038 | 0.0040 | 0.0034 |
| F06 | 27220001 | 27320000 | 0.0049 | 0.0040 | 0.0045 |
| F06 | 27240001 | 27340000 | 0.0051 | 0.0036 | 0.0046 |
| F06 | 27260001 | 27360000 | 0.0047 | 0.0026 | 0.0043 |
| F06 | 27280001 | 27380000 | 0.0035 | 0.0021 | 0.0030 |
| F06 | 27300001 | 27400000 | 0.0032 | 0.0010 | 0.0027 |
| F06 | 27320001 | 27420000 | 0.0021 | 0.0023 | 0.0016 |
| F06 | 27340001 | 27440000 | 0.0032 | 0.0027 | 0.0028 |
| F06 | 27360001 | 27460000 | 0.0035 | 0.0024 | 0.0031 |
| F06 | 27380001 | 27480000 | 0.0033 | 0.0026 | 0.0029 |
| F06 | 27400001 | 27500000 | 0.0033 | 0.0026 | 0.0030 |
| F06 | 27420001 | 27520000 | 0.0033 | 0.0013 | 0.0030 |
| F06 | 27440001 | 27540000 | 0.0023 | 0.0013 | 0.0019 |
| F06 | 27460001 | 27560000 | 0.0022 | 0.0015 | 0.0018 |
| F06 | 27480001 | 27580000 | 0.0022 | 0.0016 | 0.0020 |
| F06 | 27500001 | 27600000 | 0.0026 | 0.0016 | 0.0022 |
| F06 | 27520001 | 27620000 | 0.0025 | 0.0023 | 0.0021 |
| F06 | 27540001 | 27640000 | 0.0032 | 0.0025 | 0.0029 |
| F06 | 27560001 | 27660000 | 0.0039 | 0.0032 | 0.0034 |
| F06 | 27580001 | 27680000 | 0.0048 | 0.0030 | 0.0042 |

| Chr | Start | End | both | A1 | A2 |
|---|---|---|---|---|---|
| F06 | 27600001 | 27700000 | 0.0042 | 0.0032 | 0.0038 |
| F06 | 27620001 | 27720000 | 0.0044 | 0.0029 | 0.0040 |
| F06 | 27640001 | 27740000 | 0.0042 | 0.0025 | 0.0038 |
| F06 | 27660001 | 27760000 | 0.0036 | 0.0026 | 0.0033 |
| F06 | 27680001 | 27780000 | 0.0036 | 0.0031 | 0.0033 |
| F06 | 27700001 | 27800000 | 0.0043 | 0.0029 | 0.0039 |
| F06 | 27720001 | 27820000 | 0.0040 | 0.0030 | 0.0036 |
| F06 | 27740001 | 27840000 | 0.0040 | 0.0031 | 0.0036 |
| F06 | 27760001 | 27860000 | 0.0038 | 0.0021 | 0.0035 |
| F06 | 27780001 | 27880000 | 0.0028 | 0.0019 | 0.0026 |
| F06 | 27800001 | 27900000 | 0.0026 | 0.0022 | 0.0025 |
| F06 | 27820001 | 27920000 | 0.0031 | 0.0017 | 0.0028 |
| F06 | 27840001 | 27940000 | 0.0026 | 0.0016 | 0.0023 |
| F06 | 27860001 | 27960000 | 0.0024 | 0.0015 | 0.0021 |
| F06 | 27880001 | 27980000 | 0.0022 | 0.0008 | 0.0019 |
| F06 | 27900001 | 28000000 | 0.0013 | 0.0006 | 0.0010 |
| F06 | 27920001 | 28020000 | 0.0009 | 0.0005 | 0.0008 |
| F06 | 27940001 | 28040000 | 0.0007 | 0.0003 | 0.0006 |
| F06 | 27960001 | 28060000 | 0.0004 | 0.0003 | 0.0003 |
| F06 | 27980001 | 28080000 | 0.0005 | 0.0008 | 0.0004 |
| F06 | 28000001 | 28100000 | 0.0009 | 0.0013 | 0.0008 |
| F06 | 28020001 | 28120000 | 0.0018 | 0.0021 | 0.0016 |
| F06 | 28040001 | 28140000 | 0.0027 | 0.0023 | 0.0024 |
| F06 | 28060001 | 28160000 | 0.0030 | 0.0026 | 0.0026 |
| F06 | 28080001 | 28180000 | 0.0037 | 0.0032 | 0.0031 |
| F06 | 28100001 | 28200000 | 0.0047 | 0.0026 | 0.0038 |
| F06 | 28120001 | 28220000 | 0.0039 | 0.0023 | 0.0031 |
| F06 | 28140001 | 28240000 | 0.0035 | 0.0028 | 0.0028 |
| F06 | 28160001 | 28260000 | 0.0040 | 0.0029 | 0.0033 |
| F06 | 28180001 | 28280000 | 0.0038 | 0.0026 | 0.0033 |
| F06 | 28200001 | 28300000 | 0.0031 | 0.0033 | 0.0029 |
| F06 | 28220001 | 28320000 | 0.0038 | 0.0037 | 0.0037 |
| F06 | 28240001 | 28340000 | 0.0043 | 0.0031 | 0.0041 |
| F06 | 28260001 | 28360000 | 0.0039 | 0.0033 | 0.0035 |
| F06 | 28280001 | 28380000 | 0.0042 | 0.0026 | 0.0037 |
| F06 | 28300001 | 28400000 | 0.0039 | 0.0024 | 0.0031 |
| F06 | 28320001 | 28420000 | 0.0036 | 0.0018 | 0.0028 |
| F06 | 28340001 | 28440000 | 0.0030 | 0.0019 | 0.0023 |
| F06 | 28360001 | 28460000 | 0.0030 | 0.0018 | 0.0024 |
| F06 | 28380001 | 28480000 | 0.0029 | 0.0022 | 0.0022 |
| F06 | 28400001 | 28500000 | 0.0030 | 0.0018 | 0.0025 |
| F06 | 28420001 | 28520000 | 0.0028 | 0.0021 | 0.0022 |
| F06 | 28440001 | 28540000 | 0.0031 | 0.0020 | 0.0025 |
| F06 | 28460001 | 28560000 | 0.0030 | 0.0016 | 0.0024 |
| F06 | 28480001 | 28580000 | 0.0026 | 0.0013 | 0.0021 |
| F06 | 28500001 | 28600000 | 0.0022 | 0.0010 | 0.0016 |
| F06 | 28520001 | 28620000 | 0.0017 | 0.0006 | 0.0013 |
| F06 | 28540001 | 28640000 | 0.0012 | 0.0015 | 0.0009 |
| F06 | 28560001 | 28660000 | 0.0019 | 0.0013 | 0.0017 |
| F06 | 28580001 | 28680000 | 0.0019 | 0.0015 | 0.0015 |
| F06 | 28600001 | 28700000 | 0.0023 | 0.0018 | 0.0017 |
| F06 | 28620001 | 28720000 | 0.0028 | 0.0023 | 0.0021 |
| F06 | 28640001 | 28740000 | 0.0032 | 0.0022 | 0.0025 |
| F06 | 28660001 | 28760000 | 0.0034 | 0.0027 | 0.0026 |
| F06 | 28680001 | 28780000 | 0.0040 | 0.0028 | 0.0031 |
| F06 | 28700001 | 28800000 | 0.0038 | 0.0032 | 0.0030 |
| F06 | 28720001 | 28820000 | 0.0042 | 0.0027 | 0.0034 |
| F06 | 28740001 | 28840000 | 0.0037 | 0.0020 | 0.0029 |

| Chr | Start | End | both | A1 | A2 |
|---|---|---|---|---|---|
| F06 | 28760001 | 28860000 | 0.0031 | 0.0015 | 0.0023 |
| F06 | 28780001 | 28880000 | 0.0034 | 0.0018 | 0.0026 |
| F06 | 28800001 | 28900000 | 0.0038 | 0.0011 | 0.0030 |
| F06 | 28820001 | 28920000 | 0.0030 | 0.0014 | 0.0022 |
| F06 | 28840001 | 28940000 | 0.0035 | 0.0015 | 0.0025 |
| F06 | 28860001 | 28960000 | 0.0034 | 0.0017 | 0.0025 |
| F06 | 28880001 | 28980000 | 0.0027 | 0.0013 | 0.0019 |
| F06 | 28900001 | 29000000 | 0.0023 | 0.0016 | 0.0016 |
| F06 | 28920001 | 29020000 | 0.0028 | 0.0017 | 0.0020 |
| F06 | 28940001 | 29040000 | 0.0026 | 0.0019 | 0.0020 |
| F06 | 28960001 | 29060000 | 0.0027 | 0.0035 | 0.0022 |
| F06 | 28980001 | 29080000 | 0.0044 | 0.0034 | 0.0038 |
| F06 | 29000001 | 29100000 | 0.0043 | 0.0035 | 0.0036 |
| F06 | 29020001 | 29120000 | 0.0043 | 0.0036 | 0.0037 |
| F06 | 29040001 | 29140000 | 0.0045 | 0.0035 | 0.0038 |
| F06 | 29060001 | 29160000 | 0.0045 | 0.0022 | 0.0037 |
| F06 | 29080001 | 29180000 | 0.0032 | 0.0026 | 0.0024 |
| F06 | 29100001 | 29200000 | 0.0037 | 0.0024 | 0.0029 |
| F06 | 29120001 | 29220000 | 0.0037 | 0.0027 | 0.0027 |
| F06 | 29140001 | 29240000 | 0.0043 | 0.0028 | 0.0031 |
| F06 | 29160001 | 29260000 | 0.0046 | 0.0027 | 0.0033 |
| F06 | 29180001 | 29280000 | 0.0044 | 0.0028 | 0.0031 |
| F06 | 29200001 | 29300000 | 0.0044 | 0.0026 | 0.0031 |
| F06 | 29220001 | 29320000 | 0.0039 | 0.0021 | 0.0029 |
| F06 | 29240001 | 29340000 | 0.0031 | 0.0017 | 0.0023 |
| F06 | 29260001 | 29360000 | 0.0031 | 0.0026 | 0.0021 |
| F06 | 29280001 | 29380000 | 0.0040 | 0.0024 | 0.0030 |
| F06 | 29300001 | 29400000 | 0.0041 | 0.0028 | 0.0029 |
| F06 | 29320001 | 29420000 | 0.0048 | 0.0029 | 0.0033 |
| F06 | 29340001 | 29440000 | 0.0048 | 0.0033 | 0.0035 |
| F06 | 29360001 | 29460000 | 0.0050 | 0.0027 | 0.0039 |
| F06 | 29380001 | 29480000 | 0.0042 | 0.0024 | 0.0033 |
| F06 | 29400001 | 29500000 | 0.0037 | 0.0025 | 0.0029 |
| F06 | 29420001 | 29520000 | 0.0037 | 0.0023 | 0.0031 |
| F06 | 29440001 | 29540000 | 0.0036 | 0.0023 | 0.0029 |
| F06 | 29460001 | 29560000 | 0.0035 | 0.0021 | 0.0027 |
| F06 | 29480001 | 29580000 | 0.0033 | 0.0022 | 0.0024 |
| F06 | 29500001 | 29600000 | 0.0035 | 0.0018 | 0.0025 |
| F06 | 29520001 | 29620000 | 0.0031 | 0.0024 | 0.0021 |
| F06 | 29540001 | 29640000 | 0.0036 | 0.0022 | 0.0027 |
| F06 | 29560001 | 29660000 | 0.0035 | 0.0024 | 0.0026 |
| F06 | 29580001 | 29680000 | 0.0039 | 0.0029 | 0.0029 |
| F06 | 29600001 | 29700000 | 0.0042 | 0.0038 | 0.0033 |
| F06 | 29620001 | 29720000 | 0.0052 | 0.0038 | 0.0042 |
| F06 | 29640001 | 29740000 | 0.0052 | 0.0034 | 0.0043 |
| F06 | 29660001 | 29760000 | 0.0047 | 0.0029 | 0.0037 |
| F06 | 29680001 | 29780000 | 0.0040 | 0.0025 | 0.0031 |
| F06 | 29700001 | 29800000 | 0.0039 | 0.0019 | 0.0030 |
| F06 | 29720001 | 29820000 | 0.0030 | 0.0017 | 0.0023 |
| F06 | 29740001 | 29840000 | 0.0026 | 0.0016 | 0.0019 |
| F06 | 29760001 | 29860000 | 0.0024 | 0.0021 | 0.0019 |
| F06 | 29780001 | 29880000 | 0.0028 | 0.0021 | 0.0024 |
| F06 | 29800001 | 29900000 | 0.0025 | 0.0023 | 0.0022 |
| F06 | 29820001 | 29920000 | 0.0029 | 0.0020 | 0.0024 |
| F06 | 29840001 | 29940000 | 0.0027 | 0.0019 | 0.0022 |
| F06 | 29860001 | 29960000 | 0.0028 | 0.0017 | 0.0020 |
| F06 | 29880001 | 29980000 | 0.0027 | 0.0017 | 0.0020 |
| F06 | 29900001 | 30000000 | 0.0030 | 0.0012 | 0.0021 |

| Chr | Start | End | both | A1 | A2 |
|---|---|---|---|---|---|
| F06 | 29920001 | 30020000 | 0.0022 | 0.0022 | 0.0014 |
| F06 | 29940001 | 30040000 | 0.0034 | 0.0026 | 0.0026 |
| F06 | 29960001 | 30060000 | 0.0038 | 0.0031 | 0.0031 |
| F06 | 29980001 | 30080000 | 0.0043 | 0.0031 | 0.0037 |
| F06 | 30000001 | 30100000 | 0.0038 | 0.0032 | 0.0035 |
| F06 | 30020001 | 30120000 | 0.0043 | 0.0027 | 0.0037 |
| F06 | 30040001 | 30140000 | 0.0033 | 0.0025 | 0.0029 |
| F06 | 30060001 | 30160000 | 0.0031 | 0.0027 | 0.0028 |
| F06 | 30080001 | 30180000 | 0.0031 | 0.0031 | 0.0028 |
| F06 | 30100001 | 30200000 | 0.0034 | 0.0028 | 0.0031 |
| F06 | 30120001 | 30220000 | 0.0029 | 0.0034 | 0.0028 |
| F06 | 30140001 | 30240000 | 0.0038 | 0.0035 | 0.0037 |
| F06 | 30160001 | 30260000 | 0.0038 | 0.0028 | 0.0037 |
| F06 | 30180001 | 30280000 | 0.0030 | 0.0023 | 0.0030 |
| F06 | 30200001 | 30300000 | 0.0026 | 0.0030 | 0.0025 |
| F06 | 30220001 | 30320000 | 0.0034 | 0.0020 | 0.0033 |
| F06 | 30240001 | 30340000 | 0.0023 | 0.0025 | 0.0021 |
| F06 | 30260001 | 30360000 | 0.0029 | 0.0026 | 0.0027 |
| F06 | 30280001 | 30380000 | 0.0029 | 0.0027 | 0.0028 |
| F06 | 30300001 | 30400000 | 0.0030 | 0.0027 | 0.0029 |
| F06 | 30320001 | 30420000 | 0.0031 | 0.0030 | 0.0028 |
| F06 | 30340001 | 30440000 | 0.0035 | 0.0029 | 0.0031 |
| F06 | 30360001 | 30460000 | 0.0036 | 0.0032 | 0.0032 |
| F06 | 30380001 | 30480000 | 0.0042 | 0.0034 | 0.0035 |
| F06 | 30400001 | 30500000 | 0.0044 | 0.0034 | 0.0037 |
| F06 | 30420001 | 30520000 | 0.0043 | 0.0033 | 0.0036 |
| F06 | 30440001 | 30540000 | 0.0043 | 0.0024 | 0.0035 |
| F06 | 30460001 | 30560000 | 0.0033 | 0.0019 | 0.0025 |
| F06 | 30480001 | 30580000 | 0.0031 | 0.0017 | 0.0021 |
| F06 | 30500001 | 30600000 | 0.0034 | 0.0010 | 0.0019 |
| F06 | 30520001 | 30620000 | 0.0025 | 0.0009 | 0.0012 |
| F06 | 30540001 | 30640000 | 0.0025 | 0.0011 | 0.0011 |
| F06 | 30560001 | 30660000 | 0.0026 | 0.0013 | 0.0012 |
| F06 | 30580001 | 30680000 | 0.0024 | 0.0016 | 0.0013 |
| F06 | 30600001 | 30700000 | 0.0023 | 0.0018 | 0.0015 |
| F06 | 30620001 | 30720000 | 0.0026 | 0.0027 | 0.0017 |
| F06 | 30640001 | 30740000 | 0.0036 | 0.0045 | 0.0027 |
| F06 | 30660001 | 30760000 | 0.0056 | 0.0048 | 0.0045 |
| F06 | 30680001 | 30780000 | 0.0059 | 0.0051 | 0.0050 |
| F06 | 30700001 | 30800000 | 0.0066 | 0.0056 | 0.0055 |
| F06 | 30720001 | 30820000 | 0.0072 | 0.0054 | 0.0059 |
| F06 | 30740001 | 30840000 | 0.0070 | 0.0036 | 0.0058 |
| F06 | 30760001 | 30860000 | 0.0052 | 0.0032 | 0.0039 |
| F06 | 30780001 | 30880000 | 0.0046 | 0.0034 | 0.0034 |
| F06 | 30800001 | 30900000 | 0.0047 | 0.0034 | 0.0036 |
| F06 | 30820001 | 30920000 | 0.0046 | 0.0030 | 0.0037 |
| F06 | 30840001 | 30940000 | 0.0044 | 0.0031 | 0.0033 |
| F06 | 30860001 | 30960000 | 0.0044 | 0.0032 | 0.0035 |
| F06 | 30880001 | 30980000 | 0.0047 | 0.0029 | 0.0038 |
| F06 | 30900001 | 31000000 | 0.0044 | 0.0027 | 0.0035 |
| F06 | 30920001 | 31020000 | 0.0041 | 0.0027 | 0.0032 |
| F06 | 30940001 | 31040000 | 0.0038 | 0.0038 | 0.0030 |
| F06 | 30960001 | 31060000 | 0.0049 | 0.0048 | 0.0040 |
| F06 | 30980001 | 31080000 | 0.0059 | 0.0053 | 0.0050 |
| F06 | 31000001 | 31100000 | 0.0064 | 0.0064 | 0.0054 |
| F06 | 31020001 | 31120000 | 0.0074 | 0.0077 | 0.0064 |
| F06 | 31040001 | 31140000 | 0.0088 | 0.0072 | 0.0079 |
| F06 | 31060001 | 31160000 | 0.0083 | 0.0065 | 0.0074 |

| Chr | Start | End | both | A1 | A2 |
|---|---|---|---|---|---|
| F06 | 31080001 | 31180000 | 0.0078 | 0.0054 | 0.0070 |
| F06 | 31100001 | 31200000 | 0.0065 | 0.0043 | 0.0060 |
| F06 | 31120001 | 31220000 | 0.0056 | 0.0024 | 0.0050 |
| F06 | 31140001 | 31240000 | 0.0034 | 0.0026 | 0.0029 |
| F06 | 31160001 | 31260000 | 0.0038 | 0.0029 | 0.0034 |
| F06 | 31180001 | 31280000 | 0.0039 | 0.0036 | 0.0032 |
| F06 | 31200001 | 31300000 | 0.0051 | 0.0038 | 0.0038 |
| F06 | 31220001 | 31320000 | 0.0052 | 0.0043 | 0.0041 |
| F06 | 31240001 | 31340000 | 0.0061 | 0.0040 | 0.0047 |
| F06 | 31260001 | 31360000 | 0.0054 | 0.0037 | 0.0040 |
| F06 | 31280001 | 31380000 | 0.0053 | 0.0046 | 0.0038 |
| F06 | 31300001 | 31400000 | 0.0058 | 0.0047 | 0.0047 |
| F06 | 31320001 | 31420000 | 0.0059 | 0.0053 | 0.0049 |
| F06 | 31340001 | 31440000 | 0.0062 | 0.0053 | 0.0054 |
| F06 | 31360001 | 31460000 | 0.0065 | 0.0053 | 0.0058 |
| F06 | 31380001 | 31480000 | 0.0066 | 0.0037 | 0.0061 |
| F06 | 31400001 | 31500000 | 0.0049 | 0.0031 | 0.0044 |
| F06 | 31420001 | 31520000 | 0.0045 | 0.0026 | 0.0037 |
| F06 | 31440001 | 31540000 | 0.0039 | 0.0032 | 0.0031 |
| F06 | 31460001 | 31560000 | 0.0043 | 0.0042 | 0.0036 |
| F06 | 31480001 | 31580000 | 0.0052 | 0.0044 | 0.0044 |
| F06 | 31500001 | 31600000 | 0.0054 | 0.0045 | 0.0046 |
| F06 | 31520001 | 31620000 | 0.0052 | 0.0044 | 0.0047 |
| F06 | 31540001 | 31640000 | 0.0052 | 0.0037 | 0.0047 |
| F06 | 31560001 | 31660000 | 0.0046 | 0.0019 | 0.0038 |
| F06 | 31580001 | 31680000 | 0.0028 | 0.0024 | 0.0020 |
| F06 | 31600001 | 31700000 | 0.0029 | 0.0024 | 0.0023 |
| F06 | 31620001 | 31720000 | 0.0029 | 0.0022 | 0.0023 |
| F06 | 31640001 | 31740000 | 0.0028 | 0.0022 | 0.0022 |
| F06 | 31660001 | 31760000 | 0.0029 | 0.0024 | 0.0024 |
| F06 | 31680001 | 31780000 | 0.0034 | 0.0023 | 0.0028 |
| F06 | 31700001 | 31800000 | 0.0037 | 0.0029 | 0.0030 |
| F06 | 31720001 | 31820000 | 0.0045 | 0.0039 | 0.0038 |
| F06 | 31740001 | 31840000 | 0.0052 | 0.0041 | 0.0045 |
| F06 | 31760001 | 31860000 | 0.0051 | 0.0041 | 0.0043 |
| F06 | 31780001 | 31880000 | 0.0049 | 0.0038 | 0.0041 |
| F06 | 31800001 | 31900000 | 0.0044 | 0.0035 | 0.0037 |
| F06 | 31820001 | 31920000 | 0.0039 | 0.0044 | 0.0032 |
| F06 | 31840001 | 31940000 | 0.0047 | 0.0039 | 0.0040 |
| F06 | 31860001 | 31960000 | 0.0041 | 0.0040 | 0.0037 |
| F06 | 31880001 | 31980000 | 0.0046 | 0.0038 | 0.0040 |
| F06 | 31900001 | 32000000 | 0.0046 | 0.0038 | 0.0037 |
| F06 | 31920001 | 32020000 | 0.0046 | 0.0019 | 0.0037 |
| F06 | 31940001 | 32040000 | 0.0030 | 0.0022 | 0.0023 |
| F06 | 31960001 | 32060000 | 0.0032 | 0.0021 | 0.0026 |
| F06 | 31980001 | 32080000 | 0.0029 | 0.0022 | 0.0025 |
| F06 | 32000001 | 32100000 | 0.0027 | 0.0017 | 0.0026 |
| F06 | 32020001 | 32120000 | 0.0021 | 0.0017 | 0.0021 |
| F06 | 32040001 | 32140000 | 0.0020 | 0.0015 | 0.0020 |
| F06 | 32060001 | 32160000 | 0.0017 | 0.0017 | 0.0017 |
| F06 | 32080001 | 32180000 | 0.0020 | 0.0016 | 0.0020 |
| F06 | 32100001 | 32200000 | 0.0019 | 0.0019 | 0.0019 |
| F06 | 32120001 | 32220000 | 0.0022 | 0.0018 | 0.0022 |
| F06 | 32140001 | 32240000 | 0.0021 | 0.0017 | 0.0020 |
| F06 | 32160001 | 32260000 | 0.0020 | 0.0013 | 0.0019 |
| F06 | 32180001 | 32280000 | 0.0015 | 0.0013 | 0.0014 |
| F06 | 32200001 | 32300000 | 0.0014 | 0.0016 | 0.0014 |
| F06 | 32220001 | 32320000 | 0.0019 | 0.0019 | 0.0017 |

| Chr | Start | End | both | A1 | A2 |
|---|---|---|---|---|---|
| F06 | 32240001 | 32340000 | 0.0023 | 0.0025 | 0.0021 |
| F06 | 32260001 | 32360000 | 0.0030 | 0.0024 | 0.0025 |
| F06 | 32280001 | 32380000 | 0.0032 | 0.0031 | 0.0024 |
| F06 | 32300001 | 32400000 | 0.0040 | 0.0030 | 0.0031 |
| F06 | 32320001 | 32420000 | 0.0039 | 0.0028 | 0.0032 |
| F06 | 32340001 | 32440000 | 0.0037 | 0.0028 | 0.0030 |
| F06 | 32360001 | 32460000 | 0.0038 | 0.0036 | 0.0033 |
| F06 | 32380001 | 32480000 | 0.0046 | 0.0032 | 0.0041 |
| F06 | 32400001 | 32500000 | 0.0041 | 0.0033 | 0.0037 |
| F06 | 32420001 | 32520000 | 0.0042 | 0.0039 | 0.0036 |
| F06 | 32440001 | 32540000 | 0.0049 | 0.0040 | 0.0043 |
| F06 | 32460001 | 32560000 | 0.0049 | 0.0032 | 0.0044 |
| F06 | 32480001 | 32580000 | 0.0040 | 0.0033 | 0.0036 |
| F06 | 32500001 | 32600000 | 0.0041 | 0.0026 | 0.0037 |
| F06 | 32520001 | 32620000 | 0.0032 | 0.0020 | 0.0029 |
| F06 | 32540001 | 32640000 | 0.0025 | 0.0021 | 0.0021 |
| F06 | 32560001 | 32660000 | 0.0026 | 0.0026 | 0.0022 |
| F06 | 32580001 | 32680000 | 0.0031 | 0.0025 | 0.0028 |
| F06 | 32600001 | 32700000 | 0.0033 | 0.0031 | 0.0029 |
| F06 | 32620001 | 32720000 | 0.0043 | 0.0033 | 0.0036 |
| F06 | 32640001 | 32740000 | 0.0045 | 0.0031 | 0.0039 |
| F06 | 32660001 | 32760000 | 0.0045 | 0.0025 | 0.0037 |
| F06 | 32680001 | 32780000 | 0.0039 | 0.0028 | 0.0030 |
| F06 | 32700001 | 32800000 | 0.0041 | 0.0024 | 0.0032 |
| F06 | 32720001 | 32820000 | 0.0035 | 0.0033 | 0.0027 |
| F06 | 32740001 | 32840000 | 0.0046 | 0.0026 | 0.0036 |
| F06 | 32760001 | 32860000 | 0.0038 | 0.0035 | 0.0028 |
| F06 | 32780001 | 32880000 | 0.0050 | 0.0040 | 0.0040 |
| F06 | 32800001 | 32900000 | 0.0054 | 0.0047 | 0.0045 |
| F06 | 32820001 | 32920000 | 0.0060 | 0.0038 | 0.0052 |
| F06 | 32840001 | 32940000 | 0.0050 | 0.0042 | 0.0044 |
| F06 | 32860001 | 32960000 | 0.0052 | 0.0033 | 0.0048 |
| F06 | 32880001 | 32980000 | 0.0042 | 0.0026 | 0.0038 |
| F06 | 32900001 | 33000000 | 0.0035 | 0.0033 | 0.0032 |
| F06 | 32920001 | 33020000 | 0.0042 | 0.0037 | 0.0039 |
| F06 | 32940001 | 33040000 | 0.0046 | 0.0035 | 0.0043 |
| F06 | 32960001 | 33060000 | 0.0045 | 0.0034 | 0.0041 |
| F06 | 32980001 | 33080000 | 0.0044 | 0.0037 | 0.0040 |
| F06 | 33000001 | 33100000 | 0.0047 | 0.0028 | 0.0043 |
| F06 | 33020001 | 33120000 | 0.0036 | 0.0027 | 0.0033 |
| F06 | 33040001 | 33140000 | 0.0033 | 0.0027 | 0.0029 |
| F06 | 33060001 | 33160000 | 0.0032 | 0.0028 | 0.0028 |
| F06 | 33080001 | 33180000 | 0.0031 | 0.0026 | 0.0028 |
| F06 | 33100001 | 33200000 | 0.0027 | 0.0020 | 0.0025 |
| F06 | 33120001 | 33220000 | 0.0022 | 0.0019 | 0.0019 |
| F06 | 33140001 | 33240000 | 0.0025 | 0.0017 | 0.0020 |
| F06 | 33160001 | 33260000 | 0.0023 | 0.0017 | 0.0019 |
| F06 | 33180001 | 33280000 | 0.0027 | 0.0036 | 0.0020 |
| F06 | 33200001 | 33300000 | 0.0046 | 0.0048 | 0.0038 |
| F06 | 33220001 | 33320000 | 0.0058 | 0.0043 | 0.0051 |
| F06 | 33240001 | 33340000 | 0.0051 | 0.0048 | 0.0044 |
| F06 | 33260001 | 33360000 | 0.0057 | 0.0054 | 0.0049 |
| F06 | 33280001 | 33380000 | 0.0061 | 0.0038 | 0.0056 |
| F06 | 33300001 | 33400000 | 0.0046 | 0.0036 | 0.0042 |
| F06 | 33320001 | 33420000 | 0.0045 | 0.0042 | 0.0039 |
| F06 | 33340001 | 33440000 | 0.0053 | 0.0038 | 0.0047 |
| F06 | 33360001 | 33460000 | 0.0049 | 0.0037 | 0.0042 |
| F06 | 33380001 | 33480000 | 0.0047 | 0.0032 | 0.0041 |

| Chr | Start | End | both | A1 | A2 |
|---|---|---|---|---|---|
| F06 | 33400001 | 33500000 | 0.0040 | 0.0027 | 0.0033 |
| F06 | 33420001 | 33520000 | 0.0036 | 0.0030 | 0.0029 |
| F06 | 33440001 | 33540000 | 0.0039 | 0.0030 | 0.0032 |
| F06 | 33460001 | 33560000 | 0.0039 | 0.0026 | 0.0033 |
| F06 | 33480001 | 33580000 | 0.0035 | 0.0025 | 0.0028 |
| F06 | 33500001 | 33600000 | 0.0037 | 0.0023 | 0.0029 |
| F06 | 33520001 | 33620000 | 0.0033 | 0.0018 | 0.0027 |
| F06 | 33540001 | 33640000 | 0.0026 | 0.0018 | 0.0021 |
| F06 | 33560001 | 33660000 | 0.0028 | 0.0021 | 0.0020 |
| F06 | 33580001 | 33680000 | 0.0031 | 0.0021 | 0.0022 |
| F06 | 33600001 | 33700000 | 0.0032 | 0.0024 | 0.0021 |
| F06 | 33620001 | 33720000 | 0.0035 | 0.0019 | 0.0024 |
| F06 | 33640001 | 33740000 | 0.0030 | 0.0017 | 0.0019 |
| F06 | 33660001 | 33760000 | 0.0026 | 0.0014 | 0.0017 |
| F06 | 33680001 | 33780000 | 0.0022 | 0.0013 | 0.0016 |
| F06 | 33700001 | 33800000 | 0.0018 | 0.0009 | 0.0015 |
| F06 | 33720001 | 33820000 | 0.0013 | 0.0010 | 0.0011 |
| F06 | 33740001 | 33840000 | 0.0012 | 0.0012 | 0.0011 |
| F06 | 33760001 | 33860000 | 0.0015 | 0.0009 | 0.0014 |
| F06 | 33780001 | 33880000 | 0.0011 | 0.0016 | 0.0010 |
| F06 | 33800001 | 33900000 | 0.0019 | 0.0017 | 0.0018 |
| F06 | 33820001 | 33920000 | 0.0022 | 0.0022 | 0.0020 |
| F06 | 33840001 | 33940000 | 0.0027 | 0.0019 | 0.0025 |
| F06 | 33860001 | 33960000 | 0.0023 | 0.0035 | 0.0023 |
| F06 | 33880001 | 33980000 | 0.0039 | 0.0033 | 0.0038 |
| F06 | 33900001 | 34000000 | 0.0037 | 0.0038 | 0.0035 |
| F06 | 33920001 | 34020000 | 0.0044 | 0.0037 | 0.0040 |
| F06 | 33940001 | 34040000 | 0.0044 | 0.0054 | 0.0039 |
| F06 | 33960001 | 34060000 | 0.0064 | 0.0042 | 0.0058 |
| F06 | 33980001 | 34080000 | 0.0054 | 0.0042 | 0.0047 |
| F06 | 34000001 | 34100000 | 0.0053 | 0.0043 | 0.0047 |
| F06 | 34020001 | 34120000 | 0.0052 | 0.0050 | 0.0048 |
| F06 | 34040001 | 34140000 | 0.0061 | 0.0039 | 0.0055 |
| F06 | 34060001 | 34160000 | 0.0049 | 0.0036 | 0.0044 |
| F06 | 34080001 | 34180000 | 0.0046 | 0.0031 | 0.0040 |
| F06 | 34100001 | 34200000 | 0.0042 | 0.0027 | 0.0035 |
| F06 | 34120001 | 34220000 | 0.0039 | 0.0021 | 0.0031 |
| F06 | 34140001 | 34240000 | 0.0033 | 0.0016 | 0.0026 |
| F06 | 34160001 | 34260000 | 0.0028 | 0.0020 | 0.0020 |
| F06 | 34180001 | 34280000 | 0.0033 | 0.0030 | 0.0023 |
| F06 | 34200001 | 34300000 | 0.0043 | 0.0034 | 0.0034 |
| F06 | 34220001 | 34320000 | 0.0046 | 0.0034 | 0.0038 |
| F06 | 34240001 | 34340000 | 0.0046 | 0.0035 | 0.0038 |
| F06 | 34260001 | 34360000 | 0.0045 | 0.0037 | 0.0038 |
| F06 | 34280001 | 34380000 | 0.0046 | 0.0029 | 0.0040 |
| F06 | 34300001 | 34400000 | 0.0039 | 0.0024 | 0.0033 |
| F06 | 34320001 | 34420000 | 0.0034 | 0.0027 | 0.0028 |
| F06 | 34340001 | 34440000 | 0.0038 | 0.0027 | 0.0030 |
| F06 | 34360001 | 34460000 | 0.0039 | 0.0020 | 0.0030 |
| F06 | 34380001 | 34480000 | 0.0032 | 0.0026 | 0.0022 |
| F06 | 34400001 | 34500000 | 0.0036 | 0.0027 | 0.0027 |
| F06 | 34420001 | 34520000 | 0.0039 | 0.0019 | 0.0027 |
| F06 | 34440001 | 34540000 | 0.0030 | 0.0028 | 0.0019 |
| F06 | 34460001 | 34560000 | 0.0037 | 0.0032 | 0.0027 |
| F06 | 34480001 | 34580000 | 0.0042 | 0.0025 | 0.0032 |
| F06 | 34500001 | 34600000 | 0.0036 | 0.0021 | 0.0026 |
| F06 | 34520001 | 34620000 | 0.0034 | 0.0022 | 0.0023 |
| F06 | 34540001 | 34640000 | 0.0034 | 0.0011 | 0.0026 |

| Chr | Start | End | both | A1 | A2 |
|---|---|---|---|---|---|
| F06 | 34560001 | 34660000 | 0.0022 | 0.0009 | 0.0015 |
| F06 | 34580001 | 34680000 | 0.0017 | 0.0008 | 0.0013 |
| F06 | 34600001 | 34700000 | 0.0014 | 0.0009 | 0.0010 |
| F06 | 34620001 | 34720000 | 0.0010 | 0.0010 | 0.0009 |
| F06 | 34640001 | 34740000 | 0.0012 | 0.0014 | 0.0011 |
| F06 | 34660001 | 34760000 | 0.0015 | 0.0014 | 0.0014 |
| F06 | 34680001 | 34780000 | 0.0015 | 0.0016 | 0.0013 |
| F06 | 34700001 | 34800000 | 0.0020 | 0.0014 | 0.0016 |
| F06 | 34720001 | 34820000 | 0.0019 | 0.0019 | 0.0015 |
| F06 | 34740001 | 34840000 | 0.0028 | 0.0015 | 0.0021 |
| F06 | 34760001 | 34860000 | 0.0026 | 0.0017 | 0.0018 |
| F06 | 34780001 | 34880000 | 0.0030 | 0.0016 | 0.0019 |
| F06 | 34800001 | 34900000 | 0.0027 | 0.0017 | 0.0017 |
| F06 | 34820001 | 34920000 | 0.0027 | 0.0013 | 0.0018 |
| F06 | 34840001 | 34940000 | 0.0022 | 0.0014 | 0.0012 |
| F06 | 34860001 | 34960000 | 0.0024 | 0.0020 | 0.0013 |
| F06 | 34880001 | 34980000 | 0.0030 | 0.0026 | 0.0021 |
| F06 | 34900001 | 35000000 | 0.0036 | 0.0032 | 0.0026 |
| F06 | 34920001 | 35020000 | 0.0044 | 0.0027 | 0.0032 |
| F06 | 34940001 | 35040000 | 0.0036 | 0.0030 | 0.0028 |
| F06 | 34960001 | 35060000 | 0.0038 | 0.0023 | 0.0032 |
| F06 | 34980001 | 35080000 | 0.0029 | 0.0018 | 0.0023 |
| F06 | 35000001 | 35100000 | 0.0030 | 0.0014 | 0.0022 |
| F06 | 35020001 | 35120000 | 0.0026 | 0.0015 | 0.0019 |
| F06 | 35040001 | 35140000 | 0.0027 | 0.0015 | 0.0019 |
| F06 | 35060001 | 35160000 | 0.0027 | 0.0015 | 0.0019 |
| F06 | 35080001 | 35180000 | 0.0026 | 0.0015 | 0.0019 |
| F06 | 35100001 | 35200000 | 0.0021 | 0.0020 | 0.0017 |
| F06 | 35120001 | 35220000 | 0.0026 | 0.0028 | 0.0023 |
| F06 | 35140001 | 35240000 | 0.0033 | 0.0027 | 0.0032 |
| F06 | 35160001 | 35260000 | 0.0034 | 0.0035 | 0.0032 |
| F06 | 35180001 | 35280000 | 0.0043 | 0.0051 | 0.0040 |
| F06 | 35200001 | 35300000 | 0.0062 | 0.0049 | 0.0058 |
| F06 | 35220001 | 35320000 | 0.0059 | 0.0044 | 0.0054 |
| F06 | 35240001 | 35340000 | 0.0054 | 0.0046 | 0.0049 |
| F06 | 35260001 | 35360000 | 0.0055 | 0.0044 | 0.0051 |
| F06 | 35280001 | 35380000 | 0.0050 | 0.0027 | 0.0047 |
| F06 | 35300001 | 35400000 | 0.0030 | 0.0021 | 0.0027 |
| F06 | 35320001 | 35420000 | 0.0024 | 0.0023 | 0.0021 |
| F06 | 35340001 | 35440000 | 0.0025 | 0.0018 | 0.0024 |
| F06 | 35360001 | 35460000 | 0.0020 | 0.0012 | 0.0019 |
| F06 | 35380001 | 35480000 | 0.0014 | 0.0012 | 0.0013 |
| F06 | 35400001 | 35500000 | 0.0014 | 0.0019 | 0.0013 |
| F06 | 35420001 | 35520000 | 0.0021 | 0.0014 | 0.0020 |
| F06 | 35440001 | 35540000 | 0.0017 | 0.0013 | 0.0015 |
| F06 | 35460001 | 35560000 | 0.0017 | 0.0018 | 0.0014 |
| F06 | 35480001 | 35580000 | 0.0024 | 0.0018 | 0.0019 |
| F06 | 35500001 | 35600000 | 0.0026 | 0.0013 | 0.0020 |
| F06 | 35520001 | 35620000 | 0.0022 | 0.0012 | 0.0016 |
| F06 | 35540001 | 35640000 | 0.0020 | 0.0013 | 0.0015 |
| F06 | 35560001 | 35660000 | 0.0022 | 0.0021 | 0.0016 |
| F06 | 35580001 | 35680000 | 0.0028 | 0.0024 | 0.0023 |
| F06 | 35600001 | 35700000 | 0.0031 | 0.0025 | 0.0025 |
| F06 | 35620001 | 35720000 | 0.0032 | 0.0036 | 0.0026 |
| F06 | 35640001 | 35740000 | 0.0046 | 0.0040 | 0.0039 |
| F06 | 35660001 | 35760000 | 0.0050 | 0.0044 | 0.0042 |
| F06 | 35680001 | 35780000 | 0.0056 | 0.0047 | 0.0049 |
| F06 | 35700001 | 35800000 | 0.0060 | 0.0047 | 0.0054 |

| Chr | Start | End | both | A1 | A2 |
|---|---|---|---|---|---|
| F06 | 35720001 | 35820000 | 0.0062 | 0.0037 | 0.0056 |
| F06 | 35740001 | 35840000 | 0.0049 | 0.0044 | 0.0045 |
| F06 | 35760001 | 35860000 | 0.0061 | 0.0034 | 0.0056 |
| F06 | 35780001 | 35880000 | 0.0048 | 0.0024 | 0.0044 |
| F06 | 35800001 | 35900000 | 0.0039 | 0.0027 | 0.0032 |
| F06 | 35820001 | 35920000 | 0.0042 | 0.0028 | 0.0034 |
| F06 | 35840001 | 35940000 | 0.0042 | 0.0017 | 0.0035 |
| F06 | 35860001 | 35960000 | 0.0027 | 0.0013 | 0.0020 |
| F06 | 35880001 | 35980000 | 0.0023 | 0.0014 | 0.0015 |
| F06 | 35900001 | 36000000 | 0.0022 | 0.0009 | 0.0016 |
| F06 | 35920001 | 36020000 | 0.0016 | 0.0013 | 0.0010 |
| F06 | 35940001 | 36040000 | 0.0022 | 0.0018 | 0.0015 |
| F06 | 35960001 | 36060000 | 0.0029 | 0.0021 | 0.0021 |
| F06 | 35980001 | 36080000 | 0.0033 | 0.0021 | 0.0024 |
| F06 | 36000001 | 36100000 | 0.0032 | 0.0022 | 0.0025 |
| F06 | 36020001 | 36120000 | 0.0031 | 0.0021 | 0.0025 |
| F06 | 36040001 | 36140000 | 0.0029 | 0.0015 | 0.0024 |
| F06 | 36060001 | 36160000 | 0.0019 | 0.0015 | 0.0016 |
| F06 | 36080001 | 36180000 | 0.0017 | 0.0020 | 0.0017 |
| F06 | 36100001 | 36200000 | 0.0023 | 0.0020 | 0.0022 |
| F06 | 36120001 | 36220000 | 0.0026 | 0.0015 | 0.0022 |
| F06 | 36140001 | 36240000 | 0.0023 | 0.0018 | 0.0017 |
| F06 | 36160001 | 36260000 | 0.0026 | 0.0016 | 0.0020 |
| F06 | 36180001 | 36280000 | 0.0024 | 0.0019 | 0.0018 |
| F06 | 36200001 | 36300000 | 0.0028 | 0.0019 | 0.0021 |
| F06 | 36220001 | 36320000 | 0.0026 | 0.0026 | 0.0021 |
| F06 | 36240001 | 36340000 | 0.0032 | 0.0026 | 0.0029 |
| F06 | 36260001 | 36360000 | 0.0033 | 0.0036 | 0.0028 |
| F06 | 36280001 | 36380000 | 0.0044 | 0.0026 | 0.0038 |
| F06 | 36300001 | 36400000 | 0.0036 | 0.0028 | 0.0029 |
| F06 | 36320001 | 36420000 | 0.0039 | 0.0035 | 0.0031 |
| F06 | 36340001 | 36440000 | 0.0047 | 0.0047 | 0.0037 |
| F06 | 36360001 | 36460000 | 0.0062 | 0.0038 | 0.0051 |
| F06 | 36380001 | 36480000 | 0.0052 | 0.0050 | 0.0041 |
| F06 | 36400001 | 36500000 | 0.0063 | 0.0050 | 0.0054 |
| F06 | 36420001 | 36520000 | 0.0064 | 0.0042 | 0.0056 |
| F06 | 36440001 | 36540000 | 0.0059 | 0.0031 | 0.0050 |
| F06 | 36460001 | 36560000 | 0.0045 | 0.0034 | 0.0037 |
| F06 | 36480001 | 36580000 | 0.0049 | 0.0034 | 0.0040 |
| F06 | 36500001 | 36600000 | 0.0047 | 0.0035 | 0.0038 |
| F06 | 36520001 | 36620000 | 0.0046 | 0.0034 | 0.0038 |
| F06 | 36540001 | 36640000 | 0.0044 | 0.0032 | 0.0036 |
| F06 | 36560001 | 36660000 | 0.0041 | 0.0029 | 0.0035 |
| F06 | 36580001 | 36680000 | 0.0038 | 0.0028 | 0.0033 |
| F06 | 36600001 | 36700000 | 0.0038 | 0.0024 | 0.0031 |
| F06 | 36620001 | 36720000 | 0.0034 | 0.0018 | 0.0027 |
| F06 | 36640001 | 36740000 | 0.0026 | 0.0017 | 0.0020 |
| F06 | 36660001 | 36760000 | 0.0027 | 0.0023 | 0.0019 |
| F06 | 36680001 | 36780000 | 0.0033 | 0.0022 | 0.0024 |
| F06 | 36700001 | 36800000 | 0.0032 | 0.0025 | 0.0022 |
| F06 | 36720001 | 36820000 | 0.0039 | 0.0032 | 0.0026 |
| F06 | 36740001 | 36840000 | 0.0046 | 0.0030 | 0.0033 |
| F06 | 36760001 | 36860000 | 0.0044 | 0.0024 | 0.0032 |
| F06 | 36780001 | 36880000 | 0.0036 | 0.0031 | 0.0025 |
| F06 | 36800001 | 36900000 | 0.0043 | 0.0028 | 0.0032 |
| F06 | 36820001 | 36920000 | 0.0036 | 0.0026 | 0.0028 |
| F06 | 36840001 | 36940000 | 0.0032 | 0.0030 | 0.0025 |
| F06 | 36860001 | 36960000 | 0.0036 | 0.0033 | 0.0030 |

| Chr | Start | End | both | A1 | A2 |
|---|---|---|---|---|---|
| F06 | 36880001 | 36980000 | 0.0040 | 0.0020 | 0.0034 |
| F06 | 36900001 | 37000000 | 0.0026 | 0.0025 | 0.0023 |
| F06 | 36920001 | 37020000 | 0.0030 | 0.0022 | 0.0027 |
| F06 | 36940001 | 37040000 | 0.0026 | 0.0019 | 0.0026 |
| F06 | 36960001 | 37060000 | 0.0027 | 0.0027 | 0.0022 |
| F06 | 36980001 | 37080000 | 0.0037 | 0.0030 | 0.0030 |
| F06 | 37000001 | 37100000 | 0.0043 | 0.0025 | 0.0032 |
| F06 | 37020001 | 37120000 | 0.0039 | 0.0046 | 0.0028 |
| F06 | 37040001 | 37140000 | 0.0061 | 0.0048 | 0.0046 |
| F06 | 37060001 | 37160000 | 0.0058 | 0.0043 | 0.0047 |
| F06 | 37080001 | 37180000 | 0.0051 | 0.0042 | 0.0041 |
| F06 | 37100001 | 37200000 | 0.0052 | 0.0049 | 0.0042 |
| F06 | 37120001 | 37220000 | 0.0057 | 0.0025 | 0.0048 |
| F06 | 37140001 | 37240000 | 0.0032 | 0.0023 | 0.0027 |
| F06 | 37160001 | 37260000 | 0.0034 | 0.0018 | 0.0026 |
| F06 | 37180001 | 37280000 | 0.0030 | 0.0022 | 0.0023 |
| F06 | 37200001 | 37300000 | 0.0029 | 0.0021 | 0.0025 |
| F06 | 37220001 | 37320000 | 0.0029 | 0.0031 | 0.0026 |
| F06 | 37240001 | 37340000 | 0.0042 | 0.0034 | 0.0036 |
| F06 | 37260001 | 37360000 | 0.0043 | 0.0040 | 0.0037 |
| F06 | 37280001 | 37380000 | 0.0051 | 0.0038 | 0.0042 |
| F06 | 37300001 | 37400000 | 0.0050 | 0.0033 | 0.0039 |
| F06 | 37320001 | 37420000 | 0.0044 | 0.0024 | 0.0034 |
| F06 | 37340001 | 37440000 | 0.0034 | 0.0022 | 0.0025 |
| F06 | 37360001 | 37460000 | 0.0030 | 0.0012 | 0.0023 |
| F06 | 37380001 | 37480000 | 0.0016 | 0.0012 | 0.0012 |
| F06 | 37400001 | 37500000 | 0.0015 | 0.0015 | 0.0012 |
| F06 | 37420001 | 37520000 | 0.0019 | 0.0016 | 0.0016 |
| F06 | 37440001 | 37540000 | 0.0020 | 0.0027 | 0.0016 |
| F06 | 37460001 | 37560000 | 0.0032 | 0.0034 | 0.0027 |
| F06 | 37480001 | 37580000 | 0.0041 | 0.0031 | 0.0034 |
| F06 | 37500001 | 37600000 | 0.0040 | 0.0027 | 0.0032 |
| F06 | 37520001 | 37620000 | 0.0037 | 0.0034 | 0.0027 |
| F06 | 37540001 | 37640000 | 0.0043 | 0.0025 | 0.0036 |
| F06 | 37560001 | 37660000 | 0.0034 | 0.0022 | 0.0028 |
| F06 | 37580001 | 37680000 | 0.0031 | 0.0022 | 0.0024 |
| F06 | 37600001 | 37700000 | 0.0030 | 0.0024 | 0.0024 |
| F06 | 37620001 | 37720000 | 0.0033 | 0.0019 | 0.0026 |
| F06 | 37640001 | 37740000 | 0.0028 | 0.0018 | 0.0019 |
| F06 | 37660001 | 37760000 | 0.0028 | 0.0016 | 0.0017 |
| F06 | 37680001 | 37780000 | 0.0027 | 0.0022 | 0.0018 |
| F06 | 37700001 | 37800000 | 0.0033 | 0.0021 | 0.0025 |
| F06 | 37720001 | 37820000 | 0.0031 | 0.0018 | 0.0023 |
| F06 | 37740001 | 37840000 | 0.0030 | 0.0018 | 0.0021 |
| F06 | 37760001 | 37860000 | 0.0030 | 0.0015 | 0.0021 |
| F06 | 37780001 | 37880000 | 0.0028 | 0.0007 | 0.0019 |
| F06 | 37800001 | 37900000 | 0.0021 | 0.0005 | 0.0010 |
| F06 | 37820001 | 37920000 | 0.0021 | 0.0005 | 0.0008 |
| F06 | 37840001 | 37940000 | 0.0020 | 0.0006 | 0.0007 |
| F06 | 37860001 | 37960000 | 0.0021 | 0.0007 | 0.0008 |
| F06 | 37880001 | 37980000 | 0.0020 | 0.0010 | 0.0008 |
| F06 | 37900001 | 38000000 | 0.0021 | 0.0017 | 0.0011 |
| F06 | 37920001 | 38020000 | 0.0026 | 0.0022 | 0.0018 |
| F06 | 37940001 | 38040000 | 0.0031 | 0.0025 | 0.0024 |
| F06 | 37960001 | 38060000 | 0.0033 | 0.0027 | 0.0025 |
| F06 | 37980001 | 38080000 | 0.0033 | 0.0029 | 0.0026 |
| F06 | 38000001 | 38100000 | 0.0039 | 0.0027 | 0.0029 |
| F06 | 38020001 | 38120000 | 0.0037 | 0.0026 | 0.0028 |

| Chr | Start | End | both | A1 | A2 |
|---|---|---|---|---|---|
| F06 | 38040001 | 38140000 | 0.0034 | 0.0027 | 0.0026 |
| F06 | 38060001 | 38160000 | 0.0036 | 0.0026 | 0.0029 |
| F06 | 38080001 | 38180000 | 0.0035 | 0.0025 | 0.0030 |
| F06 | 38100001 | 38200000 | 0.0032 | 0.0023 | 0.0028 |
| F06 | 38120001 | 38220000 | 0.0031 | 0.0023 | 0.0027 |
| F06 | 38140001 | 38240000 | 0.0029 | 0.0017 | 0.0027 |
| F06 | 38160001 | 38260000 | 0.0022 | 0.0015 | 0.0020 |
| F06 | 38180001 | 38280000 | 0.0019 | 0.0013 | 0.0017 |
| F06 | 38200001 | 38300000 | 0.0016 | 0.0016 | 0.0015 |
| F06 | 38220001 | 38320000 | 0.0019 | 0.0014 | 0.0017 |
| F06 | 38240001 | 38340000 | 0.0018 | 0.0021 | 0.0015 |
| F06 | 38260001 | 38360000 | 0.0030 | 0.0030 | 0.0025 |
| F06 | 38280001 | 38380000 | 0.0042 | 0.0028 | 0.0033 |
| F06 | 38300001 | 38400000 | 0.0043 | 0.0023 | 0.0031 |
| F06 | 38320001 | 38420000 | 0.0037 | 0.0023 | 0.0025 |
| F06 | 38340001 | 38440000 | 0.0036 | 0.0015 | 0.0025 |
| F06 | 38360001 | 38460000 | 0.0024 | 0.0014 | 0.0015 |
| F06 | 38380001 | 38480000 | 0.0021 | 0.0018 | 0.0013 |
| F06 | 38400001 | 38500000 | 0.0023 | 0.0018 | 0.0018 |
| F06 | 38420001 | 38520000 | 0.0023 | 0.0017 | 0.0018 |
| F06 | 38440001 | 38540000 | 0.0023 | 0.0018 | 0.0017 |
| F06 | 38460001 | 38560000 | 0.0025 | 0.0023 | 0.0019 |
| F06 | 38480001 | 38580000 | 0.0031 | 0.0024 | 0.0027 |
| F06 | 38500001 | 38600000 | 0.0032 | 0.0024 | 0.0028 |
| F06 | 38520001 | 38620000 | 0.0033 | 0.0025 | 0.0029 |
| F06 | 38540001 | 38640000 | 0.0034 | 0.0032 | 0.0030 |
| F06 | 38560001 | 38660000 | 0.0041 | 0.0021 | 0.0035 |
| F06 | 38580001 | 38680000 | 0.0027 | 0.0016 | 0.0023 |
| F06 | 38600001 | 38700000 | 0.0021 | 0.0019 | 0.0017 |
| F06 | 38620001 | 38720000 | 0.0025 | 0.0024 | 0.0022 |
| F06 | 38640001 | 38740000 | 0.0031 | 0.0021 | 0.0025 |
| F06 | 38660001 | 38760000 | 0.0032 | 0.0028 | 0.0026 |
| F06 | 38680001 | 38780000 | 0.0041 | 0.0028 | 0.0033 |
| F06 | 38700001 | 38800000 | 0.0042 | 0.0031 | 0.0033 |
| F06 | 38720001 | 38820000 | 0.0045 | 0.0027 | 0.0036 |
| F06 | 38740001 | 38840000 | 0.0043 | 0.0023 | 0.0033 |
| F06 | 38760001 | 38860000 | 0.0036 | 0.0022 | 0.0025 |
| F06 | 38780001 | 38880000 | 0.0035 | 0.0025 | 0.0024 |
| F06 | 38800001 | 38900000 | 0.0038 | 0.0019 | 0.0028 |
| F06 | 38820001 | 38920000 | 0.0032 | 0.0021 | 0.0022 |
| F06 | 38840001 | 38940000 | 0.0030 | 0.0019 | 0.0024 |
| F06 | 38860001 | 38960000 | 0.0026 | 0.0012 | 0.0022 |
| F06 | 38880001 | 38980000 | 0.0017 | 0.0013 | 0.0015 |
| F06 | 38900001 | 39000000 | 0.0016 | 0.0014 | 0.0014 |
| F06 | 38920001 | 39020000 | 0.0016 | 0.0013 | 0.0015 |
| F06 | 38940001 | 39040000 | 0.0019 | 0.0025 | 0.0014 |
| F06 | 38960001 | 39060000 | 0.0035 | 0.0030 | 0.0028 |
| F06 | 38980001 | 39080000 | 0.0041 | 0.0025 | 0.0032 |
| F06 | 39000001 | 39100000 | 0.0039 | 0.0024 | 0.0028 |
| F06 | 39020001 | 39120000 | 0.0038 | 0.0034 | 0.0027 |
| F06 | 39040001 | 39140000 | 0.0045 | 0.0022 | 0.0037 |
| F06 | 39060001 | 39160000 | 0.0034 | 0.0020 | 0.0023 |
| F06 | 39080001 | 39180000 | 0.0032 | 0.0024 | 0.0021 |
| F06 | 39100001 | 39200000 | 0.0037 | 0.0023 | 0.0026 |
| F06 | 39120001 | 39220000 | 0.0037 | 0.0016 | 0.0025 |
| F06 | 39140001 | 39240000 | 0.0029 | 0.0019 | 0.0017 |
| F06 | 39160001 | 39260000 | 0.0029 | 0.0021 | 0.0021 |
| F06 | 39180001 | 39280000 | 0.0030 | 0.0020 | 0.0023 |

| Chr | Start | End | both | A1 | A2 |
|---|---|---|---|---|---|
| F06 | 39200001 | 39300000 | 0.0029 | 0.0020 | 0.0022 |
| F06 | 39220001 | 39320000 | 0.0026 | 0.0020 | 0.0020 |
| F06 | 39240001 | 39340000 | 0.0026 | 0.0019 | 0.0022 |
| F06 | 39260001 | 39360000 | 0.0027 | 0.0019 | 0.0020 |
| F06 | 39280001 | 39380000 | 0.0026 | 0.0030 | 0.0019 |
| F06 | 39300001 | 39400000 | 0.0036 | 0.0032 | 0.0029 |
| F06 | 39320001 | 39420000 | 0.0039 | 0.0041 | 0.0031 |
| F06 | 39340001 | 39440000 | 0.0048 | 0.0042 | 0.0039 |
| F06 | 39360001 | 39460000 | 0.0052 | 0.0042 | 0.0042 |
| F06 | 39380001 | 39480000 | 0.0052 | 0.0029 | 0.0042 |
| F06 | 39400001 | 39500000 | 0.0042 | 0.0027 | 0.0029 |
| F06 | 39420001 | 39520000 | 0.0040 | 0.0020 | 0.0028 |
| F06 | 39440001 | 39540000 | 0.0034 | 0.0019 | 0.0021 |
| F06 | 39460001 | 39560000 | 0.0028 | 0.0023 | 0.0018 |
| F06 | 39480001 | 39580000 | 0.0031 | 0.0027 | 0.0022 |
| F06 | 39500001 | 39600000 | 0.0035 | 0.0029 | 0.0028 |
| F06 | 39520001 | 39620000 | 0.0040 | 0.0028 | 0.0031 |
| F06 | 39540001 | 39640000 | 0.0038 | 0.0027 | 0.0030 |
| F06 | 39560001 | 39660000 | 0.0038 | 0.0021 | 0.0031 |
| F06 | 39580001 | 39680000 | 0.0037 | 0.0015 | 0.0027 |
| F06 | 39600001 | 39700000 | 0.0028 | 0.0012 | 0.0019 |
| F06 | 39620001 | 39720000 | 0.0023 | 0.0009 | 0.0016 |
| F06 | 39640001 | 39740000 | 0.0020 | 0.0008 | 0.0013 |
| F06 | 39660001 | 39760000 | 0.0018 | 0.0024 | 0.0011 |
| F06 | 39680001 | 39780000 | 0.0038 | 0.0033 | 0.0033 |
| F06 | 39700001 | 39800000 | 0.0049 | 0.0033 | 0.0044 |
| F06 | 39720001 | 39820000 | 0.0047 | 0.0034 | 0.0044 |
| F06 | 39740001 | 39840000 | 0.0047 | 0.0037 | 0.0044 |
| F06 | 39760001 | 39860000 | 0.0050 | 0.0020 | 0.0047 |
| F06 | 39780001 | 39880000 | 0.0028 | 0.0014 | 0.0026 |
| F06 | 39800001 | 39900000 | 0.0020 | 0.0015 | 0.0019 |
| F06 | 39820001 | 39920000 | 0.0021 | 0.0017 | 0.0020 |
| F06 | 39840001 | 39940000 | 0.0022 | 0.0015 | 0.0020 |
| F06 | 39860001 | 39960000 | 0.0022 | 0.0016 | 0.0018 |
| F06 | 39880001 | 39980000 | 0.0022 | 0.0020 | 0.0018 |
| F06 | 39900001 | 40000000 | 0.0027 | 0.0024 | 0.0022 |
| F06 | 39920001 | 40020000 | 0.0034 | 0.0025 | 0.0027 |
| F06 | 39940001 | 40040000 | 0.0036 | 0.0033 | 0.0028 |
| F06 | 39960001 | 40060000 | 0.0043 | 0.0034 | 0.0037 |
| F06 | 39980001 | 40080000 | 0.0045 | 0.0032 | 0.0039 |
| F06 | 40000001 | 40100000 | 0.0041 | 0.0030 | 0.0037 |
| F06 | 40020001 | 40120000 | 0.0038 | 0.0029 | 0.0034 |
| F06 | 40040001 | 40140000 | 0.0035 | 0.0020 | 0.0033 |
| F06 | 40060001 | 40160000 | 0.0026 | 0.0019 | 0.0022 |
| F06 | 40080001 | 40180000 | 0.0025 | 0.0021 | 0.0020 |
| F06 | 40100001 | 40200000 | 0.0028 | 0.0022 | 0.0024 |
| F06 | 40120001 | 40220000 | 0.0029 | 0.0024 | 0.0024 |
| F06 | 40140001 | 40240000 | 0.0032 | 0.0029 | 0.0026 |
| F06 | 40160001 | 40260000 | 0.0040 | 0.0027 | 0.0035 |
| F06 | 40180001 | 40280000 | 0.0038 | 0.0025 | 0.0033 |
| F06 | 40200001 | 40300000 | 0.0035 | 0.0033 | 0.0030 |
| F06 | 40220001 | 40320000 | 0.0046 | 0.0032 | 0.0040 |
| F06 | 40240001 | 40340000 | 0.0044 | 0.0025 | 0.0041 |
| F06 | 40260001 | 40360000 | 0.0035 | 0.0036 | 0.0031 |
| F06 | 40280001 | 40380000 | 0.0048 | 0.0039 | 0.0043 |
| F06 | 40300001 | 40400000 | 0.0051 | 0.0031 | 0.0047 |
| F06 | 40320001 | 40420000 | 0.0042 | 0.0030 | 0.0036 |
| F06 | 40340001 | 40440000 | 0.0040 | 0.0032 | 0.0033 |

| Chr | Start | End | both | A1 | A2 |
|---|---|---|---|---|---|
| F06 | 40360001 | 40460000 | 0.0041 | 0.0026 | 0.0034 |
| F06 | 40380001 | 40480000 | 0.0032 | 0.0018 | 0.0026 |
| F06 | 40400001 | 40500000 | 0.0022 | 0.0017 | 0.0017 |
| F06 | 40420001 | 40520000 | 0.0019 | 0.0014 | 0.0016 |
| F06 | 40440001 | 40540000 | 0.0015 | 0.0014 | 0.0012 |
| F06 | 40460001 | 40560000 | 0.0016 | 0.0015 | 0.0014 |
| F06 | 40480001 | 40580000 | 0.0018 | 0.0014 | 0.0017 |
| F06 | 40500001 | 40600000 | 0.0020 | 0.0018 | 0.0017 |
| F06 | 40520001 | 40620000 | 0.0028 | 0.0024 | 0.0022 |
| F06 | 40540001 | 40640000 | 0.0038 | 0.0039 | 0.0028 |
| F06 | 40560001 | 40660000 | 0.0054 | 0.0033 | 0.0041 |
| F06 | 40580001 | 40680000 | 0.0047 | 0.0041 | 0.0031 |
| F06 | 40600001 | 40700000 | 0.0054 | 0.0043 | 0.0040 |
| F06 | 40620001 | 40720000 | 0.0054 | 0.0047 | 0.0041 |
| F06 | 40640001 | 40740000 | 0.0057 | 0.0034 | 0.0046 |
| F06 | 40660001 | 40760000 | 0.0043 | 0.0040 | 0.0036 |
| F06 | 40680001 | 40780000 | 0.0049 | 0.0036 | 0.0042 |
| F06 | 40700001 | 40800000 | 0.0044 | 0.0038 | 0.0037 |
| F06 | 40720001 | 40820000 | 0.0049 | 0.0040 | 0.0039 |
| F06 | 40740001 | 40840000 | 0.0052 | 0.0040 | 0.0043 |
| F06 | 40760001 | 40860000 | 0.0054 | 0.0043 | 0.0043 |
| F06 | 40780001 | 40880000 | 0.0056 | 0.0048 | 0.0047 |
| F06 | 40800001 | 40900000 | 0.0064 | 0.0048 | 0.0055 |
| F06 | 40820001 | 40920000 | 0.0059 | 0.0042 | 0.0054 |
| F06 | 40840001 | 40940000 | 0.0051 | 0.0043 | 0.0045 |
| F06 | 40860001 | 40960000 | 0.0050 | 0.0035 | 0.0044 |
| F06 | 40880001 | 40980000 | 0.0042 | 0.0025 | 0.0034 |
| F06 | 40900001 | 41000000 | 0.0033 | 0.0020 | 0.0024 |
| F06 | 40920001 | 41020000 | 0.0030 | 0.0027 | 0.0019 |
| F06 | 40940001 | 41040000 | 0.0038 | 0.0028 | 0.0024 |
| F06 | 40960001 | 41060000 | 0.0039 | 0.0028 | 0.0025 |
| F06 | 40980001 | 41080000 | 0.0037 | 0.0028 | 0.0024 |
| F06 | 41000001 | 41100000 | 0.0036 | 0.0027 | 0.0025 |
| F06 | 41020001 | 41120000 | 0.0034 | 0.0014 | 0.0023 |
| F06 | 41040001 | 41140000 | 0.0023 | 0.0012 | 0.0015 |
| F06 | 41060001 | 41160000 | 0.0021 | 0.0019 | 0.0013 |
| F06 | 41080001 | 41180000 | 0.0029 | 0.0024 | 0.0021 |
| F06 | 41100001 | 41200000 | 0.0034 | 0.0024 | 0.0025 |
| F06 | 41120001 | 41220000 | 0.0036 | 0.0028 | 0.0027 |
| F06 | 41140001 | 41240000 | 0.0038 | 0.0035 | 0.0029 |
| F06 | 41160001 | 41260000 | 0.0045 | 0.0036 | 0.0037 |
| F06 | 41180001 | 41280000 | 0.0046 | 0.0032 | 0.0039 |
| F06 | 41200001 | 41300000 | 0.0043 | 0.0037 | 0.0035 |
| F06 | 41220001 | 41320000 | 0.0048 | 0.0036 | 0.0039 |
| F06 | 41240001 | 41340000 | 0.0047 | 0.0028 | 0.0039 |
| F06 | 41260001 | 41360000 | 0.0040 | 0.0027 | 0.0030 |
| F06 | 41280001 | 41380000 | 0.0038 | 0.0029 | 0.0028 |
| F06 | 41300001 | 41400000 | 0.0041 | 0.0024 | 0.0031 |
| F06 | 41320001 | 41420000 | 0.0035 | 0.0022 | 0.0026 |
| F06 | 41340001 | 41440000 | 0.0037 | 0.0040 | 0.0025 |
| F06 | 41360001 | 41460000 | 0.0053 | 0.0034 | 0.0043 |
| F06 | 41380001 | 41480000 | 0.0049 | 0.0040 | 0.0037 |
| F06 | 41400001 | 41500000 | 0.0056 | 0.0044 | 0.0043 |
| F06 | 41420001 | 41520000 | 0.0061 | 0.0048 | 0.0046 |
| F06 | 41440001 | 41540000 | 0.0063 | 0.0041 | 0.0050 |
| F06 | 41460001 | 41560000 | 0.0055 | 0.0045 | 0.0043 |
| F06 | 41480001 | 41580000 | 0.0057 | 0.0045 | 0.0048 |
| F06 | 41500001 | 41600000 | 0.0054 | 0.0056 | 0.0048 |

| Chr | Start | End | both | A1 | A2 |
|---|---|---|---|---|---|
| F06 | 41520001 | 41620000 | 0.0065 | 0.0051 | 0.0059 |
| F06 | 41540001 | 41640000 | 0.0057 | 0.0039 | 0.0053 |
| F06 | 41560001 | 41660000 | 0.0046 | 0.0053 | 0.0042 |
| F06 | 41580001 | 41680000 | 0.0058 | 0.0051 | 0.0053 |
| F06 | 41600001 | 41700000 | 0.0056 | 0.0034 | 0.0051 |
| F06 | 41620001 | 41720000 | 0.0038 | 0.0040 | 0.0034 |
| F06 | 41640001 | 41740000 | 0.0050 | 0.0040 | 0.0043 |
| F06 | 41660001 | 41760000 | 0.0052 | 0.0025 | 0.0044 |
| F06 | 41680001 | 41780000 | 0.0037 | 0.0024 | 0.0030 |
| F06 | 41700001 | 41800000 | 0.0037 | 0.0028 | 0.0030 |
| F06 | 41720001 | 41820000 | 0.0041 | 0.0020 | 0.0036 |
| F06 | 41740001 | 41840000 | 0.0027 | 0.0018 | 0.0024 |
| F06 | 41760001 | 41860000 | 0.0024 | 0.0016 | 0.0022 |
| F06 | 41780001 | 41880000 | 0.0022 | 0.0011 | 0.0021 |
| F06 | 41800001 | 41900000 | 0.0013 | 0.0008 | 0.0013 |
| F06 | 41820001 | 41920000 | 0.0009 | 0.0011 | 0.0009 |
| F06 | 41840001 | 41940000 | 0.0013 | 0.0012 | 0.0013 |
| F06 | 41860001 | 41960000 | 0.0014 | 0.0011 | 0.0014 |
| F06 | 41880001 | 41980000 | 0.0012 | 0.0017 | 0.0012 |
| F06 | 41900001 | 42000000 | 0.0019 | 0.0014 | 0.0019 |
| F06 | 41920001 | 42020000 | 0.0016 | 0.0013 | 0.0016 |
| F06 | 41940001 | 42040000 | 0.0014 | 0.0017 | 0.0014 |
| F06 | 41960001 | 42060000 | 0.0019 | 0.0017 | 0.0017 |
| F06 | 41980001 | 42080000 | 0.0020 | 0.0012 | 0.0018 |
| F06 | 42000001 | 42100000 | 0.0014 | 0.0021 | 0.0011 |
| F06 | 42020001 | 42120000 | 0.0027 | 0.0026 | 0.0023 |
| F06 | 42040001 | 42140000 | 0.0034 | 0.0042 | 0.0029 |
| F06 | 42060001 | 42160000 | 0.0055 | 0.0043 | 0.0051 |
| F06 | 42080001 | 42180000 | 0.0059 | 0.0044 | 0.0051 |
| F06 | 42100001 | 42200000 | 0.0063 | 0.0038 | 0.0054 |
| F06 | 42120001 | 42220000 | 0.0056 | 0.0041 | 0.0046 |
| F06 | 42140001 | 42240000 | 0.0059 | 0.0021 | 0.0049 |
| F06 | 42160001 | 42260000 | 0.0034 | 0.0023 | 0.0024 |
| F06 | 42180001 | 42280000 | 0.0032 | 0.0022 | 0.0027 |
| F06 | 42200001 | 42300000 | 0.0031 | 0.0023 | 0.0026 |
| F06 | 42220001 | 42320000 | 0.0030 | 0.0014 | 0.0028 |
| F06 | 42240001 | 42340000 | 0.0020 | 0.0013 | 0.0018 |
| F06 | 42260001 | 42360000 | 0.0019 | 0.0015 | 0.0017 |
| F06 | 42280001 | 42380000 | 0.0021 | 0.0013 | 0.0020 |
| F06 | 42300001 | 42400000 | 0.0017 | 0.0010 | 0.0017 |
| F06 | 42320001 | 42420000 | 0.0013 | 0.0010 | 0.0012 |
| F06 | 42340001 | 42440000 | 0.0011 | 0.0012 | 0.0011 |
| F06 | 42360001 | 42460000 | 0.0014 | 0.0006 | 0.0013 |
| F06 | 42380001 | 42480000 | 0.0006 | 0.0006 | 0.0006 |
| F06 | 42400001 | 42500000 | 0.0006 | 0.0006 | 0.0006 |
| F06 | 42420001 | 42520000 | 0.0007 | 0.0008 | 0.0006 |
| F06 | 42440001 | 42540000 | 0.0009 | 0.0007 | 0.0009 |
| F06 | 42460001 | 42560000 | 0.0008 | 0.0008 | 0.0008 |
| F06 | 42480001 | 42580000 | 0.0010 | 0.0012 | 0.0009 |
| F06 | 42500001 | 42600000 | 0.0015 | 0.0014 | 0.0015 |
| F06 | 42520001 | 42620000 | 0.0016 | 0.0011 | 0.0016 |
| F06 | 42540001 | 42640000 | 0.0013 | 0.0010 | 0.0013 |
| F06 | 42560001 | 42660000 | 0.0013 | 0.0016 | 0.0013 |
| F06 | 42580001 | 42680000 | 0.0018 | 0.0017 | 0.0017 |
| F06 | 42600001 | 42700000 | 0.0018 | 0.0024 | 0.0018 |
| F06 | 42620001 | 42720000 | 0.0029 | 0.0031 | 0.0027 |
| F06 | 42640001 | 42740000 | 0.0038 | 0.0034 | 0.0034 |
| F06 | 42660001 | 42760000 | 0.0040 | 0.0033 | 0.0036 |

| Chr | Start | End | both | A1 | A2 |
|---|---|---|---|---|---|
| F06 | 42680001 | 42780000 | 0.0040 | 0.0030 | 0.0036 |
| F06 | 42700001 | 42800000 | 0.0037 | 0.0030 | 0.0033 |
| F06 | 42720001 | 42820000 | 0.0035 | 0.0029 | 0.0032 |
| F06 | 42740001 | 42840000 | 0.0033 | 0.0025 | 0.0030 |
| F06 | 42760001 | 42860000 | 0.0031 | 0.0020 | 0.0027 |
| F06 | 42780001 | 42880000 | 0.0026 | 0.0019 | 0.0021 |
| F06 | 42800001 | 42900000 | 0.0026 | 0.0011 | 0.0020 |
| F06 | 42820001 | 42920000 | 0.0020 | 0.0005 | 0.0012 |
| F06 | 42840001 | 42940000 | 0.0013 | 0.0009 | 0.0007 |
| F06 | 42860001 | 42960000 | 0.0016 | 0.0012 | 0.0011 |
| F06 | 42880001 | 42980000 | 0.0018 | 0.0012 | 0.0014 |
| F06 | 42900001 | 43000000 | 0.0017 | 0.0015 | 0.0014 |
| F06 | 42920001 | 43020000 | 0.0018 | 0.0026 | 0.0017 |
| F06 | 42940001 | 43040000 | 0.0029 | 0.0026 | 0.0028 |
| F06 | 42960001 | 43060000 | 0.0029 | 0.0026 | 0.0027 |
| F06 | 42980001 | 43080000 | 0.0029 | 0.0031 | 0.0026 |
| F06 | 43000001 | 43100000 | 0.0034 | 0.0028 | 0.0031 |
| F06 | 43020001 | 43120000 | 0.0031 | 0.0019 | 0.0028 |
| F06 | 43040001 | 43140000 | 0.0022 | 0.0019 | 0.0019 |
| F06 | 43060001 | 43160000 | 0.0022 | 0.0023 | 0.0020 |
| F06 | 43080001 | 43180000 | 0.0026 | 0.0022 | 0.0023 |
| F06 | 43100001 | 43200000 | 0.0026 | 0.0029 | 0.0023 |
| F06 | 43120001 | 43220000 | 0.0031 | 0.0027 | 0.0028 |
| F06 | 43140001 | 43240000 | 0.0028 | 0.0026 | 0.0025 |
| F06 | 43160001 | 43260000 | 0.0029 | 0.0025 | 0.0025 |
| F06 | 43180001 | 43280000 | 0.0029 | 0.0021 | 0.0025 |
| F06 | 43200001 | 43300000 | 0.0024 | 0.0015 | 0.0020 |
| F06 | 43220001 | 43320000 | 0.0021 | 0.0018 | 0.0017 |
| F06 | 43240001 | 43340000 | 0.0025 | 0.0015 | 0.0021 |
| F06 | 43260001 | 43360000 | 0.0020 | 0.0016 | 0.0018 |
| F06 | 43280001 | 43380000 | 0.0022 | 0.0020 | 0.0019 |
| F06 | 43300001 | 43400000 | 0.0024 | 0.0018 | 0.0021 |
| F06 | 43320001 | 43420000 | 0.0023 | 0.0020 | 0.0020 |
| F06 | 43340001 | 43440000 | 0.0024 | 0.0019 | 0.0021 |
| F06 | 43360001 | 43460000 | 0.0023 | 0.0015 | 0.0020 |
| F06 | 43380001 | 43480000 | 0.0017 | 0.0014 | 0.0016 |
| F06 | 43400001 | 43500000 | 0.0016 | 0.0016 | 0.0016 |
| F06 | 43420001 | 43520000 | 0.0017 | 0.0019 | 0.0016 |
| F06 | 43440001 | 43540000 | 0.0021 | 0.0020 | 0.0020 |
| F06 | 43460001 | 43560000 | 0.0022 | 0.0020 | 0.0022 |
| F06 | 43480001 | 43580000 | 0.0023 | 0.0021 | 0.0021 |
| F06 | 43500001 | 43600000 | 0.0024 | 0.0018 | 0.0022 |
| F06 | 43520001 | 43620000 | 0.0021 | 0.0011 | 0.0020 |
| F06 | 43540001 | 43640000 | 0.0014 | 0.0012 | 0.0013 |
| F06 | 43560001 | 43660000 | 0.0014 | 0.0011 | 0.0013 |
| F06 | 43580001 | 43680000 | 0.0014 | 0.0009 | 0.0014 |
| F06 | 43600001 | 43700000 | 0.0012 | 0.0013 | 0.0012 |
| F06 | 43620001 | 43720000 | 0.0017 | 0.0023 | 0.0017 |
| F06 | 43640001 | 43740000 | 0.0027 | 0.0025 | 0.0026 |
| F06 | 43660001 | 43760000 | 0.0031 | 0.0022 | 0.0029 |
| F06 | 43680001 | 43780000 | 0.0028 | 0.0024 | 0.0025 |
| F06 | 43700001 | 43800000 | 0.0030 | 0.0027 | 0.0026 |
| F06 | 43720001 | 43820000 | 0.0036 | 0.0023 | 0.0030 |
| F06 | 43740001 | 43840000 | 0.0033 | 0.0023 | 0.0026 |
| F06 | 43760001 | 43860000 | 0.0035 | 0.0031 | 0.0026 |
| F06 | 43780001 | 43880000 | 0.0045 | 0.0034 | 0.0036 |
| F06 | 43800001 | 43900000 | 0.0051 | 0.0026 | 0.0041 |
| F06 | 43820001 | 43920000 | 0.0039 | 0.0021 | 0.0032 |

| Chr | Start | End | both | A1 | A2 |
|---|---|---|---|---|---|
| F06 | 43840001 | 43940000 | 0.0032 | 0.0019 | 0.0026 |
| F06 | 43860001 | 43960000 | 0.0027 | 0.0014 | 0.0023 |
| F06 | 43880001 | 43980000 | 0.0018 | 0.0008 | 0.0017 |
| F06 | 43900001 | 44000000 | 0.0010 | 0.0009 | 0.0010 |
| F06 | 43920001 | 44020000 | 0.0012 | 0.0012 | 0.0012 |
| F06 | 43940001 | 44040000 | 0.0016 | 0.0013 | 0.0016 |
| F06 | 43960001 | 44060000 | 0.0018 | 0.0020 | 0.0016 |
| F06 | 43980001 | 44080000 | 0.0025 | 0.0022 | 0.0022 |
| F06 | 44000001 | 44100000 | 0.0027 | 0.0023 | 0.0024 |
| F06 | 44020001 | 44120000 | 0.0029 | 0.0021 | 0.0024 |
| F06 | 44040001 | 44140000 | 0.0027 | 0.0021 | 0.0022 |
| F06 | 44060001 | 44160000 | 0.0025 | 0.0017 | 0.0021 |
| F06 | 44080001 | 44180000 | 0.0021 | 0.0017 | 0.0017 |
| F06 | 44100001 | 44200000 | 0.0022 | 0.0018 | 0.0016 |
| F06 | 44120001 | 44220000 | 0.0022 | 0.0019 | 0.0018 |
| F06 | 44140001 | 44240000 | 0.0022 | 0.0020 | 0.0018 |
| F06 | 44160001 | 44260000 | 0.0024 | 0.0025 | 0.0021 |
| F06 | 44180001 | 44280000 | 0.0029 | 0.0024 | 0.0026 |
| F06 | 44200001 | 44300000 | 0.0028 | 0.0022 | 0.0026 |
| F06 | 44220001 | 44320000 | 0.0026 | 0.0029 | 0.0023 |
| F06 | 44240001 | 44340000 | 0.0033 | 0.0042 | 0.0030 |
| F06 | 44260001 | 44360000 | 0.0050 | 0.0041 | 0.0045 |
| F06 | 44280001 | 44380000 | 0.0049 | 0.0050 | 0.0045 |
| F06 | 44300001 | 44400000 | 0.0061 | 0.0050 | 0.0054 |
| F06 | 44320001 | 44420000 | 0.0064 | 0.0049 | 0.0055 |
| F06 | 44340001 | 44440000 | 0.0063 | 0.0040 | 0.0051 |
| F06 | 44360001 | 44460000 | 0.0052 | 0.0032 | 0.0043 |
| F06 | 44380001 | 44480000 | 0.0045 | 0.0030 | 0.0034 |
| F06 | 44400001 | 44500000 | 0.0041 | 0.0033 | 0.0031 |
| F06 | 44420001 | 44520000 | 0.0042 | 0.0030 | 0.0035 |
| F06 | 44440001 | 44540000 | 0.0040 | 0.0028 | 0.0034 |
| F06 | 44460001 | 44560000 | 0.0037 | 0.0041 | 0.0032 |
| F06 | 44480001 | 44580000 | 0.0048 | 0.0040 | 0.0044 |
| F06 | 44500001 | 44600000 | 0.0045 | 0.0044 | 0.0044 |
| F06 | 44520001 | 44620000 | 0.0049 | 0.0045 | 0.0047 |
| F06 | 44540001 | 44640000 | 0.0050 | 0.0039 | 0.0049 |
| F06 | 44560001 | 44660000 | 0.0044 | 0.0031 | 0.0043 |
| F06 | 44580001 | 44680000 | 0.0036 | 0.0039 | 0.0035 |
| F06 | 44600001 | 44700000 | 0.0046 | 0.0031 | 0.0042 |
| F06 | 44620001 | 44720000 | 0.0037 | 0.0034 | 0.0033 |
| F06 | 44640001 | 44740000 | 0.0040 | 0.0051 | 0.0035 |
| F06 | 44660001 | 44760000 | 0.0061 | 0.0057 | 0.0054 |
| F06 | 44680001 | 44780000 | 0.0071 | 0.0047 | 0.0063 |
| F06 | 44700001 | 44800000 | 0.0059 | 0.0054 | 0.0053 |
| F06 | 44720001 | 44820000 | 0.0067 | 0.0049 | 0.0059 |
| F06 | 44740001 | 44840000 | 0.0061 | 0.0033 | 0.0056 |
| F06 | 44760001 | 44860000 | 0.0041 | 0.0026 | 0.0037 |
| F06 | 44780001 | 44880000 | 0.0031 | 0.0024 | 0.0027 |
| F06 | 44800001 | 44900000 | 0.0030 | 0.0021 | 0.0025 |
| F06 | 44820001 | 44920000 | 0.0026 | 0.0017 | 0.0022 |
| F06 | 44840001 | 44940000 | 0.0022 | 0.0017 | 0.0018 |
| F06 | 44860001 | 44960000 | 0.0022 | 0.0012 | 0.0018 |
| F06 | 44880001 | 44980000 | 0.0015 | 0.0011 | 0.0014 |
| F06 | 44900001 | 45000000 | 0.0012 | 0.0010 | 0.0012 |
| F06 | 44920001 | 45020000 | 0.0012 | 0.0011 | 0.0011 |
| F06 | 44940001 | 45040000 | 0.0012 | 0.0013 | 0.0012 |
| F06 | 44960001 | 45060000 | 0.0014 | 0.0014 | 0.0014 |
| F06 | 44980001 | 45080000 | 0.0015 | 0.0013 | 0.0015 |

| Chr | Start | End | both | A1 | A2 |
|---|---|---|---|---|---|
| F06 | 45000001 | 45100000 | 0.0015 | 0.0010 | 0.0014 |
| F06 | 45020001 | 45120000 | 0.0011 | 0.0008 | 0.0011 |
| F06 | 45040001 | 45140000 | 0.0009 | 0.0011 | 0.0009 |
| F06 | 45060001 | 45160000 | 0.0012 | 0.0012 | 0.0012 |
| F06 | 45080001 | 45180000 | 0.0014 | 0.0013 | 0.0013 |
| F06 | 45100001 | 45200000 | 0.0015 | 0.0015 | 0.0014 |
| F06 | 45120001 | 45220000 | 0.0016 | 0.0015 | 0.0016 |
| F06 | 45140001 | 45240000 | 0.0017 | 0.0010 | 0.0016 |
| F06 | 45160001 | 45260000 | 0.0011 | 0.0009 | 0.0011 |
| F06 | 45180001 | 45280000 | 0.0010 | 0.0007 | 0.0010 |
| F06 | 45200001 | 45300000 | 0.0008 | 0.0008 | 0.0007 |
| F06 | 45220001 | 45320000 | 0.0009 | 0.0006 | 0.0008 |
| F06 | 45240001 | 45340000 | 0.0007 | 0.0006 | 0.0007 |
| F06 | 45260001 | 45360000 | 0.0006 | 0.0004 | 0.0006 |
| F06 | 45280001 | 45380000 | 0.0005 | 0.0004 | 0.0004 |
| F06 | 45300001 | 45400000 | 0.0005 | 0.0004 | 0.0005 |
| F06 | 45320001 | 45420000 | 0.0005 | 0.0006 | 0.0005 |
| F06 | 45340001 | 45440000 | 0.0007 | 0.0015 | 0.0007 |
| F06 | 45360001 | 45460000 | 0.0016 | 0.0014 | 0.0015 |
| F06 | 45380001 | 45480000 | 0.0015 | 0.0015 | 0.0014 |
| F06 | 45400001 | 45500000 | 0.0016 | 0.0014 | 0.0015 |
| F06 | 45420001 | 45520000 | 0.0015 | 0.0014 | 0.0014 |
| F06 | 45440001 | 45540000 | 0.0015 | 0.0008 | 0.0014 |
| F06 | 45460001 | 45560000 | 0.0009 | 0.0011 | 0.0009 |
| F06 | 45480001 | 45580000 | 0.0012 | 0.0010 | 0.0012 |
| F06 | 45500001 | 45600000 | 0.0012 | 0.0008 | 0.0012 |
| F06 | 45520001 | 45620000 | 0.0010 | 0.0008 | 0.0010 |
| F06 | 45540001 | 45640000 | 0.0009 | 0.0006 | 0.0009 |
| F06 | 45560001 | 45660000 | 0.0008 | 0.0009 | 0.0008 |
| F06 | 45580001 | 45680000 | 0.0011 | 0.0013 | 0.0010 |
| F06 | 45600001 | 45700000 | 0.0016 | 0.0019 | 0.0016 |
| F06 | 45620001 | 45720000 | 0.0022 | 0.0021 | 0.0022 |
| F06 | 45640001 | 45740000 | 0.0025 | 0.0026 | 0.0025 |
| F06 | 45660001 | 45760000 | 0.0030 | 0.0026 | 0.0029 |
| F06 | 45680001 | 45780000 | 0.0030 | 0.0022 | 0.0030 |
| F06 | 45700001 | 45800000 | 0.0025 | 0.0020 | 0.0024 |
| F06 | 45720001 | 45820000 | 0.0024 | 0.0022 | 0.0023 |
| F06 | 45740001 | 45840000 | 0.0025 | 0.0020 | 0.0024 |
| F06 | 45760001 | 45860000 | 0.0024 | 0.0018 | 0.0023 |
| F06 | 45780001 | 45880000 | 0.0021 | 0.0019 | 0.0020 |
| F06 | 45800001 | 45900000 | 0.0022 | 0.0019 | 0.0021 |
| F06 | 45820001 | 45920000 | 0.0021 | 0.0022 | 0.0020 |
| F06 | 45840001 | 45940000 | 0.0024 | 0.0027 | 0.0023 |
| F06 | 45860001 | 45960000 | 0.0029 | 0.0043 | 0.0028 |
| F06 | 45880001 | 45980000 | 0.0049 | 0.0046 | 0.0048 |
| F06 | 45900001 | 46000000 | 0.0053 | 0.0048 | 0.0052 |
| F06 | 45920001 | 46020000 | 0.0055 | 0.0045 | 0.0054 |
| F06 | 45940001 | 46040000 | 0.0053 | 0.0039 | 0.0052 |
| F06 | 45960001 | 46060000 | 0.0047 | 0.0030 | 0.0045 |
| F06 | 45980001 | 46080000 | 0.0035 | 0.0030 | 0.0032 |
| F06 | 46000001 | 46100000 | 0.0037 | 0.0027 | 0.0031 |
| F06 | 46020001 | 46120000 | 0.0035 | 0.0027 | 0.0029 |
| F06 | 46040001 | 46140000 | 0.0034 | 0.0031 | 0.0029 |
| F06 | 46060001 | 46160000 | 0.0039 | 0.0030 | 0.0034 |
| F06 | 46080001 | 46180000 | 0.0038 | 0.0030 | 0.0034 |
| F06 | 46100001 | 46200000 | 0.0036 | 0.0030 | 0.0034 |
| F06 | 46120001 | 46220000 | 0.0036 | 0.0030 | 0.0034 |
| F06 | 46140001 | 46240000 | 0.0034 | 0.0024 | 0.0032 |

| Chr | Start | End | both | A1 | A2 |
|---|---|---|---|---|---|
| F06 | 46160001 | 46260000 | 0.0028 | 0.0020 | 0.0027 |
| F06 | 46180001 | 46280000 | 0.0024 | 0.0021 | 0.0022 |
| F06 | 46200001 | 46300000 | 0.0024 | 0.0028 | 0.0023 |
| F06 | 46220001 | 46320000 | 0.0034 | 0.0031 | 0.0031 |
| F06 | 46240001 | 46340000 | 0.0040 | 0.0032 | 0.0037 |
| F06 | 46260001 | 46360000 | 0.0041 | 0.0036 | 0.0038 |
| F06 | 46280001 | 46380000 | 0.0048 | 0.0035 | 0.0044 |
| F06 | 46300001 | 46400000 | 0.0047 | 0.0027 | 0.0044 |
| F06 | 46320001 | 46420000 | 0.0037 | 0.0046 | 0.0036 |
| F06 | 46340001 | 46440000 | 0.0052 | 0.0048 | 0.0047 |
| F06 | 46360001 | 46460000 | 0.0052 | 0.0042 | 0.0047 |
| F06 | 46380001 | 46480000 | 0.0044 | 0.0041 | 0.0038 |
| F06 | 46400001 | 46500000 | 0.0042 | 0.0043 | 0.0036 |
| F06 | 46420001 | 46520000 | 0.0043 | 0.0019 | 0.0038 |
| F06 | 46440001 | 46540000 | 0.0021 | 0.0015 | 0.0019 |
| F06 | 46460001 | 46560000 | 0.0018 | 0.0012 | 0.0016 |
| F06 | 46480001 | 46580000 | 0.0014 | 0.0010 | 0.0014 |
| F06 | 46500001 | 46600000 | 0.0011 | 0.0005 | 0.0011 |
| F06 | 46520001 | 46620000 | 0.0006 | 0.0005 | 0.0006 |
| F06 | 46540001 | 46640000 | 0.0005 | 0.0006 | 0.0005 |
| F06 | 46560001 | 46660000 | 0.0006 | 0.0008 | 0.0006 |
| F06 | 46580001 | 46680000 | 0.0008 | 0.0012 | 0.0008 |
| F06 | 46600001 | 46700000 | 0.0014 | 0.0012 | 0.0013 |
| F06 | 46620001 | 46720000 | 0.0014 | 0.0014 | 0.0014 |
| F06 | 46640001 | 46740000 | 0.0016 | 0.0012 | 0.0016 |
| F06 | 46660001 | 46760000 | 0.0015 | 0.0010 | 0.0015 |
| F06 | 46680001 | 46780000 | 0.0013 | 0.0017 | 0.0013 |
| F06 | 46700001 | 46800000 | 0.0023 | 0.0026 | 0.0023 |
| F06 | 46720001 | 46820000 | 0.0034 | 0.0028 | 0.0034 |
| F06 | 46740001 | 46840000 | 0.0036 | 0.0029 | 0.0034 |
| F06 | 46760001 | 46860000 | 0.0038 | 0.0031 | 0.0035 |
| F06 | 46780001 | 46880000 | 0.0043 | 0.0026 | 0.0038 |
| F06 | 46800001 | 46900000 | 0.0034 | 0.0019 | 0.0028 |
| F06 | 46820001 | 46920000 | 0.0026 | 0.0016 | 0.0020 |
| F06 | 46840001 | 46940000 | 0.0024 | 0.0020 | 0.0018 |
| F06 | 46860001 | 46960000 | 0.0028 | 0.0019 | 0.0023 |
| F06 | 46880001 | 46980000 | 0.0024 | 0.0018 | 0.0021 |
| F06 | 46900001 | 47000000 | 0.0023 | 0.0019 | 0.0021 |
| F06 | 46920001 | 47020000 | 0.0023 | 0.0020 | 0.0022 |
| F06 | 46940001 | 47040000 | 0.0025 | 0.0020 | 0.0023 |
| F06 | 46960001 | 47060000 | 0.0026 | 0.0022 | 0.0024 |
| F06 | 46980001 | 47080000 | 0.0028 | 0.0016 | 0.0026 |
| F06 | 47000001 | 47100000 | 0.0022 | 0.0015 | 0.0020 |
| F06 | 47020001 | 47120000 | 0.0020 | 0.0016 | 0.0019 |
| F06 | 47040001 | 47140000 | 0.0019 | 0.0011 | 0.0018 |
| F06 | 47060001 | 47160000 | 0.0013 | 0.0010 | 0.0012 |
| F06 | 47080001 | 47180000 | 0.0012 | 0.0015 | 0.0012 |
| F06 | 47100001 | 47200000 | 0.0019 | 0.0018 | 0.0019 |
| F06 | 47120001 | 47220000 | 0.0022 | 0.0017 | 0.0022 |
| F06 | 47140001 | 47240000 | 0.0022 | 0.0020 | 0.0021 |
| F06 | 47160001 | 47260000 | 0.0025 | 0.0019 | 0.0023 |
| F06 | 47180001 | 47280000 | 0.0023 | 0.0017 | 0.0021 |
| F06 | 47200001 | 47300000 | 0.0020 | 0.0016 | 0.0018 |
| F06 | 47220001 | 47320000 | 0.0019 | 0.0020 | 0.0017 |
| F06 | 47240001 | 47340000 | 0.0024 | 0.0020 | 0.0022 |
| F06 | 47260001 | 47360000 | 0.0026 | 0.0019 | 0.0023 |
| F06 | 47280001 | 47380000 | 0.0026 | 0.0016 | 0.0022 |
| F06 | 47300001 | 47400000 | 0.0022 | 0.0012 | 0.0019 |

| Chr | Start | End | both | A1 | A2 |
|---|---|---|---|---|---|
| F06 | 47320001 | 47420000 | 0.0019 | 0.0013 | 0.0015 |
| F06 | 47340001 | 47440000 | 0.0019 | 0.0018 | 0.0015 |
| F06 | 47360001 | 47460000 | 0.0024 | 0.0022 | 0.0020 |
| F06 | 47380001 | 47480000 | 0.0029 | 0.0029 | 0.0025 |
| F06 | 47400001 | 47500000 | 0.0040 | 0.0031 | 0.0034 |
| F06 | 47420001 | 47520000 | 0.0041 | 0.0025 | 0.0035 |
| F06 | 47440001 | 47540000 | 0.0034 | 0.0022 | 0.0029 |
| F06 | 47460001 | 47560000 | 0.0031 | 0.0027 | 0.0026 |
| F06 | 47480001 | 47580000 | 0.0038 | 0.0024 | 0.0031 |
| F06 | 47500001 | 47600000 | 0.0033 | 0.0038 | 0.0026 |
| F06 | 47520001 | 47620000 | 0.0048 | 0.0042 | 0.0040 |
| F06 | 47540001 | 47640000 | 0.0055 | 0.0044 | 0.0043 |
| F06 | 47560001 | 47660000 | 0.0056 | 0.0048 | 0.0045 |
| F06 | 47580001 | 47680000 | 0.0060 | 0.0048 | 0.0048 |
| F06 | 47600001 | 47700000 | 0.0060 | 0.0036 | 0.0050 |
| F06 | 47620001 | 47720000 | 0.0045 | 0.0045 | 0.0036 |
| F06 | 47640001 | 47740000 | 0.0053 | 0.0045 | 0.0047 |
| F06 | 47660001 | 47760000 | 0.0052 | 0.0034 | 0.0046 |
| F06 | 47680001 | 47780000 | 0.0043 | 0.0036 | 0.0035 |
| F06 | 47700001 | 47800000 | 0.0045 | 0.0044 | 0.0038 |
| F06 | 47720001 | 47820000 | 0.0056 | 0.0033 | 0.0048 |
| F06 | 47740001 | 47840000 | 0.0047 | 0.0026 | 0.0037 |
| F06 | 47760001 | 47860000 | 0.0041 | 0.0025 | 0.0030 |
| F06 | 47780001 | 47880000 | 0.0036 | 0.0025 | 0.0027 |
| F06 | 47800001 | 47900000 | 0.0035 | 0.0018 | 0.0026 |
| F06 | 47820001 | 47920000 | 0.0028 | 0.0026 | 0.0019 |
| F06 | 47840001 | 47940000 | 0.0034 | 0.0035 | 0.0026 |
| F06 | 47860001 | 47960000 | 0.0041 | 0.0042 | 0.0036 |
| F06 | 47880001 | 47980000 | 0.0049 | 0.0044 | 0.0043 |
| F06 | 47900001 | 48000000 | 0.0049 | 0.0046 | 0.0043 |
| F06 | 47920001 | 48020000 | 0.0050 | 0.0038 | 0.0044 |
| F06 | 47940001 | 48040000 | 0.0043 | 0.0040 | 0.0038 |
| F06 | 47960001 | 48060000 | 0.0046 | 0.0047 | 0.0041 |
| F06 | 47980001 | 48080000 | 0.0055 | 0.0046 | 0.0051 |
| F06 | 48000001 | 48100000 | 0.0056 | 0.0047 | 0.0050 |
| F06 | 48020001 | 48120000 | 0.0059 | 0.0047 | 0.0052 |
| F06 | 48040001 | 48140000 | 0.0063 | 0.0039 | 0.0053 |
| F06 | 48060001 | 48160000 | 0.0055 | 0.0030 | 0.0044 |
| F06 | 48080001 | 48180000 | 0.0045 | 0.0031 | 0.0034 |
| F06 | 48100001 | 48200000 | 0.0042 | 0.0032 | 0.0036 |
| F06 | 48120001 | 48220000 | 0.0042 | 0.0031 | 0.0037 |
| F06 | 48140001 | 48240000 | 0.0038 | 0.0043 | 0.0034 |
| F06 | 48160001 | 48260000 | 0.0048 | 0.0044 | 0.0046 |
| F06 | 48180001 | 48280000 | 0.0051 | 0.0043 | 0.0046 |
| F06 | 48200001 | 48300000 | 0.0049 | 0.0038 | 0.0044 |
| F06 | 48220001 | 48320000 | 0.0045 | 0.0046 | 0.0039 |
| F06 | 48240001 | 48340000 | 0.0052 | 0.0035 | 0.0046 |
| F06 | 48260001 | 48360000 | 0.0042 | 0.0037 | 0.0036 |
| F06 | 48280001 | 48380000 | 0.0042 | 0.0035 | 0.0038 |
| F06 | 48300001 | 48400000 | 0.0040 | 0.0038 | 0.0038 |
| F06 | 48320001 | 48420000 | 0.0042 | 0.0029 | 0.0040 |
| F06 | 48340001 | 48440000 | 0.0034 | 0.0031 | 0.0032 |
| F06 | 48360001 | 48460000 | 0.0036 | 0.0027 | 0.0033 |
| F06 | 48380001 | 48480000 | 0.0033 | 0.0041 | 0.0029 |
| F06 | 48400001 | 48500000 | 0.0048 | 0.0045 | 0.0042 |
| F06 | 48420001 | 48520000 | 0.0056 | 0.0048 | 0.0047 |
| F06 | 48440001 | 48540000 | 0.0059 | 0.0046 | 0.0051 |
| F06 | 48460001 | 48560000 | 0.0057 | 0.0042 | 0.0048 |

| Chr | Start | End | both | A1 | A2 |
|---|---|---|---|---|---|
| F06 | 48480001 | 48580000 | 0.0051 | 0.0032 | 0.0044 |
| F06 | 48500001 | 48600000 | 0.0042 | 0.0027 | 0.0037 |
| F06 | 48520001 | 48620000 | 0.0034 | 0.0026 | 0.0031 |
| F06 | 48540001 | 48640000 | 0.0032 | 0.0024 | 0.0030 |
| F06 | 48560001 | 48660000 | 0.0031 | 0.0023 | 0.0029 |
| F06 | 48580001 | 48680000 | 0.0030 | 0.0029 | 0.0027 |
| F06 | 48600001 | 48700000 | 0.0036 | 0.0037 | 0.0033 |
| F06 | 48620001 | 48720000 | 0.0045 | 0.0038 | 0.0041 |
| F06 | 48640001 | 48740000 | 0.0046 | 0.0044 | 0.0041 |
| F06 | 48660001 | 48760000 | 0.0050 | 0.0047 | 0.0046 |
| F06 | 48680001 | 48780000 | 0.0051 | 0.0043 | 0.0048 |
| F06 | 48700001 | 48800000 | 0.0047 | 0.0039 | 0.0044 |
| F06 | 48720001 | 48820000 | 0.0042 | 0.0036 | 0.0040 |
| F06 | 48740001 | 48840000 | 0.0041 | 0.0036 | 0.0038 |
| F06 | 48760001 | 48860000 | 0.0043 | 0.0035 | 0.0040 |
| F06 | 48780001 | 48880000 | 0.0043 | 0.0028 | 0.0039 |
| F06 | 48800001 | 48900000 | 0.0033 | 0.0018 | 0.0030 |
| F06 | 48820001 | 48920000 | 0.0023 | 0.0018 | 0.0020 |
| F06 | 48840001 | 48940000 | 0.0022 | 0.0025 | 0.0019 |
| F06 | 48860001 | 48960000 | 0.0029 | 0.0026 | 0.0026 |
| F06 | 48880001 | 48980000 | 0.0032 | 0.0028 | 0.0028 |
| F06 | 48900001 | 49000000 | 0.0034 | 0.0029 | 0.0029 |
| F06 | 48920001 | 49020000 | 0.0037 | 0.0042 | 0.0032 |
| F06 | 48940001 | 49040000 | 0.0051 | 0.0033 | 0.0047 |
| F06 | 48960001 | 49060000 | 0.0040 | 0.0037 | 0.0037 |
| F06 | 48980001 | 49080000 | 0.0043 | 0.0041 | 0.0041 |
| F06 | 49000001 | 49100000 | 0.0047 | 0.0046 | 0.0046 |
| F06 | 49020001 | 49120000 | 0.0050 | 0.0036 | 0.0048 |
| F06 | 49040001 | 49140000 | 0.0039 | 0.0032 | 0.0037 |
| F06 | 49060001 | 49160000 | 0.0035 | 0.0027 | 0.0034 |
| F06 | 49080001 | 49180000 | 0.0030 | 0.0031 | 0.0030 |
| F06 | 49100001 | 49200000 | 0.0035 | 0.0025 | 0.0034 |
| F06 | 49120001 | 49220000 | 0.0030 | 0.0022 | 0.0030 |
| F06 | 49140001 | 49240000 | 0.0027 | 0.0021 | 0.0027 |
| F06 | 49160001 | 49260000 | 0.0026 | 0.0022 | 0.0026 |
| F06 | 49180001 | 49280000 | 0.0028 | 0.0012 | 0.0027 |
| F06 | 49200001 | 49300000 | 0.0017 | 0.0025 | 0.0016 |
| F06 | 49220001 | 49320000 | 0.0033 | 0.0023 | 0.0031 |
| F06 | 49240001 | 49340000 | 0.0030 | 0.0023 | 0.0028 |
| F06 | 49260001 | 49360000 | 0.0031 | 0.0019 | 0.0029 |
| F06 | 49280001 | 49380000 | 0.0024 | 0.0026 | 0.0023 |
| F06 | 49300001 | 49400000 | 0.0029 | 0.0015 | 0.0028 |
| F06 | 49320001 | 49420000 | 0.0016 | 0.0016 | 0.0015 |
| F06 | 49340001 | 49440000 | 0.0018 | 0.0016 | 0.0017 |
| F06 | 49360001 | 49460000 | 0.0019 | 0.0016 | 0.0018 |
| F06 | 49380001 | 49480000 | 0.0021 | 0.0012 | 0.0020 |
| F06 | 49400001 | 49500000 | 0.0017 | 0.0014 | 0.0017 |
| F06 | 49420001 | 49520000 | 0.0019 | 0.0013 | 0.0019 |
| F06 | 49440001 | 49540000 | 0.0019 | 0.0013 | 0.0019 |
| F06 | 49460001 | 49560000 | 0.0016 | 0.0016 | 0.0016 |
| F06 | 49480001 | 49580000 | 0.0019 | 0.0023 | 0.0019 |
| F06 | 49500001 | 49600000 | 0.0030 | 0.0025 | 0.0029 |
| F06 | 49520001 | 49620000 | 0.0031 | 0.0025 | 0.0030 |
| F06 | 49540001 | 49640000 | 0.0031 | 0.0045 | 0.0030 |
| F06 | 49560001 | 49660000 | 0.0051 | 0.0042 | 0.0050 |
| F06 | 49580001 | 49680000 | 0.0048 | 0.0031 | 0.0047 |
| F06 | 49600001 | 49700000 | 0.0033 | 0.0026 | 0.0033 |
| F06 | 49620001 | 49720000 | 0.0029 | 0.0030 | 0.0028 |

| Chr | Start | End | both | A1 | A2 |
|---|---|---|---|---|---|
| F06 | 49640001 | 49740000 | 0.0033 | 0.0011 | 0.0033 |
| F06 | 49660001 | 49760000 | 0.0012 | 0.0014 | 0.0012 |
| F06 | 49680001 | 49780000 | 0.0016 | 0.0013 | 0.0015 |
| F06 | 49700001 | 49800000 | 0.0015 | 0.0012 | 0.0015 |
| F06 | 49720001 | 49820000 | 0.0014 | 0.0006 | 0.0013 |
| F06 | 49740001 | 49840000 | 0.0008 | 0.0008 | 0.0008 |
| F06 | 49760001 | 49860000 | 0.0010 | 0.0006 | 0.0010 |
| F06 | 49780001 | 49880000 | 0.0008 | 0.0007 | 0.0008 |
| F06 | 49800001 | 49900000 | 0.0010 | 0.0007 | 0.0010 |
| F06 | 49820001 | 49920000 | 0.0010 | 0.0009 | 0.0010 |
| F06 | 49840001 | 49940000 | 0.0012 | 0.0007 | 0.0012 |
| F06 | 49860001 | 49960000 | 0.0009 | 0.0005 | 0.0009 |
| F06 | 49880001 | 49980000 | 0.0006 | 0.0002 | 0.0006 |
| F06 | 49900001 | 50000000 | 0.0004 | 0.0002 | 0.0004 |
| F06 | 49920001 | 50020000 | 0.0003 | 0.0001 | 0.0004 |
| F06 | 49940001 | 50040000 | 0.0002 | 0.0002 | 0.0002 |
| F06 | 49960001 | 50060000 | 0.0002 | 0.0004 | 0.0002 |
| F06 | 49980001 | 50080000 | 0.0006 | 0.0008 | 0.0006 |
| F06 | 50000001 | 50100000 | 0.0010 | 0.0010 | 0.0010 |
| F06 | 50020001 | 50120000 | 0.0013 | 0.0010 | 0.0013 |
| F06 | 50040001 | 50140000 | 0.0013 | 0.0010 | 0.0013 |
| F06 | 50060001 | 50160000 | 0.0012 | 0.0010 | 0.0012 |
| F06 | 50080001 | 50180000 | 0.0011 | 0.0006 | 0.0011 |
| F06 | 50100001 | 50200000 | 0.0007 | 0.0014 | 0.0007 |
| F06 | 50120001 | 50220000 | 0.0018 | 0.0015 | 0.0018 |
| F06 | 50140001 | 50240000 | 0.0019 | 0.0015 | 0.0019 |
| F06 | 50160001 | 50260000 | 0.0019 | 0.0013 | 0.0019 |
| F06 | 50180001 | 50280000 | 0.0017 | 0.0014 | 0.0017 |
| F06 | 50200001 | 50300000 | 0.0019 | 0.0010 | 0.0018 |
| F06 | 50220001 | 50320000 | 0.0013 | 0.0010 | 0.0013 |
| F06 | 50240001 | 50340000 | 0.0012 | 0.0015 | 0.0012 |
| F06 | 50260001 | 50360000 | 0.0018 | 0.0017 | 0.0017 |
| F06 | 50280001 | 50380000 | 0.0020 | 0.0018 | 0.0020 |
| F06 | 50300001 | 50400000 | 0.0022 | 0.0016 | 0.0021 |
| F06 | 50320001 | 50420000 | 0.0020 | 0.0019 | 0.0020 |
| F06 | 50340001 | 50440000 | 0.0024 | 0.0022 | 0.0024 |
| F06 | 50360001 | 50460000 | 0.0026 | 0.0022 | 0.0025 |
| F06 | 50380001 | 50480000 | 0.0026 | 0.0021 | 0.0025 |
| F06 | 50400001 | 50500000 | 0.0024 | 0.0020 | 0.0023 |
| F06 | 50420001 | 50520000 | 0.0022 | 0.0015 | 0.0021 |
| F06 | 50440001 | 50540000 | 0.0016 | 0.0007 | 0.0015 |
| F06 | 50460001 | 50560000 | 0.0009 | 0.0009 | 0.0009 |
| F06 | 50480001 | 50580000 | 0.0011 | 0.0009 | 0.0011 |
| F06 | 50500001 | 50600000 | 0.0012 | 0.0007 | 0.0012 |
| F06 | 50520001 | 50620000 | 0.0009 | 0.0014 | 0.0010 |
| F06 | 50540001 | 50640000 | 0.0017 | 0.0020 | 0.0017 |
| F06 | 50560001 | 50660000 | 0.0025 | 0.0016 | 0.0025 |
| F06 | 50580001 | 50680000 | 0.0020 | 0.0020 | 0.0020 |
| F06 | 50600001 | 50700000 | 0.0023 | 0.0023 | 0.0023 |
| F06 | 50620001 | 50720000 | 0.0027 | 0.0024 | 0.0026 |
| F06 | 50640001 | 50740000 | 0.0028 | 0.0023 | 0.0028 |
| F06 | 50660001 | 50760000 | 0.0025 | 0.0029 | 0.0025 |
| F06 | 50680001 | 50780000 | 0.0033 | 0.0028 | 0.0032 |
| F06 | 50700001 | 50800000 | 0.0033 | 0.0035 | 0.0033 |
| F06 | 50720001 | 50820000 | 0.0039 | 0.0033 | 0.0037 |
| F06 | 50740001 | 50840000 | 0.0038 | 0.0039 | 0.0034 |
| F06 | 50760001 | 50860000 | 0.0046 | 0.0048 | 0.0040 |
| F06 | 50780001 | 50880000 | 0.0056 | 0.0053 | 0.0048 |

| Chr | Start | End | both | A1 | A2 |
|---|---|---|---|---|---|
| F06 | 50800001 | 50900000 | 0.0063 | 0.0051 | 0.0053 |
| F06 | 50820001 | 50920000 | 0.0065 | 0.0062 | 0.0052 |
| F06 | 50840001 | 50940000 | 0.0075 | 0.0052 | 0.0063 |
| F06 | 50860001 | 50960000 | 0.0067 | 0.0042 | 0.0052 |
| F06 | 50880001 | 50980000 | 0.0053 | 0.0037 | 0.0040 |
| F06 | 50900001 | 51000000 | 0.0047 | 0.0036 | 0.0037 |
| F06 | 50920001 | 51020000 | 0.0045 | 0.0021 | 0.0038 |
| F06 | 50940001 | 51040000 | 0.0028 | 0.0024 | 0.0022 |
| F06 | 50960001 | 51060000 | 0.0028 | 0.0021 | 0.0025 |
| F06 | 50980001 | 51080000 | 0.0025 | 0.0024 | 0.0024 |
| F06 | 51000001 | 51100000 | 0.0027 | 0.0019 | 0.0024 |
| F06 | 51020001 | 51120000 | 0.0025 | 0.0022 | 0.0020 |
| F06 | 51040001 | 51140000 | 0.0029 | 0.0021 | 0.0024 |
| F06 | 51060001 | 51160000 | 0.0028 | 0.0025 | 0.0023 |
| F06 | 51080001 | 51180000 | 0.0033 | 0.0022 | 0.0026 |
| F06 | 51100001 | 51200000 | 0.0032 | 0.0020 | 0.0024 |
| F06 | 51120001 | 51220000 | 0.0029 | 0.0021 | 0.0021 |
| F06 | 51140001 | 51240000 | 0.0030 | 0.0034 | 0.0022 |
| F06 | 51160001 | 51260000 | 0.0045 | 0.0032 | 0.0036 |
| F06 | 51180001 | 51280000 | 0.0047 | 0.0037 | 0.0036 |
| F06 | 51200001 | 51300000 | 0.0049 | 0.0041 | 0.0040 |
| F06 | 51220001 | 51320000 | 0.0053 | 0.0051 | 0.0043 |
| F06 | 51240001 | 51340000 | 0.0063 | 0.0048 | 0.0052 |
| F06 | 51260001 | 51360000 | 0.0057 | 0.0049 | 0.0048 |
| F06 | 51280001 | 51380000 | 0.0057 | 0.0049 | 0.0048 |
| F06 | 51300001 | 51400000 | 0.0057 | 0.0044 | 0.0050 |
| F06 | 51320001 | 51420000 | 0.0051 | 0.0028 | 0.0046 |
| F06 | 51340001 | 51440000 | 0.0036 | 0.0028 | 0.0032 |
| F06 | 51360001 | 51460000 | 0.0036 | 0.0031 | 0.0031 |
| F06 | 51380001 | 51480000 | 0.0037 | 0.0027 | 0.0033 |
| F06 | 51400001 | 51500000 | 0.0030 | 0.0035 | 0.0027 |
| F06 | 51420001 | 51520000 | 0.0041 | 0.0039 | 0.0036 |
| F06 | 51440001 | 51540000 | 0.0045 | 0.0030 | 0.0041 |
| F06 | 51460001 | 51560000 | 0.0035 | 0.0020 | 0.0032 |
| F06 | 51480001 | 51580000 | 0.0026 | 0.0020 | 0.0024 |
| F06 | 51500001 | 51600000 | 0.0025 | 0.0011 | 0.0023 |
| F06 | 51520001 | 51620000 | 0.0015 | 0.0010 | 0.0015 |
| F06 | 51540001 | 51640000 | 0.0013 | 0.0010 | 0.0013 |
| F06 | 51560001 | 51660000 | 0.0012 | 0.0014 | 0.0012 |
| F06 | 51580001 | 51680000 | 0.0017 | 0.0016 | 0.0016 |
| F06 | 51600001 | 51700000 | 0.0019 | 0.0016 | 0.0018 |
| F06 | 51620001 | 51720000 | 0.0019 | 0.0015 | 0.0018 |
| F06 | 51640001 | 51740000 | 0.0017 | 0.0017 | 0.0016 |
| F06 | 51660001 | 51760000 | 0.0019 | 0.0014 | 0.0018 |
| F06 | 51680001 | 51780000 | 0.0017 | 0.0012 | 0.0015 |
| F06 | 51700001 | 51800000 | 0.0015 | 0.0010 | 0.0014 |
| F06 | 51720001 | 51820000 | 0.0013 | 0.0011 | 0.0012 |
| F06 | 51740001 | 51840000 | 0.0014 | 0.0011 | 0.0013 |
| F06 | 51760001 | 51860000 | 0.0015 | 0.0013 | 0.0014 |
| F06 | 51780001 | 51880000 | 0.0015 | 0.0013 | 0.0014 |
| F06 | 51800001 | 51900000 | 0.0016 | 0.0017 | 0.0014 |
| F06 | 51820001 | 51920000 | 0.0020 | 0.0022 | 0.0019 |
| F06 | 51840001 | 51940000 | 0.0026 | 0.0028 | 0.0023 |
| F06 | 51860001 | 51960000 | 0.0031 | 0.0029 | 0.0029 |
| F06 | 51880001 | 51980000 | 0.0033 | 0.0034 | 0.0029 |
| F06 | 51900001 | 52000000 | 0.0040 | 0.0035 | 0.0035 |
| F06 | 51920001 | 52020000 | 0.0042 | 0.0031 | 0.0035 |
| F06 | 51940001 | 52040000 | 0.0037 | 0.0034 | 0.0032 |

| Chr | Start | End | both | A1 | A2 |
|---|---|---|---|---|---|
| F06 | 51960001 | 52060000 | 0.0042 | 0.0035 | 0.0035 |
| F06 | 51980001 | 52080000 | 0.0042 | 0.0028 | 0.0037 |
| F06 | 52000001 | 52100000 | 0.0036 | 0.0030 | 0.0031 |
| F06 | 52020001 | 52120000 | 0.0040 | 0.0034 | 0.0033 |
| F06 | 52040001 | 52140000 | 0.0047 | 0.0023 | 0.0038 |
| F06 | 52060001 | 52160000 | 0.0037 | 0.0024 | 0.0028 |
| F06 | 52080001 | 52180000 | 0.0038 | 0.0027 | 0.0028 |
| F06 | 52100001 | 52200000 | 0.0044 | 0.0020 | 0.0031 |
| F06 | 52120001 | 52220000 | 0.0036 | 0.0027 | 0.0025 |
| F06 | 52140001 | 52240000 | 0.0041 | 0.0027 | 0.0028 |
| F06 | 52160001 | 52260000 | 0.0040 | 0.0025 | 0.0028 |
| F06 | 52180001 | 52280000 | 0.0037 | 0.0020 | 0.0026 |
| F06 | 52200001 | 52300000 | 0.0027 | 0.0024 | 0.0021 |
| F06 | 52220001 | 52320000 | 0.0029 | 0.0013 | 0.0025 |
| F06 | 52240001 | 52340000 | 0.0017 | 0.0013 | 0.0016 |
| F06 | 52260001 | 52360000 | 0.0016 | 0.0019 | 0.0016 |
| F06 | 52280001 | 52380000 | 0.0020 | 0.0021 | 0.0020 |
| F06 | 52300001 | 52400000 | 0.0023 | 0.0018 | 0.0022 |
| F06 | 52320001 | 52420000 | 0.0020 | 0.0016 | 0.0019 |
| F06 | 52340001 | 52440000 | 0.0017 | 0.0016 | 0.0017 |
| F06 | 52360001 | 52460000 | 0.0018 | 0.0011 | 0.0017 |
| F06 | 52380001 | 52480000 | 0.0014 | 0.0013 | 0.0012 |
| F06 | 52400001 | 52500000 | 0.0017 | 0.0016 | 0.0014 |
| F06 | 52420001 | 52520000 | 0.0021 | 0.0017 | 0.0017 |
| F06 | 52440001 | 52540000 | 0.0022 | 0.0015 | 0.0018 |
| F06 | 52460001 | 52560000 | 0.0020 | 0.0014 | 0.0017 |
| F06 | 52480001 | 52580000 | 0.0018 | 0.0012 | 0.0016 |
| F06 | 52500001 | 52600000 | 0.0016 | 0.0011 | 0.0015 |
| F06 | 52520001 | 52620000 | 0.0014 | 0.0014 | 0.0014 |
| F06 | 52540001 | 52640000 | 0.0018 | 0.0013 | 0.0016 |
| F06 | 52560001 | 52660000 | 0.0016 | 0.0013 | 0.0014 |
| F06 | 52580001 | 52680000 | 0.0017 | 0.0016 | 0.0015 |
| F06 | 52600001 | 52700000 | 0.0021 | 0.0016 | 0.0017 |
| F06 | 52620001 | 52720000 | 0.0021 | 0.0016 | 0.0018 |
| F06 | 52640001 | 52740000 | 0.0020 | 0.0016 | 0.0019 |
| F06 | 52660001 | 52760000 | 0.0021 | 0.0015 | 0.0019 |
| F06 | 52680001 | 52780000 | 0.0019 | 0.0015 | 0.0018 |
| F06 | 52700001 | 52800000 | 0.0017 | 0.0013 | 0.0016 |
| F06 | 52720001 | 52820000 | 0.0015 | 0.0011 | 0.0014 |
| F06 | 52740001 | 52840000 | 0.0013 | 0.0013 | 0.0012 |
| F06 | 52760001 | 52860000 | 0.0015 | 0.0013 | 0.0013 |
| F06 | 52780001 | 52880000 | 0.0016 | 0.0013 | 0.0015 |
| F06 | 52800001 | 52900000 | 0.0017 | 0.0015 | 0.0016 |
| F06 | 52820001 | 52920000 | 0.0019 | 0.0013 | 0.0019 |
| F06 | 52840001 | 52940000 | 0.0016 | 0.0016 | 0.0016 |
| F06 | 52860001 | 52960000 | 0.0020 | 0.0028 | 0.0020 |
| F06 | 52880001 | 52980000 | 0.0035 | 0.0026 | 0.0035 |
| F06 | 52900001 | 53000000 | 0.0032 | 0.0030 | 0.0032 |
| F06 | 52920001 | 53020000 | 0.0038 | 0.0033 | 0.0037 |
| F06 | 52940001 | 53040000 | 0.0042 | 0.0034 | 0.0041 |
| F06 | 52960001 | 53060000 | 0.0043 | 0.0026 | 0.0041 |
| F06 | 52980001 | 53080000 | 0.0034 | 0.0025 | 0.0031 |
| F06 | 53000001 | 53100000 | 0.0032 | 0.0020 | 0.0028 |
| F06 | 53020001 | 53120000 | 0.0027 | 0.0031 | 0.0022 |
| F06 | 53040001 | 53140000 | 0.0039 | 0.0025 | 0.0034 |
| F06 | 53060001 | 53160000 | 0.0035 | 0.0025 | 0.0029 |
| F06 | 53080001 | 53180000 | 0.0036 | 0.0026 | 0.0029 |
| F06 | 53100001 | 53200000 | 0.0040 | 0.0034 | 0.0030 |

| Chr | Start | End | both | A1 | A2 |
|---|---|---|---|---|---|
| F06 | 53120001 | 53220000 | 0.0049 | 0.0022 | 0.0040 |
| F06 | 53140001 | 53240000 | 0.0035 | 0.0029 | 0.0025 |
| F06 | 53160001 | 53260000 | 0.0042 | 0.0034 | 0.0033 |
| F06 | 53180001 | 53280000 | 0.0047 | 0.0042 | 0.0037 |
| F06 | 53200001 | 53300000 | 0.0055 | 0.0039 | 0.0047 |
| F06 | 53220001 | 53320000 | 0.0051 | 0.0040 | 0.0044 |
| F06 | 53240001 | 53340000 | 0.0053 | 0.0038 | 0.0046 |
| F06 | 53260001 | 53360000 | 0.0050 | 0.0036 | 0.0043 |
| F06 | 53280001 | 53380000 | 0.0046 | 0.0032 | 0.0041 |
| F06 | 53300001 | 53400000 | 0.0040 | 0.0028 | 0.0034 |
| F06 | 53320001 | 53420000 | 0.0039 | 0.0025 | 0.0031 |
| F06 | 53340001 | 53440000 | 0.0034 | 0.0024 | 0.0027 |
| F06 | 53360001 | 53460000 | 0.0036 | 0.0021 | 0.0026 |
| F06 | 53380001 | 53480000 | 0.0035 | 0.0017 | 0.0024 |
| F06 | 53400001 | 53500000 | 0.0031 | 0.0021 | 0.0019 |
| F06 | 53420001 | 53520000 | 0.0033 | 0.0029 | 0.0022 |
| F06 | 53440001 | 53540000 | 0.0043 | 0.0035 | 0.0031 |
| F06 | 53460001 | 53560000 | 0.0046 | 0.0035 | 0.0038 |
| F06 | 53480001 | 53580000 | 0.0043 | 0.0039 | 0.0037 |
| F06 | 53500001 | 53600000 | 0.0047 | 0.0034 | 0.0043 |
| F06 | 53520001 | 53620000 | 0.0041 | 0.0031 | 0.0038 |
| F06 | 53540001 | 53640000 | 0.0037 | 0.0035 | 0.0036 |
| F06 | 53560001 | 53660000 | 0.0038 | 0.0045 | 0.0035 |
| F06 | 53580001 | 53680000 | 0.0051 | 0.0042 | 0.0046 |
| F06 | 53600001 | 53700000 | 0.0052 | 0.0040 | 0.0044 |
| F06 | 53620001 | 53720000 | 0.0052 | 0.0036 | 0.0041 |
| F06 | 53640001 | 53740000 | 0.0048 | 0.0028 | 0.0036 |
| F06 | 53660001 | 53760000 | 0.0041 | 0.0013 | 0.0031 |
| F06 | 53680001 | 53780000 | 0.0023 | 0.0017 | 0.0014 |
| F06 | 53700001 | 53800000 | 0.0024 | 0.0022 | 0.0018 |
| F06 | 53720001 | 53820000 | 0.0031 | 0.0022 | 0.0023 |
| F06 | 53740001 | 53840000 | 0.0033 | 0.0022 | 0.0022 |
| F06 | 53760001 | 53860000 | 0.0032 | 0.0035 | 0.0022 |
| F06 | 53780001 | 53880000 | 0.0046 | 0.0030 | 0.0035 |
| F06 | 53800001 | 53900000 | 0.0040 | 0.0031 | 0.0029 |
| F06 | 53820001 | 53920000 | 0.0038 | 0.0036 | 0.0030 |
| F06 | 53840001 | 53940000 | 0.0040 | 0.0040 | 0.0036 |
| F06 | 53860001 | 53960000 | 0.0044 | 0.0028 | 0.0040 |
| F06 | 53880001 | 53980000 | 0.0033 | 0.0026 | 0.0031 |
| F06 | 53900001 | 54000000 | 0.0030 | 0.0019 | 0.0029 |
| F06 | 53920001 | 54020000 | 0.0025 | 0.0018 | 0.0023 |
| F06 | 53940001 | 54040000 | 0.0024 | 0.0019 | 0.0021 |
| F06 | 53960001 | 54060000 | 0.0028 | 0.0030 | 0.0025 |
| F06 | 53980001 | 54080000 | 0.0038 | 0.0040 | 0.0034 |
| F06 | 54000001 | 54100000 | 0.0050 | 0.0044 | 0.0045 |
| F06 | 54020001 | 54120000 | 0.0055 | 0.0047 | 0.0048 |
| F06 | 54040001 | 54140000 | 0.0059 | 0.0044 | 0.0051 |
| F06 | 54060001 | 54160000 | 0.0054 | 0.0044 | 0.0046 |
| F06 | 54080001 | 54180000 | 0.0053 | 0.0038 | 0.0045 |
| F06 | 54100001 | 54200000 | 0.0048 | 0.0047 | 0.0041 |
| F06 | 54120001 | 54220000 | 0.0058 | 0.0045 | 0.0052 |
| F06 | 54140001 | 54240000 | 0.0056 | 0.0048 | 0.0050 |
| F06 | 54160001 | 54260000 | 0.0060 | 0.0038 | 0.0053 |
| F06 | 54180001 | 54280000 | 0.0050 | 0.0039 | 0.0042 |
| F06 | 54200001 | 54300000 | 0.0053 | 0.0028 | 0.0042 |
| F06 | 54220001 | 54320000 | 0.0040 | 0.0027 | 0.0029 |
| F06 | 54240001 | 54340000 | 0.0038 | 0.0027 | 0.0029 |
| F06 | 54260001 | 54360000 | 0.0038 | 0.0032 | 0.0029 |

| Chr | Start | End | both | A1 | A2 |
|---|---|---|---|---|---|
| F06 | 54280001 | 54380000 | 0.0044 | 0.0031 | 0.0035 |
| F06 | 54300001 | 54400000 | 0.0041 | 0.0041 | 0.0034 |
| F06 | 54320001 | 54420000 | 0.0051 | 0.0039 | 0.0044 |
| F06 | 54340001 | 54440000 | 0.0050 | 0.0033 | 0.0043 |
| F06 | 54360001 | 54460000 | 0.0041 | 0.0029 | 0.0036 |
| F06 | 54380001 | 54480000 | 0.0036 | 0.0030 | 0.0033 |
| F06 | 54400001 | 54500000 | 0.0037 | 0.0024 | 0.0034 |
| F06 | 54420001 | 54520000 | 0.0031 | 0.0028 | 0.0029 |
| F06 | 54440001 | 54540000 | 0.0035 | 0.0028 | 0.0032 |
| F06 | 54460001 | 54560000 | 0.0034 | 0.0030 | 0.0031 |
| F06 | 54480001 | 54580000 | 0.0036 | 0.0027 | 0.0033 |
| F06 | 54500001 | 54600000 | 0.0033 | 0.0022 | 0.0030 |
| F06 | 54520001 | 54620000 | 0.0026 | 0.0018 | 0.0023 |
| F06 | 54540001 | 54640000 | 0.0024 | 0.0024 | 0.0020 |
| F06 | 54560001 | 54660000 | 0.0032 | 0.0027 | 0.0025 |
| F06 | 54580001 | 54680000 | 0.0036 | 0.0027 | 0.0029 |
| F06 | 54600001 | 54700000 | 0.0035 | 0.0031 | 0.0028 |
| F06 | 54620001 | 54720000 | 0.0040 | 0.0032 | 0.0033 |
| F06 | 54640001 | 54740000 | 0.0039 | 0.0027 | 0.0034 |
| F06 | 54660001 | 54760000 | 0.0034 | 0.0024 | 0.0030 |
| F06 | 54680001 | 54780000 | 0.0031 | 0.0025 | 0.0027 |
| F06 | 54700001 | 54800000 | 0.0033 | 0.0025 | 0.0027 |
| F06 | 54720001 | 54820000 | 0.0032 | 0.0030 | 0.0026 |
| F06 | 54740001 | 54840000 | 0.0039 | 0.0030 | 0.0033 |
| F06 | 54760001 | 54860000 | 0.0038 | 0.0028 | 0.0033 |
| F06 | 54780001 | 54880000 | 0.0036 | 0.0028 | 0.0030 |
| F06 | 54800001 | 54900000 | 0.0034 | 0.0027 | 0.0030 |
| F06 | 54820001 | 54920000 | 0.0036 | 0.0018 | 0.0028 |
| F06 | 54840001 | 54940000 | 0.0026 | 0.0022 | 0.0018 |
| F06 | 54860001 | 54960000 | 0.0032 | 0.0025 | 0.0022 |
| F06 | 54880001 | 54980000 | 0.0037 | 0.0023 | 0.0027 |
| F06 | 54900001 | 55000000 | 0.0034 | 0.0027 | 0.0025 |
| F06 | 54920001 | 55020000 | 0.0038 | 0.0026 | 0.0031 |
| F06 | 54940001 | 55040000 | 0.0038 | 0.0019 | 0.0029 |
| F06 | 54960001 | 55060000 | 0.0030 | 0.0014 | 0.0021 |
| F06 | 54980001 | 55080000 | 0.0026 | 0.0022 | 0.0016 |
| F06 | 55000001 | 55100000 | 0.0036 | 0.0021 | 0.0025 |
| F06 | 55020001 | 55120000 | 0.0033 | 0.0024 | 0.0024 |
| F06 | 55040001 | 55140000 | 0.0035 | 0.0027 | 0.0028 |
| F06 | 55060001 | 55160000 | 0.0036 | 0.0028 | 0.0031 |
| F06 | 55080001 | 55180000 | 0.0035 | 0.0022 | 0.0033 |
| F06 | 55100001 | 55200000 | 0.0027 | 0.0022 | 0.0026 |
| F06 | 55120001 | 55220000 | 0.0027 | 0.0025 | 0.0025 |
| F06 | 55140001 | 55240000 | 0.0031 | 0.0026 | 0.0029 |
| F06 | 55160001 | 55260000 | 0.0032 | 0.0024 | 0.0030 |
| F06 | 55180001 | 55280000 | 0.0031 | 0.0022 | 0.0028 |
| F06 | 55200001 | 55300000 | 0.0028 | 0.0020 | 0.0025 |
| F06 | 55220001 | 55320000 | 0.0025 | 0.0014 | 0.0022 |
| F06 | 55240001 | 55340000 | 0.0022 | 0.0020 | 0.0015 |
| F06 | 55260001 | 55360000 | 0.0029 | 0.0028 | 0.0021 |
| F06 | 55280001 | 55380000 | 0.0039 | 0.0038 | 0.0030 |
| F06 | 55300001 | 55400000 | 0.0050 | 0.0041 | 0.0038 |
| F06 | 55320001 | 55420000 | 0.0056 | 0.0042 | 0.0043 |
| F06 | 55340001 | 55440000 | 0.0055 | 0.0035 | 0.0044 |
| F06 | 55360001 | 55460000 | 0.0049 | 0.0043 | 0.0035 |
| F06 | 55380001 | 55480000 | 0.0057 | 0.0040 | 0.0042 |
| F06 | 55400001 | 55500000 | 0.0054 | 0.0037 | 0.0040 |
| F06 | 55420001 | 55520000 | 0.0049 | 0.0039 | 0.0036 |

| Chr | Start | End | both | A1 | A2 |
|---|---|---|---|---|---|
| F06 | 55440001 | 55540000 | 0.0050 | 0.0040 | 0.0040 |
| F06 | 55460001 | 55560000 | 0.0047 | 0.0021 | 0.0040 |
| F06 | 55480001 | 55580000 | 0.0027 | 0.0025 | 0.0022 |
| F06 | 55500001 | 55600000 | 0.0031 | 0.0029 | 0.0027 |
| F06 | 55520001 | 55620000 | 0.0035 | 0.0026 | 0.0030 |
| F06 | 55540001 | 55640000 | 0.0033 | 0.0034 | 0.0027 |
| F06 | 55560001 | 55660000 | 0.0043 | 0.0035 | 0.0036 |
| F06 | 55580001 | 55680000 | 0.0042 | 0.0033 | 0.0037 |
| F06 | 55600001 | 55700000 | 0.0040 | 0.0038 | 0.0036 |
| F06 | 55620001 | 55720000 | 0.0046 | 0.0044 | 0.0043 |
| F06 | 55640001 | 55740000 | 0.0054 | 0.0039 | 0.0048 |
| F06 | 55660001 | 55760000 | 0.0050 | 0.0038 | 0.0042 |
| F06 | 55680001 | 55780000 | 0.0051 | 0.0029 | 0.0041 |
| F06 | 55700001 | 55800000 | 0.0041 | 0.0019 | 0.0031 |
| F06 | 55720001 | 55820000 | 0.0031 | 0.0024 | 0.0019 |
| F06 | 55740001 | 55840000 | 0.0034 | 0.0023 | 0.0025 |
| F06 | 55760001 | 55860000 | 0.0031 | 0.0029 | 0.0024 |
| F06 | 55780001 | 55880000 | 0.0038 | 0.0030 | 0.0031 |
| F06 | 55800001 | 55900000 | 0.0038 | 0.0033 | 0.0033 |
| F06 | 55820001 | 55920000 | 0.0039 | 0.0022 | 0.0036 |
| F06 | 55840001 | 55940000 | 0.0027 | 0.0022 | 0.0024 |
| F06 | 55860001 | 55960000 | 0.0028 | 0.0022 | 0.0025 |
| F06 | 55880001 | 55980000 | 0.0029 | 0.0026 | 0.0025 |
| F06 | 55900001 | 56000000 | 0.0034 | 0.0027 | 0.0028 |
| F06 | 55920001 | 56020000 | 0.0036 | 0.0036 | 0.0029 |
| F06 | 55940001 | 56040000 | 0.0047 | 0.0033 | 0.0039 |
| F06 | 55960001 | 56060000 | 0.0042 | 0.0030 | 0.0035 |
| F06 | 55980001 | 56080000 | 0.0038 | 0.0025 | 0.0032 |
| F06 | 56000001 | 56100000 | 0.0031 | 0.0021 | 0.0027 |
| F06 | 56020001 | 56120000 | 0.0025 | 0.0021 | 0.0024 |
| F06 | 56040001 | 56140000 | 0.0025 | 0.0024 | 0.0025 |
| F06 | 56060001 | 56160000 | 0.0029 | 0.0021 | 0.0029 |
| F06 | 56080001 | 56180000 | 0.0026 | 0.0023 | 0.0025 |
| F06 | 56100001 | 56200000 | 0.0029 | 0.0024 | 0.0027 |
| F06 | 56120001 | 56220000 | 0.0030 | 0.0016 | 0.0027 |
| F06 | 56140001 | 56240000 | 0.0022 | 0.0014 | 0.0018 |
| F06 | 56160001 | 56260000 | 0.0021 | 0.0027 | 0.0016 |
| F06 | 56180001 | 56280000 | 0.0035 | 0.0035 | 0.0032 |
| F06 | 56200001 | 56300000 | 0.0047 | 0.0040 | 0.0043 |
| F06 | 56220001 | 56320000 | 0.0053 | 0.0039 | 0.0049 |
| F06 | 56240001 | 56340000 | 0.0050 | 0.0040 | 0.0048 |
| F06 | 56260001 | 56360000 | 0.0051 | 0.0028 | 0.0050 |
| F06 | 56280001 | 56380000 | 0.0037 | 0.0022 | 0.0036 |
| F06 | 56300001 | 56400000 | 0.0027 | 0.0018 | 0.0027 |
| F06 | 56320001 | 56420000 | 0.0021 | 0.0019 | 0.0021 |
| F06 | 56340001 | 56440000 | 0.0023 | 0.0022 | 0.0022 |
| F06 | 56360001 | 56460000 | 0.0027 | 0.0023 | 0.0025 |
| F06 | 56380001 | 56480000 | 0.0028 | 0.0031 | 0.0026 |
| F06 | 56400001 | 56500000 | 0.0035 | 0.0034 | 0.0033 |
| F06 | 56420001 | 56520000 | 0.0040 | 0.0033 | 0.0037 |
| F06 | 56440001 | 56540000 | 0.0039 | 0.0036 | 0.0036 |
| F06 | 56460001 | 56560000 | 0.0042 | 0.0038 | 0.0040 |
| F06 | 56480001 | 56580000 | 0.0047 | 0.0030 | 0.0042 |
| F06 | 56500001 | 56600000 | 0.0041 | 0.0028 | 0.0036 |
| F06 | 56520001 | 56620000 | 0.0038 | 0.0036 | 0.0033 |
| F06 | 56540001 | 56640000 | 0.0051 | 0.0027 | 0.0042 |
| F06 | 56560001 | 56660000 | 0.0044 | 0.0023 | 0.0034 |
| F06 | 56580001 | 56680000 | 0.0039 | 0.0027 | 0.0029 |

| Chr | Start | End | both | A1 | A2 |
|---|---|---|---|---|---|
| F06 | 56600001 | 56700000 | 0.0041 | 0.0027 | 0.0033 |
| F06 | 56620001 | 56720000 | 0.0041 | 0.0031 | 0.0034 |
| F06 | 56640001 | 56740000 | 0.0041 | 0.0036 | 0.0037 |
| F06 | 56660001 | 56760000 | 0.0044 | 0.0038 | 0.0041 |
| F06 | 56680001 | 56780000 | 0.0048 | 0.0037 | 0.0044 |
| F06 | 56700001 | 56800000 | 0.0048 | 0.0037 | 0.0043 |
| F06 | 56720001 | 56820000 | 0.0047 | 0.0023 | 0.0042 |
| F06 | 56740001 | 56840000 | 0.0033 | 0.0015 | 0.0028 |
| F06 | 56760001 | 56860000 | 0.0025 | 0.0018 | 0.0020 |
| F06 | 56780001 | 56880000 | 0.0027 | 0.0016 | 0.0022 |
| F06 | 56800001 | 56900000 | 0.0025 | 0.0016 | 0.0019 |
| F06 | 56820001 | 56920000 | 0.0026 | 0.0020 | 0.0020 |
| F06 | 56840001 | 56940000 | 0.0033 | 0.0030 | 0.0025 |
| F06 | 56860001 | 56960000 | 0.0044 | 0.0034 | 0.0036 |
| F06 | 56880001 | 56980000 | 0.0047 | 0.0041 | 0.0040 |
| F06 | 56900001 | 57000000 | 0.0053 | 0.0039 | 0.0048 |
| F06 | 56920001 | 57020000 | 0.0050 | 0.0048 | 0.0045 |
| F06 | 56940001 | 57040000 | 0.0057 | 0.0047 | 0.0052 |
| F06 | 56960001 | 57060000 | 0.0057 | 0.0053 | 0.0053 |
| F06 | 56980001 | 57080000 | 0.0061 | 0.0045 | 0.0057 |
| F06 | 57000001 | 57100000 | 0.0051 | 0.0043 | 0.0048 |
| F06 | 57020001 | 57120000 | 0.0049 | 0.0030 | 0.0046 |
| F06 | 57040001 | 57140000 | 0.0035 | 0.0028 | 0.0033 |
| F06 | 57060001 | 57160000 | 0.0031 | 0.0018 | 0.0030 |
| F06 | 57080001 | 57180000 | 0.0021 | 0.0017 | 0.0021 |
| F06 | 57100001 | 57200000 | 0.0021 | 0.0030 | 0.0021 |
| F06 | 57120001 | 57220000 | 0.0034 | 0.0036 | 0.0033 |
| F06 | 57140001 | 57240000 | 0.0041 | 0.0039 | 0.0040 |
| F06 | 57160001 | 57260000 | 0.0048 | 0.0034 | 0.0044 |
| F06 | 57180001 | 57280000 | 0.0043 | 0.0035 | 0.0039 |
| F06 | 57200001 | 57300000 | 0.0045 | 0.0025 | 0.0038 |
| F06 | 57220001 | 57320000 | 0.0038 | 0.0025 | 0.0029 |
| F06 | 57240001 | 57340000 | 0.0040 | 0.0018 | 0.0027 |
| F06 | 57260001 | 57360000 | 0.0030 | 0.0026 | 0.0019 |
| F06 | 57280001 | 57380000 | 0.0040 | 0.0027 | 0.0028 |
| F06 | 57300001 | 57400000 | 0.0041 | 0.0024 | 0.0030 |
| F06 | 57320001 | 57420000 | 0.0035 | 0.0024 | 0.0027 |
| F06 | 57340001 | 57440000 | 0.0031 | 0.0022 | 0.0026 |
| F06 | 57360001 | 57460000 | 0.0026 | 0.0017 | 0.0023 |
| F06 | 57380001 | 57480000 | 0.0020 | 0.0019 | 0.0018 |
| F06 | 57400001 | 57500000 | 0.0021 | 0.0021 | 0.0020 |
| F06 | 57420001 | 57520000 | 0.0024 | 0.0020 | 0.0023 |
| F06 | 57440001 | 57540000 | 0.0023 | 0.0032 | 0.0021 |
| F06 | 57460001 | 57560000 | 0.0037 | 0.0027 | 0.0033 |
| F06 | 57480001 | 57580000 | 0.0032 | 0.0031 | 0.0029 |
| F06 | 57500001 | 57600000 | 0.0043 | 0.0031 | 0.0034 |
| F06 | 57520001 | 57620000 | 0.0044 | 0.0030 | 0.0033 |
| F06 | 57540001 | 57640000 | 0.0042 | 0.0022 | 0.0033 |
| F06 | 57560001 | 57660000 | 0.0034 | 0.0034 | 0.0025 |
| F06 | 57580001 | 57680000 | 0.0047 | 0.0029 | 0.0037 |
| F06 | 57600001 | 57700000 | 0.0037 | 0.0029 | 0.0030 |
| F06 | 57620001 | 57720000 | 0.0037 | 0.0029 | 0.0031 |
| F06 | 57640001 | 57740000 | 0.0037 | 0.0026 | 0.0032 |
| F06 | 57660001 | 57760000 | 0.0038 | 0.0018 | 0.0029 |
| F06 | 57680001 | 57780000 | 0.0030 | 0.0015 | 0.0021 |
| F06 | 57700001 | 57800000 | 0.0027 | 0.0016 | 0.0017 |
| F06 | 57720001 | 57820000 | 0.0031 | 0.0016 | 0.0018 |
| F06 | 57740001 | 57840000 | 0.0033 | 0.0019 | 0.0018 |

| Chr | Start | End | both | A1 | A2 |
|---|---|---|---|---|---|
| F06 | 57760001 | 57860000 | 0.0034 | 0.0026 | 0.0022 |
| F06 | 57780001 | 57880000 | 0.0041 | 0.0032 | 0.0028 |
| F06 | 57800001 | 57900000 | 0.0050 | 0.0034 | 0.0037 |
| F06 | 57820001 | 57920000 | 0.0048 | 0.0032 | 0.0039 |
| F06 | 57840001 | 57940000 | 0.0043 | 0.0030 | 0.0037 |
| F06 | 57860001 | 57960000 | 0.0039 | 0.0034 | 0.0034 |
| F06 | 57880001 | 57980000 | 0.0041 | 0.0028 | 0.0039 |
| F06 | 57900001 | 58000000 | 0.0032 | 0.0029 | 0.0032 |
| F06 | 57920001 | 58020000 | 0.0035 | 0.0031 | 0.0031 |
| F06 | 57940001 | 58040000 | 0.0040 | 0.0031 | 0.0034 |
| F06 | 57960001 | 58060000 | 0.0043 | 0.0020 | 0.0035 |
| F06 | 57980001 | 58080000 | 0.0032 | 0.0020 | 0.0023 |
| F06 | 58000001 | 58100000 | 0.0034 | 0.0022 | 0.0022 |
| F06 | 58020001 | 58120000 | 0.0035 | 0.0029 | 0.0024 |
| F06 | 58040001 | 58140000 | 0.0040 | 0.0026 | 0.0030 |
| F06 | 58060001 | 58160000 | 0.0037 | 0.0027 | 0.0027 |
| F06 | 58080001 | 58180000 | 0.0038 | 0.0033 | 0.0028 |
| F06 | 58100001 | 58200000 | 0.0044 | 0.0034 | 0.0034 |
| F06 | 58120001 | 58220000 | 0.0046 | 0.0033 | 0.0038 |
| F06 | 58140001 | 58240000 | 0.0045 | 0.0037 | 0.0038 |
| F06 | 58160001 | 58260000 | 0.0048 | 0.0036 | 0.0043 |
| F06 | 58180001 | 58280000 | 0.0045 | 0.0029 | 0.0042 |
| F06 | 58200001 | 58300000 | 0.0038 | 0.0023 | 0.0035 |
| F06 | 58220001 | 58320000 | 0.0029 | 0.0014 | 0.0027 |
| F06 | 58240001 | 58340000 | 0.0021 | 0.0012 | 0.0017 |
| F06 | 58260001 | 58360000 | 0.0020 | 0.0016 | 0.0016 |
| F06 | 58280001 | 58380000 | 0.0022 | 0.0016 | 0.0019 |
| F06 | 58300001 | 58400000 | 0.0025 | 0.0021 | 0.0020 |
| F06 | 58320001 | 58420000 | 0.0033 | 0.0022 | 0.0025 |
| F06 | 58340001 | 58440000 | 0.0031 | 0.0021 | 0.0025 |
| F06 | 58360001 | 58460000 | 0.0031 | 0.0015 | 0.0024 |
| F06 | 58380001 | 58480000 | 0.0025 | 0.0021 | 0.0018 |
| F06 | 58400001 | 58500000 | 0.0026 | 0.0021 | 0.0022 |
| F06 | 58420001 | 58520000 | 0.0028 | 0.0032 | 0.0022 |
| F06 | 58440001 | 58540000 | 0.0043 | 0.0037 | 0.0037 |
| F06 | 58460001 | 58560000 | 0.0048 | 0.0042 | 0.0042 |
| F06 | 58480001 | 58580000 | 0.0055 | 0.0044 | 0.0050 |
| F06 | 58500001 | 58600000 | 0.0057 | 0.0048 | 0.0051 |
| F06 | 58520001 | 58620000 | 0.0058 | 0.0046 | 0.0053 |
| F06 | 58540001 | 58640000 | 0.0057 | 0.0043 | 0.0050 |
| F06 | 58560001 | 58660000 | 0.0053 | 0.0044 | 0.0047 |
| F06 | 58580001 | 58680000 | 0.0055 | 0.0042 | 0.0047 |
| F06 | 58600001 | 58700000 | 0.0052 | 0.0037 | 0.0046 |
| F06 | 58620001 | 58720000 | 0.0048 | 0.0041 | 0.0043 |
| F06 | 58640001 | 58740000 | 0.0049 | 0.0051 | 0.0045 |
| F06 | 58660001 | 58760000 | 0.0061 | 0.0049 | 0.0056 |
| F06 | 58680001 | 58780000 | 0.0058 | 0.0049 | 0.0053 |
| F06 | 58700001 | 58800000 | 0.0060 | 0.0045 | 0.0053 |
| F06 | 58720001 | 58820000 | 0.0056 | 0.0034 | 0.0049 |
| F06 | 58740001 | 58840000 | 0.0043 | 0.0020 | 0.0038 |
| F06 | 58760001 | 58860000 | 0.0025 | 0.0017 | 0.0022 |
| F06 | 58780001 | 58880000 | 0.0022 | 0.0011 | 0.0020 |
| F06 | 58800001 | 58900000 | 0.0014 | 0.0013 | 0.0013 |
| F06 | 58820001 | 58920000 | 0.0015 | 0.0018 | 0.0014 |
| F06 | 58840001 | 58940000 | 0.0024 | 0.0025 | 0.0020 |
| F06 | 58860001 | 58960000 | 0.0033 | 0.0027 | 0.0028 |
| F06 | 58880001 | 58980000 | 0.0037 | 0.0028 | 0.0031 |
| F06 | 58900001 | 59000000 | 0.0039 | 0.0031 | 0.0033 |

| Chr | Start | End | both | A1 | A2 |
|---|---|---|---|---|---|
| F06 | 58920001 | 59020000 | 0.0042 | 0.0032 | 0.0036 |
| F06 | 58940001 | 59040000 | 0.0040 | 0.0028 | 0.0036 |
| F06 | 58960001 | 59060000 | 0.0037 | 0.0024 | 0.0033 |
| F06 | 58980001 | 59080000 | 0.0030 | 0.0032 | 0.0027 |
| F06 | 59000001 | 59100000 | 0.0039 | 0.0027 | 0.0035 |
| F06 | 59020001 | 59120000 | 0.0032 | 0.0022 | 0.0029 |
| F06 | 59040001 | 59140000 | 0.0027 | 0.0022 | 0.0024 |
| F06 | 59060001 | 59160000 | 0.0027 | 0.0023 | 0.0022 |
| F06 | 59080001 | 59180000 | 0.0029 | 0.0019 | 0.0024 |
| F06 | 59100001 | 59200000 | 0.0025 | 0.0025 | 0.0018 |
| F06 | 59120001 | 59220000 | 0.0033 | 0.0024 | 0.0025 |
| F06 | 59140001 | 59240000 | 0.0032 | 0.0020 | 0.0023 |
| F06 | 59160001 | 59260000 | 0.0029 | 0.0022 | 0.0019 |
| F06 | 59180001 | 59280000 | 0.0031 | 0.0019 | 0.0021 |
| F06 | 59200001 | 59300000 | 0.0028 | 0.0016 | 0.0018 |
| F06 | 59220001 | 59320000 | 0.0027 | 0.0015 | 0.0016 |
| F06 | 59240001 | 59340000 | 0.0026 | 0.0016 | 0.0017 |
| F06 | 59260001 | 59360000 | 0.0026 | 0.0018 | 0.0018 |
| F06 | 59280001 | 59380000 | 0.0030 | 0.0017 | 0.0020 |
| F06 | 59300001 | 59400000 | 0.0027 | 0.0018 | 0.0018 |
| F06 | 59320001 | 59420000 | 0.0029 | 0.0025 | 0.0021 |
| F06 | 59340001 | 59440000 | 0.0035 | 0.0024 | 0.0026 |
| F06 | 59360001 | 59460000 | 0.0033 | 0.0019 | 0.0026 |
| F06 | 59380001 | 59480000 | 0.0026 | 0.0026 | 0.0021 |
| F06 | 59400001 | 59500000 | 0.0034 | 0.0027 | 0.0029 |
| F06 | 59420001 | 59520000 | 0.0031 | 0.0025 | 0.0028 |
| F06 | 59440001 | 59540000 | 0.0030 | 0.0028 | 0.0028 |
| F06 | 59460001 | 59560000 | 0.0032 | 0.0035 | 0.0030 |
| F06 | 59480001 | 59580000 | 0.0040 | 0.0026 | 0.0039 |
| F06 | 59500001 | 59600000 | 0.0029 | 0.0021 | 0.0029 |
| F06 | 59520001 | 59620000 | 0.0025 | 0.0023 | 0.0024 |
| F06 | 59540001 | 59640000 | 0.0029 | 0.0021 | 0.0025 |
| F06 | 59560001 | 59660000 | 0.0027 | 0.0017 | 0.0023 |
| F06 | 59580001 | 59680000 | 0.0023 | 0.0024 | 0.0018 |
| F06 | 59600001 | 59700000 | 0.0033 | 0.0024 | 0.0028 |
| F06 | 59620001 | 59720000 | 0.0032 | 0.0015 | 0.0028 |
| F06 | 59640001 | 59740000 | 0.0022 | 0.0014 | 0.0019 |
| F06 | 59660001 | 59760000 | 0.0022 | 0.0016 | 0.0018 |
| F06 | 59680001 | 59780000 | 0.0022 | 0.0013 | 0.0020 |
| F06 | 59700001 | 59800000 | 0.0018 | 0.0014 | 0.0013 |
| F06 | 59720001 | 59820000 | 0.0019 | 0.0020 | 0.0014 |
| F06 | 59740001 | 59840000 | 0.0025 | 0.0023 | 0.0019 |
| F06 | 59760001 | 59860000 | 0.0029 | 0.0020 | 0.0024 |
| F06 | 59780001 | 59880000 | 0.0029 | 0.0016 | 0.0021 |
| F06 | 59800001 | 59900000 | 0.0024 | 0.0019 | 0.0018 |
| F06 | 59820001 | 59920000 | 0.0026 | 0.0025 | 0.0020 |
| F06 | 59840001 | 59940000 | 0.0032 | 0.0026 | 0.0024 |
| F06 | 59860001 | 59960000 | 0.0031 | 0.0029 | 0.0024 |
| F06 | 59880001 | 59980000 | 0.0031 | 0.0032 | 0.0025 |
| F06 | 59900001 | 60000000 | 0.0033 | 0.0034 | 0.0027 |
| F06 | 59920001 | 60020000 | 0.0037 | 0.0026 | 0.0031 |
| F06 | 59940001 | 60040000 | 0.0028 | 0.0028 | 0.0023 |
| F06 | 59960001 | 60060000 | 0.0030 | 0.0024 | 0.0023 |
| F06 | 59980001 | 60080000 | 0.0025 | 0.0025 | 0.0018 |
| F06 | 60000001 | 60100000 | 0.0026 | 0.0027 | 0.0020 |
| F06 | 60020001 | 60120000 | 0.0027 | 0.0027 | 0.0020 |
| F06 | 60040001 | 60140000 | 0.0027 | 0.0022 | 0.0022 |
| F06 | 60060001 | 60160000 | 0.0023 | 0.0021 | 0.0019 |

| Chr | Start | End | both | A1 | A2 |
|---|---|---|---|---|---|
| F06 | 60080001 | 60180000 | 0.0022 | 0.0022 | 0.0019 |
| F06 | 60100001 | 60200000 | 0.0022 | 0.0022 | 0.0018 |
| F06 | 60120001 | 60220000 | 0.0023 | 0.0039 | 0.0019 |
| F06 | 60140001 | 60240000 | 0.0040 | 0.0045 | 0.0036 |
| F06 | 60160001 | 60260000 | 0.0047 | 0.0052 | 0.0043 |
| F06 | 60180001 | 60280000 | 0.0054 | 0.0054 | 0.0050 |
| F06 | 60200001 | 60300000 | 0.0058 | 0.0046 | 0.0054 |
| F06 | 60220001 | 60320000 | 0.0050 | 0.0033 | 0.0046 |
| F06 | 60240001 | 60340000 | 0.0037 | 0.0034 | 0.0033 |
| F06 | 60260001 | 60360000 | 0.0036 | 0.0035 | 0.0031 |
| F06 | 60280001 | 60380000 | 0.0034 | 0.0030 | 0.0029 |
| F06 | 60300001 | 60400000 | 0.0028 | 0.0028 | 0.0024 |
| F06 | 60320001 | 60420000 | 0.0026 | 0.0025 | 0.0022 |
| F06 | 60340001 | 60440000 | 0.0023 | 0.0020 | 0.0019 |
| F06 | 60360001 | 60460000 | 0.0021 | 0.0036 | 0.0019 |
| F06 | 60380001 | 60480000 | 0.0039 | 0.0036 | 0.0036 |
| F06 | 60400001 | 60500000 | 0.0039 | 0.0041 | 0.0037 |
| F06 | 60420001 | 60520000 | 0.0045 | 0.0040 | 0.0043 |
| F06 | 60440001 | 60540000 | 0.0043 | 0.0038 | 0.0042 |
| F06 | 60460001 | 60560000 | 0.0040 | 0.0017 | 0.0039 |
| F06 | 60480001 | 60580000 | 0.0020 | 0.0025 | 0.0019 |
| F06 | 60500001 | 60600000 | 0.0027 | 0.0021 | 0.0025 |
| F06 | 60520001 | 60620000 | 0.0022 | 0.0024 | 0.0020 |
| F06 | 60540001 | 60640000 | 0.0025 | 0.0026 | 0.0022 |
| F06 | 60560001 | 60660000 | 0.0028 | 0.0023 | 0.0024 |
| F06 | 60580001 | 60680000 | 0.0025 | 0.0020 | 0.0023 |
| F06 | 60600001 | 60700000 | 0.0024 | 0.0025 | 0.0021 |
| F06 | 60620001 | 60720000 | 0.0027 | 0.0024 | 0.0025 |
| F06 | 60640001 | 60740000 | 0.0028 | 0.0021 | 0.0026 |
| F06 | 60660001 | 60760000 | 0.0023 | 0.0023 | 0.0022 |
| F06 | 60680001 | 60780000 | 0.0025 | 0.0024 | 0.0022 |
| F06 | 60700001 | 60800000 | 0.0024 | 0.0025 | 0.0020 |
| F06 | 60720001 | 60820000 | 0.0028 | 0.0024 | 0.0024 |
| F06 | 60740001 | 60840000 | 0.0026 | 0.0030 | 0.0022 |
| F06 | 60760001 | 60860000 | 0.0033 | 0.0032 | 0.0028 |
| F06 | 60780001 | 60880000 | 0.0035 | 0.0025 | 0.0031 |
| F06 | 60800001 | 60900000 | 0.0028 | 0.0030 | 0.0027 |
| F06 | 60820001 | 60920000 | 0.0032 | 0.0031 | 0.0030 |
| F06 | 60840001 | 60940000 | 0.0034 | 0.0029 | 0.0032 |
| F06 | 60860001 | 60960000 | 0.0031 | 0.0030 | 0.0029 |
| F06 | 60880001 | 60980000 | 0.0031 | 0.0039 | 0.0027 |
| F06 | 60900001 | 61000000 | 0.0039 | 0.0037 | 0.0034 |
| F06 | 60920001 | 61020000 | 0.0038 | 0.0036 | 0.0034 |
| F06 | 60940001 | 61040000 | 0.0037 | 0.0034 | 0.0033 |
| F06 | 60960001 | 61060000 | 0.0036 | 0.0035 | 0.0032 |
| F06 | 60980001 | 61080000 | 0.0039 | 0.0027 | 0.0038 |
| F06 | 61000001 | 61100000 | 0.0032 | 0.0016 | 0.0031 |
| F06 | 61020001 | 61120000 | 0.0019 | 0.0019 | 0.0019 |
| F06 | 61040001 | 61140000 | 0.0022 | 0.0023 | 0.0021 |
| F06 | 61060001 | 61160000 | 0.0025 | 0.0020 | 0.0022 |
| F06 | 61080001 | 61180000 | 0.0021 | 0.0026 | 0.0019 |
| F06 | 61100001 | 61200000 | 0.0028 | 0.0028 | 0.0024 |
| F06 | 61120001 | 61220000 | 0.0030 | 0.0026 | 0.0026 |
| F06 | 61140001 | 61240000 | 0.0026 | 0.0026 | 0.0021 |
| F06 | 61160001 | 61260000 | 0.0028 | 0.0030 | 0.0024 |
| F06 | 61180001 | 61280000 | 0.0033 | 0.0037 | 0.0029 |
| F06 | 61200001 | 61300000 | 0.0042 | 0.0039 | 0.0038 |
| F06 | 61220001 | 61320000 | 0.0044 | 0.0036 | 0.0041 |

| Chr | Start | End | both | A1 | A2 |
|---|---|---|---|---|---|
| F06 | 61240001 | 61340000 | 0.0042 | 0.0031 | 0.0041 |
| F06 | 61260001 | 61360000 | 0.0036 | 0.0026 | 0.0035 |
| F06 | 61280001 | 61380000 | 0.0030 | 0.0013 | 0.0029 |
| F06 | 61300001 | 61400000 | 0.0015 | 0.0013 | 0.0014 |
| F06 | 61320001 | 61420000 | 0.0015 | 0.0028 | 0.0015 |
| F06 | 61340001 | 61440000 | 0.0030 | 0.0035 | 0.0028 |
| F06 | 61360001 | 61460000 | 0.0037 | 0.0038 | 0.0034 |
| F06 | 61380001 | 61480000 | 0.0041 | 0.0042 | 0.0038 |
| F06 | 61400001 | 61500000 | 0.0046 | 0.0039 | 0.0043 |
| F06 | 61420001 | 61520000 | 0.0041 | 0.0025 | 0.0038 |
| F06 | 61440001 | 61540000 | 0.0028 | 0.0023 | 0.0026 |
| F06 | 61460001 | 61560000 | 0.0026 | 0.0023 | 0.0025 |
| F06 | 61480001 | 61580000 | 0.0025 | 0.0023 | 0.0022 |
| F06 | 61500001 | 61600000 | 0.0025 | 0.0024 | 0.0022 |
| F06 | 61520001 | 61620000 | 0.0027 | 0.0028 | 0.0024 |
| F06 | 61540001 | 61640000 | 0.0031 | 0.0024 | 0.0028 |
| F06 | 61560001 | 61660000 | 0.0028 | 0.0023 | 0.0025 |
| F06 | 61580001 | 61680000 | 0.0027 | 0.0023 | 0.0025 |
| F06 | 61600001 | 61700000 | 0.0026 | 0.0023 | 0.0023 |
| F06 | 61620001 | 61720000 | 0.0026 | 0.0017 | 0.0024 |
| F06 | 61640001 | 61740000 | 0.0019 | 0.0018 | 0.0017 |
| F06 | 61660001 | 61760000 | 0.0019 | 0.0015 | 0.0016 |
| F06 | 61680001 | 61780000 | 0.0015 | 0.0016 | 0.0013 |
| F06 | 61700001 | 61800000 | 0.0017 | 0.0017 | 0.0015 |
| F06 | 61720001 | 61820000 | 0.0019 | 0.0018 | 0.0017 |
| F06 | 61740001 | 61840000 | 0.0020 | 0.0013 | 0.0018 |
| F06 | 61760001 | 61860000 | 0.0015 | 0.0011 | 0.0014 |
| F06 | 61780001 | 61880000 | 0.0012 | 0.0008 | 0.0012 |
| F06 | 61800001 | 61900000 | 0.0010 | 0.0009 | 0.0010 |
| F06 | 61820001 | 61920000 | 0.0012 | 0.0011 | 0.0011 |
| F06 | 61840001 | 61940000 | 0.0013 | 0.0013 | 0.0013 |
| F06 | 61860001 | 61960000 | 0.0015 | 0.0014 | 0.0015 |
| F06 | 61880001 | 61980000 | 0.0017 | 0.0012 | 0.0016 |
| F06 | 61900001 | 62000000 | 0.0015 | 0.0009 | 0.0014 |
| F06 | 61920001 | 62020000 | 0.0010 | 0.0010 | 0.0009 |
| F06 | 61940001 | 62040000 | 0.0012 | 0.0012 | 0.0010 |
| F06 | 61960001 | 62060000 | 0.0013 | 0.0013 | 0.0011 |
| F06 | 61980001 | 62080000 | 0.0014 | 0.0017 | 0.0013 |
| F06 | 62000001 | 62100000 | 0.0018 | 0.0018 | 0.0016 |
| F06 | 62020001 | 62120000 | 0.0019 | 0.0023 | 0.0017 |
| F06 | 62040001 | 62140000 | 0.0024 | 0.0036 | 0.0022 |
| F06 | 62060001 | 62160000 | 0.0041 | 0.0039 | 0.0039 |
| F06 | 62080001 | 62180000 | 0.0043 | 0.0037 | 0.0041 |
| F06 | 62100001 | 62200000 | 0.0042 | 0.0035 | 0.0041 |
| F06 | 62120001 | 62220000 | 0.0040 | 0.0029 | 0.0040 |
| F06 | 62140001 | 62240000 | 0.0033 | 0.0015 | 0.0032 |
| F06 | 62160001 | 62260000 | 0.0016 | 0.0013 | 0.0015 |
| F06 | 62180001 | 62280000 | 0.0013 | 0.0013 | 0.0010 |
| F06 | 62200001 | 62300000 | 0.0012 | 0.0022 | 0.0008 |
| F06 | 62220001 | 62320000 | 0.0023 | 0.0031 | 0.0019 |
| F06 | 62240001 | 62340000 | 0.0032 | 0.0035 | 0.0027 |
| F06 | 62260001 | 62360000 | 0.0035 | 0.0037 | 0.0029 |
| F06 | 62280001 | 62380000 | 0.0038 | 0.0034 | 0.0034 |
| F06 | 62300001 | 62400000 | 0.0037 | 0.0023 | 0.0033 |
| F06 | 62320001 | 62420000 | 0.0024 | 0.0014 | 0.0021 |
| F06 | 62340001 | 62440000 | 0.0015 | 0.0008 | 0.0012 |
| F06 | 62360001 | 62460000 | 0.0009 | 0.0007 | 0.0009 |
| F06 | 62380001 | 62480000 | 0.0007 | 0.0012 | 0.0007 |

| Chr | Start | End | both | A1 | A2 |
|---|---|---|---|---|---|
| F06 | 62400001 | 62500000 | 0.0014 | 0.0016 | 0.0012 |
| F06 | 62420001 | 62520000 | 0.0020 | 0.0015 | 0.0017 |
| F06 | 62440001 | 62540000 | 0.0020 | 0.0022 | 0.0016 |
| F06 | 62460001 | 62560000 | 0.0029 | 0.0022 | 0.0025 |
| F06 | 62480001 | 62580000 | 0.0029 | 0.0024 | 0.0025 |
| F06 | 62500001 | 62600000 | 0.0031 | 0.0020 | 0.0028 |
| F06 | 62520001 | 62620000 | 0.0027 | 0.0026 | 0.0024 |
| F06 | 62540001 | 62640000 | 0.0033 | 0.0019 | 0.0030 |
| F06 | 62560001 | 62660000 | 0.0023 | 0.0017 | 0.0022 |
| F06 | 62580001 | 62680000 | 0.0021 | 0.0011 | 0.0020 |
| F06 | 62600001 | 62700000 | 0.0014 | 0.0012 | 0.0013 |
| F06 | 62620001 | 62720000 | 0.0014 | 0.0012 | 0.0014 |
| F06 | 62640001 | 62740000 | 0.0014 | 0.0014 | 0.0013 |
| F06 | 62660001 | 62760000 | 0.0017 | 0.0016 | 0.0016 |
| F06 | 62680001 | 62780000 | 0.0021 | 0.0021 | 0.0019 |
| F06 | 62700001 | 62800000 | 0.0026 | 0.0023 | 0.0023 |
| F06 | 62720001 | 62820000 | 0.0031 | 0.0022 | 0.0025 |
| F06 | 62740001 | 62840000 | 0.0032 | 0.0020 | 0.0024 |
| F06 | 62760001 | 62860000 | 0.0029 | 0.0021 | 0.0022 |
| F06 | 62780001 | 62880000 | 0.0027 | 0.0016 | 0.0021 |
| F06 | 62800001 | 62900000 | 0.0022 | 0.0016 | 0.0017 |
| F06 | 62820001 | 62920000 | 0.0020 | 0.0013 | 0.0017 |
| F06 | 62840001 | 62940000 | 0.0014 | 0.0015 | 0.0013 |
| F06 | 62860001 | 62960000 | 0.0016 | 0.0015 | 0.0015 |
| F06 | 62880001 | 62980000 | 0.0017 | 0.0013 | 0.0014 |
| F06 | 62900001 | 63000000 | 0.0015 | 0.0014 | 0.0012 |
| F06 | 62920001 | 63020000 | 0.0016 | 0.0016 | 0.0012 |
| F06 | 62940001 | 63040000 | 0.0020 | 0.0015 | 0.0016 |
| F06 | 62960001 | 63060000 | 0.0018 | 0.0017 | 0.0015 |
| F06 | 62980001 | 63080000 | 0.0021 | 0.0018 | 0.0020 |
| F06 | 63000001 | 63100000 | 0.0022 | 0.0015 | 0.0020 |
| F06 | 63020001 | 63120000 | 0.0019 | 0.0013 | 0.0018 |
| F06 | 63040001 | 63140000 | 0.0017 | 0.0013 | 0.0015 |
| F06 | 63060001 | 63160000 | 0.0018 | 0.0009 | 0.0015 |
| F06 | 63080001 | 63180000 | 0.0015 | 0.0015 | 0.0010 |
| F06 | 63100001 | 63200000 | 0.0022 | 0.0016 | 0.0016 |
| F06 | 63120001 | 63220000 | 0.0023 | 0.0019 | 0.0016 |
| F06 | 63140001 | 63240000 | 0.0026 | 0.0018 | 0.0019 |
| F06 | 63160001 | 63260000 | 0.0025 | 0.0021 | 0.0018 |
| F06 | 63180001 | 63280000 | 0.0027 | 0.0016 | 0.0022 |
| F06 | 63200001 | 63300000 | 0.0022 | 0.0015 | 0.0016 |
| F06 | 63220001 | 63320000 | 0.0021 | 0.0012 | 0.0015 |
| F06 | 63240001 | 63340000 | 0.0017 | 0.0011 | 0.0012 |
| F06 | 63260001 | 63360000 | 0.0016 | 0.0007 | 0.0012 |
| F06 | 63280001 | 63380000 | 0.0010 | 0.0008 | 0.0009 |
| F06 | 63300001 | 63400000 | 0.0009 | 0.0006 | 0.0009 |
| F06 | 63320001 | 63420000 | 0.0007 | 0.0005 | 0.0007 |
| F06 | 63340001 | 63440000 | 0.0007 | 0.0010 | 0.0007 |
| F06 | 63360001 | 63460000 | 0.0011 | 0.0011 | 0.0010 |
| F06 | 63380001 | 63480000 | 0.0012 | 0.0010 | 0.0012 |
| F06 | 63400001 | 63500000 | 0.0012 | 0.0011 | 0.0011 |
| F06 | 63420001 | 63520000 | 0.0013 | 0.0009 | 0.0013 |
| F06 | 63440001 | 63540000 | 0.0011 | 0.0012 | 0.0011 |
| F06 | 63460001 | 63560000 | 0.0014 | 0.0012 | 0.0014 |
| F06 | 63480001 | 63580000 | 0.0014 | 0.0022 | 0.0013 |
| F06 | 63500001 | 63600000 | 0.0026 | 0.0032 | 0.0024 |
| F06 | 63520001 | 63620000 | 0.0039 | 0.0032 | 0.0035 |
| F06 | 63540001 | 63640000 | 0.0040 | 0.0027 | 0.0036 |

| Chr | Start | End | both | A1 | A2 |
|---|---|---|---|---|---|
| F06 | 63560001 | 63660000 | 0.0035 | 0.0027 | 0.0030 |
| F06 | 63580001 | 63680000 | 0.0033 | 0.0018 | 0.0029 |
| F06 | 63600001 | 63700000 | 0.0023 | 0.0009 | 0.0021 |
| F06 | 63620001 | 63720000 | 0.0012 | 0.0011 | 0.0010 |
| F06 | 63640001 | 63740000 | 0.0013 | 0.0017 | 0.0012 |
| F06 | 63660001 | 63760000 | 0.0019 | 0.0025 | 0.0017 |
| F06 | 63680001 | 63780000 | 0.0027 | 0.0026 | 0.0024 |
| F06 | 63700001 | 63800000 | 0.0029 | 0.0026 | 0.0027 |
| F06 | 63720001 | 63820000 | 0.0031 | 0.0024 | 0.0027 |
| F06 | 63740001 | 63840000 | 0.0029 | 0.0022 | 0.0025 |
| F06 | 63760001 | 63860000 | 0.0028 | 0.0016 | 0.0024 |
| F06 | 63780001 | 63880000 | 0.0025 | 0.0018 | 0.0020 |
| F06 | 63800001 | 63900000 | 0.0025 | 0.0017 | 0.0020 |
| F06 | 63820001 | 63920000 | 0.0023 | 0.0017 | 0.0019 |
| F06 | 63840001 | 63940000 | 0.0023 | 0.0013 | 0.0019 |
| F06 | 63860001 | 63960000 | 0.0017 | 0.0013 | 0.0015 |
| F06 | 63880001 | 63980000 | 0.0014 | 0.0008 | 0.0012 |
| F06 | 63900001 | 64000000 | 0.0010 | 0.0010 | 0.0010 |
| F06 | 63920001 | 64020000 | 0.0012 | 0.0013 | 0.0011 |
| F06 | 63940001 | 64040000 | 0.0016 | 0.0013 | 0.0015 |
| F06 | 63960001 | 64060000 | 0.0015 | 0.0018 | 0.0014 |
| F06 | 63980001 | 64080000 | 0.0020 | 0.0019 | 0.0019 |
| F06 | 64000001 | 64100000 | 0.0023 | 0.0016 | 0.0019 |
| F06 | 64020001 | 64120000 | 0.0020 | 0.0018 | 0.0016 |
| F06 | 64040001 | 64140000 | 0.0021 | 0.0021 | 0.0018 |
| F06 | 64060001 | 64160000 | 0.0025 | 0.0016 | 0.0020 |
| F06 | 64080001 | 64180000 | 0.0022 | 0.0017 | 0.0017 |
| F06 | 64100001 | 64200000 | 0.0022 | 0.0021 | 0.0018 |
| F06 | 64120001 | 64220000 | 0.0027 | 0.0016 | 0.0022 |
| F06 | 64140001 | 64240000 | 0.0022 | 0.0012 | 0.0017 |
| F06 | 64160001 | 64260000 | 0.0017 | 0.0013 | 0.0013 |
| F06 | 64180001 | 64280000 | 0.0016 | 0.0014 | 0.0014 |
| F06 | 64200001 | 64300000 | 0.0017 | 0.0017 | 0.0014 |
| F06 | 64220001 | 64320000 | 0.0021 | 0.0016 | 0.0018 |
| F06 | 64240001 | 64340000 | 0.0020 | 0.0019 | 0.0017 |
| F06 | 64260001 | 64360000 | 0.0023 | 0.0025 | 0.0020 |
| F06 | 64280001 | 64380000 | 0.0030 | 0.0022 | 0.0026 |
| F06 | 64300001 | 64400000 | 0.0028 | 0.0017 | 0.0023 |
| F06 | 64320001 | 64420000 | 0.0022 | 0.0019 | 0.0016 |
| F06 | 64340001 | 64440000 | 0.0025 | 0.0021 | 0.0019 |
| F06 | 64360001 | 64460000 | 0.0028 | 0.0012 | 0.0021 |
| F06 | 64380001 | 64480000 | 0.0018 | 0.0011 | 0.0013 |
| F06 | 64400001 | 64500000 | 0.0015 | 0.0013 | 0.0012 |
| F06 | 64420001 | 64520000 | 0.0015 | 0.0012 | 0.0014 |
| F06 | 64440001 | 64540000 | 0.0014 | 0.0009 | 0.0013 |
| F06 | 64460001 | 64560000 | 0.0010 | 0.0012 | 0.0010 |
| F06 | 64480001 | 64580000 | 0.0015 | 0.0014 | 0.0013 |
| F06 | 64500001 | 64600000 | 0.0018 | 0.0012 | 0.0015 |
| F06 | 64520001 | 64620000 | 0.0018 | 0.0016 | 0.0014 |
| F06 | 64540001 | 64640000 | 0.0022 | 0.0021 | 0.0017 |
| F06 | 64560001 | 64660000 | 0.0029 | 0.0019 | 0.0022 |
| F06 | 64580001 | 64680000 | 0.0027 | 0.0020 | 0.0021 |
| F06 | 64600001 | 64700000 | 0.0030 | 0.0026 | 0.0022 |
| F06 | 64620001 | 64720000 | 0.0034 | 0.0023 | 0.0027 |
| F06 | 64640001 | 64740000 | 0.0032 | 0.0023 | 0.0026 |
| F06 | 64660001 | 64760000 | 0.0031 | 0.0029 | 0.0025 |
| F06 | 64680001 | 64780000 | 0.0038 | 0.0037 | 0.0032 |
| F06 | 64700001 | 64800000 | 0.0047 | 0.0033 | 0.0042 |

| Chr | Start | End | both | A1 | A2 |
|---|---|---|---|---|---|
| F06 | 64720001 | 64820000 | 0.0043 | 0.0033 | 0.0040 |
| F06 | 64740001 | 64840000 | 0.0043 | 0.0029 | 0.0038 |
| F06 | 64760001 | 64860000 | 0.0039 | 0.0022 | 0.0034 |
| F06 | 64780001 | 64880000 | 0.0030 | 0.0020 | 0.0027 |
| F06 | 64800001 | 64900000 | 0.0026 | 0.0021 | 0.0022 |
| F06 | 64820001 | 64920000 | 0.0026 | 0.0020 | 0.0022 |
| F06 | 64840001 | 64940000 | 0.0025 | 0.0019 | 0.0021 |
| F06 | 64860001 | 64960000 | 0.0023 | 0.0028 | 0.0020 |
| F06 | 64880001 | 64980000 | 0.0034 | 0.0024 | 0.0031 |
| F06 | 64900001 | 65000000 | 0.0029 | 0.0021 | 0.0027 |
| F06 | 64920001 | 65020000 | 0.0027 | 0.0022 | 0.0024 |
| F06 | 64940001 | 65040000 | 0.0031 | 0.0024 | 0.0026 |
| F06 | 64960001 | 65060000 | 0.0034 | 0.0022 | 0.0029 |
| F06 | 64980001 | 65080000 | 0.0032 | 0.0020 | 0.0026 |
| F06 | 65000001 | 65100000 | 0.0033 | 0.0023 | 0.0025 |
| F06 | 65020001 | 65120000 | 0.0034 | 0.0023 | 0.0027 |
| F06 | 65040001 | 65140000 | 0.0030 | 0.0026 | 0.0025 |
| F06 | 65060001 | 65160000 | 0.0033 | 0.0024 | 0.0028 |
| F06 | 65080001 | 65180000 | 0.0033 | 0.0028 | 0.0029 |
| F06 | 65100001 | 65200000 | 0.0035 | 0.0025 | 0.0033 |
| F06 | 65120001 | 65220000 | 0.0039 | 0.0027 | 0.0035 |
| F06 | 65140001 | 65240000 | 0.0041 | 0.0026 | 0.0036 |
| F06 | 65160001 | 65260000 | 0.0042 | 0.0021 | 0.0036 |
| F06 | 65180001 | 65280000 | 0.0038 | 0.0020 | 0.0028 |
| F06 | 65200001 | 65300000 | 0.0041 | 0.0027 | 0.0027 |
| F06 | 65220001 | 65320000 | 0.0044 | 0.0028 | 0.0029 |
| F06 | 65240001 | 65340000 | 0.0047 | 0.0027 | 0.0031 |
| F06 | 65260001 | 65360000 | 0.0044 | 0.0029 | 0.0029 |
| F06 | 65280001 | 65380000 | 0.0043 | 0.0028 | 0.0031 |
| F06 | 65300001 | 65400000 | 0.0039 | 0.0027 | 0.0030 |
| F06 | 65320001 | 65420000 | 0.0036 | 0.0029 | 0.0029 |
| F06 | 65340001 | 65440000 | 0.0037 | 0.0030 | 0.0030 |
| F06 | 65360001 | 65460000 | 0.0036 | 0.0030 | 0.0031 |
| F06 | 65380001 | 65480000 | 0.0039 | 0.0035 | 0.0032 |
| F06 | 65400001 | 65500000 | 0.0046 | 0.0031 | 0.0038 |
| F06 | 65420001 | 65520000 | 0.0042 | 0.0032 | 0.0033 |
| F06 | 65440001 | 65540000 | 0.0046 | 0.0031 | 0.0035 |
| F06 | 65460001 | 65560000 | 0.0045 | 0.0030 | 0.0034 |
| F06 | 65480001 | 65580000 | 0.0045 | 0.0033 | 0.0033 |
| F06 | 65500001 | 65600000 | 0.0047 | 0.0035 | 0.0035 |
| F06 | 65520001 | 65620000 | 0.0049 | 0.0031 | 0.0038 |
| F06 | 65540001 | 65640000 | 0.0042 | 0.0029 | 0.0035 |
| F06 | 65560001 | 65660000 | 0.0042 | 0.0035 | 0.0033 |
| F06 | 65580001 | 65680000 | 0.0048 | 0.0023 | 0.0038 |
| F06 | 65600001 | 65700000 | 0.0034 | 0.0025 | 0.0026 |
| F06 | 65620001 | 65720000 | 0.0036 | 0.0031 | 0.0028 |
| F06 | 65640001 | 65740000 | 0.0042 | 0.0043 | 0.0034 |
| F06 | 65660001 | 65760000 | 0.0050 | 0.0035 | 0.0044 |
| F06 | 65680001 | 65780000 | 0.0040 | 0.0038 | 0.0036 |
| F06 | 65700001 | 65800000 | 0.0045 | 0.0042 | 0.0041 |
| F06 | 65720001 | 65820000 | 0.0050 | 0.0031 | 0.0046 |
| F06 | 65740001 | 65840000 | 0.0041 | 0.0024 | 0.0036 |
| F06 | 65760001 | 65860000 | 0.0038 | 0.0027 | 0.0033 |
| F06 | 65780001 | 65880000 | 0.0039 | 0.0032 | 0.0035 |
| F06 | 65800001 | 65900000 | 0.0045 | 0.0032 | 0.0039 |
| F06 | 65820001 | 65920000 | 0.0045 | 0.0035 | 0.0037 |
| F06 | 65840001 | 65940000 | 0.0045 | 0.0032 | 0.0040 |
| F06 | 65860001 | 65960000 | 0.0042 | 0.0029 | 0.0036 |

| Chr | Start | End | both | A1 | A2 |
|---|---|---|---|---|---|
| F06 | 65880001 | 65980000 | 0.0038 | 0.0024 | 0.0032 |
| F06 | 65900001 | 66000000 | 0.0032 | 0.0022 | 0.0027 |
| F06 | 65920001 | 66020000 | 0.0030 | 0.0018 | 0.0025 |
| F06 | 65940001 | 66040000 | 0.0025 | 0.0013 | 0.0020 |
| F06 | 65960001 | 66060000 | 0.0018 | 0.0016 | 0.0015 |
| F06 | 65980001 | 66080000 | 0.0021 | 0.0015 | 0.0017 |
| F06 | 66000001 | 66100000 | 0.0019 | 0.0016 | 0.0015 |
| F06 | 66020001 | 66120000 | 0.0017 | 0.0021 | 0.0014 |
| F06 | 66040001 | 66140000 | 0.0023 | 0.0024 | 0.0020 |
| F06 | 66060001 | 66160000 | 0.0027 | 0.0024 | 0.0024 |
| F06 | 66080001 | 66180000 | 0.0025 | 0.0021 | 0.0023 |
| F06 | 66100001 | 66200000 | 0.0022 | 0.0014 | 0.0020 |
| F06 | 66120001 | 66220000 | 0.0017 | 0.0012 | 0.0016 |
| F06 | 66140001 | 66240000 | 0.0013 | 0.0014 | 0.0012 |
| F06 | 66160001 | 66260000 | 0.0015 | 0.0014 | 0.0014 |
| F06 | 66180001 | 66280000 | 0.0016 | 0.0014 | 0.0016 |
| F06 | 66200001 | 66300000 | 0.0017 | 0.0017 | 0.0017 |
| F06 | 66220001 | 66320000 | 0.0018 | 0.0020 | 0.0017 |
| F06 | 66240001 | 66340000 | 0.0022 | 0.0018 | 0.0020 |
| F06 | 66260001 | 66360000 | 0.0022 | 0.0021 | 0.0017 |
| F06 | 66280001 | 66380000 | 0.0026 | 0.0028 | 0.0020 |
| F06 | 66300001 | 66400000 | 0.0036 | 0.0036 | 0.0028 |
| F06 | 66320001 | 66420000 | 0.0047 | 0.0034 | 0.0039 |
| F06 | 66340001 | 66440000 | 0.0043 | 0.0032 | 0.0036 |
| F06 | 66360001 | 66460000 | 0.0039 | 0.0031 | 0.0035 |
| F06 | 66380001 | 66480000 | 0.0039 | 0.0029 | 0.0034 |
| F06 | 66400001 | 66500000 | 0.0036 | 0.0026 | 0.0032 |
| F06 | 66420001 | 66520000 | 0.0032 | 0.0035 | 0.0028 |
| F06 | 66440001 | 66540000 | 0.0046 | 0.0036 | 0.0041 |
| F06 | 66460001 | 66560000 | 0.0047 | 0.0035 | 0.0042 |
| F06 | 66480001 | 66580000 | 0.0047 | 0.0030 | 0.0041 |
| F06 | 66500001 | 66600000 | 0.0041 | 0.0037 | 0.0035 |
| F06 | 66520001 | 66620000 | 0.0047 | 0.0029 | 0.0041 |
| F06 | 66540001 | 66640000 | 0.0039 | 0.0028 | 0.0029 |
| F06 | 66560001 | 66660000 | 0.0037 | 0.0026 | 0.0028 |
| F06 | 66580001 | 66680000 | 0.0034 | 0.0036 | 0.0026 |
| F06 | 66600001 | 66700000 | 0.0045 | 0.0028 | 0.0037 |
| F06 | 66620001 | 66720000 | 0.0037 | 0.0025 | 0.0028 |
| F06 | 66640001 | 66740000 | 0.0033 | 0.0036 | 0.0025 |
| F06 | 66660001 | 66760000 | 0.0048 | 0.0038 | 0.0038 |
| F06 | 66680001 | 66780000 | 0.0049 | 0.0026 | 0.0040 |
| F06 | 66700001 | 66800000 | 0.0035 | 0.0028 | 0.0028 |
| F06 | 66720001 | 66820000 | 0.0036 | 0.0029 | 0.0030 |
| F06 | 66740001 | 66840000 | 0.0034 | 0.0033 | 0.0031 |
| F06 | 66760001 | 66860000 | 0.0037 | 0.0039 | 0.0035 |
| F06 | 66780001 | 66880000 | 0.0046 | 0.0043 | 0.0043 |
| F06 | 66800001 | 66900000 | 0.0052 | 0.0040 | 0.0048 |
| F06 | 66820001 | 66920000 | 0.0049 | 0.0047 | 0.0044 |
| F06 | 66840001 | 66940000 | 0.0058 | 0.0032 | 0.0052 |
| F06 | 66860001 | 66960000 | 0.0042 | 0.0021 | 0.0036 |
| F06 | 66880001 | 66980000 | 0.0029 | 0.0019 | 0.0024 |
| F06 | 66900001 | 67000000 | 0.0025 | 0.0025 | 0.0020 |
| F06 | 66920001 | 67020000 | 0.0031 | 0.0033 | 0.0028 |
| F06 | 66940001 | 67040000 | 0.0039 | 0.0032 | 0.0038 |
| F06 | 66960001 | 67060000 | 0.0039 | 0.0037 | 0.0037 |
| F06 | 66980001 | 67080000 | 0.0047 | 0.0047 | 0.0041 |
| F06 | 67000001 | 67100000 | 0.0059 | 0.0048 | 0.0051 |
| F06 | 67020001 | 67120000 | 0.0058 | 0.0032 | 0.0051 |

| Chr | Start | End | both | A1 | A2 |
|---|---|---|---|---|---|
| F06 | 67040001 | 67140000 | 0.0040 | 0.0037 | 0.0033 |
| F06 | 67060001 | 67160000 | 0.0046 | 0.0041 | 0.0038 |
| F06 | 67080001 | 67180000 | 0.0049 | 0.0035 | 0.0045 |
| F06 | 67100001 | 67200000 | 0.0041 | 0.0023 | 0.0038 |
| F06 | 67120001 | 67220000 | 0.0029 | 0.0024 | 0.0026 |
| F06 | 67140001 | 67240000 | 0.0032 | 0.0030 | 0.0028 |
| F06 | 67160001 | 67260000 | 0.0039 | 0.0023 | 0.0033 |
| F06 | 67180001 | 67280000 | 0.0032 | 0.0018 | 0.0024 |
| F06 | 67200001 | 67300000 | 0.0028 | 0.0018 | 0.0019 |
| F06 | 67220001 | 67320000 | 0.0028 | 0.0016 | 0.0019 |
| F06 | 67240001 | 67340000 | 0.0022 | 0.0010 | 0.0015 |
| F06 | 67260001 | 67360000 | 0.0015 | 0.0008 | 0.0010 |
| F06 | 67280001 | 67380000 | 0.0011 | 0.0014 | 0.0009 |
| F06 | 67300001 | 67400000 | 0.0018 | 0.0018 | 0.0015 |
| F06 | 67320001 | 67420000 | 0.0021 | 0.0027 | 0.0019 |
| F06 | 67340001 | 67440000 | 0.0032 | 0.0027 | 0.0028 |
| F06 | 67360001 | 67460000 | 0.0033 | 0.0032 | 0.0029 |
| F06 | 67380001 | 67480000 | 0.0039 | 0.0031 | 0.0034 |
| F06 | 67400001 | 67500000 | 0.0036 | 0.0038 | 0.0033 |
| F06 | 67420001 | 67520000 | 0.0044 | 0.0029 | 0.0040 |
| F06 | 67440001 | 67540000 | 0.0035 | 0.0028 | 0.0032 |
| F06 | 67460001 | 67560000 | 0.0037 | 0.0027 | 0.0033 |
| F06 | 67480001 | 67580000 | 0.0037 | 0.0024 | 0.0033 |
| F06 | 67500001 | 67600000 | 0.0036 | 0.0017 | 0.0030 |
| F06 | 67520001 | 67620000 | 0.0027 | 0.0019 | 0.0022 |
| F06 | 67540001 | 67640000 | 0.0029 | 0.0018 | 0.0025 |
| F06 | 67560001 | 67660000 | 0.0025 | 0.0014 | 0.0020 |
| F06 | 67580001 | 67680000 | 0.0023 | 0.0014 | 0.0016 |
| F06 | 67600001 | 67700000 | 0.0022 | 0.0020 | 0.0016 |
| F06 | 67620001 | 67720000 | 0.0028 | 0.0027 | 0.0021 |
| F06 | 67640001 | 67740000 | 0.0037 | 0.0034 | 0.0030 |
| F06 | 67660001 | 67760000 | 0.0046 | 0.0046 | 0.0038 |
| F06 | 67680001 | 67780000 | 0.0056 | 0.0050 | 0.0051 |
| F06 | 67700001 | 67800000 | 0.0059 | 0.0044 | 0.0054 |
| F06 | 67720001 | 67820000 | 0.0052 | 0.0042 | 0.0049 |
| F06 | 67740001 | 67840000 | 0.0051 | 0.0037 | 0.0045 |
| F06 | 67760001 | 67860000 | 0.0044 | 0.0025 | 0.0039 |
| F06 | 67780001 | 67880000 | 0.0030 | 0.0029 | 0.0026 |
| F06 | 67800001 | 67900000 | 0.0035 | 0.0036 | 0.0029 |
| F06 | 67820001 | 67920000 | 0.0043 | 0.0035 | 0.0036 |
| F06 | 67840001 | 67940000 | 0.0040 | 0.0036 | 0.0035 |
| F06 | 67860001 | 67960000 | 0.0044 | 0.0051 | 0.0037 |
| F06 | 67880001 | 67980000 | 0.0059 | 0.0051 | 0.0052 |
| F06 | 67900001 | 68000000 | 0.0061 | 0.0045 | 0.0054 |
| F06 | 67920001 | 68020000 | 0.0054 | 0.0041 | 0.0049 |
| F06 | 67940001 | 68040000 | 0.0052 | 0.0042 | 0.0045 |
| F06 | 67960001 | 68060000 | 0.0051 | 0.0027 | 0.0045 |
| F06 | 67980001 | 68080000 | 0.0036 | 0.0019 | 0.0031 |
| F06 | 68000001 | 68100000 | 0.0029 | 0.0021 | 0.0022 |
| F06 | 68020001 | 68120000 | 0.0034 | 0.0030 | 0.0025 |
| F06 | 68040001 | 68140000 | 0.0043 | 0.0035 | 0.0035 |
| F06 | 68060001 | 68160000 | 0.0047 | 0.0042 | 0.0040 |
| F06 | 68080001 | 68180000 | 0.0057 | 0.0042 | 0.0048 |
| F06 | 68100001 | 68200000 | 0.0056 | 0.0041 | 0.0049 |
| F06 | 68120001 | 68220000 | 0.0055 | 0.0031 | 0.0047 |
| F06 | 68140001 | 68240000 | 0.0043 | 0.0021 | 0.0035 |
| F06 | 68160001 | 68260000 | 0.0032 | 0.0016 | 0.0025 |
| F06 | 68180001 | 68280000 | 0.0024 | 0.0016 | 0.0017 |

| Chr | Start | End | both | A1 | A2 |
|---|---|---|---|---|---|
| F06 | 68200001 | 68300000 | 0.0023 | 0.0014 | 0.0017 |
| F06 | 68220001 | 68320000 | 0.0021 | 0.0018 | 0.0016 |
| F06 | 68240001 | 68340000 | 0.0028 | 0.0018 | 0.0020 |
| F06 | 68260001 | 68360000 | 0.0030 | 0.0016 | 0.0020 |
| F06 | 68280001 | 68380000 | 0.0028 | 0.0018 | 0.0019 |
| F06 | 68300001 | 68400000 | 0.0030 | 0.0021 | 0.0020 |
| F06 | 68320001 | 68420000 | 0.0033 | 0.0022 | 0.0024 |
| F06 | 68340001 | 68440000 | 0.0031 | 0.0021 | 0.0023 |
| F06 | 68360001 | 68460000 | 0.0031 | 0.0020 | 0.0022 |
| F06 | 68380001 | 68480000 | 0.0030 | 0.0017 | 0.0021 |
| F06 | 68400001 | 68500000 | 0.0025 | 0.0030 | 0.0019 |
| F06 | 68420001 | 68520000 | 0.0039 | 0.0028 | 0.0032 |
| F06 | 68440001 | 68540000 | 0.0039 | 0.0033 | 0.0030 |
| F06 | 68460001 | 68560000 | 0.0043 | 0.0033 | 0.0035 |
| F06 | 68480001 | 68580000 | 0.0046 | 0.0039 | 0.0035 |
| F06 | 68500001 | 68600000 | 0.0052 | 0.0024 | 0.0038 |
| F06 | 68520001 | 68620000 | 0.0037 | 0.0026 | 0.0022 |
| F06 | 68540001 | 68640000 | 0.0037 | 0.0022 | 0.0024 |
| F06 | 68560001 | 68660000 | 0.0032 | 0.0024 | 0.0020 |
| F06 | 68580001 | 68680000 | 0.0034 | 0.0019 | 0.0022 |
| F06 | 68600001 | 68700000 | 0.0029 | 0.0016 | 0.0020 |
| F06 | 68620001 | 68720000 | 0.0023 | 0.0013 | 0.0016 |
| F06 | 68640001 | 68740000 | 0.0019 | 0.0012 | 0.0012 |
| F06 | 68660001 | 68760000 | 0.0016 | 0.0011 | 0.0011 |
| F06 | 68680001 | 68780000 | 0.0013 | 0.0007 | 0.0010 |
| F06 | 68700001 | 68800000 | 0.0009 | 0.0009 | 0.0006 |
| F06 | 68720001 | 68820000 | 0.0011 | 0.0007 | 0.0009 |
| F06 | 68740001 | 68840000 | 0.0008 | 0.0008 | 0.0008 |
| F06 | 68760001 | 68860000 | 0.0009 | 0.0008 | 0.0008 |
| F06 | 68780001 | 68880000 | 0.0010 | 0.0017 | 0.0009 |
| F06 | 68800001 | 68900000 | 0.0020 | 0.0018 | 0.0018 |
| F06 | 68820001 | 68920000 | 0.0024 | 0.0029 | 0.0019 |
| F06 | 68840001 | 68940000 | 0.0036 | 0.0031 | 0.0031 |
| F06 | 68860001 | 68960000 | 0.0039 | 0.0034 | 0.0034 |
| F06 | 68880001 | 68980000 | 0.0041 | 0.0027 | 0.0035 |
| F06 | 68900001 | 69000000 | 0.0033 | 0.0029 | 0.0028 |
| F06 | 68920001 | 69020000 | 0.0034 | 0.0020 | 0.0031 |
| F06 | 68940001 | 69040000 | 0.0023 | 0.0021 | 0.0021 |
| F06 | 68960001 | 69060000 | 0.0026 | 0.0016 | 0.0022 |
| F06 | 68980001 | 69080000 | 0.0022 | 0.0016 | 0.0018 |
| F06 | 69000001 | 69100000 | 0.0023 | 0.0010 | 0.0018 |
| F06 | 69020001 | 69120000 | 0.0017 | 0.0010 | 0.0011 |
| F06 | 69040001 | 69140000 | 0.0018 | 0.0007 | 0.0011 |
| F06 | 69060001 | 69160000 | 0.0014 | 0.0013 | 0.0008 |
| F06 | 69080001 | 69180000 | 0.0019 | 0.0012 | 0.0014 |
| F06 | 69100001 | 69200000 | 0.0016 | 0.0015 | 0.0012 |
| F06 | 69120001 | 69220000 | 0.0018 | 0.0017 | 0.0015 |
| F06 | 69140001 | 69240000 | 0.0021 | 0.0018 | 0.0018 |
| F06 | 69160001 | 69260000 | 0.0024 | 0.0020 | 0.0020 |
| F06 | 69180001 | 69280000 | 0.0027 | 0.0020 | 0.0023 |
| F06 | 69200001 | 69300000 | 0.0027 | 0.0016 | 0.0024 |
| F06 | 69220001 | 69320000 | 0.0023 | 0.0017 | 0.0020 |
| F06 | 69240001 | 69340000 | 0.0022 | 0.0014 | 0.0021 |
| F06 | 69260001 | 69360000 | 0.0018 | 0.0009 | 0.0016 |
| F06 | 69280001 | 69380000 | 0.0010 | 0.0010 | 0.0009 |
| F06 | 69300001 | 69400000 | 0.0013 | 0.0012 | 0.0010 |
| F06 | 69320001 | 69420000 | 0.0017 | 0.0009 | 0.0012 |
| F06 | 69340001 | 69440000 | 0.0016 | 0.0011 | 0.0009 |

| Chr | Start | End | both | A1 | A2 |
|---|---|---|---|---|---|
| F06 | 69360001 | 69460000 | 0.0018 | 0.0011 | 0.0012 |
| F06 | 69380001 | 69480000 | 0.0019 | 0.0011 | 0.0013 |
| F06 | 69400001 | 69500000 | 0.0017 | 0.0009 | 0.0012 |
| F06 | 69420001 | 69520000 | 0.0014 | 0.0008 | 0.0011 |
| F06 | 69440001 | 69540000 | 0.0010 | 0.0005 | 0.0010 |
| F06 | 69460001 | 69560000 | 0.0006 | 0.0003 | 0.0006 |
| F06 | 69480001 | 69580000 | 0.0004 | 0.0005 | 0.0004 |
| F06 | 69500001 | 69600000 | 0.0005 | 0.0006 | 0.0005 |
| F06 | 69520001 | 69620000 | 0.0006 | 0.0008 | 0.0006 |
| F06 | 69540001 | 69640000 | 0.0009 | 0.0008 | 0.0009 |
| F06 | 69560001 | 69660000 | 0.0009 | 0.0008 | 0.0009 |
| F06 | 69580001 | 69680000 | 0.0010 | 0.0006 | 0.0010 |
| F06 | 69600001 | 69700000 | 0.0007 | 0.0006 | 0.0007 |
| F06 | 69620001 | 69720000 | 0.0008 | 0.0005 | 0.0008 |
| F06 | 69640001 | 69740000 | 0.0006 | 0.0005 | 0.0006 |
| F06 | 69660001 | 69760000 | 0.0006 | 0.0004 | 0.0006 |
| F06 | 69680001 | 69780000 | 0.0005 | 0.0006 | 0.0005 |
| F06 | 69700001 | 69800000 | 0.0007 | 0.0003 | 0.0007 |
| F06 | 69720001 | 69820000 | 0.0004 | 0.0003 | 0.0004 |
| F06 | 69740001 | 69840000 | 0.0004 | 0.0008 | 0.0004 |
| F06 | 69760001 | 69860000 | 0.0010 | 0.0008 | 0.0010 |
| F06 | 69780001 | 69880000 | 0.0010 | 0.0007 | 0.0010 |
| F06 | 69800001 | 69900000 | 0.0009 | 0.0010 | 0.0009 |
| F06 | 69820001 | 69920000 | 0.0011 | 0.0009 | 0.0011 |
| F06 | 69840001 | 69940000 | 0.0010 | 0.0004 | 0.0010 |
| F06 | 69860001 | 69960000 | 0.0005 | 0.0008 | 0.0005 |
| F06 | 69880001 | 69980000 | 0.0009 | 0.0010 | 0.0009 |
| F06 | 69900001 | 70000000 | 0.0012 | 0.0009 | 0.0011 |
| F06 | 69920001 | 70020000 | 0.0012 | 0.0011 | 0.0010 |
| F06 | 69940001 | 70040000 | 0.0015 | 0.0011 | 0.0013 |
| F06 | 69960001 | 70060000 | 0.0014 | 0.0012 | 0.0013 |
| F06 | 69980001 | 70080000 | 0.0017 | 0.0012 | 0.0015 |
| F06 | 70000001 | 70100000 | 0.0017 | 0.0016 | 0.0016 |
| F06 | 70020001 | 70120000 | 0.0021 | 0.0016 | 0.0018 |
| F06 | 70040001 | 70140000 | 0.0020 | 0.0016 | 0.0017 |
| F06 | 70060001 | 70160000 | 0.0022 | 0.0013 | 0.0018 |
| F06 | 70080001 | 70180000 | 0.0019 | 0.0015 | 0.0015 |
| F06 | 70100001 | 70200000 | 0.0020 | 0.0010 | 0.0016 |
| F06 | 70120001 | 70220000 | 0.0014 | 0.0014 | 0.0012 |
| F06 | 70140001 | 70240000 | 0.0017 | 0.0016 | 0.0015 |
| F06 | 70160001 | 70260000 | 0.0020 | 0.0023 | 0.0017 |
| F06 | 70180001 | 70280000 | 0.0028 | 0.0020 | 0.0023 |
| F06 | 70200001 | 70300000 | 0.0027 | 0.0022 | 0.0021 |
| F06 | 70220001 | 70320000 | 0.0029 | 0.0021 | 0.0023 |
| F06 | 70240001 | 70340000 | 0.0031 | 0.0018 | 0.0022 |
| F06 | 70260001 | 70360000 | 0.0028 | 0.0010 | 0.0019 |
| F06 | 70280001 | 70380000 | 0.0019 | 0.0014 | 0.0011 |
| F06 | 70300001 | 70400000 | 0.0021 | 0.0012 | 0.0015 |
| F06 | 70320001 | 70420000 | 0.0019 | 0.0015 | 0.0014 |
| F06 | 70340001 | 70440000 | 0.0019 | 0.0017 | 0.0016 |
| F06 | 70360001 | 70460000 | 0.0019 | 0.0020 | 0.0019 |
| F06 | 70380001 | 70480000 | 0.0023 | 0.0020 | 0.0022 |
| F06 | 70400001 | 70500000 | 0.0023 | 0.0025 | 0.0022 |
| F06 | 70420001 | 70520000 | 0.0029 | 0.0020 | 0.0028 |
| F06 | 70440001 | 70540000 | 0.0024 | 0.0024 | 0.0023 |
| F06 | 70460001 | 70560000 | 0.0030 | 0.0025 | 0.0025 |
| F06 | 70480001 | 70580000 | 0.0033 | 0.0023 | 0.0027 |
| F06 | 70500001 | 70600000 | 0.0031 | 0.0020 | 0.0025 |

| Chr | Start | End | both | A1 | A2 |
|---|---|---|---|---|---|
| F06 | 70520001 | 70620000 | 0.0027 | 0.0022 | 0.0020 |
| F06 | 70540001 | 70640000 | 0.0029 | 0.0017 | 0.0022 |
| F06 | 70560001 | 70660000 | 0.0023 | 0.0012 | 0.0018 |
| F06 | 70580001 | 70680000 | 0.0017 | 0.0009 | 0.0014 |
| F06 | 70600001 | 70700000 | 0.0013 | 0.0008 | 0.0010 |
| F06 | 70620001 | 70720000 | 0.0011 | 0.0004 | 0.0009 |
| F06 | 70640001 | 70740000 | 0.0008 | 0.0004 | 0.0006 |
| F06 | 70660001 | 70760000 | 0.0005 | 0.0005 | 0.0005 |
| F06 | 70680001 | 70780000 | 0.0007 | 0.0007 | 0.0007 |
| F06 | 70700001 | 70800000 | 0.0009 | 0.0007 | 0.0009 |
| F06 | 70720001 | 70820000 | 0.0008 | 0.0014 | 0.0008 |
| F06 | 70740001 | 70840000 | 0.0017 | 0.0014 | 0.0017 |
| F06 | 70760001 | 70860000 | 0.0018 | 0.0011 | 0.0018 |
| F06 | 70780001 | 70880000 | 0.0014 | 0.0010 | 0.0014 |
| F06 | 70800001 | 70900000 | 0.0013 | 0.0014 | 0.0013 |
| F06 | 70820001 | 70920000 | 0.0016 | 0.0009 | 0.0016 |
| F06 | 70840001 | 70940000 | 0.0010 | 0.0014 | 0.0010 |
| F06 | 70860001 | 70960000 | 0.0017 | 0.0016 | 0.0017 |
| F06 | 70880001 | 70980000 | 0.0019 | 0.0015 | 0.0019 |
| F06 | 70900001 | 71000000 | 0.0017 | 0.0013 | 0.0017 |
| F06 | 70920001 | 71020000 | 0.0016 | 0.0015 | 0.0016 |
| F06 | 70940001 | 71040000 | 0.0017 | 0.0009 | 0.0017 |
| F06 | 70960001 | 71060000 | 0.0010 | 0.0009 | 0.0010 |
| F06 | 70980001 | 71080000 | 0.0011 | 0.0015 | 0.0010 |
| F06 | 71000001 | 71100000 | 0.0017 | 0.0015 | 0.0016 |
| F06 | 71020001 | 71120000 | 0.0016 | 0.0012 | 0.0016 |
| F06 | 71040001 | 71140000 | 0.0014 | 0.0012 | 0.0014 |
| F06 | 71060001 | 71160000 | 0.0014 | 0.0011 | 0.0014 |
| F06 | 71080001 | 71180000 | 0.0013 | 0.0006 | 0.0013 |
| F06 | 71100001 | 71200000 | 0.0008 | 0.0003 | 0.0007 |
| F06 | 71120001 | 71220000 | 0.0006 | 0.0002 | 0.0004 |
| F06 | 71140001 | 71240000 | 0.0005 | 0.0003 | 0.0003 |
| F06 | 71160001 | 71260000 | 0.0009 | 0.0004 | 0.0007 |
| F06 | 71180001 | 71280000 | 0.0010 | 0.0005 | 0.0008 |
| F06 | 71200001 | 71300000 | 0.0011 | 0.0007 | 0.0009 |
| F06 | 71220001 | 71320000 | 0.0014 | 0.0015 | 0.0011 |
| F06 | 71240001 | 71340000 | 0.0025 | 0.0020 | 0.0021 |
| F06 | 71260001 | 71360000 | 0.0027 | 0.0021 | 0.0024 |
| F06 | 71280001 | 71380000 | 0.0027 | 0.0021 | 0.0024 |
| F06 | 71300001 | 71400000 | 0.0027 | 0.0021 | 0.0025 |
| F06 | 71320001 | 71420000 | 0.0026 | 0.0015 | 0.0024 |
| F06 | 71340001 | 71440000 | 0.0017 | 0.0009 | 0.0016 |
| F06 | 71360001 | 71460000 | 0.0010 | 0.0008 | 0.0010 |
| F06 | 71380001 | 71480000 | 0.0010 | 0.0007 | 0.0010 |
| F06 | 71400001 | 71500000 | 0.0008 | 0.0010 | 0.0008 |
| F06 | 71420001 | 71520000 | 0.0012 | 0.0008 | 0.0011 |
| F06 | 71440001 | 71540000 | 0.0010 | 0.0008 | 0.0010 |
| F06 | 71460001 | 71560000 | 0.0009 | 0.0014 | 0.0009 |
| F06 | 71480001 | 71580000 | 0.0016 | 0.0014 | 0.0016 |
| F06 | 71500001 | 71600000 | 0.0016 | 0.0014 | 0.0016 |
| F06 | 71520001 | 71620000 | 0.0015 | 0.0016 | 0.0015 |
| F06 | 71540001 | 71640000 | 0.0019 | 0.0017 | 0.0019 |
| F06 | 71560001 | 71660000 | 0.0020 | 0.0009 | 0.0020 |
| F06 | 71580001 | 71680000 | 0.0011 | 0.0009 | 0.0011 |
| F06 | 71600001 | 71700000 | 0.0011 | 0.0008 | 0.0011 |
| F06 | 71620001 | 71720000 | 0.0010 | 0.0007 | 0.0010 |
| F06 | 71640001 | 71740000 | 0.0008 | 0.0006 | 0.0008 |
| F06 | 71660001 | 71760000 | 0.0006 | 0.0007 | 0.0006 |

| Chr | Start | End | both | A1 | A2 |
|---|---|---|---|---|---|
| F06 | 71680001 | 71780000 | 0.0007 | 0.0007 | 0.0007 |
| F06 | 71700001 | 71800000 | 0.0008 | 0.0004 | 0.0007 |
| F06 | 71720001 | 71820000 | 0.0004 | 0.0004 | 0.0003 |
| F06 | 71740001 | 71840000 | 0.0005 | 0.0008 | 0.0004 |
| F06 | 71760001 | 71860000 | 0.0009 | 0.0012 | 0.0008 |
| F06 | 71780001 | 71880000 | 0.0014 | 0.0014 | 0.0011 |
| F06 | 71800001 | 71900000 | 0.0018 | 0.0019 | 0.0012 |
| F06 | 71820001 | 71920000 | 0.0025 | 0.0020 | 0.0017 |
| F06 | 71840001 | 71940000 | 0.0026 | 0.0021 | 0.0018 |
| F06 | 71860001 | 71960000 | 0.0027 | 0.0016 | 0.0017 |
| F06 | 71880001 | 71980000 | 0.0021 | 0.0015 | 0.0014 |
| F06 | 71900001 | 72000000 | 0.0018 | 0.0011 | 0.0014 |
| F06 | 71920001 | 72020000 | 0.0012 | 0.0007 | 0.0010 |
| F06 | 71940001 | 72040000 | 0.0010 | 0.0004 | 0.0008 |
| F06 | 71960001 | 72060000 | 0.0007 | 0.0006 | 0.0005 |
| F06 | 71980001 | 72080000 | 0.0008 | 0.0005 | 0.0007 |
| F06 | 72000001 | 72100000 | 0.0008 | 0.0006 | 0.0007 |
| F06 | 72020001 | 72120000 | 0.0008 | 0.0008 | 0.0007 |
| F06 | 72040001 | 72140000 | 0.0010 | 0.0007 | 0.0010 |
| F06 | 72060001 | 72160000 | 0.0009 | 0.0007 | 0.0009 |
| F06 | 72080001 | 72180000 | 0.0009 | 0.0010 | 0.0009 |
| F06 | 72100001 | 72200000 | 0.0012 | 0.0009 | 0.0012 |
| F06 | 72120001 | 72220000 | 0.0012 | 0.0014 | 0.0012 |
| F06 | 72140001 | 72240000 | 0.0016 | 0.0024 | 0.0016 |
| F06 | 72160001 | 72260000 | 0.0026 | 0.0025 | 0.0025 |
| F06 | 72180001 | 72280000 | 0.0028 | 0.0023 | 0.0026 |
| F06 | 72200001 | 72300000 | 0.0029 | 0.0022 | 0.0024 |
| F06 | 72220001 | 72320000 | 0.0028 | 0.0018 | 0.0023 |
| F06 | 72240001 | 72340000 | 0.0022 | 0.0009 | 0.0018 |
| F06 | 72260001 | 72360000 | 0.0014 | 0.0006 | 0.0010 |
| F06 | 72280001 | 72380000 | 0.0011 | 0.0011 | 0.0009 |
| F06 | 72300001 | 72400000 | 0.0012 | 0.0016 | 0.0012 |
| F06 | 72320001 | 72420000 | 0.0018 | 0.0013 | 0.0016 |
| F06 | 72340001 | 72440000 | 0.0017 | 0.0014 | 0.0014 |
| F06 | 72360001 | 72460000 | 0.0018 | 0.0016 | 0.0015 |
| F06 | 72380001 | 72480000 | 0.0020 | 0.0011 | 0.0016 |
| F06 | 72400001 | 72500000 | 0.0018 | 0.0013 | 0.0011 |
| F06 | 72420001 | 72520000 | 0.0018 | 0.0013 | 0.0013 |
| F06 | 72440001 | 72540000 | 0.0017 | 0.0012 | 0.0013 |
| F06 | 72460001 | 72560000 | 0.0016 | 0.0012 | 0.0012 |
| F06 | 72480001 | 72580000 | 0.0016 | 0.0012 | 0.0013 |
| F06 | 72500001 | 72600000 | 0.0013 | 0.0012 | 0.0012 |
| F06 | 72520001 | 72620000 | 0.0014 | 0.0018 | 0.0013 |
| F06 | 72540001 | 72640000 | 0.0022 | 0.0020 | 0.0020 |
| F06 | 72560001 | 72660000 | 0.0024 | 0.0023 | 0.0022 |
| F06 | 72580001 | 72680000 | 0.0031 | 0.0024 | 0.0028 |
| F06 | 72600001 | 72700000 | 0.0034 | 0.0018 | 0.0029 |
| F06 | 72620001 | 72720000 | 0.0027 | 0.0015 | 0.0022 |
| F06 | 72640001 | 72740000 | 0.0024 | 0.0014 | 0.0019 |
| F06 | 72660001 | 72760000 | 0.0023 | 0.0010 | 0.0018 |
| F06 | 72680001 | 72780000 | 0.0017 | 0.0008 | 0.0011 |
| F06 | 72700001 | 72800000 | 0.0013 | 0.0008 | 0.0009 |
| F06 | 72720001 | 72820000 | 0.0013 | 0.0006 | 0.0009 |
| F06 | 72740001 | 72840000 | 0.0011 | 0.0007 | 0.0007 |
| F06 | 72760001 | 72860000 | 0.0011 | 0.0005 | 0.0007 |
| F06 | 72780001 | 72880000 | 0.0010 | 0.0007 | 0.0007 |
| F06 | 72800001 | 72900000 | 0.0013 | 0.0007 | 0.0009 |
| F06 | 72820001 | 72920000 | 0.0012 | 0.0005 | 0.0009 |

| Chr | Start | End | both | A1 | A2 |
|---|---|---|---|---|---|
| F06 | 72840001 | 72940000 | 0.0010 | 0.0006 | 0.0008 |
| F06 | 72860001 | 72960000 | 0.0011 | 0.0008 | 0.0009 |
| F06 | 72880001 | 72980000 | 0.0012 | 0.0011 | 0.0009 |
| F06 | 72900001 | 73000000 | 0.0016 | 0.0011 | 0.0013 |
| F06 | 72920001 | 73020000 | 0.0017 | 0.0013 | 0.0013 |
| F06 | 72940001 | 73040000 | 0.0018 | 0.0014 | 0.0015 |
| F06 | 72960001 | 73060000 | 0.0020 | 0.0015 | 0.0016 |
| F06 | 72980001 | 73080000 | 0.0020 | 0.0011 | 0.0016 |
| F06 | 73000001 | 73100000 | 0.0015 | 0.0010 | 0.0011 |
| F06 | 73020001 | 73120000 | 0.0015 | 0.0009 | 0.0010 |
| F06 | 73040001 | 73140000 | 0.0014 | 0.0011 | 0.0010 |
| F06 | 73060001 | 73160000 | 0.0018 | 0.0011 | 0.0012 |
| F06 | 73080001 | 73180000 | 0.0017 | 0.0012 | 0.0012 |
| F06 | 73100001 | 73200000 | 0.0016 | 0.0019 | 0.0014 |
| F06 | 73120001 | 73220000 | 0.0023 | 0.0020 | 0.0019 |
| F06 | 73140001 | 73240000 | 0.0024 | 0.0019 | 0.0020 |
| F06 | 73160001 | 73260000 | 0.0020 | 0.0019 | 0.0018 |
| F06 | 73180001 | 73280000 | 0.0020 | 0.0022 | 0.0018 |
| F06 | 73200001 | 73300000 | 0.0023 | 0.0020 | 0.0020 |
| F06 | 73220001 | 73320000 | 0.0020 | 0.0018 | 0.0019 |
| F06 | 73240001 | 73340000 | 0.0019 | 0.0015 | 0.0017 |
| F06 | 73260001 | 73360000 | 0.0015 | 0.0014 | 0.0014 |
| F06 | 73280001 | 73380000 | 0.0014 | 0.0014 | 0.0012 |
| F06 | 73300001 | 73400000 | 0.0014 | 0.0012 | 0.0012 |
| F06 | 73320001 | 73420000 | 0.0013 | 0.0016 | 0.0011 |
| F06 | 73340001 | 73440000 | 0.0016 | 0.0019 | 0.0014 |
| F06 | 73360001 | 73460000 | 0.0019 | 0.0019 | 0.0016 |
| F06 | 73380001 | 73480000 | 0.0019 | 0.0016 | 0.0017 |
| F06 | 73400001 | 73500000 | 0.0016 | 0.0014 | 0.0015 |
| F06 | 73420001 | 73520000 | 0.0015 | 0.0014 | 0.0013 |
| F06 | 73440001 | 73540000 | 0.0014 | 0.0009 | 0.0012 |
| F06 | 73460001 | 73560000 | 0.0010 | 0.0009 | 0.0009 |
| F06 | 73480001 | 73580000 | 0.0010 | 0.0010 | 0.0009 |
| F06 | 73500001 | 73600000 | 0.0010 | 0.0009 | 0.0009 |
| F06 | 73520001 | 73620000 | 0.0009 | 0.0007 | 0.0009 |
| F06 | 73540001 | 73640000 | 0.0008 | 0.0008 | 0.0008 |
| F06 | 73560001 | 73660000 | 0.0008 | 0.0006 | 0.0008 |
| F06 | 73580001 | 73680000 | 0.0007 | 0.0007 | 0.0007 |
| F06 | 73600001 | 73700000 | 0.0008 | 0.0008 | 0.0007 |
| F06 | 73620001 | 73720000 | 0.0010 | 0.0010 | 0.0009 |
| F06 | 73640001 | 73740000 | 0.0011 | 0.0016 | 0.0010 |
| F06 | 73660001 | 73760000 | 0.0017 | 0.0022 | 0.0015 |
| F06 | 73680001 | 73780000 | 0.0022 | 0.0022 | 0.0018 |
| F06 | 73700001 | 73800000 | 0.0023 | 0.0021 | 0.0019 |
| F06 | 73720001 | 73820000 | 0.0021 | 0.0022 | 0.0017 |
| F06 | 73740001 | 73840000 | 0.0022 | 0.0018 | 0.0018 |
| F06 | 73760001 | 73860000 | 0.0018 | 0.0013 | 0.0015 |
| F06 | 73780001 | 73880000 | 0.0014 | 0.0015 | 0.0012 |
| F06 | 73800001 | 73900000 | 0.0016 | 0.0016 | 0.0013 |
| F06 | 73820001 | 73920000 | 0.0016 | 0.0012 | 0.0013 |
| F06 | 73840001 | 73940000 | 0.0012 | 0.0013 | 0.0010 |
| F06 | 73860001 | 73960000 | 0.0013 | 0.0023 | 0.0011 |
| F06 | 73880001 | 73980000 | 0.0022 | 0.0017 | 0.0019 |
| F06 | 73900001 | 74000000 | 0.0016 | 0.0016 | 0.0014 |
| F06 | 73920001 | 74020000 | 0.0016 | 0.0018 | 0.0014 |
| F06 | 73940001 | 74040000 | 0.0017 | 0.0017 | 0.0015 |
| F06 | 73960001 | 74060000 | 0.0017 | 0.0010 | 0.0015 |
| F06 | 73980001 | 74080000 | 0.0011 | 0.0014 | 0.0009 |

| Chr | Start | End | both | A1 | A2 |
|---|---|---|---|---|---|
| F06 | 74000001 | 74100000 | 0.0014 | 0.0014 | 0.0012 |
| F06 | 74020001 | 74120000 | 0.0015 | 0.0013 | 0.0012 |
| F06 | 74040001 | 74140000 | 0.0014 | 0.0010 | 0.0012 |
| F06 | 74060001 | 74160000 | 0.0010 | 0.0007 | 0.0009 |
| F06 | 74080001 | 74180000 | 0.0008 | 0.0005 | 0.0007 |
| F06 | 74100001 | 74200000 | 0.0006 | 0.0006 | 0.0005 |
| F06 | 74120001 | 74220000 | 0.0006 | 0.0006 | 0.0005 |
| F06 | 74140001 | 74240000 | 0.0006 | 0.0006 | 0.0006 |
| F06 | 74160001 | 74260000 | 0.0008 | 0.0009 | 0.0008 |
| F06 | 74180001 | 74280000 | 0.0011 | 0.0013 | 0.0011 |
| F06 | 74200001 | 74300000 | 0.0016 | 0.0012 | 0.0015 |
| F06 | 74220001 | 74320000 | 0.0015 | 0.0015 | 0.0013 |
| F06 | 74240001 | 74340000 | 0.0017 | 0.0015 | 0.0015 |
| F06 | 74260001 | 74360000 | 0.0018 | 0.0015 | 0.0015 |
| F06 | 74280001 | 74380000 | 0.0016 | 0.0011 | 0.0014 |
| F06 | 74300001 | 74400000 | 0.0012 | 0.0019 | 0.0011 |
| F06 | 74320001 | 74420000 | 0.0020 | 0.0021 | 0.0019 |
| F06 | 74340001 | 74440000 | 0.0023 | 0.0020 | 0.0022 |
| F06 | 74360001 | 74460000 | 0.0021 | 0.0019 | 0.0020 |
| F06 | 74380001 | 74480000 | 0.0020 | 0.0019 | 0.0019 |
| F06 | 74400001 | 74500000 | 0.0020 | 0.0012 | 0.0019 |
| F06 | 74420001 | 74520000 | 0.0014 | 0.0007 | 0.0013 |
| F06 | 74440001 | 74540000 | 0.0008 | 0.0006 | 0.0008 |
| F06 | 74460001 | 74560000 | 0.0007 | 0.0009 | 0.0007 |
| F06 | 74480001 | 74580000 | 0.0011 | 0.0010 | 0.0010 |
| F06 | 74500001 | 74600000 | 0.0012 | 0.0009 | 0.0011 |
| F06 | 74520001 | 74620000 | 0.0011 | 0.0012 | 0.0010 |
| F06 | 74540001 | 74640000 | 0.0014 | 0.0017 | 0.0012 |
| F06 | 74560001 | 74660000 | 0.0018 | 0.0014 | 0.0017 |
| F06 | 74580001 | 74680000 | 0.0016 | 0.0016 | 0.0015 |
| F06 | 74600001 | 74700000 | 0.0018 | 0.0017 | 0.0017 |
| F06 | 74620001 | 74720000 | 0.0019 | 0.0017 | 0.0018 |
| F06 | 74640001 | 74740000 | 0.0019 | 0.0016 | 0.0018 |
| F06 | 74660001 | 74760000 | 0.0018 | 0.0018 | 0.0015 |
| F06 | 74680001 | 74780000 | 0.0018 | 0.0016 | 0.0015 |
| F06 | 74700001 | 74800000 | 0.0016 | 0.0021 | 0.0013 |
| F06 | 74720001 | 74820000 | 0.0021 | 0.0024 | 0.0016 |
| F06 | 74740001 | 74840000 | 0.0023 | 0.0023 | 0.0018 |
| F06 | 74760001 | 74860000 | 0.0022 | 0.0021 | 0.0018 |
| F06 | 74780001 | 74880000 | 0.0021 | 0.0023 | 0.0018 |
| F06 | 74800001 | 74900000 | 0.0023 | 0.0016 | 0.0020 |
| F06 | 74820001 | 74920000 | 0.0017 | 0.0010 | 0.0015 |
| F06 | 74840001 | 74940000 | 0.0011 | 0.0008 | 0.0010 |
| F06 | 74860001 | 74960000 | 0.0010 | 0.0009 | 0.0010 |
| F06 | 74880001 | 74980000 | 0.0010 | 0.0005 | 0.0010 |
| F06 | 74900001 | 75000000 | 0.0007 | 0.0006 | 0.0007 |
| F06 | 74920001 | 75020000 | 0.0007 | 0.0008 | 0.0007 |
| F06 | 74940001 | 75040000 | 0.0009 | 0.0010 | 0.0008 |
| F06 | 74960001 | 75060000 | 0.0011 | 0.0013 | 0.0008 |
| F06 | 74980001 | 75080000 | 0.0013 | 0.0012 | 0.0010 |
| F06 | 75000001 | 75100000 | 0.0011 | 0.0011 | 0.0008 |
| F06 | 75020001 | 75120000 | 0.0011 | 0.0015 | 0.0007 |
| F06 | 75040001 | 75140000 | 0.0017 | 0.0014 | 0.0014 |
| F06 | 75060001 | 75160000 | 0.0016 | 0.0012 | 0.0013 |
| F06 | 75080001 | 75180000 | 0.0014 | 0.0013 | 0.0011 |
| F06 | 75100001 | 75200000 | 0.0015 | 0.0018 | 0.0012 |
| F06 | 75120001 | 75220000 | 0.0020 | 0.0016 | 0.0016 |
| F06 | 75140001 | 75240000 | 0.0015 | 0.0016 | 0.0010 |

| Chr | Start | End | both | A1 | A2 |
|---|---|---|---|---|---|
| F06 | 75160001 | 75260000 | 0.0014 | 0.0015 | 0.0010 |
| F06 | 75180001 | 75280000 | 0.0014 | 0.0014 | 0.0010 |
| F06 | 75200001 | 75300000 | 0.0014 | 0.0010 | 0.0010 |
| F06 | 75220001 | 75320000 | 0.0010 | 0.0009 | 0.0007 |
| F06 | 75240001 | 75340000 | 0.0009 | 0.0014 | 0.0007 |
| F06 | 75260001 | 75360000 | 0.0016 | 0.0013 | 0.0013 |
| F06 | 75280001 | 75380000 | 0.0016 | 0.0021 | 0.0013 |
| F06 | 75300001 | 75400000 | 0.0025 | 0.0026 | 0.0022 |
| F06 | 75320001 | 75420000 | 0.0030 | 0.0029 | 0.0026 |
| F06 | 75340001 | 75440000 | 0.0034 | 0.0024 | 0.0031 |
| F06 | 75360001 | 75460000 | 0.0028 | 0.0027 | 0.0026 |
| F06 | 75380001 | 75480000 | 0.0031 | 0.0020 | 0.0029 |
| F06 | 75400001 | 75500000 | 0.0022 | 0.0015 | 0.0019 |
| F06 | 75420001 | 75520000 | 0.0017 | 0.0014 | 0.0014 |
| F06 | 75440001 | 75540000 | 0.0015 | 0.0014 | 0.0013 |
| F06 | 75460001 | 75560000 | 0.0016 | 0.0014 | 0.0013 |
| F06 | 75480001 | 75580000 | 0.0015 | 0.0014 | 0.0014 |
| F06 | 75500001 | 75600000 | 0.0016 | 0.0014 | 0.0015 |
| F06 | 75520001 | 75620000 | 0.0015 | 0.0012 | 0.0015 |
| F06 | 75540001 | 75640000 | 0.0013 | 0.0011 | 0.0013 |
| F06 | 75560001 | 75660000 | 0.0013 | 0.0010 | 0.0012 |
| F06 | 75580001 | 75680000 | 0.0011 | 0.0010 | 0.0010 |
| F06 | 75600001 | 75700000 | 0.0011 | 0.0010 | 0.0011 |
| F06 | 75620001 | 75720000 | 0.0011 | 0.0010 | 0.0011 |
| F06 | 75640001 | 75740000 | 0.0011 | 0.0011 | 0.0011 |
| F06 | 75660001 | 75760000 | 0.0012 | 0.0010 | 0.0011 |
| F06 | 75680001 | 75780000 | 0.0011 | 0.0010 | 0.0011 |
| F06 | 75700001 | 75800000 | 0.0011 | 0.0013 | 0.0010 |
| F06 | 75720001 | 75820000 | 0.0014 | 0.0010 | 0.0013 |
| F06 | 75740001 | 75840000 | 0.0011 | 0.0008 | 0.0011 |
| F06 | 75760001 | 75860000 | 0.0009 | 0.0007 | 0.0009 |
| F06 | 75780001 | 75880000 | 0.0008 | 0.0007 | 0.0008 |
| F06 | 75800001 | 75900000 | 0.0008 | 0.0006 | 0.0008 |
| F06 | 75820001 | 75920000 | 0.0007 | 0.0008 | 0.0006 |
| F06 | 75840001 | 75940000 | 0.0010 | 0.0009 | 0.0008 |
| F06 | 75860001 | 75960000 | 0.0010 | 0.0010 | 0.0009 |
| F06 | 75880001 | 75980000 | 0.0011 | 0.0011 | 0.0009 |
| F06 | 75900001 | 76000000 | 0.0011 | 0.0012 | 0.0008 |
| F06 | 75920001 | 76020000 | 0.0012 | 0.0010 | 0.0009 |
| F06 | 75940001 | 76040000 | 0.0011 | 0.0012 | 0.0008 |
| F06 | 75960001 | 76060000 | 0.0013 | 0.0011 | 0.0010 |
| F06 | 75980001 | 76080000 | 0.0013 | 0.0009 | 0.0011 |
| F06 | 76000001 | 76100000 | 0.0011 | 0.0009 | 0.0011 |
| F06 | 76020001 | 76120000 | 0.0012 | 0.0010 | 0.0012 |
| F06 | 76040001 | 76140000 | 0.0013 | 0.0008 | 0.0014 |
| F06 | 76060001 | 76160000 | 0.0011 | 0.0007 | 0.0012 |
| F06 | 76080001 | 76180000 | 0.0009 | 0.0008 | 0.0010 |
| F06 | 76100001 | 76200000 | 0.0010 | 0.0007 | 0.0010 |
| F06 | 76120001 | 76220000 | 0.0009 | 0.0012 | 0.0008 |
| F06 | 76140001 | 76240000 | 0.0013 | 0.0014 | 0.0012 |
| F06 | 76160001 | 76260000 | 0.0015 | 0.0014 | 0.0014 |
| F06 | 76180001 | 76280000 | 0.0015 | 0.0017 | 0.0015 |
| F06 | 76200001 | 76300000 | 0.0018 | 0.0016 | 0.0018 |
| F06 | 76220001 | 76320000 | 0.0018 | 0.0011 | 0.0017 |
| F06 | 76240001 | 76340000 | 0.0012 | 0.0015 | 0.0011 |
| F06 | 76260001 | 76360000 | 0.0016 | 0.0014 | 0.0015 |
| F06 | 76280001 | 76380000 | 0.0014 | 0.0011 | 0.0014 |
| F06 | 76300001 | 76400000 | 0.0011 | 0.0011 | 0.0010 |

| Chr | Start | End | both | A1 | A2 |
|---|---|---|---|---|---|
| F06 | 76320001 | 76420000 | 0.0010 | 0.0012 | 0.0010 |
| F06 | 76340001 | 76440000 | 0.0012 | 0.0006 | 0.0011 |
| F06 | 76360001 | 76460000 | 0.0007 | 0.0008 | 0.0007 |
| F06 | 76380001 | 76480000 | 0.0010 | 0.0008 | 0.0010 |
| F06 | 76400001 | 76500000 | 0.0011 | 0.0011 | 0.0010 |
| F06 | 76420001 | 76520000 | 0.0012 | 0.0016 | 0.0011 |
| F06 | 76440001 | 76540000 | 0.0018 | 0.0018 | 0.0014 |
| F06 | 76460001 | 76560000 | 0.0020 | 0.0020 | 0.0016 |
| F06 | 76480001 | 76580000 | 0.0022 | 0.0023 | 0.0017 |
| F06 | 76500001 | 76600000 | 0.0026 | 0.0027 | 0.0020 |
| F06 | 76520001 | 76620000 | 0.0030 | 0.0023 | 0.0022 |
| F06 | 76540001 | 76640000 | 0.0025 | 0.0024 | 0.0019 |
| F06 | 76560001 | 76660000 | 0.0026 | 0.0029 | 0.0020 |
| F06 | 76580001 | 76680000 | 0.0029 | 0.0029 | 0.0023 |
| F06 | 76600001 | 76700000 | 0.0030 | 0.0023 | 0.0023 |
| F06 | 76620001 | 76720000 | 0.0025 | 0.0022 | 0.0020 |
| F06 | 76640001 | 76740000 | 0.0023 | 0.0020 | 0.0018 |
| F06 | 76660001 | 76760000 | 0.0020 | 0.0011 | 0.0015 |
| F06 | 76680001 | 76780000 | 0.0012 | 0.0008 | 0.0008 |
| F06 | 76700001 | 76800000 | 0.0008 | 0.0009 | 0.0006 |
| F06 | 76720001 | 76820000 | 0.0009 | 0.0007 | 0.0007 |
| F06 | 76740001 | 76840000 | 0.0007 | 0.0005 | 0.0007 |
| F06 | 76760001 | 76860000 | 0.0006 | 0.0009 | 0.0005 |
| F06 | 76780001 | 76880000 | 0.0009 | 0.0009 | 0.0007 |
| F06 | 76800001 | 76900000 | 0.0010 | 0.0013 | 0.0009 |
| F06 | 76820001 | 76920000 | 0.0014 | 0.0012 | 0.0011 |
| F06 | 76840001 | 76940000 | 0.0013 | 0.0016 | 0.0010 |
| F06 | 76860001 | 76960000 | 0.0017 | 0.0014 | 0.0015 |
| F06 | 76880001 | 76980000 | 0.0016 | 0.0017 | 0.0014 |
| F06 | 76900001 | 77000000 | 0.0019 | 0.0019 | 0.0017 |
| F06 | 76920001 | 77020000 | 0.0020 | 0.0024 | 0.0019 |
| F06 | 76940001 | 77040000 | 0.0026 | 0.0024 | 0.0024 |
| F06 | 76960001 | 77060000 | 0.0028 | 0.0028 | 0.0025 |
| F06 | 76980001 | 77080000 | 0.0031 | 0.0027 | 0.0026 |
| F06 | 77000001 | 77100000 | 0.0029 | 0.0021 | 0.0023 |
| F06 | 77020001 | 77120000 | 0.0022 | 0.0023 | 0.0018 |
| F06 | 77040001 | 77140000 | 0.0023 | 0.0022 | 0.0018 |
| F06 | 77060001 | 77160000 | 0.0019 | 0.0023 | 0.0014 |
| F06 | 77080001 | 77180000 | 0.0022 | 0.0024 | 0.0018 |
| F06 | 77100001 | 77200000 | 0.0023 | 0.0040 | 0.0019 |
| F06 | 77120001 | 77220000 | 0.0039 | 0.0036 | 0.0034 |
| F06 | 77140001 | 77240000 | 0.0037 | 0.0035 | 0.0033 |
| F06 | 77160001 | 77260000 | 0.0037 | 0.0031 | 0.0033 |
| F06 | 77180001 | 77280000 | 0.0031 | 0.0030 | 0.0027 |
| F06 | 77200001 | 77300000 | 0.0032 | 0.0018 | 0.0029 |
| F06 | 77220001 | 77320000 | 0.0019 | 0.0016 | 0.0017 |
| F06 | 77240001 | 77340000 | 0.0017 | 0.0020 | 0.0016 |
| F06 | 77260001 | 77360000 | 0.0021 | 0.0018 | 0.0019 |
| F06 | 77280001 | 77380000 | 0.0020 | 0.0017 | 0.0018 |
| F06 | 77300001 | 77400000 | 0.0018 | 0.0015 | 0.0015 |
| F06 | 77320001 | 77420000 | 0.0014 | 0.0017 | 0.0011 |
| F06 | 77340001 | 77440000 | 0.0016 | 0.0011 | 0.0013 |
| F06 | 77360001 | 77460000 | 0.0011 | 0.0012 | 0.0008 |
| F06 | 77380001 | 77480000 | 0.0012 | 0.0017 | 0.0009 |
| F06 | 77400001 | 77500000 | 0.0017 | 0.0017 | 0.0014 |
| F06 | 77420001 | 77520000 | 0.0017 | 0.0022 | 0.0014 |
| F06 | 77440001 | 77540000 | 0.0022 | 0.0021 | 0.0017 |
| F06 | 77460001 | 77560000 | 0.0022 | 0.0021 | 0.0017 |

| Chr | Start | End | both | A1 | A2 |
|---|---|---|---|---|---|
| F06 | 77480001 | 77580000 | 0.0022 | 0.0022 | 0.0019 |
| F06 | 77500001 | 77600000 | 0.0023 | 0.0024 | 0.0018 |
| F06 | 77520001 | 77620000 | 0.0025 | 0.0028 | 0.0021 |
| F06 | 77540001 | 77640000 | 0.0029 | 0.0034 | 0.0023 |
| F06 | 77560001 | 77660000 | 0.0035 | 0.0040 | 0.0028 |
| F06 | 77580001 | 77680000 | 0.0041 | 0.0036 | 0.0031 |
| F06 | 77600001 | 77700000 | 0.0037 | 0.0037 | 0.0030 |
| F06 | 77620001 | 77720000 | 0.0038 | 0.0028 | 0.0030 |
| F06 | 77640001 | 77740000 | 0.0029 | 0.0027 | 0.0023 |
| F06 | 77660001 | 77760000 | 0.0028 | 0.0023 | 0.0022 |
| F06 | 77680001 | 77780000 | 0.0024 | 0.0020 | 0.0020 |
| F06 | 77700001 | 77800000 | 0.0020 | 0.0018 | 0.0017 |
| F06 | 77720001 | 77820000 | 0.0018 | 0.0027 | 0.0015 |
| F06 | 77740001 | 77840000 | 0.0027 | 0.0026 | 0.0025 |
| F06 | 77760001 | 77860000 | 0.0028 | 0.0026 | 0.0027 |
| F06 | 77780001 | 77880000 | 0.0028 | 0.0032 | 0.0027 |
| F06 | 77800001 | 77900000 | 0.0033 | 0.0029 | 0.0032 |
| F06 | 77820001 | 77920000 | 0.0031 | 0.0021 | 0.0030 |
| F06 | 77840001 | 77940000 | 0.0022 | 0.0019 | 0.0020 |
| F06 | 77860001 | 77960000 | 0.0021 | 0.0031 | 0.0019 |
| F06 | 77880001 | 77980000 | 0.0033 | 0.0033 | 0.0031 |
| F06 | 77900001 | 78000000 | 0.0037 | 0.0033 | 0.0035 |
| F06 | 77920001 | 78020000 | 0.0037 | 0.0033 | 0.0035 |
| F06 | 77940001 | 78040000 | 0.0038 | 0.0034 | 0.0037 |
| F06 | 77960001 | 78060000 | 0.0039 | 0.0023 | 0.0037 |
| F06 | 77980001 | 78080000 | 0.0028 | 0.0016 | 0.0025 |
| F06 | 78000001 | 78100000 | 0.0019 | 0.0019 | 0.0017 |
| F06 | 78020001 | 78120000 | 0.0022 | 0.0017 | 0.0019 |
| F06 | 78040001 | 78140000 | 0.0020 | 0.0013 | 0.0016 |
| F06 | 78060001 | 78160000 | 0.0014 | 0.0009 | 0.0011 |
| F06 | 78080001 | 78180000 | 0.0009 | 0.0008 | 0.0007 |
| F06 | 78100001 | 78200000 | 0.0009 | 0.0012 | 0.0008 |
| F06 | 78120001 | 78220000 | 0.0013 | 0.0012 | 0.0011 |
| F06 | 78140001 | 78240000 | 0.0013 | 0.0015 | 0.0012 |
| F06 | 78160001 | 78260000 | 0.0016 | 0.0022 | 0.0014 |
| F06 | 78180001 | 78280000 | 0.0023 | 0.0026 | 0.0020 |
| F06 | 78200001 | 78300000 | 0.0026 | 0.0023 | 0.0020 |
| F06 | 78220001 | 78320000 | 0.0023 | 0.0024 | 0.0016 |
| F06 | 78240001 | 78340000 | 0.0024 | 0.0023 | 0.0016 |
| F06 | 78260001 | 78360000 | 0.0023 | 0.0017 | 0.0017 |
| F06 | 78280001 | 78380000 | 0.0017 | 0.0016 | 0.0010 |
| F06 | 78300001 | 78400000 | 0.0016 | 0.0021 | 0.0010 |
| F06 | 78320001 | 78420000 | 0.0021 | 0.0019 | 0.0015 |
| F06 | 78340001 | 78440000 | 0.0020 | 0.0024 | 0.0014 |
| F06 | 78360001 | 78460000 | 0.0024 | 0.0027 | 0.0017 |
| F06 | 78380001 | 78480000 | 0.0026 | 0.0030 | 0.0020 |
| F06 | 78400001 | 78500000 | 0.0030 | 0.0022 | 0.0024 |
| F06 | 78420001 | 78520000 | 0.0023 | 0.0026 | 0.0018 |
| F06 | 78440001 | 78540000 | 0.0028 | 0.0023 | 0.0023 |
| F06 | 78460001 | 78560000 | 0.0025 | 0.0024 | 0.0021 |
| F06 | 78480001 | 78580000 | 0.0026 | 0.0018 | 0.0022 |
| F06 | 78500001 | 78600000 | 0.0021 | 0.0016 | 0.0019 |
| F06 | 78520001 | 78620000 | 0.0019 | 0.0014 | 0.0017 |
| F06 | 78540001 | 78640000 | 0.0016 | 0.0017 | 0.0013 |
| F06 | 78560001 | 78660000 | 0.0018 | 0.0018 | 0.0014 |
| F06 | 78580001 | 78680000 | 0.0020 | 0.0018 | 0.0017 |
| F06 | 78600001 | 78700000 | 0.0019 | 0.0020 | 0.0015 |
| F06 | 78620001 | 78720000 | 0.0021 | 0.0022 | 0.0016 |

| Chr | Start | End | both | A1 | A2 |
|---|---|---|---|---|---|
| F06 | 78640001 | 78740000 | 0.0023 | 0.0019 | 0.0018 |
| F06 | 78660001 | 78760000 | 0.0019 | 0.0017 | 0.0015 |
| F06 | 78680001 | 78780000 | 0.0017 | 0.0018 | 0.0012 |
| F06 | 78700001 | 78800000 | 0.0017 | 0.0019 | 0.0013 |
| F06 | 78720001 | 78820000 | 0.0019 | 0.0016 | 0.0013 |
| F06 | 78740001 | 78840000 | 0.0015 | 0.0014 | 0.0010 |
| F06 | 78760001 | 78860000 | 0.0014 | 0.0012 | 0.0010 |
| F06 | 78780001 | 78880000 | 0.0012 | 0.0011 | 0.0008 |
| F06 | 78800001 | 78900000 | 0.0012 | 0.0010 | 0.0009 |
| F06 | 78820001 | 78920000 | 0.0011 | 0.0012 | 0.0009 |
| F06 | 78840001 | 78940000 | 0.0013 | 0.0019 | 0.0012 |
| F06 | 78860001 | 78960000 | 0.0020 | 0.0017 | 0.0017 |
| F06 | 78880001 | 78980000 | 0.0018 | 0.0016 | 0.0015 |
| F06 | 78900001 | 79000000 | 0.0017 | 0.0028 | 0.0016 |
| F06 | 78920001 | 79020000 | 0.0030 | 0.0030 | 0.0028 |
| F06 | 78940001 | 79040000 | 0.0031 | 0.0024 | 0.0029 |
| F06 | 78960001 | 79060000 | 0.0025 | 0.0025 | 0.0024 |
| F06 | 78980001 | 79080000 | 0.0027 | 0.0025 | 0.0026 |
| F06 | 79000001 | 79100000 | 0.0026 | 0.0010 | 0.0025 |
| F06 | 79020001 | 79120000 | 0.0011 | 0.0012 | 0.0010 |
| F06 | 79040001 | 79140000 | 0.0014 | 0.0015 | 0.0014 |
| F06 | 79060001 | 79160000 | 0.0017 | 0.0017 | 0.0016 |
| F06 | 79080001 | 79180000 | 0.0020 | 0.0019 | 0.0018 |
| F06 | 79100001 | 79200000 | 0.0020 | 0.0021 | 0.0018 |
| F06 | 79120001 | 79220000 | 0.0022 | 0.0018 | 0.0020 |
| F06 | 79140001 | 79240000 | 0.0018 | 0.0014 | 0.0015 |
| F06 | 79160001 | 79260000 | 0.0014 | 0.0010 | 0.0012 |
| F06 | 79180001 | 79280000 | 0.0010 | 0.0008 | 0.0009 |
| F06 | 79200001 | 79300000 | 0.0009 | 0.0008 | 0.0008 |
| F06 | 79220001 | 79320000 | 0.0010 | 0.0007 | 0.0010 |
| F06 | 79240001 | 79340000 | 0.0009 | 0.0008 | 0.0009 |
| F06 | 79260001 | 79360000 | 0.0010 | 0.0008 | 0.0010 |
| F06 | 79280001 | 79380000 | 0.0011 | 0.0008 | 0.0011 |
| F06 | 79300001 | 79400000 | 0.0010 | 0.0006 | 0.0010 |
| F06 | 79320001 | 79420000 | 0.0007 | 0.0006 | 0.0007 |
| F06 | 79340001 | 79440000 | 0.0007 | 0.0005 | 0.0007 |
| F06 | 79360001 | 79460000 | 0.0006 | 0.0009 | 0.0006 |
| F06 | 79380001 | 79480000 | 0.0011 | 0.0010 | 0.0011 |
| F06 | 79400001 | 79500000 | 0.0012 | 0.0021 | 0.0012 |
| F06 | 79420001 | 79520000 | 0.0024 | 0.0019 | 0.0024 |
| F06 | 79440001 | 79540000 | 0.0022 | 0.0022 | 0.0022 |
| F06 | 79460001 | 79560000 | 0.0025 | 0.0017 | 0.0025 |
| F06 | 79480001 | 79580000 | 0.0020 | 0.0017 | 0.0019 |
| F06 | 79500001 | 79600000 | 0.0020 | 0.0006 | 0.0019 |
| F06 | 79520001 | 79620000 | 0.0007 | 0.0006 | 0.0007 |
| F06 | 79540001 | 79640000 | 0.0008 | 0.0006 | 0.0008 |
| F06 | 79560001 | 79660000 | 0.0007 | 0.0010 | 0.0007 |
| F06 | 79580001 | 79680000 | 0.0011 | 0.0009 | 0.0011 |
| F06 | 79600001 | 79700000 | 0.0010 | 0.0012 | 0.0010 |
| F06 | 79620001 | 79720000 | 0.0014 | 0.0011 | 0.0014 |
| F06 | 79640001 | 79740000 | 0.0014 | 0.0008 | 0.0013 |
| F06 | 79660001 | 79760000 | 0.0010 | 0.0005 | 0.0010 |
| F06 | 79680001 | 79780000 | 0.0007 | 0.0005 | 0.0007 |
| F06 | 79700001 | 79800000 | 0.0007 | 0.0002 | 0.0007 |
| F06 | 79720001 | 79820000 | 0.0002 | 0.0002 | 0.0002 |
| F06 | 79740001 | 79840000 | 0.0003 | 0.0006 | 0.0003 |
| F06 | 79760001 | 79860000 | 0.0007 | 0.0005 | 0.0007 |
| F06 | 79780001 | 79880000 | 0.0006 | 0.0008 | 0.0006 |

| Chr | Start | End | both | A1 | A2 |
|---|---|---|---|---|---|
| F06 | 79800001 | 79900000 | 0.0010 | 0.0010 | 0.0009 |
| F06 | 79820001 | 79920000 | 0.0011 | 0.0010 | 0.0010 |
| F06 | 79840001 | 79940000 | 0.0011 | 0.0007 | 0.0009 |
| F06 | 79860001 | 79960000 | 0.0007 | 0.0010 | 0.0006 |
| F06 | 79880001 | 79980000 | 0.0010 | 0.0007 | 0.0008 |
| F06 | 79900001 | 80000000 | 0.0007 | 0.0006 | 0.0006 |
| F06 | 79920001 | 80020000 | 0.0006 | 0.0007 | 0.0005 |
| F06 | 79940001 | 80040000 | 0.0007 | 0.0009 | 0.0007 |
| F06 | 79960001 | 80060000 | 0.0009 | 0.0009 | 0.0009 |
| F06 | 79980001 | 80080000 | 0.0010 | 0.0009 | 0.0010 |
| F06 | 80000001 | 80100000 | 0.0011 | 0.0008 | 0.0010 |
| F06 | 80020001 | 80120000 | 0.0010 | 0.0008 | 0.0009 |
| F06 | 80040001 | 80140000 | 0.0009 | 0.0007 | 0.0009 |
| F06 | 80060001 | 80160000 | 0.0008 | 0.0005 | 0.0008 |
| F06 | 80080001 | 80180000 | 0.0006 | 0.0005 | 0.0005 |
| F06 | 80100001 | 80200000 | 0.0006 | 0.0007 | 0.0006 |
| F06 | 80120001 | 80220000 | 0.0008 | 0.0011 | 0.0008 |
| F06 | 80140001 | 80240000 | 0.0012 | 0.0012 | 0.0011 |
| F06 | 80160001 | 80260000 | 0.0013 | 0.0020 | 0.0011 |
| F06 | 80180001 | 80280000 | 0.0021 | 0.0024 | 0.0017 |
| F06 | 80200001 | 80300000 | 0.0025 | 0.0026 | 0.0022 |
| F06 | 80220001 | 80320000 | 0.0029 | 0.0022 | 0.0025 |
| F06 | 80240001 | 80340000 | 0.0025 | 0.0021 | 0.0022 |
| F06 | 80260001 | 80360000 | 0.0023 | 0.0014 | 0.0021 |
| F06 | 80280001 | 80380000 | 0.0016 | 0.0009 | 0.0015 |
| F06 | 80300001 | 80400000 | 0.0012 | 0.0009 | 0.0009 |
| F06 | 80320001 | 80420000 | 0.0011 | 0.0017 | 0.0009 |
| F06 | 80340001 | 80440000 | 0.0019 | 0.0019 | 0.0017 |
| F06 | 80360001 | 80460000 | 0.0022 | 0.0029 | 0.0019 |
| F06 | 80380001 | 80480000 | 0.0035 | 0.0031 | 0.0031 |
| F06 | 80400001 | 80500000 | 0.0039 | 0.0030 | 0.0033 |
| F06 | 80420001 | 80520000 | 0.0038 | 0.0028 | 0.0032 |
| F06 | 80440001 | 80540000 | 0.0037 | 0.0025 | 0.0030 |
| F06 | 80460001 | 80560000 | 0.0033 | 0.0014 | 0.0027 |
| F06 | 80480001 | 80580000 | 0.0020 | 0.0014 | 0.0014 |
| F06 | 80500001 | 80600000 | 0.0020 | 0.0010 | 0.0014 |
| F06 | 80520001 | 80620000 | 0.0016 | 0.0008 | 0.0011 |
| F06 | 80540001 | 80640000 | 0.0013 | 0.0011 | 0.0008 |
| F06 | 80560001 | 80660000 | 0.0018 | 0.0015 | 0.0012 |
| F06 | 80580001 | 80680000 | 0.0022 | 0.0015 | 0.0016 |
| F06 | 80600001 | 80700000 | 0.0021 | 0.0016 | 0.0016 |
| F06 | 80620001 | 80720000 | 0.0022 | 0.0019 | 0.0016 |
| F06 | 80640001 | 80740000 | 0.0024 | 0.0018 | 0.0018 |
| F06 | 80660001 | 80760000 | 0.0021 | 0.0022 | 0.0016 |
| F06 | 80680001 | 80780000 | 0.0024 | 0.0024 | 0.0018 |
| F06 | 80700001 | 80800000 | 0.0027 | 0.0028 | 0.0021 |
| F06 | 80720001 | 80820000 | 0.0031 | 0.0024 | 0.0024 |
| F06 | 80740001 | 80840000 | 0.0028 | 0.0024 | 0.0022 |
| F06 | 80760001 | 80860000 | 0.0027 | 0.0025 | 0.0022 |
| F06 | 80780001 | 80880000 | 0.0029 | 0.0029 | 0.0023 |
| F06 | 80800001 | 80900000 | 0.0032 | 0.0028 | 0.0025 |
| F06 | 80820001 | 80920000 | 0.0031 | 0.0029 | 0.0024 |
| F06 | 80840001 | 80940000 | 0.0032 | 0.0028 | 0.0025 |
| F06 | 80860001 | 80960000 | 0.0031 | 0.0023 | 0.0026 |
| F06 | 80880001 | 80980000 | 0.0026 | 0.0019 | 0.0020 |
| F06 | 80900001 | 81000000 | 0.0022 | 0.0022 | 0.0018 |
| F06 | 80920001 | 81020000 | 0.0024 | 0.0024 | 0.0019 |
| F06 | 80940001 | 81040000 | 0.0025 | 0.0025 | 0.0019 |

| Chr | Start | End | both | A1 | A2 |
|---|---|---|---|---|---|
| F06 | 80960001 | 81060000 | 0.0027 | 0.0025 | 0.0021 |
| F06 | 80980001 | 81080000 | 0.0027 | 0.0021 | 0.0021 |
| F06 | 81000001 | 81100000 | 0.0023 | 0.0018 | 0.0018 |
| F06 | 81020001 | 81120000 | 0.0020 | 0.0015 | 0.0016 |
| F06 | 81040001 | 81140000 | 0.0017 | 0.0015 | 0.0015 |
| F06 | 81060001 | 81160000 | 0.0016 | 0.0018 | 0.0014 |
| F06 | 81080001 | 81180000 | 0.0019 | 0.0019 | 0.0017 |
| F06 | 81100001 | 81200000 | 0.0020 | 0.0016 | 0.0016 |
| F06 | 81120001 | 81220000 | 0.0017 | 0.0016 | 0.0014 |
| F06 | 81140001 | 81240000 | 0.0017 | 0.0016 | 0.0013 |
| F06 | 81160001 | 81260000 | 0.0019 | 0.0014 | 0.0015 |
| F06 | 81180001 | 81280000 | 0.0017 | 0.0015 | 0.0012 |
| F06 | 81200001 | 81300000 | 0.0017 | 0.0022 | 0.0013 |
| F06 | 81220001 | 81320000 | 0.0027 | 0.0026 | 0.0022 |
| F06 | 81240001 | 81340000 | 0.0031 | 0.0023 | 0.0028 |
| F06 | 81260001 | 81360000 | 0.0027 | 0.0022 | 0.0025 |
| F06 | 81280001 | 81380000 | 0.0026 | 0.0025 | 0.0026 |
| F06 | 81300001 | 81400000 | 0.0030 | 0.0019 | 0.0030 |
| F06 | 81320001 | 81420000 | 0.0022 | 0.0015 | 0.0022 |
| F06 | 81340001 | 81440000 | 0.0017 | 0.0016 | 0.0016 |
| F06 | 81360001 | 81460000 | 0.0019 | 0.0016 | 0.0017 |
| F06 | 81380001 | 81480000 | 0.0018 | 0.0016 | 0.0016 |
| F06 | 81400001 | 81500000 | 0.0018 | 0.0017 | 0.0013 |
| F06 | 81420001 | 81520000 | 0.0018 | 0.0016 | 0.0014 |
| F06 | 81440001 | 81540000 | 0.0018 | 0.0014 | 0.0012 |
| F06 | 81460001 | 81560000 | 0.0015 | 0.0016 | 0.0011 |
| F06 | 81480001 | 81580000 | 0.0017 | 0.0012 | 0.0011 |
| F06 | 81500001 | 81600000 | 0.0012 | 0.0016 | 0.0007 |
| F06 | 81520001 | 81620000 | 0.0018 | 0.0015 | 0.0012 |
| F06 | 81540001 | 81640000 | 0.0016 | 0.0017 | 0.0011 |
| F06 | 81560001 | 81660000 | 0.0019 | 0.0023 | 0.0014 |
| F06 | 81580001 | 81680000 | 0.0026 | 0.0026 | 0.0022 |
| F06 | 81600001 | 81700000 | 0.0029 | 0.0024 | 0.0026 |
| F06 | 81620001 | 81720000 | 0.0026 | 0.0031 | 0.0023 |
| F06 | 81640001 | 81740000 | 0.0034 | 0.0032 | 0.0029 |
| F06 | 81660001 | 81760000 | 0.0035 | 0.0022 | 0.0030 |
| F06 | 81680001 | 81780000 | 0.0024 | 0.0021 | 0.0019 |
| F06 | 81700001 | 81800000 | 0.0023 | 0.0019 | 0.0019 |
| F06 | 81720001 | 81820000 | 0.0021 | 0.0015 | 0.0016 |
| F06 | 81740001 | 81840000 | 0.0017 | 0.0016 | 0.0013 |
| F06 | 81760001 | 81860000 | 0.0018 | 0.0016 | 0.0013 |
| F06 | 81780001 | 81880000 | 0.0018 | 0.0013 | 0.0014 |
| F06 | 81800001 | 81900000 | 0.0014 | 0.0015 | 0.0010 |
| F06 | 81820001 | 81920000 | 0.0016 | 0.0016 | 0.0013 |
| F06 | 81840001 | 81940000 | 0.0021 | 0.0025 | 0.0016 |
| F06 | 81860001 | 81960000 | 0.0031 | 0.0026 | 0.0026 |
| F06 | 81880001 | 81980000 | 0.0035 | 0.0026 | 0.0029 |
| F06 | 81900001 | 82000000 | 0.0034 | 0.0023 | 0.0029 |
| F06 | 81920001 | 82020000 | 0.0032 | 0.0022 | 0.0025 |
| F06 | 81940001 | 82040000 | 0.0031 | 0.0013 | 0.0024 |
| F06 | 81960001 | 82060000 | 0.0019 | 0.0014 | 0.0015 |
| F06 | 81980001 | 82080000 | 0.0019 | 0.0018 | 0.0017 |
| F06 | 82000001 | 82100000 | 0.0025 | 0.0019 | 0.0020 |
| F06 | 82020001 | 82120000 | 0.0028 | 0.0022 | 0.0022 |
| F06 | 82040001 | 82140000 | 0.0030 | 0.0021 | 0.0023 |
| F06 | 82060001 | 82160000 | 0.0031 | 0.0022 | 0.0022 |
| F06 | 82080001 | 82180000 | 0.0032 | 0.0024 | 0.0022 |
| F06 | 82100001 | 82200000 | 0.0034 | 0.0023 | 0.0026 |

| Chr | Start | End | both | A1 | A2 |
|---|---|---|---|---|---|
| F06 | 82120001 | 82220000 | 0.0030 | 0.0021 | 0.0024 |
| F06 | 82140001 | 82240000 | 0.0029 | 0.0023 | 0.0022 |
| F06 | 82160001 | 82260000 | 0.0030 | 0.0023 | 0.0025 |
| F06 | 82180001 | 82280000 | 0.0031 | 0.0023 | 0.0025 |
| F06 | 82200001 | 82300000 | 0.0033 | 0.0027 | 0.0024 |
| F06 | 82220001 | 82320000 | 0.0040 | 0.0024 | 0.0027 |
| F06 | 82240001 | 82340000 | 0.0037 | 0.0024 | 0.0025 |
| F06 | 82260001 | 82360000 | 0.0038 | 0.0022 | 0.0025 |
| F06 | 82280001 | 82380000 | 0.0034 | 0.0017 | 0.0024 |
| F06 | 82300001 | 82400000 | 0.0027 | 0.0016 | 0.0020 |
| F06 | 82320001 | 82420000 | 0.0023 | 0.0015 | 0.0018 |
| F06 | 82340001 | 82440000 | 0.0020 | 0.0014 | 0.0017 |
| F06 | 82360001 | 82460000 | 0.0017 | 0.0013 | 0.0017 |
| F06 | 82380001 | 82480000 | 0.0016 | 0.0024 | 0.0017 |
| F06 | 82400001 | 82500000 | 0.0028 | 0.0023 | 0.0027 |
| F06 | 82420001 | 82520000 | 0.0028 | 0.0022 | 0.0027 |
| F06 | 82440001 | 82540000 | 0.0025 | 0.0018 | 0.0024 |
| F06 | 82460001 | 82560000 | 0.0020 | 0.0019 | 0.0019 |
| F06 | 82480001 | 82580000 | 0.0022 | 0.0009 | 0.0020 |
| F06 | 82500001 | 82600000 | 0.0013 | 0.0007 | 0.0010 |
| F06 | 82520001 | 82620000 | 0.0011 | 0.0007 | 0.0008 |
| F06 | 82540001 | 82640000 | 0.0012 | 0.0010 | 0.0009 |
| F06 | 82560001 | 82660000 | 0.0015 | 0.0007 | 0.0011 |
| F06 | 82580001 | 82680000 | 0.0011 | 0.0008 | 0.0007 |
| F06 | 82600001 | 82700000 | 0.0012 | 0.0009 | 0.0010 |
| F06 | 82620001 | 82720000 | 0.0014 | 0.0016 | 0.0011 |
| F06 | 82640001 | 82740000 | 0.0022 | 0.0014 | 0.0019 |
| F06 | 82660001 | 82760000 | 0.0020 | 0.0014 | 0.0017 |
| F06 | 82680001 | 82780000 | 0.0020 | 0.0013 | 0.0017 |
| F06 | 82700001 | 82800000 | 0.0018 | 0.0013 | 0.0015 |
| F06 | 82720001 | 82820000 | 0.0018 | 0.0010 | 0.0016 |
| F06 | 82740001 | 82840000 | 0.0015 | 0.0011 | 0.0011 |
| F06 | 82760001 | 82860000 | 0.0018 | 0.0011 | 0.0012 |
| F06 | 82780001 | 82880000 | 0.0018 | 0.0008 | 0.0012 |
| F06 | 82800001 | 82900000 | 0.0014 | 0.0008 | 0.0010 |
| F06 | 82820001 | 82920000 | 0.0013 | 0.0005 | 0.0008 |
| F06 | 82840001 | 82940000 | 0.0008 | 0.0013 | 0.0006 |
| F06 | 82860001 | 82960000 | 0.0016 | 0.0018 | 0.0015 |
| F06 | 82880001 | 82980000 | 0.0021 | 0.0030 | 0.0019 |
| F06 | 82900001 | 83000000 | 0.0035 | 0.0036 | 0.0031 |
| F06 | 82920001 | 83020000 | 0.0043 | 0.0039 | 0.0038 |
| F06 | 82940001 | 83040000 | 0.0047 | 0.0030 | 0.0041 |
| F06 | 82960001 | 83060000 | 0.0039 | 0.0027 | 0.0034 |
| F06 | 82980001 | 83080000 | 0.0035 | 0.0019 | 0.0030 |
| F06 | 83000001 | 83100000 | 0.0028 | 0.0018 | 0.0022 |
| F06 | 83020001 | 83120000 | 0.0025 | 0.0019 | 0.0021 |
| F06 | 83040001 | 83140000 | 0.0026 | 0.0021 | 0.0022 |
| F06 | 83060001 | 83160000 | 0.0030 | 0.0022 | 0.0024 |
| F06 | 83080001 | 83180000 | 0.0033 | 0.0023 | 0.0026 |
| F06 | 83100001 | 83200000 | 0.0035 | 0.0019 | 0.0026 |
| F06 | 83120001 | 83220000 | 0.0030 | 0.0017 | 0.0021 |
| F06 | 83140001 | 83240000 | 0.0028 | 0.0018 | 0.0020 |
| F06 | 83160001 | 83260000 | 0.0028 | 0.0020 | 0.0020 |
| F06 | 83180001 | 83280000 | 0.0028 | 0.0016 | 0.0022 |
| F06 | 83200001 | 83300000 | 0.0022 | 0.0016 | 0.0018 |
| F06 | 83220001 | 83320000 | 0.0024 | 0.0016 | 0.0018 |
| F06 | 83240001 | 83340000 | 0.0023 | 0.0016 | 0.0016 |
| F06 | 83260001 | 83360000 | 0.0023 | 0.0012 | 0.0015 |

| Chr | Start | End | both | A1 | A2 |
|---|---|---|---|---|---|
| F06 | 83280001 | 83380000 | 0.0018 | 0.0017 | 0.0011 |
| F06 | 83300001 | 83400000 | 0.0024 | 0.0022 | 0.0016 |
| F06 | 83320001 | 83420000 | 0.0027 | 0.0020 | 0.0022 |
| F06 | 83340001 | 83440000 | 0.0025 | 0.0017 | 0.0021 |
| F06 | 83360001 | 83460000 | 0.0020 | 0.0018 | 0.0018 |
| F06 | 83380001 | 83480000 | 0.0023 | 0.0015 | 0.0020 |
| F06 | 83400001 | 83500000 | 0.0018 | 0.0011 | 0.0017 |
| F06 | 83420001 | 83520000 | 0.0016 | 0.0026 | 0.0013 |
| F06 | 83440001 | 83540000 | 0.0032 | 0.0029 | 0.0026 |
| F06 | 83460001 | 83560000 | 0.0036 | 0.0028 | 0.0029 |
| F06 | 83480001 | 83580000 | 0.0035 | 0.0029 | 0.0028 |
| F06 | 83500001 | 83600000 | 0.0038 | 0.0032 | 0.0028 |
| F06 | 83520001 | 83620000 | 0.0040 | 0.0016 | 0.0032 |
| F06 | 83540001 | 83640000 | 0.0025 | 0.0014 | 0.0018 |
| F06 | 83560001 | 83660000 | 0.0025 | 0.0017 | 0.0017 |
| F06 | 83580001 | 83680000 | 0.0029 | 0.0015 | 0.0019 |
| F06 | 83600001 | 83700000 | 0.0027 | 0.0010 | 0.0018 |
| F06 | 83620001 | 83720000 | 0.0022 | 0.0011 | 0.0011 |
| F06 | 83640001 | 83740000 | 0.0022 | 0.0013 | 0.0012 |
| F06 | 83660001 | 83760000 | 0.0024 | 0.0009 | 0.0014 |
| F06 | 83680001 | 83780000 | 0.0019 | 0.0012 | 0.0009 |
| F06 | 83700001 | 83800000 | 0.0021 | 0.0014 | 0.0012 |
| F06 | 83720001 | 83820000 | 0.0023 | 0.0015 | 0.0014 |
| F06 | 83740001 | 83840000 | 0.0024 | 0.0017 | 0.0015 |
| F06 | 83760001 | 83860000 | 0.0025 | 0.0020 | 0.0017 |
| F06 | 83780001 | 83880000 | 0.0028 | 0.0016 | 0.0020 |
| F06 | 83800001 | 83900000 | 0.0025 | 0.0015 | 0.0017 |
| F06 | 83820001 | 83920000 | 0.0023 | 0.0014 | 0.0017 |
| F06 | 83840001 | 83940000 | 0.0021 | 0.0009 | 0.0016 |
| F06 | 83860001 | 83960000 | 0.0018 | 0.0011 | 0.0012 |
| F06 | 83880001 | 83980000 | 0.0018 | 0.0016 | 0.0013 |
| F06 | 83900001 | 84000000 | 0.0022 | 0.0017 | 0.0018 |
| F06 | 83920001 | 84020000 | 0.0025 | 0.0017 | 0.0019 |
| F06 | 83940001 | 84040000 | 0.0029 | 0.0021 | 0.0020 |
| F06 | 83960001 | 84060000 | 0.0034 | 0.0018 | 0.0024 |
| F06 | 83980001 | 84080000 | 0.0034 | 0.0014 | 0.0022 |
| F06 | 84000001 | 84100000 | 0.0031 | 0.0011 | 0.0017 |
| F06 | 84020001 | 84120000 | 0.0025 | 0.0009 | 0.0014 |
| F06 | 84040001 | 84140000 | 0.0022 | 0.0009 | 0.0012 |
| F06 | 84060001 | 84160000 | 0.0020 | 0.0012 | 0.0011 |
| F06 | 84080001 | 84180000 | 0.0021 | 0.0012 | 0.0015 |
| F06 | 84100001 | 84200000 | 0.0018 | 0.0015 | 0.0014 |
| F06 | 84120001 | 84220000 | 0.0023 | 0.0016 | 0.0019 |
| F06 | 84140001 | 84240000 | 0.0020 | 0.0014 | 0.0018 |
| F06 | 84160001 | 84260000 | 0.0017 | 0.0012 | 0.0016 |
| F06 | 84180001 | 84280000 | 0.0015 | 0.0015 | 0.0014 |
| F06 | 84200001 | 84300000 | 0.0019 | 0.0013 | 0.0018 |
| F06 | 84220001 | 84320000 | 0.0017 | 0.0017 | 0.0015 |
| F06 | 84240001 | 84340000 | 0.0025 | 0.0017 | 0.0018 |
| F06 | 84260001 | 84360000 | 0.0026 | 0.0017 | 0.0019 |
| F06 | 84280001 | 84380000 | 0.0027 | 0.0016 | 0.0019 |
| F06 | 84300001 | 84400000 | 0.0027 | 0.0015 | 0.0018 |
| F06 | 84320001 | 84420000 | 0.0026 | 0.0012 | 0.0017 |
| F06 | 84340001 | 84440000 | 0.0020 | 0.0012 | 0.0014 |
| F06 | 84360001 | 84460000 | 0.0018 | 0.0014 | 0.0011 |
| F06 | 84380001 | 84480000 | 0.0022 | 0.0015 | 0.0012 |
| F06 | 84400001 | 84500000 | 0.0023 | 0.0020 | 0.0015 |
| F06 | 84420001 | 84520000 | 0.0028 | 0.0023 | 0.0021 |

| Chr | Start | End | both | A1 | A2 |
|---|---|---|---|---|---|
| F06 | 84440001 | 84540000 | 0.0034 | 0.0022 | 0.0025 |
| F06 | 84460001 | 84560000 | 0.0035 | 0.0020 | 0.0027 |
| F06 | 84480001 | 84580000 | 0.0030 | 0.0019 | 0.0024 |
| F06 | 84500001 | 84600000 | 0.0030 | 0.0018 | 0.0023 |
| F06 | 84520001 | 84620000 | 0.0027 | 0.0017 | 0.0022 |
| F06 | 84540001 | 84640000 | 0.0025 | 0.0019 | 0.0020 |
| F06 | 84560001 | 84660000 | 0.0026 | 0.0021 | 0.0021 |
| F06 | 84580001 | 84680000 | 0.0029 | 0.0019 | 0.0023 |
| F06 | 84600001 | 84700000 | 0.0026 | 0.0016 | 0.0020 |
| F06 | 84620001 | 84720000 | 0.0023 | 0.0014 | 0.0017 |
| F06 | 84640001 | 84740000 | 0.0019 | 0.0016 | 0.0015 |
| F06 | 84660001 | 84760000 | 0.0023 | 0.0014 | 0.0018 |
| F06 | 84680001 | 84780000 | 0.0022 | 0.0014 | 0.0017 |
| F06 | 84700001 | 84800000 | 0.0022 | 0.0017 | 0.0018 |
| F06 | 84720001 | 84820000 | 0.0023 | 0.0017 | 0.0020 |
| F06 | 84740001 | 84840000 | 0.0025 | 0.0012 | 0.0020 |
| F06 | 84760001 | 84860000 | 0.0020 | 0.0016 | 0.0015 |
| F06 | 84780001 | 84880000 | 0.0025 | 0.0017 | 0.0018 |
| F06 | 84800001 | 84900000 | 0.0026 | 0.0018 | 0.0019 |
| F06 | 84820001 | 84920000 | 0.0030 | 0.0017 | 0.0021 |
| F06 | 84840001 | 84940000 | 0.0029 | 0.0018 | 0.0022 |
| F06 | 84860001 | 84960000 | 0.0028 | 0.0019 | 0.0022 |
| F06 | 84880001 | 84980000 | 0.0030 | 0.0026 | 0.0023 |
| F06 | 84900001 | 85000000 | 0.0037 | 0.0023 | 0.0032 |
| F06 | 84920001 | 85020000 | 0.0034 | 0.0027 | 0.0027 |
| F06 | 84940001 | 85040000 | 0.0040 | 0.0026 | 0.0030 |
| F06 | 84960001 | 85060000 | 0.0039 | 0.0025 | 0.0029 |
| F06 | 84980001 | 85080000 | 0.0039 | 0.0017 | 0.0028 |
| F06 | 85000001 | 85100000 | 0.0032 | 0.0022 | 0.0019 |
| F06 | 85020001 | 85120000 | 0.0038 | 0.0019 | 0.0025 |
| F06 | 85040001 | 85140000 | 0.0033 | 0.0022 | 0.0022 |
| F06 | 85060001 | 85160000 | 0.0035 | 0.0021 | 0.0025 |
| F06 | 85080001 | 85180000 | 0.0032 | 0.0028 | 0.0024 |
| F06 | 85100001 | 85200000 | 0.0039 | 0.0021 | 0.0031 |
| F06 | 85120001 | 85220000 | 0.0032 | 0.0027 | 0.0025 |
| F06 | 85140001 | 85240000 | 0.0038 | 0.0045 | 0.0030 |
| F06 | 85160001 | 85260000 | 0.0058 | 0.0040 | 0.0047 |
| F06 | 85180001 | 85280000 | 0.0054 | 0.0039 | 0.0041 |
| F06 | 85200001 | 85300000 | 0.0051 | 0.0044 | 0.0040 |
| F06 | 85220001 | 85320000 | 0.0056 | 0.0043 | 0.0045 |
| F06 | 85240001 | 85340000 | 0.0056 | 0.0025 | 0.0044 |
| F06 | 85260001 | 85360000 | 0.0038 | 0.0034 | 0.0029 |
| F06 | 85280001 | 85380000 | 0.0046 | 0.0029 | 0.0038 |
| F06 | 85300001 | 85400000 | 0.0040 | 0.0028 | 0.0034 |
| F06 | 85320001 | 85420000 | 0.0035 | 0.0025 | 0.0030 |
| F06 | 85340001 | 85440000 | 0.0031 | 0.0024 | 0.0028 |
| F06 | 85360001 | 85460000 | 0.0030 | 0.0019 | 0.0026 |
| F06 | 85380001 | 85480000 | 0.0024 | 0.0019 | 0.0021 |
| F06 | 85400001 | 85500000 | 0.0024 | 0.0015 | 0.0021 |
| F06 | 85420001 | 85520000 | 0.0022 | 0.0017 | 0.0018 |
| F06 | 85440001 | 85540000 | 0.0023 | 0.0020 | 0.0020 |
| F06 | 85460001 | 85560000 | 0.0027 | 0.0028 | 0.0022 |
| F06 | 85480001 | 85580000 | 0.0038 | 0.0026 | 0.0032 |
| F06 | 85500001 | 85600000 | 0.0035 | 0.0028 | 0.0029 |
| F06 | 85520001 | 85620000 | 0.0035 | 0.0024 | 0.0031 |
| F06 | 85540001 | 85640000 | 0.0032 | 0.0020 | 0.0027 |
| F06 | 85560001 | 85660000 | 0.0031 | 0.0017 | 0.0025 |
| F06 | 85580001 | 85680000 | 0.0029 | 0.0022 | 0.0022 |

| Chr | Start | End | both | A1 | A2 |
|---|---|---|---|---|---|
| F06 | 85600001 | 85700000 | 0.0035 | 0.0020 | 0.0028 |
| F06 | 85620001 | 85720000 | 0.0036 | 0.0020 | 0.0026 |
| F06 | 85640001 | 85740000 | 0.0035 | 0.0021 | 0.0026 |
| F06 | 85660001 | 85760000 | 0.0034 | 0.0017 | 0.0025 |
| F06 | 85680001 | 85780000 | 0.0026 | 0.0015 | 0.0019 |
| F06 | 85700001 | 85800000 | 0.0025 | 0.0022 | 0.0017 |
| F06 | 85720001 | 85820000 | 0.0029 | 0.0023 | 0.0022 |
| F06 | 85740001 | 85840000 | 0.0029 | 0.0027 | 0.0023 |
| F06 | 85760001 | 85860000 | 0.0032 | 0.0027 | 0.0028 |
| F06 | 85780001 | 85880000 | 0.0032 | 0.0025 | 0.0028 |
| F06 | 85800001 | 85900000 | 0.0030 | 0.0018 | 0.0027 |
| F06 | 85820001 | 85920000 | 0.0022 | 0.0028 | 0.0020 |
| F06 | 85840001 | 85940000 | 0.0033 | 0.0023 | 0.0029 |
| F06 | 85860001 | 85960000 | 0.0027 | 0.0016 | 0.0024 |
| F06 | 85880001 | 85980000 | 0.0021 | 0.0016 | 0.0017 |
| F06 | 85900001 | 86000000 | 0.0019 | 0.0014 | 0.0016 |
| F06 | 85920001 | 86020000 | 0.0019 | 0.0010 | 0.0015 |
| F06 | 85940001 | 86040000 | 0.0016 | 0.0027 | 0.0013 |
| F06 | 85960001 | 86060000 | 0.0037 | 0.0037 | 0.0031 |
| F06 | 85980001 | 86080000 | 0.0048 | 0.0044 | 0.0041 |
| F06 | 86000001 | 86100000 | 0.0058 | 0.0045 | 0.0049 |
| F06 | 86020001 | 86120000 | 0.0060 | 0.0042 | 0.0050 |
| F06 | 86040001 | 86140000 | 0.0056 | 0.0029 | 0.0046 |
| F06 | 86060001 | 86160000 | 0.0040 | 0.0020 | 0.0032 |
| F06 | 86080001 | 86180000 | 0.0030 | 0.0014 | 0.0023 |
| F06 | 86100001 | 86200000 | 0.0024 | 0.0019 | 0.0017 |
| F06 | 86120001 | 86220000 | 0.0031 | 0.0015 | 0.0024 |
| F06 | 86140001 | 86240000 | 0.0026 | 0.0011 | 0.0020 |
| F06 | 86160001 | 86260000 | 0.0021 | 0.0015 | 0.0016 |
| F06 | 86180001 | 86280000 | 0.0027 | 0.0020 | 0.0021 |
| F06 | 86200001 | 86300000 | 0.0031 | 0.0023 | 0.0027 |
| F06 | 86220001 | 86320000 | 0.0033 | 0.0031 | 0.0028 |
| F06 | 86240001 | 86340000 | 0.0041 | 0.0032 | 0.0036 |
| F06 | 86260001 | 86360000 | 0.0042 | 0.0028 | 0.0037 |
| F06 | 86280001 | 86380000 | 0.0033 | 0.0021 | 0.0030 |
| F06 | 86300001 | 86400000 | 0.0026 | 0.0018 | 0.0023 |
| F06 | 86320001 | 86420000 | 0.0022 | 0.0013 | 0.0019 |
| F06 | 86340001 | 86440000 | 0.0017 | 0.0011 | 0.0015 |
| F06 | 86360001 | 86460000 | 0.0016 | 0.0016 | 0.0013 |
| F06 | 86380001 | 86480000 | 0.0022 | 0.0019 | 0.0018 |
| F06 | 86400001 | 86500000 | 0.0026 | 0.0020 | 0.0021 |
| F06 | 86420001 | 86520000 | 0.0027 | 0.0021 | 0.0024 |
| F06 | 86440001 | 86540000 | 0.0028 | 0.0023 | 0.0025 |
| F06 | 86460001 | 86560000 | 0.0033 | 0.0020 | 0.0029 |
| F06 | 86480001 | 86580000 | 0.0029 | 0.0022 | 0.0026 |
| F06 | 86500001 | 86600000 | 0.0031 | 0.0019 | 0.0027 |
| F06 | 86520001 | 86620000 | 0.0028 | 0.0019 | 0.0024 |
| F06 | 86540001 | 86640000 | 0.0028 | 0.0023 | 0.0022 |
| F06 | 86560001 | 86660000 | 0.0031 | 0.0022 | 0.0025 |
| F06 | 86580001 | 86680000 | 0.0032 | 0.0025 | 0.0025 |
| F06 | 86600001 | 86700000 | 0.0037 | 0.0025 | 0.0028 |
| F06 | 86620001 | 86720000 | 0.0038 | 0.0028 | 0.0029 |
| F06 | 86640001 | 86740000 | 0.0040 | 0.0023 | 0.0031 |
| F06 | 86660001 | 86760000 | 0.0033 | 0.0024 | 0.0025 |
| F06 | 86680001 | 86780000 | 0.0034 | 0.0026 | 0.0026 |
| F06 | 86700001 | 86800000 | 0.0034 | 0.0024 | 0.0028 |
| F06 | 86720001 | 86820000 | 0.0034 | 0.0021 | 0.0027 |
| F06 | 86740001 | 86840000 | 0.0029 | 0.0024 | 0.0023 |

| Chr | Start | End | both | A1 | A2 |
|---|---|---|---|---|---|
| F06 | 86760001 | 86860000 | 0.0033 | 0.0022 | 0.0027 |
| F06 | 86780001 | 86880000 | 0.0030 | 0.0014 | 0.0025 |
| F06 | 86800001 | 86900000 | 0.0020 | 0.0015 | 0.0016 |
| F06 | 86820001 | 86920000 | 0.0021 | 0.0017 | 0.0017 |
| F06 | 86840001 | 86940000 | 0.0026 | 0.0018 | 0.0020 |
| F06 | 86860001 | 86960000 | 0.0029 | 0.0022 | 0.0021 |
| F06 | 86880001 | 86980000 | 0.0034 | 0.0026 | 0.0026 |
| F06 | 86900001 | 87000000 | 0.0042 | 0.0023 | 0.0032 |
| F06 | 86920001 | 87020000 | 0.0036 | 0.0018 | 0.0028 |
| F06 | 86940001 | 87040000 | 0.0031 | 0.0018 | 0.0023 |
| F06 | 86960001 | 87060000 | 0.0030 | 0.0023 | 0.0023 |
| F06 | 86980001 | 87080000 | 0.0038 | 0.0019 | 0.0029 |
| F06 | 87000001 | 87100000 | 0.0031 | 0.0024 | 0.0023 |
| F06 | 87020001 | 87120000 | 0.0036 | 0.0027 | 0.0028 |
| F06 | 87040001 | 87140000 | 0.0037 | 0.0026 | 0.0031 |
| F06 | 87060001 | 87160000 | 0.0035 | 0.0020 | 0.0030 |
| F06 | 87080001 | 87180000 | 0.0025 | 0.0029 | 0.0022 |
| F06 | 87100001 | 87200000 | 0.0032 | 0.0026 | 0.0031 |
| F06 | 87120001 | 87220000 | 0.0030 | 0.0024 | 0.0029 |
| F06 | 87140001 | 87240000 | 0.0029 | 0.0031 | 0.0026 |
| F06 | 87160001 | 87260000 | 0.0037 | 0.0028 | 0.0033 |
| F06 | 87180001 | 87280000 | 0.0034 | 0.0022 | 0.0030 |
| F06 | 87200001 | 87300000 | 0.0030 | 0.0021 | 0.0026 |
| F06 | 87220001 | 87320000 | 0.0029 | 0.0021 | 0.0024 |
| F06 | 87240001 | 87340000 | 0.0028 | 0.0018 | 0.0024 |
| F06 | 87260001 | 87360000 | 0.0024 | 0.0018 | 0.0021 |
| F06 | 87280001 | 87380000 | 0.0025 | 0.0015 | 0.0021 |
| F06 | 87300001 | 87400000 | 0.0020 | 0.0015 | 0.0015 |
| F06 | 87320001 | 87420000 | 0.0020 | 0.0016 | 0.0016 |
| F06 | 87340001 | 87440000 | 0.0022 | 0.0005 | 0.0017 |
| F06 | 87360001 | 87460000 | 0.0012 | 0.0005 | 0.0007 |
| F06 | 87380001 | 87480000 | 0.0013 | 0.0009 | 0.0008 |
| F06 | 87400001 | 87500000 | 0.0018 | 0.0010 | 0.0011 |
| F06 | 87420001 | 87520000 | 0.0021 | 0.0013 | 0.0011 |
| F06 | 87440001 | 87540000 | 0.0024 | 0.0016 | 0.0015 |
| F06 | 87460001 | 87560000 | 0.0026 | 0.0015 | 0.0018 |
| F06 | 87480001 | 87580000 | 0.0023 | 0.0015 | 0.0017 |
| F06 | 87500001 | 87600000 | 0.0023 | 0.0014 | 0.0017 |
| F06 | 87520001 | 87620000 | 0.0019 | 0.0010 | 0.0016 |
| F06 | 87540001 | 87640000 | 0.0012 | 0.0013 | 0.0011 |
| F06 | 87560001 | 87660000 | 0.0015 | 0.0014 | 0.0014 |
| F06 | 87580001 | 87680000 | 0.0016 | 0.0015 | 0.0016 |
| F06 | 87600001 | 87700000 | 0.0018 | 0.0017 | 0.0016 |
| F06 | 87620001 | 87720000 | 0.0020 | 0.0035 | 0.0017 |
| F06 | 87640001 | 87740000 | 0.0043 | 0.0030 | 0.0036 |
| F06 | 87660001 | 87760000 | 0.0038 | 0.0035 | 0.0031 |
| F06 | 87680001 | 87780000 | 0.0045 | 0.0038 | 0.0037 |
| F06 | 87700001 | 87800000 | 0.0050 | 0.0038 | 0.0042 |
| F06 | 87720001 | 87820000 | 0.0050 | 0.0024 | 0.0042 |
| F06 | 87740001 | 87840000 | 0.0033 | 0.0026 | 0.0028 |
| F06 | 87760001 | 87860000 | 0.0034 | 0.0026 | 0.0029 |
| F06 | 87780001 | 87880000 | 0.0032 | 0.0021 | 0.0026 |
| F06 | 87800001 | 87900000 | 0.0027 | 0.0025 | 0.0020 |
| F06 | 87820001 | 87920000 | 0.0031 | 0.0024 | 0.0024 |
| F06 | 87840001 | 87940000 | 0.0030 | 0.0025 | 0.0022 |
| F06 | 87860001 | 87960000 | 0.0034 | 0.0022 | 0.0025 |
| F06 | 87880001 | 87980000 | 0.0032 | 0.0029 | 0.0024 |
| F06 | 87900001 | 88000000 | 0.0039 | 0.0024 | 0.0030 |

| Chr | Start | End | both | A1 | A2 |
|---|---|---|---|---|---|
| F06 | 87920001 | 88020000 | 0.0034 | 0.0023 | 0.0025 |
| F06 | 87940001 | 88040000 | 0.0032 | 0.0028 | 0.0022 |
| F06 | 87960001 | 88060000 | 0.0037 | 0.0032 | 0.0028 |
| F06 | 87980001 | 88080000 | 0.0044 | 0.0024 | 0.0033 |
| F06 | 88000001 | 88100000 | 0.0035 | 0.0023 | 0.0026 |
| F06 | 88020001 | 88120000 | 0.0035 | 0.0028 | 0.0026 |
| F06 | 88040001 | 88140000 | 0.0040 | 0.0022 | 0.0031 |
| F06 | 88060001 | 88160000 | 0.0033 | 0.0018 | 0.0026 |
| F06 | 88080001 | 88180000 | 0.0025 | 0.0017 | 0.0021 |
| F06 | 88100001 | 88200000 | 0.0025 | 0.0044 | 0.0019 |
| F06 | 88120001 | 88220000 | 0.0052 | 0.0040 | 0.0047 |
| F06 | 88140001 | 88240000 | 0.0050 | 0.0045 | 0.0044 |
| F06 | 88160001 | 88260000 | 0.0057 | 0.0045 | 0.0050 |
| F06 | 88180001 | 88280000 | 0.0059 | 0.0044 | 0.0049 |
| F06 | 88200001 | 88300000 | 0.0056 | 0.0017 | 0.0048 |
| F06 | 88220001 | 88320000 | 0.0026 | 0.0016 | 0.0020 |
| F06 | 88240001 | 88340000 | 0.0022 | 0.0010 | 0.0018 |
| F06 | 88260001 | 88360000 | 0.0014 | 0.0013 | 0.0011 |
| F06 | 88280001 | 88380000 | 0.0016 | 0.0013 | 0.0015 |
| F06 | 88300001 | 88400000 | 0.0015 | 0.0012 | 0.0014 |
| F06 | 88320001 | 88420000 | 0.0014 | 0.0011 | 0.0014 |
| F06 | 88340001 | 88440000 | 0.0013 | 0.0011 | 0.0012 |
| F06 | 88360001 | 88460000 | 0.0013 | 0.0010 | 0.0012 |
| F06 | 88380001 | 88480000 | 0.0011 | 0.0010 | 0.0009 |
| F06 | 88400001 | 88500000 | 0.0014 | 0.0011 | 0.0009 |
| F06 | 88420001 | 88520000 | 0.0016 | 0.0015 | 0.0010 |
| F06 | 88440001 | 88540000 | 0.0021 | 0.0019 | 0.0014 |
| F06 | 88460001 | 88560000 | 0.0027 | 0.0018 | 0.0018 |
| F06 | 88480001 | 88580000 | 0.0028 | 0.0019 | 0.0019 |
| F06 | 88500001 | 88600000 | 0.0027 | 0.0021 | 0.0020 |
| F06 | 88520001 | 88620000 | 0.0029 | 0.0028 | 0.0023 |
| F06 | 88540001 | 88640000 | 0.0034 | 0.0026 | 0.0027 |
| F06 | 88560001 | 88660000 | 0.0030 | 0.0031 | 0.0025 |
| F06 | 88580001 | 88680000 | 0.0035 | 0.0031 | 0.0030 |
| F06 | 88600001 | 88700000 | 0.0036 | 0.0034 | 0.0029 |
| F06 | 88620001 | 88720000 | 0.0039 | 0.0022 | 0.0031 |
| F06 | 88640001 | 88740000 | 0.0027 | 0.0024 | 0.0021 |
| F06 | 88660001 | 88760000 | 0.0031 | 0.0022 | 0.0023 |
| F06 | 88680001 | 88780000 | 0.0029 | 0.0023 | 0.0020 |
| F06 | 88700001 | 88800000 | 0.0032 | 0.0025 | 0.0023 |
| F06 | 88720001 | 88820000 | 0.0034 | 0.0027 | 0.0025 |
| F06 | 88740001 | 88840000 | 0.0037 | 0.0021 | 0.0027 |
| F06 | 88760001 | 88860000 | 0.0032 | 0.0015 | 0.0022 |
| F06 | 88780001 | 88880000 | 0.0025 | 0.0017 | 0.0017 |
| F06 | 88800001 | 88900000 | 0.0023 | 0.0009 | 0.0016 |
| F06 | 88820001 | 88920000 | 0.0016 | 0.0008 | 0.0008 |
| F06 | 88840001 | 88940000 | 0.0017 | 0.0007 | 0.0008 |
| F06 | 88860001 | 88960000 | 0.0014 | 0.0006 | 0.0008 |
| F06 | 88880001 | 88980000 | 0.0015 | 0.0008 | 0.0007 |
| F06 | 88900001 | 89000000 | 0.0018 | 0.0012 | 0.0009 |
| F06 | 88920001 | 89020000 | 0.0021 | 0.0013 | 0.0013 |
| F06 | 88940001 | 89040000 | 0.0020 | 0.0019 | 0.0012 |
| F06 | 88960001 | 89060000 | 0.0028 | 0.0019 | 0.0018 |
| F06 | 88980001 | 89080000 | 0.0026 | 0.0014 | 0.0018 |
| F06 | 89000001 | 89100000 | 0.0020 | 0.0013 | 0.0014 |
| F06 | 89020001 | 89120000 | 0.0019 | 0.0015 | 0.0014 |
| F06 | 89040001 | 89140000 | 0.0020 | 0.0011 | 0.0016 |
| F06 | 89060001 | 89160000 | 0.0016 | 0.0014 | 0.0012 |

| Chr | Start | End | both | A1 | A2 |
|---|---|---|---|---|---|
| F06 | 89080001 | 89180000 | 0.0020 | 0.0019 | 0.0016 |
| F06 | 89100001 | 89200000 | 0.0024 | 0.0017 | 0.0019 |
| F06 | 89120001 | 89220000 | 0.0026 | 0.0016 | 0.0018 |
| F06 | 89140001 | 89240000 | 0.0026 | 0.0018 | 0.0016 |
| F06 | 89160001 | 89260000 | 0.0027 | 0.0017 | 0.0018 |
| F06 | 89180001 | 89280000 | 0.0026 | 0.0014 | 0.0016 |
| F06 | 89200001 | 89300000 | 0.0024 | 0.0016 | 0.0014 |
| F06 | 89220001 | 89320000 | 0.0024 | 0.0023 | 0.0015 |
| F06 | 89240001 | 89340000 | 0.0031 | 0.0023 | 0.0023 |
| F06 | 89260001 | 89360000 | 0.0032 | 0.0026 | 0.0024 |
| F06 | 89280001 | 89380000 | 0.0034 | 0.0027 | 0.0025 |
| F06 | 89300001 | 89400000 | 0.0036 | 0.0023 | 0.0026 |
| F06 | 89320001 | 89420000 | 0.0031 | 0.0019 | 0.0023 |
| F06 | 89340001 | 89440000 | 0.0025 | 0.0015 | 0.0019 |
| F06 | 89360001 | 89460000 | 0.0020 | 0.0011 | 0.0014 |
| F06 | 89380001 | 89480000 | 0.0016 | 0.0012 | 0.0011 |
| F06 | 89400001 | 89500000 | 0.0014 | 0.0012 | 0.0011 |
| F06 | 89420001 | 89520000 | 0.0016 | 0.0008 | 0.0012 |
| F06 | 89440001 | 89540000 | 0.0013 | 0.0013 | 0.0008 |
| F06 | 89460001 | 89560000 | 0.0019 | 0.0018 | 0.0012 |
| F06 | 89480001 | 89580000 | 0.0024 | 0.0016 | 0.0017 |
| F06 | 89500001 | 89600000 | 0.0024 | 0.0020 | 0.0017 |
| F06 | 89520001 | 89620000 | 0.0027 | 0.0021 | 0.0019 |
| F06 | 89540001 | 89640000 | 0.0027 | 0.0024 | 0.0021 |
| F06 | 89560001 | 89660000 | 0.0031 | 0.0021 | 0.0026 |
| F06 | 89580001 | 89680000 | 0.0028 | 0.0020 | 0.0022 |
| F06 | 89600001 | 89700000 | 0.0027 | 0.0016 | 0.0020 |
| F06 | 89620001 | 89720000 | 0.0025 | 0.0019 | 0.0018 |
| F06 | 89640001 | 89740000 | 0.0030 | 0.0012 | 0.0021 |
| F06 | 89660001 | 89760000 | 0.0022 | 0.0017 | 0.0012 |
| F06 | 89680001 | 89780000 | 0.0028 | 0.0022 | 0.0018 |
| F06 | 89700001 | 89800000 | 0.0033 | 0.0021 | 0.0024 |
| F06 | 89720001 | 89820000 | 0.0030 | 0.0015 | 0.0022 |
| F06 | 89740001 | 89840000 | 0.0025 | 0.0019 | 0.0017 |
| F06 | 89760001 | 89860000 | 0.0029 | 0.0014 | 0.0023 |
| F06 | 89780001 | 89880000 | 0.0023 | 0.0014 | 0.0016 |
| F06 | 89800001 | 89900000 | 0.0024 | 0.0015 | 0.0017 |
| F06 | 89820001 | 89920000 | 0.0026 | 0.0016 | 0.0017 |
| F06 | 89840001 | 89940000 | 0.0032 | 0.0017 | 0.0020 |
| F06 | 89860001 | 89960000 | 0.0032 | 0.0015 | 0.0019 |
| F06 | 89880001 | 89980000 | 0.0030 | 0.0011 | 0.0017 |
| F06 | 89900001 | 90000000 | 0.0025 | 0.0011 | 0.0013 |
| F06 | 89920001 | 90020000 | 0.0022 | 0.0010 | 0.0013 |
| F06 | 89940001 | 90040000 | 0.0016 | 0.0009 | 0.0011 |
| F06 | 89960001 | 90060000 | 0.0013 | 0.0011 | 0.0010 |
| F06 | 89980001 | 90080000 | 0.0015 | 0.0009 | 0.0012 |
| F06 | 90000001 | 90100000 | 0.0016 | 0.0010 | 0.0011 |
| F06 | 90020001 | 90120000 | 0.0020 | 0.0015 | 0.0013 |
| F06 | 90040001 | 90140000 | 0.0026 | 0.0014 | 0.0018 |
| F06 | 90060001 | 90160000 | 0.0027 | 0.0012 | 0.0017 |
| F06 | 90080001 | 90180000 | 0.0025 | 0.0012 | 0.0015 |
| F06 | 90100001 | 90200000 | 0.0022 | 0.0012 | 0.0013 |
| F06 | 90120001 | 90220000 | 0.0019 | 0.0008 | 0.0012 |
| F06 | 90140001 | 90240000 | 0.0018 | 0.0006 | 0.0009 |
| F06 | 90160001 | 90260000 | 0.0016 | 0.0014 | 0.0006 |
| F06 | 90180001 | 90280000 | 0.0024 | 0.0014 | 0.0013 |
| F06 | 90200001 | 90300000 | 0.0025 | 0.0019 | 0.0014 |
| F06 | 90220001 | 90320000 | 0.0031 | 0.0022 | 0.0019 |

| Chr | Start | End | both | A1 | A2 |
|---|---|---|---|---|---|
| F06 | 90240001 | 90340000 | 0.0031 | 0.0030 | 0.0021 |
| F06 | 90260001 | 90360000 | 0.0040 | 0.0023 | 0.0031 |
| F06 | 90280001 | 90380000 | 0.0033 | 0.0023 | 0.0024 |
| F06 | 90300001 | 90400000 | 0.0033 | 0.0024 | 0.0024 |
| F06 | 90320001 | 90420000 | 0.0034 | 0.0021 | 0.0026 |
| F06 | 90340001 | 90440000 | 0.0030 | 0.0014 | 0.0023 |
| F06 | 90360001 | 90460000 | 0.0023 | 0.0014 | 0.0015 |
| F06 | 90380001 | 90480000 | 0.0023 | 0.0015 | 0.0015 |
| F06 | 90400001 | 90500000 | 0.0023 | 0.0011 | 0.0016 |
| F06 | 90420001 | 90520000 | 0.0021 | 0.0012 | 0.0013 |
| F06 | 90440001 | 90540000 | 0.0024 | 0.0020 | 0.0014 |
| F06 | 90460001 | 90560000 | 0.0030 | 0.0022 | 0.0022 |
| F06 | 90480001 | 90580000 | 0.0033 | 0.0022 | 0.0025 |
| F06 | 90500001 | 90600000 | 0.0031 | 0.0019 | 0.0025 |
| F06 | 90520001 | 90620000 | 0.0029 | 0.0018 | 0.0022 |
| F06 | 90540001 | 90640000 | 0.0026 | 0.0012 | 0.0021 |
| F06 | 90560001 | 90660000 | 0.0023 | 0.0014 | 0.0016 |
| F06 | 90580001 | 90680000 | 0.0026 | 0.0014 | 0.0018 |
| F06 | 90600001 | 90700000 | 0.0027 | 0.0013 | 0.0018 |
| F06 | 90620001 | 90720000 | 0.0027 | 0.0017 | 0.0017 |
| F06 | 90640001 | 90740000 | 0.0033 | 0.0016 | 0.0023 |
| F06 | 90660001 | 90760000 | 0.0029 | 0.0013 | 0.0021 |
| F06 | 90680001 | 90780000 | 0.0026 | 0.0021 | 0.0017 |
| F06 | 90700001 | 90800000 | 0.0034 | 0.0021 | 0.0024 |
| F06 | 90720001 | 90820000 | 0.0033 | 0.0019 | 0.0023 |
| F06 | 90740001 | 90840000 | 0.0030 | 0.0018 | 0.0020 |
| F06 | 90760001 | 90860000 | 0.0026 | 0.0015 | 0.0017 |
| F06 | 90780001 | 90880000 | 0.0021 | 0.0008 | 0.0015 |
| F06 | 90800001 | 90900000 | 0.0014 | 0.0010 | 0.0009 |
| F06 | 90820001 | 90920000 | 0.0015 | 0.0008 | 0.0010 |
| F06 | 90840001 | 90940000 | 0.0013 | 0.0020 | 0.0008 |
| F06 | 90860001 | 90960000 | 0.0027 | 0.0027 | 0.0023 |
| F06 | 90880001 | 90980000 | 0.0037 | 0.0028 | 0.0030 |
| F06 | 90900001 | 91000000 | 0.0038 | 0.0028 | 0.0030 |
| F06 | 90920001 | 91020000 | 0.0037 | 0.0029 | 0.0031 |
| F06 | 90940001 | 91040000 | 0.0037 | 0.0020 | 0.0031 |
| F06 | 90960001 | 91060000 | 0.0026 | 0.0014 | 0.0017 |
| F06 | 90980001 | 91080000 | 0.0018 | 0.0011 | 0.0010 |
| F06 | 91000001 | 91100000 | 0.0014 | 0.0009 | 0.0007 |
| F06 | 91020001 | 91120000 | 0.0013 | 0.0008 | 0.0005 |
| F06 | 91040001 | 91140000 | 0.0010 | 0.0006 | 0.0003 |
| F06 | 91060001 | 91160000 | 0.0010 | 0.0008 | 0.0004 |
| F06 | 91080001 | 91180000 | 0.0012 | 0.0011 | 0.0007 |
| F06 | 91100001 | 91200000 | 0.0017 | 0.0017 | 0.0011 |
| F06 | 91120001 | 91220000 | 0.0025 | 0.0020 | 0.0017 |
| F06 | 91140001 | 91240000 | 0.0027 | 0.0019 | 0.0020 |
| F06 | 91160001 | 91260000 | 0.0028 | 0.0017 | 0.0021 |
| F06 | 91180001 | 91280000 | 0.0024 | 0.0014 | 0.0018 |
| F06 | 91200001 | 91300000 | 0.0020 | 0.0013 | 0.0016 |
| F06 | 91220001 | 91320000 | 0.0018 | 0.0011 | 0.0014 |
| F06 | 91240001 | 91340000 | 0.0015 | 0.0011 | 0.0012 |
| F06 | 91260001 | 91360000 | 0.0014 | 0.0017 | 0.0010 |
| F06 | 91280001 | 91380000 | 0.0022 | 0.0021 | 0.0016 |
| F06 | 91300001 | 91400000 | 0.0026 | 0.0018 | 0.0021 |
| F06 | 91320001 | 91420000 | 0.0023 | 0.0026 | 0.0019 |
| F06 | 91340001 | 91440000 | 0.0033 | 0.0032 | 0.0027 |
| F06 | 91360001 | 91460000 | 0.0038 | 0.0028 | 0.0033 |
| F06 | 91380001 | 91480000 | 0.0032 | 0.0025 | 0.0027 |

| Chr | Start | End | both | A1 | A2 |
|---|---|---|---|---|---|
| F06 | 91400001 | 91500000 | 0.0027 | 0.0034 | 0.0023 |
| F06 | 91420001 | 91520000 | 0.0038 | 0.0030 | 0.0033 |
| F06 | 91440001 | 91540000 | 0.0032 | 0.0024 | 0.0030 |
| F06 | 91460001 | 91560000 | 0.0026 | 0.0029 | 0.0024 |
| F06 | 91480001 | 91580000 | 0.0032 | 0.0032 | 0.0030 |
| F06 | 91500001 | 91600000 | 0.0035 | 0.0024 | 0.0034 |
| F06 | 91520001 | 91620000 | 0.0027 | 0.0030 | 0.0026 |
| F06 | 91540001 | 91640000 | 0.0033 | 0.0029 | 0.0032 |
| F06 | 91560001 | 91660000 | 0.0031 | 0.0027 | 0.0031 |
| F06 | 91580001 | 91680000 | 0.0029 | 0.0028 | 0.0027 |
| F06 | 91600001 | 91700000 | 0.0033 | 0.0028 | 0.0028 |
| F06 | 91620001 | 91720000 | 0.0034 | 0.0019 | 0.0025 |
| F06 | 91640001 | 91740000 | 0.0025 | 0.0021 | 0.0015 |
| F06 | 91660001 | 91760000 | 0.0029 | 0.0023 | 0.0018 |
| F06 | 91680001 | 91780000 | 0.0031 | 0.0023 | 0.0021 |
| F06 | 91700001 | 91800000 | 0.0028 | 0.0024 | 0.0021 |
| F06 | 91720001 | 91820000 | 0.0029 | 0.0024 | 0.0023 |
| F06 | 91740001 | 91840000 | 0.0029 | 0.0022 | 0.0023 |
| F06 | 91760001 | 91860000 | 0.0027 | 0.0021 | 0.0020 |
| F06 | 91780001 | 91880000 | 0.0025 | 0.0018 | 0.0018 |
| F06 | 91800001 | 91900000 | 0.0023 | 0.0015 | 0.0015 |
| F06 | 91820001 | 91920000 | 0.0020 | 0.0014 | 0.0013 |
| F06 | 91840001 | 91940000 | 0.0020 | 0.0017 | 0.0012 |
| F06 | 91860001 | 91960000 | 0.0023 | 0.0012 | 0.0014 |
| F06 | 91880001 | 91980000 | 0.0018 | 0.0013 | 0.0010 |
| F06 | 91900001 | 92000000 | 0.0019 | 0.0011 | 0.0010 |
| F06 | 91920001 | 92020000 | 0.0016 | 0.0011 | 0.0008 |
| F06 | 91940001 | 92040000 | 0.0016 | 0.0008 | 0.0009 |
| F06 | 91960001 | 92060000 | 0.0012 | 0.0012 | 0.0006 |
| F06 | 91980001 | 92080000 | 0.0018 | 0.0012 | 0.0010 |
| F06 | 92000001 | 92100000 | 0.0017 | 0.0016 | 0.0009 |
| F06 | 92020001 | 92120000 | 0.0023 | 0.0023 | 0.0014 |
| F06 | 92040001 | 92140000 | 0.0031 | 0.0026 | 0.0022 |
| F06 | 92060001 | 92160000 | 0.0034 | 0.0025 | 0.0027 |
| F06 | 92080001 | 92180000 | 0.0032 | 0.0024 | 0.0026 |
| F06 | 92100001 | 92200000 | 0.0029 | 0.0021 | 0.0026 |
| F06 | 92120001 | 92220000 | 0.0027 | 0.0027 | 0.0024 |
| F06 | 92140001 | 92240000 | 0.0032 | 0.0027 | 0.0028 |
| F06 | 92160001 | 92260000 | 0.0032 | 0.0028 | 0.0028 |
| F06 | 92180001 | 92280000 | 0.0037 | 0.0028 | 0.0032 |
| F06 | 92200001 | 92300000 | 0.0036 | 0.0028 | 0.0031 |
| F06 | 92220001 | 92320000 | 0.0036 | 0.0020 | 0.0030 |
| F06 | 92240001 | 92340000 | 0.0027 | 0.0043 | 0.0024 |
| F06 | 92260001 | 92360000 | 0.0051 | 0.0042 | 0.0048 |
| F06 | 92280001 | 92380000 | 0.0047 | 0.0042 | 0.0046 |
| F06 | 92300001 | 92400000 | 0.0048 | 0.0039 | 0.0047 |
| F06 | 92320001 | 92420000 | 0.0044 | 0.0036 | 0.0043 |
| F06 | 92340001 | 92440000 | 0.0041 | 0.0009 | 0.0039 |
| F06 | 92360001 | 92460000 | 0.0013 | 0.0005 | 0.0010 |
| F06 | 92380001 | 92480000 | 0.0009 | 0.0007 | 0.0006 |
| F06 | 92400001 | 92500000 | 0.0012 | 0.0009 | 0.0008 |
| F06 | 92420001 | 92520000 | 0.0014 | 0.0008 | 0.0009 |
| F06 | 92440001 | 92540000 | 0.0015 | 0.0009 | 0.0009 |
| F06 | 92460001 | 92560000 | 0.0015 | 0.0009 | 0.0010 |
| F06 | 92480001 | 92580000 | 0.0016 | 0.0011 | 0.0010 |
| F06 | 92500001 | 92600000 | 0.0019 | 0.0017 | 0.0012 |
| F06 | 92520001 | 92620000 | 0.0026 | 0.0018 | 0.0018 |
| F06 | 92540001 | 92640000 | 0.0025 | 0.0018 | 0.0018 |

| Chr | Start | End | both | A1 | A2 |
|---|---|---|---|---|---|
| F06 | 92560001 | 92660000 | 0.0026 | 0.0018 | 0.0019 |
| F06 | 92580001 | 92680000 | 0.0025 | 0.0012 | 0.0018 |
| F06 | 92600001 | 92700000 | 0.0020 | 0.0011 | 0.0013 |
| F06 | 92620001 | 92720000 | 0.0019 | 0.0012 | 0.0013 |
| F06 | 92640001 | 92740000 | 0.0019 | 0.0012 | 0.0014 |
| F06 | 92660001 | 92760000 | 0.0019 | 0.0013 | 0.0014 |
| F06 | 92680001 | 92780000 | 0.0019 | 0.0014 | 0.0016 |
| F06 | 92700001 | 92800000 | 0.0017 | 0.0009 | 0.0016 |
| F06 | 92720001 | 92820000 | 0.0012 | 0.0010 | 0.0011 |
| F06 | 92740001 | 92840000 | 0.0013 | 0.0009 | 0.0012 |
| F06 | 92760001 | 92860000 | 0.0015 | 0.0008 | 0.0011 |
| F06 | 92780001 | 92880000 | 0.0016 | 0.0017 | 0.0009 |
| F06 | 92800001 | 92900000 | 0.0026 | 0.0014 | 0.0017 |
| F06 | 92820001 | 92920000 | 0.0024 | 0.0012 | 0.0015 |
| F06 | 92840001 | 92940000 | 0.0023 | 0.0016 | 0.0011 |
| F06 | 92860001 | 92960000 | 0.0024 | 0.0019 | 0.0015 |
| F06 | 92880001 | 92980000 | 0.0035 | 0.0013 | 0.0027 |
| F06 | 92900001 | 93000000 | 0.0027 | 0.0015 | 0.0020 |
| F06 | 92920001 | 93020000 | 0.0028 | 0.0020 | 0.0022 |
| F06 | 92940001 | 93040000 | 0.0030 | 0.0017 | 0.0027 |
| F06 | 92960001 | 93060000 | 0.0026 | 0.0016 | 0.0023 |
| F06 | 92980001 | 93080000 | 0.0014 | 0.0015 | 0.0013 |
| F06 | 93000001 | 93100000 | 0.0012 | 0.0015 | 0.0011 |
| F06 | 93020001 | 93120000 | 0.0010 | 0.0014 | 0.0009 |
| F06 | 93040001 | 93140000 | 0.0009 | 0.0014 | 0.0006 |
| F06 | 93060001 | 93160000 | 0.0010 | 0.0013 | 0.0007 |
| F06 | 93080001 | 93180000 | 0.0009 | 0.0012 | 0.0006 |
| F06 | 93100001 | 93200000 | 0.0010 | 0.0016 | 0.0007 |
| F06 | 93120001 | 93220000 | 0.0015 | 0.0014 | 0.0011 |
| F06 | 93140001 | 93240000 | 0.0014 | 0.0016 | 0.0009 |
| F06 | 93160001 | 93260000 | 0.0015 | 0.0020 | 0.0009 |
| F06 | 93180001 | 93280000 | 0.0019 | 0.0022 | 0.0011 |
| F06 | 93200001 | 93300000 | 0.0021 | 0.0016 | 0.0013 |
| F06 | 93220001 | 93320000 | 0.0017 | 0.0018 | 0.0008 |
| F06 | 93240001 | 93340000 | 0.0019 | 0.0015 | 0.0012 |
| F06 | 93260001 | 93360000 | 0.0019 | 0.0011 | 0.0011 |
| F06 | 93280001 | 93380000 | 0.0016 | 0.0011 | 0.0009 |
| F06 | 93300001 | 93400000 | 0.0016 | 0.0020 | 0.0009 |
| F06 | 93320001 | 93420000 | 0.0025 | 0.0023 | 0.0018 |
| F06 | 93340001 | 93440000 | 0.0027 | 0.0035 | 0.0022 |
| F06 | 93360001 | 93460000 | 0.0038 | 0.0038 | 0.0034 |
| F06 | 93380001 | 93480000 | 0.0041 | 0.0048 | 0.0039 |
| F06 | 93400001 | 93500000 | 0.0050 | 0.0043 | 0.0047 |
| F06 | 93420001 | 93520000 | 0.0043 | 0.0038 | 0.0039 |
| F06 | 93440001 | 93540000 | 0.0038 | 0.0030 | 0.0035 |
| F06 | 93460001 | 93560000 | 0.0032 | 0.0030 | 0.0030 |
| F06 | 93480001 | 93580000 | 0.0033 | 0.0018 | 0.0028 |
| F06 | 93500001 | 93600000 | 0.0023 | 0.0016 | 0.0018 |
| F06 | 93520001 | 93620000 | 0.0023 | 0.0024 | 0.0017 |
| F06 | 93540001 | 93640000 | 0.0031 | 0.0023 | 0.0023 |
| F06 | 93560001 | 93660000 | 0.0031 | 0.0020 | 0.0022 |
| F06 | 93580001 | 93680000 | 0.0027 | 0.0020 | 0.0018 |
| F06 | 93600001 | 93700000 | 0.0027 | 0.0024 | 0.0019 |
| F06 | 93620001 | 93720000 | 0.0034 | 0.0019 | 0.0026 |
| F06 | 93640001 | 93740000 | 0.0029 | 0.0018 | 0.0022 |
| F06 | 93660001 | 93760000 | 0.0028 | 0.0020 | 0.0021 |
| F06 | 93680001 | 93780000 | 0.0032 | 0.0020 | 0.0025 |
| F06 | 93700001 | 93800000 | 0.0032 | 0.0017 | 0.0025 |

| Chr | Start | End | both | A1 | A2 |
|---|---|---|---|---|---|
| F06 | 93720001 | 93820000 | 0.0027 | 0.0015 | 0.0021 |
| F06 | 93740001 | 93840000 | 0.0025 | 0.0019 | 0.0019 |
| F06 | 93760001 | 93860000 | 0.0027 | 0.0022 | 0.0023 |
| F06 | 93780001 | 93880000 | 0.0027 | 0.0049 | 0.0024 |
| F06 | 93800001 | 93900000 | 0.0054 | 0.0057 | 0.0049 |
| F06 | 93820001 | 93920000 | 0.0057 | 0.0055 | 0.0050 |
| F06 | 93840001 | 93940000 | 0.0054 | 0.0051 | 0.0046 |
| F06 | 93860001 | 93960000 | 0.0048 | 0.0049 | 0.0038 |
| F06 | 93880001 | 93980000 | 0.0044 | 0.0030 | 0.0033 |
| F06 | 93900001 | 94000000 | 0.0025 | 0.0047 | 0.0015 |
| F06 | 93920001 | 94020000 | 0.0047 | 0.0049 | 0.0039 |
| F06 | 93940001 | 94040000 | 0.0051 | 0.0053 | 0.0044 |
| F06 | 93960001 | 94060000 | 0.0056 | 0.0055 | 0.0050 |
| F06 | 93980001 | 94080000 | 0.0059 | 0.0053 | 0.0054 |
| F06 | 94000001 | 94100000 | 0.0056 | 0.0030 | 0.0051 |
| F06 | 94020001 | 94120000 | 0.0030 | 0.0033 | 0.0024 |
| F06 | 94040001 | 94140000 | 0.0030 | 0.0029 | 0.0022 |
| F06 | 94060001 | 94160000 | 0.0027 | 0.0027 | 0.0018 |
| F06 | 94080001 | 94180000 | 0.0027 | 0.0022 | 0.0017 |
| F06 | 94100001 | 94200000 | 0.0024 | 0.0018 | 0.0015 |
| F06 | 94120001 | 94220000 | 0.0023 | 0.0020 | 0.0013 |
| F06 | 94140001 | 94240000 | 0.0027 | 0.0017 | 0.0017 |
| F06 | 94160001 | 94260000 | 0.0025 | 0.0015 | 0.0015 |
| F06 | 94180001 | 94280000 | 0.0022 | 0.0013 | 0.0012 |
| F06 | 94200001 | 94300000 | 0.0020 | 0.0013 | 0.0010 |
| F06 | 94220001 | 94320000 | 0.0020 | 0.0004 | 0.0011 |
| F06 | 94240001 | 94340000 | 0.0013 | 0.0006 | 0.0004 |
| F06 | 94260001 | 94360000 | 0.0015 | 0.0008 | 0.0007 |
| F06 | 94280001 | 94380000 | 0.0016 | 0.0010 | 0.0010 |
| F06 | 94300001 | 94400000 | 0.0016 | 0.0013 | 0.0011 |
| F06 | 94320001 | 94420000 | 0.0016 | 0.0014 | 0.0013 |
| F06 | 94340001 | 94440000 | 0.0015 | 0.0017 | 0.0012 |
| F06 | 94360001 | 94460000 | 0.0018 | 0.0019 | 0.0013 |
| F06 | 94380001 | 94480000 | 0.0017 | 0.0021 | 0.0011 |
| F06 | 94400001 | 94500000 | 0.0020 | 0.0022 | 0.0013 |
| F06 | 94420001 | 94520000 | 0.0020 | 0.0027 | 0.0012 |
| F06 | 94440001 | 94540000 | 0.0026 | 0.0026 | 0.0017 |
| F06 | 94460001 | 94560000 | 0.0024 | 0.0025 | 0.0014 |
| F06 | 94480001 | 94580000 | 0.0023 | 0.0035 | 0.0013 |
| F06 | 94500001 | 94600000 | 0.0033 | 0.0046 | 0.0022 |
| F06 | 94520001 | 94620000 | 0.0045 | 0.0045 | 0.0034 |
| F06 | 94540001 | 94640000 | 0.0040 | 0.0043 | 0.0030 |
| F06 | 94560001 | 94660000 | 0.0038 | 0.0044 | 0.0029 |
| F06 | 94580001 | 94680000 | 0.0039 | 0.0032 | 0.0033 |
| F06 | 94600001 | 94700000 | 0.0027 | 0.0023 | 0.0021 |
| F06 | 94620001 | 94720000 | 0.0016 | 0.0021 | 0.0010 |
| F06 | 94640001 | 94740000 | 0.0016 | 0.0021 | 0.0008 |
| F06 | 94660001 | 94760000 | 0.0017 | 0.0017 | 0.0010 |
| F06 | 94680001 | 94780000 | 0.0014 | 0.0019 | 0.0006 |
| F06 | 94700001 | 94800000 | 0.0014 | 0.0022 | 0.0007 |
| F06 | 94720001 | 94820000 | 0.0017 | 0.0027 | 0.0010 |
| F06 | 94740001 | 94840000 | 0.0022 | 0.0031 | 0.0013 |
| F06 | 94760001 | 94860000 | 0.0025 | 0.0036 | 0.0014 |
| F06 | 94780001 | 94880000 | 0.0031 | 0.0038 | 0.0020 |
| F06 | 94800001 | 94900000 | 0.0036 | 0.0032 | 0.0026 |
| F06 | 94820001 | 94920000 | 0.0032 | 0.0025 | 0.0022 |
| F06 | 94840001 | 94940000 | 0.0027 | 0.0023 | 0.0019 |
| F06 | 94860001 | 94960000 | 0.0026 | 0.0024 | 0.0018 |

| Chr | Start | End | both | A1 | A2 |
|---|---|---|---|---|---|
| F06 | 94880001 | 94980000 | 0.0025 | 0.0022 | 0.0015 |
| F06 | 94900001 | 95000000 | 0.0021 | 0.0025 | 0.0011 |
| F06 | 94920001 | 95020000 | 0.0024 | 0.0024 | 0.0014 |
| F06 | 94940001 | 95040000 | 0.0026 | 0.0020 | 0.0014 |
| F06 | 94960001 | 95060000 | 0.0025 | 0.0018 | 0.0014 |
| F06 | 94980001 | 95080000 | 0.0026 | 0.0016 | 0.0017 |
| F06 | 95000001 | 95100000 | 0.0024 | 0.0017 | 0.0017 |
| F06 | 95020001 | 95120000 | 0.0026 | 0.0019 | 0.0019 |
| F06 | 95040001 | 95140000 | 0.0027 | 0.0019 | 0.0023 |
| F06 | 95060001 | 95160000 | 0.0024 | 0.0026 | 0.0022 |
| F06 | 95080001 | 95180000 | 0.0030 | 0.0026 | 0.0027 |
| F06 | 95100001 | 95200000 | 0.0031 | 0.0020 | 0.0027 |
| F06 | 95120001 | 95220000 | 0.0024 | 0.0018 | 0.0021 |
| F06 | 95140001 | 95240000 | 0.0023 | 0.0023 | 0.0020 |
| F06 | 95160001 | 95260000 | 0.0028 | 0.0014 | 0.0025 |
| F06 | 95180001 | 95280000 | 0.0018 | 0.0016 | 0.0015 |
| F06 | 95200001 | 95300000 | 0.0020 | 0.0017 | 0.0018 |
| F06 | 95220001 | 95320000 | 0.0025 | 0.0018 | 0.0021 |
| F06 | 95240001 | 95340000 | 0.0027 | 0.0027 | 0.0024 |
| F06 | 95260001 | 95360000 | 0.0039 | 0.0026 | 0.0036 |
| F06 | 95280001 | 95380000 | 0.0038 | 0.0032 | 0.0036 |
| F06 | 95300001 | 95400000 | 0.0044 | 0.0032 | 0.0042 |
| F06 | 95320001 | 95420000 | 0.0049 | 0.0033 | 0.0048 |
| F06 | 95340001 | 95440000 | 0.0055 | 0.0023 | 0.0054 |
| F06 | 95360001 | 95460000 | 0.0047 | 0.0023 | 0.0046 |
| F06 | 95380001 | 95480000 | 0.0053 | 0.0015 | 0.0052 |
| F06 | 95400001 | 95500000 | 0.0046 | 0.0018 | 0.0045 |
| F06 | 95420001 | 95520000 | 0.0043 | 0.0017 | 0.0042 |
| F06 | 95440001 | 95540000 | 0.0035 | 0.0016 | 0.0034 |
| F06 | 95460001 | 95560000 | 0.0034 | 0.0015 | 0.0032 |
| F06 | 95480001 | 95580000 | 0.0029 | 0.0014 | 0.0025 |
| F06 | 95500001 | 95600000 | 0.0026 | 0.0012 | 0.0021 |
| F06 | 95520001 | 95620000 | 0.0023 | 0.0028 | 0.0016 |
| F06 | 95540001 | 95640000 | 0.0038 | 0.0029 | 0.0031 |
| F06 | 95560001 | 95660000 | 0.0032 | 0.0034 | 0.0024 |
| F06 | 95580001 | 95680000 | 0.0031 | 0.0040 | 0.0024 |
| F06 | 95600001 | 95700000 | 0.0034 | 0.0040 | 0.0027 |
| F06 | 95620001 | 95720000 | 0.0031 | 0.0028 | 0.0026 |
| F06 | 95640001 | 95740000 | 0.0019 | 0.0027 | 0.0013 |
| F06 | 95660001 | 95760000 | 0.0026 | 0.0022 | 0.0020 |
| F06 | 95680001 | 95780000 | 0.0026 | 0.0019 | 0.0021 |
| F06 | 95700001 | 95800000 | 0.0025 | 0.0017 | 0.0020 |
| F06 | 95720001 | 95820000 | 0.0028 | 0.0011 | 0.0023 |
| F06 | 95740001 | 95840000 | 0.0023 | 0.0008 | 0.0018 |
| F06 | 95760001 | 95860000 | 0.0019 | 0.0008 | 0.0014 |
| F06 | 95780001 | 95880000 | 0.0021 | 0.0009 | 0.0015 |
| F06 | 95800001 | 95900000 | 0.0023 | 0.0006 | 0.0018 |
| F06 | 95820001 | 95920000 | 0.0020 | 0.0006 | 0.0016 |
| F06 | 95840001 | 95940000 | 0.0021 | 0.0007 | 0.0016 |
| F06 | 95860001 | 95960000 | 0.0021 | 0.0026 | 0.0014 |
| F06 | 95880001 | 95980000 | 0.0036 | 0.0026 | 0.0029 |
| F06 | 95900001 | 96000000 | 0.0035 | 0.0027 | 0.0028 |
| F06 | 95920001 | 96020000 | 0.0033 | 0.0027 | 0.0027 |
| F06 | 95940001 | 96040000 | 0.0030 | 0.0026 | 0.0025 |
| F06 | 95960001 | 96060000 | 0.0027 | 0.0024 | 0.0025 |
| F06 | 95980001 | 96080000 | 0.0026 | 0.0043 | 0.0025 |
| F06 | 96000001 | 96100000 | 0.0043 | 0.0050 | 0.0041 |
| F06 | 96020001 | 96120000 | 0.0050 | 0.0059 | 0.0047 |

| Chr | Start | End | both | A1 | A2 |
|---|---|---|---|---|---|
| F06 | 96040001 | 96140000 | 0.0060 | 0.0074 | 0.0056 |
| F06 | 96060001 | 96160000 | 0.0076 | 0.0064 | 0.0072 |
| F06 | 96080001 | 96180000 | 0.0065 | 0.0053 | 0.0061 |
| F06 | 96100001 | 96200000 | 0.0054 | 0.0048 | 0.0049 |
| F06 | 96120001 | 96220000 | 0.0051 | 0.0050 | 0.0046 |
| F06 | 96140001 | 96240000 | 0.0051 | 0.0034 | 0.0042 |
| F06 | 96160001 | 96260000 | 0.0034 | 0.0033 | 0.0025 |
| F06 | 96180001 | 96280000 | 0.0034 | 0.0030 | 0.0026 |
| F06 | 96200001 | 96300000 | 0.0030 | 0.0035 | 0.0021 |
| F06 | 96220001 | 96320000 | 0.0034 | 0.0031 | 0.0024 |
| F06 | 96240001 | 96340000 | 0.0031 | 0.0033 | 0.0024 |
| F06 | 96260001 | 96360000 | 0.0032 | 0.0027 | 0.0026 |
| F06 | 96280001 | 96380000 | 0.0030 | 0.0022 | 0.0023 |
| F06 | 96300001 | 96400000 | 0.0028 | 0.0015 | 0.0023 |
| F06 | 96320001 | 96420000 | 0.0028 | 0.0009 | 0.0021 |
| F06 | 96340001 | 96440000 | 0.0023 | 0.0006 | 0.0016 |
| F06 | 96360001 | 96460000 | 0.0024 | 0.0005 | 0.0017 |
| F06 | 96380001 | 96480000 | 0.0025 | 0.0004 | 0.0018 |
| F06 | 96400001 | 96500000 | 0.0029 | 0.0004 | 0.0021 |
| F06 | 96420001 | 96520000 | 0.0028 | 0.0020 | 0.0022 |
| F06 | 96440001 | 96540000 | 0.0038 | 0.0027 | 0.0031 |
| F06 | 96460001 | 96560000 | 0.0040 | 0.0037 | 0.0033 |
| F06 | 96480001 | 96580000 | 0.0048 | 0.0038 | 0.0042 |
| F06 | 96500001 | 96600000 | 0.0039 | 0.0037 | 0.0035 |
| F06 | 96520001 | 96620000 | 0.0034 | 0.0025 | 0.0031 |
| F06 | 96540001 | 96640000 | 0.0026 | 0.0023 | 0.0025 |
| F06 | 96560001 | 96660000 | 0.0022 | 0.0018 | 0.0020 |
| F06 | 96580001 | 96680000 | 0.0014 | 0.0029 | 0.0011 |
| F06 | 96600001 | 96700000 | 0.0022 | 0.0033 | 0.0017 |
| F06 | 96620001 | 96720000 | 0.0024 | 0.0032 | 0.0016 |
| F06 | 96640001 | 96740000 | 0.0023 | 0.0034 | 0.0013 |
| F06 | 96660001 | 96760000 | 0.0027 | 0.0034 | 0.0016 |
| F06 | 96680001 | 96780000 | 0.0026 | 0.0026 | 0.0015 |
| F06 | 96700001 | 96800000 | 0.0019 | 0.0028 | 0.0009 |
| F06 | 96720001 | 96820000 | 0.0023 | 0.0033 | 0.0014 |
| F06 | 96740001 | 96840000 | 0.0025 | 0.0031 | 0.0016 |
| F06 | 96760001 | 96860000 | 0.0022 | 0.0032 | 0.0013 |
| F06 | 96780001 | 96880000 | 0.0023 | 0.0030 | 0.0012 |
| F06 | 96800001 | 96900000 | 0.0023 | 0.0026 | 0.0013 |
| F06 | 96820001 | 96920000 | 0.0019 | 0.0022 | 0.0007 |
| F06 | 96840001 | 96940000 | 0.0018 | 0.0021 | 0.0006 |
| F06 | 96860001 | 96960000 | 0.0018 | 0.0014 | 0.0007 |
| F06 | 96880001 | 96980000 | 0.0017 | 0.0012 | 0.0006 |
| F06 | 96900001 | 97000000 | 0.0016 | 0.0010 | 0.0006 |
| F06 | 96920001 | 97020000 | 0.0017 | 0.0006 | 0.0006 |
| F06 | 96940001 | 97040000 | 0.0019 | 0.0005 | 0.0008 |
| F06 | 96960001 | 97060000 | 0.0021 | 0.0009 | 0.0010 |
| F06 | 96980001 | 97080000 | 0.0026 | 0.0009 | 0.0014 |
| F06 | 97000001 | 97100000 | 0.0029 | 0.0014 | 0.0015 |
| F06 | 97020001 | 97120000 | 0.0031 | 0.0017 | 0.0014 |
| F06 | 97040001 | 97140000 | 0.0029 | 0.0018 | 0.0012 |
| F06 | 97060001 | 97160000 | 0.0028 | 0.0017 | 0.0010 |
| F06 | 97080001 | 97180000 | 0.0024 | 0.0019 | 0.0008 |
| F06 | 97100001 | 97200000 | 0.0022 | 0.0016 | 0.0008 |
| F06 | 97120001 | 97220000 | 0.0019 | 0.0017 | 0.0009 |
| F06 | 97140001 | 97240000 | 0.0019 | 0.0014 | 0.0010 |
| F06 | 97160001 | 97260000 | 0.0019 | 0.0012 | 0.0009 |
| F06 | 97180001 | 97280000 | 0.0017 | 0.0009 | 0.0007 |

| Chr | Start | End | both | A1 | A2 |
|---|---|---|---|---|---|
| F06 | 97200001 | 97300000 | 0.0018 | 0.0013 | 0.0008 |
| F06 | 97220001 | 97320000 | 0.0030 | 0.0011 | 0.0017 |
| F06 | 97240001 | 97340000 | 0.0031 | 0.0009 | 0.0018 |
| F06 | 97260001 | 97360000 | 0.0029 | 0.0014 | 0.0018 |
| F06 | 97280001 | 97380000 | 0.0030 | 0.0018 | 0.0018 |
| F06 | 97300001 | 97400000 | 0.0027 | 0.0014 | 0.0016 |
| F06 | 97320001 | 97420000 | 0.0015 | 0.0017 | 0.0008 |
| F06 | 97340001 | 97440000 | 0.0014 | 0.0019 | 0.0007 |
| F06 | 97360001 | 97460000 | 0.0016 | 0.0017 | 0.0009 |
| F06 | 97380001 | 97480000 | 0.0016 | 0.0015 | 0.0008 |
| F06 | 97400001 | 97500000 | 0.0018 | 0.0014 | 0.0009 |
| F06 | 97420001 | 97520000 | 0.0017 | 0.0014 | 0.0008 |
| F06 | 97440001 | 97540000 | 0.0016 | 0.0017 | 0.0007 |
| F06 | 97460001 | 97560000 | 0.0014 | 0.0020 | 0.0006 |
| F06 | 97480001 | 97580000 | 0.0013 | 0.0022 | 0.0005 |
| F06 | 97500001 | 97600000 | 0.0013 | 0.0029 | 0.0007 |
| F06 | 97520001 | 97620000 | 0.0017 | 0.0032 | 0.0010 |
| F06 | 97540001 | 97640000 | 0.0020 | 0.0027 | 0.0013 |
| F06 | 97560001 | 97660000 | 0.0020 | 0.0024 | 0.0013 |
| F06 | 97580001 | 97680000 | 0.0022 | 0.0022 | 0.0013 |
| F06 | 97600001 | 97700000 | 0.0022 | 0.0020 | 0.0014 |
| F06 | 97620001 | 97720000 | 0.0025 | 0.0019 | 0.0017 |
| F06 | 97640001 | 97740000 | 0.0022 | 0.0024 | 0.0014 |
| F06 | 97660001 | 97760000 | 0.0023 | 0.0024 | 0.0014 |
| F06 | 97680001 | 97780000 | 0.0020 | 0.0024 | 0.0012 |
| F06 | 97700001 | 97800000 | 0.0017 | 0.0021 | 0.0009 |
| F06 | 97720001 | 97820000 | 0.0011 | 0.0019 | 0.0004 |
| F06 | 97740001 | 97840000 | 0.0011 | 0.0015 | 0.0004 |
| F06 | 97760001 | 97860000 | 0.0011 | 0.0013 | 0.0004 |
| F06 | 97780001 | 97880000 | 0.0012 | 0.0013 | 0.0004 |
| F06 | 97800001 | 97900000 | 0.0012 | 0.0013 | 0.0004 |
| F06 | 97820001 | 97920000 | 0.0011 | 0.0013 | 0.0003 |
| F06 | 97840001 | 97940000 | 0.0012 | 0.0014 | 0.0004 |
| F06 | 97860001 | 97960000 | 0.0011 | 0.0018 | 0.0004 |
| F06 | 97880001 | 97980000 | 0.0012 | 0.0019 | 0.0004 |
| F06 | 97900001 | 98000000 | 0.0014 | 0.0036 | 0.0008 |
| F06 | 97920001 | 98020000 | 0.0033 | 0.0037 | 0.0027 |
| F06 | 97940001 | 98040000 | 0.0035 | 0.0046 | 0.0031 |
| F06 | 97960001 | 98060000 | 0.0045 | 0.0042 | 0.0042 |
| F06 | 97980001 | 98080000 | 0.0043 | 0.0039 | 0.0042 |
| F06 | 98000001 | 98100000 | 0.0040 | 0.0019 | 0.0039 |
| F06 | 98020001 | 98120000 | 0.0020 | 0.0014 | 0.0019 |
| F06 | 98040001 | 98140000 | 0.0014 | 0.0002 | 0.0013 |
| F06 | 98060001 | 98160000 | 0.0002 | 0.0001 | 0.0002 |
| F06 | 98080001 | 98180000 | 0.0001 | 0.0010 | 0.0001 |
| F07 | 1 | 100000 | 0.0027 | 0.0025 | 0.0026 |
| F07 | 20001 | 120000 | 0.0034 | 0.0032 | 0.0033 |
| F07 | 40001 | 140000 | 0.0034 | 0.0032 | 0.0034 |
| F07 | 60001 | 160000 | 0.0039 | 0.0040 | 0.0037 |
| F07 | 80001 | 180000 | 0.0037 | 0.0040 | 0.0031 |
| F07 | 100001 | 200000 | 0.0023 | 0.0029 | 0.0016 |
| F07 | 120001 | 220000 | 0.0018 | 0.0026 | 0.0008 |
| F07 | 140001 | 240000 | 0.0016 | 0.0026 | 0.0005 |
| F07 | 160001 | 260000 | 0.0014 | 0.0024 | 0.0002 |
| F07 | 180001 | 280000 | 0.0013 | 0.0023 | 0.0002 |
| F07 | 200001 | 300000 | 0.0012 | 0.0021 | 0.0002 |
| F07 | 220001 | 320000 | 0.0018 | 0.0022 | 0.0007 |
| F07 | 240001 | 340000 | 0.0019 | 0.0020 | 0.0008 |

| Chr | Start | End | both | A1 | A2 |
|---|---|---|---|---|---|
| F07 | 260001 | 360000 | 0.0018 | 0.0016 | 0.0007 |
| F07 | 280001 | 380000 | 0.0019 | 0.0015 | 0.0009 |
| F07 | 300001 | 400000 | 0.0020 | 0.0014 | 0.0009 |
| F07 | 320001 | 420000 | 0.0016 | 0.0013 | 0.0006 |
| F07 | 340001 | 440000 | 0.0020 | 0.0020 | 0.0011 |
| F07 | 360001 | 460000 | 0.0024 | 0.0028 | 0.0015 |
| F07 | 380001 | 480000 | 0.0024 | 0.0030 | 0.0016 |
| F07 | 400001 | 500000 | 0.0025 | 0.0034 | 0.0015 |
| F07 | 420001 | 520000 | 0.0025 | 0.0033 | 0.0016 |
| F07 | 440001 | 540000 | 0.0019 | 0.0028 | 0.0010 |
| F07 | 460001 | 560000 | 0.0014 | 0.0024 | 0.0006 |
| F07 | 480001 | 580000 | 0.0015 | 0.0023 | 0.0004 |
| F07 | 500001 | 600000 | 0.0015 | 0.0017 | 0.0005 |
| F07 | 520001 | 620000 | 0.0016 | 0.0013 | 0.0008 |
| F07 | 540001 | 640000 | 0.0020 | 0.0009 | 0.0013 |
| F07 | 560001 | 660000 | 0.0023 | 0.0006 | 0.0018 |
| F07 | 580001 | 680000 | 0.0034 | 0.0012 | 0.0030 |
| F07 | 600001 | 700000 | 0.0035 | 0.0012 | 0.0031 |
| F07 | 620001 | 720000 | 0.0034 | 0.0013 | 0.0030 |
| F07 | 640001 | 740000 | 0.0034 | 0.0013 | 0.0028 |
| F07 | 660001 | 760000 | 0.0040 | 0.0025 | 0.0031 |
| F07 | 680001 | 780000 | 0.0030 | 0.0015 | 0.0021 |
| F07 | 700001 | 800000 | 0.0033 | 0.0018 | 0.0023 |
| F07 | 720001 | 820000 | 0.0039 | 0.0026 | 0.0029 |
| F07 | 740001 | 840000 | 0.0038 | 0.0027 | 0.0030 |
| F07 | 760001 | 860000 | 0.0032 | 0.0015 | 0.0026 |
| F07 | 780001 | 880000 | 0.0031 | 0.0015 | 0.0026 |
| F07 | 800001 | 900000 | 0.0028 | 0.0015 | 0.0025 |
| F07 | 820001 | 920000 | 0.0020 | 0.0006 | 0.0017 |
| F07 | 840001 | 940000 | 0.0017 | 0.0005 | 0.0013 |
| F07 | 860001 | 960000 | 0.0015 | 0.0005 | 0.0009 |
| F07 | 880001 | 980000 | 0.0016 | 0.0005 | 0.0007 |
| F07 | 900001 | 1000000 | 0.0016 | 0.0002 | 0.0004 |
| F07 | 920001 | 1020000 | 0.0016 | 0.0003 | 0.0003 |
| F07 | 940001 | 1040000 | 0.0016 | 0.0008 | 0.0003 |
| F07 | 960001 | 1060000 | 0.0015 | 0.0008 | 0.0004 |
| F07 | 980001 | 1080000 | 0.0014 | 0.0009 | 0.0005 |
| F07 | 1000001 | 1100000 | 0.0013 | 0.0008 | 0.0005 |
| F07 | 1020001 | 1120000 | 0.0014 | 0.0007 | 0.0006 |
| F07 | 1040001 | 1140000 | 0.0015 | 0.0003 | 0.0005 |
| F07 | 1060001 | 1160000 | 0.0015 | 0.0003 | 0.0005 |
| F07 | 1080001 | 1180000 | 0.0015 | 0.0002 | 0.0004 |
| F07 | 1100001 | 1200000 | 0.0014 | 0.0004 | 0.0004 |
| F07 | 1120001 | 1220000 | 0.0013 | 0.0006 | 0.0003 |
| F07 | 1140001 | 1240000 | 0.0012 | 0.0008 | 0.0003 |
| F07 | 1160001 | 1260000 | 0.0013 | 0.0012 | 0.0003 |
| F07 | 1180001 | 1280000 | 0.0013 | 0.0015 | 0.0002 |
| F07 | 1200001 | 1300000 | 0.0013 | 0.0016 | 0.0004 |
| F07 | 1220001 | 1320000 | 0.0014 | 0.0015 | 0.0004 |
| F07 | 1240001 | 1340000 | 0.0016 | 0.0013 | 0.0005 |
| F07 | 1260001 | 1360000 | 0.0016 | 0.0009 | 0.0007 |
| F07 | 1280001 | 1380000 | 0.0016 | 0.0006 | 0.0006 |
| F07 | 1300001 | 1400000 | 0.0017 | 0.0005 | 0.0008 |
| F07 | 1320001 | 1420000 | 0.0017 | 0.0004 | 0.0009 |
| F07 | 1340001 | 1440000 | 0.0019 | 0.0003 | 0.0012 |
| F07 | 1360001 | 1460000 | 0.0018 | 0.0003 | 0.0013 |
| F07 | 1380001 | 1480000 | 0.0020 | 0.0004 | 0.0017 |
| F07 | 1400001 | 1500000 | 0.0020 | 0.0003 | 0.0018 |

| Chr | Start | End | both | A1 | A2 |
|---|---|---|---|---|---|
| F07 | 1420001 | 1520000 | 0.0021 | 0.0004 | 0.0021 |
| F07 | 1440001 | 1540000 | 0.0022 | 0.0005 | 0.0023 |
| F07 | 1460001 | 1560000 | 0.0024 | 0.0005 | 0.0024 |
| F07 | 1480001 | 1580000 | 0.0023 | 0.0005 | 0.0023 |
| F07 | 1500001 | 1600000 | 0.0023 | 0.0003 | 0.0023 |
| F07 | 1520001 | 1620000 | 0.0022 | 0.0004 | 0.0023 |
| F07 | 1540001 | 1640000 | 0.0020 | 0.0004 | 0.0020 |
| F07 | 1560001 | 1660000 | 0.0019 | 0.0004 | 0.0019 |
| F07 | 1580001 | 1680000 | 0.0018 | 0.0007 | 0.0017 |
| F07 | 1600001 | 1700000 | 0.0017 | 0.0009 | 0.0016 |
| F07 | 1620001 | 1720000 | 0.0019 | 0.0011 | 0.0016 |
| F07 | 1640001 | 1740000 | 0.0020 | 0.0013 | 0.0017 |
| F07 | 1660001 | 1760000 | 0.0021 | 0.0015 | 0.0017 |
| F07 | 1680001 | 1780000 | 0.0021 | 0.0015 | 0.0018 |
| F07 | 1700001 | 1800000 | 0.0020 | 0.0015 | 0.0015 |
| F07 | 1720001 | 1820000 | 0.0016 | 0.0013 | 0.0012 |
| F07 | 1740001 | 1840000 | 0.0015 | 0.0014 | 0.0010 |
| F07 | 1760001 | 1860000 | 0.0014 | 0.0012 | 0.0007 |
| F07 | 1780001 | 1880000 | 0.0013 | 0.0009 | 0.0006 |
| F07 | 1800001 | 1900000 | 0.0013 | 0.0007 | 0.0005 |
| F07 | 1820001 | 1920000 | 0.0015 | 0.0006 | 0.0007 |
| F07 | 1840001 | 1940000 | 0.0016 | 0.0003 | 0.0008 |
| F07 | 1860001 | 1960000 | 0.0017 | 0.0002 | 0.0011 |
| F07 | 1880001 | 1980000 | 0.0021 | 0.0003 | 0.0017 |
| F07 | 1900001 | 2000000 | 0.0025 | 0.0006 | 0.0021 |
| F07 | 1920001 | 2020000 | 0.0023 | 0.0005 | 0.0019 |
| F07 | 1940001 | 2040000 | 0.0020 | 0.0005 | 0.0017 |
| F07 | 1960001 | 2060000 | 0.0021 | 0.0009 | 0.0017 |
| F07 | 1980001 | 2080000 | 0.0018 | 0.0013 | 0.0013 |
| F07 | 2000001 | 2100000 | 0.0017 | 0.0011 | 0.0012 |
| F07 | 2020001 | 2120000 | 0.0017 | 0.0011 | 0.0012 |
| F07 | 2040001 | 2140000 | 0.0017 | 0.0011 | 0.0011 |
| F07 | 2060001 | 2160000 | 0.0015 | 0.0006 | 0.0008 |
| F07 | 2080001 | 2180000 | 0.0017 | 0.0006 | 0.0009 |
| F07 | 2100001 | 2200000 | 0.0013 | 0.0004 | 0.0006 |
| F07 | 2120001 | 2220000 | 0.0013 | 0.0004 | 0.0005 |
| F07 | 2140001 | 2240000 | 0.0013 | 0.0004 | 0.0005 |
| F07 | 2160001 | 2260000 | 0.0014 | 0.0009 | 0.0008 |
| F07 | 2180001 | 2280000 | 0.0016 | 0.0013 | 0.0011 |
| F07 | 2200001 | 2300000 | 0.0017 | 0.0016 | 0.0011 |
| F07 | 2220001 | 2320000 | 0.0017 | 0.0018 | 0.0013 |
| F07 | 2240001 | 2340000 | 0.0016 | 0.0020 | 0.0013 |
| F07 | 2260001 | 2360000 | 0.0017 | 0.0019 | 0.0014 |
| F07 | 2280001 | 2380000 | 0.0014 | 0.0018 | 0.0011 |
| F07 | 2300001 | 2400000 | 0.0015 | 0.0020 | 0.0011 |
| F07 | 2320001 | 2420000 | 0.0013 | 0.0022 | 0.0009 |
| F07 | 2340001 | 2440000 | 0.0013 | 0.0021 | 0.0009 |
| F07 | 2360001 | 2460000 | 0.0009 | 0.0018 | 0.0005 |
| F07 | 2380001 | 2480000 | 0.0010 | 0.0016 | 0.0007 |
| F07 | 2400001 | 2500000 | 0.0013 | 0.0013 | 0.0011 |
| F07 | 2420001 | 2520000 | 0.0019 | 0.0017 | 0.0018 |
| F07 | 2440001 | 2540000 | 0.0023 | 0.0020 | 0.0022 |
| F07 | 2460001 | 2560000 | 0.0024 | 0.0022 | 0.0023 |
| F07 | 2480001 | 2580000 | 0.0028 | 0.0028 | 0.0025 |
| F07 | 2500001 | 2600000 | 0.0024 | 0.0030 | 0.0020 |
| F07 | 2520001 | 2620000 | 0.0018 | 0.0024 | 0.0014 |
| F07 | 2540001 | 2640000 | 0.0018 | 0.0024 | 0.0014 |
| F07 | 2560001 | 2660000 | 0.0018 | 0.0026 | 0.0013 |

| Chr | Start | End | both | A1 | A2 |
|---|---|---|---|---|---|
| F07 | 2580001 | 2680000 | 0.0013 | 0.0018 | 0.0007 |
| F07 | 2600001 | 2700000 | 0.0015 | 0.0017 | 0.0007 |
| F07 | 2620001 | 2720000 | 0.0015 | 0.0015 | 0.0006 |
| F07 | 2640001 | 2740000 | 0.0013 | 0.0011 | 0.0004 |
| F07 | 2660001 | 2760000 | 0.0014 | 0.0007 | 0.0005 |
| F07 | 2680001 | 2780000 | 0.0020 | 0.0012 | 0.0011 |
| F07 | 2700001 | 2800000 | 0.0020 | 0.0010 | 0.0011 |
| F07 | 2720001 | 2820000 | 0.0024 | 0.0014 | 0.0016 |
| F07 | 2740001 | 2840000 | 0.0030 | 0.0019 | 0.0021 |
| F07 | 2760001 | 2860000 | 0.0030 | 0.0020 | 0.0022 |
| F07 | 2780001 | 2880000 | 0.0026 | 0.0013 | 0.0016 |
| F07 | 2800001 | 2900000 | 0.0027 | 0.0014 | 0.0017 |
| F07 | 2820001 | 2920000 | 0.0023 | 0.0010 | 0.0012 |
| F07 | 2840001 | 2940000 | 0.0018 | 0.0006 | 0.0006 |
| F07 | 2860001 | 2960000 | 0.0020 | 0.0007 | 0.0008 |
| F07 | 2880001 | 2980000 | 0.0020 | 0.0010 | 0.0011 |
| F07 | 2900001 | 3000000 | 0.0018 | 0.0010 | 0.0011 |
| F07 | 2920001 | 3020000 | 0.0019 | 0.0011 | 0.0012 |
| F07 | 2940001 | 3040000 | 0.0019 | 0.0009 | 0.0013 |
| F07 | 2960001 | 3060000 | 0.0016 | 0.0007 | 0.0010 |
| F07 | 2980001 | 3080000 | 0.0016 | 0.0006 | 0.0009 |
| F07 | 3000001 | 3100000 | 0.0019 | 0.0005 | 0.0013 |
| F07 | 3020001 | 3120000 | 0.0017 | 0.0004 | 0.0014 |
| F07 | 3040001 | 3140000 | 0.0017 | 0.0004 | 0.0015 |
| F07 | 3060001 | 3160000 | 0.0017 | 0.0005 | 0.0016 |
| F07 | 3080001 | 3180000 | 0.0013 | 0.0003 | 0.0014 |
| F07 | 3100001 | 3200000 | 0.0010 | 0.0003 | 0.0011 |
| F07 | 3120001 | 3220000 | 0.0012 | 0.0003 | 0.0012 |
| F07 | 3140001 | 3240000 | 0.0010 | 0.0003 | 0.0011 |
| F07 | 3160001 | 3260000 | 0.0008 | 0.0002 | 0.0009 |
| F07 | 3180001 | 3280000 | 0.0009 | 0.0003 | 0.0009 |
| F07 | 3200001 | 3300000 | 0.0009 | 0.0004 | 0.0009 |
| F07 | 3220001 | 3320000 | 0.0010 | 0.0006 | 0.0010 |
| F07 | 3240001 | 3340000 | 0.0016 | 0.0009 | 0.0015 |
| F07 | 3260001 | 3360000 | 0.0020 | 0.0010 | 0.0019 |
| F07 | 3280001 | 3380000 | 0.0021 | 0.0010 | 0.0020 |
| F07 | 3300001 | 3400000 | 0.0021 | 0.0010 | 0.0020 |
| F07 | 3320001 | 3420000 | 0.0022 | 0.0010 | 0.0018 |
| F07 | 3340001 | 3440000 | 0.0019 | 0.0008 | 0.0014 |
| F07 | 3360001 | 3460000 | 0.0018 | 0.0007 | 0.0012 |
| F07 | 3380001 | 3480000 | 0.0019 | 0.0007 | 0.0013 |
| F07 | 3400001 | 3500000 | 0.0020 | 0.0006 | 0.0014 |
| F07 | 3420001 | 3520000 | 0.0019 | 0.0008 | 0.0016 |
| F07 | 3440001 | 3540000 | 0.0021 | 0.0011 | 0.0018 |
| F07 | 3460001 | 3560000 | 0.0021 | 0.0015 | 0.0020 |
| F07 | 3480001 | 3580000 | 0.0022 | 0.0020 | 0.0020 |
| F07 | 3500001 | 3600000 | 0.0022 | 0.0025 | 0.0019 |
| F07 | 3520001 | 3620000 | 0.0020 | 0.0023 | 0.0017 |
| F07 | 3540001 | 3640000 | 0.0022 | 0.0023 | 0.0019 |
| F07 | 3560001 | 3660000 | 0.0021 | 0.0018 | 0.0018 |
| F07 | 3580001 | 3680000 | 0.0020 | 0.0014 | 0.0017 |
| F07 | 3600001 | 3700000 | 0.0017 | 0.0009 | 0.0015 |
| F07 | 3620001 | 3720000 | 0.0016 | 0.0006 | 0.0015 |
| F07 | 3640001 | 3740000 | 0.0011 | 0.0002 | 0.0011 |
| F07 | 3660001 | 3760000 | 0.0009 | 0.0004 | 0.0009 |
| F07 | 3680001 | 3780000 | 0.0010 | 0.0009 | 0.0010 |
| F07 | 3700001 | 3800000 | 0.0012 | 0.0013 | 0.0011 |
| F07 | 3720001 | 3820000 | 0.0013 | 0.0018 | 0.0011 |

| Chr | Start | End | both | A1 | A2 |
|---|---|---|---|---|---|
| F07 | 3740001 | 3840000 | 0.0013 | 0.0020 | 0.0010 |
| F07 | 3760001 | 3860000 | 0.0013 | 0.0021 | 0.0009 |
| F07 | 3780001 | 3880000 | 0.0016 | 0.0023 | 0.0013 |
| F07 | 3800001 | 3900000 | 0.0017 | 0.0023 | 0.0014 |
| F07 | 3820001 | 3920000 | 0.0017 | 0.0022 | 0.0015 |
| F07 | 3840001 | 3940000 | 0.0016 | 0.0020 | 0.0014 |
| F07 | 3860001 | 3960000 | 0.0018 | 0.0022 | 0.0015 |
| F07 | 3880001 | 3980000 | 0.0019 | 0.0023 | 0.0016 |
| F07 | 3900001 | 4000000 | 0.0016 | 0.0020 | 0.0014 |
| F07 | 3920001 | 4020000 | 0.0017 | 0.0020 | 0.0015 |
| F07 | 3940001 | 4040000 | 0.0017 | 0.0019 | 0.0015 |
| F07 | 3960001 | 4060000 | 0.0015 | 0.0016 | 0.0014 |
| F07 | 3980001 | 4080000 | 0.0008 | 0.0010 | 0.0007 |
| F07 | 4000001 | 4100000 | 0.0007 | 0.0009 | 0.0006 |
| F07 | 4020001 | 4120000 | 0.0005 | 0.0005 | 0.0004 |
| F07 | 4040001 | 4140000 | 0.0007 | 0.0006 | 0.0006 |
| F07 | 4060001 | 4160000 | 0.0014 | 0.0009 | 0.0014 |
| F07 | 4080001 | 4180000 | 0.0016 | 0.0009 | 0.0016 |
| F07 | 4100001 | 4200000 | 0.0018 | 0.0009 | 0.0019 |
| F07 | 4120001 | 4220000 | 0.0021 | 0.0009 | 0.0022 |
| F07 | 4140001 | 4240000 | 0.0028 | 0.0014 | 0.0029 |
| F07 | 4160001 | 4260000 | 0.0022 | 0.0010 | 0.0023 |
| F07 | 4180001 | 4280000 | 0.0021 | 0.0009 | 0.0021 |
| F07 | 4200001 | 4300000 | 0.0022 | 0.0011 | 0.0022 |
| F07 | 4220001 | 4320000 | 0.0021 | 0.0012 | 0.0021 |
| F07 | 4240001 | 4340000 | 0.0015 | 0.0008 | 0.0013 |
| F07 | 4260001 | 4360000 | 0.0019 | 0.0014 | 0.0017 |
| F07 | 4280001 | 4380000 | 0.0022 | 0.0018 | 0.0019 |
| F07 | 4300001 | 4400000 | 0.0021 | 0.0018 | 0.0018 |
| F07 | 4320001 | 4420000 | 0.0022 | 0.0019 | 0.0019 |
| F07 | 4340001 | 4440000 | 0.0027 | 0.0023 | 0.0025 |
| F07 | 4360001 | 4460000 | 0.0026 | 0.0018 | 0.0024 |
| F07 | 4380001 | 4480000 | 0.0024 | 0.0014 | 0.0022 |
| F07 | 4400001 | 4500000 | 0.0023 | 0.0012 | 0.0022 |
| F07 | 4420001 | 4520000 | 0.0024 | 0.0010 | 0.0024 |
| F07 | 4440001 | 4540000 | 0.0019 | 0.0004 | 0.0020 |
| F07 | 4460001 | 4560000 | 0.0016 | 0.0004 | 0.0018 |
| F07 | 4480001 | 4580000 | 0.0017 | 0.0005 | 0.0018 |
| F07 | 4500001 | 4600000 | 0.0019 | 0.0007 | 0.0020 |
| F07 | 4520001 | 4620000 | 0.0017 | 0.0008 | 0.0018 |
| F07 | 4540001 | 4640000 | 0.0019 | 0.0011 | 0.0020 |
| F07 | 4560001 | 4660000 | 0.0022 | 0.0015 | 0.0023 |
| F07 | 4580001 | 4680000 | 0.0019 | 0.0014 | 0.0019 |
| F07 | 4600001 | 4700000 | 0.0019 | 0.0012 | 0.0019 |
| F07 | 4620001 | 4720000 | 0.0019 | 0.0011 | 0.0019 |
| F07 | 4640001 | 4740000 | 0.0016 | 0.0008 | 0.0014 |
| F07 | 4660001 | 4760000 | 0.0013 | 0.0005 | 0.0010 |
| F07 | 4680001 | 4780000 | 0.0018 | 0.0009 | 0.0015 |
| F07 | 4700001 | 4800000 | 0.0015 | 0.0009 | 0.0011 |
| F07 | 4720001 | 4820000 | 0.0014 | 0.0011 | 0.0010 |
| F07 | 4740001 | 4840000 | 0.0014 | 0.0010 | 0.0009 |
| F07 | 4760001 | 4860000 | 0.0016 | 0.0012 | 0.0011 |
| F07 | 4780001 | 4880000 | 0.0015 | 0.0012 | 0.0009 |
| F07 | 4800001 | 4900000 | 0.0016 | 0.0014 | 0.0009 |
| F07 | 4820001 | 4920000 | 0.0015 | 0.0016 | 0.0009 |
| F07 | 4840001 | 4940000 | 0.0015 | 0.0020 | 0.0010 |
| F07 | 4860001 | 4960000 | 0.0013 | 0.0021 | 0.0008 |
| F07 | 4880001 | 4980000 | 0.0014 | 0.0022 | 0.0010 |

| Chr | Start | End | both | A1 | A2 |
|---|---|---|---|---|---|
| F07 | 4900001 | 5000000 | 0.0012 | 0.0020 | 0.0009 |
| F07 | 4920001 | 5020000 | 0.0012 | 0.0017 | 0.0009 |
| F07 | 4940001 | 5040000 | 0.0010 | 0.0013 | 0.0008 |
| F07 | 4960001 | 5060000 | 0.0013 | 0.0015 | 0.0012 |
| F07 | 4980001 | 5080000 | 0.0008 | 0.0010 | 0.0007 |
| F07 | 5000001 | 5100000 | 0.0009 | 0.0012 | 0.0008 |
| F07 | 5020001 | 5120000 | 0.0011 | 0.0015 | 0.0009 |
| F07 | 5040001 | 5140000 | 0.0013 | 0.0018 | 0.0010 |
| F07 | 5060001 | 5160000 | 0.0011 | 0.0015 | 0.0008 |
| F07 | 5080001 | 5180000 | 0.0012 | 0.0017 | 0.0009 |
| F07 | 5100001 | 5200000 | 0.0020 | 0.0024 | 0.0018 |
| F07 | 5120001 | 5220000 | 0.0025 | 0.0027 | 0.0024 |
| F07 | 5140001 | 5240000 | 0.0024 | 0.0025 | 0.0022 |
| F07 | 5160001 | 5260000 | 0.0021 | 0.0023 | 0.0020 |
| F07 | 5180001 | 5280000 | 0.0022 | 0.0022 | 0.0021 |
| F07 | 5200001 | 5300000 | 0.0016 | 0.0016 | 0.0015 |
| F07 | 5220001 | 5320000 | 0.0012 | 0.0012 | 0.0011 |
| F07 | 5240001 | 5340000 | 0.0020 | 0.0020 | 0.0019 |
| F07 | 5260001 | 5360000 | 0.0024 | 0.0025 | 0.0024 |
| F07 | 5280001 | 5380000 | 0.0023 | 0.0025 | 0.0022 |
| F07 | 5300001 | 5400000 | 0.0020 | 0.0023 | 0.0019 |
| F07 | 5320001 | 5420000 | 0.0018 | 0.0023 | 0.0016 |
| F07 | 5340001 | 5440000 | 0.0014 | 0.0021 | 0.0012 |
| F07 | 5360001 | 5460000 | 0.0010 | 0.0016 | 0.0008 |
| F07 | 5380001 | 5480000 | 0.0011 | 0.0015 | 0.0009 |
| F07 | 5400001 | 5500000 | 0.0012 | 0.0015 | 0.0011 |
| F07 | 5420001 | 5520000 | 0.0017 | 0.0018 | 0.0017 |
| F07 | 5440001 | 5540000 | 0.0021 | 0.0019 | 0.0021 |
| F07 | 5460001 | 5560000 | 0.0027 | 0.0025 | 0.0027 |
| F07 | 5480001 | 5580000 | 0.0030 | 0.0028 | 0.0030 |
| F07 | 5500001 | 5600000 | 0.0032 | 0.0029 | 0.0031 |
| F07 | 5520001 | 5620000 | 0.0031 | 0.0029 | 0.0031 |
| F07 | 5540001 | 5640000 | 0.0027 | 0.0022 | 0.0027 |
| F07 | 5560001 | 5660000 | 0.0026 | 0.0016 | 0.0026 |
| F07 | 5580001 | 5680000 | 0.0027 | 0.0012 | 0.0026 |
| F07 | 5600001 | 5700000 | 0.0034 | 0.0014 | 0.0033 |
| F07 | 5620001 | 5720000 | 0.0035 | 0.0010 | 0.0035 |
| F07 | 5640001 | 5740000 | 0.0035 | 0.0010 | 0.0035 |
| F07 | 5660001 | 5760000 | 0.0040 | 0.0014 | 0.0040 |
| F07 | 5680001 | 5780000 | 0.0037 | 0.0013 | 0.0036 |
| F07 | 5700001 | 5800000 | 0.0028 | 0.0008 | 0.0027 |
| F07 | 5720001 | 5820000 | 0.0024 | 0.0006 | 0.0022 |
| F07 | 5740001 | 5840000 | 0.0020 | 0.0006 | 0.0017 |
| F07 | 5760001 | 5860000 | 0.0013 | 0.0002 | 0.0007 |
| F07 | 5780001 | 5880000 | 0.0013 | 0.0003 | 0.0006 |
| F07 | 5800001 | 5900000 | 0.0011 | 0.0003 | 0.0004 |
| F07 | 5820001 | 5920000 | 0.0016 | 0.0010 | 0.0010 |
| F07 | 5840001 | 5940000 | 0.0017 | 0.0010 | 0.0010 |
| F07 | 5860001 | 5960000 | 0.0017 | 0.0009 | 0.0013 |
| F07 | 5880001 | 5980000 | 0.0021 | 0.0009 | 0.0018 |
| F07 | 5900001 | 6000000 | 0.0024 | 0.0009 | 0.0021 |
| F07 | 5920001 | 6020000 | 0.0022 | 0.0007 | 0.0022 |
| F07 | 5940001 | 6040000 | 0.0027 | 0.0009 | 0.0026 |
| F07 | 5960001 | 6060000 | 0.0026 | 0.0009 | 0.0024 |
| F07 | 5980001 | 6080000 | 0.0028 | 0.0013 | 0.0024 |
| F07 | 6000001 | 6100000 | 0.0026 | 0.0013 | 0.0022 |
| F07 | 6020001 | 6120000 | 0.0021 | 0.0009 | 0.0016 |
| F07 | 6040001 | 6140000 | 0.0019 | 0.0008 | 0.0013 |

| Chr | Start | End | both | A1 | A2 |
|---|---|---|---|---|---|
| F07 | 6060001 | 6160000 | 0.0018 | 0.0008 | 0.0013 |
| F07 | 6080001 | 6180000 | 0.0012 | 0.0005 | 0.0008 |
| F07 | 6100001 | 6200000 | 0.0012 | 0.0005 | 0.0008 |
| F07 | 6120001 | 6220000 | 0.0014 | 0.0007 | 0.0010 |
| F07 | 6140001 | 6240000 | 0.0012 | 0.0006 | 0.0008 |
| F07 | 6160001 | 6260000 | 0.0014 | 0.0008 | 0.0008 |
| F07 | 6180001 | 6280000 | 0.0016 | 0.0008 | 0.0007 |
| F07 | 6200001 | 6300000 | 0.0019 | 0.0009 | 0.0008 |
| F07 | 6220001 | 6320000 | 0.0017 | 0.0006 | 0.0006 |
| F07 | 6240001 | 6340000 | 0.0018 | 0.0006 | 0.0006 |
| F07 | 6260001 | 6360000 | 0.0021 | 0.0010 | 0.0011 |
| F07 | 6280001 | 6380000 | 0.0025 | 0.0015 | 0.0015 |
| F07 | 6300001 | 6400000 | 0.0022 | 0.0013 | 0.0013 |
| F07 | 6320001 | 6420000 | 0.0027 | 0.0018 | 0.0018 |
| F07 | 6340001 | 6440000 | 0.0026 | 0.0017 | 0.0017 |
| F07 | 6360001 | 6460000 | 0.0023 | 0.0013 | 0.0011 |
| F07 | 6380001 | 6480000 | 0.0018 | 0.0007 | 0.0007 |
| F07 | 6400001 | 6500000 | 0.0018 | 0.0007 | 0.0008 |
| F07 | 6420001 | 6520000 | 0.0017 | 0.0004 | 0.0007 |
| F07 | 6440001 | 6540000 | 0.0017 | 0.0004 | 0.0007 |
| F07 | 6460001 | 6560000 | 0.0029 | 0.0019 | 0.0022 |
| F07 | 6480001 | 6580000 | 0.0033 | 0.0024 | 0.0027 |
| F07 | 6500001 | 6600000 | 0.0042 | 0.0034 | 0.0037 |
| F07 | 6520001 | 6620000 | 0.0042 | 0.0036 | 0.0038 |
| F07 | 6540001 | 6640000 | 0.0041 | 0.0036 | 0.0039 |
| F07 | 6560001 | 6660000 | 0.0027 | 0.0023 | 0.0026 |
| F07 | 6580001 | 6680000 | 0.0029 | 0.0024 | 0.0027 |
| F07 | 6600001 | 6700000 | 0.0023 | 0.0016 | 0.0019 |
| F07 | 6620001 | 6720000 | 0.0024 | 0.0017 | 0.0021 |
| F07 | 6640001 | 6740000 | 0.0032 | 0.0024 | 0.0027 |
| F07 | 6660001 | 6760000 | 0.0039 | 0.0030 | 0.0032 |
| F07 | 6680001 | 6780000 | 0.0039 | 0.0030 | 0.0031 |
| F07 | 6700001 | 6800000 | 0.0037 | 0.0029 | 0.0029 |
| F07 | 6720001 | 6820000 | 0.0033 | 0.0024 | 0.0023 |
| F07 | 6740001 | 6840000 | 0.0027 | 0.0016 | 0.0016 |
| F07 | 6760001 | 6860000 | 0.0022 | 0.0012 | 0.0013 |
| F07 | 6780001 | 6880000 | 0.0016 | 0.0006 | 0.0007 |
| F07 | 6800001 | 6900000 | 0.0025 | 0.0014 | 0.0016 |
| F07 | 6820001 | 6920000 | 0.0029 | 0.0018 | 0.0021 |
| F07 | 6840001 | 6940000 | 0.0035 | 0.0023 | 0.0026 |
| F07 | 6860001 | 6960000 | 0.0037 | 0.0024 | 0.0027 |
| F07 | 6880001 | 6980000 | 0.0041 | 0.0028 | 0.0032 |
| F07 | 6900001 | 7000000 | 0.0036 | 0.0024 | 0.0027 |
| F07 | 6920001 | 7020000 | 0.0033 | 0.0022 | 0.0026 |
| F07 | 6940001 | 7040000 | 0.0028 | 0.0018 | 0.0021 |
| F07 | 6960001 | 7060000 | 0.0025 | 0.0014 | 0.0017 |
| F07 | 6980001 | 7080000 | 0.0022 | 0.0011 | 0.0013 |
| F07 | 7000001 | 7100000 | 0.0021 | 0.0011 | 0.0012 |
| F07 | 7020001 | 7120000 | 0.0019 | 0.0009 | 0.0010 |
| F07 | 7040001 | 7140000 | 0.0019 | 0.0009 | 0.0010 |
| F07 | 7060001 | 7160000 | 0.0017 | 0.0009 | 0.0009 |
| F07 | 7080001 | 7180000 | 0.0016 | 0.0008 | 0.0009 |
| F07 | 7100001 | 7200000 | 0.0012 | 0.0004 | 0.0004 |
| F07 | 7120001 | 7220000 | 0.0016 | 0.0007 | 0.0008 |
| F07 | 7140001 | 7240000 | 0.0019 | 0.0011 | 0.0011 |
| F07 | 7160001 | 7260000 | 0.0018 | 0.0011 | 0.0011 |
| F07 | 7180001 | 7280000 | 0.0018 | 0.0011 | 0.0012 |
| F07 | 7200001 | 7300000 | 0.0021 | 0.0015 | 0.0016 |

| Chr | Start | End | both | A1 | A2 |
|---|---|---|---|---|---|
| F07 | 7220001 | 7320000 | 0.0021 | 0.0014 | 0.0017 |
| F07 | 7240001 | 7340000 | 0.0021 | 0.0014 | 0.0017 |
| F07 | 7260001 | 7360000 | 0.0024 | 0.0017 | 0.0020 |
| F07 | 7280001 | 7380000 | 0.0027 | 0.0018 | 0.0021 |
| F07 | 7300001 | 7400000 | 0.0027 | 0.0019 | 0.0020 |
| F07 | 7320001 | 7420000 | 0.0022 | 0.0014 | 0.0014 |
| F07 | 7340001 | 7440000 | 0.0021 | 0.0014 | 0.0014 |
| F07 | 7360001 | 7460000 | 0.0018 | 0.0010 | 0.0010 |
| F07 | 7380001 | 7480000 | 0.0017 | 0.0010 | 0.0009 |
| F07 | 7400001 | 7500000 | 0.0018 | 0.0012 | 0.0011 |
| F07 | 7420001 | 7520000 | 0.0024 | 0.0016 | 0.0017 |
| F07 | 7440001 | 7540000 | 0.0024 | 0.0018 | 0.0017 |
| F07 | 7460001 | 7560000 | 0.0025 | 0.0018 | 0.0017 |
| F07 | 7480001 | 7580000 | 0.0021 | 0.0016 | 0.0015 |
| F07 | 7500001 | 7600000 | 0.0016 | 0.0012 | 0.0011 |
| F07 | 7520001 | 7620000 | 0.0017 | 0.0015 | 0.0013 |
| F07 | 7540001 | 7640000 | 0.0013 | 0.0012 | 0.0009 |
| F07 | 7560001 | 7660000 | 0.0017 | 0.0017 | 0.0014 |
| F07 | 7580001 | 7680000 | 0.0025 | 0.0024 | 0.0021 |
| F07 | 7600001 | 7700000 | 0.0032 | 0.0029 | 0.0029 |
| F07 | 7620001 | 7720000 | 0.0042 | 0.0038 | 0.0039 |
| F07 | 7640001 | 7740000 | 0.0046 | 0.0043 | 0.0042 |
| F07 | 7660001 | 7760000 | 0.0044 | 0.0041 | 0.0039 |
| F07 | 7680001 | 7780000 | 0.0045 | 0.0040 | 0.0040 |
| F07 | 7700001 | 7800000 | 0.0040 | 0.0037 | 0.0033 |
| F07 | 7720001 | 7820000 | 0.0043 | 0.0040 | 0.0037 |
| F07 | 7740001 | 7840000 | 0.0045 | 0.0039 | 0.0039 |
| F07 | 7760001 | 7860000 | 0.0060 | 0.0052 | 0.0055 |
| F07 | 7780001 | 7880000 | 0.0073 | 0.0064 | 0.0067 |
| F07 | 7800001 | 7900000 | 0.0073 | 0.0064 | 0.0068 |
| F07 | 7820001 | 7920000 | 0.0054 | 0.0047 | 0.0047 |
| F07 | 7840001 | 7940000 | 0.0050 | 0.0043 | 0.0044 |
| F07 | 7860001 | 7960000 | 0.0032 | 0.0027 | 0.0026 |
| F07 | 7880001 | 7980000 | 0.0017 | 0.0013 | 0.0008 |
| F07 | 7900001 | 8000000 | 0.0020 | 0.0016 | 0.0010 |
| F07 | 7920001 | 8020000 | 0.0021 | 0.0016 | 0.0010 |
| F07 | 7940001 | 8040000 | 0.0025 | 0.0018 | 0.0013 |
| F07 | 7960001 | 8060000 | 0.0026 | 0.0018 | 0.0012 |
| F07 | 7980001 | 8080000 | 0.0022 | 0.0013 | 0.0011 |
| F07 | 8000001 | 8100000 | 0.0020 | 0.0009 | 0.0008 |
| F07 | 8020001 | 8120000 | 0.0021 | 0.0008 | 0.0008 |
| F07 | 8040001 | 8140000 | 0.0020 | 0.0007 | 0.0009 |
| F07 | 8060001 | 8160000 | 0.0025 | 0.0014 | 0.0014 |
| F07 | 8080001 | 8180000 | 0.0030 | 0.0022 | 0.0018 |
| F07 | 8100001 | 8200000 | 0.0028 | 0.0022 | 0.0018 |
| F07 | 8120001 | 8220000 | 0.0026 | 0.0025 | 0.0017 |
| F07 | 8140001 | 8240000 | 0.0024 | 0.0026 | 0.0014 |
| F07 | 8160001 | 8260000 | 0.0021 | 0.0023 | 0.0013 |
| F07 | 8180001 | 8280000 | 0.0015 | 0.0017 | 0.0008 |
| F07 | 8200001 | 8300000 | 0.0021 | 0.0022 | 0.0013 |
| F07 | 8220001 | 8320000 | 0.0020 | 0.0021 | 0.0012 |
| F07 | 8240001 | 8340000 | 0.0021 | 0.0021 | 0.0014 |
| F07 | 8260001 | 8360000 | 0.0018 | 0.0018 | 0.0011 |
| F07 | 8280001 | 8380000 | 0.0021 | 0.0020 | 0.0016 |
| F07 | 8300001 | 8400000 | 0.0031 | 0.0030 | 0.0025 |
| F07 | 8320001 | 8420000 | 0.0035 | 0.0033 | 0.0027 |
| F07 | 8340001 | 8440000 | 0.0038 | 0.0036 | 0.0030 |
| F07 | 8360001 | 8460000 | 0.0040 | 0.0038 | 0.0032 |

| Chr | Start | End | both | A1 | A2 |
| --- | --- | --- | --- | --- | --- |
| F07 | 8380001 | 8480000 | 0.0040 | 0.0039 | 0.0032 |
| F07 | 8400001 | 8500000 | 0.0028 | 0.0027 | 0.0022 |
| F07 | 8420001 | 8520000 | 0.0029 | 0.0027 | 0.0025 |
| F07 | 8440001 | 8540000 | 0.0031 | 0.0030 | 0.0027 |
| F07 | 8460001 | 8560000 | 0.0029 | 0.0030 | 0.0024 |
| F07 | 8480001 | 8580000 | 0.0027 | 0.0028 | 0.0023 |
| F07 | 8500001 | 8600000 | 0.0024 | 0.0025 | 0.0020 |
| F07 | 8520001 | 8620000 | 0.0024 | 0.0023 | 0.0018 |
| F07 | 8540001 | 8640000 | 0.0018 | 0.0012 | 0.0009 |
| F07 | 8560001 | 8660000 | 0.0017 | 0.0009 | 0.0009 |
| F07 | 8580001 | 8680000 | 0.0018 | 0.0009 | 0.0008 |
| F07 | 8600001 | 8700000 | 0.0017 | 0.0010 | 0.0007 |
| F07 | 8620001 | 8720000 | 0.0012 | 0.0009 | 0.0004 |
| F07 | 8640001 | 8740000 | 0.0021 | 0.0021 | 0.0015 |
| F07 | 8660001 | 8760000 | 0.0022 | 0.0025 | 0.0016 |
| F07 | 8680001 | 8780000 | 0.0021 | 0.0029 | 0.0016 |
| F07 | 8700001 | 8800000 | 0.0022 | 0.0032 | 0.0017 |
| F07 | 8720001 | 8820000 | 0.0023 | 0.0031 | 0.0017 |
| F07 | 8740001 | 8840000 | 0.0012 | 0.0021 | 0.0007 |
| F07 | 8760001 | 8860000 | 0.0016 | 0.0023 | 0.0010 |
| F07 | 8780001 | 8880000 | 0.0014 | 0.0020 | 0.0009 |
| F07 | 8800001 | 8900000 | 0.0015 | 0.0017 | 0.0010 |
| F07 | 8820001 | 8920000 | 0.0017 | 0.0021 | 0.0011 |
| F07 | 8840001 | 8940000 | 0.0018 | 0.0022 | 0.0012 |
| F07 | 8860001 | 8960000 | 0.0016 | 0.0019 | 0.0010 |
| F07 | 8880001 | 8980000 | 0.0018 | 0.0021 | 0.0011 |
| F07 | 8900001 | 9000000 | 0.0023 | 0.0025 | 0.0016 |
| F07 | 8920001 | 9020000 | 0.0021 | 0.0025 | 0.0013 |
| F07 | 8940001 | 9040000 | 0.0022 | 0.0024 | 0.0014 |
| F07 | 8960001 | 9060000 | 0.0021 | 0.0023 | 0.0012 |
| F07 | 8980001 | 9080000 | 0.0019 | 0.0018 | 0.0011 |
| F07 | 9000001 | 9100000 | 0.0015 | 0.0013 | 0.0006 |
| F07 | 9020001 | 9120000 | 0.0029 | 0.0025 | 0.0021 |
| F07 | 9040001 | 9140000 | 0.0044 | 0.0038 | 0.0036 |
| F07 | 9060001 | 9160000 | 0.0048 | 0.0041 | 0.0039 |
| F07 | 9080001 | 9180000 | 0.0049 | 0.0042 | 0.0040 |
| F07 | 9100001 | 9200000 | 0.0047 | 0.0041 | 0.0038 |
| F07 | 9120001 | 9220000 | 0.0032 | 0.0028 | 0.0024 |
| F07 | 9140001 | 9240000 | 0.0021 | 0.0017 | 0.0013 |
| F07 | 9160001 | 9260000 | 0.0021 | 0.0016 | 0.0016 |
| F07 | 9180001 | 9280000 | 0.0021 | 0.0017 | 0.0017 |
| F07 | 9200001 | 9300000 | 0.0021 | 0.0017 | 0.0018 |
| F07 | 9220001 | 9320000 | 0.0021 | 0.0016 | 0.0018 |
| F07 | 9240001 | 9340000 | 0.0017 | 0.0012 | 0.0012 |
| F07 | 9260001 | 9360000 | 0.0015 | 0.0011 | 0.0009 |
| F07 | 9280001 | 9380000 | 0.0021 | 0.0017 | 0.0014 |
| F07 | 9300001 | 9400000 | 0.0029 | 0.0023 | 0.0022 |
| F07 | 9320001 | 9420000 | 0.0028 | 0.0023 | 0.0023 |
| F07 | 9340001 | 9440000 | 0.0038 | 0.0033 | 0.0036 |
| F07 | 9360001 | 9460000 | 0.0039 | 0.0035 | 0.0038 |
| F07 | 9380001 | 9480000 | 0.0053 | 0.0044 | 0.0049 |
| F07 | 9400001 | 9500000 | 0.0064 | 0.0054 | 0.0060 |
| F07 | 9420001 | 9520000 | 0.0067 | 0.0057 | 0.0063 |
| F07 | 9440001 | 9540000 | 0.0057 | 0.0047 | 0.0051 |
| F07 | 9460001 | 9560000 | 0.0056 | 0.0045 | 0.0050 |
| F07 | 9480001 | 9580000 | 0.0040 | 0.0033 | 0.0036 |
| F07 | 9500001 | 9600000 | 0.0020 | 0.0016 | 0.0015 |
| F07 | 9520001 | 9620000 | 0.0022 | 0.0017 | 0.0016 |

| Chr | Start | End | both | A1 | A2 |
|---|---|---|---|---|---|
| F07 | 9540001 | 9640000 | 0.0022 | 0.0018 | 0.0018 |
| F07 | 9560001 | 9660000 | 0.0028 | 0.0023 | 0.0025 |
| F07 | 9580001 | 9680000 | 0.0029 | 0.0023 | 0.0027 |
| F07 | 9600001 | 9700000 | 0.0028 | 0.0023 | 0.0028 |
| F07 | 9620001 | 9720000 | 0.0036 | 0.0030 | 0.0036 |
| F07 | 9640001 | 9740000 | 0.0039 | 0.0032 | 0.0038 |
| F07 | 9660001 | 9760000 | 0.0040 | 0.0034 | 0.0038 |
| F07 | 9680001 | 9780000 | 0.0035 | 0.0028 | 0.0031 |
| F07 | 9700001 | 9800000 | 0.0038 | 0.0029 | 0.0032 |
| F07 | 9720001 | 9820000 | 0.0030 | 0.0022 | 0.0023 |
| F07 | 9740001 | 9840000 | 0.0040 | 0.0029 | 0.0031 |
| F07 | 9760001 | 9860000 | 0.0042 | 0.0029 | 0.0033 |
| F07 | 9780001 | 9880000 | 0.0043 | 0.0030 | 0.0033 |
| F07 | 9800001 | 9900000 | 0.0042 | 0.0028 | 0.0031 |
| F07 | 9820001 | 9920000 | 0.0038 | 0.0024 | 0.0027 |
| F07 | 9840001 | 9940000 | 0.0026 | 0.0015 | 0.0016 |
| F07 | 9860001 | 9960000 | 0.0015 | 0.0009 | 0.0006 |
| F07 | 9880001 | 9980000 | 0.0016 | 0.0013 | 0.0008 |
| F07 | 9900001 | 10000000 | 0.0018 | 0.0017 | 0.0012 |
| F07 | 9920001 | 10020000 | 0.0023 | 0.0025 | 0.0018 |
| F07 | 9940001 | 10040000 | 0.0026 | 0.0031 | 0.0021 |
| F07 | 9960001 | 10060000 | 0.0032 | 0.0036 | 0.0027 |
| F07 | 9980001 | 10080000 | 0.0031 | 0.0036 | 0.0026 |
| F07 | 10000001 | 10100000 | 0.0028 | 0.0034 | 0.0023 |
| F07 | 10020001 | 10120000 | 0.0023 | 0.0029 | 0.0018 |
| F07 | 10040001 | 10140000 | 0.0019 | 0.0024 | 0.0014 |
| F07 | 10060001 | 10160000 | 0.0017 | 0.0022 | 0.0013 |
| F07 | 10080001 | 10180000 | 0.0019 | 0.0023 | 0.0015 |
| F07 | 10100001 | 10200000 | 0.0029 | 0.0033 | 0.0025 |
| F07 | 10120001 | 10220000 | 0.0030 | 0.0035 | 0.0026 |
| F07 | 10140001 | 10240000 | 0.0035 | 0.0040 | 0.0031 |
| F07 | 10160001 | 10260000 | 0.0035 | 0.0038 | 0.0031 |
| F07 | 10180001 | 10280000 | 0.0039 | 0.0040 | 0.0036 |
| F07 | 10200001 | 10300000 | 0.0031 | 0.0033 | 0.0028 |
| F07 | 10220001 | 10320000 | 0.0031 | 0.0035 | 0.0027 |
| F07 | 10240001 | 10340000 | 0.0025 | 0.0029 | 0.0022 |
| F07 | 10260001 | 10360000 | 0.0027 | 0.0035 | 0.0022 |
| F07 | 10280001 | 10380000 | 0.0017 | 0.0028 | 0.0012 |
| F07 | 10300001 | 10400000 | 0.0025 | 0.0032 | 0.0020 |
| F07 | 10320001 | 10420000 | 0.0039 | 0.0040 | 0.0035 |
| F07 | 10340001 | 10440000 | 0.0050 | 0.0048 | 0.0046 |
| F07 | 10360001 | 10460000 | 0.0049 | 0.0045 | 0.0046 |
| F07 | 10380001 | 10480000 | 0.0051 | 0.0047 | 0.0049 |
| F07 | 10400001 | 10500000 | 0.0043 | 0.0042 | 0.0041 |
| F07 | 10420001 | 10520000 | 0.0033 | 0.0033 | 0.0030 |
| F07 | 10440001 | 10540000 | 0.0027 | 0.0027 | 0.0026 |
| F07 | 10460001 | 10560000 | 0.0027 | 0.0026 | 0.0026 |
| F07 | 10480001 | 10580000 | 0.0028 | 0.0027 | 0.0027 |
| F07 | 10500001 | 10600000 | 0.0031 | 0.0029 | 0.0030 |
| F07 | 10520001 | 10620000 | 0.0029 | 0.0026 | 0.0028 |
| F07 | 10540001 | 10640000 | 0.0029 | 0.0027 | 0.0028 |
| F07 | 10560001 | 10660000 | 0.0037 | 0.0033 | 0.0036 |
| F07 | 10580001 | 10680000 | 0.0040 | 0.0035 | 0.0039 |
| F07 | 10600001 | 10700000 | 0.0036 | 0.0033 | 0.0035 |
| F07 | 10620001 | 10720000 | 0.0032 | 0.0030 | 0.0030 |
| F07 | 10640001 | 10740000 | 0.0026 | 0.0024 | 0.0025 |
| F07 | 10660001 | 10760000 | 0.0022 | 0.0020 | 0.0021 |
| F07 | 10680001 | 10780000 | 0.0021 | 0.0019 | 0.0020 |

| Chr | Start | End | both | A1 | A2 |
|---|---|---|---|---|---|
| F07 | 10700001 | 10800000 | 0.0029 | 0.0026 | 0.0029 |
| F07 | 10720001 | 10820000 | 0.0044 | 0.0041 | 0.0043 |
| F07 | 10740001 | 10840000 | 0.0052 | 0.0048 | 0.0050 |
| F07 | 10760001 | 10860000 | 0.0057 | 0.0055 | 0.0055 |
| F07 | 10780001 | 10880000 | 0.0055 | 0.0054 | 0.0052 |
| F07 | 10800001 | 10900000 | 0.0051 | 0.0051 | 0.0048 |
| F07 | 10820001 | 10920000 | 0.0050 | 0.0051 | 0.0047 |
| F07 | 10840001 | 10940000 | 0.0046 | 0.0047 | 0.0043 |
| F07 | 10860001 | 10960000 | 0.0033 | 0.0033 | 0.0032 |
| F07 | 10880001 | 10980000 | 0.0035 | 0.0036 | 0.0033 |
| F07 | 10900001 | 11000000 | 0.0035 | 0.0036 | 0.0033 |
| F07 | 10920001 | 11020000 | 0.0023 | 0.0024 | 0.0022 |
| F07 | 10940001 | 11040000 | 0.0022 | 0.0023 | 0.0020 |
| F07 | 10960001 | 11060000 | 0.0029 | 0.0032 | 0.0026 |
| F07 | 10980001 | 11080000 | 0.0034 | 0.0034 | 0.0032 |
| F07 | 11000001 | 11100000 | 0.0034 | 0.0033 | 0.0033 |
| F07 | 11020001 | 11120000 | 0.0040 | 0.0038 | 0.0039 |
| F07 | 11040001 | 11140000 | 0.0049 | 0.0048 | 0.0047 |
| F07 | 11060001 | 11160000 | 0.0047 | 0.0043 | 0.0046 |
| F07 | 11080001 | 11180000 | 0.0038 | 0.0035 | 0.0037 |
| F07 | 11100001 | 11200000 | 0.0032 | 0.0030 | 0.0030 |
| F07 | 11120001 | 11220000 | 0.0031 | 0.0031 | 0.0030 |
| F07 | 11140001 | 11240000 | 0.0023 | 0.0023 | 0.0021 |
| F07 | 11160001 | 11260000 | 0.0022 | 0.0025 | 0.0020 |
| F07 | 11180001 | 11280000 | 0.0028 | 0.0031 | 0.0026 |
| F07 | 11200001 | 11300000 | 0.0036 | 0.0040 | 0.0034 |
| F07 | 11220001 | 11320000 | 0.0029 | 0.0032 | 0.0026 |
| F07 | 11240001 | 11340000 | 0.0036 | 0.0039 | 0.0034 |
| F07 | 11260001 | 11360000 | 0.0041 | 0.0042 | 0.0039 |
| F07 | 11280001 | 11380000 | 0.0041 | 0.0042 | 0.0040 |
| F07 | 11300001 | 11400000 | 0.0043 | 0.0044 | 0.0041 |
| F07 | 11320001 | 11420000 | 0.0044 | 0.0046 | 0.0042 |
| F07 | 11340001 | 11440000 | 0.0046 | 0.0047 | 0.0044 |
| F07 | 11360001 | 11460000 | 0.0039 | 0.0041 | 0.0037 |
| F07 | 11380001 | 11480000 | 0.0035 | 0.0037 | 0.0033 |
| F07 | 11400001 | 11500000 | 0.0030 | 0.0031 | 0.0028 |
| F07 | 11420001 | 11520000 | 0.0036 | 0.0037 | 0.0035 |
| F07 | 11440001 | 11540000 | 0.0029 | 0.0031 | 0.0028 |
| F07 | 11460001 | 11560000 | 0.0032 | 0.0033 | 0.0030 |
| F07 | 11480001 | 11580000 | 0.0032 | 0.0033 | 0.0031 |
| F07 | 11500001 | 11600000 | 0.0034 | 0.0035 | 0.0032 |
| F07 | 11520001 | 11620000 | 0.0025 | 0.0026 | 0.0024 |
| F07 | 11540001 | 11640000 | 0.0032 | 0.0032 | 0.0031 |
| F07 | 11560001 | 11660000 | 0.0032 | 0.0030 | 0.0031 |
| F07 | 11580001 | 11680000 | 0.0029 | 0.0027 | 0.0028 |
| F07 | 11600001 | 11700000 | 0.0024 | 0.0022 | 0.0023 |
| F07 | 11620001 | 11720000 | 0.0030 | 0.0027 | 0.0030 |
| F07 | 11640001 | 11740000 | 0.0026 | 0.0025 | 0.0025 |
| F07 | 11660001 | 11760000 | 0.0033 | 0.0033 | 0.0032 |
| F07 | 11680001 | 11780000 | 0.0037 | 0.0035 | 0.0036 |
| F07 | 11700001 | 11800000 | 0.0043 | 0.0041 | 0.0042 |
| F07 | 11720001 | 11820000 | 0.0040 | 0.0039 | 0.0039 |
| F07 | 11740001 | 11840000 | 0.0039 | 0.0036 | 0.0038 |
| F07 | 11760001 | 11860000 | 0.0029 | 0.0026 | 0.0028 |
| F07 | 11780001 | 11880000 | 0.0031 | 0.0029 | 0.0030 |
| F07 | 11800001 | 11900000 | 0.0028 | 0.0026 | 0.0028 |
| F07 | 11820001 | 11920000 | 0.0027 | 0.0024 | 0.0027 |
| F07 | 11840001 | 11940000 | 0.0029 | 0.0025 | 0.0028 |

| Chr | Start | End | both | A1 | A2 |
|---|---|---|---|---|---|
| F07 | 11860001 | 11960000 | 0.0029 | 0.0025 | 0.0028 |
| F07 | 11880001 | 11980000 | 0.0024 | 0.0021 | 0.0024 |
| F07 | 11900001 | 12000000 | 0.0021 | 0.0019 | 0.0020 |
| F07 | 11920001 | 12020000 | 0.0020 | 0.0018 | 0.0019 |
| F07 | 11940001 | 12040000 | 0.0025 | 0.0023 | 0.0024 |
| F07 | 11960001 | 12060000 | 0.0031 | 0.0029 | 0.0030 |
| F07 | 11980001 | 12080000 | 0.0039 | 0.0034 | 0.0039 |
| F07 | 12000001 | 12100000 | 0.0043 | 0.0037 | 0.0044 |
| F07 | 12020001 | 12120000 | 0.0047 | 0.0040 | 0.0048 |
| F07 | 12040001 | 12140000 | 0.0074 | 0.0060 | 0.0074 |
| F07 | 12060001 | 12160000 | 0.0069 | 0.0056 | 0.0069 |
| F07 | 12080001 | 12180000 | 0.0061 | 0.0050 | 0.0061 |
| F07 | 12100001 | 12200000 | 0.0062 | 0.0052 | 0.0062 |
| F07 | 12120001 | 12220000 | 0.0062 | 0.0052 | 0.0062 |
| F07 | 12140001 | 12240000 | 0.0029 | 0.0026 | 0.0029 |
| F07 | 12160001 | 12260000 | 0.0033 | 0.0030 | 0.0033 |
| F07 | 12180001 | 12280000 | 0.0033 | 0.0030 | 0.0033 |
| F07 | 12200001 | 12300000 | 0.0031 | 0.0028 | 0.0030 |
| F07 | 12220001 | 12320000 | 0.0028 | 0.0025 | 0.0027 |
| F07 | 12240001 | 12340000 | 0.0029 | 0.0026 | 0.0028 |
| F07 | 12260001 | 12360000 | 0.0029 | 0.0026 | 0.0026 |
| F07 | 12280001 | 12380000 | 0.0027 | 0.0026 | 0.0024 |
| F07 | 12300001 | 12400000 | 0.0032 | 0.0033 | 0.0027 |
| F07 | 12320001 | 12420000 | 0.0032 | 0.0033 | 0.0026 |
| F07 | 12340001 | 12440000 | 0.0030 | 0.0031 | 0.0024 |
| F07 | 12360001 | 12460000 | 0.0032 | 0.0032 | 0.0026 |
| F07 | 12380001 | 12480000 | 0.0045 | 0.0044 | 0.0041 |
| F07 | 12400001 | 12500000 | 0.0039 | 0.0038 | 0.0036 |
| F07 | 12420001 | 12520000 | 0.0048 | 0.0046 | 0.0045 |
| F07 | 12440001 | 12540000 | 0.0051 | 0.0050 | 0.0047 |
| F07 | 12460001 | 12560000 | 0.0048 | 0.0047 | 0.0043 |
| F07 | 12480001 | 12580000 | 0.0035 | 0.0034 | 0.0031 |
| F07 | 12500001 | 12600000 | 0.0041 | 0.0039 | 0.0036 |
| F07 | 12520001 | 12620000 | 0.0042 | 0.0042 | 0.0036 |
| F07 | 12540001 | 12640000 | 0.0039 | 0.0037 | 0.0035 |
| F07 | 12560001 | 12660000 | 0.0052 | 0.0050 | 0.0049 |
| F07 | 12580001 | 12680000 | 0.0054 | 0.0051 | 0.0052 |
| F07 | 12600001 | 12700000 | 0.0045 | 0.0042 | 0.0044 |
| F07 | 12620001 | 12720000 | 0.0038 | 0.0034 | 0.0038 |
| F07 | 12640001 | 12740000 | 0.0042 | 0.0039 | 0.0041 |
| F07 | 12660001 | 12760000 | 0.0029 | 0.0028 | 0.0027 |
| F07 | 12680001 | 12780000 | 0.0037 | 0.0038 | 0.0032 |
| F07 | 12700001 | 12800000 | 0.0040 | 0.0041 | 0.0034 |
| F07 | 12720001 | 12820000 | 0.0037 | 0.0040 | 0.0029 |
| F07 | 12740001 | 12840000 | 0.0035 | 0.0038 | 0.0027 |
| F07 | 12760001 | 12860000 | 0.0033 | 0.0035 | 0.0025 |
| F07 | 12780001 | 12880000 | 0.0021 | 0.0023 | 0.0014 |
| F07 | 12800001 | 12900000 | 0.0021 | 0.0026 | 0.0013 |
| F07 | 12820001 | 12920000 | 0.0021 | 0.0026 | 0.0016 |
| F07 | 12840001 | 12940000 | 0.0019 | 0.0025 | 0.0013 |
| F07 | 12860001 | 12960000 | 0.0017 | 0.0021 | 0.0012 |
| F07 | 12880001 | 12980000 | 0.0022 | 0.0026 | 0.0018 |
| F07 | 12900001 | 13000000 | 0.0039 | 0.0039 | 0.0037 |
| F07 | 12920001 | 13020000 | 0.0039 | 0.0038 | 0.0037 |
| F07 | 12940001 | 13040000 | 0.0041 | 0.0041 | 0.0037 |
| F07 | 12960001 | 13060000 | 0.0044 | 0.0047 | 0.0039 |
| F07 | 12980001 | 13080000 | 0.0042 | 0.0046 | 0.0035 |
| F07 | 13000001 | 13100000 | 0.0023 | 0.0032 | 0.0014 |

| Chr | Start | End | both | A1 | A2 |
|---|---|---|---|---|---|
| F07 | 13020001 | 13120000 | 0.0023 | 0.0034 | 0.0014 |
| F07 | 13040001 | 13140000 | 0.0022 | 0.0032 | 0.0015 |
| F07 | 13060001 | 13160000 | 0.0036 | 0.0045 | 0.0029 |
| F07 | 13080001 | 13180000 | 0.0034 | 0.0043 | 0.0028 |
| F07 | 13100001 | 13200000 | 0.0036 | 0.0041 | 0.0032 |
| F07 | 13120001 | 13220000 | 0.0037 | 0.0041 | 0.0033 |
| F07 | 13140001 | 13240000 | 0.0052 | 0.0055 | 0.0047 |
| F07 | 13160001 | 13260000 | 0.0041 | 0.0042 | 0.0038 |
| F07 | 13180001 | 13280000 | 0.0044 | 0.0044 | 0.0041 |
| F07 | 13200001 | 13300000 | 0.0040 | 0.0041 | 0.0038 |
| F07 | 13220001 | 13320000 | 0.0050 | 0.0051 | 0.0047 |
| F07 | 13240001 | 13340000 | 0.0042 | 0.0043 | 0.0038 |
| F07 | 13260001 | 13360000 | 0.0036 | 0.0039 | 0.0031 |
| F07 | 13280001 | 13380000 | 0.0033 | 0.0038 | 0.0028 |
| F07 | 13300001 | 13400000 | 0.0032 | 0.0039 | 0.0026 |
| F07 | 13320001 | 13420000 | 0.0028 | 0.0039 | 0.0020 |
| F07 | 13340001 | 13440000 | 0.0021 | 0.0034 | 0.0013 |
| F07 | 13360001 | 13460000 | 0.0021 | 0.0033 | 0.0012 |
| F07 | 13380001 | 13480000 | 0.0019 | 0.0033 | 0.0010 |
| F07 | 13400001 | 13500000 | 0.0021 | 0.0035 | 0.0011 |
| F07 | 13420001 | 13520000 | 0.0013 | 0.0025 | 0.0007 |
| F07 | 13440001 | 13540000 | 0.0016 | 0.0026 | 0.0010 |
| F07 | 13460001 | 13560000 | 0.0023 | 0.0032 | 0.0017 |
| F07 | 13480001 | 13580000 | 0.0026 | 0.0034 | 0.0021 |
| F07 | 13500001 | 13600000 | 0.0025 | 0.0033 | 0.0021 |
| F07 | 13520001 | 13620000 | 0.0026 | 0.0032 | 0.0021 |
| F07 | 13540001 | 13640000 | 0.0021 | 0.0027 | 0.0018 |
| F07 | 13560001 | 13660000 | 0.0014 | 0.0018 | 0.0012 |
| F07 | 13580001 | 13680000 | 0.0020 | 0.0021 | 0.0018 |
| F07 | 13600001 | 13700000 | 0.0024 | 0.0025 | 0.0022 |
| F07 | 13620001 | 13720000 | 0.0026 | 0.0027 | 0.0024 |
| F07 | 13640001 | 13740000 | 0.0026 | 0.0028 | 0.0024 |
| F07 | 13660001 | 13760000 | 0.0028 | 0.0031 | 0.0024 |
| F07 | 13680001 | 13780000 | 0.0019 | 0.0025 | 0.0015 |
| F07 | 13700001 | 13800000 | 0.0020 | 0.0026 | 0.0017 |
| F07 | 13720001 | 13820000 | 0.0018 | 0.0024 | 0.0015 |
| F07 | 13740001 | 13840000 | 0.0019 | 0.0026 | 0.0015 |
| F07 | 13760001 | 13860000 | 0.0017 | 0.0024 | 0.0014 |
| F07 | 13780001 | 13880000 | 0.0019 | 0.0026 | 0.0016 |
| F07 | 13800001 | 13900000 | 0.0021 | 0.0030 | 0.0017 |
| F07 | 13820001 | 13920000 | 0.0019 | 0.0027 | 0.0014 |
| F07 | 13840001 | 13940000 | 0.0020 | 0.0028 | 0.0016 |
| F07 | 13860001 | 13960000 | 0.0022 | 0.0030 | 0.0018 |
| F07 | 13880001 | 13980000 | 0.0031 | 0.0038 | 0.0027 |
| F07 | 13900001 | 14000000 | 0.0036 | 0.0043 | 0.0032 |
| F07 | 13920001 | 14020000 | 0.0056 | 0.0061 | 0.0052 |
| F07 | 13940001 | 14040000 | 0.0103 | 0.0102 | 0.0098 |
| F07 | 13960001 | 14060000 | 0.0101 | 0.0100 | 0.0096 |
| F07 | 13980001 | 14080000 | 0.0092 | 0.0089 | 0.0088 |
| F07 | 14000001 | 14100000 | 0.0082 | 0.0077 | 0.0080 |
| F07 | 14020001 | 14120000 | 0.0061 | 0.0057 | 0.0059 |
| F07 | 14040001 | 14140000 | 0.0012 | 0.0012 | 0.0012 |
| F07 | 14060001 | 14160000 | 0.0012 | 0.0010 | 0.0011 |
| F07 | 14080001 | 14180000 | 0.0012 | 0.0009 | 0.0012 |
| F07 | 14100001 | 14200000 | 0.0010 | 0.0008 | 0.0010 |
| F07 | 14120001 | 14220000 | 0.0012 | 0.0010 | 0.0012 |
| F07 | 14140001 | 14240000 | 0.0014 | 0.0011 | 0.0014 |
| F07 | 14160001 | 14260000 | 0.0019 | 0.0018 | 0.0019 |

| Chr | Start | End | both | A1 | A2 |
|---|---|---|---|---|---|
| F07 | 14180001 | 14280000 | 0.0025 | 0.0024 | 0.0023 |
| F07 | 14200001 | 14300000 | 0.0026 | 0.0023 | 0.0022 |
| F07 | 14220001 | 14320000 | 0.0030 | 0.0026 | 0.0025 |
| F07 | 14240001 | 14340000 | 0.0031 | 0.0027 | 0.0026 |
| F07 | 14260001 | 14360000 | 0.0048 | 0.0042 | 0.0043 |
| F07 | 14280001 | 14380000 | 0.0050 | 0.0042 | 0.0044 |
| F07 | 14300001 | 14400000 | 0.0056 | 0.0047 | 0.0049 |
| F07 | 14320001 | 14420000 | 0.0064 | 0.0051 | 0.0056 |
| F07 | 14340001 | 14440000 | 0.0069 | 0.0055 | 0.0060 |
| F07 | 14360001 | 14460000 | 0.0051 | 0.0037 | 0.0043 |
| F07 | 14380001 | 14480000 | 0.0048 | 0.0035 | 0.0041 |
| F07 | 14400001 | 14500000 | 0.0042 | 0.0030 | 0.0037 |
| F07 | 14420001 | 14520000 | 0.0031 | 0.0023 | 0.0027 |
| F07 | 14440001 | 14540000 | 0.0031 | 0.0022 | 0.0028 |
| F07 | 14460001 | 14560000 | 0.0029 | 0.0019 | 0.0025 |
| F07 | 14480001 | 14580000 | 0.0026 | 0.0018 | 0.0023 |
| F07 | 14500001 | 14600000 | 0.0035 | 0.0026 | 0.0031 |
| F07 | 14520001 | 14620000 | 0.0038 | 0.0027 | 0.0032 |
| F07 | 14540001 | 14640000 | 0.0033 | 0.0023 | 0.0026 |
| F07 | 14560001 | 14660000 | 0.0036 | 0.0025 | 0.0028 |
| F07 | 14580001 | 14680000 | 0.0034 | 0.0023 | 0.0026 |
| F07 | 14600001 | 14700000 | 0.0033 | 0.0022 | 0.0024 |
| F07 | 14620001 | 14720000 | 0.0036 | 0.0026 | 0.0028 |
| F07 | 14640001 | 14740000 | 0.0035 | 0.0025 | 0.0026 |
| F07 | 14660001 | 14760000 | 0.0033 | 0.0023 | 0.0025 |
| F07 | 14680001 | 14780000 | 0.0038 | 0.0025 | 0.0027 |
| F07 | 14700001 | 14800000 | 0.0029 | 0.0017 | 0.0019 |
| F07 | 14720001 | 14820000 | 0.0022 | 0.0011 | 0.0013 |
| F07 | 14740001 | 14840000 | 0.0024 | 0.0013 | 0.0016 |
| F07 | 14760001 | 14860000 | 0.0023 | 0.0012 | 0.0016 |
| F07 | 14780001 | 14880000 | 0.0028 | 0.0019 | 0.0023 |
| F07 | 14800001 | 14900000 | 0.0029 | 0.0019 | 0.0024 |
| F07 | 14820001 | 14920000 | 0.0033 | 0.0023 | 0.0028 |
| F07 | 14840001 | 14940000 | 0.0033 | 0.0023 | 0.0027 |
| F07 | 14860001 | 14960000 | 0.0036 | 0.0029 | 0.0032 |
| F07 | 14880001 | 14980000 | 0.0025 | 0.0020 | 0.0020 |
| F07 | 14900001 | 15000000 | 0.0026 | 0.0021 | 0.0021 |
| F07 | 14920001 | 15020000 | 0.0027 | 0.0024 | 0.0022 |
| F07 | 14940001 | 15040000 | 0.0028 | 0.0026 | 0.0022 |
| F07 | 14960001 | 15060000 | 0.0023 | 0.0022 | 0.0016 |
| F07 | 14980001 | 15080000 | 0.0030 | 0.0028 | 0.0022 |
| F07 | 15000001 | 15100000 | 0.0034 | 0.0032 | 0.0025 |
| F07 | 15020001 | 15120000 | 0.0036 | 0.0032 | 0.0027 |
| F07 | 15040001 | 15140000 | 0.0031 | 0.0027 | 0.0024 |
| F07 | 15060001 | 15160000 | 0.0028 | 0.0024 | 0.0023 |
| F07 | 15080001 | 15180000 | 0.0022 | 0.0019 | 0.0017 |
| F07 | 15100001 | 15200000 | 0.0020 | 0.0015 | 0.0016 |
| F07 | 15120001 | 15220000 | 0.0012 | 0.0010 | 0.0010 |
| F07 | 15140001 | 15240000 | 0.0016 | 0.0014 | 0.0014 |
| F07 | 15160001 | 15260000 | 0.0030 | 0.0028 | 0.0026 |
| F07 | 15180001 | 15280000 | 0.0028 | 0.0027 | 0.0026 |
| F07 | 15200001 | 15300000 | 0.0025 | 0.0024 | 0.0021 |
| F07 | 15220001 | 15320000 | 0.0030 | 0.0029 | 0.0024 |
| F07 | 15240001 | 15340000 | 0.0028 | 0.0026 | 0.0021 |
| F07 | 15260001 | 15360000 | 0.0019 | 0.0017 | 0.0014 |
| F07 | 15280001 | 15380000 | 0.0022 | 0.0020 | 0.0016 |
| F07 | 15300001 | 15400000 | 0.0024 | 0.0021 | 0.0018 |
| F07 | 15320001 | 15420000 | 0.0018 | 0.0015 | 0.0013 |

| Chr | Start | End | both | A1 | A2 |
|---|---|---|---|---|---|
| F07 | 15340001 | 15440000 | 0.0022 | 0.0019 | 0.0018 |
| F07 | 15360001 | 15460000 | 0.0020 | 0.0017 | 0.0014 |
| F07 | 15380001 | 15480000 | 0.0021 | 0.0019 | 0.0015 |
| F07 | 15400001 | 15500000 | 0.0029 | 0.0027 | 0.0022 |
| F07 | 15420001 | 15520000 | 0.0030 | 0.0027 | 0.0022 |
| F07 | 15440001 | 15540000 | 0.0025 | 0.0021 | 0.0017 |
| F07 | 15460001 | 15560000 | 0.0026 | 0.0022 | 0.0019 |
| F07 | 15480001 | 15580000 | 0.0033 | 0.0024 | 0.0025 |
| F07 | 15500001 | 15600000 | 0.0023 | 0.0013 | 0.0017 |
| F07 | 15520001 | 15620000 | 0.0034 | 0.0022 | 0.0028 |
| F07 | 15540001 | 15640000 | 0.0035 | 0.0021 | 0.0028 |
| F07 | 15560001 | 15660000 | 0.0034 | 0.0019 | 0.0027 |
| F07 | 15580001 | 15680000 | 0.0029 | 0.0015 | 0.0022 |
| F07 | 15600001 | 15700000 | 0.0032 | 0.0017 | 0.0024 |
| F07 | 15620001 | 15720000 | 0.0024 | 0.0008 | 0.0015 |
| F07 | 15640001 | 15740000 | 0.0025 | 0.0010 | 0.0017 |
| F07 | 15660001 | 15760000 | 0.0023 | 0.0010 | 0.0016 |
| F07 | 15680001 | 15780000 | 0.0019 | 0.0008 | 0.0013 |
| F07 | 15700001 | 15800000 | 0.0018 | 0.0007 | 0.0012 |
| F07 | 15720001 | 15820000 | 0.0016 | 0.0006 | 0.0011 |
| F07 | 15740001 | 15840000 | 0.0019 | 0.0008 | 0.0012 |
| F07 | 15760001 | 15860000 | 0.0023 | 0.0011 | 0.0016 |
| F07 | 15780001 | 15880000 | 0.0022 | 0.0010 | 0.0015 |
| F07 | 15800001 | 15900000 | 0.0018 | 0.0008 | 0.0011 |
| F07 | 15820001 | 15920000 | 0.0021 | 0.0011 | 0.0014 |
| F07 | 15840001 | 15940000 | 0.0017 | 0.0009 | 0.0012 |
| F07 | 15860001 | 15960000 | 0.0016 | 0.0007 | 0.0011 |
| F07 | 15880001 | 15980000 | 0.0017 | 0.0009 | 0.0013 |
| F07 | 15900001 | 16000000 | 0.0023 | 0.0014 | 0.0019 |
| F07 | 15920001 | 16020000 | 0.0026 | 0.0017 | 0.0023 |
| F07 | 15940001 | 16040000 | 0.0025 | 0.0016 | 0.0022 |
| F07 | 15960001 | 16060000 | 0.0024 | 0.0015 | 0.0021 |
| F07 | 15980001 | 16080000 | 0.0027 | 0.0017 | 0.0023 |
| F07 | 16000001 | 16100000 | 0.0026 | 0.0016 | 0.0022 |
| F07 | 16020001 | 16120000 | 0.0024 | 0.0016 | 0.0019 |
| F07 | 16040001 | 16140000 | 0.0028 | 0.0017 | 0.0022 |
| F07 | 16060001 | 16160000 | 0.0030 | 0.0018 | 0.0024 |
| F07 | 16080001 | 16180000 | 0.0032 | 0.0021 | 0.0027 |
| F07 | 16100001 | 16200000 | 0.0035 | 0.0023 | 0.0028 |
| F07 | 16120001 | 16220000 | 0.0033 | 0.0019 | 0.0025 |
| F07 | 16140001 | 16240000 | 0.0038 | 0.0027 | 0.0031 |
| F07 | 16160001 | 16260000 | 0.0036 | 0.0026 | 0.0030 |
| F07 | 16180001 | 16280000 | 0.0035 | 0.0025 | 0.0028 |
| F07 | 16200001 | 16300000 | 0.0033 | 0.0022 | 0.0026 |
| F07 | 16220001 | 16320000 | 0.0031 | 0.0021 | 0.0024 |
| F07 | 16240001 | 16340000 | 0.0027 | 0.0015 | 0.0020 |
| F07 | 16260001 | 16360000 | 0.0025 | 0.0014 | 0.0019 |
| F07 | 16280001 | 16380000 | 0.0023 | 0.0012 | 0.0017 |
| F07 | 16300001 | 16400000 | 0.0019 | 0.0009 | 0.0014 |
| F07 | 16320001 | 16420000 | 0.0023 | 0.0011 | 0.0018 |
| F07 | 16340001 | 16440000 | 0.0025 | 0.0014 | 0.0020 |
| F07 | 16360001 | 16460000 | 0.0032 | 0.0019 | 0.0025 |
| F07 | 16380001 | 16480000 | 0.0033 | 0.0022 | 0.0027 |
| F07 | 16400001 | 16500000 | 0.0037 | 0.0023 | 0.0029 |
| F07 | 16420001 | 16520000 | 0.0044 | 0.0029 | 0.0036 |
| F07 | 16440001 | 16540000 | 0.0040 | 0.0022 | 0.0030 |
| F07 | 16460001 | 16560000 | 0.0035 | 0.0016 | 0.0025 |
| F07 | 16480001 | 16580000 | 0.0032 | 0.0013 | 0.0022 |

| Chr | Start | End | both | A1 | A2 |
|---|---|---|---|---|---|
| F07 | 16500001 | 16600000 | 0.0028 | 0.0013 | 0.0021 |
| F07 | 16520001 | 16620000 | 0.0016 | 0.0004 | 0.0010 |
| F07 | 16540001 | 16640000 | 0.0016 | 0.0007 | 0.0013 |
| F07 | 16560001 | 16660000 | 0.0034 | 0.0029 | 0.0032 |
| F07 | 16580001 | 16680000 | 0.0035 | 0.0029 | 0.0032 |
| F07 | 16600001 | 16700000 | 0.0036 | 0.0029 | 0.0032 |
| F07 | 16620001 | 16720000 | 0.0046 | 0.0036 | 0.0040 |
| F07 | 16640001 | 16740000 | 0.0047 | 0.0037 | 0.0041 |
| F07 | 16660001 | 16760000 | 0.0032 | 0.0023 | 0.0027 |
| F07 | 16680001 | 16780000 | 0.0040 | 0.0031 | 0.0034 |
| F07 | 16700001 | 16800000 | 0.0042 | 0.0034 | 0.0037 |
| F07 | 16720001 | 16820000 | 0.0036 | 0.0031 | 0.0032 |
| F07 | 16740001 | 16840000 | 0.0040 | 0.0034 | 0.0035 |
| F07 | 16760001 | 16860000 | 0.0034 | 0.0029 | 0.0029 |
| F07 | 16780001 | 16880000 | 0.0024 | 0.0022 | 0.0020 |
| F07 | 16800001 | 16900000 | 0.0020 | 0.0018 | 0.0016 |
| F07 | 16820001 | 16920000 | 0.0016 | 0.0017 | 0.0012 |
| F07 | 16840001 | 16940000 | 0.0007 | 0.0011 | 0.0003 |
| F07 | 16860001 | 16960000 | 0.0008 | 0.0015 | 0.0003 |
| F07 | 16880001 | 16980000 | 0.0014 | 0.0018 | 0.0010 |
| F07 | 16900001 | 17000000 | 0.0016 | 0.0022 | 0.0010 |
| F07 | 16920001 | 17020000 | 0.0017 | 0.0023 | 0.0011 |
| F07 | 16940001 | 17040000 | 0.0017 | 0.0024 | 0.0012 |
| F07 | 16960001 | 17060000 | 0.0015 | 0.0018 | 0.0011 |
| F07 | 16980001 | 17080000 | 0.0010 | 0.0016 | 0.0006 |
| F07 | 17000001 | 17100000 | 0.0010 | 0.0015 | 0.0006 |
| F07 | 17020001 | 17120000 | 0.0014 | 0.0019 | 0.0010 |
| F07 | 17040001 | 17140000 | 0.0015 | 0.0019 | 0.0010 |
| F07 | 17060001 | 17160000 | 0.0016 | 0.0023 | 0.0011 |
| F07 | 17080001 | 17180000 | 0.0017 | 0.0024 | 0.0012 |
| F07 | 17100001 | 17200000 | 0.0019 | 0.0027 | 0.0013 |
| F07 | 17120001 | 17220000 | 0.0020 | 0.0029 | 0.0012 |
| F07 | 17140001 | 17240000 | 0.0023 | 0.0034 | 0.0014 |
| F07 | 17160001 | 17260000 | 0.0022 | 0.0033 | 0.0013 |
| F07 | 17180001 | 17280000 | 0.0022 | 0.0032 | 0.0015 |
| F07 | 17200001 | 17300000 | 0.0020 | 0.0030 | 0.0013 |
| F07 | 17220001 | 17320000 | 0.0020 | 0.0028 | 0.0013 |
| F07 | 17240001 | 17340000 | 0.0020 | 0.0029 | 0.0012 |
| F07 | 17260001 | 17360000 | 0.0022 | 0.0031 | 0.0014 |
| F07 | 17280001 | 17380000 | 0.0020 | 0.0033 | 0.0010 |
| F07 | 17300001 | 17400000 | 0.0025 | 0.0037 | 0.0016 |
| F07 | 17320001 | 17420000 | 0.0026 | 0.0037 | 0.0018 |
| F07 | 17340001 | 17440000 | 0.0026 | 0.0034 | 0.0021 |
| F07 | 17360001 | 17460000 | 0.0033 | 0.0039 | 0.0030 |
| F07 | 17380001 | 17480000 | 0.0035 | 0.0039 | 0.0032 |
| F07 | 17400001 | 17500000 | 0.0030 | 0.0033 | 0.0028 |
| F07 | 17420001 | 17520000 | 0.0028 | 0.0030 | 0.0026 |
| F07 | 17440001 | 17540000 | 0.0028 | 0.0031 | 0.0026 |
| F07 | 17460001 | 17560000 | 0.0028 | 0.0031 | 0.0026 |
| F07 | 17480001 | 17580000 | 0.0027 | 0.0030 | 0.0025 |
| F07 | 17500001 | 17600000 | 0.0028 | 0.0032 | 0.0026 |
| F07 | 17520001 | 17620000 | 0.0026 | 0.0031 | 0.0024 |
| F07 | 17540001 | 17640000 | 0.0022 | 0.0028 | 0.0020 |
| F07 | 17560001 | 17660000 | 0.0023 | 0.0028 | 0.0021 |
| F07 | 17580001 | 17680000 | 0.0021 | 0.0025 | 0.0019 |
| F07 | 17600001 | 17700000 | 0.0021 | 0.0023 | 0.0019 |
| F07 | 17620001 | 17720000 | 0.0022 | 0.0025 | 0.0021 |
| F07 | 17640001 | 17740000 | 0.0030 | 0.0031 | 0.0028 |

| Chr | Start | End | both | A1 | A2 |
|---|---|---|---|---|---|
| F07 | 17660001 | 17760000 | 0.0022 | 0.0025 | 0.0020 |
| F07 | 17680001 | 17780000 | 0.0022 | 0.0026 | 0.0020 |
| F07 | 17700001 | 17800000 | 0.0032 | 0.0037 | 0.0030 |
| F07 | 17720001 | 17820000 | 0.0032 | 0.0037 | 0.0029 |
| F07 | 17740001 | 17840000 | 0.0032 | 0.0036 | 0.0030 |
| F07 | 17760001 | 17860000 | 0.0031 | 0.0037 | 0.0028 |
| F07 | 17780001 | 17880000 | 0.0037 | 0.0041 | 0.0034 |
| F07 | 17800001 | 17900000 | 0.0031 | 0.0033 | 0.0029 |
| F07 | 17820001 | 17920000 | 0.0035 | 0.0035 | 0.0033 |
| F07 | 17840001 | 17940000 | 0.0033 | 0.0037 | 0.0030 |
| F07 | 17860001 | 17960000 | 0.0030 | 0.0034 | 0.0027 |
| F07 | 17880001 | 17980000 | 0.0027 | 0.0033 | 0.0024 |
| F07 | 17900001 | 18000000 | 0.0021 | 0.0027 | 0.0018 |
| F07 | 17920001 | 18020000 | 0.0021 | 0.0027 | 0.0017 |
| F07 | 17940001 | 18040000 | 0.0015 | 0.0018 | 0.0012 |
| F07 | 17960001 | 18060000 | 0.0021 | 0.0022 | 0.0019 |
| F07 | 17980001 | 18080000 | 0.0022 | 0.0022 | 0.0019 |
| F07 | 18000001 | 18100000 | 0.0028 | 0.0029 | 0.0023 |
| F07 | 18020001 | 18120000 | 0.0028 | 0.0029 | 0.0023 |
| F07 | 18040001 | 18140000 | 0.0034 | 0.0035 | 0.0028 |
| F07 | 18060001 | 18160000 | 0.0032 | 0.0033 | 0.0026 |
| F07 | 18080001 | 18180000 | 0.0037 | 0.0035 | 0.0032 |
| F07 | 18100001 | 18200000 | 0.0031 | 0.0029 | 0.0029 |
| F07 | 18120001 | 18220000 | 0.0035 | 0.0033 | 0.0031 |
| F07 | 18140001 | 18240000 | 0.0031 | 0.0029 | 0.0028 |
| F07 | 18160001 | 18260000 | 0.0027 | 0.0027 | 0.0023 |
| F07 | 18180001 | 18280000 | 0.0022 | 0.0023 | 0.0017 |
| F07 | 18200001 | 18300000 | 0.0024 | 0.0025 | 0.0018 |
| F07 | 18220001 | 18320000 | 0.0014 | 0.0014 | 0.0011 |
| F07 | 18240001 | 18340000 | 0.0019 | 0.0019 | 0.0014 |
| F07 | 18260001 | 18360000 | 0.0024 | 0.0025 | 0.0019 |
| F07 | 18280001 | 18380000 | 0.0021 | 0.0021 | 0.0018 |
| F07 | 18300001 | 18400000 | 0.0026 | 0.0026 | 0.0021 |
| F07 | 18320001 | 18420000 | 0.0030 | 0.0030 | 0.0025 |
| F07 | 18340001 | 18440000 | 0.0024 | 0.0026 | 0.0020 |
| F07 | 18360001 | 18460000 | 0.0028 | 0.0030 | 0.0023 |
| F07 | 18380001 | 18480000 | 0.0032 | 0.0033 | 0.0026 |
| F07 | 18400001 | 18500000 | 0.0026 | 0.0028 | 0.0021 |
| F07 | 18420001 | 18520000 | 0.0024 | 0.0026 | 0.0019 |
| F07 | 18440001 | 18540000 | 0.0030 | 0.0032 | 0.0024 |
| F07 | 18460001 | 18560000 | 0.0023 | 0.0025 | 0.0017 |
| F07 | 18480001 | 18580000 | 0.0021 | 0.0024 | 0.0015 |
| F07 | 18500001 | 18600000 | 0.0024 | 0.0026 | 0.0017 |
| F07 | 18520001 | 18620000 | 0.0024 | 0.0026 | 0.0017 |
| F07 | 18540001 | 18640000 | 0.0020 | 0.0021 | 0.0015 |
| F07 | 18560001 | 18660000 | 0.0017 | 0.0018 | 0.0013 |
| F07 | 18580001 | 18680000 | 0.0016 | 0.0017 | 0.0012 |
| F07 | 18600001 | 18700000 | 0.0017 | 0.0018 | 0.0012 |
| F07 | 18620001 | 18720000 | 0.0019 | 0.0021 | 0.0015 |
| F07 | 18640001 | 18740000 | 0.0024 | 0.0026 | 0.0019 |
| F07 | 18660001 | 18760000 | 0.0031 | 0.0032 | 0.0026 |
| F07 | 18680001 | 18780000 | 0.0035 | 0.0038 | 0.0027 |
| F07 | 18700001 | 18800000 | 0.0033 | 0.0037 | 0.0026 |
| F07 | 18720001 | 18820000 | 0.0033 | 0.0036 | 0.0024 |
| F07 | 18740001 | 18840000 | 0.0025 | 0.0031 | 0.0015 |
| F07 | 18760001 | 18860000 | 0.0021 | 0.0028 | 0.0011 |
| F07 | 18780001 | 18880000 | 0.0017 | 0.0022 | 0.0009 |
| F07 | 18800001 | 18900000 | 0.0020 | 0.0024 | 0.0012 |

| Chr | Start | End | both | A1 | A2 |
|---|---|---|---|---|---|
| F07 | 18820001 | 18920000 | 0.0032 | 0.0035 | 0.0024 |
| F07 | 18840001 | 18940000 | 0.0039 | 0.0040 | 0.0031 |
| F07 | 18860001 | 18960000 | 0.0041 | 0.0042 | 0.0032 |
| F07 | 18880001 | 18980000 | 0.0042 | 0.0042 | 0.0033 |
| F07 | 18900001 | 19000000 | 0.0038 | 0.0037 | 0.0030 |
| F07 | 18920001 | 19020000 | 0.0027 | 0.0027 | 0.0019 |
| F07 | 18940001 | 19040000 | 0.0020 | 0.0019 | 0.0012 |
| F07 | 18960001 | 19060000 | 0.0022 | 0.0020 | 0.0014 |
| F07 | 18980001 | 19080000 | 0.0031 | 0.0028 | 0.0023 |
| F07 | 19000001 | 19100000 | 0.0037 | 0.0034 | 0.0028 |
| F07 | 19020001 | 19120000 | 0.0043 | 0.0040 | 0.0035 |
| F07 | 19040001 | 19140000 | 0.0046 | 0.0044 | 0.0037 |
| F07 | 19060001 | 19160000 | 0.0042 | 0.0041 | 0.0034 |
| F07 | 19080001 | 19180000 | 0.0033 | 0.0035 | 0.0024 |
| F07 | 19100001 | 19200000 | 0.0036 | 0.0037 | 0.0028 |
| F07 | 19120001 | 19220000 | 0.0034 | 0.0035 | 0.0026 |
| F07 | 19140001 | 19240000 | 0.0030 | 0.0031 | 0.0023 |
| F07 | 19160001 | 19260000 | 0.0029 | 0.0032 | 0.0021 |
| F07 | 19180001 | 19280000 | 0.0029 | 0.0031 | 0.0021 |
| F07 | 19200001 | 19300000 | 0.0023 | 0.0026 | 0.0015 |
| F07 | 19220001 | 19320000 | 0.0018 | 0.0021 | 0.0012 |
| F07 | 19240001 | 19340000 | 0.0021 | 0.0024 | 0.0015 |
| F07 | 19260001 | 19360000 | 0.0021 | 0.0022 | 0.0015 |
| F07 | 19280001 | 19380000 | 0.0030 | 0.0031 | 0.0022 |
| F07 | 19300001 | 19400000 | 0.0032 | 0.0032 | 0.0025 |
| F07 | 19320001 | 19420000 | 0.0029 | 0.0029 | 0.0020 |
| F07 | 19340001 | 19440000 | 0.0034 | 0.0033 | 0.0024 |
| F07 | 19360001 | 19460000 | 0.0037 | 0.0036 | 0.0029 |
| F07 | 19380001 | 19480000 | 0.0027 | 0.0025 | 0.0021 |
| F07 | 19400001 | 19500000 | 0.0025 | 0.0024 | 0.0018 |
| F07 | 19420001 | 19520000 | 0.0024 | 0.0023 | 0.0018 |
| F07 | 19440001 | 19540000 | 0.0021 | 0.0020 | 0.0016 |
| F07 | 19460001 | 19560000 | 0.0022 | 0.0019 | 0.0018 |
| F07 | 19480001 | 19580000 | 0.0022 | 0.0019 | 0.0018 |
| F07 | 19500001 | 19600000 | 0.0024 | 0.0020 | 0.0019 |
| F07 | 19520001 | 19620000 | 0.0024 | 0.0020 | 0.0019 |
| F07 | 19540001 | 19640000 | 0.0019 | 0.0016 | 0.0015 |
| F07 | 19560001 | 19660000 | 0.0025 | 0.0023 | 0.0020 |
| F07 | 19580001 | 19680000 | 0.0034 | 0.0031 | 0.0027 |
| F07 | 19600001 | 19700000 | 0.0033 | 0.0031 | 0.0027 |
| F07 | 19620001 | 19720000 | 0.0038 | 0.0036 | 0.0031 |
| F07 | 19640001 | 19740000 | 0.0043 | 0.0040 | 0.0032 |
| F07 | 19660001 | 19760000 | 0.0042 | 0.0040 | 0.0025 |
| F07 | 19680001 | 19780000 | 0.0038 | 0.0036 | 0.0020 |
| F07 | 19700001 | 19800000 | 0.0048 | 0.0046 | 0.0030 |
| F07 | 19720001 | 19820000 | 0.0052 | 0.0049 | 0.0034 |
| F07 | 19740001 | 19840000 | 0.0048 | 0.0046 | 0.0032 |
| F07 | 19760001 | 19860000 | 0.0038 | 0.0036 | 0.0030 |
| F07 | 19780001 | 19880000 | 0.0037 | 0.0035 | 0.0030 |
| F07 | 19800001 | 19900000 | 0.0029 | 0.0027 | 0.0024 |
| F07 | 19820001 | 19920000 | 0.0020 | 0.0019 | 0.0016 |
| F07 | 19840001 | 19940000 | 0.0024 | 0.0021 | 0.0020 |
| F07 | 19860001 | 19960000 | 0.0030 | 0.0028 | 0.0026 |
| F07 | 19880001 | 19980000 | 0.0030 | 0.0028 | 0.0026 |
| F07 | 19900001 | 20000000 | 0.0026 | 0.0024 | 0.0020 |
| F07 | 19920001 | 20020000 | 0.0027 | 0.0026 | 0.0022 |
| F07 | 19940001 | 20040000 | 0.0026 | 0.0027 | 0.0018 |
| F07 | 19960001 | 20060000 | 0.0020 | 0.0020 | 0.0013 |

| Chr | Start | End | both | A1 | A2 |
|---|---|---|---|---|---|
| F07 | 19980001 | 20080000 | 0.0019 | 0.0018 | 0.0011 |
| F07 | 20000001 | 20100000 | 0.0022 | 0.0020 | 0.0014 |
| F07 | 20020001 | 20120000 | 0.0021 | 0.0019 | 0.0012 |
| F07 | 20040001 | 20140000 | 0.0025 | 0.0020 | 0.0017 |
| F07 | 20060001 | 20160000 | 0.0030 | 0.0026 | 0.0021 |
| F07 | 20080001 | 20180000 | 0.0030 | 0.0025 | 0.0022 |
| F07 | 20100001 | 20200000 | 0.0031 | 0.0025 | 0.0025 |
| F07 | 20120001 | 20220000 | 0.0032 | 0.0025 | 0.0025 |
| F07 | 20140001 | 20240000 | 0.0029 | 0.0021 | 0.0023 |
| F07 | 20160001 | 20260000 | 0.0027 | 0.0017 | 0.0020 |
| F07 | 20180001 | 20280000 | 0.0032 | 0.0020 | 0.0023 |
| F07 | 20200001 | 20300000 | 0.0032 | 0.0018 | 0.0021 |
| F07 | 20220001 | 20320000 | 0.0033 | 0.0020 | 0.0021 |
| F07 | 20240001 | 20340000 | 0.0032 | 0.0019 | 0.0022 |
| F07 | 20260001 | 20360000 | 0.0029 | 0.0018 | 0.0020 |
| F07 | 20280001 | 20380000 | 0.0026 | 0.0015 | 0.0018 |
| F07 | 20300001 | 20400000 | 0.0024 | 0.0015 | 0.0017 |
| F07 | 20320001 | 20420000 | 0.0030 | 0.0019 | 0.0022 |
| F07 | 20340001 | 20440000 | 0.0038 | 0.0026 | 0.0028 |
| F07 | 20360001 | 20460000 | 0.0041 | 0.0026 | 0.0027 |
| F07 | 20380001 | 20480000 | 0.0038 | 0.0025 | 0.0026 |
| F07 | 20400001 | 20500000 | 0.0039 | 0.0027 | 0.0028 |
| F07 | 20420001 | 20520000 | 0.0037 | 0.0025 | 0.0026 |
| F07 | 20440001 | 20540000 | 0.0031 | 0.0019 | 0.0020 |
| F07 | 20460001 | 20560000 | 0.0028 | 0.0021 | 0.0021 |
| F07 | 20480001 | 20580000 | 0.0032 | 0.0024 | 0.0024 |
| F07 | 20500001 | 20600000 | 0.0031 | 0.0022 | 0.0021 |
| F07 | 20520001 | 20620000 | 0.0029 | 0.0022 | 0.0022 |
| F07 | 20540001 | 20640000 | 0.0025 | 0.0019 | 0.0020 |
| F07 | 20560001 | 20660000 | 0.0028 | 0.0021 | 0.0022 |
| F07 | 20580001 | 20680000 | 0.0030 | 0.0023 | 0.0024 |
| F07 | 20600001 | 20700000 | 0.0031 | 0.0023 | 0.0024 |
| F07 | 20620001 | 20720000 | 0.0032 | 0.0020 | 0.0021 |
| F07 | 20640001 | 20740000 | 0.0032 | 0.0018 | 0.0020 |
| F07 | 20660001 | 20760000 | 0.0032 | 0.0018 | 0.0021 |
| F07 | 20680001 | 20780000 | 0.0027 | 0.0013 | 0.0017 |
| F07 | 20700001 | 20800000 | 0.0023 | 0.0011 | 0.0015 |
| F07 | 20720001 | 20820000 | 0.0028 | 0.0018 | 0.0023 |
| F07 | 20740001 | 20840000 | 0.0031 | 0.0022 | 0.0027 |
| F07 | 20760001 | 20860000 | 0.0026 | 0.0018 | 0.0022 |
| F07 | 20780001 | 20880000 | 0.0027 | 0.0018 | 0.0021 |
| F07 | 20800001 | 20900000 | 0.0036 | 0.0024 | 0.0028 |
| F07 | 20820001 | 20920000 | 0.0030 | 0.0017 | 0.0021 |
| F07 | 20840001 | 20940000 | 0.0024 | 0.0012 | 0.0016 |
| F07 | 20860001 | 20960000 | 0.0029 | 0.0017 | 0.0020 |
| F07 | 20880001 | 20980000 | 0.0043 | 0.0029 | 0.0034 |
| F07 | 20900001 | 21000000 | 0.0034 | 0.0021 | 0.0026 |
| F07 | 20920001 | 21020000 | 0.0039 | 0.0023 | 0.0031 |
| F07 | 20940001 | 21040000 | 0.0042 | 0.0024 | 0.0031 |
| F07 | 20960001 | 21060000 | 0.0042 | 0.0021 | 0.0029 |
| F07 | 20980001 | 21080000 | 0.0028 | 0.0010 | 0.0015 |
| F07 | 21000001 | 21100000 | 0.0037 | 0.0020 | 0.0024 |
| F07 | 21020001 | 21120000 | 0.0030 | 0.0017 | 0.0020 |
| F07 | 21040001 | 21140000 | 0.0030 | 0.0017 | 0.0021 |
| F07 | 21060001 | 21160000 | 0.0027 | 0.0018 | 0.0022 |
| F07 | 21080001 | 21180000 | 0.0030 | 0.0019 | 0.0025 |
| F07 | 21100001 | 21200000 | 0.0024 | 0.0012 | 0.0018 |
| F07 | 21120001 | 21220000 | 0.0026 | 0.0012 | 0.0018 |

| Chr | Start | End | both | A1 | A2 |
|---|---|---|---|---|---|
| F07 | 21140001 | 21240000 | 0.0033 | 0.0020 | 0.0024 |
| F07 | 21160001 | 21260000 | 0.0034 | 0.0023 | 0.0026 |
| F07 | 21180001 | 21280000 | 0.0038 | 0.0028 | 0.0030 |
| F07 | 21200001 | 21300000 | 0.0040 | 0.0031 | 0.0033 |
| F07 | 21220001 | 21320000 | 0.0037 | 0.0029 | 0.0031 |
| F07 | 21240001 | 21340000 | 0.0034 | 0.0023 | 0.0028 |
| F07 | 21260001 | 21360000 | 0.0043 | 0.0029 | 0.0035 |
| F07 | 21280001 | 21380000 | 0.0046 | 0.0033 | 0.0038 |
| F07 | 21300001 | 21400000 | 0.0045 | 0.0032 | 0.0038 |
| F07 | 21320001 | 21420000 | 0.0047 | 0.0033 | 0.0040 |
| F07 | 21340001 | 21440000 | 0.0053 | 0.0040 | 0.0047 |
| F07 | 21360001 | 21460000 | 0.0043 | 0.0032 | 0.0037 |
| F07 | 21380001 | 21480000 | 0.0031 | 0.0024 | 0.0025 |
| F07 | 21400001 | 21500000 | 0.0031 | 0.0029 | 0.0025 |
| F07 | 21420001 | 21520000 | 0.0034 | 0.0034 | 0.0028 |
| F07 | 21440001 | 21540000 | 0.0028 | 0.0032 | 0.0022 |
| F07 | 21460001 | 21560000 | 0.0027 | 0.0035 | 0.0022 |
| F07 | 21480001 | 21580000 | 0.0028 | 0.0035 | 0.0023 |
| F07 | 21500001 | 21600000 | 0.0027 | 0.0032 | 0.0023 |
| F07 | 21520001 | 21620000 | 0.0031 | 0.0037 | 0.0027 |
| F07 | 21540001 | 21640000 | 0.0028 | 0.0032 | 0.0025 |
| F07 | 21560001 | 21660000 | 0.0027 | 0.0029 | 0.0024 |
| F07 | 21580001 | 21680000 | 0.0030 | 0.0032 | 0.0028 |
| F07 | 21600001 | 21700000 | 0.0029 | 0.0033 | 0.0027 |
| F07 | 21620001 | 21720000 | 0.0026 | 0.0031 | 0.0023 |
| F07 | 21640001 | 21740000 | 0.0027 | 0.0030 | 0.0023 |
| F07 | 21660001 | 21760000 | 0.0028 | 0.0035 | 0.0022 |
| F07 | 21680001 | 21780000 | 0.0028 | 0.0041 | 0.0020 |
| F07 | 21700001 | 21800000 | 0.0025 | 0.0037 | 0.0017 |
| F07 | 21720001 | 21820000 | 0.0022 | 0.0032 | 0.0014 |
| F07 | 21740001 | 21840000 | 0.0021 | 0.0034 | 0.0013 |
| F07 | 21760001 | 21860000 | 0.0021 | 0.0031 | 0.0015 |
| F07 | 21780001 | 21880000 | 0.0018 | 0.0021 | 0.0013 |
| F07 | 21800001 | 21900000 | 0.0021 | 0.0020 | 0.0013 |
| F07 | 21820001 | 21920000 | 0.0021 | 0.0016 | 0.0012 |
| F07 | 21840001 | 21940000 | 0.0021 | 0.0013 | 0.0011 |
| F07 | 21860001 | 21960000 | 0.0023 | 0.0014 | 0.0013 |
| F07 | 21880001 | 21980000 | 0.0028 | 0.0018 | 0.0017 |
| F07 | 21900001 | 22000000 | 0.0026 | 0.0017 | 0.0017 |
| F07 | 21920001 | 22020000 | 0.0023 | 0.0016 | 0.0017 |
| F07 | 21940001 | 22040000 | 0.0023 | 0.0015 | 0.0015 |
| F07 | 21960001 | 22060000 | 0.0023 | 0.0014 | 0.0014 |
| F07 | 21980001 | 22080000 | 0.0036 | 0.0027 | 0.0029 |
| F07 | 22000001 | 22100000 | 0.0041 | 0.0030 | 0.0032 |
| F07 | 22020001 | 22120000 | 0.0052 | 0.0037 | 0.0038 |
| F07 | 22040001 | 22140000 | 0.0053 | 0.0037 | 0.0039 |
| F07 | 22060001 | 22160000 | 0.0056 | 0.0038 | 0.0039 |
| F07 | 22080001 | 22180000 | 0.0042 | 0.0025 | 0.0023 |
| F07 | 22100001 | 22200000 | 0.0044 | 0.0029 | 0.0027 |
| F07 | 22120001 | 22220000 | 0.0035 | 0.0023 | 0.0022 |
| F07 | 22140001 | 22240000 | 0.0037 | 0.0025 | 0.0026 |
| F07 | 22160001 | 22260000 | 0.0042 | 0.0031 | 0.0033 |
| F07 | 22180001 | 22280000 | 0.0045 | 0.0034 | 0.0035 |
| F07 | 22200001 | 22300000 | 0.0047 | 0.0035 | 0.0035 |
| F07 | 22220001 | 22320000 | 0.0050 | 0.0037 | 0.0038 |
| F07 | 22240001 | 22340000 | 0.0052 | 0.0040 | 0.0039 |
| F07 | 22260001 | 22360000 | 0.0044 | 0.0033 | 0.0031 |
| F07 | 22280001 | 22380000 | 0.0041 | 0.0032 | 0.0029 |

| Chr | Start | End | both | A1 | A2 |
|---|---|---|---|---|---|
| F07 | 22300001 | 22400000 | 0.0036 | 0.0027 | 0.0025 |
| F07 | 22320001 | 22420000 | 0.0040 | 0.0030 | 0.0029 |
| F07 | 22340001 | 22440000 | 0.0033 | 0.0025 | 0.0024 |
| F07 | 22360001 | 22460000 | 0.0035 | 0.0026 | 0.0025 |
| F07 | 22380001 | 22480000 | 0.0034 | 0.0026 | 0.0027 |
| F07 | 22400001 | 22500000 | 0.0031 | 0.0024 | 0.0026 |
| F07 | 22420001 | 22520000 | 0.0029 | 0.0024 | 0.0025 |
| F07 | 22440001 | 22540000 | 0.0030 | 0.0026 | 0.0028 |
| F07 | 22460001 | 22560000 | 0.0028 | 0.0022 | 0.0024 |
| F07 | 22480001 | 22580000 | 0.0022 | 0.0016 | 0.0018 |
| F07 | 22500001 | 22600000 | 0.0027 | 0.0017 | 0.0022 |
| F07 | 22520001 | 22620000 | 0.0025 | 0.0015 | 0.0017 |
| F07 | 22540001 | 22640000 | 0.0026 | 0.0016 | 0.0019 |
| F07 | 22560001 | 22660000 | 0.0024 | 0.0015 | 0.0018 |
| F07 | 22580001 | 22680000 | 0.0024 | 0.0017 | 0.0019 |
| F07 | 22600001 | 22700000 | 0.0021 | 0.0017 | 0.0016 |
| F07 | 22620001 | 22720000 | 0.0027 | 0.0022 | 0.0022 |
| F07 | 22640001 | 22740000 | 0.0031 | 0.0026 | 0.0025 |
| F07 | 22660001 | 22760000 | 0.0036 | 0.0029 | 0.0029 |
| F07 | 22680001 | 22780000 | 0.0043 | 0.0033 | 0.0035 |
| F07 | 22700001 | 22800000 | 0.0043 | 0.0033 | 0.0035 |
| F07 | 22720001 | 22820000 | 0.0043 | 0.0031 | 0.0035 |
| F07 | 22740001 | 22840000 | 0.0039 | 0.0025 | 0.0030 |
| F07 | 22760001 | 22860000 | 0.0037 | 0.0024 | 0.0030 |
| F07 | 22780001 | 22880000 | 0.0043 | 0.0031 | 0.0037 |
| F07 | 22800001 | 22900000 | 0.0045 | 0.0033 | 0.0039 |
| F07 | 22820001 | 22920000 | 0.0043 | 0.0033 | 0.0039 |
| F07 | 22840001 | 22940000 | 0.0044 | 0.0034 | 0.0039 |
| F07 | 22860001 | 22960000 | 0.0048 | 0.0037 | 0.0042 |
| F07 | 22880001 | 22980000 | 0.0041 | 0.0030 | 0.0034 |
| F07 | 22900001 | 23000000 | 0.0055 | 0.0043 | 0.0048 |
| F07 | 22920001 | 23020000 | 0.0058 | 0.0046 | 0.0051 |
| F07 | 22940001 | 23040000 | 0.0059 | 0.0048 | 0.0051 |
| F07 | 22960001 | 23060000 | 0.0055 | 0.0044 | 0.0048 |
| F07 | 22980001 | 23080000 | 0.0055 | 0.0043 | 0.0048 |
| F07 | 23000001 | 23100000 | 0.0039 | 0.0027 | 0.0030 |
| F07 | 23020001 | 23120000 | 0.0029 | 0.0019 | 0.0020 |
| F07 | 23040001 | 23140000 | 0.0024 | 0.0015 | 0.0017 |
| F07 | 23060001 | 23160000 | 0.0021 | 0.0013 | 0.0014 |
| F07 | 23080001 | 23180000 | 0.0016 | 0.0009 | 0.0010 |
| F07 | 23100001 | 23200000 | 0.0013 | 0.0007 | 0.0008 |
| F07 | 23120001 | 23220000 | 0.0011 | 0.0005 | 0.0007 |
| F07 | 23140001 | 23240000 | 0.0014 | 0.0008 | 0.0010 |
| F07 | 23160001 | 23260000 | 0.0025 | 0.0015 | 0.0018 |
| F07 | 23180001 | 23280000 | 0.0026 | 0.0017 | 0.0020 |
| F07 | 23200001 | 23300000 | 0.0030 | 0.0019 | 0.0021 |
| F07 | 23220001 | 23320000 | 0.0031 | 0.0021 | 0.0023 |
| F07 | 23240001 | 23340000 | 0.0039 | 0.0027 | 0.0030 |
| F07 | 23260001 | 23360000 | 0.0032 | 0.0023 | 0.0024 |
| F07 | 23280001 | 23380000 | 0.0040 | 0.0029 | 0.0031 |
| F07 | 23300001 | 23400000 | 0.0041 | 0.0030 | 0.0033 |
| F07 | 23320001 | 23420000 | 0.0042 | 0.0031 | 0.0032 |
| F07 | 23340001 | 23440000 | 0.0040 | 0.0028 | 0.0028 |
| F07 | 23360001 | 23460000 | 0.0047 | 0.0034 | 0.0035 |
| F07 | 23380001 | 23480000 | 0.0038 | 0.0026 | 0.0026 |
| F07 | 23400001 | 23500000 | 0.0053 | 0.0043 | 0.0043 |
| F07 | 23420001 | 23520000 | 0.0055 | 0.0044 | 0.0045 |
| F07 | 23440001 | 23540000 | 0.0055 | 0.0047 | 0.0049 |

| Chr | Start | End | both | A1 | A2 |
|---|---|---|---|---|---|
| F07 | 23460001 | 23560000 | 0.0050 | 0.0042 | 0.0042 |
| F07 | 23480001 | 23580000 | 0.0049 | 0.0041 | 0.0041 |
| F07 | 23500001 | 23600000 | 0.0031 | 0.0024 | 0.0023 |
| F07 | 23520001 | 23620000 | 0.0026 | 0.0020 | 0.0018 |
| F07 | 23540001 | 23640000 | 0.0021 | 0.0014 | 0.0011 |
| F07 | 23560001 | 23660000 | 0.0018 | 0.0011 | 0.0009 |
| F07 | 23580001 | 23680000 | 0.0018 | 0.0011 | 0.0008 |
| F07 | 23600001 | 23700000 | 0.0015 | 0.0008 | 0.0006 |
| F07 | 23620001 | 23720000 | 0.0020 | 0.0013 | 0.0010 |
| F07 | 23640001 | 23740000 | 0.0024 | 0.0017 | 0.0015 |
| F07 | 23660001 | 23760000 | 0.0025 | 0.0020 | 0.0017 |
| F07 | 23680001 | 23780000 | 0.0026 | 0.0021 | 0.0019 |
| F07 | 23700001 | 23800000 | 0.0029 | 0.0024 | 0.0023 |
| F07 | 23720001 | 23820000 | 0.0029 | 0.0024 | 0.0024 |
| F07 | 23740001 | 23840000 | 0.0032 | 0.0027 | 0.0028 |
| F07 | 23760001 | 23860000 | 0.0062 | 0.0054 | 0.0055 |
| F07 | 23780001 | 23880000 | 0.0067 | 0.0058 | 0.0058 |
| F07 | 23800001 | 23900000 | 0.0082 | 0.0069 | 0.0071 |
| F07 | 23820001 | 23920000 | 0.0078 | 0.0065 | 0.0066 |
| F07 | 23840001 | 23940000 | 0.0072 | 0.0057 | 0.0059 |
| F07 | 23860001 | 23960000 | 0.0042 | 0.0029 | 0.0029 |
| F07 | 23880001 | 23980000 | 0.0042 | 0.0027 | 0.0027 |
| F07 | 23900001 | 24000000 | 0.0026 | 0.0014 | 0.0013 |
| F07 | 23920001 | 24020000 | 0.0040 | 0.0029 | 0.0028 |
| F07 | 23940001 | 24040000 | 0.0037 | 0.0028 | 0.0025 |
| F07 | 23960001 | 24060000 | 0.0038 | 0.0028 | 0.0024 |
| F07 | 23980001 | 24080000 | 0.0040 | 0.0030 | 0.0028 |
| F07 | 24000001 | 24100000 | 0.0039 | 0.0029 | 0.0026 |
| F07 | 24020001 | 24120000 | 0.0029 | 0.0019 | 0.0017 |
| F07 | 24040001 | 24140000 | 0.0029 | 0.0020 | 0.0018 |
| F07 | 24060001 | 24160000 | 0.0035 | 0.0026 | 0.0023 |
| F07 | 24080001 | 24180000 | 0.0034 | 0.0025 | 0.0022 |
| F07 | 24100001 | 24200000 | 0.0035 | 0.0024 | 0.0022 |
| F07 | 24120001 | 24220000 | 0.0030 | 0.0018 | 0.0015 |
| F07 | 24140001 | 24240000 | 0.0031 | 0.0018 | 0.0016 |
| F07 | 24160001 | 24260000 | 0.0027 | 0.0016 | 0.0014 |
| F07 | 24180001 | 24280000 | 0.0021 | 0.0012 | 0.0011 |
| F07 | 24200001 | 24300000 | 0.0025 | 0.0017 | 0.0014 |
| F07 | 24220001 | 24320000 | 0.0025 | 0.0017 | 0.0014 |
| F07 | 24240001 | 24340000 | 0.0027 | 0.0018 | 0.0015 |
| F07 | 24260001 | 24360000 | 0.0027 | 0.0018 | 0.0016 |
| F07 | 24280001 | 24380000 | 0.0033 | 0.0022 | 0.0022 |
| F07 | 24300001 | 24400000 | 0.0029 | 0.0019 | 0.0019 |
| F07 | 24320001 | 24420000 | 0.0034 | 0.0022 | 0.0023 |
| F07 | 24340001 | 24440000 | 0.0031 | 0.0020 | 0.0021 |
| F07 | 24360001 | 24460000 | 0.0029 | 0.0018 | 0.0020 |
| F07 | 24380001 | 24480000 | 0.0027 | 0.0018 | 0.0018 |
| F07 | 24400001 | 24500000 | 0.0032 | 0.0023 | 0.0023 |
| F07 | 24420001 | 24520000 | 0.0037 | 0.0030 | 0.0028 |
| F07 | 24440001 | 24540000 | 0.0056 | 0.0045 | 0.0048 |
| F07 | 24460001 | 24560000 | 0.0055 | 0.0045 | 0.0047 |
| F07 | 24480001 | 24580000 | 0.0054 | 0.0045 | 0.0048 |
| F07 | 24500001 | 24600000 | 0.0049 | 0.0041 | 0.0043 |
| F07 | 24520001 | 24620000 | 0.0040 | 0.0030 | 0.0035 |
| F07 | 24540001 | 24640000 | 0.0019 | 0.0014 | 0.0013 |
| F07 | 24560001 | 24660000 | 0.0022 | 0.0016 | 0.0014 |
| F07 | 24580001 | 24680000 | 0.0030 | 0.0021 | 0.0019 |
| F07 | 24600001 | 24700000 | 0.0027 | 0.0018 | 0.0016 |

| Chr | Start | End | both | A1 | A2 |
|---|---|---|---|---|---|
| F07 | 24620001 | 24720000 | 0.0031 | 0.0023 | 0.0022 |
| F07 | 24640001 | 24740000 | 0.0032 | 0.0023 | 0.0023 |
| F07 | 24660001 | 24760000 | 0.0029 | 0.0020 | 0.0021 |
| F07 | 24680001 | 24780000 | 0.0017 | 0.0013 | 0.0010 |
| F07 | 24700001 | 24800000 | 0.0020 | 0.0015 | 0.0013 |
| F07 | 24720001 | 24820000 | 0.0016 | 0.0012 | 0.0009 |
| F07 | 24740001 | 24840000 | 0.0020 | 0.0016 | 0.0014 |
| F07 | 24760001 | 24860000 | 0.0022 | 0.0020 | 0.0016 |
| F07 | 24780001 | 24880000 | 0.0024 | 0.0022 | 0.0020 |
| F07 | 24800001 | 24900000 | 0.0027 | 0.0026 | 0.0020 |
| F07 | 24820001 | 24920000 | 0.0029 | 0.0028 | 0.0022 |
| F07 | 24840001 | 24940000 | 0.0027 | 0.0028 | 0.0019 |
| F07 | 24860001 | 24960000 | 0.0030 | 0.0031 | 0.0021 |
| F07 | 24880001 | 24980000 | 0.0026 | 0.0029 | 0.0017 |
| F07 | 24900001 | 25000000 | 0.0025 | 0.0026 | 0.0017 |
| F07 | 24920001 | 25020000 | 0.0020 | 0.0022 | 0.0013 |
| F07 | 24940001 | 25040000 | 0.0019 | 0.0020 | 0.0014 |
| F07 | 24960001 | 25060000 | 0.0015 | 0.0015 | 0.0011 |
| F07 | 24980001 | 25080000 | 0.0019 | 0.0019 | 0.0014 |
| F07 | 25000001 | 25100000 | 0.0016 | 0.0016 | 0.0011 |
| F07 | 25020001 | 25120000 | 0.0016 | 0.0017 | 0.0010 |
| F07 | 25040001 | 25140000 | 0.0016 | 0.0018 | 0.0007 |
| F07 | 25060001 | 25160000 | 0.0015 | 0.0017 | 0.0005 |
| F07 | 25080001 | 25180000 | 0.0014 | 0.0016 | 0.0005 |
| F07 | 25100001 | 25200000 | 0.0014 | 0.0016 | 0.0005 |
| F07 | 25120001 | 25220000 | 0.0016 | 0.0018 | 0.0009 |
| F07 | 25140001 | 25240000 | 0.0014 | 0.0017 | 0.0009 |
| F07 | 25160001 | 25260000 | 0.0016 | 0.0022 | 0.0011 |
| F07 | 25180001 | 25280000 | 0.0017 | 0.0023 | 0.0013 |
| F07 | 25200001 | 25300000 | 0.0022 | 0.0029 | 0.0017 |
| F07 | 25220001 | 25320000 | 0.0022 | 0.0029 | 0.0018 |
| F07 | 25240001 | 25340000 | 0.0028 | 0.0033 | 0.0024 |
| F07 | 25260001 | 25360000 | 0.0024 | 0.0025 | 0.0022 |
| F07 | 25280001 | 25380000 | 0.0024 | 0.0025 | 0.0022 |
| F07 | 25300001 | 25400000 | 0.0019 | 0.0020 | 0.0018 |
| F07 | 25320001 | 25420000 | 0.0022 | 0.0024 | 0.0020 |
| F07 | 25340001 | 25440000 | 0.0020 | 0.0022 | 0.0018 |
| F07 | 25360001 | 25460000 | 0.0025 | 0.0028 | 0.0023 |
| F07 | 25380001 | 25480000 | 0.0022 | 0.0026 | 0.0020 |
| F07 | 25400001 | 25500000 | 0.0023 | 0.0026 | 0.0021 |
| F07 | 25420001 | 25520000 | 0.0018 | 0.0021 | 0.0016 |
| F07 | 25440001 | 25540000 | 0.0018 | 0.0019 | 0.0016 |
| F07 | 25460001 | 25560000 | 0.0015 | 0.0014 | 0.0014 |
| F07 | 25480001 | 25580000 | 0.0051 | 0.0049 | 0.0050 |
| F07 | 25500001 | 25600000 | 0.0050 | 0.0047 | 0.0049 |
| F07 | 25520001 | 25620000 | 0.0049 | 0.0046 | 0.0048 |
| F07 | 25540001 | 25640000 | 0.0046 | 0.0045 | 0.0045 |
| F07 | 25560001 | 25660000 | 0.0046 | 0.0045 | 0.0045 |
| F07 | 25580001 | 25680000 | 0.0012 | 0.0013 | 0.0011 |
| F07 | 25600001 | 25700000 | 0.0014 | 0.0015 | 0.0013 |
| F07 | 25620001 | 25720000 | 0.0021 | 0.0022 | 0.0020 |
| F07 | 25640001 | 25740000 | 0.0020 | 0.0022 | 0.0019 |
| F07 | 25660001 | 25760000 | 0.0018 | 0.0020 | 0.0017 |
| F07 | 25680001 | 25780000 | 0.0015 | 0.0016 | 0.0014 |
| F07 | 25700001 | 25800000 | 0.0015 | 0.0015 | 0.0014 |
| F07 | 25720001 | 25820000 | 0.0009 | 0.0009 | 0.0009 |
| F07 | 25740001 | 25840000 | 0.0008 | 0.0007 | 0.0008 |
| F07 | 25760001 | 25860000 | 0.0010 | 0.0009 | 0.0010 |

| Chr | Start | End | both | A1 | A2 |
|---|---|---|---|---|---|
| F07 | 25780001 | 25880000 | 0.0009 | 0.0008 | 0.0009 |
| F07 | 25800001 | 25900000 | 0.0008 | 0.0007 | 0.0008 |
| F07 | 25820001 | 25920000 | 0.0008 | 0.0006 | 0.0008 |
| F07 | 25840001 | 25940000 | 0.0008 | 0.0007 | 0.0008 |
| F07 | 25860001 | 25960000 | 0.0014 | 0.0015 | 0.0014 |
| F07 | 25880001 | 25980000 | 0.0019 | 0.0019 | 0.0018 |
| F07 | 25900001 | 26000000 | 0.0017 | 0.0019 | 0.0016 |
| F07 | 25920001 | 26020000 | 0.0016 | 0.0018 | 0.0015 |
| F07 | 25940001 | 26040000 | 0.0022 | 0.0022 | 0.0021 |
| F07 | 25960001 | 26060000 | 0.0014 | 0.0014 | 0.0014 |
| F07 | 25980001 | 26080000 | 0.0021 | 0.0019 | 0.0020 |
| F07 | 26000001 | 26100000 | 0.0033 | 0.0030 | 0.0032 |
| F07 | 26020001 | 26120000 | 0.0042 | 0.0037 | 0.0041 |
| F07 | 26040001 | 26140000 | 0.0043 | 0.0039 | 0.0042 |
| F07 | 26060001 | 26160000 | 0.0051 | 0.0046 | 0.0050 |
| F07 | 26080001 | 26180000 | 0.0047 | 0.0044 | 0.0046 |
| F07 | 26100001 | 26200000 | 0.0041 | 0.0039 | 0.0040 |
| F07 | 26120001 | 26220000 | 0.0038 | 0.0038 | 0.0036 |
| F07 | 26140001 | 26240000 | 0.0034 | 0.0035 | 0.0033 |
| F07 | 26160001 | 26260000 | 0.0033 | 0.0033 | 0.0031 |
| F07 | 26180001 | 26280000 | 0.0029 | 0.0031 | 0.0027 |
| F07 | 26200001 | 26300000 | 0.0024 | 0.0026 | 0.0023 |
| F07 | 26220001 | 26320000 | 0.0020 | 0.0022 | 0.0019 |
| F07 | 26240001 | 26340000 | 0.0020 | 0.0023 | 0.0018 |
| F07 | 26260001 | 26360000 | 0.0016 | 0.0021 | 0.0015 |
| F07 | 26280001 | 26380000 | 0.0013 | 0.0017 | 0.0011 |
| F07 | 26300001 | 26400000 | 0.0019 | 0.0022 | 0.0018 |
| F07 | 26320001 | 26420000 | 0.0027 | 0.0029 | 0.0026 |
| F07 | 26340001 | 26440000 | 0.0032 | 0.0034 | 0.0031 |
| F07 | 26360001 | 26460000 | 0.0044 | 0.0042 | 0.0043 |
| F07 | 26380001 | 26480000 | 0.0052 | 0.0051 | 0.0051 |
| F07 | 26400001 | 26500000 | 0.0045 | 0.0045 | 0.0044 |
| F07 | 26420001 | 26520000 | 0.0034 | 0.0035 | 0.0033 |
| F07 | 26440001 | 26540000 | 0.0028 | 0.0030 | 0.0027 |
| F07 | 26460001 | 26560000 | 0.0014 | 0.0018 | 0.0013 |
| F07 | 26480001 | 26580000 | 0.0007 | 0.0011 | 0.0006 |
| F07 | 26500001 | 26600000 | 0.0008 | 0.0011 | 0.0007 |
| F07 | 26520001 | 26620000 | 0.0008 | 0.0010 | 0.0007 |
| F07 | 26540001 | 26640000 | 0.0012 | 0.0014 | 0.0011 |
| F07 | 26560001 | 26660000 | 0.0013 | 0.0014 | 0.0013 |
| F07 | 26580001 | 26680000 | 0.0021 | 0.0020 | 0.0021 |
| F07 | 26600001 | 26700000 | 0.0033 | 0.0030 | 0.0033 |
| F07 | 26620001 | 26720000 | 0.0041 | 0.0038 | 0.0041 |
| F07 | 26640001 | 26740000 | 0.0035 | 0.0033 | 0.0035 |
| F07 | 26660001 | 26760000 | 0.0041 | 0.0039 | 0.0041 |
| F07 | 26680001 | 26780000 | 0.0037 | 0.0037 | 0.0036 |
| F07 | 26700001 | 26800000 | 0.0029 | 0.0032 | 0.0027 |
| F07 | 26720001 | 26820000 | 0.0030 | 0.0032 | 0.0028 |
| F07 | 26740001 | 26840000 | 0.0034 | 0.0037 | 0.0030 |
| F07 | 26760001 | 26860000 | 0.0032 | 0.0034 | 0.0028 |
| F07 | 26780001 | 26880000 | 0.0035 | 0.0038 | 0.0031 |
| F07 | 26800001 | 26900000 | 0.0034 | 0.0038 | 0.0029 |
| F07 | 26820001 | 26920000 | 0.0028 | 0.0032 | 0.0023 |
| F07 | 26840001 | 26940000 | 0.0027 | 0.0027 | 0.0020 |
| F07 | 26860001 | 26960000 | 0.0024 | 0.0022 | 0.0016 |
| F07 | 26880001 | 26980000 | 0.0021 | 0.0015 | 0.0011 |
| F07 | 26900001 | 27000000 | 0.0023 | 0.0014 | 0.0013 |
| F07 | 26920001 | 27020000 | 0.0025 | 0.0015 | 0.0014 |

| Chr | Start | End | both | A1 | A2 |
|---|---|---|---|---|---|
| F07 | 26940001 | 27040000 | 0.0035 | 0.0026 | 0.0026 |
| F07 | 26960001 | 27060000 | 0.0035 | 0.0026 | 0.0026 |
| F07 | 26980001 | 27080000 | 0.0033 | 0.0025 | 0.0025 |
| F07 | 27000001 | 27100000 | 0.0027 | 0.0021 | 0.0020 |
| F07 | 27020001 | 27120000 | 0.0029 | 0.0022 | 0.0022 |
| F07 | 27040001 | 27140000 | 0.0019 | 0.0011 | 0.0012 |
| F07 | 27060001 | 27160000 | 0.0022 | 0.0013 | 0.0014 |
| F07 | 27080001 | 27180000 | 0.0021 | 0.0013 | 0.0013 |
| F07 | 27100001 | 27200000 | 0.0025 | 0.0014 | 0.0015 |
| F07 | 27120001 | 27220000 | 0.0024 | 0.0014 | 0.0015 |
| F07 | 27140001 | 27240000 | 0.0023 | 0.0015 | 0.0016 |
| F07 | 27160001 | 27260000 | 0.0025 | 0.0017 | 0.0018 |
| F07 | 27180001 | 27280000 | 0.0028 | 0.0021 | 0.0023 |
| F07 | 27200001 | 27300000 | 0.0024 | 0.0020 | 0.0021 |
| F07 | 27220001 | 27320000 | 0.0022 | 0.0018 | 0.0020 |
| F07 | 27240001 | 27340000 | 0.0028 | 0.0023 | 0.0026 |
| F07 | 27260001 | 27360000 | 0.0035 | 0.0025 | 0.0033 |
| F07 | 27280001 | 27380000 | 0.0035 | 0.0025 | 0.0033 |
| F07 | 27300001 | 27400000 | 0.0043 | 0.0030 | 0.0038 |
| F07 | 27320001 | 27420000 | 0.0043 | 0.0030 | 0.0038 |
| F07 | 27340001 | 27440000 | 0.0036 | 0.0023 | 0.0030 |
| F07 | 27360001 | 27460000 | 0.0034 | 0.0025 | 0.0028 |
| F07 | 27380001 | 27480000 | 0.0034 | 0.0024 | 0.0025 |
| F07 | 27400001 | 27500000 | 0.0029 | 0.0021 | 0.0023 |
| F07 | 27420001 | 27520000 | 0.0030 | 0.0023 | 0.0025 |
| F07 | 27440001 | 27540000 | 0.0032 | 0.0024 | 0.0027 |
| F07 | 27460001 | 27560000 | 0.0023 | 0.0017 | 0.0019 |
| F07 | 27480001 | 27580000 | 0.0021 | 0.0016 | 0.0019 |
| F07 | 27500001 | 27600000 | 0.0019 | 0.0014 | 0.0016 |
| F07 | 27520001 | 27620000 | 0.0014 | 0.0009 | 0.0011 |
| F07 | 27540001 | 27640000 | 0.0012 | 0.0008 | 0.0010 |
| F07 | 27560001 | 27660000 | 0.0013 | 0.0008 | 0.0009 |
| F07 | 27580001 | 27680000 | 0.0014 | 0.0009 | 0.0010 |
| F07 | 27600001 | 27700000 | 0.0017 | 0.0010 | 0.0011 |
| F07 | 27620001 | 27720000 | 0.0021 | 0.0013 | 0.0014 |
| F07 | 27640001 | 27740000 | 0.0029 | 0.0021 | 0.0021 |
| F07 | 27660001 | 27760000 | 0.0057 | 0.0046 | 0.0048 |
| F07 | 27680001 | 27780000 | 0.0059 | 0.0046 | 0.0049 |
| F07 | 27700001 | 27800000 | 0.0060 | 0.0047 | 0.0050 |
| F07 | 27720001 | 27820000 | 0.0058 | 0.0045 | 0.0048 |
| F07 | 27740001 | 27840000 | 0.0051 | 0.0038 | 0.0042 |
| F07 | 27760001 | 27860000 | 0.0024 | 0.0015 | 0.0016 |
| F07 | 27780001 | 27880000 | 0.0021 | 0.0013 | 0.0013 |
| F07 | 27800001 | 27900000 | 0.0021 | 0.0016 | 0.0017 |
| F07 | 27820001 | 27920000 | 0.0021 | 0.0016 | 0.0017 |
| F07 | 27840001 | 27940000 | 0.0027 | 0.0017 | 0.0022 |
| F07 | 27860001 | 27960000 | 0.0025 | 0.0016 | 0.0021 |
| F07 | 27880001 | 27980000 | 0.0025 | 0.0016 | 0.0022 |
| F07 | 27900001 | 28000000 | 0.0026 | 0.0015 | 0.0021 |
| F07 | 27920001 | 28020000 | 0.0037 | 0.0023 | 0.0031 |
| F07 | 27940001 | 28040000 | 0.0036 | 0.0026 | 0.0033 |
| F07 | 27960001 | 28060000 | 0.0039 | 0.0026 | 0.0034 |
| F07 | 27980001 | 28080000 | 0.0040 | 0.0027 | 0.0034 |
| F07 | 28000001 | 28100000 | 0.0035 | 0.0025 | 0.0031 |
| F07 | 28020001 | 28120000 | 0.0026 | 0.0019 | 0.0022 |
| F07 | 28040001 | 28140000 | 0.0025 | 0.0020 | 0.0021 |
| F07 | 28060001 | 28160000 | 0.0034 | 0.0030 | 0.0031 |
| F07 | 28080001 | 28180000 | 0.0044 | 0.0039 | 0.0042 |

| Chr | Start | End | both | A1 | A2 |
|---|---|---|---|---|---|
| F07 | 28100001 | 28200000 | 0.0054 | 0.0047 | 0.0049 |
| F07 | 28120001 | 28220000 | 0.0062 | 0.0050 | 0.0055 |
| F07 | 28140001 | 28240000 | 0.0058 | 0.0045 | 0.0050 |
| F07 | 28160001 | 28260000 | 0.0053 | 0.0038 | 0.0043 |
| F07 | 28180001 | 28280000 | 0.0040 | 0.0027 | 0.0031 |
| F07 | 28200001 | 28300000 | 0.0030 | 0.0020 | 0.0025 |
| F07 | 28220001 | 28320000 | 0.0018 | 0.0013 | 0.0016 |
| F07 | 28240001 | 28340000 | 0.0020 | 0.0015 | 0.0018 |
| F07 | 28260001 | 28360000 | 0.0013 | 0.0011 | 0.0012 |
| F07 | 28280001 | 28380000 | 0.0011 | 0.0010 | 0.0011 |
| F07 | 28300001 | 28400000 | 0.0010 | 0.0009 | 0.0010 |
| F07 | 28320001 | 28420000 | 0.0011 | 0.0009 | 0.0011 |
| F07 | 28340001 | 28440000 | 0.0009 | 0.0008 | 0.0009 |
| F07 | 28360001 | 28460000 | 0.0007 | 0.0006 | 0.0007 |
| F07 | 28380001 | 28480000 | 0.0009 | 0.0007 | 0.0009 |
| F07 | 28400001 | 28500000 | 0.0018 | 0.0010 | 0.0016 |
| F07 | 28420001 | 28520000 | 0.0029 | 0.0014 | 0.0026 |
| F07 | 28440001 | 28540000 | 0.0032 | 0.0013 | 0.0029 |
| F07 | 28460001 | 28560000 | 0.0046 | 0.0024 | 0.0041 |
| F07 | 28480001 | 28580000 | 0.0051 | 0.0029 | 0.0046 |
| F07 | 28500001 | 28600000 | 0.0047 | 0.0027 | 0.0043 |
| F07 | 28520001 | 28620000 | 0.0045 | 0.0029 | 0.0042 |
| F07 | 28540001 | 28640000 | 0.0043 | 0.0029 | 0.0040 |
| F07 | 28560001 | 28660000 | 0.0041 | 0.0027 | 0.0038 |
| F07 | 28580001 | 28680000 | 0.0039 | 0.0024 | 0.0035 |
| F07 | 28600001 | 28700000 | 0.0040 | 0.0028 | 0.0037 |
| F07 | 28620001 | 28720000 | 0.0038 | 0.0025 | 0.0033 |
| F07 | 28640001 | 28740000 | 0.0042 | 0.0028 | 0.0036 |
| F07 | 28660001 | 28760000 | 0.0036 | 0.0020 | 0.0029 |
| F07 | 28680001 | 28780000 | 0.0035 | 0.0018 | 0.0028 |
| F07 | 28700001 | 28800000 | 0.0032 | 0.0012 | 0.0025 |
| F07 | 28720001 | 28820000 | 0.0029 | 0.0009 | 0.0022 |
| F07 | 28740001 | 28840000 | 0.0027 | 0.0009 | 0.0020 |
| F07 | 28760001 | 28860000 | 0.0030 | 0.0013 | 0.0025 |
| F07 | 28780001 | 28880000 | 0.0035 | 0.0019 | 0.0030 |
| F07 | 28800001 | 28900000 | 0.0055 | 0.0036 | 0.0047 |
| F07 | 28820001 | 28920000 | 0.0063 | 0.0044 | 0.0056 |
| F07 | 28840001 | 28940000 | 0.0068 | 0.0049 | 0.0061 |
| F07 | 28860001 | 28960000 | 0.0072 | 0.0051 | 0.0064 |
| F07 | 28880001 | 28980000 | 0.0073 | 0.0051 | 0.0063 |
| F07 | 28900001 | 29000000 | 0.0061 | 0.0040 | 0.0052 |
| F07 | 28920001 | 29020000 | 0.0051 | 0.0033 | 0.0043 |
| F07 | 28940001 | 29040000 | 0.0049 | 0.0030 | 0.0041 |
| F07 | 28960001 | 29060000 | 0.0041 | 0.0026 | 0.0034 |
| F07 | 28980001 | 29080000 | 0.0039 | 0.0027 | 0.0034 |
| F07 | 29000001 | 29100000 | 0.0034 | 0.0021 | 0.0028 |
| F07 | 29020001 | 29120000 | 0.0043 | 0.0029 | 0.0037 |
| F07 | 29040001 | 29140000 | 0.0042 | 0.0029 | 0.0036 |
| F07 | 29060001 | 29160000 | 0.0041 | 0.0027 | 0.0035 |
| F07 | 29080001 | 29180000 | 0.0037 | 0.0023 | 0.0031 |
| F07 | 29100001 | 29200000 | 0.0032 | 0.0023 | 0.0028 |
| F07 | 29120001 | 29220000 | 0.0029 | 0.0018 | 0.0025 |
| F07 | 29140001 | 29240000 | 0.0024 | 0.0013 | 0.0020 |
| F07 | 29160001 | 29260000 | 0.0046 | 0.0031 | 0.0042 |
| F07 | 29180001 | 29280000 | 0.0050 | 0.0033 | 0.0046 |
| F07 | 29200001 | 29300000 | 0.0054 | 0.0037 | 0.0050 |
| F07 | 29220001 | 29320000 | 0.0053 | 0.0035 | 0.0050 |
| F07 | 29240001 | 29340000 | 0.0055 | 0.0037 | 0.0053 |

| Chr | Start | End | both | A1 | A2 |
|---|---|---|---|---|---|
| F07 | 29260001 | 29360000 | 0.0034 | 0.0020 | 0.0033 |
| F07 | 29280001 | 29380000 | 0.0028 | 0.0016 | 0.0027 |
| F07 | 29300001 | 29400000 | 0.0025 | 0.0013 | 0.0024 |
| F07 | 29320001 | 29420000 | 0.0029 | 0.0017 | 0.0028 |
| F07 | 29340001 | 29440000 | 0.0024 | 0.0014 | 0.0024 |
| F07 | 29360001 | 29460000 | 0.0024 | 0.0015 | 0.0023 |
| F07 | 29380001 | 29480000 | 0.0025 | 0.0015 | 0.0024 |
| F07 | 29400001 | 29500000 | 0.0027 | 0.0017 | 0.0026 |
| F07 | 29420001 | 29520000 | 0.0021 | 0.0015 | 0.0020 |
| F07 | 29440001 | 29540000 | 0.0030 | 0.0023 | 0.0029 |
| F07 | 29460001 | 29560000 | 0.0027 | 0.0022 | 0.0026 |
| F07 | 29480001 | 29580000 | 0.0027 | 0.0023 | 0.0027 |
| F07 | 29500001 | 29600000 | 0.0022 | 0.0019 | 0.0022 |
| F07 | 29520001 | 29620000 | 0.0021 | 0.0016 | 0.0020 |
| F07 | 29540001 | 29640000 | 0.0015 | 0.0009 | 0.0014 |
| F07 | 29560001 | 29660000 | 0.0027 | 0.0019 | 0.0026 |
| F07 | 29580001 | 29680000 | 0.0026 | 0.0018 | 0.0026 |
| F07 | 29600001 | 29700000 | 0.0030 | 0.0021 | 0.0029 |
| F07 | 29620001 | 29720000 | 0.0032 | 0.0022 | 0.0031 |
| F07 | 29640001 | 29740000 | 0.0032 | 0.0023 | 0.0031 |
| F07 | 29660001 | 29760000 | 0.0018 | 0.0012 | 0.0018 |
| F07 | 29680001 | 29780000 | 0.0019 | 0.0011 | 0.0019 |
| F07 | 29700001 | 29800000 | 0.0025 | 0.0015 | 0.0025 |
| F07 | 29720001 | 29820000 | 0.0032 | 0.0022 | 0.0031 |
| F07 | 29740001 | 29840000 | 0.0031 | 0.0020 | 0.0030 |
| F07 | 29760001 | 29860000 | 0.0034 | 0.0021 | 0.0033 |
| F07 | 29780001 | 29880000 | 0.0035 | 0.0023 | 0.0034 |
| F07 | 29800001 | 29900000 | 0.0032 | 0.0021 | 0.0031 |
| F07 | 29820001 | 29920000 | 0.0025 | 0.0015 | 0.0024 |
| F07 | 29840001 | 29940000 | 0.0030 | 0.0020 | 0.0029 |
| F07 | 29860001 | 29960000 | 0.0039 | 0.0028 | 0.0038 |
| F07 | 29880001 | 29980000 | 0.0047 | 0.0031 | 0.0046 |
| F07 | 29900001 | 30000000 | 0.0044 | 0.0030 | 0.0043 |
| F07 | 29920001 | 30020000 | 0.0051 | 0.0034 | 0.0050 |
| F07 | 29940001 | 30040000 | 0.0053 | 0.0034 | 0.0052 |
| F07 | 29960001 | 30060000 | 0.0051 | 0.0033 | 0.0050 |
| F07 | 29980001 | 30080000 | 0.0041 | 0.0027 | 0.0040 |
| F07 | 30000001 | 30100000 | 0.0048 | 0.0034 | 0.0047 |
| F07 | 30020001 | 30120000 | 0.0039 | 0.0028 | 0.0039 |
| F07 | 30040001 | 30140000 | 0.0035 | 0.0024 | 0.0035 |
| F07 | 30060001 | 30160000 | 0.0030 | 0.0017 | 0.0029 |
| F07 | 30080001 | 30180000 | 0.0036 | 0.0023 | 0.0034 |
| F07 | 30100001 | 30200000 | 0.0033 | 0.0019 | 0.0032 |
| F07 | 30120001 | 30220000 | 0.0043 | 0.0028 | 0.0041 |
| F07 | 30140001 | 30240000 | 0.0050 | 0.0032 | 0.0048 |
| F07 | 30160001 | 30260000 | 0.0054 | 0.0035 | 0.0052 |
| F07 | 30180001 | 30280000 | 0.0057 | 0.0037 | 0.0055 |
| F07 | 30200001 | 30300000 | 0.0058 | 0.0038 | 0.0056 |
| F07 | 30220001 | 30320000 | 0.0049 | 0.0031 | 0.0047 |
| F07 | 30240001 | 30340000 | 0.0049 | 0.0032 | 0.0047 |
| F07 | 30260001 | 30360000 | 0.0044 | 0.0030 | 0.0043 |
| F07 | 30280001 | 30380000 | 0.0035 | 0.0021 | 0.0034 |
| F07 | 30300001 | 30400000 | 0.0029 | 0.0017 | 0.0028 |
| F07 | 30320001 | 30420000 | 0.0030 | 0.0018 | 0.0029 |
| F07 | 30340001 | 30440000 | 0.0021 | 0.0010 | 0.0021 |
| F07 | 30360001 | 30460000 | 0.0027 | 0.0016 | 0.0027 |
| F07 | 30380001 | 30480000 | 0.0028 | 0.0017 | 0.0028 |
| F07 | 30400001 | 30500000 | 0.0034 | 0.0024 | 0.0034 |

| Chr | Start | End | both | A1 | A2 |
|---|---|---|---|---|---|
| F07 | 30420001 | 30520000 | 0.0039 | 0.0025 | 0.0039 |
| F07 | 30440001 | 30540000 | 0.0040 | 0.0026 | 0.0040 |
| F07 | 30460001 | 30560000 | 0.0039 | 0.0025 | 0.0038 |
| F07 | 30480001 | 30580000 | 0.0044 | 0.0028 | 0.0042 |
| F07 | 30500001 | 30600000 | 0.0042 | 0.0024 | 0.0039 |
| F07 | 30520001 | 30620000 | 0.0041 | 0.0026 | 0.0035 |
| F07 | 30540001 | 30640000 | 0.0036 | 0.0024 | 0.0031 |
| F07 | 30560001 | 30660000 | 0.0031 | 0.0021 | 0.0026 |
| F07 | 30580001 | 30680000 | 0.0028 | 0.0022 | 0.0024 |
| F07 | 30600001 | 30700000 | 0.0024 | 0.0020 | 0.0021 |
| F07 | 30620001 | 30720000 | 0.0020 | 0.0018 | 0.0018 |
| F07 | 30640001 | 30740000 | 0.0031 | 0.0027 | 0.0028 |
| F07 | 30660001 | 30760000 | 0.0031 | 0.0027 | 0.0026 |
| F07 | 30680001 | 30780000 | 0.0038 | 0.0031 | 0.0030 |
| F07 | 30700001 | 30800000 | 0.0037 | 0.0029 | 0.0027 |
| F07 | 30720001 | 30820000 | 0.0035 | 0.0028 | 0.0026 |
| F07 | 30740001 | 30840000 | 0.0031 | 0.0023 | 0.0020 |
| F07 | 30760001 | 30860000 | 0.0032 | 0.0022 | 0.0020 |
| F07 | 30780001 | 30880000 | 0.0025 | 0.0016 | 0.0016 |
| F07 | 30800001 | 30900000 | 0.0025 | 0.0017 | 0.0017 |
| F07 | 30820001 | 30920000 | 0.0028 | 0.0019 | 0.0020 |
| F07 | 30840001 | 30940000 | 0.0023 | 0.0017 | 0.0019 |
| F07 | 30860001 | 30960000 | 0.0021 | 0.0016 | 0.0018 |
| F07 | 30880001 | 30980000 | 0.0017 | 0.0015 | 0.0016 |
| F07 | 30900001 | 31000000 | 0.0026 | 0.0023 | 0.0025 |
| F07 | 30920001 | 31020000 | 0.0026 | 0.0023 | 0.0024 |
| F07 | 30940001 | 31040000 | 0.0035 | 0.0030 | 0.0031 |
| F07 | 30960001 | 31060000 | 0.0039 | 0.0033 | 0.0034 |
| F07 | 30980001 | 31080000 | 0.0037 | 0.0031 | 0.0032 |
| F07 | 31000001 | 31100000 | 0.0026 | 0.0020 | 0.0020 |
| F07 | 31020001 | 31120000 | 0.0025 | 0.0019 | 0.0019 |
| F07 | 31040001 | 31140000 | 0.0016 | 0.0012 | 0.0012 |
| F07 | 31060001 | 31160000 | 0.0017 | 0.0014 | 0.0013 |
| F07 | 31080001 | 31180000 | 0.0022 | 0.0019 | 0.0018 |
| F07 | 31100001 | 31200000 | 0.0024 | 0.0021 | 0.0020 |
| F07 | 31120001 | 31220000 | 0.0026 | 0.0022 | 0.0022 |
| F07 | 31140001 | 31240000 | 0.0028 | 0.0024 | 0.0021 |
| F07 | 31160001 | 31260000 | 0.0031 | 0.0024 | 0.0022 |
| F07 | 31180001 | 31280000 | 0.0030 | 0.0022 | 0.0020 |
| F07 | 31200001 | 31300000 | 0.0029 | 0.0022 | 0.0020 |
| F07 | 31220001 | 31320000 | 0.0030 | 0.0024 | 0.0021 |
| F07 | 31240001 | 31340000 | 0.0038 | 0.0032 | 0.0030 |
| F07 | 31260001 | 31360000 | 0.0037 | 0.0029 | 0.0026 |
| F07 | 31280001 | 31380000 | 0.0038 | 0.0031 | 0.0027 |
| F07 | 31300001 | 31400000 | 0.0041 | 0.0032 | 0.0028 |
| F07 | 31320001 | 31420000 | 0.0034 | 0.0025 | 0.0021 |
| F07 | 31340001 | 31440000 | 0.0023 | 0.0016 | 0.0013 |
| F07 | 31360001 | 31460000 | 0.0016 | 0.0013 | 0.0011 |
| F07 | 31380001 | 31480000 | 0.0015 | 0.0012 | 0.0011 |
| F07 | 31400001 | 31500000 | 0.0019 | 0.0017 | 0.0016 |
| F07 | 31420001 | 31520000 | 0.0021 | 0.0019 | 0.0017 |
| F07 | 31440001 | 31540000 | 0.0021 | 0.0018 | 0.0015 |
| F07 | 31460001 | 31560000 | 0.0024 | 0.0019 | 0.0017 |
| F07 | 31480001 | 31580000 | 0.0023 | 0.0017 | 0.0014 |
| F07 | 31500001 | 31600000 | 0.0017 | 0.0011 | 0.0008 |
| F07 | 31520001 | 31620000 | 0.0020 | 0.0013 | 0.0013 |
| F07 | 31540001 | 31640000 | 0.0021 | 0.0013 | 0.0012 |
| F07 | 31560001 | 31660000 | 0.0026 | 0.0017 | 0.0016 |

| Chr | Start | End | both | A1 | A2 |
|---|---|---|---|---|---|
| F07 | 31580001 | 31680000 | 0.0032 | 0.0024 | 0.0021 |
| F07 | 31600001 | 31700000 | 0.0038 | 0.0029 | 0.0026 |
| F07 | 31620001 | 31720000 | 0.0037 | 0.0030 | 0.0026 |
| F07 | 31640001 | 31740000 | 0.0052 | 0.0044 | 0.0040 |
| F07 | 31660001 | 31760000 | 0.0052 | 0.0045 | 0.0043 |
| F07 | 31680001 | 31780000 | 0.0059 | 0.0052 | 0.0052 |
| F07 | 31700001 | 31800000 | 0.0053 | 0.0048 | 0.0047 |
| F07 | 31720001 | 31820000 | 0.0051 | 0.0045 | 0.0044 |
| F07 | 31740001 | 31840000 | 0.0038 | 0.0033 | 0.0031 |
| F07 | 31760001 | 31860000 | 0.0040 | 0.0032 | 0.0030 |
| F07 | 31780001 | 31880000 | 0.0028 | 0.0019 | 0.0017 |
| F07 | 31800001 | 31900000 | 0.0029 | 0.0019 | 0.0017 |
| F07 | 31820001 | 31920000 | 0.0032 | 0.0021 | 0.0020 |
| F07 | 31840001 | 31940000 | 0.0031 | 0.0022 | 0.0022 |
| F07 | 31860001 | 31960000 | 0.0022 | 0.0016 | 0.0016 |
| F07 | 31880001 | 31980000 | 0.0024 | 0.0019 | 0.0020 |
| F07 | 31900001 | 32000000 | 0.0024 | 0.0019 | 0.0021 |
| F07 | 31920001 | 32020000 | 0.0019 | 0.0016 | 0.0018 |
| F07 | 31940001 | 32040000 | 0.0021 | 0.0017 | 0.0018 |
| F07 | 31960001 | 32060000 | 0.0028 | 0.0021 | 0.0022 |
| F07 | 31980001 | 32080000 | 0.0033 | 0.0027 | 0.0025 |
| F07 | 32000001 | 32100000 | 0.0033 | 0.0025 | 0.0023 |
| F07 | 32020001 | 32120000 | 0.0035 | 0.0025 | 0.0024 |
| F07 | 32040001 | 32140000 | 0.0033 | 0.0026 | 0.0024 |
| F07 | 32060001 | 32160000 | 0.0034 | 0.0027 | 0.0025 |
| F07 | 32080001 | 32180000 | 0.0026 | 0.0017 | 0.0017 |
| F07 | 32100001 | 32200000 | 0.0024 | 0.0016 | 0.0017 |
| F07 | 32120001 | 32220000 | 0.0030 | 0.0022 | 0.0022 |
| F07 | 32140001 | 32240000 | 0.0033 | 0.0026 | 0.0025 |
| F07 | 32160001 | 32260000 | 0.0030 | 0.0025 | 0.0024 |
| F07 | 32180001 | 32280000 | 0.0029 | 0.0024 | 0.0025 |
| F07 | 32200001 | 32300000 | 0.0034 | 0.0028 | 0.0029 |
| F07 | 32220001 | 32320000 | 0.0027 | 0.0021 | 0.0022 |
| F07 | 32240001 | 32340000 | 0.0023 | 0.0018 | 0.0018 |
| F07 | 32260001 | 32360000 | 0.0023 | 0.0018 | 0.0018 |
| F07 | 32280001 | 32380000 | 0.0024 | 0.0018 | 0.0017 |
| F07 | 32300001 | 32400000 | 0.0027 | 0.0021 | 0.0020 |
| F07 | 32320001 | 32420000 | 0.0026 | 0.0020 | 0.0019 |
| F07 | 32340001 | 32440000 | 0.0023 | 0.0016 | 0.0016 |
| F07 | 32360001 | 32460000 | 0.0024 | 0.0016 | 0.0015 |
| F07 | 32380001 | 32480000 | 0.0021 | 0.0015 | 0.0014 |
| F07 | 32400001 | 32500000 | 0.0019 | 0.0011 | 0.0011 |
| F07 | 32420001 | 32520000 | 0.0021 | 0.0014 | 0.0013 |
| F07 | 32440001 | 32540000 | 0.0021 | 0.0015 | 0.0015 |
| F07 | 32460001 | 32560000 | 0.0026 | 0.0020 | 0.0020 |
| F07 | 32480001 | 32580000 | 0.0029 | 0.0023 | 0.0022 |
| F07 | 32500001 | 32600000 | 0.0031 | 0.0025 | 0.0025 |
| F07 | 32520001 | 32620000 | 0.0033 | 0.0027 | 0.0027 |
| F07 | 32540001 | 32640000 | 0.0031 | 0.0025 | 0.0025 |
| F07 | 32560001 | 32660000 | 0.0029 | 0.0023 | 0.0021 |
| F07 | 32580001 | 32680000 | 0.0026 | 0.0020 | 0.0018 |
| F07 | 32600001 | 32700000 | 0.0027 | 0.0023 | 0.0021 |
| F07 | 32620001 | 32720000 | 0.0029 | 0.0024 | 0.0022 |
| F07 | 32640001 | 32740000 | 0.0042 | 0.0032 | 0.0033 |
| F07 | 32660001 | 32760000 | 0.0039 | 0.0028 | 0.0031 |
| F07 | 32680001 | 32780000 | 0.0041 | 0.0031 | 0.0034 |
| F07 | 32700001 | 32800000 | 0.0038 | 0.0028 | 0.0030 |
| F07 | 32720001 | 32820000 | 0.0036 | 0.0026 | 0.0028 |

| Chr | Start | End | both | A1 | A2 |
|---|---|---|---|---|---|
| F07 | 32740001 | 32840000 | 0.0026 | 0.0021 | 0.0020 |
| F07 | 32760001 | 32860000 | 0.0027 | 0.0020 | 0.0021 |
| F07 | 32780001 | 32880000 | 0.0028 | 0.0020 | 0.0020 |
| F07 | 32800001 | 32900000 | 0.0024 | 0.0016 | 0.0016 |
| F07 | 32820001 | 32920000 | 0.0024 | 0.0017 | 0.0017 |
| F07 | 32840001 | 32940000 | 0.0026 | 0.0017 | 0.0016 |
| F07 | 32860001 | 32960000 | 0.0022 | 0.0015 | 0.0012 |
| F07 | 32880001 | 32980000 | 0.0024 | 0.0019 | 0.0016 |
| F07 | 32900001 | 33000000 | 0.0026 | 0.0020 | 0.0018 |
| F07 | 32920001 | 33020000 | 0.0025 | 0.0019 | 0.0017 |
| F07 | 32940001 | 33040000 | 0.0018 | 0.0015 | 0.0013 |
| F07 | 32960001 | 33060000 | 0.0021 | 0.0017 | 0.0017 |
| F07 | 32980001 | 33080000 | 0.0016 | 0.0012 | 0.0012 |
| F07 | 33000001 | 33100000 | 0.0018 | 0.0012 | 0.0014 |
| F07 | 33020001 | 33120000 | 0.0025 | 0.0019 | 0.0022 |
| F07 | 33040001 | 33140000 | 0.0027 | 0.0020 | 0.0022 |
| F07 | 33060001 | 33160000 | 0.0027 | 0.0020 | 0.0022 |
| F07 | 33080001 | 33180000 | 0.0029 | 0.0023 | 0.0023 |
| F07 | 33100001 | 33200000 | 0.0027 | 0.0022 | 0.0021 |
| F07 | 33120001 | 33220000 | 0.0021 | 0.0016 | 0.0014 |
| F07 | 33140001 | 33240000 | 0.0021 | 0.0016 | 0.0013 |
| F07 | 33160001 | 33260000 | 0.0017 | 0.0012 | 0.0010 |
| F07 | 33180001 | 33280000 | 0.0022 | 0.0015 | 0.0015 |
| F07 | 33200001 | 33300000 | 0.0027 | 0.0020 | 0.0020 |
| F07 | 33220001 | 33320000 | 0.0034 | 0.0027 | 0.0026 |
| F07 | 33240001 | 33340000 | 0.0046 | 0.0038 | 0.0039 |
| F07 | 33260001 | 33360000 | 0.0052 | 0.0043 | 0.0043 |
| F07 | 33280001 | 33380000 | 0.0051 | 0.0042 | 0.0043 |
| F07 | 33300001 | 33400000 | 0.0046 | 0.0036 | 0.0038 |
| F07 | 33320001 | 33420000 | 0.0035 | 0.0027 | 0.0029 |
| F07 | 33340001 | 33440000 | 0.0026 | 0.0019 | 0.0021 |
| F07 | 33360001 | 33460000 | 0.0021 | 0.0015 | 0.0018 |
| F07 | 33380001 | 33480000 | 0.0020 | 0.0015 | 0.0017 |
| F07 | 33400001 | 33500000 | 0.0019 | 0.0016 | 0.0018 |
| F07 | 33420001 | 33520000 | 0.0023 | 0.0019 | 0.0021 |
| F07 | 33440001 | 33540000 | 0.0029 | 0.0024 | 0.0027 |
| F07 | 33460001 | 33560000 | 0.0036 | 0.0030 | 0.0033 |
| F07 | 33480001 | 33580000 | 0.0035 | 0.0027 | 0.0029 |
| F07 | 33500001 | 33600000 | 0.0037 | 0.0027 | 0.0030 |
| F07 | 33520001 | 33620000 | 0.0035 | 0.0025 | 0.0027 |
| F07 | 33540001 | 33640000 | 0.0026 | 0.0017 | 0.0018 |
| F07 | 33560001 | 33660000 | 0.0019 | 0.0012 | 0.0013 |
| F07 | 33580001 | 33680000 | 0.0014 | 0.0009 | 0.0010 |
| F07 | 33600001 | 33700000 | 0.0008 | 0.0006 | 0.0006 |
| F07 | 33620001 | 33720000 | 0.0012 | 0.0009 | 0.0008 |
| F07 | 33640001 | 33740000 | 0.0015 | 0.0011 | 0.0010 |
| F07 | 33660001 | 33760000 | 0.0020 | 0.0013 | 0.0012 |
| F07 | 33680001 | 33780000 | 0.0021 | 0.0013 | 0.0012 |
| F07 | 33700001 | 33800000 | 0.0028 | 0.0018 | 0.0016 |
| F07 | 33720001 | 33820000 | 0.0029 | 0.0016 | 0.0016 |
| F07 | 33740001 | 33840000 | 0.0033 | 0.0020 | 0.0020 |
| F07 | 33760001 | 33860000 | 0.0035 | 0.0024 | 0.0025 |
| F07 | 33780001 | 33880000 | 0.0037 | 0.0025 | 0.0026 |
| F07 | 33800001 | 33900000 | 0.0041 | 0.0032 | 0.0032 |
| F07 | 33820001 | 33920000 | 0.0043 | 0.0033 | 0.0034 |
| F07 | 33840001 | 33940000 | 0.0047 | 0.0036 | 0.0035 |
| F07 | 33860001 | 33960000 | 0.0045 | 0.0031 | 0.0029 |
| F07 | 33880001 | 33980000 | 0.0055 | 0.0042 | 0.0039 |

| Chr | Start | End | both | A1 | A2 |
|---|---|---|---|---|---|
| F07 | 33900001 | 34000000 | 0.0047 | 0.0034 | 0.0033 |
| F07 | 33920001 | 34020000 | 0.0042 | 0.0031 | 0.0032 |
| F07 | 33940001 | 34040000 | 0.0031 | 0.0023 | 0.0025 |
| F07 | 33960001 | 34060000 | 0.0030 | 0.0024 | 0.0025 |
| F07 | 33980001 | 34080000 | 0.0022 | 0.0018 | 0.0018 |
| F07 | 34000001 | 34100000 | 0.0021 | 0.0019 | 0.0017 |
| F07 | 34020001 | 34120000 | 0.0022 | 0.0019 | 0.0017 |
| F07 | 34040001 | 34140000 | 0.0023 | 0.0021 | 0.0019 |
| F07 | 34060001 | 34160000 | 0.0024 | 0.0022 | 0.0020 |
| F07 | 34080001 | 34180000 | 0.0025 | 0.0022 | 0.0022 |
| F07 | 34100001 | 34200000 | 0.0028 | 0.0025 | 0.0025 |
| F07 | 34120001 | 34220000 | 0.0032 | 0.0027 | 0.0026 |
| F07 | 34140001 | 34240000 | 0.0033 | 0.0028 | 0.0027 |
| F07 | 34160001 | 34260000 | 0.0029 | 0.0025 | 0.0025 |
| F07 | 34180001 | 34280000 | 0.0036 | 0.0030 | 0.0031 |
| F07 | 34200001 | 34300000 | 0.0036 | 0.0028 | 0.0030 |
| F07 | 34220001 | 34320000 | 0.0036 | 0.0028 | 0.0031 |
| F07 | 34240001 | 34340000 | 0.0036 | 0.0027 | 0.0032 |
| F07 | 34260001 | 34360000 | 0.0040 | 0.0031 | 0.0036 |
| F07 | 34280001 | 34380000 | 0.0032 | 0.0025 | 0.0029 |
| F07 | 34300001 | 34400000 | 0.0041 | 0.0034 | 0.0039 |
| F07 | 34320001 | 34420000 | 0.0043 | 0.0034 | 0.0039 |
| F07 | 34340001 | 34440000 | 0.0046 | 0.0037 | 0.0042 |
| F07 | 34360001 | 34460000 | 0.0049 | 0.0041 | 0.0045 |
| F07 | 34380001 | 34480000 | 0.0049 | 0.0041 | 0.0045 |
| F07 | 34400001 | 34500000 | 0.0036 | 0.0029 | 0.0033 |
| F07 | 34420001 | 34520000 | 0.0033 | 0.0028 | 0.0031 |
| F07 | 34440001 | 34540000 | 0.0034 | 0.0027 | 0.0032 |
| F07 | 34460001 | 34560000 | 0.0041 | 0.0031 | 0.0037 |
| F07 | 34480001 | 34580000 | 0.0038 | 0.0028 | 0.0035 |
| F07 | 34500001 | 34600000 | 0.0039 | 0.0030 | 0.0036 |
| F07 | 34520001 | 34620000 | 0.0034 | 0.0026 | 0.0032 |
| F07 | 34540001 | 34640000 | 0.0034 | 0.0028 | 0.0033 |
| F07 | 34560001 | 34660000 | 0.0018 | 0.0016 | 0.0018 |
| F07 | 34580001 | 34680000 | 0.0019 | 0.0016 | 0.0019 |
| F07 | 34600001 | 34700000 | 0.0019 | 0.0016 | 0.0019 |
| F07 | 34620001 | 34720000 | 0.0021 | 0.0018 | 0.0021 |
| F07 | 34640001 | 34740000 | 0.0016 | 0.0013 | 0.0016 |
| F07 | 34660001 | 34760000 | 0.0016 | 0.0013 | 0.0017 |
| F07 | 34680001 | 34780000 | 0.0017 | 0.0014 | 0.0016 |
| F07 | 34700001 | 34800000 | 0.0019 | 0.0016 | 0.0017 |
| F07 | 34720001 | 34820000 | 0.0020 | 0.0016 | 0.0018 |
| F07 | 34740001 | 34840000 | 0.0020 | 0.0016 | 0.0017 |
| F07 | 34760001 | 34860000 | 0.0023 | 0.0018 | 0.0018 |
| F07 | 34780001 | 34880000 | 0.0021 | 0.0016 | 0.0016 |
| F07 | 34800001 | 34900000 | 0.0019 | 0.0015 | 0.0015 |
| F07 | 34820001 | 34920000 | 0.0017 | 0.0013 | 0.0013 |
| F07 | 34840001 | 34940000 | 0.0013 | 0.0009 | 0.0009 |
| F07 | 34860001 | 34960000 | 0.0012 | 0.0008 | 0.0009 |
| F07 | 34880001 | 34980000 | 0.0012 | 0.0008 | 0.0008 |
| F07 | 34900001 | 35000000 | 0.0011 | 0.0007 | 0.0008 |
| F07 | 34920001 | 35020000 | 0.0014 | 0.0010 | 0.0011 |
| F07 | 34940001 | 35040000 | 0.0016 | 0.0011 | 0.0014 |
| F07 | 34960001 | 35060000 | 0.0022 | 0.0018 | 0.0018 |
| F07 | 34980001 | 35080000 | 0.0028 | 0.0024 | 0.0024 |
| F07 | 35000001 | 35100000 | 0.0028 | 0.0023 | 0.0024 |
| F07 | 35020001 | 35120000 | 0.0029 | 0.0023 | 0.0023 |
| F07 | 35040001 | 35140000 | 0.0029 | 0.0023 | 0.0022 |

| Chr | Start | End | both | A1 | A2 |
|---|---|---|---|---|---|
| F07 | 35060001 | 35160000 | 0.0025 | 0.0017 | 0.0019 |
| F07 | 35080001 | 35180000 | 0.0023 | 0.0013 | 0.0014 |
| F07 | 35100001 | 35200000 | 0.0024 | 0.0014 | 0.0014 |
| F07 | 35120001 | 35220000 | 0.0025 | 0.0014 | 0.0015 |
| F07 | 35140001 | 35240000 | 0.0023 | 0.0013 | 0.0014 |
| F07 | 35160001 | 35260000 | 0.0031 | 0.0020 | 0.0020 |
| F07 | 35180001 | 35280000 | 0.0036 | 0.0025 | 0.0028 |
| F07 | 35200001 | 35300000 | 0.0036 | 0.0027 | 0.0030 |
| F07 | 35220001 | 35320000 | 0.0042 | 0.0034 | 0.0036 |
| F07 | 35240001 | 35340000 | 0.0045 | 0.0037 | 0.0039 |
| F07 | 35260001 | 35360000 | 0.0036 | 0.0028 | 0.0030 |
| F07 | 35280001 | 35380000 | 0.0033 | 0.0025 | 0.0025 |
| F07 | 35300001 | 35400000 | 0.0040 | 0.0029 | 0.0030 |
| F07 | 35320001 | 35420000 | 0.0032 | 0.0021 | 0.0021 |
| F07 | 35340001 | 35440000 | 0.0038 | 0.0026 | 0.0027 |
| F07 | 35360001 | 35460000 | 0.0041 | 0.0030 | 0.0031 |
| F07 | 35380001 | 35480000 | 0.0043 | 0.0031 | 0.0034 |
| F07 | 35400001 | 35500000 | 0.0041 | 0.0030 | 0.0032 |
| F07 | 35420001 | 35520000 | 0.0047 | 0.0035 | 0.0037 |
| F07 | 35440001 | 35540000 | 0.0039 | 0.0028 | 0.0031 |
| F07 | 35460001 | 35560000 | 0.0038 | 0.0028 | 0.0029 |
| F07 | 35480001 | 35580000 | 0.0037 | 0.0027 | 0.0026 |
| F07 | 35500001 | 35600000 | 0.0037 | 0.0025 | 0.0024 |
| F07 | 35520001 | 35620000 | 0.0036 | 0.0026 | 0.0025 |
| F07 | 35540001 | 35640000 | 0.0036 | 0.0025 | 0.0024 |
| F07 | 35560001 | 35660000 | 0.0032 | 0.0022 | 0.0021 |
| F07 | 35580001 | 35680000 | 0.0029 | 0.0021 | 0.0021 |
| F07 | 35600001 | 35700000 | 0.0021 | 0.0016 | 0.0016 |
| F07 | 35620001 | 35720000 | 0.0018 | 0.0013 | 0.0012 |
| F07 | 35640001 | 35740000 | 0.0022 | 0.0017 | 0.0016 |
| F07 | 35660001 | 35760000 | 0.0026 | 0.0021 | 0.0018 |
| F07 | 35680001 | 35780000 | 0.0031 | 0.0025 | 0.0022 |
| F07 | 35700001 | 35800000 | 0.0030 | 0.0024 | 0.0021 |
| F07 | 35720001 | 35820000 | 0.0029 | 0.0022 | 0.0019 |
| F07 | 35740001 | 35840000 | 0.0026 | 0.0019 | 0.0014 |
| F07 | 35760001 | 35860000 | 0.0026 | 0.0018 | 0.0016 |
| F07 | 35780001 | 35880000 | 0.0018 | 0.0011 | 0.0008 |
| F07 | 35800001 | 35900000 | 0.0027 | 0.0021 | 0.0018 |
| F07 | 35820001 | 35920000 | 0.0026 | 0.0019 | 0.0015 |
| F07 | 35840001 | 35940000 | 0.0026 | 0.0019 | 0.0016 |
| F07 | 35860001 | 35960000 | 0.0035 | 0.0031 | 0.0025 |
| F07 | 35880001 | 35980000 | 0.0036 | 0.0031 | 0.0025 |
| F07 | 35900001 | 36000000 | 0.0029 | 0.0024 | 0.0018 |
| F07 | 35920001 | 36020000 | 0.0029 | 0.0025 | 0.0019 |
| F07 | 35940001 | 36040000 | 0.0040 | 0.0036 | 0.0031 |
| F07 | 35960001 | 36060000 | 0.0036 | 0.0031 | 0.0027 |
| F07 | 35980001 | 36080000 | 0.0039 | 0.0035 | 0.0032 |
| F07 | 36000001 | 36100000 | 0.0044 | 0.0038 | 0.0036 |
| F07 | 36020001 | 36120000 | 0.0048 | 0.0043 | 0.0041 |
| F07 | 36040001 | 36140000 | 0.0042 | 0.0037 | 0.0036 |
| F07 | 36060001 | 36160000 | 0.0038 | 0.0032 | 0.0033 |
| F07 | 36080001 | 36180000 | 0.0042 | 0.0033 | 0.0036 |
| F07 | 36100001 | 36200000 | 0.0040 | 0.0031 | 0.0035 |
| F07 | 36120001 | 36220000 | 0.0035 | 0.0027 | 0.0031 |
| F07 | 36140001 | 36240000 | 0.0027 | 0.0021 | 0.0025 |
| F07 | 36160001 | 36260000 | 0.0030 | 0.0023 | 0.0028 |
| F07 | 36180001 | 36280000 | 0.0031 | 0.0024 | 0.0029 |
| F07 | 36200001 | 36300000 | 0.0028 | 0.0022 | 0.0025 |

| Chr | Start | End | both | A1 | A2 |
|---|---|---|---|---|---|
| F07 | 36220001 | 36320000 | 0.0031 | 0.0023 | 0.0026 |
| F07 | 36240001 | 36340000 | 0.0032 | 0.0024 | 0.0025 |
| F07 | 36260001 | 36360000 | 0.0027 | 0.0020 | 0.0021 |
| F07 | 36280001 | 36380000 | 0.0020 | 0.0013 | 0.0013 |
| F07 | 36300001 | 36400000 | 0.0019 | 0.0014 | 0.0013 |
| F07 | 36320001 | 36420000 | 0.0018 | 0.0013 | 0.0013 |
| F07 | 36340001 | 36440000 | 0.0014 | 0.0010 | 0.0012 |
| F07 | 36360001 | 36460000 | 0.0010 | 0.0007 | 0.0008 |
| F07 | 36380001 | 36480000 | 0.0015 | 0.0013 | 0.0013 |
| F07 | 36400001 | 36500000 | 0.0017 | 0.0015 | 0.0014 |
| F07 | 36420001 | 36520000 | 0.0023 | 0.0022 | 0.0019 |
| F07 | 36440001 | 36540000 | 0.0026 | 0.0025 | 0.0022 |
| F07 | 36460001 | 36560000 | 0.0028 | 0.0026 | 0.0023 |
| F07 | 36480001 | 36580000 | 0.0026 | 0.0023 | 0.0023 |
| F07 | 36500001 | 36600000 | 0.0031 | 0.0026 | 0.0026 |
| F07 | 36520001 | 36620000 | 0.0024 | 0.0019 | 0.0020 |
| F07 | 36540001 | 36640000 | 0.0026 | 0.0020 | 0.0023 |
| F07 | 36560001 | 36660000 | 0.0027 | 0.0021 | 0.0024 |
| F07 | 36580001 | 36680000 | 0.0022 | 0.0017 | 0.0019 |
| F07 | 36600001 | 36700000 | 0.0012 | 0.0010 | 0.0012 |
| F07 | 36620001 | 36720000 | 0.0009 | 0.0008 | 0.0009 |
| F07 | 36640001 | 36740000 | 0.0005 | 0.0004 | 0.0005 |
| F07 | 36660001 | 36760000 | 0.0005 | 0.0004 | 0.0005 |
| F07 | 36680001 | 36780000 | 0.0006 | 0.0005 | 0.0006 |
| F07 | 36700001 | 36800000 | 0.0007 | 0.0006 | 0.0007 |
| F07 | 36720001 | 36820000 | 0.0011 | 0.0010 | 0.0010 |
| F07 | 36740001 | 36840000 | 0.0016 | 0.0014 | 0.0015 |
| F07 | 36760001 | 36860000 | 0.0016 | 0.0014 | 0.0016 |
| F07 | 36780001 | 36880000 | 0.0017 | 0.0015 | 0.0016 |
| F07 | 36800001 | 36900000 | 0.0016 | 0.0015 | 0.0016 |
| F07 | 36820001 | 36920000 | 0.0013 | 0.0012 | 0.0012 |
| F07 | 36840001 | 36940000 | 0.0015 | 0.0013 | 0.0014 |
| F07 | 36860001 | 36960000 | 0.0018 | 0.0016 | 0.0018 |
| F07 | 36880001 | 36980000 | 0.0023 | 0.0020 | 0.0022 |
| F07 | 36900001 | 37000000 | 0.0025 | 0.0021 | 0.0023 |
| F07 | 36920001 | 37020000 | 0.0027 | 0.0022 | 0.0024 |
| F07 | 36940001 | 37040000 | 0.0019 | 0.0017 | 0.0017 |
| F07 | 36960001 | 37060000 | 0.0015 | 0.0013 | 0.0013 |
| F07 | 36980001 | 37080000 | 0.0015 | 0.0014 | 0.0012 |
| F07 | 37000001 | 37100000 | 0.0021 | 0.0021 | 0.0020 |
| F07 | 37020001 | 37120000 | 0.0019 | 0.0018 | 0.0018 |
| F07 | 37040001 | 37140000 | 0.0022 | 0.0020 | 0.0020 |
| F07 | 37060001 | 37160000 | 0.0023 | 0.0020 | 0.0019 |
| F07 | 37080001 | 37180000 | 0.0018 | 0.0014 | 0.0013 |
| F07 | 37100001 | 37200000 | 0.0014 | 0.0009 | 0.0007 |
| F07 | 37120001 | 37220000 | 0.0019 | 0.0012 | 0.0010 |
| F07 | 37140001 | 37240000 | 0.0015 | 0.0009 | 0.0007 |
| F07 | 37160001 | 37260000 | 0.0013 | 0.0008 | 0.0007 |
| F07 | 37180001 | 37280000 | 0.0013 | 0.0010 | 0.0010 |
| F07 | 37200001 | 37300000 | 0.0012 | 0.0009 | 0.0010 |
| F07 | 37220001 | 37320000 | 0.0018 | 0.0016 | 0.0017 |
| F07 | 37240001 | 37340000 | 0.0028 | 0.0026 | 0.0027 |
| F07 | 37260001 | 37360000 | 0.0039 | 0.0034 | 0.0036 |
| F07 | 37280001 | 37380000 | 0.0045 | 0.0039 | 0.0041 |
| F07 | 37300001 | 37400000 | 0.0044 | 0.0038 | 0.0040 |
| F07 | 37320001 | 37420000 | 0.0043 | 0.0037 | 0.0039 |
| F07 | 37340001 | 37440000 | 0.0042 | 0.0033 | 0.0035 |
| F07 | 37360001 | 37460000 | 0.0038 | 0.0031 | 0.0032 |

| Chr | Start | End | both | A1 | A2 |
|---|---|---|---|---|---|
| F07 | 37380001 | 37480000 | 0.0032 | 0.0025 | 0.0025 |
| F07 | 37400001 | 37500000 | 0.0039 | 0.0028 | 0.0030 |
| F07 | 37420001 | 37520000 | 0.0034 | 0.0025 | 0.0025 |
| F07 | 37440001 | 37540000 | 0.0029 | 0.0023 | 0.0023 |
| F07 | 37460001 | 37560000 | 0.0024 | 0.0017 | 0.0017 |
| F07 | 37480001 | 37580000 | 0.0024 | 0.0018 | 0.0019 |
| F07 | 37500001 | 37600000 | 0.0017 | 0.0014 | 0.0014 |
| F07 | 37520001 | 37620000 | 0.0013 | 0.0010 | 0.0010 |
| F07 | 37540001 | 37640000 | 0.0012 | 0.0009 | 0.0009 |
| F07 | 37560001 | 37660000 | 0.0014 | 0.0011 | 0.0012 |
| F07 | 37580001 | 37680000 | 0.0017 | 0.0015 | 0.0014 |
| F07 | 37600001 | 37700000 | 0.0020 | 0.0018 | 0.0017 |
| F07 | 37620001 | 37720000 | 0.0021 | 0.0019 | 0.0018 |
| F07 | 37640001 | 37740000 | 0.0019 | 0.0017 | 0.0018 |
| F07 | 37660001 | 37760000 | 0.0016 | 0.0014 | 0.0014 |
| F07 | 37680001 | 37780000 | 0.0013 | 0.0011 | 0.0013 |
| F07 | 37700001 | 37800000 | 0.0009 | 0.0008 | 0.0009 |
| F07 | 37720001 | 37820000 | 0.0009 | 0.0007 | 0.0009 |
| F07 | 37740001 | 37840000 | 0.0009 | 0.0007 | 0.0009 |
| F07 | 37760001 | 37860000 | 0.0014 | 0.0010 | 0.0014 |
| F07 | 37780001 | 37880000 | 0.0015 | 0.0011 | 0.0015 |
| F07 | 37800001 | 37900000 | 0.0020 | 0.0016 | 0.0019 |
| F07 | 37820001 | 37920000 | 0.0028 | 0.0021 | 0.0024 |
| F07 | 37840001 | 37940000 | 0.0036 | 0.0027 | 0.0030 |
| F07 | 37860001 | 37960000 | 0.0033 | 0.0025 | 0.0026 |
| F07 | 37880001 | 37980000 | 0.0034 | 0.0024 | 0.0024 |
| F07 | 37900001 | 38000000 | 0.0038 | 0.0026 | 0.0025 |
| F07 | 37920001 | 38020000 | 0.0034 | 0.0025 | 0.0023 |
| F07 | 37940001 | 38040000 | 0.0035 | 0.0025 | 0.0026 |
| F07 | 37960001 | 38060000 | 0.0037 | 0.0030 | 0.0029 |
| F07 | 37980001 | 38080000 | 0.0033 | 0.0028 | 0.0028 |
| F07 | 38000001 | 38100000 | 0.0027 | 0.0021 | 0.0023 |
| F07 | 38020001 | 38120000 | 0.0026 | 0.0020 | 0.0021 |
| F07 | 38040001 | 38140000 | 0.0022 | 0.0018 | 0.0016 |
| F07 | 38060001 | 38160000 | 0.0023 | 0.0017 | 0.0017 |
| F07 | 38080001 | 38180000 | 0.0028 | 0.0021 | 0.0019 |
| F07 | 38100001 | 38200000 | 0.0037 | 0.0029 | 0.0027 |
| F07 | 38120001 | 38220000 | 0.0037 | 0.0031 | 0.0028 |
| F07 | 38140001 | 38240000 | 0.0037 | 0.0031 | 0.0029 |
| F07 | 38160001 | 38260000 | 0.0038 | 0.0032 | 0.0030 |
| F07 | 38180001 | 38280000 | 0.0044 | 0.0035 | 0.0037 |
| F07 | 38200001 | 38300000 | 0.0045 | 0.0036 | 0.0038 |
| F07 | 38220001 | 38320000 | 0.0053 | 0.0040 | 0.0044 |
| F07 | 38240001 | 38340000 | 0.0050 | 0.0036 | 0.0040 |
| F07 | 38260001 | 38360000 | 0.0050 | 0.0037 | 0.0042 |
| F07 | 38280001 | 38380000 | 0.0043 | 0.0035 | 0.0037 |
| F07 | 38300001 | 38400000 | 0.0034 | 0.0027 | 0.0029 |
| F07 | 38320001 | 38420000 | 0.0029 | 0.0022 | 0.0024 |
| F07 | 38340001 | 38440000 | 0.0034 | 0.0028 | 0.0029 |
| F07 | 38360001 | 38460000 | 0.0028 | 0.0021 | 0.0020 |
| F07 | 38380001 | 38480000 | 0.0027 | 0.0020 | 0.0018 |
| F07 | 38400001 | 38500000 | 0.0034 | 0.0027 | 0.0026 |
| F07 | 38420001 | 38520000 | 0.0034 | 0.0026 | 0.0025 |
| F07 | 38440001 | 38540000 | 0.0037 | 0.0028 | 0.0026 |
| F07 | 38460001 | 38560000 | 0.0036 | 0.0028 | 0.0027 |
| F07 | 38480001 | 38580000 | 0.0035 | 0.0027 | 0.0027 |
| F07 | 38500001 | 38600000 | 0.0033 | 0.0025 | 0.0024 |
| F07 | 38520001 | 38620000 | 0.0031 | 0.0025 | 0.0025 |

| Chr | Start | End | both | A1 | A2 |
|---|---|---|---|---|---|
| F07 | 38540001 | 38640000 | 0.0024 | 0.0020 | 0.0019 |
| F07 | 38560001 | 38660000 | 0.0024 | 0.0020 | 0.0020 |
| F07 | 38580001 | 38680000 | 0.0028 | 0.0024 | 0.0024 |
| F07 | 38600001 | 38700000 | 0.0030 | 0.0027 | 0.0027 |
| F07 | 38620001 | 38720000 | 0.0035 | 0.0031 | 0.0029 |
| F07 | 38640001 | 38740000 | 0.0036 | 0.0031 | 0.0029 |
| F07 | 38660001 | 38760000 | 0.0036 | 0.0032 | 0.0028 |
| F07 | 38680001 | 38780000 | 0.0035 | 0.0029 | 0.0025 |
| F07 | 38700001 | 38800000 | 0.0029 | 0.0022 | 0.0017 |
| F07 | 38720001 | 38820000 | 0.0022 | 0.0015 | 0.0011 |
| F07 | 38740001 | 38840000 | 0.0023 | 0.0017 | 0.0014 |
| F07 | 38760001 | 38860000 | 0.0021 | 0.0016 | 0.0014 |
| F07 | 38780001 | 38880000 | 0.0017 | 0.0012 | 0.0011 |
| F07 | 38800001 | 38900000 | 0.0019 | 0.0014 | 0.0011 |
| F07 | 38820001 | 38920000 | 0.0022 | 0.0017 | 0.0015 |
| F07 | 38840001 | 38940000 | 0.0020 | 0.0013 | 0.0011 |
| F07 | 38860001 | 38960000 | 0.0026 | 0.0018 | 0.0014 |
| F07 | 38880001 | 38980000 | 0.0027 | 0.0018 | 0.0015 |
| F07 | 38900001 | 39000000 | 0.0027 | 0.0018 | 0.0017 |
| F07 | 38920001 | 39020000 | 0.0027 | 0.0016 | 0.0015 |
| F07 | 38940001 | 39040000 | 0.0027 | 0.0017 | 0.0016 |
| F07 | 38960001 | 39060000 | 0.0023 | 0.0013 | 0.0013 |
| F07 | 38980001 | 39080000 | 0.0025 | 0.0017 | 0.0017 |
| F07 | 39000001 | 39100000 | 0.0022 | 0.0015 | 0.0015 |
| F07 | 39020001 | 39120000 | 0.0018 | 0.0013 | 0.0013 |
| F07 | 39040001 | 39140000 | 0.0017 | 0.0011 | 0.0012 |
| F07 | 39060001 | 39160000 | 0.0019 | 0.0013 | 0.0015 |
| F07 | 39080001 | 39180000 | 0.0019 | 0.0013 | 0.0014 |
| F07 | 39100001 | 39200000 | 0.0025 | 0.0018 | 0.0018 |
| F07 | 39120001 | 39220000 | 0.0035 | 0.0027 | 0.0027 |
| F07 | 39140001 | 39240000 | 0.0036 | 0.0029 | 0.0029 |
| F07 | 39160001 | 39260000 | 0.0049 | 0.0041 | 0.0040 |
| F07 | 39180001 | 39280000 | 0.0049 | 0.0040 | 0.0040 |
| F07 | 39200001 | 39300000 | 0.0041 | 0.0034 | 0.0034 |
| F07 | 39220001 | 39320000 | 0.0029 | 0.0024 | 0.0024 |
| F07 | 39240001 | 39340000 | 0.0030 | 0.0023 | 0.0022 |
| F07 | 39260001 | 39360000 | 0.0015 | 0.0010 | 0.0009 |
| F07 | 39280001 | 39380000 | 0.0014 | 0.0008 | 0.0007 |
| F07 | 39300001 | 39400000 | 0.0021 | 0.0015 | 0.0011 |
| F07 | 39320001 | 39420000 | 0.0034 | 0.0026 | 0.0025 |
| F07 | 39340001 | 39440000 | 0.0040 | 0.0033 | 0.0032 |
| F07 | 39360001 | 39460000 | 0.0052 | 0.0044 | 0.0044 |
| F07 | 39380001 | 39480000 | 0.0057 | 0.0048 | 0.0048 |
| F07 | 39400001 | 39500000 | 0.0062 | 0.0052 | 0.0052 |
| F07 | 39420001 | 39520000 | 0.0056 | 0.0046 | 0.0045 |
| F07 | 39440001 | 39540000 | 0.0053 | 0.0043 | 0.0040 |
| F07 | 39460001 | 39560000 | 0.0044 | 0.0034 | 0.0030 |
| F07 | 39480001 | 39580000 | 0.0037 | 0.0028 | 0.0024 |
| F07 | 39500001 | 39600000 | 0.0034 | 0.0026 | 0.0024 |
| F07 | 39520001 | 39620000 | 0.0032 | 0.0024 | 0.0021 |
| F07 | 39540001 | 39640000 | 0.0034 | 0.0026 | 0.0024 |
| F07 | 39560001 | 39660000 | 0.0030 | 0.0022 | 0.0021 |
| F07 | 39580001 | 39680000 | 0.0030 | 0.0023 | 0.0023 |
| F07 | 39600001 | 39700000 | 0.0026 | 0.0020 | 0.0018 |
| F07 | 39620001 | 39720000 | 0.0026 | 0.0021 | 0.0019 |
| F07 | 39640001 | 39740000 | 0.0025 | 0.0020 | 0.0018 |
| F07 | 39660001 | 39760000 | 0.0028 | 0.0021 | 0.0018 |
| F07 | 39680001 | 39780000 | 0.0036 | 0.0028 | 0.0026 |

| Chr | Start | End | both | A1 | A2 |
|---|---|---|---|---|---|
| F07 | 39700001 | 39800000 | 0.0041 | 0.0033 | 0.0031 |
| F07 | 39720001 | 39820000 | 0.0043 | 0.0034 | 0.0034 |
| F07 | 39740001 | 39840000 | 0.0040 | 0.0032 | 0.0032 |
| F07 | 39760001 | 39860000 | 0.0043 | 0.0034 | 0.0034 |
| F07 | 39780001 | 39880000 | 0.0040 | 0.0032 | 0.0033 |
| F07 | 39800001 | 39900000 | 0.0034 | 0.0026 | 0.0028 |
| F07 | 39820001 | 39920000 | 0.0035 | 0.0027 | 0.0029 |
| F07 | 39840001 | 39940000 | 0.0034 | 0.0026 | 0.0028 |
| F07 | 39860001 | 39960000 | 0.0032 | 0.0027 | 0.0028 |
| F07 | 39880001 | 39980000 | 0.0028 | 0.0024 | 0.0025 |
| F07 | 39900001 | 40000000 | 0.0035 | 0.0029 | 0.0028 |
| F07 | 39920001 | 40020000 | 0.0031 | 0.0025 | 0.0023 |
| F07 | 39940001 | 40040000 | 0.0031 | 0.0025 | 0.0022 |
| F07 | 39960001 | 40060000 | 0.0031 | 0.0024 | 0.0023 |
| F07 | 39980001 | 40080000 | 0.0028 | 0.0021 | 0.0020 |
| F07 | 40000001 | 40100000 | 0.0021 | 0.0016 | 0.0015 |
| F07 | 40020001 | 40120000 | 0.0022 | 0.0018 | 0.0017 |
| F07 | 40040001 | 40140000 | 0.0026 | 0.0021 | 0.0021 |
| F07 | 40060001 | 40160000 | 0.0039 | 0.0034 | 0.0033 |
| F07 | 40080001 | 40180000 | 0.0044 | 0.0039 | 0.0037 |
| F07 | 40100001 | 40200000 | 0.0045 | 0.0038 | 0.0036 |
| F07 | 40120001 | 40220000 | 0.0047 | 0.0040 | 0.0040 |
| F07 | 40140001 | 40240000 | 0.0046 | 0.0039 | 0.0040 |
| F07 | 40160001 | 40260000 | 0.0044 | 0.0037 | 0.0036 |
| F07 | 40180001 | 40280000 | 0.0041 | 0.0032 | 0.0031 |
| F07 | 40200001 | 40300000 | 0.0046 | 0.0039 | 0.0039 |
| F07 | 40220001 | 40320000 | 0.0042 | 0.0035 | 0.0034 |
| F07 | 40240001 | 40340000 | 0.0044 | 0.0036 | 0.0035 |
| F07 | 40260001 | 40360000 | 0.0029 | 0.0023 | 0.0022 |
| F07 | 40280001 | 40380000 | 0.0027 | 0.0022 | 0.0022 |
| F07 | 40300001 | 40400000 | 0.0022 | 0.0018 | 0.0018 |
| F07 | 40320001 | 40420000 | 0.0017 | 0.0014 | 0.0015 |
| F07 | 40340001 | 40440000 | 0.0014 | 0.0011 | 0.0013 |
| F07 | 40360001 | 40460000 | 0.0014 | 0.0011 | 0.0014 |
| F07 | 40380001 | 40480000 | 0.0018 | 0.0017 | 0.0018 |
| F07 | 40400001 | 40500000 | 0.0017 | 0.0016 | 0.0016 |
| F07 | 40420001 | 40520000 | 0.0020 | 0.0019 | 0.0020 |
| F07 | 40440001 | 40540000 | 0.0017 | 0.0017 | 0.0016 |
| F07 | 40460001 | 40560000 | 0.0017 | 0.0018 | 0.0016 |
| F07 | 40480001 | 40580000 | 0.0013 | 0.0012 | 0.0012 |
| F07 | 40500001 | 40600000 | 0.0011 | 0.0010 | 0.0010 |
| F07 | 40520001 | 40620000 | 0.0009 | 0.0008 | 0.0008 |
| F07 | 40540001 | 40640000 | 0.0011 | 0.0009 | 0.0011 |
| F07 | 40560001 | 40660000 | 0.0011 | 0.0010 | 0.0011 |
| F07 | 40580001 | 40680000 | 0.0012 | 0.0010 | 0.0012 |
| F07 | 40600001 | 40700000 | 0.0013 | 0.0010 | 0.0013 |
| F07 | 40620001 | 40720000 | 0.0010 | 0.0008 | 0.0010 |
| F07 | 40640001 | 40740000 | 0.0008 | 0.0006 | 0.0008 |
| F07 | 40660001 | 40760000 | 0.0017 | 0.0013 | 0.0017 |
| F07 | 40680001 | 40780000 | 0.0024 | 0.0022 | 0.0023 |
| F07 | 40700001 | 40800000 | 0.0023 | 0.0021 | 0.0022 |
| F07 | 40720001 | 40820000 | 0.0031 | 0.0029 | 0.0031 |
| F07 | 40740001 | 40840000 | 0.0030 | 0.0029 | 0.0030 |
| F07 | 40760001 | 40860000 | 0.0019 | 0.0020 | 0.0019 |
| F07 | 40780001 | 40880000 | 0.0014 | 0.0013 | 0.0014 |
| F07 | 40800001 | 40900000 | 0.0027 | 0.0024 | 0.0025 |
| F07 | 40820001 | 40920000 | 0.0019 | 0.0016 | 0.0017 |
| F07 | 40840001 | 40940000 | 0.0026 | 0.0021 | 0.0022 |

| Chr | Start | End | both | A1 | A2 |
|---|---|---|---|---|---|
| F07 | 40860001 | 40960000 | 0.0032 | 0.0027 | 0.0027 |
| F07 | 40880001 | 40980000 | 0.0036 | 0.0029 | 0.0028 |
| F07 | 40900001 | 41000000 | 0.0031 | 0.0025 | 0.0025 |
| F07 | 40920001 | 41020000 | 0.0039 | 0.0031 | 0.0031 |
| F07 | 40940001 | 41040000 | 0.0034 | 0.0029 | 0.0029 |
| F07 | 40960001 | 41060000 | 0.0038 | 0.0030 | 0.0031 |
| F07 | 40980001 | 41080000 | 0.0034 | 0.0027 | 0.0027 |
| F07 | 41000001 | 41100000 | 0.0032 | 0.0024 | 0.0023 |
| F07 | 41020001 | 41120000 | 0.0028 | 0.0021 | 0.0018 |
| F07 | 41040001 | 41140000 | 0.0030 | 0.0022 | 0.0021 |
| F07 | 41060001 | 41160000 | 0.0025 | 0.0018 | 0.0016 |
| F07 | 41080001 | 41180000 | 0.0026 | 0.0021 | 0.0020 |
| F07 | 41100001 | 41200000 | 0.0021 | 0.0017 | 0.0018 |
| F07 | 41120001 | 41220000 | 0.0019 | 0.0016 | 0.0017 |
| F07 | 41140001 | 41240000 | 0.0021 | 0.0019 | 0.0019 |
| F07 | 41160001 | 41260000 | 0.0023 | 0.0021 | 0.0022 |
| F07 | 41180001 | 41280000 | 0.0026 | 0.0022 | 0.0024 |
| F07 | 41200001 | 41300000 | 0.0032 | 0.0026 | 0.0028 |
| F07 | 41220001 | 41320000 | 0.0040 | 0.0033 | 0.0037 |
| F07 | 41240001 | 41340000 | 0.0033 | 0.0027 | 0.0030 |
| F07 | 41260001 | 41360000 | 0.0031 | 0.0025 | 0.0028 |
| F07 | 41280001 | 41380000 | 0.0025 | 0.0020 | 0.0023 |
| F07 | 41300001 | 41400000 | 0.0025 | 0.0021 | 0.0025 |
| F07 | 41320001 | 41420000 | 0.0020 | 0.0016 | 0.0019 |
| F07 | 41340001 | 41440000 | 0.0021 | 0.0017 | 0.0020 |
| F07 | 41360001 | 41460000 | 0.0024 | 0.0020 | 0.0021 |
| F07 | 41380001 | 41480000 | 0.0032 | 0.0026 | 0.0026 |
| F07 | 41400001 | 41500000 | 0.0031 | 0.0025 | 0.0024 |
| F07 | 41420001 | 41520000 | 0.0032 | 0.0027 | 0.0025 |
| F07 | 41440001 | 41540000 | 0.0038 | 0.0032 | 0.0029 |
| F07 | 41460001 | 41560000 | 0.0031 | 0.0026 | 0.0025 |
| F07 | 41480001 | 41580000 | 0.0032 | 0.0027 | 0.0026 |
| F07 | 41500001 | 41600000 | 0.0029 | 0.0023 | 0.0022 |
| F07 | 41520001 | 41620000 | 0.0031 | 0.0023 | 0.0022 |
| F07 | 41540001 | 41640000 | 0.0030 | 0.0023 | 0.0021 |
| F07 | 41560001 | 41660000 | 0.0035 | 0.0027 | 0.0025 |
| F07 | 41580001 | 41680000 | 0.0039 | 0.0031 | 0.0031 |
| F07 | 41600001 | 41700000 | 0.0047 | 0.0040 | 0.0041 |
| F07 | 41620001 | 41720000 | 0.0039 | 0.0033 | 0.0035 |
| F07 | 41640001 | 41740000 | 0.0037 | 0.0031 | 0.0033 |
| F07 | 41660001 | 41760000 | 0.0040 | 0.0034 | 0.0037 |
| F07 | 41680001 | 41780000 | 0.0032 | 0.0026 | 0.0027 |
| F07 | 41700001 | 41800000 | 0.0024 | 0.0019 | 0.0018 |
| F07 | 41720001 | 41820000 | 0.0031 | 0.0023 | 0.0024 |
| F07 | 41740001 | 41840000 | 0.0035 | 0.0025 | 0.0025 |
| F07 | 41760001 | 41860000 | 0.0035 | 0.0024 | 0.0024 |
| F07 | 41780001 | 41880000 | 0.0028 | 0.0019 | 0.0021 |
| F07 | 41800001 | 41900000 | 0.0027 | 0.0019 | 0.0020 |
| F07 | 41820001 | 41920000 | 0.0028 | 0.0019 | 0.0020 |
| F07 | 41840001 | 41940000 | 0.0028 | 0.0021 | 0.0022 |
| F07 | 41860001 | 41960000 | 0.0026 | 0.0020 | 0.0019 |
| F07 | 41880001 | 41980000 | 0.0031 | 0.0023 | 0.0022 |
| F07 | 41900001 | 42000000 | 0.0034 | 0.0024 | 0.0022 |
| F07 | 41920001 | 42020000 | 0.0029 | 0.0021 | 0.0017 |
| F07 | 41940001 | 42040000 | 0.0035 | 0.0027 | 0.0025 |
| F07 | 41960001 | 42060000 | 0.0035 | 0.0027 | 0.0025 |
| F07 | 41980001 | 42080000 | 0.0037 | 0.0030 | 0.0028 |
| F07 | 42000001 | 42100000 | 0.0034 | 0.0028 | 0.0027 |

| Chr | Start | End | both | A1 | A2 |
|---|---|---|---|---|---|
| F07 | 42020001 | 42120000 | 0.0040 | 0.0032 | 0.0032 |
| F07 | 42040001 | 42140000 | 0.0042 | 0.0039 | 0.0030 |
| F07 | 42060001 | 42160000 | 0.0039 | 0.0035 | 0.0026 |
| F07 | 42080001 | 42180000 | 0.0035 | 0.0032 | 0.0023 |
| F07 | 42100001 | 42200000 | 0.0039 | 0.0033 | 0.0027 |
| F07 | 42120001 | 42220000 | 0.0038 | 0.0032 | 0.0026 |
| F07 | 42140001 | 42240000 | 0.0031 | 0.0022 | 0.0021 |
| F07 | 42160001 | 42260000 | 0.0031 | 0.0023 | 0.0022 |
| F07 | 42180001 | 42280000 | 0.0030 | 0.0021 | 0.0019 |
| F07 | 42200001 | 42300000 | 0.0028 | 0.0020 | 0.0015 |
| F07 | 42220001 | 42320000 | 0.0026 | 0.0019 | 0.0013 |
| F07 | 42240001 | 42340000 | 0.0021 | 0.0013 | 0.0010 |
| F07 | 42260001 | 42360000 | 0.0028 | 0.0022 | 0.0020 |
| F07 | 42280001 | 42380000 | 0.0029 | 0.0025 | 0.0021 |
| F07 | 42300001 | 42400000 | 0.0035 | 0.0029 | 0.0028 |
| F07 | 42320001 | 42420000 | 0.0030 | 0.0026 | 0.0025 |
| F07 | 42340001 | 42440000 | 0.0029 | 0.0026 | 0.0024 |
| F07 | 42360001 | 42460000 | 0.0020 | 0.0017 | 0.0014 |
| F07 | 42380001 | 42480000 | 0.0020 | 0.0014 | 0.0014 |
| F07 | 42400001 | 42500000 | 0.0015 | 0.0012 | 0.0010 |
| F07 | 42420001 | 42520000 | 0.0021 | 0.0016 | 0.0016 |
| F07 | 42440001 | 42540000 | 0.0018 | 0.0014 | 0.0015 |
| F07 | 42460001 | 42560000 | 0.0021 | 0.0016 | 0.0018 |
| F07 | 42480001 | 42580000 | 0.0020 | 0.0016 | 0.0019 |
| F07 | 42500001 | 42600000 | 0.0016 | 0.0013 | 0.0016 |
| F07 | 42520001 | 42620000 | 0.0020 | 0.0016 | 0.0016 |
| F07 | 42540001 | 42640000 | 0.0027 | 0.0021 | 0.0023 |
| F07 | 42560001 | 42660000 | 0.0033 | 0.0026 | 0.0027 |
| F07 | 42580001 | 42680000 | 0.0032 | 0.0025 | 0.0024 |
| F07 | 42600001 | 42700000 | 0.0037 | 0.0029 | 0.0029 |
| F07 | 42620001 | 42720000 | 0.0033 | 0.0026 | 0.0028 |
| F07 | 42640001 | 42740000 | 0.0026 | 0.0021 | 0.0021 |
| F07 | 42660001 | 42760000 | 0.0025 | 0.0021 | 0.0022 |
| F07 | 42680001 | 42780000 | 0.0039 | 0.0035 | 0.0037 |
| F07 | 42700001 | 42800000 | 0.0034 | 0.0030 | 0.0032 |
| F07 | 42720001 | 42820000 | 0.0032 | 0.0030 | 0.0030 |
| F07 | 42740001 | 42840000 | 0.0036 | 0.0032 | 0.0034 |
| F07 | 42760001 | 42860000 | 0.0031 | 0.0027 | 0.0028 |
| F07 | 42780001 | 42880000 | 0.0020 | 0.0014 | 0.0017 |
| F07 | 42800001 | 42900000 | 0.0021 | 0.0016 | 0.0018 |
| F07 | 42820001 | 42920000 | 0.0021 | 0.0015 | 0.0019 |
| F07 | 42840001 | 42940000 | 0.0020 | 0.0014 | 0.0017 |
| F07 | 42860001 | 42960000 | 0.0022 | 0.0016 | 0.0017 |
| F07 | 42880001 | 42980000 | 0.0019 | 0.0015 | 0.0015 |
| F07 | 42900001 | 43000000 | 0.0029 | 0.0024 | 0.0025 |
| F07 | 42920001 | 43020000 | 0.0030 | 0.0025 | 0.0024 |
| F07 | 42940001 | 43040000 | 0.0035 | 0.0030 | 0.0030 |
| F07 | 42960001 | 43060000 | 0.0031 | 0.0026 | 0.0029 |
| F07 | 42980001 | 43080000 | 0.0039 | 0.0035 | 0.0036 |
| F07 | 43000001 | 43100000 | 0.0034 | 0.0031 | 0.0030 |
| F07 | 43020001 | 43120000 | 0.0029 | 0.0027 | 0.0027 |
| F07 | 43040001 | 43140000 | 0.0024 | 0.0023 | 0.0022 |
| F07 | 43060001 | 43160000 | 0.0022 | 0.0022 | 0.0020 |
| F07 | 43080001 | 43180000 | 0.0014 | 0.0013 | 0.0013 |
| F07 | 43100001 | 43200000 | 0.0009 | 0.0009 | 0.0009 |
| F07 | 43120001 | 43220000 | 0.0009 | 0.0008 | 0.0009 |
| F07 | 43140001 | 43240000 | 0.0006 | 0.0006 | 0.0006 |
| F07 | 43160001 | 43260000 | 0.0006 | 0.0006 | 0.0006 |

| Chr | Start | End | both | A1 | A2 |
|---|---|---|---|---|---|
| F07 | 43180001 | 43280000 | 0.0005 | 0.0004 | 0.0005 |
| F07 | 43200001 | 43300000 | 0.0004 | 0.0003 | 0.0004 |
| F07 | 43220001 | 43320000 | 0.0002 | 0.0002 | 0.0002 |
| F07 | 43240001 | 43340000 | 0.0003 | 0.0002 | 0.0003 |
| F07 | 43260001 | 43360000 | 0.0005 | 0.0004 | 0.0005 |
| F07 | 43280001 | 43380000 | 0.0005 | 0.0004 | 0.0005 |
| F07 | 43300001 | 43400000 | 0.0006 | 0.0004 | 0.0006 |
| F07 | 43320001 | 43420000 | 0.0006 | 0.0005 | 0.0006 |
| F07 | 43340001 | 43440000 | 0.0006 | 0.0004 | 0.0006 |
| F07 | 43360001 | 43460000 | 0.0004 | 0.0003 | 0.0004 |
| F07 | 43380001 | 43480000 | 0.0004 | 0.0003 | 0.0004 |
| F07 | 43400001 | 43500000 | 0.0007 | 0.0006 | 0.0006 |
| F07 | 43420001 | 43520000 | 0.0014 | 0.0012 | 0.0012 |
| F07 | 43440001 | 43540000 | 0.0018 | 0.0015 | 0.0016 |
| F07 | 43460001 | 43560000 | 0.0022 | 0.0018 | 0.0019 |
| F07 | 43480001 | 43580000 | 0.0023 | 0.0018 | 0.0020 |
| F07 | 43500001 | 43600000 | 0.0019 | 0.0015 | 0.0017 |
| F07 | 43520001 | 43620000 | 0.0016 | 0.0012 | 0.0015 |
| F07 | 43540001 | 43640000 | 0.0013 | 0.0010 | 0.0012 |
| F07 | 43560001 | 43660000 | 0.0017 | 0.0012 | 0.0016 |
| F07 | 43580001 | 43680000 | 0.0016 | 0.0012 | 0.0014 |
| F07 | 43600001 | 43700000 | 0.0016 | 0.0012 | 0.0014 |
| F07 | 43620001 | 43720000 | 0.0013 | 0.0009 | 0.0011 |
| F07 | 43640001 | 43740000 | 0.0014 | 0.0011 | 0.0013 |
| F07 | 43660001 | 43760000 | 0.0009 | 0.0007 | 0.0007 |
| F07 | 43680001 | 43780000 | 0.0015 | 0.0013 | 0.0012 |
| F07 | 43700001 | 43800000 | 0.0020 | 0.0017 | 0.0017 |
| F07 | 43720001 | 43820000 | 0.0022 | 0.0018 | 0.0020 |
| F07 | 43740001 | 43840000 | 0.0020 | 0.0016 | 0.0018 |
| F07 | 43760001 | 43860000 | 0.0021 | 0.0016 | 0.0018 |
| F07 | 43780001 | 43880000 | 0.0025 | 0.0019 | 0.0023 |
| F07 | 43800001 | 43900000 | 0.0021 | 0.0017 | 0.0019 |
| F07 | 43820001 | 43920000 | 0.0018 | 0.0015 | 0.0016 |
| F07 | 43840001 | 43940000 | 0.0021 | 0.0019 | 0.0020 |
| F07 | 43860001 | 43960000 | 0.0019 | 0.0017 | 0.0017 |
| F07 | 43880001 | 43980000 | 0.0010 | 0.0009 | 0.0009 |
| F07 | 43900001 | 44000000 | 0.0009 | 0.0008 | 0.0008 |
| F07 | 43920001 | 44020000 | 0.0015 | 0.0013 | 0.0013 |
| F07 | 43940001 | 44040000 | 0.0011 | 0.0009 | 0.0010 |
| F07 | 43960001 | 44060000 | 0.0011 | 0.0009 | 0.0010 |
| F07 | 43980001 | 44080000 | 0.0010 | 0.0008 | 0.0008 |
| F07 | 44000001 | 44100000 | 0.0016 | 0.0013 | 0.0014 |
| F07 | 44020001 | 44120000 | 0.0012 | 0.0009 | 0.0011 |
| F07 | 44040001 | 44140000 | 0.0019 | 0.0015 | 0.0016 |
| F07 | 44060001 | 44160000 | 0.0027 | 0.0022 | 0.0023 |
| F07 | 44080001 | 44180000 | 0.0034 | 0.0029 | 0.0030 |
| F07 | 44100001 | 44200000 | 0.0031 | 0.0027 | 0.0027 |
| F07 | 44120001 | 44220000 | 0.0033 | 0.0029 | 0.0027 |
| F07 | 44140001 | 44240000 | 0.0030 | 0.0026 | 0.0024 |
| F07 | 44160001 | 44260000 | 0.0025 | 0.0019 | 0.0018 |
| F07 | 44180001 | 44280000 | 0.0030 | 0.0022 | 0.0023 |
| F07 | 44200001 | 44300000 | 0.0034 | 0.0025 | 0.0024 |
| F07 | 44220001 | 44320000 | 0.0032 | 0.0024 | 0.0023 |
| F07 | 44240001 | 44340000 | 0.0028 | 0.0022 | 0.0020 |
| F07 | 44260001 | 44360000 | 0.0026 | 0.0021 | 0.0020 |
| F07 | 44280001 | 44380000 | 0.0016 | 0.0012 | 0.0011 |
| F07 | 44300001 | 44400000 | 0.0008 | 0.0006 | 0.0006 |
| F07 | 44320001 | 44420000 | 0.0008 | 0.0006 | 0.0006 |

| Chr | Start | End | both | A1 | A2 |
|---|---|---|---|---|---|
| F07 | 44340001 | 44440000 | 0.0011 | 0.0008 | 0.0009 |
| F07 | 44360001 | 44460000 | 0.0014 | 0.0011 | 0.0012 |
| F07 | 44380001 | 44480000 | 0.0024 | 0.0019 | 0.0021 |
| F07 | 44400001 | 44500000 | 0.0025 | 0.0019 | 0.0021 |
| F07 | 44420001 | 44520000 | 0.0026 | 0.0020 | 0.0023 |
| F07 | 44440001 | 44540000 | 0.0030 | 0.0023 | 0.0024 |
| F07 | 44460001 | 44560000 | 0.0034 | 0.0026 | 0.0026 |
| F07 | 44480001 | 44580000 | 0.0026 | 0.0020 | 0.0020 |
| F07 | 44500001 | 44600000 | 0.0027 | 0.0022 | 0.0021 |
| F07 | 44520001 | 44620000 | 0.0022 | 0.0019 | 0.0017 |
| F07 | 44540001 | 44640000 | 0.0017 | 0.0015 | 0.0014 |
| F07 | 44560001 | 44660000 | 0.0019 | 0.0016 | 0.0018 |
| F07 | 44580001 | 44680000 | 0.0023 | 0.0020 | 0.0022 |
| F07 | 44600001 | 44700000 | 0.0022 | 0.0019 | 0.0021 |
| F07 | 44620001 | 44720000 | 0.0022 | 0.0019 | 0.0022 |
| F07 | 44640001 | 44740000 | 0.0023 | 0.0018 | 0.0021 |
| F07 | 44660001 | 44760000 | 0.0020 | 0.0016 | 0.0017 |
| F07 | 44680001 | 44780000 | 0.0013 | 0.0010 | 0.0010 |
| F07 | 44700001 | 44800000 | 0.0017 | 0.0012 | 0.0014 |
| F07 | 44720001 | 44820000 | 0.0019 | 0.0014 | 0.0015 |
| F07 | 44740001 | 44840000 | 0.0019 | 0.0015 | 0.0016 |
| F07 | 44760001 | 44860000 | 0.0015 | 0.0012 | 0.0014 |
| F07 | 44780001 | 44880000 | 0.0018 | 0.0013 | 0.0014 |
| F07 | 44800001 | 44900000 | 0.0018 | 0.0014 | 0.0014 |
| F07 | 44820001 | 44920000 | 0.0017 | 0.0013 | 0.0013 |
| F07 | 44840001 | 44940000 | 0.0018 | 0.0014 | 0.0015 |
| F07 | 44860001 | 44960000 | 0.0017 | 0.0014 | 0.0014 |
| F07 | 44880001 | 44980000 | 0.0018 | 0.0014 | 0.0014 |
| F07 | 44900001 | 45000000 | 0.0016 | 0.0011 | 0.0012 |
| F07 | 44920001 | 45020000 | 0.0017 | 0.0013 | 0.0014 |
| F07 | 44940001 | 45040000 | 0.0017 | 0.0013 | 0.0015 |
| F07 | 44960001 | 45060000 | 0.0019 | 0.0015 | 0.0015 |
| F07 | 44980001 | 45080000 | 0.0021 | 0.0017 | 0.0019 |
| F07 | 45000001 | 45100000 | 0.0022 | 0.0018 | 0.0020 |
| F07 | 45020001 | 45120000 | 0.0024 | 0.0019 | 0.0020 |
| F07 | 45040001 | 45140000 | 0.0025 | 0.0020 | 0.0021 |
| F07 | 45060001 | 45160000 | 0.0026 | 0.0020 | 0.0020 |
| F07 | 45080001 | 45180000 | 0.0023 | 0.0019 | 0.0017 |
| F07 | 45100001 | 45200000 | 0.0033 | 0.0028 | 0.0023 |
| F07 | 45120001 | 45220000 | 0.0030 | 0.0027 | 0.0022 |
| F07 | 45140001 | 45240000 | 0.0032 | 0.0028 | 0.0024 |
| F07 | 45160001 | 45260000 | 0.0039 | 0.0038 | 0.0032 |
| F07 | 45180001 | 45280000 | 0.0041 | 0.0039 | 0.0033 |
| F07 | 45200001 | 45300000 | 0.0033 | 0.0030 | 0.0029 |
| F07 | 45220001 | 45320000 | 0.0038 | 0.0033 | 0.0032 |
| F07 | 45240001 | 45340000 | 0.0033 | 0.0028 | 0.0027 |
| F07 | 45260001 | 45360000 | 0.0025 | 0.0019 | 0.0021 |
| F07 | 45280001 | 45380000 | 0.0024 | 0.0017 | 0.0020 |
| F07 | 45300001 | 45400000 | 0.0022 | 0.0016 | 0.0020 |
| F07 | 45320001 | 45420000 | 0.0015 | 0.0012 | 0.0014 |
| F07 | 45340001 | 45440000 | 0.0016 | 0.0011 | 0.0012 |
| F07 | 45360001 | 45460000 | 0.0023 | 0.0018 | 0.0018 |
| F07 | 45380001 | 45480000 | 0.0017 | 0.0013 | 0.0013 |
| F07 | 45400001 | 45500000 | 0.0018 | 0.0014 | 0.0014 |
| F07 | 45420001 | 45520000 | 0.0018 | 0.0015 | 0.0014 |
| F07 | 45440001 | 45540000 | 0.0017 | 0.0015 | 0.0015 |
| F07 | 45460001 | 45560000 | 0.0010 | 0.0009 | 0.0009 |
| F07 | 45480001 | 45580000 | 0.0016 | 0.0013 | 0.0015 |

| Chr | Start | End | both | A1 | A2 |
|---|---|---|---|---|---|
| F07 | 45500001 | 45600000 | 0.0013 | 0.0011 | 0.0012 |
| F07 | 45520001 | 45620000 | 0.0014 | 0.0012 | 0.0013 |
| F07 | 45540001 | 45640000 | 0.0013 | 0.0011 | 0.0012 |
| F07 | 45560001 | 45660000 | 0.0013 | 0.0011 | 0.0012 |
| F07 | 45580001 | 45680000 | 0.0008 | 0.0007 | 0.0007 |
| F07 | 45600001 | 45700000 | 0.0007 | 0.0006 | 0.0007 |
| F07 | 45620001 | 45720000 | 0.0006 | 0.0006 | 0.0006 |
| F07 | 45640001 | 45740000 | 0.0007 | 0.0006 | 0.0006 |
| F07 | 45660001 | 45760000 | 0.0014 | 0.0012 | 0.0013 |
| F07 | 45680001 | 45780000 | 0.0021 | 0.0018 | 0.0019 |
| F07 | 45700001 | 45800000 | 0.0030 | 0.0024 | 0.0026 |
| F07 | 45720001 | 45820000 | 0.0031 | 0.0025 | 0.0027 |
| F07 | 45740001 | 45840000 | 0.0034 | 0.0028 | 0.0031 |
| F07 | 45760001 | 45860000 | 0.0025 | 0.0020 | 0.0022 |
| F07 | 45780001 | 45880000 | 0.0019 | 0.0016 | 0.0017 |
| F07 | 45800001 | 45900000 | 0.0011 | 0.0010 | 0.0011 |
| F07 | 45820001 | 45920000 | 0.0013 | 0.0012 | 0.0013 |
| F07 | 45840001 | 45940000 | 0.0012 | 0.0011 | 0.0011 |
| F07 | 45860001 | 45960000 | 0.0013 | 0.0012 | 0.0012 |
| F07 | 45880001 | 45980000 | 0.0016 | 0.0014 | 0.0016 |
| F07 | 45900001 | 46000000 | 0.0017 | 0.0014 | 0.0016 |
| F07 | 45920001 | 46020000 | 0.0014 | 0.0010 | 0.0013 |
| F07 | 45940001 | 46040000 | 0.0012 | 0.0009 | 0.0011 |
| F07 | 45960001 | 46060000 | 0.0013 | 0.0010 | 0.0012 |
| F07 | 45980001 | 46080000 | 0.0008 | 0.0007 | 0.0008 |
| F07 | 46000001 | 46100000 | 0.0007 | 0.0007 | 0.0007 |
| F07 | 46020001 | 46120000 | 0.0010 | 0.0009 | 0.0009 |
| F07 | 46040001 | 46140000 | 0.0009 | 0.0008 | 0.0008 |
| F07 | 46060001 | 46160000 | 0.0011 | 0.0011 | 0.0010 |
| F07 | 46080001 | 46180000 | 0.0015 | 0.0015 | 0.0013 |
| F07 | 46100001 | 46200000 | 0.0018 | 0.0017 | 0.0016 |
| F07 | 46120001 | 46220000 | 0.0022 | 0.0020 | 0.0019 |
| F07 | 46140001 | 46240000 | 0.0022 | 0.0020 | 0.0019 |
| F07 | 46160001 | 46260000 | 0.0017 | 0.0014 | 0.0015 |
| F07 | 46180001 | 46280000 | 0.0019 | 0.0016 | 0.0018 |
| F07 | 46200001 | 46300000 | 0.0016 | 0.0014 | 0.0015 |
| F07 | 46220001 | 46320000 | 0.0017 | 0.0014 | 0.0015 |
| F07 | 46240001 | 46340000 | 0.0022 | 0.0018 | 0.0020 |
| F07 | 46260001 | 46360000 | 0.0029 | 0.0024 | 0.0027 |
| F07 | 46280001 | 46380000 | 0.0029 | 0.0024 | 0.0027 |
| F07 | 46300001 | 46400000 | 0.0036 | 0.0031 | 0.0033 |
| F07 | 46320001 | 46420000 | 0.0032 | 0.0028 | 0.0030 |
| F07 | 46340001 | 46440000 | 0.0036 | 0.0030 | 0.0034 |
| F07 | 46360001 | 46460000 | 0.0029 | 0.0025 | 0.0028 |
| F07 | 46380001 | 46480000 | 0.0026 | 0.0022 | 0.0025 |
| F07 | 46400001 | 46500000 | 0.0021 | 0.0017 | 0.0020 |
| F07 | 46420001 | 46520000 | 0.0020 | 0.0017 | 0.0019 |
| F07 | 46440001 | 46540000 | 0.0018 | 0.0016 | 0.0016 |
| F07 | 46460001 | 46560000 | 0.0019 | 0.0017 | 0.0017 |
| F07 | 46480001 | 46580000 | 0.0022 | 0.0020 | 0.0020 |
| F07 | 46500001 | 46600000 | 0.0019 | 0.0017 | 0.0017 |
| F07 | 46520001 | 46620000 | 0.0025 | 0.0022 | 0.0024 |
| F07 | 46540001 | 46640000 | 0.0020 | 0.0018 | 0.0019 |
| F07 | 46560001 | 46660000 | 0.0019 | 0.0017 | 0.0018 |
| F07 | 46580001 | 46680000 | 0.0011 | 0.0010 | 0.0011 |
| F07 | 46600001 | 46700000 | 0.0015 | 0.0014 | 0.0014 |
| F07 | 46620001 | 46720000 | 0.0010 | 0.0008 | 0.0007 |
| F07 | 46640001 | 46740000 | 0.0014 | 0.0012 | 0.0010 |

| Chr | Start | End | both | A1 | A2 |
|---|---|---|---|---|---|
| F07 | 46660001 | 46760000 | 0.0021 | 0.0017 | 0.0016 |
| F07 | 46680001 | 46780000 | 0.0027 | 0.0022 | 0.0022 |
| F07 | 46700001 | 46800000 | 0.0027 | 0.0022 | 0.0023 |
| F07 | 46720001 | 46820000 | 0.0030 | 0.0023 | 0.0025 |
| F07 | 46740001 | 46840000 | 0.0035 | 0.0027 | 0.0031 |
| F07 | 46760001 | 46860000 | 0.0035 | 0.0029 | 0.0031 |
| F07 | 46780001 | 46880000 | 0.0033 | 0.0027 | 0.0027 |
| F07 | 46800001 | 46900000 | 0.0030 | 0.0024 | 0.0025 |
| F07 | 46820001 | 46920000 | 0.0030 | 0.0026 | 0.0025 |
| F07 | 46840001 | 46940000 | 0.0026 | 0.0020 | 0.0018 |
| F07 | 46860001 | 46960000 | 0.0019 | 0.0015 | 0.0013 |
| F07 | 46880001 | 46980000 | 0.0022 | 0.0019 | 0.0018 |
| F07 | 46900001 | 47000000 | 0.0040 | 0.0036 | 0.0035 |
| F07 | 46920001 | 47020000 | 0.0045 | 0.0039 | 0.0038 |
| F07 | 46940001 | 47040000 | 0.0055 | 0.0049 | 0.0049 |
| F07 | 46960001 | 47060000 | 0.0059 | 0.0052 | 0.0053 |
| F07 | 46980001 | 47080000 | 0.0057 | 0.0050 | 0.0051 |
| F07 | 47000001 | 47100000 | 0.0041 | 0.0036 | 0.0037 |
| F07 | 47020001 | 47120000 | 0.0038 | 0.0034 | 0.0036 |
| F07 | 47040001 | 47140000 | 0.0023 | 0.0020 | 0.0022 |
| F07 | 47060001 | 47160000 | 0.0027 | 0.0023 | 0.0027 |
| F07 | 47080001 | 47180000 | 0.0024 | 0.0020 | 0.0024 |
| F07 | 47100001 | 47200000 | 0.0024 | 0.0019 | 0.0024 |
| F07 | 47120001 | 47220000 | 0.0021 | 0.0016 | 0.0021 |
| F07 | 47140001 | 47240000 | 0.0022 | 0.0017 | 0.0022 |
| F07 | 47160001 | 47260000 | 0.0025 | 0.0022 | 0.0024 |
| F07 | 47180001 | 47280000 | 0.0027 | 0.0025 | 0.0027 |
| F07 | 47200001 | 47300000 | 0.0029 | 0.0026 | 0.0028 |
| F07 | 47220001 | 47320000 | 0.0031 | 0.0028 | 0.0030 |
| F07 | 47240001 | 47340000 | 0.0034 | 0.0030 | 0.0032 |
| F07 | 47260001 | 47360000 | 0.0024 | 0.0021 | 0.0023 |
| F07 | 47280001 | 47380000 | 0.0021 | 0.0017 | 0.0019 |
| F07 | 47300001 | 47400000 | 0.0021 | 0.0017 | 0.0019 |
| F07 | 47320001 | 47420000 | 0.0023 | 0.0019 | 0.0019 |
| F07 | 47340001 | 47440000 | 0.0025 | 0.0020 | 0.0020 |
| F07 | 47360001 | 47460000 | 0.0023 | 0.0019 | 0.0019 |
| F07 | 47380001 | 47480000 | 0.0025 | 0.0022 | 0.0021 |
| F07 | 47400001 | 47500000 | 0.0027 | 0.0022 | 0.0020 |
| F07 | 47420001 | 47520000 | 0.0021 | 0.0017 | 0.0017 |
| F07 | 47440001 | 47540000 | 0.0032 | 0.0029 | 0.0027 |
| F07 | 47460001 | 47560000 | 0.0037 | 0.0034 | 0.0031 |
| F07 | 47480001 | 47580000 | 0.0048 | 0.0042 | 0.0038 |
| F07 | 47500001 | 47600000 | 0.0044 | 0.0041 | 0.0037 |
| F07 | 47520001 | 47620000 | 0.0046 | 0.0040 | 0.0038 |
| F07 | 47540001 | 47640000 | 0.0043 | 0.0037 | 0.0036 |
| F07 | 47560001 | 47660000 | 0.0041 | 0.0035 | 0.0035 |
| F07 | 47580001 | 47680000 | 0.0029 | 0.0025 | 0.0027 |
| F07 | 47600001 | 47700000 | 0.0041 | 0.0033 | 0.0039 |
| F07 | 47620001 | 47720000 | 0.0045 | 0.0038 | 0.0043 |
| F07 | 47640001 | 47740000 | 0.0036 | 0.0028 | 0.0036 |
| F07 | 47660001 | 47760000 | 0.0038 | 0.0028 | 0.0036 |
| F07 | 47680001 | 47780000 | 0.0039 | 0.0030 | 0.0037 |
| F07 | 47700001 | 47800000 | 0.0030 | 0.0023 | 0.0028 |
| F07 | 47720001 | 47820000 | 0.0023 | 0.0018 | 0.0021 |
| F07 | 47740001 | 47840000 | 0.0021 | 0.0016 | 0.0020 |
| F07 | 47760001 | 47860000 | 0.0018 | 0.0015 | 0.0018 |
| F07 | 47780001 | 47880000 | 0.0021 | 0.0018 | 0.0020 |
| F07 | 47800001 | 47900000 | 0.0017 | 0.0016 | 0.0017 |

| Chr | Start | End | both | A1 | A2 |
|---|---|---|---|---|---|
| F07 | 47820001 | 47920000 | 0.0020 | 0.0018 | 0.0019 |
| F07 | 47840001 | 47940000 | 0.0017 | 0.0015 | 0.0017 |
| F07 | 47860001 | 47960000 | 0.0018 | 0.0016 | 0.0018 |
| F07 | 47880001 | 47980000 | 0.0018 | 0.0015 | 0.0018 |
| F07 | 47900001 | 48000000 | 0.0017 | 0.0015 | 0.0017 |
| F07 | 47920001 | 48020000 | 0.0018 | 0.0016 | 0.0017 |
| F07 | 47940001 | 48040000 | 0.0020 | 0.0018 | 0.0020 |
| F07 | 47960001 | 48060000 | 0.0024 | 0.0021 | 0.0024 |
| F07 | 47980001 | 48080000 | 0.0026 | 0.0023 | 0.0025 |
| F07 | 48000001 | 48100000 | 0.0024 | 0.0021 | 0.0024 |
| F07 | 48020001 | 48120000 | 0.0022 | 0.0020 | 0.0022 |
| F07 | 48040001 | 48140000 | 0.0023 | 0.0021 | 0.0022 |
| F07 | 48060001 | 48160000 | 0.0018 | 0.0016 | 0.0017 |
| F07 | 48080001 | 48180000 | 0.0019 | 0.0018 | 0.0018 |
| F07 | 48100001 | 48200000 | 0.0027 | 0.0026 | 0.0026 |
| F07 | 48120001 | 48220000 | 0.0028 | 0.0026 | 0.0026 |
| F07 | 48140001 | 48240000 | 0.0024 | 0.0023 | 0.0023 |
| F07 | 48160001 | 48260000 | 0.0023 | 0.0021 | 0.0022 |
| F07 | 48180001 | 48280000 | 0.0015 | 0.0013 | 0.0014 |
| F07 | 48200001 | 48300000 | 0.0009 | 0.0008 | 0.0009 |
| F07 | 48220001 | 48320000 | 0.0013 | 0.0012 | 0.0011 |
| F07 | 48240001 | 48340000 | 0.0016 | 0.0013 | 0.0013 |
| F07 | 48260001 | 48360000 | 0.0029 | 0.0026 | 0.0024 |
| F07 | 48280001 | 48380000 | 0.0048 | 0.0042 | 0.0041 |
| F07 | 48300001 | 48400000 | 0.0046 | 0.0040 | 0.0038 |
| F07 | 48320001 | 48420000 | 0.0044 | 0.0039 | 0.0039 |
| F07 | 48340001 | 48440000 | 0.0043 | 0.0037 | 0.0038 |
| F07 | 48360001 | 48460000 | 0.0032 | 0.0025 | 0.0027 |
| F07 | 48380001 | 48480000 | 0.0026 | 0.0020 | 0.0022 |
| F07 | 48400001 | 48500000 | 0.0026 | 0.0021 | 0.0023 |
| F07 | 48420001 | 48520000 | 0.0025 | 0.0019 | 0.0022 |
| F07 | 48440001 | 48540000 | 0.0031 | 0.0024 | 0.0028 |
| F07 | 48460001 | 48560000 | 0.0036 | 0.0029 | 0.0032 |
| F07 | 48480001 | 48580000 | 0.0026 | 0.0020 | 0.0022 |
| F07 | 48500001 | 48600000 | 0.0035 | 0.0026 | 0.0030 |
| F07 | 48520001 | 48620000 | 0.0038 | 0.0030 | 0.0033 |
| F07 | 48540001 | 48640000 | 0.0048 | 0.0040 | 0.0041 |
| F07 | 48560001 | 48660000 | 0.0055 | 0.0047 | 0.0050 |
| F07 | 48580001 | 48680000 | 0.0063 | 0.0054 | 0.0058 |
| F07 | 48600001 | 48700000 | 0.0058 | 0.0053 | 0.0054 |
| F07 | 48620001 | 48720000 | 0.0055 | 0.0050 | 0.0051 |
| F07 | 48640001 | 48740000 | 0.0045 | 0.0042 | 0.0043 |
| F07 | 48660001 | 48760000 | 0.0036 | 0.0034 | 0.0035 |
| F07 | 48680001 | 48780000 | 0.0030 | 0.0028 | 0.0028 |
| F07 | 48700001 | 48800000 | 0.0028 | 0.0025 | 0.0027 |
| F07 | 48720001 | 48820000 | 0.0029 | 0.0026 | 0.0029 |
| F07 | 48740001 | 48840000 | 0.0027 | 0.0023 | 0.0026 |
| F07 | 48760001 | 48860000 | 0.0023 | 0.0019 | 0.0022 |
| F07 | 48780001 | 48880000 | 0.0026 | 0.0022 | 0.0025 |
| F07 | 48800001 | 48900000 | 0.0031 | 0.0026 | 0.0030 |
| F07 | 48820001 | 48920000 | 0.0027 | 0.0021 | 0.0026 |
| F07 | 48840001 | 48940000 | 0.0032 | 0.0028 | 0.0030 |
| F07 | 48860001 | 48960000 | 0.0031 | 0.0028 | 0.0029 |
| F07 | 48880001 | 48980000 | 0.0028 | 0.0024 | 0.0026 |
| F07 | 48900001 | 49000000 | 0.0027 | 0.0023 | 0.0025 |
| F07 | 48920001 | 49020000 | 0.0028 | 0.0024 | 0.0026 |
| F07 | 48940001 | 49040000 | 0.0022 | 0.0017 | 0.0021 |
| F07 | 48960001 | 49060000 | 0.0023 | 0.0018 | 0.0023 |

| Chr | Start | End | both | A1 | A2 |
|---|---|---|---|---|---|
| F07 | 48980001 | 49080000 | 0.0021 | 0.0017 | 0.0022 |
| F07 | 49000001 | 49100000 | 0.0020 | 0.0016 | 0.0020 |
| F07 | 49020001 | 49120000 | 0.0023 | 0.0018 | 0.0023 |
| F07 | 49040001 | 49140000 | 0.0020 | 0.0016 | 0.0020 |
| F07 | 49060001 | 49160000 | 0.0023 | 0.0019 | 0.0023 |
| F07 | 49080001 | 49180000 | 0.0024 | 0.0019 | 0.0024 |
| F07 | 49100001 | 49200000 | 0.0023 | 0.0019 | 0.0022 |
| F07 | 49120001 | 49220000 | 0.0020 | 0.0016 | 0.0019 |
| F07 | 49140001 | 49240000 | 0.0025 | 0.0021 | 0.0023 |
| F07 | 49160001 | 49260000 | 0.0023 | 0.0019 | 0.0020 |
| F07 | 49180001 | 49280000 | 0.0025 | 0.0020 | 0.0022 |
| F07 | 49200001 | 49300000 | 0.0026 | 0.0021 | 0.0023 |
| F07 | 49220001 | 49320000 | 0.0027 | 0.0023 | 0.0025 |
| F07 | 49240001 | 49340000 | 0.0029 | 0.0024 | 0.0027 |
| F07 | 49260001 | 49360000 | 0.0026 | 0.0023 | 0.0025 |
| F07 | 49280001 | 49380000 | 0.0023 | 0.0020 | 0.0021 |
| F07 | 49300001 | 49400000 | 0.0021 | 0.0018 | 0.0020 |
| F07 | 49320001 | 49420000 | 0.0019 | 0.0015 | 0.0018 |
| F07 | 49340001 | 49440000 | 0.0014 | 0.0011 | 0.0013 |
| F07 | 49360001 | 49460000 | 0.0018 | 0.0014 | 0.0016 |
| F07 | 49380001 | 49480000 | 0.0021 | 0.0017 | 0.0018 |
| F07 | 49400001 | 49500000 | 0.0025 | 0.0021 | 0.0022 |
| F07 | 49420001 | 49520000 | 0.0034 | 0.0030 | 0.0028 |
| F07 | 49440001 | 49540000 | 0.0037 | 0.0032 | 0.0030 |
| F07 | 49460001 | 49560000 | 0.0037 | 0.0031 | 0.0031 |
| F07 | 49480001 | 49580000 | 0.0037 | 0.0031 | 0.0033 |
| F07 | 49500001 | 49600000 | 0.0036 | 0.0029 | 0.0031 |
| F07 | 49520001 | 49620000 | 0.0027 | 0.0022 | 0.0026 |
| F07 | 49540001 | 49640000 | 0.0023 | 0.0018 | 0.0022 |
| F07 | 49560001 | 49660000 | 0.0018 | 0.0016 | 0.0018 |
| F07 | 49580001 | 49680000 | 0.0018 | 0.0017 | 0.0016 |
| F07 | 49600001 | 49700000 | 0.0018 | 0.0015 | 0.0015 |
| F07 | 49620001 | 49720000 | 0.0018 | 0.0014 | 0.0013 |
| F07 | 49640001 | 49740000 | 0.0029 | 0.0024 | 0.0023 |
| F07 | 49660001 | 49760000 | 0.0032 | 0.0025 | 0.0025 |
| F07 | 49680001 | 49780000 | 0.0031 | 0.0024 | 0.0025 |
| F07 | 49700001 | 49800000 | 0.0028 | 0.0023 | 0.0024 |
| F07 | 49720001 | 49820000 | 0.0034 | 0.0029 | 0.0030 |
| F07 | 49740001 | 49840000 | 0.0028 | 0.0023 | 0.0025 |
| F07 | 49760001 | 49860000 | 0.0030 | 0.0025 | 0.0027 |
| F07 | 49780001 | 49880000 | 0.0033 | 0.0027 | 0.0029 |
| F07 | 49800001 | 49900000 | 0.0033 | 0.0027 | 0.0027 |
| F07 | 49820001 | 49920000 | 0.0037 | 0.0031 | 0.0032 |
| F07 | 49840001 | 49940000 | 0.0038 | 0.0031 | 0.0032 |
| F07 | 49860001 | 49960000 | 0.0036 | 0.0028 | 0.0028 |
| F07 | 49880001 | 49980000 | 0.0039 | 0.0032 | 0.0032 |
| F07 | 49900001 | 50000000 | 0.0038 | 0.0031 | 0.0031 |
| F07 | 49920001 | 50020000 | 0.0029 | 0.0021 | 0.0021 |
| F07 | 49940001 | 50040000 | 0.0026 | 0.0019 | 0.0018 |
| F07 | 49960001 | 50060000 | 0.0026 | 0.0017 | 0.0015 |
| F07 | 49980001 | 50080000 | 0.0019 | 0.0011 | 0.0008 |
| F07 | 50000001 | 50100000 | 0.0028 | 0.0018 | 0.0016 |
| F07 | 50020001 | 50120000 | 0.0044 | 0.0035 | 0.0032 |
| F07 | 50040001 | 50140000 | 0.0056 | 0.0046 | 0.0045 |
| F07 | 50060001 | 50160000 | 0.0053 | 0.0046 | 0.0046 |
| F07 | 50080001 | 50180000 | 0.0054 | 0.0047 | 0.0047 |
| F07 | 50100001 | 50200000 | 0.0050 | 0.0045 | 0.0045 |
| F07 | 50120001 | 50220000 | 0.0033 | 0.0027 | 0.0028 |

| Chr | Start | End | both | A1 | A2 |
|---|---|---|---|---|---|
| F07 | 50140001 | 50240000 | 0.0024 | 0.0017 | 0.0016 |
| F07 | 50160001 | 50260000 | 0.0040 | 0.0032 | 0.0031 |
| F07 | 50180001 | 50280000 | 0.0042 | 0.0037 | 0.0034 |
| F07 | 50200001 | 50300000 | 0.0052 | 0.0044 | 0.0042 |
| F07 | 50220001 | 50320000 | 0.0059 | 0.0050 | 0.0049 |
| F07 | 50240001 | 50340000 | 0.0063 | 0.0055 | 0.0055 |
| F07 | 50260001 | 50360000 | 0.0055 | 0.0046 | 0.0045 |
| F07 | 50280001 | 50380000 | 0.0053 | 0.0042 | 0.0045 |
| F07 | 50300001 | 50400000 | 0.0059 | 0.0049 | 0.0052 |
| F07 | 50320001 | 50420000 | 0.0061 | 0.0052 | 0.0052 |
| F07 | 50340001 | 50440000 | 0.0066 | 0.0055 | 0.0055 |
| F07 | 50360001 | 50460000 | 0.0060 | 0.0050 | 0.0052 |
| F07 | 50380001 | 50480000 | 0.0059 | 0.0048 | 0.0050 |
| F07 | 50400001 | 50500000 | 0.0041 | 0.0031 | 0.0030 |
| F07 | 50420001 | 50520000 | 0.0039 | 0.0029 | 0.0029 |
| F07 | 50440001 | 50540000 | 0.0029 | 0.0020 | 0.0019 |
| F07 | 50460001 | 50560000 | 0.0042 | 0.0032 | 0.0030 |
| F07 | 50480001 | 50580000 | 0.0041 | 0.0032 | 0.0030 |
| F07 | 50500001 | 50600000 | 0.0048 | 0.0040 | 0.0040 |
| F07 | 50520001 | 50620000 | 0.0042 | 0.0035 | 0.0035 |
| F07 | 50540001 | 50640000 | 0.0045 | 0.0039 | 0.0039 |
| F07 | 50560001 | 50660000 | 0.0029 | 0.0025 | 0.0025 |
| F07 | 50580001 | 50680000 | 0.0031 | 0.0027 | 0.0028 |
| F07 | 50600001 | 50700000 | 0.0028 | 0.0024 | 0.0024 |
| F07 | 50620001 | 50720000 | 0.0031 | 0.0028 | 0.0028 |
| F07 | 50640001 | 50740000 | 0.0027 | 0.0023 | 0.0023 |
| F07 | 50660001 | 50760000 | 0.0030 | 0.0027 | 0.0027 |
| F07 | 50680001 | 50780000 | 0.0028 | 0.0024 | 0.0026 |
| F07 | 50700001 | 50800000 | 0.0019 | 0.0016 | 0.0017 |
| F07 | 50720001 | 50820000 | 0.0017 | 0.0014 | 0.0015 |
| F07 | 50740001 | 50840000 | 0.0015 | 0.0013 | 0.0013 |
| F07 | 50760001 | 50860000 | 0.0012 | 0.0010 | 0.0011 |
| F07 | 50780001 | 50880000 | 0.0016 | 0.0014 | 0.0014 |
| F07 | 50800001 | 50900000 | 0.0016 | 0.0014 | 0.0015 |
| F07 | 50820001 | 50920000 | 0.0013 | 0.0011 | 0.0013 |
| F07 | 50840001 | 50940000 | 0.0015 | 0.0013 | 0.0015 |
| F07 | 50860001 | 50960000 | 0.0013 | 0.0012 | 0.0013 |
| F07 | 50880001 | 50980000 | 0.0010 | 0.0008 | 0.0010 |
| F07 | 50900001 | 51000000 | 0.0012 | 0.0010 | 0.0012 |
| F07 | 50920001 | 51020000 | 0.0017 | 0.0015 | 0.0016 |
| F07 | 50940001 | 51040000 | 0.0018 | 0.0016 | 0.0017 |
| F07 | 50960001 | 51060000 | 0.0022 | 0.0019 | 0.0020 |
| F07 | 50980001 | 51080000 | 0.0024 | 0.0021 | 0.0022 |
| F07 | 51000001 | 51100000 | 0.0027 | 0.0022 | 0.0025 |
| F07 | 51020001 | 51120000 | 0.0020 | 0.0016 | 0.0018 |
| F07 | 51040001 | 51140000 | 0.0026 | 0.0022 | 0.0024 |
| F07 | 51060001 | 51160000 | 0.0028 | 0.0023 | 0.0026 |
| F07 | 51080001 | 51180000 | 0.0026 | 0.0021 | 0.0024 |
| F07 | 51100001 | 51200000 | 0.0024 | 0.0020 | 0.0022 |
| F07 | 51120001 | 51220000 | 0.0025 | 0.0021 | 0.0023 |
| F07 | 51140001 | 51240000 | 0.0016 | 0.0012 | 0.0015 |
| F07 | 51160001 | 51260000 | 0.0012 | 0.0009 | 0.0011 |
| F07 | 51180001 | 51280000 | 0.0012 | 0.0011 | 0.0012 |
| F07 | 51200001 | 51300000 | 0.0009 | 0.0008 | 0.0009 |
| F07 | 51220001 | 51320000 | 0.0016 | 0.0014 | 0.0016 |
| F07 | 51240001 | 51340000 | 0.0017 | 0.0014 | 0.0016 |
| F07 | 51260001 | 51360000 | 0.0015 | 0.0013 | 0.0015 |
| F07 | 51280001 | 51380000 | 0.0013 | 0.0011 | 0.0013 |

| Chr | Start | End | both | A1 | A2 |
|---|---|---|---|---|---|
| F07 | 51300001 | 51400000 | 0.0014 | 0.0012 | 0.0014 |
| F07 | 51320001 | 51420000 | 0.0009 | 0.0007 | 0.0009 |
| F07 | 51340001 | 51440000 | 0.0013 | 0.0012 | 0.0013 |
| F07 | 51360001 | 51460000 | 0.0019 | 0.0017 | 0.0017 |
| F07 | 51380001 | 51480000 | 0.0019 | 0.0017 | 0.0018 |
| F07 | 51400001 | 51500000 | 0.0018 | 0.0016 | 0.0017 |
| F07 | 51420001 | 51520000 | 0.0018 | 0.0018 | 0.0017 |
| F07 | 51440001 | 51540000 | 0.0024 | 0.0022 | 0.0022 |
| F07 | 51460001 | 51560000 | 0.0020 | 0.0019 | 0.0020 |
| F07 | 51480001 | 51580000 | 0.0019 | 0.0017 | 0.0018 |
| F07 | 51500001 | 51600000 | 0.0028 | 0.0025 | 0.0026 |
| F07 | 51520001 | 51620000 | 0.0031 | 0.0027 | 0.0028 |
| F07 | 51540001 | 51640000 | 0.0024 | 0.0021 | 0.0022 |
| F07 | 51560001 | 51660000 | 0.0036 | 0.0030 | 0.0033 |
| F07 | 51580001 | 51680000 | 0.0039 | 0.0034 | 0.0037 |
| F07 | 51600001 | 51700000 | 0.0030 | 0.0025 | 0.0030 |
| F07 | 51620001 | 51720000 | 0.0030 | 0.0024 | 0.0029 |
| F07 | 51640001 | 51740000 | 0.0029 | 0.0024 | 0.0029 |
| F07 | 51660001 | 51760000 | 0.0017 | 0.0014 | 0.0017 |
| F07 | 51680001 | 51780000 | 0.0014 | 0.0012 | 0.0014 |
| F07 | 51700001 | 51800000 | 0.0016 | 0.0013 | 0.0016 |
| F07 | 51720001 | 51820000 | 0.0018 | 0.0016 | 0.0018 |
| F07 | 51740001 | 51840000 | 0.0023 | 0.0020 | 0.0021 |
| F07 | 51760001 | 51860000 | 0.0028 | 0.0024 | 0.0026 |
| F07 | 51780001 | 51880000 | 0.0035 | 0.0030 | 0.0033 |
| F07 | 51800001 | 51900000 | 0.0037 | 0.0031 | 0.0034 |
| F07 | 51820001 | 51920000 | 0.0029 | 0.0024 | 0.0026 |
| F07 | 51840001 | 51940000 | 0.0024 | 0.0020 | 0.0022 |
| F07 | 51860001 | 51960000 | 0.0022 | 0.0018 | 0.0021 |
| F07 | 51880001 | 51980000 | 0.0023 | 0.0019 | 0.0022 |
| F07 | 51900001 | 52000000 | 0.0027 | 0.0023 | 0.0027 |
| F07 | 51920001 | 52020000 | 0.0037 | 0.0032 | 0.0034 |
| F07 | 51940001 | 52040000 | 0.0037 | 0.0032 | 0.0035 |
| F07 | 51960001 | 52060000 | 0.0036 | 0.0030 | 0.0032 |
| F07 | 51980001 | 52080000 | 0.0035 | 0.0030 | 0.0029 |
| F07 | 52000001 | 52100000 | 0.0030 | 0.0025 | 0.0024 |
| F07 | 52020001 | 52120000 | 0.0027 | 0.0021 | 0.0023 |
| F07 | 52040001 | 52140000 | 0.0030 | 0.0023 | 0.0025 |
| F07 | 52060001 | 52160000 | 0.0026 | 0.0020 | 0.0022 |
| F07 | 52080001 | 52180000 | 0.0022 | 0.0017 | 0.0021 |
| F07 | 52100001 | 52200000 | 0.0031 | 0.0025 | 0.0029 |
| F07 | 52120001 | 52220000 | 0.0027 | 0.0022 | 0.0024 |
| F07 | 52140001 | 52240000 | 0.0032 | 0.0028 | 0.0027 |
| F07 | 52160001 | 52260000 | 0.0039 | 0.0034 | 0.0033 |
| F07 | 52180001 | 52280000 | 0.0037 | 0.0032 | 0.0031 |
| F07 | 52200001 | 52300000 | 0.0030 | 0.0026 | 0.0026 |
| F07 | 52220001 | 52320000 | 0.0029 | 0.0026 | 0.0026 |
| F07 | 52240001 | 52340000 | 0.0024 | 0.0020 | 0.0022 |
| F07 | 52260001 | 52360000 | 0.0021 | 0.0018 | 0.0018 |
| F07 | 52280001 | 52380000 | 0.0018 | 0.0016 | 0.0017 |
| F07 | 52300001 | 52400000 | 0.0022 | 0.0021 | 0.0020 |
| F07 | 52320001 | 52420000 | 0.0025 | 0.0024 | 0.0023 |
| F07 | 52340001 | 52440000 | 0.0022 | 0.0022 | 0.0020 |
| F07 | 52360001 | 52460000 | 0.0031 | 0.0030 | 0.0029 |
| F07 | 52380001 | 52480000 | 0.0032 | 0.0031 | 0.0031 |
| F07 | 52400001 | 52500000 | 0.0028 | 0.0026 | 0.0026 |
| F07 | 52420001 | 52520000 | 0.0023 | 0.0022 | 0.0022 |
| F07 | 52440001 | 52540000 | 0.0024 | 0.0022 | 0.0022 |

| Chr | Start | End | both | A1 | A2 |
|---|---|---|---|---|---|
| F07 | 52460001 | 52560000 | 0.0018 | 0.0013 | 0.0015 |
| F07 | 52480001 | 52580000 | 0.0026 | 0.0021 | 0.0022 |
| F07 | 52500001 | 52600000 | 0.0027 | 0.0021 | 0.0022 |
| F07 | 52520001 | 52620000 | 0.0038 | 0.0028 | 0.0030 |
| F07 | 52540001 | 52640000 | 0.0038 | 0.0028 | 0.0029 |
| F07 | 52560001 | 52660000 | 0.0038 | 0.0027 | 0.0027 |
| F07 | 52580001 | 52680000 | 0.0038 | 0.0027 | 0.0027 |
| F07 | 52600001 | 52700000 | 0.0049 | 0.0034 | 0.0036 |
| F07 | 52620001 | 52720000 | 0.0053 | 0.0038 | 0.0038 |
| F07 | 52640001 | 52740000 | 0.0056 | 0.0041 | 0.0042 |
| F07 | 52660001 | 52760000 | 0.0057 | 0.0045 | 0.0045 |
| F07 | 52680001 | 52780000 | 0.0051 | 0.0040 | 0.0040 |
| F07 | 52700001 | 52800000 | 0.0043 | 0.0035 | 0.0032 |
| F07 | 52720001 | 52820000 | 0.0031 | 0.0024 | 0.0024 |
| F07 | 52740001 | 52840000 | 0.0028 | 0.0021 | 0.0019 |
| F07 | 52760001 | 52860000 | 0.0030 | 0.0022 | 0.0022 |
| F07 | 52780001 | 52880000 | 0.0030 | 0.0022 | 0.0022 |
| F07 | 52800001 | 52900000 | 0.0023 | 0.0019 | 0.0018 |
| F07 | 52820001 | 52920000 | 0.0021 | 0.0017 | 0.0017 |
| F07 | 52840001 | 52940000 | 0.0020 | 0.0018 | 0.0018 |
| F07 | 52860001 | 52960000 | 0.0016 | 0.0015 | 0.0015 |
| F07 | 52880001 | 52980000 | 0.0014 | 0.0013 | 0.0014 |
| F07 | 52900001 | 53000000 | 0.0021 | 0.0019 | 0.0020 |
| F07 | 52920001 | 53020000 | 0.0024 | 0.0022 | 0.0024 |
| F07 | 52940001 | 53040000 | 0.0025 | 0.0023 | 0.0025 |
| F07 | 52960001 | 53060000 | 0.0021 | 0.0019 | 0.0021 |
| F07 | 52980001 | 53080000 | 0.0022 | 0.0020 | 0.0020 |
| F07 | 53000001 | 53100000 | 0.0015 | 0.0013 | 0.0013 |
| F07 | 53020001 | 53120000 | 0.0012 | 0.0011 | 0.0011 |
| F07 | 53040001 | 53140000 | 0.0012 | 0.0011 | 0.0010 |
| F07 | 53060001 | 53160000 | 0.0012 | 0.0010 | 0.0010 |
| F07 | 53080001 | 53180000 | 0.0011 | 0.0009 | 0.0010 |
| F07 | 53100001 | 53200000 | 0.0011 | 0.0009 | 0.0010 |
| F07 | 53120001 | 53220000 | 0.0013 | 0.0010 | 0.0013 |
| F07 | 53140001 | 53240000 | 0.0013 | 0.0010 | 0.0013 |
| F07 | 53160001 | 53260000 | 0.0013 | 0.0010 | 0.0013 |
| F07 | 53180001 | 53280000 | 0.0021 | 0.0018 | 0.0020 |
| F07 | 53200001 | 53300000 | 0.0022 | 0.0018 | 0.0021 |
| F07 | 53220001 | 53320000 | 0.0023 | 0.0021 | 0.0021 |
| F07 | 53240001 | 53340000 | 0.0022 | 0.0020 | 0.0020 |
| F07 | 53260001 | 53360000 | 0.0022 | 0.0020 | 0.0020 |
| F07 | 53280001 | 53380000 | 0.0024 | 0.0020 | 0.0022 |
| F07 | 53300001 | 53400000 | 0.0027 | 0.0023 | 0.0024 |
| F07 | 53320001 | 53420000 | 0.0030 | 0.0024 | 0.0027 |
| F07 | 53340001 | 53440000 | 0.0033 | 0.0026 | 0.0030 |
| F07 | 53360001 | 53460000 | 0.0038 | 0.0031 | 0.0034 |
| F07 | 53380001 | 53480000 | 0.0030 | 0.0023 | 0.0025 |
| F07 | 53400001 | 53500000 | 0.0030 | 0.0023 | 0.0023 |
| F07 | 53420001 | 53520000 | 0.0041 | 0.0034 | 0.0034 |
| F07 | 53440001 | 53540000 | 0.0043 | 0.0036 | 0.0036 |
| F07 | 53460001 | 53560000 | 0.0043 | 0.0034 | 0.0036 |
| F07 | 53480001 | 53580000 | 0.0046 | 0.0038 | 0.0040 |
| F07 | 53500001 | 53600000 | 0.0056 | 0.0048 | 0.0050 |
| F07 | 53520001 | 53620000 | 0.0039 | 0.0033 | 0.0035 |
| F07 | 53540001 | 53640000 | 0.0043 | 0.0035 | 0.0038 |
| F07 | 53560001 | 53660000 | 0.0050 | 0.0043 | 0.0045 |
| F07 | 53580001 | 53680000 | 0.0052 | 0.0042 | 0.0048 |
| F07 | 53600001 | 53700000 | 0.0039 | 0.0030 | 0.0036 |

| Chr | Start | End | both | A1 | A2 |
|---|---|---|---|---|---|
| F07 | 53620001 | 53720000 | 0.0040 | 0.0032 | 0.0038 |
| F07 | 53640001 | 53740000 | 0.0038 | 0.0030 | 0.0035 |
| F07 | 53660001 | 53760000 | 0.0034 | 0.0026 | 0.0033 |
| F07 | 53680001 | 53780000 | 0.0035 | 0.0028 | 0.0035 |
| F07 | 53700001 | 53800000 | 0.0038 | 0.0031 | 0.0037 |
| F07 | 53720001 | 53820000 | 0.0035 | 0.0029 | 0.0035 |
| F07 | 53740001 | 53840000 | 0.0039 | 0.0033 | 0.0037 |
| F07 | 53760001 | 53860000 | 0.0039 | 0.0034 | 0.0036 |
| F07 | 53780001 | 53880000 | 0.0035 | 0.0030 | 0.0031 |
| F07 | 53800001 | 53900000 | 0.0033 | 0.0027 | 0.0028 |
| F07 | 53820001 | 53920000 | 0.0038 | 0.0033 | 0.0033 |
| F07 | 53840001 | 53940000 | 0.0032 | 0.0029 | 0.0029 |
| F07 | 53860001 | 53960000 | 0.0026 | 0.0024 | 0.0023 |
| F07 | 53880001 | 53980000 | 0.0019 | 0.0019 | 0.0018 |
| F07 | 53900001 | 54000000 | 0.0025 | 0.0025 | 0.0023 |
| F07 | 53920001 | 54020000 | 0.0026 | 0.0025 | 0.0025 |
| F07 | 53940001 | 54040000 | 0.0028 | 0.0026 | 0.0026 |
| F07 | 53960001 | 54060000 | 0.0032 | 0.0027 | 0.0028 |
| F07 | 53980001 | 54080000 | 0.0039 | 0.0033 | 0.0034 |
| F07 | 54000001 | 54100000 | 0.0043 | 0.0036 | 0.0036 |
| F07 | 54020001 | 54120000 | 0.0038 | 0.0030 | 0.0029 |
| F07 | 54040001 | 54140000 | 0.0046 | 0.0036 | 0.0036 |
| F07 | 54060001 | 54160000 | 0.0051 | 0.0042 | 0.0042 |
| F07 | 54080001 | 54180000 | 0.0066 | 0.0056 | 0.0056 |
| F07 | 54100001 | 54200000 | 0.0064 | 0.0054 | 0.0054 |
| F07 | 54120001 | 54220000 | 0.0065 | 0.0056 | 0.0057 |
| F07 | 54140001 | 54240000 | 0.0057 | 0.0048 | 0.0048 |
| F07 | 54160001 | 54260000 | 0.0050 | 0.0042 | 0.0040 |
| F07 | 54180001 | 54280000 | 0.0033 | 0.0024 | 0.0022 |
| F07 | 54200001 | 54300000 | 0.0029 | 0.0020 | 0.0020 |
| F07 | 54220001 | 54320000 | 0.0032 | 0.0023 | 0.0022 |
| F07 | 54240001 | 54340000 | 0.0030 | 0.0023 | 0.0022 |
| F07 | 54260001 | 54360000 | 0.0040 | 0.0031 | 0.0032 |
| F07 | 54280001 | 54380000 | 0.0041 | 0.0035 | 0.0035 |
| F07 | 54300001 | 54400000 | 0.0042 | 0.0034 | 0.0036 |
| F07 | 54320001 | 54420000 | 0.0045 | 0.0036 | 0.0037 |
| F07 | 54340001 | 54440000 | 0.0045 | 0.0035 | 0.0037 |
| F07 | 54360001 | 54460000 | 0.0032 | 0.0025 | 0.0026 |
| F07 | 54380001 | 54480000 | 0.0032 | 0.0024 | 0.0025 |
| F07 | 54400001 | 54500000 | 0.0040 | 0.0031 | 0.0030 |
| F07 | 54420001 | 54520000 | 0.0042 | 0.0032 | 0.0031 |
| F07 | 54440001 | 54540000 | 0.0040 | 0.0031 | 0.0030 |
| F07 | 54460001 | 54560000 | 0.0048 | 0.0037 | 0.0033 |
| F07 | 54480001 | 54580000 | 0.0052 | 0.0041 | 0.0038 |
| F07 | 54500001 | 54600000 | 0.0042 | 0.0033 | 0.0031 |
| F07 | 54520001 | 54620000 | 0.0034 | 0.0026 | 0.0023 |
| F07 | 54540001 | 54640000 | 0.0036 | 0.0029 | 0.0024 |
| F07 | 54560001 | 54660000 | 0.0043 | 0.0038 | 0.0034 |
| F07 | 54580001 | 54680000 | 0.0038 | 0.0033 | 0.0027 |
| F07 | 54600001 | 54700000 | 0.0039 | 0.0032 | 0.0027 |
| F07 | 54620001 | 54720000 | 0.0039 | 0.0031 | 0.0026 |
| F07 | 54640001 | 54740000 | 0.0043 | 0.0034 | 0.0030 |
| F07 | 54660001 | 54760000 | 0.0031 | 0.0020 | 0.0019 |
| F07 | 54680001 | 54780000 | 0.0036 | 0.0026 | 0.0024 |
| F07 | 54700001 | 54800000 | 0.0044 | 0.0035 | 0.0032 |
| F07 | 54720001 | 54820000 | 0.0053 | 0.0044 | 0.0043 |
| F07 | 54740001 | 54840000 | 0.0050 | 0.0043 | 0.0042 |
| F07 | 54760001 | 54860000 | 0.0052 | 0.0045 | 0.0044 |

| Chr | Start | End | both | A1 | A2 |
|---|---|---|---|---|---|
| F07 | 54780001 | 54880000 | 0.0052 | 0.0046 | 0.0045 |
| F07 | 54800001 | 54900000 | 0.0046 | 0.0040 | 0.0040 |
| F07 | 54820001 | 54920000 | 0.0043 | 0.0038 | 0.0037 |
| F07 | 54840001 | 54940000 | 0.0045 | 0.0039 | 0.0038 |
| F07 | 54860001 | 54960000 | 0.0044 | 0.0039 | 0.0036 |
| F07 | 54880001 | 54980000 | 0.0033 | 0.0030 | 0.0027 |
| F07 | 54900001 | 55000000 | 0.0029 | 0.0025 | 0.0023 |
| F07 | 54920001 | 55020000 | 0.0031 | 0.0027 | 0.0027 |
| F07 | 54940001 | 55040000 | 0.0022 | 0.0020 | 0.0020 |
| F07 | 54960001 | 55060000 | 0.0018 | 0.0016 | 0.0016 |
| F07 | 54980001 | 55080000 | 0.0017 | 0.0015 | 0.0016 |
| F07 | 55000001 | 55100000 | 0.0015 | 0.0014 | 0.0015 |
| F07 | 55020001 | 55120000 | 0.0006 | 0.0005 | 0.0005 |
| F07 | 55040001 | 55140000 | 0.0007 | 0.0006 | 0.0007 |
| F07 | 55060001 | 55160000 | 0.0011 | 0.0009 | 0.0011 |
| F07 | 55080001 | 55180000 | 0.0015 | 0.0012 | 0.0015 |
| F07 | 55100001 | 55200000 | 0.0015 | 0.0011 | 0.0015 |
| F07 | 55120001 | 55220000 | 0.0020 | 0.0018 | 0.0020 |
| F07 | 55140001 | 55240000 | 0.0019 | 0.0017 | 0.0018 |
| F07 | 55160001 | 55260000 | 0.0020 | 0.0017 | 0.0019 |
| F07 | 55180001 | 55280000 | 0.0018 | 0.0017 | 0.0018 |
| F07 | 55200001 | 55300000 | 0.0021 | 0.0019 | 0.0020 |
| F07 | 55220001 | 55320000 | 0.0034 | 0.0029 | 0.0033 |
| F07 | 55240001 | 55340000 | 0.0042 | 0.0036 | 0.0041 |
| F07 | 55260001 | 55360000 | 0.0041 | 0.0034 | 0.0040 |
| F07 | 55280001 | 55380000 | 0.0041 | 0.0032 | 0.0038 |
| F07 | 55300001 | 55400000 | 0.0049 | 0.0039 | 0.0044 |
| F07 | 55320001 | 55420000 | 0.0046 | 0.0037 | 0.0041 |
| F07 | 55340001 | 55440000 | 0.0043 | 0.0034 | 0.0039 |
| F07 | 55360001 | 55460000 | 0.0049 | 0.0039 | 0.0043 |
| F07 | 55380001 | 55480000 | 0.0048 | 0.0039 | 0.0043 |
| F07 | 55400001 | 55500000 | 0.0040 | 0.0032 | 0.0037 |
| F07 | 55420001 | 55520000 | 0.0025 | 0.0018 | 0.0021 |
| F07 | 55440001 | 55540000 | 0.0025 | 0.0019 | 0.0022 |
| F07 | 55460001 | 55560000 | 0.0027 | 0.0021 | 0.0023 |
| F07 | 55480001 | 55580000 | 0.0031 | 0.0024 | 0.0028 |
| F07 | 55500001 | 55600000 | 0.0037 | 0.0030 | 0.0033 |
| F07 | 55520001 | 55620000 | 0.0042 | 0.0033 | 0.0037 |
| F07 | 55540001 | 55640000 | 0.0038 | 0.0029 | 0.0033 |
| F07 | 55560001 | 55660000 | 0.0037 | 0.0028 | 0.0032 |
| F07 | 55580001 | 55680000 | 0.0042 | 0.0034 | 0.0037 |
| F07 | 55600001 | 55700000 | 0.0037 | 0.0030 | 0.0033 |
| F07 | 55620001 | 55720000 | 0.0048 | 0.0038 | 0.0045 |
| F07 | 55640001 | 55740000 | 0.0052 | 0.0042 | 0.0048 |
| F07 | 55660001 | 55760000 | 0.0050 | 0.0041 | 0.0048 |
| F07 | 55680001 | 55780000 | 0.0040 | 0.0031 | 0.0038 |
| F07 | 55700001 | 55800000 | 0.0039 | 0.0031 | 0.0037 |
| F07 | 55720001 | 55820000 | 0.0024 | 0.0020 | 0.0021 |
| F07 | 55740001 | 55840000 | 0.0019 | 0.0016 | 0.0017 |
| F07 | 55760001 | 55860000 | 0.0013 | 0.0011 | 0.0011 |
| F07 | 55780001 | 55880000 | 0.0025 | 0.0021 | 0.0023 |
| F07 | 55800001 | 55900000 | 0.0026 | 0.0024 | 0.0023 |
| F07 | 55820001 | 55920000 | 0.0026 | 0.0024 | 0.0024 |
| F07 | 55840001 | 55940000 | 0.0031 | 0.0029 | 0.0028 |
| F07 | 55860001 | 55960000 | 0.0035 | 0.0032 | 0.0032 |
| F07 | 55880001 | 55980000 | 0.0025 | 0.0023 | 0.0022 |
| F07 | 55900001 | 56000000 | 0.0025 | 0.0021 | 0.0021 |
| F07 | 55920001 | 56020000 | 0.0029 | 0.0025 | 0.0024 |

| Chr | Start | End | both | A1 | A2 |
|---|---|---|---|---|---|
| F07 | 55940001 | 56040000 | 0.0035 | 0.0029 | 0.0028 |
| F07 | 55960001 | 56060000 | 0.0036 | 0.0030 | 0.0029 |
| F07 | 55980001 | 56080000 | 0.0043 | 0.0038 | 0.0036 |
| F07 | 56000001 | 56100000 | 0.0043 | 0.0038 | 0.0037 |
| F07 | 56020001 | 56120000 | 0.0036 | 0.0032 | 0.0033 |
| F07 | 56040001 | 56140000 | 0.0025 | 0.0023 | 0.0023 |
| F07 | 56060001 | 56160000 | 0.0024 | 0.0021 | 0.0023 |
| F07 | 56080001 | 56180000 | 0.0028 | 0.0025 | 0.0026 |
| F07 | 56100001 | 56200000 | 0.0032 | 0.0029 | 0.0031 |
| F07 | 56120001 | 56220000 | 0.0035 | 0.0031 | 0.0033 |
| F07 | 56140001 | 56240000 | 0.0039 | 0.0034 | 0.0037 |
| F07 | 56160001 | 56260000 | 0.0041 | 0.0038 | 0.0039 |
| F07 | 56180001 | 56280000 | 0.0029 | 0.0026 | 0.0029 |
| F07 | 56200001 | 56300000 | 0.0022 | 0.0020 | 0.0021 |
| F07 | 56220001 | 56320000 | 0.0023 | 0.0020 | 0.0020 |
| F07 | 56240001 | 56340000 | 0.0017 | 0.0015 | 0.0015 |
| F07 | 56260001 | 56360000 | 0.0009 | 0.0006 | 0.0007 |
| F07 | 56280001 | 56380000 | 0.0013 | 0.0010 | 0.0010 |
| F07 | 56300001 | 56400000 | 0.0018 | 0.0015 | 0.0015 |
| F07 | 56320001 | 56420000 | 0.0024 | 0.0021 | 0.0023 |
| F07 | 56340001 | 56440000 | 0.0034 | 0.0030 | 0.0033 |
| F07 | 56360001 | 56460000 | 0.0038 | 0.0033 | 0.0037 |
| F07 | 56380001 | 56480000 | 0.0032 | 0.0028 | 0.0031 |
| F07 | 56400001 | 56500000 | 0.0029 | 0.0025 | 0.0028 |
| F07 | 56420001 | 56520000 | 0.0026 | 0.0023 | 0.0024 |
| F07 | 56440001 | 56540000 | 0.0018 | 0.0015 | 0.0016 |
| F07 | 56460001 | 56560000 | 0.0028 | 0.0022 | 0.0026 |
| F07 | 56480001 | 56580000 | 0.0032 | 0.0026 | 0.0030 |
| F07 | 56500001 | 56600000 | 0.0037 | 0.0030 | 0.0035 |
| F07 | 56520001 | 56620000 | 0.0032 | 0.0026 | 0.0032 |
| F07 | 56540001 | 56640000 | 0.0032 | 0.0027 | 0.0032 |
| F07 | 56560001 | 56660000 | 0.0022 | 0.0020 | 0.0022 |
| F07 | 56580001 | 56680000 | 0.0022 | 0.0020 | 0.0022 |
| F07 | 56600001 | 56700000 | 0.0023 | 0.0020 | 0.0021 |
| F07 | 56620001 | 56720000 | 0.0026 | 0.0021 | 0.0023 |
| F07 | 56640001 | 56740000 | 0.0031 | 0.0027 | 0.0027 |
| F07 | 56660001 | 56760000 | 0.0040 | 0.0036 | 0.0035 |
| F07 | 56680001 | 56780000 | 0.0047 | 0.0042 | 0.0041 |
| F07 | 56700001 | 56800000 | 0.0046 | 0.0040 | 0.0039 |
| F07 | 56720001 | 56820000 | 0.0056 | 0.0051 | 0.0050 |
| F07 | 56740001 | 56840000 | 0.0051 | 0.0046 | 0.0046 |
| F07 | 56760001 | 56860000 | 0.0043 | 0.0039 | 0.0038 |
| F07 | 56780001 | 56880000 | 0.0041 | 0.0038 | 0.0036 |
| F07 | 56800001 | 56900000 | 0.0037 | 0.0034 | 0.0033 |
| F07 | 56820001 | 56920000 | 0.0029 | 0.0026 | 0.0024 |
| F07 | 56840001 | 56940000 | 0.0033 | 0.0028 | 0.0027 |
| F07 | 56860001 | 56960000 | 0.0031 | 0.0025 | 0.0025 |
| F07 | 56880001 | 56980000 | 0.0024 | 0.0018 | 0.0018 |
| F07 | 56900001 | 57000000 | 0.0026 | 0.0020 | 0.0020 |
| F07 | 56920001 | 57020000 | 0.0020 | 0.0014 | 0.0014 |
| F07 | 56940001 | 57040000 | 0.0018 | 0.0014 | 0.0013 |
| F07 | 56960001 | 57060000 | 0.0016 | 0.0013 | 0.0012 |
| F07 | 56980001 | 57080000 | 0.0018 | 0.0015 | 0.0014 |
| F07 | 57000001 | 57100000 | 0.0013 | 0.0011 | 0.0010 |
| F07 | 57020001 | 57120000 | 0.0020 | 0.0018 | 0.0018 |
| F07 | 57040001 | 57140000 | 0.0025 | 0.0022 | 0.0023 |
| F07 | 57060001 | 57160000 | 0.0029 | 0.0024 | 0.0025 |
| F07 | 57080001 | 57180000 | 0.0031 | 0.0026 | 0.0027 |

| Chr | Start | End | both | A1 | A2 |
|---|---|---|---|---|---|
| F07 | 57100001 | 57200000 | 0.0037 | 0.0030 | 0.0032 |
| F07 | 57120001 | 57220000 | 0.0033 | 0.0027 | 0.0027 |
| F07 | 57140001 | 57240000 | 0.0038 | 0.0030 | 0.0031 |
| F07 | 57160001 | 57260000 | 0.0049 | 0.0040 | 0.0043 |
| F07 | 57180001 | 57280000 | 0.0047 | 0.0038 | 0.0041 |
| F07 | 57200001 | 57300000 | 0.0043 | 0.0035 | 0.0038 |
| F07 | 57220001 | 57320000 | 0.0047 | 0.0039 | 0.0043 |
| F07 | 57240001 | 57340000 | 0.0044 | 0.0035 | 0.0041 |
| F07 | 57260001 | 57360000 | 0.0035 | 0.0028 | 0.0032 |
| F07 | 57280001 | 57380000 | 0.0034 | 0.0028 | 0.0032 |
| F07 | 57300001 | 57400000 | 0.0038 | 0.0031 | 0.0034 |
| F07 | 57320001 | 57420000 | 0.0031 | 0.0024 | 0.0027 |
| F07 | 57340001 | 57440000 | 0.0024 | 0.0020 | 0.0021 |
| F07 | 57360001 | 57460000 | 0.0022 | 0.0018 | 0.0019 |
| F07 | 57380001 | 57480000 | 0.0023 | 0.0018 | 0.0018 |
| F07 | 57400001 | 57500000 | 0.0019 | 0.0014 | 0.0015 |
| F07 | 57420001 | 57520000 | 0.0026 | 0.0022 | 0.0021 |
| F07 | 57440001 | 57540000 | 0.0034 | 0.0027 | 0.0027 |
| F07 | 57460001 | 57560000 | 0.0039 | 0.0030 | 0.0030 |
| F07 | 57480001 | 57580000 | 0.0041 | 0.0031 | 0.0031 |
| F07 | 57500001 | 57600000 | 0.0048 | 0.0039 | 0.0039 |
| F07 | 57520001 | 57620000 | 0.0046 | 0.0036 | 0.0037 |
| F07 | 57540001 | 57640000 | 0.0038 | 0.0030 | 0.0030 |
| F07 | 57560001 | 57660000 | 0.0033 | 0.0028 | 0.0027 |
| F07 | 57580001 | 57680000 | 0.0031 | 0.0028 | 0.0027 |
| F07 | 57600001 | 57700000 | 0.0020 | 0.0018 | 0.0018 |
| F07 | 57620001 | 57720000 | 0.0013 | 0.0013 | 0.0013 |
| F07 | 57640001 | 57740000 | 0.0012 | 0.0011 | 0.0012 |
| F07 | 57660001 | 57760000 | 0.0014 | 0.0011 | 0.0014 |
| F07 | 57680001 | 57780000 | 0.0015 | 0.0013 | 0.0015 |
| F07 | 57700001 | 57800000 | 0.0020 | 0.0018 | 0.0020 |
| F07 | 57720001 | 57820000 | 0.0018 | 0.0016 | 0.0018 |
| F07 | 57740001 | 57840000 | 0.0019 | 0.0017 | 0.0019 |
| F07 | 57760001 | 57860000 | 0.0016 | 0.0014 | 0.0015 |
| F07 | 57780001 | 57880000 | 0.0013 | 0.0013 | 0.0012 |
| F07 | 57800001 | 57900000 | 0.0011 | 0.0011 | 0.0010 |
| F07 | 57820001 | 57920000 | 0.0012 | 0.0012 | 0.0011 |
| F07 | 57840001 | 57940000 | 0.0011 | 0.0011 | 0.0010 |
| F07 | 57860001 | 57960000 | 0.0013 | 0.0014 | 0.0012 |
| F07 | 57880001 | 57980000 | 0.0012 | 0.0011 | 0.0011 |
| F07 | 57900001 | 58000000 | 0.0009 | 0.0009 | 0.0008 |
| F07 | 57920001 | 58020000 | 0.0011 | 0.0011 | 0.0010 |
| F07 | 57940001 | 58040000 | 0.0012 | 0.0012 | 0.0012 |
| F07 | 57960001 | 58060000 | 0.0014 | 0.0013 | 0.0014 |
| F07 | 57980001 | 58080000 | 0.0015 | 0.0014 | 0.0014 |
| F07 | 58000001 | 58100000 | 0.0015 | 0.0015 | 0.0015 |
| F07 | 58020001 | 58120000 | 0.0015 | 0.0014 | 0.0015 |
| F07 | 58040001 | 58140000 | 0.0015 | 0.0014 | 0.0015 |
| F07 | 58060001 | 58160000 | 0.0009 | 0.0008 | 0.0008 |
| F07 | 58080001 | 58180000 | 0.0009 | 0.0008 | 0.0009 |
| F07 | 58100001 | 58200000 | 0.0009 | 0.0008 | 0.0009 |
| F07 | 58120001 | 58220000 | 0.0007 | 0.0005 | 0.0007 |
| F07 | 58140001 | 58240000 | 0.0008 | 0.0007 | 0.0008 |
| F07 | 58160001 | 58260000 | 0.0012 | 0.0010 | 0.0012 |
| F07 | 58180001 | 58280000 | 0.0020 | 0.0018 | 0.0019 |
| F07 | 58200001 | 58300000 | 0.0021 | 0.0019 | 0.0021 |
| F07 | 58220001 | 58320000 | 0.0028 | 0.0025 | 0.0026 |
| F07 | 58240001 | 58340000 | 0.0031 | 0.0027 | 0.0027 |

| Chr | Start | End | both | A1 | A2 |
|---|---|---|---|---|---|
| F07 | 58260001 | 58360000 | 0.0051 | 0.0044 | 0.0047 |
| F07 | 58280001 | 58380000 | 0.0049 | 0.0042 | 0.0044 |
| F07 | 58300001 | 58400000 | 0.0049 | 0.0042 | 0.0043 |
| F07 | 58320001 | 58420000 | 0.0047 | 0.0038 | 0.0039 |
| F07 | 58340001 | 58440000 | 0.0046 | 0.0037 | 0.0039 |
| F07 | 58360001 | 58460000 | 0.0030 | 0.0024 | 0.0024 |
| F07 | 58380001 | 58480000 | 0.0026 | 0.0020 | 0.0020 |
| F07 | 58400001 | 58500000 | 0.0031 | 0.0025 | 0.0027 |
| F07 | 58420001 | 58520000 | 0.0029 | 0.0025 | 0.0028 |
| F07 | 58440001 | 58540000 | 0.0027 | 0.0024 | 0.0026 |
| F07 | 58460001 | 58560000 | 0.0023 | 0.0021 | 0.0023 |
| F07 | 58480001 | 58580000 | 0.0030 | 0.0028 | 0.0029 |
| F07 | 58500001 | 58600000 | 0.0028 | 0.0027 | 0.0027 |
| F07 | 58520001 | 58620000 | 0.0034 | 0.0031 | 0.0033 |
| F07 | 58540001 | 58640000 | 0.0036 | 0.0033 | 0.0035 |
| F07 | 58560001 | 58660000 | 0.0037 | 0.0034 | 0.0036 |
| F07 | 58580001 | 58680000 | 0.0037 | 0.0032 | 0.0035 |
| F07 | 58600001 | 58700000 | 0.0041 | 0.0034 | 0.0037 |
| F07 | 58620001 | 58720000 | 0.0040 | 0.0033 | 0.0036 |
| F07 | 58640001 | 58740000 | 0.0040 | 0.0031 | 0.0036 |
| F07 | 58660001 | 58760000 | 0.0037 | 0.0029 | 0.0033 |
| F07 | 58680001 | 58780000 | 0.0033 | 0.0027 | 0.0029 |
| F07 | 58700001 | 58800000 | 0.0026 | 0.0022 | 0.0024 |
| F07 | 58720001 | 58820000 | 0.0019 | 0.0017 | 0.0018 |
| F07 | 58740001 | 58840000 | 0.0014 | 0.0012 | 0.0012 |
| F07 | 58760001 | 58860000 | 0.0011 | 0.0009 | 0.0010 |
| F07 | 58780001 | 58880000 | 0.0009 | 0.0006 | 0.0008 |
| F07 | 58800001 | 58900000 | 0.0006 | 0.0004 | 0.0006 |
| F07 | 58820001 | 58920000 | 0.0006 | 0.0004 | 0.0006 |
| F07 | 58840001 | 58940000 | 0.0007 | 0.0005 | 0.0007 |
| F07 | 58860001 | 58960000 | 0.0010 | 0.0008 | 0.0010 |
| F07 | 58880001 | 58980000 | 0.0011 | 0.0010 | 0.0011 |
| F07 | 58900001 | 59000000 | 0.0009 | 0.0008 | 0.0009 |
| F07 | 58920001 | 59020000 | 0.0011 | 0.0010 | 0.0011 |
| F07 | 58940001 | 59040000 | 0.0013 | 0.0011 | 0.0013 |
| F07 | 58960001 | 59060000 | 0.0012 | 0.0011 | 0.0012 |
| F07 | 58980001 | 59080000 | 0.0011 | 0.0010 | 0.0011 |
| F07 | 59000001 | 59100000 | 0.0011 | 0.0010 | 0.0011 |
| F07 | 59020001 | 59120000 | 0.0010 | 0.0009 | 0.0010 |
| F07 | 59040001 | 59140000 | 0.0008 | 0.0008 | 0.0008 |
| F07 | 59060001 | 59160000 | 0.0008 | 0.0007 | 0.0008 |
| F07 | 59080001 | 59180000 | 0.0008 | 0.0007 | 0.0008 |
| F07 | 59100001 | 59200000 | 0.0012 | 0.0010 | 0.0012 |
| F07 | 59120001 | 59220000 | 0.0014 | 0.0011 | 0.0014 |
| F07 | 59140001 | 59240000 | 0.0017 | 0.0015 | 0.0017 |
| F07 | 59160001 | 59260000 | 0.0018 | 0.0015 | 0.0018 |
| F07 | 59180001 | 59280000 | 0.0021 | 0.0018 | 0.0021 |
| F07 | 59200001 | 59300000 | 0.0018 | 0.0015 | 0.0018 |
| F07 | 59220001 | 59320000 | 0.0015 | 0.0014 | 0.0015 |
| F07 | 59240001 | 59340000 | 0.0011 | 0.0010 | 0.0011 |
| F07 | 59260001 | 59360000 | 0.0011 | 0.0010 | 0.0011 |
| F07 | 59280001 | 59380000 | 0.0008 | 0.0007 | 0.0008 |
| F07 | 59300001 | 59400000 | 0.0008 | 0.0007 | 0.0008 |
| F07 | 59320001 | 59420000 | 0.0013 | 0.0011 | 0.0013 |
| F07 | 59340001 | 59440000 | 0.0015 | 0.0012 | 0.0015 |
| F07 | 59360001 | 59460000 | 0.0018 | 0.0015 | 0.0018 |
| F07 | 59380001 | 59480000 | 0.0018 | 0.0014 | 0.0018 |
| F07 | 59400001 | 59500000 | 0.0019 | 0.0015 | 0.0019 |

| Chr | Start | End | both | A1 | A2 |
|---|---|---|---|---|---|
| F07 | 59420001 | 59520000 | 0.0015 | 0.0012 | 0.0015 |
| F07 | 59440001 | 59540000 | 0.0015 | 0.0012 | 0.0015 |
| F07 | 59460001 | 59560000 | 0.0010 | 0.0008 | 0.0010 |
| F07 | 59480001 | 59580000 | 0.0008 | 0.0007 | 0.0008 |
| F07 | 59500001 | 59600000 | 0.0009 | 0.0008 | 0.0009 |
| F07 | 59520001 | 59620000 | 0.0015 | 0.0014 | 0.0015 |
| F07 | 59540001 | 59640000 | 0.0013 | 0.0013 | 0.0013 |
| F07 | 59560001 | 59660000 | 0.0012 | 0.0011 | 0.0011 |
| F07 | 59580001 | 59680000 | 0.0011 | 0.0010 | 0.0010 |
| F07 | 59600001 | 59700000 | 0.0010 | 0.0009 | 0.0010 |
| F07 | 59620001 | 59720000 | 0.0004 | 0.0004 | 0.0004 |
| F07 | 59640001 | 59740000 | 0.0013 | 0.0011 | 0.0013 |
| F07 | 59660001 | 59760000 | 0.0014 | 0.0012 | 0.0014 |
| F07 | 59680001 | 59780000 | 0.0017 | 0.0015 | 0.0017 |
| F07 | 59700001 | 59800000 | 0.0016 | 0.0014 | 0.0016 |
| F07 | 59720001 | 59820000 | 0.0015 | 0.0014 | 0.0015 |
| F07 | 59740001 | 59840000 | 0.0008 | 0.0008 | 0.0007 |
| F07 | 59760001 | 59860000 | 0.0009 | 0.0009 | 0.0009 |
| F07 | 59780001 | 59880000 | 0.0009 | 0.0008 | 0.0009 |
| F07 | 59800001 | 59900000 | 0.0009 | 0.0009 | 0.0009 |
| F07 | 59820001 | 59920000 | 0.0008 | 0.0007 | 0.0008 |
| F07 | 59840001 | 59940000 | 0.0013 | 0.0010 | 0.0013 |
| F07 | 59860001 | 59960000 | 0.0012 | 0.0010 | 0.0012 |
| F07 | 59880001 | 59980000 | 0.0011 | 0.0009 | 0.0010 |
| F07 | 59900001 | 60000000 | 0.0013 | 0.0009 | 0.0012 |
| F07 | 59920001 | 60020000 | 0.0018 | 0.0014 | 0.0018 |
| F07 | 59940001 | 60040000 | 0.0020 | 0.0016 | 0.0018 |
| F07 | 59960001 | 60060000 | 0.0020 | 0.0016 | 0.0018 |
| F07 | 59980001 | 60080000 | 0.0024 | 0.0019 | 0.0022 |
| F07 | 60000001 | 60100000 | 0.0023 | 0.0019 | 0.0022 |
| F07 | 60020001 | 60120000 | 0.0021 | 0.0017 | 0.0019 |
| F07 | 60040001 | 60140000 | 0.0013 | 0.0011 | 0.0013 |
| F07 | 60060001 | 60160000 | 0.0013 | 0.0010 | 0.0013 |
| F07 | 60080001 | 60180000 | 0.0008 | 0.0006 | 0.0008 |
| F07 | 60100001 | 60200000 | 0.0005 | 0.0005 | 0.0005 |
| F07 | 60120001 | 60220000 | 0.0005 | 0.0004 | 0.0005 |
| F07 | 60140001 | 60240000 | 0.0008 | 0.0008 | 0.0008 |
| F07 | 60160001 | 60260000 | 0.0008 | 0.0007 | 0.0007 |
| F07 | 60180001 | 60280000 | 0.0009 | 0.0008 | 0.0008 |
| F07 | 60200001 | 60300000 | 0.0014 | 0.0012 | 0.0013 |
| F07 | 60220001 | 60320000 | 0.0017 | 0.0014 | 0.0013 |
| F07 | 60240001 | 60340000 | 0.0015 | 0.0012 | 0.0012 |
| F07 | 60260001 | 60360000 | 0.0018 | 0.0016 | 0.0014 |
| F07 | 60280001 | 60380000 | 0.0025 | 0.0021 | 0.0019 |
| F07 | 60300001 | 60400000 | 0.0027 | 0.0021 | 0.0018 |
| F07 | 60320001 | 60420000 | 0.0032 | 0.0027 | 0.0023 |
| F07 | 60340001 | 60440000 | 0.0035 | 0.0029 | 0.0026 |
| F07 | 60360001 | 60460000 | 0.0034 | 0.0028 | 0.0027 |
| F07 | 60380001 | 60480000 | 0.0031 | 0.0025 | 0.0025 |
| F07 | 60400001 | 60500000 | 0.0025 | 0.0022 | 0.0022 |
| F07 | 60420001 | 60520000 | 0.0020 | 0.0018 | 0.0020 |
| F07 | 60440001 | 60540000 | 0.0018 | 0.0016 | 0.0018 |
| F07 | 60460001 | 60560000 | 0.0015 | 0.0013 | 0.0014 |
| F07 | 60480001 | 60580000 | 0.0011 | 0.0009 | 0.0011 |
| F07 | 60500001 | 60600000 | 0.0013 | 0.0012 | 0.0013 |
| F07 | 60520001 | 60620000 | 0.0013 | 0.0011 | 0.0013 |
| F07 | 60540001 | 60640000 | 0.0010 | 0.0009 | 0.0010 |
| F07 | 60560001 | 60660000 | 0.0013 | 0.0011 | 0.0013 |

| Chr | Start | End | both | A1 | A2 |
|---|---|---|---|---|---|
| F07 | 60580001 | 60680000 | 0.0013 | 0.0011 | 0.0013 |
| F07 | 60600001 | 60700000 | 0.0011 | 0.0010 | 0.0011 |
| F07 | 60620001 | 60720000 | 0.0006 | 0.0006 | 0.0006 |
| F07 | 60640001 | 60740000 | 0.0007 | 0.0006 | 0.0007 |
| F07 | 60660001 | 60760000 | 0.0008 | 0.0005 | 0.0008 |
| F07 | 60680001 | 60780000 | 0.0008 | 0.0005 | 0.0008 |
| F07 | 60700001 | 60800000 | 0.0008 | 0.0005 | 0.0008 |
| F07 | 60720001 | 60820000 | 0.0007 | 0.0004 | 0.0007 |
| F07 | 60740001 | 60840000 | 0.0007 | 0.0004 | 0.0007 |
| F07 | 60760001 | 60860000 | 0.0002 | 0.0002 | 0.0002 |
| F07 | 60780001 | 60880000 | 0.0005 | 0.0003 | 0.0005 |
| F07 | 60800001 | 60900000 | 0.0005 | 0.0004 | 0.0005 |
| F07 | 60820001 | 60920000 | 0.0005 | 0.0004 | 0.0005 |
| F07 | 60840001 | 60940000 | 0.0005 | 0.0004 | 0.0005 |
| F07 | 60860001 | 60960000 | 0.0005 | 0.0004 | 0.0005 |
| F07 | 60880001 | 60980000 | 0.0002 | 0.0002 | 0.0002 |
| F07 | 60900001 | 61000000 | 0.0002 | 0.0001 | 0.0002 |
| F07 | 60920001 | 61020000 | 0.0004 | 0.0003 | 0.0004 |
| F07 | 60940001 | 61040000 | 0.0005 | 0.0003 | 0.0005 |
| F07 | 60960001 | 61060000 | 0.0009 | 0.0006 | 0.0009 |
| F07 | 60980001 | 61080000 | 0.0012 | 0.0008 | 0.0012 |
| F07 | 61000001 | 61100000 | 0.0012 | 0.0009 | 0.0012 |
| F07 | 61020001 | 61120000 | 0.0014 | 0.0010 | 0.0014 |
| F07 | 61040001 | 61140000 | 0.0014 | 0.0011 | 0.0014 |
| F07 | 61060001 | 61160000 | 0.0016 | 0.0011 | 0.0016 |
| F07 | 61080001 | 61180000 | 0.0016 | 0.0012 | 0.0016 |
| F07 | 61100001 | 61200000 | 0.0015 | 0.0011 | 0.0015 |
| F07 | 61120001 | 61220000 | 0.0013 | 0.0009 | 0.0013 |
| F07 | 61140001 | 61240000 | 0.0012 | 0.0009 | 0.0012 |
| F07 | 61160001 | 61260000 | 0.0008 | 0.0006 | 0.0008 |
| F07 | 61180001 | 61280000 | 0.0009 | 0.0007 | 0.0009 |
| F07 | 61200001 | 61300000 | 0.0009 | 0.0007 | 0.0009 |
| F07 | 61220001 | 61320000 | 0.0010 | 0.0008 | 0.0010 |
| F07 | 61240001 | 61340000 | 0.0011 | 0.0009 | 0.0010 |
| F07 | 61260001 | 61360000 | 0.0010 | 0.0008 | 0.0010 |
| F07 | 61280001 | 61380000 | 0.0008 | 0.0006 | 0.0008 |
| F07 | 61300001 | 61400000 | 0.0011 | 0.0008 | 0.0010 |
| F07 | 61320001 | 61420000 | 0.0011 | 0.0009 | 0.0011 |
| F07 | 61340001 | 61440000 | 0.0012 | 0.0009 | 0.0012 |
| F07 | 61360001 | 61460000 | 0.0020 | 0.0017 | 0.0020 |
| F07 | 61380001 | 61480000 | 0.0028 | 0.0023 | 0.0028 |
| F07 | 61400001 | 61500000 | 0.0025 | 0.0021 | 0.0024 |
| F07 | 61420001 | 61520000 | 0.0027 | 0.0022 | 0.0026 |
| F07 | 61440001 | 61540000 | 0.0028 | 0.0024 | 0.0028 |
| F07 | 61460001 | 61560000 | 0.0021 | 0.0017 | 0.0021 |
| F07 | 61480001 | 61580000 | 0.0016 | 0.0012 | 0.0016 |
| F07 | 61500001 | 61600000 | 0.0019 | 0.0015 | 0.0019 |
| F07 | 61520001 | 61620000 | 0.0015 | 0.0012 | 0.0015 |
| F07 | 61540001 | 61640000 | 0.0015 | 0.0011 | 0.0015 |
| F07 | 61560001 | 61660000 | 0.0019 | 0.0015 | 0.0019 |
| F07 | 61580001 | 61680000 | 0.0016 | 0.0012 | 0.0016 |
| F07 | 61600001 | 61700000 | 0.0013 | 0.0010 | 0.0013 |
| F07 | 61620001 | 61720000 | 0.0012 | 0.0010 | 0.0013 |
| F07 | 61640001 | 61740000 | 0.0009 | 0.0008 | 0.0009 |
| F07 | 61660001 | 61760000 | 0.0004 | 0.0004 | 0.0004 |
| F07 | 61680001 | 61780000 | 0.0004 | 0.0004 | 0.0004 |
| F07 | 61700001 | 61800000 | 0.0003 | 0.0003 | 0.0003 |
| F07 | 61720001 | 61820000 | 0.0012 | 0.0010 | 0.0012 |

| Chr | Start | End | both | A1 | A2 |
|---|---|---|---|---|---|
| F07 | 61740001 | 61840000 | 0.0016 | 0.0013 | 0.0016 |
| F07 | 61760001 | 61860000 | 0.0023 | 0.0020 | 0.0023 |
| F07 | 61780001 | 61880000 | 0.0023 | 0.0020 | 0.0023 |
| F07 | 61800001 | 61900000 | 0.0026 | 0.0023 | 0.0026 |
| F07 | 61820001 | 61920000 | 0.0024 | 0.0021 | 0.0023 |
| F07 | 61840001 | 61940000 | 0.0025 | 0.0021 | 0.0024 |
| F07 | 61860001 | 61960000 | 0.0020 | 0.0016 | 0.0019 |
| F07 | 61880001 | 61980000 | 0.0021 | 0.0017 | 0.0019 |
| F07 | 61900001 | 62000000 | 0.0023 | 0.0018 | 0.0022 |
| F07 | 61920001 | 62020000 | 0.0021 | 0.0016 | 0.0020 |
| F07 | 61940001 | 62040000 | 0.0017 | 0.0014 | 0.0017 |
| F07 | 61960001 | 62060000 | 0.0017 | 0.0015 | 0.0017 |
| F07 | 61980001 | 62080000 | 0.0022 | 0.0019 | 0.0021 |
| F07 | 62000001 | 62100000 | 0.0021 | 0.0019 | 0.0021 |
| F07 | 62020001 | 62120000 | 0.0023 | 0.0021 | 0.0022 |
| F07 | 62040001 | 62140000 | 0.0027 | 0.0025 | 0.0026 |
| F07 | 62060001 | 62160000 | 0.0026 | 0.0025 | 0.0026 |
| F07 | 62080001 | 62180000 | 0.0025 | 0.0023 | 0.0024 |
| F07 | 62100001 | 62200000 | 0.0026 | 0.0023 | 0.0025 |
| F07 | 62120001 | 62220000 | 0.0023 | 0.0021 | 0.0022 |
| F07 | 62140001 | 62240000 | 0.0020 | 0.0017 | 0.0019 |
| F07 | 62160001 | 62260000 | 0.0020 | 0.0017 | 0.0020 |
| F07 | 62180001 | 62280000 | 0.0016 | 0.0013 | 0.0016 |
| F07 | 62200001 | 62300000 | 0.0014 | 0.0012 | 0.0014 |
| F07 | 62220001 | 62320000 | 0.0019 | 0.0017 | 0.0019 |
| F07 | 62240001 | 62340000 | 0.0020 | 0.0018 | 0.0020 |
| F07 | 62260001 | 62360000 | 0.0021 | 0.0018 | 0.0020 |
| F07 | 62280001 | 62380000 | 0.0023 | 0.0020 | 0.0022 |
| F07 | 62300001 | 62400000 | 0.0022 | 0.0021 | 0.0021 |
| F07 | 62320001 | 62420000 | 0.0019 | 0.0018 | 0.0018 |
| F07 | 62340001 | 62440000 | 0.0018 | 0.0017 | 0.0017 |
| F07 | 62360001 | 62460000 | 0.0016 | 0.0015 | 0.0016 |
| F07 | 62380001 | 62480000 | 0.0017 | 0.0015 | 0.0017 |
| F07 | 62400001 | 62500000 | 0.0017 | 0.0015 | 0.0017 |
| F07 | 62420001 | 62520000 | 0.0022 | 0.0019 | 0.0020 |
| F07 | 62440001 | 62540000 | 0.0021 | 0.0020 | 0.0020 |
| F07 | 62460001 | 62560000 | 0.0028 | 0.0025 | 0.0027 |
| F07 | 62480001 | 62580000 | 0.0029 | 0.0027 | 0.0027 |
| F07 | 62500001 | 62600000 | 0.0026 | 0.0024 | 0.0024 |
| F07 | 62520001 | 62620000 | 0.0020 | 0.0018 | 0.0020 |
| F07 | 62540001 | 62640000 | 0.0019 | 0.0017 | 0.0019 |
| F07 | 62560001 | 62660000 | 0.0018 | 0.0017 | 0.0017 |
| F07 | 62580001 | 62680000 | 0.0018 | 0.0017 | 0.0017 |
| F07 | 62600001 | 62700000 | 0.0025 | 0.0023 | 0.0024 |
| F07 | 62620001 | 62720000 | 0.0027 | 0.0024 | 0.0026 |
| F07 | 62640001 | 62740000 | 0.0032 | 0.0030 | 0.0031 |
| F07 | 62660001 | 62760000 | 0.0028 | 0.0025 | 0.0026 |
| F07 | 62680001 | 62780000 | 0.0033 | 0.0029 | 0.0031 |
| F07 | 62700001 | 62800000 | 0.0032 | 0.0027 | 0.0030 |
| F07 | 62720001 | 62820000 | 0.0032 | 0.0027 | 0.0030 |
| F07 | 62740001 | 62840000 | 0.0026 | 0.0021 | 0.0025 |
| F07 | 62760001 | 62860000 | 0.0027 | 0.0023 | 0.0027 |
| F07 | 62780001 | 62880000 | 0.0022 | 0.0018 | 0.0021 |
| F07 | 62800001 | 62900000 | 0.0019 | 0.0016 | 0.0019 |
| F07 | 62820001 | 62920000 | 0.0020 | 0.0017 | 0.0019 |
| F07 | 62840001 | 62940000 | 0.0019 | 0.0016 | 0.0018 |
| F07 | 62860001 | 62960000 | 0.0019 | 0.0016 | 0.0019 |
| F07 | 62880001 | 62980000 | 0.0023 | 0.0019 | 0.0023 |

| Chr | Start | End | both | A1 | A2 |
|---|---|---|---|---|---|
| F07 | 62900001 | 63000000 | 0.0025 | 0.0022 | 0.0025 |
| F07 | 62920001 | 63020000 | 0.0024 | 0.0021 | 0.0024 |
| F07 | 62940001 | 63040000 | 0.0025 | 0.0022 | 0.0025 |
| F07 | 62960001 | 63060000 | 0.0021 | 0.0019 | 0.0021 |
| F07 | 62980001 | 63080000 | 0.0015 | 0.0014 | 0.0015 |
| F07 | 63000001 | 63100000 | 0.0011 | 0.0010 | 0.0011 |
| F07 | 63020001 | 63120000 | 0.0009 | 0.0008 | 0.0009 |
| F07 | 63040001 | 63140000 | 0.0011 | 0.0009 | 0.0011 |
| F07 | 63060001 | 63160000 | 0.0026 | 0.0021 | 0.0026 |
| F07 | 63080001 | 63180000 | 0.0029 | 0.0024 | 0.0028 |
| F07 | 63100001 | 63200000 | 0.0027 | 0.0023 | 0.0027 |
| F07 | 63120001 | 63220000 | 0.0026 | 0.0022 | 0.0026 |
| F07 | 63140001 | 63240000 | 0.0026 | 0.0022 | 0.0026 |
| F07 | 63160001 | 63260000 | 0.0014 | 0.0012 | 0.0014 |
| F07 | 63180001 | 63280000 | 0.0011 | 0.0010 | 0.0011 |
| F07 | 63200001 | 63300000 | 0.0021 | 0.0019 | 0.0021 |
| F07 | 63220001 | 63320000 | 0.0022 | 0.0019 | 0.0022 |
| F07 | 63240001 | 63340000 | 0.0029 | 0.0026 | 0.0029 |
| F07 | 63260001 | 63360000 | 0.0030 | 0.0027 | 0.0030 |
| F07 | 63280001 | 63380000 | 0.0033 | 0.0029 | 0.0032 |
| F07 | 63300001 | 63400000 | 0.0025 | 0.0022 | 0.0024 |
| F07 | 63320001 | 63420000 | 0.0024 | 0.0022 | 0.0024 |
| F07 | 63340001 | 63440000 | 0.0017 | 0.0015 | 0.0017 |
| F07 | 63360001 | 63460000 | 0.0014 | 0.0012 | 0.0014 |
| F07 | 63380001 | 63480000 | 0.0013 | 0.0011 | 0.0012 |
| F07 | 63400001 | 63500000 | 0.0018 | 0.0017 | 0.0018 |
| F07 | 63420001 | 63520000 | 0.0029 | 0.0026 | 0.0028 |
| F07 | 63440001 | 63540000 | 0.0029 | 0.0026 | 0.0027 |
| F07 | 63460001 | 63560000 | 0.0029 | 0.0026 | 0.0027 |
| F07 | 63480001 | 63580000 | 0.0026 | 0.0024 | 0.0024 |
| F07 | 63500001 | 63600000 | 0.0020 | 0.0017 | 0.0019 |
| F07 | 63520001 | 63620000 | 0.0014 | 0.0011 | 0.0014 |
| F07 | 63540001 | 63640000 | 0.0015 | 0.0012 | 0.0015 |
| F07 | 63560001 | 63660000 | 0.0016 | 0.0013 | 0.0017 |
| F07 | 63580001 | 63680000 | 0.0025 | 0.0019 | 0.0025 |
| F07 | 63600001 | 63700000 | 0.0022 | 0.0017 | 0.0023 |
| F07 | 63620001 | 63720000 | 0.0018 | 0.0014 | 0.0019 |
| F07 | 63640001 | 63740000 | 0.0019 | 0.0015 | 0.0019 |
| F07 | 63660001 | 63760000 | 0.0017 | 0.0013 | 0.0017 |
| F07 | 63680001 | 63780000 | 0.0018 | 0.0016 | 0.0018 |
| F07 | 63700001 | 63800000 | 0.0021 | 0.0017 | 0.0021 |
| F07 | 63720001 | 63820000 | 0.0029 | 0.0027 | 0.0029 |
| F07 | 63740001 | 63840000 | 0.0032 | 0.0030 | 0.0032 |
| F07 | 63760001 | 63860000 | 0.0038 | 0.0035 | 0.0038 |
| F07 | 63780001 | 63880000 | 0.0030 | 0.0027 | 0.0029 |
| F07 | 63800001 | 63900000 | 0.0029 | 0.0026 | 0.0029 |
| F07 | 63820001 | 63920000 | 0.0025 | 0.0021 | 0.0025 |
| F07 | 63840001 | 63940000 | 0.0022 | 0.0018 | 0.0021 |
| F07 | 63860001 | 63960000 | 0.0015 | 0.0012 | 0.0015 |
| F07 | 63880001 | 63980000 | 0.0015 | 0.0013 | 0.0014 |
| F07 | 63900001 | 64000000 | 0.0015 | 0.0014 | 0.0015 |
| F07 | 63920001 | 64020000 | 0.0011 | 0.0010 | 0.0011 |
| F07 | 63940001 | 64040000 | 0.0009 | 0.0008 | 0.0008 |
| F07 | 63960001 | 64060000 | 0.0012 | 0.0010 | 0.0011 |
| F07 | 63980001 | 64080000 | 0.0013 | 0.0011 | 0.0012 |
| F07 | 64000001 | 64100000 | 0.0017 | 0.0014 | 0.0015 |
| F07 | 64020001 | 64120000 | 0.0021 | 0.0017 | 0.0019 |
| F07 | 64040001 | 64140000 | 0.0022 | 0.0018 | 0.0021 |

| Chr | Start | End | both | A1 | A2 |
|---|---|---|---|---|---|
| F07 | 64060001 | 64160000 | 0.0022 | 0.0018 | 0.0021 |
| F07 | 64080001 | 64180000 | 0.0021 | 0.0018 | 0.0021 |
| F07 | 64100001 | 64200000 | 0.0018 | 0.0016 | 0.0017 |
| F07 | 64120001 | 64220000 | 0.0013 | 0.0012 | 0.0012 |
| F07 | 64140001 | 64240000 | 0.0015 | 0.0014 | 0.0014 |
| F07 | 64160001 | 64260000 | 0.0015 | 0.0014 | 0.0014 |
| F07 | 64180001 | 64280000 | 0.0015 | 0.0015 | 0.0014 |
| F07 | 64200001 | 64300000 | 0.0016 | 0.0014 | 0.0016 |
| F07 | 64220001 | 64320000 | 0.0017 | 0.0016 | 0.0016 |
| F07 | 64240001 | 64340000 | 0.0015 | 0.0014 | 0.0015 |
| F07 | 64260001 | 64360000 | 0.0014 | 0.0012 | 0.0013 |
| F07 | 64280001 | 64380000 | 0.0015 | 0.0013 | 0.0016 |
| F07 | 64300001 | 64400000 | 0.0012 | 0.0010 | 0.0012 |
| F07 | 64320001 | 64420000 | 0.0011 | 0.0008 | 0.0010 |
| F07 | 64340001 | 64440000 | 0.0011 | 0.0009 | 0.0010 |
| F07 | 64360001 | 64460000 | 0.0009 | 0.0008 | 0.0009 |
| F07 | 64380001 | 64480000 | 0.0007 | 0.0006 | 0.0006 |
| F07 | 64400001 | 64500000 | 0.0013 | 0.0012 | 0.0012 |
| F07 | 64420001 | 64520000 | 0.0013 | 0.0012 | 0.0012 |
| F07 | 64440001 | 64540000 | 0.0014 | 0.0012 | 0.0014 |
| F07 | 64460001 | 64560000 | 0.0018 | 0.0016 | 0.0018 |
| F07 | 64480001 | 64580000 | 0.0019 | 0.0016 | 0.0019 |
| F07 | 64500001 | 64600000 | 0.0012 | 0.0009 | 0.0012 |
| F07 | 64520001 | 64620000 | 0.0011 | 0.0009 | 0.0011 |
| F07 | 64540001 | 64640000 | 0.0016 | 0.0015 | 0.0016 |
| F07 | 64560001 | 64660000 | 0.0012 | 0.0011 | 0.0012 |
| F07 | 64580001 | 64680000 | 0.0023 | 0.0021 | 0.0022 |
| F07 | 64600001 | 64700000 | 0.0029 | 0.0026 | 0.0028 |
| F07 | 64620001 | 64720000 | 0.0038 | 0.0033 | 0.0036 |
| F07 | 64640001 | 64740000 | 0.0030 | 0.0025 | 0.0029 |
| F07 | 64660001 | 64760000 | 0.0033 | 0.0028 | 0.0031 |
| F07 | 64680001 | 64780000 | 0.0023 | 0.0019 | 0.0022 |
| F07 | 64700001 | 64800000 | 0.0020 | 0.0017 | 0.0019 |
| F07 | 64720001 | 64820000 | 0.0018 | 0.0015 | 0.0018 |
| F07 | 64740001 | 64840000 | 0.0019 | 0.0015 | 0.0019 |
| F07 | 64760001 | 64860000 | 0.0017 | 0.0014 | 0.0017 |
| F07 | 64780001 | 64880000 | 0.0024 | 0.0020 | 0.0023 |
| F07 | 64800001 | 64900000 | 0.0029 | 0.0024 | 0.0027 |
| F07 | 64820001 | 64920000 | 0.0029 | 0.0024 | 0.0025 |
| F07 | 64840001 | 64940000 | 0.0034 | 0.0029 | 0.0030 |
| F07 | 64860001 | 64960000 | 0.0039 | 0.0032 | 0.0035 |
| F07 | 64880001 | 64980000 | 0.0036 | 0.0030 | 0.0032 |
| F07 | 64900001 | 65000000 | 0.0032 | 0.0028 | 0.0029 |
| F07 | 64920001 | 65020000 | 0.0030 | 0.0025 | 0.0028 |
| F07 | 64940001 | 65040000 | 0.0024 | 0.0021 | 0.0024 |
| F07 | 64960001 | 65060000 | 0.0020 | 0.0018 | 0.0019 |
| F07 | 64980001 | 65080000 | 0.0017 | 0.0015 | 0.0017 |
| F07 | 65000001 | 65100000 | 0.0018 | 0.0015 | 0.0016 |
| F07 | 65020001 | 65120000 | 0.0023 | 0.0020 | 0.0020 |
| F07 | 65040001 | 65140000 | 0.0027 | 0.0023 | 0.0022 |
| F07 | 65060001 | 65160000 | 0.0028 | 0.0022 | 0.0023 |
| F07 | 65080001 | 65180000 | 0.0026 | 0.0021 | 0.0021 |
| F07 | 65100001 | 65200000 | 0.0029 | 0.0024 | 0.0025 |
| F07 | 65120001 | 65220000 | 0.0027 | 0.0021 | 0.0022 |
| F07 | 65140001 | 65240000 | 0.0022 | 0.0017 | 0.0018 |
| F07 | 65160001 | 65260000 | 0.0027 | 0.0023 | 0.0024 |
| F07 | 65180001 | 65280000 | 0.0032 | 0.0026 | 0.0027 |
| F07 | 65200001 | 65300000 | 0.0032 | 0.0024 | 0.0024 |

| Chr | Start | End | both | A1 | A2 |
|---|---|---|---|---|---|
| F07 | 65220001 | 65320000 | 0.0037 | 0.0028 | 0.0026 |
| F07 | 65240001 | 65340000 | 0.0044 | 0.0034 | 0.0033 |
| F07 | 65260001 | 65360000 | 0.0040 | 0.0030 | 0.0027 |
| F07 | 65280001 | 65380000 | 0.0036 | 0.0028 | 0.0024 |
| F07 | 65300001 | 65400000 | 0.0033 | 0.0028 | 0.0025 |
| F07 | 65320001 | 65420000 | 0.0035 | 0.0029 | 0.0028 |
| F07 | 65340001 | 65440000 | 0.0038 | 0.0029 | 0.0031 |
| F07 | 65360001 | 65460000 | 0.0038 | 0.0029 | 0.0032 |
| F07 | 65380001 | 65480000 | 0.0037 | 0.0029 | 0.0032 |
| F07 | 65400001 | 65500000 | 0.0040 | 0.0031 | 0.0035 |
| F07 | 65420001 | 65520000 | 0.0042 | 0.0032 | 0.0038 |
| F07 | 65440001 | 65540000 | 0.0034 | 0.0027 | 0.0031 |
| F07 | 65460001 | 65560000 | 0.0037 | 0.0030 | 0.0034 |
| F07 | 65480001 | 65580000 | 0.0037 | 0.0030 | 0.0034 |
| F07 | 65500001 | 65600000 | 0.0039 | 0.0032 | 0.0036 |
| F07 | 65520001 | 65620000 | 0.0027 | 0.0023 | 0.0024 |
| F07 | 65540001 | 65640000 | 0.0026 | 0.0023 | 0.0024 |
| F07 | 65560001 | 65660000 | 0.0023 | 0.0019 | 0.0020 |
| F07 | 65580001 | 65680000 | 0.0030 | 0.0025 | 0.0025 |
| F07 | 65600001 | 65700000 | 0.0024 | 0.0020 | 0.0019 |
| F07 | 65620001 | 65720000 | 0.0025 | 0.0019 | 0.0018 |
| F07 | 65640001 | 65740000 | 0.0038 | 0.0029 | 0.0030 |
| F07 | 65660001 | 65760000 | 0.0038 | 0.0031 | 0.0032 |
| F07 | 65680001 | 65780000 | 0.0042 | 0.0034 | 0.0035 |
| F07 | 65700001 | 65800000 | 0.0037 | 0.0030 | 0.0032 |
| F07 | 65720001 | 65820000 | 0.0036 | 0.0031 | 0.0032 |
| F07 | 65740001 | 65840000 | 0.0036 | 0.0031 | 0.0033 |
| F07 | 65760001 | 65860000 | 0.0035 | 0.0029 | 0.0032 |
| F07 | 65780001 | 65880000 | 0.0026 | 0.0022 | 0.0024 |
| F07 | 65800001 | 65900000 | 0.0026 | 0.0022 | 0.0024 |
| F07 | 65820001 | 65920000 | 0.0027 | 0.0021 | 0.0024 |
| F07 | 65840001 | 65940000 | 0.0017 | 0.0014 | 0.0014 |
| F07 | 65860001 | 65960000 | 0.0016 | 0.0013 | 0.0014 |
| F07 | 65880001 | 65980000 | 0.0021 | 0.0018 | 0.0018 |
| F07 | 65900001 | 66000000 | 0.0026 | 0.0022 | 0.0022 |
| F07 | 65920001 | 66020000 | 0.0027 | 0.0023 | 0.0023 |
| F07 | 65940001 | 66040000 | 0.0025 | 0.0021 | 0.0021 |
| F07 | 65960001 | 66060000 | 0.0028 | 0.0023 | 0.0023 |
| F07 | 65980001 | 66080000 | 0.0023 | 0.0017 | 0.0018 |
| F07 | 66000001 | 66100000 | 0.0027 | 0.0020 | 0.0022 |
| F07 | 66020001 | 66120000 | 0.0031 | 0.0025 | 0.0025 |
| F07 | 66040001 | 66140000 | 0.0029 | 0.0022 | 0.0022 |
| F07 | 66060001 | 66160000 | 0.0026 | 0.0022 | 0.0021 |
| F07 | 66080001 | 66180000 | 0.0034 | 0.0029 | 0.0029 |
| F07 | 66100001 | 66200000 | 0.0031 | 0.0027 | 0.0028 |
| F07 | 66120001 | 66220000 | 0.0034 | 0.0029 | 0.0033 |
| F07 | 66140001 | 66240000 | 0.0037 | 0.0032 | 0.0036 |
| F07 | 66160001 | 66260000 | 0.0035 | 0.0029 | 0.0034 |
| F07 | 66180001 | 66280000 | 0.0035 | 0.0030 | 0.0034 |
| F07 | 66200001 | 66300000 | 0.0031 | 0.0026 | 0.0029 |
| F07 | 66220001 | 66320000 | 0.0024 | 0.0021 | 0.0023 |
| F07 | 66240001 | 66340000 | 0.0021 | 0.0018 | 0.0020 |
| F07 | 66260001 | 66360000 | 0.0025 | 0.0021 | 0.0023 |
| F07 | 66280001 | 66380000 | 0.0019 | 0.0016 | 0.0015 |
| F07 | 66300001 | 66400000 | 0.0020 | 0.0018 | 0.0016 |
| F07 | 66320001 | 66420000 | 0.0019 | 0.0016 | 0.0015 |
| F07 | 66340001 | 66440000 | 0.0022 | 0.0018 | 0.0016 |
| F07 | 66360001 | 66460000 | 0.0019 | 0.0015 | 0.0014 |

| Chr | Start | End | both | A1 | A2 |
|---|---|---|---|---|---|
| F07 | 66380001 | 66480000 | 0.0021 | 0.0017 | 0.0017 |
| F07 | 66400001 | 66500000 | 0.0019 | 0.0015 | 0.0014 |
| F07 | 66420001 | 66520000 | 0.0021 | 0.0015 | 0.0015 |
| F07 | 66440001 | 66540000 | 0.0023 | 0.0017 | 0.0018 |
| F07 | 66460001 | 66560000 | 0.0027 | 0.0021 | 0.0022 |
| F07 | 66480001 | 66580000 | 0.0028 | 0.0020 | 0.0024 |
| F07 | 66500001 | 66600000 | 0.0030 | 0.0021 | 0.0027 |
| F07 | 66520001 | 66620000 | 0.0030 | 0.0022 | 0.0026 |
| F07 | 66540001 | 66640000 | 0.0025 | 0.0018 | 0.0021 |
| F07 | 66560001 | 66660000 | 0.0020 | 0.0014 | 0.0016 |
| F07 | 66580001 | 66680000 | 0.0021 | 0.0016 | 0.0016 |
| F07 | 66600001 | 66700000 | 0.0020 | 0.0015 | 0.0014 |
| F07 | 66620001 | 66720000 | 0.0017 | 0.0012 | 0.0011 |
| F07 | 66640001 | 66740000 | 0.0019 | 0.0013 | 0.0013 |
| F07 | 66660001 | 66760000 | 0.0021 | 0.0014 | 0.0013 |
| F07 | 66680001 | 66780000 | 0.0021 | 0.0015 | 0.0014 |
| F07 | 66700001 | 66800000 | 0.0021 | 0.0015 | 0.0017 |
| F07 | 66720001 | 66820000 | 0.0022 | 0.0018 | 0.0017 |
| F07 | 66740001 | 66840000 | 0.0022 | 0.0019 | 0.0017 |
| F07 | 66760001 | 66860000 | 0.0024 | 0.0022 | 0.0020 |
| F07 | 66780001 | 66880000 | 0.0022 | 0.0019 | 0.0020 |
| F07 | 66800001 | 66900000 | 0.0027 | 0.0027 | 0.0023 |
| F07 | 66820001 | 66920000 | 0.0027 | 0.0029 | 0.0024 |
| F07 | 66840001 | 66940000 | 0.0026 | 0.0028 | 0.0024 |
| F07 | 66860001 | 66960000 | 0.0040 | 0.0041 | 0.0038 |
| F07 | 66880001 | 66980000 | 0.0037 | 0.0039 | 0.0035 |
| F07 | 66900001 | 67000000 | 0.0029 | 0.0030 | 0.0028 |
| F07 | 66920001 | 67020000 | 0.0033 | 0.0033 | 0.0032 |
| F07 | 66940001 | 67040000 | 0.0032 | 0.0032 | 0.0030 |
| F07 | 66960001 | 67060000 | 0.0024 | 0.0026 | 0.0023 |
| F07 | 66980001 | 67080000 | 0.0024 | 0.0026 | 0.0023 |
| F07 | 67000001 | 67100000 | 0.0027 | 0.0029 | 0.0025 |
| F07 | 67020001 | 67120000 | 0.0023 | 0.0026 | 0.0022 |
| F07 | 67040001 | 67140000 | 0.0029 | 0.0031 | 0.0027 |
| F07 | 67060001 | 67160000 | 0.0023 | 0.0025 | 0.0022 |
| F07 | 67080001 | 67180000 | 0.0035 | 0.0037 | 0.0032 |
| F07 | 67100001 | 67200000 | 0.0041 | 0.0046 | 0.0038 |
| F07 | 67120001 | 67220000 | 0.0045 | 0.0046 | 0.0043 |
| F07 | 67140001 | 67240000 | 0.0040 | 0.0041 | 0.0037 |
| F07 | 67160001 | 67260000 | 0.0035 | 0.0038 | 0.0033 |
| F07 | 67180001 | 67280000 | 0.0028 | 0.0032 | 0.0026 |
| F07 | 67200001 | 67300000 | 0.0023 | 0.0026 | 0.0020 |
| F07 | 67220001 | 67320000 | 0.0017 | 0.0023 | 0.0014 |
| F07 | 67240001 | 67340000 | 0.0021 | 0.0025 | 0.0018 |
| F07 | 67260001 | 67360000 | 0.0020 | 0.0022 | 0.0017 |
| F07 | 67280001 | 67380000 | 0.0013 | 0.0014 | 0.0011 |
| F07 | 67300001 | 67400000 | 0.0011 | 0.0011 | 0.0011 |
| F07 | 67320001 | 67420000 | 0.0017 | 0.0016 | 0.0016 |
| F07 | 67340001 | 67440000 | 0.0021 | 0.0022 | 0.0019 |
| F07 | 67360001 | 67460000 | 0.0025 | 0.0027 | 0.0022 |
| F07 | 67380001 | 67480000 | 0.0026 | 0.0032 | 0.0021 |
| F07 | 67400001 | 67500000 | 0.0034 | 0.0040 | 0.0029 |
| F07 | 67420001 | 67520000 | 0.0030 | 0.0037 | 0.0025 |
| F07 | 67440001 | 67540000 | 0.0028 | 0.0037 | 0.0022 |
| F07 | 67460001 | 67560000 | 0.0031 | 0.0038 | 0.0025 |
| F07 | 67480001 | 67580000 | 0.0031 | 0.0033 | 0.0025 |
| F07 | 67500001 | 67600000 | 0.0023 | 0.0025 | 0.0017 |
| F07 | 67520001 | 67620000 | 0.0020 | 0.0019 | 0.0015 |

| Chr | Start | End | both | A1 | A2 |
|---|---|---|---|---|---|
| F07 | 67540001 | 67640000 | 0.0019 | 0.0016 | 0.0016 |
| F07 | 67560001 | 67660000 | 0.0028 | 0.0022 | 0.0022 |
| F07 | 67580001 | 67680000 | 0.0032 | 0.0024 | 0.0026 |
| F07 | 67600001 | 67700000 | 0.0036 | 0.0027 | 0.0029 |
| F07 | 67620001 | 67720000 | 0.0045 | 0.0034 | 0.0035 |
| F07 | 67640001 | 67740000 | 0.0054 | 0.0041 | 0.0041 |
| F07 | 67660001 | 67760000 | 0.0056 | 0.0043 | 0.0044 |
| F07 | 67680001 | 67780000 | 0.0059 | 0.0048 | 0.0047 |
| F07 | 67700001 | 67800000 | 0.0055 | 0.0045 | 0.0045 |
| F07 | 67720001 | 67820000 | 0.0056 | 0.0046 | 0.0047 |
| F07 | 67740001 | 67840000 | 0.0047 | 0.0039 | 0.0040 |
| F07 | 67760001 | 67860000 | 0.0032 | 0.0026 | 0.0027 |
| F07 | 67780001 | 67880000 | 0.0033 | 0.0026 | 0.0028 |
| F07 | 67800001 | 67900000 | 0.0035 | 0.0027 | 0.0027 |
| F07 | 67820001 | 67920000 | 0.0029 | 0.0022 | 0.0020 |
| F07 | 67840001 | 67940000 | 0.0032 | 0.0025 | 0.0023 |
| F07 | 67860001 | 67960000 | 0.0031 | 0.0026 | 0.0021 |
| F07 | 67880001 | 67980000 | 0.0022 | 0.0019 | 0.0013 |
| F07 | 67900001 | 68000000 | 0.0018 | 0.0018 | 0.0012 |
| F07 | 67920001 | 68020000 | 0.0020 | 0.0019 | 0.0014 |
| F07 | 67940001 | 68040000 | 0.0012 | 0.0012 | 0.0008 |
| F07 | 67960001 | 68060000 | 0.0014 | 0.0012 | 0.0010 |
| F07 | 67980001 | 68080000 | 0.0017 | 0.0012 | 0.0010 |
| F07 | 68000001 | 68100000 | 0.0018 | 0.0011 | 0.0009 |
| F07 | 68020001 | 68120000 | 0.0016 | 0.0010 | 0.0006 |
| F07 | 68040001 | 68140000 | 0.0024 | 0.0017 | 0.0014 |
| F07 | 68060001 | 68160000 | 0.0023 | 0.0017 | 0.0013 |
| F07 | 68080001 | 68180000 | 0.0023 | 0.0018 | 0.0015 |
| F07 | 68100001 | 68200000 | 0.0038 | 0.0031 | 0.0033 |
| F07 | 68120001 | 68220000 | 0.0039 | 0.0032 | 0.0034 |
| F07 | 68140001 | 68240000 | 0.0038 | 0.0030 | 0.0033 |
| F07 | 68160001 | 68260000 | 0.0046 | 0.0037 | 0.0040 |
| F07 | 68180001 | 68280000 | 0.0055 | 0.0045 | 0.0047 |
| F07 | 68200001 | 68300000 | 0.0040 | 0.0033 | 0.0029 |
| F07 | 68220001 | 68320000 | 0.0042 | 0.0033 | 0.0031 |
| F07 | 68240001 | 68340000 | 0.0044 | 0.0032 | 0.0032 |
| F07 | 68260001 | 68360000 | 0.0041 | 0.0033 | 0.0030 |
| F07 | 68280001 | 68380000 | 0.0032 | 0.0024 | 0.0022 |
| F07 | 68300001 | 68400000 | 0.0034 | 0.0026 | 0.0023 |
| F07 | 68320001 | 68420000 | 0.0035 | 0.0026 | 0.0023 |
| F07 | 68340001 | 68440000 | 0.0033 | 0.0025 | 0.0020 |
| F07 | 68360001 | 68460000 | 0.0031 | 0.0023 | 0.0019 |
| F07 | 68380001 | 68480000 | 0.0035 | 0.0028 | 0.0025 |
| F07 | 68400001 | 68500000 | 0.0040 | 0.0034 | 0.0031 |
| F07 | 68420001 | 68520000 | 0.0044 | 0.0039 | 0.0036 |
| F07 | 68440001 | 68540000 | 0.0043 | 0.0039 | 0.0038 |
| F07 | 68460001 | 68560000 | 0.0046 | 0.0040 | 0.0042 |
| F07 | 68480001 | 68580000 | 0.0047 | 0.0038 | 0.0043 |
| F07 | 68500001 | 68600000 | 0.0040 | 0.0032 | 0.0038 |
| F07 | 68520001 | 68620000 | 0.0035 | 0.0026 | 0.0032 |
| F07 | 68540001 | 68640000 | 0.0040 | 0.0027 | 0.0029 |
| F07 | 68560001 | 68660000 | 0.0047 | 0.0032 | 0.0029 |
| F07 | 68580001 | 68680000 | 0.0051 | 0.0034 | 0.0028 |
| F07 | 68600001 | 68700000 | 0.0057 | 0.0039 | 0.0032 |
| F07 | 68620001 | 68720000 | 0.0063 | 0.0046 | 0.0038 |
| F07 | 68640001 | 68740000 | 0.0064 | 0.0047 | 0.0041 |
| F07 | 68660001 | 68760000 | 0.0049 | 0.0036 | 0.0033 |
| F07 | 68680001 | 68780000 | 0.0040 | 0.0030 | 0.0029 |

| Chr | Start | End | both | A1 | A2 |
|---|---|---|---|---|---|
| F07 | 68700001 | 68800000 | 0.0031 | 0.0023 | 0.0022 |
| F07 | 68720001 | 68820000 | 0.0022 | 0.0016 | 0.0015 |
| F07 | 68740001 | 68840000 | 0.0017 | 0.0014 | 0.0014 |
| F07 | 68760001 | 68860000 | 0.0014 | 0.0012 | 0.0013 |
| F07 | 68780001 | 68880000 | 0.0020 | 0.0017 | 0.0019 |
| F07 | 68800001 | 68900000 | 0.0022 | 0.0019 | 0.0020 |
| F07 | 68820001 | 68920000 | 0.0027 | 0.0024 | 0.0024 |
| F07 | 68840001 | 68940000 | 0.0025 | 0.0021 | 0.0022 |
| F07 | 68860001 | 68960000 | 0.0030 | 0.0025 | 0.0026 |
| F07 | 68880001 | 68980000 | 0.0023 | 0.0020 | 0.0018 |
| F07 | 68900001 | 69000000 | 0.0024 | 0.0020 | 0.0019 |
| F07 | 68920001 | 69020000 | 0.0022 | 0.0017 | 0.0018 |
| F07 | 68940001 | 69040000 | 0.0023 | 0.0019 | 0.0019 |
| F07 | 68960001 | 69060000 | 0.0024 | 0.0019 | 0.0020 |
| F07 | 68980001 | 69080000 | 0.0023 | 0.0018 | 0.0019 |
| F07 | 69000001 | 69100000 | 0.0026 | 0.0021 | 0.0020 |
| F07 | 69020001 | 69120000 | 0.0027 | 0.0022 | 0.0020 |
| F07 | 69040001 | 69140000 | 0.0059 | 0.0048 | 0.0053 |
| F07 | 69060001 | 69160000 | 0.0062 | 0.0052 | 0.0056 |
| F07 | 69080001 | 69180000 | 0.0074 | 0.0059 | 0.0062 |
| F07 | 69100001 | 69200000 | 0.0085 | 0.0066 | 0.0068 |
| F07 | 69120001 | 69220000 | 0.0082 | 0.0064 | 0.0064 |
| F07 | 69140001 | 69240000 | 0.0065 | 0.0050 | 0.0046 |
| F07 | 69160001 | 69260000 | 0.0057 | 0.0043 | 0.0039 |
| F07 | 69180001 | 69280000 | 0.0052 | 0.0041 | 0.0037 |
| F07 | 69200001 | 69300000 | 0.0044 | 0.0036 | 0.0037 |
| F07 | 69220001 | 69320000 | 0.0043 | 0.0035 | 0.0037 |
| F07 | 69240001 | 69340000 | 0.0034 | 0.0025 | 0.0026 |
| F07 | 69260001 | 69360000 | 0.0040 | 0.0029 | 0.0028 |
| F07 | 69280001 | 69380000 | 0.0040 | 0.0029 | 0.0030 |
| F07 | 69300001 | 69400000 | 0.0040 | 0.0031 | 0.0030 |
| F07 | 69320001 | 69420000 | 0.0042 | 0.0032 | 0.0033 |
| F07 | 69340001 | 69440000 | 0.0043 | 0.0035 | 0.0034 |
| F07 | 69360001 | 69460000 | 0.0042 | 0.0034 | 0.0033 |
| F07 | 69380001 | 69480000 | 0.0039 | 0.0032 | 0.0028 |
| F07 | 69400001 | 69500000 | 0.0032 | 0.0025 | 0.0020 |
| F07 | 69420001 | 69520000 | 0.0026 | 0.0020 | 0.0014 |
| F07 | 69440001 | 69540000 | 0.0019 | 0.0013 | 0.0008 |
| F07 | 69460001 | 69560000 | 0.0019 | 0.0014 | 0.0010 |
| F07 | 69480001 | 69580000 | 0.0024 | 0.0017 | 0.0014 |
| F07 | 69500001 | 69600000 | 0.0033 | 0.0022 | 0.0020 |
| F07 | 69520001 | 69620000 | 0.0047 | 0.0029 | 0.0024 |
| F07 | 69540001 | 69640000 | 0.0055 | 0.0032 | 0.0023 |
| F07 | 69560001 | 69660000 | 0.0049 | 0.0027 | 0.0019 |
| F07 | 69580001 | 69680000 | 0.0049 | 0.0026 | 0.0016 |
| F07 | 69600001 | 69700000 | 0.0043 | 0.0021 | 0.0011 |
| F07 | 69620001 | 69720000 | 0.0030 | 0.0016 | 0.0009 |
| F07 | 69640001 | 69740000 | 0.0025 | 0.0015 | 0.0010 |
| F07 | 69660001 | 69760000 | 0.0029 | 0.0018 | 0.0012 |
| F07 | 69680001 | 69780000 | 0.0026 | 0.0017 | 0.0014 |
| F07 | 69700001 | 69800000 | 0.0026 | 0.0017 | 0.0016 |
| F07 | 69720001 | 69820000 | 0.0023 | 0.0015 | 0.0013 |
| F07 | 69740001 | 69840000 | 0.0021 | 0.0013 | 0.0014 |
| F07 | 69760001 | 69860000 | 0.0027 | 0.0018 | 0.0019 |
| F07 | 69780001 | 69880000 | 0.0028 | 0.0017 | 0.0019 |
| F07 | 69800001 | 69900000 | 0.0026 | 0.0017 | 0.0017 |
| F07 | 69820001 | 69920000 | 0.0039 | 0.0027 | 0.0028 |
| F07 | 69840001 | 69940000 | 0.0043 | 0.0032 | 0.0031 |

| Chr | Start | End | both | A1 | A2 |
|---|---|---|---|---|---|
| F07 | 69860001 | 69960000 | 0.0038 | 0.0029 | 0.0028 |
| F07 | 69880001 | 69980000 | 0.0043 | 0.0035 | 0.0034 |
| F07 | 69900001 | 70000000 | 0.0054 | 0.0044 | 0.0043 |
| F07 | 69920001 | 70020000 | 0.0043 | 0.0035 | 0.0034 |
| F07 | 69940001 | 70040000 | 0.0043 | 0.0033 | 0.0035 |
| F07 | 69960001 | 70060000 | 0.0043 | 0.0033 | 0.0035 |
| F07 | 69980001 | 70080000 | 0.0035 | 0.0025 | 0.0025 |
| F07 | 70000001 | 70100000 | 0.0046 | 0.0032 | 0.0040 |
| F07 | 70020001 | 70120000 | 0.0051 | 0.0035 | 0.0045 |
| F07 | 70040001 | 70140000 | 0.0050 | 0.0035 | 0.0044 |
| F07 | 70060001 | 70160000 | 0.0055 | 0.0040 | 0.0051 |
| F07 | 70080001 | 70180000 | 0.0055 | 0.0039 | 0.0053 |
| F07 | 70100001 | 70200000 | 0.0037 | 0.0026 | 0.0035 |
| F07 | 70120001 | 70220000 | 0.0035 | 0.0025 | 0.0034 |
| F07 | 70140001 | 70240000 | 0.0042 | 0.0023 | 0.0039 |
| F07 | 70160001 | 70260000 | 0.0034 | 0.0016 | 0.0030 |
| F07 | 70180001 | 70280000 | 0.0046 | 0.0019 | 0.0041 |
| F07 | 70200001 | 70300000 | 0.0044 | 0.0018 | 0.0039 |
| F07 | 70220001 | 70320000 | 0.0047 | 0.0016 | 0.0042 |
| F07 | 70240001 | 70340000 | 0.0039 | 0.0013 | 0.0035 |
| F07 | 70260001 | 70360000 | 0.0038 | 0.0012 | 0.0035 |
| F07 | 70280001 | 70380000 | 0.0032 | 0.0014 | 0.0030 |
| F07 | 70300001 | 70400000 | 0.0032 | 0.0014 | 0.0030 |
| F07 | 70320001 | 70420000 | 0.0032 | 0.0015 | 0.0030 |
| F07 | 70340001 | 70440000 | 0.0038 | 0.0015 | 0.0037 |
| F07 | 70360001 | 70460000 | 0.0036 | 0.0015 | 0.0035 |
| F07 | 70380001 | 70480000 | 0.0034 | 0.0010 | 0.0033 |
| F07 | 70400001 | 70500000 | 0.0040 | 0.0010 | 0.0039 |
| F07 | 70420001 | 70520000 | 0.0043 | 0.0017 | 0.0041 |
| F07 | 70440001 | 70540000 | 0.0041 | 0.0025 | 0.0037 |
| F07 | 70460001 | 70560000 | 0.0043 | 0.0027 | 0.0039 |
| F07 | 70480001 | 70580000 | 0.0048 | 0.0031 | 0.0038 |
| F07 | 70500001 | 70600000 | 0.0042 | 0.0032 | 0.0032 |
| F07 | 70520001 | 70620000 | 0.0049 | 0.0038 | 0.0036 |
| F07 | 70540001 | 70640000 | 0.0044 | 0.0033 | 0.0032 |
| F07 | 70560001 | 70660000 | 0.0050 | 0.0037 | 0.0037 |
| F07 | 70580001 | 70680000 | 0.0046 | 0.0040 | 0.0039 |
| F07 | 70600001 | 70700000 | 0.0063 | 0.0054 | 0.0053 |
| F07 | 70620001 | 70720000 | 0.0048 | 0.0039 | 0.0040 |
| F07 | 70640001 | 70740000 | 0.0060 | 0.0050 | 0.0049 |
| F07 | 70660001 | 70760000 | 0.0064 | 0.0053 | 0.0052 |
| F07 | 70680001 | 70780000 | 0.0070 | 0.0057 | 0.0057 |
| F07 | 70700001 | 70800000 | 0.0068 | 0.0053 | 0.0056 |
| F07 | 70720001 | 70820000 | 0.0076 | 0.0057 | 0.0063 |
| F07 | 70740001 | 70840000 | 0.0063 | 0.0046 | 0.0053 |
| F07 | 70760001 | 70860000 | 0.0060 | 0.0044 | 0.0048 |
| F07 | 70780001 | 70880000 | 0.0054 | 0.0040 | 0.0040 |
| F07 | 70800001 | 70900000 | 0.0045 | 0.0036 | 0.0033 |
| F07 | 70820001 | 70920000 | 0.0038 | 0.0032 | 0.0027 |
| F07 | 70840001 | 70940000 | 0.0042 | 0.0033 | 0.0031 |
| F07 | 70860001 | 70960000 | 0.0038 | 0.0029 | 0.0030 |
| F07 | 70880001 | 70980000 | 0.0030 | 0.0023 | 0.0026 |
| F07 | 70900001 | 71000000 | 0.0022 | 0.0015 | 0.0020 |
| F07 | 70920001 | 71020000 | 0.0021 | 0.0016 | 0.0020 |
| F07 | 70940001 | 71040000 | 0.0026 | 0.0020 | 0.0024 |
| F07 | 70960001 | 71060000 | 0.0026 | 0.0019 | 0.0023 |
| F07 | 70980001 | 71080000 | 0.0029 | 0.0020 | 0.0025 |
| F07 | 71000001 | 71100000 | 0.0034 | 0.0022 | 0.0029 |

| Chr | Start | End | both | A1 | A2 |
|---|---|---|---|---|---|
| F07 | 71020001 | 71120000 | 0.0037 | 0.0024 | 0.0032 |
| F07 | 71040001 | 71140000 | 0.0033 | 0.0021 | 0.0029 |
| F07 | 71060001 | 71160000 | 0.0031 | 0.0021 | 0.0028 |
| F07 | 71080001 | 71180000 | 0.0031 | 0.0021 | 0.0027 |
| F07 | 71100001 | 71200000 | 0.0029 | 0.0019 | 0.0025 |
| F07 | 71120001 | 71220000 | 0.0027 | 0.0020 | 0.0022 |
| F07 | 71140001 | 71240000 | 0.0024 | 0.0017 | 0.0019 |
| F07 | 71160001 | 71260000 | 0.0031 | 0.0023 | 0.0025 |
| F07 | 71180001 | 71280000 | 0.0030 | 0.0023 | 0.0024 |
| F07 | 71200001 | 71300000 | 0.0032 | 0.0025 | 0.0024 |
| F07 | 71220001 | 71320000 | 0.0030 | 0.0024 | 0.0022 |
| F07 | 71240001 | 71340000 | 0.0031 | 0.0024 | 0.0022 |
| F07 | 71260001 | 71360000 | 0.0025 | 0.0019 | 0.0014 |
| F07 | 71280001 | 71380000 | 0.0026 | 0.0021 | 0.0015 |
| F07 | 71300001 | 71400000 | 0.0028 | 0.0022 | 0.0019 |
| F07 | 71320001 | 71420000 | 0.0034 | 0.0028 | 0.0025 |
| F07 | 71340001 | 71440000 | 0.0034 | 0.0029 | 0.0026 |
| F07 | 71360001 | 71460000 | 0.0038 | 0.0033 | 0.0032 |
| F07 | 71380001 | 71480000 | 0.0034 | 0.0029 | 0.0029 |
| F07 | 71400001 | 71500000 | 0.0035 | 0.0030 | 0.0029 |
| F07 | 71420001 | 71520000 | 0.0032 | 0.0025 | 0.0025 |
| F07 | 71440001 | 71540000 | 0.0029 | 0.0022 | 0.0022 |
| F07 | 71460001 | 71560000 | 0.0029 | 0.0022 | 0.0021 |
| F07 | 71480001 | 71580000 | 0.0027 | 0.0020 | 0.0018 |
| F07 | 71500001 | 71600000 | 0.0026 | 0.0019 | 0.0019 |
| F07 | 71520001 | 71620000 | 0.0022 | 0.0017 | 0.0018 |
| F07 | 71540001 | 71640000 | 0.0028 | 0.0024 | 0.0025 |
| F07 | 71560001 | 71660000 | 0.0022 | 0.0019 | 0.0021 |
| F07 | 71580001 | 71680000 | 0.0046 | 0.0042 | 0.0044 |
| F07 | 71600001 | 71700000 | 0.0048 | 0.0043 | 0.0044 |
| F07 | 71620001 | 71720000 | 0.0054 | 0.0049 | 0.0048 |
| F07 | 71640001 | 71740000 | 0.0049 | 0.0044 | 0.0044 |
| F07 | 71660001 | 71760000 | 0.0053 | 0.0047 | 0.0047 |
| F07 | 71680001 | 71780000 | 0.0033 | 0.0028 | 0.0027 |
| F07 | 71700001 | 71800000 | 0.0025 | 0.0021 | 0.0021 |
| F07 | 71720001 | 71820000 | 0.0018 | 0.0015 | 0.0015 |
| F07 | 71740001 | 71840000 | 0.0017 | 0.0014 | 0.0014 |
| F07 | 71760001 | 71860000 | 0.0014 | 0.0012 | 0.0012 |
| F07 | 71780001 | 71880000 | 0.0015 | 0.0012 | 0.0014 |
| F07 | 71800001 | 71900000 | 0.0015 | 0.0013 | 0.0014 |
| F07 | 71820001 | 71920000 | 0.0022 | 0.0019 | 0.0021 |
| F07 | 71840001 | 71940000 | 0.0023 | 0.0019 | 0.0022 |
| F07 | 71860001 | 71960000 | 0.0030 | 0.0025 | 0.0028 |
| F07 | 71880001 | 71980000 | 0.0026 | 0.0022 | 0.0024 |
| F07 | 71900001 | 72000000 | 0.0031 | 0.0027 | 0.0030 |
| F07 | 71920001 | 72020000 | 0.0023 | 0.0020 | 0.0022 |
| F07 | 71940001 | 72040000 | 0.0022 | 0.0019 | 0.0020 |
| F07 | 71960001 | 72060000 | 0.0016 | 0.0015 | 0.0015 |
| F07 | 71980001 | 72080000 | 0.0024 | 0.0023 | 0.0022 |
| F07 | 72000001 | 72100000 | 0.0021 | 0.0020 | 0.0020 |
| F07 | 72020001 | 72120000 | 0.0023 | 0.0022 | 0.0022 |
| F07 | 72040001 | 72140000 | 0.0035 | 0.0032 | 0.0034 |
| F07 | 72060001 | 72160000 | 0.0037 | 0.0033 | 0.0036 |
| F07 | 72080001 | 72180000 | 0.0030 | 0.0026 | 0.0029 |
| F07 | 72100001 | 72200000 | 0.0029 | 0.0025 | 0.0029 |
| F07 | 72120001 | 72220000 | 0.0027 | 0.0022 | 0.0026 |
| F07 | 72140001 | 72240000 | 0.0022 | 0.0019 | 0.0022 |
| F07 | 72160001 | 72260000 | 0.0017 | 0.0016 | 0.0017 |

| Chr | Start | End | both | A1 | A2 |
|---|---|---|---|---|---|
| F07 | 72180001 | 72280000 | 0.0016 | 0.0014 | 0.0015 |
| F07 | 72200001 | 72300000 | 0.0014 | 0.0012 | 0.0013 |
| F07 | 72220001 | 72320000 | 0.0013 | 0.0011 | 0.0012 |
| F07 | 72240001 | 72340000 | 0.0003 | 0.0002 | 0.0003 |
| F07 | 72260001 | 72360000 | 0.0007 | 0.0006 | 0.0007 |
| F07 | 72280001 | 72380000 | 0.0011 | 0.0010 | 0.0010 |
| F07 | 72300001 | 72400000 | 0.0010 | 0.0009 | 0.0010 |
| F07 | 72320001 | 72420000 | 0.0015 | 0.0014 | 0.0015 |
| F07 | 72340001 | 72440000 | 0.0021 | 0.0019 | 0.0019 |
| F07 | 72360001 | 72460000 | 0.0030 | 0.0026 | 0.0028 |
| F07 | 72380001 | 72480000 | 0.0043 | 0.0037 | 0.0040 |
| F07 | 72400001 | 72500000 | 0.0050 | 0.0043 | 0.0047 |
| F07 | 72420001 | 72520000 | 0.0057 | 0.0049 | 0.0054 |
| F07 | 72440001 | 72540000 | 0.0060 | 0.0051 | 0.0058 |
| F07 | 72460001 | 72560000 | 0.0061 | 0.0051 | 0.0057 |
| F07 | 72480001 | 72580000 | 0.0046 | 0.0038 | 0.0042 |
| F07 | 72500001 | 72600000 | 0.0044 | 0.0036 | 0.0040 |
| F07 | 72520001 | 72620000 | 0.0035 | 0.0027 | 0.0030 |
| F07 | 72540001 | 72640000 | 0.0037 | 0.0030 | 0.0034 |
| F07 | 72560001 | 72660000 | 0.0027 | 0.0021 | 0.0025 |
| F07 | 72580001 | 72680000 | 0.0033 | 0.0027 | 0.0029 |
| F07 | 72600001 | 72700000 | 0.0032 | 0.0026 | 0.0025 |
| F07 | 72620001 | 72720000 | 0.0034 | 0.0028 | 0.0028 |
| F07 | 72640001 | 72740000 | 0.0027 | 0.0023 | 0.0020 |
| F07 | 72660001 | 72760000 | 0.0030 | 0.0026 | 0.0023 |
| F07 | 72680001 | 72780000 | 0.0030 | 0.0027 | 0.0025 |
| F07 | 72700001 | 72800000 | 0.0035 | 0.0032 | 0.0032 |
| F07 | 72720001 | 72820000 | 0.0038 | 0.0034 | 0.0036 |
| F07 | 72740001 | 72840000 | 0.0041 | 0.0036 | 0.0039 |
| F07 | 72760001 | 72860000 | 0.0038 | 0.0033 | 0.0037 |
| F07 | 72780001 | 72880000 | 0.0039 | 0.0032 | 0.0035 |
| F07 | 72800001 | 72900000 | 0.0038 | 0.0031 | 0.0034 |
| F07 | 72820001 | 72920000 | 0.0037 | 0.0030 | 0.0032 |
| F07 | 72840001 | 72940000 | 0.0040 | 0.0033 | 0.0035 |
| F07 | 72860001 | 72960000 | 0.0040 | 0.0033 | 0.0034 |
| F07 | 72880001 | 72980000 | 0.0040 | 0.0034 | 0.0035 |
| F07 | 72900001 | 73000000 | 0.0043 | 0.0033 | 0.0032 |
| F07 | 72920001 | 73020000 | 0.0048 | 0.0035 | 0.0034 |
| F07 | 72940001 | 73040000 | 0.0058 | 0.0044 | 0.0041 |
| F07 | 72960001 | 73060000 | 0.0068 | 0.0051 | 0.0050 |
| F07 | 72980001 | 73080000 | 0.0071 | 0.0051 | 0.0051 |
| F07 | 73000001 | 73100000 | 0.0066 | 0.0049 | 0.0049 |
| F07 | 73020001 | 73120000 | 0.0058 | 0.0044 | 0.0045 |
| F07 | 73040001 | 73140000 | 0.0038 | 0.0028 | 0.0029 |
| F07 | 73060001 | 73160000 | 0.0026 | 0.0020 | 0.0019 |
| F07 | 73080001 | 73180000 | 0.0018 | 0.0015 | 0.0014 |
| F07 | 73100001 | 73200000 | 0.0017 | 0.0016 | 0.0015 |
| F07 | 73120001 | 73220000 | 0.0023 | 0.0021 | 0.0019 |
| F07 | 73140001 | 73240000 | 0.0029 | 0.0024 | 0.0023 |
| F07 | 73160001 | 73260000 | 0.0033 | 0.0028 | 0.0027 |
| F07 | 73180001 | 73280000 | 0.0034 | 0.0027 | 0.0027 |
| F07 | 73200001 | 73300000 | 0.0033 | 0.0027 | 0.0026 |
| F07 | 73220001 | 73320000 | 0.0033 | 0.0027 | 0.0027 |
| F07 | 73240001 | 73340000 | 0.0030 | 0.0026 | 0.0025 |
| F07 | 73260001 | 73360000 | 0.0026 | 0.0022 | 0.0021 |
| F07 | 73280001 | 73380000 | 0.0037 | 0.0034 | 0.0033 |
| F07 | 73300001 | 73400000 | 0.0040 | 0.0036 | 0.0037 |
| F07 | 73320001 | 73420000 | 0.0032 | 0.0029 | 0.0029 |

| Chr | Start | End | both | A1 | A2 |
|---|---|---|---|---|---|
| F07 | 73340001 | 73440000 | 0.0032 | 0.0029 | 0.0029 |
| F07 | 73360001 | 73460000 | 0.0037 | 0.0032 | 0.0032 |
| F07 | 73380001 | 73480000 | 0.0025 | 0.0020 | 0.0019 |
| F07 | 73400001 | 73500000 | 0.0020 | 0.0015 | 0.0014 |
| F07 | 73420001 | 73520000 | 0.0023 | 0.0018 | 0.0017 |
| F07 | 73440001 | 73540000 | 0.0023 | 0.0018 | 0.0016 |
| F07 | 73460001 | 73560000 | 0.0022 | 0.0019 | 0.0018 |
| F07 | 73480001 | 73580000 | 0.0030 | 0.0027 | 0.0027 |
| F07 | 73500001 | 73600000 | 0.0034 | 0.0031 | 0.0030 |
| F07 | 73520001 | 73620000 | 0.0040 | 0.0035 | 0.0033 |
| F07 | 73540001 | 73640000 | 0.0047 | 0.0042 | 0.0039 |
| F07 | 73560001 | 73660000 | 0.0046 | 0.0040 | 0.0036 |
| F07 | 73580001 | 73680000 | 0.0041 | 0.0034 | 0.0031 |
| F07 | 73600001 | 73700000 | 0.0059 | 0.0051 | 0.0046 |
| F07 | 73620001 | 73720000 | 0.0056 | 0.0048 | 0.0043 |
| F07 | 73640001 | 73740000 | 0.0060 | 0.0051 | 0.0047 |
| F07 | 73660001 | 73760000 | 0.0059 | 0.0050 | 0.0046 |
| F07 | 73680001 | 73780000 | 0.0066 | 0.0057 | 0.0050 |
| F07 | 73700001 | 73800000 | 0.0053 | 0.0045 | 0.0039 |
| F07 | 73720001 | 73820000 | 0.0058 | 0.0050 | 0.0045 |
| F07 | 73740001 | 73840000 | 0.0071 | 0.0061 | 0.0059 |
| F07 | 73760001 | 73860000 | 0.0069 | 0.0060 | 0.0058 |
| F07 | 73780001 | 73880000 | 0.0061 | 0.0052 | 0.0051 |
| F07 | 73800001 | 73900000 | 0.0055 | 0.0046 | 0.0044 |
| F07 | 73820001 | 73920000 | 0.0049 | 0.0041 | 0.0039 |
| F07 | 73840001 | 73940000 | 0.0023 | 0.0018 | 0.0015 |
| F07 | 73860001 | 73960000 | 0.0026 | 0.0021 | 0.0018 |
| F07 | 73880001 | 73980000 | 0.0024 | 0.0019 | 0.0018 |
| F07 | 73900001 | 74000000 | 0.0023 | 0.0018 | 0.0020 |
| F07 | 73920001 | 74020000 | 0.0018 | 0.0014 | 0.0015 |
| F07 | 73940001 | 74040000 | 0.0019 | 0.0015 | 0.0015 |
| F07 | 73960001 | 74060000 | 0.0017 | 0.0013 | 0.0015 |
| F07 | 73980001 | 74080000 | 0.0019 | 0.0015 | 0.0016 |
| F07 | 74000001 | 74100000 | 0.0020 | 0.0017 | 0.0013 |
| F07 | 74020001 | 74120000 | 0.0019 | 0.0016 | 0.0013 |
| F07 | 74040001 | 74140000 | 0.0021 | 0.0017 | 0.0014 |
| F07 | 74060001 | 74160000 | 0.0034 | 0.0029 | 0.0025 |
| F07 | 74080001 | 74180000 | 0.0034 | 0.0030 | 0.0024 |
| F07 | 74100001 | 74200000 | 0.0032 | 0.0027 | 0.0023 |
| F07 | 74120001 | 74220000 | 0.0035 | 0.0030 | 0.0027 |
| F07 | 74140001 | 74240000 | 0.0036 | 0.0031 | 0.0027 |
| F07 | 74160001 | 74260000 | 0.0022 | 0.0019 | 0.0014 |
| F07 | 74180001 | 74280000 | 0.0021 | 0.0018 | 0.0014 |
| F07 | 74200001 | 74300000 | 0.0023 | 0.0020 | 0.0018 |
| F07 | 74220001 | 74320000 | 0.0035 | 0.0031 | 0.0028 |
| F07 | 74240001 | 74340000 | 0.0039 | 0.0035 | 0.0031 |
| F07 | 74260001 | 74360000 | 0.0047 | 0.0041 | 0.0038 |
| F07 | 74280001 | 74380000 | 0.0050 | 0.0042 | 0.0039 |
| F07 | 74300001 | 74400000 | 0.0052 | 0.0045 | 0.0042 |
| F07 | 74320001 | 74420000 | 0.0041 | 0.0034 | 0.0034 |
| F07 | 74340001 | 74440000 | 0.0043 | 0.0034 | 0.0036 |
| F07 | 74360001 | 74460000 | 0.0033 | 0.0026 | 0.0028 |
| F07 | 74380001 | 74480000 | 0.0037 | 0.0030 | 0.0031 |
| F07 | 74400001 | 74500000 | 0.0038 | 0.0031 | 0.0031 |
| F07 | 74420001 | 74520000 | 0.0037 | 0.0033 | 0.0029 |
| F07 | 74440001 | 74540000 | 0.0031 | 0.0029 | 0.0024 |
| F07 | 74460001 | 74560000 | 0.0034 | 0.0032 | 0.0026 |
| F07 | 74480001 | 74580000 | 0.0032 | 0.0033 | 0.0026 |

| Chr | Start | End | both | A1 | A2 |
|---|---|---|---|---|---|
| F07 | 74500001 | 74600000 | 0.0030 | 0.0031 | 0.0024 |
| F07 | 74520001 | 74620000 | 0.0025 | 0.0026 | 0.0020 |
| F07 | 74540001 | 74640000 | 0.0032 | 0.0035 | 0.0025 |
| F07 | 74560001 | 74660000 | 0.0041 | 0.0041 | 0.0035 |
| F07 | 74580001 | 74680000 | 0.0040 | 0.0041 | 0.0035 |
| F07 | 74600001 | 74700000 | 0.0037 | 0.0039 | 0.0031 |
| F07 | 74620001 | 74720000 | 0.0038 | 0.0039 | 0.0032 |
| F07 | 74640001 | 74740000 | 0.0029 | 0.0030 | 0.0025 |
| F07 | 74660001 | 74760000 | 0.0040 | 0.0038 | 0.0035 |
| F07 | 74680001 | 74780000 | 0.0045 | 0.0042 | 0.0042 |
| F07 | 74700001 | 74800000 | 0.0046 | 0.0041 | 0.0044 |
| F07 | 74720001 | 74820000 | 0.0050 | 0.0043 | 0.0046 |
| F07 | 74740001 | 74840000 | 0.0051 | 0.0044 | 0.0047 |
| F07 | 74760001 | 74860000 | 0.0034 | 0.0030 | 0.0029 |
| F07 | 74780001 | 74880000 | 0.0024 | 0.0021 | 0.0018 |
| F07 | 74800001 | 74900000 | 0.0022 | 0.0018 | 0.0016 |
| F07 | 74820001 | 74920000 | 0.0016 | 0.0014 | 0.0013 |
| F07 | 74840001 | 74940000 | 0.0014 | 0.0012 | 0.0011 |
| F07 | 74860001 | 74960000 | 0.0018 | 0.0014 | 0.0014 |
| F07 | 74880001 | 74980000 | 0.0015 | 0.0012 | 0.0013 |
| F07 | 74900001 | 75000000 | 0.0016 | 0.0013 | 0.0013 |
| F07 | 74920001 | 75020000 | 0.0022 | 0.0017 | 0.0016 |
| F07 | 74940001 | 75040000 | 0.0028 | 0.0022 | 0.0021 |
| F07 | 74960001 | 75060000 | 0.0032 | 0.0025 | 0.0024 |
| F07 | 74980001 | 75080000 | 0.0036 | 0.0028 | 0.0027 |
| F07 | 75000001 | 75100000 | 0.0037 | 0.0028 | 0.0027 |
| F07 | 75020001 | 75120000 | 0.0038 | 0.0029 | 0.0029 |
| F07 | 75040001 | 75140000 | 0.0041 | 0.0032 | 0.0032 |
| F07 | 75060001 | 75160000 | 0.0037 | 0.0028 | 0.0028 |
| F07 | 75080001 | 75180000 | 0.0042 | 0.0031 | 0.0035 |
| F07 | 75100001 | 75200000 | 0.0049 | 0.0035 | 0.0043 |
| F07 | 75120001 | 75220000 | 0.0046 | 0.0033 | 0.0041 |
| F07 | 75140001 | 75240000 | 0.0036 | 0.0023 | 0.0032 |
| F07 | 75160001 | 75260000 | 0.0032 | 0.0019 | 0.0027 |
| F07 | 75180001 | 75280000 | 0.0024 | 0.0015 | 0.0020 |
| F07 | 75200001 | 75300000 | 0.0020 | 0.0013 | 0.0014 |
| F07 | 75220001 | 75320000 | 0.0022 | 0.0014 | 0.0014 |
| F07 | 75240001 | 75340000 | 0.0027 | 0.0019 | 0.0019 |
| F07 | 75260001 | 75360000 | 0.0033 | 0.0025 | 0.0025 |
| F07 | 75280001 | 75380000 | 0.0034 | 0.0025 | 0.0025 |
| F07 | 75300001 | 75400000 | 0.0033 | 0.0025 | 0.0023 |
| F07 | 75320001 | 75420000 | 0.0036 | 0.0028 | 0.0026 |
| F07 | 75340001 | 75440000 | 0.0040 | 0.0029 | 0.0027 |
| F07 | 75360001 | 75460000 | 0.0035 | 0.0023 | 0.0020 |
| F07 | 75380001 | 75480000 | 0.0035 | 0.0023 | 0.0019 |
| F07 | 75400001 | 75500000 | 0.0031 | 0.0019 | 0.0016 |
| F07 | 75420001 | 75520000 | 0.0027 | 0.0016 | 0.0015 |
| F07 | 75440001 | 75540000 | 0.0019 | 0.0012 | 0.0011 |
| F07 | 75460001 | 75560000 | 0.0028 | 0.0020 | 0.0021 |
| F07 | 75480001 | 75580000 | 0.0031 | 0.0023 | 0.0024 |
| F07 | 75500001 | 75600000 | 0.0038 | 0.0028 | 0.0028 |
| F07 | 75520001 | 75620000 | 0.0035 | 0.0025 | 0.0026 |
| F07 | 75540001 | 75640000 | 0.0046 | 0.0032 | 0.0035 |
| F07 | 75560001 | 75660000 | 0.0045 | 0.0028 | 0.0031 |
| F07 | 75580001 | 75680000 | 0.0042 | 0.0025 | 0.0030 |
| F07 | 75600001 | 75700000 | 0.0041 | 0.0025 | 0.0029 |
| F07 | 75620001 | 75720000 | 0.0049 | 0.0031 | 0.0037 |
| F07 | 75640001 | 75740000 | 0.0042 | 0.0027 | 0.0031 |

| Chr | Start | End | both | A1 | A2 |
|---|---|---|---|---|---|
| F07 | 75660001 | 75760000 | 0.0040 | 0.0027 | 0.0030 |
| F07 | 75680001 | 75780000 | 0.0044 | 0.0029 | 0.0033 |
| F07 | 75700001 | 75800000 | 0.0051 | 0.0036 | 0.0043 |
| F07 | 75720001 | 75820000 | 0.0042 | 0.0029 | 0.0034 |
| F07 | 75740001 | 75840000 | 0.0041 | 0.0028 | 0.0035 |
| F07 | 75760001 | 75860000 | 0.0035 | 0.0025 | 0.0032 |
| F07 | 75780001 | 75880000 | 0.0029 | 0.0022 | 0.0028 |
| F07 | 75800001 | 75900000 | 0.0019 | 0.0013 | 0.0017 |
| F07 | 75820001 | 75920000 | 0.0021 | 0.0013 | 0.0017 |
| F07 | 75840001 | 75940000 | 0.0021 | 0.0015 | 0.0016 |
| F07 | 75860001 | 75960000 | 0.0033 | 0.0025 | 0.0027 |
| F07 | 75880001 | 75980000 | 0.0035 | 0.0027 | 0.0027 |
| F07 | 75900001 | 76000000 | 0.0034 | 0.0026 | 0.0026 |
| F07 | 75920001 | 76020000 | 0.0034 | 0.0028 | 0.0027 |
| F07 | 75940001 | 76040000 | 0.0028 | 0.0024 | 0.0022 |
| F07 | 75960001 | 76060000 | 0.0016 | 0.0013 | 0.0012 |
| F07 | 75980001 | 76080000 | 0.0018 | 0.0015 | 0.0014 |
| F07 | 76000001 | 76100000 | 0.0016 | 0.0014 | 0.0013 |
| F07 | 76020001 | 76120000 | 0.0013 | 0.0011 | 0.0012 |
| F07 | 76040001 | 76140000 | 0.0014 | 0.0012 | 0.0013 |
| F07 | 76060001 | 76160000 | 0.0018 | 0.0014 | 0.0013 |
| F07 | 76080001 | 76180000 | 0.0018 | 0.0013 | 0.0011 |
| F07 | 76100001 | 76200000 | 0.0019 | 0.0013 | 0.0011 |
| F07 | 76120001 | 76220000 | 0.0025 | 0.0016 | 0.0014 |
| F07 | 76140001 | 76240000 | 0.0027 | 0.0018 | 0.0014 |
| F07 | 76160001 | 76260000 | 0.0030 | 0.0022 | 0.0018 |
| F07 | 76180001 | 76280000 | 0.0029 | 0.0023 | 0.0019 |
| F07 | 76200001 | 76300000 | 0.0032 | 0.0025 | 0.0020 |
| F07 | 76220001 | 76320000 | 0.0027 | 0.0022 | 0.0018 |
| F07 | 76240001 | 76340000 | 0.0025 | 0.0021 | 0.0018 |
| F07 | 76260001 | 76360000 | 0.0019 | 0.0016 | 0.0013 |
| F07 | 76280001 | 76380000 | 0.0019 | 0.0016 | 0.0013 |
| F07 | 76300001 | 76400000 | 0.0017 | 0.0013 | 0.0011 |
| F07 | 76320001 | 76420000 | 0.0019 | 0.0015 | 0.0013 |
| F07 | 76340001 | 76440000 | 0.0019 | 0.0014 | 0.0012 |
| F07 | 76360001 | 76460000 | 0.0019 | 0.0014 | 0.0013 |
| F07 | 76380001 | 76480000 | 0.0019 | 0.0014 | 0.0015 |
| F07 | 76400001 | 76500000 | 0.0022 | 0.0017 | 0.0019 |
| F07 | 76420001 | 76520000 | 0.0022 | 0.0017 | 0.0018 |
| F07 | 76440001 | 76540000 | 0.0025 | 0.0018 | 0.0019 |
| F07 | 76460001 | 76560000 | 0.0025 | 0.0017 | 0.0018 |
| F07 | 76480001 | 76580000 | 0.0021 | 0.0014 | 0.0014 |
| F07 | 76500001 | 76600000 | 0.0021 | 0.0013 | 0.0011 |
| F07 | 76520001 | 76620000 | 0.0021 | 0.0014 | 0.0011 |
| F07 | 76540001 | 76640000 | 0.0018 | 0.0013 | 0.0009 |
| F07 | 76560001 | 76660000 | 0.0019 | 0.0014 | 0.0010 |
| F07 | 76580001 | 76680000 | 0.0023 | 0.0015 | 0.0011 |
| F07 | 76600001 | 76700000 | 0.0024 | 0.0015 | 0.0012 |
| F07 | 76620001 | 76720000 | 0.0022 | 0.0015 | 0.0013 |
| F07 | 76640001 | 76740000 | 0.0023 | 0.0016 | 0.0014 |
| F07 | 76660001 | 76760000 | 0.0029 | 0.0022 | 0.0021 |
| F07 | 76680001 | 76780000 | 0.0031 | 0.0023 | 0.0025 |
| F07 | 76700001 | 76800000 | 0.0025 | 0.0019 | 0.0022 |
| F07 | 76720001 | 76820000 | 0.0025 | 0.0019 | 0.0023 |
| F07 | 76740001 | 76840000 | 0.0029 | 0.0024 | 0.0028 |
| F07 | 76760001 | 76860000 | 0.0021 | 0.0017 | 0.0020 |
| F07 | 76780001 | 76880000 | 0.0018 | 0.0017 | 0.0018 |
| F07 | 76800001 | 76900000 | 0.0020 | 0.0018 | 0.0019 |

| Chr | Start | End | both | A1 | A2 |
|---|---|---|---|---|---|
| F07 | 76820001 | 76920000 | 0.0019 | 0.0018 | 0.0018 |
| F07 | 76840001 | 76940000 | 0.0017 | 0.0014 | 0.0016 |
| F07 | 76860001 | 76960000 | 0.0017 | 0.0014 | 0.0017 |
| F07 | 76880001 | 76980000 | 0.0016 | 0.0012 | 0.0015 |
| F07 | 76900001 | 77000000 | 0.0017 | 0.0012 | 0.0014 |
| F07 | 76920001 | 77020000 | 0.0017 | 0.0011 | 0.0013 |
| F07 | 76940001 | 77040000 | 0.0014 | 0.0009 | 0.0008 |
| F07 | 76960001 | 77060000 | 0.0015 | 0.0009 | 0.0007 |
| F07 | 76980001 | 77080000 | 0.0021 | 0.0016 | 0.0012 |
| F07 | 77000001 | 77100000 | 0.0022 | 0.0017 | 0.0013 |
| F07 | 77020001 | 77120000 | 0.0023 | 0.0021 | 0.0014 |
| F07 | 77040001 | 77140000 | 0.0022 | 0.0023 | 0.0014 |
| F07 | 77060001 | 77160000 | 0.0024 | 0.0028 | 0.0018 |
| F07 | 77080001 | 77180000 | 0.0015 | 0.0022 | 0.0011 |
| F07 | 77100001 | 77200000 | 0.0016 | 0.0025 | 0.0010 |
| F07 | 77120001 | 77220000 | 0.0014 | 0.0025 | 0.0009 |
| F07 | 77140001 | 77240000 | 0.0014 | 0.0023 | 0.0009 |
| F07 | 77160001 | 77260000 | 0.0014 | 0.0025 | 0.0009 |
| F07 | 77180001 | 77280000 | 0.0018 | 0.0030 | 0.0012 |
| F07 | 77200001 | 77300000 | 0.0020 | 0.0030 | 0.0015 |
| F07 | 77220001 | 77320000 | 0.0019 | 0.0027 | 0.0016 |
| F07 | 77240001 | 77340000 | 0.0020 | 0.0028 | 0.0017 |
| F07 | 77260001 | 77360000 | 0.0021 | 0.0025 | 0.0018 |
| F07 | 77280001 | 77380000 | 0.0017 | 0.0019 | 0.0016 |
| F07 | 77300001 | 77400000 | 0.0014 | 0.0016 | 0.0012 |
| F07 | 77320001 | 77420000 | 0.0015 | 0.0017 | 0.0013 |
| F07 | 77340001 | 77440000 | 0.0019 | 0.0021 | 0.0016 |
| F07 | 77360001 | 77460000 | 0.0016 | 0.0017 | 0.0013 |
| F07 | 77380001 | 77480000 | 0.0017 | 0.0018 | 0.0014 |
| F07 | 77400001 | 77500000 | 0.0017 | 0.0018 | 0.0014 |
| F07 | 77420001 | 77520000 | 0.0016 | 0.0018 | 0.0013 |
| F07 | 77440001 | 77540000 | 0.0012 | 0.0016 | 0.0009 |
| F07 | 77460001 | 77560000 | 0.0013 | 0.0019 | 0.0009 |
| F07 | 77480001 | 77580000 | 0.0011 | 0.0020 | 0.0007 |
| F07 | 77500001 | 77600000 | 0.0011 | 0.0019 | 0.0007 |
| F07 | 77520001 | 77620000 | 0.0013 | 0.0018 | 0.0007 |
| F07 | 77540001 | 77640000 | 0.0016 | 0.0017 | 0.0009 |
| F07 | 77560001 | 77660000 | 0.0018 | 0.0015 | 0.0010 |
| F07 | 77580001 | 77680000 | 0.0021 | 0.0014 | 0.0012 |
| F07 | 77600001 | 77700000 | 0.0024 | 0.0016 | 0.0014 |
| F07 | 77620001 | 77720000 | 0.0038 | 0.0030 | 0.0027 |
| F07 | 77640001 | 77740000 | 0.0033 | 0.0028 | 0.0025 |
| F07 | 77660001 | 77760000 | 0.0031 | 0.0028 | 0.0023 |
| F07 | 77680001 | 77780000 | 0.0032 | 0.0034 | 0.0024 |
| F07 | 77700001 | 77800000 | 0.0034 | 0.0039 | 0.0027 |
| F07 | 77720001 | 77820000 | 0.0020 | 0.0028 | 0.0013 |
| F07 | 77740001 | 77840000 | 0.0024 | 0.0036 | 0.0013 |
| F07 | 77760001 | 77860000 | 0.0024 | 0.0036 | 0.0012 |
| F07 | 77780001 | 77880000 | 0.0021 | 0.0030 | 0.0011 |
| F07 | 77800001 | 77900000 | 0.0018 | 0.0025 | 0.0008 |
| F07 | 77820001 | 77920000 | 0.0022 | 0.0025 | 0.0009 |
| F07 | 77840001 | 77940000 | 0.0024 | 0.0023 | 0.0016 |
| F07 | 77860001 | 77960000 | 0.0025 | 0.0023 | 0.0017 |
| F07 | 77880001 | 77980000 | 0.0025 | 0.0022 | 0.0015 |
| F07 | 77900001 | 78000000 | 0.0024 | 0.0021 | 0.0015 |
| F07 | 77920001 | 78020000 | 0.0021 | 0.0019 | 0.0015 |
| F07 | 77940001 | 78040000 | 0.0017 | 0.0015 | 0.0011 |
| F07 | 77960001 | 78060000 | 0.0017 | 0.0014 | 0.0012 |

| Chr | Start | End | both | A1 | A2 |
|---|---|---|---|---|---|
| F07 | 77980001 | 78080000 | 0.0016 | 0.0012 | 0.0011 |
| F07 | 78000001 | 78100000 | 0.0015 | 0.0012 | 0.0010 |
| F07 | 78020001 | 78120000 | 0.0016 | 0.0014 | 0.0010 |
| F07 | 78040001 | 78140000 | 0.0014 | 0.0014 | 0.0009 |
| F07 | 78060001 | 78160000 | 0.0013 | 0.0013 | 0.0007 |
| F07 | 78080001 | 78180000 | 0.0019 | 0.0020 | 0.0012 |
| F07 | 78100001 | 78200000 | 0.0030 | 0.0030 | 0.0023 |
| F07 | 78120001 | 78220000 | 0.0030 | 0.0031 | 0.0020 |
| F07 | 78140001 | 78240000 | 0.0030 | 0.0032 | 0.0021 |
| F07 | 78160001 | 78260000 | 0.0032 | 0.0034 | 0.0023 |
| F07 | 78180001 | 78280000 | 0.0028 | 0.0031 | 0.0018 |
| F07 | 78200001 | 78300000 | 0.0019 | 0.0026 | 0.0009 |
| F07 | 78220001 | 78320000 | 0.0024 | 0.0031 | 0.0013 |
| F07 | 78240001 | 78340000 | 0.0026 | 0.0034 | 0.0016 |
| F07 | 78260001 | 78360000 | 0.0022 | 0.0030 | 0.0013 |
| F07 | 78280001 | 78380000 | 0.0032 | 0.0038 | 0.0024 |
| F07 | 78300001 | 78400000 | 0.0034 | 0.0039 | 0.0030 |
| F07 | 78320001 | 78420000 | 0.0030 | 0.0033 | 0.0028 |
| F07 | 78340001 | 78440000 | 0.0027 | 0.0029 | 0.0026 |
| F07 | 78360001 | 78460000 | 0.0030 | 0.0033 | 0.0028 |
| F07 | 78380001 | 78480000 | 0.0031 | 0.0036 | 0.0028 |
| F07 | 78400001 | 78500000 | 0.0022 | 0.0027 | 0.0020 |
| F07 | 78420001 | 78520000 | 0.0022 | 0.0026 | 0.0020 |
| F07 | 78440001 | 78540000 | 0.0022 | 0.0024 | 0.0020 |
| F07 | 78460001 | 78560000 | 0.0025 | 0.0025 | 0.0024 |
| F07 | 78480001 | 78580000 | 0.0013 | 0.0011 | 0.0013 |
| F07 | 78500001 | 78600000 | 0.0026 | 0.0020 | 0.0026 |
| F07 | 78520001 | 78620000 | 0.0027 | 0.0024 | 0.0026 |
| F07 | 78540001 | 78640000 | 0.0031 | 0.0032 | 0.0030 |
| F07 | 78560001 | 78660000 | 0.0031 | 0.0033 | 0.0030 |
| F07 | 78580001 | 78680000 | 0.0035 | 0.0037 | 0.0033 |
| F07 | 78600001 | 78700000 | 0.0029 | 0.0033 | 0.0027 |
| F07 | 78620001 | 78720000 | 0.0028 | 0.0030 | 0.0027 |
| F07 | 78640001 | 78740000 | 0.0025 | 0.0025 | 0.0025 |
| F07 | 78660001 | 78760000 | 0.0027 | 0.0028 | 0.0026 |
| F07 | 78680001 | 78780000 | 0.0030 | 0.0031 | 0.0028 |
| F07 | 78700001 | 78800000 | 0.0029 | 0.0031 | 0.0027 |
| F07 | 78720001 | 78820000 | 0.0029 | 0.0031 | 0.0026 |
| F07 | 78740001 | 78840000 | 0.0030 | 0.0034 | 0.0027 |
| F07 | 78760001 | 78860000 | 0.0025 | 0.0029 | 0.0023 |
| F07 | 78780001 | 78880000 | 0.0027 | 0.0030 | 0.0025 |
| F07 | 78800001 | 78900000 | 0.0029 | 0.0026 | 0.0025 |
| F07 | 78820001 | 78920000 | 0.0028 | 0.0023 | 0.0025 |
| F07 | 78840001 | 78940000 | 0.0028 | 0.0018 | 0.0024 |
| F07 | 78860001 | 78960000 | 0.0027 | 0.0015 | 0.0023 |
| F07 | 78880001 | 78980000 | 0.0022 | 0.0010 | 0.0017 |
| F07 | 78900001 | 79000000 | 0.0018 | 0.0008 | 0.0014 |
| F07 | 78920001 | 79020000 | 0.0018 | 0.0008 | 0.0013 |
| F07 | 78940001 | 79040000 | 0.0017 | 0.0011 | 0.0011 |
| F07 | 78960001 | 79060000 | 0.0016 | 0.0015 | 0.0009 |
| F07 | 78980001 | 79080000 | 0.0017 | 0.0018 | 0.0010 |
| F07 | 79000001 | 79100000 | 0.0019 | 0.0022 | 0.0010 |
| F07 | 79020001 | 79120000 | 0.0018 | 0.0022 | 0.0010 |
| F07 | 79040001 | 79140000 | 0.0019 | 0.0019 | 0.0011 |
| F07 | 79060001 | 79160000 | 0.0018 | 0.0015 | 0.0010 |
| F07 | 79080001 | 79180000 | 0.0017 | 0.0011 | 0.0008 |
| F07 | 79100001 | 79200000 | 0.0016 | 0.0010 | 0.0008 |
| F07 | 79120001 | 79220000 | 0.0020 | 0.0014 | 0.0012 |

| Chr | Start | End | both | A1 | A2 |
|---|---|---|---|---|---|
| F07 | 79140001 | 79240000 | 0.0019 | 0.0014 | 0.0011 |
| F07 | 79160001 | 79260000 | 0.0020 | 0.0015 | 0.0013 |
| F07 | 79180001 | 79280000 | 0.0024 | 0.0017 | 0.0014 |
| F07 | 79200001 | 79300000 | 0.0026 | 0.0015 | 0.0014 |
| F07 | 79220001 | 79320000 | 0.0027 | 0.0015 | 0.0014 |
| F07 | 79240001 | 79340000 | 0.0028 | 0.0016 | 0.0015 |
| F07 | 79260001 | 79360000 | 0.0027 | 0.0015 | 0.0013 |
| F07 | 79280001 | 79380000 | 0.0024 | 0.0013 | 0.0014 |
| F07 | 79300001 | 79400000 | 0.0027 | 0.0017 | 0.0020 |
| F07 | 79320001 | 79420000 | 0.0026 | 0.0016 | 0.0020 |
| F07 | 79340001 | 79440000 | 0.0026 | 0.0017 | 0.0020 |
| F07 | 79360001 | 79460000 | 0.0028 | 0.0022 | 0.0024 |
| F07 | 79380001 | 79480000 | 0.0027 | 0.0025 | 0.0022 |
| F07 | 79400001 | 79500000 | 0.0023 | 0.0029 | 0.0015 |
| F07 | 79420001 | 79520000 | 0.0020 | 0.0030 | 0.0012 |
| F07 | 79440001 | 79540000 | 0.0017 | 0.0029 | 0.0009 |
| F07 | 79460001 | 79560000 | 0.0012 | 0.0025 | 0.0005 |
| F07 | 79480001 | 79580000 | 0.0017 | 0.0029 | 0.0010 |
| F07 | 79500001 | 79600000 | 0.0015 | 0.0023 | 0.0012 |
| F07 | 79520001 | 79620000 | 0.0014 | 0.0020 | 0.0011 |
| F07 | 79540001 | 79640000 | 0.0016 | 0.0022 | 0.0012 |
| F07 | 79560001 | 79660000 | 0.0021 | 0.0027 | 0.0016 |
| F07 | 79580001 | 79680000 | 0.0020 | 0.0025 | 0.0015 |
| F07 | 79600001 | 79700000 | 0.0020 | 0.0024 | 0.0013 |
| F07 | 79620001 | 79720000 | 0.0023 | 0.0023 | 0.0016 |
| F07 | 79640001 | 79740000 | 0.0023 | 0.0020 | 0.0017 |
| F07 | 79660001 | 79760000 | 0.0021 | 0.0014 | 0.0015 |
| F07 | 79680001 | 79780000 | 0.0020 | 0.0010 | 0.0014 |
| F07 | 79700001 | 79800000 | 0.0028 | 0.0016 | 0.0022 |
| F07 | 79720001 | 79820000 | 0.0033 | 0.0020 | 0.0024 |
| F07 | 79740001 | 79840000 | 0.0035 | 0.0021 | 0.0026 |
| F07 | 79760001 | 79860000 | 0.0036 | 0.0023 | 0.0027 |
| F07 | 79780001 | 79880000 | 0.0038 | 0.0024 | 0.0028 |
| F07 | 79800001 | 79900000 | 0.0034 | 0.0020 | 0.0022 |
| F07 | 79820001 | 79920000 | 0.0033 | 0.0021 | 0.0022 |
| F07 | 79840001 | 79940000 | 0.0035 | 0.0025 | 0.0022 |
| F07 | 79860001 | 79960000 | 0.0038 | 0.0025 | 0.0023 |
| F07 | 79880001 | 79980000 | 0.0038 | 0.0027 | 0.0023 |
| F07 | 79900001 | 80000000 | 0.0035 | 0.0026 | 0.0023 |
| F07 | 79920001 | 80020000 | 0.0036 | 0.0028 | 0.0026 |
| F07 | 79940001 | 80040000 | 0.0033 | 0.0024 | 0.0024 |
| F07 | 79960001 | 80060000 | 0.0030 | 0.0022 | 0.0023 |
| F07 | 79980001 | 80080000 | 0.0025 | 0.0017 | 0.0019 |
| F07 | 80000001 | 80100000 | 0.0024 | 0.0018 | 0.0019 |
| F07 | 80020001 | 80120000 | 0.0018 | 0.0015 | 0.0013 |
| F07 | 80040001 | 80140000 | 0.0025 | 0.0022 | 0.0020 |
| F07 | 80060001 | 80160000 | 0.0025 | 0.0024 | 0.0020 |
| F07 | 80080001 | 80180000 | 0.0027 | 0.0025 | 0.0021 |
| F07 | 80100001 | 80200000 | 0.0029 | 0.0023 | 0.0022 |
| F07 | 80120001 | 80220000 | 0.0029 | 0.0020 | 0.0021 |
| F07 | 80140001 | 80240000 | 0.0022 | 0.0012 | 0.0015 |
| F07 | 80160001 | 80260000 | 0.0021 | 0.0008 | 0.0015 |
| F07 | 80180001 | 80280000 | 0.0024 | 0.0008 | 0.0017 |
| F07 | 80200001 | 80300000 | 0.0024 | 0.0009 | 0.0017 |
| F07 | 80220001 | 80320000 | 0.0029 | 0.0018 | 0.0022 |
| F07 | 80240001 | 80340000 | 0.0032 | 0.0026 | 0.0024 |
| F07 | 80260001 | 80360000 | 0.0031 | 0.0028 | 0.0021 |
| F07 | 80280001 | 80380000 | 0.0028 | 0.0032 | 0.0019 |

| Chr | Start | End | both | A1 | A2 |
|---|---|---|---|---|---|
| F07 | 80300001 | 80400000 | 0.0036 | 0.0045 | 0.0026 |
| F07 | 80320001 | 80420000 | 0.0030 | 0.0039 | 0.0021 |
| F07 | 80340001 | 80440000 | 0.0027 | 0.0036 | 0.0020 |
| F07 | 80360001 | 80460000 | 0.0028 | 0.0036 | 0.0022 |
| F07 | 80380001 | 80480000 | 0.0027 | 0.0034 | 0.0022 |
| F07 | 80400001 | 80500000 | 0.0026 | 0.0025 | 0.0024 |
| F07 | 80420001 | 80520000 | 0.0027 | 0.0021 | 0.0026 |
| F07 | 80440001 | 80540000 | 0.0031 | 0.0021 | 0.0030 |
| F07 | 80460001 | 80560000 | 0.0035 | 0.0021 | 0.0033 |
| F07 | 80480001 | 80580000 | 0.0040 | 0.0025 | 0.0039 |
| F07 | 80500001 | 80600000 | 0.0031 | 0.0021 | 0.0030 |
| F07 | 80520001 | 80620000 | 0.0034 | 0.0025 | 0.0033 |
| F07 | 80540001 | 80640000 | 0.0029 | 0.0021 | 0.0027 |
| F07 | 80560001 | 80660000 | 0.0026 | 0.0020 | 0.0025 |
| F07 | 80580001 | 80680000 | 0.0021 | 0.0016 | 0.0020 |
| F07 | 80600001 | 80700000 | 0.0022 | 0.0017 | 0.0022 |
| F07 | 80620001 | 80720000 | 0.0023 | 0.0016 | 0.0022 |
| F07 | 80640001 | 80740000 | 0.0022 | 0.0017 | 0.0022 |
| F07 | 80660001 | 80760000 | 0.0022 | 0.0017 | 0.0021 |
| F07 | 80680001 | 80780000 | 0.0022 | 0.0016 | 0.0021 |
| F07 | 80700001 | 80800000 | 0.0020 | 0.0014 | 0.0018 |
| F07 | 80720001 | 80820000 | 0.0017 | 0.0012 | 0.0013 |
| F07 | 80740001 | 80840000 | 0.0020 | 0.0013 | 0.0014 |
| F07 | 80760001 | 80860000 | 0.0022 | 0.0013 | 0.0015 |
| F07 | 80780001 | 80880000 | 0.0028 | 0.0018 | 0.0019 |
| F07 | 80800001 | 80900000 | 0.0034 | 0.0024 | 0.0025 |
| F07 | 80820001 | 80920000 | 0.0032 | 0.0024 | 0.0024 |
| F07 | 80840001 | 80940000 | 0.0032 | 0.0025 | 0.0025 |
| F07 | 80860001 | 80960000 | 0.0036 | 0.0029 | 0.0029 |
| F07 | 80880001 | 80980000 | 0.0029 | 0.0022 | 0.0021 |
| F07 | 80900001 | 81000000 | 0.0025 | 0.0017 | 0.0017 |
| F07 | 80920001 | 81020000 | 0.0027 | 0.0016 | 0.0018 |
| F07 | 80940001 | 81040000 | 0.0025 | 0.0013 | 0.0015 |
| F07 | 80960001 | 81060000 | 0.0018 | 0.0006 | 0.0009 |
| F07 | 80980001 | 81080000 | 0.0018 | 0.0006 | 0.0010 |
| F07 | 81000001 | 81100000 | 0.0015 | 0.0004 | 0.0009 |
| F07 | 81020001 | 81120000 | 0.0013 | 0.0003 | 0.0006 |
| F07 | 81040001 | 81140000 | 0.0014 | 0.0004 | 0.0007 |
| F07 | 81060001 | 81160000 | 0.0031 | 0.0019 | 0.0020 |
| F07 | 81080001 | 81180000 | 0.0034 | 0.0022 | 0.0021 |
| F07 | 81100001 | 81200000 | 0.0032 | 0.0022 | 0.0021 |
| F07 | 81120001 | 81220000 | 0.0036 | 0.0028 | 0.0026 |
| F07 | 81140001 | 81240000 | 0.0036 | 0.0029 | 0.0027 |
| F07 | 81160001 | 81260000 | 0.0022 | 0.0019 | 0.0017 |
| F07 | 81180001 | 81280000 | 0.0021 | 0.0018 | 0.0016 |
| F07 | 81200001 | 81300000 | 0.0024 | 0.0018 | 0.0017 |
| F07 | 81220001 | 81320000 | 0.0033 | 0.0023 | 0.0025 |
| F07 | 81240001 | 81340000 | 0.0041 | 0.0029 | 0.0033 |
| F07 | 81260001 | 81360000 | 0.0040 | 0.0026 | 0.0033 |
| F07 | 81280001 | 81380000 | 0.0041 | 0.0026 | 0.0035 |
| F07 | 81300001 | 81400000 | 0.0040 | 0.0026 | 0.0034 |
| F07 | 81320001 | 81420000 | 0.0030 | 0.0015 | 0.0022 |
| F07 | 81340001 | 81440000 | 0.0018 | 0.0004 | 0.0009 |
| F07 | 81360001 | 81460000 | 0.0016 | 0.0003 | 0.0007 |
| F07 | 81380001 | 81480000 | 0.0016 | 0.0002 | 0.0005 |
| F07 | 81400001 | 81500000 | 0.0017 | 0.0007 | 0.0007 |
| F07 | 81420001 | 81520000 | 0.0016 | 0.0009 | 0.0006 |
| F07 | 81440001 | 81540000 | 0.0020 | 0.0016 | 0.0009 |

| Chr | Start | End | both | A1 | A2 |
|---|---|---|---|---|---|
| F07 | 81460001 | 81560000 | 0.0022 | 0.0020 | 0.0010 |
| F07 | 81480001 | 81580000 | 0.0019 | 0.0021 | 0.0010 |
| F07 | 81500001 | 81600000 | 0.0020 | 0.0019 | 0.0010 |
| F07 | 81520001 | 81620000 | 0.0024 | 0.0022 | 0.0015 |
| F07 | 81540001 | 81640000 | 0.0021 | 0.0016 | 0.0014 |
| F07 | 81560001 | 81660000 | 0.0022 | 0.0015 | 0.0015 |
| F07 | 81580001 | 81680000 | 0.0026 | 0.0014 | 0.0017 |
| F07 | 81600001 | 81700000 | 0.0026 | 0.0012 | 0.0016 |
| F07 | 81620001 | 81720000 | 0.0021 | 0.0007 | 0.0012 |
| F07 | 81640001 | 81740000 | 0.0032 | 0.0018 | 0.0021 |
| F07 | 81660001 | 81760000 | 0.0038 | 0.0024 | 0.0028 |
| F07 | 81680001 | 81780000 | 0.0033 | 0.0023 | 0.0026 |
| F07 | 81700001 | 81800000 | 0.0036 | 0.0026 | 0.0029 |
| F07 | 81720001 | 81820000 | 0.0036 | 0.0026 | 0.0028 |
| F07 | 81740001 | 81840000 | 0.0026 | 0.0014 | 0.0018 |
| F07 | 81760001 | 81860000 | 0.0019 | 0.0007 | 0.0011 |
| F07 | 81780001 | 81880000 | 0.0022 | 0.0007 | 0.0012 |
| F07 | 81800001 | 81900000 | 0.0018 | 0.0003 | 0.0008 |
| F07 | 81820001 | 81920000 | 0.0021 | 0.0004 | 0.0011 |
| F07 | 81840001 | 81940000 | 0.0024 | 0.0007 | 0.0015 |
| F07 | 81860001 | 81960000 | 0.0025 | 0.0011 | 0.0016 |
| F07 | 81880001 | 81980000 | 0.0023 | 0.0011 | 0.0015 |
| F07 | 81900001 | 82000000 | 0.0022 | 0.0012 | 0.0014 |
| F07 | 81920001 | 82020000 | 0.0022 | 0.0014 | 0.0014 |
| F07 | 81940001 | 82040000 | 0.0022 | 0.0013 | 0.0012 |
| F07 | 81960001 | 82060000 | 0.0022 | 0.0012 | 0.0012 |
| F07 | 81980001 | 82080000 | 0.0023 | 0.0012 | 0.0013 |
| F07 | 82000001 | 82100000 | 0.0023 | 0.0013 | 0.0014 |
| F07 | 82020001 | 82120000 | 0.0025 | 0.0015 | 0.0017 |
| F07 | 82040001 | 82140000 | 0.0023 | 0.0014 | 0.0016 |
| F07 | 82060001 | 82160000 | 0.0019 | 0.0013 | 0.0014 |
| F07 | 82080001 | 82180000 | 0.0032 | 0.0024 | 0.0027 |
| F07 | 82100001 | 82200000 | 0.0036 | 0.0027 | 0.0030 |
| F07 | 82120001 | 82220000 | 0.0033 | 0.0025 | 0.0028 |
| F07 | 82140001 | 82240000 | 0.0029 | 0.0023 | 0.0026 |
| F07 | 82160001 | 82260000 | 0.0026 | 0.0020 | 0.0023 |
| F07 | 82180001 | 82280000 | 0.0010 | 0.0009 | 0.0009 |
| F07 | 82200001 | 82300000 | 0.0004 | 0.0004 | 0.0004 |
| F07 | 82220001 | 82320000 | 0.0000 | 0.0000 | 0.0000 |
| F08 | 1 | 100000 | 0.0019 | 0.0017 | 0.0014 |
| F08 | 20001 | 120000 | 0.0019 | 0.0016 | 0.0015 |
| F08 | 40001 | 140000 | 0.0020 | 0.0015 | 0.0015 |
| F08 | 60001 | 160000 | 0.0024 | 0.0021 | 0.0019 |
| F08 | 80001 | 180000 | 0.0025 | 0.0025 | 0.0020 |
| F08 | 100001 | 200000 | 0.0022 | 0.0024 | 0.0017 |
| F08 | 120001 | 220000 | 0.0020 | 0.0025 | 0.0014 |
| F08 | 140001 | 240000 | 0.0019 | 0.0025 | 0.0012 |
| F08 | 160001 | 260000 | 0.0028 | 0.0032 | 0.0021 |
| F08 | 180001 | 280000 | 0.0029 | 0.0031 | 0.0023 |
| F08 | 200001 | 300000 | 0.0034 | 0.0036 | 0.0027 |
| F08 | 220001 | 320000 | 0.0034 | 0.0035 | 0.0028 |
| F08 | 240001 | 340000 | 0.0036 | 0.0038 | 0.0030 |
| F08 | 260001 | 360000 | 0.0025 | 0.0026 | 0.0019 |
| F08 | 280001 | 380000 | 0.0029 | 0.0028 | 0.0023 |
| F08 | 300001 | 400000 | 0.0026 | 0.0021 | 0.0021 |
| F08 | 320001 | 420000 | 0.0031 | 0.0019 | 0.0026 |
| F08 | 340001 | 440000 | 0.0030 | 0.0015 | 0.0024 |
| F08 | 360001 | 460000 | 0.0030 | 0.0014 | 0.0024 |

| Chr | Start | End | both | A1 | A2 |
|---|---|---|---|---|---|
| F08 | 380001 | 480000 | 0.0022 | 0.0005 | 0.0016 |
| F08 | 400001 | 500000 | 0.0021 | 0.0005 | 0.0017 |
| F08 | 420001 | 520000 | 0.0017 | 0.0004 | 0.0013 |
| F08 | 440001 | 540000 | 0.0018 | 0.0009 | 0.0015 |
| F08 | 460001 | 560000 | 0.0021 | 0.0012 | 0.0018 |
| F08 | 480001 | 580000 | 0.0022 | 0.0018 | 0.0019 |
| F08 | 500001 | 600000 | 0.0020 | 0.0023 | 0.0016 |
| F08 | 520001 | 620000 | 0.0019 | 0.0027 | 0.0014 |
| F08 | 540001 | 640000 | 0.0015 | 0.0023 | 0.0012 |
| F08 | 560001 | 660000 | 0.0010 | 0.0019 | 0.0006 |
| F08 | 580001 | 680000 | 0.0007 | 0.0019 | 0.0002 |
| F08 | 600001 | 700000 | 0.0008 | 0.0020 | 0.0003 |
| F08 | 620001 | 720000 | 0.0009 | 0.0020 | 0.0004 |
| F08 | 640001 | 740000 | 0.0011 | 0.0020 | 0.0005 |
| F08 | 660001 | 760000 | 0.0014 | 0.0025 | 0.0007 |
| F08 | 680001 | 780000 | 0.0013 | 0.0023 | 0.0007 |
| F08 | 700001 | 800000 | 0.0014 | 0.0022 | 0.0008 |
| F08 | 720001 | 820000 | 0.0015 | 0.0024 | 0.0008 |
| F08 | 740001 | 840000 | 0.0015 | 0.0025 | 0.0008 |
| F08 | 760001 | 860000 | 0.0012 | 0.0024 | 0.0007 |
| F08 | 780001 | 880000 | 0.0012 | 0.0023 | 0.0006 |
| F08 | 800001 | 900000 | 0.0014 | 0.0025 | 0.0007 |
| F08 | 820001 | 920000 | 0.0012 | 0.0022 | 0.0006 |
| F08 | 840001 | 940000 | 0.0014 | 0.0020 | 0.0007 |
| F08 | 860001 | 960000 | 0.0017 | 0.0016 | 0.0009 |
| F08 | 880001 | 980000 | 0.0020 | 0.0015 | 0.0012 |
| F08 | 900001 | 1000000 | 0.0019 | 0.0008 | 0.0011 |
| F08 | 920001 | 1020000 | 0.0020 | 0.0011 | 0.0011 |
| F08 | 940001 | 1040000 | 0.0018 | 0.0013 | 0.0011 |
| F08 | 960001 | 1060000 | 0.0014 | 0.0013 | 0.0009 |
| F08 | 980001 | 1080000 | 0.0012 | 0.0016 | 0.0008 |
| F08 | 1000001 | 1100000 | 0.0009 | 0.0017 | 0.0005 |
| F08 | 1020001 | 1120000 | 0.0009 | 0.0017 | 0.0004 |
| F08 | 1040001 | 1140000 | 0.0011 | 0.0019 | 0.0005 |
| F08 | 1060001 | 1160000 | 0.0013 | 0.0020 | 0.0004 |
| F08 | 1080001 | 1180000 | 0.0014 | 0.0020 | 0.0004 |
| F08 | 1100001 | 1200000 | 0.0016 | 0.0020 | 0.0005 |
| F08 | 1120001 | 1220000 | 0.0016 | 0.0015 | 0.0005 |
| F08 | 1140001 | 1240000 | 0.0016 | 0.0014 | 0.0006 |
| F08 | 1160001 | 1260000 | 0.0019 | 0.0017 | 0.0010 |
| F08 | 1180001 | 1280000 | 0.0018 | 0.0015 | 0.0010 |
| F08 | 1200001 | 1300000 | 0.0016 | 0.0014 | 0.0010 |
| F08 | 1220001 | 1320000 | 0.0017 | 0.0017 | 0.0013 |
| F08 | 1240001 | 1340000 | 0.0023 | 0.0020 | 0.0018 |
| F08 | 1260001 | 1360000 | 0.0023 | 0.0018 | 0.0019 |
| F08 | 1280001 | 1380000 | 0.0028 | 0.0021 | 0.0024 |
| F08 | 1300001 | 1400000 | 0.0035 | 0.0027 | 0.0030 |
| F08 | 1320001 | 1420000 | 0.0037 | 0.0028 | 0.0032 |
| F08 | 1340001 | 1440000 | 0.0032 | 0.0027 | 0.0028 |
| F08 | 1360001 | 1460000 | 0.0033 | 0.0031 | 0.0028 |
| F08 | 1380001 | 1480000 | 0.0030 | 0.0030 | 0.0026 |
| F08 | 1400001 | 1500000 | 0.0027 | 0.0028 | 0.0021 |
| F08 | 1420001 | 1520000 | 0.0024 | 0.0029 | 0.0016 |
| F08 | 1440001 | 1540000 | 0.0028 | 0.0032 | 0.0021 |
| F08 | 1460001 | 1560000 | 0.0031 | 0.0033 | 0.0025 |
| F08 | 1480001 | 1580000 | 0.0032 | 0.0032 | 0.0026 |
| F08 | 1500001 | 1600000 | 0.0032 | 0.0031 | 0.0029 |
| F08 | 1520001 | 1620000 | 0.0042 | 0.0037 | 0.0039 |

| Chr | Start | End | both | A1 | A2 |
|---|---|---|---|---|---|
| F08 | 1540001 | 1640000 | 0.0036 | 0.0032 | 0.0033 |
| F08 | 1560001 | 1660000 | 0.0029 | 0.0028 | 0.0025 |
| F08 | 1580001 | 1680000 | 0.0027 | 0.0027 | 0.0021 |
| F08 | 1600001 | 1700000 | 0.0024 | 0.0025 | 0.0016 |
| F08 | 1620001 | 1720000 | 0.0013 | 0.0015 | 0.0005 |
| F08 | 1640001 | 1740000 | 0.0010 | 0.0013 | 0.0002 |
| F08 | 1660001 | 1760000 | 0.0010 | 0.0012 | 0.0003 |
| F08 | 1680001 | 1780000 | 0.0011 | 0.0010 | 0.0003 |
| F08 | 1700001 | 1800000 | 0.0012 | 0.0008 | 0.0004 |
| F08 | 1720001 | 1820000 | 0.0014 | 0.0006 | 0.0004 |
| F08 | 1740001 | 1840000 | 0.0015 | 0.0006 | 0.0005 |
| F08 | 1760001 | 1860000 | 0.0016 | 0.0004 | 0.0005 |
| F08 | 1780001 | 1880000 | 0.0016 | 0.0005 | 0.0005 |
| F08 | 1800001 | 1900000 | 0.0015 | 0.0005 | 0.0006 |
| F08 | 1820001 | 1920000 | 0.0013 | 0.0006 | 0.0005 |
| F08 | 1840001 | 1940000 | 0.0012 | 0.0008 | 0.0005 |
| F08 | 1860001 | 1960000 | 0.0011 | 0.0011 | 0.0006 |
| F08 | 1880001 | 1980000 | 0.0012 | 0.0015 | 0.0008 |
| F08 | 1900001 | 2000000 | 0.0015 | 0.0020 | 0.0010 |
| F08 | 1920001 | 2020000 | 0.0024 | 0.0034 | 0.0018 |
| F08 | 1940001 | 2040000 | 0.0039 | 0.0051 | 0.0033 |
| F08 | 1960001 | 2060000 | 0.0040 | 0.0052 | 0.0033 |
| F08 | 1980001 | 2080000 | 0.0037 | 0.0048 | 0.0032 |
| F08 | 2000001 | 2100000 | 0.0032 | 0.0042 | 0.0028 |
| F08 | 2020001 | 2120000 | 0.0024 | 0.0031 | 0.0021 |
| F08 | 2040001 | 2140000 | 0.0009 | 0.0013 | 0.0007 |
| F08 | 2060001 | 2160000 | 0.0008 | 0.0008 | 0.0007 |
| F08 | 2080001 | 2180000 | 0.0010 | 0.0008 | 0.0009 |
| F08 | 2100001 | 2200000 | 0.0013 | 0.0007 | 0.0013 |
| F08 | 2120001 | 2220000 | 0.0014 | 0.0003 | 0.0014 |
| F08 | 2140001 | 2240000 | 0.0014 | 0.0002 | 0.0015 |
| F08 | 2160001 | 2260000 | 0.0014 | 0.0004 | 0.0016 |
| F08 | 2180001 | 2280000 | 0.0016 | 0.0007 | 0.0017 |
| F08 | 2200001 | 2300000 | 0.0016 | 0.0008 | 0.0016 |
| F08 | 2220001 | 2320000 | 0.0016 | 0.0010 | 0.0015 |
| F08 | 2240001 | 2340000 | 0.0019 | 0.0011 | 0.0015 |
| F08 | 2260001 | 2360000 | 0.0019 | 0.0010 | 0.0014 |
| F08 | 2280001 | 2380000 | 0.0018 | 0.0007 | 0.0012 |
| F08 | 2300001 | 2400000 | 0.0020 | 0.0009 | 0.0011 |
| F08 | 2320001 | 2420000 | 0.0026 | 0.0013 | 0.0016 |
| F08 | 2340001 | 2440000 | 0.0026 | 0.0013 | 0.0018 |
| F08 | 2360001 | 2460000 | 0.0026 | 0.0012 | 0.0018 |
| F08 | 2380001 | 2480000 | 0.0028 | 0.0013 | 0.0020 |
| F08 | 2400001 | 2500000 | 0.0034 | 0.0017 | 0.0026 |
| F08 | 2420001 | 2520000 | 0.0028 | 0.0012 | 0.0021 |
| F08 | 2440001 | 2540000 | 0.0026 | 0.0011 | 0.0017 |
| F08 | 2460001 | 2560000 | 0.0025 | 0.0011 | 0.0017 |
| F08 | 2480001 | 2580000 | 0.0020 | 0.0010 | 0.0014 |
| F08 | 2500001 | 2600000 | 0.0011 | 0.0004 | 0.0007 |
| F08 | 2520001 | 2620000 | 0.0011 | 0.0003 | 0.0010 |
| F08 | 2540001 | 2640000 | 0.0014 | 0.0003 | 0.0013 |
| F08 | 2560001 | 2660000 | 0.0015 | 0.0004 | 0.0014 |
| F08 | 2580001 | 2680000 | 0.0017 | 0.0004 | 0.0015 |
| F08 | 2600001 | 2700000 | 0.0017 | 0.0007 | 0.0013 |
| F08 | 2620001 | 2720000 | 0.0014 | 0.0009 | 0.0010 |
| F08 | 2640001 | 2740000 | 0.0010 | 0.0010 | 0.0006 |
| F08 | 2660001 | 2760000 | 0.0007 | 0.0011 | 0.0004 |
| F08 | 2680001 | 2780000 | 0.0005 | 0.0012 | 0.0002 |

| Chr | Start | End | both | A1 | A2 |
|---|---|---|---|---|---|
| F08 | 2700001 | 2800000 | 0.0007 | 0.0008 | 0.0003 |
| F08 | 2720001 | 2820000 | 0.0009 | 0.0006 | 0.0003 |
| F08 | 2740001 | 2840000 | 0.0011 | 0.0005 | 0.0003 |
| F08 | 2760001 | 2860000 | 0.0013 | 0.0003 | 0.0003 |
| F08 | 2780001 | 2880000 | 0.0014 | 0.0001 | 0.0003 |
| F08 | 2800001 | 2900000 | 0.0014 | 0.0002 | 0.0004 |
| F08 | 2820001 | 2920000 | 0.0013 | 0.0002 | 0.0003 |
| F08 | 2840001 | 2940000 | 0.0015 | 0.0005 | 0.0005 |
| F08 | 2860001 | 2960000 | 0.0013 | 0.0005 | 0.0005 |
| F08 | 2880001 | 2980000 | 0.0012 | 0.0005 | 0.0005 |
| F08 | 2900001 | 3000000 | 0.0012 | 0.0005 | 0.0005 |
| F08 | 2920001 | 3020000 | 0.0013 | 0.0007 | 0.0008 |
| F08 | 2940001 | 3040000 | 0.0010 | 0.0004 | 0.0006 |
| F08 | 2960001 | 3060000 | 0.0012 | 0.0004 | 0.0006 |
| F08 | 2980001 | 3080000 | 0.0015 | 0.0007 | 0.0010 |
| F08 | 3000001 | 3100000 | 0.0018 | 0.0011 | 0.0012 |
| F08 | 3020001 | 3120000 | 0.0022 | 0.0014 | 0.0015 |
| F08 | 3040001 | 3140000 | 0.0031 | 0.0024 | 0.0026 |
| F08 | 3060001 | 3160000 | 0.0039 | 0.0031 | 0.0032 |
| F08 | 3080001 | 3180000 | 0.0035 | 0.0028 | 0.0029 |
| F08 | 3100001 | 3200000 | 0.0032 | 0.0026 | 0.0027 |
| F08 | 3120001 | 3220000 | 0.0027 | 0.0022 | 0.0022 |
| F08 | 3140001 | 3240000 | 0.0018 | 0.0012 | 0.0011 |
| F08 | 3160001 | 3260000 | 0.0010 | 0.0005 | 0.0004 |
| F08 | 3180001 | 3280000 | 0.0011 | 0.0006 | 0.0004 |
| F08 | 3200001 | 3300000 | 0.0009 | 0.0003 | 0.0002 |
| F08 | 3220001 | 3320000 | 0.0009 | 0.0002 | 0.0001 |
| F08 | 3240001 | 3340000 | 0.0014 | 0.0006 | 0.0007 |
| F08 | 3260001 | 3360000 | 0.0015 | 0.0007 | 0.0008 |
| F08 | 3280001 | 3380000 | 0.0014 | 0.0007 | 0.0008 |
| F08 | 3300001 | 3400000 | 0.0017 | 0.0008 | 0.0009 |
| F08 | 3320001 | 3420000 | 0.0019 | 0.0009 | 0.0010 |
| F08 | 3340001 | 3440000 | 0.0017 | 0.0005 | 0.0006 |
| F08 | 3360001 | 3460000 | 0.0019 | 0.0004 | 0.0005 |
| F08 | 3380001 | 3480000 | 0.0018 | 0.0005 | 0.0007 |
| F08 | 3400001 | 3500000 | 0.0017 | 0.0005 | 0.0006 |
| F08 | 3420001 | 3520000 | 0.0019 | 0.0005 | 0.0010 |
| F08 | 3440001 | 3540000 | 0.0018 | 0.0005 | 0.0013 |
| F08 | 3460001 | 3560000 | 0.0017 | 0.0004 | 0.0014 |
| F08 | 3480001 | 3580000 | 0.0020 | 0.0006 | 0.0017 |
| F08 | 3500001 | 3600000 | 0.0022 | 0.0006 | 0.0019 |
| F08 | 3520001 | 3620000 | 0.0020 | 0.0006 | 0.0018 |
| F08 | 3540001 | 3640000 | 0.0018 | 0.0006 | 0.0017 |
| F08 | 3560001 | 3660000 | 0.0018 | 0.0006 | 0.0016 |
| F08 | 3580001 | 3680000 | 0.0021 | 0.0008 | 0.0018 |
| F08 | 3600001 | 3700000 | 0.0026 | 0.0016 | 0.0023 |
| F08 | 3620001 | 3720000 | 0.0028 | 0.0022 | 0.0026 |
| F08 | 3640001 | 3740000 | 0.0030 | 0.0024 | 0.0028 |
| F08 | 3660001 | 3760000 | 0.0031 | 0.0024 | 0.0028 |
| F08 | 3680001 | 3780000 | 0.0026 | 0.0019 | 0.0023 |
| F08 | 3700001 | 3800000 | 0.0019 | 0.0011 | 0.0015 |
| F08 | 3720001 | 3820000 | 0.0016 | 0.0005 | 0.0011 |
| F08 | 3740001 | 3840000 | 0.0023 | 0.0013 | 0.0017 |
| F08 | 3760001 | 3860000 | 0.0022 | 0.0012 | 0.0015 |
| F08 | 3780001 | 3880000 | 0.0021 | 0.0011 | 0.0013 |
| F08 | 3800001 | 3900000 | 0.0026 | 0.0015 | 0.0016 |
| F08 | 3820001 | 3920000 | 0.0028 | 0.0018 | 0.0019 |
| F08 | 3840001 | 3940000 | 0.0018 | 0.0009 | 0.0010 |

| Chr | Start | End | both | A1 | A2 |
|---|---|---|---|---|---|
| F08 | 3860001 | 3960000 | 0.0021 | 0.0009 | 0.0010 |
| F08 | 3880001 | 3980000 | 0.0021 | 0.0009 | 0.0010 |
| F08 | 3900001 | 4000000 | 0.0018 | 0.0007 | 0.0009 |
| F08 | 3920001 | 4020000 | 0.0015 | 0.0004 | 0.0006 |
| F08 | 3940001 | 4040000 | 0.0014 | 0.0003 | 0.0005 |
| F08 | 3960001 | 4060000 | 0.0013 | 0.0004 | 0.0006 |
| F08 | 3980001 | 4080000 | 0.0012 | 0.0004 | 0.0006 |
| F08 | 4000001 | 4100000 | 0.0012 | 0.0003 | 0.0005 |
| F08 | 4020001 | 4120000 | 0.0015 | 0.0006 | 0.0007 |
| F08 | 4040001 | 4140000 | 0.0015 | 0.0008 | 0.0009 |
| F08 | 4060001 | 4160000 | 0.0019 | 0.0012 | 0.0013 |
| F08 | 4080001 | 4180000 | 0.0020 | 0.0014 | 0.0015 |
| F08 | 4100001 | 4200000 | 0.0019 | 0.0014 | 0.0015 |
| F08 | 4120001 | 4220000 | 0.0016 | 0.0010 | 0.0011 |
| F08 | 4140001 | 4240000 | 0.0017 | 0.0011 | 0.0012 |
| F08 | 4160001 | 4260000 | 0.0013 | 0.0007 | 0.0008 |
| F08 | 4180001 | 4280000 | 0.0017 | 0.0011 | 0.0010 |
| F08 | 4200001 | 4300000 | 0.0018 | 0.0013 | 0.0012 |
| F08 | 4220001 | 4320000 | 0.0023 | 0.0019 | 0.0018 |
| F08 | 4240001 | 4340000 | 0.0032 | 0.0028 | 0.0026 |
| F08 | 4260001 | 4360000 | 0.0031 | 0.0027 | 0.0027 |
| F08 | 4280001 | 4380000 | 0.0031 | 0.0025 | 0.0025 |
| F08 | 4300001 | 4400000 | 0.0035 | 0.0029 | 0.0029 |
| F08 | 4320001 | 4420000 | 0.0035 | 0.0026 | 0.0028 |
| F08 | 4340001 | 4440000 | 0.0028 | 0.0017 | 0.0020 |
| F08 | 4360001 | 4460000 | 0.0028 | 0.0016 | 0.0019 |
| F08 | 4380001 | 4480000 | 0.0024 | 0.0013 | 0.0016 |
| F08 | 4400001 | 4500000 | 0.0018 | 0.0008 | 0.0011 |
| F08 | 4420001 | 4520000 | 0.0016 | 0.0008 | 0.0010 |
| F08 | 4440001 | 4540000 | 0.0014 | 0.0007 | 0.0009 |
| F08 | 4460001 | 4560000 | 0.0020 | 0.0013 | 0.0015 |
| F08 | 4480001 | 4580000 | 0.0029 | 0.0022 | 0.0024 |
| F08 | 4500001 | 4600000 | 0.0037 | 0.0031 | 0.0032 |
| F08 | 4520001 | 4620000 | 0.0035 | 0.0027 | 0.0029 |
| F08 | 4540001 | 4640000 | 0.0032 | 0.0026 | 0.0027 |
| F08 | 4560001 | 4660000 | 0.0026 | 0.0019 | 0.0022 |
| F08 | 4580001 | 4680000 | 0.0016 | 0.0011 | 0.0013 |
| F08 | 4600001 | 4700000 | 0.0021 | 0.0016 | 0.0018 |
| F08 | 4620001 | 4720000 | 0.0025 | 0.0021 | 0.0023 |
| F08 | 4640001 | 4740000 | 0.0055 | 0.0052 | 0.0051 |
| F08 | 4660001 | 4760000 | 0.0056 | 0.0052 | 0.0052 |
| F08 | 4680001 | 4780000 | 0.0059 | 0.0053 | 0.0054 |
| F08 | 4700001 | 4800000 | 0.0047 | 0.0039 | 0.0042 |
| F08 | 4720001 | 4820000 | 0.0043 | 0.0033 | 0.0037 |
| F08 | 4740001 | 4840000 | 0.0015 | 0.0002 | 0.0011 |
| F08 | 4760001 | 4860000 | 0.0015 | 0.0002 | 0.0011 |
| F08 | 4780001 | 4880000 | 0.0017 | 0.0004 | 0.0013 |
| F08 | 4800001 | 4900000 | 0.0016 | 0.0005 | 0.0013 |
| F08 | 4820001 | 4920000 | 0.0018 | 0.0009 | 0.0015 |
| F08 | 4840001 | 4940000 | 0.0018 | 0.0009 | 0.0015 |
| F08 | 4860001 | 4960000 | 0.0017 | 0.0009 | 0.0014 |
| F08 | 4880001 | 4980000 | 0.0016 | 0.0010 | 0.0014 |
| F08 | 4900001 | 5000000 | 0.0017 | 0.0011 | 0.0015 |
| F08 | 4920001 | 5020000 | 0.0043 | 0.0030 | 0.0039 |
| F08 | 4940001 | 5040000 | 0.0047 | 0.0032 | 0.0043 |
| F08 | 4960001 | 5060000 | 0.0045 | 0.0033 | 0.0042 |
| F08 | 4980001 | 5080000 | 0.0045 | 0.0033 | 0.0042 |
| F08 | 5000001 | 5100000 | 0.0050 | 0.0037 | 0.0047 |

| Chr | Start | End | both | A1 | A2 |
|---|---|---|---|---|---|
| F08 | 5020001 | 5120000 | 0.0024 | 0.0018 | 0.0023 |
| F08 | 5040001 | 5140000 | 0.0019 | 0.0015 | 0.0018 |
| F08 | 5060001 | 5160000 | 0.0021 | 0.0016 | 0.0020 |
| F08 | 5080001 | 5180000 | 0.0029 | 0.0024 | 0.0028 |
| F08 | 5100001 | 5200000 | 0.0020 | 0.0017 | 0.0020 |
| F08 | 5120001 | 5220000 | 0.0019 | 0.0017 | 0.0019 |
| F08 | 5140001 | 5240000 | 0.0020 | 0.0018 | 0.0020 |
| F08 | 5160001 | 5260000 | 0.0021 | 0.0019 | 0.0020 |
| F08 | 5180001 | 5280000 | 0.0016 | 0.0014 | 0.0015 |
| F08 | 5200001 | 5300000 | 0.0018 | 0.0013 | 0.0016 |
| F08 | 5220001 | 5320000 | 0.0017 | 0.0011 | 0.0014 |
| F08 | 5240001 | 5340000 | 0.0020 | 0.0010 | 0.0014 |
| F08 | 5260001 | 5360000 | 0.0018 | 0.0006 | 0.0009 |
| F08 | 5280001 | 5380000 | 0.0014 | 0.0003 | 0.0003 |
| F08 | 5300001 | 5400000 | 0.0012 | 0.0003 | 0.0002 |
| F08 | 5320001 | 5420000 | 0.0014 | 0.0005 | 0.0004 |
| F08 | 5340001 | 5440000 | 0.0017 | 0.0010 | 0.0009 |
| F08 | 5360001 | 5460000 | 0.0016 | 0.0011 | 0.0010 |
| F08 | 5380001 | 5480000 | 0.0014 | 0.0010 | 0.0010 |
| F08 | 5400001 | 5500000 | 0.0018 | 0.0014 | 0.0014 |
| F08 | 5420001 | 5520000 | 0.0015 | 0.0011 | 0.0012 |
| F08 | 5440001 | 5540000 | 0.0012 | 0.0008 | 0.0008 |
| F08 | 5460001 | 5560000 | 0.0024 | 0.0018 | 0.0019 |
| F08 | 5480001 | 5580000 | 0.0030 | 0.0023 | 0.0024 |
| F08 | 5500001 | 5600000 | 0.0028 | 0.0020 | 0.0020 |
| F08 | 5520001 | 5620000 | 0.0031 | 0.0020 | 0.0021 |
| F08 | 5540001 | 5640000 | 0.0032 | 0.0020 | 0.0021 |
| F08 | 5560001 | 5660000 | 0.0019 | 0.0009 | 0.0008 |
| F08 | 5580001 | 5680000 | 0.0015 | 0.0004 | 0.0004 |
| F08 | 5600001 | 5700000 | 0.0016 | 0.0006 | 0.0006 |
| F08 | 5620001 | 5720000 | 0.0015 | 0.0006 | 0.0007 |
| F08 | 5640001 | 5740000 | 0.0014 | 0.0005 | 0.0006 |
| F08 | 5660001 | 5760000 | 0.0016 | 0.0005 | 0.0006 |
| F08 | 5680001 | 5780000 | 0.0014 | 0.0004 | 0.0005 |
| F08 | 5700001 | 5800000 | 0.0014 | 0.0004 | 0.0005 |
| F08 | 5720001 | 5820000 | 0.0012 | 0.0003 | 0.0004 |
| F08 | 5740001 | 5840000 | 0.0014 | 0.0006 | 0.0007 |
| F08 | 5760001 | 5860000 | 0.0012 | 0.0005 | 0.0007 |
| F08 | 5780001 | 5880000 | 0.0022 | 0.0015 | 0.0018 |
| F08 | 5800001 | 5900000 | 0.0021 | 0.0016 | 0.0017 |
| F08 | 5820001 | 5920000 | 0.0028 | 0.0022 | 0.0025 |
| F08 | 5840001 | 5940000 | 0.0037 | 0.0031 | 0.0032 |
| F08 | 5860001 | 5960000 | 0.0040 | 0.0034 | 0.0036 |
| F08 | 5880001 | 5980000 | 0.0035 | 0.0028 | 0.0029 |
| F08 | 5900001 | 6000000 | 0.0035 | 0.0027 | 0.0029 |
| F08 | 5920001 | 6020000 | 0.0030 | 0.0021 | 0.0022 |
| F08 | 5940001 | 6040000 | 0.0022 | 0.0014 | 0.0014 |
| F08 | 5960001 | 6060000 | 0.0020 | 0.0011 | 0.0012 |
| F08 | 5980001 | 6080000 | 0.0016 | 0.0007 | 0.0009 |
| F08 | 6000001 | 6100000 | 0.0018 | 0.0010 | 0.0012 |
| F08 | 6020001 | 6120000 | 0.0019 | 0.0010 | 0.0013 |
| F08 | 6040001 | 6140000 | 0.0019 | 0.0009 | 0.0013 |
| F08 | 6060001 | 6160000 | 0.0025 | 0.0014 | 0.0019 |
| F08 | 6080001 | 6180000 | 0.0034 | 0.0023 | 0.0030 |
| F08 | 6100001 | 6200000 | 0.0035 | 0.0023 | 0.0030 |
| F08 | 6120001 | 6220000 | 0.0035 | 0.0023 | 0.0030 |
| F08 | 6140001 | 6240000 | 0.0035 | 0.0022 | 0.0030 |
| F08 | 6160001 | 6260000 | 0.0030 | 0.0018 | 0.0024 |

| Chr | Start | End | both | A1 | A2 |
|---|---|---|---|---|---|
| F08 | 6180001 | 6280000 | 0.0023 | 0.0013 | 0.0017 |
| F08 | 6200001 | 6300000 | 0.0020 | 0.0012 | 0.0016 |
| F08 | 6220001 | 6320000 | 0.0021 | 0.0012 | 0.0016 |
| F08 | 6240001 | 6340000 | 0.0019 | 0.0012 | 0.0014 |
| F08 | 6260001 | 6360000 | 0.0021 | 0.0014 | 0.0017 |
| F08 | 6280001 | 6380000 | 0.0022 | 0.0013 | 0.0016 |
| F08 | 6300001 | 6400000 | 0.0022 | 0.0012 | 0.0015 |
| F08 | 6320001 | 6420000 | 0.0025 | 0.0016 | 0.0020 |
| F08 | 6340001 | 6440000 | 0.0025 | 0.0016 | 0.0019 |
| F08 | 6360001 | 6460000 | 0.0028 | 0.0017 | 0.0022 |
| F08 | 6380001 | 6480000 | 0.0039 | 0.0030 | 0.0035 |
| F08 | 6400001 | 6500000 | 0.0042 | 0.0034 | 0.0039 |
| F08 | 6420001 | 6520000 | 0.0047 | 0.0038 | 0.0041 |
| F08 | 6440001 | 6540000 | 0.0047 | 0.0038 | 0.0041 |
| F08 | 6460001 | 6560000 | 0.0042 | 0.0036 | 0.0038 |
| F08 | 6480001 | 6580000 | 0.0034 | 0.0028 | 0.0028 |
| F08 | 6500001 | 6600000 | 0.0031 | 0.0023 | 0.0023 |
| F08 | 6520001 | 6620000 | 0.0021 | 0.0014 | 0.0014 |
| F08 | 6540001 | 6640000 | 0.0040 | 0.0033 | 0.0031 |
| F08 | 6560001 | 6660000 | 0.0051 | 0.0042 | 0.0039 |
| F08 | 6580001 | 6680000 | 0.0053 | 0.0043 | 0.0042 |
| F08 | 6600001 | 6700000 | 0.0058 | 0.0047 | 0.0047 |
| F08 | 6620001 | 6720000 | 0.0064 | 0.0053 | 0.0052 |
| F08 | 6640001 | 6740000 | 0.0044 | 0.0032 | 0.0034 |
| F08 | 6660001 | 6760000 | 0.0033 | 0.0021 | 0.0025 |
| F08 | 6680001 | 6780000 | 0.0026 | 0.0016 | 0.0019 |
| F08 | 6700001 | 6800000 | 0.0019 | 0.0013 | 0.0014 |
| F08 | 6720001 | 6820000 | 0.0015 | 0.0007 | 0.0009 |
| F08 | 6740001 | 6840000 | 0.0022 | 0.0014 | 0.0016 |
| F08 | 6760001 | 6860000 | 0.0027 | 0.0019 | 0.0020 |
| F08 | 6780001 | 6880000 | 0.0028 | 0.0019 | 0.0021 |
| F08 | 6800001 | 6900000 | 0.0032 | 0.0020 | 0.0022 |
| F08 | 6820001 | 6920000 | 0.0031 | 0.0020 | 0.0022 |
| F08 | 6840001 | 6940000 | 0.0024 | 0.0014 | 0.0015 |
| F08 | 6860001 | 6960000 | 0.0019 | 0.0010 | 0.0011 |
| F08 | 6880001 | 6980000 | 0.0019 | 0.0009 | 0.0011 |
| F08 | 6900001 | 7000000 | 0.0017 | 0.0010 | 0.0011 |
| F08 | 6920001 | 7020000 | 0.0017 | 0.0011 | 0.0013 |
| F08 | 6940001 | 7040000 | 0.0023 | 0.0017 | 0.0017 |
| F08 | 6960001 | 7060000 | 0.0028 | 0.0022 | 0.0022 |
| F08 | 6980001 | 7080000 | 0.0024 | 0.0019 | 0.0018 |
| F08 | 7000001 | 7100000 | 0.0028 | 0.0021 | 0.0022 |
| F08 | 7020001 | 7120000 | 0.0029 | 0.0022 | 0.0023 |
| F08 | 7040001 | 7140000 | 0.0026 | 0.0019 | 0.0022 |
| F08 | 7060001 | 7160000 | 0.0024 | 0.0017 | 0.0021 |
| F08 | 7080001 | 7180000 | 0.0029 | 0.0021 | 0.0025 |
| F08 | 7100001 | 7200000 | 0.0033 | 0.0022 | 0.0026 |
| F08 | 7120001 | 7220000 | 0.0031 | 0.0020 | 0.0024 |
| F08 | 7140001 | 7240000 | 0.0031 | 0.0017 | 0.0022 |
| F08 | 7160001 | 7260000 | 0.0040 | 0.0026 | 0.0030 |
| F08 | 7180001 | 7280000 | 0.0037 | 0.0022 | 0.0027 |
| F08 | 7200001 | 7300000 | 0.0029 | 0.0017 | 0.0020 |
| F08 | 7220001 | 7320000 | 0.0038 | 0.0026 | 0.0029 |
| F08 | 7240001 | 7340000 | 0.0039 | 0.0029 | 0.0031 |
| F08 | 7260001 | 7360000 | 0.0026 | 0.0019 | 0.0019 |
| F08 | 7280001 | 7380000 | 0.0028 | 0.0020 | 0.0021 |
| F08 | 7300001 | 7400000 | 0.0034 | 0.0025 | 0.0026 |
| F08 | 7320001 | 7420000 | 0.0033 | 0.0023 | 0.0026 |

| Chr | Start | End | both | A1 | A2 |
|---|---|---|---|---|---|
| F08 | 7340001 | 7440000 | 0.0031 | 0.0022 | 0.0025 |
| F08 | 7360001 | 7460000 | 0.0030 | 0.0019 | 0.0023 |
| F08 | 7380001 | 7480000 | 0.0028 | 0.0017 | 0.0022 |
| F08 | 7400001 | 7500000 | 0.0032 | 0.0022 | 0.0026 |
| F08 | 7420001 | 7520000 | 0.0029 | 0.0021 | 0.0023 |
| F08 | 7440001 | 7540000 | 0.0031 | 0.0021 | 0.0024 |
| F08 | 7460001 | 7560000 | 0.0032 | 0.0021 | 0.0024 |
| F08 | 7480001 | 7580000 | 0.0037 | 0.0023 | 0.0027 |
| F08 | 7500001 | 7600000 | 0.0030 | 0.0014 | 0.0017 |
| F08 | 7520001 | 7620000 | 0.0025 | 0.0010 | 0.0013 |
| F08 | 7540001 | 7640000 | 0.0024 | 0.0010 | 0.0012 |
| F08 | 7560001 | 7660000 | 0.0023 | 0.0011 | 0.0013 |
| F08 | 7580001 | 7680000 | 0.0020 | 0.0009 | 0.0011 |
| F08 | 7600001 | 7700000 | 0.0018 | 0.0008 | 0.0010 |
| F08 | 7620001 | 7720000 | 0.0026 | 0.0014 | 0.0016 |
| F08 | 7640001 | 7740000 | 0.0026 | 0.0016 | 0.0017 |
| F08 | 7660001 | 7760000 | 0.0025 | 0.0015 | 0.0016 |
| F08 | 7680001 | 7780000 | 0.0022 | 0.0013 | 0.0015 |
| F08 | 7700001 | 7800000 | 0.0020 | 0.0013 | 0.0015 |
| F08 | 7720001 | 7820000 | 0.0019 | 0.0012 | 0.0014 |
| F08 | 7740001 | 7840000 | 0.0020 | 0.0012 | 0.0015 |
| F08 | 7760001 | 7860000 | 0.0025 | 0.0017 | 0.0020 |
| F08 | 7780001 | 7880000 | 0.0034 | 0.0025 | 0.0029 |
| F08 | 7800001 | 7900000 | 0.0038 | 0.0028 | 0.0033 |
| F08 | 7820001 | 7920000 | 0.0039 | 0.0029 | 0.0033 |
| F08 | 7840001 | 7940000 | 0.0036 | 0.0027 | 0.0031 |
| F08 | 7860001 | 7960000 | 0.0034 | 0.0022 | 0.0027 |
| F08 | 7880001 | 7980000 | 0.0031 | 0.0017 | 0.0021 |
| F08 | 7900001 | 8000000 | 0.0032 | 0.0017 | 0.0022 |
| F08 | 7920001 | 8020000 | 0.0023 | 0.0010 | 0.0015 |
| F08 | 7940001 | 8040000 | 0.0022 | 0.0009 | 0.0014 |
| F08 | 7960001 | 8060000 | 0.0024 | 0.0011 | 0.0015 |
| F08 | 7980001 | 8080000 | 0.0022 | 0.0013 | 0.0015 |
| F08 | 8000001 | 8100000 | 0.0023 | 0.0014 | 0.0015 |
| F08 | 8020001 | 8120000 | 0.0028 | 0.0020 | 0.0021 |
| F08 | 8040001 | 8140000 | 0.0036 | 0.0027 | 0.0030 |
| F08 | 8060001 | 8160000 | 0.0039 | 0.0031 | 0.0034 |
| F08 | 8080001 | 8180000 | 0.0040 | 0.0028 | 0.0034 |
| F08 | 8100001 | 8200000 | 0.0047 | 0.0034 | 0.0038 |
| F08 | 8120001 | 8220000 | 0.0044 | 0.0028 | 0.0033 |
| F08 | 8140001 | 8240000 | 0.0040 | 0.0025 | 0.0028 |
| F08 | 8160001 | 8260000 | 0.0038 | 0.0021 | 0.0025 |
| F08 | 8180001 | 8280000 | 0.0036 | 0.0021 | 0.0023 |
| F08 | 8200001 | 8300000 | 0.0027 | 0.0012 | 0.0015 |
| F08 | 8220001 | 8320000 | 0.0024 | 0.0011 | 0.0014 |
| F08 | 8240001 | 8340000 | 0.0024 | 0.0009 | 0.0013 |
| F08 | 8260001 | 8360000 | 0.0034 | 0.0020 | 0.0023 |
| F08 | 8280001 | 8380000 | 0.0037 | 0.0024 | 0.0027 |
| F08 | 8300001 | 8400000 | 0.0035 | 0.0024 | 0.0026 |
| F08 | 8320001 | 8420000 | 0.0053 | 0.0034 | 0.0044 |
| F08 | 8340001 | 8440000 | 0.0050 | 0.0033 | 0.0044 |
| F08 | 8360001 | 8460000 | 0.0037 | 0.0021 | 0.0032 |
| F08 | 8380001 | 8480000 | 0.0036 | 0.0018 | 0.0030 |
| F08 | 8400001 | 8500000 | 0.0041 | 0.0021 | 0.0034 |
| F08 | 8420001 | 8520000 | 0.0026 | 0.0013 | 0.0017 |
| F08 | 8440001 | 8540000 | 0.0034 | 0.0019 | 0.0025 |
| F08 | 8460001 | 8560000 | 0.0047 | 0.0031 | 0.0036 |
| F08 | 8480001 | 8580000 | 0.0042 | 0.0030 | 0.0031 |

| Chr | Start | End | both | A1 | A2 |
|---|---|---|---|---|---|
| F08 | 8500001 | 8600000 | 0.0041 | 0.0030 | 0.0031 |
| F08 | 8520001 | 8620000 | 0.0044 | 0.0034 | 0.0036 |
| F08 | 8540001 | 8640000 | 0.0033 | 0.0026 | 0.0026 |
| F08 | 8560001 | 8660000 | 0.0021 | 0.0018 | 0.0017 |
| F08 | 8580001 | 8680000 | 0.0019 | 0.0017 | 0.0017 |
| F08 | 8600001 | 8700000 | 0.0014 | 0.0014 | 0.0014 |
| F08 | 8620001 | 8720000 | 0.0009 | 0.0009 | 0.0008 |
| F08 | 8640001 | 8740000 | 0.0011 | 0.0009 | 0.0008 |
| F08 | 8660001 | 8760000 | 0.0011 | 0.0007 | 0.0007 |
| F08 | 8680001 | 8780000 | 0.0015 | 0.0010 | 0.0011 |
| F08 | 8700001 | 8800000 | 0.0022 | 0.0013 | 0.0015 |
| F08 | 8720001 | 8820000 | 0.0023 | 0.0013 | 0.0016 |
| F08 | 8740001 | 8840000 | 0.0026 | 0.0016 | 0.0019 |
| F08 | 8760001 | 8860000 | 0.0038 | 0.0026 | 0.0032 |
| F08 | 8780001 | 8880000 | 0.0038 | 0.0026 | 0.0030 |
| F08 | 8800001 | 8900000 | 0.0035 | 0.0024 | 0.0028 |
| F08 | 8820001 | 8920000 | 0.0044 | 0.0031 | 0.0036 |
| F08 | 8840001 | 8940000 | 0.0050 | 0.0037 | 0.0042 |
| F08 | 8860001 | 8960000 | 0.0038 | 0.0027 | 0.0030 |
| F08 | 8880001 | 8980000 | 0.0038 | 0.0027 | 0.0032 |
| F08 | 8900001 | 9000000 | 0.0035 | 0.0025 | 0.0031 |
| F08 | 8920001 | 9020000 | 0.0033 | 0.0024 | 0.0029 |
| F08 | 8940001 | 9040000 | 0.0030 | 0.0021 | 0.0025 |
| F08 | 8960001 | 9060000 | 0.0035 | 0.0026 | 0.0030 |
| F08 | 8980001 | 9080000 | 0.0034 | 0.0025 | 0.0028 |
| F08 | 9000001 | 9100000 | 0.0035 | 0.0026 | 0.0028 |
| F08 | 9020001 | 9120000 | 0.0030 | 0.0020 | 0.0023 |
| F08 | 9040001 | 9140000 | 0.0035 | 0.0024 | 0.0025 |
| F08 | 9060001 | 9160000 | 0.0035 | 0.0024 | 0.0024 |
| F08 | 9080001 | 9180000 | 0.0037 | 0.0028 | 0.0027 |
| F08 | 9100001 | 9200000 | 0.0040 | 0.0031 | 0.0031 |
| F08 | 9120001 | 9220000 | 0.0046 | 0.0039 | 0.0039 |
| F08 | 9140001 | 9240000 | 0.0040 | 0.0034 | 0.0035 |
| F08 | 9160001 | 9260000 | 0.0030 | 0.0026 | 0.0028 |
| F08 | 9180001 | 9280000 | 0.0033 | 0.0026 | 0.0027 |
| F08 | 9200001 | 9300000 | 0.0027 | 0.0021 | 0.0021 |
| F08 | 9220001 | 9320000 | 0.0020 | 0.0013 | 0.0013 |
| F08 | 9240001 | 9340000 | 0.0022 | 0.0012 | 0.0013 |
| F08 | 9260001 | 9360000 | 0.0031 | 0.0019 | 0.0020 |
| F08 | 9280001 | 9380000 | 0.0030 | 0.0017 | 0.0018 |
| F08 | 9300001 | 9400000 | 0.0032 | 0.0019 | 0.0021 |
| F08 | 9320001 | 9420000 | 0.0036 | 0.0021 | 0.0022 |
| F08 | 9340001 | 9440000 | 0.0030 | 0.0017 | 0.0018 |
| F08 | 9360001 | 9460000 | 0.0027 | 0.0013 | 0.0015 |
| F08 | 9380001 | 9480000 | 0.0028 | 0.0016 | 0.0018 |
| F08 | 9400001 | 9500000 | 0.0027 | 0.0014 | 0.0017 |
| F08 | 9420001 | 9520000 | 0.0026 | 0.0013 | 0.0016 |
| F08 | 9440001 | 9540000 | 0.0026 | 0.0013 | 0.0016 |
| F08 | 9460001 | 9560000 | 0.0024 | 0.0014 | 0.0017 |
| F08 | 9480001 | 9580000 | 0.0021 | 0.0011 | 0.0013 |
| F08 | 9500001 | 9600000 | 0.0021 | 0.0013 | 0.0014 |
| F08 | 9520001 | 9620000 | 0.0020 | 0.0012 | 0.0012 |
| F08 | 9540001 | 9640000 | 0.0026 | 0.0017 | 0.0018 |
| F08 | 9560001 | 9660000 | 0.0027 | 0.0017 | 0.0018 |
| F08 | 9580001 | 9680000 | 0.0039 | 0.0029 | 0.0030 |
| F08 | 9600001 | 9700000 | 0.0044 | 0.0033 | 0.0034 |
| F08 | 9620001 | 9720000 | 0.0045 | 0.0033 | 0.0035 |
| F08 | 9640001 | 9740000 | 0.0038 | 0.0030 | 0.0030 |

| Chr | Start | End | both | A1 | A2 |
|---|---|---|---|---|---|
| F08 | 9660001 | 9760000 | 0.0044 | 0.0034 | 0.0034 |
| F08 | 9680001 | 9780000 | 0.0029 | 0.0020 | 0.0019 |
| F08 | 9700001 | 9800000 | 0.0029 | 0.0016 | 0.0018 |
| F08 | 9720001 | 9820000 | 0.0033 | 0.0021 | 0.0024 |
| F08 | 9740001 | 9840000 | 0.0038 | 0.0025 | 0.0029 |
| F08 | 9760001 | 9860000 | 0.0041 | 0.0028 | 0.0032 |
| F08 | 9780001 | 9880000 | 0.0042 | 0.0028 | 0.0031 |
| F08 | 9800001 | 9900000 | 0.0037 | 0.0026 | 0.0028 |
| F08 | 9820001 | 9920000 | 0.0034 | 0.0023 | 0.0025 |
| F08 | 9840001 | 9940000 | 0.0032 | 0.0021 | 0.0021 |
| F08 | 9860001 | 9960000 | 0.0026 | 0.0017 | 0.0017 |
| F08 | 9880001 | 9980000 | 0.0033 | 0.0021 | 0.0022 |
| F08 | 9900001 | 10000000 | 0.0034 | 0.0022 | 0.0023 |
| F08 | 9920001 | 10020000 | 0.0030 | 0.0018 | 0.0019 |
| F08 | 9940001 | 10040000 | 0.0029 | 0.0018 | 0.0018 |
| F08 | 9960001 | 10060000 | 0.0028 | 0.0014 | 0.0015 |
| F08 | 9980001 | 10080000 | 0.0021 | 0.0010 | 0.0010 |
| F08 | 10000001 | 10100000 | 0.0019 | 0.0008 | 0.0008 |
| F08 | 10020001 | 10120000 | 0.0024 | 0.0012 | 0.0013 |
| F08 | 10040001 | 10140000 | 0.0024 | 0.0013 | 0.0013 |
| F08 | 10060001 | 10160000 | 0.0020 | 0.0010 | 0.0010 |
| F08 | 10080001 | 10180000 | 0.0020 | 0.0012 | 0.0012 |
| F08 | 10100001 | 10200000 | 0.0026 | 0.0017 | 0.0018 |
| F08 | 10120001 | 10220000 | 0.0028 | 0.0017 | 0.0019 |
| F08 | 10140001 | 10240000 | 0.0032 | 0.0018 | 0.0023 |
| F08 | 10160001 | 10260000 | 0.0035 | 0.0020 | 0.0024 |
| F08 | 10180001 | 10280000 | 0.0035 | 0.0020 | 0.0024 |
| F08 | 10200001 | 10300000 | 0.0031 | 0.0016 | 0.0020 |
| F08 | 10220001 | 10320000 | 0.0030 | 0.0015 | 0.0018 |
| F08 | 10240001 | 10340000 | 0.0029 | 0.0014 | 0.0015 |
| F08 | 10260001 | 10360000 | 0.0026 | 0.0013 | 0.0013 |
| F08 | 10280001 | 10380000 | 0.0026 | 0.0013 | 0.0013 |
| F08 | 10300001 | 10400000 | 0.0026 | 0.0014 | 0.0014 |
| F08 | 10320001 | 10420000 | 0.0024 | 0.0014 | 0.0015 |
| F08 | 10340001 | 10440000 | 0.0024 | 0.0015 | 0.0017 |
| F08 | 10360001 | 10460000 | 0.0028 | 0.0019 | 0.0019 |
| F08 | 10380001 | 10480000 | 0.0025 | 0.0017 | 0.0018 |
| F08 | 10400001 | 10500000 | 0.0028 | 0.0016 | 0.0017 |
| F08 | 10420001 | 10520000 | 0.0037 | 0.0022 | 0.0023 |
| F08 | 10440001 | 10540000 | 0.0032 | 0.0020 | 0.0021 |
| F08 | 10460001 | 10560000 | 0.0028 | 0.0016 | 0.0018 |
| F08 | 10480001 | 10580000 | 0.0032 | 0.0016 | 0.0019 |
| F08 | 10500001 | 10600000 | 0.0031 | 0.0019 | 0.0022 |
| F08 | 10520001 | 10620000 | 0.0020 | 0.0010 | 0.0012 |
| F08 | 10540001 | 10640000 | 0.0020 | 0.0009 | 0.0010 |
| F08 | 10560001 | 10660000 | 0.0035 | 0.0023 | 0.0024 |
| F08 | 10580001 | 10680000 | 0.0032 | 0.0024 | 0.0024 |
| F08 | 10600001 | 10700000 | 0.0030 | 0.0020 | 0.0021 |
| F08 | 10620001 | 10720000 | 0.0030 | 0.0020 | 0.0021 |
| F08 | 10640001 | 10740000 | 0.0029 | 0.0021 | 0.0022 |
| F08 | 10660001 | 10760000 | 0.0012 | 0.0006 | 0.0008 |
| F08 | 10680001 | 10780000 | 0.0008 | 0.0005 | 0.0006 |
| F08 | 10700001 | 10800000 | 0.0005 | 0.0004 | 0.0004 |
| F08 | 10720001 | 10820000 | 0.0006 | 0.0005 | 0.0006 |
| F08 | 10740001 | 10840000 | 0.0008 | 0.0007 | 0.0006 |
| F08 | 10760001 | 10860000 | 0.0011 | 0.0008 | 0.0007 |
| F08 | 10780001 | 10880000 | 0.0019 | 0.0014 | 0.0013 |
| F08 | 10800001 | 10900000 | 0.0021 | 0.0015 | 0.0016 |

| Chr | Start | End | both | A1 | A2 |
|---|---|---|---|---|---|
| F08 | 10820001 | 10920000 | 0.0024 | 0.0017 | 0.0016 |
| F08 | 10840001 | 10940000 | 0.0024 | 0.0016 | 0.0016 |
| F08 | 10860001 | 10960000 | 0.0023 | 0.0017 | 0.0017 |
| F08 | 10880001 | 10980000 | 0.0024 | 0.0019 | 0.0020 |
| F08 | 10900001 | 11000000 | 0.0029 | 0.0023 | 0.0022 |
| F08 | 10920001 | 11020000 | 0.0027 | 0.0022 | 0.0022 |
| F08 | 10940001 | 11040000 | 0.0025 | 0.0022 | 0.0022 |
| F08 | 10960001 | 11060000 | 0.0027 | 0.0022 | 0.0023 |
| F08 | 10980001 | 11080000 | 0.0021 | 0.0015 | 0.0016 |
| F08 | 11000001 | 11100000 | 0.0014 | 0.0009 | 0.0010 |
| F08 | 11020001 | 11120000 | 0.0012 | 0.0006 | 0.0007 |
| F08 | 11040001 | 11140000 | 0.0012 | 0.0005 | 0.0007 |
| F08 | 11060001 | 11160000 | 0.0010 | 0.0005 | 0.0006 |
| F08 | 11080001 | 11180000 | 0.0009 | 0.0006 | 0.0007 |
| F08 | 11100001 | 11200000 | 0.0010 | 0.0007 | 0.0008 |
| F08 | 11120001 | 11220000 | 0.0008 | 0.0007 | 0.0008 |
| F08 | 11140001 | 11240000 | 0.0012 | 0.0011 | 0.0012 |
| F08 | 11160001 | 11260000 | 0.0013 | 0.0012 | 0.0013 |
| F08 | 11180001 | 11280000 | 0.0017 | 0.0014 | 0.0017 |
| F08 | 11200001 | 11300000 | 0.0019 | 0.0016 | 0.0019 |
| F08 | 11220001 | 11320000 | 0.0023 | 0.0019 | 0.0023 |
| F08 | 11240001 | 11340000 | 0.0022 | 0.0018 | 0.0022 |
| F08 | 11260001 | 11360000 | 0.0022 | 0.0018 | 0.0022 |
| F08 | 11280001 | 11380000 | 0.0021 | 0.0016 | 0.0019 |
| F08 | 11300001 | 11400000 | 0.0022 | 0.0017 | 0.0020 |
| F08 | 11320001 | 11420000 | 0.0020 | 0.0014 | 0.0018 |
| F08 | 11340001 | 11440000 | 0.0023 | 0.0016 | 0.0018 |
| F08 | 11360001 | 11460000 | 0.0030 | 0.0021 | 0.0023 |
| F08 | 11380001 | 11480000 | 0.0032 | 0.0021 | 0.0024 |
| F08 | 11400001 | 11500000 | 0.0032 | 0.0020 | 0.0023 |
| F08 | 11420001 | 11520000 | 0.0039 | 0.0024 | 0.0027 |
| F08 | 11440001 | 11540000 | 0.0037 | 0.0023 | 0.0026 |
| F08 | 11460001 | 11560000 | 0.0032 | 0.0017 | 0.0020 |
| F08 | 11480001 | 11580000 | 0.0032 | 0.0016 | 0.0019 |
| F08 | 11500001 | 11600000 | 0.0028 | 0.0015 | 0.0016 |
| F08 | 11520001 | 11620000 | 0.0025 | 0.0014 | 0.0015 |
| F08 | 11540001 | 11640000 | 0.0022 | 0.0011 | 0.0012 |
| F08 | 11560001 | 11660000 | 0.0023 | 0.0014 | 0.0015 |
| F08 | 11580001 | 11680000 | 0.0024 | 0.0016 | 0.0017 |
| F08 | 11600001 | 11700000 | 0.0025 | 0.0015 | 0.0016 |
| F08 | 11620001 | 11720000 | 0.0026 | 0.0013 | 0.0015 |
| F08 | 11640001 | 11740000 | 0.0027 | 0.0014 | 0.0016 |
| F08 | 11660001 | 11760000 | 0.0040 | 0.0024 | 0.0031 |
| F08 | 11680001 | 11780000 | 0.0038 | 0.0024 | 0.0030 |
| F08 | 11700001 | 11800000 | 0.0042 | 0.0028 | 0.0033 |
| F08 | 11720001 | 11820000 | 0.0040 | 0.0029 | 0.0033 |
| F08 | 11740001 | 11840000 | 0.0044 | 0.0033 | 0.0037 |
| F08 | 11760001 | 11860000 | 0.0035 | 0.0025 | 0.0026 |
| F08 | 11780001 | 11880000 | 0.0034 | 0.0022 | 0.0023 |
| F08 | 11800001 | 11900000 | 0.0033 | 0.0023 | 0.0024 |
| F08 | 11820001 | 11920000 | 0.0040 | 0.0030 | 0.0033 |
| F08 | 11840001 | 11940000 | 0.0041 | 0.0031 | 0.0034 |
| F08 | 11860001 | 11960000 | 0.0035 | 0.0027 | 0.0029 |
| F08 | 11880001 | 11980000 | 0.0036 | 0.0028 | 0.0030 |
| F08 | 11900001 | 12000000 | 0.0033 | 0.0023 | 0.0026 |
| F08 | 11920001 | 12020000 | 0.0025 | 0.0015 | 0.0017 |
| F08 | 11940001 | 12040000 | 0.0020 | 0.0011 | 0.0013 |
| F08 | 11960001 | 12060000 | 0.0022 | 0.0011 | 0.0013 |

| Chr | Start | End | both | A1 | A2 |
|---|---|---|---|---|---|
| F08 | 11980001 | 12080000 | 0.0022 | 0.0010 | 0.0011 |
| F08 | 12000001 | 12100000 | 0.0027 | 0.0013 | 0.0015 |
| F08 | 12020001 | 12120000 | 0.0029 | 0.0016 | 0.0016 |
| F08 | 12040001 | 12140000 | 0.0030 | 0.0015 | 0.0016 |
| F08 | 12060001 | 12160000 | 0.0032 | 0.0020 | 0.0020 |
| F08 | 12080001 | 12180000 | 0.0039 | 0.0027 | 0.0029 |
| F08 | 12100001 | 12200000 | 0.0033 | 0.0024 | 0.0026 |
| F08 | 12120001 | 12220000 | 0.0031 | 0.0022 | 0.0023 |
| F08 | 12140001 | 12240000 | 0.0030 | 0.0021 | 0.0023 |
| F08 | 12160001 | 12260000 | 0.0025 | 0.0014 | 0.0017 |
| F08 | 12180001 | 12280000 | 0.0016 | 0.0007 | 0.0009 |
| F08 | 12200001 | 12300000 | 0.0017 | 0.0008 | 0.0009 |
| F08 | 12220001 | 12320000 | 0.0016 | 0.0010 | 0.0010 |
| F08 | 12240001 | 12340000 | 0.0018 | 0.0010 | 0.0010 |
| F08 | 12260001 | 12360000 | 0.0024 | 0.0014 | 0.0014 |
| F08 | 12280001 | 12380000 | 0.0026 | 0.0015 | 0.0016 |
| F08 | 12300001 | 12400000 | 0.0024 | 0.0014 | 0.0016 |
| F08 | 12320001 | 12420000 | 0.0022 | 0.0012 | 0.0015 |
| F08 | 12340001 | 12440000 | 0.0018 | 0.0011 | 0.0013 |
| F08 | 12360001 | 12460000 | 0.0010 | 0.0007 | 0.0008 |
| F08 | 12380001 | 12480000 | 0.0009 | 0.0007 | 0.0008 |
| F08 | 12400001 | 12500000 | 0.0016 | 0.0013 | 0.0014 |
| F08 | 12420001 | 12520000 | 0.0015 | 0.0012 | 0.0014 |
| F08 | 12440001 | 12540000 | 0.0015 | 0.0012 | 0.0013 |
| F08 | 12460001 | 12560000 | 0.0023 | 0.0018 | 0.0021 |
| F08 | 12480001 | 12580000 | 0.0025 | 0.0019 | 0.0023 |
| F08 | 12500001 | 12600000 | 0.0017 | 0.0013 | 0.0016 |
| F08 | 12520001 | 12620000 | 0.0027 | 0.0019 | 0.0023 |
| F08 | 12540001 | 12640000 | 0.0032 | 0.0022 | 0.0027 |
| F08 | 12560001 | 12660000 | 0.0035 | 0.0025 | 0.0028 |
| F08 | 12580001 | 12680000 | 0.0031 | 0.0021 | 0.0023 |
| F08 | 12600001 | 12700000 | 0.0039 | 0.0029 | 0.0031 |
| F08 | 12620001 | 12720000 | 0.0037 | 0.0029 | 0.0031 |
| F08 | 12640001 | 12740000 | 0.0043 | 0.0033 | 0.0036 |
| F08 | 12660001 | 12760000 | 0.0037 | 0.0027 | 0.0031 |
| F08 | 12680001 | 12780000 | 0.0039 | 0.0028 | 0.0032 |
| F08 | 12700001 | 12800000 | 0.0039 | 0.0027 | 0.0030 |
| F08 | 12720001 | 12820000 | 0.0045 | 0.0032 | 0.0034 |
| F08 | 12740001 | 12840000 | 0.0040 | 0.0030 | 0.0030 |
| F08 | 12760001 | 12860000 | 0.0041 | 0.0032 | 0.0031 |
| F08 | 12780001 | 12880000 | 0.0043 | 0.0034 | 0.0035 |
| F08 | 12800001 | 12900000 | 0.0033 | 0.0027 | 0.0028 |
| F08 | 12820001 | 12920000 | 0.0026 | 0.0022 | 0.0023 |
| F08 | 12840001 | 12940000 | 0.0025 | 0.0016 | 0.0020 |
| F08 | 12860001 | 12960000 | 0.0022 | 0.0012 | 0.0017 |
| F08 | 12880001 | 12980000 | 0.0020 | 0.0011 | 0.0015 |
| F08 | 12900001 | 13000000 | 0.0023 | 0.0015 | 0.0018 |
| F08 | 12920001 | 13020000 | 0.0022 | 0.0015 | 0.0017 |
| F08 | 12940001 | 13040000 | 0.0019 | 0.0015 | 0.0015 |
| F08 | 12960001 | 13060000 | 0.0018 | 0.0015 | 0.0015 |
| F08 | 12980001 | 13080000 | 0.0019 | 0.0017 | 0.0016 |
| F08 | 13000001 | 13100000 | 0.0019 | 0.0015 | 0.0016 |
| F08 | 13020001 | 13120000 | 0.0023 | 0.0020 | 0.0021 |
| F08 | 13040001 | 13140000 | 0.0031 | 0.0027 | 0.0028 |
| F08 | 13060001 | 13160000 | 0.0034 | 0.0029 | 0.0031 |
| F08 | 13080001 | 13180000 | 0.0036 | 0.0030 | 0.0031 |
| F08 | 13100001 | 13200000 | 0.0046 | 0.0040 | 0.0040 |
| F08 | 13120001 | 13220000 | 0.0042 | 0.0036 | 0.0037 |

| Chr | Start | End | both | A1 | A2 |
|---|---|---|---|---|---|
| F08 | 13140001 | 13240000 | 0.0036 | 0.0030 | 0.0031 |
| F08 | 13160001 | 13260000 | 0.0042 | 0.0037 | 0.0038 |
| F08 | 13180001 | 13280000 | 0.0042 | 0.0037 | 0.0040 |
| F08 | 13200001 | 13300000 | 0.0033 | 0.0028 | 0.0031 |
| F08 | 13220001 | 13320000 | 0.0031 | 0.0027 | 0.0029 |
| F08 | 13240001 | 13340000 | 0.0038 | 0.0031 | 0.0033 |
| F08 | 13260001 | 13360000 | 0.0030 | 0.0023 | 0.0025 |
| F08 | 13280001 | 13380000 | 0.0028 | 0.0020 | 0.0022 |
| F08 | 13300001 | 13400000 | 0.0037 | 0.0029 | 0.0031 |
| F08 | 13320001 | 13420000 | 0.0042 | 0.0031 | 0.0036 |
| F08 | 13340001 | 13440000 | 0.0036 | 0.0026 | 0.0032 |
| F08 | 13360001 | 13460000 | 0.0036 | 0.0028 | 0.0033 |
| F08 | 13380001 | 13480000 | 0.0038 | 0.0028 | 0.0033 |
| F08 | 13400001 | 13500000 | 0.0029 | 0.0019 | 0.0023 |
| F08 | 13420001 | 13520000 | 0.0030 | 0.0020 | 0.0023 |
| F08 | 13440001 | 13540000 | 0.0038 | 0.0027 | 0.0030 |
| F08 | 13460001 | 13560000 | 0.0037 | 0.0027 | 0.0029 |
| F08 | 13480001 | 13580000 | 0.0043 | 0.0033 | 0.0037 |
| F08 | 13500001 | 13600000 | 0.0043 | 0.0037 | 0.0038 |
| F08 | 13520001 | 13620000 | 0.0043 | 0.0038 | 0.0039 |
| F08 | 13540001 | 13640000 | 0.0036 | 0.0032 | 0.0033 |
| F08 | 13560001 | 13660000 | 0.0046 | 0.0039 | 0.0041 |
| F08 | 13580001 | 13680000 | 0.0040 | 0.0032 | 0.0033 |
| F08 | 13600001 | 13700000 | 0.0038 | 0.0027 | 0.0031 |
| F08 | 13620001 | 13720000 | 0.0028 | 0.0018 | 0.0021 |
| F08 | 13640001 | 13740000 | 0.0034 | 0.0022 | 0.0025 |
| F08 | 13660001 | 13760000 | 0.0031 | 0.0022 | 0.0024 |
| F08 | 13680001 | 13780000 | 0.0029 | 0.0022 | 0.0025 |
| F08 | 13700001 | 13800000 | 0.0032 | 0.0024 | 0.0027 |
| F08 | 13720001 | 13820000 | 0.0042 | 0.0034 | 0.0036 |
| F08 | 13740001 | 13840000 | 0.0036 | 0.0029 | 0.0032 |
| F08 | 13760001 | 13860000 | 0.0027 | 0.0020 | 0.0022 |
| F08 | 13780001 | 13880000 | 0.0038 | 0.0029 | 0.0031 |
| F08 | 13800001 | 13900000 | 0.0043 | 0.0032 | 0.0035 |
| F08 | 13820001 | 13920000 | 0.0033 | 0.0022 | 0.0026 |
| F08 | 13840001 | 13940000 | 0.0032 | 0.0021 | 0.0025 |
| F08 | 13860001 | 13960000 | 0.0039 | 0.0028 | 0.0033 |
| F08 | 13880001 | 13980000 | 0.0032 | 0.0023 | 0.0025 |
| F08 | 13900001 | 14000000 | 0.0027 | 0.0023 | 0.0023 |
| F08 | 13920001 | 14020000 | 0.0039 | 0.0033 | 0.0034 |
| F08 | 13940001 | 14040000 | 0.0038 | 0.0031 | 0.0031 |
| F08 | 13960001 | 14060000 | 0.0049 | 0.0040 | 0.0040 |
| F08 | 13980001 | 14080000 | 0.0047 | 0.0037 | 0.0041 |
| F08 | 14000001 | 14100000 | 0.0041 | 0.0032 | 0.0035 |
| F08 | 14020001 | 14120000 | 0.0033 | 0.0025 | 0.0027 |
| F08 | 14040001 | 14140000 | 0.0036 | 0.0029 | 0.0031 |
| F08 | 14060001 | 14160000 | 0.0019 | 0.0015 | 0.0016 |
| F08 | 14080001 | 14180000 | 0.0017 | 0.0011 | 0.0011 |
| F08 | 14100001 | 14200000 | 0.0018 | 0.0011 | 0.0011 |
| F08 | 14120001 | 14220000 | 0.0016 | 0.0009 | 0.0009 |
| F08 | 14140001 | 14240000 | 0.0021 | 0.0015 | 0.0014 |
| F08 | 14160001 | 14260000 | 0.0022 | 0.0013 | 0.0013 |
| F08 | 14180001 | 14280000 | 0.0026 | 0.0017 | 0.0018 |
| F08 | 14200001 | 14300000 | 0.0030 | 0.0020 | 0.0022 |
| F08 | 14220001 | 14320000 | 0.0039 | 0.0026 | 0.0032 |
| F08 | 14240001 | 14340000 | 0.0036 | 0.0021 | 0.0028 |
| F08 | 14260001 | 14360000 | 0.0041 | 0.0025 | 0.0033 |
| F08 | 14280001 | 14380000 | 0.0036 | 0.0023 | 0.0030 |

| Chr | Start | End | both | A1 | A2 |
|---|---|---|---|---|---|
| F08 | 14300001 | 14400000 | 0.0040 | 0.0027 | 0.0034 |
| F08 | 14320001 | 14420000 | 0.0035 | 0.0022 | 0.0028 |
| F08 | 14340001 | 14440000 | 0.0038 | 0.0028 | 0.0032 |
| F08 | 14360001 | 14460000 | 0.0037 | 0.0027 | 0.0031 |
| F08 | 14380001 | 14480000 | 0.0045 | 0.0033 | 0.0037 |
| F08 | 14400001 | 14500000 | 0.0039 | 0.0028 | 0.0032 |
| F08 | 14420001 | 14520000 | 0.0035 | 0.0028 | 0.0029 |
| F08 | 14440001 | 14540000 | 0.0028 | 0.0022 | 0.0024 |
| F08 | 14460001 | 14560000 | 0.0023 | 0.0020 | 0.0021 |
| F08 | 14480001 | 14580000 | 0.0015 | 0.0012 | 0.0012 |
| F08 | 14500001 | 14600000 | 0.0018 | 0.0016 | 0.0017 |
| F08 | 14520001 | 14620000 | 0.0021 | 0.0018 | 0.0020 |
| F08 | 14540001 | 14640000 | 0.0021 | 0.0017 | 0.0018 |
| F08 | 14560001 | 14660000 | 0.0023 | 0.0017 | 0.0017 |
| F08 | 14580001 | 14680000 | 0.0028 | 0.0022 | 0.0023 |
| F08 | 14600001 | 14700000 | 0.0023 | 0.0016 | 0.0017 |
| F08 | 14620001 | 14720000 | 0.0021 | 0.0012 | 0.0012 |
| F08 | 14640001 | 14740000 | 0.0027 | 0.0018 | 0.0018 |
| F08 | 14660001 | 14760000 | 0.0031 | 0.0022 | 0.0024 |
| F08 | 14680001 | 14780000 | 0.0032 | 0.0022 | 0.0024 |
| F08 | 14700001 | 14800000 | 0.0033 | 0.0023 | 0.0025 |
| F08 | 14720001 | 14820000 | 0.0030 | 0.0021 | 0.0022 |
| F08 | 14740001 | 14840000 | 0.0023 | 0.0013 | 0.0015 |
| F08 | 14760001 | 14860000 | 0.0019 | 0.0009 | 0.0009 |
| F08 | 14780001 | 14880000 | 0.0017 | 0.0008 | 0.0008 |
| F08 | 14800001 | 14900000 | 0.0015 | 0.0007 | 0.0007 |
| F08 | 14820001 | 14920000 | 0.0025 | 0.0016 | 0.0017 |
| F08 | 14840001 | 14940000 | 0.0032 | 0.0022 | 0.0023 |
| F08 | 14860001 | 14960000 | 0.0043 | 0.0032 | 0.0035 |
| F08 | 14880001 | 14980000 | 0.0045 | 0.0035 | 0.0037 |
| F08 | 14900001 | 15000000 | 0.0051 | 0.0041 | 0.0043 |
| F08 | 14920001 | 15020000 | 0.0040 | 0.0031 | 0.0033 |
| F08 | 14940001 | 15040000 | 0.0039 | 0.0033 | 0.0035 |
| F08 | 14960001 | 15060000 | 0.0030 | 0.0025 | 0.0027 |
| F08 | 14980001 | 15080000 | 0.0025 | 0.0020 | 0.0022 |
| F08 | 15000001 | 15100000 | 0.0020 | 0.0014 | 0.0015 |
| F08 | 15020001 | 15120000 | 0.0044 | 0.0036 | 0.0037 |
| F08 | 15040001 | 15140000 | 0.0037 | 0.0030 | 0.0030 |
| F08 | 15060001 | 15160000 | 0.0037 | 0.0031 | 0.0031 |
| F08 | 15080001 | 15180000 | 0.0039 | 0.0032 | 0.0033 |
| F08 | 15100001 | 15200000 | 0.0040 | 0.0034 | 0.0036 |
| F08 | 15120001 | 15220000 | 0.0019 | 0.0015 | 0.0018 |
| F08 | 15140001 | 15240000 | 0.0020 | 0.0015 | 0.0018 |
| F08 | 15160001 | 15260000 | 0.0019 | 0.0014 | 0.0017 |
| F08 | 15180001 | 15280000 | 0.0026 | 0.0020 | 0.0023 |
| F08 | 15200001 | 15300000 | 0.0023 | 0.0018 | 0.0020 |
| F08 | 15220001 | 15320000 | 0.0027 | 0.0021 | 0.0022 |
| F08 | 15240001 | 15340000 | 0.0034 | 0.0025 | 0.0026 |
| F08 | 15260001 | 15360000 | 0.0035 | 0.0026 | 0.0026 |
| F08 | 15280001 | 15380000 | 0.0029 | 0.0022 | 0.0022 |
| F08 | 15300001 | 15400000 | 0.0030 | 0.0023 | 0.0023 |
| F08 | 15320001 | 15420000 | 0.0027 | 0.0020 | 0.0021 |
| F08 | 15340001 | 15440000 | 0.0020 | 0.0018 | 0.0018 |
| F08 | 15360001 | 15460000 | 0.0023 | 0.0022 | 0.0023 |
| F08 | 15380001 | 15480000 | 0.0030 | 0.0027 | 0.0029 |
| F08 | 15400001 | 15500000 | 0.0033 | 0.0029 | 0.0032 |
| F08 | 15420001 | 15520000 | 0.0034 | 0.0027 | 0.0031 |
| F08 | 15440001 | 15540000 | 0.0038 | 0.0028 | 0.0032 |

| Chr | Start | End | both | A1 | A2 |
|---|---|---|---|---|---|
| F08 | 15460001 | 15560000 | 0.0034 | 0.0022 | 0.0025 |
| F08 | 15480001 | 15580000 | 0.0023 | 0.0012 | 0.0013 |
| F08 | 15500001 | 15600000 | 0.0025 | 0.0012 | 0.0013 |
| F08 | 15520001 | 15620000 | 0.0025 | 0.0013 | 0.0015 |
| F08 | 15540001 | 15640000 | 0.0019 | 0.0009 | 0.0011 |
| F08 | 15560001 | 15660000 | 0.0021 | 0.0014 | 0.0016 |
| F08 | 15580001 | 15680000 | 0.0021 | 0.0015 | 0.0017 |
| F08 | 15600001 | 15700000 | 0.0024 | 0.0017 | 0.0022 |
| F08 | 15620001 | 15720000 | 0.0023 | 0.0015 | 0.0021 |
| F08 | 15640001 | 15740000 | 0.0030 | 0.0021 | 0.0028 |
| F08 | 15660001 | 15760000 | 0.0026 | 0.0017 | 0.0023 |
| F08 | 15680001 | 15780000 | 0.0027 | 0.0018 | 0.0023 |
| F08 | 15700001 | 15800000 | 0.0026 | 0.0020 | 0.0021 |
| F08 | 15720001 | 15820000 | 0.0025 | 0.0019 | 0.0020 |
| F08 | 15740001 | 15840000 | 0.0022 | 0.0017 | 0.0017 |
| F08 | 15760001 | 15860000 | 0.0022 | 0.0016 | 0.0017 |
| F08 | 15780001 | 15880000 | 0.0023 | 0.0017 | 0.0018 |
| F08 | 15800001 | 15900000 | 0.0019 | 0.0013 | 0.0015 |
| F08 | 15820001 | 15920000 | 0.0024 | 0.0017 | 0.0019 |
| F08 | 15840001 | 15940000 | 0.0031 | 0.0024 | 0.0024 |
| F08 | 15860001 | 15960000 | 0.0042 | 0.0034 | 0.0034 |
| F08 | 15880001 | 15980000 | 0.0044 | 0.0035 | 0.0035 |
| F08 | 15900001 | 16000000 | 0.0056 | 0.0046 | 0.0046 |
| F08 | 15920001 | 16020000 | 0.0058 | 0.0050 | 0.0049 |
| F08 | 15940001 | 16040000 | 0.0053 | 0.0043 | 0.0044 |
| F08 | 15960001 | 16060000 | 0.0042 | 0.0035 | 0.0035 |
| F08 | 15980001 | 16080000 | 0.0040 | 0.0034 | 0.0034 |
| F08 | 16000001 | 16100000 | 0.0035 | 0.0025 | 0.0025 |
| F08 | 16020001 | 16120000 | 0.0029 | 0.0019 | 0.0021 |
| F08 | 16040001 | 16140000 | 0.0030 | 0.0020 | 0.0022 |
| F08 | 16060001 | 16160000 | 0.0033 | 0.0021 | 0.0024 |
| F08 | 16080001 | 16180000 | 0.0035 | 0.0022 | 0.0025 |
| F08 | 16100001 | 16200000 | 0.0026 | 0.0017 | 0.0020 |
| F08 | 16120001 | 16220000 | 0.0029 | 0.0021 | 0.0023 |
| F08 | 16140001 | 16240000 | 0.0029 | 0.0024 | 0.0024 |
| F08 | 16160001 | 16260000 | 0.0032 | 0.0027 | 0.0028 |
| F08 | 16180001 | 16280000 | 0.0027 | 0.0025 | 0.0025 |
| F08 | 16200001 | 16300000 | 0.0027 | 0.0025 | 0.0026 |
| F08 | 16220001 | 16320000 | 0.0023 | 0.0022 | 0.0022 |
| F08 | 16240001 | 16340000 | 0.0019 | 0.0018 | 0.0019 |
| F08 | 16260001 | 16360000 | 0.0015 | 0.0014 | 0.0015 |
| F08 | 16280001 | 16380000 | 0.0019 | 0.0016 | 0.0017 |
| F08 | 16300001 | 16400000 | 0.0033 | 0.0030 | 0.0029 |
| F08 | 16320001 | 16420000 | 0.0035 | 0.0029 | 0.0028 |
| F08 | 16340001 | 16440000 | 0.0036 | 0.0028 | 0.0026 |
| F08 | 16360001 | 16460000 | 0.0041 | 0.0033 | 0.0030 |
| F08 | 16380001 | 16480000 | 0.0044 | 0.0034 | 0.0030 |
| F08 | 16400001 | 16500000 | 0.0038 | 0.0026 | 0.0025 |
| F08 | 16420001 | 16520000 | 0.0036 | 0.0025 | 0.0024 |
| F08 | 16440001 | 16540000 | 0.0040 | 0.0029 | 0.0029 |
| F08 | 16460001 | 16560000 | 0.0046 | 0.0035 | 0.0037 |
| F08 | 16480001 | 16580000 | 0.0039 | 0.0031 | 0.0033 |
| F08 | 16500001 | 16600000 | 0.0034 | 0.0029 | 0.0030 |
| F08 | 16520001 | 16620000 | 0.0034 | 0.0029 | 0.0032 |
| F08 | 16540001 | 16640000 | 0.0025 | 0.0023 | 0.0025 |
| F08 | 16560001 | 16660000 | 0.0009 | 0.0009 | 0.0009 |
| F08 | 16580001 | 16680000 | 0.0010 | 0.0010 | 0.0010 |
| F08 | 16600001 | 16700000 | 0.0006 | 0.0005 | 0.0006 |

| Chr | Start | End | both | A1 | A2 |
|---|---|---|---|---|---|
| F08 | 16620001 | 16720000 | 0.0007 | 0.0006 | 0.0007 |
| F08 | 16640001 | 16740000 | 0.0009 | 0.0008 | 0.0009 |
| F08 | 16660001 | 16760000 | 0.0016 | 0.0014 | 0.0016 |
| F08 | 16680001 | 16780000 | 0.0019 | 0.0018 | 0.0019 |
| F08 | 16700001 | 16800000 | 0.0023 | 0.0020 | 0.0022 |
| F08 | 16720001 | 16820000 | 0.0029 | 0.0024 | 0.0028 |
| F08 | 16740001 | 16840000 | 0.0030 | 0.0025 | 0.0029 |
| F08 | 16760001 | 16860000 | 0.0025 | 0.0021 | 0.0024 |
| F08 | 16780001 | 16880000 | 0.0023 | 0.0018 | 0.0022 |
| F08 | 16800001 | 16900000 | 0.0020 | 0.0018 | 0.0019 |
| F08 | 16820001 | 16920000 | 0.0012 | 0.0012 | 0.0012 |
| F08 | 16840001 | 16940000 | 0.0016 | 0.0015 | 0.0016 |
| F08 | 16860001 | 16960000 | 0.0022 | 0.0020 | 0.0021 |
| F08 | 16880001 | 16980000 | 0.0023 | 0.0022 | 0.0022 |
| F08 | 16900001 | 17000000 | 0.0023 | 0.0021 | 0.0023 |
| F08 | 16920001 | 17020000 | 0.0025 | 0.0023 | 0.0024 |
| F08 | 16940001 | 17040000 | 0.0021 | 0.0019 | 0.0020 |
| F08 | 16960001 | 17060000 | 0.0017 | 0.0014 | 0.0016 |
| F08 | 16980001 | 17080000 | 0.0016 | 0.0011 | 0.0013 |
| F08 | 17000001 | 17100000 | 0.0017 | 0.0011 | 0.0012 |
| F08 | 17020001 | 17120000 | 0.0015 | 0.0007 | 0.0009 |
| F08 | 17040001 | 17140000 | 0.0018 | 0.0012 | 0.0012 |
| F08 | 17060001 | 17160000 | 0.0021 | 0.0014 | 0.0016 |
| F08 | 17080001 | 17180000 | 0.0025 | 0.0019 | 0.0021 |
| F08 | 17100001 | 17200000 | 0.0031 | 0.0024 | 0.0026 |
| F08 | 17120001 | 17220000 | 0.0037 | 0.0029 | 0.0033 |
| F08 | 17140001 | 17240000 | 0.0032 | 0.0025 | 0.0028 |
| F08 | 17160001 | 17260000 | 0.0035 | 0.0028 | 0.0031 |
| F08 | 17180001 | 17280000 | 0.0034 | 0.0027 | 0.0031 |
| F08 | 17200001 | 17300000 | 0.0026 | 0.0021 | 0.0025 |
| F08 | 17220001 | 17320000 | 0.0020 | 0.0017 | 0.0019 |
| F08 | 17240001 | 17340000 | 0.0026 | 0.0021 | 0.0024 |
| F08 | 17260001 | 17360000 | 0.0024 | 0.0018 | 0.0020 |
| F08 | 17280001 | 17380000 | 0.0027 | 0.0020 | 0.0021 |
| F08 | 17300001 | 17400000 | 0.0034 | 0.0024 | 0.0025 |
| F08 | 17320001 | 17420000 | 0.0033 | 0.0024 | 0.0025 |
| F08 | 17340001 | 17440000 | 0.0026 | 0.0018 | 0.0019 |
| F08 | 17360001 | 17460000 | 0.0020 | 0.0014 | 0.0015 |
| F08 | 17380001 | 17480000 | 0.0013 | 0.0008 | 0.0009 |
| F08 | 17400001 | 17500000 | 0.0011 | 0.0006 | 0.0008 |
| F08 | 17420001 | 17520000 | 0.0014 | 0.0008 | 0.0011 |
| F08 | 17440001 | 17540000 | 0.0024 | 0.0016 | 0.0019 |
| F08 | 17460001 | 17560000 | 0.0030 | 0.0020 | 0.0022 |
| F08 | 17480001 | 17580000 | 0.0030 | 0.0021 | 0.0023 |
| F08 | 17500001 | 17600000 | 0.0027 | 0.0020 | 0.0021 |
| F08 | 17520001 | 17620000 | 0.0024 | 0.0017 | 0.0018 |
| F08 | 17540001 | 17640000 | 0.0019 | 0.0014 | 0.0013 |
| F08 | 17560001 | 17660000 | 0.0024 | 0.0020 | 0.0019 |
| F08 | 17580001 | 17680000 | 0.0025 | 0.0019 | 0.0018 |
| F08 | 17600001 | 17700000 | 0.0040 | 0.0031 | 0.0031 |
| F08 | 17620001 | 17720000 | 0.0051 | 0.0041 | 0.0041 |
| F08 | 17640001 | 17740000 | 0.0053 | 0.0039 | 0.0040 |
| F08 | 17660001 | 17760000 | 0.0051 | 0.0035 | 0.0037 |
| F08 | 17680001 | 17780000 | 0.0059 | 0.0045 | 0.0047 |
| F08 | 17700001 | 17800000 | 0.0052 | 0.0036 | 0.0040 |
| F08 | 17720001 | 17820000 | 0.0048 | 0.0029 | 0.0034 |
| F08 | 17740001 | 17840000 | 0.0050 | 0.0031 | 0.0037 |
| F08 | 17760001 | 17860000 | 0.0043 | 0.0024 | 0.0030 |

| Chr | Start | End | both | A1 | A2 |
|---|---|---|---|---|---|
| F08 | 17780001 | 17880000 | 0.0034 | 0.0015 | 0.0020 |
| F08 | 17800001 | 17900000 | 0.0029 | 0.0011 | 0.0013 |
| F08 | 17820001 | 17920000 | 0.0027 | 0.0011 | 0.0012 |
| F08 | 17840001 | 17940000 | 0.0027 | 0.0014 | 0.0013 |
| F08 | 17860001 | 17960000 | 0.0027 | 0.0015 | 0.0014 |
| F08 | 17880001 | 17980000 | 0.0029 | 0.0019 | 0.0018 |
| F08 | 17900001 | 18000000 | 0.0032 | 0.0024 | 0.0023 |
| F08 | 17920001 | 18020000 | 0.0032 | 0.0024 | 0.0024 |
| F08 | 17940001 | 18040000 | 0.0028 | 0.0020 | 0.0022 |
| F08 | 17960001 | 18060000 | 0.0030 | 0.0021 | 0.0024 |
| F08 | 17980001 | 18080000 | 0.0037 | 0.0025 | 0.0029 |
| F08 | 18000001 | 18100000 | 0.0033 | 0.0022 | 0.0026 |
| F08 | 18020001 | 18120000 | 0.0033 | 0.0022 | 0.0028 |
| F08 | 18040001 | 18140000 | 0.0030 | 0.0020 | 0.0024 |
| F08 | 18060001 | 18160000 | 0.0034 | 0.0022 | 0.0027 |
| F08 | 18080001 | 18180000 | 0.0033 | 0.0023 | 0.0028 |
| F08 | 18100001 | 18200000 | 0.0037 | 0.0025 | 0.0029 |
| F08 | 18120001 | 18220000 | 0.0042 | 0.0030 | 0.0033 |
| F08 | 18140001 | 18240000 | 0.0047 | 0.0033 | 0.0037 |
| F08 | 18160001 | 18260000 | 0.0042 | 0.0030 | 0.0033 |
| F08 | 18180001 | 18280000 | 0.0043 | 0.0033 | 0.0036 |
| F08 | 18200001 | 18300000 | 0.0050 | 0.0041 | 0.0046 |
| F08 | 18220001 | 18320000 | 0.0050 | 0.0040 | 0.0044 |
| F08 | 18240001 | 18340000 | 0.0064 | 0.0054 | 0.0057 |
| F08 | 18260001 | 18360000 | 0.0070 | 0.0061 | 0.0062 |
| F08 | 18280001 | 18380000 | 0.0057 | 0.0050 | 0.0050 |
| F08 | 18300001 | 18400000 | 0.0046 | 0.0040 | 0.0040 |
| F08 | 18320001 | 18420000 | 0.0042 | 0.0038 | 0.0037 |
| F08 | 18340001 | 18440000 | 0.0023 | 0.0021 | 0.0021 |
| F08 | 18360001 | 18460000 | 0.0015 | 0.0013 | 0.0014 |
| F08 | 18380001 | 18480000 | 0.0018 | 0.0016 | 0.0018 |
| F08 | 18400001 | 18500000 | 0.0015 | 0.0013 | 0.0015 |
| F08 | 18420001 | 18520000 | 0.0011 | 0.0009 | 0.0011 |
| F08 | 18440001 | 18540000 | 0.0011 | 0.0010 | 0.0011 |
| F08 | 18460001 | 18560000 | 0.0012 | 0.0010 | 0.0011 |
| F08 | 18480001 | 18580000 | 0.0016 | 0.0012 | 0.0013 |
| F08 | 18500001 | 18600000 | 0.0029 | 0.0023 | 0.0024 |
| F08 | 18520001 | 18620000 | 0.0031 | 0.0024 | 0.0026 |
| F08 | 18540001 | 18640000 | 0.0029 | 0.0023 | 0.0025 |
| F08 | 18560001 | 18660000 | 0.0031 | 0.0024 | 0.0026 |
| F08 | 18580001 | 18680000 | 0.0033 | 0.0026 | 0.0028 |
| F08 | 18600001 | 18700000 | 0.0027 | 0.0018 | 0.0021 |
| F08 | 18620001 | 18720000 | 0.0033 | 0.0022 | 0.0024 |
| F08 | 18640001 | 18740000 | 0.0042 | 0.0030 | 0.0032 |
| F08 | 18660001 | 18760000 | 0.0039 | 0.0026 | 0.0027 |
| F08 | 18680001 | 18780000 | 0.0034 | 0.0023 | 0.0024 |
| F08 | 18700001 | 18800000 | 0.0034 | 0.0025 | 0.0026 |
| F08 | 18720001 | 18820000 | 0.0036 | 0.0030 | 0.0030 |
| F08 | 18740001 | 18840000 | 0.0031 | 0.0025 | 0.0027 |
| F08 | 18760001 | 18860000 | 0.0035 | 0.0030 | 0.0033 |
| F08 | 18780001 | 18880000 | 0.0035 | 0.0031 | 0.0033 |
| F08 | 18800001 | 18900000 | 0.0030 | 0.0027 | 0.0029 |
| F08 | 18820001 | 18920000 | 0.0026 | 0.0021 | 0.0023 |
| F08 | 18840001 | 18940000 | 0.0026 | 0.0021 | 0.0022 |
| F08 | 18860001 | 18960000 | 0.0022 | 0.0017 | 0.0018 |
| F08 | 18880001 | 18980000 | 0.0020 | 0.0015 | 0.0016 |
| F08 | 18900001 | 19000000 | 0.0017 | 0.0013 | 0.0013 |
| F08 | 18920001 | 19020000 | 0.0015 | 0.0013 | 0.0012 |

| Chr | Start | End | both | A1 | A2 |
|---|---|---|---|---|---|
| F08 | 18940001 | 19040000 | 0.0011 | 0.0011 | 0.0011 |
| F08 | 18960001 | 19060000 | 0.0012 | 0.0012 | 0.0012 |
| F08 | 18980001 | 19080000 | 0.0025 | 0.0020 | 0.0024 |
| F08 | 19000001 | 19100000 | 0.0030 | 0.0022 | 0.0028 |
| F08 | 19020001 | 19120000 | 0.0028 | 0.0020 | 0.0024 |
| F08 | 19040001 | 19140000 | 0.0031 | 0.0022 | 0.0027 |
| F08 | 19060001 | 19160000 | 0.0035 | 0.0027 | 0.0031 |
| F08 | 19080001 | 19180000 | 0.0026 | 0.0021 | 0.0022 |
| F08 | 19100001 | 19200000 | 0.0025 | 0.0021 | 0.0021 |
| F08 | 19120001 | 19220000 | 0.0035 | 0.0030 | 0.0033 |
| F08 | 19140001 | 19240000 | 0.0031 | 0.0027 | 0.0029 |
| F08 | 19160001 | 19260000 | 0.0027 | 0.0023 | 0.0025 |
| F08 | 19180001 | 19280000 | 0.0022 | 0.0020 | 0.0022 |
| F08 | 19200001 | 19300000 | 0.0019 | 0.0017 | 0.0019 |
| F08 | 19220001 | 19320000 | 0.0007 | 0.0006 | 0.0007 |
| F08 | 19240001 | 19340000 | 0.0007 | 0.0007 | 0.0007 |
| F08 | 19260001 | 19360000 | 0.0012 | 0.0009 | 0.0010 |
| F08 | 19280001 | 19380000 | 0.0018 | 0.0013 | 0.0015 |
| F08 | 19300001 | 19400000 | 0.0028 | 0.0023 | 0.0024 |
| F08 | 19320001 | 19420000 | 0.0039 | 0.0031 | 0.0034 |
| F08 | 19340001 | 19440000 | 0.0042 | 0.0033 | 0.0036 |
| F08 | 19360001 | 19460000 | 0.0036 | 0.0028 | 0.0030 |
| F08 | 19380001 | 19480000 | 0.0033 | 0.0026 | 0.0027 |
| F08 | 19400001 | 19500000 | 0.0028 | 0.0018 | 0.0020 |
| F08 | 19420001 | 19520000 | 0.0018 | 0.0009 | 0.0010 |
| F08 | 19440001 | 19540000 | 0.0014 | 0.0006 | 0.0007 |
| F08 | 19460001 | 19560000 | 0.0014 | 0.0006 | 0.0007 |
| F08 | 19480001 | 19580000 | 0.0016 | 0.0007 | 0.0008 |
| F08 | 19500001 | 19600000 | 0.0015 | 0.0008 | 0.0010 |
| F08 | 19520001 | 19620000 | 0.0021 | 0.0014 | 0.0014 |
| F08 | 19540001 | 19640000 | 0.0035 | 0.0026 | 0.0029 |
| F08 | 19560001 | 19660000 | 0.0046 | 0.0035 | 0.0040 |
| F08 | 19580001 | 19680000 | 0.0055 | 0.0043 | 0.0048 |
| F08 | 19600001 | 19700000 | 0.0059 | 0.0045 | 0.0050 |
| F08 | 19620001 | 19720000 | 0.0059 | 0.0045 | 0.0050 |
| F08 | 19640001 | 19740000 | 0.0052 | 0.0040 | 0.0044 |
| F08 | 19660001 | 19760000 | 0.0045 | 0.0033 | 0.0035 |
| F08 | 19680001 | 19780000 | 0.0038 | 0.0026 | 0.0027 |
| F08 | 19700001 | 19800000 | 0.0034 | 0.0022 | 0.0022 |
| F08 | 19720001 | 19820000 | 0.0044 | 0.0032 | 0.0033 |
| F08 | 19740001 | 19840000 | 0.0042 | 0.0028 | 0.0029 |
| F08 | 19760001 | 19860000 | 0.0040 | 0.0027 | 0.0029 |
| F08 | 19780001 | 19880000 | 0.0038 | 0.0028 | 0.0030 |
| F08 | 19800001 | 19900000 | 0.0049 | 0.0039 | 0.0041 |
| F08 | 19820001 | 19920000 | 0.0034 | 0.0025 | 0.0027 |
| F08 | 19840001 | 19940000 | 0.0031 | 0.0025 | 0.0027 |
| F08 | 19860001 | 19960000 | 0.0032 | 0.0024 | 0.0025 |
| F08 | 19880001 | 19980000 | 0.0041 | 0.0033 | 0.0032 |
| F08 | 19900001 | 20000000 | 0.0043 | 0.0033 | 0.0035 |
| F08 | 19920001 | 20020000 | 0.0054 | 0.0041 | 0.0044 |
| F08 | 19940001 | 20040000 | 0.0056 | 0.0042 | 0.0044 |
| F08 | 19960001 | 20060000 | 0.0058 | 0.0044 | 0.0046 |
| F08 | 19980001 | 20080000 | 0.0048 | 0.0036 | 0.0038 |
| F08 | 20000001 | 20100000 | 0.0039 | 0.0029 | 0.0030 |
| F08 | 20020001 | 20120000 | 0.0038 | 0.0027 | 0.0029 |
| F08 | 20040001 | 20140000 | 0.0038 | 0.0028 | 0.0031 |
| F08 | 20060001 | 20160000 | 0.0031 | 0.0024 | 0.0027 |
| F08 | 20080001 | 20180000 | 0.0038 | 0.0030 | 0.0034 |

| Chr | Start | End | both | A1 | A2 |
|---|---|---|---|---|---|
| F08 | 20100001 | 20200000 | 0.0047 | 0.0039 | 0.0044 |
| F08 | 20120001 | 20220000 | 0.0035 | 0.0030 | 0.0034 |
| F08 | 20140001 | 20240000 | 0.0044 | 0.0039 | 0.0042 |
| F08 | 20160001 | 20260000 | 0.0050 | 0.0044 | 0.0047 |
| F08 | 20180001 | 20280000 | 0.0041 | 0.0036 | 0.0038 |
| F08 | 20200001 | 20300000 | 0.0026 | 0.0023 | 0.0024 |
| F08 | 20220001 | 20320000 | 0.0034 | 0.0029 | 0.0030 |
| F08 | 20240001 | 20340000 | 0.0026 | 0.0023 | 0.0023 |
| F08 | 20260001 | 20360000 | 0.0032 | 0.0028 | 0.0029 |
| F08 | 20280001 | 20380000 | 0.0043 | 0.0037 | 0.0039 |
| F08 | 20300001 | 20400000 | 0.0046 | 0.0039 | 0.0042 |
| F08 | 20320001 | 20420000 | 0.0045 | 0.0040 | 0.0043 |
| F08 | 20340001 | 20440000 | 0.0040 | 0.0034 | 0.0038 |
| F08 | 20360001 | 20460000 | 0.0031 | 0.0024 | 0.0028 |
| F08 | 20380001 | 20480000 | 0.0021 | 0.0016 | 0.0019 |
| F08 | 20400001 | 20500000 | 0.0021 | 0.0015 | 0.0017 |
| F08 | 20420001 | 20520000 | 0.0017 | 0.0011 | 0.0012 |
| F08 | 20440001 | 20540000 | 0.0021 | 0.0012 | 0.0013 |
| F08 | 20460001 | 20560000 | 0.0023 | 0.0015 | 0.0016 |
| F08 | 20480001 | 20580000 | 0.0025 | 0.0017 | 0.0018 |
| F08 | 20500001 | 20600000 | 0.0029 | 0.0019 | 0.0020 |
| F08 | 20520001 | 20620000 | 0.0038 | 0.0028 | 0.0027 |
| F08 | 20540001 | 20640000 | 0.0042 | 0.0032 | 0.0032 |
| F08 | 20560001 | 20660000 | 0.0043 | 0.0032 | 0.0032 |
| F08 | 20580001 | 20680000 | 0.0063 | 0.0052 | 0.0050 |
| F08 | 20600001 | 20700000 | 0.0063 | 0.0054 | 0.0050 |
| F08 | 20620001 | 20720000 | 0.0054 | 0.0044 | 0.0042 |
| F08 | 20640001 | 20740000 | 0.0052 | 0.0043 | 0.0038 |
| F08 | 20660001 | 20760000 | 0.0052 | 0.0045 | 0.0041 |
| F08 | 20680001 | 20780000 | 0.0032 | 0.0024 | 0.0023 |
| F08 | 20700001 | 20800000 | 0.0029 | 0.0021 | 0.0021 |
| F08 | 20720001 | 20820000 | 0.0026 | 0.0020 | 0.0020 |
| F08 | 20740001 | 20840000 | 0.0026 | 0.0019 | 0.0021 |
| F08 | 20760001 | 20860000 | 0.0031 | 0.0022 | 0.0023 |
| F08 | 20780001 | 20880000 | 0.0030 | 0.0019 | 0.0021 |
| F08 | 20800001 | 20900000 | 0.0030 | 0.0021 | 0.0022 |
| F08 | 20820001 | 20920000 | 0.0033 | 0.0024 | 0.0027 |
| F08 | 20840001 | 20940000 | 0.0039 | 0.0029 | 0.0034 |
| F08 | 20860001 | 20960000 | 0.0039 | 0.0031 | 0.0035 |
| F08 | 20880001 | 20980000 | 0.0043 | 0.0036 | 0.0041 |
| F08 | 20900001 | 21000000 | 0.0039 | 0.0033 | 0.0038 |
| F08 | 20920001 | 21020000 | 0.0037 | 0.0032 | 0.0036 |
| F08 | 20940001 | 21040000 | 0.0030 | 0.0026 | 0.0029 |
| F08 | 20960001 | 21060000 | 0.0026 | 0.0023 | 0.0026 |
| F08 | 20980001 | 21080000 | 0.0021 | 0.0019 | 0.0020 |
| F08 | 21000001 | 21100000 | 0.0034 | 0.0029 | 0.0033 |
| F08 | 21020001 | 21120000 | 0.0035 | 0.0030 | 0.0035 |
| F08 | 21040001 | 21140000 | 0.0032 | 0.0028 | 0.0032 |
| F08 | 21060001 | 21160000 | 0.0028 | 0.0023 | 0.0026 |
| F08 | 21080001 | 21180000 | 0.0031 | 0.0025 | 0.0028 |
| F08 | 21100001 | 21200000 | 0.0019 | 0.0016 | 0.0016 |
| F08 | 21120001 | 21220000 | 0.0025 | 0.0019 | 0.0020 |
| F08 | 21140001 | 21240000 | 0.0027 | 0.0019 | 0.0020 |
| F08 | 21160001 | 21260000 | 0.0029 | 0.0019 | 0.0021 |
| F08 | 21180001 | 21280000 | 0.0032 | 0.0019 | 0.0021 |
| F08 | 21200001 | 21300000 | 0.0037 | 0.0020 | 0.0023 |
| F08 | 21220001 | 21320000 | 0.0029 | 0.0014 | 0.0015 |
| F08 | 21240001 | 21340000 | 0.0036 | 0.0020 | 0.0022 |

| Chr | Start | End | both | A1 | A2 |
|---|---|---|---|---|---|
| F08 | 21260001 | 21360000 | 0.0039 | 0.0025 | 0.0026 |
| F08 | 21280001 | 21380000 | 0.0040 | 0.0030 | 0.0031 |
| F08 | 21300001 | 21400000 | 0.0038 | 0.0029 | 0.0030 |
| F08 | 21320001 | 21420000 | 0.0035 | 0.0030 | 0.0030 |
| F08 | 21340001 | 21440000 | 0.0031 | 0.0027 | 0.0027 |
| F08 | 21360001 | 21460000 | 0.0030 | 0.0025 | 0.0026 |
| F08 | 21380001 | 21480000 | 0.0037 | 0.0028 | 0.0031 |
| F08 | 21400001 | 21500000 | 0.0037 | 0.0028 | 0.0031 |
| F08 | 21420001 | 21520000 | 0.0040 | 0.0030 | 0.0033 |
| F08 | 21440001 | 21540000 | 0.0035 | 0.0026 | 0.0029 |
| F08 | 21460001 | 21560000 | 0.0033 | 0.0025 | 0.0028 |
| F08 | 21480001 | 21580000 | 0.0018 | 0.0015 | 0.0015 |
| F08 | 21500001 | 21600000 | 0.0022 | 0.0020 | 0.0021 |
| F08 | 21520001 | 21620000 | 0.0019 | 0.0018 | 0.0018 |
| F08 | 21540001 | 21640000 | 0.0022 | 0.0022 | 0.0020 |
| F08 | 21560001 | 21660000 | 0.0027 | 0.0023 | 0.0022 |
| F08 | 21580001 | 21680000 | 0.0033 | 0.0026 | 0.0026 |
| F08 | 21600001 | 21700000 | 0.0033 | 0.0025 | 0.0026 |
| F08 | 21620001 | 21720000 | 0.0039 | 0.0029 | 0.0031 |
| F08 | 21640001 | 21740000 | 0.0040 | 0.0028 | 0.0031 |
| F08 | 21660001 | 21760000 | 0.0044 | 0.0035 | 0.0038 |
| F08 | 21680001 | 21780000 | 0.0049 | 0.0042 | 0.0044 |
| F08 | 21700001 | 21800000 | 0.0042 | 0.0037 | 0.0039 |
| F08 | 21720001 | 21820000 | 0.0038 | 0.0034 | 0.0035 |
| F08 | 21740001 | 21840000 | 0.0035 | 0.0032 | 0.0035 |
| F08 | 21760001 | 21860000 | 0.0025 | 0.0023 | 0.0025 |
| F08 | 21780001 | 21880000 | 0.0014 | 0.0012 | 0.0014 |
| F08 | 21800001 | 21900000 | 0.0012 | 0.0010 | 0.0012 |
| F08 | 21820001 | 21920000 | 0.0013 | 0.0011 | 0.0013 |
| F08 | 21840001 | 21940000 | 0.0011 | 0.0009 | 0.0011 |
| F08 | 21860001 | 21960000 | 0.0010 | 0.0008 | 0.0010 |
| F08 | 21880001 | 21980000 | 0.0012 | 0.0010 | 0.0012 |
| F08 | 21900001 | 22000000 | 0.0013 | 0.0011 | 0.0013 |
| F08 | 21920001 | 22020000 | 0.0013 | 0.0010 | 0.0011 |
| F08 | 21940001 | 22040000 | 0.0032 | 0.0027 | 0.0030 |
| F08 | 21960001 | 22060000 | 0.0036 | 0.0032 | 0.0033 |
| F08 | 21980001 | 22080000 | 0.0038 | 0.0032 | 0.0031 |
| F08 | 22000001 | 22100000 | 0.0043 | 0.0036 | 0.0036 |
| F08 | 22020001 | 22120000 | 0.0040 | 0.0035 | 0.0035 |
| F08 | 22040001 | 22140000 | 0.0021 | 0.0016 | 0.0015 |
| F08 | 22060001 | 22160000 | 0.0014 | 0.0009 | 0.0009 |
| F08 | 22080001 | 22180000 | 0.0013 | 0.0011 | 0.0012 |
| F08 | 22100001 | 22200000 | 0.0010 | 0.0008 | 0.0009 |
| F08 | 22120001 | 22220000 | 0.0020 | 0.0017 | 0.0018 |
| F08 | 22140001 | 22240000 | 0.0022 | 0.0019 | 0.0021 |
| F08 | 22160001 | 22260000 | 0.0023 | 0.0020 | 0.0022 |
| F08 | 22180001 | 22280000 | 0.0027 | 0.0021 | 0.0026 |
| F08 | 22200001 | 22300000 | 0.0030 | 0.0023 | 0.0027 |
| F08 | 22220001 | 22320000 | 0.0024 | 0.0016 | 0.0021 |
| F08 | 22240001 | 22340000 | 0.0033 | 0.0022 | 0.0028 |
| F08 | 22260001 | 22360000 | 0.0037 | 0.0026 | 0.0033 |
| F08 | 22280001 | 22380000 | 0.0035 | 0.0024 | 0.0029 |
| F08 | 22300001 | 22400000 | 0.0037 | 0.0025 | 0.0029 |
| F08 | 22320001 | 22420000 | 0.0041 | 0.0029 | 0.0032 |
| F08 | 22340001 | 22440000 | 0.0036 | 0.0025 | 0.0028 |
| F08 | 22360001 | 22460000 | 0.0045 | 0.0032 | 0.0037 |
| F08 | 22380001 | 22480000 | 0.0048 | 0.0035 | 0.0041 |
| F08 | 22400001 | 22500000 | 0.0050 | 0.0039 | 0.0046 |

| Chr | Start | End | both | A1 | A2 |
|---|---|---|---|---|---|
| F08 | 22420001 | 22520000 | 0.0042 | 0.0034 | 0.0040 |
| F08 | 22440001 | 22540000 | 0.0040 | 0.0033 | 0.0038 |
| F08 | 22460001 | 22560000 | 0.0030 | 0.0026 | 0.0029 |
| F08 | 22480001 | 22580000 | 0.0028 | 0.0024 | 0.0025 |
| F08 | 22500001 | 22600000 | 0.0024 | 0.0020 | 0.0022 |
| F08 | 22520001 | 22620000 | 0.0027 | 0.0022 | 0.0024 |
| F08 | 22540001 | 22640000 | 0.0030 | 0.0024 | 0.0027 |
| F08 | 22560001 | 22660000 | 0.0033 | 0.0027 | 0.0030 |
| F08 | 22580001 | 22680000 | 0.0037 | 0.0031 | 0.0035 |
| F08 | 22600001 | 22700000 | 0.0035 | 0.0029 | 0.0033 |
| F08 | 22620001 | 22720000 | 0.0032 | 0.0028 | 0.0031 |
| F08 | 22640001 | 22740000 | 0.0027 | 0.0023 | 0.0027 |
| F08 | 22660001 | 22760000 | 0.0022 | 0.0020 | 0.0022 |
| F08 | 22680001 | 22780000 | 0.0015 | 0.0014 | 0.0015 |
| F08 | 22700001 | 22800000 | 0.0017 | 0.0016 | 0.0017 |
| F08 | 22720001 | 22820000 | 0.0022 | 0.0017 | 0.0019 |
| F08 | 22740001 | 22840000 | 0.0024 | 0.0019 | 0.0020 |
| F08 | 22760001 | 22860000 | 0.0030 | 0.0021 | 0.0025 |
| F08 | 22780001 | 22880000 | 0.0039 | 0.0027 | 0.0033 |
| F08 | 22800001 | 22900000 | 0.0038 | 0.0025 | 0.0031 |
| F08 | 22820001 | 22920000 | 0.0033 | 0.0023 | 0.0029 |
| F08 | 22840001 | 22940000 | 0.0030 | 0.0020 | 0.0026 |
| F08 | 22860001 | 22960000 | 0.0029 | 0.0022 | 0.0025 |
| F08 | 22880001 | 22980000 | 0.0021 | 0.0015 | 0.0018 |
| F08 | 22900001 | 23000000 | 0.0023 | 0.0019 | 0.0020 |
| F08 | 22920001 | 23020000 | 0.0034 | 0.0029 | 0.0031 |
| F08 | 22940001 | 23040000 | 0.0034 | 0.0030 | 0.0031 |
| F08 | 22960001 | 23060000 | 0.0026 | 0.0023 | 0.0023 |
| F08 | 22980001 | 23080000 | 0.0033 | 0.0030 | 0.0030 |
| F08 | 23000001 | 23100000 | 0.0034 | 0.0029 | 0.0032 |
| F08 | 23020001 | 23120000 | 0.0029 | 0.0023 | 0.0026 |
| F08 | 23040001 | 23140000 | 0.0033 | 0.0025 | 0.0030 |
| F08 | 23060001 | 23160000 | 0.0035 | 0.0027 | 0.0032 |
| F08 | 23080001 | 23180000 | 0.0028 | 0.0020 | 0.0026 |
| F08 | 23100001 | 23200000 | 0.0022 | 0.0016 | 0.0021 |
| F08 | 23120001 | 23220000 | 0.0024 | 0.0018 | 0.0022 |
| F08 | 23140001 | 23240000 | 0.0034 | 0.0025 | 0.0030 |
| F08 | 23160001 | 23260000 | 0.0042 | 0.0032 | 0.0037 |
| F08 | 23180001 | 23280000 | 0.0048 | 0.0037 | 0.0040 |
| F08 | 23200001 | 23300000 | 0.0058 | 0.0046 | 0.0050 |
| F08 | 23220001 | 23320000 | 0.0059 | 0.0047 | 0.0051 |
| F08 | 23240001 | 23340000 | 0.0050 | 0.0040 | 0.0042 |
| F08 | 23260001 | 23360000 | 0.0054 | 0.0048 | 0.0042 |
| F08 | 23280001 | 23380000 | 0.0049 | 0.0045 | 0.0038 |
| F08 | 23300001 | 23400000 | 0.0038 | 0.0036 | 0.0028 |
| F08 | 23320001 | 23420000 | 0.0031 | 0.0028 | 0.0018 |
| F08 | 23340001 | 23440000 | 0.0033 | 0.0029 | 0.0021 |
| F08 | 23360001 | 23460000 | 0.0023 | 0.0015 | 0.0016 |
| F08 | 23380001 | 23480000 | 0.0032 | 0.0023 | 0.0025 |
| F08 | 23400001 | 23500000 | 0.0039 | 0.0028 | 0.0031 |
| F08 | 23420001 | 23520000 | 0.0039 | 0.0030 | 0.0034 |
| F08 | 23440001 | 23540000 | 0.0036 | 0.0029 | 0.0031 |
| F08 | 23460001 | 23560000 | 0.0038 | 0.0031 | 0.0033 |
| F08 | 23480001 | 23580000 | 0.0030 | 0.0023 | 0.0024 |
| F08 | 23500001 | 23600000 | 0.0025 | 0.0018 | 0.0018 |
| F08 | 23520001 | 23620000 | 0.0025 | 0.0018 | 0.0018 |
| F08 | 23540001 | 23640000 | 0.0026 | 0.0017 | 0.0020 |
| F08 | 23560001 | 23660000 | 0.0023 | 0.0016 | 0.0018 |

| Chr | Start | End | both | A1 | A2 |
|---|---|---|---|---|---|
| F08 | 23580001 | 23680000 | 0.0024 | 0.0016 | 0.0019 |
| F08 | 23600001 | 23700000 | 0.0029 | 0.0022 | 0.0025 |
| F08 | 23620001 | 23720000 | 0.0026 | 0.0020 | 0.0023 |
| F08 | 23640001 | 23740000 | 0.0026 | 0.0022 | 0.0025 |
| F08 | 23660001 | 23760000 | 0.0021 | 0.0017 | 0.0020 |
| F08 | 23680001 | 23780000 | 0.0020 | 0.0018 | 0.0020 |
| F08 | 23700001 | 23800000 | 0.0015 | 0.0014 | 0.0015 |
| F08 | 23720001 | 23820000 | 0.0017 | 0.0015 | 0.0017 |
| F08 | 23740001 | 23840000 | 0.0019 | 0.0017 | 0.0019 |
| F08 | 23760001 | 23860000 | 0.0023 | 0.0019 | 0.0021 |
| F08 | 23780001 | 23880000 | 0.0029 | 0.0023 | 0.0025 |
| F08 | 23800001 | 23900000 | 0.0035 | 0.0028 | 0.0029 |
| F08 | 23820001 | 23920000 | 0.0034 | 0.0027 | 0.0027 |
| F08 | 23840001 | 23940000 | 0.0033 | 0.0023 | 0.0025 |
| F08 | 23860001 | 23960000 | 0.0042 | 0.0029 | 0.0036 |
| F08 | 23880001 | 23980000 | 0.0043 | 0.0029 | 0.0036 |
| F08 | 23900001 | 24000000 | 0.0034 | 0.0020 | 0.0029 |
| F08 | 23920001 | 24020000 | 0.0042 | 0.0027 | 0.0037 |
| F08 | 23940001 | 24040000 | 0.0043 | 0.0032 | 0.0040 |
| F08 | 23960001 | 24060000 | 0.0044 | 0.0036 | 0.0040 |
| F08 | 23980001 | 24080000 | 0.0041 | 0.0034 | 0.0036 |
| F08 | 24000001 | 24100000 | 0.0046 | 0.0036 | 0.0038 |
| F08 | 24020001 | 24120000 | 0.0043 | 0.0035 | 0.0035 |
| F08 | 24040001 | 24140000 | 0.0044 | 0.0035 | 0.0037 |
| F08 | 24060001 | 24160000 | 0.0039 | 0.0029 | 0.0032 |
| F08 | 24080001 | 24180000 | 0.0039 | 0.0029 | 0.0033 |
| F08 | 24100001 | 24200000 | 0.0042 | 0.0033 | 0.0038 |
| F08 | 24120001 | 24220000 | 0.0043 | 0.0033 | 0.0039 |
| F08 | 24140001 | 24240000 | 0.0040 | 0.0031 | 0.0036 |
| F08 | 24160001 | 24260000 | 0.0033 | 0.0024 | 0.0028 |
| F08 | 24180001 | 24280000 | 0.0035 | 0.0027 | 0.0029 |
| F08 | 24200001 | 24300000 | 0.0031 | 0.0023 | 0.0025 |
| F08 | 24220001 | 24320000 | 0.0034 | 0.0025 | 0.0026 |
| F08 | 24240001 | 24340000 | 0.0032 | 0.0021 | 0.0023 |
| F08 | 24260001 | 24360000 | 0.0044 | 0.0033 | 0.0035 |
| F08 | 24280001 | 24380000 | 0.0042 | 0.0030 | 0.0033 |
| F08 | 24300001 | 24400000 | 0.0044 | 0.0032 | 0.0035 |
| F08 | 24320001 | 24420000 | 0.0050 | 0.0039 | 0.0041 |
| F08 | 24340001 | 24440000 | 0.0052 | 0.0043 | 0.0045 |
| F08 | 24360001 | 24460000 | 0.0058 | 0.0048 | 0.0053 |
| F08 | 24380001 | 24480000 | 0.0067 | 0.0057 | 0.0063 |
| F08 | 24400001 | 24500000 | 0.0070 | 0.0058 | 0.0064 |
| F08 | 24420001 | 24520000 | 0.0060 | 0.0049 | 0.0055 |
| F08 | 24440001 | 24540000 | 0.0062 | 0.0048 | 0.0054 |
| F08 | 24460001 | 24560000 | 0.0044 | 0.0035 | 0.0036 |
| F08 | 24480001 | 24580000 | 0.0044 | 0.0036 | 0.0036 |
| F08 | 24500001 | 24600000 | 0.0038 | 0.0034 | 0.0033 |
| F08 | 24520001 | 24620000 | 0.0045 | 0.0039 | 0.0040 |
| F08 | 24540001 | 24640000 | 0.0041 | 0.0036 | 0.0035 |
| F08 | 24560001 | 24660000 | 0.0047 | 0.0041 | 0.0040 |
| F08 | 24580001 | 24680000 | 0.0046 | 0.0040 | 0.0039 |
| F08 | 24600001 | 24700000 | 0.0062 | 0.0052 | 0.0053 |
| F08 | 24620001 | 24720000 | 0.0055 | 0.0044 | 0.0045 |
| F08 | 24640001 | 24740000 | 0.0060 | 0.0049 | 0.0050 |
| F08 | 24660001 | 24760000 | 0.0059 | 0.0047 | 0.0049 |
| F08 | 24680001 | 24780000 | 0.0047 | 0.0033 | 0.0036 |
| F08 | 24700001 | 24800000 | 0.0040 | 0.0029 | 0.0031 |
| F08 | 24720001 | 24820000 | 0.0041 | 0.0030 | 0.0034 |

| Chr | Start | End | both | A1 | A2 |
|---|---|---|---|---|---|
| F08 | 24740001 | 24840000 | 0.0033 | 0.0023 | 0.0027 |
| F08 | 24760001 | 24860000 | 0.0028 | 0.0020 | 0.0025 |
| F08 | 24780001 | 24880000 | 0.0038 | 0.0032 | 0.0036 |
| F08 | 24800001 | 24900000 | 0.0047 | 0.0039 | 0.0044 |
| F08 | 24820001 | 24920000 | 0.0043 | 0.0036 | 0.0040 |
| F08 | 24840001 | 24940000 | 0.0044 | 0.0037 | 0.0041 |
| F08 | 24860001 | 24960000 | 0.0048 | 0.0042 | 0.0045 |
| F08 | 24880001 | 24980000 | 0.0039 | 0.0033 | 0.0036 |
| F08 | 24900001 | 25000000 | 0.0022 | 0.0020 | 0.0021 |
| F08 | 24920001 | 25020000 | 0.0021 | 0.0019 | 0.0021 |
| F08 | 24940001 | 25040000 | 0.0023 | 0.0021 | 0.0022 |
| F08 | 24960001 | 25060000 | 0.0017 | 0.0015 | 0.0017 |
| F08 | 24980001 | 25080000 | 0.0017 | 0.0015 | 0.0017 |
| F08 | 25000001 | 25100000 | 0.0011 | 0.0009 | 0.0011 |
| F08 | 25020001 | 25120000 | 0.0013 | 0.0010 | 0.0011 |
| F08 | 25040001 | 25140000 | 0.0010 | 0.0008 | 0.0008 |
| F08 | 25060001 | 25160000 | 0.0015 | 0.0013 | 0.0013 |
| F08 | 25080001 | 25180000 | 0.0016 | 0.0013 | 0.0014 |
| F08 | 25100001 | 25200000 | 0.0022 | 0.0018 | 0.0020 |
| F08 | 25120001 | 25220000 | 0.0021 | 0.0017 | 0.0021 |
| F08 | 25140001 | 25240000 | 0.0024 | 0.0019 | 0.0023 |
| F08 | 25160001 | 25260000 | 0.0025 | 0.0019 | 0.0023 |
| F08 | 25180001 | 25280000 | 0.0029 | 0.0022 | 0.0026 |
| F08 | 25200001 | 25300000 | 0.0028 | 0.0021 | 0.0025 |
| F08 | 25220001 | 25320000 | 0.0035 | 0.0027 | 0.0032 |
| F08 | 25240001 | 25340000 | 0.0032 | 0.0025 | 0.0029 |
| F08 | 25260001 | 25360000 | 0.0030 | 0.0025 | 0.0028 |
| F08 | 25280001 | 25380000 | 0.0025 | 0.0020 | 0.0022 |
| F08 | 25300001 | 25400000 | 0.0022 | 0.0018 | 0.0019 |
| F08 | 25320001 | 25420000 | 0.0020 | 0.0016 | 0.0017 |
| F08 | 25340001 | 25440000 | 0.0025 | 0.0020 | 0.0021 |
| F08 | 25360001 | 25460000 | 0.0024 | 0.0017 | 0.0020 |
| F08 | 25380001 | 25480000 | 0.0030 | 0.0021 | 0.0026 |
| F08 | 25400001 | 25500000 | 0.0032 | 0.0021 | 0.0026 |
| F08 | 25420001 | 25520000 | 0.0032 | 0.0016 | 0.0022 |
| F08 | 25440001 | 25540000 | 0.0028 | 0.0013 | 0.0018 |
| F08 | 25460001 | 25560000 | 0.0029 | 0.0014 | 0.0018 |
| F08 | 25480001 | 25580000 | 0.0035 | 0.0021 | 0.0023 |
| F08 | 25500001 | 25600000 | 0.0033 | 0.0022 | 0.0023 |
| F08 | 25520001 | 25620000 | 0.0027 | 0.0022 | 0.0022 |
| F08 | 25540001 | 25640000 | 0.0031 | 0.0025 | 0.0026 |
| F08 | 25560001 | 25660000 | 0.0024 | 0.0020 | 0.0022 |
| F08 | 25580001 | 25680000 | 0.0011 | 0.0008 | 0.0010 |
| F08 | 25600001 | 25700000 | 0.0011 | 0.0008 | 0.0011 |
| F08 | 25620001 | 25720000 | 0.0011 | 0.0008 | 0.0011 |
| F08 | 25640001 | 25740000 | 0.0011 | 0.0008 | 0.0010 |
| F08 | 25660001 | 25760000 | 0.0015 | 0.0009 | 0.0011 |
| F08 | 25680001 | 25780000 | 0.0020 | 0.0013 | 0.0014 |
| F08 | 25700001 | 25800000 | 0.0023 | 0.0015 | 0.0015 |
| F08 | 25720001 | 25820000 | 0.0028 | 0.0018 | 0.0017 |
| F08 | 25740001 | 25840000 | 0.0034 | 0.0023 | 0.0024 |
| F08 | 25760001 | 25860000 | 0.0030 | 0.0024 | 0.0024 |
| F08 | 25780001 | 25880000 | 0.0032 | 0.0026 | 0.0028 |
| F08 | 25800001 | 25900000 | 0.0035 | 0.0028 | 0.0032 |
| F08 | 25820001 | 25920000 | 0.0037 | 0.0030 | 0.0035 |
| F08 | 25840001 | 25940000 | 0.0030 | 0.0023 | 0.0026 |
| F08 | 25860001 | 25960000 | 0.0034 | 0.0025 | 0.0029 |
| F08 | 25880001 | 25980000 | 0.0034 | 0.0024 | 0.0027 |

| Chr | Start | End | both | A1 | A2 |
|---|---|---|---|---|---|
| F08 | 25900001 | 26000000 | 0.0028 | 0.0020 | 0.0022 |
| F08 | 25920001 | 26020000 | 0.0027 | 0.0020 | 0.0020 |
| F08 | 25940001 | 26040000 | 0.0024 | 0.0019 | 0.0020 |
| F08 | 25960001 | 26060000 | 0.0021 | 0.0017 | 0.0018 |
| F08 | 25980001 | 26080000 | 0.0014 | 0.0012 | 0.0013 |
| F08 | 26000001 | 26100000 | 0.0016 | 0.0013 | 0.0014 |
| F08 | 26020001 | 26120000 | 0.0019 | 0.0014 | 0.0016 |
| F08 | 26040001 | 26140000 | 0.0022 | 0.0016 | 0.0018 |
| F08 | 26060001 | 26160000 | 0.0026 | 0.0019 | 0.0021 |
| F08 | 26080001 | 26180000 | 0.0032 | 0.0023 | 0.0026 |
| F08 | 26100001 | 26200000 | 0.0031 | 0.0021 | 0.0023 |
| F08 | 26120001 | 26220000 | 0.0025 | 0.0015 | 0.0017 |
| F08 | 26140001 | 26240000 | 0.0032 | 0.0023 | 0.0026 |
| F08 | 26160001 | 26260000 | 0.0030 | 0.0022 | 0.0025 |
| F08 | 26180001 | 26280000 | 0.0028 | 0.0019 | 0.0023 |
| F08 | 26200001 | 26300000 | 0.0029 | 0.0021 | 0.0026 |
| F08 | 26220001 | 26320000 | 0.0026 | 0.0021 | 0.0025 |
| F08 | 26240001 | 26340000 | 0.0022 | 0.0015 | 0.0021 |
| F08 | 26260001 | 26360000 | 0.0019 | 0.0013 | 0.0018 |
| F08 | 26280001 | 26380000 | 0.0019 | 0.0014 | 0.0018 |
| F08 | 26300001 | 26400000 | 0.0025 | 0.0019 | 0.0020 |
| F08 | 26320001 | 26420000 | 0.0031 | 0.0025 | 0.0026 |
| F08 | 26340001 | 26440000 | 0.0031 | 0.0023 | 0.0022 |
| F08 | 26360001 | 26460000 | 0.0040 | 0.0028 | 0.0027 |
| F08 | 26380001 | 26480000 | 0.0043 | 0.0029 | 0.0028 |
| F08 | 26400001 | 26500000 | 0.0035 | 0.0023 | 0.0024 |
| F08 | 26420001 | 26520000 | 0.0032 | 0.0018 | 0.0020 |
| F08 | 26440001 | 26540000 | 0.0034 | 0.0021 | 0.0023 |
| F08 | 26460001 | 26560000 | 0.0032 | 0.0020 | 0.0023 |
| F08 | 26480001 | 26580000 | 0.0024 | 0.0016 | 0.0019 |
| F08 | 26500001 | 26600000 | 0.0023 | 0.0014 | 0.0017 |
| F08 | 26520001 | 26620000 | 0.0022 | 0.0012 | 0.0015 |
| F08 | 26540001 | 26640000 | 0.0018 | 0.0010 | 0.0010 |
| F08 | 26560001 | 26660000 | 0.0018 | 0.0011 | 0.0011 |
| F08 | 26580001 | 26680000 | 0.0021 | 0.0014 | 0.0013 |
| F08 | 26600001 | 26700000 | 0.0029 | 0.0023 | 0.0023 |
| F08 | 26620001 | 26720000 | 0.0043 | 0.0036 | 0.0036 |
| F08 | 26640001 | 26740000 | 0.0043 | 0.0037 | 0.0038 |
| F08 | 26660001 | 26760000 | 0.0043 | 0.0039 | 0.0038 |
| F08 | 26680001 | 26780000 | 0.0045 | 0.0039 | 0.0039 |
| F08 | 26700001 | 26800000 | 0.0040 | 0.0033 | 0.0033 |
| F08 | 26720001 | 26820000 | 0.0035 | 0.0030 | 0.0029 |
| F08 | 26740001 | 26840000 | 0.0032 | 0.0028 | 0.0027 |
| F08 | 26760001 | 26860000 | 0.0024 | 0.0020 | 0.0021 |
| F08 | 26780001 | 26880000 | 0.0020 | 0.0019 | 0.0019 |
| F08 | 26800001 | 26900000 | 0.0021 | 0.0020 | 0.0020 |
| F08 | 26820001 | 26920000 | 0.0011 | 0.0010 | 0.0011 |
| F08 | 26840001 | 26940000 | 0.0020 | 0.0019 | 0.0020 |
| F08 | 26860001 | 26960000 | 0.0042 | 0.0038 | 0.0041 |
| F08 | 26880001 | 26980000 | 0.0045 | 0.0041 | 0.0045 |
| F08 | 26900001 | 27000000 | 0.0050 | 0.0043 | 0.0048 |
| F08 | 26920001 | 27020000 | 0.0055 | 0.0047 | 0.0052 |
| F08 | 26940001 | 27040000 | 0.0049 | 0.0040 | 0.0044 |
| F08 | 26960001 | 27060000 | 0.0033 | 0.0024 | 0.0026 |
| F08 | 26980001 | 27080000 | 0.0030 | 0.0022 | 0.0023 |
| F08 | 27000001 | 27100000 | 0.0019 | 0.0014 | 0.0014 |
| F08 | 27020001 | 27120000 | 0.0017 | 0.0011 | 0.0013 |
| F08 | 27040001 | 27140000 | 0.0020 | 0.0014 | 0.0017 |

| Chr | Start | End | both | A1 | A2 |
|---|---|---|---|---|---|
| F08 | 27060001 | 27160000 | 0.0033 | 0.0027 | 0.0028 |
| F08 | 27080001 | 27180000 | 0.0039 | 0.0033 | 0.0034 |
| F08 | 27100001 | 27200000 | 0.0055 | 0.0047 | 0.0049 |
| F08 | 27120001 | 27220000 | 0.0055 | 0.0049 | 0.0050 |
| F08 | 27140001 | 27240000 | 0.0047 | 0.0043 | 0.0042 |
| F08 | 27160001 | 27260000 | 0.0033 | 0.0030 | 0.0032 |
| F08 | 27180001 | 27280000 | 0.0027 | 0.0023 | 0.0027 |
| F08 | 27200001 | 27300000 | 0.0017 | 0.0014 | 0.0017 |
| F08 | 27220001 | 27320000 | 0.0023 | 0.0018 | 0.0023 |
| F08 | 27240001 | 27340000 | 0.0023 | 0.0018 | 0.0023 |
| F08 | 27260001 | 27360000 | 0.0019 | 0.0014 | 0.0019 |
| F08 | 27280001 | 27380000 | 0.0019 | 0.0016 | 0.0019 |
| F08 | 27300001 | 27400000 | 0.0015 | 0.0012 | 0.0015 |
| F08 | 27320001 | 27420000 | 0.0007 | 0.0006 | 0.0007 |
| F08 | 27340001 | 27440000 | 0.0009 | 0.0007 | 0.0009 |
| F08 | 27360001 | 27460000 | 0.0011 | 0.0009 | 0.0011 |
| F08 | 27380001 | 27480000 | 0.0010 | 0.0009 | 0.0010 |
| F08 | 27400001 | 27500000 | 0.0014 | 0.0011 | 0.0014 |
| F08 | 27420001 | 27520000 | 0.0013 | 0.0011 | 0.0013 |
| F08 | 27440001 | 27540000 | 0.0016 | 0.0013 | 0.0016 |
| F08 | 27460001 | 27560000 | 0.0014 | 0.0012 | 0.0014 |
| F08 | 27480001 | 27580000 | 0.0016 | 0.0014 | 0.0015 |
| F08 | 27500001 | 27600000 | 0.0011 | 0.0010 | 0.0010 |
| F08 | 27520001 | 27620000 | 0.0015 | 0.0012 | 0.0013 |
| F08 | 27540001 | 27640000 | 0.0011 | 0.0009 | 0.0009 |
| F08 | 27560001 | 27660000 | 0.0014 | 0.0011 | 0.0011 |
| F08 | 27580001 | 27680000 | 0.0011 | 0.0008 | 0.0009 |
| F08 | 27600001 | 27700000 | 0.0012 | 0.0008 | 0.0011 |
| F08 | 27620001 | 27720000 | 0.0013 | 0.0011 | 0.0012 |
| F08 | 27640001 | 27740000 | 0.0021 | 0.0017 | 0.0018 |
| F08 | 27660001 | 27760000 | 0.0037 | 0.0030 | 0.0034 |
| F08 | 27680001 | 27780000 | 0.0050 | 0.0041 | 0.0044 |
| F08 | 27700001 | 27800000 | 0.0054 | 0.0043 | 0.0046 |
| F08 | 27720001 | 27820000 | 0.0057 | 0.0043 | 0.0048 |
| F08 | 27740001 | 27840000 | 0.0053 | 0.0041 | 0.0046 |
| F08 | 27760001 | 27860000 | 0.0040 | 0.0028 | 0.0032 |
| F08 | 27780001 | 27880000 | 0.0034 | 0.0023 | 0.0027 |
| F08 | 27800001 | 27900000 | 0.0034 | 0.0022 | 0.0025 |
| F08 | 27820001 | 27920000 | 0.0035 | 0.0022 | 0.0025 |
| F08 | 27840001 | 27940000 | 0.0041 | 0.0029 | 0.0032 |
| F08 | 27860001 | 27960000 | 0.0038 | 0.0027 | 0.0030 |
| F08 | 27880001 | 27980000 | 0.0030 | 0.0022 | 0.0023 |
| F08 | 27900001 | 28000000 | 0.0028 | 0.0022 | 0.0023 |
| F08 | 27920001 | 28020000 | 0.0019 | 0.0017 | 0.0018 |
| F08 | 27940001 | 28040000 | 0.0019 | 0.0018 | 0.0018 |
| F08 | 27960001 | 28060000 | 0.0021 | 0.0018 | 0.0018 |
| F08 | 27980001 | 28080000 | 0.0019 | 0.0016 | 0.0017 |
| F08 | 28000001 | 28100000 | 0.0021 | 0.0017 | 0.0018 |
| F08 | 28020001 | 28120000 | 0.0027 | 0.0021 | 0.0023 |
| F08 | 28040001 | 28140000 | 0.0018 | 0.0012 | 0.0015 |
| F08 | 28060001 | 28160000 | 0.0018 | 0.0013 | 0.0016 |
| F08 | 28080001 | 28180000 | 0.0023 | 0.0019 | 0.0021 |
| F08 | 28100001 | 28200000 | 0.0042 | 0.0036 | 0.0036 |
| F08 | 28120001 | 28220000 | 0.0039 | 0.0035 | 0.0033 |
| F08 | 28140001 | 28240000 | 0.0045 | 0.0038 | 0.0038 |
| F08 | 28160001 | 28260000 | 0.0041 | 0.0035 | 0.0035 |
| F08 | 28180001 | 28280000 | 0.0041 | 0.0033 | 0.0035 |
| F08 | 28200001 | 28300000 | 0.0030 | 0.0022 | 0.0027 |

| Chr | Start | End | both | A1 | A2 |
|---|---|---|---|---|---|
| F08 | 28220001 | 28320000 | 0.0026 | 0.0018 | 0.0023 |
| F08 | 28240001 | 28340000 | 0.0024 | 0.0019 | 0.0022 |
| F08 | 28260001 | 28360000 | 0.0027 | 0.0021 | 0.0025 |
| F08 | 28280001 | 28380000 | 0.0021 | 0.0017 | 0.0019 |
| F08 | 28300001 | 28400000 | 0.0013 | 0.0011 | 0.0012 |
| F08 | 28320001 | 28420000 | 0.0022 | 0.0018 | 0.0021 |
| F08 | 28340001 | 28440000 | 0.0018 | 0.0013 | 0.0016 |
| F08 | 28360001 | 28460000 | 0.0019 | 0.0014 | 0.0017 |
| F08 | 28380001 | 28480000 | 0.0019 | 0.0013 | 0.0017 |
| F08 | 28400001 | 28500000 | 0.0017 | 0.0012 | 0.0015 |
| F08 | 28420001 | 28520000 | 0.0010 | 0.0007 | 0.0008 |
| F08 | 28440001 | 28540000 | 0.0009 | 0.0008 | 0.0008 |
| F08 | 28460001 | 28560000 | 0.0006 | 0.0007 | 0.0005 |
| F08 | 28480001 | 28580000 | 0.0008 | 0.0009 | 0.0007 |
| F08 | 28500001 | 28600000 | 0.0007 | 0.0008 | 0.0006 |
| F08 | 28520001 | 28620000 | 0.0006 | 0.0007 | 0.0005 |
| F08 | 28540001 | 28640000 | 0.0005 | 0.0006 | 0.0005 |
| F08 | 28560001 | 28660000 | 0.0005 | 0.0004 | 0.0004 |
| F08 | 28580001 | 28680000 | 0.0005 | 0.0004 | 0.0005 |
| F08 | 28600001 | 28700000 | 0.0005 | 0.0004 | 0.0005 |
| F08 | 28620001 | 28720000 | 0.0005 | 0.0004 | 0.0005 |
| F08 | 28640001 | 28740000 | 0.0006 | 0.0005 | 0.0005 |
| F08 | 28660001 | 28760000 | 0.0008 | 0.0007 | 0.0008 |
| F08 | 28680001 | 28780000 | 0.0011 | 0.0008 | 0.0011 |
| F08 | 28700001 | 28800000 | 0.0016 | 0.0012 | 0.0015 |
| F08 | 28720001 | 28820000 | 0.0015 | 0.0011 | 0.0013 |
| F08 | 28740001 | 28840000 | 0.0016 | 0.0012 | 0.0014 |
| F08 | 28760001 | 28860000 | 0.0016 | 0.0011 | 0.0014 |
| F08 | 28780001 | 28880000 | 0.0013 | 0.0009 | 0.0011 |
| F08 | 28800001 | 28900000 | 0.0010 | 0.0007 | 0.0009 |
| F08 | 28820001 | 28920000 | 0.0009 | 0.0006 | 0.0009 |
| F08 | 28840001 | 28940000 | 0.0007 | 0.0005 | 0.0007 |
| F08 | 28860001 | 28960000 | 0.0006 | 0.0004 | 0.0006 |
| F08 | 28880001 | 28980000 | 0.0006 | 0.0005 | 0.0006 |
| F08 | 28900001 | 29000000 | 0.0006 | 0.0005 | 0.0006 |
| F08 | 28920001 | 29020000 | 0.0019 | 0.0016 | 0.0019 |
| F08 | 28940001 | 29040000 | 0.0023 | 0.0019 | 0.0023 |
| F08 | 28960001 | 29060000 | 0.0022 | 0.0019 | 0.0022 |
| F08 | 28980001 | 29080000 | 0.0029 | 0.0026 | 0.0028 |
| F08 | 29000001 | 29100000 | 0.0033 | 0.0029 | 0.0032 |
| F08 | 29020001 | 29120000 | 0.0024 | 0.0022 | 0.0023 |
| F08 | 29040001 | 29140000 | 0.0020 | 0.0018 | 0.0019 |
| F08 | 29060001 | 29160000 | 0.0026 | 0.0023 | 0.0025 |
| F08 | 29080001 | 29180000 | 0.0026 | 0.0023 | 0.0025 |
| F08 | 29100001 | 29200000 | 0.0025 | 0.0024 | 0.0022 |
| F08 | 29120001 | 29220000 | 0.0025 | 0.0022 | 0.0020 |
| F08 | 29140001 | 29240000 | 0.0031 | 0.0025 | 0.0024 |
| F08 | 29160001 | 29260000 | 0.0033 | 0.0027 | 0.0025 |
| F08 | 29180001 | 29280000 | 0.0026 | 0.0020 | 0.0018 |
| F08 | 29200001 | 29300000 | 0.0025 | 0.0018 | 0.0019 |
| F08 | 29220001 | 29320000 | 0.0022 | 0.0017 | 0.0019 |
| F08 | 29240001 | 29340000 | 0.0015 | 0.0013 | 0.0014 |
| F08 | 29260001 | 29360000 | 0.0011 | 0.0008 | 0.0011 |
| F08 | 29280001 | 29380000 | 0.0010 | 0.0008 | 0.0010 |
| F08 | 29300001 | 29400000 | 0.0008 | 0.0006 | 0.0008 |
| F08 | 29320001 | 29420000 | 0.0011 | 0.0009 | 0.0011 |
| F08 | 29340001 | 29440000 | 0.0012 | 0.0009 | 0.0012 |
| F08 | 29360001 | 29460000 | 0.0009 | 0.0007 | 0.0009 |

| Chr | Start | End | both | A1 | A2 |
|---|---|---|---|---|---|
| F08 | 29380001 | 29480000 | 0.0010 | 0.0008 | 0.0010 |
| F08 | 29400001 | 29500000 | 0.0009 | 0.0007 | 0.0009 |
| F08 | 29420001 | 29520000 | 0.0009 | 0.0007 | 0.0009 |
| F08 | 29440001 | 29540000 | 0.0017 | 0.0015 | 0.0017 |
| F08 | 29460001 | 29560000 | 0.0018 | 0.0015 | 0.0017 |
| F08 | 29480001 | 29580000 | 0.0016 | 0.0014 | 0.0016 |
| F08 | 29500001 | 29600000 | 0.0015 | 0.0013 | 0.0015 |
| F08 | 29520001 | 29620000 | 0.0014 | 0.0012 | 0.0013 |
| F08 | 29540001 | 29640000 | 0.0024 | 0.0020 | 0.0024 |
| F08 | 29560001 | 29660000 | 0.0033 | 0.0028 | 0.0033 |
| F08 | 29580001 | 29680000 | 0.0035 | 0.0030 | 0.0035 |
| F08 | 29600001 | 29700000 | 0.0037 | 0.0032 | 0.0037 |
| F08 | 29620001 | 29720000 | 0.0049 | 0.0034 | 0.0046 |
| F08 | 29640001 | 29740000 | 0.0032 | 0.0019 | 0.0029 |
| F08 | 29660001 | 29760000 | 0.0024 | 0.0012 | 0.0021 |
| F08 | 29680001 | 29780000 | 0.0027 | 0.0014 | 0.0024 |
| F08 | 29700001 | 29800000 | 0.0025 | 0.0013 | 0.0022 |
| F08 | 29720001 | 29820000 | 0.0015 | 0.0011 | 0.0014 |
| F08 | 29740001 | 29840000 | 0.0012 | 0.0009 | 0.0012 |
| F08 | 29760001 | 29860000 | 0.0012 | 0.0010 | 0.0012 |
| F08 | 29780001 | 29880000 | 0.0010 | 0.0008 | 0.0010 |
| F08 | 29800001 | 29900000 | 0.0023 | 0.0017 | 0.0023 |
| F08 | 29820001 | 29920000 | 0.0026 | 0.0021 | 0.0026 |
| F08 | 29840001 | 29940000 | 0.0030 | 0.0024 | 0.0029 |
| F08 | 29860001 | 29960000 | 0.0028 | 0.0022 | 0.0028 |
| F08 | 29880001 | 29980000 | 0.0027 | 0.0022 | 0.0026 |
| F08 | 29900001 | 30000000 | 0.0015 | 0.0013 | 0.0015 |
| F08 | 29920001 | 30020000 | 0.0009 | 0.0008 | 0.0009 |
| F08 | 29940001 | 30040000 | 0.0017 | 0.0015 | 0.0017 |
| F08 | 29960001 | 30060000 | 0.0026 | 0.0023 | 0.0026 |
| F08 | 29980001 | 30080000 | 0.0030 | 0.0025 | 0.0028 |
| F08 | 30000001 | 30100000 | 0.0044 | 0.0039 | 0.0040 |
| F08 | 30020001 | 30120000 | 0.0045 | 0.0040 | 0.0041 |
| F08 | 30040001 | 30140000 | 0.0035 | 0.0030 | 0.0030 |
| F08 | 30060001 | 30160000 | 0.0028 | 0.0024 | 0.0023 |
| F08 | 30080001 | 30180000 | 0.0030 | 0.0027 | 0.0027 |
| F08 | 30100001 | 30200000 | 0.0016 | 0.0013 | 0.0016 |
| F08 | 30120001 | 30220000 | 0.0017 | 0.0015 | 0.0017 |
| F08 | 30140001 | 30240000 | 0.0021 | 0.0017 | 0.0020 |
| F08 | 30160001 | 30260000 | 0.0027 | 0.0022 | 0.0026 |
| F08 | 30180001 | 30280000 | 0.0023 | 0.0019 | 0.0023 |
| F08 | 30200001 | 30300000 | 0.0027 | 0.0022 | 0.0027 |
| F08 | 30220001 | 30320000 | 0.0028 | 0.0023 | 0.0028 |
| F08 | 30240001 | 30340000 | 0.0033 | 0.0028 | 0.0032 |
| F08 | 30260001 | 30360000 | 0.0029 | 0.0022 | 0.0025 |
| F08 | 30280001 | 30380000 | 0.0029 | 0.0022 | 0.0024 |
| F08 | 30300001 | 30400000 | 0.0027 | 0.0019 | 0.0021 |
| F08 | 30320001 | 30420000 | 0.0031 | 0.0022 | 0.0023 |
| F08 | 30340001 | 30440000 | 0.0030 | 0.0023 | 0.0022 |
| F08 | 30360001 | 30460000 | 0.0028 | 0.0022 | 0.0021 |
| F08 | 30380001 | 30480000 | 0.0024 | 0.0018 | 0.0018 |
| F08 | 30400001 | 30500000 | 0.0025 | 0.0019 | 0.0019 |
| F08 | 30420001 | 30520000 | 0.0033 | 0.0026 | 0.0026 |
| F08 | 30440001 | 30540000 | 0.0031 | 0.0022 | 0.0023 |
| F08 | 30460001 | 30560000 | 0.0037 | 0.0028 | 0.0030 |
| F08 | 30480001 | 30580000 | 0.0043 | 0.0033 | 0.0034 |
| F08 | 30500001 | 30600000 | 0.0042 | 0.0033 | 0.0034 |
| F08 | 30520001 | 30620000 | 0.0039 | 0.0028 | 0.0029 |

| Chr | Start | End | both | A1 | A2 |
|---|---|---|---|---|---|
| F08 | 30540001 | 30640000 | 0.0039 | 0.0027 | 0.0028 |
| F08 | 30560001 | 30660000 | 0.0042 | 0.0029 | 0.0030 |
| F08 | 30580001 | 30680000 | 0.0045 | 0.0030 | 0.0033 |
| F08 | 30600001 | 30700000 | 0.0055 | 0.0039 | 0.0045 |
| F08 | 30620001 | 30720000 | 0.0049 | 0.0034 | 0.0040 |
| F08 | 30640001 | 30740000 | 0.0043 | 0.0031 | 0.0037 |
| F08 | 30660001 | 30760000 | 0.0031 | 0.0023 | 0.0028 |
| F08 | 30680001 | 30780000 | 0.0022 | 0.0017 | 0.0020 |
| F08 | 30700001 | 30800000 | 0.0011 | 0.0007 | 0.0009 |
| F08 | 30720001 | 30820000 | 0.0012 | 0.0009 | 0.0012 |
| F08 | 30740001 | 30840000 | 0.0020 | 0.0017 | 0.0020 |
| F08 | 30760001 | 30860000 | 0.0024 | 0.0020 | 0.0024 |
| F08 | 30780001 | 30880000 | 0.0025 | 0.0021 | 0.0024 |
| F08 | 30800001 | 30900000 | 0.0026 | 0.0023 | 0.0026 |
| F08 | 30820001 | 30920000 | 0.0021 | 0.0019 | 0.0020 |
| F08 | 30840001 | 30940000 | 0.0013 | 0.0012 | 0.0013 |
| F08 | 30860001 | 30960000 | 0.0011 | 0.0010 | 0.0010 |
| F08 | 30880001 | 30980000 | 0.0010 | 0.0009 | 0.0009 |
| F08 | 30900001 | 31000000 | 0.0005 | 0.0005 | 0.0005 |
| F08 | 30920001 | 31020000 | 0.0007 | 0.0006 | 0.0006 |
| F08 | 30940001 | 31040000 | 0.0005 | 0.0005 | 0.0005 |
| F08 | 30960001 | 31060000 | 0.0005 | 0.0004 | 0.0005 |
| F08 | 30980001 | 31080000 | 0.0011 | 0.0010 | 0.0011 |
| F08 | 31000001 | 31100000 | 0.0017 | 0.0016 | 0.0017 |
| F08 | 31020001 | 31120000 | 0.0022 | 0.0019 | 0.0022 |
| F08 | 31040001 | 31140000 | 0.0024 | 0.0021 | 0.0024 |
| F08 | 31060001 | 31160000 | 0.0032 | 0.0029 | 0.0031 |
| F08 | 31080001 | 31180000 | 0.0029 | 0.0026 | 0.0028 |
| F08 | 31100001 | 31200000 | 0.0030 | 0.0027 | 0.0030 |
| F08 | 31120001 | 31220000 | 0.0035 | 0.0029 | 0.0034 |
| F08 | 31140001 | 31240000 | 0.0039 | 0.0031 | 0.0035 |
| F08 | 31160001 | 31260000 | 0.0051 | 0.0041 | 0.0047 |
| F08 | 31180001 | 31280000 | 0.0058 | 0.0047 | 0.0053 |
| F08 | 31200001 | 31300000 | 0.0058 | 0.0048 | 0.0054 |
| F08 | 31220001 | 31320000 | 0.0056 | 0.0046 | 0.0051 |
| F08 | 31240001 | 31340000 | 0.0056 | 0.0048 | 0.0054 |
| F08 | 31260001 | 31360000 | 0.0040 | 0.0032 | 0.0037 |
| F08 | 31280001 | 31380000 | 0.0035 | 0.0027 | 0.0031 |
| F08 | 31300001 | 31400000 | 0.0030 | 0.0020 | 0.0025 |
| F08 | 31320001 | 31420000 | 0.0026 | 0.0017 | 0.0020 |
| F08 | 31340001 | 31440000 | 0.0030 | 0.0018 | 0.0023 |
| F08 | 31360001 | 31460000 | 0.0032 | 0.0022 | 0.0027 |
| F08 | 31380001 | 31480000 | 0.0029 | 0.0019 | 0.0025 |
| F08 | 31400001 | 31500000 | 0.0030 | 0.0022 | 0.0027 |
| F08 | 31420001 | 31520000 | 0.0026 | 0.0021 | 0.0025 |
| F08 | 31440001 | 31540000 | 0.0016 | 0.0014 | 0.0016 |
| F08 | 31460001 | 31560000 | 0.0012 | 0.0010 | 0.0011 |
| F08 | 31480001 | 31580000 | 0.0016 | 0.0014 | 0.0016 |
| F08 | 31500001 | 31600000 | 0.0012 | 0.0010 | 0.0012 |
| F08 | 31520001 | 31620000 | 0.0016 | 0.0014 | 0.0016 |
| F08 | 31540001 | 31640000 | 0.0015 | 0.0014 | 0.0015 |
| F08 | 31560001 | 31660000 | 0.0013 | 0.0011 | 0.0013 |
| F08 | 31580001 | 31680000 | 0.0008 | 0.0007 | 0.0008 |
| F08 | 31600001 | 31700000 | 0.0012 | 0.0012 | 0.0012 |
| F08 | 31620001 | 31720000 | 0.0008 | 0.0007 | 0.0008 |
| F08 | 31640001 | 31740000 | 0.0008 | 0.0007 | 0.0008 |
| F08 | 31660001 | 31760000 | 0.0008 | 0.0007 | 0.0008 |
| F08 | 31680001 | 31780000 | 0.0011 | 0.0010 | 0.0011 |

| Chr | Start | End | both | A1 | A2 |
|---|---|---|---|---|---|
| F08 | 31700001 | 31800000 | 0.0010 | 0.0007 | 0.0010 |
| F08 | 31720001 | 31820000 | 0.0010 | 0.0007 | 0.0010 |
| F08 | 31740001 | 31840000 | 0.0009 | 0.0006 | 0.0009 |
| F08 | 31760001 | 31860000 | 0.0012 | 0.0009 | 0.0012 |
| F08 | 31780001 | 31880000 | 0.0013 | 0.0010 | 0.0013 |
| F08 | 31800001 | 31900000 | 0.0019 | 0.0015 | 0.0017 |
| F08 | 31820001 | 31920000 | 0.0028 | 0.0022 | 0.0026 |
| F08 | 31840001 | 31940000 | 0.0030 | 0.0024 | 0.0029 |
| F08 | 31860001 | 31960000 | 0.0032 | 0.0026 | 0.0032 |
| F08 | 31880001 | 31980000 | 0.0030 | 0.0025 | 0.0030 |
| F08 | 31900001 | 32000000 | 0.0027 | 0.0022 | 0.0026 |
| F08 | 31920001 | 32020000 | 0.0019 | 0.0015 | 0.0018 |
| F08 | 31940001 | 32040000 | 0.0018 | 0.0014 | 0.0017 |
| F08 | 31960001 | 32060000 | 0.0013 | 0.0010 | 0.0012 |
| F08 | 31980001 | 32080000 | 0.0010 | 0.0008 | 0.0009 |
| F08 | 32000001 | 32100000 | 0.0006 | 0.0005 | 0.0006 |
| F08 | 32020001 | 32120000 | 0.0006 | 0.0005 | 0.0006 |
| F08 | 32040001 | 32140000 | 0.0008 | 0.0007 | 0.0008 |
| F08 | 32060001 | 32160000 | 0.0008 | 0.0007 | 0.0008 |
| F08 | 32080001 | 32180000 | 0.0010 | 0.0008 | 0.0010 |
| F08 | 32100001 | 32200000 | 0.0010 | 0.0008 | 0.0009 |
| F08 | 32120001 | 32220000 | 0.0012 | 0.0010 | 0.0011 |
| F08 | 32140001 | 32240000 | 0.0022 | 0.0020 | 0.0020 |
| F08 | 32160001 | 32260000 | 0.0023 | 0.0020 | 0.0021 |
| F08 | 32180001 | 32280000 | 0.0022 | 0.0019 | 0.0020 |
| F08 | 32200001 | 32300000 | 0.0025 | 0.0022 | 0.0023 |
| F08 | 32220001 | 32320000 | 0.0025 | 0.0022 | 0.0023 |
| F08 | 32240001 | 32340000 | 0.0023 | 0.0017 | 0.0020 |
| F08 | 32260001 | 32360000 | 0.0023 | 0.0017 | 0.0019 |
| F08 | 32280001 | 32380000 | 0.0031 | 0.0024 | 0.0027 |
| F08 | 32300001 | 32400000 | 0.0029 | 0.0022 | 0.0025 |
| F08 | 32320001 | 32420000 | 0.0031 | 0.0023 | 0.0027 |
| F08 | 32340001 | 32440000 | 0.0026 | 0.0022 | 0.0024 |
| F08 | 32360001 | 32460000 | 0.0025 | 0.0023 | 0.0024 |
| F08 | 32380001 | 32480000 | 0.0019 | 0.0017 | 0.0019 |
| F08 | 32400001 | 32500000 | 0.0023 | 0.0021 | 0.0023 |
| F08 | 32420001 | 32520000 | 0.0021 | 0.0019 | 0.0021 |
| F08 | 32440001 | 32540000 | 0.0023 | 0.0020 | 0.0022 |
| F08 | 32460001 | 32560000 | 0.0029 | 0.0025 | 0.0028 |
| F08 | 32480001 | 32580000 | 0.0029 | 0.0025 | 0.0028 |
| F08 | 32500001 | 32600000 | 0.0025 | 0.0021 | 0.0024 |
| F08 | 32520001 | 32620000 | 0.0024 | 0.0020 | 0.0023 |
| F08 | 32540001 | 32640000 | 0.0018 | 0.0015 | 0.0018 |
| F08 | 32560001 | 32660000 | 0.0011 | 0.0009 | 0.0011 |
| F08 | 32580001 | 32680000 | 0.0009 | 0.0007 | 0.0009 |
| F08 | 32600001 | 32700000 | 0.0012 | 0.0010 | 0.0012 |
| F08 | 32620001 | 32720000 | 0.0014 | 0.0011 | 0.0014 |
| F08 | 32640001 | 32740000 | 0.0023 | 0.0017 | 0.0023 |
| F08 | 32660001 | 32760000 | 0.0023 | 0.0018 | 0.0023 |
| F08 | 32680001 | 32780000 | 0.0026 | 0.0021 | 0.0026 |
| F08 | 32700001 | 32800000 | 0.0025 | 0.0020 | 0.0024 |
| F08 | 32720001 | 32820000 | 0.0024 | 0.0021 | 0.0024 |
| F08 | 32740001 | 32840000 | 0.0015 | 0.0014 | 0.0015 |
| F08 | 32760001 | 32860000 | 0.0014 | 0.0014 | 0.0014 |
| F08 | 32780001 | 32880000 | 0.0019 | 0.0016 | 0.0017 |
| F08 | 32800001 | 32900000 | 0.0020 | 0.0016 | 0.0017 |
| F08 | 32820001 | 32920000 | 0.0024 | 0.0017 | 0.0019 |
| F08 | 32840001 | 32940000 | 0.0027 | 0.0020 | 0.0022 |

| Chr | Start | End | both | A1 | A2 |
|---|---|---|---|---|---|
| F08 | 32860001 | 32960000 | 0.0038 | 0.0029 | 0.0033 |
| F08 | 32880001 | 32980000 | 0.0044 | 0.0035 | 0.0037 |
| F08 | 32900001 | 33000000 | 0.0053 | 0.0043 | 0.0044 |
| F08 | 32920001 | 33020000 | 0.0058 | 0.0050 | 0.0052 |
| F08 | 32940001 | 33040000 | 0.0059 | 0.0049 | 0.0051 |
| F08 | 32960001 | 33060000 | 0.0058 | 0.0049 | 0.0048 |
| F08 | 32980001 | 33080000 | 0.0050 | 0.0040 | 0.0043 |
| F08 | 33000001 | 33100000 | 0.0042 | 0.0033 | 0.0036 |
| F08 | 33020001 | 33120000 | 0.0036 | 0.0026 | 0.0028 |
| F08 | 33040001 | 33140000 | 0.0036 | 0.0026 | 0.0029 |
| F08 | 33060001 | 33160000 | 0.0030 | 0.0022 | 0.0025 |
| F08 | 33080001 | 33180000 | 0.0033 | 0.0025 | 0.0027 |
| F08 | 33100001 | 33200000 | 0.0037 | 0.0029 | 0.0032 |
| F08 | 33120001 | 33220000 | 0.0040 | 0.0034 | 0.0037 |
| F08 | 33140001 | 33240000 | 0.0041 | 0.0035 | 0.0037 |
| F08 | 33160001 | 33260000 | 0.0039 | 0.0032 | 0.0033 |
| F08 | 33180001 | 33280000 | 0.0038 | 0.0030 | 0.0033 |
| F08 | 33200001 | 33300000 | 0.0040 | 0.0032 | 0.0034 |
| F08 | 33220001 | 33320000 | 0.0037 | 0.0028 | 0.0028 |
| F08 | 33240001 | 33340000 | 0.0033 | 0.0022 | 0.0023 |
| F08 | 33260001 | 33360000 | 0.0033 | 0.0023 | 0.0024 |
| F08 | 33280001 | 33380000 | 0.0031 | 0.0021 | 0.0021 |
| F08 | 33300001 | 33400000 | 0.0029 | 0.0018 | 0.0018 |
| F08 | 33320001 | 33420000 | 0.0024 | 0.0014 | 0.0014 |
| F08 | 33340001 | 33440000 | 0.0022 | 0.0014 | 0.0014 |
| F08 | 33360001 | 33460000 | 0.0020 | 0.0012 | 0.0013 |
| F08 | 33380001 | 33480000 | 0.0021 | 0.0014 | 0.0015 |
| F08 | 33400001 | 33500000 | 0.0012 | 0.0009 | 0.0010 |
| F08 | 33420001 | 33520000 | 0.0013 | 0.0009 | 0.0011 |
| F08 | 33440001 | 33540000 | 0.0015 | 0.0012 | 0.0013 |
| F08 | 33460001 | 33560000 | 0.0020 | 0.0017 | 0.0018 |
| F08 | 33480001 | 33580000 | 0.0017 | 0.0014 | 0.0015 |
| F08 | 33500001 | 33600000 | 0.0016 | 0.0013 | 0.0014 |
| F08 | 33520001 | 33620000 | 0.0021 | 0.0017 | 0.0019 |
| F08 | 33540001 | 33640000 | 0.0025 | 0.0021 | 0.0023 |
| F08 | 33560001 | 33660000 | 0.0023 | 0.0018 | 0.0021 |
| F08 | 33580001 | 33680000 | 0.0020 | 0.0018 | 0.0020 |
| F08 | 33600001 | 33700000 | 0.0021 | 0.0019 | 0.0021 |
| F08 | 33620001 | 33720000 | 0.0028 | 0.0022 | 0.0025 |
| F08 | 33640001 | 33740000 | 0.0032 | 0.0026 | 0.0028 |
| F08 | 33660001 | 33760000 | 0.0029 | 0.0022 | 0.0024 |
| F08 | 33680001 | 33780000 | 0.0032 | 0.0023 | 0.0025 |
| F08 | 33700001 | 33800000 | 0.0037 | 0.0028 | 0.0029 |
| F08 | 33720001 | 33820000 | 0.0028 | 0.0022 | 0.0022 |
| F08 | 33740001 | 33840000 | 0.0026 | 0.0019 | 0.0019 |
| F08 | 33760001 | 33860000 | 0.0029 | 0.0023 | 0.0024 |
| F08 | 33780001 | 33880000 | 0.0029 | 0.0024 | 0.0024 |
| F08 | 33800001 | 33900000 | 0.0027 | 0.0020 | 0.0020 |
| F08 | 33820001 | 33920000 | 0.0034 | 0.0025 | 0.0026 |
| F08 | 33840001 | 33940000 | 0.0032 | 0.0023 | 0.0025 |
| F08 | 33860001 | 33960000 | 0.0028 | 0.0021 | 0.0022 |
| F08 | 33880001 | 33980000 | 0.0030 | 0.0022 | 0.0022 |
| F08 | 33900001 | 34000000 | 0.0028 | 0.0021 | 0.0022 |
| F08 | 33920001 | 34020000 | 0.0028 | 0.0022 | 0.0023 |
| F08 | 33940001 | 34040000 | 0.0023 | 0.0019 | 0.0018 |
| F08 | 33960001 | 34060000 | 0.0023 | 0.0017 | 0.0017 |
| F08 | 33980001 | 34080000 | 0.0022 | 0.0016 | 0.0017 |
| F08 | 34000001 | 34100000 | 0.0032 | 0.0025 | 0.0026 |

| Chr | Start | End | both | A1 | A2 |
|---|---|---|---|---|---|
| F08 | 34020001 | 34120000 | 0.0024 | 0.0018 | 0.0018 |
| F08 | 34040001 | 34140000 | 0.0026 | 0.0018 | 0.0019 |
| F08 | 34060001 | 34160000 | 0.0038 | 0.0029 | 0.0030 |
| F08 | 34080001 | 34180000 | 0.0041 | 0.0032 | 0.0033 |
| F08 | 34100001 | 34200000 | 0.0035 | 0.0026 | 0.0026 |
| F08 | 34120001 | 34220000 | 0.0035 | 0.0027 | 0.0027 |
| F08 | 34140001 | 34240000 | 0.0040 | 0.0032 | 0.0033 |
| F08 | 34160001 | 34260000 | 0.0026 | 0.0020 | 0.0021 |
| F08 | 34180001 | 34280000 | 0.0024 | 0.0020 | 0.0021 |
| F08 | 34200001 | 34300000 | 0.0025 | 0.0021 | 0.0023 |
| F08 | 34220001 | 34320000 | 0.0028 | 0.0023 | 0.0026 |
| F08 | 34240001 | 34340000 | 0.0026 | 0.0022 | 0.0025 |
| F08 | 34260001 | 34360000 | 0.0034 | 0.0028 | 0.0032 |
| F08 | 34280001 | 34380000 | 0.0033 | 0.0028 | 0.0032 |
| F08 | 34300001 | 34400000 | 0.0037 | 0.0031 | 0.0035 |
| F08 | 34320001 | 34420000 | 0.0034 | 0.0028 | 0.0032 |
| F08 | 34340001 | 34440000 | 0.0029 | 0.0023 | 0.0028 |
| F08 | 34360001 | 34460000 | 0.0032 | 0.0026 | 0.0031 |
| F08 | 34380001 | 34480000 | 0.0028 | 0.0022 | 0.0027 |
| F08 | 34400001 | 34500000 | 0.0027 | 0.0022 | 0.0026 |
| F08 | 34420001 | 34520000 | 0.0032 | 0.0026 | 0.0030 |
| F08 | 34440001 | 34540000 | 0.0035 | 0.0029 | 0.0033 |
| F08 | 34460001 | 34560000 | 0.0030 | 0.0025 | 0.0028 |
| F08 | 34480001 | 34580000 | 0.0036 | 0.0029 | 0.0033 |
| F08 | 34500001 | 34600000 | 0.0033 | 0.0026 | 0.0030 |
| F08 | 34520001 | 34620000 | 0.0033 | 0.0027 | 0.0030 |
| F08 | 34540001 | 34640000 | 0.0033 | 0.0025 | 0.0027 |
| F08 | 34560001 | 34660000 | 0.0031 | 0.0022 | 0.0023 |
| F08 | 34580001 | 34680000 | 0.0029 | 0.0020 | 0.0020 |
| F08 | 34600001 | 34700000 | 0.0028 | 0.0019 | 0.0019 |
| F08 | 34620001 | 34720000 | 0.0032 | 0.0022 | 0.0022 |
| F08 | 34640001 | 34740000 | 0.0033 | 0.0025 | 0.0025 |
| F08 | 34660001 | 34760000 | 0.0029 | 0.0023 | 0.0023 |
| F08 | 34680001 | 34780000 | 0.0029 | 0.0023 | 0.0022 |
| F08 | 34700001 | 34800000 | 0.0027 | 0.0019 | 0.0018 |
| F08 | 34720001 | 34820000 | 0.0018 | 0.0011 | 0.0011 |
| F08 | 34740001 | 34840000 | 0.0016 | 0.0008 | 0.0008 |
| F08 | 34760001 | 34860000 | 0.0021 | 0.0010 | 0.0009 |
| F08 | 34780001 | 34880000 | 0.0024 | 0.0014 | 0.0012 |
| F08 | 34800001 | 34900000 | 0.0024 | 0.0015 | 0.0015 |
| F08 | 34820001 | 34920000 | 0.0024 | 0.0015 | 0.0014 |
| F08 | 34840001 | 34940000 | 0.0032 | 0.0021 | 0.0021 |
| F08 | 34860001 | 34960000 | 0.0037 | 0.0028 | 0.0029 |
| F08 | 34880001 | 34980000 | 0.0037 | 0.0028 | 0.0031 |
| F08 | 34900001 | 35000000 | 0.0043 | 0.0033 | 0.0035 |
| F08 | 34920001 | 35020000 | 0.0050 | 0.0040 | 0.0042 |
| F08 | 34940001 | 35040000 | 0.0043 | 0.0036 | 0.0038 |
| F08 | 34960001 | 35060000 | 0.0034 | 0.0029 | 0.0030 |
| F08 | 34980001 | 35080000 | 0.0029 | 0.0025 | 0.0024 |
| F08 | 35000001 | 35100000 | 0.0023 | 0.0020 | 0.0021 |
| F08 | 35020001 | 35120000 | 0.0023 | 0.0019 | 0.0020 |
| F08 | 35040001 | 35140000 | 0.0023 | 0.0020 | 0.0020 |
| F08 | 35060001 | 35160000 | 0.0026 | 0.0022 | 0.0023 |
| F08 | 35080001 | 35180000 | 0.0035 | 0.0029 | 0.0031 |
| F08 | 35100001 | 35200000 | 0.0039 | 0.0034 | 0.0034 |
| F08 | 35120001 | 35220000 | 0.0040 | 0.0032 | 0.0034 |
| F08 | 35140001 | 35240000 | 0.0036 | 0.0027 | 0.0031 |
| F08 | 35160001 | 35260000 | 0.0036 | 0.0028 | 0.0031 |

| Chr | Start | End | both | A1 | A2 |
|---|---|---|---|---|---|
| F08 | 35180001 | 35280000 | 0.0031 | 0.0022 | 0.0024 |
| F08 | 35200001 | 35300000 | 0.0025 | 0.0017 | 0.0020 |
| F08 | 35220001 | 35320000 | 0.0016 | 0.0011 | 0.0013 |
| F08 | 35240001 | 35340000 | 0.0022 | 0.0016 | 0.0019 |
| F08 | 35260001 | 35360000 | 0.0021 | 0.0017 | 0.0017 |
| F08 | 35280001 | 35380000 | 0.0023 | 0.0019 | 0.0020 |
| F08 | 35300001 | 35400000 | 0.0023 | 0.0019 | 0.0020 |
| F08 | 35320001 | 35420000 | 0.0024 | 0.0021 | 0.0021 |
| F08 | 35340001 | 35440000 | 0.0021 | 0.0019 | 0.0019 |
| F08 | 35360001 | 35460000 | 0.0025 | 0.0022 | 0.0023 |
| F08 | 35380001 | 35480000 | 0.0022 | 0.0020 | 0.0020 |
| F08 | 35400001 | 35500000 | 0.0023 | 0.0020 | 0.0021 |
| F08 | 35420001 | 35520000 | 0.0022 | 0.0020 | 0.0020 |
| F08 | 35440001 | 35540000 | 0.0026 | 0.0023 | 0.0024 |
| F08 | 35460001 | 35560000 | 0.0024 | 0.0022 | 0.0023 |
| F08 | 35480001 | 35580000 | 0.0026 | 0.0021 | 0.0022 |
| F08 | 35500001 | 35600000 | 0.0028 | 0.0021 | 0.0024 |
| F08 | 35520001 | 35620000 | 0.0033 | 0.0026 | 0.0029 |
| F08 | 35540001 | 35640000 | 0.0028 | 0.0021 | 0.0023 |
| F08 | 35560001 | 35660000 | 0.0022 | 0.0016 | 0.0018 |
| F08 | 35580001 | 35680000 | 0.0022 | 0.0017 | 0.0020 |
| F08 | 35600001 | 35700000 | 0.0016 | 0.0014 | 0.0015 |
| F08 | 35620001 | 35720000 | 0.0022 | 0.0017 | 0.0018 |
| F08 | 35640001 | 35740000 | 0.0028 | 0.0023 | 0.0025 |
| F08 | 35660001 | 35760000 | 0.0037 | 0.0027 | 0.0032 |
| F08 | 35680001 | 35780000 | 0.0043 | 0.0031 | 0.0037 |
| F08 | 35700001 | 35800000 | 0.0051 | 0.0038 | 0.0044 |
| F08 | 35720001 | 35820000 | 0.0043 | 0.0032 | 0.0038 |
| F08 | 35740001 | 35840000 | 0.0037 | 0.0025 | 0.0031 |
| F08 | 35760001 | 35860000 | 0.0032 | 0.0023 | 0.0026 |
| F08 | 35780001 | 35880000 | 0.0027 | 0.0019 | 0.0021 |
| F08 | 35800001 | 35900000 | 0.0020 | 0.0012 | 0.0014 |
| F08 | 35820001 | 35920000 | 0.0021 | 0.0014 | 0.0016 |
| F08 | 35840001 | 35940000 | 0.0028 | 0.0022 | 0.0024 |
| F08 | 35860001 | 35960000 | 0.0025 | 0.0022 | 0.0023 |
| F08 | 35880001 | 35980000 | 0.0020 | 0.0019 | 0.0019 |
| F08 | 35900001 | 36000000 | 0.0019 | 0.0018 | 0.0018 |
| F08 | 35920001 | 36020000 | 0.0019 | 0.0017 | 0.0018 |
| F08 | 35940001 | 36040000 | 0.0012 | 0.0011 | 0.0011 |
| F08 | 35960001 | 36060000 | 0.0011 | 0.0009 | 0.0011 |
| F08 | 35980001 | 36080000 | 0.0013 | 0.0011 | 0.0013 |
| F08 | 36000001 | 36100000 | 0.0015 | 0.0012 | 0.0014 |
| F08 | 36020001 | 36120000 | 0.0017 | 0.0013 | 0.0013 |
| F08 | 36040001 | 36140000 | 0.0023 | 0.0019 | 0.0019 |
| F08 | 36060001 | 36160000 | 0.0033 | 0.0028 | 0.0029 |
| F08 | 36080001 | 36180000 | 0.0037 | 0.0030 | 0.0032 |
| F08 | 36100001 | 36200000 | 0.0045 | 0.0039 | 0.0040 |
| F08 | 36120001 | 36220000 | 0.0049 | 0.0042 | 0.0046 |
| F08 | 36140001 | 36240000 | 0.0045 | 0.0036 | 0.0040 |
| F08 | 36160001 | 36260000 | 0.0042 | 0.0032 | 0.0036 |
| F08 | 36180001 | 36280000 | 0.0044 | 0.0034 | 0.0036 |
| F08 | 36200001 | 36300000 | 0.0042 | 0.0032 | 0.0034 |
| F08 | 36220001 | 36320000 | 0.0034 | 0.0025 | 0.0027 |
| F08 | 36240001 | 36340000 | 0.0036 | 0.0028 | 0.0029 |
| F08 | 36260001 | 36360000 | 0.0031 | 0.0022 | 0.0023 |
| F08 | 36280001 | 36380000 | 0.0030 | 0.0022 | 0.0024 |
| F08 | 36300001 | 36400000 | 0.0034 | 0.0026 | 0.0028 |
| F08 | 36320001 | 36420000 | 0.0039 | 0.0030 | 0.0033 |

| Chr | Start | End | both | A1 | A2 |
|---|---|---|---|---|---|
| F08 | 36340001 | 36440000 | 0.0032 | 0.0025 | 0.0027 |
| F08 | 36360001 | 36460000 | 0.0033 | 0.0027 | 0.0030 |
| F08 | 36380001 | 36480000 | 0.0025 | 0.0020 | 0.0023 |
| F08 | 36400001 | 36500000 | 0.0015 | 0.0012 | 0.0014 |
| F08 | 36420001 | 36520000 | 0.0009 | 0.0006 | 0.0008 |
| F08 | 36440001 | 36540000 | 0.0013 | 0.0011 | 0.0013 |
| F08 | 36460001 | 36560000 | 0.0012 | 0.0010 | 0.0011 |
| F08 | 36480001 | 36580000 | 0.0014 | 0.0012 | 0.0013 |
| F08 | 36500001 | 36600000 | 0.0021 | 0.0017 | 0.0019 |
| F08 | 36520001 | 36620000 | 0.0022 | 0.0018 | 0.0020 |
| F08 | 36540001 | 36640000 | 0.0023 | 0.0019 | 0.0021 |
| F08 | 36560001 | 36660000 | 0.0030 | 0.0024 | 0.0029 |
| F08 | 36580001 | 36680000 | 0.0028 | 0.0022 | 0.0027 |
| F08 | 36600001 | 36700000 | 0.0019 | 0.0015 | 0.0019 |
| F08 | 36620001 | 36720000 | 0.0021 | 0.0017 | 0.0021 |
| F08 | 36640001 | 36740000 | 0.0022 | 0.0017 | 0.0021 |
| F08 | 36660001 | 36760000 | 0.0015 | 0.0013 | 0.0015 |
| F08 | 36680001 | 36780000 | 0.0024 | 0.0019 | 0.0021 |
| F08 | 36700001 | 36800000 | 0.0028 | 0.0022 | 0.0023 |
| F08 | 36720001 | 36820000 | 0.0028 | 0.0022 | 0.0023 |
| F08 | 36740001 | 36840000 | 0.0036 | 0.0027 | 0.0029 |
| F08 | 36760001 | 36860000 | 0.0044 | 0.0034 | 0.0036 |
| F08 | 36780001 | 36880000 | 0.0042 | 0.0034 | 0.0036 |
| F08 | 36800001 | 36900000 | 0.0041 | 0.0034 | 0.0036 |
| F08 | 36820001 | 36920000 | 0.0043 | 0.0036 | 0.0038 |
| F08 | 36840001 | 36940000 | 0.0029 | 0.0025 | 0.0025 |
| F08 | 36860001 | 36960000 | 0.0021 | 0.0017 | 0.0017 |
| F08 | 36880001 | 36980000 | 0.0018 | 0.0015 | 0.0015 |
| F08 | 36900001 | 37000000 | 0.0028 | 0.0024 | 0.0026 |
| F08 | 36920001 | 37020000 | 0.0026 | 0.0021 | 0.0022 |
| F08 | 36940001 | 37040000 | 0.0030 | 0.0024 | 0.0026 |
| F08 | 36960001 | 37060000 | 0.0029 | 0.0023 | 0.0025 |
| F08 | 36980001 | 37080000 | 0.0027 | 0.0019 | 0.0021 |
| F08 | 37000001 | 37100000 | 0.0017 | 0.0008 | 0.0009 |
| F08 | 37020001 | 37120000 | 0.0018 | 0.0009 | 0.0010 |
| F08 | 37040001 | 37140000 | 0.0025 | 0.0014 | 0.0016 |
| F08 | 37060001 | 37160000 | 0.0035 | 0.0022 | 0.0026 |
| F08 | 37080001 | 37180000 | 0.0037 | 0.0025 | 0.0030 |
| F08 | 37100001 | 37200000 | 0.0035 | 0.0026 | 0.0031 |
| F08 | 37120001 | 37220000 | 0.0040 | 0.0028 | 0.0035 |
| F08 | 37140001 | 37240000 | 0.0032 | 0.0022 | 0.0027 |
| F08 | 37160001 | 37260000 | 0.0023 | 0.0015 | 0.0019 |
| F08 | 37180001 | 37280000 | 0.0030 | 0.0021 | 0.0026 |
| F08 | 37200001 | 37300000 | 0.0029 | 0.0020 | 0.0025 |
| F08 | 37220001 | 37320000 | 0.0020 | 0.0015 | 0.0018 |
| F08 | 37240001 | 37340000 | 0.0023 | 0.0019 | 0.0022 |
| F08 | 37260001 | 37360000 | 0.0027 | 0.0023 | 0.0026 |
| F08 | 37280001 | 37380000 | 0.0016 | 0.0014 | 0.0015 |
| F08 | 37300001 | 37400000 | 0.0019 | 0.0017 | 0.0017 |
| F08 | 37320001 | 37420000 | 0.0023 | 0.0020 | 0.0021 |
| F08 | 37340001 | 37440000 | 0.0027 | 0.0024 | 0.0025 |
| F08 | 37360001 | 37460000 | 0.0022 | 0.0018 | 0.0020 |
| F08 | 37380001 | 37480000 | 0.0026 | 0.0021 | 0.0023 |
| F08 | 37400001 | 37500000 | 0.0029 | 0.0023 | 0.0025 |
| F08 | 37420001 | 37520000 | 0.0031 | 0.0025 | 0.0026 |
| F08 | 37440001 | 37540000 | 0.0029 | 0.0023 | 0.0024 |
| F08 | 37460001 | 37560000 | 0.0033 | 0.0028 | 0.0030 |
| F08 | 37480001 | 37580000 | 0.0028 | 0.0024 | 0.0025 |

| Chr | Start | End | both | A1 | A2 |
|---|---|---|---|---|---|
| F08 | 37500001 | 37600000 | 0.0022 | 0.0019 | 0.0021 |
| F08 | 37520001 | 37620000 | 0.0016 | 0.0014 | 0.0015 |
| F08 | 37540001 | 37640000 | 0.0007 | 0.0006 | 0.0007 |
| F08 | 37560001 | 37660000 | 0.0002 | 0.0002 | 0.0001 |
| F08 | 37580001 | 37680000 | 0.0004 | 0.0003 | 0.0004 |
| F08 | 37600001 | 37700000 | 0.0006 | 0.0005 | 0.0006 |
| F08 | 37620001 | 37720000 | 0.0008 | 0.0006 | 0.0007 |
| F08 | 37640001 | 37740000 | 0.0011 | 0.0010 | 0.0011 |
| F08 | 37660001 | 37760000 | 0.0010 | 0.0008 | 0.0010 |
| F08 | 37680001 | 37780000 | 0.0016 | 0.0014 | 0.0016 |
| F08 | 37700001 | 37800000 | 0.0016 | 0.0014 | 0.0015 |
| F08 | 37720001 | 37820000 | 0.0018 | 0.0015 | 0.0017 |
| F08 | 37740001 | 37840000 | 0.0016 | 0.0013 | 0.0015 |
| F08 | 37760001 | 37860000 | 0.0019 | 0.0015 | 0.0017 |
| F08 | 37780001 | 37880000 | 0.0021 | 0.0017 | 0.0017 |
| F08 | 37800001 | 37900000 | 0.0021 | 0.0017 | 0.0017 |
| F08 | 37820001 | 37920000 | 0.0020 | 0.0017 | 0.0017 |
| F08 | 37840001 | 37940000 | 0.0021 | 0.0018 | 0.0018 |
| F08 | 37860001 | 37960000 | 0.0034 | 0.0029 | 0.0029 |
| F08 | 37880001 | 37980000 | 0.0029 | 0.0022 | 0.0025 |
| F08 | 37900001 | 38000000 | 0.0034 | 0.0026 | 0.0030 |
| F08 | 37920001 | 38020000 | 0.0031 | 0.0024 | 0.0027 |
| F08 | 37940001 | 38040000 | 0.0029 | 0.0022 | 0.0025 |
| F08 | 37960001 | 38060000 | 0.0014 | 0.0009 | 0.0012 |
| F08 | 37980001 | 38080000 | 0.0010 | 0.0008 | 0.0009 |
| F08 | 38000001 | 38100000 | 0.0007 | 0.0005 | 0.0007 |
| F08 | 38020001 | 38120000 | 0.0007 | 0.0006 | 0.0007 |
| F08 | 38040001 | 38140000 | 0.0011 | 0.0008 | 0.0010 |
| F08 | 38060001 | 38160000 | 0.0020 | 0.0016 | 0.0018 |
| F08 | 38080001 | 38180000 | 0.0024 | 0.0020 | 0.0022 |
| F08 | 38100001 | 38200000 | 0.0023 | 0.0019 | 0.0020 |
| F08 | 38120001 | 38220000 | 0.0030 | 0.0022 | 0.0026 |
| F08 | 38140001 | 38240000 | 0.0034 | 0.0027 | 0.0030 |
| F08 | 38160001 | 38260000 | 0.0028 | 0.0021 | 0.0024 |
| F08 | 38180001 | 38280000 | 0.0034 | 0.0025 | 0.0030 |
| F08 | 38200001 | 38300000 | 0.0035 | 0.0027 | 0.0032 |
| F08 | 38220001 | 38320000 | 0.0044 | 0.0036 | 0.0040 |
| F08 | 38240001 | 38340000 | 0.0046 | 0.0037 | 0.0041 |
| F08 | 38260001 | 38360000 | 0.0050 | 0.0041 | 0.0045 |
| F08 | 38280001 | 38380000 | 0.0047 | 0.0039 | 0.0041 |
| F08 | 38300001 | 38400000 | 0.0055 | 0.0048 | 0.0049 |
| F08 | 38320001 | 38420000 | 0.0053 | 0.0046 | 0.0048 |
| F08 | 38340001 | 38440000 | 0.0056 | 0.0050 | 0.0053 |
| F08 | 38360001 | 38460000 | 0.0051 | 0.0045 | 0.0048 |
| F08 | 38380001 | 38480000 | 0.0053 | 0.0046 | 0.0051 |
| F08 | 38400001 | 38500000 | 0.0052 | 0.0042 | 0.0050 |
| F08 | 38420001 | 38520000 | 0.0046 | 0.0038 | 0.0043 |
| F08 | 38440001 | 38540000 | 0.0038 | 0.0027 | 0.0033 |
| F08 | 38460001 | 38560000 | 0.0044 | 0.0032 | 0.0037 |
| F08 | 38480001 | 38580000 | 0.0038 | 0.0028 | 0.0031 |
| F08 | 38500001 | 38600000 | 0.0035 | 0.0026 | 0.0027 |
| F08 | 38520001 | 38620000 | 0.0029 | 0.0020 | 0.0022 |
| F08 | 38540001 | 38640000 | 0.0026 | 0.0019 | 0.0021 |
| F08 | 38560001 | 38660000 | 0.0019 | 0.0014 | 0.0015 |
| F08 | 38580001 | 38680000 | 0.0022 | 0.0015 | 0.0018 |
| F08 | 38600001 | 38700000 | 0.0024 | 0.0016 | 0.0020 |
| F08 | 38620001 | 38720000 | 0.0023 | 0.0016 | 0.0020 |
| F08 | 38640001 | 38740000 | 0.0028 | 0.0020 | 0.0026 |

| Chr | Start | End | both | A1 | A2 |
|---|---|---|---|---|---|
| F08 | 38660001 | 38760000 | 0.0036 | 0.0026 | 0.0034 |
| F08 | 38680001 | 38780000 | 0.0036 | 0.0026 | 0.0034 |
| F08 | 38700001 | 38800000 | 0.0027 | 0.0020 | 0.0026 |
| F08 | 38720001 | 38820000 | 0.0030 | 0.0022 | 0.0029 |
| F08 | 38740001 | 38840000 | 0.0024 | 0.0018 | 0.0023 |
| F08 | 38760001 | 38860000 | 0.0017 | 0.0014 | 0.0016 |
| F08 | 38780001 | 38880000 | 0.0010 | 0.0008 | 0.0010 |
| F08 | 38800001 | 38900000 | 0.0017 | 0.0014 | 0.0016 |
| F08 | 38820001 | 38920000 | 0.0019 | 0.0017 | 0.0018 |
| F08 | 38840001 | 38940000 | 0.0021 | 0.0019 | 0.0019 |
| F08 | 38860001 | 38960000 | 0.0028 | 0.0023 | 0.0024 |
| F08 | 38880001 | 38980000 | 0.0031 | 0.0024 | 0.0025 |
| F08 | 38900001 | 39000000 | 0.0025 | 0.0018 | 0.0018 |
| F08 | 38920001 | 39020000 | 0.0034 | 0.0023 | 0.0025 |
| F08 | 38940001 | 39040000 | 0.0034 | 0.0024 | 0.0026 |
| F08 | 38960001 | 39060000 | 0.0035 | 0.0024 | 0.0027 |
| F08 | 38980001 | 39080000 | 0.0043 | 0.0033 | 0.0036 |
| F08 | 39000001 | 39100000 | 0.0041 | 0.0032 | 0.0036 |
| F08 | 39020001 | 39120000 | 0.0039 | 0.0028 | 0.0034 |
| F08 | 39040001 | 39140000 | 0.0038 | 0.0026 | 0.0033 |
| F08 | 39060001 | 39160000 | 0.0036 | 0.0026 | 0.0032 |
| F08 | 39080001 | 39180000 | 0.0028 | 0.0019 | 0.0023 |
| F08 | 39100001 | 39200000 | 0.0036 | 0.0026 | 0.0029 |
| F08 | 39120001 | 39220000 | 0.0037 | 0.0029 | 0.0029 |
| F08 | 39140001 | 39240000 | 0.0038 | 0.0030 | 0.0029 |
| F08 | 39160001 | 39260000 | 0.0034 | 0.0025 | 0.0025 |
| F08 | 39180001 | 39280000 | 0.0033 | 0.0025 | 0.0026 |
| F08 | 39200001 | 39300000 | 0.0026 | 0.0019 | 0.0019 |
| F08 | 39220001 | 39320000 | 0.0014 | 0.0009 | 0.0011 |
| F08 | 39240001 | 39340000 | 0.0017 | 0.0013 | 0.0015 |
| F08 | 39260001 | 39360000 | 0.0027 | 0.0022 | 0.0026 |
| F08 | 39280001 | 39380000 | 0.0027 | 0.0022 | 0.0027 |
| F08 | 39300001 | 39400000 | 0.0027 | 0.0022 | 0.0027 |
| F08 | 39320001 | 39420000 | 0.0028 | 0.0023 | 0.0028 |
| F08 | 39340001 | 39440000 | 0.0024 | 0.0020 | 0.0024 |
| F08 | 39360001 | 39460000 | 0.0016 | 0.0013 | 0.0016 |
| F08 | 39380001 | 39480000 | 0.0017 | 0.0015 | 0.0017 |
| F08 | 39400001 | 39500000 | 0.0032 | 0.0028 | 0.0031 |
| F08 | 39420001 | 39520000 | 0.0032 | 0.0028 | 0.0032 |
| F08 | 39440001 | 39540000 | 0.0030 | 0.0026 | 0.0029 |
| F08 | 39460001 | 39560000 | 0.0026 | 0.0022 | 0.0025 |
| F08 | 39480001 | 39580000 | 0.0023 | 0.0020 | 0.0022 |
| F08 | 39500001 | 39600000 | 0.0015 | 0.0012 | 0.0014 |
| F08 | 39520001 | 39620000 | 0.0018 | 0.0016 | 0.0016 |
| F08 | 39540001 | 39640000 | 0.0023 | 0.0021 | 0.0021 |
| F08 | 39560001 | 39660000 | 0.0022 | 0.0021 | 0.0020 |
| F08 | 39580001 | 39680000 | 0.0024 | 0.0022 | 0.0022 |
| F08 | 39600001 | 39700000 | 0.0021 | 0.0019 | 0.0019 |
| F08 | 39620001 | 39720000 | 0.0020 | 0.0017 | 0.0019 |
| F08 | 39640001 | 39740000 | 0.0033 | 0.0028 | 0.0030 |
| F08 | 39660001 | 39760000 | 0.0037 | 0.0031 | 0.0034 |
| F08 | 39680001 | 39780000 | 0.0049 | 0.0041 | 0.0043 |
| F08 | 39700001 | 39800000 | 0.0068 | 0.0056 | 0.0059 |
| F08 | 39720001 | 39820000 | 0.0064 | 0.0053 | 0.0056 |
| F08 | 39740001 | 39840000 | 0.0049 | 0.0039 | 0.0042 |
| F08 | 39760001 | 39860000 | 0.0050 | 0.0041 | 0.0044 |
| F08 | 39780001 | 39880000 | 0.0037 | 0.0029 | 0.0032 |
| F08 | 39800001 | 39900000 | 0.0013 | 0.0011 | 0.0012 |

| Chr | Start | End | both | A1 | A2 |
|---|---|---|---|---|---|
| F08 | 39820001 | 39920000 | 0.0012 | 0.0010 | 0.0011 |
| F08 | 39840001 | 39940000 | 0.0010 | 0.0009 | 0.0009 |
| F08 | 39860001 | 39960000 | 0.0005 | 0.0005 | 0.0005 |
| F08 | 39880001 | 39980000 | 0.0008 | 0.0007 | 0.0008 |
| F08 | 39900001 | 40000000 | 0.0009 | 0.0007 | 0.0008 |
| F08 | 39920001 | 40020000 | 0.0009 | 0.0007 | 0.0008 |
| F08 | 39940001 | 40040000 | 0.0008 | 0.0007 | 0.0008 |
| F08 | 39960001 | 40060000 | 0.0009 | 0.0008 | 0.0009 |
| F08 | 39980001 | 40080000 | 0.0006 | 0.0006 | 0.0006 |
| F08 | 40000001 | 40100000 | 0.0006 | 0.0005 | 0.0006 |
| F08 | 40020001 | 40120000 | 0.0005 | 0.0004 | 0.0005 |
| F08 | 40040001 | 40140000 | 0.0006 | 0.0005 | 0.0006 |
| F08 | 40060001 | 40160000 | 0.0007 | 0.0006 | 0.0006 |
| F08 | 40080001 | 40180000 | 0.0009 | 0.0008 | 0.0009 |
| F08 | 40100001 | 40200000 | 0.0016 | 0.0015 | 0.0016 |
| F08 | 40120001 | 40220000 | 0.0019 | 0.0017 | 0.0019 |
| F08 | 40140001 | 40240000 | 0.0026 | 0.0023 | 0.0025 |
| F08 | 40160001 | 40260000 | 0.0029 | 0.0026 | 0.0027 |
| F08 | 40180001 | 40280000 | 0.0027 | 0.0025 | 0.0026 |
| F08 | 40200001 | 40300000 | 0.0023 | 0.0021 | 0.0022 |
| F08 | 40220001 | 40320000 | 0.0024 | 0.0021 | 0.0022 |
| F08 | 40240001 | 40340000 | 0.0026 | 0.0020 | 0.0023 |
| F08 | 40260001 | 40360000 | 0.0023 | 0.0018 | 0.0022 |
| F08 | 40280001 | 40380000 | 0.0023 | 0.0018 | 0.0021 |
| F08 | 40300001 | 40400000 | 0.0023 | 0.0018 | 0.0021 |
| F08 | 40320001 | 40420000 | 0.0019 | 0.0016 | 0.0018 |
| F08 | 40340001 | 40440000 | 0.0018 | 0.0015 | 0.0017 |
| F08 | 40360001 | 40460000 | 0.0015 | 0.0012 | 0.0015 |
| F08 | 40380001 | 40480000 | 0.0025 | 0.0020 | 0.0025 |
| F08 | 40400001 | 40500000 | 0.0024 | 0.0019 | 0.0024 |
| F08 | 40420001 | 40520000 | 0.0025 | 0.0021 | 0.0025 |
| F08 | 40440001 | 40540000 | 0.0025 | 0.0021 | 0.0025 |
| F08 | 40460001 | 40560000 | 0.0025 | 0.0021 | 0.0025 |
| F08 | 40480001 | 40580000 | 0.0017 | 0.0014 | 0.0017 |
| F08 | 40500001 | 40600000 | 0.0016 | 0.0013 | 0.0016 |
| F08 | 40520001 | 40620000 | 0.0023 | 0.0020 | 0.0023 |
| F08 | 40540001 | 40640000 | 0.0020 | 0.0017 | 0.0020 |
| F08 | 40560001 | 40660000 | 0.0021 | 0.0019 | 0.0020 |
| F08 | 40580001 | 40680000 | 0.0029 | 0.0026 | 0.0028 |
| F08 | 40600001 | 40700000 | 0.0032 | 0.0029 | 0.0032 |
| F08 | 40620001 | 40720000 | 0.0023 | 0.0020 | 0.0023 |
| F08 | 40640001 | 40740000 | 0.0022 | 0.0020 | 0.0022 |
| F08 | 40660001 | 40760000 | 0.0022 | 0.0019 | 0.0022 |
| F08 | 40680001 | 40780000 | 0.0012 | 0.0011 | 0.0012 |
| F08 | 40700001 | 40800000 | 0.0010 | 0.0009 | 0.0009 |
| F08 | 40720001 | 40820000 | 0.0017 | 0.0015 | 0.0016 |
| F08 | 40740001 | 40840000 | 0.0022 | 0.0019 | 0.0021 |
| F08 | 40760001 | 40860000 | 0.0023 | 0.0020 | 0.0023 |
| F08 | 40780001 | 40880000 | 0.0027 | 0.0022 | 0.0027 |
| F08 | 40800001 | 40900000 | 0.0026 | 0.0021 | 0.0026 |
| F08 | 40820001 | 40920000 | 0.0021 | 0.0018 | 0.0021 |
| F08 | 40840001 | 40940000 | 0.0022 | 0.0018 | 0.0022 |
| F08 | 40860001 | 40960000 | 0.0020 | 0.0016 | 0.0020 |
| F08 | 40880001 | 40980000 | 0.0022 | 0.0020 | 0.0022 |
| F08 | 40900001 | 41000000 | 0.0025 | 0.0021 | 0.0025 |
| F08 | 40920001 | 41020000 | 0.0029 | 0.0025 | 0.0029 |
| F08 | 40940001 | 41040000 | 0.0033 | 0.0028 | 0.0033 |
| F08 | 40960001 | 41060000 | 0.0039 | 0.0033 | 0.0038 |

| Chr | Start | End | both | A1 | A2 |
|---|---|---|---|---|---|
| F08 | 40980001 | 41080000 | 0.0035 | 0.0029 | 0.0033 |
| F08 | 41000001 | 41100000 | 0.0031 | 0.0025 | 0.0028 |
| F08 | 41020001 | 41120000 | 0.0026 | 0.0021 | 0.0024 |
| F08 | 41040001 | 41140000 | 0.0014 | 0.0011 | 0.0011 |
| F08 | 41060001 | 41160000 | 0.0015 | 0.0012 | 0.0013 |
| F08 | 41080001 | 41180000 | 0.0012 | 0.0011 | 0.0012 |
| F08 | 41100001 | 41200000 | 0.0013 | 0.0011 | 0.0013 |
| F08 | 41120001 | 41220000 | 0.0016 | 0.0013 | 0.0016 |
| F08 | 41140001 | 41240000 | 0.0016 | 0.0013 | 0.0016 |
| F08 | 41160001 | 41260000 | 0.0011 | 0.0009 | 0.0010 |
| F08 | 41180001 | 41280000 | 0.0021 | 0.0018 | 0.0020 |
| F08 | 41200001 | 41300000 | 0.0023 | 0.0020 | 0.0022 |
| F08 | 41220001 | 41320000 | 0.0018 | 0.0016 | 0.0017 |
| F08 | 41240001 | 41340000 | 0.0022 | 0.0020 | 0.0021 |
| F08 | 41260001 | 41360000 | 0.0019 | 0.0016 | 0.0018 |
| F08 | 41280001 | 41380000 | 0.0016 | 0.0014 | 0.0015 |
| F08 | 41300001 | 41400000 | 0.0017 | 0.0015 | 0.0016 |
| F08 | 41320001 | 41420000 | 0.0023 | 0.0021 | 0.0022 |
| F08 | 41340001 | 41440000 | 0.0022 | 0.0020 | 0.0021 |
| F08 | 41360001 | 41460000 | 0.0024 | 0.0022 | 0.0023 |
| F08 | 41380001 | 41480000 | 0.0016 | 0.0014 | 0.0016 |
| F08 | 41400001 | 41500000 | 0.0019 | 0.0015 | 0.0018 |
| F08 | 41420001 | 41520000 | 0.0013 | 0.0009 | 0.0012 |
| F08 | 41440001 | 41540000 | 0.0016 | 0.0013 | 0.0015 |
| F08 | 41460001 | 41560000 | 0.0024 | 0.0019 | 0.0022 |
| F08 | 41480001 | 41580000 | 0.0025 | 0.0019 | 0.0023 |
| F08 | 41500001 | 41600000 | 0.0021 | 0.0018 | 0.0020 |
| F08 | 41520001 | 41620000 | 0.0020 | 0.0018 | 0.0020 |
| F08 | 41540001 | 41640000 | 0.0015 | 0.0012 | 0.0014 |
| F08 | 41560001 | 41660000 | 0.0005 | 0.0004 | 0.0005 |
| F08 | 41580001 | 41680000 | 0.0003 | 0.0003 | 0.0003 |
| F08 | 41600001 | 41700000 | 0.0001 | 0.0001 | 0.0001 |
| F08 | 41620001 | 41720000 | 0.0003 | 0.0002 | 0.0003 |
| F08 | 41640001 | 41740000 | 0.0003 | 0.0002 | 0.0003 |
| F08 | 41660001 | 41760000 | 0.0003 | 0.0002 | 0.0003 |
| F08 | 41680001 | 41780000 | 0.0003 | 0.0002 | 0.0003 |
| F08 | 41700001 | 41800000 | 0.0005 | 0.0004 | 0.0005 |
| F08 | 41720001 | 41820000 | 0.0005 | 0.0005 | 0.0005 |
| F08 | 41740001 | 41840000 | 0.0005 | 0.0005 | 0.0005 |
| F08 | 41760001 | 41860000 | 0.0006 | 0.0006 | 0.0006 |
| F08 | 41780001 | 41880000 | 0.0006 | 0.0006 | 0.0006 |
| F08 | 41800001 | 41900000 | 0.0005 | 0.0005 | 0.0005 |
| F08 | 41820001 | 41920000 | 0.0005 | 0.0004 | 0.0005 |
| F08 | 41840001 | 41940000 | 0.0005 | 0.0004 | 0.0005 |
| F08 | 41860001 | 41960000 | 0.0004 | 0.0003 | 0.0004 |
| F08 | 41880001 | 41980000 | 0.0004 | 0.0003 | 0.0004 |
| F08 | 41900001 | 42000000 | 0.0002 | 0.0002 | 0.0002 |
| F08 | 41920001 | 42020000 | 0.0001 | 0.0001 | 0.0001 |
| F08 | 41940001 | 42040000 | 0.0002 | 0.0002 | 0.0002 |
| F08 | 41960001 | 42060000 | 0.0002 | 0.0002 | 0.0002 |
| F08 | 41980001 | 42080000 | 0.0003 | 0.0002 | 0.0003 |
| F08 | 42000001 | 42100000 | 0.0004 | 0.0003 | 0.0004 |
| F08 | 42020001 | 42120000 | 0.0004 | 0.0003 | 0.0004 |
| F08 | 42040001 | 42140000 | 0.0003 | 0.0002 | 0.0003 |
| F08 | 42060001 | 42160000 | 0.0003 | 0.0002 | 0.0003 |
| F08 | 42080001 | 42180000 | 0.0003 | 0.0002 | 0.0004 |
| F08 | 42100001 | 42200000 | 0.0002 | 0.0002 | 0.0002 |
| F08 | 42120001 | 42220000 | 0.0003 | 0.0002 | 0.0003 |

| Chr | Start | End | both | A1 | A2 |
|---|---|---|---|---|---|
| F08 | 42140001 | 42240000 | 0.0003 | 0.0003 | 0.0003 |
| F08 | 42160001 | 42260000 | 0.0003 | 0.0002 | 0.0003 |
| F08 | 42180001 | 42280000 | 0.0004 | 0.0003 | 0.0004 |
| F08 | 42200001 | 42300000 | 0.0005 | 0.0004 | 0.0005 |
| F08 | 42220001 | 42320000 | 0.0009 | 0.0007 | 0.0009 |
| F08 | 42240001 | 42340000 | 0.0009 | 0.0007 | 0.0009 |
| F08 | 42260001 | 42360000 | 0.0012 | 0.0009 | 0.0013 |
| F08 | 42280001 | 42380000 | 0.0011 | 0.0009 | 0.0011 |
| F08 | 42300001 | 42400000 | 0.0011 | 0.0008 | 0.0011 |
| F08 | 42320001 | 42420000 | 0.0007 | 0.0005 | 0.0007 |
| F08 | 42340001 | 42440000 | 0.0012 | 0.0008 | 0.0012 |
| F08 | 42360001 | 42460000 | 0.0013 | 0.0009 | 0.0013 |
| F08 | 42380001 | 42480000 | 0.0016 | 0.0012 | 0.0016 |
| F08 | 42400001 | 42500000 | 0.0017 | 0.0013 | 0.0016 |
| F08 | 42420001 | 42520000 | 0.0016 | 0.0013 | 0.0016 |
| F08 | 42440001 | 42540000 | 0.0010 | 0.0009 | 0.0010 |
| F08 | 42460001 | 42560000 | 0.0008 | 0.0007 | 0.0007 |
| F08 | 42480001 | 42580000 | 0.0005 | 0.0004 | 0.0004 |
| F08 | 42500001 | 42600000 | 0.0004 | 0.0004 | 0.0004 |
| F08 | 42520001 | 42620000 | 0.0008 | 0.0007 | 0.0008 |
| F08 | 42540001 | 42640000 | 0.0010 | 0.0008 | 0.0009 |
| F08 | 42560001 | 42660000 | 0.0015 | 0.0011 | 0.0015 |
| F08 | 42580001 | 42680000 | 0.0014 | 0.0011 | 0.0014 |
| F08 | 42600001 | 42700000 | 0.0014 | 0.0010 | 0.0014 |
| F08 | 42620001 | 42720000 | 0.0012 | 0.0009 | 0.0012 |
| F08 | 42640001 | 42740000 | 0.0017 | 0.0014 | 0.0017 |
| F08 | 42660001 | 42760000 | 0.0012 | 0.0010 | 0.0011 |
| F08 | 42680001 | 42780000 | 0.0015 | 0.0013 | 0.0015 |
| F08 | 42700001 | 42800000 | 0.0016 | 0.0013 | 0.0015 |
| F08 | 42720001 | 42820000 | 0.0015 | 0.0012 | 0.0015 |
| F08 | 42740001 | 42840000 | 0.0009 | 0.0008 | 0.0009 |
| F08 | 42760001 | 42860000 | 0.0009 | 0.0007 | 0.0008 |
| F08 | 42780001 | 42880000 | 0.0011 | 0.0008 | 0.0010 |
| F08 | 42800001 | 42900000 | 0.0020 | 0.0016 | 0.0019 |
| F08 | 42820001 | 42920000 | 0.0028 | 0.0022 | 0.0026 |
| F08 | 42840001 | 42940000 | 0.0030 | 0.0024 | 0.0027 |
| F08 | 42860001 | 42960000 | 0.0041 | 0.0033 | 0.0037 |
| F08 | 42880001 | 42980000 | 0.0034 | 0.0028 | 0.0030 |
| F08 | 42900001 | 43000000 | 0.0027 | 0.0022 | 0.0024 |
| F08 | 42920001 | 43020000 | 0.0023 | 0.0018 | 0.0021 |
| F08 | 42940001 | 43040000 | 0.0019 | 0.0015 | 0.0018 |
| F08 | 42960001 | 43060000 | 0.0009 | 0.0006 | 0.0010 |
| F08 | 42980001 | 43080000 | 0.0013 | 0.0011 | 0.0012 |
| F08 | 43000001 | 43100000 | 0.0014 | 0.0012 | 0.0013 |
| F08 | 43020001 | 43120000 | 0.0014 | 0.0013 | 0.0012 |
| F08 | 43040001 | 43140000 | 0.0016 | 0.0014 | 0.0015 |
| F08 | 43060001 | 43160000 | 0.0018 | 0.0016 | 0.0015 |
| F08 | 43080001 | 43180000 | 0.0021 | 0.0019 | 0.0019 |
| F08 | 43100001 | 43200000 | 0.0019 | 0.0016 | 0.0017 |
| F08 | 43120001 | 43220000 | 0.0017 | 0.0014 | 0.0015 |
| F08 | 43140001 | 43240000 | 0.0021 | 0.0017 | 0.0017 |
| F08 | 43160001 | 43260000 | 0.0020 | 0.0016 | 0.0017 |
| F08 | 43180001 | 43280000 | 0.0017 | 0.0013 | 0.0014 |
| F08 | 43200001 | 43300000 | 0.0022 | 0.0018 | 0.0019 |
| F08 | 43220001 | 43320000 | 0.0022 | 0.0019 | 0.0020 |
| F08 | 43240001 | 43340000 | 0.0020 | 0.0017 | 0.0020 |
| F08 | 43260001 | 43360000 | 0.0019 | 0.0017 | 0.0018 |
| F08 | 43280001 | 43380000 | 0.0015 | 0.0013 | 0.0015 |

| Chr | Start | End | both | A1 | A2 |
|---|---|---|---|---|---|
| F08 | 43300001 | 43400000 | 0.0013 | 0.0011 | 0.0013 |
| F08 | 43320001 | 43420000 | 0.0010 | 0.0009 | 0.0010 |
| F08 | 43340001 | 43440000 | 0.0006 | 0.0005 | 0.0006 |
| F08 | 43360001 | 43460000 | 0.0009 | 0.0008 | 0.0009 |
| F08 | 43380001 | 43480000 | 0.0017 | 0.0014 | 0.0017 |
| F08 | 43400001 | 43500000 | 0.0015 | 0.0012 | 0.0015 |
| F08 | 43420001 | 43520000 | 0.0018 | 0.0014 | 0.0018 |
| F08 | 43440001 | 43540000 | 0.0018 | 0.0014 | 0.0018 |
| F08 | 43460001 | 43560000 | 0.0016 | 0.0012 | 0.0016 |
| F08 | 43480001 | 43580000 | 0.0011 | 0.0010 | 0.0011 |
| F08 | 43500001 | 43600000 | 0.0012 | 0.0012 | 0.0012 |
| F08 | 43520001 | 43620000 | 0.0011 | 0.0010 | 0.0010 |
| F08 | 43540001 | 43640000 | 0.0021 | 0.0019 | 0.0020 |
| F08 | 43560001 | 43660000 | 0.0026 | 0.0024 | 0.0024 |
| F08 | 43580001 | 43680000 | 0.0027 | 0.0025 | 0.0025 |
| F08 | 43600001 | 43700000 | 0.0036 | 0.0031 | 0.0034 |
| F08 | 43620001 | 43720000 | 0.0037 | 0.0033 | 0.0035 |
| F08 | 43640001 | 43740000 | 0.0039 | 0.0034 | 0.0036 |
| F08 | 43660001 | 43760000 | 0.0036 | 0.0031 | 0.0034 |
| F08 | 43680001 | 43780000 | 0.0033 | 0.0028 | 0.0031 |
| F08 | 43700001 | 43800000 | 0.0029 | 0.0026 | 0.0026 |
| F08 | 43720001 | 43820000 | 0.0031 | 0.0028 | 0.0029 |
| F08 | 43740001 | 43840000 | 0.0022 | 0.0018 | 0.0019 |
| F08 | 43760001 | 43860000 | 0.0019 | 0.0015 | 0.0017 |
| F08 | 43780001 | 43880000 | 0.0022 | 0.0018 | 0.0019 |
| F08 | 43800001 | 43900000 | 0.0018 | 0.0014 | 0.0016 |
| F08 | 43820001 | 43920000 | 0.0023 | 0.0020 | 0.0021 |
| F08 | 43840001 | 43940000 | 0.0025 | 0.0023 | 0.0024 |
| F08 | 43860001 | 43960000 | 0.0026 | 0.0024 | 0.0025 |
| F08 | 43880001 | 43980000 | 0.0023 | 0.0021 | 0.0022 |
| F08 | 43900001 | 44000000 | 0.0027 | 0.0024 | 0.0025 |
| F08 | 43920001 | 44020000 | 0.0024 | 0.0020 | 0.0022 |
| F08 | 43940001 | 44040000 | 0.0026 | 0.0022 | 0.0024 |
| F08 | 43960001 | 44060000 | 0.0025 | 0.0021 | 0.0023 |
| F08 | 43980001 | 44080000 | 0.0026 | 0.0023 | 0.0025 |
| F08 | 44000001 | 44100000 | 0.0024 | 0.0020 | 0.0024 |
| F08 | 44020001 | 44120000 | 0.0022 | 0.0018 | 0.0022 |
| F08 | 44040001 | 44140000 | 0.0025 | 0.0020 | 0.0024 |
| F08 | 44060001 | 44160000 | 0.0035 | 0.0029 | 0.0033 |
| F08 | 44080001 | 44180000 | 0.0045 | 0.0036 | 0.0042 |
| F08 | 44100001 | 44200000 | 0.0050 | 0.0042 | 0.0045 |
| F08 | 44120001 | 44220000 | 0.0050 | 0.0043 | 0.0044 |
| F08 | 44140001 | 44240000 | 0.0044 | 0.0038 | 0.0039 |
| F08 | 44160001 | 44260000 | 0.0033 | 0.0028 | 0.0030 |
| F08 | 44180001 | 44280000 | 0.0022 | 0.0019 | 0.0019 |
| F08 | 44200001 | 44300000 | 0.0011 | 0.0010 | 0.0010 |
| F08 | 44220001 | 44320000 | 0.0008 | 0.0007 | 0.0007 |
| F08 | 44240001 | 44340000 | 0.0006 | 0.0005 | 0.0005 |
| F08 | 44260001 | 44360000 | 0.0006 | 0.0005 | 0.0005 |
| F08 | 44280001 | 44380000 | 0.0012 | 0.0011 | 0.0012 |
| F08 | 44300001 | 44400000 | 0.0014 | 0.0013 | 0.0013 |
| F08 | 44320001 | 44420000 | 0.0013 | 0.0013 | 0.0013 |
| F08 | 44340001 | 44440000 | 0.0017 | 0.0016 | 0.0016 |
| F08 | 44360001 | 44460000 | 0.0018 | 0.0017 | 0.0017 |
| F08 | 44380001 | 44480000 | 0.0011 | 0.0011 | 0.0011 |
| F08 | 44400001 | 44500000 | 0.0010 | 0.0009 | 0.0009 |
| F08 | 44420001 | 44520000 | 0.0016 | 0.0015 | 0.0015 |
| F08 | 44440001 | 44540000 | 0.0028 | 0.0024 | 0.0027 |

| Chr | Start | End | both | A1 | A2 |
|---|---|---|---|---|---|
| F08 | 44460001 | 44560000 | 0.0031 | 0.0027 | 0.0030 |
| F08 | 44480001 | 44580000 | 0.0039 | 0.0033 | 0.0038 |
| F08 | 44500001 | 44600000 | 0.0039 | 0.0033 | 0.0038 |
| F08 | 44520001 | 44620000 | 0.0033 | 0.0028 | 0.0032 |
| F08 | 44540001 | 44640000 | 0.0022 | 0.0019 | 0.0021 |
| F08 | 44560001 | 44660000 | 0.0020 | 0.0017 | 0.0019 |
| F08 | 44580001 | 44680000 | 0.0023 | 0.0020 | 0.0022 |
| F08 | 44600001 | 44700000 | 0.0024 | 0.0020 | 0.0022 |
| F08 | 44620001 | 44720000 | 0.0023 | 0.0020 | 0.0022 |
| F08 | 44640001 | 44740000 | 0.0022 | 0.0017 | 0.0020 |
| F08 | 44660001 | 44760000 | 0.0025 | 0.0019 | 0.0022 |
| F08 | 44680001 | 44780000 | 0.0015 | 0.0011 | 0.0012 |
| F08 | 44700001 | 44800000 | 0.0014 | 0.0011 | 0.0013 |
| F08 | 44720001 | 44820000 | 0.0019 | 0.0015 | 0.0018 |
| F08 | 44740001 | 44840000 | 0.0018 | 0.0014 | 0.0017 |
| F08 | 44760001 | 44860000 | 0.0015 | 0.0012 | 0.0014 |
| F08 | 44780001 | 44880000 | 0.0015 | 0.0012 | 0.0015 |
| F08 | 44800001 | 44900000 | 0.0013 | 0.0010 | 0.0013 |
| F08 | 44820001 | 44920000 | 0.0012 | 0.0010 | 0.0012 |
| F08 | 44840001 | 44940000 | 0.0012 | 0.0010 | 0.0011 |
| F08 | 44860001 | 44960000 | 0.0012 | 0.0009 | 0.0011 |
| F08 | 44880001 | 44980000 | 0.0018 | 0.0015 | 0.0017 |
| F08 | 44900001 | 45000000 | 0.0026 | 0.0021 | 0.0024 |
| F08 | 44920001 | 45020000 | 0.0024 | 0.0018 | 0.0021 |
| F08 | 44940001 | 45040000 | 0.0025 | 0.0020 | 0.0022 |
| F08 | 44960001 | 45060000 | 0.0025 | 0.0020 | 0.0022 |
| F08 | 44980001 | 45080000 | 0.0018 | 0.0014 | 0.0015 |
| F08 | 45000001 | 45100000 | 0.0014 | 0.0012 | 0.0012 |
| F08 | 45020001 | 45120000 | 0.0022 | 0.0018 | 0.0019 |
| F08 | 45040001 | 45140000 | 0.0022 | 0.0017 | 0.0020 |
| F08 | 45060001 | 45160000 | 0.0026 | 0.0021 | 0.0025 |
| F08 | 45080001 | 45180000 | 0.0031 | 0.0025 | 0.0029 |
| F08 | 45100001 | 45200000 | 0.0039 | 0.0029 | 0.0036 |
| F08 | 45120001 | 45220000 | 0.0032 | 0.0023 | 0.0029 |
| F08 | 45140001 | 45240000 | 0.0035 | 0.0026 | 0.0032 |
| F08 | 45160001 | 45260000 | 0.0045 | 0.0037 | 0.0042 |
| F08 | 45180001 | 45280000 | 0.0048 | 0.0039 | 0.0045 |
| F08 | 45200001 | 45300000 | 0.0039 | 0.0032 | 0.0036 |
| F08 | 45220001 | 45320000 | 0.0051 | 0.0042 | 0.0048 |
| F08 | 45240001 | 45340000 | 0.0054 | 0.0044 | 0.0050 |
| F08 | 45260001 | 45360000 | 0.0041 | 0.0031 | 0.0036 |
| F08 | 45280001 | 45380000 | 0.0040 | 0.0030 | 0.0033 |
| F08 | 45300001 | 45400000 | 0.0048 | 0.0036 | 0.0040 |
| F08 | 45320001 | 45420000 | 0.0036 | 0.0028 | 0.0030 |
| F08 | 45340001 | 45440000 | 0.0031 | 0.0024 | 0.0026 |
| F08 | 45360001 | 45460000 | 0.0039 | 0.0032 | 0.0035 |
| F08 | 45380001 | 45480000 | 0.0036 | 0.0031 | 0.0033 |
| F08 | 45400001 | 45500000 | 0.0032 | 0.0028 | 0.0030 |
| F08 | 45420001 | 45520000 | 0.0039 | 0.0034 | 0.0036 |
| F08 | 45440001 | 45540000 | 0.0037 | 0.0033 | 0.0034 |
| F08 | 45460001 | 45560000 | 0.0036 | 0.0033 | 0.0031 |
| F08 | 45480001 | 45580000 | 0.0040 | 0.0035 | 0.0034 |
| F08 | 45500001 | 45600000 | 0.0036 | 0.0032 | 0.0030 |
| F08 | 45520001 | 45620000 | 0.0029 | 0.0026 | 0.0024 |
| F08 | 45540001 | 45640000 | 0.0034 | 0.0029 | 0.0027 |
| F08 | 45560001 | 45660000 | 0.0026 | 0.0021 | 0.0022 |
| F08 | 45580001 | 45680000 | 0.0020 | 0.0016 | 0.0017 |
| F08 | 45600001 | 45700000 | 0.0024 | 0.0017 | 0.0019 |

| Chr | Start | End | both | A1 | A2 |
|---|---|---|---|---|---|
| F08 | 45620001 | 45720000 | 0.0031 | 0.0020 | 0.0024 |
| F08 | 45640001 | 45740000 | 0.0033 | 0.0021 | 0.0027 |
| F08 | 45660001 | 45760000 | 0.0032 | 0.0021 | 0.0027 |
| F08 | 45680001 | 45780000 | 0.0039 | 0.0028 | 0.0034 |
| F08 | 45700001 | 45800000 | 0.0039 | 0.0028 | 0.0034 |
| F08 | 45720001 | 45820000 | 0.0044 | 0.0035 | 0.0039 |
| F08 | 45740001 | 45840000 | 0.0039 | 0.0032 | 0.0035 |
| F08 | 45760001 | 45860000 | 0.0041 | 0.0033 | 0.0037 |
| F08 | 45780001 | 45880000 | 0.0049 | 0.0040 | 0.0046 |
| F08 | 45800001 | 45900000 | 0.0053 | 0.0046 | 0.0051 |
| F08 | 45820001 | 45920000 | 0.0039 | 0.0033 | 0.0038 |
| F08 | 45840001 | 45940000 | 0.0042 | 0.0036 | 0.0041 |
| F08 | 45860001 | 45960000 | 0.0045 | 0.0038 | 0.0044 |
| F08 | 45880001 | 45980000 | 0.0034 | 0.0029 | 0.0033 |
| F08 | 45900001 | 46000000 | 0.0027 | 0.0024 | 0.0027 |
| F08 | 45920001 | 46020000 | 0.0034 | 0.0028 | 0.0034 |
| F08 | 45940001 | 46040000 | 0.0035 | 0.0027 | 0.0033 |
| F08 | 45960001 | 46060000 | 0.0035 | 0.0028 | 0.0032 |
| F08 | 45980001 | 46080000 | 0.0033 | 0.0025 | 0.0029 |
| F08 | 46000001 | 46100000 | 0.0036 | 0.0026 | 0.0031 |
| F08 | 46020001 | 46120000 | 0.0041 | 0.0032 | 0.0033 |
| F08 | 46040001 | 46140000 | 0.0039 | 0.0030 | 0.0032 |
| F08 | 46060001 | 46160000 | 0.0033 | 0.0024 | 0.0026 |
| F08 | 46080001 | 46180000 | 0.0034 | 0.0026 | 0.0028 |
| F08 | 46100001 | 46200000 | 0.0039 | 0.0031 | 0.0033 |
| F08 | 46120001 | 46220000 | 0.0039 | 0.0029 | 0.0034 |
| F08 | 46140001 | 46240000 | 0.0044 | 0.0036 | 0.0039 |
| F08 | 46160001 | 46260000 | 0.0051 | 0.0042 | 0.0045 |
| F08 | 46180001 | 46280000 | 0.0049 | 0.0038 | 0.0042 |
| F08 | 46200001 | 46300000 | 0.0041 | 0.0030 | 0.0032 |
| F08 | 46220001 | 46320000 | 0.0034 | 0.0024 | 0.0026 |
| F08 | 46240001 | 46340000 | 0.0026 | 0.0015 | 0.0017 |
| F08 | 46260001 | 46360000 | 0.0022 | 0.0010 | 0.0011 |
| F08 | 46280001 | 46380000 | 0.0024 | 0.0014 | 0.0014 |
| F08 | 46300001 | 46400000 | 0.0022 | 0.0014 | 0.0014 |
| F08 | 46320001 | 46420000 | 0.0017 | 0.0011 | 0.0011 |
| F08 | 46340001 | 46440000 | 0.0020 | 0.0013 | 0.0013 |
| F08 | 46360001 | 46460000 | 0.0026 | 0.0019 | 0.0020 |
| F08 | 46380001 | 46480000 | 0.0027 | 0.0018 | 0.0022 |
| F08 | 46400001 | 46500000 | 0.0033 | 0.0020 | 0.0026 |
| F08 | 46420001 | 46520000 | 0.0038 | 0.0025 | 0.0030 |
| F08 | 46440001 | 46540000 | 0.0040 | 0.0028 | 0.0033 |
| F08 | 46460001 | 46560000 | 0.0035 | 0.0023 | 0.0029 |
| F08 | 46480001 | 46580000 | 0.0033 | 0.0024 | 0.0028 |
| F08 | 46500001 | 46600000 | 0.0035 | 0.0027 | 0.0030 |
| F08 | 46520001 | 46620000 | 0.0035 | 0.0024 | 0.0027 |
| F08 | 46540001 | 46640000 | 0.0039 | 0.0027 | 0.0031 |
| F08 | 46560001 | 46660000 | 0.0037 | 0.0024 | 0.0027 |
| F08 | 46580001 | 46680000 | 0.0041 | 0.0025 | 0.0026 |
| F08 | 46600001 | 46700000 | 0.0044 | 0.0030 | 0.0032 |
| F08 | 46620001 | 46720000 | 0.0043 | 0.0032 | 0.0033 |
| F08 | 46640001 | 46740000 | 0.0029 | 0.0023 | 0.0023 |
| F08 | 46660001 | 46760000 | 0.0027 | 0.0021 | 0.0022 |
| F08 | 46680001 | 46780000 | 0.0023 | 0.0020 | 0.0022 |
| F08 | 46700001 | 46800000 | 0.0011 | 0.0009 | 0.0011 |
| F08 | 46720001 | 46820000 | 0.0009 | 0.0008 | 0.0009 |
| F08 | 46740001 | 46840000 | 0.0011 | 0.0009 | 0.0011 |
| F08 | 46760001 | 46860000 | 0.0012 | 0.0010 | 0.0012 |

| Chr | Start | End | both | A1 | A2 |
|---|---|---|---|---|---|
| F08 | 46780001 | 46880000 | 0.0008 | 0.0007 | 0.0008 |
| F08 | 46800001 | 46900000 | 0.0007 | 0.0006 | 0.0007 |
| F08 | 46820001 | 46920000 | 0.0011 | 0.0010 | 0.0011 |
| F08 | 46840001 | 46940000 | 0.0010 | 0.0009 | 0.0010 |
| F08 | 46860001 | 46960000 | 0.0011 | 0.0011 | 0.0011 |
| F08 | 46880001 | 46980000 | 0.0011 | 0.0010 | 0.0011 |
| F08 | 46900001 | 47000000 | 0.0012 | 0.0011 | 0.0012 |
| F08 | 46920001 | 47020000 | 0.0006 | 0.0005 | 0.0006 |
| F08 | 46940001 | 47040000 | 0.0006 | 0.0005 | 0.0006 |
| F08 | 46960001 | 47060000 | 0.0004 | 0.0003 | 0.0004 |
| F08 | 46980001 | 47080000 | 0.0006 | 0.0004 | 0.0006 |
| F08 | 47000001 | 47100000 | 0.0007 | 0.0006 | 0.0006 |
| F08 | 47020001 | 47120000 | 0.0009 | 0.0008 | 0.0009 |
| F08 | 47040001 | 47140000 | 0.0009 | 0.0008 | 0.0008 |
| F08 | 47060001 | 47160000 | 0.0021 | 0.0019 | 0.0021 |
| F08 | 47080001 | 47180000 | 0.0021 | 0.0018 | 0.0021 |
| F08 | 47100001 | 47200000 | 0.0020 | 0.0017 | 0.0020 |
| F08 | 47120001 | 47220000 | 0.0023 | 0.0019 | 0.0023 |
| F08 | 47140001 | 47240000 | 0.0023 | 0.0019 | 0.0023 |
| F08 | 47160001 | 47260000 | 0.0010 | 0.0008 | 0.0010 |
| F08 | 47180001 | 47280000 | 0.0012 | 0.0010 | 0.0012 |
| F08 | 47200001 | 47300000 | 0.0014 | 0.0011 | 0.0013 |
| F08 | 47220001 | 47320000 | 0.0012 | 0.0010 | 0.0012 |
| F08 | 47240001 | 47340000 | 0.0013 | 0.0011 | 0.0012 |
| F08 | 47260001 | 47360000 | 0.0014 | 0.0012 | 0.0014 |
| F08 | 47280001 | 47380000 | 0.0011 | 0.0009 | 0.0011 |
| F08 | 47300001 | 47400000 | 0.0009 | 0.0007 | 0.0009 |
| F08 | 47320001 | 47420000 | 0.0005 | 0.0004 | 0.0005 |
| F08 | 47340001 | 47440000 | 0.0005 | 0.0004 | 0.0005 |
| F08 | 47360001 | 47460000 | 0.0003 | 0.0003 | 0.0003 |
| F08 | 47380001 | 47480000 | 0.0003 | 0.0003 | 0.0003 |
| F08 | 47400001 | 47500000 | 0.0003 | 0.0003 | 0.0003 |
| F08 | 47420001 | 47520000 | 0.0003 | 0.0003 | 0.0003 |
| F08 | 47440001 | 47540000 | 0.0003 | 0.0002 | 0.0003 |
| F08 | 47460001 | 47560000 | 0.0003 | 0.0003 | 0.0003 |
| F08 | 47480001 | 47580000 | 0.0004 | 0.0003 | 0.0004 |
| F08 | 47500001 | 47600000 | 0.0005 | 0.0003 | 0.0005 |
| F08 | 47520001 | 47620000 | 0.0006 | 0.0005 | 0.0006 |
| F08 | 47540001 | 47640000 | 0.0007 | 0.0006 | 0.0007 |
| F08 | 47560001 | 47660000 | 0.0009 | 0.0008 | 0.0008 |
| F08 | 47580001 | 47680000 | 0.0009 | 0.0009 | 0.0009 |
| F08 | 47600001 | 47700000 | 0.0011 | 0.0010 | 0.0010 |
| F08 | 47620001 | 47720000 | 0.0014 | 0.0012 | 0.0014 |
| F08 | 47640001 | 47740000 | 0.0021 | 0.0018 | 0.0021 |
| F08 | 47660001 | 47760000 | 0.0018 | 0.0016 | 0.0018 |
| F08 | 47680001 | 47780000 | 0.0017 | 0.0015 | 0.0017 |
| F08 | 47700001 | 47800000 | 0.0017 | 0.0014 | 0.0016 |
| F08 | 47720001 | 47820000 | 0.0013 | 0.0010 | 0.0012 |
| F08 | 47740001 | 47840000 | 0.0006 | 0.0004 | 0.0006 |
| F08 | 47760001 | 47860000 | 0.0011 | 0.0008 | 0.0008 |
| F08 | 47780001 | 47880000 | 0.0014 | 0.0010 | 0.0011 |
| F08 | 47800001 | 47900000 | 0.0029 | 0.0025 | 0.0026 |
| F08 | 47820001 | 47920000 | 0.0030 | 0.0024 | 0.0025 |
| F08 | 47840001 | 47940000 | 0.0035 | 0.0027 | 0.0027 |
| F08 | 47860001 | 47960000 | 0.0033 | 0.0026 | 0.0027 |
| F08 | 47880001 | 47980000 | 0.0035 | 0.0028 | 0.0030 |
| F08 | 47900001 | 48000000 | 0.0023 | 0.0016 | 0.0018 |
| F08 | 47920001 | 48020000 | 0.0024 | 0.0018 | 0.0020 |

| Chr | Start | End | both | A1 | A2 |
|---|---|---|---|---|---|
| F08 | 47940001 | 48040000 | 0.0021 | 0.0016 | 0.0019 |
| F08 | 47960001 | 48060000 | 0.0026 | 0.0019 | 0.0023 |
| F08 | 47980001 | 48080000 | 0.0031 | 0.0021 | 0.0028 |
| F08 | 48000001 | 48100000 | 0.0034 | 0.0023 | 0.0030 |
| F08 | 48020001 | 48120000 | 0.0039 | 0.0027 | 0.0034 |
| F08 | 48040001 | 48140000 | 0.0036 | 0.0026 | 0.0033 |
| F08 | 48060001 | 48160000 | 0.0033 | 0.0026 | 0.0031 |
| F08 | 48080001 | 48180000 | 0.0028 | 0.0023 | 0.0026 |
| F08 | 48100001 | 48200000 | 0.0028 | 0.0024 | 0.0026 |
| F08 | 48120001 | 48220000 | 0.0027 | 0.0023 | 0.0024 |
| F08 | 48140001 | 48240000 | 0.0027 | 0.0023 | 0.0024 |
| F08 | 48160001 | 48260000 | 0.0026 | 0.0020 | 0.0021 |
| F08 | 48180001 | 48280000 | 0.0029 | 0.0022 | 0.0024 |
| F08 | 48200001 | 48300000 | 0.0029 | 0.0020 | 0.0022 |
| F08 | 48220001 | 48320000 | 0.0031 | 0.0022 | 0.0024 |
| F08 | 48240001 | 48340000 | 0.0036 | 0.0026 | 0.0029 |
| F08 | 48260001 | 48360000 | 0.0041 | 0.0032 | 0.0034 |
| F08 | 48280001 | 48380000 | 0.0039 | 0.0030 | 0.0033 |
| F08 | 48300001 | 48400000 | 0.0031 | 0.0025 | 0.0027 |
| F08 | 48320001 | 48420000 | 0.0029 | 0.0025 | 0.0026 |
| F08 | 48340001 | 48440000 | 0.0024 | 0.0022 | 0.0023 |
| F08 | 48360001 | 48460000 | 0.0018 | 0.0015 | 0.0017 |
| F08 | 48380001 | 48480000 | 0.0015 | 0.0013 | 0.0015 |
| F08 | 48400001 | 48500000 | 0.0021 | 0.0017 | 0.0020 |
| F08 | 48420001 | 48520000 | 0.0032 | 0.0025 | 0.0030 |
| F08 | 48440001 | 48540000 | 0.0035 | 0.0026 | 0.0030 |
| F08 | 48460001 | 48560000 | 0.0052 | 0.0037 | 0.0044 |
| F08 | 48480001 | 48580000 | 0.0058 | 0.0041 | 0.0049 |
| F08 | 48500001 | 48600000 | 0.0058 | 0.0040 | 0.0048 |
| F08 | 48520001 | 48620000 | 0.0041 | 0.0027 | 0.0033 |
| F08 | 48540001 | 48640000 | 0.0039 | 0.0028 | 0.0034 |
| F08 | 48560001 | 48660000 | 0.0025 | 0.0020 | 0.0023 |
| F08 | 48580001 | 48680000 | 0.0015 | 0.0013 | 0.0014 |
| F08 | 48600001 | 48700000 | 0.0020 | 0.0018 | 0.0019 |
| F08 | 48620001 | 48720000 | 0.0022 | 0.0019 | 0.0021 |
| F08 | 48640001 | 48740000 | 0.0022 | 0.0017 | 0.0019 |
| F08 | 48660001 | 48760000 | 0.0023 | 0.0016 | 0.0018 |
| F08 | 48680001 | 48780000 | 0.0034 | 0.0024 | 0.0029 |
| F08 | 48700001 | 48800000 | 0.0024 | 0.0016 | 0.0020 |
| F08 | 48720001 | 48820000 | 0.0022 | 0.0015 | 0.0018 |
| F08 | 48740001 | 48840000 | 0.0028 | 0.0023 | 0.0025 |
| F08 | 48760001 | 48860000 | 0.0025 | 0.0022 | 0.0023 |
| F08 | 48780001 | 48880000 | 0.0014 | 0.0014 | 0.0013 |
| F08 | 48800001 | 48900000 | 0.0015 | 0.0015 | 0.0014 |
| F08 | 48820001 | 48920000 | 0.0018 | 0.0017 | 0.0017 |
| F08 | 48840001 | 48940000 | 0.0010 | 0.0009 | 0.0010 |
| F08 | 48860001 | 48960000 | 0.0009 | 0.0008 | 0.0009 |
| F08 | 48880001 | 48980000 | 0.0011 | 0.0010 | 0.0011 |
| F08 | 48900001 | 49000000 | 0.0012 | 0.0009 | 0.0010 |
| F08 | 48920001 | 49020000 | 0.0019 | 0.0015 | 0.0017 |
| F08 | 48940001 | 49040000 | 0.0019 | 0.0015 | 0.0018 |
| F08 | 48960001 | 49060000 | 0.0018 | 0.0014 | 0.0017 |
| F08 | 48980001 | 49080000 | 0.0021 | 0.0016 | 0.0019 |
| F08 | 49000001 | 49100000 | 0.0020 | 0.0017 | 0.0019 |
| F08 | 49020001 | 49120000 | 0.0012 | 0.0010 | 0.0011 |
| F08 | 49040001 | 49140000 | 0.0015 | 0.0015 | 0.0015 |
| F08 | 49060001 | 49160000 | 0.0017 | 0.0016 | 0.0016 |
| F08 | 49080001 | 49180000 | 0.0014 | 0.0013 | 0.0013 |

| Chr | Start | End | both | A1 | A2 |
|---|---|---|---|---|---|
| F08 | 49100001 | 49200000 | 0.0018 | 0.0015 | 0.0016 |
| F08 | 49120001 | 49220000 | 0.0023 | 0.0019 | 0.0021 |
| F08 | 49140001 | 49240000 | 0.0018 | 0.0014 | 0.0016 |
| F08 | 49160001 | 49260000 | 0.0019 | 0.0014 | 0.0016 |
| F08 | 49180001 | 49280000 | 0.0029 | 0.0023 | 0.0027 |
| F08 | 49200001 | 49300000 | 0.0029 | 0.0024 | 0.0027 |
| F08 | 49220001 | 49320000 | 0.0026 | 0.0024 | 0.0025 |
| F08 | 49240001 | 49340000 | 0.0029 | 0.0026 | 0.0027 |
| F08 | 49260001 | 49360000 | 0.0030 | 0.0027 | 0.0028 |
| F08 | 49280001 | 49380000 | 0.0026 | 0.0024 | 0.0024 |
| F08 | 49300001 | 49400000 | 0.0023 | 0.0020 | 0.0021 |
| F08 | 49320001 | 49420000 | 0.0027 | 0.0022 | 0.0026 |
| F08 | 49340001 | 49440000 | 0.0030 | 0.0025 | 0.0029 |
| F08 | 49360001 | 49460000 | 0.0032 | 0.0026 | 0.0030 |
| F08 | 49380001 | 49480000 | 0.0027 | 0.0021 | 0.0025 |
| F08 | 49400001 | 49500000 | 0.0033 | 0.0024 | 0.0029 |
| F08 | 49420001 | 49520000 | 0.0026 | 0.0020 | 0.0022 |
| F08 | 49440001 | 49540000 | 0.0026 | 0.0017 | 0.0022 |
| F08 | 49460001 | 49560000 | 0.0031 | 0.0023 | 0.0027 |
| F08 | 49480001 | 49580000 | 0.0032 | 0.0023 | 0.0028 |
| F08 | 49500001 | 49600000 | 0.0028 | 0.0020 | 0.0025 |
| F08 | 49520001 | 49620000 | 0.0032 | 0.0022 | 0.0027 |
| F08 | 49540001 | 49640000 | 0.0033 | 0.0025 | 0.0029 |
| F08 | 49560001 | 49660000 | 0.0029 | 0.0021 | 0.0024 |
| F08 | 49580001 | 49680000 | 0.0033 | 0.0025 | 0.0028 |
| F08 | 49600001 | 49700000 | 0.0051 | 0.0040 | 0.0045 |
| F08 | 49620001 | 49720000 | 0.0049 | 0.0038 | 0.0042 |
| F08 | 49640001 | 49740000 | 0.0043 | 0.0032 | 0.0035 |
| F08 | 49660001 | 49760000 | 0.0040 | 0.0029 | 0.0033 |
| F08 | 49680001 | 49780000 | 0.0035 | 0.0025 | 0.0028 |
| F08 | 49700001 | 49800000 | 0.0022 | 0.0011 | 0.0014 |
| F08 | 49720001 | 49820000 | 0.0025 | 0.0015 | 0.0018 |
| F08 | 49740001 | 49840000 | 0.0025 | 0.0017 | 0.0020 |
| F08 | 49760001 | 49860000 | 0.0029 | 0.0020 | 0.0024 |
| F08 | 49780001 | 49880000 | 0.0033 | 0.0023 | 0.0027 |
| F08 | 49800001 | 49900000 | 0.0037 | 0.0029 | 0.0032 |
| F08 | 49820001 | 49920000 | 0.0038 | 0.0030 | 0.0033 |
| F08 | 49840001 | 49940000 | 0.0051 | 0.0040 | 0.0044 |
| F08 | 49860001 | 49960000 | 0.0056 | 0.0041 | 0.0046 |
| F08 | 49880001 | 49980000 | 0.0053 | 0.0038 | 0.0044 |
| F08 | 49900001 | 50000000 | 0.0059 | 0.0044 | 0.0051 |
| F08 | 49920001 | 50020000 | 0.0057 | 0.0041 | 0.0047 |
| F08 | 49940001 | 50040000 | 0.0045 | 0.0030 | 0.0034 |
| F08 | 49960001 | 50060000 | 0.0035 | 0.0024 | 0.0027 |
| F08 | 49980001 | 50080000 | 0.0034 | 0.0023 | 0.0025 |
| F08 | 50000001 | 50100000 | 0.0017 | 0.0008 | 0.0009 |
| F08 | 50020001 | 50120000 | 0.0016 | 0.0009 | 0.0010 |
| F08 | 50040001 | 50140000 | 0.0025 | 0.0018 | 0.0021 |
| F08 | 50060001 | 50160000 | 0.0031 | 0.0023 | 0.0026 |
| F08 | 50080001 | 50180000 | 0.0038 | 0.0029 | 0.0032 |
| F08 | 50100001 | 50200000 | 0.0043 | 0.0033 | 0.0035 |
| F08 | 50120001 | 50220000 | 0.0039 | 0.0031 | 0.0032 |
| F08 | 50140001 | 50240000 | 0.0034 | 0.0025 | 0.0026 |
| F08 | 50160001 | 50260000 | 0.0030 | 0.0021 | 0.0022 |
| F08 | 50180001 | 50280000 | 0.0028 | 0.0018 | 0.0021 |
| F08 | 50200001 | 50300000 | 0.0023 | 0.0015 | 0.0018 |
| F08 | 50220001 | 50320000 | 0.0030 | 0.0021 | 0.0025 |
| F08 | 50240001 | 50340000 | 0.0028 | 0.0019 | 0.0025 |

| Chr | Start | End | both | A1 | A2 |
|---|---|---|---|---|---|
| F08 | 50260001 | 50360000 | 0.0028 | 0.0021 | 0.0027 |
| F08 | 50280001 | 50380000 | 0.0028 | 0.0021 | 0.0026 |
| F08 | 50300001 | 50400000 | 0.0032 | 0.0026 | 0.0030 |
| F08 | 50320001 | 50420000 | 0.0030 | 0.0023 | 0.0028 |
| F08 | 50340001 | 50440000 | 0.0027 | 0.0021 | 0.0024 |
| F08 | 50360001 | 50460000 | 0.0028 | 0.0023 | 0.0025 |
| F08 | 50380001 | 50480000 | 0.0025 | 0.0022 | 0.0022 |
| F08 | 50400001 | 50500000 | 0.0026 | 0.0019 | 0.0023 |
| F08 | 50420001 | 50520000 | 0.0024 | 0.0017 | 0.0019 |
| F08 | 50440001 | 50540000 | 0.0033 | 0.0024 | 0.0029 |
| F08 | 50460001 | 50560000 | 0.0036 | 0.0024 | 0.0030 |
| F08 | 50480001 | 50580000 | 0.0048 | 0.0031 | 0.0040 |
| F08 | 50500001 | 50600000 | 0.0070 | 0.0052 | 0.0062 |
| F08 | 50520001 | 50620000 | 0.0067 | 0.0050 | 0.0059 |
| F08 | 50540001 | 50640000 | 0.0064 | 0.0048 | 0.0056 |
| F08 | 50560001 | 50660000 | 0.0061 | 0.0046 | 0.0053 |
| F08 | 50580001 | 50680000 | 0.0048 | 0.0036 | 0.0041 |
| F08 | 50600001 | 50700000 | 0.0025 | 0.0017 | 0.0019 |
| F08 | 50620001 | 50720000 | 0.0025 | 0.0018 | 0.0021 |
| F08 | 50640001 | 50740000 | 0.0026 | 0.0019 | 0.0022 |
| F08 | 50660001 | 50760000 | 0.0028 | 0.0021 | 0.0024 |
| F08 | 50680001 | 50780000 | 0.0029 | 0.0023 | 0.0025 |
| F08 | 50700001 | 50800000 | 0.0028 | 0.0020 | 0.0023 |
| F08 | 50720001 | 50820000 | 0.0037 | 0.0027 | 0.0031 |
| F08 | 50740001 | 50840000 | 0.0029 | 0.0019 | 0.0022 |
| F08 | 50760001 | 50860000 | 0.0024 | 0.0015 | 0.0019 |
| F08 | 50780001 | 50880000 | 0.0021 | 0.0013 | 0.0016 |
| F08 | 50800001 | 50900000 | 0.0025 | 0.0019 | 0.0021 |
| F08 | 50820001 | 50920000 | 0.0016 | 0.0012 | 0.0014 |
| F08 | 50840001 | 50940000 | 0.0018 | 0.0016 | 0.0017 |
| F08 | 50860001 | 50960000 | 0.0017 | 0.0016 | 0.0016 |
| F08 | 50880001 | 50980000 | 0.0021 | 0.0019 | 0.0020 |
| F08 | 50900001 | 51000000 | 0.0018 | 0.0017 | 0.0018 |
| F08 | 50920001 | 51020000 | 0.0022 | 0.0020 | 0.0021 |
| F08 | 50940001 | 51040000 | 0.0032 | 0.0029 | 0.0030 |
| F08 | 50960001 | 51060000 | 0.0037 | 0.0031 | 0.0033 |
| F08 | 50980001 | 51080000 | 0.0048 | 0.0041 | 0.0044 |
| F08 | 51000001 | 51100000 | 0.0052 | 0.0043 | 0.0046 |
| F08 | 51020001 | 51120000 | 0.0051 | 0.0041 | 0.0044 |
| F08 | 51040001 | 51140000 | 0.0039 | 0.0029 | 0.0032 |
| F08 | 51060001 | 51160000 | 0.0035 | 0.0026 | 0.0030 |
| F08 | 51080001 | 51180000 | 0.0030 | 0.0021 | 0.0025 |
| F08 | 51100001 | 51200000 | 0.0032 | 0.0025 | 0.0028 |
| F08 | 51120001 | 51220000 | 0.0036 | 0.0028 | 0.0030 |
| F08 | 51140001 | 51240000 | 0.0041 | 0.0033 | 0.0036 |
| F08 | 51160001 | 51260000 | 0.0050 | 0.0040 | 0.0043 |
| F08 | 51180001 | 51280000 | 0.0042 | 0.0034 | 0.0036 |
| F08 | 51200001 | 51300000 | 0.0041 | 0.0029 | 0.0031 |
| F08 | 51220001 | 51320000 | 0.0040 | 0.0028 | 0.0032 |
| F08 | 51240001 | 51340000 | 0.0036 | 0.0024 | 0.0027 |
| F08 | 51260001 | 51360000 | 0.0030 | 0.0021 | 0.0023 |
| F08 | 51280001 | 51380000 | 0.0031 | 0.0020 | 0.0021 |
| F08 | 51300001 | 51400000 | 0.0031 | 0.0022 | 0.0024 |
| F08 | 51320001 | 51420000 | 0.0038 | 0.0028 | 0.0029 |
| F08 | 51340001 | 51440000 | 0.0041 | 0.0030 | 0.0031 |
| F08 | 51360001 | 51460000 | 0.0041 | 0.0030 | 0.0031 |
| F08 | 51380001 | 51480000 | 0.0041 | 0.0032 | 0.0035 |
| F08 | 51400001 | 51500000 | 0.0040 | 0.0033 | 0.0034 |

| Chr | Start | End | both | A1 | A2 |
|---|---|---|---|---|---|
| F08 | 51420001 | 51520000 | 0.0035 | 0.0029 | 0.0030 |
| F08 | 51440001 | 51540000 | 0.0033 | 0.0027 | 0.0029 |
| F08 | 51460001 | 51560000 | 0.0033 | 0.0027 | 0.0029 |
| F08 | 51480001 | 51580000 | 0.0030 | 0.0024 | 0.0025 |
| F08 | 51500001 | 51600000 | 0.0023 | 0.0015 | 0.0016 |
| F08 | 51520001 | 51620000 | 0.0029 | 0.0020 | 0.0023 |
| F08 | 51540001 | 51640000 | 0.0029 | 0.0019 | 0.0022 |
| F08 | 51560001 | 51660000 | 0.0030 | 0.0018 | 0.0022 |
| F08 | 51580001 | 51680000 | 0.0031 | 0.0019 | 0.0024 |
| F08 | 51600001 | 51700000 | 0.0041 | 0.0028 | 0.0034 |
| F08 | 51620001 | 51720000 | 0.0033 | 0.0024 | 0.0026 |
| F08 | 51640001 | 51740000 | 0.0033 | 0.0025 | 0.0026 |
| F08 | 51660001 | 51760000 | 0.0030 | 0.0022 | 0.0024 |
| F08 | 51680001 | 51780000 | 0.0035 | 0.0026 | 0.0029 |
| F08 | 51700001 | 51800000 | 0.0034 | 0.0025 | 0.0029 |
| F08 | 51720001 | 51820000 | 0.0039 | 0.0029 | 0.0033 |
| F08 | 51740001 | 51840000 | 0.0047 | 0.0038 | 0.0043 |
| F08 | 51760001 | 51860000 | 0.0049 | 0.0040 | 0.0045 |
| F08 | 51780001 | 51880000 | 0.0051 | 0.0041 | 0.0045 |
| F08 | 51800001 | 51900000 | 0.0043 | 0.0034 | 0.0035 |
| F08 | 51820001 | 51920000 | 0.0040 | 0.0031 | 0.0035 |
| F08 | 51840001 | 51940000 | 0.0040 | 0.0030 | 0.0034 |
| F08 | 51860001 | 51960000 | 0.0043 | 0.0033 | 0.0036 |
| F08 | 51880001 | 51980000 | 0.0042 | 0.0034 | 0.0037 |
| F08 | 51900001 | 52000000 | 0.0041 | 0.0034 | 0.0037 |
| F08 | 51920001 | 52020000 | 0.0035 | 0.0029 | 0.0031 |
| F08 | 51940001 | 52040000 | 0.0030 | 0.0023 | 0.0024 |
| F08 | 51960001 | 52060000 | 0.0028 | 0.0022 | 0.0022 |
| F08 | 51980001 | 52080000 | 0.0031 | 0.0023 | 0.0024 |
| F08 | 52000001 | 52100000 | 0.0033 | 0.0026 | 0.0026 |
| F08 | 52020001 | 52120000 | 0.0039 | 0.0031 | 0.0030 |
| F08 | 52040001 | 52140000 | 0.0041 | 0.0034 | 0.0032 |
| F08 | 52060001 | 52160000 | 0.0039 | 0.0033 | 0.0031 |
| F08 | 52080001 | 52180000 | 0.0034 | 0.0026 | 0.0027 |
| F08 | 52100001 | 52200000 | 0.0039 | 0.0028 | 0.0030 |
| F08 | 52120001 | 52220000 | 0.0034 | 0.0021 | 0.0025 |
| F08 | 52140001 | 52240000 | 0.0035 | 0.0022 | 0.0026 |
| F08 | 52160001 | 52260000 | 0.0042 | 0.0029 | 0.0033 |
| F08 | 52180001 | 52280000 | 0.0044 | 0.0031 | 0.0036 |
| F08 | 52200001 | 52300000 | 0.0041 | 0.0031 | 0.0035 |
| F08 | 52220001 | 52320000 | 0.0056 | 0.0046 | 0.0050 |
| F08 | 52240001 | 52340000 | 0.0050 | 0.0041 | 0.0045 |
| F08 | 52260001 | 52360000 | 0.0048 | 0.0039 | 0.0043 |
| F08 | 52280001 | 52380000 | 0.0048 | 0.0041 | 0.0043 |
| F08 | 52300001 | 52400000 | 0.0047 | 0.0039 | 0.0041 |
| F08 | 52320001 | 52420000 | 0.0035 | 0.0026 | 0.0027 |
| F08 | 52340001 | 52440000 | 0.0039 | 0.0028 | 0.0027 |
| F08 | 52360001 | 52460000 | 0.0035 | 0.0023 | 0.0022 |
| F08 | 52380001 | 52480000 | 0.0031 | 0.0017 | 0.0019 |
| F08 | 52400001 | 52500000 | 0.0035 | 0.0023 | 0.0024 |
| F08 | 52420001 | 52520000 | 0.0031 | 0.0019 | 0.0021 |
| F08 | 52440001 | 52540000 | 0.0029 | 0.0021 | 0.0022 |
| F08 | 52460001 | 52560000 | 0.0024 | 0.0018 | 0.0019 |
| F08 | 52480001 | 52580000 | 0.0025 | 0.0020 | 0.0021 |
| F08 | 52500001 | 52600000 | 0.0022 | 0.0016 | 0.0017 |
| F08 | 52520001 | 52620000 | 0.0026 | 0.0021 | 0.0022 |
| F08 | 52540001 | 52640000 | 0.0035 | 0.0028 | 0.0030 |
| F08 | 52560001 | 52660000 | 0.0039 | 0.0033 | 0.0031 |

| Chr | Start | End | both | A1 | A2 |
|---|---|---|---|---|---|
| F08 | 52580001 | 52680000 | 0.0036 | 0.0029 | 0.0027 |
| F08 | 52600001 | 52700000 | 0.0035 | 0.0029 | 0.0027 |
| F08 | 52620001 | 52720000 | 0.0032 | 0.0026 | 0.0025 |
| F08 | 52640001 | 52740000 | 0.0025 | 0.0018 | 0.0017 |
| F08 | 52660001 | 52760000 | 0.0025 | 0.0016 | 0.0017 |
| F08 | 52680001 | 52780000 | 0.0028 | 0.0018 | 0.0019 |
| F08 | 52700001 | 52800000 | 0.0027 | 0.0016 | 0.0016 |
| F08 | 52720001 | 52820000 | 0.0034 | 0.0023 | 0.0022 |
| F08 | 52740001 | 52840000 | 0.0038 | 0.0029 | 0.0029 |
| F08 | 52760001 | 52860000 | 0.0040 | 0.0032 | 0.0033 |
| F08 | 52780001 | 52880000 | 0.0044 | 0.0036 | 0.0039 |
| F08 | 52800001 | 52900000 | 0.0045 | 0.0037 | 0.0041 |
| F08 | 52820001 | 52920000 | 0.0038 | 0.0029 | 0.0034 |
| F08 | 52840001 | 52940000 | 0.0027 | 0.0018 | 0.0023 |
| F08 | 52860001 | 52960000 | 0.0026 | 0.0016 | 0.0021 |
| F08 | 52880001 | 52980000 | 0.0020 | 0.0013 | 0.0016 |
| F08 | 52900001 | 53000000 | 0.0017 | 0.0009 | 0.0011 |
| F08 | 52920001 | 53020000 | 0.0023 | 0.0015 | 0.0018 |
| F08 | 52940001 | 53040000 | 0.0041 | 0.0031 | 0.0033 |
| F08 | 52960001 | 53060000 | 0.0041 | 0.0031 | 0.0034 |
| F08 | 52980001 | 53080000 | 0.0042 | 0.0033 | 0.0035 |
| F08 | 53000001 | 53100000 | 0.0052 | 0.0045 | 0.0046 |
| F08 | 53020001 | 53120000 | 0.0048 | 0.0043 | 0.0044 |
| F08 | 53040001 | 53140000 | 0.0030 | 0.0027 | 0.0027 |
| F08 | 53060001 | 53160000 | 0.0032 | 0.0026 | 0.0027 |
| F08 | 53080001 | 53180000 | 0.0032 | 0.0025 | 0.0026 |
| F08 | 53100001 | 53200000 | 0.0040 | 0.0030 | 0.0033 |
| F08 | 53120001 | 53220000 | 0.0048 | 0.0036 | 0.0040 |
| F08 | 53140001 | 53240000 | 0.0053 | 0.0041 | 0.0044 |
| F08 | 53160001 | 53260000 | 0.0055 | 0.0044 | 0.0046 |
| F08 | 53180001 | 53280000 | 0.0055 | 0.0045 | 0.0047 |
| F08 | 53200001 | 53300000 | 0.0038 | 0.0032 | 0.0032 |
| F08 | 53220001 | 53320000 | 0.0027 | 0.0024 | 0.0023 |
| F08 | 53240001 | 53340000 | 0.0025 | 0.0022 | 0.0022 |
| F08 | 53260001 | 53360000 | 0.0028 | 0.0025 | 0.0025 |
| F08 | 53280001 | 53380000 | 0.0029 | 0.0025 | 0.0025 |
| F08 | 53300001 | 53400000 | 0.0042 | 0.0035 | 0.0036 |
| F08 | 53320001 | 53420000 | 0.0047 | 0.0039 | 0.0040 |
| F08 | 53340001 | 53440000 | 0.0052 | 0.0044 | 0.0045 |
| F08 | 53360001 | 53460000 | 0.0053 | 0.0043 | 0.0045 |
| F08 | 53380001 | 53480000 | 0.0050 | 0.0042 | 0.0044 |
| F08 | 53400001 | 53500000 | 0.0039 | 0.0033 | 0.0033 |
| F08 | 53420001 | 53520000 | 0.0036 | 0.0031 | 0.0033 |
| F08 | 53440001 | 53540000 | 0.0032 | 0.0027 | 0.0030 |
| F08 | 53460001 | 53560000 | 0.0020 | 0.0017 | 0.0019 |
| F08 | 53480001 | 53580000 | 0.0020 | 0.0016 | 0.0019 |
| F08 | 53500001 | 53600000 | 0.0018 | 0.0013 | 0.0016 |
| F08 | 53520001 | 53620000 | 0.0013 | 0.0009 | 0.0012 |
| F08 | 53540001 | 53640000 | 0.0012 | 0.0008 | 0.0010 |
| F08 | 53560001 | 53660000 | 0.0019 | 0.0014 | 0.0017 |
| F08 | 53580001 | 53680000 | 0.0025 | 0.0018 | 0.0023 |
| F08 | 53600001 | 53700000 | 0.0030 | 0.0023 | 0.0028 |
| F08 | 53620001 | 53720000 | 0.0032 | 0.0024 | 0.0030 |
| F08 | 53640001 | 53740000 | 0.0030 | 0.0022 | 0.0028 |
| F08 | 53660001 | 53760000 | 0.0027 | 0.0020 | 0.0024 |
| F08 | 53680001 | 53780000 | 0.0020 | 0.0015 | 0.0019 |
| F08 | 53700001 | 53800000 | 0.0019 | 0.0015 | 0.0016 |
| F08 | 53720001 | 53820000 | 0.0019 | 0.0016 | 0.0017 |

| Chr | Start | End | both | A1 | A2 |
|---|---|---|---|---|---|
| F08 | 53740001 | 53840000 | 0.0024 | 0.0020 | 0.0022 |
| F08 | 53760001 | 53860000 | 0.0028 | 0.0024 | 0.0025 |
| F08 | 53780001 | 53880000 | 0.0039 | 0.0033 | 0.0034 |
| F08 | 53800001 | 53900000 | 0.0045 | 0.0038 | 0.0041 |
| F08 | 53820001 | 53920000 | 0.0046 | 0.0038 | 0.0041 |
| F08 | 53840001 | 53940000 | 0.0042 | 0.0035 | 0.0038 |
| F08 | 53860001 | 53960000 | 0.0044 | 0.0036 | 0.0039 |
| F08 | 53880001 | 53980000 | 0.0049 | 0.0041 | 0.0044 |
| F08 | 53900001 | 54000000 | 0.0047 | 0.0039 | 0.0042 |
| F08 | 53920001 | 54020000 | 0.0057 | 0.0043 | 0.0047 |
| F08 | 53940001 | 54040000 | 0.0063 | 0.0049 | 0.0053 |
| F08 | 53960001 | 54060000 | 0.0063 | 0.0049 | 0.0055 |
| F08 | 53980001 | 54080000 | 0.0066 | 0.0049 | 0.0058 |
| F08 | 54000001 | 54100000 | 0.0068 | 0.0050 | 0.0059 |
| F08 | 54020001 | 54120000 | 0.0058 | 0.0046 | 0.0053 |
| F08 | 54040001 | 54140000 | 0.0060 | 0.0046 | 0.0053 |
| F08 | 54060001 | 54160000 | 0.0063 | 0.0049 | 0.0055 |
| F08 | 54080001 | 54180000 | 0.0052 | 0.0043 | 0.0045 |
| F08 | 54100001 | 54200000 | 0.0046 | 0.0039 | 0.0041 |
| F08 | 54120001 | 54220000 | 0.0058 | 0.0051 | 0.0054 |
| F08 | 54140001 | 54240000 | 0.0053 | 0.0049 | 0.0051 |
| F08 | 54160001 | 54260000 | 0.0047 | 0.0044 | 0.0045 |
| F08 | 54180001 | 54280000 | 0.0039 | 0.0035 | 0.0035 |
| F08 | 54200001 | 54300000 | 0.0039 | 0.0034 | 0.0035 |
| F08 | 54220001 | 54320000 | 0.0039 | 0.0031 | 0.0035 |
| F08 | 54240001 | 54340000 | 0.0040 | 0.0031 | 0.0036 |
| F08 | 54260001 | 54360000 | 0.0051 | 0.0041 | 0.0046 |
| F08 | 54280001 | 54380000 | 0.0057 | 0.0048 | 0.0054 |
| F08 | 54300001 | 54400000 | 0.0054 | 0.0047 | 0.0051 |
| F08 | 54320001 | 54420000 | 0.0050 | 0.0044 | 0.0047 |
| F08 | 54340001 | 54440000 | 0.0046 | 0.0041 | 0.0043 |
| F08 | 54360001 | 54460000 | 0.0029 | 0.0025 | 0.0027 |
| F08 | 54380001 | 54480000 | 0.0019 | 0.0016 | 0.0018 |
| F08 | 54400001 | 54500000 | 0.0017 | 0.0014 | 0.0017 |
| F08 | 54420001 | 54520000 | 0.0010 | 0.0007 | 0.0010 |
| F08 | 54440001 | 54540000 | 0.0007 | 0.0006 | 0.0008 |
| F08 | 54460001 | 54560000 | 0.0012 | 0.0010 | 0.0012 |
| F08 | 54480001 | 54580000 | 0.0013 | 0.0011 | 0.0013 |
| F08 | 54500001 | 54600000 | 0.0011 | 0.0010 | 0.0011 |
| F08 | 54520001 | 54620000 | 0.0015 | 0.0014 | 0.0015 |
| F08 | 54540001 | 54640000 | 0.0021 | 0.0019 | 0.0021 |
| F08 | 54560001 | 54660000 | 0.0016 | 0.0014 | 0.0016 |
| F08 | 54580001 | 54680000 | 0.0019 | 0.0016 | 0.0018 |
| F08 | 54600001 | 54700000 | 0.0020 | 0.0017 | 0.0020 |
| F08 | 54620001 | 54720000 | 0.0023 | 0.0021 | 0.0022 |
| F08 | 54640001 | 54740000 | 0.0017 | 0.0016 | 0.0016 |
| F08 | 54660001 | 54760000 | 0.0021 | 0.0019 | 0.0020 |
| F08 | 54680001 | 54780000 | 0.0018 | 0.0018 | 0.0017 |
| F08 | 54700001 | 54800000 | 0.0019 | 0.0017 | 0.0017 |
| F08 | 54720001 | 54820000 | 0.0012 | 0.0010 | 0.0010 |
| F08 | 54740001 | 54840000 | 0.0019 | 0.0015 | 0.0017 |
| F08 | 54760001 | 54860000 | 0.0020 | 0.0016 | 0.0018 |
| F08 | 54780001 | 54880000 | 0.0026 | 0.0021 | 0.0024 |
| F08 | 54800001 | 54900000 | 0.0028 | 0.0025 | 0.0027 |
| F08 | 54820001 | 54920000 | 0.0029 | 0.0026 | 0.0028 |
| F08 | 54840001 | 54940000 | 0.0023 | 0.0021 | 0.0022 |
| F08 | 54860001 | 54960000 | 0.0024 | 0.0021 | 0.0023 |
| F08 | 54880001 | 54980000 | 0.0028 | 0.0025 | 0.0027 |

| Chr | Start | End | both | A1 | A2 |
|---|---|---|---|---|---|
| F08 | 54900001 | 55000000 | 0.0025 | 0.0021 | 0.0023 |
| F08 | 54920001 | 55020000 | 0.0021 | 0.0016 | 0.0018 |
| F08 | 54940001 | 55040000 | 0.0032 | 0.0024 | 0.0026 |
| F08 | 54960001 | 55060000 | 0.0029 | 0.0022 | 0.0023 |
| F08 | 54980001 | 55080000 | 0.0020 | 0.0014 | 0.0014 |
| F08 | 55000001 | 55100000 | 0.0021 | 0.0016 | 0.0016 |
| F08 | 55020001 | 55120000 | 0.0021 | 0.0017 | 0.0017 |
| F08 | 55040001 | 55140000 | 0.0015 | 0.0012 | 0.0014 |
| F08 | 55060001 | 55160000 | 0.0014 | 0.0011 | 0.0013 |
| F08 | 55080001 | 55180000 | 0.0017 | 0.0015 | 0.0016 |
| F08 | 55100001 | 55200000 | 0.0018 | 0.0015 | 0.0016 |
| F08 | 55120001 | 55220000 | 0.0024 | 0.0021 | 0.0021 |
| F08 | 55140001 | 55240000 | 0.0022 | 0.0019 | 0.0020 |
| F08 | 55160001 | 55260000 | 0.0021 | 0.0020 | 0.0019 |
| F08 | 55180001 | 55280000 | 0.0022 | 0.0020 | 0.0019 |
| F08 | 55200001 | 55300000 | 0.0020 | 0.0018 | 0.0018 |
| F08 | 55220001 | 55320000 | 0.0017 | 0.0015 | 0.0017 |
| F08 | 55240001 | 55340000 | 0.0015 | 0.0013 | 0.0014 |
| F08 | 55260001 | 55360000 | 0.0015 | 0.0013 | 0.0015 |
| F08 | 55280001 | 55380000 | 0.0016 | 0.0012 | 0.0016 |
| F08 | 55300001 | 55400000 | 0.0019 | 0.0014 | 0.0019 |
| F08 | 55320001 | 55420000 | 0.0017 | 0.0014 | 0.0017 |
| F08 | 55340001 | 55440000 | 0.0020 | 0.0016 | 0.0020 |
| F08 | 55360001 | 55460000 | 0.0024 | 0.0018 | 0.0024 |
| F08 | 55380001 | 55480000 | 0.0022 | 0.0018 | 0.0022 |
| F08 | 55400001 | 55500000 | 0.0021 | 0.0018 | 0.0021 |
| F08 | 55420001 | 55520000 | 0.0018 | 0.0016 | 0.0018 |
| F08 | 55440001 | 55540000 | 0.0018 | 0.0016 | 0.0018 |
| F08 | 55460001 | 55560000 | 0.0013 | 0.0012 | 0.0013 |
| F08 | 55480001 | 55580000 | 0.0010 | 0.0009 | 0.0010 |
| F08 | 55500001 | 55600000 | 0.0010 | 0.0008 | 0.0010 |
| F08 | 55520001 | 55620000 | 0.0019 | 0.0016 | 0.0019 |
| F08 | 55540001 | 55640000 | 0.0022 | 0.0019 | 0.0022 |
| F08 | 55560001 | 55660000 | 0.0033 | 0.0028 | 0.0031 |
| F08 | 55580001 | 55680000 | 0.0042 | 0.0035 | 0.0039 |
| F08 | 55600001 | 55700000 | 0.0042 | 0.0034 | 0.0038 |
| F08 | 55620001 | 55720000 | 0.0037 | 0.0029 | 0.0031 |
| F08 | 55640001 | 55740000 | 0.0033 | 0.0024 | 0.0027 |
| F08 | 55660001 | 55760000 | 0.0028 | 0.0021 | 0.0023 |
| F08 | 55680001 | 55780000 | 0.0021 | 0.0015 | 0.0016 |
| F08 | 55700001 | 55800000 | 0.0022 | 0.0017 | 0.0017 |
| F08 | 55720001 | 55820000 | 0.0027 | 0.0022 | 0.0024 |
| F08 | 55740001 | 55840000 | 0.0029 | 0.0024 | 0.0027 |
| F08 | 55760001 | 55860000 | 0.0031 | 0.0024 | 0.0028 |
| F08 | 55780001 | 55880000 | 0.0029 | 0.0024 | 0.0028 |
| F08 | 55800001 | 55900000 | 0.0031 | 0.0026 | 0.0031 |
| F08 | 55820001 | 55920000 | 0.0024 | 0.0019 | 0.0024 |
| F08 | 55840001 | 55940000 | 0.0032 | 0.0028 | 0.0031 |
| F08 | 55860001 | 55960000 | 0.0026 | 0.0023 | 0.0026 |
| F08 | 55880001 | 55980000 | 0.0029 | 0.0026 | 0.0028 |
| F08 | 55900001 | 56000000 | 0.0026 | 0.0022 | 0.0025 |
| F08 | 55920001 | 56020000 | 0.0026 | 0.0022 | 0.0026 |
| F08 | 55940001 | 56040000 | 0.0017 | 0.0013 | 0.0017 |
| F08 | 55960001 | 56060000 | 0.0016 | 0.0012 | 0.0016 |
| F08 | 55980001 | 56080000 | 0.0016 | 0.0013 | 0.0016 |
| F08 | 56000001 | 56100000 | 0.0017 | 0.0015 | 0.0017 |
| F08 | 56020001 | 56120000 | 0.0021 | 0.0019 | 0.0020 |
| F08 | 56040001 | 56140000 | 0.0023 | 0.0021 | 0.0022 |

| Chr | Start | End | both | A1 | A2 |
|---|---|---|---|---|---|
| F08 | 56060001 | 56160000 | 0.0030 | 0.0026 | 0.0028 |
| F08 | 56080001 | 56180000 | 0.0027 | 0.0022 | 0.0024 |
| F08 | 56100001 | 56200000 | 0.0029 | 0.0021 | 0.0026 |
| F08 | 56120001 | 56220000 | 0.0026 | 0.0018 | 0.0023 |
| F08 | 56140001 | 56240000 | 0.0028 | 0.0021 | 0.0025 |
| F08 | 56160001 | 56260000 | 0.0026 | 0.0018 | 0.0023 |
| F08 | 56180001 | 56280000 | 0.0025 | 0.0019 | 0.0023 |
| F08 | 56200001 | 56300000 | 0.0026 | 0.0021 | 0.0024 |
| F08 | 56220001 | 56320000 | 0.0027 | 0.0021 | 0.0025 |
| F08 | 56240001 | 56340000 | 0.0027 | 0.0021 | 0.0025 |
| F08 | 56260001 | 56360000 | 0.0030 | 0.0025 | 0.0028 |
| F08 | 56280001 | 56380000 | 0.0038 | 0.0030 | 0.0034 |
| F08 | 56300001 | 56400000 | 0.0032 | 0.0026 | 0.0028 |
| F08 | 56320001 | 56420000 | 0.0032 | 0.0027 | 0.0028 |
| F08 | 56340001 | 56440000 | 0.0025 | 0.0021 | 0.0022 |
| F08 | 56360001 | 56460000 | 0.0017 | 0.0014 | 0.0014 |
| F08 | 56380001 | 56480000 | 0.0015 | 0.0012 | 0.0013 |
| F08 | 56400001 | 56500000 | 0.0015 | 0.0013 | 0.0015 |
| F08 | 56420001 | 56520000 | 0.0016 | 0.0015 | 0.0016 |
| F08 | 56440001 | 56540000 | 0.0020 | 0.0017 | 0.0019 |
| F08 | 56460001 | 56560000 | 0.0022 | 0.0019 | 0.0022 |
| F08 | 56480001 | 56580000 | 0.0017 | 0.0015 | 0.0016 |
| F08 | 56500001 | 56600000 | 0.0015 | 0.0013 | 0.0014 |
| F08 | 56520001 | 56620000 | 0.0011 | 0.0009 | 0.0011 |
| F08 | 56540001 | 56640000 | 0.0010 | 0.0008 | 0.0009 |
| F08 | 56560001 | 56660000 | 0.0006 | 0.0005 | 0.0006 |
| F08 | 56580001 | 56680000 | 0.0006 | 0.0005 | 0.0006 |
| F08 | 56600001 | 56700000 | 0.0009 | 0.0008 | 0.0009 |
| F08 | 56620001 | 56720000 | 0.0008 | 0.0007 | 0.0008 |
| F08 | 56640001 | 56740000 | 0.0009 | 0.0008 | 0.0008 |
| F08 | 56660001 | 56760000 | 0.0010 | 0.0009 | 0.0010 |
| F08 | 56680001 | 56780000 | 0.0011 | 0.0009 | 0.0010 |
| F08 | 56700001 | 56800000 | 0.0009 | 0.0008 | 0.0009 |
| F08 | 56720001 | 56820000 | 0.0010 | 0.0009 | 0.0010 |
| F08 | 56740001 | 56840000 | 0.0011 | 0.0009 | 0.0011 |
| F08 | 56760001 | 56860000 | 0.0010 | 0.0008 | 0.0009 |
| F08 | 56780001 | 56880000 | 0.0011 | 0.0010 | 0.0011 |
| F08 | 56800001 | 56900000 | 0.0013 | 0.0011 | 0.0012 |
| F08 | 56820001 | 56920000 | 0.0015 | 0.0012 | 0.0015 |
| F08 | 56840001 | 56940000 | 0.0015 | 0.0014 | 0.0015 |
| F08 | 56860001 | 56960000 | 0.0017 | 0.0015 | 0.0016 |
| F08 | 56880001 | 56980000 | 0.0024 | 0.0019 | 0.0022 |
| F08 | 56900001 | 57000000 | 0.0030 | 0.0022 | 0.0026 |
| F08 | 56920001 | 57020000 | 0.0031 | 0.0022 | 0.0027 |
| F08 | 56940001 | 57040000 | 0.0032 | 0.0019 | 0.0025 |
| F08 | 56960001 | 57060000 | 0.0036 | 0.0022 | 0.0028 |
| F08 | 56980001 | 57080000 | 0.0032 | 0.0019 | 0.0025 |
| F08 | 57000001 | 57100000 | 0.0034 | 0.0022 | 0.0027 |
| F08 | 57020001 | 57120000 | 0.0038 | 0.0027 | 0.0032 |
| F08 | 57040001 | 57140000 | 0.0049 | 0.0040 | 0.0046 |
| F08 | 57060001 | 57160000 | 0.0049 | 0.0040 | 0.0046 |
| F08 | 57080001 | 57180000 | 0.0043 | 0.0037 | 0.0041 |
| F08 | 57100001 | 57200000 | 0.0034 | 0.0028 | 0.0032 |
| F08 | 57120001 | 57220000 | 0.0027 | 0.0022 | 0.0026 |
| F08 | 57140001 | 57240000 | 0.0023 | 0.0020 | 0.0022 |
| F08 | 57160001 | 57260000 | 0.0031 | 0.0027 | 0.0029 |
| F08 | 57180001 | 57280000 | 0.0034 | 0.0030 | 0.0033 |
| F08 | 57200001 | 57300000 | 0.0037 | 0.0031 | 0.0034 |

| Chr | Start | End | both | A1 | A2 |
|---|---|---|---|---|---|
| F08 | 57220001 | 57320000 | 0.0038 | 0.0032 | 0.0034 |
| F08 | 57240001 | 57340000 | 0.0031 | 0.0025 | 0.0026 |
| F08 | 57260001 | 57360000 | 0.0025 | 0.0019 | 0.0021 |
| F08 | 57280001 | 57380000 | 0.0023 | 0.0018 | 0.0019 |
| F08 | 57300001 | 57400000 | 0.0030 | 0.0024 | 0.0025 |
| F08 | 57320001 | 57420000 | 0.0034 | 0.0029 | 0.0031 |
| F08 | 57340001 | 57440000 | 0.0040 | 0.0034 | 0.0038 |
| F08 | 57360001 | 57460000 | 0.0040 | 0.0034 | 0.0038 |
| F08 | 57380001 | 57480000 | 0.0044 | 0.0037 | 0.0043 |
| F08 | 57400001 | 57500000 | 0.0037 | 0.0031 | 0.0037 |
| F08 | 57420001 | 57520000 | 0.0027 | 0.0021 | 0.0027 |
| F08 | 57440001 | 57540000 | 0.0020 | 0.0016 | 0.0019 |
| F08 | 57460001 | 57560000 | 0.0016 | 0.0013 | 0.0016 |
| F08 | 57480001 | 57580000 | 0.0011 | 0.0009 | 0.0010 |
| F08 | 57500001 | 57600000 | 0.0020 | 0.0016 | 0.0020 |
| F08 | 57520001 | 57620000 | 0.0023 | 0.0018 | 0.0023 |
| F08 | 57540001 | 57640000 | 0.0031 | 0.0022 | 0.0028 |
| F08 | 57560001 | 57660000 | 0.0041 | 0.0030 | 0.0036 |
| F08 | 57580001 | 57680000 | 0.0044 | 0.0032 | 0.0039 |
| F08 | 57600001 | 57700000 | 0.0041 | 0.0031 | 0.0035 |
| F08 | 57620001 | 57720000 | 0.0045 | 0.0033 | 0.0037 |
| F08 | 57640001 | 57740000 | 0.0038 | 0.0029 | 0.0031 |
| F08 | 57660001 | 57760000 | 0.0041 | 0.0032 | 0.0034 |
| F08 | 57680001 | 57780000 | 0.0042 | 0.0032 | 0.0035 |
| F08 | 57700001 | 57800000 | 0.0047 | 0.0034 | 0.0038 |
| F08 | 57720001 | 57820000 | 0.0045 | 0.0033 | 0.0037 |
| F08 | 57740001 | 57840000 | 0.0052 | 0.0036 | 0.0042 |
| F08 | 57760001 | 57860000 | 0.0046 | 0.0033 | 0.0037 |
| F08 | 57780001 | 57880000 | 0.0041 | 0.0030 | 0.0033 |
| F08 | 57800001 | 57900000 | 0.0031 | 0.0022 | 0.0026 |
| F08 | 57820001 | 57920000 | 0.0030 | 0.0022 | 0.0026 |
| F08 | 57840001 | 57940000 | 0.0024 | 0.0021 | 0.0023 |
| F08 | 57860001 | 57960000 | 0.0019 | 0.0014 | 0.0018 |
| F08 | 57880001 | 57980000 | 0.0028 | 0.0022 | 0.0025 |
| F08 | 57900001 | 58000000 | 0.0035 | 0.0028 | 0.0032 |
| F08 | 57920001 | 58020000 | 0.0039 | 0.0032 | 0.0037 |
| F08 | 57940001 | 58040000 | 0.0040 | 0.0031 | 0.0036 |
| F08 | 57960001 | 58060000 | 0.0038 | 0.0033 | 0.0036 |
| F08 | 57980001 | 58080000 | 0.0033 | 0.0027 | 0.0031 |
| F08 | 58000001 | 58100000 | 0.0028 | 0.0023 | 0.0026 |
| F08 | 58020001 | 58120000 | 0.0033 | 0.0028 | 0.0030 |
| F08 | 58040001 | 58140000 | 0.0033 | 0.0028 | 0.0028 |
| F08 | 58060001 | 58160000 | 0.0034 | 0.0027 | 0.0028 |
| F08 | 58080001 | 58180000 | 0.0038 | 0.0027 | 0.0031 |
| F08 | 58100001 | 58200000 | 0.0047 | 0.0034 | 0.0039 |
| F08 | 58120001 | 58220000 | 0.0051 | 0.0038 | 0.0042 |
| F08 | 58140001 | 58240000 | 0.0051 | 0.0038 | 0.0043 |
| F08 | 58160001 | 58260000 | 0.0049 | 0.0037 | 0.0042 |
| F08 | 58180001 | 58280000 | 0.0049 | 0.0040 | 0.0042 |
| F08 | 58200001 | 58300000 | 0.0040 | 0.0034 | 0.0034 |
| F08 | 58220001 | 58320000 | 0.0023 | 0.0019 | 0.0020 |
| F08 | 58240001 | 58340000 | 0.0023 | 0.0021 | 0.0020 |
| F08 | 58260001 | 58360000 | 0.0022 | 0.0020 | 0.0020 |
| F08 | 58280001 | 58380000 | 0.0016 | 0.0014 | 0.0014 |
| F08 | 58300001 | 58400000 | 0.0022 | 0.0019 | 0.0020 |
| F08 | 58320001 | 58420000 | 0.0020 | 0.0017 | 0.0018 |
| F08 | 58340001 | 58440000 | 0.0017 | 0.0015 | 0.0016 |
| F08 | 58360001 | 58460000 | 0.0029 | 0.0026 | 0.0027 |

| Chr | Start | End | both | A1 | A2 |
|---|---|---|---|---|---|
| F08 | 58380001 | 58480000 | 0.0031 | 0.0028 | 0.0029 |
| F08 | 58400001 | 58500000 | 0.0028 | 0.0023 | 0.0026 |
| F08 | 58420001 | 58520000 | 0.0037 | 0.0030 | 0.0035 |
| F08 | 58440001 | 58540000 | 0.0042 | 0.0034 | 0.0039 |
| F08 | 58460001 | 58560000 | 0.0036 | 0.0028 | 0.0032 |
| F08 | 58480001 | 58580000 | 0.0036 | 0.0027 | 0.0031 |
| F08 | 58500001 | 58600000 | 0.0031 | 0.0024 | 0.0028 |
| F08 | 58520001 | 58620000 | 0.0024 | 0.0018 | 0.0021 |
| F08 | 58540001 | 58640000 | 0.0031 | 0.0026 | 0.0027 |
| F08 | 58560001 | 58660000 | 0.0028 | 0.0024 | 0.0025 |
| F08 | 58580001 | 58680000 | 0.0040 | 0.0037 | 0.0038 |
| F08 | 58600001 | 58700000 | 0.0042 | 0.0038 | 0.0039 |
| F08 | 58620001 | 58720000 | 0.0047 | 0.0043 | 0.0045 |
| F08 | 58640001 | 58740000 | 0.0038 | 0.0033 | 0.0036 |
| F08 | 58660001 | 58760000 | 0.0038 | 0.0032 | 0.0035 |
| F08 | 58680001 | 58780000 | 0.0026 | 0.0018 | 0.0022 |
| F08 | 58700001 | 58800000 | 0.0027 | 0.0020 | 0.0024 |
| F08 | 58720001 | 58820000 | 0.0022 | 0.0017 | 0.0019 |
| F08 | 58740001 | 58840000 | 0.0025 | 0.0017 | 0.0018 |
| F08 | 58760001 | 58860000 | 0.0029 | 0.0023 | 0.0024 |
| F08 | 58780001 | 58880000 | 0.0028 | 0.0022 | 0.0023 |
| F08 | 58800001 | 58900000 | 0.0026 | 0.0021 | 0.0022 |
| F08 | 58820001 | 58920000 | 0.0029 | 0.0023 | 0.0024 |
| F08 | 58840001 | 58940000 | 0.0023 | 0.0020 | 0.0021 |
| F08 | 58860001 | 58960000 | 0.0017 | 0.0015 | 0.0015 |
| F08 | 58880001 | 58980000 | 0.0023 | 0.0020 | 0.0021 |
| F08 | 58900001 | 59000000 | 0.0023 | 0.0019 | 0.0021 |
| F08 | 58920001 | 59020000 | 0.0022 | 0.0016 | 0.0018 |
| F08 | 58940001 | 59040000 | 0.0027 | 0.0021 | 0.0023 |
| F08 | 58960001 | 59060000 | 0.0036 | 0.0029 | 0.0032 |
| F08 | 58980001 | 59080000 | 0.0035 | 0.0027 | 0.0031 |
| F08 | 59000001 | 59100000 | 0.0035 | 0.0027 | 0.0032 |
| F08 | 59020001 | 59120000 | 0.0033 | 0.0026 | 0.0031 |
| F08 | 59040001 | 59140000 | 0.0028 | 0.0022 | 0.0027 |
| F08 | 59060001 | 59160000 | 0.0026 | 0.0019 | 0.0023 |
| F08 | 59080001 | 59180000 | 0.0027 | 0.0018 | 0.0023 |
| F08 | 59100001 | 59200000 | 0.0026 | 0.0017 | 0.0023 |
| F08 | 59120001 | 59220000 | 0.0027 | 0.0018 | 0.0023 |
| F08 | 59140001 | 59240000 | 0.0029 | 0.0021 | 0.0025 |
| F08 | 59160001 | 59260000 | 0.0027 | 0.0020 | 0.0024 |
| F08 | 59180001 | 59280000 | 0.0027 | 0.0023 | 0.0025 |
| F08 | 59200001 | 59300000 | 0.0032 | 0.0027 | 0.0030 |
| F08 | 59220001 | 59320000 | 0.0041 | 0.0033 | 0.0038 |
| F08 | 59240001 | 59340000 | 0.0046 | 0.0035 | 0.0041 |
| F08 | 59260001 | 59360000 | 0.0038 | 0.0029 | 0.0034 |
| F08 | 59280001 | 59380000 | 0.0030 | 0.0022 | 0.0027 |
| F08 | 59300001 | 59400000 | 0.0023 | 0.0017 | 0.0020 |
| F08 | 59320001 | 59420000 | 0.0013 | 0.0010 | 0.0011 |
| F08 | 59340001 | 59440000 | 0.0012 | 0.0010 | 0.0010 |
| F08 | 59360001 | 59460000 | 0.0020 | 0.0017 | 0.0017 |
| F08 | 59380001 | 59480000 | 0.0026 | 0.0022 | 0.0024 |
| F08 | 59400001 | 59500000 | 0.0029 | 0.0023 | 0.0026 |
| F08 | 59420001 | 59520000 | 0.0030 | 0.0021 | 0.0024 |
| F08 | 59440001 | 59540000 | 0.0028 | 0.0019 | 0.0023 |
| F08 | 59460001 | 59560000 | 0.0021 | 0.0012 | 0.0015 |
| F08 | 59480001 | 59580000 | 0.0019 | 0.0009 | 0.0012 |
| F08 | 59500001 | 59600000 | 0.0016 | 0.0008 | 0.0010 |
| F08 | 59520001 | 59620000 | 0.0029 | 0.0021 | 0.0023 |

| Chr | Start | End | both | A1 | A2 |
|---|---|---|---|---|---|
| F08 | 59540001 | 59640000 | 0.0031 | 0.0023 | 0.0025 |
| F08 | 59560001 | 59660000 | 0.0039 | 0.0028 | 0.0032 |
| F08 | 59580001 | 59680000 | 0.0038 | 0.0029 | 0.0032 |
| F08 | 59600001 | 59700000 | 0.0044 | 0.0033 | 0.0035 |
| F08 | 59620001 | 59720000 | 0.0029 | 0.0021 | 0.0023 |
| F08 | 59640001 | 59740000 | 0.0025 | 0.0018 | 0.0021 |
| F08 | 59660001 | 59760000 | 0.0024 | 0.0018 | 0.0020 |
| F08 | 59680001 | 59780000 | 0.0032 | 0.0026 | 0.0029 |
| F08 | 59700001 | 59800000 | 0.0035 | 0.0029 | 0.0031 |
| F08 | 59720001 | 59820000 | 0.0043 | 0.0035 | 0.0037 |
| F08 | 59740001 | 59840000 | 0.0057 | 0.0047 | 0.0050 |
| F08 | 59760001 | 59860000 | 0.0054 | 0.0044 | 0.0046 |
| F08 | 59780001 | 59880000 | 0.0041 | 0.0033 | 0.0034 |
| F08 | 59800001 | 59900000 | 0.0041 | 0.0032 | 0.0035 |
| F08 | 59820001 | 59920000 | 0.0066 | 0.0055 | 0.0060 |
| F08 | 59840001 | 59940000 | 0.0060 | 0.0051 | 0.0056 |
| F08 | 59860001 | 59960000 | 0.0060 | 0.0050 | 0.0055 |
| F08 | 59880001 | 59980000 | 0.0065 | 0.0053 | 0.0058 |
| F08 | 59900001 | 60000000 | 0.0058 | 0.0046 | 0.0051 |
| F08 | 59920001 | 60020000 | 0.0028 | 0.0019 | 0.0022 |
| F08 | 59940001 | 60040000 | 0.0017 | 0.0010 | 0.0011 |
| F08 | 59960001 | 60060000 | 0.0022 | 0.0016 | 0.0018 |
| F08 | 59980001 | 60080000 | 0.0025 | 0.0020 | 0.0023 |
| F08 | 60000001 | 60100000 | 0.0025 | 0.0021 | 0.0024 |
| F08 | 60020001 | 60120000 | 0.0028 | 0.0024 | 0.0027 |
| F08 | 60040001 | 60140000 | 0.0031 | 0.0025 | 0.0029 |
| F08 | 60060001 | 60160000 | 0.0026 | 0.0021 | 0.0024 |
| F08 | 60080001 | 60180000 | 0.0023 | 0.0016 | 0.0020 |
| F08 | 60100001 | 60200000 | 0.0025 | 0.0020 | 0.0020 |
| F08 | 60120001 | 60220000 | 0.0026 | 0.0019 | 0.0019 |
| F08 | 60140001 | 60240000 | 0.0028 | 0.0021 | 0.0021 |
| F08 | 60160001 | 60260000 | 0.0023 | 0.0018 | 0.0017 |
| F08 | 60180001 | 60280000 | 0.0031 | 0.0025 | 0.0024 |
| F08 | 60200001 | 60300000 | 0.0028 | 0.0021 | 0.0024 |
| F08 | 60220001 | 60320000 | 0.0026 | 0.0020 | 0.0023 |
| F08 | 60240001 | 60340000 | 0.0022 | 0.0017 | 0.0021 |
| F08 | 60260001 | 60360000 | 0.0026 | 0.0018 | 0.0022 |
| F08 | 60280001 | 60380000 | 0.0016 | 0.0010 | 0.0013 |
| F08 | 60300001 | 60400000 | 0.0021 | 0.0014 | 0.0018 |
| F08 | 60320001 | 60420000 | 0.0024 | 0.0016 | 0.0021 |
| F08 | 60340001 | 60440000 | 0.0031 | 0.0022 | 0.0026 |
| F08 | 60360001 | 60460000 | 0.0033 | 0.0025 | 0.0030 |
| F08 | 60380001 | 60480000 | 0.0038 | 0.0029 | 0.0035 |
| F08 | 60400001 | 60500000 | 0.0032 | 0.0024 | 0.0030 |
| F08 | 60420001 | 60520000 | 0.0025 | 0.0019 | 0.0023 |
| F08 | 60440001 | 60540000 | 0.0021 | 0.0017 | 0.0019 |
| F08 | 60460001 | 60560000 | 0.0022 | 0.0017 | 0.0019 |
| F08 | 60480001 | 60580000 | 0.0021 | 0.0016 | 0.0015 |
| F08 | 60500001 | 60600000 | 0.0024 | 0.0018 | 0.0017 |
| F08 | 60520001 | 60620000 | 0.0028 | 0.0021 | 0.0022 |
| F08 | 60540001 | 60640000 | 0.0030 | 0.0022 | 0.0023 |
| F08 | 60560001 | 60660000 | 0.0034 | 0.0027 | 0.0028 |
| F08 | 60580001 | 60680000 | 0.0034 | 0.0026 | 0.0026 |
| F08 | 60600001 | 60700000 | 0.0036 | 0.0029 | 0.0029 |
| F08 | 60620001 | 60720000 | 0.0037 | 0.0029 | 0.0028 |
| F08 | 60640001 | 60740000 | 0.0040 | 0.0032 | 0.0031 |
| F08 | 60660001 | 60760000 | 0.0031 | 0.0023 | 0.0022 |
| F08 | 60680001 | 60780000 | 0.0031 | 0.0025 | 0.0024 |

| Chr | Start | End | both | A1 | A2 |
|---|---|---|---|---|---|
| F08 | 60700001 | 60800000 | 0.0033 | 0.0025 | 0.0025 |
| F08 | 60720001 | 60820000 | 0.0047 | 0.0039 | 0.0038 |
| F08 | 60740001 | 60840000 | 0.0047 | 0.0038 | 0.0039 |
| F08 | 60760001 | 60860000 | 0.0047 | 0.0038 | 0.0039 |
| F08 | 60780001 | 60880000 | 0.0050 | 0.0040 | 0.0041 |
| F08 | 60800001 | 60900000 | 0.0052 | 0.0044 | 0.0045 |
| F08 | 60820001 | 60920000 | 0.0042 | 0.0033 | 0.0037 |
| F08 | 60840001 | 60940000 | 0.0038 | 0.0028 | 0.0032 |
| F08 | 60860001 | 60960000 | 0.0040 | 0.0033 | 0.0035 |
| F08 | 60880001 | 60980000 | 0.0037 | 0.0030 | 0.0033 |
| F08 | 60900001 | 61000000 | 0.0045 | 0.0038 | 0.0040 |
| F08 | 60920001 | 61020000 | 0.0038 | 0.0034 | 0.0033 |
| F08 | 60940001 | 61040000 | 0.0037 | 0.0032 | 0.0032 |
| F08 | 60960001 | 61060000 | 0.0042 | 0.0032 | 0.0033 |
| F08 | 60980001 | 61080000 | 0.0041 | 0.0028 | 0.0030 |
| F08 | 61000001 | 61100000 | 0.0026 | 0.0013 | 0.0015 |
| F08 | 61020001 | 61120000 | 0.0027 | 0.0013 | 0.0016 |
| F08 | 61040001 | 61140000 | 0.0025 | 0.0012 | 0.0015 |
| F08 | 61060001 | 61160000 | 0.0018 | 0.0009 | 0.0011 |
| F08 | 61080001 | 61180000 | 0.0013 | 0.0008 | 0.0009 |
| F08 | 61100001 | 61200000 | 0.0014 | 0.0009 | 0.0009 |
| F08 | 61120001 | 61220000 | 0.0015 | 0.0009 | 0.0009 |
| F08 | 61140001 | 61240000 | 0.0020 | 0.0012 | 0.0014 |
| F08 | 61160001 | 61260000 | 0.0025 | 0.0016 | 0.0019 |
| F08 | 61180001 | 61280000 | 0.0034 | 0.0024 | 0.0028 |
| F08 | 61200001 | 61300000 | 0.0045 | 0.0031 | 0.0036 |
| F08 | 61220001 | 61320000 | 0.0057 | 0.0043 | 0.0047 |
| F08 | 61240001 | 61340000 | 0.0052 | 0.0040 | 0.0043 |
| F08 | 61260001 | 61360000 | 0.0049 | 0.0036 | 0.0038 |
| F08 | 61280001 | 61380000 | 0.0044 | 0.0032 | 0.0033 |
| F08 | 61300001 | 61400000 | 0.0038 | 0.0029 | 0.0030 |
| F08 | 61320001 | 61420000 | 0.0038 | 0.0029 | 0.0033 |
| F08 | 61340001 | 61440000 | 0.0042 | 0.0033 | 0.0037 |
| F08 | 61360001 | 61460000 | 0.0048 | 0.0040 | 0.0044 |
| F08 | 61380001 | 61480000 | 0.0046 | 0.0038 | 0.0043 |
| F08 | 61400001 | 61500000 | 0.0042 | 0.0036 | 0.0040 |
| F08 | 61420001 | 61520000 | 0.0031 | 0.0026 | 0.0029 |
| F08 | 61440001 | 61540000 | 0.0034 | 0.0027 | 0.0032 |
| F08 | 61460001 | 61560000 | 0.0028 | 0.0023 | 0.0026 |
| F08 | 61480001 | 61580000 | 0.0042 | 0.0035 | 0.0038 |
| F08 | 61500001 | 61600000 | 0.0051 | 0.0042 | 0.0047 |
| F08 | 61520001 | 61620000 | 0.0052 | 0.0040 | 0.0045 |
| F08 | 61540001 | 61640000 | 0.0048 | 0.0036 | 0.0040 |
| F08 | 61560001 | 61660000 | 0.0049 | 0.0036 | 0.0040 |
| F08 | 61580001 | 61680000 | 0.0034 | 0.0022 | 0.0026 |
| F08 | 61600001 | 61700000 | 0.0025 | 0.0016 | 0.0017 |
| F08 | 61620001 | 61720000 | 0.0027 | 0.0017 | 0.0019 |
| F08 | 61640001 | 61740000 | 0.0036 | 0.0024 | 0.0026 |
| F08 | 61660001 | 61760000 | 0.0045 | 0.0033 | 0.0036 |
| F08 | 61680001 | 61780000 | 0.0044 | 0.0032 | 0.0035 |
| F08 | 61700001 | 61800000 | 0.0040 | 0.0028 | 0.0031 |
| F08 | 61720001 | 61820000 | 0.0034 | 0.0024 | 0.0027 |
| F08 | 61740001 | 61840000 | 0.0023 | 0.0015 | 0.0017 |
| F08 | 61760001 | 61860000 | 0.0023 | 0.0012 | 0.0014 |
| F08 | 61780001 | 61880000 | 0.0031 | 0.0018 | 0.0021 |
| F08 | 61800001 | 61900000 | 0.0036 | 0.0021 | 0.0023 |
| F08 | 61820001 | 61920000 | 0.0038 | 0.0022 | 0.0025 |
| F08 | 61840001 | 61940000 | 0.0040 | 0.0025 | 0.0027 |

| Chr | Start | End | both | A1 | A2 |
|---|---|---|---|---|---|
| F08 | 61860001 | 61960000 | 0.0037 | 0.0022 | 0.0022 |
| F08 | 61880001 | 61980000 | 0.0031 | 0.0017 | 0.0017 |
| F08 | 61900001 | 62000000 | 0.0037 | 0.0024 | 0.0026 |
| F08 | 61920001 | 62020000 | 0.0037 | 0.0026 | 0.0028 |
| F08 | 61940001 | 62040000 | 0.0037 | 0.0028 | 0.0029 |
| F08 | 61960001 | 62060000 | 0.0044 | 0.0039 | 0.0039 |
| F08 | 61980001 | 62080000 | 0.0047 | 0.0041 | 0.0042 |
| F08 | 62000001 | 62100000 | 0.0041 | 0.0033 | 0.0033 |
| F08 | 62020001 | 62120000 | 0.0046 | 0.0034 | 0.0036 |
| F08 | 62040001 | 62140000 | 0.0046 | 0.0032 | 0.0034 |
| F08 | 62060001 | 62160000 | 0.0033 | 0.0016 | 0.0020 |
| F08 | 62080001 | 62180000 | 0.0037 | 0.0016 | 0.0020 |
| F08 | 62100001 | 62200000 | 0.0037 | 0.0017 | 0.0021 |
| F08 | 62120001 | 62220000 | 0.0030 | 0.0013 | 0.0015 |
| F08 | 62140001 | 62240000 | 0.0038 | 0.0022 | 0.0025 |
| F08 | 62160001 | 62260000 | 0.0044 | 0.0028 | 0.0030 |
| F08 | 62180001 | 62280000 | 0.0048 | 0.0036 | 0.0036 |
| F08 | 62200001 | 62300000 | 0.0052 | 0.0039 | 0.0039 |
| F08 | 62220001 | 62320000 | 0.0063 | 0.0046 | 0.0048 |
| F08 | 62240001 | 62340000 | 0.0057 | 0.0040 | 0.0040 |
| F08 | 62260001 | 62360000 | 0.0053 | 0.0036 | 0.0039 |
| F08 | 62280001 | 62380000 | 0.0058 | 0.0040 | 0.0045 |
| F08 | 62300001 | 62400000 | 0.0057 | 0.0041 | 0.0048 |
| F08 | 62320001 | 62420000 | 0.0052 | 0.0039 | 0.0045 |
| F08 | 62340001 | 62440000 | 0.0050 | 0.0039 | 0.0046 |
| F08 | 62360001 | 62460000 | 0.0054 | 0.0040 | 0.0047 |
| F08 | 62380001 | 62480000 | 0.0046 | 0.0032 | 0.0037 |
| F08 | 62400001 | 62500000 | 0.0047 | 0.0034 | 0.0038 |
| F08 | 62420001 | 62520000 | 0.0046 | 0.0035 | 0.0039 |
| F08 | 62440001 | 62540000 | 0.0049 | 0.0038 | 0.0041 |
| F08 | 62460001 | 62560000 | 0.0041 | 0.0033 | 0.0035 |
| F08 | 62480001 | 62580000 | 0.0040 | 0.0033 | 0.0036 |
| F08 | 62500001 | 62600000 | 0.0037 | 0.0030 | 0.0033 |
| F08 | 62520001 | 62620000 | 0.0034 | 0.0026 | 0.0029 |
| F08 | 62540001 | 62640000 | 0.0036 | 0.0025 | 0.0029 |
| F08 | 62560001 | 62660000 | 0.0040 | 0.0026 | 0.0031 |
| F08 | 62580001 | 62680000 | 0.0037 | 0.0025 | 0.0029 |
| F08 | 62600001 | 62700000 | 0.0034 | 0.0023 | 0.0027 |
| F08 | 62620001 | 62720000 | 0.0034 | 0.0024 | 0.0027 |
| F08 | 62640001 | 62740000 | 0.0033 | 0.0024 | 0.0027 |
| F08 | 62660001 | 62760000 | 0.0028 | 0.0022 | 0.0025 |
| F08 | 62680001 | 62780000 | 0.0024 | 0.0016 | 0.0018 |
| F08 | 62700001 | 62800000 | 0.0024 | 0.0016 | 0.0017 |
| F08 | 62720001 | 62820000 | 0.0024 | 0.0015 | 0.0017 |
| F08 | 62740001 | 62840000 | 0.0019 | 0.0011 | 0.0013 |
| F08 | 62760001 | 62860000 | 0.0024 | 0.0014 | 0.0017 |
| F08 | 62780001 | 62880000 | 0.0027 | 0.0019 | 0.0021 |
| F08 | 62800001 | 62900000 | 0.0027 | 0.0019 | 0.0022 |
| F08 | 62820001 | 62920000 | 0.0027 | 0.0020 | 0.0023 |
| F08 | 62840001 | 62940000 | 0.0037 | 0.0026 | 0.0029 |
| F08 | 62860001 | 62960000 | 0.0040 | 0.0027 | 0.0030 |
| F08 | 62880001 | 62980000 | 0.0040 | 0.0027 | 0.0031 |
| F08 | 62900001 | 63000000 | 0.0045 | 0.0030 | 0.0034 |
| F08 | 62920001 | 63020000 | 0.0043 | 0.0029 | 0.0032 |
| F08 | 62940001 | 63040000 | 0.0036 | 0.0025 | 0.0029 |
| F08 | 62960001 | 63060000 | 0.0031 | 0.0023 | 0.0027 |
| F08 | 62980001 | 63080000 | 0.0031 | 0.0023 | 0.0027 |
| F08 | 63000001 | 63100000 | 0.0026 | 0.0018 | 0.0021 |

| Chr | Start | End | both | A1 | A2 |
|---|---|---|---|---|---|
| F08 | 63020001 | 63120000 | 0.0025 | 0.0018 | 0.0022 |
| F08 | 63040001 | 63140000 | 0.0022 | 0.0016 | 0.0019 |
| F08 | 63060001 | 63160000 | 0.0020 | 0.0015 | 0.0017 |
| F08 | 63080001 | 63180000 | 0.0021 | 0.0016 | 0.0017 |
| F08 | 63100001 | 63200000 | 0.0022 | 0.0017 | 0.0019 |
| F08 | 63120001 | 63220000 | 0.0028 | 0.0021 | 0.0022 |
| F08 | 63140001 | 63240000 | 0.0032 | 0.0025 | 0.0026 |
| F08 | 63160001 | 63260000 | 0.0039 | 0.0030 | 0.0031 |
| F08 | 63180001 | 63280000 | 0.0041 | 0.0033 | 0.0034 |
| F08 | 63200001 | 63300000 | 0.0042 | 0.0033 | 0.0033 |
| F08 | 63220001 | 63320000 | 0.0043 | 0.0033 | 0.0034 |
| F08 | 63240001 | 63340000 | 0.0041 | 0.0029 | 0.0029 |
| F08 | 63260001 | 63360000 | 0.0037 | 0.0024 | 0.0025 |
| F08 | 63280001 | 63380000 | 0.0040 | 0.0025 | 0.0027 |
| F08 | 63300001 | 63400000 | 0.0038 | 0.0024 | 0.0027 |
| F08 | 63320001 | 63420000 | 0.0033 | 0.0021 | 0.0026 |
| F08 | 63340001 | 63440000 | 0.0035 | 0.0025 | 0.0031 |
| F08 | 63360001 | 63460000 | 0.0042 | 0.0033 | 0.0039 |
| F08 | 63380001 | 63480000 | 0.0035 | 0.0029 | 0.0033 |
| F08 | 63400001 | 63500000 | 0.0037 | 0.0031 | 0.0035 |
| F08 | 63420001 | 63520000 | 0.0034 | 0.0028 | 0.0032 |
| F08 | 63440001 | 63540000 | 0.0043 | 0.0037 | 0.0040 |
| F08 | 63460001 | 63560000 | 0.0041 | 0.0036 | 0.0038 |
| F08 | 63480001 | 63580000 | 0.0040 | 0.0035 | 0.0037 |
| F08 | 63500001 | 63600000 | 0.0042 | 0.0036 | 0.0037 |
| F08 | 63520001 | 63620000 | 0.0048 | 0.0040 | 0.0043 |
| F08 | 63540001 | 63640000 | 0.0047 | 0.0037 | 0.0041 |
| F08 | 63560001 | 63660000 | 0.0044 | 0.0033 | 0.0037 |
| F08 | 63580001 | 63680000 | 0.0043 | 0.0032 | 0.0036 |
| F08 | 63600001 | 63700000 | 0.0041 | 0.0031 | 0.0035 |
| F08 | 63620001 | 63720000 | 0.0036 | 0.0028 | 0.0031 |
| F08 | 63640001 | 63740000 | 0.0022 | 0.0017 | 0.0019 |
| F08 | 63660001 | 63760000 | 0.0028 | 0.0021 | 0.0025 |
| F08 | 63680001 | 63780000 | 0.0031 | 0.0024 | 0.0027 |
| F08 | 63700001 | 63800000 | 0.0031 | 0.0022 | 0.0024 |
| F08 | 63720001 | 63820000 | 0.0037 | 0.0027 | 0.0029 |
| F08 | 63740001 | 63840000 | 0.0043 | 0.0033 | 0.0036 |
| F08 | 63760001 | 63860000 | 0.0037 | 0.0029 | 0.0030 |
| F08 | 63780001 | 63880000 | 0.0040 | 0.0031 | 0.0034 |
| F08 | 63800001 | 63900000 | 0.0046 | 0.0038 | 0.0042 |
| F08 | 63820001 | 63920000 | 0.0047 | 0.0040 | 0.0043 |
| F08 | 63840001 | 63940000 | 0.0039 | 0.0034 | 0.0035 |
| F08 | 63860001 | 63960000 | 0.0047 | 0.0039 | 0.0041 |
| F08 | 63880001 | 63980000 | 0.0051 | 0.0042 | 0.0044 |
| F08 | 63900001 | 64000000 | 0.0042 | 0.0033 | 0.0033 |
| F08 | 63920001 | 64020000 | 0.0037 | 0.0029 | 0.0030 |
| F08 | 63940001 | 64040000 | 0.0044 | 0.0034 | 0.0037 |
| F08 | 63960001 | 64060000 | 0.0038 | 0.0030 | 0.0031 |
| F08 | 63980001 | 64080000 | 0.0031 | 0.0023 | 0.0025 |
| F08 | 64000001 | 64100000 | 0.0032 | 0.0025 | 0.0027 |
| F08 | 64020001 | 64120000 | 0.0029 | 0.0023 | 0.0025 |
| F08 | 64040001 | 64140000 | 0.0021 | 0.0017 | 0.0018 |
| F08 | 64060001 | 64160000 | 0.0024 | 0.0019 | 0.0022 |
| F08 | 64080001 | 64180000 | 0.0021 | 0.0018 | 0.0021 |
| F08 | 64100001 | 64200000 | 0.0023 | 0.0018 | 0.0021 |
| F08 | 64120001 | 64220000 | 0.0034 | 0.0027 | 0.0030 |
| F08 | 64140001 | 64240000 | 0.0035 | 0.0027 | 0.0032 |
| F08 | 64160001 | 64260000 | 0.0043 | 0.0034 | 0.0038 |

| Chr | Start | End | both | A1 | A2 |
|---|---|---|---|---|---|
| F08 | 64180001 | 64280000 | 0.0041 | 0.0032 | 0.0036 |
| F08 | 64200001 | 64300000 | 0.0044 | 0.0036 | 0.0040 |
| F08 | 64220001 | 64320000 | 0.0031 | 0.0025 | 0.0028 |
| F08 | 64240001 | 64340000 | 0.0028 | 0.0023 | 0.0026 |
| F08 | 64260001 | 64360000 | 0.0017 | 0.0013 | 0.0015 |
| F08 | 64280001 | 64380000 | 0.0019 | 0.0018 | 0.0018 |
| F08 | 64300001 | 64400000 | 0.0014 | 0.0014 | 0.0013 |
| F08 | 64320001 | 64420000 | 0.0014 | 0.0013 | 0.0013 |
| F08 | 64340001 | 64440000 | 0.0015 | 0.0015 | 0.0014 |
| F08 | 64360001 | 64460000 | 0.0014 | 0.0014 | 0.0014 |
| F08 | 64380001 | 64480000 | 0.0015 | 0.0013 | 0.0014 |
| F08 | 64400001 | 64500000 | 0.0032 | 0.0027 | 0.0029 |
| F08 | 64420001 | 64520000 | 0.0034 | 0.0029 | 0.0031 |
| F08 | 64440001 | 64540000 | 0.0039 | 0.0033 | 0.0036 |
| F08 | 64460001 | 64560000 | 0.0036 | 0.0031 | 0.0033 |
| F08 | 64480001 | 64580000 | 0.0037 | 0.0030 | 0.0032 |
| F08 | 64500001 | 64600000 | 0.0025 | 0.0017 | 0.0019 |
| F08 | 64520001 | 64620000 | 0.0038 | 0.0031 | 0.0032 |
| F08 | 64540001 | 64640000 | 0.0035 | 0.0028 | 0.0028 |
| F08 | 64560001 | 64660000 | 0.0044 | 0.0036 | 0.0038 |
| F08 | 64580001 | 64680000 | 0.0046 | 0.0041 | 0.0042 |
| F08 | 64600001 | 64700000 | 0.0042 | 0.0038 | 0.0039 |
| F08 | 64620001 | 64720000 | 0.0027 | 0.0025 | 0.0026 |
| F08 | 64640001 | 64740000 | 0.0024 | 0.0022 | 0.0023 |
| F08 | 64660001 | 64760000 | 0.0017 | 0.0015 | 0.0016 |
| F08 | 64680001 | 64780000 | 0.0013 | 0.0011 | 0.0012 |
| F08 | 64700001 | 64800000 | 0.0012 | 0.0011 | 0.0012 |
| F08 | 64720001 | 64820000 | 0.0014 | 0.0012 | 0.0013 |
| F08 | 64740001 | 64840000 | 0.0015 | 0.0013 | 0.0014 |
| F08 | 64760001 | 64860000 | 0.0019 | 0.0015 | 0.0016 |
| F08 | 64780001 | 64880000 | 0.0028 | 0.0024 | 0.0024 |
| F08 | 64800001 | 64900000 | 0.0034 | 0.0029 | 0.0030 |
| F08 | 64820001 | 64920000 | 0.0035 | 0.0030 | 0.0031 |
| F08 | 64840001 | 64940000 | 0.0040 | 0.0034 | 0.0035 |
| F08 | 64860001 | 64960000 | 0.0047 | 0.0040 | 0.0041 |
| F08 | 64880001 | 64980000 | 0.0038 | 0.0032 | 0.0033 |
| F08 | 64900001 | 65000000 | 0.0036 | 0.0030 | 0.0031 |
| F08 | 64920001 | 65020000 | 0.0034 | 0.0029 | 0.0029 |
| F08 | 64940001 | 65040000 | 0.0032 | 0.0027 | 0.0027 |
| F08 | 64960001 | 65060000 | 0.0019 | 0.0018 | 0.0016 |
| F08 | 64980001 | 65080000 | 0.0018 | 0.0016 | 0.0016 |
| F08 | 65000001 | 65100000 | 0.0024 | 0.0022 | 0.0021 |
| F08 | 65020001 | 65120000 | 0.0027 | 0.0024 | 0.0024 |
| F08 | 65040001 | 65140000 | 0.0031 | 0.0026 | 0.0029 |
| F08 | 65060001 | 65160000 | 0.0031 | 0.0025 | 0.0029 |
| F08 | 65080001 | 65180000 | 0.0032 | 0.0024 | 0.0030 |
| F08 | 65100001 | 65200000 | 0.0021 | 0.0014 | 0.0020 |
| F08 | 65120001 | 65220000 | 0.0017 | 0.0011 | 0.0016 |
| F08 | 65140001 | 65240000 | 0.0012 | 0.0010 | 0.0011 |
| F08 | 65160001 | 65260000 | 0.0018 | 0.0014 | 0.0016 |
| F08 | 65180001 | 65280000 | 0.0022 | 0.0019 | 0.0021 |
| F08 | 65200001 | 65300000 | 0.0027 | 0.0023 | 0.0025 |
| F08 | 65220001 | 65320000 | 0.0031 | 0.0025 | 0.0029 |
| F08 | 65240001 | 65340000 | 0.0032 | 0.0023 | 0.0029 |
| F08 | 65260001 | 65360000 | 0.0037 | 0.0028 | 0.0034 |
| F08 | 65280001 | 65380000 | 0.0036 | 0.0026 | 0.0031 |
| F08 | 65300001 | 65400000 | 0.0044 | 0.0033 | 0.0037 |
| F08 | 65320001 | 65420000 | 0.0043 | 0.0032 | 0.0035 |

| Chr | Start | End | both | A1 | A2 |
|---|---|---|---|---|---|
| F08 | 65340001 | 65440000 | 0.0042 | 0.0032 | 0.0034 |
| F08 | 65360001 | 65460000 | 0.0030 | 0.0022 | 0.0023 |
| F08 | 65380001 | 65480000 | 0.0030 | 0.0023 | 0.0024 |
| F08 | 65400001 | 65500000 | 0.0026 | 0.0019 | 0.0022 |
| F08 | 65420001 | 65520000 | 0.0023 | 0.0016 | 0.0019 |
| F08 | 65440001 | 65540000 | 0.0024 | 0.0019 | 0.0021 |
| F08 | 65460001 | 65560000 | 0.0026 | 0.0020 | 0.0023 |
| F08 | 65480001 | 65580000 | 0.0029 | 0.0021 | 0.0024 |
| F08 | 65500001 | 65600000 | 0.0024 | 0.0016 | 0.0018 |
| F08 | 65520001 | 65620000 | 0.0026 | 0.0019 | 0.0021 |
| F08 | 65540001 | 65640000 | 0.0022 | 0.0014 | 0.0017 |
| F08 | 65560001 | 65660000 | 0.0024 | 0.0015 | 0.0018 |
| F08 | 65580001 | 65680000 | 0.0019 | 0.0013 | 0.0016 |
| F08 | 65600001 | 65700000 | 0.0025 | 0.0018 | 0.0022 |
| F08 | 65620001 | 65720000 | 0.0028 | 0.0020 | 0.0025 |
| F08 | 65640001 | 65740000 | 0.0035 | 0.0028 | 0.0032 |
| F08 | 65660001 | 65760000 | 0.0035 | 0.0029 | 0.0033 |
| F08 | 65680001 | 65780000 | 0.0038 | 0.0033 | 0.0035 |
| F08 | 65700001 | 65800000 | 0.0039 | 0.0031 | 0.0033 |
| F08 | 65720001 | 65820000 | 0.0033 | 0.0027 | 0.0028 |
| F08 | 65740001 | 65840000 | 0.0028 | 0.0023 | 0.0024 |
| F08 | 65760001 | 65860000 | 0.0029 | 0.0024 | 0.0026 |
| F08 | 65780001 | 65880000 | 0.0027 | 0.0022 | 0.0024 |
| F08 | 65800001 | 65900000 | 0.0019 | 0.0018 | 0.0017 |
| F08 | 65820001 | 65920000 | 0.0020 | 0.0018 | 0.0018 |
| F08 | 65840001 | 65940000 | 0.0023 | 0.0020 | 0.0022 |
| F08 | 65860001 | 65960000 | 0.0023 | 0.0021 | 0.0021 |
| F08 | 65880001 | 65980000 | 0.0022 | 0.0019 | 0.0020 |
| F08 | 65900001 | 66000000 | 0.0022 | 0.0019 | 0.0020 |
| F08 | 65920001 | 66020000 | 0.0020 | 0.0017 | 0.0018 |
| F08 | 65940001 | 66040000 | 0.0017 | 0.0013 | 0.0013 |
| F08 | 65960001 | 66060000 | 0.0019 | 0.0014 | 0.0016 |
| F08 | 65980001 | 66080000 | 0.0016 | 0.0012 | 0.0014 |
| F08 | 66000001 | 66100000 | 0.0028 | 0.0021 | 0.0026 |
| F08 | 66020001 | 66120000 | 0.0037 | 0.0029 | 0.0034 |
| F08 | 66040001 | 66140000 | 0.0043 | 0.0036 | 0.0041 |
| F08 | 66060001 | 66160000 | 0.0044 | 0.0035 | 0.0041 |
| F08 | 66080001 | 66180000 | 0.0048 | 0.0037 | 0.0042 |
| F08 | 66100001 | 66200000 | 0.0041 | 0.0032 | 0.0034 |
| F08 | 66120001 | 66220000 | 0.0039 | 0.0030 | 0.0033 |
| F08 | 66140001 | 66240000 | 0.0045 | 0.0036 | 0.0037 |
| F08 | 66160001 | 66260000 | 0.0043 | 0.0035 | 0.0036 |
| F08 | 66180001 | 66280000 | 0.0042 | 0.0035 | 0.0037 |
| F08 | 66200001 | 66300000 | 0.0049 | 0.0042 | 0.0044 |
| F08 | 66220001 | 66320000 | 0.0061 | 0.0055 | 0.0055 |
| F08 | 66240001 | 66340000 | 0.0057 | 0.0050 | 0.0051 |
| F08 | 66260001 | 66360000 | 0.0054 | 0.0048 | 0.0049 |
| F08 | 66280001 | 66380000 | 0.0058 | 0.0051 | 0.0052 |
| F08 | 66300001 | 66400000 | 0.0061 | 0.0051 | 0.0054 |
| F08 | 66320001 | 66420000 | 0.0051 | 0.0040 | 0.0044 |
| F08 | 66340001 | 66440000 | 0.0043 | 0.0034 | 0.0039 |
| F08 | 66360001 | 66460000 | 0.0041 | 0.0033 | 0.0038 |
| F08 | 66380001 | 66480000 | 0.0037 | 0.0028 | 0.0034 |
| F08 | 66400001 | 66500000 | 0.0024 | 0.0020 | 0.0023 |
| F08 | 66420001 | 66520000 | 0.0020 | 0.0018 | 0.0019 |
| F08 | 66440001 | 66540000 | 0.0026 | 0.0022 | 0.0022 |
| F08 | 66460001 | 66560000 | 0.0028 | 0.0021 | 0.0022 |
| F08 | 66480001 | 66580000 | 0.0028 | 0.0021 | 0.0020 |

| Chr | Start | End | both | A1 | A2 |
|---|---|---|---|---|---|
| F08 | 66500001 | 66600000 | 0.0027 | 0.0019 | 0.0018 |
| F08 | 66520001 | 66620000 | 0.0025 | 0.0015 | 0.0015 |
| F08 | 66540001 | 66640000 | 0.0021 | 0.0011 | 0.0012 |
| F08 | 66560001 | 66660000 | 0.0023 | 0.0013 | 0.0015 |
| F08 | 66580001 | 66680000 | 0.0027 | 0.0016 | 0.0019 |
| F08 | 66600001 | 66700000 | 0.0035 | 0.0023 | 0.0028 |
| F08 | 66620001 | 66720000 | 0.0041 | 0.0028 | 0.0033 |
| F08 | 66640001 | 66740000 | 0.0043 | 0.0032 | 0.0036 |
| F08 | 66660001 | 66760000 | 0.0054 | 0.0042 | 0.0048 |
| F08 | 66680001 | 66780000 | 0.0050 | 0.0041 | 0.0046 |
| F08 | 66700001 | 66800000 | 0.0050 | 0.0040 | 0.0046 |
| F08 | 66720001 | 66820000 | 0.0051 | 0.0042 | 0.0048 |
| F08 | 66740001 | 66840000 | 0.0047 | 0.0038 | 0.0043 |
| F08 | 66760001 | 66860000 | 0.0034 | 0.0026 | 0.0028 |
| F08 | 66780001 | 66880000 | 0.0034 | 0.0025 | 0.0029 |
| F08 | 66800001 | 66900000 | 0.0027 | 0.0020 | 0.0023 |
| F08 | 66820001 | 66920000 | 0.0019 | 0.0013 | 0.0016 |
| F08 | 66840001 | 66940000 | 0.0027 | 0.0019 | 0.0022 |
| F08 | 66860001 | 66960000 | 0.0033 | 0.0025 | 0.0029 |
| F08 | 66880001 | 66980000 | 0.0029 | 0.0022 | 0.0026 |
| F08 | 66900001 | 67000000 | 0.0029 | 0.0021 | 0.0026 |
| F08 | 66920001 | 67020000 | 0.0027 | 0.0021 | 0.0024 |
| F08 | 66940001 | 67040000 | 0.0022 | 0.0018 | 0.0021 |
| F08 | 66960001 | 67060000 | 0.0022 | 0.0019 | 0.0021 |
| F08 | 66980001 | 67080000 | 0.0024 | 0.0021 | 0.0024 |
| F08 | 67000001 | 67100000 | 0.0024 | 0.0021 | 0.0024 |
| F08 | 67020001 | 67120000 | 0.0027 | 0.0023 | 0.0026 |
| F08 | 67040001 | 67140000 | 0.0033 | 0.0028 | 0.0032 |
| F08 | 67060001 | 67160000 | 0.0032 | 0.0025 | 0.0028 |
| F08 | 67080001 | 67180000 | 0.0034 | 0.0025 | 0.0029 |
| F08 | 67100001 | 67200000 | 0.0038 | 0.0028 | 0.0031 |
| F08 | 67120001 | 67220000 | 0.0046 | 0.0034 | 0.0038 |
| F08 | 67140001 | 67240000 | 0.0043 | 0.0031 | 0.0035 |
| F08 | 67160001 | 67260000 | 0.0036 | 0.0025 | 0.0030 |
| F08 | 67180001 | 67280000 | 0.0032 | 0.0022 | 0.0025 |
| F08 | 67200001 | 67300000 | 0.0041 | 0.0030 | 0.0034 |
| F08 | 67220001 | 67320000 | 0.0037 | 0.0026 | 0.0029 |
| F08 | 67240001 | 67340000 | 0.0032 | 0.0022 | 0.0024 |
| F08 | 67260001 | 67360000 | 0.0034 | 0.0024 | 0.0026 |
| F08 | 67280001 | 67380000 | 0.0039 | 0.0029 | 0.0032 |
| F08 | 67300001 | 67400000 | 0.0035 | 0.0025 | 0.0029 |
| F08 | 67320001 | 67420000 | 0.0029 | 0.0020 | 0.0023 |
| F08 | 67340001 | 67440000 | 0.0034 | 0.0026 | 0.0030 |
| F08 | 67360001 | 67460000 | 0.0035 | 0.0027 | 0.0032 |
| F08 | 67380001 | 67480000 | 0.0035 | 0.0028 | 0.0033 |
| F08 | 67400001 | 67500000 | 0.0026 | 0.0020 | 0.0024 |
| F08 | 67420001 | 67520000 | 0.0027 | 0.0019 | 0.0023 |
| F08 | 67440001 | 67540000 | 0.0026 | 0.0017 | 0.0020 |
| F08 | 67460001 | 67560000 | 0.0026 | 0.0019 | 0.0020 |
| F08 | 67480001 | 67580000 | 0.0023 | 0.0017 | 0.0018 |
| F08 | 67500001 | 67600000 | 0.0022 | 0.0016 | 0.0017 |
| F08 | 67520001 | 67620000 | 0.0023 | 0.0020 | 0.0020 |
| F08 | 67540001 | 67640000 | 0.0022 | 0.0021 | 0.0021 |
| F08 | 67560001 | 67660000 | 0.0022 | 0.0020 | 0.0020 |
| F08 | 67580001 | 67680000 | 0.0024 | 0.0021 | 0.0021 |
| F08 | 67600001 | 67700000 | 0.0028 | 0.0025 | 0.0025 |
| F08 | 67620001 | 67720000 | 0.0033 | 0.0029 | 0.0029 |
| F08 | 67640001 | 67740000 | 0.0031 | 0.0025 | 0.0026 |

| Chr | Start | End | both | A1 | A2 |
|---|---|---|---|---|---|
| F08 | 67660001 | 67760000 | 0.0032 | 0.0024 | 0.0027 |
| F08 | 67680001 | 67780000 | 0.0031 | 0.0025 | 0.0027 |
| F08 | 67700001 | 67800000 | 0.0031 | 0.0023 | 0.0026 |
| F08 | 67720001 | 67820000 | 0.0024 | 0.0017 | 0.0019 |
| F08 | 67740001 | 67840000 | 0.0024 | 0.0017 | 0.0018 |
| F08 | 67760001 | 67860000 | 0.0030 | 0.0025 | 0.0026 |
| F08 | 67780001 | 67880000 | 0.0027 | 0.0020 | 0.0020 |
| F08 | 67800001 | 67900000 | 0.0028 | 0.0021 | 0.0019 |
| F08 | 67820001 | 67920000 | 0.0031 | 0.0025 | 0.0024 |
| F08 | 67840001 | 67940000 | 0.0031 | 0.0026 | 0.0025 |
| F08 | 67860001 | 67960000 | 0.0022 | 0.0016 | 0.0016 |
| F08 | 67880001 | 67980000 | 0.0034 | 0.0026 | 0.0026 |
| F08 | 67900001 | 68000000 | 0.0036 | 0.0027 | 0.0029 |
| F08 | 67920001 | 68020000 | 0.0035 | 0.0025 | 0.0027 |
| F08 | 67940001 | 68040000 | 0.0039 | 0.0024 | 0.0027 |
| F08 | 67960001 | 68060000 | 0.0037 | 0.0022 | 0.0024 |
| F08 | 67980001 | 68080000 | 0.0031 | 0.0019 | 0.0022 |
| F08 | 68000001 | 68100000 | 0.0035 | 0.0025 | 0.0026 |
| F08 | 68020001 | 68120000 | 0.0031 | 0.0022 | 0.0023 |
| F08 | 68040001 | 68140000 | 0.0029 | 0.0025 | 0.0025 |
| F08 | 68060001 | 68160000 | 0.0028 | 0.0024 | 0.0025 |
| F08 | 68080001 | 68180000 | 0.0019 | 0.0017 | 0.0018 |
| F08 | 68100001 | 68200000 | 0.0016 | 0.0014 | 0.0016 |
| F08 | 68120001 | 68220000 | 0.0016 | 0.0013 | 0.0016 |
| F08 | 68140001 | 68240000 | 0.0010 | 0.0008 | 0.0010 |
| F08 | 68160001 | 68260000 | 0.0015 | 0.0012 | 0.0015 |
| F08 | 68180001 | 68280000 | 0.0021 | 0.0018 | 0.0021 |
| F08 | 68200001 | 68300000 | 0.0025 | 0.0021 | 0.0023 |
| F08 | 68220001 | 68320000 | 0.0035 | 0.0029 | 0.0032 |
| F08 | 68240001 | 68340000 | 0.0037 | 0.0030 | 0.0033 |
| F08 | 68260001 | 68360000 | 0.0042 | 0.0029 | 0.0033 |
| F08 | 68280001 | 68380000 | 0.0048 | 0.0034 | 0.0037 |
| F08 | 68300001 | 68400000 | 0.0043 | 0.0027 | 0.0030 |
| F08 | 68320001 | 68420000 | 0.0039 | 0.0023 | 0.0026 |
| F08 | 68340001 | 68440000 | 0.0041 | 0.0025 | 0.0028 |
| F08 | 68360001 | 68460000 | 0.0049 | 0.0035 | 0.0038 |
| F08 | 68380001 | 68480000 | 0.0047 | 0.0033 | 0.0036 |
| F08 | 68400001 | 68500000 | 0.0053 | 0.0040 | 0.0044 |
| F08 | 68420001 | 68520000 | 0.0053 | 0.0039 | 0.0043 |
| F08 | 68440001 | 68540000 | 0.0061 | 0.0043 | 0.0052 |
| F08 | 68460001 | 68560000 | 0.0053 | 0.0034 | 0.0046 |
| F08 | 68480001 | 68580000 | 0.0051 | 0.0033 | 0.0046 |
| F08 | 68500001 | 68600000 | 0.0042 | 0.0026 | 0.0038 |
| F08 | 68520001 | 68620000 | 0.0043 | 0.0028 | 0.0039 |
| F08 | 68540001 | 68640000 | 0.0036 | 0.0026 | 0.0033 |
| F08 | 68560001 | 68660000 | 0.0036 | 0.0026 | 0.0031 |
| F08 | 68580001 | 68680000 | 0.0038 | 0.0028 | 0.0032 |
| F08 | 68600001 | 68700000 | 0.0046 | 0.0034 | 0.0041 |
| F08 | 68620001 | 68720000 | 0.0039 | 0.0029 | 0.0034 |
| F08 | 68640001 | 68740000 | 0.0045 | 0.0031 | 0.0036 |
| F08 | 68660001 | 68760000 | 0.0038 | 0.0025 | 0.0029 |
| F08 | 68680001 | 68780000 | 0.0033 | 0.0021 | 0.0024 |
| F08 | 68700001 | 68800000 | 0.0030 | 0.0019 | 0.0020 |
| F08 | 68720001 | 68820000 | 0.0039 | 0.0027 | 0.0029 |
| F08 | 68740001 | 68840000 | 0.0029 | 0.0021 | 0.0022 |
| F08 | 68760001 | 68860000 | 0.0029 | 0.0023 | 0.0024 |
| F08 | 68780001 | 68880000 | 0.0038 | 0.0031 | 0.0032 |
| F08 | 68800001 | 68900000 | 0.0035 | 0.0028 | 0.0029 |

| Chr | Start | End | both | A1 | A2 |
|---|---|---|---|---|---|
| F08 | 68820001 | 68920000 | 0.0031 | 0.0023 | 0.0023 |
| F08 | 68840001 | 68940000 | 0.0034 | 0.0024 | 0.0025 |
| F08 | 68860001 | 68960000 | 0.0038 | 0.0026 | 0.0028 |
| F08 | 68880001 | 68980000 | 0.0035 | 0.0020 | 0.0022 |
| F08 | 68900001 | 69000000 | 0.0043 | 0.0027 | 0.0030 |
| F08 | 68920001 | 69020000 | 0.0048 | 0.0032 | 0.0035 |
| F08 | 68940001 | 69040000 | 0.0048 | 0.0033 | 0.0035 |
| F08 | 68960001 | 69060000 | 0.0042 | 0.0030 | 0.0032 |
| F08 | 68980001 | 69080000 | 0.0037 | 0.0027 | 0.0029 |
| F08 | 69000001 | 69100000 | 0.0028 | 0.0019 | 0.0019 |
| F08 | 69020001 | 69120000 | 0.0020 | 0.0012 | 0.0012 |
| F08 | 69040001 | 69140000 | 0.0026 | 0.0017 | 0.0019 |
| F08 | 69060001 | 69160000 | 0.0029 | 0.0020 | 0.0022 |
| F08 | 69080001 | 69180000 | 0.0037 | 0.0028 | 0.0030 |
| F08 | 69100001 | 69200000 | 0.0045 | 0.0034 | 0.0038 |
| F08 | 69120001 | 69220000 | 0.0049 | 0.0037 | 0.0041 |
| F08 | 69140001 | 69240000 | 0.0044 | 0.0032 | 0.0035 |
| F08 | 69160001 | 69260000 | 0.0046 | 0.0034 | 0.0037 |
| F08 | 69180001 | 69280000 | 0.0048 | 0.0035 | 0.0039 |
| F08 | 69200001 | 69300000 | 0.0049 | 0.0038 | 0.0040 |
| F08 | 69220001 | 69320000 | 0.0050 | 0.0040 | 0.0043 |
| F08 | 69240001 | 69340000 | 0.0050 | 0.0040 | 0.0043 |
| F08 | 69260001 | 69360000 | 0.0046 | 0.0037 | 0.0039 |
| F08 | 69280001 | 69380000 | 0.0032 | 0.0026 | 0.0027 |
| F08 | 69300001 | 69400000 | 0.0026 | 0.0016 | 0.0020 |
| F08 | 69320001 | 69420000 | 0.0024 | 0.0014 | 0.0017 |
| F08 | 69340001 | 69440000 | 0.0023 | 0.0015 | 0.0018 |
| F08 | 69360001 | 69460000 | 0.0025 | 0.0016 | 0.0021 |
| F08 | 69380001 | 69480000 | 0.0026 | 0.0019 | 0.0022 |
| F08 | 69400001 | 69500000 | 0.0020 | 0.0017 | 0.0019 |
| F08 | 69420001 | 69520000 | 0.0025 | 0.0021 | 0.0025 |
| F08 | 69440001 | 69540000 | 0.0027 | 0.0022 | 0.0027 |
| F08 | 69460001 | 69560000 | 0.0025 | 0.0020 | 0.0024 |
| F08 | 69480001 | 69580000 | 0.0031 | 0.0024 | 0.0029 |
| F08 | 69500001 | 69600000 | 0.0035 | 0.0028 | 0.0032 |
| F08 | 69520001 | 69620000 | 0.0030 | 0.0023 | 0.0027 |
| F08 | 69540001 | 69640000 | 0.0029 | 0.0023 | 0.0025 |
| F08 | 69560001 | 69660000 | 0.0028 | 0.0022 | 0.0024 |
| F08 | 69580001 | 69680000 | 0.0020 | 0.0016 | 0.0018 |
| F08 | 69600001 | 69700000 | 0.0015 | 0.0012 | 0.0014 |
| F08 | 69620001 | 69720000 | 0.0020 | 0.0015 | 0.0017 |
| F08 | 69640001 | 69740000 | 0.0015 | 0.0011 | 0.0013 |
| F08 | 69660001 | 69760000 | 0.0022 | 0.0017 | 0.0019 |
| F08 | 69680001 | 69780000 | 0.0019 | 0.0014 | 0.0017 |
| F08 | 69700001 | 69800000 | 0.0019 | 0.0014 | 0.0016 |
| F08 | 69720001 | 69820000 | 0.0010 | 0.0008 | 0.0010 |
| F08 | 69740001 | 69840000 | 0.0011 | 0.0009 | 0.0011 |
| F08 | 69760001 | 69860000 | 0.0008 | 0.0007 | 0.0008 |
| F08 | 69780001 | 69880000 | 0.0012 | 0.0010 | 0.0012 |
| F08 | 69800001 | 69900000 | 0.0012 | 0.0010 | 0.0012 |
| F08 | 69820001 | 69920000 | 0.0012 | 0.0010 | 0.0012 |
| F08 | 69840001 | 69940000 | 0.0013 | 0.0012 | 0.0013 |
| F08 | 69860001 | 69960000 | 0.0010 | 0.0009 | 0.0010 |
| F08 | 69880001 | 69980000 | 0.0006 | 0.0005 | 0.0006 |
| F08 | 69900001 | 70000000 | 0.0007 | 0.0006 | 0.0007 |
| F08 | 69920001 | 70020000 | 0.0007 | 0.0006 | 0.0007 |
| F08 | 69940001 | 70040000 | 0.0005 | 0.0004 | 0.0005 |
| F08 | 69960001 | 70060000 | 0.0004 | 0.0004 | 0.0004 |

| Chr | Start | End | both | A1 | A2 |
|---|---|---|---|---|---|
| F08 | 69980001 | 70080000 | 0.0006 | 0.0006 | 0.0006 |
| F08 | 70000001 | 70100000 | 0.0008 | 0.0007 | 0.0008 |
| F08 | 70020001 | 70120000 | 0.0011 | 0.0009 | 0.0010 |
| F08 | 70040001 | 70140000 | 0.0010 | 0.0008 | 0.0009 |
| F08 | 70060001 | 70160000 | 0.0010 | 0.0008 | 0.0009 |
| F08 | 70080001 | 70180000 | 0.0008 | 0.0006 | 0.0007 |
| F08 | 70100001 | 70200000 | 0.0005 | 0.0005 | 0.0005 |
| F08 | 70120001 | 70220000 | 0.0003 | 0.0002 | 0.0002 |
| F08 | 70140001 | 70240000 | 0.0002 | 0.0002 | 0.0002 |
| F08 | 70160001 | 70260000 | 0.0003 | 0.0003 | 0.0003 |
| F08 | 70180001 | 70280000 | 0.0004 | 0.0004 | 0.0004 |
| F08 | 70200001 | 70300000 | 0.0008 | 0.0007 | 0.0008 |
| F08 | 70220001 | 70320000 | 0.0010 | 0.0008 | 0.0010 |
| F08 | 70240001 | 70340000 | 0.0021 | 0.0019 | 0.0020 |
| F08 | 70260001 | 70360000 | 0.0021 | 0.0019 | 0.0020 |
| F08 | 70280001 | 70380000 | 0.0022 | 0.0020 | 0.0021 |
| F08 | 70300001 | 70400000 | 0.0024 | 0.0022 | 0.0023 |
| F08 | 70320001 | 70420000 | 0.0025 | 0.0023 | 0.0025 |
| F08 | 70340001 | 70440000 | 0.0013 | 0.0010 | 0.0013 |
| F08 | 70360001 | 70460000 | 0.0012 | 0.0009 | 0.0012 |
| F08 | 70380001 | 70480000 | 0.0014 | 0.0010 | 0.0012 |
| F08 | 70400001 | 70500000 | 0.0017 | 0.0009 | 0.0011 |
| F08 | 70420001 | 70520000 | 0.0026 | 0.0015 | 0.0018 |
| F08 | 70440001 | 70540000 | 0.0033 | 0.0022 | 0.0025 |
| F08 | 70460001 | 70560000 | 0.0036 | 0.0025 | 0.0028 |
| F08 | 70480001 | 70580000 | 0.0034 | 0.0024 | 0.0027 |
| F08 | 70500001 | 70600000 | 0.0031 | 0.0026 | 0.0029 |
| F08 | 70520001 | 70620000 | 0.0029 | 0.0024 | 0.0026 |
| F08 | 70540001 | 70640000 | 0.0023 | 0.0018 | 0.0020 |
| F08 | 70560001 | 70660000 | 0.0020 | 0.0015 | 0.0018 |
| F08 | 70580001 | 70680000 | 0.0023 | 0.0018 | 0.0020 |
| F08 | 70600001 | 70700000 | 0.0024 | 0.0018 | 0.0021 |
| F08 | 70620001 | 70720000 | 0.0020 | 0.0015 | 0.0018 |
| F08 | 70640001 | 70740000 | 0.0021 | 0.0015 | 0.0018 |
| F08 | 70660001 | 70760000 | 0.0021 | 0.0015 | 0.0018 |
| F08 | 70680001 | 70780000 | 0.0019 | 0.0014 | 0.0017 |
| F08 | 70700001 | 70800000 | 0.0020 | 0.0015 | 0.0017 |
| F08 | 70720001 | 70820000 | 0.0027 | 0.0020 | 0.0023 |
| F08 | 70740001 | 70840000 | 0.0029 | 0.0024 | 0.0027 |
| F08 | 70760001 | 70860000 | 0.0034 | 0.0028 | 0.0031 |
| F08 | 70780001 | 70880000 | 0.0030 | 0.0025 | 0.0027 |
| F08 | 70800001 | 70900000 | 0.0031 | 0.0027 | 0.0030 |
| F08 | 70820001 | 70920000 | 0.0026 | 0.0023 | 0.0025 |
| F08 | 70840001 | 70940000 | 0.0026 | 0.0024 | 0.0026 |
| F08 | 70860001 | 70960000 | 0.0027 | 0.0023 | 0.0025 |
| F08 | 70880001 | 70980000 | 0.0032 | 0.0026 | 0.0028 |
| F08 | 70900001 | 71000000 | 0.0023 | 0.0017 | 0.0019 |
| F08 | 70920001 | 71020000 | 0.0023 | 0.0013 | 0.0014 |
| F08 | 70940001 | 71040000 | 0.0017 | 0.0008 | 0.0009 |
| F08 | 70960001 | 71060000 | 0.0016 | 0.0008 | 0.0009 |
| F08 | 70980001 | 71080000 | 0.0014 | 0.0006 | 0.0006 |
| F08 | 71000001 | 71100000 | 0.0017 | 0.0009 | 0.0009 |
| F08 | 71020001 | 71120000 | 0.0019 | 0.0011 | 0.0012 |
| F08 | 71040001 | 71140000 | 0.0032 | 0.0021 | 0.0024 |
| F08 | 71060001 | 71160000 | 0.0030 | 0.0020 | 0.0022 |
| F08 | 71080001 | 71180000 | 0.0035 | 0.0025 | 0.0029 |
| F08 | 71100001 | 71200000 | 0.0030 | 0.0023 | 0.0026 |
| F08 | 71120001 | 71220000 | 0.0027 | 0.0022 | 0.0024 |

| Chr | Start | End | both | A1 | A2 |
|---|---|---|---|---|---|
| F08 | 71140001 | 71240000 | 0.0018 | 0.0014 | 0.0016 |
| F08 | 71160001 | 71260000 | 0.0028 | 0.0024 | 0.0025 |
| F08 | 71180001 | 71280000 | 0.0028 | 0.0025 | 0.0025 |
| F08 | 71200001 | 71300000 | 0.0033 | 0.0029 | 0.0031 |
| F08 | 71220001 | 71320000 | 0.0047 | 0.0043 | 0.0045 |
| F08 | 71240001 | 71340000 | 0.0052 | 0.0049 | 0.0050 |
| F08 | 71260001 | 71360000 | 0.0045 | 0.0040 | 0.0043 |
| F08 | 71280001 | 71380000 | 0.0041 | 0.0034 | 0.0036 |
| F08 | 71300001 | 71400000 | 0.0046 | 0.0036 | 0.0038 |
| F08 | 71320001 | 71420000 | 0.0036 | 0.0026 | 0.0028 |
| F08 | 71340001 | 71440000 | 0.0032 | 0.0022 | 0.0024 |
| F08 | 71360001 | 71460000 | 0.0029 | 0.0020 | 0.0021 |
| F08 | 71380001 | 71480000 | 0.0028 | 0.0021 | 0.0022 |
| F08 | 71400001 | 71500000 | 0.0021 | 0.0018 | 0.0019 |
| F08 | 71420001 | 71520000 | 0.0021 | 0.0019 | 0.0020 |
| F08 | 71440001 | 71540000 | 0.0021 | 0.0018 | 0.0020 |
| F08 | 71460001 | 71560000 | 0.0026 | 0.0021 | 0.0023 |
| F08 | 71480001 | 71580000 | 0.0031 | 0.0020 | 0.0024 |
| F08 | 71500001 | 71600000 | 0.0031 | 0.0018 | 0.0022 |
| F08 | 71520001 | 71620000 | 0.0025 | 0.0013 | 0.0015 |
| F08 | 71540001 | 71640000 | 0.0026 | 0.0014 | 0.0015 |
| F08 | 71560001 | 71660000 | 0.0024 | 0.0015 | 0.0017 |
| F08 | 71580001 | 71680000 | 0.0018 | 0.0014 | 0.0014 |
| F08 | 71600001 | 71700000 | 0.0017 | 0.0015 | 0.0016 |
| F08 | 71620001 | 71720000 | 0.0017 | 0.0015 | 0.0016 |
| F08 | 71640001 | 71740000 | 0.0018 | 0.0012 | 0.0013 |
| F08 | 71660001 | 71760000 | 0.0024 | 0.0015 | 0.0018 |
| F08 | 71680001 | 71780000 | 0.0036 | 0.0024 | 0.0028 |
| F08 | 71700001 | 71800000 | 0.0045 | 0.0031 | 0.0036 |
| F08 | 71720001 | 71820000 | 0.0052 | 0.0037 | 0.0044 |
| F08 | 71740001 | 71840000 | 0.0054 | 0.0043 | 0.0049 |
| F08 | 71760001 | 71860000 | 0.0043 | 0.0035 | 0.0040 |
| F08 | 71780001 | 71880000 | 0.0032 | 0.0027 | 0.0030 |
| F08 | 71800001 | 71900000 | 0.0022 | 0.0020 | 0.0022 |
| F08 | 71820001 | 71920000 | 0.0018 | 0.0016 | 0.0017 |
| F08 | 71840001 | 71940000 | 0.0016 | 0.0014 | 0.0016 |
| F08 | 71860001 | 71960000 | 0.0015 | 0.0013 | 0.0015 |
| F08 | 71880001 | 71980000 | 0.0013 | 0.0011 | 0.0013 |
| F08 | 71900001 | 72000000 | 0.0018 | 0.0017 | 0.0018 |
| F08 | 71920001 | 72020000 | 0.0016 | 0.0015 | 0.0016 |
| F08 | 71940001 | 72040000 | 0.0011 | 0.0011 | 0.0011 |
| F08 | 71960001 | 72060000 | 0.0017 | 0.0016 | 0.0017 |
| F08 | 71980001 | 72080000 | 0.0024 | 0.0020 | 0.0022 |
| F08 | 72000001 | 72100000 | 0.0024 | 0.0018 | 0.0020 |
| F08 | 72020001 | 72120000 | 0.0024 | 0.0017 | 0.0020 |
| F08 | 72040001 | 72140000 | 0.0025 | 0.0018 | 0.0020 |
| F08 | 72060001 | 72160000 | 0.0023 | 0.0014 | 0.0018 |
| F08 | 72080001 | 72180000 | 0.0027 | 0.0020 | 0.0023 |
| F08 | 72100001 | 72200000 | 0.0021 | 0.0016 | 0.0020 |
| F08 | 72120001 | 72220000 | 0.0024 | 0.0020 | 0.0023 |
| F08 | 72140001 | 72240000 | 0.0028 | 0.0024 | 0.0027 |
| F08 | 72160001 | 72260000 | 0.0029 | 0.0027 | 0.0028 |
| F08 | 72180001 | 72280000 | 0.0031 | 0.0027 | 0.0030 |
| F08 | 72200001 | 72300000 | 0.0031 | 0.0027 | 0.0029 |
| F08 | 72220001 | 72320000 | 0.0027 | 0.0023 | 0.0025 |
| F08 | 72240001 | 72340000 | 0.0021 | 0.0017 | 0.0020 |
| F08 | 72260001 | 72360000 | 0.0017 | 0.0013 | 0.0016 |
| F08 | 72280001 | 72380000 | 0.0006 | 0.0005 | 0.0006 |

| Chr | Start | End | both | A1 | A2 |
|---|---|---|---|---|---|
| F08 | 72300001 | 72400000 | 0.0013 | 0.0011 | 0.0013 |
| F08 | 72320001 | 72420000 | 0.0013 | 0.0011 | 0.0013 |
| F08 | 72340001 | 72440000 | 0.0013 | 0.0012 | 0.0012 |
| F08 | 72360001 | 72460000 | 0.0015 | 0.0013 | 0.0014 |
| F08 | 72380001 | 72480000 | 0.0015 | 0.0014 | 0.0015 |
| F08 | 72400001 | 72500000 | 0.0006 | 0.0006 | 0.0006 |
| F08 | 72420001 | 72520000 | 0.0009 | 0.0008 | 0.0008 |
| F08 | 72440001 | 72540000 | 0.0014 | 0.0011 | 0.0014 |
| F08 | 72460001 | 72560000 | 0.0013 | 0.0010 | 0.0012 |
| F08 | 72480001 | 72580000 | 0.0018 | 0.0015 | 0.0017 |
| F08 | 72500001 | 72600000 | 0.0022 | 0.0018 | 0.0020 |
| F08 | 72520001 | 72620000 | 0.0026 | 0.0021 | 0.0024 |
| F08 | 72540001 | 72640000 | 0.0022 | 0.0016 | 0.0019 |
| F08 | 72560001 | 72660000 | 0.0029 | 0.0020 | 0.0025 |
| F08 | 72580001 | 72680000 | 0.0032 | 0.0022 | 0.0027 |
| F08 | 72600001 | 72700000 | 0.0028 | 0.0019 | 0.0025 |
| F08 | 72620001 | 72720000 | 0.0026 | 0.0018 | 0.0023 |
| F08 | 72640001 | 72740000 | 0.0032 | 0.0024 | 0.0029 |
| F08 | 72660001 | 72760000 | 0.0025 | 0.0021 | 0.0025 |
| F08 | 72680001 | 72780000 | 0.0016 | 0.0013 | 0.0016 |
| F08 | 72700001 | 72800000 | 0.0018 | 0.0015 | 0.0018 |
| F08 | 72720001 | 72820000 | 0.0014 | 0.0012 | 0.0014 |
| F08 | 72740001 | 72840000 | 0.0007 | 0.0006 | 0.0007 |
| F08 | 72760001 | 72860000 | 0.0005 | 0.0004 | 0.0005 |
| F08 | 72780001 | 72880000 | 0.0006 | 0.0005 | 0.0006 |
| F08 | 72800001 | 72900000 | 0.0005 | 0.0004 | 0.0005 |
| F08 | 72820001 | 72920000 | 0.0011 | 0.0009 | 0.0010 |
| F08 | 72840001 | 72940000 | 0.0015 | 0.0013 | 0.0015 |
| F08 | 72860001 | 72960000 | 0.0019 | 0.0016 | 0.0018 |
| F08 | 72880001 | 72980000 | 0.0025 | 0.0021 | 0.0024 |
| F08 | 72900001 | 73000000 | 0.0030 | 0.0024 | 0.0027 |
| F08 | 72920001 | 73020000 | 0.0026 | 0.0019 | 0.0023 |
| F08 | 72940001 | 73040000 | 0.0025 | 0.0019 | 0.0022 |
| F08 | 72960001 | 73060000 | 0.0022 | 0.0016 | 0.0019 |
| F08 | 72980001 | 73080000 | 0.0014 | 0.0010 | 0.0012 |
| F08 | 73000001 | 73100000 | 0.0015 | 0.0011 | 0.0013 |
| F08 | 73020001 | 73120000 | 0.0026 | 0.0021 | 0.0024 |
| F08 | 73040001 | 73140000 | 0.0025 | 0.0018 | 0.0022 |
| F08 | 73060001 | 73160000 | 0.0027 | 0.0019 | 0.0024 |
| F08 | 73080001 | 73180000 | 0.0031 | 0.0023 | 0.0028 |
| F08 | 73100001 | 73200000 | 0.0036 | 0.0026 | 0.0032 |
| F08 | 73120001 | 73220000 | 0.0025 | 0.0017 | 0.0021 |
| F08 | 73140001 | 73240000 | 0.0031 | 0.0021 | 0.0026 |
| F08 | 73160001 | 73260000 | 0.0035 | 0.0022 | 0.0027 |
| F08 | 73180001 | 73280000 | 0.0037 | 0.0024 | 0.0030 |
| F08 | 73200001 | 73300000 | 0.0031 | 0.0020 | 0.0024 |
| F08 | 73220001 | 73320000 | 0.0035 | 0.0023 | 0.0027 |
| F08 | 73240001 | 73340000 | 0.0029 | 0.0018 | 0.0022 |
| F08 | 73260001 | 73360000 | 0.0034 | 0.0025 | 0.0027 |
| F08 | 73280001 | 73380000 | 0.0031 | 0.0019 | 0.0021 |
| F08 | 73300001 | 73400000 | 0.0031 | 0.0018 | 0.0021 |
| F08 | 73320001 | 73420000 | 0.0024 | 0.0015 | 0.0017 |
| F08 | 73340001 | 73440000 | 0.0022 | 0.0015 | 0.0017 |
| F08 | 73360001 | 73460000 | 0.0012 | 0.0007 | 0.0009 |
| F08 | 73380001 | 73480000 | 0.0011 | 0.0008 | 0.0010 |
| F08 | 73400001 | 73500000 | 0.0010 | 0.0008 | 0.0008 |
| F08 | 73420001 | 73520000 | 0.0019 | 0.0013 | 0.0014 |
| F08 | 73440001 | 73540000 | 0.0033 | 0.0025 | 0.0028 |

| Chr | Start | End | both | A1 | A2 |
|---|---|---|---|---|---|
| F08 | 73460001 | 73560000 | 0.0036 | 0.0027 | 0.0030 |
| F08 | 73480001 | 73580000 | 0.0050 | 0.0040 | 0.0043 |
| F08 | 73500001 | 73600000 | 0.0052 | 0.0039 | 0.0044 |
| F08 | 73520001 | 73620000 | 0.0053 | 0.0041 | 0.0044 |
| F08 | 73540001 | 73640000 | 0.0047 | 0.0037 | 0.0039 |
| F08 | 73560001 | 73660000 | 0.0050 | 0.0040 | 0.0042 |
| F08 | 73580001 | 73680000 | 0.0042 | 0.0030 | 0.0032 |
| F08 | 73600001 | 73700000 | 0.0040 | 0.0031 | 0.0033 |
| F08 | 73620001 | 73720000 | 0.0043 | 0.0031 | 0.0035 |
| F08 | 73640001 | 73740000 | 0.0039 | 0.0027 | 0.0030 |
| F08 | 73660001 | 73760000 | 0.0039 | 0.0029 | 0.0031 |
| F08 | 73680001 | 73780000 | 0.0034 | 0.0029 | 0.0028 |
| F08 | 73700001 | 73800000 | 0.0033 | 0.0029 | 0.0027 |
| F08 | 73720001 | 73820000 | 0.0022 | 0.0023 | 0.0019 |
| F08 | 73740001 | 73840000 | 0.0019 | 0.0021 | 0.0017 |
| F08 | 73760001 | 73860000 | 0.0014 | 0.0017 | 0.0013 |
| F08 | 73780001 | 73880000 | 0.0018 | 0.0020 | 0.0017 |
| F08 | 73800001 | 73900000 | 0.0021 | 0.0022 | 0.0020 |
| F08 | 73820001 | 73920000 | 0.0028 | 0.0029 | 0.0026 |
| F08 | 73840001 | 73940000 | 0.0029 | 0.0030 | 0.0027 |
| F08 | 73860001 | 73960000 | 0.0033 | 0.0031 | 0.0033 |
| F08 | 73880001 | 73980000 | 0.0032 | 0.0031 | 0.0031 |
| F08 | 73900001 | 74000000 | 0.0036 | 0.0039 | 0.0034 |
| F08 | 73920001 | 74020000 | 0.0033 | 0.0038 | 0.0030 |
| F08 | 73940001 | 74040000 | 0.0029 | 0.0036 | 0.0026 |
| F08 | 73960001 | 74060000 | 0.0029 | 0.0037 | 0.0026 |
| F08 | 73980001 | 74080000 | 0.0031 | 0.0036 | 0.0028 |
| F08 | 74000001 | 74100000 | 0.0026 | 0.0027 | 0.0024 |
| F08 | 74020001 | 74120000 | 0.0028 | 0.0027 | 0.0026 |
| F08 | 74040001 | 74140000 | 0.0036 | 0.0032 | 0.0035 |
| F08 | 74060001 | 74160000 | 0.0036 | 0.0032 | 0.0035 |
| F08 | 74080001 | 74180000 | 0.0045 | 0.0040 | 0.0043 |
| F08 | 74100001 | 74200000 | 0.0044 | 0.0040 | 0.0042 |
| F08 | 74120001 | 74220000 | 0.0047 | 0.0043 | 0.0045 |
| F08 | 74140001 | 74240000 | 0.0044 | 0.0044 | 0.0042 |
| F08 | 74160001 | 74260000 | 0.0039 | 0.0038 | 0.0037 |
| F08 | 74180001 | 74280000 | 0.0021 | 0.0022 | 0.0020 |
| F08 | 74200001 | 74300000 | 0.0024 | 0.0025 | 0.0023 |
| F08 | 74220001 | 74320000 | 0.0019 | 0.0019 | 0.0017 |
| F08 | 74240001 | 74340000 | 0.0016 | 0.0016 | 0.0015 |
| F08 | 74260001 | 74360000 | 0.0026 | 0.0027 | 0.0024 |
| F08 | 74280001 | 74380000 | 0.0033 | 0.0036 | 0.0030 |
| F08 | 74300001 | 74400000 | 0.0034 | 0.0036 | 0.0031 |
| F08 | 74320001 | 74420000 | 0.0037 | 0.0039 | 0.0034 |
| F08 | 74340001 | 74440000 | 0.0035 | 0.0037 | 0.0033 |
| F08 | 74360001 | 74460000 | 0.0025 | 0.0026 | 0.0024 |
| F08 | 74380001 | 74480000 | 0.0024 | 0.0023 | 0.0023 |
| F08 | 74400001 | 74500000 | 0.0020 | 0.0019 | 0.0019 |
| F08 | 74420001 | 74520000 | 0.0017 | 0.0017 | 0.0016 |
| F08 | 74440001 | 74540000 | 0.0017 | 0.0016 | 0.0016 |
| F08 | 74460001 | 74560000 | 0.0027 | 0.0027 | 0.0026 |
| F08 | 74480001 | 74580000 | 0.0041 | 0.0041 | 0.0039 |
| F08 | 74500001 | 74600000 | 0.0042 | 0.0043 | 0.0040 |
| F08 | 74520001 | 74620000 | 0.0047 | 0.0049 | 0.0044 |
| F08 | 74540001 | 74640000 | 0.0051 | 0.0054 | 0.0048 |
| F08 | 74560001 | 74660000 | 0.0042 | 0.0044 | 0.0039 |
| F08 | 74580001 | 74680000 | 0.0022 | 0.0024 | 0.0020 |
| F08 | 74600001 | 74700000 | 0.0020 | 0.0022 | 0.0019 |

| Chr | Start | End | both | A1 | A2 |
|---|---|---|---|---|---|
| F08 | 74620001 | 74720000 | 0.0016 | 0.0016 | 0.0014 |
| F08 | 74640001 | 74740000 | 0.0015 | 0.0016 | 0.0013 |
| F08 | 74660001 | 74760000 | 0.0019 | 0.0022 | 0.0016 |
| F08 | 74680001 | 74780000 | 0.0024 | 0.0027 | 0.0022 |
| F08 | 74700001 | 74800000 | 0.0031 | 0.0033 | 0.0028 |
| F08 | 74720001 | 74820000 | 0.0031 | 0.0034 | 0.0029 |
| F08 | 74740001 | 74840000 | 0.0031 | 0.0034 | 0.0029 |
| F08 | 74760001 | 74860000 | 0.0032 | 0.0034 | 0.0030 |
| F08 | 74780001 | 74880000 | 0.0030 | 0.0032 | 0.0028 |
| F08 | 74800001 | 74900000 | 0.0029 | 0.0032 | 0.0027 |
| F08 | 74820001 | 74920000 | 0.0032 | 0.0034 | 0.0029 |
| F08 | 74840001 | 74940000 | 0.0032 | 0.0033 | 0.0029 |
| F08 | 74860001 | 74960000 | 0.0040 | 0.0040 | 0.0038 |
| F08 | 74880001 | 74980000 | 0.0041 | 0.0041 | 0.0038 |
| F08 | 74900001 | 75000000 | 0.0048 | 0.0047 | 0.0046 |
| F08 | 74920001 | 75020000 | 0.0051 | 0.0048 | 0.0049 |
| F08 | 74940001 | 75040000 | 0.0048 | 0.0047 | 0.0046 |
| F08 | 74960001 | 75060000 | 0.0042 | 0.0038 | 0.0040 |
| F08 | 74980001 | 75080000 | 0.0038 | 0.0034 | 0.0037 |
| F08 | 75000001 | 75100000 | 0.0028 | 0.0027 | 0.0026 |
| F08 | 75020001 | 75120000 | 0.0021 | 0.0020 | 0.0019 |
| F08 | 75040001 | 75140000 | 0.0019 | 0.0018 | 0.0017 |
| F08 | 75060001 | 75160000 | 0.0012 | 0.0013 | 0.0010 |
| F08 | 75080001 | 75180000 | 0.0020 | 0.0020 | 0.0019 |
| F08 | 75100001 | 75200000 | 0.0028 | 0.0027 | 0.0026 |
| F08 | 75120001 | 75220000 | 0.0026 | 0.0025 | 0.0025 |
| F08 | 75140001 | 75240000 | 0.0025 | 0.0023 | 0.0024 |
| F08 | 75160001 | 75260000 | 0.0026 | 0.0024 | 0.0026 |
| F08 | 75180001 | 75280000 | 0.0021 | 0.0019 | 0.0020 |
| F08 | 75200001 | 75300000 | 0.0020 | 0.0019 | 0.0020 |
| F08 | 75220001 | 75320000 | 0.0027 | 0.0026 | 0.0027 |
| F08 | 75240001 | 75340000 | 0.0040 | 0.0038 | 0.0039 |
| F08 | 75260001 | 75360000 | 0.0049 | 0.0045 | 0.0048 |
| F08 | 75280001 | 75380000 | 0.0059 | 0.0053 | 0.0058 |
| F08 | 75300001 | 75400000 | 0.0058 | 0.0050 | 0.0057 |
| F08 | 75320001 | 75420000 | 0.0055 | 0.0047 | 0.0055 |
| F08 | 75340001 | 75440000 | 0.0046 | 0.0038 | 0.0045 |
| F08 | 75360001 | 75460000 | 0.0049 | 0.0043 | 0.0048 |
| F08 | 75380001 | 75480000 | 0.0039 | 0.0038 | 0.0037 |
| F08 | 75400001 | 75500000 | 0.0036 | 0.0037 | 0.0034 |
| F08 | 75420001 | 75520000 | 0.0038 | 0.0040 | 0.0036 |
| F08 | 75440001 | 75540000 | 0.0044 | 0.0046 | 0.0042 |
| F08 | 75460001 | 75560000 | 0.0030 | 0.0033 | 0.0028 |
| F08 | 75480001 | 75580000 | 0.0028 | 0.0030 | 0.0026 |
| F08 | 75500001 | 75600000 | 0.0028 | 0.0030 | 0.0026 |
| F08 | 75520001 | 75620000 | 0.0020 | 0.0024 | 0.0018 |
| F08 | 75540001 | 75640000 | 0.0013 | 0.0020 | 0.0011 |
| F08 | 75560001 | 75660000 | 0.0020 | 0.0026 | 0.0017 |
| F08 | 75580001 | 75680000 | 0.0031 | 0.0039 | 0.0028 |
| F08 | 75600001 | 75700000 | 0.0030 | 0.0038 | 0.0026 |
| F08 | 75620001 | 75720000 | 0.0034 | 0.0041 | 0.0030 |
| F08 | 75640001 | 75740000 | 0.0045 | 0.0052 | 0.0041 |
| F08 | 75660001 | 75760000 | 0.0045 | 0.0058 | 0.0039 |
| F08 | 75680001 | 75780000 | 0.0041 | 0.0054 | 0.0035 |
| F08 | 75700001 | 75800000 | 0.0041 | 0.0055 | 0.0034 |
| F08 | 75720001 | 75820000 | 0.0042 | 0.0061 | 0.0034 |
| F08 | 75740001 | 75840000 | 0.0040 | 0.0057 | 0.0032 |
| F08 | 75760001 | 75860000 | 0.0037 | 0.0051 | 0.0030 |

| Chr | Start | End | both | A1 | A2 |
|---|---|---|---|---|---|
| F08 | 75780001 | 75880000 | 0.0030 | 0.0042 | 0.0024 |
| F08 | 75800001 | 75900000 | 0.0027 | 0.0036 | 0.0023 |
| F08 | 75820001 | 75920000 | 0.0021 | 0.0025 | 0.0019 |
| F08 | 75840001 | 75940000 | 0.0011 | 0.0014 | 0.0009 |
| F08 | 75860001 | 75960000 | 0.0008 | 0.0008 | 0.0007 |
| F08 | 75880001 | 75980000 | 0.0014 | 0.0012 | 0.0014 |
| F08 | 75900001 | 76000000 | 0.0012 | 0.0010 | 0.0012 |
| F08 | 75920001 | 76020000 | 0.0012 | 0.0010 | 0.0012 |
| F08 | 75940001 | 76040000 | 0.0020 | 0.0020 | 0.0019 |
| F08 | 75960001 | 76060000 | 0.0020 | 0.0023 | 0.0018 |
| F08 | 75980001 | 76080000 | 0.0014 | 0.0021 | 0.0012 |
| F08 | 76000001 | 76100000 | 0.0026 | 0.0033 | 0.0023 |
| F08 | 76020001 | 76120000 | 0.0038 | 0.0043 | 0.0034 |
| F08 | 76040001 | 76140000 | 0.0039 | 0.0044 | 0.0036 |
| F08 | 76060001 | 76160000 | 0.0036 | 0.0039 | 0.0034 |
| F08 | 76080001 | 76180000 | 0.0036 | 0.0038 | 0.0035 |
| F08 | 76100001 | 76200000 | 0.0031 | 0.0032 | 0.0029 |
| F08 | 76120001 | 76220000 | 0.0019 | 0.0023 | 0.0017 |
| F08 | 76140001 | 76240000 | 0.0012 | 0.0015 | 0.0011 |
| F08 | 76160001 | 76260000 | 0.0014 | 0.0017 | 0.0013 |
| F08 | 76180001 | 76280000 | 0.0013 | 0.0015 | 0.0012 |
| F08 | 76200001 | 76300000 | 0.0008 | 0.0010 | 0.0007 |
| F08 | 76220001 | 76320000 | 0.0010 | 0.0011 | 0.0009 |
| F08 | 76240001 | 76340000 | 0.0014 | 0.0014 | 0.0013 |
| F08 | 76260001 | 76360000 | 0.0020 | 0.0021 | 0.0019 |
| F08 | 76280001 | 76380000 | 0.0026 | 0.0028 | 0.0025 |
| F08 | 76300001 | 76400000 | 0.0027 | 0.0031 | 0.0025 |
| F08 | 76320001 | 76420000 | 0.0027 | 0.0033 | 0.0025 |
| F08 | 76340001 | 76440000 | 0.0028 | 0.0033 | 0.0025 |
| F08 | 76360001 | 76460000 | 0.0023 | 0.0030 | 0.0020 |
| F08 | 76380001 | 76480000 | 0.0018 | 0.0023 | 0.0016 |
| F08 | 76400001 | 76500000 | 0.0020 | 0.0024 | 0.0018 |
| F08 | 76420001 | 76520000 | 0.0021 | 0.0023 | 0.0019 |
| F08 | 76440001 | 76540000 | 0.0026 | 0.0029 | 0.0025 |
| F08 | 76460001 | 76560000 | 0.0025 | 0.0028 | 0.0024 |
| F08 | 76480001 | 76580000 | 0.0028 | 0.0032 | 0.0026 |
| F08 | 76500001 | 76600000 | 0.0030 | 0.0034 | 0.0028 |
| F08 | 76520001 | 76620000 | 0.0034 | 0.0039 | 0.0031 |
| F08 | 76540001 | 76640000 | 0.0024 | 0.0031 | 0.0021 |
| F08 | 76560001 | 76660000 | 0.0024 | 0.0030 | 0.0022 |
| F08 | 76580001 | 76680000 | 0.0019 | 0.0024 | 0.0017 |
| F08 | 76600001 | 76700000 | 0.0021 | 0.0027 | 0.0019 |
| F08 | 76620001 | 76720000 | 0.0018 | 0.0026 | 0.0015 |
| F08 | 76640001 | 76740000 | 0.0022 | 0.0031 | 0.0018 |
| F08 | 76660001 | 76760000 | 0.0021 | 0.0032 | 0.0016 |
| F08 | 76680001 | 76780000 | 0.0024 | 0.0037 | 0.0017 |
| F08 | 76700001 | 76800000 | 0.0019 | 0.0033 | 0.0013 |
| F08 | 76720001 | 76820000 | 0.0025 | 0.0034 | 0.0019 |
| F08 | 76740001 | 76840000 | 0.0030 | 0.0039 | 0.0024 |
| F08 | 76760001 | 76860000 | 0.0031 | 0.0037 | 0.0027 |
| F08 | 76780001 | 76880000 | 0.0030 | 0.0034 | 0.0026 |
| F08 | 76800001 | 76900000 | 0.0036 | 0.0038 | 0.0032 |
| F08 | 76820001 | 76920000 | 0.0035 | 0.0041 | 0.0030 |
| F08 | 76840001 | 76940000 | 0.0029 | 0.0033 | 0.0024 |
| F08 | 76860001 | 76960000 | 0.0033 | 0.0038 | 0.0028 |
| F08 | 76880001 | 76980000 | 0.0034 | 0.0039 | 0.0029 |
| F08 | 76900001 | 77000000 | 0.0031 | 0.0038 | 0.0025 |
| F08 | 76920001 | 77020000 | 0.0025 | 0.0031 | 0.0020 |

| Chr | Start | End | both | A1 | A2 |
|---|---|---|---|---|---|
| F08 | 76940001 | 77040000 | 0.0023 | 0.0029 | 0.0017 |
| F08 | 76960001 | 77060000 | 0.0017 | 0.0023 | 0.0011 |
| F08 | 76980001 | 77080000 | 0.0017 | 0.0022 | 0.0011 |
| F08 | 77000001 | 77100000 | 0.0016 | 0.0018 | 0.0009 |
| F08 | 77020001 | 77120000 | 0.0015 | 0.0017 | 0.0009 |
| F08 | 77040001 | 77140000 | 0.0021 | 0.0022 | 0.0015 |
| F08 | 77060001 | 77160000 | 0.0023 | 0.0024 | 0.0016 |
| F08 | 77080001 | 77180000 | 0.0024 | 0.0025 | 0.0017 |
| F08 | 77100001 | 77200000 | 0.0023 | 0.0025 | 0.0016 |
| F08 | 77120001 | 77220000 | 0.0027 | 0.0029 | 0.0020 |
| F08 | 77140001 | 77240000 | 0.0026 | 0.0029 | 0.0018 |
| F08 | 77160001 | 77260000 | 0.0025 | 0.0028 | 0.0017 |
| F08 | 77180001 | 77280000 | 0.0024 | 0.0028 | 0.0015 |
| F08 | 77200001 | 77300000 | 0.0025 | 0.0028 | 0.0016 |
| F08 | 77220001 | 77320000 | 0.0025 | 0.0026 | 0.0018 |
| F08 | 77240001 | 77340000 | 0.0022 | 0.0022 | 0.0016 |
| F08 | 77260001 | 77360000 | 0.0025 | 0.0026 | 0.0020 |
| F08 | 77280001 | 77380000 | 0.0023 | 0.0024 | 0.0018 |
| F08 | 77300001 | 77400000 | 0.0022 | 0.0022 | 0.0018 |
| F08 | 77320001 | 77420000 | 0.0018 | 0.0019 | 0.0015 |
| F08 | 77340001 | 77440000 | 0.0018 | 0.0017 | 0.0015 |
| F08 | 77360001 | 77460000 | 0.0013 | 0.0012 | 0.0011 |
| F08 | 77380001 | 77480000 | 0.0016 | 0.0014 | 0.0015 |
| F08 | 77400001 | 77500000 | 0.0015 | 0.0014 | 0.0013 |
| F08 | 77420001 | 77520000 | 0.0011 | 0.0011 | 0.0009 |
| F08 | 77440001 | 77540000 | 0.0019 | 0.0018 | 0.0017 |
| F08 | 77460001 | 77560000 | 0.0024 | 0.0023 | 0.0022 |
| F08 | 77480001 | 77580000 | 0.0026 | 0.0025 | 0.0023 |
| F08 | 77500001 | 77600000 | 0.0028 | 0.0026 | 0.0026 |
| F08 | 77520001 | 77620000 | 0.0036 | 0.0036 | 0.0032 |
| F08 | 77540001 | 77640000 | 0.0034 | 0.0035 | 0.0028 |
| F08 | 77560001 | 77660000 | 0.0032 | 0.0035 | 0.0025 |
| F08 | 77580001 | 77680000 | 0.0029 | 0.0032 | 0.0022 |
| F08 | 77600001 | 77700000 | 0.0030 | 0.0033 | 0.0023 |
| F08 | 77620001 | 77720000 | 0.0026 | 0.0025 | 0.0020 |
| F08 | 77640001 | 77740000 | 0.0019 | 0.0017 | 0.0015 |
| F08 | 77660001 | 77760000 | 0.0017 | 0.0014 | 0.0015 |
| F08 | 77680001 | 77780000 | 0.0015 | 0.0013 | 0.0013 |
| F08 | 77700001 | 77800000 | 0.0016 | 0.0016 | 0.0014 |
| F08 | 77720001 | 77820000 | 0.0014 | 0.0014 | 0.0012 |
| F08 | 77740001 | 77840000 | 0.0015 | 0.0016 | 0.0013 |
| F08 | 77760001 | 77860000 | 0.0018 | 0.0018 | 0.0015 |
| F08 | 77780001 | 77880000 | 0.0023 | 0.0024 | 0.0020 |
| F08 | 77800001 | 77900000 | 0.0025 | 0.0026 | 0.0021 |
| F08 | 77820001 | 77920000 | 0.0028 | 0.0028 | 0.0024 |
| F08 | 77840001 | 77940000 | 0.0027 | 0.0029 | 0.0022 |
| F08 | 77860001 | 77960000 | 0.0032 | 0.0034 | 0.0026 |
| F08 | 77880001 | 77980000 | 0.0030 | 0.0031 | 0.0024 |
| F08 | 77900001 | 78000000 | 0.0029 | 0.0029 | 0.0023 |
| F08 | 77920001 | 78020000 | 0.0032 | 0.0033 | 0.0026 |
| F08 | 77940001 | 78040000 | 0.0040 | 0.0039 | 0.0034 |
| F08 | 77960001 | 78060000 | 0.0033 | 0.0032 | 0.0028 |
| F08 | 77980001 | 78080000 | 0.0029 | 0.0028 | 0.0026 |
| F08 | 78000001 | 78100000 | 0.0023 | 0.0023 | 0.0021 |
| F08 | 78020001 | 78120000 | 0.0029 | 0.0030 | 0.0025 |
| F08 | 78040001 | 78140000 | 0.0026 | 0.0031 | 0.0019 |
| F08 | 78060001 | 78160000 | 0.0025 | 0.0030 | 0.0017 |
| F08 | 78080001 | 78180000 | 0.0026 | 0.0030 | 0.0017 |

| Chr | Start | End | both | A1 | A2 |
|---|---|---|---|---|---|
| F08 | 78100001 | 78200000 | 0.0029 | 0.0035 | 0.0021 |
| F08 | 78120001 | 78220000 | 0.0020 | 0.0023 | 0.0012 |
| F08 | 78140001 | 78240000 | 0.0015 | 0.0016 | 0.0010 |
| F08 | 78160001 | 78260000 | 0.0019 | 0.0020 | 0.0015 |
| F08 | 78180001 | 78280000 | 0.0019 | 0.0019 | 0.0015 |
| F08 | 78200001 | 78300000 | 0.0019 | 0.0019 | 0.0015 |
| F08 | 78220001 | 78320000 | 0.0026 | 0.0024 | 0.0022 |
| F08 | 78240001 | 78340000 | 0.0034 | 0.0030 | 0.0030 |
| F08 | 78260001 | 78360000 | 0.0032 | 0.0029 | 0.0028 |
| F08 | 78280001 | 78380000 | 0.0032 | 0.0029 | 0.0028 |
| F08 | 78300001 | 78400000 | 0.0028 | 0.0026 | 0.0024 |
| F08 | 78320001 | 78420000 | 0.0029 | 0.0027 | 0.0024 |
| F08 | 78340001 | 78440000 | 0.0025 | 0.0023 | 0.0020 |
| F08 | 78360001 | 78460000 | 0.0021 | 0.0020 | 0.0018 |
| F08 | 78380001 | 78480000 | 0.0028 | 0.0025 | 0.0026 |
| F08 | 78400001 | 78500000 | 0.0029 | 0.0026 | 0.0028 |
| F08 | 78420001 | 78520000 | 0.0029 | 0.0025 | 0.0028 |
| F08 | 78440001 | 78540000 | 0.0032 | 0.0029 | 0.0030 |
| F08 | 78460001 | 78560000 | 0.0049 | 0.0045 | 0.0046 |
| F08 | 78480001 | 78580000 | 0.0046 | 0.0044 | 0.0043 |
| F08 | 78500001 | 78600000 | 0.0048 | 0.0046 | 0.0044 |
| F08 | 78520001 | 78620000 | 0.0041 | 0.0040 | 0.0037 |
| F08 | 78540001 | 78640000 | 0.0035 | 0.0034 | 0.0031 |
| F08 | 78560001 | 78660000 | 0.0026 | 0.0025 | 0.0022 |
| F08 | 78580001 | 78680000 | 0.0025 | 0.0024 | 0.0020 |
| F08 | 78600001 | 78700000 | 0.0030 | 0.0030 | 0.0025 |
| F08 | 78620001 | 78720000 | 0.0034 | 0.0033 | 0.0029 |
| F08 | 78640001 | 78740000 | 0.0035 | 0.0035 | 0.0031 |
| F08 | 78660001 | 78760000 | 0.0029 | 0.0030 | 0.0024 |
| F08 | 78680001 | 78780000 | 0.0039 | 0.0039 | 0.0035 |
| F08 | 78700001 | 78800000 | 0.0034 | 0.0034 | 0.0030 |
| F08 | 78720001 | 78820000 | 0.0030 | 0.0030 | 0.0025 |
| F08 | 78740001 | 78840000 | 0.0027 | 0.0027 | 0.0022 |
| F08 | 78760001 | 78860000 | 0.0030 | 0.0029 | 0.0026 |
| F08 | 78780001 | 78880000 | 0.0017 | 0.0016 | 0.0014 |
| F08 | 78800001 | 78900000 | 0.0014 | 0.0012 | 0.0012 |
| F08 | 78820001 | 78920000 | 0.0017 | 0.0014 | 0.0016 |
| F08 | 78840001 | 78940000 | 0.0019 | 0.0016 | 0.0017 |
| F08 | 78860001 | 78960000 | 0.0016 | 0.0014 | 0.0014 |
| F08 | 78880001 | 78980000 | 0.0019 | 0.0017 | 0.0016 |
| F08 | 78900001 | 79000000 | 0.0028 | 0.0028 | 0.0023 |
| F08 | 78920001 | 79020000 | 0.0025 | 0.0024 | 0.0021 |
| F08 | 78940001 | 79040000 | 0.0027 | 0.0026 | 0.0024 |
| F08 | 78960001 | 79060000 | 0.0036 | 0.0032 | 0.0034 |
| F08 | 78980001 | 79080000 | 0.0041 | 0.0034 | 0.0038 |
| F08 | 79000001 | 79100000 | 0.0030 | 0.0024 | 0.0029 |
| F08 | 79020001 | 79120000 | 0.0034 | 0.0029 | 0.0032 |
| F08 | 79040001 | 79140000 | 0.0033 | 0.0028 | 0.0031 |
| F08 | 79060001 | 79160000 | 0.0024 | 0.0021 | 0.0022 |
| F08 | 79080001 | 79180000 | 0.0021 | 0.0021 | 0.0019 |
| F08 | 79100001 | 79200000 | 0.0023 | 0.0023 | 0.0020 |
| F08 | 79120001 | 79220000 | 0.0022 | 0.0022 | 0.0018 |
| F08 | 79140001 | 79240000 | 0.0033 | 0.0030 | 0.0029 |
| F08 | 79160001 | 79260000 | 0.0041 | 0.0038 | 0.0036 |
| F08 | 79180001 | 79280000 | 0.0046 | 0.0042 | 0.0043 |
| F08 | 79200001 | 79300000 | 0.0044 | 0.0040 | 0.0040 |
| F08 | 79220001 | 79320000 | 0.0042 | 0.0039 | 0.0037 |
| F08 | 79240001 | 79340000 | 0.0032 | 0.0031 | 0.0026 |

| Chr | Start | End | both | A1 | A2 |
|---|---|---|---|---|---|
| F08 | 79260001 | 79360000 | 0.0027 | 0.0028 | 0.0022 |
| F08 | 79280001 | 79380000 | 0.0021 | 0.0022 | 0.0016 |
| F08 | 79300001 | 79400000 | 0.0028 | 0.0028 | 0.0022 |
| F08 | 79320001 | 79420000 | 0.0027 | 0.0026 | 0.0023 |
| F08 | 79340001 | 79440000 | 0.0029 | 0.0029 | 0.0027 |
| F08 | 79360001 | 79460000 | 0.0025 | 0.0024 | 0.0023 |
| F08 | 79380001 | 79480000 | 0.0022 | 0.0021 | 0.0021 |
| F08 | 79400001 | 79500000 | 0.0015 | 0.0013 | 0.0014 |
| F08 | 79420001 | 79520000 | 0.0012 | 0.0011 | 0.0012 |
| F08 | 79440001 | 79540000 | 0.0012 | 0.0010 | 0.0011 |
| F08 | 79460001 | 79560000 | 0.0011 | 0.0010 | 0.0011 |
| F08 | 79480001 | 79580000 | 0.0010 | 0.0009 | 0.0010 |
| F08 | 79500001 | 79600000 | 0.0013 | 0.0012 | 0.0013 |
| F08 | 79520001 | 79620000 | 0.0015 | 0.0014 | 0.0015 |
| F08 | 79540001 | 79640000 | 0.0012 | 0.0011 | 0.0011 |
| F08 | 79560001 | 79660000 | 0.0011 | 0.0010 | 0.0010 |
| F08 | 79580001 | 79680000 | 0.0009 | 0.0009 | 0.0009 |
| F08 | 79600001 | 79700000 | 0.0007 | 0.0007 | 0.0007 |
| F08 | 79620001 | 79720000 | 0.0017 | 0.0017 | 0.0014 |
| F08 | 79640001 | 79740000 | 0.0021 | 0.0022 | 0.0016 |
| F08 | 79660001 | 79760000 | 0.0025 | 0.0027 | 0.0018 |
| F08 | 79680001 | 79780000 | 0.0030 | 0.0031 | 0.0021 |
| F08 | 79700001 | 79800000 | 0.0041 | 0.0042 | 0.0032 |
| F08 | 79720001 | 79820000 | 0.0040 | 0.0040 | 0.0031 |
| F08 | 79740001 | 79840000 | 0.0038 | 0.0037 | 0.0030 |
| F08 | 79760001 | 79860000 | 0.0038 | 0.0037 | 0.0031 |
| F08 | 79780001 | 79880000 | 0.0039 | 0.0038 | 0.0033 |
| F08 | 79800001 | 79900000 | 0.0033 | 0.0031 | 0.0025 |
| F08 | 79820001 | 79920000 | 0.0032 | 0.0031 | 0.0023 |
| F08 | 79840001 | 79940000 | 0.0034 | 0.0034 | 0.0025 |
| F08 | 79860001 | 79960000 | 0.0042 | 0.0041 | 0.0034 |
| F08 | 79880001 | 79980000 | 0.0044 | 0.0041 | 0.0036 |
| F08 | 79900001 | 80000000 | 0.0041 | 0.0038 | 0.0035 |
| F08 | 79920001 | 80020000 | 0.0040 | 0.0036 | 0.0037 |
| F08 | 79940001 | 80040000 | 0.0037 | 0.0032 | 0.0035 |
| F08 | 79960001 | 80060000 | 0.0027 | 0.0022 | 0.0027 |
| F08 | 79980001 | 80080000 | 0.0020 | 0.0017 | 0.0020 |
| F08 | 80000001 | 80100000 | 0.0024 | 0.0021 | 0.0023 |
| F08 | 80020001 | 80120000 | 0.0014 | 0.0013 | 0.0014 |
| F08 | 80040001 | 80140000 | 0.0013 | 0.0012 | 0.0013 |
| F08 | 80060001 | 80160000 | 0.0010 | 0.0010 | 0.0009 |
| F08 | 80080001 | 80180000 | 0.0014 | 0.0013 | 0.0013 |
| F08 | 80100001 | 80200000 | 0.0009 | 0.0008 | 0.0009 |
| F08 | 80120001 | 80220000 | 0.0015 | 0.0012 | 0.0015 |
| F08 | 80140001 | 80240000 | 0.0017 | 0.0015 | 0.0017 |
| F08 | 80160001 | 80260000 | 0.0024 | 0.0021 | 0.0023 |
| F08 | 80180001 | 80280000 | 0.0031 | 0.0029 | 0.0029 |
| F08 | 80200001 | 80300000 | 0.0036 | 0.0033 | 0.0034 |
| F08 | 80220001 | 80320000 | 0.0036 | 0.0034 | 0.0034 |
| F08 | 80240001 | 80340000 | 0.0038 | 0.0035 | 0.0035 |
| F08 | 80260001 | 80360000 | 0.0039 | 0.0037 | 0.0033 |
| F08 | 80280001 | 80380000 | 0.0034 | 0.0031 | 0.0029 |
| F08 | 80300001 | 80400000 | 0.0031 | 0.0030 | 0.0023 |
| F08 | 80320001 | 80420000 | 0.0031 | 0.0030 | 0.0022 |
| F08 | 80340001 | 80440000 | 0.0034 | 0.0032 | 0.0026 |
| F08 | 80360001 | 80460000 | 0.0031 | 0.0030 | 0.0025 |
| F08 | 80380001 | 80480000 | 0.0028 | 0.0027 | 0.0022 |
| F08 | 80400001 | 80500000 | 0.0029 | 0.0029 | 0.0025 |

| Chr | Start | End | both | A1 | A2 |
|---|---|---|---|---|---|
| F08 | 80420001 | 80520000 | 0.0029 | 0.0028 | 0.0025 |
| F08 | 80440001 | 80540000 | 0.0031 | 0.0032 | 0.0025 |
| F08 | 80460001 | 80560000 | 0.0030 | 0.0031 | 0.0023 |
| F08 | 80480001 | 80580000 | 0.0031 | 0.0032 | 0.0025 |
| F08 | 80500001 | 80600000 | 0.0032 | 0.0031 | 0.0025 |
| F08 | 80520001 | 80620000 | 0.0031 | 0.0031 | 0.0025 |
| F08 | 80540001 | 80640000 | 0.0032 | 0.0031 | 0.0026 |
| F08 | 80560001 | 80660000 | 0.0039 | 0.0035 | 0.0033 |
| F08 | 80580001 | 80680000 | 0.0038 | 0.0036 | 0.0028 |
| F08 | 80600001 | 80700000 | 0.0039 | 0.0039 | 0.0028 |
| F08 | 80620001 | 80720000 | 0.0036 | 0.0037 | 0.0026 |
| F08 | 80640001 | 80740000 | 0.0026 | 0.0026 | 0.0018 |
| F08 | 80660001 | 80760000 | 0.0020 | 0.0022 | 0.0011 |
| F08 | 80680001 | 80780000 | 0.0020 | 0.0020 | 0.0013 |
| F08 | 80700001 | 80800000 | 0.0020 | 0.0019 | 0.0015 |
| F08 | 80720001 | 80820000 | 0.0018 | 0.0018 | 0.0014 |
| F08 | 80740001 | 80840000 | 0.0028 | 0.0029 | 0.0022 |
| F08 | 80760001 | 80860000 | 0.0027 | 0.0027 | 0.0021 |
| F08 | 80780001 | 80880000 | 0.0028 | 0.0028 | 0.0022 |
| F08 | 80800001 | 80900000 | 0.0024 | 0.0025 | 0.0019 |
| F08 | 80820001 | 80920000 | 0.0034 | 0.0035 | 0.0029 |
| F08 | 80840001 | 80940000 | 0.0029 | 0.0028 | 0.0025 |
| F08 | 80860001 | 80960000 | 0.0029 | 0.0029 | 0.0024 |
| F08 | 80880001 | 80980000 | 0.0028 | 0.0027 | 0.0023 |
| F08 | 80900001 | 81000000 | 0.0032 | 0.0031 | 0.0026 |
| F08 | 80920001 | 81020000 | 0.0027 | 0.0027 | 0.0020 |
| F08 | 80940001 | 81040000 | 0.0028 | 0.0029 | 0.0020 |
| F08 | 80960001 | 81060000 | 0.0038 | 0.0039 | 0.0030 |
| F08 | 80980001 | 81080000 | 0.0038 | 0.0039 | 0.0029 |
| F08 | 81000001 | 81100000 | 0.0037 | 0.0037 | 0.0029 |
| F08 | 81020001 | 81120000 | 0.0041 | 0.0040 | 0.0035 |
| F08 | 81040001 | 81140000 | 0.0043 | 0.0041 | 0.0038 |
| F08 | 81060001 | 81160000 | 0.0030 | 0.0030 | 0.0025 |
| F08 | 81080001 | 81180000 | 0.0028 | 0.0027 | 0.0023 |
| F08 | 81100001 | 81200000 | 0.0023 | 0.0022 | 0.0019 |
| F08 | 81120001 | 81220000 | 0.0019 | 0.0020 | 0.0014 |
| F08 | 81140001 | 81240000 | 0.0011 | 0.0012 | 0.0007 |
| F08 | 81160001 | 81260000 | 0.0031 | 0.0028 | 0.0029 |
| F08 | 81180001 | 81280000 | 0.0032 | 0.0028 | 0.0030 |
| F08 | 81200001 | 81300000 | 0.0036 | 0.0031 | 0.0034 |
| F08 | 81220001 | 81320000 | 0.0035 | 0.0030 | 0.0034 |
| F08 | 81240001 | 81340000 | 0.0043 | 0.0037 | 0.0040 |
| F08 | 81260001 | 81360000 | 0.0029 | 0.0029 | 0.0025 |
| F08 | 81280001 | 81380000 | 0.0031 | 0.0029 | 0.0026 |
| F08 | 81300001 | 81400000 | 0.0033 | 0.0036 | 0.0026 |
| F08 | 81320001 | 81420000 | 0.0043 | 0.0044 | 0.0036 |
| F08 | 81340001 | 81440000 | 0.0039 | 0.0041 | 0.0032 |
| F08 | 81360001 | 81460000 | 0.0045 | 0.0046 | 0.0040 |
| F08 | 81380001 | 81480000 | 0.0045 | 0.0047 | 0.0039 |
| F08 | 81400001 | 81500000 | 0.0039 | 0.0038 | 0.0035 |
| F08 | 81420001 | 81520000 | 0.0030 | 0.0031 | 0.0024 |
| F08 | 81440001 | 81540000 | 0.0032 | 0.0032 | 0.0026 |
| F08 | 81460001 | 81560000 | 0.0021 | 0.0022 | 0.0013 |
| F08 | 81480001 | 81580000 | 0.0025 | 0.0027 | 0.0017 |
| F08 | 81500001 | 81600000 | 0.0030 | 0.0032 | 0.0023 |
| F08 | 81520001 | 81620000 | 0.0034 | 0.0035 | 0.0026 |
| F08 | 81540001 | 81640000 | 0.0032 | 0.0034 | 0.0023 |
| F08 | 81560001 | 81660000 | 0.0034 | 0.0035 | 0.0026 |

| Chr | Start | End | both | A1 | A2 |
|---|---|---|---|---|---|
| F08 | 81580001 | 81680000 | 0.0030 | 0.0031 | 0.0023 |
| F08 | 81600001 | 81700000 | 0.0025 | 0.0026 | 0.0016 |
| F08 | 81620001 | 81720000 | 0.0022 | 0.0023 | 0.0014 |
| F08 | 81640001 | 81740000 | 0.0034 | 0.0037 | 0.0024 |
| F08 | 81660001 | 81760000 | 0.0034 | 0.0037 | 0.0025 |
| F08 | 81680001 | 81780000 | 0.0039 | 0.0042 | 0.0029 |
| F08 | 81700001 | 81800000 | 0.0039 | 0.0042 | 0.0029 |
| F08 | 81720001 | 81820000 | 0.0036 | 0.0039 | 0.0027 |
| F08 | 81740001 | 81840000 | 0.0026 | 0.0026 | 0.0021 |
| F08 | 81760001 | 81860000 | 0.0021 | 0.0021 | 0.0017 |
| F08 | 81780001 | 81880000 | 0.0015 | 0.0015 | 0.0013 |
| F08 | 81800001 | 81900000 | 0.0017 | 0.0016 | 0.0016 |
| F08 | 81820001 | 81920000 | 0.0017 | 0.0016 | 0.0016 |
| F08 | 81840001 | 81940000 | 0.0023 | 0.0021 | 0.0021 |
| F08 | 81860001 | 81960000 | 0.0035 | 0.0032 | 0.0033 |
| F08 | 81880001 | 81980000 | 0.0033 | 0.0032 | 0.0029 |
| F08 | 81900001 | 82000000 | 0.0034 | 0.0032 | 0.0030 |
| F08 | 81920001 | 82020000 | 0.0040 | 0.0036 | 0.0035 |
| F08 | 81940001 | 82040000 | 0.0035 | 0.0033 | 0.0029 |
| F08 | 81960001 | 82060000 | 0.0029 | 0.0027 | 0.0023 |
| F08 | 81980001 | 82080000 | 0.0030 | 0.0028 | 0.0023 |
| F08 | 82000001 | 82100000 | 0.0028 | 0.0026 | 0.0022 |
| F08 | 82020001 | 82120000 | 0.0032 | 0.0032 | 0.0024 |
| F08 | 82040001 | 82140000 | 0.0028 | 0.0028 | 0.0020 |
| F08 | 82060001 | 82160000 | 0.0028 | 0.0028 | 0.0019 |
| F08 | 82080001 | 82180000 | 0.0033 | 0.0033 | 0.0024 |
| F08 | 82100001 | 82200000 | 0.0039 | 0.0038 | 0.0027 |
| F08 | 82120001 | 82220000 | 0.0035 | 0.0033 | 0.0026 |
| F08 | 82140001 | 82240000 | 0.0035 | 0.0032 | 0.0027 |
| F08 | 82160001 | 82260000 | 0.0031 | 0.0029 | 0.0025 |
| F08 | 82180001 | 82280000 | 0.0026 | 0.0023 | 0.0022 |
| F08 | 82200001 | 82300000 | 0.0021 | 0.0018 | 0.0019 |
| F08 | 82220001 | 82320000 | 0.0019 | 0.0017 | 0.0018 |
| F08 | 82240001 | 82340000 | 0.0022 | 0.0020 | 0.0020 |
| F08 | 82260001 | 82360000 | 0.0021 | 0.0020 | 0.0018 |
| F08 | 82280001 | 82380000 | 0.0024 | 0.0026 | 0.0017 |
| F08 | 82300001 | 82400000 | 0.0027 | 0.0028 | 0.0019 |
| F08 | 82320001 | 82420000 | 0.0028 | 0.0030 | 0.0019 |
| F08 | 82340001 | 82440000 | 0.0026 | 0.0028 | 0.0016 |
| F08 | 82360001 | 82460000 | 0.0030 | 0.0033 | 0.0020 |
| F08 | 82380001 | 82480000 | 0.0034 | 0.0035 | 0.0027 |
| F08 | 82400001 | 82500000 | 0.0029 | 0.0030 | 0.0022 |
| F08 | 82420001 | 82520000 | 0.0027 | 0.0029 | 0.0018 |
| F08 | 82440001 | 82540000 | 0.0033 | 0.0037 | 0.0024 |
| F08 | 82460001 | 82560000 | 0.0038 | 0.0042 | 0.0029 |
| F08 | 82480001 | 82580000 | 0.0039 | 0.0043 | 0.0029 |
| F08 | 82500001 | 82600000 | 0.0044 | 0.0047 | 0.0034 |
| F08 | 82520001 | 82620000 | 0.0043 | 0.0045 | 0.0035 |
| F08 | 82540001 | 82640000 | 0.0034 | 0.0034 | 0.0029 |
| F08 | 82560001 | 82660000 | 0.0023 | 0.0023 | 0.0019 |
| F08 | 82580001 | 82680000 | 0.0022 | 0.0021 | 0.0019 |
| F08 | 82600001 | 82700000 | 0.0023 | 0.0022 | 0.0022 |
| F08 | 82620001 | 82720000 | 0.0022 | 0.0022 | 0.0019 |
| F08 | 82640001 | 82740000 | 0.0025 | 0.0026 | 0.0020 |
| F08 | 82660001 | 82760000 | 0.0026 | 0.0027 | 0.0020 |
| F08 | 82680001 | 82780000 | 0.0021 | 0.0022 | 0.0015 |
| F08 | 82700001 | 82800000 | 0.0023 | 0.0026 | 0.0015 |
| F08 | 82720001 | 82820000 | 0.0027 | 0.0031 | 0.0018 |

| Chr | Start | End | both | A1 | A2 |
|---|---|---|---|---|---|
| F08 | 82740001 | 82840000 | 0.0025 | 0.0028 | 0.0016 |
| F08 | 82760001 | 82860000 | 0.0024 | 0.0028 | 0.0016 |
| F08 | 82780001 | 82880000 | 0.0025 | 0.0029 | 0.0016 |
| F08 | 82800001 | 82900000 | 0.0021 | 0.0023 | 0.0014 |
| F08 | 82820001 | 82920000 | 0.0020 | 0.0022 | 0.0016 |
| F08 | 82840001 | 82940000 | 0.0031 | 0.0031 | 0.0025 |
| F08 | 82860001 | 82960000 | 0.0032 | 0.0032 | 0.0026 |
| F08 | 82880001 | 82980000 | 0.0034 | 0.0034 | 0.0028 |
| F08 | 82900001 | 83000000 | 0.0037 | 0.0037 | 0.0031 |
| F08 | 82920001 | 83020000 | 0.0036 | 0.0034 | 0.0031 |
| F08 | 82940001 | 83040000 | 0.0028 | 0.0026 | 0.0024 |
| F08 | 82960001 | 83060000 | 0.0035 | 0.0031 | 0.0032 |
| F08 | 82980001 | 83080000 | 0.0030 | 0.0026 | 0.0029 |
| F08 | 83000001 | 83100000 | 0.0028 | 0.0024 | 0.0028 |
| F08 | 83020001 | 83120000 | 0.0025 | 0.0021 | 0.0025 |
| F08 | 83040001 | 83140000 | 0.0029 | 0.0025 | 0.0028 |
| F08 | 83060001 | 83160000 | 0.0021 | 0.0019 | 0.0021 |
| F08 | 83080001 | 83180000 | 0.0022 | 0.0019 | 0.0021 |
| F08 | 83100001 | 83200000 | 0.0021 | 0.0019 | 0.0020 |
| F08 | 83120001 | 83220000 | 0.0029 | 0.0027 | 0.0026 |
| F08 | 83140001 | 83240000 | 0.0023 | 0.0022 | 0.0020 |
| F08 | 83160001 | 83260000 | 0.0023 | 0.0022 | 0.0020 |
| F08 | 83180001 | 83280000 | 0.0027 | 0.0026 | 0.0022 |
| F08 | 83200001 | 83300000 | 0.0022 | 0.0021 | 0.0018 |
| F08 | 83220001 | 83320000 | 0.0021 | 0.0020 | 0.0016 |
| F08 | 83240001 | 83340000 | 0.0029 | 0.0027 | 0.0025 |
| F08 | 83260001 | 83360000 | 0.0033 | 0.0031 | 0.0027 |
| F08 | 83280001 | 83380000 | 0.0030 | 0.0029 | 0.0024 |
| F08 | 83300001 | 83400000 | 0.0033 | 0.0033 | 0.0025 |
| F08 | 83320001 | 83420000 | 0.0028 | 0.0030 | 0.0020 |
| F08 | 83340001 | 83440000 | 0.0024 | 0.0027 | 0.0014 |
| F08 | 83360001 | 83460000 | 0.0026 | 0.0031 | 0.0016 |
| F08 | 83380001 | 83480000 | 0.0025 | 0.0029 | 0.0016 |
| F08 | 83400001 | 83500000 | 0.0020 | 0.0024 | 0.0013 |
| F08 | 83420001 | 83520000 | 0.0019 | 0.0021 | 0.0014 |
| F08 | 83440001 | 83540000 | 0.0016 | 0.0016 | 0.0013 |
| F08 | 83460001 | 83560000 | 0.0025 | 0.0024 | 0.0020 |
| F08 | 83480001 | 83580000 | 0.0038 | 0.0037 | 0.0031 |
| F08 | 83500001 | 83600000 | 0.0055 | 0.0054 | 0.0047 |
| F08 | 83520001 | 83620000 | 0.0067 | 0.0066 | 0.0055 |
| F08 | 83540001 | 83640000 | 0.0064 | 0.0066 | 0.0052 |
| F08 | 83560001 | 83660000 | 0.0051 | 0.0052 | 0.0039 |
| F08 | 83580001 | 83680000 | 0.0037 | 0.0039 | 0.0028 |
| F08 | 83600001 | 83700000 | 0.0023 | 0.0026 | 0.0015 |
| F08 | 83620001 | 83720000 | 0.0011 | 0.0014 | 0.0005 |
| F08 | 83640001 | 83740000 | 0.0013 | 0.0016 | 0.0007 |
| F08 | 83660001 | 83760000 | 0.0013 | 0.0015 | 0.0008 |
| F08 | 83680001 | 83780000 | 0.0013 | 0.0015 | 0.0009 |
| F08 | 83700001 | 83800000 | 0.0010 | 0.0011 | 0.0006 |
| F08 | 83720001 | 83820000 | 0.0016 | 0.0017 | 0.0013 |
| F08 | 83740001 | 83840000 | 0.0020 | 0.0020 | 0.0016 |
| F08 | 83760001 | 83860000 | 0.0027 | 0.0028 | 0.0021 |
| F08 | 83780001 | 83880000 | 0.0037 | 0.0039 | 0.0028 |
| F08 | 83800001 | 83900000 | 0.0039 | 0.0041 | 0.0030 |
| F08 | 83820001 | 83920000 | 0.0042 | 0.0041 | 0.0034 |
| F08 | 83840001 | 83940000 | 0.0036 | 0.0035 | 0.0029 |
| F08 | 83860001 | 83960000 | 0.0034 | 0.0030 | 0.0030 |
| F08 | 83880001 | 83980000 | 0.0026 | 0.0023 | 0.0024 |

| Chr | Start | End | both | A1 | A2 |
|---|---|---|---|---|---|
| F08 | 83900001 | 84000000 | 0.0030 | 0.0029 | 0.0026 |
| F08 | 83920001 | 84020000 | 0.0021 | 0.0023 | 0.0015 |
| F08 | 83940001 | 84040000 | 0.0022 | 0.0025 | 0.0014 |
| F08 | 83960001 | 84060000 | 0.0018 | 0.0024 | 0.0008 |
| F08 | 83980001 | 84080000 | 0.0018 | 0.0023 | 0.0006 |
| F08 | 84000001 | 84100000 | 0.0016 | 0.0018 | 0.0006 |
| F08 | 84020001 | 84120000 | 0.0020 | 0.0022 | 0.0009 |
| F08 | 84040001 | 84140000 | 0.0029 | 0.0031 | 0.0019 |
| F08 | 84060001 | 84160000 | 0.0032 | 0.0033 | 0.0022 |
| F08 | 84080001 | 84180000 | 0.0036 | 0.0037 | 0.0027 |
| F08 | 84100001 | 84200000 | 0.0034 | 0.0036 | 0.0024 |
| F08 | 84120001 | 84220000 | 0.0028 | 0.0030 | 0.0020 |
| F08 | 84140001 | 84240000 | 0.0023 | 0.0025 | 0.0013 |
| F08 | 84160001 | 84260000 | 0.0021 | 0.0024 | 0.0010 |
| F08 | 84180001 | 84280000 | 0.0021 | 0.0025 | 0.0012 |
| F08 | 84200001 | 84300000 | 0.0026 | 0.0029 | 0.0017 |
| F08 | 84220001 | 84320000 | 0.0050 | 0.0053 | 0.0040 |
| F08 | 84240001 | 84340000 | 0.0048 | 0.0049 | 0.0041 |
| F08 | 84260001 | 84360000 | 0.0050 | 0.0052 | 0.0042 |
| F08 | 84280001 | 84380000 | 0.0052 | 0.0052 | 0.0043 |
| F08 | 84300001 | 84400000 | 0.0048 | 0.0049 | 0.0039 |
| F08 | 84320001 | 84420000 | 0.0031 | 0.0032 | 0.0023 |
| F08 | 84340001 | 84440000 | 0.0028 | 0.0030 | 0.0020 |
| F08 | 84360001 | 84460000 | 0.0032 | 0.0033 | 0.0026 |
| F08 | 84380001 | 84480000 | 0.0030 | 0.0032 | 0.0025 |
| F08 | 84400001 | 84500000 | 0.0028 | 0.0030 | 0.0023 |
| F08 | 84420001 | 84520000 | 0.0026 | 0.0029 | 0.0019 |
| F08 | 84440001 | 84540000 | 0.0035 | 0.0036 | 0.0028 |
| F08 | 84460001 | 84560000 | 0.0028 | 0.0028 | 0.0022 |
| F08 | 84480001 | 84580000 | 0.0025 | 0.0025 | 0.0019 |
| F08 | 84500001 | 84600000 | 0.0028 | 0.0029 | 0.0022 |
| F08 | 84520001 | 84620000 | 0.0026 | 0.0025 | 0.0021 |
| F08 | 84540001 | 84640000 | 0.0028 | 0.0028 | 0.0022 |
| F08 | 84560001 | 84660000 | 0.0026 | 0.0027 | 0.0019 |
| F08 | 84580001 | 84680000 | 0.0026 | 0.0027 | 0.0019 |
| F08 | 84600001 | 84700000 | 0.0024 | 0.0025 | 0.0016 |
| F08 | 84620001 | 84720000 | 0.0027 | 0.0029 | 0.0019 |
| F08 | 84640001 | 84740000 | 0.0018 | 0.0021 | 0.0008 |
| F08 | 84660001 | 84760000 | 0.0026 | 0.0030 | 0.0016 |
| F08 | 84680001 | 84780000 | 0.0026 | 0.0030 | 0.0014 |
| F08 | 84700001 | 84800000 | 0.0033 | 0.0034 | 0.0022 |
| F08 | 84720001 | 84820000 | 0.0030 | 0.0029 | 0.0022 |
| F08 | 84740001 | 84840000 | 0.0032 | 0.0031 | 0.0025 |
| F08 | 84760001 | 84860000 | 0.0023 | 0.0020 | 0.0018 |
| F08 | 84780001 | 84880000 | 0.0019 | 0.0016 | 0.0016 |
| F08 | 84800001 | 84900000 | 0.0018 | 0.0017 | 0.0015 |
| F08 | 84820001 | 84920000 | 0.0025 | 0.0024 | 0.0022 |
| F08 | 84840001 | 84940000 | 0.0036 | 0.0034 | 0.0031 |
| F08 | 84860001 | 84960000 | 0.0043 | 0.0042 | 0.0037 |
| F08 | 84880001 | 84980000 | 0.0047 | 0.0047 | 0.0037 |
| F08 | 84900001 | 85000000 | 0.0048 | 0.0049 | 0.0037 |
| F08 | 84920001 | 85020000 | 0.0042 | 0.0044 | 0.0029 |
| F08 | 84940001 | 85040000 | 0.0034 | 0.0039 | 0.0021 |
| F08 | 84960001 | 85060000 | 0.0029 | 0.0033 | 0.0018 |
| F08 | 84980001 | 85080000 | 0.0029 | 0.0032 | 0.0019 |
| F08 | 85000001 | 85100000 | 0.0026 | 0.0026 | 0.0018 |
| F08 | 85020001 | 85120000 | 0.0034 | 0.0034 | 0.0024 |
| F08 | 85040001 | 85140000 | 0.0034 | 0.0033 | 0.0027 |

| Chr | Start | End | both | A1 | A2 |
|---|---|---|---|---|---|
| F08 | 85060001 | 85160000 | 0.0041 | 0.0039 | 0.0032 |
| F08 | 85080001 | 85180000 | 0.0044 | 0.0041 | 0.0034 |
| F08 | 85100001 | 85200000 | 0.0043 | 0.0042 | 0.0033 |
| F08 | 85120001 | 85220000 | 0.0038 | 0.0037 | 0.0028 |
| F08 | 85140001 | 85240000 | 0.0034 | 0.0034 | 0.0023 |
| F08 | 85160001 | 85260000 | 0.0031 | 0.0030 | 0.0022 |
| F08 | 85180001 | 85280000 | 0.0027 | 0.0027 | 0.0018 |
| F08 | 85200001 | 85300000 | 0.0026 | 0.0026 | 0.0016 |
| F08 | 85220001 | 85320000 | 0.0024 | 0.0024 | 0.0015 |
| F08 | 85240001 | 85340000 | 0.0023 | 0.0023 | 0.0016 |
| F08 | 85260001 | 85360000 | 0.0022 | 0.0024 | 0.0014 |
| F08 | 85280001 | 85380000 | 0.0023 | 0.0024 | 0.0015 |
| F08 | 85300001 | 85400000 | 0.0028 | 0.0028 | 0.0022 |
| F08 | 85320001 | 85420000 | 0.0031 | 0.0031 | 0.0024 |
| F08 | 85340001 | 85440000 | 0.0030 | 0.0030 | 0.0023 |
| F08 | 85360001 | 85460000 | 0.0034 | 0.0034 | 0.0027 |
| F08 | 85380001 | 85480000 | 0.0034 | 0.0034 | 0.0026 |
| F08 | 85400001 | 85500000 | 0.0031 | 0.0032 | 0.0022 |
| F08 | 85420001 | 85520000 | 0.0038 | 0.0038 | 0.0031 |
| F08 | 85440001 | 85540000 | 0.0043 | 0.0043 | 0.0035 |
| F08 | 85460001 | 85560000 | 0.0042 | 0.0042 | 0.0035 |
| F08 | 85480001 | 85580000 | 0.0050 | 0.0049 | 0.0045 |
| F08 | 85500001 | 85600000 | 0.0053 | 0.0049 | 0.0050 |
| F08 | 85520001 | 85620000 | 0.0039 | 0.0036 | 0.0034 |
| F08 | 85540001 | 85640000 | 0.0039 | 0.0035 | 0.0035 |
| F08 | 85560001 | 85660000 | 0.0037 | 0.0033 | 0.0032 |
| F08 | 85580001 | 85680000 | 0.0031 | 0.0028 | 0.0024 |
| F08 | 85600001 | 85700000 | 0.0027 | 0.0024 | 0.0019 |
| F08 | 85620001 | 85720000 | 0.0025 | 0.0022 | 0.0019 |
| F08 | 85640001 | 85740000 | 0.0023 | 0.0021 | 0.0016 |
| F08 | 85660001 | 85760000 | 0.0024 | 0.0020 | 0.0019 |
| F08 | 85680001 | 85780000 | 0.0026 | 0.0018 | 0.0020 |
| F08 | 85700001 | 85800000 | 0.0028 | 0.0019 | 0.0022 |
| F08 | 85720001 | 85820000 | 0.0038 | 0.0025 | 0.0029 |
| F08 | 85740001 | 85840000 | 0.0042 | 0.0026 | 0.0031 |
| F08 | 85760001 | 85860000 | 0.0035 | 0.0019 | 0.0024 |
| F08 | 85780001 | 85880000 | 0.0043 | 0.0026 | 0.0031 |
| F08 | 85800001 | 85900000 | 0.0049 | 0.0030 | 0.0036 |
| F08 | 85820001 | 85920000 | 0.0042 | 0.0027 | 0.0031 |
| F08 | 85840001 | 85940000 | 0.0037 | 0.0023 | 0.0027 |
| F08 | 85860001 | 85960000 | 0.0039 | 0.0025 | 0.0029 |
| F08 | 85880001 | 85980000 | 0.0025 | 0.0015 | 0.0017 |
| F08 | 85900001 | 86000000 | 0.0021 | 0.0012 | 0.0013 |
| F08 | 85920001 | 86020000 | 0.0024 | 0.0015 | 0.0016 |
| F08 | 85940001 | 86040000 | 0.0022 | 0.0014 | 0.0014 |
| F08 | 85960001 | 86060000 | 0.0026 | 0.0018 | 0.0018 |
| F08 | 85980001 | 86080000 | 0.0035 | 0.0028 | 0.0027 |
| F08 | 86000001 | 86100000 | 0.0029 | 0.0021 | 0.0023 |
| F08 | 86020001 | 86120000 | 0.0031 | 0.0024 | 0.0025 |
| F08 | 86040001 | 86140000 | 0.0042 | 0.0034 | 0.0035 |
| F08 | 86060001 | 86160000 | 0.0043 | 0.0033 | 0.0036 |
| F08 | 86080001 | 86180000 | 0.0038 | 0.0027 | 0.0031 |
| F08 | 86100001 | 86200000 | 0.0041 | 0.0029 | 0.0033 |
| F08 | 86120001 | 86220000 | 0.0041 | 0.0029 | 0.0034 |
| F08 | 86140001 | 86240000 | 0.0034 | 0.0024 | 0.0029 |
| F08 | 86160001 | 86260000 | 0.0035 | 0.0024 | 0.0030 |
| F08 | 86180001 | 86280000 | 0.0031 | 0.0021 | 0.0026 |
| F08 | 86200001 | 86300000 | 0.0031 | 0.0020 | 0.0025 |

| Chr | Start | End | both | A1 | A2 |
|---|---|---|---|---|---|
| F08 | 86220001 | 86320000 | 0.0045 | 0.0033 | 0.0039 |
| F08 | 86240001 | 86340000 | 0.0044 | 0.0028 | 0.0036 |
| F08 | 86260001 | 86360000 | 0.0036 | 0.0023 | 0.0028 |
| F08 | 86280001 | 86380000 | 0.0038 | 0.0024 | 0.0030 |
| F08 | 86300001 | 86400000 | 0.0036 | 0.0023 | 0.0029 |
| F08 | 86320001 | 86420000 | 0.0018 | 0.0008 | 0.0011 |
| F08 | 86340001 | 86440000 | 0.0015 | 0.0007 | 0.0010 |
| F08 | 86360001 | 86460000 | 0.0025 | 0.0015 | 0.0019 |
| F08 | 86380001 | 86480000 | 0.0028 | 0.0019 | 0.0023 |
| F08 | 86400001 | 86500000 | 0.0032 | 0.0023 | 0.0027 |
| F08 | 86420001 | 86520000 | 0.0029 | 0.0021 | 0.0024 |
| F08 | 86440001 | 86540000 | 0.0031 | 0.0022 | 0.0026 |
| F08 | 86460001 | 86560000 | 0.0028 | 0.0019 | 0.0024 |
| F08 | 86480001 | 86580000 | 0.0026 | 0.0016 | 0.0021 |
| F08 | 86500001 | 86600000 | 0.0021 | 0.0012 | 0.0017 |
| F08 | 86520001 | 86620000 | 0.0023 | 0.0013 | 0.0018 |
| F08 | 86540001 | 86640000 | 0.0032 | 0.0020 | 0.0027 |
| F08 | 86560001 | 86660000 | 0.0029 | 0.0016 | 0.0024 |
| F08 | 86580001 | 86680000 | 0.0030 | 0.0019 | 0.0027 |
| F08 | 86600001 | 86700000 | 0.0045 | 0.0033 | 0.0041 |
| F08 | 86620001 | 86720000 | 0.0048 | 0.0033 | 0.0042 |
| F08 | 86640001 | 86740000 | 0.0040 | 0.0028 | 0.0036 |
| F08 | 86660001 | 86760000 | 0.0041 | 0.0032 | 0.0037 |
| F08 | 86680001 | 86780000 | 0.0038 | 0.0029 | 0.0033 |
| F08 | 86700001 | 86800000 | 0.0027 | 0.0017 | 0.0023 |
| F08 | 86720001 | 86820000 | 0.0024 | 0.0016 | 0.0020 |
| F08 | 86740001 | 86840000 | 0.0029 | 0.0021 | 0.0025 |
| F08 | 86760001 | 86860000 | 0.0023 | 0.0016 | 0.0020 |
| F08 | 86780001 | 86880000 | 0.0023 | 0.0017 | 0.0020 |
| F08 | 86800001 | 86900000 | 0.0022 | 0.0018 | 0.0020 |
| F08 | 86820001 | 86920000 | 0.0022 | 0.0018 | 0.0020 |
| F08 | 86840001 | 86940000 | 0.0021 | 0.0015 | 0.0018 |
| F08 | 86860001 | 86960000 | 0.0023 | 0.0015 | 0.0019 |
| F08 | 86880001 | 86980000 | 0.0025 | 0.0014 | 0.0019 |
| F08 | 86900001 | 87000000 | 0.0026 | 0.0013 | 0.0017 |
| F08 | 86920001 | 87020000 | 0.0031 | 0.0018 | 0.0022 |
| F08 | 86940001 | 87040000 | 0.0033 | 0.0021 | 0.0024 |
| F08 | 86960001 | 87060000 | 0.0036 | 0.0025 | 0.0028 |
| F08 | 86980001 | 87080000 | 0.0039 | 0.0030 | 0.0031 |
| F08 | 87000001 | 87100000 | 0.0039 | 0.0030 | 0.0031 |
| F08 | 87020001 | 87120000 | 0.0034 | 0.0026 | 0.0025 |
| F08 | 87040001 | 87140000 | 0.0034 | 0.0025 | 0.0025 |
| F08 | 87060001 | 87160000 | 0.0039 | 0.0031 | 0.0028 |
| F08 | 87080001 | 87180000 | 0.0038 | 0.0030 | 0.0028 |
| F08 | 87100001 | 87200000 | 0.0050 | 0.0041 | 0.0039 |
| F08 | 87120001 | 87220000 | 0.0054 | 0.0045 | 0.0043 |
| F08 | 87140001 | 87240000 | 0.0052 | 0.0045 | 0.0042 |
| F08 | 87160001 | 87260000 | 0.0042 | 0.0035 | 0.0034 |
| F08 | 87180001 | 87280000 | 0.0041 | 0.0032 | 0.0031 |
| F08 | 87200001 | 87300000 | 0.0025 | 0.0019 | 0.0018 |
| F08 | 87220001 | 87320000 | 0.0023 | 0.0017 | 0.0016 |
| F08 | 87240001 | 87340000 | 0.0019 | 0.0013 | 0.0012 |
| F08 | 87260001 | 87360000 | 0.0025 | 0.0018 | 0.0017 |
| F08 | 87280001 | 87380000 | 0.0028 | 0.0021 | 0.0020 |
| F08 | 87300001 | 87400000 | 0.0046 | 0.0038 | 0.0035 |
| F08 | 87320001 | 87420000 | 0.0047 | 0.0040 | 0.0038 |
| F08 | 87340001 | 87440000 | 0.0054 | 0.0046 | 0.0042 |
| F08 | 87360001 | 87460000 | 0.0050 | 0.0043 | 0.0039 |

| Chr | Start | End | both | A1 | A2 |
|---|---|---|---|---|---|
| F08 | 87380001 | 87480000 | 0.0052 | 0.0043 | 0.0040 |
| F08 | 87400001 | 87500000 | 0.0034 | 0.0026 | 0.0026 |
| F08 | 87420001 | 87520000 | 0.0033 | 0.0025 | 0.0026 |
| F08 | 87440001 | 87540000 | 0.0027 | 0.0019 | 0.0022 |
| F08 | 87460001 | 87560000 | 0.0029 | 0.0021 | 0.0022 |
| F08 | 87480001 | 87580000 | 0.0027 | 0.0021 | 0.0024 |
| F08 | 87500001 | 87600000 | 0.0034 | 0.0026 | 0.0031 |
| F08 | 87520001 | 87620000 | 0.0033 | 0.0023 | 0.0027 |
| F08 | 87540001 | 87640000 | 0.0031 | 0.0023 | 0.0025 |
| F08 | 87560001 | 87660000 | 0.0031 | 0.0022 | 0.0026 |
| F08 | 87580001 | 87680000 | 0.0040 | 0.0032 | 0.0033 |
| F08 | 87600001 | 87700000 | 0.0030 | 0.0026 | 0.0023 |
| F08 | 87620001 | 87720000 | 0.0031 | 0.0026 | 0.0024 |
| F08 | 87640001 | 87740000 | 0.0054 | 0.0048 | 0.0046 |
| F08 | 87660001 | 87760000 | 0.0059 | 0.0051 | 0.0050 |
| F08 | 87680001 | 87780000 | 0.0048 | 0.0038 | 0.0038 |
| F08 | 87700001 | 87800000 | 0.0068 | 0.0057 | 0.0055 |
| F08 | 87720001 | 87820000 | 0.0066 | 0.0055 | 0.0053 |
| F08 | 87740001 | 87840000 | 0.0044 | 0.0033 | 0.0030 |
| F08 | 87760001 | 87860000 | 0.0042 | 0.0034 | 0.0030 |
| F08 | 87780001 | 87880000 | 0.0039 | 0.0032 | 0.0028 |
| F08 | 87800001 | 87900000 | 0.0022 | 0.0014 | 0.0012 |
| F08 | 87820001 | 87920000 | 0.0039 | 0.0031 | 0.0030 |
| F08 | 87840001 | 87940000 | 0.0039 | 0.0031 | 0.0032 |
| F08 | 87860001 | 87960000 | 0.0040 | 0.0032 | 0.0032 |
| F08 | 87880001 | 87980000 | 0.0047 | 0.0038 | 0.0040 |
| F08 | 87900001 | 88000000 | 0.0054 | 0.0045 | 0.0046 |
| F08 | 87920001 | 88020000 | 0.0037 | 0.0029 | 0.0029 |
| F08 | 87940001 | 88040000 | 0.0035 | 0.0028 | 0.0027 |
| F08 | 87960001 | 88060000 | 0.0030 | 0.0023 | 0.0022 |
| F08 | 87980001 | 88080000 | 0.0020 | 0.0015 | 0.0013 |
| F08 | 88000001 | 88100000 | 0.0014 | 0.0009 | 0.0009 |
| F08 | 88020001 | 88120000 | 0.0014 | 0.0011 | 0.0012 |
| F08 | 88040001 | 88140000 | 0.0023 | 0.0019 | 0.0020 |
| F08 | 88060001 | 88160000 | 0.0033 | 0.0030 | 0.0029 |
| F08 | 88080001 | 88180000 | 0.0036 | 0.0031 | 0.0030 |
| F08 | 88100001 | 88200000 | 0.0037 | 0.0033 | 0.0032 |
| F08 | 88120001 | 88220000 | 0.0036 | 0.0032 | 0.0031 |
| F08 | 88140001 | 88240000 | 0.0028 | 0.0025 | 0.0023 |
| F08 | 88160001 | 88260000 | 0.0017 | 0.0015 | 0.0014 |
| F08 | 88180001 | 88280000 | 0.0022 | 0.0018 | 0.0020 |
| F08 | 88200001 | 88300000 | 0.0020 | 0.0016 | 0.0016 |
| F08 | 88220001 | 88320000 | 0.0021 | 0.0016 | 0.0016 |
| F08 | 88240001 | 88340000 | 0.0028 | 0.0020 | 0.0020 |
| F08 | 88260001 | 88360000 | 0.0029 | 0.0019 | 0.0018 |
| F08 | 88280001 | 88380000 | 0.0025 | 0.0016 | 0.0014 |
| F08 | 88300001 | 88400000 | 0.0030 | 0.0019 | 0.0018 |
| F08 | 88320001 | 88420000 | 0.0041 | 0.0028 | 0.0028 |
| F08 | 88340001 | 88440000 | 0.0034 | 0.0023 | 0.0022 |
| F08 | 88360001 | 88460000 | 0.0039 | 0.0026 | 0.0028 |
| F08 | 88380001 | 88480000 | 0.0037 | 0.0025 | 0.0026 |
| F08 | 88400001 | 88500000 | 0.0037 | 0.0025 | 0.0026 |
| F08 | 88420001 | 88520000 | 0.0031 | 0.0020 | 0.0022 |
| F08 | 88440001 | 88540000 | 0.0031 | 0.0021 | 0.0022 |
| F08 | 88460001 | 88560000 | 0.0024 | 0.0017 | 0.0017 |
| F08 | 88480001 | 88580000 | 0.0025 | 0.0017 | 0.0018 |
| F08 | 88500001 | 88600000 | 0.0021 | 0.0015 | 0.0016 |
| F08 | 88520001 | 88620000 | 0.0021 | 0.0014 | 0.0015 |

| Chr | Start | End | both | A1 | A2 |
|---|---|---|---|---|---|
| F08 | 88540001 | 88640000 | 0.0021 | 0.0014 | 0.0017 |
| F08 | 88560001 | 88660000 | 0.0019 | 0.0013 | 0.0014 |
| F08 | 88580001 | 88680000 | 0.0018 | 0.0013 | 0.0014 |
| F08 | 88600001 | 88700000 | 0.0018 | 0.0014 | 0.0015 |
| F08 | 88620001 | 88720000 | 0.0016 | 0.0012 | 0.0014 |
| F08 | 88640001 | 88740000 | 0.0020 | 0.0016 | 0.0017 |
| F08 | 88660001 | 88760000 | 0.0022 | 0.0017 | 0.0019 |
| F08 | 88680001 | 88780000 | 0.0023 | 0.0018 | 0.0022 |
| F08 | 88700001 | 88800000 | 0.0023 | 0.0018 | 0.0022 |
| F08 | 88720001 | 88820000 | 0.0019 | 0.0015 | 0.0018 |
| F08 | 88740001 | 88840000 | 0.0015 | 0.0012 | 0.0013 |
| F08 | 88760001 | 88860000 | 0.0017 | 0.0014 | 0.0015 |
| F08 | 88780001 | 88880000 | 0.0025 | 0.0021 | 0.0021 |
| F08 | 88800001 | 88900000 | 0.0030 | 0.0026 | 0.0026 |
| F08 | 88820001 | 88920000 | 0.0030 | 0.0026 | 0.0025 |
| F08 | 88840001 | 88940000 | 0.0030 | 0.0026 | 0.0026 |
| F08 | 88860001 | 88960000 | 0.0027 | 0.0024 | 0.0023 |
| F08 | 88880001 | 88980000 | 0.0023 | 0.0019 | 0.0020 |
| F08 | 88900001 | 89000000 | 0.0019 | 0.0016 | 0.0016 |
| F08 | 88920001 | 89020000 | 0.0020 | 0.0016 | 0.0016 |
| F08 | 88940001 | 89040000 | 0.0023 | 0.0019 | 0.0020 |
| F08 | 88960001 | 89060000 | 0.0031 | 0.0025 | 0.0027 |
| F08 | 88980001 | 89080000 | 0.0029 | 0.0024 | 0.0026 |
| F08 | 89000001 | 89100000 | 0.0029 | 0.0024 | 0.0025 |
| F08 | 89020001 | 89120000 | 0.0034 | 0.0027 | 0.0029 |
| F08 | 89040001 | 89140000 | 0.0033 | 0.0024 | 0.0026 |
| F08 | 89060001 | 89160000 | 0.0025 | 0.0018 | 0.0018 |
| F08 | 89080001 | 89180000 | 0.0025 | 0.0016 | 0.0016 |
| F08 | 89100001 | 89200000 | 0.0022 | 0.0015 | 0.0014 |
| F08 | 89120001 | 89220000 | 0.0019 | 0.0013 | 0.0012 |
| F08 | 89140001 | 89240000 | 0.0017 | 0.0012 | 0.0011 |
| F08 | 89160001 | 89260000 | 0.0018 | 0.0014 | 0.0014 |
| F08 | 89180001 | 89280000 | 0.0022 | 0.0016 | 0.0016 |
| F08 | 89200001 | 89300000 | 0.0031 | 0.0024 | 0.0026 |
| F08 | 89220001 | 89320000 | 0.0033 | 0.0026 | 0.0028 |
| F08 | 89240001 | 89340000 | 0.0037 | 0.0028 | 0.0031 |
| F08 | 89260001 | 89360000 | 0.0039 | 0.0029 | 0.0032 |
| F08 | 89280001 | 89380000 | 0.0044 | 0.0035 | 0.0037 |
| F08 | 89300001 | 89400000 | 0.0038 | 0.0029 | 0.0029 |
| F08 | 89320001 | 89420000 | 0.0043 | 0.0033 | 0.0031 |
| F08 | 89340001 | 89440000 | 0.0040 | 0.0030 | 0.0027 |
| F08 | 89360001 | 89460000 | 0.0041 | 0.0030 | 0.0028 |
| F08 | 89380001 | 89480000 | 0.0044 | 0.0032 | 0.0031 |
| F08 | 89400001 | 89500000 | 0.0051 | 0.0040 | 0.0038 |
| F08 | 89420001 | 89520000 | 0.0049 | 0.0038 | 0.0038 |
| F08 | 89440001 | 89540000 | 0.0053 | 0.0043 | 0.0043 |
| F08 | 89460001 | 89560000 | 0.0047 | 0.0040 | 0.0040 |
| F08 | 89480001 | 89580000 | 0.0037 | 0.0032 | 0.0031 |
| F08 | 89500001 | 89600000 | 0.0028 | 0.0022 | 0.0023 |
| F08 | 89520001 | 89620000 | 0.0023 | 0.0016 | 0.0017 |
| F08 | 89540001 | 89640000 | 0.0026 | 0.0018 | 0.0020 |
| F08 | 89560001 | 89660000 | 0.0035 | 0.0027 | 0.0027 |
| F08 | 89580001 | 89680000 | 0.0038 | 0.0028 | 0.0028 |
| F08 | 89600001 | 89700000 | 0.0036 | 0.0026 | 0.0024 |
| F08 | 89620001 | 89720000 | 0.0036 | 0.0026 | 0.0024 |
| F08 | 89640001 | 89740000 | 0.0031 | 0.0022 | 0.0020 |
| F08 | 89660001 | 89760000 | 0.0035 | 0.0027 | 0.0025 |
| F08 | 89680001 | 89780000 | 0.0038 | 0.0031 | 0.0030 |

| Chr | Start | End | both | A1 | A2 |
|---|---|---|---|---|---|
| F08 | 89700001 | 89800000 | 0.0050 | 0.0045 | 0.0046 |
| F08 | 89720001 | 89820000 | 0.0050 | 0.0045 | 0.0046 |
| F08 | 89740001 | 89840000 | 0.0048 | 0.0043 | 0.0044 |
| F08 | 89760001 | 89860000 | 0.0046 | 0.0040 | 0.0041 |
| F08 | 89780001 | 89880000 | 0.0047 | 0.0038 | 0.0038 |
| F08 | 89800001 | 89900000 | 0.0034 | 0.0024 | 0.0024 |
| F08 | 89820001 | 89920000 | 0.0032 | 0.0024 | 0.0023 |
| F08 | 89840001 | 89940000 | 0.0034 | 0.0025 | 0.0023 |
| F08 | 89860001 | 89960000 | 0.0032 | 0.0021 | 0.0019 |
| F08 | 89880001 | 89980000 | 0.0039 | 0.0028 | 0.0027 |
| F08 | 89900001 | 90000000 | 0.0043 | 0.0030 | 0.0029 |
| F08 | 89920001 | 90020000 | 0.0046 | 0.0032 | 0.0033 |
| F08 | 89940001 | 90040000 | 0.0044 | 0.0031 | 0.0031 |
| F08 | 89960001 | 90060000 | 0.0039 | 0.0029 | 0.0028 |
| F08 | 89980001 | 90080000 | 0.0029 | 0.0021 | 0.0019 |
| F08 | 90000001 | 90100000 | 0.0025 | 0.0018 | 0.0018 |
| F08 | 90020001 | 90120000 | 0.0027 | 0.0020 | 0.0020 |
| F08 | 90040001 | 90140000 | 0.0035 | 0.0027 | 0.0026 |
| F08 | 90060001 | 90160000 | 0.0034 | 0.0027 | 0.0028 |
| F08 | 90080001 | 90180000 | 0.0034 | 0.0027 | 0.0028 |
| F08 | 90100001 | 90200000 | 0.0037 | 0.0028 | 0.0028 |
| F08 | 90120001 | 90220000 | 0.0032 | 0.0025 | 0.0023 |
| F08 | 90140001 | 90240000 | 0.0027 | 0.0020 | 0.0018 |
| F08 | 90160001 | 90260000 | 0.0023 | 0.0016 | 0.0013 |
| F08 | 90180001 | 90280000 | 0.0020 | 0.0013 | 0.0010 |
| F08 | 90200001 | 90300000 | 0.0026 | 0.0018 | 0.0017 |
| F08 | 90220001 | 90320000 | 0.0026 | 0.0019 | 0.0019 |
| F08 | 90240001 | 90340000 | 0.0027 | 0.0020 | 0.0021 |
| F08 | 90260001 | 90360000 | 0.0029 | 0.0023 | 0.0024 |
| F08 | 90280001 | 90380000 | 0.0042 | 0.0035 | 0.0036 |
| F08 | 90300001 | 90400000 | 0.0032 | 0.0026 | 0.0027 |
| F08 | 90320001 | 90420000 | 0.0033 | 0.0027 | 0.0027 |
| F08 | 90340001 | 90440000 | 0.0031 | 0.0025 | 0.0026 |
| F08 | 90360001 | 90460000 | 0.0030 | 0.0024 | 0.0025 |
| F08 | 90380001 | 90480000 | 0.0019 | 0.0014 | 0.0015 |
| F08 | 90400001 | 90500000 | 0.0023 | 0.0018 | 0.0018 |
| F08 | 90420001 | 90520000 | 0.0025 | 0.0018 | 0.0018 |
| F08 | 90440001 | 90540000 | 0.0027 | 0.0020 | 0.0019 |
| F08 | 90460001 | 90560000 | 0.0027 | 0.0019 | 0.0016 |
| F08 | 90480001 | 90580000 | 0.0028 | 0.0018 | 0.0015 |
| F08 | 90500001 | 90600000 | 0.0034 | 0.0027 | 0.0022 |
| F08 | 90520001 | 90620000 | 0.0040 | 0.0035 | 0.0030 |
| F08 | 90540001 | 90640000 | 0.0036 | 0.0031 | 0.0026 |
| F08 | 90560001 | 90660000 | 0.0039 | 0.0034 | 0.0029 |
| F08 | 90580001 | 90680000 | 0.0035 | 0.0031 | 0.0027 |
| F08 | 90600001 | 90700000 | 0.0029 | 0.0023 | 0.0023 |
| F08 | 90620001 | 90720000 | 0.0021 | 0.0015 | 0.0013 |
| F08 | 90640001 | 90740000 | 0.0022 | 0.0016 | 0.0015 |
| F08 | 90660001 | 90760000 | 0.0020 | 0.0015 | 0.0014 |
| F08 | 90680001 | 90780000 | 0.0033 | 0.0027 | 0.0027 |
| F08 | 90700001 | 90800000 | 0.0050 | 0.0042 | 0.0043 |
| F08 | 90720001 | 90820000 | 0.0051 | 0.0045 | 0.0046 |
| F08 | 90740001 | 90840000 | 0.0054 | 0.0048 | 0.0049 |
| F08 | 90760001 | 90860000 | 0.0056 | 0.0049 | 0.0052 |
| F08 | 90780001 | 90880000 | 0.0047 | 0.0039 | 0.0042 |
| F08 | 90800001 | 90900000 | 0.0029 | 0.0022 | 0.0022 |
| F08 | 90820001 | 90920000 | 0.0025 | 0.0019 | 0.0019 |
| F08 | 90840001 | 90940000 | 0.0023 | 0.0017 | 0.0017 |

| Chr | Start | End | both | A1 | A2 |
|---|---|---|---|---|---|
| F08 | 90860001 | 90960000 | 0.0022 | 0.0016 | 0.0015 |
| F08 | 90880001 | 90980000 | 0.0021 | 0.0016 | 0.0014 |
| F08 | 90900001 | 91000000 | 0.0025 | 0.0022 | 0.0019 |
| F08 | 90920001 | 91020000 | 0.0042 | 0.0036 | 0.0037 |
| F08 | 90940001 | 91040000 | 0.0041 | 0.0034 | 0.0033 |
| F08 | 90960001 | 91060000 | 0.0039 | 0.0032 | 0.0031 |
| F08 | 90980001 | 91080000 | 0.0039 | 0.0034 | 0.0032 |
| F08 | 91000001 | 91100000 | 0.0033 | 0.0028 | 0.0025 |
| F08 | 91020001 | 91120000 | 0.0023 | 0.0021 | 0.0014 |
| F08 | 91040001 | 91140000 | 0.0023 | 0.0021 | 0.0016 |
| F08 | 91060001 | 91160000 | 0.0024 | 0.0022 | 0.0017 |
| F08 | 91080001 | 91180000 | 0.0023 | 0.0018 | 0.0016 |
| F08 | 91100001 | 91200000 | 0.0030 | 0.0022 | 0.0024 |
| F08 | 91120001 | 91220000 | 0.0023 | 0.0016 | 0.0018 |
| F08 | 91140001 | 91240000 | 0.0031 | 0.0025 | 0.0027 |
| F08 | 91160001 | 91260000 | 0.0038 | 0.0033 | 0.0035 |
| F08 | 91180001 | 91280000 | 0.0039 | 0.0035 | 0.0037 |
| F08 | 91200001 | 91300000 | 0.0034 | 0.0029 | 0.0032 |
| F08 | 91220001 | 91320000 | 0.0036 | 0.0029 | 0.0034 |
| F08 | 91240001 | 91340000 | 0.0030 | 0.0024 | 0.0029 |
| F08 | 91260001 | 91360000 | 0.0024 | 0.0017 | 0.0024 |
| F08 | 91280001 | 91380000 | 0.0026 | 0.0018 | 0.0026 |
| F08 | 91300001 | 91400000 | 0.0030 | 0.0024 | 0.0030 |
| F08 | 91320001 | 91420000 | 0.0032 | 0.0025 | 0.0032 |
| F08 | 91340001 | 91440000 | 0.0031 | 0.0021 | 0.0032 |
| F08 | 91360001 | 91460000 | 0.0031 | 0.0018 | 0.0031 |
| F08 | 91380001 | 91480000 | 0.0028 | 0.0014 | 0.0029 |
| F08 | 91400001 | 91500000 | 0.0027 | 0.0010 | 0.0028 |
| F08 | 91420001 | 91520000 | 0.0025 | 0.0007 | 0.0025 |
| F08 | 91440001 | 91540000 | 0.0025 | 0.0008 | 0.0023 |
| F08 | 91460001 | 91560000 | 0.0030 | 0.0013 | 0.0027 |
| F08 | 91480001 | 91580000 | 0.0039 | 0.0018 | 0.0034 |
| F08 | 91500001 | 91600000 | 0.0045 | 0.0024 | 0.0039 |
| F08 | 91520001 | 91620000 | 0.0046 | 0.0024 | 0.0039 |
| F08 | 91540001 | 91640000 | 0.0046 | 0.0025 | 0.0040 |
| F08 | 91560001 | 91660000 | 0.0040 | 0.0022 | 0.0034 |
| F08 | 91580001 | 91680000 | 0.0031 | 0.0019 | 0.0027 |
| F08 | 91600001 | 91700000 | 0.0021 | 0.0012 | 0.0018 |
| F08 | 91620001 | 91720000 | 0.0021 | 0.0013 | 0.0017 |
| F08 | 91640001 | 91740000 | 0.0030 | 0.0022 | 0.0027 |
| F08 | 91660001 | 91760000 | 0.0035 | 0.0024 | 0.0030 |
| F08 | 91680001 | 91780000 | 0.0041 | 0.0026 | 0.0034 |
| F08 | 91700001 | 91800000 | 0.0043 | 0.0029 | 0.0036 |
| F08 | 91720001 | 91820000 | 0.0043 | 0.0030 | 0.0036 |
| F08 | 91740001 | 91840000 | 0.0031 | 0.0019 | 0.0023 |
| F08 | 91760001 | 91860000 | 0.0026 | 0.0016 | 0.0017 |
| F08 | 91780001 | 91880000 | 0.0029 | 0.0022 | 0.0021 |
| F08 | 91800001 | 91900000 | 0.0028 | 0.0021 | 0.0021 |
| F08 | 91820001 | 91920000 | 0.0027 | 0.0019 | 0.0019 |
| F08 | 91840001 | 91940000 | 0.0029 | 0.0021 | 0.0021 |
| F08 | 91860001 | 91960000 | 0.0031 | 0.0023 | 0.0023 |
| F08 | 91880001 | 91980000 | 0.0035 | 0.0028 | 0.0029 |
| F08 | 91900001 | 92000000 | 0.0041 | 0.0035 | 0.0035 |
| F08 | 91920001 | 92020000 | 0.0045 | 0.0038 | 0.0039 |
| F08 | 91940001 | 92040000 | 0.0045 | 0.0038 | 0.0040 |
| F08 | 91960001 | 92060000 | 0.0043 | 0.0037 | 0.0039 |
| F08 | 91980001 | 92080000 | 0.0031 | 0.0023 | 0.0025 |
| F08 | 92000001 | 92100000 | 0.0025 | 0.0016 | 0.0018 |

| Chr | Start | End | both | A1 | A2 |
|---|---|---|---|---|---|
| F08 | 92020001 | 92120000 | 0.0025 | 0.0016 | 0.0019 |
| F08 | 92040001 | 92140000 | 0.0029 | 0.0019 | 0.0023 |
| F08 | 92060001 | 92160000 | 0.0041 | 0.0033 | 0.0036 |
| F08 | 92080001 | 92180000 | 0.0059 | 0.0053 | 0.0054 |
| F08 | 92100001 | 92200000 | 0.0058 | 0.0053 | 0.0054 |
| F08 | 92120001 | 92220000 | 0.0054 | 0.0049 | 0.0050 |
| F08 | 92140001 | 92240000 | 0.0045 | 0.0042 | 0.0041 |
| F08 | 92160001 | 92260000 | 0.0042 | 0.0037 | 0.0038 |
| F08 | 92180001 | 92280000 | 0.0031 | 0.0024 | 0.0026 |
| F08 | 92200001 | 92300000 | 0.0030 | 0.0023 | 0.0024 |
| F08 | 92220001 | 92320000 | 0.0029 | 0.0023 | 0.0024 |
| F08 | 92240001 | 92340000 | 0.0034 | 0.0025 | 0.0027 |
| F08 | 92260001 | 92360000 | 0.0028 | 0.0018 | 0.0019 |
| F08 | 92280001 | 92380000 | 0.0026 | 0.0015 | 0.0015 |
| F08 | 92300001 | 92400000 | 0.0030 | 0.0018 | 0.0019 |
| F08 | 92320001 | 92420000 | 0.0037 | 0.0025 | 0.0027 |
| F08 | 92340001 | 92440000 | 0.0039 | 0.0027 | 0.0029 |
| F08 | 92360001 | 92460000 | 0.0040 | 0.0029 | 0.0032 |
| F08 | 92380001 | 92480000 | 0.0043 | 0.0034 | 0.0037 |
| F08 | 92400001 | 92500000 | 0.0039 | 0.0030 | 0.0032 |
| F08 | 92420001 | 92520000 | 0.0032 | 0.0023 | 0.0024 |
| F08 | 92440001 | 92540000 | 0.0027 | 0.0019 | 0.0018 |
| F08 | 92460001 | 92560000 | 0.0026 | 0.0018 | 0.0017 |
| F08 | 92480001 | 92580000 | 0.0015 | 0.0008 | 0.0007 |
| F08 | 92500001 | 92600000 | 0.0015 | 0.0009 | 0.0008 |
| F08 | 92520001 | 92620000 | 0.0018 | 0.0012 | 0.0011 |
| F08 | 92540001 | 92640000 | 0.0021 | 0.0014 | 0.0014 |
| F08 | 92560001 | 92660000 | 0.0019 | 0.0013 | 0.0013 |
| F08 | 92580001 | 92680000 | 0.0022 | 0.0015 | 0.0015 |
| F08 | 92600001 | 92700000 | 0.0023 | 0.0016 | 0.0016 |
| F08 | 92620001 | 92720000 | 0.0022 | 0.0016 | 0.0016 |
| F08 | 92640001 | 92740000 | 0.0019 | 0.0013 | 0.0012 |
| F08 | 92660001 | 92760000 | 0.0017 | 0.0012 | 0.0011 |
| F08 | 92680001 | 92780000 | 0.0018 | 0.0015 | 0.0013 |
| F08 | 92700001 | 92800000 | 0.0029 | 0.0023 | 0.0025 |
| F08 | 92720001 | 92820000 | 0.0030 | 0.0024 | 0.0025 |
| F08 | 92740001 | 92840000 | 0.0034 | 0.0028 | 0.0029 |
| F08 | 92760001 | 92860000 | 0.0035 | 0.0027 | 0.0028 |
| F08 | 92780001 | 92880000 | 0.0043 | 0.0034 | 0.0035 |
| F08 | 92800001 | 92900000 | 0.0035 | 0.0028 | 0.0025 |
| F08 | 92820001 | 92920000 | 0.0037 | 0.0029 | 0.0025 |
| F08 | 92840001 | 92940000 | 0.0034 | 0.0027 | 0.0023 |
| F08 | 92860001 | 92960000 | 0.0041 | 0.0033 | 0.0031 |
| F08 | 92880001 | 92980000 | 0.0032 | 0.0024 | 0.0022 |
| F08 | 92900001 | 93000000 | 0.0028 | 0.0021 | 0.0020 |
| F08 | 92920001 | 93020000 | 0.0027 | 0.0021 | 0.0021 |
| F08 | 92940001 | 93040000 | 0.0028 | 0.0021 | 0.0020 |
| F08 | 92960001 | 93060000 | 0.0021 | 0.0015 | 0.0012 |
| F08 | 92980001 | 93080000 | 0.0019 | 0.0014 | 0.0011 |
| F08 | 93000001 | 93100000 | 0.0022 | 0.0016 | 0.0013 |
| F08 | 93020001 | 93120000 | 0.0024 | 0.0017 | 0.0013 |
| F08 | 93040001 | 93140000 | 0.0022 | 0.0015 | 0.0011 |
| F08 | 93060001 | 93160000 | 0.0022 | 0.0015 | 0.0012 |
| F08 | 93080001 | 93180000 | 0.0023 | 0.0015 | 0.0013 |
| F08 | 93100001 | 93200000 | 0.0032 | 0.0026 | 0.0023 |
| F08 | 93120001 | 93220000 | 0.0047 | 0.0041 | 0.0037 |
| F08 | 93140001 | 93240000 | 0.0048 | 0.0041 | 0.0037 |
| F08 | 93160001 | 93260000 | 0.0052 | 0.0044 | 0.0042 |

| Chr | Start | End | both | A1 | A2 |
|---|---|---|---|---|---|
| F08 | 93180001 | 93280000 | 0.0052 | 0.0044 | 0.0041 |
| F08 | 93200001 | 93300000 | 0.0043 | 0.0034 | 0.0031 |
| F08 | 93220001 | 93320000 | 0.0024 | 0.0016 | 0.0014 |
| F08 | 93240001 | 93340000 | 0.0024 | 0.0017 | 0.0015 |
| F08 | 93260001 | 93360000 | 0.0022 | 0.0015 | 0.0013 |
| F08 | 93280001 | 93380000 | 0.0027 | 0.0022 | 0.0019 |
| F08 | 93300001 | 93400000 | 0.0028 | 0.0022 | 0.0022 |
| F08 | 93320001 | 93420000 | 0.0029 | 0.0022 | 0.0024 |
| F08 | 93340001 | 93440000 | 0.0033 | 0.0026 | 0.0029 |
| F08 | 93360001 | 93460000 | 0.0036 | 0.0029 | 0.0032 |
| F08 | 93380001 | 93480000 | 0.0035 | 0.0027 | 0.0030 |
| F08 | 93400001 | 93500000 | 0.0033 | 0.0026 | 0.0027 |
| F08 | 93420001 | 93520000 | 0.0041 | 0.0033 | 0.0034 |
| F08 | 93440001 | 93540000 | 0.0054 | 0.0044 | 0.0045 |
| F08 | 93460001 | 93560000 | 0.0058 | 0.0046 | 0.0049 |
| F08 | 93480001 | 93580000 | 0.0075 | 0.0061 | 0.0065 |
| F08 | 93500001 | 93600000 | 0.0081 | 0.0069 | 0.0071 |
| F08 | 93520001 | 93620000 | 0.0077 | 0.0064 | 0.0066 |
| F08 | 93540001 | 93640000 | 0.0061 | 0.0050 | 0.0050 |
| F08 | 93560001 | 93660000 | 0.0057 | 0.0044 | 0.0043 |
| F08 | 93580001 | 93680000 | 0.0038 | 0.0025 | 0.0022 |
| F08 | 93600001 | 93700000 | 0.0031 | 0.0018 | 0.0017 |
| F08 | 93620001 | 93720000 | 0.0034 | 0.0021 | 0.0022 |
| F08 | 93640001 | 93740000 | 0.0038 | 0.0026 | 0.0028 |
| F08 | 93660001 | 93760000 | 0.0031 | 0.0023 | 0.0026 |
| F08 | 93680001 | 93780000 | 0.0034 | 0.0027 | 0.0031 |
| F08 | 93700001 | 93800000 | 0.0040 | 0.0032 | 0.0035 |
| F08 | 93720001 | 93820000 | 0.0028 | 0.0023 | 0.0025 |
| F08 | 93740001 | 93840000 | 0.0037 | 0.0030 | 0.0031 |
| F08 | 93760001 | 93860000 | 0.0043 | 0.0036 | 0.0037 |
| F08 | 93780001 | 93880000 | 0.0040 | 0.0034 | 0.0035 |
| F08 | 93800001 | 93900000 | 0.0032 | 0.0028 | 0.0029 |
| F08 | 93820001 | 93920000 | 0.0037 | 0.0033 | 0.0034 |
| F08 | 93840001 | 93940000 | 0.0023 | 0.0021 | 0.0023 |
| F08 | 93860001 | 93960000 | 0.0020 | 0.0017 | 0.0019 |
| F08 | 93880001 | 93980000 | 0.0015 | 0.0013 | 0.0014 |
| F08 | 93900001 | 94000000 | 0.0016 | 0.0013 | 0.0016 |
| F08 | 93920001 | 94020000 | 0.0010 | 0.0008 | 0.0010 |
| F08 | 93940001 | 94040000 | 0.0012 | 0.0010 | 0.0011 |
| F08 | 93960001 | 94060000 | 0.0010 | 0.0008 | 0.0010 |
| F08 | 93980001 | 94080000 | 0.0016 | 0.0012 | 0.0014 |
| F08 | 94000001 | 94100000 | 0.0025 | 0.0020 | 0.0022 |
| F08 | 94020001 | 94120000 | 0.0029 | 0.0024 | 0.0025 |
| F08 | 94040001 | 94140000 | 0.0030 | 0.0024 | 0.0025 |
| F08 | 94060001 | 94160000 | 0.0032 | 0.0025 | 0.0025 |
| F08 | 94080001 | 94180000 | 0.0044 | 0.0033 | 0.0036 |
| F08 | 94100001 | 94200000 | 0.0051 | 0.0041 | 0.0043 |
| F08 | 94120001 | 94220000 | 0.0051 | 0.0041 | 0.0044 |
| F08 | 94140001 | 94240000 | 0.0059 | 0.0047 | 0.0053 |
| F08 | 94160001 | 94260000 | 0.0060 | 0.0047 | 0.0053 |
| F08 | 94180001 | 94280000 | 0.0048 | 0.0039 | 0.0041 |
| F08 | 94200001 | 94300000 | 0.0030 | 0.0023 | 0.0025 |
| F08 | 94220001 | 94320000 | 0.0029 | 0.0023 | 0.0025 |
| F08 | 94240001 | 94340000 | 0.0027 | 0.0021 | 0.0020 |
| F08 | 94260001 | 94360000 | 0.0038 | 0.0033 | 0.0032 |
| F08 | 94280001 | 94380000 | 0.0036 | 0.0031 | 0.0030 |
| F08 | 94300001 | 94400000 | 0.0048 | 0.0042 | 0.0039 |
| F08 | 94320001 | 94420000 | 0.0052 | 0.0044 | 0.0041 |

| Chr | Start | End | both | A1 | A2 |
|---|---|---|---|---|---|
| F08 | 94340001 | 94440000 | 0.0048 | 0.0040 | 0.0038 |
| F08 | 94360001 | 94460000 | 0.0044 | 0.0035 | 0.0033 |
| F08 | 94380001 | 94480000 | 0.0048 | 0.0038 | 0.0038 |
| F08 | 94400001 | 94500000 | 0.0039 | 0.0029 | 0.0031 |
| F08 | 94420001 | 94520000 | 0.0034 | 0.0025 | 0.0027 |
| F08 | 94440001 | 94540000 | 0.0033 | 0.0025 | 0.0026 |
| F08 | 94460001 | 94560000 | 0.0027 | 0.0020 | 0.0021 |
| F08 | 94480001 | 94580000 | 0.0024 | 0.0018 | 0.0017 |
| F08 | 94500001 | 94600000 | 0.0025 | 0.0018 | 0.0019 |
| F08 | 94520001 | 94620000 | 0.0037 | 0.0029 | 0.0032 |
| F08 | 94540001 | 94640000 | 0.0042 | 0.0032 | 0.0036 |
| F08 | 94560001 | 94660000 | 0.0042 | 0.0031 | 0.0035 |
| F08 | 94580001 | 94680000 | 0.0042 | 0.0032 | 0.0036 |
| F08 | 94600001 | 94700000 | 0.0039 | 0.0030 | 0.0032 |
| F08 | 94620001 | 94720000 | 0.0030 | 0.0021 | 0.0022 |
| F08 | 94640001 | 94740000 | 0.0025 | 0.0017 | 0.0017 |
| F08 | 94660001 | 94760000 | 0.0024 | 0.0018 | 0.0018 |
| F08 | 94680001 | 94780000 | 0.0023 | 0.0017 | 0.0018 |
| F08 | 94700001 | 94800000 | 0.0025 | 0.0019 | 0.0020 |
| F08 | 94720001 | 94820000 | 0.0023 | 0.0017 | 0.0018 |
| F08 | 94740001 | 94840000 | 0.0021 | 0.0016 | 0.0016 |
| F08 | 94760001 | 94860000 | 0.0029 | 0.0023 | 0.0023 |
| F08 | 94780001 | 94880000 | 0.0030 | 0.0025 | 0.0024 |
| F08 | 94800001 | 94900000 | 0.0027 | 0.0022 | 0.0020 |
| F08 | 94820001 | 94920000 | 0.0025 | 0.0021 | 0.0018 |
| F08 | 94840001 | 94940000 | 0.0034 | 0.0027 | 0.0027 |
| F08 | 94860001 | 94960000 | 0.0026 | 0.0019 | 0.0020 |
| F08 | 94880001 | 94980000 | 0.0025 | 0.0017 | 0.0019 |
| F08 | 94900001 | 95000000 | 0.0026 | 0.0017 | 0.0020 |
| F08 | 94920001 | 95020000 | 0.0025 | 0.0017 | 0.0019 |
| F08 | 94940001 | 95040000 | 0.0016 | 0.0010 | 0.0010 |
| F08 | 94960001 | 95060000 | 0.0021 | 0.0013 | 0.0014 |
| F08 | 94980001 | 95080000 | 0.0019 | 0.0011 | 0.0012 |
| F08 | 95000001 | 95100000 | 0.0025 | 0.0019 | 0.0020 |
| F08 | 95020001 | 95120000 | 0.0028 | 0.0022 | 0.0024 |
| F08 | 95040001 | 95140000 | 0.0031 | 0.0025 | 0.0027 |
| F08 | 95060001 | 95160000 | 0.0023 | 0.0019 | 0.0021 |
| F08 | 95080001 | 95180000 | 0.0029 | 0.0026 | 0.0028 |
| F08 | 95100001 | 95200000 | 0.0028 | 0.0022 | 0.0027 |
| F08 | 95120001 | 95220000 | 0.0029 | 0.0023 | 0.0028 |
| F08 | 95140001 | 95240000 | 0.0034 | 0.0028 | 0.0032 |
| F08 | 95160001 | 95260000 | 0.0044 | 0.0038 | 0.0042 |
| F08 | 95180001 | 95280000 | 0.0036 | 0.0030 | 0.0034 |
| F08 | 95200001 | 95300000 | 0.0044 | 0.0038 | 0.0041 |
| F08 | 95220001 | 95320000 | 0.0049 | 0.0043 | 0.0047 |
| F08 | 95240001 | 95340000 | 0.0054 | 0.0048 | 0.0052 |
| F08 | 95260001 | 95360000 | 0.0049 | 0.0044 | 0.0047 |
| F08 | 95280001 | 95380000 | 0.0057 | 0.0051 | 0.0055 |
| F08 | 95300001 | 95400000 | 0.0051 | 0.0046 | 0.0048 |
| F08 | 95320001 | 95420000 | 0.0051 | 0.0045 | 0.0048 |
| F08 | 95340001 | 95440000 | 0.0049 | 0.0043 | 0.0045 |
| F08 | 95360001 | 95460000 | 0.0056 | 0.0049 | 0.0052 |
| F08 | 95380001 | 95480000 | 0.0060 | 0.0053 | 0.0056 |
| F08 | 95400001 | 95500000 | 0.0055 | 0.0048 | 0.0050 |
| F08 | 95420001 | 95520000 | 0.0045 | 0.0040 | 0.0041 |
| F08 | 95440001 | 95540000 | 0.0039 | 0.0033 | 0.0033 |
| F08 | 95460001 | 95560000 | 0.0030 | 0.0026 | 0.0024 |
| F08 | 95480001 | 95580000 | 0.0022 | 0.0019 | 0.0017 |

| Chr | Start | End | both | A1 | A2 |
|---|---|---|---|---|---|
| F08 | 95500001 | 95600000 | 0.0025 | 0.0021 | 0.0019 |
| F08 | 95520001 | 95620000 | 0.0044 | 0.0038 | 0.0038 |
| F08 | 95540001 | 95640000 | 0.0044 | 0.0036 | 0.0038 |
| F08 | 95560001 | 95660000 | 0.0044 | 0.0036 | 0.0037 |
| F08 | 95580001 | 95680000 | 0.0047 | 0.0038 | 0.0039 |
| F08 | 95600001 | 95700000 | 0.0051 | 0.0043 | 0.0044 |
| F08 | 95620001 | 95720000 | 0.0033 | 0.0026 | 0.0025 |
| F08 | 95640001 | 95740000 | 0.0039 | 0.0029 | 0.0030 |
| F08 | 95660001 | 95760000 | 0.0041 | 0.0031 | 0.0034 |
| F08 | 95680001 | 95780000 | 0.0038 | 0.0028 | 0.0033 |
| F08 | 95700001 | 95800000 | 0.0031 | 0.0022 | 0.0026 |
| F08 | 95720001 | 95820000 | 0.0031 | 0.0021 | 0.0027 |
| F08 | 95740001 | 95840000 | 0.0026 | 0.0018 | 0.0024 |
| F08 | 95760001 | 95860000 | 0.0022 | 0.0015 | 0.0018 |
| F08 | 95780001 | 95880000 | 0.0025 | 0.0018 | 0.0021 |
| F08 | 95800001 | 95900000 | 0.0027 | 0.0020 | 0.0024 |
| F08 | 95820001 | 95920000 | 0.0025 | 0.0019 | 0.0022 |
| F08 | 95840001 | 95940000 | 0.0025 | 0.0020 | 0.0023 |
| F08 | 95860001 | 95960000 | 0.0028 | 0.0025 | 0.0028 |
| F08 | 95880001 | 95980000 | 0.0020 | 0.0017 | 0.0019 |
| F08 | 95900001 | 96000000 | 0.0027 | 0.0021 | 0.0023 |
| F08 | 95920001 | 96020000 | 0.0033 | 0.0027 | 0.0027 |
| F08 | 95940001 | 96040000 | 0.0033 | 0.0027 | 0.0026 |
| F08 | 95960001 | 96060000 | 0.0031 | 0.0025 | 0.0025 |
| F08 | 95980001 | 96080000 | 0.0041 | 0.0035 | 0.0036 |
| F08 | 96000001 | 96100000 | 0.0034 | 0.0029 | 0.0030 |
| F08 | 96020001 | 96120000 | 0.0033 | 0.0030 | 0.0031 |
| F08 | 96040001 | 96140000 | 0.0029 | 0.0025 | 0.0027 |
| F08 | 96060001 | 96160000 | 0.0028 | 0.0025 | 0.0026 |
| F08 | 96080001 | 96180000 | 0.0028 | 0.0026 | 0.0026 |
| F08 | 96100001 | 96200000 | 0.0028 | 0.0027 | 0.0026 |
| F08 | 96120001 | 96220000 | 0.0027 | 0.0024 | 0.0023 |
| F08 | 96140001 | 96240000 | 0.0031 | 0.0026 | 0.0027 |
| F08 | 96160001 | 96260000 | 0.0028 | 0.0023 | 0.0023 |
| F08 | 96180001 | 96280000 | 0.0020 | 0.0015 | 0.0015 |
| F08 | 96200001 | 96300000 | 0.0021 | 0.0012 | 0.0015 |
| F08 | 96220001 | 96320000 | 0.0022 | 0.0013 | 0.0017 |
| F08 | 96240001 | 96340000 | 0.0020 | 0.0010 | 0.0014 |
| F08 | 96260001 | 96360000 | 0.0021 | 0.0010 | 0.0014 |
| F08 | 96280001 | 96380000 | 0.0021 | 0.0009 | 0.0013 |
| F08 | 96300001 | 96400000 | 0.0017 | 0.0009 | 0.0011 |
| F08 | 96320001 | 96420000 | 0.0018 | 0.0010 | 0.0011 |
| F08 | 96340001 | 96440000 | 0.0020 | 0.0013 | 0.0014 |
| F08 | 96360001 | 96460000 | 0.0021 | 0.0016 | 0.0016 |
| F08 | 96380001 | 96480000 | 0.0021 | 0.0015 | 0.0014 |
| F08 | 96400001 | 96500000 | 0.0019 | 0.0013 | 0.0012 |
| F08 | 96420001 | 96520000 | 0.0029 | 0.0022 | 0.0025 |
| F08 | 96440001 | 96540000 | 0.0040 | 0.0033 | 0.0034 |
| F08 | 96460001 | 96560000 | 0.0062 | 0.0058 | 0.0055 |
| F08 | 96480001 | 96580000 | 0.0071 | 0.0068 | 0.0064 |
| F08 | 96500001 | 96600000 | 0.0089 | 0.0081 | 0.0082 |
| F08 | 96520001 | 96620000 | 0.0080 | 0.0073 | 0.0072 |
| F08 | 96540001 | 96640000 | 0.0066 | 0.0059 | 0.0058 |
| F08 | 96560001 | 96660000 | 0.0044 | 0.0036 | 0.0039 |
| F08 | 96580001 | 96680000 | 0.0039 | 0.0029 | 0.0034 |
| F08 | 96600001 | 96700000 | 0.0023 | 0.0016 | 0.0017 |
| F08 | 96620001 | 96720000 | 0.0021 | 0.0013 | 0.0013 |
| F08 | 96640001 | 96740000 | 0.0026 | 0.0016 | 0.0018 |

| Chr | Start | End | both | A1 | A2 |
|---|---|---|---|---|---|
| F08 | 96660001 | 96760000 | 0.0022 | 0.0012 | 0.0014 |
| F08 | 96680001 | 96780000 | 0.0016 | 0.0009 | 0.0010 |
| F08 | 96700001 | 96800000 | 0.0016 | 0.0009 | 0.0011 |
| F08 | 96720001 | 96820000 | 0.0014 | 0.0009 | 0.0011 |
| F08 | 96740001 | 96840000 | 0.0021 | 0.0017 | 0.0019 |
| F08 | 96760001 | 96860000 | 0.0023 | 0.0017 | 0.0020 |
| F08 | 96780001 | 96880000 | 0.0023 | 0.0017 | 0.0020 |
| F08 | 96800001 | 96900000 | 0.0031 | 0.0023 | 0.0027 |
| F08 | 96820001 | 96920000 | 0.0032 | 0.0021 | 0.0026 |
| F08 | 96840001 | 96940000 | 0.0022 | 0.0012 | 0.0016 |
| F08 | 96860001 | 96960000 | 0.0025 | 0.0016 | 0.0021 |
| F08 | 96880001 | 96980000 | 0.0025 | 0.0017 | 0.0021 |
| F08 | 96900001 | 97000000 | 0.0033 | 0.0025 | 0.0030 |
| F08 | 96920001 | 97020000 | 0.0036 | 0.0030 | 0.0033 |
| F08 | 96940001 | 97040000 | 0.0039 | 0.0032 | 0.0034 |
| F08 | 96960001 | 97060000 | 0.0035 | 0.0028 | 0.0030 |
| F08 | 96980001 | 97080000 | 0.0037 | 0.0030 | 0.0032 |
| F08 | 97000001 | 97100000 | 0.0024 | 0.0017 | 0.0017 |
| F08 | 97020001 | 97120000 | 0.0027 | 0.0022 | 0.0021 |
| F08 | 97040001 | 97140000 | 0.0023 | 0.0019 | 0.0018 |
| F08 | 97060001 | 97160000 | 0.0029 | 0.0025 | 0.0024 |
| F08 | 97080001 | 97180000 | 0.0034 | 0.0030 | 0.0030 |
| F08 | 97100001 | 97200000 | 0.0032 | 0.0028 | 0.0028 |
| F08 | 97120001 | 97220000 | 0.0024 | 0.0019 | 0.0020 |
| F08 | 97140001 | 97240000 | 0.0024 | 0.0019 | 0.0020 |
| F08 | 97160001 | 97260000 | 0.0021 | 0.0015 | 0.0016 |
| F08 | 97180001 | 97280000 | 0.0014 | 0.0008 | 0.0008 |
| F08 | 97200001 | 97300000 | 0.0015 | 0.0008 | 0.0009 |
| F08 | 97220001 | 97320000 | 0.0018 | 0.0012 | 0.0013 |
| F08 | 97240001 | 97340000 | 0.0022 | 0.0014 | 0.0015 |
| F08 | 97260001 | 97360000 | 0.0019 | 0.0012 | 0.0013 |
| F08 | 97280001 | 97380000 | 0.0021 | 0.0012 | 0.0015 |
| F08 | 97300001 | 97400000 | 0.0022 | 0.0013 | 0.0016 |
| F08 | 97320001 | 97420000 | 0.0017 | 0.0009 | 0.0012 |
| F08 | 97340001 | 97440000 | 0.0018 | 0.0010 | 0.0011 |
| F08 | 97360001 | 97460000 | 0.0019 | 0.0011 | 0.0011 |
| F08 | 97380001 | 97480000 | 0.0019 | 0.0011 | 0.0009 |
| F08 | 97400001 | 97500000 | 0.0019 | 0.0011 | 0.0010 |
| F08 | 97420001 | 97520000 | 0.0020 | 0.0011 | 0.0009 |
| F08 | 97440001 | 97540000 | 0.0041 | 0.0032 | 0.0032 |
| F08 | 97460001 | 97560000 | 0.0043 | 0.0035 | 0.0034 |
| F08 | 97480001 | 97580000 | 0.0042 | 0.0034 | 0.0033 |
| F08 | 97500001 | 97600000 | 0.0043 | 0.0037 | 0.0035 |
| F08 | 97520001 | 97620000 | 0.0043 | 0.0037 | 0.0035 |
| F08 | 97540001 | 97640000 | 0.0018 | 0.0013 | 0.0010 |
| F08 | 97560001 | 97660000 | 0.0016 | 0.0011 | 0.0007 |
| F08 | 97580001 | 97680000 | 0.0019 | 0.0014 | 0.0010 |
| F08 | 97600001 | 97700000 | 0.0019 | 0.0012 | 0.0010 |
| F08 | 97620001 | 97720000 | 0.0020 | 0.0011 | 0.0013 |
| F08 | 97640001 | 97740000 | 0.0020 | 0.0012 | 0.0015 |
| F08 | 97660001 | 97760000 | 0.0022 | 0.0014 | 0.0018 |
| F08 | 97680001 | 97780000 | 0.0019 | 0.0013 | 0.0017 |
| F08 | 97700001 | 97800000 | 0.0014 | 0.0012 | 0.0015 |
| F08 | 97720001 | 97820000 | 0.0011 | 0.0012 | 0.0011 |
| F08 | 97740001 | 97840000 | 0.0009 | 0.0010 | 0.0008 |
| F08 | 97760001 | 97860000 | 0.0009 | 0.0010 | 0.0009 |
| F08 | 97780001 | 97880000 | 0.0008 | 0.0008 | 0.0007 |
| F08 | 97800001 | 97900000 | 0.0010 | 0.0010 | 0.0009 |

| Chr | Start | End | both | A1 | A2 |
|---|---|---|---|---|---|
| F08 | 97820001 | 97920000 | 0.0010 | 0.0010 | 0.0010 |
| F08 | 97840001 | 97940000 | 0.0021 | 0.0019 | 0.0021 |
| F08 | 97860001 | 97960000 | 0.0016 | 0.0015 | 0.0016 |
| F08 | 97880001 | 97980000 | 0.0015 | 0.0014 | 0.0015 |
| F08 | 97900001 | 98000000 | 0.0016 | 0.0014 | 0.0016 |
| F08 | 97920001 | 98020000 | 0.0016 | 0.0015 | 0.0016 |
| F08 | 97940001 | 98040000 | 0.0005 | 0.0005 | 0.0005 |
| F08 | 97960001 | 98060000 | 0.0007 | 0.0006 | 0.0007 |
| F08 | 97980001 | 98080000 | 0.0009 | 0.0007 | 0.0009 |
| F08 | 98000001 | 98100000 | 0.0009 | 0.0005 | 0.0009 |
| F08 | 98020001 | 98120000 | 0.0012 | 0.0006 | 0.0011 |
| F08 | 98040001 | 98140000 | 0.0018 | 0.0009 | 0.0017 |
| F08 | 98060001 | 98160000 | 0.0019 | 0.0009 | 0.0017 |
| F08 | 98080001 | 98180000 | 0.0021 | 0.0012 | 0.0019 |
| F08 | 98100001 | 98200000 | 0.0022 | 0.0012 | 0.0019 |
| F08 | 98120001 | 98220000 | 0.0023 | 0.0011 | 0.0021 |
| F08 | 98140001 | 98240000 | 0.0022 | 0.0008 | 0.0020 |
| F08 | 98160001 | 98260000 | 0.0026 | 0.0011 | 0.0025 |
| F08 | 98180001 | 98280000 | 0.0026 | 0.0012 | 0.0025 |
| F08 | 98200001 | 98300000 | 0.0031 | 0.0017 | 0.0031 |
| F08 | 98220001 | 98320000 | 0.0035 | 0.0022 | 0.0035 |
| F08 | 98240001 | 98340000 | 0.0045 | 0.0033 | 0.0044 |
| F08 | 98260001 | 98360000 | 0.0056 | 0.0041 | 0.0056 |
| F08 | 98280001 | 98380000 | 0.0064 | 0.0046 | 0.0063 |
| F08 | 98300001 | 98400000 | 0.0056 | 0.0039 | 0.0056 |
| F08 | 98320001 | 98420000 | 0.0066 | 0.0047 | 0.0065 |
| F08 | 98340001 | 98440000 | 0.0056 | 0.0036 | 0.0055 |
| F08 | 98360001 | 98460000 | 0.0044 | 0.0029 | 0.0044 |
| F08 | 98380001 | 98480000 | 0.0048 | 0.0032 | 0.0048 |
| F08 | 98400001 | 98500000 | 0.0053 | 0.0036 | 0.0053 |
| F08 | 98420001 | 98520000 | 0.0038 | 0.0024 | 0.0039 |
| F08 | 98440001 | 98540000 | 0.0038 | 0.0024 | 0.0038 |
| F08 | 98460001 | 98560000 | 0.0047 | 0.0035 | 0.0045 |
| F08 | 98480001 | 98580000 | 0.0036 | 0.0029 | 0.0031 |
| F08 | 98500001 | 98600000 | 0.0039 | 0.0034 | 0.0034 |
| F08 | 98520001 | 98620000 | 0.0038 | 0.0036 | 0.0032 |
| F08 | 98540001 | 98640000 | 0.0047 | 0.0046 | 0.0041 |
| F08 | 98560001 | 98660000 | 0.0035 | 0.0036 | 0.0030 |
| F08 | 98580001 | 98680000 | 0.0036 | 0.0038 | 0.0031 |
| F08 | 98600001 | 98700000 | 0.0031 | 0.0035 | 0.0024 |
| F08 | 98620001 | 98720000 | 0.0034 | 0.0037 | 0.0027 |
| F08 | 98640001 | 98740000 | 0.0021 | 0.0028 | 0.0015 |
| F08 | 98660001 | 98760000 | 0.0019 | 0.0026 | 0.0013 |
| F08 | 98680001 | 98780000 | 0.0014 | 0.0021 | 0.0010 |
| F08 | 98700001 | 98800000 | 0.0011 | 0.0017 | 0.0009 |
| F08 | 98720001 | 98820000 | 0.0017 | 0.0022 | 0.0015 |
| F08 | 98740001 | 98840000 | 0.0035 | 0.0037 | 0.0033 |
| F08 | 98760001 | 98860000 | 0.0046 | 0.0047 | 0.0044 |
| F08 | 98780001 | 98880000 | 0.0051 | 0.0052 | 0.0050 |
| F08 | 98800001 | 98900000 | 0.0057 | 0.0058 | 0.0056 |
| F08 | 98820001 | 98920000 | 0.0059 | 0.0058 | 0.0058 |
| F08 | 98840001 | 98940000 | 0.0048 | 0.0046 | 0.0046 |
| F08 | 98860001 | 98960000 | 0.0038 | 0.0034 | 0.0037 |
| F08 | 98880001 | 98980000 | 0.0034 | 0.0027 | 0.0032 |
| F08 | 98900001 | 99000000 | 0.0042 | 0.0032 | 0.0040 |
| F08 | 98920001 | 99020000 | 0.0036 | 0.0026 | 0.0034 |
| F08 | 98940001 | 99040000 | 0.0033 | 0.0026 | 0.0031 |
| F08 | 98960001 | 99060000 | 0.0032 | 0.0026 | 0.0030 |

| Chr | Start | End | both | A1 | A2 |
|---|---|---|---|---|---|
| F08 | 98980001 | 99080000 | 0.0033 | 0.0029 | 0.0032 |
| F08 | 99000001 | 99100000 | 0.0019 | 0.0017 | 0.0018 |
| F08 | 99020001 | 99120000 | 0.0013 | 0.0012 | 0.0013 |
| F08 | 99040001 | 99140000 | 0.0010 | 0.0010 | 0.0010 |
| F08 | 99060001 | 99160000 | 0.0010 | 0.0009 | 0.0009 |
| F08 | 99080001 | 99180000 | 0.0008 | 0.0007 | 0.0007 |
| F08 | 99100001 | 99200000 | 0.0009 | 0.0008 | 0.0008 |
| F08 | 99120001 | 99220000 | 0.0016 | 0.0013 | 0.0015 |
| F08 | 99140001 | 99240000 | 0.0025 | 0.0019 | 0.0023 |
| F08 | 99160001 | 99260000 | 0.0037 | 0.0028 | 0.0035 |
| F08 | 99180001 | 99280000 | 0.0040 | 0.0029 | 0.0037 |
| F08 | 99200001 | 99300000 | 0.0049 | 0.0035 | 0.0045 |
| F08 | 99220001 | 99320000 | 0.0046 | 0.0035 | 0.0043 |
| F08 | 99240001 | 99340000 | 0.0043 | 0.0035 | 0.0040 |
| F08 | 99260001 | 99360000 | 0.0033 | 0.0027 | 0.0029 |
| F08 | 99280001 | 99380000 | 0.0028 | 0.0026 | 0.0025 |
| F08 | 99300001 | 99400000 | 0.0021 | 0.0021 | 0.0016 |
| F08 | 99320001 | 99420000 | 0.0016 | 0.0017 | 0.0011 |
| F08 | 99340001 | 99440000 | 0.0010 | 0.0012 | 0.0005 |
| F08 | 99360001 | 99460000 | 0.0010 | 0.0015 | 0.0006 |
| F08 | 99380001 | 99480000 | 0.0014 | 0.0018 | 0.0011 |
| F08 | 99400001 | 99500000 | 0.0015 | 0.0020 | 0.0012 |
| F08 | 99420001 | 99520000 | 0.0016 | 0.0021 | 0.0012 |
| F08 | 99440001 | 99540000 | 0.0028 | 0.0027 | 0.0023 |
| F08 | 99460001 | 99560000 | 0.0027 | 0.0024 | 0.0022 |
| F08 | 99480001 | 99580000 | 0.0030 | 0.0025 | 0.0025 |
| F08 | 99500001 | 99600000 | 0.0027 | 0.0021 | 0.0023 |
| F08 | 99520001 | 99620000 | 0.0028 | 0.0022 | 0.0023 |
| F08 | 99540001 | 99640000 | 0.0018 | 0.0016 | 0.0012 |
| F08 | 99560001 | 99660000 | 0.0020 | 0.0019 | 0.0016 |
| F08 | 99580001 | 99680000 | 0.0013 | 0.0012 | 0.0008 |
| F08 | 99600001 | 99700000 | 0.0018 | 0.0016 | 0.0014 |
| F08 | 99620001 | 99720000 | 0.0017 | 0.0015 | 0.0015 |
| F08 | 99640001 | 99740000 | 0.0017 | 0.0015 | 0.0016 |
| F08 | 99660001 | 99760000 | 0.0016 | 0.0013 | 0.0014 |
| F08 | 99680001 | 99780000 | 0.0019 | 0.0016 | 0.0017 |
| F08 | 99700001 | 99800000 | 0.0017 | 0.0015 | 0.0015 |
| F08 | 99720001 | 99820000 | 0.0016 | 0.0014 | 0.0014 |
| F08 | 99740001 | 99840000 | 0.0016 | 0.0016 | 0.0012 |
| F08 | 99760001 | 99860000 | 0.0019 | 0.0019 | 0.0015 |
| F08 | 99780001 | 99880000 | 0.0016 | 0.0018 | 0.0012 |
| F08 | 99800001 | 99900000 | 0.0029 | 0.0030 | 0.0024 |
| F08 | 99820001 | 99920000 | 0.0032 | 0.0034 | 0.0026 |
| F08 | 99840001 | 99940000 | 0.0032 | 0.0035 | 0.0026 |
| F08 | 99860001 | 99960000 | 0.0029 | 0.0032 | 0.0023 |
| F08 | 99880001 | 99980000 | 0.0032 | 0.0036 | 0.0027 |
| F08 | 99900001 | 1E+08 | 0.0019 | 0.0026 | 0.0013 |
| F08 | 99920001 | 1E+08 | 0.0017 | 0.0022 | 0.0011 |
| F08 | 99940001 | 1E+08 | 0.0019 | 0.0020 | 0.0012 |
| F08 | 99960001 | 1E+08 | 0.0022 | 0.0022 | 0.0015 |
| F08 | 99980001 | 1E+08 | 0.0020 | 0.0017 | 0.0012 |
| F08 | 100000001 | 1E+08 | 0.0018 | 0.0013 | 0.0010 |
| F08 | 100020001 | 1E+08 | 0.0018 | 0.0013 | 0.0010 |
| F08 | 100040001 | 1E+08 | 0.0017 | 0.0012 | 0.0009 |
| F08 | 100060001 | 1E+08 | 0.0013 | 0.0007 | 0.0004 |
| F08 | 100080001 | 1E+08 | 0.0012 | 0.0007 | 0.0004 |
| F08 | 100100001 | 1E+08 | 0.0013 | 0.0008 | 0.0006 |
| F08 | 100120001 | 1E+08 | 0.0015 | 0.0010 | 0.0009 |

| Chr | Start | End | both | A1 | A2 |
|---|---|---|---|---|---|
| F08 | 100140001 | 1E+08 | 0.0016 | 0.0010 | 0.0009 |
| F08 | 100160001 | 1E+08 | 0.0016 | 0.0011 | 0.0009 |
| F08 | 100180001 | 1E+08 | 0.0017 | 0.0012 | 0.0011 |
| F08 | 100200001 | 1E+08 | 0.0016 | 0.0011 | 0.0009 |
| F08 | 100220001 | 1E+08 | 0.0013 | 0.0007 | 0.0005 |
| F08 | 100240001 | 1E+08 | 0.0014 | 0.0008 | 0.0005 |
| F08 | 100260001 | 1E+08 | 0.0015 | 0.0009 | 0.0005 |
| F08 | 100280001 | 1E+08 | 0.0015 | 0.0008 | 0.0005 |
| F08 | 100300001 | 1E+08 | 0.0019 | 0.0010 | 0.0009 |
| F08 | 100320001 | 1E+08 | 0.0021 | 0.0012 | 0.0011 |
| F08 | 100340001 | 1E+08 | 0.0025 | 0.0015 | 0.0018 |
| F08 | 100360001 | 1E+08 | 0.0025 | 0.0015 | 0.0019 |
| F08 | 100380001 | 1E+08 | 0.0025 | 0.0013 | 0.0019 |
| F08 | 100400001 | 1.01E+08 | 0.0025 | 0.0010 | 0.0018 |
| F08 | 100420001 | 1.01E+08 | 0.0025 | 0.0009 | 0.0018 |
| F08 | 100440001 | 1.01E+08 | 0.0020 | 0.0007 | 0.0013 |
| F08 | 100460001 | 1.01E+08 | 0.0021 | 0.0007 | 0.0013 |
| F08 | 100480001 | 1.01E+08 | 0.0020 | 0.0008 | 0.0012 |
| F08 | 100500001 | 1.01E+08 | 0.0016 | 0.0009 | 0.0009 |
| F08 | 100520001 | 1.01E+08 | 0.0016 | 0.0011 | 0.0009 |
| F08 | 100540001 | 1.01E+08 | 0.0019 | 0.0014 | 0.0012 |
| F08 | 100560001 | 1.01E+08 | 0.0036 | 0.0032 | 0.0031 |
| F08 | 100580001 | 1.01E+08 | 0.0039 | 0.0036 | 0.0036 |
| F08 | 100600001 | 1.01E+08 | 0.0039 | 0.0035 | 0.0036 |
| F08 | 100620001 | 1.01E+08 | 0.0040 | 0.0038 | 0.0036 |
| F08 | 100640001 | 1.01E+08 | 0.0037 | 0.0036 | 0.0032 |
| F08 | 100660001 | 1.01E+08 | 0.0019 | 0.0019 | 0.0014 |
| F08 | 100680001 | 1.01E+08 | 0.0017 | 0.0018 | 0.0011 |
| F08 | 100700001 | 1.01E+08 | 0.0016 | 0.0018 | 0.0011 |
| F08 | 100720001 | 1.01E+08 | 0.0012 | 0.0014 | 0.0008 |
| F08 | 100740001 | 1.01E+08 | 0.0009 | 0.0012 | 0.0007 |
| F08 | 100760001 | 1.01E+08 | 0.0018 | 0.0020 | 0.0017 |
| F08 | 100780001 | 1.01E+08 | 0.0020 | 0.0022 | 0.0019 |
| F08 | 100800001 | 1.01E+08 | 0.0019 | 0.0022 | 0.0018 |
| F08 | 100820001 | 1.01E+08 | 0.0019 | 0.0022 | 0.0018 |
| F08 | 100840001 | 1.01E+08 | 0.0020 | 0.0023 | 0.0018 |
| F08 | 100860001 | 1.01E+08 | 0.0029 | 0.0034 | 0.0028 |
| F08 | 100880001 | 1.01E+08 | 0.0032 | 0.0037 | 0.0030 |
| F08 | 100900001 | 1.01E+08 | 0.0038 | 0.0043 | 0.0036 |
| F08 | 100920001 | 1.01E+08 | 0.0052 | 0.0056 | 0.0050 |
| F08 | 100940001 | 1.01E+08 | 0.0053 | 0.0057 | 0.0052 |
| F08 | 100960001 | 1.01E+08 | 0.0043 | 0.0044 | 0.0041 |
| F08 | 100980001 | 1.01E+08 | 0.0034 | 0.0035 | 0.0033 |
| F08 | 101000001 | 1.01E+08 | 0.0029 | 0.0029 | 0.0028 |
| F08 | 101020001 | 1.01E+08 | 0.0014 | 0.0015 | 0.0014 |
| F08 | 101040001 | 1.01E+08 | 0.0013 | 0.0014 | 0.0012 |
| F08 | 101060001 | 1.01E+08 | 0.0003 | 0.0006 | 0.0002 |
| F08 | 101080001 | 1.01E+08 | 0.0005 | 0.0008 | 0.0004 |
| F08 | 101100001 | 1.01E+08 | 0.0005 | 0.0009 | 0.0004 |
| F08 | 101120001 | 1.01E+08 | 0.0008 | 0.0013 | 0.0006 |
| F08 | 101140001 | 1.01E+08 | 0.0008 | 0.0014 | 0.0005 |
| F08 | 101160001 | 1.01E+08 | 0.0009 | 0.0018 | 0.0005 |
| F08 | 101180001 | 1.01E+08 | 0.0011 | 0.0022 | 0.0006 |
| F08 | 101200001 | 1.01E+08 | 0.0012 | 0.0021 | 0.0007 |
| F08 | 101220001 | 1.01E+08 | 0.0011 | 0.0019 | 0.0007 |
| F08 | 101240001 | 1.01E+08 | 0.0011 | 0.0018 | 0.0008 |
| F08 | 101260001 | 1.01E+08 | 0.0014 | 0.0018 | 0.0012 |
| F08 | 101280001 | 1.01E+08 | 0.0017 | 0.0019 | 0.0016 |

| Chr | Start | End | both | A1 | A2 |
|---|---|---|---|---|---|
| F08 | 101300001 | 1.01E+08 | 0.0016 | 0.0018 | 0.0014 |
| F08 | 101320001 | 1.01E+08 | 0.0014 | 0.0017 | 0.0012 |
| F08 | 101340001 | 1.01E+08 | 0.0013 | 0.0017 | 0.0011 |
| F08 | 101360001 | 1.01E+08 | 0.0017 | 0.0022 | 0.0014 |
| F08 | 101380001 | 1.01E+08 | 0.0011 | 0.0018 | 0.0008 |
| F08 | 101400001 | 1.02E+08 | 0.0012 | 0.0020 | 0.0007 |
| F08 | 101420001 | 1.02E+08 | 0.0015 | 0.0025 | 0.0009 |
| F08 | 101440001 | 1.02E+08 | 0.0015 | 0.0025 | 0.0008 |
| F08 | 101460001 | 1.02E+08 | 0.0009 | 0.0017 | 0.0002 |
| F08 | 101480001 | 1.02E+08 | 0.0029 | 0.0037 | 0.0022 |
| F08 | 101500001 | 1.02E+08 | 0.0032 | 0.0037 | 0.0027 |
| F08 | 101520001 | 1.02E+08 | 0.0036 | 0.0040 | 0.0031 |
| F08 | 101540001 | 1.02E+08 | 0.0042 | 0.0044 | 0.0038 |
| F08 | 101560001 | 1.02E+08 | 0.0043 | 0.0045 | 0.0038 |
| F08 | 101580001 | 1.02E+08 | 0.0024 | 0.0027 | 0.0019 |
| F08 | 101600001 | 1.02E+08 | 0.0021 | 0.0024 | 0.0014 |
| F08 | 101620001 | 1.02E+08 | 0.0017 | 0.0018 | 0.0009 |
| F08 | 101640001 | 1.02E+08 | 0.0015 | 0.0015 | 0.0005 |
| F08 | 101660001 | 1.02E+08 | 0.0015 | 0.0013 | 0.0005 |
| F08 | 101680001 | 1.02E+08 | 0.0015 | 0.0011 | 0.0004 |
| F08 | 101700001 | 1.02E+08 | 0.0020 | 0.0019 | 0.0008 |
| F08 | 101720001 | 1.02E+08 | 0.0020 | 0.0022 | 0.0008 |
| F08 | 101740001 | 1.02E+08 | 0.0021 | 0.0026 | 0.0011 |
| F08 | 101760001 | 1.02E+08 | 0.0025 | 0.0031 | 0.0016 |
| F08 | 101780001 | 1.02E+08 | 0.0024 | 0.0032 | 0.0017 |
| F08 | 101800001 | 1.02E+08 | 0.0019 | 0.0024 | 0.0013 |
| F08 | 101820001 | 1.02E+08 | 0.0020 | 0.0023 | 0.0014 |
| F08 | 101840001 | 1.02E+08 | 0.0018 | 0.0020 | 0.0011 |
| F08 | 101860001 | 1.02E+08 | 0.0031 | 0.0032 | 0.0025 |
| F08 | 101880001 | 1.02E+08 | 0.0037 | 0.0037 | 0.0031 |
| F08 | 101900001 | 1.02E+08 | 0.0039 | 0.0038 | 0.0035 |
| F08 | 101920001 | 1.02E+08 | 0.0042 | 0.0040 | 0.0039 |
| F08 | 101940001 | 1.02E+08 | 0.0040 | 0.0037 | 0.0038 |
| F08 | 101960001 | 1.02E+08 | 0.0022 | 0.0019 | 0.0021 |
| F08 | 101980001 | 1.02E+08 | 0.0019 | 0.0016 | 0.0017 |
| F08 | 102000001 | 1.02E+08 | 0.0021 | 0.0019 | 0.0018 |
| F08 | 102020001 | 1.02E+08 | 0.0017 | 0.0016 | 0.0013 |
| F08 | 102040001 | 1.02E+08 | 0.0017 | 0.0016 | 0.0013 |
| F08 | 102060001 | 1.02E+08 | 0.0018 | 0.0018 | 0.0013 |
| F08 | 102080001 | 1.02E+08 | 0.0016 | 0.0017 | 0.0011 |
| F08 | 102100001 | 1.02E+08 | 0.0013 | 0.0015 | 0.0007 |
| F08 | 102120001 | 1.02E+08 | 0.0013 | 0.0015 | 0.0007 |
| F08 | 102140001 | 1.02E+08 | 0.0015 | 0.0019 | 0.0009 |
| F08 | 102160001 | 1.02E+08 | 0.0026 | 0.0028 | 0.0020 |
| F08 | 102180001 | 1.02E+08 | 0.0025 | 0.0028 | 0.0021 |
| F08 | 102200001 | 1.02E+08 | 0.0027 | 0.0026 | 0.0024 |
| F08 | 102220001 | 1.02E+08 | 0.0031 | 0.0027 | 0.0028 |
| F08 | 102240001 | 1.02E+08 | 0.0029 | 0.0022 | 0.0024 |
| F08 | 102260001 | 1.02E+08 | 0.0018 | 0.0010 | 0.0013 |
| F08 | 102280001 | 1.02E+08 | 0.0017 | 0.0009 | 0.0012 |
| F08 | 102300001 | 1.02E+08 | 0.0014 | 0.0010 | 0.0009 |
| F08 | 102320001 | 1.02E+08 | 0.0009 | 0.0008 | 0.0004 |
| F08 | 102340001 | 1.02E+08 | 0.0007 | 0.0009 | 0.0004 |
| F08 | 102360001 | 1.02E+08 | 0.0008 | 0.0013 | 0.0005 |
| F08 | 102380001 | 1.02E+08 | 0.0011 | 0.0018 | 0.0008 |
| F08 | 102400001 | 1.03E+08 | 0.0011 | 0.0019 | 0.0009 |
| F08 | 102420001 | 1.03E+08 | 0.0011 | 0.0019 | 0.0008 |
| F08 | 102440001 | 1.03E+08 | 0.0011 | 0.0018 | 0.0008 |

| Chr | Start | End | both | A1 | A2 |
|---|---|---|---|---|---|
| F08 | 102460001 | 1.03E+08 | 0.0010 | 0.0017 | 0.0007 |
| F08 | 102480001 | 1.03E+08 | 0.0005 | 0.0012 | 0.0003 |
| F08 | 102500001 | 1.03E+08 | 0.0005 | 0.0011 | 0.0003 |
| F08 | 102520001 | 1.03E+08 | 0.0008 | 0.0013 | 0.0006 |
| F08 | 102540001 | 1.03E+08 | 0.0009 | 0.0015 | 0.0006 |
| F08 | 102560001 | 1.03E+08 | 0.0010 | 0.0014 | 0.0005 |
| F08 | 102580001 | 1.03E+08 | 0.0014 | 0.0015 | 0.0008 |
| F08 | 102600001 | 1.03E+08 | 0.0017 | 0.0014 | 0.0009 |
| F08 | 102620001 | 1.03E+08 | 0.0017 | 0.0011 | 0.0006 |
| F08 | 102640001 | 1.03E+08 | 0.0017 | 0.0008 | 0.0006 |
| F08 | 102660001 | 1.03E+08 | 0.0018 | 0.0008 | 0.0007 |
| F08 | 102680001 | 1.03E+08 | 0.0017 | 0.0007 | 0.0004 |
| F08 | 102700001 | 1.03E+08 | 0.0016 | 0.0008 | 0.0004 |
| F08 | 102720001 | 1.03E+08 | 0.0016 | 0.0007 | 0.0004 |
| F08 | 102740001 | 1.03E+08 | 0.0016 | 0.0007 | 0.0005 |
| F08 | 102760001 | 1.03E+08 | 0.0016 | 0.0008 | 0.0006 |
| F08 | 102780001 | 1.03E+08 | 0.0016 | 0.0009 | 0.0007 |
| F08 | 102800001 | 1.03E+08 | 0.0016 | 0.0010 | 0.0007 |
| F08 | 102820001 | 1.03E+08 | 0.0017 | 0.0011 | 0.0007 |
| F08 | 102840001 | 1.03E+08 | 0.0017 | 0.0010 | 0.0008 |
| F08 | 102860001 | 1.03E+08 | 0.0017 | 0.0008 | 0.0008 |
| F08 | 102880001 | 1.03E+08 | 0.0018 | 0.0007 | 0.0010 |
| F08 | 102900001 | 1.03E+08 | 0.0025 | 0.0014 | 0.0020 |
| F08 | 102920001 | 1.03E+08 | 0.0023 | 0.0013 | 0.0020 |
| F08 | 102940001 | 1.03E+08 | 0.0023 | 0.0013 | 0.0019 |
| F08 | 102960001 | 1.03E+08 | 0.0021 | 0.0013 | 0.0017 |
| F08 | 102980001 | 1.03E+08 | 0.0019 | 0.0012 | 0.0014 |
| F08 | 103000001 | 1.03E+08 | 0.0015 | 0.0007 | 0.0009 |
| F08 | 103020001 | 1.03E+08 | 0.0017 | 0.0008 | 0.0009 |
| F08 | 103040001 | 1.03E+08 | 0.0017 | 0.0008 | 0.0009 |
| F08 | 103060001 | 1.03E+08 | 0.0019 | 0.0010 | 0.0010 |
| F08 | 103080001 | 1.03E+08 | 0.0019 | 0.0010 | 0.0011 |
| F08 | 103100001 | 1.03E+08 | 0.0018 | 0.0007 | 0.0010 |
| F08 | 103120001 | 1.03E+08 | 0.0020 | 0.0007 | 0.0012 |
| F08 | 103140001 | 1.03E+08 | 0.0021 | 0.0007 | 0.0013 |
| F08 | 103160001 | 1.03E+08 | 0.0021 | 0.0006 | 0.0013 |
| F08 | 103180001 | 1.03E+08 | 0.0022 | 0.0006 | 0.0011 |
| F08 | 103200001 | 1.03E+08 | 0.0022 | 0.0007 | 0.0010 |
| F08 | 103220001 | 1.03E+08 | 0.0020 | 0.0007 | 0.0009 |
| F08 | 103240001 | 1.03E+08 | 0.0019 | 0.0007 | 0.0007 |
| F08 | 103260001 | 1.03E+08 | 0.0021 | 0.0008 | 0.0009 |
| F08 | 103280001 | 1.03E+08 | 0.0019 | 0.0008 | 0.0009 |
| F08 | 103300001 | 1.03E+08 | 0.0018 | 0.0007 | 0.0009 |
| F08 | 103320001 | 1.03E+08 | 0.0019 | 0.0008 | 0.0010 |
| F08 | 103340001 | 1.03E+08 | 0.0021 | 0.0010 | 0.0013 |
| F08 | 103360001 | 1.03E+08 | 0.0020 | 0.0009 | 0.0011 |
| F08 | 103380001 | 1.03E+08 | 0.0021 | 0.0010 | 0.0011 |
| F08 | 103400001 | 1.04E+08 | 0.0019 | 0.0011 | 0.0009 |
| F08 | 103420001 | 1.04E+08 | 0.0018 | 0.0010 | 0.0006 |
| F08 | 103440001 | 1.04E+08 | 0.0015 | 0.0009 | 0.0003 |
| F08 | 103460001 | 1.04E+08 | 0.0015 | 0.0008 | 0.0004 |
| F08 | 103480001 | 1.04E+08 | 0.0017 | 0.0007 | 0.0005 |
| F08 | 103500001 | 1.04E+08 | 0.0017 | 0.0005 | 0.0005 |
| F08 | 103520001 | 1.04E+08 | 0.0016 | 0.0004 | 0.0005 |
| F08 | 103540001 | 1.04E+08 | 0.0017 | 0.0003 | 0.0006 |
| F08 | 103560001 | 1.04E+08 | 0.0016 | 0.0002 | 0.0005 |
| F08 | 103580001 | 1.04E+08 | 0.0014 | 0.0001 | 0.0004 |
| F08 | 103600001 | 1.04E+08 | 0.0018 | 0.0004 | 0.0007 |

| Chr | Start | End | both | A1 | A2 |
|---|---|---|---|---|---|
| F08 | 103620001 | 1.04E+08 | 0.0018 | 0.0005 | 0.0007 |
| F08 | 103640001 | 1.04E+08 | 0.0017 | 0.0006 | 0.0007 |
| F08 | 103660001 | 1.04E+08 | 0.0020 | 0.0009 | 0.0011 |
| F08 | 103680001 | 1.04E+08 | 0.0021 | 0.0010 | 0.0011 |
| F08 | 103700001 | 1.04E+08 | 0.0019 | 0.0011 | 0.0009 |
| F08 | 103720001 | 1.04E+08 | 0.0020 | 0.0014 | 0.0011 |
| F08 | 103740001 | 1.04E+08 | 0.0021 | 0.0017 | 0.0013 |
| F08 | 103760001 | 1.04E+08 | 0.0017 | 0.0015 | 0.0008 |
| F08 | 103780001 | 1.04E+08 | 0.0016 | 0.0015 | 0.0009 |
| F08 | 103800001 | 1.04E+08 | 0.0014 | 0.0012 | 0.0009 |
| F08 | 103820001 | 1.04E+08 | 0.0015 | 0.0009 | 0.0009 |
| F08 | 103840001 | 1.04E+08 | 0.0014 | 0.0005 | 0.0008 |
| F08 | 103860001 | 1.04E+08 | 0.0016 | 0.0004 | 0.0010 |
| F08 | 103880001 | 1.04E+08 | 0.0017 | 0.0004 | 0.0011 |
| F08 | 103900001 | 1.04E+08 | 0.0022 | 0.0009 | 0.0016 |
| F08 | 103920001 | 1.04E+08 | 0.0019 | 0.0010 | 0.0014 |
| F08 | 103940001 | 1.04E+08 | 0.0018 | 0.0013 | 0.0013 |
| F08 | 103960001 | 1.04E+08 | 0.0016 | 0.0016 | 0.0011 |
| F08 | 103980001 | 1.04E+08 | 0.0017 | 0.0022 | 0.0011 |
| F08 | 104000001 | 1.04E+08 | 0.0013 | 0.0020 | 0.0006 |
| F08 | 104020001 | 1.04E+08 | 0.0013 | 0.0019 | 0.0006 |
| F08 | 104040001 | 1.04E+08 | 0.0016 | 0.0018 | 0.0009 |
| F08 | 104060001 | 1.04E+08 | 0.0018 | 0.0016 | 0.0011 |
| F08 | 104080001 | 1.04E+08 | 0.0017 | 0.0009 | 0.0012 |
| F08 | 104100001 | 1.04E+08 | 0.0019 | 0.0007 | 0.0015 |
| F08 | 104120001 | 1.04E+08 | 0.0019 | 0.0005 | 0.0015 |
| F08 | 104140001 | 1.04E+08 | 0.0017 | 0.0003 | 0.0013 |
| F08 | 104160001 | 1.04E+08 | 0.0018 | 0.0005 | 0.0012 |
| F08 | 104180001 | 1.04E+08 | 0.0016 | 0.0007 | 0.0009 |
| F08 | 104200001 | 1.04E+08 | 0.0015 | 0.0010 | 0.0007 |
| F08 | 104220001 | 1.04E+08 | 0.0018 | 0.0016 | 0.0009 |
| F08 | 104240001 | 1.04E+08 | 0.0017 | 0.0017 | 0.0009 |
| F08 | 104260001 | 1.04E+08 | 0.0016 | 0.0016 | 0.0007 |
| F08 | 104280001 | 1.04E+08 | 0.0015 | 0.0014 | 0.0007 |
| F08 | 104300001 | 1.04E+08 | 0.0014 | 0.0011 | 0.0006 |
| F08 | 104320001 | 1.04E+08 | 0.0011 | 0.0005 | 0.0003 |
| F08 | 104340001 | 1.04E+08 | 0.0016 | 0.0008 | 0.0006 |
| F08 | 104360001 | 1.04E+08 | 0.0014 | 0.0006 | 0.0005 |
| F08 | 104380001 | 1.04E+08 | 0.0014 | 0.0005 | 0.0006 |
| F08 | 104400001 | 1.05E+08 | 0.0013 | 0.0006 | 0.0005 |
| F08 | 104420001 | 1.05E+08 | 0.0014 | 0.0007 | 0.0007 |
| F08 | 104440001 | 1.05E+08 | 0.0012 | 0.0005 | 0.0005 |
| F08 | 104460001 | 1.05E+08 | 0.0012 | 0.0006 | 0.0007 |
| F08 | 104480001 | 1.05E+08 | 0.0023 | 0.0017 | 0.0017 |
| F08 | 104500001 | 1.05E+08 | 0.0031 | 0.0027 | 0.0026 |
| F08 | 104520001 | 1.05E+08 | 0.0036 | 0.0032 | 0.0032 |
| F08 | 104540001 | 1.05E+08 | 0.0032 | 0.0030 | 0.0030 |
| F08 | 104560001 | 1.05E+08 | 0.0030 | 0.0029 | 0.0028 |
| F08 | 104580001 | 1.05E+08 | 0.0018 | 0.0017 | 0.0017 |
| F08 | 104600001 | 1.05E+08 | 0.0008 | 0.0007 | 0.0008 |
| F09 | 1 | 100000 | 0.0020 | 0.0015 | 0.0017 |
| F09 | 20001 | 120000 | 0.0017 | 0.0012 | 0.0013 |
| F09 | 40001 | 140000 | 0.0017 | 0.0012 | 0.0012 |
| F09 | 60001 | 160000 | 0.0028 | 0.0017 | 0.0020 |
| F09 | 80001 | 180000 | 0.0031 | 0.0018 | 0.0022 |
| F09 | 100001 | 200000 | 0.0034 | 0.0021 | 0.0024 |
| F09 | 120001 | 220000 | 0.0034 | 0.0018 | 0.0026 |
| F09 | 140001 | 240000 | 0.0037 | 0.0020 | 0.0030 |

| Chr | Start | End | both | A1 | A2 |
|---|---|---|---|---|---|
| F09 | 160001 | 260000 | 0.0032 | 0.0015 | 0.0029 |
| F09 | 180001 | 280000 | 0.0036 | 0.0015 | 0.0035 |
| F09 | 200001 | 300000 | 0.0045 | 0.0022 | 0.0045 |
| F09 | 220001 | 320000 | 0.0045 | 0.0024 | 0.0045 |
| F09 | 240001 | 340000 | 0.0052 | 0.0030 | 0.0052 |
| F09 | 260001 | 360000 | 0.0047 | 0.0031 | 0.0046 |
| F09 | 280001 | 380000 | 0.0042 | 0.0031 | 0.0040 |
| F09 | 300001 | 400000 | 0.0032 | 0.0021 | 0.0028 |
| F09 | 320001 | 420000 | 0.0036 | 0.0023 | 0.0028 |
| F09 | 340001 | 440000 | 0.0024 | 0.0013 | 0.0015 |
| F09 | 360001 | 460000 | 0.0024 | 0.0013 | 0.0015 |
| F09 | 380001 | 480000 | 0.0027 | 0.0017 | 0.0018 |
| F09 | 400001 | 500000 | 0.0031 | 0.0021 | 0.0022 |
| F09 | 420001 | 520000 | 0.0025 | 0.0018 | 0.0018 |
| F09 | 440001 | 540000 | 0.0028 | 0.0021 | 0.0022 |
| F09 | 460001 | 560000 | 0.0031 | 0.0023 | 0.0026 |
| F09 | 480001 | 580000 | 0.0025 | 0.0018 | 0.0021 |
| F09 | 500001 | 600000 | 0.0021 | 0.0015 | 0.0018 |
| F09 | 520001 | 620000 | 0.0021 | 0.0015 | 0.0020 |
| F09 | 540001 | 640000 | 0.0018 | 0.0013 | 0.0018 |
| F09 | 560001 | 660000 | 0.0020 | 0.0014 | 0.0016 |
| F09 | 580001 | 680000 | 0.0020 | 0.0014 | 0.0015 |
| F09 | 600001 | 700000 | 0.0020 | 0.0014 | 0.0013 |
| F09 | 620001 | 720000 | 0.0019 | 0.0015 | 0.0011 |
| F09 | 640001 | 740000 | 0.0023 | 0.0018 | 0.0012 |
| F09 | 660001 | 760000 | 0.0018 | 0.0016 | 0.0010 |
| F09 | 680001 | 780000 | 0.0020 | 0.0016 | 0.0011 |
| F09 | 700001 | 800000 | 0.0029 | 0.0020 | 0.0019 |
| F09 | 720001 | 820000 | 0.0029 | 0.0019 | 0.0020 |
| F09 | 740001 | 840000 | 0.0026 | 0.0017 | 0.0018 |
| F09 | 760001 | 860000 | 0.0027 | 0.0017 | 0.0021 |
| F09 | 780001 | 880000 | 0.0026 | 0.0017 | 0.0021 |
| F09 | 800001 | 900000 | 0.0017 | 0.0013 | 0.0014 |
| F09 | 820001 | 920000 | 0.0016 | 0.0011 | 0.0013 |
| F09 | 840001 | 940000 | 0.0016 | 0.0010 | 0.0012 |
| F09 | 860001 | 960000 | 0.0017 | 0.0007 | 0.0011 |
| F09 | 880001 | 980000 | 0.0019 | 0.0007 | 0.0013 |
| F09 | 900001 | 1000000 | 0.0023 | 0.0009 | 0.0017 |
| F09 | 920001 | 1020000 | 0.0023 | 0.0009 | 0.0017 |
| F09 | 940001 | 1040000 | 0.0027 | 0.0012 | 0.0020 |
| F09 | 960001 | 1060000 | 0.0040 | 0.0023 | 0.0028 |
| F09 | 980001 | 1080000 | 0.0046 | 0.0032 | 0.0034 |
| F09 | 1000001 | 1100000 | 0.0045 | 0.0034 | 0.0033 |
| F09 | 1020001 | 1120000 | 0.0051 | 0.0039 | 0.0037 |
| F09 | 1040001 | 1140000 | 0.0047 | 0.0038 | 0.0036 |
| F09 | 1060001 | 1160000 | 0.0040 | 0.0031 | 0.0032 |
| F09 | 1080001 | 1180000 | 0.0046 | 0.0031 | 0.0036 |
| F09 | 1100001 | 1200000 | 0.0048 | 0.0030 | 0.0035 |
| F09 | 1120001 | 1220000 | 0.0047 | 0.0028 | 0.0034 |
| F09 | 1140001 | 1240000 | 0.0049 | 0.0030 | 0.0036 |
| F09 | 1160001 | 1260000 | 0.0039 | 0.0025 | 0.0030 |
| F09 | 1180001 | 1280000 | 0.0024 | 0.0014 | 0.0015 |
| F09 | 1200001 | 1300000 | 0.0020 | 0.0012 | 0.0013 |
| F09 | 1220001 | 1320000 | 0.0019 | 0.0013 | 0.0011 |
| F09 | 1240001 | 1340000 | 0.0018 | 0.0010 | 0.0010 |
| F09 | 1260001 | 1360000 | 0.0024 | 0.0014 | 0.0015 |
| F09 | 1280001 | 1380000 | 0.0025 | 0.0014 | 0.0018 |
| F09 | 1300001 | 1400000 | 0.0029 | 0.0017 | 0.0021 |

| Chr | Start | End | both | A1 | A2 |
|---|---|---|---|---|---|
| F09 | 1320001 | 1420000 | 0.0034 | 0.0019 | 0.0027 |
| F09 | 1340001 | 1440000 | 0.0036 | 0.0019 | 0.0027 |
| F09 | 1360001 | 1460000 | 0.0034 | 0.0016 | 0.0026 |
| F09 | 1380001 | 1480000 | 0.0039 | 0.0022 | 0.0032 |
| F09 | 1400001 | 1500000 | 0.0038 | 0.0021 | 0.0033 |
| F09 | 1420001 | 1520000 | 0.0032 | 0.0017 | 0.0029 |
| F09 | 1440001 | 1540000 | 0.0035 | 0.0021 | 0.0034 |
| F09 | 1460001 | 1560000 | 0.0039 | 0.0026 | 0.0039 |
| F09 | 1480001 | 1580000 | 0.0036 | 0.0022 | 0.0035 |
| F09 | 1500001 | 1600000 | 0.0033 | 0.0021 | 0.0032 |
| F09 | 1520001 | 1620000 | 0.0035 | 0.0022 | 0.0033 |
| F09 | 1540001 | 1640000 | 0.0036 | 0.0019 | 0.0033 |
| F09 | 1560001 | 1660000 | 0.0044 | 0.0017 | 0.0038 |
| F09 | 1580001 | 1680000 | 0.0049 | 0.0018 | 0.0042 |
| F09 | 1600001 | 1700000 | 0.0060 | 0.0025 | 0.0051 |
| F09 | 1620001 | 1720000 | 0.0060 | 0.0024 | 0.0050 |
| F09 | 1640001 | 1740000 | 0.0069 | 0.0036 | 0.0059 |
| F09 | 1660001 | 1760000 | 0.0059 | 0.0035 | 0.0050 |
| F09 | 1680001 | 1780000 | 0.0059 | 0.0039 | 0.0052 |
| F09 | 1700001 | 1800000 | 0.0059 | 0.0042 | 0.0052 |
| F09 | 1720001 | 1820000 | 0.0057 | 0.0040 | 0.0050 |
| F09 | 1740001 | 1840000 | 0.0048 | 0.0029 | 0.0041 |
| F09 | 1760001 | 1860000 | 0.0046 | 0.0027 | 0.0038 |
| F09 | 1780001 | 1880000 | 0.0040 | 0.0020 | 0.0030 |
| F09 | 1800001 | 1900000 | 0.0029 | 0.0008 | 0.0020 |
| F09 | 1820001 | 1920000 | 0.0032 | 0.0009 | 0.0023 |
| F09 | 1840001 | 1940000 | 0.0031 | 0.0011 | 0.0022 |
| F09 | 1860001 | 1960000 | 0.0035 | 0.0018 | 0.0026 |
| F09 | 1880001 | 1980000 | 0.0038 | 0.0021 | 0.0031 |
| F09 | 1900001 | 2000000 | 0.0046 | 0.0026 | 0.0039 |
| F09 | 1920001 | 2020000 | 0.0042 | 0.0023 | 0.0035 |
| F09 | 1940001 | 2040000 | 0.0034 | 0.0017 | 0.0029 |
| F09 | 1960001 | 2060000 | 0.0029 | 0.0010 | 0.0025 |
| F09 | 1980001 | 2080000 | 0.0024 | 0.0006 | 0.0020 |
| F09 | 2000001 | 2100000 | 0.0019 | 0.0004 | 0.0015 |
| F09 | 2020001 | 2120000 | 0.0021 | 0.0004 | 0.0016 |
| F09 | 2040001 | 2140000 | 0.0021 | 0.0005 | 0.0016 |
| F09 | 2060001 | 2160000 | 0.0018 | 0.0005 | 0.0012 |
| F09 | 2080001 | 2180000 | 0.0017 | 0.0005 | 0.0010 |
| F09 | 2100001 | 2200000 | 0.0013 | 0.0004 | 0.0006 |
| F09 | 2120001 | 2220000 | 0.0009 | 0.0003 | 0.0004 |
| F09 | 2140001 | 2240000 | 0.0011 | 0.0005 | 0.0006 |
| F09 | 2160001 | 2260000 | 0.0010 | 0.0006 | 0.0006 |
| F09 | 2180001 | 2280000 | 0.0010 | 0.0006 | 0.0006 |
| F09 | 2200001 | 2300000 | 0.0009 | 0.0005 | 0.0006 |
| F09 | 2220001 | 2320000 | 0.0017 | 0.0009 | 0.0010 |
| F09 | 2240001 | 2340000 | 0.0027 | 0.0017 | 0.0018 |
| F09 | 2260001 | 2360000 | 0.0038 | 0.0026 | 0.0028 |
| F09 | 2280001 | 2380000 | 0.0041 | 0.0028 | 0.0031 |
| F09 | 2300001 | 2400000 | 0.0055 | 0.0038 | 0.0044 |
| F09 | 2320001 | 2420000 | 0.0055 | 0.0042 | 0.0047 |
| F09 | 2340001 | 2440000 | 0.0045 | 0.0031 | 0.0037 |
| F09 | 2360001 | 2460000 | 0.0043 | 0.0027 | 0.0033 |
| F09 | 2380001 | 2480000 | 0.0041 | 0.0029 | 0.0033 |
| F09 | 2400001 | 2500000 | 0.0034 | 0.0020 | 0.0025 |
| F09 | 2420001 | 2520000 | 0.0035 | 0.0017 | 0.0023 |
| F09 | 2440001 | 2540000 | 0.0041 | 0.0023 | 0.0028 |
| F09 | 2460001 | 2560000 | 0.0043 | 0.0026 | 0.0030 |

| Chr | Start | End | both | A1 | A2 |
|---|---|---|---|---|---|
| F09 | 2480001 | 2580000 | 0.0047 | 0.0032 | 0.0035 |
| F09 | 2500001 | 2600000 | 0.0040 | 0.0029 | 0.0030 |
| F09 | 2520001 | 2620000 | 0.0038 | 0.0028 | 0.0028 |
| F09 | 2540001 | 2640000 | 0.0030 | 0.0022 | 0.0023 |
| F09 | 2560001 | 2660000 | 0.0027 | 0.0016 | 0.0020 |
| F09 | 2580001 | 2680000 | 0.0031 | 0.0014 | 0.0023 |
| F09 | 2600001 | 2700000 | 0.0042 | 0.0020 | 0.0033 |
| F09 | 2620001 | 2720000 | 0.0044 | 0.0022 | 0.0034 |
| F09 | 2640001 | 2740000 | 0.0049 | 0.0028 | 0.0037 |
| F09 | 2660001 | 2760000 | 0.0046 | 0.0036 | 0.0034 |
| F09 | 2680001 | 2780000 | 0.0037 | 0.0036 | 0.0024 |
| F09 | 2700001 | 2800000 | 0.0029 | 0.0036 | 0.0014 |
| F09 | 2720001 | 2820000 | 0.0028 | 0.0033 | 0.0011 |
| F09 | 2740001 | 2840000 | 0.0028 | 0.0031 | 0.0013 |
| F09 | 2760001 | 2860000 | 0.0037 | 0.0029 | 0.0022 |
| F09 | 2780001 | 2880000 | 0.0043 | 0.0028 | 0.0028 |
| F09 | 2800001 | 2900000 | 0.0046 | 0.0026 | 0.0032 |
| F09 | 2820001 | 2920000 | 0.0043 | 0.0027 | 0.0033 |
| F09 | 2840001 | 2940000 | 0.0038 | 0.0024 | 0.0029 |
| F09 | 2860001 | 2960000 | 0.0035 | 0.0023 | 0.0027 |
| F09 | 2880001 | 2980000 | 0.0029 | 0.0022 | 0.0024 |
| F09 | 2900001 | 3000000 | 0.0024 | 0.0020 | 0.0020 |
| F09 | 2920001 | 3020000 | 0.0022 | 0.0018 | 0.0018 |
| F09 | 2940001 | 3040000 | 0.0025 | 0.0022 | 0.0023 |
| F09 | 2960001 | 3060000 | 0.0020 | 0.0022 | 0.0019 |
| F09 | 2980001 | 3080000 | 0.0025 | 0.0027 | 0.0024 |
| F09 | 3000001 | 3100000 | 0.0028 | 0.0029 | 0.0027 |
| F09 | 3020001 | 3120000 | 0.0032 | 0.0034 | 0.0031 |
| F09 | 3040001 | 3140000 | 0.0029 | 0.0031 | 0.0027 |
| F09 | 3060001 | 3160000 | 0.0027 | 0.0024 | 0.0022 |
| F09 | 3080001 | 3180000 | 0.0026 | 0.0016 | 0.0016 |
| F09 | 3100001 | 3200000 | 0.0024 | 0.0015 | 0.0012 |
| F09 | 3120001 | 3220000 | 0.0022 | 0.0011 | 0.0008 |
| F09 | 3140001 | 3240000 | 0.0028 | 0.0017 | 0.0012 |
| F09 | 3160001 | 3260000 | 0.0026 | 0.0017 | 0.0013 |
| F09 | 3180001 | 3280000 | 0.0018 | 0.0014 | 0.0010 |
| F09 | 3200001 | 3300000 | 0.0020 | 0.0016 | 0.0014 |
| F09 | 3220001 | 3320000 | 0.0020 | 0.0018 | 0.0014 |
| F09 | 3240001 | 3340000 | 0.0022 | 0.0019 | 0.0016 |
| F09 | 3260001 | 3360000 | 0.0026 | 0.0024 | 0.0019 |
| F09 | 3280001 | 3380000 | 0.0033 | 0.0031 | 0.0023 |
| F09 | 3300001 | 3400000 | 0.0031 | 0.0029 | 0.0020 |
| F09 | 3320001 | 3420000 | 0.0032 | 0.0031 | 0.0020 |
| F09 | 3340001 | 3440000 | 0.0024 | 0.0024 | 0.0013 |
| F09 | 3360001 | 3460000 | 0.0023 | 0.0024 | 0.0014 |
| F09 | 3380001 | 3480000 | 0.0029 | 0.0028 | 0.0023 |
| F09 | 3400001 | 3500000 | 0.0030 | 0.0029 | 0.0026 |
| F09 | 3420001 | 3520000 | 0.0034 | 0.0033 | 0.0032 |
| F09 | 3440001 | 3540000 | 0.0047 | 0.0045 | 0.0046 |
| F09 | 3460001 | 3560000 | 0.0063 | 0.0061 | 0.0062 |
| F09 | 3480001 | 3580000 | 0.0060 | 0.0061 | 0.0058 |
| F09 | 3500001 | 3600000 | 0.0070 | 0.0073 | 0.0068 |
| F09 | 3520001 | 3620000 | 0.0092 | 0.0095 | 0.0089 |
| F09 | 3540001 | 3640000 | 0.0094 | 0.0097 | 0.0092 |
| F09 | 3560001 | 3660000 | 0.0095 | 0.0096 | 0.0093 |
| F09 | 3580001 | 3680000 | 0.0103 | 0.0102 | 0.0101 |
| F09 | 3600001 | 3700000 | 0.0114 | 0.0112 | 0.0112 |
| F09 | 3620001 | 3720000 | 0.0124 | 0.0121 | 0.0122 |

| Chr | Start | End | both | A1 | A2 |
|---|---|---|---|---|---|
| F09 | 3640001 | 3740000 | 0.0125 | 0.0122 | 0.0122 |
| F09 | 3660001 | 3760000 | 0.0123 | 0.0121 | 0.0120 |
| F09 | 3680001 | 3780000 | 0.0111 | 0.0109 | 0.0108 |
| F09 | 3700001 | 3800000 | 0.0105 | 0.0105 | 0.0102 |
| F09 | 3720001 | 3820000 | 0.0076 | 0.0076 | 0.0074 |
| F09 | 3740001 | 3840000 | 0.0065 | 0.0066 | 0.0063 |
| F09 | 3760001 | 3860000 | 0.0061 | 0.0064 | 0.0059 |
| F09 | 3780001 | 3880000 | 0.0066 | 0.0069 | 0.0064 |
| F09 | 3800001 | 3900000 | 0.0063 | 0.0058 | 0.0060 |
| F09 | 3820001 | 3920000 | 0.0061 | 0.0055 | 0.0058 |
| F09 | 3840001 | 3940000 | 0.0070 | 0.0061 | 0.0066 |
| F09 | 3860001 | 3960000 | 0.0061 | 0.0051 | 0.0058 |
| F09 | 3880001 | 3980000 | 0.0060 | 0.0051 | 0.0056 |
| F09 | 3900001 | 4000000 | 0.0058 | 0.0053 | 0.0055 |
| F09 | 3920001 | 4020000 | 0.0053 | 0.0048 | 0.0050 |
| F09 | 3940001 | 4040000 | 0.0047 | 0.0043 | 0.0042 |
| F09 | 3960001 | 4060000 | 0.0044 | 0.0040 | 0.0040 |
| F09 | 3980001 | 4080000 | 0.0037 | 0.0033 | 0.0033 |
| F09 | 4000001 | 4100000 | 0.0024 | 0.0021 | 0.0020 |
| F09 | 4020001 | 4120000 | 0.0023 | 0.0020 | 0.0019 |
| F09 | 4040001 | 4140000 | 0.0015 | 0.0010 | 0.0012 |
| F09 | 4060001 | 4160000 | 0.0013 | 0.0007 | 0.0008 |
| F09 | 4080001 | 4180000 | 0.0013 | 0.0005 | 0.0007 |
| F09 | 4100001 | 4200000 | 0.0022 | 0.0013 | 0.0015 |
| F09 | 4120001 | 4220000 | 0.0025 | 0.0015 | 0.0017 |
| F09 | 4140001 | 4240000 | 0.0033 | 0.0022 | 0.0025 |
| F09 | 4160001 | 4260000 | 0.0035 | 0.0023 | 0.0027 |
| F09 | 4180001 | 4280000 | 0.0037 | 0.0022 | 0.0025 |
| F09 | 4200001 | 4300000 | 0.0032 | 0.0015 | 0.0020 |
| F09 | 4220001 | 4320000 | 0.0039 | 0.0015 | 0.0025 |
| F09 | 4240001 | 4340000 | 0.0048 | 0.0027 | 0.0037 |
| F09 | 4260001 | 4360000 | 0.0050 | 0.0033 | 0.0039 |
| F09 | 4280001 | 4380000 | 0.0055 | 0.0042 | 0.0047 |
| F09 | 4300001 | 4400000 | 0.0058 | 0.0051 | 0.0051 |
| F09 | 4320001 | 4420000 | 0.0051 | 0.0057 | 0.0047 |
| F09 | 4340001 | 4440000 | 0.0054 | 0.0062 | 0.0049 |
| F09 | 4360001 | 4460000 | 0.0048 | 0.0054 | 0.0044 |
| F09 | 4380001 | 4480000 | 0.0047 | 0.0057 | 0.0041 |
| F09 | 4400001 | 4500000 | 0.0053 | 0.0063 | 0.0046 |
| F09 | 4420001 | 4520000 | 0.0051 | 0.0060 | 0.0045 |
| F09 | 4440001 | 4540000 | 0.0034 | 0.0046 | 0.0028 |
| F09 | 4460001 | 4560000 | 0.0039 | 0.0057 | 0.0031 |
| F09 | 4480001 | 4580000 | 0.0038 | 0.0056 | 0.0027 |
| F09 | 4500001 | 4600000 | 0.0026 | 0.0044 | 0.0016 |
| F09 | 4520001 | 4620000 | 0.0023 | 0.0040 | 0.0013 |
| F09 | 4540001 | 4640000 | 0.0023 | 0.0040 | 0.0013 |
| F09 | 4560001 | 4660000 | 0.0033 | 0.0045 | 0.0025 |
| F09 | 4580001 | 4680000 | 0.0030 | 0.0037 | 0.0026 |
| F09 | 4600001 | 4700000 | 0.0034 | 0.0039 | 0.0031 |
| F09 | 4620001 | 4720000 | 0.0037 | 0.0040 | 0.0033 |
| F09 | 4640001 | 4740000 | 0.0032 | 0.0031 | 0.0029 |
| F09 | 4660001 | 4760000 | 0.0019 | 0.0017 | 0.0016 |
| F09 | 4680001 | 4780000 | 0.0019 | 0.0017 | 0.0015 |
| F09 | 4700001 | 4800000 | 0.0017 | 0.0018 | 0.0014 |
| F09 | 4720001 | 4820000 | 0.0022 | 0.0024 | 0.0019 |
| F09 | 4740001 | 4840000 | 0.0023 | 0.0023 | 0.0018 |
| F09 | 4760001 | 4860000 | 0.0029 | 0.0027 | 0.0021 |
| F09 | 4780001 | 4880000 | 0.0043 | 0.0034 | 0.0029 |

| Chr | Start | End | both | A1 | A2 |
|---|---|---|---|---|---|
| F09 | 4800001 | 4900000 | 0.0042 | 0.0029 | 0.0027 |
| F09 | 4820001 | 4920000 | 0.0036 | 0.0022 | 0.0022 |
| F09 | 4840001 | 4940000 | 0.0047 | 0.0032 | 0.0033 |
| F09 | 4860001 | 4960000 | 0.0052 | 0.0038 | 0.0037 |
| F09 | 4880001 | 4980000 | 0.0053 | 0.0036 | 0.0036 |
| F09 | 4900001 | 5000000 | 0.0061 | 0.0046 | 0.0044 |
| F09 | 4920001 | 5020000 | 0.0061 | 0.0045 | 0.0043 |
| F09 | 4940001 | 5040000 | 0.0058 | 0.0042 | 0.0038 |
| F09 | 4960001 | 5060000 | 0.0048 | 0.0035 | 0.0033 |
| F09 | 4980001 | 5080000 | 0.0035 | 0.0028 | 0.0025 |
| F09 | 5000001 | 5100000 | 0.0030 | 0.0019 | 0.0018 |
| F09 | 5020001 | 5120000 | 0.0031 | 0.0020 | 0.0020 |
| F09 | 5040001 | 5140000 | 0.0033 | 0.0014 | 0.0016 |
| F09 | 5060001 | 5160000 | 0.0034 | 0.0014 | 0.0015 |
| F09 | 5080001 | 5180000 | 0.0038 | 0.0014 | 0.0017 |
| F09 | 5100001 | 5200000 | 0.0046 | 0.0023 | 0.0025 |
| F09 | 5120001 | 5220000 | 0.0044 | 0.0022 | 0.0023 |
| F09 | 5140001 | 5240000 | 0.0036 | 0.0023 | 0.0024 |
| F09 | 5160001 | 5260000 | 0.0037 | 0.0024 | 0.0025 |
| F09 | 5180001 | 5280000 | 0.0030 | 0.0021 | 0.0021 |
| F09 | 5200001 | 5300000 | 0.0020 | 0.0012 | 0.0014 |
| F09 | 5220001 | 5320000 | 0.0020 | 0.0012 | 0.0015 |
| F09 | 5240001 | 5340000 | 0.0021 | 0.0012 | 0.0015 |
| F09 | 5260001 | 5360000 | 0.0017 | 0.0011 | 0.0013 |
| F09 | 5280001 | 5380000 | 0.0021 | 0.0013 | 0.0016 |
| F09 | 5300001 | 5400000 | 0.0018 | 0.0012 | 0.0014 |
| F09 | 5320001 | 5420000 | 0.0023 | 0.0014 | 0.0015 |
| F09 | 5340001 | 5440000 | 0.0024 | 0.0014 | 0.0016 |
| F09 | 5360001 | 5460000 | 0.0029 | 0.0019 | 0.0020 |
| F09 | 5380001 | 5480000 | 0.0027 | 0.0017 | 0.0017 |
| F09 | 5400001 | 5500000 | 0.0032 | 0.0021 | 0.0023 |
| F09 | 5420001 | 5520000 | 0.0041 | 0.0032 | 0.0033 |
| F09 | 5440001 | 5540000 | 0.0041 | 0.0032 | 0.0032 |
| F09 | 5460001 | 5560000 | 0.0036 | 0.0028 | 0.0029 |
| F09 | 5480001 | 5580000 | 0.0038 | 0.0030 | 0.0032 |
| F09 | 5500001 | 5600000 | 0.0034 | 0.0025 | 0.0026 |
| F09 | 5520001 | 5620000 | 0.0021 | 0.0012 | 0.0014 |
| F09 | 5540001 | 5640000 | 0.0024 | 0.0015 | 0.0016 |
| F09 | 5560001 | 5660000 | 0.0026 | 0.0015 | 0.0018 |
| F09 | 5580001 | 5680000 | 0.0027 | 0.0016 | 0.0018 |
| F09 | 5600001 | 5700000 | 0.0029 | 0.0020 | 0.0021 |
| F09 | 5620001 | 5720000 | 0.0032 | 0.0021 | 0.0024 |
| F09 | 5640001 | 5740000 | 0.0031 | 0.0021 | 0.0023 |
| F09 | 5660001 | 5760000 | 0.0027 | 0.0019 | 0.0021 |
| F09 | 5680001 | 5780000 | 0.0024 | 0.0016 | 0.0018 |
| F09 | 5700001 | 5800000 | 0.0029 | 0.0017 | 0.0020 |
| F09 | 5720001 | 5820000 | 0.0027 | 0.0017 | 0.0019 |
| F09 | 5740001 | 5840000 | 0.0021 | 0.0011 | 0.0014 |
| F09 | 5760001 | 5860000 | 0.0023 | 0.0012 | 0.0014 |
| F09 | 5780001 | 5880000 | 0.0024 | 0.0012 | 0.0014 |
| F09 | 5800001 | 5900000 | 0.0017 | 0.0007 | 0.0007 |
| F09 | 5820001 | 5920000 | 0.0016 | 0.0004 | 0.0006 |
| F09 | 5840001 | 5940000 | 0.0020 | 0.0004 | 0.0006 |
| F09 | 5860001 | 5960000 | 0.0020 | 0.0003 | 0.0005 |
| F09 | 5880001 | 5980000 | 0.0030 | 0.0009 | 0.0012 |
| F09 | 5900001 | 6000000 | 0.0031 | 0.0010 | 0.0014 |
| F09 | 5920001 | 6020000 | 0.0030 | 0.0012 | 0.0015 |
| F09 | 5940001 | 6040000 | 0.0035 | 0.0018 | 0.0021 |

| Chr | Start | End | both | A1 | A2 |
|---|---|---|---|---|---|
| F09 | 5960001 | 6060000 | 0.0037 | 0.0020 | 0.0023 |
| F09 | 5980001 | 6080000 | 0.0035 | 0.0019 | 0.0024 |
| F09 | 6000001 | 6100000 | 0.0038 | 0.0019 | 0.0026 |
| F09 | 6020001 | 6120000 | 0.0041 | 0.0018 | 0.0027 |
| F09 | 6040001 | 6140000 | 0.0038 | 0.0014 | 0.0025 |
| F09 | 6060001 | 6160000 | 0.0044 | 0.0022 | 0.0033 |
| F09 | 6080001 | 6180000 | 0.0039 | 0.0021 | 0.0029 |
| F09 | 6100001 | 6200000 | 0.0036 | 0.0022 | 0.0028 |
| F09 | 6120001 | 6220000 | 0.0041 | 0.0026 | 0.0035 |
| F09 | 6140001 | 6240000 | 0.0036 | 0.0025 | 0.0030 |
| F09 | 6160001 | 6260000 | 0.0034 | 0.0021 | 0.0024 |
| F09 | 6180001 | 6280000 | 0.0038 | 0.0022 | 0.0025 |
| F09 | 6200001 | 6300000 | 0.0039 | 0.0020 | 0.0023 |
| F09 | 6220001 | 6320000 | 0.0037 | 0.0018 | 0.0020 |
| F09 | 6240001 | 6340000 | 0.0038 | 0.0020 | 0.0022 |
| F09 | 6260001 | 6360000 | 0.0031 | 0.0016 | 0.0019 |
| F09 | 6280001 | 6380000 | 0.0031 | 0.0019 | 0.0021 |
| F09 | 6300001 | 6400000 | 0.0029 | 0.0021 | 0.0022 |
| F09 | 6320001 | 6420000 | 0.0032 | 0.0028 | 0.0025 |
| F09 | 6340001 | 6440000 | 0.0030 | 0.0029 | 0.0023 |
| F09 | 6360001 | 6460000 | 0.0039 | 0.0033 | 0.0032 |
| F09 | 6380001 | 6480000 | 0.0035 | 0.0029 | 0.0028 |
| F09 | 6400001 | 6500000 | 0.0051 | 0.0043 | 0.0043 |
| F09 | 6420001 | 6520000 | 0.0061 | 0.0051 | 0.0054 |
| F09 | 6440001 | 6540000 | 0.0067 | 0.0056 | 0.0060 |
| F09 | 6460001 | 6560000 | 0.0060 | 0.0057 | 0.0051 |
| F09 | 6480001 | 6580000 | 0.0060 | 0.0058 | 0.0051 |
| F09 | 6500001 | 6600000 | 0.0047 | 0.0045 | 0.0038 |
| F09 | 6520001 | 6620000 | 0.0036 | 0.0034 | 0.0028 |
| F09 | 6540001 | 6640000 | 0.0032 | 0.0030 | 0.0026 |
| F09 | 6560001 | 6660000 | 0.0033 | 0.0026 | 0.0026 |
| F09 | 6580001 | 6680000 | 0.0036 | 0.0023 | 0.0025 |
| F09 | 6600001 | 6700000 | 0.0038 | 0.0027 | 0.0029 |
| F09 | 6620001 | 6720000 | 0.0036 | 0.0022 | 0.0025 |
| F09 | 6640001 | 6740000 | 0.0032 | 0.0019 | 0.0021 |
| F09 | 6660001 | 6760000 | 0.0029 | 0.0018 | 0.0020 |
| F09 | 6680001 | 6780000 | 0.0023 | 0.0017 | 0.0018 |
| F09 | 6700001 | 6800000 | 0.0020 | 0.0011 | 0.0013 |
| F09 | 6720001 | 6820000 | 0.0017 | 0.0009 | 0.0010 |
| F09 | 6740001 | 6840000 | 0.0031 | 0.0016 | 0.0023 |
| F09 | 6760001 | 6860000 | 0.0035 | 0.0019 | 0.0026 |
| F09 | 6780001 | 6880000 | 0.0038 | 0.0021 | 0.0027 |
| F09 | 6800001 | 6900000 | 0.0035 | 0.0021 | 0.0027 |
| F09 | 6820001 | 6920000 | 0.0032 | 0.0018 | 0.0026 |
| F09 | 6840001 | 6940000 | 0.0024 | 0.0015 | 0.0017 |
| F09 | 6860001 | 6960000 | 0.0025 | 0.0016 | 0.0019 |
| F09 | 6880001 | 6980000 | 0.0030 | 0.0019 | 0.0023 |
| F09 | 6900001 | 7000000 | 0.0041 | 0.0029 | 0.0034 |
| F09 | 6920001 | 7020000 | 0.0044 | 0.0032 | 0.0037 |
| F09 | 6940001 | 7040000 | 0.0045 | 0.0034 | 0.0039 |
| F09 | 6960001 | 7060000 | 0.0049 | 0.0039 | 0.0043 |
| F09 | 6980001 | 7080000 | 0.0053 | 0.0041 | 0.0043 |
| F09 | 7000001 | 7100000 | 0.0043 | 0.0032 | 0.0034 |
| F09 | 7020001 | 7120000 | 0.0040 | 0.0031 | 0.0031 |
| F09 | 7040001 | 7140000 | 0.0038 | 0.0029 | 0.0030 |
| F09 | 7060001 | 7160000 | 0.0036 | 0.0026 | 0.0029 |
| F09 | 7080001 | 7180000 | 0.0032 | 0.0023 | 0.0027 |
| F09 | 7100001 | 7200000 | 0.0036 | 0.0025 | 0.0029 |

| Chr | Start | End | both | A1 | A2 |
|---|---|---|---|---|---|
| F09 | 7120001 | 7220000 | 0.0050 | 0.0040 | 0.0043 |
| F09 | 7140001 | 7240000 | 0.0051 | 0.0042 | 0.0044 |
| F09 | 7160001 | 7260000 | 0.0042 | 0.0036 | 0.0035 |
| F09 | 7180001 | 7280000 | 0.0035 | 0.0030 | 0.0029 |
| F09 | 7200001 | 7300000 | 0.0033 | 0.0032 | 0.0027 |
| F09 | 7220001 | 7320000 | 0.0018 | 0.0017 | 0.0013 |
| F09 | 7240001 | 7340000 | 0.0020 | 0.0019 | 0.0014 |
| F09 | 7260001 | 7360000 | 0.0023 | 0.0024 | 0.0018 |
| F09 | 7280001 | 7380000 | 0.0025 | 0.0027 | 0.0019 |
| F09 | 7300001 | 7400000 | 0.0027 | 0.0026 | 0.0019 |
| F09 | 7320001 | 7420000 | 0.0029 | 0.0028 | 0.0020 |
| F09 | 7340001 | 7440000 | 0.0026 | 0.0027 | 0.0016 |
| F09 | 7360001 | 7460000 | 0.0028 | 0.0032 | 0.0018 |
| F09 | 7380001 | 7480000 | 0.0031 | 0.0037 | 0.0021 |
| F09 | 7400001 | 7500000 | 0.0031 | 0.0041 | 0.0022 |
| F09 | 7420001 | 7520000 | 0.0029 | 0.0041 | 0.0020 |
| F09 | 7440001 | 7540000 | 0.0029 | 0.0041 | 0.0020 |
| F09 | 7460001 | 7560000 | 0.0024 | 0.0037 | 0.0015 |
| F09 | 7480001 | 7580000 | 0.0021 | 0.0032 | 0.0012 |
| F09 | 7500001 | 7600000 | 0.0016 | 0.0024 | 0.0008 |
| F09 | 7520001 | 7620000 | 0.0019 | 0.0024 | 0.0012 |
| F09 | 7540001 | 7640000 | 0.0031 | 0.0034 | 0.0025 |
| F09 | 7560001 | 7660000 | 0.0031 | 0.0031 | 0.0027 |
| F09 | 7580001 | 7680000 | 0.0032 | 0.0031 | 0.0028 |
| F09 | 7600001 | 7700000 | 0.0032 | 0.0031 | 0.0029 |
| F09 | 7620001 | 7720000 | 0.0033 | 0.0033 | 0.0029 |
| F09 | 7640001 | 7740000 | 0.0019 | 0.0021 | 0.0014 |
| F09 | 7660001 | 7760000 | 0.0019 | 0.0022 | 0.0012 |
| F09 | 7680001 | 7780000 | 0.0021 | 0.0027 | 0.0010 |
| F09 | 7700001 | 7800000 | 0.0024 | 0.0029 | 0.0011 |
| F09 | 7720001 | 7820000 | 0.0028 | 0.0031 | 0.0015 |
| F09 | 7740001 | 7840000 | 0.0027 | 0.0029 | 0.0016 |
| F09 | 7760001 | 7860000 | 0.0030 | 0.0026 | 0.0019 |
| F09 | 7780001 | 7880000 | 0.0030 | 0.0021 | 0.0020 |
| F09 | 7800001 | 7900000 | 0.0031 | 0.0018 | 0.0020 |
| F09 | 7820001 | 7920000 | 0.0032 | 0.0016 | 0.0016 |
| F09 | 7840001 | 7940000 | 0.0034 | 0.0014 | 0.0015 |
| F09 | 7860001 | 7960000 | 0.0034 | 0.0016 | 0.0016 |
| F09 | 7880001 | 7980000 | 0.0035 | 0.0016 | 0.0017 |
| F09 | 7900001 | 8000000 | 0.0036 | 0.0017 | 0.0017 |
| F09 | 7920001 | 8020000 | 0.0043 | 0.0024 | 0.0028 |
| F09 | 7940001 | 8040000 | 0.0047 | 0.0028 | 0.0031 |
| F09 | 7960001 | 8060000 | 0.0043 | 0.0024 | 0.0028 |
| F09 | 7980001 | 8080000 | 0.0039 | 0.0021 | 0.0025 |
| F09 | 8000001 | 8100000 | 0.0042 | 0.0025 | 0.0030 |
| F09 | 8020001 | 8120000 | 0.0034 | 0.0020 | 0.0023 |
| F09 | 8040001 | 8140000 | 0.0035 | 0.0023 | 0.0026 |
| F09 | 8060001 | 8160000 | 0.0037 | 0.0023 | 0.0026 |
| F09 | 8080001 | 8180000 | 0.0036 | 0.0023 | 0.0027 |
| F09 | 8100001 | 8200000 | 0.0033 | 0.0020 | 0.0023 |
| F09 | 8120001 | 8220000 | 0.0035 | 0.0020 | 0.0026 |
| F09 | 8140001 | 8240000 | 0.0031 | 0.0016 | 0.0023 |
| F09 | 8160001 | 8260000 | 0.0027 | 0.0017 | 0.0022 |
| F09 | 8180001 | 8280000 | 0.0028 | 0.0019 | 0.0025 |
| F09 | 8200001 | 8300000 | 0.0026 | 0.0019 | 0.0024 |
| F09 | 8220001 | 8320000 | 0.0013 | 0.0010 | 0.0012 |
| F09 | 8240001 | 8340000 | 0.0009 | 0.0007 | 0.0009 |
| F09 | 8260001 | 8360000 | 0.0011 | 0.0009 | 0.0010 |

| Chr | Start | End | both | A1 | A2 |
|---|---|---|---|---|---|
| F09 | 8280001 | 8380000 | 0.0009 | 0.0007 | 0.0009 |
| F09 | 8300001 | 8400000 | 0.0014 | 0.0011 | 0.0013 |
| F09 | 8320001 | 8420000 | 0.0018 | 0.0013 | 0.0016 |
| F09 | 8340001 | 8440000 | 0.0020 | 0.0015 | 0.0017 |
| F09 | 8360001 | 8460000 | 0.0017 | 0.0012 | 0.0015 |
| F09 | 8380001 | 8480000 | 0.0018 | 0.0013 | 0.0016 |
| F09 | 8400001 | 8500000 | 0.0012 | 0.0007 | 0.0010 |
| F09 | 8420001 | 8520000 | 0.0007 | 0.0006 | 0.0007 |
| F09 | 8440001 | 8540000 | 0.0005 | 0.0004 | 0.0005 |
| F09 | 8460001 | 8560000 | 0.0008 | 0.0007 | 0.0008 |
| F09 | 8480001 | 8580000 | 0.0006 | 0.0005 | 0.0006 |
| F09 | 8500001 | 8600000 | 0.0009 | 0.0008 | 0.0009 |
| F09 | 8520001 | 8620000 | 0.0010 | 0.0009 | 0.0010 |
| F09 | 8540001 | 8640000 | 0.0016 | 0.0013 | 0.0016 |
| F09 | 8560001 | 8660000 | 0.0014 | 0.0012 | 0.0014 |
| F09 | 8580001 | 8680000 | 0.0015 | 0.0013 | 0.0014 |
| F09 | 8600001 | 8700000 | 0.0013 | 0.0009 | 0.0011 |
| F09 | 8620001 | 8720000 | 0.0018 | 0.0014 | 0.0015 |
| F09 | 8640001 | 8740000 | 0.0017 | 0.0013 | 0.0014 |
| F09 | 8660001 | 8760000 | 0.0030 | 0.0024 | 0.0027 |
| F09 | 8680001 | 8780000 | 0.0038 | 0.0030 | 0.0033 |
| F09 | 8700001 | 8800000 | 0.0040 | 0.0031 | 0.0036 |
| F09 | 8720001 | 8820000 | 0.0036 | 0.0028 | 0.0033 |
| F09 | 8740001 | 8840000 | 0.0035 | 0.0027 | 0.0032 |
| F09 | 8760001 | 8860000 | 0.0026 | 0.0019 | 0.0022 |
| F09 | 8780001 | 8880000 | 0.0026 | 0.0020 | 0.0024 |
| F09 | 8800001 | 8900000 | 0.0035 | 0.0029 | 0.0032 |
| F09 | 8820001 | 8920000 | 0.0043 | 0.0036 | 0.0040 |
| F09 | 8840001 | 8940000 | 0.0050 | 0.0040 | 0.0045 |
| F09 | 8860001 | 8960000 | 0.0055 | 0.0045 | 0.0050 |
| F09 | 8880001 | 8980000 | 0.0053 | 0.0042 | 0.0047 |
| F09 | 8900001 | 9000000 | 0.0047 | 0.0034 | 0.0038 |
| F09 | 8920001 | 9020000 | 0.0039 | 0.0027 | 0.0030 |
| F09 | 8940001 | 9040000 | 0.0033 | 0.0020 | 0.0023 |
| F09 | 8960001 | 9060000 | 0.0028 | 0.0016 | 0.0018 |
| F09 | 8980001 | 9080000 | 0.0026 | 0.0015 | 0.0017 |
| F09 | 9000001 | 9100000 | 0.0019 | 0.0012 | 0.0014 |
| F09 | 9020001 | 9120000 | 0.0015 | 0.0010 | 0.0011 |
| F09 | 9040001 | 9140000 | 0.0013 | 0.0010 | 0.0012 |
| F09 | 9060001 | 9160000 | 0.0019 | 0.0012 | 0.0017 |
| F09 | 9080001 | 9180000 | 0.0022 | 0.0016 | 0.0019 |
| F09 | 9100001 | 9200000 | 0.0031 | 0.0022 | 0.0026 |
| F09 | 9120001 | 9220000 | 0.0037 | 0.0025 | 0.0030 |
| F09 | 9140001 | 9240000 | 0.0042 | 0.0028 | 0.0032 |
| F09 | 9160001 | 9260000 | 0.0039 | 0.0028 | 0.0029 |
| F09 | 9180001 | 9280000 | 0.0043 | 0.0028 | 0.0035 |
| F09 | 9200001 | 9300000 | 0.0044 | 0.0029 | 0.0037 |
| F09 | 9220001 | 9320000 | 0.0042 | 0.0030 | 0.0036 |
| F09 | 9240001 | 9340000 | 0.0041 | 0.0029 | 0.0036 |
| F09 | 9260001 | 9360000 | 0.0047 | 0.0034 | 0.0042 |
| F09 | 9280001 | 9380000 | 0.0045 | 0.0035 | 0.0040 |
| F09 | 9300001 | 9400000 | 0.0037 | 0.0030 | 0.0033 |
| F09 | 9320001 | 9420000 | 0.0034 | 0.0027 | 0.0030 |
| F09 | 9340001 | 9440000 | 0.0029 | 0.0024 | 0.0027 |
| F09 | 9360001 | 9460000 | 0.0014 | 0.0012 | 0.0013 |
| F09 | 9380001 | 9480000 | 0.0009 | 0.0006 | 0.0008 |
| F09 | 9400001 | 9500000 | 0.0012 | 0.0007 | 0.0011 |
| F09 | 9420001 | 9520000 | 0.0015 | 0.0009 | 0.0012 |

| Chr | Start | End | both | A1 | A2 |
|---|---|---|---|---|---|
| F09 | 9440001 | 9540000 | 0.0019 | 0.0011 | 0.0015 |
| F09 | 9460001 | 9560000 | 0.0022 | 0.0012 | 0.0017 |
| F09 | 9480001 | 9580000 | 0.0022 | 0.0011 | 0.0016 |
| F09 | 9500001 | 9600000 | 0.0025 | 0.0016 | 0.0019 |
| F09 | 9520001 | 9620000 | 0.0027 | 0.0018 | 0.0020 |
| F09 | 9540001 | 9640000 | 0.0025 | 0.0015 | 0.0018 |
| F09 | 9560001 | 9660000 | 0.0034 | 0.0024 | 0.0027 |
| F09 | 9580001 | 9680000 | 0.0039 | 0.0029 | 0.0031 |
| F09 | 9600001 | 9700000 | 0.0038 | 0.0027 | 0.0029 |
| F09 | 9620001 | 9720000 | 0.0035 | 0.0025 | 0.0028 |
| F09 | 9640001 | 9740000 | 0.0037 | 0.0030 | 0.0031 |
| F09 | 9660001 | 9760000 | 0.0027 | 0.0021 | 0.0022 |
| F09 | 9680001 | 9780000 | 0.0019 | 0.0016 | 0.0017 |
| F09 | 9700001 | 9800000 | 0.0017 | 0.0013 | 0.0016 |
| F09 | 9720001 | 9820000 | 0.0016 | 0.0013 | 0.0015 |
| F09 | 9740001 | 9840000 | 0.0019 | 0.0015 | 0.0019 |
| F09 | 9760001 | 9860000 | 0.0019 | 0.0016 | 0.0020 |
| F09 | 9780001 | 9880000 | 0.0023 | 0.0019 | 0.0023 |
| F09 | 9800001 | 9900000 | 0.0020 | 0.0017 | 0.0020 |
| F09 | 9820001 | 9920000 | 0.0020 | 0.0018 | 0.0020 |
| F09 | 9840001 | 9940000 | 0.0015 | 0.0013 | 0.0015 |
| F09 | 9860001 | 9960000 | 0.0017 | 0.0013 | 0.0014 |
| F09 | 9880001 | 9980000 | 0.0020 | 0.0015 | 0.0017 |
| F09 | 9900001 | 10000000 | 0.0023 | 0.0016 | 0.0018 |
| F09 | 9920001 | 10020000 | 0.0024 | 0.0015 | 0.0019 |
| F09 | 9940001 | 10040000 | 0.0023 | 0.0015 | 0.0018 |
| F09 | 9960001 | 10060000 | 0.0026 | 0.0018 | 0.0021 |
| F09 | 9980001 | 10080000 | 0.0022 | 0.0014 | 0.0018 |
| F09 | 10000001 | 10100000 | 0.0023 | 0.0016 | 0.0021 |
| F09 | 10020001 | 10120000 | 0.0025 | 0.0017 | 0.0021 |
| F09 | 10040001 | 10140000 | 0.0026 | 0.0018 | 0.0021 |
| F09 | 10060001 | 10160000 | 0.0025 | 0.0018 | 0.0021 |
| F09 | 10080001 | 10180000 | 0.0025 | 0.0018 | 0.0021 |
| F09 | 10100001 | 10200000 | 0.0019 | 0.0013 | 0.0016 |
| F09 | 10120001 | 10220000 | 0.0020 | 0.0016 | 0.0017 |
| F09 | 10140001 | 10240000 | 0.0024 | 0.0019 | 0.0021 |
| F09 | 10160001 | 10260000 | 0.0023 | 0.0017 | 0.0019 |
| F09 | 10180001 | 10280000 | 0.0021 | 0.0014 | 0.0017 |
| F09 | 10200001 | 10300000 | 0.0021 | 0.0013 | 0.0017 |
| F09 | 10220001 | 10320000 | 0.0017 | 0.0010 | 0.0015 |
| F09 | 10240001 | 10340000 | 0.0015 | 0.0007 | 0.0011 |
| F09 | 10260001 | 10360000 | 0.0012 | 0.0006 | 0.0009 |
| F09 | 10280001 | 10380000 | 0.0015 | 0.0008 | 0.0011 |
| F09 | 10300001 | 10400000 | 0.0019 | 0.0011 | 0.0014 |
| F09 | 10320001 | 10420000 | 0.0021 | 0.0011 | 0.0014 |
| F09 | 10340001 | 10440000 | 0.0024 | 0.0013 | 0.0016 |
| F09 | 10360001 | 10460000 | 0.0023 | 0.0012 | 0.0014 |
| F09 | 10380001 | 10480000 | 0.0028 | 0.0017 | 0.0021 |
| F09 | 10400001 | 10500000 | 0.0029 | 0.0018 | 0.0022 |
| F09 | 10420001 | 10520000 | 0.0032 | 0.0020 | 0.0024 |
| F09 | 10440001 | 10540000 | 0.0029 | 0.0019 | 0.0024 |
| F09 | 10460001 | 10560000 | 0.0032 | 0.0023 | 0.0028 |
| F09 | 10480001 | 10580000 | 0.0029 | 0.0021 | 0.0024 |
| F09 | 10500001 | 10600000 | 0.0030 | 0.0022 | 0.0025 |
| F09 | 10520001 | 10620000 | 0.0032 | 0.0025 | 0.0028 |
| F09 | 10540001 | 10640000 | 0.0028 | 0.0022 | 0.0024 |
| F09 | 10560001 | 10660000 | 0.0031 | 0.0023 | 0.0026 |
| F09 | 10580001 | 10680000 | 0.0030 | 0.0024 | 0.0026 |

| Chr | Start | End | both | A1 | A2 |
|---|---|---|---|---|---|
| F09 | 10600001 | 10700000 | 0.0027 | 0.0020 | 0.0022 |
| F09 | 10620001 | 10720000 | 0.0031 | 0.0022 | 0.0026 |
| F09 | 10640001 | 10740000 | 0.0034 | 0.0024 | 0.0028 |
| F09 | 10660001 | 10760000 | 0.0035 | 0.0026 | 0.0029 |
| F09 | 10680001 | 10780000 | 0.0038 | 0.0026 | 0.0030 |
| F09 | 10700001 | 10800000 | 0.0044 | 0.0032 | 0.0036 |
| F09 | 10720001 | 10820000 | 0.0035 | 0.0025 | 0.0027 |
| F09 | 10740001 | 10840000 | 0.0038 | 0.0028 | 0.0030 |
| F09 | 10760001 | 10860000 | 0.0035 | 0.0024 | 0.0027 |
| F09 | 10780001 | 10880000 | 0.0028 | 0.0015 | 0.0019 |
| F09 | 10800001 | 10900000 | 0.0024 | 0.0013 | 0.0017 |
| F09 | 10820001 | 10920000 | 0.0032 | 0.0021 | 0.0025 |
| F09 | 10840001 | 10940000 | 0.0033 | 0.0018 | 0.0024 |
| F09 | 10860001 | 10960000 | 0.0034 | 0.0021 | 0.0025 |
| F09 | 10880001 | 10980000 | 0.0035 | 0.0022 | 0.0026 |
| F09 | 10900001 | 11000000 | 0.0033 | 0.0020 | 0.0023 |
| F09 | 10920001 | 11020000 | 0.0029 | 0.0017 | 0.0020 |
| F09 | 10940001 | 11040000 | 0.0028 | 0.0017 | 0.0019 |
| F09 | 10960001 | 11060000 | 0.0023 | 0.0013 | 0.0015 |
| F09 | 10980001 | 11080000 | 0.0022 | 0.0015 | 0.0017 |
| F09 | 11000001 | 11100000 | 0.0022 | 0.0019 | 0.0017 |
| F09 | 11020001 | 11120000 | 0.0018 | 0.0013 | 0.0012 |
| F09 | 11040001 | 11140000 | 0.0020 | 0.0014 | 0.0013 |
| F09 | 11060001 | 11160000 | 0.0026 | 0.0018 | 0.0017 |
| F09 | 11080001 | 11180000 | 0.0025 | 0.0014 | 0.0014 |
| F09 | 11100001 | 11200000 | 0.0033 | 0.0016 | 0.0020 |
| F09 | 11120001 | 11220000 | 0.0035 | 0.0019 | 0.0022 |
| F09 | 11140001 | 11240000 | 0.0031 | 0.0018 | 0.0020 |
| F09 | 11160001 | 11260000 | 0.0034 | 0.0022 | 0.0024 |
| F09 | 11180001 | 11280000 | 0.0040 | 0.0027 | 0.0032 |
| F09 | 11200001 | 11300000 | 0.0037 | 0.0026 | 0.0032 |
| F09 | 11220001 | 11320000 | 0.0040 | 0.0030 | 0.0036 |
| F09 | 11240001 | 11340000 | 0.0039 | 0.0032 | 0.0038 |
| F09 | 11260001 | 11360000 | 0.0034 | 0.0028 | 0.0033 |
| F09 | 11280001 | 11380000 | 0.0028 | 0.0024 | 0.0027 |
| F09 | 11300001 | 11400000 | 0.0023 | 0.0021 | 0.0023 |
| F09 | 11320001 | 11420000 | 0.0025 | 0.0022 | 0.0024 |
| F09 | 11340001 | 11440000 | 0.0034 | 0.0030 | 0.0033 |
| F09 | 11360001 | 11460000 | 0.0032 | 0.0028 | 0.0030 |
| F09 | 11380001 | 11480000 | 0.0030 | 0.0026 | 0.0028 |
| F09 | 11400001 | 11500000 | 0.0036 | 0.0030 | 0.0032 |
| F09 | 11420001 | 11520000 | 0.0035 | 0.0026 | 0.0029 |
| F09 | 11440001 | 11540000 | 0.0029 | 0.0022 | 0.0019 |
| F09 | 11460001 | 11560000 | 0.0027 | 0.0019 | 0.0017 |
| F09 | 11480001 | 11580000 | 0.0033 | 0.0024 | 0.0022 |
| F09 | 11500001 | 11600000 | 0.0029 | 0.0021 | 0.0020 |
| F09 | 11520001 | 11620000 | 0.0027 | 0.0020 | 0.0018 |
| F09 | 11540001 | 11640000 | 0.0029 | 0.0020 | 0.0023 |
| F09 | 11560001 | 11660000 | 0.0049 | 0.0041 | 0.0043 |
| F09 | 11580001 | 11680000 | 0.0048 | 0.0039 | 0.0041 |
| F09 | 11600001 | 11700000 | 0.0057 | 0.0047 | 0.0049 |
| F09 | 11620001 | 11720000 | 0.0058 | 0.0051 | 0.0052 |
| F09 | 11640001 | 11740000 | 0.0068 | 0.0060 | 0.0061 |
| F09 | 11660001 | 11760000 | 0.0060 | 0.0047 | 0.0050 |
| F09 | 11680001 | 11780000 | 0.0068 | 0.0054 | 0.0057 |
| F09 | 11700001 | 11800000 | 0.0059 | 0.0046 | 0.0048 |
| F09 | 11720001 | 11820000 | 0.0054 | 0.0041 | 0.0042 |
| F09 | 11740001 | 11840000 | 0.0037 | 0.0025 | 0.0027 |

| Chr | Start | End | both | A1 | A2 |
|---|---|---|---|---|---|
| F09 | 11760001 | 11860000 | 0.0028 | 0.0020 | 0.0019 |
| F09 | 11780001 | 11880000 | 0.0018 | 0.0010 | 0.0010 |
| F09 | 11800001 | 11900000 | 0.0015 | 0.0008 | 0.0007 |
| F09 | 11820001 | 11920000 | 0.0017 | 0.0009 | 0.0009 |
| F09 | 11840001 | 11940000 | 0.0017 | 0.0008 | 0.0008 |
| F09 | 11860001 | 11960000 | 0.0014 | 0.0006 | 0.0006 |
| F09 | 11880001 | 11980000 | 0.0015 | 0.0007 | 0.0009 |
| F09 | 11900001 | 12000000 | 0.0016 | 0.0008 | 0.0009 |
| F09 | 11920001 | 12020000 | 0.0014 | 0.0006 | 0.0007 |
| F09 | 11940001 | 12040000 | 0.0014 | 0.0006 | 0.0008 |
| F09 | 11960001 | 12060000 | 0.0012 | 0.0006 | 0.0007 |
| F09 | 11980001 | 12080000 | 0.0016 | 0.0009 | 0.0012 |
| F09 | 12000001 | 12100000 | 0.0015 | 0.0009 | 0.0012 |
| F09 | 12020001 | 12120000 | 0.0017 | 0.0011 | 0.0013 |
| F09 | 12040001 | 12140000 | 0.0021 | 0.0015 | 0.0018 |
| F09 | 12060001 | 12160000 | 0.0024 | 0.0015 | 0.0018 |
| F09 | 12080001 | 12180000 | 0.0024 | 0.0017 | 0.0018 |
| F09 | 12100001 | 12200000 | 0.0027 | 0.0020 | 0.0021 |
| F09 | 12120001 | 12220000 | 0.0026 | 0.0021 | 0.0023 |
| F09 | 12140001 | 12240000 | 0.0027 | 0.0021 | 0.0022 |
| F09 | 12160001 | 12260000 | 0.0030 | 0.0023 | 0.0023 |
| F09 | 12180001 | 12280000 | 0.0026 | 0.0019 | 0.0020 |
| F09 | 12200001 | 12300000 | 0.0029 | 0.0022 | 0.0023 |
| F09 | 12220001 | 12320000 | 0.0027 | 0.0019 | 0.0019 |
| F09 | 12240001 | 12340000 | 0.0024 | 0.0016 | 0.0016 |
| F09 | 12260001 | 12360000 | 0.0019 | 0.0015 | 0.0016 |
| F09 | 12280001 | 12380000 | 0.0016 | 0.0011 | 0.0012 |
| F09 | 12300001 | 12400000 | 0.0011 | 0.0006 | 0.0007 |
| F09 | 12320001 | 12420000 | 0.0012 | 0.0009 | 0.0010 |
| F09 | 12340001 | 12440000 | 0.0011 | 0.0009 | 0.0009 |
| F09 | 12360001 | 12460000 | 0.0011 | 0.0009 | 0.0009 |
| F09 | 12380001 | 12480000 | 0.0014 | 0.0012 | 0.0012 |
| F09 | 12400001 | 12500000 | 0.0012 | 0.0011 | 0.0012 |
| F09 | 12420001 | 12520000 | 0.0012 | 0.0011 | 0.0012 |
| F09 | 12440001 | 12540000 | 0.0017 | 0.0014 | 0.0017 |
| F09 | 12460001 | 12560000 | 0.0017 | 0.0014 | 0.0017 |
| F09 | 12480001 | 12580000 | 0.0016 | 0.0012 | 0.0016 |
| F09 | 12500001 | 12600000 | 0.0015 | 0.0011 | 0.0015 |
| F09 | 12520001 | 12620000 | 0.0014 | 0.0010 | 0.0014 |
| F09 | 12540001 | 12640000 | 0.0016 | 0.0011 | 0.0016 |
| F09 | 12560001 | 12660000 | 0.0018 | 0.0013 | 0.0018 |
| F09 | 12580001 | 12680000 | 0.0018 | 0.0014 | 0.0018 |
| F09 | 12600001 | 12700000 | 0.0020 | 0.0016 | 0.0020 |
| F09 | 12620001 | 12720000 | 0.0019 | 0.0015 | 0.0019 |
| F09 | 12640001 | 12740000 | 0.0014 | 0.0011 | 0.0014 |
| F09 | 12660001 | 12760000 | 0.0011 | 0.0009 | 0.0011 |
| F09 | 12680001 | 12780000 | 0.0009 | 0.0007 | 0.0009 |
| F09 | 12700001 | 12800000 | 0.0010 | 0.0008 | 0.0010 |
| F09 | 12720001 | 12820000 | 0.0009 | 0.0008 | 0.0009 |
| F09 | 12740001 | 12840000 | 0.0006 | 0.0005 | 0.0006 |
| F09 | 12760001 | 12860000 | 0.0007 | 0.0006 | 0.0007 |
| F09 | 12780001 | 12880000 | 0.0010 | 0.0008 | 0.0010 |
| F09 | 12800001 | 12900000 | 0.0006 | 0.0005 | 0.0006 |
| F09 | 12820001 | 12920000 | 0.0008 | 0.0006 | 0.0008 |
| F09 | 12840001 | 12940000 | 0.0008 | 0.0006 | 0.0008 |
| F09 | 12860001 | 12960000 | 0.0007 | 0.0006 | 0.0007 |
| F09 | 12880001 | 12980000 | 0.0004 | 0.0003 | 0.0004 |
| F09 | 12900001 | 13000000 | 0.0006 | 0.0006 | 0.0006 |

| Chr | Start | End | both | A1 | A2 |
|---|---|---|---|---|---|
| F09 | 12920001 | 13020000 | 0.0006 | 0.0006 | 0.0005 |
| F09 | 12940001 | 13040000 | 0.0007 | 0.0007 | 0.0007 |
| F09 | 12960001 | 13060000 | 0.0007 | 0.0007 | 0.0007 |
| F09 | 12980001 | 13080000 | 0.0013 | 0.0012 | 0.0013 |
| F09 | 13000001 | 13100000 | 0.0017 | 0.0016 | 0.0016 |
| F09 | 13020001 | 13120000 | 0.0016 | 0.0015 | 0.0015 |
| F09 | 13040001 | 13140000 | 0.0014 | 0.0013 | 0.0013 |
| F09 | 13060001 | 13160000 | 0.0016 | 0.0015 | 0.0015 |
| F09 | 13080001 | 13180000 | 0.0013 | 0.0012 | 0.0012 |
| F09 | 13100001 | 13200000 | 0.0016 | 0.0011 | 0.0013 |
| F09 | 13120001 | 13220000 | 0.0019 | 0.0013 | 0.0016 |
| F09 | 13140001 | 13240000 | 0.0030 | 0.0020 | 0.0024 |
| F09 | 13160001 | 13260000 | 0.0030 | 0.0019 | 0.0023 |
| F09 | 13180001 | 13280000 | 0.0038 | 0.0026 | 0.0030 |
| F09 | 13200001 | 13300000 | 0.0032 | 0.0023 | 0.0026 |
| F09 | 13220001 | 13320000 | 0.0036 | 0.0026 | 0.0030 |
| F09 | 13240001 | 13340000 | 0.0028 | 0.0020 | 0.0024 |
| F09 | 13260001 | 13360000 | 0.0033 | 0.0026 | 0.0030 |
| F09 | 13280001 | 13380000 | 0.0025 | 0.0020 | 0.0023 |
| F09 | 13300001 | 13400000 | 0.0025 | 0.0020 | 0.0023 |
| F09 | 13320001 | 13420000 | 0.0022 | 0.0018 | 0.0021 |
| F09 | 13340001 | 13440000 | 0.0023 | 0.0019 | 0.0022 |
| F09 | 13360001 | 13460000 | 0.0016 | 0.0012 | 0.0015 |
| F09 | 13380001 | 13480000 | 0.0017 | 0.0011 | 0.0016 |
| F09 | 13400001 | 13500000 | 0.0016 | 0.0010 | 0.0014 |
| F09 | 13420001 | 13520000 | 0.0011 | 0.0007 | 0.0010 |
| F09 | 13440001 | 13540000 | 0.0011 | 0.0007 | 0.0008 |
| F09 | 13460001 | 13560000 | 0.0017 | 0.0011 | 0.0013 |
| F09 | 13480001 | 13580000 | 0.0018 | 0.0011 | 0.0013 |
| F09 | 13500001 | 13600000 | 0.0018 | 0.0012 | 0.0014 |
| F09 | 13520001 | 13620000 | 0.0029 | 0.0018 | 0.0020 |
| F09 | 13540001 | 13640000 | 0.0032 | 0.0021 | 0.0024 |
| F09 | 13560001 | 13660000 | 0.0027 | 0.0018 | 0.0020 |
| F09 | 13580001 | 13680000 | 0.0035 | 0.0025 | 0.0030 |
| F09 | 13600001 | 13700000 | 0.0033 | 0.0024 | 0.0028 |
| F09 | 13620001 | 13720000 | 0.0027 | 0.0020 | 0.0025 |
| F09 | 13640001 | 13740000 | 0.0044 | 0.0032 | 0.0038 |
| F09 | 13660001 | 13760000 | 0.0044 | 0.0032 | 0.0039 |
| F09 | 13680001 | 13780000 | 0.0033 | 0.0023 | 0.0027 |
| F09 | 13700001 | 13800000 | 0.0039 | 0.0024 | 0.0029 |
| F09 | 13720001 | 13820000 | 0.0041 | 0.0025 | 0.0030 |
| F09 | 13740001 | 13840000 | 0.0019 | 0.0009 | 0.0012 |
| F09 | 13760001 | 13860000 | 0.0021 | 0.0011 | 0.0014 |
| F09 | 13780001 | 13880000 | 0.0030 | 0.0019 | 0.0022 |
| F09 | 13800001 | 13900000 | 0.0030 | 0.0021 | 0.0027 |
| F09 | 13820001 | 13920000 | 0.0026 | 0.0019 | 0.0024 |
| F09 | 13840001 | 13940000 | 0.0029 | 0.0022 | 0.0028 |
| F09 | 13860001 | 13960000 | 0.0027 | 0.0020 | 0.0026 |
| F09 | 13880001 | 13980000 | 0.0018 | 0.0013 | 0.0018 |
| F09 | 13900001 | 14000000 | 0.0015 | 0.0011 | 0.0015 |
| F09 | 13920001 | 14020000 | 0.0014 | 0.0010 | 0.0013 |
| F09 | 13940001 | 14040000 | 0.0014 | 0.0010 | 0.0012 |
| F09 | 13960001 | 14060000 | 0.0018 | 0.0014 | 0.0016 |
| F09 | 13980001 | 14080000 | 0.0028 | 0.0023 | 0.0024 |
| F09 | 14000001 | 14100000 | 0.0026 | 0.0021 | 0.0022 |
| F09 | 14020001 | 14120000 | 0.0025 | 0.0021 | 0.0022 |
| F09 | 14040001 | 14140000 | 0.0022 | 0.0019 | 0.0020 |
| F09 | 14060001 | 14160000 | 0.0016 | 0.0013 | 0.0014 |

| Chr | Start | End | both | A1 | A2 |
|---|---|---|---|---|---|
| F09 | 14080001 | 14180000 | 0.0013 | 0.0011 | 0.0013 |
| F09 | 14100001 | 14200000 | 0.0013 | 0.0012 | 0.0013 |
| F09 | 14120001 | 14220000 | 0.0015 | 0.0013 | 0.0014 |
| F09 | 14140001 | 14240000 | 0.0015 | 0.0014 | 0.0014 |
| F09 | 14160001 | 14260000 | 0.0016 | 0.0015 | 0.0015 |
| F09 | 14180001 | 14280000 | 0.0008 | 0.0007 | 0.0007 |
| F09 | 14200001 | 14300000 | 0.0005 | 0.0005 | 0.0005 |
| F09 | 14220001 | 14320000 | 0.0008 | 0.0008 | 0.0008 |
| F09 | 14240001 | 14340000 | 0.0007 | 0.0005 | 0.0006 |
| F09 | 14260001 | 14360000 | 0.0006 | 0.0005 | 0.0005 |
| F09 | 14280001 | 14380000 | 0.0006 | 0.0005 | 0.0005 |
| F09 | 14300001 | 14400000 | 0.0011 | 0.0010 | 0.0010 |
| F09 | 14320001 | 14420000 | 0.0007 | 0.0007 | 0.0007 |
| F09 | 14340001 | 14440000 | 0.0010 | 0.0009 | 0.0010 |
| F09 | 14360001 | 14460000 | 0.0021 | 0.0018 | 0.0020 |
| F09 | 14380001 | 14480000 | 0.0024 | 0.0020 | 0.0023 |
| F09 | 14400001 | 14500000 | 0.0028 | 0.0022 | 0.0026 |
| F09 | 14420001 | 14520000 | 0.0033 | 0.0024 | 0.0028 |
| F09 | 14440001 | 14540000 | 0.0036 | 0.0026 | 0.0030 |
| F09 | 14460001 | 14560000 | 0.0030 | 0.0021 | 0.0025 |
| F09 | 14480001 | 14580000 | 0.0035 | 0.0025 | 0.0030 |
| F09 | 14500001 | 14600000 | 0.0033 | 0.0022 | 0.0027 |
| F09 | 14520001 | 14620000 | 0.0028 | 0.0021 | 0.0026 |
| F09 | 14540001 | 14640000 | 0.0027 | 0.0021 | 0.0026 |
| F09 | 14560001 | 14660000 | 0.0023 | 0.0017 | 0.0022 |
| F09 | 14580001 | 14680000 | 0.0017 | 0.0012 | 0.0015 |
| F09 | 14600001 | 14700000 | 0.0021 | 0.0017 | 0.0018 |
| F09 | 14620001 | 14720000 | 0.0024 | 0.0017 | 0.0020 |
| F09 | 14640001 | 14740000 | 0.0021 | 0.0015 | 0.0017 |
| F09 | 14660001 | 14760000 | 0.0028 | 0.0021 | 0.0024 |
| F09 | 14680001 | 14780000 | 0.0028 | 0.0022 | 0.0023 |
| F09 | 14700001 | 14800000 | 0.0021 | 0.0017 | 0.0019 |
| F09 | 14720001 | 14820000 | 0.0019 | 0.0015 | 0.0015 |
| F09 | 14740001 | 14840000 | 0.0017 | 0.0014 | 0.0014 |
| F09 | 14760001 | 14860000 | 0.0015 | 0.0011 | 0.0010 |
| F09 | 14780001 | 14880000 | 0.0019 | 0.0014 | 0.0015 |
| F09 | 14800001 | 14900000 | 0.0016 | 0.0011 | 0.0013 |
| F09 | 14820001 | 14920000 | 0.0017 | 0.0013 | 0.0015 |
| F09 | 14840001 | 14940000 | 0.0019 | 0.0014 | 0.0016 |
| F09 | 14860001 | 14960000 | 0.0023 | 0.0018 | 0.0022 |
| F09 | 14880001 | 14980000 | 0.0021 | 0.0016 | 0.0020 |
| F09 | 14900001 | 15000000 | 0.0028 | 0.0022 | 0.0027 |
| F09 | 14920001 | 15020000 | 0.0032 | 0.0023 | 0.0029 |
| F09 | 14940001 | 15040000 | 0.0032 | 0.0023 | 0.0029 |
| F09 | 14960001 | 15060000 | 0.0029 | 0.0022 | 0.0026 |
| F09 | 14980001 | 15080000 | 0.0032 | 0.0024 | 0.0027 |
| F09 | 15000001 | 15100000 | 0.0027 | 0.0019 | 0.0021 |
| F09 | 15020001 | 15120000 | 0.0024 | 0.0019 | 0.0020 |
| F09 | 15040001 | 15140000 | 0.0028 | 0.0023 | 0.0024 |
| F09 | 15060001 | 15160000 | 0.0029 | 0.0022 | 0.0024 |
| F09 | 15080001 | 15180000 | 0.0029 | 0.0022 | 0.0025 |
| F09 | 15100001 | 15200000 | 0.0029 | 0.0023 | 0.0026 |
| F09 | 15120001 | 15220000 | 0.0025 | 0.0019 | 0.0022 |
| F09 | 15140001 | 15240000 | 0.0021 | 0.0015 | 0.0017 |
| F09 | 15160001 | 15260000 | 0.0021 | 0.0016 | 0.0018 |
| F09 | 15180001 | 15280000 | 0.0028 | 0.0024 | 0.0026 |
| F09 | 15200001 | 15300000 | 0.0034 | 0.0026 | 0.0031 |
| F09 | 15220001 | 15320000 | 0.0041 | 0.0032 | 0.0038 |

| Chr | Start | End | both | A1 | A2 |
|---|---|---|---|---|---|
| F09 | 15240001 | 15340000 | 0.0044 | 0.0035 | 0.0041 |
| F09 | 15260001 | 15360000 | 0.0041 | 0.0030 | 0.0037 |
| F09 | 15280001 | 15380000 | 0.0032 | 0.0020 | 0.0026 |
| F09 | 15300001 | 15400000 | 0.0027 | 0.0018 | 0.0022 |
| F09 | 15320001 | 15420000 | 0.0023 | 0.0015 | 0.0017 |
| F09 | 15340001 | 15440000 | 0.0025 | 0.0018 | 0.0020 |
| F09 | 15360001 | 15460000 | 0.0025 | 0.0019 | 0.0021 |
| F09 | 15380001 | 15480000 | 0.0026 | 0.0021 | 0.0025 |
| F09 | 15400001 | 15500000 | 0.0030 | 0.0023 | 0.0027 |
| F09 | 15420001 | 15520000 | 0.0035 | 0.0025 | 0.0031 |
| F09 | 15440001 | 15540000 | 0.0044 | 0.0031 | 0.0037 |
| F09 | 15460001 | 15560000 | 0.0049 | 0.0036 | 0.0042 |
| F09 | 15480001 | 15580000 | 0.0047 | 0.0035 | 0.0040 |
| F09 | 15500001 | 15600000 | 0.0039 | 0.0031 | 0.0035 |
| F09 | 15520001 | 15620000 | 0.0032 | 0.0027 | 0.0029 |
| F09 | 15540001 | 15640000 | 0.0017 | 0.0015 | 0.0017 |
| F09 | 15560001 | 15660000 | 0.0014 | 0.0012 | 0.0014 |
| F09 | 15580001 | 15680000 | 0.0014 | 0.0012 | 0.0014 |
| F09 | 15600001 | 15700000 | 0.0017 | 0.0014 | 0.0017 |
| F09 | 15620001 | 15720000 | 0.0025 | 0.0021 | 0.0024 |
| F09 | 15640001 | 15740000 | 0.0027 | 0.0023 | 0.0026 |
| F09 | 15660001 | 15760000 | 0.0024 | 0.0020 | 0.0022 |
| F09 | 15680001 | 15780000 | 0.0033 | 0.0030 | 0.0030 |
| F09 | 15700001 | 15800000 | 0.0040 | 0.0037 | 0.0037 |
| F09 | 15720001 | 15820000 | 0.0041 | 0.0037 | 0.0038 |
| F09 | 15740001 | 15840000 | 0.0039 | 0.0035 | 0.0035 |
| F09 | 15760001 | 15860000 | 0.0041 | 0.0037 | 0.0037 |
| F09 | 15780001 | 15880000 | 0.0029 | 0.0023 | 0.0026 |
| F09 | 15800001 | 15900000 | 0.0038 | 0.0032 | 0.0035 |
| F09 | 15820001 | 15920000 | 0.0032 | 0.0025 | 0.0029 |
| F09 | 15840001 | 15940000 | 0.0034 | 0.0028 | 0.0033 |
| F09 | 15860001 | 15960000 | 0.0032 | 0.0026 | 0.0032 |
| F09 | 15880001 | 15980000 | 0.0034 | 0.0030 | 0.0034 |
| F09 | 15900001 | 16000000 | 0.0020 | 0.0017 | 0.0019 |
| F09 | 15920001 | 16020000 | 0.0028 | 0.0026 | 0.0026 |
| F09 | 15940001 | 16040000 | 0.0029 | 0.0026 | 0.0025 |
| F09 | 15960001 | 16060000 | 0.0027 | 0.0024 | 0.0023 |
| F09 | 15980001 | 16080000 | 0.0030 | 0.0026 | 0.0026 |
| F09 | 16000001 | 16100000 | 0.0027 | 0.0022 | 0.0022 |
| F09 | 16020001 | 16120000 | 0.0018 | 0.0014 | 0.0014 |
| F09 | 16040001 | 16140000 | 0.0018 | 0.0015 | 0.0016 |
| F09 | 16060001 | 16160000 | 0.0019 | 0.0017 | 0.0018 |
| F09 | 16080001 | 16180000 | 0.0016 | 0.0014 | 0.0014 |
| F09 | 16100001 | 16200000 | 0.0015 | 0.0014 | 0.0014 |
| F09 | 16120001 | 16220000 | 0.0014 | 0.0013 | 0.0014 |
| F09 | 16140001 | 16240000 | 0.0013 | 0.0011 | 0.0012 |
| F09 | 16160001 | 16260000 | 0.0010 | 0.0009 | 0.0009 |
| F09 | 16180001 | 16280000 | 0.0019 | 0.0014 | 0.0014 |
| F09 | 16200001 | 16300000 | 0.0019 | 0.0013 | 0.0014 |
| F09 | 16220001 | 16320000 | 0.0023 | 0.0016 | 0.0016 |
| F09 | 16240001 | 16340000 | 0.0026 | 0.0019 | 0.0020 |
| F09 | 16260001 | 16360000 | 0.0034 | 0.0023 | 0.0024 |
| F09 | 16280001 | 16380000 | 0.0026 | 0.0019 | 0.0020 |
| F09 | 16300001 | 16400000 | 0.0032 | 0.0024 | 0.0025 |
| F09 | 16320001 | 16420000 | 0.0030 | 0.0023 | 0.0024 |
| F09 | 16340001 | 16440000 | 0.0031 | 0.0025 | 0.0025 |
| F09 | 16360001 | 16460000 | 0.0027 | 0.0023 | 0.0024 |
| F09 | 16380001 | 16480000 | 0.0030 | 0.0027 | 0.0028 |

| Chr | Start | End | both | A1 | A2 |
|---|---|---|---|---|---|
| F09 | 16400001 | 16500000 | 0.0033 | 0.0028 | 0.0029 |
| F09 | 16420001 | 16520000 | 0.0035 | 0.0031 | 0.0031 |
| F09 | 16440001 | 16540000 | 0.0027 | 0.0025 | 0.0024 |
| F09 | 16460001 | 16560000 | 0.0024 | 0.0022 | 0.0022 |
| F09 | 16480001 | 16580000 | 0.0022 | 0.0020 | 0.0019 |
| F09 | 16500001 | 16600000 | 0.0017 | 0.0014 | 0.0014 |
| F09 | 16520001 | 16620000 | 0.0014 | 0.0012 | 0.0012 |
| F09 | 16540001 | 16640000 | 0.0019 | 0.0014 | 0.0016 |
| F09 | 16560001 | 16660000 | 0.0021 | 0.0015 | 0.0017 |
| F09 | 16580001 | 16680000 | 0.0022 | 0.0014 | 0.0015 |
| F09 | 16600001 | 16700000 | 0.0028 | 0.0020 | 0.0021 |
| F09 | 16620001 | 16720000 | 0.0033 | 0.0022 | 0.0025 |
| F09 | 16640001 | 16740000 | 0.0028 | 0.0019 | 0.0020 |
| F09 | 16660001 | 16760000 | 0.0030 | 0.0021 | 0.0023 |
| F09 | 16680001 | 16780000 | 0.0026 | 0.0019 | 0.0021 |
| F09 | 16700001 | 16800000 | 0.0017 | 0.0012 | 0.0014 |
| F09 | 16720001 | 16820000 | 0.0011 | 0.0009 | 0.0010 |
| F09 | 16740001 | 16840000 | 0.0014 | 0.0013 | 0.0014 |
| F09 | 16760001 | 16860000 | 0.0010 | 0.0010 | 0.0010 |
| F09 | 16780001 | 16880000 | 0.0012 | 0.0011 | 0.0012 |
| F09 | 16800001 | 16900000 | 0.0012 | 0.0010 | 0.0011 |
| F09 | 16820001 | 16920000 | 0.0022 | 0.0019 | 0.0020 |
| F09 | 16840001 | 16940000 | 0.0024 | 0.0021 | 0.0022 |
| F09 | 16860001 | 16960000 | 0.0042 | 0.0037 | 0.0038 |
| F09 | 16880001 | 16980000 | 0.0046 | 0.0040 | 0.0041 |
| F09 | 16900001 | 17000000 | 0.0050 | 0.0044 | 0.0045 |
| F09 | 16920001 | 17020000 | 0.0044 | 0.0038 | 0.0040 |
| F09 | 16940001 | 17040000 | 0.0040 | 0.0035 | 0.0038 |
| F09 | 16960001 | 17060000 | 0.0035 | 0.0028 | 0.0031 |
| F09 | 16980001 | 17080000 | 0.0033 | 0.0026 | 0.0030 |
| F09 | 17000001 | 17100000 | 0.0030 | 0.0023 | 0.0027 |
| F09 | 17020001 | 17120000 | 0.0025 | 0.0019 | 0.0022 |
| F09 | 17040001 | 17140000 | 0.0027 | 0.0020 | 0.0022 |
| F09 | 17060001 | 17160000 | 0.0018 | 0.0014 | 0.0015 |
| F09 | 17080001 | 17180000 | 0.0021 | 0.0017 | 0.0018 |
| F09 | 17100001 | 17200000 | 0.0032 | 0.0027 | 0.0028 |
| F09 | 17120001 | 17220000 | 0.0045 | 0.0037 | 0.0040 |
| F09 | 17140001 | 17240000 | 0.0046 | 0.0037 | 0.0041 |
| F09 | 17160001 | 17260000 | 0.0051 | 0.0041 | 0.0044 |
| F09 | 17180001 | 17280000 | 0.0053 | 0.0044 | 0.0045 |
| F09 | 17200001 | 17300000 | 0.0045 | 0.0037 | 0.0038 |
| F09 | 17220001 | 17320000 | 0.0034 | 0.0028 | 0.0028 |
| F09 | 17240001 | 17340000 | 0.0028 | 0.0025 | 0.0024 |
| F09 | 17260001 | 17360000 | 0.0032 | 0.0029 | 0.0028 |
| F09 | 17280001 | 17380000 | 0.0029 | 0.0023 | 0.0025 |
| F09 | 17300001 | 17400000 | 0.0029 | 0.0023 | 0.0024 |
| F09 | 17320001 | 17420000 | 0.0029 | 0.0023 | 0.0023 |
| F09 | 17340001 | 17440000 | 0.0036 | 0.0027 | 0.0030 |
| F09 | 17360001 | 17460000 | 0.0026 | 0.0018 | 0.0021 |
| F09 | 17380001 | 17480000 | 0.0030 | 0.0025 | 0.0025 |
| F09 | 17400001 | 17500000 | 0.0025 | 0.0019 | 0.0020 |
| F09 | 17420001 | 17520000 | 0.0027 | 0.0021 | 0.0023 |
| F09 | 17440001 | 17540000 | 0.0020 | 0.0018 | 0.0017 |
| F09 | 17460001 | 17560000 | 0.0024 | 0.0021 | 0.0021 |
| F09 | 17480001 | 17580000 | 0.0015 | 0.0010 | 0.0013 |
| F09 | 17500001 | 17600000 | 0.0018 | 0.0013 | 0.0017 |
| F09 | 17520001 | 17620000 | 0.0016 | 0.0012 | 0.0015 |
| F09 | 17540001 | 17640000 | 0.0021 | 0.0015 | 0.0019 |

| Chr | Start | End | both | A1 | A2 |
|---|---|---|---|---|---|
| F09 | 17560001 | 17660000 | 0.0014 | 0.0010 | 0.0013 |
| F09 | 17580001 | 17680000 | 0.0022 | 0.0019 | 0.0021 |
| F09 | 17600001 | 17700000 | 0.0018 | 0.0017 | 0.0017 |
| F09 | 17620001 | 17720000 | 0.0023 | 0.0021 | 0.0022 |
| F09 | 17640001 | 17740000 | 0.0017 | 0.0016 | 0.0016 |
| F09 | 17660001 | 17760000 | 0.0020 | 0.0017 | 0.0018 |
| F09 | 17680001 | 17780000 | 0.0015 | 0.0011 | 0.0014 |
| F09 | 17700001 | 17800000 | 0.0024 | 0.0019 | 0.0022 |
| F09 | 17720001 | 17820000 | 0.0023 | 0.0018 | 0.0022 |
| F09 | 17740001 | 17840000 | 0.0024 | 0.0019 | 0.0023 |
| F09 | 17760001 | 17860000 | 0.0022 | 0.0019 | 0.0021 |
| F09 | 17780001 | 17880000 | 0.0019 | 0.0017 | 0.0018 |
| F09 | 17800001 | 17900000 | 0.0015 | 0.0012 | 0.0014 |
| F09 | 17820001 | 17920000 | 0.0019 | 0.0014 | 0.0017 |
| F09 | 17840001 | 17940000 | 0.0025 | 0.0019 | 0.0023 |
| F09 | 17860001 | 17960000 | 0.0032 | 0.0025 | 0.0030 |
| F09 | 17880001 | 17980000 | 0.0040 | 0.0033 | 0.0037 |
| F09 | 17900001 | 18000000 | 0.0042 | 0.0035 | 0.0037 |
| F09 | 17920001 | 18020000 | 0.0032 | 0.0029 | 0.0029 |
| F09 | 17940001 | 18040000 | 0.0026 | 0.0024 | 0.0023 |
| F09 | 17960001 | 18060000 | 0.0022 | 0.0021 | 0.0019 |
| F09 | 17980001 | 18080000 | 0.0017 | 0.0015 | 0.0014 |
| F09 | 18000001 | 18100000 | 0.0013 | 0.0012 | 0.0013 |
| F09 | 18020001 | 18120000 | 0.0020 | 0.0019 | 0.0019 |
| F09 | 18040001 | 18140000 | 0.0020 | 0.0019 | 0.0019 |
| F09 | 18060001 | 18160000 | 0.0024 | 0.0023 | 0.0023 |
| F09 | 18080001 | 18180000 | 0.0022 | 0.0021 | 0.0021 |
| F09 | 18100001 | 18200000 | 0.0021 | 0.0020 | 0.0020 |
| F09 | 18120001 | 18220000 | 0.0018 | 0.0016 | 0.0018 |
| F09 | 18140001 | 18240000 | 0.0018 | 0.0016 | 0.0018 |
| F09 | 18160001 | 18260000 | 0.0015 | 0.0011 | 0.0015 |
| F09 | 18180001 | 18280000 | 0.0017 | 0.0012 | 0.0017 |
| F09 | 18200001 | 18300000 | 0.0018 | 0.0012 | 0.0017 |
| F09 | 18220001 | 18320000 | 0.0016 | 0.0012 | 0.0015 |
| F09 | 18240001 | 18340000 | 0.0018 | 0.0014 | 0.0017 |
| F09 | 18260001 | 18360000 | 0.0015 | 0.0012 | 0.0015 |
| F09 | 18280001 | 18380000 | 0.0018 | 0.0013 | 0.0017 |
| F09 | 18300001 | 18400000 | 0.0016 | 0.0012 | 0.0016 |
| F09 | 18320001 | 18420000 | 0.0015 | 0.0010 | 0.0015 |
| F09 | 18340001 | 18440000 | 0.0012 | 0.0007 | 0.0012 |
| F09 | 18360001 | 18460000 | 0.0019 | 0.0013 | 0.0018 |
| F09 | 18380001 | 18480000 | 0.0015 | 0.0011 | 0.0014 |
| F09 | 18400001 | 18500000 | 0.0021 | 0.0015 | 0.0019 |
| F09 | 18420001 | 18520000 | 0.0025 | 0.0018 | 0.0023 |
| F09 | 18440001 | 18540000 | 0.0028 | 0.0019 | 0.0025 |
| F09 | 18460001 | 18560000 | 0.0025 | 0.0016 | 0.0022 |
| F09 | 18480001 | 18580000 | 0.0027 | 0.0018 | 0.0023 |
| F09 | 18500001 | 18600000 | 0.0028 | 0.0021 | 0.0024 |
| F09 | 18520001 | 18620000 | 0.0027 | 0.0020 | 0.0021 |
| F09 | 18540001 | 18640000 | 0.0023 | 0.0019 | 0.0019 |
| F09 | 18560001 | 18660000 | 0.0021 | 0.0017 | 0.0017 |
| F09 | 18580001 | 18680000 | 0.0023 | 0.0018 | 0.0018 |
| F09 | 18600001 | 18700000 | 0.0022 | 0.0017 | 0.0019 |
| F09 | 18620001 | 18720000 | 0.0024 | 0.0019 | 0.0021 |
| F09 | 18640001 | 18740000 | 0.0033 | 0.0026 | 0.0030 |
| F09 | 18660001 | 18760000 | 0.0036 | 0.0026 | 0.0029 |
| F09 | 18680001 | 18780000 | 0.0040 | 0.0029 | 0.0034 |
| F09 | 18700001 | 18800000 | 0.0036 | 0.0025 | 0.0029 |

| Chr | Start | End | both | A1 | A2 |
|---|---|---|---|---|---|
| F09 | 18720001 | 18820000 | 0.0039 | 0.0028 | 0.0032 |
| F09 | 18740001 | 18840000 | 0.0041 | 0.0029 | 0.0032 |
| F09 | 18760001 | 18860000 | 0.0043 | 0.0033 | 0.0035 |
| F09 | 18780001 | 18880000 | 0.0050 | 0.0039 | 0.0040 |
| F09 | 18800001 | 18900000 | 0.0047 | 0.0037 | 0.0037 |
| F09 | 18820001 | 18920000 | 0.0050 | 0.0038 | 0.0040 |
| F09 | 18840001 | 18940000 | 0.0053 | 0.0041 | 0.0043 |
| F09 | 18860001 | 18960000 | 0.0047 | 0.0036 | 0.0039 |
| F09 | 18880001 | 18980000 | 0.0034 | 0.0025 | 0.0028 |
| F09 | 18900001 | 19000000 | 0.0036 | 0.0028 | 0.0032 |
| F09 | 18920001 | 19020000 | 0.0025 | 0.0020 | 0.0023 |
| F09 | 18940001 | 19040000 | 0.0012 | 0.0009 | 0.0011 |
| F09 | 18960001 | 19060000 | 0.0010 | 0.0008 | 0.0010 |
| F09 | 18980001 | 19080000 | 0.0006 | 0.0005 | 0.0006 |
| F09 | 19000001 | 19100000 | 0.0003 | 0.0002 | 0.0003 |
| F09 | 19020001 | 19120000 | 0.0002 | 0.0002 | 0.0002 |
| F09 | 19040001 | 19140000 | 0.0004 | 0.0004 | 0.0004 |
| F09 | 19060001 | 19160000 | 0.0018 | 0.0011 | 0.0017 |
| F09 | 19080001 | 19180000 | 0.0025 | 0.0017 | 0.0024 |
| F09 | 19100001 | 19200000 | 0.0027 | 0.0019 | 0.0026 |
| F09 | 19120001 | 19220000 | 0.0030 | 0.0021 | 0.0028 |
| F09 | 19140001 | 19240000 | 0.0034 | 0.0023 | 0.0032 |
| F09 | 19160001 | 19260000 | 0.0023 | 0.0018 | 0.0023 |
| F09 | 19180001 | 19280000 | 0.0023 | 0.0017 | 0.0023 |
| F09 | 19200001 | 19300000 | 0.0036 | 0.0029 | 0.0035 |
| F09 | 19220001 | 19320000 | 0.0038 | 0.0030 | 0.0037 |
| F09 | 19240001 | 19340000 | 0.0040 | 0.0029 | 0.0035 |
| F09 | 19260001 | 19360000 | 0.0062 | 0.0049 | 0.0054 |
| F09 | 19280001 | 19380000 | 0.0061 | 0.0048 | 0.0053 |
| F09 | 19300001 | 19400000 | 0.0056 | 0.0044 | 0.0049 |
| F09 | 19320001 | 19420000 | 0.0056 | 0.0044 | 0.0049 |
| F09 | 19340001 | 19440000 | 0.0056 | 0.0047 | 0.0050 |
| F09 | 19360001 | 19460000 | 0.0045 | 0.0036 | 0.0041 |
| F09 | 19380001 | 19480000 | 0.0048 | 0.0037 | 0.0041 |
| F09 | 19400001 | 19500000 | 0.0039 | 0.0029 | 0.0032 |
| F09 | 19420001 | 19520000 | 0.0035 | 0.0026 | 0.0028 |
| F09 | 19440001 | 19540000 | 0.0029 | 0.0022 | 0.0024 |
| F09 | 19460001 | 19560000 | 0.0019 | 0.0016 | 0.0015 |
| F09 | 19480001 | 19580000 | 0.0023 | 0.0023 | 0.0021 |
| F09 | 19500001 | 19600000 | 0.0022 | 0.0022 | 0.0020 |
| F09 | 19520001 | 19620000 | 0.0037 | 0.0032 | 0.0035 |
| F09 | 19540001 | 19640000 | 0.0042 | 0.0033 | 0.0041 |
| F09 | 19560001 | 19660000 | 0.0042 | 0.0032 | 0.0042 |
| F09 | 19580001 | 19680000 | 0.0030 | 0.0020 | 0.0031 |
| F09 | 19600001 | 19700000 | 0.0031 | 0.0020 | 0.0031 |
| F09 | 19620001 | 19720000 | 0.0020 | 0.0012 | 0.0019 |
| F09 | 19640001 | 19740000 | 0.0011 | 0.0007 | 0.0010 |
| F09 | 19660001 | 19760000 | 0.0027 | 0.0019 | 0.0026 |
| F09 | 19680001 | 19780000 | 0.0035 | 0.0024 | 0.0034 |
| F09 | 19700001 | 19800000 | 0.0040 | 0.0029 | 0.0039 |
| F09 | 19720001 | 19820000 | 0.0036 | 0.0027 | 0.0036 |
| F09 | 19740001 | 19840000 | 0.0040 | 0.0030 | 0.0040 |
| F09 | 19760001 | 19860000 | 0.0022 | 0.0017 | 0.0022 |
| F09 | 19780001 | 19880000 | 0.0021 | 0.0017 | 0.0021 |
| F09 | 19800001 | 19900000 | 0.0023 | 0.0019 | 0.0023 |
| F09 | 19820001 | 19920000 | 0.0024 | 0.0020 | 0.0024 |
| F09 | 19840001 | 19940000 | 0.0021 | 0.0017 | 0.0021 |
| F09 | 19860001 | 19960000 | 0.0020 | 0.0016 | 0.0020 |

| Chr | Start | End | both | A1 | A2 |
|---|---|---|---|---|---|
| F09 | 19880001 | 19980000 | 0.0014 | 0.0012 | 0.0014 |
| F09 | 19900001 | 20000000 | 0.0007 | 0.0006 | 0.0007 |
| F09 | 19920001 | 20020000 | 0.0008 | 0.0007 | 0.0009 |
| F09 | 19940001 | 20040000 | 0.0011 | 0.0009 | 0.0011 |
| F09 | 19960001 | 20060000 | 0.0014 | 0.0012 | 0.0014 |
| F09 | 19980001 | 20080000 | 0.0012 | 0.0011 | 0.0012 |
| F09 | 20000001 | 20100000 | 0.0015 | 0.0013 | 0.0014 |
| F09 | 20020001 | 20120000 | 0.0015 | 0.0013 | 0.0015 |
| F09 | 20040001 | 20140000 | 0.0014 | 0.0013 | 0.0014 |
| F09 | 20060001 | 20160000 | 0.0017 | 0.0012 | 0.0015 |
| F09 | 20080001 | 20180000 | 0.0022 | 0.0017 | 0.0020 |
| F09 | 20100001 | 20200000 | 0.0020 | 0.0015 | 0.0018 |
| F09 | 20120001 | 20220000 | 0.0019 | 0.0014 | 0.0017 |
| F09 | 20140001 | 20240000 | 0.0019 | 0.0013 | 0.0015 |
| F09 | 20160001 | 20260000 | 0.0015 | 0.0011 | 0.0013 |
| F09 | 20180001 | 20280000 | 0.0013 | 0.0010 | 0.0011 |
| F09 | 20200001 | 20300000 | 0.0019 | 0.0015 | 0.0017 |
| F09 | 20220001 | 20320000 | 0.0020 | 0.0015 | 0.0018 |
| F09 | 20240001 | 20340000 | 0.0023 | 0.0018 | 0.0022 |
| F09 | 20260001 | 20360000 | 0.0022 | 0.0017 | 0.0020 |
| F09 | 20280001 | 20380000 | 0.0032 | 0.0022 | 0.0026 |
| F09 | 20300001 | 20400000 | 0.0030 | 0.0020 | 0.0024 |
| F09 | 20320001 | 20420000 | 0.0030 | 0.0021 | 0.0025 |
| F09 | 20340001 | 20440000 | 0.0029 | 0.0021 | 0.0024 |
| F09 | 20360001 | 20460000 | 0.0036 | 0.0026 | 0.0029 |
| F09 | 20380001 | 20480000 | 0.0031 | 0.0022 | 0.0026 |
| F09 | 20400001 | 20500000 | 0.0037 | 0.0028 | 0.0032 |
| F09 | 20420001 | 20520000 | 0.0040 | 0.0031 | 0.0033 |
| F09 | 20440001 | 20540000 | 0.0054 | 0.0043 | 0.0047 |
| F09 | 20460001 | 20560000 | 0.0057 | 0.0046 | 0.0053 |
| F09 | 20480001 | 20580000 | 0.0058 | 0.0046 | 0.0052 |
| F09 | 20500001 | 20600000 | 0.0055 | 0.0044 | 0.0049 |
| F09 | 20520001 | 20620000 | 0.0070 | 0.0055 | 0.0063 |
| F09 | 20540001 | 20640000 | 0.0063 | 0.0048 | 0.0054 |
| F09 | 20560001 | 20660000 | 0.0057 | 0.0042 | 0.0047 |
| F09 | 20580001 | 20680000 | 0.0056 | 0.0041 | 0.0045 |
| F09 | 20600001 | 20700000 | 0.0047 | 0.0034 | 0.0037 |
| F09 | 20620001 | 20720000 | 0.0033 | 0.0023 | 0.0025 |
| F09 | 20640001 | 20740000 | 0.0025 | 0.0017 | 0.0019 |
| F09 | 20660001 | 20760000 | 0.0021 | 0.0015 | 0.0017 |
| F09 | 20680001 | 20780000 | 0.0018 | 0.0017 | 0.0018 |
| F09 | 20700001 | 20800000 | 0.0023 | 0.0020 | 0.0023 |
| F09 | 20720001 | 20820000 | 0.0025 | 0.0021 | 0.0023 |
| F09 | 20740001 | 20840000 | 0.0024 | 0.0019 | 0.0021 |
| F09 | 20760001 | 20860000 | 0.0026 | 0.0021 | 0.0023 |
| F09 | 20780001 | 20880000 | 0.0035 | 0.0029 | 0.0031 |
| F09 | 20800001 | 20900000 | 0.0035 | 0.0030 | 0.0031 |
| F09 | 20820001 | 20920000 | 0.0036 | 0.0032 | 0.0033 |
| F09 | 20840001 | 20940000 | 0.0038 | 0.0033 | 0.0037 |
| F09 | 20860001 | 20960000 | 0.0038 | 0.0033 | 0.0036 |
| F09 | 20880001 | 20980000 | 0.0029 | 0.0024 | 0.0030 |
| F09 | 20900001 | 21000000 | 0.0035 | 0.0029 | 0.0035 |
| F09 | 20920001 | 21020000 | 0.0025 | 0.0021 | 0.0026 |
| F09 | 20940001 | 21040000 | 0.0025 | 0.0021 | 0.0024 |
| F09 | 20960001 | 21060000 | 0.0027 | 0.0021 | 0.0026 |
| F09 | 20980001 | 21080000 | 0.0031 | 0.0026 | 0.0031 |
| F09 | 21000001 | 21100000 | 0.0028 | 0.0023 | 0.0028 |
| F09 | 21020001 | 21120000 | 0.0032 | 0.0026 | 0.0031 |

| Chr | Start | End | both | A1 | A2 |
|---|---|---|---|---|---|
| F09 | 21040001 | 21140000 | 0.0040 | 0.0034 | 0.0039 |
| F09 | 21060001 | 21160000 | 0.0040 | 0.0034 | 0.0039 |
| F09 | 21080001 | 21180000 | 0.0031 | 0.0026 | 0.0030 |
| F09 | 21100001 | 21200000 | 0.0026 | 0.0020 | 0.0025 |
| F09 | 21120001 | 21220000 | 0.0025 | 0.0020 | 0.0024 |
| F09 | 21140001 | 21240000 | 0.0014 | 0.0011 | 0.0013 |
| F09 | 21160001 | 21260000 | 0.0019 | 0.0018 | 0.0019 |
| F09 | 21180001 | 21280000 | 0.0026 | 0.0021 | 0.0022 |
| F09 | 21200001 | 21300000 | 0.0027 | 0.0023 | 0.0024 |
| F09 | 21220001 | 21320000 | 0.0027 | 0.0023 | 0.0024 |
| F09 | 21240001 | 21340000 | 0.0031 | 0.0025 | 0.0027 |
| F09 | 21260001 | 21360000 | 0.0025 | 0.0020 | 0.0022 |
| F09 | 21280001 | 21380000 | 0.0025 | 0.0022 | 0.0023 |
| F09 | 21300001 | 21400000 | 0.0026 | 0.0021 | 0.0023 |
| F09 | 21320001 | 21420000 | 0.0029 | 0.0022 | 0.0024 |
| F09 | 21340001 | 21440000 | 0.0036 | 0.0023 | 0.0027 |
| F09 | 21360001 | 21460000 | 0.0035 | 0.0022 | 0.0025 |
| F09 | 21380001 | 21480000 | 0.0035 | 0.0022 | 0.0026 |
| F09 | 21400001 | 21500000 | 0.0045 | 0.0033 | 0.0036 |
| F09 | 21420001 | 21520000 | 0.0042 | 0.0030 | 0.0032 |
| F09 | 21440001 | 21540000 | 0.0038 | 0.0027 | 0.0028 |
| F09 | 21460001 | 21560000 | 0.0041 | 0.0027 | 0.0028 |
| F09 | 21480001 | 21580000 | 0.0042 | 0.0029 | 0.0029 |
| F09 | 21500001 | 21600000 | 0.0041 | 0.0027 | 0.0030 |
| F09 | 21520001 | 21620000 | 0.0049 | 0.0036 | 0.0039 |
| F09 | 21540001 | 21640000 | 0.0046 | 0.0034 | 0.0037 |
| F09 | 21560001 | 21660000 | 0.0052 | 0.0039 | 0.0043 |
| F09 | 21580001 | 21680000 | 0.0052 | 0.0037 | 0.0041 |
| F09 | 21600001 | 21700000 | 0.0049 | 0.0035 | 0.0038 |
| F09 | 21620001 | 21720000 | 0.0053 | 0.0038 | 0.0042 |
| F09 | 21640001 | 21740000 | 0.0062 | 0.0047 | 0.0053 |
| F09 | 21660001 | 21760000 | 0.0065 | 0.0051 | 0.0058 |
| F09 | 21680001 | 21780000 | 0.0065 | 0.0052 | 0.0059 |
| F09 | 21700001 | 21800000 | 0.0071 | 0.0055 | 0.0064 |
| F09 | 21720001 | 21820000 | 0.0063 | 0.0048 | 0.0056 |
| F09 | 21740001 | 21840000 | 0.0056 | 0.0044 | 0.0051 |
| F09 | 21760001 | 21860000 | 0.0044 | 0.0032 | 0.0037 |
| F09 | 21780001 | 21880000 | 0.0047 | 0.0037 | 0.0041 |
| F09 | 21800001 | 21900000 | 0.0037 | 0.0028 | 0.0031 |
| F09 | 21820001 | 21920000 | 0.0034 | 0.0025 | 0.0027 |
| F09 | 21840001 | 21940000 | 0.0034 | 0.0024 | 0.0026 |
| F09 | 21860001 | 21960000 | 0.0037 | 0.0027 | 0.0029 |
| F09 | 21880001 | 21980000 | 0.0030 | 0.0023 | 0.0022 |
| F09 | 21900001 | 22000000 | 0.0032 | 0.0027 | 0.0027 |
| F09 | 21920001 | 22020000 | 0.0035 | 0.0030 | 0.0031 |
| F09 | 21940001 | 22040000 | 0.0045 | 0.0042 | 0.0042 |
| F09 | 21960001 | 22060000 | 0.0045 | 0.0042 | 0.0043 |
| F09 | 21980001 | 22080000 | 0.0047 | 0.0041 | 0.0045 |
| F09 | 22000001 | 22100000 | 0.0047 | 0.0042 | 0.0043 |
| F09 | 22020001 | 22120000 | 0.0050 | 0.0043 | 0.0044 |
| F09 | 22040001 | 22140000 | 0.0043 | 0.0034 | 0.0037 |
| F09 | 22060001 | 22160000 | 0.0044 | 0.0036 | 0.0038 |
| F09 | 22080001 | 22180000 | 0.0043 | 0.0035 | 0.0037 |
| F09 | 22100001 | 22200000 | 0.0036 | 0.0028 | 0.0032 |
| F09 | 22120001 | 22220000 | 0.0030 | 0.0025 | 0.0028 |
| F09 | 22140001 | 22240000 | 0.0026 | 0.0022 | 0.0024 |
| F09 | 22160001 | 22260000 | 0.0035 | 0.0029 | 0.0032 |
| F09 | 22180001 | 22280000 | 0.0035 | 0.0030 | 0.0032 |

| Chr | Start | End | both | A1 | A2 |
|---|---|---|---|---|---|
| F09 | 22200001 | 22300000 | 0.0044 | 0.0035 | 0.0039 |
| F09 | 22220001 | 22320000 | 0.0051 | 0.0041 | 0.0046 |
| F09 | 22240001 | 22340000 | 0.0049 | 0.0039 | 0.0043 |
| F09 | 22260001 | 22360000 | 0.0044 | 0.0033 | 0.0036 |
| F09 | 22280001 | 22380000 | 0.0041 | 0.0031 | 0.0035 |
| F09 | 22300001 | 22400000 | 0.0032 | 0.0026 | 0.0027 |
| F09 | 22320001 | 22420000 | 0.0032 | 0.0025 | 0.0026 |
| F09 | 22340001 | 22440000 | 0.0036 | 0.0028 | 0.0029 |
| F09 | 22360001 | 22460000 | 0.0026 | 0.0020 | 0.0022 |
| F09 | 22380001 | 22480000 | 0.0030 | 0.0024 | 0.0026 |
| F09 | 22400001 | 22500000 | 0.0029 | 0.0023 | 0.0025 |
| F09 | 22420001 | 22520000 | 0.0029 | 0.0022 | 0.0023 |
| F09 | 22440001 | 22540000 | 0.0024 | 0.0018 | 0.0020 |
| F09 | 22460001 | 22560000 | 0.0029 | 0.0023 | 0.0025 |
| F09 | 22480001 | 22580000 | 0.0027 | 0.0021 | 0.0023 |
| F09 | 22500001 | 22600000 | 0.0030 | 0.0023 | 0.0023 |
| F09 | 22520001 | 22620000 | 0.0026 | 0.0021 | 0.0021 |
| F09 | 22540001 | 22640000 | 0.0026 | 0.0021 | 0.0020 |
| F09 | 22560001 | 22660000 | 0.0024 | 0.0017 | 0.0017 |
| F09 | 22580001 | 22680000 | 0.0029 | 0.0020 | 0.0022 |
| F09 | 22600001 | 22700000 | 0.0035 | 0.0028 | 0.0029 |
| F09 | 22620001 | 22720000 | 0.0034 | 0.0025 | 0.0027 |
| F09 | 22640001 | 22740000 | 0.0038 | 0.0029 | 0.0031 |
| F09 | 22660001 | 22760000 | 0.0039 | 0.0031 | 0.0032 |
| F09 | 22680001 | 22780000 | 0.0032 | 0.0024 | 0.0023 |
| F09 | 22700001 | 22800000 | 0.0024 | 0.0016 | 0.0015 |
| F09 | 22720001 | 22820000 | 0.0024 | 0.0017 | 0.0015 |
| F09 | 22740001 | 22840000 | 0.0026 | 0.0017 | 0.0017 |
| F09 | 22760001 | 22860000 | 0.0025 | 0.0016 | 0.0017 |
| F09 | 22780001 | 22880000 | 0.0030 | 0.0021 | 0.0023 |
| F09 | 22800001 | 22900000 | 0.0037 | 0.0027 | 0.0029 |
| F09 | 22820001 | 22920000 | 0.0032 | 0.0024 | 0.0026 |
| F09 | 22840001 | 22940000 | 0.0026 | 0.0020 | 0.0022 |
| F09 | 22860001 | 22960000 | 0.0026 | 0.0020 | 0.0022 |
| F09 | 22880001 | 22980000 | 0.0024 | 0.0020 | 0.0022 |
| F09 | 22900001 | 23000000 | 0.0017 | 0.0014 | 0.0017 |
| F09 | 22920001 | 23020000 | 0.0022 | 0.0019 | 0.0022 |
| F09 | 22940001 | 23040000 | 0.0027 | 0.0022 | 0.0026 |
| F09 | 22960001 | 23060000 | 0.0029 | 0.0023 | 0.0026 |
| F09 | 22980001 | 23080000 | 0.0030 | 0.0024 | 0.0027 |
| F09 | 23000001 | 23100000 | 0.0033 | 0.0027 | 0.0030 |
| F09 | 23020001 | 23120000 | 0.0033 | 0.0026 | 0.0030 |
| F09 | 23040001 | 23140000 | 0.0029 | 0.0023 | 0.0027 |
| F09 | 23060001 | 23160000 | 0.0027 | 0.0023 | 0.0027 |
| F09 | 23080001 | 23180000 | 0.0022 | 0.0020 | 0.0022 |
| F09 | 23100001 | 23200000 | 0.0020 | 0.0019 | 0.0020 |
| F09 | 23120001 | 23220000 | 0.0019 | 0.0019 | 0.0019 |
| F09 | 23140001 | 23240000 | 0.0019 | 0.0019 | 0.0018 |
| F09 | 23160001 | 23260000 | 0.0021 | 0.0021 | 0.0019 |
| F09 | 23180001 | 23280000 | 0.0021 | 0.0020 | 0.0020 |
| F09 | 23200001 | 23300000 | 0.0021 | 0.0020 | 0.0020 |
| F09 | 23220001 | 23320000 | 0.0016 | 0.0015 | 0.0015 |
| F09 | 23240001 | 23340000 | 0.0016 | 0.0014 | 0.0015 |
| F09 | 23260001 | 23360000 | 0.0014 | 0.0010 | 0.0013 |
| F09 | 23280001 | 23380000 | 0.0025 | 0.0019 | 0.0021 |
| F09 | 23300001 | 23400000 | 0.0024 | 0.0018 | 0.0020 |
| F09 | 23320001 | 23420000 | 0.0031 | 0.0025 | 0.0026 |
| F09 | 23340001 | 23440000 | 0.0037 | 0.0029 | 0.0031 |

| Chr | Start | End | both | A1 | A2 |
|---|---|---|---|---|---|
| F09 | 23360001 | 23460000 | 0.0042 | 0.0035 | 0.0035 |
| F09 | 23380001 | 23480000 | 0.0030 | 0.0027 | 0.0026 |
| F09 | 23400001 | 23500000 | 0.0029 | 0.0025 | 0.0026 |
| F09 | 23420001 | 23520000 | 0.0024 | 0.0021 | 0.0022 |
| F09 | 23440001 | 23540000 | 0.0021 | 0.0018 | 0.0019 |
| F09 | 23460001 | 23560000 | 0.0016 | 0.0013 | 0.0015 |
| F09 | 23480001 | 23580000 | 0.0016 | 0.0014 | 0.0016 |
| F09 | 23500001 | 23600000 | 0.0018 | 0.0016 | 0.0018 |
| F09 | 23520001 | 23620000 | 0.0017 | 0.0015 | 0.0017 |
| F09 | 23540001 | 23640000 | 0.0021 | 0.0019 | 0.0020 |
| F09 | 23560001 | 23660000 | 0.0020 | 0.0018 | 0.0019 |
| F09 | 23580001 | 23680000 | 0.0017 | 0.0015 | 0.0017 |
| F09 | 23600001 | 23700000 | 0.0019 | 0.0016 | 0.0018 |
| F09 | 23620001 | 23720000 | 0.0018 | 0.0015 | 0.0018 |
| F09 | 23640001 | 23740000 | 0.0010 | 0.0008 | 0.0010 |
| F09 | 23660001 | 23760000 | 0.0011 | 0.0008 | 0.0011 |
| F09 | 23680001 | 23780000 | 0.0016 | 0.0013 | 0.0016 |
| F09 | 23700001 | 23800000 | 0.0014 | 0.0012 | 0.0014 |
| F09 | 23720001 | 23820000 | 0.0015 | 0.0012 | 0.0014 |
| F09 | 23740001 | 23840000 | 0.0027 | 0.0022 | 0.0025 |
| F09 | 23760001 | 23860000 | 0.0030 | 0.0025 | 0.0026 |
| F09 | 23780001 | 23880000 | 0.0029 | 0.0025 | 0.0026 |
| F09 | 23800001 | 23900000 | 0.0034 | 0.0029 | 0.0031 |
| F09 | 23820001 | 23920000 | 0.0035 | 0.0030 | 0.0032 |
| F09 | 23840001 | 23940000 | 0.0026 | 0.0023 | 0.0025 |
| F09 | 23860001 | 23960000 | 0.0021 | 0.0018 | 0.0020 |
| F09 | 23880001 | 23980000 | 0.0016 | 0.0014 | 0.0016 |
| F09 | 23900001 | 24000000 | 0.0013 | 0.0010 | 0.0012 |
| F09 | 23920001 | 24020000 | 0.0014 | 0.0011 | 0.0014 |
| F09 | 23940001 | 24040000 | 0.0015 | 0.0012 | 0.0015 |
| F09 | 23960001 | 24060000 | 0.0019 | 0.0016 | 0.0018 |
| F09 | 23980001 | 24080000 | 0.0019 | 0.0015 | 0.0017 |
| F09 | 24000001 | 24100000 | 0.0017 | 0.0013 | 0.0015 |
| F09 | 24020001 | 24120000 | 0.0014 | 0.0011 | 0.0012 |
| F09 | 24040001 | 24140000 | 0.0010 | 0.0008 | 0.0009 |
| F09 | 24060001 | 24160000 | 0.0010 | 0.0007 | 0.0010 |
| F09 | 24080001 | 24180000 | 0.0012 | 0.0010 | 0.0012 |
| F09 | 24100001 | 24200000 | 0.0010 | 0.0009 | 0.0011 |
| F09 | 24120001 | 24220000 | 0.0015 | 0.0012 | 0.0015 |
| F09 | 24140001 | 24240000 | 0.0017 | 0.0013 | 0.0017 |
| F09 | 24160001 | 24260000 | 0.0015 | 0.0013 | 0.0015 |
| F09 | 24180001 | 24280000 | 0.0020 | 0.0014 | 0.0020 |
| F09 | 24200001 | 24300000 | 0.0021 | 0.0015 | 0.0021 |
| F09 | 24220001 | 24320000 | 0.0017 | 0.0012 | 0.0018 |
| F09 | 24240001 | 24340000 | 0.0017 | 0.0012 | 0.0017 |
| F09 | 24260001 | 24360000 | 0.0019 | 0.0014 | 0.0019 |
| F09 | 24280001 | 24380000 | 0.0011 | 0.0009 | 0.0011 |
| F09 | 24300001 | 24400000 | 0.0010 | 0.0008 | 0.0010 |
| F09 | 24320001 | 24420000 | 0.0009 | 0.0009 | 0.0009 |
| F09 | 24340001 | 24440000 | 0.0009 | 0.0008 | 0.0009 |
| F09 | 24360001 | 24460000 | 0.0004 | 0.0003 | 0.0004 |
| F09 | 24380001 | 24480000 | 0.0011 | 0.0010 | 0.0011 |
| F09 | 24400001 | 24500000 | 0.0025 | 0.0021 | 0.0025 |
| F09 | 24420001 | 24520000 | 0.0023 | 0.0019 | 0.0023 |
| F09 | 24440001 | 24540000 | 0.0024 | 0.0020 | 0.0024 |
| F09 | 24460001 | 24560000 | 0.0036 | 0.0031 | 0.0036 |
| F09 | 24480001 | 24580000 | 0.0035 | 0.0030 | 0.0035 |
| F09 | 24500001 | 24600000 | 0.0027 | 0.0022 | 0.0026 |

| Chr | Start | End | both | A1 | A2 |
|---|---|---|---|---|---|
| F09 | 24520001 | 24620000 | 0.0033 | 0.0026 | 0.0030 |
| F09 | 24540001 | 24640000 | 0.0036 | 0.0028 | 0.0031 |
| F09 | 24560001 | 24660000 | 0.0028 | 0.0020 | 0.0022 |
| F09 | 24580001 | 24680000 | 0.0025 | 0.0018 | 0.0019 |
| F09 | 24600001 | 24700000 | 0.0020 | 0.0016 | 0.0016 |
| F09 | 24620001 | 24720000 | 0.0022 | 0.0018 | 0.0019 |
| F09 | 24640001 | 24740000 | 0.0019 | 0.0017 | 0.0018 |
| F09 | 24660001 | 24760000 | 0.0018 | 0.0015 | 0.0016 |
| F09 | 24680001 | 24780000 | 0.0015 | 0.0012 | 0.0014 |
| F09 | 24700001 | 24800000 | 0.0013 | 0.0010 | 0.0012 |
| F09 | 24720001 | 24820000 | 0.0009 | 0.0005 | 0.0007 |
| F09 | 24740001 | 24840000 | 0.0010 | 0.0007 | 0.0008 |
| F09 | 24760001 | 24860000 | 0.0009 | 0.0007 | 0.0009 |
| F09 | 24780001 | 24880000 | 0.0009 | 0.0006 | 0.0008 |
| F09 | 24800001 | 24900000 | 0.0008 | 0.0006 | 0.0008 |
| F09 | 24820001 | 24920000 | 0.0010 | 0.0008 | 0.0010 |
| F09 | 24840001 | 24940000 | 0.0010 | 0.0007 | 0.0010 |
| F09 | 24860001 | 24960000 | 0.0007 | 0.0005 | 0.0007 |
| F09 | 24880001 | 24980000 | 0.0008 | 0.0005 | 0.0008 |
| F09 | 24900001 | 25000000 | 0.0008 | 0.0005 | 0.0008 |
| F09 | 24920001 | 25020000 | 0.0003 | 0.0002 | 0.0003 |
| F09 | 24940001 | 25040000 | 0.0006 | 0.0005 | 0.0006 |
| F09 | 24960001 | 25060000 | 0.0007 | 0.0006 | 0.0008 |
| F09 | 24980001 | 25080000 | 0.0008 | 0.0008 | 0.0008 |
| F09 | 25000001 | 25100000 | 0.0011 | 0.0010 | 0.0011 |
| F09 | 25020001 | 25120000 | 0.0012 | 0.0010 | 0.0011 |
| F09 | 25040001 | 25140000 | 0.0009 | 0.0008 | 0.0009 |
| F09 | 25060001 | 25160000 | 0.0010 | 0.0009 | 0.0010 |
| F09 | 25080001 | 25180000 | 0.0010 | 0.0008 | 0.0010 |
| F09 | 25100001 | 25200000 | 0.0007 | 0.0005 | 0.0007 |
| F09 | 25120001 | 25220000 | 0.0007 | 0.0006 | 0.0007 |
| F09 | 25140001 | 25240000 | 0.0006 | 0.0005 | 0.0006 |
| F09 | 25160001 | 25260000 | 0.0006 | 0.0005 | 0.0005 |
| F09 | 25180001 | 25280000 | 0.0008 | 0.0005 | 0.0007 |
| F09 | 25200001 | 25300000 | 0.0008 | 0.0005 | 0.0007 |
| F09 | 25220001 | 25320000 | 0.0011 | 0.0008 | 0.0011 |
| F09 | 25240001 | 25340000 | 0.0013 | 0.0010 | 0.0012 |
| F09 | 25260001 | 25360000 | 0.0012 | 0.0009 | 0.0012 |
| F09 | 25280001 | 25380000 | 0.0011 | 0.0008 | 0.0010 |
| F09 | 25300001 | 25400000 | 0.0018 | 0.0014 | 0.0016 |
| F09 | 25320001 | 25420000 | 0.0028 | 0.0023 | 0.0026 |
| F09 | 25340001 | 25440000 | 0.0028 | 0.0023 | 0.0026 |
| F09 | 25360001 | 25460000 | 0.0027 | 0.0022 | 0.0025 |
| F09 | 25380001 | 25480000 | 0.0027 | 0.0024 | 0.0025 |
| F09 | 25400001 | 25500000 | 0.0025 | 0.0022 | 0.0024 |
| F09 | 25420001 | 25520000 | 0.0014 | 0.0012 | 0.0014 |
| F09 | 25440001 | 25540000 | 0.0013 | 0.0011 | 0.0013 |
| F09 | 25460001 | 25560000 | 0.0015 | 0.0012 | 0.0015 |
| F09 | 25480001 | 25580000 | 0.0015 | 0.0011 | 0.0015 |
| F09 | 25500001 | 25600000 | 0.0013 | 0.0009 | 0.0013 |
| F09 | 25520001 | 25620000 | 0.0010 | 0.0007 | 0.0010 |
| F09 | 25540001 | 25640000 | 0.0013 | 0.0009 | 0.0013 |
| F09 | 25560001 | 25660000 | 0.0011 | 0.0007 | 0.0011 |
| F09 | 25580001 | 25680000 | 0.0011 | 0.0008 | 0.0011 |
| F09 | 25600001 | 25700000 | 0.0008 | 0.0006 | 0.0008 |
| F09 | 25620001 | 25720000 | 0.0009 | 0.0006 | 0.0010 |
| F09 | 25640001 | 25740000 | 0.0004 | 0.0003 | 0.0004 |
| F09 | 25660001 | 25760000 | 0.0004 | 0.0003 | 0.0004 |

| Chr | Start | End | both | A1 | A2 |
|---|---|---|---|---|---|
| F09 | 25680001 | 25780000 | 0.0002 | 0.0001 | 0.0002 |
| F09 | 25700001 | 25800000 | 0.0002 | 0.0001 | 0.0002 |
| F09 | 25720001 | 25820000 | 0.0014 | 0.0009 | 0.0015 |
| F09 | 25740001 | 25840000 | 0.0016 | 0.0010 | 0.0016 |
| F09 | 25760001 | 25860000 | 0.0016 | 0.0010 | 0.0016 |
| F09 | 25780001 | 25880000 | 0.0016 | 0.0010 | 0.0016 |
| F09 | 25800001 | 25900000 | 0.0016 | 0.0010 | 0.0016 |
| F09 | 25820001 | 25920000 | 0.0007 | 0.0004 | 0.0006 |
| F09 | 25840001 | 25940000 | 0.0006 | 0.0004 | 0.0006 |
| F09 | 25860001 | 25960000 | 0.0008 | 0.0005 | 0.0007 |
| F09 | 25880001 | 25980000 | 0.0008 | 0.0005 | 0.0007 |
| F09 | 25900001 | 26000000 | 0.0010 | 0.0007 | 0.0010 |
| F09 | 25920001 | 26020000 | 0.0005 | 0.0005 | 0.0005 |
| F09 | 25940001 | 26040000 | 0.0005 | 0.0004 | 0.0005 |
| F09 | 25960001 | 26060000 | 0.0004 | 0.0003 | 0.0004 |
| F09 | 25980001 | 26080000 | 0.0005 | 0.0004 | 0.0004 |
| F09 | 26000001 | 26100000 | 0.0003 | 0.0003 | 0.0003 |
| F09 | 26020001 | 26120000 | 0.0005 | 0.0004 | 0.0004 |
| F09 | 26040001 | 26140000 | 0.0003 | 0.0003 | 0.0003 |
| F09 | 26060001 | 26160000 | 0.0004 | 0.0004 | 0.0004 |
| F09 | 26080001 | 26180000 | 0.0004 | 0.0004 | 0.0004 |
| F09 | 26100001 | 26200000 | 0.0004 | 0.0003 | 0.0003 |
| F09 | 26120001 | 26220000 | 0.0003 | 0.0003 | 0.0003 |
| F09 | 26140001 | 26240000 | 0.0005 | 0.0004 | 0.0005 |
| F09 | 26160001 | 26260000 | 0.0006 | 0.0004 | 0.0006 |
| F09 | 26180001 | 26280000 | 0.0006 | 0.0004 | 0.0006 |
| F09 | 26200001 | 26300000 | 0.0007 | 0.0004 | 0.0007 |
| F09 | 26220001 | 26320000 | 0.0007 | 0.0004 | 0.0006 |
| F09 | 26240001 | 26340000 | 0.0005 | 0.0003 | 0.0005 |
| F09 | 26260001 | 26360000 | 0.0004 | 0.0003 | 0.0004 |
| F09 | 26280001 | 26380000 | 0.0006 | 0.0005 | 0.0006 |
| F09 | 26300001 | 26400000 | 0.0005 | 0.0004 | 0.0005 |
| F09 | 26320001 | 26420000 | 0.0005 | 0.0004 | 0.0005 |
| F09 | 26340001 | 26440000 | 0.0005 | 0.0004 | 0.0005 |
| F09 | 26360001 | 26460000 | 0.0004 | 0.0003 | 0.0004 |
| F09 | 26380001 | 26480000 | 0.0003 | 0.0002 | 0.0003 |
| F09 | 26400001 | 26500000 | 0.0003 | 0.0002 | 0.0003 |
| F09 | 26420001 | 26520000 | 0.0002 | 0.0002 | 0.0002 |
| F09 | 26440001 | 26540000 | 0.0002 | 0.0002 | 0.0002 |
| F09 | 26460001 | 26560000 | 0.0002 | 0.0001 | 0.0002 |
| F09 | 26480001 | 26580000 | 0.0003 | 0.0002 | 0.0003 |
| F09 | 26500001 | 26600000 | 0.0004 | 0.0003 | 0.0004 |
| F09 | 26520001 | 26620000 | 0.0004 | 0.0003 | 0.0004 |
| F09 | 26540001 | 26640000 | 0.0005 | 0.0003 | 0.0004 |
| F09 | 26560001 | 26660000 | 0.0005 | 0.0003 | 0.0005 |
| F09 | 26580001 | 26680000 | 0.0003 | 0.0003 | 0.0003 |
| F09 | 26600001 | 26700000 | 0.0002 | 0.0001 | 0.0002 |
| F09 | 26620001 | 26720000 | 0.0003 | 0.0002 | 0.0003 |
| F09 | 26640001 | 26740000 | 0.0003 | 0.0002 | 0.0003 |
| F09 | 26660001 | 26760000 | 0.0010 | 0.0007 | 0.0010 |
| F09 | 26680001 | 26780000 | 0.0010 | 0.0007 | 0.0010 |
| F09 | 26700001 | 26800000 | 0.0012 | 0.0008 | 0.0012 |
| F09 | 26720001 | 26820000 | 0.0010 | 0.0007 | 0.0010 |
| F09 | 26740001 | 26840000 | 0.0009 | 0.0007 | 0.0009 |
| F09 | 26760001 | 26860000 | 0.0003 | 0.0002 | 0.0003 |
| F09 | 26780001 | 26880000 | 0.0003 | 0.0002 | 0.0003 |
| F09 | 26800001 | 26900000 | 0.0002 | 0.0001 | 0.0001 |
| F09 | 26820001 | 26920000 | 0.0007 | 0.0006 | 0.0007 |

| Chr | Start | End | both | A1 | A2 |
|---|---|---|---|---|---|
| F09 | 26840001 | 26940000 | 0.0009 | 0.0008 | 0.0009 |
| F09 | 26860001 | 26960000 | 0.0009 | 0.0008 | 0.0009 |
| F09 | 26880001 | 26980000 | 0.0011 | 0.0009 | 0.0010 |
| F09 | 26900001 | 27000000 | 0.0011 | 0.0010 | 0.0011 |
| F09 | 26920001 | 27020000 | 0.0007 | 0.0006 | 0.0007 |
| F09 | 26940001 | 27040000 | 0.0009 | 0.0007 | 0.0009 |
| F09 | 26960001 | 27060000 | 0.0011 | 0.0009 | 0.0011 |
| F09 | 26980001 | 27080000 | 0.0017 | 0.0014 | 0.0016 |
| F09 | 27000001 | 27100000 | 0.0019 | 0.0016 | 0.0018 |
| F09 | 27020001 | 27120000 | 0.0031 | 0.0025 | 0.0031 |
| F09 | 27040001 | 27140000 | 0.0030 | 0.0025 | 0.0029 |
| F09 | 27060001 | 27160000 | 0.0030 | 0.0025 | 0.0029 |
| F09 | 27080001 | 27180000 | 0.0023 | 0.0019 | 0.0022 |
| F09 | 27100001 | 27200000 | 0.0021 | 0.0017 | 0.0021 |
| F09 | 27120001 | 27220000 | 0.0018 | 0.0015 | 0.0018 |
| F09 | 27140001 | 27240000 | 0.0021 | 0.0017 | 0.0021 |
| F09 | 27160001 | 27260000 | 0.0019 | 0.0015 | 0.0019 |
| F09 | 27180001 | 27280000 | 0.0019 | 0.0015 | 0.0019 |
| F09 | 27200001 | 27300000 | 0.0018 | 0.0015 | 0.0018 |
| F09 | 27220001 | 27320000 | 0.0012 | 0.0009 | 0.0012 |
| F09 | 27240001 | 27340000 | 0.0012 | 0.0009 | 0.0012 |
| F09 | 27260001 | 27360000 | 0.0015 | 0.0011 | 0.0015 |
| F09 | 27280001 | 27380000 | 0.0019 | 0.0014 | 0.0019 |
| F09 | 27300001 | 27400000 | 0.0034 | 0.0027 | 0.0034 |
| F09 | 27320001 | 27420000 | 0.0032 | 0.0026 | 0.0032 |
| F09 | 27340001 | 27440000 | 0.0032 | 0.0028 | 0.0032 |
| F09 | 27360001 | 27460000 | 0.0033 | 0.0028 | 0.0032 |
| F09 | 27380001 | 27480000 | 0.0036 | 0.0031 | 0.0035 |
| F09 | 27400001 | 27500000 | 0.0022 | 0.0018 | 0.0021 |
| F09 | 27420001 | 27520000 | 0.0025 | 0.0022 | 0.0025 |
| F09 | 27440001 | 27540000 | 0.0025 | 0.0021 | 0.0025 |
| F09 | 27460001 | 27560000 | 0.0026 | 0.0023 | 0.0026 |
| F09 | 27480001 | 27580000 | 0.0020 | 0.0018 | 0.0019 |
| F09 | 27500001 | 27600000 | 0.0024 | 0.0019 | 0.0023 |
| F09 | 27520001 | 27620000 | 0.0020 | 0.0016 | 0.0019 |
| F09 | 27540001 | 27640000 | 0.0016 | 0.0012 | 0.0015 |
| F09 | 27560001 | 27660000 | 0.0013 | 0.0010 | 0.0013 |
| F09 | 27580001 | 27680000 | 0.0016 | 0.0012 | 0.0016 |
| F09 | 27600001 | 27700000 | 0.0016 | 0.0014 | 0.0016 |
| F09 | 27620001 | 27720000 | 0.0019 | 0.0015 | 0.0019 |
| F09 | 27640001 | 27740000 | 0.0025 | 0.0020 | 0.0025 |
| F09 | 27660001 | 27760000 | 0.0031 | 0.0025 | 0.0030 |
| F09 | 27680001 | 27780000 | 0.0031 | 0.0026 | 0.0030 |
| F09 | 27700001 | 27800000 | 0.0031 | 0.0025 | 0.0030 |
| F09 | 27720001 | 27820000 | 0.0034 | 0.0028 | 0.0033 |
| F09 | 27740001 | 27840000 | 0.0033 | 0.0027 | 0.0032 |
| F09 | 27760001 | 27860000 | 0.0035 | 0.0030 | 0.0034 |
| F09 | 27780001 | 27880000 | 0.0038 | 0.0031 | 0.0035 |
| F09 | 27800001 | 27900000 | 0.0039 | 0.0032 | 0.0036 |
| F09 | 27820001 | 27920000 | 0.0036 | 0.0030 | 0.0033 |
| F09 | 27840001 | 27940000 | 0.0033 | 0.0027 | 0.0030 |
| F09 | 27860001 | 27960000 | 0.0028 | 0.0022 | 0.0025 |
| F09 | 27880001 | 27980000 | 0.0035 | 0.0030 | 0.0033 |
| F09 | 27900001 | 28000000 | 0.0031 | 0.0027 | 0.0030 |
| F09 | 27920001 | 28020000 | 0.0045 | 0.0040 | 0.0043 |
| F09 | 27940001 | 28040000 | 0.0052 | 0.0047 | 0.0050 |
| F09 | 27960001 | 28060000 | 0.0047 | 0.0042 | 0.0046 |
| F09 | 27980001 | 28080000 | 0.0037 | 0.0033 | 0.0036 |

| Chr | Start | End | both | A1 | A2 |
|---|---|---|---|---|---|
| F09 | 28000001 | 28100000 | 0.0041 | 0.0036 | 0.0039 |
| F09 | 28020001 | 28120000 | 0.0036 | 0.0031 | 0.0036 |
| F09 | 28040001 | 28140000 | 0.0029 | 0.0024 | 0.0029 |
| F09 | 28060001 | 28160000 | 0.0034 | 0.0028 | 0.0034 |
| F09 | 28080001 | 28180000 | 0.0031 | 0.0025 | 0.0032 |
| F09 | 28100001 | 28200000 | 0.0028 | 0.0023 | 0.0028 |
| F09 | 28120001 | 28220000 | 0.0015 | 0.0012 | 0.0015 |
| F09 | 28140001 | 28240000 | 0.0015 | 0.0012 | 0.0015 |
| F09 | 28160001 | 28260000 | 0.0011 | 0.0010 | 0.0011 |
| F09 | 28180001 | 28280000 | 0.0012 | 0.0010 | 0.0011 |
| F09 | 28200001 | 28300000 | 0.0013 | 0.0011 | 0.0012 |
| F09 | 28220001 | 28320000 | 0.0010 | 0.0008 | 0.0010 |
| F09 | 28240001 | 28340000 | 0.0008 | 0.0007 | 0.0007 |
| F09 | 28260001 | 28360000 | 0.0007 | 0.0006 | 0.0007 |
| F09 | 28280001 | 28380000 | 0.0006 | 0.0006 | 0.0006 |
| F09 | 28300001 | 28400000 | 0.0007 | 0.0006 | 0.0007 |
| F09 | 28320001 | 28420000 | 0.0013 | 0.0012 | 0.0013 |
| F09 | 28340001 | 28440000 | 0.0013 | 0.0010 | 0.0013 |
| F09 | 28360001 | 28460000 | 0.0012 | 0.0010 | 0.0012 |
| F09 | 28380001 | 28480000 | 0.0016 | 0.0013 | 0.0015 |
| F09 | 28400001 | 28500000 | 0.0019 | 0.0018 | 0.0019 |
| F09 | 28420001 | 28520000 | 0.0019 | 0.0017 | 0.0018 |
| F09 | 28440001 | 28540000 | 0.0022 | 0.0019 | 0.0019 |
| F09 | 28460001 | 28560000 | 0.0028 | 0.0024 | 0.0025 |
| F09 | 28480001 | 28580000 | 0.0034 | 0.0027 | 0.0031 |
| F09 | 28500001 | 28600000 | 0.0027 | 0.0020 | 0.0024 |
| F09 | 28520001 | 28620000 | 0.0023 | 0.0016 | 0.0019 |
| F09 | 28540001 | 28640000 | 0.0020 | 0.0015 | 0.0019 |
| F09 | 28560001 | 28660000 | 0.0018 | 0.0014 | 0.0017 |
| F09 | 28580001 | 28680000 | 0.0013 | 0.0011 | 0.0013 |
| F09 | 28600001 | 28700000 | 0.0013 | 0.0012 | 0.0013 |
| F09 | 28620001 | 28720000 | 0.0016 | 0.0014 | 0.0016 |
| F09 | 28640001 | 28740000 | 0.0017 | 0.0015 | 0.0016 |
| F09 | 28660001 | 28760000 | 0.0015 | 0.0013 | 0.0014 |
| F09 | 28680001 | 28780000 | 0.0020 | 0.0016 | 0.0017 |
| F09 | 28700001 | 28800000 | 0.0021 | 0.0017 | 0.0018 |
| F09 | 28720001 | 28820000 | 0.0020 | 0.0016 | 0.0017 |
| F09 | 28740001 | 28840000 | 0.0025 | 0.0017 | 0.0019 |
| F09 | 28760001 | 28860000 | 0.0025 | 0.0016 | 0.0018 |
| F09 | 28780001 | 28880000 | 0.0020 | 0.0013 | 0.0014 |
| F09 | 28800001 | 28900000 | 0.0022 | 0.0015 | 0.0016 |
| F09 | 28820001 | 28920000 | 0.0029 | 0.0021 | 0.0023 |
| F09 | 28840001 | 28940000 | 0.0025 | 0.0020 | 0.0022 |
| F09 | 28860001 | 28960000 | 0.0028 | 0.0022 | 0.0023 |
| F09 | 28880001 | 28980000 | 0.0035 | 0.0028 | 0.0030 |
| F09 | 28900001 | 29000000 | 0.0037 | 0.0029 | 0.0032 |
| F09 | 28920001 | 29020000 | 0.0029 | 0.0022 | 0.0024 |
| F09 | 28940001 | 29040000 | 0.0026 | 0.0020 | 0.0022 |
| F09 | 28960001 | 29060000 | 0.0022 | 0.0018 | 0.0020 |
| F09 | 28980001 | 29080000 | 0.0009 | 0.0008 | 0.0009 |
| F09 | 29000001 | 29100000 | 0.0004 | 0.0003 | 0.0003 |
| F09 | 29020001 | 29120000 | 0.0003 | 0.0003 | 0.0003 |
| F09 | 29040001 | 29140000 | 0.0005 | 0.0004 | 0.0005 |
| F09 | 29060001 | 29160000 | 0.0007 | 0.0006 | 0.0007 |
| F09 | 29080001 | 29180000 | 0.0007 | 0.0006 | 0.0007 |
| F09 | 29100001 | 29200000 | 0.0008 | 0.0006 | 0.0008 |
| F09 | 29120001 | 29220000 | 0.0013 | 0.0010 | 0.0013 |
| F09 | 29140001 | 29240000 | 0.0013 | 0.0011 | 0.0013 |

| Chr | Start | End | both | A1 | A2 |
|---|---|---|---|---|---|
| F09 | 29160001 | 29260000 | 0.0010 | 0.0008 | 0.0010 |
| F09 | 29180001 | 29280000 | 0.0011 | 0.0009 | 0.0011 |
| F09 | 29200001 | 29300000 | 0.0011 | 0.0009 | 0.0011 |
| F09 | 29220001 | 29320000 | 0.0005 | 0.0004 | 0.0005 |
| F09 | 29240001 | 29340000 | 0.0004 | 0.0003 | 0.0004 |
| F09 | 29260001 | 29360000 | 0.0004 | 0.0003 | 0.0004 |
| F09 | 29280001 | 29380000 | 0.0013 | 0.0012 | 0.0013 |
| F09 | 29300001 | 29400000 | 0.0015 | 0.0013 | 0.0015 |
| F09 | 29320001 | 29420000 | 0.0018 | 0.0016 | 0.0018 |
| F09 | 29340001 | 29440000 | 0.0018 | 0.0015 | 0.0017 |
| F09 | 29360001 | 29460000 | 0.0021 | 0.0017 | 0.0022 |
| F09 | 29380001 | 29480000 | 0.0011 | 0.0007 | 0.0011 |
| F09 | 29400001 | 29500000 | 0.0009 | 0.0007 | 0.0010 |
| F09 | 29420001 | 29520000 | 0.0008 | 0.0005 | 0.0008 |
| F09 | 29440001 | 29540000 | 0.0012 | 0.0008 | 0.0012 |
| F09 | 29460001 | 29560000 | 0.0015 | 0.0010 | 0.0013 |
| F09 | 29480001 | 29580000 | 0.0016 | 0.0012 | 0.0015 |
| F09 | 29500001 | 29600000 | 0.0020 | 0.0012 | 0.0015 |
| F09 | 29520001 | 29620000 | 0.0023 | 0.0011 | 0.0016 |
| F09 | 29540001 | 29640000 | 0.0021 | 0.0011 | 0.0015 |
| F09 | 29560001 | 29660000 | 0.0019 | 0.0007 | 0.0011 |
| F09 | 29580001 | 29680000 | 0.0027 | 0.0014 | 0.0018 |
| F09 | 29600001 | 29700000 | 0.0036 | 0.0026 | 0.0029 |
| F09 | 29620001 | 29720000 | 0.0040 | 0.0032 | 0.0033 |
| F09 | 29640001 | 29740000 | 0.0039 | 0.0031 | 0.0032 |
| F09 | 29660001 | 29760000 | 0.0046 | 0.0037 | 0.0038 |
| F09 | 29680001 | 29780000 | 0.0038 | 0.0029 | 0.0031 |
| F09 | 29700001 | 29800000 | 0.0025 | 0.0017 | 0.0019 |
| F09 | 29720001 | 29820000 | 0.0022 | 0.0016 | 0.0018 |
| F09 | 29740001 | 29840000 | 0.0027 | 0.0017 | 0.0021 |
| F09 | 29760001 | 29860000 | 0.0026 | 0.0016 | 0.0020 |
| F09 | 29780001 | 29880000 | 0.0029 | 0.0018 | 0.0023 |
| F09 | 29800001 | 29900000 | 0.0032 | 0.0020 | 0.0025 |
| F09 | 29820001 | 29920000 | 0.0027 | 0.0016 | 0.0021 |
| F09 | 29840001 | 29940000 | 0.0024 | 0.0018 | 0.0020 |
| F09 | 29860001 | 29960000 | 0.0020 | 0.0017 | 0.0018 |
| F09 | 29880001 | 29980000 | 0.0018 | 0.0015 | 0.0015 |
| F09 | 29900001 | 30000000 | 0.0021 | 0.0016 | 0.0016 |
| F09 | 29920001 | 30020000 | 0.0044 | 0.0038 | 0.0039 |
| F09 | 29940001 | 30040000 | 0.0043 | 0.0034 | 0.0037 |
| F09 | 29960001 | 30060000 | 0.0041 | 0.0031 | 0.0034 |
| F09 | 29980001 | 30080000 | 0.0041 | 0.0032 | 0.0036 |
| F09 | 30000001 | 30100000 | 0.0040 | 0.0032 | 0.0034 |
| F09 | 30020001 | 30120000 | 0.0018 | 0.0012 | 0.0012 |
| F09 | 30040001 | 30140000 | 0.0020 | 0.0014 | 0.0014 |
| F09 | 30060001 | 30160000 | 0.0021 | 0.0014 | 0.0014 |
| F09 | 30080001 | 30180000 | 0.0025 | 0.0016 | 0.0015 |
| F09 | 30100001 | 30200000 | 0.0028 | 0.0018 | 0.0019 |
| F09 | 30120001 | 30220000 | 0.0030 | 0.0019 | 0.0020 |
| F09 | 30140001 | 30240000 | 0.0037 | 0.0025 | 0.0027 |
| F09 | 30160001 | 30260000 | 0.0038 | 0.0027 | 0.0028 |
| F09 | 30180001 | 30280000 | 0.0044 | 0.0033 | 0.0035 |
| F09 | 30200001 | 30300000 | 0.0042 | 0.0033 | 0.0035 |
| F09 | 30220001 | 30320000 | 0.0038 | 0.0031 | 0.0033 |
| F09 | 30240001 | 30340000 | 0.0030 | 0.0025 | 0.0027 |
| F09 | 30260001 | 30360000 | 0.0034 | 0.0027 | 0.0029 |
| F09 | 30280001 | 30380000 | 0.0028 | 0.0023 | 0.0023 |
| F09 | 30300001 | 30400000 | 0.0025 | 0.0018 | 0.0018 |

| Chr | Start | End | both | A1 | A2 |
|---|---|---|---|---|---|
| F09 | 30320001 | 30420000 | 0.0030 | 0.0020 | 0.0020 |
| F09 | 30340001 | 30440000 | 0.0035 | 0.0024 | 0.0024 |
| F09 | 30360001 | 30460000 | 0.0026 | 0.0018 | 0.0019 |
| F09 | 30380001 | 30480000 | 0.0023 | 0.0015 | 0.0017 |
| F09 | 30400001 | 30500000 | 0.0030 | 0.0021 | 0.0022 |
| F09 | 30420001 | 30520000 | 0.0028 | 0.0019 | 0.0022 |
| F09 | 30440001 | 30540000 | 0.0027 | 0.0018 | 0.0022 |
| F09 | 30460001 | 30560000 | 0.0034 | 0.0024 | 0.0029 |
| F09 | 30480001 | 30580000 | 0.0031 | 0.0021 | 0.0026 |
| F09 | 30500001 | 30600000 | 0.0027 | 0.0017 | 0.0023 |
| F09 | 30520001 | 30620000 | 0.0026 | 0.0017 | 0.0022 |
| F09 | 30540001 | 30640000 | 0.0024 | 0.0016 | 0.0019 |
| F09 | 30560001 | 30660000 | 0.0020 | 0.0012 | 0.0014 |
| F09 | 30580001 | 30680000 | 0.0026 | 0.0019 | 0.0022 |
| F09 | 30600001 | 30700000 | 0.0034 | 0.0028 | 0.0030 |
| F09 | 30620001 | 30720000 | 0.0032 | 0.0028 | 0.0030 |
| F09 | 30640001 | 30740000 | 0.0028 | 0.0024 | 0.0025 |
| F09 | 30660001 | 30760000 | 0.0027 | 0.0024 | 0.0025 |
| F09 | 30680001 | 30780000 | 0.0020 | 0.0018 | 0.0018 |
| F09 | 30700001 | 30800000 | 0.0009 | 0.0007 | 0.0008 |
| F09 | 30720001 | 30820000 | 0.0009 | 0.0007 | 0.0008 |
| F09 | 30740001 | 30840000 | 0.0010 | 0.0008 | 0.0010 |
| F09 | 30760001 | 30860000 | 0.0009 | 0.0007 | 0.0009 |
| F09 | 30780001 | 30880000 | 0.0008 | 0.0006 | 0.0008 |
| F09 | 30800001 | 30900000 | 0.0016 | 0.0013 | 0.0016 |
| F09 | 30820001 | 30920000 | 0.0021 | 0.0019 | 0.0021 |
| F09 | 30840001 | 30940000 | 0.0023 | 0.0020 | 0.0019 |
| F09 | 30860001 | 30960000 | 0.0034 | 0.0031 | 0.0030 |
| F09 | 30880001 | 30980000 | 0.0035 | 0.0031 | 0.0030 |
| F09 | 30900001 | 31000000 | 0.0028 | 0.0025 | 0.0024 |
| F09 | 30920001 | 31020000 | 0.0024 | 0.0022 | 0.0020 |
| F09 | 30940001 | 31040000 | 0.0020 | 0.0019 | 0.0019 |
| F09 | 30960001 | 31060000 | 0.0007 | 0.0007 | 0.0007 |
| F09 | 30980001 | 31080000 | 0.0011 | 0.0010 | 0.0011 |
| F09 | 31000001 | 31100000 | 0.0016 | 0.0016 | 0.0016 |
| F09 | 31020001 | 31120000 | 0.0015 | 0.0015 | 0.0014 |
| F09 | 31040001 | 31140000 | 0.0019 | 0.0018 | 0.0018 |
| F09 | 31060001 | 31160000 | 0.0022 | 0.0020 | 0.0021 |
| F09 | 31080001 | 31180000 | 0.0018 | 0.0017 | 0.0018 |
| F09 | 31100001 | 31200000 | 0.0019 | 0.0018 | 0.0019 |
| F09 | 31120001 | 31220000 | 0.0028 | 0.0026 | 0.0027 |
| F09 | 31140001 | 31240000 | 0.0025 | 0.0024 | 0.0024 |
| F09 | 31160001 | 31260000 | 0.0028 | 0.0026 | 0.0028 |
| F09 | 31180001 | 31280000 | 0.0030 | 0.0028 | 0.0030 |
| F09 | 31200001 | 31300000 | 0.0028 | 0.0023 | 0.0027 |
| F09 | 31220001 | 31320000 | 0.0020 | 0.0016 | 0.0019 |
| F09 | 31240001 | 31340000 | 0.0022 | 0.0016 | 0.0020 |
| F09 | 31260001 | 31360000 | 0.0025 | 0.0018 | 0.0023 |
| F09 | 31280001 | 31380000 | 0.0028 | 0.0017 | 0.0022 |
| F09 | 31300001 | 31400000 | 0.0035 | 0.0022 | 0.0029 |
| F09 | 31320001 | 31420000 | 0.0041 | 0.0027 | 0.0034 |
| F09 | 31340001 | 31440000 | 0.0043 | 0.0029 | 0.0036 |
| F09 | 31360001 | 31460000 | 0.0034 | 0.0023 | 0.0027 |
| F09 | 31380001 | 31480000 | 0.0031 | 0.0023 | 0.0027 |
| F09 | 31400001 | 31500000 | 0.0026 | 0.0018 | 0.0021 |
| F09 | 31420001 | 31520000 | 0.0021 | 0.0014 | 0.0018 |
| F09 | 31440001 | 31540000 | 0.0017 | 0.0012 | 0.0015 |
| F09 | 31460001 | 31560000 | 0.0027 | 0.0018 | 0.0023 |

| Chr | Start | End | both | A1 | A2 |
|---|---|---|---|---|---|
| F09 | 31480001 | 31580000 | 0.0027 | 0.0019 | 0.0022 |
| F09 | 31500001 | 31600000 | 0.0027 | 0.0021 | 0.0023 |
| F09 | 31520001 | 31620000 | 0.0034 | 0.0028 | 0.0030 |
| F09 | 31540001 | 31640000 | 0.0033 | 0.0027 | 0.0029 |
| F09 | 31560001 | 31660000 | 0.0025 | 0.0023 | 0.0023 |
| F09 | 31580001 | 31680000 | 0.0027 | 0.0025 | 0.0026 |
| F09 | 31600001 | 31700000 | 0.0019 | 0.0018 | 0.0018 |
| F09 | 31620001 | 31720000 | 0.0020 | 0.0017 | 0.0020 |
| F09 | 31640001 | 31740000 | 0.0023 | 0.0019 | 0.0023 |
| F09 | 31660001 | 31760000 | 0.0030 | 0.0025 | 0.0030 |
| F09 | 31680001 | 31780000 | 0.0028 | 0.0024 | 0.0028 |
| F09 | 31700001 | 31800000 | 0.0032 | 0.0027 | 0.0032 |
| F09 | 31720001 | 31820000 | 0.0029 | 0.0025 | 0.0029 |
| F09 | 31740001 | 31840000 | 0.0029 | 0.0024 | 0.0029 |
| F09 | 31760001 | 31860000 | 0.0027 | 0.0021 | 0.0025 |
| F09 | 31780001 | 31880000 | 0.0029 | 0.0023 | 0.0027 |
| F09 | 31800001 | 31900000 | 0.0025 | 0.0020 | 0.0023 |
| F09 | 31820001 | 31920000 | 0.0021 | 0.0016 | 0.0018 |
| F09 | 31840001 | 31940000 | 0.0019 | 0.0015 | 0.0017 |
| F09 | 31860001 | 31960000 | 0.0013 | 0.0010 | 0.0013 |
| F09 | 31880001 | 31980000 | 0.0021 | 0.0019 | 0.0021 |
| F09 | 31900001 | 32000000 | 0.0038 | 0.0034 | 0.0038 |
| F09 | 31920001 | 32020000 | 0.0038 | 0.0033 | 0.0038 |
| F09 | 31940001 | 32040000 | 0.0041 | 0.0036 | 0.0040 |
| F09 | 31960001 | 32060000 | 0.0042 | 0.0037 | 0.0042 |
| F09 | 31980001 | 32080000 | 0.0033 | 0.0028 | 0.0033 |
| F09 | 32000001 | 32100000 | 0.0018 | 0.0015 | 0.0017 |
| F09 | 32020001 | 32120000 | 0.0023 | 0.0019 | 0.0021 |
| F09 | 32040001 | 32140000 | 0.0020 | 0.0017 | 0.0018 |
| F09 | 32060001 | 32160000 | 0.0023 | 0.0020 | 0.0021 |
| F09 | 32080001 | 32180000 | 0.0021 | 0.0018 | 0.0019 |
| F09 | 32100001 | 32200000 | 0.0019 | 0.0017 | 0.0018 |
| F09 | 32120001 | 32220000 | 0.0015 | 0.0014 | 0.0015 |
| F09 | 32140001 | 32240000 | 0.0018 | 0.0014 | 0.0016 |
| F09 | 32160001 | 32260000 | 0.0022 | 0.0015 | 0.0017 |
| F09 | 32180001 | 32280000 | 0.0034 | 0.0025 | 0.0027 |
| F09 | 32200001 | 32300000 | 0.0039 | 0.0029 | 0.0031 |
| F09 | 32220001 | 32320000 | 0.0043 | 0.0033 | 0.0035 |
| F09 | 32240001 | 32340000 | 0.0039 | 0.0032 | 0.0034 |
| F09 | 32260001 | 32360000 | 0.0035 | 0.0029 | 0.0031 |
| F09 | 32280001 | 32380000 | 0.0032 | 0.0025 | 0.0028 |
| F09 | 32300001 | 32400000 | 0.0033 | 0.0027 | 0.0030 |
| F09 | 32320001 | 32420000 | 0.0037 | 0.0028 | 0.0032 |
| F09 | 32340001 | 32440000 | 0.0041 | 0.0029 | 0.0034 |
| F09 | 32360001 | 32460000 | 0.0044 | 0.0032 | 0.0037 |
| F09 | 32380001 | 32480000 | 0.0039 | 0.0028 | 0.0031 |
| F09 | 32400001 | 32500000 | 0.0031 | 0.0021 | 0.0024 |
| F09 | 32420001 | 32520000 | 0.0022 | 0.0015 | 0.0016 |
| F09 | 32440001 | 32540000 | 0.0021 | 0.0017 | 0.0018 |
| F09 | 32460001 | 32560000 | 0.0013 | 0.0011 | 0.0012 |
| F09 | 32480001 | 32580000 | 0.0013 | 0.0009 | 0.0011 |
| F09 | 32500001 | 32600000 | 0.0014 | 0.0010 | 0.0011 |
| F09 | 32520001 | 32620000 | 0.0017 | 0.0012 | 0.0013 |
| F09 | 32540001 | 32640000 | 0.0020 | 0.0015 | 0.0016 |
| F09 | 32560001 | 32660000 | 0.0026 | 0.0019 | 0.0020 |
| F09 | 32580001 | 32680000 | 0.0032 | 0.0026 | 0.0028 |
| F09 | 32600001 | 32700000 | 0.0037 | 0.0029 | 0.0031 |
| F09 | 32620001 | 32720000 | 0.0041 | 0.0031 | 0.0033 |

| Chr | Start | End | both | A1 | A2 |
|---|---|---|---|---|---|
| F09 | 32640001 | 32740000 | 0.0038 | 0.0027 | 0.0029 |
| F09 | 32660001 | 32760000 | 0.0038 | 0.0026 | 0.0029 |
| F09 | 32680001 | 32780000 | 0.0039 | 0.0026 | 0.0028 |
| F09 | 32700001 | 32800000 | 0.0037 | 0.0025 | 0.0029 |
| F09 | 32720001 | 32820000 | 0.0032 | 0.0022 | 0.0026 |
| F09 | 32740001 | 32840000 | 0.0027 | 0.0020 | 0.0023 |
| F09 | 32760001 | 32860000 | 0.0025 | 0.0020 | 0.0023 |
| F09 | 32780001 | 32880000 | 0.0015 | 0.0013 | 0.0015 |
| F09 | 32800001 | 32900000 | 0.0011 | 0.0010 | 0.0011 |
| F09 | 32820001 | 32920000 | 0.0016 | 0.0014 | 0.0016 |
| F09 | 32840001 | 32940000 | 0.0017 | 0.0014 | 0.0017 |
| F09 | 32860001 | 32960000 | 0.0020 | 0.0016 | 0.0019 |
| F09 | 32880001 | 32980000 | 0.0025 | 0.0019 | 0.0022 |
| F09 | 32900001 | 33000000 | 0.0032 | 0.0024 | 0.0028 |
| F09 | 32920001 | 33020000 | 0.0038 | 0.0031 | 0.0033 |
| F09 | 32940001 | 33040000 | 0.0038 | 0.0031 | 0.0033 |
| F09 | 32960001 | 33060000 | 0.0038 | 0.0032 | 0.0034 |
| F09 | 32980001 | 33080000 | 0.0036 | 0.0030 | 0.0032 |
| F09 | 33000001 | 33100000 | 0.0039 | 0.0032 | 0.0034 |
| F09 | 33020001 | 33120000 | 0.0030 | 0.0024 | 0.0026 |
| F09 | 33040001 | 33140000 | 0.0036 | 0.0026 | 0.0031 |
| F09 | 33060001 | 33160000 | 0.0036 | 0.0027 | 0.0032 |
| F09 | 33080001 | 33180000 | 0.0035 | 0.0026 | 0.0032 |
| F09 | 33100001 | 33200000 | 0.0029 | 0.0022 | 0.0027 |
| F09 | 33120001 | 33220000 | 0.0030 | 0.0024 | 0.0028 |
| F09 | 33140001 | 33240000 | 0.0022 | 0.0020 | 0.0022 |
| F09 | 33160001 | 33260000 | 0.0020 | 0.0017 | 0.0019 |
| F09 | 33180001 | 33280000 | 0.0024 | 0.0019 | 0.0021 |
| F09 | 33200001 | 33300000 | 0.0025 | 0.0019 | 0.0022 |
| F09 | 33220001 | 33320000 | 0.0028 | 0.0020 | 0.0024 |
| F09 | 33240001 | 33340000 | 0.0035 | 0.0024 | 0.0029 |
| F09 | 33260001 | 33360000 | 0.0035 | 0.0023 | 0.0028 |
| F09 | 33280001 | 33380000 | 0.0031 | 0.0019 | 0.0022 |
| F09 | 33300001 | 33400000 | 0.0037 | 0.0025 | 0.0026 |
| F09 | 33320001 | 33420000 | 0.0041 | 0.0028 | 0.0030 |
| F09 | 33340001 | 33440000 | 0.0039 | 0.0028 | 0.0029 |
| F09 | 33360001 | 33460000 | 0.0043 | 0.0030 | 0.0031 |
| F09 | 33380001 | 33480000 | 0.0048 | 0.0037 | 0.0039 |
| F09 | 33400001 | 33500000 | 0.0045 | 0.0033 | 0.0036 |
| F09 | 33420001 | 33520000 | 0.0041 | 0.0030 | 0.0033 |
| F09 | 33440001 | 33540000 | 0.0041 | 0.0030 | 0.0032 |
| F09 | 33460001 | 33560000 | 0.0045 | 0.0034 | 0.0038 |
| F09 | 33480001 | 33580000 | 0.0040 | 0.0031 | 0.0033 |
| F09 | 33500001 | 33600000 | 0.0040 | 0.0030 | 0.0032 |
| F09 | 33520001 | 33620000 | 0.0041 | 0.0031 | 0.0033 |
| F09 | 33540001 | 33640000 | 0.0041 | 0.0030 | 0.0033 |
| F09 | 33560001 | 33660000 | 0.0030 | 0.0023 | 0.0025 |
| F09 | 33580001 | 33680000 | 0.0027 | 0.0020 | 0.0022 |
| F09 | 33600001 | 33700000 | 0.0022 | 0.0019 | 0.0020 |
| F09 | 33620001 | 33720000 | 0.0014 | 0.0011 | 0.0013 |
| F09 | 33640001 | 33740000 | 0.0010 | 0.0008 | 0.0009 |
| F09 | 33660001 | 33760000 | 0.0008 | 0.0007 | 0.0008 |
| F09 | 33680001 | 33780000 | 0.0016 | 0.0014 | 0.0016 |
| F09 | 33700001 | 33800000 | 0.0013 | 0.0011 | 0.0012 |
| F09 | 33720001 | 33820000 | 0.0016 | 0.0014 | 0.0015 |
| F09 | 33740001 | 33840000 | 0.0020 | 0.0015 | 0.0017 |
| F09 | 33760001 | 33860000 | 0.0027 | 0.0021 | 0.0023 |
| F09 | 33780001 | 33880000 | 0.0020 | 0.0014 | 0.0016 |

| Chr | Start | End | both | A1 | A2 |
|---|---|---|---|---|---|
| F09 | 33800001 | 33900000 | 0.0021 | 0.0015 | 0.0017 |
| F09 | 33820001 | 33920000 | 0.0020 | 0.0015 | 0.0016 |
| F09 | 33840001 | 33940000 | 0.0016 | 0.0012 | 0.0013 |
| F09 | 33860001 | 33960000 | 0.0009 | 0.0007 | 0.0009 |
| F09 | 33880001 | 33980000 | 0.0008 | 0.0006 | 0.0007 |
| F09 | 33900001 | 34000000 | 0.0008 | 0.0005 | 0.0007 |
| F09 | 33920001 | 34020000 | 0.0005 | 0.0003 | 0.0005 |
| F09 | 33940001 | 34040000 | 0.0005 | 0.0004 | 0.0006 |
| F09 | 33960001 | 34060000 | 0.0007 | 0.0005 | 0.0007 |
| F09 | 33980001 | 34080000 | 0.0014 | 0.0011 | 0.0014 |
| F09 | 34000001 | 34100000 | 0.0016 | 0.0014 | 0.0015 |
| F09 | 34020001 | 34120000 | 0.0015 | 0.0014 | 0.0015 |
| F09 | 34040001 | 34140000 | 0.0018 | 0.0017 | 0.0018 |
| F09 | 34060001 | 34160000 | 0.0019 | 0.0017 | 0.0018 |
| F09 | 34080001 | 34180000 | 0.0015 | 0.0013 | 0.0015 |
| F09 | 34100001 | 34200000 | 0.0014 | 0.0011 | 0.0013 |
| F09 | 34120001 | 34220000 | 0.0029 | 0.0020 | 0.0024 |
| F09 | 34140001 | 34240000 | 0.0030 | 0.0019 | 0.0022 |
| F09 | 34160001 | 34260000 | 0.0038 | 0.0024 | 0.0028 |
| F09 | 34180001 | 34280000 | 0.0039 | 0.0026 | 0.0029 |
| F09 | 34200001 | 34300000 | 0.0050 | 0.0035 | 0.0040 |
| F09 | 34220001 | 34320000 | 0.0047 | 0.0038 | 0.0043 |
| F09 | 34240001 | 34340000 | 0.0050 | 0.0041 | 0.0046 |
| F09 | 34260001 | 34360000 | 0.0041 | 0.0035 | 0.0040 |
| F09 | 34280001 | 34380000 | 0.0042 | 0.0036 | 0.0041 |
| F09 | 34300001 | 34400000 | 0.0031 | 0.0027 | 0.0031 |
| F09 | 34320001 | 34420000 | 0.0020 | 0.0017 | 0.0019 |
| F09 | 34340001 | 34440000 | 0.0015 | 0.0011 | 0.0013 |
| F09 | 34360001 | 34460000 | 0.0022 | 0.0016 | 0.0018 |
| F09 | 34380001 | 34480000 | 0.0017 | 0.0011 | 0.0012 |
| F09 | 34400001 | 34500000 | 0.0014 | 0.0009 | 0.0010 |
| F09 | 34420001 | 34520000 | 0.0014 | 0.0009 | 0.0010 |
| F09 | 34440001 | 34540000 | 0.0020 | 0.0015 | 0.0017 |
| F09 | 34460001 | 34560000 | 0.0014 | 0.0012 | 0.0014 |
| F09 | 34480001 | 34580000 | 0.0015 | 0.0013 | 0.0015 |
| F09 | 34500001 | 34600000 | 0.0014 | 0.0012 | 0.0014 |
| F09 | 34520001 | 34620000 | 0.0015 | 0.0012 | 0.0014 |
| F09 | 34540001 | 34640000 | 0.0012 | 0.0011 | 0.0012 |
| F09 | 34560001 | 34660000 | 0.0010 | 0.0009 | 0.0010 |
| F09 | 34580001 | 34680000 | 0.0015 | 0.0013 | 0.0015 |
| F09 | 34600001 | 34700000 | 0.0015 | 0.0013 | 0.0015 |
| F09 | 34620001 | 34720000 | 0.0015 | 0.0012 | 0.0015 |
| F09 | 34640001 | 34740000 | 0.0014 | 0.0011 | 0.0012 |
| F09 | 34660001 | 34760000 | 0.0016 | 0.0012 | 0.0013 |
| F09 | 34680001 | 34780000 | 0.0022 | 0.0015 | 0.0016 |
| F09 | 34700001 | 34800000 | 0.0034 | 0.0023 | 0.0024 |
| F09 | 34720001 | 34820000 | 0.0039 | 0.0025 | 0.0026 |
| F09 | 34740001 | 34840000 | 0.0042 | 0.0026 | 0.0027 |
| F09 | 34760001 | 34860000 | 0.0042 | 0.0027 | 0.0028 |
| F09 | 34780001 | 34880000 | 0.0039 | 0.0022 | 0.0024 |
| F09 | 34800001 | 34900000 | 0.0044 | 0.0027 | 0.0030 |
| F09 | 34820001 | 34920000 | 0.0044 | 0.0030 | 0.0032 |
| F09 | 34840001 | 34940000 | 0.0051 | 0.0040 | 0.0041 |
| F09 | 34860001 | 34960000 | 0.0053 | 0.0041 | 0.0042 |
| F09 | 34880001 | 34980000 | 0.0053 | 0.0045 | 0.0046 |
| F09 | 34900001 | 35000000 | 0.0049 | 0.0041 | 0.0043 |
| F09 | 34920001 | 35020000 | 0.0047 | 0.0037 | 0.0039 |
| F09 | 34940001 | 35040000 | 0.0038 | 0.0026 | 0.0030 |

| Chr | Start | End | both | A1 | A2 |
|---|---|---|---|---|---|
| F09 | 34960001 | 35060000 | 0.0037 | 0.0025 | 0.0029 |
| F09 | 34980001 | 35080000 | 0.0031 | 0.0021 | 0.0024 |
| F09 | 35000001 | 35100000 | 0.0023 | 0.0015 | 0.0017 |
| F09 | 35020001 | 35120000 | 0.0025 | 0.0016 | 0.0020 |
| F09 | 35040001 | 35140000 | 0.0033 | 0.0025 | 0.0030 |
| F09 | 35060001 | 35160000 | 0.0036 | 0.0029 | 0.0033 |
| F09 | 35080001 | 35180000 | 0.0037 | 0.0029 | 0.0034 |
| F09 | 35100001 | 35200000 | 0.0037 | 0.0030 | 0.0034 |
| F09 | 35120001 | 35220000 | 0.0040 | 0.0035 | 0.0036 |
| F09 | 35140001 | 35240000 | 0.0034 | 0.0029 | 0.0028 |
| F09 | 35160001 | 35260000 | 0.0035 | 0.0030 | 0.0029 |
| F09 | 35180001 | 35280000 | 0.0032 | 0.0025 | 0.0024 |
| F09 | 35200001 | 35300000 | 0.0037 | 0.0026 | 0.0026 |
| F09 | 35220001 | 35320000 | 0.0033 | 0.0021 | 0.0025 |
| F09 | 35240001 | 35340000 | 0.0037 | 0.0023 | 0.0031 |
| F09 | 35260001 | 35360000 | 0.0038 | 0.0024 | 0.0031 |
| F09 | 35280001 | 35380000 | 0.0044 | 0.0030 | 0.0037 |
| F09 | 35300001 | 35400000 | 0.0040 | 0.0028 | 0.0035 |
| F09 | 35320001 | 35420000 | 0.0036 | 0.0027 | 0.0031 |
| F09 | 35340001 | 35440000 | 0.0033 | 0.0025 | 0.0026 |
| F09 | 35360001 | 35460000 | 0.0038 | 0.0027 | 0.0031 |
| F09 | 35380001 | 35480000 | 0.0039 | 0.0027 | 0.0032 |
| F09 | 35400001 | 35500000 | 0.0039 | 0.0028 | 0.0033 |
| F09 | 35420001 | 35520000 | 0.0052 | 0.0037 | 0.0044 |
| F09 | 35440001 | 35540000 | 0.0051 | 0.0035 | 0.0043 |
| F09 | 35460001 | 35560000 | 0.0045 | 0.0029 | 0.0036 |
| F09 | 35480001 | 35580000 | 0.0041 | 0.0026 | 0.0032 |
| F09 | 35500001 | 35600000 | 0.0043 | 0.0025 | 0.0031 |
| F09 | 35520001 | 35620000 | 0.0030 | 0.0015 | 0.0018 |
| F09 | 35540001 | 35640000 | 0.0030 | 0.0015 | 0.0018 |
| F09 | 35560001 | 35660000 | 0.0029 | 0.0015 | 0.0018 |
| F09 | 35580001 | 35680000 | 0.0027 | 0.0015 | 0.0018 |
| F09 | 35600001 | 35700000 | 0.0065 | 0.0054 | 0.0056 |
| F09 | 35620001 | 35720000 | 0.0074 | 0.0063 | 0.0066 |
| F09 | 35640001 | 35740000 | 0.0071 | 0.0061 | 0.0064 |
| F09 | 35660001 | 35760000 | 0.0072 | 0.0064 | 0.0067 |
| F09 | 35680001 | 35780000 | 0.0074 | 0.0065 | 0.0069 |
| F09 | 35700001 | 35800000 | 0.0037 | 0.0029 | 0.0032 |
| F09 | 35720001 | 35820000 | 0.0035 | 0.0030 | 0.0031 |
| F09 | 35740001 | 35840000 | 0.0039 | 0.0031 | 0.0033 |
| F09 | 35760001 | 35860000 | 0.0038 | 0.0029 | 0.0029 |
| F09 | 35780001 | 35880000 | 0.0035 | 0.0027 | 0.0027 |
| F09 | 35800001 | 35900000 | 0.0027 | 0.0020 | 0.0019 |
| F09 | 35820001 | 35920000 | 0.0018 | 0.0011 | 0.0012 |
| F09 | 35840001 | 35940000 | 0.0012 | 0.0008 | 0.0008 |
| F09 | 35860001 | 35960000 | 0.0011 | 0.0007 | 0.0008 |
| F09 | 35880001 | 35980000 | 0.0020 | 0.0017 | 0.0017 |
| F09 | 35900001 | 36000000 | 0.0036 | 0.0031 | 0.0031 |
| F09 | 35920001 | 36020000 | 0.0042 | 0.0036 | 0.0037 |
| F09 | 35940001 | 36040000 | 0.0042 | 0.0036 | 0.0036 |
| F09 | 35960001 | 36060000 | 0.0039 | 0.0033 | 0.0033 |
| F09 | 35980001 | 36080000 | 0.0038 | 0.0029 | 0.0030 |
| F09 | 36000001 | 36100000 | 0.0025 | 0.0017 | 0.0019 |
| F09 | 36020001 | 36120000 | 0.0020 | 0.0012 | 0.0014 |
| F09 | 36040001 | 36140000 | 0.0031 | 0.0019 | 0.0025 |
| F09 | 36060001 | 36160000 | 0.0037 | 0.0024 | 0.0031 |
| F09 | 36080001 | 36180000 | 0.0030 | 0.0021 | 0.0028 |
| F09 | 36100001 | 36200000 | 0.0029 | 0.0021 | 0.0027 |

| Chr | Start | End | both | A1 | A2 |
|---|---|---|---|---|---|
| F09 | 36120001 | 36220000 | 0.0035 | 0.0025 | 0.0032 |
| F09 | 36140001 | 36240000 | 0.0029 | 0.0021 | 0.0023 |
| F09 | 36160001 | 36260000 | 0.0025 | 0.0016 | 0.0017 |
| F09 | 36180001 | 36280000 | 0.0024 | 0.0015 | 0.0016 |
| F09 | 36200001 | 36300000 | 0.0022 | 0.0013 | 0.0014 |
| F09 | 36220001 | 36320000 | 0.0018 | 0.0012 | 0.0013 |
| F09 | 36240001 | 36340000 | 0.0016 | 0.0013 | 0.0013 |
| F09 | 36260001 | 36360000 | 0.0018 | 0.0016 | 0.0017 |
| F09 | 36280001 | 36380000 | 0.0020 | 0.0019 | 0.0020 |
| F09 | 36300001 | 36400000 | 0.0029 | 0.0026 | 0.0028 |
| F09 | 36320001 | 36420000 | 0.0032 | 0.0028 | 0.0032 |
| F09 | 36340001 | 36440000 | 0.0035 | 0.0030 | 0.0034 |
| F09 | 36360001 | 36460000 | 0.0033 | 0.0028 | 0.0032 |
| F09 | 36380001 | 36480000 | 0.0032 | 0.0027 | 0.0031 |
| F09 | 36400001 | 36500000 | 0.0030 | 0.0025 | 0.0028 |
| F09 | 36420001 | 36520000 | 0.0024 | 0.0021 | 0.0022 |
| F09 | 36440001 | 36540000 | 0.0029 | 0.0024 | 0.0026 |
| F09 | 36460001 | 36560000 | 0.0030 | 0.0026 | 0.0027 |
| F09 | 36480001 | 36580000 | 0.0026 | 0.0022 | 0.0023 |
| F09 | 36500001 | 36600000 | 0.0027 | 0.0023 | 0.0025 |
| F09 | 36520001 | 36620000 | 0.0026 | 0.0021 | 0.0024 |
| F09 | 36540001 | 36640000 | 0.0022 | 0.0017 | 0.0020 |
| F09 | 36560001 | 36660000 | 0.0027 | 0.0019 | 0.0022 |
| F09 | 36580001 | 36680000 | 0.0033 | 0.0022 | 0.0026 |
| F09 | 36600001 | 36700000 | 0.0028 | 0.0017 | 0.0019 |
| F09 | 36620001 | 36720000 | 0.0039 | 0.0027 | 0.0030 |
| F09 | 36640001 | 36740000 | 0.0046 | 0.0034 | 0.0037 |
| F09 | 36660001 | 36760000 | 0.0041 | 0.0032 | 0.0033 |
| F09 | 36680001 | 36780000 | 0.0039 | 0.0029 | 0.0029 |
| F09 | 36700001 | 36800000 | 0.0038 | 0.0029 | 0.0029 |
| F09 | 36720001 | 36820000 | 0.0031 | 0.0022 | 0.0021 |
| F09 | 36740001 | 36840000 | 0.0024 | 0.0013 | 0.0012 |
| F09 | 36760001 | 36860000 | 0.0026 | 0.0015 | 0.0015 |
| F09 | 36780001 | 36880000 | 0.0023 | 0.0014 | 0.0014 |
| F09 | 36800001 | 36900000 | 0.0030 | 0.0020 | 0.0020 |
| F09 | 36820001 | 36920000 | 0.0029 | 0.0020 | 0.0019 |
| F09 | 36840001 | 36940000 | 0.0026 | 0.0017 | 0.0017 |
| F09 | 36860001 | 36960000 | 0.0025 | 0.0016 | 0.0017 |
| F09 | 36880001 | 36980000 | 0.0032 | 0.0023 | 0.0023 |
| F09 | 36900001 | 37000000 | 0.0032 | 0.0023 | 0.0024 |
| F09 | 36920001 | 37020000 | 0.0034 | 0.0028 | 0.0028 |
| F09 | 36940001 | 37040000 | 0.0037 | 0.0030 | 0.0031 |
| F09 | 36960001 | 37060000 | 0.0035 | 0.0028 | 0.0030 |
| F09 | 36980001 | 37080000 | 0.0033 | 0.0027 | 0.0029 |
| F09 | 37000001 | 37100000 | 0.0031 | 0.0025 | 0.0027 |
| F09 | 37020001 | 37120000 | 0.0031 | 0.0021 | 0.0024 |
| F09 | 37040001 | 37140000 | 0.0035 | 0.0024 | 0.0027 |
| F09 | 37060001 | 37160000 | 0.0035 | 0.0024 | 0.0026 |
| F09 | 37080001 | 37180000 | 0.0030 | 0.0019 | 0.0021 |
| F09 | 37100001 | 37200000 | 0.0034 | 0.0024 | 0.0027 |
| F09 | 37120001 | 37220000 | 0.0027 | 0.0020 | 0.0023 |
| F09 | 37140001 | 37240000 | 0.0025 | 0.0018 | 0.0022 |
| F09 | 37160001 | 37260000 | 0.0032 | 0.0018 | 0.0026 |
| F09 | 37180001 | 37280000 | 0.0043 | 0.0023 | 0.0036 |
| F09 | 37200001 | 37300000 | 0.0044 | 0.0022 | 0.0035 |
| F09 | 37220001 | 37320000 | 0.0051 | 0.0027 | 0.0041 |
| F09 | 37240001 | 37340000 | 0.0044 | 0.0024 | 0.0036 |
| F09 | 37260001 | 37360000 | 0.0038 | 0.0024 | 0.0032 |

| Chr | Start | End | both | A1 | A2 |
|---|---|---|---|---|---|
| F09 | 37280001 | 37380000 | 0.0028 | 0.0021 | 0.0026 |
| F09 | 37300001 | 37400000 | 0.0029 | 0.0023 | 0.0028 |
| F09 | 37320001 | 37420000 | 0.0030 | 0.0021 | 0.0025 |
| F09 | 37340001 | 37440000 | 0.0033 | 0.0023 | 0.0026 |
| F09 | 37360001 | 37460000 | 0.0036 | 0.0023 | 0.0027 |
| F09 | 37380001 | 37480000 | 0.0032 | 0.0020 | 0.0024 |
| F09 | 37400001 | 37500000 | 0.0020 | 0.0011 | 0.0012 |
| F09 | 37420001 | 37520000 | 0.0017 | 0.0010 | 0.0012 |
| F09 | 37440001 | 37540000 | 0.0013 | 0.0010 | 0.0011 |
| F09 | 37460001 | 37560000 | 0.0024 | 0.0022 | 0.0023 |
| F09 | 37480001 | 37580000 | 0.0025 | 0.0022 | 0.0023 |
| F09 | 37500001 | 37600000 | 0.0033 | 0.0028 | 0.0030 |
| F09 | 37520001 | 37620000 | 0.0032 | 0.0025 | 0.0027 |
| F09 | 37540001 | 37640000 | 0.0034 | 0.0026 | 0.0028 |
| F09 | 37560001 | 37660000 | 0.0025 | 0.0016 | 0.0019 |
| F09 | 37580001 | 37680000 | 0.0028 | 0.0018 | 0.0020 |
| F09 | 37600001 | 37700000 | 0.0026 | 0.0016 | 0.0017 |
| F09 | 37620001 | 37720000 | 0.0025 | 0.0017 | 0.0018 |
| F09 | 37640001 | 37740000 | 0.0022 | 0.0015 | 0.0016 |
| F09 | 37660001 | 37760000 | 0.0017 | 0.0011 | 0.0011 |
| F09 | 37680001 | 37780000 | 0.0013 | 0.0008 | 0.0008 |
| F09 | 37700001 | 37800000 | 0.0014 | 0.0010 | 0.0011 |
| F09 | 37720001 | 37820000 | 0.0016 | 0.0012 | 0.0012 |
| F09 | 37740001 | 37840000 | 0.0015 | 0.0011 | 0.0011 |
| F09 | 37760001 | 37860000 | 0.0015 | 0.0012 | 0.0012 |
| F09 | 37780001 | 37880000 | 0.0016 | 0.0013 | 0.0014 |
| F09 | 37800001 | 37900000 | 0.0015 | 0.0012 | 0.0013 |
| F09 | 37820001 | 37920000 | 0.0013 | 0.0011 | 0.0012 |
| F09 | 37840001 | 37940000 | 0.0017 | 0.0013 | 0.0015 |
| F09 | 37860001 | 37960000 | 0.0024 | 0.0018 | 0.0020 |
| F09 | 37880001 | 37980000 | 0.0032 | 0.0025 | 0.0028 |
| F09 | 37900001 | 38000000 | 0.0036 | 0.0026 | 0.0030 |
| F09 | 37920001 | 38020000 | 0.0035 | 0.0026 | 0.0030 |
| F09 | 37940001 | 38040000 | 0.0036 | 0.0027 | 0.0030 |
| F09 | 37960001 | 38060000 | 0.0041 | 0.0032 | 0.0036 |
| F09 | 37980001 | 38080000 | 0.0044 | 0.0036 | 0.0037 |
| F09 | 38000001 | 38100000 | 0.0046 | 0.0037 | 0.0038 |
| F09 | 38020001 | 38120000 | 0.0050 | 0.0040 | 0.0042 |
| F09 | 38040001 | 38140000 | 0.0052 | 0.0041 | 0.0043 |
| F09 | 38060001 | 38160000 | 0.0056 | 0.0046 | 0.0047 |
| F09 | 38080001 | 38180000 | 0.0046 | 0.0035 | 0.0037 |
| F09 | 38100001 | 38200000 | 0.0046 | 0.0037 | 0.0036 |
| F09 | 38120001 | 38220000 | 0.0047 | 0.0038 | 0.0037 |
| F09 | 38140001 | 38240000 | 0.0050 | 0.0044 | 0.0041 |
| F09 | 38160001 | 38260000 | 0.0040 | 0.0034 | 0.0032 |
| F09 | 38180001 | 38280000 | 0.0042 | 0.0036 | 0.0036 |
| F09 | 38200001 | 38300000 | 0.0036 | 0.0032 | 0.0032 |
| F09 | 38220001 | 38320000 | 0.0037 | 0.0033 | 0.0031 |
| F09 | 38240001 | 38340000 | 0.0031 | 0.0024 | 0.0026 |
| F09 | 38260001 | 38360000 | 0.0027 | 0.0023 | 0.0024 |
| F09 | 38280001 | 38380000 | 0.0028 | 0.0021 | 0.0022 |
| F09 | 38300001 | 38400000 | 0.0030 | 0.0019 | 0.0021 |
| F09 | 38320001 | 38420000 | 0.0025 | 0.0013 | 0.0016 |
| F09 | 38340001 | 38440000 | 0.0029 | 0.0015 | 0.0016 |
| F09 | 38360001 | 38460000 | 0.0040 | 0.0025 | 0.0026 |
| F09 | 38380001 | 38480000 | 0.0047 | 0.0033 | 0.0035 |
| F09 | 38400001 | 38500000 | 0.0048 | 0.0037 | 0.0038 |
| F09 | 38420001 | 38520000 | 0.0058 | 0.0047 | 0.0049 |

| Chr | Start | End | both | A1 | A2 |
|---|---|---|---|---|---|
| F09 | 38440001 | 38540000 | 0.0063 | 0.0051 | 0.0055 |
| F09 | 38460001 | 38560000 | 0.0059 | 0.0049 | 0.0051 |
| F09 | 38480001 | 38580000 | 0.0065 | 0.0050 | 0.0055 |
| F09 | 38500001 | 38600000 | 0.0066 | 0.0049 | 0.0052 |
| F09 | 38520001 | 38620000 | 0.0067 | 0.0048 | 0.0052 |
| F09 | 38540001 | 38640000 | 0.0061 | 0.0043 | 0.0047 |
| F09 | 38560001 | 38660000 | 0.0051 | 0.0031 | 0.0036 |
| F09 | 38580001 | 38680000 | 0.0040 | 0.0024 | 0.0026 |
| F09 | 38600001 | 38700000 | 0.0039 | 0.0025 | 0.0027 |
| F09 | 38620001 | 38720000 | 0.0039 | 0.0024 | 0.0028 |
| F09 | 38640001 | 38740000 | 0.0044 | 0.0028 | 0.0034 |
| F09 | 38660001 | 38760000 | 0.0049 | 0.0033 | 0.0039 |
| F09 | 38680001 | 38780000 | 0.0050 | 0.0033 | 0.0041 |
| F09 | 38700001 | 38800000 | 0.0049 | 0.0033 | 0.0042 |
| F09 | 38720001 | 38820000 | 0.0043 | 0.0029 | 0.0036 |
| F09 | 38740001 | 38840000 | 0.0046 | 0.0034 | 0.0039 |
| F09 | 38760001 | 38860000 | 0.0046 | 0.0033 | 0.0039 |
| F09 | 38780001 | 38880000 | 0.0040 | 0.0031 | 0.0035 |
| F09 | 38800001 | 38900000 | 0.0034 | 0.0027 | 0.0030 |
| F09 | 38820001 | 38920000 | 0.0035 | 0.0026 | 0.0029 |
| F09 | 38840001 | 38940000 | 0.0029 | 0.0021 | 0.0023 |
| F09 | 38860001 | 38960000 | 0.0029 | 0.0019 | 0.0020 |
| F09 | 38880001 | 38980000 | 0.0036 | 0.0023 | 0.0026 |
| F09 | 38900001 | 39000000 | 0.0050 | 0.0033 | 0.0037 |
| F09 | 38920001 | 39020000 | 0.0052 | 0.0034 | 0.0037 |
| F09 | 38940001 | 39040000 | 0.0050 | 0.0031 | 0.0037 |
| F09 | 38960001 | 39060000 | 0.0054 | 0.0038 | 0.0044 |
| F09 | 38980001 | 39080000 | 0.0053 | 0.0039 | 0.0044 |
| F09 | 39000001 | 39100000 | 0.0039 | 0.0029 | 0.0033 |
| F09 | 39020001 | 39120000 | 0.0039 | 0.0031 | 0.0035 |
| F09 | 39040001 | 39140000 | 0.0035 | 0.0029 | 0.0032 |
| F09 | 39060001 | 39160000 | 0.0029 | 0.0023 | 0.0027 |
| F09 | 39080001 | 39180000 | 0.0023 | 0.0017 | 0.0021 |
| F09 | 39100001 | 39200000 | 0.0028 | 0.0019 | 0.0023 |
| F09 | 39120001 | 39220000 | 0.0025 | 0.0015 | 0.0018 |
| F09 | 39140001 | 39240000 | 0.0025 | 0.0017 | 0.0019 |
| F09 | 39160001 | 39260000 | 0.0033 | 0.0024 | 0.0026 |
| F09 | 39180001 | 39280000 | 0.0035 | 0.0028 | 0.0030 |
| F09 | 39200001 | 39300000 | 0.0031 | 0.0028 | 0.0029 |
| F09 | 39220001 | 39320000 | 0.0032 | 0.0030 | 0.0031 |
| F09 | 39240001 | 39340000 | 0.0029 | 0.0026 | 0.0029 |
| F09 | 39260001 | 39360000 | 0.0029 | 0.0026 | 0.0029 |
| F09 | 39280001 | 39380000 | 0.0034 | 0.0029 | 0.0033 |
| F09 | 39300001 | 39400000 | 0.0040 | 0.0032 | 0.0037 |
| F09 | 39320001 | 39420000 | 0.0048 | 0.0039 | 0.0043 |
| F09 | 39340001 | 39440000 | 0.0052 | 0.0039 | 0.0043 |
| F09 | 39360001 | 39460000 | 0.0043 | 0.0032 | 0.0035 |
| F09 | 39380001 | 39480000 | 0.0043 | 0.0032 | 0.0035 |
| F09 | 39400001 | 39500000 | 0.0039 | 0.0031 | 0.0031 |
| F09 | 39420001 | 39520000 | 0.0030 | 0.0021 | 0.0021 |
| F09 | 39440001 | 39540000 | 0.0028 | 0.0020 | 0.0020 |
| F09 | 39460001 | 39560000 | 0.0039 | 0.0030 | 0.0028 |
| F09 | 39480001 | 39580000 | 0.0036 | 0.0027 | 0.0025 |
| F09 | 39500001 | 39600000 | 0.0046 | 0.0037 | 0.0036 |
| F09 | 39520001 | 39620000 | 0.0053 | 0.0040 | 0.0044 |
| F09 | 39540001 | 39640000 | 0.0062 | 0.0048 | 0.0053 |
| F09 | 39560001 | 39660000 | 0.0054 | 0.0038 | 0.0045 |
| F09 | 39580001 | 39680000 | 0.0053 | 0.0036 | 0.0044 |

| Chr | Start | End | both | A1 | A2 |
|---|---|---|---|---|---|
| F09 | 39600001 | 39700000 | 0.0044 | 0.0027 | 0.0033 |
| F09 | 39620001 | 39720000 | 0.0033 | 0.0021 | 0.0023 |
| F09 | 39640001 | 39740000 | 0.0026 | 0.0014 | 0.0016 |
| F09 | 39660001 | 39760000 | 0.0021 | 0.0012 | 0.0013 |
| F09 | 39680001 | 39780000 | 0.0019 | 0.0010 | 0.0011 |
| F09 | 39700001 | 39800000 | 0.0023 | 0.0015 | 0.0016 |
| F09 | 39720001 | 39820000 | 0.0024 | 0.0015 | 0.0016 |
| F09 | 39740001 | 39840000 | 0.0027 | 0.0018 | 0.0020 |
| F09 | 39760001 | 39860000 | 0.0029 | 0.0020 | 0.0021 |
| F09 | 39780001 | 39880000 | 0.0034 | 0.0023 | 0.0024 |
| F09 | 39800001 | 39900000 | 0.0034 | 0.0021 | 0.0023 |
| F09 | 39820001 | 39920000 | 0.0045 | 0.0032 | 0.0034 |
| F09 | 39840001 | 39940000 | 0.0045 | 0.0031 | 0.0035 |
| F09 | 39860001 | 39960000 | 0.0039 | 0.0028 | 0.0032 |
| F09 | 39880001 | 39980000 | 0.0040 | 0.0029 | 0.0034 |
| F09 | 39900001 | 40000000 | 0.0039 | 0.0029 | 0.0032 |
| F09 | 39920001 | 40020000 | 0.0033 | 0.0023 | 0.0026 |
| F09 | 39940001 | 40040000 | 0.0028 | 0.0018 | 0.0021 |
| F09 | 39960001 | 40060000 | 0.0031 | 0.0018 | 0.0022 |
| F09 | 39980001 | 40080000 | 0.0032 | 0.0019 | 0.0022 |
| F09 | 40000001 | 40100000 | 0.0037 | 0.0025 | 0.0029 |
| F09 | 40020001 | 40120000 | 0.0028 | 0.0021 | 0.0023 |
| F09 | 40040001 | 40140000 | 0.0030 | 0.0024 | 0.0026 |
| F09 | 40060001 | 40160000 | 0.0038 | 0.0032 | 0.0033 |
| F09 | 40080001 | 40180000 | 0.0038 | 0.0032 | 0.0033 |
| F09 | 40100001 | 40200000 | 0.0042 | 0.0034 | 0.0036 |
| F09 | 40120001 | 40220000 | 0.0042 | 0.0034 | 0.0037 |
| F09 | 40140001 | 40240000 | 0.0041 | 0.0031 | 0.0034 |
| F09 | 40160001 | 40260000 | 0.0034 | 0.0026 | 0.0028 |
| F09 | 40180001 | 40280000 | 0.0034 | 0.0026 | 0.0028 |
| F09 | 40200001 | 40300000 | 0.0023 | 0.0016 | 0.0016 |
| F09 | 40220001 | 40320000 | 0.0024 | 0.0015 | 0.0015 |
| F09 | 40240001 | 40340000 | 0.0024 | 0.0017 | 0.0017 |
| F09 | 40260001 | 40360000 | 0.0030 | 0.0021 | 0.0022 |
| F09 | 40280001 | 40380000 | 0.0038 | 0.0028 | 0.0029 |
| F09 | 40300001 | 40400000 | 0.0040 | 0.0030 | 0.0033 |
| F09 | 40320001 | 40420000 | 0.0042 | 0.0035 | 0.0037 |
| F09 | 40340001 | 40440000 | 0.0047 | 0.0038 | 0.0041 |
| F09 | 40360001 | 40460000 | 0.0048 | 0.0039 | 0.0044 |
| F09 | 40380001 | 40480000 | 0.0046 | 0.0036 | 0.0040 |
| F09 | 40400001 | 40500000 | 0.0046 | 0.0035 | 0.0039 |
| F09 | 40420001 | 40520000 | 0.0045 | 0.0032 | 0.0038 |
| F09 | 40440001 | 40540000 | 0.0044 | 0.0032 | 0.0036 |
| F09 | 40460001 | 40560000 | 0.0041 | 0.0028 | 0.0030 |
| F09 | 40480001 | 40580000 | 0.0033 | 0.0022 | 0.0025 |
| F09 | 40500001 | 40600000 | 0.0037 | 0.0029 | 0.0029 |
| F09 | 40520001 | 40620000 | 0.0046 | 0.0038 | 0.0037 |
| F09 | 40540001 | 40640000 | 0.0041 | 0.0034 | 0.0034 |
| F09 | 40560001 | 40660000 | 0.0037 | 0.0034 | 0.0034 |
| F09 | 40580001 | 40680000 | 0.0033 | 0.0029 | 0.0029 |
| F09 | 40600001 | 40700000 | 0.0027 | 0.0020 | 0.0021 |
| F09 | 40620001 | 40720000 | 0.0024 | 0.0014 | 0.0016 |
| F09 | 40640001 | 40740000 | 0.0026 | 0.0015 | 0.0017 |
| F09 | 40660001 | 40760000 | 0.0032 | 0.0018 | 0.0022 |
| F09 | 40680001 | 40780000 | 0.0035 | 0.0021 | 0.0025 |
| F09 | 40700001 | 40800000 | 0.0035 | 0.0023 | 0.0028 |
| F09 | 40720001 | 40820000 | 0.0034 | 0.0024 | 0.0029 |
| F09 | 40740001 | 40840000 | 0.0038 | 0.0027 | 0.0033 |

| Chr | Start | End | both | A1 | A2 |
|---|---|---|---|---|---|
| F09 | 40760001 | 40860000 | 0.0031 | 0.0022 | 0.0026 |
| F09 | 40780001 | 40880000 | 0.0033 | 0.0024 | 0.0028 |
| F09 | 40800001 | 40900000 | 0.0049 | 0.0040 | 0.0044 |
| F09 | 40820001 | 40920000 | 0.0047 | 0.0039 | 0.0042 |
| F09 | 40840001 | 40940000 | 0.0045 | 0.0037 | 0.0040 |
| F09 | 40860001 | 40960000 | 0.0067 | 0.0059 | 0.0064 |
| F09 | 40880001 | 40980000 | 0.0066 | 0.0057 | 0.0062 |
| F09 | 40900001 | 41000000 | 0.0048 | 0.0039 | 0.0042 |
| F09 | 40920001 | 41020000 | 0.0047 | 0.0037 | 0.0041 |
| F09 | 40940001 | 41040000 | 0.0050 | 0.0039 | 0.0045 |
| F09 | 40960001 | 41060000 | 0.0026 | 0.0018 | 0.0021 |
| F09 | 40980001 | 41080000 | 0.0033 | 0.0026 | 0.0029 |
| F09 | 41000001 | 41100000 | 0.0037 | 0.0030 | 0.0034 |
| F09 | 41020001 | 41120000 | 0.0047 | 0.0041 | 0.0045 |
| F09 | 41040001 | 41140000 | 0.0042 | 0.0038 | 0.0041 |
| F09 | 41060001 | 41160000 | 0.0041 | 0.0037 | 0.0041 |
| F09 | 41080001 | 41180000 | 0.0031 | 0.0028 | 0.0031 |
| F09 | 41100001 | 41200000 | 0.0032 | 0.0028 | 0.0031 |
| F09 | 41120001 | 41220000 | 0.0016 | 0.0014 | 0.0016 |
| F09 | 41140001 | 41240000 | 0.0016 | 0.0015 | 0.0016 |
| F09 | 41160001 | 41260000 | 0.0020 | 0.0019 | 0.0020 |
| F09 | 41180001 | 41280000 | 0.0018 | 0.0017 | 0.0017 |
| F09 | 41200001 | 41300000 | 0.0010 | 0.0010 | 0.0010 |
| F09 | 41220001 | 41320000 | 0.0016 | 0.0014 | 0.0015 |
| F09 | 41240001 | 41340000 | 0.0022 | 0.0020 | 0.0022 |
| F09 | 41260001 | 41360000 | 0.0030 | 0.0027 | 0.0029 |
| F09 | 41280001 | 41380000 | 0.0034 | 0.0031 | 0.0032 |
| F09 | 41300001 | 41400000 | 0.0035 | 0.0032 | 0.0034 |
| F09 | 41320001 | 41420000 | 0.0034 | 0.0032 | 0.0032 |
| F09 | 41340001 | 41440000 | 0.0035 | 0.0033 | 0.0033 |
| F09 | 41360001 | 41460000 | 0.0038 | 0.0035 | 0.0037 |
| F09 | 41380001 | 41480000 | 0.0063 | 0.0059 | 0.0062 |
| F09 | 41400001 | 41500000 | 0.0063 | 0.0058 | 0.0062 |
| F09 | 41420001 | 41520000 | 0.0059 | 0.0054 | 0.0058 |
| F09 | 41440001 | 41540000 | 0.0049 | 0.0045 | 0.0048 |
| F09 | 41460001 | 41560000 | 0.0040 | 0.0037 | 0.0039 |
| F09 | 41480001 | 41580000 | 0.0011 | 0.0009 | 0.0011 |
| F09 | 41500001 | 41600000 | 0.0010 | 0.0008 | 0.0010 |
| F09 | 41520001 | 41620000 | 0.0011 | 0.0009 | 0.0011 |
| F09 | 41540001 | 41640000 | 0.0026 | 0.0022 | 0.0025 |
| F09 | 41560001 | 41660000 | 0.0032 | 0.0027 | 0.0028 |
| F09 | 41580001 | 41680000 | 0.0038 | 0.0033 | 0.0035 |
| F09 | 41600001 | 41700000 | 0.0042 | 0.0036 | 0.0038 |
| F09 | 41620001 | 41720000 | 0.0056 | 0.0048 | 0.0049 |
| F09 | 41640001 | 41740000 | 0.0056 | 0.0046 | 0.0051 |
| F09 | 41660001 | 41760000 | 0.0063 | 0.0051 | 0.0058 |
| F09 | 41680001 | 41780000 | 0.0060 | 0.0049 | 0.0055 |
| F09 | 41700001 | 41800000 | 0.0063 | 0.0052 | 0.0058 |
| F09 | 41720001 | 41820000 | 0.0065 | 0.0053 | 0.0063 |
| F09 | 41740001 | 41840000 | 0.0057 | 0.0048 | 0.0055 |
| F09 | 41760001 | 41860000 | 0.0049 | 0.0041 | 0.0048 |
| F09 | 41780001 | 41880000 | 0.0053 | 0.0046 | 0.0052 |
| F09 | 41800001 | 41900000 | 0.0076 | 0.0066 | 0.0075 |
| F09 | 41820001 | 41920000 | 0.0075 | 0.0066 | 0.0074 |
| F09 | 41840001 | 41940000 | 0.0078 | 0.0067 | 0.0075 |
| F09 | 41860001 | 41960000 | 0.0072 | 0.0060 | 0.0068 |
| F09 | 41880001 | 41980000 | 0.0076 | 0.0063 | 0.0071 |
| F09 | 41900001 | 42000000 | 0.0057 | 0.0047 | 0.0052 |

| Chr | Start | End | both | A1 | A2 |
|---|---|---|---|---|---|
| F09 | 41920001 | 42020000 | 0.0051 | 0.0041 | 0.0046 |
| F09 | 41940001 | 42040000 | 0.0055 | 0.0047 | 0.0051 |
| F09 | 41960001 | 42060000 | 0.0062 | 0.0055 | 0.0059 |
| F09 | 41980001 | 42080000 | 0.0058 | 0.0051 | 0.0056 |
| F09 | 42000001 | 42100000 | 0.0055 | 0.0049 | 0.0053 |
| F09 | 42020001 | 42120000 | 0.0050 | 0.0044 | 0.0047 |
| F09 | 42040001 | 42140000 | 0.0047 | 0.0039 | 0.0041 |
| F09 | 42060001 | 42160000 | 0.0050 | 0.0041 | 0.0042 |
| F09 | 42080001 | 42180000 | 0.0045 | 0.0037 | 0.0038 |
| F09 | 42100001 | 42200000 | 0.0050 | 0.0039 | 0.0040 |
| F09 | 42120001 | 42220000 | 0.0050 | 0.0039 | 0.0041 |
| F09 | 42140001 | 42240000 | 0.0038 | 0.0031 | 0.0032 |
| F09 | 42160001 | 42260000 | 0.0027 | 0.0020 | 0.0021 |
| F09 | 42180001 | 42280000 | 0.0032 | 0.0023 | 0.0027 |
| F09 | 42200001 | 42300000 | 0.0022 | 0.0016 | 0.0020 |
| F09 | 42220001 | 42320000 | 0.0021 | 0.0015 | 0.0019 |
| F09 | 42240001 | 42340000 | 0.0026 | 0.0018 | 0.0023 |
| F09 | 42260001 | 42360000 | 0.0024 | 0.0019 | 0.0023 |
| F09 | 42280001 | 42380000 | 0.0023 | 0.0019 | 0.0021 |
| F09 | 42300001 | 42400000 | 0.0025 | 0.0020 | 0.0023 |
| F09 | 42320001 | 42420000 | 0.0023 | 0.0019 | 0.0021 |
| F09 | 42340001 | 42440000 | 0.0021 | 0.0017 | 0.0018 |
| F09 | 42360001 | 42460000 | 0.0028 | 0.0023 | 0.0025 |
| F09 | 42380001 | 42480000 | 0.0020 | 0.0018 | 0.0018 |
| F09 | 42400001 | 42500000 | 0.0021 | 0.0019 | 0.0020 |
| F09 | 42420001 | 42520000 | 0.0027 | 0.0024 | 0.0024 |
| F09 | 42440001 | 42540000 | 0.0032 | 0.0027 | 0.0028 |
| F09 | 42460001 | 42560000 | 0.0046 | 0.0038 | 0.0041 |
| F09 | 42480001 | 42580000 | 0.0049 | 0.0040 | 0.0042 |
| F09 | 42500001 | 42600000 | 0.0049 | 0.0038 | 0.0040 |
| F09 | 42520001 | 42620000 | 0.0043 | 0.0033 | 0.0036 |
| F09 | 42540001 | 42640000 | 0.0038 | 0.0029 | 0.0032 |
| F09 | 42560001 | 42660000 | 0.0028 | 0.0020 | 0.0020 |
| F09 | 42580001 | 42680000 | 0.0030 | 0.0018 | 0.0022 |
| F09 | 42600001 | 42700000 | 0.0034 | 0.0022 | 0.0026 |
| F09 | 42620001 | 42720000 | 0.0039 | 0.0026 | 0.0030 |
| F09 | 42640001 | 42740000 | 0.0040 | 0.0028 | 0.0032 |
| F09 | 42660001 | 42760000 | 0.0035 | 0.0024 | 0.0027 |
| F09 | 42680001 | 42780000 | 0.0035 | 0.0025 | 0.0027 |
| F09 | 42700001 | 42800000 | 0.0031 | 0.0023 | 0.0025 |
| F09 | 42720001 | 42820000 | 0.0028 | 0.0021 | 0.0023 |
| F09 | 42740001 | 42840000 | 0.0036 | 0.0027 | 0.0029 |
| F09 | 42760001 | 42860000 | 0.0046 | 0.0037 | 0.0040 |
| F09 | 42780001 | 42880000 | 0.0046 | 0.0035 | 0.0038 |
| F09 | 42800001 | 42900000 | 0.0059 | 0.0046 | 0.0051 |
| F09 | 42820001 | 42920000 | 0.0061 | 0.0047 | 0.0051 |
| F09 | 42840001 | 42940000 | 0.0059 | 0.0042 | 0.0047 |
| F09 | 42860001 | 42960000 | 0.0056 | 0.0036 | 0.0040 |
| F09 | 42880001 | 42980000 | 0.0057 | 0.0038 | 0.0042 |
| F09 | 42900001 | 43000000 | 0.0045 | 0.0028 | 0.0029 |
| F09 | 42920001 | 43020000 | 0.0044 | 0.0027 | 0.0030 |
| F09 | 42940001 | 43040000 | 0.0038 | 0.0024 | 0.0026 |
| F09 | 42960001 | 43060000 | 0.0035 | 0.0020 | 0.0024 |
| F09 | 42980001 | 43080000 | 0.0033 | 0.0019 | 0.0022 |
| F09 | 43000001 | 43100000 | 0.0034 | 0.0020 | 0.0024 |
| F09 | 43020001 | 43120000 | 0.0035 | 0.0018 | 0.0022 |
| F09 | 43040001 | 43140000 | 0.0034 | 0.0016 | 0.0020 |
| F09 | 43060001 | 43160000 | 0.0041 | 0.0027 | 0.0028 |

| Chr | Start | End | both | A1 | A2 |
|---|---|---|---|---|---|
| F09 | 43080001 | 43180000 | 0.0046 | 0.0031 | 0.0033 |
| F09 | 43100001 | 43200000 | 0.0046 | 0.0032 | 0.0033 |
| F09 | 43120001 | 43220000 | 0.0042 | 0.0032 | 0.0032 |
| F09 | 43140001 | 43240000 | 0.0051 | 0.0038 | 0.0041 |
| F09 | 43160001 | 43260000 | 0.0045 | 0.0032 | 0.0036 |
| F09 | 43180001 | 43280000 | 0.0039 | 0.0029 | 0.0033 |
| F09 | 43200001 | 43300000 | 0.0033 | 0.0024 | 0.0027 |
| F09 | 43220001 | 43320000 | 0.0039 | 0.0027 | 0.0031 |
| F09 | 43240001 | 43340000 | 0.0032 | 0.0023 | 0.0024 |
| F09 | 43260001 | 43360000 | 0.0030 | 0.0021 | 0.0021 |
| F09 | 43280001 | 43380000 | 0.0030 | 0.0021 | 0.0020 |
| F09 | 43300001 | 43400000 | 0.0034 | 0.0025 | 0.0024 |
| F09 | 43320001 | 43420000 | 0.0028 | 0.0020 | 0.0020 |
| F09 | 43340001 | 43440000 | 0.0038 | 0.0030 | 0.0030 |
| F09 | 43360001 | 43460000 | 0.0043 | 0.0036 | 0.0037 |
| F09 | 43380001 | 43480000 | 0.0042 | 0.0035 | 0.0037 |
| F09 | 43400001 | 43500000 | 0.0040 | 0.0033 | 0.0035 |
| F09 | 43420001 | 43520000 | 0.0043 | 0.0036 | 0.0039 |
| F09 | 43440001 | 43540000 | 0.0031 | 0.0025 | 0.0027 |
| F09 | 43460001 | 43560000 | 0.0019 | 0.0014 | 0.0016 |
| F09 | 43480001 | 43580000 | 0.0017 | 0.0011 | 0.0014 |
| F09 | 43500001 | 43600000 | 0.0023 | 0.0017 | 0.0020 |
| F09 | 43520001 | 43620000 | 0.0019 | 0.0015 | 0.0018 |
| F09 | 43540001 | 43640000 | 0.0022 | 0.0017 | 0.0021 |
| F09 | 43560001 | 43660000 | 0.0028 | 0.0023 | 0.0027 |
| F09 | 43580001 | 43680000 | 0.0043 | 0.0034 | 0.0040 |
| F09 | 43600001 | 43700000 | 0.0047 | 0.0037 | 0.0044 |
| F09 | 43620001 | 43720000 | 0.0051 | 0.0040 | 0.0048 |
| F09 | 43640001 | 43740000 | 0.0052 | 0.0042 | 0.0049 |
| F09 | 43660001 | 43760000 | 0.0049 | 0.0038 | 0.0046 |
| F09 | 43680001 | 43780000 | 0.0044 | 0.0034 | 0.0039 |
| F09 | 43700001 | 43800000 | 0.0046 | 0.0036 | 0.0040 |
| F09 | 43720001 | 43820000 | 0.0046 | 0.0035 | 0.0038 |
| F09 | 43740001 | 43840000 | 0.0043 | 0.0031 | 0.0034 |
| F09 | 43760001 | 43860000 | 0.0051 | 0.0035 | 0.0039 |
| F09 | 43780001 | 43880000 | 0.0048 | 0.0034 | 0.0038 |
| F09 | 43800001 | 43900000 | 0.0039 | 0.0028 | 0.0030 |
| F09 | 43820001 | 43920000 | 0.0035 | 0.0025 | 0.0028 |
| F09 | 43840001 | 43940000 | 0.0035 | 0.0027 | 0.0029 |
| F09 | 43860001 | 43960000 | 0.0026 | 0.0023 | 0.0024 |
| F09 | 43880001 | 43980000 | 0.0020 | 0.0017 | 0.0019 |
| F09 | 43900001 | 44000000 | 0.0023 | 0.0020 | 0.0022 |
| F09 | 43920001 | 44020000 | 0.0022 | 0.0019 | 0.0020 |
| F09 | 43940001 | 44040000 | 0.0026 | 0.0022 | 0.0024 |
| F09 | 43960001 | 44060000 | 0.0027 | 0.0024 | 0.0025 |
| F09 | 43980001 | 44080000 | 0.0030 | 0.0027 | 0.0028 |
| F09 | 44000001 | 44100000 | 0.0020 | 0.0018 | 0.0018 |
| F09 | 44020001 | 44120000 | 0.0022 | 0.0019 | 0.0020 |
| F09 | 44040001 | 44140000 | 0.0025 | 0.0021 | 0.0022 |
| F09 | 44060001 | 44160000 | 0.0026 | 0.0019 | 0.0022 |
| F09 | 44080001 | 44180000 | 0.0031 | 0.0022 | 0.0025 |
| F09 | 44100001 | 44200000 | 0.0049 | 0.0036 | 0.0040 |
| F09 | 44120001 | 44220000 | 0.0058 | 0.0043 | 0.0049 |
| F09 | 44140001 | 44240000 | 0.0057 | 0.0043 | 0.0048 |
| F09 | 44160001 | 44260000 | 0.0062 | 0.0047 | 0.0051 |
| F09 | 44180001 | 44280000 | 0.0064 | 0.0049 | 0.0054 |
| F09 | 44200001 | 44300000 | 0.0058 | 0.0044 | 0.0049 |
| F09 | 44220001 | 44320000 | 0.0050 | 0.0036 | 0.0040 |

| Chr | Start | End | both | A1 | A2 |
|---|---|---|---|---|---|
| F09 | 44240001 | 44340000 | 0.0048 | 0.0035 | 0.0039 |
| F09 | 44260001 | 44360000 | 0.0044 | 0.0033 | 0.0037 |
| F09 | 44280001 | 44380000 | 0.0039 | 0.0028 | 0.0031 |
| F09 | 44300001 | 44400000 | 0.0035 | 0.0025 | 0.0028 |
| F09 | 44320001 | 44420000 | 0.0035 | 0.0025 | 0.0027 |
| F09 | 44340001 | 44440000 | 0.0033 | 0.0023 | 0.0024 |
| F09 | 44360001 | 44460000 | 0.0035 | 0.0024 | 0.0025 |
| F09 | 44380001 | 44480000 | 0.0040 | 0.0027 | 0.0030 |
| F09 | 44400001 | 44500000 | 0.0043 | 0.0027 | 0.0029 |
| F09 | 44420001 | 44520000 | 0.0043 | 0.0029 | 0.0032 |
| F09 | 44440001 | 44540000 | 0.0040 | 0.0025 | 0.0029 |
| F09 | 44460001 | 44560000 | 0.0037 | 0.0021 | 0.0025 |
| F09 | 44480001 | 44580000 | 0.0028 | 0.0014 | 0.0015 |
| F09 | 44500001 | 44600000 | 0.0026 | 0.0014 | 0.0015 |
| F09 | 44520001 | 44620000 | 0.0022 | 0.0012 | 0.0013 |
| F09 | 44540001 | 44640000 | 0.0027 | 0.0016 | 0.0017 |
| F09 | 44560001 | 44660000 | 0.0025 | 0.0015 | 0.0017 |
| F09 | 44580001 | 44680000 | 0.0031 | 0.0023 | 0.0024 |
| F09 | 44600001 | 44700000 | 0.0026 | 0.0017 | 0.0018 |
| F09 | 44620001 | 44720000 | 0.0030 | 0.0018 | 0.0019 |
| F09 | 44640001 | 44740000 | 0.0030 | 0.0018 | 0.0019 |
| F09 | 44660001 | 44760000 | 0.0037 | 0.0025 | 0.0025 |
| F09 | 44680001 | 44780000 | 0.0032 | 0.0018 | 0.0018 |
| F09 | 44700001 | 44800000 | 0.0035 | 0.0020 | 0.0020 |
| F09 | 44720001 | 44820000 | 0.0037 | 0.0021 | 0.0021 |
| F09 | 44740001 | 44840000 | 0.0033 | 0.0020 | 0.0020 |
| F09 | 44760001 | 44860000 | 0.0032 | 0.0018 | 0.0019 |
| F09 | 44780001 | 44880000 | 0.0031 | 0.0016 | 0.0018 |
| F09 | 44800001 | 44900000 | 0.0027 | 0.0014 | 0.0017 |
| F09 | 44820001 | 44920000 | 0.0028 | 0.0013 | 0.0021 |
| F09 | 44840001 | 44940000 | 0.0031 | 0.0014 | 0.0025 |
| F09 | 44860001 | 44960000 | 0.0029 | 0.0011 | 0.0025 |
| F09 | 44880001 | 44980000 | 0.0029 | 0.0010 | 0.0027 |
| F09 | 44900001 | 45000000 | 0.0031 | 0.0013 | 0.0031 |
| F09 | 44920001 | 45020000 | 0.0036 | 0.0017 | 0.0035 |
| F09 | 44940001 | 45040000 | 0.0030 | 0.0014 | 0.0029 |
| F09 | 44960001 | 45060000 | 0.0032 | 0.0019 | 0.0032 |
| F09 | 44980001 | 45080000 | 0.0036 | 0.0025 | 0.0036 |
| F09 | 45000001 | 45100000 | 0.0033 | 0.0023 | 0.0032 |
| F09 | 45020001 | 45120000 | 0.0023 | 0.0020 | 0.0023 |
| F09 | 45040001 | 45140000 | 0.0036 | 0.0029 | 0.0036 |
| F09 | 45060001 | 45160000 | 0.0029 | 0.0020 | 0.0028 |
| F09 | 45080001 | 45180000 | 0.0028 | 0.0020 | 0.0027 |
| F09 | 45100001 | 45200000 | 0.0034 | 0.0023 | 0.0033 |
| F09 | 45120001 | 45220000 | 0.0035 | 0.0022 | 0.0034 |
| F09 | 45140001 | 45240000 | 0.0023 | 0.0013 | 0.0023 |
| F09 | 45160001 | 45260000 | 0.0032 | 0.0017 | 0.0031 |
| F09 | 45180001 | 45280000 | 0.0033 | 0.0017 | 0.0032 |
| F09 | 45200001 | 45300000 | 0.0031 | 0.0015 | 0.0031 |
| F09 | 45220001 | 45320000 | 0.0040 | 0.0021 | 0.0040 |
| F09 | 45240001 | 45340000 | 0.0049 | 0.0029 | 0.0048 |
| F09 | 45260001 | 45360000 | 0.0045 | 0.0030 | 0.0045 |
| F09 | 45280001 | 45380000 | 0.0046 | 0.0028 | 0.0045 |
| F09 | 45300001 | 45400000 | 0.0042 | 0.0027 | 0.0041 |
| F09 | 45320001 | 45420000 | 0.0048 | 0.0035 | 0.0048 |
| F09 | 45340001 | 45440000 | 0.0048 | 0.0034 | 0.0047 |
| F09 | 45360001 | 45460000 | 0.0047 | 0.0035 | 0.0046 |
| F09 | 45380001 | 45480000 | 0.0041 | 0.0032 | 0.0041 |

| Chr | Start | End | both | A1 | A2 |
|---|---|---|---|---|---|
| F09 | 45400001 | 45500000 | 0.0044 | 0.0033 | 0.0044 |
| F09 | 45420001 | 45520000 | 0.0034 | 0.0019 | 0.0033 |
| F09 | 45440001 | 45540000 | 0.0031 | 0.0013 | 0.0030 |
| F09 | 45460001 | 45560000 | 0.0040 | 0.0017 | 0.0039 |
| F09 | 45480001 | 45580000 | 0.0056 | 0.0027 | 0.0054 |
| F09 | 45500001 | 45600000 | 0.0069 | 0.0037 | 0.0066 |
| F09 | 45520001 | 45620000 | 0.0075 | 0.0042 | 0.0073 |
| F09 | 45540001 | 45640000 | 0.0075 | 0.0045 | 0.0073 |
| F09 | 45560001 | 45660000 | 0.0069 | 0.0041 | 0.0067 |
| F09 | 45580001 | 45680000 | 0.0054 | 0.0031 | 0.0052 |
| F09 | 45600001 | 45700000 | 0.0042 | 0.0022 | 0.0041 |
| F09 | 45620001 | 45720000 | 0.0032 | 0.0019 | 0.0031 |
| F09 | 45640001 | 45740000 | 0.0029 | 0.0019 | 0.0029 |
| F09 | 45660001 | 45760000 | 0.0036 | 0.0022 | 0.0035 |
| F09 | 45680001 | 45780000 | 0.0042 | 0.0028 | 0.0041 |
| F09 | 45700001 | 45800000 | 0.0045 | 0.0031 | 0.0043 |
| F09 | 45720001 | 45820000 | 0.0050 | 0.0032 | 0.0048 |
| F09 | 45740001 | 45840000 | 0.0051 | 0.0032 | 0.0049 |
| F09 | 45760001 | 45860000 | 0.0043 | 0.0028 | 0.0041 |
| F09 | 45780001 | 45880000 | 0.0036 | 0.0023 | 0.0034 |
| F09 | 45800001 | 45900000 | 0.0036 | 0.0022 | 0.0034 |
| F09 | 45820001 | 45920000 | 0.0029 | 0.0020 | 0.0028 |
| F09 | 45840001 | 45940000 | 0.0025 | 0.0016 | 0.0024 |
| F09 | 45860001 | 45960000 | 0.0025 | 0.0016 | 0.0024 |
| F09 | 45880001 | 45980000 | 0.0024 | 0.0016 | 0.0023 |
| F09 | 45900001 | 46000000 | 0.0019 | 0.0010 | 0.0018 |
| F09 | 45920001 | 46020000 | 0.0018 | 0.0009 | 0.0016 |
| F09 | 45940001 | 46040000 | 0.0017 | 0.0013 | 0.0015 |
| F09 | 45960001 | 46060000 | 0.0020 | 0.0018 | 0.0018 |
| F09 | 45980001 | 46080000 | 0.0024 | 0.0022 | 0.0022 |
| F09 | 46000001 | 46100000 | 0.0032 | 0.0034 | 0.0030 |
| F09 | 46020001 | 46120000 | 0.0034 | 0.0038 | 0.0032 |
| F09 | 46040001 | 46140000 | 0.0034 | 0.0039 | 0.0032 |
| F09 | 46060001 | 46160000 | 0.0029 | 0.0035 | 0.0026 |
| F09 | 46080001 | 46180000 | 0.0025 | 0.0031 | 0.0022 |
| F09 | 46100001 | 46200000 | 0.0016 | 0.0022 | 0.0014 |
| F09 | 46120001 | 46220000 | 0.0014 | 0.0019 | 0.0013 |
| F09 | 46140001 | 46240000 | 0.0017 | 0.0019 | 0.0016 |
| F09 | 46160001 | 46260000 | 0.0013 | 0.0014 | 0.0012 |
| F09 | 46180001 | 46280000 | 0.0016 | 0.0017 | 0.0015 |
| F09 | 46200001 | 46300000 | 0.0016 | 0.0018 | 0.0015 |
| F09 | 46220001 | 46320000 | 0.0019 | 0.0020 | 0.0018 |
| F09 | 46240001 | 46340000 | 0.0017 | 0.0019 | 0.0015 |
| F09 | 46260001 | 46360000 | 0.0023 | 0.0026 | 0.0022 |
| F09 | 46280001 | 46380000 | 0.0025 | 0.0028 | 0.0023 |
| F09 | 46300001 | 46400000 | 0.0036 | 0.0037 | 0.0034 |
| F09 | 46320001 | 46420000 | 0.0033 | 0.0034 | 0.0031 |
| F09 | 46340001 | 46440000 | 0.0035 | 0.0037 | 0.0034 |
| F09 | 46360001 | 46460000 | 0.0037 | 0.0039 | 0.0035 |
| F09 | 46380001 | 46480000 | 0.0040 | 0.0040 | 0.0039 |
| F09 | 46400001 | 46500000 | 0.0029 | 0.0030 | 0.0027 |
| F09 | 46420001 | 46520000 | 0.0029 | 0.0032 | 0.0028 |
| F09 | 46440001 | 46540000 | 0.0024 | 0.0026 | 0.0022 |
| F09 | 46460001 | 46560000 | 0.0017 | 0.0021 | 0.0016 |
| F09 | 46480001 | 46580000 | 0.0017 | 0.0022 | 0.0015 |
| F09 | 46500001 | 46600000 | 0.0016 | 0.0022 | 0.0014 |
| F09 | 46520001 | 46620000 | 0.0024 | 0.0028 | 0.0022 |
| F09 | 46540001 | 46640000 | 0.0032 | 0.0037 | 0.0029 |

| Chr | Start | End | both | A1 | A2 |
|---|---|---|---|---|---|
| F09 | 46560001 | 46660000 | 0.0030 | 0.0034 | 0.0027 |
| F09 | 46580001 | 46680000 | 0.0021 | 0.0027 | 0.0018 |
| F09 | 46600001 | 46700000 | 0.0032 | 0.0029 | 0.0030 |
| F09 | 46620001 | 46720000 | 0.0023 | 0.0022 | 0.0021 |
| F09 | 46640001 | 46740000 | 0.0019 | 0.0017 | 0.0018 |
| F09 | 46660001 | 46760000 | 0.0036 | 0.0029 | 0.0035 |
| F09 | 46680001 | 46780000 | 0.0040 | 0.0034 | 0.0039 |
| F09 | 46700001 | 46800000 | 0.0038 | 0.0041 | 0.0036 |
| F09 | 46720001 | 46820000 | 0.0046 | 0.0049 | 0.0044 |
| F09 | 46740001 | 46840000 | 0.0051 | 0.0055 | 0.0048 |
| F09 | 46760001 | 46860000 | 0.0037 | 0.0044 | 0.0034 |
| F09 | 46780001 | 46880000 | 0.0036 | 0.0043 | 0.0033 |
| F09 | 46800001 | 46900000 | 0.0033 | 0.0037 | 0.0031 |
| F09 | 46820001 | 46920000 | 0.0030 | 0.0033 | 0.0028 |
| F09 | 46840001 | 46940000 | 0.0027 | 0.0027 | 0.0026 |
| F09 | 46860001 | 46960000 | 0.0028 | 0.0030 | 0.0026 |
| F09 | 46880001 | 46980000 | 0.0029 | 0.0030 | 0.0028 |
| F09 | 46900001 | 47000000 | 0.0032 | 0.0033 | 0.0031 |
| F09 | 46920001 | 47020000 | 0.0027 | 0.0029 | 0.0026 |
| F09 | 46940001 | 47040000 | 0.0029 | 0.0030 | 0.0028 |
| F09 | 46960001 | 47060000 | 0.0030 | 0.0028 | 0.0029 |
| F09 | 46980001 | 47080000 | 0.0026 | 0.0025 | 0.0026 |
| F09 | 47000001 | 47100000 | 0.0017 | 0.0015 | 0.0017 |
| F09 | 47020001 | 47120000 | 0.0018 | 0.0016 | 0.0018 |
| F09 | 47040001 | 47140000 | 0.0023 | 0.0020 | 0.0023 |
| F09 | 47060001 | 47160000 | 0.0022 | 0.0020 | 0.0022 |
| F09 | 47080001 | 47180000 | 0.0027 | 0.0024 | 0.0027 |
| F09 | 47100001 | 47200000 | 0.0035 | 0.0032 | 0.0034 |
| F09 | 47120001 | 47220000 | 0.0043 | 0.0038 | 0.0043 |
| F09 | 47140001 | 47240000 | 0.0044 | 0.0041 | 0.0043 |
| F09 | 47160001 | 47260000 | 0.0043 | 0.0041 | 0.0042 |
| F09 | 47180001 | 47280000 | 0.0040 | 0.0038 | 0.0039 |
| F09 | 47200001 | 47300000 | 0.0031 | 0.0031 | 0.0030 |
| F09 | 47220001 | 47320000 | 0.0024 | 0.0026 | 0.0022 |
| F09 | 47240001 | 47340000 | 0.0021 | 0.0022 | 0.0020 |
| F09 | 47260001 | 47360000 | 0.0024 | 0.0023 | 0.0023 |
| F09 | 47280001 | 47380000 | 0.0023 | 0.0023 | 0.0022 |
| F09 | 47300001 | 47400000 | 0.0040 | 0.0037 | 0.0039 |
| F09 | 47320001 | 47420000 | 0.0043 | 0.0039 | 0.0043 |
| F09 | 47340001 | 47440000 | 0.0035 | 0.0032 | 0.0034 |
| F09 | 47360001 | 47460000 | 0.0041 | 0.0039 | 0.0040 |
| F09 | 47380001 | 47480000 | 0.0042 | 0.0039 | 0.0041 |
| F09 | 47400001 | 47500000 | 0.0029 | 0.0027 | 0.0028 |
| F09 | 47420001 | 47520000 | 0.0028 | 0.0028 | 0.0028 |
| F09 | 47440001 | 47540000 | 0.0030 | 0.0029 | 0.0029 |
| F09 | 47460001 | 47560000 | 0.0023 | 0.0022 | 0.0022 |
| F09 | 47480001 | 47580000 | 0.0034 | 0.0034 | 0.0032 |
| F09 | 47500001 | 47600000 | 0.0039 | 0.0041 | 0.0037 |
| F09 | 47520001 | 47620000 | 0.0043 | 0.0045 | 0.0041 |
| F09 | 47540001 | 47640000 | 0.0053 | 0.0054 | 0.0051 |
| F09 | 47560001 | 47660000 | 0.0066 | 0.0069 | 0.0063 |
| F09 | 47580001 | 47680000 | 0.0054 | 0.0058 | 0.0052 |
| F09 | 47600001 | 47700000 | 0.0050 | 0.0054 | 0.0047 |
| F09 | 47620001 | 47720000 | 0.0045 | 0.0049 | 0.0043 |
| F09 | 47640001 | 47740000 | 0.0046 | 0.0049 | 0.0043 |
| F09 | 47660001 | 47760000 | 0.0042 | 0.0043 | 0.0040 |
| F09 | 47680001 | 47780000 | 0.0047 | 0.0048 | 0.0045 |
| F09 | 47700001 | 47800000 | 0.0057 | 0.0057 | 0.0055 |

| Chr | Start | End | both | A1 | A2 |
|---|---|---|---|---|---|
| F09 | 47720001 | 47820000 | 0.0060 | 0.0059 | 0.0058 |
| F09 | 47740001 | 47840000 | 0.0062 | 0.0060 | 0.0060 |
| F09 | 47760001 | 47860000 | 0.0050 | 0.0052 | 0.0048 |
| F09 | 47780001 | 47880000 | 0.0048 | 0.0047 | 0.0046 |
| F09 | 47800001 | 47900000 | 0.0048 | 0.0047 | 0.0046 |
| F09 | 47820001 | 47920000 | 0.0053 | 0.0054 | 0.0051 |
| F09 | 47840001 | 47940000 | 0.0051 | 0.0051 | 0.0049 |
| F09 | 47860001 | 47960000 | 0.0051 | 0.0050 | 0.0049 |
| F09 | 47880001 | 47980000 | 0.0046 | 0.0047 | 0.0044 |
| F09 | 47900001 | 48000000 | 0.0032 | 0.0036 | 0.0030 |
| F09 | 47920001 | 48020000 | 0.0027 | 0.0030 | 0.0024 |
| F09 | 47940001 | 48040000 | 0.0022 | 0.0028 | 0.0019 |
| F09 | 47960001 | 48060000 | 0.0020 | 0.0027 | 0.0018 |
| F09 | 47980001 | 48080000 | 0.0023 | 0.0032 | 0.0020 |
| F09 | 48000001 | 48100000 | 0.0029 | 0.0035 | 0.0026 |
| F09 | 48020001 | 48120000 | 0.0022 | 0.0029 | 0.0019 |
| F09 | 48040001 | 48140000 | 0.0032 | 0.0036 | 0.0029 |
| F09 | 48060001 | 48160000 | 0.0036 | 0.0040 | 0.0034 |
| F09 | 48080001 | 48180000 | 0.0035 | 0.0037 | 0.0032 |
| F09 | 48100001 | 48200000 | 0.0031 | 0.0034 | 0.0029 |
| F09 | 48120001 | 48220000 | 0.0040 | 0.0043 | 0.0037 |
| F09 | 48140001 | 48240000 | 0.0030 | 0.0032 | 0.0027 |
| F09 | 48160001 | 48260000 | 0.0026 | 0.0028 | 0.0024 |
| F09 | 48180001 | 48280000 | 0.0030 | 0.0032 | 0.0028 |
| F09 | 48200001 | 48300000 | 0.0032 | 0.0029 | 0.0028 |
| F09 | 48220001 | 48320000 | 0.0027 | 0.0023 | 0.0024 |
| F09 | 48240001 | 48340000 | 0.0033 | 0.0025 | 0.0028 |
| F09 | 48260001 | 48360000 | 0.0032 | 0.0025 | 0.0028 |
| F09 | 48280001 | 48380000 | 0.0029 | 0.0020 | 0.0024 |
| F09 | 48300001 | 48400000 | 0.0035 | 0.0027 | 0.0032 |
| F09 | 48320001 | 48420000 | 0.0039 | 0.0030 | 0.0036 |
| F09 | 48340001 | 48440000 | 0.0038 | 0.0029 | 0.0035 |
| F09 | 48360001 | 48460000 | 0.0047 | 0.0037 | 0.0043 |
| F09 | 48380001 | 48480000 | 0.0051 | 0.0039 | 0.0047 |
| F09 | 48400001 | 48500000 | 0.0043 | 0.0030 | 0.0038 |
| F09 | 48420001 | 48520000 | 0.0035 | 0.0023 | 0.0031 |
| F09 | 48440001 | 48540000 | 0.0028 | 0.0016 | 0.0024 |
| F09 | 48460001 | 48560000 | 0.0020 | 0.0009 | 0.0017 |
| F09 | 48480001 | 48580000 | 0.0018 | 0.0010 | 0.0016 |
| F09 | 48500001 | 48600000 | 0.0021 | 0.0012 | 0.0018 |
| F09 | 48520001 | 48620000 | 0.0029 | 0.0018 | 0.0026 |
| F09 | 48540001 | 48640000 | 0.0029 | 0.0018 | 0.0026 |
| F09 | 48560001 | 48660000 | 0.0031 | 0.0019 | 0.0027 |
| F09 | 48580001 | 48680000 | 0.0034 | 0.0022 | 0.0030 |
| F09 | 48600001 | 48700000 | 0.0029 | 0.0020 | 0.0026 |
| F09 | 48620001 | 48720000 | 0.0026 | 0.0015 | 0.0022 |
| F09 | 48640001 | 48740000 | 0.0028 | 0.0019 | 0.0025 |
| F09 | 48660001 | 48760000 | 0.0031 | 0.0020 | 0.0027 |
| F09 | 48680001 | 48780000 | 0.0025 | 0.0013 | 0.0022 |
| F09 | 48700001 | 48800000 | 0.0027 | 0.0013 | 0.0023 |
| F09 | 48720001 | 48820000 | 0.0033 | 0.0019 | 0.0029 |
| F09 | 48740001 | 48840000 | 0.0030 | 0.0015 | 0.0025 |
| F09 | 48760001 | 48860000 | 0.0035 | 0.0020 | 0.0029 |
| F09 | 48780001 | 48880000 | 0.0037 | 0.0022 | 0.0031 |
| F09 | 48800001 | 48900000 | 0.0050 | 0.0035 | 0.0043 |
| F09 | 48820001 | 48920000 | 0.0040 | 0.0027 | 0.0034 |
| F09 | 48840001 | 48940000 | 0.0049 | 0.0032 | 0.0040 |
| F09 | 48860001 | 48960000 | 0.0042 | 0.0029 | 0.0034 |

| Chr | Start | End | both | A1 | A2 |
|---|---|---|---|---|---|
| F09 | 48880001 | 48980000 | 0.0041 | 0.0029 | 0.0032 |
| F09 | 48900001 | 49000000 | 0.0029 | 0.0016 | 0.0018 |
| F09 | 48920001 | 49020000 | 0.0028 | 0.0019 | 0.0015 |
| F09 | 48940001 | 49040000 | 0.0024 | 0.0022 | 0.0013 |
| F09 | 48960001 | 49060000 | 0.0021 | 0.0022 | 0.0011 |
| F09 | 48980001 | 49080000 | 0.0022 | 0.0027 | 0.0012 |
| F09 | 49000001 | 49100000 | 0.0023 | 0.0033 | 0.0015 |
| F09 | 49020001 | 49120000 | 0.0027 | 0.0038 | 0.0021 |
| F09 | 49040001 | 49140000 | 0.0025 | 0.0037 | 0.0019 |
| F09 | 49060001 | 49160000 | 0.0026 | 0.0039 | 0.0019 |
| F09 | 49080001 | 49180000 | 0.0030 | 0.0040 | 0.0024 |
| F09 | 49100001 | 49200000 | 0.0028 | 0.0038 | 0.0021 |
| F09 | 49120001 | 49220000 | 0.0024 | 0.0033 | 0.0017 |
| F09 | 49140001 | 49240000 | 0.0028 | 0.0032 | 0.0023 |
| F09 | 49160001 | 49260000 | 0.0026 | 0.0029 | 0.0022 |
| F09 | 49180001 | 49280000 | 0.0021 | 0.0027 | 0.0016 |
| F09 | 49200001 | 49300000 | 0.0024 | 0.0029 | 0.0018 |
| F09 | 49220001 | 49320000 | 0.0023 | 0.0029 | 0.0016 |
| F09 | 49240001 | 49340000 | 0.0021 | 0.0028 | 0.0013 |
| F09 | 49260001 | 49360000 | 0.0033 | 0.0038 | 0.0025 |
| F09 | 49280001 | 49380000 | 0.0039 | 0.0042 | 0.0031 |
| F09 | 49300001 | 49400000 | 0.0039 | 0.0039 | 0.0032 |
| F09 | 49320001 | 49420000 | 0.0038 | 0.0037 | 0.0032 |
| F09 | 49340001 | 49440000 | 0.0034 | 0.0033 | 0.0028 |
| F09 | 49360001 | 49460000 | 0.0023 | 0.0024 | 0.0017 |
| F09 | 49380001 | 49480000 | 0.0016 | 0.0017 | 0.0010 |
| F09 | 49400001 | 49500000 | 0.0012 | 0.0016 | 0.0006 |
| F09 | 49420001 | 49520000 | 0.0019 | 0.0022 | 0.0013 |
| F09 | 49440001 | 49540000 | 0.0028 | 0.0030 | 0.0022 |
| F09 | 49460001 | 49560000 | 0.0033 | 0.0037 | 0.0028 |
| F09 | 49480001 | 49580000 | 0.0033 | 0.0038 | 0.0029 |
| F09 | 49500001 | 49600000 | 0.0033 | 0.0039 | 0.0030 |
| F09 | 49520001 | 49620000 | 0.0029 | 0.0034 | 0.0026 |
| F09 | 49540001 | 49640000 | 0.0031 | 0.0036 | 0.0028 |
| F09 | 49560001 | 49660000 | 0.0024 | 0.0028 | 0.0021 |
| F09 | 49580001 | 49680000 | 0.0033 | 0.0034 | 0.0031 |
| F09 | 49600001 | 49700000 | 0.0032 | 0.0035 | 0.0030 |
| F09 | 49620001 | 49720000 | 0.0030 | 0.0032 | 0.0028 |
| F09 | 49640001 | 49740000 | 0.0019 | 0.0024 | 0.0017 |
| F09 | 49660001 | 49760000 | 0.0021 | 0.0026 | 0.0019 |
| F09 | 49680001 | 49780000 | 0.0011 | 0.0018 | 0.0009 |
| F09 | 49700001 | 49800000 | 0.0013 | 0.0017 | 0.0011 |
| F09 | 49720001 | 49820000 | 0.0023 | 0.0026 | 0.0022 |
| F09 | 49740001 | 49840000 | 0.0036 | 0.0039 | 0.0033 |
| F09 | 49760001 | 49860000 | 0.0037 | 0.0041 | 0.0035 |
| F09 | 49780001 | 49880000 | 0.0041 | 0.0042 | 0.0039 |
| F09 | 49800001 | 49900000 | 0.0042 | 0.0044 | 0.0040 |
| F09 | 49820001 | 49920000 | 0.0036 | 0.0039 | 0.0033 |
| F09 | 49840001 | 49940000 | 0.0021 | 0.0025 | 0.0019 |
| F09 | 49860001 | 49960000 | 0.0018 | 0.0023 | 0.0015 |
| F09 | 49880001 | 49980000 | 0.0024 | 0.0029 | 0.0021 |
| F09 | 49900001 | 50000000 | 0.0028 | 0.0032 | 0.0026 |
| F09 | 49920001 | 50020000 | 0.0024 | 0.0028 | 0.0021 |
| F09 | 49940001 | 50040000 | 0.0034 | 0.0037 | 0.0031 |
| F09 | 49960001 | 50060000 | 0.0039 | 0.0040 | 0.0037 |
| F09 | 49980001 | 50080000 | 0.0040 | 0.0043 | 0.0038 |
| F09 | 50000001 | 50100000 | 0.0034 | 0.0039 | 0.0032 |
| F09 | 50020001 | 50120000 | 0.0034 | 0.0040 | 0.0032 |

| Chr | Start | End | both | A1 | A2 |
|---|---|---|---|---|---|
| F09 | 50040001 | 50140000 | 0.0026 | 0.0033 | 0.0023 |
| F09 | 50060001 | 50160000 | 0.0046 | 0.0049 | 0.0044 |
| F09 | 50080001 | 50180000 | 0.0045 | 0.0046 | 0.0042 |
| F09 | 50100001 | 50200000 | 0.0042 | 0.0043 | 0.0040 |
| F09 | 50120001 | 50220000 | 0.0042 | 0.0042 | 0.0040 |
| F09 | 50140001 | 50240000 | 0.0042 | 0.0041 | 0.0040 |
| F09 | 50160001 | 50260000 | 0.0015 | 0.0020 | 0.0013 |
| F09 | 50180001 | 50280000 | 0.0008 | 0.0013 | 0.0006 |
| F09 | 50200001 | 50300000 | 0.0009 | 0.0014 | 0.0007 |
| F09 | 50220001 | 50320000 | 0.0009 | 0.0014 | 0.0007 |
| F09 | 50240001 | 50340000 | 0.0010 | 0.0015 | 0.0009 |
| F09 | 50260001 | 50360000 | 0.0018 | 0.0020 | 0.0017 |
| F09 | 50280001 | 50380000 | 0.0026 | 0.0026 | 0.0024 |
| F09 | 50300001 | 50400000 | 0.0027 | 0.0028 | 0.0025 |
| F09 | 50320001 | 50420000 | 0.0033 | 0.0035 | 0.0031 |
| F09 | 50340001 | 50440000 | 0.0034 | 0.0037 | 0.0031 |
| F09 | 50360001 | 50460000 | 0.0028 | 0.0033 | 0.0026 |
| F09 | 50380001 | 50480000 | 0.0025 | 0.0030 | 0.0021 |
| F09 | 50400001 | 50500000 | 0.0036 | 0.0036 | 0.0033 |
| F09 | 50420001 | 50520000 | 0.0041 | 0.0039 | 0.0039 |
| F09 | 50440001 | 50540000 | 0.0043 | 0.0039 | 0.0042 |
| F09 | 50460001 | 50560000 | 0.0041 | 0.0037 | 0.0040 |
| F09 | 50480001 | 50580000 | 0.0045 | 0.0042 | 0.0044 |
| F09 | 50500001 | 50600000 | 0.0044 | 0.0044 | 0.0043 |
| F09 | 50520001 | 50620000 | 0.0036 | 0.0037 | 0.0034 |
| F09 | 50540001 | 50640000 | 0.0032 | 0.0035 | 0.0029 |
| F09 | 50560001 | 50660000 | 0.0037 | 0.0041 | 0.0035 |
| F09 | 50580001 | 50680000 | 0.0030 | 0.0034 | 0.0028 |
| F09 | 50600001 | 50700000 | 0.0019 | 0.0024 | 0.0016 |
| F09 | 50620001 | 50720000 | 0.0021 | 0.0025 | 0.0017 |
| F09 | 50640001 | 50740000 | 0.0019 | 0.0023 | 0.0016 |
| F09 | 50660001 | 50760000 | 0.0014 | 0.0018 | 0.0011 |
| F09 | 50680001 | 50780000 | 0.0014 | 0.0018 | 0.0012 |
| F09 | 50700001 | 50800000 | 0.0015 | 0.0019 | 0.0013 |
| F09 | 50720001 | 50820000 | 0.0018 | 0.0020 | 0.0016 |
| F09 | 50740001 | 50840000 | 0.0025 | 0.0024 | 0.0023 |
| F09 | 50760001 | 50860000 | 0.0030 | 0.0029 | 0.0029 |
| F09 | 50780001 | 50880000 | 0.0027 | 0.0026 | 0.0025 |
| F09 | 50800001 | 50900000 | 0.0024 | 0.0023 | 0.0023 |
| F09 | 50820001 | 50920000 | 0.0026 | 0.0024 | 0.0025 |
| F09 | 50840001 | 50940000 | 0.0022 | 0.0023 | 0.0020 |
| F09 | 50860001 | 50960000 | 0.0030 | 0.0031 | 0.0028 |
| F09 | 50880001 | 50980000 | 0.0039 | 0.0041 | 0.0036 |
| F09 | 50900001 | 51000000 | 0.0044 | 0.0048 | 0.0040 |
| F09 | 50920001 | 51020000 | 0.0039 | 0.0046 | 0.0034 |
| F09 | 50940001 | 51040000 | 0.0038 | 0.0045 | 0.0032 |
| F09 | 50960001 | 51060000 | 0.0031 | 0.0037 | 0.0024 |
| F09 | 50980001 | 51080000 | 0.0022 | 0.0029 | 0.0016 |
| F09 | 51000001 | 51100000 | 0.0021 | 0.0027 | 0.0014 |
| F09 | 51020001 | 51120000 | 0.0026 | 0.0030 | 0.0020 |
| F09 | 51040001 | 51140000 | 0.0028 | 0.0030 | 0.0023 |
| F09 | 51060001 | 51160000 | 0.0025 | 0.0026 | 0.0020 |
| F09 | 51080001 | 51180000 | 0.0025 | 0.0022 | 0.0021 |
| F09 | 51100001 | 51200000 | 0.0025 | 0.0018 | 0.0021 |
| F09 | 51120001 | 51220000 | 0.0018 | 0.0011 | 0.0013 |
| F09 | 51140001 | 51240000 | 0.0020 | 0.0012 | 0.0014 |
| F09 | 51160001 | 51260000 | 0.0020 | 0.0012 | 0.0014 |
| F09 | 51180001 | 51280000 | 0.0029 | 0.0021 | 0.0022 |

| Chr | Start | End | both | A1 | A2 |
|---|---|---|---|---|---|
| F09 | 51200001 | 51300000 | 0.0031 | 0.0025 | 0.0022 |
| F09 | 51220001 | 51320000 | 0.0032 | 0.0026 | 0.0024 |
| F09 | 51240001 | 51340000 | 0.0036 | 0.0030 | 0.0029 |
| F09 | 51260001 | 51360000 | 0.0033 | 0.0028 | 0.0026 |
| F09 | 51280001 | 51380000 | 0.0032 | 0.0026 | 0.0026 |
| F09 | 51300001 | 51400000 | 0.0038 | 0.0032 | 0.0034 |
| F09 | 51320001 | 51420000 | 0.0038 | 0.0032 | 0.0035 |
| F09 | 51340001 | 51440000 | 0.0033 | 0.0027 | 0.0030 |
| F09 | 51360001 | 51460000 | 0.0033 | 0.0028 | 0.0031 |
| F09 | 51380001 | 51480000 | 0.0023 | 0.0020 | 0.0022 |
| F09 | 51400001 | 51500000 | 0.0056 | 0.0052 | 0.0054 |
| F09 | 51420001 | 51520000 | 0.0059 | 0.0054 | 0.0056 |
| F09 | 51440001 | 51540000 | 0.0057 | 0.0052 | 0.0054 |
| F09 | 51460001 | 51560000 | 0.0062 | 0.0056 | 0.0057 |
| F09 | 51480001 | 51580000 | 0.0070 | 0.0060 | 0.0061 |
| F09 | 51500001 | 51600000 | 0.0031 | 0.0022 | 0.0021 |
| F09 | 51520001 | 51620000 | 0.0027 | 0.0018 | 0.0017 |
| F09 | 51540001 | 51640000 | 0.0026 | 0.0017 | 0.0017 |
| F09 | 51560001 | 51660000 | 0.0025 | 0.0016 | 0.0017 |
| F09 | 51580001 | 51680000 | 0.0018 | 0.0011 | 0.0013 |
| F09 | 51600001 | 51700000 | 0.0015 | 0.0010 | 0.0011 |
| F09 | 51620001 | 51720000 | 0.0026 | 0.0018 | 0.0022 |
| F09 | 51640001 | 51740000 | 0.0029 | 0.0021 | 0.0024 |
| F09 | 51660001 | 51760000 | 0.0033 | 0.0024 | 0.0025 |
| F09 | 51680001 | 51780000 | 0.0034 | 0.0025 | 0.0027 |
| F09 | 51700001 | 51800000 | 0.0035 | 0.0026 | 0.0027 |
| F09 | 51720001 | 51820000 | 0.0024 | 0.0015 | 0.0015 |
| F09 | 51740001 | 51840000 | 0.0024 | 0.0016 | 0.0016 |
| F09 | 51760001 | 51860000 | 0.0019 | 0.0012 | 0.0013 |
| F09 | 51780001 | 51880000 | 0.0019 | 0.0012 | 0.0014 |
| F09 | 51800001 | 51900000 | 0.0022 | 0.0016 | 0.0018 |
| F09 | 51820001 | 51920000 | 0.0023 | 0.0019 | 0.0020 |
| F09 | 51840001 | 51940000 | 0.0022 | 0.0020 | 0.0018 |
| F09 | 51860001 | 51960000 | 0.0024 | 0.0023 | 0.0021 |
| F09 | 51880001 | 51980000 | 0.0025 | 0.0023 | 0.0021 |
| F09 | 51900001 | 52000000 | 0.0026 | 0.0024 | 0.0024 |
| F09 | 51920001 | 52020000 | 0.0026 | 0.0024 | 0.0023 |
| F09 | 51940001 | 52040000 | 0.0021 | 0.0019 | 0.0019 |
| F09 | 51960001 | 52060000 | 0.0019 | 0.0018 | 0.0016 |
| F09 | 51980001 | 52080000 | 0.0029 | 0.0028 | 0.0026 |
| F09 | 52000001 | 52100000 | 0.0030 | 0.0031 | 0.0026 |
| F09 | 52020001 | 52120000 | 0.0036 | 0.0036 | 0.0031 |
| F09 | 52040001 | 52140000 | 0.0037 | 0.0038 | 0.0033 |
| F09 | 52060001 | 52160000 | 0.0041 | 0.0045 | 0.0037 |
| F09 | 52080001 | 52180000 | 0.0033 | 0.0039 | 0.0029 |
| F09 | 52100001 | 52200000 | 0.0028 | 0.0032 | 0.0025 |
| F09 | 52120001 | 52220000 | 0.0025 | 0.0030 | 0.0022 |
| F09 | 52140001 | 52240000 | 0.0028 | 0.0033 | 0.0026 |
| F09 | 52160001 | 52260000 | 0.0027 | 0.0030 | 0.0025 |
| F09 | 52180001 | 52280000 | 0.0025 | 0.0028 | 0.0023 |
| F09 | 52200001 | 52300000 | 0.0024 | 0.0031 | 0.0021 |
| F09 | 52220001 | 52320000 | 0.0029 | 0.0036 | 0.0026 |
| F09 | 52240001 | 52340000 | 0.0031 | 0.0039 | 0.0028 |
| F09 | 52260001 | 52360000 | 0.0030 | 0.0037 | 0.0025 |
| F09 | 52280001 | 52380000 | 0.0032 | 0.0035 | 0.0027 |
| F09 | 52300001 | 52400000 | 0.0030 | 0.0028 | 0.0025 |
| F09 | 52320001 | 52420000 | 0.0029 | 0.0025 | 0.0025 |
| F09 | 52340001 | 52440000 | 0.0025 | 0.0018 | 0.0020 |

| Chr | Start | End | both | A1 | A2 |
|---|---|---|---|---|---|
| F09 | 52360001 | 52460000 | 0.0022 | 0.0015 | 0.0017 |
| F09 | 52380001 | 52480000 | 0.0019 | 0.0013 | 0.0014 |
| F09 | 52400001 | 52500000 | 0.0025 | 0.0016 | 0.0017 |
| F09 | 52420001 | 52520000 | 0.0021 | 0.0009 | 0.0010 |
| F09 | 52440001 | 52540000 | 0.0024 | 0.0012 | 0.0012 |
| F09 | 52460001 | 52560000 | 0.0026 | 0.0014 | 0.0013 |
| F09 | 52480001 | 52580000 | 0.0031 | 0.0016 | 0.0015 |
| F09 | 52500001 | 52600000 | 0.0026 | 0.0014 | 0.0012 |
| F09 | 52520001 | 52620000 | 0.0025 | 0.0015 | 0.0013 |
| F09 | 52540001 | 52640000 | 0.0021 | 0.0010 | 0.0009 |
| F09 | 52560001 | 52660000 | 0.0021 | 0.0011 | 0.0009 |
| F09 | 52580001 | 52680000 | 0.0027 | 0.0019 | 0.0019 |
| F09 | 52600001 | 52700000 | 0.0028 | 0.0020 | 0.0020 |
| F09 | 52620001 | 52720000 | 0.0035 | 0.0026 | 0.0026 |
| F09 | 52640001 | 52740000 | 0.0037 | 0.0029 | 0.0030 |
| F09 | 52660001 | 52760000 | 0.0033 | 0.0026 | 0.0027 |
| F09 | 52680001 | 52780000 | 0.0025 | 0.0020 | 0.0019 |
| F09 | 52700001 | 52800000 | 0.0026 | 0.0021 | 0.0020 |
| F09 | 52720001 | 52820000 | 0.0025 | 0.0020 | 0.0022 |
| F09 | 52740001 | 52840000 | 0.0023 | 0.0019 | 0.0020 |
| F09 | 52760001 | 52860000 | 0.0023 | 0.0019 | 0.0021 |
| F09 | 52780001 | 52880000 | 0.0019 | 0.0016 | 0.0018 |
| F09 | 52800001 | 52900000 | 0.0022 | 0.0019 | 0.0021 |
| F09 | 52820001 | 52920000 | 0.0026 | 0.0024 | 0.0025 |
| F09 | 52840001 | 52940000 | 0.0032 | 0.0029 | 0.0031 |
| F09 | 52860001 | 52960000 | 0.0032 | 0.0029 | 0.0031 |
| F09 | 52880001 | 52980000 | 0.0032 | 0.0029 | 0.0032 |
| F09 | 52900001 | 53000000 | 0.0030 | 0.0028 | 0.0030 |
| F09 | 52920001 | 53020000 | 0.0028 | 0.0023 | 0.0027 |
| F09 | 52940001 | 53040000 | 0.0038 | 0.0030 | 0.0037 |
| F09 | 52960001 | 53060000 | 0.0063 | 0.0053 | 0.0062 |
| F09 | 52980001 | 53080000 | 0.0078 | 0.0067 | 0.0076 |
| F09 | 53000001 | 53100000 | 0.0078 | 0.0067 | 0.0076 |
| F09 | 53020001 | 53120000 | 0.0080 | 0.0068 | 0.0077 |
| F09 | 53040001 | 53140000 | 0.0066 | 0.0056 | 0.0062 |
| F09 | 53060001 | 53160000 | 0.0044 | 0.0034 | 0.0040 |
| F09 | 53080001 | 53180000 | 0.0033 | 0.0022 | 0.0027 |
| F09 | 53100001 | 53200000 | 0.0037 | 0.0025 | 0.0030 |
| F09 | 53120001 | 53220000 | 0.0031 | 0.0021 | 0.0024 |
| F09 | 53140001 | 53240000 | 0.0026 | 0.0017 | 0.0020 |
| F09 | 53160001 | 53260000 | 0.0025 | 0.0016 | 0.0018 |
| F09 | 53180001 | 53280000 | 0.0024 | 0.0016 | 0.0017 |
| F09 | 53200001 | 53300000 | 0.0017 | 0.0009 | 0.0010 |
| F09 | 53220001 | 53320000 | 0.0012 | 0.0006 | 0.0007 |
| F09 | 53240001 | 53340000 | 0.0014 | 0.0007 | 0.0008 |
| F09 | 53260001 | 53360000 | 0.0019 | 0.0010 | 0.0012 |
| F09 | 53280001 | 53380000 | 0.0017 | 0.0010 | 0.0010 |
| F09 | 53300001 | 53400000 | 0.0018 | 0.0011 | 0.0012 |
| F09 | 53320001 | 53420000 | 0.0019 | 0.0011 | 0.0012 |
| F09 | 53340001 | 53440000 | 0.0021 | 0.0011 | 0.0013 |
| F09 | 53360001 | 53460000 | 0.0021 | 0.0010 | 0.0010 |
| F09 | 53380001 | 53480000 | 0.0023 | 0.0012 | 0.0012 |
| F09 | 53400001 | 53500000 | 0.0022 | 0.0011 | 0.0011 |
| F09 | 53420001 | 53520000 | 0.0022 | 0.0013 | 0.0013 |
| F09 | 53440001 | 53540000 | 0.0032 | 0.0024 | 0.0023 |
| F09 | 53460001 | 53560000 | 0.0030 | 0.0024 | 0.0023 |
| F09 | 53480001 | 53580000 | 0.0026 | 0.0020 | 0.0020 |
| F09 | 53500001 | 53600000 | 0.0028 | 0.0022 | 0.0022 |

| Chr | Start | End | both | A1 | A2 |
|---|---|---|---|---|---|
| F09 | 53520001 | 53620000 | 0.0032 | 0.0024 | 0.0025 |
| F09 | 53540001 | 53640000 | 0.0021 | 0.0011 | 0.0012 |
| F09 | 53560001 | 53660000 | 0.0018 | 0.0010 | 0.0011 |
| F09 | 53580001 | 53680000 | 0.0019 | 0.0011 | 0.0013 |
| F09 | 53600001 | 53700000 | 0.0018 | 0.0008 | 0.0010 |
| F09 | 53620001 | 53720000 | 0.0014 | 0.0003 | 0.0004 |
| F09 | 53640001 | 53740000 | 0.0015 | 0.0006 | 0.0007 |
| F09 | 53660001 | 53760000 | 0.0022 | 0.0009 | 0.0011 |
| F09 | 53680001 | 53780000 | 0.0022 | 0.0008 | 0.0011 |
| F09 | 53700001 | 53800000 | 0.0022 | 0.0008 | 0.0011 |
| F09 | 53720001 | 53820000 | 0.0022 | 0.0010 | 0.0013 |
| F09 | 53740001 | 53840000 | 0.0024 | 0.0012 | 0.0015 |
| F09 | 53760001 | 53860000 | 0.0026 | 0.0014 | 0.0017 |
| F09 | 53780001 | 53880000 | 0.0025 | 0.0015 | 0.0016 |
| F09 | 53800001 | 53900000 | 0.0027 | 0.0017 | 0.0017 |
| F09 | 53820001 | 53920000 | 0.0033 | 0.0022 | 0.0023 |
| F09 | 53840001 | 53940000 | 0.0031 | 0.0020 | 0.0022 |
| F09 | 53860001 | 53960000 | 0.0027 | 0.0016 | 0.0018 |
| F09 | 53880001 | 53980000 | 0.0027 | 0.0017 | 0.0019 |
| F09 | 53900001 | 54000000 | 0.0028 | 0.0018 | 0.0021 |
| F09 | 53920001 | 54020000 | 0.0026 | 0.0017 | 0.0018 |
| F09 | 53940001 | 54040000 | 0.0025 | 0.0016 | 0.0018 |
| F09 | 53960001 | 54060000 | 0.0024 | 0.0016 | 0.0018 |
| F09 | 53980001 | 54080000 | 0.0024 | 0.0016 | 0.0018 |
| F09 | 54000001 | 54100000 | 0.0029 | 0.0021 | 0.0023 |
| F09 | 54020001 | 54120000 | 0.0026 | 0.0020 | 0.0020 |
| F09 | 54040001 | 54140000 | 0.0027 | 0.0023 | 0.0019 |
| F09 | 54060001 | 54160000 | 0.0025 | 0.0023 | 0.0016 |
| F09 | 54080001 | 54180000 | 0.0052 | 0.0050 | 0.0044 |
| F09 | 54100001 | 54200000 | 0.0046 | 0.0047 | 0.0037 |
| F09 | 54120001 | 54220000 | 0.0054 | 0.0055 | 0.0043 |
| F09 | 54140001 | 54240000 | 0.0052 | 0.0053 | 0.0043 |
| F09 | 54160001 | 54260000 | 0.0051 | 0.0051 | 0.0043 |
| F09 | 54180001 | 54280000 | 0.0023 | 0.0025 | 0.0015 |
| F09 | 54200001 | 54300000 | 0.0021 | 0.0023 | 0.0014 |
| F09 | 54220001 | 54320000 | 0.0014 | 0.0016 | 0.0009 |
| F09 | 54240001 | 54340000 | 0.0014 | 0.0016 | 0.0008 |
| F09 | 54260001 | 54360000 | 0.0015 | 0.0017 | 0.0009 |
| F09 | 54280001 | 54380000 | 0.0016 | 0.0017 | 0.0009 |
| F09 | 54300001 | 54400000 | 0.0019 | 0.0020 | 0.0013 |
| F09 | 54320001 | 54420000 | 0.0018 | 0.0019 | 0.0012 |
| F09 | 54340001 | 54440000 | 0.0022 | 0.0022 | 0.0016 |
| F09 | 54360001 | 54460000 | 0.0025 | 0.0025 | 0.0019 |
| F09 | 54380001 | 54480000 | 0.0030 | 0.0031 | 0.0026 |
| F09 | 54400001 | 54500000 | 0.0038 | 0.0038 | 0.0035 |
| F09 | 54420001 | 54520000 | 0.0044 | 0.0044 | 0.0039 |
| F09 | 54440001 | 54540000 | 0.0040 | 0.0041 | 0.0033 |
| F09 | 54460001 | 54560000 | 0.0035 | 0.0036 | 0.0029 |
| F09 | 54480001 | 54580000 | 0.0028 | 0.0030 | 0.0022 |
| F09 | 54500001 | 54600000 | 0.0018 | 0.0020 | 0.0012 |
| F09 | 54520001 | 54620000 | 0.0013 | 0.0015 | 0.0008 |
| F09 | 54540001 | 54640000 | 0.0015 | 0.0017 | 0.0009 |
| F09 | 54560001 | 54660000 | 0.0015 | 0.0017 | 0.0009 |
| F09 | 54580001 | 54680000 | 0.0019 | 0.0020 | 0.0012 |
| F09 | 54600001 | 54700000 | 0.0024 | 0.0026 | 0.0016 |
| F09 | 54620001 | 54720000 | 0.0023 | 0.0024 | 0.0014 |
| F09 | 54640001 | 54740000 | 0.0022 | 0.0022 | 0.0014 |
| F09 | 54660001 | 54760000 | 0.0024 | 0.0023 | 0.0017 |

| Chr | Start | End | both | A1 | A2 |
|---|---|---|---|---|---|
| F09 | 54680001 | 54780000 | 0.0021 | 0.0021 | 0.0014 |
| F09 | 54700001 | 54800000 | 0.0015 | 0.0015 | 0.0009 |
| F09 | 54720001 | 54820000 | 0.0017 | 0.0017 | 0.0011 |
| F09 | 54740001 | 54840000 | 0.0018 | 0.0019 | 0.0012 |
| F09 | 54760001 | 54860000 | 0.0018 | 0.0019 | 0.0012 |
| F09 | 54780001 | 54880000 | 0.0020 | 0.0020 | 0.0013 |
| F09 | 54800001 | 54900000 | 0.0019 | 0.0019 | 0.0012 |
| F09 | 54820001 | 54920000 | 0.0021 | 0.0020 | 0.0013 |
| F09 | 54840001 | 54940000 | 0.0020 | 0.0020 | 0.0011 |
| F09 | 54860001 | 54960000 | 0.0021 | 0.0024 | 0.0011 |
| F09 | 54880001 | 54980000 | 0.0021 | 0.0026 | 0.0013 |
| F09 | 54900001 | 55000000 | 0.0025 | 0.0031 | 0.0015 |
| F09 | 54920001 | 55020000 | 0.0025 | 0.0033 | 0.0018 |
| F09 | 54940001 | 55040000 | 0.0033 | 0.0040 | 0.0027 |
| F09 | 54960001 | 55060000 | 0.0032 | 0.0038 | 0.0026 |
| F09 | 54980001 | 55080000 | 0.0040 | 0.0046 | 0.0034 |
| F09 | 55000001 | 55100000 | 0.0040 | 0.0043 | 0.0035 |
| F09 | 55020001 | 55120000 | 0.0036 | 0.0040 | 0.0030 |
| F09 | 55040001 | 55140000 | 0.0027 | 0.0033 | 0.0021 |
| F09 | 55060001 | 55160000 | 0.0027 | 0.0032 | 0.0021 |
| F09 | 55080001 | 55180000 | 0.0017 | 0.0021 | 0.0011 |
| F09 | 55100001 | 55200000 | 0.0015 | 0.0020 | 0.0008 |
| F09 | 55120001 | 55220000 | 0.0015 | 0.0018 | 0.0007 |
| F09 | 55140001 | 55240000 | 0.0014 | 0.0016 | 0.0007 |
| F09 | 55160001 | 55260000 | 0.0013 | 0.0013 | 0.0006 |
| F09 | 55180001 | 55280000 | 0.0019 | 0.0016 | 0.0013 |
| F09 | 55200001 | 55300000 | 0.0020 | 0.0016 | 0.0014 |
| F09 | 55220001 | 55320000 | 0.0022 | 0.0020 | 0.0016 |
| F09 | 55240001 | 55340000 | 0.0023 | 0.0021 | 0.0016 |
| F09 | 55260001 | 55360000 | 0.0023 | 0.0021 | 0.0017 |
| F09 | 55280001 | 55380000 | 0.0022 | 0.0023 | 0.0014 |
| F09 | 55300001 | 55400000 | 0.0026 | 0.0027 | 0.0020 |
| F09 | 55320001 | 55420000 | 0.0034 | 0.0035 | 0.0027 |
| F09 | 55340001 | 55440000 | 0.0035 | 0.0037 | 0.0027 |
| F09 | 55360001 | 55460000 | 0.0035 | 0.0038 | 0.0028 |
| F09 | 55380001 | 55480000 | 0.0032 | 0.0034 | 0.0024 |
| F09 | 55400001 | 55500000 | 0.0025 | 0.0028 | 0.0018 |
| F09 | 55420001 | 55520000 | 0.0019 | 0.0019 | 0.0013 |
| F09 | 55440001 | 55540000 | 0.0016 | 0.0016 | 0.0012 |
| F09 | 55460001 | 55560000 | 0.0017 | 0.0016 | 0.0012 |
| F09 | 55480001 | 55580000 | 0.0019 | 0.0018 | 0.0014 |
| F09 | 55500001 | 55600000 | 0.0020 | 0.0019 | 0.0015 |
| F09 | 55520001 | 55620000 | 0.0015 | 0.0016 | 0.0009 |
| F09 | 55540001 | 55640000 | 0.0016 | 0.0016 | 0.0011 |
| F09 | 55560001 | 55660000 | 0.0015 | 0.0015 | 0.0010 |
| F09 | 55580001 | 55680000 | 0.0017 | 0.0014 | 0.0011 |
| F09 | 55600001 | 55700000 | 0.0018 | 0.0015 | 0.0012 |
| F09 | 55620001 | 55720000 | 0.0021 | 0.0016 | 0.0014 |
| F09 | 55640001 | 55740000 | 0.0020 | 0.0015 | 0.0012 |
| F09 | 55660001 | 55760000 | 0.0019 | 0.0012 | 0.0011 |
| F09 | 55680001 | 55780000 | 0.0014 | 0.0009 | 0.0007 |
| F09 | 55700001 | 55800000 | 0.0013 | 0.0007 | 0.0005 |
| F09 | 55720001 | 55820000 | 0.0018 | 0.0012 | 0.0010 |
| F09 | 55740001 | 55840000 | 0.0019 | 0.0011 | 0.0011 |
| F09 | 55760001 | 55860000 | 0.0021 | 0.0013 | 0.0015 |
| F09 | 55780001 | 55880000 | 0.0020 | 0.0013 | 0.0015 |
| F09 | 55800001 | 55900000 | 0.0023 | 0.0015 | 0.0018 |
| F09 | 55820001 | 55920000 | 0.0022 | 0.0015 | 0.0019 |

| Chr | Start | End | both | A1 | A2 |
|---|---|---|---|---|---|
| F09 | 55840001 | 55940000 | 0.0023 | 0.0015 | 0.0019 |
| F09 | 55860001 | 55960000 | 0.0029 | 0.0020 | 0.0025 |
| F09 | 55880001 | 55980000 | 0.0040 | 0.0023 | 0.0034 |
| F09 | 55900001 | 56000000 | 0.0040 | 0.0022 | 0.0036 |
| F09 | 55920001 | 56020000 | 0.0035 | 0.0015 | 0.0031 |
| F09 | 55940001 | 56040000 | 0.0039 | 0.0018 | 0.0036 |
| F09 | 55960001 | 56060000 | 0.0035 | 0.0014 | 0.0033 |
| F09 | 55980001 | 56080000 | 0.0029 | 0.0015 | 0.0029 |
| F09 | 56000001 | 56100000 | 0.0034 | 0.0021 | 0.0033 |
| F09 | 56020001 | 56120000 | 0.0037 | 0.0025 | 0.0037 |
| F09 | 56040001 | 56140000 | 0.0035 | 0.0024 | 0.0035 |
| F09 | 56060001 | 56160000 | 0.0035 | 0.0021 | 0.0035 |
| F09 | 56080001 | 56180000 | 0.0030 | 0.0016 | 0.0029 |
| F09 | 56100001 | 56200000 | 0.0023 | 0.0008 | 0.0022 |
| F09 | 56120001 | 56220000 | 0.0020 | 0.0006 | 0.0019 |
| F09 | 56140001 | 56240000 | 0.0021 | 0.0008 | 0.0020 |
| F09 | 56160001 | 56260000 | 0.0022 | 0.0012 | 0.0020 |
| F09 | 56180001 | 56280000 | 0.0023 | 0.0013 | 0.0022 |
| F09 | 56200001 | 56300000 | 0.0027 | 0.0017 | 0.0026 |
| F09 | 56220001 | 56320000 | 0.0024 | 0.0016 | 0.0023 |
| F09 | 56240001 | 56340000 | 0.0023 | 0.0014 | 0.0022 |
| F09 | 56260001 | 56360000 | 0.0026 | 0.0016 | 0.0024 |
| F09 | 56280001 | 56380000 | 0.0026 | 0.0016 | 0.0024 |
| F09 | 56300001 | 56400000 | 0.0023 | 0.0013 | 0.0020 |
| F09 | 56320001 | 56420000 | 0.0024 | 0.0015 | 0.0021 |
| F09 | 56340001 | 56440000 | 0.0021 | 0.0013 | 0.0018 |
| F09 | 56360001 | 56460000 | 0.0021 | 0.0012 | 0.0017 |
| F09 | 56380001 | 56480000 | 0.0024 | 0.0014 | 0.0019 |
| F09 | 56400001 | 56500000 | 0.0025 | 0.0013 | 0.0019 |
| F09 | 56420001 | 56520000 | 0.0035 | 0.0021 | 0.0028 |
| F09 | 56440001 | 56540000 | 0.0035 | 0.0020 | 0.0027 |
| F09 | 56460001 | 56560000 | 0.0030 | 0.0016 | 0.0023 |
| F09 | 56480001 | 56580000 | 0.0028 | 0.0016 | 0.0022 |
| F09 | 56500001 | 56600000 | 0.0029 | 0.0020 | 0.0024 |
| F09 | 56520001 | 56620000 | 0.0019 | 0.0012 | 0.0014 |
| F09 | 56540001 | 56640000 | 0.0022 | 0.0014 | 0.0017 |
| F09 | 56560001 | 56660000 | 0.0020 | 0.0012 | 0.0014 |
| F09 | 56580001 | 56680000 | 0.0019 | 0.0012 | 0.0012 |
| F09 | 56600001 | 56700000 | 0.0025 | 0.0017 | 0.0017 |
| F09 | 56620001 | 56720000 | 0.0029 | 0.0022 | 0.0022 |
| F09 | 56640001 | 56740000 | 0.0025 | 0.0020 | 0.0018 |
| F09 | 56660001 | 56760000 | 0.0028 | 0.0023 | 0.0022 |
| F09 | 56680001 | 56780000 | 0.0032 | 0.0027 | 0.0026 |
| F09 | 56700001 | 56800000 | 0.0021 | 0.0016 | 0.0015 |
| F09 | 56720001 | 56820000 | 0.0021 | 0.0016 | 0.0016 |
| F09 | 56740001 | 56840000 | 0.0023 | 0.0016 | 0.0017 |
| F09 | 56760001 | 56860000 | 0.0024 | 0.0015 | 0.0016 |
| F09 | 56780001 | 56880000 | 0.0019 | 0.0011 | 0.0012 |
| F09 | 56800001 | 56900000 | 0.0023 | 0.0014 | 0.0016 |
| F09 | 56820001 | 56920000 | 0.0021 | 0.0014 | 0.0014 |
| F09 | 56840001 | 56940000 | 0.0024 | 0.0017 | 0.0017 |
| F09 | 56860001 | 56960000 | 0.0023 | 0.0017 | 0.0017 |
| F09 | 56880001 | 56980000 | 0.0033 | 0.0025 | 0.0027 |
| F09 | 56900001 | 57000000 | 0.0034 | 0.0025 | 0.0027 |
| F09 | 56920001 | 57020000 | 0.0030 | 0.0021 | 0.0024 |
| F09 | 56940001 | 57040000 | 0.0028 | 0.0018 | 0.0020 |
| F09 | 56960001 | 57060000 | 0.0035 | 0.0024 | 0.0029 |
| F09 | 56980001 | 57080000 | 0.0034 | 0.0024 | 0.0028 |

| Chr | Start | End | both | A1 | A2 |
|---|---|---|---|---|---|
| F09 | 57000001 | 57100000 | 0.0041 | 0.0031 | 0.0036 |
| F09 | 57020001 | 57120000 | 0.0046 | 0.0035 | 0.0041 |
| F09 | 57040001 | 57140000 | 0.0045 | 0.0036 | 0.0043 |
| F09 | 57060001 | 57160000 | 0.0032 | 0.0027 | 0.0031 |
| F09 | 57080001 | 57180000 | 0.0030 | 0.0024 | 0.0029 |
| F09 | 57100001 | 57200000 | 0.0024 | 0.0019 | 0.0022 |
| F09 | 57120001 | 57220000 | 0.0026 | 0.0020 | 0.0022 |
| F09 | 57140001 | 57240000 | 0.0025 | 0.0018 | 0.0020 |
| F09 | 57160001 | 57260000 | 0.0033 | 0.0022 | 0.0025 |
| F09 | 57180001 | 57280000 | 0.0038 | 0.0027 | 0.0029 |
| F09 | 57200001 | 57300000 | 0.0033 | 0.0021 | 0.0023 |
| F09 | 57220001 | 57320000 | 0.0026 | 0.0015 | 0.0016 |
| F09 | 57240001 | 57340000 | 0.0027 | 0.0015 | 0.0017 |
| F09 | 57260001 | 57360000 | 0.0024 | 0.0014 | 0.0014 |
| F09 | 57280001 | 57380000 | 0.0019 | 0.0010 | 0.0010 |
| F09 | 57300001 | 57400000 | 0.0024 | 0.0014 | 0.0017 |
| F09 | 57320001 | 57420000 | 0.0029 | 0.0017 | 0.0020 |
| F09 | 57340001 | 57440000 | 0.0033 | 0.0022 | 0.0026 |
| F09 | 57360001 | 57460000 | 0.0030 | 0.0019 | 0.0024 |
| F09 | 57380001 | 57480000 | 0.0026 | 0.0015 | 0.0020 |
| F09 | 57400001 | 57500000 | 0.0025 | 0.0015 | 0.0018 |
| F09 | 57420001 | 57520000 | 0.0023 | 0.0015 | 0.0017 |
| F09 | 57440001 | 57540000 | 0.0017 | 0.0011 | 0.0013 |
| F09 | 57460001 | 57560000 | 0.0019 | 0.0012 | 0.0013 |
| F09 | 57480001 | 57580000 | 0.0018 | 0.0011 | 0.0012 |
| F09 | 57500001 | 57600000 | 0.0022 | 0.0014 | 0.0016 |
| F09 | 57520001 | 57620000 | 0.0021 | 0.0011 | 0.0012 |
| F09 | 57540001 | 57640000 | 0.0023 | 0.0012 | 0.0013 |
| F09 | 57560001 | 57660000 | 0.0023 | 0.0012 | 0.0013 |
| F09 | 57580001 | 57680000 | 0.0026 | 0.0014 | 0.0016 |
| F09 | 57600001 | 57700000 | 0.0022 | 0.0011 | 0.0014 |
| F09 | 57620001 | 57720000 | 0.0029 | 0.0019 | 0.0021 |
| F09 | 57640001 | 57740000 | 0.0032 | 0.0022 | 0.0023 |
| F09 | 57660001 | 57760000 | 0.0031 | 0.0021 | 0.0023 |
| F09 | 57680001 | 57780000 | 0.0027 | 0.0020 | 0.0020 |
| F09 | 57700001 | 57800000 | 0.0022 | 0.0018 | 0.0016 |
| F09 | 57720001 | 57820000 | 0.0014 | 0.0011 | 0.0009 |
| F09 | 57740001 | 57840000 | 0.0008 | 0.0007 | 0.0005 |
| F09 | 57760001 | 57860000 | 0.0014 | 0.0017 | 0.0012 |
| F09 | 57780001 | 57880000 | 0.0016 | 0.0020 | 0.0014 |
| F09 | 57800001 | 57900000 | 0.0019 | 0.0023 | 0.0017 |
| F09 | 57820001 | 57920000 | 0.0027 | 0.0029 | 0.0024 |
| F09 | 57840001 | 57940000 | 0.0029 | 0.0032 | 0.0027 |
| F09 | 57860001 | 57960000 | 0.0023 | 0.0026 | 0.0021 |
| F09 | 57880001 | 57980000 | 0.0023 | 0.0025 | 0.0021 |
| F09 | 57900001 | 58000000 | 0.0018 | 0.0022 | 0.0016 |
| F09 | 57920001 | 58020000 | 0.0018 | 0.0022 | 0.0016 |
| F09 | 57940001 | 58040000 | 0.0015 | 0.0020 | 0.0013 |
| F09 | 57960001 | 58060000 | 0.0015 | 0.0020 | 0.0013 |
| F09 | 57980001 | 58080000 | 0.0021 | 0.0026 | 0.0019 |
| F09 | 58000001 | 58100000 | 0.0024 | 0.0029 | 0.0022 |
| F09 | 58020001 | 58120000 | 0.0017 | 0.0022 | 0.0015 |
| F09 | 58040001 | 58140000 | 0.0019 | 0.0023 | 0.0017 |
| F09 | 58060001 | 58160000 | 0.0017 | 0.0021 | 0.0015 |
| F09 | 58080001 | 58180000 | 0.0011 | 0.0017 | 0.0009 |
| F09 | 58100001 | 58200000 | 0.0019 | 0.0024 | 0.0016 |
| F09 | 58120001 | 58220000 | 0.0023 | 0.0027 | 0.0021 |
| F09 | 58140001 | 58240000 | 0.0023 | 0.0030 | 0.0020 |

| Chr | Start | End | both | A1 | A2 |
|---|---|---|---|---|---|
| F09 | 58160001 | 58260000 | 0.0025 | 0.0032 | 0.0021 |
| F09 | 58180001 | 58280000 | 0.0024 | 0.0029 | 0.0020 |
| F09 | 58200001 | 58300000 | 0.0018 | 0.0021 | 0.0013 |
| F09 | 58220001 | 58320000 | 0.0015 | 0.0016 | 0.0009 |
| F09 | 58240001 | 58340000 | 0.0015 | 0.0011 | 0.0008 |
| F09 | 58260001 | 58360000 | 0.0016 | 0.0008 | 0.0009 |
| F09 | 58280001 | 58380000 | 0.0017 | 0.0007 | 0.0010 |
| F09 | 58300001 | 58400000 | 0.0013 | 0.0005 | 0.0008 |
| F09 | 58320001 | 58420000 | 0.0013 | 0.0007 | 0.0009 |
| F09 | 58340001 | 58440000 | 0.0015 | 0.0010 | 0.0012 |
| F09 | 58360001 | 58460000 | 0.0031 | 0.0023 | 0.0027 |
| F09 | 58380001 | 58480000 | 0.0035 | 0.0026 | 0.0031 |
| F09 | 58400001 | 58500000 | 0.0037 | 0.0030 | 0.0033 |
| F09 | 58420001 | 58520000 | 0.0042 | 0.0034 | 0.0038 |
| F09 | 58440001 | 58540000 | 0.0041 | 0.0034 | 0.0037 |
| F09 | 58460001 | 58560000 | 0.0028 | 0.0023 | 0.0023 |
| F09 | 58480001 | 58580000 | 0.0029 | 0.0023 | 0.0024 |
| F09 | 58500001 | 58600000 | 0.0028 | 0.0019 | 0.0022 |
| F09 | 58520001 | 58620000 | 0.0025 | 0.0013 | 0.0016 |
| F09 | 58540001 | 58640000 | 0.0028 | 0.0011 | 0.0018 |
| F09 | 58560001 | 58660000 | 0.0033 | 0.0015 | 0.0024 |
| F09 | 58580001 | 58680000 | 0.0028 | 0.0012 | 0.0020 |
| F09 | 58600001 | 58700000 | 0.0036 | 0.0021 | 0.0030 |
| F09 | 58620001 | 58720000 | 0.0038 | 0.0024 | 0.0032 |
| F09 | 58640001 | 58740000 | 0.0033 | 0.0023 | 0.0029 |
| F09 | 58660001 | 58760000 | 0.0024 | 0.0015 | 0.0021 |
| F09 | 58680001 | 58780000 | 0.0028 | 0.0016 | 0.0024 |
| F09 | 58700001 | 58800000 | 0.0022 | 0.0007 | 0.0017 |
| F09 | 58720001 | 58820000 | 0.0019 | 0.0007 | 0.0015 |
| F09 | 58740001 | 58840000 | 0.0021 | 0.0005 | 0.0015 |
| F09 | 58760001 | 58860000 | 0.0022 | 0.0006 | 0.0016 |
| F09 | 58780001 | 58880000 | 0.0021 | 0.0007 | 0.0016 |
| F09 | 58800001 | 58900000 | 0.0023 | 0.0009 | 0.0018 |
| F09 | 58820001 | 58920000 | 0.0023 | 0.0007 | 0.0017 |
| F09 | 58840001 | 58940000 | 0.0030 | 0.0015 | 0.0024 |
| F09 | 58860001 | 58960000 | 0.0028 | 0.0015 | 0.0022 |
| F09 | 58880001 | 58980000 | 0.0025 | 0.0014 | 0.0020 |
| F09 | 58900001 | 59000000 | 0.0020 | 0.0013 | 0.0017 |
| F09 | 58920001 | 59020000 | 0.0018 | 0.0012 | 0.0015 |
| F09 | 58940001 | 59040000 | 0.0015 | 0.0010 | 0.0012 |
| F09 | 58960001 | 59060000 | 0.0018 | 0.0013 | 0.0016 |
| F09 | 58980001 | 59080000 | 0.0016 | 0.0012 | 0.0014 |
| F09 | 59000001 | 59100000 | 0.0016 | 0.0013 | 0.0014 |
| F09 | 59020001 | 59120000 | 0.0019 | 0.0015 | 0.0017 |
| F09 | 59040001 | 59140000 | 0.0015 | 0.0012 | 0.0014 |
| F09 | 59060001 | 59160000 | 0.0015 | 0.0010 | 0.0013 |
| F09 | 59080001 | 59180000 | 0.0016 | 0.0010 | 0.0014 |
| F09 | 59100001 | 59200000 | 0.0019 | 0.0012 | 0.0017 |
| F09 | 59120001 | 59220000 | 0.0021 | 0.0011 | 0.0017 |
| F09 | 59140001 | 59240000 | 0.0020 | 0.0010 | 0.0014 |
| F09 | 59160001 | 59260000 | 0.0022 | 0.0013 | 0.0016 |
| F09 | 59180001 | 59280000 | 0.0022 | 0.0012 | 0.0015 |
| F09 | 59200001 | 59300000 | 0.0020 | 0.0010 | 0.0012 |
| F09 | 59220001 | 59320000 | 0.0017 | 0.0009 | 0.0010 |
| F09 | 59240001 | 59340000 | 0.0020 | 0.0011 | 0.0012 |
| F09 | 59260001 | 59360000 | 0.0018 | 0.0009 | 0.0010 |
| F09 | 59280001 | 59380000 | 0.0022 | 0.0011 | 0.0012 |
| F09 | 59300001 | 59400000 | 0.0030 | 0.0016 | 0.0017 |

| Chr | Start | End | both | A1 | A2 |
|---|---|---|---|---|---|
| F09 | 59320001 | 59420000 | 0.0030 | 0.0016 | 0.0018 |
| F09 | 59340001 | 59440000 | 0.0030 | 0.0015 | 0.0017 |
| F09 | 59360001 | 59460000 | 0.0027 | 0.0014 | 0.0015 |
| F09 | 59380001 | 59480000 | 0.0032 | 0.0021 | 0.0023 |
| F09 | 59400001 | 59500000 | 0.0023 | 0.0013 | 0.0015 |
| F09 | 59420001 | 59520000 | 0.0022 | 0.0013 | 0.0014 |
| F09 | 59440001 | 59540000 | 0.0023 | 0.0014 | 0.0016 |
| F09 | 59460001 | 59560000 | 0.0032 | 0.0022 | 0.0025 |
| F09 | 59480001 | 59580000 | 0.0029 | 0.0018 | 0.0021 |
| F09 | 59500001 | 59600000 | 0.0030 | 0.0018 | 0.0021 |
| F09 | 59520001 | 59620000 | 0.0030 | 0.0018 | 0.0022 |
| F09 | 59540001 | 59640000 | 0.0028 | 0.0018 | 0.0021 |
| F09 | 59560001 | 59660000 | 0.0018 | 0.0009 | 0.0011 |
| F09 | 59580001 | 59680000 | 0.0014 | 0.0005 | 0.0008 |
| F09 | 59600001 | 59700000 | 0.0016 | 0.0008 | 0.0011 |
| F09 | 59620001 | 59720000 | 0.0015 | 0.0009 | 0.0012 |
| F09 | 59640001 | 59740000 | 0.0017 | 0.0008 | 0.0013 |
| F09 | 59660001 | 59760000 | 0.0026 | 0.0016 | 0.0022 |
| F09 | 59680001 | 59780000 | 0.0029 | 0.0018 | 0.0025 |
| F09 | 59700001 | 59800000 | 0.0027 | 0.0015 | 0.0022 |
| F09 | 59720001 | 59820000 | 0.0032 | 0.0018 | 0.0025 |
| F09 | 59740001 | 59840000 | 0.0032 | 0.0019 | 0.0023 |
| F09 | 59760001 | 59860000 | 0.0024 | 0.0011 | 0.0014 |
| F09 | 59780001 | 59880000 | 0.0021 | 0.0009 | 0.0010 |
| F09 | 59800001 | 59900000 | 0.0019 | 0.0009 | 0.0010 |
| F09 | 59820001 | 59920000 | 0.0015 | 0.0005 | 0.0006 |
| F09 | 59840001 | 59940000 | 0.0013 | 0.0004 | 0.0005 |
| F09 | 59860001 | 59960000 | 0.0012 | 0.0003 | 0.0004 |
| F09 | 59880001 | 59980000 | 0.0012 | 0.0005 | 0.0005 |
| F09 | 59900001 | 60000000 | 0.0013 | 0.0005 | 0.0006 |
| F09 | 59920001 | 60020000 | 0.0015 | 0.0005 | 0.0006 |
| F09 | 59940001 | 60040000 | 0.0016 | 0.0009 | 0.0009 |
| F09 | 59960001 | 60060000 | 0.0016 | 0.0008 | 0.0009 |
| F09 | 59980001 | 60080000 | 0.0020 | 0.0010 | 0.0012 |
| F09 | 60000001 | 60100000 | 0.0022 | 0.0012 | 0.0013 |
| F09 | 60020001 | 60120000 | 0.0025 | 0.0017 | 0.0018 |
| F09 | 60040001 | 60140000 | 0.0023 | 0.0013 | 0.0015 |
| F09 | 60060001 | 60160000 | 0.0024 | 0.0015 | 0.0016 |
| F09 | 60080001 | 60180000 | 0.0024 | 0.0016 | 0.0017 |
| F09 | 60100001 | 60200000 | 0.0023 | 0.0015 | 0.0015 |
| F09 | 60120001 | 60220000 | 0.0017 | 0.0009 | 0.0009 |
| F09 | 60140001 | 60240000 | 0.0022 | 0.0012 | 0.0013 |
| F09 | 60160001 | 60260000 | 0.0021 | 0.0012 | 0.0013 |
| F09 | 60180001 | 60280000 | 0.0019 | 0.0011 | 0.0011 |
| F09 | 60200001 | 60300000 | 0.0021 | 0.0013 | 0.0014 |
| F09 | 60220001 | 60320000 | 0.0028 | 0.0020 | 0.0022 |
| F09 | 60240001 | 60340000 | 0.0027 | 0.0018 | 0.0021 |
| F09 | 60260001 | 60360000 | 0.0031 | 0.0022 | 0.0025 |
| F09 | 60280001 | 60380000 | 0.0030 | 0.0020 | 0.0024 |
| F09 | 60300001 | 60400000 | 0.0031 | 0.0023 | 0.0027 |
| F09 | 60320001 | 60420000 | 0.0025 | 0.0017 | 0.0020 |
| F09 | 60340001 | 60440000 | 0.0023 | 0.0016 | 0.0019 |
| F09 | 60360001 | 60460000 | 0.0019 | 0.0012 | 0.0015 |
| F09 | 60380001 | 60480000 | 0.0019 | 0.0011 | 0.0013 |
| F09 | 60400001 | 60500000 | 0.0016 | 0.0007 | 0.0008 |
| F09 | 60420001 | 60520000 | 0.0022 | 0.0011 | 0.0013 |
| F09 | 60440001 | 60540000 | 0.0025 | 0.0013 | 0.0015 |
| F09 | 60460001 | 60560000 | 0.0025 | 0.0013 | 0.0014 |

| Chr | Start | End | both | A1 | A2 |
|---|---|---|---|---|---|
| F09 | 60480001 | 60580000 | 0.0029 | 0.0017 | 0.0020 |
| F09 | 60500001 | 60600000 | 0.0037 | 0.0027 | 0.0029 |
| F09 | 60520001 | 60620000 | 0.0033 | 0.0023 | 0.0025 |
| F09 | 60540001 | 60640000 | 0.0029 | 0.0019 | 0.0020 |
| F09 | 60560001 | 60660000 | 0.0032 | 0.0022 | 0.0023 |
| F09 | 60580001 | 60680000 | 0.0030 | 0.0018 | 0.0018 |
| F09 | 60600001 | 60700000 | 0.0021 | 0.0008 | 0.0008 |
| F09 | 60620001 | 60720000 | 0.0019 | 0.0007 | 0.0007 |
| F09 | 60640001 | 60740000 | 0.0018 | 0.0007 | 0.0007 |
| F09 | 60660001 | 60760000 | 0.0016 | 0.0006 | 0.0005 |
| F09 | 60680001 | 60780000 | 0.0013 | 0.0004 | 0.0004 |
| F09 | 60700001 | 60800000 | 0.0013 | 0.0004 | 0.0004 |
| F09 | 60720001 | 60820000 | 0.0015 | 0.0004 | 0.0005 |
| F09 | 60740001 | 60840000 | 0.0018 | 0.0007 | 0.0008 |
| F09 | 60760001 | 60860000 | 0.0018 | 0.0006 | 0.0009 |
| F09 | 60780001 | 60880000 | 0.0020 | 0.0008 | 0.0011 |
| F09 | 60800001 | 60900000 | 0.0025 | 0.0015 | 0.0017 |
| F09 | 60820001 | 60920000 | 0.0025 | 0.0015 | 0.0017 |
| F09 | 60840001 | 60940000 | 0.0023 | 0.0013 | 0.0014 |
| F09 | 60860001 | 60960000 | 0.0021 | 0.0012 | 0.0012 |
| F09 | 60880001 | 60980000 | 0.0020 | 0.0012 | 0.0012 |
| F09 | 60900001 | 61000000 | 0.0014 | 0.0005 | 0.0006 |
| F09 | 60920001 | 61020000 | 0.0016 | 0.0007 | 0.0008 |
| F09 | 60940001 | 61040000 | 0.0026 | 0.0015 | 0.0020 |
| F09 | 60960001 | 61060000 | 0.0027 | 0.0017 | 0.0022 |
| F09 | 60980001 | 61080000 | 0.0026 | 0.0014 | 0.0019 |
| F09 | 61000001 | 61100000 | 0.0035 | 0.0024 | 0.0029 |
| F09 | 61020001 | 61120000 | 0.0034 | 0.0025 | 0.0030 |
| F09 | 61040001 | 61140000 | 0.0025 | 0.0018 | 0.0019 |
| F09 | 61060001 | 61160000 | 0.0026 | 0.0019 | 0.0020 |
| F09 | 61080001 | 61180000 | 0.0027 | 0.0022 | 0.0023 |
| F09 | 61100001 | 61200000 | 0.0017 | 0.0012 | 0.0012 |
| F09 | 61120001 | 61220000 | 0.0019 | 0.0012 | 0.0014 |
| F09 | 61140001 | 61240000 | 0.0025 | 0.0018 | 0.0021 |
| F09 | 61160001 | 61260000 | 0.0037 | 0.0026 | 0.0033 |
| F09 | 61180001 | 61280000 | 0.0035 | 0.0023 | 0.0030 |
| F09 | 61200001 | 61300000 | 0.0036 | 0.0024 | 0.0031 |
| F09 | 61220001 | 61320000 | 0.0042 | 0.0031 | 0.0038 |
| F09 | 61240001 | 61340000 | 0.0036 | 0.0025 | 0.0031 |
| F09 | 61260001 | 61360000 | 0.0024 | 0.0016 | 0.0018 |
| F09 | 61280001 | 61380000 | 0.0023 | 0.0016 | 0.0019 |
| F09 | 61300001 | 61400000 | 0.0024 | 0.0018 | 0.0020 |
| F09 | 61320001 | 61420000 | 0.0014 | 0.0007 | 0.0008 |
| F09 | 61340001 | 61440000 | 0.0016 | 0.0008 | 0.0010 |
| F09 | 61360001 | 61460000 | 0.0015 | 0.0007 | 0.0009 |
| F09 | 61380001 | 61480000 | 0.0017 | 0.0007 | 0.0008 |
| F09 | 61400001 | 61500000 | 0.0015 | 0.0005 | 0.0006 |
| F09 | 61420001 | 61520000 | 0.0022 | 0.0012 | 0.0014 |
| F09 | 61440001 | 61540000 | 0.0020 | 0.0012 | 0.0012 |
| F09 | 61460001 | 61560000 | 0.0024 | 0.0016 | 0.0016 |
| F09 | 61480001 | 61580000 | 0.0023 | 0.0017 | 0.0017 |
| F09 | 61500001 | 61600000 | 0.0022 | 0.0018 | 0.0018 |
| F09 | 61520001 | 61620000 | 0.0014 | 0.0011 | 0.0011 |
| F09 | 61540001 | 61640000 | 0.0013 | 0.0010 | 0.0011 |
| F09 | 61560001 | 61660000 | 0.0008 | 0.0007 | 0.0006 |
| F09 | 61580001 | 61680000 | 0.0008 | 0.0008 | 0.0005 |
| F09 | 61600001 | 61700000 | 0.0009 | 0.0010 | 0.0004 |
| F09 | 61620001 | 61720000 | 0.0011 | 0.0012 | 0.0005 |

| Chr | Start | End | both | A1 | A2 |
|---|---|---|---|---|---|
| F09 | 61640001 | 61740000 | 0.0013 | 0.0015 | 0.0006 |
| F09 | 61660001 | 61760000 | 0.0017 | 0.0019 | 0.0010 |
| F09 | 61680001 | 61780000 | 0.0019 | 0.0020 | 0.0013 |
| F09 | 61700001 | 61800000 | 0.0018 | 0.0019 | 0.0012 |
| F09 | 61720001 | 61820000 | 0.0017 | 0.0018 | 0.0011 |
| F09 | 61740001 | 61840000 | 0.0016 | 0.0017 | 0.0010 |
| F09 | 61760001 | 61860000 | 0.0012 | 0.0013 | 0.0005 |
| F09 | 61780001 | 61880000 | 0.0012 | 0.0013 | 0.0004 |
| F09 | 61800001 | 61900000 | 0.0016 | 0.0017 | 0.0009 |
| F09 | 61820001 | 61920000 | 0.0019 | 0.0019 | 0.0012 |
| F09 | 61840001 | 61940000 | 0.0018 | 0.0018 | 0.0011 |
| F09 | 61860001 | 61960000 | 0.0019 | 0.0019 | 0.0012 |
| F09 | 61880001 | 61980000 | 0.0021 | 0.0021 | 0.0015 |
| F09 | 61900001 | 62000000 | 0.0016 | 0.0016 | 0.0011 |
| F09 | 61920001 | 62020000 | 0.0016 | 0.0017 | 0.0008 |
| F09 | 61940001 | 62040000 | 0.0015 | 0.0017 | 0.0008 |
| F09 | 61960001 | 62060000 | 0.0019 | 0.0020 | 0.0012 |
| F09 | 61980001 | 62080000 | 0.0016 | 0.0017 | 0.0008 |
| F09 | 62000001 | 62100000 | 0.0029 | 0.0030 | 0.0020 |
| F09 | 62020001 | 62120000 | 0.0037 | 0.0037 | 0.0029 |
| F09 | 62040001 | 62140000 | 0.0039 | 0.0039 | 0.0031 |
| F09 | 62060001 | 62160000 | 0.0042 | 0.0043 | 0.0034 |
| F09 | 62080001 | 62180000 | 0.0045 | 0.0044 | 0.0037 |
| F09 | 62100001 | 62200000 | 0.0033 | 0.0032 | 0.0026 |
| F09 | 62120001 | 62220000 | 0.0028 | 0.0027 | 0.0020 |
| F09 | 62140001 | 62240000 | 0.0027 | 0.0027 | 0.0020 |
| F09 | 62160001 | 62260000 | 0.0022 | 0.0022 | 0.0015 |
| F09 | 62180001 | 62280000 | 0.0020 | 0.0020 | 0.0013 |
| F09 | 62200001 | 62300000 | 0.0019 | 0.0020 | 0.0012 |
| F09 | 62220001 | 62320000 | 0.0015 | 0.0016 | 0.0007 |
| F09 | 62240001 | 62340000 | 0.0013 | 0.0014 | 0.0006 |
| F09 | 62260001 | 62360000 | 0.0015 | 0.0016 | 0.0008 |
| F09 | 62280001 | 62380000 | 0.0018 | 0.0020 | 0.0012 |
| F09 | 62300001 | 62400000 | 0.0018 | 0.0020 | 0.0011 |
| F09 | 62320001 | 62420000 | 0.0017 | 0.0019 | 0.0010 |
| F09 | 62340001 | 62440000 | 0.0019 | 0.0020 | 0.0011 |
| F09 | 62360001 | 62460000 | 0.0015 | 0.0017 | 0.0007 |
| F09 | 62380001 | 62480000 | 0.0012 | 0.0014 | 0.0003 |
| F09 | 62400001 | 62500000 | 0.0012 | 0.0014 | 0.0004 |
| F09 | 62420001 | 62520000 | 0.0013 | 0.0012 | 0.0005 |
| F09 | 62440001 | 62540000 | 0.0013 | 0.0009 | 0.0004 |
| F09 | 62460001 | 62560000 | 0.0013 | 0.0006 | 0.0004 |
| F09 | 62480001 | 62580000 | 0.0012 | 0.0003 | 0.0004 |
| F09 | 62500001 | 62600000 | 0.0013 | 0.0003 | 0.0004 |
| F09 | 62520001 | 62620000 | 0.0013 | 0.0004 | 0.0003 |
| F09 | 62540001 | 62640000 | 0.0015 | 0.0005 | 0.0005 |
| F09 | 62560001 | 62660000 | 0.0014 | 0.0007 | 0.0008 |
| F09 | 62580001 | 62680000 | 0.0016 | 0.0010 | 0.0010 |
| F09 | 62600001 | 62700000 | 0.0016 | 0.0009 | 0.0010 |
| F09 | 62620001 | 62720000 | 0.0018 | 0.0010 | 0.0010 |
| F09 | 62640001 | 62740000 | 0.0018 | 0.0010 | 0.0010 |
| F09 | 62660001 | 62760000 | 0.0017 | 0.0008 | 0.0008 |
| F09 | 62680001 | 62780000 | 0.0015 | 0.0005 | 0.0005 |
| F09 | 62700001 | 62800000 | 0.0015 | 0.0005 | 0.0005 |
| F09 | 62720001 | 62820000 | 0.0016 | 0.0005 | 0.0007 |
| F09 | 62740001 | 62840000 | 0.0015 | 0.0007 | 0.0009 |
| F09 | 62760001 | 62860000 | 0.0016 | 0.0006 | 0.0008 |
| F09 | 62780001 | 62880000 | 0.0020 | 0.0009 | 0.0011 |

| Chr | Start | End | both | A1 | A2 |
|---|---|---|---|---|---|
| F09 | 62800001 | 62900000 | 0.0023 | 0.0014 | 0.0015 |
| F09 | 62820001 | 62920000 | 0.0020 | 0.0013 | 0.0013 |
| F09 | 62840001 | 62940000 | 0.0023 | 0.0014 | 0.0015 |
| F09 | 62860001 | 62960000 | 0.0031 | 0.0022 | 0.0025 |
| F09 | 62880001 | 62980000 | 0.0029 | 0.0020 | 0.0023 |
| F09 | 62900001 | 63000000 | 0.0035 | 0.0026 | 0.0029 |
| F09 | 62920001 | 63020000 | 0.0042 | 0.0032 | 0.0036 |
| F09 | 62940001 | 63040000 | 0.0041 | 0.0032 | 0.0037 |
| F09 | 62960001 | 63060000 | 0.0043 | 0.0035 | 0.0039 |
| F09 | 62980001 | 63080000 | 0.0045 | 0.0037 | 0.0041 |
| F09 | 63000001 | 63100000 | 0.0036 | 0.0027 | 0.0032 |
| F09 | 63020001 | 63120000 | 0.0030 | 0.0021 | 0.0025 |
| F09 | 63040001 | 63140000 | 0.0029 | 0.0018 | 0.0021 |
| F09 | 63060001 | 63160000 | 0.0018 | 0.0009 | 0.0010 |
| F09 | 63080001 | 63180000 | 0.0017 | 0.0007 | 0.0008 |
| F09 | 63100001 | 63200000 | 0.0015 | 0.0006 | 0.0006 |
| F09 | 63120001 | 63220000 | 0.0020 | 0.0009 | 0.0011 |
| F09 | 63140001 | 63240000 | 0.0019 | 0.0008 | 0.0009 |
| F09 | 63160001 | 63260000 | 0.0019 | 0.0007 | 0.0008 |
| F09 | 63180001 | 63280000 | 0.0022 | 0.0012 | 0.0014 |
| F09 | 63200001 | 63300000 | 0.0022 | 0.0012 | 0.0014 |
| F09 | 63220001 | 63320000 | 0.0018 | 0.0008 | 0.0010 |
| F09 | 63240001 | 63340000 | 0.0019 | 0.0008 | 0.0010 |
| F09 | 63260001 | 63360000 | 0.0019 | 0.0007 | 0.0010 |
| F09 | 63280001 | 63380000 | 0.0014 | 0.0001 | 0.0003 |
| F09 | 63300001 | 63400000 | 0.0014 | 0.0002 | 0.0004 |
| F09 | 63320001 | 63420000 | 0.0012 | 0.0002 | 0.0003 |
| F09 | 63340001 | 63440000 | 0.0012 | 0.0003 | 0.0004 |
| F09 | 63360001 | 63460000 | 0.0013 | 0.0003 | 0.0005 |
| F09 | 63380001 | 63480000 | 0.0013 | 0.0003 | 0.0005 |
| F09 | 63400001 | 63500000 | 0.0013 | 0.0002 | 0.0004 |
| F09 | 63420001 | 63520000 | 0.0015 | 0.0002 | 0.0005 |
| F09 | 63440001 | 63540000 | 0.0016 | 0.0003 | 0.0005 |
| F09 | 63460001 | 63560000 | 0.0015 | 0.0003 | 0.0005 |
| F09 | 63480001 | 63580000 | 0.0015 | 0.0003 | 0.0005 |
| F09 | 63500001 | 63600000 | 0.0014 | 0.0003 | 0.0005 |
| F09 | 63520001 | 63620000 | 0.0013 | 0.0003 | 0.0006 |
| F09 | 63540001 | 63640000 | 0.0013 | 0.0002 | 0.0005 |
| F09 | 63560001 | 63660000 | 0.0013 | 0.0002 | 0.0005 |
| F09 | 63580001 | 63680000 | 0.0013 | 0.0002 | 0.0005 |
| F09 | 63600001 | 63700000 | 0.0013 | 0.0002 | 0.0004 |
| F09 | 63620001 | 63720000 | 0.0014 | 0.0002 | 0.0004 |
| F09 | 63640001 | 63740000 | 0.0013 | 0.0002 | 0.0003 |
| F09 | 63660001 | 63760000 | 0.0014 | 0.0002 | 0.0003 |
| F09 | 63680001 | 63780000 | 0.0014 | 0.0003 | 0.0005 |
| F09 | 63700001 | 63800000 | 0.0013 | 0.0004 | 0.0006 |
| F09 | 63720001 | 63820000 | 0.0014 | 0.0005 | 0.0007 |
| F09 | 63740001 | 63840000 | 0.0023 | 0.0014 | 0.0016 |
| F09 | 63760001 | 63860000 | 0.0025 | 0.0017 | 0.0018 |
| F09 | 63780001 | 63880000 | 0.0037 | 0.0026 | 0.0028 |
| F09 | 63800001 | 63900000 | 0.0038 | 0.0027 | 0.0029 |
| F09 | 63820001 | 63920000 | 0.0039 | 0.0029 | 0.0030 |
| F09 | 63840001 | 63940000 | 0.0031 | 0.0020 | 0.0021 |
| F09 | 63860001 | 63960000 | 0.0029 | 0.0017 | 0.0018 |
| F09 | 63880001 | 63980000 | 0.0017 | 0.0006 | 0.0007 |
| F09 | 63900001 | 64000000 | 0.0015 | 0.0004 | 0.0006 |
| F09 | 63920001 | 64020000 | 0.0013 | 0.0001 | 0.0003 |
| F09 | 63940001 | 64040000 | 0.0013 | 0.0001 | 0.0003 |

| Chr | Start | End | both | A1 | A2 |
|---|---|---|---|---|---|
| F09 | 63960001 | 64060000 | 0.0014 | 0.0003 | 0.0006 |
| F09 | 63980001 | 64080000 | 0.0017 | 0.0006 | 0.0009 |
| F09 | 64000001 | 64100000 | 0.0016 | 0.0006 | 0.0008 |
| F09 | 64020001 | 64120000 | 0.0016 | 0.0007 | 0.0009 |
| F09 | 64040001 | 64140000 | 0.0021 | 0.0013 | 0.0015 |
| F09 | 64060001 | 64160000 | 0.0021 | 0.0015 | 0.0014 |
| F09 | 64080001 | 64180000 | 0.0018 | 0.0013 | 0.0011 |
| F09 | 64100001 | 64200000 | 0.0019 | 0.0014 | 0.0012 |
| F09 | 64120001 | 64220000 | 0.0020 | 0.0015 | 0.0013 |
| F09 | 64140001 | 64240000 | 0.0015 | 0.0009 | 0.0008 |
| F09 | 64160001 | 64260000 | 0.0014 | 0.0005 | 0.0006 |
| F09 | 64180001 | 64280000 | 0.0015 | 0.0005 | 0.0007 |
| F09 | 64200001 | 64300000 | 0.0014 | 0.0004 | 0.0005 |
| F09 | 64220001 | 64320000 | 0.0015 | 0.0006 | 0.0007 |
| F09 | 64240001 | 64340000 | 0.0016 | 0.0006 | 0.0008 |
| F09 | 64260001 | 64360000 | 0.0017 | 0.0006 | 0.0008 |
| F09 | 64280001 | 64380000 | 0.0016 | 0.0006 | 0.0008 |
| F09 | 64300001 | 64400000 | 0.0017 | 0.0008 | 0.0009 |
| F09 | 64320001 | 64420000 | 0.0018 | 0.0006 | 0.0008 |
| F09 | 64340001 | 64440000 | 0.0018 | 0.0005 | 0.0006 |
| F09 | 64360001 | 64460000 | 0.0019 | 0.0005 | 0.0006 |
| F09 | 64380001 | 64480000 | 0.0026 | 0.0006 | 0.0011 |
| F09 | 64400001 | 64500000 | 0.0026 | 0.0005 | 0.0011 |
| F09 | 64420001 | 64520000 | 0.0024 | 0.0005 | 0.0009 |
| F09 | 64440001 | 64540000 | 0.0022 | 0.0006 | 0.0009 |
| F09 | 64460001 | 64560000 | 0.0020 | 0.0006 | 0.0009 |
| F09 | 64480001 | 64580000 | 0.0013 | 0.0005 | 0.0004 |
| F09 | 64500001 | 64600000 | 0.0010 | 0.0005 | 0.0002 |
| F09 | 64520001 | 64620000 | 0.0012 | 0.0005 | 0.0004 |
| F09 | 64540001 | 64640000 | 0.0013 | 0.0007 | 0.0005 |
| F09 | 64560001 | 64660000 | 0.0012 | 0.0009 | 0.0006 |
| F09 | 64580001 | 64680000 | 0.0014 | 0.0014 | 0.0008 |
| F09 | 64600001 | 64700000 | 0.0016 | 0.0018 | 0.0008 |
| F09 | 64620001 | 64720000 | 0.0013 | 0.0018 | 0.0005 |
| F09 | 64640001 | 64740000 | 0.0012 | 0.0016 | 0.0004 |
| F09 | 64660001 | 64760000 | 0.0017 | 0.0018 | 0.0010 |
| F09 | 64680001 | 64780000 | 0.0018 | 0.0017 | 0.0011 |
| F09 | 64700001 | 64800000 | 0.0023 | 0.0017 | 0.0015 |
| F09 | 64720001 | 64820000 | 0.0026 | 0.0018 | 0.0019 |
| F09 | 64740001 | 64840000 | 0.0029 | 0.0020 | 0.0021 |
| F09 | 64760001 | 64860000 | 0.0025 | 0.0017 | 0.0016 |
| F09 | 64780001 | 64880000 | 0.0022 | 0.0014 | 0.0013 |
| F09 | 64800001 | 64900000 | 0.0017 | 0.0010 | 0.0009 |
| F09 | 64820001 | 64920000 | 0.0015 | 0.0008 | 0.0007 |
| F09 | 64840001 | 64940000 | 0.0013 | 0.0006 | 0.0006 |
| F09 | 64860001 | 64960000 | 0.0012 | 0.0007 | 0.0006 |
| F09 | 64880001 | 64980000 | 0.0012 | 0.0010 | 0.0006 |
| F09 | 64900001 | 65000000 | 0.0012 | 0.0013 | 0.0006 |
| F09 | 64920001 | 65020000 | 0.0011 | 0.0015 | 0.0006 |
| F09 | 64940001 | 65040000 | 0.0023 | 0.0027 | 0.0017 |
| F09 | 64960001 | 65060000 | 0.0023 | 0.0026 | 0.0018 |
| F09 | 64980001 | 65080000 | 0.0023 | 0.0025 | 0.0019 |
| F09 | 65000001 | 65100000 | 0.0026 | 0.0024 | 0.0022 |
| F09 | 65020001 | 65120000 | 0.0031 | 0.0027 | 0.0028 |
| F09 | 65040001 | 65140000 | 0.0019 | 0.0014 | 0.0016 |
| F09 | 65060001 | 65160000 | 0.0027 | 0.0020 | 0.0023 |
| F09 | 65080001 | 65180000 | 0.0027 | 0.0019 | 0.0023 |
| F09 | 65100001 | 65200000 | 0.0023 | 0.0015 | 0.0019 |

| Chr | Start | End | both | A1 | A2 |
|---|---|---|---|---|---|
| F09 | 65120001 | 65220000 | 0.0019 | 0.0011 | 0.0013 |
| F09 | 65140001 | 65240000 | 0.0018 | 0.0009 | 0.0010 |
| F09 | 65160001 | 65260000 | 0.0012 | 0.0003 | 0.0004 |
| F09 | 65180001 | 65280000 | 0.0012 | 0.0002 | 0.0003 |
| F09 | 65200001 | 65300000 | 0.0013 | 0.0002 | 0.0004 |
| F09 | 65220001 | 65320000 | 0.0013 | 0.0002 | 0.0003 |
| F09 | 65240001 | 65340000 | 0.0013 | 0.0003 | 0.0004 |
| F09 | 65260001 | 65360000 | 0.0012 | 0.0001 | 0.0002 |
| F09 | 65280001 | 65380000 | 0.0012 | 0.0002 | 0.0002 |
| F09 | 65300001 | 65400000 | 0.0016 | 0.0007 | 0.0007 |
| F09 | 65320001 | 65420000 | 0.0017 | 0.0007 | 0.0007 |
| F09 | 65340001 | 65440000 | 0.0019 | 0.0009 | 0.0010 |
| F09 | 65360001 | 65460000 | 0.0020 | 0.0009 | 0.0010 |
| F09 | 65380001 | 65480000 | 0.0020 | 0.0009 | 0.0011 |
| F09 | 65400001 | 65500000 | 0.0016 | 0.0004 | 0.0006 |
| F09 | 65420001 | 65520000 | 0.0016 | 0.0004 | 0.0006 |
| F09 | 65440001 | 65540000 | 0.0015 | 0.0002 | 0.0004 |
| F09 | 65460001 | 65560000 | 0.0014 | 0.0003 | 0.0005 |
| F09 | 65480001 | 65580000 | 0.0014 | 0.0003 | 0.0004 |
| F09 | 65500001 | 65600000 | 0.0015 | 0.0004 | 0.0005 |
| F09 | 65520001 | 65620000 | 0.0026 | 0.0016 | 0.0019 |
| F09 | 65540001 | 65640000 | 0.0038 | 0.0026 | 0.0031 |
| F09 | 65560001 | 65660000 | 0.0042 | 0.0031 | 0.0035 |
| F09 | 65580001 | 65680000 | 0.0041 | 0.0031 | 0.0034 |
| F09 | 65600001 | 65700000 | 0.0043 | 0.0034 | 0.0037 |
| F09 | 65620001 | 65720000 | 0.0031 | 0.0021 | 0.0024 |
| F09 | 65640001 | 65740000 | 0.0018 | 0.0011 | 0.0011 |
| F09 | 65660001 | 65760000 | 0.0016 | 0.0005 | 0.0008 |
| F09 | 65680001 | 65780000 | 0.0018 | 0.0006 | 0.0009 |
| F09 | 65700001 | 65800000 | 0.0016 | 0.0003 | 0.0006 |
| F09 | 65720001 | 65820000 | 0.0015 | 0.0007 | 0.0006 |
| F09 | 65740001 | 65840000 | 0.0013 | 0.0011 | 0.0006 |
| F09 | 65760001 | 65860000 | 0.0012 | 0.0014 | 0.0005 |
| F09 | 65780001 | 65880000 | 0.0010 | 0.0014 | 0.0004 |
| F09 | 65800001 | 65900000 | 0.0008 | 0.0016 | 0.0003 |
| F09 | 65820001 | 65920000 | 0.0010 | 0.0014 | 0.0005 |
| F09 | 65840001 | 65940000 | 0.0012 | 0.0011 | 0.0005 |
| F09 | 65860001 | 65960000 | 0.0015 | 0.0011 | 0.0007 |
| F09 | 65880001 | 65980000 | 0.0017 | 0.0008 | 0.0007 |
| F09 | 65900001 | 66000000 | 0.0019 | 0.0008 | 0.0009 |
| F09 | 65920001 | 66020000 | 0.0019 | 0.0006 | 0.0008 |
| F09 | 65940001 | 66040000 | 0.0018 | 0.0005 | 0.0008 |
| F09 | 65960001 | 66060000 | 0.0016 | 0.0003 | 0.0006 |
| F09 | 65980001 | 66080000 | 0.0020 | 0.0009 | 0.0011 |
| F09 | 66000001 | 66100000 | 0.0027 | 0.0018 | 0.0020 |
| F09 | 66020001 | 66120000 | 0.0029 | 0.0022 | 0.0023 |
| F09 | 66040001 | 66140000 | 0.0029 | 0.0022 | 0.0023 |
| F09 | 66060001 | 66160000 | 0.0029 | 0.0022 | 0.0022 |
| F09 | 66080001 | 66180000 | 0.0027 | 0.0019 | 0.0019 |
| F09 | 66100001 | 66200000 | 0.0018 | 0.0008 | 0.0008 |
| F09 | 66120001 | 66220000 | 0.0016 | 0.0004 | 0.0005 |
| F09 | 66140001 | 66240000 | 0.0018 | 0.0005 | 0.0006 |
| F09 | 66160001 | 66260000 | 0.0019 | 0.0005 | 0.0006 |
| F09 | 66180001 | 66280000 | 0.0017 | 0.0003 | 0.0004 |
| F09 | 66200001 | 66300000 | 0.0017 | 0.0003 | 0.0004 |
| F09 | 66220001 | 66320000 | 0.0017 | 0.0003 | 0.0004 |
| F09 | 66240001 | 66340000 | 0.0015 | 0.0002 | 0.0002 |
| F09 | 66260001 | 66360000 | 0.0015 | 0.0002 | 0.0002 |

| Chr | Start | End | both | A1 | A2 |
|---|---|---|---|---|---|
| F09 | 66280001 | 66380000 | 0.0014 | 0.0001 | 0.0001 |
| F09 | 66300001 | 66400000 | 0.0021 | 0.0004 | 0.0009 |
| F09 | 66320001 | 66420000 | 0.0022 | 0.0008 | 0.0012 |
| F09 | 66340001 | 66440000 | 0.0025 | 0.0011 | 0.0016 |
| F09 | 66360001 | 66460000 | 0.0025 | 0.0011 | 0.0016 |
| F09 | 66380001 | 66480000 | 0.0026 | 0.0011 | 0.0016 |
| F09 | 66400001 | 66500000 | 0.0020 | 0.0009 | 0.0010 |
| F09 | 66420001 | 66520000 | 0.0022 | 0.0007 | 0.0014 |
| F09 | 66440001 | 66540000 | 0.0020 | 0.0004 | 0.0014 |
| F09 | 66460001 | 66560000 | 0.0021 | 0.0004 | 0.0017 |
| F09 | 66480001 | 66580000 | 0.0024 | 0.0008 | 0.0022 |
| F09 | 66500001 | 66600000 | 0.0026 | 0.0010 | 0.0026 |
| F09 | 66520001 | 66620000 | 0.0023 | 0.0008 | 0.0023 |
| F09 | 66540001 | 66640000 | 0.0023 | 0.0008 | 0.0022 |
| F09 | 66560001 | 66660000 | 0.0026 | 0.0009 | 0.0025 |
| F09 | 66580001 | 66680000 | 0.0026 | 0.0007 | 0.0023 |
| F09 | 66600001 | 66700000 | 0.0024 | 0.0003 | 0.0022 |
| F09 | 66620001 | 66720000 | 0.0023 | 0.0003 | 0.0021 |
| F09 | 66640001 | 66740000 | 0.0023 | 0.0003 | 0.0020 |
| F09 | 66660001 | 66760000 | 0.0020 | 0.0003 | 0.0015 |
| F09 | 66680001 | 66780000 | 0.0018 | 0.0002 | 0.0012 |
| F09 | 66700001 | 66800000 | 0.0021 | 0.0008 | 0.0014 |
| F09 | 66720001 | 66820000 | 0.0022 | 0.0009 | 0.0014 |
| F09 | 66740001 | 66840000 | 0.0024 | 0.0009 | 0.0017 |
| F09 | 66760001 | 66860000 | 0.0024 | 0.0009 | 0.0018 |
| F09 | 66780001 | 66880000 | 0.0022 | 0.0009 | 0.0018 |
| F09 | 66800001 | 66900000 | 0.0019 | 0.0003 | 0.0013 |
| F09 | 66820001 | 66920000 | 0.0018 | 0.0002 | 0.0011 |
| F09 | 66840001 | 66940000 | 0.0017 | 0.0003 | 0.0008 |
| F09 | 66860001 | 66960000 | 0.0015 | 0.0003 | 0.0006 |
| F09 | 66880001 | 66980000 | 0.0015 | 0.0003 | 0.0006 |
| F09 | 66900001 | 67000000 | 0.0013 | 0.0003 | 0.0006 |
| F09 | 66920001 | 67020000 | 0.0014 | 0.0006 | 0.0008 |
| F09 | 66940001 | 67040000 | 0.0015 | 0.0006 | 0.0010 |
| F09 | 66960001 | 67060000 | 0.0015 | 0.0006 | 0.0010 |
| F09 | 66980001 | 67080000 | 0.0016 | 0.0006 | 0.0010 |
| F09 | 67000001 | 67100000 | 0.0018 | 0.0006 | 0.0011 |
| F09 | 67020001 | 67120000 | 0.0016 | 0.0004 | 0.0009 |
| F09 | 67040001 | 67140000 | 0.0015 | 0.0003 | 0.0008 |
| F09 | 67060001 | 67160000 | 0.0018 | 0.0003 | 0.0010 |
| F09 | 67080001 | 67180000 | 0.0023 | 0.0005 | 0.0014 |
| F09 | 67100001 | 67200000 | 0.0025 | 0.0007 | 0.0017 |
| F09 | 67120001 | 67220000 | 0.0026 | 0.0007 | 0.0019 |
| F09 | 67140001 | 67240000 | 0.0026 | 0.0009 | 0.0021 |
| F09 | 67160001 | 67260000 | 0.0024 | 0.0009 | 0.0021 |
| F09 | 67180001 | 67280000 | 0.0021 | 0.0007 | 0.0019 |
| F09 | 67200001 | 67300000 | 0.0020 | 0.0006 | 0.0018 |
| F09 | 67220001 | 67320000 | 0.0020 | 0.0007 | 0.0019 |
| F09 | 67240001 | 67340000 | 0.0021 | 0.0004 | 0.0020 |
| F09 | 67260001 | 67360000 | 0.0024 | 0.0004 | 0.0020 |
| F09 | 67280001 | 67380000 | 0.0023 | 0.0004 | 0.0018 |
| F09 | 67300001 | 67400000 | 0.0023 | 0.0005 | 0.0016 |
| F09 | 67320001 | 67420000 | 0.0020 | 0.0007 | 0.0011 |
| F09 | 67340001 | 67440000 | 0.0018 | 0.0009 | 0.0008 |
| F09 | 67360001 | 67460000 | 0.0016 | 0.0010 | 0.0007 |
| F09 | 67380001 | 67480000 | 0.0017 | 0.0011 | 0.0010 |
| F09 | 67400001 | 67500000 | 0.0013 | 0.0008 | 0.0008 |
| F09 | 67420001 | 67520000 | 0.0013 | 0.0006 | 0.0009 |

| Chr | Start | End | both | A1 | A2 |
|---|---|---|---|---|---|
| F09 | 67440001 | 67540000 | 0.0017 | 0.0007 | 0.0012 |
| F09 | 67460001 | 67560000 | 0.0017 | 0.0008 | 0.0013 |
| F09 | 67480001 | 67580000 | 0.0019 | 0.0006 | 0.0013 |
| F09 | 67500001 | 67600000 | 0.0021 | 0.0006 | 0.0014 |
| F09 | 67520001 | 67620000 | 0.0021 | 0.0006 | 0.0014 |
| F09 | 67540001 | 67640000 | 0.0019 | 0.0006 | 0.0012 |
| F09 | 67560001 | 67660000 | 0.0016 | 0.0006 | 0.0010 |
| F09 | 67580001 | 67680000 | 0.0013 | 0.0007 | 0.0009 |
| F09 | 67600001 | 67700000 | 0.0015 | 0.0010 | 0.0011 |
| F09 | 67620001 | 67720000 | 0.0016 | 0.0013 | 0.0013 |
| F09 | 67640001 | 67740000 | 0.0014 | 0.0010 | 0.0011 |
| F09 | 67660001 | 67760000 | 0.0016 | 0.0008 | 0.0012 |
| F09 | 67680001 | 67780000 | 0.0017 | 0.0007 | 0.0012 |
| F09 | 67700001 | 67800000 | 0.0014 | 0.0004 | 0.0010 |
| F09 | 67720001 | 67820000 | 0.0014 | 0.0001 | 0.0008 |
| F09 | 67740001 | 67840000 | 0.0014 | 0.0001 | 0.0009 |
| F09 | 67760001 | 67860000 | 0.0013 | 0.0001 | 0.0008 |
| F09 | 67780001 | 67880000 | 0.0016 | 0.0004 | 0.0012 |
| F09 | 67800001 | 67900000 | 0.0017 | 0.0004 | 0.0014 |
| F09 | 67820001 | 67920000 | 0.0018 | 0.0004 | 0.0016 |
| F09 | 67840001 | 67940000 | 0.0018 | 0.0004 | 0.0016 |
| F09 | 67860001 | 67960000 | 0.0018 | 0.0004 | 0.0015 |
| F09 | 67880001 | 67980000 | 0.0014 | 0.0002 | 0.0010 |
| F09 | 67900001 | 68000000 | 0.0015 | 0.0002 | 0.0011 |
| F09 | 67920001 | 68020000 | 0.0016 | 0.0002 | 0.0010 |
| F09 | 67940001 | 68040000 | 0.0016 | 0.0002 | 0.0010 |
| F09 | 67960001 | 68060000 | 0.0018 | 0.0002 | 0.0009 |
| F09 | 67980001 | 68080000 | 0.0019 | 0.0004 | 0.0010 |
| F09 | 68000001 | 68100000 | 0.0017 | 0.0005 | 0.0007 |
| F09 | 68020001 | 68120000 | 0.0021 | 0.0010 | 0.0009 |
| F09 | 68040001 | 68140000 | 0.0024 | 0.0013 | 0.0011 |
| F09 | 68060001 | 68160000 | 0.0025 | 0.0014 | 0.0013 |
| F09 | 68080001 | 68180000 | 0.0023 | 0.0013 | 0.0012 |
| F09 | 68100001 | 68200000 | 0.0027 | 0.0017 | 0.0016 |
| F09 | 68120001 | 68220000 | 0.0021 | 0.0012 | 0.0010 |
| F09 | 68140001 | 68240000 | 0.0019 | 0.0009 | 0.0009 |
| F09 | 68160001 | 68260000 | 0.0017 | 0.0008 | 0.0008 |
| F09 | 68180001 | 68280000 | 0.0018 | 0.0008 | 0.0008 |
| F09 | 68200001 | 68300000 | 0.0013 | 0.0002 | 0.0004 |
| F09 | 68220001 | 68320000 | 0.0015 | 0.0004 | 0.0006 |
| F09 | 68240001 | 68340000 | 0.0015 | 0.0004 | 0.0006 |
| F09 | 68260001 | 68360000 | 0.0015 | 0.0004 | 0.0007 |
| F09 | 68280001 | 68380000 | 0.0016 | 0.0004 | 0.0008 |
| F09 | 68300001 | 68400000 | 0.0016 | 0.0004 | 0.0011 |
| F09 | 68320001 | 68420000 | 0.0015 | 0.0002 | 0.0011 |
| F09 | 68340001 | 68440000 | 0.0018 | 0.0006 | 0.0016 |
| F09 | 68360001 | 68460000 | 0.0019 | 0.0006 | 0.0018 |
| F09 | 68380001 | 68480000 | 0.0021 | 0.0008 | 0.0018 |
| F09 | 68400001 | 68500000 | 0.0020 | 0.0008 | 0.0016 |
| F09 | 68420001 | 68520000 | 0.0017 | 0.0008 | 0.0013 |
| F09 | 68440001 | 68540000 | 0.0014 | 0.0004 | 0.0008 |
| F09 | 68460001 | 68560000 | 0.0013 | 0.0008 | 0.0006 |
| F09 | 68480001 | 68580000 | 0.0011 | 0.0010 | 0.0005 |
| F09 | 68500001 | 68600000 | 0.0012 | 0.0011 | 0.0004 |
| F09 | 68520001 | 68620000 | 0.0016 | 0.0015 | 0.0008 |
| F09 | 68540001 | 68640000 | 0.0017 | 0.0015 | 0.0007 |
| F09 | 68560001 | 68660000 | 0.0017 | 0.0011 | 0.0006 |
| F09 | 68580001 | 68680000 | 0.0017 | 0.0008 | 0.0006 |

| Chr | Start | End | both | A1 | A2 |
|---|---|---|---|---|---|
| F09 | 68600001 | 68700000 | 0.0019 | 0.0009 | 0.0010 |
| F09 | 68620001 | 68720000 | 0.0017 | 0.0006 | 0.0007 |
| F09 | 68640001 | 68740000 | 0.0016 | 0.0006 | 0.0007 |
| F09 | 68660001 | 68760000 | 0.0020 | 0.0011 | 0.0012 |
| F09 | 68680001 | 68780000 | 0.0018 | 0.0010 | 0.0011 |
| F09 | 68700001 | 68800000 | 0.0016 | 0.0008 | 0.0008 |
| F09 | 68720001 | 68820000 | 0.0014 | 0.0007 | 0.0006 |
| F09 | 68740001 | 68840000 | 0.0015 | 0.0007 | 0.0008 |
| F09 | 68760001 | 68860000 | 0.0010 | 0.0002 | 0.0003 |
| F09 | 68780001 | 68880000 | 0.0012 | 0.0003 | 0.0004 |
| F09 | 68800001 | 68900000 | 0.0013 | 0.0004 | 0.0006 |
| F09 | 68820001 | 68920000 | 0.0015 | 0.0005 | 0.0009 |
| F09 | 68840001 | 68940000 | 0.0015 | 0.0004 | 0.0010 |
| F09 | 68860001 | 68960000 | 0.0017 | 0.0004 | 0.0012 |
| F09 | 68880001 | 68980000 | 0.0016 | 0.0003 | 0.0012 |
| F09 | 68900001 | 69000000 | 0.0017 | 0.0002 | 0.0012 |
| F09 | 68920001 | 69020000 | 0.0018 | 0.0002 | 0.0011 |
| F09 | 68940001 | 69040000 | 0.0018 | 0.0003 | 0.0011 |
| F09 | 68960001 | 69060000 | 0.0017 | 0.0002 | 0.0009 |
| F09 | 68980001 | 69080000 | 0.0016 | 0.0002 | 0.0008 |
| F09 | 69000001 | 69100000 | 0.0015 | 0.0002 | 0.0006 |
| F09 | 69020001 | 69120000 | 0.0014 | 0.0002 | 0.0005 |
| F09 | 69040001 | 69140000 | 0.0014 | 0.0003 | 0.0004 |
| F09 | 69060001 | 69160000 | 0.0014 | 0.0003 | 0.0004 |
| F09 | 69080001 | 69180000 | 0.0016 | 0.0004 | 0.0005 |
| F09 | 69100001 | 69200000 | 0.0017 | 0.0004 | 0.0006 |
| F09 | 69120001 | 69220000 | 0.0018 | 0.0007 | 0.0008 |
| F09 | 69140001 | 69240000 | 0.0018 | 0.0006 | 0.0008 |
| F09 | 69160001 | 69260000 | 0.0018 | 0.0006 | 0.0009 |
| F09 | 69180001 | 69280000 | 0.0018 | 0.0006 | 0.0011 |
| F09 | 69200001 | 69300000 | 0.0019 | 0.0006 | 0.0012 |
| F09 | 69220001 | 69320000 | 0.0014 | 0.0003 | 0.0008 |
| F09 | 69240001 | 69340000 | 0.0014 | 0.0002 | 0.0006 |
| F09 | 69260001 | 69360000 | 0.0015 | 0.0004 | 0.0006 |
| F09 | 69280001 | 69380000 | 0.0013 | 0.0003 | 0.0003 |
| F09 | 69300001 | 69400000 | 0.0012 | 0.0003 | 0.0002 |
| F09 | 69320001 | 69420000 | 0.0013 | 0.0003 | 0.0003 |
| F09 | 69340001 | 69440000 | 0.0012 | 0.0004 | 0.0003 |
| F09 | 69360001 | 69460000 | 0.0013 | 0.0002 | 0.0002 |
| F09 | 69380001 | 69480000 | 0.0013 | 0.0002 | 0.0003 |
| F09 | 69400001 | 69500000 | 0.0015 | 0.0002 | 0.0004 |
| F09 | 69420001 | 69520000 | 0.0015 | 0.0002 | 0.0004 |
| F09 | 69440001 | 69540000 | 0.0015 | 0.0002 | 0.0004 |
| F09 | 69460001 | 69560000 | 0.0018 | 0.0004 | 0.0008 |
| F09 | 69480001 | 69580000 | 0.0018 | 0.0004 | 0.0008 |
| F09 | 69500001 | 69600000 | 0.0016 | 0.0004 | 0.0009 |
| F09 | 69520001 | 69620000 | 0.0015 | 0.0004 | 0.0010 |
| F09 | 69540001 | 69640000 | 0.0014 | 0.0004 | 0.0012 |
| F09 | 69560001 | 69660000 | 0.0012 | 0.0003 | 0.0012 |
| F09 | 69580001 | 69680000 | 0.0025 | 0.0015 | 0.0026 |
| F09 | 69600001 | 69700000 | 0.0029 | 0.0017 | 0.0030 |
| F09 | 69620001 | 69720000 | 0.0036 | 0.0019 | 0.0037 |
| F09 | 69640001 | 69740000 | 0.0039 | 0.0021 | 0.0040 |
| F09 | 69660001 | 69760000 | 0.0037 | 0.0019 | 0.0038 |
| F09 | 69680001 | 69780000 | 0.0026 | 0.0008 | 0.0024 |
| F09 | 69700001 | 69800000 | 0.0024 | 0.0005 | 0.0019 |
| F09 | 69720001 | 69820000 | 0.0018 | 0.0003 | 0.0011 |
| F09 | 69740001 | 69840000 | 0.0024 | 0.0009 | 0.0016 |

| Chr | Start | End | both | A1 | A2 |
|---|---|---|---|---|---|
| F09 | 69760001 | 69860000 | 0.0026 | 0.0009 | 0.0018 |
| F09 | 69780001 | 69880000 | 0.0026 | 0.0009 | 0.0020 |
| F09 | 69800001 | 69900000 | 0.0026 | 0.0009 | 0.0021 |
| F09 | 69820001 | 69920000 | 0.0026 | 0.0009 | 0.0022 |
| F09 | 69840001 | 69940000 | 0.0018 | 0.0001 | 0.0012 |
| F09 | 69860001 | 69960000 | 0.0020 | 0.0004 | 0.0013 |
| F09 | 69880001 | 69980000 | 0.0019 | 0.0004 | 0.0011 |
| F09 | 69900001 | 70000000 | 0.0017 | 0.0004 | 0.0010 |
| F09 | 69920001 | 70020000 | 0.0017 | 0.0004 | 0.0011 |
| F09 | 69940001 | 70040000 | 0.0024 | 0.0004 | 0.0020 |
| F09 | 69960001 | 70060000 | 0.0023 | 0.0002 | 0.0021 |
| F09 | 69980001 | 70080000 | 0.0024 | 0.0001 | 0.0023 |
| F09 | 70000001 | 70100000 | 0.0025 | 0.0001 | 0.0025 |
| F09 | 70020001 | 70120000 | 0.0030 | 0.0006 | 0.0030 |
| F09 | 70040001 | 70140000 | 0.0024 | 0.0006 | 0.0024 |
| F09 | 70060001 | 70160000 | 0.0022 | 0.0006 | 0.0021 |
| F09 | 70080001 | 70180000 | 0.0027 | 0.0010 | 0.0025 |
| F09 | 70100001 | 70200000 | 0.0025 | 0.0010 | 0.0023 |
| F09 | 70120001 | 70220000 | 0.0020 | 0.0006 | 0.0018 |
| F09 | 70140001 | 70240000 | 0.0020 | 0.0005 | 0.0019 |
| F09 | 70160001 | 70260000 | 0.0019 | 0.0005 | 0.0020 |
| F09 | 70180001 | 70280000 | 0.0013 | 0.0001 | 0.0014 |
| F09 | 70200001 | 70300000 | 0.0014 | 0.0002 | 0.0015 |
| F09 | 70220001 | 70320000 | 0.0016 | 0.0002 | 0.0015 |
| F09 | 70240001 | 70340000 | 0.0015 | 0.0003 | 0.0013 |
| F09 | 70260001 | 70360000 | 0.0015 | 0.0003 | 0.0012 |
| F09 | 70280001 | 70380000 | 0.0023 | 0.0010 | 0.0018 |
| F09 | 70300001 | 70400000 | 0.0023 | 0.0011 | 0.0018 |
| F09 | 70320001 | 70420000 | 0.0028 | 0.0018 | 0.0022 |
| F09 | 70340001 | 70440000 | 0.0031 | 0.0019 | 0.0023 |
| F09 | 70360001 | 70460000 | 0.0040 | 0.0029 | 0.0032 |
| F09 | 70380001 | 70480000 | 0.0033 | 0.0023 | 0.0024 |
| F09 | 70400001 | 70500000 | 0.0032 | 0.0021 | 0.0022 |
| F09 | 70420001 | 70520000 | 0.0025 | 0.0014 | 0.0016 |
| F09 | 70440001 | 70540000 | 0.0022 | 0.0012 | 0.0013 |
| F09 | 70460001 | 70560000 | 0.0012 | 0.0003 | 0.0003 |
| F09 | 70480001 | 70580000 | 0.0012 | 0.0004 | 0.0004 |
| F09 | 70500001 | 70600000 | 0.0017 | 0.0009 | 0.0010 |
| F09 | 70520001 | 70620000 | 0.0015 | 0.0009 | 0.0010 |
| F09 | 70540001 | 70640000 | 0.0013 | 0.0009 | 0.0010 |
| F09 | 70560001 | 70660000 | 0.0010 | 0.0007 | 0.0008 |
| F09 | 70580001 | 70680000 | 0.0008 | 0.0005 | 0.0006 |
| F10 | 1 | 100000 | 0.0018 | 0.0014 | 0.0019 |
| F10 | 20001 | 120000 | 0.0023 | 0.0010 | 0.0024 |
| F10 | 40001 | 140000 | 0.0021 | 0.0006 | 0.0022 |
| F10 | 60001 | 160000 | 0.0017 | 0.0008 | 0.0018 |
| F10 | 80001 | 180000 | 0.0020 | 0.0009 | 0.0021 |
| F10 | 100001 | 200000 | 0.0023 | 0.0010 | 0.0024 |
| F10 | 120001 | 220000 | 0.0022 | 0.0012 | 0.0022 |
| F10 | 140001 | 240000 | 0.0022 | 0.0017 | 0.0021 |
| F10 | 160001 | 260000 | 0.0024 | 0.0018 | 0.0021 |
| F10 | 180001 | 280000 | 0.0022 | 0.0018 | 0.0018 |
| F10 | 200001 | 300000 | 0.0020 | 0.0016 | 0.0014 |
| F10 | 220001 | 320000 | 0.0019 | 0.0015 | 0.0012 |
| F10 | 240001 | 340000 | 0.0019 | 0.0011 | 0.0011 |
| F10 | 260001 | 360000 | 0.0017 | 0.0010 | 0.0008 |
| F10 | 280001 | 380000 | 0.0015 | 0.0008 | 0.0007 |
| F10 | 300001 | 400000 | 0.0014 | 0.0009 | 0.0006 |

| Chr | Start | End | both | A1 | A2 |
|---|---|---|---|---|---|
| F10 | 320001 | 420000 | 0.0014 | 0.0012 | 0.0007 |
| F10 | 340001 | 440000 | 0.0016 | 0.0012 | 0.0010 |
| F10 | 360001 | 460000 | 0.0016 | 0.0032 | 0.0012 |
| F10 | 380001 | 480000 | 0.0036 | 0.0033 | 0.0032 |
| F10 | 400001 | 500000 | 0.0038 | 0.0030 | 0.0035 |
| F10 | 420001 | 520000 | 0.0038 | 0.0025 | 0.0034 |
| F10 | 440001 | 540000 | 0.0037 | 0.0026 | 0.0031 |
| F10 | 460001 | 560000 | 0.0038 | 0.0009 | 0.0031 |
| F10 | 480001 | 580000 | 0.0022 | 0.0007 | 0.0014 |
| F10 | 500001 | 600000 | 0.0019 | 0.0008 | 0.0011 |
| F10 | 520001 | 620000 | 0.0020 | 0.0008 | 0.0010 |
| F10 | 540001 | 640000 | 0.0019 | 0.0009 | 0.0009 |
| F10 | 560001 | 660000 | 0.0020 | 0.0006 | 0.0010 |
| F10 | 580001 | 680000 | 0.0018 | 0.0007 | 0.0009 |
| F10 | 600001 | 700000 | 0.0019 | 0.0006 | 0.0010 |
| F10 | 620001 | 720000 | 0.0019 | 0.0009 | 0.0012 |
| F10 | 640001 | 740000 | 0.0021 | 0.0007 | 0.0013 |
| F10 | 660001 | 760000 | 0.0021 | 0.0011 | 0.0013 |
| F10 | 680001 | 780000 | 0.0019 | 0.0013 | 0.0010 |
| F10 | 700001 | 800000 | 0.0017 | 0.0018 | 0.0009 |
| F10 | 720001 | 820000 | 0.0016 | 0.0019 | 0.0007 |
| F10 | 740001 | 840000 | 0.0014 | 0.0023 | 0.0008 |
| F10 | 760001 | 860000 | 0.0013 | 0.0020 | 0.0008 |
| F10 | 780001 | 880000 | 0.0013 | 0.0021 | 0.0008 |
| F10 | 800001 | 900000 | 0.0013 | 0.0020 | 0.0008 |
| F10 | 820001 | 920000 | 0.0012 | 0.0022 | 0.0007 |
| F10 | 840001 | 940000 | 0.0012 | 0.0024 | 0.0007 |
| F10 | 860001 | 960000 | 0.0010 | 0.0023 | 0.0004 |
| F10 | 880001 | 980000 | 0.0011 | 0.0024 | 0.0006 |
| F10 | 900001 | 1000000 | 0.0014 | 0.0024 | 0.0009 |
| F10 | 920001 | 1020000 | 0.0014 | 0.0023 | 0.0009 |
| F10 | 940001 | 1040000 | 0.0012 | 0.0022 | 0.0008 |
| F10 | 960001 | 1060000 | 0.0012 | 0.0022 | 0.0008 |
| F10 | 980001 | 1080000 | 0.0010 | 0.0020 | 0.0006 |
| F10 | 1000001 | 1100000 | 0.0008 | 0.0020 | 0.0004 |
| F10 | 1020001 | 1120000 | 0.0008 | 0.0019 | 0.0004 |
| F10 | 1040001 | 1140000 | 0.0008 | 0.0018 | 0.0004 |
| F10 | 1060001 | 1160000 | 0.0007 | 0.0017 | 0.0004 |
| F10 | 1080001 | 1180000 | 0.0007 | 0.0021 | 0.0004 |
| F10 | 1100001 | 1200000 | 0.0012 | 0.0036 | 0.0009 |
| F10 | 1120001 | 1220000 | 0.0030 | 0.0047 | 0.0028 |
| F10 | 1140001 | 1240000 | 0.0043 | 0.0053 | 0.0040 |
| F10 | 1160001 | 1260000 | 0.0050 | 0.0059 | 0.0048 |
| F10 | 1180001 | 1280000 | 0.0058 | 0.0056 | 0.0056 |
| F10 | 1200001 | 1300000 | 0.0056 | 0.0040 | 0.0054 |
| F10 | 1220001 | 1320000 | 0.0038 | 0.0028 | 0.0036 |
| F10 | 1240001 | 1340000 | 0.0025 | 0.0022 | 0.0024 |
| F10 | 1260001 | 1360000 | 0.0020 | 0.0016 | 0.0019 |
| F10 | 1280001 | 1380000 | 0.0013 | 0.0016 | 0.0011 |
| F10 | 1300001 | 1400000 | 0.0011 | 0.0019 | 0.0008 |
| F10 | 1320001 | 1420000 | 0.0014 | 0.0017 | 0.0012 |
| F10 | 1340001 | 1440000 | 0.0013 | 0.0023 | 0.0011 |
| F10 | 1360001 | 1460000 | 0.0020 | 0.0022 | 0.0018 |
| F10 | 1380001 | 1480000 | 0.0022 | 0.0022 | 0.0020 |
| F10 | 1400001 | 1500000 | 0.0023 | 0.0017 | 0.0022 |
| F10 | 1420001 | 1520000 | 0.0019 | 0.0018 | 0.0018 |
| F10 | 1440001 | 1540000 | 0.0021 | 0.0014 | 0.0021 |
| F10 | 1460001 | 1560000 | 0.0016 | 0.0011 | 0.0015 |

| Chr | Start | End | both | A1 | A2 |
|---|---|---|---|---|---|
| F10 | 1480001 | 1580000 | 0.0013 | 0.0010 | 0.0013 |
| F10 | 1500001 | 1600000 | 0.0012 | 0.0016 | 0.0012 |
| F10 | 1520001 | 1620000 | 0.0017 | 0.0024 | 0.0017 |
| F10 | 1540001 | 1640000 | 0.0025 | 0.0024 | 0.0025 |
| F10 | 1560001 | 1660000 | 0.0024 | 0.0026 | 0.0024 |
| F10 | 1580001 | 1680000 | 0.0025 | 0.0030 | 0.0024 |
| F10 | 1600001 | 1700000 | 0.0025 | 0.0026 | 0.0023 |
| F10 | 1620001 | 1720000 | 0.0019 | 0.0021 | 0.0016 |
| F10 | 1640001 | 1740000 | 0.0012 | 0.0020 | 0.0007 |
| F10 | 1660001 | 1760000 | 0.0008 | 0.0024 | 0.0003 |
| F10 | 1680001 | 1780000 | 0.0009 | 0.0027 | 0.0004 |
| F10 | 1700001 | 1800000 | 0.0010 | 0.0026 | 0.0004 |
| F10 | 1720001 | 1820000 | 0.0010 | 0.0026 | 0.0004 |
| F10 | 1740001 | 1840000 | 0.0008 | 0.0026 | 0.0003 |
| F10 | 1760001 | 1860000 | 0.0009 | 0.0024 | 0.0003 |
| F10 | 1780001 | 1880000 | 0.0008 | 0.0022 | 0.0002 |
| F10 | 1800001 | 1900000 | 0.0007 | 0.0023 | 0.0002 |
| F10 | 1820001 | 1920000 | 0.0008 | 0.0020 | 0.0003 |
| F10 | 1840001 | 1940000 | 0.0007 | 0.0019 | 0.0003 |
| F10 | 1860001 | 1960000 | 0.0007 | 0.0019 | 0.0003 |
| F10 | 1880001 | 1980000 | 0.0007 | 0.0018 | 0.0003 |
| F10 | 1900001 | 2000000 | 0.0008 | 0.0018 | 0.0004 |
| F10 | 1920001 | 2020000 | 0.0007 | 0.0021 | 0.0004 |
| F10 | 1940001 | 2040000 | 0.0009 | 0.0019 | 0.0005 |
| F10 | 1960001 | 2060000 | 0.0009 | 0.0019 | 0.0005 |
| F10 | 1980001 | 2080000 | 0.0009 | 0.0020 | 0.0005 |
| F10 | 2000001 | 2100000 | 0.0009 | 0.0020 | 0.0004 |
| F10 | 2020001 | 2120000 | 0.0009 | 0.0019 | 0.0003 |
| F10 | 2040001 | 2140000 | 0.0009 | 0.0021 | 0.0002 |
| F10 | 2060001 | 2160000 | 0.0011 | 0.0022 | 0.0003 |
| F10 | 2080001 | 2180000 | 0.0012 | 0.0022 | 0.0003 |
| F10 | 2100001 | 2200000 | 0.0012 | 0.0023 | 0.0004 |
| F10 | 2120001 | 2220000 | 0.0013 | 0.0024 | 0.0005 |
| F10 | 2140001 | 2240000 | 0.0015 | 0.0022 | 0.0008 |
| F10 | 2160001 | 2260000 | 0.0012 | 0.0021 | 0.0006 |
| F10 | 2180001 | 2280000 | 0.0011 | 0.0020 | 0.0006 |
| F10 | 2200001 | 2300000 | 0.0010 | 0.0020 | 0.0006 |
| F10 | 2220001 | 2320000 | 0.0009 | 0.0020 | 0.0005 |
| F10 | 2240001 | 2340000 | 0.0007 | 0.0023 | 0.0003 |
| F10 | 2260001 | 2360000 | 0.0009 | 0.0022 | 0.0004 |
| F10 | 2280001 | 2380000 | 0.0009 | 0.0020 | 0.0005 |
| F10 | 2300001 | 2400000 | 0.0009 | 0.0020 | 0.0005 |
| F10 | 2320001 | 2420000 | 0.0009 | 0.0018 | 0.0006 |
| F10 | 2340001 | 2440000 | 0.0007 | 0.0016 | 0.0004 |
| F10 | 2360001 | 2460000 | 0.0007 | 0.0018 | 0.0003 |
| F10 | 2380001 | 2480000 | 0.0008 | 0.0020 | 0.0004 |
| F10 | 2400001 | 2500000 | 0.0011 | 0.0019 | 0.0007 |
| F10 | 2420001 | 2520000 | 0.0012 | 0.0018 | 0.0008 |
| F10 | 2440001 | 2540000 | 0.0013 | 0.0017 | 0.0009 |
| F10 | 2460001 | 2560000 | 0.0014 | 0.0020 | 0.0010 |
| F10 | 2480001 | 2580000 | 0.0017 | 0.0019 | 0.0014 |
| F10 | 2500001 | 2600000 | 0.0016 | 0.0019 | 0.0013 |
| F10 | 2520001 | 2620000 | 0.0014 | 0.0021 | 0.0011 |
| F10 | 2540001 | 2640000 | 0.0015 | 0.0021 | 0.0012 |
| F10 | 2560001 | 2660000 | 0.0014 | 0.0021 | 0.0011 |
| F10 | 2580001 | 2680000 | 0.0015 | 0.0023 | 0.0012 |
| F10 | 2600001 | 2700000 | 0.0017 | 0.0024 | 0.0014 |
| F10 | 2620001 | 2720000 | 0.0017 | 0.0025 | 0.0014 |

| Chr | Start | End | both | A1 | A2 |
|---|---|---|---|---|---|
| F10 | 2640001 | 2740000 | 0.0017 | 0.0027 | 0.0013 |
| F10 | 2660001 | 2760000 | 0.0020 | 0.0022 | 0.0015 |
| F10 | 2680001 | 2780000 | 0.0014 | 0.0021 | 0.0008 |
| F10 | 2700001 | 2800000 | 0.0013 | 0.0022 | 0.0005 |
| F10 | 2720001 | 2820000 | 0.0014 | 0.0020 | 0.0005 |
| F10 | 2740001 | 2840000 | 0.0013 | 0.0018 | 0.0004 |
| F10 | 2760001 | 2860000 | 0.0011 | 0.0020 | 0.0002 |
| F10 | 2780001 | 2880000 | 0.0013 | 0.0019 | 0.0004 |
| F10 | 2800001 | 2900000 | 0.0014 | 0.0016 | 0.0005 |
| F10 | 2820001 | 2920000 | 0.0014 | 0.0015 | 0.0007 |
| F10 | 2840001 | 2940000 | 0.0014 | 0.0015 | 0.0008 |
| F10 | 2860001 | 2960000 | 0.0013 | 0.0012 | 0.0008 |
| F10 | 2880001 | 2980000 | 0.0010 | 0.0011 | 0.0006 |
| F10 | 2900001 | 3000000 | 0.0008 | 0.0017 | 0.0005 |
| F10 | 2920001 | 3020000 | 0.0010 | 0.0017 | 0.0007 |
| F10 | 2940001 | 3040000 | 0.0009 | 0.0018 | 0.0006 |
| F10 | 2960001 | 3060000 | 0.0010 | 0.0019 | 0.0007 |
| F10 | 2980001 | 3080000 | 0.0010 | 0.0018 | 0.0007 |
| F10 | 3000001 | 3100000 | 0.0012 | 0.0013 | 0.0007 |
| F10 | 3020001 | 3120000 | 0.0011 | 0.0012 | 0.0005 |
| F10 | 3040001 | 3140000 | 0.0013 | 0.0009 | 0.0006 |
| F10 | 3060001 | 3160000 | 0.0014 | 0.0008 | 0.0005 |
| F10 | 3080001 | 3180000 | 0.0017 | 0.0008 | 0.0007 |
| F10 | 3100001 | 3200000 | 0.0017 | 0.0006 | 0.0007 |
| F10 | 3120001 | 3220000 | 0.0016 | 0.0006 | 0.0006 |
| F10 | 3140001 | 3240000 | 0.0016 | 0.0006 | 0.0006 |
| F10 | 3160001 | 3260000 | 0.0015 | 0.0008 | 0.0005 |
| F10 | 3180001 | 3280000 | 0.0017 | 0.0008 | 0.0008 |
| F10 | 3200001 | 3300000 | 0.0018 | 0.0011 | 0.0009 |
| F10 | 3220001 | 3320000 | 0.0021 | 0.0012 | 0.0013 |
| F10 | 3240001 | 3340000 | 0.0020 | 0.0012 | 0.0015 |
| F10 | 3260001 | 3360000 | 0.0020 | 0.0008 | 0.0015 |
| F10 | 3280001 | 3380000 | 0.0016 | 0.0011 | 0.0012 |
| F10 | 3300001 | 3400000 | 0.0018 | 0.0010 | 0.0015 |
| F10 | 3320001 | 3420000 | 0.0016 | 0.0009 | 0.0013 |
| F10 | 3340001 | 3440000 | 0.0017 | 0.0016 | 0.0013 |
| F10 | 3360001 | 3460000 | 0.0026 | 0.0017 | 0.0022 |
| F10 | 3380001 | 3480000 | 0.0028 | 0.0015 | 0.0022 |
| F10 | 3400001 | 3500000 | 0.0026 | 0.0013 | 0.0019 |
| F10 | 3420001 | 3520000 | 0.0024 | 0.0013 | 0.0017 |
| F10 | 3440001 | 3540000 | 0.0022 | 0.0007 | 0.0015 |
| F10 | 3460001 | 3560000 | 0.0015 | 0.0004 | 0.0007 |
| F10 | 3480001 | 3580000 | 0.0012 | 0.0011 | 0.0004 |
| F10 | 3500001 | 3600000 | 0.0015 | 0.0015 | 0.0008 |
| F10 | 3520001 | 3620000 | 0.0019 | 0.0023 | 0.0013 |
| F10 | 3540001 | 3640000 | 0.0023 | 0.0025 | 0.0018 |
| F10 | 3560001 | 3660000 | 0.0022 | 0.0030 | 0.0017 |
| F10 | 3580001 | 3680000 | 0.0023 | 0.0028 | 0.0018 |
| F10 | 3600001 | 3700000 | 0.0019 | 0.0029 | 0.0014 |
| F10 | 3620001 | 3720000 | 0.0020 | 0.0023 | 0.0013 |
| F10 | 3640001 | 3740000 | 0.0014 | 0.0022 | 0.0009 |
| F10 | 3660001 | 3760000 | 0.0013 | 0.0018 | 0.0008 |
| F10 | 3680001 | 3780000 | 0.0014 | 0.0014 | 0.0008 |
| F10 | 3700001 | 3800000 | 0.0014 | 0.0007 | 0.0007 |
| F10 | 3720001 | 3820000 | 0.0010 | 0.0007 | 0.0004 |
| F10 | 3740001 | 3840000 | 0.0013 | 0.0008 | 0.0004 |
| F10 | 3760001 | 3860000 | 0.0017 | 0.0010 | 0.0006 |
| F10 | 3780001 | 3880000 | 0.0016 | 0.0011 | 0.0006 |

| Chr | Start | End | both | A1 | A2 |
|---|---|---|---|---|---|
| F10 | 3800001 | 3900000 | 0.0013 | 0.0013 | 0.0004 |
| F10 | 3820001 | 3920000 | 0.0012 | 0.0015 | 0.0004 |
| F10 | 3840001 | 3940000 | 0.0010 | 0.0018 | 0.0004 |
| F10 | 3860001 | 3960000 | 0.0014 | 0.0015 | 0.0009 |
| F10 | 3880001 | 3980000 | 0.0015 | 0.0014 | 0.0008 |
| F10 | 3900001 | 4000000 | 0.0017 | 0.0012 | 0.0009 |
| F10 | 3920001 | 4020000 | 0.0019 | 0.0009 | 0.0010 |
| F10 | 3940001 | 4040000 | 0.0019 | 0.0005 | 0.0011 |
| F10 | 3960001 | 4060000 | 0.0015 | 0.0005 | 0.0008 |
| F10 | 3980001 | 4080000 | 0.0015 | 0.0008 | 0.0008 |
| F10 | 4000001 | 4100000 | 0.0017 | 0.0014 | 0.0011 |
| F10 | 4020001 | 4120000 | 0.0024 | 0.0015 | 0.0018 |
| F10 | 4040001 | 4140000 | 0.0025 | 0.0017 | 0.0018 |
| F10 | 4060001 | 4160000 | 0.0028 | 0.0017 | 0.0021 |
| F10 | 4080001 | 4180000 | 0.0026 | 0.0014 | 0.0022 |
| F10 | 4100001 | 4200000 | 0.0025 | 0.0009 | 0.0022 |
| F10 | 4120001 | 4220000 | 0.0019 | 0.0011 | 0.0018 |
| F10 | 4140001 | 4240000 | 0.0021 | 0.0007 | 0.0019 |
| F10 | 4160001 | 4260000 | 0.0017 | 0.0006 | 0.0013 |
| F10 | 4180001 | 4280000 | 0.0018 | 0.0008 | 0.0012 |
| F10 | 4200001 | 4300000 | 0.0018 | 0.0007 | 0.0010 |
| F10 | 4220001 | 4320000 | 0.0017 | 0.0005 | 0.0007 |
| F10 | 4240001 | 4340000 | 0.0014 | 0.0004 | 0.0006 |
| F10 | 4260001 | 4360000 | 0.0012 | 0.0005 | 0.0006 |
| F10 | 4280001 | 4380000 | 0.0016 | 0.0005 | 0.0011 |
| F10 | 4300001 | 4400000 | 0.0015 | 0.0004 | 0.0011 |
| F10 | 4320001 | 4420000 | 0.0015 | 0.0005 | 0.0012 |
| F10 | 4340001 | 4440000 | 0.0015 | 0.0005 | 0.0014 |
| F10 | 4360001 | 4460000 | 0.0015 | 0.0009 | 0.0014 |
| F10 | 4380001 | 4480000 | 0.0017 | 0.0008 | 0.0017 |
| F10 | 4400001 | 4500000 | 0.0019 | 0.0009 | 0.0019 |
| F10 | 4420001 | 4520000 | 0.0022 | 0.0008 | 0.0021 |
| F10 | 4440001 | 4540000 | 0.0024 | 0.0008 | 0.0022 |
| F10 | 4460001 | 4560000 | 0.0027 | 0.0003 | 0.0025 |
| F10 | 4480001 | 4580000 | 0.0023 | 0.0006 | 0.0021 |
| F10 | 4500001 | 4600000 | 0.0026 | 0.0009 | 0.0021 |
| F10 | 4520001 | 4620000 | 0.0027 | 0.0011 | 0.0021 |
| F10 | 4540001 | 4640000 | 0.0025 | 0.0011 | 0.0018 |
| F10 | 4560001 | 4660000 | 0.0023 | 0.0015 | 0.0015 |
| F10 | 4580001 | 4680000 | 0.0025 | 0.0020 | 0.0017 |
| F10 | 4600001 | 4700000 | 0.0029 | 0.0017 | 0.0023 |
| F10 | 4620001 | 4720000 | 0.0026 | 0.0015 | 0.0019 |
| F10 | 4640001 | 4740000 | 0.0026 | 0.0016 | 0.0018 |
| F10 | 4660001 | 4760000 | 0.0026 | 0.0014 | 0.0018 |
| F10 | 4680001 | 4780000 | 0.0024 | 0.0010 | 0.0015 |
| F10 | 4700001 | 4800000 | 0.0020 | 0.0015 | 0.0009 |
| F10 | 4720001 | 4820000 | 0.0023 | 0.0017 | 0.0014 |
| F10 | 4740001 | 4840000 | 0.0024 | 0.0027 | 0.0015 |
| F10 | 4760001 | 4860000 | 0.0036 | 0.0026 | 0.0026 |
| F10 | 4780001 | 4880000 | 0.0035 | 0.0021 | 0.0024 |
| F10 | 4800001 | 4900000 | 0.0031 | 0.0018 | 0.0021 |
| F10 | 4820001 | 4920000 | 0.0028 | 0.0018 | 0.0017 |
| F10 | 4840001 | 4940000 | 0.0027 | 0.0011 | 0.0017 |
| F10 | 4860001 | 4960000 | 0.0019 | 0.0012 | 0.0011 |
| F10 | 4880001 | 4980000 | 0.0022 | 0.0021 | 0.0014 |
| F10 | 4900001 | 5000000 | 0.0032 | 0.0021 | 0.0025 |
| F10 | 4920001 | 5020000 | 0.0033 | 0.0020 | 0.0026 |
| F10 | 4940001 | 5040000 | 0.0032 | 0.0017 | 0.0025 |

| Chr | Start | End | both | A1 | A2 |
|---|---|---|---|---|---|
| F10 | 4960001 | 5060000 | 0.0029 | 0.0018 | 0.0022 |
| F10 | 4980001 | 5080000 | 0.0027 | 0.0009 | 0.0021 |
| F10 | 5000001 | 5100000 | 0.0018 | 0.0017 | 0.0012 |
| F10 | 5020001 | 5120000 | 0.0025 | 0.0027 | 0.0020 |
| F10 | 5040001 | 5140000 | 0.0036 | 0.0034 | 0.0031 |
| F10 | 5060001 | 5160000 | 0.0041 | 0.0039 | 0.0037 |
| F10 | 5080001 | 5180000 | 0.0045 | 0.0049 | 0.0042 |
| F10 | 5100001 | 5200000 | 0.0055 | 0.0039 | 0.0053 |
| F10 | 5120001 | 5220000 | 0.0046 | 0.0031 | 0.0042 |
| F10 | 5140001 | 5240000 | 0.0038 | 0.0024 | 0.0032 |
| F10 | 5160001 | 5260000 | 0.0032 | 0.0021 | 0.0025 |
| F10 | 5180001 | 5280000 | 0.0031 | 0.0010 | 0.0022 |
| F10 | 5200001 | 5300000 | 0.0020 | 0.0011 | 0.0010 |
| F10 | 5220001 | 5320000 | 0.0021 | 0.0009 | 0.0011 |
| F10 | 5240001 | 5340000 | 0.0019 | 0.0012 | 0.0009 |
| F10 | 5260001 | 5360000 | 0.0021 | 0.0010 | 0.0013 |
| F10 | 5280001 | 5380000 | 0.0019 | 0.0010 | 0.0011 |
| F10 | 5300001 | 5400000 | 0.0017 | 0.0013 | 0.0011 |
| F10 | 5320001 | 5420000 | 0.0020 | 0.0014 | 0.0014 |
| F10 | 5340001 | 5440000 | 0.0021 | 0.0011 | 0.0015 |
| F10 | 5360001 | 5460000 | 0.0019 | 0.0010 | 0.0011 |
| F10 | 5380001 | 5480000 | 0.0018 | 0.0012 | 0.0011 |
| F10 | 5400001 | 5500000 | 0.0019 | 0.0008 | 0.0012 |
| F10 | 5420001 | 5520000 | 0.0014 | 0.0010 | 0.0007 |
| F10 | 5440001 | 5540000 | 0.0019 | 0.0015 | 0.0011 |
| F10 | 5460001 | 5560000 | 0.0022 | 0.0019 | 0.0015 |
| F10 | 5480001 | 5580000 | 0.0029 | 0.0024 | 0.0019 |
| F10 | 5500001 | 5600000 | 0.0034 | 0.0025 | 0.0024 |
| F10 | 5520001 | 5620000 | 0.0034 | 0.0021 | 0.0024 |
| F10 | 5540001 | 5640000 | 0.0028 | 0.0015 | 0.0019 |
| F10 | 5560001 | 5660000 | 0.0024 | 0.0018 | 0.0015 |
| F10 | 5580001 | 5680000 | 0.0024 | 0.0019 | 0.0016 |
| F10 | 5600001 | 5700000 | 0.0027 | 0.0019 | 0.0021 |
| F10 | 5620001 | 5720000 | 0.0030 | 0.0022 | 0.0023 |
| F10 | 5640001 | 5740000 | 0.0033 | 0.0025 | 0.0026 |
| F10 | 5660001 | 5760000 | 0.0035 | 0.0018 | 0.0029 |
| F10 | 5680001 | 5780000 | 0.0028 | 0.0023 | 0.0022 |
| F10 | 5700001 | 5800000 | 0.0029 | 0.0029 | 0.0024 |
| F10 | 5720001 | 5820000 | 0.0031 | 0.0026 | 0.0027 |
| F10 | 5740001 | 5840000 | 0.0030 | 0.0032 | 0.0025 |
| F10 | 5760001 | 5860000 | 0.0036 | 0.0030 | 0.0030 |
| F10 | 5780001 | 5880000 | 0.0036 | 0.0021 | 0.0028 |
| F10 | 5800001 | 5900000 | 0.0027 | 0.0024 | 0.0019 |
| F10 | 5820001 | 5920000 | 0.0033 | 0.0024 | 0.0024 |
| F10 | 5840001 | 5940000 | 0.0034 | 0.0019 | 0.0025 |
| F10 | 5860001 | 5960000 | 0.0027 | 0.0020 | 0.0020 |
| F10 | 5880001 | 5980000 | 0.0026 | 0.0046 | 0.0022 |
| F10 | 5900001 | 6000000 | 0.0053 | 0.0047 | 0.0048 |
| F10 | 5920001 | 6020000 | 0.0051 | 0.0050 | 0.0047 |
| F10 | 5940001 | 6040000 | 0.0054 | 0.0051 | 0.0051 |
| F10 | 5960001 | 6060000 | 0.0057 | 0.0051 | 0.0051 |
| F10 | 5980001 | 6080000 | 0.0061 | 0.0024 | 0.0052 |
| F10 | 6000001 | 6100000 | 0.0034 | 0.0018 | 0.0025 |
| F10 | 6020001 | 6120000 | 0.0028 | 0.0014 | 0.0020 |
| F10 | 6040001 | 6140000 | 0.0022 | 0.0012 | 0.0016 |
| F10 | 6060001 | 6160000 | 0.0018 | 0.0011 | 0.0014 |
| F10 | 6080001 | 6180000 | 0.0015 | 0.0012 | 0.0013 |
| F10 | 6100001 | 6200000 | 0.0016 | 0.0011 | 0.0013 |

| Chr | Start | End | both | A1 | A2 |
|---|---|---|---|---|---|
| F10 | 6120001 | 6220000 | 0.0016 | 0.0014 | 0.0011 |
| F10 | 6140001 | 6240000 | 0.0019 | 0.0012 | 0.0014 |
| F10 | 6160001 | 6260000 | 0.0018 | 0.0018 | 0.0012 |
| F10 | 6180001 | 6280000 | 0.0024 | 0.0017 | 0.0018 |
| F10 | 6200001 | 6300000 | 0.0024 | 0.0018 | 0.0016 |
| F10 | 6220001 | 6320000 | 0.0023 | 0.0016 | 0.0017 |
| F10 | 6240001 | 6340000 | 0.0023 | 0.0018 | 0.0015 |
| F10 | 6260001 | 6360000 | 0.0025 | 0.0016 | 0.0016 |
| F10 | 6280001 | 6380000 | 0.0024 | 0.0021 | 0.0015 |
| F10 | 6300001 | 6400000 | 0.0030 | 0.0017 | 0.0021 |
| F10 | 6320001 | 6420000 | 0.0026 | 0.0017 | 0.0018 |
| F10 | 6340001 | 6440000 | 0.0024 | 0.0018 | 0.0017 |
| F10 | 6360001 | 6460000 | 0.0026 | 0.0013 | 0.0018 |
| F10 | 6380001 | 6480000 | 0.0018 | 0.0009 | 0.0012 |
| F10 | 6400001 | 6500000 | 0.0015 | 0.0014 | 0.0007 |
| F10 | 6420001 | 6520000 | 0.0020 | 0.0017 | 0.0012 |
| F10 | 6440001 | 6540000 | 0.0025 | 0.0022 | 0.0017 |
| F10 | 6460001 | 6560000 | 0.0028 | 0.0025 | 0.0019 |
| F10 | 6480001 | 6580000 | 0.0030 | 0.0026 | 0.0021 |
| F10 | 6500001 | 6600000 | 0.0030 | 0.0022 | 0.0023 |
| F10 | 6520001 | 6620000 | 0.0025 | 0.0020 | 0.0018 |
| F10 | 6540001 | 6640000 | 0.0021 | 0.0019 | 0.0013 |
| F10 | 6560001 | 6660000 | 0.0021 | 0.0019 | 0.0014 |
| F10 | 6580001 | 6680000 | 0.0021 | 0.0020 | 0.0014 |
| F10 | 6600001 | 6700000 | 0.0023 | 0.0021 | 0.0014 |
| F10 | 6620001 | 6720000 | 0.0023 | 0.0020 | 0.0014 |
| F10 | 6640001 | 6740000 | 0.0022 | 0.0018 | 0.0014 |
| F10 | 6660001 | 6760000 | 0.0020 | 0.0017 | 0.0013 |
| F10 | 6680001 | 6780000 | 0.0018 | 0.0013 | 0.0013 |
| F10 | 6700001 | 6800000 | 0.0014 | 0.0020 | 0.0010 |
| F10 | 6720001 | 6820000 | 0.0021 | 0.0024 | 0.0017 |
| F10 | 6740001 | 6840000 | 0.0025 | 0.0026 | 0.0019 |
| F10 | 6760001 | 6860000 | 0.0027 | 0.0026 | 0.0020 |
| F10 | 6780001 | 6880000 | 0.0027 | 0.0026 | 0.0019 |
| F10 | 6800001 | 6900000 | 0.0028 | 0.0023 | 0.0019 |
| F10 | 6820001 | 6920000 | 0.0025 | 0.0026 | 0.0014 |
| F10 | 6840001 | 6940000 | 0.0024 | 0.0031 | 0.0014 |
| F10 | 6860001 | 6960000 | 0.0030 | 0.0031 | 0.0020 |
| F10 | 6880001 | 6980000 | 0.0029 | 0.0033 | 0.0019 |
| F10 | 6900001 | 7000000 | 0.0031 | 0.0032 | 0.0020 |
| F10 | 6920001 | 7020000 | 0.0031 | 0.0028 | 0.0022 |
| F10 | 6940001 | 7040000 | 0.0032 | 0.0022 | 0.0022 |
| F10 | 6960001 | 7060000 | 0.0026 | 0.0029 | 0.0016 |
| F10 | 6980001 | 7080000 | 0.0034 | 0.0027 | 0.0024 |
| F10 | 7000001 | 7100000 | 0.0033 | 0.0029 | 0.0024 |
| F10 | 7020001 | 7120000 | 0.0035 | 0.0035 | 0.0028 |
| F10 | 7040001 | 7140000 | 0.0037 | 0.0033 | 0.0031 |
| F10 | 7060001 | 7160000 | 0.0035 | 0.0026 | 0.0029 |
| F10 | 7080001 | 7180000 | 0.0027 | 0.0025 | 0.0021 |
| F10 | 7100001 | 7200000 | 0.0026 | 0.0021 | 0.0019 |
| F10 | 7120001 | 7220000 | 0.0021 | 0.0013 | 0.0013 |
| F10 | 7140001 | 7240000 | 0.0014 | 0.0013 | 0.0007 |
| F10 | 7160001 | 7260000 | 0.0015 | 0.0013 | 0.0007 |
| F10 | 7180001 | 7280000 | 0.0016 | 0.0014 | 0.0008 |
| F10 | 7200001 | 7300000 | 0.0017 | 0.0015 | 0.0009 |
| F10 | 7220001 | 7320000 | 0.0018 | 0.0015 | 0.0011 |
| F10 | 7240001 | 7340000 | 0.0019 | 0.0016 | 0.0011 |
| F10 | 7260001 | 7360000 | 0.0018 | 0.0018 | 0.0010 |

| Chr | Start | End | both | A1 | A2 |
|---|---|---|---|---|---|
| F10 | 7280001 | 7380000 | 0.0020 | 0.0023 | 0.0014 |
| F10 | 7300001 | 7400000 | 0.0026 | 0.0022 | 0.0019 |
| F10 | 7320001 | 7420000 | 0.0025 | 0.0023 | 0.0016 |
| F10 | 7340001 | 7440000 | 0.0026 | 0.0020 | 0.0017 |
| F10 | 7360001 | 7460000 | 0.0023 | 0.0022 | 0.0015 |
| F10 | 7380001 | 7480000 | 0.0026 | 0.0021 | 0.0018 |
| F10 | 7400001 | 7500000 | 0.0025 | 0.0021 | 0.0018 |
| F10 | 7420001 | 7520000 | 0.0024 | 0.0029 | 0.0019 |
| F10 | 7440001 | 7540000 | 0.0032 | 0.0036 | 0.0029 |
| F10 | 7460001 | 7560000 | 0.0041 | 0.0044 | 0.0039 |
| F10 | 7480001 | 7580000 | 0.0049 | 0.0050 | 0.0046 |
| F10 | 7500001 | 7600000 | 0.0055 | 0.0053 | 0.0053 |
| F10 | 7520001 | 7620000 | 0.0059 | 0.0050 | 0.0056 |
| F10 | 7540001 | 7640000 | 0.0056 | 0.0043 | 0.0054 |
| F10 | 7560001 | 7660000 | 0.0049 | 0.0031 | 0.0045 |
| F10 | 7580001 | 7680000 | 0.0036 | 0.0023 | 0.0034 |
| F10 | 7600001 | 7700000 | 0.0026 | 0.0029 | 0.0023 |
| F10 | 7620001 | 7720000 | 0.0032 | 0.0024 | 0.0029 |
| F10 | 7640001 | 7740000 | 0.0025 | 0.0023 | 0.0023 |
| F10 | 7660001 | 7760000 | 0.0024 | 0.0030 | 0.0023 |
| F10 | 7680001 | 7780000 | 0.0030 | 0.0036 | 0.0027 |
| F10 | 7700001 | 7800000 | 0.0036 | 0.0030 | 0.0033 |
| F10 | 7720001 | 7820000 | 0.0030 | 0.0038 | 0.0027 |
| F10 | 7740001 | 7840000 | 0.0041 | 0.0045 | 0.0038 |
| F10 | 7760001 | 7860000 | 0.0048 | 0.0042 | 0.0045 |
| F10 | 7780001 | 7880000 | 0.0046 | 0.0034 | 0.0042 |
| F10 | 7800001 | 7900000 | 0.0038 | 0.0033 | 0.0034 |
| F10 | 7820001 | 7920000 | 0.0038 | 0.0021 | 0.0034 |
| F10 | 7840001 | 7940000 | 0.0025 | 0.0016 | 0.0020 |
| F10 | 7860001 | 7960000 | 0.0019 | 0.0011 | 0.0015 |
| F10 | 7880001 | 7980000 | 0.0014 | 0.0010 | 0.0011 |
| F10 | 7900001 | 8000000 | 0.0015 | 0.0008 | 0.0012 |
| F10 | 7920001 | 8020000 | 0.0011 | 0.0012 | 0.0009 |
| F10 | 7940001 | 8040000 | 0.0017 | 0.0012 | 0.0015 |
| F10 | 7960001 | 8060000 | 0.0018 | 0.0014 | 0.0016 |
| F10 | 7980001 | 8080000 | 0.0021 | 0.0017 | 0.0019 |
| F10 | 8000001 | 8100000 | 0.0024 | 0.0019 | 0.0022 |
| F10 | 8020001 | 8120000 | 0.0028 | 0.0020 | 0.0025 |
| F10 | 8040001 | 8140000 | 0.0029 | 0.0020 | 0.0027 |
| F10 | 8060001 | 8160000 | 0.0031 | 0.0020 | 0.0028 |
| F10 | 8080001 | 8180000 | 0.0031 | 0.0021 | 0.0028 |
| F10 | 8100001 | 8200000 | 0.0030 | 0.0027 | 0.0027 |
| F10 | 8120001 | 8220000 | 0.0035 | 0.0022 | 0.0032 |
| F10 | 8140001 | 8240000 | 0.0029 | 0.0022 | 0.0027 |
| F10 | 8160001 | 8260000 | 0.0029 | 0.0019 | 0.0027 |
| F10 | 8180001 | 8280000 | 0.0024 | 0.0017 | 0.0022 |
| F10 | 8200001 | 8300000 | 0.0023 | 0.0010 | 0.0021 |
| F10 | 8220001 | 8320000 | 0.0018 | 0.0013 | 0.0016 |
| F10 | 8240001 | 8340000 | 0.0021 | 0.0012 | 0.0019 |
| F10 | 8260001 | 8360000 | 0.0020 | 0.0015 | 0.0018 |
| F10 | 8280001 | 8380000 | 0.0024 | 0.0014 | 0.0022 |
| F10 | 8300001 | 8400000 | 0.0023 | 0.0012 | 0.0020 |
| F10 | 8320001 | 8420000 | 0.0021 | 0.0010 | 0.0019 |
| F10 | 8340001 | 8440000 | 0.0020 | 0.0026 | 0.0017 |
| F10 | 8360001 | 8460000 | 0.0036 | 0.0027 | 0.0034 |
| F10 | 8380001 | 8480000 | 0.0039 | 0.0040 | 0.0036 |
| F10 | 8400001 | 8500000 | 0.0053 | 0.0041 | 0.0050 |
| F10 | 8420001 | 8520000 | 0.0053 | 0.0041 | 0.0051 |

| Chr | Start | End | both | A1 | A2 |
|---|---|---|---|---|---|
| F10 | 8440001 | 8540000 | 0.0052 | 0.0028 | 0.0050 |
| F10 | 8460001 | 8560000 | 0.0036 | 0.0039 | 0.0035 |
| F10 | 8480001 | 8580000 | 0.0043 | 0.0029 | 0.0040 |
| F10 | 8500001 | 8600000 | 0.0032 | 0.0032 | 0.0029 |
| F10 | 8520001 | 8620000 | 0.0037 | 0.0031 | 0.0033 |
| F10 | 8540001 | 8640000 | 0.0037 | 0.0029 | 0.0033 |
| F10 | 8560001 | 8660000 | 0.0035 | 0.0014 | 0.0031 |
| F10 | 8580001 | 8680000 | 0.0024 | 0.0012 | 0.0022 |
| F10 | 8600001 | 8700000 | 0.0024 | 0.0006 | 0.0023 |
| F10 | 8620001 | 8720000 | 0.0018 | 0.0010 | 0.0017 |
| F10 | 8640001 | 8740000 | 0.0021 | 0.0010 | 0.0020 |
| F10 | 8660001 | 8760000 | 0.0022 | 0.0011 | 0.0022 |
| F10 | 8680001 | 8780000 | 0.0025 | 0.0016 | 0.0025 |
| F10 | 8700001 | 8800000 | 0.0029 | 0.0024 | 0.0028 |
| F10 | 8720001 | 8820000 | 0.0038 | 0.0028 | 0.0036 |
| F10 | 8740001 | 8840000 | 0.0043 | 0.0030 | 0.0041 |
| F10 | 8760001 | 8860000 | 0.0043 | 0.0030 | 0.0040 |
| F10 | 8780001 | 8880000 | 0.0038 | 0.0023 | 0.0035 |
| F10 | 8800001 | 8900000 | 0.0030 | 0.0027 | 0.0028 |
| F10 | 8820001 | 8920000 | 0.0033 | 0.0019 | 0.0032 |
| F10 | 8840001 | 8940000 | 0.0025 | 0.0015 | 0.0024 |
| F10 | 8860001 | 8960000 | 0.0024 | 0.0019 | 0.0023 |
| F10 | 8880001 | 8980000 | 0.0032 | 0.0028 | 0.0030 |
| F10 | 8900001 | 9000000 | 0.0044 | 0.0030 | 0.0041 |
| F10 | 8920001 | 9020000 | 0.0046 | 0.0030 | 0.0043 |
| F10 | 8940001 | 9040000 | 0.0049 | 0.0037 | 0.0046 |
| F10 | 8960001 | 9060000 | 0.0057 | 0.0039 | 0.0053 |
| F10 | 8980001 | 9080000 | 0.0055 | 0.0032 | 0.0051 |
| F10 | 9000001 | 9100000 | 0.0047 | 0.0027 | 0.0043 |
| F10 | 9020001 | 9120000 | 0.0044 | 0.0031 | 0.0040 |
| F10 | 9040001 | 9140000 | 0.0046 | 0.0030 | 0.0042 |
| F10 | 9060001 | 9160000 | 0.0045 | 0.0027 | 0.0041 |
| F10 | 9080001 | 9180000 | 0.0045 | 0.0031 | 0.0040 |
| F10 | 9100001 | 9200000 | 0.0050 | 0.0023 | 0.0045 |
| F10 | 9120001 | 9220000 | 0.0040 | 0.0021 | 0.0035 |
| F10 | 9140001 | 9240000 | 0.0037 | 0.0031 | 0.0032 |
| F10 | 9160001 | 9260000 | 0.0046 | 0.0028 | 0.0041 |
| F10 | 9180001 | 9280000 | 0.0040 | 0.0043 | 0.0036 |
| F10 | 9200001 | 9300000 | 0.0053 | 0.0071 | 0.0050 |
| F10 | 9220001 | 9320000 | 0.0083 | 0.0074 | 0.0081 |
| F10 | 9240001 | 9340000 | 0.0089 | 0.0060 | 0.0087 |
| F10 | 9260001 | 9360000 | 0.0078 | 0.0061 | 0.0075 |
| F10 | 9280001 | 9380000 | 0.0083 | 0.0044 | 0.0080 |
| F10 | 9300001 | 9400000 | 0.0066 | 0.0015 | 0.0063 |
| F10 | 9320001 | 9420000 | 0.0034 | 0.0012 | 0.0031 |
| F10 | 9340001 | 9440000 | 0.0027 | 0.0014 | 0.0026 |
| F10 | 9360001 | 9460000 | 0.0028 | 0.0013 | 0.0028 |
| F10 | 9380001 | 9480000 | 0.0023 | 0.0014 | 0.0023 |
| F10 | 9400001 | 9500000 | 0.0025 | 0.0019 | 0.0026 |
| F10 | 9420001 | 9520000 | 0.0030 | 0.0021 | 0.0031 |
| F10 | 9440001 | 9540000 | 0.0033 | 0.0018 | 0.0034 |
| F10 | 9460001 | 9560000 | 0.0028 | 0.0027 | 0.0029 |
| F10 | 9480001 | 9580000 | 0.0038 | 0.0023 | 0.0039 |
| F10 | 9500001 | 9600000 | 0.0033 | 0.0021 | 0.0034 |
| F10 | 9520001 | 9620000 | 0.0031 | 0.0020 | 0.0032 |
| F10 | 9540001 | 9640000 | 0.0031 | 0.0022 | 0.0032 |
| F10 | 9560001 | 9660000 | 0.0034 | 0.0014 | 0.0035 |
| F10 | 9580001 | 9680000 | 0.0025 | 0.0014 | 0.0026 |

| Chr | Start | End | both | A1 | A2 |
|---|---|---|---|---|---|
| F10 | 9600001 | 9700000 | 0.0022 | 0.0019 | 0.0023 |
| F10 | 9620001 | 9720000 | 0.0027 | 0.0020 | 0.0028 |
| F10 | 9640001 | 9740000 | 0.0028 | 0.0025 | 0.0029 |
| F10 | 9660001 | 9760000 | 0.0032 | 0.0026 | 0.0033 |
| F10 | 9680001 | 9780000 | 0.0036 | 0.0025 | 0.0037 |
| F10 | 9700001 | 9800000 | 0.0035 | 0.0018 | 0.0036 |
| F10 | 9720001 | 9820000 | 0.0029 | 0.0014 | 0.0029 |
| F10 | 9740001 | 9840000 | 0.0026 | 0.0010 | 0.0026 |
| F10 | 9760001 | 9860000 | 0.0024 | 0.0014 | 0.0023 |
| F10 | 9780001 | 9880000 | 0.0029 | 0.0020 | 0.0027 |
| F10 | 9800001 | 9900000 | 0.0036 | 0.0028 | 0.0031 |
| F10 | 9820001 | 9920000 | 0.0045 | 0.0034 | 0.0039 |
| F10 | 9840001 | 9940000 | 0.0048 | 0.0032 | 0.0042 |
| F10 | 9860001 | 9960000 | 0.0041 | 0.0028 | 0.0035 |
| F10 | 9880001 | 9980000 | 0.0036 | 0.0035 | 0.0029 |
| F10 | 9900001 | 10000000 | 0.0044 | 0.0033 | 0.0040 |
| F10 | 9920001 | 10020000 | 0.0043 | 0.0031 | 0.0037 |
| F10 | 9940001 | 10040000 | 0.0043 | 0.0032 | 0.0037 |
| F10 | 9960001 | 10060000 | 0.0045 | 0.0034 | 0.0039 |
| F10 | 9980001 | 10080000 | 0.0047 | 0.0021 | 0.0041 |
| F10 | 10000001 | 10100000 | 0.0034 | 0.0019 | 0.0027 |
| F10 | 10020001 | 10120000 | 0.0030 | 0.0017 | 0.0023 |
| F10 | 10040001 | 10140000 | 0.0027 | 0.0026 | 0.0020 |
| F10 | 10060001 | 10160000 | 0.0037 | 0.0027 | 0.0029 |
| F10 | 10080001 | 10180000 | 0.0038 | 0.0028 | 0.0031 |
| F10 | 10100001 | 10200000 | 0.0039 | 0.0025 | 0.0032 |
| F10 | 10120001 | 10220000 | 0.0036 | 0.0025 | 0.0031 |
| F10 | 10140001 | 10240000 | 0.0035 | 0.0014 | 0.0030 |
| F10 | 10160001 | 10260000 | 0.0024 | 0.0010 | 0.0018 |
| F10 | 10180001 | 10280000 | 0.0020 | 0.0008 | 0.0013 |
| F10 | 10200001 | 10300000 | 0.0017 | 0.0014 | 0.0011 |
| F10 | 10220001 | 10320000 | 0.0021 | 0.0013 | 0.0015 |
| F10 | 10240001 | 10340000 | 0.0019 | 0.0015 | 0.0014 |
| F10 | 10260001 | 10360000 | 0.0020 | 0.0026 | 0.0016 |
| F10 | 10280001 | 10380000 | 0.0032 | 0.0029 | 0.0028 |
| F10 | 10300001 | 10400000 | 0.0035 | 0.0024 | 0.0031 |
| F10 | 10320001 | 10420000 | 0.0031 | 0.0037 | 0.0028 |
| F10 | 10340001 | 10440000 | 0.0045 | 0.0045 | 0.0041 |
| F10 | 10360001 | 10460000 | 0.0054 | 0.0037 | 0.0050 |
| F10 | 10380001 | 10480000 | 0.0046 | 0.0035 | 0.0042 |
| F10 | 10400001 | 10500000 | 0.0046 | 0.0035 | 0.0039 |
| F10 | 10420001 | 10520000 | 0.0047 | 0.0026 | 0.0038 |
| F10 | 10440001 | 10540000 | 0.0038 | 0.0027 | 0.0030 |
| F10 | 10460001 | 10560000 | 0.0039 | 0.0024 | 0.0031 |
| F10 | 10480001 | 10580000 | 0.0034 | 0.0032 | 0.0028 |
| F10 | 10500001 | 10600000 | 0.0043 | 0.0035 | 0.0036 |
| F10 | 10520001 | 10620000 | 0.0045 | 0.0030 | 0.0037 |
| F10 | 10540001 | 10640000 | 0.0041 | 0.0022 | 0.0031 |
| F10 | 10560001 | 10660000 | 0.0034 | 0.0023 | 0.0022 |
| F10 | 10580001 | 10680000 | 0.0035 | 0.0015 | 0.0022 |
| F10 | 10600001 | 10700000 | 0.0024 | 0.0009 | 0.0012 |
| F10 | 10620001 | 10720000 | 0.0020 | 0.0015 | 0.0007 |
| F10 | 10640001 | 10740000 | 0.0025 | 0.0017 | 0.0013 |
| F10 | 10660001 | 10760000 | 0.0026 | 0.0015 | 0.0017 |
| F10 | 10680001 | 10780000 | 0.0025 | 0.0019 | 0.0014 |
| F10 | 10700001 | 10800000 | 0.0029 | 0.0025 | 0.0019 |
| F10 | 10720001 | 10820000 | 0.0034 | 0.0027 | 0.0026 |
| F10 | 10740001 | 10840000 | 0.0035 | 0.0025 | 0.0029 |

| Chr | Start | End | both | A1 | A2 |
|---|---|---|---|---|---|
| F10 | 10760001 | 10860000 | 0.0034 | 0.0032 | 0.0028 |
| F10 | 10780001 | 10880000 | 0.0040 | 0.0033 | 0.0035 |
| F10 | 10800001 | 10900000 | 0.0042 | 0.0033 | 0.0037 |
| F10 | 10820001 | 10920000 | 0.0042 | 0.0033 | 0.0038 |
| F10 | 10840001 | 10940000 | 0.0042 | 0.0037 | 0.0035 |
| F10 | 10860001 | 10960000 | 0.0047 | 0.0041 | 0.0038 |
| F10 | 10880001 | 10980000 | 0.0050 | 0.0042 | 0.0041 |
| F10 | 10900001 | 11000000 | 0.0050 | 0.0039 | 0.0040 |
| F10 | 10920001 | 11020000 | 0.0048 | 0.0039 | 0.0036 |
| F10 | 10940001 | 11040000 | 0.0050 | 0.0033 | 0.0039 |
| F10 | 10960001 | 11060000 | 0.0044 | 0.0023 | 0.0033 |
| F10 | 10980001 | 11080000 | 0.0034 | 0.0019 | 0.0023 |
| F10 | 11000001 | 11100000 | 0.0030 | 0.0021 | 0.0020 |
| F10 | 11020001 | 11120000 | 0.0030 | 0.0013 | 0.0022 |
| F10 | 11040001 | 11140000 | 0.0021 | 0.0020 | 0.0014 |
| F10 | 11060001 | 11160000 | 0.0029 | 0.0023 | 0.0021 |
| F10 | 11080001 | 11180000 | 0.0031 | 0.0023 | 0.0025 |
| F10 | 11100001 | 11200000 | 0.0031 | 0.0019 | 0.0025 |
| F10 | 11120001 | 11220000 | 0.0027 | 0.0020 | 0.0019 |
| F10 | 11140001 | 11240000 | 0.0028 | 0.0010 | 0.0019 |
| F10 | 11160001 | 11260000 | 0.0019 | 0.0009 | 0.0010 |
| F10 | 11180001 | 11280000 | 0.0021 | 0.0007 | 0.0009 |
| F10 | 11200001 | 11300000 | 0.0019 | 0.0009 | 0.0007 |
| F10 | 11220001 | 11320000 | 0.0022 | 0.0008 | 0.0010 |
| F10 | 11240001 | 11340000 | 0.0022 | 0.0008 | 0.0009 |
| F10 | 11260001 | 11360000 | 0.0022 | 0.0006 | 0.0009 |
| F10 | 11280001 | 11380000 | 0.0019 | 0.0008 | 0.0005 |
| F10 | 11300001 | 11400000 | 0.0021 | 0.0017 | 0.0007 |
| F10 | 11320001 | 11420000 | 0.0028 | 0.0028 | 0.0017 |
| F10 | 11340001 | 11440000 | 0.0038 | 0.0034 | 0.0027 |
| F10 | 11360001 | 11460000 | 0.0042 | 0.0039 | 0.0033 |
| F10 | 11380001 | 11480000 | 0.0047 | 0.0044 | 0.0040 |
| F10 | 11400001 | 11500000 | 0.0051 | 0.0035 | 0.0046 |
| F10 | 11420001 | 11520000 | 0.0042 | 0.0032 | 0.0035 |
| F10 | 11440001 | 11540000 | 0.0039 | 0.0028 | 0.0032 |
| F10 | 11460001 | 11560000 | 0.0037 | 0.0031 | 0.0028 |
| F10 | 11480001 | 11580000 | 0.0038 | 0.0042 | 0.0028 |
| F10 | 11500001 | 11600000 | 0.0049 | 0.0045 | 0.0038 |
| F10 | 11520001 | 11620000 | 0.0052 | 0.0037 | 0.0040 |
| F10 | 11540001 | 11640000 | 0.0044 | 0.0036 | 0.0033 |
| F10 | 11560001 | 11660000 | 0.0044 | 0.0027 | 0.0033 |
| F10 | 11580001 | 11680000 | 0.0036 | 0.0028 | 0.0026 |
| F10 | 11600001 | 11700000 | 0.0038 | 0.0025 | 0.0027 |
| F10 | 11620001 | 11720000 | 0.0035 | 0.0030 | 0.0025 |
| F10 | 11640001 | 11740000 | 0.0040 | 0.0034 | 0.0030 |
| F10 | 11660001 | 11760000 | 0.0043 | 0.0042 | 0.0034 |
| F10 | 11680001 | 11780000 | 0.0051 | 0.0027 | 0.0041 |
| F10 | 11700001 | 11800000 | 0.0034 | 0.0033 | 0.0026 |
| F10 | 11720001 | 11820000 | 0.0041 | 0.0032 | 0.0033 |
| F10 | 11740001 | 11840000 | 0.0041 | 0.0030 | 0.0034 |
| F10 | 11760001 | 11860000 | 0.0038 | 0.0027 | 0.0030 |
| F10 | 11780001 | 11880000 | 0.0034 | 0.0023 | 0.0028 |
| F10 | 11800001 | 11900000 | 0.0030 | 0.0019 | 0.0024 |
| F10 | 11820001 | 11920000 | 0.0024 | 0.0017 | 0.0018 |
| F10 | 11840001 | 11940000 | 0.0022 | 0.0025 | 0.0016 |
| F10 | 11860001 | 11960000 | 0.0034 | 0.0024 | 0.0027 |
| F10 | 11880001 | 11980000 | 0.0033 | 0.0027 | 0.0026 |
| F10 | 11900001 | 12000000 | 0.0036 | 0.0028 | 0.0028 |

| Chr | Start | End | both | A1 | A2 |
|---|---|---|---|---|---|
| F10 | 11920001 | 12020000 | 0.0038 | 0.0028 | 0.0030 |
| F10 | 11940001 | 12040000 | 0.0038 | 0.0018 | 0.0030 |
| F10 | 11960001 | 12060000 | 0.0025 | 0.0022 | 0.0017 |
| F10 | 11980001 | 12080000 | 0.0031 | 0.0024 | 0.0024 |
| F10 | 12000001 | 12100000 | 0.0033 | 0.0022 | 0.0026 |
| F10 | 12020001 | 12120000 | 0.0030 | 0.0019 | 0.0023 |
| F10 | 12040001 | 12140000 | 0.0026 | 0.0026 | 0.0020 |
| F10 | 12060001 | 12160000 | 0.0034 | 0.0020 | 0.0027 |
| F10 | 12080001 | 12180000 | 0.0027 | 0.0018 | 0.0019 |
| F10 | 12100001 | 12200000 | 0.0027 | 0.0016 | 0.0017 |
| F10 | 12120001 | 12220000 | 0.0026 | 0.0023 | 0.0015 |
| F10 | 12140001 | 12240000 | 0.0035 | 0.0021 | 0.0022 |
| F10 | 12160001 | 12260000 | 0.0033 | 0.0019 | 0.0022 |
| F10 | 12180001 | 12280000 | 0.0031 | 0.0018 | 0.0021 |
| F10 | 12200001 | 12300000 | 0.0028 | 0.0024 | 0.0019 |
| F10 | 12220001 | 12320000 | 0.0034 | 0.0024 | 0.0025 |
| F10 | 12240001 | 12340000 | 0.0032 | 0.0031 | 0.0025 |
| F10 | 12260001 | 12360000 | 0.0039 | 0.0031 | 0.0032 |
| F10 | 12280001 | 12380000 | 0.0038 | 0.0032 | 0.0032 |
| F10 | 12300001 | 12400000 | 0.0040 | 0.0032 | 0.0034 |
| F10 | 12320001 | 12420000 | 0.0041 | 0.0031 | 0.0034 |
| F10 | 12340001 | 12440000 | 0.0041 | 0.0026 | 0.0033 |
| F10 | 12360001 | 12460000 | 0.0037 | 0.0026 | 0.0027 |
| F10 | 12380001 | 12480000 | 0.0039 | 0.0027 | 0.0027 |
| F10 | 12400001 | 12500000 | 0.0039 | 0.0024 | 0.0027 |
| F10 | 12420001 | 12520000 | 0.0035 | 0.0028 | 0.0024 |
| F10 | 12440001 | 12540000 | 0.0036 | 0.0036 | 0.0027 |
| F10 | 12460001 | 12560000 | 0.0045 | 0.0042 | 0.0037 |
| F10 | 12480001 | 12580000 | 0.0048 | 0.0046 | 0.0042 |
| F10 | 12500001 | 12600000 | 0.0052 | 0.0046 | 0.0047 |
| F10 | 12520001 | 12620000 | 0.0054 | 0.0042 | 0.0047 |
| F10 | 12540001 | 12640000 | 0.0052 | 0.0028 | 0.0043 |
| F10 | 12560001 | 12660000 | 0.0036 | 0.0024 | 0.0027 |
| F10 | 12580001 | 12680000 | 0.0036 | 0.0016 | 0.0023 |
| F10 | 12600001 | 12700000 | 0.0029 | 0.0013 | 0.0016 |
| F10 | 12620001 | 12720000 | 0.0025 | 0.0009 | 0.0012 |
| F10 | 12640001 | 12740000 | 0.0020 | 0.0010 | 0.0009 |
| F10 | 12660001 | 12760000 | 0.0020 | 0.0008 | 0.0010 |
| F10 | 12680001 | 12780000 | 0.0017 | 0.0012 | 0.0008 |
| F10 | 12700001 | 12800000 | 0.0021 | 0.0014 | 0.0011 |
| F10 | 12720001 | 12820000 | 0.0024 | 0.0015 | 0.0014 |
| F10 | 12740001 | 12840000 | 0.0025 | 0.0021 | 0.0015 |
| F10 | 12760001 | 12860000 | 0.0030 | 0.0029 | 0.0020 |
| F10 | 12780001 | 12880000 | 0.0036 | 0.0030 | 0.0028 |
| F10 | 12800001 | 12900000 | 0.0037 | 0.0027 | 0.0031 |
| F10 | 12820001 | 12920000 | 0.0034 | 0.0032 | 0.0028 |
| F10 | 12840001 | 12940000 | 0.0040 | 0.0029 | 0.0031 |
| F10 | 12860001 | 12960000 | 0.0039 | 0.0028 | 0.0030 |
| F10 | 12880001 | 12980000 | 0.0040 | 0.0023 | 0.0030 |
| F10 | 12900001 | 13000000 | 0.0035 | 0.0037 | 0.0024 |
| F10 | 12920001 | 13020000 | 0.0051 | 0.0043 | 0.0040 |
| F10 | 12940001 | 13040000 | 0.0056 | 0.0040 | 0.0049 |
| F10 | 12960001 | 13060000 | 0.0051 | 0.0033 | 0.0045 |
| F10 | 12980001 | 13080000 | 0.0041 | 0.0034 | 0.0037 |
| F10 | 13000001 | 13100000 | 0.0043 | 0.0025 | 0.0039 |
| F10 | 13020001 | 13120000 | 0.0031 | 0.0018 | 0.0029 |
| F10 | 13040001 | 13140000 | 0.0023 | 0.0021 | 0.0021 |
| F10 | 13060001 | 13160000 | 0.0027 | 0.0025 | 0.0025 |

| Chr | Start | End | both | A1 | A2 |
|---|---|---|---|---|---|
| F10 | 13080001 | 13180000 | 0.0032 | 0.0027 | 0.0030 |
| F10 | 13100001 | 13200000 | 0.0034 | 0.0026 | 0.0031 |
| F10 | 13120001 | 13220000 | 0.0034 | 0.0027 | 0.0030 |
| F10 | 13140001 | 13240000 | 0.0036 | 0.0024 | 0.0031 |
| F10 | 13160001 | 13260000 | 0.0033 | 0.0018 | 0.0025 |
| F10 | 13180001 | 13280000 | 0.0028 | 0.0017 | 0.0019 |
| F10 | 13200001 | 13300000 | 0.0027 | 0.0022 | 0.0018 |
| F10 | 13220001 | 13320000 | 0.0031 | 0.0020 | 0.0021 |
| F10 | 13240001 | 13340000 | 0.0027 | 0.0020 | 0.0017 |
| F10 | 13260001 | 13360000 | 0.0026 | 0.0022 | 0.0018 |
| F10 | 13280001 | 13380000 | 0.0028 | 0.0019 | 0.0020 |
| F10 | 13300001 | 13400000 | 0.0027 | 0.0032 | 0.0018 |
| F10 | 13320001 | 13420000 | 0.0040 | 0.0029 | 0.0031 |
| F10 | 13340001 | 13440000 | 0.0038 | 0.0028 | 0.0030 |
| F10 | 13360001 | 13460000 | 0.0038 | 0.0030 | 0.0028 |
| F10 | 13380001 | 13480000 | 0.0042 | 0.0034 | 0.0031 |
| F10 | 13400001 | 13500000 | 0.0043 | 0.0015 | 0.0034 |
| F10 | 13420001 | 13520000 | 0.0023 | 0.0015 | 0.0016 |
| F10 | 13440001 | 13540000 | 0.0023 | 0.0019 | 0.0016 |
| F10 | 13460001 | 13560000 | 0.0027 | 0.0018 | 0.0021 |
| F10 | 13480001 | 13580000 | 0.0026 | 0.0017 | 0.0020 |
| F10 | 13500001 | 13600000 | 0.0025 | 0.0030 | 0.0018 |
| F10 | 13520001 | 13620000 | 0.0039 | 0.0037 | 0.0031 |
| F10 | 13540001 | 13640000 | 0.0046 | 0.0037 | 0.0037 |
| F10 | 13560001 | 13660000 | 0.0044 | 0.0037 | 0.0036 |
| F10 | 13580001 | 13680000 | 0.0044 | 0.0040 | 0.0036 |
| F10 | 13600001 | 13700000 | 0.0044 | 0.0026 | 0.0039 |
| F10 | 13620001 | 13720000 | 0.0030 | 0.0019 | 0.0024 |
| F10 | 13640001 | 13740000 | 0.0024 | 0.0021 | 0.0017 |
| F10 | 13660001 | 13760000 | 0.0030 | 0.0020 | 0.0019 |
| F10 | 13680001 | 13780000 | 0.0029 | 0.0027 | 0.0020 |
| F10 | 13700001 | 13800000 | 0.0040 | 0.0032 | 0.0028 |
| F10 | 13720001 | 13820000 | 0.0046 | 0.0037 | 0.0032 |
| F10 | 13740001 | 13840000 | 0.0052 | 0.0037 | 0.0037 |
| F10 | 13760001 | 13860000 | 0.0049 | 0.0038 | 0.0036 |
| F10 | 13780001 | 13880000 | 0.0051 | 0.0028 | 0.0038 |
| F10 | 13800001 | 13900000 | 0.0038 | 0.0025 | 0.0027 |
| F10 | 13820001 | 13920000 | 0.0034 | 0.0022 | 0.0025 |
| F10 | 13840001 | 13940000 | 0.0033 | 0.0017 | 0.0023 |
| F10 | 13860001 | 13960000 | 0.0028 | 0.0014 | 0.0017 |
| F10 | 13880001 | 13980000 | 0.0026 | 0.0014 | 0.0014 |
| F10 | 13900001 | 14000000 | 0.0026 | 0.0018 | 0.0014 |
| F10 | 13920001 | 14020000 | 0.0029 | 0.0037 | 0.0018 |
| F10 | 13940001 | 14040000 | 0.0044 | 0.0038 | 0.0034 |
| F10 | 13960001 | 14060000 | 0.0044 | 0.0040 | 0.0035 |
| F10 | 13980001 | 14080000 | 0.0044 | 0.0039 | 0.0038 |
| F10 | 14000001 | 14100000 | 0.0044 | 0.0035 | 0.0037 |
| F10 | 14020001 | 14120000 | 0.0038 | 0.0013 | 0.0033 |
| F10 | 14040001 | 14140000 | 0.0018 | 0.0012 | 0.0014 |
| F10 | 14060001 | 14160000 | 0.0017 | 0.0029 | 0.0013 |
| F10 | 14080001 | 14180000 | 0.0035 | 0.0034 | 0.0029 |
| F10 | 14100001 | 14200000 | 0.0040 | 0.0032 | 0.0032 |
| F10 | 14120001 | 14220000 | 0.0038 | 0.0038 | 0.0030 |
| F10 | 14140001 | 14240000 | 0.0047 | 0.0042 | 0.0037 |
| F10 | 14160001 | 14260000 | 0.0050 | 0.0022 | 0.0041 |
| F10 | 14180001 | 14280000 | 0.0030 | 0.0020 | 0.0022 |
| F10 | 14200001 | 14300000 | 0.0028 | 0.0021 | 0.0020 |
| F10 | 14220001 | 14320000 | 0.0030 | 0.0025 | 0.0023 |

| Chr | Start | End | both | A1 | A2 |
|---|---|---|---|---|---|
| F10 | 14240001 | 14340000 | 0.0032 | 0.0019 | 0.0024 |
| F10 | 14260001 | 14360000 | 0.0027 | 0.0021 | 0.0018 |
| F10 | 14280001 | 14380000 | 0.0029 | 0.0029 | 0.0019 |
| F10 | 14300001 | 14400000 | 0.0037 | 0.0029 | 0.0029 |
| F10 | 14320001 | 14420000 | 0.0038 | 0.0026 | 0.0030 |
| F10 | 14340001 | 14440000 | 0.0035 | 0.0039 | 0.0027 |
| F10 | 14360001 | 14460000 | 0.0049 | 0.0039 | 0.0042 |
| F10 | 14380001 | 14480000 | 0.0048 | 0.0029 | 0.0041 |
| F10 | 14400001 | 14500000 | 0.0036 | 0.0028 | 0.0029 |
| F10 | 14420001 | 14520000 | 0.0036 | 0.0025 | 0.0028 |
| F10 | 14440001 | 14540000 | 0.0034 | 0.0013 | 0.0026 |
| F10 | 14460001 | 14560000 | 0.0022 | 0.0020 | 0.0013 |
| F10 | 14480001 | 14580000 | 0.0029 | 0.0020 | 0.0020 |
| F10 | 14500001 | 14600000 | 0.0029 | 0.0020 | 0.0020 |
| F10 | 14520001 | 14620000 | 0.0029 | 0.0019 | 0.0020 |
| F10 | 14540001 | 14640000 | 0.0029 | 0.0019 | 0.0019 |
| F10 | 14560001 | 14660000 | 0.0028 | 0.0012 | 0.0019 |
| F10 | 14580001 | 14680000 | 0.0021 | 0.0016 | 0.0012 |
| F10 | 14600001 | 14700000 | 0.0025 | 0.0016 | 0.0017 |
| F10 | 14620001 | 14720000 | 0.0023 | 0.0019 | 0.0017 |
| F10 | 14640001 | 14740000 | 0.0025 | 0.0021 | 0.0021 |
| F10 | 14660001 | 14760000 | 0.0030 | 0.0024 | 0.0024 |
| F10 | 14680001 | 14780000 | 0.0033 | 0.0028 | 0.0026 |
| F10 | 14700001 | 14800000 | 0.0037 | 0.0028 | 0.0029 |
| F10 | 14720001 | 14820000 | 0.0037 | 0.0024 | 0.0030 |
| F10 | 14740001 | 14840000 | 0.0031 | 0.0029 | 0.0024 |
| F10 | 14760001 | 14860000 | 0.0038 | 0.0034 | 0.0031 |
| F10 | 14780001 | 14880000 | 0.0042 | 0.0024 | 0.0037 |
| F10 | 14800001 | 14900000 | 0.0032 | 0.0024 | 0.0028 |
| F10 | 14820001 | 14920000 | 0.0033 | 0.0025 | 0.0026 |
| F10 | 14840001 | 14940000 | 0.0036 | 0.0019 | 0.0027 |
| F10 | 14860001 | 14960000 | 0.0025 | 0.0012 | 0.0018 |
| F10 | 14880001 | 14980000 | 0.0020 | 0.0013 | 0.0010 |
| F10 | 14900001 | 15000000 | 0.0022 | 0.0014 | 0.0013 |
| F10 | 14920001 | 15020000 | 0.0021 | 0.0011 | 0.0013 |
| F10 | 14940001 | 15040000 | 0.0018 | 0.0020 | 0.0011 |
| F10 | 14960001 | 15060000 | 0.0028 | 0.0026 | 0.0022 |
| F10 | 14980001 | 15080000 | 0.0034 | 0.0029 | 0.0028 |
| F10 | 15000001 | 15100000 | 0.0038 | 0.0043 | 0.0030 |
| F10 | 15020001 | 15120000 | 0.0051 | 0.0049 | 0.0043 |
| F10 | 15040001 | 15140000 | 0.0057 | 0.0038 | 0.0048 |
| F10 | 15060001 | 15160000 | 0.0045 | 0.0031 | 0.0036 |
| F10 | 15080001 | 15180000 | 0.0036 | 0.0030 | 0.0029 |
| F10 | 15100001 | 15200000 | 0.0033 | 0.0017 | 0.0028 |
| F10 | 15120001 | 15220000 | 0.0020 | 0.0015 | 0.0016 |
| F10 | 15140001 | 15240000 | 0.0019 | 0.0017 | 0.0016 |
| F10 | 15160001 | 15260000 | 0.0022 | 0.0016 | 0.0018 |
| F10 | 15180001 | 15280000 | 0.0020 | 0.0015 | 0.0017 |
| F10 | 15200001 | 15300000 | 0.0019 | 0.0020 | 0.0014 |
| F10 | 15220001 | 15320000 | 0.0025 | 0.0020 | 0.0018 |
| F10 | 15240001 | 15340000 | 0.0025 | 0.0019 | 0.0017 |
| F10 | 15260001 | 15360000 | 0.0024 | 0.0020 | 0.0016 |
| F10 | 15280001 | 15380000 | 0.0025 | 0.0017 | 0.0017 |
| F10 | 15300001 | 15400000 | 0.0022 | 0.0012 | 0.0015 |
| F10 | 15320001 | 15420000 | 0.0019 | 0.0009 | 0.0010 |
| F10 | 15340001 | 15440000 | 0.0016 | 0.0013 | 0.0009 |
| F10 | 15360001 | 15460000 | 0.0021 | 0.0027 | 0.0012 |
| F10 | 15380001 | 15480000 | 0.0036 | 0.0035 | 0.0026 |

| Chr | Start | End | both | A1 | A2 |
|---|---|---|---|---|---|
| F10 | 15400001 | 15500000 | 0.0045 | 0.0038 | 0.0033 |
| F10 | 15420001 | 15520000 | 0.0045 | 0.0040 | 0.0037 |
| F10 | 15440001 | 15540000 | 0.0047 | 0.0039 | 0.0039 |
| F10 | 15460001 | 15560000 | 0.0044 | 0.0028 | 0.0039 |
| F10 | 15480001 | 15580000 | 0.0033 | 0.0026 | 0.0028 |
| F10 | 15500001 | 15600000 | 0.0033 | 0.0028 | 0.0028 |
| F10 | 15520001 | 15620000 | 0.0037 | 0.0026 | 0.0030 |
| F10 | 15540001 | 15640000 | 0.0035 | 0.0021 | 0.0028 |
| F10 | 15560001 | 15660000 | 0.0029 | 0.0020 | 0.0022 |
| F10 | 15580001 | 15680000 | 0.0029 | 0.0019 | 0.0023 |
| F10 | 15600001 | 15700000 | 0.0026 | 0.0018 | 0.0022 |
| F10 | 15620001 | 15720000 | 0.0023 | 0.0019 | 0.0019 |
| F10 | 15640001 | 15740000 | 0.0027 | 0.0025 | 0.0019 |
| F10 | 15660001 | 15760000 | 0.0034 | 0.0024 | 0.0026 |
| F10 | 15680001 | 15780000 | 0.0032 | 0.0020 | 0.0023 |
| F10 | 15700001 | 15800000 | 0.0030 | 0.0017 | 0.0020 |
| F10 | 15720001 | 15820000 | 0.0028 | 0.0020 | 0.0017 |
| F10 | 15740001 | 15840000 | 0.0028 | 0.0019 | 0.0019 |
| F10 | 15760001 | 15860000 | 0.0027 | 0.0017 | 0.0018 |
| F10 | 15780001 | 15880000 | 0.0025 | 0.0028 | 0.0016 |
| F10 | 15800001 | 15900000 | 0.0032 | 0.0031 | 0.0023 |
| F10 | 15820001 | 15920000 | 0.0034 | 0.0028 | 0.0026 |
| F10 | 15840001 | 15940000 | 0.0033 | 0.0023 | 0.0024 |
| F10 | 15860001 | 15960000 | 0.0027 | 0.0031 | 0.0019 |
| F10 | 15880001 | 15980000 | 0.0036 | 0.0022 | 0.0027 |
| F10 | 15900001 | 16000000 | 0.0030 | 0.0025 | 0.0023 |
| F10 | 15920001 | 16020000 | 0.0032 | 0.0023 | 0.0025 |
| F10 | 15940001 | 16040000 | 0.0031 | 0.0028 | 0.0023 |
| F10 | 15960001 | 16060000 | 0.0035 | 0.0019 | 0.0027 |
| F10 | 15980001 | 16080000 | 0.0026 | 0.0015 | 0.0018 |
| F10 | 16000001 | 16100000 | 0.0020 | 0.0014 | 0.0013 |
| F10 | 16020001 | 16120000 | 0.0021 | 0.0016 | 0.0014 |
| F10 | 16040001 | 16140000 | 0.0023 | 0.0018 | 0.0018 |
| F10 | 16060001 | 16160000 | 0.0024 | 0.0023 | 0.0020 |
| F10 | 16080001 | 16180000 | 0.0031 | 0.0027 | 0.0025 |
| F10 | 16100001 | 16200000 | 0.0036 | 0.0023 | 0.0032 |
| F10 | 16120001 | 16220000 | 0.0030 | 0.0023 | 0.0027 |
| F10 | 16140001 | 16240000 | 0.0029 | 0.0019 | 0.0026 |
| F10 | 16160001 | 16260000 | 0.0026 | 0.0018 | 0.0023 |
| F10 | 16180001 | 16280000 | 0.0024 | 0.0019 | 0.0023 |
| F10 | 16200001 | 16300000 | 0.0024 | 0.0022 | 0.0022 |
| F10 | 16220001 | 16320000 | 0.0027 | 0.0024 | 0.0025 |
| F10 | 16240001 | 16340000 | 0.0029 | 0.0025 | 0.0025 |
| F10 | 16260001 | 16360000 | 0.0030 | 0.0037 | 0.0027 |
| F10 | 16280001 | 16380000 | 0.0041 | 0.0036 | 0.0034 |
| F10 | 16300001 | 16400000 | 0.0041 | 0.0038 | 0.0033 |
| F10 | 16320001 | 16420000 | 0.0044 | 0.0038 | 0.0034 |
| F10 | 16340001 | 16440000 | 0.0046 | 0.0037 | 0.0036 |
| F10 | 16360001 | 16460000 | 0.0043 | 0.0025 | 0.0034 |
| F10 | 16380001 | 16480000 | 0.0030 | 0.0023 | 0.0024 |
| F10 | 16400001 | 16500000 | 0.0027 | 0.0018 | 0.0022 |
| F10 | 16420001 | 16520000 | 0.0022 | 0.0015 | 0.0017 |
| F10 | 16440001 | 16540000 | 0.0018 | 0.0016 | 0.0013 |
| F10 | 16460001 | 16560000 | 0.0022 | 0.0024 | 0.0015 |
| F10 | 16480001 | 16580000 | 0.0031 | 0.0021 | 0.0024 |
| F10 | 16500001 | 16600000 | 0.0029 | 0.0021 | 0.0021 |
| F10 | 16520001 | 16620000 | 0.0030 | 0.0031 | 0.0021 |
| F10 | 16540001 | 16640000 | 0.0039 | 0.0036 | 0.0030 |

| Chr | Start | End | both | A1 | A2 |
|---|---|---|---|---|---|
| F10 | 16560001 | 16660000 | 0.0044 | 0.0033 | 0.0034 |
| F10 | 16580001 | 16680000 | 0.0044 | 0.0038 | 0.0034 |
| F10 | 16600001 | 16700000 | 0.0051 | 0.0038 | 0.0040 |
| F10 | 16620001 | 16720000 | 0.0050 | 0.0033 | 0.0041 |
| F10 | 16640001 | 16740000 | 0.0045 | 0.0027 | 0.0038 |
| F10 | 16660001 | 16760000 | 0.0039 | 0.0023 | 0.0034 |
| F10 | 16680001 | 16780000 | 0.0031 | 0.0024 | 0.0028 |
| F10 | 16700001 | 16800000 | 0.0029 | 0.0024 | 0.0027 |
| F10 | 16720001 | 16820000 | 0.0030 | 0.0024 | 0.0028 |
| F10 | 16740001 | 16840000 | 0.0030 | 0.0025 | 0.0027 |
| F10 | 16760001 | 16860000 | 0.0030 | 0.0026 | 0.0026 |
| F10 | 16780001 | 16880000 | 0.0032 | 0.0024 | 0.0026 |
| F10 | 16800001 | 16900000 | 0.0032 | 0.0029 | 0.0023 |
| F10 | 16820001 | 16920000 | 0.0036 | 0.0030 | 0.0028 |
| F10 | 16840001 | 16940000 | 0.0038 | 0.0027 | 0.0032 |
| F10 | 16860001 | 16960000 | 0.0036 | 0.0029 | 0.0029 |
| F10 | 16880001 | 16980000 | 0.0037 | 0.0031 | 0.0032 |
| F10 | 16900001 | 17000000 | 0.0039 | 0.0031 | 0.0036 |
| F10 | 16920001 | 17020000 | 0.0039 | 0.0024 | 0.0037 |
| F10 | 16940001 | 17040000 | 0.0031 | 0.0033 | 0.0029 |
| F10 | 16960001 | 17060000 | 0.0042 | 0.0033 | 0.0040 |
| F10 | 16980001 | 17080000 | 0.0041 | 0.0034 | 0.0037 |
| F10 | 17000001 | 17100000 | 0.0043 | 0.0028 | 0.0037 |
| F10 | 17020001 | 17120000 | 0.0037 | 0.0029 | 0.0028 |
| F10 | 17040001 | 17140000 | 0.0040 | 0.0026 | 0.0029 |
| F10 | 17060001 | 17160000 | 0.0035 | 0.0032 | 0.0025 |
| F10 | 17080001 | 17180000 | 0.0043 | 0.0025 | 0.0032 |
| F10 | 17100001 | 17200000 | 0.0033 | 0.0025 | 0.0025 |
| F10 | 17120001 | 17220000 | 0.0032 | 0.0026 | 0.0026 |
| F10 | 17140001 | 17240000 | 0.0032 | 0.0023 | 0.0028 |
| F10 | 17160001 | 17260000 | 0.0027 | 0.0011 | 0.0024 |
| F10 | 17180001 | 17280000 | 0.0013 | 0.0015 | 0.0013 |
| F10 | 17200001 | 17300000 | 0.0018 | 0.0026 | 0.0018 |
| F10 | 17220001 | 17320000 | 0.0031 | 0.0026 | 0.0029 |
| F10 | 17240001 | 17340000 | 0.0033 | 0.0025 | 0.0029 |
| F10 | 17260001 | 17360000 | 0.0036 | 0.0024 | 0.0030 |
| F10 | 17280001 | 17380000 | 0.0036 | 0.0024 | 0.0029 |
| F10 | 17300001 | 17400000 | 0.0036 | 0.0014 | 0.0028 |
| F10 | 17320001 | 17420000 | 0.0028 | 0.0015 | 0.0019 |
| F10 | 17340001 | 17440000 | 0.0029 | 0.0013 | 0.0018 |
| F10 | 17360001 | 17460000 | 0.0027 | 0.0013 | 0.0015 |
| F10 | 17380001 | 17480000 | 0.0028 | 0.0020 | 0.0016 |
| F10 | 17400001 | 17500000 | 0.0035 | 0.0020 | 0.0022 |
| F10 | 17420001 | 17520000 | 0.0033 | 0.0021 | 0.0022 |
| F10 | 17440001 | 17540000 | 0.0033 | 0.0020 | 0.0025 |
| F10 | 17460001 | 17560000 | 0.0030 | 0.0024 | 0.0024 |
| F10 | 17480001 | 17580000 | 0.0033 | 0.0030 | 0.0026 |
| F10 | 17500001 | 17600000 | 0.0040 | 0.0030 | 0.0033 |
| F10 | 17520001 | 17620000 | 0.0039 | 0.0034 | 0.0032 |
| F10 | 17540001 | 17640000 | 0.0042 | 0.0035 | 0.0035 |
| F10 | 17560001 | 17660000 | 0.0043 | 0.0033 | 0.0035 |
| F10 | 17580001 | 17680000 | 0.0043 | 0.0023 | 0.0035 |
| F10 | 17600001 | 17700000 | 0.0033 | 0.0023 | 0.0025 |
| F10 | 17620001 | 17720000 | 0.0035 | 0.0016 | 0.0026 |
| F10 | 17640001 | 17740000 | 0.0028 | 0.0016 | 0.0019 |
| F10 | 17660001 | 17760000 | 0.0029 | 0.0017 | 0.0020 |
| F10 | 17680001 | 17780000 | 0.0030 | 0.0012 | 0.0020 |
| F10 | 17700001 | 17800000 | 0.0022 | 0.0012 | 0.0013 |

| Chr | Start | End | both | A1 | A2 |
|---|---|---|---|---|---|
| F10 | 17720001 | 17820000 | 0.0020 | 0.0017 | 0.0012 |
| F10 | 17740001 | 17840000 | 0.0024 | 0.0016 | 0.0014 |
| F10 | 17760001 | 17860000 | 0.0022 | 0.0015 | 0.0013 |
| F10 | 17780001 | 17880000 | 0.0018 | 0.0019 | 0.0012 |
| F10 | 17800001 | 17900000 | 0.0025 | 0.0016 | 0.0015 |
| F10 | 17820001 | 17920000 | 0.0023 | 0.0012 | 0.0013 |
| F10 | 17840001 | 17940000 | 0.0020 | 0.0020 | 0.0011 |
| F10 | 17860001 | 17960000 | 0.0026 | 0.0021 | 0.0017 |
| F10 | 17880001 | 17980000 | 0.0027 | 0.0022 | 0.0017 |
| F10 | 17900001 | 18000000 | 0.0027 | 0.0024 | 0.0019 |
| F10 | 17920001 | 18020000 | 0.0027 | 0.0025 | 0.0021 |
| F10 | 17940001 | 18040000 | 0.0028 | 0.0027 | 0.0023 |
| F10 | 17960001 | 18060000 | 0.0032 | 0.0025 | 0.0027 |
| F10 | 17980001 | 18080000 | 0.0032 | 0.0024 | 0.0028 |
| F10 | 18000001 | 18100000 | 0.0033 | 0.0024 | 0.0028 |
| F10 | 18020001 | 18120000 | 0.0035 | 0.0026 | 0.0027 |
| F10 | 18040001 | 18140000 | 0.0038 | 0.0017 | 0.0031 |
| F10 | 18060001 | 18160000 | 0.0029 | 0.0021 | 0.0021 |
| F10 | 18080001 | 18180000 | 0.0033 | 0.0020 | 0.0025 |
| F10 | 18100001 | 18200000 | 0.0031 | 0.0022 | 0.0024 |
| F10 | 18120001 | 18220000 | 0.0031 | 0.0017 | 0.0025 |
| F10 | 18140001 | 18240000 | 0.0026 | 0.0016 | 0.0020 |
| F10 | 18160001 | 18260000 | 0.0024 | 0.0011 | 0.0018 |
| F10 | 18180001 | 18280000 | 0.0018 | 0.0015 | 0.0013 |
| F10 | 18200001 | 18300000 | 0.0021 | 0.0016 | 0.0015 |
| F10 | 18220001 | 18320000 | 0.0023 | 0.0020 | 0.0017 |
| F10 | 18240001 | 18340000 | 0.0027 | 0.0022 | 0.0022 |
| F10 | 18260001 | 18360000 | 0.0029 | 0.0023 | 0.0023 |
| F10 | 18280001 | 18380000 | 0.0033 | 0.0020 | 0.0025 |
| F10 | 18300001 | 18400000 | 0.0028 | 0.0019 | 0.0021 |
| F10 | 18320001 | 18420000 | 0.0032 | 0.0021 | 0.0024 |
| F10 | 18340001 | 18440000 | 0.0036 | 0.0019 | 0.0026 |
| F10 | 18360001 | 18460000 | 0.0036 | 0.0024 | 0.0025 |
| F10 | 18380001 | 18480000 | 0.0040 | 0.0043 | 0.0030 |
| F10 | 18400001 | 18500000 | 0.0063 | 0.0040 | 0.0050 |
| F10 | 18420001 | 18520000 | 0.0056 | 0.0046 | 0.0043 |
| F10 | 18440001 | 18540000 | 0.0061 | 0.0053 | 0.0049 |
| F10 | 18460001 | 18560000 | 0.0067 | 0.0063 | 0.0056 |
| F10 | 18480001 | 18580000 | 0.0076 | 0.0048 | 0.0067 |
| F10 | 18500001 | 18600000 | 0.0058 | 0.0054 | 0.0051 |
| F10 | 18520001 | 18620000 | 0.0066 | 0.0045 | 0.0059 |
| F10 | 18540001 | 18640000 | 0.0056 | 0.0040 | 0.0050 |
| F10 | 18560001 | 18660000 | 0.0051 | 0.0025 | 0.0044 |
| F10 | 18580001 | 18680000 | 0.0036 | 0.0026 | 0.0028 |
| F10 | 18600001 | 18700000 | 0.0039 | 0.0029 | 0.0031 |
| F10 | 18620001 | 18720000 | 0.0041 | 0.0027 | 0.0035 |
| F10 | 18640001 | 18740000 | 0.0037 | 0.0027 | 0.0031 |
| F10 | 18660001 | 18760000 | 0.0038 | 0.0028 | 0.0031 |
| F10 | 18680001 | 18780000 | 0.0038 | 0.0021 | 0.0032 |
| F10 | 18700001 | 18800000 | 0.0030 | 0.0020 | 0.0023 |
| F10 | 18720001 | 18820000 | 0.0030 | 0.0019 | 0.0023 |
| F10 | 18740001 | 18840000 | 0.0028 | 0.0016 | 0.0021 |
| F10 | 18760001 | 18860000 | 0.0027 | 0.0013 | 0.0019 |
| F10 | 18780001 | 18880000 | 0.0023 | 0.0014 | 0.0016 |
| F10 | 18800001 | 18900000 | 0.0022 | 0.0005 | 0.0017 |
| F10 | 18820001 | 18920000 | 0.0009 | 0.0009 | 0.0006 |
| F10 | 18840001 | 18940000 | 0.0014 | 0.0008 | 0.0010 |
| F10 | 18860001 | 18960000 | 0.0011 | 0.0008 | 0.0009 |

| Chr | Start | End | both | A1 | A2 |
|---|---|---|---|---|---|
| F10 | 18880001 | 18980000 | 0.0010 | 0.0006 | 0.0009 |
| F10 | 18900001 | 19000000 | 0.0007 | 0.0007 | 0.0007 |
| F10 | 18920001 | 19020000 | 0.0009 | 0.0005 | 0.0009 |
| F10 | 18940001 | 19040000 | 0.0006 | 0.0006 | 0.0006 |
| F10 | 18960001 | 19060000 | 0.0007 | 0.0010 | 0.0007 |
| F10 | 18980001 | 19080000 | 0.0012 | 0.0019 | 0.0012 |
| F10 | 19000001 | 19100000 | 0.0022 | 0.0022 | 0.0022 |
| F10 | 19020001 | 19120000 | 0.0026 | 0.0023 | 0.0026 |
| F10 | 19040001 | 19140000 | 0.0028 | 0.0033 | 0.0027 |
| F10 | 19060001 | 19160000 | 0.0038 | 0.0030 | 0.0038 |
| F10 | 19080001 | 19180000 | 0.0037 | 0.0028 | 0.0034 |
| F10 | 19100001 | 19200000 | 0.0037 | 0.0024 | 0.0033 |
| F10 | 19120001 | 19220000 | 0.0031 | 0.0026 | 0.0027 |
| F10 | 19140001 | 19240000 | 0.0034 | 0.0019 | 0.0027 |
| F10 | 19160001 | 19260000 | 0.0029 | 0.0027 | 0.0021 |
| F10 | 19180001 | 19280000 | 0.0037 | 0.0026 | 0.0031 |
| F10 | 19200001 | 19300000 | 0.0036 | 0.0028 | 0.0030 |
| F10 | 19220001 | 19320000 | 0.0039 | 0.0028 | 0.0033 |
| F10 | 19240001 | 19340000 | 0.0037 | 0.0025 | 0.0033 |
| F10 | 19260001 | 19360000 | 0.0034 | 0.0020 | 0.0030 |
| F10 | 19280001 | 19380000 | 0.0028 | 0.0027 | 0.0024 |
| F10 | 19300001 | 19400000 | 0.0033 | 0.0033 | 0.0030 |
| F10 | 19320001 | 19420000 | 0.0041 | 0.0033 | 0.0036 |
| F10 | 19340001 | 19440000 | 0.0040 | 0.0032 | 0.0036 |
| F10 | 19360001 | 19460000 | 0.0038 | 0.0033 | 0.0034 |
| F10 | 19380001 | 19480000 | 0.0039 | 0.0031 | 0.0034 |
| F10 | 19400001 | 19500000 | 0.0036 | 0.0028 | 0.0031 |
| F10 | 19420001 | 19520000 | 0.0033 | 0.0036 | 0.0030 |
| F10 | 19440001 | 19540000 | 0.0043 | 0.0042 | 0.0038 |
| F10 | 19460001 | 19560000 | 0.0052 | 0.0046 | 0.0047 |
| F10 | 19480001 | 19580000 | 0.0058 | 0.0037 | 0.0053 |
| F10 | 19500001 | 19600000 | 0.0049 | 0.0045 | 0.0044 |
| F10 | 19520001 | 19620000 | 0.0058 | 0.0038 | 0.0054 |
| F10 | 19540001 | 19640000 | 0.0050 | 0.0031 | 0.0047 |
| F10 | 19560001 | 19660000 | 0.0040 | 0.0030 | 0.0038 |
| F10 | 19580001 | 19680000 | 0.0036 | 0.0034 | 0.0035 |
| F10 | 19600001 | 19700000 | 0.0043 | 0.0019 | 0.0039 |
| F10 | 19620001 | 19720000 | 0.0026 | 0.0024 | 0.0020 |
| F10 | 19640001 | 19740000 | 0.0032 | 0.0031 | 0.0024 |
| F10 | 19660001 | 19760000 | 0.0041 | 0.0040 | 0.0033 |
| F10 | 19680001 | 19780000 | 0.0049 | 0.0037 | 0.0041 |
| F10 | 19700001 | 19800000 | 0.0046 | 0.0046 | 0.0037 |
| F10 | 19720001 | 19820000 | 0.0054 | 0.0045 | 0.0046 |
| F10 | 19740001 | 19840000 | 0.0054 | 0.0039 | 0.0047 |
| F10 | 19760001 | 19860000 | 0.0046 | 0.0023 | 0.0038 |
| F10 | 19780001 | 19880000 | 0.0032 | 0.0024 | 0.0023 |
| F10 | 19800001 | 19900000 | 0.0034 | 0.0020 | 0.0024 |
| F10 | 19820001 | 19920000 | 0.0028 | 0.0017 | 0.0021 |
| F10 | 19840001 | 19940000 | 0.0025 | 0.0023 | 0.0017 |
| F10 | 19860001 | 19960000 | 0.0029 | 0.0025 | 0.0022 |
| F10 | 19880001 | 19980000 | 0.0032 | 0.0028 | 0.0025 |
| F10 | 19900001 | 20000000 | 0.0035 | 0.0029 | 0.0028 |
| F10 | 19920001 | 20020000 | 0.0037 | 0.0024 | 0.0030 |
| F10 | 19940001 | 20040000 | 0.0032 | 0.0018 | 0.0025 |
| F10 | 19960001 | 20060000 | 0.0028 | 0.0020 | 0.0020 |
| F10 | 19980001 | 20080000 | 0.0029 | 0.0019 | 0.0021 |
| F10 | 20000001 | 20100000 | 0.0027 | 0.0014 | 0.0020 |
| F10 | 20020001 | 20120000 | 0.0022 | 0.0016 | 0.0014 |

| Chr | Start | End | both | A1 | A2 |
|---|---|---|---|---|---|
| F10 | 20040001 | 20140000 | 0.0025 | 0.0024 | 0.0016 |
| F10 | 20060001 | 20160000 | 0.0032 | 0.0021 | 0.0024 |
| F10 | 20080001 | 20180000 | 0.0029 | 0.0016 | 0.0021 |
| F10 | 20100001 | 20200000 | 0.0024 | 0.0019 | 0.0014 |
| F10 | 20120001 | 20220000 | 0.0027 | 0.0018 | 0.0017 |
| F10 | 20140001 | 20240000 | 0.0024 | 0.0016 | 0.0016 |
| F10 | 20160001 | 20260000 | 0.0022 | 0.0019 | 0.0013 |
| F10 | 20180001 | 20280000 | 0.0026 | 0.0024 | 0.0018 |
| F10 | 20200001 | 20300000 | 0.0031 | 0.0024 | 0.0024 |
| F10 | 20220001 | 20320000 | 0.0030 | 0.0030 | 0.0027 |
| F10 | 20240001 | 20340000 | 0.0038 | 0.0031 | 0.0035 |
| F10 | 20260001 | 20360000 | 0.0042 | 0.0029 | 0.0040 |
| F10 | 20280001 | 20380000 | 0.0039 | 0.0026 | 0.0038 |
| F10 | 20300001 | 20400000 | 0.0035 | 0.0034 | 0.0035 |
| F10 | 20320001 | 20420000 | 0.0042 | 0.0031 | 0.0041 |
| F10 | 20340001 | 20440000 | 0.0038 | 0.0026 | 0.0038 |
| F10 | 20360001 | 20460000 | 0.0030 | 0.0027 | 0.0029 |
| F10 | 20380001 | 20480000 | 0.0031 | 0.0034 | 0.0030 |
| F10 | 20400001 | 20500000 | 0.0041 | 0.0025 | 0.0037 |
| F10 | 20420001 | 20520000 | 0.0033 | 0.0021 | 0.0029 |
| F10 | 20440001 | 20540000 | 0.0028 | 0.0020 | 0.0024 |
| F10 | 20460001 | 20560000 | 0.0027 | 0.0022 | 0.0021 |
| F10 | 20480001 | 20580000 | 0.0028 | 0.0020 | 0.0023 |
| F10 | 20500001 | 20600000 | 0.0023 | 0.0021 | 0.0020 |
| F10 | 20520001 | 20620000 | 0.0025 | 0.0029 | 0.0021 |
| F10 | 20540001 | 20640000 | 0.0035 | 0.0034 | 0.0030 |
| F10 | 20560001 | 20660000 | 0.0042 | 0.0033 | 0.0036 |
| F10 | 20580001 | 20680000 | 0.0042 | 0.0040 | 0.0035 |
| F10 | 20600001 | 20700000 | 0.0049 | 0.0048 | 0.0042 |
| F10 | 20620001 | 20720000 | 0.0059 | 0.0046 | 0.0051 |
| F10 | 20640001 | 20740000 | 0.0057 | 0.0043 | 0.0049 |
| F10 | 20660001 | 20760000 | 0.0052 | 0.0045 | 0.0047 |
| F10 | 20680001 | 20780000 | 0.0052 | 0.0042 | 0.0049 |
| F10 | 20700001 | 20800000 | 0.0053 | 0.0045 | 0.0048 |
| F10 | 20720001 | 20820000 | 0.0057 | 0.0042 | 0.0052 |
| F10 | 20740001 | 20840000 | 0.0053 | 0.0050 | 0.0048 |
| F10 | 20760001 | 20860000 | 0.0064 | 0.0048 | 0.0056 |
| F10 | 20780001 | 20880000 | 0.0064 | 0.0037 | 0.0054 |
| F10 | 20800001 | 20900000 | 0.0048 | 0.0025 | 0.0040 |
| F10 | 20820001 | 20920000 | 0.0033 | 0.0024 | 0.0025 |
| F10 | 20840001 | 20940000 | 0.0032 | 0.0013 | 0.0023 |
| F10 | 20860001 | 20960000 | 0.0020 | 0.0009 | 0.0012 |
| F10 | 20880001 | 20980000 | 0.0015 | 0.0010 | 0.0008 |
| F10 | 20900001 | 21000000 | 0.0017 | 0.0012 | 0.0010 |
| F10 | 20920001 | 21020000 | 0.0019 | 0.0015 | 0.0012 |
| F10 | 20940001 | 21040000 | 0.0025 | 0.0022 | 0.0018 |
| F10 | 20960001 | 21060000 | 0.0031 | 0.0025 | 0.0024 |
| F10 | 20980001 | 21080000 | 0.0034 | 0.0025 | 0.0027 |
| F10 | 21000001 | 21100000 | 0.0034 | 0.0041 | 0.0026 |
| F10 | 21020001 | 21120000 | 0.0050 | 0.0039 | 0.0043 |
| F10 | 21040001 | 21140000 | 0.0043 | 0.0032 | 0.0039 |
| F10 | 21060001 | 21160000 | 0.0035 | 0.0030 | 0.0033 |
| F10 | 21080001 | 21180000 | 0.0033 | 0.0033 | 0.0031 |
| F10 | 21100001 | 21200000 | 0.0038 | 0.0013 | 0.0036 |
| F10 | 21120001 | 21220000 | 0.0018 | 0.0013 | 0.0014 |
| F10 | 21140001 | 21240000 | 0.0020 | 0.0017 | 0.0016 |
| F10 | 21160001 | 21260000 | 0.0023 | 0.0016 | 0.0019 |
| F10 | 21180001 | 21280000 | 0.0022 | 0.0015 | 0.0018 |

| Chr | Start | End | both | A1 | A2 |
|---|---|---|---|---|---|
| F10 | 21200001 | 21300000 | 0.0023 | 0.0020 | 0.0016 |
| F10 | 21220001 | 21320000 | 0.0026 | 0.0021 | 0.0020 |
| F10 | 21240001 | 21340000 | 0.0028 | 0.0030 | 0.0020 |
| F10 | 21260001 | 21360000 | 0.0037 | 0.0033 | 0.0029 |
| F10 | 21280001 | 21380000 | 0.0042 | 0.0033 | 0.0032 |
| F10 | 21300001 | 21400000 | 0.0042 | 0.0029 | 0.0032 |
| F10 | 21320001 | 21420000 | 0.0040 | 0.0030 | 0.0030 |
| F10 | 21340001 | 21440000 | 0.0040 | 0.0021 | 0.0031 |
| F10 | 21360001 | 21460000 | 0.0029 | 0.0024 | 0.0020 |
| F10 | 21380001 | 21480000 | 0.0030 | 0.0020 | 0.0023 |
| F10 | 21400001 | 21500000 | 0.0026 | 0.0025 | 0.0019 |
| F10 | 21420001 | 21520000 | 0.0031 | 0.0025 | 0.0024 |
| F10 | 21440001 | 21540000 | 0.0031 | 0.0024 | 0.0026 |
| F10 | 21460001 | 21560000 | 0.0033 | 0.0025 | 0.0027 |
| F10 | 21480001 | 21580000 | 0.0035 | 0.0028 | 0.0027 |
| F10 | 21500001 | 21600000 | 0.0038 | 0.0020 | 0.0030 |
| F10 | 21520001 | 21620000 | 0.0029 | 0.0018 | 0.0021 |
| F10 | 21540001 | 21640000 | 0.0028 | 0.0017 | 0.0018 |
| F10 | 21560001 | 21660000 | 0.0026 | 0.0023 | 0.0018 |
| F10 | 21580001 | 21680000 | 0.0030 | 0.0029 | 0.0023 |
| F10 | 21600001 | 21700000 | 0.0036 | 0.0032 | 0.0029 |
| F10 | 21620001 | 21720000 | 0.0041 | 0.0033 | 0.0031 |
| F10 | 21640001 | 21740000 | 0.0042 | 0.0043 | 0.0032 |
| F10 | 21660001 | 21760000 | 0.0054 | 0.0032 | 0.0042 |
| F10 | 21680001 | 21780000 | 0.0044 | 0.0028 | 0.0032 |
| F10 | 21700001 | 21800000 | 0.0040 | 0.0026 | 0.0027 |
| F10 | 21720001 | 21820000 | 0.0037 | 0.0027 | 0.0026 |
| F10 | 21740001 | 21840000 | 0.0038 | 0.0016 | 0.0028 |
| F10 | 21760001 | 21860000 | 0.0023 | 0.0019 | 0.0017 |
| F10 | 21780001 | 21880000 | 0.0026 | 0.0018 | 0.0021 |
| F10 | 21800001 | 21900000 | 0.0024 | 0.0027 | 0.0020 |
| F10 | 21820001 | 21920000 | 0.0031 | 0.0028 | 0.0029 |
| F10 | 21840001 | 21940000 | 0.0032 | 0.0027 | 0.0029 |
| F10 | 21860001 | 21960000 | 0.0032 | 0.0028 | 0.0029 |
| F10 | 21880001 | 21980000 | 0.0033 | 0.0034 | 0.0027 |
| F10 | 21900001 | 22000000 | 0.0040 | 0.0025 | 0.0033 |
| F10 | 21920001 | 22020000 | 0.0033 | 0.0022 | 0.0025 |
| F10 | 21940001 | 22040000 | 0.0028 | 0.0025 | 0.0021 |
| F10 | 21960001 | 22060000 | 0.0032 | 0.0020 | 0.0025 |
| F10 | 21980001 | 22080000 | 0.0026 | 0.0022 | 0.0021 |
| F10 | 22000001 | 22100000 | 0.0031 | 0.0028 | 0.0023 |
| F10 | 22020001 | 22120000 | 0.0038 | 0.0030 | 0.0029 |
| F10 | 22040001 | 22140000 | 0.0040 | 0.0026 | 0.0032 |
| F10 | 22060001 | 22160000 | 0.0035 | 0.0033 | 0.0027 |
| F10 | 22080001 | 22180000 | 0.0043 | 0.0027 | 0.0034 |
| F10 | 22100001 | 22200000 | 0.0035 | 0.0022 | 0.0026 |
| F10 | 22120001 | 22220000 | 0.0030 | 0.0022 | 0.0021 |
| F10 | 22140001 | 22240000 | 0.0029 | 0.0025 | 0.0020 |
| F10 | 22160001 | 22260000 | 0.0034 | 0.0021 | 0.0024 |
| F10 | 22180001 | 22280000 | 0.0027 | 0.0018 | 0.0020 |
| F10 | 22200001 | 22300000 | 0.0024 | 0.0021 | 0.0018 |
| F10 | 22220001 | 22320000 | 0.0027 | 0.0023 | 0.0021 |
| F10 | 22240001 | 22340000 | 0.0027 | 0.0023 | 0.0022 |
| F10 | 22260001 | 22360000 | 0.0030 | 0.0028 | 0.0022 |
| F10 | 22280001 | 22380000 | 0.0037 | 0.0028 | 0.0027 |
| F10 | 22300001 | 22400000 | 0.0038 | 0.0027 | 0.0028 |
| F10 | 22320001 | 22420000 | 0.0035 | 0.0027 | 0.0028 |
| F10 | 22340001 | 22440000 | 0.0036 | 0.0028 | 0.0029 |

| Chr | Start | End | both | A1 | A2 |
|---|---|---|---|---|---|
| F10 | 22360001 | 22460000 | 0.0035 | 0.0025 | 0.0030 |
| F10 | 22380001 | 22480000 | 0.0031 | 0.0024 | 0.0026 |
| F10 | 22400001 | 22500000 | 0.0028 | 0.0020 | 0.0024 |
| F10 | 22420001 | 22520000 | 0.0025 | 0.0016 | 0.0020 |
| F10 | 22440001 | 22540000 | 0.0021 | 0.0011 | 0.0015 |
| F10 | 22460001 | 22560000 | 0.0016 | 0.0007 | 0.0010 |
| F10 | 22480001 | 22580000 | 0.0013 | 0.0011 | 0.0005 |
| F10 | 22500001 | 22600000 | 0.0018 | 0.0014 | 0.0011 |
| F10 | 22520001 | 22620000 | 0.0020 | 0.0020 | 0.0013 |
| F10 | 22540001 | 22640000 | 0.0027 | 0.0022 | 0.0020 |
| F10 | 22560001 | 22660000 | 0.0031 | 0.0026 | 0.0023 |
| F10 | 22580001 | 22680000 | 0.0034 | 0.0028 | 0.0030 |
| F10 | 22600001 | 22700000 | 0.0037 | 0.0031 | 0.0031 |
| F10 | 22620001 | 22720000 | 0.0042 | 0.0027 | 0.0035 |
| F10 | 22640001 | 22740000 | 0.0039 | 0.0026 | 0.0030 |
| F10 | 22660001 | 22760000 | 0.0037 | 0.0025 | 0.0029 |
| F10 | 22680001 | 22780000 | 0.0038 | 0.0019 | 0.0028 |
| F10 | 22700001 | 22800000 | 0.0030 | 0.0015 | 0.0020 |
| F10 | 22720001 | 22820000 | 0.0026 | 0.0017 | 0.0017 |
| F10 | 22740001 | 22840000 | 0.0026 | 0.0021 | 0.0018 |
| F10 | 22760001 | 22860000 | 0.0030 | 0.0025 | 0.0023 |
| F10 | 22780001 | 22880000 | 0.0035 | 0.0025 | 0.0026 |
| F10 | 22800001 | 22900000 | 0.0035 | 0.0021 | 0.0025 |
| F10 | 22820001 | 22920000 | 0.0029 | 0.0019 | 0.0021 |
| F10 | 22840001 | 22940000 | 0.0028 | 0.0023 | 0.0020 |
| F10 | 22860001 | 22960000 | 0.0031 | 0.0018 | 0.0024 |
| F10 | 22880001 | 22980000 | 0.0024 | 0.0022 | 0.0019 |
| F10 | 22900001 | 23000000 | 0.0030 | 0.0028 | 0.0024 |
| F10 | 22920001 | 23020000 | 0.0036 | 0.0028 | 0.0030 |
| F10 | 22940001 | 23040000 | 0.0037 | 0.0025 | 0.0032 |
| F10 | 22960001 | 23060000 | 0.0034 | 0.0022 | 0.0029 |
| F10 | 22980001 | 23080000 | 0.0030 | 0.0017 | 0.0025 |
| F10 | 23000001 | 23100000 | 0.0022 | 0.0016 | 0.0020 |
| F10 | 23020001 | 23120000 | 0.0021 | 0.0019 | 0.0019 |
| F10 | 23040001 | 23140000 | 0.0023 | 0.0016 | 0.0021 |
| F10 | 23060001 | 23160000 | 0.0020 | 0.0027 | 0.0018 |
| F10 | 23080001 | 23180000 | 0.0031 | 0.0031 | 0.0028 |
| F10 | 23100001 | 23200000 | 0.0039 | 0.0026 | 0.0033 |
| F10 | 23120001 | 23220000 | 0.0036 | 0.0024 | 0.0028 |
| F10 | 23140001 | 23240000 | 0.0033 | 0.0024 | 0.0024 |
| F10 | 23160001 | 23260000 | 0.0034 | 0.0014 | 0.0024 |
| F10 | 23180001 | 23280000 | 0.0024 | 0.0015 | 0.0015 |
| F10 | 23200001 | 23300000 | 0.0022 | 0.0017 | 0.0012 |
| F10 | 23220001 | 23320000 | 0.0026 | 0.0016 | 0.0016 |
| F10 | 23240001 | 23340000 | 0.0025 | 0.0014 | 0.0016 |
| F10 | 23260001 | 23360000 | 0.0022 | 0.0015 | 0.0014 |
| F10 | 23280001 | 23380000 | 0.0023 | 0.0014 | 0.0015 |
| F10 | 23300001 | 23400000 | 0.0022 | 0.0020 | 0.0016 |
| F10 | 23320001 | 23420000 | 0.0028 | 0.0020 | 0.0023 |
| F10 | 23340001 | 23440000 | 0.0029 | 0.0023 | 0.0022 |
| F10 | 23360001 | 23460000 | 0.0031 | 0.0025 | 0.0024 |
| F10 | 23380001 | 23480000 | 0.0034 | 0.0031 | 0.0026 |
| F10 | 23400001 | 23500000 | 0.0038 | 0.0029 | 0.0029 |
| F10 | 23420001 | 23520000 | 0.0037 | 0.0040 | 0.0027 |
| F10 | 23440001 | 23540000 | 0.0048 | 0.0042 | 0.0038 |
| F10 | 23460001 | 23560000 | 0.0050 | 0.0042 | 0.0041 |
| F10 | 23480001 | 23580000 | 0.0049 | 0.0036 | 0.0041 |
| F10 | 23500001 | 23600000 | 0.0045 | 0.0033 | 0.0037 |

| Chr | Start | End | both | A1 | A2 |
|---|---|---|---|---|---|
| F10 | 23520001 | 23620000 | 0.0041 | 0.0027 | 0.0034 |
| F10 | 23540001 | 23640000 | 0.0034 | 0.0023 | 0.0029 |
| F10 | 23560001 | 23660000 | 0.0031 | 0.0023 | 0.0025 |
| F10 | 23580001 | 23680000 | 0.0032 | 0.0024 | 0.0023 |
| F10 | 23600001 | 23700000 | 0.0033 | 0.0024 | 0.0025 |
| F10 | 23620001 | 23720000 | 0.0033 | 0.0019 | 0.0024 |
| F10 | 23640001 | 23740000 | 0.0028 | 0.0020 | 0.0019 |
| F10 | 23660001 | 23760000 | 0.0028 | 0.0020 | 0.0020 |
| F10 | 23680001 | 23780000 | 0.0025 | 0.0021 | 0.0021 |
| F10 | 23700001 | 23800000 | 0.0026 | 0.0019 | 0.0023 |
| F10 | 23720001 | 23820000 | 0.0023 | 0.0017 | 0.0022 |
| F10 | 23740001 | 23840000 | 0.0021 | 0.0021 | 0.0021 |
| F10 | 23760001 | 23860000 | 0.0026 | 0.0022 | 0.0025 |
| F10 | 23780001 | 23880000 | 0.0027 | 0.0024 | 0.0026 |
| F10 | 23800001 | 23900000 | 0.0027 | 0.0023 | 0.0027 |
| F10 | 23820001 | 23920000 | 0.0025 | 0.0026 | 0.0025 |
| F10 | 23840001 | 23940000 | 0.0028 | 0.0021 | 0.0028 |
| F10 | 23860001 | 23960000 | 0.0023 | 0.0019 | 0.0023 |
| F10 | 23880001 | 23980000 | 0.0021 | 0.0012 | 0.0020 |
| F10 | 23900001 | 24000000 | 0.0015 | 0.0018 | 0.0015 |
| F10 | 23920001 | 24020000 | 0.0024 | 0.0014 | 0.0023 |
| F10 | 23940001 | 24040000 | 0.0021 | 0.0018 | 0.0018 |
| F10 | 23960001 | 24060000 | 0.0027 | 0.0018 | 0.0024 |
| F10 | 23980001 | 24080000 | 0.0028 | 0.0019 | 0.0023 |
| F10 | 24000001 | 24100000 | 0.0030 | 0.0021 | 0.0023 |
| F10 | 24020001 | 24120000 | 0.0031 | 0.0022 | 0.0023 |
| F10 | 24040001 | 24140000 | 0.0033 | 0.0018 | 0.0024 |
| F10 | 24060001 | 24160000 | 0.0028 | 0.0022 | 0.0018 |
| F10 | 24080001 | 24180000 | 0.0033 | 0.0024 | 0.0022 |
| F10 | 24100001 | 24200000 | 0.0037 | 0.0017 | 0.0025 |
| F10 | 24120001 | 24220000 | 0.0030 | 0.0016 | 0.0018 |
| F10 | 24140001 | 24240000 | 0.0029 | 0.0018 | 0.0017 |
| F10 | 24160001 | 24260000 | 0.0033 | 0.0016 | 0.0019 |
| F10 | 24180001 | 24280000 | 0.0031 | 0.0020 | 0.0015 |
| F10 | 24200001 | 24300000 | 0.0032 | 0.0019 | 0.0018 |
| F10 | 24220001 | 24320000 | 0.0030 | 0.0020 | 0.0018 |
| F10 | 24240001 | 24340000 | 0.0031 | 0.0017 | 0.0019 |
| F10 | 24260001 | 24360000 | 0.0027 | 0.0016 | 0.0018 |
| F10 | 24280001 | 24380000 | 0.0024 | 0.0009 | 0.0017 |
| F10 | 24300001 | 24400000 | 0.0018 | 0.0010 | 0.0010 |
| F10 | 24320001 | 24420000 | 0.0019 | 0.0009 | 0.0010 |
| F10 | 24340001 | 24440000 | 0.0017 | 0.0018 | 0.0010 |
| F10 | 24360001 | 24460000 | 0.0024 | 0.0022 | 0.0018 |
| F10 | 24380001 | 24480000 | 0.0030 | 0.0027 | 0.0023 |
| F10 | 24400001 | 24500000 | 0.0035 | 0.0042 | 0.0027 |
| F10 | 24420001 | 24520000 | 0.0050 | 0.0044 | 0.0042 |
| F10 | 24440001 | 24540000 | 0.0053 | 0.0037 | 0.0044 |
| F10 | 24460001 | 24560000 | 0.0047 | 0.0034 | 0.0036 |
| F10 | 24480001 | 24580000 | 0.0042 | 0.0029 | 0.0032 |
| F10 | 24500001 | 24600000 | 0.0038 | 0.0016 | 0.0029 |
| F10 | 24520001 | 24620000 | 0.0026 | 0.0017 | 0.0016 |
| F10 | 24540001 | 24640000 | 0.0028 | 0.0015 | 0.0017 |
| F10 | 24560001 | 24660000 | 0.0025 | 0.0011 | 0.0016 |
| F10 | 24580001 | 24680000 | 0.0021 | 0.0011 | 0.0012 |
| F10 | 24600001 | 24700000 | 0.0019 | 0.0013 | 0.0012 |
| F10 | 24620001 | 24720000 | 0.0024 | 0.0011 | 0.0015 |
| F10 | 24640001 | 24740000 | 0.0018 | 0.0010 | 0.0012 |
| F10 | 24660001 | 24760000 | 0.0017 | 0.0011 | 0.0011 |

| Chr | Start | End | both | A1 | A2 |
|---|---|---|---|---|---|
| F10 | 24680001 | 24780000 | 0.0019 | 0.0015 | 0.0012 |
| F10 | 24700001 | 24800000 | 0.0024 | 0.0016 | 0.0015 |
| F10 | 24720001 | 24820000 | 0.0020 | 0.0015 | 0.0015 |
| F10 | 24740001 | 24840000 | 0.0018 | 0.0018 | 0.0014 |
| F10 | 24760001 | 24860000 | 0.0020 | 0.0018 | 0.0017 |
| F10 | 24780001 | 24880000 | 0.0019 | 0.0013 | 0.0017 |
| F10 | 24800001 | 24900000 | 0.0015 | 0.0008 | 0.0013 |
| F10 | 24820001 | 24920000 | 0.0010 | 0.0015 | 0.0007 |
| F10 | 24840001 | 24940000 | 0.0018 | 0.0018 | 0.0014 |
| F10 | 24860001 | 24960000 | 0.0024 | 0.0022 | 0.0021 |
| F10 | 24880001 | 24980000 | 0.0029 | 0.0032 | 0.0023 |
| F10 | 24900001 | 25000000 | 0.0039 | 0.0034 | 0.0035 |
| F10 | 24920001 | 25020000 | 0.0041 | 0.0029 | 0.0037 |
| F10 | 24940001 | 25040000 | 0.0036 | 0.0028 | 0.0033 |
| F10 | 24960001 | 25060000 | 0.0033 | 0.0028 | 0.0030 |
| F10 | 24980001 | 25080000 | 0.0033 | 0.0020 | 0.0031 |
| F10 | 25000001 | 25100000 | 0.0025 | 0.0023 | 0.0022 |
| F10 | 25020001 | 25120000 | 0.0027 | 0.0028 | 0.0024 |
| F10 | 25040001 | 25140000 | 0.0035 | 0.0029 | 0.0031 |
| F10 | 25060001 | 25160000 | 0.0036 | 0.0027 | 0.0032 |
| F10 | 25080001 | 25180000 | 0.0035 | 0.0027 | 0.0029 |
| F10 | 25100001 | 25200000 | 0.0035 | 0.0021 | 0.0029 |
| F10 | 25120001 | 25220000 | 0.0030 | 0.0018 | 0.0023 |
| F10 | 25140001 | 25240000 | 0.0025 | 0.0019 | 0.0018 |
| F10 | 25160001 | 25260000 | 0.0026 | 0.0018 | 0.0019 |
| F10 | 25180001 | 25280000 | 0.0025 | 0.0019 | 0.0018 |
| F10 | 25200001 | 25300000 | 0.0026 | 0.0018 | 0.0020 |
| F10 | 25220001 | 25320000 | 0.0024 | 0.0021 | 0.0019 |
| F10 | 25240001 | 25340000 | 0.0027 | 0.0017 | 0.0023 |
| F10 | 25260001 | 25360000 | 0.0023 | 0.0018 | 0.0019 |
| F10 | 25280001 | 25380000 | 0.0022 | 0.0018 | 0.0020 |
| F10 | 25300001 | 25400000 | 0.0021 | 0.0019 | 0.0019 |
| F10 | 25320001 | 25420000 | 0.0023 | 0.0012 | 0.0021 |
| F10 | 25340001 | 25440000 | 0.0014 | 0.0012 | 0.0012 |
| F10 | 25360001 | 25460000 | 0.0013 | 0.0011 | 0.0012 |
| F10 | 25380001 | 25480000 | 0.0012 | 0.0008 | 0.0012 |
| F10 | 25400001 | 25500000 | 0.0009 | 0.0007 | 0.0009 |
| F10 | 25420001 | 25520000 | 0.0008 | 0.0008 | 0.0008 |
| F10 | 25440001 | 25540000 | 0.0011 | 0.0010 | 0.0011 |
| F10 | 25460001 | 25560000 | 0.0013 | 0.0010 | 0.0012 |
| F10 | 25480001 | 25580000 | 0.0013 | 0.0017 | 0.0012 |
| F10 | 25500001 | 25600000 | 0.0021 | 0.0027 | 0.0020 |
| F10 | 25520001 | 25620000 | 0.0033 | 0.0030 | 0.0031 |
| F10 | 25540001 | 25640000 | 0.0035 | 0.0029 | 0.0033 |
| F10 | 25560001 | 25660000 | 0.0035 | 0.0027 | 0.0034 |
| F10 | 25580001 | 25680000 | 0.0033 | 0.0020 | 0.0031 |
| F10 | 25600001 | 25700000 | 0.0024 | 0.0009 | 0.0022 |
| F10 | 25620001 | 25720000 | 0.0010 | 0.0005 | 0.0010 |
| F10 | 25640001 | 25740000 | 0.0006 | 0.0003 | 0.0006 |
| F10 | 25660001 | 25760000 | 0.0004 | 0.0007 | 0.0004 |
| F10 | 25680001 | 25780000 | 0.0008 | 0.0009 | 0.0008 |
| F10 | 25700001 | 25800000 | 0.0010 | 0.0009 | 0.0010 |
| F10 | 25720001 | 25820000 | 0.0010 | 0.0010 | 0.0010 |
| F10 | 25740001 | 25840000 | 0.0012 | 0.0008 | 0.0011 |
| F10 | 25760001 | 25860000 | 0.0010 | 0.0008 | 0.0010 |
| F10 | 25780001 | 25880000 | 0.0010 | 0.0008 | 0.0009 |
| F10 | 25800001 | 25900000 | 0.0009 | 0.0015 | 0.0008 |
| F10 | 25820001 | 25920000 | 0.0017 | 0.0018 | 0.0016 |

| Chr | Start | End | both | A1 | A2 |
|---|---|---|---|---|---|
| F10 | 25840001 | 25940000 | 0.0020 | 0.0019 | 0.0019 |
| F10 | 25860001 | 25960000 | 0.0021 | 0.0021 | 0.0020 |
| F10 | 25880001 | 25980000 | 0.0025 | 0.0025 | 0.0023 |
| F10 | 25900001 | 26000000 | 0.0029 | 0.0021 | 0.0027 |
| F10 | 25920001 | 26020000 | 0.0024 | 0.0022 | 0.0022 |
| F10 | 25940001 | 26040000 | 0.0024 | 0.0025 | 0.0022 |
| F10 | 25960001 | 26060000 | 0.0028 | 0.0028 | 0.0025 |
| F10 | 25980001 | 26080000 | 0.0029 | 0.0033 | 0.0025 |
| F10 | 26000001 | 26100000 | 0.0033 | 0.0030 | 0.0029 |
| F10 | 26020001 | 26120000 | 0.0029 | 0.0027 | 0.0026 |
| F10 | 26040001 | 26140000 | 0.0028 | 0.0025 | 0.0025 |
| F10 | 26060001 | 26160000 | 0.0026 | 0.0016 | 0.0023 |
| F10 | 26080001 | 26180000 | 0.0017 | 0.0007 | 0.0016 |
| F10 | 26100001 | 26200000 | 0.0008 | 0.0020 | 0.0008 |
| F10 | 26120001 | 26220000 | 0.0023 | 0.0026 | 0.0022 |
| F10 | 26140001 | 26240000 | 0.0030 | 0.0035 | 0.0027 |
| F10 | 26160001 | 26260000 | 0.0041 | 0.0043 | 0.0036 |
| F10 | 26180001 | 26280000 | 0.0053 | 0.0052 | 0.0045 |
| F10 | 26200001 | 26300000 | 0.0064 | 0.0044 | 0.0056 |
| F10 | 26220001 | 26320000 | 0.0054 | 0.0037 | 0.0045 |
| F10 | 26240001 | 26340000 | 0.0047 | 0.0032 | 0.0040 |
| F10 | 26260001 | 26360000 | 0.0042 | 0.0025 | 0.0036 |
| F10 | 26280001 | 26380000 | 0.0031 | 0.0018 | 0.0028 |
| F10 | 26300001 | 26400000 | 0.0025 | 0.0015 | 0.0021 |
| F10 | 26320001 | 26420000 | 0.0020 | 0.0018 | 0.0019 |
| F10 | 26340001 | 26440000 | 0.0022 | 0.0014 | 0.0023 |
| F10 | 26360001 | 26460000 | 0.0017 | 0.0014 | 0.0017 |
| F10 | 26380001 | 26480000 | 0.0016 | 0.0011 | 0.0015 |
| F10 | 26400001 | 26500000 | 0.0012 | 0.0013 | 0.0012 |
| F10 | 26420001 | 26520000 | 0.0014 | 0.0012 | 0.0013 |
| F10 | 26440001 | 26540000 | 0.0012 | 0.0009 | 0.0012 |
| F10 | 26460001 | 26560000 | 0.0010 | 0.0010 | 0.0010 |
| F10 | 26480001 | 26580000 | 0.0012 | 0.0013 | 0.0012 |
| F10 | 26500001 | 26600000 | 0.0015 | 0.0014 | 0.0015 |
| F10 | 26520001 | 26620000 | 0.0018 | 0.0012 | 0.0016 |
| F10 | 26540001 | 26640000 | 0.0017 | 0.0019 | 0.0014 |
| F10 | 26560001 | 26660000 | 0.0024 | 0.0018 | 0.0021 |
| F10 | 26580001 | 26680000 | 0.0023 | 0.0020 | 0.0020 |
| F10 | 26600001 | 26700000 | 0.0027 | 0.0021 | 0.0023 |
| F10 | 26620001 | 26720000 | 0.0027 | 0.0024 | 0.0026 |
| F10 | 26640001 | 26740000 | 0.0031 | 0.0022 | 0.0030 |
| F10 | 26660001 | 26760000 | 0.0029 | 0.0026 | 0.0029 |
| F10 | 26680001 | 26780000 | 0.0034 | 0.0026 | 0.0033 |
| F10 | 26700001 | 26800000 | 0.0035 | 0.0028 | 0.0032 |
| F10 | 26720001 | 26820000 | 0.0038 | 0.0028 | 0.0034 |
| F10 | 26740001 | 26840000 | 0.0039 | 0.0028 | 0.0033 |
| F10 | 26760001 | 26860000 | 0.0039 | 0.0026 | 0.0032 |
| F10 | 26780001 | 26880000 | 0.0037 | 0.0045 | 0.0030 |
| F10 | 26800001 | 26900000 | 0.0054 | 0.0045 | 0.0048 |
| F10 | 26820001 | 26920000 | 0.0052 | 0.0041 | 0.0047 |
| F10 | 26840001 | 26940000 | 0.0050 | 0.0049 | 0.0044 |
| F10 | 26860001 | 26960000 | 0.0058 | 0.0052 | 0.0049 |
| F10 | 26880001 | 26980000 | 0.0061 | 0.0028 | 0.0052 |
| F10 | 26900001 | 27000000 | 0.0037 | 0.0024 | 0.0028 |
| F10 | 26920001 | 27020000 | 0.0032 | 0.0032 | 0.0024 |
| F10 | 26940001 | 27040000 | 0.0038 | 0.0034 | 0.0031 |
| F10 | 26960001 | 27060000 | 0.0043 | 0.0035 | 0.0035 |
| F10 | 26980001 | 27080000 | 0.0043 | 0.0052 | 0.0033 |

| Chr | Start | End | both | A1 | A2 |
|---|---|---|---|---|---|
| F10 | 27000001 | 27100000 | 0.0062 | 0.0079 | 0.0051 |
| F10 | 27020001 | 27120000 | 0.0092 | 0.0085 | 0.0080 |
| F10 | 27040001 | 27140000 | 0.0098 | 0.0075 | 0.0089 |
| F10 | 27060001 | 27160000 | 0.0085 | 0.0072 | 0.0079 |
| F10 | 27080001 | 27180000 | 0.0082 | 0.0056 | 0.0078 |
| F10 | 27100001 | 27200000 | 0.0063 | 0.0032 | 0.0062 |
| F10 | 27120001 | 27220000 | 0.0036 | 0.0017 | 0.0035 |
| F10 | 27140001 | 27240000 | 0.0020 | 0.0013 | 0.0019 |
| F10 | 27160001 | 27260000 | 0.0015 | 0.0010 | 0.0014 |
| F10 | 27180001 | 27280000 | 0.0012 | 0.0009 | 0.0012 |
| F10 | 27200001 | 27300000 | 0.0010 | 0.0004 | 0.0010 |
| F10 | 27220001 | 27320000 | 0.0005 | 0.0010 | 0.0005 |
| F10 | 27240001 | 27340000 | 0.0012 | 0.0011 | 0.0011 |
| F10 | 27260001 | 27360000 | 0.0012 | 0.0021 | 0.0011 |
| F10 | 27280001 | 27380000 | 0.0022 | 0.0021 | 0.0021 |
| F10 | 27300001 | 27400000 | 0.0022 | 0.0020 | 0.0022 |
| F10 | 27320001 | 27420000 | 0.0022 | 0.0019 | 0.0022 |
| F10 | 27340001 | 27440000 | 0.0020 | 0.0023 | 0.0020 |
| F10 | 27360001 | 27460000 | 0.0025 | 0.0012 | 0.0025 |
| F10 | 27380001 | 27480000 | 0.0013 | 0.0013 | 0.0013 |
| F10 | 27400001 | 27500000 | 0.0014 | 0.0014 | 0.0014 |
| F10 | 27420001 | 27520000 | 0.0015 | 0.0009 | 0.0014 |
| F10 | 27440001 | 27540000 | 0.0009 | 0.0010 | 0.0009 |
| F10 | 27460001 | 27560000 | 0.0011 | 0.0014 | 0.0010 |
| F10 | 27480001 | 27580000 | 0.0014 | 0.0022 | 0.0013 |
| F10 | 27500001 | 27600000 | 0.0024 | 0.0031 | 0.0023 |
| F10 | 27520001 | 27620000 | 0.0037 | 0.0037 | 0.0034 |
| F10 | 27540001 | 27640000 | 0.0054 | 0.0035 | 0.0038 |
| F10 | 27560001 | 27660000 | 0.0060 | 0.0042 | 0.0033 |
| F10 | 27580001 | 27680000 | 0.0071 | 0.0035 | 0.0040 |
| F10 | 27600001 | 27700000 | 0.0065 | 0.0030 | 0.0031 |
| F10 | 27620001 | 27720000 | 0.0057 | 0.0028 | 0.0024 |
| F10 | 27640001 | 27740000 | 0.0048 | 0.0043 | 0.0023 |
| F10 | 27660001 | 27760000 | 0.0063 | 0.0036 | 0.0039 |
| F10 | 27680001 | 27780000 | 0.0057 | 0.0042 | 0.0031 |
| F10 | 27700001 | 27800000 | 0.0064 | 0.0042 | 0.0037 |
| F10 | 27720001 | 27820000 | 0.0068 | 0.0038 | 0.0039 |
| F10 | 27740001 | 27840000 | 0.0062 | 0.0029 | 0.0036 |
| F10 | 27760001 | 27860000 | 0.0046 | 0.0032 | 0.0029 |
| F10 | 27780001 | 27880000 | 0.0047 | 0.0027 | 0.0034 |
| F10 | 27800001 | 27900000 | 0.0040 | 0.0023 | 0.0031 |
| F10 | 27820001 | 27920000 | 0.0033 | 0.0025 | 0.0026 |
| F10 | 27840001 | 27940000 | 0.0034 | 0.0017 | 0.0029 |
| F10 | 27860001 | 27960000 | 0.0026 | 0.0012 | 0.0021 |
| F10 | 27880001 | 27980000 | 0.0018 | 0.0009 | 0.0014 |
| F10 | 27900001 | 28000000 | 0.0013 | 0.0008 | 0.0010 |
| F10 | 27920001 | 28020000 | 0.0013 | 0.0008 | 0.0009 |
| F10 | 27940001 | 28040000 | 0.0015 | 0.0025 | 0.0009 |
| F10 | 27960001 | 28060000 | 0.0033 | 0.0031 | 0.0028 |
| F10 | 27980001 | 28080000 | 0.0041 | 0.0036 | 0.0035 |
| F10 | 28000001 | 28100000 | 0.0046 | 0.0040 | 0.0040 |
| F10 | 28020001 | 28120000 | 0.0051 | 0.0040 | 0.0045 |
| F10 | 28040001 | 28140000 | 0.0050 | 0.0027 | 0.0045 |
| F10 | 28060001 | 28160000 | 0.0033 | 0.0021 | 0.0029 |
| F10 | 28080001 | 28180000 | 0.0025 | 0.0022 | 0.0022 |
| F10 | 28100001 | 28200000 | 0.0026 | 0.0020 | 0.0024 |
| F10 | 28120001 | 28220000 | 0.0024 | 0.0018 | 0.0021 |
| F10 | 28140001 | 28240000 | 0.0023 | 0.0019 | 0.0020 |

| Chr | Start | End | both | A1 | A2 |
|---|---|---|---|---|---|
| F10 | 28160001 | 28260000 | 0.0026 | 0.0023 | 0.0021 |
| F10 | 28180001 | 28280000 | 0.0032 | 0.0021 | 0.0024 |
| F10 | 28200001 | 28300000 | 0.0031 | 0.0022 | 0.0022 |
| F10 | 28220001 | 28320000 | 0.0032 | 0.0030 | 0.0022 |
| F10 | 28240001 | 28340000 | 0.0041 | 0.0025 | 0.0031 |
| F10 | 28260001 | 28360000 | 0.0036 | 0.0024 | 0.0024 |
| F10 | 28280001 | 28380000 | 0.0034 | 0.0021 | 0.0023 |
| F10 | 28300001 | 28400000 | 0.0031 | 0.0020 | 0.0021 |
| F10 | 28320001 | 28420000 | 0.0029 | 0.0009 | 0.0021 |
| F10 | 28340001 | 28440000 | 0.0019 | 0.0011 | 0.0010 |
| F10 | 28360001 | 28460000 | 0.0021 | 0.0011 | 0.0013 |
| F10 | 28380001 | 28480000 | 0.0023 | 0.0016 | 0.0014 |
| F10 | 28400001 | 28500000 | 0.0028 | 0.0016 | 0.0018 |
| F10 | 28420001 | 28520000 | 0.0028 | 0.0023 | 0.0018 |
| F10 | 28440001 | 28540000 | 0.0034 | 0.0021 | 0.0025 |
| F10 | 28460001 | 28560000 | 0.0031 | 0.0026 | 0.0022 |
| F10 | 28480001 | 28580000 | 0.0035 | 0.0020 | 0.0027 |
| F10 | 28500001 | 28600000 | 0.0029 | 0.0020 | 0.0022 |
| F10 | 28520001 | 28620000 | 0.0029 | 0.0014 | 0.0021 |
| F10 | 28540001 | 28640000 | 0.0022 | 0.0013 | 0.0015 |
| F10 | 28560001 | 28660000 | 0.0022 | 0.0013 | 0.0014 |
| F10 | 28580001 | 28680000 | 0.0021 | 0.0024 | 0.0012 |
| F10 | 28600001 | 28700000 | 0.0034 | 0.0023 | 0.0023 |
| F10 | 28620001 | 28720000 | 0.0035 | 0.0027 | 0.0022 |
| F10 | 28640001 | 28740000 | 0.0039 | 0.0030 | 0.0026 |
| F10 | 28660001 | 28760000 | 0.0041 | 0.0028 | 0.0029 |
| F10 | 28680001 | 28780000 | 0.0039 | 0.0028 | 0.0028 |
| F10 | 28700001 | 28800000 | 0.0038 | 0.0037 | 0.0028 |
| F10 | 28720001 | 28820000 | 0.0047 | 0.0041 | 0.0038 |
| F10 | 28740001 | 28840000 | 0.0050 | 0.0040 | 0.0042 |
| F10 | 28760001 | 28860000 | 0.0048 | 0.0035 | 0.0042 |
| F10 | 28780001 | 28880000 | 0.0042 | 0.0025 | 0.0037 |
| F10 | 28800001 | 28900000 | 0.0030 | 0.0019 | 0.0028 |
| F10 | 28820001 | 28920000 | 0.0025 | 0.0018 | 0.0024 |
| F10 | 28840001 | 28940000 | 0.0027 | 0.0016 | 0.0025 |
| F10 | 28860001 | 28960000 | 0.0025 | 0.0016 | 0.0022 |
| F10 | 28880001 | 28980000 | 0.0026 | 0.0016 | 0.0021 |
| F10 | 28900001 | 29000000 | 0.0027 | 0.0014 | 0.0021 |
| F10 | 28920001 | 29020000 | 0.0025 | 0.0006 | 0.0019 |
| F10 | 28940001 | 29040000 | 0.0014 | 0.0015 | 0.0009 |
| F10 | 28960001 | 29060000 | 0.0021 | 0.0018 | 0.0016 |
| F10 | 28980001 | 29080000 | 0.0023 | 0.0022 | 0.0019 |
| F10 | 29000001 | 29100000 | 0.0025 | 0.0021 | 0.0022 |
| F10 | 29020001 | 29120000 | 0.0022 | 0.0023 | 0.0020 |
| F10 | 29040001 | 29140000 | 0.0024 | 0.0018 | 0.0021 |
| F10 | 29060001 | 29160000 | 0.0022 | 0.0019 | 0.0019 |
| F10 | 29080001 | 29180000 | 0.0024 | 0.0014 | 0.0020 |
| F10 | 29100001 | 29200000 | 0.0020 | 0.0018 | 0.0015 |
| F10 | 29120001 | 29220000 | 0.0025 | 0.0020 | 0.0019 |
| F10 | 29140001 | 29240000 | 0.0027 | 0.0020 | 0.0021 |
| F10 | 29160001 | 29260000 | 0.0024 | 0.0017 | 0.0020 |
| F10 | 29180001 | 29280000 | 0.0021 | 0.0020 | 0.0017 |
| F10 | 29200001 | 29300000 | 0.0023 | 0.0019 | 0.0020 |
| F10 | 29220001 | 29320000 | 0.0022 | 0.0022 | 0.0020 |
| F10 | 29240001 | 29340000 | 0.0026 | 0.0023 | 0.0024 |
| F10 | 29260001 | 29360000 | 0.0029 | 0.0022 | 0.0026 |
| F10 | 29280001 | 29380000 | 0.0027 | 0.0027 | 0.0025 |
| F10 | 29300001 | 29400000 | 0.0033 | 0.0023 | 0.0031 |

| Chr | Start | End | both | A1 | A2 |
|---|---|---|---|---|---|
| F10 | 29320001 | 29420000 | 0.0029 | 0.0019 | 0.0027 |
| F10 | 29340001 | 29440000 | 0.0023 | 0.0015 | 0.0022 |
| F10 | 29360001 | 29460000 | 0.0017 | 0.0019 | 0.0017 |
| F10 | 29380001 | 29480000 | 0.0022 | 0.0019 | 0.0021 |
| F10 | 29400001 | 29500000 | 0.0020 | 0.0029 | 0.0019 |
| F10 | 29420001 | 29520000 | 0.0032 | 0.0033 | 0.0030 |
| F10 | 29440001 | 29540000 | 0.0036 | 0.0039 | 0.0034 |
| F10 | 29460001 | 29560000 | 0.0043 | 0.0041 | 0.0041 |
| F10 | 29480001 | 29580000 | 0.0046 | 0.0040 | 0.0044 |
| F10 | 29500001 | 29600000 | 0.0046 | 0.0029 | 0.0043 |
| F10 | 29520001 | 29620000 | 0.0036 | 0.0026 | 0.0031 |
| F10 | 29540001 | 29640000 | 0.0034 | 0.0033 | 0.0027 |
| F10 | 29560001 | 29660000 | 0.0043 | 0.0028 | 0.0034 |
| F10 | 29580001 | 29680000 | 0.0038 | 0.0022 | 0.0028 |
| F10 | 29600001 | 29700000 | 0.0032 | 0.0027 | 0.0022 |
| F10 | 29620001 | 29720000 | 0.0038 | 0.0027 | 0.0027 |
| F10 | 29640001 | 29740000 | 0.0038 | 0.0014 | 0.0028 |
| F10 | 29660001 | 29760000 | 0.0023 | 0.0013 | 0.0014 |
| F10 | 29680001 | 29780000 | 0.0022 | 0.0012 | 0.0012 |
| F10 | 29700001 | 29800000 | 0.0021 | 0.0013 | 0.0012 |
| F10 | 29720001 | 29820000 | 0.0020 | 0.0015 | 0.0014 |
| F10 | 29740001 | 29840000 | 0.0021 | 0.0022 | 0.0015 |
| F10 | 29760001 | 29860000 | 0.0030 | 0.0022 | 0.0023 |
| F10 | 29780001 | 29880000 | 0.0030 | 0.0025 | 0.0023 |
| F10 | 29800001 | 29900000 | 0.0035 | 0.0023 | 0.0027 |
| F10 | 29820001 | 29920000 | 0.0034 | 0.0021 | 0.0025 |
| F10 | 29840001 | 29940000 | 0.0035 | 0.0024 | 0.0023 |
| F10 | 29860001 | 29960000 | 0.0040 | 0.0029 | 0.0030 |
| F10 | 29880001 | 29980000 | 0.0046 | 0.0035 | 0.0035 |
| F10 | 29900001 | 30000000 | 0.0053 | 0.0039 | 0.0039 |
| F10 | 29920001 | 30020000 | 0.0053 | 0.0042 | 0.0041 |
| F10 | 29940001 | 30040000 | 0.0055 | 0.0035 | 0.0043 |
| F10 | 29960001 | 30060000 | 0.0045 | 0.0033 | 0.0032 |
| F10 | 29980001 | 30080000 | 0.0042 | 0.0029 | 0.0031 |
| F10 | 30000001 | 30100000 | 0.0037 | 0.0024 | 0.0028 |
| F10 | 30020001 | 30120000 | 0.0034 | 0.0022 | 0.0023 |
| F10 | 30040001 | 30140000 | 0.0031 | 0.0020 | 0.0020 |
| F10 | 30060001 | 30160000 | 0.0031 | 0.0021 | 0.0019 |
| F10 | 30080001 | 30180000 | 0.0032 | 0.0024 | 0.0020 |
| F10 | 30100001 | 30200000 | 0.0035 | 0.0027 | 0.0023 |
| F10 | 30120001 | 30220000 | 0.0037 | 0.0025 | 0.0027 |
| F10 | 30140001 | 30240000 | 0.0036 | 0.0029 | 0.0026 |
| F10 | 30160001 | 30260000 | 0.0039 | 0.0033 | 0.0030 |
| F10 | 30180001 | 30280000 | 0.0044 | 0.0028 | 0.0036 |
| F10 | 30200001 | 30300000 | 0.0038 | 0.0025 | 0.0032 |
| F10 | 30220001 | 30320000 | 0.0034 | 0.0023 | 0.0029 |
| F10 | 30240001 | 30340000 | 0.0032 | 0.0023 | 0.0028 |
| F10 | 30260001 | 30360000 | 0.0033 | 0.0018 | 0.0029 |
| F10 | 30280001 | 30380000 | 0.0025 | 0.0021 | 0.0022 |
| F10 | 30300001 | 30400000 | 0.0028 | 0.0026 | 0.0024 |
| F10 | 30320001 | 30420000 | 0.0034 | 0.0030 | 0.0030 |
| F10 | 30340001 | 30440000 | 0.0037 | 0.0027 | 0.0034 |
| F10 | 30360001 | 30460000 | 0.0034 | 0.0027 | 0.0031 |
| F10 | 30380001 | 30480000 | 0.0034 | 0.0026 | 0.0030 |
| F10 | 30400001 | 30500000 | 0.0033 | 0.0023 | 0.0029 |
| F10 | 30420001 | 30520000 | 0.0029 | 0.0021 | 0.0024 |
| F10 | 30440001 | 30540000 | 0.0028 | 0.0026 | 0.0023 |
| F10 | 30460001 | 30560000 | 0.0033 | 0.0026 | 0.0028 |

| Chr | Start | End | both | A1 | A2 |
|---|---|---|---|---|---|
| F10 | 30480001 | 30580000 | 0.0032 | 0.0027 | 0.0028 |
| F10 | 30500001 | 30600000 | 0.0035 | 0.0026 | 0.0029 |
| F10 | 30520001 | 30620000 | 0.0034 | 0.0025 | 0.0029 |
| F10 | 30540001 | 30640000 | 0.0032 | 0.0019 | 0.0029 |
| F10 | 30560001 | 30660000 | 0.0023 | 0.0023 | 0.0021 |
| F10 | 30580001 | 30680000 | 0.0028 | 0.0028 | 0.0026 |
| F10 | 30600001 | 30700000 | 0.0032 | 0.0046 | 0.0031 |
| F10 | 30620001 | 30720000 | 0.0050 | 0.0048 | 0.0048 |
| F10 | 30640001 | 30740000 | 0.0053 | 0.0046 | 0.0049 |
| F10 | 30660001 | 30760000 | 0.0052 | 0.0040 | 0.0047 |
| F10 | 30680001 | 30780000 | 0.0045 | 0.0032 | 0.0041 |
| F10 | 30700001 | 30800000 | 0.0035 | 0.0014 | 0.0032 |
| F10 | 30720001 | 30820000 | 0.0018 | 0.0014 | 0.0015 |
| F10 | 30740001 | 30840000 | 0.0021 | 0.0021 | 0.0015 |
| F10 | 30760001 | 30860000 | 0.0028 | 0.0026 | 0.0020 |
| F10 | 30780001 | 30880000 | 0.0036 | 0.0031 | 0.0025 |
| F10 | 30800001 | 30900000 | 0.0042 | 0.0029 | 0.0029 |
| F10 | 30820001 | 30920000 | 0.0041 | 0.0033 | 0.0028 |
| F10 | 30840001 | 30940000 | 0.0041 | 0.0028 | 0.0032 |
| F10 | 30860001 | 30960000 | 0.0037 | 0.0023 | 0.0028 |
| F10 | 30880001 | 30980000 | 0.0030 | 0.0017 | 0.0023 |
| F10 | 30900001 | 31000000 | 0.0022 | 0.0016 | 0.0016 |
| F10 | 30920001 | 31020000 | 0.0021 | 0.0009 | 0.0016 |
| F10 | 30940001 | 31040000 | 0.0014 | 0.0009 | 0.0010 |
| F10 | 30960001 | 31060000 | 0.0014 | 0.0012 | 0.0010 |
| F10 | 30980001 | 31080000 | 0.0017 | 0.0017 | 0.0014 |
| F10 | 31000001 | 31100000 | 0.0025 | 0.0021 | 0.0020 |
| F10 | 31020001 | 31120000 | 0.0031 | 0.0026 | 0.0025 |
| F10 | 31040001 | 31140000 | 0.0037 | 0.0027 | 0.0030 |
| F10 | 31060001 | 31160000 | 0.0039 | 0.0021 | 0.0033 |
| F10 | 31080001 | 31180000 | 0.0034 | 0.0017 | 0.0027 |
| F10 | 31100001 | 31200000 | 0.0027 | 0.0013 | 0.0021 |
| F10 | 31120001 | 31220000 | 0.0022 | 0.0007 | 0.0016 |
| F10 | 31140001 | 31240000 | 0.0016 | 0.0011 | 0.0010 |
| F10 | 31160001 | 31260000 | 0.0019 | 0.0015 | 0.0012 |
| F10 | 31180001 | 31280000 | 0.0022 | 0.0019 | 0.0016 |
| F10 | 31200001 | 31300000 | 0.0029 | 0.0020 | 0.0023 |
| F10 | 31220001 | 31320000 | 0.0031 | 0.0025 | 0.0023 |
| F10 | 31240001 | 31340000 | 0.0039 | 0.0027 | 0.0029 |
| F10 | 31260001 | 31360000 | 0.0045 | 0.0025 | 0.0031 |
| F10 | 31280001 | 31380000 | 0.0043 | 0.0030 | 0.0028 |
| F10 | 31300001 | 31400000 | 0.0047 | 0.0033 | 0.0031 |
| F10 | 31320001 | 31420000 | 0.0049 | 0.0034 | 0.0033 |
| F10 | 31340001 | 31440000 | 0.0047 | 0.0043 | 0.0033 |
| F10 | 31360001 | 31460000 | 0.0053 | 0.0041 | 0.0043 |
| F10 | 31380001 | 31480000 | 0.0050 | 0.0038 | 0.0042 |
| F10 | 31400001 | 31500000 | 0.0047 | 0.0046 | 0.0039 |
| F10 | 31420001 | 31520000 | 0.0056 | 0.0050 | 0.0050 |
| F10 | 31440001 | 31540000 | 0.0060 | 0.0043 | 0.0056 |
| F10 | 31460001 | 31560000 | 0.0054 | 0.0044 | 0.0048 |
| F10 | 31480001 | 31580000 | 0.0056 | 0.0044 | 0.0049 |
| F10 | 31500001 | 31600000 | 0.0054 | 0.0034 | 0.0048 |
| F10 | 31520001 | 31620000 | 0.0042 | 0.0033 | 0.0035 |
| F10 | 31540001 | 31640000 | 0.0043 | 0.0030 | 0.0034 |
| F10 | 31560001 | 31660000 | 0.0040 | 0.0030 | 0.0032 |
| F10 | 31580001 | 31680000 | 0.0039 | 0.0031 | 0.0031 |
| F10 | 31600001 | 31700000 | 0.0041 | 0.0035 | 0.0032 |
| F10 | 31620001 | 31720000 | 0.0048 | 0.0035 | 0.0038 |

| Chr | Start | End | both | A1 | A2 |
|---|---|---|---|---|---|
| F10 | 31640001 | 31740000 | 0.0047 | 0.0033 | 0.0039 |
| F10 | 31660001 | 31760000 | 0.0047 | 0.0034 | 0.0039 |
| F10 | 31680001 | 31780000 | 0.0047 | 0.0033 | 0.0040 |
| F10 | 31700001 | 31800000 | 0.0043 | 0.0030 | 0.0037 |
| F10 | 31720001 | 31820000 | 0.0038 | 0.0027 | 0.0033 |
| F10 | 31740001 | 31840000 | 0.0032 | 0.0023 | 0.0027 |
| F10 | 31760001 | 31860000 | 0.0027 | 0.0025 | 0.0023 |
| F10 | 31780001 | 31880000 | 0.0031 | 0.0020 | 0.0025 |
| F10 | 31800001 | 31900000 | 0.0028 | 0.0022 | 0.0021 |
| F10 | 31820001 | 31920000 | 0.0028 | 0.0016 | 0.0021 |
| F10 | 31840001 | 31940000 | 0.0023 | 0.0013 | 0.0017 |
| F10 | 31860001 | 31960000 | 0.0022 | 0.0041 | 0.0015 |
| F10 | 31880001 | 31980000 | 0.0048 | 0.0041 | 0.0041 |
| F10 | 31900001 | 32000000 | 0.0050 | 0.0036 | 0.0044 |
| F10 | 31920001 | 32020000 | 0.0045 | 0.0039 | 0.0040 |
| F10 | 31940001 | 32040000 | 0.0048 | 0.0051 | 0.0045 |
| F10 | 31960001 | 32060000 | 0.0056 | 0.0025 | 0.0054 |
| F10 | 31980001 | 32080000 | 0.0030 | 0.0026 | 0.0029 |
| F10 | 32000001 | 32100000 | 0.0030 | 0.0027 | 0.0029 |
| F10 | 32020001 | 32120000 | 0.0032 | 0.0033 | 0.0030 |
| F10 | 32040001 | 32140000 | 0.0038 | 0.0025 | 0.0036 |
| F10 | 32060001 | 32160000 | 0.0032 | 0.0025 | 0.0029 |
| F10 | 32080001 | 32180000 | 0.0036 | 0.0026 | 0.0032 |
| F10 | 32100001 | 32200000 | 0.0039 | 0.0031 | 0.0034 |
| F10 | 32120001 | 32220000 | 0.0044 | 0.0023 | 0.0040 |
| F10 | 32140001 | 32240000 | 0.0034 | 0.0024 | 0.0030 |
| F10 | 32160001 | 32260000 | 0.0034 | 0.0032 | 0.0030 |
| F10 | 32180001 | 32280000 | 0.0038 | 0.0030 | 0.0035 |
| F10 | 32200001 | 32300000 | 0.0034 | 0.0036 | 0.0033 |
| F10 | 32220001 | 32320000 | 0.0040 | 0.0041 | 0.0038 |
| F10 | 32240001 | 32340000 | 0.0046 | 0.0053 | 0.0043 |
| F10 | 32260001 | 32360000 | 0.0060 | 0.0047 | 0.0058 |
| F10 | 32280001 | 32380000 | 0.0055 | 0.0045 | 0.0053 |
| F10 | 32300001 | 32400000 | 0.0053 | 0.0035 | 0.0051 |
| F10 | 32320001 | 32420000 | 0.0044 | 0.0051 | 0.0042 |
| F10 | 32340001 | 32440000 | 0.0067 | 0.0043 | 0.0063 |
| F10 | 32360001 | 32460000 | 0.0059 | 0.0045 | 0.0054 |
| F10 | 32380001 | 32480000 | 0.0062 | 0.0047 | 0.0056 |
| F10 | 32400001 | 32500000 | 0.0064 | 0.0043 | 0.0058 |
| F10 | 32420001 | 32520000 | 0.0059 | 0.0023 | 0.0054 |
| F10 | 32440001 | 32540000 | 0.0034 | 0.0020 | 0.0029 |
| F10 | 32460001 | 32560000 | 0.0032 | 0.0013 | 0.0027 |
| F10 | 32480001 | 32580000 | 0.0024 | 0.0014 | 0.0019 |
| F10 | 32500001 | 32600000 | 0.0027 | 0.0016 | 0.0021 |
| F10 | 32520001 | 32620000 | 0.0032 | 0.0024 | 0.0024 |
| F10 | 32540001 | 32640000 | 0.0044 | 0.0026 | 0.0033 |
| F10 | 32560001 | 32660000 | 0.0043 | 0.0028 | 0.0033 |
| F10 | 32580001 | 32680000 | 0.0047 | 0.0026 | 0.0036 |
| F10 | 32600001 | 32700000 | 0.0045 | 0.0032 | 0.0034 |
| F10 | 32620001 | 32720000 | 0.0051 | 0.0023 | 0.0041 |
| F10 | 32640001 | 32740000 | 0.0037 | 0.0018 | 0.0029 |
| F10 | 32660001 | 32760000 | 0.0035 | 0.0023 | 0.0027 |
| F10 | 32680001 | 32780000 | 0.0042 | 0.0027 | 0.0035 |
| F10 | 32700001 | 32800000 | 0.0049 | 0.0023 | 0.0041 |
| F10 | 32720001 | 32820000 | 0.0043 | 0.0032 | 0.0037 |
| F10 | 32740001 | 32840000 | 0.0055 | 0.0030 | 0.0049 |
| F10 | 32760001 | 32860000 | 0.0052 | 0.0028 | 0.0045 |
| F10 | 32780001 | 32880000 | 0.0048 | 0.0029 | 0.0041 |

| Chr | Start | End | both | A1 | A2 |
|---|---|---|---|---|---|
| F10 | 32800001 | 32900000 | 0.0046 | 0.0026 | 0.0040 |
| F10 | 32820001 | 32920000 | 0.0045 | 0.0017 | 0.0037 |
| F10 | 32840001 | 32940000 | 0.0035 | 0.0018 | 0.0026 |
| F10 | 32860001 | 32960000 | 0.0036 | 0.0016 | 0.0028 |
| F10 | 32880001 | 32980000 | 0.0032 | 0.0014 | 0.0024 |
| F10 | 32900001 | 33000000 | 0.0031 | 0.0016 | 0.0022 |
| F10 | 32920001 | 33020000 | 0.0033 | 0.0021 | 0.0024 |
| F10 | 32940001 | 33040000 | 0.0038 | 0.0020 | 0.0029 |
| F10 | 32960001 | 33060000 | 0.0036 | 0.0021 | 0.0028 |
| F10 | 32980001 | 33080000 | 0.0039 | 0.0022 | 0.0029 |
| F10 | 33000001 | 33100000 | 0.0037 | 0.0023 | 0.0029 |
| F10 | 33020001 | 33120000 | 0.0035 | 0.0029 | 0.0029 |
| F10 | 33040001 | 33140000 | 0.0041 | 0.0032 | 0.0035 |
| F10 | 33060001 | 33160000 | 0.0043 | 0.0031 | 0.0037 |
| F10 | 33080001 | 33180000 | 0.0039 | 0.0032 | 0.0035 |
| F10 | 33100001 | 33200000 | 0.0039 | 0.0030 | 0.0035 |
| F10 | 33120001 | 33220000 | 0.0038 | 0.0019 | 0.0033 |
| F10 | 33140001 | 33240000 | 0.0026 | 0.0018 | 0.0022 |
| F10 | 33160001 | 33260000 | 0.0026 | 0.0021 | 0.0021 |
| F10 | 33180001 | 33280000 | 0.0035 | 0.0021 | 0.0029 |
| F10 | 33200001 | 33300000 | 0.0038 | 0.0023 | 0.0030 |
| F10 | 33220001 | 33320000 | 0.0039 | 0.0022 | 0.0031 |
| F10 | 33240001 | 33340000 | 0.0039 | 0.0023 | 0.0030 |
| F10 | 33260001 | 33360000 | 0.0038 | 0.0017 | 0.0031 |
| F10 | 33280001 | 33380000 | 0.0028 | 0.0015 | 0.0023 |
| F10 | 33300001 | 33400000 | 0.0022 | 0.0018 | 0.0019 |
| F10 | 33320001 | 33420000 | 0.0026 | 0.0022 | 0.0023 |
| F10 | 33340001 | 33440000 | 0.0031 | 0.0020 | 0.0028 |
| F10 | 33360001 | 33460000 | 0.0033 | 0.0024 | 0.0030 |
| F10 | 33380001 | 33480000 | 0.0039 | 0.0023 | 0.0036 |
| F10 | 33400001 | 33500000 | 0.0039 | 0.0023 | 0.0037 |
| F10 | 33420001 | 33520000 | 0.0039 | 0.0022 | 0.0037 |
| F10 | 33440001 | 33540000 | 0.0036 | 0.0023 | 0.0034 |
| F10 | 33460001 | 33560000 | 0.0033 | 0.0019 | 0.0032 |
| F10 | 33480001 | 33580000 | 0.0028 | 0.0022 | 0.0026 |
| F10 | 33500001 | 33600000 | 0.0031 | 0.0033 | 0.0029 |
| F10 | 33520001 | 33620000 | 0.0046 | 0.0045 | 0.0044 |
| F10 | 33540001 | 33640000 | 0.0056 | 0.0047 | 0.0055 |
| F10 | 33560001 | 33660000 | 0.0059 | 0.0048 | 0.0057 |
| F10 | 33580001 | 33680000 | 0.0060 | 0.0046 | 0.0059 |
| F10 | 33600001 | 33700000 | 0.0061 | 0.0034 | 0.0060 |
| F10 | 33620001 | 33720000 | 0.0047 | 0.0023 | 0.0046 |
| F10 | 33640001 | 33740000 | 0.0036 | 0.0024 | 0.0035 |
| F10 | 33660001 | 33760000 | 0.0034 | 0.0023 | 0.0032 |
| F10 | 33680001 | 33780000 | 0.0033 | 0.0022 | 0.0032 |
| F10 | 33700001 | 33800000 | 0.0027 | 0.0018 | 0.0026 |
| F10 | 33720001 | 33820000 | 0.0020 | 0.0019 | 0.0019 |
| F10 | 33740001 | 33840000 | 0.0021 | 0.0014 | 0.0021 |
| F10 | 33760001 | 33860000 | 0.0016 | 0.0011 | 0.0016 |
| F10 | 33780001 | 33880000 | 0.0012 | 0.0010 | 0.0012 |
| F10 | 33800001 | 33900000 | 0.0011 | 0.0011 | 0.0011 |
| F10 | 33820001 | 33920000 | 0.0012 | 0.0007 | 0.0012 |
| F10 | 33840001 | 33940000 | 0.0006 | 0.0007 | 0.0006 |
| F10 | 33860001 | 33960000 | 0.0007 | 0.0016 | 0.0007 |
| F10 | 33880001 | 33980000 | 0.0018 | 0.0015 | 0.0018 |
| F10 | 33900001 | 34000000 | 0.0016 | 0.0016 | 0.0016 |
| F10 | 33920001 | 34020000 | 0.0020 | 0.0020 | 0.0020 |
| F10 | 33940001 | 34040000 | 0.0025 | 0.0022 | 0.0025 |

| Chr | Start | End | both | A1 | A2 |
|---|---|---|---|---|---|
| F10 | 33960001 | 34060000 | 0.0027 | 0.0017 | 0.0027 |
| F10 | 33980001 | 34080000 | 0.0021 | 0.0024 | 0.0021 |
| F10 | 34000001 | 34100000 | 0.0029 | 0.0030 | 0.0029 |
| F10 | 34020001 | 34120000 | 0.0036 | 0.0033 | 0.0036 |
| F10 | 34040001 | 34140000 | 0.0040 | 0.0033 | 0.0040 |
| F10 | 34060001 | 34160000 | 0.0041 | 0.0031 | 0.0041 |
| F10 | 34080001 | 34180000 | 0.0038 | 0.0032 | 0.0038 |
| F10 | 34100001 | 34200000 | 0.0037 | 0.0030 | 0.0037 |
| F10 | 34120001 | 34220000 | 0.0034 | 0.0025 | 0.0032 |
| F10 | 34140001 | 34240000 | 0.0026 | 0.0020 | 0.0025 |
| F10 | 34160001 | 34260000 | 0.0022 | 0.0022 | 0.0021 |
| F10 | 34180001 | 34280000 | 0.0029 | 0.0024 | 0.0028 |
| F10 | 34200001 | 34300000 | 0.0033 | 0.0021 | 0.0033 |
| F10 | 34220001 | 34320000 | 0.0032 | 0.0030 | 0.0033 |
| F10 | 34240001 | 34340000 | 0.0048 | 0.0037 | 0.0049 |
| F10 | 34260001 | 34360000 | 0.0058 | 0.0035 | 0.0060 |
| F10 | 34280001 | 34380000 | 0.0054 | 0.0032 | 0.0055 |
| F10 | 34300001 | 34400000 | 0.0050 | 0.0032 | 0.0051 |
| F10 | 34320001 | 34420000 | 0.0049 | 0.0032 | 0.0050 |
| F10 | 34340001 | 34440000 | 0.0044 | 0.0030 | 0.0045 |
| F10 | 34360001 | 34460000 | 0.0042 | 0.0035 | 0.0042 |
| F10 | 34380001 | 34480000 | 0.0048 | 0.0033 | 0.0049 |
| F10 | 34400001 | 34500000 | 0.0045 | 0.0035 | 0.0046 |
| F10 | 34420001 | 34520000 | 0.0046 | 0.0028 | 0.0047 |
| F10 | 34440001 | 34540000 | 0.0038 | 0.0024 | 0.0039 |
| F10 | 34460001 | 34560000 | 0.0037 | 0.0020 | 0.0039 |
| F10 | 34480001 | 34580000 | 0.0033 | 0.0020 | 0.0034 |
| F10 | 34500001 | 34600000 | 0.0036 | 0.0018 | 0.0037 |
| F10 | 34520001 | 34620000 | 0.0039 | 0.0022 | 0.0040 |
| F10 | 34540001 | 34640000 | 0.0041 | 0.0029 | 0.0042 |
| F10 | 34560001 | 34660000 | 0.0043 | 0.0029 | 0.0043 |
| F10 | 34580001 | 34680000 | 0.0041 | 0.0028 | 0.0041 |
| F10 | 34600001 | 34700000 | 0.0037 | 0.0024 | 0.0037 |
| F10 | 34620001 | 34720000 | 0.0030 | 0.0025 | 0.0029 |
| F10 | 34640001 | 34740000 | 0.0032 | 0.0024 | 0.0032 |
| F10 | 34660001 | 34760000 | 0.0032 | 0.0025 | 0.0031 |
| F10 | 34680001 | 34780000 | 0.0033 | 0.0027 | 0.0032 |
| F10 | 34700001 | 34800000 | 0.0034 | 0.0034 | 0.0033 |
| F10 | 34720001 | 34820000 | 0.0043 | 0.0029 | 0.0042 |
| F10 | 34740001 | 34840000 | 0.0041 | 0.0026 | 0.0040 |
| F10 | 34760001 | 34860000 | 0.0037 | 0.0024 | 0.0036 |
| F10 | 34780001 | 34880000 | 0.0038 | 0.0021 | 0.0038 |
| F10 | 34800001 | 34900000 | 0.0039 | 0.0020 | 0.0038 |
| F10 | 34820001 | 34920000 | 0.0038 | 0.0019 | 0.0037 |
| F10 | 34840001 | 34940000 | 0.0041 | 0.0018 | 0.0040 |
| F10 | 34860001 | 34960000 | 0.0039 | 0.0022 | 0.0039 |
| F10 | 34880001 | 34980000 | 0.0044 | 0.0026 | 0.0043 |
| F10 | 34900001 | 35000000 | 0.0047 | 0.0023 | 0.0045 |
| F10 | 34920001 | 35020000 | 0.0045 | 0.0023 | 0.0043 |
| F10 | 34940001 | 35040000 | 0.0042 | 0.0024 | 0.0040 |
| F10 | 34960001 | 35060000 | 0.0044 | 0.0019 | 0.0042 |
| F10 | 34980001 | 35080000 | 0.0041 | 0.0017 | 0.0038 |
| F10 | 35000001 | 35100000 | 0.0040 | 0.0016 | 0.0036 |
| F10 | 35020001 | 35120000 | 0.0036 | 0.0017 | 0.0033 |
| F10 | 35040001 | 35140000 | 0.0039 | 0.0015 | 0.0034 |
| F10 | 35060001 | 35160000 | 0.0036 | 0.0016 | 0.0032 |
| F10 | 35080001 | 35180000 | 0.0034 | 0.0013 | 0.0030 |
| F10 | 35100001 | 35200000 | 0.0030 | 0.0015 | 0.0026 |

| Chr | Start | End | both | A1 | A2 |
|---|---|---|---|---|---|
| F10 | 35120001 | 35220000 | 0.0031 | 0.0014 | 0.0027 |
| F10 | 35140001 | 35240000 | 0.0025 | 0.0021 | 0.0022 |
| F10 | 35160001 | 35260000 | 0.0032 | 0.0021 | 0.0028 |
| F10 | 35180001 | 35280000 | 0.0032 | 0.0028 | 0.0028 |
| F10 | 35200001 | 35300000 | 0.0040 | 0.0024 | 0.0037 |
| F10 | 35220001 | 35320000 | 0.0039 | 0.0022 | 0.0035 |
| F10 | 35240001 | 35340000 | 0.0036 | 0.0018 | 0.0033 |
| F10 | 35260001 | 35360000 | 0.0031 | 0.0024 | 0.0029 |
| F10 | 35280001 | 35380000 | 0.0035 | 0.0027 | 0.0033 |
| F10 | 35300001 | 35400000 | 0.0036 | 0.0031 | 0.0034 |
| F10 | 35320001 | 35420000 | 0.0038 | 0.0037 | 0.0036 |
| F10 | 35340001 | 35440000 | 0.0045 | 0.0038 | 0.0043 |
| F10 | 35360001 | 35460000 | 0.0049 | 0.0041 | 0.0045 |
| F10 | 35380001 | 35480000 | 0.0055 | 0.0038 | 0.0051 |
| F10 | 35400001 | 35500000 | 0.0050 | 0.0035 | 0.0043 |
| F10 | 35420001 | 35520000 | 0.0050 | 0.0046 | 0.0042 |
| F10 | 35440001 | 35540000 | 0.0061 | 0.0056 | 0.0053 |
| F10 | 35460001 | 35560000 | 0.0071 | 0.0050 | 0.0064 |
| F10 | 35480001 | 35580000 | 0.0063 | 0.0052 | 0.0058 |
| F10 | 35500001 | 35600000 | 0.0066 | 0.0055 | 0.0063 |
| F10 | 35520001 | 35620000 | 0.0065 | 0.0043 | 0.0064 |
| F10 | 35540001 | 35640000 | 0.0053 | 0.0033 | 0.0051 |
| F10 | 35560001 | 35660000 | 0.0042 | 0.0030 | 0.0039 |
| F10 | 35580001 | 35680000 | 0.0038 | 0.0025 | 0.0034 |
| F10 | 35600001 | 35700000 | 0.0034 | 0.0027 | 0.0029 |
| F10 | 35620001 | 35720000 | 0.0038 | 0.0026 | 0.0033 |
| F10 | 35640001 | 35740000 | 0.0035 | 0.0022 | 0.0031 |
| F10 | 35660001 | 35760000 | 0.0032 | 0.0022 | 0.0028 |
| F10 | 35680001 | 35780000 | 0.0032 | 0.0022 | 0.0028 |
| F10 | 35700001 | 35800000 | 0.0031 | 0.0016 | 0.0027 |
| F10 | 35720001 | 35820000 | 0.0026 | 0.0024 | 0.0022 |
| F10 | 35740001 | 35840000 | 0.0036 | 0.0029 | 0.0032 |
| F10 | 35760001 | 35860000 | 0.0043 | 0.0031 | 0.0039 |
| F10 | 35780001 | 35880000 | 0.0048 | 0.0030 | 0.0042 |
| F10 | 35800001 | 35900000 | 0.0048 | 0.0031 | 0.0043 |
| F10 | 35820001 | 35920000 | 0.0049 | 0.0027 | 0.0043 |
| F10 | 35840001 | 35940000 | 0.0045 | 0.0023 | 0.0038 |
| F10 | 35860001 | 35960000 | 0.0040 | 0.0029 | 0.0033 |
| F10 | 35880001 | 35980000 | 0.0043 | 0.0033 | 0.0037 |
| F10 | 35900001 | 36000000 | 0.0048 | 0.0031 | 0.0041 |
| F10 | 35920001 | 36020000 | 0.0045 | 0.0033 | 0.0039 |
| F10 | 35940001 | 36040000 | 0.0050 | 0.0038 | 0.0042 |
| F10 | 35960001 | 36060000 | 0.0058 | 0.0039 | 0.0050 |
| F10 | 35980001 | 36080000 | 0.0061 | 0.0038 | 0.0052 |
| F10 | 36000001 | 36100000 | 0.0057 | 0.0045 | 0.0050 |
| F10 | 36020001 | 36120000 | 0.0066 | 0.0054 | 0.0059 |
| F10 | 36040001 | 36140000 | 0.0070 | 0.0048 | 0.0065 |
| F10 | 36060001 | 36160000 | 0.0061 | 0.0039 | 0.0056 |
| F10 | 36080001 | 36180000 | 0.0053 | 0.0036 | 0.0048 |
| F10 | 36100001 | 36200000 | 0.0049 | 0.0031 | 0.0044 |
| F10 | 36120001 | 36220000 | 0.0041 | 0.0014 | 0.0037 |
| F10 | 36140001 | 36240000 | 0.0024 | 0.0021 | 0.0020 |
| F10 | 36160001 | 36260000 | 0.0031 | 0.0022 | 0.0027 |
| F10 | 36180001 | 36280000 | 0.0032 | 0.0027 | 0.0029 |
| F10 | 36200001 | 36300000 | 0.0040 | 0.0027 | 0.0036 |
| F10 | 36220001 | 36320000 | 0.0041 | 0.0033 | 0.0036 |
| F10 | 36240001 | 36340000 | 0.0047 | 0.0029 | 0.0041 |
| F10 | 36260001 | 36360000 | 0.0044 | 0.0031 | 0.0038 |

| Chr | Start | End | both | A1 | A2 |
|---|---|---|---|---|---|
| F10 | 36280001 | 36380000 | 0.0045 | 0.0026 | 0.0039 |
| F10 | 36300001 | 36400000 | 0.0039 | 0.0023 | 0.0033 |
| F10 | 36320001 | 36420000 | 0.0038 | 0.0020 | 0.0032 |
| F10 | 36340001 | 36440000 | 0.0034 | 0.0017 | 0.0028 |
| F10 | 36360001 | 36460000 | 0.0028 | 0.0011 | 0.0024 |
| F10 | 36380001 | 36480000 | 0.0019 | 0.0019 | 0.0016 |
| F10 | 36400001 | 36500000 | 0.0028 | 0.0021 | 0.0027 |
| F10 | 36420001 | 36520000 | 0.0030 | 0.0028 | 0.0029 |
| F10 | 36440001 | 36540000 | 0.0038 | 0.0034 | 0.0036 |
| F10 | 36460001 | 36560000 | 0.0048 | 0.0035 | 0.0044 |
| F10 | 36480001 | 36580000 | 0.0049 | 0.0035 | 0.0045 |
| F10 | 36500001 | 36600000 | 0.0049 | 0.0042 | 0.0043 |
| F10 | 36520001 | 36620000 | 0.0055 | 0.0039 | 0.0050 |
| F10 | 36540001 | 36640000 | 0.0051 | 0.0031 | 0.0046 |
| F10 | 36560001 | 36660000 | 0.0041 | 0.0031 | 0.0037 |
| F10 | 36580001 | 36680000 | 0.0043 | 0.0022 | 0.0037 |
| F10 | 36600001 | 36700000 | 0.0033 | 0.0012 | 0.0027 |
| F10 | 36620001 | 36720000 | 0.0025 | 0.0008 | 0.0019 |
| F10 | 36640001 | 36740000 | 0.0020 | 0.0007 | 0.0014 |
| F10 | 36660001 | 36760000 | 0.0017 | 0.0006 | 0.0013 |
| F10 | 36680001 | 36780000 | 0.0013 | 0.0013 | 0.0011 |
| F10 | 36700001 | 36800000 | 0.0018 | 0.0015 | 0.0017 |
| F10 | 36720001 | 36820000 | 0.0018 | 0.0018 | 0.0017 |
| F10 | 36740001 | 36840000 | 0.0021 | 0.0019 | 0.0020 |
| F10 | 36760001 | 36860000 | 0.0021 | 0.0031 | 0.0020 |
| F10 | 36780001 | 36880000 | 0.0032 | 0.0026 | 0.0031 |
| F10 | 36800001 | 36900000 | 0.0027 | 0.0024 | 0.0026 |
| F10 | 36820001 | 36920000 | 0.0025 | 0.0025 | 0.0024 |
| F10 | 36840001 | 36940000 | 0.0026 | 0.0024 | 0.0025 |
| F10 | 36860001 | 36960000 | 0.0026 | 0.0013 | 0.0025 |
| F10 | 36880001 | 36980000 | 0.0015 | 0.0010 | 0.0015 |
| F10 | 36900001 | 37000000 | 0.0015 | 0.0013 | 0.0014 |
| F10 | 36920001 | 37020000 | 0.0018 | 0.0009 | 0.0017 |
| F10 | 36940001 | 37040000 | 0.0014 | 0.0010 | 0.0013 |
| F10 | 36960001 | 37060000 | 0.0015 | 0.0010 | 0.0014 |
| F10 | 36980001 | 37080000 | 0.0015 | 0.0010 | 0.0013 |
| F10 | 37000001 | 37100000 | 0.0012 | 0.0007 | 0.0011 |
| F10 | 37020001 | 37120000 | 0.0009 | 0.0007 | 0.0008 |
| F10 | 37040001 | 37140000 | 0.0009 | 0.0011 | 0.0008 |
| F10 | 37060001 | 37160000 | 0.0015 | 0.0014 | 0.0014 |
| F10 | 37080001 | 37180000 | 0.0020 | 0.0017 | 0.0018 |
| F10 | 37100001 | 37200000 | 0.0026 | 0.0016 | 0.0023 |
| F10 | 37120001 | 37220000 | 0.0025 | 0.0019 | 0.0022 |
| F10 | 37140001 | 37240000 | 0.0028 | 0.0016 | 0.0025 |
| F10 | 37160001 | 37260000 | 0.0024 | 0.0017 | 0.0021 |
| F10 | 37180001 | 37280000 | 0.0023 | 0.0013 | 0.0021 |
| F10 | 37200001 | 37300000 | 0.0017 | 0.0015 | 0.0015 |
| F10 | 37220001 | 37320000 | 0.0019 | 0.0011 | 0.0017 |
| F10 | 37240001 | 37340000 | 0.0014 | 0.0012 | 0.0012 |
| F10 | 37260001 | 37360000 | 0.0016 | 0.0008 | 0.0014 |
| F10 | 37280001 | 37380000 | 0.0011 | 0.0007 | 0.0009 |
| F10 | 37300001 | 37400000 | 0.0010 | 0.0007 | 0.0008 |
| F10 | 37320001 | 37420000 | 0.0008 | 0.0013 | 0.0007 |
| F10 | 37340001 | 37440000 | 0.0015 | 0.0016 | 0.0014 |
| F10 | 37360001 | 37460000 | 0.0020 | 0.0018 | 0.0017 |
| F10 | 37380001 | 37480000 | 0.0023 | 0.0023 | 0.0019 |
| F10 | 37400001 | 37500000 | 0.0028 | 0.0025 | 0.0023 |
| F10 | 37420001 | 37520000 | 0.0030 | 0.0018 | 0.0024 |

| Chr | Start | End | both | A1 | A2 |
|---|---|---|---|---|---|
| F10 | 37440001 | 37540000 | 0.0025 | 0.0013 | 0.0018 |
| F10 | 37460001 | 37560000 | 0.0019 | 0.0014 | 0.0013 |
| F10 | 37480001 | 37580000 | 0.0020 | 0.0010 | 0.0014 |
| F10 | 37500001 | 37600000 | 0.0016 | 0.0008 | 0.0011 |
| F10 | 37520001 | 37620000 | 0.0014 | 0.0008 | 0.0009 |
| F10 | 37540001 | 37640000 | 0.0015 | 0.0012 | 0.0010 |
| F10 | 37560001 | 37660000 | 0.0018 | 0.0018 | 0.0013 |
| F10 | 37580001 | 37680000 | 0.0028 | 0.0021 | 0.0020 |
| F10 | 37600001 | 37700000 | 0.0031 | 0.0023 | 0.0023 |
| F10 | 37620001 | 37720000 | 0.0035 | 0.0022 | 0.0026 |
| F10 | 37640001 | 37740000 | 0.0033 | 0.0019 | 0.0025 |
| F10 | 37660001 | 37760000 | 0.0031 | 0.0016 | 0.0022 |
| F10 | 37680001 | 37780000 | 0.0027 | 0.0015 | 0.0018 |
| F10 | 37700001 | 37800000 | 0.0025 | 0.0019 | 0.0017 |
| F10 | 37720001 | 37820000 | 0.0028 | 0.0021 | 0.0021 |
| F10 | 37740001 | 37840000 | 0.0029 | 0.0021 | 0.0023 |
| F10 | 37760001 | 37860000 | 0.0027 | 0.0024 | 0.0023 |
| F10 | 37780001 | 37880000 | 0.0027 | 0.0033 | 0.0025 |
| F10 | 37800001 | 37900000 | 0.0040 | 0.0030 | 0.0035 |
| F10 | 37820001 | 37920000 | 0.0040 | 0.0031 | 0.0034 |
| F10 | 37840001 | 37940000 | 0.0041 | 0.0037 | 0.0034 |
| F10 | 37860001 | 37960000 | 0.0047 | 0.0037 | 0.0040 |
| F10 | 37880001 | 37980000 | 0.0049 | 0.0034 | 0.0041 |
| F10 | 37900001 | 38000000 | 0.0042 | 0.0041 | 0.0035 |
| F10 | 37920001 | 38020000 | 0.0049 | 0.0043 | 0.0041 |
| F10 | 37940001 | 38040000 | 0.0054 | 0.0038 | 0.0045 |
| F10 | 37960001 | 38060000 | 0.0053 | 0.0033 | 0.0041 |
| F10 | 37980001 | 38080000 | 0.0048 | 0.0026 | 0.0036 |
| F10 | 38000001 | 38100000 | 0.0041 | 0.0023 | 0.0030 |
| F10 | 38020001 | 38120000 | 0.0038 | 0.0020 | 0.0026 |
| F10 | 38040001 | 38140000 | 0.0031 | 0.0020 | 0.0022 |
| F10 | 38060001 | 38160000 | 0.0028 | 0.0016 | 0.0020 |
| F10 | 38080001 | 38180000 | 0.0022 | 0.0021 | 0.0016 |
| F10 | 38100001 | 38200000 | 0.0027 | 0.0014 | 0.0022 |
| F10 | 38120001 | 38220000 | 0.0019 | 0.0014 | 0.0017 |
| F10 | 38140001 | 38240000 | 0.0021 | 0.0015 | 0.0018 |
| F10 | 38160001 | 38260000 | 0.0022 | 0.0016 | 0.0018 |
| F10 | 38180001 | 38280000 | 0.0025 | 0.0017 | 0.0019 |
| F10 | 38200001 | 38300000 | 0.0025 | 0.0021 | 0.0019 |
| F10 | 38220001 | 38320000 | 0.0030 | 0.0023 | 0.0024 |
| F10 | 38240001 | 38340000 | 0.0031 | 0.0021 | 0.0026 |
| F10 | 38260001 | 38360000 | 0.0026 | 0.0024 | 0.0023 |
| F10 | 38280001 | 38380000 | 0.0028 | 0.0023 | 0.0026 |
| F10 | 38300001 | 38400000 | 0.0026 | 0.0018 | 0.0025 |
| F10 | 38320001 | 38420000 | 0.0020 | 0.0017 | 0.0019 |
| F10 | 38340001 | 38440000 | 0.0020 | 0.0018 | 0.0020 |
| F10 | 38360001 | 38460000 | 0.0022 | 0.0016 | 0.0021 |
| F10 | 38380001 | 38480000 | 0.0020 | 0.0010 | 0.0018 |
| F10 | 38400001 | 38500000 | 0.0016 | 0.0012 | 0.0014 |
| F10 | 38420001 | 38520000 | 0.0018 | 0.0013 | 0.0015 |
| F10 | 38440001 | 38540000 | 0.0017 | 0.0013 | 0.0015 |
| F10 | 38460001 | 38560000 | 0.0018 | 0.0013 | 0.0017 |
| F10 | 38480001 | 38580000 | 0.0017 | 0.0015 | 0.0016 |
| F10 | 38500001 | 38600000 | 0.0019 | 0.0018 | 0.0018 |
| F10 | 38520001 | 38620000 | 0.0026 | 0.0020 | 0.0020 |
| F10 | 38540001 | 38640000 | 0.0028 | 0.0017 | 0.0020 |
| F10 | 38560001 | 38660000 | 0.0026 | 0.0017 | 0.0017 |
| F10 | 38580001 | 38680000 | 0.0025 | 0.0015 | 0.0017 |

| Chr | Start | End | both | A1 | A2 |
|---|---|---|---|---|---|
| F10 | 38600001 | 38700000 | 0.0022 | 0.0016 | 0.0014 |
| F10 | 38620001 | 38720000 | 0.0020 | 0.0017 | 0.0017 |
| F10 | 38640001 | 38740000 | 0.0022 | 0.0021 | 0.0019 |
| F10 | 38660001 | 38760000 | 0.0029 | 0.0027 | 0.0023 |
| F10 | 38680001 | 38780000 | 0.0037 | 0.0027 | 0.0030 |
| F10 | 38700001 | 38800000 | 0.0037 | 0.0022 | 0.0030 |
| F10 | 38720001 | 38820000 | 0.0032 | 0.0018 | 0.0024 |
| F10 | 38740001 | 38840000 | 0.0029 | 0.0018 | 0.0020 |
| F10 | 38760001 | 38860000 | 0.0029 | 0.0012 | 0.0022 |
| F10 | 38780001 | 38880000 | 0.0022 | 0.0014 | 0.0016 |
| F10 | 38800001 | 38900000 | 0.0025 | 0.0018 | 0.0018 |
| F10 | 38820001 | 38920000 | 0.0029 | 0.0022 | 0.0021 |
| F10 | 38840001 | 38940000 | 0.0030 | 0.0025 | 0.0025 |
| F10 | 38860001 | 38960000 | 0.0031 | 0.0028 | 0.0027 |
| F10 | 38880001 | 38980000 | 0.0035 | 0.0027 | 0.0030 |
| F10 | 38900001 | 39000000 | 0.0034 | 0.0025 | 0.0029 |
| F10 | 38920001 | 39020000 | 0.0032 | 0.0023 | 0.0029 |
| F10 | 38940001 | 39040000 | 0.0029 | 0.0018 | 0.0026 |
| F10 | 38960001 | 39060000 | 0.0025 | 0.0017 | 0.0022 |
| F10 | 38980001 | 39080000 | 0.0023 | 0.0017 | 0.0021 |
| F10 | 39000001 | 39100000 | 0.0023 | 0.0015 | 0.0021 |
| F10 | 39020001 | 39120000 | 0.0022 | 0.0017 | 0.0019 |
| F10 | 39040001 | 39140000 | 0.0023 | 0.0021 | 0.0020 |
| F10 | 39060001 | 39160000 | 0.0031 | 0.0028 | 0.0028 |
| F10 | 39080001 | 39180000 | 0.0038 | 0.0030 | 0.0035 |
| F10 | 39100001 | 39200000 | 0.0041 | 0.0035 | 0.0038 |
| F10 | 39120001 | 39220000 | 0.0048 | 0.0030 | 0.0045 |
| F10 | 39140001 | 39240000 | 0.0042 | 0.0032 | 0.0039 |
| F10 | 39160001 | 39260000 | 0.0046 | 0.0025 | 0.0043 |
| F10 | 39180001 | 39280000 | 0.0037 | 0.0025 | 0.0034 |
| F10 | 39200001 | 39300000 | 0.0037 | 0.0019 | 0.0035 |
| F10 | 39220001 | 39320000 | 0.0028 | 0.0020 | 0.0026 |
| F10 | 39240001 | 39340000 | 0.0028 | 0.0012 | 0.0028 |
| F10 | 39260001 | 39360000 | 0.0016 | 0.0013 | 0.0016 |
| F10 | 39280001 | 39380000 | 0.0015 | 0.0012 | 0.0015 |
| F10 | 39300001 | 39400000 | 0.0014 | 0.0012 | 0.0014 |
| F10 | 39320001 | 39420000 | 0.0014 | 0.0010 | 0.0014 |
| F10 | 39340001 | 39440000 | 0.0012 | 0.0010 | 0.0012 |
| F10 | 39360001 | 39460000 | 0.0013 | 0.0010 | 0.0012 |
| F10 | 39380001 | 39480000 | 0.0012 | 0.0010 | 0.0012 |
| F10 | 39400001 | 39500000 | 0.0012 | 0.0010 | 0.0012 |
| F10 | 39420001 | 39520000 | 0.0012 | 0.0011 | 0.0012 |
| F10 | 39440001 | 39540000 | 0.0013 | 0.0010 | 0.0013 |
| F10 | 39460001 | 39560000 | 0.0011 | 0.0008 | 0.0011 |
| F10 | 39480001 | 39580000 | 0.0009 | 0.0006 | 0.0009 |
| F10 | 39500001 | 39600000 | 0.0007 | 0.0007 | 0.0007 |
| F10 | 39520001 | 39620000 | 0.0008 | 0.0008 | 0.0008 |
| F10 | 39540001 | 39640000 | 0.0009 | 0.0008 | 0.0009 |
| F10 | 39560001 | 39660000 | 0.0009 | 0.0008 | 0.0009 |
| F10 | 39580001 | 39680000 | 0.0010 | 0.0008 | 0.0010 |
| F10 | 39600001 | 39700000 | 0.0009 | 0.0007 | 0.0009 |
| F10 | 39620001 | 39720000 | 0.0007 | 0.0008 | 0.0007 |
| F10 | 39640001 | 39740000 | 0.0009 | 0.0006 | 0.0009 |
| F10 | 39660001 | 39760000 | 0.0008 | 0.0009 | 0.0007 |
| F10 | 39680001 | 39780000 | 0.0011 | 0.0011 | 0.0011 |
| F10 | 39700001 | 39800000 | 0.0013 | 0.0011 | 0.0013 |
| F10 | 39720001 | 39820000 | 0.0014 | 0.0012 | 0.0014 |
| F10 | 39740001 | 39840000 | 0.0013 | 0.0013 | 0.0013 |

| Chr | Start | End | both | A1 | A2 |
|---|---|---|---|---|---|
| F10 | 39760001 | 39860000 | 0.0015 | 0.0009 | 0.0014 |
| F10 | 39780001 | 39880000 | 0.0010 | 0.0009 | 0.0010 |
| F10 | 39800001 | 39900000 | 0.0010 | 0.0010 | 0.0010 |
| F10 | 39820001 | 39920000 | 0.0012 | 0.0009 | 0.0012 |
| F10 | 39840001 | 39940000 | 0.0010 | 0.0010 | 0.0010 |
| F10 | 39860001 | 39960000 | 0.0011 | 0.0012 | 0.0011 |
| F10 | 39880001 | 39980000 | 0.0013 | 0.0012 | 0.0012 |
| F10 | 39900001 | 40000000 | 0.0012 | 0.0011 | 0.0012 |
| F10 | 39920001 | 40020000 | 0.0011 | 0.0011 | 0.0010 |
| F10 | 39940001 | 40040000 | 0.0011 | 0.0011 | 0.0011 |
| F10 | 39960001 | 40060000 | 0.0012 | 0.0012 | 0.0011 |
| F10 | 39980001 | 40080000 | 0.0014 | 0.0012 | 0.0014 |
| F10 | 40000001 | 40100000 | 0.0013 | 0.0011 | 0.0012 |
| F10 | 40020001 | 40120000 | 0.0012 | 0.0012 | 0.0011 |
| F10 | 40040001 | 40140000 | 0.0013 | 0.0012 | 0.0013 |
| F10 | 40060001 | 40160000 | 0.0014 | 0.0012 | 0.0014 |
| F10 | 40080001 | 40180000 | 0.0015 | 0.0012 | 0.0015 |
| F10 | 40100001 | 40200000 | 0.0015 | 0.0018 | 0.0015 |
| F10 | 40120001 | 40220000 | 0.0021 | 0.0019 | 0.0021 |
| F10 | 40140001 | 40240000 | 0.0022 | 0.0020 | 0.0022 |
| F10 | 40160001 | 40260000 | 0.0023 | 0.0018 | 0.0022 |
| F10 | 40180001 | 40280000 | 0.0020 | 0.0017 | 0.0019 |
| F10 | 40200001 | 40300000 | 0.0019 | 0.0012 | 0.0018 |
| F10 | 40220001 | 40320000 | 0.0014 | 0.0015 | 0.0013 |
| F10 | 40240001 | 40340000 | 0.0017 | 0.0015 | 0.0017 |
| F10 | 40260001 | 40360000 | 0.0017 | 0.0017 | 0.0017 |
| F10 | 40280001 | 40380000 | 0.0020 | 0.0023 | 0.0019 |
| F10 | 40300001 | 40400000 | 0.0026 | 0.0024 | 0.0025 |
| F10 | 40320001 | 40420000 | 0.0030 | 0.0020 | 0.0028 |
| F10 | 40340001 | 40440000 | 0.0027 | 0.0025 | 0.0024 |
| F10 | 40360001 | 40460000 | 0.0032 | 0.0022 | 0.0029 |
| F10 | 40380001 | 40480000 | 0.0029 | 0.0020 | 0.0026 |
| F10 | 40400001 | 40500000 | 0.0026 | 0.0029 | 0.0023 |
| F10 | 40420001 | 40520000 | 0.0034 | 0.0034 | 0.0032 |
| F10 | 40440001 | 40540000 | 0.0039 | 0.0035 | 0.0037 |
| F10 | 40460001 | 40560000 | 0.0042 | 0.0036 | 0.0040 |
| F10 | 40480001 | 40580000 | 0.0046 | 0.0035 | 0.0041 |
| F10 | 40500001 | 40600000 | 0.0046 | 0.0029 | 0.0039 |
| F10 | 40520001 | 40620000 | 0.0039 | 0.0028 | 0.0032 |
| F10 | 40540001 | 40640000 | 0.0040 | 0.0025 | 0.0031 |
| F10 | 40560001 | 40660000 | 0.0035 | 0.0030 | 0.0028 |
| F10 | 40580001 | 40680000 | 0.0039 | 0.0035 | 0.0034 |
| F10 | 40600001 | 40700000 | 0.0042 | 0.0033 | 0.0038 |
| F10 | 40620001 | 40720000 | 0.0038 | 0.0030 | 0.0035 |
| F10 | 40640001 | 40740000 | 0.0034 | 0.0027 | 0.0033 |
| F10 | 40660001 | 40760000 | 0.0030 | 0.0024 | 0.0029 |
| F10 | 40680001 | 40780000 | 0.0027 | 0.0018 | 0.0027 |
| F10 | 40700001 | 40800000 | 0.0021 | 0.0021 | 0.0021 |
| F10 | 40720001 | 40820000 | 0.0026 | 0.0016 | 0.0026 |
| F10 | 40740001 | 40840000 | 0.0021 | 0.0014 | 0.0021 |
| F10 | 40760001 | 40860000 | 0.0018 | 0.0021 | 0.0018 |
| F10 | 40780001 | 40880000 | 0.0026 | 0.0023 | 0.0026 |
| F10 | 40800001 | 40900000 | 0.0027 | 0.0018 | 0.0027 |
| F10 | 40820001 | 40920000 | 0.0021 | 0.0024 | 0.0021 |
| F10 | 40840001 | 40940000 | 0.0028 | 0.0022 | 0.0028 |
| F10 | 40860001 | 40960000 | 0.0025 | 0.0013 | 0.0025 |
| F10 | 40880001 | 40980000 | 0.0015 | 0.0014 | 0.0015 |
| F10 | 40900001 | 41000000 | 0.0017 | 0.0017 | 0.0016 |

| Chr | Start | End | both | A1 | A2 |
|---|---|---|---|---|---|
| F10 | 40920001 | 41020000 | 0.0019 | 0.0013 | 0.0019 |
| F10 | 40940001 | 41040000 | 0.0014 | 0.0025 | 0.0014 |
| F10 | 40960001 | 41060000 | 0.0029 | 0.0028 | 0.0028 |
| F10 | 40980001 | 41080000 | 0.0033 | 0.0032 | 0.0032 |
| F10 | 41000001 | 41100000 | 0.0037 | 0.0037 | 0.0036 |
| F10 | 41020001 | 41120000 | 0.0046 | 0.0044 | 0.0043 |
| F10 | 41040001 | 41140000 | 0.0054 | 0.0040 | 0.0051 |
| F10 | 41060001 | 41160000 | 0.0049 | 0.0037 | 0.0045 |
| F10 | 41080001 | 41180000 | 0.0045 | 0.0039 | 0.0041 |
| F10 | 41100001 | 41200000 | 0.0047 | 0.0035 | 0.0042 |
| F10 | 41120001 | 41220000 | 0.0040 | 0.0036 | 0.0037 |
| F10 | 41140001 | 41240000 | 0.0045 | 0.0031 | 0.0039 |
| F10 | 41160001 | 41260000 | 0.0040 | 0.0034 | 0.0034 |
| F10 | 41180001 | 41280000 | 0.0043 | 0.0025 | 0.0038 |
| F10 | 41200001 | 41300000 | 0.0034 | 0.0025 | 0.0029 |
| F10 | 41220001 | 41320000 | 0.0034 | 0.0025 | 0.0030 |
| F10 | 41240001 | 41340000 | 0.0031 | 0.0029 | 0.0029 |
| F10 | 41260001 | 41360000 | 0.0036 | 0.0035 | 0.0032 |
| F10 | 41280001 | 41380000 | 0.0042 | 0.0041 | 0.0038 |
| F10 | 41300001 | 41400000 | 0.0049 | 0.0044 | 0.0043 |
| F10 | 41320001 | 41420000 | 0.0051 | 0.0036 | 0.0045 |
| F10 | 41340001 | 41440000 | 0.0042 | 0.0032 | 0.0036 |
| F10 | 41360001 | 41460000 | 0.0037 | 0.0022 | 0.0032 |
| F10 | 41380001 | 41480000 | 0.0025 | 0.0018 | 0.0022 |
| F10 | 41400001 | 41500000 | 0.0024 | 0.0021 | 0.0021 |
| F10 | 41420001 | 41520000 | 0.0027 | 0.0021 | 0.0025 |
| F10 | 41440001 | 41540000 | 0.0028 | 0.0020 | 0.0026 |
| F10 | 41460001 | 41560000 | 0.0027 | 0.0027 | 0.0025 |
| F10 | 41480001 | 41580000 | 0.0035 | 0.0024 | 0.0033 |
| F10 | 41500001 | 41600000 | 0.0030 | 0.0017 | 0.0029 |
| F10 | 41520001 | 41620000 | 0.0023 | 0.0019 | 0.0021 |
| F10 | 41540001 | 41640000 | 0.0025 | 0.0019 | 0.0022 |
| F10 | 41560001 | 41660000 | 0.0027 | 0.0020 | 0.0024 |
| F10 | 41580001 | 41680000 | 0.0026 | 0.0022 | 0.0023 |
| F10 | 41600001 | 41700000 | 0.0030 | 0.0022 | 0.0025 |
| F10 | 41620001 | 41720000 | 0.0030 | 0.0023 | 0.0025 |
| F10 | 41640001 | 41740000 | 0.0029 | 0.0022 | 0.0026 |
| F10 | 41660001 | 41760000 | 0.0027 | 0.0025 | 0.0026 |
| F10 | 41680001 | 41780000 | 0.0031 | 0.0029 | 0.0029 |
| F10 | 41700001 | 41800000 | 0.0035 | 0.0034 | 0.0033 |
| F10 | 41720001 | 41820000 | 0.0041 | 0.0036 | 0.0040 |
| F10 | 41740001 | 41840000 | 0.0045 | 0.0044 | 0.0044 |
| F10 | 41760001 | 41860000 | 0.0054 | 0.0046 | 0.0052 |
| F10 | 41780001 | 41880000 | 0.0056 | 0.0039 | 0.0053 |
| F10 | 41800001 | 41900000 | 0.0047 | 0.0032 | 0.0044 |
| F10 | 41820001 | 41920000 | 0.0042 | 0.0031 | 0.0039 |
| F10 | 41840001 | 41940000 | 0.0041 | 0.0025 | 0.0037 |
| F10 | 41860001 | 41960000 | 0.0034 | 0.0026 | 0.0030 |
| F10 | 41880001 | 41980000 | 0.0035 | 0.0030 | 0.0032 |
| F10 | 41900001 | 42000000 | 0.0040 | 0.0029 | 0.0037 |
| F10 | 41920001 | 42020000 | 0.0036 | 0.0023 | 0.0034 |
| F10 | 41940001 | 42040000 | 0.0031 | 0.0029 | 0.0028 |
| F10 | 41960001 | 42060000 | 0.0036 | 0.0019 | 0.0033 |
| F10 | 41980001 | 42080000 | 0.0024 | 0.0015 | 0.0022 |
| F10 | 42000001 | 42100000 | 0.0020 | 0.0016 | 0.0018 |
| F10 | 42020001 | 42120000 | 0.0020 | 0.0023 | 0.0018 |
| F10 | 42040001 | 42140000 | 0.0025 | 0.0016 | 0.0023 |
| F10 | 42060001 | 42160000 | 0.0018 | 0.0018 | 0.0017 |

| Chr | Start | End | both | A1 | A2 |
|---|---|---|---|---|---|
| F10 | 42080001 | 42180000 | 0.0019 | 0.0028 | 0.0018 |
| F10 | 42100001 | 42200000 | 0.0032 | 0.0033 | 0.0029 |
| F10 | 42120001 | 42220000 | 0.0037 | 0.0028 | 0.0034 |
| F10 | 42140001 | 42240000 | 0.0035 | 0.0027 | 0.0032 |
| F10 | 42160001 | 42260000 | 0.0036 | 0.0029 | 0.0031 |
| F10 | 42180001 | 42280000 | 0.0039 | 0.0024 | 0.0033 |
| F10 | 42200001 | 42300000 | 0.0036 | 0.0025 | 0.0030 |
| F10 | 42220001 | 42320000 | 0.0037 | 0.0024 | 0.0030 |
| F10 | 42240001 | 42340000 | 0.0036 | 0.0026 | 0.0030 |
| F10 | 42260001 | 42360000 | 0.0038 | 0.0023 | 0.0032 |
| F10 | 42280001 | 42380000 | 0.0034 | 0.0022 | 0.0028 |
| F10 | 42300001 | 42400000 | 0.0031 | 0.0021 | 0.0025 |
| F10 | 42320001 | 42420000 | 0.0029 | 0.0021 | 0.0024 |
| F10 | 42340001 | 42440000 | 0.0027 | 0.0026 | 0.0023 |
| F10 | 42360001 | 42460000 | 0.0031 | 0.0031 | 0.0028 |
| F10 | 42380001 | 42480000 | 0.0037 | 0.0029 | 0.0034 |
| F10 | 42400001 | 42500000 | 0.0034 | 0.0026 | 0.0032 |
| F10 | 42420001 | 42520000 | 0.0029 | 0.0023 | 0.0028 |
| F10 | 42440001 | 42540000 | 0.0026 | 0.0018 | 0.0025 |
| F10 | 42460001 | 42560000 | 0.0019 | 0.0012 | 0.0019 |
| F10 | 42480001 | 42580000 | 0.0013 | 0.0012 | 0.0013 |
| F10 | 42500001 | 42600000 | 0.0014 | 0.0011 | 0.0014 |
| F10 | 42520001 | 42620000 | 0.0012 | 0.0012 | 0.0012 |
| F10 | 42540001 | 42640000 | 0.0014 | 0.0014 | 0.0014 |
| F10 | 42560001 | 42660000 | 0.0017 | 0.0018 | 0.0017 |
| F10 | 42580001 | 42680000 | 0.0022 | 0.0018 | 0.0021 |
| F10 | 42600001 | 42700000 | 0.0022 | 0.0019 | 0.0022 |
| F10 | 42620001 | 42720000 | 0.0025 | 0.0019 | 0.0024 |
| F10 | 42640001 | 42740000 | 0.0027 | 0.0017 | 0.0023 |
| F10 | 42660001 | 42760000 | 0.0025 | 0.0014 | 0.0021 |
| F10 | 42680001 | 42780000 | 0.0021 | 0.0014 | 0.0016 |
| F10 | 42700001 | 42800000 | 0.0020 | 0.0017 | 0.0016 |
| F10 | 42720001 | 42820000 | 0.0023 | 0.0017 | 0.0018 |
| F10 | 42740001 | 42840000 | 0.0020 | 0.0016 | 0.0017 |
| F10 | 42760001 | 42860000 | 0.0018 | 0.0022 | 0.0016 |
| F10 | 42780001 | 42880000 | 0.0025 | 0.0020 | 0.0023 |
| F10 | 42800001 | 42900000 | 0.0023 | 0.0022 | 0.0020 |
| F10 | 42820001 | 42920000 | 0.0026 | 0.0027 | 0.0024 |
| F10 | 42840001 | 42940000 | 0.0031 | 0.0028 | 0.0029 |
| F10 | 42860001 | 42960000 | 0.0032 | 0.0021 | 0.0030 |
| F10 | 42880001 | 42980000 | 0.0024 | 0.0027 | 0.0022 |
| F10 | 42900001 | 43000000 | 0.0030 | 0.0030 | 0.0029 |
| F10 | 42920001 | 43020000 | 0.0035 | 0.0027 | 0.0033 |
| F10 | 42940001 | 43040000 | 0.0031 | 0.0027 | 0.0031 |
| F10 | 42960001 | 43060000 | 0.0032 | 0.0028 | 0.0031 |
| F10 | 42980001 | 43080000 | 0.0034 | 0.0026 | 0.0033 |
| F10 | 43000001 | 43100000 | 0.0031 | 0.0018 | 0.0031 |
| F10 | 43020001 | 43120000 | 0.0022 | 0.0019 | 0.0020 |
| F10 | 43040001 | 43140000 | 0.0024 | 0.0028 | 0.0020 |
| F10 | 43060001 | 43160000 | 0.0034 | 0.0028 | 0.0028 |
| F10 | 43080001 | 43180000 | 0.0034 | 0.0022 | 0.0027 |
| F10 | 43100001 | 43200000 | 0.0028 | 0.0021 | 0.0021 |
| F10 | 43120001 | 43220000 | 0.0027 | 0.0019 | 0.0021 |
| F10 | 43140001 | 43240000 | 0.0024 | 0.0014 | 0.0020 |
| F10 | 43160001 | 43260000 | 0.0019 | 0.0023 | 0.0017 |
| F10 | 43180001 | 43280000 | 0.0032 | 0.0034 | 0.0027 |
| F10 | 43200001 | 43300000 | 0.0046 | 0.0038 | 0.0039 |
| F10 | 43220001 | 43320000 | 0.0050 | 0.0045 | 0.0043 |

| Chr | Start | End | both | A1 | A2 |
|---|---|---|---|---|---|
| F10 | 43240001 | 43340000 | 0.0059 | 0.0046 | 0.0050 |
| F10 | 43260001 | 43360000 | 0.0061 | 0.0039 | 0.0051 |
| F10 | 43280001 | 43380000 | 0.0051 | 0.0027 | 0.0043 |
| F10 | 43300001 | 43400000 | 0.0035 | 0.0030 | 0.0029 |
| F10 | 43320001 | 43420000 | 0.0038 | 0.0024 | 0.0032 |
| F10 | 43340001 | 43440000 | 0.0029 | 0.0019 | 0.0026 |
| F10 | 43360001 | 43460000 | 0.0023 | 0.0023 | 0.0022 |
| F10 | 43380001 | 43480000 | 0.0026 | 0.0028 | 0.0023 |
| F10 | 43400001 | 43500000 | 0.0031 | 0.0021 | 0.0028 |
| F10 | 43420001 | 43520000 | 0.0026 | 0.0022 | 0.0021 |
| F10 | 43440001 | 43540000 | 0.0026 | 0.0026 | 0.0021 |
| F10 | 43460001 | 43560000 | 0.0032 | 0.0019 | 0.0026 |
| F10 | 43480001 | 43580000 | 0.0025 | 0.0015 | 0.0022 |
| F10 | 43500001 | 43600000 | 0.0020 | 0.0013 | 0.0017 |
| F10 | 43520001 | 43620000 | 0.0017 | 0.0010 | 0.0016 |
| F10 | 43540001 | 43640000 | 0.0012 | 0.0004 | 0.0012 |
| F10 | 43560001 | 43660000 | 0.0005 | 0.0003 | 0.0005 |
| F10 | 43580001 | 43680000 | 0.0004 | 0.0003 | 0.0004 |
| F10 | 43600001 | 43700000 | 0.0004 | 0.0009 | 0.0004 |
| F10 | 43620001 | 43720000 | 0.0013 | 0.0037 | 0.0011 |
| F10 | 43640001 | 43740000 | 0.0044 | 0.0045 | 0.0040 |
| F10 | 43660001 | 43760000 | 0.0053 | 0.0061 | 0.0047 |
| F10 | 43680001 | 43780000 | 0.0072 | 0.0071 | 0.0066 |
| F10 | 43700001 | 43800000 | 0.0088 | 0.0072 | 0.0080 |
| F10 | 43720001 | 43820000 | 0.0089 | 0.0077 | 0.0083 |
| F10 | 43740001 | 43840000 | 0.0093 | 0.0073 | 0.0086 |
| F10 | 43760001 | 43860000 | 0.0092 | 0.0058 | 0.0082 |
| F10 | 43780001 | 43880000 | 0.0074 | 0.0054 | 0.0065 |
| F10 | 43800001 | 43900000 | 0.0065 | 0.0053 | 0.0057 |
| F10 | 43820001 | 43920000 | 0.0066 | 0.0021 | 0.0056 |
| F10 | 43840001 | 43940000 | 0.0034 | 0.0018 | 0.0026 |
| F10 | 43860001 | 43960000 | 0.0029 | 0.0020 | 0.0022 |
| F10 | 43880001 | 43980000 | 0.0034 | 0.0023 | 0.0024 |
| F10 | 43900001 | 44000000 | 0.0038 | 0.0022 | 0.0027 |
| F10 | 43920001 | 44020000 | 0.0033 | 0.0021 | 0.0024 |
| F10 | 43940001 | 44040000 | 0.0030 | 0.0020 | 0.0022 |
| F10 | 43960001 | 44060000 | 0.0028 | 0.0026 | 0.0020 |
| F10 | 43980001 | 44080000 | 0.0032 | 0.0017 | 0.0026 |
| F10 | 44000001 | 44100000 | 0.0022 | 0.0019 | 0.0017 |
| F10 | 44020001 | 44120000 | 0.0023 | 0.0022 | 0.0017 |
| F10 | 44040001 | 44140000 | 0.0026 | 0.0024 | 0.0020 |
| F10 | 44060001 | 44160000 | 0.0028 | 0.0015 | 0.0023 |
| F10 | 44080001 | 44180000 | 0.0018 | 0.0017 | 0.0014 |
| F10 | 44100001 | 44200000 | 0.0019 | 0.0012 | 0.0016 |
| F10 | 44120001 | 44220000 | 0.0014 | 0.0011 | 0.0012 |
| F10 | 44140001 | 44240000 | 0.0014 | 0.0011 | 0.0013 |
| F10 | 44160001 | 44260000 | 0.0013 | 0.0017 | 0.0012 |
| F10 | 44180001 | 44280000 | 0.0020 | 0.0017 | 0.0019 |
| F10 | 44200001 | 44300000 | 0.0019 | 0.0019 | 0.0018 |
| F10 | 44220001 | 44320000 | 0.0023 | 0.0021 | 0.0021 |
| F10 | 44240001 | 44340000 | 0.0026 | 0.0020 | 0.0023 |
| F10 | 44260001 | 44360000 | 0.0026 | 0.0017 | 0.0023 |
| F10 | 44280001 | 44380000 | 0.0021 | 0.0023 | 0.0019 |
| F10 | 44300001 | 44400000 | 0.0028 | 0.0024 | 0.0025 |
| F10 | 44320001 | 44420000 | 0.0030 | 0.0023 | 0.0028 |
| F10 | 44340001 | 44440000 | 0.0028 | 0.0021 | 0.0025 |
| F10 | 44360001 | 44460000 | 0.0026 | 0.0019 | 0.0023 |
| F10 | 44380001 | 44480000 | 0.0024 | 0.0012 | 0.0021 |

| Chr | Start | End | both | A1 | A2 |
|---|---|---|---|---|---|
| F10 | 44400001 | 44500000 | 0.0017 | 0.0008 | 0.0013 |
| F10 | 44420001 | 44520000 | 0.0013 | 0.0005 | 0.0008 |
| F10 | 44440001 | 44540000 | 0.0010 | 0.0017 | 0.0005 |
| F10 | 44460001 | 44560000 | 0.0022 | 0.0020 | 0.0017 |
| F10 | 44480001 | 44580000 | 0.0026 | 0.0027 | 0.0021 |
| F10 | 44500001 | 44600000 | 0.0033 | 0.0029 | 0.0028 |
| F10 | 44520001 | 44620000 | 0.0037 | 0.0031 | 0.0032 |
| F10 | 44540001 | 44640000 | 0.0040 | 0.0024 | 0.0035 |
| F10 | 44560001 | 44660000 | 0.0032 | 0.0020 | 0.0028 |
| F10 | 44580001 | 44680000 | 0.0027 | 0.0013 | 0.0022 |
| F10 | 44600001 | 44700000 | 0.0018 | 0.0014 | 0.0015 |
| F10 | 44620001 | 44720000 | 0.0018 | 0.0021 | 0.0015 |
| F10 | 44640001 | 44740000 | 0.0026 | 0.0019 | 0.0021 |
| F10 | 44660001 | 44760000 | 0.0024 | 0.0020 | 0.0019 |
| F10 | 44680001 | 44780000 | 0.0026 | 0.0020 | 0.0021 |
| F10 | 44700001 | 44800000 | 0.0027 | 0.0018 | 0.0022 |
| F10 | 44720001 | 44820000 | 0.0023 | 0.0014 | 0.0020 |
| F10 | 44740001 | 44840000 | 0.0017 | 0.0015 | 0.0016 |
| F10 | 44760001 | 44860000 | 0.0021 | 0.0018 | 0.0019 |
| F10 | 44780001 | 44880000 | 0.0023 | 0.0021 | 0.0021 |
| F10 | 44800001 | 44900000 | 0.0026 | 0.0024 | 0.0023 |
| F10 | 44820001 | 44920000 | 0.0030 | 0.0033 | 0.0026 |
| F10 | 44840001 | 44940000 | 0.0039 | 0.0038 | 0.0035 |
| F10 | 44860001 | 44960000 | 0.0041 | 0.0037 | 0.0038 |
| F10 | 44880001 | 44980000 | 0.0042 | 0.0035 | 0.0037 |
| F10 | 44900001 | 45000000 | 0.0039 | 0.0036 | 0.0037 |
| F10 | 44920001 | 45020000 | 0.0041 | 0.0025 | 0.0038 |
| F10 | 44940001 | 45040000 | 0.0030 | 0.0017 | 0.0027 |
| F10 | 44960001 | 45060000 | 0.0023 | 0.0024 | 0.0018 |
| F10 | 44980001 | 45080000 | 0.0032 | 0.0029 | 0.0027 |
| F10 | 45000001 | 45100000 | 0.0038 | 0.0029 | 0.0032 |
| F10 | 45020001 | 45120000 | 0.0038 | 0.0029 | 0.0031 |
| F10 | 45040001 | 45140000 | 0.0038 | 0.0032 | 0.0032 |
| F10 | 45060001 | 45160000 | 0.0040 | 0.0024 | 0.0036 |
| F10 | 45080001 | 45180000 | 0.0030 | 0.0019 | 0.0027 |
| F10 | 45100001 | 45200000 | 0.0024 | 0.0015 | 0.0022 |
| F10 | 45120001 | 45220000 | 0.0017 | 0.0013 | 0.0017 |
| F10 | 45140001 | 45240000 | 0.0013 | 0.0018 | 0.0013 |
| F10 | 45160001 | 45260000 | 0.0019 | 0.0018 | 0.0018 |
| F10 | 45180001 | 45280000 | 0.0019 | 0.0018 | 0.0019 |
| F10 | 45200001 | 45300000 | 0.0020 | 0.0017 | 0.0020 |
| F10 | 45220001 | 45320000 | 0.0019 | 0.0017 | 0.0018 |
| F10 | 45240001 | 45340000 | 0.0019 | 0.0018 | 0.0018 |
| F10 | 45260001 | 45360000 | 0.0022 | 0.0018 | 0.0020 |
| F10 | 45280001 | 45380000 | 0.0021 | 0.0023 | 0.0020 |
| F10 | 45300001 | 45400000 | 0.0026 | 0.0027 | 0.0025 |
| F10 | 45320001 | 45420000 | 0.0031 | 0.0030 | 0.0030 |
| F10 | 45340001 | 45440000 | 0.0035 | 0.0027 | 0.0033 |
| F10 | 45360001 | 45460000 | 0.0032 | 0.0030 | 0.0031 |
| F10 | 45380001 | 45480000 | 0.0035 | 0.0028 | 0.0033 |
| F10 | 45400001 | 45500000 | 0.0032 | 0.0024 | 0.0030 |
| F10 | 45420001 | 45520000 | 0.0029 | 0.0023 | 0.0025 |
| F10 | 45440001 | 45540000 | 0.0028 | 0.0018 | 0.0024 |
| F10 | 45460001 | 45560000 | 0.0023 | 0.0020 | 0.0019 |
| F10 | 45480001 | 45580000 | 0.0029 | 0.0022 | 0.0024 |
| F10 | 45500001 | 45600000 | 0.0032 | 0.0024 | 0.0027 |
| F10 | 45520001 | 45620000 | 0.0033 | 0.0026 | 0.0029 |
| F10 | 45540001 | 45640000 | 0.0038 | 0.0028 | 0.0035 |

| Chr | Start | End | both | A1 | A2 |
|---|---|---|---|---|---|
| F10 | 45560001 | 45660000 | 0.0041 | 0.0026 | 0.0036 |
| F10 | 45580001 | 45680000 | 0.0037 | 0.0026 | 0.0030 |
| F10 | 45600001 | 45700000 | 0.0041 | 0.0029 | 0.0032 |
| F10 | 45620001 | 45720000 | 0.0045 | 0.0027 | 0.0036 |
| F10 | 45640001 | 45740000 | 0.0039 | 0.0025 | 0.0030 |
| F10 | 45660001 | 45760000 | 0.0038 | 0.0022 | 0.0030 |
| F10 | 45680001 | 45780000 | 0.0035 | 0.0023 | 0.0027 |
| F10 | 45700001 | 45800000 | 0.0033 | 0.0019 | 0.0026 |
| F10 | 45720001 | 45820000 | 0.0027 | 0.0019 | 0.0022 |
| F10 | 45740001 | 45840000 | 0.0027 | 0.0018 | 0.0023 |
| F10 | 45760001 | 45860000 | 0.0023 | 0.0017 | 0.0020 |
| F10 | 45780001 | 45880000 | 0.0021 | 0.0010 | 0.0020 |
| F10 | 45800001 | 45900000 | 0.0014 | 0.0019 | 0.0013 |
| F10 | 45820001 | 45920000 | 0.0025 | 0.0030 | 0.0022 |
| F10 | 45840001 | 45940000 | 0.0039 | 0.0034 | 0.0035 |
| F10 | 45860001 | 45960000 | 0.0042 | 0.0039 | 0.0038 |
| F10 | 45880001 | 45980000 | 0.0049 | 0.0040 | 0.0046 |
| F10 | 45900001 | 46000000 | 0.0050 | 0.0036 | 0.0047 |
| F10 | 45920001 | 46020000 | 0.0043 | 0.0025 | 0.0040 |
| F10 | 45940001 | 46040000 | 0.0033 | 0.0029 | 0.0027 |
| F10 | 45960001 | 46060000 | 0.0038 | 0.0028 | 0.0032 |
| F10 | 45980001 | 46080000 | 0.0037 | 0.0033 | 0.0029 |
| F10 | 46000001 | 46100000 | 0.0041 | 0.0027 | 0.0035 |
| F10 | 46020001 | 46120000 | 0.0036 | 0.0025 | 0.0030 |
| F10 | 46040001 | 46140000 | 0.0031 | 0.0020 | 0.0029 |
| F10 | 46060001 | 46160000 | 0.0024 | 0.0017 | 0.0022 |
| F10 | 46080001 | 46180000 | 0.0021 | 0.0012 | 0.0019 |
| F10 | 46100001 | 46200000 | 0.0016 | 0.0016 | 0.0012 |
| F10 | 46120001 | 46220000 | 0.0022 | 0.0019 | 0.0018 |
| F10 | 46140001 | 46240000 | 0.0025 | 0.0017 | 0.0021 |
| F10 | 46160001 | 46260000 | 0.0023 | 0.0016 | 0.0019 |
| F10 | 46180001 | 46280000 | 0.0022 | 0.0014 | 0.0018 |
| F10 | 46200001 | 46300000 | 0.0018 | 0.0017 | 0.0016 |
| F10 | 46220001 | 46320000 | 0.0019 | 0.0017 | 0.0016 |
| F10 | 46240001 | 46340000 | 0.0020 | 0.0018 | 0.0016 |
| F10 | 46260001 | 46360000 | 0.0021 | 0.0016 | 0.0015 |
| F10 | 46280001 | 46380000 | 0.0019 | 0.0018 | 0.0014 |
| F10 | 46300001 | 46400000 | 0.0021 | 0.0013 | 0.0016 |
| F10 | 46320001 | 46420000 | 0.0017 | 0.0008 | 0.0014 |
| F10 | 46340001 | 46440000 | 0.0011 | 0.0008 | 0.0010 |
| F10 | 46360001 | 46460000 | 0.0010 | 0.0009 | 0.0009 |
| F10 | 46380001 | 46480000 | 0.0012 | 0.0011 | 0.0011 |
| F10 | 46400001 | 46500000 | 0.0015 | 0.0008 | 0.0013 |
| F10 | 46420001 | 46520000 | 0.0013 | 0.0010 | 0.0009 |
| F10 | 46440001 | 46540000 | 0.0015 | 0.0010 | 0.0011 |
| F10 | 46460001 | 46560000 | 0.0015 | 0.0010 | 0.0011 |
| F10 | 46480001 | 46580000 | 0.0015 | 0.0008 | 0.0012 |
| F10 | 46500001 | 46600000 | 0.0011 | 0.0007 | 0.0008 |
| F10 | 46520001 | 46620000 | 0.0010 | 0.0007 | 0.0008 |
| F10 | 46540001 | 46640000 | 0.0010 | 0.0008 | 0.0009 |
| F10 | 46560001 | 46660000 | 0.0011 | 0.0008 | 0.0010 |
| F10 | 46580001 | 46680000 | 0.0010 | 0.0007 | 0.0010 |
| F10 | 46600001 | 46700000 | 0.0009 | 0.0007 | 0.0010 |
| F10 | 46620001 | 46720000 | 0.0009 | 0.0006 | 0.0009 |
| F10 | 46640001 | 46740000 | 0.0008 | 0.0007 | 0.0008 |
| F10 | 46660001 | 46760000 | 0.0008 | 0.0009 | 0.0008 |
| F10 | 46680001 | 46780000 | 0.0010 | 0.0014 | 0.0010 |
| F10 | 46700001 | 46800000 | 0.0017 | 0.0017 | 0.0017 |

| Chr | Start | End | both | A1 | A2 |
|---|---|---|---|---|---|
| F10 | 46720001 | 46820000 | 0.0022 | 0.0019 | 0.0021 |
| F10 | 46740001 | 46840000 | 0.0024 | 0.0020 | 0.0023 |
| F10 | 46760001 | 46860000 | 0.0026 | 0.0018 | 0.0025 |
| F10 | 46780001 | 46880000 | 0.0023 | 0.0014 | 0.0022 |
| F10 | 46800001 | 46900000 | 0.0018 | 0.0011 | 0.0017 |
| F10 | 46820001 | 46920000 | 0.0013 | 0.0010 | 0.0012 |
| F10 | 46840001 | 46940000 | 0.0012 | 0.0009 | 0.0012 |
| F10 | 46860001 | 46960000 | 0.0011 | 0.0008 | 0.0011 |
| F10 | 46880001 | 46980000 | 0.0009 | 0.0006 | 0.0009 |
| F10 | 46900001 | 47000000 | 0.0007 | 0.0006 | 0.0007 |
| F10 | 46920001 | 47020000 | 0.0007 | 0.0004 | 0.0007 |
| F10 | 46940001 | 47040000 | 0.0005 | 0.0002 | 0.0005 |
| F10 | 46960001 | 47060000 | 0.0002 | 0.0001 | 0.0002 |
| F10 | 46980001 | 47080000 | 0.0002 | 0.0001 | 0.0002 |
| F10 | 47000001 | 47100000 | 0.0002 | 0.0002 | 0.0002 |
| F10 | 47020001 | 47120000 | 0.0002 | 0.0002 | 0.0002 |
| F10 | 47040001 | 47140000 | 0.0003 | 0.0002 | 0.0003 |
| F10 | 47060001 | 47160000 | 0.0002 | 0.0003 | 0.0002 |
| F10 | 47080001 | 47180000 | 0.0003 | 0.0004 | 0.0003 |
| F10 | 47100001 | 47200000 | 0.0005 | 0.0006 | 0.0005 |
| F10 | 47120001 | 47220000 | 0.0007 | 0.0015 | 0.0007 |
| F10 | 47140001 | 47240000 | 0.0017 | 0.0017 | 0.0016 |
| F10 | 47160001 | 47260000 | 0.0020 | 0.0016 | 0.0018 |
| F10 | 47180001 | 47280000 | 0.0020 | 0.0018 | 0.0017 |
| F10 | 47200001 | 47300000 | 0.0024 | 0.0024 | 0.0019 |
| F10 | 47220001 | 47320000 | 0.0032 | 0.0018 | 0.0026 |
| F10 | 47240001 | 47340000 | 0.0026 | 0.0019 | 0.0020 |
| F10 | 47260001 | 47360000 | 0.0028 | 0.0020 | 0.0022 |
| F10 | 47280001 | 47380000 | 0.0026 | 0.0017 | 0.0022 |
| F10 | 47300001 | 47400000 | 0.0020 | 0.0010 | 0.0019 |
| F10 | 47320001 | 47420000 | 0.0011 | 0.0010 | 0.0009 |
| F10 | 47340001 | 47440000 | 0.0011 | 0.0007 | 0.0010 |
| F10 | 47360001 | 47460000 | 0.0007 | 0.0010 | 0.0007 |
| F10 | 47380001 | 47480000 | 0.0011 | 0.0011 | 0.0010 |
| F10 | 47400001 | 47500000 | 0.0012 | 0.0013 | 0.0012 |
| F10 | 47420001 | 47520000 | 0.0015 | 0.0010 | 0.0014 |
| F10 | 47440001 | 47540000 | 0.0012 | 0.0014 | 0.0012 |
| F10 | 47460001 | 47560000 | 0.0017 | 0.0016 | 0.0015 |
| F10 | 47480001 | 47580000 | 0.0019 | 0.0023 | 0.0016 |
| F10 | 47500001 | 47600000 | 0.0026 | 0.0027 | 0.0023 |
| F10 | 47520001 | 47620000 | 0.0032 | 0.0036 | 0.0027 |
| F10 | 47540001 | 47640000 | 0.0041 | 0.0040 | 0.0035 |
| F10 | 47560001 | 47660000 | 0.0044 | 0.0050 | 0.0039 |
| F10 | 47580001 | 47680000 | 0.0054 | 0.0054 | 0.0049 |
| F10 | 47600001 | 47700000 | 0.0057 | 0.0054 | 0.0050 |
| F10 | 47620001 | 47720000 | 0.0057 | 0.0049 | 0.0051 |
| F10 | 47640001 | 47740000 | 0.0056 | 0.0050 | 0.0048 |
| F10 | 47660001 | 47760000 | 0.0059 | 0.0042 | 0.0050 |
| F10 | 47680001 | 47780000 | 0.0052 | 0.0042 | 0.0043 |
| F10 | 47700001 | 47800000 | 0.0054 | 0.0037 | 0.0047 |
| F10 | 47720001 | 47820000 | 0.0048 | 0.0033 | 0.0042 |
| F10 | 47740001 | 47840000 | 0.0043 | 0.0028 | 0.0037 |
| F10 | 47760001 | 47860000 | 0.0035 | 0.0024 | 0.0031 |
| F10 | 47780001 | 47880000 | 0.0032 | 0.0024 | 0.0029 |
| F10 | 47800001 | 47900000 | 0.0031 | 0.0033 | 0.0028 |
| F10 | 47820001 | 47920000 | 0.0042 | 0.0038 | 0.0038 |
| F10 | 47840001 | 47940000 | 0.0046 | 0.0039 | 0.0043 |
| F10 | 47860001 | 47960000 | 0.0047 | 0.0037 | 0.0044 |

| Chr | Start | End | both | A1 | A2 |
|---|---|---|---|---|---|
| F10 | 47880001 | 47980000 | 0.0046 | 0.0029 | 0.0043 |
| F10 | 47900001 | 48000000 | 0.0036 | 0.0022 | 0.0033 |
| F10 | 47920001 | 48020000 | 0.0028 | 0.0029 | 0.0026 |
| F10 | 47940001 | 48040000 | 0.0034 | 0.0032 | 0.0033 |
| F10 | 47960001 | 48060000 | 0.0039 | 0.0035 | 0.0038 |
| F10 | 47980001 | 48080000 | 0.0041 | 0.0033 | 0.0041 |
| F10 | 48000001 | 48100000 | 0.0039 | 0.0032 | 0.0039 |
| F10 | 48020001 | 48120000 | 0.0040 | 0.0024 | 0.0037 |
| F10 | 48040001 | 48140000 | 0.0032 | 0.0020 | 0.0029 |
| F10 | 48060001 | 48160000 | 0.0027 | 0.0022 | 0.0024 |
| F10 | 48080001 | 48180000 | 0.0029 | 0.0021 | 0.0024 |
| F10 | 48100001 | 48200000 | 0.0027 | 0.0020 | 0.0023 |
| F10 | 48120001 | 48220000 | 0.0024 | 0.0020 | 0.0022 |
| F10 | 48140001 | 48240000 | 0.0024 | 0.0023 | 0.0022 |
| F10 | 48160001 | 48260000 | 0.0027 | 0.0025 | 0.0025 |
| F10 | 48180001 | 48280000 | 0.0030 | 0.0030 | 0.0026 |
| F10 | 48200001 | 48300000 | 0.0036 | 0.0035 | 0.0031 |
| F10 | 48220001 | 48320000 | 0.0042 | 0.0043 | 0.0037 |
| F10 | 48240001 | 48340000 | 0.0048 | 0.0041 | 0.0043 |
| F10 | 48260001 | 48360000 | 0.0046 | 0.0037 | 0.0042 |
| F10 | 48280001 | 48380000 | 0.0042 | 0.0041 | 0.0038 |
| F10 | 48300001 | 48400000 | 0.0046 | 0.0036 | 0.0042 |
| F10 | 48320001 | 48420000 | 0.0041 | 0.0027 | 0.0038 |
| F10 | 48340001 | 48440000 | 0.0033 | 0.0037 | 0.0029 |
| F10 | 48360001 | 48460000 | 0.0044 | 0.0039 | 0.0040 |
| F10 | 48380001 | 48480000 | 0.0048 | 0.0037 | 0.0043 |
| F10 | 48400001 | 48500000 | 0.0049 | 0.0044 | 0.0045 |
| F10 | 48420001 | 48520000 | 0.0058 | 0.0049 | 0.0052 |
| F10 | 48440001 | 48540000 | 0.0064 | 0.0040 | 0.0058 |
| F10 | 48460001 | 48560000 | 0.0054 | 0.0033 | 0.0047 |
| F10 | 48480001 | 48580000 | 0.0045 | 0.0035 | 0.0040 |
| F10 | 48500001 | 48600000 | 0.0044 | 0.0032 | 0.0038 |
| F10 | 48520001 | 48620000 | 0.0039 | 0.0029 | 0.0034 |
| F10 | 48540001 | 48640000 | 0.0034 | 0.0027 | 0.0030 |
| F10 | 48560001 | 48660000 | 0.0032 | 0.0031 | 0.0029 |
| F10 | 48580001 | 48680000 | 0.0038 | 0.0027 | 0.0033 |
| F10 | 48600001 | 48700000 | 0.0035 | 0.0025 | 0.0029 |
| F10 | 48620001 | 48720000 | 0.0033 | 0.0022 | 0.0027 |
| F10 | 48640001 | 48740000 | 0.0029 | 0.0024 | 0.0024 |
| F10 | 48660001 | 48760000 | 0.0033 | 0.0021 | 0.0029 |
| F10 | 48680001 | 48780000 | 0.0031 | 0.0025 | 0.0025 |
| F10 | 48700001 | 48800000 | 0.0033 | 0.0032 | 0.0029 |
| F10 | 48720001 | 48820000 | 0.0041 | 0.0038 | 0.0037 |
| F10 | 48740001 | 48840000 | 0.0047 | 0.0037 | 0.0044 |
| F10 | 48760001 | 48860000 | 0.0044 | 0.0049 | 0.0040 |
| F10 | 48780001 | 48880000 | 0.0051 | 0.0047 | 0.0046 |
| F10 | 48800001 | 48900000 | 0.0047 | 0.0044 | 0.0041 |
| F10 | 48820001 | 48920000 | 0.0043 | 0.0039 | 0.0036 |
| F10 | 48840001 | 48940000 | 0.0037 | 0.0045 | 0.0030 |
| F10 | 48860001 | 48960000 | 0.0043 | 0.0032 | 0.0037 |
| F10 | 48880001 | 48980000 | 0.0033 | 0.0030 | 0.0029 |
| F10 | 48900001 | 49000000 | 0.0034 | 0.0024 | 0.0030 |
| F10 | 48920001 | 49020000 | 0.0029 | 0.0038 | 0.0026 |
| F10 | 48940001 | 49040000 | 0.0043 | 0.0039 | 0.0038 |
| F10 | 48960001 | 49060000 | 0.0046 | 0.0040 | 0.0039 |
| F10 | 48980001 | 49080000 | 0.0048 | 0.0046 | 0.0040 |
| F10 | 49000001 | 49100000 | 0.0053 | 0.0045 | 0.0044 |
| F10 | 49020001 | 49120000 | 0.0054 | 0.0033 | 0.0046 |

| Chr | Start | End | both | A1 | A2 |
|---|---|---|---|---|---|
| F10 | 49040001 | 49140000 | 0.0042 | 0.0021 | 0.0036 |
| F10 | 49060001 | 49160000 | 0.0028 | 0.0026 | 0.0023 |
| F10 | 49080001 | 49180000 | 0.0032 | 0.0025 | 0.0028 |
| F10 | 49100001 | 49200000 | 0.0033 | 0.0035 | 0.0027 |
| F10 | 49120001 | 49220000 | 0.0043 | 0.0040 | 0.0038 |
| F10 | 49140001 | 49240000 | 0.0050 | 0.0045 | 0.0045 |
| F10 | 49160001 | 49260000 | 0.0058 | 0.0047 | 0.0051 |
| F10 | 49180001 | 49280000 | 0.0061 | 0.0044 | 0.0053 |
| F10 | 49200001 | 49300000 | 0.0056 | 0.0035 | 0.0050 |
| F10 | 49220001 | 49320000 | 0.0046 | 0.0033 | 0.0038 |
| F10 | 49240001 | 49340000 | 0.0044 | 0.0034 | 0.0036 |
| F10 | 49260001 | 49360000 | 0.0041 | 0.0026 | 0.0037 |
| F10 | 49280001 | 49380000 | 0.0031 | 0.0023 | 0.0029 |
| F10 | 49300001 | 49400000 | 0.0027 | 0.0019 | 0.0025 |
| F10 | 49320001 | 49420000 | 0.0023 | 0.0015 | 0.0023 |
| F10 | 49340001 | 49440000 | 0.0019 | 0.0013 | 0.0020 |
| F10 | 49360001 | 49460000 | 0.0015 | 0.0015 | 0.0014 |
| F10 | 49380001 | 49480000 | 0.0017 | 0.0020 | 0.0017 |
| F10 | 49400001 | 49500000 | 0.0022 | 0.0021 | 0.0022 |
| F10 | 49420001 | 49520000 | 0.0023 | 0.0023 | 0.0023 |
| F10 | 49440001 | 49540000 | 0.0025 | 0.0021 | 0.0024 |
| F10 | 49460001 | 49560000 | 0.0023 | 0.0021 | 0.0023 |
| F10 | 49480001 | 49580000 | 0.0025 | 0.0016 | 0.0025 |
| F10 | 49500001 | 49600000 | 0.0020 | 0.0015 | 0.0021 |
| F10 | 49520001 | 49620000 | 0.0019 | 0.0025 | 0.0020 |
| F10 | 49540001 | 49640000 | 0.0029 | 0.0025 | 0.0029 |
| F10 | 49560001 | 49660000 | 0.0029 | 0.0021 | 0.0029 |
| F10 | 49580001 | 49680000 | 0.0024 | 0.0023 | 0.0024 |
| F10 | 49600001 | 49700000 | 0.0026 | 0.0021 | 0.0025 |
| F10 | 49620001 | 49720000 | 0.0026 | 0.0014 | 0.0023 |
| F10 | 49640001 | 49740000 | 0.0017 | 0.0015 | 0.0014 |
| F10 | 49660001 | 49760000 | 0.0018 | 0.0019 | 0.0014 |
| F10 | 49680001 | 49780000 | 0.0022 | 0.0015 | 0.0018 |
| F10 | 49700001 | 49800000 | 0.0018 | 0.0015 | 0.0015 |
| F10 | 49720001 | 49820000 | 0.0016 | 0.0008 | 0.0015 |
| F10 | 49740001 | 49840000 | 0.0011 | 0.0009 | 0.0010 |
| F10 | 49760001 | 49860000 | 0.0011 | 0.0008 | 0.0011 |
| F10 | 49780001 | 49880000 | 0.0010 | 0.0012 | 0.0010 |
| F10 | 49800001 | 49900000 | 0.0014 | 0.0013 | 0.0014 |
| F10 | 49820001 | 49920000 | 0.0015 | 0.0015 | 0.0015 |
| F10 | 49840001 | 49940000 | 0.0018 | 0.0019 | 0.0017 |
| F10 | 49860001 | 49960000 | 0.0021 | 0.0028 | 0.0020 |
| F10 | 49880001 | 49980000 | 0.0033 | 0.0027 | 0.0030 |
| F10 | 49900001 | 50000000 | 0.0033 | 0.0028 | 0.0030 |
| F10 | 49920001 | 50020000 | 0.0034 | 0.0028 | 0.0030 |
| F10 | 49940001 | 50040000 | 0.0033 | 0.0022 | 0.0030 |
| F10 | 49960001 | 50060000 | 0.0028 | 0.0020 | 0.0024 |
| F10 | 49980001 | 50080000 | 0.0026 | 0.0021 | 0.0022 |
| F10 | 50000001 | 50100000 | 0.0029 | 0.0023 | 0.0023 |
| F10 | 50020001 | 50120000 | 0.0031 | 0.0020 | 0.0024 |
| F10 | 50040001 | 50140000 | 0.0027 | 0.0027 | 0.0022 |
| F10 | 50060001 | 50160000 | 0.0034 | 0.0019 | 0.0030 |
| F10 | 50080001 | 50180000 | 0.0024 | 0.0019 | 0.0020 |
| F10 | 50100001 | 50200000 | 0.0024 | 0.0018 | 0.0022 |
| F10 | 50120001 | 50220000 | 0.0025 | 0.0020 | 0.0022 |
| F10 | 50140001 | 50240000 | 0.0027 | 0.0014 | 0.0024 |
| F10 | 50160001 | 50260000 | 0.0021 | 0.0024 | 0.0017 |
| F10 | 50180001 | 50280000 | 0.0030 | 0.0027 | 0.0026 |

| Chr | Start | End | both | A1 | A2 |
|---|---|---|---|---|---|
| F10 | 50200001 | 50300000 | 0.0031 | 0.0025 | 0.0028 |
| F10 | 50220001 | 50320000 | 0.0029 | 0.0024 | 0.0027 |
| F10 | 50240001 | 50340000 | 0.0028 | 0.0031 | 0.0025 |
| F10 | 50260001 | 50360000 | 0.0037 | 0.0028 | 0.0031 |
| F10 | 50280001 | 50380000 | 0.0035 | 0.0026 | 0.0029 |
| F10 | 50300001 | 50400000 | 0.0032 | 0.0028 | 0.0025 |
| F10 | 50320001 | 50420000 | 0.0034 | 0.0028 | 0.0027 |
| F10 | 50340001 | 50440000 | 0.0035 | 0.0024 | 0.0029 |
| F10 | 50360001 | 50460000 | 0.0032 | 0.0016 | 0.0028 |
| F10 | 50380001 | 50480000 | 0.0024 | 0.0019 | 0.0021 |
| F10 | 50400001 | 50500000 | 0.0027 | 0.0030 | 0.0026 |
| F10 | 50420001 | 50520000 | 0.0040 | 0.0035 | 0.0039 |
| F10 | 50440001 | 50540000 | 0.0045 | 0.0034 | 0.0043 |
| F10 | 50460001 | 50560000 | 0.0042 | 0.0037 | 0.0041 |
| F10 | 50480001 | 50580000 | 0.0045 | 0.0032 | 0.0044 |
| F10 | 50500001 | 50600000 | 0.0039 | 0.0022 | 0.0038 |
| F10 | 50520001 | 50620000 | 0.0028 | 0.0019 | 0.0027 |
| F10 | 50540001 | 50640000 | 0.0022 | 0.0017 | 0.0020 |
| F10 | 50560001 | 50660000 | 0.0018 | 0.0013 | 0.0018 |
| F10 | 50580001 | 50680000 | 0.0015 | 0.0017 | 0.0014 |
| F10 | 50600001 | 50700000 | 0.0019 | 0.0014 | 0.0017 |
| F10 | 50620001 | 50720000 | 0.0016 | 0.0015 | 0.0015 |
| F10 | 50640001 | 50740000 | 0.0020 | 0.0017 | 0.0016 |
| F10 | 50660001 | 50760000 | 0.0023 | 0.0022 | 0.0019 |
| F10 | 50680001 | 50780000 | 0.0029 | 0.0018 | 0.0025 |
| F10 | 50700001 | 50800000 | 0.0026 | 0.0021 | 0.0023 |
| F10 | 50720001 | 50820000 | 0.0028 | 0.0026 | 0.0025 |
| F10 | 50740001 | 50840000 | 0.0032 | 0.0027 | 0.0032 |
| F10 | 50760001 | 50860000 | 0.0033 | 0.0028 | 0.0033 |
| F10 | 50780001 | 50880000 | 0.0033 | 0.0032 | 0.0032 |
| F10 | 50800001 | 50900000 | 0.0039 | 0.0031 | 0.0038 |
| F10 | 50820001 | 50920000 | 0.0040 | 0.0024 | 0.0038 |
| F10 | 50840001 | 50940000 | 0.0031 | 0.0023 | 0.0029 |
| F10 | 50860001 | 50960000 | 0.0030 | 0.0018 | 0.0027 |
| F10 | 50880001 | 50980000 | 0.0023 | 0.0014 | 0.0021 |
| F10 | 50900001 | 51000000 | 0.0017 | 0.0014 | 0.0015 |
| F10 | 50920001 | 51020000 | 0.0015 | 0.0015 | 0.0014 |
| F10 | 50940001 | 51040000 | 0.0015 | 0.0017 | 0.0014 |
| F10 | 50960001 | 51060000 | 0.0017 | 0.0037 | 0.0016 |
| F10 | 50980001 | 51080000 | 0.0038 | 0.0037 | 0.0036 |
| F10 | 51000001 | 51100000 | 0.0039 | 0.0035 | 0.0035 |
| F10 | 51020001 | 51120000 | 0.0040 | 0.0034 | 0.0035 |
| F10 | 51040001 | 51140000 | 0.0042 | 0.0034 | 0.0033 |
| F10 | 51060001 | 51160000 | 0.0041 | 0.0016 | 0.0031 |
| F10 | 51080001 | 51180000 | 0.0025 | 0.0017 | 0.0015 |
| F10 | 51100001 | 51200000 | 0.0026 | 0.0017 | 0.0016 |
| F10 | 51120001 | 51220000 | 0.0024 | 0.0018 | 0.0016 |
| F10 | 51140001 | 51240000 | 0.0021 | 0.0018 | 0.0016 |
| F10 | 51160001 | 51260000 | 0.0023 | 0.0020 | 0.0018 |
| F10 | 51180001 | 51280000 | 0.0026 | 0.0017 | 0.0020 |
| F10 | 51200001 | 51300000 | 0.0022 | 0.0016 | 0.0017 |
| F10 | 51220001 | 51320000 | 0.0021 | 0.0014 | 0.0015 |
| F10 | 51240001 | 51340000 | 0.0019 | 0.0012 | 0.0015 |
| F10 | 51260001 | 51360000 | 0.0018 | 0.0011 | 0.0013 |
| F10 | 51280001 | 51380000 | 0.0017 | 0.0012 | 0.0012 |
| F10 | 51300001 | 51400000 | 0.0018 | 0.0019 | 0.0013 |
| F10 | 51320001 | 51420000 | 0.0026 | 0.0025 | 0.0020 |
| F10 | 51340001 | 51440000 | 0.0036 | 0.0026 | 0.0027 |

| Chr | Start | End | both | A1 | A2 |
|---|---|---|---|---|---|
| F10 | 51360001 | 51460000 | 0.0035 | 0.0029 | 0.0028 |
| F10 | 51380001 | 51480000 | 0.0036 | 0.0033 | 0.0031 |
| F10 | 51400001 | 51500000 | 0.0042 | 0.0026 | 0.0037 |
| F10 | 51420001 | 51520000 | 0.0034 | 0.0021 | 0.0030 |
| F10 | 51440001 | 51540000 | 0.0028 | 0.0021 | 0.0023 |
| F10 | 51460001 | 51560000 | 0.0032 | 0.0017 | 0.0024 |
| F10 | 51480001 | 51580000 | 0.0028 | 0.0017 | 0.0020 |
| F10 | 51500001 | 51600000 | 0.0026 | 0.0015 | 0.0020 |
| F10 | 51520001 | 51620000 | 0.0025 | 0.0016 | 0.0018 |
| F10 | 51540001 | 51640000 | 0.0025 | 0.0015 | 0.0019 |
| F10 | 51560001 | 51660000 | 0.0021 | 0.0012 | 0.0017 |
| F10 | 51580001 | 51680000 | 0.0016 | 0.0009 | 0.0013 |
| F10 | 51600001 | 51700000 | 0.0012 | 0.0009 | 0.0009 |
| F10 | 51620001 | 51720000 | 0.0013 | 0.0007 | 0.0009 |
| F10 | 51640001 | 51740000 | 0.0009 | 0.0012 | 0.0008 |
| F10 | 51660001 | 51760000 | 0.0014 | 0.0017 | 0.0013 |
| F10 | 51680001 | 51780000 | 0.0019 | 0.0020 | 0.0018 |
| F10 | 51700001 | 51800000 | 0.0023 | 0.0023 | 0.0023 |
| F10 | 51720001 | 51820000 | 0.0027 | 0.0029 | 0.0026 |
| F10 | 51740001 | 51840000 | 0.0035 | 0.0030 | 0.0034 |
| F10 | 51760001 | 51860000 | 0.0038 | 0.0033 | 0.0035 |
| F10 | 51780001 | 51880000 | 0.0040 | 0.0035 | 0.0037 |
| F10 | 51800001 | 51900000 | 0.0041 | 0.0033 | 0.0037 |
| F10 | 51820001 | 51920000 | 0.0039 | 0.0029 | 0.0035 |
| F10 | 51840001 | 51940000 | 0.0033 | 0.0019 | 0.0029 |
| F10 | 51860001 | 51960000 | 0.0022 | 0.0016 | 0.0019 |
| F10 | 51880001 | 51980000 | 0.0019 | 0.0013 | 0.0017 |
| F10 | 51900001 | 52000000 | 0.0015 | 0.0012 | 0.0014 |
| F10 | 51920001 | 52020000 | 0.0015 | 0.0012 | 0.0012 |
| F10 | 51940001 | 52040000 | 0.0016 | 0.0013 | 0.0013 |
| F10 | 51960001 | 52060000 | 0.0017 | 0.0016 | 0.0014 |
| F10 | 51980001 | 52080000 | 0.0020 | 0.0013 | 0.0017 |
| F10 | 52000001 | 52100000 | 0.0018 | 0.0013 | 0.0014 |
| F10 | 52020001 | 52120000 | 0.0018 | 0.0012 | 0.0015 |
| F10 | 52040001 | 52140000 | 0.0016 | 0.0019 | 0.0013 |
| F10 | 52060001 | 52160000 | 0.0023 | 0.0016 | 0.0021 |
| F10 | 52080001 | 52180000 | 0.0020 | 0.0013 | 0.0018 |
| F10 | 52100001 | 52200000 | 0.0017 | 0.0015 | 0.0016 |
| F10 | 52120001 | 52220000 | 0.0019 | 0.0019 | 0.0018 |
| F10 | 52140001 | 52240000 | 0.0025 | 0.0021 | 0.0025 |
| F10 | 52160001 | 52260000 | 0.0027 | 0.0020 | 0.0027 |
| F10 | 52180001 | 52280000 | 0.0025 | 0.0028 | 0.0023 |
| F10 | 52200001 | 52300000 | 0.0031 | 0.0034 | 0.0027 |
| F10 | 52220001 | 52320000 | 0.0037 | 0.0037 | 0.0034 |
| F10 | 52240001 | 52340000 | 0.0038 | 0.0028 | 0.0035 |
| F10 | 52260001 | 52360000 | 0.0027 | 0.0029 | 0.0024 |
| F10 | 52280001 | 52380000 | 0.0028 | 0.0029 | 0.0025 |
| F10 | 52300001 | 52400000 | 0.0032 | 0.0020 | 0.0030 |
| F10 | 52320001 | 52420000 | 0.0022 | 0.0014 | 0.0020 |
| F10 | 52340001 | 52440000 | 0.0015 | 0.0018 | 0.0013 |
| F10 | 52360001 | 52460000 | 0.0020 | 0.0020 | 0.0017 |
| F10 | 52380001 | 52480000 | 0.0025 | 0.0016 | 0.0020 |
| F10 | 52400001 | 52500000 | 0.0020 | 0.0020 | 0.0015 |
| F10 | 52420001 | 52520000 | 0.0025 | 0.0024 | 0.0018 |
| F10 | 52440001 | 52540000 | 0.0033 | 0.0026 | 0.0025 |
| F10 | 52460001 | 52560000 | 0.0034 | 0.0026 | 0.0027 |
| F10 | 52480001 | 52580000 | 0.0034 | 0.0024 | 0.0028 |
| F10 | 52500001 | 52600000 | 0.0031 | 0.0024 | 0.0026 |

| Chr | Start | End | both | A1 | A2 |
|---|---|---|---|---|---|
| F10 | 52520001 | 52620000 | 0.0030 | 0.0021 | 0.0025 |
| F10 | 52540001 | 52640000 | 0.0026 | 0.0016 | 0.0022 |
| F10 | 52560001 | 52660000 | 0.0020 | 0.0015 | 0.0016 |
| F10 | 52580001 | 52680000 | 0.0018 | 0.0015 | 0.0015 |
| F10 | 52600001 | 52700000 | 0.0019 | 0.0013 | 0.0015 |
| F10 | 52620001 | 52720000 | 0.0020 | 0.0012 | 0.0014 |
| F10 | 52640001 | 52740000 | 0.0020 | 0.0013 | 0.0014 |
| F10 | 52660001 | 52760000 | 0.0021 | 0.0016 | 0.0015 |
| F10 | 52680001 | 52780000 | 0.0025 | 0.0024 | 0.0019 |
| F10 | 52700001 | 52800000 | 0.0034 | 0.0033 | 0.0027 |
| F10 | 52720001 | 52820000 | 0.0043 | 0.0030 | 0.0034 |
| F10 | 52740001 | 52840000 | 0.0039 | 0.0033 | 0.0031 |
| F10 | 52760001 | 52860000 | 0.0043 | 0.0036 | 0.0033 |
| F10 | 52780001 | 52880000 | 0.0045 | 0.0032 | 0.0035 |
| F10 | 52800001 | 52900000 | 0.0041 | 0.0032 | 0.0033 |
| F10 | 52820001 | 52920000 | 0.0042 | 0.0041 | 0.0034 |
| F10 | 52840001 | 52940000 | 0.0049 | 0.0045 | 0.0041 |
| F10 | 52860001 | 52960000 | 0.0052 | 0.0039 | 0.0044 |
| F10 | 52880001 | 52980000 | 0.0047 | 0.0036 | 0.0039 |
| F10 | 52900001 | 53000000 | 0.0044 | 0.0029 | 0.0035 |
| F10 | 52920001 | 53020000 | 0.0035 | 0.0026 | 0.0030 |
| F10 | 52940001 | 53040000 | 0.0035 | 0.0020 | 0.0029 |
| F10 | 52960001 | 53060000 | 0.0027 | 0.0029 | 0.0023 |
| F10 | 52980001 | 53080000 | 0.0035 | 0.0027 | 0.0031 |
| F10 | 53000001 | 53100000 | 0.0031 | 0.0028 | 0.0028 |
| F10 | 53020001 | 53120000 | 0.0032 | 0.0021 | 0.0028 |
| F10 | 53040001 | 53140000 | 0.0023 | 0.0026 | 0.0022 |
| F10 | 53060001 | 53160000 | 0.0028 | 0.0017 | 0.0028 |
| F10 | 53080001 | 53180000 | 0.0022 | 0.0019 | 0.0019 |
| F10 | 53100001 | 53200000 | 0.0024 | 0.0018 | 0.0019 |
| F10 | 53120001 | 53220000 | 0.0023 | 0.0020 | 0.0014 |
| F10 | 53140001 | 53240000 | 0.0026 | 0.0023 | 0.0015 |
| F10 | 53160001 | 53260000 | 0.0032 | 0.0032 | 0.0019 |
| F10 | 53180001 | 53280000 | 0.0042 | 0.0034 | 0.0030 |
| F10 | 53200001 | 53300000 | 0.0043 | 0.0035 | 0.0034 |
| F10 | 53220001 | 53320000 | 0.0045 | 0.0049 | 0.0038 |
| F10 | 53240001 | 53340000 | 0.0057 | 0.0039 | 0.0051 |
| F10 | 53260001 | 53360000 | 0.0045 | 0.0028 | 0.0040 |
| F10 | 53280001 | 53380000 | 0.0030 | 0.0024 | 0.0027 |
| F10 | 53300001 | 53400000 | 0.0026 | 0.0025 | 0.0022 |
| F10 | 53320001 | 53420000 | 0.0027 | 0.0014 | 0.0024 |
| F10 | 53340001 | 53440000 | 0.0018 | 0.0015 | 0.0016 |
| F10 | 53360001 | 53460000 | 0.0019 | 0.0021 | 0.0017 |
| F10 | 53380001 | 53480000 | 0.0026 | 0.0028 | 0.0025 |
| F10 | 53400001 | 53500000 | 0.0034 | 0.0023 | 0.0034 |
| F10 | 53420001 | 53520000 | 0.0028 | 0.0032 | 0.0028 |
| F10 | 53440001 | 53540000 | 0.0037 | 0.0033 | 0.0036 |
| F10 | 53460001 | 53560000 | 0.0038 | 0.0027 | 0.0037 |
| F10 | 53480001 | 53580000 | 0.0033 | 0.0027 | 0.0031 |
| F10 | 53500001 | 53600000 | 0.0033 | 0.0034 | 0.0030 |
| F10 | 53520001 | 53620000 | 0.0041 | 0.0023 | 0.0039 |
| F10 | 53540001 | 53640000 | 0.0029 | 0.0022 | 0.0028 |
| F10 | 53560001 | 53660000 | 0.0028 | 0.0023 | 0.0026 |
| F10 | 53580001 | 53680000 | 0.0029 | 0.0020 | 0.0027 |
| F10 | 53600001 | 53700000 | 0.0026 | 0.0021 | 0.0022 |
| F10 | 53620001 | 53720000 | 0.0026 | 0.0029 | 0.0021 |
| F10 | 53640001 | 53740000 | 0.0037 | 0.0034 | 0.0030 |
| F10 | 53660001 | 53760000 | 0.0042 | 0.0036 | 0.0036 |

| Chr | Start | End | both | A1 | A2 |
|---|---|---|---|---|---|
| F10 | 53680001 | 53780000 | 0.0042 | 0.0036 | 0.0037 |
| F10 | 53700001 | 53800000 | 0.0040 | 0.0030 | 0.0037 |
| F10 | 53720001 | 53820000 | 0.0033 | 0.0023 | 0.0031 |
| F10 | 53740001 | 53840000 | 0.0026 | 0.0022 | 0.0024 |
| F10 | 53760001 | 53860000 | 0.0025 | 0.0021 | 0.0023 |
| F10 | 53780001 | 53880000 | 0.0024 | 0.0022 | 0.0021 |
| F10 | 53800001 | 53900000 | 0.0025 | 0.0025 | 0.0023 |
| F10 | 53820001 | 53920000 | 0.0029 | 0.0022 | 0.0027 |
| F10 | 53840001 | 53940000 | 0.0024 | 0.0016 | 0.0022 |
| F10 | 53860001 | 53960000 | 0.0018 | 0.0013 | 0.0016 |
| F10 | 53880001 | 53980000 | 0.0016 | 0.0010 | 0.0015 |
| F10 | 53900001 | 54000000 | 0.0011 | 0.0005 | 0.0011 |
| F10 | 53920001 | 54020000 | 0.0005 | 0.0004 | 0.0005 |
| F10 | 53940001 | 54040000 | 0.0004 | 0.0004 | 0.0004 |
| F10 | 53960001 | 54060000 | 0.0005 | 0.0005 | 0.0005 |
| F10 | 53980001 | 54080000 | 0.0006 | 0.0003 | 0.0006 |
| F10 | 54000001 | 54100000 | 0.0005 | 0.0003 | 0.0005 |
| F10 | 54020001 | 54120000 | 0.0005 | 0.0004 | 0.0005 |
| F10 | 54040001 | 54140000 | 0.0006 | 0.0004 | 0.0006 |
| F10 | 54060001 | 54160000 | 0.0005 | 0.0002 | 0.0005 |
| F10 | 54080001 | 54180000 | 0.0003 | 0.0003 | 0.0003 |
| F10 | 54100001 | 54200000 | 0.0004 | 0.0003 | 0.0004 |
| F10 | 54120001 | 54220000 | 0.0004 | 0.0002 | 0.0004 |
| F10 | 54140001 | 54240000 | 0.0002 | 0.0002 | 0.0002 |
| F10 | 54160001 | 54260000 | 0.0002 | 0.0002 | 0.0002 |
| F10 | 54180001 | 54280000 | 0.0003 | 0.0002 | 0.0003 |
| F10 | 54200001 | 54300000 | 0.0003 | 0.0006 | 0.0003 |
| F10 | 54220001 | 54320000 | 0.0006 | 0.0005 | 0.0005 |
| F10 | 54240001 | 54340000 | 0.0006 | 0.0008 | 0.0005 |
| F10 | 54260001 | 54360000 | 0.0008 | 0.0008 | 0.0008 |
| F10 | 54280001 | 54380000 | 0.0009 | 0.0010 | 0.0009 |
| F10 | 54300001 | 54400000 | 0.0011 | 0.0007 | 0.0010 |
| F10 | 54320001 | 54420000 | 0.0009 | 0.0008 | 0.0009 |
| F10 | 54340001 | 54440000 | 0.0010 | 0.0007 | 0.0010 |
| F10 | 54360001 | 54460000 | 0.0008 | 0.0006 | 0.0008 |
| F10 | 54380001 | 54480000 | 0.0008 | 0.0007 | 0.0008 |
| F10 | 54400001 | 54500000 | 0.0010 | 0.0006 | 0.0010 |
| F10 | 54420001 | 54520000 | 0.0008 | 0.0005 | 0.0009 |
| F10 | 54440001 | 54540000 | 0.0007 | 0.0006 | 0.0007 |
| F10 | 54460001 | 54560000 | 0.0009 | 0.0007 | 0.0010 |
| F10 | 54480001 | 54580000 | 0.0010 | 0.0005 | 0.0010 |
| F10 | 54500001 | 54600000 | 0.0007 | 0.0005 | 0.0008 |
| F10 | 54520001 | 54620000 | 0.0007 | 0.0006 | 0.0007 |
| F10 | 54540001 | 54640000 | 0.0010 | 0.0005 | 0.0010 |
| F10 | 54560001 | 54660000 | 0.0007 | 0.0007 | 0.0007 |
| F10 | 54580001 | 54680000 | 0.0009 | 0.0007 | 0.0009 |
| F10 | 54600001 | 54700000 | 0.0008 | 0.0007 | 0.0008 |
| F10 | 54620001 | 54720000 | 0.0008 | 0.0004 | 0.0008 |
| F10 | 54640001 | 54740000 | 0.0005 | 0.0008 | 0.0005 |
| F10 | 54660001 | 54760000 | 0.0009 | 0.0009 | 0.0008 |
| F10 | 54680001 | 54780000 | 0.0011 | 0.0013 | 0.0011 |
| F10 | 54700001 | 54800000 | 0.0017 | 0.0016 | 0.0018 |
| F10 | 54720001 | 54820000 | 0.0020 | 0.0017 | 0.0020 |
| F10 | 54740001 | 54840000 | 0.0021 | 0.0014 | 0.0021 |
| F10 | 54760001 | 54860000 | 0.0018 | 0.0016 | 0.0019 |
| F10 | 54780001 | 54880000 | 0.0020 | 0.0011 | 0.0021 |
| F10 | 54800001 | 54900000 | 0.0014 | 0.0013 | 0.0015 |
| F10 | 54820001 | 54920000 | 0.0015 | 0.0011 | 0.0015 |

| Chr | Start | End | both | A1 | A2 |
|---|---|---|---|---|---|
| F10 | 54840001 | 54940000 | 0.0014 | 0.0020 | 0.0014 |
| F10 | 54860001 | 54960000 | 0.0025 | 0.0020 | 0.0024 |
| F10 | 54880001 | 54980000 | 0.0023 | 0.0022 | 0.0022 |
| F10 | 54900001 | 55000000 | 0.0026 | 0.0023 | 0.0025 |
| F10 | 54920001 | 55020000 | 0.0028 | 0.0024 | 0.0027 |
| F10 | 54940001 | 55040000 | 0.0028 | 0.0014 | 0.0028 |
| F10 | 54960001 | 55060000 | 0.0017 | 0.0010 | 0.0016 |
| F10 | 54980001 | 55080000 | 0.0013 | 0.0009 | 0.0013 |
| F10 | 55000001 | 55100000 | 0.0010 | 0.0006 | 0.0010 |
| F10 | 55020001 | 55120000 | 0.0008 | 0.0011 | 0.0008 |
| F10 | 55040001 | 55140000 | 0.0013 | 0.0016 | 0.0012 |
| F10 | 55060001 | 55160000 | 0.0020 | 0.0020 | 0.0019 |
| F10 | 55080001 | 55180000 | 0.0024 | 0.0022 | 0.0023 |
| F10 | 55100001 | 55200000 | 0.0028 | 0.0022 | 0.0026 |
| F10 | 55120001 | 55220000 | 0.0027 | 0.0021 | 0.0026 |
| F10 | 55140001 | 55240000 | 0.0026 | 0.0016 | 0.0025 |
| F10 | 55160001 | 55260000 | 0.0020 | 0.0013 | 0.0020 |
| F10 | 55180001 | 55280000 | 0.0016 | 0.0012 | 0.0016 |
| F10 | 55200001 | 55300000 | 0.0014 | 0.0013 | 0.0014 |
| F10 | 55220001 | 55320000 | 0.0014 | 0.0016 | 0.0014 |
| F10 | 55240001 | 55340000 | 0.0016 | 0.0016 | 0.0015 |
| F10 | 55260001 | 55360000 | 0.0018 | 0.0018 | 0.0016 |
| F10 | 55280001 | 55380000 | 0.0019 | 0.0019 | 0.0017 |
| F10 | 55300001 | 55400000 | 0.0020 | 0.0020 | 0.0018 |
| F10 | 55320001 | 55420000 | 0.0021 | 0.0017 | 0.0019 |
| F10 | 55340001 | 55440000 | 0.0021 | 0.0017 | 0.0020 |
| F10 | 55360001 | 55460000 | 0.0021 | 0.0016 | 0.0020 |
| F10 | 55380001 | 55480000 | 0.0021 | 0.0015 | 0.0021 |
| F10 | 55400001 | 55500000 | 0.0019 | 0.0014 | 0.0018 |
| F10 | 55420001 | 55520000 | 0.0019 | 0.0017 | 0.0018 |
| F10 | 55440001 | 55540000 | 0.0018 | 0.0019 | 0.0018 |
| F10 | 55460001 | 55560000 | 0.0021 | 0.0024 | 0.0021 |
| F10 | 55480001 | 55580000 | 0.0026 | 0.0031 | 0.0024 |
| F10 | 55500001 | 55600000 | 0.0036 | 0.0032 | 0.0030 |
| F10 | 55520001 | 55620000 | 0.0041 | 0.0024 | 0.0034 |
| F10 | 55540001 | 55640000 | 0.0034 | 0.0023 | 0.0027 |
| F10 | 55560001 | 55660000 | 0.0033 | 0.0018 | 0.0026 |
| F10 | 55580001 | 55680000 | 0.0027 | 0.0015 | 0.0022 |
| F10 | 55600001 | 55700000 | 0.0020 | 0.0012 | 0.0018 |
| F10 | 55620001 | 55720000 | 0.0015 | 0.0015 | 0.0013 |
| F10 | 55640001 | 55740000 | 0.0018 | 0.0014 | 0.0017 |
| F10 | 55660001 | 55760000 | 0.0017 | 0.0015 | 0.0015 |
| F10 | 55680001 | 55780000 | 0.0019 | 0.0015 | 0.0017 |
| F10 | 55700001 | 55800000 | 0.0020 | 0.0017 | 0.0018 |
| F10 | 55720001 | 55820000 | 0.0023 | 0.0024 | 0.0022 |
| F10 | 55740001 | 55840000 | 0.0028 | 0.0021 | 0.0026 |
| F10 | 55760001 | 55860000 | 0.0025 | 0.0019 | 0.0023 |
| F10 | 55780001 | 55880000 | 0.0022 | 0.0024 | 0.0021 |
| F10 | 55800001 | 55900000 | 0.0027 | 0.0020 | 0.0025 |
| F10 | 55820001 | 55920000 | 0.0022 | 0.0014 | 0.0020 |
| F10 | 55840001 | 55940000 | 0.0018 | 0.0015 | 0.0016 |
| F10 | 55860001 | 55960000 | 0.0020 | 0.0015 | 0.0017 |
| F10 | 55880001 | 55980000 | 0.0019 | 0.0011 | 0.0017 |
| F10 | 55900001 | 56000000 | 0.0013 | 0.0012 | 0.0012 |
| F10 | 55920001 | 56020000 | 0.0014 | 0.0023 | 0.0012 |
| F10 | 55940001 | 56040000 | 0.0026 | 0.0027 | 0.0025 |
| F10 | 55960001 | 56060000 | 0.0030 | 0.0029 | 0.0029 |
| F10 | 55980001 | 56080000 | 0.0032 | 0.0030 | 0.0032 |

| Chr | Start | End | both | A1 | A2 |
|---|---|---|---|---|---|
| F10 | 56000001 | 56100000 | 0.0034 | 0.0029 | 0.0033 |
| F10 | 56020001 | 56120000 | 0.0033 | 0.0015 | 0.0032 |
| F10 | 56040001 | 56140000 | 0.0017 | 0.0014 | 0.0017 |
| F10 | 56060001 | 56160000 | 0.0016 | 0.0015 | 0.0016 |
| F10 | 56080001 | 56180000 | 0.0018 | 0.0014 | 0.0018 |
| F10 | 56100001 | 56200000 | 0.0018 | 0.0019 | 0.0018 |
| F10 | 56120001 | 56220000 | 0.0025 | 0.0025 | 0.0023 |
| F10 | 56140001 | 56240000 | 0.0031 | 0.0026 | 0.0029 |
| F10 | 56160001 | 56260000 | 0.0031 | 0.0023 | 0.0029 |
| F10 | 56180001 | 56280000 | 0.0028 | 0.0019 | 0.0026 |
| F10 | 56200001 | 56300000 | 0.0022 | 0.0018 | 0.0020 |
| F10 | 56220001 | 56320000 | 0.0020 | 0.0017 | 0.0019 |
| F10 | 56240001 | 56340000 | 0.0022 | 0.0015 | 0.0019 |
| F10 | 56260001 | 56360000 | 0.0023 | 0.0014 | 0.0016 |
| F10 | 56280001 | 56380000 | 0.0021 | 0.0018 | 0.0015 |
| F10 | 56300001 | 56400000 | 0.0025 | 0.0018 | 0.0019 |
| F10 | 56320001 | 56420000 | 0.0026 | 0.0014 | 0.0020 |
| F10 | 56340001 | 56440000 | 0.0020 | 0.0012 | 0.0017 |
| F10 | 56360001 | 56460000 | 0.0015 | 0.0016 | 0.0015 |
| F10 | 56380001 | 56480000 | 0.0019 | 0.0016 | 0.0019 |
| F10 | 56400001 | 56500000 | 0.0020 | 0.0012 | 0.0019 |
| F10 | 56420001 | 56520000 | 0.0014 | 0.0012 | 0.0014 |
| F10 | 56440001 | 56540000 | 0.0014 | 0.0012 | 0.0014 |
| F10 | 56460001 | 56560000 | 0.0015 | 0.0020 | 0.0014 |
| F10 | 56480001 | 56580000 | 0.0026 | 0.0020 | 0.0021 |
| F10 | 56500001 | 56600000 | 0.0028 | 0.0028 | 0.0020 |
| F10 | 56520001 | 56620000 | 0.0039 | 0.0035 | 0.0026 |
| F10 | 56540001 | 56640000 | 0.0047 | 0.0043 | 0.0033 |
| F10 | 56560001 | 56660000 | 0.0056 | 0.0035 | 0.0042 |
| F10 | 56580001 | 56680000 | 0.0046 | 0.0031 | 0.0036 |
| F10 | 56600001 | 56700000 | 0.0039 | 0.0023 | 0.0032 |
| F10 | 56620001 | 56720000 | 0.0028 | 0.0020 | 0.0026 |
| F10 | 56640001 | 56740000 | 0.0025 | 0.0010 | 0.0024 |
| F10 | 56660001 | 56760000 | 0.0013 | 0.0007 | 0.0013 |
| F10 | 56680001 | 56780000 | 0.0008 | 0.0010 | 0.0008 |
| F10 | 56700001 | 56800000 | 0.0012 | 0.0010 | 0.0012 |
| F10 | 56720001 | 56820000 | 0.0012 | 0.0005 | 0.0012 |
| F10 | 56740001 | 56840000 | 0.0006 | 0.0007 | 0.0006 |
| F10 | 56760001 | 56860000 | 0.0009 | 0.0008 | 0.0009 |
| F10 | 56780001 | 56880000 | 0.0010 | 0.0008 | 0.0010 |
| F10 | 56800001 | 56900000 | 0.0010 | 0.0009 | 0.0010 |
| F10 | 56820001 | 56920000 | 0.0010 | 0.0007 | 0.0010 |
| F10 | 56840001 | 56940000 | 0.0009 | 0.0009 | 0.0009 |
| F10 | 56860001 | 56960000 | 0.0011 | 0.0009 | 0.0011 |
| F10 | 56880001 | 56980000 | 0.0010 | 0.0010 | 0.0009 |
| F10 | 56900001 | 57000000 | 0.0012 | 0.0013 | 0.0011 |
| F10 | 56920001 | 57020000 | 0.0016 | 0.0015 | 0.0015 |
| F10 | 56940001 | 57040000 | 0.0018 | 0.0011 | 0.0016 |
| F10 | 56960001 | 57060000 | 0.0014 | 0.0015 | 0.0013 |
| F10 | 56980001 | 57080000 | 0.0020 | 0.0015 | 0.0019 |
| F10 | 57000001 | 57100000 | 0.0020 | 0.0012 | 0.0017 |
| F10 | 57020001 | 57120000 | 0.0017 | 0.0015 | 0.0014 |
| F10 | 57040001 | 57140000 | 0.0021 | 0.0015 | 0.0016 |
| F10 | 57060001 | 57160000 | 0.0020 | 0.0014 | 0.0016 |
| F10 | 57080001 | 57180000 | 0.0018 | 0.0012 | 0.0014 |
| F10 | 57100001 | 57200000 | 0.0014 | 0.0014 | 0.0013 |
| F10 | 57120001 | 57220000 | 0.0015 | 0.0011 | 0.0014 |
| F10 | 57140001 | 57240000 | 0.0011 | 0.0011 | 0.0011 |

| Chr | Start | End | both | A1 | A2 |
|---|---|---|---|---|---|
| F10 | 57160001 | 57260000 | 0.0012 | 0.0007 | 0.0011 |
| F10 | 57180001 | 57280000 | 0.0008 | 0.0006 | 0.0008 |
| F10 | 57200001 | 57300000 | 0.0008 | 0.0006 | 0.0007 |
| F10 | 57220001 | 57320000 | 0.0008 | 0.0008 | 0.0007 |
| F10 | 57240001 | 57340000 | 0.0010 | 0.0013 | 0.0010 |
| F10 | 57260001 | 57360000 | 0.0017 | 0.0016 | 0.0016 |
| F10 | 57280001 | 57380000 | 0.0020 | 0.0015 | 0.0019 |
| F10 | 57300001 | 57400000 | 0.0018 | 0.0018 | 0.0017 |
| F10 | 57320001 | 57420000 | 0.0023 | 0.0016 | 0.0022 |
| F10 | 57340001 | 57440000 | 0.0021 | 0.0011 | 0.0020 |
| F10 | 57360001 | 57460000 | 0.0016 | 0.0008 | 0.0016 |
| F10 | 57380001 | 57480000 | 0.0012 | 0.0010 | 0.0012 |
| F10 | 57400001 | 57500000 | 0.0015 | 0.0005 | 0.0015 |
| F10 | 57420001 | 57520000 | 0.0008 | 0.0010 | 0.0008 |
| F10 | 57440001 | 57540000 | 0.0014 | 0.0019 | 0.0013 |
| F10 | 57460001 | 57560000 | 0.0023 | 0.0020 | 0.0020 |
| F10 | 57480001 | 57580000 | 0.0023 | 0.0021 | 0.0020 |
| F10 | 57500001 | 57600000 | 0.0024 | 0.0026 | 0.0021 |
| F10 | 57520001 | 57620000 | 0.0030 | 0.0022 | 0.0027 |
| F10 | 57540001 | 57640000 | 0.0026 | 0.0019 | 0.0023 |
| F10 | 57560001 | 57660000 | 0.0021 | 0.0020 | 0.0020 |
| F10 | 57580001 | 57680000 | 0.0023 | 0.0018 | 0.0021 |
| F10 | 57600001 | 57700000 | 0.0021 | 0.0016 | 0.0019 |
| F10 | 57620001 | 57720000 | 0.0019 | 0.0015 | 0.0017 |
| F10 | 57640001 | 57740000 | 0.0018 | 0.0012 | 0.0016 |
| F10 | 57660001 | 57760000 | 0.0017 | 0.0012 | 0.0014 |
| F10 | 57680001 | 57780000 | 0.0017 | 0.0012 | 0.0014 |
| F10 | 57700001 | 57800000 | 0.0016 | 0.0010 | 0.0014 |
| F10 | 57720001 | 57820000 | 0.0014 | 0.0013 | 0.0012 |
| F10 | 57740001 | 57840000 | 0.0017 | 0.0010 | 0.0014 |
| F10 | 57760001 | 57860000 | 0.0012 | 0.0012 | 0.0011 |
| F10 | 57780001 | 57880000 | 0.0015 | 0.0012 | 0.0013 |
| F10 | 57800001 | 57900000 | 0.0015 | 0.0013 | 0.0013 |
| F10 | 57820001 | 57920000 | 0.0016 | 0.0014 | 0.0014 |
| F10 | 57840001 | 57940000 | 0.0017 | 0.0016 | 0.0016 |
| F10 | 57860001 | 57960000 | 0.0019 | 0.0019 | 0.0018 |
| F10 | 57880001 | 57980000 | 0.0023 | 0.0023 | 0.0022 |
| F10 | 57900001 | 58000000 | 0.0029 | 0.0023 | 0.0027 |
| F10 | 57920001 | 58020000 | 0.0030 | 0.0018 | 0.0027 |
| F10 | 57940001 | 58040000 | 0.0025 | 0.0017 | 0.0022 |
| F10 | 57960001 | 58060000 | 0.0024 | 0.0011 | 0.0021 |
| F10 | 57980001 | 58080000 | 0.0016 | 0.0009 | 0.0013 |
| F10 | 58000001 | 58100000 | 0.0012 | 0.0010 | 0.0010 |
| F10 | 58020001 | 58120000 | 0.0013 | 0.0011 | 0.0012 |
| F10 | 58040001 | 58140000 | 0.0013 | 0.0009 | 0.0013 |
| F10 | 58060001 | 58160000 | 0.0011 | 0.0012 | 0.0011 |
| F10 | 58080001 | 58180000 | 0.0015 | 0.0014 | 0.0015 |
| F10 | 58100001 | 58200000 | 0.0017 | 0.0014 | 0.0017 |
| F10 | 58120001 | 58220000 | 0.0015 | 0.0017 | 0.0015 |
| F10 | 58140001 | 58240000 | 0.0018 | 0.0017 | 0.0018 |
| F10 | 58160001 | 58260000 | 0.0019 | 0.0019 | 0.0018 |
| F10 | 58180001 | 58280000 | 0.0020 | 0.0017 | 0.0020 |
| F10 | 58200001 | 58300000 | 0.0018 | 0.0014 | 0.0017 |
| F10 | 58220001 | 58320000 | 0.0016 | 0.0018 | 0.0016 |
| F10 | 58240001 | 58340000 | 0.0020 | 0.0025 | 0.0019 |
| F10 | 58260001 | 58360000 | 0.0029 | 0.0020 | 0.0028 |
| F10 | 58280001 | 58380000 | 0.0023 | 0.0020 | 0.0022 |
| F10 | 58300001 | 58400000 | 0.0022 | 0.0018 | 0.0021 |

| Chr | Start | End | both | A1 | A2 |
|---|---|---|---|---|---|
| F10 | 58320001 | 58420000 | 0.0021 | 0.0010 | 0.0019 |
| F10 | 58340001 | 58440000 | 0.0012 | 0.0003 | 0.0011 |
| F10 | 58360001 | 58460000 | 0.0002 | 0.0005 | 0.0002 |
| F10 | 58380001 | 58480000 | 0.0005 | 0.0007 | 0.0004 |
| F10 | 58400001 | 58500000 | 0.0008 | 0.0008 | 0.0007 |
| F10 | 58420001 | 58520000 | 0.0010 | 0.0009 | 0.0010 |
| F10 | 58440001 | 58540000 | 0.0010 | 0.0013 | 0.0010 |
| F10 | 58460001 | 58560000 | 0.0015 | 0.0030 | 0.0014 |
| F10 | 58480001 | 58580000 | 0.0036 | 0.0037 | 0.0032 |
| F10 | 58500001 | 58600000 | 0.0043 | 0.0035 | 0.0040 |
| F10 | 58520001 | 58620000 | 0.0041 | 0.0044 | 0.0037 |
| F10 | 58540001 | 58640000 | 0.0050 | 0.0041 | 0.0046 |
| F10 | 58560001 | 58660000 | 0.0047 | 0.0021 | 0.0043 |
| F10 | 58580001 | 58680000 | 0.0023 | 0.0017 | 0.0022 |
| F10 | 58600001 | 58700000 | 0.0019 | 0.0022 | 0.0018 |
| F10 | 58620001 | 58720000 | 0.0025 | 0.0013 | 0.0023 |
| F10 | 58640001 | 58740000 | 0.0016 | 0.0012 | 0.0014 |
| F10 | 58660001 | 58760000 | 0.0015 | 0.0014 | 0.0013 |
| F10 | 58680001 | 58780000 | 0.0016 | 0.0010 | 0.0014 |
| F10 | 58700001 | 58800000 | 0.0012 | 0.0008 | 0.0010 |
| F10 | 58720001 | 58820000 | 0.0009 | 0.0009 | 0.0008 |
| F10 | 58740001 | 58840000 | 0.0010 | 0.0010 | 0.0009 |
| F10 | 58760001 | 58860000 | 0.0012 | 0.0010 | 0.0011 |
| F10 | 58780001 | 58880000 | 0.0012 | 0.0012 | 0.0011 |
| F10 | 58800001 | 58900000 | 0.0014 | 0.0008 | 0.0013 |
| F10 | 58820001 | 58920000 | 0.0010 | 0.0009 | 0.0010 |
| F10 | 58840001 | 58940000 | 0.0010 | 0.0007 | 0.0010 |
| F10 | 58860001 | 58960000 | 0.0009 | 0.0007 | 0.0009 |
| F10 | 58880001 | 58980000 | 0.0009 | 0.0005 | 0.0009 |
| F10 | 58900001 | 59000000 | 0.0007 | 0.0005 | 0.0007 |
| F10 | 58920001 | 59020000 | 0.0007 | 0.0006 | 0.0008 |
| F10 | 58940001 | 59040000 | 0.0008 | 0.0006 | 0.0008 |
| F10 | 58960001 | 59060000 | 0.0008 | 0.0004 | 0.0008 |
| F10 | 58980001 | 59080000 | 0.0006 | 0.0006 | 0.0006 |
| F10 | 59000001 | 59100000 | 0.0006 | 0.0008 | 0.0006 |
| F10 | 59020001 | 59120000 | 0.0011 | 0.0007 | 0.0008 |
| F10 | 59040001 | 59140000 | 0.0010 | 0.0011 | 0.0007 |
| F10 | 59060001 | 59160000 | 0.0015 | 0.0014 | 0.0012 |
| F10 | 59080001 | 59180000 | 0.0018 | 0.0012 | 0.0015 |
| F10 | 59100001 | 59200000 | 0.0016 | 0.0012 | 0.0013 |
| F10 | 59120001 | 59220000 | 0.0014 | 0.0014 | 0.0014 |
| F10 | 59140001 | 59240000 | 0.0016 | 0.0013 | 0.0016 |
| F10 | 59160001 | 59260000 | 0.0015 | 0.0015 | 0.0015 |
| F10 | 59180001 | 59280000 | 0.0017 | 0.0016 | 0.0017 |
| F10 | 59200001 | 59300000 | 0.0019 | 0.0016 | 0.0019 |
| F10 | 59220001 | 59320000 | 0.0020 | 0.0014 | 0.0020 |
| F10 | 59240001 | 59340000 | 0.0018 | 0.0014 | 0.0018 |
| F10 | 59260001 | 59360000 | 0.0017 | 0.0011 | 0.0017 |
| F10 | 59280001 | 59380000 | 0.0013 | 0.0009 | 0.0013 |
| F10 | 59300001 | 59400000 | 0.0011 | 0.0006 | 0.0010 |
| F10 | 59320001 | 59420000 | 0.0007 | 0.0011 | 0.0007 |
| F10 | 59340001 | 59440000 | 0.0013 | 0.0010 | 0.0012 |
| F10 | 59360001 | 59460000 | 0.0011 | 0.0010 | 0.0010 |
| F10 | 59380001 | 59480000 | 0.0011 | 0.0010 | 0.0010 |
| F10 | 59400001 | 59500000 | 0.0011 | 0.0013 | 0.0010 |
| F10 | 59420001 | 59520000 | 0.0014 | 0.0009 | 0.0013 |
| F10 | 59440001 | 59540000 | 0.0011 | 0.0007 | 0.0010 |
| F10 | 59460001 | 59560000 | 0.0009 | 0.0008 | 0.0009 |

| Chr | Start | End | both | A1 | A2 |
|---|---|---|---|---|---|
| F10 | 59480001 | 59580000 | 0.0010 | 0.0011 | 0.0010 |
| F10 | 59500001 | 59600000 | 0.0013 | 0.0013 | 0.0012 |
| F10 | 59520001 | 59620000 | 0.0016 | 0.0014 | 0.0013 |
| F10 | 59540001 | 59640000 | 0.0019 | 0.0025 | 0.0014 |
| F10 | 59560001 | 59660000 | 0.0032 | 0.0030 | 0.0027 |
| F10 | 59580001 | 59680000 | 0.0038 | 0.0028 | 0.0033 |
| F10 | 59600001 | 59700000 | 0.0037 | 0.0026 | 0.0032 |
| F10 | 59620001 | 59720000 | 0.0034 | 0.0025 | 0.0031 |
| F10 | 59640001 | 59740000 | 0.0032 | 0.0017 | 0.0031 |
| F10 | 59660001 | 59760000 | 0.0022 | 0.0012 | 0.0021 |
| F10 | 59680001 | 59780000 | 0.0015 | 0.0013 | 0.0014 |
| F10 | 59700001 | 59800000 | 0.0015 | 0.0011 | 0.0014 |
| F10 | 59720001 | 59820000 | 0.0012 | 0.0010 | 0.0012 |
| F10 | 59740001 | 59840000 | 0.0010 | 0.0009 | 0.0010 |
| F10 | 59760001 | 59860000 | 0.0010 | 0.0018 | 0.0009 |
| F10 | 59780001 | 59880000 | 0.0021 | 0.0022 | 0.0020 |
| F10 | 59800001 | 59900000 | 0.0028 | 0.0023 | 0.0025 |
| F10 | 59820001 | 59920000 | 0.0029 | 0.0025 | 0.0026 |
| F10 | 59840001 | 59940000 | 0.0031 | 0.0027 | 0.0028 |
| F10 | 59860001 | 59960000 | 0.0034 | 0.0020 | 0.0031 |
| F10 | 59880001 | 59980000 | 0.0024 | 0.0021 | 0.0022 |
| F10 | 59900001 | 60000000 | 0.0027 | 0.0019 | 0.0026 |
| F10 | 59920001 | 60020000 | 0.0024 | 0.0019 | 0.0023 |
| F10 | 59940001 | 60040000 | 0.0026 | 0.0019 | 0.0025 |
| F10 | 59960001 | 60060000 | 0.0025 | 0.0019 | 0.0024 |
| F10 | 59980001 | 60080000 | 0.0025 | 0.0013 | 0.0024 |
| F10 | 60000001 | 60100000 | 0.0015 | 0.0013 | 0.0015 |
| F10 | 60020001 | 60120000 | 0.0016 | 0.0011 | 0.0015 |
| F10 | 60040001 | 60140000 | 0.0012 | 0.0007 | 0.0012 |
| F10 | 60060001 | 60160000 | 0.0008 | 0.0006 | 0.0008 |
| F10 | 60080001 | 60180000 | 0.0007 | 0.0007 | 0.0007 |
| F10 | 60100001 | 60200000 | 0.0009 | 0.0008 | 0.0009 |
| F10 | 60120001 | 60220000 | 0.0010 | 0.0006 | 0.0010 |
| F10 | 60140001 | 60240000 | 0.0008 | 0.0006 | 0.0008 |
| F10 | 60160001 | 60260000 | 0.0008 | 0.0005 | 0.0008 |
| F10 | 60180001 | 60280000 | 0.0006 | 0.0002 | 0.0006 |
| F10 | 60200001 | 60300000 | 0.0002 | 0.0001 | 0.0002 |
| F10 | 60220001 | 60320000 | 0.0001 | 0.0001 | 0.0001 |
| F10 | 60240001 | 60340000 | 0.0001 | 0.0004 | 0.0001 |
| F10 | 60260001 | 60360000 | 0.0004 | 0.0008 | 0.0004 |
| F10 | 60280001 | 60380000 | 0.0008 | 0.0010 | 0.0008 |
| F10 | 60300001 | 60400000 | 0.0011 | 0.0010 | 0.0010 |
| F10 | 60320001 | 60420000 | 0.0011 | 0.0010 | 0.0010 |
| F10 | 60340001 | 60440000 | 0.0011 | 0.0010 | 0.0011 |
| F10 | 60360001 | 60460000 | 0.0011 | 0.0007 | 0.0011 |
| F10 | 60380001 | 60480000 | 0.0008 | 0.0006 | 0.0008 |
| F10 | 60400001 | 60500000 | 0.0007 | 0.0007 | 0.0006 |
| F10 | 60420001 | 60520000 | 0.0008 | 0.0009 | 0.0007 |
| F10 | 60440001 | 60540000 | 0.0010 | 0.0014 | 0.0008 |
| F10 | 60460001 | 60560000 | 0.0015 | 0.0018 | 0.0012 |
| F10 | 60480001 | 60580000 | 0.0020 | 0.0029 | 0.0016 |
| F10 | 60500001 | 60600000 | 0.0032 | 0.0033 | 0.0028 |
| F10 | 60520001 | 60620000 | 0.0036 | 0.0035 | 0.0031 |
| F10 | 60540001 | 60640000 | 0.0039 | 0.0032 | 0.0033 |
| F10 | 60560001 | 60660000 | 0.0037 | 0.0030 | 0.0031 |
| F10 | 60580001 | 60680000 | 0.0037 | 0.0026 | 0.0029 |
| F10 | 60600001 | 60700000 | 0.0033 | 0.0029 | 0.0024 |
| F10 | 60620001 | 60720000 | 0.0038 | 0.0030 | 0.0027 |

| Chr | Start | End | both | A1 | A2 |
|---|---|---|---|---|---|
| F10 | 60640001 | 60740000 | 0.0038 | 0.0033 | 0.0029 |
| F10 | 60660001 | 60760000 | 0.0040 | 0.0031 | 0.0033 |
| F10 | 60680001 | 60780000 | 0.0037 | 0.0030 | 0.0032 |
| F10 | 60700001 | 60800000 | 0.0039 | 0.0025 | 0.0030 |
| F10 | 60720001 | 60820000 | 0.0034 | 0.0026 | 0.0026 |
| F10 | 60740001 | 60840000 | 0.0037 | 0.0018 | 0.0029 |
| F10 | 60760001 | 60860000 | 0.0029 | 0.0017 | 0.0020 |
| F10 | 60780001 | 60880000 | 0.0027 | 0.0013 | 0.0018 |
| F10 | 60800001 | 60900000 | 0.0020 | 0.0018 | 0.0016 |
| F10 | 60820001 | 60920000 | 0.0024 | 0.0017 | 0.0022 |
| F10 | 60840001 | 60940000 | 0.0021 | 0.0021 | 0.0019 |
| F10 | 60860001 | 60960000 | 0.0025 | 0.0023 | 0.0023 |
| F10 | 60880001 | 60980000 | 0.0027 | 0.0027 | 0.0026 |
| F10 | 60900001 | 61000000 | 0.0033 | 0.0025 | 0.0031 |
| F10 | 60920001 | 61020000 | 0.0030 | 0.0029 | 0.0028 |
| F10 | 60940001 | 61040000 | 0.0034 | 0.0029 | 0.0032 |
| F10 | 60960001 | 61060000 | 0.0035 | 0.0027 | 0.0032 |
| F10 | 60980001 | 61080000 | 0.0033 | 0.0020 | 0.0031 |
| F10 | 61000001 | 61100000 | 0.0024 | 0.0015 | 0.0022 |
| F10 | 61020001 | 61120000 | 0.0019 | 0.0011 | 0.0016 |
| F10 | 61040001 | 61140000 | 0.0014 | 0.0012 | 0.0011 |
| F10 | 61060001 | 61160000 | 0.0014 | 0.0013 | 0.0011 |
| F10 | 61080001 | 61180000 | 0.0016 | 0.0024 | 0.0011 |
| F10 | 61100001 | 61200000 | 0.0027 | 0.0029 | 0.0022 |
| F10 | 61120001 | 61220000 | 0.0034 | 0.0028 | 0.0029 |
| F10 | 61140001 | 61240000 | 0.0033 | 0.0027 | 0.0029 |
| F10 | 61160001 | 61260000 | 0.0033 | 0.0028 | 0.0030 |
| F10 | 61180001 | 61280000 | 0.0033 | 0.0017 | 0.0032 |
| F10 | 61200001 | 61300000 | 0.0021 | 0.0016 | 0.0021 |
| F10 | 61220001 | 61320000 | 0.0021 | 0.0014 | 0.0018 |
| F10 | 61240001 | 61340000 | 0.0018 | 0.0010 | 0.0016 |
| F10 | 61260001 | 61360000 | 0.0014 | 0.0009 | 0.0011 |
| F10 | 61280001 | 61380000 | 0.0013 | 0.0011 | 0.0010 |
| F10 | 61300001 | 61400000 | 0.0016 | 0.0014 | 0.0013 |
| F10 | 61320001 | 61420000 | 0.0018 | 0.0019 | 0.0015 |
| F10 | 61340001 | 61440000 | 0.0025 | 0.0019 | 0.0021 |
| F10 | 61360001 | 61460000 | 0.0025 | 0.0018 | 0.0022 |
| F10 | 61380001 | 61480000 | 0.0023 | 0.0020 | 0.0020 |
| F10 | 61400001 | 61500000 | 0.0026 | 0.0019 | 0.0023 |
| F10 | 61420001 | 61520000 | 0.0024 | 0.0014 | 0.0024 |
| F10 | 61440001 | 61540000 | 0.0017 | 0.0016 | 0.0017 |
| F10 | 61460001 | 61560000 | 0.0018 | 0.0022 | 0.0018 |
| F10 | 61480001 | 61580000 | 0.0024 | 0.0018 | 0.0023 |
| F10 | 61500001 | 61600000 | 0.0019 | 0.0014 | 0.0018 |
| F10 | 61520001 | 61620000 | 0.0014 | 0.0015 | 0.0012 |
| F10 | 61540001 | 61640000 | 0.0016 | 0.0016 | 0.0014 |
| F10 | 61560001 | 61660000 | 0.0018 | 0.0012 | 0.0016 |
| F10 | 61580001 | 61680000 | 0.0013 | 0.0017 | 0.0012 |
| F10 | 61600001 | 61700000 | 0.0020 | 0.0016 | 0.0018 |
| F10 | 61620001 | 61720000 | 0.0021 | 0.0016 | 0.0017 |
| F10 | 61640001 | 61740000 | 0.0022 | 0.0021 | 0.0016 |
| F10 | 61660001 | 61760000 | 0.0028 | 0.0020 | 0.0020 |
| F10 | 61680001 | 61780000 | 0.0029 | 0.0024 | 0.0019 |
| F10 | 61700001 | 61800000 | 0.0031 | 0.0022 | 0.0022 |
| F10 | 61720001 | 61820000 | 0.0028 | 0.0020 | 0.0020 |
| F10 | 61740001 | 61840000 | 0.0024 | 0.0013 | 0.0019 |
| F10 | 61760001 | 61860000 | 0.0015 | 0.0013 | 0.0011 |
| F10 | 61780001 | 61880000 | 0.0013 | 0.0005 | 0.0011 |

| Chr | Start | End | both | A1 | A2 |
|---|---|---|---|---|---|
| F10 | 61800001 | 61900000 | 0.0005 | 0.0005 | 0.0005 |
| F10 | 61820001 | 61920000 | 0.0006 | 0.0007 | 0.0006 |
| F10 | 61840001 | 61940000 | 0.0008 | 0.0019 | 0.0008 |
| F10 | 61860001 | 61960000 | 0.0020 | 0.0022 | 0.0020 |
| F10 | 61880001 | 61980000 | 0.0024 | 0.0029 | 0.0023 |
| F10 | 61900001 | 62000000 | 0.0031 | 0.0033 | 0.0030 |
| F10 | 61920001 | 62020000 | 0.0039 | 0.0042 | 0.0035 |
| F10 | 61940001 | 62040000 | 0.0049 | 0.0035 | 0.0044 |
| F10 | 61960001 | 62060000 | 0.0044 | 0.0039 | 0.0038 |
| F10 | 61980001 | 62080000 | 0.0049 | 0.0038 | 0.0042 |
| F10 | 62000001 | 62100000 | 0.0050 | 0.0038 | 0.0040 |
| F10 | 62020001 | 62120000 | 0.0050 | 0.0033 | 0.0040 |
| F10 | 62040001 | 62140000 | 0.0045 | 0.0030 | 0.0035 |
| F10 | 62060001 | 62160000 | 0.0040 | 0.0031 | 0.0030 |
| F10 | 62080001 | 62180000 | 0.0043 | 0.0034 | 0.0032 |
| F10 | 62100001 | 62200000 | 0.0043 | 0.0034 | 0.0036 |
| F10 | 62120001 | 62220000 | 0.0043 | 0.0041 | 0.0035 |
| F10 | 62140001 | 62240000 | 0.0050 | 0.0045 | 0.0042 |
| F10 | 62160001 | 62260000 | 0.0055 | 0.0043 | 0.0047 |
| F10 | 62180001 | 62280000 | 0.0052 | 0.0037 | 0.0043 |
| F10 | 62200001 | 62300000 | 0.0048 | 0.0046 | 0.0036 |
| F10 | 62220001 | 62320000 | 0.0059 | 0.0045 | 0.0049 |
| F10 | 62240001 | 62340000 | 0.0059 | 0.0043 | 0.0046 |
| F10 | 62260001 | 62360000 | 0.0058 | 0.0041 | 0.0043 |
| F10 | 62280001 | 62380000 | 0.0056 | 0.0045 | 0.0043 |
| F10 | 62300001 | 62400000 | 0.0058 | 0.0037 | 0.0047 |
| F10 | 62320001 | 62420000 | 0.0050 | 0.0030 | 0.0038 |
| F10 | 62340001 | 62440000 | 0.0045 | 0.0033 | 0.0034 |
| F10 | 62360001 | 62460000 | 0.0047 | 0.0035 | 0.0039 |
| F10 | 62380001 | 62480000 | 0.0050 | 0.0039 | 0.0041 |
| F10 | 62400001 | 62500000 | 0.0057 | 0.0038 | 0.0047 |
| F10 | 62420001 | 62520000 | 0.0054 | 0.0035 | 0.0044 |
| F10 | 62440001 | 62540000 | 0.0049 | 0.0031 | 0.0040 |
| F10 | 62460001 | 62560000 | 0.0045 | 0.0031 | 0.0035 |
| F10 | 62480001 | 62580000 | 0.0045 | 0.0025 | 0.0036 |
| F10 | 62500001 | 62600000 | 0.0037 | 0.0024 | 0.0027 |
| F10 | 62520001 | 62620000 | 0.0034 | 0.0032 | 0.0025 |
| F10 | 62540001 | 62640000 | 0.0043 | 0.0029 | 0.0034 |
| F10 | 62560001 | 62660000 | 0.0039 | 0.0028 | 0.0030 |
| F10 | 62580001 | 62680000 | 0.0040 | 0.0033 | 0.0029 |
| F10 | 62600001 | 62700000 | 0.0046 | 0.0036 | 0.0033 |
| F10 | 62620001 | 62720000 | 0.0050 | 0.0030 | 0.0035 |
| F10 | 62640001 | 62740000 | 0.0043 | 0.0031 | 0.0029 |
| F10 | 62660001 | 62760000 | 0.0044 | 0.0027 | 0.0030 |
| F10 | 62680001 | 62780000 | 0.0038 | 0.0021 | 0.0025 |
| F10 | 62700001 | 62800000 | 0.0029 | 0.0020 | 0.0020 |
| F10 | 62720001 | 62820000 | 0.0029 | 0.0017 | 0.0020 |
| F10 | 62740001 | 62840000 | 0.0027 | 0.0021 | 0.0015 |
| F10 | 62760001 | 62860000 | 0.0032 | 0.0022 | 0.0020 |
| F10 | 62780001 | 62880000 | 0.0035 | 0.0026 | 0.0023 |
| F10 | 62800001 | 62900000 | 0.0042 | 0.0028 | 0.0029 |
| F10 | 62820001 | 62920000 | 0.0045 | 0.0028 | 0.0036 |
| F10 | 62840001 | 62940000 | 0.0042 | 0.0029 | 0.0038 |
| F10 | 62860001 | 62960000 | 0.0042 | 0.0030 | 0.0036 |
| F10 | 62880001 | 62980000 | 0.0041 | 0.0026 | 0.0036 |
| F10 | 62900001 | 63000000 | 0.0036 | 0.0027 | 0.0030 |
| F10 | 62920001 | 63020000 | 0.0035 | 0.0027 | 0.0029 |
| F10 | 62940001 | 63040000 | 0.0035 | 0.0021 | 0.0027 |

| Chr | Start | End | both | A1 | A2 |
|---|---|---|---|---|---|
| F10 | 62960001 | 63060000 | 0.0028 | 0.0019 | 0.0023 |
| F10 | 62980001 | 63080000 | 0.0026 | 0.0020 | 0.0022 |
| F10 | 63000001 | 63100000 | 0.0027 | 0.0014 | 0.0022 |
| F10 | 63020001 | 63120000 | 0.0019 | 0.0019 | 0.0014 |
| F10 | 63040001 | 63140000 | 0.0024 | 0.0024 | 0.0019 |
| F10 | 63060001 | 63160000 | 0.0030 | 0.0022 | 0.0024 |
| F10 | 63080001 | 63180000 | 0.0028 | 0.0019 | 0.0022 |
| F10 | 63100001 | 63200000 | 0.0024 | 0.0017 | 0.0019 |
| F10 | 63120001 | 63220000 | 0.0022 | 0.0013 | 0.0017 |
| F10 | 63140001 | 63240000 | 0.0018 | 0.0012 | 0.0015 |
| F10 | 63160001 | 63260000 | 0.0018 | 0.0016 | 0.0015 |
| F10 | 63180001 | 63280000 | 0.0025 | 0.0018 | 0.0019 |
| F10 | 63200001 | 63300000 | 0.0027 | 0.0021 | 0.0020 |
| F10 | 63220001 | 63320000 | 0.0034 | 0.0020 | 0.0025 |
| F10 | 63240001 | 63340000 | 0.0034 | 0.0018 | 0.0022 |
| F10 | 63260001 | 63360000 | 0.0035 | 0.0016 | 0.0021 |
| F10 | 63280001 | 63380000 | 0.0030 | 0.0014 | 0.0019 |
| F10 | 63300001 | 63400000 | 0.0028 | 0.0016 | 0.0018 |
| F10 | 63320001 | 63420000 | 0.0029 | 0.0021 | 0.0018 |
| F10 | 63340001 | 63440000 | 0.0032 | 0.0022 | 0.0023 |
| F10 | 63360001 | 63460000 | 0.0032 | 0.0022 | 0.0023 |
| F10 | 63380001 | 63480000 | 0.0030 | 0.0029 | 0.0021 |
| F10 | 63400001 | 63500000 | 0.0037 | 0.0029 | 0.0029 |
| F10 | 63420001 | 63520000 | 0.0037 | 0.0024 | 0.0030 |
| F10 | 63440001 | 63540000 | 0.0035 | 0.0027 | 0.0027 |
| F10 | 63460001 | 63560000 | 0.0040 | 0.0029 | 0.0031 |
| F10 | 63480001 | 63580000 | 0.0045 | 0.0025 | 0.0033 |
| F10 | 63500001 | 63600000 | 0.0040 | 0.0024 | 0.0029 |
| F10 | 63520001 | 63620000 | 0.0038 | 0.0026 | 0.0026 |
| F10 | 63540001 | 63640000 | 0.0037 | 0.0021 | 0.0025 |
| F10 | 63560001 | 63660000 | 0.0029 | 0.0025 | 0.0020 |
| F10 | 63580001 | 63680000 | 0.0032 | 0.0026 | 0.0023 |
| F10 | 63600001 | 63700000 | 0.0034 | 0.0026 | 0.0024 |
| F10 | 63620001 | 63720000 | 0.0032 | 0.0030 | 0.0023 |
| F10 | 63640001 | 63740000 | 0.0035 | 0.0035 | 0.0028 |
| F10 | 63660001 | 63760000 | 0.0043 | 0.0029 | 0.0034 |
| F10 | 63680001 | 63780000 | 0.0036 | 0.0024 | 0.0029 |
| F10 | 63700001 | 63800000 | 0.0032 | 0.0025 | 0.0025 |
| F10 | 63720001 | 63820000 | 0.0033 | 0.0022 | 0.0026 |
| F10 | 63740001 | 63840000 | 0.0029 | 0.0032 | 0.0022 |
| F10 | 63760001 | 63860000 | 0.0039 | 0.0033 | 0.0033 |
| F10 | 63780001 | 63880000 | 0.0038 | 0.0042 | 0.0033 |
| F10 | 63800001 | 63900000 | 0.0048 | 0.0048 | 0.0041 |
| F10 | 63820001 | 63920000 | 0.0056 | 0.0048 | 0.0049 |
| F10 | 63840001 | 63940000 | 0.0056 | 0.0047 | 0.0049 |
| F10 | 63860001 | 63960000 | 0.0056 | 0.0047 | 0.0051 |
| F10 | 63880001 | 63980000 | 0.0056 | 0.0041 | 0.0051 |
| F10 | 63900001 | 64000000 | 0.0048 | 0.0040 | 0.0044 |
| F10 | 63920001 | 64020000 | 0.0048 | 0.0044 | 0.0043 |
| F10 | 63940001 | 64040000 | 0.0052 | 0.0031 | 0.0046 |
| F10 | 63960001 | 64060000 | 0.0037 | 0.0030 | 0.0032 |
| F10 | 63980001 | 64080000 | 0.0037 | 0.0031 | 0.0031 |
| F10 | 64000001 | 64100000 | 0.0039 | 0.0030 | 0.0034 |
| F10 | 64020001 | 64120000 | 0.0039 | 0.0025 | 0.0032 |
| F10 | 64040001 | 64140000 | 0.0034 | 0.0023 | 0.0026 |
| F10 | 64060001 | 64160000 | 0.0033 | 0.0022 | 0.0024 |
| F10 | 64080001 | 64180000 | 0.0030 | 0.0029 | 0.0024 |
| F10 | 64100001 | 64200000 | 0.0038 | 0.0024 | 0.0032 |

| Chr | Start | End | both | A1 | A2 |
|---|---|---|---|---|---|
| F10 | 64120001 | 64220000 | 0.0035 | 0.0025 | 0.0029 |
| F10 | 64140001 | 64240000 | 0.0035 | 0.0023 | 0.0029 |
| F10 | 64160001 | 64260000 | 0.0032 | 0.0023 | 0.0027 |
| F10 | 64180001 | 64280000 | 0.0033 | 0.0022 | 0.0027 |
| F10 | 64200001 | 64300000 | 0.0032 | 0.0022 | 0.0025 |
| F10 | 64220001 | 64320000 | 0.0029 | 0.0020 | 0.0025 |
| F10 | 64240001 | 64340000 | 0.0025 | 0.0026 | 0.0023 |
| F10 | 64260001 | 64360000 | 0.0032 | 0.0030 | 0.0029 |
| F10 | 64280001 | 64380000 | 0.0035 | 0.0021 | 0.0033 |
| F10 | 64300001 | 64400000 | 0.0026 | 0.0023 | 0.0023 |
| F10 | 64320001 | 64420000 | 0.0028 | 0.0023 | 0.0024 |
| F10 | 64340001 | 64440000 | 0.0029 | 0.0024 | 0.0025 |
| F10 | 64360001 | 64460000 | 0.0031 | 0.0026 | 0.0025 |
| F10 | 64380001 | 64480000 | 0.0031 | 0.0023 | 0.0026 |
| F10 | 64400001 | 64500000 | 0.0028 | 0.0018 | 0.0023 |
| F10 | 64420001 | 64520000 | 0.0023 | 0.0020 | 0.0019 |
| F10 | 64440001 | 64540000 | 0.0025 | 0.0018 | 0.0021 |
| F10 | 64460001 | 64560000 | 0.0021 | 0.0014 | 0.0019 |
| F10 | 64480001 | 64580000 | 0.0018 | 0.0021 | 0.0016 |
| F10 | 64500001 | 64600000 | 0.0026 | 0.0023 | 0.0023 |
| F10 | 64520001 | 64620000 | 0.0029 | 0.0026 | 0.0024 |
| F10 | 64540001 | 64640000 | 0.0034 | 0.0024 | 0.0029 |
| F10 | 64560001 | 64660000 | 0.0033 | 0.0024 | 0.0029 |
| F10 | 64580001 | 64680000 | 0.0032 | 0.0021 | 0.0029 |
| F10 | 64600001 | 64700000 | 0.0029 | 0.0024 | 0.0025 |
| F10 | 64620001 | 64720000 | 0.0033 | 0.0023 | 0.0028 |
| F10 | 64640001 | 64740000 | 0.0032 | 0.0027 | 0.0026 |
| F10 | 64660001 | 64760000 | 0.0037 | 0.0026 | 0.0029 |
| F10 | 64680001 | 64780000 | 0.0038 | 0.0024 | 0.0026 |
| F10 | 64700001 | 64800000 | 0.0036 | 0.0024 | 0.0024 |
| F10 | 64720001 | 64820000 | 0.0035 | 0.0025 | 0.0024 |
| F10 | 64740001 | 64840000 | 0.0036 | 0.0025 | 0.0025 |
| F10 | 64760001 | 64860000 | 0.0035 | 0.0029 | 0.0026 |
| F10 | 64780001 | 64880000 | 0.0038 | 0.0038 | 0.0030 |
| F10 | 64800001 | 64900000 | 0.0046 | 0.0034 | 0.0039 |
| F10 | 64820001 | 64920000 | 0.0044 | 0.0029 | 0.0038 |
| F10 | 64840001 | 64940000 | 0.0039 | 0.0025 | 0.0032 |
| F10 | 64860001 | 64960000 | 0.0037 | 0.0021 | 0.0029 |
| F10 | 64880001 | 64980000 | 0.0032 | 0.0011 | 0.0026 |
| F10 | 64900001 | 65000000 | 0.0020 | 0.0021 | 0.0016 |
| F10 | 64920001 | 65020000 | 0.0031 | 0.0029 | 0.0027 |
| F10 | 64940001 | 65040000 | 0.0039 | 0.0035 | 0.0037 |
| F10 | 64960001 | 65060000 | 0.0042 | 0.0035 | 0.0041 |
| F10 | 64980001 | 65080000 | 0.0042 | 0.0046 | 0.0042 |
| F10 | 65000001 | 65100000 | 0.0055 | 0.0037 | 0.0054 |
| F10 | 65020001 | 65120000 | 0.0045 | 0.0031 | 0.0043 |
| F10 | 65040001 | 65140000 | 0.0038 | 0.0023 | 0.0035 |
| F10 | 65060001 | 65160000 | 0.0030 | 0.0020 | 0.0026 |
| F10 | 65080001 | 65180000 | 0.0027 | 0.0015 | 0.0023 |
| F10 | 65100001 | 65200000 | 0.0021 | 0.0020 | 0.0017 |
| F10 | 65120001 | 65220000 | 0.0027 | 0.0030 | 0.0023 |
| F10 | 65140001 | 65240000 | 0.0037 | 0.0035 | 0.0033 |
| F10 | 65160001 | 65260000 | 0.0044 | 0.0044 | 0.0038 |
| F10 | 65180001 | 65280000 | 0.0054 | 0.0045 | 0.0049 |
| F10 | 65200001 | 65300000 | 0.0054 | 0.0045 | 0.0050 |
| F10 | 65220001 | 65320000 | 0.0052 | 0.0036 | 0.0047 |
| F10 | 65240001 | 65340000 | 0.0042 | 0.0034 | 0.0036 |
| F10 | 65260001 | 65360000 | 0.0039 | 0.0029 | 0.0035 |

| Chr | Start | End | both | A1 | A2 |
|---|---|---|---|---|---|
| F10 | 65280001 | 65380000 | 0.0032 | 0.0031 | 0.0027 |
| F10 | 65300001 | 65400000 | 0.0036 | 0.0024 | 0.0031 |
| F10 | 65320001 | 65420000 | 0.0030 | 0.0030 | 0.0025 |
| F10 | 65340001 | 65440000 | 0.0036 | 0.0031 | 0.0033 |
| F10 | 65360001 | 65460000 | 0.0039 | 0.0041 | 0.0035 |
| F10 | 65380001 | 65480000 | 0.0051 | 0.0037 | 0.0047 |
| F10 | 65400001 | 65500000 | 0.0043 | 0.0041 | 0.0040 |
| F10 | 65420001 | 65520000 | 0.0047 | 0.0030 | 0.0045 |
| F10 | 65440001 | 65540000 | 0.0036 | 0.0031 | 0.0033 |
| F10 | 65460001 | 65560000 | 0.0036 | 0.0021 | 0.0032 |
| F10 | 65480001 | 65580000 | 0.0026 | 0.0025 | 0.0023 |
| F10 | 65500001 | 65600000 | 0.0031 | 0.0023 | 0.0025 |
| F10 | 65520001 | 65620000 | 0.0030 | 0.0024 | 0.0022 |
| F10 | 65540001 | 65640000 | 0.0032 | 0.0021 | 0.0024 |
| F10 | 65560001 | 65660000 | 0.0029 | 0.0017 | 0.0020 |
| F10 | 65580001 | 65680000 | 0.0025 | 0.0016 | 0.0017 |
| F10 | 65600001 | 65700000 | 0.0024 | 0.0016 | 0.0017 |
| F10 | 65620001 | 65720000 | 0.0023 | 0.0015 | 0.0018 |
| F10 | 65640001 | 65740000 | 0.0023 | 0.0015 | 0.0016 |
| F10 | 65660001 | 65760000 | 0.0021 | 0.0014 | 0.0016 |
| F10 | 65680001 | 65780000 | 0.0021 | 0.0009 | 0.0015 |
| F10 | 65700001 | 65800000 | 0.0016 | 0.0008 | 0.0011 |
| F10 | 65720001 | 65820000 | 0.0014 | 0.0007 | 0.0010 |
| F10 | 65740001 | 65840000 | 0.0011 | 0.0007 | 0.0009 |
| F10 | 65760001 | 65860000 | 0.0012 | 0.0006 | 0.0010 |
| F10 | 65780001 | 65880000 | 0.0010 | 0.0005 | 0.0009 |
| F10 | 65800001 | 65900000 | 0.0007 | 0.0004 | 0.0007 |
| F10 | 65820001 | 65920000 | 0.0006 | 0.0007 | 0.0006 |
| F10 | 65840001 | 65940000 | 0.0009 | 0.0008 | 0.0009 |
| F10 | 65860001 | 65960000 | 0.0010 | 0.0013 | 0.0009 |
| F10 | 65880001 | 65980000 | 0.0016 | 0.0013 | 0.0014 |
| F10 | 65900001 | 66000000 | 0.0016 | 0.0014 | 0.0014 |
| F10 | 65920001 | 66020000 | 0.0017 | 0.0015 | 0.0016 |
| F10 | 65940001 | 66040000 | 0.0020 | 0.0013 | 0.0016 |
| F10 | 65960001 | 66060000 | 0.0019 | 0.0011 | 0.0016 |
| F10 | 65980001 | 66080000 | 0.0015 | 0.0015 | 0.0012 |
| F10 | 66000001 | 66100000 | 0.0021 | 0.0021 | 0.0016 |
| F10 | 66020001 | 66120000 | 0.0029 | 0.0021 | 0.0023 |
| F10 | 66040001 | 66140000 | 0.0028 | 0.0021 | 0.0023 |
| F10 | 66060001 | 66160000 | 0.0028 | 0.0022 | 0.0024 |
| F10 | 66080001 | 66180000 | 0.0028 | 0.0020 | 0.0025 |
| F10 | 66100001 | 66200000 | 0.0025 | 0.0014 | 0.0023 |
| F10 | 66120001 | 66220000 | 0.0017 | 0.0012 | 0.0016 |
| F10 | 66140001 | 66240000 | 0.0013 | 0.0010 | 0.0013 |
| F10 | 66160001 | 66260000 | 0.0012 | 0.0009 | 0.0012 |
| F10 | 66180001 | 66280000 | 0.0010 | 0.0007 | 0.0010 |
| F10 | 66200001 | 66300000 | 0.0008 | 0.0006 | 0.0008 |
| F10 | 66220001 | 66320000 | 0.0007 | 0.0007 | 0.0007 |
| F10 | 66240001 | 66340000 | 0.0008 | 0.0009 | 0.0008 |
| F10 | 66260001 | 66360000 | 0.0010 | 0.0010 | 0.0010 |
| F10 | 66280001 | 66380000 | 0.0011 | 0.0015 | 0.0011 |
| F10 | 66300001 | 66400000 | 0.0017 | 0.0020 | 0.0017 |
| F10 | 66320001 | 66420000 | 0.0024 | 0.0020 | 0.0023 |
| F10 | 66340001 | 66440000 | 0.0026 | 0.0019 | 0.0023 |
| F10 | 66360001 | 66460000 | 0.0025 | 0.0023 | 0.0021 |
| F10 | 66380001 | 66480000 | 0.0030 | 0.0026 | 0.0026 |
| F10 | 66400001 | 66500000 | 0.0035 | 0.0021 | 0.0028 |
| F10 | 66420001 | 66520000 | 0.0033 | 0.0022 | 0.0025 |

| Chr | Start | End | both | A1 | A2 |
|---|---|---|---|---|---|
| F10 | 66440001 | 66540000 | 0.0032 | 0.0021 | 0.0025 |
| F10 | 66460001 | 66560000 | 0.0031 | 0.0018 | 0.0025 |
| F10 | 66480001 | 66580000 | 0.0031 | 0.0014 | 0.0022 |
| F10 | 66500001 | 66600000 | 0.0025 | 0.0015 | 0.0017 |
| F10 | 66520001 | 66620000 | 0.0025 | 0.0017 | 0.0017 |
| F10 | 66540001 | 66640000 | 0.0027 | 0.0016 | 0.0020 |
| F10 | 66560001 | 66660000 | 0.0024 | 0.0023 | 0.0017 |
| F10 | 66580001 | 66680000 | 0.0030 | 0.0022 | 0.0025 |
| F10 | 66600001 | 66700000 | 0.0026 | 0.0032 | 0.0022 |
| F10 | 66620001 | 66720000 | 0.0036 | 0.0035 | 0.0032 |
| F10 | 66640001 | 66740000 | 0.0041 | 0.0039 | 0.0035 |
| F10 | 66660001 | 66760000 | 0.0045 | 0.0031 | 0.0039 |
| F10 | 66680001 | 66780000 | 0.0037 | 0.0036 | 0.0030 |
| F10 | 66700001 | 66800000 | 0.0043 | 0.0024 | 0.0035 |
| F10 | 66720001 | 66820000 | 0.0032 | 0.0021 | 0.0024 |
| F10 | 66740001 | 66840000 | 0.0027 | 0.0019 | 0.0021 |
| F10 | 66760001 | 66860000 | 0.0025 | 0.0019 | 0.0019 |
| F10 | 66780001 | 66880000 | 0.0023 | 0.0015 | 0.0020 |
| F10 | 66800001 | 66900000 | 0.0019 | 0.0020 | 0.0017 |
| F10 | 66820001 | 66920000 | 0.0023 | 0.0021 | 0.0022 |
| F10 | 66840001 | 66940000 | 0.0024 | 0.0024 | 0.0023 |
| F10 | 66860001 | 66960000 | 0.0026 | 0.0028 | 0.0026 |
| F10 | 66880001 | 66980000 | 0.0033 | 0.0029 | 0.0032 |
| F10 | 66900001 | 67000000 | 0.0036 | 0.0027 | 0.0034 |
| F10 | 66920001 | 67020000 | 0.0038 | 0.0025 | 0.0034 |
| F10 | 66940001 | 67040000 | 0.0035 | 0.0034 | 0.0031 |
| F10 | 66960001 | 67060000 | 0.0044 | 0.0036 | 0.0037 |
| F10 | 66980001 | 67080000 | 0.0045 | 0.0042 | 0.0037 |
| F10 | 67000001 | 67100000 | 0.0049 | 0.0039 | 0.0042 |
| F10 | 67020001 | 67120000 | 0.0044 | 0.0039 | 0.0036 |
| F10 | 67040001 | 67140000 | 0.0045 | 0.0028 | 0.0038 |
| F10 | 67060001 | 67160000 | 0.0034 | 0.0035 | 0.0028 |
| F10 | 67080001 | 67180000 | 0.0042 | 0.0031 | 0.0037 |
| F10 | 67100001 | 67200000 | 0.0038 | 0.0032 | 0.0035 |
| F10 | 67120001 | 67220000 | 0.0038 | 0.0035 | 0.0036 |
| F10 | 67140001 | 67240000 | 0.0041 | 0.0037 | 0.0039 |
| F10 | 67160001 | 67260000 | 0.0044 | 0.0022 | 0.0042 |
| F10 | 67180001 | 67280000 | 0.0026 | 0.0016 | 0.0026 |
| F10 | 67200001 | 67300000 | 0.0021 | 0.0014 | 0.0020 |
| F10 | 67220001 | 67320000 | 0.0019 | 0.0010 | 0.0018 |
| F10 | 67240001 | 67340000 | 0.0016 | 0.0007 | 0.0013 |
| F10 | 67260001 | 67360000 | 0.0014 | 0.0007 | 0.0009 |
| F10 | 67280001 | 67380000 | 0.0015 | 0.0008 | 0.0009 |
| F10 | 67300001 | 67400000 | 0.0015 | 0.0014 | 0.0010 |
| F10 | 67320001 | 67420000 | 0.0022 | 0.0016 | 0.0015 |
| F10 | 67340001 | 67440000 | 0.0024 | 0.0017 | 0.0019 |
| F10 | 67360001 | 67460000 | 0.0024 | 0.0021 | 0.0019 |
| F10 | 67380001 | 67480000 | 0.0029 | 0.0028 | 0.0025 |
| F10 | 67400001 | 67500000 | 0.0036 | 0.0030 | 0.0030 |
| F10 | 67420001 | 67520000 | 0.0039 | 0.0041 | 0.0033 |
| F10 | 67440001 | 67540000 | 0.0051 | 0.0039 | 0.0043 |
| F10 | 67460001 | 67560000 | 0.0048 | 0.0036 | 0.0042 |
| F10 | 67480001 | 67580000 | 0.0043 | 0.0028 | 0.0037 |
| F10 | 67500001 | 67600000 | 0.0036 | 0.0029 | 0.0031 |
| F10 | 67520001 | 67620000 | 0.0036 | 0.0019 | 0.0031 |
| F10 | 67540001 | 67640000 | 0.0025 | 0.0024 | 0.0020 |
| F10 | 67560001 | 67660000 | 0.0032 | 0.0033 | 0.0025 |
| F10 | 67580001 | 67680000 | 0.0042 | 0.0031 | 0.0034 |

| Chr | Start | End | both | A1 | A2 |
|---|---|---|---|---|---|
| F10 | 67600001 | 67700000 | 0.0039 | 0.0027 | 0.0033 |
| F10 | 67620001 | 67720000 | 0.0036 | 0.0031 | 0.0029 |
| F10 | 67640001 | 67740000 | 0.0040 | 0.0028 | 0.0032 |
| F10 | 67660001 | 67760000 | 0.0036 | 0.0021 | 0.0029 |
| F10 | 67680001 | 67780000 | 0.0026 | 0.0024 | 0.0020 |
| F10 | 67700001 | 67800000 | 0.0032 | 0.0022 | 0.0024 |
| F10 | 67720001 | 67820000 | 0.0030 | 0.0022 | 0.0022 |
| F10 | 67740001 | 67840000 | 0.0030 | 0.0021 | 0.0022 |
| F10 | 67760001 | 67860000 | 0.0028 | 0.0022 | 0.0020 |
| F10 | 67780001 | 67880000 | 0.0031 | 0.0019 | 0.0023 |
| F10 | 67800001 | 67900000 | 0.0028 | 0.0020 | 0.0020 |
| F10 | 67820001 | 67920000 | 0.0028 | 0.0016 | 0.0022 |
| F10 | 67840001 | 67940000 | 0.0023 | 0.0024 | 0.0017 |
| F10 | 67860001 | 67960000 | 0.0035 | 0.0039 | 0.0029 |
| F10 | 67880001 | 67980000 | 0.0052 | 0.0041 | 0.0046 |
| F10 | 67900001 | 68000000 | 0.0054 | 0.0042 | 0.0048 |
| F10 | 67920001 | 68020000 | 0.0056 | 0.0040 | 0.0049 |
| F10 | 67940001 | 68040000 | 0.0053 | 0.0037 | 0.0047 |
| F10 | 67960001 | 68060000 | 0.0050 | 0.0024 | 0.0042 |
| F10 | 67980001 | 68080000 | 0.0036 | 0.0020 | 0.0028 |
| F10 | 68000001 | 68100000 | 0.0030 | 0.0015 | 0.0023 |
| F10 | 68020001 | 68120000 | 0.0024 | 0.0017 | 0.0017 |
| F10 | 68040001 | 68140000 | 0.0026 | 0.0015 | 0.0020 |
| F10 | 68060001 | 68160000 | 0.0022 | 0.0011 | 0.0019 |
| F10 | 68080001 | 68180000 | 0.0017 | 0.0017 | 0.0013 |
| F10 | 68100001 | 68200000 | 0.0024 | 0.0021 | 0.0019 |
| F10 | 68120001 | 68220000 | 0.0031 | 0.0019 | 0.0023 |
| F10 | 68140001 | 68240000 | 0.0032 | 0.0018 | 0.0021 |
| F10 | 68160001 | 68260000 | 0.0033 | 0.0021 | 0.0021 |
| F10 | 68180001 | 68280000 | 0.0035 | 0.0022 | 0.0025 |
| F10 | 68200001 | 68300000 | 0.0037 | 0.0027 | 0.0027 |
| F10 | 68220001 | 68320000 | 0.0040 | 0.0032 | 0.0032 |
| F10 | 68240001 | 68340000 | 0.0045 | 0.0030 | 0.0038 |
| F10 | 68260001 | 68360000 | 0.0041 | 0.0040 | 0.0036 |
| F10 | 68280001 | 68380000 | 0.0052 | 0.0044 | 0.0046 |
| F10 | 68300001 | 68400000 | 0.0056 | 0.0044 | 0.0050 |
| F10 | 68320001 | 68420000 | 0.0057 | 0.0041 | 0.0052 |
| F10 | 68340001 | 68440000 | 0.0052 | 0.0042 | 0.0049 |
| F10 | 68360001 | 68460000 | 0.0051 | 0.0032 | 0.0048 |
| F10 | 68380001 | 68480000 | 0.0040 | 0.0024 | 0.0038 |
| F10 | 68400001 | 68500000 | 0.0034 | 0.0020 | 0.0030 |
| F10 | 68420001 | 68520000 | 0.0028 | 0.0017 | 0.0024 |
| F10 | 68440001 | 68540000 | 0.0026 | 0.0022 | 0.0021 |
| F10 | 68460001 | 68560000 | 0.0030 | 0.0022 | 0.0024 |
| F10 | 68480001 | 68580000 | 0.0030 | 0.0024 | 0.0024 |
| F10 | 68500001 | 68600000 | 0.0028 | 0.0029 | 0.0026 |
| F10 | 68520001 | 68620000 | 0.0033 | 0.0029 | 0.0030 |
| F10 | 68540001 | 68640000 | 0.0035 | 0.0022 | 0.0031 |
| F10 | 68560001 | 68660000 | 0.0028 | 0.0028 | 0.0025 |
| F10 | 68580001 | 68680000 | 0.0036 | 0.0031 | 0.0032 |
| F10 | 68600001 | 68700000 | 0.0045 | 0.0027 | 0.0039 |
| F10 | 68620001 | 68720000 | 0.0042 | 0.0026 | 0.0035 |
| F10 | 68640001 | 68740000 | 0.0041 | 0.0033 | 0.0034 |
| F10 | 68660001 | 68760000 | 0.0049 | 0.0025 | 0.0041 |
| F10 | 68680001 | 68780000 | 0.0042 | 0.0017 | 0.0032 |
| F10 | 68700001 | 68800000 | 0.0034 | 0.0016 | 0.0020 |
| F10 | 68720001 | 68820000 | 0.0034 | 0.0019 | 0.0020 |
| F10 | 68740001 | 68840000 | 0.0038 | 0.0016 | 0.0023 |

| Chr | Start | End | both | A1 | A2 |
|---|---|---|---|---|---|
| F10 | 68760001 | 68860000 | 0.0032 | 0.0020 | 0.0019 |
| F10 | 68780001 | 68880000 | 0.0033 | 0.0019 | 0.0022 |
| F10 | 68800001 | 68900000 | 0.0026 | 0.0018 | 0.0020 |
| F10 | 68820001 | 68920000 | 0.0023 | 0.0014 | 0.0018 |
| F10 | 68840001 | 68940000 | 0.0019 | 0.0014 | 0.0015 |
| F10 | 68860001 | 68960000 | 0.0020 | 0.0013 | 0.0015 |
| F10 | 68880001 | 68980000 | 0.0021 | 0.0018 | 0.0014 |
| F10 | 68900001 | 69000000 | 0.0027 | 0.0018 | 0.0019 |
| F10 | 68920001 | 69020000 | 0.0025 | 0.0025 | 0.0018 |
| F10 | 68940001 | 69040000 | 0.0033 | 0.0023 | 0.0026 |
| F10 | 68960001 | 69060000 | 0.0028 | 0.0030 | 0.0024 |
| F10 | 68980001 | 69080000 | 0.0037 | 0.0030 | 0.0033 |
| F10 | 69000001 | 69100000 | 0.0036 | 0.0033 | 0.0033 |
| F10 | 69020001 | 69120000 | 0.0040 | 0.0040 | 0.0036 |
| F10 | 69040001 | 69140000 | 0.0048 | 0.0044 | 0.0045 |
| F10 | 69060001 | 69160000 | 0.0055 | 0.0032 | 0.0050 |
| F10 | 69080001 | 69180000 | 0.0042 | 0.0029 | 0.0036 |
| F10 | 69100001 | 69200000 | 0.0040 | 0.0033 | 0.0033 |
| F10 | 69120001 | 69220000 | 0.0044 | 0.0025 | 0.0037 |
| F10 | 69140001 | 69240000 | 0.0034 | 0.0030 | 0.0026 |
| F10 | 69160001 | 69260000 | 0.0038 | 0.0030 | 0.0030 |
| F10 | 69180001 | 69280000 | 0.0039 | 0.0042 | 0.0031 |
| F10 | 69200001 | 69300000 | 0.0052 | 0.0039 | 0.0043 |
| F10 | 69220001 | 69320000 | 0.0049 | 0.0040 | 0.0039 |
| F10 | 69240001 | 69340000 | 0.0051 | 0.0031 | 0.0041 |
| F10 | 69260001 | 69360000 | 0.0040 | 0.0035 | 0.0032 |
| F10 | 69280001 | 69380000 | 0.0044 | 0.0029 | 0.0035 |
| F10 | 69300001 | 69400000 | 0.0037 | 0.0025 | 0.0030 |
| F10 | 69320001 | 69420000 | 0.0031 | 0.0019 | 0.0026 |
| F10 | 69340001 | 69440000 | 0.0025 | 0.0023 | 0.0021 |
| F10 | 69360001 | 69460000 | 0.0030 | 0.0027 | 0.0025 |
| F10 | 69380001 | 69480000 | 0.0031 | 0.0023 | 0.0027 |
| F10 | 69400001 | 69500000 | 0.0028 | 0.0022 | 0.0024 |
| F10 | 69420001 | 69520000 | 0.0027 | 0.0023 | 0.0023 |
| F10 | 69440001 | 69540000 | 0.0028 | 0.0026 | 0.0024 |
| F10 | 69460001 | 69560000 | 0.0035 | 0.0023 | 0.0029 |
| F10 | 69480001 | 69580000 | 0.0034 | 0.0029 | 0.0029 |
| F10 | 69500001 | 69600000 | 0.0041 | 0.0035 | 0.0034 |
| F10 | 69520001 | 69620000 | 0.0052 | 0.0040 | 0.0042 |
| F10 | 69540001 | 69640000 | 0.0057 | 0.0038 | 0.0047 |
| F10 | 69560001 | 69660000 | 0.0052 | 0.0035 | 0.0043 |
| F10 | 69580001 | 69680000 | 0.0050 | 0.0033 | 0.0040 |
| F10 | 69600001 | 69700000 | 0.0047 | 0.0027 | 0.0039 |
| F10 | 69620001 | 69720000 | 0.0039 | 0.0024 | 0.0031 |
| F10 | 69640001 | 69740000 | 0.0036 | 0.0029 | 0.0029 |
| F10 | 69660001 | 69760000 | 0.0041 | 0.0031 | 0.0033 |
| F10 | 69680001 | 69780000 | 0.0044 | 0.0026 | 0.0036 |
| F10 | 69700001 | 69800000 | 0.0036 | 0.0024 | 0.0030 |
| F10 | 69720001 | 69820000 | 0.0035 | 0.0020 | 0.0028 |
| F10 | 69740001 | 69840000 | 0.0030 | 0.0015 | 0.0023 |
| F10 | 69760001 | 69860000 | 0.0028 | 0.0017 | 0.0018 |
| F10 | 69780001 | 69880000 | 0.0029 | 0.0017 | 0.0020 |
| F10 | 69800001 | 69900000 | 0.0029 | 0.0023 | 0.0019 |
| F10 | 69820001 | 69920000 | 0.0034 | 0.0025 | 0.0025 |
| F10 | 69840001 | 69940000 | 0.0036 | 0.0024 | 0.0028 |
| F10 | 69860001 | 69960000 | 0.0033 | 0.0019 | 0.0027 |
| F10 | 69880001 | 69980000 | 0.0027 | 0.0028 | 0.0021 |
| F10 | 69900001 | 70000000 | 0.0036 | 0.0025 | 0.0030 |

| Chr | Start | End | both | A1 | A2 |
|---|---|---|---|---|---|
| F10 | 69920001 | 70020000 | 0.0035 | 0.0025 | 0.0026 |
| F10 | 69940001 | 70040000 | 0.0032 | 0.0034 | 0.0025 |
| F10 | 69960001 | 70060000 | 0.0042 | 0.0033 | 0.0036 |
| F10 | 69980001 | 70080000 | 0.0042 | 0.0025 | 0.0033 |
| F10 | 70000001 | 70100000 | 0.0034 | 0.0023 | 0.0026 |
| F10 | 70020001 | 70120000 | 0.0030 | 0.0024 | 0.0024 |
| F10 | 70040001 | 70140000 | 0.0034 | 0.0015 | 0.0026 |
| F10 | 70060001 | 70160000 | 0.0027 | 0.0014 | 0.0018 |
| F10 | 70080001 | 70180000 | 0.0025 | 0.0018 | 0.0018 |
| F10 | 70100001 | 70200000 | 0.0030 | 0.0020 | 0.0024 |
| F10 | 70120001 | 70220000 | 0.0033 | 0.0021 | 0.0027 |
| F10 | 70140001 | 70240000 | 0.0030 | 0.0026 | 0.0027 |
| F10 | 70160001 | 70260000 | 0.0032 | 0.0033 | 0.0027 |
| F10 | 70180001 | 70280000 | 0.0040 | 0.0036 | 0.0035 |
| F10 | 70200001 | 70300000 | 0.0043 | 0.0041 | 0.0037 |
| F10 | 70220001 | 70320000 | 0.0046 | 0.0037 | 0.0041 |
| F10 | 70240001 | 70340000 | 0.0044 | 0.0039 | 0.0037 |
| F10 | 70260001 | 70360000 | 0.0048 | 0.0034 | 0.0042 |
| F10 | 70280001 | 70380000 | 0.0043 | 0.0032 | 0.0036 |
| F10 | 70300001 | 70400000 | 0.0043 | 0.0041 | 0.0033 |
| F10 | 70320001 | 70420000 | 0.0054 | 0.0045 | 0.0042 |
| F10 | 70340001 | 70440000 | 0.0057 | 0.0042 | 0.0046 |
| F10 | 70360001 | 70460000 | 0.0055 | 0.0056 | 0.0041 |
| F10 | 70380001 | 70480000 | 0.0072 | 0.0058 | 0.0058 |
| F10 | 70400001 | 70500000 | 0.0074 | 0.0047 | 0.0062 |
| F10 | 70420001 | 70520000 | 0.0063 | 0.0058 | 0.0052 |
| F10 | 70440001 | 70540000 | 0.0075 | 0.0051 | 0.0062 |
| F10 | 70460001 | 70560000 | 0.0067 | 0.0038 | 0.0055 |
| F10 | 70480001 | 70580000 | 0.0054 | 0.0036 | 0.0040 |
| F10 | 70500001 | 70600000 | 0.0051 | 0.0036 | 0.0036 |
| F10 | 70520001 | 70620000 | 0.0050 | 0.0032 | 0.0035 |
| F10 | 70540001 | 70640000 | 0.0047 | 0.0031 | 0.0033 |
| F10 | 70560001 | 70660000 | 0.0046 | 0.0028 | 0.0032 |
| F10 | 70580001 | 70680000 | 0.0040 | 0.0026 | 0.0030 |
| F10 | 70600001 | 70700000 | 0.0037 | 0.0031 | 0.0027 |
| F10 | 70620001 | 70720000 | 0.0043 | 0.0030 | 0.0033 |
| F10 | 70640001 | 70740000 | 0.0041 | 0.0034 | 0.0032 |
| F10 | 70660001 | 70760000 | 0.0046 | 0.0038 | 0.0037 |
| F10 | 70680001 | 70780000 | 0.0050 | 0.0038 | 0.0040 |
| F10 | 70700001 | 70800000 | 0.0050 | 0.0036 | 0.0040 |
| F10 | 70720001 | 70820000 | 0.0047 | 0.0026 | 0.0037 |
| F10 | 70740001 | 70840000 | 0.0036 | 0.0027 | 0.0027 |
| F10 | 70760001 | 70860000 | 0.0037 | 0.0028 | 0.0028 |
| F10 | 70780001 | 70880000 | 0.0041 | 0.0026 | 0.0031 |
| F10 | 70800001 | 70900000 | 0.0040 | 0.0021 | 0.0030 |
| F10 | 70820001 | 70920000 | 0.0033 | 0.0028 | 0.0023 |
| F10 | 70840001 | 70940000 | 0.0040 | 0.0041 | 0.0029 |
| F10 | 70860001 | 70960000 | 0.0050 | 0.0041 | 0.0040 |
| F10 | 70880001 | 70980000 | 0.0048 | 0.0049 | 0.0041 |
| F10 | 70900001 | 71000000 | 0.0056 | 0.0054 | 0.0048 |
| F10 | 70920001 | 71020000 | 0.0063 | 0.0050 | 0.0054 |
| F10 | 70940001 | 71040000 | 0.0060 | 0.0040 | 0.0051 |
| F10 | 70960001 | 71060000 | 0.0052 | 0.0044 | 0.0041 |
| F10 | 70980001 | 71080000 | 0.0056 | 0.0038 | 0.0044 |
| F10 | 71000001 | 71100000 | 0.0050 | 0.0037 | 0.0039 |
| F10 | 71020001 | 71120000 | 0.0048 | 0.0036 | 0.0038 |
| F10 | 71040001 | 71140000 | 0.0048 | 0.0035 | 0.0036 |
| F10 | 71060001 | 71160000 | 0.0045 | 0.0029 | 0.0035 |

| Chr | Start | End | both | A1 | A2 |
|---|---|---|---|---|---|
| F10 | 71080001 | 71180000 | 0.0037 | 0.0030 | 0.0028 |
| F10 | 71100001 | 71200000 | 0.0038 | 0.0024 | 0.0028 |
| F10 | 71120001 | 71220000 | 0.0035 | 0.0026 | 0.0025 |
| F10 | 71140001 | 71240000 | 0.0036 | 0.0038 | 0.0027 |
| F10 | 71160001 | 71260000 | 0.0050 | 0.0044 | 0.0041 |
| F10 | 71180001 | 71280000 | 0.0057 | 0.0040 | 0.0046 |
| F10 | 71200001 | 71300000 | 0.0050 | 0.0039 | 0.0043 |
| F10 | 71220001 | 71320000 | 0.0049 | 0.0037 | 0.0040 |
| F10 | 71240001 | 71340000 | 0.0048 | 0.0020 | 0.0037 |
| F10 | 71260001 | 71360000 | 0.0028 | 0.0015 | 0.0019 |
| F10 | 71280001 | 71380000 | 0.0026 | 0.0019 | 0.0017 |
| F10 | 71300001 | 71400000 | 0.0033 | 0.0022 | 0.0021 |
| F10 | 71320001 | 71420000 | 0.0036 | 0.0028 | 0.0024 |
| F10 | 71340001 | 71440000 | 0.0042 | 0.0026 | 0.0031 |
| F10 | 71360001 | 71460000 | 0.0041 | 0.0024 | 0.0030 |
| F10 | 71380001 | 71480000 | 0.0035 | 0.0024 | 0.0025 |
| F10 | 71400001 | 71500000 | 0.0033 | 0.0028 | 0.0026 |
| F10 | 71420001 | 71520000 | 0.0035 | 0.0020 | 0.0030 |
| F10 | 71440001 | 71540000 | 0.0027 | 0.0023 | 0.0022 |
| F10 | 71460001 | 71560000 | 0.0031 | 0.0023 | 0.0026 |
| F10 | 71480001 | 71580000 | 0.0030 | 0.0024 | 0.0026 |
| F10 | 71500001 | 71600000 | 0.0031 | 0.0020 | 0.0025 |
| F10 | 71520001 | 71620000 | 0.0027 | 0.0027 | 0.0022 |
| F10 | 71540001 | 71640000 | 0.0035 | 0.0030 | 0.0027 |
| F10 | 71560001 | 71660000 | 0.0039 | 0.0038 | 0.0030 |
| F10 | 71580001 | 71680000 | 0.0047 | 0.0038 | 0.0040 |
| F10 | 71600001 | 71700000 | 0.0049 | 0.0042 | 0.0042 |
| F10 | 71620001 | 71720000 | 0.0052 | 0.0035 | 0.0045 |
| F10 | 71640001 | 71740000 | 0.0042 | 0.0035 | 0.0039 |
| F10 | 71660001 | 71760000 | 0.0040 | 0.0033 | 0.0039 |
| F10 | 71680001 | 71780000 | 0.0038 | 0.0029 | 0.0037 |
| F10 | 71700001 | 71800000 | 0.0033 | 0.0027 | 0.0032 |
| F10 | 71720001 | 71820000 | 0.0033 | 0.0030 | 0.0032 |
| F10 | 71740001 | 71840000 | 0.0034 | 0.0034 | 0.0033 |
| F10 | 71760001 | 71860000 | 0.0040 | 0.0029 | 0.0039 |
| F10 | 71780001 | 71880000 | 0.0036 | 0.0031 | 0.0033 |
| F10 | 71800001 | 71900000 | 0.0039 | 0.0033 | 0.0034 |
| F10 | 71820001 | 71920000 | 0.0040 | 0.0029 | 0.0036 |
| F10 | 71840001 | 71940000 | 0.0037 | 0.0020 | 0.0033 |
| F10 | 71860001 | 71960000 | 0.0027 | 0.0019 | 0.0023 |
| F10 | 71880001 | 71980000 | 0.0024 | 0.0017 | 0.0021 |
| F10 | 71900001 | 72000000 | 0.0021 | 0.0012 | 0.0021 |
| F10 | 71920001 | 72020000 | 0.0016 | 0.0016 | 0.0016 |
| F10 | 71940001 | 72040000 | 0.0020 | 0.0015 | 0.0019 |
| F10 | 71960001 | 72060000 | 0.0019 | 0.0015 | 0.0018 |
| F10 | 71980001 | 72080000 | 0.0020 | 0.0018 | 0.0018 |
| F10 | 72000001 | 72100000 | 0.0022 | 0.0017 | 0.0020 |
| F10 | 72020001 | 72120000 | 0.0022 | 0.0018 | 0.0018 |
| F10 | 72040001 | 72140000 | 0.0023 | 0.0016 | 0.0018 |
| F10 | 72060001 | 72160000 | 0.0023 | 0.0016 | 0.0016 |
| F10 | 72080001 | 72180000 | 0.0023 | 0.0012 | 0.0017 |
| F10 | 72100001 | 72200000 | 0.0020 | 0.0020 | 0.0013 |
| F10 | 72120001 | 72220000 | 0.0028 | 0.0023 | 0.0021 |
| F10 | 72140001 | 72240000 | 0.0031 | 0.0026 | 0.0025 |
| F10 | 72160001 | 72260000 | 0.0036 | 0.0026 | 0.0028 |
| F10 | 72180001 | 72280000 | 0.0037 | 0.0034 | 0.0027 |
| F10 | 72200001 | 72300000 | 0.0046 | 0.0029 | 0.0037 |
| F10 | 72220001 | 72320000 | 0.0039 | 0.0027 | 0.0031 |

| Chr | Start | End | both | A1 | A2 |
|---|---|---|---|---|---|
| F10 | 72240001 | 72340000 | 0.0035 | 0.0030 | 0.0028 |
| F10 | 72260001 | 72360000 | 0.0037 | 0.0032 | 0.0032 |
| F10 | 72280001 | 72380000 | 0.0037 | 0.0026 | 0.0034 |
| F10 | 72300001 | 72400000 | 0.0031 | 0.0025 | 0.0027 |
| F10 | 72320001 | 72420000 | 0.0033 | 0.0025 | 0.0025 |
| F10 | 72340001 | 72440000 | 0.0034 | 0.0025 | 0.0025 |
| F10 | 72360001 | 72460000 | 0.0035 | 0.0030 | 0.0025 |
| F10 | 72380001 | 72480000 | 0.0041 | 0.0030 | 0.0030 |
| F10 | 72400001 | 72500000 | 0.0041 | 0.0030 | 0.0032 |
| F10 | 72420001 | 72520000 | 0.0040 | 0.0031 | 0.0032 |
| F10 | 72440001 | 72540000 | 0.0039 | 0.0028 | 0.0031 |
| F10 | 72460001 | 72560000 | 0.0036 | 0.0022 | 0.0027 |
| F10 | 72480001 | 72580000 | 0.0029 | 0.0021 | 0.0020 |
| F10 | 72500001 | 72600000 | 0.0027 | 0.0024 | 0.0018 |
| F10 | 72520001 | 72620000 | 0.0030 | 0.0023 | 0.0022 |
| F10 | 72540001 | 72640000 | 0.0032 | 0.0028 | 0.0025 |
| F10 | 72560001 | 72660000 | 0.0036 | 0.0029 | 0.0030 |
| F10 | 72580001 | 72680000 | 0.0037 | 0.0038 | 0.0030 |
| F10 | 72600001 | 72700000 | 0.0048 | 0.0037 | 0.0038 |
| F10 | 72620001 | 72720000 | 0.0047 | 0.0034 | 0.0037 |
| F10 | 72640001 | 72740000 | 0.0043 | 0.0032 | 0.0030 |
| F10 | 72660001 | 72760000 | 0.0041 | 0.0034 | 0.0027 |
| F10 | 72680001 | 72780000 | 0.0044 | 0.0030 | 0.0030 |
| F10 | 72700001 | 72800000 | 0.0043 | 0.0028 | 0.0026 |
| F10 | 72720001 | 72820000 | 0.0040 | 0.0027 | 0.0024 |
| F10 | 72740001 | 72840000 | 0.0039 | 0.0025 | 0.0025 |
| F10 | 72760001 | 72860000 | 0.0037 | 0.0022 | 0.0024 |
| F10 | 72780001 | 72880000 | 0.0032 | 0.0017 | 0.0022 |
| F10 | 72800001 | 72900000 | 0.0025 | 0.0021 | 0.0017 |
| F10 | 72820001 | 72920000 | 0.0030 | 0.0020 | 0.0021 |
| F10 | 72840001 | 72940000 | 0.0029 | 0.0017 | 0.0021 |
| F10 | 72860001 | 72960000 | 0.0027 | 0.0020 | 0.0019 |
| F10 | 72880001 | 72980000 | 0.0031 | 0.0026 | 0.0022 |
| F10 | 72900001 | 73000000 | 0.0037 | 0.0023 | 0.0031 |
| F10 | 72920001 | 73020000 | 0.0032 | 0.0029 | 0.0026 |
| F10 | 72940001 | 73040000 | 0.0042 | 0.0032 | 0.0031 |
| F10 | 72960001 | 73060000 | 0.0043 | 0.0032 | 0.0035 |
| F10 | 72980001 | 73080000 | 0.0042 | 0.0030 | 0.0034 |
| F10 | 73000001 | 73100000 | 0.0040 | 0.0033 | 0.0032 |
| F10 | 73020001 | 73120000 | 0.0044 | 0.0034 | 0.0036 |
| F10 | 73040001 | 73140000 | 0.0043 | 0.0041 | 0.0036 |
| F10 | 73060001 | 73160000 | 0.0049 | 0.0043 | 0.0041 |
| F10 | 73080001 | 73180000 | 0.0052 | 0.0037 | 0.0042 |
| F10 | 73100001 | 73200000 | 0.0044 | 0.0034 | 0.0034 |
| F10 | 73120001 | 73220000 | 0.0040 | 0.0027 | 0.0031 |
| F10 | 73140001 | 73240000 | 0.0034 | 0.0016 | 0.0024 |
| F10 | 73160001 | 73260000 | 0.0023 | 0.0013 | 0.0016 |
| F10 | 73180001 | 73280000 | 0.0020 | 0.0015 | 0.0012 |
| F10 | 73200001 | 73300000 | 0.0023 | 0.0021 | 0.0015 |
| F10 | 73220001 | 73320000 | 0.0030 | 0.0024 | 0.0023 |
| F10 | 73240001 | 73340000 | 0.0033 | 0.0028 | 0.0026 |
| F10 | 73260001 | 73360000 | 0.0039 | 0.0030 | 0.0030 |
| F10 | 73280001 | 73380000 | 0.0039 | 0.0035 | 0.0032 |
| F10 | 73300001 | 73400000 | 0.0043 | 0.0030 | 0.0038 |
| F10 | 73320001 | 73420000 | 0.0041 | 0.0029 | 0.0034 |
| F10 | 73340001 | 73440000 | 0.0040 | 0.0024 | 0.0033 |
| F10 | 73360001 | 73460000 | 0.0034 | 0.0021 | 0.0027 |
| F10 | 73380001 | 73480000 | 0.0032 | 0.0014 | 0.0024 |

| Chr | Start | End | both | A1 | A2 |
|---|---|---|---|---|---|
| F10 | 73400001 | 73500000 | 0.0024 | 0.0016 | 0.0016 |
| F10 | 73420001 | 73520000 | 0.0024 | 0.0024 | 0.0018 |
| F10 | 73440001 | 73540000 | 0.0033 | 0.0031 | 0.0027 |
| F10 | 73460001 | 73560000 | 0.0038 | 0.0034 | 0.0034 |
| F10 | 73480001 | 73580000 | 0.0042 | 0.0045 | 0.0038 |
| F10 | 73500001 | 73600000 | 0.0054 | 0.0042 | 0.0049 |
| F10 | 73520001 | 73620000 | 0.0052 | 0.0042 | 0.0046 |
| F10 | 73540001 | 73640000 | 0.0049 | 0.0036 | 0.0043 |
| F10 | 73560001 | 73660000 | 0.0045 | 0.0034 | 0.0037 |
| F10 | 73580001 | 73680000 | 0.0044 | 0.0026 | 0.0034 |
| F10 | 73600001 | 73700000 | 0.0038 | 0.0026 | 0.0027 |
| F10 | 73620001 | 73720000 | 0.0037 | 0.0027 | 0.0028 |
| F10 | 73640001 | 73740000 | 0.0040 | 0.0032 | 0.0031 |
| F10 | 73660001 | 73760000 | 0.0043 | 0.0034 | 0.0034 |
| F10 | 73680001 | 73780000 | 0.0043 | 0.0040 | 0.0037 |
| F10 | 73700001 | 73800000 | 0.0046 | 0.0040 | 0.0042 |
| F10 | 73720001 | 73820000 | 0.0046 | 0.0028 | 0.0042 |
| F10 | 73740001 | 73840000 | 0.0033 | 0.0023 | 0.0028 |
| F10 | 73760001 | 73860000 | 0.0027 | 0.0024 | 0.0025 |
| F10 | 73780001 | 73880000 | 0.0028 | 0.0020 | 0.0026 |
| F10 | 73800001 | 73900000 | 0.0024 | 0.0018 | 0.0022 |
| F10 | 73820001 | 73920000 | 0.0022 | 0.0020 | 0.0020 |
| F10 | 73840001 | 73940000 | 0.0025 | 0.0025 | 0.0022 |
| F10 | 73860001 | 73960000 | 0.0031 | 0.0035 | 0.0027 |
| F10 | 73880001 | 73980000 | 0.0040 | 0.0036 | 0.0036 |
| F10 | 73900001 | 74000000 | 0.0042 | 0.0037 | 0.0037 |
| F10 | 73920001 | 74020000 | 0.0042 | 0.0045 | 0.0037 |
| F10 | 73940001 | 74040000 | 0.0050 | 0.0043 | 0.0046 |
| F10 | 73960001 | 74060000 | 0.0049 | 0.0026 | 0.0044 |
| F10 | 73980001 | 74080000 | 0.0031 | 0.0021 | 0.0027 |
| F10 | 74000001 | 74100000 | 0.0025 | 0.0019 | 0.0020 |
| F10 | 74020001 | 74120000 | 0.0023 | 0.0008 | 0.0019 |
| F10 | 74040001 | 74140000 | 0.0013 | 0.0007 | 0.0009 |
| F10 | 74060001 | 74160000 | 0.0012 | 0.0011 | 0.0008 |
| F10 | 74080001 | 74180000 | 0.0018 | 0.0018 | 0.0012 |
| F10 | 74100001 | 74200000 | 0.0024 | 0.0031 | 0.0019 |
| F10 | 74120001 | 74220000 | 0.0039 | 0.0035 | 0.0032 |
| F10 | 74140001 | 74240000 | 0.0041 | 0.0037 | 0.0035 |
| F10 | 74160001 | 74260000 | 0.0040 | 0.0034 | 0.0035 |
| F10 | 74180001 | 74280000 | 0.0035 | 0.0038 | 0.0031 |
| F10 | 74200001 | 74300000 | 0.0039 | 0.0033 | 0.0037 |
| F10 | 74220001 | 74320000 | 0.0033 | 0.0045 | 0.0031 |
| F10 | 74240001 | 74340000 | 0.0044 | 0.0050 | 0.0043 |
| F10 | 74260001 | 74360000 | 0.0051 | 0.0052 | 0.0049 |
| F10 | 74280001 | 74380000 | 0.0053 | 0.0046 | 0.0051 |
| F10 | 74300001 | 74400000 | 0.0045 | 0.0040 | 0.0043 |
| F10 | 74320001 | 74420000 | 0.0039 | 0.0024 | 0.0037 |
| F10 | 74340001 | 74440000 | 0.0025 | 0.0015 | 0.0024 |
| F10 | 74360001 | 74460000 | 0.0015 | 0.0020 | 0.0015 |
| F10 | 74380001 | 74480000 | 0.0020 | 0.0019 | 0.0020 |
| F10 | 74400001 | 74500000 | 0.0020 | 0.0018 | 0.0019 |
| F10 | 74420001 | 74520000 | 0.0017 | 0.0020 | 0.0016 |
| F10 | 74440001 | 74540000 | 0.0018 | 0.0027 | 0.0016 |
| F10 | 74460001 | 74560000 | 0.0025 | 0.0030 | 0.0022 |
| F10 | 74480001 | 74580000 | 0.0027 | 0.0035 | 0.0025 |
| F10 | 74500001 | 74600000 | 0.0034 | 0.0045 | 0.0032 |
| F10 | 74520001 | 74620000 | 0.0045 | 0.0044 | 0.0043 |
| F10 | 74540001 | 74640000 | 0.0044 | 0.0040 | 0.0043 |

| Chr | Start | End | both | A1 | A2 |
|---|---|---|---|---|---|
| F10 | 74560001 | 74660000 | 0.0039 | 0.0035 | 0.0037 |
| F10 | 74580001 | 74680000 | 0.0035 | 0.0032 | 0.0033 |
| F10 | 74600001 | 74700000 | 0.0030 | 0.0023 | 0.0028 |
| F10 | 74620001 | 74720000 | 0.0021 | 0.0024 | 0.0019 |
| F10 | 74640001 | 74740000 | 0.0021 | 0.0026 | 0.0020 |
| F10 | 74660001 | 74760000 | 0.0024 | 0.0026 | 0.0022 |
| F10 | 74680001 | 74780000 | 0.0021 | 0.0026 | 0.0019 |
| F10 | 74700001 | 74800000 | 0.0020 | 0.0031 | 0.0018 |
| F10 | 74720001 | 74820000 | 0.0026 | 0.0032 | 0.0023 |
| F10 | 74740001 | 74840000 | 0.0026 | 0.0033 | 0.0023 |
| F10 | 74760001 | 74860000 | 0.0026 | 0.0033 | 0.0024 |
| F10 | 74780001 | 74880000 | 0.0030 | 0.0038 | 0.0028 |
| F10 | 74800001 | 74900000 | 0.0036 | 0.0043 | 0.0035 |
| F10 | 74820001 | 74920000 | 0.0045 | 0.0045 | 0.0044 |
| F10 | 74840001 | 74940000 | 0.0048 | 0.0046 | 0.0047 |
| F10 | 74860001 | 74960000 | 0.0049 | 0.0043 | 0.0048 |
| F10 | 74880001 | 74980000 | 0.0045 | 0.0039 | 0.0043 |
| F10 | 74900001 | 75000000 | 0.0040 | 0.0026 | 0.0038 |
| F10 | 74920001 | 75020000 | 0.0024 | 0.0027 | 0.0022 |
| F10 | 74940001 | 75040000 | 0.0025 | 0.0021 | 0.0024 |
| F10 | 74960001 | 75060000 | 0.0020 | 0.0024 | 0.0018 |
| F10 | 74980001 | 75080000 | 0.0023 | 0.0027 | 0.0021 |
| F10 | 75000001 | 75100000 | 0.0028 | 0.0030 | 0.0027 |
| F10 | 75020001 | 75120000 | 0.0030 | 0.0047 | 0.0028 |
| F10 | 75040001 | 75140000 | 0.0047 | 0.0050 | 0.0045 |
| F10 | 75060001 | 75160000 | 0.0050 | 0.0046 | 0.0049 |
| F10 | 75080001 | 75180000 | 0.0047 | 0.0040 | 0.0046 |
| F10 | 75100001 | 75200000 | 0.0042 | 0.0044 | 0.0041 |
| F10 | 75120001 | 75220000 | 0.0050 | 0.0026 | 0.0049 |
| F10 | 75140001 | 75240000 | 0.0032 | 0.0024 | 0.0032 |
| F10 | 75160001 | 75260000 | 0.0029 | 0.0022 | 0.0029 |
| F10 | 75180001 | 75280000 | 0.0027 | 0.0021 | 0.0027 |
| F10 | 75200001 | 75300000 | 0.0026 | 0.0016 | 0.0026 |
| F10 | 75220001 | 75320000 | 0.0019 | 0.0014 | 0.0019 |
| F10 | 75240001 | 75340000 | 0.0016 | 0.0017 | 0.0016 |
| F10 | 75260001 | 75360000 | 0.0019 | 0.0024 | 0.0019 |
| F10 | 75280001 | 75380000 | 0.0026 | 0.0027 | 0.0026 |
| F10 | 75300001 | 75400000 | 0.0029 | 0.0026 | 0.0029 |
| F10 | 75320001 | 75420000 | 0.0027 | 0.0029 | 0.0027 |
| F10 | 75340001 | 75440000 | 0.0031 | 0.0027 | 0.0031 |
| F10 | 75360001 | 75460000 | 0.0029 | 0.0037 | 0.0029 |
| F10 | 75380001 | 75480000 | 0.0040 | 0.0047 | 0.0040 |
| F10 | 75400001 | 75500000 | 0.0049 | 0.0049 | 0.0049 |
| F10 | 75420001 | 75520000 | 0.0051 | 0.0052 | 0.0051 |
| F10 | 75440001 | 75540000 | 0.0055 | 0.0067 | 0.0055 |
| F10 | 75460001 | 75560000 | 0.0071 | 0.0054 | 0.0071 |
| F10 | 75480001 | 75580000 | 0.0058 | 0.0042 | 0.0058 |
| F10 | 75500001 | 75600000 | 0.0045 | 0.0045 | 0.0045 |
| F10 | 75520001 | 75620000 | 0.0048 | 0.0036 | 0.0048 |
| F10 | 75540001 | 75640000 | 0.0039 | 0.0027 | 0.0039 |
| F10 | 75560001 | 75660000 | 0.0029 | 0.0035 | 0.0029 |
| F10 | 75580001 | 75680000 | 0.0036 | 0.0043 | 0.0036 |
| F10 | 75600001 | 75700000 | 0.0045 | 0.0039 | 0.0045 |
| F10 | 75620001 | 75720000 | 0.0042 | 0.0043 | 0.0042 |
| F10 | 75640001 | 75740000 | 0.0045 | 0.0042 | 0.0045 |
| F10 | 75660001 | 75760000 | 0.0044 | 0.0041 | 0.0044 |
| F10 | 75680001 | 75780000 | 0.0043 | 0.0035 | 0.0043 |
| F10 | 75700001 | 75800000 | 0.0037 | 0.0037 | 0.0037 |

| Chr | Start | End | both | A1 | A2 |
|---|---|---|---|---|---|
| F10 | 75720001 | 75820000 | 0.0039 | 0.0036 | 0.0039 |
| F10 | 75740001 | 75840000 | 0.0038 | 0.0034 | 0.0038 |
| F10 | 75760001 | 75860000 | 0.0036 | 0.0029 | 0.0036 |
| F10 | 75780001 | 75880000 | 0.0031 | 0.0026 | 0.0031 |
| F10 | 75800001 | 75900000 | 0.0028 | 0.0024 | 0.0027 |
| F10 | 75820001 | 75920000 | 0.0024 | 0.0026 | 0.0024 |
| F10 | 75840001 | 75940000 | 0.0030 | 0.0027 | 0.0030 |
| F10 | 75860001 | 75960000 | 0.0031 | 0.0029 | 0.0031 |
| F10 | 75880001 | 75980000 | 0.0034 | 0.0032 | 0.0033 |
| F10 | 75900001 | 76000000 | 0.0036 | 0.0034 | 0.0036 |
| F10 | 75920001 | 76020000 | 0.0039 | 0.0028 | 0.0039 |
| F10 | 75940001 | 76040000 | 0.0030 | 0.0029 | 0.0030 |
| F10 | 75960001 | 76060000 | 0.0030 | 0.0035 | 0.0029 |
| F10 | 75980001 | 76080000 | 0.0038 | 0.0030 | 0.0037 |
| F10 | 76000001 | 76100000 | 0.0033 | 0.0027 | 0.0032 |
| F10 | 76020001 | 76120000 | 0.0029 | 0.0033 | 0.0028 |
| F10 | 76040001 | 76140000 | 0.0036 | 0.0030 | 0.0036 |
| F10 | 76060001 | 76160000 | 0.0033 | 0.0016 | 0.0032 |
| F10 | 76080001 | 76180000 | 0.0016 | 0.0019 | 0.0016 |
| F10 | 76100001 | 76200000 | 0.0019 | 0.0036 | 0.0019 |
| F10 | 76120001 | 76220000 | 0.0038 | 0.0027 | 0.0037 |
| F10 | 76140001 | 76240000 | 0.0029 | 0.0030 | 0.0028 |
| F10 | 76160001 | 76260000 | 0.0032 | 0.0045 | 0.0031 |
| F10 | 76180001 | 76280000 | 0.0049 | 0.0046 | 0.0049 |
| F10 | 76200001 | 76300000 | 0.0052 | 0.0038 | 0.0052 |
| F10 | 76220001 | 76320000 | 0.0044 | 0.0041 | 0.0044 |
| F10 | 76240001 | 76340000 | 0.0047 | 0.0041 | 0.0047 |
| F10 | 76260001 | 76360000 | 0.0047 | 0.0032 | 0.0046 |
| F10 | 76280001 | 76380000 | 0.0038 | 0.0049 | 0.0037 |
| F10 | 76300001 | 76400000 | 0.0056 | 0.0042 | 0.0053 |
| F10 | 76320001 | 76420000 | 0.0050 | 0.0042 | 0.0047 |
| F10 | 76340001 | 76440000 | 0.0055 | 0.0041 | 0.0052 |
| F10 | 76360001 | 76460000 | 0.0053 | 0.0034 | 0.0051 |
| F10 | 76380001 | 76480000 | 0.0044 | 0.0013 | 0.0043 |
| F10 | 76400001 | 76500000 | 0.0021 | 0.0017 | 0.0022 |
| F10 | 76420001 | 76520000 | 0.0023 | 0.0022 | 0.0023 |
| F10 | 76440001 | 76540000 | 0.0022 | 0.0029 | 0.0021 |
| F10 | 76460001 | 76560000 | 0.0027 | 0.0030 | 0.0026 |
| F10 | 76480001 | 76580000 | 0.0027 | 0.0033 | 0.0025 |
| F10 | 76500001 | 76600000 | 0.0029 | 0.0028 | 0.0027 |
| F10 | 76520001 | 76620000 | 0.0023 | 0.0027 | 0.0020 |
| F10 | 76540001 | 76640000 | 0.0024 | 0.0019 | 0.0021 |
| F10 | 76560001 | 76660000 | 0.0016 | 0.0023 | 0.0014 |
| F10 | 76580001 | 76680000 | 0.0022 | 0.0021 | 0.0020 |
| F10 | 76600001 | 76700000 | 0.0020 | 0.0024 | 0.0018 |
| F10 | 76620001 | 76720000 | 0.0024 | 0.0020 | 0.0022 |
| F10 | 76640001 | 76740000 | 0.0019 | 0.0021 | 0.0017 |
| F10 | 76660001 | 76760000 | 0.0020 | 0.0016 | 0.0018 |
| F10 | 76680001 | 76780000 | 0.0015 | 0.0019 | 0.0014 |
| F10 | 76700001 | 76800000 | 0.0018 | 0.0015 | 0.0017 |
| F10 | 76720001 | 76820000 | 0.0014 | 0.0016 | 0.0012 |
| F10 | 76740001 | 76840000 | 0.0015 | 0.0020 | 0.0013 |
| F10 | 76760001 | 76860000 | 0.0018 | 0.0021 | 0.0016 |
| F10 | 76780001 | 76880000 | 0.0017 | 0.0023 | 0.0015 |
| F10 | 76800001 | 76900000 | 0.0020 | 0.0024 | 0.0018 |
| F10 | 76820001 | 76920000 | 0.0022 | 0.0024 | 0.0020 |
| F10 | 76840001 | 76940000 | 0.0023 | 0.0036 | 0.0021 |
| F10 | 76860001 | 76960000 | 0.0036 | 0.0038 | 0.0035 |

| Chr | Start | End | both | A1 | A2 |
|---|---|---|---|---|---|
| F10 | 76880001 | 76980000 | 0.0039 | 0.0035 | 0.0037 |
| F10 | 76900001 | 77000000 | 0.0036 | 0.0044 | 0.0035 |
| F10 | 76920001 | 77020000 | 0.0046 | 0.0066 | 0.0045 |
| F10 | 76940001 | 77040000 | 0.0068 | 0.0054 | 0.0067 |
| F10 | 76960001 | 77060000 | 0.0055 | 0.0057 | 0.0054 |
| F10 | 76980001 | 77080000 | 0.0057 | 0.0066 | 0.0055 |
| F10 | 77000001 | 77100000 | 0.0066 | 0.0056 | 0.0064 |
| F10 | 77020001 | 77120000 | 0.0055 | 0.0034 | 0.0053 |
| F10 | 77040001 | 77140000 | 0.0031 | 0.0031 | 0.0029 |
| F10 | 77060001 | 77160000 | 0.0028 | 0.0028 | 0.0026 |
| F10 | 77080001 | 77180000 | 0.0026 | 0.0031 | 0.0024 |
| F10 | 77100001 | 77200000 | 0.0029 | 0.0051 | 0.0027 |
| F10 | 77120001 | 77220000 | 0.0051 | 0.0054 | 0.0049 |
| F10 | 77140001 | 77240000 | 0.0053 | 0.0065 | 0.0051 |
| F10 | 77160001 | 77260000 | 0.0067 | 0.0066 | 0.0065 |
| F10 | 77180001 | 77280000 | 0.0068 | 0.0057 | 0.0066 |
| F10 | 77200001 | 77300000 | 0.0057 | 0.0039 | 0.0054 |
| F10 | 77220001 | 77320000 | 0.0039 | 0.0037 | 0.0036 |
| F10 | 77240001 | 77340000 | 0.0038 | 0.0030 | 0.0036 |
| F10 | 77260001 | 77360000 | 0.0029 | 0.0037 | 0.0027 |
| F10 | 77280001 | 77380000 | 0.0037 | 0.0037 | 0.0036 |
| F10 | 77300001 | 77400000 | 0.0040 | 0.0038 | 0.0038 |
| F10 | 77320001 | 77420000 | 0.0037 | 0.0039 | 0.0035 |
| F10 | 77340001 | 77440000 | 0.0038 | 0.0034 | 0.0036 |
| F10 | 77360001 | 77460000 | 0.0033 | 0.0027 | 0.0032 |
| F10 | 77380001 | 77480000 | 0.0027 | 0.0024 | 0.0025 |
| F10 | 77400001 | 77500000 | 0.0023 | 0.0029 | 0.0021 |
| F10 | 77420001 | 77520000 | 0.0033 | 0.0029 | 0.0032 |
| F10 | 77440001 | 77540000 | 0.0035 | 0.0031 | 0.0035 |
| F10 | 77460001 | 77560000 | 0.0036 | 0.0029 | 0.0035 |
| F10 | 77480001 | 77580000 | 0.0032 | 0.0035 | 0.0031 |
| F10 | 77500001 | 77600000 | 0.0038 | 0.0026 | 0.0036 |
| F10 | 77520001 | 77620000 | 0.0026 | 0.0023 | 0.0024 |
| F10 | 77540001 | 77640000 | 0.0021 | 0.0027 | 0.0019 |
| F10 | 77560001 | 77660000 | 0.0027 | 0.0028 | 0.0025 |
| F10 | 77580001 | 77680000 | 0.0030 | 0.0022 | 0.0029 |
| F10 | 77600001 | 77700000 | 0.0025 | 0.0027 | 0.0024 |
| F10 | 77620001 | 77720000 | 0.0029 | 0.0028 | 0.0028 |
| F10 | 77640001 | 77740000 | 0.0030 | 0.0033 | 0.0028 |
| F10 | 77660001 | 77760000 | 0.0034 | 0.0039 | 0.0032 |
| F10 | 77680001 | 77780000 | 0.0041 | 0.0041 | 0.0040 |
| F10 | 77700001 | 77800000 | 0.0045 | 0.0046 | 0.0044 |
| F10 | 77720001 | 77820000 | 0.0048 | 0.0048 | 0.0047 |
| F10 | 77740001 | 77840000 | 0.0051 | 0.0039 | 0.0049 |
| F10 | 77760001 | 77860000 | 0.0041 | 0.0032 | 0.0040 |
| F10 | 77780001 | 77880000 | 0.0031 | 0.0027 | 0.0029 |
| F10 | 77800001 | 77900000 | 0.0025 | 0.0020 | 0.0023 |
| F10 | 77820001 | 77920000 | 0.0018 | 0.0021 | 0.0016 |
| F10 | 77840001 | 77940000 | 0.0018 | 0.0020 | 0.0017 |
| F10 | 77860001 | 77960000 | 0.0018 | 0.0020 | 0.0016 |
| F10 | 77880001 | 77980000 | 0.0018 | 0.0022 | 0.0017 |
| F10 | 77900001 | 78000000 | 0.0019 | 0.0024 | 0.0018 |
| F10 | 77920001 | 78020000 | 0.0019 | 0.0021 | 0.0017 |
| F10 | 77940001 | 78040000 | 0.0017 | 0.0022 | 0.0015 |
| F10 | 77960001 | 78060000 | 0.0019 | 0.0026 | 0.0018 |
| F10 | 77980001 | 78080000 | 0.0023 | 0.0030 | 0.0022 |
| F10 | 78000001 | 78100000 | 0.0027 | 0.0031 | 0.0025 |
| F10 | 78020001 | 78120000 | 0.0030 | 0.0028 | 0.0029 |

| Chr | Start | End | both | A1 | A2 |
|---|---|---|---|---|---|
| F10 | 78040001 | 78140000 | 0.0028 | 0.0029 | 0.0027 |
| F10 | 78060001 | 78160000 | 0.0029 | 0.0026 | 0.0027 |
| F10 | 78080001 | 78180000 | 0.0024 | 0.0027 | 0.0023 |
| F10 | 78100001 | 78200000 | 0.0023 | 0.0027 | 0.0020 |
| F10 | 78120001 | 78220000 | 0.0020 | 0.0032 | 0.0016 |
| F10 | 78140001 | 78240000 | 0.0022 | 0.0035 | 0.0016 |
| F10 | 78160001 | 78260000 | 0.0027 | 0.0033 | 0.0021 |
| F10 | 78180001 | 78280000 | 0.0025 | 0.0025 | 0.0019 |
| F10 | 78200001 | 78300000 | 0.0023 | 0.0024 | 0.0018 |
| F10 | 78220001 | 78320000 | 0.0024 | 0.0015 | 0.0020 |
| F10 | 78240001 | 78340000 | 0.0018 | 0.0008 | 0.0016 |
| F10 | 78260001 | 78360000 | 0.0010 | 0.0008 | 0.0009 |
| F10 | 78280001 | 78380000 | 0.0012 | 0.0007 | 0.0009 |
| F10 | 78300001 | 78400000 | 0.0015 | 0.0009 | 0.0009 |
| F10 | 78320001 | 78420000 | 0.0019 | 0.0012 | 0.0011 |
| F10 | 78340001 | 78440000 | 0.0025 | 0.0012 | 0.0016 |
| F10 | 78360001 | 78460000 | 0.0029 | 0.0016 | 0.0017 |
| F10 | 78380001 | 78480000 | 0.0033 | 0.0020 | 0.0021 |
| F10 | 78400001 | 78500000 | 0.0033 | 0.0023 | 0.0024 |
| F10 | 78420001 | 78520000 | 0.0034 | 0.0020 | 0.0027 |
| F10 | 78440001 | 78540000 | 0.0031 | 0.0021 | 0.0023 |
| F10 | 78460001 | 78560000 | 0.0030 | 0.0021 | 0.0024 |
| F10 | 78480001 | 78580000 | 0.0033 | 0.0023 | 0.0026 |
| F10 | 78500001 | 78600000 | 0.0037 | 0.0016 | 0.0029 |
| F10 | 78520001 | 78620000 | 0.0030 | 0.0016 | 0.0021 |
| F10 | 78540001 | 78640000 | 0.0032 | 0.0014 | 0.0024 |
| F10 | 78560001 | 78660000 | 0.0028 | 0.0009 | 0.0021 |
| F10 | 78580001 | 78680000 | 0.0021 | 0.0010 | 0.0015 |
| F10 | 78600001 | 78700000 | 0.0021 | 0.0013 | 0.0016 |
| F10 | 78620001 | 78720000 | 0.0025 | 0.0013 | 0.0020 |
| F10 | 78640001 | 78740000 | 0.0022 | 0.0015 | 0.0018 |
| F10 | 78660001 | 78760000 | 0.0025 | 0.0016 | 0.0019 |
| F10 | 78680001 | 78780000 | 0.0029 | 0.0009 | 0.0019 |
| F10 | 78700001 | 78800000 | 0.0022 | 0.0012 | 0.0012 |
| F10 | 78720001 | 78820000 | 0.0023 | 0.0014 | 0.0013 |
| F10 | 78740001 | 78840000 | 0.0027 | 0.0019 | 0.0015 |
| F10 | 78760001 | 78860000 | 0.0030 | 0.0019 | 0.0021 |
| F10 | 78780001 | 78880000 | 0.0027 | 0.0024 | 0.0020 |
| F10 | 78800001 | 78900000 | 0.0033 | 0.0019 | 0.0026 |
| F10 | 78820001 | 78920000 | 0.0028 | 0.0019 | 0.0020 |
| F10 | 78840001 | 78940000 | 0.0028 | 0.0014 | 0.0020 |
| F10 | 78860001 | 78960000 | 0.0023 | 0.0012 | 0.0015 |
| F10 | 78880001 | 78980000 | 0.0023 | 0.0021 | 0.0015 |
| F10 | 78900001 | 79000000 | 0.0030 | 0.0030 | 0.0022 |
| F10 | 78920001 | 79020000 | 0.0040 | 0.0027 | 0.0033 |
| F10 | 78940001 | 79040000 | 0.0037 | 0.0033 | 0.0032 |
| F10 | 78960001 | 79060000 | 0.0043 | 0.0036 | 0.0039 |
| F10 | 78980001 | 79080000 | 0.0046 | 0.0029 | 0.0043 |
| F10 | 79000001 | 79100000 | 0.0039 | 0.0023 | 0.0037 |
| F10 | 79020001 | 79120000 | 0.0034 | 0.0024 | 0.0032 |
| F10 | 79040001 | 79140000 | 0.0037 | 0.0023 | 0.0034 |
| F10 | 79060001 | 79160000 | 0.0039 | 0.0027 | 0.0035 |
| F10 | 79080001 | 79180000 | 0.0044 | 0.0024 | 0.0038 |
| F10 | 79100001 | 79200000 | 0.0043 | 0.0033 | 0.0037 |
| F10 | 79120001 | 79220000 | 0.0051 | 0.0033 | 0.0045 |
| F10 | 79140001 | 79240000 | 0.0048 | 0.0031 | 0.0042 |
| F10 | 79160001 | 79260000 | 0.0046 | 0.0030 | 0.0041 |
| F10 | 79180001 | 79280000 | 0.0043 | 0.0038 | 0.0039 |

| Chr | Start | End | both | A1 | A2 |
|---|---|---|---|---|---|
| F10 | 79200001 | 79300000 | 0.0054 | 0.0047 | 0.0048 |
| F10 | 79220001 | 79320000 | 0.0061 | 0.0050 | 0.0056 |
| F10 | 79240001 | 79340000 | 0.0068 | 0.0050 | 0.0061 |
| F10 | 79260001 | 79360000 | 0.0066 | 0.0045 | 0.0060 |
| F10 | 79280001 | 79380000 | 0.0061 | 0.0039 | 0.0056 |
| F10 | 79300001 | 79400000 | 0.0054 | 0.0022 | 0.0048 |
| F10 | 79320001 | 79420000 | 0.0038 | 0.0029 | 0.0032 |
| F10 | 79340001 | 79440000 | 0.0045 | 0.0027 | 0.0039 |
| F10 | 79360001 | 79460000 | 0.0044 | 0.0027 | 0.0037 |
| F10 | 79380001 | 79480000 | 0.0045 | 0.0027 | 0.0039 |
| F10 | 79400001 | 79500000 | 0.0041 | 0.0027 | 0.0037 |
| F10 | 79420001 | 79520000 | 0.0039 | 0.0022 | 0.0035 |
| F10 | 79440001 | 79540000 | 0.0032 | 0.0030 | 0.0029 |
| F10 | 79460001 | 79560000 | 0.0038 | 0.0030 | 0.0035 |
| F10 | 79480001 | 79580000 | 0.0038 | 0.0025 | 0.0035 |
| F10 | 79500001 | 79600000 | 0.0038 | 0.0026 | 0.0036 |
| F10 | 79520001 | 79620000 | 0.0039 | 0.0020 | 0.0037 |
| F10 | 79540001 | 79640000 | 0.0037 | 0.0014 | 0.0036 |
| F10 | 79560001 | 79660000 | 0.0035 | 0.0014 | 0.0035 |
| F10 | 79580001 | 79680000 | 0.0032 | 0.0024 | 0.0033 |
| F10 | 79600001 | 79700000 | 0.0038 | 0.0037 | 0.0038 |
| F10 | 79620001 | 79720000 | 0.0052 | 0.0048 | 0.0052 |
| F10 | 79640001 | 79740000 | 0.0060 | 0.0054 | 0.0059 |
| F10 | 79660001 | 79760000 | 0.0062 | 0.0064 | 0.0062 |
| F10 | 79680001 | 79780000 | 0.0072 | 0.0059 | 0.0071 |
| F10 | 79700001 | 79800000 | 0.0070 | 0.0043 | 0.0070 |
| F10 | 79720001 | 79820000 | 0.0056 | 0.0032 | 0.0056 |
| F10 | 79740001 | 79840000 | 0.0045 | 0.0026 | 0.0045 |
| F10 | 79760001 | 79860000 | 0.0037 | 0.0016 | 0.0037 |
| F10 | 79780001 | 79880000 | 0.0030 | 0.0010 | 0.0030 |
| F10 | 79800001 | 79900000 | 0.0022 | 0.0011 | 0.0022 |
| F10 | 79820001 | 79920000 | 0.0024 | 0.0014 | 0.0025 |
| F10 | 79840001 | 79940000 | 0.0029 | 0.0013 | 0.0030 |
| F10 | 79860001 | 79960000 | 0.0030 | 0.0016 | 0.0031 |
| F10 | 79880001 | 79980000 | 0.0034 | 0.0020 | 0.0034 |
| F10 | 79900001 | 80000000 | 0.0037 | 0.0018 | 0.0037 |
| F10 | 79920001 | 80020000 | 0.0036 | 0.0014 | 0.0036 |
| F10 | 79940001 | 80040000 | 0.0029 | 0.0017 | 0.0029 |
| F10 | 79960001 | 80060000 | 0.0032 | 0.0016 | 0.0033 |
| F10 | 79980001 | 80080000 | 0.0033 | 0.0014 | 0.0035 |
| F10 | 80000001 | 80100000 | 0.0031 | 0.0022 | 0.0033 |
| F10 | 80020001 | 80120000 | 0.0039 | 0.0026 | 0.0040 |
| F10 | 80040001 | 80140000 | 0.0042 | 0.0025 | 0.0043 |
| F10 | 80060001 | 80160000 | 0.0040 | 0.0024 | 0.0041 |
| F10 | 80080001 | 80180000 | 0.0036 | 0.0023 | 0.0037 |
| F10 | 80100001 | 80200000 | 0.0034 | 0.0016 | 0.0034 |
| F10 | 80120001 | 80220000 | 0.0026 | 0.0019 | 0.0026 |
| F10 | 80140001 | 80240000 | 0.0033 | 0.0023 | 0.0034 |
| F10 | 80160001 | 80260000 | 0.0038 | 0.0022 | 0.0040 |
| F10 | 80180001 | 80280000 | 0.0036 | 0.0022 | 0.0038 |
| F10 | 80200001 | 80300000 | 0.0036 | 0.0026 | 0.0038 |
| F10 | 80220001 | 80320000 | 0.0039 | 0.0022 | 0.0040 |
| F10 | 80240001 | 80340000 | 0.0031 | 0.0022 | 0.0032 |
| F10 | 80260001 | 80360000 | 0.0028 | 0.0027 | 0.0028 |
| F10 | 80280001 | 80380000 | 0.0033 | 0.0032 | 0.0033 |
| F10 | 80300001 | 80400000 | 0.0041 | 0.0029 | 0.0042 |
| F10 | 80320001 | 80420000 | 0.0038 | 0.0031 | 0.0038 |
| F10 | 80340001 | 80440000 | 0.0040 | 0.0030 | 0.0040 |

| Chr | Start | End | both | A1 | A2 |
|---|---|---|---|---|---|
| F10 | 80360001 | 80460000 | 0.0037 | 0.0042 | 0.0038 |
| F10 | 80380001 | 80480000 | 0.0049 | 0.0038 | 0.0048 |
| F10 | 80400001 | 80500000 | 0.0041 | 0.0034 | 0.0040 |
| F10 | 80420001 | 80520000 | 0.0038 | 0.0029 | 0.0036 |
| F10 | 80440001 | 80540000 | 0.0035 | 0.0026 | 0.0034 |
| F10 | 80460001 | 80560000 | 0.0033 | 0.0009 | 0.0032 |
| F10 | 80480001 | 80580000 | 0.0018 | 0.0009 | 0.0019 |
| F10 | 80500001 | 80600000 | 0.0021 | 0.0011 | 0.0022 |
| F10 | 80520001 | 80620000 | 0.0026 | 0.0021 | 0.0027 |
| F10 | 80540001 | 80640000 | 0.0034 | 0.0020 | 0.0034 |
| F10 | 80560001 | 80660000 | 0.0038 | 0.0021 | 0.0039 |
| F10 | 80580001 | 80680000 | 0.0041 | 0.0027 | 0.0041 |
| F10 | 80600001 | 80700000 | 0.0046 | 0.0027 | 0.0045 |
| F10 | 80620001 | 80720000 | 0.0046 | 0.0019 | 0.0045 |
| F10 | 80640001 | 80740000 | 0.0041 | 0.0033 | 0.0040 |
| F10 | 80660001 | 80760000 | 0.0052 | 0.0042 | 0.0050 |
| F10 | 80680001 | 80780000 | 0.0063 | 0.0036 | 0.0060 |
| F10 | 80700001 | 80800000 | 0.0059 | 0.0038 | 0.0055 |
| F10 | 80720001 | 80820000 | 0.0063 | 0.0047 | 0.0060 |
| F10 | 80740001 | 80840000 | 0.0070 | 0.0032 | 0.0068 |
| F10 | 80760001 | 80860000 | 0.0056 | 0.0024 | 0.0055 |
| F10 | 80780001 | 80880000 | 0.0051 | 0.0024 | 0.0051 |
| F10 | 80800001 | 80900000 | 0.0051 | 0.0022 | 0.0051 |
| F10 | 80820001 | 80920000 | 0.0044 | 0.0012 | 0.0044 |
| F10 | 80840001 | 80940000 | 0.0032 | 0.0010 | 0.0033 |
| F10 | 80860001 | 80960000 | 0.0027 | 0.0010 | 0.0028 |
| F10 | 80880001 | 80980000 | 0.0020 | 0.0014 | 0.0021 |
| F10 | 80900001 | 81000000 | 0.0021 | 0.0013 | 0.0022 |
| F10 | 80920001 | 81020000 | 0.0020 | 0.0013 | 0.0020 |
| F10 | 80940001 | 81040000 | 0.0020 | 0.0021 | 0.0021 |
| F10 | 80960001 | 81060000 | 0.0029 | 0.0018 | 0.0030 |
| F10 | 80980001 | 81080000 | 0.0027 | 0.0015 | 0.0028 |
| F10 | 81000001 | 81100000 | 0.0024 | 0.0020 | 0.0024 |
| F10 | 81020001 | 81120000 | 0.0032 | 0.0026 | 0.0033 |
| F10 | 81040001 | 81140000 | 0.0037 | 0.0018 | 0.0038 |
| F10 | 81060001 | 81160000 | 0.0029 | 0.0020 | 0.0031 |
| F10 | 81080001 | 81180000 | 0.0032 | 0.0022 | 0.0033 |
| F10 | 81100001 | 81200000 | 0.0034 | 0.0017 | 0.0036 |
| F10 | 81120001 | 81220000 | 0.0028 | 0.0017 | 0.0029 |
| F10 | 81140001 | 81240000 | 0.0027 | 0.0019 | 0.0029 |
| F10 | 81160001 | 81260000 | 0.0027 | 0.0018 | 0.0027 |
| F10 | 81180001 | 81280000 | 0.0024 | 0.0015 | 0.0024 |
| F10 | 81200001 | 81300000 | 0.0019 | 0.0017 | 0.0019 |
| F10 | 81220001 | 81320000 | 0.0019 | 0.0024 | 0.0019 |
| F10 | 81240001 | 81340000 | 0.0026 | 0.0026 | 0.0026 |
| F10 | 81260001 | 81360000 | 0.0028 | 0.0025 | 0.0028 |
| F10 | 81280001 | 81380000 | 0.0027 | 0.0040 | 0.0026 |
| F10 | 81300001 | 81400000 | 0.0044 | 0.0038 | 0.0044 |
| F10 | 81320001 | 81420000 | 0.0042 | 0.0025 | 0.0042 |
| F10 | 81340001 | 81440000 | 0.0029 | 0.0028 | 0.0029 |
| F10 | 81360001 | 81460000 | 0.0032 | 0.0033 | 0.0032 |
| F10 | 81380001 | 81480000 | 0.0036 | 0.0022 | 0.0036 |
| F10 | 81400001 | 81500000 | 0.0023 | 0.0036 | 0.0023 |
| F10 | 81420001 | 81520000 | 0.0037 | 0.0036 | 0.0036 |
| F10 | 81440001 | 81540000 | 0.0037 | 0.0034 | 0.0036 |
| F10 | 81460001 | 81560000 | 0.0035 | 0.0034 | 0.0034 |
| F10 | 81480001 | 81580000 | 0.0036 | 0.0034 | 0.0035 |
| F10 | 81500001 | 81600000 | 0.0036 | 0.0023 | 0.0035 |

| Chr | Start | End | both | A1 | A2 |
|---|---|---|---|---|---|
| F10 | 81520001 | 81620000 | 0.0025 | 0.0022 | 0.0024 |
| F10 | 81540001 | 81640000 | 0.0023 | 0.0018 | 0.0023 |
| F10 | 81560001 | 81660000 | 0.0018 | 0.0016 | 0.0018 |
| F10 | 81580001 | 81680000 | 0.0016 | 0.0023 | 0.0016 |
| F10 | 81600001 | 81700000 | 0.0022 | 0.0035 | 0.0022 |
| F10 | 81620001 | 81720000 | 0.0034 | 0.0046 | 0.0033 |
| F10 | 81640001 | 81740000 | 0.0045 | 0.0058 | 0.0044 |
| F10 | 81660001 | 81760000 | 0.0057 | 0.0057 | 0.0056 |
| F10 | 81680001 | 81780000 | 0.0057 | 0.0057 | 0.0056 |
| F10 | 81700001 | 81800000 | 0.0058 | 0.0044 | 0.0057 |
| F10 | 81720001 | 81820000 | 0.0047 | 0.0037 | 0.0047 |
| F10 | 81740001 | 81840000 | 0.0041 | 0.0027 | 0.0041 |
| F10 | 81760001 | 81860000 | 0.0033 | 0.0024 | 0.0032 |
| F10 | 81780001 | 81880000 | 0.0029 | 0.0021 | 0.0028 |
| F10 | 81800001 | 81900000 | 0.0025 | 0.0026 | 0.0024 |
| F10 | 81820001 | 81920000 | 0.0028 | 0.0034 | 0.0026 |
| F10 | 81840001 | 81940000 | 0.0036 | 0.0034 | 0.0034 |
| F10 | 81860001 | 81960000 | 0.0035 | 0.0034 | 0.0034 |
| F10 | 81880001 | 81980000 | 0.0035 | 0.0027 | 0.0034 |
| F10 | 81900001 | 82000000 | 0.0028 | 0.0020 | 0.0028 |
| F10 | 81920001 | 82020000 | 0.0022 | 0.0010 | 0.0022 |
| F10 | 81940001 | 82040000 | 0.0011 | 0.0010 | 0.0011 |
| F10 | 81960001 | 82060000 | 0.0011 | 0.0010 | 0.0011 |
| F10 | 81980001 | 82080000 | 0.0011 | 0.0011 | 0.0011 |
| F10 | 82000001 | 82100000 | 0.0013 | 0.0015 | 0.0013 |
| F10 | 82020001 | 82120000 | 0.0018 | 0.0023 | 0.0018 |
| F10 | 82040001 | 82140000 | 0.0026 | 0.0034 | 0.0026 |
| F10 | 82060001 | 82160000 | 0.0038 | 0.0046 | 0.0038 |
| F10 | 82080001 | 82180000 | 0.0053 | 0.0044 | 0.0052 |
| F10 | 82100001 | 82200000 | 0.0052 | 0.0051 | 0.0052 |
| F10 | 82120001 | 82220000 | 0.0058 | 0.0042 | 0.0057 |
| F10 | 82140001 | 82240000 | 0.0048 | 0.0038 | 0.0048 |
| F10 | 82160001 | 82260000 | 0.0043 | 0.0032 | 0.0042 |
| F10 | 82180001 | 82280000 | 0.0036 | 0.0034 | 0.0035 |
| F10 | 82200001 | 82300000 | 0.0037 | 0.0031 | 0.0035 |
| F10 | 82220001 | 82320000 | 0.0036 | 0.0038 | 0.0036 |
| F10 | 82240001 | 82340000 | 0.0044 | 0.0036 | 0.0044 |
| F10 | 82260001 | 82360000 | 0.0043 | 0.0035 | 0.0044 |
| F10 | 82280001 | 82380000 | 0.0047 | 0.0039 | 0.0048 |
| F10 | 82300001 | 82400000 | 0.0052 | 0.0030 | 0.0054 |
| F10 | 82320001 | 82420000 | 0.0041 | 0.0026 | 0.0043 |
| F10 | 82340001 | 82440000 | 0.0038 | 0.0026 | 0.0039 |
| F10 | 82360001 | 82460000 | 0.0038 | 0.0023 | 0.0039 |
| F10 | 82380001 | 82480000 | 0.0033 | 0.0025 | 0.0034 |
| F10 | 82400001 | 82500000 | 0.0035 | 0.0028 | 0.0035 |
| F10 | 82420001 | 82520000 | 0.0041 | 0.0030 | 0.0042 |
| F10 | 82440001 | 82540000 | 0.0041 | 0.0041 | 0.0042 |
| F10 | 82460001 | 82560000 | 0.0052 | 0.0078 | 0.0053 |
| F10 | 82480001 | 82580000 | 0.0091 | 0.0092 | 0.0092 |
| F10 | 82500001 | 82600000 | 0.0104 | 0.0132 | 0.0104 |
| F10 | 82520001 | 82620000 | 0.0145 | 0.0141 | 0.0144 |
| F10 | 82540001 | 82640000 | 0.0161 | 0.0147 | 0.0160 |
| F10 | 82560001 | 82660000 | 0.0170 | 0.0114 | 0.0169 |
| F10 | 82580001 | 82680000 | 0.0134 | 0.0097 | 0.0133 |
| F10 | 82600001 | 82700000 | 0.0117 | 0.0061 | 0.0116 |
| F10 | 82620001 | 82720000 | 0.0076 | 0.0054 | 0.0077 |
| F10 | 82640001 | 82740000 | 0.0063 | 0.0031 | 0.0063 |
| F10 | 82660001 | 82760000 | 0.0035 | 0.0032 | 0.0035 |

| Chr | Start | End | both | A1 | A2 |
|---|---|---|---|---|---|
| F10 | 82680001 | 82780000 | 0.0040 | 0.0034 | 0.0040 |
| F10 | 82700001 | 82800000 | 0.0042 | 0.0071 | 0.0042 |
| F10 | 82720001 | 82820000 | 0.0082 | 0.0079 | 0.0082 |
| F10 | 82740001 | 82840000 | 0.0091 | 0.0083 | 0.0090 |
| F10 | 82760001 | 82860000 | 0.0097 | 0.0083 | 0.0097 |
| F10 | 82780001 | 82880000 | 0.0097 | 0.0080 | 0.0096 |
| F10 | 82800001 | 82900000 | 0.0094 | 0.0037 | 0.0093 |
| F10 | 82820001 | 82920000 | 0.0047 | 0.0024 | 0.0046 |
| F10 | 82840001 | 82940000 | 0.0033 | 0.0021 | 0.0031 |
| F10 | 82860001 | 82960000 | 0.0031 | 0.0016 | 0.0027 |
| F10 | 82880001 | 82980000 | 0.0023 | 0.0018 | 0.0020 |
| F10 | 82900001 | 83000000 | 0.0025 | 0.0023 | 0.0021 |
| F10 | 82920001 | 83020000 | 0.0031 | 0.0029 | 0.0028 |
| F10 | 82940001 | 83040000 | 0.0035 | 0.0031 | 0.0033 |
| F10 | 82960001 | 83060000 | 0.0036 | 0.0030 | 0.0036 |
| F10 | 82980001 | 83080000 | 0.0035 | 0.0029 | 0.0034 |
| F10 | 83000001 | 83100000 | 0.0035 | 0.0025 | 0.0035 |
| F10 | 83020001 | 83120000 | 0.0029 | 0.0023 | 0.0029 |
| F10 | 83040001 | 83140000 | 0.0027 | 0.0022 | 0.0027 |
| F10 | 83060001 | 83160000 | 0.0026 | 0.0023 | 0.0026 |
| F10 | 83080001 | 83180000 | 0.0027 | 0.0026 | 0.0027 |
| F10 | 83100001 | 83200000 | 0.0029 | 0.0024 | 0.0029 |
| F10 | 83120001 | 83220000 | 0.0028 | 0.0020 | 0.0027 |
| F10 | 83140001 | 83240000 | 0.0025 | 0.0016 | 0.0023 |
| F10 | 83160001 | 83260000 | 0.0021 | 0.0011 | 0.0018 |
| F10 | 83180001 | 83280000 | 0.0017 | 0.0005 | 0.0013 |
| F10 | 83200001 | 83300000 | 0.0011 | 0.0012 | 0.0006 |
| F10 | 83220001 | 83320000 | 0.0017 | 0.0016 | 0.0011 |
| F10 | 83240001 | 83340000 | 0.0027 | 0.0023 | 0.0022 |
| F10 | 83260001 | 83360000 | 0.0037 | 0.0029 | 0.0033 |
| F10 | 83280001 | 83380000 | 0.0046 | 0.0028 | 0.0044 |
| F10 | 83300001 | 83400000 | 0.0053 | 0.0022 | 0.0049 |
| F10 | 83320001 | 83420000 | 0.0054 | 0.0020 | 0.0047 |
| F10 | 83340001 | 83440000 | 0.0057 | 0.0030 | 0.0049 |
| F10 | 83360001 | 83460000 | 0.0063 | 0.0028 | 0.0055 |
| F10 | 83380001 | 83480000 | 0.0058 | 0.0033 | 0.0050 |
| F10 | 83400001 | 83500000 | 0.0055 | 0.0043 | 0.0050 |
| F10 | 83420001 | 83520000 | 0.0068 | 0.0045 | 0.0065 |
| F10 | 83440001 | 83540000 | 0.0063 | 0.0033 | 0.0061 |
| F10 | 83460001 | 83560000 | 0.0054 | 0.0034 | 0.0051 |
| F10 | 83480001 | 83580000 | 0.0054 | 0.0033 | 0.0050 |
| F10 | 83500001 | 83600000 | 0.0054 | 0.0023 | 0.0048 |
| F10 | 83520001 | 83620000 | 0.0042 | 0.0017 | 0.0029 |
| F10 | 83540001 | 83640000 | 0.0036 | 0.0014 | 0.0020 |
| F10 | 83560001 | 83660000 | 0.0031 | 0.0018 | 0.0015 |
| F10 | 83580001 | 83680000 | 0.0037 | 0.0030 | 0.0019 |
| F10 | 83600001 | 83700000 | 0.0049 | 0.0030 | 0.0031 |
| F10 | 83620001 | 83720000 | 0.0042 | 0.0046 | 0.0032 |
| F10 | 83640001 | 83740000 | 0.0055 | 0.0055 | 0.0046 |
| F10 | 83660001 | 83760000 | 0.0061 | 0.0065 | 0.0054 |
| F10 | 83680001 | 83780000 | 0.0067 | 0.0054 | 0.0061 |
| F10 | 83700001 | 83800000 | 0.0051 | 0.0060 | 0.0046 |
| F10 | 83720001 | 83820000 | 0.0054 | 0.0051 | 0.0049 |
| F10 | 83740001 | 83840000 | 0.0045 | 0.0048 | 0.0038 |
| F10 | 83760001 | 83860000 | 0.0041 | 0.0043 | 0.0034 |
| F10 | 83780001 | 83880000 | 0.0037 | 0.0036 | 0.0030 |
| F10 | 83800001 | 83900000 | 0.0035 | 0.0036 | 0.0028 |
| F10 | 83820001 | 83920000 | 0.0038 | 0.0031 | 0.0030 |

| Chr | Start | End | both | A1 | A2 |
|---|---|---|---|---|---|
| F10 | 83840001 | 83940000 | 0.0037 | 0.0031 | 0.0029 |
| F10 | 83860001 | 83960000 | 0.0038 | 0.0028 | 0.0030 |
| F10 | 83880001 | 83980000 | 0.0037 | 0.0033 | 0.0028 |
| F10 | 83900001 | 84000000 | 0.0041 | 0.0037 | 0.0032 |
| F10 | 83920001 | 84020000 | 0.0045 | 0.0038 | 0.0037 |
| F10 | 83940001 | 84040000 | 0.0043 | 0.0037 | 0.0036 |
| F10 | 83960001 | 84060000 | 0.0043 | 0.0027 | 0.0037 |
| F10 | 83980001 | 84080000 | 0.0032 | 0.0023 | 0.0028 |
| F10 | 84000001 | 84100000 | 0.0027 | 0.0022 | 0.0025 |
| F10 | 84020001 | 84120000 | 0.0027 | 0.0019 | 0.0025 |
| F10 | 84040001 | 84140000 | 0.0024 | 0.0020 | 0.0024 |
| F10 | 84060001 | 84160000 | 0.0024 | 0.0025 | 0.0023 |
| F10 | 84080001 | 84180000 | 0.0030 | 0.0029 | 0.0029 |
| F10 | 84100001 | 84200000 | 0.0036 | 0.0037 | 0.0033 |
| F10 | 84120001 | 84220000 | 0.0044 | 0.0040 | 0.0040 |
| F10 | 84140001 | 84240000 | 0.0046 | 0.0036 | 0.0041 |
| F10 | 84160001 | 84260000 | 0.0042 | 0.0030 | 0.0036 |
| F10 | 84180001 | 84280000 | 0.0035 | 0.0029 | 0.0030 |
| F10 | 84200001 | 84300000 | 0.0033 | 0.0017 | 0.0029 |
| F10 | 84220001 | 84320000 | 0.0017 | 0.0020 | 0.0013 |
| F10 | 84240001 | 84340000 | 0.0019 | 0.0021 | 0.0015 |
| F10 | 84260001 | 84360000 | 0.0018 | 0.0028 | 0.0015 |
| F10 | 84280001 | 84380000 | 0.0024 | 0.0033 | 0.0021 |
| F10 | 84300001 | 84400000 | 0.0027 | 0.0038 | 0.0024 |
| F10 | 84320001 | 84420000 | 0.0028 | 0.0034 | 0.0024 |
| F10 | 84340001 | 84440000 | 0.0025 | 0.0039 | 0.0020 |
| F10 | 84360001 | 84460000 | 0.0029 | 0.0045 | 0.0025 |
| F10 | 84380001 | 84480000 | 0.0034 | 0.0042 | 0.0029 |
| F10 | 84400001 | 84500000 | 0.0030 | 0.0039 | 0.0025 |
| F10 | 84420001 | 84520000 | 0.0034 | 0.0039 | 0.0027 |
| F10 | 84440001 | 84540000 | 0.0036 | 0.0031 | 0.0029 |
| F10 | 84460001 | 84560000 | 0.0031 | 0.0026 | 0.0024 |
| F10 | 84480001 | 84580000 | 0.0027 | 0.0028 | 0.0020 |
| F10 | 84500001 | 84600000 | 0.0031 | 0.0033 | 0.0024 |
| F10 | 84520001 | 84620000 | 0.0033 | 0.0038 | 0.0027 |
| F10 | 84540001 | 84640000 | 0.0035 | 0.0047 | 0.0030 |
| F10 | 84560001 | 84660000 | 0.0044 | 0.0043 | 0.0038 |
| F10 | 84580001 | 84680000 | 0.0038 | 0.0047 | 0.0032 |
| F10 | 84600001 | 84700000 | 0.0044 | 0.0042 | 0.0039 |
| F10 | 84620001 | 84720000 | 0.0039 | 0.0042 | 0.0035 |
| F10 | 84640001 | 84740000 | 0.0040 | 0.0044 | 0.0036 |
| F10 | 84660001 | 84760000 | 0.0040 | 0.0048 | 0.0036 |
| F10 | 84680001 | 84780000 | 0.0040 | 0.0046 | 0.0035 |
| F10 | 84700001 | 84800000 | 0.0035 | 0.0045 | 0.0029 |
| F10 | 84720001 | 84820000 | 0.0033 | 0.0048 | 0.0027 |
| F10 | 84740001 | 84840000 | 0.0036 | 0.0042 | 0.0029 |
| F10 | 84760001 | 84860000 | 0.0032 | 0.0053 | 0.0024 |
| F10 | 84780001 | 84880000 | 0.0047 | 0.0050 | 0.0035 |
| F10 | 84800001 | 84900000 | 0.0046 | 0.0049 | 0.0035 |
| F10 | 84820001 | 84920000 | 0.0045 | 0.0041 | 0.0033 |
| F10 | 84840001 | 84940000 | 0.0039 | 0.0037 | 0.0028 |
| F10 | 84860001 | 84960000 | 0.0036 | 0.0028 | 0.0027 |
| F10 | 84880001 | 84980000 | 0.0023 | 0.0032 | 0.0018 |
| F10 | 84900001 | 85000000 | 0.0025 | 0.0033 | 0.0019 |
| F10 | 84920001 | 85020000 | 0.0026 | 0.0034 | 0.0020 |
| F10 | 84940001 | 85040000 | 0.0027 | 0.0038 | 0.0022 |
| F10 | 84960001 | 85060000 | 0.0030 | 0.0037 | 0.0024 |
| F10 | 84980001 | 85080000 | 0.0030 | 0.0032 | 0.0025 |

| Chr | Start | End | both | A1 | A2 |
|---|---|---|---|---|---|
| F10 | 85000001 | 85100000 | 0.0026 | 0.0036 | 0.0022 |
| F10 | 85020001 | 85120000 | 0.0028 | 0.0043 | 0.0023 |
| F10 | 85040001 | 85140000 | 0.0035 | 0.0042 | 0.0030 |
| F10 | 85060001 | 85160000 | 0.0036 | 0.0042 | 0.0032 |
| F10 | 85080001 | 85180000 | 0.0038 | 0.0039 | 0.0035 |
| F10 | 85100001 | 85200000 | 0.0036 | 0.0037 | 0.0033 |
| F10 | 85120001 | 85220000 | 0.0035 | 0.0030 | 0.0032 |
| F10 | 85140001 | 85240000 | 0.0026 | 0.0032 | 0.0023 |
| F10 | 85160001 | 85260000 | 0.0026 | 0.0036 | 0.0022 |
| F10 | 85180001 | 85280000 | 0.0031 | 0.0033 | 0.0027 |
| F10 | 85200001 | 85300000 | 0.0028 | 0.0033 | 0.0024 |
| F10 | 85220001 | 85320000 | 0.0033 | 0.0037 | 0.0030 |
| F10 | 85240001 | 85340000 | 0.0038 | 0.0043 | 0.0036 |
| F10 | 85260001 | 85360000 | 0.0045 | 0.0043 | 0.0043 |
| F10 | 85280001 | 85380000 | 0.0042 | 0.0054 | 0.0039 |
| F10 | 85300001 | 85400000 | 0.0052 | 0.0050 | 0.0049 |
| F10 | 85320001 | 85420000 | 0.0047 | 0.0047 | 0.0044 |
| F10 | 85340001 | 85440000 | 0.0044 | 0.0040 | 0.0040 |
| F10 | 85360001 | 85460000 | 0.0037 | 0.0030 | 0.0033 |
| F10 | 85380001 | 85480000 | 0.0028 | 0.0025 | 0.0026 |
| F10 | 85400001 | 85500000 | 0.0025 | 0.0027 | 0.0024 |
| F10 | 85420001 | 85520000 | 0.0027 | 0.0022 | 0.0026 |
| F10 | 85440001 | 85540000 | 0.0024 | 0.0025 | 0.0024 |
| F10 | 85460001 | 85560000 | 0.0028 | 0.0025 | 0.0027 |
| F10 | 85480001 | 85580000 | 0.0027 | 0.0023 | 0.0027 |
| F10 | 85500001 | 85600000 | 0.0024 | 0.0028 | 0.0023 |
| F10 | 85520001 | 85620000 | 0.0031 | 0.0037 | 0.0030 |
| F10 | 85540001 | 85640000 | 0.0039 | 0.0032 | 0.0038 |
| F10 | 85560001 | 85660000 | 0.0036 | 0.0038 | 0.0036 |
| F10 | 85580001 | 85680000 | 0.0042 | 0.0039 | 0.0042 |
| F10 | 85600001 | 85700000 | 0.0042 | 0.0033 | 0.0041 |
| F10 | 85620001 | 85720000 | 0.0032 | 0.0024 | 0.0030 |
| F10 | 85640001 | 85740000 | 0.0023 | 0.0025 | 0.0021 |
| F10 | 85660001 | 85760000 | 0.0023 | 0.0022 | 0.0021 |
| F10 | 85680001 | 85780000 | 0.0019 | 0.0021 | 0.0017 |
| F10 | 85700001 | 85800000 | 0.0020 | 0.0026 | 0.0018 |
| F10 | 85720001 | 85820000 | 0.0028 | 0.0030 | 0.0027 |
| F10 | 85740001 | 85840000 | 0.0032 | 0.0028 | 0.0030 |
| F10 | 85760001 | 85860000 | 0.0029 | 0.0027 | 0.0028 |
| F10 | 85780001 | 85880000 | 0.0029 | 0.0023 | 0.0028 |
| F10 | 85800001 | 85900000 | 0.0025 | 0.0016 | 0.0024 |
| F10 | 85820001 | 85920000 | 0.0018 | 0.0014 | 0.0017 |
| F10 | 85840001 | 85940000 | 0.0015 | 0.0013 | 0.0014 |
| F10 | 85860001 | 85960000 | 0.0014 | 0.0013 | 0.0014 |
| F10 | 85880001 | 85980000 | 0.0013 | 0.0022 | 0.0011 |
| F10 | 85900001 | 86000000 | 0.0021 | 0.0031 | 0.0019 |
| F10 | 85920001 | 86020000 | 0.0029 | 0.0031 | 0.0027 |
| F10 | 85940001 | 86040000 | 0.0028 | 0.0033 | 0.0026 |
| F10 | 85960001 | 86060000 | 0.0031 | 0.0036 | 0.0028 |
| F10 | 85980001 | 86080000 | 0.0033 | 0.0044 | 0.0029 |
| F10 | 86000001 | 86100000 | 0.0043 | 0.0047 | 0.0040 |
| F10 | 86020001 | 86120000 | 0.0045 | 0.0046 | 0.0041 |
| F10 | 86040001 | 86140000 | 0.0044 | 0.0043 | 0.0041 |
| F10 | 86060001 | 86160000 | 0.0042 | 0.0042 | 0.0039 |
| F10 | 86080001 | 86180000 | 0.0042 | 0.0026 | 0.0039 |
| F10 | 86100001 | 86200000 | 0.0023 | 0.0014 | 0.0021 |
| F10 | 86120001 | 86220000 | 0.0011 | 0.0018 | 0.0009 |
| F10 | 86140001 | 86240000 | 0.0017 | 0.0026 | 0.0013 |

| Chr | Start | End | both | A1 | A2 |
|---|---|---|---|---|---|
| F10 | 86160001 | 86260000 | 0.0023 | 0.0022 | 0.0017 |
| F10 | 86180001 | 86280000 | 0.0019 | 0.0025 | 0.0012 |
| F10 | 86200001 | 86300000 | 0.0021 | 0.0024 | 0.0013 |
| F10 | 86220001 | 86320000 | 0.0021 | 0.0018 | 0.0014 |
| F10 | 86240001 | 86340000 | 0.0014 | 0.0017 | 0.0010 |
| F10 | 86260001 | 86360000 | 0.0015 | 0.0021 | 0.0012 |
| F10 | 86280001 | 86380000 | 0.0020 | 0.0023 | 0.0017 |
| F10 | 86300001 | 86400000 | 0.0022 | 0.0026 | 0.0020 |
| F10 | 86320001 | 86420000 | 0.0024 | 0.0032 | 0.0021 |
| F10 | 86340001 | 86440000 | 0.0027 | 0.0030 | 0.0024 |
| F10 | 86360001 | 86460000 | 0.0021 | 0.0028 | 0.0017 |
| F10 | 86380001 | 86480000 | 0.0016 | 0.0026 | 0.0011 |
| F10 | 86400001 | 86500000 | 0.0013 | 0.0026 | 0.0008 |
| F10 | 86420001 | 86520000 | 0.0013 | 0.0023 | 0.0008 |
| F10 | 86440001 | 86540000 | 0.0011 | 0.0021 | 0.0006 |
| F10 | 86460001 | 86560000 | 0.0011 | 0.0023 | 0.0007 |
| F10 | 86480001 | 86580000 | 0.0014 | 0.0028 | 0.0010 |
| F10 | 86500001 | 86600000 | 0.0019 | 0.0026 | 0.0015 |
| F10 | 86520001 | 86620000 | 0.0017 | 0.0027 | 0.0013 |
| F10 | 86540001 | 86640000 | 0.0018 | 0.0036 | 0.0014 |
| F10 | 86560001 | 86660000 | 0.0029 | 0.0032 | 0.0025 |
| F10 | 86580001 | 86680000 | 0.0026 | 0.0027 | 0.0023 |
| F10 | 86600001 | 86700000 | 0.0024 | 0.0026 | 0.0021 |
| F10 | 86620001 | 86720000 | 0.0025 | 0.0031 | 0.0022 |
| F10 | 86640001 | 86740000 | 0.0036 | 0.0022 | 0.0030 |
| F10 | 86660001 | 86760000 | 0.0029 | 0.0020 | 0.0022 |
| F10 | 86680001 | 86780000 | 0.0032 | 0.0018 | 0.0023 |
| F10 | 86700001 | 86800000 | 0.0032 | 0.0017 | 0.0021 |
| F10 | 86720001 | 86820000 | 0.0033 | 0.0010 | 0.0020 |
| F10 | 86740001 | 86840000 | 0.0023 | 0.0007 | 0.0013 |
| F10 | 86760001 | 86860000 | 0.0019 | 0.0006 | 0.0009 |
| F10 | 86780001 | 86880000 | 0.0015 | 0.0008 | 0.0007 |
| F10 | 86800001 | 86900000 | 0.0015 | 0.0007 | 0.0008 |
| F10 | 86820001 | 86920000 | 0.0015 | 0.0007 | 0.0007 |
| F10 | 86840001 | 86940000 | 0.0016 | 0.0008 | 0.0009 |
| F10 | 86860001 | 86960000 | 0.0017 | 0.0008 | 0.0009 |
| F10 | 86880001 | 86980000 | 0.0019 | 0.0008 | 0.0009 |
| F10 | 86900001 | 87000000 | 0.0018 | 0.0009 | 0.0009 |
| F10 | 86920001 | 87020000 | 0.0018 | 0.0008 | 0.0011 |
| F10 | 86940001 | 87040000 | 0.0016 | 0.0025 | 0.0009 |
| F10 | 86960001 | 87060000 | 0.0039 | 0.0036 | 0.0032 |
| F10 | 86980001 | 87080000 | 0.0058 | 0.0044 | 0.0051 |
| F10 | 87000001 | 87100000 | 0.0078 | 0.0046 | 0.0070 |
| F10 | 87020001 | 87120000 | 0.0081 | 0.0046 | 0.0072 |
| F10 | 87040001 | 87140000 | 0.0081 | 0.0034 | 0.0073 |
| F10 | 87060001 | 87160000 | 0.0065 | 0.0032 | 0.0059 |
| F10 | 87080001 | 87180000 | 0.0056 | 0.0040 | 0.0051 |
| F10 | 87100001 | 87200000 | 0.0056 | 0.0043 | 0.0053 |
| F10 | 87120001 | 87220000 | 0.0061 | 0.0047 | 0.0058 |
| F10 | 87140001 | 87240000 | 0.0065 | 0.0046 | 0.0062 |
| F10 | 87160001 | 87260000 | 0.0062 | 0.0039 | 0.0059 |
| F10 | 87180001 | 87280000 | 0.0052 | 0.0030 | 0.0050 |
| F10 | 87200001 | 87300000 | 0.0038 | 0.0030 | 0.0036 |
| F10 | 87220001 | 87320000 | 0.0035 | 0.0026 | 0.0033 |
| F10 | 87240001 | 87340000 | 0.0035 | 0.0023 | 0.0030 |
| F10 | 87260001 | 87360000 | 0.0032 | 0.0026 | 0.0027 |
| F10 | 87280001 | 87380000 | 0.0038 | 0.0024 | 0.0032 |
| F10 | 87300001 | 87400000 | 0.0039 | 0.0021 | 0.0032 |

| Chr | Start | End | both | A1 | A2 |
|---|---|---|---|---|---|
| F10 | 87320001 | 87420000 | 0.0037 | 0.0025 | 0.0028 |
| F10 | 87340001 | 87440000 | 0.0041 | 0.0031 | 0.0031 |
| F10 | 87360001 | 87460000 | 0.0048 | 0.0028 | 0.0038 |
| F10 | 87380001 | 87480000 | 0.0045 | 0.0022 | 0.0033 |
| F10 | 87400001 | 87500000 | 0.0038 | 0.0019 | 0.0026 |
| F10 | 87420001 | 87520000 | 0.0035 | 0.0014 | 0.0023 |
| F10 | 87440001 | 87540000 | 0.0029 | 0.0011 | 0.0019 |
| F10 | 87460001 | 87560000 | 0.0029 | 0.0010 | 0.0017 |
| F10 | 87480001 | 87580000 | 0.0029 | 0.0013 | 0.0019 |
| F10 | 87500001 | 87600000 | 0.0031 | 0.0017 | 0.0022 |
| F10 | 87520001 | 87620000 | 0.0034 | 0.0018 | 0.0026 |
| F10 | 87540001 | 87640000 | 0.0034 | 0.0024 | 0.0028 |
| F10 | 87560001 | 87660000 | 0.0036 | 0.0028 | 0.0030 |
| F10 | 87580001 | 87680000 | 0.0037 | 0.0027 | 0.0031 |
| F10 | 87600001 | 87700000 | 0.0038 | 0.0026 | 0.0032 |
| F10 | 87620001 | 87720000 | 0.0039 | 0.0024 | 0.0031 |
| F10 | 87640001 | 87740000 | 0.0036 | 0.0015 | 0.0028 |
| F10 | 87660001 | 87760000 | 0.0030 | 0.0014 | 0.0021 |
| F10 | 87680001 | 87780000 | 0.0028 | 0.0010 | 0.0020 |
| F10 | 87700001 | 87800000 | 0.0022 | 0.0011 | 0.0014 |
| F10 | 87720001 | 87820000 | 0.0021 | 0.0013 | 0.0015 |
| F10 | 87740001 | 87840000 | 0.0024 | 0.0019 | 0.0019 |
| F10 | 87760001 | 87860000 | 0.0029 | 0.0021 | 0.0024 |
| F10 | 87780001 | 87880000 | 0.0033 | 0.0025 | 0.0027 |
| F10 | 87800001 | 87900000 | 0.0036 | 0.0024 | 0.0030 |
| F10 | 87820001 | 87920000 | 0.0033 | 0.0022 | 0.0027 |
| F10 | 87840001 | 87940000 | 0.0031 | 0.0017 | 0.0024 |
| F10 | 87860001 | 87960000 | 0.0027 | 0.0019 | 0.0018 |
| F10 | 87880001 | 87980000 | 0.0031 | 0.0017 | 0.0022 |
| F10 | 87900001 | 88000000 | 0.0030 | 0.0031 | 0.0021 |
| F10 | 87920001 | 88020000 | 0.0044 | 0.0031 | 0.0035 |
| F10 | 87940001 | 88040000 | 0.0043 | 0.0030 | 0.0035 |
| F10 | 87960001 | 88060000 | 0.0041 | 0.0022 | 0.0034 |
| F10 | 87980001 | 88080000 | 0.0032 | 0.0022 | 0.0024 |
| F10 | 88000001 | 88100000 | 0.0034 | 0.0008 | 0.0025 |
| F10 | 88020001 | 88120000 | 0.0022 | 0.0013 | 0.0012 |
| F10 | 88040001 | 88140000 | 0.0026 | 0.0014 | 0.0017 |
| F10 | 88060001 | 88160000 | 0.0026 | 0.0017 | 0.0018 |
| F10 | 88080001 | 88180000 | 0.0025 | 0.0019 | 0.0018 |
| F10 | 88100001 | 88200000 | 0.0023 | 0.0017 | 0.0016 |
| F10 | 88120001 | 88220000 | 0.0020 | 0.0012 | 0.0014 |
| F10 | 88140001 | 88240000 | 0.0017 | 0.0011 | 0.0010 |
| F10 | 88160001 | 88260000 | 0.0017 | 0.0013 | 0.0007 |
| F10 | 88180001 | 88280000 | 0.0024 | 0.0013 | 0.0015 |
| F10 | 88200001 | 88300000 | 0.0025 | 0.0013 | 0.0018 |
| F10 | 88220001 | 88320000 | 0.0024 | 0.0012 | 0.0019 |
| F10 | 88240001 | 88340000 | 0.0025 | 0.0013 | 0.0020 |
| F10 | 88260001 | 88360000 | 0.0025 | 0.0012 | 0.0022 |
| F10 | 88280001 | 88380000 | 0.0021 | 0.0012 | 0.0019 |
| F10 | 88300001 | 88400000 | 0.0019 | 0.0015 | 0.0016 |
| F10 | 88320001 | 88420000 | 0.0019 | 0.0016 | 0.0014 |
| F10 | 88340001 | 88440000 | 0.0018 | 0.0015 | 0.0011 |
| F10 | 88360001 | 88460000 | 0.0017 | 0.0011 | 0.0009 |
| F10 | 88380001 | 88480000 | 0.0015 | 0.0009 | 0.0005 |
| F10 | 88400001 | 88500000 | 0.0015 | 0.0009 | 0.0006 |
| F10 | 88420001 | 88520000 | 0.0019 | 0.0014 | 0.0009 |
| F10 | 88440001 | 88540000 | 0.0025 | 0.0018 | 0.0015 |
| F10 | 88460001 | 88560000 | 0.0031 | 0.0021 | 0.0021 |

| Chr | Start | End | both | A1 | A2 |
|---|---|---|---|---|---|
| F10 | 88480001 | 88580000 | 0.0034 | 0.0021 | 0.0025 |
| F10 | 88500001 | 88600000 | 0.0033 | 0.0022 | 0.0024 |
| F10 | 88520001 | 88620000 | 0.0033 | 0.0017 | 0.0028 |
| F10 | 88540001 | 88640000 | 0.0028 | 0.0020 | 0.0022 |
| F10 | 88560001 | 88660000 | 0.0031 | 0.0029 | 0.0025 |
| F10 | 88580001 | 88680000 | 0.0035 | 0.0035 | 0.0026 |
| F10 | 88600001 | 88700000 | 0.0040 | 0.0051 | 0.0031 |
| F10 | 88620001 | 88720000 | 0.0055 | 0.0050 | 0.0044 |
| F10 | 88640001 | 88740000 | 0.0054 | 0.0043 | 0.0043 |
| F10 | 88660001 | 88760000 | 0.0045 | 0.0031 | 0.0035 |
| F10 | 88680001 | 88780000 | 0.0038 | 0.0025 | 0.0030 |
| F10 | 88700001 | 88800000 | 0.0034 | 0.0009 | 0.0024 |
| F10 | 88720001 | 88820000 | 0.0014 | 0.0013 | 0.0005 |
| F10 | 88740001 | 88840000 | 0.0016 | 0.0015 | 0.0008 |
| F10 | 88760001 | 88860000 | 0.0016 | 0.0024 | 0.0009 |
| F10 | 88780001 | 88880000 | 0.0023 | 0.0031 | 0.0017 |
| F10 | 88800001 | 88900000 | 0.0031 | 0.0036 | 0.0025 |
| F10 | 88820001 | 88920000 | 0.0042 | 0.0040 | 0.0037 |
| F10 | 88840001 | 88940000 | 0.0052 | 0.0043 | 0.0048 |
| F10 | 88860001 | 88960000 | 0.0063 | 0.0039 | 0.0060 |
| F10 | 88880001 | 88980000 | 0.0064 | 0.0044 | 0.0061 |
| F10 | 88900001 | 89000000 | 0.0071 | 0.0045 | 0.0071 |
| F10 | 88920001 | 89020000 | 0.0071 | 0.0041 | 0.0071 |
| F10 | 88940001 | 89040000 | 0.0070 | 0.0038 | 0.0070 |
| F10 | 88960001 | 89060000 | 0.0070 | 0.0040 | 0.0066 |
| F10 | 88980001 | 89080000 | 0.0071 | 0.0030 | 0.0065 |
| F10 | 89000001 | 89100000 | 0.0056 | 0.0023 | 0.0049 |
| F10 | 89020001 | 89120000 | 0.0047 | 0.0016 | 0.0040 |
| F10 | 89040001 | 89140000 | 0.0033 | 0.0018 | 0.0027 |
| F10 | 89060001 | 89160000 | 0.0026 | 0.0016 | 0.0022 |
| F10 | 89080001 | 89180000 | 0.0021 | 0.0017 | 0.0020 |
| F10 | 89100001 | 89200000 | 0.0020 | 0.0027 | 0.0019 |
| F10 | 89120001 | 89220000 | 0.0024 | 0.0030 | 0.0021 |
| F10 | 89140001 | 89240000 | 0.0028 | 0.0030 | 0.0023 |
| F10 | 89160001 | 89260000 | 0.0032 | 0.0043 | 0.0023 |
| F10 | 89180001 | 89280000 | 0.0047 | 0.0061 | 0.0038 |
| F10 | 89200001 | 89300000 | 0.0074 | 0.0052 | 0.0059 |
| F10 | 89220001 | 89320000 | 0.0078 | 0.0067 | 0.0061 |
| F10 | 89240001 | 89340000 | 0.0099 | 0.0063 | 0.0079 |
| F10 | 89260001 | 89360000 | 0.0092 | 0.0051 | 0.0074 |
| F10 | 89280001 | 89380000 | 0.0075 | 0.0036 | 0.0057 |
| F10 | 89300001 | 89400000 | 0.0050 | 0.0043 | 0.0036 |
| F10 | 89320001 | 89420000 | 0.0051 | 0.0036 | 0.0042 |
| F10 | 89340001 | 89440000 | 0.0039 | 0.0048 | 0.0035 |
| F10 | 89360001 | 89460000 | 0.0050 | 0.0054 | 0.0047 |
| F10 | 89380001 | 89480000 | 0.0059 | 0.0051 | 0.0057 |
| F10 | 89400001 | 89500000 | 0.0057 | 0.0048 | 0.0056 |
| F10 | 89420001 | 89520000 | 0.0053 | 0.0059 | 0.0052 |
| F10 | 89440001 | 89540000 | 0.0066 | 0.0055 | 0.0065 |
| F10 | 89460001 | 89560000 | 0.0065 | 0.0060 | 0.0064 |
| F10 | 89480001 | 89580000 | 0.0068 | 0.0059 | 0.0067 |
| F10 | 89500001 | 89600000 | 0.0066 | 0.0060 | 0.0065 |
| F10 | 89520001 | 89620000 | 0.0070 | 0.0039 | 0.0069 |
| F10 | 89540001 | 89640000 | 0.0046 | 0.0042 | 0.0045 |
| F10 | 89560001 | 89660000 | 0.0046 | 0.0039 | 0.0045 |
| F10 | 89580001 | 89680000 | 0.0045 | 0.0043 | 0.0044 |
| F10 | 89600001 | 89700000 | 0.0049 | 0.0040 | 0.0048 |
| F10 | 89620001 | 89720000 | 0.0041 | 0.0040 | 0.0040 |

| Chr | Start | End | both | A1 | A2 |
|---|---|---|---|---|---|
| F10 | 89640001 | 89740000 | 0.0042 | 0.0041 | 0.0041 |
| F10 | 89660001 | 89760000 | 0.0043 | 0.0032 | 0.0042 |
| F10 | 89680001 | 89780000 | 0.0034 | 0.0027 | 0.0032 |
| F10 | 89700001 | 89800000 | 0.0029 | 0.0022 | 0.0028 |
| F10 | 89720001 | 89820000 | 0.0024 | 0.0020 | 0.0024 |
| F10 | 89740001 | 89840000 | 0.0022 | 0.0018 | 0.0022 |
| F10 | 89760001 | 89860000 | 0.0018 | 0.0017 | 0.0017 |
| F10 | 89780001 | 89880000 | 0.0016 | 0.0019 | 0.0015 |
| F10 | 89800001 | 89900000 | 0.0018 | 0.0019 | 0.0017 |
| F10 | 89820001 | 89920000 | 0.0019 | 0.0030 | 0.0018 |
| F10 | 89840001 | 89940000 | 0.0029 | 0.0018 | 0.0028 |
| F10 | 89860001 | 89960000 | 0.0018 | 0.0044 | 0.0018 |
| F10 | 89880001 | 89980000 | 0.0038 | 0.0043 | 0.0033 |
| F10 | 89900001 | 90000000 | 0.0037 | 0.0052 | 0.0032 |
| F10 | 89920001 | 90020000 | 0.0041 | 0.0047 | 0.0035 |
| F10 | 89940001 | 90040000 | 0.0035 | 0.0053 | 0.0028 |
| F10 | 89960001 | 90060000 | 0.0036 | 0.0028 | 0.0028 |
| F10 | 89980001 | 90080000 | 0.0016 | 0.0032 | 0.0011 |
| F10 | 90000001 | 90100000 | 0.0021 | 0.0023 | 0.0015 |
| F10 | 90020001 | 90120000 | 0.0016 | 0.0023 | 0.0012 |
| F10 | 90040001 | 90140000 | 0.0017 | 0.0024 | 0.0014 |
| F10 | 90060001 | 90160000 | 0.0022 | 0.0019 | 0.0020 |
| F10 | 90080001 | 90180000 | 0.0019 | 0.0029 | 0.0017 |
| F10 | 90100001 | 90200000 | 0.0028 | 0.0040 | 0.0026 |
| F10 | 90120001 | 90220000 | 0.0036 | 0.0041 | 0.0031 |
| F10 | 90140001 | 90240000 | 0.0040 | 0.0046 | 0.0033 |
| F10 | 90160001 | 90260000 | 0.0047 | 0.0055 | 0.0039 |
| F10 | 90180001 | 90280000 | 0.0062 | 0.0045 | 0.0049 |
| F10 | 90200001 | 90300000 | 0.0052 | 0.0037 | 0.0039 |
| F10 | 90220001 | 90320000 | 0.0047 | 0.0037 | 0.0038 |
| F10 | 90240001 | 90340000 | 0.0047 | 0.0034 | 0.0038 |
| F10 | 90260001 | 90360000 | 0.0045 | 0.0069 | 0.0038 |
| F10 | 90280001 | 90380000 | 0.0074 | 0.0085 | 0.0070 |
| F10 | 90300001 | 90400000 | 0.0089 | 0.0090 | 0.0085 |
| F10 | 90320001 | 90420000 | 0.0094 | 0.0088 | 0.0088 |
| F10 | 90340001 | 90440000 | 0.0093 | 0.0081 | 0.0086 |
| F10 | 90360001 | 90460000 | 0.0085 | 0.0039 | 0.0077 |
| F10 | 90380001 | 90480000 | 0.0045 | 0.0036 | 0.0039 |
| F10 | 90400001 | 90500000 | 0.0042 | 0.0029 | 0.0037 |
| F10 | 90420001 | 90520000 | 0.0035 | 0.0030 | 0.0031 |
| F10 | 90440001 | 90540000 | 0.0038 | 0.0032 | 0.0035 |
| F10 | 90460001 | 90560000 | 0.0043 | 0.0031 | 0.0041 |
| F10 | 90480001 | 90580000 | 0.0046 | 0.0013 | 0.0043 |
| F10 | 90500001 | 90600000 | 0.0032 | 0.0016 | 0.0030 |
| F10 | 90520001 | 90620000 | 0.0037 | 0.0011 | 0.0034 |
| F10 | 90540001 | 90640000 | 0.0029 | 0.0009 | 0.0026 |
| F10 | 90560001 | 90660000 | 0.0022 | 0.0014 | 0.0021 |
| F10 | 90580001 | 90680000 | 0.0025 | 0.0025 | 0.0024 |
| F10 | 90600001 | 90700000 | 0.0034 | 0.0028 | 0.0032 |
| F10 | 90620001 | 90720000 | 0.0038 | 0.0055 | 0.0036 |
| F10 | 90640001 | 90740000 | 0.0067 | 0.0061 | 0.0065 |
| F10 | 90660001 | 90760000 | 0.0073 | 0.0056 | 0.0072 |
| F10 | 90680001 | 90780000 | 0.0068 | 0.0053 | 0.0066 |
| F10 | 90700001 | 90800000 | 0.0065 | 0.0051 | 0.0062 |
| F10 | 90720001 | 90820000 | 0.0061 | 0.0023 | 0.0058 |
| F10 | 90740001 | 90840000 | 0.0032 | 0.0017 | 0.0028 |
| F10 | 90760001 | 90860000 | 0.0024 | 0.0022 | 0.0020 |
| F10 | 90780001 | 90880000 | 0.0026 | 0.0020 | 0.0023 |

| Chr | Start | End | both | A1 | A2 |
|---|---|---|---|---|---|
| F10 | 90800001 | 90900000 | 0.0022 | 0.0028 | 0.0021 |
| F10 | 90820001 | 90920000 | 0.0030 | 0.0040 | 0.0027 |
| F10 | 90840001 | 90940000 | 0.0043 | 0.0043 | 0.0038 |
| F10 | 90860001 | 90960000 | 0.0045 | 0.0041 | 0.0037 |
| F10 | 90880001 | 90980000 | 0.0043 | 0.0038 | 0.0035 |
| F10 | 90900001 | 91000000 | 0.0040 | 0.0040 | 0.0032 |
| F10 | 90920001 | 91020000 | 0.0041 | 0.0028 | 0.0035 |
| F10 | 90940001 | 91040000 | 0.0027 | 0.0053 | 0.0023 |
| F10 | 90960001 | 91060000 | 0.0052 | 0.0055 | 0.0051 |
| F10 | 90980001 | 91080000 | 0.0054 | 0.0054 | 0.0053 |
| F10 | 91000001 | 91100000 | 0.0052 | 0.0056 | 0.0051 |
| F10 | 91020001 | 91120000 | 0.0054 | 0.0062 | 0.0053 |
| F10 | 91040001 | 91140000 | 0.0063 | 0.0037 | 0.0062 |
| F10 | 91060001 | 91160000 | 0.0039 | 0.0043 | 0.0039 |
| F10 | 91080001 | 91180000 | 0.0045 | 0.0056 | 0.0044 |
| F10 | 91100001 | 91200000 | 0.0054 | 0.0061 | 0.0052 |
| F10 | 91120001 | 91220000 | 0.0059 | 0.0063 | 0.0057 |
| F10 | 91140001 | 91240000 | 0.0061 | 0.0082 | 0.0058 |
| F10 | 91160001 | 91260000 | 0.0083 | 0.0086 | 0.0080 |
| F10 | 91180001 | 91280000 | 0.0086 | 0.0092 | 0.0084 |
| F10 | 91200001 | 91300000 | 0.0096 | 0.0090 | 0.0094 |
| F10 | 91220001 | 91320000 | 0.0090 | 0.0097 | 0.0088 |
| F10 | 91240001 | 91340000 | 0.0097 | 0.0077 | 0.0095 |
| F10 | 91260001 | 91360000 | 0.0072 | 0.0068 | 0.0069 |
| F10 | 91280001 | 91380000 | 0.0059 | 0.0059 | 0.0055 |
| F10 | 91300001 | 91400000 | 0.0051 | 0.0048 | 0.0046 |
| F10 | 91320001 | 91420000 | 0.0042 | 0.0049 | 0.0037 |
| F10 | 91340001 | 91440000 | 0.0042 | 0.0051 | 0.0035 |
| F10 | 91360001 | 91460000 | 0.0045 | 0.0058 | 0.0039 |
| F10 | 91380001 | 91480000 | 0.0057 | 0.0054 | 0.0051 |
| F10 | 91400001 | 91500000 | 0.0052 | 0.0057 | 0.0047 |
| F10 | 91420001 | 91520000 | 0.0055 | 0.0047 | 0.0050 |
| F10 | 91440001 | 91540000 | 0.0044 | 0.0050 | 0.0036 |
| F10 | 91460001 | 91560000 | 0.0044 | 0.0037 | 0.0033 |
| F10 | 91480001 | 91580000 | 0.0032 | 0.0032 | 0.0021 |
| F10 | 91500001 | 91600000 | 0.0028 | 0.0045 | 0.0016 |
| F10 | 91520001 | 91620000 | 0.0044 | 0.0056 | 0.0028 |
| F10 | 91540001 | 91640000 | 0.0062 | 0.0073 | 0.0044 |
| F10 | 91560001 | 91660000 | 0.0087 | 0.0079 | 0.0069 |
| F10 | 91580001 | 91680000 | 0.0090 | 0.0082 | 0.0069 |
| F10 | 91600001 | 91700000 | 0.0092 | 0.0058 | 0.0069 |
| F10 | 91620001 | 91720000 | 0.0067 | 0.0040 | 0.0047 |
| F10 | 91640001 | 91740000 | 0.0048 | 0.0020 | 0.0029 |
| F10 | 91660001 | 91760000 | 0.0022 | 0.0015 | 0.0005 |
| F10 | 91680001 | 91780000 | 0.0019 | 0.0014 | 0.0005 |
| F10 | 91700001 | 91800000 | 0.0019 | 0.0015 | 0.0008 |
| F10 | 91720001 | 91820000 | 0.0025 | 0.0016 | 0.0015 |
| F10 | 91740001 | 91840000 | 0.0025 | 0.0016 | 0.0018 |
| F10 | 91760001 | 91860000 | 0.0026 | 0.0017 | 0.0022 |
| F10 | 91780001 | 91880000 | 0.0031 | 0.0015 | 0.0024 |
| F10 | 91800001 | 91900000 | 0.0029 | 0.0012 | 0.0021 |
| F10 | 91820001 | 91920000 | 0.0025 | 0.0010 | 0.0014 |
| F10 | 91840001 | 91940000 | 0.0022 | 0.0006 | 0.0011 |
| F10 | 91860001 | 91960000 | 0.0020 | 0.0003 | 0.0006 |
| F10 | 91880001 | 91980000 | 0.0015 | 0.0002 | 0.0003 |
| F10 | 91900001 | 92000000 | 0.0016 | 0.0003 | 0.0005 |
| F10 | 91920001 | 92020000 | 0.0018 | 0.0003 | 0.0009 |
| F10 | 91940001 | 92040000 | 0.0020 | 0.0006 | 0.0010 |

| Chr | Start | End | both | A1 | A2 |
|---|---|---|---|---|---|
| F10 | 91960001 | 92060000 | 0.0021 | 0.0007 | 0.0011 |
| F10 | 91980001 | 92080000 | 0.0023 | 0.0020 | 0.0013 |
| F10 | 92000001 | 92100000 | 0.0037 | 0.0027 | 0.0025 |
| F10 | 92020001 | 92120000 | 0.0043 | 0.0028 | 0.0032 |
| F10 | 92040001 | 92140000 | 0.0042 | 0.0035 | 0.0033 |
| F10 | 92060001 | 92160000 | 0.0058 | 0.0033 | 0.0051 |
| F10 | 92080001 | 92180000 | 0.0057 | 0.0021 | 0.0050 |
| F10 | 92100001 | 92200000 | 0.0043 | 0.0019 | 0.0037 |
| F10 | 92120001 | 92220000 | 0.0038 | 0.0044 | 0.0031 |
| F10 | 92140001 | 92240000 | 0.0065 | 0.0050 | 0.0055 |
| F10 | 92160001 | 92260000 | 0.0061 | 0.0050 | 0.0049 |
| F10 | 92180001 | 92280000 | 0.0061 | 0.0055 | 0.0048 |
| F10 | 92200001 | 92300000 | 0.0065 | 0.0058 | 0.0053 |
| F10 | 92220001 | 92320000 | 0.0068 | 0.0032 | 0.0056 |
| F10 | 92240001 | 92340000 | 0.0042 | 0.0020 | 0.0030 |
| F10 | 92260001 | 92360000 | 0.0029 | 0.0031 | 0.0020 |
| F10 | 92280001 | 92380000 | 0.0041 | 0.0033 | 0.0033 |
| F10 | 92300001 | 92400000 | 0.0043 | 0.0032 | 0.0037 |
| F10 | 92320001 | 92420000 | 0.0044 | 0.0051 | 0.0038 |
| F10 | 92340001 | 92440000 | 0.0065 | 0.0059 | 0.0058 |
| F10 | 92360001 | 92460000 | 0.0073 | 0.0051 | 0.0065 |
| F10 | 92380001 | 92480000 | 0.0064 | 0.0043 | 0.0054 |
| F10 | 92400001 | 92500000 | 0.0057 | 0.0037 | 0.0045 |
| F10 | 92420001 | 92520000 | 0.0049 | 0.0017 | 0.0037 |
| F10 | 92440001 | 92540000 | 0.0031 | 0.0011 | 0.0018 |
| F10 | 92460001 | 92560000 | 0.0031 | 0.0013 | 0.0015 |
| F10 | 92480001 | 92580000 | 0.0035 | 0.0013 | 0.0019 |
| F10 | 92500001 | 92600000 | 0.0037 | 0.0020 | 0.0020 |
| F10 | 92520001 | 92620000 | 0.0046 | 0.0029 | 0.0027 |
| F10 | 92540001 | 92640000 | 0.0053 | 0.0024 | 0.0038 |
| F10 | 92560001 | 92660000 | 0.0041 | 0.0038 | 0.0029 |
| F10 | 92580001 | 92680000 | 0.0051 | 0.0038 | 0.0042 |
| F10 | 92600001 | 92700000 | 0.0048 | 0.0033 | 0.0041 |
| F10 | 92620001 | 92720000 | 0.0039 | 0.0042 | 0.0034 |
| F10 | 92640001 | 92740000 | 0.0046 | 0.0054 | 0.0038 |
| F10 | 92660001 | 92760000 | 0.0060 | 0.0044 | 0.0051 |
| F10 | 92680001 | 92780000 | 0.0048 | 0.0043 | 0.0038 |
| F10 | 92700001 | 92800000 | 0.0048 | 0.0042 | 0.0037 |
| F10 | 92720001 | 92820000 | 0.0050 | 0.0023 | 0.0038 |
| F10 | 92740001 | 92840000 | 0.0035 | 0.0011 | 0.0027 |
| F10 | 92760001 | 92860000 | 0.0026 | 0.0003 | 0.0017 |
| F10 | 92780001 | 92880000 | 0.0021 | 0.0004 | 0.0010 |
| F10 | 92800001 | 92900000 | 0.0022 | 0.0004 | 0.0011 |
| F10 | 92820001 | 92920000 | 0.0022 | 0.0005 | 0.0013 |
| F10 | 92840001 | 92940000 | 0.0024 | 0.0015 | 0.0016 |
| F10 | 92860001 | 92960000 | 0.0030 | 0.0020 | 0.0025 |
| F10 | 92880001 | 92980000 | 0.0033 | 0.0035 | 0.0031 |
| F10 | 92900001 | 93000000 | 0.0046 | 0.0035 | 0.0044 |
| F10 | 92920001 | 93020000 | 0.0043 | 0.0038 | 0.0043 |
| F10 | 92940001 | 93040000 | 0.0040 | 0.0034 | 0.0038 |
| F10 | 92960001 | 93060000 | 0.0034 | 0.0046 | 0.0031 |
| F10 | 92980001 | 93080000 | 0.0045 | 0.0054 | 0.0041 |
| F10 | 93000001 | 93100000 | 0.0057 | 0.0061 | 0.0053 |
| F10 | 93020001 | 93120000 | 0.0067 | 0.0066 | 0.0061 |
| F10 | 93040001 | 93140000 | 0.0073 | 0.0071 | 0.0067 |
| F10 | 93060001 | 93160000 | 0.0082 | 0.0072 | 0.0076 |
| F10 | 93080001 | 93180000 | 0.0082 | 0.0074 | 0.0076 |
| F10 | 93100001 | 93200000 | 0.0080 | 0.0072 | 0.0075 |

| Chr | Start | End | both | A1 | A2 |
|---|---|---|---|---|---|
| F10 | 93120001 | 93220000 | 0.0075 | 0.0063 | 0.0071 |
| F10 | 93140001 | 93240000 | 0.0065 | 0.0054 | 0.0062 |
| F10 | 93160001 | 93260000 | 0.0054 | 0.0052 | 0.0052 |
| F10 | 93180001 | 93280000 | 0.0052 | 0.0039 | 0.0051 |
| F10 | 93200001 | 93300000 | 0.0040 | 0.0037 | 0.0039 |
| F10 | 93220001 | 93320000 | 0.0039 | 0.0041 | 0.0038 |
| F10 | 93240001 | 93340000 | 0.0042 | 0.0058 | 0.0041 |
| F10 | 93260001 | 93360000 | 0.0058 | 0.0053 | 0.0057 |
| F10 | 93280001 | 93380000 | 0.0054 | 0.0050 | 0.0052 |
| F10 | 93300001 | 93400000 | 0.0050 | 0.0058 | 0.0048 |
| F10 | 93320001 | 93420000 | 0.0057 | 0.0077 | 0.0054 |
| F10 | 93340001 | 93440000 | 0.0075 | 0.0088 | 0.0072 |
| F10 | 93360001 | 93460000 | 0.0083 | 0.0103 | 0.0078 |
| F10 | 93380001 | 93480000 | 0.0098 | 0.0109 | 0.0091 |
| F10 | 93400001 | 93500000 | 0.0096 | 0.0108 | 0.0086 |
| F10 | 93420001 | 93520000 | 0.0100 | 0.0095 | 0.0089 |
| F10 | 93440001 | 93540000 | 0.0086 | 0.0074 | 0.0075 |
| F10 | 93460001 | 93560000 | 0.0069 | 0.0054 | 0.0060 |
| F10 | 93480001 | 93580000 | 0.0050 | 0.0053 | 0.0044 |
| F10 | 93500001 | 93600000 | 0.0055 | 0.0050 | 0.0052 |
| F10 | 93520001 | 93620000 | 0.0048 | 0.0043 | 0.0045 |
| F10 | 93540001 | 93640000 | 0.0041 | 0.0038 | 0.0039 |
| F10 | 93560001 | 93660000 | 0.0035 | 0.0045 | 0.0031 |
| F10 | 93580001 | 93680000 | 0.0041 | 0.0042 | 0.0037 |
| F10 | 93600001 | 93700000 | 0.0038 | 0.0044 | 0.0035 |
| F10 | 93620001 | 93720000 | 0.0041 | 0.0064 | 0.0037 |
| F10 | 93640001 | 93740000 | 0.0055 | 0.0068 | 0.0051 |
| F10 | 93660001 | 93760000 | 0.0062 | 0.0064 | 0.0059 |
| F10 | 93680001 | 93780000 | 0.0057 | 0.0054 | 0.0054 |
| F10 | 93700001 | 93800000 | 0.0052 | 0.0043 | 0.0048 |
| F10 | 93720001 | 93820000 | 0.0044 | 0.0024 | 0.0040 |
| F10 | 93740001 | 93840000 | 0.0030 | 0.0018 | 0.0026 |
| F10 | 93760001 | 93860000 | 0.0024 | 0.0015 | 0.0019 |
| F10 | 93780001 | 93880000 | 0.0018 | 0.0020 | 0.0012 |
| F10 | 93800001 | 93900000 | 0.0015 | 0.0024 | 0.0010 |
| F10 | 93820001 | 93920000 | 0.0014 | 0.0027 | 0.0009 |
| F10 | 93840001 | 93940000 | 0.0018 | 0.0028 | 0.0013 |
| F10 | 93860001 | 93960000 | 0.0018 | 0.0046 | 0.0014 |
| F10 | 93880001 | 93980000 | 0.0039 | 0.0053 | 0.0036 |
| F10 | 93900001 | 94000000 | 0.0049 | 0.0053 | 0.0047 |
| F10 | 93920001 | 94020000 | 0.0052 | 0.0060 | 0.0050 |
| F10 | 93940001 | 94040000 | 0.0058 | 0.0064 | 0.0056 |
| F10 | 93960001 | 94060000 | 0.0062 | 0.0046 | 0.0060 |
| F10 | 93980001 | 94080000 | 0.0042 | 0.0038 | 0.0040 |
| F10 | 94000001 | 94100000 | 0.0034 | 0.0036 | 0.0030 |
| F10 | 94020001 | 94120000 | 0.0031 | 0.0027 | 0.0026 |
| F10 | 94040001 | 94140000 | 0.0020 | 0.0021 | 0.0015 |
| F10 | 94060001 | 94160000 | 0.0013 | 0.0022 | 0.0007 |
| F10 | 94080001 | 94180000 | 0.0012 | 0.0037 | 0.0005 |
| F10 | 94100001 | 94200000 | 0.0025 | 0.0054 | 0.0017 |
| F10 | 94120001 | 94220000 | 0.0041 | 0.0050 | 0.0031 |
| F10 | 94140001 | 94240000 | 0.0046 | 0.0047 | 0.0035 |
| F10 | 94160001 | 94260000 | 0.0047 | 0.0044 | 0.0036 |
| F10 | 94180001 | 94280000 | 0.0051 | 0.0031 | 0.0042 |
| F10 | 94200001 | 94300000 | 0.0042 | 0.0017 | 0.0033 |
| F10 | 94220001 | 94320000 | 0.0028 | 0.0022 | 0.0023 |
| F10 | 94240001 | 94340000 | 0.0023 | 0.0023 | 0.0019 |
| F10 | 94260001 | 94360000 | 0.0021 | 0.0024 | 0.0017 |

| Chr | Start | End | both | A1 | A2 |
|---|---|---|---|---|---|
| F10 | 94280001 | 94380000 | 0.0018 | 0.0022 | 0.0014 |
| F10 | 94300001 | 94400000 | 0.0016 | 0.0023 | 0.0012 |
| F10 | 94320001 | 94420000 | 0.0017 | 0.0020 | 0.0012 |
| F10 | 94340001 | 94440000 | 0.0021 | 0.0019 | 0.0018 |
| F10 | 94360001 | 94460000 | 0.0025 | 0.0016 | 0.0024 |
| F10 | 94380001 | 94480000 | 0.0026 | 0.0013 | 0.0025 |
| F10 | 94400001 | 94500000 | 0.0025 | 0.0010 | 0.0025 |
| F10 | 94420001 | 94520000 | 0.0022 | 0.0015 | 0.0024 |
| F10 | 94440001 | 94540000 | 0.0019 | 0.0020 | 0.0018 |
| F10 | 94460001 | 94560000 | 0.0018 | 0.0020 | 0.0016 |
| F10 | 94480001 | 94580000 | 0.0020 | 0.0019 | 0.0016 |
| F10 | 94500001 | 94600000 | 0.0023 | 0.0019 | 0.0019 |
| F10 | 94520001 | 94620000 | 0.0024 | 0.0017 | 0.0020 |
| F10 | 94540001 | 94640000 | 0.0032 | 0.0016 | 0.0030 |
| F10 | 94560001 | 94660000 | 0.0039 | 0.0023 | 0.0038 |
| F10 | 94580001 | 94680000 | 0.0037 | 0.0032 | 0.0036 |
| F10 | 94600001 | 94700000 | 0.0036 | 0.0045 | 0.0031 |
| F10 | 94620001 | 94720000 | 0.0045 | 0.0042 | 0.0039 |
| F10 | 94640001 | 94740000 | 0.0037 | 0.0047 | 0.0030 |
| F10 | 94660001 | 94760000 | 0.0032 | 0.0040 | 0.0023 |
| F10 | 94680001 | 94780000 | 0.0030 | 0.0038 | 0.0023 |
| F10 | 94700001 | 94800000 | 0.0032 | 0.0028 | 0.0028 |
| F10 | 94720001 | 94820000 | 0.0023 | 0.0031 | 0.0019 |
| F10 | 94740001 | 94840000 | 0.0023 | 0.0029 | 0.0018 |
| F10 | 94760001 | 94860000 | 0.0024 | 0.0030 | 0.0020 |
| F10 | 94780001 | 94880000 | 0.0023 | 0.0031 | 0.0018 |
| F10 | 94800001 | 94900000 | 0.0023 | 0.0034 | 0.0018 |
| F10 | 94820001 | 94920000 | 0.0029 | 0.0030 | 0.0024 |
| F10 | 94840001 | 94940000 | 0.0028 | 0.0027 | 0.0024 |
| F10 | 94860001 | 94960000 | 0.0024 | 0.0029 | 0.0017 |
| F10 | 94880001 | 94980000 | 0.0027 | 0.0025 | 0.0017 |
| F10 | 94900001 | 95000000 | 0.0024 | 0.0022 | 0.0012 |
| F10 | 94920001 | 95020000 | 0.0019 | 0.0030 | 0.0005 |
| F10 | 94940001 | 95040000 | 0.0027 | 0.0028 | 0.0010 |
| F10 | 94960001 | 95060000 | 0.0028 | 0.0025 | 0.0014 |
| F10 | 94980001 | 95080000 | 0.0026 | 0.0021 | 0.0016 |
| F10 | 95000001 | 95100000 | 0.0022 | 0.0014 | 0.0015 |
| F10 | 95020001 | 95120000 | 0.0016 | 0.0006 | 0.0013 |
| F10 | 95040001 | 95140000 | 0.0007 | 0.0003 | 0.0007 |
| F10 | 95060001 | 95160000 | 0.0003 | 0.0000 | 0.0003 |
| F10 | 95080001 | 95180000 | 0.0000 | 0.0043 | 0.0000 |
| F11 | 1 | 100000 | 0.0020 | 0.0019 | 0.0019 |
| F11 | 20001 | 120000 | 0.0011 | 0.0011 | 0.0011 |
| F11 | 40001 | 140000 | 0.0006 | 0.0006 | 0.0006 |
| F11 | 60001 | 160000 | 0.0003 | 0.0003 | 0.0003 |
| F11 | 80001 | 180000 | 0.0003 | 0.0003 | 0.0003 |
| F11 | 100001 | 200000 | 0.0003 | 0.0003 | 0.0003 |
| F11 | 120001 | 220000 | 0.0004 | 0.0004 | 0.0004 |
| F11 | 140001 | 240000 | 0.0011 | 0.0011 | 0.0011 |
| F11 | 160001 | 260000 | 0.0013 | 0.0011 | 0.0013 |
| F11 | 180001 | 280000 | 0.0013 | 0.0010 | 0.0013 |
| F11 | 200001 | 300000 | 0.0016 | 0.0014 | 0.0016 |
| F11 | 220001 | 320000 | 0.0022 | 0.0018 | 0.0022 |
| F11 | 240001 | 340000 | 0.0017 | 0.0013 | 0.0017 |
| F11 | 260001 | 360000 | 0.0019 | 0.0016 | 0.0019 |
| F11 | 280001 | 380000 | 0.0019 | 0.0016 | 0.0019 |
| F11 | 300001 | 400000 | 0.0018 | 0.0012 | 0.0018 |
| F11 | 320001 | 420000 | 0.0014 | 0.0007 | 0.0013 |

| Chr | Start | End | both | A1 | A2 |
|---|---|---|---|---|---|
| F11 | 340001 | 440000 | 0.0015 | 0.0005 | 0.0013 |
| F11 | 360001 | 460000 | 0.0013 | 0.0001 | 0.0010 |
| F11 | 380001 | 480000 | 0.0014 | 0.0001 | 0.0012 |
| F11 | 400001 | 500000 | 0.0015 | 0.0001 | 0.0012 |
| F11 | 420001 | 520000 | 0.0017 | 0.0001 | 0.0015 |
| F11 | 440001 | 540000 | 0.0015 | 0.0001 | 0.0015 |
| F11 | 460001 | 560000 | 0.0015 | 0.0001 | 0.0015 |
| F11 | 480001 | 580000 | 0.0017 | 0.0001 | 0.0017 |
| F11 | 500001 | 600000 | 0.0016 | 0.0001 | 0.0016 |
| F11 | 520001 | 620000 | 0.0014 | 0.0001 | 0.0015 |
| F11 | 540001 | 640000 | 0.0017 | 0.0002 | 0.0018 |
| F11 | 560001 | 660000 | 0.0018 | 0.0002 | 0.0017 |
| F11 | 580001 | 680000 | 0.0018 | 0.0002 | 0.0015 |
| F11 | 600001 | 700000 | 0.0020 | 0.0002 | 0.0015 |
| F11 | 620001 | 720000 | 0.0021 | 0.0004 | 0.0014 |
| F11 | 640001 | 740000 | 0.0019 | 0.0003 | 0.0012 |
| F11 | 660001 | 760000 | 0.0020 | 0.0003 | 0.0012 |
| F11 | 680001 | 780000 | 0.0018 | 0.0003 | 0.0010 |
| F11 | 700001 | 800000 | 0.0015 | 0.0003 | 0.0007 |
| F11 | 720001 | 820000 | 0.0015 | 0.0003 | 0.0007 |
| F11 | 740001 | 840000 | 0.0018 | 0.0009 | 0.0010 |
| F11 | 760001 | 860000 | 0.0020 | 0.0014 | 0.0012 |
| F11 | 780001 | 880000 | 0.0020 | 0.0015 | 0.0012 |
| F11 | 800001 | 900000 | 0.0020 | 0.0016 | 0.0012 |
| F11 | 820001 | 920000 | 0.0029 | 0.0022 | 0.0023 |
| F11 | 840001 | 940000 | 0.0026 | 0.0018 | 0.0019 |
| F11 | 860001 | 960000 | 0.0023 | 0.0014 | 0.0017 |
| F11 | 880001 | 980000 | 0.0024 | 0.0013 | 0.0018 |
| F11 | 900001 | 1000000 | 0.0026 | 0.0012 | 0.0022 |
| F11 | 920001 | 1020000 | 0.0017 | 0.0003 | 0.0013 |
| F11 | 940001 | 1040000 | 0.0018 | 0.0001 | 0.0015 |
| F11 | 960001 | 1060000 | 0.0019 | 0.0001 | 0.0018 |
| F11 | 980001 | 1080000 | 0.0021 | 0.0002 | 0.0020 |
| F11 | 1000001 | 1100000 | 0.0019 | 0.0002 | 0.0017 |
| F11 | 1020001 | 1120000 | 0.0017 | 0.0002 | 0.0014 |
| F11 | 1040001 | 1140000 | 0.0016 | 0.0003 | 0.0011 |
| F11 | 1060001 | 1160000 | 0.0014 | 0.0002 | 0.0008 |
| F11 | 1080001 | 1180000 | 0.0015 | 0.0002 | 0.0007 |
| F11 | 1100001 | 1200000 | 0.0016 | 0.0001 | 0.0009 |
| F11 | 1120001 | 1220000 | 0.0017 | 0.0002 | 0.0010 |
| F11 | 1140001 | 1240000 | 0.0027 | 0.0011 | 0.0020 |
| F11 | 1160001 | 1260000 | 0.0027 | 0.0011 | 0.0019 |
| F11 | 1180001 | 1280000 | 0.0023 | 0.0013 | 0.0016 |
| F11 | 1200001 | 1300000 | 0.0022 | 0.0013 | 0.0013 |
| F11 | 1220001 | 1320000 | 0.0022 | 0.0013 | 0.0012 |
| F11 | 1240001 | 1340000 | 0.0015 | 0.0007 | 0.0005 |
| F11 | 1260001 | 1360000 | 0.0016 | 0.0008 | 0.0005 |
| F11 | 1280001 | 1380000 | 0.0018 | 0.0008 | 0.0005 |
| F11 | 1300001 | 1400000 | 0.0021 | 0.0012 | 0.0010 |
| F11 | 1320001 | 1420000 | 0.0022 | 0.0012 | 0.0012 |
| F11 | 1340001 | 1440000 | 0.0021 | 0.0010 | 0.0011 |
| F11 | 1360001 | 1460000 | 0.0022 | 0.0010 | 0.0014 |
| F11 | 1380001 | 1480000 | 0.0022 | 0.0010 | 0.0017 |
| F11 | 1400001 | 1500000 | 0.0020 | 0.0007 | 0.0015 |
| F11 | 1420001 | 1520000 | 0.0018 | 0.0008 | 0.0014 |
| F11 | 1440001 | 1540000 | 0.0018 | 0.0010 | 0.0013 |
| F11 | 1460001 | 1560000 | 0.0016 | 0.0010 | 0.0011 |
| F11 | 1480001 | 1580000 | 0.0015 | 0.0010 | 0.0008 |

| Chr | Start | End | both | A1 | A2 |
|---|---|---|---|---|---|
| F11 | 1500001 | 1600000 | 0.0015 | 0.0009 | 0.0006 |
| F11 | 1520001 | 1620000 | 0.0015 | 0.0009 | 0.0006 |
| F11 | 1540001 | 1640000 | 0.0014 | 0.0007 | 0.0005 |
| F11 | 1560001 | 1660000 | 0.0016 | 0.0007 | 0.0008 |
| F11 | 1580001 | 1680000 | 0.0017 | 0.0007 | 0.0011 |
| F11 | 1600001 | 1700000 | 0.0019 | 0.0007 | 0.0014 |
| F11 | 1620001 | 1720000 | 0.0019 | 0.0006 | 0.0013 |
| F11 | 1640001 | 1740000 | 0.0019 | 0.0006 | 0.0013 |
| F11 | 1660001 | 1760000 | 0.0017 | 0.0006 | 0.0009 |
| F11 | 1680001 | 1780000 | 0.0016 | 0.0007 | 0.0007 |
| F11 | 1700001 | 1800000 | 0.0013 | 0.0006 | 0.0004 |
| F11 | 1720001 | 1820000 | 0.0012 | 0.0006 | 0.0003 |
| F11 | 1740001 | 1840000 | 0.0013 | 0.0006 | 0.0004 |
| F11 | 1760001 | 1860000 | 0.0014 | 0.0006 | 0.0004 |
| F11 | 1780001 | 1880000 | 0.0016 | 0.0007 | 0.0005 |
| F11 | 1800001 | 1900000 | 0.0029 | 0.0021 | 0.0018 |
| F11 | 1820001 | 1920000 | 0.0031 | 0.0023 | 0.0022 |
| F11 | 1840001 | 1940000 | 0.0030 | 0.0023 | 0.0022 |
| F11 | 1860001 | 1960000 | 0.0030 | 0.0024 | 0.0023 |
| F11 | 1880001 | 1980000 | 0.0028 | 0.0022 | 0.0020 |
| F11 | 1900001 | 2000000 | 0.0016 | 0.0009 | 0.0008 |
| F11 | 1920001 | 2020000 | 0.0014 | 0.0008 | 0.0005 |
| F11 | 1940001 | 2040000 | 0.0014 | 0.0007 | 0.0004 |
| F11 | 1960001 | 2060000 | 0.0014 | 0.0006 | 0.0003 |
| F11 | 1980001 | 2080000 | 0.0017 | 0.0008 | 0.0005 |
| F11 | 2000001 | 2100000 | 0.0016 | 0.0008 | 0.0005 |
| F11 | 2020001 | 2120000 | 0.0016 | 0.0007 | 0.0005 |
| F11 | 2040001 | 2140000 | 0.0017 | 0.0007 | 0.0005 |
| F11 | 2060001 | 2160000 | 0.0018 | 0.0007 | 0.0006 |
| F11 | 2080001 | 2180000 | 0.0015 | 0.0005 | 0.0004 |
| F11 | 2100001 | 2200000 | 0.0015 | 0.0005 | 0.0004 |
| F11 | 2120001 | 2220000 | 0.0016 | 0.0006 | 0.0005 |
| F11 | 2140001 | 2240000 | 0.0016 | 0.0009 | 0.0005 |
| F11 | 2160001 | 2260000 | 0.0014 | 0.0010 | 0.0004 |
| F11 | 2180001 | 2280000 | 0.0016 | 0.0011 | 0.0007 |
| F11 | 2200001 | 2300000 | 0.0015 | 0.0010 | 0.0006 |
| F11 | 2220001 | 2320000 | 0.0016 | 0.0012 | 0.0009 |
| F11 | 2240001 | 2340000 | 0.0016 | 0.0010 | 0.0008 |
| F11 | 2260001 | 2360000 | 0.0016 | 0.0009 | 0.0009 |
| F11 | 2280001 | 2380000 | 0.0014 | 0.0008 | 0.0006 |
| F11 | 2300001 | 2400000 | 0.0013 | 0.0009 | 0.0006 |
| F11 | 2320001 | 2420000 | 0.0012 | 0.0007 | 0.0003 |
| F11 | 2340001 | 2440000 | 0.0011 | 0.0008 | 0.0003 |
| F11 | 2360001 | 2460000 | 0.0013 | 0.0010 | 0.0004 |
| F11 | 2380001 | 2480000 | 0.0013 | 0.0010 | 0.0004 |
| F11 | 2400001 | 2500000 | 0.0017 | 0.0014 | 0.0007 |
| F11 | 2420001 | 2520000 | 0.0023 | 0.0019 | 0.0013 |
| F11 | 2440001 | 2540000 | 0.0023 | 0.0019 | 0.0012 |
| F11 | 2460001 | 2560000 | 0.0022 | 0.0018 | 0.0013 |
| F11 | 2480001 | 2580000 | 0.0022 | 0.0018 | 0.0014 |
| F11 | 2500001 | 2600000 | 0.0019 | 0.0014 | 0.0010 |
| F11 | 2520001 | 2620000 | 0.0014 | 0.0009 | 0.0005 |
| F11 | 2540001 | 2640000 | 0.0014 | 0.0010 | 0.0006 |
| F11 | 2560001 | 2660000 | 0.0015 | 0.0009 | 0.0005 |
| F11 | 2580001 | 2680000 | 0.0022 | 0.0015 | 0.0011 |
| F11 | 2600001 | 2700000 | 0.0024 | 0.0016 | 0.0012 |
| F11 | 2620001 | 2720000 | 0.0025 | 0.0016 | 0.0012 |
| F11 | 2640001 | 2740000 | 0.0025 | 0.0014 | 0.0012 |

| Chr | Start | End | both | A1 | A2 |
|---|---|---|---|---|---|
| F11 | 2660001 | 2760000 | 0.0023 | 0.0013 | 0.0015 |
| F11 | 2680001 | 2780000 | 0.0018 | 0.0007 | 0.0012 |
| F11 | 2700001 | 2800000 | 0.0016 | 0.0004 | 0.0012 |
| F11 | 2720001 | 2820000 | 0.0012 | 0.0003 | 0.0012 |
| F11 | 2740001 | 2840000 | 0.0011 | 0.0002 | 0.0012 |
| F11 | 2760001 | 2860000 | 0.0011 | 0.0001 | 0.0012 |
| F11 | 2780001 | 2880000 | 0.0009 | 0.0001 | 0.0011 |
| F11 | 2800001 | 2900000 | 0.0011 | 0.0003 | 0.0013 |
| F11 | 2820001 | 2920000 | 0.0012 | 0.0003 | 0.0013 |
| F11 | 2840001 | 2940000 | 0.0010 | 0.0003 | 0.0011 |
| F11 | 2860001 | 2960000 | 0.0012 | 0.0004 | 0.0013 |
| F11 | 2880001 | 2980000 | 0.0011 | 0.0004 | 0.0012 |
| F11 | 2900001 | 3000000 | 0.0011 | 0.0003 | 0.0009 |
| F11 | 2920001 | 3020000 | 0.0015 | 0.0006 | 0.0011 |
| F11 | 2940001 | 3040000 | 0.0017 | 0.0008 | 0.0012 |
| F11 | 2960001 | 3060000 | 0.0015 | 0.0007 | 0.0008 |
| F11 | 2980001 | 3080000 | 0.0015 | 0.0007 | 0.0006 |
| F11 | 3000001 | 3100000 | 0.0015 | 0.0007 | 0.0007 |
| F11 | 3020001 | 3120000 | 0.0012 | 0.0004 | 0.0005 |
| F11 | 3040001 | 3140000 | 0.0018 | 0.0009 | 0.0011 |
| F11 | 3060001 | 3160000 | 0.0018 | 0.0008 | 0.0012 |
| F11 | 3080001 | 3180000 | 0.0020 | 0.0008 | 0.0014 |
| F11 | 3100001 | 3200000 | 0.0019 | 0.0007 | 0.0013 |
| F11 | 3120001 | 3220000 | 0.0018 | 0.0007 | 0.0013 |
| F11 | 3140001 | 3240000 | 0.0018 | 0.0002 | 0.0010 |
| F11 | 3160001 | 3260000 | 0.0021 | 0.0003 | 0.0012 |
| F11 | 3180001 | 3280000 | 0.0021 | 0.0005 | 0.0013 |
| F11 | 3200001 | 3300000 | 0.0022 | 0.0006 | 0.0013 |
| F11 | 3220001 | 3320000 | 0.0023 | 0.0006 | 0.0012 |
| F11 | 3240001 | 3340000 | 0.0019 | 0.0007 | 0.0009 |
| F11 | 3260001 | 3360000 | 0.0017 | 0.0007 | 0.0007 |
| F11 | 3280001 | 3380000 | 0.0016 | 0.0005 | 0.0005 |
| F11 | 3300001 | 3400000 | 0.0018 | 0.0008 | 0.0007 |
| F11 | 3320001 | 3420000 | 0.0018 | 0.0008 | 0.0008 |
| F11 | 3340001 | 3440000 | 0.0017 | 0.0006 | 0.0007 |
| F11 | 3360001 | 3460000 | 0.0016 | 0.0006 | 0.0006 |
| F11 | 3380001 | 3480000 | 0.0015 | 0.0006 | 0.0007 |
| F11 | 3400001 | 3500000 | 0.0014 | 0.0004 | 0.0005 |
| F11 | 3420001 | 3520000 | 0.0014 | 0.0004 | 0.0004 |
| F11 | 3440001 | 3540000 | 0.0011 | 0.0004 | 0.0003 |
| F11 | 3460001 | 3560000 | 0.0017 | 0.0010 | 0.0011 |
| F11 | 3480001 | 3580000 | 0.0018 | 0.0010 | 0.0011 |
| F11 | 3500001 | 3600000 | 0.0021 | 0.0014 | 0.0015 |
| F11 | 3520001 | 3620000 | 0.0021 | 0.0015 | 0.0016 |
| F11 | 3540001 | 3640000 | 0.0023 | 0.0015 | 0.0016 |
| F11 | 3560001 | 3660000 | 0.0020 | 0.0012 | 0.0013 |
| F11 | 3580001 | 3680000 | 0.0021 | 0.0012 | 0.0015 |
| F11 | 3600001 | 3700000 | 0.0018 | 0.0008 | 0.0012 |
| F11 | 3620001 | 3720000 | 0.0021 | 0.0010 | 0.0015 |
| F11 | 3640001 | 3740000 | 0.0022 | 0.0009 | 0.0017 |
| F11 | 3660001 | 3760000 | 0.0019 | 0.0006 | 0.0014 |
| F11 | 3680001 | 3780000 | 0.0019 | 0.0006 | 0.0015 |
| F11 | 3700001 | 3800000 | 0.0019 | 0.0006 | 0.0016 |
| F11 | 3720001 | 3820000 | 0.0017 | 0.0003 | 0.0014 |
| F11 | 3740001 | 3840000 | 0.0018 | 0.0003 | 0.0015 |
| F11 | 3760001 | 3860000 | 0.0020 | 0.0004 | 0.0017 |
| F11 | 3780001 | 3880000 | 0.0019 | 0.0003 | 0.0016 |
| F11 | 3800001 | 3900000 | 0.0019 | 0.0006 | 0.0015 |

| Chr | Start | End | both | A1 | A2 |
|---|---|---|---|---|---|
| F11 | 3820001 | 3920000 | 0.0019 | 0.0006 | 0.0013 |
| F11 | 3840001 | 3940000 | 0.0018 | 0.0006 | 0.0011 |
| F11 | 3860001 | 3960000 | 0.0014 | 0.0006 | 0.0007 |
| F11 | 3880001 | 3980000 | 0.0017 | 0.0008 | 0.0009 |
| F11 | 3900001 | 4000000 | 0.0020 | 0.0008 | 0.0013 |
| F11 | 3920001 | 4020000 | 0.0033 | 0.0020 | 0.0027 |
| F11 | 3940001 | 4040000 | 0.0034 | 0.0020 | 0.0029 |
| F11 | 3960001 | 4060000 | 0.0035 | 0.0021 | 0.0032 |
| F11 | 3980001 | 4080000 | 0.0034 | 0.0019 | 0.0032 |
| F11 | 4000001 | 4100000 | 0.0031 | 0.0017 | 0.0030 |
| F11 | 4020001 | 4120000 | 0.0019 | 0.0004 | 0.0019 |
| F11 | 4040001 | 4140000 | 0.0017 | 0.0003 | 0.0017 |
| F11 | 4060001 | 4160000 | 0.0016 | 0.0003 | 0.0015 |
| F11 | 4080001 | 4180000 | 0.0016 | 0.0004 | 0.0015 |
| F11 | 4100001 | 4200000 | 0.0017 | 0.0004 | 0.0016 |
| F11 | 4120001 | 4220000 | 0.0019 | 0.0009 | 0.0018 |
| F11 | 4140001 | 4240000 | 0.0021 | 0.0010 | 0.0019 |
| F11 | 4160001 | 4260000 | 0.0024 | 0.0014 | 0.0023 |
| F11 | 4180001 | 4280000 | 0.0023 | 0.0014 | 0.0022 |
| F11 | 4200001 | 4300000 | 0.0022 | 0.0013 | 0.0020 |
| F11 | 4220001 | 4320000 | 0.0018 | 0.0009 | 0.0016 |
| F11 | 4240001 | 4340000 | 0.0017 | 0.0007 | 0.0014 |
| F11 | 4260001 | 4360000 | 0.0015 | 0.0004 | 0.0012 |
| F11 | 4280001 | 4380000 | 0.0018 | 0.0003 | 0.0013 |
| F11 | 4300001 | 4400000 | 0.0016 | 0.0003 | 0.0011 |
| F11 | 4320001 | 4420000 | 0.0020 | 0.0005 | 0.0014 |
| F11 | 4340001 | 4440000 | 0.0020 | 0.0004 | 0.0015 |
| F11 | 4360001 | 4460000 | 0.0031 | 0.0015 | 0.0027 |
| F11 | 4380001 | 4480000 | 0.0027 | 0.0015 | 0.0026 |
| F11 | 4400001 | 4500000 | 0.0025 | 0.0015 | 0.0025 |
| F11 | 4420001 | 4520000 | 0.0020 | 0.0013 | 0.0020 |
| F11 | 4440001 | 4540000 | 0.0018 | 0.0014 | 0.0018 |
| F11 | 4460001 | 4560000 | 0.0005 | 0.0003 | 0.0005 |
| F11 | 4480001 | 4580000 | 0.0004 | 0.0002 | 0.0004 |
| F11 | 4500001 | 4600000 | 0.0008 | 0.0002 | 0.0008 |
| F11 | 4520001 | 4620000 | 0.0010 | 0.0002 | 0.0010 |
| F11 | 4540001 | 4640000 | 0.0010 | 0.0001 | 0.0010 |
| F11 | 4560001 | 4660000 | 0.0012 | 0.0002 | 0.0010 |
| F11 | 4580001 | 4680000 | 0.0012 | 0.0002 | 0.0009 |
| F11 | 4600001 | 4700000 | 0.0011 | 0.0003 | 0.0006 |
| F11 | 4620001 | 4720000 | 0.0013 | 0.0005 | 0.0007 |
| F11 | 4640001 | 4740000 | 0.0017 | 0.0011 | 0.0008 |
| F11 | 4660001 | 4760000 | 0.0018 | 0.0011 | 0.0008 |
| F11 | 4680001 | 4780000 | 0.0018 | 0.0012 | 0.0008 |
| F11 | 4700001 | 4800000 | 0.0018 | 0.0011 | 0.0008 |
| F11 | 4720001 | 4820000 | 0.0016 | 0.0009 | 0.0005 |
| F11 | 4740001 | 4840000 | 0.0014 | 0.0004 | 0.0004 |
| F11 | 4760001 | 4860000 | 0.0014 | 0.0004 | 0.0004 |
| F11 | 4780001 | 4880000 | 0.0015 | 0.0005 | 0.0005 |
| F11 | 4800001 | 4900000 | 0.0015 | 0.0005 | 0.0005 |
| F11 | 4820001 | 4920000 | 0.0017 | 0.0008 | 0.0007 |
| F11 | 4840001 | 4940000 | 0.0015 | 0.0007 | 0.0006 |
| F11 | 4860001 | 4960000 | 0.0015 | 0.0007 | 0.0005 |
| F11 | 4880001 | 4980000 | 0.0014 | 0.0006 | 0.0005 |
| F11 | 4900001 | 5000000 | 0.0018 | 0.0010 | 0.0008 |
| F11 | 4920001 | 5020000 | 0.0025 | 0.0018 | 0.0016 |
| F11 | 4940001 | 5040000 | 0.0025 | 0.0017 | 0.0016 |
| F11 | 4960001 | 5060000 | 0.0027 | 0.0019 | 0.0019 |

| Chr | Start | End | both | A1 | A2 |
|---|---|---|---|---|---|
| F11 | 4980001 | 5080000 | 0.0028 | 0.0021 | 0.0021 |
| F11 | 5000001 | 5100000 | 0.0031 | 0.0024 | 0.0025 |
| F11 | 5020001 | 5120000 | 0.0023 | 0.0014 | 0.0016 |
| F11 | 5040001 | 5140000 | 0.0024 | 0.0015 | 0.0016 |
| F11 | 5060001 | 5160000 | 0.0023 | 0.0013 | 0.0014 |
| F11 | 5080001 | 5180000 | 0.0022 | 0.0012 | 0.0013 |
| F11 | 5100001 | 5200000 | 0.0015 | 0.0006 | 0.0008 |
| F11 | 5120001 | 5220000 | 0.0016 | 0.0009 | 0.0010 |
| F11 | 5140001 | 5240000 | 0.0015 | 0.0009 | 0.0010 |
| F11 | 5160001 | 5260000 | 0.0014 | 0.0009 | 0.0010 |
| F11 | 5180001 | 5280000 | 0.0017 | 0.0008 | 0.0012 |
| F11 | 5200001 | 5300000 | 0.0017 | 0.0007 | 0.0011 |
| F11 | 5220001 | 5320000 | 0.0016 | 0.0004 | 0.0010 |
| F11 | 5240001 | 5340000 | 0.0019 | 0.0006 | 0.0014 |
| F11 | 5260001 | 5360000 | 0.0026 | 0.0012 | 0.0021 |
| F11 | 5280001 | 5380000 | 0.0024 | 0.0012 | 0.0020 |
| F11 | 5300001 | 5400000 | 0.0028 | 0.0015 | 0.0022 |
| F11 | 5320001 | 5420000 | 0.0028 | 0.0015 | 0.0021 |
| F11 | 5340001 | 5440000 | 0.0025 | 0.0013 | 0.0018 |
| F11 | 5360001 | 5460000 | 0.0019 | 0.0007 | 0.0011 |
| F11 | 5380001 | 5480000 | 0.0019 | 0.0007 | 0.0011 |
| F11 | 5400001 | 5500000 | 0.0015 | 0.0004 | 0.0006 |
| F11 | 5420001 | 5520000 | 0.0015 | 0.0006 | 0.0007 |
| F11 | 5440001 | 5540000 | 0.0015 | 0.0005 | 0.0006 |
| F11 | 5460001 | 5560000 | 0.0015 | 0.0006 | 0.0005 |
| F11 | 5480001 | 5580000 | 0.0014 | 0.0005 | 0.0004 |
| F11 | 5500001 | 5600000 | 0.0013 | 0.0005 | 0.0003 |
| F11 | 5520001 | 5620000 | 0.0013 | 0.0004 | 0.0002 |
| F11 | 5540001 | 5640000 | 0.0019 | 0.0009 | 0.0008 |
| F11 | 5560001 | 5660000 | 0.0020 | 0.0010 | 0.0009 |
| F11 | 5580001 | 5680000 | 0.0022 | 0.0012 | 0.0012 |
| F11 | 5600001 | 5700000 | 0.0022 | 0.0012 | 0.0012 |
| F11 | 5620001 | 5720000 | 0.0022 | 0.0011 | 0.0012 |
| F11 | 5640001 | 5740000 | 0.0016 | 0.0007 | 0.0006 |
| F11 | 5660001 | 5760000 | 0.0015 | 0.0006 | 0.0005 |
| F11 | 5680001 | 5780000 | 0.0013 | 0.0006 | 0.0003 |
| F11 | 5700001 | 5800000 | 0.0013 | 0.0006 | 0.0003 |
| F11 | 5720001 | 5820000 | 0.0016 | 0.0010 | 0.0007 |
| F11 | 5740001 | 5840000 | 0.0019 | 0.0015 | 0.0011 |
| F11 | 5760001 | 5860000 | 0.0020 | 0.0015 | 0.0012 |
| F11 | 5780001 | 5880000 | 0.0023 | 0.0019 | 0.0016 |
| F11 | 5800001 | 5900000 | 0.0025 | 0.0020 | 0.0018 |
| F11 | 5820001 | 5920000 | 0.0027 | 0.0023 | 0.0020 |
| F11 | 5840001 | 5940000 | 0.0027 | 0.0024 | 0.0020 |
| F11 | 5860001 | 5960000 | 0.0028 | 0.0025 | 0.0021 |
| F11 | 5880001 | 5980000 | 0.0024 | 0.0022 | 0.0017 |
| F11 | 5900001 | 6000000 | 0.0023 | 0.0020 | 0.0016 |
| F11 | 5920001 | 6020000 | 0.0019 | 0.0019 | 0.0011 |
| F11 | 5940001 | 6040000 | 0.0016 | 0.0014 | 0.0008 |
| F11 | 5960001 | 6060000 | 0.0016 | 0.0012 | 0.0007 |
| F11 | 5980001 | 6080000 | 0.0016 | 0.0010 | 0.0006 |
| F11 | 6000001 | 6100000 | 0.0017 | 0.0012 | 0.0007 |
| F11 | 6020001 | 6120000 | 0.0015 | 0.0006 | 0.0005 |
| F11 | 6040001 | 6140000 | 0.0016 | 0.0006 | 0.0005 |
| F11 | 6060001 | 6160000 | 0.0021 | 0.0011 | 0.0011 |
| F11 | 6080001 | 6180000 | 0.0024 | 0.0014 | 0.0014 |
| F11 | 6100001 | 6200000 | 0.0025 | 0.0014 | 0.0015 |
| F11 | 6120001 | 6220000 | 0.0030 | 0.0020 | 0.0020 |

| Chr | Start | End | both | A1 | A2 |
|---|---|---|---|---|---|
| F11 | 6140001 | 6240000 | 0.0029 | 0.0020 | 0.0020 |
| F11 | 6160001 | 6260000 | 0.0023 | 0.0015 | 0.0014 |
| F11 | 6180001 | 6280000 | 0.0021 | 0.0011 | 0.0011 |
| F11 | 6200001 | 6300000 | 0.0018 | 0.0010 | 0.0009 |
| F11 | 6220001 | 6320000 | 0.0016 | 0.0008 | 0.0008 |
| F11 | 6240001 | 6340000 | 0.0014 | 0.0006 | 0.0006 |
| F11 | 6260001 | 6360000 | 0.0015 | 0.0008 | 0.0008 |
| F11 | 6280001 | 6380000 | 0.0021 | 0.0014 | 0.0014 |
| F11 | 6300001 | 6400000 | 0.0022 | 0.0014 | 0.0014 |
| F11 | 6320001 | 6420000 | 0.0018 | 0.0010 | 0.0010 |
| F11 | 6340001 | 6440000 | 0.0022 | 0.0013 | 0.0014 |
| F11 | 6360001 | 6460000 | 0.0022 | 0.0013 | 0.0015 |
| F11 | 6380001 | 6480000 | 0.0018 | 0.0011 | 0.0012 |
| F11 | 6400001 | 6500000 | 0.0017 | 0.0011 | 0.0012 |
| F11 | 6420001 | 6520000 | 0.0018 | 0.0011 | 0.0012 |
| F11 | 6440001 | 6540000 | 0.0014 | 0.0009 | 0.0008 |
| F11 | 6460001 | 6560000 | 0.0012 | 0.0007 | 0.0006 |
| F11 | 6480001 | 6580000 | 0.0013 | 0.0007 | 0.0005 |
| F11 | 6500001 | 6600000 | 0.0016 | 0.0008 | 0.0007 |
| F11 | 6520001 | 6620000 | 0.0015 | 0.0008 | 0.0007 |
| F11 | 6540001 | 6640000 | 0.0016 | 0.0009 | 0.0008 |
| F11 | 6560001 | 6660000 | 0.0015 | 0.0009 | 0.0008 |
| F11 | 6580001 | 6680000 | 0.0012 | 0.0006 | 0.0005 |
| F11 | 6600001 | 6700000 | 0.0012 | 0.0005 | 0.0004 |
| F11 | 6620001 | 6720000 | 0.0013 | 0.0006 | 0.0006 |
| F11 | 6640001 | 6740000 | 0.0017 | 0.0008 | 0.0011 |
| F11 | 6660001 | 6760000 | 0.0023 | 0.0014 | 0.0016 |
| F11 | 6680001 | 6780000 | 0.0024 | 0.0015 | 0.0017 |
| F11 | 6700001 | 6800000 | 0.0023 | 0.0014 | 0.0016 |
| F11 | 6720001 | 6820000 | 0.0021 | 0.0012 | 0.0014 |
| F11 | 6740001 | 6840000 | 0.0020 | 0.0012 | 0.0011 |
| F11 | 6760001 | 6860000 | 0.0014 | 0.0006 | 0.0006 |
| F11 | 6780001 | 6880000 | 0.0021 | 0.0013 | 0.0013 |
| F11 | 6800001 | 6900000 | 0.0021 | 0.0013 | 0.0013 |
| F11 | 6820001 | 6920000 | 0.0035 | 0.0022 | 0.0026 |
| F11 | 6840001 | 6940000 | 0.0034 | 0.0020 | 0.0026 |
| F11 | 6860001 | 6960000 | 0.0035 | 0.0020 | 0.0027 |
| F11 | 6880001 | 6980000 | 0.0030 | 0.0016 | 0.0022 |
| F11 | 6900001 | 7000000 | 0.0030 | 0.0016 | 0.0021 |
| F11 | 6920001 | 7020000 | 0.0018 | 0.0007 | 0.0009 |
| F11 | 6940001 | 7040000 | 0.0015 | 0.0007 | 0.0006 |
| F11 | 6960001 | 7060000 | 0.0016 | 0.0008 | 0.0006 |
| F11 | 6980001 | 7080000 | 0.0015 | 0.0006 | 0.0005 |
| F11 | 7000001 | 7100000 | 0.0016 | 0.0007 | 0.0005 |
| F11 | 7020001 | 7120000 | 0.0014 | 0.0005 | 0.0004 |
| F11 | 7040001 | 7140000 | 0.0018 | 0.0008 | 0.0008 |
| F11 | 7060001 | 7160000 | 0.0028 | 0.0018 | 0.0019 |
| F11 | 7080001 | 7180000 | 0.0027 | 0.0017 | 0.0020 |
| F11 | 7100001 | 7200000 | 0.0029 | 0.0018 | 0.0024 |
| F11 | 7120001 | 7220000 | 0.0031 | 0.0019 | 0.0029 |
| F11 | 7140001 | 7240000 | 0.0030 | 0.0016 | 0.0026 |
| F11 | 7160001 | 7260000 | 0.0033 | 0.0019 | 0.0030 |
| F11 | 7180001 | 7280000 | 0.0035 | 0.0022 | 0.0031 |
| F11 | 7200001 | 7300000 | 0.0037 | 0.0024 | 0.0030 |
| F11 | 7220001 | 7320000 | 0.0035 | 0.0023 | 0.0026 |
| F11 | 7240001 | 7340000 | 0.0031 | 0.0022 | 0.0024 |
| F11 | 7260001 | 7360000 | 0.0017 | 0.0009 | 0.0009 |
| F11 | 7280001 | 7380000 | 0.0015 | 0.0008 | 0.0005 |

| Chr | Start | End | both | A1 | A2 |
|---|---|---|---|---|---|
| F11 | 7300001 | 7400000 | 0.0011 | 0.0006 | 0.0002 |
| F11 | 7320001 | 7420000 | 0.0012 | 0.0009 | 0.0003 |
| F11 | 7340001 | 7440000 | 0.0012 | 0.0011 | 0.0004 |
| F11 | 7360001 | 7460000 | 0.0010 | 0.0011 | 0.0004 |
| F11 | 7380001 | 7480000 | 0.0008 | 0.0011 | 0.0004 |
| F11 | 7400001 | 7500000 | 0.0007 | 0.0012 | 0.0003 |
| F11 | 7420001 | 7520000 | 0.0011 | 0.0018 | 0.0009 |
| F11 | 7440001 | 7540000 | 0.0011 | 0.0017 | 0.0009 |
| F11 | 7460001 | 7560000 | 0.0014 | 0.0021 | 0.0011 |
| F11 | 7480001 | 7580000 | 0.0018 | 0.0024 | 0.0015 |
| F11 | 7500001 | 7600000 | 0.0022 | 0.0028 | 0.0019 |
| F11 | 7520001 | 7620000 | 0.0018 | 0.0023 | 0.0014 |
| F11 | 7540001 | 7640000 | 0.0026 | 0.0028 | 0.0022 |
| F11 | 7560001 | 7660000 | 0.0025 | 0.0023 | 0.0020 |
| F11 | 7580001 | 7680000 | 0.0028 | 0.0024 | 0.0024 |
| F11 | 7600001 | 7700000 | 0.0029 | 0.0022 | 0.0025 |
| F11 | 7620001 | 7720000 | 0.0028 | 0.0019 | 0.0026 |
| F11 | 7640001 | 7740000 | 0.0025 | 0.0014 | 0.0024 |
| F11 | 7660001 | 7760000 | 0.0026 | 0.0014 | 0.0025 |
| F11 | 7680001 | 7780000 | 0.0024 | 0.0012 | 0.0024 |
| F11 | 7700001 | 7800000 | 0.0024 | 0.0011 | 0.0024 |
| F11 | 7720001 | 7820000 | 0.0028 | 0.0015 | 0.0028 |
| F11 | 7740001 | 7840000 | 0.0028 | 0.0016 | 0.0028 |
| F11 | 7760001 | 7860000 | 0.0027 | 0.0022 | 0.0026 |
| F11 | 7780001 | 7880000 | 0.0023 | 0.0023 | 0.0020 |
| F11 | 7800001 | 7900000 | 0.0019 | 0.0024 | 0.0016 |
| F11 | 7820001 | 7920000 | 0.0014 | 0.0023 | 0.0009 |
| F11 | 7840001 | 7940000 | 0.0012 | 0.0024 | 0.0007 |
| F11 | 7860001 | 7960000 | 0.0013 | 0.0023 | 0.0008 |
| F11 | 7880001 | 7980000 | 0.0016 | 0.0026 | 0.0011 |
| F11 | 7900001 | 8000000 | 0.0020 | 0.0029 | 0.0015 |
| F11 | 7920001 | 8020000 | 0.0020 | 0.0028 | 0.0014 |
| F11 | 7940001 | 8040000 | 0.0018 | 0.0025 | 0.0012 |
| F11 | 7960001 | 8060000 | 0.0018 | 0.0024 | 0.0013 |
| F11 | 7980001 | 8080000 | 0.0018 | 0.0021 | 0.0013 |
| F11 | 8000001 | 8100000 | 0.0020 | 0.0021 | 0.0014 |
| F11 | 8020001 | 8120000 | 0.0022 | 0.0021 | 0.0017 |
| F11 | 8040001 | 8140000 | 0.0023 | 0.0019 | 0.0018 |
| F11 | 8060001 | 8160000 | 0.0023 | 0.0017 | 0.0015 |
| F11 | 8080001 | 8180000 | 0.0021 | 0.0014 | 0.0013 |
| F11 | 8100001 | 8200000 | 0.0015 | 0.0007 | 0.0007 |
| F11 | 8120001 | 8220000 | 0.0023 | 0.0013 | 0.0014 |
| F11 | 8140001 | 8240000 | 0.0027 | 0.0018 | 0.0019 |
| F11 | 8160001 | 8260000 | 0.0025 | 0.0017 | 0.0018 |
| F11 | 8180001 | 8280000 | 0.0024 | 0.0017 | 0.0018 |
| F11 | 8200001 | 8300000 | 0.0028 | 0.0020 | 0.0023 |
| F11 | 8220001 | 8320000 | 0.0024 | 0.0017 | 0.0019 |
| F11 | 8240001 | 8340000 | 0.0021 | 0.0014 | 0.0015 |
| F11 | 8260001 | 8360000 | 0.0021 | 0.0014 | 0.0015 |
| F11 | 8280001 | 8380000 | 0.0035 | 0.0026 | 0.0027 |
| F11 | 8300001 | 8400000 | 0.0031 | 0.0024 | 0.0023 |
| F11 | 8320001 | 8420000 | 0.0035 | 0.0024 | 0.0025 |
| F11 | 8340001 | 8440000 | 0.0033 | 0.0023 | 0.0024 |
| F11 | 8360001 | 8460000 | 0.0036 | 0.0027 | 0.0028 |
| F11 | 8380001 | 8480000 | 0.0026 | 0.0018 | 0.0019 |
| F11 | 8400001 | 8500000 | 0.0027 | 0.0019 | 0.0019 |
| F11 | 8420001 | 8520000 | 0.0022 | 0.0017 | 0.0016 |
| F11 | 8440001 | 8540000 | 0.0023 | 0.0018 | 0.0016 |

| Chr | Start | End | both | A1 | A2 |
|---|---|---|---|---|---|
| F11 | 8460001 | 8560000 | 0.0022 | 0.0015 | 0.0014 |
| F11 | 8480001 | 8580000 | 0.0020 | 0.0013 | 0.0013 |
| F11 | 8500001 | 8600000 | 0.0021 | 0.0014 | 0.0015 |
| F11 | 8520001 | 8620000 | 0.0019 | 0.0012 | 0.0011 |
| F11 | 8540001 | 8640000 | 0.0019 | 0.0012 | 0.0012 |
| F11 | 8560001 | 8660000 | 0.0017 | 0.0011 | 0.0009 |
| F11 | 8580001 | 8680000 | 0.0016 | 0.0009 | 0.0008 |
| F11 | 8600001 | 8700000 | 0.0015 | 0.0009 | 0.0008 |
| F11 | 8620001 | 8720000 | 0.0015 | 0.0010 | 0.0009 |
| F11 | 8640001 | 8740000 | 0.0018 | 0.0012 | 0.0012 |
| F11 | 8660001 | 8760000 | 0.0020 | 0.0014 | 0.0015 |
| F11 | 8680001 | 8780000 | 0.0023 | 0.0019 | 0.0020 |
| F11 | 8700001 | 8800000 | 0.0024 | 0.0017 | 0.0019 |
| F11 | 8720001 | 8820000 | 0.0023 | 0.0015 | 0.0017 |
| F11 | 8740001 | 8840000 | 0.0027 | 0.0018 | 0.0019 |
| F11 | 8760001 | 8860000 | 0.0035 | 0.0026 | 0.0028 |
| F11 | 8780001 | 8880000 | 0.0034 | 0.0024 | 0.0027 |
| F11 | 8800001 | 8900000 | 0.0034 | 0.0026 | 0.0028 |
| F11 | 8820001 | 8920000 | 0.0034 | 0.0026 | 0.0028 |
| F11 | 8840001 | 8940000 | 0.0026 | 0.0019 | 0.0021 |
| F11 | 8860001 | 8960000 | 0.0021 | 0.0013 | 0.0014 |
| F11 | 8880001 | 8980000 | 0.0019 | 0.0011 | 0.0012 |
| F11 | 8900001 | 9000000 | 0.0024 | 0.0016 | 0.0017 |
| F11 | 8920001 | 9020000 | 0.0024 | 0.0015 | 0.0017 |
| F11 | 8940001 | 9040000 | 0.0024 | 0.0016 | 0.0017 |
| F11 | 8960001 | 9060000 | 0.0021 | 0.0013 | 0.0013 |
| F11 | 8980001 | 9080000 | 0.0025 | 0.0016 | 0.0016 |
| F11 | 9000001 | 9100000 | 0.0017 | 0.0007 | 0.0008 |
| F11 | 9020001 | 9120000 | 0.0020 | 0.0011 | 0.0011 |
| F11 | 9040001 | 9140000 | 0.0020 | 0.0010 | 0.0011 |
| F11 | 9060001 | 9160000 | 0.0021 | 0.0010 | 0.0010 |
| F11 | 9080001 | 9180000 | 0.0018 | 0.0008 | 0.0008 |
| F11 | 9100001 | 9200000 | 0.0018 | 0.0009 | 0.0008 |
| F11 | 9120001 | 9220000 | 0.0021 | 0.0011 | 0.0011 |
| F11 | 9140001 | 9240000 | 0.0022 | 0.0012 | 0.0011 |
| F11 | 9160001 | 9260000 | 0.0021 | 0.0012 | 0.0011 |
| F11 | 9180001 | 9280000 | 0.0022 | 0.0013 | 0.0013 |
| F11 | 9200001 | 9300000 | 0.0023 | 0.0014 | 0.0014 |
| F11 | 9220001 | 9320000 | 0.0017 | 0.0008 | 0.0008 |
| F11 | 9240001 | 9340000 | 0.0017 | 0.0008 | 0.0008 |
| F11 | 9260001 | 9360000 | 0.0020 | 0.0011 | 0.0013 |
| F11 | 9280001 | 9380000 | 0.0019 | 0.0010 | 0.0012 |
| F11 | 9300001 | 9400000 | 0.0018 | 0.0009 | 0.0011 |
| F11 | 9320001 | 9420000 | 0.0024 | 0.0015 | 0.0017 |
| F11 | 9340001 | 9440000 | 0.0024 | 0.0016 | 0.0018 |
| F11 | 9360001 | 9460000 | 0.0021 | 0.0012 | 0.0014 |
| F11 | 9380001 | 9480000 | 0.0021 | 0.0011 | 0.0013 |
| F11 | 9400001 | 9500000 | 0.0022 | 0.0010 | 0.0013 |
| F11 | 9420001 | 9520000 | 0.0016 | 0.0005 | 0.0007 |
| F11 | 9440001 | 9540000 | 0.0022 | 0.0012 | 0.0013 |
| F11 | 9460001 | 9560000 | 0.0024 | 0.0014 | 0.0016 |
| F11 | 9480001 | 9580000 | 0.0024 | 0.0015 | 0.0017 |
| F11 | 9500001 | 9600000 | 0.0024 | 0.0015 | 0.0018 |
| F11 | 9520001 | 9620000 | 0.0023 | 0.0016 | 0.0018 |
| F11 | 9540001 | 9640000 | 0.0018 | 0.0010 | 0.0013 |
| F11 | 9560001 | 9660000 | 0.0016 | 0.0008 | 0.0012 |
| F11 | 9580001 | 9680000 | 0.0016 | 0.0007 | 0.0011 |
| F11 | 9600001 | 9700000 | 0.0015 | 0.0007 | 0.0009 |

| Chr | Start | End | both | A1 | A2 |
|---|---|---|---|---|---|
| F11 | 9620001 | 9720000 | 0.0019 | 0.0010 | 0.0013 |
| F11 | 9640001 | 9740000 | 0.0021 | 0.0012 | 0.0014 |
| F11 | 9660001 | 9760000 | 0.0029 | 0.0019 | 0.0021 |
| F11 | 9680001 | 9780000 | 0.0028 | 0.0019 | 0.0021 |
| F11 | 9700001 | 9800000 | 0.0034 | 0.0024 | 0.0025 |
| F11 | 9720001 | 9820000 | 0.0032 | 0.0022 | 0.0021 |
| F11 | 9740001 | 9840000 | 0.0028 | 0.0019 | 0.0018 |
| F11 | 9760001 | 9860000 | 0.0021 | 0.0014 | 0.0012 |
| F11 | 9780001 | 9880000 | 0.0023 | 0.0016 | 0.0014 |
| F11 | 9800001 | 9900000 | 0.0019 | 0.0011 | 0.0009 |
| F11 | 9820001 | 9920000 | 0.0018 | 0.0011 | 0.0010 |
| F11 | 9840001 | 9940000 | 0.0022 | 0.0015 | 0.0013 |
| F11 | 9860001 | 9960000 | 0.0021 | 0.0015 | 0.0011 |
| F11 | 9880001 | 9980000 | 0.0021 | 0.0016 | 0.0013 |
| F11 | 9900001 | 10000000 | 0.0020 | 0.0015 | 0.0012 |
| F11 | 9920001 | 10020000 | 0.0018 | 0.0014 | 0.0010 |
| F11 | 9940001 | 10040000 | 0.0016 | 0.0011 | 0.0008 |
| F11 | 9960001 | 10060000 | 0.0016 | 0.0009 | 0.0006 |
| F11 | 9980001 | 10080000 | 0.0017 | 0.0007 | 0.0005 |
| F11 | 10000001 | 10100000 | 0.0016 | 0.0007 | 0.0005 |
| F11 | 10020001 | 10120000 | 0.0016 | 0.0007 | 0.0006 |
| F11 | 10040001 | 10140000 | 0.0015 | 0.0006 | 0.0004 |
| F11 | 10060001 | 10160000 | 0.0019 | 0.0009 | 0.0009 |
| F11 | 10080001 | 10180000 | 0.0019 | 0.0011 | 0.0010 |
| F11 | 10100001 | 10200000 | 0.0019 | 0.0013 | 0.0012 |
| F11 | 10120001 | 10220000 | 0.0019 | 0.0014 | 0.0014 |
| F11 | 10140001 | 10240000 | 0.0018 | 0.0014 | 0.0015 |
| F11 | 10160001 | 10260000 | 0.0012 | 0.0010 | 0.0010 |
| F11 | 10180001 | 10280000 | 0.0011 | 0.0007 | 0.0008 |
| F11 | 10200001 | 10300000 | 0.0012 | 0.0007 | 0.0008 |
| F11 | 10220001 | 10320000 | 0.0010 | 0.0005 | 0.0006 |
| F11 | 10240001 | 10340000 | 0.0010 | 0.0004 | 0.0004 |
| F11 | 10260001 | 10360000 | 0.0014 | 0.0007 | 0.0007 |
| F11 | 10280001 | 10380000 | 0.0012 | 0.0005 | 0.0006 |
| F11 | 10300001 | 10400000 | 0.0029 | 0.0020 | 0.0022 |
| F11 | 10320001 | 10420000 | 0.0033 | 0.0023 | 0.0025 |
| F11 | 10340001 | 10440000 | 0.0038 | 0.0027 | 0.0029 |
| F11 | 10360001 | 10460000 | 0.0040 | 0.0028 | 0.0030 |
| F11 | 10380001 | 10480000 | 0.0041 | 0.0028 | 0.0032 |
| F11 | 10400001 | 10500000 | 0.0024 | 0.0013 | 0.0015 |
| F11 | 10420001 | 10520000 | 0.0023 | 0.0010 | 0.0013 |
| F11 | 10440001 | 10540000 | 0.0019 | 0.0007 | 0.0010 |
| F11 | 10460001 | 10560000 | 0.0017 | 0.0006 | 0.0009 |
| F11 | 10480001 | 10580000 | 0.0019 | 0.0008 | 0.0011 |
| F11 | 10500001 | 10600000 | 0.0023 | 0.0012 | 0.0015 |
| F11 | 10520001 | 10620000 | 0.0024 | 0.0012 | 0.0015 |
| F11 | 10540001 | 10640000 | 0.0025 | 0.0012 | 0.0016 |
| F11 | 10560001 | 10660000 | 0.0035 | 0.0019 | 0.0024 |
| F11 | 10580001 | 10680000 | 0.0034 | 0.0019 | 0.0023 |
| F11 | 10600001 | 10700000 | 0.0028 | 0.0015 | 0.0018 |
| F11 | 10620001 | 10720000 | 0.0028 | 0.0014 | 0.0017 |
| F11 | 10640001 | 10740000 | 0.0024 | 0.0013 | 0.0016 |
| F11 | 10660001 | 10760000 | 0.0017 | 0.0010 | 0.0011 |
| F11 | 10680001 | 10780000 | 0.0017 | 0.0010 | 0.0011 |
| F11 | 10700001 | 10800000 | 0.0026 | 0.0016 | 0.0020 |
| F11 | 10720001 | 10820000 | 0.0035 | 0.0022 | 0.0030 |
| F11 | 10740001 | 10840000 | 0.0040 | 0.0026 | 0.0034 |
| F11 | 10760001 | 10860000 | 0.0040 | 0.0025 | 0.0033 |

| Chr | Start | End | both | A1 | A2 |
|---|---|---|---|---|---|
| F11 | 10780001 | 10880000 | 0.0040 | 0.0027 | 0.0035 |
| F11 | 10800001 | 10900000 | 0.0036 | 0.0028 | 0.0032 |
| F11 | 10820001 | 10920000 | 0.0031 | 0.0026 | 0.0027 |
| F11 | 10840001 | 10940000 | 0.0028 | 0.0022 | 0.0024 |
| F11 | 10860001 | 10960000 | 0.0031 | 0.0022 | 0.0024 |
| F11 | 10880001 | 10980000 | 0.0032 | 0.0022 | 0.0024 |
| F11 | 10900001 | 11000000 | 0.0031 | 0.0019 | 0.0023 |
| F11 | 10920001 | 11020000 | 0.0028 | 0.0016 | 0.0019 |
| F11 | 10940001 | 11040000 | 0.0028 | 0.0017 | 0.0020 |
| F11 | 10960001 | 11060000 | 0.0021 | 0.0011 | 0.0013 |
| F11 | 10980001 | 11080000 | 0.0017 | 0.0007 | 0.0009 |
| F11 | 11000001 | 11100000 | 0.0014 | 0.0006 | 0.0007 |
| F11 | 11020001 | 11120000 | 0.0018 | 0.0010 | 0.0011 |
| F11 | 11040001 | 11140000 | 0.0035 | 0.0025 | 0.0027 |
| F11 | 11060001 | 11160000 | 0.0035 | 0.0026 | 0.0028 |
| F11 | 11080001 | 11180000 | 0.0037 | 0.0026 | 0.0029 |
| F11 | 11100001 | 11200000 | 0.0039 | 0.0027 | 0.0030 |
| F11 | 11120001 | 11220000 | 0.0036 | 0.0024 | 0.0027 |
| F11 | 11140001 | 11240000 | 0.0018 | 0.0008 | 0.0010 |
| F11 | 11160001 | 11260000 | 0.0017 | 0.0007 | 0.0009 |
| F11 | 11180001 | 11280000 | 0.0027 | 0.0017 | 0.0020 |
| F11 | 11200001 | 11300000 | 0.0028 | 0.0019 | 0.0022 |
| F11 | 11220001 | 11320000 | 0.0030 | 0.0018 | 0.0021 |
| F11 | 11240001 | 11340000 | 0.0032 | 0.0020 | 0.0023 |
| F11 | 11260001 | 11360000 | 0.0032 | 0.0021 | 0.0025 |
| F11 | 11280001 | 11380000 | 0.0023 | 0.0012 | 0.0015 |
| F11 | 11300001 | 11400000 | 0.0019 | 0.0008 | 0.0010 |
| F11 | 11320001 | 11420000 | 0.0017 | 0.0010 | 0.0011 |
| F11 | 11340001 | 11440000 | 0.0017 | 0.0009 | 0.0009 |
| F11 | 11360001 | 11460000 | 0.0018 | 0.0009 | 0.0010 |
| F11 | 11380001 | 11480000 | 0.0021 | 0.0015 | 0.0014 |
| F11 | 11400001 | 11500000 | 0.0020 | 0.0018 | 0.0012 |
| F11 | 11420001 | 11520000 | 0.0019 | 0.0024 | 0.0010 |
| F11 | 11440001 | 11540000 | 0.0022 | 0.0031 | 0.0014 |
| F11 | 11460001 | 11560000 | 0.0026 | 0.0036 | 0.0018 |
| F11 | 11480001 | 11580000 | 0.0027 | 0.0038 | 0.0020 |
| F11 | 11500001 | 11600000 | 0.0031 | 0.0041 | 0.0024 |
| F11 | 11520001 | 11620000 | 0.0031 | 0.0037 | 0.0026 |
| F11 | 11540001 | 11640000 | 0.0036 | 0.0041 | 0.0031 |
| F11 | 11560001 | 11660000 | 0.0040 | 0.0045 | 0.0035 |
| F11 | 11580001 | 11680000 | 0.0033 | 0.0037 | 0.0029 |
| F11 | 11600001 | 11700000 | 0.0027 | 0.0031 | 0.0025 |
| F11 | 11620001 | 11720000 | 0.0027 | 0.0030 | 0.0025 |
| F11 | 11640001 | 11740000 | 0.0017 | 0.0020 | 0.0015 |
| F11 | 11660001 | 11760000 | 0.0010 | 0.0014 | 0.0007 |
| F11 | 11680001 | 11780000 | 0.0010 | 0.0015 | 0.0007 |
| F11 | 11700001 | 11800000 | 0.0012 | 0.0016 | 0.0009 |
| F11 | 11720001 | 11820000 | 0.0016 | 0.0021 | 0.0013 |
| F11 | 11740001 | 11840000 | 0.0018 | 0.0024 | 0.0014 |
| F11 | 11760001 | 11860000 | 0.0025 | 0.0031 | 0.0020 |
| F11 | 11780001 | 11880000 | 0.0025 | 0.0031 | 0.0020 |
| F11 | 11800001 | 11900000 | 0.0029 | 0.0034 | 0.0024 |
| F11 | 11820001 | 11920000 | 0.0025 | 0.0028 | 0.0020 |
| F11 | 11840001 | 11940000 | 0.0025 | 0.0026 | 0.0022 |
| F11 | 11860001 | 11960000 | 0.0019 | 0.0019 | 0.0014 |
| F11 | 11880001 | 11980000 | 0.0018 | 0.0019 | 0.0013 |
| F11 | 11900001 | 12000000 | 0.0013 | 0.0015 | 0.0007 |
| F11 | 11920001 | 12020000 | 0.0014 | 0.0012 | 0.0005 |

| Chr | Start | End | both | A1 | A2 |
|---|---|---|---|---|---|
| F11 | 11940001 | 12040000 | 0.0015 | 0.0013 | 0.0007 |
| F11 | 11960001 | 12060000 | 0.0014 | 0.0011 | 0.0008 |
| F11 | 11980001 | 12080000 | 0.0016 | 0.0010 | 0.0009 |
| F11 | 12000001 | 12100000 | 0.0029 | 0.0020 | 0.0022 |
| F11 | 12020001 | 12120000 | 0.0029 | 0.0022 | 0.0025 |
| F11 | 12040001 | 12140000 | 0.0028 | 0.0021 | 0.0023 |
| F11 | 12060001 | 12160000 | 0.0031 | 0.0022 | 0.0025 |
| F11 | 12080001 | 12180000 | 0.0032 | 0.0023 | 0.0026 |
| F11 | 12100001 | 12200000 | 0.0027 | 0.0020 | 0.0022 |
| F11 | 12120001 | 12220000 | 0.0027 | 0.0020 | 0.0023 |
| F11 | 12140001 | 12240000 | 0.0025 | 0.0018 | 0.0020 |
| F11 | 12160001 | 12260000 | 0.0025 | 0.0018 | 0.0020 |
| F11 | 12180001 | 12280000 | 0.0023 | 0.0017 | 0.0017 |
| F11 | 12200001 | 12300000 | 0.0017 | 0.0011 | 0.0012 |
| F11 | 12220001 | 12320000 | 0.0016 | 0.0008 | 0.0009 |
| F11 | 12240001 | 12340000 | 0.0017 | 0.0009 | 0.0009 |
| F11 | 12260001 | 12360000 | 0.0024 | 0.0016 | 0.0016 |
| F11 | 12280001 | 12380000 | 0.0029 | 0.0021 | 0.0021 |
| F11 | 12300001 | 12400000 | 0.0030 | 0.0023 | 0.0022 |
| F11 | 12320001 | 12420000 | 0.0029 | 0.0023 | 0.0020 |
| F11 | 12340001 | 12440000 | 0.0029 | 0.0024 | 0.0022 |
| F11 | 12360001 | 12460000 | 0.0023 | 0.0019 | 0.0016 |
| F11 | 12380001 | 12480000 | 0.0020 | 0.0016 | 0.0014 |
| F11 | 12400001 | 12500000 | 0.0017 | 0.0013 | 0.0011 |
| F11 | 12420001 | 12520000 | 0.0026 | 0.0021 | 0.0022 |
| F11 | 12440001 | 12540000 | 0.0036 | 0.0030 | 0.0031 |
| F11 | 12460001 | 12560000 | 0.0035 | 0.0028 | 0.0028 |
| F11 | 12480001 | 12580000 | 0.0035 | 0.0027 | 0.0026 |
| F11 | 12500001 | 12600000 | 0.0036 | 0.0027 | 0.0027 |
| F11 | 12520001 | 12620000 | 0.0027 | 0.0018 | 0.0016 |
| F11 | 12540001 | 12640000 | 0.0016 | 0.0009 | 0.0007 |
| F11 | 12560001 | 12660000 | 0.0021 | 0.0013 | 0.0012 |
| F11 | 12580001 | 12680000 | 0.0018 | 0.0011 | 0.0011 |
| F11 | 12600001 | 12700000 | 0.0023 | 0.0017 | 0.0016 |
| F11 | 12620001 | 12720000 | 0.0023 | 0.0018 | 0.0016 |
| F11 | 12640001 | 12740000 | 0.0025 | 0.0021 | 0.0019 |
| F11 | 12660001 | 12760000 | 0.0026 | 0.0024 | 0.0019 |
| F11 | 12680001 | 12780000 | 0.0027 | 0.0027 | 0.0021 |
| F11 | 12700001 | 12800000 | 0.0022 | 0.0022 | 0.0016 |
| F11 | 12720001 | 12820000 | 0.0022 | 0.0022 | 0.0017 |
| F11 | 12740001 | 12840000 | 0.0020 | 0.0019 | 0.0014 |
| F11 | 12760001 | 12860000 | 0.0013 | 0.0013 | 0.0009 |
| F11 | 12780001 | 12880000 | 0.0014 | 0.0013 | 0.0009 |
| F11 | 12800001 | 12900000 | 0.0014 | 0.0011 | 0.0008 |
| F11 | 12820001 | 12920000 | 0.0016 | 0.0013 | 0.0009 |
| F11 | 12840001 | 12940000 | 0.0017 | 0.0013 | 0.0010 |
| F11 | 12860001 | 12960000 | 0.0018 | 0.0013 | 0.0012 |
| F11 | 12880001 | 12980000 | 0.0019 | 0.0014 | 0.0013 |
| F11 | 12900001 | 13000000 | 0.0020 | 0.0017 | 0.0015 |
| F11 | 12920001 | 13020000 | 0.0021 | 0.0019 | 0.0015 |
| F11 | 12940001 | 13040000 | 0.0024 | 0.0021 | 0.0019 |
| F11 | 12960001 | 13060000 | 0.0026 | 0.0027 | 0.0020 |
| F11 | 12980001 | 13080000 | 0.0026 | 0.0029 | 0.0020 |
| F11 | 13000001 | 13100000 | 0.0023 | 0.0029 | 0.0018 |
| F11 | 13020001 | 13120000 | 0.0020 | 0.0026 | 0.0015 |
| F11 | 13040001 | 13140000 | 0.0021 | 0.0028 | 0.0014 |
| F11 | 13060001 | 13160000 | 0.0030 | 0.0035 | 0.0022 |
| F11 | 13080001 | 13180000 | 0.0034 | 0.0035 | 0.0027 |

| Chr | Start | End | both | A1 | A2 |
|---|---|---|---|---|---|
| F11 | 13100001 | 13200000 | 0.0035 | 0.0034 | 0.0027 |
| F11 | 13120001 | 13220000 | 0.0036 | 0.0033 | 0.0028 |
| F11 | 13140001 | 13240000 | 0.0037 | 0.0029 | 0.0027 |
| F11 | 13160001 | 13260000 | 0.0023 | 0.0014 | 0.0014 |
| F11 | 13180001 | 13280000 | 0.0016 | 0.0007 | 0.0007 |
| F11 | 13200001 | 13300000 | 0.0019 | 0.0007 | 0.0010 |
| F11 | 13220001 | 13320000 | 0.0020 | 0.0008 | 0.0012 |
| F11 | 13240001 | 13340000 | 0.0019 | 0.0008 | 0.0012 |
| F11 | 13260001 | 13360000 | 0.0024 | 0.0012 | 0.0016 |
| F11 | 13280001 | 13380000 | 0.0031 | 0.0017 | 0.0022 |
| F11 | 13300001 | 13400000 | 0.0030 | 0.0017 | 0.0021 |
| F11 | 13320001 | 13420000 | 0.0033 | 0.0019 | 0.0023 |
| F11 | 13340001 | 13440000 | 0.0044 | 0.0031 | 0.0035 |
| F11 | 13360001 | 13460000 | 0.0053 | 0.0041 | 0.0046 |
| F11 | 13380001 | 13480000 | 0.0050 | 0.0037 | 0.0041 |
| F11 | 13400001 | 13500000 | 0.0049 | 0.0036 | 0.0040 |
| F11 | 13420001 | 13520000 | 0.0046 | 0.0034 | 0.0037 |
| F11 | 13440001 | 13540000 | 0.0033 | 0.0022 | 0.0024 |
| F11 | 13460001 | 13560000 | 0.0023 | 0.0009 | 0.0012 |
| F11 | 13480001 | 13580000 | 0.0019 | 0.0007 | 0.0010 |
| F11 | 13500001 | 13600000 | 0.0018 | 0.0007 | 0.0010 |
| F11 | 13520001 | 13620000 | 0.0016 | 0.0005 | 0.0008 |
| F11 | 13540001 | 13640000 | 0.0015 | 0.0003 | 0.0006 |
| F11 | 13560001 | 13660000 | 0.0012 | 0.0002 | 0.0003 |
| F11 | 13580001 | 13680000 | 0.0013 | 0.0002 | 0.0003 |
| F11 | 13600001 | 13700000 | 0.0015 | 0.0002 | 0.0003 |
| F11 | 13620001 | 13720000 | 0.0015 | 0.0002 | 0.0003 |
| F11 | 13640001 | 13740000 | 0.0015 | 0.0002 | 0.0002 |
| F11 | 13660001 | 13760000 | 0.0020 | 0.0008 | 0.0008 |
| F11 | 13680001 | 13780000 | 0.0019 | 0.0007 | 0.0007 |
| F11 | 13700001 | 13800000 | 0.0020 | 0.0007 | 0.0007 |
| F11 | 13720001 | 13820000 | 0.0020 | 0.0007 | 0.0007 |
| F11 | 13740001 | 13840000 | 0.0022 | 0.0009 | 0.0009 |
| F11 | 13760001 | 13860000 | 0.0021 | 0.0010 | 0.0009 |
| F11 | 13780001 | 13880000 | 0.0023 | 0.0013 | 0.0013 |
| F11 | 13800001 | 13900000 | 0.0025 | 0.0018 | 0.0018 |
| F11 | 13820001 | 13920000 | 0.0032 | 0.0027 | 0.0027 |
| F11 | 13840001 | 13940000 | 0.0034 | 0.0029 | 0.0030 |
| F11 | 13860001 | 13960000 | 0.0035 | 0.0029 | 0.0032 |
| F11 | 13880001 | 13980000 | 0.0035 | 0.0028 | 0.0032 |
| F11 | 13900001 | 14000000 | 0.0038 | 0.0030 | 0.0035 |
| F11 | 13920001 | 14020000 | 0.0034 | 0.0026 | 0.0030 |
| F11 | 13940001 | 14040000 | 0.0032 | 0.0023 | 0.0027 |
| F11 | 13960001 | 14060000 | 0.0031 | 0.0023 | 0.0025 |
| F11 | 13980001 | 14080000 | 0.0033 | 0.0026 | 0.0027 |
| F11 | 14000001 | 14100000 | 0.0031 | 0.0025 | 0.0024 |
| F11 | 14020001 | 14120000 | 0.0032 | 0.0026 | 0.0026 |
| F11 | 14040001 | 14140000 | 0.0042 | 0.0037 | 0.0037 |
| F11 | 14060001 | 14160000 | 0.0044 | 0.0037 | 0.0038 |
| F11 | 14080001 | 14180000 | 0.0043 | 0.0034 | 0.0036 |
| F11 | 14100001 | 14200000 | 0.0040 | 0.0029 | 0.0031 |
| F11 | 14120001 | 14220000 | 0.0037 | 0.0026 | 0.0028 |
| F11 | 14140001 | 14240000 | 0.0027 | 0.0015 | 0.0017 |
| F11 | 14160001 | 14260000 | 0.0031 | 0.0019 | 0.0023 |
| F11 | 14180001 | 14280000 | 0.0029 | 0.0018 | 0.0021 |
| F11 | 14200001 | 14300000 | 0.0034 | 0.0021 | 0.0025 |
| F11 | 14220001 | 14320000 | 0.0033 | 0.0023 | 0.0026 |
| F11 | 14240001 | 14340000 | 0.0041 | 0.0030 | 0.0033 |

| Chr | Start | End | both | A1 | A2 |
|---|---|---|---|---|---|
| F11 | 14260001 | 14360000 | 0.0031 | 0.0021 | 0.0022 |
| F11 | 14280001 | 14380000 | 0.0032 | 0.0023 | 0.0024 |
| F11 | 14300001 | 14400000 | 0.0027 | 0.0021 | 0.0022 |
| F11 | 14320001 | 14420000 | 0.0026 | 0.0020 | 0.0021 |
| F11 | 14340001 | 14440000 | 0.0020 | 0.0013 | 0.0016 |
| F11 | 14360001 | 14460000 | 0.0019 | 0.0013 | 0.0016 |
| F11 | 14380001 | 14480000 | 0.0018 | 0.0011 | 0.0014 |
| F11 | 14400001 | 14500000 | 0.0022 | 0.0016 | 0.0018 |
| F11 | 14420001 | 14520000 | 0.0023 | 0.0015 | 0.0017 |
| F11 | 14440001 | 14540000 | 0.0026 | 0.0020 | 0.0022 |
| F11 | 14460001 | 14560000 | 0.0028 | 0.0021 | 0.0023 |
| F11 | 14480001 | 14580000 | 0.0030 | 0.0022 | 0.0025 |
| F11 | 14500001 | 14600000 | 0.0029 | 0.0019 | 0.0025 |
| F11 | 14520001 | 14620000 | 0.0040 | 0.0029 | 0.0036 |
| F11 | 14540001 | 14640000 | 0.0033 | 0.0022 | 0.0029 |
| F11 | 14560001 | 14660000 | 0.0037 | 0.0026 | 0.0033 |
| F11 | 14580001 | 14680000 | 0.0038 | 0.0028 | 0.0032 |
| F11 | 14600001 | 14700000 | 0.0037 | 0.0026 | 0.0030 |
| F11 | 14620001 | 14720000 | 0.0027 | 0.0017 | 0.0020 |
| F11 | 14640001 | 14740000 | 0.0027 | 0.0018 | 0.0021 |
| F11 | 14660001 | 14760000 | 0.0019 | 0.0012 | 0.0015 |
| F11 | 14680001 | 14780000 | 0.0018 | 0.0008 | 0.0013 |
| F11 | 14700001 | 14800000 | 0.0014 | 0.0008 | 0.0010 |
| F11 | 14720001 | 14820000 | 0.0018 | 0.0011 | 0.0013 |
| F11 | 14740001 | 14840000 | 0.0020 | 0.0010 | 0.0014 |
| F11 | 14760001 | 14860000 | 0.0025 | 0.0013 | 0.0017 |
| F11 | 14780001 | 14880000 | 0.0023 | 0.0014 | 0.0016 |
| F11 | 14800001 | 14900000 | 0.0025 | 0.0012 | 0.0017 |
| F11 | 14820001 | 14920000 | 0.0025 | 0.0010 | 0.0016 |
| F11 | 14840001 | 14940000 | 0.0028 | 0.0012 | 0.0019 |
| F11 | 14860001 | 14960000 | 0.0026 | 0.0008 | 0.0017 |
| F11 | 14880001 | 14980000 | 0.0033 | 0.0014 | 0.0025 |
| F11 | 14900001 | 15000000 | 0.0031 | 0.0015 | 0.0026 |
| F11 | 14920001 | 15020000 | 0.0026 | 0.0012 | 0.0022 |
| F11 | 14940001 | 15040000 | 0.0025 | 0.0012 | 0.0020 |
| F11 | 14960001 | 15060000 | 0.0025 | 0.0013 | 0.0021 |
| F11 | 14980001 | 15080000 | 0.0022 | 0.0008 | 0.0016 |
| F11 | 15000001 | 15100000 | 0.0022 | 0.0006 | 0.0016 |
| F11 | 15020001 | 15120000 | 0.0021 | 0.0007 | 0.0016 |
| F11 | 15040001 | 15140000 | 0.0019 | 0.0004 | 0.0014 |
| F11 | 15060001 | 15160000 | 0.0023 | 0.0009 | 0.0018 |
| F11 | 15080001 | 15180000 | 0.0024 | 0.0013 | 0.0020 |
| F11 | 15100001 | 15200000 | 0.0026 | 0.0017 | 0.0023 |
| F11 | 15120001 | 15220000 | 0.0027 | 0.0020 | 0.0025 |
| F11 | 15140001 | 15240000 | 0.0030 | 0.0025 | 0.0028 |
| F11 | 15160001 | 15260000 | 0.0036 | 0.0030 | 0.0034 |
| F11 | 15180001 | 15280000 | 0.0037 | 0.0032 | 0.0036 |
| F11 | 15200001 | 15300000 | 0.0049 | 0.0043 | 0.0046 |
| F11 | 15220001 | 15320000 | 0.0050 | 0.0044 | 0.0047 |
| F11 | 15240001 | 15340000 | 0.0050 | 0.0046 | 0.0047 |
| F11 | 15260001 | 15360000 | 0.0043 | 0.0044 | 0.0039 |
| F11 | 15280001 | 15380000 | 0.0041 | 0.0043 | 0.0036 |
| F11 | 15300001 | 15400000 | 0.0026 | 0.0032 | 0.0023 |
| F11 | 15320001 | 15420000 | 0.0023 | 0.0031 | 0.0019 |
| F11 | 15340001 | 15440000 | 0.0021 | 0.0027 | 0.0017 |
| F11 | 15360001 | 15460000 | 0.0019 | 0.0025 | 0.0015 |
| F11 | 15380001 | 15480000 | 0.0015 | 0.0021 | 0.0011 |
| F11 | 15400001 | 15500000 | 0.0015 | 0.0019 | 0.0012 |

| Chr | Start | End | both | A1 | A2 |
|---|---|---|---|---|---|
| F11 | 15420001 | 15520000 | 0.0018 | 0.0021 | 0.0016 |
| F11 | 15440001 | 15540000 | 0.0028 | 0.0032 | 0.0025 |
| F11 | 15460001 | 15560000 | 0.0025 | 0.0028 | 0.0022 |
| F11 | 15480001 | 15580000 | 0.0031 | 0.0036 | 0.0026 |
| F11 | 15500001 | 15600000 | 0.0034 | 0.0041 | 0.0029 |
| F11 | 15520001 | 15620000 | 0.0035 | 0.0041 | 0.0030 |
| F11 | 15540001 | 15640000 | 0.0026 | 0.0032 | 0.0022 |
| F11 | 15560001 | 15660000 | 0.0028 | 0.0033 | 0.0023 |
| F11 | 15580001 | 15680000 | 0.0035 | 0.0035 | 0.0031 |
| F11 | 15600001 | 15700000 | 0.0035 | 0.0035 | 0.0031 |
| F11 | 15620001 | 15720000 | 0.0032 | 0.0030 | 0.0026 |
| F11 | 15640001 | 15740000 | 0.0031 | 0.0027 | 0.0025 |
| F11 | 15660001 | 15760000 | 0.0034 | 0.0025 | 0.0029 |
| F11 | 15680001 | 15780000 | 0.0024 | 0.0014 | 0.0018 |
| F11 | 15700001 | 15800000 | 0.0021 | 0.0011 | 0.0017 |
| F11 | 15720001 | 15820000 | 0.0027 | 0.0016 | 0.0023 |
| F11 | 15740001 | 15840000 | 0.0044 | 0.0029 | 0.0038 |
| F11 | 15760001 | 15860000 | 0.0041 | 0.0027 | 0.0036 |
| F11 | 15780001 | 15880000 | 0.0049 | 0.0034 | 0.0044 |
| F11 | 15800001 | 15900000 | 0.0050 | 0.0032 | 0.0044 |
| F11 | 15820001 | 15920000 | 0.0050 | 0.0031 | 0.0045 |
| F11 | 15840001 | 15940000 | 0.0040 | 0.0025 | 0.0036 |
| F11 | 15860001 | 15960000 | 0.0045 | 0.0032 | 0.0041 |
| F11 | 15880001 | 15980000 | 0.0036 | 0.0024 | 0.0032 |
| F11 | 15900001 | 16000000 | 0.0042 | 0.0030 | 0.0038 |
| F11 | 15920001 | 16020000 | 0.0039 | 0.0026 | 0.0034 |
| F11 | 15940001 | 16040000 | 0.0030 | 0.0017 | 0.0025 |
| F11 | 15960001 | 16060000 | 0.0031 | 0.0017 | 0.0024 |
| F11 | 15980001 | 16080000 | 0.0034 | 0.0019 | 0.0026 |
| F11 | 16000001 | 16100000 | 0.0030 | 0.0019 | 0.0023 |
| F11 | 16020001 | 16120000 | 0.0027 | 0.0019 | 0.0021 |
| F11 | 16040001 | 16140000 | 0.0028 | 0.0019 | 0.0021 |
| F11 | 16060001 | 16160000 | 0.0019 | 0.0013 | 0.0014 |
| F11 | 16080001 | 16180000 | 0.0020 | 0.0015 | 0.0015 |
| F11 | 16100001 | 16200000 | 0.0023 | 0.0014 | 0.0015 |
| F11 | 16120001 | 16220000 | 0.0025 | 0.0015 | 0.0016 |
| F11 | 16140001 | 16240000 | 0.0023 | 0.0016 | 0.0016 |
| F11 | 16160001 | 16260000 | 0.0028 | 0.0020 | 0.0021 |
| F11 | 16180001 | 16280000 | 0.0033 | 0.0025 | 0.0027 |
| F11 | 16200001 | 16300000 | 0.0032 | 0.0027 | 0.0027 |
| F11 | 16220001 | 16320000 | 0.0032 | 0.0027 | 0.0028 |
| F11 | 16240001 | 16340000 | 0.0033 | 0.0028 | 0.0028 |
| F11 | 16260001 | 16360000 | 0.0037 | 0.0032 | 0.0033 |
| F11 | 16280001 | 16380000 | 0.0035 | 0.0030 | 0.0030 |
| F11 | 16300001 | 16400000 | 0.0029 | 0.0023 | 0.0023 |
| F11 | 16320001 | 16420000 | 0.0032 | 0.0030 | 0.0026 |
| F11 | 16340001 | 16440000 | 0.0036 | 0.0037 | 0.0029 |
| F11 | 16360001 | 16460000 | 0.0036 | 0.0041 | 0.0028 |
| F11 | 16380001 | 16480000 | 0.0030 | 0.0038 | 0.0022 |
| F11 | 16400001 | 16500000 | 0.0029 | 0.0037 | 0.0020 |
| F11 | 16420001 | 16520000 | 0.0024 | 0.0032 | 0.0016 |
| F11 | 16440001 | 16540000 | 0.0019 | 0.0026 | 0.0012 |
| F11 | 16460001 | 16560000 | 0.0011 | 0.0018 | 0.0004 |
| F11 | 16480001 | 16580000 | 0.0011 | 0.0018 | 0.0005 |
| F11 | 16500001 | 16600000 | 0.0014 | 0.0021 | 0.0008 |
| F11 | 16520001 | 16620000 | 0.0016 | 0.0022 | 0.0010 |
| F11 | 16540001 | 16640000 | 0.0024 | 0.0028 | 0.0018 |
| F11 | 16560001 | 16660000 | 0.0024 | 0.0027 | 0.0018 |

| Chr | Start | End | both | A1 | A2 |
|---|---|---|---|---|---|
| F11 | 16580001 | 16680000 | 0.0026 | 0.0027 | 0.0021 |
| F11 | 16600001 | 16700000 | 0.0025 | 0.0026 | 0.0021 |
| F11 | 16620001 | 16720000 | 0.0024 | 0.0026 | 0.0019 |
| F11 | 16640001 | 16740000 | 0.0018 | 0.0022 | 0.0013 |
| F11 | 16660001 | 16760000 | 0.0020 | 0.0022 | 0.0015 |
| F11 | 16680001 | 16780000 | 0.0020 | 0.0023 | 0.0015 |
| F11 | 16700001 | 16800000 | 0.0022 | 0.0024 | 0.0017 |
| F11 | 16720001 | 16820000 | 0.0023 | 0.0025 | 0.0018 |
| F11 | 16740001 | 16840000 | 0.0022 | 0.0025 | 0.0017 |
| F11 | 16760001 | 16860000 | 0.0018 | 0.0022 | 0.0014 |
| F11 | 16780001 | 16880000 | 0.0018 | 0.0022 | 0.0014 |
| F11 | 16800001 | 16900000 | 0.0014 | 0.0019 | 0.0010 |
| F11 | 16820001 | 16920000 | 0.0016 | 0.0018 | 0.0013 |
| F11 | 16840001 | 16940000 | 0.0016 | 0.0017 | 0.0014 |
| F11 | 16860001 | 16960000 | 0.0016 | 0.0016 | 0.0014 |
| F11 | 16880001 | 16980000 | 0.0018 | 0.0018 | 0.0017 |
| F11 | 16900001 | 17000000 | 0.0027 | 0.0027 | 0.0025 |
| F11 | 16920001 | 17020000 | 0.0028 | 0.0029 | 0.0026 |
| F11 | 16940001 | 17040000 | 0.0030 | 0.0030 | 0.0027 |
| F11 | 16960001 | 17060000 | 0.0030 | 0.0031 | 0.0027 |
| F11 | 16980001 | 17080000 | 0.0026 | 0.0027 | 0.0021 |
| F11 | 17000001 | 17100000 | 0.0019 | 0.0021 | 0.0012 |
| F11 | 17020001 | 17120000 | 0.0017 | 0.0019 | 0.0009 |
| F11 | 17040001 | 17140000 | 0.0015 | 0.0019 | 0.0008 |
| F11 | 17060001 | 17160000 | 0.0020 | 0.0024 | 0.0012 |
| F11 | 17080001 | 17180000 | 0.0023 | 0.0028 | 0.0014 |
| F11 | 17100001 | 17200000 | 0.0020 | 0.0023 | 0.0013 |
| F11 | 17120001 | 17220000 | 0.0019 | 0.0022 | 0.0012 |
| F11 | 17140001 | 17240000 | 0.0017 | 0.0021 | 0.0010 |
| F11 | 17160001 | 17260000 | 0.0018 | 0.0022 | 0.0012 |
| F11 | 17180001 | 17280000 | 0.0026 | 0.0028 | 0.0021 |
| F11 | 17200001 | 17300000 | 0.0027 | 0.0029 | 0.0023 |
| F11 | 17220001 | 17320000 | 0.0025 | 0.0027 | 0.0021 |
| F11 | 17240001 | 17340000 | 0.0029 | 0.0031 | 0.0024 |
| F11 | 17260001 | 17360000 | 0.0023 | 0.0024 | 0.0019 |
| F11 | 17280001 | 17380000 | 0.0013 | 0.0014 | 0.0009 |
| F11 | 17300001 | 17400000 | 0.0017 | 0.0019 | 0.0012 |
| F11 | 17320001 | 17420000 | 0.0018 | 0.0019 | 0.0012 |
| F11 | 17340001 | 17440000 | 0.0017 | 0.0018 | 0.0012 |
| F11 | 17360001 | 17460000 | 0.0020 | 0.0021 | 0.0014 |
| F11 | 17380001 | 17480000 | 0.0017 | 0.0017 | 0.0012 |
| F11 | 17400001 | 17500000 | 0.0019 | 0.0019 | 0.0013 |
| F11 | 17420001 | 17520000 | 0.0027 | 0.0027 | 0.0020 |
| F11 | 17440001 | 17540000 | 0.0025 | 0.0025 | 0.0020 |
| F11 | 17460001 | 17560000 | 0.0024 | 0.0023 | 0.0020 |
| F11 | 17480001 | 17580000 | 0.0027 | 0.0026 | 0.0022 |
| F11 | 17500001 | 17600000 | 0.0023 | 0.0020 | 0.0018 |
| F11 | 17520001 | 17620000 | 0.0014 | 0.0010 | 0.0011 |
| F11 | 17540001 | 17640000 | 0.0013 | 0.0009 | 0.0010 |
| F11 | 17560001 | 17660000 | 0.0014 | 0.0008 | 0.0011 |
| F11 | 17580001 | 17680000 | 0.0013 | 0.0006 | 0.0011 |
| F11 | 17600001 | 17700000 | 0.0014 | 0.0004 | 0.0012 |
| F11 | 17620001 | 17720000 | 0.0015 | 0.0004 | 0.0013 |
| F11 | 17640001 | 17740000 | 0.0016 | 0.0005 | 0.0015 |
| F11 | 17660001 | 17760000 | 0.0019 | 0.0004 | 0.0018 |
| F11 | 17680001 | 17780000 | 0.0019 | 0.0004 | 0.0018 |
| F11 | 17700001 | 17800000 | 0.0019 | 0.0004 | 0.0018 |
| F11 | 17720001 | 17820000 | 0.0026 | 0.0008 | 0.0024 |

| Chr | Start | End | both | A1 | A2 |
|---|---|---|---|---|---|
| F11 | 17740001 | 17840000 | 0.0028 | 0.0011 | 0.0027 |
| F11 | 17760001 | 17860000 | 0.0023 | 0.0011 | 0.0023 |
| F11 | 17780001 | 17880000 | 0.0023 | 0.0013 | 0.0022 |
| F11 | 17800001 | 17900000 | 0.0025 | 0.0017 | 0.0024 |
| F11 | 17820001 | 17920000 | 0.0017 | 0.0014 | 0.0017 |
| F11 | 17840001 | 17940000 | 0.0020 | 0.0016 | 0.0019 |
| F11 | 17860001 | 17960000 | 0.0019 | 0.0016 | 0.0019 |
| F11 | 17880001 | 17980000 | 0.0019 | 0.0016 | 0.0018 |
| F11 | 17900001 | 18000000 | 0.0021 | 0.0018 | 0.0020 |
| F11 | 17920001 | 18020000 | 0.0022 | 0.0019 | 0.0021 |
| F11 | 17940001 | 18040000 | 0.0021 | 0.0020 | 0.0021 |
| F11 | 17960001 | 18060000 | 0.0021 | 0.0019 | 0.0020 |
| F11 | 17980001 | 18080000 | 0.0020 | 0.0018 | 0.0019 |
| F11 | 18000001 | 18100000 | 0.0013 | 0.0011 | 0.0014 |
| F11 | 18020001 | 18120000 | 0.0018 | 0.0014 | 0.0018 |
| F11 | 18040001 | 18140000 | 0.0017 | 0.0013 | 0.0017 |
| F11 | 18060001 | 18160000 | 0.0021 | 0.0017 | 0.0021 |
| F11 | 18080001 | 18180000 | 0.0027 | 0.0021 | 0.0027 |
| F11 | 18100001 | 18200000 | 0.0032 | 0.0024 | 0.0033 |
| F11 | 18120001 | 18220000 | 0.0029 | 0.0019 | 0.0030 |
| F11 | 18140001 | 18240000 | 0.0026 | 0.0017 | 0.0027 |
| F11 | 18160001 | 18260000 | 0.0030 | 0.0017 | 0.0031 |
| F11 | 18180001 | 18280000 | 0.0030 | 0.0015 | 0.0032 |
| F11 | 18200001 | 18300000 | 0.0027 | 0.0013 | 0.0029 |
| F11 | 18220001 | 18320000 | 0.0032 | 0.0017 | 0.0032 |
| F11 | 18240001 | 18340000 | 0.0031 | 0.0015 | 0.0031 |
| F11 | 18260001 | 18360000 | 0.0025 | 0.0012 | 0.0025 |
| F11 | 18280001 | 18380000 | 0.0022 | 0.0011 | 0.0021 |
| F11 | 18300001 | 18400000 | 0.0022 | 0.0012 | 0.0020 |
| F11 | 18320001 | 18420000 | 0.0018 | 0.0010 | 0.0016 |
| F11 | 18340001 | 18440000 | 0.0019 | 0.0012 | 0.0017 |
| F11 | 18360001 | 18460000 | 0.0023 | 0.0015 | 0.0020 |
| F11 | 18380001 | 18480000 | 0.0024 | 0.0017 | 0.0021 |
| F11 | 18400001 | 18500000 | 0.0022 | 0.0017 | 0.0020 |
| F11 | 18420001 | 18520000 | 0.0021 | 0.0017 | 0.0019 |
| F11 | 18440001 | 18540000 | 0.0019 | 0.0015 | 0.0017 |
| F11 | 18460001 | 18560000 | 0.0016 | 0.0012 | 0.0012 |
| F11 | 18480001 | 18580000 | 0.0014 | 0.0011 | 0.0009 |
| F11 | 18500001 | 18600000 | 0.0019 | 0.0017 | 0.0013 |
| F11 | 18520001 | 18620000 | 0.0018 | 0.0015 | 0.0011 |
| F11 | 18540001 | 18640000 | 0.0018 | 0.0016 | 0.0011 |
| F11 | 18560001 | 18660000 | 0.0019 | 0.0018 | 0.0013 |
| F11 | 18580001 | 18680000 | 0.0020 | 0.0019 | 0.0013 |
| F11 | 18600001 | 18700000 | 0.0018 | 0.0017 | 0.0011 |
| F11 | 18620001 | 18720000 | 0.0022 | 0.0022 | 0.0015 |
| F11 | 18640001 | 18740000 | 0.0039 | 0.0037 | 0.0031 |
| F11 | 18660001 | 18760000 | 0.0044 | 0.0041 | 0.0036 |
| F11 | 18680001 | 18780000 | 0.0045 | 0.0042 | 0.0037 |
| F11 | 18700001 | 18800000 | 0.0045 | 0.0041 | 0.0038 |
| F11 | 18720001 | 18820000 | 0.0045 | 0.0040 | 0.0037 |
| F11 | 18740001 | 18840000 | 0.0027 | 0.0022 | 0.0019 |
| F11 | 18760001 | 18860000 | 0.0028 | 0.0022 | 0.0020 |
| F11 | 18780001 | 18880000 | 0.0030 | 0.0022 | 0.0023 |
| F11 | 18800001 | 18900000 | 0.0029 | 0.0019 | 0.0020 |
| F11 | 18820001 | 18920000 | 0.0028 | 0.0018 | 0.0021 |
| F11 | 18840001 | 18940000 | 0.0031 | 0.0021 | 0.0024 |
| F11 | 18860001 | 18960000 | 0.0030 | 0.0018 | 0.0021 |
| F11 | 18880001 | 18980000 | 0.0026 | 0.0014 | 0.0017 |

| Chr | Start | End | both | A1 | A2 |
|---|---|---|---|---|---|
| F11 | 18900001 | 19000000 | 0.0023 | 0.0013 | 0.0014 |
| F11 | 18920001 | 19020000 | 0.0019 | 0.0010 | 0.0010 |
| F11 | 18940001 | 19040000 | 0.0016 | 0.0008 | 0.0008 |
| F11 | 18960001 | 19060000 | 0.0010 | 0.0005 | 0.0006 |
| F11 | 18980001 | 19080000 | 0.0017 | 0.0011 | 0.0012 |
| F11 | 19000001 | 19100000 | 0.0021 | 0.0013 | 0.0016 |
| F11 | 19020001 | 19120000 | 0.0021 | 0.0012 | 0.0016 |
| F11 | 19040001 | 19140000 | 0.0022 | 0.0011 | 0.0015 |
| F11 | 19060001 | 19160000 | 0.0027 | 0.0017 | 0.0018 |
| F11 | 19080001 | 19180000 | 0.0025 | 0.0014 | 0.0015 |
| F11 | 19100001 | 19200000 | 0.0022 | 0.0013 | 0.0013 |
| F11 | 19120001 | 19220000 | 0.0030 | 0.0021 | 0.0021 |
| F11 | 19140001 | 19240000 | 0.0033 | 0.0022 | 0.0023 |
| F11 | 19160001 | 19260000 | 0.0032 | 0.0017 | 0.0020 |
| F11 | 19180001 | 19280000 | 0.0034 | 0.0018 | 0.0021 |
| F11 | 19200001 | 19300000 | 0.0043 | 0.0024 | 0.0029 |
| F11 | 19220001 | 19320000 | 0.0037 | 0.0018 | 0.0022 |
| F11 | 19240001 | 19340000 | 0.0033 | 0.0017 | 0.0020 |
| F11 | 19260001 | 19360000 | 0.0031 | 0.0019 | 0.0022 |
| F11 | 19280001 | 19380000 | 0.0029 | 0.0018 | 0.0022 |
| F11 | 19300001 | 19400000 | 0.0023 | 0.0013 | 0.0016 |
| F11 | 19320001 | 19420000 | 0.0024 | 0.0014 | 0.0018 |
| F11 | 19340001 | 19440000 | 0.0030 | 0.0020 | 0.0024 |
| F11 | 19360001 | 19460000 | 0.0038 | 0.0026 | 0.0031 |
| F11 | 19380001 | 19480000 | 0.0044 | 0.0033 | 0.0037 |
| F11 | 19400001 | 19500000 | 0.0042 | 0.0031 | 0.0035 |
| F11 | 19420001 | 19520000 | 0.0049 | 0.0037 | 0.0040 |
| F11 | 19440001 | 19540000 | 0.0048 | 0.0034 | 0.0037 |
| F11 | 19460001 | 19560000 | 0.0048 | 0.0031 | 0.0034 |
| F11 | 19480001 | 19580000 | 0.0040 | 0.0023 | 0.0024 |
| F11 | 19500001 | 19600000 | 0.0042 | 0.0025 | 0.0027 |
| F11 | 19520001 | 19620000 | 0.0031 | 0.0017 | 0.0018 |
| F11 | 19540001 | 19640000 | 0.0026 | 0.0014 | 0.0015 |
| F11 | 19560001 | 19660000 | 0.0018 | 0.0010 | 0.0011 |
| F11 | 19580001 | 19680000 | 0.0020 | 0.0013 | 0.0015 |
| F11 | 19600001 | 19700000 | 0.0018 | 0.0011 | 0.0012 |
| F11 | 19620001 | 19720000 | 0.0020 | 0.0011 | 0.0014 |
| F11 | 19640001 | 19740000 | 0.0022 | 0.0011 | 0.0015 |
| F11 | 19660001 | 19760000 | 0.0022 | 0.0010 | 0.0013 |
| F11 | 19680001 | 19780000 | 0.0021 | 0.0008 | 0.0011 |
| F11 | 19700001 | 19800000 | 0.0037 | 0.0022 | 0.0025 |
| F11 | 19720001 | 19820000 | 0.0037 | 0.0022 | 0.0025 |
| F11 | 19740001 | 19840000 | 0.0038 | 0.0024 | 0.0027 |
| F11 | 19760001 | 19860000 | 0.0037 | 0.0024 | 0.0027 |
| F11 | 19780001 | 19880000 | 0.0038 | 0.0026 | 0.0030 |
| F11 | 19800001 | 19900000 | 0.0027 | 0.0015 | 0.0018 |
| F11 | 19820001 | 19920000 | 0.0029 | 0.0017 | 0.0018 |
| F11 | 19840001 | 19940000 | 0.0034 | 0.0019 | 0.0021 |
| F11 | 19860001 | 19960000 | 0.0037 | 0.0023 | 0.0024 |
| F11 | 19880001 | 19980000 | 0.0043 | 0.0028 | 0.0029 |
| F11 | 19900001 | 20000000 | 0.0039 | 0.0025 | 0.0026 |
| F11 | 19920001 | 20020000 | 0.0037 | 0.0023 | 0.0025 |
| F11 | 19940001 | 20040000 | 0.0029 | 0.0017 | 0.0019 |
| F11 | 19960001 | 20060000 | 0.0023 | 0.0014 | 0.0015 |
| F11 | 19980001 | 20080000 | 0.0018 | 0.0008 | 0.0011 |
| F11 | 20000001 | 20100000 | 0.0015 | 0.0008 | 0.0010 |
| F11 | 20020001 | 20120000 | 0.0012 | 0.0007 | 0.0009 |
| F11 | 20040001 | 20140000 | 0.0013 | 0.0009 | 0.0011 |

| Chr | Start | End | both | A1 | A2 |
|---|---|---|---|---|---|
| F11 | 20060001 | 20160000 | 0.0012 | 0.0008 | 0.0009 |
| F11 | 20080001 | 20180000 | 0.0016 | 0.0012 | 0.0013 |
| F11 | 20100001 | 20200000 | 0.0028 | 0.0022 | 0.0024 |
| F11 | 20120001 | 20220000 | 0.0033 | 0.0025 | 0.0026 |
| F11 | 20140001 | 20240000 | 0.0039 | 0.0029 | 0.0031 |
| F11 | 20160001 | 20260000 | 0.0042 | 0.0032 | 0.0034 |
| F11 | 20180001 | 20280000 | 0.0033 | 0.0023 | 0.0024 |
| F11 | 20200001 | 20300000 | 0.0022 | 0.0013 | 0.0014 |
| F11 | 20220001 | 20320000 | 0.0020 | 0.0012 | 0.0012 |
| F11 | 20240001 | 20340000 | 0.0019 | 0.0012 | 0.0014 |
| F11 | 20260001 | 20360000 | 0.0022 | 0.0014 | 0.0015 |
| F11 | 20280001 | 20380000 | 0.0035 | 0.0025 | 0.0027 |
| F11 | 20300001 | 20400000 | 0.0046 | 0.0037 | 0.0039 |
| F11 | 20320001 | 20420000 | 0.0048 | 0.0036 | 0.0039 |
| F11 | 20340001 | 20440000 | 0.0066 | 0.0054 | 0.0055 |
| F11 | 20360001 | 20460000 | 0.0068 | 0.0056 | 0.0059 |
| F11 | 20380001 | 20480000 | 0.0061 | 0.0050 | 0.0054 |
| F11 | 20400001 | 20500000 | 0.0050 | 0.0040 | 0.0044 |
| F11 | 20420001 | 20520000 | 0.0048 | 0.0040 | 0.0042 |
| F11 | 20440001 | 20540000 | 0.0025 | 0.0016 | 0.0020 |
| F11 | 20460001 | 20560000 | 0.0021 | 0.0014 | 0.0017 |
| F11 | 20480001 | 20580000 | 0.0020 | 0.0012 | 0.0015 |
| F11 | 20500001 | 20600000 | 0.0026 | 0.0019 | 0.0021 |
| F11 | 20520001 | 20620000 | 0.0028 | 0.0019 | 0.0021 |
| F11 | 20540001 | 20640000 | 0.0029 | 0.0020 | 0.0022 |
| F11 | 20560001 | 20660000 | 0.0028 | 0.0017 | 0.0019 |
| F11 | 20580001 | 20680000 | 0.0028 | 0.0017 | 0.0018 |
| F11 | 20600001 | 20700000 | 0.0022 | 0.0010 | 0.0010 |
| F11 | 20620001 | 20720000 | 0.0020 | 0.0009 | 0.0009 |
| F11 | 20640001 | 20740000 | 0.0018 | 0.0008 | 0.0009 |
| F11 | 20660001 | 20760000 | 0.0015 | 0.0006 | 0.0007 |
| F11 | 20680001 | 20780000 | 0.0010 | 0.0003 | 0.0003 |
| F11 | 20700001 | 20800000 | 0.0008 | 0.0002 | 0.0002 |
| F11 | 20720001 | 20820000 | 0.0016 | 0.0009 | 0.0009 |
| F11 | 20740001 | 20840000 | 0.0026 | 0.0019 | 0.0019 |
| F11 | 20760001 | 20860000 | 0.0030 | 0.0020 | 0.0022 |
| F11 | 20780001 | 20880000 | 0.0040 | 0.0030 | 0.0033 |
| F11 | 20800001 | 20900000 | 0.0044 | 0.0032 | 0.0034 |
| F11 | 20820001 | 20920000 | 0.0040 | 0.0028 | 0.0030 |
| F11 | 20840001 | 20940000 | 0.0031 | 0.0019 | 0.0020 |
| F11 | 20860001 | 20960000 | 0.0032 | 0.0022 | 0.0022 |
| F11 | 20880001 | 20980000 | 0.0030 | 0.0018 | 0.0018 |
| F11 | 20900001 | 21000000 | 0.0032 | 0.0019 | 0.0020 |
| F11 | 20920001 | 21020000 | 0.0032 | 0.0019 | 0.0020 |
| F11 | 20940001 | 21040000 | 0.0033 | 0.0022 | 0.0023 |
| F11 | 20960001 | 21060000 | 0.0033 | 0.0022 | 0.0022 |
| F11 | 20980001 | 21080000 | 0.0028 | 0.0019 | 0.0020 |
| F11 | 21000001 | 21100000 | 0.0024 | 0.0018 | 0.0019 |
| F11 | 21020001 | 21120000 | 0.0027 | 0.0021 | 0.0023 |
| F11 | 21040001 | 21140000 | 0.0023 | 0.0017 | 0.0020 |
| F11 | 21060001 | 21160000 | 0.0024 | 0.0017 | 0.0020 |
| F11 | 21080001 | 21180000 | 0.0023 | 0.0015 | 0.0017 |
| F11 | 21100001 | 21200000 | 0.0024 | 0.0015 | 0.0017 |
| F11 | 21120001 | 21220000 | 0.0024 | 0.0014 | 0.0016 |
| F11 | 21140001 | 21240000 | 0.0032 | 0.0020 | 0.0023 |
| F11 | 21160001 | 21260000 | 0.0027 | 0.0016 | 0.0018 |
| F11 | 21180001 | 21280000 | 0.0032 | 0.0017 | 0.0020 |
| F11 | 21200001 | 21300000 | 0.0028 | 0.0016 | 0.0018 |

| Chr | Start | End | both | A1 | A2 |
|---|---|---|---|---|---|
| F11 | 21220001 | 21320000 | 0.0025 | 0.0014 | 0.0016 |
| F11 | 21240001 | 21340000 | 0.0024 | 0.0014 | 0.0014 |
| F11 | 21260001 | 21360000 | 0.0027 | 0.0016 | 0.0018 |
| F11 | 21280001 | 21380000 | 0.0027 | 0.0018 | 0.0018 |
| F11 | 21300001 | 21400000 | 0.0031 | 0.0020 | 0.0022 |
| F11 | 21320001 | 21420000 | 0.0032 | 0.0021 | 0.0024 |
| F11 | 21340001 | 21440000 | 0.0033 | 0.0020 | 0.0023 |
| F11 | 21360001 | 21460000 | 0.0038 | 0.0025 | 0.0029 |
| F11 | 21380001 | 21480000 | 0.0034 | 0.0024 | 0.0027 |
| F11 | 21400001 | 21500000 | 0.0046 | 0.0034 | 0.0039 |
| F11 | 21420001 | 21520000 | 0.0046 | 0.0034 | 0.0039 |
| F11 | 21440001 | 21540000 | 0.0045 | 0.0033 | 0.0038 |
| F11 | 21460001 | 21560000 | 0.0041 | 0.0030 | 0.0034 |
| F11 | 21480001 | 21580000 | 0.0042 | 0.0029 | 0.0032 |
| F11 | 21500001 | 21600000 | 0.0029 | 0.0019 | 0.0020 |
| F11 | 21520001 | 21620000 | 0.0034 | 0.0023 | 0.0025 |
| F11 | 21540001 | 21640000 | 0.0033 | 0.0024 | 0.0026 |
| F11 | 21560001 | 21660000 | 0.0036 | 0.0027 | 0.0030 |
| F11 | 21580001 | 21680000 | 0.0033 | 0.0027 | 0.0030 |
| F11 | 21600001 | 21700000 | 0.0041 | 0.0032 | 0.0035 |
| F11 | 21620001 | 21720000 | 0.0036 | 0.0028 | 0.0030 |
| F11 | 21640001 | 21740000 | 0.0040 | 0.0029 | 0.0032 |
| F11 | 21660001 | 21760000 | 0.0043 | 0.0032 | 0.0034 |
| F11 | 21680001 | 21780000 | 0.0050 | 0.0037 | 0.0040 |
| F11 | 21700001 | 21800000 | 0.0045 | 0.0032 | 0.0037 |
| F11 | 21720001 | 21820000 | 0.0045 | 0.0031 | 0.0036 |
| F11 | 21740001 | 21840000 | 0.0037 | 0.0026 | 0.0029 |
| F11 | 21760001 | 21860000 | 0.0028 | 0.0019 | 0.0022 |
| F11 | 21780001 | 21880000 | 0.0020 | 0.0013 | 0.0015 |
| F11 | 21800001 | 21900000 | 0.0017 | 0.0012 | 0.0013 |
| F11 | 21820001 | 21920000 | 0.0011 | 0.0008 | 0.0008 |
| F11 | 21840001 | 21940000 | 0.0010 | 0.0008 | 0.0009 |
| F11 | 21860001 | 21960000 | 0.0007 | 0.0005 | 0.0006 |
| F11 | 21880001 | 21980000 | 0.0006 | 0.0005 | 0.0006 |
| F11 | 21900001 | 22000000 | 0.0004 | 0.0004 | 0.0004 |
| F11 | 21920001 | 22020000 | 0.0007 | 0.0006 | 0.0007 |
| F11 | 21940001 | 22040000 | 0.0014 | 0.0012 | 0.0013 |
| F11 | 21960001 | 22060000 | 0.0015 | 0.0012 | 0.0013 |
| F11 | 21980001 | 22080000 | 0.0017 | 0.0012 | 0.0013 |
| F11 | 22000001 | 22100000 | 0.0027 | 0.0019 | 0.0020 |
| F11 | 22020001 | 22120000 | 0.0034 | 0.0022 | 0.0023 |
| F11 | 22040001 | 22140000 | 0.0036 | 0.0025 | 0.0025 |
| F11 | 22060001 | 22160000 | 0.0037 | 0.0025 | 0.0026 |
| F11 | 22080001 | 22180000 | 0.0037 | 0.0028 | 0.0029 |
| F11 | 22100001 | 22200000 | 0.0034 | 0.0024 | 0.0027 |
| F11 | 22120001 | 22220000 | 0.0030 | 0.0024 | 0.0027 |
| F11 | 22140001 | 22240000 | 0.0024 | 0.0018 | 0.0020 |
| F11 | 22160001 | 22260000 | 0.0036 | 0.0030 | 0.0032 |
| F11 | 22180001 | 22280000 | 0.0039 | 0.0031 | 0.0034 |
| F11 | 22200001 | 22300000 | 0.0034 | 0.0027 | 0.0028 |
| F11 | 22220001 | 22320000 | 0.0037 | 0.0027 | 0.0030 |
| F11 | 22240001 | 22340000 | 0.0037 | 0.0028 | 0.0032 |
| F11 | 22260001 | 22360000 | 0.0028 | 0.0020 | 0.0024 |
| F11 | 22280001 | 22380000 | 0.0027 | 0.0019 | 0.0023 |
| F11 | 22300001 | 22400000 | 0.0030 | 0.0021 | 0.0025 |
| F11 | 22320001 | 22420000 | 0.0027 | 0.0019 | 0.0021 |
| F11 | 22340001 | 22440000 | 0.0027 | 0.0018 | 0.0019 |
| F11 | 22360001 | 22460000 | 0.0024 | 0.0014 | 0.0014 |

| Chr | Start | End | both | A1 | A2 |
|---|---|---|---|---|---|
| F11 | 22380001 | 22480000 | 0.0030 | 0.0018 | 0.0020 |
| F11 | 22400001 | 22500000 | 0.0035 | 0.0020 | 0.0023 |
| F11 | 22420001 | 22520000 | 0.0030 | 0.0018 | 0.0020 |
| F11 | 22440001 | 22540000 | 0.0040 | 0.0023 | 0.0029 |
| F11 | 22460001 | 22560000 | 0.0044 | 0.0028 | 0.0035 |
| F11 | 22480001 | 22580000 | 0.0038 | 0.0023 | 0.0029 |
| F11 | 22500001 | 22600000 | 0.0032 | 0.0019 | 0.0025 |
| F11 | 22520001 | 22620000 | 0.0036 | 0.0022 | 0.0028 |
| F11 | 22540001 | 22640000 | 0.0024 | 0.0014 | 0.0017 |
| F11 | 22560001 | 22660000 | 0.0023 | 0.0013 | 0.0014 |
| F11 | 22580001 | 22680000 | 0.0033 | 0.0022 | 0.0023 |
| F11 | 22600001 | 22700000 | 0.0038 | 0.0028 | 0.0028 |
| F11 | 22620001 | 22720000 | 0.0038 | 0.0028 | 0.0028 |
| F11 | 22640001 | 22740000 | 0.0038 | 0.0028 | 0.0028 |
| F11 | 22660001 | 22760000 | 0.0034 | 0.0026 | 0.0026 |
| F11 | 22680001 | 22780000 | 0.0025 | 0.0017 | 0.0017 |
| F11 | 22700001 | 22800000 | 0.0018 | 0.0010 | 0.0010 |
| F11 | 22720001 | 22820000 | 0.0016 | 0.0007 | 0.0007 |
| F11 | 22740001 | 22840000 | 0.0017 | 0.0007 | 0.0009 |
| F11 | 22760001 | 22860000 | 0.0016 | 0.0005 | 0.0007 |
| F11 | 22780001 | 22880000 | 0.0014 | 0.0003 | 0.0006 |
| F11 | 22800001 | 22900000 | 0.0013 | 0.0003 | 0.0005 |
| F11 | 22820001 | 22920000 | 0.0018 | 0.0008 | 0.0010 |
| F11 | 22840001 | 22940000 | 0.0029 | 0.0020 | 0.0022 |
| F11 | 22860001 | 22960000 | 0.0033 | 0.0023 | 0.0024 |
| F11 | 22880001 | 22980000 | 0.0040 | 0.0027 | 0.0029 |
| F11 | 22900001 | 23000000 | 0.0044 | 0.0031 | 0.0034 |
| F11 | 22920001 | 23020000 | 0.0040 | 0.0029 | 0.0032 |
| F11 | 22940001 | 23040000 | 0.0032 | 0.0017 | 0.0020 |
| F11 | 22960001 | 23060000 | 0.0031 | 0.0017 | 0.0020 |
| F11 | 22980001 | 23080000 | 0.0022 | 0.0013 | 0.0015 |
| F11 | 23000001 | 23100000 | 0.0019 | 0.0009 | 0.0011 |
| F11 | 23020001 | 23120000 | 0.0017 | 0.0006 | 0.0008 |
| F11 | 23040001 | 23140000 | 0.0010 | 0.0004 | 0.0005 |
| F11 | 23060001 | 23160000 | 0.0010 | 0.0005 | 0.0006 |
| F11 | 23080001 | 23180000 | 0.0022 | 0.0014 | 0.0016 |
| F11 | 23100001 | 23200000 | 0.0024 | 0.0016 | 0.0018 |
| F11 | 23120001 | 23220000 | 0.0038 | 0.0027 | 0.0032 |
| F11 | 23140001 | 23240000 | 0.0045 | 0.0033 | 0.0039 |
| F11 | 23160001 | 23260000 | 0.0044 | 0.0033 | 0.0039 |
| F11 | 23180001 | 23280000 | 0.0034 | 0.0026 | 0.0031 |
| F11 | 23200001 | 23300000 | 0.0037 | 0.0030 | 0.0036 |
| F11 | 23220001 | 23320000 | 0.0030 | 0.0025 | 0.0030 |
| F11 | 23240001 | 23340000 | 0.0028 | 0.0024 | 0.0028 |
| F11 | 23260001 | 23360000 | 0.0030 | 0.0025 | 0.0030 |
| F11 | 23280001 | 23380000 | 0.0028 | 0.0024 | 0.0028 |
| F11 | 23300001 | 23400000 | 0.0022 | 0.0018 | 0.0022 |
| F11 | 23320001 | 23420000 | 0.0022 | 0.0018 | 0.0021 |
| F11 | 23340001 | 23440000 | 0.0018 | 0.0013 | 0.0015 |
| F11 | 23360001 | 23460000 | 0.0025 | 0.0021 | 0.0021 |
| F11 | 23380001 | 23480000 | 0.0028 | 0.0023 | 0.0024 |
| F11 | 23400001 | 23500000 | 0.0034 | 0.0028 | 0.0029 |
| F11 | 23420001 | 23520000 | 0.0036 | 0.0031 | 0.0031 |
| F11 | 23440001 | 23540000 | 0.0033 | 0.0030 | 0.0031 |
| F11 | 23460001 | 23560000 | 0.0022 | 0.0020 | 0.0020 |
| F11 | 23480001 | 23580000 | 0.0027 | 0.0022 | 0.0024 |
| F11 | 23500001 | 23600000 | 0.0028 | 0.0024 | 0.0025 |
| F11 | 23520001 | 23620000 | 0.0024 | 0.0017 | 0.0019 |

| Chr | Start | End | both | A1 | A2 |
|---|---|---|---|---|---|
| F11 | 23540001 | 23640000 | 0.0029 | 0.0020 | 0.0022 |
| F11 | 23560001 | 23660000 | 0.0033 | 0.0024 | 0.0025 |
| F11 | 23580001 | 23680000 | 0.0030 | 0.0023 | 0.0024 |
| F11 | 23600001 | 23700000 | 0.0023 | 0.0016 | 0.0017 |
| F11 | 23620001 | 23720000 | 0.0023 | 0.0016 | 0.0017 |
| F11 | 23640001 | 23740000 | 0.0027 | 0.0020 | 0.0021 |
| F11 | 23660001 | 23760000 | 0.0031 | 0.0021 | 0.0024 |
| F11 | 23680001 | 23780000 | 0.0035 | 0.0021 | 0.0025 |
| F11 | 23700001 | 23800000 | 0.0039 | 0.0024 | 0.0027 |
| F11 | 23720001 | 23820000 | 0.0038 | 0.0023 | 0.0027 |
| F11 | 23740001 | 23840000 | 0.0038 | 0.0023 | 0.0028 |
| F11 | 23760001 | 23860000 | 0.0032 | 0.0018 | 0.0022 |
| F11 | 23780001 | 23880000 | 0.0027 | 0.0016 | 0.0019 |
| F11 | 23800001 | 23900000 | 0.0027 | 0.0018 | 0.0021 |
| F11 | 23820001 | 23920000 | 0.0036 | 0.0025 | 0.0029 |
| F11 | 23840001 | 23940000 | 0.0032 | 0.0021 | 0.0023 |
| F11 | 23860001 | 23960000 | 0.0040 | 0.0030 | 0.0030 |
| F11 | 23880001 | 23980000 | 0.0050 | 0.0041 | 0.0041 |
| F11 | 23900001 | 24000000 | 0.0049 | 0.0038 | 0.0037 |
| F11 | 23920001 | 24020000 | 0.0041 | 0.0032 | 0.0029 |
| F11 | 23940001 | 24040000 | 0.0048 | 0.0039 | 0.0038 |
| F11 | 23960001 | 24060000 | 0.0039 | 0.0031 | 0.0032 |
| F11 | 23980001 | 24080000 | 0.0036 | 0.0028 | 0.0029 |
| F11 | 24000001 | 24100000 | 0.0034 | 0.0028 | 0.0030 |
| F11 | 24020001 | 24120000 | 0.0035 | 0.0031 | 0.0032 |
| F11 | 24040001 | 24140000 | 0.0026 | 0.0023 | 0.0024 |
| F11 | 24060001 | 24160000 | 0.0031 | 0.0027 | 0.0029 |
| F11 | 24080001 | 24180000 | 0.0023 | 0.0018 | 0.0020 |
| F11 | 24100001 | 24200000 | 0.0025 | 0.0019 | 0.0022 |
| F11 | 24120001 | 24220000 | 0.0021 | 0.0014 | 0.0017 |
| F11 | 24140001 | 24240000 | 0.0037 | 0.0027 | 0.0032 |
| F11 | 24160001 | 24260000 | 0.0037 | 0.0029 | 0.0030 |
| F11 | 24180001 | 24280000 | 0.0038 | 0.0030 | 0.0030 |
| F11 | 24200001 | 24300000 | 0.0037 | 0.0028 | 0.0028 |
| F11 | 24220001 | 24320000 | 0.0042 | 0.0032 | 0.0033 |
| F11 | 24240001 | 24340000 | 0.0027 | 0.0019 | 0.0018 |
| F11 | 24260001 | 24360000 | 0.0022 | 0.0011 | 0.0013 |
| F11 | 24280001 | 24380000 | 0.0025 | 0.0014 | 0.0018 |
| F11 | 24300001 | 24400000 | 0.0027 | 0.0016 | 0.0020 |
| F11 | 24320001 | 24420000 | 0.0022 | 0.0013 | 0.0017 |
| F11 | 24340001 | 24440000 | 0.0024 | 0.0015 | 0.0019 |
| F11 | 24360001 | 24460000 | 0.0024 | 0.0015 | 0.0019 |
| F11 | 24380001 | 24480000 | 0.0022 | 0.0013 | 0.0017 |
| F11 | 24400001 | 24500000 | 0.0022 | 0.0014 | 0.0019 |
| F11 | 24420001 | 24520000 | 0.0020 | 0.0013 | 0.0017 |
| F11 | 24440001 | 24540000 | 0.0024 | 0.0016 | 0.0019 |
| F11 | 24460001 | 24560000 | 0.0026 | 0.0017 | 0.0019 |
| F11 | 24480001 | 24580000 | 0.0028 | 0.0017 | 0.0019 |
| F11 | 24500001 | 24600000 | 0.0034 | 0.0023 | 0.0025 |
| F11 | 24520001 | 24620000 | 0.0043 | 0.0030 | 0.0033 |
| F11 | 24540001 | 24640000 | 0.0037 | 0.0024 | 0.0028 |
| F11 | 24560001 | 24660000 | 0.0038 | 0.0026 | 0.0031 |
| F11 | 24580001 | 24680000 | 0.0040 | 0.0028 | 0.0033 |
| F11 | 24600001 | 24700000 | 0.0032 | 0.0018 | 0.0025 |
| F11 | 24620001 | 24720000 | 0.0032 | 0.0020 | 0.0026 |
| F11 | 24640001 | 24740000 | 0.0038 | 0.0024 | 0.0030 |
| F11 | 24660001 | 24760000 | 0.0049 | 0.0031 | 0.0041 |
| F11 | 24680001 | 24780000 | 0.0045 | 0.0031 | 0.0040 |

| Chr | Start | End | both | A1 | A2 |
|---|---|---|---|---|---|
| F11 | 24700001 | 24800000 | 0.0050 | 0.0035 | 0.0046 |
| F11 | 24720001 | 24820000 | 0.0048 | 0.0032 | 0.0044 |
| F11 | 24740001 | 24840000 | 0.0055 | 0.0039 | 0.0052 |
| F11 | 24760001 | 24860000 | 0.0040 | 0.0027 | 0.0036 |
| F11 | 24780001 | 24880000 | 0.0038 | 0.0025 | 0.0033 |
| F11 | 24800001 | 24900000 | 0.0032 | 0.0022 | 0.0027 |
| F11 | 24820001 | 24920000 | 0.0025 | 0.0017 | 0.0021 |
| F11 | 24840001 | 24940000 | 0.0019 | 0.0014 | 0.0016 |
| F11 | 24860001 | 24960000 | 0.0026 | 0.0021 | 0.0023 |
| F11 | 24880001 | 24980000 | 0.0027 | 0.0022 | 0.0024 |
| F11 | 24900001 | 25000000 | 0.0027 | 0.0021 | 0.0023 |
| F11 | 24920001 | 25020000 | 0.0027 | 0.0020 | 0.0023 |
| F11 | 24940001 | 25040000 | 0.0024 | 0.0015 | 0.0018 |
| F11 | 24960001 | 25060000 | 0.0023 | 0.0013 | 0.0016 |
| F11 | 24980001 | 25080000 | 0.0024 | 0.0015 | 0.0019 |
| F11 | 25000001 | 25100000 | 0.0031 | 0.0020 | 0.0025 |
| F11 | 25020001 | 25120000 | 0.0040 | 0.0028 | 0.0034 |
| F11 | 25040001 | 25140000 | 0.0042 | 0.0031 | 0.0036 |
| F11 | 25060001 | 25160000 | 0.0048 | 0.0036 | 0.0042 |
| F11 | 25080001 | 25180000 | 0.0049 | 0.0038 | 0.0043 |
| F11 | 25100001 | 25200000 | 0.0043 | 0.0034 | 0.0037 |
| F11 | 25120001 | 25220000 | 0.0044 | 0.0034 | 0.0038 |
| F11 | 25140001 | 25240000 | 0.0040 | 0.0032 | 0.0036 |
| F11 | 25160001 | 25260000 | 0.0026 | 0.0022 | 0.0024 |
| F11 | 25180001 | 25280000 | 0.0030 | 0.0023 | 0.0027 |
| F11 | 25200001 | 25300000 | 0.0035 | 0.0028 | 0.0032 |
| F11 | 25220001 | 25320000 | 0.0030 | 0.0025 | 0.0028 |
| F11 | 25240001 | 25340000 | 0.0033 | 0.0025 | 0.0029 |
| F11 | 25260001 | 25360000 | 0.0036 | 0.0028 | 0.0032 |
| F11 | 25280001 | 25380000 | 0.0028 | 0.0021 | 0.0025 |
| F11 | 25300001 | 25400000 | 0.0025 | 0.0017 | 0.0021 |
| F11 | 25320001 | 25420000 | 0.0021 | 0.0014 | 0.0018 |
| F11 | 25340001 | 25440000 | 0.0018 | 0.0012 | 0.0016 |
| F11 | 25360001 | 25460000 | 0.0020 | 0.0013 | 0.0018 |
| F11 | 25380001 | 25480000 | 0.0027 | 0.0020 | 0.0025 |
| F11 | 25400001 | 25500000 | 0.0025 | 0.0020 | 0.0024 |
| F11 | 25420001 | 25520000 | 0.0026 | 0.0021 | 0.0025 |
| F11 | 25440001 | 25540000 | 0.0032 | 0.0026 | 0.0030 |
| F11 | 25460001 | 25560000 | 0.0034 | 0.0027 | 0.0031 |
| F11 | 25480001 | 25580000 | 0.0036 | 0.0028 | 0.0032 |
| F11 | 25500001 | 25600000 | 0.0035 | 0.0025 | 0.0030 |
| F11 | 25520001 | 25620000 | 0.0034 | 0.0023 | 0.0028 |
| F11 | 25540001 | 25640000 | 0.0028 | 0.0017 | 0.0023 |
| F11 | 25560001 | 25660000 | 0.0034 | 0.0025 | 0.0029 |
| F11 | 25580001 | 25680000 | 0.0029 | 0.0018 | 0.0023 |
| F11 | 25600001 | 25700000 | 0.0032 | 0.0023 | 0.0028 |
| F11 | 25620001 | 25720000 | 0.0034 | 0.0026 | 0.0030 |
| F11 | 25640001 | 25740000 | 0.0032 | 0.0027 | 0.0030 |
| F11 | 25660001 | 25760000 | 0.0026 | 0.0022 | 0.0024 |
| F11 | 25680001 | 25780000 | 0.0031 | 0.0026 | 0.0030 |
| F11 | 25700001 | 25800000 | 0.0035 | 0.0029 | 0.0032 |
| F11 | 25720001 | 25820000 | 0.0040 | 0.0034 | 0.0037 |
| F11 | 25740001 | 25840000 | 0.0038 | 0.0033 | 0.0036 |
| F11 | 25760001 | 25860000 | 0.0051 | 0.0043 | 0.0048 |
| F11 | 25780001 | 25880000 | 0.0042 | 0.0037 | 0.0040 |
| F11 | 25800001 | 25900000 | 0.0043 | 0.0039 | 0.0042 |
| F11 | 25820001 | 25920000 | 0.0043 | 0.0037 | 0.0042 |
| F11 | 25840001 | 25940000 | 0.0043 | 0.0037 | 0.0042 |

| Chr | Start | End | both | A1 | A2 |
|---|---|---|---|---|---|
| F11 | 25860001 | 25960000 | 0.0026 | 0.0022 | 0.0025 |
| F11 | 25880001 | 25980000 | 0.0023 | 0.0019 | 0.0023 |
| F11 | 25900001 | 26000000 | 0.0015 | 0.0012 | 0.0015 |
| F11 | 25920001 | 26020000 | 0.0019 | 0.0015 | 0.0018 |
| F11 | 25940001 | 26040000 | 0.0023 | 0.0017 | 0.0020 |
| F11 | 25960001 | 26060000 | 0.0021 | 0.0017 | 0.0018 |
| F11 | 25980001 | 26080000 | 0.0030 | 0.0023 | 0.0026 |
| F11 | 26000001 | 26100000 | 0.0043 | 0.0035 | 0.0039 |
| F11 | 26020001 | 26120000 | 0.0042 | 0.0035 | 0.0037 |
| F11 | 26040001 | 26140000 | 0.0043 | 0.0034 | 0.0037 |
| F11 | 26060001 | 26160000 | 0.0044 | 0.0034 | 0.0039 |
| F11 | 26080001 | 26180000 | 0.0040 | 0.0030 | 0.0035 |
| F11 | 26100001 | 26200000 | 0.0033 | 0.0023 | 0.0028 |
| F11 | 26120001 | 26220000 | 0.0030 | 0.0018 | 0.0023 |
| F11 | 26140001 | 26240000 | 0.0031 | 0.0020 | 0.0026 |
| F11 | 26160001 | 26260000 | 0.0034 | 0.0022 | 0.0027 |
| F11 | 26180001 | 26280000 | 0.0040 | 0.0026 | 0.0031 |
| F11 | 26200001 | 26300000 | 0.0039 | 0.0026 | 0.0030 |
| F11 | 26220001 | 26320000 | 0.0033 | 0.0022 | 0.0025 |
| F11 | 26240001 | 26340000 | 0.0035 | 0.0022 | 0.0026 |
| F11 | 26260001 | 26360000 | 0.0032 | 0.0020 | 0.0025 |
| F11 | 26280001 | 26380000 | 0.0025 | 0.0014 | 0.0017 |
| F11 | 26300001 | 26400000 | 0.0024 | 0.0015 | 0.0018 |
| F11 | 26320001 | 26420000 | 0.0026 | 0.0017 | 0.0019 |
| F11 | 26340001 | 26440000 | 0.0025 | 0.0016 | 0.0018 |
| F11 | 26360001 | 26460000 | 0.0026 | 0.0018 | 0.0019 |
| F11 | 26380001 | 26480000 | 0.0030 | 0.0022 | 0.0023 |
| F11 | 26400001 | 26500000 | 0.0023 | 0.0016 | 0.0017 |
| F11 | 26420001 | 26520000 | 0.0026 | 0.0018 | 0.0019 |
| F11 | 26440001 | 26540000 | 0.0031 | 0.0024 | 0.0023 |
| F11 | 26460001 | 26560000 | 0.0030 | 0.0022 | 0.0021 |
| F11 | 26480001 | 26580000 | 0.0027 | 0.0018 | 0.0017 |
| F11 | 26500001 | 26600000 | 0.0030 | 0.0019 | 0.0018 |
| F11 | 26520001 | 26620000 | 0.0032 | 0.0020 | 0.0021 |
| F11 | 26540001 | 26640000 | 0.0028 | 0.0016 | 0.0018 |
| F11 | 26560001 | 26660000 | 0.0032 | 0.0021 | 0.0024 |
| F11 | 26580001 | 26680000 | 0.0030 | 0.0021 | 0.0024 |
| F11 | 26600001 | 26700000 | 0.0030 | 0.0023 | 0.0025 |
| F11 | 26620001 | 26720000 | 0.0023 | 0.0018 | 0.0019 |
| F11 | 26640001 | 26740000 | 0.0018 | 0.0014 | 0.0015 |
| F11 | 26660001 | 26760000 | 0.0024 | 0.0018 | 0.0019 |
| F11 | 26680001 | 26780000 | 0.0025 | 0.0019 | 0.0020 |
| F11 | 26700001 | 26800000 | 0.0030 | 0.0023 | 0.0025 |
| F11 | 26720001 | 26820000 | 0.0034 | 0.0028 | 0.0030 |
| F11 | 26740001 | 26840000 | 0.0038 | 0.0032 | 0.0035 |
| F11 | 26760001 | 26860000 | 0.0028 | 0.0025 | 0.0027 |
| F11 | 26780001 | 26880000 | 0.0030 | 0.0026 | 0.0028 |
| F11 | 26800001 | 26900000 | 0.0032 | 0.0026 | 0.0028 |
| F11 | 26820001 | 26920000 | 0.0030 | 0.0022 | 0.0024 |
| F11 | 26840001 | 26940000 | 0.0037 | 0.0028 | 0.0029 |
| F11 | 26860001 | 26960000 | 0.0038 | 0.0030 | 0.0031 |
| F11 | 26880001 | 26980000 | 0.0038 | 0.0029 | 0.0030 |
| F11 | 26900001 | 27000000 | 0.0030 | 0.0023 | 0.0025 |
| F11 | 26920001 | 27020000 | 0.0040 | 0.0035 | 0.0038 |
| F11 | 26940001 | 27040000 | 0.0032 | 0.0027 | 0.0030 |
| F11 | 26960001 | 27060000 | 0.0031 | 0.0026 | 0.0030 |
| F11 | 26980001 | 27080000 | 0.0032 | 0.0027 | 0.0029 |
| F11 | 27000001 | 27100000 | 0.0039 | 0.0033 | 0.0033 |

| Chr | Start | End | both | A1 | A2 |
|---|---|---|---|---|---|
| F11 | 27020001 | 27120000 | 0.0028 | 0.0024 | 0.0023 |
| F11 | 27040001 | 27140000 | 0.0025 | 0.0020 | 0.0019 |
| F11 | 27060001 | 27160000 | 0.0027 | 0.0021 | 0.0021 |
| F11 | 27080001 | 27180000 | 0.0024 | 0.0019 | 0.0020 |
| F11 | 27100001 | 27200000 | 0.0017 | 0.0013 | 0.0015 |
| F11 | 27120001 | 27220000 | 0.0018 | 0.0014 | 0.0016 |
| F11 | 27140001 | 27240000 | 0.0024 | 0.0019 | 0.0023 |
| F11 | 27160001 | 27260000 | 0.0026 | 0.0022 | 0.0025 |
| F11 | 27180001 | 27280000 | 0.0027 | 0.0022 | 0.0025 |
| F11 | 27200001 | 27300000 | 0.0036 | 0.0029 | 0.0032 |
| F11 | 27220001 | 27320000 | 0.0040 | 0.0033 | 0.0036 |
| F11 | 27240001 | 27340000 | 0.0035 | 0.0030 | 0.0031 |
| F11 | 27260001 | 27360000 | 0.0035 | 0.0029 | 0.0029 |
| F11 | 27280001 | 27380000 | 0.0037 | 0.0028 | 0.0029 |
| F11 | 27300001 | 27400000 | 0.0034 | 0.0024 | 0.0026 |
| F11 | 27320001 | 27420000 | 0.0049 | 0.0038 | 0.0040 |
| F11 | 27340001 | 27440000 | 0.0050 | 0.0041 | 0.0041 |
| F11 | 27360001 | 27460000 | 0.0042 | 0.0034 | 0.0034 |
| F11 | 27380001 | 27480000 | 0.0050 | 0.0046 | 0.0045 |
| F11 | 27400001 | 27500000 | 0.0044 | 0.0042 | 0.0041 |
| F11 | 27420001 | 27520000 | 0.0035 | 0.0033 | 0.0033 |
| F11 | 27440001 | 27540000 | 0.0035 | 0.0032 | 0.0032 |
| F11 | 27460001 | 27560000 | 0.0037 | 0.0033 | 0.0034 |
| F11 | 27480001 | 27580000 | 0.0031 | 0.0026 | 0.0029 |
| F11 | 27500001 | 27600000 | 0.0030 | 0.0026 | 0.0028 |
| F11 | 27520001 | 27620000 | 0.0019 | 0.0016 | 0.0017 |
| F11 | 27540001 | 27640000 | 0.0018 | 0.0014 | 0.0016 |
| F11 | 27560001 | 27660000 | 0.0019 | 0.0016 | 0.0018 |
| F11 | 27580001 | 27680000 | 0.0014 | 0.0012 | 0.0013 |
| F11 | 27600001 | 27700000 | 0.0017 | 0.0015 | 0.0017 |
| F11 | 27620001 | 27720000 | 0.0025 | 0.0022 | 0.0024 |
| F11 | 27640001 | 27740000 | 0.0027 | 0.0023 | 0.0025 |
| F11 | 27660001 | 27760000 | 0.0028 | 0.0021 | 0.0023 |
| F11 | 27680001 | 27780000 | 0.0031 | 0.0020 | 0.0022 |
| F11 | 27700001 | 27800000 | 0.0034 | 0.0022 | 0.0025 |
| F11 | 27720001 | 27820000 | 0.0023 | 0.0013 | 0.0015 |
| F11 | 27740001 | 27840000 | 0.0022 | 0.0013 | 0.0016 |
| F11 | 27760001 | 27860000 | 0.0017 | 0.0011 | 0.0013 |
| F11 | 27780001 | 27880000 | 0.0014 | 0.0012 | 0.0014 |
| F11 | 27800001 | 27900000 | 0.0010 | 0.0008 | 0.0010 |
| F11 | 27820001 | 27920000 | 0.0012 | 0.0009 | 0.0012 |
| F11 | 27840001 | 27940000 | 0.0010 | 0.0008 | 0.0009 |
| F11 | 27860001 | 27960000 | 0.0012 | 0.0010 | 0.0011 |
| F11 | 27880001 | 27980000 | 0.0013 | 0.0010 | 0.0011 |
| F11 | 27900001 | 28000000 | 0.0013 | 0.0010 | 0.0010 |
| F11 | 27920001 | 28020000 | 0.0014 | 0.0011 | 0.0012 |
| F11 | 27940001 | 28040000 | 0.0015 | 0.0011 | 0.0012 |
| F11 | 27960001 | 28060000 | 0.0020 | 0.0015 | 0.0018 |
| F11 | 27980001 | 28080000 | 0.0028 | 0.0021 | 0.0026 |
| F11 | 28000001 | 28100000 | 0.0030 | 0.0023 | 0.0028 |
| F11 | 28020001 | 28120000 | 0.0026 | 0.0020 | 0.0025 |
| F11 | 28040001 | 28140000 | 0.0028 | 0.0022 | 0.0027 |
| F11 | 28060001 | 28160000 | 0.0034 | 0.0027 | 0.0031 |
| F11 | 28080001 | 28180000 | 0.0027 | 0.0020 | 0.0022 |
| F11 | 28100001 | 28200000 | 0.0026 | 0.0019 | 0.0020 |
| F11 | 28120001 | 28220000 | 0.0026 | 0.0019 | 0.0020 |
| F11 | 28140001 | 28240000 | 0.0022 | 0.0016 | 0.0016 |
| F11 | 28160001 | 28260000 | 0.0019 | 0.0015 | 0.0015 |

| Chr | Start | End | both | A1 | A2 |
|---|---|---|---|---|---|
| F11 | 28180001 | 28280000 | 0.0025 | 0.0019 | 0.0021 |
| F11 | 28200001 | 28300000 | 0.0031 | 0.0023 | 0.0026 |
| F11 | 28220001 | 28320000 | 0.0038 | 0.0027 | 0.0031 |
| F11 | 28240001 | 28340000 | 0.0045 | 0.0030 | 0.0036 |
| F11 | 28260001 | 28360000 | 0.0046 | 0.0027 | 0.0033 |
| F11 | 28280001 | 28380000 | 0.0042 | 0.0025 | 0.0030 |
| F11 | 28300001 | 28400000 | 0.0039 | 0.0021 | 0.0027 |
| F11 | 28320001 | 28420000 | 0.0038 | 0.0022 | 0.0026 |
| F11 | 28340001 | 28440000 | 0.0034 | 0.0020 | 0.0023 |
| F11 | 28360001 | 28460000 | 0.0026 | 0.0017 | 0.0018 |
| F11 | 28380001 | 28480000 | 0.0020 | 0.0013 | 0.0014 |
| F11 | 28400001 | 28500000 | 0.0017 | 0.0013 | 0.0013 |
| F11 | 28420001 | 28520000 | 0.0012 | 0.0011 | 0.0011 |
| F11 | 28440001 | 28540000 | 0.0011 | 0.0009 | 0.0009 |
| F11 | 28460001 | 28560000 | 0.0012 | 0.0009 | 0.0010 |
| F11 | 28480001 | 28580000 | 0.0016 | 0.0010 | 0.0012 |
| F11 | 28500001 | 28600000 | 0.0018 | 0.0008 | 0.0010 |
| F11 | 28520001 | 28620000 | 0.0028 | 0.0016 | 0.0018 |
| F11 | 28540001 | 28640000 | 0.0026 | 0.0016 | 0.0018 |
| F11 | 28560001 | 28660000 | 0.0027 | 0.0016 | 0.0018 |
| F11 | 28580001 | 28680000 | 0.0025 | 0.0014 | 0.0015 |
| F11 | 28600001 | 28700000 | 0.0028 | 0.0018 | 0.0021 |
| F11 | 28620001 | 28720000 | 0.0023 | 0.0015 | 0.0018 |
| F11 | 28640001 | 28740000 | 0.0026 | 0.0017 | 0.0020 |
| F11 | 28660001 | 28760000 | 0.0029 | 0.0020 | 0.0023 |
| F11 | 28680001 | 28780000 | 0.0027 | 0.0020 | 0.0023 |
| F11 | 28700001 | 28800000 | 0.0023 | 0.0016 | 0.0019 |
| F11 | 28720001 | 28820000 | 0.0018 | 0.0011 | 0.0014 |
| F11 | 28740001 | 28840000 | 0.0021 | 0.0010 | 0.0014 |
| F11 | 28760001 | 28860000 | 0.0017 | 0.0007 | 0.0011 |
| F11 | 28780001 | 28880000 | 0.0021 | 0.0010 | 0.0014 |
| F11 | 28800001 | 28900000 | 0.0016 | 0.0007 | 0.0011 |
| F11 | 28820001 | 28920000 | 0.0014 | 0.0006 | 0.0009 |
| F11 | 28840001 | 28940000 | 0.0013 | 0.0009 | 0.0010 |
| F11 | 28860001 | 28960000 | 0.0019 | 0.0014 | 0.0016 |
| F11 | 28880001 | 28980000 | 0.0024 | 0.0018 | 0.0020 |
| F11 | 28900001 | 29000000 | 0.0029 | 0.0021 | 0.0023 |
| F11 | 28920001 | 29020000 | 0.0031 | 0.0023 | 0.0025 |
| F11 | 28940001 | 29040000 | 0.0034 | 0.0023 | 0.0026 |
| F11 | 28960001 | 29060000 | 0.0029 | 0.0018 | 0.0022 |
| F11 | 28980001 | 29080000 | 0.0039 | 0.0028 | 0.0033 |
| F11 | 29000001 | 29100000 | 0.0035 | 0.0026 | 0.0031 |
| F11 | 29020001 | 29120000 | 0.0038 | 0.0029 | 0.0035 |
| F11 | 29040001 | 29140000 | 0.0033 | 0.0027 | 0.0032 |
| F11 | 29060001 | 29160000 | 0.0032 | 0.0027 | 0.0031 |
| F11 | 29080001 | 29180000 | 0.0013 | 0.0011 | 0.0011 |
| F11 | 29100001 | 29200000 | 0.0031 | 0.0028 | 0.0027 |
| F11 | 29120001 | 29220000 | 0.0038 | 0.0035 | 0.0034 |
| F11 | 29140001 | 29240000 | 0.0055 | 0.0049 | 0.0050 |
| F11 | 29160001 | 29260000 | 0.0054 | 0.0048 | 0.0049 |
| F11 | 29180001 | 29280000 | 0.0054 | 0.0047 | 0.0049 |
| F11 | 29200001 | 29300000 | 0.0039 | 0.0031 | 0.0034 |
| F11 | 29220001 | 29320000 | 0.0027 | 0.0019 | 0.0022 |
| F11 | 29240001 | 29340000 | 0.0024 | 0.0015 | 0.0018 |
| F11 | 29260001 | 29360000 | 0.0031 | 0.0022 | 0.0024 |
| F11 | 29280001 | 29380000 | 0.0032 | 0.0024 | 0.0026 |
| F11 | 29300001 | 29400000 | 0.0031 | 0.0025 | 0.0026 |
| F11 | 29320001 | 29420000 | 0.0032 | 0.0026 | 0.0028 |

| Chr | Start | End | both | A1 | A2 |
|---|---|---|---|---|---|
| F11 | 29340001 | 29440000 | 0.0017 | 0.0015 | 0.0015 |
| F11 | 29360001 | 29460000 | 0.0014 | 0.0012 | 0.0014 |
| F11 | 29380001 | 29480000 | 0.0022 | 0.0015 | 0.0020 |
| F11 | 29400001 | 29500000 | 0.0023 | 0.0015 | 0.0019 |
| F11 | 29420001 | 29520000 | 0.0033 | 0.0024 | 0.0028 |
| F11 | 29440001 | 29540000 | 0.0036 | 0.0027 | 0.0031 |
| F11 | 29460001 | 29560000 | 0.0044 | 0.0036 | 0.0039 |
| F11 | 29480001 | 29580000 | 0.0042 | 0.0037 | 0.0037 |
| F11 | 29500001 | 29600000 | 0.0056 | 0.0051 | 0.0048 |
| F11 | 29520001 | 29620000 | 0.0047 | 0.0044 | 0.0040 |
| F11 | 29540001 | 29640000 | 0.0045 | 0.0040 | 0.0037 |
| F11 | 29560001 | 29660000 | 0.0033 | 0.0028 | 0.0026 |
| F11 | 29580001 | 29680000 | 0.0028 | 0.0023 | 0.0021 |
| F11 | 29600001 | 29700000 | 0.0020 | 0.0015 | 0.0016 |
| F11 | 29620001 | 29720000 | 0.0023 | 0.0017 | 0.0019 |
| F11 | 29640001 | 29740000 | 0.0033 | 0.0027 | 0.0026 |
| F11 | 29660001 | 29760000 | 0.0033 | 0.0025 | 0.0025 |
| F11 | 29680001 | 29780000 | 0.0039 | 0.0028 | 0.0028 |
| F11 | 29700001 | 29800000 | 0.0041 | 0.0028 | 0.0030 |
| F11 | 29720001 | 29820000 | 0.0040 | 0.0030 | 0.0031 |
| F11 | 29740001 | 29840000 | 0.0036 | 0.0025 | 0.0029 |
| F11 | 29760001 | 29860000 | 0.0044 | 0.0032 | 0.0036 |
| F11 | 29780001 | 29880000 | 0.0041 | 0.0030 | 0.0036 |
| F11 | 29800001 | 29900000 | 0.0044 | 0.0028 | 0.0037 |
| F11 | 29820001 | 29920000 | 0.0042 | 0.0026 | 0.0034 |
| F11 | 29840001 | 29940000 | 0.0036 | 0.0020 | 0.0028 |
| F11 | 29860001 | 29960000 | 0.0032 | 0.0017 | 0.0024 |
| F11 | 29880001 | 29980000 | 0.0030 | 0.0015 | 0.0019 |
| F11 | 29900001 | 30000000 | 0.0029 | 0.0018 | 0.0020 |
| F11 | 29920001 | 30020000 | 0.0029 | 0.0017 | 0.0020 |
| F11 | 29940001 | 30040000 | 0.0036 | 0.0023 | 0.0026 |
| F11 | 29960001 | 30060000 | 0.0038 | 0.0023 | 0.0027 |
| F11 | 29980001 | 30080000 | 0.0043 | 0.0030 | 0.0034 |
| F11 | 30000001 | 30100000 | 0.0040 | 0.0028 | 0.0031 |
| F11 | 30020001 | 30120000 | 0.0038 | 0.0027 | 0.0030 |
| F11 | 30040001 | 30140000 | 0.0033 | 0.0023 | 0.0026 |
| F11 | 30060001 | 30160000 | 0.0029 | 0.0023 | 0.0024 |
| F11 | 30080001 | 30180000 | 0.0027 | 0.0018 | 0.0020 |
| F11 | 30100001 | 30200000 | 0.0022 | 0.0013 | 0.0014 |
| F11 | 30120001 | 30220000 | 0.0024 | 0.0014 | 0.0015 |
| F11 | 30140001 | 30240000 | 0.0024 | 0.0013 | 0.0014 |
| F11 | 30160001 | 30260000 | 0.0023 | 0.0012 | 0.0014 |
| F11 | 30180001 | 30280000 | 0.0024 | 0.0015 | 0.0017 |
| F11 | 30200001 | 30300000 | 0.0025 | 0.0018 | 0.0019 |
| F11 | 30220001 | 30320000 | 0.0020 | 0.0015 | 0.0014 |
| F11 | 30240001 | 30340000 | 0.0024 | 0.0017 | 0.0018 |
| F11 | 30260001 | 30360000 | 0.0024 | 0.0017 | 0.0017 |
| F11 | 30280001 | 30380000 | 0.0024 | 0.0015 | 0.0016 |
| F11 | 30300001 | 30400000 | 0.0030 | 0.0018 | 0.0020 |
| F11 | 30320001 | 30420000 | 0.0036 | 0.0022 | 0.0025 |
| F11 | 30340001 | 30440000 | 0.0032 | 0.0019 | 0.0021 |
| F11 | 30360001 | 30460000 | 0.0039 | 0.0025 | 0.0027 |
| F11 | 30380001 | 30480000 | 0.0035 | 0.0023 | 0.0024 |
| F11 | 30400001 | 30500000 | 0.0027 | 0.0017 | 0.0017 |
| F11 | 30420001 | 30520000 | 0.0023 | 0.0012 | 0.0012 |
| F11 | 30440001 | 30540000 | 0.0023 | 0.0014 | 0.0014 |
| F11 | 30460001 | 30560000 | 0.0018 | 0.0010 | 0.0011 |
| F11 | 30480001 | 30580000 | 0.0023 | 0.0014 | 0.0016 |

| Chr | Start | End | both | A1 | A2 |
|---|---|---|---|---|---|
| F11 | 30500001 | 30600000 | 0.0026 | 0.0017 | 0.0020 |
| F11 | 30520001 | 30620000 | 0.0027 | 0.0019 | 0.0021 |
| F11 | 30540001 | 30640000 | 0.0035 | 0.0026 | 0.0030 |
| F11 | 30560001 | 30660000 | 0.0038 | 0.0029 | 0.0033 |
| F11 | 30580001 | 30680000 | 0.0034 | 0.0027 | 0.0029 |
| F11 | 30600001 | 30700000 | 0.0028 | 0.0022 | 0.0025 |
| F11 | 30620001 | 30720000 | 0.0026 | 0.0020 | 0.0023 |
| F11 | 30640001 | 30740000 | 0.0019 | 0.0013 | 0.0015 |
| F11 | 30660001 | 30760000 | 0.0013 | 0.0009 | 0.0010 |
| F11 | 30680001 | 30780000 | 0.0020 | 0.0014 | 0.0016 |
| F11 | 30700001 | 30800000 | 0.0025 | 0.0018 | 0.0020 |
| F11 | 30720001 | 30820000 | 0.0029 | 0.0022 | 0.0025 |
| F11 | 30740001 | 30840000 | 0.0028 | 0.0021 | 0.0023 |
| F11 | 30760001 | 30860000 | 0.0030 | 0.0023 | 0.0025 |
| F11 | 30780001 | 30880000 | 0.0022 | 0.0018 | 0.0018 |
| F11 | 30800001 | 30900000 | 0.0020 | 0.0015 | 0.0017 |
| F11 | 30820001 | 30920000 | 0.0017 | 0.0014 | 0.0015 |
| F11 | 30840001 | 30940000 | 0.0015 | 0.0013 | 0.0014 |
| F11 | 30860001 | 30960000 | 0.0022 | 0.0018 | 0.0022 |
| F11 | 30880001 | 30980000 | 0.0022 | 0.0019 | 0.0022 |
| F11 | 30900001 | 31000000 | 0.0020 | 0.0018 | 0.0020 |
| F11 | 30920001 | 31020000 | 0.0020 | 0.0018 | 0.0020 |
| F11 | 30940001 | 31040000 | 0.0023 | 0.0021 | 0.0023 |
| F11 | 30960001 | 31060000 | 0.0018 | 0.0018 | 0.0017 |
| F11 | 30980001 | 31080000 | 0.0020 | 0.0019 | 0.0019 |
| F11 | 31000001 | 31100000 | 0.0024 | 0.0021 | 0.0021 |
| F11 | 31020001 | 31120000 | 0.0022 | 0.0018 | 0.0019 |
| F11 | 31040001 | 31140000 | 0.0019 | 0.0015 | 0.0016 |
| F11 | 31060001 | 31160000 | 0.0012 | 0.0008 | 0.0009 |
| F11 | 31080001 | 31180000 | 0.0011 | 0.0006 | 0.0006 |
| F11 | 31100001 | 31200000 | 0.0008 | 0.0004 | 0.0005 |
| F11 | 31120001 | 31220000 | 0.0009 | 0.0005 | 0.0006 |
| F11 | 31140001 | 31240000 | 0.0010 | 0.0006 | 0.0006 |
| F11 | 31160001 | 31260000 | 0.0015 | 0.0010 | 0.0012 |
| F11 | 31180001 | 31280000 | 0.0014 | 0.0010 | 0.0012 |
| F11 | 31200001 | 31300000 | 0.0015 | 0.0012 | 0.0014 |
| F11 | 31220001 | 31320000 | 0.0013 | 0.0011 | 0.0013 |
| F11 | 31240001 | 31340000 | 0.0014 | 0.0010 | 0.0012 |
| F11 | 31260001 | 31360000 | 0.0014 | 0.0009 | 0.0010 |
| F11 | 31280001 | 31380000 | 0.0018 | 0.0010 | 0.0011 |
| F11 | 31300001 | 31400000 | 0.0028 | 0.0016 | 0.0019 |
| F11 | 31320001 | 31420000 | 0.0046 | 0.0033 | 0.0035 |
| F11 | 31340001 | 31440000 | 0.0050 | 0.0036 | 0.0040 |
| F11 | 31360001 | 31460000 | 0.0047 | 0.0036 | 0.0038 |
| F11 | 31380001 | 31480000 | 0.0051 | 0.0040 | 0.0043 |
| F11 | 31400001 | 31500000 | 0.0045 | 0.0034 | 0.0037 |
| F11 | 31420001 | 31520000 | 0.0029 | 0.0020 | 0.0022 |
| F11 | 31440001 | 31540000 | 0.0026 | 0.0019 | 0.0020 |
| F11 | 31460001 | 31560000 | 0.0029 | 0.0019 | 0.0022 |
| F11 | 31480001 | 31580000 | 0.0025 | 0.0015 | 0.0017 |
| F11 | 31500001 | 31600000 | 0.0023 | 0.0016 | 0.0017 |
| F11 | 31520001 | 31620000 | 0.0028 | 0.0017 | 0.0018 |
| F11 | 31540001 | 31640000 | 0.0028 | 0.0017 | 0.0019 |
| F11 | 31560001 | 31660000 | 0.0026 | 0.0016 | 0.0019 |
| F11 | 31580001 | 31680000 | 0.0022 | 0.0014 | 0.0016 |
| F11 | 31600001 | 31700000 | 0.0019 | 0.0012 | 0.0013 |
| F11 | 31620001 | 31720000 | 0.0018 | 0.0012 | 0.0014 |
| F11 | 31640001 | 31740000 | 0.0017 | 0.0010 | 0.0011 |

| Chr | Start | End | both | A1 | A2 |
|---|---|---|---|---|---|
| F11 | 31660001 | 31760000 | 0.0020 | 0.0012 | 0.0012 |
| F11 | 31680001 | 31780000 | 0.0034 | 0.0025 | 0.0026 |
| F11 | 31700001 | 31800000 | 0.0035 | 0.0025 | 0.0024 |
| F11 | 31720001 | 31820000 | 0.0032 | 0.0021 | 0.0020 |
| F11 | 31740001 | 31840000 | 0.0031 | 0.0022 | 0.0021 |
| F11 | 31760001 | 31860000 | 0.0035 | 0.0025 | 0.0026 |
| F11 | 31780001 | 31880000 | 0.0021 | 0.0012 | 0.0013 |
| F11 | 31800001 | 31900000 | 0.0019 | 0.0012 | 0.0013 |
| F11 | 31820001 | 31920000 | 0.0022 | 0.0016 | 0.0019 |
| F11 | 31840001 | 31940000 | 0.0025 | 0.0019 | 0.0022 |
| F11 | 31860001 | 31960000 | 0.0019 | 0.0015 | 0.0017 |
| F11 | 31880001 | 31980000 | 0.0022 | 0.0016 | 0.0019 |
| F11 | 31900001 | 32000000 | 0.0021 | 0.0015 | 0.0018 |
| F11 | 31920001 | 32020000 | 0.0025 | 0.0018 | 0.0019 |
| F11 | 31940001 | 32040000 | 0.0025 | 0.0015 | 0.0016 |
| F11 | 31960001 | 32060000 | 0.0025 | 0.0016 | 0.0015 |
| F11 | 31980001 | 32080000 | 0.0025 | 0.0017 | 0.0016 |
| F11 | 32000001 | 32100000 | 0.0029 | 0.0020 | 0.0019 |
| F11 | 32020001 | 32120000 | 0.0024 | 0.0016 | 0.0015 |
| F11 | 32040001 | 32140000 | 0.0021 | 0.0014 | 0.0015 |
| F11 | 32060001 | 32160000 | 0.0019 | 0.0012 | 0.0013 |
| F11 | 32080001 | 32180000 | 0.0017 | 0.0010 | 0.0011 |
| F11 | 32100001 | 32200000 | 0.0016 | 0.0009 | 0.0011 |
| F11 | 32120001 | 32220000 | 0.0017 | 0.0010 | 0.0012 |
| F11 | 32140001 | 32240000 | 0.0016 | 0.0009 | 0.0011 |
| F11 | 32160001 | 32260000 | 0.0020 | 0.0010 | 0.0013 |
| F11 | 32180001 | 32280000 | 0.0027 | 0.0016 | 0.0019 |
| F11 | 32200001 | 32300000 | 0.0028 | 0.0017 | 0.0020 |
| F11 | 32220001 | 32320000 | 0.0031 | 0.0020 | 0.0024 |
| F11 | 32240001 | 32340000 | 0.0034 | 0.0021 | 0.0026 |
| F11 | 32260001 | 32360000 | 0.0032 | 0.0021 | 0.0025 |
| F11 | 32280001 | 32380000 | 0.0026 | 0.0016 | 0.0020 |
| F11 | 32300001 | 32400000 | 0.0022 | 0.0013 | 0.0016 |
| F11 | 32320001 | 32420000 | 0.0017 | 0.0010 | 0.0012 |
| F11 | 32340001 | 32440000 | 0.0018 | 0.0011 | 0.0013 |
| F11 | 32360001 | 32460000 | 0.0025 | 0.0016 | 0.0019 |
| F11 | 32380001 | 32480000 | 0.0027 | 0.0018 | 0.0020 |
| F11 | 32400001 | 32500000 | 0.0027 | 0.0018 | 0.0019 |
| F11 | 32420001 | 32520000 | 0.0036 | 0.0021 | 0.0025 |
| F11 | 32440001 | 32540000 | 0.0038 | 0.0021 | 0.0026 |
| F11 | 32460001 | 32560000 | 0.0028 | 0.0014 | 0.0019 |
| F11 | 32480001 | 32580000 | 0.0043 | 0.0030 | 0.0036 |
| F11 | 32500001 | 32600000 | 0.0051 | 0.0037 | 0.0044 |
| F11 | 32520001 | 32620000 | 0.0060 | 0.0051 | 0.0054 |
| F11 | 32540001 | 32640000 | 0.0064 | 0.0055 | 0.0059 |
| F11 | 32560001 | 32660000 | 0.0070 | 0.0060 | 0.0064 |
| F11 | 32580001 | 32680000 | 0.0059 | 0.0048 | 0.0053 |
| F11 | 32600001 | 32700000 | 0.0053 | 0.0043 | 0.0047 |
| F11 | 32620001 | 32720000 | 0.0042 | 0.0031 | 0.0036 |
| F11 | 32640001 | 32740000 | 0.0039 | 0.0031 | 0.0034 |
| F11 | 32660001 | 32760000 | 0.0041 | 0.0031 | 0.0035 |
| F11 | 32680001 | 32780000 | 0.0039 | 0.0030 | 0.0034 |
| F11 | 32700001 | 32800000 | 0.0048 | 0.0037 | 0.0042 |
| F11 | 32720001 | 32820000 | 0.0046 | 0.0034 | 0.0039 |
| F11 | 32740001 | 32840000 | 0.0048 | 0.0037 | 0.0041 |
| F11 | 32760001 | 32860000 | 0.0050 | 0.0040 | 0.0042 |
| F11 | 32780001 | 32880000 | 0.0056 | 0.0048 | 0.0048 |
| F11 | 32800001 | 32900000 | 0.0058 | 0.0049 | 0.0049 |

| Chr | Start | End | both | A1 | A2 |
|---|---|---|---|---|---|
| F11 | 32820001 | 32920000 | 0.0060 | 0.0053 | 0.0052 |
| F11 | 32840001 | 32940000 | 0.0058 | 0.0049 | 0.0050 |
| F11 | 32860001 | 32960000 | 0.0052 | 0.0043 | 0.0045 |
| F11 | 32880001 | 32980000 | 0.0046 | 0.0034 | 0.0039 |
| F11 | 32900001 | 33000000 | 0.0036 | 0.0025 | 0.0029 |
| F11 | 32920001 | 33020000 | 0.0033 | 0.0022 | 0.0026 |
| F11 | 32940001 | 33040000 | 0.0034 | 0.0022 | 0.0026 |
| F11 | 32960001 | 33060000 | 0.0041 | 0.0028 | 0.0034 |
| F11 | 32980001 | 33080000 | 0.0035 | 0.0026 | 0.0030 |
| F11 | 33000001 | 33100000 | 0.0041 | 0.0033 | 0.0036 |
| F11 | 33020001 | 33120000 | 0.0043 | 0.0032 | 0.0037 |
| F11 | 33040001 | 33140000 | 0.0042 | 0.0031 | 0.0036 |
| F11 | 33060001 | 33160000 | 0.0043 | 0.0034 | 0.0037 |
| F11 | 33080001 | 33180000 | 0.0042 | 0.0032 | 0.0035 |
| F11 | 33100001 | 33200000 | 0.0037 | 0.0026 | 0.0030 |
| F11 | 33120001 | 33220000 | 0.0037 | 0.0027 | 0.0030 |
| F11 | 33140001 | 33240000 | 0.0033 | 0.0025 | 0.0028 |
| F11 | 33160001 | 33260000 | 0.0025 | 0.0015 | 0.0019 |
| F11 | 33180001 | 33280000 | 0.0031 | 0.0018 | 0.0022 |
| F11 | 33200001 | 33300000 | 0.0038 | 0.0023 | 0.0028 |
| F11 | 33220001 | 33320000 | 0.0031 | 0.0018 | 0.0021 |
| F11 | 33240001 | 33340000 | 0.0031 | 0.0018 | 0.0021 |
| F11 | 33260001 | 33360000 | 0.0031 | 0.0016 | 0.0019 |
| F11 | 33280001 | 33380000 | 0.0025 | 0.0013 | 0.0015 |
| F11 | 33300001 | 33400000 | 0.0024 | 0.0009 | 0.0014 |
| F11 | 33320001 | 33420000 | 0.0031 | 0.0014 | 0.0020 |
| F11 | 33340001 | 33440000 | 0.0041 | 0.0025 | 0.0029 |
| F11 | 33360001 | 33460000 | 0.0041 | 0.0028 | 0.0031 |
| F11 | 33380001 | 33480000 | 0.0046 | 0.0031 | 0.0035 |
| F11 | 33400001 | 33500000 | 0.0036 | 0.0026 | 0.0027 |
| F11 | 33420001 | 33520000 | 0.0030 | 0.0022 | 0.0022 |
| F11 | 33440001 | 33540000 | 0.0018 | 0.0010 | 0.0011 |
| F11 | 33460001 | 33560000 | 0.0015 | 0.0007 | 0.0008 |
| F11 | 33480001 | 33580000 | 0.0011 | 0.0003 | 0.0004 |
| F11 | 33500001 | 33600000 | 0.0020 | 0.0008 | 0.0008 |
| F11 | 33520001 | 33620000 | 0.0020 | 0.0007 | 0.0008 |
| F11 | 33540001 | 33640000 | 0.0021 | 0.0008 | 0.0010 |
| F11 | 33560001 | 33660000 | 0.0033 | 0.0019 | 0.0021 |
| F11 | 33580001 | 33680000 | 0.0049 | 0.0035 | 0.0037 |
| F11 | 33600001 | 33700000 | 0.0050 | 0.0038 | 0.0041 |
| F11 | 33620001 | 33720000 | 0.0053 | 0.0043 | 0.0045 |
| F11 | 33640001 | 33740000 | 0.0051 | 0.0041 | 0.0043 |
| F11 | 33660001 | 33760000 | 0.0041 | 0.0033 | 0.0034 |
| F11 | 33680001 | 33780000 | 0.0026 | 0.0019 | 0.0020 |
| F11 | 33700001 | 33800000 | 0.0024 | 0.0015 | 0.0015 |
| F11 | 33720001 | 33820000 | 0.0021 | 0.0010 | 0.0011 |
| F11 | 33740001 | 33840000 | 0.0026 | 0.0015 | 0.0015 |
| F11 | 33760001 | 33860000 | 0.0031 | 0.0017 | 0.0018 |
| F11 | 33780001 | 33880000 | 0.0030 | 0.0017 | 0.0019 |
| F11 | 33800001 | 33900000 | 0.0029 | 0.0018 | 0.0020 |
| F11 | 33820001 | 33920000 | 0.0028 | 0.0018 | 0.0020 |
| F11 | 33840001 | 33940000 | 0.0026 | 0.0015 | 0.0017 |
| F11 | 33860001 | 33960000 | 0.0026 | 0.0016 | 0.0017 |
| F11 | 33880001 | 33980000 | 0.0028 | 0.0015 | 0.0018 |
| F11 | 33900001 | 34000000 | 0.0028 | 0.0015 | 0.0017 |
| F11 | 33920001 | 34020000 | 0.0027 | 0.0014 | 0.0016 |
| F11 | 33940001 | 34040000 | 0.0026 | 0.0014 | 0.0016 |
| F11 | 33960001 | 34060000 | 0.0030 | 0.0018 | 0.0020 |

| Chr | Start | End | both | A1 | A2 |
|---|---|---|---|---|---|
| F11 | 33980001 | 34080000 | 0.0037 | 0.0026 | 0.0028 |
| F11 | 34000001 | 34100000 | 0.0044 | 0.0032 | 0.0036 |
| F11 | 34020001 | 34120000 | 0.0052 | 0.0036 | 0.0043 |
| F11 | 34040001 | 34140000 | 0.0055 | 0.0040 | 0.0047 |
| F11 | 34060001 | 34160000 | 0.0047 | 0.0032 | 0.0040 |
| F11 | 34080001 | 34180000 | 0.0041 | 0.0026 | 0.0032 |
| F11 | 34100001 | 34200000 | 0.0029 | 0.0017 | 0.0021 |
| F11 | 34120001 | 34220000 | 0.0024 | 0.0014 | 0.0016 |
| F11 | 34140001 | 34240000 | 0.0019 | 0.0010 | 0.0011 |
| F11 | 34160001 | 34260000 | 0.0018 | 0.0009 | 0.0010 |
| F11 | 34180001 | 34280000 | 0.0019 | 0.0010 | 0.0012 |
| F11 | 34200001 | 34300000 | 0.0027 | 0.0015 | 0.0018 |
| F11 | 34220001 | 34320000 | 0.0030 | 0.0019 | 0.0023 |
| F11 | 34240001 | 34340000 | 0.0039 | 0.0024 | 0.0029 |
| F11 | 34260001 | 34360000 | 0.0042 | 0.0025 | 0.0032 |
| F11 | 34280001 | 34380000 | 0.0036 | 0.0022 | 0.0029 |
| F11 | 34300001 | 34400000 | 0.0040 | 0.0027 | 0.0031 |
| F11 | 34320001 | 34420000 | 0.0038 | 0.0025 | 0.0028 |
| F11 | 34340001 | 34440000 | 0.0030 | 0.0020 | 0.0022 |
| F11 | 34360001 | 34460000 | 0.0033 | 0.0024 | 0.0024 |
| F11 | 34380001 | 34480000 | 0.0036 | 0.0026 | 0.0027 |
| F11 | 34400001 | 34500000 | 0.0028 | 0.0019 | 0.0021 |
| F11 | 34420001 | 34520000 | 0.0028 | 0.0017 | 0.0021 |
| F11 | 34440001 | 34540000 | 0.0031 | 0.0019 | 0.0023 |
| F11 | 34460001 | 34560000 | 0.0034 | 0.0022 | 0.0026 |
| F11 | 34480001 | 34580000 | 0.0031 | 0.0019 | 0.0022 |
| F11 | 34500001 | 34600000 | 0.0031 | 0.0017 | 0.0022 |
| F11 | 34520001 | 34620000 | 0.0031 | 0.0022 | 0.0022 |
| F11 | 34540001 | 34640000 | 0.0042 | 0.0032 | 0.0034 |
| F11 | 34560001 | 34660000 | 0.0040 | 0.0029 | 0.0032 |
| F11 | 34580001 | 34680000 | 0.0042 | 0.0030 | 0.0033 |
| F11 | 34600001 | 34700000 | 0.0042 | 0.0033 | 0.0035 |
| F11 | 34620001 | 34720000 | 0.0045 | 0.0031 | 0.0037 |
| F11 | 34640001 | 34740000 | 0.0044 | 0.0030 | 0.0035 |
| F11 | 34660001 | 34760000 | 0.0042 | 0.0030 | 0.0034 |
| F11 | 34680001 | 34780000 | 0.0040 | 0.0031 | 0.0035 |
| F11 | 34700001 | 34800000 | 0.0043 | 0.0031 | 0.0036 |
| F11 | 34720001 | 34820000 | 0.0040 | 0.0030 | 0.0035 |
| F11 | 34740001 | 34840000 | 0.0030 | 0.0022 | 0.0026 |
| F11 | 34760001 | 34860000 | 0.0027 | 0.0016 | 0.0021 |
| F11 | 34780001 | 34880000 | 0.0026 | 0.0015 | 0.0020 |
| F11 | 34800001 | 34900000 | 0.0021 | 0.0013 | 0.0016 |
| F11 | 34820001 | 34920000 | 0.0018 | 0.0012 | 0.0014 |
| F11 | 34840001 | 34940000 | 0.0015 | 0.0010 | 0.0012 |
| F11 | 34860001 | 34960000 | 0.0022 | 0.0018 | 0.0020 |
| F11 | 34880001 | 34980000 | 0.0024 | 0.0019 | 0.0022 |
| F11 | 34900001 | 35000000 | 0.0027 | 0.0021 | 0.0026 |
| F11 | 34920001 | 35020000 | 0.0037 | 0.0029 | 0.0035 |
| F11 | 34940001 | 35040000 | 0.0040 | 0.0031 | 0.0037 |
| F11 | 34960001 | 35060000 | 0.0036 | 0.0028 | 0.0033 |
| F11 | 34980001 | 35080000 | 0.0037 | 0.0027 | 0.0032 |
| F11 | 35000001 | 35100000 | 0.0036 | 0.0025 | 0.0028 |
| F11 | 35020001 | 35120000 | 0.0029 | 0.0017 | 0.0017 |
| F11 | 35040001 | 35140000 | 0.0026 | 0.0013 | 0.0013 |
| F11 | 35060001 | 35160000 | 0.0024 | 0.0011 | 0.0010 |
| F11 | 35080001 | 35180000 | 0.0024 | 0.0012 | 0.0011 |
| F11 | 35100001 | 35200000 | 0.0025 | 0.0013 | 0.0013 |
| F11 | 35120001 | 35220000 | 0.0023 | 0.0014 | 0.0014 |

| Chr | Start | End | both | A1 | A2 |
|---|---|---|---|---|---|
| F11 | 35140001 | 35240000 | 0.0031 | 0.0021 | 0.0023 |
| F11 | 35160001 | 35260000 | 0.0032 | 0.0021 | 0.0024 |
| F11 | 35180001 | 35280000 | 0.0037 | 0.0024 | 0.0028 |
| F11 | 35200001 | 35300000 | 0.0033 | 0.0022 | 0.0026 |
| F11 | 35220001 | 35320000 | 0.0029 | 0.0018 | 0.0022 |
| F11 | 35240001 | 35340000 | 0.0028 | 0.0017 | 0.0022 |
| F11 | 35260001 | 35360000 | 0.0033 | 0.0024 | 0.0026 |
| F11 | 35280001 | 35380000 | 0.0028 | 0.0022 | 0.0024 |
| F11 | 35300001 | 35400000 | 0.0036 | 0.0030 | 0.0031 |
| F11 | 35320001 | 35420000 | 0.0040 | 0.0032 | 0.0034 |
| F11 | 35340001 | 35440000 | 0.0035 | 0.0028 | 0.0029 |
| F11 | 35360001 | 35460000 | 0.0029 | 0.0020 | 0.0022 |
| F11 | 35380001 | 35480000 | 0.0026 | 0.0016 | 0.0019 |
| F11 | 35400001 | 35500000 | 0.0019 | 0.0011 | 0.0013 |
| F11 | 35420001 | 35520000 | 0.0016 | 0.0010 | 0.0012 |
| F11 | 35440001 | 35540000 | 0.0012 | 0.0007 | 0.0009 |
| F11 | 35460001 | 35560000 | 0.0011 | 0.0007 | 0.0007 |
| F11 | 35480001 | 35580000 | 0.0013 | 0.0007 | 0.0007 |
| F11 | 35500001 | 35600000 | 0.0021 | 0.0013 | 0.0014 |
| F11 | 35520001 | 35620000 | 0.0025 | 0.0016 | 0.0016 |
| F11 | 35540001 | 35640000 | 0.0030 | 0.0018 | 0.0019 |
| F11 | 35560001 | 35660000 | 0.0038 | 0.0025 | 0.0027 |
| F11 | 35580001 | 35680000 | 0.0039 | 0.0029 | 0.0030 |
| F11 | 35600001 | 35700000 | 0.0030 | 0.0023 | 0.0022 |
| F11 | 35620001 | 35720000 | 0.0029 | 0.0022 | 0.0023 |
| F11 | 35640001 | 35740000 | 0.0028 | 0.0022 | 0.0024 |
| F11 | 35660001 | 35760000 | 0.0027 | 0.0022 | 0.0024 |
| F11 | 35680001 | 35780000 | 0.0034 | 0.0026 | 0.0031 |
| F11 | 35700001 | 35800000 | 0.0036 | 0.0029 | 0.0033 |
| F11 | 35720001 | 35820000 | 0.0033 | 0.0026 | 0.0029 |
| F11 | 35740001 | 35840000 | 0.0034 | 0.0025 | 0.0029 |
| F11 | 35760001 | 35860000 | 0.0038 | 0.0030 | 0.0032 |
| F11 | 35780001 | 35880000 | 0.0030 | 0.0024 | 0.0025 |
| F11 | 35800001 | 35900000 | 0.0032 | 0.0026 | 0.0026 |
| F11 | 35820001 | 35920000 | 0.0040 | 0.0033 | 0.0033 |
| F11 | 35840001 | 35940000 | 0.0043 | 0.0039 | 0.0038 |
| F11 | 35860001 | 35960000 | 0.0046 | 0.0039 | 0.0039 |
| F11 | 35880001 | 35980000 | 0.0044 | 0.0036 | 0.0036 |
| F11 | 35900001 | 36000000 | 0.0047 | 0.0039 | 0.0039 |
| F11 | 35920001 | 36020000 | 0.0051 | 0.0043 | 0.0044 |
| F11 | 35940001 | 36040000 | 0.0058 | 0.0051 | 0.0050 |
| F11 | 35960001 | 36060000 | 0.0051 | 0.0046 | 0.0046 |
| F11 | 35980001 | 36080000 | 0.0051 | 0.0046 | 0.0046 |
| F11 | 36000001 | 36100000 | 0.0051 | 0.0046 | 0.0047 |
| F11 | 36020001 | 36120000 | 0.0042 | 0.0036 | 0.0036 |
| F11 | 36040001 | 36140000 | 0.0040 | 0.0029 | 0.0033 |
| F11 | 36060001 | 36160000 | 0.0040 | 0.0027 | 0.0032 |
| F11 | 36080001 | 36180000 | 0.0041 | 0.0028 | 0.0032 |
| F11 | 36100001 | 36200000 | 0.0038 | 0.0024 | 0.0029 |
| F11 | 36120001 | 36220000 | 0.0038 | 0.0026 | 0.0030 |
| F11 | 36140001 | 36240000 | 0.0032 | 0.0023 | 0.0026 |
| F11 | 36160001 | 36260000 | 0.0027 | 0.0020 | 0.0022 |
| F11 | 36180001 | 36280000 | 0.0029 | 0.0025 | 0.0026 |
| F11 | 36200001 | 36300000 | 0.0024 | 0.0022 | 0.0022 |
| F11 | 36220001 | 36320000 | 0.0025 | 0.0023 | 0.0023 |
| F11 | 36240001 | 36340000 | 0.0027 | 0.0024 | 0.0027 |
| F11 | 36260001 | 36360000 | 0.0026 | 0.0023 | 0.0026 |
| F11 | 36280001 | 36380000 | 0.0029 | 0.0025 | 0.0028 |

| Chr | Start | End | both | A1 | A2 |
|---|---|---|---|---|---|
| F11 | 36300001 | 36400000 | 0.0031 | 0.0025 | 0.0029 |
| F11 | 36320001 | 36420000 | 0.0031 | 0.0024 | 0.0028 |
| F11 | 36340001 | 36440000 | 0.0026 | 0.0018 | 0.0022 |
| F11 | 36360001 | 36460000 | 0.0031 | 0.0021 | 0.0025 |
| F11 | 36380001 | 36480000 | 0.0027 | 0.0017 | 0.0020 |
| F11 | 36400001 | 36500000 | 0.0031 | 0.0020 | 0.0025 |
| F11 | 36420001 | 36520000 | 0.0030 | 0.0019 | 0.0025 |
| F11 | 36440001 | 36540000 | 0.0033 | 0.0021 | 0.0025 |
| F11 | 36460001 | 36560000 | 0.0037 | 0.0025 | 0.0031 |
| F11 | 36480001 | 36580000 | 0.0037 | 0.0025 | 0.0031 |
| F11 | 36500001 | 36600000 | 0.0041 | 0.0027 | 0.0033 |
| F11 | 36520001 | 36620000 | 0.0049 | 0.0036 | 0.0041 |
| F11 | 36540001 | 36640000 | 0.0043 | 0.0033 | 0.0038 |
| F11 | 36560001 | 36660000 | 0.0033 | 0.0026 | 0.0029 |
| F11 | 36580001 | 36680000 | 0.0034 | 0.0028 | 0.0029 |
| F11 | 36600001 | 36700000 | 0.0028 | 0.0023 | 0.0022 |
| F11 | 36620001 | 36720000 | 0.0021 | 0.0016 | 0.0013 |
| F11 | 36640001 | 36740000 | 0.0022 | 0.0016 | 0.0013 |
| F11 | 36660001 | 36760000 | 0.0023 | 0.0017 | 0.0015 |
| F11 | 36680001 | 36780000 | 0.0022 | 0.0015 | 0.0014 |
| F11 | 36700001 | 36800000 | 0.0023 | 0.0016 | 0.0017 |
| F11 | 36720001 | 36820000 | 0.0023 | 0.0015 | 0.0017 |
| F11 | 36740001 | 36840000 | 0.0022 | 0.0013 | 0.0016 |
| F11 | 36760001 | 36860000 | 0.0026 | 0.0017 | 0.0020 |
| F11 | 36780001 | 36880000 | 0.0026 | 0.0018 | 0.0021 |
| F11 | 36800001 | 36900000 | 0.0026 | 0.0018 | 0.0020 |
| F11 | 36820001 | 36920000 | 0.0023 | 0.0016 | 0.0018 |
| F11 | 36840001 | 36940000 | 0.0022 | 0.0017 | 0.0018 |
| F11 | 36860001 | 36960000 | 0.0025 | 0.0020 | 0.0020 |
| F11 | 36880001 | 36980000 | 0.0024 | 0.0019 | 0.0020 |
| F11 | 36900001 | 37000000 | 0.0025 | 0.0018 | 0.0019 |
| F11 | 36920001 | 37020000 | 0.0025 | 0.0017 | 0.0019 |
| F11 | 36940001 | 37040000 | 0.0026 | 0.0016 | 0.0018 |
| F11 | 36960001 | 37060000 | 0.0027 | 0.0017 | 0.0018 |
| F11 | 36980001 | 37080000 | 0.0026 | 0.0016 | 0.0016 |
| F11 | 37000001 | 37100000 | 0.0036 | 0.0026 | 0.0025 |
| F11 | 37020001 | 37120000 | 0.0042 | 0.0031 | 0.0030 |
| F11 | 37040001 | 37140000 | 0.0045 | 0.0035 | 0.0036 |
| F11 | 37060001 | 37160000 | 0.0042 | 0.0032 | 0.0033 |
| F11 | 37080001 | 37180000 | 0.0047 | 0.0036 | 0.0038 |
| F11 | 37100001 | 37200000 | 0.0035 | 0.0025 | 0.0028 |
| F11 | 37120001 | 37220000 | 0.0031 | 0.0021 | 0.0025 |
| F11 | 37140001 | 37240000 | 0.0030 | 0.0019 | 0.0022 |
| F11 | 37160001 | 37260000 | 0.0025 | 0.0013 | 0.0016 |
| F11 | 37180001 | 37280000 | 0.0021 | 0.0011 | 0.0013 |
| F11 | 37200001 | 37300000 | 0.0021 | 0.0011 | 0.0014 |
| F11 | 37220001 | 37320000 | 0.0031 | 0.0020 | 0.0022 |
| F11 | 37240001 | 37340000 | 0.0034 | 0.0025 | 0.0026 |
| F11 | 37260001 | 37360000 | 0.0038 | 0.0028 | 0.0031 |
| F11 | 37280001 | 37380000 | 0.0041 | 0.0030 | 0.0033 |
| F11 | 37300001 | 37400000 | 0.0042 | 0.0033 | 0.0035 |
| F11 | 37320001 | 37420000 | 0.0030 | 0.0024 | 0.0026 |
| F11 | 37340001 | 37440000 | 0.0029 | 0.0023 | 0.0026 |
| F11 | 37360001 | 37460000 | 0.0032 | 0.0026 | 0.0029 |
| F11 | 37380001 | 37480000 | 0.0026 | 0.0023 | 0.0025 |
| F11 | 37400001 | 37500000 | 0.0027 | 0.0022 | 0.0025 |
| F11 | 37420001 | 37520000 | 0.0032 | 0.0025 | 0.0029 |
| F11 | 37440001 | 37540000 | 0.0030 | 0.0021 | 0.0025 |

| Chr | Start | End | both | A1 | A2 |
|---|---|---|---|---|---|
| F11 | 37460001 | 37560000 | 0.0025 | 0.0016 | 0.0019 |
| F11 | 37480001 | 37580000 | 0.0026 | 0.0017 | 0.0019 |
| F11 | 37500001 | 37600000 | 0.0034 | 0.0023 | 0.0027 |
| F11 | 37520001 | 37620000 | 0.0037 | 0.0026 | 0.0030 |
| F11 | 37540001 | 37640000 | 0.0037 | 0.0027 | 0.0031 |
| F11 | 37560001 | 37660000 | 0.0046 | 0.0035 | 0.0039 |
| F11 | 37580001 | 37680000 | 0.0046 | 0.0034 | 0.0038 |
| F11 | 37600001 | 37700000 | 0.0035 | 0.0026 | 0.0027 |
| F11 | 37620001 | 37720000 | 0.0034 | 0.0025 | 0.0025 |
| F11 | 37640001 | 37740000 | 0.0037 | 0.0028 | 0.0029 |
| F11 | 37660001 | 37760000 | 0.0030 | 0.0020 | 0.0021 |
| F11 | 37680001 | 37780000 | 0.0039 | 0.0028 | 0.0030 |
| F11 | 37700001 | 37800000 | 0.0040 | 0.0028 | 0.0031 |
| F11 | 37720001 | 37820000 | 0.0034 | 0.0025 | 0.0028 |
| F11 | 37740001 | 37840000 | 0.0029 | 0.0020 | 0.0023 |
| F11 | 37760001 | 37860000 | 0.0025 | 0.0019 | 0.0021 |
| F11 | 37780001 | 37880000 | 0.0016 | 0.0012 | 0.0013 |
| F11 | 37800001 | 37900000 | 0.0022 | 0.0018 | 0.0020 |
| F11 | 37820001 | 37920000 | 0.0031 | 0.0025 | 0.0028 |
| F11 | 37840001 | 37940000 | 0.0030 | 0.0025 | 0.0028 |
| F11 | 37860001 | 37960000 | 0.0036 | 0.0032 | 0.0035 |
| F11 | 37880001 | 37980000 | 0.0036 | 0.0033 | 0.0035 |
| F11 | 37900001 | 38000000 | 0.0038 | 0.0033 | 0.0037 |
| F11 | 37920001 | 38020000 | 0.0028 | 0.0024 | 0.0027 |
| F11 | 37940001 | 38040000 | 0.0027 | 0.0023 | 0.0026 |
| F11 | 37960001 | 38060000 | 0.0022 | 0.0019 | 0.0022 |
| F11 | 37980001 | 38080000 | 0.0021 | 0.0017 | 0.0021 |
| F11 | 38000001 | 38100000 | 0.0017 | 0.0015 | 0.0016 |
| F11 | 38020001 | 38120000 | 0.0020 | 0.0019 | 0.0019 |
| F11 | 38040001 | 38140000 | 0.0020 | 0.0019 | 0.0019 |
| F11 | 38060001 | 38160000 | 0.0020 | 0.0019 | 0.0019 |
| F11 | 38080001 | 38180000 | 0.0027 | 0.0025 | 0.0026 |
| F11 | 38100001 | 38200000 | 0.0030 | 0.0027 | 0.0028 |
| F11 | 38120001 | 38220000 | 0.0039 | 0.0032 | 0.0036 |
| F11 | 38140001 | 38240000 | 0.0042 | 0.0035 | 0.0040 |
| F11 | 38160001 | 38260000 | 0.0041 | 0.0033 | 0.0039 |
| F11 | 38180001 | 38280000 | 0.0038 | 0.0031 | 0.0036 |
| F11 | 38200001 | 38300000 | 0.0037 | 0.0029 | 0.0036 |
| F11 | 38220001 | 38320000 | 0.0028 | 0.0023 | 0.0027 |
| F11 | 38240001 | 38340000 | 0.0031 | 0.0028 | 0.0031 |
| F11 | 38260001 | 38360000 | 0.0030 | 0.0027 | 0.0030 |
| F11 | 38280001 | 38380000 | 0.0032 | 0.0029 | 0.0031 |
| F11 | 38300001 | 38400000 | 0.0026 | 0.0024 | 0.0025 |
| F11 | 38320001 | 38420000 | 0.0026 | 0.0024 | 0.0025 |
| F11 | 38340001 | 38440000 | 0.0024 | 0.0020 | 0.0022 |
| F11 | 38360001 | 38460000 | 0.0025 | 0.0022 | 0.0023 |
| F11 | 38380001 | 38480000 | 0.0018 | 0.0014 | 0.0016 |
| F11 | 38400001 | 38500000 | 0.0020 | 0.0016 | 0.0019 |
| F11 | 38420001 | 38520000 | 0.0017 | 0.0014 | 0.0016 |
| F11 | 38440001 | 38540000 | 0.0014 | 0.0012 | 0.0014 |
| F11 | 38460001 | 38560000 | 0.0012 | 0.0011 | 0.0012 |
| F11 | 38480001 | 38580000 | 0.0017 | 0.0015 | 0.0017 |
| F11 | 38500001 | 38600000 | 0.0014 | 0.0012 | 0.0014 |
| F11 | 38520001 | 38620000 | 0.0016 | 0.0014 | 0.0016 |
| F11 | 38540001 | 38640000 | 0.0013 | 0.0011 | 0.0013 |
| F11 | 38560001 | 38660000 | 0.0015 | 0.0012 | 0.0014 |
| F11 | 38580001 | 38680000 | 0.0014 | 0.0011 | 0.0012 |
| F11 | 38600001 | 38700000 | 0.0015 | 0.0012 | 0.0013 |

| Chr | Start | End | both | A1 | A2 |
|---|---|---|---|---|---|
| F11 | 38620001 | 38720000 | 0.0018 | 0.0015 | 0.0016 |
| F11 | 38640001 | 38740000 | 0.0024 | 0.0019 | 0.0021 |
| F11 | 38660001 | 38760000 | 0.0021 | 0.0017 | 0.0019 |
| F11 | 38680001 | 38780000 | 0.0024 | 0.0019 | 0.0022 |
| F11 | 38700001 | 38800000 | 0.0027 | 0.0021 | 0.0022 |
| F11 | 38720001 | 38820000 | 0.0032 | 0.0026 | 0.0025 |
| F11 | 38740001 | 38840000 | 0.0030 | 0.0024 | 0.0024 |
| F11 | 38760001 | 38860000 | 0.0033 | 0.0024 | 0.0025 |
| F11 | 38780001 | 38880000 | 0.0029 | 0.0021 | 0.0022 |
| F11 | 38800001 | 38900000 | 0.0026 | 0.0019 | 0.0022 |
| F11 | 38820001 | 38920000 | 0.0025 | 0.0019 | 0.0021 |
| F11 | 38840001 | 38940000 | 0.0024 | 0.0018 | 0.0019 |
| F11 | 38860001 | 38960000 | 0.0023 | 0.0018 | 0.0019 |
| F11 | 38880001 | 38980000 | 0.0023 | 0.0018 | 0.0019 |
| F11 | 38900001 | 39000000 | 0.0024 | 0.0017 | 0.0020 |
| F11 | 38920001 | 39020000 | 0.0015 | 0.0010 | 0.0013 |
| F11 | 38940001 | 39040000 | 0.0016 | 0.0012 | 0.0015 |
| F11 | 38960001 | 39060000 | 0.0017 | 0.0012 | 0.0015 |
| F11 | 38980001 | 39080000 | 0.0019 | 0.0016 | 0.0018 |
| F11 | 39000001 | 39100000 | 0.0018 | 0.0015 | 0.0016 |
| F11 | 39020001 | 39120000 | 0.0029 | 0.0025 | 0.0025 |
| F11 | 39040001 | 39140000 | 0.0032 | 0.0027 | 0.0028 |
| F11 | 39060001 | 39160000 | 0.0035 | 0.0029 | 0.0029 |
| F11 | 39080001 | 39180000 | 0.0038 | 0.0030 | 0.0032 |
| F11 | 39100001 | 39200000 | 0.0040 | 0.0032 | 0.0035 |
| F11 | 39120001 | 39220000 | 0.0031 | 0.0024 | 0.0026 |
| F11 | 39140001 | 39240000 | 0.0031 | 0.0023 | 0.0027 |
| F11 | 39160001 | 39260000 | 0.0031 | 0.0024 | 0.0028 |
| F11 | 39180001 | 39280000 | 0.0035 | 0.0029 | 0.0033 |
| F11 | 39200001 | 39300000 | 0.0037 | 0.0030 | 0.0034 |
| F11 | 39220001 | 39320000 | 0.0036 | 0.0029 | 0.0034 |
| F11 | 39240001 | 39340000 | 0.0038 | 0.0031 | 0.0034 |
| F11 | 39260001 | 39360000 | 0.0042 | 0.0035 | 0.0038 |
| F11 | 39280001 | 39380000 | 0.0032 | 0.0027 | 0.0027 |
| F11 | 39300001 | 39400000 | 0.0046 | 0.0042 | 0.0038 |
| F11 | 39320001 | 39420000 | 0.0052 | 0.0047 | 0.0043 |
| F11 | 39340001 | 39440000 | 0.0060 | 0.0053 | 0.0052 |
| F11 | 39360001 | 39460000 | 0.0061 | 0.0054 | 0.0054 |
| F11 | 39380001 | 39480000 | 0.0060 | 0.0053 | 0.0054 |
| F11 | 39400001 | 39500000 | 0.0050 | 0.0040 | 0.0047 |
| F11 | 39420001 | 39520000 | 0.0042 | 0.0034 | 0.0040 |
| F11 | 39440001 | 39540000 | 0.0034 | 0.0024 | 0.0030 |
| F11 | 39460001 | 39560000 | 0.0025 | 0.0016 | 0.0019 |
| F11 | 39480001 | 39580000 | 0.0024 | 0.0014 | 0.0018 |
| F11 | 39500001 | 39600000 | 0.0032 | 0.0023 | 0.0026 |
| F11 | 39520001 | 39620000 | 0.0044 | 0.0031 | 0.0036 |
| F11 | 39540001 | 39640000 | 0.0042 | 0.0032 | 0.0037 |
| F11 | 39560001 | 39660000 | 0.0047 | 0.0036 | 0.0043 |
| F11 | 39580001 | 39680000 | 0.0049 | 0.0040 | 0.0045 |
| F11 | 39600001 | 39700000 | 0.0037 | 0.0030 | 0.0034 |
| F11 | 39620001 | 39720000 | 0.0031 | 0.0026 | 0.0028 |
| F11 | 39640001 | 39740000 | 0.0036 | 0.0031 | 0.0032 |
| F11 | 39660001 | 39760000 | 0.0038 | 0.0034 | 0.0035 |
| F11 | 39680001 | 39780000 | 0.0037 | 0.0032 | 0.0033 |
| F11 | 39700001 | 39800000 | 0.0036 | 0.0031 | 0.0031 |
| F11 | 39720001 | 39820000 | 0.0032 | 0.0027 | 0.0027 |
| F11 | 39740001 | 39840000 | 0.0025 | 0.0021 | 0.0022 |
| F11 | 39760001 | 39860000 | 0.0029 | 0.0022 | 0.0026 |

| Chr | Start | End | both | A1 | A2 |
|---|---|---|---|---|---|
| F11 | 39780001 | 39880000 | 0.0038 | 0.0028 | 0.0036 |
| F11 | 39800001 | 39900000 | 0.0038 | 0.0029 | 0.0038 |
| F11 | 39820001 | 39920000 | 0.0042 | 0.0033 | 0.0042 |
| F11 | 39840001 | 39940000 | 0.0044 | 0.0034 | 0.0045 |
| F11 | 39860001 | 39960000 | 0.0033 | 0.0026 | 0.0033 |
| F11 | 39880001 | 39980000 | 0.0024 | 0.0019 | 0.0023 |
| F11 | 39900001 | 40000000 | 0.0019 | 0.0015 | 0.0018 |
| F11 | 39920001 | 40020000 | 0.0019 | 0.0015 | 0.0018 |
| F11 | 39940001 | 40040000 | 0.0017 | 0.0014 | 0.0016 |
| F11 | 39960001 | 40060000 | 0.0022 | 0.0019 | 0.0021 |
| F11 | 39980001 | 40080000 | 0.0023 | 0.0018 | 0.0022 |
| F11 | 40000001 | 40100000 | 0.0024 | 0.0019 | 0.0022 |
| F11 | 40020001 | 40120000 | 0.0023 | 0.0018 | 0.0020 |
| F11 | 40040001 | 40140000 | 0.0025 | 0.0019 | 0.0022 |
| F11 | 40060001 | 40160000 | 0.0021 | 0.0017 | 0.0018 |
| F11 | 40080001 | 40180000 | 0.0025 | 0.0022 | 0.0022 |
| F11 | 40100001 | 40200000 | 0.0028 | 0.0026 | 0.0026 |
| F11 | 40120001 | 40220000 | 0.0031 | 0.0027 | 0.0027 |
| F11 | 40140001 | 40240000 | 0.0037 | 0.0030 | 0.0032 |
| F11 | 40160001 | 40260000 | 0.0046 | 0.0038 | 0.0040 |
| F11 | 40180001 | 40280000 | 0.0042 | 0.0032 | 0.0034 |
| F11 | 40200001 | 40300000 | 0.0040 | 0.0030 | 0.0032 |
| F11 | 40220001 | 40320000 | 0.0043 | 0.0032 | 0.0035 |
| F11 | 40240001 | 40340000 | 0.0039 | 0.0030 | 0.0031 |
| F11 | 40260001 | 40360000 | 0.0035 | 0.0025 | 0.0026 |
| F11 | 40280001 | 40380000 | 0.0040 | 0.0029 | 0.0033 |
| F11 | 40300001 | 40400000 | 0.0045 | 0.0032 | 0.0036 |
| F11 | 40320001 | 40420000 | 0.0039 | 0.0027 | 0.0030 |
| F11 | 40340001 | 40440000 | 0.0040 | 0.0027 | 0.0030 |
| F11 | 40360001 | 40460000 | 0.0046 | 0.0033 | 0.0037 |
| F11 | 40380001 | 40480000 | 0.0044 | 0.0032 | 0.0036 |
| F11 | 40400001 | 40500000 | 0.0041 | 0.0031 | 0.0035 |
| F11 | 40420001 | 40520000 | 0.0046 | 0.0035 | 0.0041 |
| F11 | 40440001 | 40540000 | 0.0046 | 0.0033 | 0.0040 |
| F11 | 40460001 | 40560000 | 0.0038 | 0.0027 | 0.0033 |
| F11 | 40480001 | 40580000 | 0.0035 | 0.0022 | 0.0027 |
| F11 | 40500001 | 40600000 | 0.0044 | 0.0032 | 0.0037 |
| F11 | 40520001 | 40620000 | 0.0044 | 0.0033 | 0.0037 |
| F11 | 40540001 | 40640000 | 0.0039 | 0.0031 | 0.0033 |
| F11 | 40560001 | 40660000 | 0.0044 | 0.0035 | 0.0040 |
| F11 | 40580001 | 40680000 | 0.0041 | 0.0036 | 0.0040 |
| F11 | 40600001 | 40700000 | 0.0030 | 0.0026 | 0.0028 |
| F11 | 40620001 | 40720000 | 0.0024 | 0.0020 | 0.0022 |
| F11 | 40640001 | 40740000 | 0.0022 | 0.0017 | 0.0020 |
| F11 | 40660001 | 40760000 | 0.0019 | 0.0016 | 0.0017 |
| F11 | 40680001 | 40780000 | 0.0020 | 0.0017 | 0.0018 |
| F11 | 40700001 | 40800000 | 0.0018 | 0.0015 | 0.0017 |
| F11 | 40720001 | 40820000 | 0.0018 | 0.0015 | 0.0016 |
| F11 | 40740001 | 40840000 | 0.0025 | 0.0021 | 0.0022 |
| F11 | 40760001 | 40860000 | 0.0026 | 0.0021 | 0.0023 |
| F11 | 40780001 | 40880000 | 0.0029 | 0.0024 | 0.0025 |
| F11 | 40800001 | 40900000 | 0.0032 | 0.0026 | 0.0029 |
| F11 | 40820001 | 40920000 | 0.0033 | 0.0029 | 0.0031 |
| F11 | 40840001 | 40940000 | 0.0033 | 0.0029 | 0.0030 |
| F11 | 40860001 | 40960000 | 0.0039 | 0.0034 | 0.0034 |
| F11 | 40880001 | 40980000 | 0.0037 | 0.0031 | 0.0032 |
| F11 | 40900001 | 41000000 | 0.0044 | 0.0037 | 0.0040 |
| F11 | 40920001 | 41020000 | 0.0048 | 0.0040 | 0.0043 |

| Chr | Start | End | both | A1 | A2 |
|---|---|---|---|---|---|
| F11 | 40940001 | 41040000 | 0.0048 | 0.0041 | 0.0046 |
| F11 | 40960001 | 41060000 | 0.0038 | 0.0032 | 0.0036 |
| F11 | 40980001 | 41080000 | 0.0040 | 0.0034 | 0.0036 |
| F11 | 41000001 | 41100000 | 0.0042 | 0.0035 | 0.0038 |
| F11 | 41020001 | 41120000 | 0.0050 | 0.0041 | 0.0044 |
| F11 | 41040001 | 41140000 | 0.0043 | 0.0033 | 0.0036 |
| F11 | 41060001 | 41160000 | 0.0042 | 0.0032 | 0.0035 |
| F11 | 41080001 | 41180000 | 0.0042 | 0.0032 | 0.0034 |
| F11 | 41100001 | 41200000 | 0.0035 | 0.0024 | 0.0027 |
| F11 | 41120001 | 41220000 | 0.0020 | 0.0011 | 0.0013 |
| F11 | 41140001 | 41240000 | 0.0025 | 0.0018 | 0.0019 |
| F11 | 41160001 | 41260000 | 0.0026 | 0.0019 | 0.0020 |
| F11 | 41180001 | 41280000 | 0.0026 | 0.0021 | 0.0021 |
| F11 | 41200001 | 41300000 | 0.0021 | 0.0018 | 0.0017 |
| F11 | 41220001 | 41320000 | 0.0030 | 0.0023 | 0.0023 |
| F11 | 41240001 | 41340000 | 0.0031 | 0.0022 | 0.0023 |
| F11 | 41260001 | 41360000 | 0.0039 | 0.0030 | 0.0031 |
| F11 | 41280001 | 41380000 | 0.0034 | 0.0025 | 0.0027 |
| F11 | 41300001 | 41400000 | 0.0047 | 0.0037 | 0.0040 |
| F11 | 41320001 | 41420000 | 0.0044 | 0.0036 | 0.0039 |
| F11 | 41340001 | 41440000 | 0.0043 | 0.0036 | 0.0039 |
| F11 | 41360001 | 41460000 | 0.0045 | 0.0036 | 0.0042 |
| F11 | 41380001 | 41480000 | 0.0055 | 0.0044 | 0.0050 |
| F11 | 41400001 | 41500000 | 0.0049 | 0.0036 | 0.0042 |
| F11 | 41420001 | 41520000 | 0.0046 | 0.0034 | 0.0038 |
| F11 | 41440001 | 41540000 | 0.0049 | 0.0038 | 0.0039 |
| F11 | 41460001 | 41560000 | 0.0037 | 0.0029 | 0.0028 |
| F11 | 41480001 | 41580000 | 0.0039 | 0.0030 | 0.0031 |
| F11 | 41500001 | 41600000 | 0.0035 | 0.0028 | 0.0029 |
| F11 | 41520001 | 41620000 | 0.0042 | 0.0036 | 0.0038 |
| F11 | 41540001 | 41640000 | 0.0036 | 0.0029 | 0.0032 |
| F11 | 41560001 | 41660000 | 0.0036 | 0.0029 | 0.0032 |
| F11 | 41580001 | 41680000 | 0.0027 | 0.0023 | 0.0024 |
| F11 | 41600001 | 41700000 | 0.0022 | 0.0019 | 0.0021 |
| F11 | 41620001 | 41720000 | 0.0013 | 0.0010 | 0.0012 |
| F11 | 41640001 | 41740000 | 0.0016 | 0.0014 | 0.0016 |
| F11 | 41660001 | 41760000 | 0.0015 | 0.0012 | 0.0015 |
| F11 | 41680001 | 41780000 | 0.0015 | 0.0013 | 0.0015 |
| F11 | 41700001 | 41800000 | 0.0018 | 0.0016 | 0.0018 |
| F11 | 41720001 | 41820000 | 0.0018 | 0.0015 | 0.0018 |
| F11 | 41740001 | 41840000 | 0.0014 | 0.0013 | 0.0014 |
| F11 | 41760001 | 41860000 | 0.0023 | 0.0022 | 0.0022 |
| F11 | 41780001 | 41880000 | 0.0030 | 0.0026 | 0.0026 |
| F11 | 41800001 | 41900000 | 0.0031 | 0.0027 | 0.0026 |
| F11 | 41820001 | 41920000 | 0.0036 | 0.0030 | 0.0031 |
| F11 | 41840001 | 41940000 | 0.0041 | 0.0035 | 0.0036 |
| F11 | 41860001 | 41960000 | 0.0039 | 0.0033 | 0.0034 |
| F11 | 41880001 | 41980000 | 0.0041 | 0.0037 | 0.0040 |
| F11 | 41900001 | 42000000 | 0.0049 | 0.0043 | 0.0047 |
| F11 | 41920001 | 42020000 | 0.0048 | 0.0042 | 0.0046 |
| F11 | 41940001 | 42040000 | 0.0045 | 0.0037 | 0.0042 |
| F11 | 41960001 | 42060000 | 0.0044 | 0.0035 | 0.0040 |
| F11 | 41980001 | 42080000 | 0.0035 | 0.0027 | 0.0030 |
| F11 | 42000001 | 42100000 | 0.0026 | 0.0019 | 0.0021 |
| F11 | 42020001 | 42120000 | 0.0025 | 0.0019 | 0.0021 |
| F11 | 42040001 | 42140000 | 0.0026 | 0.0021 | 0.0023 |
| F11 | 42060001 | 42160000 | 0.0030 | 0.0026 | 0.0028 |
| F11 | 42080001 | 42180000 | 0.0033 | 0.0027 | 0.0031 |

| Chr | Start | End | both | A1 | A2 |
|---|---|---|---|---|---|
| F11 | 42100001 | 42200000 | 0.0040 | 0.0034 | 0.0038 |
| F11 | 42120001 | 42220000 | 0.0037 | 0.0031 | 0.0034 |
| F11 | 42140001 | 42240000 | 0.0037 | 0.0029 | 0.0033 |
| F11 | 42160001 | 42260000 | 0.0033 | 0.0025 | 0.0028 |
| F11 | 42180001 | 42280000 | 0.0035 | 0.0024 | 0.0028 |
| F11 | 42200001 | 42300000 | 0.0030 | 0.0019 | 0.0023 |
| F11 | 42220001 | 42320000 | 0.0030 | 0.0021 | 0.0024 |
| F11 | 42240001 | 42340000 | 0.0030 | 0.0022 | 0.0023 |
| F11 | 42260001 | 42360000 | 0.0031 | 0.0025 | 0.0025 |
| F11 | 42280001 | 42380000 | 0.0024 | 0.0022 | 0.0021 |
| F11 | 42300001 | 42400000 | 0.0024 | 0.0023 | 0.0021 |
| F11 | 42320001 | 42420000 | 0.0025 | 0.0023 | 0.0022 |
| F11 | 42340001 | 42440000 | 0.0020 | 0.0018 | 0.0018 |
| F11 | 42360001 | 42460000 | 0.0016 | 0.0011 | 0.0013 |
| F11 | 42380001 | 42480000 | 0.0017 | 0.0012 | 0.0014 |
| F11 | 42400001 | 42500000 | 0.0019 | 0.0012 | 0.0014 |
| F11 | 42420001 | 42520000 | 0.0021 | 0.0014 | 0.0016 |
| F11 | 42440001 | 42540000 | 0.0030 | 0.0018 | 0.0022 |
| F11 | 42460001 | 42560000 | 0.0036 | 0.0025 | 0.0028 |
| F11 | 42480001 | 42580000 | 0.0048 | 0.0037 | 0.0040 |
| F11 | 42500001 | 42600000 | 0.0049 | 0.0037 | 0.0041 |
| F11 | 42520001 | 42620000 | 0.0042 | 0.0032 | 0.0035 |
| F11 | 42540001 | 42640000 | 0.0034 | 0.0029 | 0.0031 |
| F11 | 42560001 | 42660000 | 0.0028 | 0.0025 | 0.0027 |
| F11 | 42580001 | 42680000 | 0.0021 | 0.0016 | 0.0019 |
| F11 | 42600001 | 42700000 | 0.0020 | 0.0012 | 0.0015 |
| F11 | 42620001 | 42720000 | 0.0023 | 0.0014 | 0.0017 |
| F11 | 42640001 | 42740000 | 0.0025 | 0.0014 | 0.0018 |
| F11 | 42660001 | 42760000 | 0.0031 | 0.0018 | 0.0022 |
| F11 | 42680001 | 42780000 | 0.0029 | 0.0016 | 0.0020 |
| F11 | 42700001 | 42800000 | 0.0030 | 0.0019 | 0.0023 |
| F11 | 42720001 | 42820000 | 0.0030 | 0.0020 | 0.0024 |
| F11 | 42740001 | 42840000 | 0.0032 | 0.0023 | 0.0026 |
| F11 | 42760001 | 42860000 | 0.0031 | 0.0023 | 0.0027 |
| F11 | 42780001 | 42880000 | 0.0033 | 0.0025 | 0.0030 |
| F11 | 42800001 | 42900000 | 0.0045 | 0.0034 | 0.0041 |
| F11 | 42820001 | 42920000 | 0.0050 | 0.0038 | 0.0045 |
| F11 | 42840001 | 42940000 | 0.0050 | 0.0036 | 0.0043 |
| F11 | 42860001 | 42960000 | 0.0058 | 0.0044 | 0.0051 |
| F11 | 42880001 | 42980000 | 0.0058 | 0.0046 | 0.0050 |
| F11 | 42900001 | 43000000 | 0.0059 | 0.0048 | 0.0053 |
| F11 | 42920001 | 43020000 | 0.0059 | 0.0050 | 0.0053 |
| F11 | 42940001 | 43040000 | 0.0055 | 0.0048 | 0.0052 |
| F11 | 42960001 | 43060000 | 0.0049 | 0.0042 | 0.0044 |
| F11 | 42980001 | 43080000 | 0.0045 | 0.0037 | 0.0041 |
| F11 | 43000001 | 43100000 | 0.0030 | 0.0023 | 0.0026 |
| F11 | 43020001 | 43120000 | 0.0027 | 0.0019 | 0.0022 |
| F11 | 43040001 | 43140000 | 0.0034 | 0.0023 | 0.0026 |
| F11 | 43060001 | 43160000 | 0.0049 | 0.0037 | 0.0041 |
| F11 | 43080001 | 43180000 | 0.0049 | 0.0039 | 0.0042 |
| F11 | 43100001 | 43200000 | 0.0060 | 0.0049 | 0.0053 |
| F11 | 43120001 | 43220000 | 0.0066 | 0.0054 | 0.0058 |
| F11 | 43140001 | 43240000 | 0.0070 | 0.0056 | 0.0064 |
| F11 | 43160001 | 43260000 | 0.0059 | 0.0046 | 0.0052 |
| F11 | 43180001 | 43280000 | 0.0059 | 0.0045 | 0.0052 |
| F11 | 43200001 | 43300000 | 0.0046 | 0.0033 | 0.0039 |
| F11 | 43220001 | 43320000 | 0.0041 | 0.0032 | 0.0036 |
| F11 | 43240001 | 43340000 | 0.0031 | 0.0024 | 0.0025 |

| Chr | Start | End | both | A1 | A2 |
|---|---|---|---|---|---|
| F11 | 43260001 | 43360000 | 0.0021 | 0.0015 | 0.0015 |
| F11 | 43280001 | 43380000 | 0.0018 | 0.0012 | 0.0012 |
| F11 | 43300001 | 43400000 | 0.0019 | 0.0013 | 0.0014 |
| F11 | 43320001 | 43420000 | 0.0017 | 0.0012 | 0.0012 |
| F11 | 43340001 | 43440000 | 0.0015 | 0.0013 | 0.0013 |
| F11 | 43360001 | 43460000 | 0.0012 | 0.0010 | 0.0010 |
| F11 | 43380001 | 43480000 | 0.0013 | 0.0011 | 0.0012 |
| F11 | 43400001 | 43500000 | 0.0013 | 0.0009 | 0.0011 |
| F11 | 43420001 | 43520000 | 0.0013 | 0.0008 | 0.0011 |
| F11 | 43440001 | 43540000 | 0.0017 | 0.0011 | 0.0013 |
| F11 | 43460001 | 43560000 | 0.0021 | 0.0013 | 0.0015 |
| F11 | 43480001 | 43580000 | 0.0026 | 0.0016 | 0.0017 |
| F11 | 43500001 | 43600000 | 0.0035 | 0.0022 | 0.0023 |
| F11 | 43520001 | 43620000 | 0.0035 | 0.0021 | 0.0022 |
| F11 | 43540001 | 43640000 | 0.0037 | 0.0020 | 0.0024 |
| F11 | 43560001 | 43660000 | 0.0048 | 0.0032 | 0.0036 |
| F11 | 43580001 | 43680000 | 0.0051 | 0.0035 | 0.0041 |
| F11 | 43600001 | 43700000 | 0.0045 | 0.0031 | 0.0035 |
| F11 | 43620001 | 43720000 | 0.0041 | 0.0029 | 0.0033 |
| F11 | 43640001 | 43740000 | 0.0040 | 0.0030 | 0.0031 |
| F11 | 43660001 | 43760000 | 0.0033 | 0.0023 | 0.0024 |
| F11 | 43680001 | 43780000 | 0.0034 | 0.0024 | 0.0024 |
| F11 | 43700001 | 43800000 | 0.0038 | 0.0030 | 0.0031 |
| F11 | 43720001 | 43820000 | 0.0041 | 0.0033 | 0.0035 |
| F11 | 43740001 | 43840000 | 0.0049 | 0.0042 | 0.0046 |
| F11 | 43760001 | 43860000 | 0.0042 | 0.0036 | 0.0039 |
| F11 | 43780001 | 43880000 | 0.0034 | 0.0031 | 0.0033 |
| F11 | 43800001 | 43900000 | 0.0029 | 0.0025 | 0.0029 |
| F11 | 43820001 | 43920000 | 0.0027 | 0.0025 | 0.0027 |
| F11 | 43840001 | 43940000 | 0.0016 | 0.0014 | 0.0015 |
| F11 | 43860001 | 43960000 | 0.0016 | 0.0014 | 0.0016 |
| F11 | 43880001 | 43980000 | 0.0015 | 0.0013 | 0.0015 |
| F11 | 43900001 | 44000000 | 0.0013 | 0.0012 | 0.0013 |
| F11 | 43920001 | 44020000 | 0.0012 | 0.0010 | 0.0012 |
| F11 | 43940001 | 44040000 | 0.0012 | 0.0011 | 0.0012 |
| F11 | 43960001 | 44060000 | 0.0013 | 0.0012 | 0.0013 |
| F11 | 43980001 | 44080000 | 0.0013 | 0.0011 | 0.0013 |
| F11 | 44000001 | 44100000 | 0.0017 | 0.0013 | 0.0017 |
| F11 | 44020001 | 44120000 | 0.0014 | 0.0012 | 0.0014 |
| F11 | 44040001 | 44140000 | 0.0011 | 0.0009 | 0.0012 |
| F11 | 44060001 | 44160000 | 0.0019 | 0.0016 | 0.0019 |
| F11 | 44080001 | 44180000 | 0.0017 | 0.0016 | 0.0016 |
| F11 | 44100001 | 44200000 | 0.0015 | 0.0014 | 0.0014 |
| F11 | 44120001 | 44220000 | 0.0017 | 0.0016 | 0.0016 |
| F11 | 44140001 | 44240000 | 0.0020 | 0.0019 | 0.0020 |
| F11 | 44160001 | 44260000 | 0.0015 | 0.0012 | 0.0014 |
| F11 | 44180001 | 44280000 | 0.0019 | 0.0015 | 0.0019 |
| F11 | 44200001 | 44300000 | 0.0023 | 0.0020 | 0.0023 |
| F11 | 44220001 | 44320000 | 0.0024 | 0.0020 | 0.0024 |
| F11 | 44240001 | 44340000 | 0.0021 | 0.0018 | 0.0021 |
| F11 | 44260001 | 44360000 | 0.0018 | 0.0016 | 0.0018 |
| F11 | 44280001 | 44380000 | 0.0013 | 0.0012 | 0.0013 |
| F11 | 44300001 | 44400000 | 0.0008 | 0.0007 | 0.0008 |
| F11 | 44320001 | 44420000 | 0.0008 | 0.0007 | 0.0008 |
| F11 | 44340001 | 44440000 | 0.0017 | 0.0014 | 0.0015 |
| F11 | 44360001 | 44460000 | 0.0018 | 0.0014 | 0.0015 |
| F11 | 44380001 | 44480000 | 0.0022 | 0.0016 | 0.0017 |
| F11 | 44400001 | 44500000 | 0.0024 | 0.0017 | 0.0018 |

| Chr | Start | End | both | A1 | A2 |
|---|---|---|---|---|---|
| F11 | 44420001 | 44520000 | 0.0026 | 0.0018 | 0.0018 |
| F11 | 44440001 | 44540000 | 0.0019 | 0.0012 | 0.0013 |
| F11 | 44460001 | 44560000 | 0.0025 | 0.0016 | 0.0018 |
| F11 | 44480001 | 44580000 | 0.0022 | 0.0014 | 0.0016 |
| F11 | 44500001 | 44600000 | 0.0040 | 0.0031 | 0.0034 |
| F11 | 44520001 | 44620000 | 0.0044 | 0.0036 | 0.0038 |
| F11 | 44540001 | 44640000 | 0.0059 | 0.0048 | 0.0050 |
| F11 | 44560001 | 44660000 | 0.0067 | 0.0056 | 0.0058 |
| F11 | 44580001 | 44680000 | 0.0074 | 0.0061 | 0.0063 |
| F11 | 44600001 | 44700000 | 0.0061 | 0.0048 | 0.0050 |
| F11 | 44620001 | 44720000 | 0.0054 | 0.0041 | 0.0045 |
| F11 | 44640001 | 44740000 | 0.0042 | 0.0031 | 0.0033 |
| F11 | 44660001 | 44760000 | 0.0033 | 0.0024 | 0.0028 |
| F11 | 44680001 | 44780000 | 0.0032 | 0.0023 | 0.0027 |
| F11 | 44700001 | 44800000 | 0.0023 | 0.0016 | 0.0019 |
| F11 | 44720001 | 44820000 | 0.0033 | 0.0027 | 0.0029 |
| F11 | 44740001 | 44840000 | 0.0035 | 0.0030 | 0.0032 |
| F11 | 44760001 | 44860000 | 0.0035 | 0.0029 | 0.0029 |
| F11 | 44780001 | 44880000 | 0.0034 | 0.0028 | 0.0028 |
| F11 | 44800001 | 44900000 | 0.0043 | 0.0035 | 0.0037 |
| F11 | 44820001 | 44920000 | 0.0036 | 0.0027 | 0.0029 |
| F11 | 44840001 | 44940000 | 0.0035 | 0.0027 | 0.0028 |
| F11 | 44860001 | 44960000 | 0.0035 | 0.0028 | 0.0030 |
| F11 | 44880001 | 44980000 | 0.0036 | 0.0031 | 0.0033 |
| F11 | 44900001 | 45000000 | 0.0031 | 0.0026 | 0.0028 |
| F11 | 44920001 | 45020000 | 0.0029 | 0.0024 | 0.0028 |
| F11 | 44940001 | 45040000 | 0.0028 | 0.0023 | 0.0026 |
| F11 | 44960001 | 45060000 | 0.0021 | 0.0017 | 0.0020 |
| F11 | 44980001 | 45080000 | 0.0022 | 0.0018 | 0.0020 |
| F11 | 45000001 | 45100000 | 0.0024 | 0.0020 | 0.0022 |
| F11 | 45020001 | 45120000 | 0.0025 | 0.0022 | 0.0023 |
| F11 | 45040001 | 45140000 | 0.0025 | 0.0022 | 0.0024 |
| F11 | 45060001 | 45160000 | 0.0030 | 0.0025 | 0.0028 |
| F11 | 45080001 | 45180000 | 0.0028 | 0.0022 | 0.0027 |
| F11 | 45100001 | 45200000 | 0.0029 | 0.0023 | 0.0028 |
| F11 | 45120001 | 45220000 | 0.0026 | 0.0020 | 0.0025 |
| F11 | 45140001 | 45240000 | 0.0024 | 0.0018 | 0.0023 |
| F11 | 45160001 | 45260000 | 0.0031 | 0.0024 | 0.0030 |
| F11 | 45180001 | 45280000 | 0.0029 | 0.0023 | 0.0029 |
| F11 | 45200001 | 45300000 | 0.0023 | 0.0018 | 0.0022 |
| F11 | 45220001 | 45320000 | 0.0024 | 0.0019 | 0.0023 |
| F11 | 45240001 | 45340000 | 0.0025 | 0.0021 | 0.0024 |
| F11 | 45260001 | 45360000 | 0.0015 | 0.0013 | 0.0013 |
| F11 | 45280001 | 45380000 | 0.0019 | 0.0015 | 0.0017 |
| F11 | 45300001 | 45400000 | 0.0021 | 0.0017 | 0.0019 |
| F11 | 45320001 | 45420000 | 0.0026 | 0.0023 | 0.0024 |
| F11 | 45340001 | 45440000 | 0.0031 | 0.0027 | 0.0029 |
| F11 | 45360001 | 45460000 | 0.0036 | 0.0033 | 0.0034 |
| F11 | 45380001 | 45480000 | 0.0036 | 0.0032 | 0.0033 |
| F11 | 45400001 | 45500000 | 0.0037 | 0.0033 | 0.0033 |
| F11 | 45420001 | 45520000 | 0.0033 | 0.0028 | 0.0028 |
| F11 | 45440001 | 45540000 | 0.0028 | 0.0022 | 0.0022 |
| F11 | 45460001 | 45560000 | 0.0025 | 0.0018 | 0.0020 |
| F11 | 45480001 | 45580000 | 0.0023 | 0.0015 | 0.0017 |
| F11 | 45500001 | 45600000 | 0.0039 | 0.0027 | 0.0033 |
| F11 | 45520001 | 45620000 | 0.0040 | 0.0028 | 0.0034 |
| F11 | 45540001 | 45640000 | 0.0049 | 0.0036 | 0.0041 |
| F11 | 45560001 | 45660000 | 0.0047 | 0.0035 | 0.0040 |

| Chr | Start | End | both | A1 | A2 |
|---|---|---|---|---|---|
| F11 | 45580001 | 45680000 | 0.0049 | 0.0039 | 0.0043 |
| F11 | 45600001 | 45700000 | 0.0029 | 0.0024 | 0.0025 |
| F11 | 45620001 | 45720000 | 0.0024 | 0.0020 | 0.0020 |
| F11 | 45640001 | 45740000 | 0.0015 | 0.0012 | 0.0014 |
| F11 | 45660001 | 45760000 | 0.0014 | 0.0012 | 0.0013 |
| F11 | 45680001 | 45780000 | 0.0007 | 0.0007 | 0.0007 |
| F11 | 45700001 | 45800000 | 0.0011 | 0.0010 | 0.0011 |
| F11 | 45720001 | 45820000 | 0.0013 | 0.0011 | 0.0013 |
| F11 | 45740001 | 45840000 | 0.0014 | 0.0012 | 0.0014 |
| F11 | 45760001 | 45860000 | 0.0017 | 0.0014 | 0.0017 |
| F11 | 45780001 | 45880000 | 0.0026 | 0.0022 | 0.0025 |
| F11 | 45800001 | 45900000 | 0.0024 | 0.0022 | 0.0024 |
| F11 | 45820001 | 45920000 | 0.0025 | 0.0022 | 0.0024 |
| F11 | 45840001 | 45940000 | 0.0024 | 0.0022 | 0.0024 |
| F11 | 45860001 | 45960000 | 0.0023 | 0.0021 | 0.0022 |
| F11 | 45880001 | 45980000 | 0.0016 | 0.0014 | 0.0016 |
| F11 | 45900001 | 46000000 | 0.0014 | 0.0012 | 0.0014 |
| F11 | 45920001 | 46020000 | 0.0011 | 0.0010 | 0.0011 |
| F11 | 45940001 | 46040000 | 0.0007 | 0.0006 | 0.0007 |
| F11 | 45960001 | 46060000 | 0.0007 | 0.0006 | 0.0007 |
| F11 | 45980001 | 46080000 | 0.0005 | 0.0004 | 0.0005 |
| F11 | 46000001 | 46100000 | 0.0008 | 0.0006 | 0.0007 |
| F11 | 46020001 | 46120000 | 0.0011 | 0.0009 | 0.0010 |
| F11 | 46040001 | 46140000 | 0.0012 | 0.0009 | 0.0010 |
| F11 | 46060001 | 46160000 | 0.0014 | 0.0010 | 0.0013 |
| F11 | 46080001 | 46180000 | 0.0018 | 0.0013 | 0.0017 |
| F11 | 46100001 | 46200000 | 0.0023 | 0.0018 | 0.0023 |
| F11 | 46120001 | 46220000 | 0.0032 | 0.0025 | 0.0032 |
| F11 | 46140001 | 46240000 | 0.0039 | 0.0030 | 0.0036 |
| F11 | 46160001 | 46260000 | 0.0052 | 0.0040 | 0.0048 |
| F11 | 46180001 | 46280000 | 0.0053 | 0.0040 | 0.0048 |
| F11 | 46200001 | 46300000 | 0.0050 | 0.0039 | 0.0044 |
| F11 | 46220001 | 46320000 | 0.0051 | 0.0039 | 0.0044 |
| F11 | 46240001 | 46340000 | 0.0058 | 0.0046 | 0.0051 |
| F11 | 46260001 | 46360000 | 0.0047 | 0.0037 | 0.0039 |
| F11 | 46280001 | 46380000 | 0.0043 | 0.0034 | 0.0037 |
| F11 | 46300001 | 46400000 | 0.0049 | 0.0039 | 0.0042 |
| F11 | 46320001 | 46420000 | 0.0041 | 0.0034 | 0.0036 |
| F11 | 46340001 | 46440000 | 0.0035 | 0.0029 | 0.0031 |
| F11 | 46360001 | 46460000 | 0.0036 | 0.0029 | 0.0032 |
| F11 | 46380001 | 46480000 | 0.0042 | 0.0035 | 0.0037 |
| F11 | 46400001 | 46500000 | 0.0039 | 0.0031 | 0.0034 |
| F11 | 46420001 | 46520000 | 0.0053 | 0.0042 | 0.0046 |
| F11 | 46440001 | 46540000 | 0.0046 | 0.0036 | 0.0040 |
| F11 | 46460001 | 46560000 | 0.0051 | 0.0041 | 0.0045 |
| F11 | 46480001 | 46580000 | 0.0049 | 0.0041 | 0.0045 |
| F11 | 46500001 | 46600000 | 0.0051 | 0.0045 | 0.0048 |
| F11 | 46520001 | 46620000 | 0.0037 | 0.0034 | 0.0035 |
| F11 | 46540001 | 46640000 | 0.0038 | 0.0034 | 0.0036 |
| F11 | 46560001 | 46660000 | 0.0031 | 0.0028 | 0.0029 |
| F11 | 46580001 | 46680000 | 0.0029 | 0.0025 | 0.0026 |
| F11 | 46600001 | 46700000 | 0.0018 | 0.0015 | 0.0016 |
| F11 | 46620001 | 46720000 | 0.0018 | 0.0015 | 0.0015 |
| F11 | 46640001 | 46740000 | 0.0022 | 0.0018 | 0.0018 |
| F11 | 46660001 | 46760000 | 0.0020 | 0.0017 | 0.0017 |
| F11 | 46680001 | 46780000 | 0.0021 | 0.0017 | 0.0017 |
| F11 | 46700001 | 46800000 | 0.0038 | 0.0032 | 0.0033 |
| F11 | 46720001 | 46820000 | 0.0037 | 0.0030 | 0.0032 |

| Chr | Start | End | both | A1 | A2 |
|---|---|---|---|---|---|
| F11 | 46740001 | 46840000 | 0.0031 | 0.0026 | 0.0028 |
| F11 | 46760001 | 46860000 | 0.0031 | 0.0026 | 0.0028 |
| F11 | 46780001 | 46880000 | 0.0027 | 0.0023 | 0.0025 |
| F11 | 46800001 | 46900000 | 0.0009 | 0.0008 | 0.0009 |
| F11 | 46820001 | 46920000 | 0.0006 | 0.0006 | 0.0006 |
| F11 | 46840001 | 46940000 | 0.0013 | 0.0012 | 0.0013 |
| F11 | 46860001 | 46960000 | 0.0015 | 0.0014 | 0.0015 |
| F11 | 46880001 | 46980000 | 0.0020 | 0.0018 | 0.0020 |
| F11 | 46900001 | 47000000 | 0.0022 | 0.0020 | 0.0022 |
| F11 | 46920001 | 47020000 | 0.0028 | 0.0025 | 0.0028 |
| F11 | 46940001 | 47040000 | 0.0024 | 0.0020 | 0.0023 |
| F11 | 46960001 | 47060000 | 0.0031 | 0.0026 | 0.0029 |
| F11 | 46980001 | 47080000 | 0.0039 | 0.0031 | 0.0034 |
| F11 | 47000001 | 47100000 | 0.0043 | 0.0032 | 0.0036 |
| F11 | 47020001 | 47120000 | 0.0051 | 0.0038 | 0.0042 |
| F11 | 47040001 | 47140000 | 0.0054 | 0.0040 | 0.0046 |
| F11 | 47060001 | 47160000 | 0.0050 | 0.0037 | 0.0043 |
| F11 | 47080001 | 47180000 | 0.0037 | 0.0028 | 0.0034 |
| F11 | 47100001 | 47200000 | 0.0032 | 0.0025 | 0.0030 |
| F11 | 47120001 | 47220000 | 0.0020 | 0.0016 | 0.0019 |
| F11 | 47140001 | 47240000 | 0.0012 | 0.0011 | 0.0012 |
| F11 | 47160001 | 47260000 | 0.0010 | 0.0008 | 0.0010 |
| F11 | 47180001 | 47280000 | 0.0010 | 0.0008 | 0.0010 |
| F11 | 47200001 | 47300000 | 0.0010 | 0.0008 | 0.0010 |
| F11 | 47220001 | 47320000 | 0.0013 | 0.0010 | 0.0013 |
| F11 | 47240001 | 47340000 | 0.0016 | 0.0012 | 0.0016 |
| F11 | 47260001 | 47360000 | 0.0012 | 0.0010 | 0.0012 |
| F11 | 47280001 | 47380000 | 0.0011 | 0.0010 | 0.0011 |
| F11 | 47300001 | 47400000 | 0.0012 | 0.0010 | 0.0011 |
| F11 | 47320001 | 47420000 | 0.0008 | 0.0008 | 0.0008 |
| F11 | 47340001 | 47440000 | 0.0008 | 0.0007 | 0.0008 |
| F11 | 47360001 | 47460000 | 0.0012 | 0.0010 | 0.0011 |
| F11 | 47380001 | 47480000 | 0.0012 | 0.0010 | 0.0012 |
| F11 | 47400001 | 47500000 | 0.0013 | 0.0010 | 0.0013 |
| F11 | 47420001 | 47520000 | 0.0016 | 0.0013 | 0.0016 |
| F11 | 47440001 | 47540000 | 0.0015 | 0.0012 | 0.0015 |
| F11 | 47460001 | 47560000 | 0.0012 | 0.0010 | 0.0012 |
| F11 | 47480001 | 47580000 | 0.0015 | 0.0012 | 0.0014 |
| F11 | 47500001 | 47600000 | 0.0015 | 0.0012 | 0.0014 |
| F11 | 47520001 | 47620000 | 0.0013 | 0.0010 | 0.0013 |
| F11 | 47540001 | 47640000 | 0.0016 | 0.0013 | 0.0016 |
| F11 | 47560001 | 47660000 | 0.0016 | 0.0013 | 0.0016 |
| F11 | 47580001 | 47680000 | 0.0013 | 0.0010 | 0.0013 |
| F11 | 47600001 | 47700000 | 0.0012 | 0.0010 | 0.0012 |
| F11 | 47620001 | 47720000 | 0.0011 | 0.0010 | 0.0011 |
| F11 | 47640001 | 47740000 | 0.0011 | 0.0010 | 0.0011 |
| F11 | 47660001 | 47760000 | 0.0013 | 0.0011 | 0.0012 |
| F11 | 47680001 | 47780000 | 0.0015 | 0.0011 | 0.0012 |
| F11 | 47700001 | 47800000 | 0.0018 | 0.0014 | 0.0014 |
| F11 | 47720001 | 47820000 | 0.0019 | 0.0013 | 0.0014 |
| F11 | 47740001 | 47840000 | 0.0022 | 0.0014 | 0.0015 |
| F11 | 47760001 | 47860000 | 0.0031 | 0.0022 | 0.0023 |
| F11 | 47780001 | 47880000 | 0.0036 | 0.0026 | 0.0028 |
| F11 | 47800001 | 47900000 | 0.0045 | 0.0033 | 0.0037 |
| F11 | 47820001 | 47920000 | 0.0060 | 0.0045 | 0.0051 |
| F11 | 47840001 | 47940000 | 0.0058 | 0.0044 | 0.0049 |
| F11 | 47860001 | 47960000 | 0.0047 | 0.0033 | 0.0039 |
| F11 | 47880001 | 47980000 | 0.0051 | 0.0038 | 0.0044 |

| Chr | Start | End | both | A1 | A2 |
|---|---|---|---|---|---|
| F11 | 47900001 | 48000000 | 0.0040 | 0.0030 | 0.0033 |
| F11 | 47920001 | 48020000 | 0.0030 | 0.0023 | 0.0025 |
| F11 | 47940001 | 48040000 | 0.0029 | 0.0024 | 0.0025 |
| F11 | 47960001 | 48060000 | 0.0029 | 0.0024 | 0.0025 |
| F11 | 47980001 | 48080000 | 0.0022 | 0.0019 | 0.0019 |
| F11 | 48000001 | 48100000 | 0.0027 | 0.0021 | 0.0024 |
| F11 | 48020001 | 48120000 | 0.0020 | 0.0016 | 0.0018 |
| F11 | 48040001 | 48140000 | 0.0022 | 0.0016 | 0.0020 |
| F11 | 48060001 | 48160000 | 0.0023 | 0.0018 | 0.0022 |
| F11 | 48080001 | 48180000 | 0.0029 | 0.0020 | 0.0025 |
| F11 | 48100001 | 48200000 | 0.0025 | 0.0019 | 0.0022 |
| F11 | 48120001 | 48220000 | 0.0026 | 0.0019 | 0.0022 |
| F11 | 48140001 | 48240000 | 0.0024 | 0.0017 | 0.0018 |
| F11 | 48160001 | 48260000 | 0.0023 | 0.0015 | 0.0015 |
| F11 | 48180001 | 48280000 | 0.0022 | 0.0014 | 0.0014 |
| F11 | 48200001 | 48300000 | 0.0025 | 0.0016 | 0.0016 |
| F11 | 48220001 | 48320000 | 0.0024 | 0.0016 | 0.0016 |
| F11 | 48240001 | 48340000 | 0.0025 | 0.0019 | 0.0018 |
| F11 | 48260001 | 48360000 | 0.0023 | 0.0018 | 0.0019 |
| F11 | 48280001 | 48380000 | 0.0015 | 0.0012 | 0.0012 |
| F11 | 48300001 | 48400000 | 0.0008 | 0.0008 | 0.0008 |
| F11 | 48320001 | 48420000 | 0.0009 | 0.0009 | 0.0009 |
| F11 | 48340001 | 48440000 | 0.0008 | 0.0006 | 0.0007 |
| F11 | 48360001 | 48460000 | 0.0008 | 0.0004 | 0.0006 |
| F11 | 48380001 | 48480000 | 0.0015 | 0.0010 | 0.0013 |
| F11 | 48400001 | 48500000 | 0.0022 | 0.0016 | 0.0018 |
| F11 | 48420001 | 48520000 | 0.0031 | 0.0022 | 0.0024 |
| F11 | 48440001 | 48540000 | 0.0030 | 0.0023 | 0.0023 |
| F11 | 48460001 | 48560000 | 0.0043 | 0.0033 | 0.0034 |
| F11 | 48480001 | 48580000 | 0.0041 | 0.0029 | 0.0030 |
| F11 | 48500001 | 48600000 | 0.0036 | 0.0023 | 0.0025 |
| F11 | 48520001 | 48620000 | 0.0034 | 0.0021 | 0.0023 |
| F11 | 48540001 | 48640000 | 0.0037 | 0.0021 | 0.0025 |
| F11 | 48560001 | 48660000 | 0.0031 | 0.0019 | 0.0022 |
| F11 | 48580001 | 48680000 | 0.0034 | 0.0025 | 0.0027 |
| F11 | 48600001 | 48700000 | 0.0038 | 0.0028 | 0.0031 |
| F11 | 48620001 | 48720000 | 0.0041 | 0.0032 | 0.0036 |
| F11 | 48640001 | 48740000 | 0.0039 | 0.0031 | 0.0034 |
| F11 | 48660001 | 48760000 | 0.0034 | 0.0027 | 0.0030 |
| F11 | 48680001 | 48780000 | 0.0030 | 0.0024 | 0.0026 |
| F11 | 48700001 | 48800000 | 0.0026 | 0.0021 | 0.0022 |
| F11 | 48720001 | 48820000 | 0.0024 | 0.0020 | 0.0020 |
| F11 | 48740001 | 48840000 | 0.0026 | 0.0021 | 0.0022 |
| F11 | 48760001 | 48860000 | 0.0025 | 0.0018 | 0.0020 |
| F11 | 48780001 | 48880000 | 0.0031 | 0.0020 | 0.0025 |
| F11 | 48800001 | 48900000 | 0.0037 | 0.0024 | 0.0032 |
| F11 | 48820001 | 48920000 | 0.0031 | 0.0019 | 0.0027 |
| F11 | 48840001 | 48940000 | 0.0036 | 0.0023 | 0.0029 |
| F11 | 48860001 | 48960000 | 0.0035 | 0.0023 | 0.0028 |
| F11 | 48880001 | 48980000 | 0.0029 | 0.0018 | 0.0020 |
| F11 | 48900001 | 49000000 | 0.0033 | 0.0020 | 0.0021 |
| F11 | 48920001 | 49020000 | 0.0037 | 0.0023 | 0.0026 |
| F11 | 48940001 | 49040000 | 0.0037 | 0.0023 | 0.0027 |
| F11 | 48960001 | 49060000 | 0.0043 | 0.0027 | 0.0033 |
| F11 | 48980001 | 49080000 | 0.0048 | 0.0033 | 0.0040 |
| F11 | 49000001 | 49100000 | 0.0041 | 0.0029 | 0.0034 |
| F11 | 49020001 | 49120000 | 0.0047 | 0.0036 | 0.0039 |
| F11 | 49040001 | 49140000 | 0.0052 | 0.0040 | 0.0044 |

| Chr | Start | End | both | A1 | A2 |
|---|---|---|---|---|---|
| F11 | 49060001 | 49160000 | 0.0044 | 0.0035 | 0.0038 |
| F11 | 49080001 | 49180000 | 0.0043 | 0.0036 | 0.0038 |
| F11 | 49100001 | 49200000 | 0.0049 | 0.0041 | 0.0044 |
| F11 | 49120001 | 49220000 | 0.0035 | 0.0029 | 0.0031 |
| F11 | 49140001 | 49240000 | 0.0030 | 0.0022 | 0.0026 |
| F11 | 49160001 | 49260000 | 0.0031 | 0.0024 | 0.0027 |
| F11 | 49180001 | 49280000 | 0.0027 | 0.0021 | 0.0023 |
| F11 | 49200001 | 49300000 | 0.0017 | 0.0012 | 0.0013 |
| F11 | 49220001 | 49320000 | 0.0021 | 0.0015 | 0.0018 |
| F11 | 49240001 | 49340000 | 0.0016 | 0.0013 | 0.0014 |
| F11 | 49260001 | 49360000 | 0.0018 | 0.0014 | 0.0015 |
| F11 | 49280001 | 49380000 | 0.0021 | 0.0016 | 0.0018 |
| F11 | 49300001 | 49400000 | 0.0034 | 0.0026 | 0.0029 |
| F11 | 49320001 | 49420000 | 0.0036 | 0.0028 | 0.0029 |
| F11 | 49340001 | 49440000 | 0.0041 | 0.0033 | 0.0035 |
| F11 | 49360001 | 49460000 | 0.0044 | 0.0035 | 0.0036 |
| F11 | 49380001 | 49480000 | 0.0038 | 0.0030 | 0.0031 |
| F11 | 49400001 | 49500000 | 0.0032 | 0.0023 | 0.0025 |
| F11 | 49420001 | 49520000 | 0.0028 | 0.0020 | 0.0022 |
| F11 | 49440001 | 49540000 | 0.0022 | 0.0014 | 0.0016 |
| F11 | 49460001 | 49560000 | 0.0019 | 0.0012 | 0.0014 |
| F11 | 49480001 | 49580000 | 0.0021 | 0.0012 | 0.0014 |
| F11 | 49500001 | 49600000 | 0.0021 | 0.0015 | 0.0016 |
| F11 | 49520001 | 49620000 | 0.0023 | 0.0017 | 0.0020 |
| F11 | 49540001 | 49640000 | 0.0022 | 0.0016 | 0.0018 |
| F11 | 49560001 | 49660000 | 0.0021 | 0.0016 | 0.0018 |
| F11 | 49580001 | 49680000 | 0.0028 | 0.0020 | 0.0023 |
| F11 | 49600001 | 49700000 | 0.0029 | 0.0021 | 0.0022 |
| F11 | 49620001 | 49720000 | 0.0029 | 0.0019 | 0.0020 |
| F11 | 49640001 | 49740000 | 0.0033 | 0.0019 | 0.0022 |
| F11 | 49660001 | 49760000 | 0.0042 | 0.0027 | 0.0030 |
| F11 | 49680001 | 49780000 | 0.0041 | 0.0030 | 0.0033 |
| F11 | 49700001 | 49800000 | 0.0041 | 0.0030 | 0.0033 |
| F11 | 49720001 | 49820000 | 0.0036 | 0.0028 | 0.0030 |
| F11 | 49740001 | 49840000 | 0.0029 | 0.0024 | 0.0026 |
| F11 | 49760001 | 49860000 | 0.0034 | 0.0030 | 0.0029 |
| F11 | 49780001 | 49880000 | 0.0028 | 0.0026 | 0.0025 |
| F11 | 49800001 | 49900000 | 0.0025 | 0.0025 | 0.0021 |
| F11 | 49820001 | 49920000 | 0.0028 | 0.0027 | 0.0025 |
| F11 | 49840001 | 49940000 | 0.0031 | 0.0030 | 0.0027 |
| F11 | 49860001 | 49960000 | 0.0029 | 0.0027 | 0.0027 |
| F11 | 49880001 | 49980000 | 0.0027 | 0.0025 | 0.0026 |
| F11 | 49900001 | 50000000 | 0.0029 | 0.0024 | 0.0028 |
| F11 | 49920001 | 50020000 | 0.0034 | 0.0029 | 0.0034 |
| F11 | 49940001 | 50040000 | 0.0043 | 0.0036 | 0.0042 |
| F11 | 49960001 | 50060000 | 0.0036 | 0.0029 | 0.0032 |
| F11 | 49980001 | 50080000 | 0.0038 | 0.0032 | 0.0035 |
| F11 | 50000001 | 50100000 | 0.0037 | 0.0031 | 0.0033 |
| F11 | 50020001 | 50120000 | 0.0036 | 0.0029 | 0.0031 |
| F11 | 50040001 | 50140000 | 0.0034 | 0.0029 | 0.0029 |
| F11 | 50060001 | 50160000 | 0.0034 | 0.0030 | 0.0031 |
| F11 | 50080001 | 50180000 | 0.0037 | 0.0030 | 0.0033 |
| F11 | 50100001 | 50200000 | 0.0039 | 0.0029 | 0.0033 |
| F11 | 50120001 | 50220000 | 0.0040 | 0.0029 | 0.0035 |
| F11 | 50140001 | 50240000 | 0.0039 | 0.0027 | 0.0034 |
| F11 | 50160001 | 50260000 | 0.0046 | 0.0029 | 0.0036 |
| F11 | 50180001 | 50280000 | 0.0043 | 0.0028 | 0.0033 |
| F11 | 50200001 | 50300000 | 0.0041 | 0.0027 | 0.0033 |

| Chr | Start | End | both | A1 | A2 |
|---|---|---|---|---|---|
| F11 | 50220001 | 50320000 | 0.0040 | 0.0027 | 0.0031 |
| F11 | 50240001 | 50340000 | 0.0042 | 0.0030 | 0.0032 |
| F11 | 50260001 | 50360000 | 0.0036 | 0.0025 | 0.0028 |
| F11 | 50280001 | 50380000 | 0.0035 | 0.0025 | 0.0027 |
| F11 | 50300001 | 50400000 | 0.0034 | 0.0024 | 0.0025 |
| F11 | 50320001 | 50420000 | 0.0036 | 0.0027 | 0.0026 |
| F11 | 50340001 | 50440000 | 0.0029 | 0.0021 | 0.0021 |
| F11 | 50360001 | 50460000 | 0.0030 | 0.0023 | 0.0023 |
| F11 | 50380001 | 50480000 | 0.0027 | 0.0021 | 0.0022 |
| F11 | 50400001 | 50500000 | 0.0031 | 0.0026 | 0.0028 |
| F11 | 50420001 | 50520000 | 0.0022 | 0.0019 | 0.0021 |
| F11 | 50440001 | 50540000 | 0.0019 | 0.0017 | 0.0019 |
| F11 | 50460001 | 50560000 | 0.0020 | 0.0018 | 0.0019 |
| F11 | 50480001 | 50580000 | 0.0029 | 0.0024 | 0.0027 |
| F11 | 50500001 | 50600000 | 0.0031 | 0.0025 | 0.0026 |
| F11 | 50520001 | 50620000 | 0.0036 | 0.0029 | 0.0030 |
| F11 | 50540001 | 50640000 | 0.0033 | 0.0027 | 0.0028 |
| F11 | 50560001 | 50660000 | 0.0028 | 0.0021 | 0.0023 |
| F11 | 50580001 | 50680000 | 0.0024 | 0.0018 | 0.0019 |
| F11 | 50600001 | 50700000 | 0.0020 | 0.0016 | 0.0019 |
| F11 | 50620001 | 50720000 | 0.0016 | 0.0013 | 0.0016 |
| F11 | 50640001 | 50740000 | 0.0021 | 0.0017 | 0.0021 |
| F11 | 50660001 | 50760000 | 0.0031 | 0.0027 | 0.0030 |
| F11 | 50680001 | 50780000 | 0.0034 | 0.0029 | 0.0033 |
| F11 | 50700001 | 50800000 | 0.0039 | 0.0032 | 0.0036 |
| F11 | 50720001 | 50820000 | 0.0048 | 0.0039 | 0.0043 |
| F11 | 50740001 | 50840000 | 0.0059 | 0.0048 | 0.0053 |
| F11 | 50760001 | 50860000 | 0.0053 | 0.0040 | 0.0047 |
| F11 | 50780001 | 50880000 | 0.0047 | 0.0035 | 0.0041 |
| F11 | 50800001 | 50900000 | 0.0045 | 0.0035 | 0.0041 |
| F11 | 50820001 | 50920000 | 0.0044 | 0.0036 | 0.0042 |
| F11 | 50840001 | 50940000 | 0.0028 | 0.0022 | 0.0027 |
| F11 | 50860001 | 50960000 | 0.0024 | 0.0020 | 0.0023 |
| F11 | 50880001 | 50980000 | 0.0030 | 0.0026 | 0.0030 |
| F11 | 50900001 | 51000000 | 0.0034 | 0.0023 | 0.0032 |
| F11 | 50920001 | 51020000 | 0.0038 | 0.0024 | 0.0036 |
| F11 | 50940001 | 51040000 | 0.0046 | 0.0025 | 0.0042 |
| F11 | 50960001 | 51060000 | 0.0049 | 0.0027 | 0.0044 |
| F11 | 50980001 | 51080000 | 0.0051 | 0.0026 | 0.0043 |
| F11 | 51000001 | 51100000 | 0.0047 | 0.0028 | 0.0039 |
| F11 | 51020001 | 51120000 | 0.0046 | 0.0029 | 0.0036 |
| F11 | 51040001 | 51140000 | 0.0043 | 0.0028 | 0.0032 |
| F11 | 51060001 | 51160000 | 0.0052 | 0.0038 | 0.0041 |
| F11 | 51080001 | 51180000 | 0.0043 | 0.0033 | 0.0035 |
| F11 | 51100001 | 51200000 | 0.0035 | 0.0026 | 0.0028 |
| F11 | 51120001 | 51220000 | 0.0032 | 0.0024 | 0.0026 |
| F11 | 51140001 | 51240000 | 0.0030 | 0.0024 | 0.0027 |
| F11 | 51160001 | 51260000 | 0.0015 | 0.0011 | 0.0013 |
| F11 | 51180001 | 51280000 | 0.0016 | 0.0012 | 0.0013 |
| F11 | 51200001 | 51300000 | 0.0018 | 0.0012 | 0.0014 |
| F11 | 51220001 | 51320000 | 0.0013 | 0.0008 | 0.0009 |
| F11 | 51240001 | 51340000 | 0.0022 | 0.0016 | 0.0017 |
| F11 | 51260001 | 51360000 | 0.0022 | 0.0016 | 0.0017 |
| F11 | 51280001 | 51380000 | 0.0025 | 0.0019 | 0.0021 |
| F11 | 51300001 | 51400000 | 0.0026 | 0.0021 | 0.0022 |
| F11 | 51320001 | 51420000 | 0.0028 | 0.0024 | 0.0026 |
| F11 | 51340001 | 51440000 | 0.0022 | 0.0019 | 0.0021 |
| F11 | 51360001 | 51460000 | 0.0024 | 0.0021 | 0.0022 |

| Chr | Start | End | both | A1 | A2 |
|---|---|---|---|---|---|
| F11 | 51380001 | 51480000 | 0.0019 | 0.0017 | 0.0018 |
| F11 | 51400001 | 51500000 | 0.0022 | 0.0018 | 0.0020 |
| F11 | 51420001 | 51520000 | 0.0015 | 0.0012 | 0.0014 |
| F11 | 51440001 | 51540000 | 0.0013 | 0.0010 | 0.0012 |
| F11 | 51460001 | 51560000 | 0.0015 | 0.0011 | 0.0014 |
| F11 | 51480001 | 51580000 | 0.0015 | 0.0011 | 0.0013 |
| F11 | 51500001 | 51600000 | 0.0010 | 0.0007 | 0.0009 |
| F11 | 51520001 | 51620000 | 0.0010 | 0.0008 | 0.0010 |
| F11 | 51540001 | 51640000 | 0.0011 | 0.0008 | 0.0011 |
| F11 | 51560001 | 51660000 | 0.0011 | 0.0008 | 0.0010 |
| F11 | 51580001 | 51680000 | 0.0010 | 0.0008 | 0.0010 |
| F11 | 51600001 | 51700000 | 0.0011 | 0.0009 | 0.0011 |
| F11 | 51620001 | 51720000 | 0.0010 | 0.0009 | 0.0010 |
| F11 | 51640001 | 51740000 | 0.0011 | 0.0008 | 0.0010 |
| F11 | 51660001 | 51760000 | 0.0018 | 0.0013 | 0.0016 |
| F11 | 51680001 | 51780000 | 0.0021 | 0.0014 | 0.0017 |
| F11 | 51700001 | 51800000 | 0.0030 | 0.0019 | 0.0024 |
| F11 | 51720001 | 51820000 | 0.0033 | 0.0023 | 0.0027 |
| F11 | 51740001 | 51840000 | 0.0037 | 0.0027 | 0.0030 |
| F11 | 51760001 | 51860000 | 0.0034 | 0.0023 | 0.0026 |
| F11 | 51780001 | 51880000 | 0.0041 | 0.0031 | 0.0034 |
| F11 | 51800001 | 51900000 | 0.0040 | 0.0031 | 0.0033 |
| F11 | 51820001 | 51920000 | 0.0039 | 0.0028 | 0.0032 |
| F11 | 51840001 | 51940000 | 0.0036 | 0.0025 | 0.0028 |
| F11 | 51860001 | 51960000 | 0.0039 | 0.0028 | 0.0031 |
| F11 | 51880001 | 51980000 | 0.0030 | 0.0019 | 0.0022 |
| F11 | 51900001 | 52000000 | 0.0026 | 0.0019 | 0.0019 |
| F11 | 51920001 | 52020000 | 0.0025 | 0.0019 | 0.0019 |
| F11 | 51940001 | 52040000 | 0.0023 | 0.0018 | 0.0018 |
| F11 | 51960001 | 52060000 | 0.0015 | 0.0013 | 0.0014 |
| F11 | 51980001 | 52080000 | 0.0014 | 0.0012 | 0.0013 |
| F11 | 52000001 | 52100000 | 0.0012 | 0.0010 | 0.0011 |
| F11 | 52020001 | 52120000 | 0.0017 | 0.0014 | 0.0015 |
| F11 | 52040001 | 52140000 | 0.0018 | 0.0015 | 0.0016 |
| F11 | 52060001 | 52160000 | 0.0017 | 0.0014 | 0.0015 |
| F11 | 52080001 | 52180000 | 0.0027 | 0.0020 | 0.0024 |
| F11 | 52100001 | 52200000 | 0.0034 | 0.0028 | 0.0031 |
| F11 | 52120001 | 52220000 | 0.0036 | 0.0028 | 0.0032 |
| F11 | 52140001 | 52240000 | 0.0035 | 0.0026 | 0.0030 |
| F11 | 52160001 | 52260000 | 0.0045 | 0.0033 | 0.0037 |
| F11 | 52180001 | 52280000 | 0.0039 | 0.0029 | 0.0030 |
| F11 | 52200001 | 52300000 | 0.0032 | 0.0021 | 0.0024 |
| F11 | 52220001 | 52320000 | 0.0029 | 0.0019 | 0.0022 |
| F11 | 52240001 | 52340000 | 0.0028 | 0.0018 | 0.0022 |
| F11 | 52260001 | 52360000 | 0.0018 | 0.0011 | 0.0014 |
| F11 | 52280001 | 52380000 | 0.0024 | 0.0017 | 0.0022 |
| F11 | 52300001 | 52400000 | 0.0025 | 0.0019 | 0.0023 |
| F11 | 52320001 | 52420000 | 0.0020 | 0.0016 | 0.0019 |
| F11 | 52340001 | 52440000 | 0.0019 | 0.0016 | 0.0018 |
| F11 | 52360001 | 52460000 | 0.0020 | 0.0017 | 0.0019 |
| F11 | 52380001 | 52480000 | 0.0013 | 0.0011 | 0.0012 |
| F11 | 52400001 | 52500000 | 0.0009 | 0.0007 | 0.0008 |
| F11 | 52420001 | 52520000 | 0.0020 | 0.0014 | 0.0020 |
| F11 | 52440001 | 52540000 | 0.0021 | 0.0015 | 0.0020 |
| F11 | 52460001 | 52560000 | 0.0026 | 0.0018 | 0.0024 |
| F11 | 52480001 | 52580000 | 0.0024 | 0.0016 | 0.0023 |
| F11 | 52500001 | 52600000 | 0.0044 | 0.0033 | 0.0040 |
| F11 | 52520001 | 52620000 | 0.0038 | 0.0032 | 0.0035 |

| Chr | Start | End | both | A1 | A2 |
|---|---|---|---|---|---|
| F11 | 52540001 | 52640000 | 0.0046 | 0.0038 | 0.0043 |
| F11 | 52560001 | 52660000 | 0.0054 | 0.0045 | 0.0050 |
| F11 | 52580001 | 52680000 | 0.0056 | 0.0048 | 0.0053 |
| F11 | 52600001 | 52700000 | 0.0041 | 0.0035 | 0.0039 |
| F11 | 52620001 | 52720000 | 0.0041 | 0.0033 | 0.0039 |
| F11 | 52640001 | 52740000 | 0.0033 | 0.0028 | 0.0032 |
| F11 | 52660001 | 52760000 | 0.0020 | 0.0016 | 0.0020 |
| F11 | 52680001 | 52780000 | 0.0024 | 0.0018 | 0.0023 |
| F11 | 52700001 | 52800000 | 0.0023 | 0.0017 | 0.0022 |
| F11 | 52720001 | 52820000 | 0.0022 | 0.0016 | 0.0021 |
| F11 | 52740001 | 52840000 | 0.0023 | 0.0017 | 0.0021 |
| F11 | 52760001 | 52860000 | 0.0029 | 0.0022 | 0.0028 |
| F11 | 52780001 | 52880000 | 0.0024 | 0.0019 | 0.0023 |
| F11 | 52800001 | 52900000 | 0.0024 | 0.0019 | 0.0023 |
| F11 | 52820001 | 52920000 | 0.0026 | 0.0021 | 0.0024 |
| F11 | 52840001 | 52940000 | 0.0030 | 0.0025 | 0.0029 |
| F11 | 52860001 | 52960000 | 0.0025 | 0.0020 | 0.0023 |
| F11 | 52880001 | 52980000 | 0.0024 | 0.0020 | 0.0023 |
| F11 | 52900001 | 53000000 | 0.0024 | 0.0018 | 0.0021 |
| F11 | 52920001 | 53020000 | 0.0021 | 0.0014 | 0.0016 |
| F11 | 52940001 | 53040000 | 0.0020 | 0.0013 | 0.0015 |
| F11 | 52960001 | 53060000 | 0.0024 | 0.0017 | 0.0019 |
| F11 | 52980001 | 53080000 | 0.0031 | 0.0023 | 0.0026 |
| F11 | 53000001 | 53100000 | 0.0028 | 0.0022 | 0.0025 |
| F11 | 53020001 | 53120000 | 0.0022 | 0.0019 | 0.0022 |
| F11 | 53040001 | 53140000 | 0.0017 | 0.0014 | 0.0017 |
| F11 | 53060001 | 53160000 | 0.0015 | 0.0012 | 0.0015 |
| F11 | 53080001 | 53180000 | 0.0009 | 0.0007 | 0.0009 |
| F11 | 53100001 | 53200000 | 0.0012 | 0.0009 | 0.0012 |
| F11 | 53120001 | 53220000 | 0.0018 | 0.0010 | 0.0015 |
| F11 | 53140001 | 53240000 | 0.0020 | 0.0012 | 0.0017 |
| F11 | 53160001 | 53260000 | 0.0018 | 0.0011 | 0.0015 |
| F11 | 53180001 | 53280000 | 0.0016 | 0.0009 | 0.0012 |
| F11 | 53200001 | 53300000 | 0.0017 | 0.0012 | 0.0014 |
| F11 | 53220001 | 53320000 | 0.0013 | 0.0011 | 0.0012 |
| F11 | 53240001 | 53340000 | 0.0011 | 0.0009 | 0.0010 |
| F11 | 53260001 | 53360000 | 0.0009 | 0.0008 | 0.0008 |
| F11 | 53280001 | 53380000 | 0.0013 | 0.0011 | 0.0012 |
| F11 | 53300001 | 53400000 | 0.0008 | 0.0007 | 0.0008 |
| F11 | 53320001 | 53420000 | 0.0008 | 0.0007 | 0.0008 |
| F11 | 53340001 | 53440000 | 0.0008 | 0.0007 | 0.0008 |
| F11 | 53360001 | 53460000 | 0.0009 | 0.0008 | 0.0009 |
| F11 | 53380001 | 53480000 | 0.0008 | 0.0006 | 0.0008 |
| F11 | 53400001 | 53500000 | 0.0007 | 0.0005 | 0.0007 |
| F11 | 53420001 | 53520000 | 0.0006 | 0.0004 | 0.0006 |
| F11 | 53440001 | 53540000 | 0.0009 | 0.0006 | 0.0008 |
| F11 | 53460001 | 53560000 | 0.0009 | 0.0006 | 0.0008 |
| F11 | 53480001 | 53580000 | 0.0011 | 0.0008 | 0.0010 |
| F11 | 53500001 | 53600000 | 0.0013 | 0.0010 | 0.0013 |
| F11 | 53520001 | 53620000 | 0.0021 | 0.0017 | 0.0020 |
| F11 | 53540001 | 53640000 | 0.0019 | 0.0017 | 0.0019 |
| F11 | 53560001 | 53660000 | 0.0020 | 0.0018 | 0.0020 |
| F11 | 53580001 | 53680000 | 0.0017 | 0.0015 | 0.0017 |
| F11 | 53600001 | 53700000 | 0.0016 | 0.0015 | 0.0016 |
| F11 | 53620001 | 53720000 | 0.0010 | 0.0009 | 0.0009 |
| F11 | 53640001 | 53740000 | 0.0011 | 0.0010 | 0.0011 |
| F11 | 53660001 | 53760000 | 0.0012 | 0.0010 | 0.0011 |
| F11 | 53680001 | 53780000 | 0.0011 | 0.0009 | 0.0010 |

| Chr | Start | End | both | A1 | A2 |
|---|---|---|---|---|---|
| F11 | 53700001 | 53800000 | 0.0014 | 0.0013 | 0.0014 |
| F11 | 53720001 | 53820000 | 0.0014 | 0.0012 | 0.0014 |
| F11 | 53740001 | 53840000 | 0.0011 | 0.0010 | 0.0010 |
| F11 | 53760001 | 53860000 | 0.0009 | 0.0008 | 0.0009 |
| F11 | 53780001 | 53880000 | 0.0022 | 0.0019 | 0.0019 |
| F11 | 53800001 | 53900000 | 0.0026 | 0.0021 | 0.0021 |
| F11 | 53820001 | 53920000 | 0.0030 | 0.0024 | 0.0025 |
| F11 | 53840001 | 53940000 | 0.0042 | 0.0035 | 0.0036 |
| F11 | 53860001 | 53960000 | 0.0055 | 0.0047 | 0.0048 |
| F11 | 53880001 | 53980000 | 0.0051 | 0.0043 | 0.0044 |
| F11 | 53900001 | 54000000 | 0.0046 | 0.0040 | 0.0041 |
| F11 | 53920001 | 54020000 | 0.0042 | 0.0036 | 0.0038 |
| F11 | 53940001 | 54040000 | 0.0032 | 0.0027 | 0.0029 |
| F11 | 53960001 | 54060000 | 0.0021 | 0.0017 | 0.0019 |
| F11 | 53980001 | 54080000 | 0.0016 | 0.0013 | 0.0016 |
| F11 | 54000001 | 54100000 | 0.0019 | 0.0016 | 0.0019 |
| F11 | 54020001 | 54120000 | 0.0019 | 0.0016 | 0.0018 |
| F11 | 54040001 | 54140000 | 0.0018 | 0.0015 | 0.0017 |
| F11 | 54060001 | 54160000 | 0.0018 | 0.0015 | 0.0018 |
| F11 | 54080001 | 54180000 | 0.0014 | 0.0010 | 0.0013 |
| F11 | 54100001 | 54200000 | 0.0010 | 0.0006 | 0.0009 |
| F11 | 54120001 | 54220000 | 0.0010 | 0.0007 | 0.0009 |
| F11 | 54140001 | 54240000 | 0.0009 | 0.0006 | 0.0008 |
| F11 | 54160001 | 54260000 | 0.0007 | 0.0005 | 0.0006 |
| F11 | 54180001 | 54280000 | 0.0008 | 0.0007 | 0.0008 |
| F11 | 54200001 | 54300000 | 0.0005 | 0.0005 | 0.0005 |
| F11 | 54220001 | 54320000 | 0.0007 | 0.0007 | 0.0007 |
| F11 | 54240001 | 54340000 | 0.0014 | 0.0013 | 0.0013 |
| F11 | 54260001 | 54360000 | 0.0016 | 0.0015 | 0.0016 |
| F11 | 54280001 | 54380000 | 0.0017 | 0.0015 | 0.0016 |
| F11 | 54300001 | 54400000 | 0.0022 | 0.0020 | 0.0022 |
| F11 | 54320001 | 54420000 | 0.0022 | 0.0019 | 0.0021 |
| F11 | 54340001 | 54440000 | 0.0016 | 0.0013 | 0.0015 |
| F11 | 54360001 | 54460000 | 0.0017 | 0.0014 | 0.0015 |
| F11 | 54380001 | 54480000 | 0.0021 | 0.0018 | 0.0019 |
| F11 | 54400001 | 54500000 | 0.0020 | 0.0017 | 0.0017 |
| F11 | 54420001 | 54520000 | 0.0018 | 0.0016 | 0.0015 |
| F11 | 54440001 | 54540000 | 0.0022 | 0.0018 | 0.0019 |
| F11 | 54460001 | 54560000 | 0.0026 | 0.0022 | 0.0024 |
| F11 | 54480001 | 54580000 | 0.0022 | 0.0017 | 0.0019 |
| F11 | 54500001 | 54600000 | 0.0031 | 0.0025 | 0.0029 |
| F11 | 54520001 | 54620000 | 0.0036 | 0.0029 | 0.0033 |
| F11 | 54540001 | 54640000 | 0.0034 | 0.0027 | 0.0031 |
| F11 | 54560001 | 54660000 | 0.0025 | 0.0020 | 0.0023 |
| F11 | 54580001 | 54680000 | 0.0032 | 0.0026 | 0.0030 |
| F11 | 54600001 | 54700000 | 0.0027 | 0.0019 | 0.0022 |
| F11 | 54620001 | 54720000 | 0.0026 | 0.0016 | 0.0019 |
| F11 | 54640001 | 54740000 | 0.0027 | 0.0017 | 0.0020 |
| F11 | 54660001 | 54760000 | 0.0027 | 0.0017 | 0.0021 |
| F11 | 54680001 | 54780000 | 0.0019 | 0.0011 | 0.0013 |
| F11 | 54700001 | 54800000 | 0.0015 | 0.0008 | 0.0011 |
| F11 | 54720001 | 54820000 | 0.0013 | 0.0009 | 0.0013 |
| F11 | 54740001 | 54840000 | 0.0021 | 0.0014 | 0.0021 |
| F11 | 54760001 | 54860000 | 0.0023 | 0.0016 | 0.0023 |
| F11 | 54780001 | 54880000 | 0.0022 | 0.0015 | 0.0022 |
| F11 | 54800001 | 54900000 | 0.0019 | 0.0014 | 0.0019 |
| F11 | 54820001 | 54920000 | 0.0021 | 0.0016 | 0.0021 |
| F11 | 54840001 | 54940000 | 0.0015 | 0.0013 | 0.0015 |

| Chr | Start | End | both | A1 | A2 |
|---|---|---|---|---|---|
| F11 | 54860001 | 54960000 | 0.0017 | 0.0015 | 0.0016 |
| F11 | 54880001 | 54980000 | 0.0023 | 0.0019 | 0.0022 |
| F11 | 54900001 | 55000000 | 0.0024 | 0.0020 | 0.0023 |
| F11 | 54920001 | 55020000 | 0.0025 | 0.0016 | 0.0020 |
| F11 | 54940001 | 55040000 | 0.0024 | 0.0015 | 0.0018 |
| F11 | 54960001 | 55060000 | 0.0021 | 0.0013 | 0.0016 |
| F11 | 54980001 | 55080000 | 0.0025 | 0.0014 | 0.0018 |
| F11 | 55000001 | 55100000 | 0.0025 | 0.0013 | 0.0017 |
| F11 | 55020001 | 55120000 | 0.0026 | 0.0018 | 0.0019 |
| F11 | 55040001 | 55140000 | 0.0045 | 0.0034 | 0.0036 |
| F11 | 55060001 | 55160000 | 0.0049 | 0.0037 | 0.0039 |
| F11 | 55080001 | 55180000 | 0.0047 | 0.0037 | 0.0039 |
| F11 | 55100001 | 55200000 | 0.0054 | 0.0045 | 0.0047 |
| F11 | 55120001 | 55220000 | 0.0047 | 0.0039 | 0.0042 |
| F11 | 55140001 | 55240000 | 0.0030 | 0.0025 | 0.0028 |
| F11 | 55160001 | 55260000 | 0.0031 | 0.0026 | 0.0029 |
| F11 | 55180001 | 55280000 | 0.0024 | 0.0019 | 0.0021 |
| F11 | 55200001 | 55300000 | 0.0016 | 0.0011 | 0.0014 |
| F11 | 55220001 | 55320000 | 0.0023 | 0.0017 | 0.0021 |
| F11 | 55240001 | 55340000 | 0.0027 | 0.0021 | 0.0025 |
| F11 | 55260001 | 55360000 | 0.0032 | 0.0026 | 0.0029 |
| F11 | 55280001 | 55380000 | 0.0032 | 0.0027 | 0.0031 |
| F11 | 55300001 | 55400000 | 0.0035 | 0.0029 | 0.0033 |
| F11 | 55320001 | 55420000 | 0.0037 | 0.0031 | 0.0034 |
| F11 | 55340001 | 55440000 | 0.0041 | 0.0032 | 0.0035 |
| F11 | 55360001 | 55460000 | 0.0033 | 0.0025 | 0.0027 |
| F11 | 55380001 | 55480000 | 0.0036 | 0.0027 | 0.0030 |
| F11 | 55400001 | 55500000 | 0.0036 | 0.0028 | 0.0031 |
| F11 | 55420001 | 55520000 | 0.0035 | 0.0027 | 0.0030 |
| F11 | 55440001 | 55540000 | 0.0032 | 0.0027 | 0.0030 |
| F11 | 55460001 | 55560000 | 0.0034 | 0.0028 | 0.0030 |
| F11 | 55480001 | 55580000 | 0.0032 | 0.0025 | 0.0029 |
| F11 | 55500001 | 55600000 | 0.0041 | 0.0031 | 0.0037 |
| F11 | 55520001 | 55620000 | 0.0045 | 0.0036 | 0.0040 |
| F11 | 55540001 | 55640000 | 0.0038 | 0.0028 | 0.0032 |
| F11 | 55560001 | 55660000 | 0.0036 | 0.0027 | 0.0031 |
| F11 | 55580001 | 55680000 | 0.0043 | 0.0033 | 0.0037 |
| F11 | 55600001 | 55700000 | 0.0050 | 0.0041 | 0.0044 |
| F11 | 55620001 | 55720000 | 0.0043 | 0.0035 | 0.0038 |
| F11 | 55640001 | 55740000 | 0.0044 | 0.0036 | 0.0039 |
| F11 | 55660001 | 55760000 | 0.0048 | 0.0037 | 0.0041 |
| F11 | 55680001 | 55780000 | 0.0044 | 0.0033 | 0.0037 |
| F11 | 55700001 | 55800000 | 0.0026 | 0.0017 | 0.0020 |
| F11 | 55720001 | 55820000 | 0.0031 | 0.0019 | 0.0024 |
| F11 | 55740001 | 55840000 | 0.0035 | 0.0023 | 0.0028 |
| F11 | 55760001 | 55860000 | 0.0034 | 0.0023 | 0.0027 |
| F11 | 55780001 | 55880000 | 0.0030 | 0.0021 | 0.0024 |
| F11 | 55800001 | 55900000 | 0.0033 | 0.0022 | 0.0025 |
| F11 | 55820001 | 55920000 | 0.0029 | 0.0020 | 0.0024 |
| F11 | 55840001 | 55940000 | 0.0032 | 0.0022 | 0.0026 |
| F11 | 55860001 | 55960000 | 0.0033 | 0.0023 | 0.0029 |
| F11 | 55880001 | 55980000 | 0.0040 | 0.0029 | 0.0034 |
| F11 | 55900001 | 56000000 | 0.0040 | 0.0031 | 0.0034 |
| F11 | 55920001 | 56020000 | 0.0039 | 0.0029 | 0.0031 |
| F11 | 55940001 | 56040000 | 0.0035 | 0.0026 | 0.0027 |
| F11 | 55960001 | 56060000 | 0.0030 | 0.0022 | 0.0023 |
| F11 | 55980001 | 56080000 | 0.0027 | 0.0018 | 0.0018 |
| F11 | 56000001 | 56100000 | 0.0027 | 0.0019 | 0.0019 |

| Chr | Start | End | both | A1 | A2 |
|---|---|---|---|---|---|
| F11 | 56020001 | 56120000 | 0.0046 | 0.0037 | 0.0038 |
| F11 | 56040001 | 56140000 | 0.0056 | 0.0047 | 0.0049 |
| F11 | 56060001 | 56160000 | 0.0055 | 0.0046 | 0.0048 |
| F11 | 56080001 | 56180000 | 0.0051 | 0.0044 | 0.0046 |
| F11 | 56100001 | 56200000 | 0.0050 | 0.0042 | 0.0044 |
| F11 | 56120001 | 56220000 | 0.0035 | 0.0029 | 0.0032 |
| F11 | 56140001 | 56240000 | 0.0024 | 0.0021 | 0.0022 |
| F11 | 56160001 | 56260000 | 0.0029 | 0.0025 | 0.0026 |
| F11 | 56180001 | 56280000 | 0.0036 | 0.0032 | 0.0033 |
| F11 | 56200001 | 56300000 | 0.0045 | 0.0039 | 0.0042 |
| F11 | 56220001 | 56320000 | 0.0035 | 0.0031 | 0.0032 |
| F11 | 56240001 | 56340000 | 0.0036 | 0.0031 | 0.0033 |
| F11 | 56260001 | 56360000 | 0.0030 | 0.0026 | 0.0028 |
| F11 | 56280001 | 56380000 | 0.0029 | 0.0023 | 0.0027 |
| F11 | 56300001 | 56400000 | 0.0024 | 0.0018 | 0.0021 |
| F11 | 56320001 | 56420000 | 0.0027 | 0.0021 | 0.0024 |
| F11 | 56340001 | 56440000 | 0.0023 | 0.0017 | 0.0021 |
| F11 | 56360001 | 56460000 | 0.0030 | 0.0021 | 0.0027 |
| F11 | 56380001 | 56480000 | 0.0024 | 0.0018 | 0.0021 |
| F11 | 56400001 | 56500000 | 0.0028 | 0.0022 | 0.0027 |
| F11 | 56420001 | 56520000 | 0.0029 | 0.0024 | 0.0028 |
| F11 | 56440001 | 56540000 | 0.0041 | 0.0034 | 0.0038 |
| F11 | 56460001 | 56560000 | 0.0037 | 0.0031 | 0.0035 |
| F11 | 56480001 | 56580000 | 0.0051 | 0.0044 | 0.0048 |
| F11 | 56500001 | 56600000 | 0.0044 | 0.0038 | 0.0042 |
| F11 | 56520001 | 56620000 | 0.0044 | 0.0038 | 0.0043 |
| F11 | 56540001 | 56640000 | 0.0035 | 0.0031 | 0.0034 |
| F11 | 56560001 | 56660000 | 0.0039 | 0.0033 | 0.0037 |
| F11 | 56580001 | 56680000 | 0.0034 | 0.0028 | 0.0031 |
| F11 | 56600001 | 56700000 | 0.0041 | 0.0032 | 0.0034 |
| F11 | 56620001 | 56720000 | 0.0037 | 0.0029 | 0.0030 |
| F11 | 56640001 | 56740000 | 0.0035 | 0.0027 | 0.0029 |
| F11 | 56660001 | 56760000 | 0.0030 | 0.0024 | 0.0026 |
| F11 | 56680001 | 56780000 | 0.0025 | 0.0020 | 0.0021 |
| F11 | 56700001 | 56800000 | 0.0016 | 0.0014 | 0.0015 |
| F11 | 56720001 | 56820000 | 0.0017 | 0.0015 | 0.0016 |
| F11 | 56740001 | 56840000 | 0.0018 | 0.0015 | 0.0016 |
| F11 | 56760001 | 56860000 | 0.0022 | 0.0019 | 0.0020 |
| F11 | 56780001 | 56880000 | 0.0023 | 0.0020 | 0.0022 |
| F11 | 56800001 | 56900000 | 0.0026 | 0.0023 | 0.0024 |
| F11 | 56820001 | 56920000 | 0.0026 | 0.0022 | 0.0025 |
| F11 | 56840001 | 56940000 | 0.0033 | 0.0027 | 0.0031 |
| F11 | 56860001 | 56960000 | 0.0034 | 0.0026 | 0.0030 |
| F11 | 56880001 | 56980000 | 0.0030 | 0.0021 | 0.0025 |
| F11 | 56900001 | 57000000 | 0.0035 | 0.0026 | 0.0031 |
| F11 | 56920001 | 57020000 | 0.0036 | 0.0029 | 0.0032 |
| F11 | 56940001 | 57040000 | 0.0028 | 0.0022 | 0.0024 |
| F11 | 56960001 | 57060000 | 0.0023 | 0.0020 | 0.0021 |
| F11 | 56980001 | 57080000 | 0.0023 | 0.0020 | 0.0022 |
| F11 | 57000001 | 57100000 | 0.0018 | 0.0014 | 0.0017 |
| F11 | 57020001 | 57120000 | 0.0015 | 0.0010 | 0.0014 |
| F11 | 57040001 | 57140000 | 0.0017 | 0.0013 | 0.0017 |
| F11 | 57060001 | 57160000 | 0.0017 | 0.0012 | 0.0017 |
| F11 | 57080001 | 57180000 | 0.0015 | 0.0011 | 0.0015 |
| F11 | 57100001 | 57200000 | 0.0014 | 0.0012 | 0.0014 |
| F11 | 57120001 | 57220000 | 0.0023 | 0.0021 | 0.0022 |
| F11 | 57140001 | 57240000 | 0.0023 | 0.0021 | 0.0022 |
| F11 | 57160001 | 57260000 | 0.0029 | 0.0026 | 0.0028 |

| Chr | Start | End | both | A1 | A2 |
|---|---|---|---|---|---|
| F11 | 57180001 | 57280000 | 0.0032 | 0.0029 | 0.0031 |
| F11 | 57200001 | 57300000 | 0.0040 | 0.0034 | 0.0039 |
| F11 | 57220001 | 57320000 | 0.0032 | 0.0026 | 0.0031 |
| F11 | 57240001 | 57340000 | 0.0032 | 0.0026 | 0.0031 |
| F11 | 57260001 | 57360000 | 0.0026 | 0.0022 | 0.0026 |
| F11 | 57280001 | 57380000 | 0.0024 | 0.0019 | 0.0023 |
| F11 | 57300001 | 57400000 | 0.0018 | 0.0015 | 0.0018 |
| F11 | 57320001 | 57420000 | 0.0018 | 0.0016 | 0.0018 |
| F11 | 57340001 | 57440000 | 0.0022 | 0.0019 | 0.0022 |
| F11 | 57360001 | 57460000 | 0.0034 | 0.0028 | 0.0033 |
| F11 | 57380001 | 57480000 | 0.0037 | 0.0030 | 0.0036 |
| F11 | 57400001 | 57500000 | 0.0039 | 0.0031 | 0.0038 |
| F11 | 57420001 | 57520000 | 0.0039 | 0.0029 | 0.0037 |
| F11 | 57440001 | 57540000 | 0.0037 | 0.0028 | 0.0035 |
| F11 | 57460001 | 57560000 | 0.0026 | 0.0019 | 0.0025 |
| F11 | 57480001 | 57580000 | 0.0022 | 0.0016 | 0.0021 |
| F11 | 57500001 | 57600000 | 0.0022 | 0.0018 | 0.0020 |
| F11 | 57520001 | 57620000 | 0.0032 | 0.0029 | 0.0031 |
| F11 | 57540001 | 57640000 | 0.0034 | 0.0031 | 0.0032 |
| F11 | 57560001 | 57660000 | 0.0042 | 0.0039 | 0.0040 |
| F11 | 57580001 | 57680000 | 0.0044 | 0.0040 | 0.0042 |
| F11 | 57600001 | 57700000 | 0.0040 | 0.0034 | 0.0039 |
| F11 | 57620001 | 57720000 | 0.0028 | 0.0024 | 0.0027 |
| F11 | 57640001 | 57740000 | 0.0024 | 0.0020 | 0.0023 |
| F11 | 57660001 | 57760000 | 0.0017 | 0.0013 | 0.0017 |
| F11 | 57680001 | 57780000 | 0.0016 | 0.0011 | 0.0016 |
| F11 | 57700001 | 57800000 | 0.0016 | 0.0011 | 0.0016 |
| F11 | 57720001 | 57820000 | 0.0014 | 0.0010 | 0.0015 |
| F11 | 57740001 | 57840000 | 0.0010 | 0.0006 | 0.0010 |
| F11 | 57760001 | 57860000 | 0.0006 | 0.0004 | 0.0006 |
| F11 | 57780001 | 57880000 | 0.0008 | 0.0006 | 0.0008 |
| F11 | 57800001 | 57900000 | 0.0008 | 0.0006 | 0.0008 |
| F11 | 57820001 | 57920000 | 0.0010 | 0.0008 | 0.0010 |
| F11 | 57840001 | 57940000 | 0.0010 | 0.0008 | 0.0010 |
| F11 | 57860001 | 57960000 | 0.0008 | 0.0006 | 0.0008 |
| F11 | 57880001 | 57980000 | 0.0007 | 0.0005 | 0.0007 |
| F11 | 57900001 | 58000000 | 0.0007 | 0.0006 | 0.0006 |
| F11 | 57920001 | 58020000 | 0.0009 | 0.0008 | 0.0009 |
| F11 | 57940001 | 58040000 | 0.0011 | 0.0009 | 0.0011 |
| F11 | 57960001 | 58060000 | 0.0011 | 0.0009 | 0.0011 |
| F11 | 57980001 | 58080000 | 0.0009 | 0.0008 | 0.0009 |
| F11 | 58000001 | 58100000 | 0.0008 | 0.0006 | 0.0007 |
| F11 | 58020001 | 58120000 | 0.0004 | 0.0003 | 0.0004 |
| F11 | 58040001 | 58140000 | 0.0002 | 0.0002 | 0.0002 |
| F11 | 58060001 | 58160000 | 0.0003 | 0.0002 | 0.0002 |
| F11 | 58080001 | 58180000 | 0.0003 | 0.0003 | 0.0003 |
| F11 | 58100001 | 58200000 | 0.0004 | 0.0003 | 0.0004 |
| F11 | 58120001 | 58220000 | 0.0003 | 0.0002 | 0.0002 |
| F11 | 58140001 | 58240000 | 0.0003 | 0.0002 | 0.0003 |
| F11 | 58160001 | 58260000 | 0.0004 | 0.0003 | 0.0004 |
| F11 | 58180001 | 58280000 | 0.0004 | 0.0003 | 0.0004 |
| F11 | 58200001 | 58300000 | 0.0007 | 0.0005 | 0.0006 |
| F11 | 58220001 | 58320000 | 0.0006 | 0.0005 | 0.0006 |
| F11 | 58240001 | 58340000 | 0.0009 | 0.0006 | 0.0008 |
| F11 | 58260001 | 58360000 | 0.0008 | 0.0005 | 0.0008 |
| F11 | 58280001 | 58380000 | 0.0009 | 0.0006 | 0.0009 |
| F11 | 58300001 | 58400000 | 0.0006 | 0.0004 | 0.0006 |
| F11 | 58320001 | 58420000 | 0.0024 | 0.0017 | 0.0024 |

| Chr | Start | End | both | A1 | A2 |
|---|---|---|---|---|---|
| F11 | 58340001 | 58440000 | 0.0023 | 0.0016 | 0.0023 |
| F11 | 58360001 | 58460000 | 0.0024 | 0.0017 | 0.0024 |
| F11 | 58380001 | 58480000 | 0.0025 | 0.0017 | 0.0025 |
| F11 | 58400001 | 58500000 | 0.0027 | 0.0018 | 0.0027 |
| F11 | 58420001 | 58520000 | 0.0011 | 0.0007 | 0.0012 |
| F11 | 58440001 | 58540000 | 0.0009 | 0.0005 | 0.0010 |
| F11 | 58460001 | 58560000 | 0.0008 | 0.0005 | 0.0008 |
| F11 | 58480001 | 58580000 | 0.0006 | 0.0004 | 0.0006 |
| F11 | 58500001 | 58600000 | 0.0004 | 0.0003 | 0.0004 |
| F11 | 58520001 | 58620000 | 0.0002 | 0.0002 | 0.0002 |
| F11 | 58540001 | 58640000 | 0.0004 | 0.0003 | 0.0004 |
| F11 | 58560001 | 58660000 | 0.0006 | 0.0005 | 0.0007 |
| F11 | 58580001 | 58680000 | 0.0006 | 0.0005 | 0.0006 |
| F11 | 58600001 | 58700000 | 0.0008 | 0.0006 | 0.0008 |
| F11 | 58620001 | 58720000 | 0.0011 | 0.0009 | 0.0011 |
| F11 | 58640001 | 58740000 | 0.0014 | 0.0010 | 0.0014 |
| F11 | 58660001 | 58760000 | 0.0015 | 0.0011 | 0.0015 |
| F11 | 58680001 | 58780000 | 0.0018 | 0.0014 | 0.0018 |
| F11 | 58700001 | 58800000 | 0.0021 | 0.0016 | 0.0021 |
| F11 | 58720001 | 58820000 | 0.0021 | 0.0016 | 0.0021 |
| F11 | 58740001 | 58840000 | 0.0026 | 0.0021 | 0.0026 |
| F11 | 58760001 | 58860000 | 0.0026 | 0.0022 | 0.0026 |
| F11 | 58780001 | 58880000 | 0.0025 | 0.0021 | 0.0025 |
| F11 | 58800001 | 58900000 | 0.0021 | 0.0017 | 0.0020 |
| F11 | 58820001 | 58920000 | 0.0022 | 0.0019 | 0.0022 |
| F11 | 58840001 | 58940000 | 0.0022 | 0.0018 | 0.0022 |
| F11 | 58860001 | 58960000 | 0.0021 | 0.0018 | 0.0021 |
| F11 | 58880001 | 58980000 | 0.0019 | 0.0016 | 0.0019 |
| F11 | 58900001 | 59000000 | 0.0033 | 0.0027 | 0.0033 |
| F11 | 58920001 | 59020000 | 0.0031 | 0.0025 | 0.0031 |
| F11 | 58940001 | 59040000 | 0.0068 | 0.0051 | 0.0068 |
| F11 | 58960001 | 59060000 | 0.0065 | 0.0049 | 0.0065 |
| F11 | 58980001 | 59080000 | 0.0075 | 0.0057 | 0.0074 |
| F11 | 59000001 | 59100000 | 0.0064 | 0.0048 | 0.0063 |
| F11 | 59020001 | 59120000 | 0.0060 | 0.0045 | 0.0060 |
| F11 | 59040001 | 59140000 | 0.0017 | 0.0013 | 0.0016 |
| F11 | 59060001 | 59160000 | 0.0018 | 0.0015 | 0.0018 |
| F11 | 59080001 | 59180000 | 0.0010 | 0.0007 | 0.0010 |
| F11 | 59100001 | 59200000 | 0.0007 | 0.0005 | 0.0007 |
| F11 | 59120001 | 59220000 | 0.0013 | 0.0010 | 0.0013 |
| F11 | 59140001 | 59240000 | 0.0012 | 0.0010 | 0.0012 |
| F11 | 59160001 | 59260000 | 0.0010 | 0.0008 | 0.0010 |
| F11 | 59180001 | 59280000 | 0.0013 | 0.0010 | 0.0012 |
| F11 | 59200001 | 59300000 | 0.0014 | 0.0010 | 0.0012 |
| F11 | 59220001 | 59320000 | 0.0011 | 0.0008 | 0.0010 |
| F11 | 59240001 | 59340000 | 0.0011 | 0.0008 | 0.0009 |
| F11 | 59260001 | 59360000 | 0.0018 | 0.0014 | 0.0016 |
| F11 | 59280001 | 59380000 | 0.0021 | 0.0016 | 0.0018 |
| F11 | 59300001 | 59400000 | 0.0021 | 0.0017 | 0.0019 |
| F11 | 59320001 | 59420000 | 0.0021 | 0.0017 | 0.0019 |
| F11 | 59340001 | 59440000 | 0.0024 | 0.0019 | 0.0022 |
| F11 | 59360001 | 59460000 | 0.0021 | 0.0016 | 0.0019 |
| F11 | 59380001 | 59480000 | 0.0018 | 0.0013 | 0.0016 |
| F11 | 59400001 | 59500000 | 0.0019 | 0.0014 | 0.0018 |
| F11 | 59420001 | 59520000 | 0.0018 | 0.0013 | 0.0017 |
| F11 | 59440001 | 59540000 | 0.0015 | 0.0011 | 0.0014 |
| F11 | 59460001 | 59560000 | 0.0016 | 0.0013 | 0.0015 |
| F11 | 59480001 | 59580000 | 0.0020 | 0.0017 | 0.0019 |

| Chr | Start | End | both | A1 | A2 |
|---|---|---|---|---|---|
| F11 | 59500001 | 59600000 | 0.0022 | 0.0017 | 0.0021 |
| F11 | 59520001 | 59620000 | 0.0019 | 0.0015 | 0.0018 |
| F11 | 59540001 | 59640000 | 0.0020 | 0.0016 | 0.0020 |
| F11 | 59560001 | 59660000 | 0.0019 | 0.0015 | 0.0020 |
| F11 | 59580001 | 59680000 | 0.0015 | 0.0011 | 0.0015 |
| F11 | 59600001 | 59700000 | 0.0012 | 0.0010 | 0.0012 |
| F11 | 59620001 | 59720000 | 0.0013 | 0.0010 | 0.0013 |
| F11 | 59640001 | 59740000 | 0.0018 | 0.0015 | 0.0017 |
| F11 | 59660001 | 59760000 | 0.0019 | 0.0016 | 0.0018 |
| F11 | 59680001 | 59780000 | 0.0019 | 0.0017 | 0.0018 |
| F11 | 59700001 | 59800000 | 0.0018 | 0.0016 | 0.0017 |
| F11 | 59720001 | 59820000 | 0.0019 | 0.0016 | 0.0018 |
| F11 | 59740001 | 59840000 | 0.0016 | 0.0013 | 0.0015 |
| F11 | 59760001 | 59860000 | 0.0010 | 0.0008 | 0.0010 |
| F11 | 59780001 | 59880000 | 0.0012 | 0.0009 | 0.0011 |
| F11 | 59800001 | 59900000 | 0.0011 | 0.0009 | 0.0011 |
| F11 | 59820001 | 59920000 | 0.0012 | 0.0009 | 0.0011 |
| F11 | 59840001 | 59940000 | 0.0011 | 0.0009 | 0.0011 |
| F11 | 59860001 | 59960000 | 0.0010 | 0.0009 | 0.0010 |
| F11 | 59880001 | 59980000 | 0.0010 | 0.0008 | 0.0010 |
| F11 | 59900001 | 60000000 | 0.0011 | 0.0009 | 0.0011 |
| F11 | 59920001 | 60020000 | 0.0010 | 0.0009 | 0.0010 |
| F11 | 59940001 | 60040000 | 0.0011 | 0.0009 | 0.0011 |
| F11 | 59960001 | 60060000 | 0.0018 | 0.0014 | 0.0017 |
| F11 | 59980001 | 60080000 | 0.0019 | 0.0015 | 0.0018 |
| F11 | 60000001 | 60100000 | 0.0021 | 0.0017 | 0.0021 |
| F11 | 60020001 | 60120000 | 0.0023 | 0.0018 | 0.0022 |
| F11 | 60040001 | 60140000 | 0.0025 | 0.0020 | 0.0023 |
| F11 | 60060001 | 60160000 | 0.0026 | 0.0020 | 0.0023 |
| F11 | 60080001 | 60180000 | 0.0022 | 0.0016 | 0.0019 |
| F11 | 60100001 | 60200000 | 0.0033 | 0.0027 | 0.0029 |
| F11 | 60120001 | 60220000 | 0.0037 | 0.0030 | 0.0032 |
| F11 | 60140001 | 60240000 | 0.0034 | 0.0029 | 0.0030 |
| F11 | 60160001 | 60260000 | 0.0036 | 0.0032 | 0.0032 |
| F11 | 60180001 | 60280000 | 0.0041 | 0.0036 | 0.0037 |
| F11 | 60200001 | 60300000 | 0.0029 | 0.0025 | 0.0026 |
| F11 | 60220001 | 60320000 | 0.0030 | 0.0025 | 0.0026 |
| F11 | 60240001 | 60340000 | 0.0028 | 0.0022 | 0.0024 |
| F11 | 60260001 | 60360000 | 0.0027 | 0.0021 | 0.0024 |
| F11 | 60280001 | 60380000 | 0.0025 | 0.0021 | 0.0023 |
| F11 | 60300001 | 60400000 | 0.0022 | 0.0018 | 0.0020 |
| F11 | 60320001 | 60420000 | 0.0017 | 0.0014 | 0.0016 |
| F11 | 60340001 | 60440000 | 0.0016 | 0.0014 | 0.0015 |
| F11 | 60360001 | 60460000 | 0.0008 | 0.0007 | 0.0008 |
| F11 | 60380001 | 60480000 | 0.0006 | 0.0006 | 0.0006 |
| F11 | 60400001 | 60500000 | 0.0011 | 0.0010 | 0.0011 |
| F11 | 60420001 | 60520000 | 0.0014 | 0.0012 | 0.0014 |
| F11 | 60440001 | 60540000 | 0.0022 | 0.0016 | 0.0022 |
| F11 | 60460001 | 60560000 | 0.0032 | 0.0017 | 0.0031 |
| F11 | 60480001 | 60580000 | 0.0042 | 0.0025 | 0.0040 |
| F11 | 60500001 | 60600000 | 0.0039 | 0.0023 | 0.0038 |
| F11 | 60520001 | 60620000 | 0.0036 | 0.0020 | 0.0034 |
| F11 | 60540001 | 60640000 | 0.0028 | 0.0016 | 0.0027 |
| F11 | 60560001 | 60660000 | 0.0022 | 0.0017 | 0.0021 |
| F11 | 60580001 | 60680000 | 0.0021 | 0.0016 | 0.0020 |
| F11 | 60600001 | 60700000 | 0.0022 | 0.0016 | 0.0020 |
| F11 | 60620001 | 60720000 | 0.0024 | 0.0017 | 0.0021 |
| F11 | 60640001 | 60740000 | 0.0027 | 0.0018 | 0.0024 |

| Chr | Start | End | both | A1 | A2 |
|---|---|---|---|---|---|
| F11 | 60660001 | 60760000 | 0.0025 | 0.0017 | 0.0021 |
| F11 | 60680001 | 60780000 | 0.0019 | 0.0013 | 0.0017 |
| F11 | 60700001 | 60800000 | 0.0017 | 0.0013 | 0.0017 |
| F11 | 60720001 | 60820000 | 0.0016 | 0.0012 | 0.0016 |
| F11 | 60740001 | 60840000 | 0.0012 | 0.0009 | 0.0012 |
| F11 | 60760001 | 60860000 | 0.0017 | 0.0014 | 0.0017 |
| F11 | 60780001 | 60880000 | 0.0012 | 0.0010 | 0.0012 |
| F11 | 60800001 | 60900000 | 0.0012 | 0.0010 | 0.0012 |
| F11 | 60820001 | 60920000 | 0.0014 | 0.0013 | 0.0014 |
| F11 | 60840001 | 60940000 | 0.0015 | 0.0012 | 0.0014 |
| F11 | 60860001 | 60960000 | 0.0018 | 0.0014 | 0.0017 |
| F11 | 60880001 | 60980000 | 0.0021 | 0.0017 | 0.0020 |
| F11 | 60900001 | 61000000 | 0.0027 | 0.0021 | 0.0025 |
| F11 | 60920001 | 61020000 | 0.0030 | 0.0023 | 0.0027 |
| F11 | 60940001 | 61040000 | 0.0030 | 0.0024 | 0.0027 |
| F11 | 60960001 | 61060000 | 0.0029 | 0.0022 | 0.0026 |
| F11 | 60980001 | 61080000 | 0.0026 | 0.0021 | 0.0024 |
| F11 | 61000001 | 61100000 | 0.0023 | 0.0020 | 0.0021 |
| F11 | 61020001 | 61120000 | 0.0026 | 0.0023 | 0.0025 |
| F11 | 61040001 | 61140000 | 0.0026 | 0.0023 | 0.0025 |
| F11 | 61060001 | 61160000 | 0.0028 | 0.0024 | 0.0026 |
| F11 | 61080001 | 61180000 | 0.0035 | 0.0029 | 0.0031 |
| F11 | 61100001 | 61200000 | 0.0045 | 0.0035 | 0.0040 |
| F11 | 61120001 | 61220000 | 0.0039 | 0.0029 | 0.0033 |
| F11 | 61140001 | 61240000 | 0.0043 | 0.0033 | 0.0037 |
| F11 | 61160001 | 61260000 | 0.0045 | 0.0035 | 0.0038 |
| F11 | 61180001 | 61280000 | 0.0042 | 0.0033 | 0.0037 |
| F11 | 61200001 | 61300000 | 0.0035 | 0.0028 | 0.0031 |
| F11 | 61220001 | 61320000 | 0.0035 | 0.0028 | 0.0030 |
| F11 | 61240001 | 61340000 | 0.0039 | 0.0030 | 0.0034 |
| F11 | 61260001 | 61360000 | 0.0033 | 0.0025 | 0.0028 |
| F11 | 61280001 | 61380000 | 0.0035 | 0.0025 | 0.0030 |
| F11 | 61300001 | 61400000 | 0.0037 | 0.0026 | 0.0033 |
| F11 | 61320001 | 61420000 | 0.0043 | 0.0031 | 0.0038 |
| F11 | 61340001 | 61440000 | 0.0038 | 0.0027 | 0.0034 |
| F11 | 61360001 | 61460000 | 0.0037 | 0.0027 | 0.0033 |
| F11 | 61380001 | 61480000 | 0.0037 | 0.0028 | 0.0031 |
| F11 | 61400001 | 61500000 | 0.0029 | 0.0021 | 0.0022 |
| F11 | 61420001 | 61520000 | 0.0032 | 0.0024 | 0.0025 |
| F11 | 61440001 | 61540000 | 0.0037 | 0.0027 | 0.0029 |
| F11 | 61460001 | 61560000 | 0.0060 | 0.0047 | 0.0051 |
| F11 | 61480001 | 61580000 | 0.0061 | 0.0050 | 0.0054 |
| F11 | 61500001 | 61600000 | 0.0059 | 0.0049 | 0.0055 |
| F11 | 61520001 | 61620000 | 0.0049 | 0.0041 | 0.0046 |
| F11 | 61540001 | 61640000 | 0.0039 | 0.0034 | 0.0037 |
| F11 | 61560001 | 61660000 | 0.0021 | 0.0014 | 0.0019 |
| F11 | 61580001 | 61680000 | 0.0014 | 0.0009 | 0.0012 |
| F11 | 61600001 | 61700000 | 0.0020 | 0.0012 | 0.0016 |
| F11 | 61620001 | 61720000 | 0.0025 | 0.0016 | 0.0020 |
| F11 | 61640001 | 61740000 | 0.0028 | 0.0018 | 0.0023 |
| F11 | 61660001 | 61760000 | 0.0030 | 0.0022 | 0.0024 |
| F11 | 61680001 | 61780000 | 0.0031 | 0.0022 | 0.0024 |
| F11 | 61700001 | 61800000 | 0.0029 | 0.0022 | 0.0024 |
| F11 | 61720001 | 61820000 | 0.0027 | 0.0020 | 0.0021 |
| F11 | 61740001 | 61840000 | 0.0029 | 0.0022 | 0.0023 |
| F11 | 61760001 | 61860000 | 0.0022 | 0.0016 | 0.0018 |
| F11 | 61780001 | 61880000 | 0.0023 | 0.0016 | 0.0018 |
| F11 | 61800001 | 61900000 | 0.0028 | 0.0020 | 0.0021 |

| Chr | Start | End | both | A1 | A2 |
|---|---|---|---|---|---|
| F11 | 61820001 | 61920000 | 0.0030 | 0.0019 | 0.0022 |
| F11 | 61840001 | 61940000 | 0.0035 | 0.0022 | 0.0027 |
| F11 | 61860001 | 61960000 | 0.0037 | 0.0024 | 0.0029 |
| F11 | 61880001 | 61980000 | 0.0041 | 0.0028 | 0.0033 |
| F11 | 61900001 | 62000000 | 0.0037 | 0.0027 | 0.0030 |
| F11 | 61920001 | 62020000 | 0.0035 | 0.0026 | 0.0030 |
| F11 | 61940001 | 62040000 | 0.0031 | 0.0022 | 0.0024 |
| F11 | 61960001 | 62060000 | 0.0036 | 0.0026 | 0.0028 |
| F11 | 61980001 | 62080000 | 0.0031 | 0.0022 | 0.0023 |
| F11 | 62000001 | 62100000 | 0.0031 | 0.0022 | 0.0024 |
| F11 | 62020001 | 62120000 | 0.0038 | 0.0030 | 0.0031 |
| F11 | 62040001 | 62140000 | 0.0042 | 0.0037 | 0.0036 |
| F11 | 62060001 | 62160000 | 0.0034 | 0.0031 | 0.0031 |
| F11 | 62080001 | 62180000 | 0.0035 | 0.0030 | 0.0030 |
| F11 | 62100001 | 62200000 | 0.0033 | 0.0028 | 0.0027 |
| F11 | 62120001 | 62220000 | 0.0034 | 0.0030 | 0.0027 |
| F11 | 62140001 | 62240000 | 0.0047 | 0.0038 | 0.0040 |
| F11 | 62160001 | 62260000 | 0.0052 | 0.0042 | 0.0043 |
| F11 | 62180001 | 62280000 | 0.0051 | 0.0042 | 0.0044 |
| F11 | 62200001 | 62300000 | 0.0048 | 0.0042 | 0.0043 |
| F11 | 62220001 | 62320000 | 0.0041 | 0.0034 | 0.0037 |
| F11 | 62240001 | 62340000 | 0.0026 | 0.0023 | 0.0024 |
| F11 | 62260001 | 62360000 | 0.0022 | 0.0020 | 0.0020 |
| F11 | 62280001 | 62380000 | 0.0025 | 0.0021 | 0.0020 |
| F11 | 62300001 | 62400000 | 0.0032 | 0.0023 | 0.0023 |
| F11 | 62320001 | 62420000 | 0.0033 | 0.0024 | 0.0023 |
| F11 | 62340001 | 62440000 | 0.0029 | 0.0019 | 0.0019 |
| F11 | 62360001 | 62460000 | 0.0033 | 0.0022 | 0.0023 |
| F11 | 62380001 | 62480000 | 0.0031 | 0.0021 | 0.0022 |
| F11 | 62400001 | 62500000 | 0.0024 | 0.0017 | 0.0019 |
| F11 | 62420001 | 62520000 | 0.0023 | 0.0016 | 0.0018 |
| F11 | 62440001 | 62540000 | 0.0019 | 0.0013 | 0.0014 |
| F11 | 62460001 | 62560000 | 0.0020 | 0.0015 | 0.0016 |
| F11 | 62480001 | 62580000 | 0.0020 | 0.0016 | 0.0017 |
| F11 | 62500001 | 62600000 | 0.0021 | 0.0017 | 0.0019 |
| F11 | 62520001 | 62620000 | 0.0023 | 0.0019 | 0.0022 |
| F11 | 62540001 | 62640000 | 0.0024 | 0.0020 | 0.0023 |
| F11 | 62560001 | 62660000 | 0.0023 | 0.0019 | 0.0023 |
| F11 | 62580001 | 62680000 | 0.0033 | 0.0028 | 0.0031 |
| F11 | 62600001 | 62700000 | 0.0031 | 0.0027 | 0.0029 |
| F11 | 62620001 | 62720000 | 0.0040 | 0.0032 | 0.0035 |
| F11 | 62640001 | 62740000 | 0.0041 | 0.0032 | 0.0034 |
| F11 | 62660001 | 62760000 | 0.0045 | 0.0035 | 0.0039 |
| F11 | 62680001 | 62780000 | 0.0038 | 0.0028 | 0.0032 |
| F11 | 62700001 | 62800000 | 0.0046 | 0.0037 | 0.0040 |
| F11 | 62720001 | 62820000 | 0.0037 | 0.0030 | 0.0034 |
| F11 | 62740001 | 62840000 | 0.0039 | 0.0033 | 0.0037 |
| F11 | 62760001 | 62860000 | 0.0028 | 0.0024 | 0.0026 |
| F11 | 62780001 | 62880000 | 0.0027 | 0.0023 | 0.0026 |
| F11 | 62800001 | 62900000 | 0.0033 | 0.0029 | 0.0032 |
| F11 | 62820001 | 62920000 | 0.0033 | 0.0029 | 0.0033 |
| F11 | 62840001 | 62940000 | 0.0035 | 0.0032 | 0.0034 |
| F11 | 62860001 | 62960000 | 0.0033 | 0.0031 | 0.0032 |
| F11 | 62880001 | 62980000 | 0.0036 | 0.0034 | 0.0035 |
| F11 | 62900001 | 63000000 | 0.0020 | 0.0018 | 0.0020 |
| F11 | 62920001 | 63020000 | 0.0016 | 0.0014 | 0.0016 |
| F11 | 62940001 | 63040000 | 0.0018 | 0.0015 | 0.0017 |
| F11 | 62960001 | 63060000 | 0.0022 | 0.0019 | 0.0022 |

| Chr | Start | End | both | A1 | A2 |
|---|---|---|---|---|---|
| F11 | 62980001 | 63080000 | 0.0018 | 0.0015 | 0.0017 |
| F11 | 63000001 | 63100000 | 0.0020 | 0.0017 | 0.0019 |
| F11 | 63020001 | 63120000 | 0.0021 | 0.0018 | 0.0021 |
| F11 | 63040001 | 63140000 | 0.0015 | 0.0013 | 0.0015 |
| F11 | 63060001 | 63160000 | 0.0011 | 0.0009 | 0.0011 |
| F11 | 63080001 | 63180000 | 0.0009 | 0.0008 | 0.0009 |
| F11 | 63100001 | 63200000 | 0.0008 | 0.0007 | 0.0008 |
| F11 | 63120001 | 63220000 | 0.0008 | 0.0006 | 0.0008 |
| F11 | 63140001 | 63240000 | 0.0005 | 0.0004 | 0.0006 |
| F11 | 63160001 | 63260000 | 0.0009 | 0.0008 | 0.0009 |
| F11 | 63180001 | 63280000 | 0.0010 | 0.0009 | 0.0010 |
| F11 | 63200001 | 63300000 | 0.0007 | 0.0007 | 0.0007 |
| F11 | 63220001 | 63320000 | 0.0012 | 0.0011 | 0.0012 |
| F11 | 63240001 | 63340000 | 0.0016 | 0.0014 | 0.0016 |
| F11 | 63260001 | 63360000 | 0.0023 | 0.0020 | 0.0023 |
| F11 | 63280001 | 63380000 | 0.0025 | 0.0021 | 0.0025 |
| F11 | 63300001 | 63400000 | 0.0028 | 0.0023 | 0.0028 |
| F11 | 63320001 | 63420000 | 0.0024 | 0.0021 | 0.0023 |
| F11 | 63340001 | 63440000 | 0.0021 | 0.0019 | 0.0021 |
| F11 | 63360001 | 63460000 | 0.0012 | 0.0011 | 0.0012 |
| F11 | 63380001 | 63480000 | 0.0011 | 0.0010 | 0.0011 |
| F11 | 63400001 | 63500000 | 0.0008 | 0.0008 | 0.0008 |
| F11 | 63420001 | 63520000 | 0.0005 | 0.0005 | 0.0005 |
| F11 | 63440001 | 63540000 | 0.0004 | 0.0004 | 0.0004 |
| F11 | 63460001 | 63560000 | 0.0005 | 0.0004 | 0.0005 |
| F11 | 63480001 | 63580000 | 0.0003 | 0.0003 | 0.0003 |
| F11 | 63500001 | 63600000 | 0.0004 | 0.0004 | 0.0004 |
| F11 | 63520001 | 63620000 | 0.0004 | 0.0003 | 0.0004 |
| F11 | 63540001 | 63640000 | 0.0006 | 0.0005 | 0.0007 |
| F11 | 63560001 | 63660000 | 0.0009 | 0.0007 | 0.0009 |
| F11 | 63580001 | 63680000 | 0.0013 | 0.0011 | 0.0013 |
| F11 | 63600001 | 63700000 | 0.0015 | 0.0012 | 0.0015 |
| F11 | 63620001 | 63720000 | 0.0017 | 0.0014 | 0.0017 |
| F11 | 63640001 | 63740000 | 0.0019 | 0.0018 | 0.0019 |
| F11 | 63660001 | 63760000 | 0.0018 | 0.0016 | 0.0017 |
| F11 | 63680001 | 63780000 | 0.0022 | 0.0020 | 0.0020 |
| F11 | 63700001 | 63800000 | 0.0021 | 0.0018 | 0.0018 |
| F11 | 63720001 | 63820000 | 0.0019 | 0.0016 | 0.0016 |
| F11 | 63740001 | 63840000 | 0.0014 | 0.0011 | 0.0012 |
| F11 | 63760001 | 63860000 | 0.0015 | 0.0013 | 0.0014 |
| F11 | 63780001 | 63880000 | 0.0010 | 0.0008 | 0.0010 |
| F11 | 63800001 | 63900000 | 0.0010 | 0.0008 | 0.0010 |
| F11 | 63820001 | 63920000 | 0.0013 | 0.0011 | 0.0012 |
| F11 | 63840001 | 63940000 | 0.0016 | 0.0014 | 0.0014 |
| F11 | 63860001 | 63960000 | 0.0017 | 0.0015 | 0.0015 |
| F11 | 63880001 | 63980000 | 0.0016 | 0.0013 | 0.0015 |
| F11 | 63900001 | 64000000 | 0.0017 | 0.0014 | 0.0015 |
| F11 | 63920001 | 64020000 | 0.0017 | 0.0014 | 0.0015 |
| F11 | 63940001 | 64040000 | 0.0013 | 0.0011 | 0.0013 |
| F11 | 63960001 | 64060000 | 0.0009 | 0.0007 | 0.0008 |
| F11 | 63980001 | 64080000 | 0.0008 | 0.0007 | 0.0007 |
| F11 | 64000001 | 64100000 | 0.0007 | 0.0007 | 0.0007 |
| F11 | 64020001 | 64120000 | 0.0006 | 0.0006 | 0.0006 |
| F11 | 64040001 | 64140000 | 0.0011 | 0.0009 | 0.0011 |
| F11 | 64060001 | 64160000 | 0.0013 | 0.0011 | 0.0012 |
| F11 | 64080001 | 64180000 | 0.0014 | 0.0012 | 0.0014 |
| F11 | 64100001 | 64200000 | 0.0022 | 0.0019 | 0.0022 |
| F11 | 64120001 | 64220000 | 0.0026 | 0.0023 | 0.0025 |

| Chr | Start | End | both | A1 | A2 |
|---|---|---|---|---|---|
| F11 | 64140001 | 64240000 | 0.0024 | 0.0021 | 0.0021 |
| F11 | 64160001 | 64260000 | 0.0024 | 0.0020 | 0.0020 |
| F11 | 64180001 | 64280000 | 0.0024 | 0.0020 | 0.0021 |
| F11 | 64200001 | 64300000 | 0.0016 | 0.0013 | 0.0013 |
| F11 | 64220001 | 64320000 | 0.0016 | 0.0012 | 0.0012 |
| F11 | 64240001 | 64340000 | 0.0018 | 0.0015 | 0.0015 |
| F11 | 64260001 | 64360000 | 0.0029 | 0.0025 | 0.0026 |
| F11 | 64280001 | 64380000 | 0.0036 | 0.0030 | 0.0033 |
| F11 | 64300001 | 64400000 | 0.0057 | 0.0048 | 0.0053 |
| F11 | 64320001 | 64420000 | 0.0078 | 0.0066 | 0.0075 |
| F11 | 64340001 | 64440000 | 0.0076 | 0.0065 | 0.0074 |
| F11 | 64360001 | 64460000 | 0.0071 | 0.0060 | 0.0069 |
| F11 | 64380001 | 64480000 | 0.0063 | 0.0053 | 0.0061 |
| F11 | 64400001 | 64500000 | 0.0046 | 0.0037 | 0.0044 |
| F11 | 64420001 | 64520000 | 0.0023 | 0.0018 | 0.0023 |
| F11 | 64440001 | 64540000 | 0.0021 | 0.0016 | 0.0021 |
| F11 | 64460001 | 64560000 | 0.0015 | 0.0010 | 0.0014 |
| F11 | 64480001 | 64580000 | 0.0016 | 0.0012 | 0.0015 |
| F11 | 64500001 | 64600000 | 0.0016 | 0.0014 | 0.0016 |
| F11 | 64520001 | 64620000 | 0.0015 | 0.0013 | 0.0015 |
| F11 | 64540001 | 64640000 | 0.0015 | 0.0013 | 0.0015 |
| F11 | 64560001 | 64660000 | 0.0020 | 0.0018 | 0.0019 |
| F11 | 64580001 | 64680000 | 0.0029 | 0.0025 | 0.0027 |
| F11 | 64600001 | 64700000 | 0.0032 | 0.0027 | 0.0029 |
| F11 | 64620001 | 64720000 | 0.0032 | 0.0027 | 0.0029 |
| F11 | 64640001 | 64740000 | 0.0039 | 0.0033 | 0.0035 |
| F11 | 64660001 | 64760000 | 0.0037 | 0.0031 | 0.0034 |
| F11 | 64680001 | 64780000 | 0.0028 | 0.0024 | 0.0026 |
| F11 | 64700001 | 64800000 | 0.0024 | 0.0022 | 0.0023 |
| F11 | 64720001 | 64820000 | 0.0022 | 0.0020 | 0.0021 |
| F11 | 64740001 | 64840000 | 0.0020 | 0.0017 | 0.0017 |
| F11 | 64760001 | 64860000 | 0.0027 | 0.0024 | 0.0023 |
| F11 | 64780001 | 64880000 | 0.0027 | 0.0024 | 0.0023 |
| F11 | 64800001 | 64900000 | 0.0027 | 0.0024 | 0.0022 |
| F11 | 64820001 | 64920000 | 0.0031 | 0.0027 | 0.0024 |
| F11 | 64840001 | 64940000 | 0.0029 | 0.0025 | 0.0025 |
| F11 | 64860001 | 64960000 | 0.0027 | 0.0021 | 0.0020 |
| F11 | 64880001 | 64980000 | 0.0035 | 0.0027 | 0.0027 |
| F11 | 64900001 | 65000000 | 0.0033 | 0.0025 | 0.0026 |
| F11 | 64920001 | 65020000 | 0.0037 | 0.0028 | 0.0032 |
| F11 | 64940001 | 65040000 | 0.0039 | 0.0030 | 0.0033 |
| F11 | 64960001 | 65060000 | 0.0038 | 0.0030 | 0.0034 |
| F11 | 64980001 | 65080000 | 0.0029 | 0.0022 | 0.0025 |
| F11 | 65000001 | 65100000 | 0.0034 | 0.0026 | 0.0029 |
| F11 | 65020001 | 65120000 | 0.0030 | 0.0021 | 0.0022 |
| F11 | 65040001 | 65140000 | 0.0032 | 0.0022 | 0.0024 |
| F11 | 65060001 | 65160000 | 0.0033 | 0.0021 | 0.0024 |
| F11 | 65080001 | 65180000 | 0.0038 | 0.0025 | 0.0028 |
| F11 | 65100001 | 65200000 | 0.0041 | 0.0026 | 0.0031 |
| F11 | 65120001 | 65220000 | 0.0044 | 0.0031 | 0.0035 |
| F11 | 65140001 | 65240000 | 0.0043 | 0.0032 | 0.0034 |
| F11 | 65160001 | 65260000 | 0.0055 | 0.0044 | 0.0047 |
| F11 | 65180001 | 65280000 | 0.0061 | 0.0050 | 0.0053 |
| F11 | 65200001 | 65300000 | 0.0056 | 0.0047 | 0.0050 |
| F11 | 65220001 | 65320000 | 0.0055 | 0.0046 | 0.0049 |
| F11 | 65240001 | 65340000 | 0.0048 | 0.0042 | 0.0043 |
| F11 | 65260001 | 65360000 | 0.0032 | 0.0027 | 0.0028 |
| F11 | 65280001 | 65380000 | 0.0023 | 0.0019 | 0.0021 |

| Chr | Start | End | both | A1 | A2 |
|---|---|---|---|---|---|
| F11 | 65300001 | 65400000 | 0.0022 | 0.0019 | 0.0020 |
| F11 | 65320001 | 65420000 | 0.0021 | 0.0016 | 0.0019 |
| F11 | 65340001 | 65440000 | 0.0027 | 0.0019 | 0.0021 |
| F11 | 65360001 | 65460000 | 0.0027 | 0.0016 | 0.0017 |
| F11 | 65380001 | 65480000 | 0.0032 | 0.0019 | 0.0021 |
| F11 | 65400001 | 65500000 | 0.0031 | 0.0019 | 0.0021 |
| F11 | 65420001 | 65520000 | 0.0033 | 0.0021 | 0.0020 |
| F11 | 65440001 | 65540000 | 0.0031 | 0.0019 | 0.0018 |
| F11 | 65460001 | 65560000 | 0.0041 | 0.0031 | 0.0033 |
| F11 | 65480001 | 65580000 | 0.0049 | 0.0039 | 0.0040 |
| F11 | 65500001 | 65600000 | 0.0053 | 0.0042 | 0.0042 |
| F11 | 65520001 | 65620000 | 0.0047 | 0.0037 | 0.0041 |
| F11 | 65540001 | 65640000 | 0.0051 | 0.0041 | 0.0045 |
| F11 | 65560001 | 65660000 | 0.0044 | 0.0036 | 0.0038 |
| F11 | 65580001 | 65680000 | 0.0032 | 0.0024 | 0.0027 |
| F11 | 65600001 | 65700000 | 0.0031 | 0.0022 | 0.0025 |
| F11 | 65620001 | 65720000 | 0.0033 | 0.0024 | 0.0027 |
| F11 | 65640001 | 65740000 | 0.0028 | 0.0020 | 0.0023 |
| F11 | 65660001 | 65760000 | 0.0020 | 0.0014 | 0.0016 |
| F11 | 65680001 | 65780000 | 0.0015 | 0.0011 | 0.0013 |
| F11 | 65700001 | 65800000 | 0.0012 | 0.0009 | 0.0011 |
| F11 | 65720001 | 65820000 | 0.0015 | 0.0013 | 0.0014 |
| F11 | 65740001 | 65840000 | 0.0015 | 0.0012 | 0.0014 |
| F11 | 65760001 | 65860000 | 0.0016 | 0.0012 | 0.0014 |
| F11 | 65780001 | 65880000 | 0.0018 | 0.0015 | 0.0017 |
| F11 | 65800001 | 65900000 | 0.0020 | 0.0017 | 0.0018 |
| F11 | 65820001 | 65920000 | 0.0016 | 0.0012 | 0.0013 |
| F11 | 65840001 | 65940000 | 0.0020 | 0.0017 | 0.0018 |
| F11 | 65860001 | 65960000 | 0.0024 | 0.0020 | 0.0021 |
| F11 | 65880001 | 65980000 | 0.0036 | 0.0028 | 0.0031 |
| F11 | 65900001 | 66000000 | 0.0035 | 0.0026 | 0.0029 |
| F11 | 65920001 | 66020000 | 0.0039 | 0.0030 | 0.0033 |
| F11 | 65940001 | 66040000 | 0.0040 | 0.0029 | 0.0033 |
| F11 | 65960001 | 66060000 | 0.0042 | 0.0031 | 0.0036 |
| F11 | 65980001 | 66080000 | 0.0034 | 0.0023 | 0.0026 |
| F11 | 66000001 | 66100000 | 0.0056 | 0.0044 | 0.0048 |
| F11 | 66020001 | 66120000 | 0.0054 | 0.0041 | 0.0046 |
| F11 | 66040001 | 66140000 | 0.0049 | 0.0039 | 0.0041 |
| F11 | 66060001 | 66160000 | 0.0045 | 0.0035 | 0.0037 |
| F11 | 66080001 | 66180000 | 0.0050 | 0.0041 | 0.0043 |
| F11 | 66100001 | 66200000 | 0.0035 | 0.0028 | 0.0028 |
| F11 | 66120001 | 66220000 | 0.0041 | 0.0033 | 0.0034 |
| F11 | 66140001 | 66240000 | 0.0048 | 0.0040 | 0.0041 |
| F11 | 66160001 | 66260000 | 0.0056 | 0.0045 | 0.0049 |
| F11 | 66180001 | 66280000 | 0.0056 | 0.0044 | 0.0047 |
| F11 | 66200001 | 66300000 | 0.0060 | 0.0049 | 0.0051 |
| F11 | 66220001 | 66320000 | 0.0053 | 0.0042 | 0.0044 |
| F11 | 66240001 | 66340000 | 0.0051 | 0.0038 | 0.0040 |
| F11 | 66260001 | 66360000 | 0.0047 | 0.0034 | 0.0034 |
| F11 | 66280001 | 66380000 | 0.0044 | 0.0030 | 0.0030 |
| F11 | 66300001 | 66400000 | 0.0046 | 0.0031 | 0.0033 |
| F11 | 66320001 | 66420000 | 0.0049 | 0.0035 | 0.0038 |
| F11 | 66340001 | 66440000 | 0.0062 | 0.0049 | 0.0053 |
| F11 | 66360001 | 66460000 | 0.0065 | 0.0056 | 0.0059 |
| F11 | 66380001 | 66480000 | 0.0061 | 0.0053 | 0.0057 |
| F11 | 66400001 | 66500000 | 0.0049 | 0.0041 | 0.0045 |
| F11 | 66420001 | 66520000 | 0.0045 | 0.0037 | 0.0040 |
| F11 | 66440001 | 66540000 | 0.0027 | 0.0019 | 0.0021 |

| Chr | Start | End | both | A1 | A2 |
|---|---|---|---|---|---|
| F11 | 66460001 | 66560000 | 0.0020 | 0.0010 | 0.0013 |
| F11 | 66480001 | 66580000 | 0.0035 | 0.0024 | 0.0027 |
| F11 | 66500001 | 66600000 | 0.0036 | 0.0026 | 0.0028 |
| F11 | 66520001 | 66620000 | 0.0045 | 0.0035 | 0.0036 |
| F11 | 66540001 | 66640000 | 0.0055 | 0.0041 | 0.0043 |
| F11 | 66560001 | 66660000 | 0.0054 | 0.0041 | 0.0042 |
| F11 | 66580001 | 66680000 | 0.0042 | 0.0028 | 0.0030 |
| F11 | 66600001 | 66700000 | 0.0042 | 0.0028 | 0.0031 |
| F11 | 66620001 | 66720000 | 0.0037 | 0.0024 | 0.0026 |
| F11 | 66640001 | 66740000 | 0.0037 | 0.0027 | 0.0029 |
| F11 | 66660001 | 66760000 | 0.0050 | 0.0040 | 0.0043 |
| F11 | 66680001 | 66780000 | 0.0051 | 0.0041 | 0.0044 |
| F11 | 66700001 | 66800000 | 0.0050 | 0.0040 | 0.0043 |
| F11 | 66720001 | 66820000 | 0.0049 | 0.0037 | 0.0043 |
| F11 | 66740001 | 66840000 | 0.0043 | 0.0030 | 0.0036 |
| F11 | 66760001 | 66860000 | 0.0037 | 0.0022 | 0.0029 |
| F11 | 66780001 | 66880000 | 0.0043 | 0.0028 | 0.0036 |
| F11 | 66800001 | 66900000 | 0.0045 | 0.0031 | 0.0039 |
| F11 | 66820001 | 66920000 | 0.0042 | 0.0031 | 0.0037 |
| F11 | 66840001 | 66940000 | 0.0048 | 0.0037 | 0.0043 |
| F11 | 66860001 | 66960000 | 0.0044 | 0.0034 | 0.0041 |
| F11 | 66880001 | 66980000 | 0.0035 | 0.0026 | 0.0030 |
| F11 | 66900001 | 67000000 | 0.0030 | 0.0020 | 0.0024 |
| F11 | 66920001 | 67020000 | 0.0036 | 0.0023 | 0.0027 |
| F11 | 66940001 | 67040000 | 0.0029 | 0.0017 | 0.0022 |
| F11 | 66960001 | 67060000 | 0.0044 | 0.0029 | 0.0035 |
| F11 | 66980001 | 67080000 | 0.0048 | 0.0034 | 0.0040 |
| F11 | 67000001 | 67100000 | 0.0052 | 0.0040 | 0.0046 |
| F11 | 67020001 | 67120000 | 0.0050 | 0.0041 | 0.0046 |
| F11 | 67040001 | 67140000 | 0.0055 | 0.0043 | 0.0048 |
| F11 | 67060001 | 67160000 | 0.0045 | 0.0036 | 0.0038 |
| F11 | 67080001 | 67180000 | 0.0044 | 0.0033 | 0.0036 |
| F11 | 67100001 | 67200000 | 0.0049 | 0.0037 | 0.0041 |
| F11 | 67120001 | 67220000 | 0.0055 | 0.0040 | 0.0045 |
| F11 | 67140001 | 67240000 | 0.0050 | 0.0038 | 0.0045 |
| F11 | 67160001 | 67260000 | 0.0046 | 0.0034 | 0.0039 |
| F11 | 67180001 | 67280000 | 0.0046 | 0.0037 | 0.0042 |
| F11 | 67200001 | 67300000 | 0.0038 | 0.0029 | 0.0033 |
| F11 | 67220001 | 67320000 | 0.0029 | 0.0023 | 0.0025 |
| F11 | 67240001 | 67340000 | 0.0025 | 0.0021 | 0.0021 |
| F11 | 67260001 | 67360000 | 0.0028 | 0.0025 | 0.0026 |
| F11 | 67280001 | 67380000 | 0.0024 | 0.0020 | 0.0020 |
| F11 | 67300001 | 67400000 | 0.0024 | 0.0019 | 0.0022 |
| F11 | 67320001 | 67420000 | 0.0023 | 0.0017 | 0.0019 |
| F11 | 67340001 | 67440000 | 0.0027 | 0.0019 | 0.0022 |
| F11 | 67360001 | 67460000 | 0.0030 | 0.0019 | 0.0022 |
| F11 | 67380001 | 67480000 | 0.0034 | 0.0020 | 0.0024 |
| F11 | 67400001 | 67500000 | 0.0036 | 0.0019 | 0.0022 |
| F11 | 67420001 | 67520000 | 0.0042 | 0.0023 | 0.0029 |
| F11 | 67440001 | 67540000 | 0.0044 | 0.0024 | 0.0031 |
| F11 | 67460001 | 67560000 | 0.0042 | 0.0021 | 0.0027 |
| F11 | 67480001 | 67580000 | 0.0035 | 0.0019 | 0.0023 |
| F11 | 67500001 | 67600000 | 0.0034 | 0.0021 | 0.0026 |
| F11 | 67520001 | 67620000 | 0.0036 | 0.0025 | 0.0029 |
| F11 | 67540001 | 67640000 | 0.0034 | 0.0024 | 0.0027 |
| F11 | 67560001 | 67660000 | 0.0031 | 0.0024 | 0.0027 |
| F11 | 67580001 | 67680000 | 0.0031 | 0.0024 | 0.0027 |
| F11 | 67600001 | 67700000 | 0.0043 | 0.0032 | 0.0036 |

| Chr | Start | End | both | A1 | A2 |
|---|---|---|---|---|---|
| F11 | 67620001 | 67720000 | 0.0038 | 0.0028 | 0.0030 |
| F11 | 67640001 | 67740000 | 0.0035 | 0.0026 | 0.0027 |
| F11 | 67660001 | 67760000 | 0.0033 | 0.0024 | 0.0026 |
| F11 | 67680001 | 67780000 | 0.0039 | 0.0030 | 0.0031 |
| F11 | 67700001 | 67800000 | 0.0037 | 0.0030 | 0.0030 |
| F11 | 67720001 | 67820000 | 0.0042 | 0.0035 | 0.0035 |
| F11 | 67740001 | 67840000 | 0.0046 | 0.0037 | 0.0037 |
| F11 | 67760001 | 67860000 | 0.0055 | 0.0045 | 0.0046 |
| F11 | 67780001 | 67880000 | 0.0054 | 0.0043 | 0.0045 |
| F11 | 67800001 | 67900000 | 0.0049 | 0.0039 | 0.0043 |
| F11 | 67820001 | 67920000 | 0.0043 | 0.0035 | 0.0038 |
| F11 | 67840001 | 67940000 | 0.0045 | 0.0040 | 0.0042 |
| F11 | 67860001 | 67960000 | 0.0036 | 0.0032 | 0.0034 |
| F11 | 67880001 | 67980000 | 0.0034 | 0.0029 | 0.0032 |
| F11 | 67900001 | 68000000 | 0.0029 | 0.0026 | 0.0027 |
| F11 | 67920001 | 68020000 | 0.0026 | 0.0022 | 0.0023 |
| F11 | 67940001 | 68040000 | 0.0020 | 0.0012 | 0.0015 |
| F11 | 67960001 | 68060000 | 0.0022 | 0.0015 | 0.0016 |
| F11 | 67980001 | 68080000 | 0.0032 | 0.0024 | 0.0026 |
| F11 | 68000001 | 68100000 | 0.0038 | 0.0027 | 0.0030 |
| F11 | 68020001 | 68120000 | 0.0046 | 0.0032 | 0.0036 |
| F11 | 68040001 | 68140000 | 0.0046 | 0.0033 | 0.0037 |
| F11 | 68060001 | 68160000 | 0.0046 | 0.0029 | 0.0035 |
| F11 | 68080001 | 68180000 | 0.0039 | 0.0021 | 0.0025 |
| F11 | 68100001 | 68200000 | 0.0033 | 0.0017 | 0.0019 |
| F11 | 68120001 | 68220000 | 0.0036 | 0.0023 | 0.0024 |
| F11 | 68140001 | 68240000 | 0.0038 | 0.0024 | 0.0026 |
| F11 | 68160001 | 68260000 | 0.0045 | 0.0032 | 0.0033 |
| F11 | 68180001 | 68280000 | 0.0047 | 0.0036 | 0.0036 |
| F11 | 68200001 | 68300000 | 0.0063 | 0.0053 | 0.0052 |
| F11 | 68220001 | 68320000 | 0.0067 | 0.0056 | 0.0057 |
| F11 | 68240001 | 68340000 | 0.0078 | 0.0067 | 0.0070 |
| F11 | 68260001 | 68360000 | 0.0067 | 0.0059 | 0.0061 |
| F11 | 68280001 | 68380000 | 0.0060 | 0.0053 | 0.0056 |
| F11 | 68300001 | 68400000 | 0.0046 | 0.0039 | 0.0044 |
| F11 | 68320001 | 68420000 | 0.0032 | 0.0028 | 0.0030 |
| F11 | 68340001 | 68440000 | 0.0017 | 0.0015 | 0.0016 |
| F11 | 68360001 | 68460000 | 0.0014 | 0.0013 | 0.0014 |
| F11 | 68380001 | 68480000 | 0.0018 | 0.0016 | 0.0017 |
| F11 | 68400001 | 68500000 | 0.0016 | 0.0014 | 0.0016 |
| F11 | 68420001 | 68520000 | 0.0019 | 0.0017 | 0.0018 |
| F11 | 68440001 | 68540000 | 0.0020 | 0.0017 | 0.0019 |
| F11 | 68460001 | 68560000 | 0.0027 | 0.0025 | 0.0026 |
| F11 | 68480001 | 68580000 | 0.0030 | 0.0025 | 0.0026 |
| F11 | 68500001 | 68600000 | 0.0033 | 0.0026 | 0.0029 |
| F11 | 68520001 | 68620000 | 0.0030 | 0.0021 | 0.0023 |
| F11 | 68540001 | 68640000 | 0.0032 | 0.0021 | 0.0022 |
| F11 | 68560001 | 68660000 | 0.0029 | 0.0017 | 0.0019 |
| F11 | 68580001 | 68680000 | 0.0032 | 0.0019 | 0.0022 |
| F11 | 68600001 | 68700000 | 0.0041 | 0.0028 | 0.0031 |
| F11 | 68620001 | 68720000 | 0.0044 | 0.0033 | 0.0036 |
| F11 | 68640001 | 68740000 | 0.0049 | 0.0038 | 0.0042 |
| F11 | 68660001 | 68760000 | 0.0043 | 0.0033 | 0.0037 |
| F11 | 68680001 | 68780000 | 0.0033 | 0.0027 | 0.0030 |
| F11 | 68700001 | 68800000 | 0.0023 | 0.0017 | 0.0019 |
| F11 | 68720001 | 68820000 | 0.0021 | 0.0016 | 0.0018 |
| F11 | 68740001 | 68840000 | 0.0020 | 0.0017 | 0.0018 |
| F11 | 68760001 | 68860000 | 0.0033 | 0.0027 | 0.0030 |

| Chr | Start | End | both | A1 | A2 |
|---|---|---|---|---|---|
| F11 | 68780001 | 68880000 | 0.0033 | 0.0026 | 0.0029 |
| F11 | 68800001 | 68900000 | 0.0035 | 0.0027 | 0.0030 |
| F11 | 68820001 | 68920000 | 0.0033 | 0.0026 | 0.0029 |
| F11 | 68840001 | 68940000 | 0.0034 | 0.0025 | 0.0030 |
| F11 | 68860001 | 68960000 | 0.0026 | 0.0019 | 0.0024 |
| F11 | 68880001 | 68980000 | 0.0026 | 0.0020 | 0.0025 |
| F11 | 68900001 | 69000000 | 0.0019 | 0.0015 | 0.0019 |
| F11 | 68920001 | 69020000 | 0.0017 | 0.0013 | 0.0017 |
| F11 | 68940001 | 69040000 | 0.0008 | 0.0007 | 0.0008 |
| F11 | 68960001 | 69060000 | 0.0003 | 0.0003 | 0.0003 |
| F11 | 68980001 | 69080000 | 0.0002 | 0.0002 | 0.0002 |
| F11 | 69000001 | 69100000 | 0.0002 | 0.0002 | 0.0002 |
| F11 | 69020001 | 69120000 | 0.0002 | 0.0001 | 0.0002 |
| F11 | 69040001 | 69140000 | 0.0004 | 0.0004 | 0.0004 |
| F11 | 69060001 | 69160000 | 0.0007 | 0.0006 | 0.0007 |
| F11 | 69080001 | 69180000 | 0.0014 | 0.0013 | 0.0013 |
| F11 | 69100001 | 69200000 | 0.0020 | 0.0018 | 0.0020 |
| F11 | 69120001 | 69220000 | 0.0023 | 0.0020 | 0.0023 |
| F11 | 69140001 | 69240000 | 0.0022 | 0.0019 | 0.0022 |
| F11 | 69160001 | 69260000 | 0.0027 | 0.0022 | 0.0026 |
| F11 | 69180001 | 69280000 | 0.0022 | 0.0018 | 0.0021 |
| F11 | 69200001 | 69300000 | 0.0025 | 0.0019 | 0.0022 |
| F11 | 69220001 | 69320000 | 0.0029 | 0.0022 | 0.0024 |
| F11 | 69240001 | 69340000 | 0.0043 | 0.0034 | 0.0037 |
| F11 | 69260001 | 69360000 | 0.0041 | 0.0032 | 0.0035 |
| F11 | 69280001 | 69380000 | 0.0044 | 0.0036 | 0.0039 |
| F11 | 69300001 | 69400000 | 0.0037 | 0.0030 | 0.0033 |
| F11 | 69320001 | 69420000 | 0.0036 | 0.0030 | 0.0035 |
| F11 | 69340001 | 69440000 | 0.0023 | 0.0019 | 0.0023 |
| F11 | 69360001 | 69460000 | 0.0024 | 0.0020 | 0.0024 |
| F11 | 69380001 | 69480000 | 0.0023 | 0.0019 | 0.0023 |
| F11 | 69400001 | 69500000 | 0.0022 | 0.0018 | 0.0022 |
| F11 | 69420001 | 69520000 | 0.0017 | 0.0014 | 0.0017 |
| F11 | 69440001 | 69540000 | 0.0014 | 0.0012 | 0.0014 |
| F11 | 69460001 | 69560000 | 0.0008 | 0.0007 | 0.0008 |
| F11 | 69480001 | 69580000 | 0.0006 | 0.0004 | 0.0006 |
| F11 | 69500001 | 69600000 | 0.0009 | 0.0008 | 0.0009 |
| F11 | 69520001 | 69620000 | 0.0012 | 0.0009 | 0.0012 |
| F11 | 69540001 | 69640000 | 0.0012 | 0.0010 | 0.0012 |
| F11 | 69560001 | 69660000 | 0.0012 | 0.0010 | 0.0012 |
| F11 | 69580001 | 69680000 | 0.0012 | 0.0011 | 0.0011 |
| F11 | 69600001 | 69700000 | 0.0013 | 0.0010 | 0.0012 |
| F11 | 69620001 | 69720000 | 0.0013 | 0.0010 | 0.0012 |
| F11 | 69640001 | 69740000 | 0.0015 | 0.0011 | 0.0014 |
| F11 | 69660001 | 69760000 | 0.0019 | 0.0013 | 0.0017 |
| F11 | 69680001 | 69780000 | 0.0018 | 0.0012 | 0.0016 |
| F11 | 69700001 | 69800000 | 0.0016 | 0.0011 | 0.0015 |
| F11 | 69720001 | 69820000 | 0.0013 | 0.0009 | 0.0012 |
| F11 | 69740001 | 69840000 | 0.0017 | 0.0012 | 0.0015 |
| F11 | 69760001 | 69860000 | 0.0019 | 0.0014 | 0.0015 |
| F11 | 69780001 | 69880000 | 0.0018 | 0.0014 | 0.0015 |
| F11 | 69800001 | 69900000 | 0.0028 | 0.0022 | 0.0023 |
| F11 | 69820001 | 69920000 | 0.0033 | 0.0027 | 0.0029 |
| F11 | 69840001 | 69940000 | 0.0034 | 0.0028 | 0.0030 |
| F11 | 69860001 | 69960000 | 0.0030 | 0.0025 | 0.0028 |
| F11 | 69880001 | 69980000 | 0.0032 | 0.0026 | 0.0029 |
| F11 | 69900001 | 70000000 | 0.0026 | 0.0022 | 0.0024 |
| F11 | 69920001 | 70020000 | 0.0022 | 0.0018 | 0.0020 |

| Chr | Start | End | both | A1 | A2 |
|---|---|---|---|---|---|
| F11 | 69940001 | 70040000 | 0.0021 | 0.0017 | 0.0018 |
| F11 | 69960001 | 70060000 | 0.0029 | 0.0021 | 0.0024 |
| F11 | 69980001 | 70080000 | 0.0037 | 0.0027 | 0.0030 |
| F11 | 70000001 | 70100000 | 0.0036 | 0.0024 | 0.0028 |
| F11 | 70020001 | 70120000 | 0.0041 | 0.0029 | 0.0033 |
| F11 | 70040001 | 70140000 | 0.0045 | 0.0032 | 0.0037 |
| F11 | 70060001 | 70160000 | 0.0044 | 0.0032 | 0.0038 |
| F11 | 70080001 | 70180000 | 0.0037 | 0.0028 | 0.0033 |
| F11 | 70100001 | 70200000 | 0.0034 | 0.0028 | 0.0031 |
| F11 | 70120001 | 70220000 | 0.0031 | 0.0025 | 0.0028 |
| F11 | 70140001 | 70240000 | 0.0024 | 0.0021 | 0.0023 |
| F11 | 70160001 | 70260000 | 0.0018 | 0.0014 | 0.0016 |
| F11 | 70180001 | 70280000 | 0.0023 | 0.0018 | 0.0019 |
| F11 | 70200001 | 70300000 | 0.0030 | 0.0022 | 0.0023 |
| F11 | 70220001 | 70320000 | 0.0036 | 0.0023 | 0.0026 |
| F11 | 70240001 | 70340000 | 0.0044 | 0.0029 | 0.0033 |
| F11 | 70260001 | 70360000 | 0.0043 | 0.0029 | 0.0033 |
| F11 | 70280001 | 70380000 | 0.0037 | 0.0024 | 0.0028 |
| F11 | 70300001 | 70400000 | 0.0032 | 0.0022 | 0.0023 |
| F11 | 70320001 | 70420000 | 0.0024 | 0.0018 | 0.0019 |
| F11 | 70340001 | 70440000 | 0.0019 | 0.0013 | 0.0015 |
| F11 | 70360001 | 70460000 | 0.0025 | 0.0019 | 0.0021 |
| F11 | 70380001 | 70480000 | 0.0026 | 0.0019 | 0.0021 |
| F11 | 70400001 | 70500000 | 0.0024 | 0.0017 | 0.0021 |
| F11 | 70420001 | 70520000 | 0.0023 | 0.0016 | 0.0019 |
| F11 | 70440001 | 70540000 | 0.0022 | 0.0016 | 0.0017 |
| F11 | 70460001 | 70560000 | 0.0023 | 0.0014 | 0.0016 |
| F11 | 70480001 | 70580000 | 0.0023 | 0.0015 | 0.0017 |
| F11 | 70500001 | 70600000 | 0.0044 | 0.0034 | 0.0036 |
| F11 | 70520001 | 70620000 | 0.0049 | 0.0037 | 0.0039 |
| F11 | 70540001 | 70640000 | 0.0048 | 0.0034 | 0.0038 |
| F11 | 70560001 | 70660000 | 0.0055 | 0.0041 | 0.0045 |
| F11 | 70580001 | 70680000 | 0.0060 | 0.0042 | 0.0047 |
| F11 | 70600001 | 70700000 | 0.0043 | 0.0025 | 0.0029 |
| F11 | 70620001 | 70720000 | 0.0049 | 0.0033 | 0.0037 |
| F11 | 70640001 | 70740000 | 0.0057 | 0.0040 | 0.0044 |
| F11 | 70660001 | 70760000 | 0.0049 | 0.0033 | 0.0035 |
| F11 | 70680001 | 70780000 | 0.0050 | 0.0031 | 0.0034 |
| F11 | 70700001 | 70800000 | 0.0045 | 0.0028 | 0.0031 |
| F11 | 70720001 | 70820000 | 0.0036 | 0.0021 | 0.0023 |
| F11 | 70740001 | 70840000 | 0.0030 | 0.0017 | 0.0019 |
| F11 | 70760001 | 70860000 | 0.0024 | 0.0014 | 0.0016 |
| F11 | 70780001 | 70880000 | 0.0023 | 0.0017 | 0.0019 |
| F11 | 70800001 | 70900000 | 0.0026 | 0.0019 | 0.0021 |
| F11 | 70820001 | 70920000 | 0.0031 | 0.0024 | 0.0027 |
| F11 | 70840001 | 70940000 | 0.0038 | 0.0029 | 0.0034 |
| F11 | 70860001 | 70960000 | 0.0049 | 0.0039 | 0.0045 |
| F11 | 70880001 | 70980000 | 0.0043 | 0.0035 | 0.0041 |
| F11 | 70900001 | 71000000 | 0.0045 | 0.0040 | 0.0043 |
| F11 | 70920001 | 71020000 | 0.0044 | 0.0037 | 0.0039 |
| F11 | 70940001 | 71040000 | 0.0050 | 0.0038 | 0.0043 |
| F11 | 70960001 | 71060000 | 0.0039 | 0.0028 | 0.0032 |
| F11 | 70980001 | 71080000 | 0.0042 | 0.0031 | 0.0035 |
| F11 | 71000001 | 71100000 | 0.0038 | 0.0028 | 0.0030 |
| F11 | 71020001 | 71120000 | 0.0042 | 0.0032 | 0.0035 |
| F11 | 71040001 | 71140000 | 0.0035 | 0.0028 | 0.0029 |
| F11 | 71060001 | 71160000 | 0.0040 | 0.0034 | 0.0034 |
| F11 | 71080001 | 71180000 | 0.0042 | 0.0034 | 0.0035 |

| Chr | Start | End | both | A1 | A2 |
|---|---|---|---|---|---|
| F11 | 71100001 | 71200000 | 0.0038 | 0.0029 | 0.0032 |
| F11 | 71120001 | 71220000 | 0.0032 | 0.0025 | 0.0027 |
| F11 | 71140001 | 71240000 | 0.0026 | 0.0019 | 0.0021 |
| F11 | 71160001 | 71260000 | 0.0025 | 0.0016 | 0.0018 |
| F11 | 71180001 | 71280000 | 0.0026 | 0.0015 | 0.0016 |
| F11 | 71200001 | 71300000 | 0.0036 | 0.0023 | 0.0025 |
| F11 | 71220001 | 71320000 | 0.0038 | 0.0022 | 0.0025 |
| F11 | 71240001 | 71340000 | 0.0042 | 0.0025 | 0.0029 |
| F11 | 71260001 | 71360000 | 0.0052 | 0.0035 | 0.0039 |
| F11 | 71280001 | 71380000 | 0.0050 | 0.0036 | 0.0040 |
| F11 | 71300001 | 71400000 | 0.0051 | 0.0036 | 0.0040 |
| F11 | 71320001 | 71420000 | 0.0049 | 0.0034 | 0.0038 |
| F11 | 71340001 | 71440000 | 0.0044 | 0.0032 | 0.0034 |
| F11 | 71360001 | 71460000 | 0.0037 | 0.0026 | 0.0028 |
| F11 | 71380001 | 71480000 | 0.0039 | 0.0026 | 0.0031 |
| F11 | 71400001 | 71500000 | 0.0032 | 0.0022 | 0.0025 |
| F11 | 71420001 | 71520000 | 0.0035 | 0.0027 | 0.0030 |
| F11 | 71440001 | 71540000 | 0.0043 | 0.0034 | 0.0037 |
| F11 | 71460001 | 71560000 | 0.0040 | 0.0031 | 0.0035 |
| F11 | 71480001 | 71580000 | 0.0046 | 0.0036 | 0.0038 |
| F11 | 71500001 | 71600000 | 0.0048 | 0.0037 | 0.0038 |
| F11 | 71520001 | 71620000 | 0.0047 | 0.0034 | 0.0035 |
| F11 | 71540001 | 71640000 | 0.0061 | 0.0047 | 0.0050 |
| F11 | 71560001 | 71660000 | 0.0066 | 0.0051 | 0.0055 |
| F11 | 71580001 | 71680000 | 0.0061 | 0.0046 | 0.0050 |
| F11 | 71600001 | 71700000 | 0.0061 | 0.0046 | 0.0050 |
| F11 | 71620001 | 71720000 | 0.0064 | 0.0048 | 0.0054 |
| F11 | 71640001 | 71740000 | 0.0049 | 0.0033 | 0.0038 |
| F11 | 71660001 | 71760000 | 0.0048 | 0.0032 | 0.0035 |
| F11 | 71680001 | 71780000 | 0.0043 | 0.0028 | 0.0031 |
| F11 | 71700001 | 71800000 | 0.0044 | 0.0029 | 0.0033 |
| F11 | 71720001 | 71820000 | 0.0043 | 0.0028 | 0.0030 |
| F11 | 71740001 | 71840000 | 0.0042 | 0.0029 | 0.0031 |
| F11 | 71760001 | 71860000 | 0.0048 | 0.0031 | 0.0036 |
| F11 | 71780001 | 71880000 | 0.0060 | 0.0041 | 0.0047 |
| F11 | 71800001 | 71900000 | 0.0062 | 0.0049 | 0.0050 |
| F11 | 71820001 | 71920000 | 0.0058 | 0.0046 | 0.0048 |
| F11 | 71840001 | 71940000 | 0.0064 | 0.0053 | 0.0054 |
| F11 | 71860001 | 71960000 | 0.0053 | 0.0045 | 0.0044 |
| F11 | 71880001 | 71980000 | 0.0046 | 0.0038 | 0.0036 |
| F11 | 71900001 | 72000000 | 0.0043 | 0.0033 | 0.0033 |
| F11 | 71920001 | 72020000 | 0.0047 | 0.0034 | 0.0036 |
| F11 | 71940001 | 72040000 | 0.0035 | 0.0025 | 0.0025 |
| F11 | 71960001 | 72060000 | 0.0040 | 0.0028 | 0.0029 |
| F11 | 71980001 | 72080000 | 0.0045 | 0.0032 | 0.0036 |
| F11 | 72000001 | 72100000 | 0.0043 | 0.0029 | 0.0034 |
| F11 | 72020001 | 72120000 | 0.0040 | 0.0025 | 0.0028 |
| F11 | 72040001 | 72140000 | 0.0048 | 0.0031 | 0.0036 |
| F11 | 72060001 | 72160000 | 0.0045 | 0.0029 | 0.0034 |
| F11 | 72080001 | 72180000 | 0.0036 | 0.0021 | 0.0024 |
| F11 | 72100001 | 72200000 | 0.0036 | 0.0020 | 0.0024 |
| F11 | 72120001 | 72220000 | 0.0038 | 0.0022 | 0.0027 |
| F11 | 72140001 | 72240000 | 0.0037 | 0.0022 | 0.0025 |
| F11 | 72160001 | 72260000 | 0.0035 | 0.0020 | 0.0025 |
| F11 | 72180001 | 72280000 | 0.0036 | 0.0022 | 0.0025 |
| F11 | 72200001 | 72300000 | 0.0043 | 0.0028 | 0.0033 |
| F11 | 72220001 | 72320000 | 0.0045 | 0.0033 | 0.0036 |
| F11 | 72240001 | 72340000 | 0.0038 | 0.0028 | 0.0030 |

| Chr | Start | End | both | A1 | A2 |
|---|---|---|---|---|---|
| F11 | 72260001 | 72360000 | 0.0042 | 0.0032 | 0.0033 |
| F11 | 72280001 | 72380000 | 0.0043 | 0.0033 | 0.0035 |
| F11 | 72300001 | 72400000 | 0.0039 | 0.0030 | 0.0031 |
| F11 | 72320001 | 72420000 | 0.0039 | 0.0029 | 0.0031 |
| F11 | 72340001 | 72440000 | 0.0041 | 0.0030 | 0.0032 |
| F11 | 72360001 | 72460000 | 0.0033 | 0.0024 | 0.0027 |
| F11 | 72380001 | 72480000 | 0.0030 | 0.0022 | 0.0024 |
| F11 | 72400001 | 72500000 | 0.0026 | 0.0018 | 0.0021 |
| F11 | 72420001 | 72520000 | 0.0022 | 0.0016 | 0.0018 |
| F11 | 72440001 | 72540000 | 0.0021 | 0.0015 | 0.0018 |
| F11 | 72460001 | 72560000 | 0.0025 | 0.0017 | 0.0021 |
| F11 | 72480001 | 72580000 | 0.0027 | 0.0020 | 0.0022 |
| F11 | 72500001 | 72600000 | 0.0027 | 0.0021 | 0.0024 |
| F11 | 72520001 | 72620000 | 0.0022 | 0.0018 | 0.0020 |
| F11 | 72540001 | 72640000 | 0.0017 | 0.0015 | 0.0016 |
| F11 | 72560001 | 72660000 | 0.0017 | 0.0017 | 0.0016 |
| F11 | 72580001 | 72680000 | 0.0022 | 0.0020 | 0.0022 |
| F11 | 72600001 | 72700000 | 0.0019 | 0.0016 | 0.0018 |
| F11 | 72620001 | 72720000 | 0.0028 | 0.0023 | 0.0027 |
| F11 | 72640001 | 72740000 | 0.0037 | 0.0030 | 0.0034 |
| F11 | 72660001 | 72760000 | 0.0036 | 0.0029 | 0.0034 |
| F11 | 72680001 | 72780000 | 0.0035 | 0.0027 | 0.0030 |
| F11 | 72700001 | 72800000 | 0.0039 | 0.0029 | 0.0032 |
| F11 | 72720001 | 72820000 | 0.0032 | 0.0023 | 0.0024 |
| F11 | 72740001 | 72840000 | 0.0031 | 0.0023 | 0.0023 |
| F11 | 72760001 | 72860000 | 0.0032 | 0.0022 | 0.0023 |
| F11 | 72780001 | 72880000 | 0.0030 | 0.0019 | 0.0021 |
| F11 | 72800001 | 72900000 | 0.0033 | 0.0023 | 0.0026 |
| F11 | 72820001 | 72920000 | 0.0047 | 0.0035 | 0.0040 |
| F11 | 72840001 | 72940000 | 0.0049 | 0.0035 | 0.0041 |
| F11 | 72860001 | 72960000 | 0.0053 | 0.0037 | 0.0044 |
| F11 | 72880001 | 72980000 | 0.0057 | 0.0042 | 0.0048 |
| F11 | 72900001 | 73000000 | 0.0056 | 0.0041 | 0.0044 |
| F11 | 72920001 | 73020000 | 0.0051 | 0.0037 | 0.0039 |
| F11 | 72940001 | 73040000 | 0.0047 | 0.0035 | 0.0037 |
| F11 | 72960001 | 73060000 | 0.0046 | 0.0037 | 0.0037 |
| F11 | 72980001 | 73080000 | 0.0047 | 0.0039 | 0.0040 |
| F11 | 73000001 | 73100000 | 0.0053 | 0.0044 | 0.0046 |
| F11 | 73020001 | 73120000 | 0.0050 | 0.0039 | 0.0041 |
| F11 | 73040001 | 73140000 | 0.0053 | 0.0038 | 0.0043 |
| F11 | 73060001 | 73160000 | 0.0054 | 0.0038 | 0.0044 |
| F11 | 73080001 | 73180000 | 0.0052 | 0.0036 | 0.0040 |
| F11 | 73100001 | 73200000 | 0.0044 | 0.0028 | 0.0032 |
| F11 | 73120001 | 73220000 | 0.0042 | 0.0028 | 0.0032 |
| F11 | 73140001 | 73240000 | 0.0037 | 0.0028 | 0.0029 |
| F11 | 73160001 | 73260000 | 0.0036 | 0.0027 | 0.0029 |
| F11 | 73180001 | 73280000 | 0.0033 | 0.0025 | 0.0027 |
| F11 | 73200001 | 73300000 | 0.0033 | 0.0026 | 0.0028 |
| F11 | 73220001 | 73320000 | 0.0034 | 0.0026 | 0.0029 |
| F11 | 73240001 | 73340000 | 0.0036 | 0.0027 | 0.0029 |
| F11 | 73260001 | 73360000 | 0.0033 | 0.0023 | 0.0025 |
| F11 | 73280001 | 73380000 | 0.0034 | 0.0024 | 0.0025 |
| F11 | 73300001 | 73400000 | 0.0043 | 0.0028 | 0.0034 |
| F11 | 73320001 | 73420000 | 0.0044 | 0.0028 | 0.0037 |
| F11 | 73340001 | 73440000 | 0.0043 | 0.0027 | 0.0035 |
| F11 | 73360001 | 73460000 | 0.0042 | 0.0026 | 0.0034 |
| F11 | 73380001 | 73480000 | 0.0036 | 0.0020 | 0.0029 |
| F11 | 73400001 | 73500000 | 0.0025 | 0.0014 | 0.0019 |

| Chr | Start | End | both | A1 | A2 |
|---|---|---|---|---|---|
| F11 | 73420001 | 73520000 | 0.0021 | 0.0012 | 0.0013 |
| F11 | 73440001 | 73540000 | 0.0022 | 0.0012 | 0.0014 |
| F11 | 73460001 | 73560000 | 0.0029 | 0.0018 | 0.0020 |
| F11 | 73480001 | 73580000 | 0.0035 | 0.0024 | 0.0027 |
| F11 | 73500001 | 73600000 | 0.0032 | 0.0022 | 0.0023 |
| F11 | 73520001 | 73620000 | 0.0035 | 0.0024 | 0.0026 |
| F11 | 73540001 | 73640000 | 0.0032 | 0.0024 | 0.0025 |
| F11 | 73560001 | 73660000 | 0.0028 | 0.0021 | 0.0021 |
| F11 | 73580001 | 73680000 | 0.0023 | 0.0014 | 0.0014 |
| F11 | 73600001 | 73700000 | 0.0026 | 0.0017 | 0.0019 |
| F11 | 73620001 | 73720000 | 0.0028 | 0.0019 | 0.0022 |
| F11 | 73640001 | 73740000 | 0.0025 | 0.0015 | 0.0019 |
| F11 | 73660001 | 73760000 | 0.0025 | 0.0016 | 0.0020 |
| F11 | 73680001 | 73780000 | 0.0033 | 0.0023 | 0.0027 |
| F11 | 73700001 | 73800000 | 0.0033 | 0.0023 | 0.0028 |
| F11 | 73720001 | 73820000 | 0.0032 | 0.0022 | 0.0026 |
| F11 | 73740001 | 73840000 | 0.0033 | 0.0023 | 0.0028 |
| F11 | 73760001 | 73860000 | 0.0032 | 0.0023 | 0.0027 |
| F11 | 73780001 | 73880000 | 0.0026 | 0.0018 | 0.0022 |
| F11 | 73800001 | 73900000 | 0.0025 | 0.0018 | 0.0020 |
| F11 | 73820001 | 73920000 | 0.0021 | 0.0016 | 0.0017 |
| F11 | 73840001 | 73940000 | 0.0019 | 0.0015 | 0.0017 |
| F11 | 73860001 | 73960000 | 0.0017 | 0.0014 | 0.0015 |
| F11 | 73880001 | 73980000 | 0.0015 | 0.0013 | 0.0015 |
| F11 | 73900001 | 74000000 | 0.0016 | 0.0014 | 0.0016 |
| F11 | 73920001 | 74020000 | 0.0018 | 0.0016 | 0.0018 |
| F11 | 73940001 | 74040000 | 0.0018 | 0.0014 | 0.0017 |
| F11 | 73960001 | 74060000 | 0.0028 | 0.0022 | 0.0025 |
| F11 | 73980001 | 74080000 | 0.0044 | 0.0036 | 0.0040 |
| F11 | 74000001 | 74100000 | 0.0046 | 0.0036 | 0.0040 |
| F11 | 74020001 | 74120000 | 0.0047 | 0.0035 | 0.0040 |
| F11 | 74040001 | 74140000 | 0.0053 | 0.0040 | 0.0044 |
| F11 | 74060001 | 74160000 | 0.0050 | 0.0038 | 0.0042 |
| F11 | 74080001 | 74180000 | 0.0034 | 0.0024 | 0.0028 |
| F11 | 74100001 | 74200000 | 0.0037 | 0.0027 | 0.0029 |
| F11 | 74120001 | 74220000 | 0.0040 | 0.0028 | 0.0031 |
| F11 | 74140001 | 74240000 | 0.0038 | 0.0027 | 0.0031 |
| F11 | 74160001 | 74260000 | 0.0036 | 0.0025 | 0.0029 |
| F11 | 74180001 | 74280000 | 0.0035 | 0.0024 | 0.0027 |
| F11 | 74200001 | 74300000 | 0.0033 | 0.0020 | 0.0026 |
| F11 | 74220001 | 74320000 | 0.0031 | 0.0020 | 0.0026 |
| F11 | 74240001 | 74340000 | 0.0034 | 0.0023 | 0.0029 |
| F11 | 74260001 | 74360000 | 0.0037 | 0.0027 | 0.0031 |
| F11 | 74280001 | 74380000 | 0.0047 | 0.0037 | 0.0040 |
| F11 | 74300001 | 74400000 | 0.0040 | 0.0033 | 0.0034 |
| F11 | 74320001 | 74420000 | 0.0034 | 0.0030 | 0.0030 |
| F11 | 74340001 | 74440000 | 0.0032 | 0.0026 | 0.0027 |
| F11 | 74360001 | 74460000 | 0.0029 | 0.0022 | 0.0024 |
| F11 | 74380001 | 74480000 | 0.0021 | 0.0016 | 0.0019 |
| F11 | 74400001 | 74500000 | 0.0029 | 0.0023 | 0.0026 |
| F11 | 74420001 | 74520000 | 0.0030 | 0.0023 | 0.0027 |
| F11 | 74440001 | 74540000 | 0.0025 | 0.0020 | 0.0023 |
| F11 | 74460001 | 74560000 | 0.0025 | 0.0020 | 0.0023 |
| F11 | 74480001 | 74580000 | 0.0024 | 0.0019 | 0.0022 |
| F11 | 74500001 | 74600000 | 0.0020 | 0.0015 | 0.0018 |
| F11 | 74520001 | 74620000 | 0.0028 | 0.0023 | 0.0024 |
| F11 | 74540001 | 74640000 | 0.0038 | 0.0031 | 0.0035 |
| F11 | 74560001 | 74660000 | 0.0036 | 0.0029 | 0.0033 |

| Chr | Start | End | both | A1 | A2 |
|---|---|---|---|---|---|
| F11 | 74580001 | 74680000 | 0.0039 | 0.0030 | 0.0035 |
| F11 | 74600001 | 74700000 | 0.0037 | 0.0030 | 0.0034 |
| F11 | 74620001 | 74720000 | 0.0033 | 0.0026 | 0.0030 |
| F11 | 74640001 | 74740000 | 0.0028 | 0.0022 | 0.0026 |
| F11 | 74660001 | 74760000 | 0.0026 | 0.0019 | 0.0022 |
| F11 | 74680001 | 74780000 | 0.0031 | 0.0023 | 0.0028 |
| F11 | 74700001 | 74800000 | 0.0040 | 0.0031 | 0.0035 |
| F11 | 74720001 | 74820000 | 0.0043 | 0.0031 | 0.0036 |
| F11 | 74740001 | 74840000 | 0.0042 | 0.0030 | 0.0033 |
| F11 | 74760001 | 74860000 | 0.0046 | 0.0033 | 0.0035 |
| F11 | 74780001 | 74880000 | 0.0047 | 0.0034 | 0.0035 |
| F11 | 74800001 | 74900000 | 0.0037 | 0.0025 | 0.0027 |
| F11 | 74820001 | 74920000 | 0.0032 | 0.0024 | 0.0025 |
| F11 | 74840001 | 74940000 | 0.0031 | 0.0022 | 0.0024 |
| F11 | 74860001 | 74960000 | 0.0035 | 0.0027 | 0.0030 |
| F11 | 74880001 | 74980000 | 0.0030 | 0.0024 | 0.0027 |
| F11 | 74900001 | 75000000 | 0.0030 | 0.0024 | 0.0027 |
| F11 | 74920001 | 75020000 | 0.0041 | 0.0034 | 0.0036 |
| F11 | 74940001 | 75040000 | 0.0041 | 0.0033 | 0.0035 |
| F11 | 74960001 | 75060000 | 0.0044 | 0.0032 | 0.0034 |
| F11 | 74980001 | 75080000 | 0.0038 | 0.0024 | 0.0026 |
| F11 | 75000001 | 75100000 | 0.0044 | 0.0031 | 0.0032 |
| F11 | 75020001 | 75120000 | 0.0030 | 0.0018 | 0.0019 |
| F11 | 75040001 | 75140000 | 0.0027 | 0.0015 | 0.0016 |
| F11 | 75060001 | 75160000 | 0.0021 | 0.0012 | 0.0011 |
| F11 | 75080001 | 75180000 | 0.0022 | 0.0013 | 0.0012 |
| F11 | 75100001 | 75200000 | 0.0017 | 0.0006 | 0.0007 |
| F11 | 75120001 | 75220000 | 0.0026 | 0.0014 | 0.0016 |
| F11 | 75140001 | 75240000 | 0.0029 | 0.0016 | 0.0020 |
| F11 | 75160001 | 75260000 | 0.0028 | 0.0016 | 0.0020 |
| F11 | 75180001 | 75280000 | 0.0034 | 0.0022 | 0.0026 |
| F11 | 75200001 | 75300000 | 0.0036 | 0.0023 | 0.0027 |
| F11 | 75220001 | 75320000 | 0.0029 | 0.0015 | 0.0017 |
| F11 | 75240001 | 75340000 | 0.0028 | 0.0015 | 0.0016 |
| F11 | 75260001 | 75360000 | 0.0027 | 0.0014 | 0.0015 |
| F11 | 75280001 | 75380000 | 0.0023 | 0.0009 | 0.0010 |
| F11 | 75300001 | 75400000 | 0.0028 | 0.0015 | 0.0016 |
| F11 | 75320001 | 75420000 | 0.0025 | 0.0015 | 0.0016 |
| F11 | 75340001 | 75440000 | 0.0026 | 0.0017 | 0.0019 |
| F11 | 75360001 | 75460000 | 0.0023 | 0.0015 | 0.0017 |
| F11 | 75380001 | 75480000 | 0.0021 | 0.0016 | 0.0018 |
| F11 | 75400001 | 75500000 | 0.0025 | 0.0020 | 0.0024 |
| F11 | 75420001 | 75520000 | 0.0028 | 0.0023 | 0.0028 |
| F11 | 75440001 | 75540000 | 0.0039 | 0.0033 | 0.0038 |
| F11 | 75460001 | 75560000 | 0.0048 | 0.0041 | 0.0046 |
| F11 | 75480001 | 75580000 | 0.0047 | 0.0041 | 0.0046 |
| F11 | 75500001 | 75600000 | 0.0049 | 0.0043 | 0.0047 |
| F11 | 75520001 | 75620000 | 0.0054 | 0.0046 | 0.0052 |
| F11 | 75540001 | 75640000 | 0.0050 | 0.0044 | 0.0049 |
| F11 | 75560001 | 75660000 | 0.0049 | 0.0039 | 0.0048 |
| F11 | 75580001 | 75680000 | 0.0051 | 0.0041 | 0.0050 |
| F11 | 75600001 | 75700000 | 0.0039 | 0.0029 | 0.0039 |
| F11 | 75620001 | 75720000 | 0.0033 | 0.0024 | 0.0033 |
| F11 | 75640001 | 75740000 | 0.0026 | 0.0018 | 0.0026 |
| F11 | 75660001 | 75760000 | 0.0022 | 0.0018 | 0.0022 |
| F11 | 75680001 | 75780000 | 0.0022 | 0.0018 | 0.0022 |
| F11 | 75700001 | 75800000 | 0.0022 | 0.0018 | 0.0022 |
| F11 | 75720001 | 75820000 | 0.0019 | 0.0015 | 0.0018 |

| Chr | Start | End | both | A1 | A2 |
|---|---|---|---|---|---|
| F11 | 75740001 | 75840000 | 0.0018 | 0.0014 | 0.0018 |
| F11 | 75760001 | 75860000 | 0.0024 | 0.0016 | 0.0024 |
| F11 | 75780001 | 75880000 | 0.0019 | 0.0012 | 0.0020 |
| F11 | 75800001 | 75900000 | 0.0018 | 0.0011 | 0.0018 |
| F11 | 75820001 | 75920000 | 0.0029 | 0.0020 | 0.0028 |
| F11 | 75840001 | 75940000 | 0.0025 | 0.0016 | 0.0024 |
| F11 | 75860001 | 75960000 | 0.0018 | 0.0014 | 0.0017 |
| F11 | 75880001 | 75980000 | 0.0020 | 0.0015 | 0.0020 |
| F11 | 75900001 | 76000000 | 0.0024 | 0.0019 | 0.0024 |
| F11 | 75920001 | 76020000 | 0.0020 | 0.0016 | 0.0020 |
| F11 | 75940001 | 76040000 | 0.0039 | 0.0030 | 0.0038 |
| F11 | 75960001 | 76060000 | 0.0037 | 0.0029 | 0.0036 |
| F11 | 75980001 | 76080000 | 0.0034 | 0.0027 | 0.0034 |
| F11 | 76000001 | 76100000 | 0.0029 | 0.0023 | 0.0028 |
| F11 | 76020001 | 76120000 | 0.0032 | 0.0026 | 0.0032 |
| F11 | 76040001 | 76140000 | 0.0015 | 0.0013 | 0.0015 |
| F11 | 76060001 | 76160000 | 0.0019 | 0.0014 | 0.0019 |
| F11 | 76080001 | 76180000 | 0.0020 | 0.0015 | 0.0020 |
| F11 | 76100001 | 76200000 | 0.0023 | 0.0018 | 0.0023 |
| F11 | 76120001 | 76220000 | 0.0016 | 0.0012 | 0.0016 |
| F11 | 76140001 | 76240000 | 0.0015 | 0.0010 | 0.0015 |
| F11 | 76160001 | 76260000 | 0.0018 | 0.0016 | 0.0018 |
| F11 | 76180001 | 76280000 | 0.0029 | 0.0024 | 0.0027 |
| F11 | 76200001 | 76300000 | 0.0027 | 0.0021 | 0.0025 |
| F11 | 76220001 | 76320000 | 0.0026 | 0.0020 | 0.0023 |
| F11 | 76240001 | 76340000 | 0.0036 | 0.0027 | 0.0030 |
| F11 | 76260001 | 76360000 | 0.0031 | 0.0022 | 0.0026 |
| F11 | 76280001 | 76380000 | 0.0023 | 0.0014 | 0.0017 |
| F11 | 76300001 | 76400000 | 0.0026 | 0.0016 | 0.0019 |
| F11 | 76320001 | 76420000 | 0.0030 | 0.0020 | 0.0023 |
| F11 | 76340001 | 76440000 | 0.0027 | 0.0017 | 0.0019 |
| F11 | 76360001 | 76460000 | 0.0027 | 0.0013 | 0.0016 |
| F11 | 76380001 | 76480000 | 0.0032 | 0.0018 | 0.0021 |
| F11 | 76400001 | 76500000 | 0.0035 | 0.0020 | 0.0024 |
| F11 | 76420001 | 76520000 | 0.0033 | 0.0018 | 0.0023 |
| F11 | 76440001 | 76540000 | 0.0030 | 0.0016 | 0.0021 |
| F11 | 76460001 | 76560000 | 0.0038 | 0.0026 | 0.0032 |
| F11 | 76480001 | 76580000 | 0.0031 | 0.0021 | 0.0026 |
| F11 | 76500001 | 76600000 | 0.0028 | 0.0017 | 0.0022 |
| F11 | 76520001 | 76620000 | 0.0022 | 0.0013 | 0.0018 |
| F11 | 76540001 | 76640000 | 0.0028 | 0.0018 | 0.0023 |
| F11 | 76560001 | 76660000 | 0.0019 | 0.0011 | 0.0015 |
| F11 | 76580001 | 76680000 | 0.0027 | 0.0018 | 0.0022 |
| F11 | 76600001 | 76700000 | 0.0030 | 0.0022 | 0.0025 |
| F11 | 76620001 | 76720000 | 0.0040 | 0.0029 | 0.0032 |
| F11 | 76640001 | 76740000 | 0.0055 | 0.0034 | 0.0035 |
| F11 | 76660001 | 76760000 | 0.0058 | 0.0034 | 0.0033 |
| F11 | 76680001 | 76780000 | 0.0054 | 0.0029 | 0.0028 |
| F11 | 76700001 | 76800000 | 0.0057 | 0.0030 | 0.0029 |
| F11 | 76720001 | 76820000 | 0.0059 | 0.0028 | 0.0027 |
| F11 | 76740001 | 76840000 | 0.0045 | 0.0022 | 0.0020 |
| F11 | 76760001 | 76860000 | 0.0046 | 0.0021 | 0.0022 |
| F11 | 76780001 | 76880000 | 0.0044 | 0.0021 | 0.0022 |
| F11 | 76800001 | 76900000 | 0.0043 | 0.0021 | 0.0022 |
| F11 | 76820001 | 76920000 | 0.0036 | 0.0018 | 0.0019 |
| F11 | 76840001 | 76940000 | 0.0030 | 0.0014 | 0.0016 |
| F11 | 76860001 | 76960000 | 0.0026 | 0.0014 | 0.0015 |
| F11 | 76880001 | 76980000 | 0.0031 | 0.0019 | 0.0020 |

| Chr | Start | End | both | A1 | A2 |
|---|---|---|---|---|---|
| F11 | 76900001 | 77000000 | 0.0027 | 0.0017 | 0.0019 |
| F11 | 76920001 | 77020000 | 0.0042 | 0.0032 | 0.0035 |
| F11 | 76940001 | 77040000 | 0.0051 | 0.0038 | 0.0043 |
| F11 | 76960001 | 77060000 | 0.0062 | 0.0048 | 0.0055 |
| F11 | 76980001 | 77080000 | 0.0063 | 0.0045 | 0.0054 |
| F11 | 77000001 | 77100000 | 0.0060 | 0.0043 | 0.0051 |
| F11 | 77020001 | 77120000 | 0.0043 | 0.0028 | 0.0034 |
| F11 | 77040001 | 77140000 | 0.0033 | 0.0022 | 0.0027 |
| F11 | 77060001 | 77160000 | 0.0020 | 0.0009 | 0.0013 |
| F11 | 77080001 | 77180000 | 0.0012 | 0.0005 | 0.0006 |
| F11 | 77100001 | 77200000 | 0.0009 | 0.0003 | 0.0004 |
| F11 | 77120001 | 77220000 | 0.0010 | 0.0003 | 0.0004 |
| F11 | 77140001 | 77240000 | 0.0016 | 0.0007 | 0.0008 |
| F11 | 77160001 | 77260000 | 0.0028 | 0.0015 | 0.0018 |
| F11 | 77180001 | 77280000 | 0.0029 | 0.0016 | 0.0019 |
| F11 | 77200001 | 77300000 | 0.0036 | 0.0022 | 0.0026 |
| F11 | 77220001 | 77320000 | 0.0044 | 0.0024 | 0.0036 |
| F11 | 77240001 | 77340000 | 0.0051 | 0.0031 | 0.0043 |
| F11 | 77260001 | 77360000 | 0.0048 | 0.0028 | 0.0041 |
| F11 | 77280001 | 77380000 | 0.0050 | 0.0030 | 0.0044 |
| F11 | 77300001 | 77400000 | 0.0048 | 0.0029 | 0.0041 |
| F11 | 77320001 | 77420000 | 0.0040 | 0.0027 | 0.0032 |
| F11 | 77340001 | 77440000 | 0.0035 | 0.0024 | 0.0026 |
| F11 | 77360001 | 77460000 | 0.0032 | 0.0024 | 0.0025 |
| F11 | 77380001 | 77480000 | 0.0049 | 0.0039 | 0.0042 |
| F11 | 77400001 | 77500000 | 0.0056 | 0.0044 | 0.0048 |
| F11 | 77420001 | 77520000 | 0.0057 | 0.0045 | 0.0050 |
| F11 | 77440001 | 77540000 | 0.0057 | 0.0046 | 0.0049 |
| F11 | 77460001 | 77560000 | 0.0056 | 0.0044 | 0.0047 |
| F11 | 77480001 | 77580000 | 0.0037 | 0.0025 | 0.0027 |
| F11 | 77500001 | 77600000 | 0.0038 | 0.0027 | 0.0031 |
| F11 | 77520001 | 77620000 | 0.0039 | 0.0029 | 0.0032 |
| F11 | 77540001 | 77640000 | 0.0037 | 0.0028 | 0.0032 |
| F11 | 77560001 | 77660000 | 0.0047 | 0.0038 | 0.0043 |
| F11 | 77580001 | 77680000 | 0.0054 | 0.0045 | 0.0050 |
| F11 | 77600001 | 77700000 | 0.0040 | 0.0033 | 0.0035 |
| F11 | 77620001 | 77720000 | 0.0037 | 0.0031 | 0.0033 |
| F11 | 77640001 | 77740000 | 0.0030 | 0.0023 | 0.0025 |
| F11 | 77660001 | 77760000 | 0.0013 | 0.0010 | 0.0011 |
| F11 | 77680001 | 77780000 | 0.0006 | 0.0004 | 0.0004 |
| F11 | 77700001 | 77800000 | 0.0008 | 0.0006 | 0.0007 |
| F11 | 77720001 | 77820000 | 0.0010 | 0.0004 | 0.0005 |
| F11 | 77740001 | 77840000 | 0.0023 | 0.0013 | 0.0016 |
| F11 | 77760001 | 77860000 | 0.0031 | 0.0019 | 0.0023 |
| F11 | 77780001 | 77880000 | 0.0034 | 0.0020 | 0.0023 |
| F11 | 77800001 | 77900000 | 0.0034 | 0.0021 | 0.0023 |
| F11 | 77820001 | 77920000 | 0.0035 | 0.0023 | 0.0026 |
| F11 | 77840001 | 77940000 | 0.0029 | 0.0020 | 0.0021 |
| F11 | 77860001 | 77960000 | 0.0026 | 0.0017 | 0.0018 |
| F11 | 77880001 | 77980000 | 0.0027 | 0.0020 | 0.0021 |
| F11 | 77900001 | 78000000 | 0.0034 | 0.0025 | 0.0026 |
| F11 | 77920001 | 78020000 | 0.0031 | 0.0023 | 0.0025 |
| F11 | 77940001 | 78040000 | 0.0030 | 0.0023 | 0.0025 |
| F11 | 77960001 | 78060000 | 0.0034 | 0.0027 | 0.0029 |
| F11 | 77980001 | 78080000 | 0.0031 | 0.0024 | 0.0026 |
| F11 | 78000001 | 78100000 | 0.0022 | 0.0018 | 0.0020 |
| F11 | 78020001 | 78120000 | 0.0020 | 0.0017 | 0.0018 |
| F11 | 78040001 | 78140000 | 0.0019 | 0.0016 | 0.0018 |

| Chr | Start | End | both | A1 | A2 |
|---|---|---|---|---|---|
| F11 | 78060001 | 78160000 | 0.0011 | 0.0009 | 0.0011 |
| F11 | 78080001 | 78180000 | 0.0010 | 0.0009 | 0.0010 |
| F11 | 78100001 | 78200000 | 0.0010 | 0.0008 | 0.0010 |
| F11 | 78120001 | 78220000 | 0.0012 | 0.0010 | 0.0012 |
| F11 | 78140001 | 78240000 | 0.0015 | 0.0012 | 0.0014 |
| F11 | 78160001 | 78260000 | 0.0016 | 0.0014 | 0.0015 |
| F11 | 78180001 | 78280000 | 0.0022 | 0.0018 | 0.0021 |
| F11 | 78200001 | 78300000 | 0.0038 | 0.0033 | 0.0036 |
| F11 | 78220001 | 78320000 | 0.0045 | 0.0038 | 0.0041 |
| F11 | 78240001 | 78340000 | 0.0048 | 0.0040 | 0.0043 |
| F11 | 78260001 | 78360000 | 0.0050 | 0.0040 | 0.0043 |
| F11 | 78280001 | 78380000 | 0.0046 | 0.0036 | 0.0040 |
| F11 | 78300001 | 78400000 | 0.0028 | 0.0021 | 0.0024 |
| F11 | 78320001 | 78420000 | 0.0021 | 0.0015 | 0.0018 |
| F11 | 78340001 | 78440000 | 0.0010 | 0.0007 | 0.0008 |
| F11 | 78360001 | 78460000 | 0.0009 | 0.0007 | 0.0009 |
| F11 | 78380001 | 78480000 | 0.0006 | 0.0005 | 0.0005 |
| F11 | 78400001 | 78500000 | 0.0009 | 0.0007 | 0.0009 |
| F11 | 78420001 | 78520000 | 0.0010 | 0.0008 | 0.0010 |
| F11 | 78440001 | 78540000 | 0.0010 | 0.0009 | 0.0010 |
| F11 | 78460001 | 78560000 | 0.0010 | 0.0008 | 0.0010 |
| F11 | 78480001 | 78580000 | 0.0011 | 0.0010 | 0.0011 |
| F11 | 78500001 | 78600000 | 0.0010 | 0.0009 | 0.0009 |
| F11 | 78520001 | 78620000 | 0.0010 | 0.0008 | 0.0009 |
| F11 | 78540001 | 78640000 | 0.0010 | 0.0008 | 0.0008 |
| F11 | 78560001 | 78660000 | 0.0017 | 0.0013 | 0.0014 |
| F11 | 78580001 | 78680000 | 0.0022 | 0.0017 | 0.0018 |
| F11 | 78600001 | 78700000 | 0.0031 | 0.0027 | 0.0028 |
| F11 | 78620001 | 78720000 | 0.0039 | 0.0031 | 0.0033 |
| F11 | 78640001 | 78740000 | 0.0047 | 0.0037 | 0.0040 |
| F11 | 78660001 | 78760000 | 0.0041 | 0.0031 | 0.0033 |
| F11 | 78680001 | 78780000 | 0.0039 | 0.0028 | 0.0030 |
| F11 | 78700001 | 78800000 | 0.0034 | 0.0021 | 0.0023 |
| F11 | 78720001 | 78820000 | 0.0030 | 0.0016 | 0.0017 |
| F11 | 78740001 | 78840000 | 0.0024 | 0.0011 | 0.0012 |
| F11 | 78760001 | 78860000 | 0.0028 | 0.0014 | 0.0016 |
| F11 | 78780001 | 78880000 | 0.0028 | 0.0014 | 0.0017 |
| F11 | 78800001 | 78900000 | 0.0034 | 0.0020 | 0.0023 |
| F11 | 78820001 | 78920000 | 0.0031 | 0.0020 | 0.0023 |
| F11 | 78840001 | 78940000 | 0.0029 | 0.0018 | 0.0020 |
| F11 | 78860001 | 78960000 | 0.0027 | 0.0015 | 0.0018 |
| F11 | 78880001 | 78980000 | 0.0031 | 0.0018 | 0.0020 |
| F11 | 78900001 | 79000000 | 0.0027 | 0.0014 | 0.0018 |
| F11 | 78920001 | 79020000 | 0.0036 | 0.0022 | 0.0025 |
| F11 | 78940001 | 79040000 | 0.0045 | 0.0031 | 0.0035 |
| F11 | 78960001 | 79060000 | 0.0044 | 0.0031 | 0.0034 |
| F11 | 78980001 | 79080000 | 0.0043 | 0.0032 | 0.0034 |
| F11 | 79000001 | 79100000 | 0.0043 | 0.0031 | 0.0033 |
| F11 | 79020001 | 79120000 | 0.0034 | 0.0024 | 0.0027 |
| F11 | 79040001 | 79140000 | 0.0027 | 0.0016 | 0.0019 |
| F11 | 79060001 | 79160000 | 0.0036 | 0.0022 | 0.0024 |
| F11 | 79080001 | 79180000 | 0.0035 | 0.0021 | 0.0024 |
| F11 | 79100001 | 79200000 | 0.0027 | 0.0015 | 0.0017 |
| F11 | 79120001 | 79220000 | 0.0028 | 0.0016 | 0.0018 |
| F11 | 79140001 | 79240000 | 0.0033 | 0.0021 | 0.0025 |
| F11 | 79160001 | 79260000 | 0.0027 | 0.0019 | 0.0024 |
| F11 | 79180001 | 79280000 | 0.0028 | 0.0019 | 0.0024 |
| F11 | 79200001 | 79300000 | 0.0029 | 0.0020 | 0.0026 |

| Chr | Start | End | both | A1 | A2 |
|---|---|---|---|---|---|
| F11 | 79220001 | 79320000 | 0.0030 | 0.0020 | 0.0026 |
| F11 | 79240001 | 79340000 | 0.0027 | 0.0016 | 0.0021 |
| F11 | 79260001 | 79360000 | 0.0026 | 0.0014 | 0.0019 |
| F11 | 79280001 | 79380000 | 0.0020 | 0.0010 | 0.0014 |
| F11 | 79300001 | 79400000 | 0.0022 | 0.0010 | 0.0014 |
| F11 | 79320001 | 79420000 | 0.0021 | 0.0010 | 0.0014 |
| F11 | 79340001 | 79440000 | 0.0025 | 0.0015 | 0.0020 |
| F11 | 79360001 | 79460000 | 0.0031 | 0.0021 | 0.0027 |
| F11 | 79380001 | 79480000 | 0.0035 | 0.0024 | 0.0031 |
| F11 | 79400001 | 79500000 | 0.0034 | 0.0026 | 0.0032 |
| F11 | 79420001 | 79520000 | 0.0032 | 0.0025 | 0.0030 |
| F11 | 79440001 | 79540000 | 0.0022 | 0.0017 | 0.0021 |
| F11 | 79460001 | 79560000 | 0.0016 | 0.0009 | 0.0012 |
| F11 | 79480001 | 79580000 | 0.0016 | 0.0009 | 0.0012 |
| F11 | 79500001 | 79600000 | 0.0022 | 0.0013 | 0.0016 |
| F11 | 79520001 | 79620000 | 0.0021 | 0.0013 | 0.0017 |
| F11 | 79540001 | 79640000 | 0.0024 | 0.0015 | 0.0019 |
| F11 | 79560001 | 79660000 | 0.0024 | 0.0017 | 0.0021 |
| F11 | 79580001 | 79680000 | 0.0025 | 0.0018 | 0.0023 |
| F11 | 79600001 | 79700000 | 0.0025 | 0.0019 | 0.0023 |
| F11 | 79620001 | 79720000 | 0.0027 | 0.0019 | 0.0023 |
| F11 | 79640001 | 79740000 | 0.0037 | 0.0029 | 0.0032 |
| F11 | 79660001 | 79760000 | 0.0039 | 0.0031 | 0.0034 |
| F11 | 79680001 | 79780000 | 0.0037 | 0.0029 | 0.0032 |
| F11 | 79700001 | 79800000 | 0.0032 | 0.0026 | 0.0028 |
| F11 | 79720001 | 79820000 | 0.0033 | 0.0026 | 0.0028 |
| F11 | 79740001 | 79840000 | 0.0030 | 0.0023 | 0.0026 |
| F11 | 79760001 | 79860000 | 0.0034 | 0.0028 | 0.0030 |
| F11 | 79780001 | 79880000 | 0.0039 | 0.0032 | 0.0035 |
| F11 | 79800001 | 79900000 | 0.0036 | 0.0030 | 0.0033 |
| F11 | 79820001 | 79920000 | 0.0032 | 0.0029 | 0.0030 |
| F11 | 79840001 | 79940000 | 0.0023 | 0.0020 | 0.0022 |
| F11 | 79860001 | 79960000 | 0.0015 | 0.0014 | 0.0015 |
| F11 | 79880001 | 79980000 | 0.0010 | 0.0010 | 0.0010 |
| F11 | 79900001 | 80000000 | 0.0009 | 0.0009 | 0.0009 |
| F11 | 79920001 | 80020000 | 0.0011 | 0.0010 | 0.0010 |
| F11 | 79940001 | 80040000 | 0.0011 | 0.0012 | 0.0011 |
| F11 | 79960001 | 80060000 | 0.0009 | 0.0009 | 0.0008 |
| F11 | 79980001 | 80080000 | 0.0005 | 0.0005 | 0.0004 |
| F11 | 80000001 | 80100000 | 0.0005 | 0.0005 | 0.0005 |
| F11 | 80020001 | 80120000 | 0.0009 | 0.0009 | 0.0009 |
| F11 | 80040001 | 80140000 | 0.0019 | 0.0018 | 0.0018 |
| F11 | 80060001 | 80160000 | 0.0020 | 0.0018 | 0.0019 |
| F11 | 80080001 | 80180000 | 0.0022 | 0.0020 | 0.0021 |
| F11 | 80100001 | 80200000 | 0.0030 | 0.0028 | 0.0028 |
| F11 | 80120001 | 80220000 | 0.0033 | 0.0029 | 0.0030 |
| F11 | 80140001 | 80240000 | 0.0027 | 0.0023 | 0.0024 |
| F11 | 80160001 | 80260000 | 0.0028 | 0.0022 | 0.0023 |
| F11 | 80180001 | 80280000 | 0.0035 | 0.0027 | 0.0028 |
| F11 | 80200001 | 80300000 | 0.0035 | 0.0024 | 0.0028 |
| F11 | 80220001 | 80320000 | 0.0033 | 0.0021 | 0.0023 |
| F11 | 80240001 | 80340000 | 0.0035 | 0.0021 | 0.0024 |
| F11 | 80260001 | 80360000 | 0.0041 | 0.0028 | 0.0031 |
| F11 | 80280001 | 80380000 | 0.0038 | 0.0025 | 0.0029 |
| F11 | 80300001 | 80400000 | 0.0033 | 0.0023 | 0.0025 |
| F11 | 80320001 | 80420000 | 0.0031 | 0.0020 | 0.0023 |
| F11 | 80340001 | 80440000 | 0.0030 | 0.0020 | 0.0023 |
| F11 | 80360001 | 80460000 | 0.0028 | 0.0019 | 0.0021 |

| Chr | Start | End | both | A1 | A2 |
|---|---|---|---|---|---|
| F11 | 80380001 | 80480000 | 0.0046 | 0.0035 | 0.0038 |
| F11 | 80400001 | 80500000 | 0.0049 | 0.0036 | 0.0040 |
| F11 | 80420001 | 80520000 | 0.0057 | 0.0045 | 0.0050 |
| F11 | 80440001 | 80540000 | 0.0051 | 0.0040 | 0.0044 |
| F11 | 80460001 | 80560000 | 0.0045 | 0.0035 | 0.0039 |
| F11 | 80480001 | 80580000 | 0.0025 | 0.0017 | 0.0021 |
| F11 | 80500001 | 80600000 | 0.0022 | 0.0017 | 0.0021 |
| F11 | 80520001 | 80620000 | 0.0018 | 0.0015 | 0.0018 |
| F11 | 80540001 | 80640000 | 0.0018 | 0.0014 | 0.0018 |
| F11 | 80560001 | 80660000 | 0.0020 | 0.0016 | 0.0020 |
| F11 | 80580001 | 80680000 | 0.0020 | 0.0016 | 0.0020 |
| F11 | 80600001 | 80700000 | 0.0019 | 0.0014 | 0.0016 |
| F11 | 80620001 | 80720000 | 0.0018 | 0.0013 | 0.0016 |
| F11 | 80640001 | 80740000 | 0.0027 | 0.0020 | 0.0023 |
| F11 | 80660001 | 80760000 | 0.0029 | 0.0022 | 0.0026 |
| F11 | 80680001 | 80780000 | 0.0037 | 0.0028 | 0.0033 |
| F11 | 80700001 | 80800000 | 0.0040 | 0.0030 | 0.0035 |
| F11 | 80720001 | 80820000 | 0.0036 | 0.0025 | 0.0030 |
| F11 | 80740001 | 80840000 | 0.0028 | 0.0019 | 0.0023 |
| F11 | 80760001 | 80860000 | 0.0025 | 0.0015 | 0.0020 |
| F11 | 80780001 | 80880000 | 0.0017 | 0.0011 | 0.0014 |
| F11 | 80800001 | 80900000 | 0.0011 | 0.0008 | 0.0010 |
| F11 | 80820001 | 80920000 | 0.0009 | 0.0007 | 0.0009 |
| F11 | 80840001 | 80940000 | 0.0018 | 0.0013 | 0.0017 |
| F11 | 80860001 | 80960000 | 0.0018 | 0.0013 | 0.0017 |
| F11 | 80880001 | 80980000 | 0.0019 | 0.0014 | 0.0019 |
| F11 | 80900001 | 81000000 | 0.0020 | 0.0016 | 0.0020 |
| F11 | 80920001 | 81020000 | 0.0024 | 0.0019 | 0.0023 |
| F11 | 80940001 | 81040000 | 0.0029 | 0.0024 | 0.0027 |
| F11 | 80960001 | 81060000 | 0.0033 | 0.0027 | 0.0031 |
| F11 | 80980001 | 81080000 | 0.0031 | 0.0026 | 0.0029 |
| F11 | 81000001 | 81100000 | 0.0033 | 0.0027 | 0.0030 |
| F11 | 81020001 | 81120000 | 0.0035 | 0.0028 | 0.0031 |
| F11 | 81040001 | 81140000 | 0.0023 | 0.0018 | 0.0021 |
| F11 | 81060001 | 81160000 | 0.0023 | 0.0019 | 0.0022 |
| F11 | 81080001 | 81180000 | 0.0026 | 0.0022 | 0.0024 |
| F11 | 81100001 | 81200000 | 0.0033 | 0.0029 | 0.0032 |
| F11 | 81120001 | 81220000 | 0.0031 | 0.0027 | 0.0030 |
| F11 | 81140001 | 81240000 | 0.0035 | 0.0030 | 0.0035 |
| F11 | 81160001 | 81260000 | 0.0039 | 0.0033 | 0.0039 |
| F11 | 81180001 | 81280000 | 0.0036 | 0.0030 | 0.0036 |
| F11 | 81200001 | 81300000 | 0.0027 | 0.0022 | 0.0027 |
| F11 | 81220001 | 81320000 | 0.0025 | 0.0020 | 0.0025 |
| F11 | 81240001 | 81340000 | 0.0019 | 0.0016 | 0.0019 |
| F11 | 81260001 | 81360000 | 0.0010 | 0.0010 | 0.0010 |
| F11 | 81280001 | 81380000 | 0.0014 | 0.0013 | 0.0014 |
| F11 | 81300001 | 81400000 | 0.0018 | 0.0017 | 0.0017 |
| F11 | 81320001 | 81420000 | 0.0026 | 0.0025 | 0.0025 |
| F11 | 81340001 | 81440000 | 0.0028 | 0.0027 | 0.0027 |
| F11 | 81360001 | 81460000 | 0.0039 | 0.0038 | 0.0038 |
| F11 | 81380001 | 81480000 | 0.0039 | 0.0037 | 0.0038 |
| F11 | 81400001 | 81500000 | 0.0040 | 0.0037 | 0.0039 |
| F11 | 81420001 | 81520000 | 0.0032 | 0.0029 | 0.0032 |
| F11 | 81440001 | 81540000 | 0.0030 | 0.0027 | 0.0030 |
| F11 | 81460001 | 81560000 | 0.0018 | 0.0015 | 0.0018 |
| F11 | 81480001 | 81580000 | 0.0012 | 0.0010 | 0.0012 |
| F11 | 81500001 | 81600000 | 0.0010 | 0.0007 | 0.0010 |
| F11 | 81520001 | 81620000 | 0.0014 | 0.0011 | 0.0014 |

| Chr | Start | End | both | A1 | A2 |
|---|---|---|---|---|---|
| F11 | 81540001 | 81640000 | 0.0014 | 0.0012 | 0.0014 |
| F11 | 81560001 | 81660000 | 0.0021 | 0.0016 | 0.0020 |
| F11 | 81580001 | 81680000 | 0.0023 | 0.0018 | 0.0022 |
| F11 | 81600001 | 81700000 | 0.0026 | 0.0019 | 0.0023 |
| F11 | 81620001 | 81720000 | 0.0027 | 0.0020 | 0.0024 |
| F11 | 81640001 | 81740000 | 0.0027 | 0.0019 | 0.0024 |
| F11 | 81660001 | 81760000 | 0.0030 | 0.0021 | 0.0027 |
| F11 | 81680001 | 81780000 | 0.0032 | 0.0021 | 0.0028 |
| F11 | 81700001 | 81800000 | 0.0030 | 0.0019 | 0.0026 |
| F11 | 81720001 | 81820000 | 0.0032 | 0.0020 | 0.0026 |
| F11 | 81740001 | 81840000 | 0.0040 | 0.0027 | 0.0034 |
| F11 | 81760001 | 81860000 | 0.0036 | 0.0025 | 0.0030 |
| F11 | 81780001 | 81880000 | 0.0041 | 0.0029 | 0.0034 |
| F11 | 81800001 | 81900000 | 0.0042 | 0.0031 | 0.0035 |
| F11 | 81820001 | 81920000 | 0.0038 | 0.0025 | 0.0030 |
| F11 | 81840001 | 81940000 | 0.0034 | 0.0021 | 0.0026 |
| F11 | 81860001 | 81960000 | 0.0035 | 0.0019 | 0.0025 |
| F11 | 81880001 | 81980000 | 0.0034 | 0.0019 | 0.0025 |
| F11 | 81900001 | 82000000 | 0.0028 | 0.0016 | 0.0021 |
| F11 | 81920001 | 82020000 | 0.0025 | 0.0015 | 0.0019 |
| F11 | 81940001 | 82040000 | 0.0023 | 0.0015 | 0.0018 |
| F11 | 81960001 | 82060000 | 0.0021 | 0.0017 | 0.0019 |
| F11 | 81980001 | 82080000 | 0.0012 | 0.0009 | 0.0009 |
| F11 | 82000001 | 82100000 | 0.0013 | 0.0009 | 0.0010 |
| F11 | 82020001 | 82120000 | 0.0020 | 0.0015 | 0.0016 |
| F11 | 82040001 | 82140000 | 0.0018 | 0.0013 | 0.0014 |
| F11 | 82060001 | 82160000 | 0.0018 | 0.0011 | 0.0012 |
| F11 | 82080001 | 82180000 | 0.0029 | 0.0020 | 0.0022 |
| F11 | 82100001 | 82200000 | 0.0036 | 0.0024 | 0.0026 |
| F11 | 82120001 | 82220000 | 0.0031 | 0.0020 | 0.0023 |
| F11 | 82140001 | 82240000 | 0.0031 | 0.0018 | 0.0022 |
| F11 | 82160001 | 82260000 | 0.0035 | 0.0024 | 0.0028 |
| F11 | 82180001 | 82280000 | 0.0030 | 0.0019 | 0.0024 |
| F11 | 82200001 | 82300000 | 0.0027 | 0.0016 | 0.0021 |
| F11 | 82220001 | 82320000 | 0.0027 | 0.0015 | 0.0019 |
| F11 | 82240001 | 82340000 | 0.0031 | 0.0015 | 0.0020 |
| F11 | 82260001 | 82360000 | 0.0033 | 0.0012 | 0.0017 |
| F11 | 82280001 | 82380000 | 0.0033 | 0.0011 | 0.0016 |
| F11 | 82300001 | 82400000 | 0.0034 | 0.0012 | 0.0017 |
| F11 | 82320001 | 82420000 | 0.0033 | 0.0012 | 0.0017 |
| F11 | 82340001 | 82440000 | 0.0029 | 0.0012 | 0.0015 |
| F11 | 82360001 | 82460000 | 0.0032 | 0.0019 | 0.0021 |
| F11 | 82380001 | 82480000 | 0.0032 | 0.0020 | 0.0022 |
| F11 | 82400001 | 82500000 | 0.0035 | 0.0021 | 0.0023 |
| F11 | 82420001 | 82520000 | 0.0036 | 0.0022 | 0.0025 |
| F11 | 82440001 | 82540000 | 0.0038 | 0.0024 | 0.0026 |
| F11 | 82460001 | 82560000 | 0.0035 | 0.0022 | 0.0023 |
| F11 | 82480001 | 82580000 | 0.0031 | 0.0021 | 0.0020 |
| F11 | 82500001 | 82600000 | 0.0031 | 0.0024 | 0.0023 |
| F11 | 82520001 | 82620000 | 0.0031 | 0.0025 | 0.0023 |
| F11 | 82540001 | 82640000 | 0.0030 | 0.0024 | 0.0022 |
| F11 | 82560001 | 82660000 | 0.0022 | 0.0015 | 0.0014 |
| F11 | 82580001 | 82680000 | 0.0021 | 0.0014 | 0.0013 |
| F11 | 82600001 | 82700000 | 0.0015 | 0.0007 | 0.0007 |
| F11 | 82620001 | 82720000 | 0.0015 | 0.0006 | 0.0007 |
| F11 | 82640001 | 82740000 | 0.0019 | 0.0007 | 0.0009 |
| F11 | 82660001 | 82760000 | 0.0021 | 0.0007 | 0.0010 |
| F11 | 82680001 | 82780000 | 0.0031 | 0.0017 | 0.0021 |

| Chr | Start | End | both | A1 | A2 |
|---|---|---|---|---|---|
| F11 | 82700001 | 82800000 | 0.0036 | 0.0023 | 0.0028 |
| F11 | 82720001 | 82820000 | 0.0038 | 0.0026 | 0.0031 |
| F11 | 82740001 | 82840000 | 0.0038 | 0.0025 | 0.0030 |
| F11 | 82760001 | 82860000 | 0.0047 | 0.0034 | 0.0038 |
| F11 | 82780001 | 82880000 | 0.0044 | 0.0031 | 0.0035 |
| F11 | 82800001 | 82900000 | 0.0043 | 0.0031 | 0.0034 |
| F11 | 82820001 | 82920000 | 0.0044 | 0.0031 | 0.0034 |
| F11 | 82840001 | 82940000 | 0.0042 | 0.0031 | 0.0034 |
| F11 | 82860001 | 82960000 | 0.0032 | 0.0021 | 0.0023 |
| F11 | 82880001 | 82980000 | 0.0029 | 0.0019 | 0.0019 |
| F11 | 82900001 | 83000000 | 0.0032 | 0.0017 | 0.0021 |
| F11 | 82920001 | 83020000 | 0.0029 | 0.0014 | 0.0018 |
| F11 | 82940001 | 83040000 | 0.0036 | 0.0020 | 0.0023 |
| F11 | 82960001 | 83060000 | 0.0045 | 0.0029 | 0.0032 |
| F11 | 82980001 | 83080000 | 0.0046 | 0.0030 | 0.0035 |
| F11 | 83000001 | 83100000 | 0.0047 | 0.0034 | 0.0034 |
| F11 | 83020001 | 83120000 | 0.0049 | 0.0037 | 0.0037 |
| F11 | 83040001 | 83140000 | 0.0047 | 0.0036 | 0.0037 |
| F11 | 83060001 | 83160000 | 0.0040 | 0.0031 | 0.0032 |
| F11 | 83080001 | 83180000 | 0.0036 | 0.0028 | 0.0029 |
| F11 | 83100001 | 83200000 | 0.0027 | 0.0022 | 0.0023 |
| F11 | 83120001 | 83220000 | 0.0045 | 0.0038 | 0.0041 |
| F11 | 83140001 | 83240000 | 0.0039 | 0.0033 | 0.0035 |
| F11 | 83160001 | 83260000 | 0.0035 | 0.0029 | 0.0030 |
| F11 | 83180001 | 83280000 | 0.0039 | 0.0033 | 0.0034 |
| F11 | 83200001 | 83300000 | 0.0044 | 0.0037 | 0.0039 |
| F11 | 83220001 | 83320000 | 0.0028 | 0.0023 | 0.0023 |
| F11 | 83240001 | 83340000 | 0.0036 | 0.0029 | 0.0031 |
| F11 | 83260001 | 83360000 | 0.0046 | 0.0036 | 0.0038 |
| F11 | 83280001 | 83380000 | 0.0038 | 0.0029 | 0.0031 |
| F11 | 83300001 | 83400000 | 0.0038 | 0.0028 | 0.0030 |
| F11 | 83320001 | 83420000 | 0.0039 | 0.0025 | 0.0028 |
| F11 | 83340001 | 83440000 | 0.0032 | 0.0018 | 0.0020 |
| F11 | 83360001 | 83460000 | 0.0025 | 0.0013 | 0.0015 |
| F11 | 83380001 | 83480000 | 0.0029 | 0.0014 | 0.0017 |
| F11 | 83400001 | 83500000 | 0.0027 | 0.0012 | 0.0014 |
| F11 | 83420001 | 83520000 | 0.0034 | 0.0019 | 0.0021 |
| F11 | 83440001 | 83540000 | 0.0048 | 0.0032 | 0.0035 |
| F11 | 83460001 | 83560000 | 0.0049 | 0.0033 | 0.0037 |
| F11 | 83480001 | 83580000 | 0.0049 | 0.0032 | 0.0035 |
| F11 | 83500001 | 83600000 | 0.0054 | 0.0031 | 0.0041 |
| F11 | 83520001 | 83620000 | 0.0044 | 0.0024 | 0.0033 |
| F11 | 83540001 | 83640000 | 0.0041 | 0.0022 | 0.0029 |
| F11 | 83560001 | 83660000 | 0.0044 | 0.0026 | 0.0031 |
| F11 | 83580001 | 83680000 | 0.0050 | 0.0033 | 0.0039 |
| F11 | 83600001 | 83700000 | 0.0044 | 0.0033 | 0.0033 |
| F11 | 83620001 | 83720000 | 0.0044 | 0.0032 | 0.0033 |
| F11 | 83640001 | 83740000 | 0.0033 | 0.0024 | 0.0025 |
| F11 | 83660001 | 83760000 | 0.0038 | 0.0027 | 0.0032 |
| F11 | 83680001 | 83780000 | 0.0036 | 0.0024 | 0.0030 |
| F11 | 83700001 | 83800000 | 0.0035 | 0.0024 | 0.0031 |
| F11 | 83720001 | 83820000 | 0.0032 | 0.0021 | 0.0027 |
| F11 | 83740001 | 83840000 | 0.0032 | 0.0021 | 0.0025 |
| F11 | 83760001 | 83860000 | 0.0024 | 0.0015 | 0.0013 |
| F11 | 83780001 | 83880000 | 0.0027 | 0.0020 | 0.0018 |
| F11 | 83800001 | 83900000 | 0.0031 | 0.0023 | 0.0020 |
| F11 | 83820001 | 83920000 | 0.0032 | 0.0025 | 0.0023 |
| F11 | 83840001 | 83940000 | 0.0041 | 0.0033 | 0.0031 |

| Chr | Start | End | both | A1 | A2 |
|---|---|---|---|---|---|
| F11 | 83860001 | 83960000 | 0.0041 | 0.0034 | 0.0031 |
| F11 | 83880001 | 83980000 | 0.0035 | 0.0029 | 0.0026 |
| F11 | 83900001 | 84000000 | 0.0031 | 0.0026 | 0.0024 |
| F11 | 83920001 | 84020000 | 0.0030 | 0.0025 | 0.0022 |
| F11 | 83940001 | 84040000 | 0.0025 | 0.0020 | 0.0018 |
| F11 | 83960001 | 84060000 | 0.0028 | 0.0023 | 0.0023 |
| F11 | 83980001 | 84080000 | 0.0032 | 0.0027 | 0.0027 |
| F11 | 84000001 | 84100000 | 0.0031 | 0.0026 | 0.0026 |
| F11 | 84020001 | 84120000 | 0.0035 | 0.0030 | 0.0031 |
| F11 | 84040001 | 84140000 | 0.0039 | 0.0035 | 0.0032 |
| F11 | 84060001 | 84160000 | 0.0036 | 0.0032 | 0.0030 |
| F11 | 84080001 | 84180000 | 0.0032 | 0.0028 | 0.0027 |
| F11 | 84100001 | 84200000 | 0.0036 | 0.0033 | 0.0032 |
| F11 | 84120001 | 84220000 | 0.0039 | 0.0035 | 0.0034 |
| F11 | 84140001 | 84240000 | 0.0033 | 0.0026 | 0.0028 |
| F11 | 84160001 | 84260000 | 0.0035 | 0.0029 | 0.0028 |
| F11 | 84180001 | 84280000 | 0.0039 | 0.0041 | 0.0026 |
| F11 | 84200001 | 84300000 | 0.0043 | 0.0052 | 0.0025 |
| F11 | 84220001 | 84320000 | 0.0037 | 0.0050 | 0.0016 |
| F11 | 84240001 | 84340000 | 0.0039 | 0.0053 | 0.0019 |
| F11 | 84260001 | 84360000 | 0.0035 | 0.0049 | 0.0019 |
| F11 | 84280001 | 84380000 | 0.0028 | 0.0033 | 0.0017 |
| F11 | 84300001 | 84400000 | 0.0021 | 0.0017 | 0.0013 |
| F11 | 84320001 | 84420000 | 0.0021 | 0.0011 | 0.0014 |
| F11 | 84340001 | 84440000 | 0.0021 | 0.0010 | 0.0012 |
| F11 | 84360001 | 84460000 | 0.0030 | 0.0017 | 0.0021 |
| F11 | 84380001 | 84480000 | 0.0028 | 0.0015 | 0.0018 |
| F11 | 84400001 | 84500000 | 0.0028 | 0.0015 | 0.0019 |
| F11 | 84420001 | 84520000 | 0.0029 | 0.0018 | 0.0022 |
| F11 | 84440001 | 84540000 | 0.0031 | 0.0021 | 0.0025 |
| F11 | 84460001 | 84560000 | 0.0025 | 0.0016 | 0.0018 |
| F11 | 84480001 | 84580000 | 0.0030 | 0.0021 | 0.0023 |
| F11 | 84500001 | 84600000 | 0.0032 | 0.0022 | 0.0025 |
| F11 | 84520001 | 84620000 | 0.0028 | 0.0020 | 0.0022 |
| F11 | 84540001 | 84640000 | 0.0028 | 0.0018 | 0.0022 |
| F11 | 84560001 | 84660000 | 0.0024 | 0.0014 | 0.0019 |
| F11 | 84580001 | 84680000 | 0.0025 | 0.0014 | 0.0018 |
| F11 | 84600001 | 84700000 | 0.0028 | 0.0015 | 0.0021 |
| F11 | 84620001 | 84720000 | 0.0033 | 0.0018 | 0.0025 |
| F11 | 84640001 | 84740000 | 0.0028 | 0.0015 | 0.0020 |
| F11 | 84660001 | 84760000 | 0.0033 | 0.0021 | 0.0025 |
| F11 | 84680001 | 84780000 | 0.0031 | 0.0020 | 0.0023 |
| F11 | 84700001 | 84800000 | 0.0033 | 0.0025 | 0.0028 |
| F11 | 84720001 | 84820000 | 0.0036 | 0.0030 | 0.0031 |
| F11 | 84740001 | 84840000 | 0.0041 | 0.0032 | 0.0035 |
| F11 | 84760001 | 84860000 | 0.0037 | 0.0029 | 0.0031 |
| F11 | 84780001 | 84880000 | 0.0034 | 0.0025 | 0.0028 |
| F11 | 84800001 | 84900000 | 0.0030 | 0.0021 | 0.0023 |
| F11 | 84820001 | 84920000 | 0.0022 | 0.0011 | 0.0014 |
| F11 | 84840001 | 84940000 | 0.0025 | 0.0014 | 0.0018 |
| F11 | 84860001 | 84960000 | 0.0024 | 0.0013 | 0.0017 |
| F11 | 84880001 | 84980000 | 0.0028 | 0.0019 | 0.0022 |
| F11 | 84900001 | 85000000 | 0.0035 | 0.0024 | 0.0027 |
| F11 | 84920001 | 85020000 | 0.0045 | 0.0034 | 0.0038 |
| F11 | 84940001 | 85040000 | 0.0042 | 0.0031 | 0.0033 |
| F11 | 84960001 | 85060000 | 0.0053 | 0.0041 | 0.0044 |
| F11 | 84980001 | 85080000 | 0.0059 | 0.0044 | 0.0051 |
| F11 | 85000001 | 85100000 | 0.0065 | 0.0051 | 0.0059 |

| Chr | Start | End | both | A1 | A2 |
|---|---|---|---|---|---|
| F11 | 85020001 | 85120000 | 0.0060 | 0.0046 | 0.0053 |
| F11 | 85040001 | 85140000 | 0.0062 | 0.0049 | 0.0056 |
| F11 | 85060001 | 85160000 | 0.0057 | 0.0045 | 0.0052 |
| F11 | 85080001 | 85180000 | 0.0060 | 0.0049 | 0.0053 |
| F11 | 85100001 | 85200000 | 0.0047 | 0.0036 | 0.0040 |
| F11 | 85120001 | 85220000 | 0.0052 | 0.0041 | 0.0045 |
| F11 | 85140001 | 85240000 | 0.0066 | 0.0053 | 0.0058 |
| F11 | 85160001 | 85260000 | 0.0060 | 0.0047 | 0.0052 |
| F11 | 85180001 | 85280000 | 0.0052 | 0.0040 | 0.0045 |
| F11 | 85200001 | 85300000 | 0.0055 | 0.0043 | 0.0049 |
| F11 | 85220001 | 85320000 | 0.0050 | 0.0038 | 0.0042 |
| F11 | 85240001 | 85340000 | 0.0031 | 0.0021 | 0.0023 |
| F11 | 85260001 | 85360000 | 0.0032 | 0.0021 | 0.0023 |
| F11 | 85280001 | 85380000 | 0.0029 | 0.0017 | 0.0020 |
| F11 | 85300001 | 85400000 | 0.0021 | 0.0010 | 0.0011 |
| F11 | 85320001 | 85420000 | 0.0016 | 0.0006 | 0.0007 |
| F11 | 85340001 | 85440000 | 0.0016 | 0.0005 | 0.0006 |
| F11 | 85360001 | 85460000 | 0.0015 | 0.0003 | 0.0004 |
| F11 | 85380001 | 85480000 | 0.0018 | 0.0004 | 0.0005 |
| F11 | 85400001 | 85500000 | 0.0019 | 0.0005 | 0.0006 |
| F11 | 85420001 | 85520000 | 0.0022 | 0.0008 | 0.0009 |
| F11 | 85440001 | 85540000 | 0.0023 | 0.0009 | 0.0011 |
| F11 | 85460001 | 85560000 | 0.0026 | 0.0011 | 0.0015 |
| F11 | 85480001 | 85580000 | 0.0020 | 0.0008 | 0.0012 |
| F11 | 85500001 | 85600000 | 0.0024 | 0.0011 | 0.0016 |
| F11 | 85520001 | 85620000 | 0.0025 | 0.0013 | 0.0017 |
| F11 | 85540001 | 85640000 | 0.0025 | 0.0014 | 0.0018 |
| F11 | 85560001 | 85660000 | 0.0026 | 0.0017 | 0.0020 |
| F11 | 85580001 | 85680000 | 0.0034 | 0.0023 | 0.0027 |
| F11 | 85600001 | 85700000 | 0.0037 | 0.0026 | 0.0030 |
| F11 | 85620001 | 85720000 | 0.0041 | 0.0030 | 0.0033 |
| F11 | 85640001 | 85740000 | 0.0048 | 0.0036 | 0.0040 |
| F11 | 85660001 | 85760000 | 0.0042 | 0.0031 | 0.0035 |
| F11 | 85680001 | 85780000 | 0.0050 | 0.0039 | 0.0042 |
| F11 | 85700001 | 85800000 | 0.0056 | 0.0040 | 0.0045 |
| F11 | 85720001 | 85820000 | 0.0059 | 0.0046 | 0.0051 |
| F11 | 85740001 | 85840000 | 0.0057 | 0.0044 | 0.0048 |
| F11 | 85760001 | 85860000 | 0.0082 | 0.0058 | 0.0070 |
| F11 | 85780001 | 85880000 | 0.0084 | 0.0059 | 0.0072 |
| F11 | 85800001 | 85900000 | 0.0076 | 0.0055 | 0.0066 |
| F11 | 85820001 | 85920000 | 0.0064 | 0.0042 | 0.0053 |
| F11 | 85840001 | 85940000 | 0.0060 | 0.0038 | 0.0048 |
| F11 | 85860001 | 85960000 | 0.0043 | 0.0029 | 0.0032 |
| F11 | 85880001 | 85980000 | 0.0028 | 0.0014 | 0.0017 |
| F11 | 85900001 | 86000000 | 0.0025 | 0.0015 | 0.0016 |
| F11 | 85920001 | 86020000 | 0.0026 | 0.0016 | 0.0017 |
| F11 | 85940001 | 86040000 | 0.0027 | 0.0019 | 0.0020 |
| F11 | 85960001 | 86060000 | 0.0023 | 0.0016 | 0.0018 |
| F11 | 85980001 | 86080000 | 0.0025 | 0.0018 | 0.0020 |
| F11 | 86000001 | 86100000 | 0.0024 | 0.0016 | 0.0018 |
| F11 | 86020001 | 86120000 | 0.0022 | 0.0014 | 0.0016 |
| F11 | 86040001 | 86140000 | 0.0027 | 0.0019 | 0.0021 |
| F11 | 86060001 | 86160000 | 0.0028 | 0.0021 | 0.0022 |
| F11 | 86080001 | 86180000 | 0.0029 | 0.0022 | 0.0023 |
| F11 | 86100001 | 86200000 | 0.0031 | 0.0022 | 0.0024 |
| F11 | 86120001 | 86220000 | 0.0034 | 0.0024 | 0.0027 |
| F11 | 86140001 | 86240000 | 0.0031 | 0.0019 | 0.0023 |
| F11 | 86160001 | 86260000 | 0.0052 | 0.0035 | 0.0042 |

| Chr | Start | End | both | A1 | A2 |
|---|---|---|---|---|---|
| F11 | 86180001 | 86280000 | 0.0056 | 0.0039 | 0.0046 |
| F11 | 86200001 | 86300000 | 0.0056 | 0.0040 | 0.0046 |
| F11 | 86220001 | 86320000 | 0.0064 | 0.0047 | 0.0053 |
| F11 | 86240001 | 86340000 | 0.0061 | 0.0045 | 0.0051 |
| F11 | 86260001 | 86360000 | 0.0041 | 0.0029 | 0.0032 |
| F11 | 86280001 | 86380000 | 0.0033 | 0.0024 | 0.0025 |
| F11 | 86300001 | 86400000 | 0.0033 | 0.0021 | 0.0021 |
| F11 | 86320001 | 86420000 | 0.0023 | 0.0014 | 0.0015 |
| F11 | 86340001 | 86440000 | 0.0020 | 0.0012 | 0.0013 |
| F11 | 86360001 | 86460000 | 0.0024 | 0.0015 | 0.0017 |
| F11 | 86380001 | 86480000 | 0.0026 | 0.0014 | 0.0018 |
| F11 | 86400001 | 86500000 | 0.0027 | 0.0016 | 0.0021 |
| F11 | 86420001 | 86520000 | 0.0026 | 0.0015 | 0.0019 |
| F11 | 86440001 | 86540000 | 0.0041 | 0.0027 | 0.0031 |
| F11 | 86460001 | 86560000 | 0.0037 | 0.0022 | 0.0026 |
| F11 | 86480001 | 86580000 | 0.0036 | 0.0022 | 0.0025 |
| F11 | 86500001 | 86600000 | 0.0035 | 0.0021 | 0.0025 |
| F11 | 86520001 | 86620000 | 0.0036 | 0.0022 | 0.0026 |
| F11 | 86540001 | 86640000 | 0.0028 | 0.0013 | 0.0017 |
| F11 | 86560001 | 86660000 | 0.0027 | 0.0011 | 0.0014 |
| F11 | 86580001 | 86680000 | 0.0027 | 0.0011 | 0.0014 |
| F11 | 86600001 | 86700000 | 0.0026 | 0.0011 | 0.0015 |
| F11 | 86620001 | 86720000 | 0.0023 | 0.0009 | 0.0013 |
| F11 | 86640001 | 86740000 | 0.0017 | 0.0005 | 0.0009 |
| F11 | 86660001 | 86760000 | 0.0015 | 0.0005 | 0.0009 |
| F11 | 86680001 | 86780000 | 0.0012 | 0.0004 | 0.0007 |
| F11 | 86700001 | 86800000 | 0.0012 | 0.0004 | 0.0006 |
| F11 | 86720001 | 86820000 | 0.0016 | 0.0009 | 0.0010 |
| F11 | 86740001 | 86840000 | 0.0028 | 0.0021 | 0.0024 |
| F11 | 86760001 | 86860000 | 0.0030 | 0.0021 | 0.0026 |
| F11 | 86780001 | 86880000 | 0.0033 | 0.0025 | 0.0029 |
| F11 | 86800001 | 86900000 | 0.0043 | 0.0035 | 0.0040 |
| F11 | 86820001 | 86920000 | 0.0048 | 0.0037 | 0.0044 |
| F11 | 86840001 | 86940000 | 0.0039 | 0.0027 | 0.0033 |
| F11 | 86860001 | 86960000 | 0.0041 | 0.0029 | 0.0034 |
| F11 | 86880001 | 86980000 | 0.0041 | 0.0028 | 0.0034 |
| F11 | 86900001 | 87000000 | 0.0037 | 0.0021 | 0.0026 |
| F11 | 86920001 | 87020000 | 0.0029 | 0.0014 | 0.0018 |
| F11 | 86940001 | 87040000 | 0.0026 | 0.0013 | 0.0015 |
| F11 | 86960001 | 87060000 | 0.0023 | 0.0012 | 0.0013 |
| F11 | 86980001 | 87080000 | 0.0020 | 0.0010 | 0.0010 |
| F11 | 87000001 | 87100000 | 0.0014 | 0.0005 | 0.0006 |
| F11 | 87020001 | 87120000 | 0.0018 | 0.0008 | 0.0008 |
| F11 | 87040001 | 87140000 | 0.0025 | 0.0015 | 0.0016 |
| F11 | 87060001 | 87160000 | 0.0036 | 0.0025 | 0.0028 |
| F11 | 87080001 | 87180000 | 0.0045 | 0.0034 | 0.0038 |
| F11 | 87100001 | 87200000 | 0.0044 | 0.0035 | 0.0040 |
| F11 | 87120001 | 87220000 | 0.0048 | 0.0038 | 0.0044 |
| F11 | 87140001 | 87240000 | 0.0040 | 0.0030 | 0.0036 |
| F11 | 87160001 | 87260000 | 0.0029 | 0.0022 | 0.0025 |
| F11 | 87180001 | 87280000 | 0.0022 | 0.0014 | 0.0017 |
| F11 | 87200001 | 87300000 | 0.0022 | 0.0013 | 0.0015 |
| F11 | 87220001 | 87320000 | 0.0015 | 0.0008 | 0.0009 |
| F11 | 87240001 | 87340000 | 0.0022 | 0.0011 | 0.0014 |
| F11 | 87260001 | 87360000 | 0.0022 | 0.0010 | 0.0013 |
| F11 | 87280001 | 87380000 | 0.0024 | 0.0011 | 0.0015 |
| F11 | 87300001 | 87400000 | 0.0023 | 0.0011 | 0.0015 |
| F11 | 87320001 | 87420000 | 0.0024 | 0.0012 | 0.0016 |

| Chr | Start | End | both | A1 | A2 |
|---|---|---|---|---|---|
| F11 | 87340001 | 87440000 | 0.0023 | 0.0013 | 0.0015 |
| F11 | 87360001 | 87460000 | 0.0023 | 0.0014 | 0.0016 |
| F11 | 87380001 | 87480000 | 0.0023 | 0.0014 | 0.0016 |
| F11 | 87400001 | 87500000 | 0.0023 | 0.0015 | 0.0017 |
| F11 | 87420001 | 87520000 | 0.0031 | 0.0021 | 0.0024 |
| F11 | 87440001 | 87540000 | 0.0028 | 0.0019 | 0.0021 |
| F11 | 87460001 | 87560000 | 0.0032 | 0.0024 | 0.0023 |
| F11 | 87480001 | 87580000 | 0.0032 | 0.0027 | 0.0024 |
| F11 | 87500001 | 87600000 | 0.0041 | 0.0035 | 0.0032 |
| F11 | 87520001 | 87620000 | 0.0033 | 0.0030 | 0.0024 |
| F11 | 87540001 | 87640000 | 0.0035 | 0.0034 | 0.0026 |
| F11 | 87560001 | 87660000 | 0.0029 | 0.0030 | 0.0021 |
| F11 | 87580001 | 87680000 | 0.0027 | 0.0028 | 0.0018 |
| F11 | 87600001 | 87700000 | 0.0021 | 0.0023 | 0.0013 |
| F11 | 87620001 | 87720000 | 0.0020 | 0.0020 | 0.0012 |
| F11 | 87640001 | 87740000 | 0.0018 | 0.0014 | 0.0010 |
| F11 | 87660001 | 87760000 | 0.0026 | 0.0019 | 0.0017 |
| F11 | 87680001 | 87780000 | 0.0031 | 0.0020 | 0.0021 |
| F11 | 87700001 | 87800000 | 0.0034 | 0.0021 | 0.0023 |
| F11 | 87720001 | 87820000 | 0.0040 | 0.0028 | 0.0030 |
| F11 | 87740001 | 87840000 | 0.0043 | 0.0031 | 0.0032 |
| F11 | 87760001 | 87860000 | 0.0036 | 0.0024 | 0.0025 |
| F11 | 87780001 | 87880000 | 0.0035 | 0.0024 | 0.0026 |
| F11 | 87800001 | 87900000 | 0.0028 | 0.0018 | 0.0018 |
| F11 | 87820001 | 87920000 | 0.0023 | 0.0013 | 0.0013 |
| F11 | 87840001 | 87940000 | 0.0019 | 0.0009 | 0.0010 |
| F11 | 87860001 | 87960000 | 0.0026 | 0.0017 | 0.0018 |
| F11 | 87880001 | 87980000 | 0.0027 | 0.0017 | 0.0018 |
| F11 | 87900001 | 88000000 | 0.0031 | 0.0022 | 0.0023 |
| F11 | 87920001 | 88020000 | 0.0031 | 0.0023 | 0.0023 |
| F11 | 87940001 | 88040000 | 0.0032 | 0.0026 | 0.0027 |
| F11 | 87960001 | 88060000 | 0.0032 | 0.0026 | 0.0027 |
| F11 | 87980001 | 88080000 | 0.0030 | 0.0026 | 0.0025 |
| F11 | 88000001 | 88100000 | 0.0033 | 0.0028 | 0.0026 |
| F11 | 88020001 | 88120000 | 0.0035 | 0.0029 | 0.0027 |
| F11 | 88040001 | 88140000 | 0.0031 | 0.0026 | 0.0024 |
| F11 | 88060001 | 88160000 | 0.0028 | 0.0023 | 0.0022 |
| F11 | 88080001 | 88180000 | 0.0024 | 0.0020 | 0.0020 |
| F11 | 88100001 | 88200000 | 0.0023 | 0.0018 | 0.0019 |
| F11 | 88120001 | 88220000 | 0.0027 | 0.0024 | 0.0024 |
| F11 | 88140001 | 88240000 | 0.0036 | 0.0031 | 0.0031 |
| F11 | 88160001 | 88260000 | 0.0037 | 0.0033 | 0.0031 |
| F11 | 88180001 | 88280000 | 0.0041 | 0.0037 | 0.0034 |
| F11 | 88200001 | 88300000 | 0.0042 | 0.0040 | 0.0036 |
| F11 | 88220001 | 88320000 | 0.0035 | 0.0031 | 0.0028 |
| F11 | 88240001 | 88340000 | 0.0031 | 0.0025 | 0.0023 |
| F11 | 88260001 | 88360000 | 0.0025 | 0.0018 | 0.0016 |
| F11 | 88280001 | 88380000 | 0.0024 | 0.0014 | 0.0014 |
| F11 | 88300001 | 88400000 | 0.0042 | 0.0028 | 0.0030 |
| F11 | 88320001 | 88420000 | 0.0040 | 0.0025 | 0.0029 |
| F11 | 88340001 | 88440000 | 0.0040 | 0.0026 | 0.0030 |
| F11 | 88360001 | 88460000 | 0.0041 | 0.0031 | 0.0033 |
| F11 | 88380001 | 88480000 | 0.0045 | 0.0035 | 0.0038 |
| F11 | 88400001 | 88500000 | 0.0021 | 0.0016 | 0.0016 |
| F11 | 88420001 | 88520000 | 0.0033 | 0.0032 | 0.0028 |
| F11 | 88440001 | 88540000 | 0.0036 | 0.0033 | 0.0030 |
| F11 | 88460001 | 88560000 | 0.0033 | 0.0029 | 0.0027 |
| F11 | 88480001 | 88580000 | 0.0030 | 0.0024 | 0.0021 |

| Chr | Start | End | both | A1 | A2 |
|---|---|---|---|---|---|
| F11 | 88500001 | 88600000 | 0.0029 | 0.0020 | 0.0019 |
| F11 | 88520001 | 88620000 | 0.0021 | 0.0009 | 0.0011 |
| F11 | 88540001 | 88640000 | 0.0023 | 0.0012 | 0.0015 |
| F11 | 88560001 | 88660000 | 0.0023 | 0.0013 | 0.0015 |
| F11 | 88580001 | 88680000 | 0.0022 | 0.0012 | 0.0015 |
| F11 | 88600001 | 88700000 | 0.0026 | 0.0015 | 0.0019 |
| F11 | 88620001 | 88720000 | 0.0024 | 0.0012 | 0.0017 |
| F11 | 88640001 | 88740000 | 0.0017 | 0.0005 | 0.0009 |
| F11 | 88660001 | 88760000 | 0.0022 | 0.0006 | 0.0013 |
| F11 | 88680001 | 88780000 | 0.0023 | 0.0006 | 0.0013 |
| F11 | 88700001 | 88800000 | 0.0018 | 0.0004 | 0.0009 |
| F11 | 88720001 | 88820000 | 0.0018 | 0.0003 | 0.0008 |
| F11 | 88740001 | 88840000 | 0.0020 | 0.0005 | 0.0010 |
| F11 | 88760001 | 88860000 | 0.0024 | 0.0009 | 0.0015 |
| F11 | 88780001 | 88880000 | 0.0042 | 0.0029 | 0.0034 |
| F11 | 88800001 | 88900000 | 0.0070 | 0.0052 | 0.0062 |
| F11 | 88820001 | 88920000 | 0.0084 | 0.0069 | 0.0077 |
| F11 | 88840001 | 88940000 | 0.0089 | 0.0074 | 0.0082 |
| F11 | 88860001 | 88960000 | 0.0086 | 0.0073 | 0.0080 |
| F11 | 88880001 | 88980000 | 0.0082 | 0.0071 | 0.0077 |
| F11 | 88900001 | 89000000 | 0.0063 | 0.0059 | 0.0058 |
| F11 | 88920001 | 89020000 | 0.0057 | 0.0051 | 0.0052 |
| F11 | 88940001 | 89040000 | 0.0057 | 0.0051 | 0.0052 |
| F11 | 88960001 | 89060000 | 0.0053 | 0.0046 | 0.0047 |
| F11 | 88980001 | 89080000 | 0.0038 | 0.0029 | 0.0031 |
| F11 | 89000001 | 89100000 | 0.0032 | 0.0019 | 0.0022 |
| F11 | 89020001 | 89120000 | 0.0029 | 0.0012 | 0.0018 |
| F11 | 89040001 | 89140000 | 0.0032 | 0.0015 | 0.0021 |
| F11 | 89060001 | 89160000 | 0.0033 | 0.0018 | 0.0023 |
| F11 | 89080001 | 89180000 | 0.0040 | 0.0024 | 0.0029 |
| F11 | 89100001 | 89200000 | 0.0039 | 0.0022 | 0.0028 |
| F11 | 89120001 | 89220000 | 0.0034 | 0.0020 | 0.0023 |
| F11 | 89140001 | 89240000 | 0.0026 | 0.0017 | 0.0014 |
| F11 | 89160001 | 89260000 | 0.0028 | 0.0026 | 0.0012 |
| F11 | 89180001 | 89280000 | 0.0031 | 0.0037 | 0.0013 |
| F11 | 89200001 | 89300000 | 0.0033 | 0.0044 | 0.0018 |
| F11 | 89220001 | 89320000 | 0.0039 | 0.0054 | 0.0026 |
| F11 | 89240001 | 89340000 | 0.0039 | 0.0054 | 0.0025 |
| F11 | 89260001 | 89360000 | 0.0036 | 0.0047 | 0.0027 |
| F11 | 89280001 | 89380000 | 0.0040 | 0.0050 | 0.0032 |
| F11 | 89300001 | 89400000 | 0.0040 | 0.0052 | 0.0032 |
| F11 | 89320001 | 89420000 | 0.0041 | 0.0058 | 0.0032 |
| F11 | 89340001 | 89440000 | 0.0050 | 0.0065 | 0.0043 |
| F11 | 89360001 | 89460000 | 0.0052 | 0.0066 | 0.0046 |
| F11 | 89380001 | 89480000 | 0.0039 | 0.0048 | 0.0035 |
| F11 | 89400001 | 89500000 | 0.0041 | 0.0047 | 0.0037 |
| F11 | 89420001 | 89520000 | 0.0038 | 0.0036 | 0.0036 |
| F11 | 89440001 | 89540000 | 0.0038 | 0.0038 | 0.0036 |
| F11 | 89460001 | 89560000 | 0.0034 | 0.0034 | 0.0032 |
| F11 | 89480001 | 89580000 | 0.0036 | 0.0033 | 0.0034 |
| F11 | 89500001 | 89600000 | 0.0028 | 0.0026 | 0.0027 |
| F11 | 89520001 | 89620000 | 0.0039 | 0.0036 | 0.0038 |
| F11 | 89540001 | 89640000 | 0.0040 | 0.0037 | 0.0038 |
| F11 | 89560001 | 89660000 | 0.0040 | 0.0036 | 0.0038 |
| F11 | 89580001 | 89680000 | 0.0043 | 0.0043 | 0.0042 |
| F11 | 89600001 | 89700000 | 0.0046 | 0.0047 | 0.0044 |
| F11 | 89620001 | 89720000 | 0.0031 | 0.0035 | 0.0028 |
| F11 | 89640001 | 89740000 | 0.0026 | 0.0028 | 0.0025 |

| Chr | Start | End | both | A1 | A2 |
|---|---|---|---|---|---|
| F11 | 89660001 | 89760000 | 0.0036 | 0.0036 | 0.0035 |
| F11 | 89680001 | 89780000 | 0.0044 | 0.0038 | 0.0043 |
| F11 | 89700001 | 89800000 | 0.0054 | 0.0051 | 0.0053 |
| F11 | 89720001 | 89820000 | 0.0062 | 0.0055 | 0.0060 |
| F11 | 89740001 | 89840000 | 0.0052 | 0.0046 | 0.0050 |
| F11 | 89760001 | 89860000 | 0.0043 | 0.0039 | 0.0041 |
| F11 | 89780001 | 89880000 | 0.0034 | 0.0033 | 0.0031 |
| F11 | 89800001 | 89900000 | 0.0025 | 0.0021 | 0.0024 |
| F11 | 89820001 | 89920000 | 0.0017 | 0.0016 | 0.0016 |
| F11 | 89840001 | 89940000 | 0.0018 | 0.0016 | 0.0017 |
| F11 | 89860001 | 89960000 | 0.0025 | 0.0023 | 0.0024 |
| F11 | 89880001 | 89980000 | 0.0033 | 0.0030 | 0.0032 |
| F11 | 89900001 | 90000000 | 0.0030 | 0.0028 | 0.0029 |
| F11 | 89920001 | 90020000 | 0.0040 | 0.0036 | 0.0039 |
| F11 | 89940001 | 90040000 | 0.0045 | 0.0041 | 0.0044 |
| F11 | 89960001 | 90060000 | 0.0040 | 0.0035 | 0.0039 |
| F11 | 89980001 | 90080000 | 0.0031 | 0.0027 | 0.0030 |
| F11 | 90000001 | 90100000 | 0.0033 | 0.0030 | 0.0033 |
| F11 | 90020001 | 90120000 | 0.0032 | 0.0029 | 0.0031 |
| F11 | 90040001 | 90140000 | 0.0029 | 0.0025 | 0.0028 |
| F11 | 90060001 | 90160000 | 0.0030 | 0.0028 | 0.0028 |
| F11 | 90080001 | 90180000 | 0.0031 | 0.0029 | 0.0028 |
| F11 | 90100001 | 90200000 | 0.0033 | 0.0032 | 0.0030 |
| F11 | 90120001 | 90220000 | 0.0038 | 0.0039 | 0.0033 |
| F11 | 90140001 | 90240000 | 0.0051 | 0.0051 | 0.0046 |
| F11 | 90160001 | 90260000 | 0.0079 | 0.0078 | 0.0075 |
| F11 | 90180001 | 90280000 | 0.0084 | 0.0084 | 0.0079 |
| F11 | 90200001 | 90300000 | 0.0099 | 0.0100 | 0.0092 |
| F11 | 90220001 | 90320000 | 0.0105 | 0.0109 | 0.0096 |
| F11 | 90240001 | 90340000 | 0.0098 | 0.0106 | 0.0088 |
| F11 | 90260001 | 90360000 | 0.0069 | 0.0078 | 0.0059 |
| F11 | 90280001 | 90380000 | 0.0062 | 0.0069 | 0.0052 |
| F11 | 90300001 | 90400000 | 0.0041 | 0.0047 | 0.0033 |
| F11 | 90320001 | 90420000 | 0.0023 | 0.0025 | 0.0019 |
| F11 | 90340001 | 90440000 | 0.0030 | 0.0028 | 0.0029 |
| F11 | 90360001 | 90460000 | 0.0042 | 0.0041 | 0.0038 |
| F11 | 90380001 | 90480000 | 0.0046 | 0.0044 | 0.0043 |
| F11 | 90400001 | 90500000 | 0.0046 | 0.0044 | 0.0043 |
| F11 | 90420001 | 90520000 | 0.0049 | 0.0047 | 0.0046 |
| F11 | 90440001 | 90540000 | 0.0045 | 0.0045 | 0.0042 |
| F11 | 90460001 | 90560000 | 0.0040 | 0.0037 | 0.0037 |
| F11 | 90480001 | 90580000 | 0.0031 | 0.0030 | 0.0029 |
| F11 | 90500001 | 90600000 | 0.0036 | 0.0034 | 0.0033 |
| F11 | 90520001 | 90620000 | 0.0031 | 0.0028 | 0.0028 |
| F11 | 90540001 | 90640000 | 0.0026 | 0.0022 | 0.0024 |
| F11 | 90560001 | 90660000 | 0.0020 | 0.0015 | 0.0019 |
| F11 | 90580001 | 90680000 | 0.0044 | 0.0037 | 0.0043 |
| F11 | 90600001 | 90700000 | 0.0046 | 0.0038 | 0.0046 |
| F11 | 90620001 | 90720000 | 0.0059 | 0.0051 | 0.0059 |
| F11 | 90640001 | 90740000 | 0.0063 | 0.0055 | 0.0062 |
| F11 | 90660001 | 90760000 | 0.0060 | 0.0053 | 0.0060 |
| F11 | 90680001 | 90780000 | 0.0047 | 0.0043 | 0.0047 |
| F11 | 90700001 | 90800000 | 0.0048 | 0.0044 | 0.0048 |
| F11 | 90720001 | 90820000 | 0.0047 | 0.0042 | 0.0048 |
| F11 | 90740001 | 90840000 | 0.0043 | 0.0035 | 0.0044 |
| F11 | 90760001 | 90860000 | 0.0052 | 0.0045 | 0.0053 |
| F11 | 90780001 | 90880000 | 0.0050 | 0.0043 | 0.0051 |
| F11 | 90800001 | 90900000 | 0.0045 | 0.0038 | 0.0045 |

| Chr | Start | End | both | A1 | A2 |
|---|---|---|---|---|---|
| F11 | 90820001 | 90920000 | 0.0034 | 0.0030 | 0.0034 |
| F11 | 90840001 | 90940000 | 0.0039 | 0.0037 | 0.0039 |
| F11 | 90860001 | 90960000 | 0.0044 | 0.0042 | 0.0044 |
| F11 | 90880001 | 90980000 | 0.0044 | 0.0040 | 0.0044 |
| F11 | 90900001 | 91000000 | 0.0054 | 0.0047 | 0.0054 |
| F11 | 90920001 | 91020000 | 0.0067 | 0.0059 | 0.0068 |
| F11 | 90940001 | 91040000 | 0.0055 | 0.0048 | 0.0056 |
| F11 | 90960001 | 91060000 | 0.0045 | 0.0036 | 0.0045 |
| F11 | 90980001 | 91080000 | 0.0051 | 0.0033 | 0.0045 |
| F11 | 91000001 | 91100000 | 0.0059 | 0.0040 | 0.0049 |
| F11 | 91020001 | 91120000 | 0.0051 | 0.0030 | 0.0037 |
| F11 | 91040001 | 91140000 | 0.0066 | 0.0038 | 0.0046 |
| F11 | 91060001 | 91160000 | 0.0062 | 0.0035 | 0.0043 |
| F11 | 91080001 | 91180000 | 0.0050 | 0.0033 | 0.0036 |
| F11 | 91100001 | 91200000 | 0.0034 | 0.0018 | 0.0023 |
| F11 | 91120001 | 91220000 | 0.0029 | 0.0016 | 0.0022 |
| F11 | 91140001 | 91240000 | 0.0017 | 0.0009 | 0.0014 |
| F11 | 91160001 | 91260000 | 0.0037 | 0.0026 | 0.0029 |
| F11 | 91180001 | 91280000 | 0.0043 | 0.0032 | 0.0036 |
| F11 | 91200001 | 91300000 | 0.0050 | 0.0038 | 0.0041 |
| F11 | 91220001 | 91320000 | 0.0055 | 0.0042 | 0.0045 |
| F11 | 91240001 | 91340000 | 0.0059 | 0.0051 | 0.0049 |
| F11 | 91260001 | 91360000 | 0.0064 | 0.0063 | 0.0054 |
| F11 | 91280001 | 91380000 | 0.0079 | 0.0083 | 0.0067 |
| F11 | 91300001 | 91400000 | 0.0077 | 0.0082 | 0.0066 |
| F11 | 91320001 | 91420000 | 0.0081 | 0.0086 | 0.0071 |
| F11 | 91340001 | 91440000 | 0.0086 | 0.0087 | 0.0076 |
| F11 | 91360001 | 91460000 | 0.0065 | 0.0062 | 0.0058 |
| F11 | 91380001 | 91480000 | 0.0050 | 0.0041 | 0.0044 |
| F11 | 91400001 | 91500000 | 0.0044 | 0.0035 | 0.0037 |
| F11 | 91420001 | 91520000 | 0.0032 | 0.0025 | 0.0026 |
| F11 | 91440001 | 91540000 | 0.0026 | 0.0020 | 0.0020 |
| F11 | 91460001 | 91560000 | 0.0020 | 0.0016 | 0.0015 |
| F11 | 91480001 | 91580000 | 0.0016 | 0.0013 | 0.0013 |
| F11 | 91500001 | 91600000 | 0.0020 | 0.0018 | 0.0017 |
| F11 | 91520001 | 91620000 | 0.0020 | 0.0017 | 0.0017 |
| F11 | 91540001 | 91640000 | 0.0013 | 0.0012 | 0.0011 |
| F11 | 91560001 | 91660000 | 0.0015 | 0.0014 | 0.0013 |
| F11 | 91580001 | 91680000 | 0.0014 | 0.0014 | 0.0011 |
| F11 | 91600001 | 91700000 | 0.0010 | 0.0011 | 0.0007 |
| F11 | 91620001 | 91720000 | 0.0013 | 0.0015 | 0.0007 |
| F11 | 91640001 | 91740000 | 0.0019 | 0.0021 | 0.0012 |
| F11 | 91660001 | 91760000 | 0.0021 | 0.0024 | 0.0015 |
| F11 | 91680001 | 91780000 | 0.0023 | 0.0026 | 0.0017 |
| F11 | 91700001 | 91800000 | 0.0035 | 0.0042 | 0.0025 |
| F11 | 91720001 | 91820000 | 0.0038 | 0.0046 | 0.0029 |
| F11 | 91740001 | 91840000 | 0.0034 | 0.0042 | 0.0026 |
| F11 | 91760001 | 91860000 | 0.0035 | 0.0041 | 0.0026 |
| F11 | 91780001 | 91880000 | 0.0030 | 0.0035 | 0.0022 |
| F11 | 91800001 | 91900000 | 0.0021 | 0.0025 | 0.0017 |
| F11 | 91820001 | 91920000 | 0.0023 | 0.0025 | 0.0020 |
| F11 | 91840001 | 91940000 | 0.0027 | 0.0029 | 0.0023 |
| F11 | 91860001 | 91960000 | 0.0025 | 0.0028 | 0.0021 |
| F11 | 91880001 | 91980000 | 0.0041 | 0.0042 | 0.0037 |
| F11 | 91900001 | 92000000 | 0.0044 | 0.0039 | 0.0040 |
| F11 | 91920001 | 92020000 | 0.0040 | 0.0035 | 0.0037 |
| F11 | 91940001 | 92040000 | 0.0048 | 0.0038 | 0.0044 |
| F11 | 91960001 | 92060000 | 0.0058 | 0.0044 | 0.0052 |

| Chr | Start | End | both | A1 | A2 |
|---|---|---|---|---|---|
| F11 | 91980001 | 92080000 | 0.0046 | 0.0032 | 0.0039 |
| F11 | 92000001 | 92100000 | 0.0038 | 0.0026 | 0.0031 |
| F11 | 92020001 | 92120000 | 0.0038 | 0.0025 | 0.0030 |
| F11 | 92040001 | 92140000 | 0.0025 | 0.0017 | 0.0018 |
| F11 | 92060001 | 92160000 | 0.0014 | 0.0008 | 0.0008 |
| F11 | 92080001 | 92180000 | 0.0018 | 0.0012 | 0.0010 |
| F11 | 92100001 | 92200000 | 0.0023 | 0.0016 | 0.0015 |
| F11 | 92120001 | 92220000 | 0.0027 | 0.0024 | 0.0014 |
| F11 | 92140001 | 92240000 | 0.0032 | 0.0029 | 0.0019 |
| F11 | 92160001 | 92260000 | 0.0031 | 0.0028 | 0.0020 |
| F11 | 92180001 | 92280000 | 0.0024 | 0.0023 | 0.0018 |
| F11 | 92200001 | 92300000 | 0.0020 | 0.0019 | 0.0014 |
| F11 | 92220001 | 92320000 | 0.0033 | 0.0022 | 0.0030 |
| F11 | 92240001 | 92340000 | 0.0028 | 0.0018 | 0.0026 |
| F11 | 92260001 | 92360000 | 0.0041 | 0.0034 | 0.0036 |
| F11 | 92280001 | 92380000 | 0.0045 | 0.0043 | 0.0039 |
| F11 | 92300001 | 92400000 | 0.0050 | 0.0049 | 0.0044 |
| F11 | 92320001 | 92420000 | 0.0035 | 0.0043 | 0.0030 |
| F11 | 92340001 | 92440000 | 0.0049 | 0.0063 | 0.0042 |
| F11 | 92360001 | 92460000 | 0.0051 | 0.0069 | 0.0044 |
| F11 | 92380001 | 92480000 | 0.0048 | 0.0066 | 0.0041 |
| F11 | 92400001 | 92500000 | 0.0043 | 0.0063 | 0.0036 |
| F11 | 92420001 | 92520000 | 0.0038 | 0.0059 | 0.0031 |
| F11 | 92440001 | 92540000 | 0.0025 | 0.0041 | 0.0020 |
| F11 | 92460001 | 92560000 | 0.0011 | 0.0021 | 0.0008 |
| F11 | 92480001 | 92580000 | 0.0014 | 0.0023 | 0.0010 |
| F11 | 92500001 | 92600000 | 0.0019 | 0.0025 | 0.0014 |
| F11 | 92520001 | 92620000 | 0.0020 | 0.0025 | 0.0015 |
| F11 | 92540001 | 92640000 | 0.0025 | 0.0028 | 0.0020 |
| F11 | 92560001 | 92660000 | 0.0024 | 0.0026 | 0.0020 |
| F11 | 92580001 | 92680000 | 0.0025 | 0.0023 | 0.0023 |
| F11 | 92600001 | 92700000 | 0.0059 | 0.0056 | 0.0057 |
| F11 | 92620001 | 92720000 | 0.0065 | 0.0060 | 0.0063 |
| F11 | 92640001 | 92740000 | 0.0063 | 0.0059 | 0.0061 |
| F11 | 92660001 | 92760000 | 0.0075 | 0.0070 | 0.0073 |
| F11 | 92680001 | 92780000 | 0.0106 | 0.0094 | 0.0103 |
| F11 | 92700001 | 92800000 | 0.0070 | 0.0059 | 0.0068 |
| F11 | 92720001 | 92820000 | 0.0077 | 0.0066 | 0.0075 |
| F11 | 92740001 | 92840000 | 0.0092 | 0.0074 | 0.0091 |
| F11 | 92760001 | 92860000 | 0.0098 | 0.0077 | 0.0096 |
| F11 | 92780001 | 92880000 | 0.0066 | 0.0052 | 0.0064 |
| F11 | 92800001 | 92900000 | 0.0078 | 0.0063 | 0.0076 |
| F11 | 92820001 | 92920000 | 0.0071 | 0.0055 | 0.0069 |
| F11 | 92840001 | 92940000 | 0.0063 | 0.0050 | 0.0060 |
| F11 | 92860001 | 92960000 | 0.0048 | 0.0041 | 0.0044 |
| F11 | 92880001 | 92980000 | 0.0072 | 0.0064 | 0.0066 |
| F11 | 92900001 | 93000000 | 0.0069 | 0.0062 | 0.0062 |
| F11 | 92920001 | 93020000 | 0.0066 | 0.0059 | 0.0060 |
| F11 | 92940001 | 93040000 | 0.0064 | 0.0057 | 0.0057 |
| F11 | 92960001 | 93060000 | 0.0070 | 0.0061 | 0.0065 |
| F11 | 92980001 | 93080000 | 0.0062 | 0.0064 | 0.0055 |
| F11 | 93000001 | 93100000 | 0.0073 | 0.0076 | 0.0065 |
| F11 | 93020001 | 93120000 | 0.0072 | 0.0076 | 0.0063 |
| F11 | 93040001 | 93140000 | 0.0069 | 0.0073 | 0.0060 |
| F11 | 93060001 | 93160000 | 0.0066 | 0.0071 | 0.0057 |
| F11 | 93080001 | 93180000 | 0.0046 | 0.0044 | 0.0041 |
| F11 | 93100001 | 93200000 | 0.0043 | 0.0043 | 0.0035 |
| F11 | 93120001 | 93220000 | 0.0048 | 0.0050 | 0.0040 |

| Chr | Start | End | both | A1 | A2 |
|---|---|---|---|---|---|
| F11 | 93140001 | 93240000 | 0.0057 | 0.0061 | 0.0047 |
| F11 | 93160001 | 93260000 | 0.0051 | 0.0057 | 0.0040 |
| F11 | 93180001 | 93280000 | 0.0053 | 0.0062 | 0.0040 |
| F11 | 93200001 | 93300000 | 0.0043 | 0.0049 | 0.0033 |
| F11 | 93220001 | 93320000 | 0.0037 | 0.0042 | 0.0028 |
| F11 | 93240001 | 93340000 | 0.0030 | 0.0034 | 0.0021 |
| F11 | 93260001 | 93360000 | 0.0046 | 0.0047 | 0.0037 |
| F11 | 93280001 | 93380000 | 0.0046 | 0.0044 | 0.0038 |
| F11 | 93300001 | 93400000 | 0.0043 | 0.0043 | 0.0034 |
| F11 | 93320001 | 93420000 | 0.0056 | 0.0060 | 0.0039 |
| F11 | 93340001 | 93440000 | 0.0060 | 0.0064 | 0.0043 |
| F11 | 93360001 | 93460000 | 0.0048 | 0.0054 | 0.0029 |
| F11 | 93380001 | 93480000 | 0.0043 | 0.0049 | 0.0026 |
| F11 | 93400001 | 93500000 | 0.0037 | 0.0043 | 0.0021 |
| F11 | 93420001 | 93520000 | 0.0024 | 0.0024 | 0.0014 |
| F11 | 93440001 | 93540000 | 0.0015 | 0.0016 | 0.0006 |
| F11 | 93460001 | 93560000 | 0.0019 | 0.0025 | 0.0010 |
| F11 | 93480001 | 93580000 | 0.0034 | 0.0045 | 0.0022 |
| F11 | 93500001 | 93600000 | 0.0039 | 0.0054 | 0.0024 |
| F11 | 93520001 | 93620000 | 0.0059 | 0.0075 | 0.0044 |
| F11 | 93540001 | 93640000 | 0.0061 | 0.0076 | 0.0048 |
| F11 | 93560001 | 93660000 | 0.0057 | 0.0070 | 0.0046 |
| F11 | 93580001 | 93680000 | 0.0050 | 0.0058 | 0.0042 |
| F11 | 93600001 | 93700000 | 0.0046 | 0.0050 | 0.0040 |
| F11 | 93620001 | 93720000 | 0.0027 | 0.0030 | 0.0022 |
| F11 | 93640001 | 93740000 | 0.0029 | 0.0032 | 0.0022 |
| F11 | 93660001 | 93760000 | 0.0026 | 0.0026 | 0.0019 |
| F11 | 93680001 | 93780000 | 0.0019 | 0.0018 | 0.0012 |
| F11 | 93700001 | 93800000 | 0.0020 | 0.0018 | 0.0013 |
| F11 | 93720001 | 93820000 | 0.0022 | 0.0022 | 0.0013 |
| F11 | 93740001 | 93840000 | 0.0023 | 0.0024 | 0.0014 |
| F11 | 93760001 | 93860000 | 0.0040 | 0.0042 | 0.0030 |
| F11 | 93780001 | 93880000 | 0.0042 | 0.0044 | 0.0033 |
| F11 | 93800001 | 93900000 | 0.0045 | 0.0046 | 0.0038 |
| F11 | 93820001 | 93920000 | 0.0046 | 0.0046 | 0.0042 |
| F11 | 93840001 | 93940000 | 0.0047 | 0.0047 | 0.0044 |
| F11 | 93860001 | 93960000 | 0.0041 | 0.0040 | 0.0039 |
| F11 | 93880001 | 93980000 | 0.0045 | 0.0044 | 0.0041 |
| F11 | 93900001 | 94000000 | 0.0041 | 0.0041 | 0.0036 |
| F11 | 93920001 | 94020000 | 0.0052 | 0.0048 | 0.0043 |
| F11 | 93940001 | 94040000 | 0.0060 | 0.0055 | 0.0049 |
| F11 | 93960001 | 94060000 | 0.0051 | 0.0045 | 0.0041 |
| F11 | 93980001 | 94080000 | 0.0042 | 0.0038 | 0.0035 |
| F11 | 94000001 | 94100000 | 0.0050 | 0.0046 | 0.0042 |
| F11 | 94020001 | 94120000 | 0.0040 | 0.0040 | 0.0035 |
| F11 | 94040001 | 94140000 | 0.0029 | 0.0030 | 0.0024 |
| F11 | 94060001 | 94160000 | 0.0027 | 0.0029 | 0.0021 |
| F11 | 94080001 | 94180000 | 0.0027 | 0.0029 | 0.0021 |
| F11 | 94100001 | 94200000 | 0.0021 | 0.0022 | 0.0015 |
| F11 | 94120001 | 94220000 | 0.0015 | 0.0015 | 0.0009 |
| F11 | 94140001 | 94240000 | 0.0020 | 0.0019 | 0.0015 |
| F11 | 94160001 | 94260000 | 0.0020 | 0.0019 | 0.0015 |
| F11 | 94180001 | 94280000 | 0.0030 | 0.0028 | 0.0025 |
| F11 | 94200001 | 94300000 | 0.0037 | 0.0033 | 0.0033 |
| F11 | 94220001 | 94320000 | 0.0042 | 0.0039 | 0.0037 |
| F11 | 94240001 | 94340000 | 0.0032 | 0.0030 | 0.0028 |
| F11 | 94260001 | 94360000 | 0.0033 | 0.0030 | 0.0029 |
| F11 | 94280001 | 94380000 | 0.0032 | 0.0029 | 0.0028 |

| Chr | Start | End | both | A1 | A2 |
|---|---|---|---|---|---|
| F11 | 94300001 | 94400000 | 0.0031 | 0.0029 | 0.0026 |
| F11 | 94320001 | 94420000 | 0.0027 | 0.0025 | 0.0023 |
| F11 | 94340001 | 94440000 | 0.0028 | 0.0026 | 0.0023 |
| F11 | 94360001 | 94460000 | 0.0027 | 0.0026 | 0.0022 |
| F11 | 94380001 | 94480000 | 0.0024 | 0.0025 | 0.0019 |
| F11 | 94400001 | 94500000 | 0.0023 | 0.0023 | 0.0017 |
| F11 | 94420001 | 94520000 | 0.0034 | 0.0040 | 0.0023 |
| F11 | 94440001 | 94540000 | 0.0037 | 0.0046 | 0.0026 |
| F11 | 94460001 | 94560000 | 0.0040 | 0.0048 | 0.0028 |
| F11 | 94480001 | 94580000 | 0.0041 | 0.0048 | 0.0029 |
| F11 | 94500001 | 94600000 | 0.0038 | 0.0045 | 0.0027 |
| F11 | 94520001 | 94620000 | 0.0028 | 0.0028 | 0.0021 |
| F11 | 94540001 | 94640000 | 0.0028 | 0.0026 | 0.0021 |
| F11 | 94560001 | 94660000 | 0.0025 | 0.0024 | 0.0017 |
| F11 | 94580001 | 94680000 | 0.0019 | 0.0019 | 0.0011 |
| F11 | 94600001 | 94700000 | 0.0021 | 0.0022 | 0.0010 |
| F11 | 94620001 | 94720000 | 0.0021 | 0.0023 | 0.0008 |
| F11 | 94640001 | 94740000 | 0.0019 | 0.0021 | 0.0007 |
| F11 | 94660001 | 94760000 | 0.0021 | 0.0021 | 0.0008 |
| F11 | 94680001 | 94780000 | 0.0024 | 0.0025 | 0.0010 |
| F11 | 94700001 | 94800000 | 0.0026 | 0.0027 | 0.0012 |
| F11 | 94720001 | 94820000 | 0.0024 | 0.0025 | 0.0013 |
| F11 | 94740001 | 94840000 | 0.0038 | 0.0037 | 0.0028 |
| F11 | 94760001 | 94860000 | 0.0050 | 0.0050 | 0.0042 |
| F11 | 94780001 | 94880000 | 0.0047 | 0.0047 | 0.0043 |
| F11 | 94800001 | 94900000 | 0.0043 | 0.0043 | 0.0041 |
| F11 | 94820001 | 94920000 | 0.0050 | 0.0050 | 0.0048 |
| F11 | 94840001 | 94940000 | 0.0043 | 0.0042 | 0.0041 |
| F11 | 94860001 | 94960000 | 0.0037 | 0.0040 | 0.0034 |
| F11 | 94880001 | 94980000 | 0.0035 | 0.0038 | 0.0032 |
| F11 | 94900001 | 95000000 | 0.0034 | 0.0037 | 0.0031 |
| F11 | 94920001 | 95020000 | 0.0035 | 0.0036 | 0.0033 |
| F11 | 94940001 | 95040000 | 0.0025 | 0.0027 | 0.0023 |
| F11 | 94960001 | 95060000 | 0.0018 | 0.0015 | 0.0018 |
| F11 | 94980001 | 95080000 | 0.0020 | 0.0017 | 0.0019 |
| F11 | 95000001 | 95100000 | 0.0026 | 0.0027 | 0.0024 |
| F11 | 95020001 | 95120000 | 0.0019 | 0.0022 | 0.0017 |
| F11 | 95040001 | 95140000 | 0.0019 | 0.0022 | 0.0016 |
| F11 | 95060001 | 95160000 | 0.0022 | 0.0025 | 0.0020 |
| F11 | 95080001 | 95180000 | 0.0021 | 0.0024 | 0.0019 |
| F11 | 95100001 | 95200000 | 0.0027 | 0.0024 | 0.0025 |
| F11 | 95120001 | 95220000 | 0.0026 | 0.0022 | 0.0024 |
| F11 | 95140001 | 95240000 | 0.0032 | 0.0028 | 0.0029 |
| F11 | 95160001 | 95260000 | 0.0029 | 0.0025 | 0.0026 |
| F11 | 95180001 | 95280000 | 0.0034 | 0.0030 | 0.0031 |
| F11 | 95200001 | 95300000 | 0.0027 | 0.0024 | 0.0024 |
| F11 | 95220001 | 95320000 | 0.0029 | 0.0027 | 0.0026 |
| F11 | 95240001 | 95340000 | 0.0026 | 0.0023 | 0.0024 |
| F11 | 95260001 | 95360000 | 0.0046 | 0.0041 | 0.0042 |
| F11 | 95280001 | 95380000 | 0.0052 | 0.0046 | 0.0048 |
| F11 | 95300001 | 95400000 | 0.0049 | 0.0043 | 0.0045 |
| F11 | 95320001 | 95420000 | 0.0053 | 0.0046 | 0.0049 |
| F11 | 95340001 | 95440000 | 0.0056 | 0.0048 | 0.0050 |
| F11 | 95360001 | 95460000 | 0.0037 | 0.0030 | 0.0032 |
| F11 | 95380001 | 95480000 | 0.0035 | 0.0030 | 0.0031 |
| F11 | 95400001 | 95500000 | 0.0040 | 0.0039 | 0.0035 |
| F11 | 95420001 | 95520000 | 0.0043 | 0.0040 | 0.0037 |
| F11 | 95440001 | 95540000 | 0.0052 | 0.0050 | 0.0046 |

| Chr | Start | End | both | A1 | A2 |
|---|---|---|---|---|---|
| F11 | 95460001 | 95560000 | 0.0048 | 0.0047 | 0.0043 |
| F11 | 95480001 | 95580000 | 0.0036 | 0.0036 | 0.0032 |
| F11 | 95500001 | 95600000 | 0.0038 | 0.0030 | 0.0026 |
| F11 | 95520001 | 95620000 | 0.0035 | 0.0024 | 0.0017 |
| F11 | 95540001 | 95640000 | 0.0022 | 0.0012 | 0.0004 |
| F11 | 95560001 | 95660000 | 0.0024 | 0.0016 | 0.0004 |
| F11 | 95580001 | 95680000 | 0.0025 | 0.0019 | 0.0005 |
| F11 | 95600001 | 95700000 | 0.0017 | 0.0017 | 0.0005 |
| F11 | 95620001 | 95720000 | 0.0011 | 0.0017 | 0.0005 |
| F11 | 95640001 | 95740000 | 0.0013 | 0.0023 | 0.0008 |
| F11 | 95660001 | 95760000 | 0.0016 | 0.0025 | 0.0011 |
| F11 | 95680001 | 95780000 | 0.0017 | 0.0025 | 0.0013 |
| F11 | 95700001 | 95800000 | 0.0016 | 0.0024 | 0.0013 |
| F11 | 95720001 | 95820000 | 0.0018 | 0.0025 | 0.0015 |
| F11 | 95740001 | 95840000 | 0.0017 | 0.0023 | 0.0015 |
| F11 | 95760001 | 95860000 | 0.0014 | 0.0018 | 0.0013 |
| F11 | 95780001 | 95880000 | 0.0012 | 0.0015 | 0.0011 |
| F11 | 95800001 | 95900000 | 0.0029 | 0.0034 | 0.0027 |
| F11 | 95820001 | 95920000 | 0.0044 | 0.0049 | 0.0042 |
| F11 | 95840001 | 95940000 | 0.0050 | 0.0058 | 0.0048 |
| F11 | 95860001 | 95960000 | 0.0063 | 0.0072 | 0.0060 |
| F11 | 95880001 | 95980000 | 0.0068 | 0.0077 | 0.0064 |
| F11 | 95900001 | 96000000 | 0.0057 | 0.0066 | 0.0054 |
| F11 | 95920001 | 96020000 | 0.0041 | 0.0048 | 0.0038 |
| F11 | 95940001 | 96040000 | 0.0034 | 0.0039 | 0.0032 |
| F11 | 95960001 | 96060000 | 0.0022 | 0.0026 | 0.0021 |
| F11 | 95980001 | 96080000 | 0.0023 | 0.0027 | 0.0021 |
| F11 | 96000001 | 96100000 | 0.0025 | 0.0030 | 0.0024 |
| F11 | 96020001 | 96120000 | 0.0025 | 0.0032 | 0.0023 |
| F11 | 96040001 | 96140000 | 0.0023 | 0.0032 | 0.0021 |
| F11 | 96060001 | 96160000 | 0.0025 | 0.0035 | 0.0022 |
| F11 | 96080001 | 96180000 | 0.0021 | 0.0031 | 0.0018 |
| F11 | 96100001 | 96200000 | 0.0012 | 0.0021 | 0.0008 |
| F11 | 96120001 | 96220000 | 0.0013 | 0.0020 | 0.0008 |
| F11 | 96140001 | 96240000 | 0.0015 | 0.0018 | 0.0008 |
| F11 | 96160001 | 96260000 | 0.0015 | 0.0016 | 0.0005 |
| F11 | 96180001 | 96280000 | 0.0018 | 0.0015 | 0.0006 |
| F11 | 96200001 | 96300000 | 0.0019 | 0.0014 | 0.0007 |
| F11 | 96220001 | 96320000 | 0.0021 | 0.0013 | 0.0010 |
| F11 | 96240001 | 96340000 | 0.0022 | 0.0011 | 0.0014 |
| F11 | 96260001 | 96360000 | 0.0023 | 0.0009 | 0.0018 |
| F11 | 96280001 | 96380000 | 0.0026 | 0.0009 | 0.0023 |
| F11 | 96300001 | 96400000 | 0.0027 | 0.0007 | 0.0023 |
| F11 | 96320001 | 96420000 | 0.0025 | 0.0007 | 0.0023 |
| F11 | 96340001 | 96440000 | 0.0024 | 0.0010 | 0.0020 |
| F11 | 96360001 | 96460000 | 0.0022 | 0.0009 | 0.0016 |
| F11 | 96380001 | 96480000 | 0.0018 | 0.0007 | 0.0010 |
| F11 | 96400001 | 96500000 | 0.0019 | 0.0007 | 0.0009 |
| F11 | 96420001 | 96520000 | 0.0019 | 0.0006 | 0.0008 |
| F11 | 96440001 | 96540000 | 0.0018 | 0.0004 | 0.0006 |
| F11 | 96460001 | 96560000 | 0.0017 | 0.0006 | 0.0006 |
| F11 | 96480001 | 96580000 | 0.0016 | 0.0006 | 0.0005 |
| F11 | 96500001 | 96600000 | 0.0015 | 0.0008 | 0.0007 |
| F11 | 96520001 | 96620000 | 0.0012 | 0.0008 | 0.0006 |
| F11 | 96540001 | 96640000 | 0.0012 | 0.0008 | 0.0008 |
| F11 | 96560001 | 96660000 | 0.0015 | 0.0007 | 0.0012 |
| F11 | 96580001 | 96680000 | 0.0018 | 0.0006 | 0.0015 |
| F11 | 96600001 | 96700000 | 0.0018 | 0.0005 | 0.0014 |

| Chr | Start | End | both | A1 | A2 |
|---|---|---|---|---|---|
| F11 | 96620001 | 96720000 | 0.0030 | 0.0015 | 0.0025 |
| F11 | 96640001 | 96740000 | 0.0039 | 0.0024 | 0.0033 |
| F11 | 96660001 | 96760000 | 0.0038 | 0.0025 | 0.0033 |
| F11 | 96680001 | 96780000 | 0.0037 | 0.0025 | 0.0034 |
| F11 | 96700001 | 96800000 | 0.0036 | 0.0025 | 0.0035 |
| F11 | 96720001 | 96820000 | 0.0025 | 0.0015 | 0.0027 |
| F11 | 96740001 | 96840000 | 0.0015 | 0.0006 | 0.0017 |
| F11 | 96760001 | 96860000 | 0.0020 | 0.0012 | 0.0021 |
| F11 | 96780001 | 96880000 | 0.0023 | 0.0016 | 0.0024 |
| F11 | 96800001 | 96900000 | 0.0039 | 0.0032 | 0.0040 |
| F11 | 96820001 | 96920000 | 0.0070 | 0.0064 | 0.0070 |
| F11 | 96840001 | 96940000 | 0.0078 | 0.0072 | 0.0078 |
| F11 | 96860001 | 96960000 | 0.0095 | 0.0089 | 0.0096 |
| F11 | 96880001 | 96980000 | 0.0104 | 0.0093 | 0.0102 |
| F11 | 96900001 | 97000000 | 0.0089 | 0.0076 | 0.0087 |
| F11 | 96920001 | 97020000 | 0.0061 | 0.0045 | 0.0058 |
| F11 | 96940001 | 97040000 | 0.0057 | 0.0043 | 0.0054 |
| F11 | 96960001 | 97060000 | 0.0043 | 0.0029 | 0.0039 |
| F11 | 96980001 | 97080000 | 0.0029 | 0.0020 | 0.0026 |
| F11 | 97000001 | 97100000 | 0.0024 | 0.0019 | 0.0022 |
| F11 | 97020001 | 97120000 | 0.0025 | 0.0022 | 0.0024 |
| F11 | 97040001 | 97140000 | 0.0031 | 0.0028 | 0.0030 |
| F11 | 97060001 | 97160000 | 0.0022 | 0.0019 | 0.0021 |
| F11 | 97080001 | 97180000 | 0.0037 | 0.0030 | 0.0036 |
| F11 | 97100001 | 97200000 | 0.0038 | 0.0031 | 0.0037 |
| F11 | 97120001 | 97220000 | 0.0039 | 0.0031 | 0.0039 |
| F11 | 97140001 | 97240000 | 0.0029 | 0.0022 | 0.0029 |
| F11 | 97160001 | 97260000 | 0.0028 | 0.0022 | 0.0028 |
| F11 | 97180001 | 97280000 | 0.0022 | 0.0015 | 0.0022 |
| F11 | 97200001 | 97300000 | 0.0035 | 0.0026 | 0.0034 |
| F11 | 97220001 | 97320000 | 0.0036 | 0.0025 | 0.0035 |
| F11 | 97240001 | 97340000 | 0.0042 | 0.0031 | 0.0042 |
| F11 | 97260001 | 97360000 | 0.0043 | 0.0032 | 0.0042 |
| F11 | 97280001 | 97380000 | 0.0041 | 0.0034 | 0.0040 |
| F11 | 97300001 | 97400000 | 0.0030 | 0.0024 | 0.0030 |
| F11 | 97320001 | 97420000 | 0.0034 | 0.0028 | 0.0034 |
| F11 | 97340001 | 97440000 | 0.0030 | 0.0024 | 0.0030 |
| F11 | 97360001 | 97460000 | 0.0034 | 0.0027 | 0.0034 |
| F11 | 97380001 | 97480000 | 0.0039 | 0.0033 | 0.0039 |
| F11 | 97400001 | 97500000 | 0.0043 | 0.0037 | 0.0043 |
| F11 | 97420001 | 97520000 | 0.0035 | 0.0031 | 0.0033 |
| F11 | 97440001 | 97540000 | 0.0035 | 0.0030 | 0.0032 |
| F11 | 97460001 | 97560000 | 0.0036 | 0.0031 | 0.0031 |
| F11 | 97480001 | 97580000 | 0.0029 | 0.0022 | 0.0022 |
| F11 | 97500001 | 97600000 | 0.0027 | 0.0018 | 0.0016 |
| F11 | 97520001 | 97620000 | 0.0028 | 0.0019 | 0.0015 |
| F11 | 97540001 | 97640000 | 0.0045 | 0.0033 | 0.0027 |
| F11 | 97560001 | 97660000 | 0.0046 | 0.0032 | 0.0024 |
| F11 | 97580001 | 97680000 | 0.0046 | 0.0035 | 0.0025 |
| F11 | 97600001 | 97700000 | 0.0042 | 0.0035 | 0.0024 |
| F11 | 97620001 | 97720000 | 0.0038 | 0.0036 | 0.0023 |
| F11 | 97640001 | 97740000 | 0.0025 | 0.0029 | 0.0015 |
| F11 | 97660001 | 97760000 | 0.0041 | 0.0049 | 0.0037 |
| F11 | 97680001 | 97780000 | 0.0062 | 0.0070 | 0.0059 |
| F11 | 97700001 | 97800000 | 0.0072 | 0.0081 | 0.0070 |
| F11 | 97720001 | 97820000 | 0.0094 | 0.0098 | 0.0092 |
| F11 | 97740001 | 97840000 | 0.0090 | 0.0093 | 0.0088 |
| F11 | 97760001 | 97860000 | 0.0080 | 0.0084 | 0.0078 |

| Chr | Start | End | both | A1 | A2 |
|---|---|---|---|---|---|
| F11 | 97780001 | 97880000 | 0.0067 | 0.0072 | 0.0065 |
| F11 | 97800001 | 97900000 | 0.0070 | 0.0076 | 0.0068 |
| F11 | 97820001 | 97920000 | 0.0048 | 0.0055 | 0.0046 |
| F11 | 97840001 | 97940000 | 0.0044 | 0.0050 | 0.0042 |
| F11 | 97860001 | 97960000 | 0.0028 | 0.0035 | 0.0026 |
| F11 | 97880001 | 97980000 | 0.0021 | 0.0021 | 0.0020 |
| F11 | 97900001 | 98000000 | 0.0010 | 0.0006 | 0.0010 |
| F11 | 97920001 | 98020000 | 0.0013 | 0.0005 | 0.0012 |
| F11 | 97940001 | 98040000 | 0.0019 | 0.0005 | 0.0016 |
| F11 | 97960001 | 98060000 | 0.0031 | 0.0013 | 0.0027 |
| F11 | 97980001 | 98080000 | 0.0030 | 0.0013 | 0.0024 |
| F11 | 98000001 | 98100000 | 0.0041 | 0.0025 | 0.0033 |
| F11 | 98020001 | 98120000 | 0.0042 | 0.0024 | 0.0034 |
| F11 | 98040001 | 98140000 | 0.0039 | 0.0022 | 0.0032 |
| F11 | 98060001 | 98160000 | 0.0040 | 0.0024 | 0.0034 |
| F11 | 98080001 | 98180000 | 0.0040 | 0.0026 | 0.0036 |
| F11 | 98100001 | 98200000 | 0.0051 | 0.0037 | 0.0048 |
| F11 | 98120001 | 98220000 | 0.0068 | 0.0046 | 0.0067 |
| F11 | 98140001 | 98240000 | 0.0076 | 0.0055 | 0.0075 |
| F11 | 98160001 | 98260000 | 0.0069 | 0.0049 | 0.0069 |
| F11 | 98180001 | 98280000 | 0.0073 | 0.0052 | 0.0072 |
| F11 | 98200001 | 98300000 | 0.0057 | 0.0038 | 0.0056 |
| F11 | 98220001 | 98320000 | 0.0037 | 0.0030 | 0.0036 |
| F11 | 98240001 | 98340000 | 0.0031 | 0.0025 | 0.0030 |
| F11 | 98260001 | 98360000 | 0.0035 | 0.0030 | 0.0035 |
| F11 | 98280001 | 98380000 | 0.0045 | 0.0040 | 0.0045 |
| F11 | 98300001 | 98400000 | 0.0061 | 0.0054 | 0.0061 |
| F11 | 98320001 | 98420000 | 0.0069 | 0.0059 | 0.0069 |
| F11 | 98340001 | 98440000 | 0.0075 | 0.0065 | 0.0075 |
| F11 | 98360001 | 98460000 | 0.0076 | 0.0066 | 0.0076 |
| F11 | 98380001 | 98480000 | 0.0063 | 0.0052 | 0.0063 |
| F11 | 98400001 | 98500000 | 0.0042 | 0.0033 | 0.0042 |
| F11 | 98420001 | 98520000 | 0.0048 | 0.0037 | 0.0048 |
| F11 | 98440001 | 98540000 | 0.0043 | 0.0030 | 0.0043 |
| F11 | 98460001 | 98560000 | 0.0038 | 0.0025 | 0.0038 |
| F11 | 98480001 | 98580000 | 0.0039 | 0.0024 | 0.0038 |
| F11 | 98500001 | 98600000 | 0.0059 | 0.0040 | 0.0054 |
| F11 | 98520001 | 98620000 | 0.0045 | 0.0031 | 0.0039 |
| F11 | 98540001 | 98640000 | 0.0047 | 0.0032 | 0.0039 |
| F11 | 98560001 | 98660000 | 0.0052 | 0.0034 | 0.0042 |
| F11 | 98580001 | 98680000 | 0.0062 | 0.0035 | 0.0050 |
| F11 | 98600001 | 98700000 | 0.0041 | 0.0017 | 0.0033 |
| F11 | 98620001 | 98720000 | 0.0045 | 0.0018 | 0.0038 |
| F11 | 98640001 | 98740000 | 0.0052 | 0.0026 | 0.0045 |
| F11 | 98660001 | 98760000 | 0.0052 | 0.0029 | 0.0046 |
| F11 | 98680001 | 98780000 | 0.0048 | 0.0035 | 0.0043 |
| F11 | 98700001 | 98800000 | 0.0054 | 0.0040 | 0.0048 |
| F11 | 98720001 | 98820000 | 0.0061 | 0.0048 | 0.0054 |
| F11 | 98740001 | 98840000 | 0.0057 | 0.0051 | 0.0048 |
| F11 | 98760001 | 98860000 | 0.0051 | 0.0051 | 0.0044 |
| F11 | 98780001 | 98880000 | 0.0049 | 0.0047 | 0.0043 |
| F11 | 98800001 | 98900000 | 0.0045 | 0.0050 | 0.0036 |
| F11 | 98820001 | 98920000 | 0.0049 | 0.0067 | 0.0034 |
| F11 | 98840001 | 98940000 | 0.0040 | 0.0057 | 0.0027 |
| F11 | 98860001 | 98960000 | 0.0034 | 0.0053 | 0.0021 |
| F11 | 98880001 | 98980000 | 0.0028 | 0.0051 | 0.0014 |
| F11 | 98900001 | 99000000 | 0.0026 | 0.0044 | 0.0015 |
| F11 | 98920001 | 99020000 | 0.0011 | 0.0019 | 0.0008 |

| Chr | Start | End | both | A1 | A2 |
|---|---|---|---|---|---|
| F11 | 98940001 | 99040000 | 0.0019 | 0.0024 | 0.0017 |
| F11 | 98960001 | 99060000 | 0.0020 | 0.0026 | 0.0018 |
| F11 | 98980001 | 99080000 | 0.0022 | 0.0027 | 0.0020 |
| F11 | 99000001 | 99100000 | 0.0025 | 0.0030 | 0.0023 |
| F11 | 99020001 | 99120000 | 0.0025 | 0.0033 | 0.0021 |
| F11 | 99040001 | 99140000 | 0.0016 | 0.0024 | 0.0012 |
| F11 | 99060001 | 99160000 | 0.0014 | 0.0022 | 0.0009 |
| F11 | 99080001 | 99180000 | 0.0016 | 0.0022 | 0.0011 |
| F11 | 99100001 | 99200000 | 0.0012 | 0.0017 | 0.0008 |
| F11 | 99120001 | 99220000 | 0.0015 | 0.0013 | 0.0010 |
| F11 | 99140001 | 99240000 | 0.0017 | 0.0010 | 0.0013 |
| F11 | 99160001 | 99260000 | 0.0033 | 0.0020 | 0.0030 |
| F11 | 99180001 | 99280000 | 0.0045 | 0.0027 | 0.0041 |
| F11 | 99200001 | 99300000 | 0.0044 | 0.0026 | 0.0041 |
| F11 | 99220001 | 99320000 | 0.0045 | 0.0028 | 0.0045 |
| F11 | 99240001 | 99340000 | 0.0047 | 0.0029 | 0.0044 |
| F11 | 99260001 | 99360000 | 0.0032 | 0.0016 | 0.0029 |
| F11 | 99280001 | 99380000 | 0.0023 | 0.0006 | 0.0020 |
| F11 | 99300001 | 99400000 | 0.0023 | 0.0006 | 0.0020 |
| F11 | 99320001 | 99420000 | 0.0019 | 0.0004 | 0.0016 |
| F11 | 99340001 | 99440000 | 0.0015 | 0.0003 | 0.0015 |
| F11 | 99360001 | 99460000 | 0.0013 | 0.0002 | 0.0013 |
| F11 | 99380001 | 99480000 | 0.0013 | 0.0005 | 0.0014 |
| F11 | 99400001 | 99500000 | 0.0012 | 0.0005 | 0.0013 |
| F11 | 99420001 | 99520000 | 0.0012 | 0.0005 | 0.0013 |
| F11 | 99440001 | 99540000 | 0.0012 | 0.0005 | 0.0012 |
| F11 | 99460001 | 99560000 | 0.0015 | 0.0008 | 0.0015 |
| F11 | 99480001 | 99580000 | 0.0013 | 0.0006 | 0.0012 |
| F11 | 99500001 | 99600000 | 0.0012 | 0.0006 | 0.0010 |
| F11 | 99520001 | 99620000 | 0.0009 | 0.0004 | 0.0008 |
| F11 | 99540001 | 99640000 | 0.0008 | 0.0004 | 0.0008 |
| F11 | 99560001 | 99660000 | 0.0003 | 0.0001 | 0.0003 |
| F11 | 99580001 | 99680000 | 0.0000 | | 0.0000 |
| F12 | 1 | 100000 | 0.0013 | 0.0008 | 0.0007 |
| F12 | 20001 | 120000 | 0.0011 | 0.0007 | 0.0004 |
| F12 | 40001 | 140000 | 0.0009 | 0.0006 | 0.0004 |
| F12 | 60001 | 160000 | 0.0011 | 0.0005 | 0.0006 |
| F12 | 80001 | 180000 | 0.0011 | 0.0003 | 0.0004 |
| F12 | 100001 | 200000 | 0.0012 | 0.0003 | 0.0005 |
| F12 | 120001 | 220000 | 0.0014 | 0.0003 | 0.0007 |
| F12 | 140001 | 240000 | 0.0015 | 0.0003 | 0.0007 |
| F12 | 160001 | 260000 | 0.0015 | 0.0004 | 0.0008 |
| F12 | 180001 | 280000 | 0.0015 | 0.0005 | 0.0010 |
| F12 | 200001 | 300000 | 0.0016 | 0.0005 | 0.0012 |
| F12 | 220001 | 320000 | 0.0017 | 0.0004 | 0.0014 |
| F12 | 240001 | 340000 | 0.0018 | 0.0004 | 0.0015 |
| F12 | 260001 | 360000 | 0.0018 | 0.0004 | 0.0017 |
| F12 | 280001 | 380000 | 0.0023 | 0.0006 | 0.0021 |
| F12 | 300001 | 400000 | 0.0023 | 0.0006 | 0.0021 |
| F12 | 320001 | 420000 | 0.0025 | 0.0009 | 0.0024 |
| F12 | 340001 | 440000 | 0.0028 | 0.0011 | 0.0028 |
| F12 | 360001 | 460000 | 0.0027 | 0.0011 | 0.0027 |
| F12 | 380001 | 480000 | 0.0023 | 0.0008 | 0.0022 |
| F12 | 400001 | 500000 | 0.0024 | 0.0012 | 0.0022 |
| F12 | 420001 | 520000 | 0.0022 | 0.0012 | 0.0018 |
| F12 | 440001 | 540000 | 0.0019 | 0.0011 | 0.0015 |
| F12 | 460001 | 560000 | 0.0019 | 0.0011 | 0.0014 |
| F12 | 480001 | 580000 | 0.0018 | 0.0011 | 0.0015 |

| Chr | Start | End | both | A1 | A2 |
|---|---|---|---|---|---|
| F12 | 500001 | 600000 | 0.0024 | 0.0008 | 0.0022 |
| F12 | 520001 | 620000 | 0.0027 | 0.0007 | 0.0026 |
| F12 | 540001 | 640000 | 0.0028 | 0.0006 | 0.0026 |
| F12 | 560001 | 660000 | 0.0033 | 0.0009 | 0.0028 |
| F12 | 580001 | 680000 | 0.0040 | 0.0017 | 0.0034 |
| F12 | 600001 | 700000 | 0.0033 | 0.0015 | 0.0025 |
| F12 | 620001 | 720000 | 0.0028 | 0.0014 | 0.0020 |
| F12 | 640001 | 740000 | 0.0028 | 0.0014 | 0.0020 |
| F12 | 660001 | 760000 | 0.0027 | 0.0012 | 0.0023 |
| F12 | 680001 | 780000 | 0.0024 | 0.0008 | 0.0020 |
| F12 | 700001 | 800000 | 0.0024 | 0.0007 | 0.0021 |
| F12 | 720001 | 820000 | 0.0027 | 0.0009 | 0.0025 |
| F12 | 740001 | 840000 | 0.0026 | 0.0009 | 0.0025 |
| F12 | 760001 | 860000 | 0.0021 | 0.0009 | 0.0020 |
| F12 | 780001 | 880000 | 0.0019 | 0.0005 | 0.0018 |
| F12 | 800001 | 900000 | 0.0019 | 0.0005 | 0.0018 |
| F12 | 820001 | 920000 | 0.0016 | 0.0003 | 0.0016 |
| F12 | 840001 | 940000 | 0.0016 | 0.0003 | 0.0015 |
| F12 | 860001 | 960000 | 0.0017 | 0.0004 | 0.0016 |
| F12 | 880001 | 980000 | 0.0024 | 0.0015 | 0.0024 |
| F12 | 900001 | 1000000 | 0.0023 | 0.0016 | 0.0023 |
| F12 | 920001 | 1020000 | 0.0036 | 0.0028 | 0.0034 |
| F12 | 940001 | 1040000 | 0.0040 | 0.0030 | 0.0037 |
| F12 | 960001 | 1060000 | 0.0039 | 0.0027 | 0.0034 |
| F12 | 980001 | 1080000 | 0.0034 | 0.0022 | 0.0027 |
| F12 | 1000001 | 1100000 | 0.0035 | 0.0022 | 0.0026 |
| F12 | 1020001 | 1120000 | 0.0022 | 0.0011 | 0.0014 |
| F12 | 1040001 | 1140000 | 0.0021 | 0.0013 | 0.0012 |
| F12 | 1060001 | 1160000 | 0.0023 | 0.0017 | 0.0013 |
| F12 | 1080001 | 1180000 | 0.0021 | 0.0013 | 0.0009 |
| F12 | 1100001 | 1200000 | 0.0020 | 0.0012 | 0.0009 |
| F12 | 1120001 | 1220000 | 0.0019 | 0.0011 | 0.0009 |
| F12 | 1140001 | 1240000 | 0.0017 | 0.0009 | 0.0009 |
| F12 | 1160001 | 1260000 | 0.0017 | 0.0007 | 0.0011 |
| F12 | 1180001 | 1280000 | 0.0015 | 0.0006 | 0.0012 |
| F12 | 1200001 | 1300000 | 0.0022 | 0.0011 | 0.0020 |
| F12 | 1220001 | 1320000 | 0.0022 | 0.0011 | 0.0022 |
| F12 | 1240001 | 1340000 | 0.0022 | 0.0011 | 0.0021 |
| F12 | 1260001 | 1360000 | 0.0020 | 0.0009 | 0.0020 |
| F12 | 1280001 | 1380000 | 0.0021 | 0.0011 | 0.0021 |
| F12 | 1300001 | 1400000 | 0.0018 | 0.0009 | 0.0018 |
| F12 | 1320001 | 1420000 | 0.0025 | 0.0014 | 0.0024 |
| F12 | 1340001 | 1440000 | 0.0029 | 0.0017 | 0.0028 |
| F12 | 1360001 | 1460000 | 0.0033 | 0.0019 | 0.0031 |
| F12 | 1380001 | 1480000 | 0.0034 | 0.0018 | 0.0031 |
| F12 | 1400001 | 1500000 | 0.0033 | 0.0014 | 0.0029 |
| F12 | 1420001 | 1520000 | 0.0028 | 0.0009 | 0.0024 |
| F12 | 1440001 | 1540000 | 0.0029 | 0.0010 | 0.0026 |
| F12 | 1460001 | 1560000 | 0.0026 | 0.0009 | 0.0023 |
| F12 | 1480001 | 1580000 | 0.0024 | 0.0011 | 0.0023 |
| F12 | 1500001 | 1600000 | 0.0026 | 0.0016 | 0.0025 |
| F12 | 1520001 | 1620000 | 0.0024 | 0.0017 | 0.0023 |
| F12 | 1540001 | 1640000 | 0.0019 | 0.0014 | 0.0019 |
| F12 | 1560001 | 1660000 | 0.0020 | 0.0018 | 0.0020 |
| F12 | 1580001 | 1680000 | 0.0019 | 0.0018 | 0.0019 |
| F12 | 1600001 | 1700000 | 0.0016 | 0.0015 | 0.0015 |
| F12 | 1620001 | 1720000 | 0.0017 | 0.0018 | 0.0016 |
| F12 | 1640001 | 1740000 | 0.0018 | 0.0019 | 0.0017 |

| Chr | Start | End | both | A1 | A2 |
|---|---|---|---|---|---|
| F12 | 1660001 | 1760000 | 0.0018 | 0.0020 | 0.0017 |
| F12 | 1680001 | 1780000 | 0.0017 | 0.0019 | 0.0016 |
| F12 | 1700001 | 1800000 | 0.0019 | 0.0022 | 0.0018 |
| F12 | 1720001 | 1820000 | 0.0019 | 0.0022 | 0.0017 |
| F12 | 1740001 | 1840000 | 0.0016 | 0.0020 | 0.0014 |
| F12 | 1760001 | 1860000 | 0.0015 | 0.0018 | 0.0013 |
| F12 | 1780001 | 1880000 | 0.0019 | 0.0022 | 0.0017 |
| F12 | 1800001 | 1900000 | 0.0031 | 0.0034 | 0.0028 |
| F12 | 1820001 | 1920000 | 0.0031 | 0.0034 | 0.0028 |
| F12 | 1840001 | 1940000 | 0.0033 | 0.0036 | 0.0031 |
| F12 | 1860001 | 1960000 | 0.0035 | 0.0038 | 0.0032 |
| F12 | 1880001 | 1980000 | 0.0033 | 0.0038 | 0.0030 |
| F12 | 1900001 | 2000000 | 0.0019 | 0.0025 | 0.0016 |
| F12 | 1920001 | 2020000 | 0.0022 | 0.0028 | 0.0019 |
| F12 | 1940001 | 2040000 | 0.0024 | 0.0031 | 0.0021 |
| F12 | 1960001 | 2060000 | 0.0026 | 0.0031 | 0.0023 |
| F12 | 1980001 | 2080000 | 0.0027 | 0.0029 | 0.0024 |
| F12 | 2000001 | 2100000 | 0.0029 | 0.0025 | 0.0025 |
| F12 | 2020001 | 2120000 | 0.0030 | 0.0024 | 0.0024 |
| F12 | 2040001 | 2140000 | 0.0025 | 0.0017 | 0.0018 |
| F12 | 2060001 | 2160000 | 0.0023 | 0.0012 | 0.0015 |
| F12 | 2080001 | 2180000 | 0.0020 | 0.0009 | 0.0012 |
| F12 | 2100001 | 2200000 | 0.0018 | 0.0009 | 0.0012 |
| F12 | 2120001 | 2220000 | 0.0013 | 0.0005 | 0.0008 |
| F12 | 2140001 | 2240000 | 0.0013 | 0.0004 | 0.0008 |
| F12 | 2160001 | 2260000 | 0.0019 | 0.0010 | 0.0015 |
| F12 | 2180001 | 2280000 | 0.0017 | 0.0008 | 0.0013 |
| F12 | 2200001 | 2300000 | 0.0028 | 0.0019 | 0.0023 |
| F12 | 2220001 | 2320000 | 0.0041 | 0.0030 | 0.0035 |
| F12 | 2240001 | 2340000 | 0.0043 | 0.0033 | 0.0036 |
| F12 | 2260001 | 2360000 | 0.0036 | 0.0028 | 0.0030 |
| F12 | 2280001 | 2380000 | 0.0037 | 0.0031 | 0.0032 |
| F12 | 2300001 | 2400000 | 0.0033 | 0.0025 | 0.0028 |
| F12 | 2320001 | 2420000 | 0.0021 | 0.0014 | 0.0016 |
| F12 | 2340001 | 2440000 | 0.0018 | 0.0012 | 0.0014 |
| F12 | 2360001 | 2460000 | 0.0017 | 0.0009 | 0.0012 |
| F12 | 2380001 | 2480000 | 0.0018 | 0.0008 | 0.0012 |
| F12 | 2400001 | 2500000 | 0.0016 | 0.0004 | 0.0009 |
| F12 | 2420001 | 2520000 | 0.0020 | 0.0008 | 0.0013 |
| F12 | 2440001 | 2540000 | 0.0021 | 0.0008 | 0.0013 |
| F12 | 2460001 | 2560000 | 0.0029 | 0.0014 | 0.0019 |
| F12 | 2480001 | 2580000 | 0.0039 | 0.0024 | 0.0029 |
| F12 | 2500001 | 2600000 | 0.0035 | 0.0024 | 0.0027 |
| F12 | 2520001 | 2620000 | 0.0034 | 0.0023 | 0.0026 |
| F12 | 2540001 | 2640000 | 0.0052 | 0.0040 | 0.0043 |
| F12 | 2560001 | 2660000 | 0.0050 | 0.0042 | 0.0044 |
| F12 | 2580001 | 2680000 | 0.0043 | 0.0035 | 0.0038 |
| F12 | 2600001 | 2700000 | 0.0060 | 0.0047 | 0.0052 |
| F12 | 2620001 | 2720000 | 0.0061 | 0.0046 | 0.0051 |
| F12 | 2640001 | 2740000 | 0.0054 | 0.0039 | 0.0046 |
| F12 | 2660001 | 2760000 | 0.0049 | 0.0033 | 0.0040 |
| F12 | 2680001 | 2780000 | 0.0044 | 0.0028 | 0.0034 |
| F12 | 2700001 | 2800000 | 0.0026 | 0.0015 | 0.0019 |
| F12 | 2720001 | 2820000 | 0.0027 | 0.0017 | 0.0021 |
| F12 | 2740001 | 2840000 | 0.0020 | 0.0008 | 0.0013 |
| F12 | 2760001 | 2860000 | 0.0033 | 0.0019 | 0.0024 |
| F12 | 2780001 | 2880000 | 0.0038 | 0.0026 | 0.0030 |
| F12 | 2800001 | 2900000 | 0.0039 | 0.0026 | 0.0030 |

| Chr | Start | End | both | A1 | A2 |
|---|---|---|---|---|---|
| F12 | 2820001 | 2920000 | 0.0041 | 0.0029 | 0.0032 |
| F12 | 2840001 | 2940000 | 0.0039 | 0.0028 | 0.0031 |
| F12 | 2860001 | 2960000 | 0.0029 | 0.0018 | 0.0021 |
| F12 | 2880001 | 2980000 | 0.0024 | 0.0013 | 0.0016 |
| F12 | 2900001 | 3000000 | 0.0023 | 0.0014 | 0.0016 |
| F12 | 2920001 | 3020000 | 0.0020 | 0.0011 | 0.0013 |
| F12 | 2940001 | 3040000 | 0.0023 | 0.0014 | 0.0017 |
| F12 | 2960001 | 3060000 | 0.0025 | 0.0015 | 0.0019 |
| F12 | 2980001 | 3080000 | 0.0047 | 0.0034 | 0.0043 |
| F12 | 3000001 | 3100000 | 0.0053 | 0.0039 | 0.0049 |
| F12 | 3020001 | 3120000 | 0.0055 | 0.0041 | 0.0051 |
| F12 | 3040001 | 3140000 | 0.0049 | 0.0038 | 0.0046 |
| F12 | 3060001 | 3160000 | 0.0046 | 0.0036 | 0.0044 |
| F12 | 3080001 | 3180000 | 0.0022 | 0.0015 | 0.0021 |
| F12 | 3100001 | 3200000 | 0.0018 | 0.0011 | 0.0018 |
| F12 | 3120001 | 3220000 | 0.0017 | 0.0011 | 0.0017 |
| F12 | 3140001 | 3240000 | 0.0019 | 0.0013 | 0.0019 |
| F12 | 3160001 | 3260000 | 0.0020 | 0.0016 | 0.0020 |
| F12 | 3180001 | 3280000 | 0.0026 | 0.0025 | 0.0026 |
| F12 | 3200001 | 3300000 | 0.0032 | 0.0031 | 0.0031 |
| F12 | 3220001 | 3320000 | 0.0027 | 0.0026 | 0.0027 |
| F12 | 3240001 | 3340000 | 0.0029 | 0.0027 | 0.0029 |
| F12 | 3260001 | 3360000 | 0.0029 | 0.0022 | 0.0027 |
| F12 | 3280001 | 3380000 | 0.0024 | 0.0014 | 0.0021 |
| F12 | 3300001 | 3400000 | 0.0018 | 0.0005 | 0.0013 |
| F12 | 3320001 | 3420000 | 0.0020 | 0.0006 | 0.0013 |
| F12 | 3340001 | 3440000 | 0.0019 | 0.0004 | 0.0010 |
| F12 | 3360001 | 3460000 | 0.0020 | 0.0006 | 0.0011 |
| F12 | 3380001 | 3480000 | 0.0027 | 0.0014 | 0.0018 |
| F12 | 3400001 | 3500000 | 0.0028 | 0.0017 | 0.0020 |
| F12 | 3420001 | 3520000 | 0.0033 | 0.0021 | 0.0027 |
| F12 | 3440001 | 3540000 | 0.0033 | 0.0021 | 0.0027 |
| F12 | 3460001 | 3560000 | 0.0033 | 0.0021 | 0.0026 |
| F12 | 3480001 | 3580000 | 0.0028 | 0.0016 | 0.0022 |
| F12 | 3500001 | 3600000 | 0.0026 | 0.0013 | 0.0019 |
| F12 | 3520001 | 3620000 | 0.0024 | 0.0010 | 0.0012 |
| F12 | 3540001 | 3640000 | 0.0031 | 0.0017 | 0.0019 |
| F12 | 3560001 | 3660000 | 0.0028 | 0.0015 | 0.0016 |
| F12 | 3580001 | 3680000 | 0.0028 | 0.0015 | 0.0017 |
| F12 | 3600001 | 3700000 | 0.0030 | 0.0016 | 0.0020 |
| F12 | 3620001 | 3720000 | 0.0026 | 0.0015 | 0.0019 |
| F12 | 3640001 | 3740000 | 0.0017 | 0.0007 | 0.0012 |
| F12 | 3660001 | 3760000 | 0.0023 | 0.0009 | 0.0019 |
| F12 | 3680001 | 3780000 | 0.0028 | 0.0012 | 0.0025 |
| F12 | 3700001 | 3800000 | 0.0029 | 0.0015 | 0.0028 |
| F12 | 3720001 | 3820000 | 0.0031 | 0.0016 | 0.0030 |
| F12 | 3740001 | 3840000 | 0.0034 | 0.0018 | 0.0034 |
| F12 | 3760001 | 3860000 | 0.0033 | 0.0020 | 0.0033 |
| F12 | 3780001 | 3880000 | 0.0030 | 0.0017 | 0.0030 |
| F12 | 3800001 | 3900000 | 0.0032 | 0.0017 | 0.0033 |
| F12 | 3820001 | 3920000 | 0.0034 | 0.0016 | 0.0034 |
| F12 | 3840001 | 3940000 | 0.0033 | 0.0016 | 0.0034 |
| F12 | 3860001 | 3960000 | 0.0030 | 0.0013 | 0.0030 |
| F12 | 3880001 | 3980000 | 0.0029 | 0.0013 | 0.0026 |
| F12 | 3900001 | 4000000 | 0.0024 | 0.0009 | 0.0020 |
| F12 | 3920001 | 4020000 | 0.0025 | 0.0011 | 0.0019 |
| F12 | 3940001 | 4040000 | 0.0022 | 0.0011 | 0.0015 |
| F12 | 3960001 | 4060000 | 0.0025 | 0.0014 | 0.0018 |

| Chr | Start | End | both | A1 | A2 |
|---|---|---|---|---|---|
| F12 | 3980001 | 4080000 | 0.0030 | 0.0020 | 0.0026 |
| F12 | 4000001 | 4100000 | 0.0032 | 0.0019 | 0.0028 |
| F12 | 4020001 | 4120000 | 0.0033 | 0.0021 | 0.0030 |
| F12 | 4040001 | 4140000 | 0.0032 | 0.0019 | 0.0029 |
| F12 | 4060001 | 4160000 | 0.0035 | 0.0019 | 0.0028 |
| F12 | 4080001 | 4180000 | 0.0029 | 0.0012 | 0.0020 |
| F12 | 4100001 | 4200000 | 0.0029 | 0.0015 | 0.0018 |
| F12 | 4120001 | 4220000 | 0.0026 | 0.0012 | 0.0015 |
| F12 | 4140001 | 4240000 | 0.0028 | 0.0013 | 0.0016 |
| F12 | 4160001 | 4260000 | 0.0022 | 0.0010 | 0.0012 |
| F12 | 4180001 | 4280000 | 0.0022 | 0.0010 | 0.0012 |
| F12 | 4200001 | 4300000 | 0.0019 | 0.0007 | 0.0009 |
| F12 | 4220001 | 4320000 | 0.0022 | 0.0009 | 0.0011 |
| F12 | 4240001 | 4340000 | 0.0022 | 0.0008 | 0.0010 |
| F12 | 4260001 | 4360000 | 0.0024 | 0.0010 | 0.0011 |
| F12 | 4280001 | 4380000 | 0.0024 | 0.0008 | 0.0010 |
| F12 | 4300001 | 4400000 | 0.0032 | 0.0016 | 0.0018 |
| F12 | 4320001 | 4420000 | 0.0029 | 0.0014 | 0.0015 |
| F12 | 4340001 | 4440000 | 0.0031 | 0.0018 | 0.0020 |
| F12 | 4360001 | 4460000 | 0.0029 | 0.0016 | 0.0018 |
| F12 | 4380001 | 4480000 | 0.0032 | 0.0019 | 0.0022 |
| F12 | 4400001 | 4500000 | 0.0027 | 0.0016 | 0.0018 |
| F12 | 4420001 | 4520000 | 0.0030 | 0.0020 | 0.0022 |
| F12 | 4440001 | 4540000 | 0.0027 | 0.0016 | 0.0019 |
| F12 | 4460001 | 4560000 | 0.0027 | 0.0018 | 0.0021 |
| F12 | 4480001 | 4580000 | 0.0029 | 0.0017 | 0.0020 |
| F12 | 4500001 | 4600000 | 0.0025 | 0.0011 | 0.0015 |
| F12 | 4520001 | 4620000 | 0.0024 | 0.0011 | 0.0014 |
| F12 | 4540001 | 4640000 | 0.0024 | 0.0010 | 0.0014 |
| F12 | 4560001 | 4660000 | 0.0028 | 0.0011 | 0.0015 |
| F12 | 4580001 | 4680000 | 0.0023 | 0.0009 | 0.0012 |
| F12 | 4600001 | 4700000 | 0.0027 | 0.0011 | 0.0016 |
| F12 | 4620001 | 4720000 | 0.0026 | 0.0010 | 0.0015 |
| F12 | 4640001 | 4740000 | 0.0028 | 0.0015 | 0.0019 |
| F12 | 4660001 | 4760000 | 0.0042 | 0.0030 | 0.0035 |
| F12 | 4680001 | 4780000 | 0.0046 | 0.0035 | 0.0039 |
| F12 | 4700001 | 4800000 | 0.0047 | 0.0039 | 0.0040 |
| F12 | 4720001 | 4820000 | 0.0045 | 0.0036 | 0.0039 |
| F12 | 4740001 | 4840000 | 0.0043 | 0.0033 | 0.0037 |
| F12 | 4760001 | 4860000 | 0.0026 | 0.0017 | 0.0020 |
| F12 | 4780001 | 4880000 | 0.0020 | 0.0013 | 0.0015 |
| F12 | 4800001 | 4900000 | 0.0023 | 0.0017 | 0.0019 |
| F12 | 4820001 | 4920000 | 0.0028 | 0.0021 | 0.0024 |
| F12 | 4840001 | 4940000 | 0.0026 | 0.0022 | 0.0024 |
| F12 | 4860001 | 4960000 | 0.0030 | 0.0024 | 0.0027 |
| F12 | 4880001 | 4980000 | 0.0034 | 0.0026 | 0.0030 |
| F12 | 4900001 | 5000000 | 0.0030 | 0.0021 | 0.0026 |
| F12 | 4920001 | 5020000 | 0.0023 | 0.0016 | 0.0019 |
| F12 | 4940001 | 5040000 | 0.0021 | 0.0013 | 0.0016 |
| F12 | 4960001 | 5060000 | 0.0017 | 0.0012 | 0.0014 |
| F12 | 4980001 | 5080000 | 0.0014 | 0.0011 | 0.0013 |
| F12 | 5000001 | 5100000 | 0.0012 | 0.0006 | 0.0009 |
| F12 | 5020001 | 5120000 | 0.0020 | 0.0010 | 0.0014 |
| F12 | 5040001 | 5140000 | 0.0024 | 0.0014 | 0.0017 |
| F12 | 5060001 | 5160000 | 0.0046 | 0.0034 | 0.0036 |
| F12 | 5080001 | 5180000 | 0.0047 | 0.0036 | 0.0036 |
| F12 | 5100001 | 5200000 | 0.0048 | 0.0037 | 0.0037 |
| F12 | 5120001 | 5220000 | 0.0042 | 0.0033 | 0.0032 |

| Chr | Start | End | both | A1 | A2 |
|---|---|---|---|---|---|
| F12 | 5140001 | 5240000 | 0.0060 | 0.0050 | 0.0051 |
| F12 | 5160001 | 5260000 | 0.0052 | 0.0044 | 0.0045 |
| F12 | 5180001 | 5280000 | 0.0055 | 0.0046 | 0.0046 |
| F12 | 5200001 | 5300000 | 0.0052 | 0.0045 | 0.0045 |
| F12 | 5220001 | 5320000 | 0.0057 | 0.0049 | 0.0049 |
| F12 | 5240001 | 5340000 | 0.0037 | 0.0029 | 0.0029 |
| F12 | 5260001 | 5360000 | 0.0024 | 0.0016 | 0.0016 |
| F12 | 5280001 | 5380000 | 0.0017 | 0.0010 | 0.0011 |
| F12 | 5300001 | 5400000 | 0.0016 | 0.0010 | 0.0012 |
| F12 | 5320001 | 5420000 | 0.0015 | 0.0008 | 0.0011 |
| F12 | 5340001 | 5440000 | 0.0020 | 0.0013 | 0.0016 |
| F12 | 5360001 | 5460000 | 0.0026 | 0.0015 | 0.0021 |
| F12 | 5380001 | 5480000 | 0.0028 | 0.0016 | 0.0022 |
| F12 | 5400001 | 5500000 | 0.0031 | 0.0016 | 0.0022 |
| F12 | 5420001 | 5520000 | 0.0027 | 0.0015 | 0.0019 |
| F12 | 5440001 | 5540000 | 0.0025 | 0.0011 | 0.0016 |
| F12 | 5460001 | 5560000 | 0.0018 | 0.0008 | 0.0010 |
| F12 | 5480001 | 5580000 | 0.0028 | 0.0016 | 0.0020 |
| F12 | 5500001 | 5600000 | 0.0036 | 0.0026 | 0.0030 |
| F12 | 5520001 | 5620000 | 0.0037 | 0.0029 | 0.0032 |
| F12 | 5540001 | 5640000 | 0.0038 | 0.0030 | 0.0035 |
| F12 | 5560001 | 5660000 | 0.0036 | 0.0030 | 0.0034 |
| F12 | 5580001 | 5680000 | 0.0032 | 0.0028 | 0.0031 |
| F12 | 5600001 | 5700000 | 0.0022 | 0.0018 | 0.0021 |
| F12 | 5620001 | 5720000 | 0.0028 | 0.0023 | 0.0027 |
| F12 | 5640001 | 5740000 | 0.0026 | 0.0021 | 0.0026 |
| F12 | 5660001 | 5760000 | 0.0036 | 0.0030 | 0.0036 |
| F12 | 5680001 | 5780000 | 0.0031 | 0.0026 | 0.0030 |
| F12 | 5700001 | 5800000 | 0.0037 | 0.0031 | 0.0036 |
| F12 | 5720001 | 5820000 | 0.0033 | 0.0027 | 0.0033 |
| F12 | 5740001 | 5840000 | 0.0036 | 0.0030 | 0.0035 |
| F12 | 5760001 | 5860000 | 0.0031 | 0.0026 | 0.0031 |
| F12 | 5780001 | 5880000 | 0.0031 | 0.0026 | 0.0031 |
| F12 | 5800001 | 5900000 | 0.0035 | 0.0030 | 0.0035 |
| F12 | 5820001 | 5920000 | 0.0035 | 0.0030 | 0.0035 |
| F12 | 5840001 | 5940000 | 0.0029 | 0.0024 | 0.0028 |
| F12 | 5860001 | 5960000 | 0.0031 | 0.0025 | 0.0029 |
| F12 | 5880001 | 5980000 | 0.0034 | 0.0027 | 0.0033 |
| F12 | 5900001 | 6000000 | 0.0027 | 0.0022 | 0.0025 |
| F12 | 5920001 | 6020000 | 0.0025 | 0.0020 | 0.0022 |
| F12 | 5940001 | 6040000 | 0.0028 | 0.0020 | 0.0023 |
| F12 | 5960001 | 6060000 | 0.0024 | 0.0018 | 0.0019 |
| F12 | 5980001 | 6080000 | 0.0023 | 0.0015 | 0.0017 |
| F12 | 6000001 | 6100000 | 0.0022 | 0.0012 | 0.0014 |
| F12 | 6020001 | 6120000 | 0.0021 | 0.0011 | 0.0013 |
| F12 | 6040001 | 6140000 | 0.0021 | 0.0015 | 0.0013 |
| F12 | 6060001 | 6160000 | 0.0021 | 0.0016 | 0.0013 |
| F12 | 6080001 | 6180000 | 0.0019 | 0.0017 | 0.0010 |
| F12 | 6100001 | 6200000 | 0.0018 | 0.0021 | 0.0008 |
| F12 | 6120001 | 6220000 | 0.0022 | 0.0026 | 0.0014 |
| F12 | 6140001 | 6240000 | 0.0022 | 0.0024 | 0.0015 |
| F12 | 6160001 | 6260000 | 0.0028 | 0.0028 | 0.0022 |
| F12 | 6180001 | 6280000 | 0.0038 | 0.0037 | 0.0033 |
| F12 | 6200001 | 6300000 | 0.0050 | 0.0045 | 0.0047 |
| F12 | 6220001 | 6320000 | 0.0045 | 0.0041 | 0.0041 |
| F12 | 6240001 | 6340000 | 0.0043 | 0.0039 | 0.0037 |
| F12 | 6260001 | 6360000 | 0.0039 | 0.0036 | 0.0029 |
| F12 | 6280001 | 6380000 | 0.0036 | 0.0033 | 0.0026 |

| Chr | Start | End | both | A1 | A2 |
|---|---|---|---|---|---|
| F12 | 6300001 | 6400000 | 0.0023 | 0.0022 | 0.0012 |
| F12 | 6320001 | 6420000 | 0.0024 | 0.0024 | 0.0013 |
| F12 | 6340001 | 6440000 | 0.0024 | 0.0024 | 0.0014 |
| F12 | 6360001 | 6460000 | 0.0025 | 0.0025 | 0.0017 |
| F12 | 6380001 | 6480000 | 0.0022 | 0.0019 | 0.0014 |
| F12 | 6400001 | 6500000 | 0.0029 | 0.0024 | 0.0018 |
| F12 | 6420001 | 6520000 | 0.0027 | 0.0020 | 0.0017 |
| F12 | 6440001 | 6540000 | 0.0031 | 0.0024 | 0.0019 |
| F12 | 6460001 | 6560000 | 0.0029 | 0.0020 | 0.0016 |
| F12 | 6480001 | 6580000 | 0.0031 | 0.0021 | 0.0018 |
| F12 | 6500001 | 6600000 | 0.0026 | 0.0017 | 0.0015 |
| F12 | 6520001 | 6620000 | 0.0034 | 0.0024 | 0.0021 |
| F12 | 6540001 | 6640000 | 0.0033 | 0.0023 | 0.0022 |
| F12 | 6560001 | 6660000 | 0.0031 | 0.0021 | 0.0023 |
| F12 | 6580001 | 6680000 | 0.0023 | 0.0016 | 0.0017 |
| F12 | 6600001 | 6700000 | 0.0026 | 0.0017 | 0.0019 |
| F12 | 6620001 | 6720000 | 0.0027 | 0.0019 | 0.0022 |
| F12 | 6640001 | 6740000 | 0.0029 | 0.0020 | 0.0024 |
| F12 | 6660001 | 6760000 | 0.0029 | 0.0021 | 0.0023 |
| F12 | 6680001 | 6780000 | 0.0029 | 0.0021 | 0.0023 |
| F12 | 6700001 | 6800000 | 0.0028 | 0.0022 | 0.0024 |
| F12 | 6720001 | 6820000 | 0.0025 | 0.0017 | 0.0019 |
| F12 | 6740001 | 6840000 | 0.0028 | 0.0021 | 0.0021 |
| F12 | 6760001 | 6860000 | 0.0027 | 0.0018 | 0.0019 |
| F12 | 6780001 | 6880000 | 0.0031 | 0.0021 | 0.0022 |
| F12 | 6800001 | 6900000 | 0.0037 | 0.0026 | 0.0027 |
| F12 | 6820001 | 6920000 | 0.0033 | 0.0026 | 0.0026 |
| F12 | 6840001 | 6940000 | 0.0034 | 0.0029 | 0.0029 |
| F12 | 6860001 | 6960000 | 0.0037 | 0.0033 | 0.0033 |
| F12 | 6880001 | 6980000 | 0.0039 | 0.0037 | 0.0037 |
| F12 | 6900001 | 7000000 | 0.0035 | 0.0034 | 0.0033 |
| F12 | 6920001 | 7020000 | 0.0033 | 0.0032 | 0.0032 |
| F12 | 6940001 | 7040000 | 0.0024 | 0.0023 | 0.0022 |
| F12 | 6960001 | 7060000 | 0.0023 | 0.0022 | 0.0021 |
| F12 | 6980001 | 7080000 | 0.0017 | 0.0014 | 0.0013 |
| F12 | 7000001 | 7100000 | 0.0013 | 0.0008 | 0.0008 |
| F12 | 7020001 | 7120000 | 0.0020 | 0.0014 | 0.0015 |
| F12 | 7040001 | 7140000 | 0.0030 | 0.0022 | 0.0022 |
| F12 | 7060001 | 7160000 | 0.0027 | 0.0019 | 0.0020 |
| F12 | 7080001 | 7180000 | 0.0030 | 0.0024 | 0.0024 |
| F12 | 7100001 | 7200000 | 0.0030 | 0.0026 | 0.0026 |
| F12 | 7120001 | 7220000 | 0.0028 | 0.0024 | 0.0024 |
| F12 | 7140001 | 7240000 | 0.0025 | 0.0022 | 0.0023 |
| F12 | 7160001 | 7260000 | 0.0027 | 0.0025 | 0.0026 |
| F12 | 7180001 | 7280000 | 0.0040 | 0.0034 | 0.0037 |
| F12 | 7200001 | 7300000 | 0.0039 | 0.0032 | 0.0035 |
| F12 | 7220001 | 7320000 | 0.0041 | 0.0033 | 0.0036 |
| F12 | 7240001 | 7340000 | 0.0038 | 0.0028 | 0.0031 |
| F12 | 7260001 | 7360000 | 0.0037 | 0.0027 | 0.0030 |
| F12 | 7280001 | 7380000 | 0.0029 | 0.0021 | 0.0020 |
| F12 | 7300001 | 7400000 | 0.0028 | 0.0020 | 0.0019 |
| F12 | 7320001 | 7420000 | 0.0032 | 0.0022 | 0.0022 |
| F12 | 7340001 | 7440000 | 0.0032 | 0.0023 | 0.0023 |
| F12 | 7360001 | 7460000 | 0.0032 | 0.0021 | 0.0021 |
| F12 | 7380001 | 7480000 | 0.0031 | 0.0018 | 0.0021 |
| F12 | 7400001 | 7500000 | 0.0034 | 0.0020 | 0.0024 |
| F12 | 7420001 | 7520000 | 0.0026 | 0.0014 | 0.0016 |
| F12 | 7440001 | 7540000 | 0.0023 | 0.0012 | 0.0014 |

| Chr | Start | End | both | A1 | A2 |
|---|---|---|---|---|---|
| F12 | 7460001 | 7560000 | 0.0025 | 0.0014 | 0.0015 |
| F12 | 7480001 | 7580000 | 0.0022 | 0.0011 | 0.0013 |
| F12 | 7500001 | 7600000 | 0.0022 | 0.0012 | 0.0013 |
| F12 | 7520001 | 7620000 | 0.0020 | 0.0010 | 0.0011 |
| F12 | 7540001 | 7640000 | 0.0028 | 0.0016 | 0.0018 |
| F12 | 7560001 | 7660000 | 0.0032 | 0.0018 | 0.0024 |
| F12 | 7580001 | 7680000 | 0.0036 | 0.0021 | 0.0028 |
| F12 | 7600001 | 7700000 | 0.0036 | 0.0021 | 0.0027 |
| F12 | 7620001 | 7720000 | 0.0037 | 0.0024 | 0.0030 |
| F12 | 7640001 | 7740000 | 0.0029 | 0.0019 | 0.0022 |
| F12 | 7660001 | 7760000 | 0.0020 | 0.0013 | 0.0013 |
| F12 | 7680001 | 7780000 | 0.0015 | 0.0012 | 0.0011 |
| F12 | 7700001 | 7800000 | 0.0013 | 0.0011 | 0.0008 |
| F12 | 7720001 | 7820000 | 0.0013 | 0.0012 | 0.0006 |
| F12 | 7740001 | 7840000 | 0.0015 | 0.0014 | 0.0008 |
| F12 | 7760001 | 7860000 | 0.0020 | 0.0020 | 0.0012 |
| F12 | 7780001 | 7880000 | 0.0027 | 0.0028 | 0.0017 |
| F12 | 7800001 | 7900000 | 0.0029 | 0.0030 | 0.0019 |
| F12 | 7820001 | 7920000 | 0.0033 | 0.0033 | 0.0023 |
| F12 | 7840001 | 7940000 | 0.0032 | 0.0032 | 0.0022 |
| F12 | 7860001 | 7960000 | 0.0031 | 0.0032 | 0.0022 |
| F12 | 7880001 | 7980000 | 0.0024 | 0.0024 | 0.0017 |
| F12 | 7900001 | 8000000 | 0.0026 | 0.0025 | 0.0022 |
| F12 | 7920001 | 8020000 | 0.0018 | 0.0017 | 0.0015 |
| F12 | 7940001 | 8040000 | 0.0020 | 0.0018 | 0.0019 |
| F12 | 7960001 | 8060000 | 0.0021 | 0.0019 | 0.0021 |
| F12 | 7980001 | 8080000 | 0.0020 | 0.0019 | 0.0017 |
| F12 | 8000001 | 8100000 | 0.0014 | 0.0014 | 0.0011 |
| F12 | 8020001 | 8120000 | 0.0016 | 0.0017 | 0.0013 |
| F12 | 8040001 | 8140000 | 0.0018 | 0.0019 | 0.0013 |
| F12 | 8060001 | 8160000 | 0.0017 | 0.0015 | 0.0010 |
| F12 | 8080001 | 8180000 | 0.0019 | 0.0014 | 0.0012 |
| F12 | 8100001 | 8200000 | 0.0024 | 0.0015 | 0.0014 |
| F12 | 8120001 | 8220000 | 0.0025 | 0.0015 | 0.0016 |
| F12 | 8140001 | 8240000 | 0.0023 | 0.0011 | 0.0013 |
| F12 | 8160001 | 8260000 | 0.0025 | 0.0014 | 0.0016 |
| F12 | 8180001 | 8280000 | 0.0033 | 0.0022 | 0.0025 |
| F12 | 8200001 | 8300000 | 0.0034 | 0.0028 | 0.0029 |
| F12 | 8220001 | 8320000 | 0.0045 | 0.0035 | 0.0040 |
| F12 | 8240001 | 8340000 | 0.0046 | 0.0036 | 0.0042 |
| F12 | 8260001 | 8360000 | 0.0046 | 0.0035 | 0.0041 |
| F12 | 8280001 | 8380000 | 0.0038 | 0.0027 | 0.0032 |
| F12 | 8300001 | 8400000 | 0.0047 | 0.0034 | 0.0040 |
| F12 | 8320001 | 8420000 | 0.0041 | 0.0030 | 0.0032 |
| F12 | 8340001 | 8440000 | 0.0040 | 0.0029 | 0.0031 |
| F12 | 8360001 | 8460000 | 0.0037 | 0.0026 | 0.0028 |
| F12 | 8380001 | 8480000 | 0.0038 | 0.0029 | 0.0029 |
| F12 | 8400001 | 8500000 | 0.0032 | 0.0024 | 0.0024 |
| F12 | 8420001 | 8520000 | 0.0047 | 0.0038 | 0.0040 |
| F12 | 8440001 | 8540000 | 0.0055 | 0.0045 | 0.0045 |
| F12 | 8460001 | 8560000 | 0.0057 | 0.0048 | 0.0047 |
| F12 | 8480001 | 8580000 | 0.0058 | 0.0048 | 0.0049 |
| F12 | 8500001 | 8600000 | 0.0057 | 0.0045 | 0.0047 |
| F12 | 8520001 | 8620000 | 0.0041 | 0.0028 | 0.0029 |
| F12 | 8540001 | 8640000 | 0.0034 | 0.0021 | 0.0022 |
| F12 | 8560001 | 8660000 | 0.0031 | 0.0017 | 0.0018 |
| F12 | 8580001 | 8680000 | 0.0031 | 0.0016 | 0.0018 |
| F12 | 8600001 | 8700000 | 0.0027 | 0.0014 | 0.0015 |

| Chr | Start | End | both | A1 | A2 |
|---|---|---|---|---|---|
| F12 | 8620001 | 8720000 | 0.0021 | 0.0010 | 0.0010 |
| F12 | 8640001 | 8740000 | 0.0019 | 0.0010 | 0.0011 |
| F12 | 8660001 | 8760000 | 0.0030 | 0.0023 | 0.0022 |
| F12 | 8680001 | 8780000 | 0.0028 | 0.0020 | 0.0018 |
| F12 | 8700001 | 8800000 | 0.0027 | 0.0018 | 0.0015 |
| F12 | 8720001 | 8820000 | 0.0032 | 0.0023 | 0.0020 |
| F12 | 8740001 | 8840000 | 0.0031 | 0.0023 | 0.0019 |
| F12 | 8760001 | 8860000 | 0.0026 | 0.0018 | 0.0015 |
| F12 | 8780001 | 8880000 | 0.0024 | 0.0018 | 0.0015 |
| F12 | 8800001 | 8900000 | 0.0036 | 0.0029 | 0.0027 |
| F12 | 8820001 | 8920000 | 0.0033 | 0.0024 | 0.0023 |
| F12 | 8840001 | 8940000 | 0.0036 | 0.0027 | 0.0028 |
| F12 | 8860001 | 8960000 | 0.0036 | 0.0025 | 0.0026 |
| F12 | 8880001 | 8980000 | 0.0040 | 0.0028 | 0.0030 |
| F12 | 8900001 | 9000000 | 0.0029 | 0.0019 | 0.0021 |
| F12 | 8920001 | 9020000 | 0.0031 | 0.0020 | 0.0022 |
| F12 | 8940001 | 9040000 | 0.0028 | 0.0017 | 0.0019 |
| F12 | 8960001 | 9060000 | 0.0025 | 0.0015 | 0.0017 |
| F12 | 8980001 | 9080000 | 0.0027 | 0.0015 | 0.0018 |
| F12 | 9000001 | 9100000 | 0.0032 | 0.0020 | 0.0023 |
| F12 | 9020001 | 9120000 | 0.0035 | 0.0025 | 0.0027 |
| F12 | 9040001 | 9140000 | 0.0038 | 0.0027 | 0.0030 |
| F12 | 9060001 | 9160000 | 0.0042 | 0.0029 | 0.0030 |
| F12 | 9080001 | 9180000 | 0.0041 | 0.0028 | 0.0027 |
| F12 | 9100001 | 9200000 | 0.0046 | 0.0032 | 0.0031 |
| F12 | 9120001 | 9220000 | 0.0052 | 0.0037 | 0.0037 |
| F12 | 9140001 | 9240000 | 0.0064 | 0.0049 | 0.0049 |
| F12 | 9160001 | 9260000 | 0.0066 | 0.0053 | 0.0053 |
| F12 | 9180001 | 9280000 | 0.0060 | 0.0049 | 0.0051 |
| F12 | 9200001 | 9300000 | 0.0057 | 0.0045 | 0.0046 |
| F12 | 9220001 | 9320000 | 0.0045 | 0.0035 | 0.0035 |
| F12 | 9240001 | 9340000 | 0.0030 | 0.0020 | 0.0020 |
| F12 | 9260001 | 9360000 | 0.0026 | 0.0013 | 0.0015 |
| F12 | 9280001 | 9380000 | 0.0034 | 0.0019 | 0.0022 |
| F12 | 9300001 | 9400000 | 0.0031 | 0.0015 | 0.0018 |
| F12 | 9320001 | 9420000 | 0.0037 | 0.0018 | 0.0022 |
| F12 | 9340001 | 9440000 | 0.0036 | 0.0019 | 0.0022 |
| F12 | 9360001 | 9460000 | 0.0031 | 0.0018 | 0.0020 |
| F12 | 9380001 | 9480000 | 0.0026 | 0.0013 | 0.0015 |
| F12 | 9400001 | 9500000 | 0.0026 | 0.0014 | 0.0014 |
| F12 | 9420001 | 9520000 | 0.0024 | 0.0015 | 0.0014 |
| F12 | 9440001 | 9540000 | 0.0024 | 0.0015 | 0.0015 |
| F12 | 9460001 | 9560000 | 0.0032 | 0.0020 | 0.0021 |
| F12 | 9480001 | 9580000 | 0.0039 | 0.0027 | 0.0027 |
| F12 | 9500001 | 9600000 | 0.0034 | 0.0024 | 0.0024 |
| F12 | 9520001 | 9620000 | 0.0030 | 0.0024 | 0.0019 |
| F12 | 9540001 | 9640000 | 0.0035 | 0.0030 | 0.0023 |
| F12 | 9560001 | 9660000 | 0.0034 | 0.0032 | 0.0021 |
| F12 | 9580001 | 9680000 | 0.0031 | 0.0031 | 0.0017 |
| F12 | 9600001 | 9700000 | 0.0033 | 0.0031 | 0.0019 |
| F12 | 9620001 | 9720000 | 0.0036 | 0.0031 | 0.0022 |
| F12 | 9640001 | 9740000 | 0.0034 | 0.0029 | 0.0020 |
| F12 | 9660001 | 9760000 | 0.0031 | 0.0025 | 0.0019 |
| F12 | 9680001 | 9780000 | 0.0029 | 0.0023 | 0.0019 |
| F12 | 9700001 | 9800000 | 0.0026 | 0.0020 | 0.0017 |
| F12 | 9720001 | 9820000 | 0.0025 | 0.0016 | 0.0016 |
| F12 | 9740001 | 9840000 | 0.0026 | 0.0015 | 0.0018 |
| F12 | 9760001 | 9860000 | 0.0029 | 0.0016 | 0.0021 |

| Chr | Start | End | both | A1 | A2 |
|---|---|---|---|---|---|
| F12 | 9780001 | 9880000 | 0.0031 | 0.0017 | 0.0023 |
| F12 | 9800001 | 9900000 | 0.0036 | 0.0023 | 0.0028 |
| F12 | 9820001 | 9920000 | 0.0037 | 0.0024 | 0.0029 |
| F12 | 9840001 | 9940000 | 0.0035 | 0.0024 | 0.0029 |
| F12 | 9860001 | 9960000 | 0.0028 | 0.0019 | 0.0023 |
| F12 | 9880001 | 9980000 | 0.0020 | 0.0012 | 0.0015 |
| F12 | 9900001 | 10000000 | 0.0015 | 0.0006 | 0.0010 |
| F12 | 9920001 | 10020000 | 0.0026 | 0.0016 | 0.0020 |
| F12 | 9940001 | 10040000 | 0.0027 | 0.0015 | 0.0018 |
| F12 | 9960001 | 10060000 | 0.0037 | 0.0023 | 0.0027 |
| F12 | 9980001 | 10080000 | 0.0041 | 0.0027 | 0.0031 |
| F12 | 10000001 | 10100000 | 0.0046 | 0.0030 | 0.0036 |
| F12 | 10020001 | 10120000 | 0.0037 | 0.0021 | 0.0028 |
| F12 | 10040001 | 10140000 | 0.0037 | 0.0024 | 0.0030 |
| F12 | 10060001 | 10160000 | 0.0032 | 0.0019 | 0.0024 |
| F12 | 10080001 | 10180000 | 0.0031 | 0.0017 | 0.0023 |
| F12 | 10100001 | 10200000 | 0.0039 | 0.0026 | 0.0031 |
| F12 | 10120001 | 10220000 | 0.0042 | 0.0031 | 0.0035 |
| F12 | 10140001 | 10240000 | 0.0038 | 0.0029 | 0.0034 |
| F12 | 10160001 | 10260000 | 0.0039 | 0.0031 | 0.0036 |
| F12 | 10180001 | 10280000 | 0.0041 | 0.0032 | 0.0038 |
| F12 | 10200001 | 10300000 | 0.0030 | 0.0022 | 0.0027 |
| F12 | 10220001 | 10320000 | 0.0035 | 0.0021 | 0.0025 |
| F12 | 10240001 | 10340000 | 0.0034 | 0.0020 | 0.0022 |
| F12 | 10260001 | 10360000 | 0.0035 | 0.0020 | 0.0023 |
| F12 | 10280001 | 10380000 | 0.0032 | 0.0019 | 0.0021 |
| F12 | 10300001 | 10400000 | 0.0037 | 0.0021 | 0.0024 |
| F12 | 10320001 | 10420000 | 0.0026 | 0.0017 | 0.0020 |
| F12 | 10340001 | 10440000 | 0.0024 | 0.0017 | 0.0020 |
| F12 | 10360001 | 10460000 | 0.0022 | 0.0014 | 0.0016 |
| F12 | 10380001 | 10480000 | 0.0026 | 0.0016 | 0.0018 |
| F12 | 10400001 | 10500000 | 0.0025 | 0.0016 | 0.0018 |
| F12 | 10420001 | 10520000 | 0.0032 | 0.0021 | 0.0024 |
| F12 | 10440001 | 10540000 | 0.0041 | 0.0027 | 0.0032 |
| F12 | 10460001 | 10560000 | 0.0049 | 0.0036 | 0.0040 |
| F12 | 10480001 | 10580000 | 0.0045 | 0.0033 | 0.0037 |
| F12 | 10500001 | 10600000 | 0.0042 | 0.0030 | 0.0033 |
| F12 | 10520001 | 10620000 | 0.0039 | 0.0027 | 0.0030 |
| F12 | 10540001 | 10640000 | 0.0036 | 0.0024 | 0.0026 |
| F12 | 10560001 | 10660000 | 0.0042 | 0.0028 | 0.0029 |
| F12 | 10580001 | 10680000 | 0.0042 | 0.0029 | 0.0030 |
| F12 | 10600001 | 10700000 | 0.0042 | 0.0030 | 0.0030 |
| F12 | 10620001 | 10720000 | 0.0039 | 0.0025 | 0.0024 |
| F12 | 10640001 | 10740000 | 0.0033 | 0.0022 | 0.0021 |
| F12 | 10660001 | 10760000 | 0.0021 | 0.0011 | 0.0012 |
| F12 | 10680001 | 10780000 | 0.0022 | 0.0011 | 0.0013 |
| F12 | 10700001 | 10800000 | 0.0026 | 0.0017 | 0.0018 |
| F12 | 10720001 | 10820000 | 0.0027 | 0.0019 | 0.0021 |
| F12 | 10740001 | 10840000 | 0.0033 | 0.0023 | 0.0026 |
| F12 | 10760001 | 10860000 | 0.0039 | 0.0024 | 0.0028 |
| F12 | 10780001 | 10880000 | 0.0042 | 0.0025 | 0.0030 |
| F12 | 10800001 | 10900000 | 0.0047 | 0.0028 | 0.0035 |
| F12 | 10820001 | 10920000 | 0.0046 | 0.0027 | 0.0034 |
| F12 | 10840001 | 10940000 | 0.0040 | 0.0022 | 0.0027 |
| F12 | 10860001 | 10960000 | 0.0040 | 0.0026 | 0.0030 |
| F12 | 10880001 | 10980000 | 0.0036 | 0.0023 | 0.0025 |
| F12 | 10900001 | 11000000 | 0.0027 | 0.0016 | 0.0016 |
| F12 | 10920001 | 11020000 | 0.0030 | 0.0020 | 0.0020 |

| Chr | Start | End | both | A1 | A2 |
|---|---|---|---|---|---|
| F12 | 10940001 | 11040000 | 0.0035 | 0.0027 | 0.0027 |
| F12 | 10960001 | 11060000 | 0.0035 | 0.0028 | 0.0027 |
| F12 | 10980001 | 11080000 | 0.0037 | 0.0031 | 0.0030 |
| F12 | 11000001 | 11100000 | 0.0037 | 0.0031 | 0.0029 |
| F12 | 11020001 | 11120000 | 0.0038 | 0.0028 | 0.0028 |
| F12 | 11040001 | 11140000 | 0.0034 | 0.0024 | 0.0025 |
| F12 | 11060001 | 11160000 | 0.0023 | 0.0015 | 0.0017 |
| F12 | 11080001 | 11180000 | 0.0033 | 0.0024 | 0.0027 |
| F12 | 11100001 | 11200000 | 0.0032 | 0.0024 | 0.0028 |
| F12 | 11120001 | 11220000 | 0.0029 | 0.0021 | 0.0024 |
| F12 | 11140001 | 11240000 | 0.0033 | 0.0023 | 0.0027 |
| F12 | 11160001 | 11260000 | 0.0036 | 0.0024 | 0.0028 |
| F12 | 11180001 | 11280000 | 0.0025 | 0.0012 | 0.0016 |
| F12 | 11200001 | 11300000 | 0.0026 | 0.0012 | 0.0016 |
| F12 | 11220001 | 11320000 | 0.0029 | 0.0016 | 0.0020 |
| F12 | 11240001 | 11340000 | 0.0026 | 0.0013 | 0.0016 |
| F12 | 11260001 | 11360000 | 0.0030 | 0.0015 | 0.0020 |
| F12 | 11280001 | 11380000 | 0.0032 | 0.0019 | 0.0022 |
| F12 | 11300001 | 11400000 | 0.0029 | 0.0016 | 0.0019 |
| F12 | 11320001 | 11420000 | 0.0026 | 0.0015 | 0.0017 |
| F12 | 11340001 | 11440000 | 0.0024 | 0.0015 | 0.0016 |
| F12 | 11360001 | 11460000 | 0.0022 | 0.0012 | 0.0013 |
| F12 | 11380001 | 11480000 | 0.0025 | 0.0015 | 0.0016 |
| F12 | 11400001 | 11500000 | 0.0032 | 0.0022 | 0.0022 |
| F12 | 11420001 | 11520000 | 0.0033 | 0.0022 | 0.0023 |
| F12 | 11440001 | 11540000 | 0.0041 | 0.0028 | 0.0031 |
| F12 | 11460001 | 11560000 | 0.0040 | 0.0028 | 0.0031 |
| F12 | 11480001 | 11580000 | 0.0038 | 0.0021 | 0.0026 |
| F12 | 11500001 | 11600000 | 0.0040 | 0.0019 | 0.0026 |
| F12 | 11520001 | 11620000 | 0.0042 | 0.0021 | 0.0028 |
| F12 | 11540001 | 11640000 | 0.0038 | 0.0018 | 0.0022 |
| F12 | 11560001 | 11660000 | 0.0040 | 0.0019 | 0.0023 |
| F12 | 11580001 | 11680000 | 0.0039 | 0.0023 | 0.0025 |
| F12 | 11600001 | 11700000 | 0.0032 | 0.0020 | 0.0021 |
| F12 | 11620001 | 11720000 | 0.0028 | 0.0016 | 0.0016 |
| F12 | 11640001 | 11740000 | 0.0031 | 0.0020 | 0.0021 |
| F12 | 11660001 | 11760000 | 0.0030 | 0.0022 | 0.0022 |
| F12 | 11680001 | 11780000 | 0.0031 | 0.0021 | 0.0021 |
| F12 | 11700001 | 11800000 | 0.0036 | 0.0026 | 0.0027 |
| F12 | 11720001 | 11820000 | 0.0043 | 0.0031 | 0.0034 |
| F12 | 11740001 | 11840000 | 0.0044 | 0.0029 | 0.0034 |
| F12 | 11760001 | 11860000 | 0.0040 | 0.0025 | 0.0031 |
| F12 | 11780001 | 11880000 | 0.0039 | 0.0025 | 0.0031 |
| F12 | 11800001 | 11900000 | 0.0036 | 0.0023 | 0.0029 |
| F12 | 11820001 | 11920000 | 0.0028 | 0.0017 | 0.0022 |
| F12 | 11840001 | 11940000 | 0.0020 | 0.0012 | 0.0014 |
| F12 | 11860001 | 11960000 | 0.0021 | 0.0013 | 0.0016 |
| F12 | 11880001 | 11980000 | 0.0019 | 0.0010 | 0.0013 |
| F12 | 11900001 | 12000000 | 0.0016 | 0.0006 | 0.0010 |
| F12 | 11920001 | 12020000 | 0.0018 | 0.0006 | 0.0009 |
| F12 | 11940001 | 12040000 | 0.0019 | 0.0010 | 0.0012 |
| F12 | 11960001 | 12060000 | 0.0021 | 0.0009 | 0.0012 |
| F12 | 11980001 | 12080000 | 0.0037 | 0.0021 | 0.0026 |
| F12 | 12000001 | 12100000 | 0.0037 | 0.0021 | 0.0026 |
| F12 | 12020001 | 12120000 | 0.0037 | 0.0021 | 0.0026 |
| F12 | 12040001 | 12140000 | 0.0037 | 0.0019 | 0.0024 |
| F12 | 12060001 | 12160000 | 0.0040 | 0.0025 | 0.0028 |
| F12 | 12080001 | 12180000 | 0.0028 | 0.0015 | 0.0017 |

| Chr | Start | End | both | A1 | A2 |
|---|---|---|---|---|---|
| F12 | 12100001 | 12200000 | 0.0032 | 0.0019 | 0.0020 |
| F12 | 12120001 | 12220000 | 0.0033 | 0.0019 | 0.0021 |
| F12 | 12140001 | 12240000 | 0.0029 | 0.0018 | 0.0020 |
| F12 | 12160001 | 12260000 | 0.0027 | 0.0015 | 0.0020 |
| F12 | 12180001 | 12280000 | 0.0024 | 0.0015 | 0.0018 |
| F12 | 12200001 | 12300000 | 0.0021 | 0.0012 | 0.0016 |
| F12 | 12220001 | 12320000 | 0.0019 | 0.0011 | 0.0014 |
| F12 | 12240001 | 12340000 | 0.0023 | 0.0015 | 0.0018 |
| F12 | 12260001 | 12360000 | 0.0021 | 0.0013 | 0.0014 |
| F12 | 12280001 | 12380000 | 0.0022 | 0.0014 | 0.0016 |
| F12 | 12300001 | 12400000 | 0.0032 | 0.0023 | 0.0025 |
| F12 | 12320001 | 12420000 | 0.0043 | 0.0031 | 0.0036 |
| F12 | 12340001 | 12440000 | 0.0043 | 0.0030 | 0.0034 |
| F12 | 12360001 | 12460000 | 0.0047 | 0.0036 | 0.0037 |
| F12 | 12380001 | 12480000 | 0.0049 | 0.0036 | 0.0037 |
| F12 | 12400001 | 12500000 | 0.0043 | 0.0029 | 0.0030 |
| F12 | 12420001 | 12520000 | 0.0035 | 0.0023 | 0.0022 |
| F12 | 12440001 | 12540000 | 0.0035 | 0.0023 | 0.0023 |
| F12 | 12460001 | 12560000 | 0.0028 | 0.0016 | 0.0018 |
| F12 | 12480001 | 12580000 | 0.0030 | 0.0018 | 0.0020 |
| F12 | 12500001 | 12600000 | 0.0030 | 0.0019 | 0.0023 |
| F12 | 12520001 | 12620000 | 0.0036 | 0.0025 | 0.0030 |
| F12 | 12540001 | 12640000 | 0.0039 | 0.0027 | 0.0033 |
| F12 | 12560001 | 12660000 | 0.0041 | 0.0028 | 0.0035 |
| F12 | 12580001 | 12680000 | 0.0040 | 0.0027 | 0.0033 |
| F12 | 12600001 | 12700000 | 0.0039 | 0.0028 | 0.0032 |
| F12 | 12620001 | 12720000 | 0.0034 | 0.0023 | 0.0027 |
| F12 | 12640001 | 12740000 | 0.0028 | 0.0018 | 0.0020 |
| F12 | 12660001 | 12760000 | 0.0027 | 0.0018 | 0.0019 |
| F12 | 12680001 | 12780000 | 0.0028 | 0.0020 | 0.0021 |
| F12 | 12700001 | 12800000 | 0.0027 | 0.0019 | 0.0020 |
| F12 | 12720001 | 12820000 | 0.0028 | 0.0019 | 0.0021 |
| F12 | 12740001 | 12840000 | 0.0032 | 0.0023 | 0.0026 |
| F12 | 12760001 | 12860000 | 0.0035 | 0.0026 | 0.0029 |
| F12 | 12780001 | 12880000 | 0.0032 | 0.0023 | 0.0027 |
| F12 | 12800001 | 12900000 | 0.0035 | 0.0026 | 0.0031 |
| F12 | 12820001 | 12920000 | 0.0039 | 0.0031 | 0.0036 |
| F12 | 12840001 | 12940000 | 0.0037 | 0.0029 | 0.0033 |
| F12 | 12860001 | 12960000 | 0.0040 | 0.0032 | 0.0037 |
| F12 | 12880001 | 12980000 | 0.0040 | 0.0033 | 0.0037 |
| F12 | 12900001 | 13000000 | 0.0036 | 0.0029 | 0.0032 |
| F12 | 12920001 | 13020000 | 0.0026 | 0.0020 | 0.0022 |
| F12 | 12940001 | 13040000 | 0.0029 | 0.0023 | 0.0026 |
| F12 | 12960001 | 13060000 | 0.0023 | 0.0017 | 0.0019 |
| F12 | 12980001 | 13080000 | 0.0019 | 0.0014 | 0.0015 |
| F12 | 13000001 | 13100000 | 0.0021 | 0.0015 | 0.0017 |
| F12 | 13020001 | 13120000 | 0.0035 | 0.0025 | 0.0031 |
| F12 | 13040001 | 13140000 | 0.0038 | 0.0029 | 0.0034 |
| F12 | 13060001 | 13160000 | 0.0041 | 0.0031 | 0.0037 |
| F12 | 13080001 | 13180000 | 0.0045 | 0.0033 | 0.0039 |
| F12 | 13100001 | 13200000 | 0.0042 | 0.0032 | 0.0037 |
| F12 | 13120001 | 13220000 | 0.0038 | 0.0031 | 0.0034 |
| F12 | 13140001 | 13240000 | 0.0042 | 0.0034 | 0.0036 |
| F12 | 13160001 | 13260000 | 0.0053 | 0.0044 | 0.0048 |
| F12 | 13180001 | 13280000 | 0.0054 | 0.0047 | 0.0051 |
| F12 | 13200001 | 13300000 | 0.0054 | 0.0044 | 0.0048 |
| F12 | 13220001 | 13320000 | 0.0048 | 0.0040 | 0.0040 |
| F12 | 13240001 | 13340000 | 0.0037 | 0.0027 | 0.0028 |

| Chr | Start | End | both | A1 | A2 |
|---|---|---|---|---|---|
| F12 | 13260001 | 13360000 | 0.0023 | 0.0015 | 0.0013 |
| F12 | 13280001 | 13380000 | 0.0027 | 0.0017 | 0.0015 |
| F12 | 13300001 | 13400000 | 0.0029 | 0.0020 | 0.0018 |
| F12 | 13320001 | 13420000 | 0.0027 | 0.0019 | 0.0017 |
| F12 | 13340001 | 13440000 | 0.0028 | 0.0020 | 0.0020 |
| F12 | 13360001 | 13460000 | 0.0027 | 0.0020 | 0.0020 |
| F12 | 13380001 | 13480000 | 0.0021 | 0.0015 | 0.0014 |
| F12 | 13400001 | 13500000 | 0.0016 | 0.0011 | 0.0011 |
| F12 | 13420001 | 13520000 | 0.0018 | 0.0011 | 0.0012 |
| F12 | 13440001 | 13540000 | 0.0020 | 0.0011 | 0.0013 |
| F12 | 13460001 | 13560000 | 0.0029 | 0.0020 | 0.0022 |
| F12 | 13480001 | 13580000 | 0.0029 | 0.0020 | 0.0023 |
| F12 | 13500001 | 13600000 | 0.0039 | 0.0028 | 0.0030 |
| F12 | 13520001 | 13620000 | 0.0043 | 0.0032 | 0.0035 |
| F12 | 13540001 | 13640000 | 0.0042 | 0.0032 | 0.0035 |
| F12 | 13560001 | 13660000 | 0.0035 | 0.0023 | 0.0027 |
| F12 | 13580001 | 13680000 | 0.0043 | 0.0031 | 0.0034 |
| F12 | 13600001 | 13700000 | 0.0038 | 0.0028 | 0.0031 |
| F12 | 13620001 | 13720000 | 0.0045 | 0.0031 | 0.0037 |
| F12 | 13640001 | 13740000 | 0.0040 | 0.0029 | 0.0034 |
| F12 | 13660001 | 13760000 | 0.0044 | 0.0033 | 0.0039 |
| F12 | 13680001 | 13780000 | 0.0037 | 0.0024 | 0.0030 |
| F12 | 13700001 | 13800000 | 0.0036 | 0.0021 | 0.0027 |
| F12 | 13720001 | 13820000 | 0.0026 | 0.0013 | 0.0016 |
| F12 | 13740001 | 13840000 | 0.0028 | 0.0015 | 0.0017 |
| F12 | 13760001 | 13860000 | 0.0024 | 0.0010 | 0.0012 |
| F12 | 13780001 | 13880000 | 0.0023 | 0.0014 | 0.0014 |
| F12 | 13800001 | 13900000 | 0.0025 | 0.0014 | 0.0016 |
| F12 | 13820001 | 13920000 | 0.0026 | 0.0016 | 0.0018 |
| F12 | 13840001 | 13940000 | 0.0028 | 0.0017 | 0.0020 |
| F12 | 13860001 | 13960000 | 0.0029 | 0.0019 | 0.0022 |
| F12 | 13880001 | 13980000 | 0.0026 | 0.0015 | 0.0018 |
| F12 | 13900001 | 14000000 | 0.0023 | 0.0015 | 0.0016 |
| F12 | 13920001 | 14020000 | 0.0025 | 0.0016 | 0.0019 |
| F12 | 13940001 | 14040000 | 0.0022 | 0.0013 | 0.0015 |
| F12 | 13960001 | 14060000 | 0.0024 | 0.0014 | 0.0015 |
| F12 | 13980001 | 14080000 | 0.0029 | 0.0021 | 0.0022 |
| F12 | 14000001 | 14100000 | 0.0029 | 0.0020 | 0.0022 |
| F12 | 14020001 | 14120000 | 0.0023 | 0.0015 | 0.0014 |
| F12 | 14040001 | 14140000 | 0.0027 | 0.0017 | 0.0018 |
| F12 | 14060001 | 14160000 | 0.0024 | 0.0014 | 0.0015 |
| F12 | 14080001 | 14180000 | 0.0027 | 0.0016 | 0.0016 |
| F12 | 14100001 | 14200000 | 0.0031 | 0.0020 | 0.0021 |
| F12 | 14120001 | 14220000 | 0.0030 | 0.0021 | 0.0021 |
| F12 | 14140001 | 14240000 | 0.0027 | 0.0021 | 0.0019 |
| F12 | 14160001 | 14260000 | 0.0037 | 0.0031 | 0.0030 |
| F12 | 14180001 | 14280000 | 0.0032 | 0.0026 | 0.0028 |
| F12 | 14200001 | 14300000 | 0.0029 | 0.0025 | 0.0027 |
| F12 | 14220001 | 14320000 | 0.0039 | 0.0032 | 0.0035 |
| F12 | 14240001 | 14340000 | 0.0037 | 0.0029 | 0.0032 |
| F12 | 14260001 | 14360000 | 0.0027 | 0.0021 | 0.0023 |
| F12 | 14280001 | 14380000 | 0.0035 | 0.0028 | 0.0029 |
| F12 | 14300001 | 14400000 | 0.0032 | 0.0025 | 0.0025 |
| F12 | 14320001 | 14420000 | 0.0021 | 0.0017 | 0.0016 |
| F12 | 14340001 | 14440000 | 0.0029 | 0.0024 | 0.0024 |
| F12 | 14360001 | 14460000 | 0.0031 | 0.0028 | 0.0025 |
| F12 | 14380001 | 14480000 | 0.0026 | 0.0022 | 0.0020 |
| F12 | 14400001 | 14500000 | 0.0031 | 0.0024 | 0.0023 |

| Chr | Start | End | both | A1 | A2 |
|---|---|---|---|---|---|
| F12 | 14420001 | 14520000 | 0.0033 | 0.0024 | 0.0024 |
| F12 | 14440001 | 14540000 | 0.0031 | 0.0021 | 0.0021 |
| F12 | 14460001 | 14560000 | 0.0029 | 0.0016 | 0.0019 |
| F12 | 14480001 | 14580000 | 0.0022 | 0.0011 | 0.0012 |
| F12 | 14500001 | 14600000 | 0.0015 | 0.0009 | 0.0009 |
| F12 | 14520001 | 14620000 | 0.0013 | 0.0007 | 0.0007 |
| F12 | 14540001 | 14640000 | 0.0009 | 0.0008 | 0.0005 |
| F12 | 14560001 | 14660000 | 0.0009 | 0.0011 | 0.0008 |
| F12 | 14580001 | 14680000 | 0.0010 | 0.0011 | 0.0009 |
| F12 | 14600001 | 14700000 | 0.0010 | 0.0011 | 0.0009 |
| F12 | 14620001 | 14720000 | 0.0013 | 0.0013 | 0.0012 |
| F12 | 14640001 | 14740000 | 0.0011 | 0.0010 | 0.0011 |
| F12 | 14660001 | 14760000 | 0.0010 | 0.0009 | 0.0010 |
| F12 | 14680001 | 14780000 | 0.0012 | 0.0011 | 0.0011 |
| F12 | 14700001 | 14800000 | 0.0020 | 0.0017 | 0.0017 |
| F12 | 14720001 | 14820000 | 0.0030 | 0.0025 | 0.0025 |
| F12 | 14740001 | 14840000 | 0.0031 | 0.0026 | 0.0026 |
| F12 | 14760001 | 14860000 | 0.0034 | 0.0027 | 0.0028 |
| F12 | 14780001 | 14880000 | 0.0034 | 0.0026 | 0.0030 |
| F12 | 14800001 | 14900000 | 0.0030 | 0.0024 | 0.0027 |
| F12 | 14820001 | 14920000 | 0.0020 | 0.0015 | 0.0018 |
| F12 | 14840001 | 14940000 | 0.0017 | 0.0013 | 0.0017 |
| F12 | 14860001 | 14960000 | 0.0016 | 0.0013 | 0.0016 |
| F12 | 14880001 | 14980000 | 0.0015 | 0.0012 | 0.0014 |
| F12 | 14900001 | 15000000 | 0.0013 | 0.0012 | 0.0012 |
| F12 | 14920001 | 15020000 | 0.0015 | 0.0013 | 0.0014 |
| F12 | 14940001 | 15040000 | 0.0021 | 0.0018 | 0.0020 |
| F12 | 14960001 | 15060000 | 0.0020 | 0.0017 | 0.0019 |
| F12 | 14980001 | 15080000 | 0.0020 | 0.0017 | 0.0020 |
| F12 | 15000001 | 15100000 | 0.0022 | 0.0018 | 0.0020 |
| F12 | 15020001 | 15120000 | 0.0022 | 0.0018 | 0.0021 |
| F12 | 15040001 | 15140000 | 0.0030 | 0.0024 | 0.0028 |
| F12 | 15060001 | 15160000 | 0.0033 | 0.0026 | 0.0031 |
| F12 | 15080001 | 15180000 | 0.0033 | 0.0026 | 0.0031 |
| F12 | 15100001 | 15200000 | 0.0031 | 0.0025 | 0.0030 |
| F12 | 15120001 | 15220000 | 0.0032 | 0.0026 | 0.0031 |
| F12 | 15140001 | 15240000 | 0.0030 | 0.0024 | 0.0027 |
| F12 | 15160001 | 15260000 | 0.0032 | 0.0026 | 0.0029 |
| F12 | 15180001 | 15280000 | 0.0041 | 0.0033 | 0.0037 |
| F12 | 15200001 | 15300000 | 0.0041 | 0.0033 | 0.0036 |
| F12 | 15220001 | 15320000 | 0.0040 | 0.0029 | 0.0033 |
| F12 | 15240001 | 15340000 | 0.0034 | 0.0023 | 0.0027 |
| F12 | 15260001 | 15360000 | 0.0028 | 0.0018 | 0.0023 |
| F12 | 15280001 | 15380000 | 0.0020 | 0.0010 | 0.0013 |
| F12 | 15300001 | 15400000 | 0.0021 | 0.0012 | 0.0014 |
| F12 | 15320001 | 15420000 | 0.0024 | 0.0015 | 0.0017 |
| F12 | 15340001 | 15440000 | 0.0025 | 0.0018 | 0.0020 |
| F12 | 15360001 | 15460000 | 0.0027 | 0.0018 | 0.0018 |
| F12 | 15380001 | 15480000 | 0.0031 | 0.0022 | 0.0023 |
| F12 | 15400001 | 15500000 | 0.0031 | 0.0022 | 0.0022 |
| F12 | 15420001 | 15520000 | 0.0040 | 0.0032 | 0.0031 |
| F12 | 15440001 | 15540000 | 0.0045 | 0.0036 | 0.0036 |
| F12 | 15460001 | 15560000 | 0.0045 | 0.0035 | 0.0036 |
| F12 | 15480001 | 15580000 | 0.0043 | 0.0032 | 0.0035 |
| F12 | 15500001 | 15600000 | 0.0049 | 0.0031 | 0.0042 |
| F12 | 15520001 | 15620000 | 0.0042 | 0.0023 | 0.0035 |
| F12 | 15540001 | 15640000 | 0.0035 | 0.0017 | 0.0028 |
| F12 | 15560001 | 15660000 | 0.0044 | 0.0026 | 0.0039 |

| Chr | Start | End | both | A1 | A2 |
|---|---|---|---|---|---|
| F12 | 15580001 | 15680000 | 0.0043 | 0.0028 | 0.0038 |
| F12 | 15600001 | 15700000 | 0.0043 | 0.0032 | 0.0036 |
| F12 | 15620001 | 15720000 | 0.0043 | 0.0031 | 0.0034 |
| F12 | 15640001 | 15740000 | 0.0038 | 0.0028 | 0.0031 |
| F12 | 15660001 | 15760000 | 0.0026 | 0.0018 | 0.0019 |
| F12 | 15680001 | 15780000 | 0.0024 | 0.0016 | 0.0017 |
| F12 | 15700001 | 15800000 | 0.0020 | 0.0013 | 0.0014 |
| F12 | 15720001 | 15820000 | 0.0020 | 0.0013 | 0.0013 |
| F12 | 15740001 | 15840000 | 0.0030 | 0.0020 | 0.0021 |
| F12 | 15760001 | 15860000 | 0.0036 | 0.0023 | 0.0024 |
| F12 | 15780001 | 15880000 | 0.0048 | 0.0036 | 0.0038 |
| F12 | 15800001 | 15900000 | 0.0054 | 0.0041 | 0.0043 |
| F12 | 15820001 | 15920000 | 0.0049 | 0.0037 | 0.0039 |
| F12 | 15840001 | 15940000 | 0.0044 | 0.0033 | 0.0036 |
| F12 | 15860001 | 15960000 | 0.0042 | 0.0030 | 0.0034 |
| F12 | 15880001 | 15980000 | 0.0033 | 0.0020 | 0.0024 |
| F12 | 15900001 | 16000000 | 0.0027 | 0.0015 | 0.0018 |
| F12 | 15920001 | 16020000 | 0.0030 | 0.0016 | 0.0019 |
| F12 | 15940001 | 16040000 | 0.0030 | 0.0016 | 0.0018 |
| F12 | 15960001 | 16060000 | 0.0029 | 0.0016 | 0.0017 |
| F12 | 15980001 | 16080000 | 0.0028 | 0.0014 | 0.0015 |
| F12 | 16000001 | 16100000 | 0.0028 | 0.0015 | 0.0017 |
| F12 | 16020001 | 16120000 | 0.0033 | 0.0021 | 0.0023 |
| F12 | 16040001 | 16140000 | 0.0029 | 0.0020 | 0.0021 |
| F12 | 16060001 | 16160000 | 0.0035 | 0.0025 | 0.0029 |
| F12 | 16080001 | 16180000 | 0.0043 | 0.0033 | 0.0038 |
| F12 | 16100001 | 16200000 | 0.0047 | 0.0036 | 0.0042 |
| F12 | 16120001 | 16220000 | 0.0052 | 0.0038 | 0.0045 |
| F12 | 16140001 | 16240000 | 0.0051 | 0.0036 | 0.0043 |
| F12 | 16160001 | 16260000 | 0.0046 | 0.0031 | 0.0037 |
| F12 | 16180001 | 16280000 | 0.0046 | 0.0030 | 0.0037 |
| F12 | 16200001 | 16300000 | 0.0045 | 0.0031 | 0.0038 |
| F12 | 16220001 | 16320000 | 0.0044 | 0.0031 | 0.0037 |
| F12 | 16240001 | 16340000 | 0.0044 | 0.0032 | 0.0038 |
| F12 | 16260001 | 16360000 | 0.0040 | 0.0031 | 0.0036 |
| F12 | 16280001 | 16380000 | 0.0034 | 0.0028 | 0.0031 |
| F12 | 16300001 | 16400000 | 0.0030 | 0.0024 | 0.0027 |
| F12 | 16320001 | 16420000 | 0.0023 | 0.0018 | 0.0021 |
| F12 | 16340001 | 16440000 | 0.0022 | 0.0018 | 0.0020 |
| F12 | 16360001 | 16460000 | 0.0029 | 0.0022 | 0.0025 |
| F12 | 16380001 | 16480000 | 0.0029 | 0.0022 | 0.0024 |
| F12 | 16400001 | 16500000 | 0.0032 | 0.0024 | 0.0027 |
| F12 | 16420001 | 16520000 | 0.0032 | 0.0025 | 0.0027 |
| F12 | 16440001 | 16540000 | 0.0035 | 0.0026 | 0.0029 |
| F12 | 16460001 | 16560000 | 0.0042 | 0.0033 | 0.0035 |
| F12 | 16480001 | 16580000 | 0.0045 | 0.0036 | 0.0041 |
| F12 | 16500001 | 16600000 | 0.0043 | 0.0035 | 0.0039 |
| F12 | 16520001 | 16620000 | 0.0047 | 0.0037 | 0.0041 |
| F12 | 16540001 | 16640000 | 0.0055 | 0.0044 | 0.0048 |
| F12 | 16560001 | 16660000 | 0.0046 | 0.0037 | 0.0041 |
| F12 | 16580001 | 16680000 | 0.0044 | 0.0033 | 0.0036 |
| F12 | 16600001 | 16700000 | 0.0046 | 0.0035 | 0.0039 |
| F12 | 16620001 | 16720000 | 0.0043 | 0.0031 | 0.0038 |
| F12 | 16640001 | 16740000 | 0.0047 | 0.0036 | 0.0042 |
| F12 | 16660001 | 16760000 | 0.0049 | 0.0037 | 0.0047 |
| F12 | 16680001 | 16780000 | 0.0047 | 0.0038 | 0.0046 |
| F12 | 16700001 | 16800000 | 0.0043 | 0.0033 | 0.0042 |
| F12 | 16720001 | 16820000 | 0.0048 | 0.0035 | 0.0044 |

| Chr | Start | End | both | A1 | A2 |
|---|---|---|---|---|---|
| F12 | 16740001 | 16840000 | 0.0037 | 0.0025 | 0.0033 |
| F12 | 16760001 | 16860000 | 0.0036 | 0.0026 | 0.0029 |
| F12 | 16780001 | 16880000 | 0.0034 | 0.0024 | 0.0026 |
| F12 | 16800001 | 16900000 | 0.0036 | 0.0024 | 0.0026 |
| F12 | 16820001 | 16920000 | 0.0031 | 0.0021 | 0.0022 |
| F12 | 16840001 | 16940000 | 0.0033 | 0.0021 | 0.0025 |
| F12 | 16860001 | 16960000 | 0.0027 | 0.0016 | 0.0021 |
| F12 | 16880001 | 16980000 | 0.0026 | 0.0015 | 0.0020 |
| F12 | 16900001 | 17000000 | 0.0018 | 0.0011 | 0.0014 |
| F12 | 16920001 | 17020000 | 0.0015 | 0.0010 | 0.0011 |
| F12 | 16940001 | 17040000 | 0.0015 | 0.0010 | 0.0010 |
| F12 | 16960001 | 17060000 | 0.0020 | 0.0011 | 0.0013 |
| F12 | 16980001 | 17080000 | 0.0024 | 0.0016 | 0.0017 |
| F12 | 17000001 | 17100000 | 0.0030 | 0.0020 | 0.0023 |
| F12 | 17020001 | 17120000 | 0.0037 | 0.0026 | 0.0030 |
| F12 | 17040001 | 17140000 | 0.0033 | 0.0026 | 0.0030 |
| F12 | 17060001 | 17160000 | 0.0042 | 0.0038 | 0.0041 |
| F12 | 17080001 | 17180000 | 0.0036 | 0.0032 | 0.0036 |
| F12 | 17100001 | 17200000 | 0.0038 | 0.0034 | 0.0037 |
| F12 | 17120001 | 17220000 | 0.0041 | 0.0038 | 0.0039 |
| F12 | 17140001 | 17240000 | 0.0043 | 0.0040 | 0.0040 |
| F12 | 17160001 | 17260000 | 0.0042 | 0.0036 | 0.0038 |
| F12 | 17180001 | 17280000 | 0.0053 | 0.0045 | 0.0047 |
| F12 | 17200001 | 17300000 | 0.0063 | 0.0053 | 0.0056 |
| F12 | 17220001 | 17320000 | 0.0062 | 0.0051 | 0.0055 |
| F12 | 17240001 | 17340000 | 0.0065 | 0.0050 | 0.0056 |
| F12 | 17260001 | 17360000 | 0.0056 | 0.0041 | 0.0047 |
| F12 | 17280001 | 17380000 | 0.0045 | 0.0032 | 0.0037 |
| F12 | 17300001 | 17400000 | 0.0041 | 0.0026 | 0.0031 |
| F12 | 17320001 | 17420000 | 0.0044 | 0.0027 | 0.0032 |
| F12 | 17340001 | 17440000 | 0.0060 | 0.0043 | 0.0049 |
| F12 | 17360001 | 17460000 | 0.0062 | 0.0046 | 0.0052 |
| F12 | 17380001 | 17480000 | 0.0064 | 0.0048 | 0.0056 |
| F12 | 17400001 | 17500000 | 0.0054 | 0.0043 | 0.0050 |
| F12 | 17420001 | 17520000 | 0.0046 | 0.0039 | 0.0044 |
| F12 | 17440001 | 17540000 | 0.0026 | 0.0021 | 0.0026 |
| F12 | 17460001 | 17560000 | 0.0021 | 0.0016 | 0.0019 |
| F12 | 17480001 | 17580000 | 0.0018 | 0.0013 | 0.0015 |
| F12 | 17500001 | 17600000 | 0.0032 | 0.0025 | 0.0028 |
| F12 | 17520001 | 17620000 | 0.0036 | 0.0028 | 0.0031 |
| F12 | 17540001 | 17640000 | 0.0038 | 0.0028 | 0.0030 |
| F12 | 17560001 | 17660000 | 0.0038 | 0.0027 | 0.0029 |
| F12 | 17580001 | 17680000 | 0.0044 | 0.0031 | 0.0034 |
| F12 | 17600001 | 17700000 | 0.0037 | 0.0024 | 0.0027 |
| F12 | 17620001 | 17720000 | 0.0039 | 0.0025 | 0.0027 |
| F12 | 17640001 | 17740000 | 0.0038 | 0.0024 | 0.0027 |
| F12 | 17660001 | 17760000 | 0.0045 | 0.0031 | 0.0036 |
| F12 | 17680001 | 17780000 | 0.0041 | 0.0028 | 0.0032 |
| F12 | 17700001 | 17800000 | 0.0040 | 0.0026 | 0.0030 |
| F12 | 17720001 | 17820000 | 0.0032 | 0.0020 | 0.0024 |
| F12 | 17740001 | 17840000 | 0.0036 | 0.0024 | 0.0029 |
| F12 | 17760001 | 17860000 | 0.0036 | 0.0024 | 0.0029 |
| F12 | 17780001 | 17880000 | 0.0052 | 0.0039 | 0.0046 |
| F12 | 17800001 | 17900000 | 0.0049 | 0.0038 | 0.0043 |
| F12 | 17820001 | 17920000 | 0.0052 | 0.0042 | 0.0047 |
| F12 | 17840001 | 17940000 | 0.0044 | 0.0035 | 0.0039 |
| F12 | 17860001 | 17960000 | 0.0037 | 0.0030 | 0.0032 |
| F12 | 17880001 | 17980000 | 0.0029 | 0.0024 | 0.0025 |

| Chr | Start | End | both | A1 | A2 |
|---|---|---|---|---|---|
| F12 | 17900001 | 18000000 | 0.0026 | 0.0021 | 0.0024 |
| F12 | 17920001 | 18020000 | 0.0026 | 0.0023 | 0.0025 |
| F12 | 17940001 | 18040000 | 0.0034 | 0.0028 | 0.0031 |
| F12 | 17960001 | 18060000 | 0.0033 | 0.0027 | 0.0030 |
| F12 | 17980001 | 18080000 | 0.0026 | 0.0019 | 0.0022 |
| F12 | 18000001 | 18100000 | 0.0032 | 0.0022 | 0.0025 |
| F12 | 18020001 | 18120000 | 0.0033 | 0.0021 | 0.0024 |
| F12 | 18040001 | 18140000 | 0.0034 | 0.0024 | 0.0026 |
| F12 | 18060001 | 18160000 | 0.0039 | 0.0030 | 0.0031 |
| F12 | 18080001 | 18180000 | 0.0062 | 0.0049 | 0.0052 |
| F12 | 18100001 | 18200000 | 0.0064 | 0.0053 | 0.0056 |
| F12 | 18120001 | 18220000 | 0.0062 | 0.0052 | 0.0056 |
| F12 | 18140001 | 18240000 | 0.0057 | 0.0049 | 0.0054 |
| F12 | 18160001 | 18260000 | 0.0058 | 0.0050 | 0.0053 |
| F12 | 18180001 | 18280000 | 0.0041 | 0.0034 | 0.0037 |
| F12 | 18200001 | 18300000 | 0.0040 | 0.0033 | 0.0034 |
| F12 | 18220001 | 18320000 | 0.0040 | 0.0033 | 0.0034 |
| F12 | 18240001 | 18340000 | 0.0043 | 0.0034 | 0.0034 |
| F12 | 18260001 | 18360000 | 0.0038 | 0.0028 | 0.0030 |
| F12 | 18280001 | 18380000 | 0.0034 | 0.0024 | 0.0025 |
| F12 | 18300001 | 18400000 | 0.0031 | 0.0021 | 0.0022 |
| F12 | 18320001 | 18420000 | 0.0039 | 0.0026 | 0.0028 |
| F12 | 18340001 | 18440000 | 0.0041 | 0.0027 | 0.0032 |
| F12 | 18360001 | 18460000 | 0.0040 | 0.0026 | 0.0030 |
| F12 | 18380001 | 18480000 | 0.0054 | 0.0038 | 0.0042 |
| F12 | 18400001 | 18500000 | 0.0065 | 0.0048 | 0.0052 |
| F12 | 18420001 | 18520000 | 0.0067 | 0.0050 | 0.0054 |
| F12 | 18440001 | 18540000 | 0.0069 | 0.0053 | 0.0055 |
| F12 | 18460001 | 18560000 | 0.0072 | 0.0058 | 0.0060 |
| F12 | 18480001 | 18580000 | 0.0087 | 0.0072 | 0.0074 |
| F12 | 18500001 | 18600000 | 0.0071 | 0.0057 | 0.0060 |
| F12 | 18520001 | 18620000 | 0.0072 | 0.0058 | 0.0064 |
| F12 | 18540001 | 18640000 | 0.0065 | 0.0052 | 0.0058 |
| F12 | 18560001 | 18660000 | 0.0083 | 0.0070 | 0.0074 |
| F12 | 18580001 | 18680000 | 0.0068 | 0.0056 | 0.0061 |
| F12 | 18600001 | 18700000 | 0.0072 | 0.0060 | 0.0064 |
| F12 | 18620001 | 18720000 | 0.0059 | 0.0049 | 0.0050 |
| F12 | 18640001 | 18740000 | 0.0065 | 0.0056 | 0.0055 |
| F12 | 18660001 | 18760000 | 0.0045 | 0.0037 | 0.0038 |
| F12 | 18680001 | 18780000 | 0.0037 | 0.0029 | 0.0030 |
| F12 | 18700001 | 18800000 | 0.0035 | 0.0026 | 0.0028 |
| F12 | 18720001 | 18820000 | 0.0040 | 0.0030 | 0.0033 |
| F12 | 18740001 | 18840000 | 0.0042 | 0.0030 | 0.0034 |
| F12 | 18760001 | 18860000 | 0.0051 | 0.0035 | 0.0039 |
| F12 | 18780001 | 18880000 | 0.0044 | 0.0030 | 0.0033 |
| F12 | 18800001 | 18900000 | 0.0043 | 0.0031 | 0.0035 |
| F12 | 18820001 | 18920000 | 0.0037 | 0.0025 | 0.0028 |
| F12 | 18840001 | 18940000 | 0.0028 | 0.0016 | 0.0019 |
| F12 | 18860001 | 18960000 | 0.0027 | 0.0017 | 0.0019 |
| F12 | 18880001 | 18980000 | 0.0035 | 0.0025 | 0.0026 |
| F12 | 18900001 | 19000000 | 0.0037 | 0.0028 | 0.0027 |
| F12 | 18920001 | 19020000 | 0.0043 | 0.0032 | 0.0033 |
| F12 | 18940001 | 19040000 | 0.0040 | 0.0030 | 0.0031 |
| F12 | 18960001 | 19060000 | 0.0031 | 0.0022 | 0.0021 |
| F12 | 18980001 | 19080000 | 0.0026 | 0.0014 | 0.0015 |
| F12 | 19000001 | 19100000 | 0.0023 | 0.0012 | 0.0013 |
| F12 | 19020001 | 19120000 | 0.0016 | 0.0008 | 0.0007 |
| F12 | 19040001 | 19140000 | 0.0017 | 0.0009 | 0.0009 |

| Chr | Start | End | both | A1 | A2 |
|---|---|---|---|---|---|
| F12 | 19060001 | 19160000 | 0.0019 | 0.0013 | 0.0014 |
| F12 | 19080001 | 19180000 | 0.0019 | 0.0018 | 0.0018 |
| F12 | 19100001 | 19200000 | 0.0017 | 0.0015 | 0.0016 |
| F12 | 19120001 | 19220000 | 0.0022 | 0.0020 | 0.0021 |
| F12 | 19140001 | 19240000 | 0.0023 | 0.0021 | 0.0022 |
| F12 | 19160001 | 19260000 | 0.0024 | 0.0021 | 0.0023 |
| F12 | 19180001 | 19280000 | 0.0020 | 0.0016 | 0.0018 |
| F12 | 19200001 | 19300000 | 0.0019 | 0.0015 | 0.0018 |
| F12 | 19220001 | 19320000 | 0.0028 | 0.0021 | 0.0024 |
| F12 | 19240001 | 19340000 | 0.0032 | 0.0024 | 0.0029 |
| F12 | 19260001 | 19360000 | 0.0044 | 0.0034 | 0.0039 |
| F12 | 19280001 | 19380000 | 0.0045 | 0.0036 | 0.0041 |
| F12 | 19300001 | 19400000 | 0.0057 | 0.0045 | 0.0051 |
| F12 | 19320001 | 19420000 | 0.0060 | 0.0049 | 0.0055 |
| F12 | 19340001 | 19440000 | 0.0061 | 0.0049 | 0.0055 |
| F12 | 19360001 | 19460000 | 0.0051 | 0.0042 | 0.0046 |
| F12 | 19380001 | 19480000 | 0.0051 | 0.0041 | 0.0044 |
| F12 | 19400001 | 19500000 | 0.0046 | 0.0037 | 0.0040 |
| F12 | 19420001 | 19520000 | 0.0038 | 0.0028 | 0.0031 |
| F12 | 19440001 | 19540000 | 0.0038 | 0.0027 | 0.0030 |
| F12 | 19460001 | 19560000 | 0.0038 | 0.0026 | 0.0030 |
| F12 | 19480001 | 19580000 | 0.0038 | 0.0028 | 0.0031 |
| F12 | 19500001 | 19600000 | 0.0037 | 0.0029 | 0.0030 |
| F12 | 19520001 | 19620000 | 0.0037 | 0.0027 | 0.0028 |
| F12 | 19540001 | 19640000 | 0.0035 | 0.0026 | 0.0028 |
| F12 | 19560001 | 19660000 | 0.0031 | 0.0023 | 0.0025 |
| F12 | 19580001 | 19680000 | 0.0038 | 0.0030 | 0.0032 |
| F12 | 19600001 | 19700000 | 0.0042 | 0.0030 | 0.0033 |
| F12 | 19620001 | 19720000 | 0.0039 | 0.0027 | 0.0031 |
| F12 | 19640001 | 19740000 | 0.0042 | 0.0031 | 0.0034 |
| F12 | 19660001 | 19760000 | 0.0045 | 0.0033 | 0.0035 |
| F12 | 19680001 | 19780000 | 0.0038 | 0.0024 | 0.0026 |
| F12 | 19700001 | 19800000 | 0.0034 | 0.0022 | 0.0024 |
| F12 | 19720001 | 19820000 | 0.0034 | 0.0023 | 0.0024 |
| F12 | 19740001 | 19840000 | 0.0033 | 0.0021 | 0.0024 |
| F12 | 19760001 | 19860000 | 0.0030 | 0.0021 | 0.0023 |
| F12 | 19780001 | 19880000 | 0.0036 | 0.0030 | 0.0031 |
| F12 | 19800001 | 19900000 | 0.0032 | 0.0028 | 0.0028 |
| F12 | 19820001 | 19920000 | 0.0035 | 0.0030 | 0.0032 |
| F12 | 19840001 | 19940000 | 0.0031 | 0.0027 | 0.0029 |
| F12 | 19860001 | 19960000 | 0.0034 | 0.0029 | 0.0031 |
| F12 | 19880001 | 19980000 | 0.0037 | 0.0029 | 0.0034 |
| F12 | 19900001 | 20000000 | 0.0038 | 0.0029 | 0.0036 |
| F12 | 19920001 | 20020000 | 0.0036 | 0.0029 | 0.0034 |
| F12 | 19940001 | 20040000 | 0.0033 | 0.0027 | 0.0031 |
| F12 | 19960001 | 20060000 | 0.0030 | 0.0023 | 0.0028 |
| F12 | 19980001 | 20080000 | 0.0024 | 0.0018 | 0.0022 |
| F12 | 20000001 | 20100000 | 0.0024 | 0.0017 | 0.0020 |
| F12 | 20020001 | 20120000 | 0.0028 | 0.0018 | 0.0024 |
| F12 | 20040001 | 20140000 | 0.0029 | 0.0022 | 0.0025 |
| F12 | 20060001 | 20160000 | 0.0029 | 0.0023 | 0.0026 |
| F12 | 20080001 | 20180000 | 0.0030 | 0.0023 | 0.0026 |
| F12 | 20100001 | 20200000 | 0.0038 | 0.0031 | 0.0034 |
| F12 | 20120001 | 20220000 | 0.0034 | 0.0026 | 0.0029 |
| F12 | 20140001 | 20240000 | 0.0040 | 0.0029 | 0.0035 |
| F12 | 20160001 | 20260000 | 0.0046 | 0.0034 | 0.0040 |
| F12 | 20180001 | 20280000 | 0.0046 | 0.0034 | 0.0039 |
| F12 | 20200001 | 20300000 | 0.0036 | 0.0026 | 0.0030 |

| Chr | Start | End | both | A1 | A2 |
|---|---|---|---|---|---|
| F12 | 20220001 | 20320000 | 0.0040 | 0.0031 | 0.0035 |
| F12 | 20240001 | 20340000 | 0.0044 | 0.0034 | 0.0038 |
| F12 | 20260001 | 20360000 | 0.0042 | 0.0032 | 0.0036 |
| F12 | 20280001 | 20380000 | 0.0038 | 0.0031 | 0.0034 |
| F12 | 20300001 | 20400000 | 0.0044 | 0.0036 | 0.0040 |
| F12 | 20320001 | 20420000 | 0.0044 | 0.0034 | 0.0041 |
| F12 | 20340001 | 20440000 | 0.0035 | 0.0028 | 0.0033 |
| F12 | 20360001 | 20460000 | 0.0029 | 0.0023 | 0.0027 |
| F12 | 20380001 | 20480000 | 0.0030 | 0.0024 | 0.0027 |
| F12 | 20400001 | 20500000 | 0.0023 | 0.0018 | 0.0020 |
| F12 | 20420001 | 20520000 | 0.0016 | 0.0012 | 0.0012 |
| F12 | 20440001 | 20540000 | 0.0018 | 0.0014 | 0.0013 |
| F12 | 20460001 | 20560000 | 0.0031 | 0.0027 | 0.0027 |
| F12 | 20480001 | 20580000 | 0.0031 | 0.0027 | 0.0028 |
| F12 | 20500001 | 20600000 | 0.0030 | 0.0027 | 0.0027 |
| F12 | 20520001 | 20620000 | 0.0035 | 0.0032 | 0.0034 |
| F12 | 20540001 | 20640000 | 0.0039 | 0.0033 | 0.0037 |
| F12 | 20560001 | 20660000 | 0.0028 | 0.0021 | 0.0026 |
| F12 | 20580001 | 20680000 | 0.0031 | 0.0022 | 0.0027 |
| F12 | 20600001 | 20700000 | 0.0040 | 0.0029 | 0.0036 |
| F12 | 20620001 | 20720000 | 0.0035 | 0.0023 | 0.0029 |
| F12 | 20640001 | 20740000 | 0.0031 | 0.0021 | 0.0025 |
| F12 | 20660001 | 20760000 | 0.0026 | 0.0018 | 0.0022 |
| F12 | 20680001 | 20780000 | 0.0024 | 0.0018 | 0.0021 |
| F12 | 20700001 | 20800000 | 0.0015 | 0.0011 | 0.0013 |
| F12 | 20720001 | 20820000 | 0.0018 | 0.0016 | 0.0016 |
| F12 | 20740001 | 20840000 | 0.0016 | 0.0015 | 0.0016 |
| F12 | 20760001 | 20860000 | 0.0016 | 0.0015 | 0.0015 |
| F12 | 20780001 | 20880000 | 0.0019 | 0.0016 | 0.0019 |
| F12 | 20800001 | 20900000 | 0.0019 | 0.0017 | 0.0019 |
| F12 | 20820001 | 20920000 | 0.0016 | 0.0013 | 0.0016 |
| F12 | 20840001 | 20940000 | 0.0017 | 0.0013 | 0.0016 |
| F12 | 20860001 | 20960000 | 0.0018 | 0.0014 | 0.0017 |
| F12 | 20880001 | 20980000 | 0.0010 | 0.0009 | 0.0010 |
| F12 | 20900001 | 21000000 | 0.0012 | 0.0010 | 0.0011 |
| F12 | 20920001 | 21020000 | 0.0014 | 0.0012 | 0.0013 |
| F12 | 20940001 | 21040000 | 0.0010 | 0.0009 | 0.0010 |
| F12 | 20960001 | 21060000 | 0.0010 | 0.0009 | 0.0009 |
| F12 | 20980001 | 21080000 | 0.0012 | 0.0012 | 0.0012 |
| F12 | 21000001 | 21100000 | 0.0011 | 0.0010 | 0.0010 |
| F12 | 21020001 | 21120000 | 0.0014 | 0.0012 | 0.0014 |
| F12 | 21040001 | 21140000 | 0.0015 | 0.0013 | 0.0015 |
| F12 | 21060001 | 21160000 | 0.0014 | 0.0011 | 0.0014 |
| F12 | 21080001 | 21180000 | 0.0012 | 0.0009 | 0.0012 |
| F12 | 21100001 | 21200000 | 0.0014 | 0.0010 | 0.0014 |
| F12 | 21120001 | 21220000 | 0.0008 | 0.0007 | 0.0008 |
| F12 | 21140001 | 21240000 | 0.0012 | 0.0010 | 0.0012 |
| F12 | 21160001 | 21260000 | 0.0019 | 0.0015 | 0.0019 |
| F12 | 21180001 | 21280000 | 0.0018 | 0.0014 | 0.0018 |
| F12 | 21200001 | 21300000 | 0.0015 | 0.0012 | 0.0015 |
| F12 | 21220001 | 21320000 | 0.0015 | 0.0012 | 0.0015 |
| F12 | 21240001 | 21340000 | 0.0011 | 0.0009 | 0.0011 |
| F12 | 21260001 | 21360000 | 0.0006 | 0.0006 | 0.0006 |
| F12 | 21280001 | 21380000 | 0.0006 | 0.0006 | 0.0006 |
| F12 | 21300001 | 21400000 | 0.0011 | 0.0010 | 0.0011 |
| F12 | 21320001 | 21420000 | 0.0023 | 0.0020 | 0.0023 |
| F12 | 21340001 | 21440000 | 0.0026 | 0.0021 | 0.0026 |
| F12 | 21360001 | 21460000 | 0.0027 | 0.0022 | 0.0027 |

| Chr | Start | End | both | A1 | A2 |
|---|---|---|---|---|---|
| F12 | 21380001 | 21480000 | 0.0029 | 0.0025 | 0.0029 |
| F12 | 21400001 | 21500000 | 0.0025 | 0.0022 | 0.0025 |
| F12 | 21420001 | 21520000 | 0.0014 | 0.0012 | 0.0013 |
| F12 | 21440001 | 21540000 | 0.0013 | 0.0012 | 0.0012 |
| F12 | 21460001 | 21560000 | 0.0012 | 0.0011 | 0.0011 |
| F12 | 21480001 | 21580000 | 0.0011 | 0.0008 | 0.0010 |
| F12 | 21500001 | 21600000 | 0.0014 | 0.0011 | 0.0013 |
| F12 | 21520001 | 21620000 | 0.0023 | 0.0017 | 0.0019 |
| F12 | 21540001 | 21640000 | 0.0025 | 0.0019 | 0.0021 |
| F12 | 21560001 | 21660000 | 0.0024 | 0.0018 | 0.0020 |
| F12 | 21580001 | 21680000 | 0.0031 | 0.0024 | 0.0026 |
| F12 | 21600001 | 21700000 | 0.0034 | 0.0024 | 0.0028 |
| F12 | 21620001 | 21720000 | 0.0028 | 0.0021 | 0.0024 |
| F12 | 21640001 | 21740000 | 0.0026 | 0.0018 | 0.0022 |
| F12 | 21660001 | 21760000 | 0.0042 | 0.0033 | 0.0038 |
| F12 | 21680001 | 21780000 | 0.0037 | 0.0030 | 0.0034 |
| F12 | 21700001 | 21800000 | 0.0034 | 0.0028 | 0.0031 |
| F12 | 21720001 | 21820000 | 0.0033 | 0.0026 | 0.0029 |
| F12 | 21740001 | 21840000 | 0.0040 | 0.0032 | 0.0036 |
| F12 | 21760001 | 21860000 | 0.0041 | 0.0033 | 0.0034 |
| F12 | 21780001 | 21880000 | 0.0048 | 0.0037 | 0.0041 |
| F12 | 21800001 | 21900000 | 0.0054 | 0.0041 | 0.0046 |
| F12 | 21820001 | 21920000 | 0.0054 | 0.0043 | 0.0048 |
| F12 | 21840001 | 21940000 | 0.0054 | 0.0042 | 0.0048 |
| F12 | 21860001 | 21960000 | 0.0040 | 0.0030 | 0.0038 |
| F12 | 21880001 | 21980000 | 0.0035 | 0.0028 | 0.0033 |
| F12 | 21900001 | 22000000 | 0.0033 | 0.0027 | 0.0031 |
| F12 | 21920001 | 22020000 | 0.0037 | 0.0031 | 0.0035 |
| F12 | 21940001 | 22040000 | 0.0033 | 0.0025 | 0.0031 |
| F12 | 21960001 | 22060000 | 0.0036 | 0.0029 | 0.0035 |
| F12 | 21980001 | 22080000 | 0.0037 | 0.0029 | 0.0036 |
| F12 | 22000001 | 22100000 | 0.0043 | 0.0035 | 0.0041 |
| F12 | 22020001 | 22120000 | 0.0042 | 0.0033 | 0.0039 |
| F12 | 22040001 | 22140000 | 0.0048 | 0.0039 | 0.0044 |
| F12 | 22060001 | 22160000 | 0.0051 | 0.0041 | 0.0046 |
| F12 | 22080001 | 22180000 | 0.0046 | 0.0035 | 0.0040 |
| F12 | 22100001 | 22200000 | 0.0035 | 0.0026 | 0.0029 |
| F12 | 22120001 | 22220000 | 0.0031 | 0.0024 | 0.0027 |
| F12 | 22140001 | 22240000 | 0.0038 | 0.0029 | 0.0034 |
| F12 | 22160001 | 22260000 | 0.0030 | 0.0021 | 0.0025 |
| F12 | 22180001 | 22280000 | 0.0032 | 0.0022 | 0.0027 |
| F12 | 22200001 | 22300000 | 0.0035 | 0.0026 | 0.0029 |
| F12 | 22220001 | 22320000 | 0.0037 | 0.0026 | 0.0029 |
| F12 | 22240001 | 22340000 | 0.0026 | 0.0015 | 0.0019 |
| F12 | 22260001 | 22360000 | 0.0030 | 0.0019 | 0.0023 |
| F12 | 22280001 | 22380000 | 0.0036 | 0.0024 | 0.0030 |
| F12 | 22300001 | 22400000 | 0.0040 | 0.0026 | 0.0035 |
| F12 | 22320001 | 22420000 | 0.0044 | 0.0028 | 0.0039 |
| F12 | 22340001 | 22440000 | 0.0041 | 0.0027 | 0.0035 |
| F12 | 22360001 | 22460000 | 0.0042 | 0.0028 | 0.0036 |
| F12 | 22380001 | 22480000 | 0.0044 | 0.0031 | 0.0038 |
| F12 | 22400001 | 22500000 | 0.0043 | 0.0032 | 0.0038 |
| F12 | 22420001 | 22520000 | 0.0040 | 0.0032 | 0.0034 |
| F12 | 22440001 | 22540000 | 0.0039 | 0.0032 | 0.0035 |
| F12 | 22460001 | 22560000 | 0.0037 | 0.0030 | 0.0033 |
| F12 | 22480001 | 22580000 | 0.0033 | 0.0027 | 0.0029 |
| F12 | 22500001 | 22600000 | 0.0026 | 0.0020 | 0.0023 |
| F12 | 22520001 | 22620000 | 0.0027 | 0.0020 | 0.0023 |

| Chr | Start | End | both | A1 | A2 |
|---|---|---|---|---|---|
| F12 | 22540001 | 22640000 | 0.0028 | 0.0021 | 0.0024 |
| F12 | 22560001 | 22660000 | 0.0025 | 0.0019 | 0.0022 |
| F12 | 22580001 | 22680000 | 0.0022 | 0.0017 | 0.0019 |
| F12 | 22600001 | 22700000 | 0.0023 | 0.0018 | 0.0021 |
| F12 | 22620001 | 22720000 | 0.0024 | 0.0020 | 0.0023 |
| F12 | 22640001 | 22740000 | 0.0025 | 0.0022 | 0.0025 |
| F12 | 22660001 | 22760000 | 0.0032 | 0.0028 | 0.0030 |
| F12 | 22680001 | 22780000 | 0.0035 | 0.0030 | 0.0034 |
| F12 | 22700001 | 22800000 | 0.0039 | 0.0034 | 0.0036 |
| F12 | 22720001 | 22820000 | 0.0045 | 0.0037 | 0.0039 |
| F12 | 22740001 | 22840000 | 0.0046 | 0.0037 | 0.0040 |
| F12 | 22760001 | 22860000 | 0.0038 | 0.0030 | 0.0033 |
| F12 | 22780001 | 22880000 | 0.0036 | 0.0028 | 0.0031 |
| F12 | 22800001 | 22900000 | 0.0030 | 0.0024 | 0.0027 |
| F12 | 22820001 | 22920000 | 0.0030 | 0.0024 | 0.0028 |
| F12 | 22840001 | 22940000 | 0.0027 | 0.0020 | 0.0024 |
| F12 | 22860001 | 22960000 | 0.0033 | 0.0025 | 0.0031 |
| F12 | 22880001 | 22980000 | 0.0031 | 0.0022 | 0.0028 |
| F12 | 22900001 | 23000000 | 0.0042 | 0.0032 | 0.0037 |
| F12 | 22920001 | 23020000 | 0.0031 | 0.0023 | 0.0027 |
| F12 | 22940001 | 23040000 | 0.0030 | 0.0023 | 0.0027 |
| F12 | 22960001 | 23060000 | 0.0028 | 0.0021 | 0.0023 |
| F12 | 22980001 | 23080000 | 0.0026 | 0.0020 | 0.0023 |
| F12 | 23000001 | 23100000 | 0.0018 | 0.0013 | 0.0015 |
| F12 | 23020001 | 23120000 | 0.0022 | 0.0017 | 0.0019 |
| F12 | 23040001 | 23140000 | 0.0023 | 0.0019 | 0.0021 |
| F12 | 23060001 | 23160000 | 0.0028 | 0.0023 | 0.0026 |
| F12 | 23080001 | 23180000 | 0.0031 | 0.0025 | 0.0027 |
| F12 | 23100001 | 23200000 | 0.0029 | 0.0022 | 0.0025 |
| F12 | 23120001 | 23220000 | 0.0025 | 0.0018 | 0.0021 |
| F12 | 23140001 | 23240000 | 0.0021 | 0.0015 | 0.0017 |
| F12 | 23160001 | 23260000 | 0.0016 | 0.0012 | 0.0013 |
| F12 | 23180001 | 23280000 | 0.0014 | 0.0011 | 0.0012 |
| F12 | 23200001 | 23300000 | 0.0016 | 0.0013 | 0.0015 |
| F12 | 23220001 | 23320000 | 0.0022 | 0.0017 | 0.0020 |
| F12 | 23240001 | 23340000 | 0.0026 | 0.0019 | 0.0022 |
| F12 | 23260001 | 23360000 | 0.0026 | 0.0018 | 0.0023 |
| F12 | 23280001 | 23380000 | 0.0025 | 0.0017 | 0.0022 |
| F12 | 23300001 | 23400000 | 0.0023 | 0.0016 | 0.0020 |
| F12 | 23320001 | 23420000 | 0.0022 | 0.0013 | 0.0017 |
| F12 | 23340001 | 23440000 | 0.0023 | 0.0017 | 0.0020 |
| F12 | 23360001 | 23460000 | 0.0027 | 0.0020 | 0.0023 |
| F12 | 23380001 | 23480000 | 0.0028 | 0.0020 | 0.0024 |
| F12 | 23400001 | 23500000 | 0.0026 | 0.0019 | 0.0022 |
| F12 | 23420001 | 23520000 | 0.0023 | 0.0019 | 0.0021 |
| F12 | 23440001 | 23540000 | 0.0017 | 0.0013 | 0.0015 |
| F12 | 23460001 | 23560000 | 0.0009 | 0.0007 | 0.0009 |
| F12 | 23480001 | 23580000 | 0.0016 | 0.0013 | 0.0015 |
| F12 | 23500001 | 23600000 | 0.0021 | 0.0017 | 0.0019 |
| F12 | 23520001 | 23620000 | 0.0028 | 0.0023 | 0.0026 |
| F12 | 23540001 | 23640000 | 0.0038 | 0.0032 | 0.0036 |
| F12 | 23560001 | 23660000 | 0.0045 | 0.0037 | 0.0043 |
| F12 | 23580001 | 23680000 | 0.0044 | 0.0035 | 0.0041 |
| F12 | 23600001 | 23700000 | 0.0043 | 0.0036 | 0.0041 |
| F12 | 23620001 | 23720000 | 0.0043 | 0.0035 | 0.0039 |
| F12 | 23640001 | 23740000 | 0.0041 | 0.0031 | 0.0035 |
| F12 | 23660001 | 23760000 | 0.0036 | 0.0028 | 0.0031 |
| F12 | 23680001 | 23780000 | 0.0035 | 0.0028 | 0.0030 |

| Chr | Start | End | both | A1 | A2 |
|---|---|---|---|---|---|
| F12 | 23700001 | 23800000 | 0.0032 | 0.0023 | 0.0025 |
| F12 | 23720001 | 23820000 | 0.0029 | 0.0020 | 0.0023 |
| F12 | 23740001 | 23840000 | 0.0026 | 0.0016 | 0.0019 |
| F12 | 23760001 | 23860000 | 0.0028 | 0.0017 | 0.0021 |
| F12 | 23780001 | 23880000 | 0.0039 | 0.0026 | 0.0032 |
| F12 | 23800001 | 23900000 | 0.0044 | 0.0031 | 0.0037 |
| F12 | 23820001 | 23920000 | 0.0051 | 0.0038 | 0.0043 |
| F12 | 23840001 | 23940000 | 0.0054 | 0.0040 | 0.0046 |
| F12 | 23860001 | 23960000 | 0.0062 | 0.0044 | 0.0050 |
| F12 | 23880001 | 23980000 | 0.0051 | 0.0036 | 0.0039 |
| F12 | 23900001 | 24000000 | 0.0050 | 0.0034 | 0.0039 |
| F12 | 23920001 | 24020000 | 0.0047 | 0.0031 | 0.0037 |
| F12 | 23940001 | 24040000 | 0.0043 | 0.0031 | 0.0036 |
| F12 | 23960001 | 24060000 | 0.0037 | 0.0028 | 0.0034 |
| F12 | 23980001 | 24080000 | 0.0037 | 0.0029 | 0.0034 |
| F12 | 24000001 | 24100000 | 0.0044 | 0.0038 | 0.0041 |
| F12 | 24020001 | 24120000 | 0.0036 | 0.0031 | 0.0033 |
| F12 | 24040001 | 24140000 | 0.0038 | 0.0032 | 0.0034 |
| F12 | 24060001 | 24160000 | 0.0043 | 0.0035 | 0.0038 |
| F12 | 24080001 | 24180000 | 0.0041 | 0.0034 | 0.0036 |
| F12 | 24100001 | 24200000 | 0.0030 | 0.0022 | 0.0024 |
| F12 | 24120001 | 24220000 | 0.0036 | 0.0026 | 0.0029 |
| F12 | 24140001 | 24240000 | 0.0035 | 0.0027 | 0.0030 |
| F12 | 24160001 | 24260000 | 0.0034 | 0.0027 | 0.0030 |
| F12 | 24180001 | 24280000 | 0.0042 | 0.0031 | 0.0034 |
| F12 | 24200001 | 24300000 | 0.0048 | 0.0038 | 0.0042 |
| F12 | 24220001 | 24320000 | 0.0051 | 0.0042 | 0.0045 |
| F12 | 24240001 | 24340000 | 0.0045 | 0.0037 | 0.0039 |
| F12 | 24260001 | 24360000 | 0.0035 | 0.0029 | 0.0030 |
| F12 | 24280001 | 24380000 | 0.0026 | 0.0023 | 0.0023 |
| F12 | 24300001 | 24400000 | 0.0019 | 0.0017 | 0.0017 |
| F12 | 24320001 | 24420000 | 0.0021 | 0.0019 | 0.0020 |
| F12 | 24340001 | 24440000 | 0.0023 | 0.0020 | 0.0022 |
| F12 | 24360001 | 24460000 | 0.0033 | 0.0028 | 0.0031 |
| F12 | 24380001 | 24480000 | 0.0034 | 0.0029 | 0.0032 |
| F12 | 24400001 | 24500000 | 0.0037 | 0.0030 | 0.0033 |
| F12 | 24420001 | 24520000 | 0.0034 | 0.0027 | 0.0030 |
| F12 | 24440001 | 24540000 | 0.0041 | 0.0032 | 0.0035 |
| F12 | 24460001 | 24560000 | 0.0045 | 0.0033 | 0.0038 |
| F12 | 24480001 | 24580000 | 0.0048 | 0.0035 | 0.0040 |
| F12 | 24500001 | 24600000 | 0.0048 | 0.0036 | 0.0042 |
| F12 | 24520001 | 24620000 | 0.0047 | 0.0035 | 0.0041 |
| F12 | 24540001 | 24640000 | 0.0047 | 0.0035 | 0.0040 |
| F12 | 24560001 | 24660000 | 0.0035 | 0.0026 | 0.0030 |
| F12 | 24580001 | 24680000 | 0.0032 | 0.0024 | 0.0028 |
| F12 | 24600001 | 24700000 | 0.0036 | 0.0028 | 0.0031 |
| F12 | 24620001 | 24720000 | 0.0032 | 0.0026 | 0.0028 |
| F12 | 24640001 | 24740000 | 0.0026 | 0.0022 | 0.0023 |
| F12 | 24660001 | 24760000 | 0.0025 | 0.0022 | 0.0023 |
| F12 | 24680001 | 24780000 | 0.0023 | 0.0021 | 0.0022 |
| F12 | 24700001 | 24800000 | 0.0015 | 0.0013 | 0.0015 |
| F12 | 24720001 | 24820000 | 0.0012 | 0.0009 | 0.0012 |
| F12 | 24740001 | 24840000 | 0.0008 | 0.0006 | 0.0008 |
| F12 | 24760001 | 24860000 | 0.0016 | 0.0011 | 0.0015 |
| F12 | 24780001 | 24880000 | 0.0023 | 0.0018 | 0.0021 |
| F12 | 24800001 | 24900000 | 0.0024 | 0.0019 | 0.0021 |
| F12 | 24820001 | 24920000 | 0.0026 | 0.0021 | 0.0023 |
| F12 | 24840001 | 24940000 | 0.0028 | 0.0022 | 0.0024 |

| Chr | Start | End | both | A1 | A2 |
|---|---|---|---|---|---|
| F12 | 24860001 | 24960000 | 0.0022 | 0.0019 | 0.0020 |
| F12 | 24880001 | 24980000 | 0.0030 | 0.0025 | 0.0028 |
| F12 | 24900001 | 25000000 | 0.0038 | 0.0028 | 0.0036 |
| F12 | 24920001 | 25020000 | 0.0037 | 0.0027 | 0.0035 |
| F12 | 24940001 | 25040000 | 0.0036 | 0.0027 | 0.0035 |
| F12 | 24960001 | 25060000 | 0.0032 | 0.0023 | 0.0030 |
| F12 | 24980001 | 25080000 | 0.0019 | 0.0013 | 0.0018 |
| F12 | 25000001 | 25100000 | 0.0015 | 0.0013 | 0.0014 |
| F12 | 25020001 | 25120000 | 0.0022 | 0.0018 | 0.0020 |
| F12 | 25040001 | 25140000 | 0.0027 | 0.0021 | 0.0024 |
| F12 | 25060001 | 25160000 | 0.0036 | 0.0030 | 0.0033 |
| F12 | 25080001 | 25180000 | 0.0039 | 0.0033 | 0.0035 |
| F12 | 25100001 | 25200000 | 0.0036 | 0.0031 | 0.0034 |
| F12 | 25120001 | 25220000 | 0.0033 | 0.0028 | 0.0031 |
| F12 | 25140001 | 25240000 | 0.0026 | 0.0023 | 0.0025 |
| F12 | 25160001 | 25260000 | 0.0019 | 0.0017 | 0.0018 |
| F12 | 25180001 | 25280000 | 0.0012 | 0.0010 | 0.0012 |
| F12 | 25200001 | 25300000 | 0.0011 | 0.0010 | 0.0011 |
| F12 | 25220001 | 25320000 | 0.0010 | 0.0009 | 0.0010 |
| F12 | 25240001 | 25340000 | 0.0012 | 0.0011 | 0.0012 |
| F12 | 25260001 | 25360000 | 0.0014 | 0.0012 | 0.0014 |
| F12 | 25280001 | 25380000 | 0.0015 | 0.0013 | 0.0015 |
| F12 | 25300001 | 25400000 | 0.0012 | 0.0010 | 0.0012 |
| F12 | 25320001 | 25420000 | 0.0010 | 0.0007 | 0.0010 |
| F12 | 25340001 | 25440000 | 0.0013 | 0.0011 | 0.0013 |
| F12 | 25360001 | 25460000 | 0.0012 | 0.0011 | 0.0012 |
| F12 | 25380001 | 25480000 | 0.0022 | 0.0019 | 0.0021 |
| F12 | 25400001 | 25500000 | 0.0028 | 0.0023 | 0.0027 |
| F12 | 25420001 | 25520000 | 0.0027 | 0.0023 | 0.0027 |
| F12 | 25440001 | 25540000 | 0.0025 | 0.0021 | 0.0025 |
| F12 | 25460001 | 25560000 | 0.0022 | 0.0017 | 0.0022 |
| F12 | 25480001 | 25580000 | 0.0012 | 0.0010 | 0.0012 |
| F12 | 25500001 | 25600000 | 0.0007 | 0.0006 | 0.0007 |
| F12 | 25520001 | 25620000 | 0.0013 | 0.0012 | 0.0012 |
| F12 | 25540001 | 25640000 | 0.0011 | 0.0010 | 0.0010 |
| F12 | 25560001 | 25660000 | 0.0013 | 0.0010 | 0.0010 |
| F12 | 25580001 | 25680000 | 0.0017 | 0.0012 | 0.0014 |
| F12 | 25600001 | 25700000 | 0.0017 | 0.0013 | 0.0014 |
| F12 | 25620001 | 25720000 | 0.0019 | 0.0013 | 0.0017 |
| F12 | 25640001 | 25740000 | 0.0020 | 0.0014 | 0.0018 |
| F12 | 25660001 | 25760000 | 0.0021 | 0.0016 | 0.0020 |
| F12 | 25680001 | 25780000 | 0.0017 | 0.0014 | 0.0017 |
| F12 | 25700001 | 25800000 | 0.0016 | 0.0014 | 0.0016 |
| F12 | 25720001 | 25820000 | 0.0015 | 0.0012 | 0.0014 |
| F12 | 25740001 | 25840000 | 0.0013 | 0.0010 | 0.0013 |
| F12 | 25760001 | 25860000 | 0.0011 | 0.0009 | 0.0011 |
| F12 | 25780001 | 25880000 | 0.0013 | 0.0010 | 0.0013 |
| F12 | 25800001 | 25900000 | 0.0012 | 0.0010 | 0.0012 |
| F12 | 25820001 | 25920000 | 0.0012 | 0.0010 | 0.0011 |
| F12 | 25840001 | 25940000 | 0.0013 | 0.0012 | 0.0013 |
| F12 | 25860001 | 25960000 | 0.0015 | 0.0014 | 0.0015 |
| F12 | 25880001 | 25980000 | 0.0018 | 0.0015 | 0.0018 |
| F12 | 25900001 | 26000000 | 0.0023 | 0.0020 | 0.0023 |
| F12 | 25920001 | 26020000 | 0.0025 | 0.0018 | 0.0023 |
| F12 | 25940001 | 26040000 | 0.0024 | 0.0017 | 0.0021 |
| F12 | 25960001 | 26060000 | 0.0024 | 0.0016 | 0.0021 |
| F12 | 25980001 | 26080000 | 0.0027 | 0.0019 | 0.0024 |
| F12 | 26000001 | 26100000 | 0.0024 | 0.0016 | 0.0021 |

| Chr | Start | End | both | A1 | A2 |
|---|---|---|---|---|---|
| F12 | 26020001 | 26120000 | 0.0020 | 0.0014 | 0.0019 |
| F12 | 26040001 | 26140000 | 0.0025 | 0.0018 | 0.0024 |
| F12 | 26060001 | 26160000 | 0.0024 | 0.0018 | 0.0023 |
| F12 | 26080001 | 26180000 | 0.0015 | 0.0011 | 0.0014 |
| F12 | 26100001 | 26200000 | 0.0019 | 0.0015 | 0.0018 |
| F12 | 26120001 | 26220000 | 0.0020 | 0.0016 | 0.0019 |
| F12 | 26140001 | 26240000 | 0.0015 | 0.0013 | 0.0015 |
| F12 | 26160001 | 26260000 | 0.0015 | 0.0013 | 0.0015 |
| F12 | 26180001 | 26280000 | 0.0024 | 0.0021 | 0.0023 |
| F12 | 26200001 | 26300000 | 0.0017 | 0.0015 | 0.0017 |
| F12 | 26220001 | 26320000 | 0.0019 | 0.0016 | 0.0019 |
| F12 | 26240001 | 26340000 | 0.0019 | 0.0017 | 0.0019 |
| F12 | 26260001 | 26360000 | 0.0018 | 0.0016 | 0.0018 |
| F12 | 26280001 | 26380000 | 0.0014 | 0.0012 | 0.0014 |
| F12 | 26300001 | 26400000 | 0.0025 | 0.0021 | 0.0025 |
| F12 | 26320001 | 26420000 | 0.0022 | 0.0019 | 0.0022 |
| F12 | 26340001 | 26440000 | 0.0023 | 0.0019 | 0.0023 |
| F12 | 26360001 | 26460000 | 0.0038 | 0.0033 | 0.0038 |
| F12 | 26380001 | 26480000 | 0.0043 | 0.0038 | 0.0042 |
| F12 | 26400001 | 26500000 | 0.0038 | 0.0034 | 0.0038 |
| F12 | 26420001 | 26520000 | 0.0044 | 0.0039 | 0.0043 |
| F12 | 26440001 | 26540000 | 0.0048 | 0.0043 | 0.0046 |
| F12 | 26460001 | 26560000 | 0.0043 | 0.0037 | 0.0041 |
| F12 | 26480001 | 26580000 | 0.0041 | 0.0035 | 0.0038 |
| F12 | 26500001 | 26600000 | 0.0037 | 0.0031 | 0.0034 |
| F12 | 26520001 | 26620000 | 0.0034 | 0.0029 | 0.0032 |
| F12 | 26540001 | 26640000 | 0.0040 | 0.0034 | 0.0038 |
| F12 | 26560001 | 26660000 | 0.0035 | 0.0030 | 0.0034 |
| F12 | 26580001 | 26680000 | 0.0036 | 0.0032 | 0.0035 |
| F12 | 26600001 | 26700000 | 0.0036 | 0.0032 | 0.0035 |
| F12 | 26620001 | 26720000 | 0.0033 | 0.0029 | 0.0032 |
| F12 | 26640001 | 26740000 | 0.0024 | 0.0021 | 0.0023 |
| F12 | 26660001 | 26760000 | 0.0020 | 0.0017 | 0.0020 |
| F12 | 26680001 | 26780000 | 0.0016 | 0.0013 | 0.0016 |
| F12 | 26700001 | 26800000 | 0.0015 | 0.0011 | 0.0015 |
| F12 | 26720001 | 26820000 | 0.0014 | 0.0010 | 0.0014 |
| F12 | 26740001 | 26840000 | 0.0011 | 0.0009 | 0.0011 |
| F12 | 26760001 | 26860000 | 0.0012 | 0.0009 | 0.0012 |
| F12 | 26780001 | 26880000 | 0.0010 | 0.0008 | 0.0009 |
| F12 | 26800001 | 26900000 | 0.0009 | 0.0008 | 0.0009 |
| F12 | 26820001 | 26920000 | 0.0008 | 0.0007 | 0.0008 |
| F12 | 26840001 | 26940000 | 0.0011 | 0.0009 | 0.0011 |
| F12 | 26860001 | 26960000 | 0.0016 | 0.0013 | 0.0015 |
| F12 | 26880001 | 26980000 | 0.0021 | 0.0018 | 0.0020 |
| F12 | 26900001 | 27000000 | 0.0024 | 0.0021 | 0.0023 |
| F12 | 26920001 | 27020000 | 0.0029 | 0.0024 | 0.0027 |
| F12 | 26940001 | 27040000 | 0.0028 | 0.0025 | 0.0028 |
| F12 | 26960001 | 27060000 | 0.0027 | 0.0024 | 0.0027 |
| F12 | 26980001 | 27080000 | 0.0034 | 0.0029 | 0.0031 |
| F12 | 27000001 | 27100000 | 0.0043 | 0.0037 | 0.0040 |
| F12 | 27020001 | 27120000 | 0.0043 | 0.0036 | 0.0039 |
| F12 | 27040001 | 27140000 | 0.0044 | 0.0037 | 0.0039 |
| F12 | 27060001 | 27160000 | 0.0042 | 0.0035 | 0.0037 |
| F12 | 27080001 | 27180000 | 0.0034 | 0.0027 | 0.0032 |
| F12 | 27100001 | 27200000 | 0.0031 | 0.0023 | 0.0029 |
| F12 | 27120001 | 27220000 | 0.0029 | 0.0024 | 0.0028 |
| F12 | 27140001 | 27240000 | 0.0026 | 0.0022 | 0.0026 |
| F12 | 27160001 | 27260000 | 0.0021 | 0.0017 | 0.0021 |

| Chr | Start | End | both | A1 | A2 |
|---|---|---|---|---|---|
| F12 | 27180001 | 27280000 | 0.0017 | 0.0014 | 0.0017 |
| F12 | 27200001 | 27300000 | 0.0008 | 0.0008 | 0.0008 |
| F12 | 27220001 | 27320000 | 0.0013 | 0.0010 | 0.0013 |
| F12 | 27240001 | 27340000 | 0.0013 | 0.0011 | 0.0013 |
| F12 | 27260001 | 27360000 | 0.0013 | 0.0011 | 0.0013 |
| F12 | 27280001 | 27380000 | 0.0021 | 0.0017 | 0.0021 |
| F12 | 27300001 | 27400000 | 0.0021 | 0.0017 | 0.0021 |
| F12 | 27320001 | 27420000 | 0.0016 | 0.0015 | 0.0016 |
| F12 | 27340001 | 27440000 | 0.0017 | 0.0015 | 0.0016 |
| F12 | 27360001 | 27460000 | 0.0021 | 0.0019 | 0.0021 |
| F12 | 27380001 | 27480000 | 0.0014 | 0.0012 | 0.0014 |
| F12 | 27400001 | 27500000 | 0.0015 | 0.0012 | 0.0014 |
| F12 | 27420001 | 27520000 | 0.0012 | 0.0010 | 0.0012 |
| F12 | 27440001 | 27540000 | 0.0022 | 0.0019 | 0.0022 |
| F12 | 27460001 | 27560000 | 0.0022 | 0.0020 | 0.0022 |
| F12 | 27480001 | 27580000 | 0.0023 | 0.0020 | 0.0022 |
| F12 | 27500001 | 27600000 | 0.0030 | 0.0028 | 0.0029 |
| F12 | 27520001 | 27620000 | 0.0031 | 0.0028 | 0.0030 |
| F12 | 27540001 | 27640000 | 0.0022 | 0.0020 | 0.0021 |
| F12 | 27560001 | 27660000 | 0.0019 | 0.0017 | 0.0019 |
| F12 | 27580001 | 27680000 | 0.0019 | 0.0017 | 0.0019 |
| F12 | 27600001 | 27700000 | 0.0018 | 0.0015 | 0.0018 |
| F12 | 27620001 | 27720000 | 0.0020 | 0.0015 | 0.0020 |
| F12 | 27640001 | 27740000 | 0.0024 | 0.0019 | 0.0023 |
| F12 | 27660001 | 27760000 | 0.0025 | 0.0020 | 0.0025 |
| F12 | 27680001 | 27780000 | 0.0030 | 0.0024 | 0.0030 |
| F12 | 27700001 | 27800000 | 0.0022 | 0.0017 | 0.0021 |
| F12 | 27720001 | 27820000 | 0.0028 | 0.0024 | 0.0027 |
| F12 | 27740001 | 27840000 | 0.0025 | 0.0021 | 0.0025 |
| F12 | 27760001 | 27860000 | 0.0022 | 0.0019 | 0.0021 |
| F12 | 27780001 | 27880000 | 0.0017 | 0.0014 | 0.0016 |
| F12 | 27800001 | 27900000 | 0.0022 | 0.0020 | 0.0021 |
| F12 | 27820001 | 27920000 | 0.0012 | 0.0010 | 0.0011 |
| F12 | 27840001 | 27940000 | 0.0009 | 0.0008 | 0.0008 |
| F12 | 27860001 | 27960000 | 0.0010 | 0.0009 | 0.0009 |
| F12 | 27880001 | 27980000 | 0.0010 | 0.0009 | 0.0009 |
| F12 | 27900001 | 28000000 | 0.0004 | 0.0003 | 0.0004 |
| F12 | 27920001 | 28020000 | 0.0006 | 0.0005 | 0.0006 |
| F12 | 27940001 | 28040000 | 0.0008 | 0.0007 | 0.0008 |
| F12 | 27960001 | 28060000 | 0.0007 | 0.0006 | 0.0007 |
| F12 | 27980001 | 28080000 | 0.0005 | 0.0005 | 0.0005 |
| F12 | 28000001 | 28100000 | 0.0007 | 0.0006 | 0.0006 |
| F12 | 28020001 | 28120000 | 0.0007 | 0.0005 | 0.0007 |
| F12 | 28040001 | 28140000 | 0.0006 | 0.0005 | 0.0005 |
| F12 | 28060001 | 28160000 | 0.0006 | 0.0005 | 0.0005 |
| F12 | 28080001 | 28180000 | 0.0006 | 0.0005 | 0.0006 |
| F12 | 28100001 | 28200000 | 0.0012 | 0.0009 | 0.0012 |
| F12 | 28120001 | 28220000 | 0.0012 | 0.0010 | 0.0012 |
| F12 | 28140001 | 28240000 | 0.0013 | 0.0009 | 0.0013 |
| F12 | 28160001 | 28260000 | 0.0013 | 0.0010 | 0.0013 |
| F12 | 28180001 | 28280000 | 0.0013 | 0.0009 | 0.0013 |
| F12 | 28200001 | 28300000 | 0.0008 | 0.0006 | 0.0008 |
| F12 | 28220001 | 28320000 | 0.0010 | 0.0007 | 0.0010 |
| F12 | 28240001 | 28340000 | 0.0010 | 0.0008 | 0.0010 |
| F12 | 28260001 | 28360000 | 0.0011 | 0.0008 | 0.0011 |
| F12 | 28280001 | 28380000 | 0.0012 | 0.0009 | 0.0012 |
| F12 | 28300001 | 28400000 | 0.0015 | 0.0011 | 0.0015 |
| F12 | 28320001 | 28420000 | 0.0013 | 0.0010 | 0.0013 |

| Chr | Start | End | both | A1 | A2 |
|---|---|---|---|---|---|
| F12 | 28340001 | 28440000 | 0.0013 | 0.0010 | 0.0012 |
| F12 | 28360001 | 28460000 | 0.0013 | 0.0010 | 0.0013 |
| F12 | 28380001 | 28480000 | 0.0014 | 0.0011 | 0.0014 |
| F12 | 28400001 | 28500000 | 0.0011 | 0.0009 | 0.0011 |
| F12 | 28420001 | 28520000 | 0.0015 | 0.0012 | 0.0015 |
| F12 | 28440001 | 28540000 | 0.0014 | 0.0011 | 0.0014 |
| F12 | 28460001 | 28560000 | 0.0020 | 0.0016 | 0.0019 |
| F12 | 28480001 | 28580000 | 0.0020 | 0.0017 | 0.0020 |
| F12 | 28500001 | 28600000 | 0.0018 | 0.0015 | 0.0017 |
| F12 | 28520001 | 28620000 | 0.0012 | 0.0010 | 0.0012 |
| F12 | 28540001 | 28640000 | 0.0013 | 0.0011 | 0.0013 |
| F12 | 28560001 | 28660000 | 0.0010 | 0.0008 | 0.0010 |
| F12 | 28580001 | 28680000 | 0.0015 | 0.0012 | 0.0015 |
| F12 | 28600001 | 28700000 | 0.0026 | 0.0021 | 0.0026 |
| F12 | 28620001 | 28720000 | 0.0028 | 0.0023 | 0.0028 |
| F12 | 28640001 | 28740000 | 0.0028 | 0.0022 | 0.0028 |
| F12 | 28660001 | 28760000 | 0.0026 | 0.0021 | 0.0026 |
| F12 | 28680001 | 28780000 | 0.0025 | 0.0019 | 0.0023 |
| F12 | 28700001 | 28800000 | 0.0020 | 0.0015 | 0.0017 |
| F12 | 28720001 | 28820000 | 0.0022 | 0.0016 | 0.0019 |
| F12 | 28740001 | 28840000 | 0.0033 | 0.0026 | 0.0029 |
| F12 | 28760001 | 28860000 | 0.0040 | 0.0033 | 0.0036 |
| F12 | 28780001 | 28880000 | 0.0049 | 0.0041 | 0.0045 |
| F12 | 28800001 | 28900000 | 0.0048 | 0.0041 | 0.0045 |
| F12 | 28820001 | 28920000 | 0.0053 | 0.0045 | 0.0049 |
| F12 | 28840001 | 28940000 | 0.0046 | 0.0041 | 0.0043 |
| F12 | 28860001 | 28960000 | 0.0045 | 0.0038 | 0.0041 |
| F12 | 28880001 | 28980000 | 0.0037 | 0.0032 | 0.0034 |
| F12 | 28900001 | 29000000 | 0.0032 | 0.0027 | 0.0030 |
| F12 | 28920001 | 29020000 | 0.0031 | 0.0024 | 0.0027 |
| F12 | 28940001 | 29040000 | 0.0033 | 0.0024 | 0.0028 |
| F12 | 28960001 | 29060000 | 0.0030 | 0.0022 | 0.0025 |
| F12 | 28980001 | 29080000 | 0.0031 | 0.0023 | 0.0026 |
| F12 | 29000001 | 29100000 | 0.0035 | 0.0026 | 0.0030 |
| F12 | 29020001 | 29120000 | 0.0028 | 0.0023 | 0.0026 |
| F12 | 29040001 | 29140000 | 0.0025 | 0.0020 | 0.0023 |
| F12 | 29060001 | 29160000 | 0.0026 | 0.0022 | 0.0025 |
| F12 | 29080001 | 29180000 | 0.0028 | 0.0022 | 0.0027 |
| F12 | 29100001 | 29200000 | 0.0029 | 0.0022 | 0.0026 |
| F12 | 29120001 | 29220000 | 0.0035 | 0.0026 | 0.0030 |
| F12 | 29140001 | 29240000 | 0.0037 | 0.0026 | 0.0030 |
| F12 | 29160001 | 29260000 | 0.0032 | 0.0023 | 0.0025 |
| F12 | 29180001 | 29280000 | 0.0028 | 0.0020 | 0.0021 |
| F12 | 29200001 | 29300000 | 0.0021 | 0.0016 | 0.0017 |
| F12 | 29220001 | 29320000 | 0.0019 | 0.0015 | 0.0015 |
| F12 | 29240001 | 29340000 | 0.0015 | 0.0012 | 0.0013 |
| F12 | 29260001 | 29360000 | 0.0016 | 0.0013 | 0.0016 |
| F12 | 29280001 | 29380000 | 0.0018 | 0.0014 | 0.0017 |
| F12 | 29300001 | 29400000 | 0.0020 | 0.0016 | 0.0019 |
| F12 | 29320001 | 29420000 | 0.0017 | 0.0013 | 0.0017 |
| F12 | 29340001 | 29440000 | 0.0019 | 0.0015 | 0.0018 |
| F12 | 29360001 | 29460000 | 0.0017 | 0.0012 | 0.0016 |
| F12 | 29380001 | 29480000 | 0.0018 | 0.0014 | 0.0017 |
| F12 | 29400001 | 29500000 | 0.0028 | 0.0022 | 0.0027 |
| F12 | 29420001 | 29520000 | 0.0036 | 0.0029 | 0.0035 |
| F12 | 29440001 | 29540000 | 0.0034 | 0.0027 | 0.0033 |
| F12 | 29460001 | 29560000 | 0.0039 | 0.0033 | 0.0038 |
| F12 | 29480001 | 29580000 | 0.0044 | 0.0038 | 0.0043 |

| Chr | Start | End | both | A1 | A2 |
|---|---|---|---|---|---|
| F12 | 29500001 | 29600000 | 0.0039 | 0.0035 | 0.0038 |
| F12 | 29520001 | 29620000 | 0.0030 | 0.0027 | 0.0030 |
| F12 | 29540001 | 29640000 | 0.0034 | 0.0031 | 0.0034 |
| F12 | 29560001 | 29660000 | 0.0029 | 0.0025 | 0.0027 |
| F12 | 29580001 | 29680000 | 0.0020 | 0.0014 | 0.0017 |
| F12 | 29600001 | 29700000 | 0.0018 | 0.0012 | 0.0014 |
| F12 | 29620001 | 29720000 | 0.0025 | 0.0017 | 0.0019 |
| F12 | 29640001 | 29740000 | 0.0021 | 0.0014 | 0.0015 |
| F12 | 29660001 | 29760000 | 0.0023 | 0.0017 | 0.0018 |
| F12 | 29680001 | 29780000 | 0.0029 | 0.0024 | 0.0025 |
| F12 | 29700001 | 29800000 | 0.0029 | 0.0025 | 0.0026 |
| F12 | 29720001 | 29820000 | 0.0025 | 0.0021 | 0.0021 |
| F12 | 29740001 | 29840000 | 0.0028 | 0.0022 | 0.0024 |
| F12 | 29760001 | 29860000 | 0.0027 | 0.0021 | 0.0022 |
| F12 | 29780001 | 29880000 | 0.0019 | 0.0014 | 0.0015 |
| F12 | 29800001 | 29900000 | 0.0017 | 0.0012 | 0.0014 |
| F12 | 29820001 | 29920000 | 0.0016 | 0.0012 | 0.0015 |
| F12 | 29840001 | 29940000 | 0.0017 | 0.0013 | 0.0017 |
| F12 | 29860001 | 29960000 | 0.0022 | 0.0018 | 0.0021 |
| F12 | 29880001 | 29980000 | 0.0033 | 0.0028 | 0.0032 |
| F12 | 29900001 | 30000000 | 0.0029 | 0.0025 | 0.0029 |
| F12 | 29920001 | 30020000 | 0.0033 | 0.0029 | 0.0032 |
| F12 | 29940001 | 30040000 | 0.0037 | 0.0034 | 0.0036 |
| F12 | 29960001 | 30060000 | 0.0028 | 0.0026 | 0.0028 |
| F12 | 29980001 | 30080000 | 0.0017 | 0.0015 | 0.0016 |
| F12 | 30000001 | 30100000 | 0.0024 | 0.0021 | 0.0023 |
| F12 | 30020001 | 30120000 | 0.0021 | 0.0018 | 0.0020 |
| F12 | 30040001 | 30140000 | 0.0016 | 0.0012 | 0.0015 |
| F12 | 30060001 | 30160000 | 0.0020 | 0.0016 | 0.0019 |
| F12 | 30080001 | 30180000 | 0.0022 | 0.0017 | 0.0021 |
| F12 | 30100001 | 30200000 | 0.0015 | 0.0011 | 0.0015 |
| F12 | 30120001 | 30220000 | 0.0019 | 0.0015 | 0.0018 |
| F12 | 30140001 | 30240000 | 0.0018 | 0.0015 | 0.0017 |
| F12 | 30160001 | 30260000 | 0.0015 | 0.0013 | 0.0015 |
| F12 | 30180001 | 30280000 | 0.0015 | 0.0013 | 0.0015 |
| F12 | 30200001 | 30300000 | 0.0016 | 0.0013 | 0.0016 |
| F12 | 30220001 | 30320000 | 0.0010 | 0.0009 | 0.0010 |
| F12 | 30240001 | 30340000 | 0.0009 | 0.0007 | 0.0009 |
| F12 | 30260001 | 30360000 | 0.0014 | 0.0012 | 0.0014 |
| F12 | 30280001 | 30380000 | 0.0020 | 0.0019 | 0.0019 |
| F12 | 30300001 | 30400000 | 0.0029 | 0.0027 | 0.0029 |
| F12 | 30320001 | 30420000 | 0.0035 | 0.0031 | 0.0035 |
| F12 | 30340001 | 30440000 | 0.0034 | 0.0030 | 0.0033 |
| F12 | 30360001 | 30460000 | 0.0030 | 0.0027 | 0.0030 |
| F12 | 30380001 | 30480000 | 0.0031 | 0.0025 | 0.0028 |
| F12 | 30400001 | 30500000 | 0.0023 | 0.0017 | 0.0020 |
| F12 | 30420001 | 30520000 | 0.0016 | 0.0012 | 0.0013 |
| F12 | 30440001 | 30540000 | 0.0028 | 0.0023 | 0.0024 |
| F12 | 30460001 | 30560000 | 0.0030 | 0.0022 | 0.0026 |
| F12 | 30480001 | 30580000 | 0.0025 | 0.0019 | 0.0022 |
| F12 | 30500001 | 30600000 | 0.0028 | 0.0023 | 0.0026 |
| F12 | 30520001 | 30620000 | 0.0032 | 0.0026 | 0.0030 |
| F12 | 30540001 | 30640000 | 0.0020 | 0.0015 | 0.0018 |
| F12 | 30560001 | 30660000 | 0.0019 | 0.0016 | 0.0018 |
| F12 | 30580001 | 30680000 | 0.0020 | 0.0017 | 0.0020 |
| F12 | 30600001 | 30700000 | 0.0031 | 0.0025 | 0.0029 |
| F12 | 30620001 | 30720000 | 0.0030 | 0.0024 | 0.0028 |
| F12 | 30640001 | 30740000 | 0.0040 | 0.0032 | 0.0036 |

| Chr | Start | End | both | A1 | A2 |
|---|---|---|---|---|---|
| F12 | 30660001 | 30760000 | 0.0049 | 0.0037 | 0.0042 |
| F12 | 30680001 | 30780000 | 0.0049 | 0.0035 | 0.0041 |
| F12 | 30700001 | 30800000 | 0.0036 | 0.0024 | 0.0029 |
| F12 | 30720001 | 30820000 | 0.0037 | 0.0025 | 0.0031 |
| F12 | 30740001 | 30840000 | 0.0027 | 0.0017 | 0.0023 |
| F12 | 30760001 | 30860000 | 0.0019 | 0.0012 | 0.0018 |
| F12 | 30780001 | 30880000 | 0.0024 | 0.0018 | 0.0022 |
| F12 | 30800001 | 30900000 | 0.0027 | 0.0019 | 0.0023 |
| F12 | 30820001 | 30920000 | 0.0028 | 0.0021 | 0.0022 |
| F12 | 30840001 | 30940000 | 0.0034 | 0.0026 | 0.0027 |
| F12 | 30860001 | 30960000 | 0.0030 | 0.0023 | 0.0024 |
| F12 | 30880001 | 30980000 | 0.0035 | 0.0030 | 0.0030 |
| F12 | 30900001 | 31000000 | 0.0045 | 0.0040 | 0.0041 |
| F12 | 30920001 | 31020000 | 0.0046 | 0.0041 | 0.0043 |
| F12 | 30940001 | 31040000 | 0.0050 | 0.0045 | 0.0048 |
| F12 | 30960001 | 31060000 | 0.0051 | 0.0045 | 0.0049 |
| F12 | 30980001 | 31080000 | 0.0039 | 0.0034 | 0.0037 |
| F12 | 31000001 | 31100000 | 0.0032 | 0.0029 | 0.0031 |
| F12 | 31020001 | 31120000 | 0.0034 | 0.0029 | 0.0033 |
| F12 | 31040001 | 31140000 | 0.0024 | 0.0021 | 0.0024 |
| F12 | 31060001 | 31160000 | 0.0025 | 0.0022 | 0.0024 |
| F12 | 31080001 | 31180000 | 0.0022 | 0.0020 | 0.0022 |
| F12 | 31100001 | 31200000 | 0.0018 | 0.0016 | 0.0018 |
| F12 | 31120001 | 31220000 | 0.0013 | 0.0012 | 0.0013 |
| F12 | 31140001 | 31240000 | 0.0021 | 0.0019 | 0.0021 |
| F12 | 31160001 | 31260000 | 0.0027 | 0.0024 | 0.0027 |
| F12 | 31180001 | 31280000 | 0.0034 | 0.0029 | 0.0033 |
| F12 | 31200001 | 31300000 | 0.0034 | 0.0027 | 0.0033 |
| F12 | 31220001 | 31320000 | 0.0039 | 0.0033 | 0.0037 |
| F12 | 31240001 | 31340000 | 0.0039 | 0.0031 | 0.0037 |
| F12 | 31260001 | 31360000 | 0.0033 | 0.0027 | 0.0031 |
| F12 | 31280001 | 31380000 | 0.0028 | 0.0023 | 0.0026 |
| F12 | 31300001 | 31400000 | 0.0026 | 0.0021 | 0.0024 |
| F12 | 31320001 | 31420000 | 0.0017 | 0.0012 | 0.0015 |
| F12 | 31340001 | 31440000 | 0.0012 | 0.0010 | 0.0011 |
| F12 | 31360001 | 31460000 | 0.0017 | 0.0014 | 0.0015 |
| F12 | 31380001 | 31480000 | 0.0014 | 0.0013 | 0.0014 |
| F12 | 31400001 | 31500000 | 0.0018 | 0.0016 | 0.0018 |
| F12 | 31420001 | 31520000 | 0.0021 | 0.0018 | 0.0020 |
| F12 | 31440001 | 31540000 | 0.0019 | 0.0016 | 0.0019 |
| F12 | 31460001 | 31560000 | 0.0013 | 0.0010 | 0.0013 |
| F12 | 31480001 | 31580000 | 0.0014 | 0.0011 | 0.0014 |
| F12 | 31500001 | 31600000 | 0.0011 | 0.0010 | 0.0011 |
| F12 | 31520001 | 31620000 | 0.0013 | 0.0011 | 0.0013 |
| F12 | 31540001 | 31640000 | 0.0016 | 0.0014 | 0.0016 |
| F12 | 31560001 | 31660000 | 0.0015 | 0.0014 | 0.0015 |
| F12 | 31580001 | 31680000 | 0.0014 | 0.0013 | 0.0014 |
| F12 | 31600001 | 31700000 | 0.0016 | 0.0014 | 0.0016 |
| F12 | 31620001 | 31720000 | 0.0014 | 0.0011 | 0.0014 |
| F12 | 31640001 | 31740000 | 0.0012 | 0.0010 | 0.0012 |
| F12 | 31660001 | 31760000 | 0.0010 | 0.0008 | 0.0010 |
| F12 | 31680001 | 31780000 | 0.0013 | 0.0010 | 0.0012 |
| F12 | 31700001 | 31800000 | 0.0016 | 0.0013 | 0.0014 |
| F12 | 31720001 | 31820000 | 0.0017 | 0.0015 | 0.0015 |
| F12 | 31740001 | 31840000 | 0.0026 | 0.0023 | 0.0023 |
| F12 | 31760001 | 31860000 | 0.0038 | 0.0032 | 0.0035 |
| F12 | 31780001 | 31880000 | 0.0047 | 0.0037 | 0.0041 |
| F12 | 31800001 | 31900000 | 0.0043 | 0.0034 | 0.0038 |

| Chr | Start | End | both | A1 | A2 |
|---|---|---|---|---|---|
| F12 | 31820001 | 31920000 | 0.0043 | 0.0033 | 0.0037 |
| F12 | 31840001 | 31940000 | 0.0031 | 0.0022 | 0.0026 |
| F12 | 31860001 | 31960000 | 0.0021 | 0.0015 | 0.0017 |
| F12 | 31880001 | 31980000 | 0.0022 | 0.0017 | 0.0019 |
| F12 | 31900001 | 32000000 | 0.0024 | 0.0020 | 0.0021 |
| F12 | 31920001 | 32020000 | 0.0023 | 0.0019 | 0.0021 |
| F12 | 31940001 | 32040000 | 0.0023 | 0.0019 | 0.0021 |
| F12 | 31960001 | 32060000 | 0.0031 | 0.0024 | 0.0028 |
| F12 | 31980001 | 32080000 | 0.0025 | 0.0020 | 0.0023 |
| F12 | 32000001 | 32100000 | 0.0023 | 0.0017 | 0.0021 |
| F12 | 32020001 | 32120000 | 0.0028 | 0.0022 | 0.0026 |
| F12 | 32040001 | 32140000 | 0.0034 | 0.0028 | 0.0033 |
| F12 | 32060001 | 32160000 | 0.0026 | 0.0022 | 0.0025 |
| F12 | 32080001 | 32180000 | 0.0034 | 0.0029 | 0.0032 |
| F12 | 32100001 | 32200000 | 0.0033 | 0.0029 | 0.0031 |
| F12 | 32120001 | 32220000 | 0.0035 | 0.0030 | 0.0034 |
| F12 | 32140001 | 32240000 | 0.0032 | 0.0027 | 0.0031 |
| F12 | 32160001 | 32260000 | 0.0042 | 0.0036 | 0.0040 |
| F12 | 32180001 | 32280000 | 0.0029 | 0.0025 | 0.0027 |
| F12 | 32200001 | 32300000 | 0.0031 | 0.0028 | 0.0030 |
| F12 | 32220001 | 32320000 | 0.0035 | 0.0031 | 0.0032 |
| F12 | 32240001 | 32340000 | 0.0036 | 0.0032 | 0.0032 |
| F12 | 32260001 | 32360000 | 0.0030 | 0.0026 | 0.0025 |
| F12 | 32280001 | 32380000 | 0.0037 | 0.0031 | 0.0032 |
| F12 | 32300001 | 32400000 | 0.0037 | 0.0031 | 0.0031 |
| F12 | 32320001 | 32420000 | 0.0039 | 0.0031 | 0.0034 |
| F12 | 32340001 | 32440000 | 0.0047 | 0.0040 | 0.0042 |
| F12 | 32360001 | 32460000 | 0.0047 | 0.0039 | 0.0042 |
| F12 | 32380001 | 32480000 | 0.0046 | 0.0038 | 0.0040 |
| F12 | 32400001 | 32500000 | 0.0044 | 0.0037 | 0.0040 |
| F12 | 32420001 | 32520000 | 0.0032 | 0.0028 | 0.0029 |
| F12 | 32440001 | 32540000 | 0.0021 | 0.0017 | 0.0019 |
| F12 | 32460001 | 32560000 | 0.0018 | 0.0016 | 0.0017 |
| F12 | 32480001 | 32580000 | 0.0017 | 0.0015 | 0.0016 |
| F12 | 32500001 | 32600000 | 0.0019 | 0.0016 | 0.0018 |
| F12 | 32520001 | 32620000 | 0.0016 | 0.0014 | 0.0015 |
| F12 | 32540001 | 32640000 | 0.0020 | 0.0017 | 0.0019 |
| F12 | 32560001 | 32660000 | 0.0021 | 0.0019 | 0.0020 |
| F12 | 32580001 | 32680000 | 0.0023 | 0.0020 | 0.0022 |
| F12 | 32600001 | 32700000 | 0.0021 | 0.0018 | 0.0020 |
| F12 | 32620001 | 32720000 | 0.0029 | 0.0026 | 0.0029 |
| F12 | 32640001 | 32740000 | 0.0027 | 0.0023 | 0.0026 |
| F12 | 32660001 | 32760000 | 0.0029 | 0.0026 | 0.0027 |
| F12 | 32680001 | 32780000 | 0.0024 | 0.0022 | 0.0022 |
| F12 | 32700001 | 32800000 | 0.0022 | 0.0020 | 0.0020 |
| F12 | 32720001 | 32820000 | 0.0019 | 0.0016 | 0.0017 |
| F12 | 32740001 | 32840000 | 0.0024 | 0.0021 | 0.0021 |
| F12 | 32760001 | 32860000 | 0.0020 | 0.0016 | 0.0019 |
| F12 | 32780001 | 32880000 | 0.0022 | 0.0018 | 0.0021 |
| F12 | 32800001 | 32900000 | 0.0021 | 0.0018 | 0.0020 |
| F12 | 32820001 | 32920000 | 0.0023 | 0.0020 | 0.0022 |
| F12 | 32840001 | 32940000 | 0.0017 | 0.0014 | 0.0016 |
| F12 | 32860001 | 32960000 | 0.0020 | 0.0016 | 0.0017 |
| F12 | 32880001 | 32980000 | 0.0019 | 0.0015 | 0.0017 |
| F12 | 32900001 | 33000000 | 0.0021 | 0.0016 | 0.0019 |
| F12 | 32920001 | 33020000 | 0.0020 | 0.0013 | 0.0016 |
| F12 | 32940001 | 33040000 | 0.0021 | 0.0015 | 0.0018 |
| F12 | 32960001 | 33060000 | 0.0030 | 0.0022 | 0.0026 |

| Chr | Start | End | both | A1 | A2 |
|---|---|---|---|---|---|
| F12 | 32980001 | 33080000 | 0.0031 | 0.0023 | 0.0026 |
| F12 | 33000001 | 33100000 | 0.0034 | 0.0025 | 0.0028 |
| F12 | 33020001 | 33120000 | 0.0028 | 0.0020 | 0.0024 |
| F12 | 33040001 | 33140000 | 0.0034 | 0.0026 | 0.0028 |
| F12 | 33060001 | 33160000 | 0.0024 | 0.0019 | 0.0020 |
| F12 | 33080001 | 33180000 | 0.0022 | 0.0018 | 0.0018 |
| F12 | 33100001 | 33200000 | 0.0029 | 0.0025 | 0.0024 |
| F12 | 33120001 | 33220000 | 0.0044 | 0.0039 | 0.0039 |
| F12 | 33140001 | 33240000 | 0.0037 | 0.0032 | 0.0033 |
| F12 | 33160001 | 33260000 | 0.0038 | 0.0033 | 0.0035 |
| F12 | 33180001 | 33280000 | 0.0039 | 0.0035 | 0.0036 |
| F12 | 33200001 | 33300000 | 0.0033 | 0.0030 | 0.0031 |
| F12 | 33220001 | 33320000 | 0.0023 | 0.0021 | 0.0020 |
| F12 | 33240001 | 33340000 | 0.0032 | 0.0027 | 0.0028 |
| F12 | 33260001 | 33360000 | 0.0032 | 0.0026 | 0.0026 |
| F12 | 33280001 | 33380000 | 0.0034 | 0.0027 | 0.0028 |
| F12 | 33300001 | 33400000 | 0.0032 | 0.0025 | 0.0026 |
| F12 | 33320001 | 33420000 | 0.0029 | 0.0022 | 0.0024 |
| F12 | 33340001 | 33440000 | 0.0021 | 0.0017 | 0.0018 |
| F12 | 33360001 | 33460000 | 0.0019 | 0.0017 | 0.0017 |
| F12 | 33380001 | 33480000 | 0.0017 | 0.0016 | 0.0015 |
| F12 | 33400001 | 33500000 | 0.0013 | 0.0013 | 0.0012 |
| F12 | 33420001 | 33520000 | 0.0010 | 0.0010 | 0.0010 |
| F12 | 33440001 | 33540000 | 0.0011 | 0.0011 | 0.0011 |
| F12 | 33460001 | 33560000 | 0.0010 | 0.0009 | 0.0009 |
| F12 | 33480001 | 33580000 | 0.0007 | 0.0007 | 0.0007 |
| F12 | 33500001 | 33600000 | 0.0009 | 0.0008 | 0.0009 |
| F12 | 33520001 | 33620000 | 0.0012 | 0.0010 | 0.0012 |
| F12 | 33540001 | 33640000 | 0.0011 | 0.0009 | 0.0010 |
| F12 | 33560001 | 33660000 | 0.0012 | 0.0009 | 0.0011 |
| F12 | 33580001 | 33680000 | 0.0020 | 0.0015 | 0.0018 |
| F12 | 33600001 | 33700000 | 0.0036 | 0.0031 | 0.0034 |
| F12 | 33620001 | 33720000 | 0.0040 | 0.0035 | 0.0037 |
| F12 | 33640001 | 33740000 | 0.0039 | 0.0032 | 0.0035 |
| F12 | 33660001 | 33760000 | 0.0040 | 0.0034 | 0.0036 |
| F12 | 33680001 | 33780000 | 0.0036 | 0.0029 | 0.0031 |
| F12 | 33700001 | 33800000 | 0.0025 | 0.0019 | 0.0019 |
| F12 | 33720001 | 33820000 | 0.0027 | 0.0019 | 0.0019 |
| F12 | 33740001 | 33840000 | 0.0036 | 0.0028 | 0.0028 |
| F12 | 33760001 | 33860000 | 0.0038 | 0.0028 | 0.0029 |
| F12 | 33780001 | 33880000 | 0.0044 | 0.0035 | 0.0037 |
| F12 | 33800001 | 33900000 | 0.0048 | 0.0039 | 0.0040 |
| F12 | 33820001 | 33920000 | 0.0045 | 0.0037 | 0.0037 |
| F12 | 33840001 | 33940000 | 0.0036 | 0.0027 | 0.0027 |
| F12 | 33860001 | 33960000 | 0.0042 | 0.0031 | 0.0033 |
| F12 | 33880001 | 33980000 | 0.0037 | 0.0025 | 0.0027 |
| F12 | 33900001 | 34000000 | 0.0025 | 0.0015 | 0.0017 |
| F12 | 33920001 | 34020000 | 0.0023 | 0.0013 | 0.0016 |
| F12 | 33940001 | 34040000 | 0.0021 | 0.0014 | 0.0017 |
| F12 | 33960001 | 34060000 | 0.0023 | 0.0018 | 0.0021 |
| F12 | 33980001 | 34080000 | 0.0020 | 0.0017 | 0.0020 |
| F12 | 34000001 | 34100000 | 0.0027 | 0.0021 | 0.0025 |
| F12 | 34020001 | 34120000 | 0.0031 | 0.0023 | 0.0027 |
| F12 | 34040001 | 34140000 | 0.0031 | 0.0022 | 0.0026 |
| F12 | 34060001 | 34160000 | 0.0024 | 0.0015 | 0.0018 |
| F12 | 34080001 | 34180000 | 0.0034 | 0.0023 | 0.0026 |
| F12 | 34100001 | 34200000 | 0.0029 | 0.0019 | 0.0023 |
| F12 | 34120001 | 34220000 | 0.0026 | 0.0016 | 0.0019 |

| Chr | Start | End | both | A1 | A2 |
|---|---|---|---|---|---|
| F12 | 34140001 | 34240000 | 0.0032 | 0.0021 | 0.0024 |
| F12 | 34160001 | 34260000 | 0.0042 | 0.0031 | 0.0034 |
| F12 | 34180001 | 34280000 | 0.0038 | 0.0028 | 0.0031 |
| F12 | 34200001 | 34300000 | 0.0048 | 0.0035 | 0.0037 |
| F12 | 34220001 | 34320000 | 0.0050 | 0.0038 | 0.0040 |
| F12 | 34240001 | 34340000 | 0.0051 | 0.0039 | 0.0041 |
| F12 | 34260001 | 34360000 | 0.0046 | 0.0035 | 0.0038 |
| F12 | 34280001 | 34380000 | 0.0041 | 0.0031 | 0.0033 |
| F12 | 34300001 | 34400000 | 0.0035 | 0.0027 | 0.0030 |
| F12 | 34320001 | 34420000 | 0.0033 | 0.0026 | 0.0029 |
| F12 | 34340001 | 34440000 | 0.0024 | 0.0020 | 0.0022 |
| F12 | 34360001 | 34460000 | 0.0022 | 0.0017 | 0.0019 |
| F12 | 34380001 | 34480000 | 0.0034 | 0.0026 | 0.0028 |
| F12 | 34400001 | 34500000 | 0.0040 | 0.0032 | 0.0035 |
| F12 | 34420001 | 34520000 | 0.0043 | 0.0033 | 0.0036 |
| F12 | 34440001 | 34540000 | 0.0050 | 0.0038 | 0.0040 |
| F12 | 34460001 | 34560000 | 0.0048 | 0.0036 | 0.0037 |
| F12 | 34480001 | 34580000 | 0.0045 | 0.0034 | 0.0033 |
| F12 | 34500001 | 34600000 | 0.0052 | 0.0039 | 0.0040 |
| F12 | 34520001 | 34620000 | 0.0052 | 0.0039 | 0.0040 |
| F12 | 34540001 | 34640000 | 0.0046 | 0.0034 | 0.0035 |
| F12 | 34560001 | 34660000 | 0.0043 | 0.0032 | 0.0035 |
| F12 | 34580001 | 34680000 | 0.0043 | 0.0033 | 0.0037 |
| F12 | 34600001 | 34700000 | 0.0028 | 0.0021 | 0.0024 |
| F12 | 34620001 | 34720000 | 0.0025 | 0.0019 | 0.0021 |
| F12 | 34640001 | 34740000 | 0.0029 | 0.0023 | 0.0026 |
| F12 | 34660001 | 34760000 | 0.0028 | 0.0023 | 0.0026 |
| F12 | 34680001 | 34780000 | 0.0018 | 0.0014 | 0.0016 |
| F12 | 34700001 | 34800000 | 0.0024 | 0.0020 | 0.0022 |
| F12 | 34720001 | 34820000 | 0.0024 | 0.0020 | 0.0021 |
| F12 | 34740001 | 34840000 | 0.0027 | 0.0021 | 0.0023 |
| F12 | 34760001 | 34860000 | 0.0026 | 0.0020 | 0.0021 |
| F12 | 34780001 | 34880000 | 0.0038 | 0.0029 | 0.0031 |
| F12 | 34800001 | 34900000 | 0.0029 | 0.0021 | 0.0023 |
| F12 | 34820001 | 34920000 | 0.0034 | 0.0025 | 0.0027 |
| F12 | 34840001 | 34940000 | 0.0028 | 0.0022 | 0.0024 |
| F12 | 34860001 | 34960000 | 0.0026 | 0.0021 | 0.0023 |
| F12 | 34880001 | 34980000 | 0.0017 | 0.0013 | 0.0015 |
| F12 | 34900001 | 35000000 | 0.0029 | 0.0025 | 0.0025 |
| F12 | 34920001 | 35020000 | 0.0025 | 0.0020 | 0.0021 |
| F12 | 34940001 | 35040000 | 0.0026 | 0.0019 | 0.0019 |
| F12 | 34960001 | 35060000 | 0.0036 | 0.0024 | 0.0027 |
| F12 | 34980001 | 35080000 | 0.0036 | 0.0023 | 0.0025 |
| F12 | 35000001 | 35100000 | 0.0034 | 0.0020 | 0.0024 |
| F12 | 35020001 | 35120000 | 0.0039 | 0.0024 | 0.0029 |
| F12 | 35040001 | 35140000 | 0.0045 | 0.0033 | 0.0037 |
| F12 | 35060001 | 35160000 | 0.0040 | 0.0031 | 0.0034 |
| F12 | 35080001 | 35180000 | 0.0042 | 0.0034 | 0.0037 |
| F12 | 35100001 | 35200000 | 0.0033 | 0.0027 | 0.0030 |
| F12 | 35120001 | 35220000 | 0.0025 | 0.0022 | 0.0023 |
| F12 | 35140001 | 35240000 | 0.0017 | 0.0015 | 0.0016 |
| F12 | 35160001 | 35260000 | 0.0020 | 0.0019 | 0.0020 |
| F12 | 35180001 | 35280000 | 0.0023 | 0.0020 | 0.0023 |
| F12 | 35200001 | 35300000 | 0.0027 | 0.0025 | 0.0027 |
| F12 | 35220001 | 35320000 | 0.0029 | 0.0026 | 0.0029 |
| F12 | 35240001 | 35340000 | 0.0033 | 0.0028 | 0.0030 |
| F12 | 35260001 | 35360000 | 0.0044 | 0.0037 | 0.0040 |
| F12 | 35280001 | 35380000 | 0.0041 | 0.0035 | 0.0036 |

| Chr | Start | End | both | A1 | A2 |
|---|---|---|---|---|---|
| F12 | 35300001 | 35400000 | 0.0038 | 0.0032 | 0.0033 |
| F12 | 35320001 | 35420000 | 0.0049 | 0.0041 | 0.0043 |
| F12 | 35340001 | 35440000 | 0.0060 | 0.0052 | 0.0055 |
| F12 | 35360001 | 35460000 | 0.0049 | 0.0042 | 0.0043 |
| F12 | 35380001 | 35480000 | 0.0048 | 0.0039 | 0.0041 |
| F12 | 35400001 | 35500000 | 0.0052 | 0.0040 | 0.0042 |
| F12 | 35420001 | 35520000 | 0.0048 | 0.0035 | 0.0038 |
| F12 | 35440001 | 35540000 | 0.0039 | 0.0026 | 0.0029 |
| F12 | 35460001 | 35560000 | 0.0036 | 0.0024 | 0.0027 |
| F12 | 35480001 | 35580000 | 0.0035 | 0.0022 | 0.0026 |
| F12 | 35500001 | 35600000 | 0.0041 | 0.0029 | 0.0031 |
| F12 | 35520001 | 35620000 | 0.0042 | 0.0030 | 0.0031 |
| F12 | 35540001 | 35640000 | 0.0038 | 0.0024 | 0.0026 |
| F12 | 35560001 | 35660000 | 0.0037 | 0.0023 | 0.0025 |
| F12 | 35580001 | 35680000 | 0.0039 | 0.0027 | 0.0029 |
| F12 | 35600001 | 35700000 | 0.0036 | 0.0021 | 0.0026 |
| F12 | 35620001 | 35720000 | 0.0033 | 0.0017 | 0.0022 |
| F12 | 35640001 | 35740000 | 0.0036 | 0.0019 | 0.0024 |
| F12 | 35660001 | 35760000 | 0.0031 | 0.0016 | 0.0021 |
| F12 | 35680001 | 35780000 | 0.0035 | 0.0018 | 0.0023 |
| F12 | 35700001 | 35800000 | 0.0032 | 0.0018 | 0.0022 |
| F12 | 35720001 | 35820000 | 0.0033 | 0.0022 | 0.0025 |
| F12 | 35740001 | 35840000 | 0.0038 | 0.0026 | 0.0029 |
| F12 | 35760001 | 35860000 | 0.0043 | 0.0029 | 0.0032 |
| F12 | 35780001 | 35880000 | 0.0059 | 0.0046 | 0.0049 |
| F12 | 35800001 | 35900000 | 0.0067 | 0.0051 | 0.0054 |
| F12 | 35820001 | 35920000 | 0.0065 | 0.0049 | 0.0052 |
| F12 | 35840001 | 35940000 | 0.0064 | 0.0050 | 0.0052 |
| F12 | 35860001 | 35960000 | 0.0062 | 0.0048 | 0.0051 |
| F12 | 35880001 | 35980000 | 0.0038 | 0.0028 | 0.0030 |
| F12 | 35900001 | 36000000 | 0.0027 | 0.0020 | 0.0021 |
| F12 | 35920001 | 36020000 | 0.0031 | 0.0025 | 0.0027 |
| F12 | 35940001 | 36040000 | 0.0023 | 0.0018 | 0.0021 |
| F12 | 35960001 | 36060000 | 0.0027 | 0.0023 | 0.0026 |
| F12 | 35980001 | 36080000 | 0.0025 | 0.0021 | 0.0024 |
| F12 | 36000001 | 36100000 | 0.0029 | 0.0024 | 0.0026 |
| F12 | 36020001 | 36120000 | 0.0028 | 0.0023 | 0.0023 |
| F12 | 36040001 | 36140000 | 0.0042 | 0.0035 | 0.0034 |
| F12 | 36060001 | 36160000 | 0.0045 | 0.0036 | 0.0036 |
| F12 | 36080001 | 36180000 | 0.0049 | 0.0036 | 0.0036 |
| F12 | 36100001 | 36200000 | 0.0050 | 0.0036 | 0.0038 |
| F12 | 36120001 | 36220000 | 0.0049 | 0.0036 | 0.0038 |
| F12 | 36140001 | 36240000 | 0.0039 | 0.0028 | 0.0029 |
| F12 | 36160001 | 36260000 | 0.0041 | 0.0030 | 0.0031 |
| F12 | 36180001 | 36280000 | 0.0043 | 0.0034 | 0.0035 |
| F12 | 36200001 | 36300000 | 0.0035 | 0.0028 | 0.0029 |
| F12 | 36220001 | 36320000 | 0.0032 | 0.0024 | 0.0025 |
| F12 | 36240001 | 36340000 | 0.0034 | 0.0027 | 0.0028 |
| F12 | 36260001 | 36360000 | 0.0024 | 0.0019 | 0.0020 |
| F12 | 36280001 | 36380000 | 0.0018 | 0.0014 | 0.0015 |
| F12 | 36300001 | 36400000 | 0.0020 | 0.0016 | 0.0017 |
| F12 | 36320001 | 36420000 | 0.0019 | 0.0017 | 0.0017 |
| F12 | 36340001 | 36440000 | 0.0011 | 0.0010 | 0.0011 |
| F12 | 36360001 | 36460000 | 0.0014 | 0.0012 | 0.0013 |
| F12 | 36380001 | 36480000 | 0.0018 | 0.0015 | 0.0017 |
| F12 | 36400001 | 36500000 | 0.0021 | 0.0017 | 0.0019 |
| F12 | 36420001 | 36520000 | 0.0021 | 0.0016 | 0.0018 |
| F12 | 36440001 | 36540000 | 0.0022 | 0.0017 | 0.0018 |

| Chr | Start | End | both | A1 | A2 |
|---|---|---|---|---|---|
| F12 | 36460001 | 36560000 | 0.0019 | 0.0014 | 0.0016 |
| F12 | 36480001 | 36580000 | 0.0027 | 0.0023 | 0.0025 |
| F12 | 36500001 | 36600000 | 0.0024 | 0.0021 | 0.0022 |
| F12 | 36520001 | 36620000 | 0.0027 | 0.0021 | 0.0024 |
| F12 | 36540001 | 36640000 | 0.0034 | 0.0028 | 0.0032 |
| F12 | 36560001 | 36660000 | 0.0047 | 0.0038 | 0.0041 |
| F12 | 36580001 | 36680000 | 0.0036 | 0.0028 | 0.0031 |
| F12 | 36600001 | 36700000 | 0.0034 | 0.0027 | 0.0029 |
| F12 | 36620001 | 36720000 | 0.0040 | 0.0032 | 0.0036 |
| F12 | 36640001 | 36740000 | 0.0037 | 0.0027 | 0.0032 |
| F12 | 36660001 | 36760000 | 0.0027 | 0.0020 | 0.0026 |
| F12 | 36680001 | 36780000 | 0.0027 | 0.0021 | 0.0025 |
| F12 | 36700001 | 36800000 | 0.0034 | 0.0027 | 0.0031 |
| F12 | 36720001 | 36820000 | 0.0031 | 0.0027 | 0.0028 |
| F12 | 36740001 | 36840000 | 0.0029 | 0.0027 | 0.0026 |
| F12 | 36760001 | 36860000 | 0.0025 | 0.0022 | 0.0022 |
| F12 | 36780001 | 36880000 | 0.0029 | 0.0023 | 0.0024 |
| F12 | 36800001 | 36900000 | 0.0026 | 0.0019 | 0.0020 |
| F12 | 36820001 | 36920000 | 0.0022 | 0.0013 | 0.0017 |
| F12 | 36840001 | 36940000 | 0.0021 | 0.0010 | 0.0014 |
| F12 | 36860001 | 36960000 | 0.0023 | 0.0012 | 0.0016 |
| F12 | 36880001 | 36980000 | 0.0030 | 0.0018 | 0.0024 |
| F12 | 36900001 | 37000000 | 0.0040 | 0.0027 | 0.0034 |
| F12 | 36920001 | 37020000 | 0.0043 | 0.0031 | 0.0037 |
| F12 | 36940001 | 37040000 | 0.0045 | 0.0034 | 0.0040 |
| F12 | 36960001 | 37060000 | 0.0048 | 0.0037 | 0.0043 |
| F12 | 36980001 | 37080000 | 0.0039 | 0.0031 | 0.0036 |
| F12 | 37000001 | 37100000 | 0.0027 | 0.0022 | 0.0025 |
| F12 | 37020001 | 37120000 | 0.0024 | 0.0019 | 0.0021 |
| F12 | 37040001 | 37140000 | 0.0028 | 0.0019 | 0.0022 |
| F12 | 37060001 | 37160000 | 0.0025 | 0.0017 | 0.0019 |
| F12 | 37080001 | 37180000 | 0.0030 | 0.0018 | 0.0021 |
| F12 | 37100001 | 37200000 | 0.0033 | 0.0019 | 0.0023 |
| F12 | 37120001 | 37220000 | 0.0030 | 0.0017 | 0.0021 |
| F12 | 37140001 | 37240000 | 0.0025 | 0.0015 | 0.0020 |
| F12 | 37160001 | 37260000 | 0.0026 | 0.0017 | 0.0021 |
| F12 | 37180001 | 37280000 | 0.0023 | 0.0017 | 0.0021 |
| F12 | 37200001 | 37300000 | 0.0021 | 0.0015 | 0.0017 |
| F12 | 37220001 | 37320000 | 0.0032 | 0.0025 | 0.0028 |
| F12 | 37240001 | 37340000 | 0.0038 | 0.0028 | 0.0030 |
| F12 | 37260001 | 37360000 | 0.0038 | 0.0027 | 0.0029 |
| F12 | 37280001 | 37380000 | 0.0039 | 0.0027 | 0.0030 |
| F12 | 37300001 | 37400000 | 0.0052 | 0.0038 | 0.0043 |
| F12 | 37320001 | 37420000 | 0.0039 | 0.0028 | 0.0032 |
| F12 | 37340001 | 37440000 | 0.0029 | 0.0022 | 0.0027 |
| F12 | 37360001 | 37460000 | 0.0029 | 0.0022 | 0.0028 |
| F12 | 37380001 | 37480000 | 0.0028 | 0.0021 | 0.0027 |
| F12 | 37400001 | 37500000 | 0.0015 | 0.0010 | 0.0015 |
| F12 | 37420001 | 37520000 | 0.0022 | 0.0015 | 0.0022 |
| F12 | 37440001 | 37540000 | 0.0023 | 0.0014 | 0.0020 |
| F12 | 37460001 | 37560000 | 0.0021 | 0.0013 | 0.0017 |
| F12 | 37480001 | 37580000 | 0.0016 | 0.0009 | 0.0012 |
| F12 | 37500001 | 37600000 | 0.0018 | 0.0010 | 0.0014 |
| F12 | 37520001 | 37620000 | 0.0012 | 0.0006 | 0.0008 |
| F12 | 37540001 | 37640000 | 0.0009 | 0.0006 | 0.0008 |
| F12 | 37560001 | 37660000 | 0.0011 | 0.0009 | 0.0011 |
| F12 | 37580001 | 37680000 | 0.0010 | 0.0009 | 0.0010 |
| F12 | 37600001 | 37700000 | 0.0008 | 0.0007 | 0.0008 |

| Chr | Start | End | both | A1 | A2 |
|---|---|---|---|---|---|
| F12 | 37620001 | 37720000 | 0.0008 | 0.0006 | 0.0007 |
| F12 | 37640001 | 37740000 | 0.0014 | 0.0012 | 0.0014 |
| F12 | 37660001 | 37760000 | 0.0012 | 0.0010 | 0.0012 |
| F12 | 37680001 | 37780000 | 0.0013 | 0.0011 | 0.0013 |
| F12 | 37700001 | 37800000 | 0.0012 | 0.0010 | 0.0012 |
| F12 | 37720001 | 37820000 | 0.0012 | 0.0010 | 0.0012 |
| F12 | 37740001 | 37840000 | 0.0007 | 0.0006 | 0.0007 |
| F12 | 37760001 | 37860000 | 0.0014 | 0.0012 | 0.0014 |
| F12 | 37780001 | 37880000 | 0.0014 | 0.0012 | 0.0014 |
| F12 | 37800001 | 37900000 | 0.0021 | 0.0018 | 0.0021 |
| F12 | 37820001 | 37920000 | 0.0028 | 0.0024 | 0.0027 |
| F12 | 37840001 | 37940000 | 0.0030 | 0.0025 | 0.0028 |
| F12 | 37860001 | 37960000 | 0.0030 | 0.0026 | 0.0028 |
| F12 | 37880001 | 37980000 | 0.0028 | 0.0024 | 0.0027 |
| F12 | 37900001 | 38000000 | 0.0023 | 0.0019 | 0.0022 |
| F12 | 37920001 | 38020000 | 0.0022 | 0.0018 | 0.0021 |
| F12 | 37940001 | 38040000 | 0.0020 | 0.0016 | 0.0019 |
| F12 | 37960001 | 38060000 | 0.0023 | 0.0019 | 0.0021 |
| F12 | 37980001 | 38080000 | 0.0027 | 0.0020 | 0.0022 |
| F12 | 38000001 | 38100000 | 0.0028 | 0.0019 | 0.0022 |
| F12 | 38020001 | 38120000 | 0.0033 | 0.0023 | 0.0025 |
| F12 | 38040001 | 38140000 | 0.0033 | 0.0025 | 0.0025 |
| F12 | 38060001 | 38160000 | 0.0027 | 0.0018 | 0.0019 |
| F12 | 38080001 | 38180000 | 0.0029 | 0.0022 | 0.0021 |
| F12 | 38100001 | 38200000 | 0.0031 | 0.0023 | 0.0023 |
| F12 | 38120001 | 38220000 | 0.0025 | 0.0017 | 0.0018 |
| F12 | 38140001 | 38240000 | 0.0031 | 0.0020 | 0.0024 |
| F12 | 38160001 | 38260000 | 0.0033 | 0.0023 | 0.0027 |
| F12 | 38180001 | 38280000 | 0.0032 | 0.0022 | 0.0026 |
| F12 | 38200001 | 38300000 | 0.0030 | 0.0022 | 0.0026 |
| F12 | 38220001 | 38320000 | 0.0034 | 0.0028 | 0.0031 |
| F12 | 38240001 | 38340000 | 0.0033 | 0.0028 | 0.0031 |
| F12 | 38260001 | 38360000 | 0.0030 | 0.0026 | 0.0029 |
| F12 | 38280001 | 38380000 | 0.0038 | 0.0032 | 0.0036 |
| F12 | 38300001 | 38400000 | 0.0040 | 0.0033 | 0.0038 |
| F12 | 38320001 | 38420000 | 0.0032 | 0.0025 | 0.0029 |
| F12 | 38340001 | 38440000 | 0.0031 | 0.0023 | 0.0028 |
| F12 | 38360001 | 38460000 | 0.0032 | 0.0023 | 0.0027 |
| F12 | 38380001 | 38480000 | 0.0021 | 0.0014 | 0.0017 |
| F12 | 38400001 | 38500000 | 0.0018 | 0.0012 | 0.0015 |
| F12 | 38420001 | 38520000 | 0.0018 | 0.0013 | 0.0015 |
| F12 | 38440001 | 38540000 | 0.0016 | 0.0013 | 0.0015 |
| F12 | 38460001 | 38560000 | 0.0018 | 0.0014 | 0.0017 |
| F12 | 38480001 | 38580000 | 0.0024 | 0.0020 | 0.0023 |
| F12 | 38500001 | 38600000 | 0.0029 | 0.0025 | 0.0027 |
| F12 | 38520001 | 38620000 | 0.0031 | 0.0027 | 0.0028 |
| F12 | 38540001 | 38640000 | 0.0030 | 0.0024 | 0.0026 |
| F12 | 38560001 | 38660000 | 0.0030 | 0.0025 | 0.0026 |
| F12 | 38580001 | 38680000 | 0.0029 | 0.0024 | 0.0025 |
| F12 | 38600001 | 38700000 | 0.0026 | 0.0020 | 0.0022 |
| F12 | 38620001 | 38720000 | 0.0033 | 0.0023 | 0.0026 |
| F12 | 38640001 | 38740000 | 0.0037 | 0.0028 | 0.0032 |
| F12 | 38660001 | 38760000 | 0.0038 | 0.0028 | 0.0032 |
| F12 | 38680001 | 38780000 | 0.0035 | 0.0023 | 0.0027 |
| F12 | 38700001 | 38800000 | 0.0034 | 0.0023 | 0.0026 |
| F12 | 38720001 | 38820000 | 0.0027 | 0.0020 | 0.0022 |
| F12 | 38740001 | 38840000 | 0.0025 | 0.0019 | 0.0020 |
| F12 | 38760001 | 38860000 | 0.0019 | 0.0015 | 0.0017 |

| Chr | Start | End | both | A1 | A2 |
|---|---|---|---|---|---|
| F12 | 38780001 | 38880000 | 0.0019 | 0.0017 | 0.0018 |
| F12 | 38800001 | 38900000 | 0.0018 | 0.0016 | 0.0018 |
| F12 | 38820001 | 38920000 | 0.0018 | 0.0016 | 0.0018 |
| F12 | 38840001 | 38940000 | 0.0021 | 0.0018 | 0.0021 |
| F12 | 38860001 | 38960000 | 0.0019 | 0.0017 | 0.0019 |
| F12 | 38880001 | 38980000 | 0.0017 | 0.0015 | 0.0017 |
| F12 | 38900001 | 39000000 | 0.0020 | 0.0016 | 0.0020 |
| F12 | 38920001 | 39020000 | 0.0022 | 0.0017 | 0.0022 |
| F12 | 38940001 | 39040000 | 0.0015 | 0.0011 | 0.0015 |
| F12 | 38960001 | 39060000 | 0.0017 | 0.0013 | 0.0017 |
| F12 | 38980001 | 39080000 | 0.0024 | 0.0020 | 0.0024 |
| F12 | 39000001 | 39100000 | 0.0025 | 0.0021 | 0.0024 |
| F12 | 39020001 | 39120000 | 0.0024 | 0.0021 | 0.0024 |
| F12 | 39040001 | 39140000 | 0.0024 | 0.0021 | 0.0023 |
| F12 | 39060001 | 39160000 | 0.0031 | 0.0023 | 0.0029 |
| F12 | 39080001 | 39180000 | 0.0028 | 0.0020 | 0.0025 |
| F12 | 39100001 | 39200000 | 0.0023 | 0.0015 | 0.0019 |
| F12 | 39120001 | 39220000 | 0.0024 | 0.0015 | 0.0019 |
| F12 | 39140001 | 39240000 | 0.0027 | 0.0017 | 0.0021 |
| F12 | 39160001 | 39260000 | 0.0032 | 0.0025 | 0.0027 |
| F12 | 39180001 | 39280000 | 0.0036 | 0.0029 | 0.0032 |
| F12 | 39200001 | 39300000 | 0.0045 | 0.0038 | 0.0042 |
| F12 | 39220001 | 39320000 | 0.0055 | 0.0048 | 0.0052 |
| F12 | 39240001 | 39340000 | 0.0054 | 0.0048 | 0.0052 |
| F12 | 39260001 | 39360000 | 0.0042 | 0.0038 | 0.0041 |
| F12 | 39280001 | 39380000 | 0.0036 | 0.0033 | 0.0034 |
| F12 | 39300001 | 39400000 | 0.0026 | 0.0024 | 0.0025 |
| F12 | 39320001 | 39420000 | 0.0014 | 0.0013 | 0.0014 |
| F12 | 39340001 | 39440000 | 0.0014 | 0.0012 | 0.0014 |
| F12 | 39360001 | 39460000 | 0.0017 | 0.0014 | 0.0017 |
| F12 | 39380001 | 39480000 | 0.0021 | 0.0017 | 0.0021 |
| F12 | 39400001 | 39500000 | 0.0033 | 0.0027 | 0.0031 |
| F12 | 39420001 | 39520000 | 0.0038 | 0.0031 | 0.0035 |
| F12 | 39440001 | 39540000 | 0.0042 | 0.0034 | 0.0036 |
| F12 | 39460001 | 39560000 | 0.0047 | 0.0037 | 0.0038 |
| F12 | 39480001 | 39580000 | 0.0042 | 0.0032 | 0.0033 |
| F12 | 39500001 | 39600000 | 0.0034 | 0.0023 | 0.0024 |
| F12 | 39520001 | 39620000 | 0.0033 | 0.0019 | 0.0020 |
| F12 | 39540001 | 39640000 | 0.0031 | 0.0016 | 0.0017 |
| F12 | 39560001 | 39660000 | 0.0023 | 0.0010 | 0.0011 |
| F12 | 39580001 | 39680000 | 0.0029 | 0.0014 | 0.0015 |
| F12 | 39600001 | 39700000 | 0.0032 | 0.0014 | 0.0014 |
| F12 | 39620001 | 39720000 | 0.0032 | 0.0017 | 0.0018 |
| F12 | 39640001 | 39740000 | 0.0035 | 0.0021 | 0.0023 |
| F12 | 39660001 | 39760000 | 0.0039 | 0.0024 | 0.0026 |
| F12 | 39680001 | 39780000 | 0.0045 | 0.0027 | 0.0029 |
| F12 | 39700001 | 39800000 | 0.0050 | 0.0032 | 0.0035 |
| F12 | 39720001 | 39820000 | 0.0048 | 0.0030 | 0.0033 |
| F12 | 39740001 | 39840000 | 0.0042 | 0.0023 | 0.0025 |
| F12 | 39760001 | 39860000 | 0.0040 | 0.0024 | 0.0026 |
| F12 | 39780001 | 39880000 | 0.0035 | 0.0023 | 0.0025 |
| F12 | 39800001 | 39900000 | 0.0027 | 0.0018 | 0.0019 |
| F12 | 39820001 | 39920000 | 0.0028 | 0.0021 | 0.0022 |
| F12 | 39840001 | 39940000 | 0.0031 | 0.0024 | 0.0025 |
| F12 | 39860001 | 39960000 | 0.0032 | 0.0024 | 0.0025 |
| F12 | 39880001 | 39980000 | 0.0031 | 0.0022 | 0.0023 |
| F12 | 39900001 | 40000000 | 0.0036 | 0.0028 | 0.0029 |
| F12 | 39920001 | 40020000 | 0.0034 | 0.0027 | 0.0029 |

| Chr | Start | End | both | A1 | A2 |
|---|---|---|---|---|---|
| F12 | 39940001 | 40040000 | 0.0032 | 0.0026 | 0.0029 |
| F12 | 39960001 | 40060000 | 0.0035 | 0.0031 | 0.0033 |
| F12 | 39980001 | 40080000 | 0.0032 | 0.0028 | 0.0030 |
| F12 | 40000001 | 40100000 | 0.0030 | 0.0026 | 0.0026 |
| F12 | 40020001 | 40120000 | 0.0032 | 0.0026 | 0.0025 |
| F12 | 40040001 | 40140000 | 0.0043 | 0.0036 | 0.0034 |
| F12 | 40060001 | 40160000 | 0.0036 | 0.0028 | 0.0026 |
| F12 | 40080001 | 40180000 | 0.0037 | 0.0030 | 0.0027 |
| F12 | 40100001 | 40200000 | 0.0041 | 0.0033 | 0.0034 |
| F12 | 40120001 | 40220000 | 0.0036 | 0.0028 | 0.0030 |
| F12 | 40140001 | 40240000 | 0.0028 | 0.0020 | 0.0022 |
| F12 | 40160001 | 40260000 | 0.0042 | 0.0031 | 0.0035 |
| F12 | 40180001 | 40280000 | 0.0037 | 0.0028 | 0.0031 |
| F12 | 40200001 | 40300000 | 0.0033 | 0.0022 | 0.0026 |
| F12 | 40220001 | 40320000 | 0.0044 | 0.0035 | 0.0037 |
| F12 | 40240001 | 40340000 | 0.0043 | 0.0035 | 0.0038 |
| F12 | 40260001 | 40360000 | 0.0035 | 0.0026 | 0.0028 |
| F12 | 40280001 | 40380000 | 0.0044 | 0.0032 | 0.0036 |
| F12 | 40300001 | 40400000 | 0.0044 | 0.0035 | 0.0036 |
| F12 | 40320001 | 40420000 | 0.0036 | 0.0026 | 0.0028 |
| F12 | 40340001 | 40440000 | 0.0038 | 0.0027 | 0.0030 |
| F12 | 40360001 | 40460000 | 0.0038 | 0.0031 | 0.0033 |
| F12 | 40380001 | 40480000 | 0.0030 | 0.0025 | 0.0027 |
| F12 | 40400001 | 40500000 | 0.0023 | 0.0018 | 0.0021 |
| F12 | 40420001 | 40520000 | 0.0018 | 0.0015 | 0.0017 |
| F12 | 40440001 | 40540000 | 0.0015 | 0.0013 | 0.0014 |
| F12 | 40460001 | 40560000 | 0.0015 | 0.0012 | 0.0013 |
| F12 | 40480001 | 40580000 | 0.0017 | 0.0014 | 0.0015 |
| F12 | 40500001 | 40600000 | 0.0027 | 0.0022 | 0.0023 |
| F12 | 40520001 | 40620000 | 0.0035 | 0.0028 | 0.0032 |
| F12 | 40540001 | 40640000 | 0.0031 | 0.0024 | 0.0028 |
| F12 | 40560001 | 40660000 | 0.0030 | 0.0023 | 0.0027 |
| F12 | 40580001 | 40680000 | 0.0029 | 0.0023 | 0.0025 |
| F12 | 40600001 | 40700000 | 0.0031 | 0.0024 | 0.0028 |
| F12 | 40620001 | 40720000 | 0.0030 | 0.0023 | 0.0026 |
| F12 | 40640001 | 40740000 | 0.0038 | 0.0029 | 0.0033 |
| F12 | 40660001 | 40760000 | 0.0037 | 0.0027 | 0.0032 |
| F12 | 40680001 | 40780000 | 0.0042 | 0.0033 | 0.0038 |
| F12 | 40700001 | 40800000 | 0.0042 | 0.0033 | 0.0037 |
| F12 | 40720001 | 40820000 | 0.0039 | 0.0032 | 0.0035 |
| F12 | 40740001 | 40840000 | 0.0033 | 0.0028 | 0.0031 |
| F12 | 40760001 | 40860000 | 0.0034 | 0.0029 | 0.0032 |
| F12 | 40780001 | 40880000 | 0.0026 | 0.0021 | 0.0025 |
| F12 | 40800001 | 40900000 | 0.0017 | 0.0014 | 0.0016 |
| F12 | 40820001 | 40920000 | 0.0013 | 0.0009 | 0.0011 |
| F12 | 40840001 | 40940000 | 0.0013 | 0.0010 | 0.0011 |
| F12 | 40860001 | 40960000 | 0.0013 | 0.0010 | 0.0011 |
| F12 | 40880001 | 40980000 | 0.0013 | 0.0009 | 0.0011 |
| F12 | 40900001 | 41000000 | 0.0013 | 0.0008 | 0.0012 |
| F12 | 40920001 | 41020000 | 0.0021 | 0.0014 | 0.0020 |
| F12 | 40940001 | 41040000 | 0.0020 | 0.0013 | 0.0019 |
| F12 | 40960001 | 41060000 | 0.0015 | 0.0009 | 0.0015 |
| F12 | 40980001 | 41080000 | 0.0016 | 0.0010 | 0.0015 |
| F12 | 41000001 | 41100000 | 0.0020 | 0.0015 | 0.0020 |
| F12 | 41020001 | 41120000 | 0.0019 | 0.0016 | 0.0017 |
| F12 | 41040001 | 41140000 | 0.0021 | 0.0018 | 0.0019 |
| F12 | 41060001 | 41160000 | 0.0036 | 0.0031 | 0.0034 |
| F12 | 41080001 | 41180000 | 0.0056 | 0.0050 | 0.0052 |

| Chr | Start | End | both | A1 | A2 |
|---|---|---|---|---|---|
| F12 | 41100001 | 41200000 | 0.0050 | 0.0046 | 0.0047 |
| F12 | 41120001 | 41220000 | 0.0057 | 0.0051 | 0.0054 |
| F12 | 41140001 | 41240000 | 0.0056 | 0.0050 | 0.0054 |
| F12 | 41160001 | 41260000 | 0.0042 | 0.0038 | 0.0040 |
| F12 | 41180001 | 41280000 | 0.0022 | 0.0018 | 0.0021 |
| F12 | 41200001 | 41300000 | 0.0022 | 0.0018 | 0.0021 |
| F12 | 41220001 | 41320000 | 0.0013 | 0.0010 | 0.0013 |
| F12 | 41240001 | 41340000 | 0.0011 | 0.0008 | 0.0010 |
| F12 | 41260001 | 41360000 | 0.0016 | 0.0012 | 0.0015 |
| F12 | 41280001 | 41380000 | 0.0021 | 0.0016 | 0.0020 |
| F12 | 41300001 | 41400000 | 0.0030 | 0.0026 | 0.0029 |
| F12 | 41320001 | 41420000 | 0.0033 | 0.0027 | 0.0029 |
| F12 | 41340001 | 41440000 | 0.0039 | 0.0030 | 0.0034 |
| F12 | 41360001 | 41460000 | 0.0045 | 0.0036 | 0.0040 |
| F12 | 41380001 | 41480000 | 0.0046 | 0.0037 | 0.0041 |
| F12 | 41400001 | 41500000 | 0.0041 | 0.0031 | 0.0036 |
| F12 | 41420001 | 41520000 | 0.0033 | 0.0026 | 0.0031 |
| F12 | 41440001 | 41540000 | 0.0027 | 0.0022 | 0.0026 |
| F12 | 41460001 | 41560000 | 0.0018 | 0.0013 | 0.0017 |
| F12 | 41480001 | 41580000 | 0.0016 | 0.0013 | 0.0016 |
| F12 | 41500001 | 41600000 | 0.0012 | 0.0011 | 0.0012 |
| F12 | 41520001 | 41620000 | 0.0014 | 0.0012 | 0.0014 |
| F12 | 41540001 | 41640000 | 0.0016 | 0.0015 | 0.0016 |
| F12 | 41560001 | 41660000 | 0.0014 | 0.0013 | 0.0014 |
| F12 | 41580001 | 41680000 | 0.0009 | 0.0007 | 0.0009 |
| F12 | 41600001 | 41700000 | 0.0010 | 0.0008 | 0.0009 |
| F12 | 41620001 | 41720000 | 0.0015 | 0.0014 | 0.0015 |
| F12 | 41640001 | 41740000 | 0.0017 | 0.0016 | 0.0017 |
| F12 | 41660001 | 41760000 | 0.0019 | 0.0017 | 0.0018 |
| F12 | 41680001 | 41780000 | 0.0023 | 0.0020 | 0.0022 |
| F12 | 41700001 | 41800000 | 0.0029 | 0.0025 | 0.0027 |
| F12 | 41720001 | 41820000 | 0.0021 | 0.0018 | 0.0020 |
| F12 | 41740001 | 41840000 | 0.0027 | 0.0023 | 0.0025 |
| F12 | 41760001 | 41860000 | 0.0044 | 0.0037 | 0.0040 |
| F12 | 41780001 | 41880000 | 0.0046 | 0.0038 | 0.0040 |
| F12 | 41800001 | 41900000 | 0.0047 | 0.0037 | 0.0038 |
| F12 | 41820001 | 41920000 | 0.0056 | 0.0044 | 0.0045 |
| F12 | 41840001 | 41940000 | 0.0057 | 0.0044 | 0.0046 |
| F12 | 41860001 | 41960000 | 0.0045 | 0.0035 | 0.0036 |
| F12 | 41880001 | 41980000 | 0.0044 | 0.0035 | 0.0037 |
| F12 | 41900001 | 42000000 | 0.0045 | 0.0036 | 0.0037 |
| F12 | 41920001 | 42020000 | 0.0051 | 0.0040 | 0.0042 |
| F12 | 41940001 | 42040000 | 0.0051 | 0.0039 | 0.0040 |
| F12 | 41960001 | 42060000 | 0.0052 | 0.0040 | 0.0041 |
| F12 | 41980001 | 42080000 | 0.0057 | 0.0045 | 0.0046 |
| F12 | 42000001 | 42100000 | 0.0056 | 0.0045 | 0.0048 |
| F12 | 42020001 | 42120000 | 0.0052 | 0.0043 | 0.0046 |
| F12 | 42040001 | 42140000 | 0.0051 | 0.0042 | 0.0045 |
| F12 | 42060001 | 42160000 | 0.0047 | 0.0039 | 0.0043 |
| F12 | 42080001 | 42180000 | 0.0042 | 0.0034 | 0.0038 |
| F12 | 42100001 | 42200000 | 0.0033 | 0.0028 | 0.0031 |
| F12 | 42120001 | 42220000 | 0.0030 | 0.0025 | 0.0028 |
| F12 | 42140001 | 42240000 | 0.0025 | 0.0021 | 0.0025 |
| F12 | 42160001 | 42260000 | 0.0030 | 0.0024 | 0.0029 |
| F12 | 42180001 | 42280000 | 0.0038 | 0.0032 | 0.0036 |
| F12 | 42200001 | 42300000 | 0.0039 | 0.0033 | 0.0037 |
| F12 | 42220001 | 42320000 | 0.0035 | 0.0029 | 0.0033 |
| F12 | 42240001 | 42340000 | 0.0039 | 0.0035 | 0.0037 |

| Chr | Start | End | both | A1 | A2 |
|---|---|---|---|---|---|
| F12 | 42260001 | 42360000 | 0.0038 | 0.0033 | 0.0037 |
| F12 | 42280001 | 42380000 | 0.0030 | 0.0025 | 0.0028 |
| F12 | 42300001 | 42400000 | 0.0034 | 0.0028 | 0.0032 |
| F12 | 42320001 | 42420000 | 0.0038 | 0.0030 | 0.0035 |
| F12 | 42340001 | 42440000 | 0.0036 | 0.0026 | 0.0030 |
| F12 | 42360001 | 42460000 | 0.0037 | 0.0026 | 0.0030 |
| F12 | 42380001 | 42480000 | 0.0035 | 0.0023 | 0.0027 |
| F12 | 42400001 | 42500000 | 0.0045 | 0.0031 | 0.0035 |
| F12 | 42420001 | 42520000 | 0.0056 | 0.0041 | 0.0047 |
| F12 | 42440001 | 42540000 | 0.0057 | 0.0043 | 0.0048 |
| F12 | 42460001 | 42560000 | 0.0059 | 0.0046 | 0.0051 |
| F12 | 42480001 | 42580000 | 0.0065 | 0.0053 | 0.0058 |
| F12 | 42500001 | 42600000 | 0.0055 | 0.0044 | 0.0048 |
| F12 | 42520001 | 42620000 | 0.0040 | 0.0032 | 0.0034 |
| F12 | 42540001 | 42640000 | 0.0038 | 0.0029 | 0.0033 |
| F12 | 42560001 | 42660000 | 0.0027 | 0.0019 | 0.0022 |
| F12 | 42580001 | 42680000 | 0.0020 | 0.0013 | 0.0016 |
| F12 | 42600001 | 42700000 | 0.0018 | 0.0013 | 0.0014 |
| F12 | 42620001 | 42720000 | 0.0019 | 0.0014 | 0.0016 |
| F12 | 42640001 | 42740000 | 0.0028 | 0.0023 | 0.0028 |
| F12 | 42660001 | 42760000 | 0.0045 | 0.0040 | 0.0044 |
| F12 | 42680001 | 42780000 | 0.0051 | 0.0045 | 0.0049 |
| F12 | 42700001 | 42800000 | 0.0053 | 0.0044 | 0.0049 |
| F12 | 42720001 | 42820000 | 0.0054 | 0.0045 | 0.0047 |
| F12 | 42740001 | 42840000 | 0.0043 | 0.0035 | 0.0036 |
| F12 | 42760001 | 42860000 | 0.0033 | 0.0023 | 0.0026 |
| F12 | 42780001 | 42880000 | 0.0030 | 0.0020 | 0.0022 |
| F12 | 42800001 | 42900000 | 0.0029 | 0.0020 | 0.0022 |
| F12 | 42820001 | 42920000 | 0.0030 | 0.0021 | 0.0025 |
| F12 | 42840001 | 42940000 | 0.0031 | 0.0021 | 0.0025 |
| F12 | 42860001 | 42960000 | 0.0028 | 0.0016 | 0.0021 |
| F12 | 42880001 | 42980000 | 0.0025 | 0.0014 | 0.0019 |
| F12 | 42900001 | 43000000 | 0.0031 | 0.0019 | 0.0024 |
| F12 | 42920001 | 43020000 | 0.0028 | 0.0017 | 0.0021 |
| F12 | 42940001 | 43040000 | 0.0025 | 0.0015 | 0.0019 |
| F12 | 42960001 | 43060000 | 0.0026 | 0.0021 | 0.0021 |
| F12 | 42980001 | 43080000 | 0.0029 | 0.0025 | 0.0024 |
| F12 | 43000001 | 43100000 | 0.0029 | 0.0025 | 0.0024 |
| F12 | 43020001 | 43120000 | 0.0033 | 0.0028 | 0.0027 |
| F12 | 43040001 | 43140000 | 0.0035 | 0.0029 | 0.0028 |
| F12 | 43060001 | 43160000 | 0.0030 | 0.0022 | 0.0024 |
| F12 | 43080001 | 43180000 | 0.0029 | 0.0020 | 0.0022 |
| F12 | 43100001 | 43200000 | 0.0030 | 0.0020 | 0.0024 |
| F12 | 43120001 | 43220000 | 0.0028 | 0.0020 | 0.0023 |
| F12 | 43140001 | 43240000 | 0.0032 | 0.0023 | 0.0026 |
| F12 | 43160001 | 43260000 | 0.0040 | 0.0029 | 0.0033 |
| F12 | 43180001 | 43280000 | 0.0039 | 0.0030 | 0.0034 |
| F12 | 43200001 | 43300000 | 0.0032 | 0.0026 | 0.0029 |
| F12 | 43220001 | 43320000 | 0.0029 | 0.0023 | 0.0026 |
| F12 | 43240001 | 43340000 | 0.0028 | 0.0024 | 0.0027 |
| F12 | 43260001 | 43360000 | 0.0024 | 0.0019 | 0.0022 |
| F12 | 43280001 | 43380000 | 0.0034 | 0.0024 | 0.0028 |
| F12 | 43300001 | 43400000 | 0.0040 | 0.0027 | 0.0032 |
| F12 | 43320001 | 43420000 | 0.0041 | 0.0028 | 0.0032 |
| F12 | 43340001 | 43440000 | 0.0039 | 0.0026 | 0.0028 |
| F12 | 43360001 | 43460000 | 0.0044 | 0.0030 | 0.0033 |
| F12 | 43380001 | 43480000 | 0.0036 | 0.0025 | 0.0028 |
| F12 | 43400001 | 43500000 | 0.0030 | 0.0023 | 0.0024 |

| Chr | Start | End | both | A1 | A2 |
|---|---|---|---|---|---|
| F12 | 43420001 | 43520000 | 0.0024 | 0.0018 | 0.0020 |
| F12 | 43440001 | 43540000 | 0.0022 | 0.0016 | 0.0019 |
| F12 | 43460001 | 43560000 | 0.0015 | 0.0012 | 0.0013 |
| F12 | 43480001 | 43580000 | 0.0013 | 0.0010 | 0.0012 |
| F12 | 43500001 | 43600000 | 0.0013 | 0.0010 | 0.0012 |
| F12 | 43520001 | 43620000 | 0.0017 | 0.0013 | 0.0016 |
| F12 | 43540001 | 43640000 | 0.0016 | 0.0013 | 0.0016 |
| F12 | 43560001 | 43660000 | 0.0015 | 0.0012 | 0.0015 |
| F12 | 43580001 | 43680000 | 0.0016 | 0.0013 | 0.0016 |
| F12 | 43600001 | 43700000 | 0.0011 | 0.0009 | 0.0011 |
| F12 | 43620001 | 43720000 | 0.0007 | 0.0006 | 0.0007 |
| F12 | 43640001 | 43740000 | 0.0016 | 0.0014 | 0.0016 |
| F12 | 43660001 | 43760000 | 0.0015 | 0.0014 | 0.0015 |
| F12 | 43680001 | 43780000 | 0.0013 | 0.0011 | 0.0012 |
| F12 | 43700001 | 43800000 | 0.0015 | 0.0013 | 0.0015 |
| F12 | 43720001 | 43820000 | 0.0015 | 0.0013 | 0.0015 |
| F12 | 43740001 | 43840000 | 0.0008 | 0.0006 | 0.0008 |
| F12 | 43760001 | 43860000 | 0.0008 | 0.0006 | 0.0008 |
| F12 | 43780001 | 43880000 | 0.0008 | 0.0006 | 0.0008 |
| F12 | 43800001 | 43900000 | 0.0006 | 0.0005 | 0.0006 |
| F12 | 43820001 | 43920000 | 0.0007 | 0.0005 | 0.0006 |
| F12 | 43840001 | 43940000 | 0.0006 | 0.0004 | 0.0004 |
| F12 | 43860001 | 43960000 | 0.0007 | 0.0005 | 0.0005 |
| F12 | 43880001 | 43980000 | 0.0013 | 0.0011 | 0.0011 |
| F12 | 43900001 | 44000000 | 0.0018 | 0.0015 | 0.0015 |
| F12 | 43920001 | 44020000 | 0.0017 | 0.0014 | 0.0015 |
| F12 | 43940001 | 44040000 | 0.0015 | 0.0013 | 0.0014 |
| F12 | 43960001 | 44060000 | 0.0015 | 0.0013 | 0.0015 |
| F12 | 43980001 | 44080000 | 0.0014 | 0.0012 | 0.0013 |
| F12 | 44000001 | 44100000 | 0.0014 | 0.0012 | 0.0014 |
| F12 | 44020001 | 44120000 | 0.0033 | 0.0027 | 0.0031 |
| F12 | 44040001 | 44140000 | 0.0038 | 0.0032 | 0.0036 |
| F12 | 44060001 | 44160000 | 0.0039 | 0.0032 | 0.0037 |
| F12 | 44080001 | 44180000 | 0.0047 | 0.0039 | 0.0044 |
| F12 | 44100001 | 44200000 | 0.0047 | 0.0040 | 0.0044 |
| F12 | 44120001 | 44220000 | 0.0044 | 0.0038 | 0.0042 |
| F12 | 44140001 | 44240000 | 0.0040 | 0.0034 | 0.0037 |
| F12 | 44160001 | 44260000 | 0.0044 | 0.0038 | 0.0040 |
| F12 | 44180001 | 44280000 | 0.0036 | 0.0030 | 0.0032 |
| F12 | 44200001 | 44300000 | 0.0035 | 0.0029 | 0.0031 |
| F12 | 44220001 | 44320000 | 0.0029 | 0.0024 | 0.0025 |
| F12 | 44240001 | 44340000 | 0.0031 | 0.0026 | 0.0028 |
| F12 | 44260001 | 44360000 | 0.0028 | 0.0024 | 0.0026 |
| F12 | 44280001 | 44380000 | 0.0026 | 0.0022 | 0.0024 |
| F12 | 44300001 | 44400000 | 0.0034 | 0.0030 | 0.0031 |
| F12 | 44320001 | 44420000 | 0.0030 | 0.0027 | 0.0027 |
| F12 | 44340001 | 44440000 | 0.0027 | 0.0024 | 0.0024 |
| F12 | 44360001 | 44460000 | 0.0027 | 0.0024 | 0.0025 |
| F12 | 44380001 | 44480000 | 0.0026 | 0.0024 | 0.0024 |
| F12 | 44400001 | 44500000 | 0.0017 | 0.0016 | 0.0017 |
| F12 | 44420001 | 44520000 | 0.0017 | 0.0014 | 0.0015 |
| F12 | 44440001 | 44540000 | 0.0016 | 0.0014 | 0.0014 |
| F12 | 44460001 | 44560000 | 0.0019 | 0.0017 | 0.0017 |
| F12 | 44480001 | 44580000 | 0.0023 | 0.0018 | 0.0020 |
| F12 | 44500001 | 44600000 | 0.0022 | 0.0018 | 0.0020 |
| F12 | 44520001 | 44620000 | 0.0019 | 0.0016 | 0.0017 |
| F12 | 44540001 | 44640000 | 0.0023 | 0.0017 | 0.0019 |
| F12 | 44560001 | 44660000 | 0.0020 | 0.0012 | 0.0016 |

| Chr | Start | End | both | A1 | A2 |
|---|---|---|---|---|---|
| F12 | 44580001 | 44680000 | 0.0015 | 0.0009 | 0.0012 |
| F12 | 44600001 | 44700000 | 0.0028 | 0.0020 | 0.0025 |
| F12 | 44620001 | 44720000 | 0.0034 | 0.0027 | 0.0031 |
| F12 | 44640001 | 44740000 | 0.0032 | 0.0026 | 0.0030 |
| F12 | 44660001 | 44760000 | 0.0027 | 0.0024 | 0.0027 |
| F12 | 44680001 | 44780000 | 0.0032 | 0.0029 | 0.0032 |
| F12 | 44700001 | 44800000 | 0.0018 | 0.0017 | 0.0018 |
| F12 | 44720001 | 44820000 | 0.0013 | 0.0011 | 0.0013 |
| F12 | 44740001 | 44840000 | 0.0012 | 0.0010 | 0.0012 |
| F12 | 44760001 | 44860000 | 0.0012 | 0.0010 | 0.0012 |
| F12 | 44780001 | 44880000 | 0.0007 | 0.0006 | 0.0007 |
| F12 | 44800001 | 44900000 | 0.0005 | 0.0005 | 0.0005 |
| F12 | 44820001 | 44920000 | 0.0009 | 0.0007 | 0.0009 |
| F12 | 44840001 | 44940000 | 0.0012 | 0.0010 | 0.0012 |
| F12 | 44860001 | 44960000 | 0.0016 | 0.0013 | 0.0016 |
| F12 | 44880001 | 44980000 | 0.0019 | 0.0015 | 0.0019 |
| F12 | 44900001 | 45000000 | 0.0019 | 0.0015 | 0.0019 |
| F12 | 44920001 | 45020000 | 0.0014 | 0.0011 | 0.0014 |
| F12 | 44940001 | 45040000 | 0.0012 | 0.0010 | 0.0012 |
| F12 | 44960001 | 45060000 | 0.0009 | 0.0007 | 0.0009 |
| F12 | 44980001 | 45080000 | 0.0006 | 0.0005 | 0.0006 |
| F12 | 45000001 | 45100000 | 0.0007 | 0.0006 | 0.0007 |
| F12 | 45020001 | 45120000 | 0.0007 | 0.0006 | 0.0007 |
| F12 | 45040001 | 45140000 | 0.0010 | 0.0008 | 0.0010 |
| F12 | 45060001 | 45160000 | 0.0023 | 0.0020 | 0.0023 |
| F12 | 45080001 | 45180000 | 0.0026 | 0.0022 | 0.0025 |
| F12 | 45100001 | 45200000 | 0.0037 | 0.0031 | 0.0036 |
| F12 | 45120001 | 45220000 | 0.0042 | 0.0037 | 0.0041 |
| F12 | 45140001 | 45240000 | 0.0042 | 0.0037 | 0.0041 |
| F12 | 45160001 | 45260000 | 0.0032 | 0.0029 | 0.0032 |
| F12 | 45180001 | 45280000 | 0.0036 | 0.0032 | 0.0035 |
| F12 | 45200001 | 45300000 | 0.0027 | 0.0023 | 0.0026 |
| F12 | 45220001 | 45320000 | 0.0024 | 0.0020 | 0.0023 |
| F12 | 45240001 | 45340000 | 0.0051 | 0.0045 | 0.0048 |
| F12 | 45260001 | 45360000 | 0.0053 | 0.0046 | 0.0049 |
| F12 | 45280001 | 45380000 | 0.0053 | 0.0046 | 0.0049 |
| F12 | 45300001 | 45400000 | 0.0056 | 0.0047 | 0.0050 |
| F12 | 45320001 | 45420000 | 0.0058 | 0.0049 | 0.0052 |
| F12 | 45340001 | 45440000 | 0.0026 | 0.0019 | 0.0021 |
| F12 | 45360001 | 45460000 | 0.0024 | 0.0017 | 0.0021 |
| F12 | 45380001 | 45480000 | 0.0025 | 0.0019 | 0.0023 |
| F12 | 45400001 | 45500000 | 0.0027 | 0.0023 | 0.0026 |
| F12 | 45420001 | 45520000 | 0.0023 | 0.0020 | 0.0023 |
| F12 | 45440001 | 45540000 | 0.0025 | 0.0022 | 0.0024 |
| F12 | 45460001 | 45560000 | 0.0022 | 0.0021 | 0.0021 |
| F12 | 45480001 | 45580000 | 0.0018 | 0.0017 | 0.0017 |
| F12 | 45500001 | 45600000 | 0.0015 | 0.0014 | 0.0015 |
| F12 | 45520001 | 45620000 | 0.0013 | 0.0012 | 0.0012 |
| F12 | 45540001 | 45640000 | 0.0011 | 0.0011 | 0.0011 |
| F12 | 45560001 | 45660000 | 0.0010 | 0.0009 | 0.0009 |
| F12 | 45580001 | 45680000 | 0.0006 | 0.0005 | 0.0005 |
| F12 | 45600001 | 45700000 | 0.0004 | 0.0003 | 0.0004 |
| F12 | 45620001 | 45720000 | 0.0010 | 0.0008 | 0.0009 |
| F12 | 45640001 | 45740000 | 0.0013 | 0.0011 | 0.0012 |
| F12 | 45660001 | 45760000 | 0.0021 | 0.0018 | 0.0020 |
| F12 | 45680001 | 45780000 | 0.0022 | 0.0018 | 0.0021 |
| F12 | 45700001 | 45800000 | 0.0022 | 0.0018 | 0.0021 |
| F12 | 45720001 | 45820000 | 0.0022 | 0.0017 | 0.0021 |

| Chr | Start | End | both | A1 | A2 |
|---|---|---|---|---|---|
| F12 | 45740001 | 45840000 | 0.0026 | 0.0020 | 0.0025 |
| F12 | 45760001 | 45860000 | 0.0024 | 0.0019 | 0.0022 |
| F12 | 45780001 | 45880000 | 0.0027 | 0.0021 | 0.0024 |
| F12 | 45800001 | 45900000 | 0.0031 | 0.0022 | 0.0027 |
| F12 | 45820001 | 45920000 | 0.0030 | 0.0022 | 0.0026 |
| F12 | 45840001 | 45940000 | 0.0030 | 0.0023 | 0.0027 |
| F12 | 45860001 | 45960000 | 0.0028 | 0.0022 | 0.0027 |
| F12 | 45880001 | 45980000 | 0.0033 | 0.0028 | 0.0031 |
| F12 | 45900001 | 46000000 | 0.0035 | 0.0032 | 0.0034 |
| F12 | 45920001 | 46020000 | 0.0034 | 0.0030 | 0.0032 |
| F12 | 45940001 | 46040000 | 0.0032 | 0.0027 | 0.0029 |
| F12 | 45960001 | 46060000 | 0.0031 | 0.0026 | 0.0028 |
| F12 | 45980001 | 46080000 | 0.0028 | 0.0022 | 0.0026 |
| F12 | 46000001 | 46100000 | 0.0025 | 0.0020 | 0.0022 |
| F12 | 46020001 | 46120000 | 0.0027 | 0.0022 | 0.0025 |
| F12 | 46040001 | 46140000 | 0.0038 | 0.0032 | 0.0036 |
| F12 | 46060001 | 46160000 | 0.0046 | 0.0038 | 0.0043 |
| F12 | 46080001 | 46180000 | 0.0053 | 0.0042 | 0.0049 |
| F12 | 46100001 | 46200000 | 0.0050 | 0.0039 | 0.0046 |
| F12 | 46120001 | 46220000 | 0.0053 | 0.0043 | 0.0048 |
| F12 | 46140001 | 46240000 | 0.0043 | 0.0035 | 0.0039 |
| F12 | 46160001 | 46260000 | 0.0041 | 0.0034 | 0.0037 |
| F12 | 46180001 | 46280000 | 0.0035 | 0.0031 | 0.0031 |
| F12 | 46200001 | 46300000 | 0.0036 | 0.0033 | 0.0033 |
| F12 | 46220001 | 46320000 | 0.0034 | 0.0030 | 0.0031 |
| F12 | 46240001 | 46340000 | 0.0036 | 0.0030 | 0.0032 |
| F12 | 46260001 | 46360000 | 0.0045 | 0.0040 | 0.0042 |
| F12 | 46280001 | 46380000 | 0.0040 | 0.0035 | 0.0038 |
| F12 | 46300001 | 46400000 | 0.0036 | 0.0032 | 0.0034 |
| F12 | 46320001 | 46420000 | 0.0031 | 0.0026 | 0.0029 |
| F12 | 46340001 | 46440000 | 0.0030 | 0.0027 | 0.0028 |
| F12 | 46360001 | 46460000 | 0.0020 | 0.0017 | 0.0017 |
| F12 | 46380001 | 46480000 | 0.0026 | 0.0021 | 0.0021 |
| F12 | 46400001 | 46500000 | 0.0041 | 0.0034 | 0.0035 |
| F12 | 46420001 | 46520000 | 0.0051 | 0.0041 | 0.0045 |
| F12 | 46440001 | 46540000 | 0.0049 | 0.0038 | 0.0042 |
| F12 | 46460001 | 46560000 | 0.0050 | 0.0040 | 0.0044 |
| F12 | 46480001 | 46580000 | 0.0045 | 0.0037 | 0.0040 |
| F12 | 46500001 | 46600000 | 0.0032 | 0.0025 | 0.0028 |
| F12 | 46520001 | 46620000 | 0.0028 | 0.0023 | 0.0024 |
| F12 | 46540001 | 46640000 | 0.0028 | 0.0025 | 0.0026 |
| F12 | 46560001 | 46660000 | 0.0023 | 0.0019 | 0.0021 |
| F12 | 46580001 | 46680000 | 0.0024 | 0.0019 | 0.0022 |
| F12 | 46600001 | 46700000 | 0.0024 | 0.0019 | 0.0023 |
| F12 | 46620001 | 46720000 | 0.0026 | 0.0020 | 0.0025 |
| F12 | 46640001 | 46740000 | 0.0024 | 0.0019 | 0.0024 |
| F12 | 46660001 | 46760000 | 0.0022 | 0.0016 | 0.0021 |
| F12 | 46680001 | 46780000 | 0.0019 | 0.0014 | 0.0018 |
| F12 | 46700001 | 46800000 | 0.0018 | 0.0014 | 0.0017 |
| F12 | 46720001 | 46820000 | 0.0011 | 0.0010 | 0.0011 |
| F12 | 46740001 | 46840000 | 0.0012 | 0.0011 | 0.0011 |
| F12 | 46760001 | 46860000 | 0.0010 | 0.0009 | 0.0009 |
| F12 | 46780001 | 46880000 | 0.0014 | 0.0014 | 0.0014 |
| F12 | 46800001 | 46900000 | 0.0013 | 0.0012 | 0.0012 |
| F12 | 46820001 | 46920000 | 0.0015 | 0.0014 | 0.0015 |
| F12 | 46840001 | 46940000 | 0.0017 | 0.0015 | 0.0016 |
| F12 | 46860001 | 46960000 | 0.0020 | 0.0018 | 0.0020 |
| F12 | 46880001 | 46980000 | 0.0025 | 0.0022 | 0.0025 |

| Chr | Start | End | both | A1 | A2 |
|---|---|---|---|---|---|
| F12 | 46900001 | 47000000 | 0.0029 | 0.0025 | 0.0028 |
| F12 | 46920001 | 47020000 | 0.0028 | 0.0025 | 0.0028 |
| F12 | 46940001 | 47040000 | 0.0027 | 0.0022 | 0.0025 |
| F12 | 46960001 | 47060000 | 0.0036 | 0.0029 | 0.0032 |
| F12 | 46980001 | 47080000 | 0.0031 | 0.0024 | 0.0027 |
| F12 | 47000001 | 47100000 | 0.0030 | 0.0023 | 0.0026 |
| F12 | 47020001 | 47120000 | 0.0030 | 0.0023 | 0.0025 |
| F12 | 47040001 | 47140000 | 0.0026 | 0.0022 | 0.0022 |
| F12 | 47060001 | 47160000 | 0.0012 | 0.0011 | 0.0010 |
| F12 | 47080001 | 47180000 | 0.0008 | 0.0007 | 0.0006 |
| F12 | 47100001 | 47200000 | 0.0012 | 0.0010 | 0.0009 |
| F12 | 47120001 | 47220000 | 0.0013 | 0.0009 | 0.0010 |
| F12 | 47140001 | 47240000 | 0.0018 | 0.0013 | 0.0015 |
| F12 | 47160001 | 47260000 | 0.0021 | 0.0015 | 0.0018 |
| F12 | 47180001 | 47280000 | 0.0023 | 0.0018 | 0.0020 |
| F12 | 47200001 | 47300000 | 0.0022 | 0.0018 | 0.0020 |
| F12 | 47220001 | 47320000 | 0.0016 | 0.0015 | 0.0015 |
| F12 | 47240001 | 47340000 | 0.0013 | 0.0012 | 0.0012 |
| F12 | 47260001 | 47360000 | 0.0012 | 0.0012 | 0.0012 |
| F12 | 47280001 | 47380000 | 0.0010 | 0.0009 | 0.0010 |
| F12 | 47300001 | 47400000 | 0.0007 | 0.0005 | 0.0006 |
| F12 | 47320001 | 47420000 | 0.0012 | 0.0009 | 0.0011 |
| F12 | 47340001 | 47440000 | 0.0014 | 0.0011 | 0.0013 |
| F12 | 47360001 | 47460000 | 0.0014 | 0.0011 | 0.0013 |
| F12 | 47380001 | 47480000 | 0.0014 | 0.0011 | 0.0013 |
| F12 | 47400001 | 47500000 | 0.0020 | 0.0018 | 0.0019 |
| F12 | 47420001 | 47520000 | 0.0018 | 0.0017 | 0.0018 |
| F12 | 47440001 | 47540000 | 0.0018 | 0.0016 | 0.0018 |
| F12 | 47460001 | 47560000 | 0.0018 | 0.0016 | 0.0017 |
| F12 | 47480001 | 47580000 | 0.0021 | 0.0018 | 0.0020 |
| F12 | 47500001 | 47600000 | 0.0013 | 0.0011 | 0.0012 |
| F12 | 47520001 | 47620000 | 0.0010 | 0.0008 | 0.0009 |
| F12 | 47540001 | 47640000 | 0.0009 | 0.0007 | 0.0008 |
| F12 | 47560001 | 47660000 | 0.0008 | 0.0005 | 0.0007 |
| F12 | 47580001 | 47680000 | 0.0005 | 0.0004 | 0.0005 |
| F12 | 47600001 | 47700000 | 0.0005 | 0.0004 | 0.0004 |
| F12 | 47620001 | 47720000 | 0.0007 | 0.0005 | 0.0007 |
| F12 | 47640001 | 47740000 | 0.0009 | 0.0007 | 0.0009 |
| F12 | 47660001 | 47760000 | 0.0007 | 0.0005 | 0.0007 |
| F12 | 47680001 | 47780000 | 0.0009 | 0.0007 | 0.0009 |
| F12 | 47700001 | 47800000 | 0.0015 | 0.0011 | 0.0015 |
| F12 | 47720001 | 47820000 | 0.0015 | 0.0011 | 0.0014 |
| F12 | 47740001 | 47840000 | 0.0014 | 0.0010 | 0.0013 |
| F12 | 47760001 | 47860000 | 0.0014 | 0.0009 | 0.0013 |
| F12 | 47780001 | 47880000 | 0.0020 | 0.0015 | 0.0018 |
| F12 | 47800001 | 47900000 | 0.0017 | 0.0012 | 0.0015 |
| F12 | 47820001 | 47920000 | 0.0020 | 0.0017 | 0.0018 |
| F12 | 47840001 | 47940000 | 0.0026 | 0.0023 | 0.0023 |
| F12 | 47860001 | 47960000 | 0.0026 | 0.0023 | 0.0024 |
| F12 | 47880001 | 47980000 | 0.0018 | 0.0016 | 0.0017 |
| F12 | 47900001 | 48000000 | 0.0016 | 0.0014 | 0.0015 |
| F12 | 47920001 | 48020000 | 0.0009 | 0.0008 | 0.0009 |
| F12 | 47940001 | 48040000 | 0.0005 | 0.0004 | 0.0005 |
| F12 | 47960001 | 48060000 | 0.0006 | 0.0005 | 0.0006 |
| F12 | 47980001 | 48080000 | 0.0006 | 0.0005 | 0.0006 |
| F12 | 48000001 | 48100000 | 0.0010 | 0.0008 | 0.0010 |
| F12 | 48020001 | 48120000 | 0.0012 | 0.0010 | 0.0012 |
| F12 | 48040001 | 48140000 | 0.0011 | 0.0009 | 0.0011 |

| Chr | Start | End | both | A1 | A2 |
|---|---|---|---|---|---|
| F12 | 48060001 | 48160000 | 0.0010 | 0.0009 | 0.0010 |
| F12 | 48080001 | 48180000 | 0.0010 | 0.0009 | 0.0010 |
| F12 | 48100001 | 48200000 | 0.0006 | 0.0006 | 0.0006 |
| F12 | 48120001 | 48220000 | 0.0011 | 0.0011 | 0.0011 |
| F12 | 48140001 | 48240000 | 0.0023 | 0.0022 | 0.0022 |
| F12 | 48160001 | 48260000 | 0.0025 | 0.0023 | 0.0024 |
| F12 | 48180001 | 48280000 | 0.0029 | 0.0025 | 0.0026 |
| F12 | 48200001 | 48300000 | 0.0029 | 0.0025 | 0.0027 |
| F12 | 48220001 | 48320000 | 0.0026 | 0.0021 | 0.0024 |
| F12 | 48240001 | 48340000 | 0.0017 | 0.0012 | 0.0014 |
| F12 | 48260001 | 48360000 | 0.0015 | 0.0010 | 0.0012 |
| F12 | 48280001 | 48380000 | 0.0012 | 0.0009 | 0.0011 |
| F12 | 48300001 | 48400000 | 0.0014 | 0.0011 | 0.0013 |
| F12 | 48320001 | 48420000 | 0.0014 | 0.0011 | 0.0012 |
| F12 | 48340001 | 48440000 | 0.0019 | 0.0018 | 0.0018 |
| F12 | 48360001 | 48460000 | 0.0023 | 0.0021 | 0.0022 |
| F12 | 48380001 | 48480000 | 0.0024 | 0.0023 | 0.0023 |
| F12 | 48400001 | 48500000 | 0.0022 | 0.0021 | 0.0021 |
| F12 | 48420001 | 48520000 | 0.0025 | 0.0023 | 0.0024 |
| F12 | 48440001 | 48540000 | 0.0023 | 0.0020 | 0.0023 |
| F12 | 48460001 | 48560000 | 0.0021 | 0.0018 | 0.0021 |
| F12 | 48480001 | 48580000 | 0.0023 | 0.0019 | 0.0023 |
| F12 | 48500001 | 48600000 | 0.0029 | 0.0024 | 0.0029 |
| F12 | 48520001 | 48620000 | 0.0027 | 0.0022 | 0.0026 |
| F12 | 48540001 | 48640000 | 0.0025 | 0.0021 | 0.0023 |
| F12 | 48560001 | 48660000 | 0.0033 | 0.0027 | 0.0030 |
| F12 | 48580001 | 48680000 | 0.0036 | 0.0027 | 0.0030 |
| F12 | 48600001 | 48700000 | 0.0029 | 0.0022 | 0.0023 |
| F12 | 48620001 | 48720000 | 0.0028 | 0.0021 | 0.0024 |
| F12 | 48640001 | 48740000 | 0.0032 | 0.0024 | 0.0027 |
| F12 | 48660001 | 48760000 | 0.0026 | 0.0019 | 0.0021 |
| F12 | 48680001 | 48780000 | 0.0019 | 0.0016 | 0.0018 |
| F12 | 48700001 | 48800000 | 0.0019 | 0.0016 | 0.0018 |
| F12 | 48720001 | 48820000 | 0.0019 | 0.0015 | 0.0017 |
| F12 | 48740001 | 48840000 | 0.0009 | 0.0008 | 0.0009 |
| F12 | 48760001 | 48860000 | 0.0008 | 0.0007 | 0.0007 |
| F12 | 48780001 | 48880000 | 0.0009 | 0.0008 | 0.0009 |
| F12 | 48800001 | 48900000 | 0.0010 | 0.0009 | 0.0010 |
| F12 | 48820001 | 48920000 | 0.0010 | 0.0010 | 0.0010 |
| F12 | 48840001 | 48940000 | 0.0010 | 0.0010 | 0.0010 |
| F12 | 48860001 | 48960000 | 0.0010 | 0.0009 | 0.0009 |
| F12 | 48880001 | 48980000 | 0.0012 | 0.0011 | 0.0012 |
| F12 | 48900001 | 49000000 | 0.0017 | 0.0015 | 0.0017 |
| F12 | 48920001 | 49020000 | 0.0017 | 0.0014 | 0.0017 |
| F12 | 48940001 | 49040000 | 0.0021 | 0.0018 | 0.0021 |
| F12 | 48960001 | 49060000 | 0.0023 | 0.0019 | 0.0023 |
| F12 | 48980001 | 49080000 | 0.0031 | 0.0022 | 0.0030 |
| F12 | 49000001 | 49100000 | 0.0030 | 0.0022 | 0.0029 |
| F12 | 49020001 | 49120000 | 0.0034 | 0.0026 | 0.0033 |
| F12 | 49040001 | 49140000 | 0.0047 | 0.0036 | 0.0046 |
| F12 | 49060001 | 49160000 | 0.0062 | 0.0048 | 0.0061 |
| F12 | 49080001 | 49180000 | 0.0055 | 0.0046 | 0.0054 |
| F12 | 49100001 | 49200000 | 0.0047 | 0.0039 | 0.0047 |
| F12 | 49120001 | 49220000 | 0.0047 | 0.0039 | 0.0046 |
| F12 | 49140001 | 49240000 | 0.0037 | 0.0032 | 0.0035 |
| F12 | 49160001 | 49260000 | 0.0028 | 0.0024 | 0.0026 |
| F12 | 49180001 | 49280000 | 0.0031 | 0.0028 | 0.0028 |
| F12 | 49200001 | 49300000 | 0.0041 | 0.0037 | 0.0038 |

| Chr | Start | End | both | A1 | A2 |
|---|---|---|---|---|---|
| F12 | 49220001 | 49320000 | 0.0039 | 0.0035 | 0.0036 |
| F12 | 49240001 | 49340000 | 0.0032 | 0.0029 | 0.0030 |
| F12 | 49260001 | 49360000 | 0.0031 | 0.0027 | 0.0030 |
| F12 | 49280001 | 49380000 | 0.0035 | 0.0029 | 0.0032 |
| F12 | 49300001 | 49400000 | 0.0031 | 0.0025 | 0.0028 |
| F12 | 49320001 | 49420000 | 0.0029 | 0.0023 | 0.0027 |
| F12 | 49340001 | 49440000 | 0.0029 | 0.0023 | 0.0027 |
| F12 | 49360001 | 49460000 | 0.0024 | 0.0017 | 0.0020 |
| F12 | 49380001 | 49480000 | 0.0014 | 0.0010 | 0.0012 |
| F12 | 49400001 | 49500000 | 0.0014 | 0.0008 | 0.0010 |
| F12 | 49420001 | 49520000 | 0.0019 | 0.0011 | 0.0014 |
| F12 | 49440001 | 49540000 | 0.0026 | 0.0015 | 0.0019 |
| F12 | 49460001 | 49560000 | 0.0025 | 0.0014 | 0.0019 |
| F12 | 49480001 | 49580000 | 0.0031 | 0.0017 | 0.0024 |
| F12 | 49500001 | 49600000 | 0.0034 | 0.0021 | 0.0026 |
| F12 | 49520001 | 49620000 | 0.0040 | 0.0027 | 0.0031 |
| F12 | 49540001 | 49640000 | 0.0045 | 0.0033 | 0.0036 |
| F12 | 49560001 | 49660000 | 0.0049 | 0.0039 | 0.0041 |
| F12 | 49580001 | 49680000 | 0.0049 | 0.0040 | 0.0040 |
| F12 | 49600001 | 49700000 | 0.0051 | 0.0042 | 0.0043 |
| F12 | 49620001 | 49720000 | 0.0039 | 0.0033 | 0.0035 |
| F12 | 49640001 | 49740000 | 0.0032 | 0.0027 | 0.0029 |
| F12 | 49660001 | 49760000 | 0.0027 | 0.0022 | 0.0025 |
| F12 | 49680001 | 49780000 | 0.0032 | 0.0029 | 0.0031 |
| F12 | 49700001 | 49800000 | 0.0029 | 0.0025 | 0.0029 |
| F12 | 49720001 | 49820000 | 0.0029 | 0.0025 | 0.0029 |
| F12 | 49740001 | 49840000 | 0.0029 | 0.0025 | 0.0029 |
| F12 | 49760001 | 49860000 | 0.0034 | 0.0029 | 0.0033 |
| F12 | 49780001 | 49880000 | 0.0027 | 0.0021 | 0.0027 |
| F12 | 49800001 | 49900000 | 0.0022 | 0.0018 | 0.0022 |
| F12 | 49820001 | 49920000 | 0.0029 | 0.0024 | 0.0029 |
| F12 | 49840001 | 49940000 | 0.0030 | 0.0025 | 0.0030 |
| F12 | 49860001 | 49960000 | 0.0029 | 0.0022 | 0.0026 |
| F12 | 49880001 | 49980000 | 0.0032 | 0.0025 | 0.0028 |
| F12 | 49900001 | 50000000 | 0.0032 | 0.0025 | 0.0028 |
| F12 | 49920001 | 50020000 | 0.0032 | 0.0026 | 0.0028 |
| F12 | 49940001 | 50040000 | 0.0029 | 0.0023 | 0.0025 |
| F12 | 49960001 | 50060000 | 0.0032 | 0.0027 | 0.0029 |
| F12 | 49980001 | 50080000 | 0.0034 | 0.0028 | 0.0031 |
| F12 | 50000001 | 50100000 | 0.0043 | 0.0035 | 0.0038 |
| F12 | 50020001 | 50120000 | 0.0036 | 0.0029 | 0.0032 |
| F12 | 50040001 | 50140000 | 0.0039 | 0.0031 | 0.0033 |
| F12 | 50060001 | 50160000 | 0.0038 | 0.0028 | 0.0031 |
| F12 | 50080001 | 50180000 | 0.0029 | 0.0020 | 0.0022 |
| F12 | 50100001 | 50200000 | 0.0026 | 0.0017 | 0.0020 |
| F12 | 50120001 | 50220000 | 0.0031 | 0.0020 | 0.0023 |
| F12 | 50140001 | 50240000 | 0.0031 | 0.0020 | 0.0023 |
| F12 | 50160001 | 50260000 | 0.0030 | 0.0018 | 0.0021 |
| F12 | 50180001 | 50280000 | 0.0034 | 0.0022 | 0.0025 |
| F12 | 50200001 | 50300000 | 0.0038 | 0.0022 | 0.0028 |
| F12 | 50220001 | 50320000 | 0.0037 | 0.0021 | 0.0028 |
| F12 | 50240001 | 50340000 | 0.0033 | 0.0017 | 0.0025 |
| F12 | 50260001 | 50360000 | 0.0038 | 0.0021 | 0.0030 |
| F12 | 50280001 | 50380000 | 0.0040 | 0.0022 | 0.0031 |
| F12 | 50300001 | 50400000 | 0.0033 | 0.0020 | 0.0026 |
| F12 | 50320001 | 50420000 | 0.0031 | 0.0021 | 0.0025 |
| F12 | 50340001 | 50440000 | 0.0036 | 0.0025 | 0.0028 |
| F12 | 50360001 | 50460000 | 0.0034 | 0.0026 | 0.0028 |

| Chr | Start | End | both | A1 | A2 |
|---|---|---|---|---|---|
| F12 | 50380001 | 50480000 | 0.0044 | 0.0034 | 0.0037 |
| F12 | 50400001 | 50500000 | 0.0047 | 0.0034 | 0.0039 |
| F12 | 50420001 | 50520000 | 0.0050 | 0.0036 | 0.0042 |
| F12 | 50440001 | 50540000 | 0.0054 | 0.0041 | 0.0048 |
| F12 | 50460001 | 50560000 | 0.0057 | 0.0044 | 0.0051 |
| F12 | 50480001 | 50580000 | 0.0052 | 0.0042 | 0.0046 |
| F12 | 50500001 | 50600000 | 0.0056 | 0.0045 | 0.0051 |
| F12 | 50520001 | 50620000 | 0.0066 | 0.0054 | 0.0059 |
| F12 | 50540001 | 50640000 | 0.0056 | 0.0046 | 0.0049 |
| F12 | 50560001 | 50660000 | 0.0050 | 0.0040 | 0.0042 |
| F12 | 50580001 | 50680000 | 0.0042 | 0.0033 | 0.0037 |
| F12 | 50600001 | 50700000 | 0.0035 | 0.0029 | 0.0030 |
| F12 | 50620001 | 50720000 | 0.0018 | 0.0014 | 0.0015 |
| F12 | 50640001 | 50740000 | 0.0020 | 0.0015 | 0.0016 |
| F12 | 50660001 | 50760000 | 0.0031 | 0.0024 | 0.0023 |
| F12 | 50680001 | 50780000 | 0.0047 | 0.0039 | 0.0037 |
| F12 | 50700001 | 50800000 | 0.0046 | 0.0037 | 0.0036 |
| F12 | 50720001 | 50820000 | 0.0056 | 0.0044 | 0.0045 |
| F12 | 50740001 | 50840000 | 0.0055 | 0.0045 | 0.0045 |
| F12 | 50760001 | 50860000 | 0.0047 | 0.0038 | 0.0039 |
| F12 | 50780001 | 50880000 | 0.0038 | 0.0029 | 0.0032 |
| F12 | 50800001 | 50900000 | 0.0042 | 0.0032 | 0.0036 |
| F12 | 50820001 | 50920000 | 0.0045 | 0.0035 | 0.0038 |
| F12 | 50840001 | 50940000 | 0.0049 | 0.0037 | 0.0041 |
| F12 | 50860001 | 50960000 | 0.0049 | 0.0038 | 0.0043 |
| F12 | 50880001 | 50980000 | 0.0040 | 0.0031 | 0.0034 |
| F12 | 50900001 | 51000000 | 0.0041 | 0.0031 | 0.0035 |
| F12 | 50920001 | 51020000 | 0.0037 | 0.0027 | 0.0031 |
| F12 | 50940001 | 51040000 | 0.0045 | 0.0035 | 0.0039 |
| F12 | 50960001 | 51060000 | 0.0047 | 0.0038 | 0.0041 |
| F12 | 50980001 | 51080000 | 0.0050 | 0.0040 | 0.0044 |
| F12 | 51000001 | 51100000 | 0.0049 | 0.0039 | 0.0043 |
| F12 | 51020001 | 51120000 | 0.0043 | 0.0037 | 0.0040 |
| F12 | 51040001 | 51140000 | 0.0029 | 0.0025 | 0.0027 |
| F12 | 51060001 | 51160000 | 0.0028 | 0.0024 | 0.0028 |
| F12 | 51080001 | 51180000 | 0.0027 | 0.0024 | 0.0026 |
| F12 | 51100001 | 51200000 | 0.0025 | 0.0021 | 0.0023 |
| F12 | 51120001 | 51220000 | 0.0031 | 0.0027 | 0.0028 |
| F12 | 51140001 | 51240000 | 0.0037 | 0.0030 | 0.0031 |
| F12 | 51160001 | 51260000 | 0.0032 | 0.0025 | 0.0024 |
| F12 | 51180001 | 51280000 | 0.0034 | 0.0026 | 0.0027 |
| F12 | 51200001 | 51300000 | 0.0035 | 0.0027 | 0.0027 |
| F12 | 51220001 | 51320000 | 0.0030 | 0.0021 | 0.0023 |
| F12 | 51240001 | 51340000 | 0.0023 | 0.0017 | 0.0018 |
| F12 | 51260001 | 51360000 | 0.0019 | 0.0014 | 0.0016 |
| F12 | 51280001 | 51380000 | 0.0014 | 0.0008 | 0.0009 |
| F12 | 51300001 | 51400000 | 0.0012 | 0.0008 | 0.0009 |
| F12 | 51320001 | 51420000 | 0.0019 | 0.0013 | 0.0014 |
| F12 | 51340001 | 51440000 | 0.0028 | 0.0021 | 0.0023 |
| F12 | 51360001 | 51460000 | 0.0039 | 0.0030 | 0.0034 |
| F12 | 51380001 | 51480000 | 0.0038 | 0.0031 | 0.0034 |
| F12 | 51400001 | 51500000 | 0.0040 | 0.0033 | 0.0037 |
| F12 | 51420001 | 51520000 | 0.0038 | 0.0033 | 0.0036 |
| F12 | 51440001 | 51540000 | 0.0051 | 0.0044 | 0.0048 |
| F12 | 51460001 | 51560000 | 0.0041 | 0.0036 | 0.0038 |
| F12 | 51480001 | 51580000 | 0.0042 | 0.0036 | 0.0040 |
| F12 | 51500001 | 51600000 | 0.0038 | 0.0033 | 0.0036 |
| F12 | 51520001 | 51620000 | 0.0038 | 0.0033 | 0.0036 |

| Chr | Start | End | both | A1 | A2 |
|---|---|---|---|---|---|
| F12 | 51540001 | 51640000 | 0.0017 | 0.0014 | 0.0017 |
| F12 | 51560001 | 51660000 | 0.0018 | 0.0015 | 0.0018 |
| F12 | 51580001 | 51680000 | 0.0026 | 0.0023 | 0.0026 |
| F12 | 51600001 | 51700000 | 0.0025 | 0.0021 | 0.0024 |
| F12 | 51620001 | 51720000 | 0.0020 | 0.0018 | 0.0019 |
| F12 | 51640001 | 51740000 | 0.0022 | 0.0019 | 0.0021 |
| F12 | 51660001 | 51760000 | 0.0019 | 0.0017 | 0.0019 |
| F12 | 51680001 | 51780000 | 0.0011 | 0.0010 | 0.0011 |
| F12 | 51700001 | 51800000 | 0.0012 | 0.0010 | 0.0011 |
| F12 | 51720001 | 51820000 | 0.0013 | 0.0010 | 0.0012 |
| F12 | 51740001 | 51840000 | 0.0011 | 0.0009 | 0.0010 |
| F12 | 51760001 | 51860000 | 0.0016 | 0.0014 | 0.0015 |
| F12 | 51780001 | 51880000 | 0.0016 | 0.0013 | 0.0014 |
| F12 | 51800001 | 51900000 | 0.0023 | 0.0018 | 0.0019 |
| F12 | 51820001 | 51920000 | 0.0021 | 0.0017 | 0.0018 |
| F12 | 51840001 | 51940000 | 0.0034 | 0.0027 | 0.0029 |
| F12 | 51860001 | 51960000 | 0.0033 | 0.0025 | 0.0028 |
| F12 | 51880001 | 51980000 | 0.0038 | 0.0032 | 0.0034 |
| F12 | 51900001 | 52000000 | 0.0041 | 0.0034 | 0.0036 |
| F12 | 51920001 | 52020000 | 0.0041 | 0.0035 | 0.0037 |
| F12 | 51940001 | 52040000 | 0.0033 | 0.0028 | 0.0031 |
| F12 | 51960001 | 52060000 | 0.0031 | 0.0027 | 0.0029 |
| F12 | 51980001 | 52080000 | 0.0027 | 0.0022 | 0.0025 |
| F12 | 52000001 | 52100000 | 0.0023 | 0.0020 | 0.0022 |
| F12 | 52020001 | 52120000 | 0.0027 | 0.0024 | 0.0027 |
| F12 | 52040001 | 52140000 | 0.0024 | 0.0021 | 0.0023 |
| F12 | 52060001 | 52160000 | 0.0022 | 0.0020 | 0.0022 |
| F12 | 52080001 | 52180000 | 0.0023 | 0.0021 | 0.0023 |
| F12 | 52100001 | 52200000 | 0.0021 | 0.0017 | 0.0020 |
| F12 | 52120001 | 52220000 | 0.0015 | 0.0012 | 0.0014 |
| F12 | 52140001 | 52240000 | 0.0018 | 0.0015 | 0.0017 |
| F12 | 52160001 | 52260000 | 0.0025 | 0.0020 | 0.0022 |
| F12 | 52180001 | 52280000 | 0.0022 | 0.0019 | 0.0020 |
| F12 | 52200001 | 52300000 | 0.0021 | 0.0019 | 0.0020 |
| F12 | 52220001 | 52320000 | 0.0026 | 0.0024 | 0.0024 |
| F12 | 52240001 | 52340000 | 0.0023 | 0.0021 | 0.0022 |
| F12 | 52260001 | 52360000 | 0.0022 | 0.0019 | 0.0020 |
| F12 | 52280001 | 52380000 | 0.0030 | 0.0027 | 0.0029 |
| F12 | 52300001 | 52400000 | 0.0042 | 0.0035 | 0.0039 |
| F12 | 52320001 | 52420000 | 0.0039 | 0.0031 | 0.0036 |
| F12 | 52340001 | 52440000 | 0.0038 | 0.0030 | 0.0035 |
| F12 | 52360001 | 52460000 | 0.0047 | 0.0040 | 0.0044 |
| F12 | 52380001 | 52480000 | 0.0037 | 0.0030 | 0.0033 |
| F12 | 52400001 | 52500000 | 0.0038 | 0.0033 | 0.0035 |
| F12 | 52420001 | 52520000 | 0.0034 | 0.0031 | 0.0032 |
| F12 | 52440001 | 52540000 | 0.0039 | 0.0035 | 0.0035 |
| F12 | 52460001 | 52560000 | 0.0028 | 0.0024 | 0.0026 |
| F12 | 52480001 | 52580000 | 0.0030 | 0.0025 | 0.0027 |
| F12 | 52500001 | 52600000 | 0.0021 | 0.0017 | 0.0019 |
| F12 | 52520001 | 52620000 | 0.0024 | 0.0020 | 0.0022 |
| F12 | 52540001 | 52640000 | 0.0021 | 0.0018 | 0.0020 |
| F12 | 52560001 | 52660000 | 0.0019 | 0.0018 | 0.0018 |
| F12 | 52580001 | 52680000 | 0.0018 | 0.0017 | 0.0017 |
| F12 | 52600001 | 52700000 | 0.0015 | 0.0014 | 0.0014 |
| F12 | 52620001 | 52720000 | 0.0018 | 0.0017 | 0.0018 |
| F12 | 52640001 | 52740000 | 0.0018 | 0.0016 | 0.0018 |
| F12 | 52660001 | 52760000 | 0.0016 | 0.0014 | 0.0016 |
| F12 | 52680001 | 52780000 | 0.0020 | 0.0018 | 0.0020 |

| Chr | Start | End | both | A1 | A2 |
|---|---|---|---|---|---|
| F12 | 52700001 | 52800000 | 0.0025 | 0.0023 | 0.0025 |
| F12 | 52720001 | 52820000 | 0.0033 | 0.0029 | 0.0032 |
| F12 | 52740001 | 52840000 | 0.0043 | 0.0038 | 0.0041 |
| F12 | 52760001 | 52860000 | 0.0050 | 0.0042 | 0.0046 |
| F12 | 52780001 | 52880000 | 0.0061 | 0.0050 | 0.0056 |
| F12 | 52800001 | 52900000 | 0.0064 | 0.0052 | 0.0059 |
| F12 | 52820001 | 52920000 | 0.0056 | 0.0044 | 0.0050 |
| F12 | 52840001 | 52940000 | 0.0049 | 0.0038 | 0.0044 |
| F12 | 52860001 | 52960000 | 0.0049 | 0.0039 | 0.0045 |
| F12 | 52880001 | 52980000 | 0.0039 | 0.0030 | 0.0034 |
| F12 | 52900001 | 53000000 | 0.0030 | 0.0021 | 0.0023 |
| F12 | 52920001 | 53020000 | 0.0025 | 0.0018 | 0.0019 |
| F12 | 52940001 | 53040000 | 0.0020 | 0.0013 | 0.0014 |
| F12 | 52960001 | 53060000 | 0.0025 | 0.0018 | 0.0018 |
| F12 | 52980001 | 53080000 | 0.0027 | 0.0021 | 0.0022 |
| F12 | 53000001 | 53100000 | 0.0036 | 0.0027 | 0.0032 |
| F12 | 53020001 | 53120000 | 0.0047 | 0.0035 | 0.0042 |
| F12 | 53040001 | 53140000 | 0.0051 | 0.0040 | 0.0047 |
| F12 | 53060001 | 53160000 | 0.0046 | 0.0034 | 0.0042 |
| F12 | 53080001 | 53180000 | 0.0058 | 0.0045 | 0.0053 |
| F12 | 53100001 | 53200000 | 0.0051 | 0.0040 | 0.0044 |
| F12 | 53120001 | 53220000 | 0.0041 | 0.0032 | 0.0034 |
| F12 | 53140001 | 53240000 | 0.0047 | 0.0038 | 0.0040 |
| F12 | 53160001 | 53260000 | 0.0046 | 0.0038 | 0.0039 |
| F12 | 53180001 | 53280000 | 0.0032 | 0.0026 | 0.0026 |
| F12 | 53200001 | 53300000 | 0.0032 | 0.0026 | 0.0027 |
| F12 | 53220001 | 53320000 | 0.0031 | 0.0027 | 0.0028 |
| F12 | 53240001 | 53340000 | 0.0030 | 0.0026 | 0.0026 |
| F12 | 53260001 | 53360000 | 0.0029 | 0.0024 | 0.0024 |
| F12 | 53280001 | 53380000 | 0.0038 | 0.0032 | 0.0032 |
| F12 | 53300001 | 53400000 | 0.0039 | 0.0033 | 0.0033 |
| F12 | 53320001 | 53420000 | 0.0044 | 0.0038 | 0.0037 |
| F12 | 53340001 | 53440000 | 0.0039 | 0.0034 | 0.0032 |
| F12 | 53360001 | 53460000 | 0.0042 | 0.0038 | 0.0038 |
| F12 | 53380001 | 53480000 | 0.0035 | 0.0031 | 0.0032 |
| F12 | 53400001 | 53500000 | 0.0044 | 0.0037 | 0.0041 |
| F12 | 53420001 | 53520000 | 0.0041 | 0.0034 | 0.0040 |
| F12 | 53440001 | 53540000 | 0.0050 | 0.0041 | 0.0046 |
| F12 | 53460001 | 53560000 | 0.0057 | 0.0046 | 0.0052 |
| F12 | 53480001 | 53580000 | 0.0056 | 0.0045 | 0.0049 |
| F12 | 53500001 | 53600000 | 0.0046 | 0.0038 | 0.0040 |
| F12 | 53520001 | 53620000 | 0.0044 | 0.0037 | 0.0038 |
| F12 | 53540001 | 53640000 | 0.0041 | 0.0034 | 0.0036 |
| F12 | 53560001 | 53660000 | 0.0028 | 0.0023 | 0.0023 |
| F12 | 53580001 | 53680000 | 0.0035 | 0.0028 | 0.0029 |
| F12 | 53600001 | 53700000 | 0.0034 | 0.0027 | 0.0029 |
| F12 | 53620001 | 53720000 | 0.0039 | 0.0033 | 0.0031 |
| F12 | 53640001 | 53740000 | 0.0042 | 0.0036 | 0.0035 |
| F12 | 53660001 | 53760000 | 0.0041 | 0.0036 | 0.0035 |
| F12 | 53680001 | 53780000 | 0.0032 | 0.0029 | 0.0028 |
| F12 | 53700001 | 53800000 | 0.0029 | 0.0026 | 0.0025 |
| F12 | 53720001 | 53820000 | 0.0030 | 0.0025 | 0.0029 |
| F12 | 53740001 | 53840000 | 0.0021 | 0.0016 | 0.0020 |
| F12 | 53760001 | 53860000 | 0.0020 | 0.0016 | 0.0019 |
| F12 | 53780001 | 53880000 | 0.0026 | 0.0021 | 0.0025 |
| F12 | 53800001 | 53900000 | 0.0029 | 0.0023 | 0.0028 |
| F12 | 53820001 | 53920000 | 0.0022 | 0.0017 | 0.0021 |
| F12 | 53840001 | 53940000 | 0.0022 | 0.0017 | 0.0021 |

| Chr | Start | End | both | A1 | A2 |
|---|---|---|---|---|---|
| F12 | 53860001 | 53960000 | 0.0021 | 0.0016 | 0.0020 |
| F12 | 53880001 | 53980000 | 0.0014 | 0.0010 | 0.0013 |
| F12 | 53900001 | 54000000 | 0.0011 | 0.0007 | 0.0010 |
| F12 | 53920001 | 54020000 | 0.0009 | 0.0007 | 0.0009 |
| F12 | 53940001 | 54040000 | 0.0006 | 0.0005 | 0.0006 |
| F12 | 53960001 | 54060000 | 0.0007 | 0.0006 | 0.0007 |
| F12 | 53980001 | 54080000 | 0.0008 | 0.0006 | 0.0007 |
| F12 | 54000001 | 54100000 | 0.0009 | 0.0006 | 0.0008 |
| F12 | 54020001 | 54120000 | 0.0011 | 0.0007 | 0.0010 |
| F12 | 54040001 | 54140000 | 0.0011 | 0.0008 | 0.0010 |
| F12 | 54060001 | 54160000 | 0.0011 | 0.0007 | 0.0010 |
| F12 | 54080001 | 54180000 | 0.0009 | 0.0007 | 0.0008 |
| F12 | 54100001 | 54200000 | 0.0010 | 0.0007 | 0.0008 |
| F12 | 54120001 | 54220000 | 0.0012 | 0.0008 | 0.0009 |
| F12 | 54140001 | 54240000 | 0.0022 | 0.0016 | 0.0020 |
| F12 | 54160001 | 54260000 | 0.0030 | 0.0021 | 0.0026 |
| F12 | 54180001 | 54280000 | 0.0040 | 0.0029 | 0.0035 |
| F12 | 54200001 | 54300000 | 0.0058 | 0.0049 | 0.0052 |
| F12 | 54220001 | 54320000 | 0.0056 | 0.0047 | 0.0050 |
| F12 | 54240001 | 54340000 | 0.0046 | 0.0040 | 0.0041 |
| F12 | 54260001 | 54360000 | 0.0045 | 0.0042 | 0.0042 |
| F12 | 54280001 | 54380000 | 0.0038 | 0.0036 | 0.0036 |
| F12 | 54300001 | 54400000 | 0.0020 | 0.0019 | 0.0020 |
| F12 | 54320001 | 54420000 | 0.0020 | 0.0018 | 0.0020 |
| F12 | 54340001 | 54440000 | 0.0021 | 0.0019 | 0.0021 |
| F12 | 54360001 | 54460000 | 0.0013 | 0.0011 | 0.0013 |
| F12 | 54380001 | 54480000 | 0.0010 | 0.0009 | 0.0010 |
| F12 | 54400001 | 54500000 | 0.0010 | 0.0008 | 0.0010 |
| F12 | 54420001 | 54520000 | 0.0012 | 0.0009 | 0.0012 |
| F12 | 54440001 | 54540000 | 0.0012 | 0.0009 | 0.0012 |
| F12 | 54460001 | 54560000 | 0.0013 | 0.0011 | 0.0013 |
| F12 | 54480001 | 54580000 | 0.0020 | 0.0017 | 0.0020 |
| F12 | 54500001 | 54600000 | 0.0021 | 0.0019 | 0.0021 |
| F12 | 54520001 | 54620000 | 0.0019 | 0.0018 | 0.0019 |
| F12 | 54540001 | 54640000 | 0.0019 | 0.0016 | 0.0017 |
| F12 | 54560001 | 54660000 | 0.0020 | 0.0017 | 0.0018 |
| F12 | 54580001 | 54680000 | 0.0019 | 0.0015 | 0.0017 |
| F12 | 54600001 | 54700000 | 0.0024 | 0.0016 | 0.0019 |
| F12 | 54620001 | 54720000 | 0.0026 | 0.0016 | 0.0020 |
| F12 | 54640001 | 54740000 | 0.0026 | 0.0017 | 0.0020 |
| F12 | 54660001 | 54760000 | 0.0033 | 0.0024 | 0.0027 |
| F12 | 54680001 | 54780000 | 0.0032 | 0.0023 | 0.0026 |
| F12 | 54700001 | 54800000 | 0.0034 | 0.0027 | 0.0029 |
| F12 | 54720001 | 54820000 | 0.0031 | 0.0026 | 0.0028 |
| F12 | 54740001 | 54840000 | 0.0034 | 0.0030 | 0.0031 |
| F12 | 54760001 | 54860000 | 0.0031 | 0.0026 | 0.0028 |
| F12 | 54780001 | 54880000 | 0.0030 | 0.0026 | 0.0026 |
| F12 | 54800001 | 54900000 | 0.0028 | 0.0025 | 0.0024 |
| F12 | 54820001 | 54920000 | 0.0027 | 0.0025 | 0.0024 |
| F12 | 54840001 | 54940000 | 0.0028 | 0.0023 | 0.0024 |
| F12 | 54860001 | 54960000 | 0.0022 | 0.0018 | 0.0018 |
| F12 | 54880001 | 54980000 | 0.0026 | 0.0021 | 0.0024 |
| F12 | 54900001 | 55000000 | 0.0023 | 0.0016 | 0.0020 |
| F12 | 54920001 | 55020000 | 0.0026 | 0.0017 | 0.0021 |
| F12 | 54940001 | 55040000 | 0.0025 | 0.0018 | 0.0021 |
| F12 | 54960001 | 55060000 | 0.0035 | 0.0026 | 0.0031 |
| F12 | 54980001 | 55080000 | 0.0033 | 0.0023 | 0.0028 |
| F12 | 55000001 | 55100000 | 0.0035 | 0.0026 | 0.0030 |

| Chr | Start | End | both | A1 | A2 |
|---|---|---|---|---|---|
| F12 | 55020001 | 55120000 | 0.0040 | 0.0032 | 0.0035 |
| F12 | 55040001 | 55140000 | 0.0038 | 0.0029 | 0.0032 |
| F12 | 55060001 | 55160000 | 0.0027 | 0.0019 | 0.0020 |
| F12 | 55080001 | 55180000 | 0.0027 | 0.0018 | 0.0020 |
| F12 | 55100001 | 55200000 | 0.0034 | 0.0025 | 0.0026 |
| F12 | 55120001 | 55220000 | 0.0031 | 0.0022 | 0.0023 |
| F12 | 55140001 | 55240000 | 0.0032 | 0.0024 | 0.0025 |
| F12 | 55160001 | 55260000 | 0.0042 | 0.0033 | 0.0034 |
| F12 | 55180001 | 55280000 | 0.0048 | 0.0041 | 0.0039 |
| F12 | 55200001 | 55300000 | 0.0040 | 0.0035 | 0.0032 |
| F12 | 55220001 | 55320000 | 0.0036 | 0.0031 | 0.0030 |
| F12 | 55240001 | 55340000 | 0.0038 | 0.0032 | 0.0033 |
| F12 | 55260001 | 55360000 | 0.0036 | 0.0032 | 0.0032 |
| F12 | 55280001 | 55380000 | 0.0029 | 0.0026 | 0.0027 |
| F12 | 55300001 | 55400000 | 0.0025 | 0.0022 | 0.0024 |
| F12 | 55320001 | 55420000 | 0.0027 | 0.0023 | 0.0025 |
| F12 | 55340001 | 55440000 | 0.0045 | 0.0039 | 0.0042 |
| F12 | 55360001 | 55460000 | 0.0055 | 0.0056 | 0.0049 |
| F12 | 55380001 | 55480000 | 0.0063 | 0.0062 | 0.0056 |
| F12 | 55400001 | 55500000 | 0.0067 | 0.0064 | 0.0059 |
| F12 | 55420001 | 55520000 | 0.0068 | 0.0066 | 0.0060 |
| F12 | 55440001 | 55540000 | 0.0058 | 0.0059 | 0.0050 |
| F12 | 55460001 | 55560000 | 0.0045 | 0.0035 | 0.0040 |
| F12 | 55480001 | 55580000 | 0.0039 | 0.0029 | 0.0035 |
| F12 | 55500001 | 55600000 | 0.0035 | 0.0026 | 0.0032 |
| F12 | 55520001 | 55620000 | 0.0041 | 0.0029 | 0.0039 |
| F12 | 55540001 | 55640000 | 0.0037 | 0.0024 | 0.0035 |
| F12 | 55560001 | 55660000 | 0.0033 | 0.0027 | 0.0030 |
| F12 | 55580001 | 55680000 | 0.0032 | 0.0026 | 0.0028 |
| F12 | 55600001 | 55700000 | 0.0034 | 0.0028 | 0.0030 |
| F12 | 55620001 | 55720000 | 0.0027 | 0.0023 | 0.0023 |
| F12 | 55640001 | 55740000 | 0.0025 | 0.0023 | 0.0020 |
| F12 | 55660001 | 55760000 | 0.0026 | 0.0023 | 0.0023 |
| F12 | 55680001 | 55780000 | 0.0034 | 0.0031 | 0.0030 |
| F12 | 55700001 | 55800000 | 0.0032 | 0.0029 | 0.0028 |
| F12 | 55720001 | 55820000 | 0.0032 | 0.0029 | 0.0028 |
| F12 | 55740001 | 55840000 | 0.0025 | 0.0022 | 0.0021 |
| F12 | 55760001 | 55860000 | 0.0022 | 0.0018 | 0.0017 |
| F12 | 55780001 | 55880000 | 0.0014 | 0.0009 | 0.0011 |
| F12 | 55800001 | 55900000 | 0.0014 | 0.0010 | 0.0011 |
| F12 | 55820001 | 55920000 | 0.0016 | 0.0012 | 0.0013 |
| F12 | 55840001 | 55940000 | 0.0021 | 0.0015 | 0.0016 |
| F12 | 55860001 | 55960000 | 0.0024 | 0.0018 | 0.0019 |
| F12 | 55880001 | 55980000 | 0.0027 | 0.0020 | 0.0022 |
| F12 | 55900001 | 56000000 | 0.0038 | 0.0028 | 0.0030 |
| F12 | 55920001 | 56020000 | 0.0038 | 0.0029 | 0.0030 |
| F12 | 55940001 | 56040000 | 0.0043 | 0.0034 | 0.0037 |
| F12 | 55960001 | 56060000 | 0.0048 | 0.0036 | 0.0040 |
| F12 | 55980001 | 56080000 | 0.0044 | 0.0034 | 0.0037 |
| F12 | 56000001 | 56100000 | 0.0035 | 0.0026 | 0.0030 |
| F12 | 56020001 | 56120000 | 0.0029 | 0.0020 | 0.0024 |
| F12 | 56040001 | 56140000 | 0.0023 | 0.0015 | 0.0017 |
| F12 | 56060001 | 56160000 | 0.0016 | 0.0010 | 0.0011 |
| F12 | 56080001 | 56180000 | 0.0020 | 0.0013 | 0.0014 |
| F12 | 56100001 | 56200000 | 0.0019 | 0.0014 | 0.0014 |
| F12 | 56120001 | 56220000 | 0.0019 | 0.0013 | 0.0014 |
| F12 | 56140001 | 56240000 | 0.0016 | 0.0011 | 0.0012 |
| F12 | 56160001 | 56260000 | 0.0032 | 0.0026 | 0.0029 |

| Chr | Start | End | both | A1 | A2 |
|---|---|---|---|---|---|
| F12 | 56180001 | 56280000 | 0.0024 | 0.0020 | 0.0022 |
| F12 | 56200001 | 56300000 | 0.0024 | 0.0020 | 0.0022 |
| F12 | 56220001 | 56320000 | 0.0023 | 0.0020 | 0.0022 |
| F12 | 56240001 | 56340000 | 0.0028 | 0.0022 | 0.0025 |
| F12 | 56260001 | 56360000 | 0.0016 | 0.0013 | 0.0014 |
| F12 | 56280001 | 56380000 | 0.0028 | 0.0023 | 0.0026 |
| F12 | 56300001 | 56400000 | 0.0029 | 0.0024 | 0.0026 |
| F12 | 56320001 | 56420000 | 0.0032 | 0.0025 | 0.0028 |
| F12 | 56340001 | 56440000 | 0.0031 | 0.0025 | 0.0029 |
| F12 | 56360001 | 56460000 | 0.0034 | 0.0025 | 0.0032 |
| F12 | 56380001 | 56480000 | 0.0022 | 0.0014 | 0.0020 |
| F12 | 56400001 | 56500000 | 0.0018 | 0.0012 | 0.0017 |
| F12 | 56420001 | 56520000 | 0.0015 | 0.0010 | 0.0014 |
| F12 | 56440001 | 56540000 | 0.0014 | 0.0009 | 0.0013 |
| F12 | 56460001 | 56560000 | 0.0011 | 0.0008 | 0.0010 |
| F12 | 56480001 | 56580000 | 0.0014 | 0.0009 | 0.0010 |
| F12 | 56500001 | 56600000 | 0.0014 | 0.0008 | 0.0010 |
| F12 | 56520001 | 56620000 | 0.0015 | 0.0010 | 0.0012 |
| F12 | 56540001 | 56640000 | 0.0011 | 0.0007 | 0.0008 |
| F12 | 56560001 | 56660000 | 0.0008 | 0.0006 | 0.0006 |
| F12 | 56580001 | 56680000 | 0.0007 | 0.0006 | 0.0006 |
| F12 | 56600001 | 56700000 | 0.0013 | 0.0011 | 0.0012 |
| F12 | 56620001 | 56720000 | 0.0021 | 0.0017 | 0.0019 |
| F12 | 56640001 | 56740000 | 0.0023 | 0.0019 | 0.0021 |
| F12 | 56660001 | 56760000 | 0.0025 | 0.0019 | 0.0022 |
| F12 | 56680001 | 56780000 | 0.0028 | 0.0023 | 0.0025 |
| F12 | 56700001 | 56800000 | 0.0024 | 0.0020 | 0.0022 |
| F12 | 56720001 | 56820000 | 0.0017 | 0.0013 | 0.0015 |
| F12 | 56740001 | 56840000 | 0.0020 | 0.0017 | 0.0017 |
| F12 | 56760001 | 56860000 | 0.0018 | 0.0016 | 0.0015 |
| F12 | 56780001 | 56880000 | 0.0017 | 0.0014 | 0.0014 |
| F12 | 56800001 | 56900000 | 0.0024 | 0.0020 | 0.0021 |
| F12 | 56820001 | 56920000 | 0.0029 | 0.0025 | 0.0025 |
| F12 | 56840001 | 56940000 | 0.0028 | 0.0023 | 0.0023 |
| F12 | 56860001 | 56960000 | 0.0029 | 0.0024 | 0.0023 |
| F12 | 56880001 | 56980000 | 0.0032 | 0.0024 | 0.0025 |
| F12 | 56900001 | 57000000 | 0.0028 | 0.0021 | 0.0020 |
| F12 | 56920001 | 57020000 | 0.0024 | 0.0016 | 0.0018 |
| F12 | 56940001 | 57040000 | 0.0029 | 0.0020 | 0.0023 |
| F12 | 56960001 | 57060000 | 0.0031 | 0.0022 | 0.0025 |
| F12 | 56980001 | 57080000 | 0.0036 | 0.0027 | 0.0028 |
| F12 | 57000001 | 57100000 | 0.0042 | 0.0031 | 0.0034 |
| F12 | 57020001 | 57120000 | 0.0047 | 0.0034 | 0.0037 |
| F12 | 57040001 | 57140000 | 0.0047 | 0.0035 | 0.0038 |
| F12 | 57060001 | 57160000 | 0.0042 | 0.0032 | 0.0036 |
| F12 | 57080001 | 57180000 | 0.0034 | 0.0024 | 0.0028 |
| F12 | 57100001 | 57200000 | 0.0031 | 0.0021 | 0.0025 |
| F12 | 57120001 | 57220000 | 0.0022 | 0.0016 | 0.0018 |
| F12 | 57140001 | 57240000 | 0.0012 | 0.0008 | 0.0009 |
| F12 | 57160001 | 57260000 | 0.0014 | 0.0008 | 0.0008 |
| F12 | 57180001 | 57280000 | 0.0020 | 0.0013 | 0.0013 |
| F12 | 57200001 | 57300000 | 0.0013 | 0.0008 | 0.0008 |
| F12 | 57220001 | 57320000 | 0.0014 | 0.0010 | 0.0010 |
| F12 | 57240001 | 57340000 | 0.0016 | 0.0012 | 0.0012 |
| F12 | 57260001 | 57360000 | 0.0016 | 0.0013 | 0.0014 |
| F12 | 57280001 | 57380000 | 0.0013 | 0.0011 | 0.0012 |
| F12 | 57300001 | 57400000 | 0.0015 | 0.0014 | 0.0014 |
| F12 | 57320001 | 57420000 | 0.0018 | 0.0017 | 0.0016 |

| Chr | Start | End | both | A1 | A2 |
|---|---|---|---|---|---|
| F12 | 57340001 | 57440000 | 0.0018 | 0.0017 | 0.0016 |
| F12 | 57360001 | 57460000 | 0.0021 | 0.0020 | 0.0019 |
| F12 | 57380001 | 57480000 | 0.0032 | 0.0028 | 0.0030 |
| F12 | 57400001 | 57500000 | 0.0031 | 0.0027 | 0.0029 |
| F12 | 57420001 | 57520000 | 0.0031 | 0.0025 | 0.0027 |
| F12 | 57440001 | 57540000 | 0.0033 | 0.0026 | 0.0027 |
| F12 | 57460001 | 57560000 | 0.0032 | 0.0023 | 0.0025 |
| F12 | 57480001 | 57580000 | 0.0021 | 0.0013 | 0.0014 |
| F12 | 57500001 | 57600000 | 0.0020 | 0.0011 | 0.0012 |
| F12 | 57520001 | 57620000 | 0.0019 | 0.0009 | 0.0011 |
| F12 | 57540001 | 57640000 | 0.0023 | 0.0013 | 0.0015 |
| F12 | 57560001 | 57660000 | 0.0023 | 0.0014 | 0.0014 |
| F12 | 57580001 | 57680000 | 0.0018 | 0.0011 | 0.0011 |
| F12 | 57600001 | 57700000 | 0.0025 | 0.0017 | 0.0017 |
| F12 | 57620001 | 57720000 | 0.0032 | 0.0023 | 0.0023 |
| F12 | 57640001 | 57740000 | 0.0027 | 0.0018 | 0.0020 |
| F12 | 57660001 | 57760000 | 0.0027 | 0.0019 | 0.0021 |
| F12 | 57680001 | 57780000 | 0.0030 | 0.0022 | 0.0025 |
| F12 | 57700001 | 57800000 | 0.0027 | 0.0021 | 0.0024 |
| F12 | 57720001 | 57820000 | 0.0020 | 0.0015 | 0.0018 |
| F12 | 57740001 | 57840000 | 0.0020 | 0.0014 | 0.0017 |
| F12 | 57760001 | 57860000 | 0.0023 | 0.0017 | 0.0020 |
| F12 | 57780001 | 57880000 | 0.0031 | 0.0023 | 0.0027 |
| F12 | 57800001 | 57900000 | 0.0029 | 0.0022 | 0.0025 |
| F12 | 57820001 | 57920000 | 0.0027 | 0.0021 | 0.0023 |
| F12 | 57840001 | 57940000 | 0.0026 | 0.0022 | 0.0024 |
| F12 | 57860001 | 57960000 | 0.0021 | 0.0018 | 0.0020 |
| F12 | 57880001 | 57980000 | 0.0015 | 0.0013 | 0.0013 |
| F12 | 57900001 | 58000000 | 0.0016 | 0.0013 | 0.0014 |
| F12 | 57920001 | 58020000 | 0.0020 | 0.0014 | 0.0016 |
| F12 | 57940001 | 58040000 | 0.0023 | 0.0017 | 0.0019 |
| F12 | 57960001 | 58060000 | 0.0026 | 0.0019 | 0.0022 |
| F12 | 57980001 | 58080000 | 0.0023 | 0.0017 | 0.0019 |
| F12 | 58000001 | 58100000 | 0.0021 | 0.0016 | 0.0019 |
| F12 | 58020001 | 58120000 | 0.0019 | 0.0016 | 0.0018 |
| F12 | 58040001 | 58140000 | 0.0016 | 0.0012 | 0.0014 |
| F12 | 58060001 | 58160000 | 0.0016 | 0.0012 | 0.0014 |
| F12 | 58080001 | 58180000 | 0.0020 | 0.0016 | 0.0018 |
| F12 | 58100001 | 58200000 | 0.0018 | 0.0015 | 0.0016 |
| F12 | 58120001 | 58220000 | 0.0017 | 0.0014 | 0.0015 |
| F12 | 58140001 | 58240000 | 0.0021 | 0.0018 | 0.0020 |
| F12 | 58160001 | 58260000 | 0.0021 | 0.0018 | 0.0020 |
| F12 | 58180001 | 58280000 | 0.0018 | 0.0015 | 0.0017 |
| F12 | 58200001 | 58300000 | 0.0022 | 0.0019 | 0.0021 |
| F12 | 58220001 | 58320000 | 0.0023 | 0.0020 | 0.0022 |
| F12 | 58240001 | 58340000 | 0.0017 | 0.0016 | 0.0017 |
| F12 | 58260001 | 58360000 | 0.0014 | 0.0012 | 0.0013 |
| F12 | 58280001 | 58380000 | 0.0013 | 0.0011 | 0.0013 |
| F12 | 58300001 | 58400000 | 0.0008 | 0.0006 | 0.0008 |
| F12 | 58320001 | 58420000 | 0.0011 | 0.0008 | 0.0011 |
| F12 | 58340001 | 58440000 | 0.0011 | 0.0008 | 0.0011 |
| F12 | 58360001 | 58460000 | 0.0019 | 0.0015 | 0.0017 |
| F12 | 58380001 | 58480000 | 0.0022 | 0.0017 | 0.0020 |
| F12 | 58400001 | 58500000 | 0.0025 | 0.0019 | 0.0023 |
| F12 | 58420001 | 58520000 | 0.0027 | 0.0021 | 0.0026 |
| F12 | 58440001 | 58540000 | 0.0028 | 0.0022 | 0.0026 |
| F12 | 58460001 | 58560000 | 0.0019 | 0.0015 | 0.0019 |
| F12 | 58480001 | 58580000 | 0.0015 | 0.0012 | 0.0015 |

| Chr | Start | End | both | A1 | A2 |
|---|---|---|---|---|---|
| F12 | 58500001 | 58600000 | 0.0011 | 0.0009 | 0.0011 |
| F12 | 58520001 | 58620000 | 0.0005 | 0.0004 | 0.0005 |
| F12 | 58540001 | 58640000 | 0.0004 | 0.0004 | 0.0004 |
| F12 | 58560001 | 58660000 | 0.0005 | 0.0005 | 0.0005 |
| F12 | 58580001 | 58680000 | 0.0008 | 0.0007 | 0.0008 |
| F12 | 58600001 | 58700000 | 0.0009 | 0.0008 | 0.0009 |
| F12 | 58620001 | 58720000 | 0.0012 | 0.0009 | 0.0010 |
| F12 | 58640001 | 58740000 | 0.0014 | 0.0010 | 0.0012 |
| F12 | 58660001 | 58760000 | 0.0014 | 0.0009 | 0.0012 |
| F12 | 58680001 | 58780000 | 0.0011 | 0.0007 | 0.0008 |
| F12 | 58700001 | 58800000 | 0.0014 | 0.0009 | 0.0011 |
| F12 | 58720001 | 58820000 | 0.0018 | 0.0013 | 0.0015 |
| F12 | 58740001 | 58840000 | 0.0020 | 0.0014 | 0.0016 |
| F12 | 58760001 | 58860000 | 0.0029 | 0.0021 | 0.0024 |
| F12 | 58780001 | 58880000 | 0.0034 | 0.0025 | 0.0028 |
| F12 | 58800001 | 58900000 | 0.0035 | 0.0025 | 0.0028 |
| F12 | 58820001 | 58920000 | 0.0035 | 0.0027 | 0.0028 |
| F12 | 58840001 | 58940000 | 0.0033 | 0.0027 | 0.0028 |
| F12 | 58860001 | 58960000 | 0.0025 | 0.0021 | 0.0022 |
| F12 | 58880001 | 58980000 | 0.0021 | 0.0018 | 0.0017 |
| F12 | 58900001 | 59000000 | 0.0020 | 0.0016 | 0.0015 |
| F12 | 58920001 | 59020000 | 0.0014 | 0.0009 | 0.0010 |
| F12 | 58940001 | 59040000 | 0.0023 | 0.0017 | 0.0019 |
| F12 | 58960001 | 59060000 | 0.0024 | 0.0017 | 0.0018 |
| F12 | 58980001 | 59080000 | 0.0035 | 0.0028 | 0.0029 |
| F12 | 59000001 | 59100000 | 0.0034 | 0.0029 | 0.0030 |
| F12 | 59020001 | 59120000 | 0.0039 | 0.0030 | 0.0033 |
| F12 | 59040001 | 59140000 | 0.0028 | 0.0020 | 0.0022 |
| F12 | 59060001 | 59160000 | 0.0029 | 0.0022 | 0.0023 |
| F12 | 59080001 | 59180000 | 0.0026 | 0.0017 | 0.0018 |
| F12 | 59100001 | 59200000 | 0.0027 | 0.0019 | 0.0018 |
| F12 | 59120001 | 59220000 | 0.0032 | 0.0026 | 0.0025 |
| F12 | 59140001 | 59240000 | 0.0033 | 0.0028 | 0.0026 |
| F12 | 59160001 | 59260000 | 0.0032 | 0.0027 | 0.0027 |
| F12 | 59180001 | 59280000 | 0.0025 | 0.0023 | 0.0023 |
| F12 | 59200001 | 59300000 | 0.0024 | 0.0020 | 0.0022 |
| F12 | 59220001 | 59320000 | 0.0015 | 0.0011 | 0.0014 |
| F12 | 59240001 | 59340000 | 0.0022 | 0.0016 | 0.0020 |
| F12 | 59260001 | 59360000 | 0.0022 | 0.0016 | 0.0020 |
| F12 | 59280001 | 59380000 | 0.0021 | 0.0014 | 0.0018 |
| F12 | 59300001 | 59400000 | 0.0022 | 0.0016 | 0.0019 |
| F12 | 59320001 | 59420000 | 0.0021 | 0.0016 | 0.0019 |
| F12 | 59340001 | 59440000 | 0.0013 | 0.0010 | 0.0011 |
| F12 | 59360001 | 59460000 | 0.0008 | 0.0006 | 0.0008 |
| F12 | 59380001 | 59480000 | 0.0008 | 0.0006 | 0.0008 |
| F12 | 59400001 | 59500000 | 0.0013 | 0.0010 | 0.0013 |
| F12 | 59420001 | 59520000 | 0.0018 | 0.0014 | 0.0018 |
| F12 | 59440001 | 59540000 | 0.0019 | 0.0015 | 0.0019 |
| F12 | 59460001 | 59560000 | 0.0020 | 0.0017 | 0.0020 |
| F12 | 59480001 | 59580000 | 0.0021 | 0.0017 | 0.0020 |
| F12 | 59500001 | 59600000 | 0.0011 | 0.0010 | 0.0011 |
| F12 | 59520001 | 59620000 | 0.0009 | 0.0009 | 0.0009 |
| F12 | 59540001 | 59640000 | 0.0010 | 0.0009 | 0.0010 |
| F12 | 59560001 | 59660000 | 0.0009 | 0.0008 | 0.0009 |
| F12 | 59580001 | 59680000 | 0.0010 | 0.0008 | 0.0009 |
| F12 | 59600001 | 59700000 | 0.0009 | 0.0007 | 0.0008 |
| F12 | 59620001 | 59720000 | 0.0006 | 0.0004 | 0.0006 |
| F12 | 59640001 | 59740000 | 0.0005 | 0.0004 | 0.0005 |

| Chr | Start | End | both | A1 | A2 |
|---|---|---|---|---|---|
| F12 | 59660001 | 59760000 | 0.0004 | 0.0003 | 0.0004 |
| F12 | 59680001 | 59780000 | 0.0004 | 0.0004 | 0.0004 |
| F12 | 59700001 | 59800000 | 0.0009 | 0.0007 | 0.0009 |
| F12 | 59720001 | 59820000 | 0.0009 | 0.0009 | 0.0009 |
| F12 | 59740001 | 59840000 | 0.0008 | 0.0008 | 0.0008 |
| F12 | 59760001 | 59860000 | 0.0012 | 0.0012 | 0.0012 |
| F12 | 59780001 | 59880000 | 0.0012 | 0.0012 | 0.0011 |
| F12 | 59800001 | 59900000 | 0.0008 | 0.0008 | 0.0007 |
| F12 | 59820001 | 59920000 | 0.0008 | 0.0008 | 0.0008 |
| F12 | 59840001 | 59940000 | 0.0011 | 0.0010 | 0.0010 |
| F12 | 59860001 | 59960000 | 0.0015 | 0.0011 | 0.0015 |
| F12 | 59880001 | 59980000 | 0.0019 | 0.0013 | 0.0018 |
| F12 | 59900001 | 60000000 | 0.0026 | 0.0020 | 0.0025 |
| F12 | 59920001 | 60020000 | 0.0029 | 0.0023 | 0.0027 |
| F12 | 59940001 | 60040000 | 0.0032 | 0.0025 | 0.0030 |
| F12 | 59960001 | 60060000 | 0.0025 | 0.0021 | 0.0023 |
| F12 | 59980001 | 60080000 | 0.0025 | 0.0021 | 0.0022 |
| F12 | 60000001 | 60100000 | 0.0023 | 0.0017 | 0.0018 |
| F12 | 60020001 | 60120000 | 0.0025 | 0.0018 | 0.0020 |
| F12 | 60040001 | 60140000 | 0.0018 | 0.0013 | 0.0014 |
| F12 | 60060001 | 60160000 | 0.0017 | 0.0012 | 0.0013 |
| F12 | 60080001 | 60180000 | 0.0018 | 0.0013 | 0.0014 |
| F12 | 60100001 | 60200000 | 0.0019 | 0.0015 | 0.0016 |
| F12 | 60120001 | 60220000 | 0.0016 | 0.0013 | 0.0013 |
| F12 | 60140001 | 60240000 | 0.0026 | 0.0022 | 0.0023 |
| F12 | 60160001 | 60260000 | 0.0031 | 0.0027 | 0.0027 |
| F12 | 60180001 | 60280000 | 0.0029 | 0.0025 | 0.0026 |
| F12 | 60200001 | 60300000 | 0.0027 | 0.0023 | 0.0025 |
| F12 | 60220001 | 60320000 | 0.0025 | 0.0021 | 0.0023 |
| F12 | 60240001 | 60340000 | 0.0016 | 0.0014 | 0.0015 |
| F12 | 60260001 | 60360000 | 0.0012 | 0.0011 | 0.0012 |
| F12 | 60280001 | 60380000 | 0.0019 | 0.0017 | 0.0018 |
| F12 | 60300001 | 60400000 | 0.0020 | 0.0018 | 0.0020 |
| F12 | 60320001 | 60420000 | 0.0018 | 0.0016 | 0.0018 |
| F12 | 60340001 | 60440000 | 0.0019 | 0.0015 | 0.0018 |
| F12 | 60360001 | 60460000 | 0.0023 | 0.0018 | 0.0020 |
| F12 | 60380001 | 60480000 | 0.0024 | 0.0018 | 0.0020 |
| F12 | 60400001 | 60500000 | 0.0026 | 0.0020 | 0.0022 |
| F12 | 60420001 | 60520000 | 0.0033 | 0.0025 | 0.0027 |
| F12 | 60440001 | 60540000 | 0.0033 | 0.0025 | 0.0028 |
| F12 | 60460001 | 60560000 | 0.0032 | 0.0026 | 0.0028 |
| F12 | 60480001 | 60580000 | 0.0034 | 0.0025 | 0.0031 |
| F12 | 60500001 | 60600000 | 0.0030 | 0.0021 | 0.0027 |
| F12 | 60520001 | 60620000 | 0.0025 | 0.0018 | 0.0024 |
| F12 | 60540001 | 60640000 | 0.0023 | 0.0017 | 0.0023 |
| F12 | 60560001 | 60660000 | 0.0028 | 0.0021 | 0.0027 |
| F12 | 60580001 | 60680000 | 0.0025 | 0.0021 | 0.0023 |
| F12 | 60600001 | 60700000 | 0.0030 | 0.0025 | 0.0027 |
| F12 | 60620001 | 60720000 | 0.0031 | 0.0026 | 0.0029 |
| F12 | 60640001 | 60740000 | 0.0032 | 0.0027 | 0.0029 |
| F12 | 60660001 | 60760000 | 0.0024 | 0.0020 | 0.0022 |
| F12 | 60680001 | 60780000 | 0.0018 | 0.0014 | 0.0017 |
| F12 | 60700001 | 60800000 | 0.0008 | 0.0006 | 0.0008 |
| F12 | 60720001 | 60820000 | 0.0006 | 0.0005 | 0.0006 |
| F12 | 60740001 | 60840000 | 0.0015 | 0.0014 | 0.0014 |
| F12 | 60760001 | 60860000 | 0.0015 | 0.0013 | 0.0014 |
| F12 | 60780001 | 60880000 | 0.0013 | 0.0012 | 0.0013 |
| F12 | 60800001 | 60900000 | 0.0014 | 0.0013 | 0.0013 |

| Chr | Start | End | both | A1 | A2 |
|---|---|---|---|---|---|
| F12 | 60820001 | 60920000 | 0.0013 | 0.0013 | 0.0013 |
| F12 | 60840001 | 60940000 | 0.0006 | 0.0004 | 0.0006 |
| F12 | 60860001 | 60960000 | 0.0006 | 0.0005 | 0.0006 |
| F12 | 60880001 | 60980000 | 0.0008 | 0.0007 | 0.0008 |
| F12 | 60900001 | 61000000 | 0.0012 | 0.0010 | 0.0012 |
| F12 | 60920001 | 61020000 | 0.0013 | 0.0011 | 0.0013 |
| F12 | 60940001 | 61040000 | 0.0012 | 0.0010 | 0.0012 |
| F12 | 60960001 | 61060000 | 0.0011 | 0.0009 | 0.0010 |
| F12 | 60980001 | 61080000 | 0.0011 | 0.0010 | 0.0011 |
| F12 | 61000001 | 61100000 | 0.0011 | 0.0011 | 0.0011 |
| F12 | 61020001 | 61120000 | 0.0010 | 0.0010 | 0.0009 |
| F12 | 61040001 | 61140000 | 0.0028 | 0.0027 | 0.0027 |
| F12 | 61060001 | 61160000 | 0.0044 | 0.0038 | 0.0041 |
| F12 | 61080001 | 61180000 | 0.0048 | 0.0042 | 0.0044 |
| F12 | 61100001 | 61200000 | 0.0045 | 0.0037 | 0.0041 |
| F12 | 61120001 | 61220000 | 0.0046 | 0.0038 | 0.0041 |
| F12 | 61140001 | 61240000 | 0.0032 | 0.0026 | 0.0028 |
| F12 | 61160001 | 61260000 | 0.0018 | 0.0015 | 0.0014 |
| F12 | 61180001 | 61280000 | 0.0014 | 0.0011 | 0.0011 |
| F12 | 61200001 | 61300000 | 0.0025 | 0.0021 | 0.0022 |
| F12 | 61220001 | 61320000 | 0.0028 | 0.0023 | 0.0024 |
| F12 | 61240001 | 61340000 | 0.0029 | 0.0022 | 0.0023 |
| F12 | 61260001 | 61360000 | 0.0027 | 0.0020 | 0.0022 |
| F12 | 61280001 | 61380000 | 0.0030 | 0.0021 | 0.0025 |
| F12 | 61300001 | 61400000 | 0.0021 | 0.0013 | 0.0016 |
| F12 | 61320001 | 61420000 | 0.0028 | 0.0021 | 0.0022 |
| F12 | 61340001 | 61440000 | 0.0031 | 0.0027 | 0.0025 |
| F12 | 61360001 | 61460000 | 0.0034 | 0.0030 | 0.0028 |
| F12 | 61380001 | 61480000 | 0.0032 | 0.0028 | 0.0026 |
| F12 | 61400001 | 61500000 | 0.0031 | 0.0028 | 0.0026 |
| F12 | 61420001 | 61520000 | 0.0029 | 0.0023 | 0.0023 |
| F12 | 61440001 | 61540000 | 0.0022 | 0.0015 | 0.0016 |
| F12 | 61460001 | 61560000 | 0.0022 | 0.0015 | 0.0017 |
| F12 | 61480001 | 61580000 | 0.0022 | 0.0015 | 0.0017 |
| F12 | 61500001 | 61600000 | 0.0024 | 0.0017 | 0.0019 |
| F12 | 61520001 | 61620000 | 0.0025 | 0.0020 | 0.0022 |
| F12 | 61540001 | 61640000 | 0.0033 | 0.0027 | 0.0032 |
| F12 | 61560001 | 61660000 | 0.0034 | 0.0028 | 0.0033 |
| F12 | 61580001 | 61680000 | 0.0038 | 0.0031 | 0.0037 |
| F12 | 61600001 | 61700000 | 0.0037 | 0.0030 | 0.0036 |
| F12 | 61620001 | 61720000 | 0.0034 | 0.0025 | 0.0031 |
| F12 | 61640001 | 61740000 | 0.0032 | 0.0025 | 0.0029 |
| F12 | 61660001 | 61760000 | 0.0028 | 0.0022 | 0.0025 |
| F12 | 61680001 | 61780000 | 0.0022 | 0.0016 | 0.0018 |
| F12 | 61700001 | 61800000 | 0.0018 | 0.0015 | 0.0016 |
| F12 | 61720001 | 61820000 | 0.0013 | 0.0011 | 0.0012 |
| F12 | 61740001 | 61840000 | 0.0012 | 0.0011 | 0.0011 |
| F12 | 61760001 | 61860000 | 0.0013 | 0.0012 | 0.0012 |
| F12 | 61780001 | 61880000 | 0.0012 | 0.0012 | 0.0012 |
| F12 | 61800001 | 61900000 | 0.0017 | 0.0015 | 0.0016 |
| F12 | 61820001 | 61920000 | 0.0018 | 0.0016 | 0.0015 |
| F12 | 61840001 | 61940000 | 0.0015 | 0.0011 | 0.0010 |
| F12 | 61860001 | 61960000 | 0.0028 | 0.0023 | 0.0023 |
| F12 | 61880001 | 61980000 | 0.0033 | 0.0026 | 0.0027 |
| F12 | 61900001 | 62000000 | 0.0030 | 0.0024 | 0.0024 |
| F12 | 61920001 | 62020000 | 0.0029 | 0.0022 | 0.0024 |
| F12 | 61940001 | 62040000 | 0.0028 | 0.0022 | 0.0024 |
| F12 | 61960001 | 62060000 | 0.0017 | 0.0011 | 0.0014 |

| Chr | Start | End | both | A1 | A2 |
|---|---|---|---|---|---|
| F12 | 61980001 | 62080000 | 0.0019 | 0.0014 | 0.0018 |
| F12 | 62000001 | 62100000 | 0.0021 | 0.0015 | 0.0019 |
| F12 | 62020001 | 62120000 | 0.0027 | 0.0021 | 0.0026 |
| F12 | 62040001 | 62140000 | 0.0026 | 0.0020 | 0.0024 |
| F12 | 62060001 | 62160000 | 0.0028 | 0.0022 | 0.0026 |
| F12 | 62080001 | 62180000 | 0.0023 | 0.0017 | 0.0019 |
| F12 | 62100001 | 62200000 | 0.0024 | 0.0018 | 0.0019 |
| F12 | 62120001 | 62220000 | 0.0020 | 0.0014 | 0.0013 |
| F12 | 62140001 | 62240000 | 0.0021 | 0.0013 | 0.0012 |
| F12 | 62160001 | 62260000 | 0.0020 | 0.0011 | 0.0010 |
| F12 | 62180001 | 62280000 | 0.0021 | 0.0012 | 0.0012 |
| F12 | 62200001 | 62300000 | 0.0020 | 0.0012 | 0.0013 |
| F12 | 62220001 | 62320000 | 0.0026 | 0.0018 | 0.0019 |
| F12 | 62240001 | 62340000 | 0.0023 | 0.0017 | 0.0019 |
| F12 | 62260001 | 62360000 | 0.0020 | 0.0014 | 0.0016 |
| F12 | 62280001 | 62380000 | 0.0019 | 0.0014 | 0.0015 |
| F12 | 62300001 | 62400000 | 0.0021 | 0.0017 | 0.0016 |
| F12 | 62320001 | 62420000 | 0.0014 | 0.0009 | 0.0008 |
| F12 | 62340001 | 62440000 | 0.0019 | 0.0012 | 0.0012 |
| F12 | 62360001 | 62460000 | 0.0032 | 0.0024 | 0.0024 |
| F12 | 62380001 | 62480000 | 0.0036 | 0.0027 | 0.0027 |
| F12 | 62400001 | 62500000 | 0.0035 | 0.0026 | 0.0028 |
| F12 | 62420001 | 62520000 | 0.0039 | 0.0029 | 0.0031 |
| F12 | 62440001 | 62540000 | 0.0034 | 0.0027 | 0.0029 |
| F12 | 62460001 | 62560000 | 0.0023 | 0.0015 | 0.0018 |
| F12 | 62480001 | 62580000 | 0.0019 | 0.0011 | 0.0013 |
| F12 | 62500001 | 62600000 | 0.0017 | 0.0008 | 0.0011 |
| F12 | 62520001 | 62620000 | 0.0025 | 0.0016 | 0.0020 |
| F12 | 62540001 | 62640000 | 0.0029 | 0.0021 | 0.0024 |
| F12 | 62560001 | 62660000 | 0.0032 | 0.0024 | 0.0027 |
| F12 | 62580001 | 62680000 | 0.0035 | 0.0027 | 0.0030 |
| F12 | 62600001 | 62700000 | 0.0037 | 0.0029 | 0.0030 |
| F12 | 62620001 | 62720000 | 0.0030 | 0.0024 | 0.0024 |
| F12 | 62640001 | 62740000 | 0.0026 | 0.0019 | 0.0019 |
| F12 | 62660001 | 62760000 | 0.0031 | 0.0024 | 0.0022 |
| F12 | 62680001 | 62780000 | 0.0029 | 0.0022 | 0.0021 |
| F12 | 62700001 | 62800000 | 0.0025 | 0.0017 | 0.0017 |
| F12 | 62720001 | 62820000 | 0.0021 | 0.0013 | 0.0013 |
| F12 | 62740001 | 62840000 | 0.0026 | 0.0017 | 0.0018 |
| F12 | 62760001 | 62860000 | 0.0019 | 0.0012 | 0.0014 |
| F12 | 62780001 | 62880000 | 0.0022 | 0.0015 | 0.0017 |
| F12 | 62800001 | 62900000 | 0.0024 | 0.0015 | 0.0018 |
| F12 | 62820001 | 62920000 | 0.0024 | 0.0014 | 0.0016 |
| F12 | 62840001 | 62940000 | 0.0029 | 0.0017 | 0.0020 |
| F12 | 62860001 | 62960000 | 0.0028 | 0.0014 | 0.0017 |
| F12 | 62880001 | 62980000 | 0.0026 | 0.0014 | 0.0017 |
| F12 | 62900001 | 63000000 | 0.0025 | 0.0014 | 0.0016 |
| F12 | 62920001 | 63020000 | 0.0035 | 0.0023 | 0.0026 |
| F12 | 62940001 | 63040000 | 0.0027 | 0.0019 | 0.0021 |
| F12 | 62960001 | 63060000 | 0.0034 | 0.0027 | 0.0029 |
| F12 | 62980001 | 63080000 | 0.0040 | 0.0033 | 0.0033 |
| F12 | 63000001 | 63100000 | 0.0042 | 0.0035 | 0.0035 |
| F12 | 63020001 | 63120000 | 0.0032 | 0.0025 | 0.0024 |
| F12 | 63040001 | 63140000 | 0.0029 | 0.0021 | 0.0021 |
| F12 | 63060001 | 63160000 | 0.0026 | 0.0018 | 0.0017 |
| F12 | 63080001 | 63180000 | 0.0026 | 0.0018 | 0.0018 |
| F12 | 63100001 | 63200000 | 0.0033 | 0.0025 | 0.0025 |
| F12 | 63120001 | 63220000 | 0.0035 | 0.0025 | 0.0026 |

| Chr | Start | End | both | A1 | A2 |
|---|---|---|---|---|---|
| F12 | 63140001 | 63240000 | 0.0038 | 0.0030 | 0.0029 |
| F12 | 63160001 | 63260000 | 0.0038 | 0.0032 | 0.0032 |
| F12 | 63180001 | 63280000 | 0.0029 | 0.0025 | 0.0023 |
| F12 | 63200001 | 63300000 | 0.0020 | 0.0017 | 0.0015 |
| F12 | 63220001 | 63320000 | 0.0018 | 0.0019 | 0.0016 |
| F12 | 63240001 | 63340000 | 0.0021 | 0.0019 | 0.0019 |
| F12 | 63260001 | 63360000 | 0.0018 | 0.0014 | 0.0014 |
| F12 | 63280001 | 63380000 | 0.0020 | 0.0014 | 0.0014 |
| F12 | 63300001 | 63400000 | 0.0024 | 0.0015 | 0.0016 |
| F12 | 63320001 | 63420000 | 0.0027 | 0.0015 | 0.0017 |
| F12 | 63340001 | 63440000 | 0.0028 | 0.0014 | 0.0016 |
| F12 | 63360001 | 63460000 | 0.0032 | 0.0019 | 0.0022 |
| F12 | 63380001 | 63480000 | 0.0040 | 0.0027 | 0.0030 |
| F12 | 63400001 | 63500000 | 0.0038 | 0.0025 | 0.0028 |
| F12 | 63420001 | 63520000 | 0.0037 | 0.0025 | 0.0027 |
| F12 | 63440001 | 63540000 | 0.0030 | 0.0022 | 0.0023 |
| F12 | 63460001 | 63560000 | 0.0025 | 0.0018 | 0.0019 |
| F12 | 63480001 | 63580000 | 0.0020 | 0.0014 | 0.0015 |
| F12 | 63500001 | 63600000 | 0.0021 | 0.0017 | 0.0017 |
| F12 | 63520001 | 63620000 | 0.0021 | 0.0016 | 0.0017 |
| F12 | 63540001 | 63640000 | 0.0034 | 0.0027 | 0.0028 |
| F12 | 63560001 | 63660000 | 0.0037 | 0.0028 | 0.0031 |
| F12 | 63580001 | 63680000 | 0.0035 | 0.0027 | 0.0030 |
| F12 | 63600001 | 63700000 | 0.0040 | 0.0031 | 0.0034 |
| F12 | 63620001 | 63720000 | 0.0039 | 0.0031 | 0.0033 |
| F12 | 63640001 | 63740000 | 0.0026 | 0.0018 | 0.0021 |
| F12 | 63660001 | 63760000 | 0.0027 | 0.0019 | 0.0021 |
| F12 | 63680001 | 63780000 | 0.0030 | 0.0019 | 0.0021 |
| F12 | 63700001 | 63800000 | 0.0027 | 0.0017 | 0.0020 |
| F12 | 63720001 | 63820000 | 0.0027 | 0.0017 | 0.0019 |
| F12 | 63740001 | 63840000 | 0.0032 | 0.0021 | 0.0023 |
| F12 | 63760001 | 63860000 | 0.0028 | 0.0018 | 0.0020 |
| F12 | 63780001 | 63880000 | 0.0027 | 0.0019 | 0.0019 |
| F12 | 63800001 | 63900000 | 0.0026 | 0.0017 | 0.0017 |
| F12 | 63820001 | 63920000 | 0.0024 | 0.0017 | 0.0017 |
| F12 | 63840001 | 63940000 | 0.0030 | 0.0023 | 0.0024 |
| F12 | 63860001 | 63960000 | 0.0032 | 0.0022 | 0.0023 |
| F12 | 63880001 | 63980000 | 0.0028 | 0.0019 | 0.0019 |
| F12 | 63900001 | 64000000 | 0.0033 | 0.0023 | 0.0025 |
| F12 | 63920001 | 64020000 | 0.0040 | 0.0029 | 0.0030 |
| F12 | 63940001 | 64040000 | 0.0034 | 0.0023 | 0.0026 |
| F12 | 63960001 | 64060000 | 0.0030 | 0.0021 | 0.0023 |
| F12 | 63980001 | 64080000 | 0.0036 | 0.0028 | 0.0028 |
| F12 | 64000001 | 64100000 | 0.0030 | 0.0024 | 0.0023 |
| F12 | 64020001 | 64120000 | 0.0025 | 0.0016 | 0.0016 |
| F12 | 64040001 | 64140000 | 0.0021 | 0.0013 | 0.0012 |
| F12 | 64060001 | 64160000 | 0.0026 | 0.0016 | 0.0014 |
| F12 | 64080001 | 64180000 | 0.0021 | 0.0008 | 0.0007 |
| F12 | 64100001 | 64200000 | 0.0021 | 0.0007 | 0.0006 |
| F12 | 64120001 | 64220000 | 0.0024 | 0.0010 | 0.0010 |
| F12 | 64140001 | 64240000 | 0.0026 | 0.0010 | 0.0009 |
| F12 | 64160001 | 64260000 | 0.0025 | 0.0012 | 0.0013 |
| F12 | 64180001 | 64280000 | 0.0025 | 0.0014 | 0.0014 |
| F12 | 64200001 | 64300000 | 0.0021 | 0.0013 | 0.0012 |
| F12 | 64220001 | 64320000 | 0.0017 | 0.0010 | 0.0009 |
| F12 | 64240001 | 64340000 | 0.0015 | 0.0010 | 0.0009 |
| F12 | 64260001 | 64360000 | 0.0013 | 0.0005 | 0.0005 |
| F12 | 64280001 | 64380000 | 0.0012 | 0.0004 | 0.0004 |

| Chr | Start | End | both | A1 | A2 |
|---|---|---|---|---|---|
| F12 | 64300001 | 64400000 | 0.0014 | 0.0005 | 0.0005 |
| F12 | 64320001 | 64420000 | 0.0019 | 0.0009 | 0.0010 |
| F12 | 64340001 | 64440000 | 0.0026 | 0.0015 | 0.0017 |
| F12 | 64360001 | 64460000 | 0.0026 | 0.0018 | 0.0019 |
| F12 | 64380001 | 64480000 | 0.0027 | 0.0020 | 0.0021 |
| F12 | 64400001 | 64500000 | 0.0027 | 0.0019 | 0.0021 |
| F12 | 64420001 | 64520000 | 0.0027 | 0.0018 | 0.0020 |
| F12 | 64440001 | 64540000 | 0.0021 | 0.0011 | 0.0013 |
| F12 | 64460001 | 64560000 | 0.0020 | 0.0010 | 0.0011 |
| F12 | 64480001 | 64580000 | 0.0018 | 0.0007 | 0.0009 |
| F12 | 64500001 | 64600000 | 0.0027 | 0.0016 | 0.0018 |
| F12 | 64520001 | 64620000 | 0.0025 | 0.0016 | 0.0017 |
| F12 | 64540001 | 64640000 | 0.0035 | 0.0024 | 0.0025 |
| F12 | 64560001 | 64660000 | 0.0040 | 0.0028 | 0.0030 |
| F12 | 64580001 | 64680000 | 0.0042 | 0.0032 | 0.0033 |
| F12 | 64600001 | 64700000 | 0.0035 | 0.0025 | 0.0026 |
| F12 | 64620001 | 64720000 | 0.0036 | 0.0024 | 0.0025 |
| F12 | 64640001 | 64740000 | 0.0027 | 0.0018 | 0.0018 |
| F12 | 64660001 | 64760000 | 0.0021 | 0.0012 | 0.0012 |
| F12 | 64680001 | 64780000 | 0.0018 | 0.0009 | 0.0009 |
| F12 | 64700001 | 64800000 | 0.0015 | 0.0006 | 0.0006 |
| F12 | 64720001 | 64820000 | 0.0015 | 0.0004 | 0.0005 |
| F12 | 64740001 | 64840000 | 0.0022 | 0.0009 | 0.0011 |
| F12 | 64760001 | 64860000 | 0.0023 | 0.0013 | 0.0014 |
| F12 | 64780001 | 64880000 | 0.0025 | 0.0015 | 0.0017 |
| F12 | 64800001 | 64900000 | 0.0024 | 0.0015 | 0.0016 |
| F12 | 64820001 | 64920000 | 0.0024 | 0.0016 | 0.0017 |
| F12 | 64840001 | 64940000 | 0.0015 | 0.0011 | 0.0011 |
| F12 | 64860001 | 64960000 | 0.0013 | 0.0009 | 0.0010 |
| F12 | 64880001 | 64980000 | 0.0010 | 0.0007 | 0.0007 |
| F12 | 64900001 | 65000000 | 0.0011 | 0.0008 | 0.0009 |
| F12 | 64920001 | 65020000 | 0.0015 | 0.0012 | 0.0014 |
| F12 | 64940001 | 65040000 | 0.0020 | 0.0017 | 0.0019 |
| F12 | 64960001 | 65060000 | 0.0030 | 0.0026 | 0.0028 |
| F12 | 64980001 | 65080000 | 0.0030 | 0.0025 | 0.0027 |
| F12 | 65000001 | 65100000 | 0.0033 | 0.0026 | 0.0027 |
| F12 | 65020001 | 65120000 | 0.0033 | 0.0025 | 0.0027 |
| F12 | 65040001 | 65140000 | 0.0034 | 0.0023 | 0.0027 |
| F12 | 65060001 | 65160000 | 0.0025 | 0.0014 | 0.0018 |
| F12 | 65080001 | 65180000 | 0.0027 | 0.0016 | 0.0020 |
| F12 | 65100001 | 65200000 | 0.0027 | 0.0015 | 0.0018 |
| F12 | 65120001 | 65220000 | 0.0023 | 0.0012 | 0.0014 |
| F12 | 65140001 | 65240000 | 0.0019 | 0.0010 | 0.0011 |
| F12 | 65160001 | 65260000 | 0.0021 | 0.0013 | 0.0014 |
| F12 | 65180001 | 65280000 | 0.0025 | 0.0017 | 0.0019 |
| F12 | 65200001 | 65300000 | 0.0029 | 0.0020 | 0.0025 |
| F12 | 65220001 | 65320000 | 0.0032 | 0.0024 | 0.0029 |
| F12 | 65240001 | 65340000 | 0.0032 | 0.0023 | 0.0029 |
| F12 | 65260001 | 65360000 | 0.0028 | 0.0020 | 0.0026 |
| F12 | 65280001 | 65380000 | 0.0025 | 0.0014 | 0.0024 |
| F12 | 65300001 | 65400000 | 0.0026 | 0.0014 | 0.0023 |
| F12 | 65320001 | 65420000 | 0.0024 | 0.0013 | 0.0021 |
| F12 | 65340001 | 65440000 | 0.0023 | 0.0011 | 0.0017 |
| F12 | 65360001 | 65460000 | 0.0036 | 0.0022 | 0.0028 |
| F12 | 65380001 | 65480000 | 0.0041 | 0.0029 | 0.0031 |
| F12 | 65400001 | 65500000 | 0.0041 | 0.0031 | 0.0033 |
| F12 | 65420001 | 65520000 | 0.0044 | 0.0035 | 0.0037 |
| F12 | 65440001 | 65540000 | 0.0044 | 0.0036 | 0.0039 |

| Chr | Start | End | both | A1 | A2 |
|---|---|---|---|---|---|
| F12 | 65460001 | 65560000 | 0.0040 | 0.0034 | 0.0037 |
| F12 | 65480001 | 65580000 | 0.0037 | 0.0031 | 0.0035 |
| F12 | 65500001 | 65600000 | 0.0038 | 0.0031 | 0.0035 |
| F12 | 65520001 | 65620000 | 0.0044 | 0.0032 | 0.0038 |
| F12 | 65540001 | 65640000 | 0.0047 | 0.0034 | 0.0039 |
| F12 | 65560001 | 65660000 | 0.0043 | 0.0030 | 0.0034 |
| F12 | 65580001 | 65680000 | 0.0042 | 0.0028 | 0.0032 |
| F12 | 65600001 | 65700000 | 0.0039 | 0.0027 | 0.0030 |
| F12 | 65620001 | 65720000 | 0.0029 | 0.0019 | 0.0022 |
| F12 | 65640001 | 65740000 | 0.0035 | 0.0026 | 0.0027 |
| F12 | 65660001 | 65760000 | 0.0030 | 0.0021 | 0.0023 |
| F12 | 65680001 | 65780000 | 0.0034 | 0.0024 | 0.0025 |
| F12 | 65700001 | 65800000 | 0.0036 | 0.0025 | 0.0026 |
| F12 | 65720001 | 65820000 | 0.0034 | 0.0025 | 0.0025 |
| F12 | 65740001 | 65840000 | 0.0025 | 0.0015 | 0.0016 |
| F12 | 65760001 | 65860000 | 0.0025 | 0.0015 | 0.0016 |
| F12 | 65780001 | 65880000 | 0.0025 | 0.0013 | 0.0015 |
| F12 | 65800001 | 65900000 | 0.0018 | 0.0007 | 0.0008 |
| F12 | 65820001 | 65920000 | 0.0020 | 0.0008 | 0.0008 |
| F12 | 65840001 | 65940000 | 0.0022 | 0.0010 | 0.0011 |
| F12 | 65860001 | 65960000 | 0.0023 | 0.0010 | 0.0012 |
| F12 | 65880001 | 65980000 | 0.0018 | 0.0007 | 0.0009 |
| F12 | 65900001 | 66000000 | 0.0019 | 0.0007 | 0.0009 |
| F12 | 65920001 | 66020000 | 0.0018 | 0.0006 | 0.0008 |
| F12 | 65940001 | 66040000 | 0.0016 | 0.0004 | 0.0005 |
| F12 | 65960001 | 66060000 | 0.0016 | 0.0006 | 0.0007 |
| F12 | 65980001 | 66080000 | 0.0018 | 0.0008 | 0.0008 |
| F12 | 66000001 | 66100000 | 0.0017 | 0.0009 | 0.0009 |
| F12 | 66020001 | 66120000 | 0.0023 | 0.0017 | 0.0017 |
| F12 | 66040001 | 66140000 | 0.0021 | 0.0019 | 0.0017 |
| F12 | 66060001 | 66160000 | 0.0024 | 0.0023 | 0.0020 |
| F12 | 66080001 | 66180000 | 0.0021 | 0.0021 | 0.0019 |
| F12 | 66100001 | 66200000 | 0.0026 | 0.0029 | 0.0024 |
| F12 | 66120001 | 66220000 | 0.0019 | 0.0023 | 0.0017 |
| F12 | 66140001 | 66240000 | 0.0019 | 0.0023 | 0.0017 |
| F12 | 66160001 | 66260000 | 0.0015 | 0.0019 | 0.0013 |
| F12 | 66180001 | 66280000 | 0.0012 | 0.0017 | 0.0010 |
| F12 | 66200001 | 66300000 | 0.0006 | 0.0010 | 0.0005 |
| F12 | 66220001 | 66320000 | 0.0005 | 0.0009 | 0.0003 |
| F12 | 66240001 | 66340000 | 0.0012 | 0.0014 | 0.0011 |
| F12 | 66260001 | 66360000 | 0.0013 | 0.0017 | 0.0012 |
| F12 | 66280001 | 66380000 | 0.0022 | 0.0024 | 0.0020 |
| F12 | 66300001 | 66400000 | 0.0025 | 0.0025 | 0.0023 |
| F12 | 66320001 | 66420000 | 0.0029 | 0.0028 | 0.0028 |
| F12 | 66340001 | 66440000 | 0.0027 | 0.0029 | 0.0026 |
| F12 | 66360001 | 66460000 | 0.0038 | 0.0038 | 0.0037 |
| F12 | 66380001 | 66480000 | 0.0035 | 0.0038 | 0.0033 |
| F12 | 66400001 | 66500000 | 0.0039 | 0.0040 | 0.0037 |
| F12 | 66420001 | 66520000 | 0.0039 | 0.0043 | 0.0037 |
| F12 | 66440001 | 66540000 | 0.0033 | 0.0036 | 0.0031 |
| F12 | 66460001 | 66560000 | 0.0023 | 0.0026 | 0.0022 |
| F12 | 66480001 | 66580000 | 0.0018 | 0.0020 | 0.0017 |
| F12 | 66500001 | 66600000 | 0.0020 | 0.0024 | 0.0019 |
| F12 | 66520001 | 66620000 | 0.0017 | 0.0021 | 0.0016 |
| F12 | 66540001 | 66640000 | 0.0022 | 0.0024 | 0.0020 |
| F12 | 66560001 | 66660000 | 0.0025 | 0.0029 | 0.0023 |
| F12 | 66580001 | 66680000 | 0.0026 | 0.0030 | 0.0024 |
| F12 | 66600001 | 66700000 | 0.0020 | 0.0025 | 0.0018 |

| Chr | Start | End | both | A1 | A2 |
|---|---|---|---|---|---|
| F12 | 66620001 | 66720000 | 0.0020 | 0.0024 | 0.0018 |
| F12 | 66640001 | 66740000 | 0.0017 | 0.0022 | 0.0015 |
| F12 | 66660001 | 66760000 | 0.0013 | 0.0018 | 0.0011 |
| F12 | 66680001 | 66780000 | 0.0017 | 0.0021 | 0.0015 |
| F12 | 66700001 | 66800000 | 0.0015 | 0.0019 | 0.0014 |
| F12 | 66720001 | 66820000 | 0.0015 | 0.0018 | 0.0014 |
| F12 | 66740001 | 66840000 | 0.0016 | 0.0020 | 0.0014 |
| F12 | 66760001 | 66860000 | 0.0014 | 0.0018 | 0.0013 |
| F12 | 66780001 | 66880000 | 0.0010 | 0.0016 | 0.0008 |
| F12 | 66800001 | 66900000 | 0.0011 | 0.0019 | 0.0008 |
| F12 | 66820001 | 66920000 | 0.0019 | 0.0029 | 0.0016 |
| F12 | 66840001 | 66940000 | 0.0019 | 0.0029 | 0.0016 |
| F12 | 66860001 | 66960000 | 0.0020 | 0.0028 | 0.0016 |
| F12 | 66880001 | 66980000 | 0.0019 | 0.0026 | 0.0017 |
| F12 | 66900001 | 67000000 | 0.0026 | 0.0028 | 0.0024 |
| F12 | 66920001 | 67020000 | 0.0026 | 0.0027 | 0.0024 |
| F12 | 66940001 | 67040000 | 0.0031 | 0.0029 | 0.0030 |
| F12 | 66960001 | 67060000 | 0.0033 | 0.0032 | 0.0031 |
| F12 | 66980001 | 67080000 | 0.0039 | 0.0040 | 0.0038 |
| F12 | 67000001 | 67100000 | 0.0036 | 0.0038 | 0.0034 |
| F12 | 67020001 | 67120000 | 0.0035 | 0.0035 | 0.0033 |
| F12 | 67040001 | 67140000 | 0.0041 | 0.0041 | 0.0039 |
| F12 | 67060001 | 67160000 | 0.0042 | 0.0040 | 0.0041 |
| F12 | 67080001 | 67180000 | 0.0040 | 0.0034 | 0.0039 |
| F12 | 67100001 | 67200000 | 0.0037 | 0.0033 | 0.0036 |
| F12 | 67120001 | 67220000 | 0.0030 | 0.0030 | 0.0029 |
| F12 | 67140001 | 67240000 | 0.0022 | 0.0024 | 0.0021 |
| F12 | 67160001 | 67260000 | 0.0027 | 0.0029 | 0.0025 |
| F12 | 67180001 | 67280000 | 0.0029 | 0.0032 | 0.0027 |
| F12 | 67200001 | 67300000 | 0.0030 | 0.0033 | 0.0028 |
| F12 | 67220001 | 67320000 | 0.0032 | 0.0033 | 0.0031 |
| F12 | 67240001 | 67340000 | 0.0029 | 0.0029 | 0.0028 |
| F12 | 67260001 | 67360000 | 0.0030 | 0.0030 | 0.0029 |
| F12 | 67280001 | 67380000 | 0.0027 | 0.0027 | 0.0026 |
| F12 | 67300001 | 67400000 | 0.0032 | 0.0031 | 0.0031 |
| F12 | 67320001 | 67420000 | 0.0036 | 0.0034 | 0.0035 |
| F12 | 67340001 | 67440000 | 0.0038 | 0.0035 | 0.0037 |
| F12 | 67360001 | 67460000 | 0.0036 | 0.0033 | 0.0035 |
| F12 | 67380001 | 67480000 | 0.0035 | 0.0031 | 0.0035 |
| F12 | 67400001 | 67500000 | 0.0033 | 0.0026 | 0.0033 |
| F12 | 67420001 | 67520000 | 0.0033 | 0.0029 | 0.0033 |
| F12 | 67440001 | 67540000 | 0.0029 | 0.0027 | 0.0028 |
| F12 | 67460001 | 67560000 | 0.0021 | 0.0020 | 0.0021 |
| F12 | 67480001 | 67580000 | 0.0021 | 0.0020 | 0.0020 |
| F12 | 67500001 | 67600000 | 0.0025 | 0.0027 | 0.0024 |
| F12 | 67520001 | 67620000 | 0.0020 | 0.0023 | 0.0019 |
| F12 | 67540001 | 67640000 | 0.0022 | 0.0023 | 0.0021 |
| F12 | 67560001 | 67660000 | 0.0024 | 0.0027 | 0.0023 |
| F12 | 67580001 | 67680000 | 0.0030 | 0.0033 | 0.0029 |
| F12 | 67600001 | 67700000 | 0.0033 | 0.0034 | 0.0032 |
| F12 | 67620001 | 67720000 | 0.0043 | 0.0043 | 0.0041 |
| F12 | 67640001 | 67740000 | 0.0046 | 0.0047 | 0.0044 |
| F12 | 67660001 | 67760000 | 0.0049 | 0.0049 | 0.0047 |
| F12 | 67680001 | 67780000 | 0.0039 | 0.0040 | 0.0037 |
| F12 | 67700001 | 67800000 | 0.0045 | 0.0046 | 0.0043 |
| F12 | 67720001 | 67820000 | 0.0044 | 0.0045 | 0.0042 |
| F12 | 67740001 | 67840000 | 0.0038 | 0.0040 | 0.0037 |
| F12 | 67760001 | 67860000 | 0.0043 | 0.0044 | 0.0041 |

| Chr | Start | End | both | A1 | A2 |
|---|---|---|---|---|---|
| F12 | 67780001 | 67880000 | 0.0050 | 0.0050 | 0.0048 |
| F12 | 67800001 | 67900000 | 0.0044 | 0.0046 | 0.0042 |
| F12 | 67820001 | 67920000 | 0.0037 | 0.0040 | 0.0035 |
| F12 | 67840001 | 67940000 | 0.0045 | 0.0046 | 0.0043 |
| F12 | 67860001 | 67960000 | 0.0036 | 0.0039 | 0.0033 |
| F12 | 67880001 | 67980000 | 0.0031 | 0.0035 | 0.0029 |
| F12 | 67900001 | 68000000 | 0.0026 | 0.0030 | 0.0024 |
| F12 | 67920001 | 68020000 | 0.0028 | 0.0031 | 0.0026 |
| F12 | 67940001 | 68040000 | 0.0021 | 0.0024 | 0.0020 |
| F12 | 67960001 | 68060000 | 0.0023 | 0.0025 | 0.0022 |
| F12 | 67980001 | 68080000 | 0.0025 | 0.0026 | 0.0024 |
| F12 | 68000001 | 68100000 | 0.0018 | 0.0021 | 0.0016 |
| F12 | 68020001 | 68120000 | 0.0012 | 0.0017 | 0.0010 |
| F12 | 68040001 | 68140000 | 0.0012 | 0.0017 | 0.0009 |
| F12 | 68060001 | 68160000 | 0.0011 | 0.0016 | 0.0008 |
| F12 | 68080001 | 68180000 | 0.0008 | 0.0012 | 0.0004 |
| F12 | 68100001 | 68200000 | 0.0010 | 0.0010 | 0.0004 |
| F12 | 68120001 | 68220000 | 0.0016 | 0.0011 | 0.0008 |
| F12 | 68140001 | 68240000 | 0.0023 | 0.0013 | 0.0014 |
| F12 | 68160001 | 68260000 | 0.0023 | 0.0013 | 0.0014 |
| F12 | 68180001 | 68280000 | 0.0023 | 0.0012 | 0.0014 |
| F12 | 68200001 | 68300000 | 0.0026 | 0.0015 | 0.0018 |
| F12 | 68220001 | 68320000 | 0.0027 | 0.0017 | 0.0020 |
| F12 | 68240001 | 68340000 | 0.0022 | 0.0012 | 0.0014 |
| F12 | 68260001 | 68360000 | 0.0025 | 0.0015 | 0.0017 |
| F12 | 68280001 | 68380000 | 0.0031 | 0.0020 | 0.0022 |
| F12 | 68300001 | 68400000 | 0.0029 | 0.0019 | 0.0020 |
| F12 | 68320001 | 68420000 | 0.0032 | 0.0021 | 0.0021 |
| F12 | 68340001 | 68440000 | 0.0039 | 0.0028 | 0.0028 |
| F12 | 68360001 | 68460000 | 0.0037 | 0.0027 | 0.0026 |
| F12 | 68380001 | 68480000 | 0.0036 | 0.0026 | 0.0025 |
| F12 | 68400001 | 68500000 | 0.0034 | 0.0024 | 0.0024 |
| F12 | 68420001 | 68520000 | 0.0028 | 0.0018 | 0.0018 |
| F12 | 68440001 | 68540000 | 0.0020 | 0.0011 | 0.0010 |
| F12 | 68460001 | 68560000 | 0.0017 | 0.0009 | 0.0009 |
| F12 | 68480001 | 68580000 | 0.0013 | 0.0006 | 0.0006 |
| F12 | 68500001 | 68600000 | 0.0014 | 0.0007 | 0.0007 |
| F12 | 68520001 | 68620000 | 0.0017 | 0.0009 | 0.0010 |
| F12 | 68540001 | 68640000 | 0.0017 | 0.0009 | 0.0010 |
| F12 | 68560001 | 68660000 | 0.0019 | 0.0011 | 0.0011 |
| F12 | 68580001 | 68680000 | 0.0022 | 0.0012 | 0.0013 |
| F12 | 68600001 | 68700000 | 0.0020 | 0.0010 | 0.0012 |
| F12 | 68620001 | 68720000 | 0.0018 | 0.0010 | 0.0011 |
| F12 | 68640001 | 68740000 | 0.0035 | 0.0013 | 0.0028 |
| F12 | 68660001 | 68760000 | 0.0036 | 0.0014 | 0.0028 |
| F12 | 68680001 | 68780000 | 0.0037 | 0.0016 | 0.0028 |
| F12 | 68700001 | 68800000 | 0.0037 | 0.0017 | 0.0029 |
| F12 | 68720001 | 68820000 | 0.0033 | 0.0015 | 0.0026 |
| F12 | 68740001 | 68840000 | 0.0022 | 0.0014 | 0.0013 |
| F12 | 68760001 | 68860000 | 0.0021 | 0.0012 | 0.0012 |
| F12 | 68780001 | 68880000 | 0.0021 | 0.0009 | 0.0013 |
| F12 | 68800001 | 68900000 | 0.0026 | 0.0011 | 0.0016 |
| F12 | 68820001 | 68920000 | 0.0030 | 0.0013 | 0.0019 |
| F12 | 68840001 | 68940000 | 0.0026 | 0.0013 | 0.0018 |
| F12 | 68860001 | 68960000 | 0.0027 | 0.0012 | 0.0019 |
| F12 | 68880001 | 68980000 | 0.0026 | 0.0012 | 0.0018 |
| F12 | 68900001 | 69000000 | 0.0024 | 0.0010 | 0.0016 |
| F12 | 68920001 | 69020000 | 0.0036 | 0.0023 | 0.0028 |

| Chr | Start | End | both | A1 | A2 |
|---|---|---|---|---|---|
| F12 | 68940001 | 69040000 | 0.0039 | 0.0025 | 0.0029 |
| F12 | 68960001 | 69060000 | 0.0041 | 0.0028 | 0.0032 |
| F12 | 68980001 | 69080000 | 0.0043 | 0.0030 | 0.0034 |
| F12 | 69000001 | 69100000 | 0.0049 | 0.0037 | 0.0041 |
| F12 | 69020001 | 69120000 | 0.0035 | 0.0023 | 0.0028 |
| F12 | 69040001 | 69140000 | 0.0039 | 0.0028 | 0.0033 |
| F12 | 69060001 | 69160000 | 0.0038 | 0.0029 | 0.0034 |
| F12 | 69080001 | 69180000 | 0.0035 | 0.0027 | 0.0032 |
| F12 | 69100001 | 69200000 | 0.0029 | 0.0022 | 0.0024 |
| F12 | 69120001 | 69220000 | 0.0039 | 0.0028 | 0.0032 |
| F12 | 69140001 | 69240000 | 0.0037 | 0.0024 | 0.0030 |
| F12 | 69160001 | 69260000 | 0.0033 | 0.0021 | 0.0025 |
| F12 | 69180001 | 69280000 | 0.0036 | 0.0023 | 0.0028 |
| F12 | 69200001 | 69300000 | 0.0035 | 0.0019 | 0.0026 |
| F12 | 69220001 | 69320000 | 0.0021 | 0.0010 | 0.0015 |
| F12 | 69240001 | 69340000 | 0.0023 | 0.0013 | 0.0015 |
| F12 | 69260001 | 69360000 | 0.0025 | 0.0013 | 0.0015 |
| F12 | 69280001 | 69380000 | 0.0031 | 0.0019 | 0.0022 |
| F12 | 69300001 | 69400000 | 0.0033 | 0.0022 | 0.0024 |
| F12 | 69320001 | 69420000 | 0.0038 | 0.0028 | 0.0030 |
| F12 | 69340001 | 69440000 | 0.0030 | 0.0023 | 0.0024 |
| F12 | 69360001 | 69460000 | 0.0028 | 0.0024 | 0.0025 |
| F12 | 69380001 | 69480000 | 0.0022 | 0.0020 | 0.0018 |
| F12 | 69400001 | 69500000 | 0.0027 | 0.0024 | 0.0024 |
| F12 | 69420001 | 69520000 | 0.0037 | 0.0030 | 0.0033 |
| F12 | 69440001 | 69540000 | 0.0037 | 0.0029 | 0.0034 |
| F12 | 69460001 | 69560000 | 0.0042 | 0.0032 | 0.0038 |
| F12 | 69480001 | 69580000 | 0.0042 | 0.0032 | 0.0039 |
| F12 | 69500001 | 69600000 | 0.0035 | 0.0024 | 0.0032 |
| F12 | 69520001 | 69620000 | 0.0023 | 0.0012 | 0.0019 |
| F12 | 69540001 | 69640000 | 0.0023 | 0.0012 | 0.0018 |
| F12 | 69560001 | 69660000 | 0.0022 | 0.0010 | 0.0016 |
| F12 | 69580001 | 69680000 | 0.0020 | 0.0008 | 0.0014 |
| F12 | 69600001 | 69700000 | 0.0026 | 0.0016 | 0.0021 |
| F12 | 69620001 | 69720000 | 0.0028 | 0.0016 | 0.0022 |
| F12 | 69640001 | 69740000 | 0.0031 | 0.0020 | 0.0026 |
| F12 | 69660001 | 69760000 | 0.0033 | 0.0022 | 0.0028 |
| F12 | 69680001 | 69780000 | 0.0042 | 0.0031 | 0.0036 |
| F12 | 69700001 | 69800000 | 0.0041 | 0.0026 | 0.0033 |
| F12 | 69720001 | 69820000 | 0.0038 | 0.0026 | 0.0032 |
| F12 | 69740001 | 69840000 | 0.0038 | 0.0022 | 0.0029 |
| F12 | 69760001 | 69860000 | 0.0035 | 0.0018 | 0.0025 |
| F12 | 69780001 | 69880000 | 0.0031 | 0.0014 | 0.0022 |
| F12 | 69800001 | 69900000 | 0.0025 | 0.0011 | 0.0017 |
| F12 | 69820001 | 69920000 | 0.0026 | 0.0013 | 0.0019 |
| F12 | 69840001 | 69940000 | 0.0024 | 0.0013 | 0.0018 |
| F12 | 69860001 | 69960000 | 0.0024 | 0.0013 | 0.0018 |
| F12 | 69880001 | 69980000 | 0.0018 | 0.0008 | 0.0011 |
| F12 | 69900001 | 70000000 | 0.0019 | 0.0008 | 0.0011 |
| F12 | 69920001 | 70020000 | 0.0018 | 0.0007 | 0.0010 |
| F12 | 69940001 | 70040000 | 0.0017 | 0.0006 | 0.0009 |
| F12 | 69960001 | 70060000 | 0.0018 | 0.0009 | 0.0011 |
| F12 | 69980001 | 70080000 | 0.0019 | 0.0009 | 0.0012 |
| F12 | 70000001 | 70100000 | 0.0018 | 0.0009 | 0.0011 |
| F12 | 70020001 | 70120000 | 0.0025 | 0.0015 | 0.0019 |
| F12 | 70040001 | 70140000 | 0.0037 | 0.0029 | 0.0032 |
| F12 | 70060001 | 70160000 | 0.0034 | 0.0024 | 0.0029 |
| F12 | 70080001 | 70180000 | 0.0042 | 0.0029 | 0.0034 |

| Chr | Start | End | both | A1 | A2 |
|---|---|---|---|---|---|
| F12 | 70100001 | 70200000 | 0.0046 | 0.0032 | 0.0037 |
| F12 | 70120001 | 70220000 | 0.0042 | 0.0028 | 0.0030 |
| F12 | 70140001 | 70240000 | 0.0036 | 0.0019 | 0.0024 |
| F12 | 70160001 | 70260000 | 0.0043 | 0.0026 | 0.0031 |
| F12 | 70180001 | 70280000 | 0.0037 | 0.0021 | 0.0026 |
| F12 | 70200001 | 70300000 | 0.0041 | 0.0027 | 0.0031 |
| F12 | 70220001 | 70320000 | 0.0038 | 0.0025 | 0.0029 |
| F12 | 70240001 | 70340000 | 0.0033 | 0.0021 | 0.0025 |
| F12 | 70260001 | 70360000 | 0.0034 | 0.0021 | 0.0026 |
| F12 | 70280001 | 70380000 | 0.0035 | 0.0023 | 0.0028 |
| F12 | 70300001 | 70400000 | 0.0035 | 0.0022 | 0.0028 |
| F12 | 70320001 | 70420000 | 0.0039 | 0.0026 | 0.0031 |
| F12 | 70340001 | 70440000 | 0.0050 | 0.0039 | 0.0044 |
| F12 | 70360001 | 70460000 | 0.0043 | 0.0034 | 0.0038 |
| F12 | 70380001 | 70480000 | 0.0043 | 0.0035 | 0.0038 |
| F12 | 70400001 | 70500000 | 0.0047 | 0.0040 | 0.0042 |
| F12 | 70420001 | 70520000 | 0.0044 | 0.0035 | 0.0038 |
| F12 | 70440001 | 70540000 | 0.0031 | 0.0021 | 0.0024 |
| F12 | 70460001 | 70560000 | 0.0031 | 0.0020 | 0.0023 |
| F12 | 70480001 | 70580000 | 0.0029 | 0.0016 | 0.0020 |
| F12 | 70500001 | 70600000 | 0.0017 | 0.0003 | 0.0006 |
| F12 | 70520001 | 70620000 | 0.0019 | 0.0006 | 0.0008 |
| F12 | 70540001 | 70640000 | 0.0024 | 0.0011 | 0.0012 |
| F12 | 70560001 | 70660000 | 0.0043 | 0.0035 | 0.0031 |
| F12 | 70580001 | 70680000 | 0.0040 | 0.0035 | 0.0031 |
| F12 | 70600001 | 70700000 | 0.0038 | 0.0035 | 0.0030 |
| F12 | 70620001 | 70720000 | 0.0038 | 0.0033 | 0.0028 |
| F12 | 70640001 | 70740000 | 0.0039 | 0.0033 | 0.0029 |
| F12 | 70660001 | 70760000 | 0.0024 | 0.0016 | 0.0015 |
| F12 | 70680001 | 70780000 | 0.0027 | 0.0017 | 0.0017 |
| F12 | 70700001 | 70800000 | 0.0042 | 0.0034 | 0.0033 |
| F12 | 70720001 | 70820000 | 0.0045 | 0.0037 | 0.0035 |
| F12 | 70740001 | 70840000 | 0.0038 | 0.0031 | 0.0029 |
| F12 | 70760001 | 70860000 | 0.0038 | 0.0032 | 0.0029 |
| F12 | 70780001 | 70880000 | 0.0047 | 0.0039 | 0.0037 |
| F12 | 70800001 | 70900000 | 0.0040 | 0.0033 | 0.0031 |
| F12 | 70820001 | 70920000 | 0.0036 | 0.0028 | 0.0028 |
| F12 | 70840001 | 70940000 | 0.0041 | 0.0032 | 0.0032 |
| F12 | 70860001 | 70960000 | 0.0039 | 0.0025 | 0.0028 |
| F12 | 70880001 | 70980000 | 0.0033 | 0.0023 | 0.0024 |
| F12 | 70900001 | 71000000 | 0.0032 | 0.0018 | 0.0020 |
| F12 | 70920001 | 71020000 | 0.0034 | 0.0020 | 0.0022 |
| F12 | 70940001 | 71040000 | 0.0031 | 0.0016 | 0.0018 |
| F12 | 70960001 | 71060000 | 0.0029 | 0.0017 | 0.0018 |
| F12 | 70980001 | 71080000 | 0.0025 | 0.0011 | 0.0013 |
| F12 | 71000001 | 71100000 | 0.0019 | 0.0007 | 0.0009 |
| F12 | 71020001 | 71120000 | 0.0019 | 0.0007 | 0.0010 |
| F12 | 71040001 | 71140000 | 0.0019 | 0.0008 | 0.0011 |
| F12 | 71060001 | 71160000 | 0.0021 | 0.0010 | 0.0012 |
| F12 | 71080001 | 71180000 | 0.0022 | 0.0013 | 0.0015 |
| F12 | 71100001 | 71200000 | 0.0030 | 0.0023 | 0.0024 |
| F12 | 71120001 | 71220000 | 0.0035 | 0.0028 | 0.0028 |
| F12 | 71140001 | 71240000 | 0.0035 | 0.0027 | 0.0026 |
| F12 | 71160001 | 71260000 | 0.0033 | 0.0024 | 0.0024 |
| F12 | 71180001 | 71280000 | 0.0035 | 0.0024 | 0.0025 |
| F12 | 71200001 | 71300000 | 0.0026 | 0.0014 | 0.0015 |
| F12 | 71220001 | 71320000 | 0.0019 | 0.0007 | 0.0009 |
| F12 | 71240001 | 71340000 | 0.0018 | 0.0007 | 0.0008 |

| Chr | Start | End | both | A1 | A2 |
|---|---|---|---|---|---|
| F12 | 71260001 | 71360000 | 0.0017 | 0.0009 | 0.0009 |
| F12 | 71280001 | 71380000 | 0.0015 | 0.0008 | 0.0007 |
| F12 | 71300001 | 71400000 | 0.0016 | 0.0010 | 0.0010 |
| F12 | 71320001 | 71420000 | 0.0021 | 0.0015 | 0.0015 |
| F12 | 71340001 | 71440000 | 0.0026 | 0.0018 | 0.0019 |
| F12 | 71360001 | 71460000 | 0.0026 | 0.0016 | 0.0018 |
| F12 | 71380001 | 71480000 | 0.0023 | 0.0014 | 0.0017 |
| F12 | 71400001 | 71500000 | 0.0021 | 0.0013 | 0.0015 |
| F12 | 71420001 | 71520000 | 0.0015 | 0.0007 | 0.0009 |
| F12 | 71440001 | 71540000 | 0.0010 | 0.0004 | 0.0006 |
| F12 | 71460001 | 71560000 | 0.0014 | 0.0004 | 0.0008 |
| F12 | 71480001 | 71580000 | 0.0016 | 0.0007 | 0.0010 |
| F12 | 71500001 | 71600000 | 0.0021 | 0.0012 | 0.0015 |
| F12 | 71520001 | 71620000 | 0.0028 | 0.0019 | 0.0022 |
| F12 | 71540001 | 71640000 | 0.0028 | 0.0020 | 0.0023 |
| F12 | 71560001 | 71660000 | 0.0032 | 0.0026 | 0.0028 |
| F12 | 71580001 | 71680000 | 0.0031 | 0.0023 | 0.0026 |
| F12 | 71600001 | 71700000 | 0.0029 | 0.0021 | 0.0024 |
| F12 | 71620001 | 71720000 | 0.0025 | 0.0015 | 0.0018 |
| F12 | 71640001 | 71740000 | 0.0030 | 0.0019 | 0.0023 |
| F12 | 71660001 | 71760000 | 0.0024 | 0.0014 | 0.0018 |
| F12 | 71680001 | 71780000 | 0.0020 | 0.0012 | 0.0015 |
| F12 | 71700001 | 71800000 | 0.0024 | 0.0016 | 0.0020 |
| F12 | 71720001 | 71820000 | 0.0021 | 0.0015 | 0.0018 |
| F12 | 71740001 | 71840000 | 0.0014 | 0.0011 | 0.0012 |
| F12 | 71760001 | 71860000 | 0.0030 | 0.0027 | 0.0028 |
| F12 | 71780001 | 71880000 | 0.0037 | 0.0033 | 0.0035 |
| F12 | 71800001 | 71900000 | 0.0032 | 0.0027 | 0.0029 |
| F12 | 71820001 | 71920000 | 0.0036 | 0.0030 | 0.0033 |
| F12 | 71840001 | 71940000 | 0.0040 | 0.0033 | 0.0036 |
| F12 | 71860001 | 71960000 | 0.0031 | 0.0024 | 0.0028 |
| F12 | 71880001 | 71980000 | 0.0028 | 0.0021 | 0.0025 |
| F12 | 71900001 | 72000000 | 0.0029 | 0.0023 | 0.0027 |
| F12 | 71920001 | 72020000 | 0.0027 | 0.0022 | 0.0025 |
| F12 | 71940001 | 72040000 | 0.0027 | 0.0019 | 0.0024 |
| F12 | 71960001 | 72060000 | 0.0028 | 0.0018 | 0.0023 |
| F12 | 71980001 | 72080000 | 0.0034 | 0.0021 | 0.0028 |
| F12 | 72000001 | 72100000 | 0.0035 | 0.0020 | 0.0026 |
| F12 | 72020001 | 72120000 | 0.0034 | 0.0018 | 0.0025 |
| F12 | 72040001 | 72140000 | 0.0035 | 0.0023 | 0.0028 |
| F12 | 72060001 | 72160000 | 0.0038 | 0.0023 | 0.0032 |
| F12 | 72080001 | 72180000 | 0.0033 | 0.0021 | 0.0027 |
| F12 | 72100001 | 72200000 | 0.0033 | 0.0023 | 0.0030 |
| F12 | 72120001 | 72220000 | 0.0036 | 0.0026 | 0.0033 |
| F12 | 72140001 | 72240000 | 0.0037 | 0.0024 | 0.0032 |
| F12 | 72160001 | 72260000 | 0.0026 | 0.0018 | 0.0022 |
| F12 | 72180001 | 72280000 | 0.0027 | 0.0016 | 0.0019 |
| F12 | 72200001 | 72300000 | 0.0029 | 0.0016 | 0.0019 |
| F12 | 72220001 | 72320000 | 0.0030 | 0.0018 | 0.0020 |
| F12 | 72240001 | 72340000 | 0.0025 | 0.0014 | 0.0016 |
| F12 | 72260001 | 72360000 | 0.0023 | 0.0015 | 0.0016 |
| F12 | 72280001 | 72380000 | 0.0024 | 0.0017 | 0.0020 |
| F12 | 72300001 | 72400000 | 0.0022 | 0.0016 | 0.0019 |
| F12 | 72320001 | 72420000 | 0.0020 | 0.0015 | 0.0018 |
| F12 | 72340001 | 72440000 | 0.0020 | 0.0014 | 0.0017 |
| F12 | 72360001 | 72460000 | 0.0025 | 0.0017 | 0.0022 |
| F12 | 72380001 | 72480000 | 0.0025 | 0.0016 | 0.0020 |
| F12 | 72400001 | 72500000 | 0.0023 | 0.0012 | 0.0017 |

| Chr | Start | End | both | A1 | A2 |
|---|---|---|---|---|---|
| F12 | 72420001 | 72520000 | 0.0021 | 0.0008 | 0.0012 |
| F12 | 72440001 | 72540000 | 0.0019 | 0.0008 | 0.0011 |
| F12 | 72460001 | 72560000 | 0.0017 | 0.0008 | 0.0011 |
| F12 | 72480001 | 72580000 | 0.0019 | 0.0012 | 0.0015 |
| F12 | 72500001 | 72600000 | 0.0025 | 0.0016 | 0.0020 |
| F12 | 72520001 | 72620000 | 0.0022 | 0.0016 | 0.0020 |
| F12 | 72540001 | 72640000 | 0.0022 | 0.0016 | 0.0020 |
| F12 | 72560001 | 72660000 | 0.0020 | 0.0013 | 0.0016 |
| F12 | 72580001 | 72680000 | 0.0013 | 0.0006 | 0.0008 |
| F12 | 72600001 | 72700000 | 0.0009 | 0.0002 | 0.0003 |
| F12 | 72620001 | 72720000 | 0.0029 | 0.0020 | 0.0022 |
| F12 | 72640001 | 72740000 | 0.0030 | 0.0020 | 0.0022 |
| F12 | 72660001 | 72760000 | 0.0045 | 0.0035 | 0.0036 |
| F12 | 72680001 | 72780000 | 0.0054 | 0.0044 | 0.0047 |
| F12 | 72700001 | 72800000 | 0.0061 | 0.0051 | 0.0054 |
| F12 | 72720001 | 72820000 | 0.0047 | 0.0039 | 0.0041 |
| F12 | 72740001 | 72840000 | 0.0048 | 0.0040 | 0.0042 |
| F12 | 72760001 | 72860000 | 0.0040 | 0.0031 | 0.0032 |
| F12 | 72780001 | 72880000 | 0.0029 | 0.0020 | 0.0021 |
| F12 | 72800001 | 72900000 | 0.0022 | 0.0013 | 0.0014 |
| F12 | 72820001 | 72920000 | 0.0020 | 0.0011 | 0.0011 |
| F12 | 72840001 | 72940000 | 0.0024 | 0.0014 | 0.0017 |
| F12 | 72860001 | 72960000 | 0.0022 | 0.0012 | 0.0016 |
| F12 | 72880001 | 72980000 | 0.0029 | 0.0020 | 0.0024 |
| F12 | 72900001 | 73000000 | 0.0034 | 0.0025 | 0.0029 |
| F12 | 72920001 | 73020000 | 0.0033 | 0.0022 | 0.0027 |
| F12 | 72940001 | 73040000 | 0.0029 | 0.0019 | 0.0022 |
| F12 | 72960001 | 73060000 | 0.0026 | 0.0017 | 0.0019 |
| F12 | 72980001 | 73080000 | 0.0022 | 0.0011 | 0.0013 |
| F12 | 73000001 | 73100000 | 0.0016 | 0.0006 | 0.0008 |
| F12 | 73020001 | 73120000 | 0.0018 | 0.0007 | 0.0010 |
| F12 | 73040001 | 73140000 | 0.0017 | 0.0005 | 0.0008 |
| F12 | 73060001 | 73160000 | 0.0016 | 0.0005 | 0.0008 |
| F12 | 73080001 | 73180000 | 0.0018 | 0.0005 | 0.0009 |
| F12 | 73100001 | 73200000 | 0.0019 | 0.0006 | 0.0010 |
| F12 | 73120001 | 73220000 | 0.0020 | 0.0006 | 0.0009 |
| F12 | 73140001 | 73240000 | 0.0025 | 0.0013 | 0.0017 |
| F12 | 73160001 | 73260000 | 0.0023 | 0.0013 | 0.0016 |
| F12 | 73180001 | 73280000 | 0.0019 | 0.0012 | 0.0015 |
| F12 | 73200001 | 73300000 | 0.0019 | 0.0013 | 0.0016 |
| F12 | 73220001 | 73320000 | 0.0014 | 0.0012 | 0.0014 |
| F12 | 73240001 | 73340000 | 0.0007 | 0.0006 | 0.0007 |
| F12 | 73260001 | 73360000 | 0.0018 | 0.0015 | 0.0017 |
| F12 | 73280001 | 73380000 | 0.0018 | 0.0015 | 0.0018 |
| F12 | 73300001 | 73400000 | 0.0016 | 0.0014 | 0.0016 |
| F12 | 73320001 | 73420000 | 0.0018 | 0.0016 | 0.0018 |
| F12 | 73340001 | 73440000 | 0.0019 | 0.0016 | 0.0019 |
| F12 | 73360001 | 73460000 | 0.0016 | 0.0014 | 0.0015 |
| F12 | 73380001 | 73480000 | 0.0017 | 0.0015 | 0.0016 |
| F12 | 73400001 | 73500000 | 0.0020 | 0.0016 | 0.0018 |
| F12 | 73420001 | 73520000 | 0.0024 | 0.0020 | 0.0022 |
| F12 | 73440001 | 73540000 | 0.0029 | 0.0025 | 0.0027 |
| F12 | 73460001 | 73560000 | 0.0026 | 0.0021 | 0.0022 |
| F12 | 73480001 | 73580000 | 0.0029 | 0.0021 | 0.0023 |
| F12 | 73500001 | 73600000 | 0.0035 | 0.0026 | 0.0028 |
| F12 | 73520001 | 73620000 | 0.0036 | 0.0025 | 0.0027 |
| F12 | 73540001 | 73640000 | 0.0034 | 0.0022 | 0.0024 |
| F12 | 73560001 | 73660000 | 0.0034 | 0.0023 | 0.0025 |

| Chr | Start | End | both | A1 | A2 |
|---|---|---|---|---|---|
| F12 | 73580001 | 73680000 | 0.0033 | 0.0022 | 0.0024 |
| F12 | 73600001 | 73700000 | 0.0024 | 0.0016 | 0.0016 |
| F12 | 73620001 | 73720000 | 0.0022 | 0.0014 | 0.0015 |
| F12 | 73640001 | 73740000 | 0.0026 | 0.0018 | 0.0018 |
| F12 | 73660001 | 73760000 | 0.0023 | 0.0016 | 0.0016 |
| F12 | 73680001 | 73780000 | 0.0024 | 0.0016 | 0.0017 |
| F12 | 73700001 | 73800000 | 0.0030 | 0.0016 | 0.0019 |
| F12 | 73720001 | 73820000 | 0.0027 | 0.0013 | 0.0017 |
| F12 | 73740001 | 73840000 | 0.0026 | 0.0011 | 0.0016 |
| F12 | 73760001 | 73860000 | 0.0029 | 0.0012 | 0.0017 |
| F12 | 73780001 | 73880000 | 0.0034 | 0.0014 | 0.0022 |
| F12 | 73800001 | 73900000 | 0.0035 | 0.0019 | 0.0025 |
| F12 | 73820001 | 73920000 | 0.0041 | 0.0023 | 0.0030 |
| F12 | 73840001 | 73940000 | 0.0043 | 0.0026 | 0.0032 |
| F12 | 73860001 | 73960000 | 0.0046 | 0.0028 | 0.0036 |
| F12 | 73880001 | 73980000 | 0.0040 | 0.0027 | 0.0031 |
| F12 | 73900001 | 74000000 | 0.0037 | 0.0026 | 0.0030 |
| F12 | 73920001 | 74020000 | 0.0033 | 0.0024 | 0.0027 |
| F12 | 73940001 | 74040000 | 0.0023 | 0.0016 | 0.0018 |
| F12 | 73960001 | 74060000 | 0.0023 | 0.0013 | 0.0017 |
| F12 | 73980001 | 74080000 | 0.0028 | 0.0015 | 0.0019 |
| F12 | 74000001 | 74100000 | 0.0027 | 0.0014 | 0.0018 |
| F12 | 74020001 | 74120000 | 0.0029 | 0.0014 | 0.0017 |
| F12 | 74040001 | 74140000 | 0.0037 | 0.0022 | 0.0026 |
| F12 | 74060001 | 74160000 | 0.0035 | 0.0022 | 0.0024 |
| F12 | 74080001 | 74180000 | 0.0030 | 0.0018 | 0.0021 |
| F12 | 74100001 | 74200000 | 0.0029 | 0.0019 | 0.0021 |
| F12 | 74120001 | 74220000 | 0.0027 | 0.0018 | 0.0022 |
| F12 | 74140001 | 74240000 | 0.0022 | 0.0013 | 0.0015 |
| F12 | 74160001 | 74260000 | 0.0027 | 0.0018 | 0.0021 |
| F12 | 74180001 | 74280000 | 0.0030 | 0.0020 | 0.0022 |
| F12 | 74200001 | 74300000 | 0.0034 | 0.0023 | 0.0026 |
| F12 | 74220001 | 74320000 | 0.0031 | 0.0019 | 0.0023 |
| F12 | 74240001 | 74340000 | 0.0028 | 0.0017 | 0.0020 |
| F12 | 74260001 | 74360000 | 0.0022 | 0.0011 | 0.0014 |
| F12 | 74280001 | 74380000 | 0.0023 | 0.0010 | 0.0014 |
| F12 | 74300001 | 74400000 | 0.0021 | 0.0011 | 0.0013 |
| F12 | 74320001 | 74420000 | 0.0028 | 0.0018 | 0.0021 |
| F12 | 74340001 | 74440000 | 0.0039 | 0.0028 | 0.0032 |
| F12 | 74360001 | 74460000 | 0.0038 | 0.0029 | 0.0032 |
| F12 | 74380001 | 74480000 | 0.0034 | 0.0027 | 0.0029 |
| F12 | 74400001 | 74500000 | 0.0029 | 0.0022 | 0.0025 |
| F12 | 74420001 | 74520000 | 0.0022 | 0.0015 | 0.0018 |
| F12 | 74440001 | 74540000 | 0.0019 | 0.0012 | 0.0015 |
| F12 | 74460001 | 74560000 | 0.0022 | 0.0013 | 0.0017 |
| F12 | 74480001 | 74580000 | 0.0024 | 0.0013 | 0.0017 |
| F12 | 74500001 | 74600000 | 0.0030 | 0.0018 | 0.0022 |
| F12 | 74520001 | 74620000 | 0.0037 | 0.0025 | 0.0029 |
| F12 | 74540001 | 74640000 | 0.0030 | 0.0020 | 0.0023 |
| F12 | 74560001 | 74660000 | 0.0028 | 0.0021 | 0.0023 |
| F12 | 74580001 | 74680000 | 0.0028 | 0.0022 | 0.0024 |
| F12 | 74600001 | 74700000 | 0.0022 | 0.0017 | 0.0018 |
| F12 | 74620001 | 74720000 | 0.0014 | 0.0009 | 0.0010 |
| F12 | 74640001 | 74740000 | 0.0021 | 0.0015 | 0.0017 |
| F12 | 74660001 | 74760000 | 0.0020 | 0.0013 | 0.0016 |
| F12 | 74680001 | 74780000 | 0.0022 | 0.0014 | 0.0016 |
| F12 | 74700001 | 74800000 | 0.0021 | 0.0014 | 0.0016 |
| F12 | 74720001 | 74820000 | 0.0022 | 0.0014 | 0.0016 |

| Chr | Start | End | both | A1 | A2 |
|---|---|---|---|---|---|
| F12 | 74740001 | 74840000 | 0.0026 | 0.0017 | 0.0018 |
| F12 | 74760001 | 74860000 | 0.0025 | 0.0015 | 0.0016 |
| F12 | 74780001 | 74880000 | 0.0030 | 0.0020 | 0.0019 |
| F12 | 74800001 | 74900000 | 0.0032 | 0.0022 | 0.0020 |
| F12 | 74820001 | 74920000 | 0.0034 | 0.0023 | 0.0022 |
| F12 | 74840001 | 74940000 | 0.0025 | 0.0014 | 0.0014 |
| F12 | 74860001 | 74960000 | 0.0026 | 0.0018 | 0.0018 |
| F12 | 74880001 | 74980000 | 0.0020 | 0.0011 | 0.0012 |
| F12 | 74900001 | 75000000 | 0.0018 | 0.0009 | 0.0011 |
| F12 | 74920001 | 75020000 | 0.0020 | 0.0010 | 0.0013 |
| F12 | 74940001 | 75040000 | 0.0018 | 0.0009 | 0.0012 |
| F12 | 74960001 | 75060000 | 0.0016 | 0.0007 | 0.0010 |
| F12 | 74980001 | 75080000 | 0.0016 | 0.0006 | 0.0009 |
| F12 | 75000001 | 75100000 | 0.0018 | 0.0009 | 0.0012 |
| F12 | 75020001 | 75120000 | 0.0021 | 0.0013 | 0.0014 |
| F12 | 75040001 | 75140000 | 0.0022 | 0.0016 | 0.0016 |
| F12 | 75060001 | 75160000 | 0.0027 | 0.0020 | 0.0022 |
| F12 | 75080001 | 75180000 | 0.0026 | 0.0022 | 0.0023 |
| F12 | 75100001 | 75200000 | 0.0026 | 0.0024 | 0.0025 |
| F12 | 75120001 | 75220000 | 0.0024 | 0.0021 | 0.0023 |
| F12 | 75140001 | 75240000 | 0.0027 | 0.0023 | 0.0027 |
| F12 | 75160001 | 75260000 | 0.0020 | 0.0017 | 0.0019 |
| F12 | 75180001 | 75280000 | 0.0020 | 0.0017 | 0.0019 |
| F12 | 75200001 | 75300000 | 0.0015 | 0.0013 | 0.0015 |
| F12 | 75220001 | 75320000 | 0.0010 | 0.0008 | 0.0010 |
| F12 | 75240001 | 75340000 | 0.0003 | 0.0002 | 0.0003 |
| F12 | 75260001 | 75360000 | 0.0004 | 0.0003 | 0.0004 |
| F12 | 75280001 | 75380000 | 0.0004 | 0.0003 | 0.0004 |
| F12 | 75300001 | 75400000 | 0.0009 | 0.0006 | 0.0008 |
| F12 | 75320001 | 75420000 | 0.0015 | 0.0010 | 0.0013 |
| F12 | 75340001 | 75440000 | 0.0022 | 0.0015 | 0.0019 |
| F12 | 75360001 | 75460000 | 0.0027 | 0.0018 | 0.0022 |
| F12 | 75380001 | 75480000 | 0.0030 | 0.0021 | 0.0024 |
| F12 | 75400001 | 75500000 | 0.0027 | 0.0018 | 0.0020 |
| F12 | 75420001 | 75520000 | 0.0024 | 0.0014 | 0.0015 |
| F12 | 75440001 | 75540000 | 0.0019 | 0.0010 | 0.0010 |
| F12 | 75460001 | 75560000 | 0.0016 | 0.0006 | 0.0007 |
| F12 | 75480001 | 75580000 | 0.0013 | 0.0002 | 0.0003 |
| F12 | 75500001 | 75600000 | 0.0014 | 0.0002 | 0.0004 |
| F12 | 75520001 | 75620000 | 0.0015 | 0.0004 | 0.0006 |
| F12 | 75540001 | 75640000 | 0.0015 | 0.0003 | 0.0006 |
| F12 | 75560001 | 75660000 | 0.0020 | 0.0008 | 0.0010 |
| F12 | 75580001 | 75680000 | 0.0024 | 0.0011 | 0.0014 |
| F12 | 75600001 | 75700000 | 0.0024 | 0.0014 | 0.0016 |
| F12 | 75620001 | 75720000 | 0.0024 | 0.0012 | 0.0014 |
| F12 | 75640001 | 75740000 | 0.0025 | 0.0012 | 0.0014 |
| F12 | 75660001 | 75760000 | 0.0020 | 0.0009 | 0.0010 |
| F12 | 75680001 | 75780000 | 0.0022 | 0.0010 | 0.0011 |
| F12 | 75700001 | 75800000 | 0.0026 | 0.0012 | 0.0014 |
| F12 | 75720001 | 75820000 | 0.0024 | 0.0013 | 0.0015 |
| F12 | 75740001 | 75840000 | 0.0024 | 0.0013 | 0.0016 |
| F12 | 75760001 | 75860000 | 0.0027 | 0.0016 | 0.0019 |
| F12 | 75780001 | 75880000 | 0.0029 | 0.0021 | 0.0025 |
| F12 | 75800001 | 75900000 | 0.0025 | 0.0017 | 0.0020 |
| F12 | 75820001 | 75920000 | 0.0037 | 0.0029 | 0.0032 |
| F12 | 75840001 | 75940000 | 0.0035 | 0.0028 | 0.0031 |
| F12 | 75860001 | 75960000 | 0.0031 | 0.0025 | 0.0026 |
| F12 | 75880001 | 75980000 | 0.0022 | 0.0015 | 0.0015 |

| Chr | Start | End | both | A1 | A2 |
|---|---|---|---|---|---|
| F12 | 75900001 | 76000000 | 0.0024 | 0.0015 | 0.0017 |
| F12 | 75920001 | 76020000 | 0.0011 | 0.0003 | 0.0004 |
| F12 | 75940001 | 76040000 | 0.0018 | 0.0008 | 0.0010 |
| F12 | 75960001 | 76060000 | 0.0022 | 0.0012 | 0.0013 |
| F12 | 75980001 | 76080000 | 0.0025 | 0.0014 | 0.0016 |
| F12 | 76000001 | 76100000 | 0.0025 | 0.0014 | 0.0015 |
| F12 | 76020001 | 76120000 | 0.0027 | 0.0016 | 0.0017 |
| F12 | 76040001 | 76140000 | 0.0022 | 0.0013 | 0.0014 |
| F12 | 76060001 | 76160000 | 0.0016 | 0.0009 | 0.0010 |
| F12 | 76080001 | 76180000 | 0.0014 | 0.0008 | 0.0009 |
| F12 | 76100001 | 76200000 | 0.0012 | 0.0007 | 0.0009 |
| F12 | 76120001 | 76220000 | 0.0012 | 0.0007 | 0.0007 |
| F12 | 76140001 | 76240000 | 0.0019 | 0.0013 | 0.0013 |
| F12 | 76160001 | 76260000 | 0.0027 | 0.0020 | 0.0021 |
| F12 | 76180001 | 76280000 | 0.0031 | 0.0022 | 0.0024 |
| F12 | 76200001 | 76300000 | 0.0031 | 0.0023 | 0.0025 |
| F12 | 76220001 | 76320000 | 0.0033 | 0.0025 | 0.0028 |
| F12 | 76240001 | 76340000 | 0.0028 | 0.0021 | 0.0023 |
| F12 | 76260001 | 76360000 | 0.0025 | 0.0017 | 0.0020 |
| F12 | 76280001 | 76380000 | 0.0022 | 0.0013 | 0.0015 |
| F12 | 76300001 | 76400000 | 0.0025 | 0.0015 | 0.0018 |
| F12 | 76320001 | 76420000 | 0.0021 | 0.0011 | 0.0013 |
| F12 | 76340001 | 76440000 | 0.0020 | 0.0009 | 0.0010 |
| F12 | 76360001 | 76460000 | 0.0018 | 0.0008 | 0.0008 |
| F12 | 76380001 | 76480000 | 0.0020 | 0.0011 | 0.0013 |
| F12 | 76400001 | 76500000 | 0.0020 | 0.0010 | 0.0011 |
| F12 | 76420001 | 76520000 | 0.0020 | 0.0010 | 0.0011 |
| F12 | 76440001 | 76540000 | 0.0022 | 0.0014 | 0.0014 |
| F12 | 76460001 | 76560000 | 0.0020 | 0.0013 | 0.0013 |
| F12 | 76480001 | 76580000 | 0.0018 | 0.0011 | 0.0011 |
| F12 | 76500001 | 76600000 | 0.0021 | 0.0017 | 0.0017 |
| F12 | 76520001 | 76620000 | 0.0028 | 0.0023 | 0.0023 |
| F12 | 76540001 | 76640000 | 0.0024 | 0.0020 | 0.0019 |
| F12 | 76560001 | 76660000 | 0.0029 | 0.0023 | 0.0024 |
| F12 | 76580001 | 76680000 | 0.0039 | 0.0032 | 0.0033 |
| F12 | 76600001 | 76700000 | 0.0035 | 0.0026 | 0.0027 |
| F12 | 76620001 | 76720000 | 0.0030 | 0.0021 | 0.0023 |
| F12 | 76640001 | 76740000 | 0.0036 | 0.0027 | 0.0028 |
| F12 | 76660001 | 76760000 | 0.0039 | 0.0031 | 0.0032 |
| F12 | 76680001 | 76780000 | 0.0030 | 0.0023 | 0.0024 |
| F12 | 76700001 | 76800000 | 0.0027 | 0.0022 | 0.0023 |
| F12 | 76720001 | 76820000 | 0.0024 | 0.0019 | 0.0020 |
| F12 | 76740001 | 76840000 | 0.0018 | 0.0012 | 0.0013 |
| F12 | 76760001 | 76860000 | 0.0012 | 0.0004 | 0.0005 |
| F12 | 76780001 | 76880000 | 0.0009 | 0.0001 | 0.0002 |
| F12 | 76800001 | 76900000 | 0.0012 | 0.0003 | 0.0004 |
| F12 | 76820001 | 76920000 | 0.0019 | 0.0010 | 0.0011 |
| F12 | 76840001 | 76940000 | 0.0018 | 0.0010 | 0.0011 |
| F12 | 76860001 | 76960000 | 0.0018 | 0.0010 | 0.0011 |
| F12 | 76880001 | 76980000 | 0.0018 | 0.0011 | 0.0011 |
| F12 | 76900001 | 77000000 | 0.0021 | 0.0014 | 0.0014 |
| F12 | 76920001 | 77020000 | 0.0017 | 0.0009 | 0.0009 |
| F12 | 76940001 | 77040000 | 0.0021 | 0.0013 | 0.0013 |
| F12 | 76960001 | 77060000 | 0.0025 | 0.0017 | 0.0017 |
| F12 | 76980001 | 77080000 | 0.0027 | 0.0017 | 0.0017 |
| F12 | 77000001 | 77100000 | 0.0026 | 0.0016 | 0.0016 |
| F12 | 77020001 | 77120000 | 0.0025 | 0.0015 | 0.0015 |
| F12 | 77040001 | 77140000 | 0.0025 | 0.0015 | 0.0016 |

| Chr | Start | End | both | A1 | A2 |
|---|---|---|---|---|---|
| F12 | 77060001 | 77160000 | 0.0020 | 0.0011 | 0.0013 |
| F12 | 77080001 | 77180000 | 0.0018 | 0.0011 | 0.0012 |
| F12 | 77100001 | 77200000 | 0.0020 | 0.0015 | 0.0015 |
| F12 | 77120001 | 77220000 | 0.0020 | 0.0015 | 0.0016 |
| F12 | 77140001 | 77240000 | 0.0019 | 0.0015 | 0.0014 |
| F12 | 77160001 | 77260000 | 0.0022 | 0.0018 | 0.0015 |
| F12 | 77180001 | 77280000 | 0.0024 | 0.0019 | 0.0017 |
| F12 | 77200001 | 77300000 | 0.0019 | 0.0013 | 0.0012 |
| F12 | 77220001 | 77320000 | 0.0020 | 0.0015 | 0.0013 |
| F12 | 77240001 | 77340000 | 0.0017 | 0.0013 | 0.0013 |
| F12 | 77260001 | 77360000 | 0.0015 | 0.0010 | 0.0011 |
| F12 | 77280001 | 77380000 | 0.0013 | 0.0009 | 0.0010 |
| F12 | 77300001 | 77400000 | 0.0014 | 0.0010 | 0.0011 |
| F12 | 77320001 | 77420000 | 0.0014 | 0.0009 | 0.0010 |
| F12 | 77340001 | 77440000 | 0.0019 | 0.0013 | 0.0015 |
| F12 | 77360001 | 77460000 | 0.0028 | 0.0022 | 0.0025 |
| F12 | 77380001 | 77480000 | 0.0031 | 0.0024 | 0.0027 |
| F12 | 77400001 | 77500000 | 0.0031 | 0.0023 | 0.0027 |
| F12 | 77420001 | 77520000 | 0.0032 | 0.0023 | 0.0027 |
| F12 | 77440001 | 77540000 | 0.0028 | 0.0020 | 0.0023 |
| F12 | 77460001 | 77560000 | 0.0031 | 0.0023 | 0.0025 |
| F12 | 77480001 | 77580000 | 0.0041 | 0.0033 | 0.0035 |
| F12 | 77500001 | 77600000 | 0.0046 | 0.0037 | 0.0040 |
| F12 | 77520001 | 77620000 | 0.0050 | 0.0041 | 0.0043 |
| F12 | 77540001 | 77640000 | 0.0049 | 0.0041 | 0.0041 |
| F12 | 77560001 | 77660000 | 0.0044 | 0.0036 | 0.0036 |
| F12 | 77580001 | 77680000 | 0.0033 | 0.0024 | 0.0025 |
| F12 | 77600001 | 77700000 | 0.0030 | 0.0021 | 0.0021 |
| F12 | 77620001 | 77720000 | 0.0025 | 0.0017 | 0.0017 |
| F12 | 77640001 | 77740000 | 0.0024 | 0.0013 | 0.0015 |
| F12 | 77660001 | 77760000 | 0.0020 | 0.0008 | 0.0010 |
| F12 | 77680001 | 77780000 | 0.0023 | 0.0012 | 0.0013 |
| F12 | 77700001 | 77800000 | 0.0022 | 0.0013 | 0.0014 |
| F12 | 77720001 | 77820000 | 0.0027 | 0.0018 | 0.0019 |
| F12 | 77740001 | 77840000 | 0.0030 | 0.0022 | 0.0023 |
| F12 | 77760001 | 77860000 | 0.0028 | 0.0021 | 0.0022 |
| F12 | 77780001 | 77880000 | 0.0024 | 0.0016 | 0.0017 |
| F12 | 77800001 | 77900000 | 0.0022 | 0.0012 | 0.0012 |
| F12 | 77820001 | 77920000 | 0.0021 | 0.0010 | 0.0011 |
| F12 | 77840001 | 77940000 | 0.0017 | 0.0007 | 0.0008 |
| F12 | 77860001 | 77960000 | 0.0025 | 0.0015 | 0.0016 |
| F12 | 77880001 | 77980000 | 0.0025 | 0.0018 | 0.0019 |
| F12 | 77900001 | 78000000 | 0.0028 | 0.0021 | 0.0022 |
| F12 | 77920001 | 78020000 | 0.0026 | 0.0020 | 0.0020 |
| F12 | 77940001 | 78040000 | 0.0034 | 0.0027 | 0.0028 |
| F12 | 77960001 | 78060000 | 0.0030 | 0.0022 | 0.0023 |
| F12 | 77980001 | 78080000 | 0.0033 | 0.0023 | 0.0025 |
| F12 | 78000001 | 78100000 | 0.0032 | 0.0022 | 0.0024 |
| F12 | 78020001 | 78120000 | 0.0031 | 0.0021 | 0.0023 |
| F12 | 78040001 | 78140000 | 0.0023 | 0.0013 | 0.0015 |
| F12 | 78060001 | 78160000 | 0.0019 | 0.0010 | 0.0011 |
| F12 | 78080001 | 78180000 | 0.0015 | 0.0007 | 0.0007 |
| F12 | 78100001 | 78200000 | 0.0009 | 0.0004 | 0.0004 |
| F12 | 78120001 | 78220000 | 0.0008 | 0.0004 | 0.0002 |
| F12 | 78140001 | 78240000 | 0.0007 | 0.0006 | 0.0002 |
| F12 | 78160001 | 78260000 | 0.0006 | 0.0007 | 0.0001 |
| F12 | 78180001 | 78280000 | 0.0004 | 0.0008 | 0.0001 |
| F12 | 78200001 | 78300000 | 0.0004 | 0.0009 | 0.0002 |

| Chr | Start | End | both | A1 | A2 |
|---|---|---|---|---|---|
| F12 | 78220001 | 78320000 | 0.0010 | 0.0012 | 0.0008 |
| F12 | 78240001 | 78340000 | 0.0012 | 0.0013 | 0.0010 |
| F12 | 78260001 | 78360000 | 0.0013 | 0.0014 | 0.0012 |
| F12 | 78280001 | 78380000 | 0.0019 | 0.0019 | 0.0018 |
| F12 | 78300001 | 78400000 | 0.0020 | 0.0020 | 0.0020 |
| F12 | 78320001 | 78420000 | 0.0020 | 0.0024 | 0.0018 |
| F12 | 78340001 | 78440000 | 0.0023 | 0.0025 | 0.0021 |
| F12 | 78360001 | 78460000 | 0.0022 | 0.0026 | 0.0020 |
| F12 | 78380001 | 78480000 | 0.0018 | 0.0023 | 0.0015 |
| F12 | 78400001 | 78500000 | 0.0018 | 0.0023 | 0.0015 |
| F12 | 78420001 | 78520000 | 0.0014 | 0.0014 | 0.0009 |
| F12 | 78440001 | 78540000 | 0.0020 | 0.0019 | 0.0014 |
| F12 | 78460001 | 78560000 | 0.0027 | 0.0022 | 0.0020 |
| F12 | 78480001 | 78580000 | 0.0028 | 0.0020 | 0.0022 |
| F12 | 78500001 | 78600000 | 0.0042 | 0.0033 | 0.0035 |
| F12 | 78520001 | 78620000 | 0.0043 | 0.0035 | 0.0037 |
| F12 | 78540001 | 78640000 | 0.0039 | 0.0031 | 0.0034 |
| F12 | 78560001 | 78660000 | 0.0032 | 0.0027 | 0.0028 |
| F12 | 78580001 | 78680000 | 0.0032 | 0.0027 | 0.0028 |
| F12 | 78600001 | 78700000 | 0.0036 | 0.0028 | 0.0031 |
| F12 | 78620001 | 78720000 | 0.0034 | 0.0027 | 0.0030 |
| F12 | 78640001 | 78740000 | 0.0030 | 0.0022 | 0.0024 |
| F12 | 78660001 | 78760000 | 0.0037 | 0.0027 | 0.0029 |
| F12 | 78680001 | 78780000 | 0.0034 | 0.0025 | 0.0027 |
| F12 | 78700001 | 78800000 | 0.0022 | 0.0015 | 0.0016 |
| F12 | 78720001 | 78820000 | 0.0025 | 0.0016 | 0.0018 |
| F12 | 78740001 | 78840000 | 0.0025 | 0.0018 | 0.0020 |
| F12 | 78760001 | 78860000 | 0.0018 | 0.0012 | 0.0014 |
| F12 | 78780001 | 78880000 | 0.0028 | 0.0019 | 0.0024 |
| F12 | 78800001 | 78900000 | 0.0024 | 0.0014 | 0.0018 |
| F12 | 78820001 | 78920000 | 0.0021 | 0.0011 | 0.0014 |
| F12 | 78840001 | 78940000 | 0.0026 | 0.0015 | 0.0017 |
| F12 | 78860001 | 78960000 | 0.0029 | 0.0017 | 0.0019 |
| F12 | 78880001 | 78980000 | 0.0028 | 0.0019 | 0.0018 |
| F12 | 78900001 | 79000000 | 0.0030 | 0.0021 | 0.0021 |
| F12 | 78920001 | 79020000 | 0.0032 | 0.0023 | 0.0023 |
| F12 | 78940001 | 79040000 | 0.0028 | 0.0021 | 0.0022 |
| F12 | 78960001 | 79060000 | 0.0031 | 0.0025 | 0.0025 |
| F12 | 78980001 | 79080000 | 0.0023 | 0.0015 | 0.0016 |
| F12 | 79000001 | 79100000 | 0.0021 | 0.0015 | 0.0015 |
| F12 | 79020001 | 79120000 | 0.0020 | 0.0016 | 0.0016 |
| F12 | 79040001 | 79140000 | 0.0021 | 0.0015 | 0.0015 |
| F12 | 79060001 | 79160000 | 0.0017 | 0.0011 | 0.0012 |
| F12 | 79080001 | 79180000 | 0.0021 | 0.0013 | 0.0015 |
| F12 | 79100001 | 79200000 | 0.0022 | 0.0011 | 0.0017 |
| F12 | 79120001 | 79220000 | 0.0020 | 0.0008 | 0.0015 |
| F12 | 79140001 | 79240000 | 0.0018 | 0.0005 | 0.0014 |
| F12 | 79160001 | 79260000 | 0.0027 | 0.0015 | 0.0025 |
| F12 | 79180001 | 79280000 | 0.0035 | 0.0021 | 0.0034 |
| F12 | 79200001 | 79300000 | 0.0031 | 0.0022 | 0.0031 |
| F12 | 79220001 | 79320000 | 0.0035 | 0.0025 | 0.0035 |
| F12 | 79240001 | 79340000 | 0.0039 | 0.0029 | 0.0039 |
| F12 | 79260001 | 79360000 | 0.0028 | 0.0019 | 0.0027 |
| F12 | 79280001 | 79380000 | 0.0017 | 0.0011 | 0.0017 |
| F12 | 79300001 | 79400000 | 0.0020 | 0.0013 | 0.0020 |
| F12 | 79320001 | 79420000 | 0.0025 | 0.0018 | 0.0024 |
| F12 | 79340001 | 79440000 | 0.0037 | 0.0029 | 0.0036 |
| F12 | 79360001 | 79460000 | 0.0050 | 0.0040 | 0.0049 |

| Chr | Start | End | both | A1 | A2 |
|---|---|---|---|---|---|
| F12 | 79380001 | 79480000 | 0.0057 | 0.0046 | 0.0056 |
| F12 | 79400001 | 79500000 | 0.0056 | 0.0045 | 0.0055 |
| F12 | 79420001 | 79520000 | 0.0064 | 0.0052 | 0.0063 |
| F12 | 79440001 | 79540000 | 0.0050 | 0.0036 | 0.0049 |
| F12 | 79460001 | 79560000 | 0.0045 | 0.0030 | 0.0045 |
| F12 | 79480001 | 79580000 | 0.0042 | 0.0026 | 0.0042 |
| F12 | 79500001 | 79600000 | 0.0044 | 0.0027 | 0.0045 |
| F12 | 79520001 | 79620000 | 0.0032 | 0.0017 | 0.0033 |
| F12 | 79540001 | 79640000 | 0.0034 | 0.0021 | 0.0035 |
| F12 | 79560001 | 79660000 | 0.0031 | 0.0022 | 0.0031 |
| F12 | 79580001 | 79680000 | 0.0028 | 0.0019 | 0.0028 |
| F12 | 79600001 | 79700000 | 0.0027 | 0.0018 | 0.0027 |
| F12 | 79620001 | 79720000 | 0.0023 | 0.0014 | 0.0024 |
| F12 | 79640001 | 79740000 | 0.0022 | 0.0014 | 0.0023 |
| F12 | 79660001 | 79760000 | 0.0024 | 0.0015 | 0.0025 |
| F12 | 79680001 | 79780000 | 0.0024 | 0.0015 | 0.0025 |
| F12 | 79700001 | 79800000 | 0.0027 | 0.0018 | 0.0027 |
| F12 | 79720001 | 79820000 | 0.0028 | 0.0019 | 0.0029 |
| F12 | 79740001 | 79840000 | 0.0027 | 0.0016 | 0.0028 |
| F12 | 79760001 | 79860000 | 0.0030 | 0.0020 | 0.0031 |
| F12 | 79780001 | 79880000 | 0.0034 | 0.0023 | 0.0036 |
| F12 | 79800001 | 79900000 | 0.0031 | 0.0021 | 0.0033 |
| F12 | 79820001 | 79920000 | 0.0038 | 0.0028 | 0.0039 |
| F12 | 79840001 | 79940000 | 0.0039 | 0.0030 | 0.0039 |
| F12 | 79860001 | 79960000 | 0.0028 | 0.0019 | 0.0029 |
| F12 | 79880001 | 79980000 | 0.0024 | 0.0018 | 0.0025 |
| F12 | 79900001 | 80000000 | 0.0026 | 0.0019 | 0.0026 |
| F12 | 79920001 | 80020000 | 0.0023 | 0.0014 | 0.0024 |
| F12 | 79940001 | 80040000 | 0.0024 | 0.0015 | 0.0025 |
| F12 | 79960001 | 80060000 | 0.0025 | 0.0015 | 0.0027 |
| F12 | 79980001 | 80080000 | 0.0025 | 0.0014 | 0.0026 |
| F12 | 80000001 | 80100000 | 0.0029 | 0.0019 | 0.0030 |
| F12 | 80020001 | 80120000 | 0.0023 | 0.0016 | 0.0024 |
| F12 | 80040001 | 80140000 | 0.0027 | 0.0021 | 0.0027 |
| F12 | 80060001 | 80160000 | 0.0029 | 0.0023 | 0.0030 |
| F12 | 80080001 | 80180000 | 0.0034 | 0.0027 | 0.0034 |
| F12 | 80100001 | 80200000 | 0.0035 | 0.0026 | 0.0036 |
| F12 | 80120001 | 80220000 | 0.0038 | 0.0027 | 0.0039 |
| F12 | 80140001 | 80240000 | 0.0029 | 0.0018 | 0.0030 |
| F12 | 80160001 | 80260000 | 0.0030 | 0.0019 | 0.0032 |
| F12 | 80180001 | 80280000 | 0.0028 | 0.0018 | 0.0029 |
| F12 | 80200001 | 80300000 | 0.0028 | 0.0022 | 0.0028 |
| F12 | 80220001 | 80320000 | 0.0026 | 0.0020 | 0.0026 |
| F12 | 80240001 | 80340000 | 0.0028 | 0.0020 | 0.0028 |
| F12 | 80260001 | 80360000 | 0.0023 | 0.0015 | 0.0024 |
| F12 | 80280001 | 80380000 | 0.0019 | 0.0012 | 0.0019 |
| F12 | 80300001 | 80400000 | 0.0015 | 0.0007 | 0.0016 |
| F12 | 80320001 | 80420000 | 0.0017 | 0.0006 | 0.0017 |
| F12 | 80340001 | 80440000 | 0.0018 | 0.0006 | 0.0018 |
| F12 | 80360001 | 80460000 | 0.0020 | 0.0006 | 0.0020 |
| F12 | 80380001 | 80480000 | 0.0027 | 0.0007 | 0.0027 |
| F12 | 80400001 | 80500000 | 0.0025 | 0.0005 | 0.0025 |
| F12 | 80420001 | 80520000 | 0.0022 | 0.0005 | 0.0022 |
| F12 | 80440001 | 80540000 | 0.0021 | 0.0005 | 0.0021 |
| F12 | 80460001 | 80560000 | 0.0022 | 0.0009 | 0.0021 |
| F12 | 80480001 | 80580000 | 0.0014 | 0.0008 | 0.0014 |
| F12 | 80500001 | 80600000 | 0.0015 | 0.0008 | 0.0015 |
| F12 | 80520001 | 80620000 | 0.0017 | 0.0010 | 0.0017 |

| Chr | Start | End | both | A1 | A2 |
|---|---|---|---|---|---|
| F12 | 80540001 | 80640000 | 0.0035 | 0.0025 | 0.0035 |
| F12 | 80560001 | 80660000 | 0.0034 | 0.0023 | 0.0034 |
| F12 | 80580001 | 80680000 | 0.0035 | 0.0024 | 0.0033 |
| F12 | 80600001 | 80700000 | 0.0037 | 0.0026 | 0.0034 |
| F12 | 80620001 | 80720000 | 0.0046 | 0.0034 | 0.0043 |
| F12 | 80640001 | 80740000 | 0.0036 | 0.0028 | 0.0033 |
| F12 | 80660001 | 80760000 | 0.0038 | 0.0030 | 0.0034 |
| F12 | 80680001 | 80780000 | 0.0041 | 0.0034 | 0.0038 |
| F12 | 80700001 | 80800000 | 0.0049 | 0.0041 | 0.0044 |
| F12 | 80720001 | 80820000 | 0.0047 | 0.0038 | 0.0040 |
| F12 | 80740001 | 80840000 | 0.0044 | 0.0035 | 0.0036 |
| F12 | 80760001 | 80860000 | 0.0043 | 0.0035 | 0.0036 |
| F12 | 80780001 | 80880000 | 0.0040 | 0.0032 | 0.0033 |
| F12 | 80800001 | 80900000 | 0.0029 | 0.0023 | 0.0025 |
| F12 | 80820001 | 80920000 | 0.0032 | 0.0028 | 0.0028 |
| F12 | 80840001 | 80940000 | 0.0030 | 0.0026 | 0.0026 |
| F12 | 80860001 | 80960000 | 0.0038 | 0.0035 | 0.0033 |
| F12 | 80880001 | 80980000 | 0.0042 | 0.0039 | 0.0037 |
| F12 | 80900001 | 81000000 | 0.0042 | 0.0038 | 0.0036 |
| F12 | 80920001 | 81020000 | 0.0038 | 0.0034 | 0.0035 |
| F12 | 80940001 | 81040000 | 0.0059 | 0.0053 | 0.0054 |
| F12 | 80960001 | 81060000 | 0.0062 | 0.0055 | 0.0056 |
| F12 | 80980001 | 81080000 | 0.0058 | 0.0050 | 0.0051 |
| F12 | 81000001 | 81100000 | 0.0058 | 0.0049 | 0.0050 |
| F12 | 81020001 | 81120000 | 0.0058 | 0.0049 | 0.0049 |
| F12 | 81040001 | 81140000 | 0.0034 | 0.0025 | 0.0025 |
| F12 | 81060001 | 81160000 | 0.0022 | 0.0015 | 0.0015 |
| F12 | 81080001 | 81180000 | 0.0027 | 0.0019 | 0.0020 |
| F12 | 81100001 | 81200000 | 0.0027 | 0.0019 | 0.0020 |
| F12 | 81120001 | 81220000 | 0.0021 | 0.0015 | 0.0015 |
| F12 | 81140001 | 81240000 | 0.0029 | 0.0023 | 0.0024 |
| F12 | 81160001 | 81260000 | 0.0037 | 0.0032 | 0.0031 |
| F12 | 81180001 | 81280000 | 0.0033 | 0.0029 | 0.0028 |
| F12 | 81200001 | 81300000 | 0.0031 | 0.0027 | 0.0026 |
| F12 | 81220001 | 81320000 | 0.0033 | 0.0029 | 0.0028 |
| F12 | 81240001 | 81340000 | 0.0028 | 0.0025 | 0.0022 |
| F12 | 81260001 | 81360000 | 0.0021 | 0.0018 | 0.0015 |
| F12 | 81280001 | 81380000 | 0.0026 | 0.0024 | 0.0021 |
| F12 | 81300001 | 81400000 | 0.0048 | 0.0041 | 0.0042 |
| F12 | 81320001 | 81420000 | 0.0046 | 0.0038 | 0.0039 |
| F12 | 81340001 | 81440000 | 0.0043 | 0.0033 | 0.0036 |
| F12 | 81360001 | 81460000 | 0.0046 | 0.0034 | 0.0039 |
| F12 | 81380001 | 81480000 | 0.0046 | 0.0033 | 0.0038 |
| F12 | 81400001 | 81500000 | 0.0028 | 0.0018 | 0.0019 |
| F12 | 81420001 | 81520000 | 0.0027 | 0.0017 | 0.0018 |
| F12 | 81440001 | 81540000 | 0.0026 | 0.0017 | 0.0017 |
| F12 | 81460001 | 81560000 | 0.0018 | 0.0011 | 0.0009 |
| F12 | 81480001 | 81580000 | 0.0018 | 0.0012 | 0.0010 |
| F12 | 81500001 | 81600000 | 0.0041 | 0.0034 | 0.0036 |
| F12 | 81520001 | 81620000 | 0.0051 | 0.0042 | 0.0046 |
| F12 | 81540001 | 81640000 | 0.0052 | 0.0044 | 0.0047 |
| F12 | 81560001 | 81660000 | 0.0054 | 0.0045 | 0.0050 |
| F12 | 81580001 | 81680000 | 0.0046 | 0.0038 | 0.0042 |
| F12 | 81600001 | 81700000 | 0.0020 | 0.0014 | 0.0015 |
| F12 | 81620001 | 81720000 | 0.0012 | 0.0006 | 0.0005 |
| F12 | 81640001 | 81740000 | 0.0012 | 0.0006 | 0.0006 |
| F12 | 81660001 | 81760000 | 0.0016 | 0.0009 | 0.0009 |
| F12 | 81680001 | 81780000 | 0.0017 | 0.0009 | 0.0010 |

| Chr | Start | End | both | A1 | A2 |
|---|---|---|---|---|---|
| F12 | 81700001 | 81800000 | 0.0018 | 0.0009 | 0.0011 |
| F12 | 81720001 | 81820000 | 0.0018 | 0.0009 | 0.0011 |
| F12 | 81740001 | 81840000 | 0.0019 | 0.0008 | 0.0010 |
| F12 | 81760001 | 81860000 | 0.0015 | 0.0004 | 0.0008 |
| F12 | 81780001 | 81880000 | 0.0023 | 0.0012 | 0.0016 |
| F12 | 81800001 | 81900000 | 0.0027 | 0.0017 | 0.0021 |
| F12 | 81820001 | 81920000 | 0.0025 | 0.0017 | 0.0020 |
| F12 | 81840001 | 81940000 | 0.0033 | 0.0026 | 0.0030 |
| F12 | 81860001 | 81960000 | 0.0038 | 0.0032 | 0.0035 |
| F12 | 81880001 | 81980000 | 0.0033 | 0.0025 | 0.0028 |
| F12 | 81900001 | 82000000 | 0.0030 | 0.0022 | 0.0025 |
| F12 | 81920001 | 82020000 | 0.0033 | 0.0023 | 0.0027 |
| F12 | 81940001 | 82040000 | 0.0024 | 0.0015 | 0.0018 |
| F12 | 81960001 | 82060000 | 0.0020 | 0.0011 | 0.0015 |
| F12 | 81980001 | 82080000 | 0.0020 | 0.0012 | 0.0014 |
| F12 | 82000001 | 82100000 | 0.0015 | 0.0009 | 0.0010 |
| F12 | 82020001 | 82120000 | 0.0014 | 0.0009 | 0.0009 |
| F12 | 82040001 | 82140000 | 0.0015 | 0.0008 | 0.0010 |
| F12 | 82060001 | 82160000 | 0.0017 | 0.0009 | 0.0012 |
| F12 | 82080001 | 82180000 | 0.0021 | 0.0010 | 0.0016 |
| F12 | 82100001 | 82200000 | 0.0027 | 0.0011 | 0.0023 |
| F12 | 82120001 | 82220000 | 0.0030 | 0.0011 | 0.0027 |
| F12 | 82140001 | 82240000 | 0.0030 | 0.0010 | 0.0027 |
| F12 | 82160001 | 82260000 | 0.0045 | 0.0024 | 0.0042 |
| F12 | 82180001 | 82280000 | 0.0041 | 0.0023 | 0.0039 |
| F12 | 82200001 | 82300000 | 0.0039 | 0.0023 | 0.0036 |
| F12 | 82220001 | 82320000 | 0.0038 | 0.0024 | 0.0036 |
| F12 | 82240001 | 82340000 | 0.0038 | 0.0026 | 0.0036 |
| F12 | 82260001 | 82360000 | 0.0020 | 0.0012 | 0.0020 |
| F12 | 82280001 | 82380000 | 0.0018 | 0.0009 | 0.0018 |
| F12 | 82300001 | 82400000 | 0.0019 | 0.0010 | 0.0019 |
| F12 | 82320001 | 82420000 | 0.0020 | 0.0012 | 0.0020 |
| F12 | 82340001 | 82440000 | 0.0022 | 0.0011 | 0.0021 |
| F12 | 82360001 | 82460000 | 0.0021 | 0.0009 | 0.0019 |
| F12 | 82380001 | 82480000 | 0.0022 | 0.0012 | 0.0020 |
| F12 | 82400001 | 82500000 | 0.0019 | 0.0012 | 0.0017 |
| F12 | 82420001 | 82520000 | 0.0016 | 0.0011 | 0.0014 |
| F12 | 82440001 | 82540000 | 0.0019 | 0.0016 | 0.0017 |
| F12 | 82460001 | 82560000 | 0.0019 | 0.0016 | 0.0017 |
| F12 | 82480001 | 82580000 | 0.0023 | 0.0019 | 0.0021 |
| F12 | 82500001 | 82600000 | 0.0024 | 0.0018 | 0.0019 |
| F12 | 82520001 | 82620000 | 0.0025 | 0.0016 | 0.0018 |
| F12 | 82540001 | 82640000 | 0.0020 | 0.0010 | 0.0012 |
| F12 | 82560001 | 82660000 | 0.0018 | 0.0007 | 0.0010 |
| F12 | 82580001 | 82680000 | 0.0014 | 0.0002 | 0.0005 |
| F12 | 82600001 | 82700000 | 0.0014 | 0.0003 | 0.0006 |
| F12 | 82620001 | 82720000 | 0.0014 | 0.0002 | 0.0005 |
| F12 | 82640001 | 82740000 | 0.0017 | 0.0005 | 0.0008 |
| F12 | 82660001 | 82760000 | 0.0023 | 0.0010 | 0.0013 |
| F12 | 82680001 | 82780000 | 0.0021 | 0.0010 | 0.0012 |
| F12 | 82700001 | 82800000 | 0.0023 | 0.0012 | 0.0014 |
| F12 | 82720001 | 82820000 | 0.0023 | 0.0013 | 0.0015 |
| F12 | 82740001 | 82840000 | 0.0021 | 0.0010 | 0.0013 |
| F12 | 82760001 | 82860000 | 0.0015 | 0.0005 | 0.0007 |
| F12 | 82780001 | 82880000 | 0.0021 | 0.0008 | 0.0012 |
| F12 | 82800001 | 82900000 | 0.0029 | 0.0016 | 0.0021 |
| F12 | 82820001 | 82920000 | 0.0038 | 0.0026 | 0.0030 |
| F12 | 82840001 | 82940000 | 0.0039 | 0.0026 | 0.0030 |

| Chr | Start | End | both | A1 | A2 |
|---|---|---|---|---|---|
| F12 | 82860001 | 82960000 | 0.0041 | 0.0028 | 0.0032 |
| F12 | 82880001 | 82980000 | 0.0037 | 0.0027 | 0.0030 |
| F12 | 82900001 | 83000000 | 0.0030 | 0.0019 | 0.0022 |
| F12 | 82920001 | 83020000 | 0.0019 | 0.0009 | 0.0011 |
| F12 | 82940001 | 83040000 | 0.0026 | 0.0017 | 0.0020 |
| F12 | 82960001 | 83060000 | 0.0026 | 0.0017 | 0.0020 |
| F12 | 82980001 | 83080000 | 0.0028 | 0.0020 | 0.0023 |
| F12 | 83000001 | 83100000 | 0.0025 | 0.0017 | 0.0020 |
| F12 | 83020001 | 83120000 | 0.0026 | 0.0017 | 0.0020 |
| F12 | 83040001 | 83140000 | 0.0016 | 0.0008 | 0.0011 |
| F12 | 83060001 | 83160000 | 0.0019 | 0.0010 | 0.0014 |
| F12 | 83080001 | 83180000 | 0.0018 | 0.0008 | 0.0014 |
| F12 | 83100001 | 83200000 | 0.0018 | 0.0008 | 0.0015 |
| F12 | 83120001 | 83220000 | 0.0019 | 0.0009 | 0.0019 |
| F12 | 83140001 | 83240000 | 0.0024 | 0.0013 | 0.0024 |
| F12 | 83160001 | 83260000 | 0.0022 | 0.0010 | 0.0021 |
| F12 | 83180001 | 83280000 | 0.0030 | 0.0018 | 0.0029 |
| F12 | 83200001 | 83300000 | 0.0039 | 0.0027 | 0.0036 |
| F12 | 83220001 | 83320000 | 0.0037 | 0.0026 | 0.0033 |
| F12 | 83240001 | 83340000 | 0.0031 | 0.0022 | 0.0026 |
| F12 | 83260001 | 83360000 | 0.0027 | 0.0020 | 0.0023 |
| F12 | 83280001 | 83380000 | 0.0021 | 0.0016 | 0.0016 |
| F12 | 83300001 | 83400000 | 0.0019 | 0.0014 | 0.0014 |
| F12 | 83320001 | 83420000 | 0.0037 | 0.0034 | 0.0032 |
| F12 | 83340001 | 83440000 | 0.0041 | 0.0036 | 0.0034 |
| F12 | 83360001 | 83460000 | 0.0043 | 0.0037 | 0.0035 |
| F12 | 83380001 | 83480000 | 0.0042 | 0.0034 | 0.0034 |
| F12 | 83400001 | 83500000 | 0.0035 | 0.0028 | 0.0027 |
| F12 | 83420001 | 83520000 | 0.0018 | 0.0010 | 0.0011 |
| F12 | 83440001 | 83540000 | 0.0024 | 0.0016 | 0.0018 |
| F12 | 83460001 | 83560000 | 0.0024 | 0.0018 | 0.0019 |
| F12 | 83480001 | 83580000 | 0.0027 | 0.0022 | 0.0022 |
| F12 | 83500001 | 83600000 | 0.0028 | 0.0024 | 0.0023 |
| F12 | 83520001 | 83620000 | 0.0030 | 0.0025 | 0.0026 |
| F12 | 83540001 | 83640000 | 0.0024 | 0.0019 | 0.0019 |
| F12 | 83560001 | 83660000 | 0.0026 | 0.0019 | 0.0020 |
| F12 | 83580001 | 83680000 | 0.0018 | 0.0011 | 0.0013 |
| F12 | 83600001 | 83700000 | 0.0018 | 0.0010 | 0.0012 |
| F12 | 83620001 | 83720000 | 0.0014 | 0.0007 | 0.0008 |
| F12 | 83640001 | 83740000 | 0.0015 | 0.0008 | 0.0010 |
| F12 | 83660001 | 83760000 | 0.0017 | 0.0010 | 0.0013 |
| F12 | 83680001 | 83780000 | 0.0027 | 0.0019 | 0.0023 |
| F12 | 83700001 | 83800000 | 0.0034 | 0.0027 | 0.0032 |
| F12 | 83720001 | 83820000 | 0.0036 | 0.0028 | 0.0035 |
| F12 | 83740001 | 83840000 | 0.0034 | 0.0028 | 0.0034 |
| F12 | 83760001 | 83860000 | 0.0029 | 0.0025 | 0.0029 |
| F12 | 83780001 | 83880000 | 0.0020 | 0.0017 | 0.0020 |
| F12 | 83800001 | 83900000 | 0.0015 | 0.0012 | 0.0014 |
| F12 | 83820001 | 83920000 | 0.0013 | 0.0011 | 0.0013 |
| F12 | 83840001 | 83940000 | 0.0011 | 0.0008 | 0.0011 |
| F12 | 83860001 | 83960000 | 0.0011 | 0.0008 | 0.0011 |
| F12 | 83880001 | 83980000 | 0.0014 | 0.0011 | 0.0014 |
| F12 | 83900001 | 84000000 | 0.0014 | 0.0011 | 0.0014 |
| F12 | 83920001 | 84020000 | 0.0014 | 0.0011 | 0.0014 |
| F12 | 83940001 | 84040000 | 0.0013 | 0.0010 | 0.0013 |
| F12 | 83960001 | 84060000 | 0.0013 | 0.0011 | 0.0013 |
| F12 | 83980001 | 84080000 | 0.0014 | 0.0012 | 0.0014 |
| F12 | 84000001 | 84100000 | 0.0014 | 0.0012 | 0.0014 |

| Chr | Start | End | both | A1 | A2 |
|---|---|---|---|---|---|
| F12 | 84020001 | 84120000 | 0.0018 | 0.0015 | 0.0018 |
| F12 | 84040001 | 84140000 | 0.0020 | 0.0017 | 0.0020 |
| F12 | 84060001 | 84160000 | 0.0026 | 0.0022 | 0.0026 |
| F12 | 84080001 | 84180000 | 0.0033 | 0.0029 | 0.0033 |
| F12 | 84100001 | 84200000 | 0.0030 | 0.0027 | 0.0029 |
| F12 | 84120001 | 84220000 | 0.0029 | 0.0026 | 0.0028 |
| F12 | 84140001 | 84240000 | 0.0032 | 0.0028 | 0.0030 |
| F12 | 84160001 | 84260000 | 0.0030 | 0.0026 | 0.0029 |
| F12 | 84180001 | 84280000 | 0.0021 | 0.0015 | 0.0019 |
| F12 | 84200001 | 84300000 | 0.0021 | 0.0015 | 0.0018 |
| F12 | 84220001 | 84320000 | 0.0024 | 0.0016 | 0.0019 |
| F12 | 84240001 | 84340000 | 0.0021 | 0.0013 | 0.0015 |
| F12 | 84260001 | 84360000 | 0.0023 | 0.0015 | 0.0015 |
| F12 | 84280001 | 84380000 | 0.0024 | 0.0016 | 0.0017 |
| F12 | 84300001 | 84400000 | 0.0025 | 0.0018 | 0.0019 |
| F12 | 84320001 | 84420000 | 0.0028 | 0.0021 | 0.0022 |
| F12 | 84340001 | 84440000 | 0.0030 | 0.0022 | 0.0023 |
| F12 | 84360001 | 84460000 | 0.0024 | 0.0017 | 0.0018 |
| F12 | 84380001 | 84480000 | 0.0029 | 0.0020 | 0.0021 |
| F12 | 84400001 | 84500000 | 0.0031 | 0.0021 | 0.0023 |
| F12 | 84420001 | 84520000 | 0.0022 | 0.0013 | 0.0013 |
| F12 | 84440001 | 84540000 | 0.0020 | 0.0012 | 0.0011 |
| F12 | 84460001 | 84560000 | 0.0021 | 0.0013 | 0.0013 |
| F12 | 84480001 | 84580000 | 0.0014 | 0.0006 | 0.0005 |
| F12 | 84500001 | 84600000 | 0.0011 | 0.0003 | 0.0003 |
| F12 | 84520001 | 84620000 | 0.0011 | 0.0002 | 0.0003 |
| F12 | 84540001 | 84640000 | 0.0026 | 0.0014 | 0.0018 |
| F12 | 84560001 | 84660000 | 0.0025 | 0.0012 | 0.0017 |
| F12 | 84580001 | 84680000 | 0.0027 | 0.0012 | 0.0017 |
| F12 | 84600001 | 84700000 | 0.0025 | 0.0013 | 0.0017 |
| F12 | 84620001 | 84720000 | 0.0026 | 0.0014 | 0.0018 |
| F12 | 84640001 | 84740000 | 0.0011 | 0.0002 | 0.0004 |
| F12 | 84660001 | 84760000 | 0.0009 | 0.0003 | 0.0004 |
| F12 | 84680001 | 84780000 | 0.0009 | 0.0003 | 0.0004 |
| F12 | 84700001 | 84800000 | 0.0010 | 0.0003 | 0.0004 |
| F12 | 84720001 | 84820000 | 0.0009 | 0.0001 | 0.0003 |
| F12 | 84740001 | 84840000 | 0.0011 | 0.0002 | 0.0004 |
| F12 | 84760001 | 84860000 | 0.0015 | 0.0003 | 0.0006 |
| F12 | 84780001 | 84880000 | 0.0013 | 0.0003 | 0.0006 |
| F12 | 84800001 | 84900000 | 0.0019 | 0.0009 | 0.0014 |
| F12 | 84820001 | 84920000 | 0.0021 | 0.0012 | 0.0016 |
| F12 | 84840001 | 84940000 | 0.0020 | 0.0012 | 0.0016 |
| F12 | 84860001 | 84960000 | 0.0019 | 0.0012 | 0.0016 |
| F12 | 84880001 | 84980000 | 0.0026 | 0.0017 | 0.0023 |
| F12 | 84900001 | 85000000 | 0.0021 | 0.0014 | 0.0019 |
| F12 | 84920001 | 85020000 | 0.0021 | 0.0011 | 0.0018 |
| F12 | 84940001 | 85040000 | 0.0022 | 0.0011 | 0.0019 |
| F12 | 84960001 | 85060000 | 0.0029 | 0.0016 | 0.0026 |
| F12 | 84980001 | 85080000 | 0.0025 | 0.0011 | 0.0022 |
| F12 | 85000001 | 85100000 | 0.0025 | 0.0009 | 0.0023 |
| F12 | 85020001 | 85120000 | 0.0026 | 0.0009 | 0.0024 |
| F12 | 85040001 | 85140000 | 0.0028 | 0.0011 | 0.0027 |
| F12 | 85060001 | 85160000 | 0.0021 | 0.0006 | 0.0021 |
| F12 | 85080001 | 85180000 | 0.0019 | 0.0005 | 0.0018 |
| F12 | 85100001 | 85200000 | 0.0020 | 0.0005 | 0.0019 |
| F12 | 85120001 | 85220000 | 0.0021 | 0.0008 | 0.0019 |
| F12 | 85140001 | 85240000 | 0.0019 | 0.0008 | 0.0015 |
| F12 | 85160001 | 85260000 | 0.0018 | 0.0009 | 0.0013 |

| Chr | Start | End | both | A1 | A2 |
|---|---|---|---|---|---|
| F12 | 85180001 | 85280000 | 0.0020 | 0.0013 | 0.0013 |
| F12 | 85200001 | 85300000 | 0.0020 | 0.0015 | 0.0012 |
| F12 | 85220001 | 85320000 | 0.0017 | 0.0013 | 0.0009 |
| F12 | 85240001 | 85340000 | 0.0016 | 0.0011 | 0.0009 |
| F12 | 85260001 | 85360000 | 0.0017 | 0.0008 | 0.0010 |
| F12 | 85280001 | 85380000 | 0.0017 | 0.0005 | 0.0010 |
| F12 | 85300001 | 85400000 | 0.0019 | 0.0005 | 0.0012 |
| F12 | 85320001 | 85420000 | 0.0019 | 0.0006 | 0.0011 |
| F12 | 85340001 | 85440000 | 0.0019 | 0.0005 | 0.0010 |
| F12 | 85360001 | 85460000 | 0.0022 | 0.0008 | 0.0013 |
| F12 | 85380001 | 85480000 | 0.0024 | 0.0009 | 0.0015 |
| F12 | 85400001 | 85500000 | 0.0019 | 0.0007 | 0.0010 |
| F12 | 85420001 | 85520000 | 0.0021 | 0.0012 | 0.0013 |
| F12 | 85440001 | 85540000 | 0.0020 | 0.0014 | 0.0013 |
| F12 | 85460001 | 85560000 | 0.0019 | 0.0012 | 0.0013 |
| F12 | 85480001 | 85580000 | 0.0018 | 0.0011 | 0.0013 |
| F12 | 85500001 | 85600000 | 0.0018 | 0.0010 | 0.0015 |
| F12 | 85520001 | 85620000 | 0.0016 | 0.0005 | 0.0015 |
| F12 | 85540001 | 85640000 | 0.0031 | 0.0016 | 0.0030 |
| F12 | 85560001 | 85660000 | 0.0032 | 0.0019 | 0.0031 |
| F12 | 85580001 | 85680000 | 0.0036 | 0.0025 | 0.0036 |
| F12 | 85600001 | 85700000 | 0.0050 | 0.0040 | 0.0050 |
| F12 | 85620001 | 85720000 | 0.0052 | 0.0044 | 0.0052 |
| F12 | 85640001 | 85740000 | 0.0044 | 0.0036 | 0.0044 |
| F12 | 85660001 | 85760000 | 0.0041 | 0.0031 | 0.0041 |
| F12 | 85680001 | 85780000 | 0.0037 | 0.0029 | 0.0037 |
| F12 | 85700001 | 85800000 | 0.0029 | 0.0022 | 0.0028 |
| F12 | 85720001 | 85820000 | 0.0027 | 0.0021 | 0.0026 |
| F12 | 85740001 | 85840000 | 0.0022 | 0.0020 | 0.0021 |
| F12 | 85760001 | 85860000 | 0.0022 | 0.0024 | 0.0019 |
| F12 | 85780001 | 85880000 | 0.0019 | 0.0025 | 0.0016 |
| F12 | 85800001 | 85900000 | 0.0012 | 0.0019 | 0.0009 |
| F12 | 85820001 | 85920000 | 0.0014 | 0.0023 | 0.0011 |
| F12 | 85840001 | 85940000 | 0.0011 | 0.0019 | 0.0008 |
| F12 | 85860001 | 85960000 | 0.0014 | 0.0020 | 0.0011 |
| F12 | 85880001 | 85980000 | 0.0015 | 0.0020 | 0.0013 |
| F12 | 85900001 | 86000000 | 0.0014 | 0.0020 | 0.0012 |
| F12 | 85920001 | 86020000 | 0.0010 | 0.0017 | 0.0008 |
| F12 | 85940001 | 86040000 | 0.0012 | 0.0022 | 0.0008 |
| F12 | 85960001 | 86060000 | 0.0010 | 0.0020 | 0.0005 |
| F12 | 85980001 | 86080000 | 0.0009 | 0.0019 | 0.0004 |
| F12 | 86000001 | 86100000 | 0.0010 | 0.0019 | 0.0005 |
| F12 | 86020001 | 86120000 | 0.0011 | 0.0018 | 0.0005 |
| F12 | 86040001 | 86140000 | 0.0010 | 0.0014 | 0.0006 |
| F12 | 86060001 | 86160000 | 0.0010 | 0.0012 | 0.0007 |
| F12 | 86080001 | 86180000 | 0.0009 | 0.0010 | 0.0007 |
| F12 | 86100001 | 86200000 | 0.0011 | 0.0010 | 0.0010 |
| F12 | 86120001 | 86220000 | 0.0011 | 0.0008 | 0.0011 |
| F12 | 86140001 | 86240000 | 0.0014 | 0.0007 | 0.0015 |
| F12 | 86160001 | 86260000 | 0.0013 | 0.0004 | 0.0014 |
| F12 | 86180001 | 86280000 | 0.0015 | 0.0005 | 0.0016 |
| F12 | 86200001 | 86300000 | 0.0013 | 0.0002 | 0.0014 |
| F12 | 86220001 | 86320000 | 0.0015 | 0.0002 | 0.0016 |
| F12 | 86240001 | 86340000 | 0.0014 | 0.0005 | 0.0015 |
| F12 | 86260001 | 86360000 | 0.0015 | 0.0005 | 0.0017 |
| F12 | 86280001 | 86380000 | 0.0021 | 0.0008 | 0.0022 |
| F12 | 86300001 | 86400000 | 0.0024 | 0.0012 | 0.0025 |
| F12 | 86320001 | 86420000 | 0.0028 | 0.0017 | 0.0029 |

| Chr | Start | End | both | A1 | A2 |
|---|---|---|---|---|---|
| F12 | 86340001 | 86440000 | 0.0031 | 0.0019 | 0.0033 |
| F12 | 86360001 | 86460000 | 0.0029 | 0.0020 | 0.0030 |
| F12 | 86380001 | 86480000 | 0.0022 | 0.0018 | 0.0022 |
| F12 | 86400001 | 86500000 | 0.0025 | 0.0021 | 0.0025 |
| F12 | 86420001 | 86520000 | 0.0021 | 0.0020 | 0.0021 |
| F12 | 86440001 | 86540000 | 0.0017 | 0.0018 | 0.0016 |
| F12 | 86460001 | 86560000 | 0.0018 | 0.0021 | 0.0017 |
| F12 | 86480001 | 86580000 | 0.0018 | 0.0021 | 0.0016 |
| F12 | 86500001 | 86600000 | 0.0010 | 0.0016 | 0.0008 |
| F12 | 86520001 | 86620000 | 0.0008 | 0.0015 | 0.0006 |
| F12 | 86540001 | 86640000 | 0.0031 | 0.0036 | 0.0028 |
| F12 | 86560001 | 86660000 | 0.0030 | 0.0034 | 0.0027 |
| F12 | 86580001 | 86680000 | 0.0035 | 0.0037 | 0.0033 |
| F12 | 86600001 | 86700000 | 0.0036 | 0.0038 | 0.0033 |
| F12 | 86620001 | 86720000 | 0.0036 | 0.0038 | 0.0033 |
| F12 | 86640001 | 86740000 | 0.0011 | 0.0016 | 0.0010 |
| F12 | 86660001 | 86760000 | 0.0017 | 0.0021 | 0.0015 |
| F12 | 86680001 | 86780000 | 0.0013 | 0.0017 | 0.0011 |
| F12 | 86700001 | 86800000 | 0.0018 | 0.0022 | 0.0016 |
| F12 | 86720001 | 86820000 | 0.0017 | 0.0021 | 0.0016 |
| F12 | 86740001 | 86840000 | 0.0017 | 0.0020 | 0.0015 |
| F12 | 86760001 | 86860000 | 0.0014 | 0.0019 | 0.0012 |
| F12 | 86780001 | 86880000 | 0.0014 | 0.0019 | 0.0011 |
| F12 | 86800001 | 86900000 | 0.0008 | 0.0015 | 0.0006 |
| F12 | 86820001 | 86920000 | 0.0008 | 0.0014 | 0.0005 |
| F12 | 86840001 | 86940000 | 0.0008 | 0.0015 | 0.0005 |
| F12 | 86860001 | 86960000 | 0.0004 | 0.0012 | 0.0002 |
| F12 | 86880001 | 86980000 | 0.0004 | 0.0012 | 0.0002 |
| F12 | 86900001 | 87000000 | 0.0004 | 0.0011 | 0.0002 |
| F12 | 86920001 | 87020000 | 0.0004 | 0.0012 | 0.0002 |
| F12 | 86940001 | 87040000 | 0.0004 | 0.0012 | 0.0002 |
| F12 | 86960001 | 87060000 | 0.0006 | 0.0012 | 0.0003 |
| F12 | 86980001 | 87080000 | 0.0006 | 0.0011 | 0.0002 |
| F12 | 87000001 | 87100000 | 0.0008 | 0.0011 | 0.0002 |
| F12 | 87020001 | 87120000 | 0.0015 | 0.0016 | 0.0007 |
| F12 | 87040001 | 87140000 | 0.0017 | 0.0015 | 0.0008 |
| F12 | 87060001 | 87160000 | 0.0016 | 0.0013 | 0.0007 |
| F12 | 87080001 | 87180000 | 0.0019 | 0.0013 | 0.0009 |
| F12 | 87100001 | 87200000 | 0.0043 | 0.0035 | 0.0035 |
| F12 | 87120001 | 87220000 | 0.0041 | 0.0031 | 0.0034 |
| F12 | 87140001 | 87240000 | 0.0040 | 0.0031 | 0.0034 |
| F12 | 87160001 | 87260000 | 0.0041 | 0.0031 | 0.0034 |
| F12 | 87180001 | 87280000 | 0.0039 | 0.0029 | 0.0032 |
| F12 | 87200001 | 87300000 | 0.0013 | 0.0006 | 0.0007 |
| F12 | 87220001 | 87320000 | 0.0014 | 0.0005 | 0.0006 |
| F12 | 87240001 | 87340000 | 0.0016 | 0.0007 | 0.0007 |
| F12 | 87260001 | 87360000 | 0.0019 | 0.0010 | 0.0011 |
| F12 | 87280001 | 87380000 | 0.0020 | 0.0013 | 0.0013 |
| F12 | 87300001 | 87400000 | 0.0023 | 0.0014 | 0.0015 |
| F12 | 87320001 | 87420000 | 0.0021 | 0.0012 | 0.0012 |
| F12 | 87340001 | 87440000 | 0.0019 | 0.0010 | 0.0010 |
| F12 | 87360001 | 87460000 | 0.0016 | 0.0007 | 0.0007 |
| F12 | 87380001 | 87480000 | 0.0016 | 0.0006 | 0.0006 |
| F12 | 87400001 | 87500000 | 0.0013 | 0.0004 | 0.0004 |
| F12 | 87420001 | 87520000 | 0.0012 | 0.0004 | 0.0005 |
| F12 | 87440001 | 87540000 | 0.0015 | 0.0006 | 0.0007 |
| F12 | 87460001 | 87560000 | 0.0016 | 0.0007 | 0.0008 |
| F12 | 87480001 | 87580000 | 0.0014 | 0.0005 | 0.0007 |

| Chr | Start | End | both | A1 | A2 |
|---|---|---|---|---|---|
| F12 | 87500001 | 87600000 | 0.0014 | 0.0005 | 0.0007 |
| F12 | 87520001 | 87620000 | 0.0014 | 0.0005 | 0.0007 |
| F12 | 87540001 | 87640000 | 0.0011 | 0.0002 | 0.0004 |
| F12 | 87560001 | 87660000 | 0.0013 | 0.0004 | 0.0006 |
| F12 | 87580001 | 87680000 | 0.0044 | 0.0034 | 0.0037 |
| F12 | 87600001 | 87700000 | 0.0044 | 0.0034 | 0.0037 |
| F12 | 87620001 | 87720000 | 0.0045 | 0.0035 | 0.0038 |
| F12 | 87640001 | 87740000 | 0.0044 | 0.0035 | 0.0037 |
| F12 | 87660001 | 87760000 | 0.0041 | 0.0032 | 0.0034 |
| F12 | 87680001 | 87780000 | 0.0012 | 0.0002 | 0.0003 |
| F12 | 87700001 | 87800000 | 0.0012 | 0.0004 | 0.0002 |
| F12 | 87720001 | 87820000 | 0.0013 | 0.0004 | 0.0003 |
| F12 | 87740001 | 87840000 | 0.0015 | 0.0006 | 0.0004 |
| F12 | 87760001 | 87860000 | 0.0016 | 0.0006 | 0.0004 |
| F12 | 87780001 | 87880000 | 0.0019 | 0.0010 | 0.0007 |
| F12 | 87800001 | 87900000 | 0.0019 | 0.0008 | 0.0007 |
| F12 | 87820001 | 87920000 | 0.0018 | 0.0008 | 0.0008 |
| F12 | 87840001 | 87940000 | 0.0017 | 0.0006 | 0.0007 |
| F12 | 87860001 | 87960000 | 0.0017 | 0.0006 | 0.0007 |
| F12 | 87880001 | 87980000 | 0.0014 | 0.0002 | 0.0005 |
| F12 | 87900001 | 88000000 | 0.0017 | 0.0004 | 0.0008 |
| F12 | 87920001 | 88020000 | 0.0016 | 0.0004 | 0.0007 |
| F12 | 87940001 | 88040000 | 0.0018 | 0.0004 | 0.0008 |
| F12 | 87960001 | 88060000 | 0.0016 | 0.0004 | 0.0008 |
| F12 | 87980001 | 88080000 | 0.0015 | 0.0004 | 0.0007 |
| F12 | 88000001 | 88100000 | 0.0014 | 0.0002 | 0.0006 |
| F12 | 88020001 | 88120000 | 0.0015 | 0.0002 | 0.0007 |
| F12 | 88040001 | 88140000 | 0.0015 | 0.0004 | 0.0008 |
| F12 | 88060001 | 88160000 | 0.0016 | 0.0004 | 0.0009 |
| F12 | 88080001 | 88180000 | 0.0018 | 0.0005 | 0.0010 |
| F12 | 88100001 | 88200000 | 0.0022 | 0.0009 | 0.0015 |
| F12 | 88120001 | 88220000 | 0.0022 | 0.0008 | 0.0016 |
| F12 | 88140001 | 88240000 | 0.0021 | 0.0006 | 0.0016 |
| F12 | 88160001 | 88260000 | 0.0022 | 0.0008 | 0.0019 |
| F12 | 88180001 | 88280000 | 0.0025 | 0.0012 | 0.0023 |
| F12 | 88200001 | 88300000 | 0.0023 | 0.0009 | 0.0023 |
| F12 | 88220001 | 88320000 | 0.0030 | 0.0012 | 0.0029 |
| F12 | 88240001 | 88340000 | 0.0032 | 0.0011 | 0.0031 |
| F12 | 88260001 | 88360000 | 0.0029 | 0.0010 | 0.0027 |
| F12 | 88280001 | 88380000 | 0.0026 | 0.0007 | 0.0022 |
| F12 | 88300001 | 88400000 | 0.0023 | 0.0010 | 0.0018 |
| F12 | 88320001 | 88420000 | 0.0016 | 0.0010 | 0.0009 |
| F12 | 88340001 | 88440000 | 0.0014 | 0.0011 | 0.0006 |
| F12 | 88360001 | 88460000 | 0.0014 | 0.0014 | 0.0005 |
| F12 | 88380001 | 88480000 | 0.0012 | 0.0014 | 0.0004 |
| F12 | 88400001 | 88500000 | 0.0011 | 0.0011 | 0.0002 |
| F12 | 88420001 | 88520000 | 0.0010 | 0.0009 | 0.0002 |
| F12 | 88440001 | 88540000 | 0.0011 | 0.0008 | 0.0002 |
| F12 | 88460001 | 88560000 | 0.0010 | 0.0007 | 0.0002 |
| F12 | 88480001 | 88580000 | 0.0010 | 0.0008 | 0.0003 |
| F12 | 88500001 | 88600000 | 0.0010 | 0.0010 | 0.0004 |
| F12 | 88520001 | 88620000 | 0.0009 | 0.0011 | 0.0003 |
| F12 | 88540001 | 88640000 | 0.0008 | 0.0014 | 0.0005 |
| F12 | 88560001 | 88660000 | 0.0008 | 0.0014 | 0.0005 |
| F12 | 88580001 | 88680000 | 0.0008 | 0.0012 | 0.0006 |
| F12 | 88600001 | 88700000 | 0.0016 | 0.0020 | 0.0013 |
| F12 | 88620001 | 88720000 | 0.0017 | 0.0022 | 0.0013 |
| F12 | 88640001 | 88740000 | 0.0016 | 0.0020 | 0.0013 |

| Chr | Start | End | both | A1 | A2 |
|---|---|---|---|---|---|
| F12 | 88660001 | 88760000 | 0.0016 | 0.0018 | 0.0013 |
| F12 | 88680001 | 88780000 | 0.0018 | 0.0018 | 0.0014 |
| F12 | 88700001 | 88800000 | 0.0014 | 0.0009 | 0.0012 |
| F12 | 88720001 | 88820000 | 0.0014 | 0.0006 | 0.0013 |
| F12 | 88740001 | 88840000 | 0.0015 | 0.0004 | 0.0016 |
| F12 | 88760001 | 88860000 | 0.0018 | 0.0004 | 0.0018 |
| F12 | 88780001 | 88880000 | 0.0018 | 0.0003 | 0.0018 |
| F12 | 88800001 | 88900000 | 0.0016 | 0.0002 | 0.0015 |
| F12 | 88820001 | 88920000 | 0.0016 | 0.0002 | 0.0012 |
| F12 | 88840001 | 88940000 | 0.0016 | 0.0002 | 0.0010 |
| F12 | 88860001 | 88960000 | 0.0017 | 0.0003 | 0.0010 |
| F12 | 88880001 | 88980000 | 0.0016 | 0.0003 | 0.0009 |
| F12 | 88900001 | 89000000 | 0.0015 | 0.0004 | 0.0009 |
| F12 | 88920001 | 89020000 | 0.0019 | 0.0004 | 0.0012 |
| F12 | 88940001 | 89040000 | 0.0020 | 0.0005 | 0.0014 |
| F12 | 88960001 | 89060000 | 0.0019 | 0.0007 | 0.0012 |
| F12 | 88980001 | 89080000 | 0.0019 | 0.0007 | 0.0012 |
| F12 | 89000001 | 89100000 | 0.0020 | 0.0010 | 0.0012 |
| F12 | 89020001 | 89120000 | 0.0024 | 0.0019 | 0.0018 |
| F12 | 89040001 | 89140000 | 0.0022 | 0.0018 | 0.0016 |
| F12 | 89060001 | 89160000 | 0.0021 | 0.0016 | 0.0015 |
| F12 | 89080001 | 89180000 | 0.0021 | 0.0017 | 0.0016 |
| F12 | 89100001 | 89200000 | 0.0022 | 0.0018 | 0.0018 |
| F12 | 89120001 | 89220000 | 0.0014 | 0.0010 | 0.0010 |
| F12 | 89140001 | 89240000 | 0.0017 | 0.0012 | 0.0012 |
| F12 | 89160001 | 89260000 | 0.0022 | 0.0014 | 0.0017 |
| F12 | 89180001 | 89280000 | 0.0021 | 0.0013 | 0.0016 |
| F12 | 89200001 | 89300000 | 0.0019 | 0.0009 | 0.0013 |
| F12 | 89220001 | 89320000 | 0.0022 | 0.0011 | 0.0015 |
| F12 | 89240001 | 89340000 | 0.0021 | 0.0009 | 0.0014 |
| F12 | 89260001 | 89360000 | 0.0020 | 0.0008 | 0.0011 |
| F12 | 89280001 | 89380000 | 0.0018 | 0.0008 | 0.0010 |
| F12 | 89300001 | 89400000 | 0.0017 | 0.0009 | 0.0010 |
| F12 | 89320001 | 89420000 | 0.0016 | 0.0006 | 0.0007 |
| F12 | 89340001 | 89440000 | 0.0016 | 0.0006 | 0.0006 |
| F12 | 89360001 | 89460000 | 0.0013 | 0.0003 | 0.0002 |
| F12 | 89380001 | 89480000 | 0.0015 | 0.0003 | 0.0003 |
| F12 | 89400001 | 89500000 | 0.0016 | 0.0003 | 0.0004 |
| F12 | 89420001 | 89520000 | 0.0016 | 0.0002 | 0.0006 |
| F12 | 89440001 | 89540000 | 0.0017 | 0.0001 | 0.0007 |
| F12 | 89460001 | 89560000 | 0.0016 | 0.0002 | 0.0007 |
| F12 | 89480001 | 89580000 | 0.0015 | 0.0003 | 0.0006 |
| F12 | 89500001 | 89600000 | 0.0014 | 0.0003 | 0.0005 |
| F12 | 89520001 | 89620000 | 0.0015 | 0.0004 | 0.0005 |
| F12 | 89540001 | 89640000 | 0.0015 | 0.0004 | 0.0006 |
| F12 | 89560001 | 89660000 | 0.0018 | 0.0004 | 0.0009 |
| F12 | 89580001 | 89680000 | 0.0020 | 0.0002 | 0.0011 |
| F12 | 89600001 | 89700000 | 0.0022 | 0.0003 | 0.0014 |
| F12 | 89620001 | 89720000 | 0.0022 | 0.0002 | 0.0015 |
| F12 | 89640001 | 89740000 | 0.0023 | 0.0003 | 0.0015 |
| F12 | 89660001 | 89760000 | 0.0022 | 0.0004 | 0.0015 |
| F12 | 89680001 | 89780000 | 0.0021 | 0.0004 | 0.0014 |
| F12 | 89700001 | 89800000 | 0.0020 | 0.0004 | 0.0012 |
| F12 | 89720001 | 89820000 | 0.0018 | 0.0004 | 0.0010 |
| F12 | 89740001 | 89840000 | 0.0017 | 0.0005 | 0.0010 |
| F12 | 89760001 | 89860000 | 0.0016 | 0.0004 | 0.0008 |
| F12 | 89780001 | 89880000 | 0.0018 | 0.0007 | 0.0011 |
| F12 | 89800001 | 89900000 | 0.0019 | 0.0006 | 0.0011 |

| Chr | Start | End | both | A1 | A2 |
|---|---|---|---|---|---|
| F12 | 89820001 | 89920000 | 0.0019 | 0.0006 | 0.0010 |
| F12 | 89840001 | 89940000 | 0.0018 | 0.0005 | 0.0008 |
| F12 | 89860001 | 89960000 | 0.0017 | 0.0004 | 0.0008 |
| F12 | 89880001 | 89980000 | 0.0014 | 0.0002 | 0.0004 |
| F12 | 89900001 | 90000000 | 0.0017 | 0.0006 | 0.0007 |
| F12 | 89920001 | 90020000 | 0.0018 | 0.0010 | 0.0009 |
| F12 | 89940001 | 90040000 | 0.0017 | 0.0013 | 0.0009 |
| F12 | 89960001 | 90060000 | 0.0016 | 0.0016 | 0.0009 |
| F12 | 89980001 | 90080000 | 0.0017 | 0.0021 | 0.0012 |
| F12 | 90000001 | 90100000 | 0.0016 | 0.0019 | 0.0012 |
| F12 | 90020001 | 90120000 | 0.0017 | 0.0017 | 0.0012 |
| F12 | 90040001 | 90140000 | 0.0021 | 0.0016 | 0.0013 |
| F12 | 90060001 | 90160000 | 0.0024 | 0.0013 | 0.0014 |
| F12 | 90080001 | 90180000 | 0.0022 | 0.0010 | 0.0012 |
| F12 | 90100001 | 90200000 | 0.0020 | 0.0008 | 0.0008 |
| F12 | 90120001 | 90220000 | 0.0018 | 0.0007 | 0.0007 |
| F12 | 90140001 | 90240000 | 0.0025 | 0.0014 | 0.0014 |
| F12 | 90160001 | 90260000 | 0.0028 | 0.0015 | 0.0016 |
| F12 | 90180001 | 90280000 | 0.0029 | 0.0014 | 0.0016 |
| F12 | 90200001 | 90300000 | 0.0029 | 0.0013 | 0.0015 |
| F12 | 90220001 | 90320000 | 0.0030 | 0.0013 | 0.0017 |
| F12 | 90240001 | 90340000 | 0.0024 | 0.0006 | 0.0013 |
| F12 | 90260001 | 90360000 | 0.0021 | 0.0004 | 0.0014 |
| F12 | 90280001 | 90380000 | 0.0021 | 0.0004 | 0.0016 |
| F12 | 90300001 | 90400000 | 0.0023 | 0.0005 | 0.0019 |
| F12 | 90320001 | 90420000 | 0.0026 | 0.0003 | 0.0023 |
| F12 | 90340001 | 90440000 | 0.0025 | 0.0001 | 0.0022 |
| F12 | 90360001 | 90460000 | 0.0027 | 0.0001 | 0.0024 |
| F12 | 90380001 | 90480000 | 0.0027 | 0.0001 | 0.0024 |
| F12 | 90400001 | 90500000 | 0.0027 | 0.0003 | 0.0025 |
| F12 | 90420001 | 90520000 | 0.0024 | 0.0003 | 0.0023 |
| F12 | 90440001 | 90540000 | 0.0025 | 0.0006 | 0.0025 |
| F12 | 90460001 | 90560000 | 0.0025 | 0.0011 | 0.0025 |
| F12 | 90480001 | 90580000 | 0.0026 | 0.0012 | 0.0025 |
| F12 | 90500001 | 90600000 | 0.0026 | 0.0014 | 0.0026 |
| F12 | 90520001 | 90620000 | 0.0025 | 0.0015 | 0.0024 |
| F12 | 90540001 | 90640000 | 0.0022 | 0.0014 | 0.0019 |
| F12 | 90560001 | 90660000 | 0.0018 | 0.0011 | 0.0014 |
| F12 | 90580001 | 90680000 | 0.0017 | 0.0012 | 0.0011 |
| F12 | 90600001 | 90700000 | 0.0016 | 0.0012 | 0.0008 |
| F12 | 90620001 | 90720000 | 0.0016 | 0.0013 | 0.0006 |
| F12 | 90640001 | 90740000 | 0.0017 | 0.0014 | 0.0008 |
| F12 | 90660001 | 90760000 | 0.0018 | 0.0015 | 0.0010 |
| F12 | 90680001 | 90780000 | 0.0019 | 0.0014 | 0.0012 |
| F12 | 90700001 | 90800000 | 0.0018 | 0.0012 | 0.0012 |
| F12 | 90720001 | 90820000 | 0.0016 | 0.0009 | 0.0011 |
| F12 | 90740001 | 90840000 | 0.0018 | 0.0009 | 0.0011 |
| F12 | 90760001 | 90860000 | 0.0019 | 0.0007 | 0.0011 |
| F12 | 90780001 | 90880000 | 0.0017 | 0.0005 | 0.0010 |
| F12 | 90800001 | 90900000 | 0.0018 | 0.0004 | 0.0012 |
| F12 | 90820001 | 90920000 | 0.0021 | 0.0005 | 0.0014 |
| F12 | 90840001 | 90940000 | 0.0019 | 0.0002 | 0.0014 |
| F12 | 90860001 | 90960000 | 0.0019 | 0.0002 | 0.0015 |
| F12 | 90880001 | 90980000 | 0.0019 | 0.0002 | 0.0015 |
| F12 | 90900001 | 91000000 | 0.0020 | 0.0002 | 0.0015 |
| F12 | 90920001 | 91020000 | 0.0026 | 0.0010 | 0.0021 |
| F12 | 90940001 | 91040000 | 0.0027 | 0.0013 | 0.0022 |
| F12 | 90960001 | 91060000 | 0.0029 | 0.0017 | 0.0024 |

| Chr | Start | End | both | A1 | A2 |
|---|---|---|---|---|---|
| F12 | 90980001 | 91080000 | 0.0028 | 0.0017 | 0.0024 |
| F12 | 91000001 | 91100000 | 0.0029 | 0.0020 | 0.0025 |
| F12 | 91020001 | 91120000 | 0.0022 | 0.0013 | 0.0019 |
| F12 | 91040001 | 91140000 | 0.0018 | 0.0010 | 0.0016 |
| F12 | 91060001 | 91160000 | 0.0015 | 0.0007 | 0.0013 |
| F12 | 91080001 | 91180000 | 0.0014 | 0.0006 | 0.0011 |
| F12 | 91100001 | 91200000 | 0.0012 | 0.0003 | 0.0009 |
| F12 | 91120001 | 91220000 | 0.0009 | 0.0002 | 0.0008 |
| F12 | 91140001 | 91240000 | 0.0012 | 0.0004 | 0.0010 |
| F12 | 91160001 | 91260000 | 0.0015 | 0.0007 | 0.0013 |
| F12 | 91180001 | 91280000 | 0.0025 | 0.0015 | 0.0022 |
| F12 | 91200001 | 91300000 | 0.0024 | 0.0017 | 0.0022 |
| F12 | 91220001 | 91320000 | 0.0023 | 0.0017 | 0.0021 |
| F12 | 91240001 | 91340000 | 0.0018 | 0.0014 | 0.0017 |
| F12 | 91260001 | 91360000 | 0.0013 | 0.0011 | 0.0012 |
| F12 | 91280001 | 91380000 | 0.0002 | 0.0002 | 0.0002 |
| F13 | 1 | 100000 | 0.0040 | 0.0014 | 0.0038 |
| F13 | 20001 | 120000 | 0.0044 | 0.0037 | 0.0040 |
| F13 | 40001 | 140000 | 0.0046 | 0.0038 | 0.0041 |
| F13 | 60001 | 160000 | 0.0038 | 0.0028 | 0.0030 |
| F13 | 80001 | 180000 | 0.0029 | 0.0016 | 0.0019 |
| F13 | 100001 | 200000 | 0.0020 | 0.0006 | 0.0010 |
| F13 | 120001 | 220000 | 0.0019 | 0.0006 | 0.0009 |
| F13 | 140001 | 240000 | 0.0026 | 0.0011 | 0.0012 |
| F13 | 160001 | 260000 | 0.0031 | 0.0017 | 0.0019 |
| F13 | 180001 | 280000 | 0.0030 | 0.0018 | 0.0018 |
| F13 | 200001 | 300000 | 0.0028 | 0.0017 | 0.0017 |
| F13 | 220001 | 320000 | 0.0029 | 0.0019 | 0.0020 |
| F13 | 240001 | 340000 | 0.0024 | 0.0017 | 0.0020 |
| F13 | 260001 | 360000 | 0.0020 | 0.0014 | 0.0018 |
| F13 | 280001 | 380000 | 0.0021 | 0.0013 | 0.0018 |
| F13 | 300001 | 400000 | 0.0021 | 0.0012 | 0.0017 |
| F13 | 320001 | 420000 | 0.0023 | 0.0011 | 0.0019 |
| F13 | 340001 | 440000 | 0.0023 | 0.0009 | 0.0018 |
| F13 | 360001 | 460000 | 0.0024 | 0.0006 | 0.0019 |
| F13 | 380001 | 480000 | 0.0023 | 0.0006 | 0.0020 |
| F13 | 400001 | 500000 | 0.0025 | 0.0008 | 0.0024 |
| F13 | 420001 | 520000 | 0.0024 | 0.0012 | 0.0024 |
| F13 | 440001 | 540000 | 0.0023 | 0.0014 | 0.0024 |
| F13 | 460001 | 560000 | 0.0030 | 0.0026 | 0.0031 |
| F13 | 480001 | 580000 | 0.0046 | 0.0044 | 0.0047 |
| F13 | 500001 | 600000 | 0.0043 | 0.0043 | 0.0043 |
| F13 | 520001 | 620000 | 0.0042 | 0.0040 | 0.0042 |
| F13 | 540001 | 640000 | 0.0051 | 0.0050 | 0.0051 |
| F13 | 560001 | 660000 | 0.0046 | 0.0037 | 0.0047 |
| F13 | 580001 | 680000 | 0.0032 | 0.0023 | 0.0033 |
| F13 | 600001 | 700000 | 0.0036 | 0.0023 | 0.0038 |
| F13 | 620001 | 720000 | 0.0035 | 0.0023 | 0.0037 |
| F13 | 640001 | 740000 | 0.0026 | 0.0013 | 0.0028 |
| F13 | 660001 | 760000 | 0.0020 | 0.0012 | 0.0021 |
| F13 | 680001 | 780000 | 0.0017 | 0.0008 | 0.0019 |
| F13 | 700001 | 800000 | 0.0016 | 0.0006 | 0.0017 |
| F13 | 720001 | 820000 | 0.0017 | 0.0005 | 0.0018 |
| F13 | 740001 | 840000 | 0.0022 | 0.0004 | 0.0022 |
| F13 | 760001 | 860000 | 0.0031 | 0.0010 | 0.0028 |
| F13 | 780001 | 880000 | 0.0033 | 0.0010 | 0.0028 |
| F13 | 800001 | 900000 | 0.0038 | 0.0017 | 0.0028 |
| F13 | 820001 | 920000 | 0.0042 | 0.0023 | 0.0030 |

| Chr | Start | End | both | A1 | A2 |
|---|---|---|---|---|---|
| F13 | 840001 | 940000 | 0.0054 | 0.0039 | 0.0036 |
| F13 | 860001 | 960000 | 0.0050 | 0.0041 | 0.0031 |
| F13 | 880001 | 980000 | 0.0047 | 0.0042 | 0.0031 |
| F13 | 900001 | 1000000 | 0.0042 | 0.0035 | 0.0030 |
| F13 | 920001 | 1020000 | 0.0036 | 0.0028 | 0.0026 |
| F13 | 940001 | 1040000 | 0.0019 | 0.0009 | 0.0016 |
| F13 | 960001 | 1060000 | 0.0016 | 0.0004 | 0.0017 |
| F13 | 980001 | 1080000 | 0.0019 | 0.0004 | 0.0021 |
| F13 | 1000001 | 1100000 | 0.0019 | 0.0006 | 0.0019 |
| F13 | 1020001 | 1120000 | 0.0019 | 0.0008 | 0.0017 |
| F13 | 1040001 | 1140000 | 0.0020 | 0.0012 | 0.0019 |
| F13 | 1060001 | 1160000 | 0.0022 | 0.0016 | 0.0020 |
| F13 | 1080001 | 1180000 | 0.0022 | 0.0021 | 0.0020 |
| F13 | 1100001 | 1200000 | 0.0022 | 0.0021 | 0.0021 |
| F13 | 1120001 | 1220000 | 0.0023 | 0.0022 | 0.0022 |
| F13 | 1140001 | 1240000 | 0.0025 | 0.0024 | 0.0023 |
| F13 | 1160001 | 1260000 | 0.0030 | 0.0030 | 0.0026 |
| F13 | 1180001 | 1280000 | 0.0028 | 0.0027 | 0.0022 |
| F13 | 1200001 | 1300000 | 0.0034 | 0.0032 | 0.0026 |
| F13 | 1220001 | 1320000 | 0.0042 | 0.0040 | 0.0034 |
| F13 | 1240001 | 1340000 | 0.0039 | 0.0036 | 0.0031 |
| F13 | 1260001 | 1360000 | 0.0032 | 0.0027 | 0.0023 |
| F13 | 1280001 | 1380000 | 0.0032 | 0.0024 | 0.0023 |
| F13 | 1300001 | 1400000 | 0.0028 | 0.0019 | 0.0021 |
| F13 | 1320001 | 1420000 | 0.0021 | 0.0010 | 0.0016 |
| F13 | 1340001 | 1440000 | 0.0024 | 0.0011 | 0.0021 |
| F13 | 1360001 | 1460000 | 0.0026 | 0.0012 | 0.0024 |
| F13 | 1380001 | 1480000 | 0.0026 | 0.0014 | 0.0025 |
| F13 | 1400001 | 1500000 | 0.0026 | 0.0014 | 0.0024 |
| F13 | 1420001 | 1520000 | 0.0029 | 0.0016 | 0.0025 |
| F13 | 1440001 | 1540000 | 0.0029 | 0.0015 | 0.0023 |
| F13 | 1460001 | 1560000 | 0.0029 | 0.0015 | 0.0023 |
| F13 | 1480001 | 1580000 | 0.0029 | 0.0014 | 0.0024 |
| F13 | 1500001 | 1600000 | 0.0026 | 0.0014 | 0.0021 |
| F13 | 1520001 | 1620000 | 0.0031 | 0.0023 | 0.0026 |
| F13 | 1540001 | 1640000 | 0.0035 | 0.0030 | 0.0030 |
| F13 | 1560001 | 1660000 | 0.0032 | 0.0031 | 0.0027 |
| F13 | 1580001 | 1680000 | 0.0031 | 0.0034 | 0.0026 |
| F13 | 1600001 | 1700000 | 0.0029 | 0.0035 | 0.0024 |
| F13 | 1620001 | 1720000 | 0.0022 | 0.0029 | 0.0016 |
| F13 | 1640001 | 1740000 | 0.0014 | 0.0025 | 0.0008 |
| F13 | 1660001 | 1760000 | 0.0015 | 0.0027 | 0.0008 |
| F13 | 1680001 | 1780000 | 0.0014 | 0.0024 | 0.0007 |
| F13 | 1700001 | 1800000 | 0.0018 | 0.0023 | 0.0011 |
| F13 | 1720001 | 1820000 | 0.0017 | 0.0018 | 0.0011 |
| F13 | 1740001 | 1840000 | 0.0018 | 0.0014 | 0.0011 |
| F13 | 1760001 | 1860000 | 0.0021 | 0.0012 | 0.0014 |
| F13 | 1780001 | 1880000 | 0.0023 | 0.0012 | 0.0014 |
| F13 | 1800001 | 1900000 | 0.0020 | 0.0008 | 0.0011 |
| F13 | 1820001 | 1920000 | 0.0022 | 0.0008 | 0.0012 |
| F13 | 1840001 | 1940000 | 0.0022 | 0.0009 | 0.0012 |
| F13 | 1860001 | 1960000 | 0.0018 | 0.0006 | 0.0009 |
| F13 | 1880001 | 1980000 | 0.0017 | 0.0005 | 0.0008 |
| F13 | 1900001 | 2000000 | 0.0016 | 0.0004 | 0.0007 |
| F13 | 1920001 | 2020000 | 0.0013 | 0.0004 | 0.0005 |
| F13 | 1940001 | 2040000 | 0.0014 | 0.0004 | 0.0004 |
| F13 | 1960001 | 2060000 | 0.0029 | 0.0020 | 0.0020 |
| F13 | 1980001 | 2080000 | 0.0028 | 0.0022 | 0.0021 |

| Chr | Start | End | both | A1 | A2 |
|---|---|---|---|---|---|
| F13 | 2000001 | 2100000 | 0.0033 | 0.0027 | 0.0026 |
| F13 | 2020001 | 2120000 | 0.0034 | 0.0028 | 0.0027 |
| F13 | 2040001 | 2140000 | 0.0037 | 0.0033 | 0.0033 |
| F13 | 2060001 | 2160000 | 0.0022 | 0.0018 | 0.0019 |
| F13 | 2080001 | 2180000 | 0.0023 | 0.0019 | 0.0021 |
| F13 | 2100001 | 2200000 | 0.0020 | 0.0017 | 0.0019 |
| F13 | 2120001 | 2220000 | 0.0023 | 0.0019 | 0.0022 |
| F13 | 2140001 | 2240000 | 0.0022 | 0.0019 | 0.0020 |
| F13 | 2160001 | 2260000 | 0.0025 | 0.0021 | 0.0021 |
| F13 | 2180001 | 2280000 | 0.0024 | 0.0019 | 0.0019 |
| F13 | 2200001 | 2300000 | 0.0025 | 0.0021 | 0.0017 |
| F13 | 2220001 | 2320000 | 0.0021 | 0.0019 | 0.0013 |
| F13 | 2240001 | 2340000 | 0.0017 | 0.0017 | 0.0007 |
| F13 | 2260001 | 2360000 | 0.0017 | 0.0020 | 0.0007 |
| F13 | 2280001 | 2380000 | 0.0019 | 0.0023 | 0.0009 |
| F13 | 2300001 | 2400000 | 0.0016 | 0.0017 | 0.0007 |
| F13 | 2320001 | 2420000 | 0.0016 | 0.0015 | 0.0008 |
| F13 | 2340001 | 2440000 | 0.0015 | 0.0011 | 0.0009 |
| F13 | 2360001 | 2460000 | 0.0021 | 0.0012 | 0.0011 |
| F13 | 2380001 | 2480000 | 0.0019 | 0.0010 | 0.0008 |
| F13 | 2400001 | 2500000 | 0.0025 | 0.0017 | 0.0014 |
| F13 | 2420001 | 2520000 | 0.0024 | 0.0017 | 0.0013 |
| F13 | 2440001 | 2540000 | 0.0024 | 0.0017 | 0.0012 |
| F13 | 2460001 | 2560000 | 0.0017 | 0.0014 | 0.0008 |
| F13 | 2480001 | 2580000 | 0.0015 | 0.0013 | 0.0008 |
| F13 | 2500001 | 2600000 | 0.0012 | 0.0008 | 0.0006 |
| F13 | 2520001 | 2620000 | 0.0016 | 0.0011 | 0.0010 |
| F13 | 2540001 | 2640000 | 0.0020 | 0.0013 | 0.0013 |
| F13 | 2560001 | 2660000 | 0.0030 | 0.0021 | 0.0023 |
| F13 | 2580001 | 2680000 | 0.0036 | 0.0024 | 0.0029 |
| F13 | 2600001 | 2700000 | 0.0042 | 0.0031 | 0.0037 |
| F13 | 2620001 | 2720000 | 0.0044 | 0.0030 | 0.0037 |
| F13 | 2640001 | 2740000 | 0.0042 | 0.0029 | 0.0036 |
| F13 | 2660001 | 2760000 | 0.0031 | 0.0019 | 0.0026 |
| F13 | 2680001 | 2780000 | 0.0026 | 0.0016 | 0.0022 |
| F13 | 2700001 | 2800000 | 0.0018 | 0.0007 | 0.0011 |
| F13 | 2720001 | 2820000 | 0.0015 | 0.0008 | 0.0010 |
| F13 | 2740001 | 2840000 | 0.0021 | 0.0010 | 0.0012 |
| F13 | 2760001 | 2860000 | 0.0028 | 0.0013 | 0.0015 |
| F13 | 2780001 | 2880000 | 0.0041 | 0.0022 | 0.0023 |
| F13 | 2800001 | 2900000 | 0.0042 | 0.0023 | 0.0024 |
| F13 | 2820001 | 2920000 | 0.0042 | 0.0021 | 0.0022 |
| F13 | 2840001 | 2940000 | 0.0037 | 0.0017 | 0.0018 |
| F13 | 2860001 | 2960000 | 0.0030 | 0.0013 | 0.0015 |
| F13 | 2880001 | 2980000 | 0.0017 | 0.0004 | 0.0005 |
| F13 | 2900001 | 3000000 | 0.0015 | 0.0003 | 0.0004 |
| F13 | 2920001 | 3020000 | 0.0015 | 0.0003 | 0.0004 |
| F13 | 2940001 | 3040000 | 0.0017 | 0.0003 | 0.0004 |
| F13 | 2960001 | 3060000 | 0.0015 | 0.0003 | 0.0004 |
| F13 | 2980001 | 3080000 | 0.0017 | 0.0005 | 0.0007 |
| F13 | 3000001 | 3100000 | 0.0019 | 0.0007 | 0.0009 |
| F13 | 3020001 | 3120000 | 0.0022 | 0.0015 | 0.0011 |
| F13 | 3040001 | 3140000 | 0.0030 | 0.0027 | 0.0022 |
| F13 | 3060001 | 3160000 | 0.0033 | 0.0028 | 0.0023 |
| F13 | 3080001 | 3180000 | 0.0033 | 0.0025 | 0.0020 |
| F13 | 3100001 | 3200000 | 0.0032 | 0.0023 | 0.0018 |
| F13 | 3120001 | 3220000 | 0.0029 | 0.0017 | 0.0018 |
| F13 | 3140001 | 3240000 | 0.0017 | 0.0005 | 0.0007 |

| Chr | Start | End | both | A1 | A2 |
|---|---|---|---|---|---|
| F13 | 3160001 | 3260000 | 0.0016 | 0.0004 | 0.0005 |
| F13 | 3180001 | 3280000 | 0.0015 | 0.0006 | 0.0007 |
| F13 | 3200001 | 3300000 | 0.0016 | 0.0007 | 0.0008 |
| F13 | 3220001 | 3320000 | 0.0015 | 0.0005 | 0.0007 |
| F13 | 3240001 | 3340000 | 0.0017 | 0.0007 | 0.0007 |
| F13 | 3260001 | 3360000 | 0.0043 | 0.0032 | 0.0031 |
| F13 | 3280001 | 3380000 | 0.0043 | 0.0032 | 0.0030 |
| F13 | 3300001 | 3400000 | 0.0045 | 0.0034 | 0.0032 |
| F13 | 3320001 | 3420000 | 0.0044 | 0.0032 | 0.0031 |
| F13 | 3340001 | 3440000 | 0.0043 | 0.0031 | 0.0030 |
| F13 | 3360001 | 3460000 | 0.0018 | 0.0005 | 0.0006 |
| F13 | 3380001 | 3480000 | 0.0017 | 0.0003 | 0.0004 |
| F13 | 3400001 | 3500000 | 0.0015 | 0.0002 | 0.0003 |
| F13 | 3420001 | 3520000 | 0.0017 | 0.0004 | 0.0004 |
| F13 | 3440001 | 3540000 | 0.0016 | 0.0004 | 0.0005 |
| F13 | 3460001 | 3560000 | 0.0020 | 0.0008 | 0.0010 |
| F13 | 3480001 | 3580000 | 0.0020 | 0.0008 | 0.0010 |
| F13 | 3500001 | 3600000 | 0.0020 | 0.0007 | 0.0008 |
| F13 | 3520001 | 3620000 | 0.0017 | 0.0005 | 0.0006 |
| F13 | 3540001 | 3640000 | 0.0019 | 0.0007 | 0.0007 |
| F13 | 3560001 | 3660000 | 0.0015 | 0.0004 | 0.0004 |
| F13 | 3580001 | 3680000 | 0.0016 | 0.0005 | 0.0005 |
| F13 | 3600001 | 3700000 | 0.0019 | 0.0008 | 0.0008 |
| F13 | 3620001 | 3720000 | 0.0022 | 0.0010 | 0.0011 |
| F13 | 3640001 | 3740000 | 0.0022 | 0.0011 | 0.0012 |
| F13 | 3660001 | 3760000 | 0.0025 | 0.0013 | 0.0015 |
| F13 | 3680001 | 3780000 | 0.0031 | 0.0021 | 0.0021 |
| F13 | 3700001 | 3800000 | 0.0040 | 0.0030 | 0.0030 |
| F13 | 3720001 | 3820000 | 0.0039 | 0.0029 | 0.0029 |
| F13 | 3740001 | 3840000 | 0.0048 | 0.0037 | 0.0036 |
| F13 | 3760001 | 3860000 | 0.0046 | 0.0037 | 0.0036 |
| F13 | 3780001 | 3880000 | 0.0057 | 0.0047 | 0.0048 |
| F13 | 3800001 | 3900000 | 0.0067 | 0.0054 | 0.0059 |
| F13 | 3820001 | 3920000 | 0.0072 | 0.0060 | 0.0064 |
| F13 | 3840001 | 3940000 | 0.0065 | 0.0055 | 0.0057 |
| F13 | 3860001 | 3960000 | 0.0063 | 0.0054 | 0.0055 |
| F13 | 3880001 | 3980000 | 0.0046 | 0.0036 | 0.0038 |
| F13 | 3900001 | 4000000 | 0.0022 | 0.0016 | 0.0014 |
| F13 | 3920001 | 4020000 | 0.0019 | 0.0013 | 0.0012 |
| F13 | 3940001 | 4040000 | 0.0021 | 0.0014 | 0.0015 |
| F13 | 3960001 | 4060000 | 0.0024 | 0.0015 | 0.0018 |
| F13 | 3980001 | 4080000 | 0.0026 | 0.0017 | 0.0020 |
| F13 | 4000001 | 4100000 | 0.0030 | 0.0020 | 0.0023 |
| F13 | 4020001 | 4120000 | 0.0032 | 0.0022 | 0.0024 |
| F13 | 4040001 | 4140000 | 0.0029 | 0.0019 | 0.0020 |
| F13 | 4060001 | 4160000 | 0.0031 | 0.0021 | 0.0021 |
| F13 | 4080001 | 4180000 | 0.0029 | 0.0023 | 0.0020 |
| F13 | 4100001 | 4200000 | 0.0033 | 0.0026 | 0.0025 |
| F13 | 4120001 | 4220000 | 0.0033 | 0.0026 | 0.0025 |
| F13 | 4140001 | 4240000 | 0.0032 | 0.0024 | 0.0022 |
| F13 | 4160001 | 4260000 | 0.0027 | 0.0019 | 0.0017 |
| F13 | 4180001 | 4280000 | 0.0031 | 0.0021 | 0.0021 |
| F13 | 4200001 | 4300000 | 0.0027 | 0.0015 | 0.0015 |
| F13 | 4220001 | 4320000 | 0.0024 | 0.0012 | 0.0012 |
| F13 | 4240001 | 4340000 | 0.0024 | 0.0011 | 0.0012 |
| F13 | 4260001 | 4360000 | 0.0025 | 0.0011 | 0.0012 |
| F13 | 4280001 | 4380000 | 0.0020 | 0.0007 | 0.0007 |
| F13 | 4300001 | 4400000 | 0.0020 | 0.0009 | 0.0009 |

| Chr | Start | End | both | A1 | A2 |
|---|---|---|---|---|---|
| F13 | 4320001 | 4420000 | 0.0020 | 0.0010 | 0.0009 |
| F13 | 4340001 | 4440000 | 0.0017 | 0.0010 | 0.0008 |
| F13 | 4360001 | 4460000 | 0.0019 | 0.0013 | 0.0012 |
| F13 | 4380001 | 4480000 | 0.0031 | 0.0025 | 0.0025 |
| F13 | 4400001 | 4500000 | 0.0029 | 0.0025 | 0.0025 |
| F13 | 4420001 | 4520000 | 0.0026 | 0.0023 | 0.0024 |
| F13 | 4440001 | 4540000 | 0.0028 | 0.0024 | 0.0025 |
| F13 | 4460001 | 4560000 | 0.0030 | 0.0025 | 0.0026 |
| F13 | 4480001 | 4580000 | 0.0027 | 0.0020 | 0.0022 |
| F13 | 4500001 | 4600000 | 0.0032 | 0.0023 | 0.0027 |
| F13 | 4520001 | 4620000 | 0.0034 | 0.0024 | 0.0028 |
| F13 | 4540001 | 4640000 | 0.0039 | 0.0029 | 0.0033 |
| F13 | 4560001 | 4660000 | 0.0035 | 0.0025 | 0.0029 |
| F13 | 4580001 | 4680000 | 0.0030 | 0.0018 | 0.0023 |
| F13 | 4600001 | 4700000 | 0.0027 | 0.0013 | 0.0016 |
| F13 | 4620001 | 4720000 | 0.0030 | 0.0013 | 0.0016 |
| F13 | 4640001 | 4740000 | 0.0025 | 0.0008 | 0.0010 |
| F13 | 4660001 | 4760000 | 0.0025 | 0.0009 | 0.0012 |
| F13 | 4680001 | 4780000 | 0.0019 | 0.0007 | 0.0007 |
| F13 | 4700001 | 4800000 | 0.0017 | 0.0008 | 0.0007 |
| F13 | 4720001 | 4820000 | 0.0014 | 0.0006 | 0.0006 |
| F13 | 4740001 | 4840000 | 0.0016 | 0.0010 | 0.0010 |
| F13 | 4760001 | 4860000 | 0.0017 | 0.0010 | 0.0009 |
| F13 | 4780001 | 4880000 | 0.0023 | 0.0014 | 0.0013 |
| F13 | 4800001 | 4900000 | 0.0032 | 0.0021 | 0.0022 |
| F13 | 4820001 | 4920000 | 0.0035 | 0.0024 | 0.0025 |
| F13 | 4840001 | 4940000 | 0.0032 | 0.0020 | 0.0021 |
| F13 | 4860001 | 4960000 | 0.0032 | 0.0021 | 0.0023 |
| F13 | 4880001 | 4980000 | 0.0028 | 0.0016 | 0.0019 |
| F13 | 4900001 | 5000000 | 0.0027 | 0.0018 | 0.0019 |
| F13 | 4920001 | 5020000 | 0.0029 | 0.0018 | 0.0022 |
| F13 | 4940001 | 5040000 | 0.0030 | 0.0020 | 0.0024 |
| F13 | 4960001 | 5060000 | 0.0029 | 0.0017 | 0.0022 |
| F13 | 4980001 | 5080000 | 0.0027 | 0.0017 | 0.0021 |
| F13 | 5000001 | 5100000 | 0.0018 | 0.0010 | 0.0013 |
| F13 | 5020001 | 5120000 | 0.0013 | 0.0005 | 0.0006 |
| F13 | 5040001 | 5140000 | 0.0012 | 0.0004 | 0.0006 |
| F13 | 5060001 | 5160000 | 0.0021 | 0.0014 | 0.0014 |
| F13 | 5080001 | 5180000 | 0.0028 | 0.0020 | 0.0021 |
| F13 | 5100001 | 5200000 | 0.0029 | 0.0019 | 0.0021 |
| F13 | 5120001 | 5220000 | 0.0030 | 0.0021 | 0.0024 |
| F13 | 5140001 | 5240000 | 0.0029 | 0.0022 | 0.0024 |
| F13 | 5160001 | 5260000 | 0.0025 | 0.0020 | 0.0021 |
| F13 | 5180001 | 5280000 | 0.0026 | 0.0023 | 0.0021 |
| F13 | 5200001 | 5300000 | 0.0036 | 0.0035 | 0.0031 |
| F13 | 5220001 | 5320000 | 0.0038 | 0.0038 | 0.0034 |
| F13 | 5240001 | 5340000 | 0.0036 | 0.0037 | 0.0032 |
| F13 | 5260001 | 5360000 | 0.0037 | 0.0036 | 0.0034 |
| F13 | 5280001 | 5380000 | 0.0033 | 0.0031 | 0.0030 |
| F13 | 5300001 | 5400000 | 0.0027 | 0.0020 | 0.0024 |
| F13 | 5320001 | 5420000 | 0.0031 | 0.0020 | 0.0027 |
| F13 | 5340001 | 5440000 | 0.0039 | 0.0026 | 0.0034 |
| F13 | 5360001 | 5460000 | 0.0035 | 0.0021 | 0.0029 |
| F13 | 5380001 | 5480000 | 0.0038 | 0.0021 | 0.0030 |
| F13 | 5400001 | 5500000 | 0.0033 | 0.0020 | 0.0026 |
| F13 | 5420001 | 5520000 | 0.0027 | 0.0019 | 0.0019 |
| F13 | 5440001 | 5540000 | 0.0026 | 0.0018 | 0.0017 |
| F13 | 5460001 | 5560000 | 0.0026 | 0.0019 | 0.0017 |

| Chr | Start | End | both | A1 | A2 |
|---|---|---|---|---|---|
| F13 | 5480001 | 5580000 | 0.0025 | 0.0019 | 0.0015 |
| F13 | 5500001 | 5600000 | 0.0028 | 0.0021 | 0.0019 |
| F13 | 5520001 | 5620000 | 0.0033 | 0.0024 | 0.0026 |
| F13 | 5540001 | 5640000 | 0.0040 | 0.0030 | 0.0032 |
| F13 | 5560001 | 5660000 | 0.0040 | 0.0029 | 0.0032 |
| F13 | 5580001 | 5680000 | 0.0037 | 0.0026 | 0.0028 |
| F13 | 5600001 | 5700000 | 0.0038 | 0.0026 | 0.0029 |
| F13 | 5620001 | 5720000 | 0.0032 | 0.0022 | 0.0023 |
| F13 | 5640001 | 5740000 | 0.0036 | 0.0025 | 0.0027 |
| F13 | 5660001 | 5760000 | 0.0035 | 0.0022 | 0.0026 |
| F13 | 5680001 | 5780000 | 0.0035 | 0.0022 | 0.0028 |
| F13 | 5700001 | 5800000 | 0.0035 | 0.0020 | 0.0028 |
| F13 | 5720001 | 5820000 | 0.0038 | 0.0020 | 0.0031 |
| F13 | 5740001 | 5840000 | 0.0044 | 0.0025 | 0.0038 |
| F13 | 5760001 | 5860000 | 0.0048 | 0.0029 | 0.0041 |
| F13 | 5780001 | 5880000 | 0.0054 | 0.0033 | 0.0047 |
| F13 | 5800001 | 5900000 | 0.0055 | 0.0034 | 0.0048 |
| F13 | 5820001 | 5920000 | 0.0054 | 0.0035 | 0.0048 |
| F13 | 5840001 | 5940000 | 0.0035 | 0.0021 | 0.0030 |
| F13 | 5860001 | 5960000 | 0.0035 | 0.0021 | 0.0031 |
| F13 | 5880001 | 5980000 | 0.0044 | 0.0035 | 0.0042 |
| F13 | 5900001 | 6000000 | 0.0043 | 0.0034 | 0.0040 |
| F13 | 5920001 | 6020000 | 0.0043 | 0.0032 | 0.0037 |
| F13 | 5940001 | 6040000 | 0.0042 | 0.0031 | 0.0036 |
| F13 | 5960001 | 6060000 | 0.0039 | 0.0029 | 0.0033 |
| F13 | 5980001 | 6080000 | 0.0024 | 0.0014 | 0.0017 |
| F13 | 6000001 | 6100000 | 0.0021 | 0.0012 | 0.0014 |
| F13 | 6020001 | 6120000 | 0.0018 | 0.0012 | 0.0012 |
| F13 | 6040001 | 6140000 | 0.0015 | 0.0009 | 0.0009 |
| F13 | 6060001 | 6160000 | 0.0015 | 0.0008 | 0.0008 |
| F13 | 6080001 | 6180000 | 0.0015 | 0.0007 | 0.0007 |
| F13 | 6100001 | 6200000 | 0.0015 | 0.0007 | 0.0008 |
| F13 | 6120001 | 6220000 | 0.0019 | 0.0008 | 0.0010 |
| F13 | 6140001 | 6240000 | 0.0024 | 0.0011 | 0.0012 |
| F13 | 6160001 | 6260000 | 0.0032 | 0.0020 | 0.0022 |
| F13 | 6180001 | 6280000 | 0.0037 | 0.0025 | 0.0027 |
| F13 | 6200001 | 6300000 | 0.0043 | 0.0031 | 0.0034 |
| F13 | 6220001 | 6320000 | 0.0042 | 0.0033 | 0.0035 |
| F13 | 6240001 | 6340000 | 0.0041 | 0.0033 | 0.0035 |
| F13 | 6260001 | 6360000 | 0.0039 | 0.0033 | 0.0033 |
| F13 | 6280001 | 6380000 | 0.0033 | 0.0027 | 0.0028 |
| F13 | 6300001 | 6400000 | 0.0025 | 0.0021 | 0.0022 |
| F13 | 6320001 | 6420000 | 0.0030 | 0.0025 | 0.0026 |
| F13 | 6340001 | 6440000 | 0.0031 | 0.0026 | 0.0027 |
| F13 | 6360001 | 6460000 | 0.0028 | 0.0021 | 0.0023 |
| F13 | 6380001 | 6480000 | 0.0030 | 0.0020 | 0.0023 |
| F13 | 6400001 | 6500000 | 0.0034 | 0.0024 | 0.0026 |
| F13 | 6420001 | 6520000 | 0.0027 | 0.0016 | 0.0018 |
| F13 | 6440001 | 6540000 | 0.0024 | 0.0013 | 0.0015 |
| F13 | 6460001 | 6560000 | 0.0025 | 0.0016 | 0.0018 |
| F13 | 6480001 | 6580000 | 0.0034 | 0.0026 | 0.0029 |
| F13 | 6500001 | 6600000 | 0.0034 | 0.0026 | 0.0029 |
| F13 | 6520001 | 6620000 | 0.0036 | 0.0028 | 0.0031 |
| F13 | 6540001 | 6640000 | 0.0038 | 0.0032 | 0.0034 |
| F13 | 6560001 | 6660000 | 0.0034 | 0.0028 | 0.0030 |
| F13 | 6580001 | 6680000 | 0.0032 | 0.0027 | 0.0027 |
| F13 | 6600001 | 6700000 | 0.0031 | 0.0027 | 0.0026 |
| F13 | 6620001 | 6720000 | 0.0036 | 0.0032 | 0.0033 |

| Chr | Start | End | both | A1 | A2 |
|---|---|---|---|---|---|
| F13 | 6640001 | 6740000 | 0.0042 | 0.0037 | 0.0038 |
| F13 | 6660001 | 6760000 | 0.0066 | 0.0058 | 0.0062 |
| F13 | 6680001 | 6780000 | 0.0069 | 0.0059 | 0.0064 |
| F13 | 6700001 | 6800000 | 0.0069 | 0.0059 | 0.0064 |
| F13 | 6720001 | 6820000 | 0.0067 | 0.0057 | 0.0062 |
| F13 | 6740001 | 6840000 | 0.0060 | 0.0049 | 0.0054 |
| F13 | 6760001 | 6860000 | 0.0041 | 0.0031 | 0.0033 |
| F13 | 6780001 | 6880000 | 0.0033 | 0.0023 | 0.0025 |
| F13 | 6800001 | 6900000 | 0.0033 | 0.0022 | 0.0025 |
| F13 | 6820001 | 6920000 | 0.0033 | 0.0021 | 0.0023 |
| F13 | 6840001 | 6940000 | 0.0036 | 0.0024 | 0.0027 |
| F13 | 6860001 | 6960000 | 0.0034 | 0.0024 | 0.0027 |
| F13 | 6880001 | 6980000 | 0.0029 | 0.0021 | 0.0024 |
| F13 | 6900001 | 7000000 | 0.0027 | 0.0020 | 0.0023 |
| F13 | 6920001 | 7020000 | 0.0023 | 0.0018 | 0.0020 |
| F13 | 6940001 | 7040000 | 0.0017 | 0.0014 | 0.0016 |
| F13 | 6960001 | 7060000 | 0.0011 | 0.0009 | 0.0010 |
| F13 | 6980001 | 7080000 | 0.0009 | 0.0008 | 0.0008 |
| F13 | 7000001 | 7100000 | 0.0017 | 0.0016 | 0.0015 |
| F13 | 7020001 | 7120000 | 0.0025 | 0.0021 | 0.0024 |
| F13 | 7040001 | 7140000 | 0.0036 | 0.0031 | 0.0034 |
| F13 | 7060001 | 7160000 | 0.0041 | 0.0036 | 0.0039 |
| F13 | 7080001 | 7180000 | 0.0045 | 0.0040 | 0.0043 |
| F13 | 7100001 | 7200000 | 0.0038 | 0.0032 | 0.0035 |
| F13 | 7120001 | 7220000 | 0.0034 | 0.0028 | 0.0029 |
| F13 | 7140001 | 7240000 | 0.0030 | 0.0022 | 0.0024 |
| F13 | 7160001 | 7260000 | 0.0028 | 0.0019 | 0.0021 |
| F13 | 7180001 | 7280000 | 0.0028 | 0.0019 | 0.0022 |
| F13 | 7200001 | 7300000 | 0.0030 | 0.0021 | 0.0023 |
| F13 | 7220001 | 7320000 | 0.0026 | 0.0017 | 0.0018 |
| F13 | 7240001 | 7340000 | 0.0023 | 0.0014 | 0.0014 |
| F13 | 7260001 | 7360000 | 0.0025 | 0.0013 | 0.0012 |
| F13 | 7280001 | 7380000 | 0.0022 | 0.0011 | 0.0010 |
| F13 | 7300001 | 7400000 | 0.0022 | 0.0011 | 0.0011 |
| F13 | 7320001 | 7420000 | 0.0028 | 0.0017 | 0.0018 |
| F13 | 7340001 | 7440000 | 0.0025 | 0.0016 | 0.0017 |
| F13 | 7360001 | 7460000 | 0.0025 | 0.0016 | 0.0017 |
| F13 | 7380001 | 7480000 | 0.0029 | 0.0020 | 0.0021 |
| F13 | 7400001 | 7500000 | 0.0029 | 0.0021 | 0.0024 |
| F13 | 7420001 | 7520000 | 0.0037 | 0.0026 | 0.0027 |
| F13 | 7440001 | 7540000 | 0.0042 | 0.0028 | 0.0029 |
| F13 | 7460001 | 7560000 | 0.0052 | 0.0039 | 0.0041 |
| F13 | 7480001 | 7580000 | 0.0048 | 0.0034 | 0.0037 |
| F13 | 7500001 | 7600000 | 0.0046 | 0.0034 | 0.0036 |
| F13 | 7520001 | 7620000 | 0.0034 | 0.0027 | 0.0028 |
| F13 | 7540001 | 7640000 | 0.0035 | 0.0029 | 0.0031 |
| F13 | 7560001 | 7660000 | 0.0029 | 0.0024 | 0.0025 |
| F13 | 7580001 | 7680000 | 0.0030 | 0.0023 | 0.0024 |
| F13 | 7600001 | 7700000 | 0.0027 | 0.0019 | 0.0019 |
| F13 | 7620001 | 7720000 | 0.0028 | 0.0019 | 0.0019 |
| F13 | 7640001 | 7740000 | 0.0036 | 0.0022 | 0.0027 |
| F13 | 7660001 | 7760000 | 0.0035 | 0.0018 | 0.0027 |
| F13 | 7680001 | 7780000 | 0.0040 | 0.0022 | 0.0033 |
| F13 | 7700001 | 7800000 | 0.0040 | 0.0022 | 0.0033 |
| F13 | 7720001 | 7820000 | 0.0043 | 0.0027 | 0.0038 |
| F13 | 7740001 | 7840000 | 0.0030 | 0.0019 | 0.0026 |
| F13 | 7760001 | 7860000 | 0.0036 | 0.0026 | 0.0033 |
| F13 | 7780001 | 7880000 | 0.0039 | 0.0027 | 0.0036 |

| Chr | Start | End | both | A1 | A2 |
|---|---|---|---|---|---|
| F13 | 7800001 | 7900000 | 0.0043 | 0.0029 | 0.0041 |
| F13 | 7820001 | 7920000 | 0.0036 | 0.0021 | 0.0034 |
| F13 | 7840001 | 7940000 | 0.0036 | 0.0020 | 0.0034 |
| F13 | 7860001 | 7960000 | 0.0035 | 0.0017 | 0.0032 |
| F13 | 7880001 | 7980000 | 0.0029 | 0.0013 | 0.0026 |
| F13 | 7900001 | 8000000 | 0.0030 | 0.0014 | 0.0025 |
| F13 | 7920001 | 8020000 | 0.0031 | 0.0014 | 0.0025 |
| F13 | 7940001 | 8040000 | 0.0033 | 0.0014 | 0.0024 |
| F13 | 7960001 | 8060000 | 0.0023 | 0.0007 | 0.0013 |
| F13 | 7980001 | 8080000 | 0.0018 | 0.0004 | 0.0008 |
| F13 | 8000001 | 8100000 | 0.0018 | 0.0005 | 0.0008 |
| F13 | 8020001 | 8120000 | 0.0025 | 0.0010 | 0.0013 |
| F13 | 8040001 | 8140000 | 0.0027 | 0.0015 | 0.0019 |
| F13 | 8060001 | 8160000 | 0.0028 | 0.0015 | 0.0018 |
| F13 | 8080001 | 8180000 | 0.0029 | 0.0016 | 0.0020 |
| F13 | 8100001 | 8200000 | 0.0027 | 0.0015 | 0.0019 |
| F13 | 8120001 | 8220000 | 0.0025 | 0.0016 | 0.0018 |
| F13 | 8140001 | 8240000 | 0.0020 | 0.0010 | 0.0011 |
| F13 | 8160001 | 8260000 | 0.0020 | 0.0011 | 0.0013 |
| F13 | 8180001 | 8280000 | 0.0018 | 0.0010 | 0.0011 |
| F13 | 8200001 | 8300000 | 0.0017 | 0.0010 | 0.0012 |
| F13 | 8220001 | 8320000 | 0.0014 | 0.0006 | 0.0007 |
| F13 | 8240001 | 8340000 | 0.0013 | 0.0006 | 0.0007 |
| F13 | 8260001 | 8360000 | 0.0014 | 0.0008 | 0.0009 |
| F13 | 8280001 | 8380000 | 0.0019 | 0.0011 | 0.0010 |
| F13 | 8300001 | 8400000 | 0.0039 | 0.0028 | 0.0029 |
| F13 | 8320001 | 8420000 | 0.0063 | 0.0051 | 0.0052 |
| F13 | 8340001 | 8440000 | 0.0075 | 0.0061 | 0.0064 |
| F13 | 8360001 | 8460000 | 0.0081 | 0.0066 | 0.0068 |
| F13 | 8380001 | 8480000 | 0.0098 | 0.0082 | 0.0088 |
| F13 | 8400001 | 8500000 | 0.0086 | 0.0072 | 0.0078 |
| F13 | 8420001 | 8520000 | 0.0063 | 0.0051 | 0.0055 |
| F13 | 8440001 | 8540000 | 0.0053 | 0.0042 | 0.0045 |
| F13 | 8460001 | 8560000 | 0.0049 | 0.0037 | 0.0041 |
| F13 | 8480001 | 8580000 | 0.0028 | 0.0018 | 0.0020 |
| F13 | 8500001 | 8600000 | 0.0019 | 0.0011 | 0.0012 |
| F13 | 8520001 | 8620000 | 0.0021 | 0.0014 | 0.0015 |
| F13 | 8540001 | 8640000 | 0.0023 | 0.0017 | 0.0018 |
| F13 | 8560001 | 8660000 | 0.0026 | 0.0019 | 0.0020 |
| F13 | 8580001 | 8680000 | 0.0031 | 0.0024 | 0.0025 |
| F13 | 8600001 | 8700000 | 0.0037 | 0.0028 | 0.0030 |
| F13 | 8620001 | 8720000 | 0.0034 | 0.0024 | 0.0026 |
| F13 | 8640001 | 8740000 | 0.0032 | 0.0022 | 0.0024 |
| F13 | 8660001 | 8760000 | 0.0033 | 0.0024 | 0.0026 |
| F13 | 8680001 | 8780000 | 0.0033 | 0.0022 | 0.0025 |
| F13 | 8700001 | 8800000 | 0.0033 | 0.0018 | 0.0022 |
| F13 | 8720001 | 8820000 | 0.0038 | 0.0021 | 0.0025 |
| F13 | 8740001 | 8840000 | 0.0043 | 0.0025 | 0.0030 |
| F13 | 8760001 | 8860000 | 0.0045 | 0.0027 | 0.0032 |
| F13 | 8780001 | 8880000 | 0.0042 | 0.0025 | 0.0030 |
| F13 | 8800001 | 8900000 | 0.0038 | 0.0027 | 0.0030 |
| F13 | 8820001 | 8920000 | 0.0032 | 0.0025 | 0.0028 |
| F13 | 8840001 | 8940000 | 0.0027 | 0.0020 | 0.0023 |
| F13 | 8860001 | 8960000 | 0.0017 | 0.0012 | 0.0015 |
| F13 | 8880001 | 8980000 | 0.0017 | 0.0015 | 0.0017 |
| F13 | 8900001 | 9000000 | 0.0016 | 0.0013 | 0.0016 |
| F13 | 8920001 | 9020000 | 0.0019 | 0.0016 | 0.0019 |
| F13 | 8940001 | 9040000 | 0.0022 | 0.0019 | 0.0022 |

| Chr | Start | End | both | A1 | A2 |
|---|---|---|---|---|---|
| F13 | 8960001 | 9060000 | 0.0022 | 0.0018 | 0.0021 |
| F13 | 8980001 | 9080000 | 0.0020 | 0.0016 | 0.0019 |
| F13 | 9000001 | 9100000 | 0.0028 | 0.0022 | 0.0026 |
| F13 | 9020001 | 9120000 | 0.0031 | 0.0021 | 0.0027 |
| F13 | 9040001 | 9140000 | 0.0031 | 0.0022 | 0.0027 |
| F13 | 9060001 | 9160000 | 0.0035 | 0.0024 | 0.0030 |
| F13 | 9080001 | 9180000 | 0.0037 | 0.0024 | 0.0031 |
| F13 | 9100001 | 9200000 | 0.0031 | 0.0018 | 0.0024 |
| F13 | 9120001 | 9220000 | 0.0028 | 0.0016 | 0.0022 |
| F13 | 9140001 | 9240000 | 0.0024 | 0.0011 | 0.0016 |
| F13 | 9160001 | 9260000 | 0.0025 | 0.0013 | 0.0018 |
| F13 | 9180001 | 9280000 | 0.0028 | 0.0017 | 0.0022 |
| F13 | 9200001 | 9300000 | 0.0028 | 0.0015 | 0.0020 |
| F13 | 9220001 | 9320000 | 0.0036 | 0.0027 | 0.0030 |
| F13 | 9240001 | 9340000 | 0.0036 | 0.0029 | 0.0030 |
| F13 | 9260001 | 9360000 | 0.0039 | 0.0032 | 0.0032 |
| F13 | 9280001 | 9380000 | 0.0044 | 0.0035 | 0.0035 |
| F13 | 9300001 | 9400000 | 0.0041 | 0.0034 | 0.0034 |
| F13 | 9320001 | 9420000 | 0.0036 | 0.0028 | 0.0029 |
| F13 | 9340001 | 9440000 | 0.0049 | 0.0036 | 0.0040 |
| F13 | 9360001 | 9460000 | 0.0045 | 0.0032 | 0.0035 |
| F13 | 9380001 | 9480000 | 0.0041 | 0.0031 | 0.0034 |
| F13 | 9400001 | 9500000 | 0.0045 | 0.0035 | 0.0037 |
| F13 | 9420001 | 9520000 | 0.0040 | 0.0028 | 0.0029 |
| F13 | 9440001 | 9540000 | 0.0031 | 0.0022 | 0.0022 |
| F13 | 9460001 | 9560000 | 0.0033 | 0.0023 | 0.0024 |
| F13 | 9480001 | 9580000 | 0.0028 | 0.0017 | 0.0017 |
| F13 | 9500001 | 9600000 | 0.0028 | 0.0014 | 0.0016 |
| F13 | 9520001 | 9620000 | 0.0028 | 0.0015 | 0.0017 |
| F13 | 9540001 | 9640000 | 0.0030 | 0.0017 | 0.0019 |
| F13 | 9560001 | 9660000 | 0.0027 | 0.0014 | 0.0016 |
| F13 | 9580001 | 9680000 | 0.0027 | 0.0016 | 0.0019 |
| F13 | 9600001 | 9700000 | 0.0023 | 0.0013 | 0.0016 |
| F13 | 9620001 | 9720000 | 0.0021 | 0.0014 | 0.0017 |
| F13 | 9640001 | 9740000 | 0.0017 | 0.0010 | 0.0014 |
| F13 | 9660001 | 9760000 | 0.0028 | 0.0020 | 0.0023 |
| F13 | 9680001 | 9780000 | 0.0032 | 0.0025 | 0.0027 |
| F13 | 9700001 | 9800000 | 0.0033 | 0.0026 | 0.0028 |
| F13 | 9720001 | 9820000 | 0.0034 | 0.0028 | 0.0028 |
| F13 | 9740001 | 9840000 | 0.0037 | 0.0031 | 0.0032 |
| F13 | 9760001 | 9860000 | 0.0031 | 0.0026 | 0.0028 |
| F13 | 9780001 | 9880000 | 0.0033 | 0.0026 | 0.0029 |
| F13 | 9800001 | 9900000 | 0.0047 | 0.0037 | 0.0041 |
| F13 | 9820001 | 9920000 | 0.0048 | 0.0037 | 0.0042 |
| F13 | 9840001 | 9940000 | 0.0054 | 0.0044 | 0.0047 |
| F13 | 9860001 | 9960000 | 0.0053 | 0.0043 | 0.0045 |
| F13 | 9880001 | 9980000 | 0.0056 | 0.0049 | 0.0048 |
| F13 | 9900001 | 10000000 | 0.0047 | 0.0041 | 0.0041 |
| F13 | 9920001 | 10020000 | 0.0054 | 0.0047 | 0.0047 |
| F13 | 9940001 | 10040000 | 0.0050 | 0.0040 | 0.0042 |
| F13 | 9960001 | 10060000 | 0.0060 | 0.0047 | 0.0053 |
| F13 | 9980001 | 10080000 | 0.0056 | 0.0041 | 0.0049 |
| F13 | 10000001 | 10100000 | 0.0067 | 0.0049 | 0.0059 |
| F13 | 10020001 | 10120000 | 0.0058 | 0.0043 | 0.0050 |
| F13 | 10040001 | 10140000 | 0.0054 | 0.0041 | 0.0048 |
| F13 | 10060001 | 10160000 | 0.0043 | 0.0034 | 0.0036 |
| F13 | 10080001 | 10180000 | 0.0040 | 0.0032 | 0.0034 |
| F13 | 10100001 | 10200000 | 0.0032 | 0.0026 | 0.0024 |

| Chr | Start | End | both | A1 | A2 |
|---|---|---|---|---|---|
| F13 | 10120001 | 10220000 | 0.0034 | 0.0027 | 0.0025 |
| F13 | 10140001 | 10240000 | 0.0040 | 0.0030 | 0.0026 |
| F13 | 10160001 | 10260000 | 0.0058 | 0.0045 | 0.0040 |
| F13 | 10180001 | 10280000 | 0.0077 | 0.0059 | 0.0046 |
| F13 | 10200001 | 10300000 | 0.0078 | 0.0061 | 0.0049 |
| F13 | 10220001 | 10320000 | 0.0080 | 0.0062 | 0.0049 |
| F13 | 10240001 | 10340000 | 0.0092 | 0.0072 | 0.0063 |
| F13 | 10260001 | 10360000 | 0.0075 | 0.0057 | 0.0051 |
| F13 | 10280001 | 10380000 | 0.0052 | 0.0042 | 0.0043 |
| F13 | 10300001 | 10400000 | 0.0042 | 0.0033 | 0.0034 |
| F13 | 10320001 | 10420000 | 0.0038 | 0.0032 | 0.0034 |
| F13 | 10340001 | 10440000 | 0.0025 | 0.0022 | 0.0022 |
| F13 | 10360001 | 10460000 | 0.0025 | 0.0022 | 0.0021 |
| F13 | 10380001 | 10480000 | 0.0029 | 0.0025 | 0.0023 |
| F13 | 10400001 | 10500000 | 0.0034 | 0.0030 | 0.0024 |
| F13 | 10420001 | 10520000 | 0.0040 | 0.0035 | 0.0030 |
| F13 | 10440001 | 10540000 | 0.0041 | 0.0038 | 0.0030 |
| F13 | 10460001 | 10560000 | 0.0040 | 0.0037 | 0.0029 |
| F13 | 10480001 | 10580000 | 0.0035 | 0.0033 | 0.0026 |
| F13 | 10500001 | 10600000 | 0.0039 | 0.0036 | 0.0032 |
| F13 | 10520001 | 10620000 | 0.0040 | 0.0034 | 0.0032 |
| F13 | 10540001 | 10640000 | 0.0035 | 0.0027 | 0.0026 |
| F13 | 10560001 | 10660000 | 0.0032 | 0.0025 | 0.0024 |
| F13 | 10580001 | 10680000 | 0.0032 | 0.0024 | 0.0025 |
| F13 | 10600001 | 10700000 | 0.0029 | 0.0021 | 0.0022 |
| F13 | 10620001 | 10720000 | 0.0025 | 0.0019 | 0.0019 |
| F13 | 10640001 | 10740000 | 0.0024 | 0.0019 | 0.0019 |
| F13 | 10660001 | 10760000 | 0.0036 | 0.0029 | 0.0029 |
| F13 | 10680001 | 10780000 | 0.0040 | 0.0032 | 0.0031 |
| F13 | 10700001 | 10800000 | 0.0037 | 0.0031 | 0.0030 |
| F13 | 10720001 | 10820000 | 0.0039 | 0.0030 | 0.0030 |
| F13 | 10740001 | 10840000 | 0.0046 | 0.0036 | 0.0038 |
| F13 | 10760001 | 10860000 | 0.0035 | 0.0027 | 0.0029 |
| F13 | 10780001 | 10880000 | 0.0032 | 0.0023 | 0.0025 |
| F13 | 10800001 | 10900000 | 0.0040 | 0.0027 | 0.0031 |
| F13 | 10820001 | 10920000 | 0.0053 | 0.0043 | 0.0044 |
| F13 | 10840001 | 10940000 | 0.0045 | 0.0036 | 0.0036 |
| F13 | 10860001 | 10960000 | 0.0054 | 0.0043 | 0.0044 |
| F13 | 10880001 | 10980000 | 0.0063 | 0.0054 | 0.0055 |
| F13 | 10900001 | 11000000 | 0.0063 | 0.0053 | 0.0054 |
| F13 | 10920001 | 11020000 | 0.0042 | 0.0032 | 0.0035 |
| F13 | 10940001 | 11040000 | 0.0053 | 0.0041 | 0.0045 |
| F13 | 10960001 | 11060000 | 0.0053 | 0.0038 | 0.0043 |
| F13 | 10980001 | 11080000 | 0.0045 | 0.0029 | 0.0034 |
| F13 | 11000001 | 11100000 | 0.0042 | 0.0026 | 0.0031 |
| F13 | 11020001 | 11120000 | 0.0044 | 0.0028 | 0.0033 |
| F13 | 11040001 | 11140000 | 0.0041 | 0.0026 | 0.0031 |
| F13 | 11060001 | 11160000 | 0.0037 | 0.0022 | 0.0028 |
| F13 | 11080001 | 11180000 | 0.0037 | 0.0024 | 0.0030 |
| F13 | 11100001 | 11200000 | 0.0037 | 0.0027 | 0.0032 |
| F13 | 11120001 | 11220000 | 0.0038 | 0.0027 | 0.0033 |
| F13 | 11140001 | 11240000 | 0.0031 | 0.0023 | 0.0027 |
| F13 | 11160001 | 11260000 | 0.0029 | 0.0022 | 0.0025 |
| F13 | 11180001 | 11280000 | 0.0029 | 0.0022 | 0.0024 |
| F13 | 11200001 | 11300000 | 0.0029 | 0.0022 | 0.0024 |
| F13 | 11220001 | 11320000 | 0.0026 | 0.0019 | 0.0021 |
| F13 | 11240001 | 11340000 | 0.0031 | 0.0022 | 0.0026 |
| F13 | 11260001 | 11360000 | 0.0029 | 0.0021 | 0.0025 |

| Chr | Start | End | both | A1 | A2 |
|---|---|---|---|---|---|
| F13 | 11280001 | 11380000 | 0.0023 | 0.0016 | 0.0021 |
| F13 | 11300001 | 11400000 | 0.0017 | 0.0012 | 0.0015 |
| F13 | 11320001 | 11420000 | 0.0017 | 0.0013 | 0.0016 |
| F13 | 11340001 | 11440000 | 0.0008 | 0.0007 | 0.0008 |
| F13 | 11360001 | 11460000 | 0.0010 | 0.0008 | 0.0009 |
| F13 | 11380001 | 11480000 | 0.0013 | 0.0010 | 0.0011 |
| F13 | 11400001 | 11500000 | 0.0021 | 0.0016 | 0.0019 |
| F13 | 11420001 | 11520000 | 0.0019 | 0.0015 | 0.0016 |
| F13 | 11440001 | 11540000 | 0.0020 | 0.0016 | 0.0017 |
| F13 | 11460001 | 11560000 | 0.0018 | 0.0013 | 0.0014 |
| F13 | 11480001 | 11580000 | 0.0021 | 0.0014 | 0.0015 |
| F13 | 11500001 | 11600000 | 0.0015 | 0.0010 | 0.0010 |
| F13 | 11520001 | 11620000 | 0.0020 | 0.0012 | 0.0013 |
| F13 | 11540001 | 11640000 | 0.0026 | 0.0016 | 0.0019 |
| F13 | 11560001 | 11660000 | 0.0027 | 0.0017 | 0.0021 |
| F13 | 11580001 | 11680000 | 0.0024 | 0.0016 | 0.0020 |
| F13 | 11600001 | 11700000 | 0.0035 | 0.0029 | 0.0031 |
| F13 | 11620001 | 11720000 | 0.0033 | 0.0028 | 0.0029 |
| F13 | 11640001 | 11740000 | 0.0035 | 0.0031 | 0.0032 |
| F13 | 11660001 | 11760000 | 0.0035 | 0.0033 | 0.0032 |
| F13 | 11680001 | 11780000 | 0.0039 | 0.0037 | 0.0035 |
| F13 | 11700001 | 11800000 | 0.0029 | 0.0027 | 0.0025 |
| F13 | 11720001 | 11820000 | 0.0036 | 0.0031 | 0.0031 |
| F13 | 11740001 | 11840000 | 0.0032 | 0.0024 | 0.0025 |
| F13 | 11760001 | 11860000 | 0.0032 | 0.0021 | 0.0023 |
| F13 | 11780001 | 11880000 | 0.0030 | 0.0018 | 0.0021 |
| F13 | 11800001 | 11900000 | 0.0029 | 0.0015 | 0.0021 |
| F13 | 11820001 | 11920000 | 0.0022 | 0.0011 | 0.0015 |
| F13 | 11840001 | 11940000 | 0.0016 | 0.0009 | 0.0011 |
| F13 | 11860001 | 11960000 | 0.0015 | 0.0010 | 0.0012 |
| F13 | 11880001 | 11980000 | 0.0014 | 0.0009 | 0.0010 |
| F13 | 11900001 | 12000000 | 0.0012 | 0.0008 | 0.0008 |
| F13 | 11920001 | 12020000 | 0.0023 | 0.0013 | 0.0018 |
| F13 | 11940001 | 12040000 | 0.0024 | 0.0014 | 0.0019 |
| F13 | 11960001 | 12060000 | 0.0027 | 0.0016 | 0.0020 |
| F13 | 11980001 | 12080000 | 0.0026 | 0.0016 | 0.0020 |
| F13 | 12000001 | 12100000 | 0.0027 | 0.0016 | 0.0020 |
| F13 | 12020001 | 12120000 | 0.0018 | 0.0009 | 0.0010 |
| F13 | 12040001 | 12140000 | 0.0022 | 0.0011 | 0.0012 |
| F13 | 12060001 | 12160000 | 0.0022 | 0.0010 | 0.0011 |
| F13 | 12080001 | 12180000 | 0.0024 | 0.0012 | 0.0012 |
| F13 | 12100001 | 12200000 | 0.0029 | 0.0017 | 0.0016 |
| F13 | 12120001 | 12220000 | 0.0029 | 0.0019 | 0.0018 |
| F13 | 12140001 | 12240000 | 0.0032 | 0.0022 | 0.0020 |
| F13 | 12160001 | 12260000 | 0.0027 | 0.0019 | 0.0017 |
| F13 | 12180001 | 12280000 | 0.0025 | 0.0016 | 0.0016 |
| F13 | 12200001 | 12300000 | 0.0020 | 0.0012 | 0.0013 |
| F13 | 12220001 | 12320000 | 0.0026 | 0.0017 | 0.0019 |
| F13 | 12240001 | 12340000 | 0.0027 | 0.0017 | 0.0020 |
| F13 | 12260001 | 12360000 | 0.0032 | 0.0022 | 0.0024 |
| F13 | 12280001 | 12380000 | 0.0036 | 0.0027 | 0.0028 |
| F13 | 12300001 | 12400000 | 0.0040 | 0.0030 | 0.0031 |
| F13 | 12320001 | 12420000 | 0.0044 | 0.0031 | 0.0034 |
| F13 | 12340001 | 12440000 | 0.0044 | 0.0030 | 0.0034 |
| F13 | 12360001 | 12460000 | 0.0040 | 0.0028 | 0.0032 |
| F13 | 12380001 | 12480000 | 0.0044 | 0.0031 | 0.0037 |
| F13 | 12400001 | 12500000 | 0.0040 | 0.0028 | 0.0034 |
| F13 | 12420001 | 12520000 | 0.0031 | 0.0020 | 0.0024 |

| Chr | Start | End | both | A1 | A2 |
|---|---|---|---|---|---|
| F13 | 12440001 | 12540000 | 0.0022 | 0.0015 | 0.0017 |
| F13 | 12460001 | 12560000 | 0.0024 | 0.0015 | 0.0018 |
| F13 | 12480001 | 12580000 | 0.0014 | 0.0006 | 0.0007 |
| F13 | 12500001 | 12600000 | 0.0017 | 0.0008 | 0.0009 |
| F13 | 12520001 | 12620000 | 0.0016 | 0.0007 | 0.0009 |
| F13 | 12540001 | 12640000 | 0.0025 | 0.0012 | 0.0016 |
| F13 | 12560001 | 12660000 | 0.0023 | 0.0012 | 0.0015 |
| F13 | 12580001 | 12680000 | 0.0030 | 0.0018 | 0.0021 |
| F13 | 12600001 | 12700000 | 0.0031 | 0.0021 | 0.0024 |
| F13 | 12620001 | 12720000 | 0.0034 | 0.0025 | 0.0027 |
| F13 | 12640001 | 12740000 | 0.0031 | 0.0022 | 0.0024 |
| F13 | 12660001 | 12760000 | 0.0029 | 0.0021 | 0.0023 |
| F13 | 12680001 | 12780000 | 0.0026 | 0.0018 | 0.0019 |
| F13 | 12700001 | 12800000 | 0.0029 | 0.0021 | 0.0023 |
| F13 | 12720001 | 12820000 | 0.0026 | 0.0017 | 0.0020 |
| F13 | 12740001 | 12840000 | 0.0027 | 0.0018 | 0.0022 |
| F13 | 12760001 | 12860000 | 0.0030 | 0.0020 | 0.0025 |
| F13 | 12780001 | 12880000 | 0.0026 | 0.0018 | 0.0023 |
| F13 | 12800001 | 12900000 | 0.0023 | 0.0013 | 0.0018 |
| F13 | 12820001 | 12920000 | 0.0031 | 0.0022 | 0.0025 |
| F13 | 12840001 | 12940000 | 0.0037 | 0.0030 | 0.0031 |
| F13 | 12860001 | 12960000 | 0.0036 | 0.0030 | 0.0030 |
| F13 | 12880001 | 12980000 | 0.0036 | 0.0030 | 0.0030 |
| F13 | 12900001 | 13000000 | 0.0035 | 0.0033 | 0.0031 |
| F13 | 12920001 | 13020000 | 0.0029 | 0.0027 | 0.0026 |
| F13 | 12940001 | 13040000 | 0.0023 | 0.0019 | 0.0019 |
| F13 | 12960001 | 13060000 | 0.0030 | 0.0023 | 0.0025 |
| F13 | 12980001 | 13080000 | 0.0033 | 0.0025 | 0.0028 |
| F13 | 13000001 | 13100000 | 0.0031 | 0.0021 | 0.0025 |
| F13 | 13020001 | 13120000 | 0.0028 | 0.0017 | 0.0022 |
| F13 | 13040001 | 13140000 | 0.0029 | 0.0016 | 0.0021 |
| F13 | 13060001 | 13160000 | 0.0020 | 0.0009 | 0.0014 |
| F13 | 13080001 | 13180000 | 0.0015 | 0.0006 | 0.0009 |
| F13 | 13100001 | 13200000 | 0.0015 | 0.0007 | 0.0010 |
| F13 | 13120001 | 13220000 | 0.0017 | 0.0009 | 0.0013 |
| F13 | 13140001 | 13240000 | 0.0021 | 0.0014 | 0.0018 |
| F13 | 13160001 | 13260000 | 0.0036 | 0.0029 | 0.0031 |
| F13 | 13180001 | 13280000 | 0.0040 | 0.0032 | 0.0034 |
| F13 | 13200001 | 13300000 | 0.0040 | 0.0032 | 0.0034 |
| F13 | 13220001 | 13320000 | 0.0035 | 0.0029 | 0.0030 |
| F13 | 13240001 | 13340000 | 0.0026 | 0.0021 | 0.0022 |
| F13 | 13260001 | 13360000 | 0.0013 | 0.0009 | 0.0011 |
| F13 | 13280001 | 13380000 | 0.0018 | 0.0016 | 0.0018 |
| F13 | 13300001 | 13400000 | 0.0023 | 0.0020 | 0.0023 |
| F13 | 13320001 | 13420000 | 0.0035 | 0.0028 | 0.0032 |
| F13 | 13340001 | 13440000 | 0.0036 | 0.0029 | 0.0033 |
| F13 | 13360001 | 13460000 | 0.0033 | 0.0027 | 0.0030 |
| F13 | 13380001 | 13480000 | 0.0023 | 0.0018 | 0.0020 |
| F13 | 13400001 | 13500000 | 0.0019 | 0.0014 | 0.0017 |
| F13 | 13420001 | 13520000 | 0.0012 | 0.0009 | 0.0011 |
| F13 | 13440001 | 13540000 | 0.0011 | 0.0009 | 0.0010 |
| F13 | 13460001 | 13560000 | 0.0013 | 0.0011 | 0.0012 |
| F13 | 13480001 | 13580000 | 0.0012 | 0.0010 | 0.0012 |
| F13 | 13500001 | 13600000 | 0.0013 | 0.0010 | 0.0013 |
| F13 | 13520001 | 13620000 | 0.0012 | 0.0010 | 0.0011 |
| F13 | 13540001 | 13640000 | 0.0012 | 0.0010 | 0.0011 |
| F13 | 13560001 | 13660000 | 0.0016 | 0.0012 | 0.0015 |
| F13 | 13580001 | 13680000 | 0.0025 | 0.0019 | 0.0022 |

| Chr | Start | End | both | A1 | A2 |
|---|---|---|---|---|---|
| F13 | 13600001 | 13700000 | 0.0028 | 0.0022 | 0.0025 |
| F13 | 13620001 | 13720000 | 0.0024 | 0.0019 | 0.0022 |
| F13 | 13640001 | 13740000 | 0.0028 | 0.0021 | 0.0026 |
| F13 | 13660001 | 13760000 | 0.0026 | 0.0020 | 0.0023 |
| F13 | 13680001 | 13780000 | 0.0022 | 0.0016 | 0.0018 |
| F13 | 13700001 | 13800000 | 0.0015 | 0.0010 | 0.0011 |
| F13 | 13720001 | 13820000 | 0.0019 | 0.0014 | 0.0016 |
| F13 | 13740001 | 13840000 | 0.0017 | 0.0012 | 0.0012 |
| F13 | 13760001 | 13860000 | 0.0016 | 0.0012 | 0.0012 |
| F13 | 13780001 | 13880000 | 0.0014 | 0.0012 | 0.0012 |
| F13 | 13800001 | 13900000 | 0.0019 | 0.0016 | 0.0017 |
| F13 | 13820001 | 13920000 | 0.0019 | 0.0013 | 0.0016 |
| F13 | 13840001 | 13940000 | 0.0028 | 0.0019 | 0.0025 |
| F13 | 13860001 | 13960000 | 0.0030 | 0.0021 | 0.0027 |
| F13 | 13880001 | 13980000 | 0.0029 | 0.0021 | 0.0027 |
| F13 | 13900001 | 14000000 | 0.0028 | 0.0018 | 0.0025 |
| F13 | 13920001 | 14020000 | 0.0032 | 0.0022 | 0.0028 |
| F13 | 13940001 | 14040000 | 0.0024 | 0.0018 | 0.0021 |
| F13 | 13960001 | 14060000 | 0.0023 | 0.0017 | 0.0020 |
| F13 | 13980001 | 14080000 | 0.0026 | 0.0019 | 0.0023 |
| F13 | 14000001 | 14100000 | 0.0034 | 0.0026 | 0.0030 |
| F13 | 14020001 | 14120000 | 0.0035 | 0.0025 | 0.0031 |
| F13 | 14040001 | 14140000 | 0.0036 | 0.0026 | 0.0032 |
| F13 | 14060001 | 14160000 | 0.0040 | 0.0029 | 0.0035 |
| F13 | 14080001 | 14180000 | 0.0042 | 0.0030 | 0.0035 |
| F13 | 14100001 | 14200000 | 0.0033 | 0.0023 | 0.0026 |
| F13 | 14120001 | 14220000 | 0.0027 | 0.0021 | 0.0023 |
| F13 | 14140001 | 14240000 | 0.0025 | 0.0018 | 0.0021 |
| F13 | 14160001 | 14260000 | 0.0018 | 0.0013 | 0.0015 |
| F13 | 14180001 | 14280000 | 0.0015 | 0.0010 | 0.0013 |
| F13 | 14200001 | 14300000 | 0.0025 | 0.0010 | 0.0022 |
| F13 | 14220001 | 14320000 | 0.0031 | 0.0014 | 0.0025 |
| F13 | 14240001 | 14340000 | 0.0034 | 0.0018 | 0.0028 |
| F13 | 14260001 | 14360000 | 0.0046 | 0.0028 | 0.0039 |
| F13 | 14280001 | 14380000 | 0.0049 | 0.0033 | 0.0042 |
| F13 | 14300001 | 14400000 | 0.0040 | 0.0033 | 0.0032 |
| F13 | 14320001 | 14420000 | 0.0032 | 0.0026 | 0.0025 |
| F13 | 14340001 | 14440000 | 0.0029 | 0.0024 | 0.0024 |
| F13 | 14360001 | 14460000 | 0.0027 | 0.0020 | 0.0020 |
| F13 | 14380001 | 14480000 | 0.0024 | 0.0015 | 0.0016 |
| F13 | 14400001 | 14500000 | 0.0028 | 0.0019 | 0.0021 |
| F13 | 14420001 | 14520000 | 0.0031 | 0.0022 | 0.0024 |
| F13 | 14440001 | 14540000 | 0.0035 | 0.0026 | 0.0029 |
| F13 | 14460001 | 14560000 | 0.0028 | 0.0022 | 0.0023 |
| F13 | 14480001 | 14580000 | 0.0028 | 0.0022 | 0.0022 |
| F13 | 14500001 | 14600000 | 0.0025 | 0.0019 | 0.0020 |
| F13 | 14520001 | 14620000 | 0.0026 | 0.0018 | 0.0020 |
| F13 | 14540001 | 14640000 | 0.0025 | 0.0017 | 0.0017 |
| F13 | 14560001 | 14660000 | 0.0028 | 0.0019 | 0.0020 |
| F13 | 14580001 | 14680000 | 0.0030 | 0.0021 | 0.0023 |
| F13 | 14600001 | 14700000 | 0.0035 | 0.0024 | 0.0026 |
| F13 | 14620001 | 14720000 | 0.0039 | 0.0028 | 0.0029 |
| F13 | 14640001 | 14740000 | 0.0035 | 0.0025 | 0.0027 |
| F13 | 14660001 | 14760000 | 0.0030 | 0.0021 | 0.0024 |
| F13 | 14680001 | 14780000 | 0.0030 | 0.0019 | 0.0022 |
| F13 | 14700001 | 14800000 | 0.0037 | 0.0027 | 0.0031 |
| F13 | 14720001 | 14820000 | 0.0033 | 0.0023 | 0.0028 |
| F13 | 14740001 | 14840000 | 0.0037 | 0.0025 | 0.0031 |

| Chr | Start | End | both | A1 | A2 |
|---|---|---|---|---|---|
| F13 | 14760001 | 14860000 | 0.0041 | 0.0027 | 0.0033 |
| F13 | 14780001 | 14880000 | 0.0046 | 0.0033 | 0.0039 |
| F13 | 14800001 | 14900000 | 0.0031 | 0.0022 | 0.0025 |
| F13 | 14820001 | 14920000 | 0.0035 | 0.0024 | 0.0028 |
| F13 | 14840001 | 14940000 | 0.0035 | 0.0025 | 0.0030 |
| F13 | 14860001 | 14960000 | 0.0030 | 0.0023 | 0.0027 |
| F13 | 14880001 | 14980000 | 0.0022 | 0.0017 | 0.0020 |
| F13 | 14900001 | 15000000 | 0.0023 | 0.0019 | 0.0022 |
| F13 | 14920001 | 15020000 | 0.0030 | 0.0027 | 0.0029 |
| F13 | 14940001 | 15040000 | 0.0033 | 0.0030 | 0.0032 |
| F13 | 14960001 | 15060000 | 0.0036 | 0.0032 | 0.0035 |
| F13 | 14980001 | 15080000 | 0.0035 | 0.0031 | 0.0034 |
| F13 | 15000001 | 15100000 | 0.0036 | 0.0031 | 0.0034 |
| F13 | 15020001 | 15120000 | 0.0035 | 0.0028 | 0.0032 |
| F13 | 15040001 | 15140000 | 0.0055 | 0.0046 | 0.0050 |
| F13 | 15060001 | 15160000 | 0.0060 | 0.0048 | 0.0054 |
| F13 | 15080001 | 15180000 | 0.0065 | 0.0052 | 0.0057 |
| F13 | 15100001 | 15200000 | 0.0063 | 0.0050 | 0.0056 |
| F13 | 15120001 | 15220000 | 0.0052 | 0.0042 | 0.0047 |
| F13 | 15140001 | 15240000 | 0.0024 | 0.0017 | 0.0021 |
| F13 | 15160001 | 15260000 | 0.0017 | 0.0014 | 0.0016 |
| F13 | 15180001 | 15280000 | 0.0012 | 0.0010 | 0.0011 |
| F13 | 15200001 | 15300000 | 0.0009 | 0.0008 | 0.0009 |
| F13 | 15220001 | 15320000 | 0.0008 | 0.0007 | 0.0007 |
| F13 | 15240001 | 15340000 | 0.0007 | 0.0005 | 0.0006 |
| F13 | 15260001 | 15360000 | 0.0013 | 0.0009 | 0.0011 |
| F13 | 15280001 | 15380000 | 0.0024 | 0.0017 | 0.0021 |
| F13 | 15300001 | 15400000 | 0.0024 | 0.0017 | 0.0021 |
| F13 | 15320001 | 15420000 | 0.0026 | 0.0019 | 0.0023 |
| F13 | 15340001 | 15440000 | 0.0052 | 0.0044 | 0.0048 |
| F13 | 15360001 | 15460000 | 0.0046 | 0.0041 | 0.0043 |
| F13 | 15380001 | 15480000 | 0.0038 | 0.0035 | 0.0037 |
| F13 | 15400001 | 15500000 | 0.0044 | 0.0040 | 0.0042 |
| F13 | 15420001 | 15520000 | 0.0049 | 0.0044 | 0.0046 |
| F13 | 15440001 | 15540000 | 0.0024 | 0.0020 | 0.0021 |
| F13 | 15460001 | 15560000 | 0.0022 | 0.0017 | 0.0019 |
| F13 | 15480001 | 15580000 | 0.0023 | 0.0018 | 0.0020 |
| F13 | 15500001 | 15600000 | 0.0020 | 0.0014 | 0.0017 |
| F13 | 15520001 | 15620000 | 0.0012 | 0.0008 | 0.0010 |
| F13 | 15540001 | 15640000 | 0.0013 | 0.0010 | 0.0012 |
| F13 | 15560001 | 15660000 | 0.0014 | 0.0011 | 0.0012 |
| F13 | 15580001 | 15680000 | 0.0013 | 0.0010 | 0.0011 |
| F13 | 15600001 | 15700000 | 0.0012 | 0.0009 | 0.0009 |
| F13 | 15620001 | 15720000 | 0.0018 | 0.0014 | 0.0015 |
| F13 | 15640001 | 15740000 | 0.0020 | 0.0014 | 0.0016 |
| F13 | 15660001 | 15760000 | 0.0023 | 0.0015 | 0.0018 |
| F13 | 15680001 | 15780000 | 0.0025 | 0.0017 | 0.0021 |
| F13 | 15700001 | 15800000 | 0.0028 | 0.0018 | 0.0022 |
| F13 | 15720001 | 15820000 | 0.0023 | 0.0014 | 0.0017 |
| F13 | 15740001 | 15840000 | 0.0022 | 0.0011 | 0.0014 |
| F13 | 15760001 | 15860000 | 0.0020 | 0.0008 | 0.0010 |
| F13 | 15780001 | 15880000 | 0.0017 | 0.0003 | 0.0005 |
| F13 | 15800001 | 15900000 | 0.0023 | 0.0009 | 0.0012 |
| F13 | 15820001 | 15920000 | 0.0026 | 0.0012 | 0.0015 |
| F13 | 15840001 | 15940000 | 0.0032 | 0.0019 | 0.0024 |
| F13 | 15860001 | 15960000 | 0.0034 | 0.0025 | 0.0029 |
| F13 | 15880001 | 15980000 | 0.0038 | 0.0030 | 0.0034 |
| F13 | 15900001 | 16000000 | 0.0034 | 0.0028 | 0.0031 |

| Chr | Start | End | both | A1 | A2 |
|---|---|---|---|---|---|
| F13 | 15920001 | 16020000 | 0.0040 | 0.0034 | 0.0037 |
| F13 | 15940001 | 16040000 | 0.0038 | 0.0031 | 0.0033 |
| F13 | 15960001 | 16060000 | 0.0041 | 0.0032 | 0.0035 |
| F13 | 15980001 | 16080000 | 0.0040 | 0.0031 | 0.0034 |
| F13 | 16000001 | 16100000 | 0.0039 | 0.0030 | 0.0033 |
| F13 | 16020001 | 16120000 | 0.0037 | 0.0027 | 0.0032 |
| F13 | 16040001 | 16140000 | 0.0032 | 0.0024 | 0.0027 |
| F13 | 16060001 | 16160000 | 0.0028 | 0.0019 | 0.0022 |
| F13 | 16080001 | 16180000 | 0.0027 | 0.0019 | 0.0021 |
| F13 | 16100001 | 16200000 | 0.0025 | 0.0018 | 0.0021 |
| F13 | 16120001 | 16220000 | 0.0023 | 0.0017 | 0.0019 |
| F13 | 16140001 | 16240000 | 0.0025 | 0.0020 | 0.0022 |
| F13 | 16160001 | 16260000 | 0.0021 | 0.0018 | 0.0020 |
| F13 | 16180001 | 16280000 | 0.0020 | 0.0017 | 0.0019 |
| F13 | 16200001 | 16300000 | 0.0017 | 0.0013 | 0.0016 |
| F13 | 16220001 | 16320000 | 0.0014 | 0.0012 | 0.0014 |
| F13 | 16240001 | 16340000 | 0.0014 | 0.0011 | 0.0013 |
| F13 | 16260001 | 16360000 | 0.0015 | 0.0012 | 0.0015 |
| F13 | 16280001 | 16380000 | 0.0016 | 0.0012 | 0.0015 |
| F13 | 16300001 | 16400000 | 0.0017 | 0.0011 | 0.0014 |
| F13 | 16320001 | 16420000 | 0.0016 | 0.0008 | 0.0011 |
| F13 | 16340001 | 16440000 | 0.0019 | 0.0010 | 0.0012 |
| F13 | 16360001 | 16460000 | 0.0024 | 0.0015 | 0.0017 |
| F13 | 16380001 | 16480000 | 0.0023 | 0.0014 | 0.0016 |
| F13 | 16400001 | 16500000 | 0.0022 | 0.0016 | 0.0018 |
| F13 | 16420001 | 16520000 | 0.0030 | 0.0024 | 0.0026 |
| F13 | 16440001 | 16540000 | 0.0026 | 0.0020 | 0.0023 |
| F13 | 16460001 | 16560000 | 0.0025 | 0.0019 | 0.0022 |
| F13 | 16480001 | 16580000 | 0.0027 | 0.0021 | 0.0023 |
| F13 | 16500001 | 16600000 | 0.0029 | 0.0023 | 0.0025 |
| F13 | 16520001 | 16620000 | 0.0023 | 0.0017 | 0.0020 |
| F13 | 16540001 | 16640000 | 0.0035 | 0.0030 | 0.0031 |
| F13 | 16560001 | 16660000 | 0.0031 | 0.0025 | 0.0026 |
| F13 | 16580001 | 16680000 | 0.0034 | 0.0027 | 0.0027 |
| F13 | 16600001 | 16700000 | 0.0039 | 0.0031 | 0.0032 |
| F13 | 16620001 | 16720000 | 0.0037 | 0.0030 | 0.0031 |
| F13 | 16640001 | 16740000 | 0.0021 | 0.0015 | 0.0017 |
| F13 | 16660001 | 16760000 | 0.0021 | 0.0016 | 0.0019 |
| F13 | 16680001 | 16780000 | 0.0018 | 0.0014 | 0.0017 |
| F13 | 16700001 | 16800000 | 0.0012 | 0.0008 | 0.0010 |
| F13 | 16720001 | 16820000 | 0.0015 | 0.0009 | 0.0010 |
| F13 | 16740001 | 16840000 | 0.0019 | 0.0010 | 0.0012 |
| F13 | 16760001 | 16860000 | 0.0022 | 0.0012 | 0.0013 |
| F13 | 16780001 | 16880000 | 0.0022 | 0.0009 | 0.0010 |
| F13 | 16800001 | 16900000 | 0.0024 | 0.0012 | 0.0013 |
| F13 | 16820001 | 16920000 | 0.0022 | 0.0012 | 0.0013 |
| F13 | 16840001 | 16940000 | 0.0022 | 0.0012 | 0.0014 |
| F13 | 16860001 | 16960000 | 0.0025 | 0.0016 | 0.0019 |
| F13 | 16880001 | 16980000 | 0.0023 | 0.0015 | 0.0018 |
| F13 | 16900001 | 17000000 | 0.0020 | 0.0014 | 0.0015 |
| F13 | 16920001 | 17020000 | 0.0024 | 0.0018 | 0.0019 |
| F13 | 16940001 | 17040000 | 0.0029 | 0.0021 | 0.0024 |
| F13 | 16960001 | 17060000 | 0.0030 | 0.0022 | 0.0024 |
| F13 | 16980001 | 17080000 | 0.0032 | 0.0023 | 0.0024 |
| F13 | 17000001 | 17100000 | 0.0040 | 0.0026 | 0.0030 |
| F13 | 17020001 | 17120000 | 0.0037 | 0.0024 | 0.0028 |
| F13 | 17040001 | 17140000 | 0.0038 | 0.0024 | 0.0029 |
| F13 | 17060001 | 17160000 | 0.0030 | 0.0017 | 0.0022 |

| Chr | Start | End | both | A1 | A2 |
|---|---|---|---|---|---|
| F13 | 17080001 | 17180000 | 0.0035 | 0.0024 | 0.0028 |
| F13 | 17100001 | 17200000 | 0.0031 | 0.0024 | 0.0026 |
| F13 | 17120001 | 17220000 | 0.0037 | 0.0028 | 0.0032 |
| F13 | 17140001 | 17240000 | 0.0033 | 0.0025 | 0.0028 |
| F13 | 17160001 | 17260000 | 0.0040 | 0.0029 | 0.0034 |
| F13 | 17180001 | 17280000 | 0.0037 | 0.0025 | 0.0029 |
| F13 | 17200001 | 17300000 | 0.0034 | 0.0023 | 0.0028 |
| F13 | 17220001 | 17320000 | 0.0030 | 0.0021 | 0.0025 |
| F13 | 17240001 | 17340000 | 0.0033 | 0.0026 | 0.0029 |
| F13 | 17260001 | 17360000 | 0.0028 | 0.0025 | 0.0025 |
| F13 | 17280001 | 17380000 | 0.0026 | 0.0022 | 0.0024 |
| F13 | 17300001 | 17400000 | 0.0029 | 0.0025 | 0.0027 |
| F13 | 17320001 | 17420000 | 0.0025 | 0.0022 | 0.0023 |
| F13 | 17340001 | 17440000 | 0.0018 | 0.0014 | 0.0016 |
| F13 | 17360001 | 17460000 | 0.0016 | 0.0011 | 0.0015 |
| F13 | 17380001 | 17480000 | 0.0015 | 0.0012 | 0.0015 |
| F13 | 17400001 | 17500000 | 0.0012 | 0.0011 | 0.0011 |
| F13 | 17420001 | 17520000 | 0.0012 | 0.0011 | 0.0011 |
| F13 | 17440001 | 17540000 | 0.0017 | 0.0016 | 0.0016 |
| F13 | 17460001 | 17560000 | 0.0019 | 0.0018 | 0.0018 |
| F13 | 17480001 | 17580000 | 0.0020 | 0.0018 | 0.0019 |
| F13 | 17500001 | 17600000 | 0.0015 | 0.0013 | 0.0015 |
| F13 | 17520001 | 17620000 | 0.0022 | 0.0018 | 0.0020 |
| F13 | 17540001 | 17640000 | 0.0019 | 0.0014 | 0.0017 |
| F13 | 17560001 | 17660000 | 0.0019 | 0.0014 | 0.0016 |
| F13 | 17580001 | 17680000 | 0.0025 | 0.0017 | 0.0021 |
| F13 | 17600001 | 17700000 | 0.0035 | 0.0025 | 0.0030 |
| F13 | 17620001 | 17720000 | 0.0029 | 0.0021 | 0.0026 |
| F13 | 17640001 | 17740000 | 0.0028 | 0.0020 | 0.0024 |
| F13 | 17660001 | 17760000 | 0.0025 | 0.0018 | 0.0022 |
| F13 | 17680001 | 17780000 | 0.0017 | 0.0014 | 0.0016 |
| F13 | 17700001 | 17800000 | 0.0014 | 0.0011 | 0.0012 |
| F13 | 17720001 | 17820000 | 0.0018 | 0.0013 | 0.0015 |
| F13 | 17740001 | 17840000 | 0.0038 | 0.0032 | 0.0035 |
| F13 | 17760001 | 17860000 | 0.0041 | 0.0033 | 0.0035 |
| F13 | 17780001 | 17880000 | 0.0044 | 0.0034 | 0.0037 |
| F13 | 17800001 | 17900000 | 0.0039 | 0.0029 | 0.0032 |
| F13 | 17820001 | 17920000 | 0.0036 | 0.0026 | 0.0029 |
| F13 | 17840001 | 17940000 | 0.0018 | 0.0008 | 0.0011 |
| F13 | 17860001 | 17960000 | 0.0020 | 0.0010 | 0.0013 |
| F13 | 17880001 | 17980000 | 0.0020 | 0.0010 | 0.0013 |
| F13 | 17900001 | 18000000 | 0.0019 | 0.0010 | 0.0012 |
| F13 | 17920001 | 18020000 | 0.0021 | 0.0013 | 0.0015 |
| F13 | 17940001 | 18040000 | 0.0025 | 0.0017 | 0.0018 |
| F13 | 17960001 | 18060000 | 0.0027 | 0.0022 | 0.0021 |
| F13 | 17980001 | 18080000 | 0.0023 | 0.0020 | 0.0019 |
| F13 | 18000001 | 18100000 | 0.0024 | 0.0021 | 0.0021 |
| F13 | 18020001 | 18120000 | 0.0019 | 0.0016 | 0.0017 |
| F13 | 18040001 | 18140000 | 0.0019 | 0.0019 | 0.0018 |
| F13 | 18060001 | 18160000 | 0.0021 | 0.0020 | 0.0020 |
| F13 | 18080001 | 18180000 | 0.0026 | 0.0024 | 0.0024 |
| F13 | 18100001 | 18200000 | 0.0029 | 0.0027 | 0.0026 |
| F13 | 18120001 | 18220000 | 0.0031 | 0.0029 | 0.0028 |
| F13 | 18140001 | 18240000 | 0.0030 | 0.0026 | 0.0027 |
| F13 | 18160001 | 18260000 | 0.0028 | 0.0024 | 0.0026 |
| F13 | 18180001 | 18280000 | 0.0023 | 0.0019 | 0.0022 |
| F13 | 18200001 | 18300000 | 0.0023 | 0.0019 | 0.0022 |
| F13 | 18220001 | 18320000 | 0.0023 | 0.0019 | 0.0021 |

| Chr | Start | End | both | A1 | A2 |
|---|---|---|---|---|---|
| F13 | 18240001 | 18340000 | 0.0020 | 0.0017 | 0.0018 |
| F13 | 18260001 | 18360000 | 0.0018 | 0.0014 | 0.0015 |
| F13 | 18280001 | 18380000 | 0.0019 | 0.0015 | 0.0015 |
| F13 | 18300001 | 18400000 | 0.0018 | 0.0012 | 0.0013 |
| F13 | 18320001 | 18420000 | 0.0019 | 0.0013 | 0.0015 |
| F13 | 18340001 | 18440000 | 0.0019 | 0.0011 | 0.0014 |
| F13 | 18360001 | 18460000 | 0.0018 | 0.0011 | 0.0013 |
| F13 | 18380001 | 18480000 | 0.0027 | 0.0020 | 0.0023 |
| F13 | 18400001 | 18500000 | 0.0024 | 0.0018 | 0.0020 |
| F13 | 18420001 | 18520000 | 0.0022 | 0.0018 | 0.0020 |
| F13 | 18440001 | 18540000 | 0.0029 | 0.0022 | 0.0026 |
| F13 | 18460001 | 18560000 | 0.0036 | 0.0029 | 0.0034 |
| F13 | 18480001 | 18580000 | 0.0029 | 0.0023 | 0.0027 |
| F13 | 18500001 | 18600000 | 0.0030 | 0.0023 | 0.0027 |
| F13 | 18520001 | 18620000 | 0.0030 | 0.0023 | 0.0028 |
| F13 | 18540001 | 18640000 | 0.0020 | 0.0018 | 0.0019 |
| F13 | 18560001 | 18660000 | 0.0010 | 0.0009 | 0.0010 |
| F13 | 18580001 | 18680000 | 0.0008 | 0.0008 | 0.0008 |
| F13 | 18600001 | 18700000 | 0.0011 | 0.0010 | 0.0010 |
| F13 | 18620001 | 18720000 | 0.0008 | 0.0007 | 0.0008 |
| F13 | 18640001 | 18740000 | 0.0008 | 0.0007 | 0.0008 |
| F13 | 18660001 | 18760000 | 0.0009 | 0.0008 | 0.0009 |
| F13 | 18680001 | 18780000 | 0.0008 | 0.0006 | 0.0008 |
| F13 | 18700001 | 18800000 | 0.0014 | 0.0009 | 0.0012 |
| F13 | 18720001 | 18820000 | 0.0024 | 0.0017 | 0.0019 |
| F13 | 18740001 | 18840000 | 0.0029 | 0.0020 | 0.0023 |
| F13 | 18760001 | 18860000 | 0.0032 | 0.0021 | 0.0024 |
| F13 | 18780001 | 18880000 | 0.0033 | 0.0020 | 0.0024 |
| F13 | 18800001 | 18900000 | 0.0029 | 0.0017 | 0.0021 |
| F13 | 18820001 | 18920000 | 0.0026 | 0.0014 | 0.0018 |
| F13 | 18840001 | 18940000 | 0.0036 | 0.0026 | 0.0030 |
| F13 | 18860001 | 18960000 | 0.0039 | 0.0028 | 0.0034 |
| F13 | 18880001 | 18980000 | 0.0036 | 0.0028 | 0.0033 |
| F13 | 18900001 | 19000000 | 0.0035 | 0.0029 | 0.0034 |
| F13 | 18920001 | 19020000 | 0.0031 | 0.0026 | 0.0030 |
| F13 | 18940001 | 19040000 | 0.0031 | 0.0022 | 0.0029 |
| F13 | 18960001 | 19060000 | 0.0033 | 0.0026 | 0.0029 |
| F13 | 18980001 | 19080000 | 0.0041 | 0.0033 | 0.0037 |
| F13 | 19000001 | 19100000 | 0.0041 | 0.0033 | 0.0036 |
| F13 | 19020001 | 19120000 | 0.0044 | 0.0035 | 0.0038 |
| F13 | 19040001 | 19140000 | 0.0047 | 0.0038 | 0.0041 |
| F13 | 19060001 | 19160000 | 0.0047 | 0.0038 | 0.0043 |
| F13 | 19080001 | 19180000 | 0.0042 | 0.0034 | 0.0038 |
| F13 | 19100001 | 19200000 | 0.0045 | 0.0036 | 0.0040 |
| F13 | 19120001 | 19220000 | 0.0046 | 0.0037 | 0.0041 |
| F13 | 19140001 | 19240000 | 0.0034 | 0.0025 | 0.0028 |
| F13 | 19160001 | 19260000 | 0.0026 | 0.0019 | 0.0020 |
| F13 | 19180001 | 19280000 | 0.0024 | 0.0017 | 0.0018 |
| F13 | 19200001 | 19300000 | 0.0021 | 0.0015 | 0.0015 |
| F13 | 19220001 | 19320000 | 0.0017 | 0.0010 | 0.0010 |
| F13 | 19240001 | 19340000 | 0.0014 | 0.0007 | 0.0008 |
| F13 | 19260001 | 19360000 | 0.0022 | 0.0013 | 0.0014 |
| F13 | 19280001 | 19380000 | 0.0028 | 0.0019 | 0.0019 |
| F13 | 19300001 | 19400000 | 0.0033 | 0.0022 | 0.0021 |
| F13 | 19320001 | 19420000 | 0.0032 | 0.0022 | 0.0021 |
| F13 | 19340001 | 19440000 | 0.0041 | 0.0030 | 0.0031 |
| F13 | 19360001 | 19460000 | 0.0039 | 0.0028 | 0.0030 |
| F13 | 19380001 | 19480000 | 0.0033 | 0.0023 | 0.0026 |

| Chr | Start | End | both | A1 | A2 |
|---|---|---|---|---|---|
| F13 | 19400001 | 19500000 | 0.0032 | 0.0022 | 0.0027 |
| F13 | 19420001 | 19520000 | 0.0032 | 0.0022 | 0.0027 |
| F13 | 19440001 | 19540000 | 0.0023 | 0.0016 | 0.0019 |
| F13 | 19460001 | 19560000 | 0.0018 | 0.0013 | 0.0016 |
| F13 | 19480001 | 19580000 | 0.0018 | 0.0012 | 0.0015 |
| F13 | 19500001 | 19600000 | 0.0016 | 0.0010 | 0.0012 |
| F13 | 19520001 | 19620000 | 0.0020 | 0.0012 | 0.0014 |
| F13 | 19540001 | 19640000 | 0.0022 | 0.0015 | 0.0016 |
| F13 | 19560001 | 19660000 | 0.0030 | 0.0021 | 0.0021 |
| F13 | 19580001 | 19680000 | 0.0033 | 0.0025 | 0.0025 |
| F13 | 19600001 | 19700000 | 0.0036 | 0.0026 | 0.0027 |
| F13 | 19620001 | 19720000 | 0.0042 | 0.0032 | 0.0035 |
| F13 | 19640001 | 19740000 | 0.0043 | 0.0033 | 0.0036 |
| F13 | 19660001 | 19760000 | 0.0035 | 0.0026 | 0.0030 |
| F13 | 19680001 | 19780000 | 0.0032 | 0.0022 | 0.0025 |
| F13 | 19700001 | 19800000 | 0.0027 | 0.0020 | 0.0022 |
| F13 | 19720001 | 19820000 | 0.0021 | 0.0015 | 0.0016 |
| F13 | 19740001 | 19840000 | 0.0022 | 0.0015 | 0.0017 |
| F13 | 19760001 | 19860000 | 0.0023 | 0.0017 | 0.0019 |
| F13 | 19780001 | 19880000 | 0.0023 | 0.0017 | 0.0020 |
| F13 | 19800001 | 19900000 | 0.0026 | 0.0018 | 0.0024 |
| F13 | 19820001 | 19920000 | 0.0026 | 0.0019 | 0.0024 |
| F13 | 19840001 | 19940000 | 0.0024 | 0.0017 | 0.0022 |
| F13 | 19860001 | 19960000 | 0.0026 | 0.0018 | 0.0022 |
| F13 | 19880001 | 19980000 | 0.0027 | 0.0019 | 0.0023 |
| F13 | 19900001 | 20000000 | 0.0030 | 0.0021 | 0.0022 |
| F13 | 19920001 | 20020000 | 0.0031 | 0.0021 | 0.0022 |
| F13 | 19940001 | 20040000 | 0.0030 | 0.0020 | 0.0021 |
| F13 | 19960001 | 20060000 | 0.0036 | 0.0027 | 0.0028 |
| F13 | 19980001 | 20080000 | 0.0038 | 0.0030 | 0.0030 |
| F13 | 20000001 | 20100000 | 0.0037 | 0.0030 | 0.0032 |
| F13 | 20020001 | 20120000 | 0.0034 | 0.0028 | 0.0030 |
| F13 | 20040001 | 20140000 | 0.0033 | 0.0028 | 0.0029 |
| F13 | 20060001 | 20160000 | 0.0029 | 0.0024 | 0.0025 |
| F13 | 20080001 | 20180000 | 0.0027 | 0.0021 | 0.0023 |
| F13 | 20100001 | 20200000 | 0.0023 | 0.0017 | 0.0017 |
| F13 | 20120001 | 20220000 | 0.0027 | 0.0020 | 0.0020 |
| F13 | 20140001 | 20240000 | 0.0026 | 0.0017 | 0.0018 |
| F13 | 20160001 | 20260000 | 0.0027 | 0.0018 | 0.0018 |
| F13 | 20180001 | 20280000 | 0.0024 | 0.0016 | 0.0016 |
| F13 | 20200001 | 20300000 | 0.0025 | 0.0018 | 0.0018 |
| F13 | 20220001 | 20320000 | 0.0020 | 0.0014 | 0.0014 |
| F13 | 20240001 | 20340000 | 0.0026 | 0.0021 | 0.0019 |
| F13 | 20260001 | 20360000 | 0.0024 | 0.0019 | 0.0018 |
| F13 | 20280001 | 20380000 | 0.0023 | 0.0018 | 0.0018 |
| F13 | 20300001 | 20400000 | 0.0024 | 0.0020 | 0.0020 |
| F13 | 20320001 | 20420000 | 0.0024 | 0.0020 | 0.0020 |
| F13 | 20340001 | 20440000 | 0.0020 | 0.0018 | 0.0020 |
| F13 | 20360001 | 20460000 | 0.0017 | 0.0017 | 0.0015 |
| F13 | 20380001 | 20480000 | 0.0016 | 0.0016 | 0.0014 |
| F13 | 20400001 | 20500000 | 0.0012 | 0.0012 | 0.0011 |
| F13 | 20420001 | 20520000 | 0.0011 | 0.0011 | 0.0010 |
| F13 | 20440001 | 20540000 | 0.0009 | 0.0007 | 0.0006 |
| F13 | 20460001 | 20560000 | 0.0012 | 0.0009 | 0.0010 |
| F13 | 20480001 | 20580000 | 0.0018 | 0.0013 | 0.0015 |
| F13 | 20500001 | 20600000 | 0.0026 | 0.0019 | 0.0020 |
| F13 | 20520001 | 20620000 | 0.0029 | 0.0019 | 0.0021 |
| F13 | 20540001 | 20640000 | 0.0034 | 0.0025 | 0.0025 |

| Chr | Start | End | both | A1 | A2 |
|---|---|---|---|---|---|
| F13 | 20560001 | 20660000 | 0.0031 | 0.0022 | 0.0023 |
| F13 | 20580001 | 20680000 | 0.0030 | 0.0023 | 0.0023 |
| F13 | 20600001 | 20700000 | 0.0025 | 0.0020 | 0.0019 |
| F13 | 20620001 | 20720000 | 0.0027 | 0.0022 | 0.0022 |
| F13 | 20640001 | 20740000 | 0.0018 | 0.0015 | 0.0016 |
| F13 | 20660001 | 20760000 | 0.0017 | 0.0014 | 0.0015 |
| F13 | 20680001 | 20780000 | 0.0012 | 0.0009 | 0.0010 |
| F13 | 20700001 | 20800000 | 0.0012 | 0.0007 | 0.0009 |
| F13 | 20720001 | 20820000 | 0.0016 | 0.0011 | 0.0013 |
| F13 | 20740001 | 20840000 | 0.0022 | 0.0016 | 0.0018 |
| F13 | 20760001 | 20860000 | 0.0032 | 0.0027 | 0.0027 |
| F13 | 20780001 | 20880000 | 0.0035 | 0.0029 | 0.0030 |
| F13 | 20800001 | 20900000 | 0.0037 | 0.0033 | 0.0034 |
| F13 | 20820001 | 20920000 | 0.0034 | 0.0030 | 0.0030 |
| F13 | 20840001 | 20940000 | 0.0035 | 0.0030 | 0.0031 |
| F13 | 20860001 | 20960000 | 0.0025 | 0.0020 | 0.0022 |
| F13 | 20880001 | 20980000 | 0.0022 | 0.0018 | 0.0020 |
| F13 | 20900001 | 21000000 | 0.0021 | 0.0016 | 0.0017 |
| F13 | 20920001 | 21020000 | 0.0020 | 0.0016 | 0.0018 |
| F13 | 20940001 | 21040000 | 0.0024 | 0.0019 | 0.0021 |
| F13 | 20960001 | 21060000 | 0.0029 | 0.0025 | 0.0026 |
| F13 | 20980001 | 21080000 | 0.0028 | 0.0024 | 0.0024 |
| F13 | 21000001 | 21100000 | 0.0025 | 0.0022 | 0.0023 |
| F13 | 21020001 | 21120000 | 0.0022 | 0.0020 | 0.0020 |
| F13 | 21040001 | 21140000 | 0.0014 | 0.0014 | 0.0012 |
| F13 | 21060001 | 21160000 | 0.0007 | 0.0007 | 0.0007 |
| F13 | 21080001 | 21180000 | 0.0010 | 0.0009 | 0.0010 |
| F13 | 21100001 | 21200000 | 0.0009 | 0.0008 | 0.0008 |
| F13 | 21120001 | 21220000 | 0.0008 | 0.0007 | 0.0008 |
| F13 | 21140001 | 21240000 | 0.0009 | 0.0008 | 0.0009 |
| F13 | 21160001 | 21260000 | 0.0017 | 0.0015 | 0.0016 |
| F13 | 21180001 | 21280000 | 0.0014 | 0.0013 | 0.0014 |
| F13 | 21200001 | 21300000 | 0.0013 | 0.0012 | 0.0012 |
| F13 | 21220001 | 21320000 | 0.0013 | 0.0012 | 0.0013 |
| F13 | 21240001 | 21340000 | 0.0018 | 0.0016 | 0.0017 |
| F13 | 21260001 | 21360000 | 0.0012 | 0.0010 | 0.0010 |
| F13 | 21280001 | 21380000 | 0.0014 | 0.0011 | 0.0012 |
| F13 | 21300001 | 21400000 | 0.0015 | 0.0012 | 0.0012 |
| F13 | 21320001 | 21420000 | 0.0015 | 0.0012 | 0.0012 |
| F13 | 21340001 | 21440000 | 0.0008 | 0.0006 | 0.0006 |
| F13 | 21360001 | 21460000 | 0.0018 | 0.0014 | 0.0015 |
| F13 | 21380001 | 21480000 | 0.0019 | 0.0015 | 0.0016 |
| F13 | 21400001 | 21500000 | 0.0028 | 0.0020 | 0.0022 |
| F13 | 21420001 | 21520000 | 0.0035 | 0.0027 | 0.0029 |
| F13 | 21440001 | 21540000 | 0.0042 | 0.0033 | 0.0034 |
| F13 | 21460001 | 21560000 | 0.0033 | 0.0026 | 0.0025 |
| F13 | 21480001 | 21580000 | 0.0032 | 0.0024 | 0.0025 |
| F13 | 21500001 | 21600000 | 0.0030 | 0.0022 | 0.0024 |
| F13 | 21520001 | 21620000 | 0.0031 | 0.0019 | 0.0022 |
| F13 | 21540001 | 21640000 | 0.0024 | 0.0013 | 0.0017 |
| F13 | 21560001 | 21660000 | 0.0025 | 0.0012 | 0.0017 |
| F13 | 21580001 | 21680000 | 0.0026 | 0.0013 | 0.0018 |
| F13 | 21600001 | 21700000 | 0.0022 | 0.0011 | 0.0014 |
| F13 | 21620001 | 21720000 | 0.0022 | 0.0008 | 0.0010 |
| F13 | 21640001 | 21740000 | 0.0034 | 0.0011 | 0.0014 |
| F13 | 21660001 | 21760000 | 0.0039 | 0.0017 | 0.0021 |
| F13 | 21680001 | 21780000 | 0.0043 | 0.0018 | 0.0021 |
| F13 | 21700001 | 21800000 | 0.0043 | 0.0017 | 0.0019 |

| Chr | Start | End | both | A1 | A2 |
|---|---|---|---|---|---|
| F13 | 21720001 | 21820000 | 0.0036 | 0.0016 | 0.0017 |
| F13 | 21740001 | 21840000 | 0.0026 | 0.0013 | 0.0016 |
| F13 | 21760001 | 21860000 | 0.0019 | 0.0008 | 0.0009 |
| F13 | 21780001 | 21880000 | 0.0014 | 0.0006 | 0.0008 |
| F13 | 21800001 | 21900000 | 0.0014 | 0.0007 | 0.0010 |
| F13 | 21820001 | 21920000 | 0.0020 | 0.0014 | 0.0016 |
| F13 | 21840001 | 21940000 | 0.0018 | 0.0012 | 0.0013 |
| F13 | 21860001 | 21960000 | 0.0016 | 0.0012 | 0.0012 |
| F13 | 21880001 | 21980000 | 0.0019 | 0.0015 | 0.0015 |
| F13 | 21900001 | 22000000 | 0.0018 | 0.0014 | 0.0014 |
| F13 | 21920001 | 22020000 | 0.0010 | 0.0006 | 0.0007 |
| F13 | 21940001 | 22040000 | 0.0013 | 0.0007 | 0.0008 |
| F13 | 21960001 | 22060000 | 0.0018 | 0.0010 | 0.0011 |
| F13 | 21980001 | 22080000 | 0.0019 | 0.0010 | 0.0011 |
| F13 | 22000001 | 22100000 | 0.0023 | 0.0014 | 0.0016 |
| F13 | 22020001 | 22120000 | 0.0031 | 0.0020 | 0.0022 |
| F13 | 22040001 | 22140000 | 0.0030 | 0.0019 | 0.0021 |
| F13 | 22060001 | 22160000 | 0.0027 | 0.0017 | 0.0018 |
| F13 | 22080001 | 22180000 | 0.0024 | 0.0016 | 0.0017 |
| F13 | 22100001 | 22200000 | 0.0029 | 0.0018 | 0.0020 |
| F13 | 22120001 | 22220000 | 0.0030 | 0.0018 | 0.0022 |
| F13 | 22140001 | 22240000 | 0.0041 | 0.0026 | 0.0032 |
| F13 | 22160001 | 22260000 | 0.0047 | 0.0030 | 0.0036 |
| F13 | 22180001 | 22280000 | 0.0048 | 0.0032 | 0.0038 |
| F13 | 22200001 | 22300000 | 0.0040 | 0.0026 | 0.0030 |
| F13 | 22220001 | 22320000 | 0.0036 | 0.0024 | 0.0028 |
| F13 | 22240001 | 22340000 | 0.0031 | 0.0020 | 0.0021 |
| F13 | 22260001 | 22360000 | 0.0032 | 0.0019 | 0.0021 |
| F13 | 22280001 | 22380000 | 0.0031 | 0.0017 | 0.0019 |
| F13 | 22300001 | 22400000 | 0.0033 | 0.0017 | 0.0019 |
| F13 | 22320001 | 22420000 | 0.0040 | 0.0020 | 0.0023 |
| F13 | 22340001 | 22440000 | 0.0039 | 0.0021 | 0.0024 |
| F13 | 22360001 | 22460000 | 0.0034 | 0.0019 | 0.0021 |
| F13 | 22380001 | 22480000 | 0.0033 | 0.0018 | 0.0019 |
| F13 | 22400001 | 22500000 | 0.0036 | 0.0025 | 0.0026 |
| F13 | 22420001 | 22520000 | 0.0031 | 0.0023 | 0.0022 |
| F13 | 22440001 | 22540000 | 0.0028 | 0.0021 | 0.0020 |
| F13 | 22460001 | 22560000 | 0.0030 | 0.0025 | 0.0024 |
| F13 | 22480001 | 22580000 | 0.0040 | 0.0035 | 0.0034 |
| F13 | 22500001 | 22600000 | 0.0038 | 0.0032 | 0.0031 |
| F13 | 22520001 | 22620000 | 0.0035 | 0.0028 | 0.0029 |
| F13 | 22540001 | 22640000 | 0.0033 | 0.0025 | 0.0027 |
| F13 | 22560001 | 22660000 | 0.0036 | 0.0024 | 0.0026 |
| F13 | 22580001 | 22680000 | 0.0026 | 0.0016 | 0.0017 |
| F13 | 22600001 | 22700000 | 0.0030 | 0.0019 | 0.0021 |
| F13 | 22620001 | 22720000 | 0.0032 | 0.0020 | 0.0022 |
| F13 | 22640001 | 22740000 | 0.0032 | 0.0022 | 0.0023 |
| F13 | 22660001 | 22760000 | 0.0026 | 0.0020 | 0.0021 |
| F13 | 22680001 | 22780000 | 0.0027 | 0.0021 | 0.0023 |
| F13 | 22700001 | 22800000 | 0.0019 | 0.0014 | 0.0016 |
| F13 | 22720001 | 22820000 | 0.0016 | 0.0013 | 0.0014 |
| F13 | 22740001 | 22840000 | 0.0018 | 0.0011 | 0.0013 |
| F13 | 22760001 | 22860000 | 0.0022 | 0.0014 | 0.0016 |
| F13 | 22780001 | 22880000 | 0.0019 | 0.0012 | 0.0014 |
| F13 | 22800001 | 22900000 | 0.0019 | 0.0012 | 0.0014 |
| F13 | 22820001 | 22920000 | 0.0016 | 0.0011 | 0.0013 |
| F13 | 22840001 | 22940000 | 0.0014 | 0.0012 | 0.0014 |
| F13 | 22860001 | 22960000 | 0.0022 | 0.0020 | 0.0021 |

| Chr | Start | End | both | A1 | A2 |
|---|---|---|---|---|---|
| F13 | 22880001 | 22980000 | 0.0030 | 0.0022 | 0.0025 |
| F13 | 22900001 | 23000000 | 0.0030 | 0.0022 | 0.0025 |
| F13 | 22920001 | 23020000 | 0.0034 | 0.0023 | 0.0026 |
| F13 | 22940001 | 23040000 | 0.0037 | 0.0024 | 0.0029 |
| F13 | 22960001 | 23060000 | 0.0031 | 0.0015 | 0.0020 |
| F13 | 22980001 | 23080000 | 0.0026 | 0.0014 | 0.0018 |
| F13 | 23000001 | 23100000 | 0.0030 | 0.0017 | 0.0020 |
| F13 | 23020001 | 23120000 | 0.0031 | 0.0018 | 0.0022 |
| F13 | 23040001 | 23140000 | 0.0036 | 0.0021 | 0.0026 |
| F13 | 23060001 | 23160000 | 0.0034 | 0.0021 | 0.0025 |
| F13 | 23080001 | 23180000 | 0.0035 | 0.0023 | 0.0027 |
| F13 | 23100001 | 23200000 | 0.0031 | 0.0020 | 0.0024 |
| F13 | 23120001 | 23220000 | 0.0029 | 0.0019 | 0.0021 |
| F13 | 23140001 | 23240000 | 0.0025 | 0.0016 | 0.0016 |
| F13 | 23160001 | 23260000 | 0.0023 | 0.0015 | 0.0015 |
| F13 | 23180001 | 23280000 | 0.0022 | 0.0015 | 0.0015 |
| F13 | 23200001 | 23300000 | 0.0027 | 0.0020 | 0.0019 |
| F13 | 23220001 | 23320000 | 0.0025 | 0.0019 | 0.0019 |
| F13 | 23240001 | 23340000 | 0.0023 | 0.0016 | 0.0019 |
| F13 | 23260001 | 23360000 | 0.0024 | 0.0018 | 0.0019 |
| F13 | 23280001 | 23380000 | 0.0029 | 0.0023 | 0.0024 |
| F13 | 23300001 | 23400000 | 0.0026 | 0.0021 | 0.0022 |
| F13 | 23320001 | 23420000 | 0.0035 | 0.0030 | 0.0031 |
| F13 | 23340001 | 23440000 | 0.0040 | 0.0034 | 0.0034 |
| F13 | 23360001 | 23460000 | 0.0042 | 0.0035 | 0.0037 |
| F13 | 23380001 | 23480000 | 0.0037 | 0.0030 | 0.0033 |
| F13 | 23400001 | 23500000 | 0.0043 | 0.0034 | 0.0038 |
| F13 | 23420001 | 23520000 | 0.0036 | 0.0027 | 0.0032 |
| F13 | 23440001 | 23540000 | 0.0037 | 0.0029 | 0.0034 |
| F13 | 23460001 | 23560000 | 0.0036 | 0.0028 | 0.0032 |
| F13 | 23480001 | 23580000 | 0.0039 | 0.0029 | 0.0033 |
| F13 | 23500001 | 23600000 | 0.0033 | 0.0024 | 0.0029 |
| F13 | 23520001 | 23620000 | 0.0034 | 0.0024 | 0.0028 |
| F13 | 23540001 | 23640000 | 0.0028 | 0.0016 | 0.0019 |
| F13 | 23560001 | 23660000 | 0.0024 | 0.0012 | 0.0015 |
| F13 | 23580001 | 23680000 | 0.0016 | 0.0006 | 0.0007 |
| F13 | 23600001 | 23700000 | 0.0017 | 0.0005 | 0.0006 |
| F13 | 23620001 | 23720000 | 0.0014 | 0.0005 | 0.0005 |
| F13 | 23640001 | 23740000 | 0.0015 | 0.0007 | 0.0007 |
| F13 | 23660001 | 23760000 | 0.0014 | 0.0008 | 0.0009 |
| F13 | 23680001 | 23780000 | 0.0016 | 0.0010 | 0.0011 |
| F13 | 23700001 | 23800000 | 0.0018 | 0.0013 | 0.0014 |
| F13 | 23720001 | 23820000 | 0.0016 | 0.0013 | 0.0013 |
| F13 | 23740001 | 23840000 | 0.0019 | 0.0016 | 0.0017 |
| F13 | 23760001 | 23860000 | 0.0022 | 0.0017 | 0.0019 |
| F13 | 23780001 | 23880000 | 0.0025 | 0.0019 | 0.0020 |
| F13 | 23800001 | 23900000 | 0.0030 | 0.0019 | 0.0021 |
| F13 | 23820001 | 23920000 | 0.0038 | 0.0026 | 0.0028 |
| F13 | 23840001 | 23940000 | 0.0038 | 0.0025 | 0.0026 |
| F13 | 23860001 | 23960000 | 0.0036 | 0.0025 | 0.0026 |
| F13 | 23880001 | 23980000 | 0.0035 | 0.0025 | 0.0028 |
| F13 | 23900001 | 24000000 | 0.0029 | 0.0023 | 0.0024 |
| F13 | 23920001 | 24020000 | 0.0021 | 0.0016 | 0.0018 |
| F13 | 23940001 | 24040000 | 0.0018 | 0.0014 | 0.0016 |
| F13 | 23960001 | 24060000 | 0.0015 | 0.0011 | 0.0013 |
| F13 | 23980001 | 24080000 | 0.0015 | 0.0011 | 0.0012 |
| F13 | 24000001 | 24100000 | 0.0014 | 0.0011 | 0.0012 |
| F13 | 24020001 | 24120000 | 0.0012 | 0.0010 | 0.0010 |

| Chr | Start | End | both | A1 | A2 |
|---|---|---|---|---|---|
| F13 | 24040001 | 24140000 | 0.0013 | 0.0011 | 0.0011 |
| F13 | 24060001 | 24160000 | 0.0015 | 0.0013 | 0.0014 |
| F13 | 24080001 | 24180000 | 0.0018 | 0.0016 | 0.0016 |
| F13 | 24100001 | 24200000 | 0.0021 | 0.0016 | 0.0016 |
| F13 | 24120001 | 24220000 | 0.0031 | 0.0023 | 0.0025 |
| F13 | 24140001 | 24240000 | 0.0032 | 0.0023 | 0.0026 |
| F13 | 24160001 | 24260000 | 0.0037 | 0.0026 | 0.0030 |
| F13 | 24180001 | 24280000 | 0.0036 | 0.0023 | 0.0028 |
| F13 | 24200001 | 24300000 | 0.0032 | 0.0022 | 0.0026 |
| F13 | 24220001 | 24320000 | 0.0027 | 0.0018 | 0.0021 |
| F13 | 24240001 | 24340000 | 0.0031 | 0.0019 | 0.0022 |
| F13 | 24260001 | 24360000 | 0.0027 | 0.0016 | 0.0019 |
| F13 | 24280001 | 24380000 | 0.0024 | 0.0016 | 0.0017 |
| F13 | 24300001 | 24400000 | 0.0025 | 0.0016 | 0.0017 |
| F13 | 24320001 | 24420000 | 0.0029 | 0.0020 | 0.0022 |
| F13 | 24340001 | 24440000 | 0.0037 | 0.0030 | 0.0033 |
| F13 | 24360001 | 24460000 | 0.0037 | 0.0030 | 0.0033 |
| F13 | 24380001 | 24480000 | 0.0037 | 0.0030 | 0.0032 |
| F13 | 24400001 | 24500000 | 0.0040 | 0.0032 | 0.0034 |
| F13 | 24420001 | 24520000 | 0.0033 | 0.0026 | 0.0027 |
| F13 | 24440001 | 24540000 | 0.0018 | 0.0012 | 0.0012 |
| F13 | 24460001 | 24560000 | 0.0013 | 0.0009 | 0.0010 |
| F13 | 24480001 | 24580000 | 0.0014 | 0.0009 | 0.0009 |
| F13 | 24500001 | 24600000 | 0.0015 | 0.0011 | 0.0012 |
| F13 | 24520001 | 24620000 | 0.0018 | 0.0013 | 0.0015 |
| F13 | 24540001 | 24640000 | 0.0018 | 0.0013 | 0.0016 |
| F13 | 24560001 | 24660000 | 0.0019 | 0.0014 | 0.0017 |
| F13 | 24580001 | 24680000 | 0.0016 | 0.0010 | 0.0015 |
| F13 | 24600001 | 24700000 | 0.0015 | 0.0010 | 0.0014 |
| F13 | 24620001 | 24720000 | 0.0010 | 0.0008 | 0.0010 |
| F13 | 24640001 | 24740000 | 0.0013 | 0.0011 | 0.0012 |
| F13 | 24660001 | 24760000 | 0.0013 | 0.0013 | 0.0013 |
| F13 | 24680001 | 24780000 | 0.0015 | 0.0014 | 0.0015 |
| F13 | 24700001 | 24800000 | 0.0014 | 0.0012 | 0.0014 |
| F13 | 24720001 | 24820000 | 0.0014 | 0.0012 | 0.0014 |
| F13 | 24740001 | 24840000 | 0.0014 | 0.0011 | 0.0013 |
| F13 | 24760001 | 24860000 | 0.0017 | 0.0014 | 0.0017 |
| F13 | 24780001 | 24880000 | 0.0017 | 0.0013 | 0.0016 |
| F13 | 24800001 | 24900000 | 0.0019 | 0.0014 | 0.0018 |
| F13 | 24820001 | 24920000 | 0.0021 | 0.0016 | 0.0020 |
| F13 | 24840001 | 24940000 | 0.0021 | 0.0015 | 0.0020 |
| F13 | 24860001 | 24960000 | 0.0024 | 0.0016 | 0.0024 |
| F13 | 24880001 | 24980000 | 0.0029 | 0.0018 | 0.0026 |
| F13 | 24900001 | 25000000 | 0.0027 | 0.0016 | 0.0021 |
| F13 | 24920001 | 25020000 | 0.0029 | 0.0017 | 0.0022 |
| F13 | 24940001 | 25040000 | 0.0030 | 0.0018 | 0.0023 |
| F13 | 24960001 | 25060000 | 0.0027 | 0.0016 | 0.0018 |
| F13 | 24980001 | 25080000 | 0.0027 | 0.0018 | 0.0020 |
| F13 | 25000001 | 25100000 | 0.0027 | 0.0018 | 0.0021 |
| F13 | 25020001 | 25120000 | 0.0025 | 0.0017 | 0.0019 |
| F13 | 25040001 | 25140000 | 0.0024 | 0.0016 | 0.0017 |
| F13 | 25060001 | 25160000 | 0.0028 | 0.0017 | 0.0019 |
| F13 | 25080001 | 25180000 | 0.0028 | 0.0017 | 0.0019 |
| F13 | 25100001 | 25200000 | 0.0030 | 0.0018 | 0.0019 |
| F13 | 25120001 | 25220000 | 0.0035 | 0.0021 | 0.0022 |
| F13 | 25140001 | 25240000 | 0.0034 | 0.0021 | 0.0023 |
| F13 | 25160001 | 25260000 | 0.0030 | 0.0021 | 0.0021 |
| F13 | 25180001 | 25280000 | 0.0024 | 0.0017 | 0.0016 |

| Chr | Start | End | both | A1 | A2 |
|---|---|---|---|---|---|
| F13 | 25200001 | 25300000 | 0.0021 | 0.0014 | 0.0015 |
| F13 | 25220001 | 25320000 | 0.0016 | 0.0010 | 0.0011 |
| F13 | 25240001 | 25340000 | 0.0022 | 0.0016 | 0.0017 |
| F13 | 25260001 | 25360000 | 0.0020 | 0.0015 | 0.0016 |
| F13 | 25280001 | 25380000 | 0.0022 | 0.0016 | 0.0017 |
| F13 | 25300001 | 25400000 | 0.0021 | 0.0017 | 0.0018 |
| F13 | 25320001 | 25420000 | 0.0024 | 0.0021 | 0.0019 |
| F13 | 25340001 | 25440000 | 0.0020 | 0.0016 | 0.0013 |
| F13 | 25360001 | 25460000 | 0.0021 | 0.0016 | 0.0015 |
| F13 | 25380001 | 25480000 | 0.0023 | 0.0018 | 0.0017 |
| F13 | 25400001 | 25500000 | 0.0022 | 0.0017 | 0.0016 |
| F13 | 25420001 | 25520000 | 0.0018 | 0.0013 | 0.0014 |
| F13 | 25440001 | 25540000 | 0.0021 | 0.0017 | 0.0019 |
| F13 | 25460001 | 25560000 | 0.0027 | 0.0024 | 0.0025 |
| F13 | 25480001 | 25580000 | 0.0023 | 0.0020 | 0.0021 |
| F13 | 25500001 | 25600000 | 0.0029 | 0.0026 | 0.0026 |
| F13 | 25520001 | 25620000 | 0.0039 | 0.0033 | 0.0036 |
| F13 | 25540001 | 25640000 | 0.0037 | 0.0031 | 0.0034 |
| F13 | 25560001 | 25660000 | 0.0029 | 0.0022 | 0.0025 |
| F13 | 25580001 | 25680000 | 0.0031 | 0.0022 | 0.0025 |
| F13 | 25600001 | 25700000 | 0.0030 | 0.0019 | 0.0021 |
| F13 | 25620001 | 25720000 | 0.0033 | 0.0024 | 0.0024 |
| F13 | 25640001 | 25740000 | 0.0036 | 0.0026 | 0.0027 |
| F13 | 25660001 | 25760000 | 0.0036 | 0.0027 | 0.0028 |
| F13 | 25680001 | 25780000 | 0.0037 | 0.0031 | 0.0031 |
| F13 | 25700001 | 25800000 | 0.0032 | 0.0028 | 0.0029 |
| F13 | 25720001 | 25820000 | 0.0021 | 0.0017 | 0.0017 |
| F13 | 25740001 | 25840000 | 0.0013 | 0.0011 | 0.0010 |
| F13 | 25760001 | 25860000 | 0.0009 | 0.0008 | 0.0008 |
| F13 | 25780001 | 25880000 | 0.0012 | 0.0010 | 0.0010 |
| F13 | 25800001 | 25900000 | 0.0013 | 0.0011 | 0.0011 |
| F13 | 25820001 | 25920000 | 0.0012 | 0.0010 | 0.0010 |
| F13 | 25840001 | 25940000 | 0.0020 | 0.0017 | 0.0018 |
| F13 | 25860001 | 25960000 | 0.0025 | 0.0021 | 0.0023 |
| F13 | 25880001 | 25980000 | 0.0021 | 0.0018 | 0.0019 |
| F13 | 25900001 | 26000000 | 0.0022 | 0.0019 | 0.0021 |
| F13 | 25920001 | 26020000 | 0.0022 | 0.0018 | 0.0020 |
| F13 | 25940001 | 26040000 | 0.0012 | 0.0010 | 0.0011 |
| F13 | 25960001 | 26060000 | 0.0010 | 0.0008 | 0.0009 |
| F13 | 25980001 | 26080000 | 0.0011 | 0.0009 | 0.0011 |
| F13 | 26000001 | 26100000 | 0.0015 | 0.0013 | 0.0014 |
| F13 | 26020001 | 26120000 | 0.0019 | 0.0015 | 0.0017 |
| F13 | 26040001 | 26140000 | 0.0024 | 0.0018 | 0.0020 |
| F13 | 26060001 | 26160000 | 0.0032 | 0.0024 | 0.0027 |
| F13 | 26080001 | 26180000 | 0.0029 | 0.0021 | 0.0023 |
| F13 | 26100001 | 26200000 | 0.0025 | 0.0017 | 0.0019 |
| F13 | 26120001 | 26220000 | 0.0023 | 0.0015 | 0.0019 |
| F13 | 26140001 | 26240000 | 0.0019 | 0.0014 | 0.0016 |
| F13 | 26160001 | 26260000 | 0.0010 | 0.0007 | 0.0009 |
| F13 | 26180001 | 26280000 | 0.0019 | 0.0016 | 0.0016 |
| F13 | 26200001 | 26300000 | 0.0030 | 0.0027 | 0.0028 |
| F13 | 26220001 | 26320000 | 0.0032 | 0.0029 | 0.0030 |
| F13 | 26240001 | 26340000 | 0.0035 | 0.0031 | 0.0032 |
| F13 | 26260001 | 26360000 | 0.0035 | 0.0032 | 0.0033 |
| F13 | 26280001 | 26380000 | 0.0028 | 0.0024 | 0.0026 |
| F13 | 26300001 | 26400000 | 0.0019 | 0.0015 | 0.0017 |
| F13 | 26320001 | 26420000 | 0.0018 | 0.0013 | 0.0016 |
| F13 | 26340001 | 26440000 | 0.0016 | 0.0012 | 0.0015 |

| Chr | Start | End | both | A1 | A2 |
|---|---|---|---|---|---|
| F13 | 26360001 | 26460000 | 0.0014 | 0.0010 | 0.0013 |
| F13 | 26380001 | 26480000 | 0.0013 | 0.0009 | 0.0012 |
| F13 | 26400001 | 26500000 | 0.0016 | 0.0013 | 0.0015 |
| F13 | 26420001 | 26520000 | 0.0021 | 0.0016 | 0.0020 |
| F13 | 26440001 | 26540000 | 0.0021 | 0.0017 | 0.0021 |
| F13 | 26460001 | 26560000 | 0.0021 | 0.0016 | 0.0021 |
| F13 | 26480001 | 26580000 | 0.0025 | 0.0018 | 0.0023 |
| F13 | 26500001 | 26600000 | 0.0022 | 0.0016 | 0.0019 |
| F13 | 26520001 | 26620000 | 0.0016 | 0.0012 | 0.0013 |
| F13 | 26540001 | 26640000 | 0.0017 | 0.0012 | 0.0013 |
| F13 | 26560001 | 26660000 | 0.0021 | 0.0016 | 0.0017 |
| F13 | 26580001 | 26680000 | 0.0020 | 0.0016 | 0.0017 |
| F13 | 26600001 | 26700000 | 0.0021 | 0.0016 | 0.0018 |
| F13 | 26620001 | 26720000 | 0.0026 | 0.0019 | 0.0022 |
| F13 | 26640001 | 26740000 | 0.0027 | 0.0019 | 0.0021 |
| F13 | 26660001 | 26760000 | 0.0024 | 0.0017 | 0.0019 |
| F13 | 26680001 | 26780000 | 0.0029 | 0.0022 | 0.0023 |
| F13 | 26700001 | 26800000 | 0.0023 | 0.0018 | 0.0018 |
| F13 | 26720001 | 26820000 | 0.0019 | 0.0016 | 0.0016 |
| F13 | 26740001 | 26840000 | 0.0019 | 0.0018 | 0.0017 |
| F13 | 26760001 | 26860000 | 0.0017 | 0.0015 | 0.0015 |
| F13 | 26780001 | 26880000 | 0.0011 | 0.0010 | 0.0010 |
| F13 | 26800001 | 26900000 | 0.0016 | 0.0015 | 0.0015 |
| F13 | 26820001 | 26920000 | 0.0015 | 0.0011 | 0.0014 |
| F13 | 26840001 | 26940000 | 0.0012 | 0.0008 | 0.0011 |
| F13 | 26860001 | 26960000 | 0.0019 | 0.0012 | 0.0018 |
| F13 | 26880001 | 26980000 | 0.0030 | 0.0021 | 0.0028 |
| F13 | 26900001 | 27000000 | 0.0033 | 0.0022 | 0.0031 |
| F13 | 26920001 | 27020000 | 0.0035 | 0.0027 | 0.0033 |
| F13 | 26940001 | 27040000 | 0.0036 | 0.0028 | 0.0035 |
| F13 | 26960001 | 27060000 | 0.0030 | 0.0024 | 0.0028 |
| F13 | 26980001 | 27080000 | 0.0021 | 0.0016 | 0.0020 |
| F13 | 27000001 | 27100000 | 0.0020 | 0.0015 | 0.0020 |
| F13 | 27020001 | 27120000 | 0.0018 | 0.0012 | 0.0018 |
| F13 | 27040001 | 27140000 | 0.0018 | 0.0011 | 0.0018 |
| F13 | 27060001 | 27160000 | 0.0025 | 0.0018 | 0.0024 |
| F13 | 27080001 | 27180000 | 0.0023 | 0.0016 | 0.0022 |
| F13 | 27100001 | 27200000 | 0.0020 | 0.0014 | 0.0019 |
| F13 | 27120001 | 27220000 | 0.0017 | 0.0012 | 0.0017 |
| F13 | 27140001 | 27240000 | 0.0024 | 0.0016 | 0.0023 |
| F13 | 27160001 | 27260000 | 0.0022 | 0.0011 | 0.0020 |
| F13 | 27180001 | 27280000 | 0.0026 | 0.0014 | 0.0022 |
| F13 | 27200001 | 27300000 | 0.0023 | 0.0013 | 0.0019 |
| F13 | 27220001 | 27320000 | 0.0035 | 0.0020 | 0.0030 |
| F13 | 27240001 | 27340000 | 0.0031 | 0.0018 | 0.0026 |
| F13 | 27260001 | 27360000 | 0.0031 | 0.0022 | 0.0027 |
| F13 | 27280001 | 27380000 | 0.0029 | 0.0020 | 0.0027 |
| F13 | 27300001 | 27400000 | 0.0038 | 0.0024 | 0.0036 |
| F13 | 27320001 | 27420000 | 0.0024 | 0.0015 | 0.0023 |
| F13 | 27340001 | 27440000 | 0.0020 | 0.0013 | 0.0019 |
| F13 | 27360001 | 27460000 | 0.0015 | 0.0009 | 0.0015 |
| F13 | 27380001 | 27480000 | 0.0011 | 0.0005 | 0.0011 |
| F13 | 27400001 | 27500000 | 0.0004 | 0.0003 | 0.0003 |
| F13 | 27420001 | 27520000 | 0.0004 | 0.0004 | 0.0004 |
| F13 | 27440001 | 27540000 | 0.0005 | 0.0005 | 0.0005 |
| F13 | 27460001 | 27560000 | 0.0009 | 0.0009 | 0.0009 |
| F13 | 27480001 | 27580000 | 0.0009 | 0.0009 | 0.0009 |
| F13 | 27500001 | 27600000 | 0.0012 | 0.0011 | 0.0011 |

| Chr | Start | End | both | A1 | A2 |
|---|---|---|---|---|---|
| F13 | 27520001 | 27620000 | 0.0013 | 0.0013 | 0.0013 |
| F13 | 27540001 | 27640000 | 0.0012 | 0.0011 | 0.0011 |
| F13 | 27560001 | 27660000 | 0.0009 | 0.0008 | 0.0008 |
| F13 | 27580001 | 27680000 | 0.0007 | 0.0007 | 0.0007 |
| F13 | 27600001 | 27700000 | 0.0006 | 0.0005 | 0.0006 |
| F13 | 27620001 | 27720000 | 0.0009 | 0.0007 | 0.0009 |
| F13 | 27640001 | 27740000 | 0.0013 | 0.0009 | 0.0011 |
| F13 | 27660001 | 27760000 | 0.0016 | 0.0011 | 0.0014 |
| F13 | 27680001 | 27780000 | 0.0024 | 0.0017 | 0.0020 |
| F13 | 27700001 | 27800000 | 0.0024 | 0.0018 | 0.0021 |
| F13 | 27720001 | 27820000 | 0.0021 | 0.0015 | 0.0018 |
| F13 | 27740001 | 27840000 | 0.0023 | 0.0014 | 0.0018 |
| F13 | 27760001 | 27860000 | 0.0028 | 0.0020 | 0.0023 |
| F13 | 27780001 | 27880000 | 0.0030 | 0.0022 | 0.0026 |
| F13 | 27800001 | 27900000 | 0.0028 | 0.0021 | 0.0024 |
| F13 | 27820001 | 27920000 | 0.0049 | 0.0040 | 0.0044 |
| F13 | 27840001 | 27940000 | 0.0047 | 0.0042 | 0.0045 |
| F13 | 27860001 | 27960000 | 0.0045 | 0.0040 | 0.0041 |
| F13 | 27880001 | 27980000 | 0.0038 | 0.0034 | 0.0035 |
| F13 | 27900001 | 28000000 | 0.0037 | 0.0032 | 0.0033 |
| F13 | 27920001 | 28020000 | 0.0020 | 0.0016 | 0.0017 |
| F13 | 27940001 | 28040000 | 0.0019 | 0.0015 | 0.0016 |
| F13 | 27960001 | 28060000 | 0.0017 | 0.0011 | 0.0014 |
| F13 | 27980001 | 28080000 | 0.0021 | 0.0014 | 0.0017 |
| F13 | 28000001 | 28100000 | 0.0024 | 0.0017 | 0.0019 |
| F13 | 28020001 | 28120000 | 0.0024 | 0.0017 | 0.0018 |
| F13 | 28040001 | 28140000 | 0.0025 | 0.0019 | 0.0020 |
| F13 | 28060001 | 28160000 | 0.0027 | 0.0019 | 0.0021 |
| F13 | 28080001 | 28180000 | 0.0024 | 0.0016 | 0.0018 |
| F13 | 28100001 | 28200000 | 0.0023 | 0.0014 | 0.0015 |
| F13 | 28120001 | 28220000 | 0.0024 | 0.0014 | 0.0017 |
| F13 | 28140001 | 28240000 | 0.0026 | 0.0016 | 0.0018 |
| F13 | 28160001 | 28260000 | 0.0030 | 0.0022 | 0.0023 |
| F13 | 28180001 | 28280000 | 0.0028 | 0.0020 | 0.0021 |
| F13 | 28200001 | 28300000 | 0.0028 | 0.0021 | 0.0022 |
| F13 | 28220001 | 28320000 | 0.0028 | 0.0022 | 0.0022 |
| F13 | 28240001 | 28340000 | 0.0027 | 0.0020 | 0.0021 |
| F13 | 28260001 | 28360000 | 0.0027 | 0.0019 | 0.0022 |
| F13 | 28280001 | 28380000 | 0.0030 | 0.0022 | 0.0025 |
| F13 | 28300001 | 28400000 | 0.0031 | 0.0024 | 0.0026 |
| F13 | 28320001 | 28420000 | 0.0026 | 0.0019 | 0.0022 |
| F13 | 28340001 | 28440000 | 0.0022 | 0.0017 | 0.0020 |
| F13 | 28360001 | 28460000 | 0.0016 | 0.0013 | 0.0014 |
| F13 | 28380001 | 28480000 | 0.0019 | 0.0016 | 0.0018 |
| F13 | 28400001 | 28500000 | 0.0023 | 0.0019 | 0.0022 |
| F13 | 28420001 | 28520000 | 0.0029 | 0.0024 | 0.0027 |
| F13 | 28440001 | 28540000 | 0.0030 | 0.0024 | 0.0027 |
| F13 | 28460001 | 28560000 | 0.0030 | 0.0024 | 0.0027 |
| F13 | 28480001 | 28580000 | 0.0031 | 0.0025 | 0.0027 |
| F13 | 28500001 | 28600000 | 0.0030 | 0.0024 | 0.0025 |
| F13 | 28520001 | 28620000 | 0.0040 | 0.0034 | 0.0036 |
| F13 | 28540001 | 28640000 | 0.0042 | 0.0037 | 0.0038 |
| F13 | 28560001 | 28660000 | 0.0040 | 0.0035 | 0.0036 |
| F13 | 28580001 | 28680000 | 0.0041 | 0.0036 | 0.0038 |
| F13 | 28600001 | 28700000 | 0.0036 | 0.0031 | 0.0033 |
| F13 | 28620001 | 28720000 | 0.0022 | 0.0018 | 0.0021 |
| F13 | 28640001 | 28740000 | 0.0020 | 0.0016 | 0.0019 |
| F13 | 28660001 | 28760000 | 0.0022 | 0.0019 | 0.0022 |

| Chr | Start | End | both | A1 | A2 |
|---|---|---|---|---|---|
| F13 | 28680001 | 28780000 | 0.0017 | 0.0014 | 0.0017 |
| F13 | 28700001 | 28800000 | 0.0022 | 0.0018 | 0.0021 |
| F13 | 28720001 | 28820000 | 0.0027 | 0.0022 | 0.0025 |
| F13 | 28740001 | 28840000 | 0.0032 | 0.0025 | 0.0029 |
| F13 | 28760001 | 28860000 | 0.0034 | 0.0027 | 0.0030 |
| F13 | 28780001 | 28880000 | 0.0043 | 0.0036 | 0.0039 |
| F13 | 28800001 | 28900000 | 0.0046 | 0.0039 | 0.0043 |
| F13 | 28820001 | 28920000 | 0.0041 | 0.0034 | 0.0038 |
| F13 | 28840001 | 28940000 | 0.0038 | 0.0033 | 0.0037 |
| F13 | 28860001 | 28960000 | 0.0037 | 0.0031 | 0.0035 |
| F13 | 28880001 | 28980000 | 0.0040 | 0.0031 | 0.0038 |
| F13 | 28900001 | 29000000 | 0.0035 | 0.0026 | 0.0032 |
| F13 | 28920001 | 29020000 | 0.0037 | 0.0029 | 0.0034 |
| F13 | 28940001 | 29040000 | 0.0035 | 0.0026 | 0.0031 |
| F13 | 28960001 | 29060000 | 0.0030 | 0.0022 | 0.0026 |
| F13 | 28980001 | 29080000 | 0.0014 | 0.0010 | 0.0012 |
| F13 | 29000001 | 29100000 | 0.0012 | 0.0010 | 0.0011 |
| F13 | 29020001 | 29120000 | 0.0011 | 0.0009 | 0.0011 |
| F13 | 29040001 | 29140000 | 0.0012 | 0.0008 | 0.0010 |
| F13 | 29060001 | 29160000 | 0.0016 | 0.0012 | 0.0015 |
| F13 | 29080001 | 29180000 | 0.0020 | 0.0015 | 0.0019 |
| F13 | 29100001 | 29200000 | 0.0027 | 0.0022 | 0.0026 |
| F13 | 29120001 | 29220000 | 0.0025 | 0.0020 | 0.0024 |
| F13 | 29140001 | 29240000 | 0.0022 | 0.0019 | 0.0022 |
| F13 | 29160001 | 29260000 | 0.0022 | 0.0019 | 0.0022 |
| F13 | 29180001 | 29280000 | 0.0028 | 0.0021 | 0.0026 |
| F13 | 29200001 | 29300000 | 0.0022 | 0.0015 | 0.0019 |
| F13 | 29220001 | 29320000 | 0.0020 | 0.0013 | 0.0016 |
| F13 | 29240001 | 29340000 | 0.0023 | 0.0015 | 0.0018 |
| F13 | 29260001 | 29360000 | 0.0022 | 0.0014 | 0.0017 |
| F13 | 29280001 | 29380000 | 0.0013 | 0.0008 | 0.0009 |
| F13 | 29300001 | 29400000 | 0.0012 | 0.0009 | 0.0010 |
| F13 | 29320001 | 29420000 | 0.0019 | 0.0015 | 0.0016 |
| F13 | 29340001 | 29440000 | 0.0018 | 0.0014 | 0.0016 |
| F13 | 29360001 | 29460000 | 0.0020 | 0.0015 | 0.0017 |
| F13 | 29380001 | 29480000 | 0.0021 | 0.0016 | 0.0018 |
| F13 | 29400001 | 29500000 | 0.0027 | 0.0021 | 0.0023 |
| F13 | 29420001 | 29520000 | 0.0030 | 0.0022 | 0.0024 |
| F13 | 29440001 | 29540000 | 0.0030 | 0.0022 | 0.0024 |
| F13 | 29460001 | 29560000 | 0.0030 | 0.0023 | 0.0024 |
| F13 | 29480001 | 29580000 | 0.0034 | 0.0028 | 0.0029 |
| F13 | 29500001 | 29600000 | 0.0034 | 0.0027 | 0.0029 |
| F13 | 29520001 | 29620000 | 0.0024 | 0.0021 | 0.0021 |
| F13 | 29540001 | 29640000 | 0.0023 | 0.0021 | 0.0020 |
| F13 | 29560001 | 29660000 | 0.0019 | 0.0018 | 0.0017 |
| F13 | 29580001 | 29680000 | 0.0015 | 0.0013 | 0.0013 |
| F13 | 29600001 | 29700000 | 0.0008 | 0.0007 | 0.0007 |
| F13 | 29620001 | 29720000 | 0.0006 | 0.0006 | 0.0006 |
| F13 | 29640001 | 29740000 | 0.0005 | 0.0004 | 0.0005 |
| F13 | 29660001 | 29760000 | 0.0009 | 0.0008 | 0.0009 |
| F13 | 29680001 | 29780000 | 0.0007 | 0.0007 | 0.0007 |
| F13 | 29700001 | 29800000 | 0.0008 | 0.0008 | 0.0007 |
| F13 | 29720001 | 29820000 | 0.0009 | 0.0009 | 0.0008 |
| F13 | 29740001 | 29840000 | 0.0008 | 0.0009 | 0.0008 |
| F13 | 29760001 | 29860000 | 0.0004 | 0.0004 | 0.0004 |
| F13 | 29780001 | 29880000 | 0.0006 | 0.0006 | 0.0005 |
| F13 | 29800001 | 29900000 | 0.0005 | 0.0005 | 0.0004 |
| F13 | 29820001 | 29920000 | 0.0005 | 0.0004 | 0.0004 |

| Chr | Start | End | both | A1 | A2 |
|---|---|---|---|---|---|
| F13 | 29840001 | 29940000 | 0.0009 | 0.0008 | 0.0008 |
| F13 | 29860001 | 29960000 | 0.0010 | 0.0008 | 0.0009 |
| F13 | 29880001 | 29980000 | 0.0008 | 0.0007 | 0.0007 |
| F13 | 29900001 | 30000000 | 0.0007 | 0.0006 | 0.0007 |
| F13 | 29920001 | 30020000 | 0.0008 | 0.0007 | 0.0008 |
| F13 | 29940001 | 30040000 | 0.0004 | 0.0003 | 0.0004 |
| F13 | 29960001 | 30060000 | 0.0004 | 0.0003 | 0.0004 |
| F13 | 29980001 | 30080000 | 0.0007 | 0.0004 | 0.0007 |
| F13 | 30000001 | 30100000 | 0.0007 | 0.0005 | 0.0007 |
| F13 | 30020001 | 30120000 | 0.0006 | 0.0004 | 0.0006 |
| F13 | 30040001 | 30140000 | 0.0006 | 0.0003 | 0.0006 |
| F13 | 30060001 | 30160000 | 0.0005 | 0.0003 | 0.0005 |
| F13 | 30080001 | 30180000 | 0.0002 | 0.0001 | 0.0002 |
| F13 | 30120001 | 30220000 | 0.0001 | 0.0001 | 0.0001 |
| F13 | 30140001 | 30240000 | 0.0002 | 0.0001 | 0.0001 |
| F13 | 30160001 | 30260000 | 0.0002 | 0.0001 | 0.0002 |
| F13 | 30180001 | 30280000 | 0.0008 | 0.0006 | 0.0007 |
| F13 | 30200001 | 30300000 | 0.0010 | 0.0007 | 0.0009 |
| F13 | 30220001 | 30320000 | 0.0009 | 0.0006 | 0.0008 |
| F13 | 30240001 | 30340000 | 0.0010 | 0.0006 | 0.0008 |
| F13 | 30260001 | 30360000 | 0.0010 | 0.0007 | 0.0009 |
| F13 | 30280001 | 30380000 | 0.0005 | 0.0003 | 0.0004 |
| F13 | 30300001 | 30400000 | 0.0003 | 0.0001 | 0.0002 |
| F13 | 30320001 | 30420000 | 0.0003 | 0.0002 | 0.0003 |
| F13 | 30340001 | 30440000 | 0.0003 | 0.0003 | 0.0003 |
| F13 | 30360001 | 30460000 | 0.0003 | 0.0002 | 0.0003 |
| F13 | 30380001 | 30480000 | 0.0002 | 0.0002 | 0.0002 |
| F13 | 30400001 | 30500000 | 0.0002 | 0.0002 | 0.0002 |
| F13 | 30420001 | 30520000 | 0.0002 | 0.0001 | 0.0002 |
| F13 | 30440001 | 30540000 | 0.0001 | 0.0001 | 0.0001 |
| F13 | 30460001 | 30560000 | 0.0003 | 0.0002 | 0.0003 |
| F13 | 30480001 | 30580000 | 0.0003 | 0.0003 | 0.0003 |
| F13 | 30500001 | 30600000 | 0.0008 | 0.0007 | 0.0007 |
| F13 | 30520001 | 30620000 | 0.0009 | 0.0008 | 0.0008 |
| F13 | 30540001 | 30640000 | 0.0008 | 0.0008 | 0.0008 |
| F13 | 30560001 | 30660000 | 0.0012 | 0.0011 | 0.0011 |
| F13 | 30580001 | 30680000 | 0.0013 | 0.0012 | 0.0013 |
| F13 | 30600001 | 30700000 | 0.0011 | 0.0009 | 0.0010 |
| F13 | 30620001 | 30720000 | 0.0009 | 0.0007 | 0.0008 |
| F13 | 30640001 | 30740000 | 0.0013 | 0.0010 | 0.0012 |
| F13 | 30660001 | 30760000 | 0.0009 | 0.0007 | 0.0009 |
| F13 | 30680001 | 30780000 | 0.0012 | 0.0009 | 0.0011 |
| F13 | 30700001 | 30800000 | 0.0014 | 0.0012 | 0.0014 |
| F13 | 30720001 | 30820000 | 0.0017 | 0.0013 | 0.0017 |
| F13 | 30740001 | 30840000 | 0.0015 | 0.0013 | 0.0015 |
| F13 | 30760001 | 30860000 | 0.0015 | 0.0013 | 0.0014 |
| F13 | 30780001 | 30880000 | 0.0020 | 0.0016 | 0.0019 |
| F13 | 30800001 | 30900000 | 0.0018 | 0.0014 | 0.0018 |
| F13 | 30820001 | 30920000 | 0.0021 | 0.0015 | 0.0020 |
| F13 | 30840001 | 30940000 | 0.0020 | 0.0014 | 0.0019 |
| F13 | 30860001 | 30960000 | 0.0028 | 0.0022 | 0.0028 |
| F13 | 30880001 | 30980000 | 0.0021 | 0.0016 | 0.0020 |
| F13 | 30900001 | 31000000 | 0.0027 | 0.0022 | 0.0027 |
| F13 | 30920001 | 31020000 | 0.0035 | 0.0032 | 0.0034 |
| F13 | 30940001 | 31040000 | 0.0037 | 0.0032 | 0.0036 |
| F13 | 30960001 | 31060000 | 0.0030 | 0.0026 | 0.0029 |
| F13 | 30980001 | 31080000 | 0.0032 | 0.0027 | 0.0030 |
| F13 | 31000001 | 31100000 | 0.0030 | 0.0023 | 0.0025 |

| Chr | Start | End | both | A1 | A2 |
|---|---|---|---|---|---|
| F13 | 31020001 | 31120000 | 0.0023 | 0.0016 | 0.0018 |
| F13 | 31040001 | 31140000 | 0.0027 | 0.0019 | 0.0021 |
| F13 | 31060001 | 31160000 | 0.0029 | 0.0021 | 0.0023 |
| F13 | 31080001 | 31180000 | 0.0027 | 0.0020 | 0.0022 |
| F13 | 31100001 | 31200000 | 0.0022 | 0.0018 | 0.0020 |
| F13 | 31120001 | 31220000 | 0.0025 | 0.0021 | 0.0024 |
| F13 | 31140001 | 31240000 | 0.0021 | 0.0018 | 0.0021 |
| F13 | 31160001 | 31260000 | 0.0018 | 0.0015 | 0.0018 |
| F13 | 31180001 | 31280000 | 0.0022 | 0.0018 | 0.0022 |
| F13 | 31200001 | 31300000 | 0.0024 | 0.0021 | 0.0024 |
| F13 | 31220001 | 31320000 | 0.0019 | 0.0015 | 0.0018 |
| F13 | 31240001 | 31340000 | 0.0019 | 0.0014 | 0.0017 |
| F13 | 31260001 | 31360000 | 0.0021 | 0.0015 | 0.0018 |
| F13 | 31280001 | 31380000 | 0.0021 | 0.0016 | 0.0017 |
| F13 | 31300001 | 31400000 | 0.0027 | 0.0021 | 0.0023 |
| F13 | 31320001 | 31420000 | 0.0028 | 0.0024 | 0.0025 |
| F13 | 31340001 | 31440000 | 0.0026 | 0.0022 | 0.0023 |
| F13 | 31360001 | 31460000 | 0.0025 | 0.0021 | 0.0022 |
| F13 | 31380001 | 31480000 | 0.0020 | 0.0017 | 0.0019 |
| F13 | 31400001 | 31500000 | 0.0021 | 0.0014 | 0.0018 |
| F13 | 31420001 | 31520000 | 0.0020 | 0.0013 | 0.0017 |
| F13 | 31440001 | 31540000 | 0.0024 | 0.0017 | 0.0021 |
| F13 | 31460001 | 31560000 | 0.0025 | 0.0017 | 0.0022 |
| F13 | 31480001 | 31580000 | 0.0029 | 0.0021 | 0.0026 |
| F13 | 31500001 | 31600000 | 0.0023 | 0.0018 | 0.0021 |
| F13 | 31520001 | 31620000 | 0.0019 | 0.0014 | 0.0017 |
| F13 | 31540001 | 31640000 | 0.0016 | 0.0011 | 0.0013 |
| F13 | 31560001 | 31660000 | 0.0016 | 0.0010 | 0.0012 |
| F13 | 31580001 | 31680000 | 0.0017 | 0.0011 | 0.0013 |
| F13 | 31600001 | 31700000 | 0.0016 | 0.0011 | 0.0012 |
| F13 | 31620001 | 31720000 | 0.0014 | 0.0011 | 0.0011 |
| F13 | 31640001 | 31740000 | 0.0016 | 0.0014 | 0.0014 |
| F13 | 31660001 | 31760000 | 0.0015 | 0.0013 | 0.0013 |
| F13 | 31680001 | 31780000 | 0.0014 | 0.0012 | 0.0012 |
| F13 | 31700001 | 31800000 | 0.0018 | 0.0015 | 0.0015 |
| F13 | 31720001 | 31820000 | 0.0027 | 0.0022 | 0.0022 |
| F13 | 31740001 | 31840000 | 0.0030 | 0.0023 | 0.0024 |
| F13 | 31760001 | 31860000 | 0.0035 | 0.0025 | 0.0028 |
| F13 | 31780001 | 31880000 | 0.0039 | 0.0028 | 0.0032 |
| F13 | 31800001 | 31900000 | 0.0032 | 0.0022 | 0.0026 |
| F13 | 31820001 | 31920000 | 0.0029 | 0.0017 | 0.0022 |
| F13 | 31840001 | 31940000 | 0.0028 | 0.0016 | 0.0020 |
| F13 | 31860001 | 31960000 | 0.0026 | 0.0016 | 0.0019 |
| F13 | 31880001 | 31980000 | 0.0022 | 0.0013 | 0.0013 |
| F13 | 31900001 | 32000000 | 0.0029 | 0.0018 | 0.0019 |
| F13 | 31920001 | 32020000 | 0.0027 | 0.0018 | 0.0017 |
| F13 | 31940001 | 32040000 | 0.0026 | 0.0017 | 0.0017 |
| F13 | 31960001 | 32060000 | 0.0026 | 0.0017 | 0.0018 |
| F13 | 31980001 | 32080000 | 0.0022 | 0.0014 | 0.0016 |
| F13 | 32000001 | 32100000 | 0.0016 | 0.0011 | 0.0012 |
| F13 | 32020001 | 32120000 | 0.0016 | 0.0012 | 0.0014 |
| F13 | 32040001 | 32140000 | 0.0014 | 0.0012 | 0.0014 |
| F13 | 32060001 | 32160000 | 0.0009 | 0.0008 | 0.0009 |
| F13 | 32080001 | 32180000 | 0.0009 | 0.0008 | 0.0009 |
| F13 | 32100001 | 32200000 | 0.0013 | 0.0009 | 0.0010 |
| F13 | 32120001 | 32220000 | 0.0013 | 0.0008 | 0.0010 |
| F13 | 32140001 | 32240000 | 0.0014 | 0.0010 | 0.0011 |
| F13 | 32160001 | 32260000 | 0.0018 | 0.0013 | 0.0014 |

| Chr | Start | End | both | A1 | A2 |
|---|---|---|---|---|---|
| F13 | 32180001 | 32280000 | 0.0019 | 0.0014 | 0.0016 |
| F13 | 32200001 | 32300000 | 0.0020 | 0.0018 | 0.0018 |
| F13 | 32220001 | 32320000 | 0.0031 | 0.0029 | 0.0029 |
| F13 | 32240001 | 32340000 | 0.0033 | 0.0029 | 0.0031 |
| F13 | 32260001 | 32360000 | 0.0044 | 0.0038 | 0.0041 |
| F13 | 32280001 | 32380000 | 0.0053 | 0.0047 | 0.0049 |
| F13 | 32300001 | 32400000 | 0.0052 | 0.0045 | 0.0048 |
| F13 | 32320001 | 32420000 | 0.0040 | 0.0033 | 0.0036 |
| F13 | 32340001 | 32440000 | 0.0037 | 0.0030 | 0.0032 |
| F13 | 32360001 | 32460000 | 0.0028 | 0.0019 | 0.0022 |
| F13 | 32380001 | 32480000 | 0.0021 | 0.0012 | 0.0017 |
| F13 | 32400001 | 32500000 | 0.0022 | 0.0012 | 0.0016 |
| F13 | 32420001 | 32520000 | 0.0028 | 0.0016 | 0.0022 |
| F13 | 32440001 | 32540000 | 0.0033 | 0.0022 | 0.0028 |
| F13 | 32460001 | 32560000 | 0.0032 | 0.0023 | 0.0030 |
| F13 | 32480001 | 32580000 | 0.0036 | 0.0027 | 0.0033 |
| F13 | 32500001 | 32600000 | 0.0036 | 0.0028 | 0.0033 |
| F13 | 32520001 | 32620000 | 0.0035 | 0.0027 | 0.0030 |
| F13 | 32540001 | 32640000 | 0.0039 | 0.0029 | 0.0032 |
| F13 | 32560001 | 32660000 | 0.0036 | 0.0027 | 0.0029 |
| F13 | 32580001 | 32680000 | 0.0036 | 0.0027 | 0.0028 |
| F13 | 32600001 | 32700000 | 0.0030 | 0.0021 | 0.0023 |
| F13 | 32620001 | 32720000 | 0.0024 | 0.0019 | 0.0020 |
| F13 | 32640001 | 32740000 | 0.0015 | 0.0012 | 0.0013 |
| F13 | 32660001 | 32760000 | 0.0017 | 0.0014 | 0.0016 |
| F13 | 32680001 | 32780000 | 0.0014 | 0.0010 | 0.0012 |
| F13 | 32700001 | 32800000 | 0.0015 | 0.0011 | 0.0013 |
| F13 | 32720001 | 32820000 | 0.0022 | 0.0019 | 0.0019 |
| F13 | 32740001 | 32840000 | 0.0025 | 0.0022 | 0.0021 |
| F13 | 32760001 | 32860000 | 0.0023 | 0.0021 | 0.0019 |
| F13 | 32780001 | 32880000 | 0.0024 | 0.0023 | 0.0020 |
| F13 | 32800001 | 32900000 | 0.0027 | 0.0027 | 0.0021 |
| F13 | 32820001 | 32920000 | 0.0019 | 0.0017 | 0.0015 |
| F13 | 32840001 | 32940000 | 0.0018 | 0.0016 | 0.0016 |
| F13 | 32860001 | 32960000 | 0.0017 | 0.0016 | 0.0016 |
| F13 | 32880001 | 32980000 | 0.0012 | 0.0011 | 0.0011 |
| F13 | 32900001 | 33000000 | 0.0013 | 0.0009 | 0.0011 |
| F13 | 32920001 | 33020000 | 0.0013 | 0.0009 | 0.0011 |
| F13 | 32940001 | 33040000 | 0.0011 | 0.0007 | 0.0009 |
| F13 | 32960001 | 33060000 | 0.0013 | 0.0007 | 0.0008 |
| F13 | 32980001 | 33080000 | 0.0020 | 0.0012 | 0.0014 |
| F13 | 33000001 | 33100000 | 0.0017 | 0.0011 | 0.0013 |
| F13 | 33020001 | 33120000 | 0.0018 | 0.0011 | 0.0013 |
| F13 | 33040001 | 33140000 | 0.0023 | 0.0016 | 0.0018 |
| F13 | 33060001 | 33160000 | 0.0029 | 0.0022 | 0.0024 |
| F13 | 33080001 | 33180000 | 0.0028 | 0.0022 | 0.0024 |
| F13 | 33100001 | 33200000 | 0.0027 | 0.0021 | 0.0023 |
| F13 | 33120001 | 33220000 | 0.0030 | 0.0023 | 0.0024 |
| F13 | 33140001 | 33240000 | 0.0025 | 0.0018 | 0.0019 |
| F13 | 33160001 | 33260000 | 0.0021 | 0.0013 | 0.0014 |
| F13 | 33180001 | 33280000 | 0.0022 | 0.0012 | 0.0014 |
| F13 | 33200001 | 33300000 | 0.0029 | 0.0017 | 0.0018 |
| F13 | 33220001 | 33320000 | 0.0033 | 0.0018 | 0.0020 |
| F13 | 33240001 | 33340000 | 0.0036 | 0.0020 | 0.0023 |
| F13 | 33260001 | 33360000 | 0.0038 | 0.0022 | 0.0026 |
| F13 | 33280001 | 33380000 | 0.0041 | 0.0024 | 0.0029 |
| F13 | 33300001 | 33400000 | 0.0039 | 0.0023 | 0.0029 |
| F13 | 33320001 | 33420000 | 0.0036 | 0.0023 | 0.0029 |

| Chr | Start | End | both | A1 | A2 |
|---|---|---|---|---|---|
| F13 | 33340001 | 33440000 | 0.0031 | 0.0021 | 0.0026 |
| F13 | 33360001 | 33460000 | 0.0026 | 0.0020 | 0.0023 |
| F13 | 33380001 | 33480000 | 0.0026 | 0.0020 | 0.0024 |
| F13 | 33400001 | 33500000 | 0.0020 | 0.0016 | 0.0019 |
| F13 | 33420001 | 33520000 | 0.0021 | 0.0017 | 0.0020 |
| F13 | 33440001 | 33540000 | 0.0026 | 0.0021 | 0.0024 |
| F13 | 33460001 | 33560000 | 0.0026 | 0.0021 | 0.0025 |
| F13 | 33480001 | 33580000 | 0.0019 | 0.0016 | 0.0018 |
| F13 | 33500001 | 33600000 | 0.0024 | 0.0019 | 0.0023 |
| F13 | 33520001 | 33620000 | 0.0019 | 0.0015 | 0.0019 |
| F13 | 33540001 | 33640000 | 0.0015 | 0.0011 | 0.0015 |
| F13 | 33560001 | 33660000 | 0.0014 | 0.0010 | 0.0013 |
| F13 | 33580001 | 33680000 | 0.0025 | 0.0020 | 0.0024 |
| F13 | 33600001 | 33700000 | 0.0027 | 0.0021 | 0.0025 |
| F13 | 33620001 | 33720000 | 0.0027 | 0.0021 | 0.0026 |
| F13 | 33640001 | 33740000 | 0.0037 | 0.0026 | 0.0032 |
| F13 | 33660001 | 33760000 | 0.0040 | 0.0029 | 0.0036 |
| F13 | 33680001 | 33780000 | 0.0028 | 0.0018 | 0.0024 |
| F13 | 33700001 | 33800000 | 0.0024 | 0.0016 | 0.0021 |
| F13 | 33720001 | 33820000 | 0.0024 | 0.0016 | 0.0020 |
| F13 | 33740001 | 33840000 | 0.0014 | 0.0011 | 0.0013 |
| F13 | 33760001 | 33860000 | 0.0010 | 0.0008 | 0.0009 |
| F13 | 33780001 | 33880000 | 0.0012 | 0.0011 | 0.0012 |
| F13 | 33800001 | 33900000 | 0.0010 | 0.0008 | 0.0009 |
| F13 | 33820001 | 33920000 | 0.0012 | 0.0010 | 0.0011 |
| F13 | 33840001 | 33940000 | 0.0011 | 0.0010 | 0.0010 |
| F13 | 33860001 | 33960000 | 0.0014 | 0.0012 | 0.0014 |
| F13 | 33880001 | 33980000 | 0.0011 | 0.0009 | 0.0011 |
| F13 | 33900001 | 34000000 | 0.0010 | 0.0008 | 0.0010 |
| F13 | 33920001 | 34020000 | 0.0014 | 0.0011 | 0.0013 |
| F13 | 33940001 | 34040000 | 0.0014 | 0.0011 | 0.0013 |
| F13 | 33960001 | 34060000 | 0.0012 | 0.0010 | 0.0011 |
| F13 | 33980001 | 34080000 | 0.0013 | 0.0011 | 0.0012 |
| F13 | 34000001 | 34100000 | 0.0016 | 0.0012 | 0.0013 |
| F13 | 34020001 | 34120000 | 0.0014 | 0.0009 | 0.0011 |
| F13 | 34040001 | 34140000 | 0.0023 | 0.0016 | 0.0019 |
| F13 | 34060001 | 34160000 | 0.0024 | 0.0016 | 0.0019 |
| F13 | 34080001 | 34180000 | 0.0025 | 0.0015 | 0.0019 |
| F13 | 34100001 | 34200000 | 0.0028 | 0.0020 | 0.0023 |
| F13 | 34120001 | 34220000 | 0.0031 | 0.0022 | 0.0026 |
| F13 | 34140001 | 34240000 | 0.0025 | 0.0018 | 0.0022 |
| F13 | 34160001 | 34260000 | 0.0025 | 0.0021 | 0.0024 |
| F13 | 34180001 | 34280000 | 0.0031 | 0.0027 | 0.0030 |
| F13 | 34200001 | 34300000 | 0.0028 | 0.0025 | 0.0028 |
| F13 | 34220001 | 34320000 | 0.0021 | 0.0019 | 0.0021 |
| F13 | 34240001 | 34340000 | 0.0023 | 0.0021 | 0.0023 |
| F13 | 34260001 | 34360000 | 0.0020 | 0.0018 | 0.0019 |
| F13 | 34280001 | 34380000 | 0.0016 | 0.0015 | 0.0016 |
| F13 | 34300001 | 34400000 | 0.0014 | 0.0013 | 0.0014 |
| F13 | 34320001 | 34420000 | 0.0016 | 0.0014 | 0.0015 |
| F13 | 34340001 | 34440000 | 0.0011 | 0.0010 | 0.0011 |
| F13 | 34360001 | 34460000 | 0.0010 | 0.0008 | 0.0010 |
| F13 | 34380001 | 34480000 | 0.0010 | 0.0008 | 0.0009 |
| F13 | 34400001 | 34500000 | 0.0012 | 0.0010 | 0.0011 |
| F13 | 34420001 | 34520000 | 0.0017 | 0.0014 | 0.0014 |
| F13 | 34440001 | 34540000 | 0.0020 | 0.0017 | 0.0017 |
| F13 | 34460001 | 34560000 | 0.0026 | 0.0022 | 0.0021 |
| F13 | 34480001 | 34580000 | 0.0033 | 0.0029 | 0.0028 |

| Chr | Start | End | both | A1 | A2 |
|---|---|---|---|---|---|
| F13 | 34500001 | 34600000 | 0.0038 | 0.0033 | 0.0033 |
| F13 | 34520001 | 34620000 | 0.0032 | 0.0029 | 0.0029 |
| F13 | 34540001 | 34640000 | 0.0033 | 0.0030 | 0.0030 |
| F13 | 34560001 | 34660000 | 0.0029 | 0.0027 | 0.0028 |
| F13 | 34580001 | 34680000 | 0.0027 | 0.0026 | 0.0026 |
| F13 | 34600001 | 34700000 | 0.0026 | 0.0024 | 0.0024 |
| F13 | 34620001 | 34720000 | 0.0032 | 0.0027 | 0.0027 |
| F13 | 34640001 | 34740000 | 0.0033 | 0.0027 | 0.0027 |
| F13 | 34660001 | 34760000 | 0.0041 | 0.0033 | 0.0033 |
| F13 | 34680001 | 34780000 | 0.0035 | 0.0025 | 0.0026 |
| F13 | 34700001 | 34800000 | 0.0030 | 0.0021 | 0.0022 |
| F13 | 34720001 | 34820000 | 0.0025 | 0.0017 | 0.0019 |
| F13 | 34740001 | 34840000 | 0.0023 | 0.0016 | 0.0018 |
| F13 | 34760001 | 34860000 | 0.0016 | 0.0010 | 0.0011 |
| F13 | 34780001 | 34880000 | 0.0020 | 0.0015 | 0.0017 |
| F13 | 34800001 | 34900000 | 0.0021 | 0.0017 | 0.0018 |
| F13 | 34820001 | 34920000 | 0.0022 | 0.0017 | 0.0020 |
| F13 | 34840001 | 34940000 | 0.0023 | 0.0018 | 0.0021 |
| F13 | 34860001 | 34960000 | 0.0024 | 0.0020 | 0.0023 |
| F13 | 34880001 | 34980000 | 0.0018 | 0.0014 | 0.0016 |
| F13 | 34900001 | 35000000 | 0.0018 | 0.0013 | 0.0015 |
| F13 | 34920001 | 35020000 | 0.0021 | 0.0015 | 0.0017 |
| F13 | 34940001 | 35040000 | 0.0016 | 0.0011 | 0.0013 |
| F13 | 34960001 | 35060000 | 0.0020 | 0.0016 | 0.0017 |
| F13 | 34980001 | 35080000 | 0.0020 | 0.0016 | 0.0017 |
| F13 | 35000001 | 35100000 | 0.0020 | 0.0017 | 0.0017 |
| F13 | 35020001 | 35120000 | 0.0015 | 0.0014 | 0.0014 |
| F13 | 35040001 | 35140000 | 0.0014 | 0.0013 | 0.0013 |
| F13 | 35060001 | 35160000 | 0.0009 | 0.0007 | 0.0008 |
| F13 | 35080001 | 35180000 | 0.0008 | 0.0006 | 0.0008 |
| F13 | 35100001 | 35200000 | 0.0008 | 0.0005 | 0.0007 |
| F13 | 35120001 | 35220000 | 0.0009 | 0.0006 | 0.0009 |
| F13 | 35140001 | 35240000 | 0.0026 | 0.0023 | 0.0025 |
| F13 | 35160001 | 35260000 | 0.0035 | 0.0030 | 0.0035 |
| F13 | 35180001 | 35280000 | 0.0038 | 0.0033 | 0.0037 |
| F13 | 35200001 | 35300000 | 0.0040 | 0.0036 | 0.0039 |
| F13 | 35220001 | 35320000 | 0.0040 | 0.0036 | 0.0040 |
| F13 | 35240001 | 35340000 | 0.0026 | 0.0021 | 0.0025 |
| F13 | 35260001 | 35360000 | 0.0028 | 0.0023 | 0.0028 |
| F13 | 35280001 | 35380000 | 0.0035 | 0.0027 | 0.0032 |
| F13 | 35300001 | 35400000 | 0.0034 | 0.0027 | 0.0032 |
| F13 | 35320001 | 35420000 | 0.0036 | 0.0029 | 0.0034 |
| F13 | 35340001 | 35440000 | 0.0038 | 0.0030 | 0.0034 |
| F13 | 35360001 | 35460000 | 0.0028 | 0.0020 | 0.0023 |
| F13 | 35380001 | 35480000 | 0.0026 | 0.0021 | 0.0022 |
| F13 | 35400001 | 35500000 | 0.0029 | 0.0024 | 0.0024 |
| F13 | 35420001 | 35520000 | 0.0027 | 0.0022 | 0.0021 |
| F13 | 35440001 | 35540000 | 0.0030 | 0.0026 | 0.0026 |
| F13 | 35460001 | 35560000 | 0.0029 | 0.0027 | 0.0025 |
| F13 | 35480001 | 35580000 | 0.0028 | 0.0025 | 0.0024 |
| F13 | 35500001 | 35600000 | 0.0025 | 0.0021 | 0.0023 |
| F13 | 35520001 | 35620000 | 0.0027 | 0.0024 | 0.0025 |
| F13 | 35540001 | 35640000 | 0.0024 | 0.0021 | 0.0021 |
| F13 | 35560001 | 35660000 | 0.0022 | 0.0021 | 0.0021 |
| F13 | 35580001 | 35680000 | 0.0017 | 0.0015 | 0.0015 |
| F13 | 35600001 | 35700000 | 0.0014 | 0.0014 | 0.0013 |
| F13 | 35620001 | 35720000 | 0.0011 | 0.0010 | 0.0010 |
| F13 | 35640001 | 35740000 | 0.0007 | 0.0006 | 0.0007 |

| Chr | Start | End | both | A1 | A2 |
|---|---|---|---|---|---|
| F13 | 35660001 | 35760000 | 0.0007 | 0.0006 | 0.0006 |
| F13 | 35680001 | 35780000 | 0.0007 | 0.0006 | 0.0007 |
| F13 | 35700001 | 35800000 | 0.0011 | 0.0009 | 0.0010 |
| F13 | 35720001 | 35820000 | 0.0017 | 0.0014 | 0.0016 |
| F13 | 35740001 | 35840000 | 0.0020 | 0.0015 | 0.0017 |
| F13 | 35760001 | 35860000 | 0.0021 | 0.0015 | 0.0017 |
| F13 | 35780001 | 35880000 | 0.0031 | 0.0023 | 0.0027 |
| F13 | 35800001 | 35900000 | 0.0033 | 0.0024 | 0.0028 |
| F13 | 35820001 | 35920000 | 0.0029 | 0.0020 | 0.0024 |
| F13 | 35840001 | 35940000 | 0.0032 | 0.0024 | 0.0029 |
| F13 | 35860001 | 35960000 | 0.0030 | 0.0023 | 0.0028 |
| F13 | 35880001 | 35980000 | 0.0021 | 0.0016 | 0.0018 |
| F13 | 35900001 | 36000000 | 0.0022 | 0.0016 | 0.0019 |
| F13 | 35920001 | 36020000 | 0.0034 | 0.0028 | 0.0032 |
| F13 | 35940001 | 36040000 | 0.0031 | 0.0025 | 0.0028 |
| F13 | 35960001 | 36060000 | 0.0031 | 0.0025 | 0.0028 |
| F13 | 35980001 | 36080000 | 0.0029 | 0.0025 | 0.0027 |
| F13 | 36000001 | 36100000 | 0.0030 | 0.0026 | 0.0028 |
| F13 | 36020001 | 36120000 | 0.0016 | 0.0012 | 0.0014 |
| F13 | 36040001 | 36140000 | 0.0019 | 0.0017 | 0.0019 |
| F13 | 36060001 | 36160000 | 0.0020 | 0.0018 | 0.0020 |
| F13 | 36080001 | 36180000 | 0.0026 | 0.0023 | 0.0025 |
| F13 | 36100001 | 36200000 | 0.0025 | 0.0021 | 0.0023 |
| F13 | 36120001 | 36220000 | 0.0031 | 0.0025 | 0.0029 |
| F13 | 36140001 | 36240000 | 0.0031 | 0.0026 | 0.0029 |
| F13 | 36160001 | 36260000 | 0.0029 | 0.0024 | 0.0027 |
| F13 | 36180001 | 36280000 | 0.0028 | 0.0020 | 0.0024 |
| F13 | 36200001 | 36300000 | 0.0032 | 0.0025 | 0.0028 |
| F13 | 36220001 | 36320000 | 0.0031 | 0.0024 | 0.0026 |
| F13 | 36240001 | 36340000 | 0.0027 | 0.0019 | 0.0023 |
| F13 | 36260001 | 36360000 | 0.0028 | 0.0019 | 0.0023 |
| F13 | 36280001 | 36380000 | 0.0027 | 0.0020 | 0.0023 |
| F13 | 36300001 | 36400000 | 0.0017 | 0.0011 | 0.0014 |
| F13 | 36320001 | 36420000 | 0.0015 | 0.0008 | 0.0011 |
| F13 | 36340001 | 36440000 | 0.0016 | 0.0010 | 0.0013 |
| F13 | 36360001 | 36460000 | 0.0015 | 0.0011 | 0.0013 |
| F13 | 36380001 | 36480000 | 0.0011 | 0.0008 | 0.0010 |
| F13 | 36400001 | 36500000 | 0.0011 | 0.0008 | 0.0010 |
| F13 | 36420001 | 36520000 | 0.0011 | 0.0009 | 0.0011 |
| F13 | 36440001 | 36540000 | 0.0006 | 0.0005 | 0.0005 |
| F13 | 36460001 | 36560000 | 0.0007 | 0.0006 | 0.0007 |
| F13 | 36480001 | 36580000 | 0.0006 | 0.0005 | 0.0006 |
| F13 | 36500001 | 36600000 | 0.0006 | 0.0005 | 0.0006 |
| F13 | 36520001 | 36620000 | 0.0006 | 0.0005 | 0.0006 |
| F13 | 36540001 | 36640000 | 0.0007 | 0.0006 | 0.0007 |
| F13 | 36560001 | 36660000 | 0.0007 | 0.0006 | 0.0007 |
| F13 | 36580001 | 36680000 | 0.0012 | 0.0010 | 0.0012 |
| F13 | 36600001 | 36700000 | 0.0013 | 0.0011 | 0.0013 |
| F13 | 36620001 | 36720000 | 0.0009 | 0.0007 | 0.0009 |
| F13 | 36640001 | 36740000 | 0.0012 | 0.0009 | 0.0012 |
| F13 | 36660001 | 36760000 | 0.0013 | 0.0010 | 0.0013 |
| F13 | 36680001 | 36780000 | 0.0009 | 0.0007 | 0.0009 |
| F13 | 36700001 | 36800000 | 0.0013 | 0.0009 | 0.0012 |
| F13 | 36720001 | 36820000 | 0.0012 | 0.0009 | 0.0012 |
| F13 | 36740001 | 36840000 | 0.0009 | 0.0007 | 0.0009 |
| F13 | 36760001 | 36860000 | 0.0009 | 0.0007 | 0.0009 |
| F13 | 36780001 | 36880000 | 0.0009 | 0.0006 | 0.0009 |
| F13 | 36800001 | 36900000 | 0.0009 | 0.0007 | 0.0009 |

| Chr | Start | End | both | A1 | A2 |
|---|---|---|---|---|---|
| F13 | 36820001 | 36920000 | 0.0014 | 0.0010 | 0.0014 |
| F13 | 36840001 | 36940000 | 0.0017 | 0.0013 | 0.0017 |
| F13 | 36860001 | 36960000 | 0.0018 | 0.0014 | 0.0018 |
| F13 | 36880001 | 36980000 | 0.0020 | 0.0016 | 0.0020 |
| F13 | 36900001 | 37000000 | 0.0021 | 0.0018 | 0.0021 |
| F13 | 36920001 | 37020000 | 0.0023 | 0.0018 | 0.0022 |
| F13 | 36940001 | 37040000 | 0.0022 | 0.0017 | 0.0021 |
| F13 | 36960001 | 37060000 | 0.0025 | 0.0021 | 0.0024 |
| F13 | 36980001 | 37080000 | 0.0032 | 0.0027 | 0.0030 |
| F13 | 37000001 | 37100000 | 0.0028 | 0.0023 | 0.0026 |
| F13 | 37020001 | 37120000 | 0.0035 | 0.0032 | 0.0032 |
| F13 | 37040001 | 37140000 | 0.0033 | 0.0031 | 0.0030 |
| F13 | 37060001 | 37160000 | 0.0029 | 0.0026 | 0.0027 |
| F13 | 37080001 | 37180000 | 0.0019 | 0.0017 | 0.0018 |
| F13 | 37100001 | 37200000 | 0.0023 | 0.0018 | 0.0022 |
| F13 | 37120001 | 37220000 | 0.0015 | 0.0007 | 0.0015 |
| F13 | 37140001 | 37240000 | 0.0016 | 0.0008 | 0.0015 |
| F13 | 37160001 | 37260000 | 0.0016 | 0.0007 | 0.0015 |
| F13 | 37180001 | 37280000 | 0.0020 | 0.0010 | 0.0019 |
| F13 | 37200001 | 37300000 | 0.0017 | 0.0009 | 0.0017 |
| F13 | 37220001 | 37320000 | 0.0013 | 0.0008 | 0.0013 |
| F13 | 37240001 | 37340000 | 0.0012 | 0.0008 | 0.0011 |
| F13 | 37260001 | 37360000 | 0.0014 | 0.0011 | 0.0014 |
| F13 | 37280001 | 37380000 | 0.0015 | 0.0012 | 0.0014 |
| F13 | 37300001 | 37400000 | 0.0014 | 0.0012 | 0.0014 |
| F13 | 37320001 | 37420000 | 0.0017 | 0.0015 | 0.0017 |
| F13 | 37340001 | 37440000 | 0.0017 | 0.0014 | 0.0016 |
| F13 | 37360001 | 37460000 | 0.0013 | 0.0011 | 0.0013 |
| F13 | 37380001 | 37480000 | 0.0011 | 0.0008 | 0.0009 |
| F13 | 37400001 | 37500000 | 0.0009 | 0.0007 | 0.0008 |
| F13 | 37420001 | 37520000 | 0.0007 | 0.0005 | 0.0005 |
| F13 | 37440001 | 37540000 | 0.0014 | 0.0011 | 0.0012 |
| F13 | 37460001 | 37560000 | 0.0014 | 0.0011 | 0.0013 |
| F13 | 37480001 | 37580000 | 0.0019 | 0.0014 | 0.0017 |
| F13 | 37500001 | 37600000 | 0.0024 | 0.0018 | 0.0022 |
| F13 | 37520001 | 37620000 | 0.0026 | 0.0019 | 0.0024 |
| F13 | 37540001 | 37640000 | 0.0022 | 0.0017 | 0.0019 |
| F13 | 37560001 | 37660000 | 0.0025 | 0.0019 | 0.0022 |
| F13 | 37580001 | 37680000 | 0.0024 | 0.0020 | 0.0021 |
| F13 | 37600001 | 37700000 | 0.0022 | 0.0017 | 0.0017 |
| F13 | 37620001 | 37720000 | 0.0021 | 0.0015 | 0.0016 |
| F13 | 37640001 | 37740000 | 0.0021 | 0.0015 | 0.0017 |
| F13 | 37660001 | 37760000 | 0.0019 | 0.0014 | 0.0016 |
| F13 | 37680001 | 37780000 | 0.0016 | 0.0012 | 0.0014 |
| F13 | 37700001 | 37800000 | 0.0017 | 0.0014 | 0.0016 |
| F13 | 37720001 | 37820000 | 0.0021 | 0.0018 | 0.0020 |
| F13 | 37740001 | 37840000 | 0.0020 | 0.0016 | 0.0017 |
| F13 | 37760001 | 37860000 | 0.0019 | 0.0014 | 0.0015 |
| F13 | 37780001 | 37880000 | 0.0029 | 0.0022 | 0.0023 |
| F13 | 37800001 | 37900000 | 0.0026 | 0.0018 | 0.0019 |
| F13 | 37820001 | 37920000 | 0.0026 | 0.0017 | 0.0017 |
| F13 | 37840001 | 37940000 | 0.0027 | 0.0018 | 0.0019 |
| F13 | 37860001 | 37960000 | 0.0033 | 0.0023 | 0.0026 |
| F13 | 37880001 | 37980000 | 0.0022 | 0.0013 | 0.0017 |
| F13 | 37900001 | 38000000 | 0.0025 | 0.0017 | 0.0020 |
| F13 | 37920001 | 38020000 | 0.0027 | 0.0018 | 0.0022 |
| F13 | 37940001 | 38040000 | 0.0031 | 0.0022 | 0.0026 |
| F13 | 37960001 | 38060000 | 0.0024 | 0.0017 | 0.0020 |

| Chr | Start | End | both | A1 | A2 |
|---|---|---|---|---|---|
| F13 | 37980001 | 38080000 | 0.0029 | 0.0020 | 0.0023 |
| F13 | 38000001 | 38100000 | 0.0027 | 0.0018 | 0.0023 |
| F13 | 38020001 | 38120000 | 0.0023 | 0.0017 | 0.0021 |
| F13 | 38040001 | 38140000 | 0.0022 | 0.0015 | 0.0019 |
| F13 | 38060001 | 38160000 | 0.0027 | 0.0019 | 0.0023 |
| F13 | 38080001 | 38180000 | 0.0024 | 0.0016 | 0.0021 |
| F13 | 38100001 | 38200000 | 0.0025 | 0.0018 | 0.0020 |
| F13 | 38120001 | 38220000 | 0.0024 | 0.0018 | 0.0019 |
| F13 | 38140001 | 38240000 | 0.0020 | 0.0015 | 0.0017 |
| F13 | 38160001 | 38260000 | 0.0033 | 0.0027 | 0.0028 |
| F13 | 38180001 | 38280000 | 0.0035 | 0.0028 | 0.0029 |
| F13 | 38200001 | 38300000 | 0.0038 | 0.0029 | 0.0032 |
| F13 | 38220001 | 38320000 | 0.0039 | 0.0031 | 0.0034 |
| F13 | 38240001 | 38340000 | 0.0049 | 0.0037 | 0.0040 |
| F13 | 38260001 | 38360000 | 0.0031 | 0.0021 | 0.0023 |
| F13 | 38280001 | 38380000 | 0.0025 | 0.0019 | 0.0019 |
| F13 | 38300001 | 38400000 | 0.0020 | 0.0014 | 0.0015 |
| F13 | 38320001 | 38420000 | 0.0019 | 0.0013 | 0.0014 |
| F13 | 38340001 | 38440000 | 0.0009 | 0.0006 | 0.0007 |
| F13 | 38360001 | 38460000 | 0.0012 | 0.0008 | 0.0010 |
| F13 | 38380001 | 38480000 | 0.0015 | 0.0011 | 0.0013 |
| F13 | 38400001 | 38500000 | 0.0016 | 0.0013 | 0.0014 |
| F13 | 38420001 | 38520000 | 0.0014 | 0.0011 | 0.0011 |
| F13 | 38440001 | 38540000 | 0.0018 | 0.0012 | 0.0015 |
| F13 | 38460001 | 38560000 | 0.0032 | 0.0026 | 0.0028 |
| F13 | 38480001 | 38580000 | 0.0035 | 0.0028 | 0.0031 |
| F13 | 38500001 | 38600000 | 0.0034 | 0.0028 | 0.0031 |
| F13 | 38520001 | 38620000 | 0.0031 | 0.0026 | 0.0028 |
| F13 | 38540001 | 38640000 | 0.0035 | 0.0032 | 0.0033 |
| F13 | 38560001 | 38660000 | 0.0024 | 0.0022 | 0.0022 |
| F13 | 38580001 | 38680000 | 0.0023 | 0.0021 | 0.0022 |
| F13 | 38600001 | 38700000 | 0.0025 | 0.0022 | 0.0024 |
| F13 | 38620001 | 38720000 | 0.0030 | 0.0028 | 0.0030 |
| F13 | 38640001 | 38740000 | 0.0029 | 0.0024 | 0.0027 |
| F13 | 38660001 | 38760000 | 0.0029 | 0.0020 | 0.0025 |
| F13 | 38680001 | 38780000 | 0.0037 | 0.0027 | 0.0033 |
| F13 | 38700001 | 38800000 | 0.0039 | 0.0028 | 0.0035 |
| F13 | 38720001 | 38820000 | 0.0035 | 0.0025 | 0.0031 |
| F13 | 38740001 | 38840000 | 0.0036 | 0.0027 | 0.0034 |
| F13 | 38760001 | 38860000 | 0.0032 | 0.0027 | 0.0032 |
| F13 | 38780001 | 38880000 | 0.0026 | 0.0022 | 0.0025 |
| F13 | 38800001 | 38900000 | 0.0024 | 0.0020 | 0.0022 |
| F13 | 38820001 | 38920000 | 0.0023 | 0.0019 | 0.0022 |
| F13 | 38840001 | 38940000 | 0.0020 | 0.0017 | 0.0017 |
| F13 | 38860001 | 38960000 | 0.0020 | 0.0017 | 0.0016 |
| F13 | 38880001 | 38980000 | 0.0014 | 0.0012 | 0.0011 |
| F13 | 38900001 | 39000000 | 0.0015 | 0.0012 | 0.0011 |
| F13 | 38920001 | 39020000 | 0.0014 | 0.0011 | 0.0011 |
| F13 | 38940001 | 39040000 | 0.0023 | 0.0019 | 0.0020 |
| F13 | 38960001 | 39060000 | 0.0027 | 0.0024 | 0.0025 |
| F13 | 38980001 | 39080000 | 0.0028 | 0.0025 | 0.0026 |
| F13 | 39000001 | 39100000 | 0.0025 | 0.0024 | 0.0025 |
| F13 | 39020001 | 39120000 | 0.0025 | 0.0024 | 0.0024 |
| F13 | 39040001 | 39140000 | 0.0023 | 0.0022 | 0.0022 |
| F13 | 39060001 | 39160000 | 0.0018 | 0.0016 | 0.0017 |
| F13 | 39080001 | 39180000 | 0.0018 | 0.0016 | 0.0018 |
| F13 | 39100001 | 39200000 | 0.0023 | 0.0019 | 0.0022 |
| F13 | 39120001 | 39220000 | 0.0029 | 0.0023 | 0.0028 |

| Chr | Start | End | both | A1 | A2 |
|---|---|---|---|---|---|
| F13 | 39140001 | 39240000 | 0.0019 | 0.0014 | 0.0019 |
| F13 | 39160001 | 39260000 | 0.0019 | 0.0014 | 0.0018 |
| F13 | 39180001 | 39280000 | 0.0022 | 0.0017 | 0.0021 |
| F13 | 39200001 | 39300000 | 0.0021 | 0.0016 | 0.0020 |
| F13 | 39220001 | 39320000 | 0.0015 | 0.0011 | 0.0014 |
| F13 | 39240001 | 39340000 | 0.0016 | 0.0012 | 0.0014 |
| F13 | 39260001 | 39360000 | 0.0016 | 0.0013 | 0.0015 |
| F13 | 39280001 | 39380000 | 0.0013 | 0.0010 | 0.0012 |
| F13 | 39300001 | 39400000 | 0.0011 | 0.0009 | 0.0010 |
| F13 | 39320001 | 39420000 | 0.0010 | 0.0008 | 0.0010 |
| F13 | 39340001 | 39440000 | 0.0011 | 0.0010 | 0.0011 |
| F13 | 39360001 | 39460000 | 0.0010 | 0.0008 | 0.0010 |
| F13 | 39380001 | 39480000 | 0.0012 | 0.0011 | 0.0012 |
| F13 | 39400001 | 39500000 | 0.0011 | 0.0009 | 0.0011 |
| F13 | 39420001 | 39520000 | 0.0010 | 0.0009 | 0.0010 |
| F13 | 39440001 | 39540000 | 0.0006 | 0.0005 | 0.0006 |
| F13 | 39460001 | 39560000 | 0.0007 | 0.0007 | 0.0007 |
| F13 | 39480001 | 39580000 | 0.0007 | 0.0007 | 0.0007 |
| F13 | 39500001 | 39600000 | 0.0007 | 0.0007 | 0.0007 |
| F13 | 39520001 | 39620000 | 0.0007 | 0.0007 | 0.0007 |
| F13 | 39540001 | 39640000 | 0.0008 | 0.0008 | 0.0008 |
| F13 | 39560001 | 39660000 | 0.0011 | 0.0010 | 0.0011 |
| F13 | 39580001 | 39680000 | 0.0010 | 0.0009 | 0.0010 |
| F13 | 39600001 | 39700000 | 0.0011 | 0.0010 | 0.0011 |
| F13 | 39620001 | 39720000 | 0.0013 | 0.0012 | 0.0012 |
| F13 | 39640001 | 39740000 | 0.0012 | 0.0011 | 0.0012 |
| F13 | 39660001 | 39760000 | 0.0006 | 0.0005 | 0.0006 |
| F13 | 39680001 | 39780000 | 0.0005 | 0.0004 | 0.0004 |
| F13 | 39700001 | 39800000 | 0.0012 | 0.0009 | 0.0012 |
| F13 | 39720001 | 39820000 | 0.0012 | 0.0008 | 0.0012 |
| F13 | 39740001 | 39840000 | 0.0016 | 0.0012 | 0.0016 |
| F13 | 39760001 | 39860000 | 0.0017 | 0.0013 | 0.0017 |
| F13 | 39780001 | 39880000 | 0.0019 | 0.0015 | 0.0019 |
| F13 | 39800001 | 39900000 | 0.0014 | 0.0012 | 0.0014 |
| F13 | 39820001 | 39920000 | 0.0023 | 0.0017 | 0.0021 |
| F13 | 39840001 | 39940000 | 0.0024 | 0.0016 | 0.0020 |
| F13 | 39860001 | 39960000 | 0.0031 | 0.0022 | 0.0026 |
| F13 | 39880001 | 39980000 | 0.0035 | 0.0023 | 0.0027 |
| F13 | 39900001 | 40000000 | 0.0033 | 0.0021 | 0.0025 |
| F13 | 39920001 | 40020000 | 0.0027 | 0.0019 | 0.0021 |
| F13 | 39940001 | 40040000 | 0.0025 | 0.0017 | 0.0019 |
| F13 | 39960001 | 40060000 | 0.0022 | 0.0014 | 0.0016 |
| F13 | 39980001 | 40080000 | 0.0016 | 0.0012 | 0.0013 |
| F13 | 40000001 | 40100000 | 0.0025 | 0.0020 | 0.0021 |
| F13 | 40020001 | 40120000 | 0.0036 | 0.0030 | 0.0030 |
| F13 | 40040001 | 40140000 | 0.0037 | 0.0031 | 0.0032 |
| F13 | 40060001 | 40160000 | 0.0046 | 0.0040 | 0.0042 |
| F13 | 40080001 | 40180000 | 0.0046 | 0.0041 | 0.0042 |
| F13 | 40100001 | 40200000 | 0.0037 | 0.0032 | 0.0034 |
| F13 | 40120001 | 40220000 | 0.0028 | 0.0024 | 0.0026 |
| F13 | 40140001 | 40240000 | 0.0032 | 0.0028 | 0.0029 |
| F13 | 40160001 | 40260000 | 0.0024 | 0.0021 | 0.0019 |
| F13 | 40180001 | 40280000 | 0.0028 | 0.0025 | 0.0023 |
| F13 | 40200001 | 40300000 | 0.0036 | 0.0033 | 0.0030 |
| F13 | 40220001 | 40320000 | 0.0034 | 0.0030 | 0.0029 |
| F13 | 40240001 | 40340000 | 0.0040 | 0.0035 | 0.0035 |
| F13 | 40260001 | 40360000 | 0.0045 | 0.0039 | 0.0040 |
| F13 | 40280001 | 40380000 | 0.0042 | 0.0034 | 0.0035 |

| Chr | Start | End | both | A1 | A2 |
|---|---|---|---|---|---|
| F13 | 40300001 | 40400000 | 0.0043 | 0.0031 | 0.0034 |
| F13 | 40320001 | 40420000 | 0.0038 | 0.0028 | 0.0030 |
| F13 | 40340001 | 40440000 | 0.0031 | 0.0022 | 0.0023 |
| F13 | 40360001 | 40460000 | 0.0027 | 0.0020 | 0.0019 |
| F13 | 40380001 | 40480000 | 0.0034 | 0.0028 | 0.0027 |
| F13 | 40400001 | 40500000 | 0.0026 | 0.0025 | 0.0022 |
| F13 | 40420001 | 40520000 | 0.0031 | 0.0028 | 0.0026 |
| F13 | 40440001 | 40540000 | 0.0031 | 0.0028 | 0.0026 |
| F13 | 40460001 | 40560000 | 0.0027 | 0.0024 | 0.0023 |
| F13 | 40480001 | 40580000 | 0.0020 | 0.0017 | 0.0017 |
| F13 | 40500001 | 40600000 | 0.0019 | 0.0015 | 0.0017 |
| F13 | 40520001 | 40620000 | 0.0016 | 0.0013 | 0.0014 |
| F13 | 40540001 | 40640000 | 0.0015 | 0.0012 | 0.0013 |
| F13 | 40560001 | 40660000 | 0.0016 | 0.0012 | 0.0013 |
| F13 | 40580001 | 40680000 | 0.0021 | 0.0016 | 0.0015 |
| F13 | 40600001 | 40700000 | 0.0028 | 0.0022 | 0.0021 |
| F13 | 40620001 | 40720000 | 0.0030 | 0.0025 | 0.0024 |
| F13 | 40640001 | 40740000 | 0.0028 | 0.0023 | 0.0022 |
| F13 | 40660001 | 40760000 | 0.0031 | 0.0025 | 0.0026 |
| F13 | 40680001 | 40780000 | 0.0028 | 0.0023 | 0.0024 |
| F13 | 40700001 | 40800000 | 0.0025 | 0.0020 | 0.0023 |
| F13 | 40720001 | 40820000 | 0.0024 | 0.0016 | 0.0019 |
| F13 | 40740001 | 40840000 | 0.0024 | 0.0016 | 0.0019 |
| F13 | 40760001 | 40860000 | 0.0022 | 0.0013 | 0.0017 |
| F13 | 40780001 | 40880000 | 0.0026 | 0.0016 | 0.0020 |
| F13 | 40800001 | 40900000 | 0.0021 | 0.0013 | 0.0015 |
| F13 | 40820001 | 40920000 | 0.0029 | 0.0021 | 0.0023 |
| F13 | 40840001 | 40940000 | 0.0033 | 0.0023 | 0.0026 |
| F13 | 40860001 | 40960000 | 0.0031 | 0.0022 | 0.0024 |
| F13 | 40880001 | 40980000 | 0.0028 | 0.0018 | 0.0020 |
| F13 | 40900001 | 41000000 | 0.0030 | 0.0021 | 0.0021 |
| F13 | 40920001 | 41020000 | 0.0030 | 0.0021 | 0.0023 |
| F13 | 40940001 | 41040000 | 0.0027 | 0.0018 | 0.0020 |
| F13 | 40960001 | 41060000 | 0.0030 | 0.0021 | 0.0021 |
| F13 | 40980001 | 41080000 | 0.0031 | 0.0021 | 0.0023 |
| F13 | 41000001 | 41100000 | 0.0030 | 0.0019 | 0.0021 |
| F13 | 41020001 | 41120000 | 0.0024 | 0.0015 | 0.0015 |
| F13 | 41040001 | 41140000 | 0.0029 | 0.0021 | 0.0021 |
| F13 | 41060001 | 41160000 | 0.0032 | 0.0023 | 0.0024 |
| F13 | 41080001 | 41180000 | 0.0033 | 0.0025 | 0.0026 |
| F13 | 41100001 | 41200000 | 0.0035 | 0.0025 | 0.0026 |
| F13 | 41120001 | 41220000 | 0.0033 | 0.0024 | 0.0024 |
| F13 | 41140001 | 41240000 | 0.0026 | 0.0016 | 0.0016 |
| F13 | 41160001 | 41260000 | 0.0018 | 0.0010 | 0.0010 |
| F13 | 41180001 | 41280000 | 0.0019 | 0.0008 | 0.0010 |
| F13 | 41200001 | 41300000 | 0.0018 | 0.0009 | 0.0010 |
| F13 | 41220001 | 41320000 | 0.0019 | 0.0010 | 0.0013 |
| F13 | 41240001 | 41340000 | 0.0019 | 0.0011 | 0.0013 |
| F13 | 41260001 | 41360000 | 0.0027 | 0.0016 | 0.0019 |
| F13 | 41280001 | 41380000 | 0.0027 | 0.0019 | 0.0021 |
| F13 | 41300001 | 41400000 | 0.0037 | 0.0027 | 0.0030 |
| F13 | 41320001 | 41420000 | 0.0036 | 0.0023 | 0.0026 |
| F13 | 41340001 | 41440000 | 0.0044 | 0.0028 | 0.0032 |
| F13 | 41360001 | 41460000 | 0.0045 | 0.0030 | 0.0033 |
| F13 | 41380001 | 41480000 | 0.0043 | 0.0028 | 0.0032 |
| F13 | 41400001 | 41500000 | 0.0039 | 0.0025 | 0.0029 |
| F13 | 41420001 | 41520000 | 0.0045 | 0.0033 | 0.0037 |
| F13 | 41440001 | 41540000 | 0.0037 | 0.0028 | 0.0031 |

| Chr | Start | End | both | A1 | A2 |
|---|---|---|---|---|---|
| F13 | 41460001 | 41560000 | 0.0031 | 0.0022 | 0.0026 |
| F13 | 41480001 | 41580000 | 0.0026 | 0.0020 | 0.0021 |
| F13 | 41500001 | 41600000 | 0.0023 | 0.0018 | 0.0018 |
| F13 | 41520001 | 41620000 | 0.0015 | 0.0011 | 0.0011 |
| F13 | 41540001 | 41640000 | 0.0017 | 0.0014 | 0.0013 |
| F13 | 41560001 | 41660000 | 0.0021 | 0.0017 | 0.0015 |
| F13 | 41580001 | 41680000 | 0.0022 | 0.0017 | 0.0016 |
| F13 | 41600001 | 41700000 | 0.0017 | 0.0013 | 0.0013 |
| F13 | 41620001 | 41720000 | 0.0027 | 0.0021 | 0.0022 |
| F13 | 41640001 | 41740000 | 0.0030 | 0.0022 | 0.0023 |
| F13 | 41660001 | 41760000 | 0.0025 | 0.0018 | 0.0021 |
| F13 | 41680001 | 41780000 | 0.0025 | 0.0017 | 0.0020 |
| F13 | 41700001 | 41800000 | 0.0025 | 0.0018 | 0.0021 |
| F13 | 41720001 | 41820000 | 0.0022 | 0.0016 | 0.0018 |
| F13 | 41740001 | 41840000 | 0.0015 | 0.0012 | 0.0014 |
| F13 | 41760001 | 41860000 | 0.0015 | 0.0013 | 0.0014 |
| F13 | 41780001 | 41880000 | 0.0016 | 0.0013 | 0.0014 |
| F13 | 41800001 | 41900000 | 0.0028 | 0.0023 | 0.0024 |
| F13 | 41820001 | 41920000 | 0.0028 | 0.0022 | 0.0023 |
| F13 | 41840001 | 41940000 | 0.0038 | 0.0031 | 0.0032 |
| F13 | 41860001 | 41960000 | 0.0041 | 0.0034 | 0.0036 |
| F13 | 41880001 | 41980000 | 0.0042 | 0.0036 | 0.0037 |
| F13 | 41900001 | 42000000 | 0.0028 | 0.0023 | 0.0025 |
| F13 | 41920001 | 42020000 | 0.0026 | 0.0022 | 0.0024 |
| F13 | 41940001 | 42040000 | 0.0019 | 0.0016 | 0.0018 |
| F13 | 41960001 | 42060000 | 0.0014 | 0.0012 | 0.0014 |
| F13 | 41980001 | 42080000 | 0.0013 | 0.0011 | 0.0013 |
| F13 | 42000001 | 42100000 | 0.0019 | 0.0016 | 0.0019 |
| F13 | 42020001 | 42120000 | 0.0014 | 0.0012 | 0.0014 |
| F13 | 42040001 | 42140000 | 0.0014 | 0.0012 | 0.0014 |
| F13 | 42060001 | 42160000 | 0.0013 | 0.0012 | 0.0013 |
| F13 | 42080001 | 42180000 | 0.0015 | 0.0013 | 0.0015 |
| F13 | 42100001 | 42200000 | 0.0014 | 0.0012 | 0.0013 |
| F13 | 42120001 | 42220000 | 0.0015 | 0.0012 | 0.0013 |
| F13 | 42140001 | 42240000 | 0.0014 | 0.0010 | 0.0011 |
| F13 | 42160001 | 42260000 | 0.0016 | 0.0012 | 0.0013 |
| F13 | 42180001 | 42280000 | 0.0018 | 0.0013 | 0.0015 |
| F13 | 42200001 | 42300000 | 0.0017 | 0.0014 | 0.0014 |
| F13 | 42220001 | 42320000 | 0.0019 | 0.0018 | 0.0018 |
| F13 | 42240001 | 42340000 | 0.0022 | 0.0019 | 0.0020 |
| F13 | 42260001 | 42360000 | 0.0027 | 0.0022 | 0.0025 |
| F13 | 42280001 | 42380000 | 0.0029 | 0.0024 | 0.0026 |
| F13 | 42300001 | 42400000 | 0.0028 | 0.0022 | 0.0026 |
| F13 | 42320001 | 42420000 | 0.0023 | 0.0018 | 0.0022 |
| F13 | 42340001 | 42440000 | 0.0019 | 0.0015 | 0.0018 |
| F13 | 42360001 | 42460000 | 0.0011 | 0.0009 | 0.0011 |
| F13 | 42380001 | 42480000 | 0.0004 | 0.0003 | 0.0004 |
| F13 | 42400001 | 42500000 | 0.0008 | 0.0006 | 0.0008 |
| F13 | 42420001 | 42520000 | 0.0009 | 0.0007 | 0.0010 |
| F13 | 42440001 | 42540000 | 0.0011 | 0.0008 | 0.0011 |
| F13 | 42460001 | 42560000 | 0.0013 | 0.0011 | 0.0013 |
| F13 | 42480001 | 42580000 | 0.0014 | 0.0011 | 0.0014 |
| F13 | 42500001 | 42600000 | 0.0010 | 0.0009 | 0.0010 |
| F13 | 42520001 | 42620000 | 0.0010 | 0.0010 | 0.0010 |
| F13 | 42540001 | 42640000 | 0.0011 | 0.0011 | 0.0010 |
| F13 | 42560001 | 42660000 | 0.0010 | 0.0010 | 0.0010 |
| F13 | 42580001 | 42680000 | 0.0018 | 0.0014 | 0.0017 |
| F13 | 42600001 | 42700000 | 0.0017 | 0.0013 | 0.0016 |

| Chr | Start | End | both | A1 | A2 |
|---|---|---|---|---|---|
| F13 | 42620001 | 42720000 | 0.0016 | 0.0012 | 0.0015 |
| F13 | 42640001 | 42740000 | 0.0018 | 0.0014 | 0.0018 |
| F13 | 42660001 | 42760000 | 0.0018 | 0.0014 | 0.0017 |
| F13 | 42680001 | 42780000 | 0.0015 | 0.0014 | 0.0015 |
| F13 | 42700001 | 42800000 | 0.0014 | 0.0013 | 0.0014 |
| F13 | 42720001 | 42820000 | 0.0017 | 0.0015 | 0.0016 |
| F13 | 42740001 | 42840000 | 0.0013 | 0.0012 | 0.0013 |
| F13 | 42760001 | 42860000 | 0.0013 | 0.0012 | 0.0013 |
| F13 | 42780001 | 42880000 | 0.0007 | 0.0007 | 0.0007 |
| F13 | 42800001 | 42900000 | 0.0006 | 0.0006 | 0.0006 |
| F13 | 42820001 | 42920000 | 0.0005 | 0.0004 | 0.0005 |
| F13 | 42840001 | 42940000 | 0.0009 | 0.0007 | 0.0009 |
| F13 | 42860001 | 42960000 | 0.0011 | 0.0008 | 0.0011 |
| F13 | 42880001 | 42980000 | 0.0012 | 0.0009 | 0.0012 |
| F13 | 42900001 | 43000000 | 0.0013 | 0.0010 | 0.0013 |
| F13 | 42920001 | 43020000 | 0.0020 | 0.0016 | 0.0019 |
| F13 | 42940001 | 43040000 | 0.0017 | 0.0014 | 0.0016 |
| F13 | 42960001 | 43060000 | 0.0014 | 0.0011 | 0.0014 |
| F13 | 42980001 | 43080000 | 0.0017 | 0.0013 | 0.0016 |
| F13 | 43000001 | 43100000 | 0.0018 | 0.0014 | 0.0016 |
| F13 | 43020001 | 43120000 | 0.0016 | 0.0011 | 0.0014 |
| F13 | 43040001 | 43140000 | 0.0015 | 0.0011 | 0.0014 |
| F13 | 43060001 | 43160000 | 0.0016 | 0.0011 | 0.0014 |
| F13 | 43080001 | 43180000 | 0.0014 | 0.0010 | 0.0013 |
| F13 | 43100001 | 43200000 | 0.0014 | 0.0011 | 0.0013 |
| F13 | 43120001 | 43220000 | 0.0008 | 0.0006 | 0.0007 |
| F13 | 43140001 | 43240000 | 0.0009 | 0.0007 | 0.0008 |
| F13 | 43160001 | 43260000 | 0.0007 | 0.0006 | 0.0007 |
| F13 | 43180001 | 43280000 | 0.0011 | 0.0008 | 0.0011 |
| F13 | 43200001 | 43300000 | 0.0017 | 0.0012 | 0.0017 |
| F13 | 43220001 | 43320000 | 0.0019 | 0.0015 | 0.0019 |
| F13 | 43240001 | 43340000 | 0.0017 | 0.0014 | 0.0017 |
| F13 | 43260001 | 43360000 | 0.0017 | 0.0013 | 0.0017 |
| F13 | 43280001 | 43380000 | 0.0014 | 0.0011 | 0.0014 |
| F13 | 43300001 | 43400000 | 0.0007 | 0.0006 | 0.0007 |
| F13 | 43320001 | 43420000 | 0.0004 | 0.0004 | 0.0004 |
| F13 | 43340001 | 43440000 | 0.0005 | 0.0004 | 0.0005 |
| F13 | 43360001 | 43460000 | 0.0006 | 0.0005 | 0.0006 |
| F13 | 43380001 | 43480000 | 0.0006 | 0.0006 | 0.0006 |
| F13 | 43400001 | 43500000 | 0.0008 | 0.0007 | 0.0008 |
| F13 | 43420001 | 43520000 | 0.0008 | 0.0007 | 0.0008 |
| F13 | 43440001 | 43540000 | 0.0010 | 0.0008 | 0.0010 |
| F13 | 43460001 | 43560000 | 0.0012 | 0.0010 | 0.0012 |
| F13 | 43480001 | 43580000 | 0.0017 | 0.0015 | 0.0017 |
| F13 | 43500001 | 43600000 | 0.0020 | 0.0018 | 0.0020 |
| F13 | 43520001 | 43620000 | 0.0026 | 0.0023 | 0.0026 |
| F13 | 43540001 | 43640000 | 0.0023 | 0.0020 | 0.0022 |
| F13 | 43560001 | 43660000 | 0.0022 | 0.0019 | 0.0021 |
| F13 | 43580001 | 43680000 | 0.0018 | 0.0015 | 0.0018 |
| F13 | 43600001 | 43700000 | 0.0016 | 0.0013 | 0.0015 |
| F13 | 43620001 | 43720000 | 0.0012 | 0.0010 | 0.0012 |
| F13 | 43640001 | 43740000 | 0.0016 | 0.0014 | 0.0016 |
| F13 | 43660001 | 43760000 | 0.0017 | 0.0015 | 0.0017 |
| F13 | 43680001 | 43780000 | 0.0017 | 0.0016 | 0.0017 |
| F13 | 43700001 | 43800000 | 0.0013 | 0.0012 | 0.0013 |
| F13 | 43720001 | 43820000 | 0.0011 | 0.0010 | 0.0011 |
| F13 | 43740001 | 43840000 | 0.0008 | 0.0007 | 0.0007 |
| F13 | 43760001 | 43860000 | 0.0008 | 0.0007 | 0.0008 |

| Chr | Start | End | both | A1 | A2 |
|---|---|---|---|---|---|
| F13 | 43780001 | 43880000 | 0.0009 | 0.0008 | 0.0009 |
| F13 | 43800001 | 43900000 | 0.0010 | 0.0008 | 0.0010 |
| F13 | 43820001 | 43920000 | 0.0010 | 0.0009 | 0.0009 |
| F13 | 43840001 | 43940000 | 0.0012 | 0.0010 | 0.0011 |
| F13 | 43860001 | 43960000 | 0.0009 | 0.0008 | 0.0009 |
| F13 | 43880001 | 43980000 | 0.0006 | 0.0006 | 0.0006 |
| F13 | 43900001 | 44000000 | 0.0006 | 0.0006 | 0.0006 |
| F13 | 43920001 | 44020000 | 0.0006 | 0.0006 | 0.0006 |
| F13 | 43940001 | 44040000 | 0.0004 | 0.0004 | 0.0004 |
| F13 | 43960001 | 44060000 | 0.0004 | 0.0004 | 0.0004 |
| F13 | 43980001 | 44080000 | 0.0004 | 0.0003 | 0.0004 |
| F13 | 44000001 | 44100000 | 0.0006 | 0.0005 | 0.0006 |
| F13 | 44020001 | 44120000 | 0.0008 | 0.0006 | 0.0008 |
| F13 | 44040001 | 44140000 | 0.0008 | 0.0007 | 0.0008 |
| F13 | 44060001 | 44160000 | 0.0011 | 0.0010 | 0.0011 |
| F13 | 44080001 | 44180000 | 0.0009 | 0.0008 | 0.0009 |
| F13 | 44100001 | 44200000 | 0.0010 | 0.0009 | 0.0010 |
| F13 | 44120001 | 44220000 | 0.0012 | 0.0011 | 0.0012 |
| F13 | 44140001 | 44240000 | 0.0012 | 0.0011 | 0.0012 |
| F13 | 44160001 | 44260000 | 0.0009 | 0.0008 | 0.0008 |
| F13 | 44180001 | 44280000 | 0.0009 | 0.0008 | 0.0008 |
| F13 | 44200001 | 44300000 | 0.0006 | 0.0006 | 0.0006 |
| F13 | 44220001 | 44320000 | 0.0003 | 0.0002 | 0.0002 |
| F13 | 44240001 | 44340000 | 0.0003 | 0.0002 | 0.0003 |
| F13 | 44260001 | 44360000 | 0.0006 | 0.0005 | 0.0006 |
| F13 | 44280001 | 44380000 | 0.0007 | 0.0006 | 0.0007 |
| F13 | 44300001 | 44400000 | 0.0008 | 0.0007 | 0.0007 |
| F13 | 44320001 | 44420000 | 0.0009 | 0.0009 | 0.0009 |
| F13 | 44340001 | 44440000 | 0.0010 | 0.0010 | 0.0009 |
| F13 | 44360001 | 44460000 | 0.0009 | 0.0008 | 0.0008 |
| F13 | 44380001 | 44480000 | 0.0012 | 0.0010 | 0.0010 |
| F13 | 44400001 | 44500000 | 0.0014 | 0.0012 | 0.0012 |
| F13 | 44420001 | 44520000 | 0.0023 | 0.0018 | 0.0020 |
| F13 | 44440001 | 44540000 | 0.0024 | 0.0019 | 0.0021 |
| F13 | 44460001 | 44560000 | 0.0022 | 0.0017 | 0.0019 |
| F13 | 44480001 | 44580000 | 0.0028 | 0.0023 | 0.0025 |
| F13 | 44500001 | 44600000 | 0.0028 | 0.0024 | 0.0026 |
| F13 | 44520001 | 44620000 | 0.0019 | 0.0016 | 0.0018 |
| F13 | 44540001 | 44640000 | 0.0020 | 0.0018 | 0.0020 |
| F13 | 44560001 | 44660000 | 0.0021 | 0.0018 | 0.0020 |
| F13 | 44580001 | 44680000 | 0.0011 | 0.0009 | 0.0011 |
| F13 | 44600001 | 44700000 | 0.0007 | 0.0006 | 0.0007 |
| F13 | 44620001 | 44720000 | 0.0006 | 0.0005 | 0.0006 |
| F13 | 44640001 | 44740000 | 0.0007 | 0.0006 | 0.0006 |
| F13 | 44660001 | 44760000 | 0.0012 | 0.0008 | 0.0009 |
| F13 | 44680001 | 44780000 | 0.0023 | 0.0017 | 0.0019 |
| F13 | 44700001 | 44800000 | 0.0024 | 0.0017 | 0.0021 |
| F13 | 44720001 | 44820000 | 0.0031 | 0.0023 | 0.0027 |
| F13 | 44740001 | 44840000 | 0.0032 | 0.0023 | 0.0029 |
| F13 | 44760001 | 44860000 | 0.0026 | 0.0020 | 0.0026 |
| F13 | 44780001 | 44880000 | 0.0017 | 0.0014 | 0.0017 |
| F13 | 44800001 | 44900000 | 0.0017 | 0.0014 | 0.0017 |
| F13 | 44820001 | 44920000 | 0.0012 | 0.0009 | 0.0012 |
| F13 | 44840001 | 44940000 | 0.0008 | 0.0007 | 0.0008 |
| F13 | 44860001 | 44960000 | 0.0008 | 0.0007 | 0.0008 |
| F13 | 44880001 | 44980000 | 0.0010 | 0.0009 | 0.0010 |
| F13 | 44900001 | 45000000 | 0.0010 | 0.0009 | 0.0010 |
| F13 | 44920001 | 45020000 | 0.0010 | 0.0008 | 0.0009 |

| Chr | Start | End | both | A1 | A2 |
|---|---|---|---|---|---|
| F13 | 44940001 | 45040000 | 0.0008 | 0.0007 | 0.0008 |
| F13 | 44960001 | 45060000 | 0.0007 | 0.0007 | 0.0007 |
| F13 | 44980001 | 45080000 | 0.0017 | 0.0016 | 0.0016 |
| F13 | 45000001 | 45100000 | 0.0017 | 0.0014 | 0.0014 |
| F13 | 45020001 | 45120000 | 0.0016 | 0.0014 | 0.0014 |
| F13 | 45040001 | 45140000 | 0.0019 | 0.0017 | 0.0017 |
| F13 | 45060001 | 45160000 | 0.0030 | 0.0025 | 0.0025 |
| F13 | 45080001 | 45180000 | 0.0020 | 0.0016 | 0.0016 |
| F13 | 45100001 | 45200000 | 0.0022 | 0.0018 | 0.0017 |
| F13 | 45120001 | 45220000 | 0.0030 | 0.0025 | 0.0026 |
| F13 | 45140001 | 45240000 | 0.0038 | 0.0032 | 0.0034 |
| F13 | 45160001 | 45260000 | 0.0034 | 0.0028 | 0.0031 |
| F13 | 45180001 | 45280000 | 0.0043 | 0.0035 | 0.0040 |
| F13 | 45200001 | 45300000 | 0.0043 | 0.0034 | 0.0040 |
| F13 | 45220001 | 45320000 | 0.0035 | 0.0027 | 0.0031 |
| F13 | 45240001 | 45340000 | 0.0024 | 0.0018 | 0.0020 |
| F13 | 45260001 | 45360000 | 0.0021 | 0.0016 | 0.0017 |
| F13 | 45280001 | 45380000 | 0.0010 | 0.0008 | 0.0007 |
| F13 | 45300001 | 45400000 | 0.0009 | 0.0007 | 0.0007 |
| F13 | 45320001 | 45420000 | 0.0008 | 0.0006 | 0.0006 |
| F13 | 45340001 | 45440000 | 0.0010 | 0.0008 | 0.0009 |
| F13 | 45360001 | 45460000 | 0.0008 | 0.0008 | 0.0008 |
| F13 | 45380001 | 45480000 | 0.0010 | 0.0009 | 0.0010 |
| F13 | 45400001 | 45500000 | 0.0011 | 0.0009 | 0.0010 |
| F13 | 45420001 | 45520000 | 0.0015 | 0.0013 | 0.0015 |
| F13 | 45440001 | 45540000 | 0.0016 | 0.0013 | 0.0016 |
| F13 | 45460001 | 45560000 | 0.0015 | 0.0012 | 0.0015 |
| F13 | 45480001 | 45580000 | 0.0013 | 0.0011 | 0.0013 |
| F13 | 45500001 | 45600000 | 0.0012 | 0.0010 | 0.0012 |
| F13 | 45520001 | 45620000 | 0.0009 | 0.0008 | 0.0009 |
| F13 | 45540001 | 45640000 | 0.0007 | 0.0007 | 0.0007 |
| F13 | 45560001 | 45660000 | 0.0010 | 0.0009 | 0.0010 |
| F13 | 45580001 | 45680000 | 0.0009 | 0.0008 | 0.0009 |
| F13 | 45600001 | 45700000 | 0.0010 | 0.0009 | 0.0010 |
| F13 | 45620001 | 45720000 | 0.0016 | 0.0014 | 0.0015 |
| F13 | 45640001 | 45740000 | 0.0017 | 0.0016 | 0.0016 |
| F13 | 45660001 | 45760000 | 0.0018 | 0.0017 | 0.0017 |
| F13 | 45680001 | 45780000 | 0.0025 | 0.0025 | 0.0023 |
| F13 | 45700001 | 45800000 | 0.0024 | 0.0024 | 0.0023 |
| F13 | 45720001 | 45820000 | 0.0022 | 0.0022 | 0.0021 |
| F13 | 45740001 | 45840000 | 0.0020 | 0.0019 | 0.0019 |
| F13 | 45760001 | 45860000 | 0.0019 | 0.0018 | 0.0019 |
| F13 | 45780001 | 45880000 | 0.0014 | 0.0011 | 0.0013 |
| F13 | 45800001 | 45900000 | 0.0020 | 0.0018 | 0.0019 |
| F13 | 45820001 | 45920000 | 0.0019 | 0.0015 | 0.0016 |
| F13 | 45840001 | 45940000 | 0.0023 | 0.0017 | 0.0018 |
| F13 | 45860001 | 45960000 | 0.0020 | 0.0014 | 0.0015 |
| F13 | 45880001 | 45980000 | 0.0022 | 0.0015 | 0.0017 |
| F13 | 45900001 | 46000000 | 0.0017 | 0.0010 | 0.0012 |
| F13 | 45920001 | 46020000 | 0.0021 | 0.0015 | 0.0018 |
| F13 | 45940001 | 46040000 | 0.0019 | 0.0015 | 0.0016 |
| F13 | 45960001 | 46060000 | 0.0028 | 0.0022 | 0.0026 |
| F13 | 45980001 | 46080000 | 0.0028 | 0.0023 | 0.0026 |
| F13 | 46000001 | 46100000 | 0.0024 | 0.0021 | 0.0024 |
| F13 | 46020001 | 46120000 | 0.0019 | 0.0016 | 0.0019 |
| F13 | 46040001 | 46140000 | 0.0017 | 0.0014 | 0.0016 |
| F13 | 46060001 | 46160000 | 0.0010 | 0.0009 | 0.0010 |
| F13 | 46080001 | 46180000 | 0.0012 | 0.0011 | 0.0012 |

| Chr | Start | End | both | A1 | A2 |
|---|---|---|---|---|---|
| F13 | 46100001 | 46200000 | 0.0012 | 0.0011 | 0.0012 |
| F13 | 46120001 | 46220000 | 0.0014 | 0.0012 | 0.0014 |
| F13 | 46140001 | 46240000 | 0.0021 | 0.0018 | 0.0020 |
| F13 | 46160001 | 46260000 | 0.0029 | 0.0026 | 0.0028 |
| F13 | 46180001 | 46280000 | 0.0026 | 0.0023 | 0.0025 |
| F13 | 46200001 | 46300000 | 0.0030 | 0.0026 | 0.0029 |
| F13 | 46220001 | 46320000 | 0.0029 | 0.0025 | 0.0027 |
| F13 | 46240001 | 46340000 | 0.0023 | 0.0019 | 0.0021 |
| F13 | 46260001 | 46360000 | 0.0011 | 0.0009 | 0.0011 |
| F13 | 46280001 | 46380000 | 0.0012 | 0.0009 | 0.0011 |
| F13 | 46300001 | 46400000 | 0.0010 | 0.0007 | 0.0009 |
| F13 | 46320001 | 46420000 | 0.0011 | 0.0009 | 0.0011 |
| F13 | 46340001 | 46440000 | 0.0014 | 0.0012 | 0.0013 |
| F13 | 46360001 | 46460000 | 0.0018 | 0.0015 | 0.0017 |
| F13 | 46380001 | 46480000 | 0.0019 | 0.0016 | 0.0019 |
| F13 | 46400001 | 46500000 | 0.0022 | 0.0020 | 0.0021 |
| F13 | 46420001 | 46520000 | 0.0024 | 0.0022 | 0.0023 |
| F13 | 46440001 | 46540000 | 0.0020 | 0.0018 | 0.0019 |
| F13 | 46460001 | 46560000 | 0.0017 | 0.0016 | 0.0016 |
| F13 | 46480001 | 46580000 | 0.0013 | 0.0012 | 0.0013 |
| F13 | 46500001 | 46600000 | 0.0010 | 0.0009 | 0.0010 |
| F13 | 46520001 | 46620000 | 0.0008 | 0.0007 | 0.0008 |
| F13 | 46540001 | 46640000 | 0.0014 | 0.0012 | 0.0014 |
| F13 | 46560001 | 46660000 | 0.0014 | 0.0012 | 0.0014 |
| F13 | 46580001 | 46680000 | 0.0014 | 0.0012 | 0.0014 |
| F13 | 46600001 | 46700000 | 0.0018 | 0.0015 | 0.0019 |
| F13 | 46620001 | 46720000 | 0.0018 | 0.0014 | 0.0018 |
| F13 | 46640001 | 46740000 | 0.0023 | 0.0019 | 0.0020 |
| F13 | 46660001 | 46760000 | 0.0025 | 0.0021 | 0.0022 |
| F13 | 46680001 | 46780000 | 0.0032 | 0.0025 | 0.0026 |
| F13 | 46700001 | 46800000 | 0.0028 | 0.0022 | 0.0021 |
| F13 | 46720001 | 46820000 | 0.0032 | 0.0025 | 0.0023 |
| F13 | 46740001 | 46840000 | 0.0021 | 0.0015 | 0.0016 |
| F13 | 46760001 | 46860000 | 0.0025 | 0.0015 | 0.0019 |
| F13 | 46780001 | 46880000 | 0.0021 | 0.0014 | 0.0016 |
| F13 | 46800001 | 46900000 | 0.0024 | 0.0017 | 0.0020 |
| F13 | 46820001 | 46920000 | 0.0026 | 0.0020 | 0.0023 |
| F13 | 46840001 | 46940000 | 0.0031 | 0.0024 | 0.0028 |
| F13 | 46860001 | 46960000 | 0.0024 | 0.0021 | 0.0023 |
| F13 | 46880001 | 46980000 | 0.0024 | 0.0022 | 0.0023 |
| F13 | 46900001 | 47000000 | 0.0021 | 0.0019 | 0.0020 |
| F13 | 46920001 | 47020000 | 0.0015 | 0.0014 | 0.0014 |
| F13 | 46940001 | 47040000 | 0.0015 | 0.0013 | 0.0013 |
| F13 | 46960001 | 47060000 | 0.0018 | 0.0015 | 0.0015 |
| F13 | 46980001 | 47080000 | 0.0022 | 0.0018 | 0.0020 |
| F13 | 47000001 | 47100000 | 0.0028 | 0.0023 | 0.0026 |
| F13 | 47020001 | 47120000 | 0.0028 | 0.0022 | 0.0026 |
| F13 | 47040001 | 47140000 | 0.0026 | 0.0020 | 0.0025 |
| F13 | 47060001 | 47160000 | 0.0026 | 0.0021 | 0.0025 |
| F13 | 47080001 | 47180000 | 0.0021 | 0.0015 | 0.0020 |
| F13 | 47100001 | 47200000 | 0.0017 | 0.0012 | 0.0016 |
| F13 | 47120001 | 47220000 | 0.0035 | 0.0028 | 0.0033 |
| F13 | 47140001 | 47240000 | 0.0047 | 0.0040 | 0.0045 |
| F13 | 47160001 | 47260000 | 0.0044 | 0.0037 | 0.0041 |
| F13 | 47180001 | 47280000 | 0.0042 | 0.0037 | 0.0039 |
| F13 | 47200001 | 47300000 | 0.0043 | 0.0038 | 0.0039 |
| F13 | 47220001 | 47320000 | 0.0027 | 0.0023 | 0.0025 |
| F13 | 47240001 | 47340000 | 0.0013 | 0.0010 | 0.0011 |

| Chr | Start | End | both | A1 | A2 |
|---|---|---|---|---|---|
| F13 | 47260001 | 47360000 | 0.0013 | 0.0010 | 0.0012 |
| F13 | 47280001 | 47380000 | 0.0013 | 0.0010 | 0.0012 |
| F13 | 47300001 | 47400000 | 0.0009 | 0.0007 | 0.0009 |
| F13 | 47320001 | 47420000 | 0.0005 | 0.0003 | 0.0005 |
| F13 | 47340001 | 47440000 | 0.0013 | 0.0010 | 0.0012 |
| F13 | 47360001 | 47460000 | 0.0025 | 0.0020 | 0.0022 |
| F13 | 47380001 | 47480000 | 0.0028 | 0.0023 | 0.0025 |
| F13 | 47400001 | 47500000 | 0.0029 | 0.0024 | 0.0026 |
| F13 | 47420001 | 47520000 | 0.0030 | 0.0025 | 0.0027 |
| F13 | 47440001 | 47540000 | 0.0022 | 0.0018 | 0.0020 |
| F13 | 47460001 | 47560000 | 0.0013 | 0.0012 | 0.0012 |
| F13 | 47480001 | 47580000 | 0.0010 | 0.0009 | 0.0010 |
| F13 | 47500001 | 47600000 | 0.0009 | 0.0008 | 0.0009 |
| F13 | 47520001 | 47620000 | 0.0009 | 0.0008 | 0.0009 |
| F13 | 47540001 | 47640000 | 0.0011 | 0.0010 | 0.0010 |
| F13 | 47560001 | 47660000 | 0.0010 | 0.0009 | 0.0010 |
| F13 | 47580001 | 47680000 | 0.0014 | 0.0011 | 0.0014 |
| F13 | 47600001 | 47700000 | 0.0017 | 0.0012 | 0.0016 |
| F13 | 47620001 | 47720000 | 0.0023 | 0.0018 | 0.0022 |
| F13 | 47640001 | 47740000 | 0.0026 | 0.0020 | 0.0025 |
| F13 | 47660001 | 47760000 | 0.0028 | 0.0022 | 0.0026 |
| F13 | 47680001 | 47780000 | 0.0027 | 0.0020 | 0.0023 |
| F13 | 47700001 | 47800000 | 0.0028 | 0.0022 | 0.0023 |
| F13 | 47720001 | 47820000 | 0.0027 | 0.0019 | 0.0022 |
| F13 | 47740001 | 47840000 | 0.0024 | 0.0018 | 0.0019 |
| F13 | 47760001 | 47860000 | 0.0022 | 0.0016 | 0.0018 |
| F13 | 47780001 | 47880000 | 0.0022 | 0.0018 | 0.0020 |
| F13 | 47800001 | 47900000 | 0.0018 | 0.0015 | 0.0017 |
| F13 | 47820001 | 47920000 | 0.0015 | 0.0013 | 0.0015 |
| F13 | 47840001 | 47940000 | 0.0022 | 0.0017 | 0.0020 |
| F13 | 47860001 | 47960000 | 0.0026 | 0.0021 | 0.0022 |
| F13 | 47880001 | 47980000 | 0.0037 | 0.0030 | 0.0032 |
| F13 | 47900001 | 48000000 | 0.0042 | 0.0033 | 0.0036 |
| F13 | 47920001 | 48020000 | 0.0042 | 0.0032 | 0.0035 |
| F13 | 47940001 | 48040000 | 0.0036 | 0.0028 | 0.0030 |
| F13 | 47960001 | 48060000 | 0.0034 | 0.0027 | 0.0028 |
| F13 | 47980001 | 48080000 | 0.0021 | 0.0016 | 0.0017 |
| F13 | 48000001 | 48100000 | 0.0026 | 0.0021 | 0.0022 |
| F13 | 48020001 | 48120000 | 0.0032 | 0.0025 | 0.0027 |
| F13 | 48040001 | 48140000 | 0.0033 | 0.0026 | 0.0028 |
| F13 | 48060001 | 48160000 | 0.0033 | 0.0025 | 0.0027 |
| F13 | 48080001 | 48180000 | 0.0036 | 0.0026 | 0.0029 |
| F13 | 48100001 | 48200000 | 0.0028 | 0.0018 | 0.0021 |
| F13 | 48120001 | 48220000 | 0.0023 | 0.0016 | 0.0019 |
| F13 | 48140001 | 48240000 | 0.0025 | 0.0017 | 0.0020 |
| F13 | 48160001 | 48260000 | 0.0021 | 0.0013 | 0.0018 |
| F13 | 48180001 | 48280000 | 0.0020 | 0.0014 | 0.0017 |
| F13 | 48200001 | 48300000 | 0.0017 | 0.0013 | 0.0016 |
| F13 | 48220001 | 48320000 | 0.0014 | 0.0010 | 0.0013 |
| F13 | 48240001 | 48340000 | 0.0014 | 0.0014 | 0.0013 |
| F13 | 48260001 | 48360000 | 0.0014 | 0.0014 | 0.0013 |
| F13 | 48280001 | 48380000 | 0.0012 | 0.0013 | 0.0011 |
| F13 | 48300001 | 48400000 | 0.0014 | 0.0014 | 0.0014 |
| F13 | 48320001 | 48420000 | 0.0021 | 0.0020 | 0.0020 |
| F13 | 48340001 | 48440000 | 0.0016 | 0.0014 | 0.0015 |
| F13 | 48360001 | 48460000 | 0.0016 | 0.0013 | 0.0015 |
| F13 | 48380001 | 48480000 | 0.0013 | 0.0011 | 0.0013 |
| F13 | 48400001 | 48500000 | 0.0014 | 0.0011 | 0.0014 |

| Chr | Start | End | both | A1 | A2 |
|---|---|---|---|---|---|
| F13 | 48420001 | 48520000 | 0.0008 | 0.0006 | 0.0008 |
| F13 | 48440001 | 48540000 | 0.0007 | 0.0006 | 0.0007 |
| F13 | 48460001 | 48560000 | 0.0011 | 0.0009 | 0.0011 |
| F13 | 48480001 | 48580000 | 0.0012 | 0.0010 | 0.0012 |
| F13 | 48500001 | 48600000 | 0.0009 | 0.0008 | 0.0009 |
| F13 | 48520001 | 48620000 | 0.0009 | 0.0008 | 0.0009 |
| F13 | 48540001 | 48640000 | 0.0011 | 0.0009 | 0.0010 |
| F13 | 48560001 | 48660000 | 0.0017 | 0.0015 | 0.0017 |
| F13 | 48580001 | 48680000 | 0.0019 | 0.0016 | 0.0018 |
| F13 | 48600001 | 48700000 | 0.0021 | 0.0018 | 0.0021 |
| F13 | 48620001 | 48720000 | 0.0025 | 0.0021 | 0.0023 |
| F13 | 48640001 | 48740000 | 0.0023 | 0.0020 | 0.0021 |
| F13 | 48660001 | 48760000 | 0.0022 | 0.0018 | 0.0020 |
| F13 | 48680001 | 48780000 | 0.0021 | 0.0018 | 0.0019 |
| F13 | 48700001 | 48800000 | 0.0023 | 0.0018 | 0.0019 |
| F13 | 48720001 | 48820000 | 0.0029 | 0.0024 | 0.0026 |
| F13 | 48740001 | 48840000 | 0.0030 | 0.0025 | 0.0027 |
| F13 | 48760001 | 48860000 | 0.0024 | 0.0019 | 0.0019 |
| F13 | 48780001 | 48880000 | 0.0022 | 0.0017 | 0.0018 |
| F13 | 48800001 | 48900000 | 0.0022 | 0.0017 | 0.0019 |
| F13 | 48820001 | 48920000 | 0.0010 | 0.0007 | 0.0008 |
| F13 | 48840001 | 48940000 | 0.0010 | 0.0007 | 0.0009 |
| F13 | 48860001 | 48960000 | 0.0009 | 0.0008 | 0.0009 |
| F13 | 48880001 | 48980000 | 0.0016 | 0.0013 | 0.0016 |
| F13 | 48900001 | 49000000 | 0.0015 | 0.0012 | 0.0014 |
| F13 | 48920001 | 49020000 | 0.0017 | 0.0013 | 0.0016 |
| F13 | 48940001 | 49040000 | 0.0016 | 0.0013 | 0.0016 |
| F13 | 48960001 | 49060000 | 0.0016 | 0.0012 | 0.0015 |
| F13 | 48980001 | 49080000 | 0.0011 | 0.0008 | 0.0010 |
| F13 | 49000001 | 49100000 | 0.0012 | 0.0010 | 0.0012 |
| F13 | 49020001 | 49120000 | 0.0017 | 0.0014 | 0.0017 |
| F13 | 49040001 | 49140000 | 0.0015 | 0.0013 | 0.0015 |
| F13 | 49060001 | 49160000 | 0.0014 | 0.0012 | 0.0014 |
| F13 | 49080001 | 49180000 | 0.0019 | 0.0016 | 0.0019 |
| F13 | 49100001 | 49200000 | 0.0020 | 0.0016 | 0.0020 |
| F13 | 49120001 | 49220000 | 0.0012 | 0.0010 | 0.0012 |
| F13 | 49140001 | 49240000 | 0.0022 | 0.0018 | 0.0021 |
| F13 | 49160001 | 49260000 | 0.0026 | 0.0021 | 0.0024 |
| F13 | 49180001 | 49280000 | 0.0021 | 0.0016 | 0.0019 |
| F13 | 49200001 | 49300000 | 0.0020 | 0.0016 | 0.0017 |
| F13 | 49220001 | 49320000 | 0.0024 | 0.0019 | 0.0021 |
| F13 | 49240001 | 49340000 | 0.0015 | 0.0013 | 0.0014 |
| F13 | 49260001 | 49360000 | 0.0015 | 0.0013 | 0.0014 |
| F13 | 49280001 | 49380000 | 0.0015 | 0.0013 | 0.0014 |
| F13 | 49300001 | 49400000 | 0.0012 | 0.0010 | 0.0012 |
| F13 | 49320001 | 49420000 | 0.0013 | 0.0011 | 0.0013 |
| F13 | 49340001 | 49440000 | 0.0016 | 0.0013 | 0.0014 |
| F13 | 49360001 | 49460000 | 0.0017 | 0.0014 | 0.0016 |
| F13 | 49380001 | 49480000 | 0.0019 | 0.0015 | 0.0017 |
| F13 | 49400001 | 49500000 | 0.0028 | 0.0023 | 0.0025 |
| F13 | 49420001 | 49520000 | 0.0031 | 0.0025 | 0.0029 |
| F13 | 49440001 | 49540000 | 0.0031 | 0.0025 | 0.0030 |
| F13 | 49460001 | 49560000 | 0.0029 | 0.0023 | 0.0028 |
| F13 | 49480001 | 49580000 | 0.0026 | 0.0021 | 0.0025 |
| F13 | 49500001 | 49600000 | 0.0020 | 0.0016 | 0.0020 |
| F13 | 49520001 | 49620000 | 0.0013 | 0.0010 | 0.0013 |
| F13 | 49540001 | 49640000 | 0.0010 | 0.0008 | 0.0010 |
| F13 | 49560001 | 49660000 | 0.0007 | 0.0006 | 0.0007 |

| Chr | Start | End | both | A1 | A2 |
|---|---|---|---|---|---|
| F13 | 49580001 | 49680000 | 0.0007 | 0.0006 | 0.0007 |
| F13 | 49600001 | 49700000 | 0.0005 | 0.0003 | 0.0004 |
| F13 | 49620001 | 49720000 | 0.0007 | 0.0004 | 0.0004 |
| F13 | 49640001 | 49740000 | 0.0005 | 0.0003 | 0.0003 |
| F13 | 49660001 | 49760000 | 0.0007 | 0.0004 | 0.0005 |
| F13 | 49680001 | 49780000 | 0.0009 | 0.0005 | 0.0005 |
| F13 | 49700001 | 49800000 | 0.0009 | 0.0005 | 0.0006 |
| F13 | 49720001 | 49820000 | 0.0020 | 0.0017 | 0.0017 |
| F13 | 49740001 | 49840000 | 0.0026 | 0.0023 | 0.0023 |
| F13 | 49760001 | 49860000 | 0.0023 | 0.0021 | 0.0021 |
| F13 | 49780001 | 49880000 | 0.0021 | 0.0020 | 0.0020 |
| F13 | 49800001 | 49900000 | 0.0022 | 0.0020 | 0.0020 |
| F13 | 49820001 | 49920000 | 0.0008 | 0.0007 | 0.0008 |
| F13 | 49840001 | 49940000 | 0.0007 | 0.0005 | 0.0006 |
| F13 | 49860001 | 49960000 | 0.0013 | 0.0009 | 0.0012 |
| F13 | 49880001 | 49980000 | 0.0023 | 0.0016 | 0.0019 |
| F13 | 49900001 | 50000000 | 0.0029 | 0.0021 | 0.0024 |
| F13 | 49920001 | 50020000 | 0.0034 | 0.0023 | 0.0027 |
| F13 | 49940001 | 50040000 | 0.0030 | 0.0020 | 0.0023 |
| F13 | 49960001 | 50060000 | 0.0028 | 0.0018 | 0.0021 |
| F13 | 49980001 | 50080000 | 0.0019 | 0.0013 | 0.0016 |
| F13 | 50000001 | 50100000 | 0.0014 | 0.0009 | 0.0012 |
| F13 | 50020001 | 50120000 | 0.0019 | 0.0014 | 0.0018 |
| F13 | 50040001 | 50140000 | 0.0030 | 0.0024 | 0.0028 |
| F13 | 50060001 | 50160000 | 0.0028 | 0.0022 | 0.0027 |
| F13 | 50080001 | 50180000 | 0.0029 | 0.0022 | 0.0027 |
| F13 | 50100001 | 50200000 | 0.0032 | 0.0023 | 0.0028 |
| F13 | 50120001 | 50220000 | 0.0029 | 0.0021 | 0.0025 |
| F13 | 50140001 | 50240000 | 0.0022 | 0.0015 | 0.0018 |
| F13 | 50160001 | 50260000 | 0.0027 | 0.0019 | 0.0022 |
| F13 | 50180001 | 50280000 | 0.0031 | 0.0024 | 0.0028 |
| F13 | 50200001 | 50300000 | 0.0040 | 0.0031 | 0.0036 |
| F13 | 50220001 | 50320000 | 0.0035 | 0.0028 | 0.0032 |
| F13 | 50240001 | 50340000 | 0.0032 | 0.0025 | 0.0029 |
| F13 | 50260001 | 50360000 | 0.0031 | 0.0025 | 0.0028 |
| F13 | 50280001 | 50380000 | 0.0028 | 0.0021 | 0.0024 |
| F13 | 50300001 | 50400000 | 0.0018 | 0.0012 | 0.0015 |
| F13 | 50320001 | 50420000 | 0.0023 | 0.0016 | 0.0020 |
| F13 | 50340001 | 50440000 | 0.0025 | 0.0018 | 0.0022 |
| F13 | 50360001 | 50460000 | 0.0027 | 0.0020 | 0.0024 |
| F13 | 50380001 | 50480000 | 0.0028 | 0.0022 | 0.0026 |
| F13 | 50400001 | 50500000 | 0.0028 | 0.0022 | 0.0025 |
| F13 | 50420001 | 50520000 | 0.0026 | 0.0021 | 0.0023 |
| F13 | 50440001 | 50540000 | 0.0029 | 0.0023 | 0.0025 |
| F13 | 50460001 | 50560000 | 0.0032 | 0.0026 | 0.0029 |
| F13 | 50480001 | 50580000 | 0.0028 | 0.0022 | 0.0025 |
| F13 | 50500001 | 50600000 | 0.0028 | 0.0022 | 0.0026 |
| F13 | 50520001 | 50620000 | 0.0029 | 0.0024 | 0.0028 |
| F13 | 50540001 | 50640000 | 0.0026 | 0.0022 | 0.0026 |
| F13 | 50560001 | 50660000 | 0.0020 | 0.0016 | 0.0019 |
| F13 | 50580001 | 50680000 | 0.0021 | 0.0016 | 0.0020 |
| F13 | 50600001 | 50700000 | 0.0032 | 0.0025 | 0.0030 |
| F13 | 50620001 | 50720000 | 0.0028 | 0.0022 | 0.0027 |
| F13 | 50640001 | 50740000 | 0.0027 | 0.0021 | 0.0026 |
| F13 | 50660001 | 50760000 | 0.0025 | 0.0020 | 0.0023 |
| F13 | 50680001 | 50780000 | 0.0024 | 0.0020 | 0.0022 |
| F13 | 50700001 | 50800000 | 0.0012 | 0.0010 | 0.0011 |
| F13 | 50720001 | 50820000 | 0.0015 | 0.0011 | 0.0013 |

| Chr | Start | End | both | A1 | A2 |
|---|---|---|---|---|---|
| F13 | 50740001 | 50840000 | 0.0015 | 0.0012 | 0.0013 |
| F13 | 50760001 | 50860000 | 0.0014 | 0.0011 | 0.0013 |
| F13 | 50780001 | 50880000 | 0.0017 | 0.0014 | 0.0016 |
| F13 | 50800001 | 50900000 | 0.0020 | 0.0017 | 0.0018 |
| F13 | 50820001 | 50920000 | 0.0016 | 0.0014 | 0.0015 |
| F13 | 50840001 | 50940000 | 0.0014 | 0.0012 | 0.0013 |
| F13 | 50860001 | 50960000 | 0.0012 | 0.0011 | 0.0011 |
| F13 | 50880001 | 50980000 | 0.0010 | 0.0009 | 0.0010 |
| F13 | 50900001 | 51000000 | 0.0008 | 0.0008 | 0.0008 |
| F13 | 50920001 | 51020000 | 0.0010 | 0.0009 | 0.0010 |
| F13 | 50940001 | 51040000 | 0.0013 | 0.0011 | 0.0012 |
| F13 | 50960001 | 51060000 | 0.0019 | 0.0015 | 0.0017 |
| F13 | 50980001 | 51080000 | 0.0019 | 0.0014 | 0.0017 |
| F13 | 51000001 | 51100000 | 0.0020 | 0.0015 | 0.0018 |
| F13 | 51020001 | 51120000 | 0.0022 | 0.0016 | 0.0019 |
| F13 | 51040001 | 51140000 | 0.0021 | 0.0015 | 0.0019 |
| F13 | 51060001 | 51160000 | 0.0015 | 0.0012 | 0.0014 |
| F13 | 51080001 | 51180000 | 0.0022 | 0.0018 | 0.0021 |
| F13 | 51100001 | 51200000 | 0.0021 | 0.0016 | 0.0019 |
| F13 | 51120001 | 51220000 | 0.0020 | 0.0015 | 0.0018 |
| F13 | 51140001 | 51240000 | 0.0022 | 0.0016 | 0.0021 |
| F13 | 51160001 | 51260000 | 0.0029 | 0.0021 | 0.0027 |
| F13 | 51180001 | 51280000 | 0.0017 | 0.0012 | 0.0017 |
| F13 | 51200001 | 51300000 | 0.0019 | 0.0015 | 0.0020 |
| F13 | 51220001 | 51320000 | 0.0019 | 0.0016 | 0.0019 |
| F13 | 51240001 | 51340000 | 0.0015 | 0.0013 | 0.0015 |
| F13 | 51260001 | 51360000 | 0.0011 | 0.0010 | 0.0011 |
| F13 | 51280001 | 51380000 | 0.0015 | 0.0013 | 0.0014 |
| F13 | 51300001 | 51400000 | 0.0012 | 0.0011 | 0.0011 |
| F13 | 51320001 | 51420000 | 0.0010 | 0.0009 | 0.0010 |
| F13 | 51340001 | 51440000 | 0.0016 | 0.0014 | 0.0016 |
| F13 | 51360001 | 51460000 | 0.0016 | 0.0014 | 0.0015 |
| F13 | 51380001 | 51480000 | 0.0012 | 0.0011 | 0.0012 |
| F13 | 51400001 | 51500000 | 0.0013 | 0.0011 | 0.0013 |
| F13 | 51420001 | 51520000 | 0.0025 | 0.0022 | 0.0025 |
| F13 | 51440001 | 51540000 | 0.0022 | 0.0019 | 0.0022 |
| F13 | 51460001 | 51560000 | 0.0022 | 0.0019 | 0.0022 |
| F13 | 51480001 | 51580000 | 0.0022 | 0.0019 | 0.0022 |
| F13 | 51500001 | 51600000 | 0.0021 | 0.0019 | 0.0021 |
| F13 | 51520001 | 51620000 | 0.0007 | 0.0006 | 0.0007 |
| F13 | 51540001 | 51640000 | 0.0009 | 0.0006 | 0.0009 |
| F13 | 51560001 | 51660000 | 0.0014 | 0.0010 | 0.0014 |
| F13 | 51580001 | 51680000 | 0.0014 | 0.0010 | 0.0014 |
| F13 | 51600001 | 51700000 | 0.0017 | 0.0013 | 0.0017 |
| F13 | 51620001 | 51720000 | 0.0022 | 0.0017 | 0.0021 |
| F13 | 51640001 | 51740000 | 0.0026 | 0.0021 | 0.0022 |
| F13 | 51660001 | 51760000 | 0.0027 | 0.0021 | 0.0022 |
| F13 | 51680001 | 51780000 | 0.0040 | 0.0032 | 0.0033 |
| F13 | 51700001 | 51800000 | 0.0043 | 0.0034 | 0.0036 |
| F13 | 51720001 | 51820000 | 0.0044 | 0.0034 | 0.0038 |
| F13 | 51740001 | 51840000 | 0.0043 | 0.0034 | 0.0040 |
| F13 | 51760001 | 51860000 | 0.0041 | 0.0032 | 0.0037 |
| F13 | 51780001 | 51880000 | 0.0034 | 0.0024 | 0.0030 |
| F13 | 51800001 | 51900000 | 0.0042 | 0.0035 | 0.0037 |
| F13 | 51820001 | 51920000 | 0.0050 | 0.0041 | 0.0044 |
| F13 | 51840001 | 51940000 | 0.0055 | 0.0044 | 0.0049 |
| F13 | 51860001 | 51960000 | 0.0056 | 0.0044 | 0.0049 |
| F13 | 51880001 | 51980000 | 0.0059 | 0.0045 | 0.0051 |

| Chr | Start | End | both | A1 | A2 |
|---|---|---|---|---|---|
| F13 | 51900001 | 52000000 | 0.0054 | 0.0039 | 0.0047 |
| F13 | 51920001 | 52020000 | 0.0045 | 0.0031 | 0.0037 |
| F13 | 51940001 | 52040000 | 0.0044 | 0.0032 | 0.0036 |
| F13 | 51960001 | 52060000 | 0.0054 | 0.0041 | 0.0047 |
| F13 | 51980001 | 52080000 | 0.0050 | 0.0039 | 0.0043 |
| F13 | 52000001 | 52100000 | 0.0051 | 0.0038 | 0.0043 |
| F13 | 52020001 | 52120000 | 0.0061 | 0.0047 | 0.0052 |
| F13 | 52040001 | 52140000 | 0.0059 | 0.0045 | 0.0050 |
| F13 | 52060001 | 52160000 | 0.0049 | 0.0037 | 0.0041 |
| F13 | 52080001 | 52180000 | 0.0045 | 0.0036 | 0.0039 |
| F13 | 52100001 | 52200000 | 0.0051 | 0.0039 | 0.0044 |
| F13 | 52120001 | 52220000 | 0.0046 | 0.0036 | 0.0040 |
| F13 | 52140001 | 52240000 | 0.0044 | 0.0032 | 0.0036 |
| F13 | 52160001 | 52260000 | 0.0045 | 0.0032 | 0.0036 |
| F13 | 52180001 | 52280000 | 0.0055 | 0.0036 | 0.0043 |
| F13 | 52200001 | 52300000 | 0.0039 | 0.0025 | 0.0030 |
| F13 | 52220001 | 52320000 | 0.0036 | 0.0021 | 0.0026 |
| F13 | 52240001 | 52340000 | 0.0040 | 0.0024 | 0.0031 |
| F13 | 52260001 | 52360000 | 0.0042 | 0.0025 | 0.0031 |
| F13 | 52280001 | 52380000 | 0.0038 | 0.0026 | 0.0029 |
| F13 | 52300001 | 52400000 | 0.0038 | 0.0026 | 0.0030 |
| F13 | 52320001 | 52420000 | 0.0033 | 0.0024 | 0.0026 |
| F13 | 52340001 | 52440000 | 0.0023 | 0.0017 | 0.0016 |
| F13 | 52360001 | 52460000 | 0.0020 | 0.0015 | 0.0014 |
| F13 | 52380001 | 52480000 | 0.0015 | 0.0010 | 0.0010 |
| F13 | 52400001 | 52500000 | 0.0018 | 0.0012 | 0.0013 |
| F13 | 52420001 | 52520000 | 0.0026 | 0.0017 | 0.0018 |
| F13 | 52440001 | 52540000 | 0.0025 | 0.0016 | 0.0017 |
| F13 | 52460001 | 52560000 | 0.0026 | 0.0017 | 0.0019 |
| F13 | 52480001 | 52580000 | 0.0030 | 0.0020 | 0.0020 |
| F13 | 52500001 | 52600000 | 0.0029 | 0.0019 | 0.0020 |
| F13 | 52520001 | 52620000 | 0.0022 | 0.0015 | 0.0015 |
| F13 | 52540001 | 52640000 | 0.0028 | 0.0020 | 0.0021 |
| F13 | 52560001 | 52660000 | 0.0041 | 0.0033 | 0.0036 |
| F13 | 52580001 | 52680000 | 0.0039 | 0.0030 | 0.0034 |
| F13 | 52600001 | 52700000 | 0.0046 | 0.0038 | 0.0042 |
| F13 | 52620001 | 52720000 | 0.0045 | 0.0037 | 0.0041 |
| F13 | 52640001 | 52740000 | 0.0049 | 0.0041 | 0.0043 |
| F13 | 52660001 | 52760000 | 0.0038 | 0.0029 | 0.0031 |
| F13 | 52680001 | 52780000 | 0.0038 | 0.0030 | 0.0032 |
| F13 | 52700001 | 52800000 | 0.0027 | 0.0021 | 0.0021 |
| F13 | 52720001 | 52820000 | 0.0031 | 0.0023 | 0.0025 |
| F13 | 52740001 | 52840000 | 0.0027 | 0.0021 | 0.0024 |
| F13 | 52760001 | 52860000 | 0.0030 | 0.0023 | 0.0029 |
| F13 | 52780001 | 52880000 | 0.0032 | 0.0026 | 0.0032 |
| F13 | 52800001 | 52900000 | 0.0038 | 0.0029 | 0.0035 |
| F13 | 52820001 | 52920000 | 0.0045 | 0.0035 | 0.0040 |
| F13 | 52840001 | 52940000 | 0.0040 | 0.0032 | 0.0035 |
| F13 | 52860001 | 52960000 | 0.0029 | 0.0023 | 0.0024 |
| F13 | 52880001 | 52980000 | 0.0029 | 0.0023 | 0.0024 |
| F13 | 52900001 | 53000000 | 0.0026 | 0.0022 | 0.0022 |
| F13 | 52920001 | 53020000 | 0.0020 | 0.0016 | 0.0018 |
| F13 | 52940001 | 53040000 | 0.0026 | 0.0020 | 0.0019 |
| F13 | 52960001 | 53060000 | 0.0031 | 0.0024 | 0.0023 |
| F13 | 52980001 | 53080000 | 0.0025 | 0.0019 | 0.0018 |
| F13 | 53000001 | 53100000 | 0.0032 | 0.0026 | 0.0025 |
| F13 | 53020001 | 53120000 | 0.0039 | 0.0029 | 0.0030 |
| F13 | 53040001 | 53140000 | 0.0042 | 0.0031 | 0.0037 |

| Chr | Start | End | both | A1 | A2 |
|---|---|---|---|---|---|
| F13 | 53060001 | 53160000 | 0.0042 | 0.0032 | 0.0037 |
| F13 | 53080001 | 53180000 | 0.0051 | 0.0038 | 0.0043 |
| F13 | 53100001 | 53200000 | 0.0059 | 0.0044 | 0.0048 |
| F13 | 53120001 | 53220000 | 0.0053 | 0.0039 | 0.0043 |
| F13 | 53140001 | 53240000 | 0.0056 | 0.0040 | 0.0046 |
| F13 | 53160001 | 53260000 | 0.0056 | 0.0041 | 0.0046 |
| F13 | 53180001 | 53280000 | 0.0047 | 0.0035 | 0.0040 |
| F13 | 53200001 | 53300000 | 0.0040 | 0.0029 | 0.0034 |
| F13 | 53220001 | 53320000 | 0.0039 | 0.0032 | 0.0035 |
| F13 | 53240001 | 53340000 | 0.0031 | 0.0028 | 0.0028 |
| F13 | 53260001 | 53360000 | 0.0031 | 0.0027 | 0.0029 |
| F13 | 53280001 | 53380000 | 0.0041 | 0.0036 | 0.0038 |
| F13 | 53300001 | 53400000 | 0.0042 | 0.0037 | 0.0040 |
| F13 | 53320001 | 53420000 | 0.0036 | 0.0030 | 0.0034 |
| F13 | 53340001 | 53440000 | 0.0033 | 0.0028 | 0.0032 |
| F13 | 53360001 | 53460000 | 0.0037 | 0.0033 | 0.0036 |
| F13 | 53380001 | 53480000 | 0.0027 | 0.0022 | 0.0026 |
| F13 | 53400001 | 53500000 | 0.0030 | 0.0023 | 0.0027 |
| F13 | 53420001 | 53520000 | 0.0035 | 0.0027 | 0.0031 |
| F13 | 53440001 | 53540000 | 0.0043 | 0.0032 | 0.0037 |
| F13 | 53460001 | 53560000 | 0.0045 | 0.0031 | 0.0037 |
| F13 | 53480001 | 53580000 | 0.0058 | 0.0043 | 0.0050 |
| F13 | 53500001 | 53600000 | 0.0051 | 0.0037 | 0.0042 |
| F13 | 53520001 | 53620000 | 0.0048 | 0.0035 | 0.0040 |
| F13 | 53540001 | 53640000 | 0.0040 | 0.0030 | 0.0033 |
| F13 | 53560001 | 53660000 | 0.0038 | 0.0030 | 0.0030 |
| F13 | 53580001 | 53680000 | 0.0027 | 0.0020 | 0.0020 |
| F13 | 53600001 | 53700000 | 0.0027 | 0.0020 | 0.0021 |
| F13 | 53620001 | 53720000 | 0.0042 | 0.0033 | 0.0035 |
| F13 | 53640001 | 53740000 | 0.0044 | 0.0034 | 0.0037 |
| F13 | 53660001 | 53760000 | 0.0037 | 0.0030 | 0.0034 |
| F13 | 53680001 | 53780000 | 0.0035 | 0.0029 | 0.0033 |
| F13 | 53700001 | 53800000 | 0.0036 | 0.0030 | 0.0034 |
| F13 | 53720001 | 53820000 | 0.0022 | 0.0018 | 0.0021 |
| F13 | 53740001 | 53840000 | 0.0021 | 0.0019 | 0.0021 |
| F13 | 53760001 | 53860000 | 0.0018 | 0.0016 | 0.0018 |
| F13 | 53780001 | 53880000 | 0.0022 | 0.0019 | 0.0021 |
| F13 | 53800001 | 53900000 | 0.0017 | 0.0015 | 0.0016 |
| F13 | 53820001 | 53920000 | 0.0025 | 0.0022 | 0.0025 |
| F13 | 53840001 | 53940000 | 0.0032 | 0.0028 | 0.0031 |
| F13 | 53860001 | 53960000 | 0.0032 | 0.0028 | 0.0031 |
| F13 | 53880001 | 53980000 | 0.0030 | 0.0025 | 0.0028 |
| F13 | 53900001 | 54000000 | 0.0032 | 0.0026 | 0.0030 |
| F13 | 53920001 | 54020000 | 0.0026 | 0.0018 | 0.0022 |
| F13 | 53940001 | 54040000 | 0.0026 | 0.0019 | 0.0022 |
| F13 | 53960001 | 54060000 | 0.0038 | 0.0025 | 0.0032 |
| F13 | 53980001 | 54080000 | 0.0037 | 0.0025 | 0.0031 |
| F13 | 54000001 | 54100000 | 0.0039 | 0.0028 | 0.0033 |
| F13 | 54020001 | 54120000 | 0.0033 | 0.0025 | 0.0030 |
| F13 | 54040001 | 54140000 | 0.0031 | 0.0022 | 0.0027 |
| F13 | 54060001 | 54160000 | 0.0026 | 0.0021 | 0.0024 |
| F13 | 54080001 | 54180000 | 0.0025 | 0.0021 | 0.0023 |
| F13 | 54100001 | 54200000 | 0.0020 | 0.0016 | 0.0019 |
| F13 | 54120001 | 54220000 | 0.0020 | 0.0015 | 0.0019 |
| F13 | 54140001 | 54240000 | 0.0023 | 0.0019 | 0.0023 |
| F13 | 54160001 | 54260000 | 0.0017 | 0.0014 | 0.0017 |
| F13 | 54180001 | 54280000 | 0.0025 | 0.0020 | 0.0023 |
| F13 | 54200001 | 54300000 | 0.0031 | 0.0025 | 0.0030 |

| Chr | Start | End | both | A1 | A2 |
|---|---|---|---|---|---|
| F13 | 54220001 | 54320000 | 0.0032 | 0.0026 | 0.0030 |
| F13 | 54240001 | 54340000 | 0.0020 | 0.0016 | 0.0018 |
| F13 | 54260001 | 54360000 | 0.0023 | 0.0020 | 0.0022 |
| F13 | 54280001 | 54380000 | 0.0017 | 0.0015 | 0.0017 |
| F13 | 54300001 | 54400000 | 0.0016 | 0.0014 | 0.0016 |
| F13 | 54320001 | 54420000 | 0.0016 | 0.0014 | 0.0016 |
| F13 | 54340001 | 54440000 | 0.0020 | 0.0017 | 0.0020 |
| F13 | 54360001 | 54460000 | 0.0017 | 0.0014 | 0.0016 |
| F13 | 54380001 | 54480000 | 0.0017 | 0.0015 | 0.0017 |
| F13 | 54400001 | 54500000 | 0.0025 | 0.0020 | 0.0023 |
| F13 | 54420001 | 54520000 | 0.0025 | 0.0021 | 0.0023 |
| F13 | 54440001 | 54540000 | 0.0030 | 0.0024 | 0.0028 |
| F13 | 54460001 | 54560000 | 0.0040 | 0.0033 | 0.0038 |
| F13 | 54480001 | 54580000 | 0.0048 | 0.0039 | 0.0044 |
| F13 | 54500001 | 54600000 | 0.0038 | 0.0031 | 0.0035 |
| F13 | 54520001 | 54620000 | 0.0050 | 0.0042 | 0.0046 |
| F13 | 54540001 | 54640000 | 0.0052 | 0.0043 | 0.0046 |
| F13 | 54560001 | 54660000 | 0.0052 | 0.0045 | 0.0044 |
| F13 | 54580001 | 54680000 | 0.0047 | 0.0039 | 0.0038 |
| F13 | 54600001 | 54700000 | 0.0055 | 0.0047 | 0.0045 |
| F13 | 54620001 | 54720000 | 0.0048 | 0.0040 | 0.0039 |
| F13 | 54640001 | 54740000 | 0.0043 | 0.0035 | 0.0034 |
| F13 | 54660001 | 54760000 | 0.0039 | 0.0030 | 0.0032 |
| F13 | 54680001 | 54780000 | 0.0044 | 0.0031 | 0.0035 |
| F13 | 54700001 | 54800000 | 0.0034 | 0.0022 | 0.0027 |
| F13 | 54720001 | 54820000 | 0.0037 | 0.0024 | 0.0029 |
| F13 | 54740001 | 54840000 | 0.0042 | 0.0028 | 0.0033 |
| F13 | 54760001 | 54860000 | 0.0045 | 0.0030 | 0.0035 |
| F13 | 54780001 | 54880000 | 0.0040 | 0.0030 | 0.0032 |
| F13 | 54800001 | 54900000 | 0.0053 | 0.0040 | 0.0043 |
| F13 | 54820001 | 54920000 | 0.0051 | 0.0038 | 0.0042 |
| F13 | 54840001 | 54940000 | 0.0055 | 0.0043 | 0.0046 |
| F13 | 54860001 | 54960000 | 0.0050 | 0.0036 | 0.0040 |
| F13 | 54880001 | 54980000 | 0.0067 | 0.0053 | 0.0058 |
| F13 | 54900001 | 55000000 | 0.0054 | 0.0044 | 0.0048 |
| F13 | 54920001 | 55020000 | 0.0048 | 0.0039 | 0.0043 |
| F13 | 54940001 | 55040000 | 0.0045 | 0.0038 | 0.0041 |
| F13 | 54960001 | 55060000 | 0.0048 | 0.0043 | 0.0044 |
| F13 | 54980001 | 55080000 | 0.0027 | 0.0024 | 0.0024 |
| F13 | 55000001 | 55100000 | 0.0038 | 0.0034 | 0.0034 |
| F13 | 55020001 | 55120000 | 0.0042 | 0.0035 | 0.0034 |
| F13 | 55040001 | 55140000 | 0.0034 | 0.0027 | 0.0025 |
| F13 | 55060001 | 55160000 | 0.0032 | 0.0024 | 0.0022 |
| F13 | 55080001 | 55180000 | 0.0041 | 0.0032 | 0.0029 |
| F13 | 55100001 | 55200000 | 0.0031 | 0.0022 | 0.0019 |
| F13 | 55120001 | 55220000 | 0.0030 | 0.0021 | 0.0021 |
| F13 | 55140001 | 55240000 | 0.0032 | 0.0023 | 0.0023 |
| F13 | 55160001 | 55260000 | 0.0028 | 0.0020 | 0.0021 |
| F13 | 55180001 | 55280000 | 0.0019 | 0.0012 | 0.0014 |
| F13 | 55200001 | 55300000 | 0.0021 | 0.0014 | 0.0016 |
| F13 | 55220001 | 55320000 | 0.0029 | 0.0021 | 0.0025 |
| F13 | 55240001 | 55340000 | 0.0027 | 0.0020 | 0.0023 |
| F13 | 55260001 | 55360000 | 0.0036 | 0.0029 | 0.0031 |
| F13 | 55280001 | 55380000 | 0.0041 | 0.0032 | 0.0034 |
| F13 | 55300001 | 55400000 | 0.0055 | 0.0044 | 0.0047 |
| F13 | 55320001 | 55420000 | 0.0047 | 0.0039 | 0.0040 |
| F13 | 55340001 | 55440000 | 0.0051 | 0.0043 | 0.0045 |
| F13 | 55360001 | 55460000 | 0.0049 | 0.0043 | 0.0043 |

| Chr | Start | End | both | A1 | A2 |
|---|---|---|---|---|---|
| F13 | 55380001 | 55480000 | 0.0052 | 0.0047 | 0.0047 |
| F13 | 55400001 | 55500000 | 0.0042 | 0.0039 | 0.0037 |
| F13 | 55420001 | 55520000 | 0.0042 | 0.0038 | 0.0036 |
| F13 | 55440001 | 55540000 | 0.0042 | 0.0036 | 0.0035 |
| F13 | 55460001 | 55560000 | 0.0041 | 0.0029 | 0.0031 |
| F13 | 55480001 | 55580000 | 0.0036 | 0.0022 | 0.0023 |
| F13 | 55500001 | 55600000 | 0.0035 | 0.0022 | 0.0023 |
| F13 | 55520001 | 55620000 | 0.0036 | 0.0025 | 0.0025 |
| F13 | 55540001 | 55640000 | 0.0036 | 0.0027 | 0.0027 |
| F13 | 55560001 | 55660000 | 0.0041 | 0.0034 | 0.0035 |
| F13 | 55580001 | 55680000 | 0.0038 | 0.0032 | 0.0034 |
| F13 | 55600001 | 55700000 | 0.0040 | 0.0030 | 0.0033 |
| F13 | 55620001 | 55720000 | 0.0045 | 0.0033 | 0.0037 |
| F13 | 55640001 | 55740000 | 0.0053 | 0.0040 | 0.0044 |
| F13 | 55660001 | 55760000 | 0.0047 | 0.0036 | 0.0040 |
| F13 | 55680001 | 55780000 | 0.0049 | 0.0037 | 0.0042 |
| F13 | 55700001 | 55800000 | 0.0043 | 0.0033 | 0.0038 |
| F13 | 55720001 | 55820000 | 0.0037 | 0.0029 | 0.0033 |
| F13 | 55740001 | 55840000 | 0.0028 | 0.0022 | 0.0026 |
| F13 | 55760001 | 55860000 | 0.0022 | 0.0016 | 0.0019 |
| F13 | 55780001 | 55880000 | 0.0036 | 0.0027 | 0.0031 |
| F13 | 55800001 | 55900000 | 0.0036 | 0.0029 | 0.0033 |
| F13 | 55820001 | 55920000 | 0.0037 | 0.0030 | 0.0034 |
| F13 | 55840001 | 55940000 | 0.0037 | 0.0027 | 0.0034 |
| F13 | 55860001 | 55960000 | 0.0046 | 0.0035 | 0.0041 |
| F13 | 55880001 | 55980000 | 0.0034 | 0.0023 | 0.0028 |
| F13 | 55900001 | 56000000 | 0.0045 | 0.0031 | 0.0036 |
| F13 | 55920001 | 56020000 | 0.0045 | 0.0030 | 0.0035 |
| F13 | 55940001 | 56040000 | 0.0045 | 0.0034 | 0.0034 |
| F13 | 55960001 | 56060000 | 0.0037 | 0.0030 | 0.0029 |
| F13 | 55980001 | 56080000 | 0.0038 | 0.0032 | 0.0032 |
| F13 | 56000001 | 56100000 | 0.0031 | 0.0028 | 0.0026 |
| F13 | 56020001 | 56120000 | 0.0026 | 0.0024 | 0.0023 |
| F13 | 56040001 | 56140000 | 0.0028 | 0.0025 | 0.0027 |
| F13 | 56060001 | 56160000 | 0.0024 | 0.0021 | 0.0023 |
| F13 | 56080001 | 56180000 | 0.0019 | 0.0017 | 0.0019 |
| F13 | 56100001 | 56200000 | 0.0012 | 0.0010 | 0.0012 |
| F13 | 56120001 | 56220000 | 0.0017 | 0.0013 | 0.0015 |
| F13 | 56140001 | 56240000 | 0.0012 | 0.0009 | 0.0010 |
| F13 | 56160001 | 56260000 | 0.0020 | 0.0014 | 0.0017 |
| F13 | 56180001 | 56280000 | 0.0031 | 0.0023 | 0.0026 |
| F13 | 56200001 | 56300000 | 0.0033 | 0.0024 | 0.0027 |
| F13 | 56220001 | 56320000 | 0.0030 | 0.0022 | 0.0024 |
| F13 | 56240001 | 56340000 | 0.0030 | 0.0022 | 0.0024 |
| F13 | 56260001 | 56360000 | 0.0032 | 0.0024 | 0.0024 |
| F13 | 56280001 | 56380000 | 0.0026 | 0.0020 | 0.0020 |
| F13 | 56300001 | 56400000 | 0.0024 | 0.0019 | 0.0019 |
| F13 | 56320001 | 56420000 | 0.0028 | 0.0023 | 0.0023 |
| F13 | 56340001 | 56440000 | 0.0028 | 0.0023 | 0.0021 |
| F13 | 56360001 | 56460000 | 0.0025 | 0.0018 | 0.0016 |
| F13 | 56380001 | 56480000 | 0.0025 | 0.0018 | 0.0016 |
| F13 | 56400001 | 56500000 | 0.0029 | 0.0022 | 0.0019 |
| F13 | 56420001 | 56520000 | 0.0027 | 0.0018 | 0.0017 |
| F13 | 56440001 | 56540000 | 0.0023 | 0.0015 | 0.0015 |
| F13 | 56460001 | 56560000 | 0.0026 | 0.0020 | 0.0021 |
| F13 | 56480001 | 56580000 | 0.0027 | 0.0018 | 0.0020 |
| F13 | 56500001 | 56600000 | 0.0024 | 0.0015 | 0.0018 |
| F13 | 56520001 | 56620000 | 0.0023 | 0.0015 | 0.0015 |

| Chr | Start | End | both | A1 | A2 |
|---|---|---|---|---|---|
| F13 | 56540001 | 56640000 | 0.0035 | 0.0024 | 0.0024 |
| F13 | 56560001 | 56660000 | 0.0031 | 0.0021 | 0.0019 |
| F13 | 56580001 | 56680000 | 0.0031 | 0.0023 | 0.0022 |
| F13 | 56600001 | 56700000 | 0.0034 | 0.0025 | 0.0025 |
| F13 | 56620001 | 56720000 | 0.0034 | 0.0025 | 0.0025 |
| F13 | 56640001 | 56740000 | 0.0026 | 0.0018 | 0.0019 |
| F13 | 56660001 | 56760000 | 0.0028 | 0.0020 | 0.0022 |
| F13 | 56680001 | 56780000 | 0.0023 | 0.0015 | 0.0016 |
| F13 | 56700001 | 56800000 | 0.0026 | 0.0020 | 0.0018 |
| F13 | 56720001 | 56820000 | 0.0025 | 0.0019 | 0.0018 |
| F13 | 56740001 | 56840000 | 0.0026 | 0.0019 | 0.0017 |
| F13 | 56760001 | 56860000 | 0.0024 | 0.0015 | 0.0013 |
| F13 | 56780001 | 56880000 | 0.0038 | 0.0027 | 0.0026 |
| F13 | 56800001 | 56900000 | 0.0035 | 0.0022 | 0.0023 |
| F13 | 56820001 | 56920000 | 0.0035 | 0.0022 | 0.0022 |
| F13 | 56840001 | 56940000 | 0.0038 | 0.0024 | 0.0025 |
| F13 | 56860001 | 56960000 | 0.0040 | 0.0026 | 0.0027 |
| F13 | 56880001 | 56980000 | 0.0031 | 0.0017 | 0.0018 |
| F13 | 56900001 | 57000000 | 0.0029 | 0.0015 | 0.0016 |
| F13 | 56920001 | 57020000 | 0.0035 | 0.0022 | 0.0022 |
| F13 | 56940001 | 57040000 | 0.0028 | 0.0019 | 0.0018 |
| F13 | 56960001 | 57060000 | 0.0029 | 0.0021 | 0.0020 |
| F13 | 56980001 | 57080000 | 0.0035 | 0.0028 | 0.0029 |
| F13 | 57000001 | 57100000 | 0.0035 | 0.0028 | 0.0029 |
| F13 | 57020001 | 57120000 | 0.0036 | 0.0030 | 0.0032 |
| F13 | 57040001 | 57140000 | 0.0038 | 0.0032 | 0.0034 |
| F13 | 57060001 | 57160000 | 0.0033 | 0.0027 | 0.0029 |
| F13 | 57080001 | 57180000 | 0.0020 | 0.0017 | 0.0017 |
| F13 | 57100001 | 57200000 | 0.0041 | 0.0035 | 0.0036 |
| F13 | 57120001 | 57220000 | 0.0039 | 0.0032 | 0.0032 |
| F13 | 57140001 | 57240000 | 0.0045 | 0.0036 | 0.0036 |
| F13 | 57160001 | 57260000 | 0.0054 | 0.0043 | 0.0045 |
| F13 | 57180001 | 57280000 | 0.0063 | 0.0049 | 0.0052 |
| F13 | 57200001 | 57300000 | 0.0048 | 0.0034 | 0.0036 |
| F13 | 57220001 | 57320000 | 0.0045 | 0.0032 | 0.0036 |
| F13 | 57240001 | 57340000 | 0.0044 | 0.0033 | 0.0037 |
| F13 | 57260001 | 57360000 | 0.0037 | 0.0029 | 0.0030 |
| F13 | 57280001 | 57380000 | 0.0033 | 0.0025 | 0.0025 |
| F13 | 57300001 | 57400000 | 0.0034 | 0.0026 | 0.0025 |
| F13 | 57320001 | 57420000 | 0.0034 | 0.0024 | 0.0023 |
| F13 | 57340001 | 57440000 | 0.0030 | 0.0018 | 0.0017 |
| F13 | 57360001 | 57460000 | 0.0029 | 0.0016 | 0.0016 |
| F13 | 57380001 | 57480000 | 0.0026 | 0.0014 | 0.0014 |
| F13 | 57400001 | 57500000 | 0.0024 | 0.0014 | 0.0014 |
| F13 | 57420001 | 57520000 | 0.0027 | 0.0016 | 0.0016 |
| F13 | 57440001 | 57540000 | 0.0026 | 0.0016 | 0.0017 |
| F13 | 57460001 | 57560000 | 0.0028 | 0.0019 | 0.0019 |
| F13 | 57480001 | 57580000 | 0.0026 | 0.0019 | 0.0019 |
| F13 | 57500001 | 57600000 | 0.0024 | 0.0019 | 0.0019 |
| F13 | 57520001 | 57620000 | 0.0020 | 0.0016 | 0.0017 |
| F13 | 57540001 | 57640000 | 0.0019 | 0.0016 | 0.0017 |
| F13 | 57560001 | 57660000 | 0.0015 | 0.0012 | 0.0013 |
| F13 | 57580001 | 57680000 | 0.0017 | 0.0012 | 0.0013 |
| F13 | 57600001 | 57700000 | 0.0019 | 0.0012 | 0.0012 |
| F13 | 57620001 | 57720000 | 0.0025 | 0.0015 | 0.0015 |
| F13 | 57640001 | 57740000 | 0.0030 | 0.0014 | 0.0016 |
| F13 | 57660001 | 57760000 | 0.0043 | 0.0023 | 0.0027 |
| F13 | 57680001 | 57780000 | 0.0046 | 0.0027 | 0.0031 |

| Chr | Start | End | both | A1 | A2 |
|---|---|---|---|---|---|
| F13 | 57700001 | 57800000 | 0.0043 | 0.0026 | 0.0032 |
| F13 | 57720001 | 57820000 | 0.0048 | 0.0032 | 0.0038 |
| F13 | 57740001 | 57840000 | 0.0042 | 0.0032 | 0.0036 |
| F13 | 57760001 | 57860000 | 0.0036 | 0.0029 | 0.0031 |
| F13 | 57780001 | 57880000 | 0.0034 | 0.0027 | 0.0028 |
| F13 | 57800001 | 57900000 | 0.0035 | 0.0026 | 0.0027 |
| F13 | 57820001 | 57920000 | 0.0040 | 0.0032 | 0.0034 |
| F13 | 57840001 | 57940000 | 0.0041 | 0.0032 | 0.0035 |
| F13 | 57860001 | 57960000 | 0.0035 | 0.0026 | 0.0029 |
| F13 | 57880001 | 57980000 | 0.0037 | 0.0029 | 0.0031 |
| F13 | 57900001 | 58000000 | 0.0041 | 0.0033 | 0.0034 |
| F13 | 57920001 | 58020000 | 0.0024 | 0.0016 | 0.0015 |
| F13 | 57940001 | 58040000 | 0.0028 | 0.0018 | 0.0016 |
| F13 | 57960001 | 58060000 | 0.0037 | 0.0026 | 0.0024 |
| F13 | 57980001 | 58080000 | 0.0038 | 0.0024 | 0.0023 |
| F13 | 58000001 | 58100000 | 0.0034 | 0.0020 | 0.0020 |
| F13 | 58020001 | 58120000 | 0.0037 | 0.0023 | 0.0022 |
| F13 | 58040001 | 58140000 | 0.0047 | 0.0032 | 0.0031 |
| F13 | 58060001 | 58160000 | 0.0049 | 0.0036 | 0.0036 |
| F13 | 58080001 | 58180000 | 0.0049 | 0.0037 | 0.0036 |
| F13 | 58100001 | 58200000 | 0.0054 | 0.0042 | 0.0044 |
| F13 | 58120001 | 58220000 | 0.0056 | 0.0043 | 0.0047 |
| F13 | 58140001 | 58240000 | 0.0049 | 0.0039 | 0.0042 |
| F13 | 58160001 | 58260000 | 0.0046 | 0.0034 | 0.0036 |
| F13 | 58180001 | 58280000 | 0.0049 | 0.0039 | 0.0042 |
| F13 | 58200001 | 58300000 | 0.0042 | 0.0033 | 0.0035 |
| F13 | 58220001 | 58320000 | 0.0048 | 0.0041 | 0.0044 |
| F13 | 58240001 | 58340000 | 0.0048 | 0.0041 | 0.0043 |
| F13 | 58260001 | 58360000 | 0.0049 | 0.0043 | 0.0045 |
| F13 | 58280001 | 58380000 | 0.0041 | 0.0034 | 0.0036 |
| F13 | 58300001 | 58400000 | 0.0042 | 0.0036 | 0.0037 |
| F13 | 58320001 | 58420000 | 0.0038 | 0.0030 | 0.0031 |
| F13 | 58340001 | 58440000 | 0.0049 | 0.0036 | 0.0041 |
| F13 | 58360001 | 58460000 | 0.0048 | 0.0037 | 0.0043 |
| F13 | 58380001 | 58480000 | 0.0054 | 0.0042 | 0.0049 |
| F13 | 58400001 | 58500000 | 0.0058 | 0.0047 | 0.0054 |
| F13 | 58420001 | 58520000 | 0.0052 | 0.0042 | 0.0049 |
| F13 | 58440001 | 58540000 | 0.0036 | 0.0030 | 0.0034 |
| F13 | 58460001 | 58560000 | 0.0041 | 0.0035 | 0.0038 |
| F13 | 58480001 | 58580000 | 0.0044 | 0.0041 | 0.0041 |
| F13 | 58500001 | 58600000 | 0.0042 | 0.0039 | 0.0039 |
| F13 | 58520001 | 58620000 | 0.0044 | 0.0040 | 0.0040 |
| F13 | 58540001 | 58640000 | 0.0044 | 0.0041 | 0.0040 |
| F13 | 58560001 | 58660000 | 0.0033 | 0.0029 | 0.0029 |
| F13 | 58580001 | 58680000 | 0.0028 | 0.0023 | 0.0024 |
| F13 | 58600001 | 58700000 | 0.0030 | 0.0024 | 0.0026 |
| F13 | 58620001 | 58720000 | 0.0029 | 0.0024 | 0.0026 |
| F13 | 58640001 | 58740000 | 0.0037 | 0.0032 | 0.0033 |
| F13 | 58660001 | 58760000 | 0.0037 | 0.0034 | 0.0034 |
| F13 | 58680001 | 58780000 | 0.0037 | 0.0035 | 0.0036 |
| F13 | 58700001 | 58800000 | 0.0030 | 0.0029 | 0.0029 |
| F13 | 58720001 | 58820000 | 0.0032 | 0.0031 | 0.0030 |
| F13 | 58740001 | 58840000 | 0.0030 | 0.0028 | 0.0029 |
| F13 | 58760001 | 58860000 | 0.0042 | 0.0037 | 0.0040 |
| F13 | 58780001 | 58880000 | 0.0057 | 0.0050 | 0.0053 |
| F13 | 58800001 | 58900000 | 0.0062 | 0.0052 | 0.0054 |
| F13 | 58820001 | 58920000 | 0.0081 | 0.0068 | 0.0071 |
| F13 | 58840001 | 58940000 | 0.0088 | 0.0074 | 0.0076 |

| Chr | Start | End | both | A1 | A2 |
|---|---|---|---|---|---|
| F13 | 58860001 | 58960000 | 0.0072 | 0.0060 | 0.0060 |
| F13 | 58880001 | 58980000 | 0.0065 | 0.0053 | 0.0052 |
| F13 | 58900001 | 59000000 | 0.0066 | 0.0055 | 0.0055 |
| F13 | 58920001 | 59020000 | 0.0048 | 0.0035 | 0.0034 |
| F13 | 58940001 | 59040000 | 0.0036 | 0.0024 | 0.0023 |
| F13 | 58960001 | 59060000 | 0.0038 | 0.0027 | 0.0026 |
| F13 | 58980001 | 59080000 | 0.0036 | 0.0024 | 0.0025 |
| F13 | 59000001 | 59100000 | 0.0038 | 0.0027 | 0.0026 |
| F13 | 59020001 | 59120000 | 0.0045 | 0.0034 | 0.0037 |
| F13 | 59040001 | 59140000 | 0.0048 | 0.0037 | 0.0040 |
| F13 | 59060001 | 59160000 | 0.0045 | 0.0034 | 0.0038 |
| F13 | 59080001 | 59180000 | 0.0040 | 0.0032 | 0.0033 |
| F13 | 59100001 | 59200000 | 0.0038 | 0.0031 | 0.0031 |
| F13 | 59120001 | 59220000 | 0.0037 | 0.0030 | 0.0031 |
| F13 | 59140001 | 59240000 | 0.0032 | 0.0027 | 0.0026 |
| F13 | 59160001 | 59260000 | 0.0041 | 0.0034 | 0.0035 |
| F13 | 59180001 | 59280000 | 0.0040 | 0.0034 | 0.0036 |
| F13 | 59200001 | 59300000 | 0.0036 | 0.0031 | 0.0033 |
| F13 | 59220001 | 59320000 | 0.0033 | 0.0025 | 0.0028 |
| F13 | 59240001 | 59340000 | 0.0037 | 0.0030 | 0.0033 |
| F13 | 59260001 | 59360000 | 0.0032 | 0.0025 | 0.0027 |
| F13 | 59280001 | 59380000 | 0.0034 | 0.0025 | 0.0027 |
| F13 | 59300001 | 59400000 | 0.0037 | 0.0026 | 0.0029 |
| F13 | 59320001 | 59420000 | 0.0032 | 0.0022 | 0.0023 |
| F13 | 59340001 | 59440000 | 0.0035 | 0.0023 | 0.0025 |
| F13 | 59360001 | 59460000 | 0.0035 | 0.0022 | 0.0025 |
| F13 | 59380001 | 59480000 | 0.0039 | 0.0027 | 0.0028 |
| F13 | 59400001 | 59500000 | 0.0039 | 0.0027 | 0.0028 |
| F13 | 59420001 | 59520000 | 0.0050 | 0.0038 | 0.0040 |
| F13 | 59440001 | 59540000 | 0.0050 | 0.0040 | 0.0041 |
| F13 | 59460001 | 59560000 | 0.0054 | 0.0043 | 0.0046 |
| F13 | 59480001 | 59580000 | 0.0041 | 0.0033 | 0.0036 |
| F13 | 59500001 | 59600000 | 0.0039 | 0.0031 | 0.0035 |
| F13 | 59520001 | 59620000 | 0.0029 | 0.0023 | 0.0026 |
| F13 | 59540001 | 59640000 | 0.0027 | 0.0021 | 0.0024 |
| F13 | 59560001 | 59660000 | 0.0022 | 0.0017 | 0.0016 |
| F13 | 59580001 | 59680000 | 0.0024 | 0.0018 | 0.0017 |
| F13 | 59600001 | 59700000 | 0.0036 | 0.0029 | 0.0031 |
| F13 | 59620001 | 59720000 | 0.0040 | 0.0033 | 0.0034 |
| F13 | 59640001 | 59740000 | 0.0036 | 0.0031 | 0.0030 |
| F13 | 59660001 | 59760000 | 0.0037 | 0.0033 | 0.0032 |
| F13 | 59680001 | 59780000 | 0.0046 | 0.0042 | 0.0041 |
| F13 | 59700001 | 59800000 | 0.0042 | 0.0041 | 0.0036 |
| F13 | 59720001 | 59820000 | 0.0042 | 0.0039 | 0.0035 |
| F13 | 59740001 | 59840000 | 0.0039 | 0.0036 | 0.0033 |
| F13 | 59760001 | 59860000 | 0.0036 | 0.0032 | 0.0030 |
| F13 | 59780001 | 59880000 | 0.0031 | 0.0027 | 0.0024 |
| F13 | 59800001 | 59900000 | 0.0026 | 0.0020 | 0.0021 |
| F13 | 59820001 | 59920000 | 0.0019 | 0.0015 | 0.0016 |
| F13 | 59840001 | 59940000 | 0.0027 | 0.0022 | 0.0023 |
| F13 | 59860001 | 59960000 | 0.0032 | 0.0027 | 0.0027 |
| F13 | 59880001 | 59980000 | 0.0028 | 0.0023 | 0.0024 |
| F13 | 59900001 | 60000000 | 0.0027 | 0.0025 | 0.0023 |
| F13 | 59920001 | 60020000 | 0.0031 | 0.0027 | 0.0026 |
| F13 | 59940001 | 60040000 | 0.0024 | 0.0022 | 0.0019 |
| F13 | 59960001 | 60060000 | 0.0018 | 0.0016 | 0.0015 |
| F13 | 59980001 | 60080000 | 0.0020 | 0.0018 | 0.0017 |
| F13 | 60000001 | 60100000 | 0.0023 | 0.0020 | 0.0020 |

| Chr | Start | End | both | A1 | A2 |
|---|---|---|---|---|---|
| F13 | 60020001 | 60120000 | 0.0019 | 0.0018 | 0.0018 |
| F13 | 60040001 | 60140000 | 0.0017 | 0.0016 | 0.0016 |
| F13 | 60060001 | 60160000 | 0.0021 | 0.0019 | 0.0021 |
| F13 | 60080001 | 60180000 | 0.0026 | 0.0022 | 0.0025 |
| F13 | 60100001 | 60200000 | 0.0020 | 0.0017 | 0.0020 |
| F13 | 60120001 | 60220000 | 0.0024 | 0.0020 | 0.0023 |
| F13 | 60140001 | 60240000 | 0.0025 | 0.0019 | 0.0024 |
| F13 | 60160001 | 60260000 | 0.0026 | 0.0019 | 0.0023 |
| F13 | 60180001 | 60280000 | 0.0026 | 0.0023 | 0.0021 |
| F13 | 60200001 | 60300000 | 0.0026 | 0.0023 | 0.0020 |
| F13 | 60220001 | 60320000 | 0.0031 | 0.0028 | 0.0024 |
| F13 | 60240001 | 60340000 | 0.0044 | 0.0038 | 0.0035 |
| F13 | 60260001 | 60360000 | 0.0045 | 0.0040 | 0.0036 |
| F13 | 60280001 | 60380000 | 0.0055 | 0.0047 | 0.0048 |
| F13 | 60300001 | 60400000 | 0.0065 | 0.0056 | 0.0057 |
| F13 | 60320001 | 60420000 | 0.0071 | 0.0062 | 0.0063 |
| F13 | 60340001 | 60440000 | 0.0061 | 0.0057 | 0.0056 |
| F13 | 60360001 | 60460000 | 0.0061 | 0.0056 | 0.0056 |
| F13 | 60380001 | 60480000 | 0.0052 | 0.0045 | 0.0046 |
| F13 | 60400001 | 60500000 | 0.0049 | 0.0038 | 0.0042 |
| F13 | 60420001 | 60520000 | 0.0044 | 0.0030 | 0.0036 |
| F13 | 60440001 | 60540000 | 0.0043 | 0.0029 | 0.0034 |
| F13 | 60460001 | 60560000 | 0.0042 | 0.0028 | 0.0032 |
| F13 | 60480001 | 60580000 | 0.0038 | 0.0024 | 0.0028 |
| F13 | 60500001 | 60600000 | 0.0036 | 0.0024 | 0.0027 |
| F13 | 60520001 | 60620000 | 0.0043 | 0.0032 | 0.0035 |
| F13 | 60540001 | 60640000 | 0.0048 | 0.0034 | 0.0039 |
| F13 | 60560001 | 60660000 | 0.0044 | 0.0031 | 0.0036 |
| F13 | 60580001 | 60680000 | 0.0046 | 0.0034 | 0.0040 |
| F13 | 60600001 | 60700000 | 0.0050 | 0.0038 | 0.0041 |
| F13 | 60620001 | 60720000 | 0.0038 | 0.0026 | 0.0028 |
| F13 | 60640001 | 60740000 | 0.0041 | 0.0026 | 0.0029 |
| F13 | 60660001 | 60760000 | 0.0049 | 0.0032 | 0.0038 |
| F13 | 60680001 | 60780000 | 0.0046 | 0.0030 | 0.0034 |
| F13 | 60700001 | 60800000 | 0.0036 | 0.0021 | 0.0028 |
| F13 | 60720001 | 60820000 | 0.0037 | 0.0023 | 0.0030 |
| F13 | 60740001 | 60840000 | 0.0040 | 0.0029 | 0.0035 |
| F13 | 60760001 | 60860000 | 0.0035 | 0.0028 | 0.0032 |
| F13 | 60780001 | 60880000 | 0.0032 | 0.0026 | 0.0030 |
| F13 | 60800001 | 60900000 | 0.0037 | 0.0031 | 0.0034 |
| F13 | 60820001 | 60920000 | 0.0033 | 0.0029 | 0.0031 |
| F13 | 60840001 | 60940000 | 0.0026 | 0.0023 | 0.0023 |
| F13 | 60860001 | 60960000 | 0.0021 | 0.0018 | 0.0018 |
| F13 | 60880001 | 60980000 | 0.0021 | 0.0018 | 0.0018 |
| F13 | 60900001 | 61000000 | 0.0023 | 0.0020 | 0.0020 |
| F13 | 60920001 | 61020000 | 0.0033 | 0.0027 | 0.0029 |
| F13 | 60940001 | 61040000 | 0.0028 | 0.0024 | 0.0026 |
| F13 | 60960001 | 61060000 | 0.0030 | 0.0026 | 0.0028 |
| F13 | 60980001 | 61080000 | 0.0031 | 0.0028 | 0.0029 |
| F13 | 61000001 | 61100000 | 0.0024 | 0.0021 | 0.0022 |
| F13 | 61020001 | 61120000 | 0.0017 | 0.0016 | 0.0015 |
| F13 | 61040001 | 61140000 | 0.0015 | 0.0014 | 0.0014 |
| F13 | 61060001 | 61160000 | 0.0014 | 0.0012 | 0.0012 |
| F13 | 61080001 | 61180000 | 0.0026 | 0.0017 | 0.0024 |
| F13 | 61100001 | 61200000 | 0.0034 | 0.0021 | 0.0030 |
| F13 | 61120001 | 61220000 | 0.0033 | 0.0019 | 0.0028 |
| F13 | 61140001 | 61240000 | 0.0041 | 0.0024 | 0.0034 |
| F13 | 61160001 | 61260000 | 0.0043 | 0.0025 | 0.0036 |

| Chr | Start | End | both | A1 | A2 |
|---|---|---|---|---|---|
| F13 | 61180001 | 61280000 | 0.0040 | 0.0027 | 0.0033 |
| F13 | 61200001 | 61300000 | 0.0040 | 0.0029 | 0.0035 |
| F13 | 61220001 | 61320000 | 0.0046 | 0.0035 | 0.0040 |
| F13 | 61240001 | 61340000 | 0.0044 | 0.0034 | 0.0038 |
| F13 | 61260001 | 61360000 | 0.0050 | 0.0039 | 0.0042 |
| F13 | 61280001 | 61380000 | 0.0040 | 0.0032 | 0.0033 |
| F13 | 61300001 | 61400000 | 0.0036 | 0.0028 | 0.0030 |
| F13 | 61320001 | 61420000 | 0.0031 | 0.0024 | 0.0026 |
| F13 | 61340001 | 61440000 | 0.0029 | 0.0022 | 0.0024 |
| F13 | 61360001 | 61460000 | 0.0024 | 0.0019 | 0.0021 |
| F13 | 61380001 | 61480000 | 0.0033 | 0.0024 | 0.0027 |
| F13 | 61400001 | 61500000 | 0.0031 | 0.0021 | 0.0024 |
| F13 | 61420001 | 61520000 | 0.0033 | 0.0021 | 0.0023 |
| F13 | 61440001 | 61540000 | 0.0037 | 0.0024 | 0.0026 |
| F13 | 61460001 | 61560000 | 0.0047 | 0.0029 | 0.0034 |
| F13 | 61480001 | 61580000 | 0.0050 | 0.0033 | 0.0040 |
| F13 | 61500001 | 61600000 | 0.0053 | 0.0036 | 0.0042 |
| F13 | 61520001 | 61620000 | 0.0053 | 0.0038 | 0.0045 |
| F13 | 61540001 | 61640000 | 0.0052 | 0.0038 | 0.0044 |
| F13 | 61560001 | 61660000 | 0.0042 | 0.0034 | 0.0036 |
| F13 | 61580001 | 61680000 | 0.0034 | 0.0027 | 0.0028 |
| F13 | 61600001 | 61700000 | 0.0031 | 0.0024 | 0.0024 |
| F13 | 61620001 | 61720000 | 0.0036 | 0.0029 | 0.0027 |
| F13 | 61640001 | 61740000 | 0.0039 | 0.0032 | 0.0029 |
| F13 | 61660001 | 61760000 | 0.0037 | 0.0029 | 0.0028 |
| F13 | 61680001 | 61780000 | 0.0043 | 0.0035 | 0.0034 |
| F13 | 61700001 | 61800000 | 0.0050 | 0.0043 | 0.0044 |
| F13 | 61720001 | 61820000 | 0.0046 | 0.0039 | 0.0042 |
| F13 | 61740001 | 61840000 | 0.0037 | 0.0031 | 0.0034 |
| F13 | 61760001 | 61860000 | 0.0039 | 0.0032 | 0.0036 |
| F13 | 61780001 | 61880000 | 0.0033 | 0.0025 | 0.0028 |
| F13 | 61800001 | 61900000 | 0.0030 | 0.0022 | 0.0024 |
| F13 | 61820001 | 61920000 | 0.0028 | 0.0021 | 0.0023 |
| F13 | 61840001 | 61940000 | 0.0035 | 0.0027 | 0.0029 |
| F13 | 61860001 | 61960000 | 0.0037 | 0.0028 | 0.0029 |
| F13 | 61880001 | 61980000 | 0.0039 | 0.0030 | 0.0032 |
| F13 | 61900001 | 62000000 | 0.0038 | 0.0029 | 0.0032 |
| F13 | 61920001 | 62020000 | 0.0033 | 0.0023 | 0.0025 |
| F13 | 61940001 | 62040000 | 0.0029 | 0.0019 | 0.0020 |
| F13 | 61960001 | 62060000 | 0.0027 | 0.0016 | 0.0018 |
| F13 | 61980001 | 62080000 | 0.0024 | 0.0014 | 0.0016 |
| F13 | 62000001 | 62100000 | 0.0023 | 0.0012 | 0.0014 |
| F13 | 62020001 | 62120000 | 0.0030 | 0.0019 | 0.0021 |
| F13 | 62040001 | 62140000 | 0.0029 | 0.0020 | 0.0023 |
| F13 | 62060001 | 62160000 | 0.0035 | 0.0026 | 0.0029 |
| F13 | 62080001 | 62180000 | 0.0045 | 0.0035 | 0.0036 |
| F13 | 62100001 | 62200000 | 0.0046 | 0.0036 | 0.0037 |
| F13 | 62120001 | 62220000 | 0.0045 | 0.0036 | 0.0035 |
| F13 | 62140001 | 62240000 | 0.0046 | 0.0038 | 0.0037 |
| F13 | 62160001 | 62260000 | 0.0044 | 0.0037 | 0.0037 |
| F13 | 62180001 | 62280000 | 0.0039 | 0.0029 | 0.0031 |
| F13 | 62200001 | 62300000 | 0.0042 | 0.0032 | 0.0035 |
| F13 | 62220001 | 62320000 | 0.0038 | 0.0029 | 0.0032 |
| F13 | 62240001 | 62340000 | 0.0034 | 0.0025 | 0.0028 |
| F13 | 62260001 | 62360000 | 0.0028 | 0.0020 | 0.0022 |
| F13 | 62280001 | 62380000 | 0.0029 | 0.0025 | 0.0027 |
| F13 | 62300001 | 62400000 | 0.0019 | 0.0017 | 0.0018 |
| F13 | 62320001 | 62420000 | 0.0018 | 0.0016 | 0.0017 |

| Chr | Start | End | both | A1 | A2 |
|---|---|---|---|---|---|
| F13 | 62340001 | 62440000 | 0.0019 | 0.0017 | 0.0019 |
| F13 | 62360001 | 62460000 | 0.0044 | 0.0041 | 0.0043 |
| F13 | 62380001 | 62480000 | 0.0050 | 0.0047 | 0.0047 |
| F13 | 62400001 | 62500000 | 0.0051 | 0.0047 | 0.0047 |
| F13 | 62420001 | 62520000 | 0.0058 | 0.0052 | 0.0052 |
| F13 | 62440001 | 62540000 | 0.0062 | 0.0054 | 0.0054 |
| F13 | 62460001 | 62560000 | 0.0043 | 0.0037 | 0.0036 |
| F13 | 62480001 | 62580000 | 0.0039 | 0.0030 | 0.0031 |
| F13 | 62500001 | 62600000 | 0.0048 | 0.0039 | 0.0039 |
| F13 | 62520001 | 62620000 | 0.0046 | 0.0038 | 0.0040 |
| F13 | 62540001 | 62640000 | 0.0052 | 0.0041 | 0.0043 |
| F13 | 62560001 | 62660000 | 0.0049 | 0.0035 | 0.0038 |
| F13 | 62580001 | 62680000 | 0.0040 | 0.0029 | 0.0031 |
| F13 | 62600001 | 62700000 | 0.0032 | 0.0020 | 0.0023 |
| F13 | 62620001 | 62720000 | 0.0033 | 0.0021 | 0.0024 |
| F13 | 62640001 | 62740000 | 0.0038 | 0.0027 | 0.0031 |
| F13 | 62660001 | 62760000 | 0.0043 | 0.0033 | 0.0037 |
| F13 | 62680001 | 62780000 | 0.0060 | 0.0049 | 0.0054 |
| F13 | 62700001 | 62800000 | 0.0060 | 0.0049 | 0.0052 |
| F13 | 62720001 | 62820000 | 0.0060 | 0.0047 | 0.0050 |
| F13 | 62740001 | 62840000 | 0.0050 | 0.0039 | 0.0041 |
| F13 | 62760001 | 62860000 | 0.0056 | 0.0044 | 0.0046 |
| F13 | 62780001 | 62880000 | 0.0045 | 0.0033 | 0.0036 |
| F13 | 62800001 | 62900000 | 0.0043 | 0.0032 | 0.0035 |
| F13 | 62820001 | 62920000 | 0.0039 | 0.0030 | 0.0032 |
| F13 | 62840001 | 62940000 | 0.0042 | 0.0034 | 0.0035 |
| F13 | 62860001 | 62960000 | 0.0050 | 0.0045 | 0.0044 |
| F13 | 62880001 | 62980000 | 0.0046 | 0.0042 | 0.0039 |
| F13 | 62900001 | 63000000 | 0.0052 | 0.0048 | 0.0047 |
| F13 | 62920001 | 63020000 | 0.0061 | 0.0057 | 0.0056 |
| F13 | 62940001 | 63040000 | 0.0056 | 0.0050 | 0.0050 |
| F13 | 62960001 | 63060000 | 0.0039 | 0.0031 | 0.0032 |
| F13 | 62980001 | 63080000 | 0.0049 | 0.0040 | 0.0042 |
| F13 | 63000001 | 63100000 | 0.0048 | 0.0038 | 0.0040 |
| F13 | 63020001 | 63120000 | 0.0039 | 0.0027 | 0.0030 |
| F13 | 63040001 | 63140000 | 0.0045 | 0.0032 | 0.0034 |
| F13 | 63060001 | 63160000 | 0.0046 | 0.0031 | 0.0035 |
| F13 | 63080001 | 63180000 | 0.0039 | 0.0025 | 0.0028 |
| F13 | 63100001 | 63200000 | 0.0039 | 0.0024 | 0.0027 |
| F13 | 63120001 | 63220000 | 0.0042 | 0.0027 | 0.0031 |
| F13 | 63140001 | 63240000 | 0.0042 | 0.0029 | 0.0032 |
| F13 | 63160001 | 63260000 | 0.0051 | 0.0039 | 0.0042 |
| F13 | 63180001 | 63280000 | 0.0053 | 0.0040 | 0.0045 |
| F13 | 63200001 | 63300000 | 0.0056 | 0.0045 | 0.0051 |
| F13 | 63220001 | 63320000 | 0.0058 | 0.0045 | 0.0051 |
| F13 | 63240001 | 63340000 | 0.0056 | 0.0041 | 0.0047 |
| F13 | 63260001 | 63360000 | 0.0056 | 0.0038 | 0.0043 |
| F13 | 63280001 | 63380000 | 0.0050 | 0.0032 | 0.0036 |
| F13 | 63300001 | 63400000 | 0.0042 | 0.0026 | 0.0029 |
| F13 | 63320001 | 63420000 | 0.0039 | 0.0025 | 0.0027 |
| F13 | 63340001 | 63440000 | 0.0033 | 0.0020 | 0.0023 |
| F13 | 63360001 | 63460000 | 0.0026 | 0.0012 | 0.0013 |
| F13 | 63380001 | 63480000 | 0.0035 | 0.0021 | 0.0022 |
| F13 | 63400001 | 63500000 | 0.0037 | 0.0021 | 0.0022 |
| F13 | 63420001 | 63520000 | 0.0042 | 0.0024 | 0.0026 |
| F13 | 63440001 | 63540000 | 0.0043 | 0.0027 | 0.0029 |
| F13 | 63460001 | 63560000 | 0.0039 | 0.0028 | 0.0032 |
| F13 | 63480001 | 63580000 | 0.0028 | 0.0019 | 0.0022 |

| Chr | Start | End | both | A1 | A2 |
|---|---|---|---|---|---|
| F13 | 63500001 | 63600000 | 0.0030 | 0.0021 | 0.0024 |
| F13 | 63520001 | 63620000 | 0.0032 | 0.0024 | 0.0026 |
| F13 | 63540001 | 63640000 | 0.0030 | 0.0021 | 0.0024 |
| F13 | 63560001 | 63660000 | 0.0027 | 0.0019 | 0.0021 |
| F13 | 63580001 | 63680000 | 0.0026 | 0.0019 | 0.0020 |
| F13 | 63600001 | 63700000 | 0.0020 | 0.0014 | 0.0016 |
| F13 | 63620001 | 63720000 | 0.0011 | 0.0007 | 0.0008 |
| F13 | 63640001 | 63740000 | 0.0010 | 0.0008 | 0.0009 |
| F13 | 63660001 | 63760000 | 0.0009 | 0.0007 | 0.0007 |
| F13 | 63680001 | 63780000 | 0.0011 | 0.0009 | 0.0010 |
| F13 | 63700001 | 63800000 | 0.0015 | 0.0012 | 0.0013 |
| F13 | 63720001 | 63820000 | 0.0027 | 0.0022 | 0.0025 |
| F13 | 63740001 | 63840000 | 0.0030 | 0.0022 | 0.0026 |
| F13 | 63760001 | 63860000 | 0.0036 | 0.0026 | 0.0031 |
| F13 | 63780001 | 63880000 | 0.0036 | 0.0026 | 0.0031 |
| F13 | 63800001 | 63900000 | 0.0033 | 0.0022 | 0.0027 |
| F13 | 63820001 | 63920000 | 0.0026 | 0.0014 | 0.0019 |
| F13 | 63840001 | 63940000 | 0.0026 | 0.0017 | 0.0021 |
| F13 | 63860001 | 63960000 | 0.0026 | 0.0015 | 0.0020 |
| F13 | 63880001 | 63980000 | 0.0033 | 0.0022 | 0.0026 |
| F13 | 63900001 | 64000000 | 0.0037 | 0.0025 | 0.0028 |
| F13 | 63920001 | 64020000 | 0.0043 | 0.0034 | 0.0035 |
| F13 | 63940001 | 64040000 | 0.0051 | 0.0038 | 0.0039 |
| F13 | 63960001 | 64060000 | 0.0052 | 0.0040 | 0.0042 |
| F13 | 63980001 | 64080000 | 0.0050 | 0.0038 | 0.0040 |
| F13 | 64000001 | 64100000 | 0.0046 | 0.0035 | 0.0040 |
| F13 | 64020001 | 64120000 | 0.0034 | 0.0024 | 0.0028 |
| F13 | 64040001 | 64140000 | 0.0026 | 0.0021 | 0.0024 |
| F13 | 64060001 | 64160000 | 0.0023 | 0.0019 | 0.0022 |
| F13 | 64080001 | 64180000 | 0.0015 | 0.0012 | 0.0014 |
| F13 | 64100001 | 64200000 | 0.0015 | 0.0013 | 0.0014 |
| F13 | 64120001 | 64220000 | 0.0019 | 0.0017 | 0.0019 |
| F13 | 64140001 | 64240000 | 0.0022 | 0.0018 | 0.0022 |
| F13 | 64160001 | 64260000 | 0.0024 | 0.0018 | 0.0021 |
| F13 | 64180001 | 64280000 | 0.0031 | 0.0024 | 0.0028 |
| F13 | 64200001 | 64300000 | 0.0035 | 0.0026 | 0.0030 |
| F13 | 64220001 | 64320000 | 0.0033 | 0.0024 | 0.0027 |
| F13 | 64240001 | 64340000 | 0.0030 | 0.0022 | 0.0024 |
| F13 | 64260001 | 64360000 | 0.0031 | 0.0021 | 0.0024 |
| F13 | 64280001 | 64380000 | 0.0028 | 0.0017 | 0.0019 |
| F13 | 64300001 | 64400000 | 0.0028 | 0.0018 | 0.0022 |
| F13 | 64320001 | 64420000 | 0.0032 | 0.0020 | 0.0024 |
| F13 | 64340001 | 64440000 | 0.0038 | 0.0025 | 0.0031 |
| F13 | 64360001 | 64460000 | 0.0040 | 0.0029 | 0.0033 |
| F13 | 64380001 | 64480000 | 0.0048 | 0.0038 | 0.0041 |
| F13 | 64400001 | 64500000 | 0.0069 | 0.0055 | 0.0059 |
| F13 | 64420001 | 64520000 | 0.0073 | 0.0061 | 0.0064 |
| F13 | 64440001 | 64540000 | 0.0066 | 0.0054 | 0.0056 |
| F13 | 64460001 | 64560000 | 0.0066 | 0.0052 | 0.0055 |
| F13 | 64480001 | 64580000 | 0.0058 | 0.0043 | 0.0045 |
| F13 | 64500001 | 64600000 | 0.0042 | 0.0030 | 0.0030 |
| F13 | 64520001 | 64620000 | 0.0042 | 0.0027 | 0.0027 |
| F13 | 64540001 | 64640000 | 0.0046 | 0.0028 | 0.0030 |
| F13 | 64560001 | 64660000 | 0.0046 | 0.0028 | 0.0030 |
| F13 | 64580001 | 64680000 | 0.0049 | 0.0032 | 0.0035 |
| F13 | 64600001 | 64700000 | 0.0047 | 0.0029 | 0.0035 |
| F13 | 64620001 | 64720000 | 0.0036 | 0.0024 | 0.0028 |
| F13 | 64640001 | 64740000 | 0.0038 | 0.0030 | 0.0034 |

| Chr | Start | End | both | A1 | A2 |
|---|---|---|---|---|---|
| F13 | 64660001 | 64760000 | 0.0033 | 0.0028 | 0.0032 |
| F13 | 64680001 | 64780000 | 0.0031 | 0.0026 | 0.0031 |
| F13 | 64700001 | 64800000 | 0.0027 | 0.0021 | 0.0026 |
| F13 | 64720001 | 64820000 | 0.0027 | 0.0022 | 0.0026 |
| F13 | 64740001 | 64840000 | 0.0021 | 0.0017 | 0.0020 |
| F13 | 64760001 | 64860000 | 0.0023 | 0.0019 | 0.0022 |
| F13 | 64780001 | 64880000 | 0.0024 | 0.0020 | 0.0023 |
| F13 | 64800001 | 64900000 | 0.0034 | 0.0030 | 0.0034 |
| F13 | 64820001 | 64920000 | 0.0035 | 0.0030 | 0.0034 |
| F13 | 64840001 | 64940000 | 0.0033 | 0.0029 | 0.0032 |
| F13 | 64860001 | 64960000 | 0.0032 | 0.0027 | 0.0031 |
| F13 | 64880001 | 64980000 | 0.0032 | 0.0027 | 0.0031 |
| F13 | 64900001 | 65000000 | 0.0019 | 0.0016 | 0.0019 |
| F13 | 64920001 | 65020000 | 0.0018 | 0.0016 | 0.0018 |
| F13 | 64940001 | 65040000 | 0.0018 | 0.0014 | 0.0016 |
| F13 | 64960001 | 65060000 | 0.0017 | 0.0012 | 0.0015 |
| F13 | 64980001 | 65080000 | 0.0017 | 0.0010 | 0.0011 |
| F13 | 65000001 | 65100000 | 0.0016 | 0.0009 | 0.0009 |
| F13 | 65020001 | 65120000 | 0.0016 | 0.0009 | 0.0009 |
| F13 | 65040001 | 65140000 | 0.0017 | 0.0013 | 0.0011 |
| F13 | 65060001 | 65160000 | 0.0020 | 0.0016 | 0.0015 |
| F13 | 65080001 | 65180000 | 0.0018 | 0.0016 | 0.0016 |
| F13 | 65100001 | 65200000 | 0.0020 | 0.0019 | 0.0019 |
| F13 | 65120001 | 65220000 | 0.0025 | 0.0020 | 0.0022 |
| F13 | 65140001 | 65240000 | 0.0026 | 0.0020 | 0.0023 |
| F13 | 65160001 | 65260000 | 0.0026 | 0.0018 | 0.0023 |
| F13 | 65180001 | 65280000 | 0.0026 | 0.0019 | 0.0022 |
| F13 | 65200001 | 65300000 | 0.0028 | 0.0020 | 0.0023 |
| F13 | 65220001 | 65320000 | 0.0033 | 0.0025 | 0.0028 |
| F13 | 65240001 | 65340000 | 0.0036 | 0.0027 | 0.0031 |
| F13 | 65260001 | 65360000 | 0.0038 | 0.0030 | 0.0032 |
| F13 | 65280001 | 65380000 | 0.0046 | 0.0037 | 0.0040 |
| F13 | 65300001 | 65400000 | 0.0048 | 0.0038 | 0.0041 |
| F13 | 65320001 | 65420000 | 0.0048 | 0.0039 | 0.0041 |
| F13 | 65340001 | 65440000 | 0.0042 | 0.0033 | 0.0035 |
| F13 | 65360001 | 65460000 | 0.0040 | 0.0031 | 0.0031 |
| F13 | 65380001 | 65480000 | 0.0038 | 0.0031 | 0.0031 |
| F13 | 65400001 | 65500000 | 0.0038 | 0.0032 | 0.0033 |
| F13 | 65420001 | 65520000 | 0.0042 | 0.0036 | 0.0036 |
| F13 | 65440001 | 65540000 | 0.0044 | 0.0039 | 0.0039 |
| F13 | 65460001 | 65560000 | 0.0043 | 0.0038 | 0.0039 |
| F13 | 65480001 | 65580000 | 0.0032 | 0.0028 | 0.0029 |
| F13 | 65500001 | 65600000 | 0.0027 | 0.0023 | 0.0024 |
| F13 | 65520001 | 65620000 | 0.0014 | 0.0010 | 0.0012 |
| F13 | 65540001 | 65640000 | 0.0015 | 0.0011 | 0.0013 |
| F13 | 65560001 | 65660000 | 0.0022 | 0.0016 | 0.0018 |
| F13 | 65580001 | 65680000 | 0.0032 | 0.0025 | 0.0029 |
| F13 | 65600001 | 65700000 | 0.0036 | 0.0025 | 0.0028 |
| F13 | 65620001 | 65720000 | 0.0039 | 0.0028 | 0.0032 |
| F13 | 65640001 | 65740000 | 0.0040 | 0.0028 | 0.0031 |
| F13 | 65660001 | 65760000 | 0.0034 | 0.0021 | 0.0023 |
| F13 | 65680001 | 65780000 | 0.0028 | 0.0013 | 0.0014 |
| F13 | 65700001 | 65800000 | 0.0024 | 0.0014 | 0.0015 |
| F13 | 65720001 | 65820000 | 0.0023 | 0.0013 | 0.0014 |
| F13 | 65740001 | 65840000 | 0.0025 | 0.0016 | 0.0017 |
| F13 | 65760001 | 65860000 | 0.0024 | 0.0016 | 0.0019 |
| F13 | 65780001 | 65880000 | 0.0033 | 0.0027 | 0.0031 |
| F13 | 65800001 | 65900000 | 0.0033 | 0.0027 | 0.0031 |

| Chr | Start | End | both | A1 | A2 |
|---|---|---|---|---|---|
| F13 | 65820001 | 65920000 | 0.0042 | 0.0035 | 0.0039 |
| F13 | 65840001 | 65940000 | 0.0035 | 0.0029 | 0.0032 |
| F13 | 65860001 | 65960000 | 0.0041 | 0.0036 | 0.0038 |
| F13 | 65880001 | 65980000 | 0.0031 | 0.0027 | 0.0029 |
| F13 | 65900001 | 66000000 | 0.0033 | 0.0029 | 0.0031 |
| F13 | 65920001 | 66020000 | 0.0024 | 0.0022 | 0.0023 |
| F13 | 65940001 | 66040000 | 0.0045 | 0.0040 | 0.0043 |
| F13 | 65960001 | 66060000 | 0.0057 | 0.0049 | 0.0053 |
| F13 | 65980001 | 66080000 | 0.0063 | 0.0054 | 0.0058 |
| F13 | 66000001 | 66100000 | 0.0061 | 0.0050 | 0.0054 |
| F13 | 66020001 | 66120000 | 0.0069 | 0.0056 | 0.0061 |
| F13 | 66040001 | 66140000 | 0.0053 | 0.0039 | 0.0043 |
| F13 | 66060001 | 66160000 | 0.0041 | 0.0028 | 0.0030 |
| F13 | 66080001 | 66180000 | 0.0038 | 0.0025 | 0.0027 |
| F13 | 66100001 | 66200000 | 0.0046 | 0.0031 | 0.0032 |
| F13 | 66120001 | 66220000 | 0.0047 | 0.0034 | 0.0031 |
| F13 | 66140001 | 66240000 | 0.0048 | 0.0038 | 0.0033 |
| F13 | 66160001 | 66260000 | 0.0042 | 0.0032 | 0.0029 |
| F13 | 66180001 | 66280000 | 0.0037 | 0.0029 | 0.0026 |
| F13 | 66200001 | 66300000 | 0.0035 | 0.0030 | 0.0028 |
| F13 | 66220001 | 66320000 | 0.0025 | 0.0020 | 0.0021 |
| F13 | 66240001 | 66340000 | 0.0019 | 0.0015 | 0.0017 |
| F13 | 66260001 | 66360000 | 0.0025 | 0.0021 | 0.0024 |
| F13 | 66280001 | 66380000 | 0.0031 | 0.0023 | 0.0026 |
| F13 | 66300001 | 66400000 | 0.0025 | 0.0016 | 0.0018 |
| F13 | 66320001 | 66420000 | 0.0031 | 0.0021 | 0.0023 |
| F13 | 66340001 | 66440000 | 0.0042 | 0.0030 | 0.0032 |
| F13 | 66360001 | 66460000 | 0.0035 | 0.0024 | 0.0024 |
| F13 | 66380001 | 66480000 | 0.0037 | 0.0028 | 0.0027 |
| F13 | 66400001 | 66500000 | 0.0037 | 0.0029 | 0.0029 |
| F13 | 66420001 | 66520000 | 0.0035 | 0.0024 | 0.0023 |
| F13 | 66440001 | 66540000 | 0.0029 | 0.0017 | 0.0017 |
| F13 | 66460001 | 66560000 | 0.0035 | 0.0022 | 0.0023 |
| F13 | 66480001 | 66580000 | 0.0036 | 0.0022 | 0.0024 |
| F13 | 66500001 | 66600000 | 0.0037 | 0.0024 | 0.0026 |
| F13 | 66520001 | 66620000 | 0.0032 | 0.0024 | 0.0026 |
| F13 | 66540001 | 66640000 | 0.0028 | 0.0021 | 0.0023 |
| F13 | 66560001 | 66660000 | 0.0020 | 0.0016 | 0.0017 |
| F13 | 66580001 | 66680000 | 0.0013 | 0.0010 | 0.0012 |
| F13 | 66600001 | 66700000 | 0.0014 | 0.0009 | 0.0009 |
| F13 | 66620001 | 66720000 | 0.0018 | 0.0011 | 0.0012 |
| F13 | 66640001 | 66740000 | 0.0021 | 0.0013 | 0.0012 |
| F13 | 66660001 | 66760000 | 0.0036 | 0.0024 | 0.0025 |
| F13 | 66680001 | 66780000 | 0.0047 | 0.0032 | 0.0032 |
| F13 | 66700001 | 66800000 | 0.0048 | 0.0033 | 0.0035 |
| F13 | 66720001 | 66820000 | 0.0049 | 0.0034 | 0.0035 |
| F13 | 66740001 | 66840000 | 0.0046 | 0.0032 | 0.0033 |
| F13 | 66760001 | 66860000 | 0.0034 | 0.0024 | 0.0025 |
| F13 | 66780001 | 66880000 | 0.0037 | 0.0028 | 0.0029 |
| F13 | 66800001 | 66900000 | 0.0035 | 0.0026 | 0.0027 |
| F13 | 66820001 | 66920000 | 0.0035 | 0.0029 | 0.0030 |
| F13 | 66840001 | 66940000 | 0.0049 | 0.0042 | 0.0040 |
| F13 | 66860001 | 66960000 | 0.0055 | 0.0047 | 0.0044 |
| F13 | 66880001 | 66980000 | 0.0054 | 0.0045 | 0.0042 |
| F13 | 66900001 | 67000000 | 0.0052 | 0.0041 | 0.0039 |
| F13 | 66920001 | 67020000 | 0.0056 | 0.0043 | 0.0040 |
| F13 | 66940001 | 67040000 | 0.0049 | 0.0036 | 0.0036 |
| F13 | 66960001 | 67060000 | 0.0048 | 0.0033 | 0.0035 |

| Chr | Start | End | both | A1 | A2 |
|---|---|---|---|---|---|
| F13 | 66980001 | 67080000 | 0.0041 | 0.0027 | 0.0030 |
| F13 | 67000001 | 67100000 | 0.0054 | 0.0037 | 0.0041 |
| F13 | 67020001 | 67120000 | 0.0047 | 0.0030 | 0.0034 |
| F13 | 67040001 | 67140000 | 0.0045 | 0.0028 | 0.0031 |
| F13 | 67060001 | 67160000 | 0.0044 | 0.0028 | 0.0030 |
| F13 | 67080001 | 67180000 | 0.0040 | 0.0025 | 0.0026 |
| F13 | 67100001 | 67200000 | 0.0026 | 0.0016 | 0.0017 |
| F13 | 67120001 | 67220000 | 0.0031 | 0.0022 | 0.0023 |
| F13 | 67140001 | 67240000 | 0.0029 | 0.0020 | 0.0020 |
| F13 | 67160001 | 67260000 | 0.0023 | 0.0014 | 0.0015 |
| F13 | 67180001 | 67280000 | 0.0025 | 0.0016 | 0.0018 |
| F13 | 67200001 | 67300000 | 0.0031 | 0.0022 | 0.0023 |
| F13 | 67220001 | 67320000 | 0.0026 | 0.0017 | 0.0019 |
| F13 | 67240001 | 67340000 | 0.0030 | 0.0023 | 0.0025 |
| F13 | 67260001 | 67360000 | 0.0031 | 0.0025 | 0.0028 |
| F13 | 67280001 | 67380000 | 0.0028 | 0.0021 | 0.0023 |
| F13 | 67300001 | 67400000 | 0.0020 | 0.0014 | 0.0016 |
| F13 | 67320001 | 67420000 | 0.0019 | 0.0014 | 0.0016 |
| F13 | 67340001 | 67440000 | 0.0012 | 0.0008 | 0.0008 |
| F13 | 67360001 | 67460000 | 0.0015 | 0.0010 | 0.0010 |
| F13 | 67380001 | 67480000 | 0.0014 | 0.0010 | 0.0010 |
| F13 | 67400001 | 67500000 | 0.0016 | 0.0010 | 0.0011 |
| F13 | 67420001 | 67520000 | 0.0023 | 0.0012 | 0.0015 |
| F13 | 67440001 | 67540000 | 0.0025 | 0.0014 | 0.0017 |
| F13 | 67460001 | 67560000 | 0.0027 | 0.0016 | 0.0019 |
| F13 | 67480001 | 67580000 | 0.0035 | 0.0021 | 0.0024 |
| F13 | 67500001 | 67600000 | 0.0037 | 0.0025 | 0.0026 |
| F13 | 67520001 | 67620000 | 0.0039 | 0.0025 | 0.0025 |
| F13 | 67540001 | 67640000 | 0.0049 | 0.0033 | 0.0032 |
| F13 | 67560001 | 67660000 | 0.0051 | 0.0035 | 0.0037 |
| F13 | 67580001 | 67680000 | 0.0047 | 0.0032 | 0.0033 |
| F13 | 67600001 | 67700000 | 0.0058 | 0.0038 | 0.0041 |
| F13 | 67620001 | 67720000 | 0.0050 | 0.0035 | 0.0038 |
| F13 | 67640001 | 67740000 | 0.0050 | 0.0034 | 0.0038 |
| F13 | 67660001 | 67760000 | 0.0049 | 0.0031 | 0.0035 |
| F13 | 67680001 | 67780000 | 0.0048 | 0.0030 | 0.0033 |
| F13 | 67700001 | 67800000 | 0.0039 | 0.0022 | 0.0025 |
| F13 | 67720001 | 67820000 | 0.0037 | 0.0020 | 0.0022 |
| F13 | 67740001 | 67840000 | 0.0042 | 0.0025 | 0.0027 |
| F13 | 67760001 | 67860000 | 0.0052 | 0.0037 | 0.0037 |
| F13 | 67780001 | 67880000 | 0.0056 | 0.0042 | 0.0042 |
| F13 | 67800001 | 67900000 | 0.0060 | 0.0048 | 0.0048 |
| F13 | 67820001 | 67920000 | 0.0065 | 0.0052 | 0.0053 |
| F13 | 67840001 | 67940000 | 0.0057 | 0.0044 | 0.0045 |
| F13 | 67860001 | 67960000 | 0.0048 | 0.0032 | 0.0035 |
| F13 | 67880001 | 67980000 | 0.0042 | 0.0028 | 0.0031 |
| F13 | 67900001 | 68000000 | 0.0053 | 0.0035 | 0.0041 |
| F13 | 67920001 | 68020000 | 0.0051 | 0.0036 | 0.0041 |
| F13 | 67940001 | 68040000 | 0.0043 | 0.0031 | 0.0036 |
| F13 | 67960001 | 68060000 | 0.0032 | 0.0024 | 0.0028 |
| F13 | 67980001 | 68080000 | 0.0041 | 0.0031 | 0.0035 |
| F13 | 68000001 | 68100000 | 0.0032 | 0.0027 | 0.0027 |
| F13 | 68020001 | 68120000 | 0.0036 | 0.0030 | 0.0031 |
| F13 | 68040001 | 68140000 | 0.0045 | 0.0036 | 0.0034 |
| F13 | 68060001 | 68160000 | 0.0046 | 0.0036 | 0.0034 |
| F13 | 68080001 | 68180000 | 0.0038 | 0.0030 | 0.0027 |
| F13 | 68100001 | 68200000 | 0.0039 | 0.0030 | 0.0030 |
| F13 | 68120001 | 68220000 | 0.0049 | 0.0040 | 0.0039 |

| Chr | Start | End | both | A1 | A2 |
|---|---|---|---|---|---|
| F13 | 68140001 | 68240000 | 0.0050 | 0.0041 | 0.0043 |
| F13 | 68160001 | 68260000 | 0.0055 | 0.0045 | 0.0047 |
| F13 | 68180001 | 68280000 | 0.0056 | 0.0043 | 0.0046 |
| F13 | 68200001 | 68300000 | 0.0044 | 0.0033 | 0.0035 |
| F13 | 68220001 | 68320000 | 0.0032 | 0.0020 | 0.0022 |
| F13 | 68240001 | 68340000 | 0.0026 | 0.0016 | 0.0016 |
| F13 | 68260001 | 68360000 | 0.0027 | 0.0017 | 0.0017 |
| F13 | 68280001 | 68380000 | 0.0030 | 0.0021 | 0.0020 |
| F13 | 68300001 | 68400000 | 0.0039 | 0.0028 | 0.0027 |
| F13 | 68320001 | 68420000 | 0.0045 | 0.0034 | 0.0035 |
| F13 | 68340001 | 68440000 | 0.0076 | 0.0063 | 0.0065 |
| F13 | 68360001 | 68460000 | 0.0086 | 0.0073 | 0.0076 |
| F13 | 68380001 | 68480000 | 0.0112 | 0.0096 | 0.0103 |
| F13 | 68400001 | 68500000 | 0.0102 | 0.0089 | 0.0096 |
| F13 | 68420001 | 68520000 | 0.0092 | 0.0080 | 0.0087 |
| F13 | 68440001 | 68540000 | 0.0070 | 0.0057 | 0.0066 |
| F13 | 68460001 | 68560000 | 0.0056 | 0.0043 | 0.0053 |
| F13 | 68480001 | 68580000 | 0.0034 | 0.0017 | 0.0030 |
| F13 | 68500001 | 68600000 | 0.0042 | 0.0025 | 0.0038 |
| F13 | 68520001 | 68620000 | 0.0041 | 0.0025 | 0.0037 |
| F13 | 68540001 | 68640000 | 0.0031 | 0.0019 | 0.0027 |
| F13 | 68560001 | 68660000 | 0.0039 | 0.0022 | 0.0035 |
| F13 | 68580001 | 68680000 | 0.0033 | 0.0022 | 0.0030 |
| F13 | 68600001 | 68700000 | 0.0032 | 0.0022 | 0.0029 |
| F13 | 68620001 | 68720000 | 0.0036 | 0.0026 | 0.0033 |
| F13 | 68640001 | 68740000 | 0.0053 | 0.0040 | 0.0048 |
| F13 | 68660001 | 68760000 | 0.0048 | 0.0041 | 0.0044 |
| F13 | 68680001 | 68780000 | 0.0045 | 0.0039 | 0.0042 |
| F13 | 68700001 | 68800000 | 0.0044 | 0.0036 | 0.0040 |
| F13 | 68720001 | 68820000 | 0.0044 | 0.0034 | 0.0038 |
| F13 | 68740001 | 68840000 | 0.0028 | 0.0019 | 0.0023 |
| F13 | 68760001 | 68860000 | 0.0021 | 0.0014 | 0.0017 |
| F13 | 68780001 | 68880000 | 0.0023 | 0.0015 | 0.0018 |
| F13 | 68800001 | 68900000 | 0.0025 | 0.0015 | 0.0019 |
| F13 | 68820001 | 68920000 | 0.0030 | 0.0021 | 0.0024 |
| F13 | 68840001 | 68940000 | 0.0029 | 0.0022 | 0.0025 |
| F13 | 68860001 | 68960000 | 0.0041 | 0.0033 | 0.0035 |
| F13 | 68880001 | 68980000 | 0.0046 | 0.0037 | 0.0040 |
| F13 | 68900001 | 69000000 | 0.0043 | 0.0036 | 0.0038 |
| F13 | 68920001 | 69020000 | 0.0036 | 0.0032 | 0.0033 |
| F13 | 68940001 | 69040000 | 0.0032 | 0.0027 | 0.0029 |
| F13 | 68960001 | 69060000 | 0.0020 | 0.0016 | 0.0018 |
| F13 | 68980001 | 69080000 | 0.0014 | 0.0011 | 0.0013 |
| F13 | 69000001 | 69100000 | 0.0014 | 0.0012 | 0.0014 |
| F13 | 69020001 | 69120000 | 0.0014 | 0.0011 | 0.0013 |
| F13 | 69040001 | 69140000 | 0.0020 | 0.0016 | 0.0020 |
| F13 | 69060001 | 69160000 | 0.0027 | 0.0022 | 0.0024 |
| F13 | 69080001 | 69180000 | 0.0033 | 0.0025 | 0.0028 |
| F13 | 69100001 | 69200000 | 0.0038 | 0.0029 | 0.0032 |
| F13 | 69120001 | 69220000 | 0.0044 | 0.0035 | 0.0038 |
| F13 | 69140001 | 69240000 | 0.0045 | 0.0035 | 0.0038 |
| F13 | 69160001 | 69260000 | 0.0041 | 0.0031 | 0.0036 |
| F13 | 69180001 | 69280000 | 0.0037 | 0.0030 | 0.0034 |
| F13 | 69200001 | 69300000 | 0.0028 | 0.0021 | 0.0024 |
| F13 | 69220001 | 69320000 | 0.0020 | 0.0014 | 0.0016 |
| F13 | 69240001 | 69340000 | 0.0013 | 0.0008 | 0.0009 |
| F13 | 69260001 | 69360000 | 0.0014 | 0.0010 | 0.0012 |
| F13 | 69280001 | 69380000 | 0.0015 | 0.0011 | 0.0014 |

| Chr | Start | End | both | A1 | A2 |
|---|---|---|---|---|---|
| F13 | 69300001 | 69400000 | 0.0021 | 0.0015 | 0.0019 |
| F13 | 69320001 | 69420000 | 0.0026 | 0.0018 | 0.0021 |
| F13 | 69340001 | 69440000 | 0.0030 | 0.0020 | 0.0023 |
| F13 | 69360001 | 69460000 | 0.0032 | 0.0026 | 0.0025 |
| F13 | 69380001 | 69480000 | 0.0026 | 0.0021 | 0.0019 |
| F13 | 69400001 | 69500000 | 0.0026 | 0.0029 | 0.0018 |
| F13 | 69420001 | 69520000 | 0.0022 | 0.0027 | 0.0017 |
| F13 | 69440001 | 69540000 | 0.0021 | 0.0027 | 0.0016 |
| F13 | 69460001 | 69560000 | 0.0015 | 0.0020 | 0.0012 |
| F13 | 69480001 | 69580000 | 0.0017 | 0.0023 | 0.0014 |
| F13 | 69500001 | 69600000 | 0.0020 | 0.0023 | 0.0017 |
| F13 | 69520001 | 69620000 | 0.0021 | 0.0024 | 0.0018 |
| F13 | 69540001 | 69640000 | 0.0021 | 0.0024 | 0.0018 |
| F13 | 69560001 | 69660000 | 0.0021 | 0.0024 | 0.0019 |
| F13 | 69580001 | 69680000 | 0.0021 | 0.0025 | 0.0018 |
| F13 | 69600001 | 69700000 | 0.0012 | 0.0014 | 0.0010 |
| F13 | 69620001 | 69720000 | 0.0016 | 0.0019 | 0.0014 |
| F13 | 69640001 | 69740000 | 0.0017 | 0.0019 | 0.0015 |
| F13 | 69660001 | 69760000 | 0.0015 | 0.0017 | 0.0014 |
| F13 | 69680001 | 69780000 | 0.0014 | 0.0015 | 0.0013 |
| F13 | 69700001 | 69800000 | 0.0015 | 0.0015 | 0.0015 |
| F13 | 69720001 | 69820000 | 0.0012 | 0.0011 | 0.0012 |
| F13 | 69740001 | 69840000 | 0.0009 | 0.0008 | 0.0009 |
| F13 | 69760001 | 69860000 | 0.0012 | 0.0010 | 0.0012 |
| F13 | 69780001 | 69880000 | 0.0011 | 0.0010 | 0.0011 |
| F13 | 69800001 | 69900000 | 0.0009 | 0.0008 | 0.0009 |
| F13 | 69820001 | 69920000 | 0.0012 | 0.0012 | 0.0012 |
| F13 | 69840001 | 69940000 | 0.0021 | 0.0019 | 0.0019 |
| F13 | 69860001 | 69960000 | 0.0020 | 0.0018 | 0.0016 |
| F13 | 69880001 | 69980000 | 0.0022 | 0.0019 | 0.0017 |
| F13 | 69900001 | 70000000 | 0.0026 | 0.0020 | 0.0019 |
| F13 | 69920001 | 70020000 | 0.0027 | 0.0018 | 0.0016 |
| F13 | 69940001 | 70040000 | 0.0025 | 0.0015 | 0.0015 |
| F13 | 69960001 | 70060000 | 0.0027 | 0.0017 | 0.0017 |
| F13 | 69980001 | 70080000 | 0.0034 | 0.0024 | 0.0023 |
| F13 | 70000001 | 70100000 | 0.0034 | 0.0025 | 0.0025 |
| F13 | 70020001 | 70120000 | 0.0035 | 0.0026 | 0.0025 |
| F13 | 70040001 | 70140000 | 0.0035 | 0.0023 | 0.0022 |
| F13 | 70060001 | 70160000 | 0.0033 | 0.0022 | 0.0022 |
| F13 | 70080001 | 70180000 | 0.0026 | 0.0015 | 0.0017 |
| F13 | 70100001 | 70200000 | 0.0024 | 0.0013 | 0.0014 |
| F13 | 70120001 | 70220000 | 0.0026 | 0.0017 | 0.0018 |
| F13 | 70140001 | 70240000 | 0.0023 | 0.0017 | 0.0017 |
| F13 | 70160001 | 70260000 | 0.0029 | 0.0021 | 0.0022 |
| F13 | 70180001 | 70280000 | 0.0042 | 0.0033 | 0.0034 |
| F13 | 70200001 | 70300000 | 0.0053 | 0.0042 | 0.0044 |
| F13 | 70220001 | 70320000 | 0.0046 | 0.0036 | 0.0038 |
| F13 | 70240001 | 70340000 | 0.0052 | 0.0041 | 0.0044 |
| F13 | 70260001 | 70360000 | 0.0053 | 0.0041 | 0.0045 |
| F13 | 70280001 | 70380000 | 0.0043 | 0.0031 | 0.0036 |
| F13 | 70300001 | 70400000 | 0.0040 | 0.0029 | 0.0034 |
| F13 | 70320001 | 70420000 | 0.0042 | 0.0030 | 0.0034 |
| F13 | 70340001 | 70440000 | 0.0040 | 0.0028 | 0.0031 |
| F13 | 70360001 | 70460000 | 0.0038 | 0.0028 | 0.0030 |
| F13 | 70380001 | 70480000 | 0.0040 | 0.0031 | 0.0031 |
| F13 | 70400001 | 70500000 | 0.0045 | 0.0032 | 0.0036 |
| F13 | 70420001 | 70520000 | 0.0044 | 0.0031 | 0.0037 |
| F13 | 70440001 | 70540000 | 0.0039 | 0.0027 | 0.0033 |

| Chr | Start | End | both | A1 | A2 |
|---|---|---|---|---|---|
| F13 | 70460001 | 70560000 | 0.0038 | 0.0026 | 0.0034 |
| F13 | 70480001 | 70580000 | 0.0036 | 0.0025 | 0.0032 |
| F13 | 70500001 | 70600000 | 0.0029 | 0.0020 | 0.0025 |
| F13 | 70520001 | 70620000 | 0.0036 | 0.0025 | 0.0031 |
| F13 | 70540001 | 70640000 | 0.0037 | 0.0026 | 0.0032 |
| F13 | 70560001 | 70660000 | 0.0037 | 0.0027 | 0.0030 |
| F13 | 70580001 | 70680000 | 0.0037 | 0.0027 | 0.0030 |
| F13 | 70600001 | 70700000 | 0.0049 | 0.0039 | 0.0042 |
| F13 | 70620001 | 70720000 | 0.0039 | 0.0031 | 0.0033 |
| F13 | 70640001 | 70740000 | 0.0037 | 0.0031 | 0.0032 |
| F13 | 70660001 | 70760000 | 0.0035 | 0.0030 | 0.0032 |
| F13 | 70680001 | 70780000 | 0.0028 | 0.0024 | 0.0026 |
| F13 | 70700001 | 70800000 | 0.0009 | 0.0007 | 0.0009 |
| F13 | 70720001 | 70820000 | 0.0010 | 0.0008 | 0.0010 |
| F13 | 70740001 | 70840000 | 0.0010 | 0.0008 | 0.0010 |
| F13 | 70760001 | 70860000 | 0.0012 | 0.0010 | 0.0012 |
| F13 | 70780001 | 70880000 | 0.0014 | 0.0012 | 0.0014 |
| F13 | 70800001 | 70900000 | 0.0016 | 0.0012 | 0.0015 |
| F13 | 70820001 | 70920000 | 0.0018 | 0.0015 | 0.0018 |
| F13 | 70840001 | 70940000 | 0.0016 | 0.0013 | 0.0016 |
| F13 | 70860001 | 70960000 | 0.0012 | 0.0010 | 0.0011 |
| F13 | 70880001 | 70980000 | 0.0011 | 0.0008 | 0.0010 |
| F13 | 70900001 | 71000000 | 0.0012 | 0.0009 | 0.0011 |
| F13 | 70920001 | 71020000 | 0.0015 | 0.0011 | 0.0014 |
| F13 | 70940001 | 71040000 | 0.0019 | 0.0015 | 0.0018 |
| F13 | 70960001 | 71060000 | 0.0016 | 0.0013 | 0.0016 |
| F13 | 70980001 | 71080000 | 0.0019 | 0.0016 | 0.0019 |
| F13 | 71000001 | 71100000 | 0.0016 | 0.0013 | 0.0016 |
| F13 | 71020001 | 71120000 | 0.0012 | 0.0010 | 0.0011 |
| F13 | 71040001 | 71140000 | 0.0012 | 0.0009 | 0.0011 |
| F13 | 71060001 | 71160000 | 0.0015 | 0.0012 | 0.0015 |
| F13 | 71080001 | 71180000 | 0.0015 | 0.0012 | 0.0014 |
| F13 | 71100001 | 71200000 | 0.0017 | 0.0014 | 0.0016 |
| F13 | 71120001 | 71220000 | 0.0026 | 0.0023 | 0.0026 |
| F13 | 71140001 | 71240000 | 0.0029 | 0.0025 | 0.0028 |
| F13 | 71160001 | 71260000 | 0.0029 | 0.0025 | 0.0029 |
| F13 | 71180001 | 71280000 | 0.0028 | 0.0025 | 0.0028 |
| F13 | 71200001 | 71300000 | 0.0031 | 0.0026 | 0.0030 |
| F13 | 71220001 | 71320000 | 0.0020 | 0.0017 | 0.0020 |
| F13 | 71240001 | 71340000 | 0.0018 | 0.0016 | 0.0018 |
| F13 | 71260001 | 71360000 | 0.0023 | 0.0020 | 0.0022 |
| F13 | 71280001 | 71380000 | 0.0028 | 0.0022 | 0.0025 |
| F13 | 71300001 | 71400000 | 0.0031 | 0.0023 | 0.0025 |
| F13 | 71320001 | 71420000 | 0.0038 | 0.0030 | 0.0032 |
| F13 | 71340001 | 71440000 | 0.0038 | 0.0029 | 0.0031 |
| F13 | 71360001 | 71460000 | 0.0030 | 0.0022 | 0.0024 |
| F13 | 71380001 | 71480000 | 0.0026 | 0.0020 | 0.0021 |
| F13 | 71400001 | 71500000 | 0.0023 | 0.0018 | 0.0018 |
| F13 | 71420001 | 71520000 | 0.0023 | 0.0017 | 0.0017 |
| F13 | 71440001 | 71540000 | 0.0026 | 0.0019 | 0.0019 |
| F13 | 71460001 | 71560000 | 0.0038 | 0.0028 | 0.0029 |
| F13 | 71480001 | 71580000 | 0.0041 | 0.0030 | 0.0031 |
| F13 | 71500001 | 71600000 | 0.0044 | 0.0030 | 0.0033 |
| F13 | 71520001 | 71620000 | 0.0047 | 0.0032 | 0.0035 |
| F13 | 71540001 | 71640000 | 0.0044 | 0.0029 | 0.0033 |
| F13 | 71560001 | 71660000 | 0.0037 | 0.0023 | 0.0025 |
| F13 | 71580001 | 71680000 | 0.0035 | 0.0020 | 0.0022 |
| F13 | 71600001 | 71700000 | 0.0035 | 0.0022 | 0.0023 |

| Chr | Start | End | both | A1 | A2 |
|---|---|---|---|---|---|
| F13 | 71620001 | 71720000 | 0.0027 | 0.0014 | 0.0016 |
| F13 | 71640001 | 71740000 | 0.0043 | 0.0027 | 0.0031 |
| F13 | 71660001 | 71760000 | 0.0043 | 0.0026 | 0.0030 |
| F13 | 71680001 | 71780000 | 0.0042 | 0.0026 | 0.0030 |
| F13 | 71700001 | 71800000 | 0.0050 | 0.0034 | 0.0037 |
| F13 | 71720001 | 71820000 | 0.0048 | 0.0034 | 0.0037 |
| F13 | 71740001 | 71840000 | 0.0036 | 0.0026 | 0.0029 |
| F13 | 71760001 | 71860000 | 0.0039 | 0.0029 | 0.0035 |
| F13 | 71780001 | 71880000 | 0.0038 | 0.0030 | 0.0036 |
| F13 | 71800001 | 71900000 | 0.0030 | 0.0022 | 0.0028 |
| F13 | 71820001 | 71920000 | 0.0036 | 0.0023 | 0.0030 |
| F13 | 71840001 | 71940000 | 0.0040 | 0.0027 | 0.0033 |
| F13 | 71860001 | 71960000 | 0.0036 | 0.0024 | 0.0027 |
| F13 | 71880001 | 71980000 | 0.0048 | 0.0034 | 0.0039 |
| F13 | 71900001 | 72000000 | 0.0051 | 0.0036 | 0.0042 |
| F13 | 71920001 | 72020000 | 0.0057 | 0.0042 | 0.0049 |
| F13 | 71940001 | 72040000 | 0.0049 | 0.0036 | 0.0043 |
| F13 | 71960001 | 72060000 | 0.0048 | 0.0037 | 0.0044 |
| F13 | 71980001 | 72080000 | 0.0040 | 0.0029 | 0.0034 |
| F13 | 72000001 | 72100000 | 0.0033 | 0.0025 | 0.0028 |
| F13 | 72020001 | 72120000 | 0.0028 | 0.0020 | 0.0022 |
| F13 | 72040001 | 72140000 | 0.0038 | 0.0026 | 0.0029 |
| F13 | 72060001 | 72160000 | 0.0040 | 0.0027 | 0.0029 |
| F13 | 72080001 | 72180000 | 0.0043 | 0.0030 | 0.0032 |
| F13 | 72100001 | 72200000 | 0.0047 | 0.0032 | 0.0035 |
| F13 | 72120001 | 72220000 | 0.0041 | 0.0031 | 0.0033 |
| F13 | 72140001 | 72240000 | 0.0047 | 0.0038 | 0.0041 |
| F13 | 72160001 | 72260000 | 0.0043 | 0.0035 | 0.0040 |
| F13 | 72180001 | 72280000 | 0.0042 | 0.0037 | 0.0038 |
| F13 | 72200001 | 72300000 | 0.0039 | 0.0034 | 0.0034 |
| F13 | 72220001 | 72320000 | 0.0045 | 0.0039 | 0.0040 |
| F13 | 72240001 | 72340000 | 0.0031 | 0.0028 | 0.0025 |
| F13 | 72260001 | 72360000 | 0.0035 | 0.0029 | 0.0028 |
| F13 | 72280001 | 72380000 | 0.0030 | 0.0024 | 0.0025 |
| F13 | 72300001 | 72400000 | 0.0029 | 0.0024 | 0.0026 |
| F13 | 72320001 | 72420000 | 0.0022 | 0.0017 | 0.0019 |
| F13 | 72340001 | 72440000 | 0.0025 | 0.0020 | 0.0023 |
| F13 | 72360001 | 72460000 | 0.0019 | 0.0016 | 0.0019 |
| F13 | 72380001 | 72480000 | 0.0017 | 0.0015 | 0.0017 |
| F13 | 72400001 | 72500000 | 0.0017 | 0.0015 | 0.0016 |
| F13 | 72420001 | 72520000 | 0.0025 | 0.0021 | 0.0025 |
| F13 | 72440001 | 72540000 | 0.0027 | 0.0022 | 0.0027 |
| F13 | 72460001 | 72560000 | 0.0028 | 0.0022 | 0.0027 |
| F13 | 72480001 | 72580000 | 0.0035 | 0.0029 | 0.0033 |
| F13 | 72500001 | 72600000 | 0.0036 | 0.0030 | 0.0034 |
| F13 | 72520001 | 72620000 | 0.0032 | 0.0024 | 0.0026 |
| F13 | 72540001 | 72640000 | 0.0026 | 0.0017 | 0.0017 |
| F13 | 72560001 | 72660000 | 0.0026 | 0.0019 | 0.0018 |
| F13 | 72580001 | 72680000 | 0.0018 | 0.0010 | 0.0010 |
| F13 | 72600001 | 72700000 | 0.0022 | 0.0011 | 0.0012 |
| F13 | 72620001 | 72720000 | 0.0023 | 0.0012 | 0.0015 |
| F13 | 72640001 | 72740000 | 0.0018 | 0.0011 | 0.0012 |
| F13 | 72660001 | 72760000 | 0.0019 | 0.0011 | 0.0013 |
| F13 | 72680001 | 72780000 | 0.0025 | 0.0014 | 0.0017 |
| F13 | 72700001 | 72800000 | 0.0024 | 0.0016 | 0.0019 |
| F13 | 72720001 | 72820000 | 0.0025 | 0.0017 | 0.0020 |
| F13 | 72740001 | 72840000 | 0.0029 | 0.0021 | 0.0024 |
| F13 | 72760001 | 72860000 | 0.0030 | 0.0020 | 0.0023 |

| Chr | Start | End | both | A1 | A2 |
|---|---|---|---|---|---|
| F13 | 72780001 | 72880000 | 0.0029 | 0.0018 | 0.0021 |
| F13 | 72800001 | 72900000 | 0.0027 | 0.0016 | 0.0018 |
| F13 | 72820001 | 72920000 | 0.0022 | 0.0012 | 0.0012 |
| F13 | 72840001 | 72940000 | 0.0026 | 0.0014 | 0.0015 |
| F13 | 72860001 | 72960000 | 0.0031 | 0.0020 | 0.0020 |
| F13 | 72880001 | 72980000 | 0.0037 | 0.0028 | 0.0027 |
| F13 | 72900001 | 73000000 | 0.0040 | 0.0031 | 0.0031 |
| F13 | 72920001 | 73020000 | 0.0047 | 0.0039 | 0.0040 |
| F13 | 72940001 | 73040000 | 0.0044 | 0.0037 | 0.0038 |
| F13 | 72960001 | 73060000 | 0.0043 | 0.0037 | 0.0039 |
| F13 | 72980001 | 73080000 | 0.0039 | 0.0032 | 0.0035 |
| F13 | 73000001 | 73100000 | 0.0034 | 0.0028 | 0.0030 |
| F13 | 73020001 | 73120000 | 0.0037 | 0.0028 | 0.0030 |
| F13 | 73040001 | 73140000 | 0.0035 | 0.0025 | 0.0027 |
| F13 | 73060001 | 73160000 | 0.0033 | 0.0022 | 0.0024 |
| F13 | 73080001 | 73180000 | 0.0036 | 0.0023 | 0.0026 |
| F13 | 73100001 | 73200000 | 0.0042 | 0.0029 | 0.0033 |
| F13 | 73120001 | 73220000 | 0.0035 | 0.0025 | 0.0028 |
| F13 | 73140001 | 73240000 | 0.0038 | 0.0028 | 0.0032 |
| F13 | 73160001 | 73260000 | 0.0041 | 0.0030 | 0.0035 |
| F13 | 73180001 | 73280000 | 0.0040 | 0.0033 | 0.0036 |
| F13 | 73200001 | 73300000 | 0.0034 | 0.0027 | 0.0030 |
| F13 | 73220001 | 73320000 | 0.0036 | 0.0027 | 0.0031 |
| F13 | 73240001 | 73340000 | 0.0037 | 0.0027 | 0.0031 |
| F13 | 73260001 | 73360000 | 0.0028 | 0.0021 | 0.0023 |
| F13 | 73280001 | 73380000 | 0.0030 | 0.0020 | 0.0024 |
| F13 | 73300001 | 73400000 | 0.0041 | 0.0030 | 0.0034 |
| F13 | 73320001 | 73420000 | 0.0043 | 0.0031 | 0.0036 |
| F13 | 73340001 | 73440000 | 0.0040 | 0.0029 | 0.0033 |
| F13 | 73360001 | 73460000 | 0.0042 | 0.0031 | 0.0035 |
| F13 | 73380001 | 73480000 | 0.0037 | 0.0026 | 0.0029 |
| F13 | 73400001 | 73500000 | 0.0032 | 0.0019 | 0.0023 |
| F13 | 73420001 | 73520000 | 0.0035 | 0.0023 | 0.0026 |
| F13 | 73440001 | 73540000 | 0.0052 | 0.0038 | 0.0042 |
| F13 | 73460001 | 73560000 | 0.0056 | 0.0041 | 0.0046 |
| F13 | 73480001 | 73580000 | 0.0063 | 0.0047 | 0.0053 |
| F13 | 73500001 | 73600000 | 0.0059 | 0.0046 | 0.0052 |
| F13 | 73520001 | 73620000 | 0.0050 | 0.0039 | 0.0045 |
| F13 | 73540001 | 73640000 | 0.0035 | 0.0027 | 0.0032 |
| F13 | 73560001 | 73660000 | 0.0038 | 0.0029 | 0.0036 |
| F13 | 73580001 | 73680000 | 0.0035 | 0.0026 | 0.0032 |
| F13 | 73600001 | 73700000 | 0.0031 | 0.0021 | 0.0026 |
| F13 | 73620001 | 73720000 | 0.0034 | 0.0024 | 0.0028 |
| F13 | 73640001 | 73740000 | 0.0040 | 0.0030 | 0.0033 |
| F13 | 73660001 | 73760000 | 0.0044 | 0.0031 | 0.0033 |
| F13 | 73680001 | 73780000 | 0.0045 | 0.0034 | 0.0035 |
| F13 | 73700001 | 73800000 | 0.0049 | 0.0036 | 0.0036 |
| F13 | 73720001 | 73820000 | 0.0047 | 0.0034 | 0.0034 |
| F13 | 73740001 | 73840000 | 0.0037 | 0.0025 | 0.0025 |
| F13 | 73760001 | 73860000 | 0.0027 | 0.0016 | 0.0016 |
| F13 | 73780001 | 73880000 | 0.0022 | 0.0011 | 0.0011 |
| F13 | 73800001 | 73900000 | 0.0017 | 0.0008 | 0.0008 |
| F13 | 73820001 | 73920000 | 0.0017 | 0.0010 | 0.0009 |
| F13 | 73840001 | 73940000 | 0.0018 | 0.0012 | 0.0011 |
| F13 | 73860001 | 73960000 | 0.0024 | 0.0020 | 0.0018 |
| F13 | 73880001 | 73980000 | 0.0029 | 0.0024 | 0.0025 |
| F13 | 73900001 | 74000000 | 0.0034 | 0.0029 | 0.0030 |
| F13 | 73920001 | 74020000 | 0.0035 | 0.0028 | 0.0031 |

| Chr | Start | End | both | A1 | A2 |
|---|---|---|---|---|---|
| F13 | 73940001 | 74040000 | 0.0036 | 0.0028 | 0.0033 |
| F13 | 73960001 | 74060000 | 0.0037 | 0.0028 | 0.0034 |
| F13 | 73980001 | 74080000 | 0.0044 | 0.0031 | 0.0037 |
| F13 | 74000001 | 74100000 | 0.0042 | 0.0029 | 0.0034 |
| F13 | 74020001 | 74120000 | 0.0049 | 0.0038 | 0.0042 |
| F13 | 74040001 | 74140000 | 0.0057 | 0.0043 | 0.0048 |
| F13 | 74060001 | 74160000 | 0.0057 | 0.0041 | 0.0048 |
| F13 | 74080001 | 74180000 | 0.0052 | 0.0038 | 0.0041 |
| F13 | 74100001 | 74200000 | 0.0054 | 0.0040 | 0.0043 |
| F13 | 74120001 | 74220000 | 0.0046 | 0.0031 | 0.0035 |
| F13 | 74140001 | 74240000 | 0.0041 | 0.0026 | 0.0030 |
| F13 | 74160001 | 74260000 | 0.0036 | 0.0024 | 0.0026 |
| F13 | 74180001 | 74280000 | 0.0031 | 0.0020 | 0.0024 |
| F13 | 74200001 | 74300000 | 0.0029 | 0.0019 | 0.0023 |
| F13 | 74220001 | 74320000 | 0.0032 | 0.0022 | 0.0026 |
| F13 | 74240001 | 74340000 | 0.0036 | 0.0030 | 0.0030 |
| F13 | 74260001 | 74360000 | 0.0032 | 0.0026 | 0.0027 |
| F13 | 74280001 | 74380000 | 0.0031 | 0.0023 | 0.0025 |
| F13 | 74300001 | 74400000 | 0.0035 | 0.0027 | 0.0028 |
| F13 | 74320001 | 74420000 | 0.0027 | 0.0022 | 0.0023 |
| F13 | 74340001 | 74440000 | 0.0016 | 0.0010 | 0.0013 |
| F13 | 74360001 | 74460000 | 0.0020 | 0.0014 | 0.0017 |
| F13 | 74380001 | 74480000 | 0.0021 | 0.0016 | 0.0020 |
| F13 | 74400001 | 74500000 | 0.0019 | 0.0015 | 0.0018 |
| F13 | 74420001 | 74520000 | 0.0024 | 0.0019 | 0.0023 |
| F13 | 74440001 | 74540000 | 0.0026 | 0.0020 | 0.0024 |
| F13 | 74460001 | 74560000 | 0.0028 | 0.0019 | 0.0023 |
| F13 | 74480001 | 74580000 | 0.0033 | 0.0022 | 0.0026 |
| F13 | 74500001 | 74600000 | 0.0028 | 0.0017 | 0.0021 |
| F13 | 74520001 | 74620000 | 0.0034 | 0.0018 | 0.0025 |
| F13 | 74540001 | 74640000 | 0.0040 | 0.0023 | 0.0030 |
| F13 | 74560001 | 74660000 | 0.0036 | 0.0021 | 0.0028 |
| F13 | 74580001 | 74680000 | 0.0038 | 0.0022 | 0.0032 |
| F13 | 74600001 | 74700000 | 0.0040 | 0.0024 | 0.0035 |
| F13 | 74620001 | 74720000 | 0.0033 | 0.0023 | 0.0029 |
| F13 | 74640001 | 74740000 | 0.0031 | 0.0019 | 0.0027 |
| F13 | 74660001 | 74760000 | 0.0031 | 0.0019 | 0.0026 |
| F13 | 74680001 | 74780000 | 0.0030 | 0.0020 | 0.0024 |
| F13 | 74700001 | 74800000 | 0.0036 | 0.0024 | 0.0029 |
| F13 | 74720001 | 74820000 | 0.0031 | 0.0019 | 0.0025 |
| F13 | 74740001 | 74840000 | 0.0025 | 0.0017 | 0.0020 |
| F13 | 74760001 | 74860000 | 0.0040 | 0.0029 | 0.0033 |
| F13 | 74780001 | 74880000 | 0.0034 | 0.0023 | 0.0028 |
| F13 | 74800001 | 74900000 | 0.0032 | 0.0020 | 0.0025 |
| F13 | 74820001 | 74920000 | 0.0036 | 0.0022 | 0.0027 |
| F13 | 74840001 | 74940000 | 0.0040 | 0.0025 | 0.0030 |
| F13 | 74860001 | 74960000 | 0.0031 | 0.0018 | 0.0023 |
| F13 | 74880001 | 74980000 | 0.0032 | 0.0019 | 0.0024 |
| F13 | 74900001 | 75000000 | 0.0035 | 0.0024 | 0.0029 |
| F13 | 74920001 | 75020000 | 0.0039 | 0.0028 | 0.0034 |
| F13 | 74940001 | 75040000 | 0.0042 | 0.0029 | 0.0037 |
| F13 | 74960001 | 75060000 | 0.0036 | 0.0025 | 0.0033 |
| F13 | 74980001 | 75080000 | 0.0039 | 0.0028 | 0.0036 |
| F13 | 75000001 | 75100000 | 0.0033 | 0.0023 | 0.0030 |
| F13 | 75020001 | 75120000 | 0.0026 | 0.0017 | 0.0022 |
| F13 | 75040001 | 75140000 | 0.0026 | 0.0016 | 0.0020 |
| F13 | 75060001 | 75160000 | 0.0024 | 0.0015 | 0.0019 |
| F13 | 75080001 | 75180000 | 0.0023 | 0.0010 | 0.0014 |

| Chr | Start | End | both | A1 | A2 |
|---|---|---|---|---|---|
| F13 | 75100001 | 75200000 | 0.0026 | 0.0009 | 0.0014 |
| F13 | 75120001 | 75220000 | 0.0035 | 0.0014 | 0.0021 |
| F13 | 75140001 | 75240000 | 0.0038 | 0.0019 | 0.0024 |
| F13 | 75160001 | 75260000 | 0.0044 | 0.0024 | 0.0030 |
| F13 | 75180001 | 75280000 | 0.0045 | 0.0030 | 0.0035 |
| F13 | 75200001 | 75300000 | 0.0043 | 0.0029 | 0.0034 |
| F13 | 75220001 | 75320000 | 0.0043 | 0.0029 | 0.0033 |
| F13 | 75240001 | 75340000 | 0.0042 | 0.0029 | 0.0033 |
| F13 | 75260001 | 75360000 | 0.0036 | 0.0023 | 0.0027 |
| F13 | 75280001 | 75380000 | 0.0033 | 0.0018 | 0.0023 |
| F13 | 75300001 | 75400000 | 0.0037 | 0.0024 | 0.0029 |
| F13 | 75320001 | 75420000 | 0.0029 | 0.0018 | 0.0024 |
| F13 | 75340001 | 75440000 | 0.0033 | 0.0020 | 0.0025 |
| F13 | 75360001 | 75460000 | 0.0045 | 0.0033 | 0.0039 |
| F13 | 75380001 | 75480000 | 0.0047 | 0.0039 | 0.0043 |
| F13 | 75400001 | 75500000 | 0.0044 | 0.0034 | 0.0039 |
| F13 | 75420001 | 75520000 | 0.0044 | 0.0036 | 0.0040 |
| F13 | 75440001 | 75540000 | 0.0036 | 0.0027 | 0.0032 |
| F13 | 75460001 | 75560000 | 0.0030 | 0.0022 | 0.0026 |
| F13 | 75480001 | 75580000 | 0.0026 | 0.0017 | 0.0022 |
| F13 | 75500001 | 75600000 | 0.0029 | 0.0020 | 0.0023 |
| F13 | 75520001 | 75620000 | 0.0043 | 0.0034 | 0.0034 |
| F13 | 75540001 | 75640000 | 0.0047 | 0.0037 | 0.0037 |
| F13 | 75560001 | 75660000 | 0.0044 | 0.0032 | 0.0031 |
| F13 | 75580001 | 75680000 | 0.0051 | 0.0035 | 0.0033 |
| F13 | 75600001 | 75700000 | 0.0053 | 0.0036 | 0.0036 |
| F13 | 75620001 | 75720000 | 0.0046 | 0.0028 | 0.0030 |
| F13 | 75640001 | 75740000 | 0.0045 | 0.0028 | 0.0030 |
| F13 | 75660001 | 75760000 | 0.0046 | 0.0032 | 0.0035 |
| F13 | 75680001 | 75780000 | 0.0043 | 0.0030 | 0.0034 |
| F13 | 75700001 | 75800000 | 0.0039 | 0.0027 | 0.0030 |
| F13 | 75720001 | 75820000 | 0.0032 | 0.0019 | 0.0023 |
| F13 | 75740001 | 75840000 | 0.0036 | 0.0025 | 0.0028 |
| F13 | 75760001 | 75860000 | 0.0034 | 0.0022 | 0.0024 |
| F13 | 75780001 | 75880000 | 0.0036 | 0.0022 | 0.0024 |
| F13 | 75800001 | 75900000 | 0.0033 | 0.0020 | 0.0022 |
| F13 | 75820001 | 75920000 | 0.0034 | 0.0020 | 0.0022 |
| F13 | 75840001 | 75940000 | 0.0026 | 0.0010 | 0.0012 |
| F13 | 75860001 | 75960000 | 0.0030 | 0.0015 | 0.0017 |
| F13 | 75880001 | 75980000 | 0.0024 | 0.0013 | 0.0014 |
| F13 | 75900001 | 76000000 | 0.0026 | 0.0017 | 0.0018 |
| F13 | 75920001 | 76020000 | 0.0030 | 0.0019 | 0.0020 |
| F13 | 75940001 | 76040000 | 0.0036 | 0.0028 | 0.0027 |
| F13 | 75960001 | 76060000 | 0.0036 | 0.0028 | 0.0026 |
| F13 | 75980001 | 76080000 | 0.0038 | 0.0026 | 0.0025 |
| F13 | 76000001 | 76100000 | 0.0037 | 0.0024 | 0.0023 |
| F13 | 76020001 | 76120000 | 0.0035 | 0.0025 | 0.0023 |
| F13 | 76040001 | 76140000 | 0.0031 | 0.0018 | 0.0018 |
| F13 | 76060001 | 76160000 | 0.0029 | 0.0014 | 0.0017 |
| F13 | 76080001 | 76180000 | 0.0031 | 0.0015 | 0.0017 |
| F13 | 76100001 | 76200000 | 0.0027 | 0.0012 | 0.0014 |
| F13 | 76120001 | 76220000 | 0.0026 | 0.0010 | 0.0012 |
| F13 | 76140001 | 76240000 | 0.0027 | 0.0010 | 0.0012 |
| F13 | 76160001 | 76260000 | 0.0025 | 0.0010 | 0.0010 |
| F13 | 76180001 | 76280000 | 0.0026 | 0.0014 | 0.0014 |
| F13 | 76200001 | 76300000 | 0.0026 | 0.0014 | 0.0014 |
| F13 | 76220001 | 76320000 | 0.0032 | 0.0022 | 0.0021 |
| F13 | 76240001 | 76340000 | 0.0030 | 0.0022 | 0.0021 |

| Chr | Start | End | both | A1 | A2 |
|---|---|---|---|---|---|
| F13 | 76260001 | 76360000 | 0.0031 | 0.0023 | 0.0024 |
| F13 | 76280001 | 76380000 | 0.0026 | 0.0019 | 0.0020 |
| F13 | 76300001 | 76400000 | 0.0031 | 0.0024 | 0.0026 |
| F13 | 76320001 | 76420000 | 0.0028 | 0.0019 | 0.0024 |
| F13 | 76340001 | 76440000 | 0.0033 | 0.0024 | 0.0029 |
| F13 | 76360001 | 76460000 | 0.0033 | 0.0018 | 0.0028 |
| F13 | 76380001 | 76480000 | 0.0040 | 0.0019 | 0.0034 |
| F13 | 76400001 | 76500000 | 0.0038 | 0.0015 | 0.0032 |
| F13 | 76420001 | 76520000 | 0.0039 | 0.0015 | 0.0031 |
| F13 | 76440001 | 76540000 | 0.0040 | 0.0017 | 0.0032 |
| F13 | 76460001 | 76560000 | 0.0036 | 0.0019 | 0.0030 |
| F13 | 76480001 | 76580000 | 0.0030 | 0.0019 | 0.0024 |
| F13 | 76500001 | 76600000 | 0.0031 | 0.0022 | 0.0025 |
| F13 | 76520001 | 76620000 | 0.0032 | 0.0023 | 0.0026 |
| F13 | 76540001 | 76640000 | 0.0028 | 0.0018 | 0.0021 |
| F13 | 76560001 | 76660000 | 0.0026 | 0.0016 | 0.0018 |
| F13 | 76580001 | 76680000 | 0.0032 | 0.0021 | 0.0023 |
| F13 | 76600001 | 76700000 | 0.0031 | 0.0018 | 0.0019 |
| F13 | 76620001 | 76720000 | 0.0027 | 0.0015 | 0.0016 |
| F13 | 76640001 | 76740000 | 0.0028 | 0.0017 | 0.0019 |
| F13 | 76660001 | 76760000 | 0.0029 | 0.0019 | 0.0022 |
| F13 | 76680001 | 76780000 | 0.0019 | 0.0010 | 0.0014 |
| F13 | 76700001 | 76800000 | 0.0021 | 0.0012 | 0.0016 |
| F13 | 76720001 | 76820000 | 0.0027 | 0.0016 | 0.0020 |
| F13 | 76740001 | 76840000 | 0.0040 | 0.0028 | 0.0030 |
| F13 | 76760001 | 76860000 | 0.0051 | 0.0036 | 0.0037 |
| F13 | 76780001 | 76880000 | 0.0054 | 0.0039 | 0.0040 |
| F13 | 76800001 | 76900000 | 0.0057 | 0.0042 | 0.0043 |
| F13 | 76820001 | 76920000 | 0.0059 | 0.0046 | 0.0046 |
| F13 | 76840001 | 76940000 | 0.0046 | 0.0034 | 0.0034 |
| F13 | 76860001 | 76960000 | 0.0036 | 0.0025 | 0.0026 |
| F13 | 76880001 | 76980000 | 0.0049 | 0.0035 | 0.0038 |
| F13 | 76900001 | 77000000 | 0.0048 | 0.0033 | 0.0036 |
| F13 | 76920001 | 77020000 | 0.0046 | 0.0034 | 0.0037 |
| F13 | 76940001 | 77040000 | 0.0049 | 0.0038 | 0.0041 |
| F13 | 76960001 | 77060000 | 0.0050 | 0.0038 | 0.0041 |
| F13 | 76980001 | 77080000 | 0.0041 | 0.0029 | 0.0032 |
| F13 | 77000001 | 77100000 | 0.0047 | 0.0032 | 0.0040 |
| F13 | 77020001 | 77120000 | 0.0036 | 0.0023 | 0.0029 |
| F13 | 77040001 | 77140000 | 0.0030 | 0.0018 | 0.0024 |
| F13 | 77060001 | 77160000 | 0.0031 | 0.0019 | 0.0025 |
| F13 | 77080001 | 77180000 | 0.0034 | 0.0024 | 0.0031 |
| F13 | 77100001 | 77200000 | 0.0029 | 0.0023 | 0.0028 |
| F13 | 77120001 | 77220000 | 0.0032 | 0.0026 | 0.0031 |
| F13 | 77140001 | 77240000 | 0.0029 | 0.0023 | 0.0028 |
| F13 | 77160001 | 77260000 | 0.0024 | 0.0020 | 0.0024 |
| F13 | 77180001 | 77280000 | 0.0013 | 0.0010 | 0.0013 |
| F13 | 77200001 | 77300000 | 0.0008 | 0.0007 | 0.0008 |
| F13 | 77220001 | 77320000 | 0.0004 | 0.0004 | 0.0004 |
| F13 | 77240001 | 77340000 | 0.0008 | 0.0006 | 0.0007 |
| F13 | 77260001 | 77360000 | 0.0013 | 0.0010 | 0.0013 |
| F13 | 77280001 | 77380000 | 0.0014 | 0.0011 | 0.0014 |
| F13 | 77300001 | 77400000 | 0.0017 | 0.0014 | 0.0017 |
| F13 | 77320001 | 77420000 | 0.0022 | 0.0019 | 0.0022 |
| F13 | 77340001 | 77440000 | 0.0026 | 0.0020 | 0.0026 |
| F13 | 77360001 | 77460000 | 0.0023 | 0.0017 | 0.0022 |
| F13 | 77380001 | 77480000 | 0.0033 | 0.0027 | 0.0032 |
| F13 | 77400001 | 77500000 | 0.0029 | 0.0024 | 0.0029 |

| Chr | Start | End | both | A1 | A2 |
|---|---|---|---|---|---|
| F13 | 77420001 | 77520000 | 0.0035 | 0.0029 | 0.0034 |
| F13 | 77440001 | 77540000 | 0.0035 | 0.0025 | 0.0033 |
| F13 | 77460001 | 77560000 | 0.0035 | 0.0025 | 0.0033 |
| F13 | 77480001 | 77580000 | 0.0027 | 0.0017 | 0.0025 |
| F13 | 77500001 | 77600000 | 0.0033 | 0.0023 | 0.0030 |
| F13 | 77520001 | 77620000 | 0.0029 | 0.0017 | 0.0025 |
| F13 | 77540001 | 77640000 | 0.0024 | 0.0017 | 0.0019 |
| F13 | 77560001 | 77660000 | 0.0025 | 0.0017 | 0.0018 |
| F13 | 77580001 | 77680000 | 0.0025 | 0.0015 | 0.0018 |
| F13 | 77600001 | 77700000 | 0.0016 | 0.0007 | 0.0009 |
| F13 | 77620001 | 77720000 | 0.0010 | 0.0003 | 0.0005 |
| F13 | 77640001 | 77740000 | 0.0011 | 0.0005 | 0.0006 |
| F13 | 77660001 | 77760000 | 0.0011 | 0.0006 | 0.0008 |
| F13 | 77680001 | 77780000 | 0.0017 | 0.0010 | 0.0011 |
| F13 | 77700001 | 77800000 | 0.0019 | 0.0012 | 0.0013 |
| F13 | 77720001 | 77820000 | 0.0025 | 0.0016 | 0.0018 |
| F13 | 77740001 | 77840000 | 0.0028 | 0.0018 | 0.0021 |
| F13 | 77760001 | 77860000 | 0.0028 | 0.0018 | 0.0021 |
| F13 | 77780001 | 77880000 | 0.0027 | 0.0020 | 0.0024 |
| F13 | 77800001 | 77900000 | 0.0034 | 0.0025 | 0.0030 |
| F13 | 77820001 | 77920000 | 0.0036 | 0.0027 | 0.0031 |
| F13 | 77840001 | 77940000 | 0.0037 | 0.0028 | 0.0032 |
| F13 | 77860001 | 77960000 | 0.0044 | 0.0036 | 0.0039 |
| F13 | 77880001 | 77980000 | 0.0042 | 0.0033 | 0.0036 |
| F13 | 77900001 | 78000000 | 0.0039 | 0.0029 | 0.0034 |
| F13 | 77920001 | 78020000 | 0.0037 | 0.0027 | 0.0032 |
| F13 | 77940001 | 78040000 | 0.0042 | 0.0031 | 0.0034 |
| F13 | 77960001 | 78060000 | 0.0042 | 0.0027 | 0.0032 |
| F13 | 77980001 | 78080000 | 0.0042 | 0.0026 | 0.0030 |
| F13 | 78000001 | 78100000 | 0.0038 | 0.0025 | 0.0027 |
| F13 | 78020001 | 78120000 | 0.0036 | 0.0022 | 0.0025 |
| F13 | 78040001 | 78140000 | 0.0026 | 0.0013 | 0.0016 |
| F13 | 78060001 | 78160000 | 0.0021 | 0.0012 | 0.0013 |
| F13 | 78080001 | 78180000 | 0.0019 | 0.0012 | 0.0013 |
| F13 | 78100001 | 78200000 | 0.0026 | 0.0017 | 0.0020 |
| F13 | 78120001 | 78220000 | 0.0027 | 0.0018 | 0.0023 |
| F13 | 78140001 | 78240000 | 0.0032 | 0.0023 | 0.0028 |
| F13 | 78160001 | 78260000 | 0.0035 | 0.0027 | 0.0032 |
| F13 | 78180001 | 78280000 | 0.0039 | 0.0031 | 0.0037 |
| F13 | 78200001 | 78300000 | 0.0032 | 0.0028 | 0.0030 |
| F13 | 78220001 | 78320000 | 0.0032 | 0.0030 | 0.0030 |
| F13 | 78240001 | 78340000 | 0.0029 | 0.0027 | 0.0027 |
| F13 | 78260001 | 78360000 | 0.0024 | 0.0021 | 0.0022 |
| F13 | 78280001 | 78380000 | 0.0020 | 0.0017 | 0.0018 |
| F13 | 78300001 | 78400000 | 0.0019 | 0.0014 | 0.0016 |
| F13 | 78320001 | 78420000 | 0.0016 | 0.0012 | 0.0014 |
| F13 | 78340001 | 78440000 | 0.0029 | 0.0024 | 0.0026 |
| F13 | 78360001 | 78460000 | 0.0037 | 0.0032 | 0.0034 |
| F13 | 78380001 | 78480000 | 0.0038 | 0.0034 | 0.0036 |
| F13 | 78400001 | 78500000 | 0.0047 | 0.0042 | 0.0046 |
| F13 | 78420001 | 78520000 | 0.0054 | 0.0048 | 0.0051 |
| F13 | 78440001 | 78540000 | 0.0047 | 0.0040 | 0.0043 |
| F13 | 78460001 | 78560000 | 0.0039 | 0.0033 | 0.0036 |
| F13 | 78480001 | 78580000 | 0.0037 | 0.0032 | 0.0033 |
| F13 | 78500001 | 78600000 | 0.0043 | 0.0037 | 0.0040 |
| F13 | 78520001 | 78620000 | 0.0053 | 0.0047 | 0.0051 |
| F13 | 78540001 | 78640000 | 0.0052 | 0.0048 | 0.0051 |
| F13 | 78560001 | 78660000 | 0.0056 | 0.0052 | 0.0055 |

| Chr | Start | End | both | A1 | A2 |
|---|---|---|---|---|---|
| F13 | 78580001 | 78680000 | 0.0059 | 0.0054 | 0.0058 |
| F13 | 78600001 | 78700000 | 0.0056 | 0.0051 | 0.0055 |
| F13 | 78620001 | 78720000 | 0.0039 | 0.0035 | 0.0039 |
| F13 | 78640001 | 78740000 | 0.0033 | 0.0030 | 0.0033 |
| F13 | 78660001 | 78760000 | 0.0029 | 0.0025 | 0.0029 |
| F13 | 78680001 | 78780000 | 0.0031 | 0.0027 | 0.0031 |
| F13 | 78700001 | 78800000 | 0.0025 | 0.0023 | 0.0025 |
| F13 | 78720001 | 78820000 | 0.0035 | 0.0029 | 0.0032 |
| F13 | 78740001 | 78840000 | 0.0037 | 0.0030 | 0.0033 |
| F13 | 78760001 | 78860000 | 0.0040 | 0.0031 | 0.0035 |
| F13 | 78780001 | 78880000 | 0.0044 | 0.0033 | 0.0037 |
| F13 | 78800001 | 78900000 | 0.0048 | 0.0036 | 0.0040 |
| F13 | 78820001 | 78920000 | 0.0038 | 0.0029 | 0.0032 |
| F13 | 78840001 | 78940000 | 0.0038 | 0.0030 | 0.0032 |
| F13 | 78860001 | 78960000 | 0.0034 | 0.0028 | 0.0029 |
| F13 | 78880001 | 78980000 | 0.0024 | 0.0020 | 0.0020 |
| F13 | 78900001 | 79000000 | 0.0020 | 0.0017 | 0.0017 |
| F13 | 78920001 | 79020000 | 0.0019 | 0.0016 | 0.0016 |
| F13 | 78940001 | 79040000 | 0.0014 | 0.0012 | 0.0013 |
| F13 | 78960001 | 79060000 | 0.0018 | 0.0015 | 0.0017 |
| F13 | 78980001 | 79080000 | 0.0022 | 0.0018 | 0.0021 |
| F13 | 79000001 | 79100000 | 0.0019 | 0.0016 | 0.0019 |
| F13 | 79020001 | 79120000 | 0.0019 | 0.0015 | 0.0019 |
| F13 | 79040001 | 79140000 | 0.0020 | 0.0017 | 0.0020 |
| F13 | 79060001 | 79160000 | 0.0018 | 0.0016 | 0.0018 |
| F13 | 79080001 | 79180000 | 0.0014 | 0.0012 | 0.0014 |
| F13 | 79100001 | 79200000 | 0.0007 | 0.0007 | 0.0007 |
| F13 | 79120001 | 79220000 | 0.0006 | 0.0006 | 0.0006 |
| F13 | 79140001 | 79240000 | 0.0006 | 0.0005 | 0.0005 |
| F13 | 79160001 | 79260000 | 0.0010 | 0.0008 | 0.0009 |
| F13 | 79180001 | 79280000 | 0.0010 | 0.0009 | 0.0010 |
| F13 | 79200001 | 79300000 | 0.0014 | 0.0013 | 0.0013 |
| F13 | 79220001 | 79320000 | 0.0013 | 0.0012 | 0.0013 |
| F13 | 79240001 | 79340000 | 0.0024 | 0.0022 | 0.0024 |
| F13 | 79260001 | 79360000 | 0.0022 | 0.0019 | 0.0020 |
| F13 | 79280001 | 79380000 | 0.0025 | 0.0021 | 0.0022 |
| F13 | 79300001 | 79400000 | 0.0023 | 0.0018 | 0.0021 |
| F13 | 79320001 | 79420000 | 0.0028 | 0.0022 | 0.0026 |
| F13 | 79340001 | 79440000 | 0.0021 | 0.0014 | 0.0017 |
| F13 | 79360001 | 79460000 | 0.0022 | 0.0016 | 0.0019 |
| F13 | 79380001 | 79480000 | 0.0021 | 0.0016 | 0.0018 |
| F13 | 79400001 | 79500000 | 0.0024 | 0.0016 | 0.0019 |
| F13 | 79420001 | 79520000 | 0.0027 | 0.0016 | 0.0018 |
| F13 | 79440001 | 79540000 | 0.0028 | 0.0019 | 0.0021 |
| F13 | 79460001 | 79560000 | 0.0030 | 0.0019 | 0.0020 |
| F13 | 79480001 | 79580000 | 0.0030 | 0.0018 | 0.0019 |
| F13 | 79500001 | 79600000 | 0.0032 | 0.0021 | 0.0021 |
| F13 | 79520001 | 79620000 | 0.0035 | 0.0025 | 0.0025 |
| F13 | 79540001 | 79640000 | 0.0031 | 0.0021 | 0.0021 |
| F13 | 79560001 | 79660000 | 0.0026 | 0.0018 | 0.0018 |
| F13 | 79580001 | 79680000 | 0.0028 | 0.0017 | 0.0018 |
| F13 | 79600001 | 79700000 | 0.0026 | 0.0015 | 0.0017 |
| F13 | 79620001 | 79720000 | 0.0021 | 0.0011 | 0.0014 |
| F13 | 79640001 | 79740000 | 0.0023 | 0.0011 | 0.0014 |
| F13 | 79660001 | 79760000 | 0.0024 | 0.0011 | 0.0014 |
| F13 | 79680001 | 79780000 | 0.0031 | 0.0018 | 0.0021 |
| F13 | 79700001 | 79800000 | 0.0029 | 0.0018 | 0.0020 |
| F13 | 79720001 | 79820000 | 0.0031 | 0.0019 | 0.0022 |

| Chr | Start | End | both | A1 | A2 |
|---|---|---|---|---|---|
| F13 | 79740001 | 79840000 | 0.0032 | 0.0021 | 0.0023 |
| F13 | 79760001 | 79860000 | 0.0031 | 0.0021 | 0.0023 |
| F13 | 79780001 | 79880000 | 0.0025 | 0.0016 | 0.0018 |
| F13 | 79800001 | 79900000 | 0.0025 | 0.0016 | 0.0018 |
| F13 | 79820001 | 79920000 | 0.0023 | 0.0015 | 0.0016 |
| F13 | 79840001 | 79940000 | 0.0024 | 0.0016 | 0.0017 |
| F13 | 79860001 | 79960000 | 0.0027 | 0.0018 | 0.0019 |
| F13 | 79880001 | 79980000 | 0.0027 | 0.0018 | 0.0020 |
| F13 | 79900001 | 80000000 | 0.0034 | 0.0022 | 0.0027 |
| F13 | 79920001 | 80020000 | 0.0034 | 0.0021 | 0.0026 |
| F13 | 79940001 | 80040000 | 0.0037 | 0.0022 | 0.0026 |
| F13 | 79960001 | 80060000 | 0.0039 | 0.0025 | 0.0029 |
| F13 | 79980001 | 80080000 | 0.0035 | 0.0022 | 0.0025 |
| F13 | 80000001 | 80100000 | 0.0027 | 0.0015 | 0.0017 |
| F13 | 80020001 | 80120000 | 0.0023 | 0.0014 | 0.0014 |
| F13 | 80040001 | 80140000 | 0.0016 | 0.0009 | 0.0011 |
| F13 | 80060001 | 80160000 | 0.0009 | 0.0004 | 0.0004 |
| F13 | 80080001 | 80180000 | 0.0012 | 0.0007 | 0.0007 |
| F13 | 80100001 | 80200000 | 0.0018 | 0.0013 | 0.0015 |
| F13 | 80120001 | 80220000 | 0.0021 | 0.0014 | 0.0015 |
| F13 | 80140001 | 80240000 | 0.0021 | 0.0014 | 0.0014 |
| F13 | 80160001 | 80260000 | 0.0033 | 0.0022 | 0.0025 |
| F13 | 80180001 | 80280000 | 0.0041 | 0.0028 | 0.0031 |
| F13 | 80200001 | 80300000 | 0.0043 | 0.0024 | 0.0029 |
| F13 | 80220001 | 80320000 | 0.0042 | 0.0023 | 0.0028 |
| F13 | 80240001 | 80340000 | 0.0044 | 0.0026 | 0.0030 |
| F13 | 80260001 | 80360000 | 0.0037 | 0.0022 | 0.0025 |
| F13 | 80280001 | 80380000 | 0.0026 | 0.0013 | 0.0016 |
| F13 | 80300001 | 80400000 | 0.0022 | 0.0015 | 0.0014 |
| F13 | 80320001 | 80420000 | 0.0022 | 0.0014 | 0.0014 |
| F13 | 80340001 | 80440000 | 0.0021 | 0.0012 | 0.0012 |
| F13 | 80360001 | 80460000 | 0.0027 | 0.0015 | 0.0017 |
| F13 | 80380001 | 80480000 | 0.0037 | 0.0025 | 0.0029 |
| F13 | 80400001 | 80500000 | 0.0036 | 0.0025 | 0.0030 |
| F13 | 80420001 | 80520000 | 0.0040 | 0.0028 | 0.0033 |
| F13 | 80440001 | 80540000 | 0.0042 | 0.0030 | 0.0035 |
| F13 | 80460001 | 80560000 | 0.0035 | 0.0023 | 0.0026 |
| F13 | 80480001 | 80580000 | 0.0030 | 0.0018 | 0.0019 |
| F13 | 80500001 | 80600000 | 0.0033 | 0.0020 | 0.0021 |
| F13 | 80520001 | 80620000 | 0.0031 | 0.0019 | 0.0020 |
| F13 | 80540001 | 80640000 | 0.0029 | 0.0020 | 0.0020 |
| F13 | 80560001 | 80660000 | 0.0032 | 0.0023 | 0.0023 |
| F13 | 80580001 | 80680000 | 0.0039 | 0.0028 | 0.0028 |
| F13 | 80600001 | 80700000 | 0.0035 | 0.0024 | 0.0023 |
| F13 | 80620001 | 80720000 | 0.0036 | 0.0025 | 0.0024 |
| F13 | 80640001 | 80740000 | 0.0044 | 0.0029 | 0.0031 |
| F13 | 80660001 | 80760000 | 0.0047 | 0.0028 | 0.0036 |
| F13 | 80680001 | 80780000 | 0.0035 | 0.0019 | 0.0026 |
| F13 | 80700001 | 80800000 | 0.0039 | 0.0026 | 0.0031 |
| F13 | 80720001 | 80820000 | 0.0045 | 0.0033 | 0.0034 |
| F13 | 80740001 | 80840000 | 0.0039 | 0.0028 | 0.0025 |
| F13 | 80760001 | 80860000 | 0.0036 | 0.0027 | 0.0018 |
| F13 | 80780001 | 80880000 | 0.0048 | 0.0038 | 0.0030 |
| F13 | 80800001 | 80900000 | 0.0055 | 0.0041 | 0.0039 |
| F13 | 80820001 | 80920000 | 0.0049 | 0.0034 | 0.0035 |
| F13 | 80840001 | 80940000 | 0.0055 | 0.0038 | 0.0041 |
| F13 | 80860001 | 80960000 | 0.0053 | 0.0042 | 0.0045 |
| F13 | 80880001 | 80980000 | 0.0040 | 0.0030 | 0.0032 |

| Chr | Start | End | both | A1 | A2 |
|---|---|---|---|---|---|
| F13 | 80900001 | 81000000 | 0.0027 | 0.0018 | 0.0019 |
| F13 | 80920001 | 81020000 | 0.0027 | 0.0018 | 0.0019 |
| F13 | 80940001 | 81040000 | 0.0022 | 0.0015 | 0.0015 |
| F13 | 80960001 | 81060000 | 0.0024 | 0.0017 | 0.0017 |
| F13 | 80980001 | 81080000 | 0.0029 | 0.0022 | 0.0022 |
| F13 | 81000001 | 81100000 | 0.0030 | 0.0022 | 0.0023 |
| F13 | 81020001 | 81120000 | 0.0032 | 0.0023 | 0.0025 |
| F13 | 81040001 | 81140000 | 0.0035 | 0.0025 | 0.0029 |
| F13 | 81060001 | 81160000 | 0.0042 | 0.0030 | 0.0035 |
| F13 | 81080001 | 81180000 | 0.0045 | 0.0032 | 0.0037 |
| F13 | 81100001 | 81200000 | 0.0041 | 0.0029 | 0.0033 |
| F13 | 81120001 | 81220000 | 0.0039 | 0.0028 | 0.0031 |
| F13 | 81140001 | 81240000 | 0.0037 | 0.0024 | 0.0025 |
| F13 | 81160001 | 81260000 | 0.0029 | 0.0017 | 0.0018 |
| F13 | 81180001 | 81280000 | 0.0023 | 0.0011 | 0.0013 |
| F13 | 81200001 | 81300000 | 0.0029 | 0.0013 | 0.0017 |
| F13 | 81220001 | 81320000 | 0.0027 | 0.0013 | 0.0016 |
| F13 | 81240001 | 81340000 | 0.0026 | 0.0015 | 0.0018 |
| F13 | 81260001 | 81360000 | 0.0026 | 0.0019 | 0.0018 |
| F13 | 81280001 | 81380000 | 0.0031 | 0.0024 | 0.0024 |
| F13 | 81300001 | 81400000 | 0.0030 | 0.0026 | 0.0025 |
| F13 | 81320001 | 81420000 | 0.0041 | 0.0035 | 0.0036 |
| F13 | 81340001 | 81440000 | 0.0051 | 0.0043 | 0.0047 |
| F13 | 81360001 | 81460000 | 0.0046 | 0.0038 | 0.0044 |
| F13 | 81380001 | 81480000 | 0.0043 | 0.0035 | 0.0040 |
| F13 | 81400001 | 81500000 | 0.0039 | 0.0034 | 0.0038 |
| F13 | 81420001 | 81520000 | 0.0029 | 0.0024 | 0.0027 |
| F13 | 81440001 | 81540000 | 0.0020 | 0.0015 | 0.0017 |
| F13 | 81460001 | 81560000 | 0.0020 | 0.0014 | 0.0016 |
| F13 | 81480001 | 81580000 | 0.0018 | 0.0011 | 0.0013 |
| F13 | 81500001 | 81600000 | 0.0020 | 0.0011 | 0.0013 |
| F13 | 81520001 | 81620000 | 0.0026 | 0.0013 | 0.0016 |
| F13 | 81540001 | 81640000 | 0.0026 | 0.0012 | 0.0015 |
| F13 | 81560001 | 81660000 | 0.0034 | 0.0019 | 0.0024 |
| F13 | 81580001 | 81680000 | 0.0038 | 0.0021 | 0.0027 |
| F13 | 81600001 | 81700000 | 0.0036 | 0.0020 | 0.0025 |
| F13 | 81620001 | 81720000 | 0.0033 | 0.0019 | 0.0024 |
| F13 | 81640001 | 81740000 | 0.0036 | 0.0020 | 0.0025 |
| F13 | 81660001 | 81760000 | 0.0039 | 0.0022 | 0.0026 |
| F13 | 81680001 | 81780000 | 0.0038 | 0.0023 | 0.0025 |
| F13 | 81700001 | 81800000 | 0.0042 | 0.0026 | 0.0030 |
| F13 | 81720001 | 81820000 | 0.0045 | 0.0029 | 0.0033 |
| F13 | 81740001 | 81840000 | 0.0038 | 0.0027 | 0.0030 |
| F13 | 81760001 | 81860000 | 0.0029 | 0.0020 | 0.0023 |
| F13 | 81780001 | 81880000 | 0.0037 | 0.0026 | 0.0032 |
| F13 | 81800001 | 81900000 | 0.0042 | 0.0033 | 0.0039 |
| F13 | 81820001 | 81920000 | 0.0036 | 0.0028 | 0.0033 |
| F13 | 81840001 | 81940000 | 0.0037 | 0.0028 | 0.0033 |
| F13 | 81860001 | 81960000 | 0.0036 | 0.0026 | 0.0031 |
| F13 | 81880001 | 81980000 | 0.0029 | 0.0020 | 0.0022 |
| F13 | 81900001 | 82000000 | 0.0017 | 0.0009 | 0.0010 |
| F13 | 81920001 | 82020000 | 0.0024 | 0.0013 | 0.0015 |
| F13 | 81940001 | 82040000 | 0.0029 | 0.0014 | 0.0020 |
| F13 | 81960001 | 82060000 | 0.0031 | 0.0016 | 0.0022 |
| F13 | 81980001 | 82080000 | 0.0026 | 0.0013 | 0.0018 |
| F13 | 82000001 | 82100000 | 0.0028 | 0.0016 | 0.0021 |
| F13 | 82020001 | 82120000 | 0.0024 | 0.0013 | 0.0018 |
| F13 | 82040001 | 82140000 | 0.0021 | 0.0015 | 0.0016 |

| Chr | Start | End | both | A1 | A2 |
|---|---|---|---|---|---|
| F13 | 82060001 | 82160000 | 0.0021 | 0.0012 | 0.0013 |
| F13 | 82080001 | 82180000 | 0.0021 | 0.0011 | 0.0014 |
| F13 | 82100001 | 82200000 | 0.0023 | 0.0009 | 0.0013 |
| F13 | 82120001 | 82220000 | 0.0027 | 0.0011 | 0.0013 |
| F13 | 82140001 | 82240000 | 0.0030 | 0.0014 | 0.0015 |
| F13 | 82160001 | 82260000 | 0.0042 | 0.0024 | 0.0025 |
| F13 | 82180001 | 82280000 | 0.0047 | 0.0027 | 0.0029 |
| F13 | 82200001 | 82300000 | 0.0050 | 0.0030 | 0.0032 |
| F13 | 82220001 | 82320000 | 0.0047 | 0.0029 | 0.0031 |
| F13 | 82240001 | 82340000 | 0.0050 | 0.0030 | 0.0033 |
| F13 | 82260001 | 82360000 | 0.0038 | 0.0022 | 0.0025 |
| F13 | 82280001 | 82380000 | 0.0041 | 0.0025 | 0.0027 |
| F13 | 82300001 | 82400000 | 0.0038 | 0.0024 | 0.0026 |
| F13 | 82320001 | 82420000 | 0.0037 | 0.0023 | 0.0025 |
| F13 | 82340001 | 82440000 | 0.0030 | 0.0019 | 0.0020 |
| F13 | 82360001 | 82460000 | 0.0031 | 0.0019 | 0.0020 |
| F13 | 82380001 | 82480000 | 0.0028 | 0.0016 | 0.0015 |
| F13 | 82400001 | 82500000 | 0.0029 | 0.0015 | 0.0015 |
| F13 | 82420001 | 82520000 | 0.0027 | 0.0016 | 0.0015 |
| F13 | 82440001 | 82540000 | 0.0025 | 0.0013 | 0.0013 |
| F13 | 82460001 | 82560000 | 0.0030 | 0.0019 | 0.0019 |
| F13 | 82480001 | 82580000 | 0.0029 | 0.0018 | 0.0019 |
| F13 | 82500001 | 82600000 | 0.0028 | 0.0018 | 0.0019 |
| F13 | 82520001 | 82620000 | 0.0029 | 0.0017 | 0.0017 |
| F13 | 82540001 | 82640000 | 0.0029 | 0.0017 | 0.0017 |
| F13 | 82560001 | 82660000 | 0.0024 | 0.0012 | 0.0013 |
| F13 | 82580001 | 82680000 | 0.0025 | 0.0011 | 0.0013 |
| F13 | 82600001 | 82700000 | 0.0021 | 0.0008 | 0.0010 |
| F13 | 82620001 | 82720000 | 0.0022 | 0.0010 | 0.0013 |
| F13 | 82640001 | 82740000 | 0.0023 | 0.0012 | 0.0015 |
| F13 | 82660001 | 82760000 | 0.0022 | 0.0012 | 0.0016 |
| F13 | 82680001 | 82780000 | 0.0019 | 0.0013 | 0.0016 |
| F13 | 82700001 | 82800000 | 0.0020 | 0.0013 | 0.0016 |
| F13 | 82720001 | 82820000 | 0.0022 | 0.0014 | 0.0018 |
| F13 | 82740001 | 82840000 | 0.0028 | 0.0019 | 0.0023 |
| F13 | 82760001 | 82860000 | 0.0026 | 0.0018 | 0.0021 |
| F13 | 82780001 | 82880000 | 0.0025 | 0.0016 | 0.0018 |
| F13 | 82800001 | 82900000 | 0.0027 | 0.0017 | 0.0020 |
| F13 | 82820001 | 82920000 | 0.0027 | 0.0019 | 0.0020 |
| F13 | 82840001 | 82940000 | 0.0029 | 0.0021 | 0.0019 |
| F13 | 82860001 | 82960000 | 0.0030 | 0.0021 | 0.0021 |
| F13 | 82880001 | 82980000 | 0.0032 | 0.0022 | 0.0022 |
| F13 | 82900001 | 83000000 | 0.0032 | 0.0024 | 0.0024 |
| F13 | 82920001 | 83020000 | 0.0041 | 0.0031 | 0.0032 |
| F13 | 82940001 | 83040000 | 0.0035 | 0.0026 | 0.0028 |
| F13 | 82960001 | 83060000 | 0.0034 | 0.0026 | 0.0027 |
| F13 | 82980001 | 83080000 | 0.0034 | 0.0025 | 0.0028 |
| F13 | 83000001 | 83100000 | 0.0033 | 0.0022 | 0.0024 |
| F13 | 83020001 | 83120000 | 0.0027 | 0.0019 | 0.0021 |
| F13 | 83040001 | 83140000 | 0.0044 | 0.0036 | 0.0036 |
| F13 | 83060001 | 83160000 | 0.0043 | 0.0036 | 0.0036 |
| F13 | 83080001 | 83180000 | 0.0046 | 0.0039 | 0.0039 |
| F13 | 83100001 | 83200000 | 0.0050 | 0.0043 | 0.0043 |
| F13 | 83120001 | 83220000 | 0.0044 | 0.0035 | 0.0036 |
| F13 | 83140001 | 83240000 | 0.0028 | 0.0018 | 0.0018 |
| F13 | 83160001 | 83260000 | 0.0027 | 0.0015 | 0.0015 |
| F13 | 83180001 | 83280000 | 0.0027 | 0.0015 | 0.0015 |
| F13 | 83200001 | 83300000 | 0.0027 | 0.0013 | 0.0013 |

| Chr | Start | End | both | A1 | A2 |
|---|---|---|---|---|---|
| F13 | 83220001 | 83320000 | 0.0028 | 0.0014 | 0.0014 |
| F13 | 83240001 | 83340000 | 0.0028 | 0.0012 | 0.0013 |
| F13 | 83260001 | 83360000 | 0.0050 | 0.0024 | 0.0037 |
| F13 | 83280001 | 83380000 | 0.0051 | 0.0024 | 0.0039 |
| F13 | 83300001 | 83400000 | 0.0049 | 0.0025 | 0.0040 |
| F13 | 83320001 | 83420000 | 0.0061 | 0.0039 | 0.0054 |
| F13 | 83340001 | 83440000 | 0.0064 | 0.0043 | 0.0057 |
| F13 | 83360001 | 83460000 | 0.0044 | 0.0035 | 0.0038 |
| F13 | 83380001 | 83480000 | 0.0050 | 0.0037 | 0.0044 |
| F13 | 83400001 | 83500000 | 0.0053 | 0.0037 | 0.0047 |
| F13 | 83420001 | 83520000 | 0.0042 | 0.0023 | 0.0036 |
| F13 | 83440001 | 83540000 | 0.0041 | 0.0022 | 0.0037 |
| F13 | 83460001 | 83560000 | 0.0037 | 0.0020 | 0.0035 |
| F13 | 83480001 | 83580000 | 0.0025 | 0.0013 | 0.0024 |
| F13 | 83500001 | 83600000 | 0.0019 | 0.0011 | 0.0020 |
| F13 | 83520001 | 83620000 | 0.0027 | 0.0020 | 0.0028 |
| F13 | 83540001 | 83640000 | 0.0026 | 0.0019 | 0.0027 |
| F13 | 83560001 | 83660000 | 0.0024 | 0.0018 | 0.0025 |
| F13 | 83580001 | 83680000 | 0.0024 | 0.0018 | 0.0025 |
| F13 | 83600001 | 83700000 | 0.0024 | 0.0018 | 0.0024 |
| F13 | 83620001 | 83720000 | 0.0017 | 0.0012 | 0.0018 |
| F13 | 83640001 | 83740000 | 0.0016 | 0.0012 | 0.0016 |
| F13 | 83660001 | 83760000 | 0.0018 | 0.0013 | 0.0018 |
| F13 | 83680001 | 83780000 | 0.0021 | 0.0015 | 0.0021 |
| F13 | 83700001 | 83800000 | 0.0021 | 0.0015 | 0.0022 |
| F13 | 83720001 | 83820000 | 0.0017 | 0.0011 | 0.0017 |
| F13 | 83740001 | 83840000 | 0.0020 | 0.0012 | 0.0021 |
| F13 | 83760001 | 83860000 | 0.0019 | 0.0011 | 0.0020 |
| F13 | 83780001 | 83880000 | 0.0023 | 0.0015 | 0.0023 |
| F13 | 83800001 | 83900000 | 0.0027 | 0.0020 | 0.0027 |
| F13 | 83820001 | 83920000 | 0.0032 | 0.0026 | 0.0032 |
| F13 | 83840001 | 83940000 | 0.0031 | 0.0027 | 0.0031 |
| F13 | 83860001 | 83960000 | 0.0038 | 0.0045 | 0.0034 |
| F13 | 83880001 | 83980000 | 0.0035 | 0.0043 | 0.0031 |
| F13 | 83900001 | 84000000 | 0.0034 | 0.0042 | 0.0029 |
| F13 | 83920001 | 84020000 | 0.0028 | 0.0036 | 0.0024 |
| F13 | 83940001 | 84040000 | 0.0036 | 0.0044 | 0.0032 |
| F13 | 83960001 | 84060000 | 0.0031 | 0.0026 | 0.0031 |
| F13 | 83980001 | 84080000 | 0.0029 | 0.0023 | 0.0029 |
| F13 | 84000001 | 84100000 | 0.0034 | 0.0027 | 0.0034 |
| F13 | 84020001 | 84120000 | 0.0034 | 0.0027 | 0.0034 |
| F13 | 84040001 | 84140000 | 0.0023 | 0.0017 | 0.0023 |
| F13 | 84060001 | 84160000 | 0.0021 | 0.0017 | 0.0021 |
| F13 | 84080001 | 84180000 | 0.0022 | 0.0019 | 0.0022 |
| F13 | 84100001 | 84200000 | 0.0017 | 0.0015 | 0.0017 |
| F13 | 84120001 | 84220000 | 0.0023 | 0.0021 | 0.0023 |
| F13 | 84140001 | 84240000 | 0.0024 | 0.0022 | 0.0024 |
| F13 | 84160001 | 84260000 | 0.0025 | 0.0022 | 0.0024 |
| F13 | 84180001 | 84280000 | 0.0025 | 0.0021 | 0.0024 |
| F13 | 84200001 | 84300000 | 0.0038 | 0.0030 | 0.0036 |
| F13 | 84220001 | 84320000 | 0.0040 | 0.0030 | 0.0037 |
| F13 | 84240001 | 84340000 | 0.0042 | 0.0030 | 0.0039 |
| F13 | 84260001 | 84360000 | 0.0042 | 0.0031 | 0.0038 |
| F13 | 84280001 | 84380000 | 0.0043 | 0.0032 | 0.0038 |
| F13 | 84300001 | 84400000 | 0.0031 | 0.0026 | 0.0026 |
| F13 | 84320001 | 84420000 | 0.0027 | 0.0024 | 0.0023 |
| F13 | 84340001 | 84440000 | 0.0024 | 0.0027 | 0.0018 |
| F13 | 84360001 | 84460000 | 0.0023 | 0.0029 | 0.0017 |

| Chr | Start | End | both | A1 | A2 |
|---|---|---|---|---|---|
| F13 | 84380001 | 84480000 | 0.0023 | 0.0029 | 0.0017 |
| F13 | 84400001 | 84500000 | 0.0021 | 0.0025 | 0.0015 |
| F13 | 84420001 | 84520000 | 0.0015 | 0.0019 | 0.0009 |
| F13 | 84440001 | 84540000 | 0.0020 | 0.0018 | 0.0015 |
| F13 | 84460001 | 84560000 | 0.0025 | 0.0021 | 0.0020 |
| F13 | 84480001 | 84580000 | 0.0030 | 0.0026 | 0.0027 |
| F13 | 84500001 | 84600000 | 0.0035 | 0.0031 | 0.0031 |
| F13 | 84520001 | 84620000 | 0.0047 | 0.0043 | 0.0041 |
| F13 | 84540001 | 84640000 | 0.0045 | 0.0042 | 0.0040 |
| F13 | 84560001 | 84660000 | 0.0043 | 0.0039 | 0.0039 |
| F13 | 84580001 | 84680000 | 0.0050 | 0.0047 | 0.0045 |
| F13 | 84600001 | 84700000 | 0.0056 | 0.0049 | 0.0050 |
| F13 | 84620001 | 84720000 | 0.0055 | 0.0048 | 0.0052 |
| F13 | 84640001 | 84740000 | 0.0054 | 0.0048 | 0.0049 |
| F13 | 84660001 | 84760000 | 0.0051 | 0.0046 | 0.0046 |
| F13 | 84680001 | 84780000 | 0.0038 | 0.0032 | 0.0034 |
| F13 | 84700001 | 84800000 | 0.0041 | 0.0038 | 0.0036 |
| F13 | 84720001 | 84820000 | 0.0044 | 0.0040 | 0.0038 |
| F13 | 84740001 | 84840000 | 0.0045 | 0.0040 | 0.0040 |
| F13 | 84760001 | 84860000 | 0.0046 | 0.0041 | 0.0041 |
| F13 | 84780001 | 84880000 | 0.0053 | 0.0048 | 0.0047 |
| F13 | 84800001 | 84900000 | 0.0042 | 0.0038 | 0.0037 |
| F13 | 84820001 | 84920000 | 0.0041 | 0.0038 | 0.0036 |
| F13 | 84840001 | 84940000 | 0.0045 | 0.0043 | 0.0038 |
| F13 | 84860001 | 84960000 | 0.0051 | 0.0050 | 0.0038 |
| F13 | 84880001 | 84980000 | 0.0040 | 0.0039 | 0.0029 |
| F13 | 84900001 | 85000000 | 0.0040 | 0.0039 | 0.0029 |
| F13 | 84920001 | 85020000 | 0.0029 | 0.0029 | 0.0018 |
| F13 | 84940001 | 85040000 | 0.0025 | 0.0025 | 0.0018 |
| F13 | 84960001 | 85060000 | 0.0019 | 0.0018 | 0.0019 |
| F13 | 84980001 | 85080000 | 0.0029 | 0.0025 | 0.0027 |
| F13 | 85000001 | 85100000 | 0.0030 | 0.0026 | 0.0027 |
| F13 | 85020001 | 85120000 | 0.0028 | 0.0024 | 0.0024 |
| F13 | 85040001 | 85140000 | 0.0024 | 0.0019 | 0.0021 |
| F13 | 85060001 | 85160000 | 0.0027 | 0.0023 | 0.0023 |
| F13 | 85080001 | 85180000 | 0.0025 | 0.0023 | 0.0024 |
| F13 | 85100001 | 85200000 | 0.0031 | 0.0029 | 0.0030 |
| F13 | 85120001 | 85220000 | 0.0035 | 0.0031 | 0.0033 |
| F13 | 85140001 | 85240000 | 0.0039 | 0.0036 | 0.0037 |
| F13 | 85160001 | 85260000 | 0.0056 | 0.0052 | 0.0049 |
| F13 | 85180001 | 85280000 | 0.0071 | 0.0064 | 0.0063 |
| F13 | 85200001 | 85300000 | 0.0065 | 0.0059 | 0.0055 |
| F13 | 85220001 | 85320000 | 0.0065 | 0.0061 | 0.0056 |
| F13 | 85240001 | 85340000 | 0.0064 | 0.0060 | 0.0055 |
| F13 | 85260001 | 85360000 | 0.0043 | 0.0039 | 0.0037 |
| F13 | 85280001 | 85380000 | 0.0026 | 0.0024 | 0.0019 |
| F13 | 85300001 | 85400000 | 0.0024 | 0.0023 | 0.0018 |
| F13 | 85320001 | 85420000 | 0.0023 | 0.0022 | 0.0016 |
| F13 | 85340001 | 85440000 | 0.0025 | 0.0024 | 0.0018 |
| F13 | 85360001 | 85460000 | 0.0030 | 0.0028 | 0.0023 |
| F13 | 85380001 | 85480000 | 0.0029 | 0.0027 | 0.0021 |
| F13 | 85400001 | 85500000 | 0.0029 | 0.0026 | 0.0021 |
| F13 | 85420001 | 85520000 | 0.0026 | 0.0023 | 0.0018 |
| F13 | 85440001 | 85540000 | 0.0049 | 0.0044 | 0.0042 |
| F13 | 85460001 | 85560000 | 0.0043 | 0.0039 | 0.0037 |
| F13 | 85480001 | 85580000 | 0.0053 | 0.0049 | 0.0048 |
| F13 | 85500001 | 85600000 | 0.0052 | 0.0047 | 0.0047 |
| F13 | 85520001 | 85620000 | 0.0051 | 0.0045 | 0.0045 |

| Chr | Start | End | both | A1 | A2 |
|---|---|---|---|---|---|
| F13 | 85540001 | 85640000 | 0.0023 | 0.0019 | 0.0018 |
| F13 | 85560001 | 85660000 | 0.0032 | 0.0027 | 0.0027 |
| F13 | 85580001 | 85680000 | 0.0033 | 0.0026 | 0.0029 |
| F13 | 85600001 | 85700000 | 0.0037 | 0.0029 | 0.0033 |
| F13 | 85620001 | 85720000 | 0.0046 | 0.0035 | 0.0041 |
| F13 | 85640001 | 85740000 | 0.0047 | 0.0035 | 0.0043 |
| F13 | 85660001 | 85760000 | 0.0044 | 0.0031 | 0.0039 |
| F13 | 85680001 | 85780000 | 0.0034 | 0.0018 | 0.0028 |
| F13 | 85700001 | 85800000 | 0.0029 | 0.0017 | 0.0025 |
| F13 | 85720001 | 85820000 | 0.0026 | 0.0016 | 0.0023 |
| F13 | 85740001 | 85840000 | 0.0022 | 0.0014 | 0.0018 |
| F13 | 85760001 | 85860000 | 0.0016 | 0.0012 | 0.0014 |
| F13 | 85780001 | 85880000 | 0.0022 | 0.0020 | 0.0021 |
| F13 | 85800001 | 85900000 | 0.0021 | 0.0019 | 0.0020 |
| F13 | 85820001 | 85920000 | 0.0019 | 0.0017 | 0.0018 |
| F13 | 85840001 | 85940000 | 0.0023 | 0.0019 | 0.0020 |
| F13 | 85860001 | 85960000 | 0.0025 | 0.0020 | 0.0023 |
| F13 | 85880001 | 85980000 | 0.0022 | 0.0016 | 0.0018 |
| F13 | 85900001 | 86000000 | 0.0027 | 0.0019 | 0.0022 |
| F13 | 85920001 | 86020000 | 0.0028 | 0.0018 | 0.0021 |
| F13 | 85940001 | 86040000 | 0.0026 | 0.0017 | 0.0020 |
| F13 | 85960001 | 86060000 | 0.0026 | 0.0019 | 0.0017 |
| F13 | 85980001 | 86080000 | 0.0024 | 0.0019 | 0.0017 |
| F13 | 86000001 | 86100000 | 0.0026 | 0.0021 | 0.0019 |
| F13 | 86020001 | 86120000 | 0.0024 | 0.0019 | 0.0017 |
| F13 | 86040001 | 86140000 | 0.0027 | 0.0024 | 0.0021 |
| F13 | 86060001 | 86160000 | 0.0026 | 0.0020 | 0.0021 |
| F13 | 86080001 | 86180000 | 0.0034 | 0.0025 | 0.0028 |
| F13 | 86100001 | 86200000 | 0.0035 | 0.0026 | 0.0029 |
| F13 | 86120001 | 86220000 | 0.0036 | 0.0027 | 0.0030 |
| F13 | 86140001 | 86240000 | 0.0032 | 0.0023 | 0.0026 |
| F13 | 86160001 | 86260000 | 0.0032 | 0.0023 | 0.0028 |
| F13 | 86180001 | 86280000 | 0.0022 | 0.0017 | 0.0018 |
| F13 | 86200001 | 86300000 | 0.0022 | 0.0015 | 0.0018 |
| F13 | 86220001 | 86320000 | 0.0029 | 0.0023 | 0.0026 |
| F13 | 86240001 | 86340000 | 0.0032 | 0.0024 | 0.0028 |
| F13 | 86260001 | 86360000 | 0.0031 | 0.0022 | 0.0026 |
| F13 | 86280001 | 86380000 | 0.0028 | 0.0019 | 0.0023 |
| F13 | 86300001 | 86400000 | 0.0024 | 0.0017 | 0.0019 |
| F13 | 86320001 | 86420000 | 0.0024 | 0.0016 | 0.0018 |
| F13 | 86340001 | 86440000 | 0.0021 | 0.0014 | 0.0015 |
| F13 | 86360001 | 86460000 | 0.0019 | 0.0014 | 0.0014 |
| F13 | 86380001 | 86480000 | 0.0019 | 0.0014 | 0.0014 |
| F13 | 86400001 | 86500000 | 0.0018 | 0.0013 | 0.0013 |
| F13 | 86420001 | 86520000 | 0.0017 | 0.0012 | 0.0011 |
| F13 | 86440001 | 86540000 | 0.0022 | 0.0019 | 0.0017 |
| F13 | 86460001 | 86560000 | 0.0026 | 0.0020 | 0.0019 |
| F13 | 86480001 | 86580000 | 0.0031 | 0.0024 | 0.0025 |
| F13 | 86500001 | 86600000 | 0.0034 | 0.0024 | 0.0029 |
| F13 | 86520001 | 86620000 | 0.0028 | 0.0018 | 0.0024 |
| F13 | 86540001 | 86640000 | 0.0026 | 0.0011 | 0.0020 |
| F13 | 86560001 | 86660000 | 0.0022 | 0.0010 | 0.0019 |
| F13 | 86580001 | 86680000 | 0.0017 | 0.0008 | 0.0014 |
| F13 | 86600001 | 86700000 | 0.0014 | 0.0007 | 0.0012 |
| F13 | 86620001 | 86720000 | 0.0018 | 0.0010 | 0.0016 |
| F13 | 86640001 | 86740000 | 0.0020 | 0.0013 | 0.0017 |
| F13 | 86660001 | 86760000 | 0.0022 | 0.0014 | 0.0020 |
| F13 | 86680001 | 86780000 | 0.0028 | 0.0019 | 0.0026 |

| Chr | Start | End | both | A1 | A2 |
|---|---|---|---|---|---|
| F13 | 86700001 | 86800000 | 0.0031 | 0.0021 | 0.0029 |
| F13 | 86720001 | 86820000 | 0.0026 | 0.0017 | 0.0025 |
| F13 | 86740001 | 86840000 | 0.0022 | 0.0015 | 0.0022 |
| F13 | 86760001 | 86860000 | 0.0021 | 0.0014 | 0.0020 |
| F13 | 86780001 | 86880000 | 0.0016 | 0.0008 | 0.0016 |
| F13 | 86800001 | 86900000 | 0.0018 | 0.0012 | 0.0019 |
| F13 | 86820001 | 86920000 | 0.0031 | 0.0024 | 0.0031 |
| F13 | 86840001 | 86940000 | 0.0031 | 0.0025 | 0.0031 |
| F13 | 86860001 | 86960000 | 0.0033 | 0.0026 | 0.0033 |
| F13 | 86880001 | 86980000 | 0.0031 | 0.0025 | 0.0031 |
| F13 | 86900001 | 87000000 | 0.0027 | 0.0020 | 0.0027 |
| F13 | 86920001 | 87020000 | 0.0016 | 0.0010 | 0.0017 |
| F13 | 86940001 | 87040000 | 0.0026 | 0.0017 | 0.0027 |
| F13 | 86960001 | 87060000 | 0.0029 | 0.0018 | 0.0030 |
| F13 | 86980001 | 87080000 | 0.0032 | 0.0019 | 0.0033 |
| F13 | 87000001 | 87100000 | 0.0031 | 0.0021 | 0.0032 |
| F13 | 87020001 | 87120000 | 0.0043 | 0.0031 | 0.0043 |
| F13 | 87040001 | 87140000 | 0.0033 | 0.0024 | 0.0033 |
| F13 | 87060001 | 87160000 | 0.0035 | 0.0025 | 0.0035 |
| F13 | 87080001 | 87180000 | 0.0035 | 0.0026 | 0.0035 |
| F13 | 87100001 | 87200000 | 0.0043 | 0.0032 | 0.0043 |
| F13 | 87120001 | 87220000 | 0.0036 | 0.0026 | 0.0037 |
| F13 | 87140001 | 87240000 | 0.0050 | 0.0039 | 0.0050 |
| F13 | 87160001 | 87260000 | 0.0060 | 0.0050 | 0.0060 |
| F13 | 87180001 | 87280000 | 0.0064 | 0.0055 | 0.0063 |
| F13 | 87200001 | 87300000 | 0.0063 | 0.0054 | 0.0063 |
| F13 | 87220001 | 87320000 | 0.0062 | 0.0054 | 0.0062 |
| F13 | 87240001 | 87340000 | 0.0051 | 0.0045 | 0.0050 |
| F13 | 87260001 | 87360000 | 0.0044 | 0.0036 | 0.0044 |
| F13 | 87280001 | 87380000 | 0.0039 | 0.0032 | 0.0039 |
| F13 | 87300001 | 87400000 | 0.0035 | 0.0028 | 0.0034 |
| F13 | 87320001 | 87420000 | 0.0038 | 0.0029 | 0.0037 |
| F13 | 87340001 | 87440000 | 0.0043 | 0.0031 | 0.0040 |
| F13 | 87360001 | 87460000 | 0.0038 | 0.0027 | 0.0034 |
| F13 | 87380001 | 87480000 | 0.0038 | 0.0027 | 0.0033 |
| F13 | 87400001 | 87500000 | 0.0034 | 0.0023 | 0.0029 |
| F13 | 87420001 | 87520000 | 0.0029 | 0.0019 | 0.0024 |
| F13 | 87440001 | 87540000 | 0.0025 | 0.0016 | 0.0020 |
| F13 | 87460001 | 87560000 | 0.0024 | 0.0015 | 0.0019 |
| F13 | 87480001 | 87580000 | 0.0022 | 0.0013 | 0.0017 |
| F13 | 87500001 | 87600000 | 0.0025 | 0.0017 | 0.0021 |
| F13 | 87520001 | 87620000 | 0.0020 | 0.0013 | 0.0017 |
| F13 | 87540001 | 87640000 | 0.0022 | 0.0016 | 0.0020 |
| F13 | 87560001 | 87660000 | 0.0022 | 0.0014 | 0.0020 |
| F13 | 87580001 | 87680000 | 0.0022 | 0.0014 | 0.0019 |
| F13 | 87600001 | 87700000 | 0.0020 | 0.0010 | 0.0016 |
| F13 | 87620001 | 87720000 | 0.0022 | 0.0011 | 0.0017 |
| F13 | 87640001 | 87740000 | 0.0022 | 0.0009 | 0.0016 |
| F13 | 87660001 | 87760000 | 0.0030 | 0.0018 | 0.0023 |
| F13 | 87680001 | 87780000 | 0.0038 | 0.0026 | 0.0033 |
| F13 | 87700001 | 87800000 | 0.0059 | 0.0045 | 0.0053 |
| F13 | 87720001 | 87820000 | 0.0089 | 0.0072 | 0.0082 |
| F13 | 87740001 | 87840000 | 0.0111 | 0.0092 | 0.0105 |
| F13 | 87760001 | 87860000 | 0.0103 | 0.0085 | 0.0098 |
| F13 | 87780001 | 87880000 | 0.0119 | 0.0102 | 0.0113 |
| F13 | 87800001 | 87900000 | 0.0108 | 0.0097 | 0.0101 |
| F13 | 87820001 | 87920000 | 0.0095 | 0.0087 | 0.0088 |
| F13 | 87840001 | 87940000 | 0.0083 | 0.0079 | 0.0076 |

| Chr | Start | End | both | A1 | A2 |
|---|---|---|---|---|---|
| F13 | 87860001 | 87960000 | 0.0083 | 0.0079 | 0.0075 |
| F13 | 87880001 | 87980000 | 0.0059 | 0.0057 | 0.0051 |
| F13 | 87900001 | 88000000 | 0.0047 | 0.0042 | 0.0042 |
| F13 | 87920001 | 88020000 | 0.0030 | 0.0024 | 0.0026 |
| F13 | 87940001 | 88040000 | 0.0012 | 0.0009 | 0.0010 |
| F13 | 87960001 | 88060000 | 0.0016 | 0.0012 | 0.0014 |
| F13 | 87980001 | 88080000 | 0.0014 | 0.0009 | 0.0012 |
| F13 | 88000001 | 88100000 | 0.0013 | 0.0009 | 0.0012 |
| F13 | 88020001 | 88120000 | 0.0016 | 0.0012 | 0.0014 |
| F13 | 88040001 | 88140000 | 0.0019 | 0.0014 | 0.0017 |
| F13 | 88060001 | 88160000 | 0.0017 | 0.0012 | 0.0016 |
| F13 | 88080001 | 88180000 | 0.0020 | 0.0014 | 0.0018 |
| F13 | 88100001 | 88200000 | 0.0024 | 0.0018 | 0.0023 |
| F13 | 88120001 | 88220000 | 0.0022 | 0.0015 | 0.0020 |
| F13 | 88140001 | 88240000 | 0.0020 | 0.0012 | 0.0017 |
| F13 | 88160001 | 88260000 | 0.0016 | 0.0010 | 0.0014 |
| F13 | 88180001 | 88280000 | 0.0021 | 0.0014 | 0.0018 |
| F13 | 88200001 | 88300000 | 0.0019 | 0.0010 | 0.0014 |
| F13 | 88220001 | 88320000 | 0.0023 | 0.0015 | 0.0019 |
| F13 | 88240001 | 88340000 | 0.0033 | 0.0026 | 0.0029 |
| F13 | 88260001 | 88360000 | 0.0034 | 0.0024 | 0.0028 |
| F13 | 88280001 | 88380000 | 0.0027 | 0.0018 | 0.0022 |
| F13 | 88300001 | 88400000 | 0.0027 | 0.0018 | 0.0021 |
| F13 | 88320001 | 88420000 | 0.0026 | 0.0016 | 0.0020 |
| F13 | 88340001 | 88440000 | 0.0022 | 0.0010 | 0.0015 |
| F13 | 88360001 | 88460000 | 0.0027 | 0.0015 | 0.0019 |
| F13 | 88380001 | 88480000 | 0.0034 | 0.0018 | 0.0026 |
| F13 | 88400001 | 88500000 | 0.0036 | 0.0020 | 0.0028 |
| F13 | 88420001 | 88520000 | 0.0039 | 0.0021 | 0.0031 |
| F13 | 88440001 | 88540000 | 0.0035 | 0.0018 | 0.0026 |
| F13 | 88460001 | 88560000 | 0.0031 | 0.0015 | 0.0024 |
| F13 | 88480001 | 88580000 | 0.0023 | 0.0013 | 0.0017 |
| F13 | 88500001 | 88600000 | 0.0022 | 0.0011 | 0.0016 |
| F13 | 88520001 | 88620000 | 0.0021 | 0.0010 | 0.0016 |
| F13 | 88540001 | 88640000 | 0.0021 | 0.0009 | 0.0016 |
| F13 | 88560001 | 88660000 | 0.0023 | 0.0011 | 0.0017 |
| F13 | 88580001 | 88680000 | 0.0025 | 0.0012 | 0.0017 |
| F13 | 88600001 | 88700000 | 0.0025 | 0.0012 | 0.0017 |
| F13 | 88620001 | 88720000 | 0.0020 | 0.0008 | 0.0012 |
| F13 | 88640001 | 88740000 | 0.0020 | 0.0009 | 0.0013 |
| F13 | 88660001 | 88760000 | 0.0015 | 0.0005 | 0.0010 |
| F13 | 88680001 | 88780000 | 0.0018 | 0.0008 | 0.0015 |
| F13 | 88700001 | 88800000 | 0.0018 | 0.0007 | 0.0017 |
| F13 | 88720001 | 88820000 | 0.0016 | 0.0007 | 0.0016 |
| F13 | 88740001 | 88840000 | 0.0017 | 0.0010 | 0.0018 |
| F13 | 88760001 | 88860000 | 0.0027 | 0.0019 | 0.0027 |
| F13 | 88780001 | 88880000 | 0.0026 | 0.0017 | 0.0026 |
| F13 | 88800001 | 88900000 | 0.0027 | 0.0019 | 0.0027 |
| F13 | 88820001 | 88920000 | 0.0029 | 0.0021 | 0.0029 |
| F13 | 88840001 | 88940000 | 0.0027 | 0.0018 | 0.0027 |
| F13 | 88860001 | 88960000 | 0.0019 | 0.0009 | 0.0020 |
| F13 | 88880001 | 88980000 | 0.0016 | 0.0007 | 0.0017 |
| F13 | 88900001 | 89000000 | 0.0017 | 0.0005 | 0.0016 |
| F13 | 88920001 | 89020000 | 0.0015 | 0.0004 | 0.0013 |
| F13 | 88940001 | 89040000 | 0.0018 | 0.0005 | 0.0013 |
| F13 | 88960001 | 89060000 | 0.0016 | 0.0005 | 0.0011 |
| F13 | 88980001 | 89080000 | 0.0018 | 0.0006 | 0.0012 |
| F13 | 89000001 | 89100000 | 0.0017 | 0.0005 | 0.0010 |

| Chr | Start | End | both | A1 | A2 |
|---|---|---|---|---|---|
| F13 | 89020001 | 89120000 | 0.0020 | 0.0007 | 0.0014 |
| F13 | 89040001 | 89140000 | 0.0025 | 0.0008 | 0.0021 |
| F13 | 89060001 | 89160000 | 0.0036 | 0.0015 | 0.0031 |
| F13 | 89080001 | 89180000 | 0.0037 | 0.0014 | 0.0031 |
| F13 | 89100001 | 89200000 | 0.0037 | 0.0014 | 0.0032 |
| F13 | 89120001 | 89220000 | 0.0033 | 0.0011 | 0.0028 |
| F13 | 89140001 | 89240000 | 0.0025 | 0.0009 | 0.0019 |
| F13 | 89160001 | 89260000 | 0.0018 | 0.0005 | 0.0012 |
| F13 | 89180001 | 89280000 | 0.0016 | 0.0004 | 0.0011 |
| F13 | 89200001 | 89300000 | 0.0019 | 0.0007 | 0.0015 |
| F13 | 89220001 | 89320000 | 0.0022 | 0.0011 | 0.0019 |
| F13 | 89240001 | 89340000 | 0.0025 | 0.0015 | 0.0023 |
| F13 | 89260001 | 89360000 | 0.0022 | 0.0014 | 0.0020 |
| F13 | 89280001 | 89380000 | 0.0021 | 0.0016 | 0.0019 |
| F13 | 89300001 | 89400000 | 0.0029 | 0.0027 | 0.0028 |
| F13 | 89320001 | 89420000 | 0.0049 | 0.0048 | 0.0047 |
| F13 | 89340001 | 89440000 | 0.0048 | 0.0047 | 0.0047 |
| F13 | 89360001 | 89460000 | 0.0048 | 0.0047 | 0.0048 |
| F13 | 89380001 | 89480000 | 0.0053 | 0.0051 | 0.0052 |
| F13 | 89400001 | 89500000 | 0.0051 | 0.0050 | 0.0051 |
| F13 | 89420001 | 89520000 | 0.0033 | 0.0032 | 0.0032 |
| F13 | 89440001 | 89540000 | 0.0045 | 0.0044 | 0.0044 |
| F13 | 89460001 | 89560000 | 0.0042 | 0.0043 | 0.0041 |
| F13 | 89480001 | 89580000 | 0.0039 | 0.0039 | 0.0038 |
| F13 | 89500001 | 89600000 | 0.0027 | 0.0027 | 0.0026 |
| F13 | 89520001 | 89620000 | 0.0020 | 0.0021 | 0.0020 |
| F13 | 89540001 | 89640000 | 0.0006 | 0.0006 | 0.0005 |
| F13 | 89560001 | 89660000 | 0.0007 | 0.0008 | 0.0007 |
| F13 | 89580001 | 89680000 | 0.0007 | 0.0007 | 0.0007 |
| F13 | 89600001 | 89700000 | 0.0006 | 0.0006 | 0.0006 |
| F13 | 89620001 | 89720000 | 0.0007 | 0.0007 | 0.0007 |
| F13 | 89640001 | 89740000 | 0.0009 | 0.0008 | 0.0009 |
| F13 | 89660001 | 89760000 | 0.0010 | 0.0009 | 0.0010 |
| F13 | 89680001 | 89780000 | 0.0013 | 0.0012 | 0.0013 |
| F13 | 89700001 | 89800000 | 0.0022 | 0.0021 | 0.0021 |
| F13 | 89720001 | 89820000 | 0.0023 | 0.0022 | 0.0022 |
| F13 | 89740001 | 89840000 | 0.0027 | 0.0026 | 0.0026 |
| F13 | 89760001 | 89860000 | 0.0033 | 0.0032 | 0.0032 |
| F13 | 89780001 | 89880000 | 0.0037 | 0.0037 | 0.0036 |
| F13 | 89800001 | 89900000 | 0.0031 | 0.0030 | 0.0031 |
| F13 | 89820001 | 89920000 | 0.0046 | 0.0045 | 0.0045 |
| F13 | 89840001 | 89940000 | 0.0043 | 0.0041 | 0.0042 |
| F13 | 89860001 | 89960000 | 0.0037 | 0.0032 | 0.0036 |
| F13 | 89880001 | 89980000 | 0.0030 | 0.0024 | 0.0029 |
| F13 | 89900001 | 90000000 | 0.0030 | 0.0021 | 0.0029 |
| F13 | 89920001 | 90020000 | 0.0019 | 0.0006 | 0.0020 |
| F13 | 89940001 | 90040000 | 0.0021 | 0.0005 | 0.0022 |
| F13 | 89960001 | 90060000 | 0.0023 | 0.0007 | 0.0022 |
| F13 | 89980001 | 90080000 | 0.0023 | 0.0007 | 0.0021 |
| F13 | 90000001 | 90100000 | 0.0025 | 0.0007 | 0.0020 |
| F13 | 90020001 | 90120000 | 0.0020 | 0.0005 | 0.0013 |
| F13 | 90040001 | 90140000 | 0.0020 | 0.0007 | 0.0011 |
| F13 | 90060001 | 90160000 | 0.0018 | 0.0005 | 0.0008 |
| F13 | 90080001 | 90180000 | 0.0017 | 0.0005 | 0.0006 |
| F13 | 90100001 | 90200000 | 0.0015 | 0.0007 | 0.0006 |
| F13 | 90120001 | 90220000 | 0.0017 | 0.0008 | 0.0007 |
| F13 | 90140001 | 90240000 | 0.0023 | 0.0010 | 0.0014 |
| F13 | 90160001 | 90260000 | 0.0023 | 0.0011 | 0.0015 |

| Chr | Start | End | both | A1 | A2 |
|---|---|---|---|---|---|
| F13 | 90180001 | 90280000 | 0.0026 | 0.0011 | 0.0018 |
| F13 | 90200001 | 90300000 | 0.0027 | 0.0009 | 0.0018 |
| F13 | 90220001 | 90320000 | 0.0030 | 0.0011 | 0.0020 |
| F13 | 90240001 | 90340000 | 0.0023 | 0.0008 | 0.0014 |
| F13 | 90260001 | 90360000 | 0.0023 | 0.0008 | 0.0012 |
| F13 | 90280001 | 90380000 | 0.0021 | 0.0007 | 0.0010 |
| F13 | 90300001 | 90400000 | 0.0020 | 0.0007 | 0.0009 |
| F13 | 90320001 | 90420000 | 0.0019 | 0.0007 | 0.0008 |
| F13 | 90340001 | 90440000 | 0.0019 | 0.0009 | 0.0008 |
| F13 | 90360001 | 90460000 | 0.0020 | 0.0011 | 0.0010 |
| F13 | 90380001 | 90480000 | 0.0023 | 0.0014 | 0.0014 |
| F13 | 90400001 | 90500000 | 0.0030 | 0.0022 | 0.0022 |
| F13 | 90420001 | 90520000 | 0.0028 | 0.0019 | 0.0020 |
| F13 | 90440001 | 90540000 | 0.0028 | 0.0018 | 0.0020 |
| F13 | 90460001 | 90560000 | 0.0028 | 0.0016 | 0.0019 |
| F13 | 90480001 | 90580000 | 0.0047 | 0.0035 | 0.0038 |
| F13 | 90500001 | 90600000 | 0.0044 | 0.0032 | 0.0034 |
| F13 | 90520001 | 90620000 | 0.0055 | 0.0043 | 0.0046 |
| F13 | 90540001 | 90640000 | 0.0055 | 0.0045 | 0.0048 |
| F13 | 90560001 | 90660000 | 0.0059 | 0.0050 | 0.0052 |
| F13 | 90580001 | 90680000 | 0.0038 | 0.0031 | 0.0031 |
| F13 | 90600001 | 90700000 | 0.0035 | 0.0027 | 0.0026 |
| F13 | 90620001 | 90720000 | 0.0023 | 0.0015 | 0.0014 |
| F13 | 90640001 | 90740000 | 0.0020 | 0.0012 | 0.0010 |
| F13 | 90660001 | 90760000 | 0.0015 | 0.0006 | 0.0006 |
| F13 | 90680001 | 90780000 | 0.0013 | 0.0004 | 0.0003 |
| F13 | 90700001 | 90800000 | 0.0011 | 0.0002 | 0.0003 |
| F13 | 90720001 | 90820000 | 0.0011 | 0.0002 | 0.0002 |
| F13 | 90740001 | 90840000 | 0.0013 | 0.0003 | 0.0004 |
| F13 | 90760001 | 90860000 | 0.0014 | 0.0005 | 0.0006 |
| F13 | 90780001 | 90880000 | 0.0015 | 0.0006 | 0.0007 |
| F13 | 90800001 | 90900000 | 0.0017 | 0.0008 | 0.0010 |
| F13 | 90820001 | 90920000 | 0.0019 | 0.0009 | 0.0011 |
| F13 | 90840001 | 90940000 | 0.0018 | 0.0009 | 0.0010 |
| F13 | 90860001 | 90960000 | 0.0019 | 0.0007 | 0.0009 |
| F13 | 90880001 | 90980000 | 0.0020 | 0.0007 | 0.0009 |
| F13 | 90900001 | 91000000 | 0.0017 | 0.0004 | 0.0007 |
| F13 | 90920001 | 91020000 | 0.0017 | 0.0003 | 0.0005 |
| F13 | 90940001 | 91040000 | 0.0019 | 0.0006 | 0.0008 |
| F13 | 90960001 | 91060000 | 0.0018 | 0.0008 | 0.0009 |
| F13 | 90980001 | 91080000 | 0.0017 | 0.0007 | 0.0008 |
| F13 | 91000001 | 91100000 | 0.0018 | 0.0009 | 0.0010 |
| F13 | 91020001 | 91120000 | 0.0021 | 0.0013 | 0.0013 |
| F13 | 91040001 | 91140000 | 0.0020 | 0.0011 | 0.0012 |
| F13 | 91060001 | 91160000 | 0.0027 | 0.0020 | 0.0020 |
| F13 | 91080001 | 91180000 | 0.0025 | 0.0020 | 0.0019 |
| F13 | 91100001 | 91200000 | 0.0024 | 0.0018 | 0.0018 |
| F13 | 91120001 | 91220000 | 0.0025 | 0.0014 | 0.0019 |
| F13 | 91140001 | 91240000 | 0.0024 | 0.0012 | 0.0018 |
| F13 | 91160001 | 91260000 | 0.0014 | 0.0001 | 0.0009 |
| F13 | 91180001 | 91280000 | 0.0019 | 0.0004 | 0.0014 |
| F13 | 91200001 | 91300000 | 0.0019 | 0.0004 | 0.0014 |
| F13 | 91220001 | 91320000 | 0.0015 | 0.0003 | 0.0012 |
| F13 | 91240001 | 91340000 | 0.0017 | 0.0003 | 0.0016 |
| F13 | 91260001 | 91360000 | 0.0024 | 0.0009 | 0.0023 |
| F13 | 91280001 | 91380000 | 0.0024 | 0.0009 | 0.0023 |
| F13 | 91300001 | 91400000 | 0.0025 | 0.0009 | 0.0024 |
| F13 | 91320001 | 91420000 | 0.0028 | 0.0011 | 0.0025 |

| Chr | Start | End | both | A1 | A2 |
|---|---|---|---|---|---|
| F13 | 91340001 | 91440000 | 0.0027 | 0.0013 | 0.0022 |
| F13 | 91360001 | 91460000 | 0.0022 | 0.0009 | 0.0016 |
| F13 | 91380001 | 91480000 | 0.0021 | 0.0008 | 0.0013 |
| F13 | 91400001 | 91500000 | 0.0022 | 0.0009 | 0.0014 |
| F13 | 91420001 | 91520000 | 0.0021 | 0.0008 | 0.0012 |
| F13 | 91440001 | 91540000 | 0.0018 | 0.0005 | 0.0010 |
| F13 | 91460001 | 91560000 | 0.0019 | 0.0007 | 0.0009 |
| F13 | 91480001 | 91580000 | 0.0022 | 0.0011 | 0.0011 |
| F13 | 91500001 | 91600000 | 0.0021 | 0.0010 | 0.0009 |
| F13 | 91520001 | 91620000 | 0.0019 | 0.0010 | 0.0008 |
| F13 | 91540001 | 91640000 | 0.0019 | 0.0010 | 0.0007 |
| F13 | 91560001 | 91660000 | 0.0019 | 0.0008 | 0.0006 |
| F13 | 91580001 | 91680000 | 0.0014 | 0.0003 | 0.0002 |
| F13 | 91600001 | 91700000 | 0.0014 | 0.0003 | 0.0001 |
| F13 | 91620001 | 91720000 | 0.0019 | 0.0008 | 0.0005 |
| F13 | 91640001 | 91740000 | 0.0020 | 0.0008 | 0.0006 |
| F13 | 91660001 | 91760000 | 0.0020 | 0.0009 | 0.0010 |
| F13 | 91680001 | 91780000 | 0.0033 | 0.0017 | 0.0025 |
| F13 | 91700001 | 91800000 | 0.0036 | 0.0020 | 0.0030 |
| F13 | 91720001 | 91820000 | 0.0044 | 0.0025 | 0.0041 |
| F13 | 91740001 | 91840000 | 0.0050 | 0.0031 | 0.0049 |
| F13 | 91760001 | 91860000 | 0.0049 | 0.0032 | 0.0048 |
| F13 | 91780001 | 91880000 | 0.0043 | 0.0027 | 0.0040 |
| F13 | 91800001 | 91900000 | 0.0043 | 0.0026 | 0.0039 |
| F13 | 91820001 | 91920000 | 0.0034 | 0.0020 | 0.0029 |
| F13 | 91840001 | 91940000 | 0.0029 | 0.0015 | 0.0022 |
| F13 | 91860001 | 91960000 | 0.0028 | 0.0015 | 0.0022 |
| F13 | 91880001 | 91980000 | 0.0021 | 0.0012 | 0.0015 |
| F13 | 91900001 | 92000000 | 0.0020 | 0.0009 | 0.0012 |
| F13 | 91920001 | 92020000 | 0.0017 | 0.0005 | 0.0009 |
| F13 | 91940001 | 92040000 | 0.0023 | 0.0013 | 0.0016 |
| F13 | 91960001 | 92060000 | 0.0027 | 0.0017 | 0.0019 |
| F13 | 91980001 | 92080000 | 0.0031 | 0.0021 | 0.0023 |
| F13 | 92000001 | 92100000 | 0.0036 | 0.0029 | 0.0031 |
| F13 | 92020001 | 92120000 | 0.0037 | 0.0033 | 0.0034 |
| F13 | 92040001 | 92140000 | 0.0034 | 0.0027 | 0.0029 |
| F13 | 92060001 | 92160000 | 0.0031 | 0.0025 | 0.0025 |
| F13 | 92080001 | 92180000 | 0.0033 | 0.0025 | 0.0026 |
| F13 | 92100001 | 92200000 | 0.0029 | 0.0017 | 0.0020 |
| F13 | 92120001 | 92220000 | 0.0029 | 0.0015 | 0.0020 |
| F13 | 92140001 | 92240000 | 0.0028 | 0.0011 | 0.0017 |
| F13 | 92160001 | 92260000 | 0.0027 | 0.0008 | 0.0015 |
| F13 | 92180001 | 92280000 | 0.0024 | 0.0005 | 0.0012 |
| F13 | 92200001 | 92300000 | 0.0023 | 0.0006 | 0.0010 |
| F13 | 92220001 | 92320000 | 0.0020 | 0.0005 | 0.0007 |
| F13 | 92240001 | 92340000 | 0.0018 | 0.0005 | 0.0005 |
| F13 | 92260001 | 92360000 | 0.0016 | 0.0005 | 0.0005 |
| F13 | 92280001 | 92380000 | 0.0016 | 0.0010 | 0.0005 |
| F13 | 92300001 | 92400000 | 0.0018 | 0.0015 | 0.0009 |
| F13 | 92320001 | 92420000 | 0.0016 | 0.0016 | 0.0008 |
| F13 | 92340001 | 92440000 | 0.0015 | 0.0016 | 0.0008 |
| F13 | 92360001 | 92460000 | 0.0018 | 0.0016 | 0.0008 |
| F13 | 92380001 | 92480000 | 0.0015 | 0.0010 | 0.0005 |
| F13 | 92400001 | 92500000 | 0.0012 | 0.0003 | 0.0001 |
| F13 | 92420001 | 92520000 | 0.0013 | 0.0001 | 0.0002 |
| F13 | 92440001 | 92540000 | 0.0015 | 0.0001 | 0.0002 |
| F13 | 92460001 | 92560000 | 0.0015 | 0.0003 | 0.0004 |
| F13 | 92480001 | 92580000 | 0.0015 | 0.0003 | 0.0005 |

| Chr | Start | End | both | A1 | A2 |
|---|---|---|---|---|---|
| F13 | 92500001 | 92600000 | 0.0016 | 0.0004 | 0.0006 |
| F13 | 92520001 | 92620000 | 0.0018 | 0.0005 | 0.0008 |
| F13 | 92540001 | 92640000 | 0.0023 | 0.0012 | 0.0014 |
| F13 | 92560001 | 92660000 | 0.0018 | 0.0010 | 0.0012 |
| F13 | 92580001 | 92680000 | 0.0020 | 0.0013 | 0.0014 |
| F13 | 92600001 | 92700000 | 0.0024 | 0.0019 | 0.0021 |
| F13 | 92620001 | 92720000 | 0.0028 | 0.0022 | 0.0025 |
| F13 | 92640001 | 92740000 | 0.0027 | 0.0023 | 0.0026 |
| F13 | 92660001 | 92760000 | 0.0031 | 0.0024 | 0.0027 |
| F13 | 92680001 | 92780000 | 0.0038 | 0.0032 | 0.0034 |
| F13 | 92700001 | 92800000 | 0.0034 | 0.0028 | 0.0031 |
| F13 | 92720001 | 92820000 | 0.0029 | 0.0024 | 0.0026 |
| F13 | 92740001 | 92840000 | 0.0032 | 0.0025 | 0.0028 |
| F13 | 92760001 | 92860000 | 0.0029 | 0.0025 | 0.0027 |
| F13 | 92780001 | 92880000 | 0.0022 | 0.0017 | 0.0020 |
| F13 | 92800001 | 92900000 | 0.0031 | 0.0023 | 0.0028 |
| F13 | 92820001 | 92920000 | 0.0032 | 0.0024 | 0.0029 |
| F13 | 92840001 | 92940000 | 0.0023 | 0.0016 | 0.0020 |
| F13 | 92860001 | 92960000 | 0.0024 | 0.0017 | 0.0022 |
| F13 | 92880001 | 92980000 | 0.0029 | 0.0021 | 0.0026 |
| F13 | 92900001 | 93000000 | 0.0023 | 0.0017 | 0.0020 |
| F13 | 92920001 | 93020000 | 0.0027 | 0.0020 | 0.0024 |
| F13 | 92940001 | 93040000 | 0.0036 | 0.0030 | 0.0033 |
| F13 | 92960001 | 93060000 | 0.0039 | 0.0034 | 0.0037 |
| F13 | 92980001 | 93080000 | 0.0041 | 0.0036 | 0.0038 |
| F13 | 93000001 | 93100000 | 0.0046 | 0.0041 | 0.0043 |
| F13 | 93020001 | 93120000 | 0.0042 | 0.0038 | 0.0040 |
| F13 | 93040001 | 93140000 | 0.0041 | 0.0036 | 0.0039 |
| F13 | 93060001 | 93160000 | 0.0058 | 0.0052 | 0.0056 |
| F13 | 93080001 | 93180000 | 0.0070 | 0.0062 | 0.0068 |
| F13 | 93100001 | 93200000 | 0.0064 | 0.0058 | 0.0062 |
| F13 | 93120001 | 93220000 | 0.0063 | 0.0057 | 0.0061 |
| F13 | 93140001 | 93240000 | 0.0061 | 0.0055 | 0.0059 |
| F13 | 93160001 | 93260000 | 0.0082 | 0.0077 | 0.0079 |
| F13 | 93180001 | 93280000 | 0.0062 | 0.0058 | 0.0059 |
| F13 | 93200001 | 93300000 | 0.0064 | 0.0060 | 0.0061 |
| F13 | 93220001 | 93320000 | 0.0068 | 0.0063 | 0.0064 |
| F13 | 93240001 | 93340000 | 0.0072 | 0.0067 | 0.0068 |
| F13 | 93260001 | 93360000 | 0.0032 | 0.0025 | 0.0026 |
| F13 | 93280001 | 93380000 | 0.0035 | 0.0026 | 0.0026 |
| F13 | 93300001 | 93400000 | 0.0031 | 0.0022 | 0.0022 |
| F13 | 93320001 | 93420000 | 0.0028 | 0.0017 | 0.0017 |
| F13 | 93340001 | 93440000 | 0.0019 | 0.0006 | 0.0006 |
| F13 | 93360001 | 93460000 | 0.0016 | 0.0006 | 0.0006 |
| F13 | 93380001 | 93480000 | 0.0015 | 0.0005 | 0.0006 |
| F13 | 93400001 | 93500000 | 0.0014 | 0.0003 | 0.0005 |
| F13 | 93420001 | 93520000 | 0.0016 | 0.0004 | 0.0007 |
| F13 | 93440001 | 93540000 | 0.0017 | 0.0006 | 0.0008 |
| F13 | 93460001 | 93560000 | 0.0020 | 0.0006 | 0.0008 |
| F13 | 93480001 | 93580000 | 0.0021 | 0.0007 | 0.0009 |
| F13 | 93500001 | 93600000 | 0.0022 | 0.0008 | 0.0011 |
| F13 | 93520001 | 93620000 | 0.0018 | 0.0007 | 0.0009 |
| F13 | 93540001 | 93640000 | 0.0018 | 0.0005 | 0.0010 |
| F13 | 93560001 | 93660000 | 0.0017 | 0.0005 | 0.0009 |
| F13 | 93580001 | 93680000 | 0.0015 | 0.0004 | 0.0009 |
| F13 | 93600001 | 93700000 | 0.0012 | 0.0002 | 0.0004 |
| F13 | 93620001 | 93720000 | 0.0015 | 0.0004 | 0.0005 |
| F13 | 93640001 | 93740000 | 0.0012 | 0.0004 | 0.0004 |

| Chr | Start | End | both | A1 | A2 |
|---|---|---|---|---|---|
| F13 | 93660001 | 93760000 | 0.0011 | 0.0005 | 0.0004 |
| F13 | 93680001 | 93780000 | 0.0009 | 0.0005 | 0.0005 |
| F13 | 93700001 | 93800000 | 0.0007 | 0.0005 | 0.0006 |
| F13 | 93720001 | 93820000 | 0.0012 | 0.0009 | 0.0011 |
| F13 | 93740001 | 93840000 | 0.0023 | 0.0018 | 0.0022 |
| F13 | 93760001 | 93860000 | 0.0044 | 0.0036 | 0.0042 |
| F13 | 93780001 | 93880000 | 0.0048 | 0.0038 | 0.0045 |
| F13 | 93800001 | 93900000 | 0.0052 | 0.0040 | 0.0049 |
| F13 | 93820001 | 93920000 | 0.0050 | 0.0037 | 0.0048 |
| F13 | 93840001 | 93940000 | 0.0042 | 0.0027 | 0.0040 |
| F13 | 93860001 | 93960000 | 0.0025 | 0.0008 | 0.0024 |
| F13 | 93880001 | 93980000 | 0.0026 | 0.0006 | 0.0026 |
| F13 | 93900001 | 94000000 | 0.0024 | 0.0003 | 0.0024 |
| F13 | 93920001 | 94020000 | 0.0023 | 0.0001 | 0.0022 |
| F13 | 93940001 | 94040000 | 0.0025 | 0.0003 | 0.0023 |
| F13 | 93960001 | 94060000 | 0.0022 | 0.0003 | 0.0018 |
| F13 | 93980001 | 94080000 | 0.0020 | 0.0006 | 0.0014 |
| F13 | 94000001 | 94100000 | 0.0029 | 0.0020 | 0.0022 |
| F13 | 94020001 | 94120000 | 0.0027 | 0.0024 | 0.0019 |
| F13 | 94040001 | 94140000 | 0.0030 | 0.0030 | 0.0021 |
| F13 | 94060001 | 94160000 | 0.0040 | 0.0038 | 0.0027 |
| F13 | 94080001 | 94180000 | 0.0055 | 0.0047 | 0.0042 |
| F13 | 94100001 | 94200000 | 0.0045 | 0.0034 | 0.0031 |
| F13 | 94120001 | 94220000 | 0.0046 | 0.0030 | 0.0032 |
| F13 | 94140001 | 94240000 | 0.0042 | 0.0023 | 0.0028 |
| F13 | 94160001 | 94260000 | 0.0042 | 0.0024 | 0.0032 |
| F13 | 94180001 | 94280000 | 0.0027 | 0.0012 | 0.0018 |
| F13 | 94200001 | 94300000 | 0.0028 | 0.0013 | 0.0018 |
| F13 | 94220001 | 94320000 | 0.0030 | 0.0017 | 0.0021 |
| F13 | 94240001 | 94340000 | 0.0028 | 0.0016 | 0.0019 |
| F13 | 94260001 | 94360000 | 0.0016 | 0.0007 | 0.0008 |
| F13 | 94280001 | 94380000 | 0.0015 | 0.0007 | 0.0006 |
| F13 | 94300001 | 94400000 | 0.0015 | 0.0006 | 0.0006 |
| F13 | 94320001 | 94420000 | 0.0012 | 0.0003 | 0.0004 |
| F13 | 94340001 | 94440000 | 0.0012 | 0.0003 | 0.0004 |
| F13 | 94360001 | 94460000 | 0.0014 | 0.0003 | 0.0006 |
| F13 | 94380001 | 94480000 | 0.0014 | 0.0002 | 0.0006 |
| F13 | 94400001 | 94500000 | 0.0012 | 0.0008 | 0.0005 |
| F13 | 94420001 | 94520000 | 0.0012 | 0.0010 | 0.0004 |
| F13 | 94440001 | 94540000 | 0.0012 | 0.0011 | 0.0004 |
| F13 | 94460001 | 94560000 | 0.0014 | 0.0012 | 0.0003 |
| F13 | 94480001 | 94580000 | 0.0014 | 0.0014 | 0.0002 |
| F13 | 94500001 | 94600000 | 0.0013 | 0.0010 | 0.0002 |
| F13 | 94520001 | 94620000 | 0.0012 | 0.0008 | 0.0002 |
| F13 | 94540001 | 94640000 | 0.0012 | 0.0010 | 0.0002 |
| F13 | 94560001 | 94660000 | 0.0013 | 0.0014 | 0.0002 |
| F13 | 94580001 | 94680000 | 0.0012 | 0.0016 | 0.0002 |
| F13 | 94600001 | 94700000 | 0.0012 | 0.0019 | 0.0002 |
| F13 | 94620001 | 94720000 | 0.0015 | 0.0026 | 0.0004 |
| F13 | 94640001 | 94740000 | 0.0015 | 0.0029 | 0.0004 |
| F13 | 94660001 | 94760000 | 0.0013 | 0.0024 | 0.0004 |
| F13 | 94680001 | 94780000 | 0.0013 | 0.0020 | 0.0004 |
| F13 | 94700001 | 94800000 | 0.0014 | 0.0016 | 0.0004 |
| F13 | 94720001 | 94820000 | 0.0018 | 0.0009 | 0.0003 |
| F13 | 94740001 | 94840000 | 0.0021 | 0.0003 | 0.0003 |
| F13 | 94760001 | 94860000 | 0.0022 | 0.0005 | 0.0004 |
| F13 | 94780001 | 94880000 | 0.0033 | 0.0016 | 0.0016 |
| F13 | 94800001 | 94900000 | 0.0031 | 0.0016 | 0.0017 |

| Chr | Start | End | both | A1 | A2 |
|---|---|---|---|---|---|
| F13 | 94820001 | 94920000 | 0.0030 | 0.0019 | 0.0019 |
| F13 | 94840001 | 94940000 | 0.0032 | 0.0023 | 0.0022 |
| F13 | 94860001 | 94960000 | 0.0032 | 0.0023 | 0.0024 |
| F13 | 94880001 | 94980000 | 0.0023 | 0.0014 | 0.0013 |
| F13 | 94900001 | 95000000 | 0.0024 | 0.0013 | 0.0012 |
| F13 | 94920001 | 95020000 | 0.0022 | 0.0009 | 0.0010 |
| F13 | 94940001 | 95040000 | 0.0021 | 0.0005 | 0.0008 |
| F13 | 94960001 | 95060000 | 0.0021 | 0.0003 | 0.0007 |
| F13 | 94980001 | 95080000 | 0.0034 | 0.0015 | 0.0020 |
| F13 | 95000001 | 95100000 | 0.0035 | 0.0015 | 0.0024 |
| F13 | 95020001 | 95120000 | 0.0035 | 0.0015 | 0.0026 |
| F13 | 95040001 | 95140000 | 0.0037 | 0.0023 | 0.0031 |
| F13 | 95060001 | 95160000 | 0.0043 | 0.0037 | 0.0036 |
| F13 | 95080001 | 95180000 | 0.0032 | 0.0031 | 0.0028 |
| F13 | 95100001 | 95200000 | 0.0039 | 0.0044 | 0.0034 |
| F13 | 95120001 | 95220000 | 0.0041 | 0.0047 | 0.0035 |
| F13 | 95140001 | 95240000 | 0.0038 | 0.0042 | 0.0030 |
| F13 | 95160001 | 95260000 | 0.0030 | 0.0029 | 0.0023 |
| F13 | 95180001 | 95280000 | 0.0028 | 0.0025 | 0.0021 |
| F13 | 95200001 | 95300000 | 0.0020 | 0.0012 | 0.0012 |
| F13 | 95220001 | 95320000 | 0.0014 | 0.0008 | 0.0007 |
| F13 | 95240001 | 95340000 | 0.0013 | 0.0006 | 0.0007 |
| F13 | 95260001 | 95360000 | 0.0015 | 0.0007 | 0.0009 |
| F13 | 95280001 | 95380000 | 0.0018 | 0.0009 | 0.0012 |
| F13 | 95300001 | 95400000 | 0.0020 | 0.0010 | 0.0016 |
| F13 | 95320001 | 95420000 | 0.0025 | 0.0013 | 0.0021 |
| F13 | 95340001 | 95440000 | 0.0035 | 0.0021 | 0.0030 |
| F13 | 95360001 | 95460000 | 0.0038 | 0.0023 | 0.0034 |
| F13 | 95380001 | 95480000 | 0.0037 | 0.0023 | 0.0032 |
| F13 | 95400001 | 95500000 | 0.0034 | 0.0023 | 0.0028 |
| F13 | 95420001 | 95520000 | 0.0032 | 0.0021 | 0.0024 |
| F13 | 95440001 | 95540000 | 0.0028 | 0.0017 | 0.0020 |
| F13 | 95460001 | 95560000 | 0.0023 | 0.0013 | 0.0014 |
| F13 | 95480001 | 95580000 | 0.0020 | 0.0012 | 0.0011 |
| F13 | 95500001 | 95600000 | 0.0019 | 0.0011 | 0.0012 |
| F13 | 95520001 | 95620000 | 0.0018 | 0.0011 | 0.0013 |
| F13 | 95540001 | 95640000 | 0.0013 | 0.0007 | 0.0009 |
| F13 | 95560001 | 95660000 | 0.0014 | 0.0009 | 0.0007 |
| F13 | 95580001 | 95680000 | 0.0013 | 0.0008 | 0.0006 |
| F13 | 95600001 | 95700000 | 0.0012 | 0.0007 | 0.0004 |
| F13 | 95620001 | 95720000 | 0.0012 | 0.0007 | 0.0003 |
| F13 | 95640001 | 95740000 | 0.0013 | 0.0007 | 0.0003 |
| F13 | 95660001 | 95760000 | 0.0012 | 0.0006 | 0.0004 |
| F13 | 95680001 | 95780000 | 0.0013 | 0.0006 | 0.0005 |
| F13 | 95700001 | 95800000 | 0.0014 | 0.0007 | 0.0008 |
| F13 | 95720001 | 95820000 | 0.0023 | 0.0016 | 0.0018 |
| F13 | 95740001 | 95840000 | 0.0026 | 0.0020 | 0.0023 |
| F13 | 95760001 | 95860000 | 0.0024 | 0.0021 | 0.0023 |
| F13 | 95780001 | 95880000 | 0.0025 | 0.0022 | 0.0024 |
| F13 | 95800001 | 95900000 | 0.0022 | 0.0019 | 0.0021 |
| F13 | 95820001 | 95920000 | 0.0011 | 0.0009 | 0.0010 |
| F13 | 95840001 | 95940000 | 0.0004 | 0.0003 | 0.0004 |
| F13 | 95860001 | 95960000 | 0.0003 | 0.0002 | 0.0003 |

**Supplementary Table 7**. Pi by chromosome and genomic compartment for *G. herbaceum* (A1), *G. arboreum* (A2), and all samples combined (both).

| | | Total | | | Chr01 | | | Chr02 | | | Chr03 | | | Chr04 | | | Chr05 | | | Chr06 | | |
|---|---|---|---|---|---|---|---|---|---|---|---|---|---|---|---|---|---|---|---|---|---|---|
| | | both | A1 | A2 | both | A1 | A2 | both | A1 | A2 | both | A1 | A2 | both | A1 | A2 | both | A1 | A2 | both | A1 | A2 |
| **Intergenic** | Mean | 0.0026 | 0.0020 | 0.0022 | 0.0025 | 0.0019 | 0.0021 | 0.0029 | 0.0022 | 0.0024 | 0.0027 | 0.0021 | 0.0023 | 0.0025 | 0.0019 | 0.0021 | 0.0024 | 0.0018 | 0.0020 | 0.0027 | 0.0021 | 0.0023 |
| | Stdev | 0.0013 | 0.0011 | 0.0012 | 0.0012 | 0.0010 | 0.0011 | 0.0013 | 0.0011 | 0.0011 | 0.0012 | 0.0010 | 0.0011 | 0.0012 | 0.0010 | 0.0011 | 0.0015 | 0.0013 | 0.0015 | 0.0013 | 0.0011 | 0.0011 |
| | Median | 0.0025 | 0.0019 | 0.0021 | 0.0024 | 0.0018 | 0.0019 | 0.0028 | 0.0021 | 0.0023 | 0.0026 | 0.0019 | 0.0021 | 0.0024 | 0.0017 | 0.0019 | 0.0021 | 0.0016 | 0.0017 | 0.0026 | 0.0020 | 0.0021 |
| | Minimum | 0.0000 | 0.0000 | 0.0000 | 0.0000 | 0.0000 | 0.0000 | 0.0001 | 0.0001 | 0.0001 | 0.0001 | 0.0000 | 0.0001 | 0.0000 | 0.0000 | 0.0000 | 0.0000 | 0.0000 | 0.0000 | 0.0001 | 0.0001 | 0.0001 |
| | Maximum | 0.0148 | 0.0125 | 0.0147 | 0.0133 | 0.0108 | 0.0130 | 0.0081 | 0.0068 | 0.0072 | 0.0088 | 0.0078 | 0.0081 | 0.0084 | 0.0077 | 0.0070 | 0.0117 | 0.0114 | 0.0108 | 0.0088 | 0.0077 | 0.0079 |
| | 1. Quartile | 0.0017 | 0.0013 | 0.0014 | 0.0017 | 0.0012 | 0.0014 | 0.0019 | 0.0014 | 0.0016 | 0.0018 | 0.0013 | 0.0015 | 0.0016 | 0.0012 | 0.0013 | 0.0012 | 0.0008 | 0.0008 | 0.0017 | 0.0013 | 0.0014 |
| | 3. Quartile | 0.0034 | 0.0027 | 0.0029 | 0.0032 | 0.0025 | 0.0027 | 0.0037 | 0.0029 | 0.0031 | 0.0035 | 0.0027 | 0.0029 | 0.0032 | 0.0024 | 0.0027 | 0.0033 | 0.0026 | 0.0028 | 0.0035 | 0.0027 | 0.0029 |
| **Exon** | Mean | 0.0002 | 0.0001 | 0.0001 | 0.0002 | 0.0001 | 0.0002 | 0.0002 | 0.0001 | 0.0001 | 0.0001 | 0.0001 | 0.0001 | 0.0002 | 0.0001 | 0.0001 | 0.0003 | 0.0002 | 0.0002 | 0.0001 | 0.0001 | 0.0001 |
| | Stdev | 0.0003 | 0.0003 | 0.0003 | 0.0003 | 0.0003 | 0.0003 | 0.0003 | 0.0003 | 0.0003 | 0.0003 | 0.0003 | 0.0003 | 0.0003 | 0.0003 | 0.0003 | 0.0003 | 0.0003 | 0.0003 | 0.0002 | 0.0002 | 0.0002 |
| | Median | 0.0001 | 0.0000 | 0.0000 | 0.0001 | 0.0000 | 0.0000 | 0.0001 | 0.0000 | 0.0000 | 0.0000 | 0.0000 | 0.0000 | 0.0001 | 0.0000 | 0.0000 | 0.0002 | 0.0001 | 0.0001 | 0.0001 | 0.0000 | 0.0000 |
| | Minimum | 0.0000 | 0.0000 | 0.0000 | 0.0000 | 0.0000 | 0.0000 | 0.0000 | 0.0000 | 0.0000 | 0.0000 | 0.0000 | 0.0000 | 0.0000 | 0.0000 | 0.0000 | 0.0000 | 0.0000 | 0.0000 | 0.0000 | 0.0000 | 0.0000 |
| | Maximum | 0.0041 | 0.0043 | 0.0040 | 0.0041 | 0.0043 | 0.0040 | 0.0033 | 0.0021 | 0.0031 | 0.0032 | 0.0040 | 0.0029 | 0.0032 | 0.0032 | 0.0031 | 0.0034 | 0.0031 | 0.0031 | 0.0018 | 0.0016 | 0.0016 |
| | 1. Quartile | 0.0000 | 0.0000 | 0.0000 | 0.0000 | 0.0000 | 0.0000 | 0.0000 | 0.0000 | 0.0000 | 0.0000 | 0.0000 | 0.0000 | 0.0000 | 0.0000 | 0.0000 | 0.0001 | 0.0000 | 0.0000 | 0.0000 | 0.0000 | 0.0000 |
| | 3. Quartile | 0.0002 | 0.0002 | 0.0002 | 0.0002 | 0.0002 | 0.0002 | 0.0002 | 0.0002 | 0.0002 | 0.0001 | 0.0001 | 0.0001 | 0.0002 | 0.0001 | 0.0001 | 0.0003 | 0.0003 | 0.0002 | 0.0001 | 0.0001 | 0.0001 |
| **Intron** | Mean | 0.0002 | 0.0001 | 0.0001 | 0.0002 | 0.0001 | 0.0002 | 0.0002 | 0.0001 | 0.0001 | 0.0002 | 0.0001 | 0.0001 | 0.0002 | 0.0001 | 0.0001 | 0.0002 | 0.0002 | 0.0002 | 0.0001 | 0.0001 | 0.0001 |
| | Stdev | 0.0003 | 0.0002 | 0.0002 | 0.0003 | 0.0003 | 0.0003 | 0.0003 | 0.0002 | 0.0002 | 0.0002 | 0.0002 | 0.0002 | 0.0003 | 0.0002 | 0.0002 | 0.0003 | 0.0003 | 0.0003 | 0.0002 | 0.0002 | 0.0002 |
| | Median | 0.0001 | 0.0000 | 0.0000 | 0.0001 | 0.0000 | 0.0000 | 0.0001 | 0.0000 | 0.0000 | 0.0001 | 0.0000 | 0.0000 | 0.0001 | 0.0000 | 0.0000 | 0.0002 | 0.0001 | 0.0001 | 0.0001 | 0.0000 | 0.0000 |
| | Minimum | 0.0000 | 0.0000 | 0.0000 | 0.0000 | 0.0000 | 0.0000 | 0.0000 | 0.0000 | 0.0000 | 0.0000 | 0.0000 | 0.0000 | 0.0000 | 0.0000 | 0.0000 | 0.0000 | 0.0000 | 0.0000 | 0.0000 | 0.0000 | 0.0000 |
| | Maximum | 0.0037 | 0.0035 | 0.0036 | 0.0033 | 0.0030 | 0.0025 | 0.0028 | 0.0024 | 0.0023 | 0.0020 | 0.0018 | 0.0018 | 0.0026 | 0.0023 | 0.0026 | 0.0028 | 0.0026 | 0.0025 | 0.0014 | 0.0014 | 0.0013 |
| | 1. Quartile | 0.0000 | 0.0000 | 0.0000 | 0.0000 | 0.0000 | 0.0000 | 0.0000 | 0.0000 | 0.0000 | 0.0000 | 0.0000 | 0.0000 | 0.0000 | 0.0000 | 0.0000 | 0.0001 | 0.0000 | 0.0000 | 0.0000 | 0.0000 | 0.0000 |
| | 3. Quartile | 0.0002 | 0.0002 | 0.0002 | 0.0003 | 0.0002 | 0.0002 | 0.0002 | 0.0002 | 0.0001 | 0.0002 | 0.0001 | 0.0001 | 0.0002 | 0.0001 | 0.0002 | 0.0003 | 0.0002 | 0.0002 | 0.0002 | 0.0001 | 0.0001 |

| | | Chr07 | | | Chr08 | | | Chr09 | | | Chr10 | | | Chr11 | | | Chr12 | | | Chr13 | | |
|---|---|---|---|---|---|---|---|---|---|---|---|---|---|---|---|---|---|---|---|---|---|---|
| | | both | A1 | A2 | both | A1 | A2 | both | A1 | A2 | both | A1 | A2 | both | A1 | A2 | both | A1 | A2 | both | A1 | A2 |
| **Intergenic** | Mean | 0.0026 | 0.0021 | 0.0021 | 0.0026 | 0.0021 | 0.0022 | 0.0025 | 0.0019 | 0.0021 | 0.0029 | 0.0023 | 0.0025 | 0.0027 | 0.0020 | 0.0022 | 0.0026 | 0.0019 | 0.0022 | 0.0026 | 0.0020 | 0.0022 |
| | Stdev | 0.0012 | 0.0010 | 0.0011 | 0.0012 | 0.0010 | 0.0011 | 0.0013 | 0.0012 | 0.0012 | 0.0014 | 0.0012 | 0.0013 | 0.0013 | 0.0011 | 0.0012 | 0.0012 | 0.0010 | 0.0011 | 0.0012 | 0.0011 | 0.0011 |
| | Median | 0.0024 | 0.0020 | 0.0020 | 0.0026 | 0.0021 | 0.0022 | 0.0024 | 0.0018 | 0.0020 | 0.0028 | 0.0022 | 0.0024 | 0.0025 | 0.0019 | 0.0021 | 0.0025 | 0.0018 | 0.0021 | 0.0025 | 0.0018 | 0.0021 |
| | Minimum | 0.0000 | 0.0000 | 0.0000 | 0.0001 | 0.0000 | 0.0001 | 0.0002 | 0.0000 | 0.0001 | 0.0001 | 0.0001 | 0.0001 | 0.0000 | 0.0000 | 0.0000 | 0.0002 | 0.0000 | 0.0001 | 0.0001 | 0.0000 | 0.0001 |
| | Maximum | 0.0090 | 0.0089 | 0.0086 | 0.0078 | 0.0066 | 0.0071 | 0.0096 | 0.0093 | 0.0094 | 0.0148 | 0.0125 | 0.0147 | 0.0104 | 0.0101 | 0.0101 | 0.0087 | 0.0072 | 0.0074 | 0.0110 | 0.0094 | 0.0104 |
| | 1. Quartile | 0.0017 | 0.0014 | 0.0013 | 0.0018 | 0.0014 | 0.0015 | 0.0015 | 0.0011 | 0.0012 | 0.0020 | 0.0015 | 0.0016 | 0.0017 | 0.0013 | 0.0014 | 0.0017 | 0.0012 | 0.0014 | 0.0017 | 0.0012 | 0.0014 |
| | 3. Quartile | 0.0033 | 0.0027 | 0.0027 | 0.0034 | 0.0028 | 0.0029 | 0.0033 | 0.0026 | 0.0028 | 0.0037 | 0.0029 | 0.0032 | 0.0035 | 0.0027 | 0.0029 | 0.0033 | 0.0026 | 0.0028 | 0.0034 | 0.0025 | 0.0028 |
| **Exon** | Mean | 0.0002 | 0.0001 | 0.0001 | 0.0001 | 0.0001 | 0.0001 | 0.0002 | 0.0001 | 0.0001 | 0.0002 | 0.0002 | 0.0002 | 0.0002 | 0.0002 | 0.0002 | 0.0001 | 0.0001 | 0.0001 | 0.0002 | 0.0001 | 0.0001 |
| | Stdev | 0.0003 | 0.0002 | 0.0003 | 0.0002 | 0.0002 | 0.0002 | 0.0003 | 0.0002 | 0.0002 | 0.0003 | 0.0003 | 0.0003 | 0.0003 | 0.0003 | 0.0003 | 0.0002 | 0.0001 | 0.0002 | 0.0002 | 0.0002 | 0.0002 |
| | Median | 0.0001 | 0.0001 | 0.0000 | 0.0001 | 0.0000 | 0.0000 | 0.0001 | 0.0000 | 0.0001 | 0.0001 | 0.0000 | 0.0000 | 0.0001 | 0.0001 | 0.0001 | 0.0001 | 0.0000 | 0.0000 | 0.0001 | 0.0000 | 0.0000 |
| | Minimum | 0.0000 | 0.0000 | 0.0000 | 0.0000 | 0.0000 | 0.0000 | 0.0000 | 0.0000 | 0.0000 | 0.0000 | 0.0000 | 0.0000 | 0.0000 | 0.0000 | 0.0000 | 0.0000 | 0.0000 | 0.0000 | 0.0000 | 0.0000 | 0.0000 |
| | Maximum | 0.0032 | 0.0026 | 0.0029 | 0.0017 | 0.0018 | 0.0016 | 0.0023 | 0.0025 | 0.0022 | 0.0035 | 0.0030 | 0.0031 | 0.0032 | 0.0029 | 0.0030 | 0.0012 | 0.0010 | 0.0012 | 0.0025 | 0.0015 | 0.0024 |
| | 1. Quartile | 0.0000 | 0.0000 | 0.0000 | 0.0000 | 0.0000 | 0.0000 | 0.0000 | 0.0000 | 0.0000 | 0.0000 | 0.0000 | 0.0000 | 0.0000 | 0.0000 | 0.0000 | 0.0000 | 0.0000 | 0.0000 | 0.0000 | 0.0000 | 0.0000 |
| | 3. Quartile | 0.0002 | 0.0002 | 0.0002 | 0.0002 | 0.0001 | 0.0001 | 0.0002 | 0.0002 | 0.0002 | 0.0002 | 0.0002 | 0.0002 | 0.0003 | 0.0002 | 0.0002 | 0.0002 | 0.0001 | 0.0001 | 0.0002 | 0.0002 | 0.0002 |
| **Intron** | Mean | 0.0002 | 0.0002 | 0.0001 | 0.0002 | 0.0001 | 0.0001 | 0.0002 | 0.0001 | 0.0001 | 0.0002 | 0.0001 | 0.0001 | 0.0002 | 0.0002 | 0.0002 | 0.0002 | 0.0001 | 0.0001 | 0.0002 | 0.0001 | 0.0001 |
| | Stdev | 0.0003 | 0.0002 | 0.0002 | 0.0002 | 0.0002 | 0.0002 | 0.0002 | 0.0002 | 0.0002 | 0.0003 | 0.0002 | 0.0002 | 0.0003 | 0.0003 | 0.0002 | 0.0002 | 0.0002 | 0.0002 | 0.0003 | 0.0003 | 0.0003 |
| | Median | 0.0001 | 0.0001 | 0.0000 | 0.0001 | 0.0000 | 0.0000 | 0.0001 | 0.0001 | 0.0001 | 0.0001 | 0.0000 | 0.0000 | 0.0001 | 0.0001 | 0.0001 | 0.0001 | 0.0000 | 0.0000 | 0.0001 | 0.0000 | 0.0000 |
| | Minimum | 0.0000 | 0.0000 | 0.0000 | 0.0000 | 0.0000 | 0.0000 | 0.0000 | 0.0000 | 0.0000 | 0.0000 | 0.0000 | 0.0000 | 0.0000 | 0.0000 | 0.0000 | 0.0000 | 0.0000 | 0.0000 | 0.0000 | 0.0000 | 0.0000 |
| | Maximum | 0.0021 | 0.0021 | 0.0021 | 0.0017 | 0.0016 | 0.0017 | 0.0019 | 0.0017 | 0.0016 | 0.0021 | 0.0020 | 0.0017 | 0.0024 | 0.0023 | 0.0022 | 0.0017 | 0.0016 | 0.0016 | 0.0037 | 0.0035 | 0.0036 |
| | 1. Quartile | 0.0000 | 0.0000 | 0.0000 | 0.0000 | 0.0000 | 0.0000 | 0.0000 | 0.0000 | 0.0000 | 0.0000 | 0.0000 | 0.0000 | 0.0000 | 0.0000 | 0.0000 | 0.0000 | 0.0000 | 0.0000 | 0.0000 | 0.0000 | 0.0000 |
| | 3. Quartile | 0.0002 | 0.0002 | 0.0002 | 0.0002 | 0.0002 | 0.0002 | 0.0003 | 0.0002 | 0.0002 | 0.0002 | 0.0002 | 0.0002 | 0.0003 | 0.0002 | 0.0002 | 0.0002 | 0.0001 | 0.0002 | 0.0003 | 0.0002 | 0.0002 |

**Supplementary Table 8:** Diagnosis of introgression from windowed phylogenetics. In some cases, tree toplogy had limited resolution or had substantial intercalation between species (complex). In cases of the latter, we note the accessions that were unexpectedly place, but do not diagnose this as introgression due to other potentially confounding forces (e.g., lineage sorting).

| Window | Chr | Window on Chr | Start | End | Diagnosis |
|---|---|---|---|---|---|
| 1 | 1 | 0 | 12,474 | 413,190 | No introgression |
| 2 | 1 | 1 | 452,802 | 1,003,958 | No introgression |
| 3 | 1 | 2 | 1,007,708 | 1,651,215 | No introgression |
| 4 | 1 | 3 | 1,668,745 | 2,330,240 | No introgression |
| 5 | 1 | 4 | 2,346,706 | 3,115,138 | No introgression |
| 6 | 1 | 5 | 3,192,350 | 4,062,757 | A1_097, A2_044 |
| 7 | 1 | 6 | 4,076,584 | 5,001,338 | No introgression |
| 8 | 1 | 7 | 5,014,268 | 5,825,507 | No introgression |
| 9 | 1 | 8 | 5,923,867 | 6,748,102 | No introgression |
| 10 | 1 | 9 | 6,749,462 | 7,707,111 | No introgression |
| 11 | 1 | 10 | 7,804,999 | 9,164,105 | No introgression |
| 12 | 1 | 11 | 9,214,566 | 10,112,653 | No introgression |
| 13 | 1 | 12 | 10,113,607 | 10,995,573 | No introgression |
| 14 | 1 | 13 | 11,009,886 | 11,935,445 | No introgression |
| 15 | 1 | 14 | 11,999,669 | 13,477,278 | No introgression |
| 16 | 1 | 15 | 13,598,857 | 16,438,326 | No introgression |
| 17 | 1 | 16 | 16,492,498 | 17,661,130 | No introgression |
| 18 | 1 | 17 | 17,696,994 | 19,969,243 | No introgression |
| 19 | 1 | 18 | 20,137,804 | 22,321,097 | No introgression |
| 20 | 1 | 19 | 22,322,759 | 27,529,381 | No introgression |
| 21 | 1 | 20 | 27,638,557 | 35,114,345 | No introgression |
| 22 | 1 | 21 | 35,120,499 | 40,011,021 | No introgression |
| 23 | 1 | 22 | 41,677,924 | 50,108,534 | No introgression |
| 24 | 1 | 23 | 50,197,011 | 60,487,723 | No introgression |
| 25 | 1 | 24 | 60,741,911 | 70,038,967 | No introgression |
| 26 | 1 | 25 | 70,056,344 | 79,419,652 | No introgression |
| 27 | 1 | 26 | 79,523,177 | 84,284,487 | No introgression |

| Window | Chr | Window on Chr | Start | End | Diagnosis |
|---|---|---|---|---|---|
| 28 | 1 | 27 | 84,322,983 | 87,731,556 | No introgression |
| 29 | 1 | 28 | 87,975,468 | 89,696,356 | No introgression |
| 30 | 1 | 29 | 89,777,037 | 90,739,241 | No introgression |
| 31 | 1 | 30 | 90,740,985 | 91,874,805 | A1_155 |
| 32 | 1 | 31 | 91,911,737 | 93,308,636 | No introgression |
| 33 | 1 | 32 | 93,328,648 | 94,368,528 | A1_155 |
| 34 | 1 | 33 | 94,427,390 | 94,843,485 | No introgression |
| 35 | 1 | 34 | 95,189,749 | 96,332,423 | No introgression |
| 36 | 1 | 35 | 96,347,519 | 97,000,140 | No introgression |
| 37 | 1 | 36 | 97,006,048 | 97,448,212 | No introgression |
| 38 | 1 | 37 | 97,474,419 | 98,008,323 | No introgression |
| 39 | 1 | 38 | 98,022,208 | 98,412,493 | No introgression |
| 40 | 1 | 39 | 98,421,174 | 98,439,501 | No introgression |
| 41 | 2 | 0 | 13,002 | 287,947 | A1_155 |
| 42 | 2 | 1 | 310,892 | 783,096 | No introgression |
| 43 | 2 | 2 | 784,227 | 1,473,012 | No introgression |
| 44 | 2 | 3 | 1,485,252 | 2,260,542 | No introgression |
| 45 | 2 | 4 | 2,283,102 | 3,157,960 | No introgression |
| 46 | 2 | 5 | 3,167,358 | 4,242,207 | No introgression |
| 47 | 2 | 6 | 4,278,567 | 5,915,875 | A2_061 |
| 48 | 2 | 7 | 5,982,523 | 7,476,123 | No introgression |
| 49 | 2 | 8 | 7,623,321 | 10,697,145 | No introgression |
| 50 | 2 | 9 | 10,825,814 | 14,864,251 | No introgression |
| 51 | 2 | 10 | 14,987,903 | 19,463,596 | No introgression |
| 52 | 2 | 11 | 19,532,551 | 25,751,692 | No introgression |
| 53 | 2 | 12 | 25,791,299 | 36,360,915 | No introgression |
| 54 | 2 | 13 | 36,435,704 | 48,476,030 | No introgression |
| 55 | 2 | 14 | 48,483,931 | 51,980,821 | No introgression |
| 56 | 2 | 15 | 51,994,767 | 56,781,018 | No introgression |
| 57 | 2 | 16 | 56,991,776 | 59,442,929 | No introgression |

| Window | Chr | Window on Chr | Start | End | Diagnosis |
|---|---|---|---|---|---|
| 58 | 2 | 17 | 59,480,134 | 62,231,133 | No introgression |
| 59 | 2 | 18 | 62,268,493 | 64,262,678 | No introgression |
| 60 | 2 | 19 | 64,300,791 | 65,828,433 | No introgression |
| 61 | 2 | 20 | 65,886,912 | 67,561,044 | No introgression |
| 62 | 2 | 21 | 67,575,989 | 68,838,176 | No introgression |
| 63 | 2 | 22 | 68,851,200 | 70,385,954 | No introgression |
| 64 | 2 | 23 | 70,389,738 | 71,565,849 | No introgression |
| 65 | 2 | 24 | 71,574,075 | 72,393,878 | No introgression |
| 66 | 2 | 25 | 72,411,484 | 73,212,302 | No introgression |
| 67 | 2 | 26 | 73,221,865 | 74,035,738 | No introgression |
| 68 | 2 | 27 | 74,044,868 | 74,899,502 | No introgression |
| 69 | 2 | 28 | 74,931,154 | 75,526,940 | No introgression |
| 70 | 2 | 29 | 75,542,583 | 76,093,305 | No introgression |
| 71 | 2 | 30 | 76,097,421 | 76,289,229 | No introgression |
| 72 | 3 | 0 | 93 | 451,850 | No introgression |
| 73 | 3 | 1 | 457,685 | 1,124,111 | No introgression |
| 74 | 3 | 2 | 1,129,385 | 1,837,035 | A1_155 |
| 75 | 3 | 3 | 1,851,196 | 2,505,639 | A1_155 |
| 76 | 3 | 4 | 2,516,965 | 3,145,288 | A1_155 |
| 77 | 3 | 5 | 3,150,576 | 3,776,637 | No introgression |
| 78 | 3 | 6 | 3,778,038 | 4,466,556 | No introgression |
| 79 | 3 | 7 | 4,467,291 | 5,614,445 | No introgression |
| 80 | 3 | 8 | 5,721,076 | 7,034,063 | No introgression |
| 81 | 3 | 9 | 7,009,767 | 8,065,888 | No introgression |
| 82 | 3 | 10 | 8,073,593 | 9,450,727 | No introgression |
| 83 | 3 | 11 | 9,485,018 | 11,198,151 | No introgression |
| 84 | 3 | 12 | 11,291,035 | 13,111,777 | No introgression |
| 85 | 3 | 13 | 13,117,351 | 15,285,519 | No introgression |
| 86 | 3 | 14 | 15,830,544 | 22,425,814 | No introgression |
| 87 | 3 | 15 | 23,146,039 | 30,897,453 | No introgression |

| Window | Chr | Window on Chr | Start | End | Diagnosis |
|---|---|---|---|---|---|
| 88 | 3 | 16 | 31,328,385 | 38,969,697 | No introgression |
| 89 | 3 | 17 | 39,064,829 | 46,552,808 | No introgression |
| 90 | 3 | 18 | 46,845,611 | 55,900,892 | No introgression |
| 91 | 3 | 19 | 56,728,132 | 61,944,681 | No introgression |
| 92 | 3 | 20 | 61,988,859 | 72,758,660 | No introgression |
| 93 | 3 | 21 | 73,094,075 | 75,825,523 | No introgression |
| 94 | 3 | 22 | 75,829,070 | 79,542,731 | No introgression |
| 95 | 3 | 23 | 80,452,000 | 87,350,791 | No introgression |
| 96 | 3 | 24 | 87,365,188 | 92,062,236 | No introgression |
| 97 | 3 | 25 | 92,090,191 | 94,558,136 | No introgression |
| 98 | 3 | 26 | 94,627,586 | 97,017,935 | No introgression |
| 99 | 3 | 27 | 97,023,031 | 99,758,951 | No introgression |
| 100 | 3 | 28 | 99,876,345 | 101,979,910 | No introgression |
| 101 | 3 | 29 | 101,982,943 | 103,102,234 | No introgression |
| 102 | 3 | 30 | 103,133,823 | 104,708,473 | No introgression |
| 103 | 3 | 31 | 104,720,900 | 105,571,022 | No introgression |
| 104 | 3 | 32 | 105,579,320 | 106,395,953 | No introgression |
| 105 | 3 | 33 | 106,407,899 | 107,463,230 | No introgression |
| 106 | 3 | 34 | 107,483,257 | 108,108,727 | No introgression |
| 107 | 3 | 35 | 108,120,658 | 108,641,743 | No introgression |
| 108 | 3 | 36 | 108,644,897 | 109,111,322 | No introgression |
| 109 | 3 | 37 | 109,173,342 | 109,620,038 | No introgression |
| 110 | 3 | 38 | 109,621,461 | 109,980,036 | A1_051, A1_133, A1_132, A1_054 |
| 111 | 3 | 39 | 109,993,696 | 110,201,909 | No introgression |
| 112 | 4 | 0 | 31,507 | 690,285 | No introgression |
| 113 | 4 | 1 | 698,024 | 1,294,933 | No introgression |
| 114 | 4 | 2 | 1,327,053 | 2,153,434 | No introgression |
| 115 | 4 | 3 | 2,172,553 | 3,148,782 | A2_101 |
| 116 | 4 | 4 | 3,160,963 | 3,973,633 | No introgression |
| 117 | 4 | 5 | 3,977,559 | 4,753,274 | No introgression |

| Window | Chr | Window on Chr | Start | End | Diagnosis |
|---|---|---|---|---|---|
| 118 | 4 | 6 | 4,755,357 | 5,777,552 | No introgression |
| 119 | 4 | 7 | 5,855,092 | 7,045,223 | No introgression |
| 120 | 4 | 8 | 7,075,626 | 7,921,338 | No introgression |
| 121 | 4 | 9 | 7,991,457 | 9,798,768 | No introgression |
| 122 | 4 | 10 | 9,823,220 | 11,019,048 | No introgression |
| 123 | 4 | 11 | 10,755,926 | 12,581,033 | No introgression |
| 124 | 4 | 12 | 12,657,510 | 15,559,777 | No introgression |
| 125 | 4 | 13 | 15,595,911 | 18,481,151 | No introgression |
| 126 | 4 | 14 | 18,487,687 | 20,442,382 | No introgression |
| 127 | 4 | 15 | 20,525,466 | 31,150,252 | No introgression |
| 128 | 4 | 16 | 31,163,360 | 45,187,777 | No introgression |
| 129 | 4 | 17 | 45,632,126 | 58,804,564 | No introgression |
| 130 | 4 | 18 | 58,815,316 | 66,210,252 | No introgression |
| 131 | 4 | 19 | 66,228,215 | 71,379,012 | No introgression |
| 132 | 4 | 20 | 70,625,551 | 73,760,343 | No introgression |
| 133 | 4 | 21 | 73,769,588 | 77,221,539 | No introgression |
| 134 | 4 | 22 | 77,305,900 | 80,359,096 | No introgression |
| 135 | 4 | 23 | 80,500,841 | 82,038,235 | No introgression |
| 136 | 4 | 24 | 82,167,157 | 84,358,484 | No introgression |
| 137 | 4 | 25 | 84,366,821 | 85,870,576 | Gar04 |
| 138 | 4 | 26 | 85,947,128 | 86,876,305 | No introgression |
| 139 | 4 | 27 | 86,887,522 | 88,056,755 | A2_065 |
| 140 | 4 | 28 | 88,076,573 | 88,700,733 | A1_074 |
| 141 | 4 | 29 | 88,709,297 | 89,139,378 | Complex |
| 142 | 4 | 30 | 89,139,569 | 89,802,659 | No introgression |
| 143 | 4 | 31 | 89,894,293 | 90,526,505 | No introgression |
| 144 | 4 | 32 | 90,544,719 | 90,978,759 | No introgression |
| 145 | 5 | 0 | 45,790 | 567,417 | No introgression |
| 146 | 5 | 1 | 582,741 | 1,039,386 | No introgression |
| 147 | 5 | 2 | 1,064,217 | 1,515,686 | No introgression |

| Window | Chr | Window on Chr | Start | End | Diagnosis |
|---|---|---|---|---|---|
| 148 | 5 | 3 | 1,520,905 | 2,067,289 | No introgression |
| 149 | 5 | 4 | 2,073,804 | 2,462,361 | No introgression |
| 150 | 5 | 5 | 2,517,849 | 2,916,403 | No introgression |
| 151 | 5 | 6 | 2,959,254 | 3,354,288 | No introgression |
| 152 | 5 | 7 | 3,356,746 | 3,803,400 | No introgression |
| 153 | 5 | 8 | 3,808,978 | 4,254,266 | No introgression |
| 154 | 5 | 9 | 4,262,551 | 4,627,931 | A1_155 |
| 155 | 5 | 10 | 4,637,158 | 4,993,128 | A1_155 |
| 156 | 5 | 11 | 5,003,090 | 5,449,381 | A1_155 |
| 157 | 5 | 12 | 5,456,299 | 5,843,709 | No introgression |
| 158 | 5 | 13 | 5,865,211 | 6,408,432 | No introgression |
| 159 | 5 | 14 | 6,399,847 | 6,889,211 | A1_155 |
| 160 | 5 | 15 | 6,901,739 | 7,321,900 | A1_155 |
| 161 | 5 | 16 | 7,329,262 | 7,689,364 | A1_155 |
| 162 | 5 | 17 | 7,698,999 | 8,152,965 | A1_155 |
| 163 | 5 | 18 | 8,168,745 | 8,689,603 | A1_155 |
| 164 | 5 | 19 | 8,690,592 | 9,257,717 | No introgression |
| 165 | 5 | 20 | 9,268,997 | 9,890,445 | No introgression |
| 166 | 5 | 21 | 9,892,820 | 10,433,072 | No introgression |
| 167 | 5 | 22 | 10,450,225 | 10,896,209 | No introgression |
| 168 | 5 | 23 | 10,913,395 | 11,544,154 | No introgression |
| 169 | 5 | 24 | 11,557,260 | 12,049,441 | No introgression |
| 170 | 5 | 25 | 12,060,856 | 12,582,038 | Complex |
| 171 | 5 | 26 | 12,602,478 | 13,157,354 | A1_155 |
| 172 | 5 | 27 | 13,159,027 | 13,774,198 | No introgression |
| 173 | 5 | 28 | 13,775,746 | 14,236,864 | A1_155 |
| 174 | 5 | 29 | 14,244,852 | 14,695,848 | No introgression |
| 175 | 5 | 30 | 14,704,737 | 15,201,120 | No introgression |
| 176 | 5 | 31 | 15,212,001 | 15,722,451 | No introgression |
| 177 | 5 | 32 | 15,723,430 | 16,264,573 | No introgression |

| Window | Chr | Window on Chr | Start | End | Diagnosis |
|---|---|---|---|---|---|
| 178 | 5 | 33 | 16,272,133 | 16,846,268 | No introgression |
| 179 | 5 | 34 | 16,958,169 | 17,566,423 | No introgression |
| 180 | 5 | 35 | 17,571,765 | 18,130,582 | No introgression |
| 181 | 5 | 36 | 18,167,686 | 18,934,467 | No introgression |
| 182 | 5 | 37 | 18,967,866 | 19,690,312 | No introgression |
| 183 | 5 | 38 | 19,711,979 | 20,420,718 | No introgression |
| 184 | 5 | 39 | 20,432,431 | 21,187,573 | No introgression |
| 185 | 5 | 40 | 21,222,993 | 22,612,355 | No introgression |
| 186 | 5 | 41 | 22,628,184 | 23,323,944 | No introgression |
| 187 | 5 | 42 | 23,348,430 | 24,293,808 | No introgression |
| 188 | 5 | 43 | 24,294,687 | 25,288,043 | No introgression |
| 189 | 5 | 44 | 25,309,629 | 26,152,758 | No introgression |
| 190 | 5 | 45 | 26,207,826 | 27,883,428 | No introgression |
| 191 | 5 | 46 | 27,958,816 | 28,830,068 | No introgression |
| 192 | 5 | 47 | 28,884,000 | 31,270,837 | No introgression |
| 193 | 5 | 48 | 31,307,780 | 33,674,888 | No introgression |
| 194 | 5 | 49 | 33,710,410 | 34,966,507 | A1_051, A1_133, A1_132, A1_054 |
| 195 | 5 | 50 | 34,990,524 | 37,778,760 | No introgression |
| 196 | 5 | 51 | 38,089,076 | 41,190,771 | No introgression |
| 197 | 5 | 52 | 41,594,780 | 49,274,219 | No introgression |
| 198 | 5 | 53 | 49,522,425 | 53,625,088 | No introgression |
| 199 | 5 | 54 | 53,983,664 | 61,306,297 | No introgression |
| 200 | 5 | 55 | 61,660,868 | 65,441,723 | No introgression |
| 201 | 5 | 56 | 65,450,970 | 67,211,237 | No introgression |
| 202 | 5 | 57 | 67,381,604 | 68,979,299 | No introgression |
| 203 | 5 | 58 | 69,002,983 | 69,984,292 | A1_155 |
| 204 | 5 | 59 | 69,996,695 | 71,057,243 | No introgression |
| 205 | 5 | 60 | 71,097,734 | 72,609,302 | No introgression |
| 206 | 5 | 61 | 72,653,627 | 74,248,239 | No introgression |
| 207 | 5 | 62 | 74,325,051 | 75,103,056 | No introgression |

| Window | Chr | Window on Chr | Start | End | Diagnosis |
|---|---|---|---|---|---|
| 208 | 5 | 63 | 75,173,470 | 75,787,573 | No introgression |
| 209 | 5 | 64 | 75,801,974 | 76,318,549 | No introgression |
| 210 | 6 | 0 | 23,761 | 584,267 | No introgression |
| 211 | 6 | 1 | 618,449 | 1,339,718 | No introgression |
| 212 | 6 | 2 | 1,343,196 | 1,823,536 | No introgression |
| 213 | 6 | 3 | 2,019,858 | 2,749,228 | No introgression |
| 214 | 6 | 4 | 2,758,691 | 3,602,605 | No introgression |
| 215 | 6 | 5 | 3,806,736 | 5,394,868 | No introgression |
| 216 | 6 | 6 | 5,444,528 | 6,572,720 | No introgression |
| 217 | 6 | 7 | 6,596,987 | 7,401,525 | No introgression |
| 218 | 6 | 8 | 7,471,622 | 8,834,538 | No introgression |
| 219 | 6 | 9 | 8,885,035 | 10,315,833 | No introgression |
| 220 | 6 | 10 | 10,336,694 | 14,098,754 | No introgression |
| 221 | 6 | 11 | 14,144,268 | 15,419,507 | No introgression |
| 222 | 6 | 12 | 15,604,513 | 17,594,377 | No introgression |
| 223 | 6 | 13 | 17,668,974 | 22,515,760 | No introgression |
| 224 | 6 | 14 | 22,953,924 | 27,213,216 | No introgression |
| 225 | 6 | 15 | 27,228,538 | 30,554,610 | No introgression |
| 226 | 6 | 16 | 30,629,803 | 34,466,698 | No introgression |
| 227 | 6 | 17 | 34,471,777 | 37,899,424 | No introgression |
| 228 | 6 | 18 | 37,906,183 | 44,409,183 | No introgression |
| 229 | 6 | 19 | 44,475,363 | 53,194,093 | No introgression |
| 230 | 6 | 20 | 53,424,311 | 59,278,779 | No introgression |
| 231 | 6 | 21 | 59,291,443 | 65,560,236 | No introgression |
| 232 | 6 | 22 | 65,651,694 | 73,766,070 | No introgression |
| 233 | 6 | 23 | 74,147,023 | 81,631,338 | No introgression |
| 234 | 6 | 24 | 81,855,968 | 85,504,140 | No introgression |
| 235 | 6 | 25 | 85,550,521 | 89,021,518 | No introgression |
| 236 | 6 | 26 | 89,025,599 | 90,447,284 | No introgression |
| 237 | 6 | 27 | 90,449,013 | 91,994,724 | A1_054 |

| Window | Chr | Window on Chr | Start | End | Diagnosis |
|---|---|---|---|---|---|
| 238 | 6 | 28 | 92,034,470 | 93,717,730 | No introgression |
| 239 | 6 | 29 | 93,768,396 | 94,698,362 | No introgression |
| 240 | 6 | 30 | 94,724,656 | 95,841,418 | No introgression |
| 241 | 6 | 31 | 95,851,863 | 96,712,000 | No introgression |
| 242 | 6 | 32 | 96,717,967 | 97,271,981 | No introgression |
| 243 | 6 | 33 | 97,284,058 | 97,764,590 | No introgression |
| 244 | 6 | 34 | 97,772,488 | 98,049,306 | No introgression |
| 245 | 7 | 0 | 62,203 | 609,103 | No introgression |
| 246 | 7 | 1 | 611,336 | 1,297,193 | No introgression |
| 247 | 7 | 2 | 1,299,792 | 1,862,892 | No introgression |
| 248 | 7 | 3 | 1,876,573 | 2,534,218 | No introgression |
| 249 | 7 | 4 | 2,535,894 | 3,054,676 | No introgression |
| 250 | 7 | 5 | 3,065,856 | 3,596,298 | No introgression |
| 251 | 7 | 6 | 3,602,775 | 4,256,790 | No introgression |
| 252 | 7 | 7 | 4,309,563 | 4,963,252 | No introgression |
| 253 | 7 | 8 | 4,984,169 | 5,752,883 | No introgression |
| 254 | 7 | 9 | 5,764,757 | 6,685,604 | No introgression |
| 255 | 7 | 10 | 6,740,859 | 7,452,999 | No introgression |
| 256 | 7 | 11 | 7,484,442 | 8,388,205 | No introgression |
| 257 | 7 | 12 | 8,404,791 | 9,766,516 | No introgression |
| 258 | 7 | 13 | 9,805,597 | 10,430,017 | Complex |
| 259 | 7 | 14 | 10,451,023 | 12,561,882 | No introgression |
| 260 | 7 | 15 | 12,934,019 | 13,570,737 | No introgression |
| 261 | 7 | 16 | 13,572,141 | 14,846,214 | No introgression |
| 262 | 7 | 17 | 14,939,489 | 16,102,782 | A2_057, A2_043 |
| 263 | 7 | 18 | 16,296,016 | 17,800,195 | No introgression |
| 264 | 7 | 19 | 17,805,506 | 19,818,251 | A1_051, A1_133, A1_132 |
| 265 | 7 | 20 | 19,838,042 | 21,619,152 | No introgression |
| 266 | 7 | 21 | 21,676,569 | 23,653,787 | A2_085 |
| 267 | 7 | 22 | 23,695,275 | 25,375,527 | A1_155 |

| Window | Chr | Window on Chr | Start | End | Diagnosis |
|---|---|---|---|---|---|
| 268 | 7 | 23 | 25,437,938 | 27,785,523 | Complex |
| 269 | 7 | 24 | 27,856,832 | 31,085,418 | No introgression |
| 270 | 7 | 25 | 31,229,438 | 35,480,735 | A1_155 |
| 271 | 7 | 26 | 35,573,636 | 39,022,440 | No introgression |
| 272 | 7 | 27 | 39,024,353 | 44,499,931 | A1_155 |
| 273 | 7 | 28 | 44,724,501 | 52,560,561 | A1_155 |
| 274 | 7 | 29 | 52,598,135 | 60,404,127 | A1_155 |
| 275 | 7 | 30 | 62,744,397 | 68,280,179 | No introgression |
| 276 | 7 | 31 | 68,402,937 | 71,332,473 | No introgression |
| 277 | 7 | 32 | 71,342,618 | 74,435,620 | No introgression |
| 278 | 7 | 33 | 74,465,022 | 76,834,961 | A1_155 |
| 279 | 7 | 34 | 76,860,337 | 77,733,220 | No introgression |
| 280 | 7 | 35 | 77,815,777 | 78,820,493 | A1_155 |
| 281 | 7 | 36 | 78,848,140 | 79,683,327 | A1_155 |
| 282 | 7 | 37 | 79,691,715 | 80,461,465 | No introgression |
| 283 | 7 | 38 | 80,491,011 | 81,389,145 | No introgression |
| 284 | 7 | 39 | 81,400,263 | 81,885,277 | No introgression |
| 285 | 7 | 40 | 81,893,913 | 82,181,728 | No introgression |
| 286 | 8 | 0 | 890 | 464,555 | No introgression |
| 287 | 8 | 1 | 484,336 | 1,146,948 | A1_155 |
| 288 | 8 | 2 | 1,166,441 | 1,791,132 | No introgression |
| 289 | 8 | 3 | 1,808,824 | 2,366,561 | No introgression |
| 290 | 8 | 4 | 2,378,318 | 2,883,902 | No introgression |
| 291 | 8 | 5 | 2,925,710 | 3,797,441 | No introgression |
| 292 | 8 | 6 | 3,810,081 | 4,843,500 | No introgression |
| 293 | 8 | 7 | 4,949,442 | 5,182,378 | No introgression |
| 294 | 8 | 8 | 6,016,419 | 7,038,560 | No introgression |
| 295 | 8 | 9 | 7,087,019 | 8,719,044 | No introgression |
| 296 | 8 | 10 | 8,721,621 | 10,331,524 | No introgression |
| 297 | 8 | 11 | 10,377,313 | 13,058,853 | No introgression |

| Window | Chr | Window on Chr | Start | End | Diagnosis |
|---|---|---|---|---|---|
| 298 | 8 | 12 | 13,399,180 | 16,393,581 | No introgression |
| 299 | 8 | 13 | 16,417,233 | 23,319,523 | No introgression |
| 300 | 8 | 14 | 24,376,924 | 34,658,092 | No introgression |
| 301 | 8 | 15 | 34,800,511 | 46,349,763 | No introgression |
| 302 | 8 | 16 | 46,449,260 | 53,166,304 | No introgression |
| 303 | 8 | 17 | 53,172,926 | 62,202,913 | No introgression |
| 304 | 8 | 18 | 62,289,915 | 68,837,049 | No introgression |
| 305 | 8 | 19 | 68,977,409 | 75,471,755 | No introgression |
| 306 | 8 | 20 | 75,584,384 | 78,114,713 | A1_051, A1_054, A1_133, A1_132, A1_077 |
| 307 | 8 | 21 | 78,142,423 | 82,492,546 | A1_051, A1_077, A1_132, A1_133 |
| 308 | 8 | 22 | 82,676,448 | 84,461,950 | A1_051, A1_077, A1_132, A1_133 |
| 309 | 8 | 23 | 84,506,388 | 86,270,986 | No introgression |
| 310 | 8 | 24 | 86,283,976 | 87,814,971 | No introgression |
| 311 | 8 | 25 | 88,007,560 | 90,167,669 | A2_062, Gar02 |
| 312 | 8 | 26 | 90,173,101 | 91,457,348 | A1_155 |
| 313 | 8 | 27 | 91,463,426 | 92,519,620 | A1_155 |
| 314 | 8 | 28 | 92,589,716 | 94,267,303 | No introgression |
| 315 | 8 | 29 | 94,326,473 | 95,700,222 | A1_155 |
| 316 | 8 | 30 | 95,713,256 | 97,196,591 | A1_155 |
| 317 | 8 | 31 | 97,198,098 | 98,101,375 | No introgression |
| 318 | 8 | 32 | 98,111,505 | 98,793,447 | No introgression |
| 319 | 8 | 33 | 98,804,895 | 99,831,364 | No introgression |
| 320 | 8 | 34 | 99,873,679 | 100,460,294 | No introgression |
| 321 | 8 | 35 | 100,473,338 | 100,945,907 | A1_155 |
| 322 | 8 | 36 | 100,951,479 | 101,547,096 | No introgression |
| 323 | 8 | 37 | 101,553,620 | 102,088,786 | No introgression |
| 324 | 8 | 38 | 102,095,318 | 102,736,230 | A1_155 |
| 325 | 8 | 39 | 102,749,920 | 103,379,606 | No introgression |
| 326 | 8 | 40 | 103,389,356 | 103,987,552 | No introgression |
| 327 | 8 | 41 | 104,022,474 | 104,459,473 | No introgression |

| Window | Chr | Window on Chr | Start | End | Diagnosis |
|---|---|---|---|---|---|
| 328 | 8 | 42 | 104,473,184 | 104,586,489 | No introgression |
| 329 | 9 | 0 | 36,170 | 817,027 | A1_155 |
| 330 | 9 | 1 | 902,870 | 2,000,882 | Complex |
| 331 | 9 | 2 | 2,010,238 | 3,131,613 | No introgression |
| 332 | 9 | 3 | 3,177,992 | 4,603,074 | Complex |
| 333 | 9 | 4 | 4,611,668 | 6,087,215 | No introgression |
| 334 | 9 | 5 | 5,982,918 | 8,106,033 | No introgression |
| 335 | 9 | 6 | 8,366,353 | 11,866,207 | No introgression |
| 336 | 9 | 7 | 11,925,386 | 21,505,956 | No introgression |
| 337 | 9 | 8 | 21,625,714 | 35,205,490 | No introgression |
| 338 | 9 | 9 | 35,219,054 | 38,218,204 | No introgression |
| 339 | 9 | 10 | 38,228,819 | 43,967,174 | No introgression |
| 340 | 9 | 11 | 43,971,462 | 45,928,116 | No introgression |
| 341 | 9 | 12 | 45,933,750 | 48,164,997 | Complex |
| 342 | 9 | 13 | 48,210,694 | 49,478,175 | No introgression |
| 343 | 9 | 14 | 49,510,254 | 50,748,640 | Complex |
| 344 | 9 | 15 | 50,802,636 | 52,044,801 | No introgression |
| 345 | 9 | 16 | 52,089,736 | 53,771,836 | No introgression |
| 346 | 9 | 17 | 53,783,884 | 54,919,509 | No introgression |
| 347 | 9 | 18 | 55,056,617 | 55,760,475 | A1_051, A1_054, A1_132, A1_133 |
| 348 | 9 | 19 | 55,762,981 | 57,023,277 | No introgression |
| 349 | 9 | 20 | 57,026,679 | 58,038,330 | No introgression |
| 350 | 9 | 21 | 58,056,011 | 59,203,439 | No introgression |
| 351 | 9 | 22 | 59,212,137 | 59,942,757 | No introgression |
| 352 | 9 | 23 | 59,984,864 | 60,754,903 | No introgression |
| 353 | 9 | 24 | 60,804,494 | 61,815,448 | No introgression |
| 354 | 9 | 25 | 61,844,943 | 62,448,568 | A1_051, A1_054, A1_133, A1_132 |
| 355 | 9 | 26 | 62,449,091 | 62,988,241 | No introgression |
| 356 | 9 | 27 | 62,993,095 | 63,596,841 | No introgression |
| 357 | 9 | 28 | 63,614,403 | 64,172,402 | No introgression |

| Window | Chr | Window on Chr | Start | End | Diagnosis |
|---|---|---|---|---|---|
| 358 | 9 | 29 | 64,238,365 | 64,946,933 | No introgression |
| 359 | 9 | 30 | 64,957,232 | 65,538,536 | No introgression |
| 360 | 9 | 31 | 65,545,352 | 66,034,195 | No introgression |
| 361 | 9 | 32 | 66,041,121 | 66,402,704 | No introgression |
| 362 | 9 | 33 | 66,477,366 | 66,966,371 | No introgression |
| 363 | 9 | 34 | 66,997,870 | 67,434,035 | No introgression |
| 364 | 9 | 35 | 67,440,687 | 67,953,247 | No introgression |
| 365 | 9 | 36 | 67,971,933 | 68,465,827 | No introgression |
| 366 | 9 | 37 | 68,270,744 | 69,056,916 | No introgression |
| 367 | 9 | 38 | 69,066,504 | 69,524,086 | No introgression |
| 368 | 9 | 39 | 69,528,447 | 70,014,929 | A2_047, A2_074, A2_141, A2_154 |
| 369 | 9 | 40 | 70,041,901 | 70,468,049 | No introgression |
| 370 | 9 | 41 | 70,493,696 | 70,589,496 | A2_074 |
| 371 | 10 | 0 | 50,135 | 511,682 | Gar04 |
| 372 | 10 | 1 | 520,592 | 922,315 | No introgression |
| 373 | 10 | 2 | 926,126 | 1,330,820 | Complex |
| 374 | 10 | 3 | 1,352,909 | 2,087,186 | Complex |
| 375 | 10 | 4 | 2,089,862 | 2,536,973 | No resolution |
| 376 | 10 | 5 | 2,548,602 | 3,098,600 | Gar04, A2_041 |
| 377 | 10 | 6 | 3,106,020 | 3,697,758 | Gar04 |
| 378 | 10 | 7 | 3,700,467 | 4,282,981 | No introgression |
| 379 | 10 | 8 | 4,287,612 | 5,100,561 | No introgression |
| 380 | 10 | 9 | 5,199,789 | 6,032,869 | No introgression |
| 381 | 10 | 10 | 6,034,810 | 7,134,577 | No introgression |
| 382 | 10 | 11 | 7,148,838 | 8,297,137 | No introgression |
| 383 | 10 | 12 | 8,302,698 | 9,613,579 | A1_132, A1_133 |
| 384 | 10 | 13 | 9,625,786 | 11,045,001 | No introgression |
| 385 | 10 | 14 | 11,064,056 | 12,243,491 | No introgression |
| 386 | 10 | 15 | 12,294,495 | 13,859,754 | No introgression |
| 387 | 10 | 16 | 14,173,537 | 17,563,051 | No introgression |

| Window | Chr | Window on Chr | Start | End | Diagnosis |
|---|---|---|---|---|---|
| 388 | 10 | 17 | 16,536,253 | 19,868,480 | No introgression |
| 389 | 10 | 18 | 19,978,840 | 22,483,273 | A1_155 |
| 390 | 10 | 19 | 22,501,595 | 26,988,976 | No introgression |
| 391 | 10 | 20 | 27,027,685 | 29,547,836 | No introgression |
| 392 | 10 | 21 | 29,605,139 | 31,666,676 | No introgression |
| 393 | 10 | 22 | 31,760,019 | 34,370,708 | A2_067, A2_079, Gar61, Gar37 |
| 394 | 10 | 23 | 34,381,415 | 37,971,275 | No introgression |
| 395 | 10 | 24 | 38,007,925 | 44,403,454 | A2_008, A2_096 |
| 396 | 10 | 25 | 45,026,439 | 52,898,312 | No introgression |
| 397 | 10 | 26 | 53,385,679 | 65,651,549 | No introgression |
| 398 | 10 | 27 | 65,653,388 | 69,226,177 | No introgression |
| 399 | 10 | 28 | 69,271,403 | 72,127,909 | No introgression |
| 400 | 10 | 29 | 72,273,729 | 75,643,180 | No introgression |
| 401 | 10 | 30 | 75,644,875 | 78,599,809 | No introgression |
| 402 | 10 | 31 | 78,675,019 | 80,031,438 | Gar04 |
| 403 | 10 | 32 | 80,034,765 | 81,504,650 | No introgression |
| 404 | 10 | 33 | 81,656,801 | 83,545,359 | No introgression |
| 405 | 10 | 34 | 83,608,345 | 84,397,330 | No introgression |
| 406 | 10 | 35 | 84,469,569 | 86,269,586 | No introgression |
| 407 | 10 | 36 | 86,352,823 | 87,690,171 | No introgression |
| 408 | 10 | 37 | 87,694,229 | 88,834,692 | No introgression |
| 409 | 10 | 38 | 88,857,223 | 90,218,905 | No introgression |
| 410 | 10 | 39 | 90,258,072 | 91,593,760 | No introgression |
| 411 | 10 | 40 | 91,600,221 | 92,289,749 | No introgression |
| 412 | 10 | 41 | 92,347,425 | 92,973,889 | No introgression |
| 413 | 10 | 42 | 92,986,842 | 94,163,975 | Complex |
| 414 | 10 | 43 | 94,179,655 | 94,959,048 | No introgression |
| 415 | 10 | 44 | 94,967,981 | 95,053,312 | A1_051, A1_132, A1_133 |
| 416 | 11 | 0 | 25,362 | 483,015 | No introgression |
| 417 | 11 | 1 | 488,084 | 1,050,474 | No introgression |

| Window | Chr | Window on Chr | Start | End | Diagnosis |
|---|---|---|---|---|---|
| 418 | 11 | 2 | 1,056,772 | 1,490,670 | No introgression |
| 419 | 11 | 3 | 1,499,101 | 1,966,306 | A1_155 |
| 420 | 11 | 4 | 1,962,283 | 2,384,631 | A1_155 |
| 421 | 11 | 5 | 2,400,487 | 2,882,632 | No introgression |
| 422 | 11 | 6 | 2,910,132 | 3,331,471 | No introgression |
| 423 | 11 | 7 | 3,360,561 | 3,813,956 | No introgression |
| 424 | 11 | 8 | 3,821,941 | 4,305,605 | No introgression |
| 425 | 11 | 9 | 4,310,369 | 4,833,830 | No introgression |
| 426 | 11 | 10 | 4,844,309 | 5,286,419 | No introgression |
| 427 | 11 | 11 | 5,297,495 | 5,731,429 | No introgression |
| 428 | 11 | 12 | 5,762,901 | 6,251,572 | No introgression |
| 429 | 11 | 13 | 6,270,684 | 6,864,764 | No introgression |
| 430 | 11 | 14 | 6,873,005 | 7,289,693 | No introgression |
| 431 | 11 | 15 | 7,313,316 | 7,868,246 | No introgression |
| 432 | 11 | 16 | 7,876,200 | 8,641,869 | No introgression |
| 433 | 11 | 17 | 8,674,657 | 9,319,038 | No introgression |
| 434 | 11 | 18 | 9,331,032 | 9,989,744 | A1_155 |
| 435 | 11 | 19 | 10,073,172 | 10,666,672 | No introgression |
| 436 | 11 | 20 | 10,690,876 | 11,444,082 | No introgression |
| 437 | 11 | 21 | 11,501,430 | 12,466,668 | No introgression |
| 438 | 11 | 22 | 12,473,161 | 13,336,281 | No introgression |
| 439 | 11 | 23 | 13,380,397 | 14,591,447 | No introgression |
| 440 | 11 | 24 | 14,609,032 | 15,292,150 | No introgression |
| 441 | 11 | 25 | 15,304,505 | 16,739,284 | No introgression |
| 442 | 11 | 26 | 16,783,202 | 17,800,763 | No introgression |
| 443 | 11 | 27 | 17,849,231 | 19,104,041 | No introgression |
| 444 | 11 | 28 | 19,139,821 | 20,539,595 | No introgression |
| 445 | 11 | 29 | 20,692,973 | 22,903,412 | No introgression |
| 446 | 11 | 30 | 22,924,909 | 26,349,743 | No introgression |
| 447 | 11 | 31 | 26,484,081 | 29,899,579 | No introgression |

| Window | Chr | Window on Chr | Start | End | Diagnosis |
|---|---|---|---|---|---|
| 448 | 11 | 32 | 29,959,399 | 33,545,383 | No introgression |
| 449 | 11 | 33 | 33,566,281 | 37,031,914 | No introgression |
| 450 | 11 | 34 | 37,181,559 | 43,324,611 | No introgression |
| 451 | 11 | 35 | 43,358,732 | 51,126,125 | No introgression |
| 452 | 11 | 36 | 51,279,925 | 61,753,088 | No introgression |
| 453 | 11 | 37 | 61,881,074 | 66,882,369 | No introgression |
| 454 | 11 | 38 | 67,005,772 | 72,728,983 | No introgression |
| 455 | 11 | 39 | 72,680,815 | 75,348,799 | No introgression |
| 456 | 11 | 40 | 75,354,519 | 78,303,660 | No introgression |
| 457 | 11 | 41 | 78,740,588 | 82,200,578 | No introgression |
| 458 | 11 | 42 | 82,274,534 | 83,259,778 | No introgression |
| 459 | 11 | 43 | 83,298,842 | 84,744,738 | No introgression |
| 460 | 11 | 44 | 84,858,057 | 86,010,703 | No introgression |
| 461 | 11 | 45 | 86,052,643 | 86,973,434 | No introgression |
| 462 | 11 | 46 | 87,073,145 | 88,055,982 | No introgression |
| 463 | 11 | 47 | 88,065,634 | 88,865,094 | No introgression |
| 464 | 11 | 48 | 89,336,271 | 89,887,270 | No introgression |
| 465 | 11 | 49 | 90,201,905 | 91,247,515 | Complex |
| 466 | 11 | 50 | 91,294,305 | 92,515,402 | No introgression |
| 467 | 11 | 51 | 92,535,569 | 93,655,663 | Ghe01, A1_051, A1_054, A1_132, A1_133 |
| 468 | 11 | 52 | 93,781,903 | 94,702,575 | Complex |
| 469 | 11 | 53 | 94,713,216 | 96,096,394 | A1_054 |
| 470 | 11 | 54 | 96,112,107 | 96,858,024 | No introgression |
| 471 | 11 | 55 | 96,866,284 | 97,997,428 | No introgression |
| 472 | 11 | 56 | 97,958,058 | 98,647,945 | No introgression |
| 473 | 11 | 57 | 98,675,755 | 99,354,429 | A1_051, A1_132, A1_133 |
| 474 | 11 | 58 | 99,356,034 | 99,568,638 | A2_074 |
| 475 | 12 | 0 | 25,910 | 512,002 | No introgression |
| 476 | 12 | 1 | 516,029 | 1,180,579 | No introgression |
| 477 | 12 | 2 | 1,216,325 | 1,909,232 | No introgression |

| Window | Chr | Window on Chr | Start | End | Diagnosis |
|---|---|---|---|---|---|
| 478 | 12 | 3 | 1,971,540 | 2,931,671 | No introgression |
| 479 | 12 | 4 | 2,942,816 | 4,014,718 | No introgression |
| 480 | 12 | 5 | 4,015,945 | 5,418,365 | No introgression |
| 481 | 12 | 6 | 5,468,857 | 6,613,264 | No introgression |
| 482 | 12 | 7 | 6,627,925 | 7,878,010 | No introgression |
| 483 | 12 | 8 | 7,883,746 | 9,344,188 | No introgression |
| 484 | 12 | 9 | 9,371,706 | 11,766,191 | No introgression |
| 485 | 12 | 10 | 11,767,801 | 13,352,370 | No introgression |
| 486 | 12 | 11 | 13,365,105 | 16,025,388 | No introgression |
| 487 | 12 | 12 | 16,088,520 | 23,410,503 | No introgression |
| 488 | 12 | 13 | 23,413,887 | 33,909,832 | No introgression |
| 489 | 12 | 14 | 33,943,176 | 40,723,575 | No introgression |
| 490 | 12 | 15 | 40,786,393 | 51,180,456 | No introgression |
| 491 | 12 | 16 | 51,234,118 | 60,679,169 | No introgression |
| 492 | 12 | 17 | 60,681,462 | 63,477,064 | No introgression |
| 493 | 12 | 18 | 63,495,549 | 64,840,519 | No introgression |
| 494 | 12 | 19 | 65,083,804 | 66,139,503 | No introgression |
| 495 | 12 | 20 | 66,208,266 | 68,038,417 | Complex |
| 496 | 12 | 21 | 68,116,073 | 69,257,418 | No introgression |
| 497 | 12 | 22 | 69,298,375 | 70,525,212 | No introgression |
| 498 | 12 | 23 | 70,563,475 | 72,246,000 | No introgression |
| 499 | 12 | 24 | 72,297,081 | 74,349,208 | No introgression |
| 500 | 12 | 25 | 74,405,570 | 76,863,153 | No introgression |
| 501 | 12 | 26 | 76,948,417 | 77,859,906 | No introgression |
| 502 | 12 | 27 | 77,867,769 | 78,734,283 | No introgression |
| 503 | 12 | 28 | 78,751,806 | 80,310,546 | No introgression |
| 504 | 12 | 29 | 80,321,120 | 81,388,593 | No introgression |
| 505 | 12 | 30 | 81,407,298 | 82,298,913 | No introgression |
| 506 | 12 | 31 | 82,383,870 | 82,986,531 | No introgression |
| 507 | 12 | 32 | 82,990,716 | 84,172,498 | No introgression |

| Window | Chr | Window on Chr | Start | End | Diagnosis |
|---|---|---|---|---|---|
| 508 | 12 | 33 | 84,296,042 | 84,836,307 | No introgression |
| 509 | 12 | 34 | 84,851,425 | 85,436,193 | No introgression |
| 510 | 12 | 35 | 85,445,389 | 85,970,013 | No introgression |
| 511 | 12 | 36 | 85,973,147 | 86,669,672 | No introgression |
| 512 | 12 | 37 | 86,693,449 | 87,158,499 | No introgression |
| 513 | 12 | 38 | 87,269,143 | 87,721,001 | No introgression |
| 514 | 12 | 39 | 87,760,350 | 88,185,296 | No introgression |
| 515 | 12 | 40 | 88,195,566 | 88,686,535 | No introgression |
| 516 | 12 | 41 | 88,697,920 | 89,203,531 | No introgression |
| 517 | 12 | 42 | 89,241,748 | 89,345,523 | No introgression |
| 518 | 12 | 43 | 89,775,254 | 90,166,749 | No introgression |
| 519 | 12 | 44 | 90,182,464 | 90,757,281 | No introgression |
| 520 | 12 | 45 | 90,778,616 | 91,242,533 | No introgression |
| 521 | 13 | 0 | 200,413 | 695,257 | No introgression |
| 522 | 13 | 1 | 699,634 | 1,159,022 | No introgression |
| 523 | 13 | 2 | 1,167,054 | 1,708,434 | No introgression |
| 524 | 13 | 3 | 1,718,917 | 2,504,159 | No introgression |
| 525 | 13 | 4 | 2,517,170 | 3,278,436 | No introgression |
| 526 | 13 | 5 | 3,293,062 | 4,223,188 | No introgression |
| 527 | 13 | 6 | 4,253,253 | 5,193,780 | No introgression |
| 528 | 13 | 7 | 5,200,000 | 7,038,181 | No introgression |
| 529 | 13 | 8 | 6,835,278 | 8,262,220 | No introgression |
| 530 | 13 | 9 | 8,311,329 | 9,589,521 | No introgression |
| 531 | 13 | 10 | 9,600,464 | 11,548,873 | No introgression |
| 532 | 13 | 11 | 11,635,977 | 16,858,876 | No introgression |
| 533 | 13 | 12 | 16,863,995 | 21,786,932 | No introgression |
| 534 | 13 | 13 | 21,800,511 | 25,865,447 | No introgression |
| 535 | 13 | 14 | 26,571,947 | 34,500,136 | No introgression |
| 536 | 13 | 15 | 34,712,717 | 47,446,837 | No introgression |
| 537 | 13 | 16 | 48,022,668 | 55,358,122 | No introgression |

| Window | Chr | Window on Chr | Start | End | Diagnosis |
|---|---|---|---|---|---|
| 538 | 13 | 17 | 55,405,365 | 58,674,891 | No introgression |
| 539 | 13 | 18 | 59,011,374 | 62,897,778 | No introgression |
| 540 | 13 | 19 | 62,907,281 | 66,209,071 | No introgression |
| 541 | 13 | 20 | 66,246,240 | 70,429,187 | No introgression |
| 542 | 13 | 21 | 70,624,701 | 74,425,242 | No introgression |
| 543 | 13 | 22 | 74,522,622 | 76,491,605 | No introgression |
| 544 | 13 | 23 | 76,497,131 | 79,681,431 | No introgression |
| 545 | 13 | 24 | 79,722,878 | 81,039,542 | No introgression |
| 546 | 13 | 25 | 81,044,133 | 82,223,132 | No introgression |
| 547 | 13 | 26 | 82,295,542 | 83,407,224 | No introgression |
| 548 | 13 | 27 | 83,418,882 | 84,856,059 | No introgression |
| 549 | 13 | 28 | 84,883,164 | 86,486,967 | No introgression |
| 550 | 13 | 29 | 86,547,489 | 87,809,134 | No introgression |
| 551 | 13 | 30 | 87,828,801 | 88,940,808 | No introgression |
| 552 | 13 | 31 | 88,947,886 | 89,628,280 | No introgression |
| 553 | 13 | 32 | 89,331,044 | 90,398,101 | No introgression |
| 554 | 13 | 33 | 90,402,775 | 90,958,138 | No introgression |
| 555 | 13 | 34 | 90,972,196 | 91,602,508 | No introgression |
| 556 | 13 | 35 | 91,617,632 | 92,358,013 | No introgression |
| 557 | 13 | 36 | 92,362,995 | 93,685,962 | No introgression |
| 558 | 13 | 37 | 93,694,802 | 94,362,121 | No introgression |
| 559 | 13 | 38 | 94,369,422 | 94,963,450 | No introgression |
| 560 | 13 | 39 | 94,972,380 | 95,459,175 | No introgression |
| 561 | 13 | 40 | 95,463,642 | 95,809,270 | No introgression |
| 562 | 13 | 41 | 95,812,061 | 95,831,174 | No introgression |

| Window | Complex Placement for these accessions; Introgression cannot be accepted or ruled out |
|---|---|
| 141 | Ghe02, A1_019, A1_028, A1_030, A1_051, A1_054, A1_077, A1_079, A1_084, A1_088, A1_097, A1_108, A1_125 |
| 170 | A1_019, A1_028, A1_051, A1_054, A1_079, A1_084, A1_097, A1_132, A1_133, A1_155 |
| 258 | Ghe02, A1_019, A1_028, A1_030, A1_079, A1_084, A1_088, A1_097, A1_125, A1_108 |
| 268 | Ghe02, A1_019, A1_028, A1_030, A1_051, A1_054, A1_077, A1_079, A1_084, A1_088, A1_097, A1_108, A1_125, A1_132, A1_133, A1_155 |
| 332 | Ghe01, A1_Af, A1_Nisa, A1_028, A1_051, A1_073, A1_074, A1_155 |
| 341 | Ghe02, A1_019, A1_028, A1_030, A1_051, A1_054, A1_077, A1_079, A1_084, A1_097, A1_088, A1_108, A1_125, A1_132, A1_133 |
| 343 | Ghe02, A1_019, A1_028, A1_030, A1_051, A1_054, A1_077, A1_079, A1_084, A1_088, A1_097, A1_108, A1_125, A1_132, A1_133 |
| 373 | Ghe02, A1_019, A1_028, A1_030, A1_051, A1_054, A1_079, A1_084, A1_088, A1_097, A1_108, A1_125, A1_132, A1_133 |
| 374 | Ghe02, A1_019, A1_028, A1_030, A1_051, A1_054, A1_079, A1_084, A1_088, A1_097, A1_108, A1_125, A1_132, A1_133 |
| 413 | Ghe02, A1_019, A1_051, A1_054, A1_077, A1_079, A1_097, A1_108, A1_132, A1_133 |
| 465 | Gar45, Gar51, Gar58, Gar60, A2_021, A2_031, A2_131, A2_156, A2_158, A2_164 |
| 495 | Ghe02, A1_019, A1_028, A1_030, A1_051, A1_054, A1_077, A1_079, A1_084, A1_088, A1_097, A1_108, A1_125, A1_132, A1_133 |

| Accession | # of windows |
|---|---|
| No resolution | 1 |
| No introgression | 478 |
| Complex | 14 |
| A1_051 | 11 |
| A1_054 | 8 |
| A1_074 | 1 |
| A1_077 | 3 |
| A1_097 | 1 |
| A1_132 | 12 |
| A1_133 | 12 |
| A1_155 | 37 |
| Ghe01 | 1 |
| A2_008 | 1 |
| A2_041 | 1 |
| A2_043 | 1 |
| A2_044 | 1 |
| A2_047 | 1 |
| A2_057 | 1 |
| A2_061 | 1 |
| A2_062 | 1 |
| A2_065 | 1 |
| A2_067 | 1 |
| A2_074 | 3 |
| A2_079 | 1 |
| A2_085 | 1 |
| A2_096 | 1 |
| A2_101 | 1 |
| A2_141 | 1 |
| A2_154 | 1 |
| Gar02 | 1 |
| Gar04 | 5 |
| Gar37 | 1 |
| Gar61 | 1 |

| | # of windows | % of windows | Complex |
|---|---|---|---|
| Chromosome 1 | 3 | 7.5 | 0 |
| Chromosome 2 | 2 | 6.5 | 0 |
| Chromosome 3 | 4 | 10.0 | 0 |
| Chromosome 4 | 4 | 12.1 | 1 |
| Chromosome 5 | 12 | 18.5 | 1 |
| Chromosome 6 | 1 | 2.9 | 0 |
| Chromosome 7 | 11 | 26.8 | 2 |
| Chromosome 8 | 11 | 25.6 | 0 |
| Chromosome 9 | 5 | 11.9 | 4 |
| Chromosome 10 | 9 | 20.0 | 3 |
| Chromosome 11 | 7 | 11.9 | 2 |
| Chromosome 12 | 0 | 0.0 | 1 |
| Chromosome 13 | 0 | 0.0 | 0 |

Supplementary Table 9. Clusters that distinguish *G. herbaceum* (A1) from *G. arboreum* (A2), the number of other clusters significantly correlated with that cluster, and the read counts for each accession.

| | annotation | p.value | padj | Distinguish A1 and A2 | # signicantly correlated | A1_019 | A1_028 | A1_030 | A1_051 | A1_054 | A1_073 | A1_074 | A1_077 | A1_079 | A1_084 | A1_088 |
|---|---|---|---|---|---|---|---|---|---|---|---|---|---|---|---|---|
| CL002 | LTR/Gypsy | 0.0000 | 0.0000 | yes | 112 | 3846 | 3934 | 3609 | 4107 | 3966 | 3957 | 3909 | 3941 | 3870 | 3717 | 3687 |
| CL003 | LTR/Gypsy | 0.0000 | 0.0000 | yes | 128 | 2380 | 2295 | 2217 | 2539 | 2363 | 2092 | 2111 | 2467 | 2445 | 2261 | 2110 |
| CL004 | LTR/Gypsy | 0.0000 | 0.0000 | yes | 144 | 2140 | 2217 | 2165 | 2301 | 2279 | 1923 | 2208 | 2310 | 2082 | 2181 | 2148 |
| CL005 | LTR/Gypsy | 0.0000 | 0.0000 | yes | 118 | 2370 | 2262 | 2168 | 2271 | 2329 | 2547 | 2358 | 2397 | 2371 | 2232 | 2299 |
| CL008 | LTR/Gypsy | 0.0000 | 0.0000 | yes | 138 | 1761 | 1743 | 1768 | 1902 | 1784 | 1545 | 1688 | 1827 | 1684 | 1749 | 1682 |
| CL009 | LTR/Gypsy | 0.0000 | 0.0000 | yes | 127 | 2004 | 1887 | 1923 | 2114 | 2049 | 2063 | 2076 | 2176 | 2012 | 1914 | 1928 |
| CL010 | LTR/Gypsy | 0.0000 | 0.0000 | yes | 116 | 1910 | 1850 | 1815 | 1994 | 1902 | 1641 | 1782 | 1872 | 1884 | 1868 | 1804 |
| CL011 | LTR/Gypsy | 0.0000 | 0.0000 | yes | 85 | 1678 | 1597 | 1579 | 1685 | 1689 | 1537 | 1522 | 1666 | 1582 | 1545 | 1449 |
| CL012 | LTR/Gypsy | 0.0000 | 0.0000 | yes | 124 | 1881 | 1905 | 1698 | 1911 | 1764 | 2088 | 1936 | 1886 | 1970 | 1742 | 1950 |
| CL013 | LTR/Gypsy | 0.0000 | 0.0000 | yes | 135 | 1570 | 1604 | 1436 | 1702 | 1589 | 1379 | 1572 | 1619 | 1571 | 1571 | 1521 |
| CL014 | LTR/Gypsy | 0.0000 | 0.0000 | yes | 93 | 1656 | 1553 | 1542 | 1705 | 1748 | 1577 | 1486 | 1637 | 1528 | 1430 | 1529 |
| CL015 | LTR/Gypsy | 0.0000 | 0.0000 | yes | 137 | 1549 | 1581 | 1526 | 1719 | 1654 | 1407 | 1554 | 1626 | 1539 | 1552 | 1494 |
| CL016 | LTR/Gypsy | 0.0000 | 0.0000 | yes | 154 | 1829 | 1964 | 1949 | 1945 | 1918 | 2319 | 1897 | 2204 | 2193 | 1827 | 1918 |
| CL017 | LTR/Gypsy | 0.0000 | 0.0000 | yes | 118 | 1429 | 1410 | 1381 | 1542 | 1372 | 1006 | 1149 | 1460 | 1423 | 1326 | 1094 |
| CL020 | LTR/Gypsy | 0.0000 | 0.0000 | yes | 122 | 1439 | 1356 | 1347 | 1390 | 1368 | 1352 | 1420 | 1495 | 1474 | 1301 | 1290 |
| CL021 | LTR/Gypsy | 0.0000 | 0.0000 | yes | 124 | 1333 | 1379 | 1262 | 1471 | 1398 | 1248 | 1286 | 1413 | 1315 | 1395 | 1258 |
| CL023 | LTR/Gypsy | 0.0000 | 0.0000 | yes | 102 | 1212 | 1168 | 1186 | 1336 | 1257 | 1107 | 1144 | 1253 | 1263 | 1144 | 1177 |
| CL024 | LTR/Gypsy | 0.0000 | 0.0000 | yes | 115 | 1353 | 1346 | 1337 | 1498 | 1295 | 1269 | 1306 | 1473 | 1304 | 1311 | 1278 |
| CL025 | LTR/Gypsy | 0.0000 | 0.0000 | yes | 145 | 1547 | 1596 | 1527 | 1568 | 1539 | 1923 | 1638 | 1604 | 1677 | 1539 | 1541 |
| CL026 | LTR/Gypsy | 0.0004 | 0.0012 | yes | 156 | 1451 | 1472 | 1611 | 1490 | 1519 | 1868 | 1408 | 1595 | 1604 | 1419 | 1489 |
| CL027 | LTR/Gypsy | 0.0000 | 0.0000 | yes | 130 | 1286 | 1258 | 1212 | 1337 | 1297 | 1243 | 1256 | 1375 | 1310 | 1275 | 1238 |
| CL030 | LTR/Gypsy | 0.0000 | 0.0001 | yes | 98 | 1221 | 1211 | 1187 | 1331 | 1230 | 1235 | 1217 | 1336 | 1308 | 1250 | 1125 |
| CL032 | LTR/Gypsy | 0.0000 | 0.0000 | yes | 126 | 1130 | 1065 | 1041 | 1220 | 1100 | 1093 | 1012 | 1159 | 1100 | 1107 | 1009 |
| CL033 | LTR/Gypsy | 0.0000 | 0.0000 | yes | 116 | 1263 | 1294 | 1216 | 1379 | 1367 | 1364 | 1356 | 1331 | 1356 | 1307 | 1177 |
| CL034 | LTR/Gypsy | 0.0000 | 0.0000 | yes | 92 | 942 | 841 | 912 | 955 | 924 | 897 | 887 | 991 | 931 | 797 | 805 |
| CL037 | LTR/Gypsy | 0.0000 | 0.0000 | yes | 123 | 1010 | 1000 | 972 | 1089 | 1005 | 921 | 1017 | 1049 | 1059 | 980 | 982 |
| CL038 | LTR/Gypsy | 0.0005 | 0.0015 | yes | 130 | 1105 | 1135 | 1064 | 1216 | 1091 | 1125 | 1087 | 1252 | 1176 | 1091 | 1055 |
| CL040 | LTR/Gypsy | 0.0001 | 0.0004 | yes | 103 | 955 | 952 | 910 | 1078 | 982 | 929 | 970 | 994 | 982 | 895 | 945 |
| CL046 | LTR/Gypsy | 0.0000 | 0.0000 | yes | 98 | 701 | 653 | 685 | 753 | 731 | 484 | 675 | 693 | 680 | 709 | 623 |
| CL047 | LTR/Gypsy | 0.0000 | 0.0000 | yes | 58 | 788 | 791 | 786 | 854 | 773 | 826 | 756 | 819 | 795 | 723 | 749 |
| CL050 | LTR/Gypsy | 0.0000 | 0.0000 | yes | 55 | 1232 | 1206 | 1140 | 1205 | 1236 | 1156 | 1037 | 1226 | 1164 | 1171 | 984 |
| CL052 | LTR/Gypsy | 0.0000 | 0.0000 | yes | 94 | 879 | 803 | 884 | 884 | 842 | 914 | 825 | 853 | 845 | 871 | 830 |
| CL053 | LTR/Gypsy | 0.0000 | 0.0000 | yes | 74 | 589 | 555 | 497 | 550 | 606 | 651 | 592 | 562 | 566 | 550 | 544 |
| CL057 | LTR/Gypsy | 0.0000 | 0.0000 | yes | 88 | 540 | 550 | 498 | 573 | 532 | 438 | 552 | 563 | 513 | 554 | 520 |
| CL058 | LTR/Gypsy | 0.0000 | 0.0000 | yes | 63 | 517 | 500 | 532 | 492 | 587 | 609 | 584 | 500 | 497 | 458 | 506 |
| CL065 | LTR/Gypsy | 0.0023 | 0.0064 | yes | 76 | 579 | 567 | 545 | 574 | 509 | 491 | 523 | 578 | 548 | 565 | 527 |
| CL066 | LTR/Copia | 0.0000 | 0.0000 | yes | 84 | 404 | 457 | 401 | 468 | 461 | 534 | 449 | 473 | 456 | 423 | 446 |
| CL067 | LTR/Gypsy | 0.0000 | 0.0000 | yes | 68 | 440 | 430 | 402 | 444 | 367 | 390 | 383 | 407 | 378 | 438 | 405 |
| CL072 | LTR/Gypsy | 0.0000 | 0.0000 | yes | 80 | 416 | 419 | 409 | 430 | 419 | 381 | 421 | 466 | 407 | 390 | 386 |
| CL081 | LTR/Copia | 0.0000 | 0.0000 | yes | 82 | 344 | 402 | 345 | 389 | 364 | 384 | 326 | 375 | 366 | 333 | 372 |
| CL093 | LTR/Gypsy | 0.0000 | 0.0000 | yes | 63 | 537 | 597 | 616 | 623 | 560 | 677 | 559 | 584 | 640 | 517 | 511 |
| CL099 | LTR/Gypsy | 0.0001 | 0.0004 | yes | 90 | 389 | 443 | 441 | 409 | 407 | 512 | 432 | 410 | 431 | 397 | 416 |

| | annotation | p.value | padj | Distinguish A1 and A2 | # signicantly correlated | A1_019 | A1_028 | A1_030 | A1_051 | A1_054 | A1_073 | A1_074 | A1_077 | A1_079 | A1_084 | A1_088 |
|---|---|---|---|---|---|---|---|---|---|---|---|---|---|---|---|---|
| CL100 | LTR/Gypsy | 0.0002 | 0.0006 | yes | 24 | 493 | 386 | 493 | 406 | 435 | 336 | 426 | 432 | 465 | 557 | 533 |
| CL101 | LTR/Gypsy | 0.0001 | 0.0004 | yes | 52 | 394 | 413 | 399 | 406 | 449 | 482 | 461 | 421 | 427 | 370 | 447 |
| CL113 | LTR/Gypsy | 0.0000 | 0.0000 | yes | 61 | 345 | 382 | 420 | 420 | 426 | 452 | 369 | 398 | 480 | 375 | 333 |
| CL118 | LTR/Copia | 0.0008 | 0.0024 | yes | 75 | 335 | 350 | 312 | 324 | 304 | 363 | 284 | 343 | 333 | 356 | 330 |
| CL128 | LTR/Copia | 0.0000 | 0.0000 | yes | 46 | 220 | 253 | 232 | 237 | 274 | 245 | 265 | 238 | 258 | 257 | 234 |
| CL133 | LTR | 0.0030 | 0.0083 | yes | 79 | 267 | 255 | 228 | 297 | 267 | 363 | 266 | 304 | 331 | 257 | 285 |
| CL137 | LTR/Gypsy | 0.0005 | 0.0016 | yes | 59 | 268 | 253 | 247 | 259 | 234 | 346 | 294 | 276 | 290 | 258 | 301 |
| CL146 | LTR/Gypsy | 0.0005 | 0.0016 | yes | 61 | 216 | 201 | 185 | 188 | 209 | 240 | 205 | 193 | 200 | 172 | 213 |
| CL148 | LTR/Gypsy | 0.0000 | 0.0000 | yes | 21 | 163 | 119 | 113 | 119 | 156 | 152 | 137 | 127 | 150 | 102 | 116 |
| CL149 | LTR/Gypsy | 0.0020 | 0.0059 | yes | 25 | 210 | 207 | 204 | 231 | 235 | 238 | 200 | 232 | 241 | 191 | 222 |
| CL152 | LTR/Gypsy | 0.0000 | 0.0000 | yes | 30 | 172 | 143 | 137 | 183 | 140 | 138 | 156 | 156 | 166 | 145 | 179 |
| CL155 | LTR/Gypsy | 0.0003 | 0.0011 | yes | 48 | 175 | 190 | 174 | 186 | 226 | 241 | 175 | 213 | 214 | 191 | 194 |
| CL168 | LTR/Gypsy | 0.0000 | 0.0002 | yes | 34 | 177 | 160 | 189 | 169 | 151 | 199 | 178 | 154 | 152 | 147 | 158 |
| CL170 | LTR/Gypsy | 0.0000 | 0.0000 | yes | 4 | 118 | 102 | 114 | 119 | 116 | 96 | 109 | 120 | 116 | 99 | 108 |
| CL174 | LTR | 0.0000 | 0.0000 | yes | 36 | 124 | 129 | 133 | 141 | 136 | 161 | 139 | 145 | 146 | 117 | 128 |
| CL176 | LTR/Gypsy | 0.0000 | 0.0000 | yes | 16 | 139 | 133 | 122 | 113 | 135 | 136 | 111 | 138 | 105 | 128 | 124 |
| CL184 | LTR/Gypsy | 0.0000 | 0.0001 | yes | 24 | 76 | 92 | 79 | 100 | 85 | 101 | 93 | 98 | 84 | 77 | 95 |
| CL196 | LTR/Copia | 0.0026 | 0.0071 | yes | 3 | 37 | 45 | 35 | 51 | 36 | 42 | 42 | 28 | 30 | 38 | 32 |
| CL198 | LTR/Gypsy | 0.0000 | 0.0000 | yes | 2 | 28 | 34 | 29 | 29 | 36 | 35 | 22 | 41 | 34 | 33 | 31 |
| CL210 | LTR/Copia | 0.0000 | 0.0000 | yes | 1 | 12 | 18 | 7 | 19 | 10 | 12 | 14 | 13 | 13 | 12 | 17 |
| CL219 | LTR/Copia | 0.0007 | 0.0021 | yes | 2 | 7 | 10 | 15 | 20 | 25 | 17 | 10 | 14 | 12 | 10 | 15 |
| CL238 | LTR | 0.0001 | 0.0003 | yes | 0 | 12 | 8 | 10 | 13 | 7 | 8 | 6 | 7 | 3 | 8 | 6 |
| CL001 | LTR/Gypsy | 0.8109 | 0.8549 | no | 111 | 4991 | 4898 | 4712 | 5075 | 4947 | 4715 | 4537 | 5121 | 4983 | 4853 | 4302 |
| CL006 | LTR/Gypsy | 0.0085 | 0.0203 | no | 155 | 2546 | 2560 | 2609 | 2580 | 2619 | 3012 | 2499 | 2582 | 2570 | 2381 | 2429 |
| CL007 | LTR/Gypsy | 0.5404 | 0.6221 | no | 145 | 2279 | 2248 | 2220 | 2298 | 2245 | 2537 | 2298 | 2324 | 2423 | 2092 | 2192 |
| CL018 | LTR/Gypsy | 0.2065 | 0.2851 | no | 152 | 1639 | 1697 | 1640 | 1763 | 1648 | 1950 | 1727 | 1749 | 1826 | 1667 | 1658 |
| CL019 | LTR/Gypsy | 0.0559 | 0.0978 | no | 96 | 1701 | 1682 | 1577 | 1779 | 1732 | 1747 | 1747 | 1848 | 1779 | 1698 | 1624 |
| CL022 | LTR/Gypsy | 0.1459 | 0.2110 | no | 116 | 1574 | 1475 | 1456 | 1610 | 1478 | 1418 | 1484 | 1537 | 1511 | 1481 | 1438 |
| CL028 | LTR/Gypsy | 0.0042 | 0.0112 | no | 127 | 1456 | 1424 | 1469 | 1449 | 1418 | 1610 | 1431 | 1451 | 1411 | 1382 | 1352 |
| CL029 | LTR/Gypsy | 0.0309 | 0.0605 | no | 108 | 1349 | 1302 | 1328 | 1286 | 1406 | 1452 | 1399 | 1345 | 1291 | 1288 | 1346 |
| CL031 | LTR/Gypsy | 0.1107 | 0.1722 | no | 92 | 1290 | 1182 | 1199 | 1381 | 1242 | 1232 | 1272 | 1431 | 1302 | 1308 | 1247 |
| CL035 | LTR/Gypsy | 0.5251 | 0.6085 | no | 111 | 1177 | 1089 | 1095 | 1245 | 1130 | 1319 | 1168 | 1128 | 1070 | 1069 | 1079 |
| CL036 | LTR | 0.0730 | 0.1205 | no | 155 | 1063 | 1089 | 1109 | 1100 | 1074 | 1276 | 1027 | 1126 | 1168 | 1085 | 1106 |
| CL039 | LTR/Gypsy | 0.1019 | 0.1607 | no | 91 | 1006 | 1069 | 993 | 1142 | 1021 | 1020 | 982 | 1118 | 1042 | 1058 | 988 |
| CL041 | LTR/Gypsy | 0.0056 | 0.0142 | no | 152 | 1012 | 1072 | 973 | 1092 | 1048 | 1343 | 1083 | 1073 | 1162 | 975 | 1126 |
| CL042 | LTR/Gypsy | 0.0867 | 0.1392 | no | 94 | 940 | 917 | 909 | 994 | 958 | 1080 | 951 | 1026 | 1069 | 904 | 893 |
| CL043 | LTR/Gypsy | 0.0853 | 0.1382 | no | 90 | 1065 | 989 | 868 | 1087 | 936 | 880 | 879 | 1023 | 961 | 954 | 878 |
| CL044 | LTR/Gypsy | 0.3754 | 0.4563 | no | 70 | 941 | 925 | 980 | 1004 | 910 | 867 | 971 | 999 | 1004 | 1019 | 913 |
| CL045 | LTR/Gypsy | 0.6140 | 0.6800 | no | 118 | 1012 | 963 | 941 | 1024 | 981 | 1070 | 940 | 970 | 965 | 892 | 929 |
| CL048 | LTR/Gypsy | 0.2901 | 0.3733 | no | 84 | 935 | 896 | 889 | 977 | 939 | 1152 | 927 | 952 | 897 | 838 | 839 |
| CL049 | LTR/Gypsy | 0.9573 | 0.9629 | no | 101 | 853 | 869 | 807 | 916 | 933 | 997 | 900 | 980 | 946 | 843 | 874 |
| CL051 | LTR/Gypsy | 0.0542 | 0.0957 | no | 123 | 898 | 885 | 814 | 887 | 884 | 916 | 872 | 903 | 887 | 811 | 837 |
| CL055 | LTR/Gypsy | 0.0580 | 0.1001 | no | 102 | 703 | 719 | 708 | 729 | 727 | 798 | 788 | 745 | 775 | 682 | 694 |
| CL056 | LTR/Gypsy | 0.0457 | 0.0842 | no | 144 | 718 | 724 | 691 | 767 | 705 | 810 | 732 | 729 | 777 | 712 | 708 |

| | annotation | p.value | padj | Distinguish A1 and A2 | # signicantly correlated | A1_019 | A1_028 | A1_030 | A1_051 | A1_054 | A1_073 | A1_074 | A1_077 | A1_079 | A1_084 | A1_088 |
|---|---|---|---|---|---|---|---|---|---|---|---|---|---|---|---|---|
| CL059 | LTR/Gypsy | 0.1189 | 0.1825 | no | 67 | 691 | 711 | 627 | 651 | 694 | 735 | 700 | 731 | 680 | 591 | 704 |
| CL060 | LTR/Gypsy | 0.0141 | 0.0309 | no | 129 | 696 | 639 | 674 | 763 | 737 | 810 | 674 | 770 | 731 | 706 | 691 |
| CL061 | LTR | 0.6546 | 0.7115 | no | 135 | 660 | 659 | 629 | 724 | 683 | 800 | 657 | 668 | 682 | 704 | 669 |
| CL062 | LTR | 0.0486 | 0.0877 | no | 94 | 711 | 681 | 701 | 713 | 660 | 727 | 618 | 739 | 666 | 695 | 647 |
| CL063 | LTR/Gypsy | 0.0083 | 0.0201 | no | 129 | 716 | 742 | 699 | 781 | 660 | 819 | 687 | 753 | 814 | 670 | 676 |
| CL064 | LTR/Gypsy | 0.1372 | 0.2035 | no | 56 | 577 | 601 | 599 | 651 | 582 | 523 | 586 | 664 | 580 | 583 | 526 |
| CL068 | LTR | 0.2958 | 0.3778 | no | 128 | 566 | 553 | 580 | 643 | 516 | 660 | 587 | 609 | 595 | 541 | 559 |
| CL070 | LTR/Gypsy | 0.2139 | 0.2902 | no | 30 | 547 | 575 | 554 | 621 | 572 | 573 | 571 | 555 | 613 | 536 | 565 |
| CL071 | LTR/Gypsy | 0.0120 | 0.0265 | no | 120 | 590 | 521 | 550 | 553 | 549 | 682 | 619 | 596 | 536 | 553 | 554 |
| CL074 | LTR/Gypsy | 0.0814 | 0.1331 | no | 110 | 503 | 504 | 550 | 525 | 524 | 643 | 552 | 545 | 601 | 507 | 509 |
| CL075 | LTR/Gypsy | 0.0583 | 0.1001 | no | 53 | 513 | 527 | 464 | 548 | 478 | 455 | 522 | 552 | 520 | 541 | 473 |
| CL076 | LTR | 0.0045 | 0.0116 | no | 126 | 482 | 550 | 538 | 611 | 559 | 681 | 538 | 576 | 591 | 508 | 530 |
| CL078 | LTR/Gypsy | 0.7097 | 0.7573 | no | 93 | 498 | 482 | 510 | 532 | 488 | 566 | 501 | 576 | 525 | 479 | 491 |
| CL079 | LTR/Gypsy | 0.0481 | 0.0877 | no | 110 | 506 | 528 | 506 | 556 | 539 | 563 | 502 | 518 | 571 | 477 | 485 |
| CL080 | LTR/Copia | 0.0364 | 0.0685 | no | 106 | 532 | 466 | 475 | 506 | 513 | 537 | 490 | 571 | 532 | 477 | 485 |
| CL082 | LTR/Gypsy | 0.0221 | 0.0456 | no | 47 | 496 | 447 | 464 | 513 | 464 | 398 | 404 | 524 | 391 | 486 | 381 |
| CL084 | LTR/Gypsy | 0.1864 | 0.2652 | no | 65 | 478 | 473 | 454 | 500 | 470 | 520 | 434 | 435 | 484 | 452 | 441 |
| CL085 | LTR/Gypsy | 0.6180 | 0.6802 | no | 52 | 494 | 465 | 462 | 507 | 465 | 545 | 499 | 473 | 463 | 490 | 476 |
| CL091 | LTR/Gypsy | 0.2118 | 0.2896 | no | 60 | 439 | 416 | 406 | 470 | 384 | 409 | 422 | 482 | 444 | 414 | 408 |
| CL092 | LTR/Gypsy | 0.0208 | 0.0438 | no | 103 | 397 | 462 | 427 | 471 | 490 | 502 | 439 | 462 | 447 | 394 | 433 |
| CL094 | DNA/MULE-MuDR | 0.3512 | 0.4389 | no | 101 | 380 | 397 | 391 | 449 | 450 | 463 | 416 | 421 | 444 | 407 | 460 |
| CL095 | LTR/Gypsy | 0.9629 | 0.9629 | no | 97 | 507 | 369 | 433 | 382 | 420 | 110 | 374 | 406 | 489 | 722 | 432 |
| CL102 | DNA/MULE-MuDR | 0.0321 | 0.0617 | no | 92 | 363 | 386 | 388 | 432 | 423 | 404 | 392 | 468 | 413 | 372 | 383 |
| CL104 | LTR/Gypsy | 0.0311 | 0.0605 | no | 82 | 360 | 403 | 383 | 415 | 399 | 512 | 389 | 451 | 429 | 403 | 383 |
| CL106 | LTR/Copia | 0.0112 | 0.0251 | no | 93 | 354 | 330 | 318 | 406 | 369 | 477 | 345 | 363 | 434 | 354 | 355 |
| CL107 | LTR/Gypsy | 0.0244 | 0.0496 | no | 93 | 335 | 334 | 337 | 363 | 353 | 429 | 359 | 379 | 369 | 340 | 363 |
| CL109 | LTR/Copia | 0.4096 | 0.4910 | no | 86 | 339 | 306 | 324 | 372 | 327 | 326 | 311 | 332 | 323 | 312 | 323 |
| CL112 | LTR/Gypsy | 0.0163 | 0.0350 | no | 88 | 333 | 346 | 325 | 348 | 353 | 405 | 341 | 343 | 334 | 305 | 306 |
| CL117 | LTR/Gypsy | 0.2187 | 0.2922 | no | 64 | 325 | 319 | 308 | 355 | 315 | 340 | 309 | 325 | 307 | 291 | 323 |
| CL119 | LTR/Gypsy | 0.0535 | 0.0956 | no | 24 | 299 | 301 | 307 | 323 | 309 | 339 | 314 | 372 | 287 | 288 | 310 |
| CL120 | LTR/Gypsy | 0.7780 | 0.8251 | no | 6 | 286 | 286 | 315 | 336 | 314 | 286 | 313 | 316 | 295 | 289 | 302 |
| CL121 | LTR/Gypsy | 0.0108 | 0.0248 | no | 40 | 314 | 286 | 293 | 356 | 314 | 325 | 339 | 356 | 328 | 294 | 276 |
| CL122 | LTR | 0.0673 | 0.1143 | no | 84 | 298 | 334 | 302 | 319 | 300 | 383 | 304 | 317 | 358 | 287 | 318 |
| CL129 | LTR/Gypsy | 0.2832 | 0.3671 | no | 74 | 253 | 268 | 301 | 314 | 267 | 284 | 317 | 305 | 303 | 260 | 299 |
| CL130 | LTR/Copia | 0.1277 | 0.1943 | no | 33 | 299 | 266 | 270 | 230 | 266 | 303 | 347 | 276 | 256 | 263 | 350 |
| CL131 | LTR/Copia | 0.0042 | 0.0112 | no | 73 | 269 | 278 | 256 | 299 | 271 | 357 | 234 | 294 | 308 | 297 | 274 |
| CL139 | LTR/Copia | 0.0218 | 0.0454 | no | 66 | 253 | 240 | 247 | 272 | 266 | 334 | 240 | 249 | 248 | 269 | 214 |
| CL143 | LTR/Gypsy | 0.1967 | 0.2753 | no | 30 | 218 | 200 | 210 | 241 | 243 | 192 | 240 | 244 | 198 | 201 | 228 |
| CL147 | LTR/Gypsy | 0.1732 | 0.2484 | no | 42 | 216 | 209 | 217 | 196 | 196 | 226 | 244 | 217 | 230 | 214 | 197 |
| CL151 | LTR/Gypsy | 0.0257 | 0.0517 | no | 54 | 206 | 201 | 204 | 208 | 206 | 237 | 202 | 196 | 228 | 202 | 214 |
| CL157 | LTR/Gypsy | 0.1419 | 0.2070 | no | 53 | 173 | 186 | 157 | 211 | 151 | 228 | 178 | 201 | 194 | 167 | 157 |
| CL158 | LTR/Gypsy | 0.3647 | 0.4494 | no | 34 | 170 | 137 | 146 | 194 | 161 | 193 | 174 | 167 | 171 | 181 | 169 |
| CL159 | LTR/Gypsy | 0.1405 | 0.2066 | no | 25 | 186 | 160 | 167 | 188 | 167 | 192 | 177 | 153 | 178 | 146 | 173 |
| CL160 | LTR/Gypsy | 0.3731 | 0.4563 | no | 8 | 161 | 163 | 139 | 176 | 159 | 152 | 181 | 153 | 167 | 157 | 168 |

| | annotation | p.value | padj | Distinguish A1 and A2 | # signicantly correlated | A1_019 | A1_028 | A1_030 | A1_051 | A1_054 | A1_073 | A1_074 | A1_077 | A1_079 | A1_084 | A1_088 |
|---|---|---|---|---|---|---|---|---|---|---|---|---|---|---|---|---|
| CL162 | LTR/Gypsy | 0.0309 | 0.0605 | no | 20 | 180 | 155 | 156 | 177 | 179 | 150 | 175 | 175 | 181 | 178 | 170 |
| CL163 | LTR | 0.2799 | 0.3655 | no | 30 | 156 | 171 | 137 | 172 | 149 | 195 | 147 | 156 | 167 | 147 | 172 |
| CL164 | LTR/Gypsy | 0.2791 | 0.3655 | no | 10 | 134 | 162 | 159 | 145 | 143 | 185 | 141 | 148 | 162 | 144 | 158 |
| CL166 | LTR | 0.0058 | 0.0145 | no | 36 | 166 | 144 | 169 | 159 | 139 | 190 | 164 | 168 | 182 | 159 | 169 |
| CL171 | LTR/Gypsy | 0.9538 | 0.9629 | no | 10 | 125 | 139 | 107 | 123 | 126 | 142 | 114 | 146 | 113 | 116 | 103 |
| CL173 | LTR/Gypsy | 0.0907 | 0.1443 | no | 34 | 95 | 105 | 106 | 129 | 121 | 148 | 120 | 139 | 120 | 131 | 98 |
| CL175 | LTR/Gypsy | 0.0729 | 0.1205 | no | 8 | 87 | 117 | 118 | 135 | 101 | 115 | 115 | 131 | 112 | 115 | 136 |
| CL177 | LTR/Gypsy | 0.2691 | 0.3567 | no | 11 | 105 | 118 | 90 | 89 | 102 | 96 | 92 | 119 | 109 | 95 | 92 |
| CL182 | LTR/Gypsy | 0.0045 | 0.0116 | no | 27 | 78 | 75 | 75 | 86 | 105 | 105 | 91 | 89 | 97 | 78 | 84 |
| CL186 | LTR/Gypsy | 0.0109 | 0.0248 | no | 24 | 80 | 71 | 74 | 72 | 86 | 99 | 66 | 67 | 87 | 76 | 85 |
| CL187 | LTR/Gypsy | 0.0164 | 0.0350 | no | 5 | 84 | 61 | 82 | 89 | 66 | 74 | 82 | 84 | 77 | 62 | 82 |
| CL189 | LTR/Gypsy | 0.6225 | 0.6808 | no | 4 | 51 | 73 | 75 | 59 | 55 | 65 | 55 | 64 | 59 | 55 | 65 |
| CL191 | LTR/Gypsy | 0.8420 | 0.8771 | no | 1 | 55 | 63 | 65 | 55 | 58 | 52 | 62 | 57 | 55 | 72 | 60 |
| CL194 | LTR/Gypsy | 0.0101 | 0.0240 | no | 9 | 45 | 36 | 51 | 48 | 44 | 53 | 45 | 50 | 48 | 43 | 38 |
| CL195 | LTR/Gypsy | 0.3388 | 0.4265 | no | 17 | 50 | 47 | 37 | 48 | 49 | 74 | 52 | 33 | 60 | 32 | 53 |
| CL197 | LTR/Gypsy | 0.5801 | 0.6466 | no | 4 | 44 | 49 | 33 | 39 | 47 | 51 | 34 | 52 | 34 | 46 | 42 |
| CL199 | LTR/Copia | 0.4189 | 0.4987 | no | 2 | 33 | 35 | 33 | 49 | 38 | 52 | 42 | 59 | 55 | 38 | 37 |
| CL200 | LTR/Gypsy | 0.5782 | 0.6466 | no | 1 | 49 | 39 | 32 | 37 | 44 | 46 | 38 | 33 | 45 | 40 | 38 |
| CL202 | LTR/Gypsy | 0.5737 | 0.6466 | no | 0 | 37 | 37 | 30 | 28 | 34 | 30 | 27 | 28 | 39 | 31 | 17 |
| CL203 | LTR/Gypsy | 0.8248 | 0.8644 | no | 13 | 31 | 39 | 27 | 41 | 39 | 35 | 25 | 35 | 38 | 22 | 41 |
| CL204 | LTR/Gypsy | 0.4485 | 0.5303 | no | 10 | 28 | 26 | 37 | 35 | 33 | 45 | 39 | 35 | 32 | 24 | 22 |
| CL205 | LTR/Gypsy | 0.7069 | 0.7573 | no | 5 | 34 | 36 | 29 | 37 | 32 | 40 | 26 | 43 | 34 | 28 | 28 |
| CL206 | LTR/Gypsy | 0.1334 | 0.1995 | no | 2 | 27 | 27 | 29 | 24 | 27 | 37 | 34 | 37 | 24 | 35 | 30 |
| CL209 | LTR/Gypsy | 0.0725 | 0.1205 | no | 2 | 19 | 30 | 30 | 36 | 30 | 34 | 28 | 19 | 40 | 29 | 32 |
| CL211 | LTR/Gypsy | 0.5782 | 0.6466 | no | 7 | 30 | 23 | 19 | 33 | 33 | 30 | 33 | 23 | 27 | 12 | 31 |
| CL212 | LTR/Gypsy | 0.2186 | 0.2922 | no | 1 | 26 | 33 | 21 | 30 | 25 | 25 | 31 | 28 | 26 | 38 | 20 |
| CL213 | LTR/Gypsy | 0.2069 | 0.2851 | no | 5 | 31 | 28 | 37 | 25 | 20 | 31 | 20 | 28 | 32 | 23 | 24 |
| CL214 | LTR/Gypsy | 0.1914 | 0.2701 | no | 5 | 26 | 32 | 28 | 24 | 25 | 28 | 30 | 23 | 25 | 26 | 20 |
| CL215 | LTR/Copia | 0.3152 | 0.3997 | no | 0 | 26 | 17 | 16 | 25 | 22 | 28 | 26 | 23 | 24 | 27 | 25 |
| CL216 | LTR | 0.5210 | 0.6078 | no | 1 | 19 | 14 | 17 | 36 | 24 | 24 | 26 | 19 | 26 | 24 | 23 |
| CL218 | LTR/Copia | 0.5683 | 0.6466 | no | 0 | 20 | 29 | 13 | 19 | 25 | 21 | 22 | 16 | 15 | 15 | 19 |
| CL220 | LTR/Gypsy | 0.9596 | 0.9629 | no | 0 | 10 | 15 | 21 | 19 | 26 | 19 | 16 | 16 | 33 | 19 | 25 |
| CL221 | LTR/Copia | 0.5030 | 0.5908 | no | 0 | 18 | 11 | 19 | 15 | 18 | 24 | 22 | 21 | 16 | 20 | 19 |
| CL222 | LTR/Gypsy | 0.3536 | 0.4389 | no | 0 | 14 | 18 | 10 | 15 | 18 | 20 | 13 | 19 | 22 | 16 | 15 |
| CL223 | LTR/Gypsy | 0.1112 | 0.1722 | no | 0 | 27 | 14 | 23 | 20 | 25 | 15 | 20 | 23 | 17 | 13 | 13 |
| CL226 | LTR/Gypsy | 0.9200 | 0.9471 | no | 0 | 15 | 9 | 17 | 14 | 16 | 21 | 17 | 16 | 23 | 21 | 19 |
| CL228 | LTR/Gypsy | 0.9046 | 0.9368 | no | 0 | 10 | 24 | 18 | 7 | 18 | 18 | 24 | 18 | 17 | 15 | 18 |
| CL229 | LTR | 0.9326 | 0.9544 | no | 1 | 16 | 17 | 23 | 20 | 19 | 25 | 14 | 23 | 9 | 7 | 21 |
| CL231 | LTR/Gypsy | 0.3904 | 0.4712 | no | 0 | 20 | 14 | 18 | 18 | 15 | 12 | 13 | 16 | 17 | 15 | 10 |
| CL232 | LTR/Gypsy | 0.0077 | 0.0190 | no | 1 | 17 | 7 | 21 | 16 | 12 | 19 | 14 | 19 | 18 | 14 | 14 |
| CL234 | LTR | 0.0421 | 0.0784 | no | 0 | 13 | 16 | 10 | 12 | 12 | 20 | 14 | 8 | 12 | 13 | 9 |
| CL235 | LTR | 0.0105 | 0.0244 | no | 0 | 13 | 9 | 8 | 18 | 18 | 16 | 15 | 16 | 17 | 11 | 12 |
| CL236 | LTR/Gypsy | 0.0350 | 0.0665 | no | 1 | 18 | 7 | 13 | 14 | 20 | 15 | 11 | 10 | 14 | 13 | 9 |
| CL237 | LTR/Gypsy | 0.6760 | 0.7303 | no | 0 | 14 | 9 | 10 | 11 | 10 | 12 | 7 | 14 | 9 | 15 | 13 |
| CL239 | LTR/Gypsy | 0.1306 | 0.1971 | no | 1 | 10 | 6 | 11 | 17 | 13 | 15 | 9 | 8 | 4 | 8 | 12 |

| | A1_097 | A1_108 | A1_125 | A1_132 | A1_133 | A1_155 | A1_Afr | A1_Nis | A2_001 | A2_004 | A2_008 | A2_016 | A2_019 | A2_021 | A2_026 | A2_029 | A2_030 | A2_031 | A2_034 |
|---|---|---|---|---|---|---|---|---|---|---|---|---|---|---|---|---|---|---|---|
| CL002 | 3802 | 3963 | 3812 | 4336 | 4176 | 4258 | 3748 | 4047 | 4842 | 4845 | 4782 | 4964 | 4631 | 4708 | 4632 | 4630 | 5039 | 4815 | 5328 |
| CL003 | 2307 | 2525 | 2100 | 2467 | 2484 | 2187 | 2318 | 2321 | 2982 | 2349 | 2990 | 2951 | 2698 | 2850 | 2954 | 2719 | 2968 | 2941 | 3051 |
| CL004 | 2262 | 2319 | 2205 | 2590 | 2410 | 1970 | 2274 | 2266 | 2646 | 1859 | 2607 | 2762 | 2480 | 2617 | 2672 | 2575 | 2614 | 2655 | 2572 |
| CL005 | 2349 | 2405 | 2354 | 2506 | 2413 | 2608 | 2279 | 2471 | 2476 | 2827 | 2491 | 2705 | 2554 | 2501 | 2493 | 2505 | 2698 | 2478 | 2816 |
| CL008 | 1772 | 1820 | 1798 | 2027 | 1943 | 1476 | 1854 | 1715 | 2115 | 1358 | 2232 | 2224 | 2083 | 2216 | 2217 | 1965 | 2189 | 2149 | 2257 |
| CL009 | 2049 | 2111 | 1997 | 2309 | 2211 | 2141 | 2178 | 2123 | 2003 | 2125 | 2117 | 2120 | 2118 | 2183 | 2145 | 2047 | 2174 | 2141 | 2348 |
| CL010 | 1790 | 1906 | 1886 | 2197 | 1978 | 1670 | 1879 | 1877 | 2053 | 1494 | 2106 | 2161 | 1861 | 2067 | 2161 | 2001 | 2072 | 2098 | 1984 |
| CL011 | 1665 | 1670 | 1536 | 1912 | 1832 | 1620 | 1556 | 1644 | 2198 | 2324 | 2080 | 2218 | 2030 | 2172 | 2158 | 2124 | 2265 | 2309 | 2726 |
| CL012 | 1831 | 1927 | 1900 | 1959 | 1941 | 2064 | 2049 | 1878 | 1918 | 2158 | 2033 | 2036 | 1998 | 1919 | 1875 | 2011 | 2053 | 2023 | 2182 |
| CL013 | 1635 | 1658 | 1609 | 1763 | 1704 | 1450 | 1553 | 1609 | 1908 | 1431 | 1900 | 2016 | 1903 | 1930 | 1986 | 1817 | 1912 | 1911 | 1891 |
| CL014 | 1574 | 1721 | 1491 | 1808 | 1776 | 1603 | 1464 | 1545 | 2012 | 2425 | 1975 | 2194 | 1935 | 2092 | 2103 | 2034 | 2103 | 2055 | 2352 |
| CL015 | 1538 | 1687 | 1610 | 1867 | 1744 | 1397 | 1612 | 1683 | 1977 | 1398 | 1914 | 2085 | 1787 | 1948 | 2068 | 1821 | 1838 | 1951 | 1900 |
| CL016 | 1926 | 1923 | 1921 | 2030 | 1881 | 2241 | 2051 | 2084 | 1743 | 2437 | 1825 | 1724 | 1762 | 1916 | 1708 | 1860 | 1697 | 1736 | 1836 |
| CL017 | 1472 | 1420 | 1212 | 1519 | 1581 | 1003 | 1374 | 1353 | 1855 | 1228 | 1814 | 1911 | 1597 | 1876 | 1905 | 1736 | 1653 | 1781 | 1515 |
| CL020 | 1390 | 1489 | 1325 | 1459 | 1413 | 1392 | 1425 | 1504 | 1508 | 1279 | 1544 | 1636 | 1393 | 1513 | 1515 | 1472 | 1522 | 1474 | 1487 |
| CL021 | 1392 | 1392 | 1403 | 1516 | 1388 | 1147 | 1469 | 1428 | 1457 | 1047 | 1535 | 1496 | 1408 | 1581 | 1582 | 1436 | 1469 | 1533 | 1555 |
| CL023 | 1162 | 1250 | 1184 | 1392 | 1399 | 1144 | 1205 | 1172 | 1457 | 1480 | 1443 | 1669 | 1498 | 1507 | 1438 | 1452 | 1527 | 1497 | 1535 |
| CL024 | 1356 | 1332 | 1377 | 1400 | 1374 | 1294 | 1432 | 1379 | 1431 | 1079 | 1467 | 1435 | 1426 | 1454 | 1501 | 1318 | 1424 | 1484 | 1418 |
| CL025 | 1573 | 1599 | 1534 | 1628 | 1638 | 1872 | 1640 | 1599 | 1420 | 1899 | 1521 | 1460 | 1495 | 1474 | 1393 | 1426 | 1482 | 1459 | 1693 |
| CL026 | 1484 | 1536 | 1479 | 1681 | 1539 | 1765 | 1485 | 1624 | 1400 | 1976 | 1364 | 1429 | 1416 | 1379 | 1425 | 1461 | 1437 | 1357 | 1650 |
| CL027 | 1363 | 1383 | 1340 | 1394 | 1362 | 1073 | 1332 | 1371 | 1378 | 940 | 1337 | 1344 | 1302 | 1364 | 1407 | 1297 | 1315 | 1361 | 1376 |
| CL030 | 1133 | 1275 | 1191 | 1303 | 1275 | 1232 | 1348 | 1304 | 1290 | 1278 | 1259 | 1465 | 1264 | 1391 | 1357 | 1280 | 1284 | 1334 | 1373 |
| CL032 | 1101 | 1160 | 1039 | 1227 | 1226 | 880 | 1197 | 1188 | 1292 | 902 | 1251 | 1310 | 1069 | 1311 | 1288 | 1219 | 1263 | 1292 | 1023 |
| CL033 | 1281 | 1369 | 1243 | 1374 | 1385 | 1523 | 1282 | 1298 | 1140 | 1383 | 1154 | 1233 | 1164 | 1146 | 1140 | 1180 | 1180 | 1176 | 1304 |
| CL034 | 867 | 940 | 854 | 1107 | 1031 | 830 | 981 | 971 | 1198 | 1069 | 1205 | 1348 | 1120 | 1180 | 1241 | 1117 | 1239 | 1212 | 1228 |
| CL037 | 1039 | 967 | 968 | 1005 | 1048 | 921 | 1056 | 1094 | 1087 | 1014 | 1078 | 1057 | 1028 | 1075 | 1073 | 1116 | 1063 | 1126 | 1203 |
| CL038 | 1102 | 1144 | 1107 | 1143 | 1153 | 1033 | 1207 | 1170 | 953 | 904 | 988 | 981 | 1014 | 1075 | 1035 | 981 | 975 | 1037 | 1060 |
| CL040 | 997 | 998 | 1043 | 1120 | 1058 | 958 | 1060 | 996 | 942 | 915 | 979 | 1062 | 992 | 1036 | 1038 | 1009 | 975 | 1035 | 1059 |
| CL046 | 728 | 651 | 665 | 782 | 774 | 476 | 654 | 639 | 930 | 422 | 958 | 998 | 877 | 994 | 1029 | 906 | 870 | 954 | 778 |
| CL047 | 761 | 831 | 712 | 927 | 902 | 877 | 806 | 830 | 1016 | 1171 | 1031 | 1136 | 955 | 1025 | 1057 | 961 | 1041 | 1054 | 1117 |
| CL050 | 1092 | 1200 | 1010 | 1316 | 1281 | 1086 | 1220 | 1327 | 817 | 911 | 845 | 912 | 737 | 911 | 845 | 866 | 803 | 888 | 860 |
| CL052 | 806 | 860 | 825 | 1027 | 909 | 919 | 886 | 910 | 800 | 1027 | 748 | 874 | 759 | 830 | 767 | 765 | 776 | 793 | 859 |
| CL053 | 545 | 614 | 533 | 612 | 615 | 709 | 567 | 596 | 757 | 1024 | 753 | 849 | 756 | 740 | 698 | 758 | 683 | 825 | 982 |
| CL057 | 597 | 622 | 501 | 616 | 574 | 466 | 571 | 574 | 665 | 427 | 710 | 714 | 622 | 711 | 705 | 709 | 719 | 707 | 603 |
| CL058 | 464 | 532 | 509 | 593 | 582 | 601 | 531 | 578 | 681 | 844 | 718 | 798 | 660 | 708 | 723 | 651 | 752 | 714 | 895 |
| CL065 | 508 | 591 | 517 | 620 | 592 | 490 | 610 | 570 | 570 | 400 | 546 | 597 | 485 | 554 | 614 | 501 | 534 | 552 | 519 |
| CL066 | 443 | 493 | 474 | 457 | 501 | 502 | 470 | 480 | 594 | 679 | 576 | 562 | 595 | 584 | 572 | 592 | 559 | 604 | 697 |
| CL067 | 388 | 378 | 429 | 455 | 432 | 384 | 422 | 443 | 606 | 356 | 661 | 602 | 571 | 567 | 549 | 570 | 594 | 573 | 591 |
| CL072 | 364 | 402 | 407 | 456 | 442 | 431 | 443 | 450 | 550 | 441 | 502 | 543 | 511 | 557 | 529 | 519 | 504 | 559 | 573 |
| CL081 | 380 | 387 | 369 | 385 | 418 | 364 | 378 | 339 | 493 | 489 | 445 | 476 | 459 | 538 | 499 | 450 | 496 | 497 | 510 |
| CL093 | 509 | 534 | 491 | 558 | 481 | 618 | 590 | 592 | 488 | 804 | 475 | 415 | 438 | 541 | 498 | 485 | 483 | 505 | 323 |
| CL099 | 437 | 400 | 380 | 453 | 426 | 503 | 459 | 431 | 387 | 438 | 410 | 414 | 419 | 400 | 400 | 415 | 402 | 368 | 434 |

| | A1_097 | A1_108 | A1_125 | A1_132 | A1_133 | A1_155 | A1_Afr | A1_Nis | A2_001 | A2_004 | A2_008 | A2_016 | A2_019 | A2_021 | A2_026 | A2_029 | A2_030 | A2_031 | A2_034 |
|---|---|---|---|---|---|---|---|---|---|---|---|---|---|---|---|---|---|---|---|
| CL100 | 546 | 433 | 462 | 399 | 446 | 404 | 465 | 419 | 407 | 345 | 347 | 312 | 381 | 331 | 416 | 382 | 427 | 392 | 380 |
| CL101 | 440 | 419 | 396 | 430 | 436 | 499 | 408 | 466 | 417 | 549 | 358 | 451 | 413 | 397 | 376 | 406 | 404 | 400 | 420 |
| CL113 | 368 | 400 | 338 | 340 | 350 | 518 | 370 | 380 | 382 | 597 | 323 | 314 | 331 | 347 | 377 | 356 | 344 | 318 | 302 |
| CL118 | 306 | 304 | 304 | 296 | 354 | 372 | 314 | 366 | 302 | 425 | 324 | 323 | 332 | 306 | 311 | 301 | 324 | 337 | 363 |
| CL128 | 228 | 255 | 221 | 278 | 267 | 257 | 251 | 246 | 268 | 275 | 294 | 286 | 264 | 284 | 269 | 285 | 264 | 287 | 319 |
| CL133 | 238 | 262 | 266 | 243 | 253 | 369 | 278 | 298 | 252 | 402 | 273 | 218 | 235 | 251 | 216 | 253 | 256 | 256 | 272 |
| CL137 | 252 | 263 | 241 | 262 | 252 | 328 | 275 | 269 | 280 | 379 | 246 | 245 | 279 | 268 | 220 | 259 | 229 | 262 | 284 |
| CL146 | 216 | 187 | 177 | 205 | 198 | 218 | 194 | 201 | 210 | 277 | 230 | 217 | 218 | 224 | 200 | 227 | 213 | 254 | 235 |
| CL148 | 123 | 146 | 121 | 155 | 135 | 164 | 123 | 130 | 254 | 253 | 220 | 217 | 206 | 235 | 209 | 220 | 232 | 242 | 268 |
| CL149 | 227 | 214 | 231 | 184 | 200 | 218 | 234 | 241 | 200 | 220 | 236 | 227 | 207 | 198 | 200 | 195 | 198 | 203 | 115 |
| CL152 | 160 | 171 | 167 | 169 | 171 | 109 | 188 | 149 | 151 | 114 | 170 | 204 | 179 | 191 | 183 | 161 | 173 | 172 | 195 |
| CL155 | 197 | 180 | 185 | 168 | 153 | 192 | 177 | 218 | 174 | 256 | 177 | 179 | 167 | 159 | 173 | 191 | 157 | 174 | 176 |
| CL168 | 138 | 150 | 142 | 153 | 119 | 209 | 167 | 163 | 185 | 217 | 145 | 150 | 154 | 143 | 147 | 153 | 127 | 147 | 135 |
| CL170 | 94 | 122 | 108 | 118 | 112 | 109 | 110 | 111 | 155 | 132 | 115 | 130 | 121 | 133 | 115 | 169 | 159 | 138 | 118 |
| CL174 | 122 | 110 | 124 | 118 | 103 | 173 | 139 | 135 | 117 | 188 | 99 | 94 | 127 | 106 | 114 | 132 | 113 | 104 | 112 |
| CL176 | 134 | 142 | 123 | 141 | 116 | 139 | 140 | 150 | 94 | 138 | 103 | 108 | 93 | 115 | 90 | 105 | 102 | 83 | 117 |
| CL184 | 77 | 78 | 81 | 72 | 81 | 96 | 93 | 98 | 75 | 125 | 82 | 71 | 84 | 101 | 86 | 99 | 69 | 76 | 81 |
| CL196 | 47 | 38 | 35 | 47 | 46 | 42 | 44 | 36 | 49 | 39 | 36 | 41 | 41 | 34 | 36 | 45 | 55 | 45 | 52 |
| CL198 | 42 | 32 | 31 | 21 | 38 | 33 | 31 | 18 | 36 | 42 | 50 | 46 | 41 | 41 | 44 | 26 | 43 | 38 | 37 |
| CL210 | 23 | 15 | 14 | 9 | 10 | 14 | 18 | 12 | 33 | 22 | 32 | 27 | 31 | 29 | 28 | 19 | 33 | 32 | 34 |
| CL219 | 8 | 8 | 18 | 19 | 15 | 21 | 19 | 24 | 19 | 25 | 24 | 10 | 25 | 16 | 30 | 20 | 22 | 20 | 24 |
| CL238 | 10 | 9 | 5 | 7 | 1 | 11 | 10 | 7 | 8 | 6 | 13 | 7 | 15 | 10 | 14 | 6 | 13 | 18 | 12 |
| CL001 | 4817 | 4919 | 4491 | 5414 | 5220 | 4400 | 5208 | 4969 | 4996 | 3960 | 4746 | 5179 | 4445 | 4808 | 5031 | 4597 | 4769 | 4654 | 4750 |
| CL006 | 2441 | 2593 | 2465 | 3011 | 2734 | 3050 | 2572 | 2789 | 2644 | 3550 | 2434 | 2782 | 2471 | 2548 | 2451 | 2539 | 2504 | 2577 | 2965 |
| CL007 | 2289 | 2515 | 2177 | 2465 | 2429 | 2651 | 2334 | 2406 | 2303 | 2882 | 2208 | 2354 | 2258 | 2235 | 2120 | 2220 | 2381 | 2243 | 2533 |
| CL018 | 1714 | 1693 | 1630 | 1832 | 1728 | 2046 | 1706 | 1806 | 1774 | 2190 | 1729 | 1720 | 1698 | 1756 | 1610 | 1653 | 1698 | 1727 | 1913 |
| CL019 | 1649 | 1755 | 1788 | 1874 | 1888 | 1589 | 1816 | 1804 | 1529 | 1424 | 1651 | 1571 | 1516 | 1720 | 1595 | 1519 | 1574 | 1615 | 1712 |
| CL022 | 1492 | 1549 | 1474 | 1677 | 1602 | 1320 | 1604 | 1615 | 1464 | 1156 | 1508 | 1493 | 1405 | 1479 | 1435 | 1385 | 1514 | 1472 | 1547 |
| CL028 | 1373 | 1577 | 1367 | 1681 | 1532 | 1603 | 1519 | 1598 | 1408 | 1819 | 1479 | 1495 | 1406 | 1460 | 1341 | 1429 | 1409 | 1477 | 1642 |
| CL029 | 1334 | 1389 | 1298 | 1548 | 1421 | 1437 | 1373 | 1355 | 1267 | 1505 | 1239 | 1368 | 1270 | 1281 | 1267 | 1302 | 1279 | 1322 | 1455 |
| CL031 | 1264 | 1294 | 1279 | 1359 | 1368 | 1125 | 1399 | 1384 | 1210 | 934 | 1272 | 1129 | 1098 | 1222 | 1304 | 1238 | 1194 | 1127 | 1301 |
| CL035 | 1135 | 1188 | 1095 | 1254 | 1216 | 1287 | 1109 | 1162 | 1168 | 1351 | 1129 | 1165 | 1135 | 1130 | 1002 | 1099 | 1170 | 1204 | 1240 |
| CL036 | 1088 | 1060 | 1005 | 1182 | 1113 | 1301 | 1117 | 1145 | 1056 | 1509 | 1060 | 1123 | 1080 | 1081 | 1081 | 1047 | 1067 | 1068 | 1320 |
| CL039 | 1022 | 1095 | 1054 | 1103 | 987 | 1035 | 1048 | 1072 | 987 | 948 | 1082 | 1046 | 1022 | 1060 | 996 | 974 | 1030 | 1059 | 1084 |
| CL041 | 1068 | 1119 | 1043 | 1101 | 1064 | 1271 | 1045 | 1121 | 989 | 1408 | 1016 | 1046 | 1000 | 1036 | 994 | 1040 | 1071 | 1028 | 1109 |
| CL042 | 982 | 1018 | 913 | 1062 | 963 | 968 | 947 | 1007 | 1027 | 950 | 939 | 976 | 962 | 947 | 928 | 912 | 963 | 953 | 1032 |
| CL043 | 960 | 1030 | 975 | 971 | 1026 | 793 | 1057 | 999 | 942 | 650 | 988 | 977 | 848 | 979 | 1061 | 868 | 906 | 917 | 872 |
| CL044 | 990 | 930 | 901 | 1049 | 985 | 962 | 966 | 990 | 940 | 764 | 950 | 989 | 942 | 925 | 929 | 885 | 907 | 944 | 987 |
| CL045 | 976 | 923 | 887 | 1060 | 1040 | 1039 | 916 | 961 | 1008 | 1167 | 990 | 1045 | 878 | 938 | 936 | 948 | 1002 | 1015 | 1188 |
| CL048 | 908 | 1019 | 875 | 1079 | 938 | 1033 | 964 | 1018 | 939 | 1256 | 868 | 966 | 900 | 937 | 859 | 878 | 990 | 936 | 1089 |
| CL049 | 868 | 968 | 896 | 1044 | 930 | 932 | 987 | 973 | 881 | 1012 | 880 | 983 | 899 | 967 | 898 | 939 | 890 | 971 | 976 |
| CL051 | 865 | 795 | 879 | 845 | 907 | 907 | 919 | 955 | 789 | 895 | 892 | 897 | 843 | 881 | 805 | 825 | 828 | 871 | 995 |
| CL055 | 718 | 763 | 701 | 784 | 709 | 821 | 749 | 742 | 682 | 859 | 706 | 727 | 639 | 701 | 705 | 744 | 714 | 734 | 779 |
| CL056 | 724 | 699 | 704 | 738 | 753 | 831 | 763 | 697 | 716 | 904 | 678 | 742 | 701 | 681 | 685 | 705 | 714 | 704 | 857 |

| | A1_097 | A1_108 | A1_125 | A1_132 | A1_133 | A1_155 | A1_Afr | A1_Nis | A2_001 | A2_004 | A2_008 | A2_016 | A2_019 | A2_021 | A2_026 | A2_029 | A2_030 | A2_031 | A2_034 |
|---|---|---|---|---|---|---|---|---|---|---|---|---|---|---|---|---|---|---|---|
| CL059 | 673 | 633 | 678 | 707 | 682 | 686 | 675 | 719 | 709 | 618 | 693 | 684 | 657 | 716 | 661 | 655 | 708 | 683 | 719 |
| CL060 | 675 | 736 | 663 | 806 | 729 | 858 | 685 | 724 | 682 | 954 | 670 | 702 | 648 | 620 | 638 | 683 | 667 | 712 | 817 |
| CL061 | 655 | 697 | 711 | 622 | 672 | 724 | 696 | 739 | 687 | 915 | 711 | 643 | 702 | 715 | 666 | 670 | 680 | 661 | 703 |
| CL062 | 710 | 684 | 629 | 751 | 747 | 670 | 717 | 713 | 678 | 851 | 639 | 688 | 624 | 716 | 717 | 674 | 661 | 686 | 725 |
| CL063 | 684 | 702 | 662 | 570 | 615 | 823 | 684 | 717 | 670 | 928 | 658 | 629 | 695 | 679 | 673 | 696 | 648 | 629 | 666 |
| CL064 | 553 | 614 | 601 | 644 | 625 | 448 | 644 | 680 | 580 | 322 | 549 | 669 | 485 | 654 | 637 | 571 | 539 | 593 | 523 |
| CL068 | 550 | 547 | 568 | 557 | 629 | 602 | 648 | 588 | 617 | 674 | 561 | 576 | 567 | 565 | 519 | 558 | 539 | 544 | 644 |
| CL070 | 551 | 572 | 568 | 670 | 597 | 534 | 582 | 566 | 558 | 542 | 579 | 543 | 564 | 561 | 509 | 543 | 559 | 554 | 644 |
| CL071 | 565 | 564 | 520 | 597 | 576 | 605 | 535 | 584 | 569 | 698 | 576 | 562 | 569 | 575 | 511 | 577 | 557 | 535 | 634 |
| CL074 | 542 | 558 | 534 | 554 | 504 | 591 | 537 | 579 | 500 | 651 | 523 | 518 | 479 | 546 | 483 | 491 | 554 | 517 | 636 |
| CL075 | 528 | 530 | 486 | 524 | 559 | 355 | 553 | 508 | 497 | 243 | 484 | 514 | 464 | 546 | 491 | 471 | 509 | 531 | 428 |
| CL076 | 532 | 523 | 517 | 538 | 490 | 654 | 597 | 588 | 541 | 791 | 532 | 493 | 526 | 517 | 505 | 529 | 564 | 491 | 601 |
| CL078 | 537 | 473 | 534 | 520 | 515 | 500 | 528 | 526 | 495 | 470 | 508 | 516 | 489 | 516 | 494 | 494 | 479 | 518 | 514 |
| CL079 | 518 | 567 | 471 | 538 | 522 | 572 | 542 | 563 | 515 | 601 | 535 | 533 | 501 | 498 | 477 | 513 | 468 | 551 | 505 |
| CL080 | 464 | 487 | 508 | 478 | 495 | 549 | 554 | 558 | 484 | 531 | 476 | 498 | 437 | 498 | 484 | 488 | 443 | 473 | 606 |
| CL082 | 479 | 486 | 426 | 525 | 507 | 369 | 517 | 487 | 492 | 378 | 498 | 511 | 409 | 459 | 518 | 466 | 424 | 474 | 405 |
| CL084 | 482 | 473 | 420 | 555 | 486 | 488 | 447 | 484 | 441 | 577 | 461 | 477 | 487 | 455 | 445 | 491 | 452 | 480 | 521 |
| CL085 | 429 | 437 | 496 | 544 | 472 | 475 | 459 | 486 | 481 | 488 | 474 | 491 | 473 | 458 | 469 | 429 | 447 | 467 | 568 |
| CL091 | 458 | 447 | 446 | 450 | 452 | 364 | 438 | 495 | 414 | 288 | 425 | 416 | 413 | 457 | 421 | 420 | 366 | 417 | 426 |
| CL092 | 464 | 453 | 434 | 489 | 487 | 516 | 442 | 463 | 422 | 558 | 430 | 458 | 413 | 451 | 401 | 420 | 400 | 438 | 515 |
| CL094 | 405 | 399 | 448 | 414 | 413 | 430 | 428 | 439 | 415 | 435 | 428 | 411 | 426 | 423 | 436 | 393 | 425 | 430 | 406 |
| CL095 | 602 | 378 | 435 | 395 | 446 | 52 | 513 | 437 | 483 | 45 | 430 | 421 | 302 | 428 | 773 | 430 | 401 | 360 | 165 |
| CL102 | 426 | 380 | 375 | 411 | 381 | 401 | 392 | 411 | 351 | 384 | 418 | 330 | 382 | 364 | 400 | 369 | 322 | 396 | 333 |
| CL104 | 409 | 300 | 355 | 308 | 296 | 511 | 360 | 363 | 418 | 611 | 376 | 336 | 399 | 372 | 406 | 418 | 339 | 324 | 387 |
| CL106 | 338 | 375 | 367 | 378 | 338 | 414 | 406 | 380 | 364 | 444 | 359 | 358 | 384 | 350 | 333 | 334 | 344 | 368 | 408 |
| CL107 | 354 | 352 | 390 | 355 | 382 | 403 | 366 | 351 | 361 | 447 | 319 | 358 | 385 | 322 | 320 | 390 | 358 | 347 | 373 |
| CL109 | 349 | 339 | 331 | 373 | 352 | 355 | 300 | 340 | 327 | 346 | 319 | 349 | 328 | 354 | 289 | 317 | 327 | 324 | 347 |
| CL112 | 343 | 339 | 299 | 392 | 330 | 425 | 334 | 367 | 349 | 460 | 363 | 342 | 310 | 352 | 287 | 361 | 323 | 325 | 382 |
| CL117 | 289 | 307 | 305 | 307 | 346 | 328 | 312 | 359 | 270 | 309 | 301 | 310 | 297 | 283 | 323 | 323 | 314 | 307 | 350 |
| CL119 | 322 | 295 | 268 | 336 | 352 | 373 | 329 | 359 | 327 | 373 | 343 | 308 | 299 | 317 | 327 | 305 | 318 | 320 | 356 |
| CL120 | 298 | 301 | 275 | 322 | 278 | 290 | 301 | 276 | 285 | 236 | 297 | 270 | 315 | 304 | 286 | 300 | 293 | 260 | 349 |
| CL121 | 312 | 328 | 335 | 333 | 338 | 276 | 336 | 333 | 284 | 225 | 310 | 287 | 285 | 318 | 294 | 275 | 306 | 265 | 290 |
| CL122 | 260 | 295 | 320 | 254 | 297 | 385 | 339 | 326 | 326 | 382 | 274 | 271 | 305 | 293 | 262 | 309 | 260 | 306 | 299 |
| CL129 | 291 | 290 | 285 | 274 | 296 | 293 | 321 | 287 | 315 | 305 | 277 | 266 | 284 | 288 | 271 | 301 | 273 | 273 | 286 |
| CL130 | 280 | 250 | 316 | 253 | 272 | 256 | 284 | 280 | 285 | 229 | 222 | 263 | 340 | 265 | 231 | 250 | 246 | 251 | 261 |
| CL131 | 256 | 269 | 291 | 278 | 287 | 317 | 264 | 320 | 266 | 423 | 261 | 257 | 296 | 250 | 271 | 264 | 250 | 309 | 347 |
| CL139 | 225 | 266 | 221 | 252 | 249 | 315 | 247 | 283 | 253 | 295 | 240 | 230 | 238 | 247 | 242 | 249 | 232 | 251 | 281 |
| CL143 | 223 | 201 | 233 | 219 | 228 | 205 | 250 | 238 | 215 | 129 | 191 | 185 | 174 | 224 | 224 | 218 | 222 | 208 | 224 |
| CL147 | 214 | 206 | 212 | 217 | 237 | 225 | 217 | 231 | 221 | 269 | 188 | 224 | 218 | 209 | 256 | 206 | 212 | 262 | 237 |
| CL151 | 215 | 202 | 200 | 185 | 182 | 266 | 214 | 233 | 216 | 265 | 185 | 189 | 218 | 198 | 208 | 195 | 182 | 203 | 228 |
| CL157 | 168 | 164 | 143 | 161 | 178 | 196 | 186 | 158 | 169 | 239 | 163 | 168 | 160 | 201 | 171 | 167 | 150 | 189 | 175 |
| CL158 | 165 | 177 | 159 | 181 | 167 | 209 | 189 | 167 | 161 | 202 | 194 | 171 | 178 | 166 | 156 | 148 | 171 | 170 | 201 |
| CL159 | 156 | 181 | 170 | 179 | 170 | 216 | 169 | 156 | 192 | 280 | 155 | 166 | 136 | 192 | 184 | 185 | 180 | 179 | 181 |
| CL160 | 158 | 142 | 160 | 162 | 169 | 150 | 182 | 196 | 165 | 127 | 166 | 143 | 136 | 160 | 158 | 149 | 138 | 140 | 143 |

| | A1_097 | A1_108 | A1_125 | A1_132 | A1_133 | A1_155 | A1_Afr | A1_Nis | A2_001 | A2_004 | A2_008 | A2_016 | A2_019 | A2_021 | A2_026 | A2_029 | A2_030 | A2_031 | A2_034 |
|---|---|---|---|---|---|---|---|---|---|---|---|---|---|---|---|---|---|---|---|
| CL162 | 158 | 158 | 152 | 171 | 162 | 181 | 179 | 194 | 186 | 170 | 159 | 164 | 152 | 149 | 175 | 153 | 168 | 164 | 189 |
| CL163 | 153 | 141 | 169 | 169 | 156 | 181 | 176 | 162 | 151 | 187 | 164 | 173 | 174 | 174 | 142 | 143 | 158 | 164 | 179 |
| CL164 | 168 | 150 | 148 | 180 | 156 | 173 | 165 | 161 | 136 | 181 | 163 | 183 | 158 | 161 | 132 | 164 | 162 | 136 | 179 |
| CL166 | 156 | 142 | 152 | 119 | 132 | 197 | 172 | 156 | 131 | 220 | 166 | 128 | 131 | 168 | 122 | 162 | 138 | 136 | 117 |
| CL171 | 119 | 137 | 122 | 156 | 133 | 152 | 115 | 167 | 118 | 89 | 134 | 121 | 115 | 152 | 120 | 128 | 92 | 132 | 131 |
| CL173 | 107 | 115 | 140 | 108 | 99 | 123 | 128 | 132 | 104 | 136 | 131 | 117 | 103 | 115 | 100 | 107 | 119 | 126 | 134 |
| CL175 | 123 | 100 | 108 | 105 | 112 | 100 | 121 | 106 | 92 | 110 | 102 | 92 | 104 | 108 | 129 | 97 | 95 | 124 | 112 |
| CL177 | 109 | 109 | 94 | 108 | 101 | 96 | 98 | 108 | 107 | 100 | 116 | 92 | 82 | 116 | 95 | 97 | 98 | 118 | 62 |
| CL182 | 75 | 59 | 93 | 78 | 76 | 122 | 106 | 93 | 84 | 105 | 86 | 61 | 79 | 81 | 75 | 99 | 74 | 65 | 87 |
| CL186 | 79 | 64 | 91 | 73 | 77 | 87 | 89 | 93 | 62 | 112 | 71 | 70 | 74 | 72 | 77 | 70 | 67 | 71 | 104 |
| CL187 | 84 | 83 | 89 | 86 | 75 | 69 | 82 | 80 | 73 | 86 | 69 | 71 | 66 | 75 | 74 | 77 | 68 | 66 | 84 |
| CL189 | 54 | 64 | 59 | 66 | 73 | 76 | 62 | 59 | 49 | 75 | 61 | 67 | 68 | 54 | 46 | 56 | 71 | 53 | 81 |
| CL191 | 56 | 43 | 41 | 58 | 74 | 61 | 68 | 57 | 57 | 91 | 50 | 51 | 53 | 55 | 55 | 44 | 69 | 53 | 71 |
| CL194 | 36 | 46 | 35 | 42 | 38 | 49 | 45 | 44 | 49 | 56 | 37 | 53 | 39 | 47 | 41 | 45 | 33 | 41 | 50 |
| CL195 | 43 | 42 | 48 | 29 | 36 | 59 | 46 | 43 | 44 | 81 | 48 | 41 | 41 | 47 | 49 | 42 | 39 | 51 | 52 |
| CL197 | 42 | 37 | 46 | 36 | 37 | 60 | 52 | 39 | 44 | 60 | 46 | 35 | 52 | 53 | 34 | 25 | 46 | 46 | 49 |
| CL199 | 33 | 46 | 35 | 43 | 46 | 36 | 44 | 41 | 40 | 27 | 39 | 38 | 39 | 30 | 42 | 53 | 36 | 32 | 33 |
| CL200 | 43 | 33 | 30 | 46 | 42 | 40 | 31 | 39 | 27 | 36 | 49 | 35 | 42 | 42 | 33 | 28 | 38 | 37 | 52 |
| CL202 | 33 | 33 | 30 | 29 | 37 | 37 | 40 | 42 | 42 | 34 | 29 | 40 | 29 | 34 | 33 | 38 | 35 | 34 | 36 |
| CL203 | 32 | 39 | 25 | 28 | 23 | 42 | 41 | 31 | 31 | 56 | 36 | 28 | 37 | 39 | 36 | 30 | 35 | 40 | 27 |
| CL204 | 34 | 27 | 41 | 42 | 38 | 38 | 31 | 34 | 31 | 47 | 42 | 23 | 26 | 27 | 23 | 36 | 27 | 29 | 36 |
| CL205 | 29 | 16 | 34 | 28 | 28 | 41 | 34 | 20 | 35 | 37 | 31 | 27 | 33 | 26 | 40 | 39 | 36 | 21 | 31 |
| CL206 | 29 | 25 | 30 | 22 | 29 | 33 | 30 | 29 | 30 | 42 | 29 | 28 | 31 | 30 | 35 | 37 | 37 | 31 | 27 |
| CL209 | 24 | 33 | 31 | 29 | 25 | 27 | 28 | 40 | 23 | 41 | 19 | 29 | 21 | 31 | 25 | 19 | 33 | 29 | 22 |
| CL211 | 23 | 24 | 16 | 28 | 18 | 41 | 26 | 29 | 21 | 35 | 25 | 28 | 30 | 27 | 22 | 39 | 24 | 24 | 35 |
| CL212 | 30 | 27 | 20 | 29 | 23 | 29 | 24 | 27 | 30 | 23 | 25 | 22 | 23 | 17 | 21 | 22 | 21 | 28 | 19 |
| CL213 | 21 | 19 | 20 | 20 | 29 | 28 | 29 | 19 | 29 | 35 | 27 | 26 | 26 | 20 | 22 | 33 | 24 | 18 | 29 |
| CL214 | 21 | 11 | 20 | 25 | 27 | 39 | 19 | 34 | 34 | 41 | 25 | 31 | 30 | 29 | 18 | 32 | 26 | 17 | 33 |
| CL215 | 24 | 19 | 22 | 23 | 27 | 20 | 27 | 19 | 17 | 25 | 25 | 24 | 27 | 23 | 29 | 19 | 24 | 20 | 24 |
| CL216 | 23 | 26 | 21 | 16 | 26 | 23 | 18 | 25 | 21 | 25 | 13 | 28 | 20 | 24 | 19 | 16 | 21 | 20 | 24 |
| CL218 | 19 | 18 | 24 | 13 | 13 | 17 | 20 | 25 | 28 | 30 | 20 | 21 | 20 | 17 | 14 | 16 | 16 | 15 | 25 |
| CL220 | 21 | 14 | 15 | 19 | 17 | 23 | 15 | 16 | 17 | 16 | 15 | 16 | 10 | 21 | 19 | 28 | 10 | 18 | 15 |
| CL221 | 23 | 15 | 14 | 24 | 11 | 14 | 19 | 21 | 16 | 19 | 16 | 18 | 11 | 17 | 20 | 17 | 18 | 17 | 22 |
| CL222 | 14 | 17 | 17 | 18 | 16 | 23 | 15 | 30 | 16 | 16 | 25 | 19 | 16 | 19 | 20 | 17 | 13 | 14 | 11 |
| CL223 | 20 | 21 | 15 | 26 | 18 | 15 | 17 | 25 | 18 | 25 | 17 | 15 | 16 | 20 | 10 | 11 | 9 | 23 | 15 |
| CL226 | 16 | 17 | 20 | 15 | 20 | 18 | 12 | 26 | 17 | 17 | 12 | 17 | 19 | 14 | 15 | 14 | 18 | 20 | 16 |
| CL228 | 9 | 23 | 25 | 22 | 16 | 8 | 16 | 18 | 20 | 21 | 15 | 24 | 20 | 10 | 21 | 12 | 10 | 17 | 25 |
| CL229 | 16 | 10 | 15 | 20 | 13 | 21 | 13 | 13 | 13 | 36 | 23 | 20 | 24 | 13 | 17 | 26 | 17 | 13 | 15 |
| CL231 | 14 | 13 | 7 | 15 | 19 | 20 | 16 | 18 | 14 | 19 | 11 | 19 | 11 | 11 | 15 | 22 | 18 | 15 | 15 |
| CL232 | 20 | 19 | 17 | 11 | 11 | 19 | 12 | 17 | 21 | 16 | 17 | 11 | 18 | 10 | 14 | 7 | 12 | 13 | 17 |
| CL234 | 12 | 19 | 15 | 13 | 11 | 10 | 16 | 11 | 8 | 11 | 8 | 7 | 7 | 8 | 13 | 12 | 17 | 11 | 13 |
| CL235 | 12 | 10 | 12 | 13 | 14 | 13 | 12 | 14 | 21 | 12 | 10 | 14 | 10 | 11 | 10 | 12 | 15 | 10 | 10 |
| CL236 | 13 | 10 | 11 | 7 | 13 | 14 | 11 | 10 | 13 | 14 | 16 | 9 | 12 | 17 | 11 | 15 | 7 | 8 | 12 |
| CL237 | 8 | 9 | 6 | 13 | 14 | 13 | 7 | 13 | 10 | 9 | 11 | 10 | 11 | 7 | 9 | 8 | 14 | 14 | 14 |
| CL239 | 11 | 15 | 19 | 11 | 12 | 10 | 12 | 16 | 15 | 24 | 9 | 16 | 10 | 13 | 11 | 11 | 8 | 11 | 8 |

| | A2_040 | A2_041 | A2_043 | A2_044 | A2_045 | A2_046 | A2_047 | A2_056 | A2_057 | A2_059 | A2_060 | A2_061 | A2_062 | A2_064 | A2_065 | A2_066 | A2_067 | A2_069 | A2_073 |
|---|---|---|---|---|---|---|---|---|---|---|---|---|---|---|---|---|---|---|---|
| CL002 | 5004 | 5235 | 4781 | 4769 | 5745 | 4952 | 4632 | 4663 | 4775 | 5116 | 5102 | 4766 | 4613 | 5342 | 5349 | 5327 | 4954 | 4741 | 4693 |
| CL003 | 3008 | 3255 | 2853 | 3175 | 3432 | 2946 | 3166 | 2745 | 2958 | 3013 | 2989 | 2841 | 2746 | 3156 | 3136 | 3202 | 2971 | 2973 | 2643 |
| CL004 | 2853 | 2912 | 2772 | 2926 | 3839 | 2709 | 2760 | 2609 | 2684 | 2863 | 2964 | 2766 | 2599 | 2864 | 2991 | 2981 | 2855 | 2684 | 2591 |
| CL005 | 2627 | 2755 | 2649 | 2622 | 2168 | 2444 | 2568 | 2518 | 2493 | 2804 | 2676 | 2583 | 2538 | 2948 | 2869 | 2682 | 2539 | 2603 | 2568 |
| CL008 | 2281 | 2269 | 2274 | 2353 | 3128 | 2205 | 2224 | 2119 | 2190 | 2309 | 2351 | 2317 | 2131 | 2366 | 2391 | 2425 | 2273 | 2296 | 2043 |
| CL009 | 2258 | 2266 | 2225 | 2230 | 2306 | 2163 | 2108 | 2149 | 2152 | 2238 | 2271 | 2150 | 2095 | 2313 | 2523 | 2391 | 2240 | 2099 | 2083 |
| CL010 | 2225 | 2195 | 1952 | 2196 | 2373 | 2152 | 2149 | 2036 | 2156 | 2210 | 2305 | 2106 | 2039 | 2301 | 2339 | 2273 | 2226 | 2150 | 1913 |
| CL011 | 2283 | 2274 | 2134 | 2183 | 2918 | 2171 | 2147 | 2193 | 2282 | 2320 | 2311 | 2156 | 2040 | 2414 | 2355 | 2446 | 2254 | 2200 | 2128 |
| CL012 | 1994 | 2137 | 2112 | 2050 | 1701 | 2005 | 2000 | 1944 | 1945 | 2121 | 2027 | 1977 | 1995 | 2197 | 2213 | 2110 | 1995 | 2043 | 2018 |
| CL013 | 2048 | 2055 | 2105 | 2116 | 2700 | 1951 | 1885 | 1874 | 1937 | 2073 | 2029 | 2011 | 1916 | 2141 | 2089 | 2072 | 2044 | 2038 | 1902 |
| CL014 | 2046 | 2188 | 2022 | 2020 | 2592 | 2013 | 2057 | 2096 | 2127 | 2171 | 2145 | 2071 | 1929 | 2153 | 2186 | 2159 | 2151 | 2020 | 1969 |
| CL015 | 1983 | 2008 | 1907 | 2063 | 2711 | 2043 | 1941 | 1884 | 1959 | 1941 | 2122 | 1831 | 1811 | 2102 | 2095 | 2166 | 2010 | 1995 | 1844 |
| CL016 | 1709 | 1656 | 1589 | 1692 | 628 | 1754 | 1759 | 1844 | 1612 | 1720 | 1675 | 1684 | 1855 | 1797 | 1685 | 1545 | 1732 | 1752 | 1620 |
| CL017 | 2053 | 1943 | 1686 | 1955 | 2143 | 1840 | 1841 | 1799 | 1918 | 1955 | 2041 | 1640 | 1600 | 1920 | 1959 | 1907 | 1926 | 1757 | 1573 |
| CL020 | 1538 | 1691 | 1557 | 1634 | 1363 | 1570 | 1690 | 1480 | 1567 | 1597 | 1610 | 1523 | 1481 | 1618 | 1590 | 1750 | 1562 | 1544 | 1424 |
| CL021 | 1605 | 1629 | 1512 | 1628 | 1867 | 1485 | 1578 | 1556 | 1570 | 1610 | 1666 | 1509 | 1457 | 1581 | 1575 | 1662 | 1597 | 1544 | 1445 |
| CL023 | 1582 | 1626 | 1536 | 1581 | 2027 | 1492 | 1517 | 1550 | 1514 | 1655 | 1537 | 1594 | 1536 | 1642 | 1694 | 1689 | 1510 | 1466 | 1441 |
| CL024 | 1535 | 1612 | 1405 | 1544 | 1523 | 1572 | 1505 | 1444 | 1459 | 1499 | 1566 | 1469 | 1380 | 1440 | 1521 | 1571 | 1576 | 1401 | 1325 |
| CL025 | 1461 | 1508 | 1446 | 1406 | 963 | 1469 | 1372 | 1402 | 1391 | 1519 | 1425 | 1476 | 1439 | 1647 | 1585 | 1423 | 1414 | 1457 | 1393 |
| CL026 | 1462 | 1397 | 1274 | 1280 | 443 | 1466 | 1449 | 1425 | 1333 | 1443 | 1317 | 1254 | 1334 | 1427 | 1478 | 1381 | 1395 | 1371 | 1249 |
| CL027 | 1457 | 1500 | 1398 | 1477 | 1792 | 1476 | 1425 | 1337 | 1364 | 1438 | 1447 | 1385 | 1333 | 1443 | 1419 | 1512 | 1442 | 1381 | 1359 |
| CL030 | 1382 | 1463 | 1259 | 1250 | 1039 | 1337 | 1303 | 1314 | 1325 | 1302 | 1419 | 1258 | 1207 | 1441 | 1358 | 1390 | 1340 | 1204 | 1228 |
| CL032 | 1319 | 1332 | 1160 | 1260 | 1113 | 1313 | 1368 | 1276 | 1353 | 1340 | 1348 | 1146 | 1134 | 1294 | 1361 | 1434 | 1355 | 1267 | 1091 |
| CL033 | 1213 | 1260 | 1159 | 1116 | 820 | 1113 | 1119 | 1108 | 1137 | 1246 | 1195 | 1103 | 1107 | 1299 | 1225 | 1257 | 1117 | 1108 | 1119 |
| CL034 | 1267 | 1278 | 1224 | 1269 | 1621 | 1263 | 1121 | 1134 | 1185 | 1271 | 1324 | 1226 | 1170 | 1263 | 1269 | 1279 | 1214 | 1140 | 1166 |
| CL037 | 1156 | 1104 | 983 | 1081 | 602 | 1167 | 1112 | 1063 | 1108 | 1069 | 1080 | 984 | 1037 | 1033 | 1081 | 1092 | 1104 | 997 | 1027 |
| CL038 | 1096 | 1120 | 1004 | 1020 | 599 | 1017 | 1110 | 1061 | 968 | 1056 | 1099 | 960 | 956 | 1081 | 1039 | 1043 | 1036 | 1007 | 864 |
| CL040 | 1041 | 1033 | 1051 | 1049 | 1007 | 1086 | 1028 | 972 | 933 | 1014 | 1088 | 1031 | 971 | 1053 | 1085 | 1133 | 997 | 1002 | 976 |
| CL046 | 1007 | 1022 | 939 | 1061 | 1449 | 952 | 1025 | 1002 | 979 | 1072 | 1071 | 977 | 894 | 1022 | 1046 | 1142 | 967 | 1010 | 911 |
| CL047 | 988 | 1074 | 998 | 1013 | 1208 | 974 | 963 | 988 | 1121 | 1026 | 1013 | 984 | 921 | 1032 | 992 | 975 | 991 | 965 | 1011 |
| CL050 | 847 | 924 | 739 | 850 | 657 | 877 | 846 | 877 | 835 | 897 | 850 | 732 | 736 | 857 | 893 | 864 | 909 | 772 | 692 |
| CL052 | 778 | 816 | 734 | 774 | 596 | 765 | 813 | 804 | 823 | 851 | 786 | 707 | 772 | 836 | 781 | 786 | 821 | 734 | 766 |
| CL053 | 812 | 818 | 783 | 725 | 674 | 749 | 776 | 761 | 748 | 800 | 745 | 775 | 700 | 744 | 809 | 790 | 789 | 714 | 712 |
| CL057 | 754 | 785 | 713 | 795 | 1066 | 704 | 720 | 696 | 746 | 813 | 759 | 741 | 683 | 730 | 795 | 795 | 750 | 741 | 653 |
| CL058 | 756 | 793 | 716 | 684 | 726 | 734 | 721 | 731 | 735 | 747 | 755 | 753 | 752 | 792 | 787 | 796 | 674 | 741 | 702 |
| CL065 | 564 | 654 | 535 | 637 | 584 | 569 | 598 | 577 | 572 | 577 | 597 | 524 | 544 | 561 | 562 | 621 | 591 | 574 | 529 |
| CL066 | 572 | 606 | 575 | 555 | 294 | 602 | 597 | 570 | 572 | 589 | 591 | 530 | 598 | 596 | 610 | 624 | 578 | 599 | 528 |
| CL067 | 625 | 636 | 624 | 653 | 755 | 600 | 647 | 603 | 624 | 641 | 636 | 586 | 564 | 590 | 613 | 687 | 627 | 633 | 545 |
| CL072 | 537 | 533 | 524 | 595 | 612 | 586 | 575 | 514 | 513 | 579 | 603 | 517 | 546 | 602 | 585 | 590 | 519 | 550 | 503 |
| CL081 | 516 | 527 | 496 | 433 | 327 | 500 | 525 | 479 | 484 | 495 | 523 | 441 | 503 | 529 | 509 | 523 | 458 | 469 | 448 |
| CL093 | 470 | 392 | 412 | 434 | 128 | 516 | 457 | 526 | 473 | 451 | 437 | 432 | 442 | 502 | 421 | 410 | 459 | 500 | 381 |
| CL099 | 405 | 403 | 370 | 331 | 166 | 425 | 361 | 409 | 375 | 381 | 380 | 352 | 357 | 378 | 385 | 348 | 405 | 385 | 357 |

| | A2_040 | A2_041 | A2_043 | A2_044 | A2_045 | A2_046 | A2_047 | A2_056 | A2_057 | A2_059 | A2_060 | A2_061 | A2_062 | A2_064 | A2_065 | A2_066 | A2_067 | A2_069 | A2_073 |
|---|---|---|---|---|---|---|---|---|---|---|---|---|---|---|---|---|---|---|---|
| CL100 | 356 | 419 | 325 | 410 | 563 | 383 | 336 | 413 | 356 | 356 | 357 | 327 | 376 | 344 | 415 | 432 | 395 | 370 | 298 |
| CL101 | 406 | 362 | 397 | 416 | 286 | 429 | 393 | 380 | 390 | 387 | 387 | 364 | 385 | 430 | 405 | 401 | 381 | 361 | 386 |
| CL113 | 344 | 312 | 320 | 326 | 87 | 350 | 354 | 363 | 333 | 330 | 338 | 324 | 338 | 369 | 326 | 313 | 339 | 361 | 289 |
| CL118 | 280 | 291 | 290 | 282 | 209 | 319 | 328 | 316 | 332 | 331 | 289 | 267 | 288 | 293 | 258 | 282 | 307 | 330 | 287 |
| CL128 | 282 | 295 | 266 | 262 | 201 | 294 | 331 | 274 | 278 | 308 | 295 | 257 | 249 | 286 | 274 | 298 | 292 | 305 | 247 |
| CL133 | 239 | 211 | 256 | 231 | 71 | 245 | 240 | 249 | 250 | 264 | 244 | 226 | 241 | 219 | 206 | 207 | 252 | 266 | 243 |
| CL137 | 253 | 250 | 259 | 222 | 106 | 246 | 228 | 244 | 233 | 270 | 236 | 222 | 243 | 248 | 246 | 217 | 272 | 233 | 221 |
| CL146 | 208 | 225 | 229 | 209 | 116 | 251 | 209 | 228 | 181 | 201 | 192 | 195 | 191 | 219 | 195 | 208 | 203 | 212 | 191 |
| CL148 | 231 | 244 | 219 | 203 | 223 | 231 | 230 | 219 | 212 | 242 | 247 | 232 | 195 | 236 | 245 | 235 | 236 | 219 | 232 |
| CL149 | 211 | 202 | 197 | 197 | 53 | 204 | 217 | 182 | 200 | 244 | 193 | 195 | 195 | 213 | 251 | 224 | 214 | 215 | 178 |
| CL152 | 188 | 210 | 174 | 181 | 98 | 184 | 218 | 178 | 167 | 166 | 205 | 164 | 165 | 170 | 193 | 144 | 178 | 214 | 149 |
| CL155 | 181 | 147 | 169 | 177 | 68 | 156 | 148 | 163 | 179 | 154 | 188 | 158 | 171 | 166 | 179 | 158 | 182 | 168 | 157 |
| CL168 | 138 | 134 | 135 | 155 | 48 | 127 | 138 | 153 | 157 | 139 | 136 | 128 | 143 | 107 | 131 | 131 | 145 | 110 | 135 |
| CL170 | 143 | 128 | 152 | 126 | 104 | 151 | 140 | 128 | 127 | 130 | 125 | 136 | 130 | 141 | 141 | 133 | 162 | 167 | 143 |
| CL174 | 95 | 103 | 123 | 113 | 35 | 103 | 121 | 109 | 109 | 96 | 107 | 98 | 114 | 106 | 84 | 101 | 125 | 115 | 124 |
| CL176 | 101 | 97 | 83 | 91 | 69 | 86 | 108 | 96 | 94 | 108 | 83 | 96 | 100 | 87 | 110 | 96 | 111 | 101 | 88 |
| CL184 | 85 | 60 | 63 | 74 | 18 | 92 | 96 | 91 | 99 | 79 | 77 | 76 | 88 | 72 | 57 | 55 | 79 | 68 | 69 |
| CL196 | 39 | 59 | 48 | 49 | 35 | 36 | 36 | 52 | 60 | 49 | 54 | 43 | 38 | 50 | 43 | 37 | 59 | 41 | 44 |
| CL198 | 38 | 46 | 47 | 46 | 45 | 42 | 47 | 45 | 42 | 39 | 33 | 45 | 57 | 26 | 26 | 43 | 33 | 40 | 53 |
| CL210 | 31 | 35 | 34 | 24 | 21 | 24 | 31 | 36 | 25 | 24 | 37 | 27 | 28 | 19 | 19 | 35 | 27 | 19 | 18 |
| CL219 | 18 | 11 | 36 | 12 | 12 | 7 | 12 | 20 | 22 | 13 | 11 | 12 | 13 | 15 | 24 | 13 | 11 | 23 | 14 |
| CL238 | 12 | 13 | 8 | 14 | 5 | 16 | 8 | 12 | 13 | 6 | 7 | 12 | 11 | 10 | 11 | 11 | 11 | 12 | 10 |
| CL001 | 4986 | 5151 | 4605 | 4987 | 3815 | 4930 | 4929 | 4643 | 4841 | 5013 | 5058 | 4387 | 4412 | 5008 | 5013 | 4992 | 5097 | 4841 | 4421 |
| CL006 | 2497 | 2401 | 2365 | 2467 | 1717 | 2455 | 2391 | 2507 | 2638 | 2654 | 2376 | 2342 | 2356 | 2465 | 2479 | 2245 | 2508 | 2461 | 2410 |
| CL007 | 2215 | 2374 | 2193 | 2215 | 1495 | 2236 | 2259 | 2190 | 2329 | 2378 | 2315 | 2187 | 2229 | 2499 | 2438 | 2442 | 2229 | 2323 | 2192 |
| CL018 | 1633 | 1698 | 1623 | 1631 | 1175 | 1681 | 1729 | 1597 | 1645 | 1755 | 1745 | 1644 | 1605 | 1768 | 1778 | 1643 | 1718 | 1712 | 1686 |
| CL019 | 1780 | 1762 | 1558 | 1712 | 1498 | 1693 | 1718 | 1623 | 1486 | 1648 | 1767 | 1640 | 1553 | 1648 | 1687 | 1739 | 1633 | 1585 | 1546 |
| CL022 | 1594 | 1578 | 1530 | 1531 | 1492 | 1576 | 1490 | 1453 | 1496 | 1582 | 1543 | 1469 | 1500 | 1528 | 1494 | 1642 | 1530 | 1441 | 1424 |
| CL028 | 1463 | 1518 | 1360 | 1361 | 1021 | 1361 | 1390 | 1426 | 1417 | 1383 | 1361 | 1350 | 1358 | 1395 | 1415 | 1379 | 1387 | 1393 | 1397 |
| CL029 | 1305 | 1349 | 1309 | 1353 | 1133 | 1294 | 1268 | 1249 | 1355 | 1322 | 1277 | 1264 | 1282 | 1377 | 1383 | 1399 | 1296 | 1316 | 1228 |
| CL031 | 1250 | 1281 | 1174 | 1174 | 1072 | 1310 | 1297 | 1166 | 1126 | 1286 | 1283 | 1146 | 1204 | 1267 | 1240 | 1316 | 1244 | 1233 | 1078 |
| CL035 | 1176 | 1144 | 1111 | 1025 | 778 | 1190 | 1136 | 1134 | 1236 | 1159 | 1206 | 1101 | 1152 | 1227 | 1237 | 1226 | 1128 | 1197 | 1072 |
| CL036 | 1058 | 1103 | 1001 | 1005 | 575 | 1090 | 1107 | 1108 | 1071 | 1063 | 990 | 989 | 1078 | 1026 | 1031 | 1025 | 1043 | 1050 | 980 |
| CL039 | 1052 | 1158 | 996 | 1077 | 765 | 1155 | 1073 | 1034 | 958 | 1077 | 1027 | 1027 | 957 | 1115 | 1136 | 1095 | 1083 | 1052 | 957 |
| CL041 | 982 | 1063 | 1039 | 980 | 499 | 982 | 1056 | 977 | 1021 | 1063 | 1008 | 930 | 1060 | 1094 | 1063 | 998 | 937 | 1070 | 932 |
| CL042 | 961 | 1032 | 902 | 1036 | 830 | 1040 | 987 | 960 | 967 | 935 | 978 | 990 | 958 | 981 | 1005 | 1027 | 1008 | 947 | 903 |
| CL043 | 998 | 1030 | 894 | 996 | 852 | 973 | 1043 | 948 | 993 | 981 | 1034 | 922 | 889 | 996 | 943 | 1097 | 1028 | 992 | 903 |
| CL044 | 970 | 1032 | 945 | 973 | 915 | 963 | 958 | 898 | 874 | 972 | 998 | 958 | 900 | 1025 | 1046 | 1011 | 956 | 947 | 932 |
| CL045 | 994 | 1017 | 971 | 993 | 893 | 957 | 948 | 914 | 998 | 989 | 979 | 933 | 994 | 927 | 943 | 925 | 1002 | 1029 | 972 |
| CL048 | 973 | 995 | 883 | 862 | 782 | 980 | 898 | 962 | 958 | 944 | 944 | 902 | 954 | 1043 | 971 | 989 | 983 | 921 | 948 |
| CL049 | 942 | 917 | 849 | 849 | 658 | 946 | 870 | 942 | 916 | 903 | 947 | 856 | 841 | 1008 | 957 | 942 | 916 | 947 | 865 |
| CL051 | 868 | 915 | 852 | 929 | 643 | 908 | 795 | 860 | 825 | 801 | 853 | 835 | 880 | 787 | 833 | 899 | 878 | 889 | 794 |
| CL055 | 696 | 783 | 707 | 697 | 515 | 785 | 731 | 648 | 707 | 753 | 731 | 713 | 713 | 727 | 781 | 736 | 691 | 728 | 686 |
| CL056 | 691 | 777 | 710 | 726 | 406 | 735 | 694 | 753 | 707 | 691 | 648 | 713 | 667 | 696 | 593 | 662 | 709 | 711 | 672 |

| | A2_040 | A2_041 | A2_043 | A2_044 | A2_045 | A2_046 | A2_047 | A2_056 | A2_057 | A2_059 | A2_060 | A2_061 | A2_062 | A2_064 | A2_065 | A2_066 | A2_067 | A2_069 | A2_073 |
|---|---|---|---|---|---|---|---|---|---|---|---|---|---|---|---|---|---|---|---|
| CL059 | 732 | 770 | 662 | 680 | 591 | 737 | 758 | 664 | 681 | 703 | 673 | 719 | 677 | 719 | 736 | 689 | 719 | 663 | 645 |
| CL060 | 705 | 740 | 660 | 660 | 371 | 720 | 687 | 705 | 718 | 670 | 701 | 653 | 653 | 694 | 699 | 702 | 631 | 685 | 640 |
| CL061 | 719 | 689 | 559 | 600 | 223 | 672 | 695 | 712 | 648 | 685 | 712 | 578 | 707 | 697 | 624 | 698 | 668 | 712 | 606 |
| CL062 | 659 | 664 | 610 | 668 | 420 | 689 | 672 | 713 | 672 | 744 | 703 | 605 | 588 | 666 | 680 | 655 | 720 | 673 | 611 |
| CL063 | 701 | 613 | 631 | 578 | 219 | 718 | 674 | 690 | 627 | 656 | 630 | 610 | 650 | 669 | 595 | 594 | 631 | 740 | 627 |
| CL064 | 628 | 652 | 601 | 630 | 513 | 602 | 550 | 614 | 611 | 646 | 661 | 597 | 585 | 614 | 627 | 636 | 626 | 590 | 506 |
| CL068 | 559 | 546 | 537 | 520 | 270 | 567 | 566 | 562 | 573 | 528 | 535 | 473 | 524 | 529 | 541 | 521 | 600 | 626 | 494 |
| CL070 | 568 | 561 | 568 | 556 | 517 | 574 | 578 | 559 | 535 | 578 | 554 | 548 | 576 | 559 | 549 | 599 | 567 | 544 | 507 |
| CL071 | 513 | 519 | 589 | 584 | 348 | 562 | 545 | 519 | 561 | 507 | 513 | 547 | 523 | 480 | 459 | 490 | 552 | 579 | 497 |
| CL074 | 507 | 573 | 544 | 516 | 377 | 525 | 520 | 508 | 535 | 526 | 517 | 506 | 542 | 522 | 569 | 486 | 502 | 519 | 510 |
| CL075 | 566 | 554 | 469 | 558 | 461 | 516 | 487 | 501 | 532 | 546 | 596 | 499 | 484 | 538 | 551 | 566 | 517 | 557 | 461 |
| CL076 | 501 | 534 | 526 | 442 | 201 | 545 | 532 | 561 | 536 | 492 | 473 | 481 | 547 | 454 | 444 | 460 | 483 | 536 | 461 |
| CL078 | 531 | 554 | 501 | 473 | 295 | 533 | 490 | 511 | 442 | 522 | 520 | 426 | 473 | 519 | 522 | 559 | 482 | 516 | 447 |
| CL079 | 520 | 542 | 525 | 458 | 242 | 498 | 511 | 501 | 475 | 546 | 512 | 503 | 503 | 519 | 531 | 482 | 523 | 518 | 449 |
| CL080 | 464 | 506 | 490 | 517 | 293 | 526 | 484 | 468 | 489 | 475 | 494 | 476 | 492 | 426 | 437 | 483 | 526 | 456 | 488 |
| CL082 | 530 | 520 | 436 | 519 | 447 | 487 | 515 | 456 | 501 | 515 | 517 | 420 | 423 | 480 | 522 | 559 | 532 | 502 | 409 |
| CL084 | 498 | 515 | 491 | 466 | 364 | 501 | 399 | 465 | 488 | 463 | 466 | 422 | 433 | 488 | 511 | 499 | 452 | 444 | 451 |
| CL085 | 453 | 503 | 460 | 441 | 490 | 465 | 482 | 412 | 466 | 459 | 467 | 482 | 465 | 469 | 445 | 486 | 475 | 490 | 455 |
| CL091 | 457 | 452 | 412 | 455 | 346 | 422 | 495 | 446 | 385 | 464 | 440 | 413 | 426 | 441 | 462 | 429 | 428 | 405 | 397 |
| CL092 | 410 | 413 | 400 | 439 | 319 | 399 | 444 | 440 | 464 | 407 | 390 | 429 | 418 | 379 | 396 | 354 | 421 | 408 | 475 |
| CL094 | 376 | 410 | 379 | 345 | 151 | 439 | 436 | 429 | 375 | 438 | 391 | 363 | 419 | 395 | 387 | 369 | 420 | 430 | 355 |
| CL095 | 393 | 342 | 284 | 457 | 1886 | 413 | 556 | 481 | 355 | 383 | 429 | 299 | 246 | 390 | 545 | 523 | 504 | 470 | 213 |
| CL102 | 425 | 417 | 363 | 330 | 155 | 378 | 393 | 398 | 365 | 410 | 384 | 346 | 348 | 392 | 371 | 416 | 435 | 354 | 313 |
| CL104 | 321 | 334 | 392 | 331 | 173 | 405 | 395 | 413 | 389 | 363 | 331 | 351 | 399 | 306 | 253 | 266 | 362 | 390 | 340 |
| CL106 | 330 | 367 | 371 | 336 | 194 | 376 | 359 | 338 | 356 | 344 | 309 | 339 | 351 | 308 | 302 | 361 | 365 | 340 | 361 |
| CL107 | 342 | 362 | 358 | 365 | 177 | 348 | 357 | 362 | 338 | 344 | 380 | 339 | 345 | 306 | 314 | 338 | 340 | 346 | 327 |
| CL109 | 313 | 333 | 335 | 295 | 164 | 353 | 360 | 322 | 291 | 352 | 328 | 314 | 323 | 352 | 322 | 333 | 338 | 322 | 271 |
| CL112 | 299 | 309 | 301 | 293 | 185 | 322 | 343 | 337 | 329 | 335 | 303 | 305 | 339 | 304 | 309 | 285 | 306 | 346 | 318 |
| CL117 | 273 | 321 | 262 | 302 | 220 | 283 | 286 | 312 | 297 | 304 | 312 | 272 | 320 | 299 | 293 | 349 | 307 | 301 | 285 |
| CL119 | 286 | 327 | 314 | 327 | 317 | 354 | 342 | 323 | 327 | 306 | 303 | 272 | 315 | 296 | 262 | 307 | 311 | 351 | 274 |
| CL120 | 294 | 322 | 266 | 318 | 310 | 288 | 320 | 308 | 301 | 342 | 306 | 295 | 301 | 306 | 299 | 280 | 295 | 288 | 308 |
| CL121 | 276 | 321 | 281 | 274 | 215 | 261 | 280 | 302 | 249 | 299 | 288 | 268 | 284 | 294 | 309 | 292 | 292 | 291 | 249 |
| CL122 | 287 | 292 | 293 | 249 | 120 | 287 | 315 | 318 | 302 | 286 | 241 | 263 | 276 | 288 | 258 | 239 | 280 | 308 | 293 |
| CL129 | 284 | 252 | 250 | 273 | 159 | 294 | 289 | 291 | 267 | 312 | 250 | 259 | 261 | 254 | 248 | 232 | 281 | 295 | 266 |
| CL130 | 261 | 262 | 294 | 260 | 675 | 235 | 272 | 250 | 283 | 248 | 228 | 362 | 269 | 229 | 235 | 285 | 273 | 251 | 382 |
| CL131 | 266 | 272 | 258 | 257 | 87 | 276 | 259 | 298 | 280 | 252 | 252 | 224 | 275 | 228 | 236 | 240 | 246 | 264 | 253 |
| CL139 | 232 | 225 | 241 | 231 | 86 | 249 | 265 | 239 | 242 | 225 | 223 | 229 | 255 | 207 | 198 | 199 | 243 | 248 | 217 |
| CL143 | 230 | 238 | 224 | 251 | 273 | 218 | 219 | 237 | 234 | 236 | 242 | 223 | 224 | 244 | 245 | 287 | 248 | 259 | 233 |
| CL147 | 216 | 212 | 221 | 217 | 128 | 203 | 214 | 216 | 212 | 216 | 164 | 228 | 208 | 198 | 215 | 200 | 223 | 219 | 219 |
| CL151 | 189 | 183 | 200 | 183 | 116 | 189 | 208 | 204 | 196 | 196 | 174 | 213 | 210 | 159 | 139 | 151 | 191 | 199 | 193 |
| CL157 | 164 | 151 | 147 | 141 | 41 | 174 | 169 | 172 | 157 | 160 | 164 | 179 | 166 | 155 | 149 | 140 | 163 | 160 | 141 |
| CL158 | 153 | 181 | 165 | 178 | 115 | 173 | 175 | 177 | 155 | 139 | 162 | 172 | 173 | 157 | 169 | 133 | 184 | 163 | 166 |
| CL159 | 176 | 157 | 168 | 168 | 101 | 163 | 150 | 173 | 197 | 167 | 184 | 154 | 145 | 178 | 153 | 176 | 183 | 168 | 172 |
| CL160 | 159 | 144 | 157 | 171 | 157 | 151 | 164 | 155 | 166 | 174 | 177 | 159 | 164 | 157 | 172 | 151 | 165 | 176 | 161 |

| | A2_040 | A2_041 | A2_043 | A2_044 | A2_045 | A2_046 | A2_047 | A2_056 | A2_057 | A2_059 | A2_060 | A2_061 | A2_062 | A2_064 | A2_065 | A2_066 | A2_067 | A2_069 | A2_073 |
|---|---|---|---|---|---|---|---|---|---|---|---|---|---|---|---|---|---|---|---|
| CL162 | 184 | 185 | 158 | 160 | 130 | 158 | 154 | 160 | 165 | 150 | 150 | 157 | 164 | 152 | 160 | 178 | 168 | 174 | 168 |
| CL163 | 163 | 181 | 178 | 168 | 108 | 167 | 163 | 147 | 169 | 160 | 162 | 156 | 147 | 157 | 114 | 135 | 162 | 158 | 157 |
| CL164 | 149 | 141 | 150 | 121 | 122 | 163 | 171 | 147 | 160 | 157 | 168 | 160 | 143 | 173 | 178 | 152 | 148 | 144 | 136 |
| CL166 | 133 | 134 | 148 | 138 | 46 | 161 | 146 | 142 | 144 | 157 | 135 | 145 | 165 | 146 | 128 | 151 | 156 | 148 | 133 |
| CL171 | 132 | 125 | 138 | 145 | 175 | 137 | 151 | 118 | 137 | 147 | 124 | 131 | 116 | 124 | 130 | 144 | 139 | 123 | 130 |
| CL173 | 102 | 98 | 95 | 111 | 50 | 117 | 122 | 118 | 116 | 116 | 101 | 118 | 102 | 110 | 102 | 101 | 99 | 124 | 88 |
| CL175 | 108 | 126 | 126 | 115 | 69 | 107 | 100 | 92 | 96 | 106 | 107 | 110 | 97 | 103 | 101 | 111 | 103 | 103 | 91 |
| CL177 | 101 | 100 | 75 | 81 | 23 | 92 | 101 | 110 | 101 | 112 | 107 | 88 | 98 | 94 | 95 | 103 | 113 | 110 | 77 |
| CL182 | 63 | 59 | 75 | 68 | 31 | 83 | 67 | 75 | 74 | 66 | 61 | 87 | 74 | 67 | 71 | 60 | 78 | 80 | 87 |
| CL186 | 72 | 64 | 68 | 69 | 32 | 72 | 69 | 86 | 67 | 71 | 58 | 63 | 66 | 81 | 60 | 62 | 71 | 83 | 65 |
| CL187 | 65 | 82 | 72 | 70 | 52 | 78 | 76 | 69 | 61 | 63 | 71 | 85 | 83 | 66 | 62 | 70 | 74 | 62 | 75 |
| CL189 | 50 | 68 | 62 | 61 | 28 | 59 | 58 | 57 | 48 | 65 | 61 | 64 | 64 | 61 | 56 | 59 | 55 | 56 | 46 |
| CL191 | 59 | 55 | 75 | 58 | 50 | 57 | 61 | 52 | 70 | 62 | 56 | 72 | 55 | 53 | 66 | 52 | 38 | 62 | 53 |
| CL194 | 43 | 45 | 44 | 46 | 36 | 49 | 57 | 41 | 28 | 41 | 53 | 49 | 46 | 45 | 31 | 44 | 41 | 48 | 44 |
| CL195 | 47 | 36 | 36 | 36 | 13 | 53 | 52 | 49 | 42 | 37 | 39 | 40 | 45 | 28 | 41 | 37 | 47 | 46 | 42 |
| CL197 | 37 | 46 | 49 | 30 | 12 | 46 | 43 | 45 | 27 | 44 | 38 | 36 | 38 | 36 | 39 | 43 | 45 | 38 | 53 |
| CL199 | 46 | 36 | 40 | 39 | 24 | 39 | 51 | 50 | 41 | 41 | 44 | 44 | 42 | 48 | 36 | 43 | 33 | 34 | 33 |
| CL200 | 38 | 38 | 37 | 43 | 34 | 35 | 30 | 28 | 45 | 28 | 42 | 34 | 34 | 31 | 27 | 41 | 52 | 37 | 45 |
| CL202 | 29 | 38 | 29 | 48 | 27 | 40 | 38 | 21 | 41 | 26 | 44 | 25 | 38 | 31 | 23 | 37 | 40 | 32 | 30 |
| CL203 | 34 | 30 | 29 | 27 | 5 | 30 | 31 | 43 | 38 | 34 | 31 | 24 | 23 | 25 | 30 | 29 | 36 | 34 | 26 |
| CL204 | 31 | 24 | 33 | 37 | 11 | 43 | 31 | 23 | 26 | 23 | 27 | 33 | 32 | 26 | 31 | 28 | 31 | 41 | 31 |
| CL205 | 34 | 23 | 19 | 26 | 15 | 37 | 31 | 26 | 22 | 47 | 36 | 20 | 36 | 20 | 28 | 26 | 29 | 33 | 34 |
| CL206 | 23 | 30 | 43 | 26 | 28 | 35 | 23 | 25 | 39 | 34 | 31 | 38 | 24 | 28 | 38 | 38 | 34 | 34 | 23 |
| CL209 | 23 | 28 | 26 | 25 | 18 | 27 | 22 | 30 | 22 | 36 | 21 | 21 | 25 | 24 | 35 | 34 | 29 | 25 | 26 |
| CL211 | 27 | 22 | 21 | 25 | 8 | 30 | 23 | 24 | 25 | 19 | 24 | 20 | 26 | 15 | 23 | 28 | 30 | 21 | 35 |
| CL212 | 21 | 27 | 19 | 27 | 11 | 28 | 20 | 20 | 24 | 35 | 21 | 19 | 27 | 27 | 24 | 34 | 20 | 31 | 21 |
| CL213 | 16 | 24 | 20 | 25 | 12 | 24 | 18 | 20 | 24 | 31 | 23 | 20 | 27 | 20 | 25 | 16 | 14 | 29 | 32 |
| CL214 | 22 | 13 | 26 | 31 | 11 | 20 | 19 | 28 | 15 | 21 | 32 | 24 | 31 | 23 | 19 | 17 | 25 | 23 | 21 |
| CL215 | 22 | 23 | 23 | 15 | 11 | 21 | 20 | 25 | 16 | 17 | 25 | 23 | 22 | 24 | 16 | 34 | 18 | 26 | 20 |
| CL216 | 22 | 16 | 22 | 19 | 10 | 18 | 23 | 20 | 15 | 22 | 22 | 14 | 20 | 26 | 15 | 18 | 25 | 20 | 24 |
| CL218 | 25 | 19 | 22 | 20 | 16 | 25 | 21 | 14 | 24 | 17 | 18 | 19 | 30 | 22 | 18 | 12 | 17 | 23 | 16 |
| CL220 | 12 | 18 | 10 | 20 | 8 | 18 | 20 | 30 | 16 | 13 | 22 | 11 | 24 | 22 | 15 | 20 | 18 | 7 | 20 |
| CL221 | 20 | 22 | 15 | 15 | 11 | 14 | 12 | 12 | 15 | 24 | 18 | 16 | 23 | 19 | 15 | 34 | 18 | 14 | 14 |
| CL222 | 18 | 14 | 24 | 16 | 14 | 14 | 26 | 13 | 20 | 17 | 25 | 19 | 20 | 14 | 11 | 14 | 14 | 16 | 24 |
| CL223 | 17 | 25 | 14 | 23 | 12 | 16 | 26 | 17 | 17 | 15 | 16 | 11 | 15 | 13 | 18 | 21 | 19 | 20 | 24 |
| CL226 | 17 | 16 | 22 | 13 | 9 | 14 | 12 | 14 | 21 | 16 | 15 | 14 | 15 | 9 | 19 | 16 | 21 | 17 | 18 |
| CL228 | 17 | 15 | 24 | 14 | 13 | 24 | 18 | 17 | 18 | 17 | 13 | 25 | 19 | 19 | 10 | 19 | 13 | 16 | 21 |
| CL229 | 15 | 11 | 19 | 16 | 6 | 14 | 11 | 20 | 11 | 13 | 15 | 14 | 17 | 13 | 14 | 14 | 8 | 22 | 12 |
| CL231 | 14 | 18 | 7 | 11 | 13 | 13 | 13 | 17 | 10 | 13 | 15 | 14 | 9 | 8 | 14 | 13 | 24 | 23 | 15 |
| CL232 | 16 | 11 | 9 | 12 | 2 | 21 | 22 | 16 | 11 | 11 | 14 | 11 | 21 | 10 | 8 | 12 | 10 | 12 | 9 |
| CL234 | 10 | 14 | 11 | 11 | 0 | 9 | 17 | 11 | 18 | 13 | 6 | 13 | 14 | 9 | 9 | 9 | 16 | 9 | 9 |
| CL235 | 12 | 3 | 6 | 15 | 1 | 12 | 22 | 9 | 14 | 10 | 13 | 10 | 16 | 7 | 9 | 8 | 11 | 13 | 9 |
| CL236 | 7 | 13 | 9 | 8 | 3 | 11 | 12 | 12 | 8 | 13 | 10 | 9 | 6 | 8 | 5 | 4 | 7 | 9 | 14 |
| CL237 | 16 | 12 | 14 | 5 | 7 | 14 | 7 | 9 | 5 | 10 | 13 | 8 | 7 | 10 | 7 | 11 | 9 | 9 | 7 |
| CL239 | 14 | 12 | 11 | 12 | 7 | 6 | 8 | 6 | 10 | 9 | 6 | 9 | 9 | 8 | 6 | 8 | 11 | 8 | 10 |

| | A2_074 | A2_076 | A2_079 | A2_080 | A2_084 | A2_085 | A2_087 | A2_091 | A2_096 | A2_100 | A2_101 | A2_113 | A2_117 | A2_118 | A2_119 | A2_123 | A2_124 | A2_131 | A2_141 |
|---|---|---|---|---|---|---|---|---|---|---|---|---|---|---|---|---|---|---|---|
| CL002 | 4770 | 6026 | 4795 | 4721 | 4755 | 5740 | 5521 | 4748 | 5701 | 5477 | 5047 | 4540 | 4828 | 4815 | 4758 | 5387 | 5610 | 5061 | 5571 |
| CL003 | 3121 | 3519 | 3046 | 2867 | 2868 | 3319 | 3233 | 3006 | 3279 | 3291 | 3110 | 2955 | 2904 | 2971 | 3140 | 3298 | 3377 | 3013 | 3057 |
| CL004 | 2731 | 3571 | 2694 | 2637 | 2599 | 3222 | 3022 | 2591 | 3277 | 3160 | 2163 | 2665 | 2667 | 2745 | 2778 | 3084 | 3164 | 2874 | 2897 |
| CL005 | 2445 | 2839 | 2693 | 2552 | 2475 | 2867 | 2739 | 2449 | 2852 | 2783 | 3130 | 2531 | 2586 | 2573 | 2717 | 2716 | 2827 | 2670 | 2878 |
| CL008 | 2316 | 2799 | 2180 | 2092 | 2090 | 2511 | 2524 | 2140 | 2529 | 2519 | 2102 | 2208 | 2114 | 2207 | 2309 | 2490 | 2520 | 2339 | 2526 |
| CL009 | 2092 | 2711 | 2155 | 2022 | 2163 | 2593 | 2438 | 2158 | 2549 | 2505 | 2160 | 2080 | 2113 | 2162 | 2176 | 2370 | 2417 | 2201 | 2348 |
| CL010 | 2099 | 2537 | 2223 | 2051 | 2025 | 2502 | 2264 | 2040 | 2332 | 2512 | 2323 | 2009 | 2146 | 2243 | 2118 | 2355 | 2462 | 2183 | 2434 |
| CL011 | 2149 | 2470 | 2198 | 2123 | 2126 | 2476 | 2342 | 2245 | 2553 | 2499 | 3513 | 2179 | 2183 | 2143 | 2199 | 2498 | 2436 | 2282 | 2463 |
| CL012 | 2009 | 2227 | 2042 | 1915 | 1968 | 2308 | 2151 | 2007 | 2156 | 2092 | 1602 | 1957 | 2016 | 2008 | 2048 | 2093 | 2225 | 2016 | 2130 |
| CL013 | 1954 | 2637 | 2009 | 1862 | 1923 | 2381 | 2178 | 1981 | 2279 | 2292 | 1716 | 1977 | 1970 | 2023 | 2061 | 2303 | 2374 | 1979 | 2108 |
| CL014 | 2026 | 2461 | 2029 | 1926 | 1984 | 2324 | 2246 | 2027 | 2267 | 2299 | 3519 | 1942 | 2109 | 2048 | 2091 | 2244 | 2357 | 2139 | 2302 |
| CL015 | 1947 | 2588 | 1916 | 1919 | 1834 | 2257 | 2192 | 1760 | 2351 | 2265 | 1886 | 1960 | 1958 | 1970 | 2039 | 2201 | 2286 | 2082 | 2125 |
| CL016 | 1755 | 1697 | 1783 | 1729 | 1683 | 1683 | 1639 | 1841 | 1592 | 1502 | 1551 | 1817 | 1687 | 1639 | 1507 | 1536 | 1633 | 1552 | 1775 |
| CL017 | 1837 | 2365 | 1855 | 1722 | 1803 | 2147 | 2115 | 1847 | 2072 | 2144 | 1867 | 1859 | 1765 | 1971 | 1807 | 2143 | 2140 | 1771 | 1985 |
| CL020 | 1629 | 1894 | 1550 | 1407 | 1548 | 1790 | 1725 | 1659 | 1725 | 1795 | 1282 | 1582 | 1557 | 1551 | 1677 | 1739 | 1785 | 1612 | 1723 |
| CL021 | 1593 | 2012 | 1544 | 1419 | 1486 | 1750 | 1600 | 1548 | 1804 | 1768 | 1098 | 1591 | 1490 | 1568 | 1582 | 1792 | 1869 | 1603 | 1595 |
| CL023 | 1501 | 1938 | 1551 | 1483 | 1564 | 1850 | 1783 | 1458 | 1835 | 1760 | 2076 | 1484 | 1516 | 1600 | 1517 | 1718 | 1799 | 1661 | 1735 |
| CL024 | 1477 | 1706 | 1552 | 1410 | 1429 | 1650 | 1588 | 1527 | 1646 | 1757 | 1241 | 1473 | 1521 | 1498 | 1544 | 1594 | 1560 | 1545 | 1487 |
| CL025 | 1495 | 1442 | 1419 | 1433 | 1460 | 1497 | 1532 | 1408 | 1560 | 1419 | 1849 | 1463 | 1480 | 1432 | 1540 | 1371 | 1569 | 1473 | 1516 |
| CL026 | 1403 | 1336 | 1349 | 1355 | 1392 | 1411 | 1389 | 1447 | 1346 | 1332 | 2058 | 1435 | 1470 | 1382 | 1340 | 1276 | 1408 | 1303 | 1532 |
| CL027 | 1400 | 1726 | 1348 | 1281 | 1307 | 1651 | 1460 | 1422 | 1660 | 1587 | 1146 | 1443 | 1414 | 1434 | 1494 | 1593 | 1610 | 1382 | 1456 |
| CL030 | 1277 | 1580 | 1374 | 1243 | 1352 | 1469 | 1398 | 1315 | 1416 | 1483 | 1935 | 1328 | 1340 | 1321 | 1390 | 1422 | 1525 | 1371 | 1451 |
| CL032 | 1326 | 1632 | 1287 | 1228 | 1284 | 1486 | 1386 | 1249 | 1474 | 1454 | 1334 | 1279 | 1337 | 1302 | 1302 | 1417 | 1509 | 1344 | 1455 |
| CL033 | 1064 | 1310 | 1157 | 1080 | 1150 | 1310 | 1312 | 1135 | 1274 | 1274 | 1150 | 1105 | 1129 | 1145 | 1206 | 1176 | 1293 | 1177 | 1271 |
| CL034 | 1150 | 1443 | 1207 | 1096 | 1165 | 1442 | 1333 | 1163 | 1379 | 1313 | 1641 | 1153 | 1230 | 1261 | 1237 | 1383 | 1396 | 1264 | 1352 |
| CL037 | 1151 | 1004 | 1089 | 939 | 1049 | 1109 | 981 | 1087 | 1066 | 1087 | 1147 | 1104 | 1086 | 1062 | 1091 | 1049 | 1073 | 1014 | 1042 |
| CL038 | 1066 | 1153 | 1026 | 974 | 984 | 1063 | 1046 | 1067 | 1080 | 1042 | 877 | 1097 | 981 | 960 | 1050 | 1000 | 1092 | 1059 | 1046 |
| CL040 | 964 | 1255 | 1029 | 1010 | 971 | 1191 | 1094 | 1017 | 1173 | 1076 | 827 | 979 | 1000 | 1054 | 1088 | 1135 | 1127 | 1038 | 1061 |
| CL046 | 1014 | 1261 | 1020 | 936 | 932 | 1089 | 1117 | 1008 | 1162 | 1081 | 551 | 953 | 984 | 1039 | 1006 | 1120 | 1193 | 962 | 1015 |
| CL047 | 989 | 1124 | 1048 | 966 | 921 | 1213 | 1019 | 966 | 1163 | 1072 | 1737 | 958 | 1017 | 936 | 996 | 1138 | 1070 | 955 | 1132 |
| CL050 | 819 | 935 | 863 | 826 | 832 | 864 | 909 | 842 | 899 | 936 | 1469 | 839 | 821 | 824 | 840 | 934 | 943 | 835 | 912 |
| CL052 | 767 | 838 | 765 | 750 | 741 | 857 | 838 | 782 | 866 | 821 | 1249 | 771 | 751 | 757 | 739 | 846 | 870 | 790 | 794 |
| CL053 | 749 | 690 | 771 | 709 | 757 | 739 | 762 | 790 | 750 | 768 | 1109 | 724 | 749 | 731 | 811 | 741 | 729 | 763 | 810 |
| CL057 | 724 | 890 | 748 | 703 | 749 | 868 | 822 | 748 | 849 | 796 | 376 | 726 | 749 | 733 | 774 | 849 | 910 | 738 | 765 |
| CL058 | 713 | 773 | 748 | 708 | 745 | 836 | 820 | 759 | 806 | 791 | 1021 | 749 | 696 | 778 | 793 | 768 | 783 | 731 | 805 |
| CL065 | 585 | 736 | 550 | 508 | 567 | 611 | 616 | 612 | 671 | 625 | 392 | 552 | 634 | 591 | 617 | 668 | 680 | 591 | 566 |
| CL066 | 606 | 572 | 612 | 566 | 549 | 636 | 587 | 596 | 613 | 619 | 451 | 585 | 603 | 538 | 609 | 627 | 616 | 602 | 627 |
| CL067 | 671 | 742 | 608 | 559 | 640 | 617 | 568 | 615 | 663 | 667 | 385 | 621 | 625 | 626 | 607 | 710 | 676 | 643 | 622 |
| CL072 | 524 | 716 | 516 | 543 | 528 | 639 | 594 | 518 | 648 | 653 | 486 | 512 | 540 | 508 | 567 | 569 | 641 | 591 | 620 |
| CL081 | 508 | 574 | 438 | 448 | 489 | 517 | 538 | 500 | 561 | 523 | 408 | 500 | 481 | 466 | 510 | 501 | 556 | 482 | 508 |
| CL093 | 494 | 416 | 467 | 505 | 480 | 449 | 430 | 520 | 368 | 323 | 242 | 519 | 487 | 423 | 337 | 388 | 411 | 374 | 453 |
| CL099 | 330 | 360 | 368 | 389 | 406 | 362 | 351 | 365 | 395 | 359 | 575 | 403 | 415 | 369 | 446 | 379 | 364 | 357 | 397 |

| | A2_074 | A2_076 | A2_079 | A2_080 | A2_084 | A2_085 | A2_087 | A2_091 | A2_096 | A2_100 | A2_101 | A2_113 | A2_117 | A2_118 | A2_119 | A2_123 | A2_124 | A2_131 | A2_141 |
|---|---|---|---|---|---|---|---|---|---|---|---|---|---|---|---|---|---|---|---|
| CL100 | 317 | 357 | 426 | 362 | 398 | 390 | 449 | 438 | 421 | 343 | 239 | 336 | 388 | 438 | 417 | 493 | 408 | 433 | 301 |
| CL101 | 400 | 400 | 359 | 410 | 401 | 412 | 406 | 362 | 421 | 388 | 635 | 361 | 373 | 396 | 388 | 371 | 442 | 375 | 412 |
| CL113 | 359 | 264 | 352 | 342 | 331 | 321 | 348 | 350 | 276 | 280 | 247 | 391 | 349 | 314 | 235 | 274 | 285 | 286 | 345 |
| CL118 | 317 | 243 | 344 | 311 | 295 | 256 | 287 | 303 | 259 | 275 | 454 | 353 | 334 | 273 | 304 | 251 | 287 | 281 | 310 |
| CL128 | 264 | 283 | 280 | 283 | 288 | 255 | 296 | 252 | 287 | 277 | 274 | 298 | 309 | 316 | 292 | 282 | 292 | 268 | 305 |
| CL133 | 232 | 155 | 262 | 264 | 242 | 198 | 222 | 257 | 158 | 205 | 233 | 273 | 274 | 221 | 218 | 174 | 179 | 198 | 248 |
| CL137 | 230 | 205 | 258 | 283 | 250 | 224 | 244 | 231 | 204 | 234 | 279 | 244 | 229 | 227 | 252 | 185 | 237 | 256 | 225 |
| CL146 | 218 | 148 | 213 | 227 | 218 | 197 | 196 | 233 | 199 | 191 | 156 | 208 | 217 | 213 | 222 | 185 | 182 | 187 | 203 |
| CL148 | 205 | 241 | 223 | 233 | 216 | 258 | 271 | 224 | 246 | 243 | 379 | 224 | 216 | 205 | 234 | 232 | 256 | 210 | 252 |
| CL149 | 194 | 227 | 220 | 184 | 195 | 225 | 246 | 209 | 190 | 190 | 88 | 205 | 195 | 221 | 179 | 226 | 203 | 225 | 195 |
| CL152 | 198 | 186 | 190 | 161 | 203 | 187 | 195 | 182 | 192 | 196 | 113 | 188 | 189 | 199 | 163 | 191 | 206 | 168 | 176 |
| CL155 | 147 | 99 | 156 | 170 | 162 | 160 | 161 | 192 | 129 | 130 | 202 | 180 | 180 | 161 | 168 | 142 | 130 | 132 | 177 |
| CL168 | 153 | 78 | 129 | 133 | 163 | 108 | 128 | 140 | 91 | 101 | 121 | 157 | 151 | 128 | 124 | 110 | 114 | 123 | 127 |
| CL170 | 150 | 151 | 157 | 137 | 134 | 147 | 127 | 175 | 124 | 146 | 71 | 140 | 130 | 124 | 128 | 152 | 155 | 125 | 146 |
| CL174 | 110 | 54 | 124 | 105 | 121 | 75 | 87 | 109 | 55 | 77 | 88 | 138 | 138 | 104 | 107 | 88 | 80 | 89 | 94 |
| CL176 | 89 | 98 | 96 | 100 | 86 | 101 | 101 | 98 | 104 | 88 | 192 | 110 | 95 | 93 | 115 | 90 | 104 | 95 | 114 |
| CL184 | 77 | 48 | 82 | 66 | 87 | 55 | 55 | 73 | 61 | 56 | 64 | 90 | 83 | 85 | 63 | 52 | 54 | 70 | 68 |
| CL196 | 51 | 51 | 50 | 40 | 35 | 41 | 32 | 42 | 50 | 39 | 36 | 39 | 56 | 44 | 53 | 34 | 43 | 60 | 29 |
| CL198 | 65 | 33 | 50 | 52 | 35 | 54 | 34 | 38 | 36 | 47 | 66 | 58 | 50 | 43 | 38 | 34 | 46 | 41 | 40 |
| CL210 | 31 | 24 | 26 | 28 | 31 | 20 | 23 | 35 | 22 | 34 | 29 | 32 | 28 | 36 | 22 | 29 | 27 | 39 | 20 |
| CL219 | 5 | 12 | 21 | 34 | 24 | 34 | 27 | 27 | 31 | 5 | 24 | 11 | 17 | 25 | 15 | 14 | 17 | 34 | 13 |
| CL238 | 12 | 4 | 7 | 6 | 11 | 16 | 7 | 9 | 19 | 6 | 13 | 5 | 13 | 15 | 16 | 13 | 16 | 7 | 11 |
| CL001 | 4831 | 5705 | 4815 | 4438 | 4695 | 5313 | 5216 | 4817 | 5485 | 5432 | 6673 | 4744 | 4951 | 4939 | 4872 | 5484 | 5416 | 5025 | 5270 |
| CL006 | 2342 | 2221 | 2444 | 2463 | 2470 | 2446 | 2445 | 2402 | 2404 | 2432 | 5105 | 2501 | 2440 | 2456 | 2502 | 2322 | 2383 | 2366 | 2756 |
| CL007 | 2300 | 2332 | 2279 | 2205 | 2192 | 2444 | 2422 | 2190 | 2474 | 2265 | 3189 | 2256 | 2250 | 2261 | 2302 | 2376 | 2458 | 2237 | 2413 |
| CL018 | 1690 | 1582 | 1631 | 1671 | 1666 | 1811 | 1817 | 1697 | 1764 | 1693 | 1835 | 1735 | 1713 | 1614 | 1706 | 1598 | 1720 | 1718 | 1879 |
| CL019 | 1577 | 1972 | 1636 | 1574 | 1562 | 1664 | 1692 | 1604 | 1774 | 1640 | 1224 | 1668 | 1662 | 1507 | 1687 | 1711 | 1807 | 1683 | 1682 |
| CL022 | 1544 | 1808 | 1441 | 1416 | 1422 | 1681 | 1606 | 1462 | 1714 | 1676 | 1074 | 1549 | 1474 | 1464 | 1611 | 1589 | 1729 | 1580 | 1547 |
| CL028 | 1433 | 1363 | 1336 | 1362 | 1368 | 1449 | 1412 | 1439 | 1420 | 1395 | 2399 | 1387 | 1391 | 1309 | 1396 | 1334 | 1454 | 1376 | 1555 |
| CL029 | 1287 | 1332 | 1228 | 1275 | 1294 | 1419 | 1398 | 1296 | 1401 | 1386 | 1732 | 1291 | 1351 | 1367 | 1332 | 1349 | 1442 | 1370 | 1415 |
| CL031 | 1227 | 1477 | 1231 | 1141 | 1116 | 1240 | 1271 | 1157 | 1280 | 1281 | 858 | 1255 | 1251 | 1222 | 1267 | 1386 | 1379 | 1261 | 1206 |
| CL035 | 1096 | 1314 | 1129 | 1146 | 1035 | 1238 | 1265 | 1058 | 1253 | 1149 | 1135 | 1144 | 1130 | 1033 | 1177 | 1153 | 1264 | 1148 | 1220 |
| CL036 | 1062 | 872 | 1090 | 1025 | 1107 | 997 | 967 | 1105 | 964 | 962 | 1503 | 1065 | 1068 | 1076 | 1038 | 966 | 1000 | 1022 | 1075 |
| CL039 | 1017 | 1234 | 1018 | 974 | 982 | 1207 | 1100 | 1034 | 1209 | 1118 | 1077 | 1082 | 1077 | 1076 | 1118 | 1214 | 1157 | 1000 | 1109 |
| CL041 | 1013 | 943 | 1036 | 971 | 919 | 1076 | 1073 | 986 | 993 | 947 | 983 | 1016 | 1040 | 1051 | 1021 | 992 | 1019 | 996 | 1091 |
| CL042 | 1030 | 1097 | 934 | 986 | 957 | 1079 | 1060 | 950 | 1027 | 1087 | 927 | 955 | 980 | 932 | 978 | 969 | 1066 | 993 | 1021 |
| CL043 | 1030 | 1101 | 933 | 838 | 973 | 1028 | 987 | 1052 | 1083 | 1117 | 956 | 1037 | 948 | 971 | 1024 | 1124 | 1155 | 995 | 948 |
| CL044 | 990 | 1197 | 975 | 873 | 902 | 1015 | 993 | 964 | 1077 | 1032 | 768 | 958 | 962 | 965 | 1040 | 1013 | 1058 | 1032 | 1030 |
| CL045 | 972 | 780 | 981 | 957 | 974 | 897 | 890 | 959 | 918 | 857 | 1377 | 978 | 985 | 952 | 1075 | 892 | 932 | 944 | 1029 |
| CL048 | 948 | 1013 | 930 | 946 | 955 | 1006 | 958 | 989 | 999 | 965 | 1492 | 927 | 970 | 925 | 896 | 926 | 974 | 959 | 1066 |
| CL049 | 900 | 1004 | 887 | 874 | 824 | 935 | 901 | 912 | 1032 | 989 | 1210 | 890 | 878 | 923 | 967 | 934 | 1001 | 918 | 930 |
| CL051 | 841 | 794 | 831 | 845 | 823 | 798 | 803 | 867 | 835 | 861 | 843 | 878 | 843 | 812 | 903 | 826 | 851 | 879 | 863 |
| CL055 | 697 | 786 | 738 | 663 | 705 | 772 | 785 | 701 | 754 | 760 | 754 | 709 | 718 | 717 | 736 | 767 | 748 | 704 | 817 |
| CL056 | 741 | 569 | 724 | 709 | 676 | 660 | 626 | 730 | 607 | 651 | 773 | 710 | 741 | 689 | 764 | 640 | 635 | 698 | 676 |

| | A2_074 | A2_076 | A2_079 | A2_080 | A2_084 | A2_085 | A2_087 | A2_091 | A2_096 | A2_100 | A2_101 | A2_113 | A2_117 | A2_118 | A2_119 | A2_123 | A2_124 | A2_131 | A2_141 |
|---|---|---|---|---|---|---|---|---|---|---|---|---|---|---|---|---|---|---|---|
| CL059 | 662 | 742 | 718 | 654 | 672 | 701 | 717 | 648 | 769 | 778 | 636 | 752 | 681 | 679 | 690 | 681 | 780 | 659 | 729 |
| CL060 | 709 | 632 | 675 | 699 | 647 | 673 | 679 | 706 | 690 | 624 | 933 | 673 | 665 | 683 | 696 | 695 | 723 | 706 | 788 |
| CL061 | 672 | 639 | 670 | 674 | 661 | 661 | 674 | 697 | 638 | 684 | 675 | 761 | 682 | 680 | 688 | 611 | 661 | 677 | 675 |
| CL062 | 626 | 654 | 703 | 655 | 664 | 661 | 667 | 658 | 718 | 737 | 1024 | 686 | 652 | 704 | 667 | 694 | 752 | 702 | 726 |
| CL063 | 725 | 473 | 692 | 666 | 663 | 517 | 595 | 685 | 526 | 572 | 552 | 698 | 661 | 625 | 624 | 485 | 542 | 556 | 583 |
| CL064 | 605 | 710 | 661 | 585 | 602 | 629 | 638 | 594 | 705 | 639 | 518 | 617 | 548 | 559 | 633 | 662 | 670 | 625 | 594 |
| CL068 | 566 | 478 | 552 | 541 | 566 | 480 | 527 | 548 | 495 | 523 | 730 | 587 | 597 | 550 | 612 | 490 | 521 | 542 | 559 |
| CL070 | 573 | 632 | 530 | 550 | 540 | 581 | 594 | 594 | 591 | 614 | 493 | 562 | 543 | 558 | 578 | 593 | 625 | 579 | 613 |
| CL071 | 547 | 387 | 545 | 595 | 544 | 479 | 494 | 532 | 491 | 477 | 829 | 579 | 567 | 510 | 542 | 456 | 454 | 517 | 523 |
| CL074 | 538 | 519 | 561 | 518 | 440 | 537 | 566 | 563 | 545 | 565 | 644 | 523 | 516 | 502 | 507 | 473 | 545 | 469 | 541 |
| CL075 | 529 | 651 | 509 | 471 | 510 | 587 | 551 | 502 | 595 | 610 | 393 | 516 | 543 | 521 | 497 | 568 | 664 | 534 | 536 |
| CL076 | 466 | 330 | 504 | 512 | 484 | 417 | 436 | 526 | 372 | 414 | 599 | 531 | 510 | 504 | 498 | 392 | 413 | 468 | 479 |
| CL078 | 520 | 581 | 462 | 472 | 475 | 508 | 522 | 512 | 507 | 474 | 325 | 547 | 493 | 502 | 552 | 496 | 536 | 488 | 458 |
| CL079 | 476 | 512 | 527 | 504 | 499 | 499 | 487 | 502 | 482 | 517 | 713 | 470 | 477 | 497 | 543 | 473 | 537 | 518 | 551 |
| CL080 | 527 | 453 | 479 | 492 | 474 | 415 | 429 | 457 | 471 | 456 | 489 | 534 | 470 | 503 | 508 | 488 | 502 | 559 | 464 |
| CL082 | 500 | 595 | 513 | 474 | 492 | 562 | 525 | 521 | 556 | 560 | 557 | 479 | 476 | 490 | 461 | 574 | 585 | 483 | 529 |
| CL084 | 456 | 520 | 548 | 421 | 476 | 538 | 510 | 414 | 519 | 519 | 458 | 471 | 468 | 441 | 471 | 490 | 535 | 504 | 529 |
| CL085 | 486 | 454 | 458 | 431 | 435 | 453 | 474 | 443 | 467 | 440 | 476 | 468 | 472 | 487 | 508 | 421 | 496 | 469 | 449 |
| CL091 | 408 | 539 | 396 | 405 | 393 | 466 | 443 | 405 | 480 | 470 | 261 | 381 | 412 | 430 | 414 | 463 | 487 | 434 | 402 |
| CL092 | 389 | 288 | 405 | 411 | 435 | 371 | 369 | 455 | 367 | 379 | 701 | 451 | 439 | 428 | 428 | 326 | 358 | 409 | 422 |
| CL094 | 387 | 408 | 431 | 392 | 402 | 422 | 435 | 432 | 363 | 378 | 372 | 430 | 407 | 356 | 350 | 388 | 387 | 373 | 387 |
| CL095 | 340 | 502 | 491 | 325 | 401 | 475 | 665 | 461 | 613 | 405 | 66 | 376 | 415 | 611 | 389 | 906 | 418 | 643 | 290 |
| CL102 | 338 | 429 | 353 | 346 | 339 | 370 | 332 | 378 | 376 | 366 | 357 | 409 | 372 | 377 | 373 | 370 | 381 | 359 | 369 |
| CL104 | 375 | 187 | 399 | 418 | 386 | 292 | 311 | 390 | 245 | 267 | 312 | 404 | 372 | 346 | 304 | 231 | 281 | 281 | 305 |
| CL106 | 357 | 286 | 362 | 331 | 349 | 300 | 345 | 350 | 307 | 330 | 358 | 381 | 356 | 340 | 339 | 295 | 317 | 327 | 329 |
| CL107 | 382 | 294 | 358 | 371 | 320 | 332 | 331 | 347 | 329 | 319 | 450 | 359 | 367 | 338 | 313 | 328 | 349 | 332 | 340 |
| CL109 | 315 | 265 | 353 | 317 | 289 | 300 | 324 | 317 | 337 | 306 | 301 | 343 | 323 | 350 | 331 | 304 | 353 | 342 | 299 |
| CL112 | 329 | 238 | 334 | 297 | 313 | 295 | 302 | 343 | 255 | 287 | 500 | 329 | 309 | 332 | 295 | 242 | 280 | 338 | 295 |
| CL117 | 272 | 312 | 298 | 242 | 269 | 309 | 291 | 309 | 333 | 355 | 297 | 289 | 288 | 286 | 335 | 289 | 322 | 309 | 327 |
| CL119 | 295 | 241 | 331 | 312 | 306 | 296 | 302 | 286 | 287 | 301 | 526 | 300 | 301 | 315 | 332 | 279 | 290 | 314 | 308 |
| CL120 | 298 | 306 | 306 | 307 | 277 | 301 | 294 | 294 | 300 | 310 | 210 | 318 | 295 | 310 | 314 | 279 | 265 | 290 | 271 |
| CL121 | 233 | 375 | 292 | 276 | 266 | 303 | 285 | 288 | 348 | 353 | 212 | 286 | 266 | 275 | 277 | 270 | 346 | 254 | 261 |
| CL122 | 295 | 221 | 311 | 292 | 295 | 216 | 288 | 292 | 206 | 253 | 240 | 301 | 320 | 253 | 299 | 198 | 247 | 254 | 277 |
| CL129 | 333 | 266 | 304 | 253 | 319 | 245 | 253 | 283 | 247 | 234 | 248 | 267 | 317 | 265 | 273 | 237 | 276 | 294 | 268 |
| CL130 | 223 | 190 | 256 | 259 | 293 | 200 | 218 | 232 | 215 | 226 | 191 | 255 | 289 | 246 | 233 | 222 | 240 | 254 | 249 |
| CL131 | 263 | 159 | 267 | 295 | 282 | 222 | 244 | 248 | 223 | 219 | 303 | 292 | 289 | 247 | 248 | 214 | 210 | 260 | 236 |
| CL139 | 225 | 149 | 229 | 207 | 217 | 209 | 224 | 246 | 189 | 211 | 270 | 255 | 245 | 242 | 198 | 189 | 188 | 243 | 237 |
| CL143 | 225 | 272 | 224 | 197 | 200 | 223 | 248 | 197 | 235 | 256 | 186 | 207 | 232 | 230 | 238 | 243 | 262 | 231 | 223 |
| CL147 | 236 | 175 | 215 | 201 | 191 | 160 | 179 | 220 | 174 | 229 | 263 | 232 | 222 | 211 | 212 | 198 | 199 | 223 | 204 |
| CL151 | 193 | 119 | 209 | 190 | 202 | 150 | 150 | 199 | 116 | 173 | 229 | 204 | 195 | 178 | 177 | 145 | 151 | 186 | 197 |
| CL157 | 178 | 126 | 150 | 161 | 154 | 153 | 159 | 157 | 157 | 140 | 140 | 191 | 151 | 152 | 143 | 131 | 135 | 159 | 158 |
| CL158 | 189 | 129 | 178 | 157 | 168 | 156 | 153 | 182 | 138 | 163 | 178 | 177 | 173 | 154 | 165 | 164 | 139 | 193 | 163 |
| CL159 | 166 | 160 | 182 | 161 | 171 | 194 | 167 | 162 | 162 | 148 | 349 | 160 | 165 | 155 | 136 | 137 | 183 | 172 | 203 |
| CL160 | 193 | 185 | 157 | 145 | 156 | 163 | 162 | 159 | 193 | 190 | 105 | 159 | 150 | 152 | 184 | 173 | 193 | 149 | 150 |

| | A2_074 | A2_076 | A2_079 | A2_080 | A2_084 | A2_085 | A2_087 | A2_091 | A2_096 | A2_100 | A2_101 | A2_113 | A2_117 | A2_118 | A2_119 | A2_123 | A2_124 | A2_131 | A2_141 |
|---|---|---|---|---|---|---|---|---|---|---|---|---|---|---|---|---|---|---|---|
| CL162 | 159 | 129 | 167 | 151 | 164 | 137 | 141 | 143 | 147 | 161 | 192 | 144 | 170 | 153 | 151 | 146 | 156 | 172 | 151 |
| CL163 | 144 | 120 | 161 | 134 | 168 | 127 | 148 | 141 | 142 | 155 | 122 | 150 | 182 | 159 | 141 | 149 | 147 | 150 | 138 |
| CL164 | 138 | 171 | 177 | 148 | 169 | 162 | 170 | 141 | 182 | 155 | 203 | 149 | 161 | 168 | 173 | 143 | 164 | 164 | 175 |
| CL166 | 149 | 110 | 159 | 136 | 157 | 124 | 153 | 142 | 108 | 100 | 54 | 145 | 138 | 158 | 162 | 116 | 110 | 134 | 135 |
| CL171 | 124 | 172 | 118 | 119 | 111 | 135 | 126 | 145 | 150 | 139 | 144 | 133 | 113 | 125 | 141 | 120 | 153 | 115 | 114 |
| CL173 | 127 | 83 | 122 | 114 | 105 | 100 | 93 | 96 | 85 | 104 | 119 | 114 | 113 | 118 | 105 | 100 | 105 | 92 | 105 |
| CL175 | 121 | 95 | 98 | 139 | 115 | 98 | 114 | 122 | 104 | 101 | 87 | 104 | 110 | 104 | 117 | 88 | 98 | 92 | 93 |
| CL177 | 89 | 106 | 99 | 81 | 100 | 88 | 90 | 130 | 92 | 109 | 41 | 114 | 101 | 94 | 103 | 98 | 124 | 89 | 95 |
| CL182 | 73 | 42 | 84 | 94 | 84 | 58 | 54 | 70 | 49 | 41 | 71 | 83 | 69 | 62 | 61 | 31 | 45 | 54 | 62 |
| CL186 | 62 | 55 | 66 | 69 | 81 | 44 | 57 | 81 | 68 | 51 | 72 | 83 | 74 | 76 | 75 | 57 | 61 | 70 | 64 |
| CL187 | 74 | 64 | 81 | 85 | 67 | 71 | 56 | 76 | 62 | 57 | 91 | 71 | 65 | 80 | 72 | 51 | 72 | 80 | 69 |
| CL189 | 73 | 58 | 57 | 56 | 62 | 53 | 64 | 57 | 56 | 59 | 66 | 49 | 61 | 53 | 75 | 57 | 62 | 55 | 75 |
| CL191 | 58 | 34 | 74 | 60 | 54 | 48 | 37 | 49 | 58 | 59 | 59 | 64 | 57 | 62 | 58 | 56 | 57 | 67 | 59 |
| CL194 | 46 | 35 | 39 | 42 | 35 | 39 | 35 | 54 | 38 | 40 | 20 | 50 | 59 | 55 | 42 | 41 | 51 | 41 | 38 |
| CL195 | 53 | 22 | 37 | 39 | 43 | 24 | 40 | 44 | 35 | 23 | 47 | 43 | 43 | 35 | 30 | 30 | 30 | 39 | 46 |
| CL197 | 30 | 42 | 36 | 38 | 34 | 24 | 42 | 42 | 57 | 39 | 51 | 34 | 39 | 43 | 48 | 34 | 46 | 50 | 47 |
| CL199 | 35 | 39 | 36 | 36 | 40 | 35 | 43 | 32 | 47 | 38 | 44 | 50 | 45 | 49 | 44 | 28 | 37 | 37 | 38 |
| CL200 | 33 | 37 | 45 | 50 | 44 | 27 | 36 | 42 | 36 | 40 | 58 | 43 | 44 | 42 | 38 | 32 | 24 | 36 | 34 |
| CL202 | 36 | 30 | 29 | 29 | 20 | 29 | 31 | 33 | 30 | 35 | 16 | 32 | 41 | 17 | 22 | 34 | 27 | 34 | 23 |
| CL203 | 21 | 24 | 34 | 30 | 38 | 31 | 36 | 49 | 14 | 29 | 14 | 30 | 23 | 38 | 24 | 18 | 21 | 27 | 20 |
| CL204 | 28 | 25 | 32 | 39 | 43 | 14 | 31 | 37 | 23 | 23 | 33 | 36 | 28 | 20 | 20 | 31 | 26 | 28 | 37 |
| CL205 | 24 | 31 | 29 | 36 | 37 | 19 | 23 | 34 | 33 | 24 | 25 | 25 | 30 | 34 | 43 | 25 | 17 | 27 | 29 |
| CL206 | 29 | 25 | 31 | 36 | 22 | 24 | 24 | 20 | 20 | 25 | 35 | 32 | 33 | 29 | 29 | 25 | 28 | 32 | 27 |
| CL209 | 26 | 29 | 24 | 20 | 23 | 23 | 21 | 32 | 17 | 23 | 47 | 29 | 22 | 28 | 23 | 25 | 15 | 24 | 25 |
| CL211 | 21 | 14 | 17 | 20 | 22 | 23 | 21 | 26 | 15 | 27 | 28 | 28 | 27 | 29 | 24 | 20 | 22 | 20 | 28 |
| CL212 | 17 | 25 | 27 | 23 | 10 | 27 | 21 | 24 | 19 | 17 | 8 | 23 | 27 | 25 | 26 | 27 | 25 | 28 | 23 |
| CL213 | 27 | 19 | 30 | 36 | 30 | 19 | 28 | 32 | 12 | 12 | 23 | 25 | 26 | 18 | 23 | 19 | 13 | 16 | 20 |
| CL214 | 20 | 13 | 41 | 29 | 25 | 19 | 19 | 23 | 14 | 21 | 51 | 29 | 21 | 15 | 20 | 17 | 17 | 13 | 23 |
| CL215 | 17 | 37 | 25 | 21 | 18 | 25 | 31 | 26 | 22 | 21 | 17 | 21 | 24 | 23 | 25 | 24 | 24 | 18 | 26 |
| CL216 | 21 | 17 | 21 | 10 | 29 | 30 | 16 | 24 | 12 | 19 | 21 | 19 | 33 | 21 | 26 | 20 | 14 | 11 | 25 |
| CL218 | 24 | 18 | 16 | 15 | 22 | 14 | 17 | 20 | 13 | 22 | 8 | 18 | 19 | 24 | 19 | 17 | 17 | 18 | 19 |
| CL220 | 18 | 17 | 26 | 14 | 24 | 18 | 13 | 27 | 13 | 21 | 16 | 13 | 17 | 18 | 19 | 14 | 17 | 25 | 13 |
| CL221 | 15 | 33 | 25 | 23 | 16 | 15 | 16 | 18 | 18 | 18 | 23 | 30 | 15 | 19 | 25 | 20 | 15 | 20 | 11 |
| CL222 | 17 | 15 | 17 | 9 | 15 | 14 | 14 | 17 | 14 | 15 | 19 | 14 | 22 | 22 | 20 | 17 | 12 | 20 | 16 |
| CL223 | 20 | 15 | 26 | 14 | 21 | 9 | 9 | 17 | 17 | 11 | 26 | 16 | 11 | 20 | 19 | 22 | 24 | 23 | 29 |
| CL226 | 20 | 12 | 16 | 18 | 11 | 12 | 13 | 17 | 24 | 15 | 14 | 19 | 22 | 13 | 16 | 15 | 14 | 23 | 17 |
| CL228 | 20 | 21 | 19 | 19 | 24 | 17 | 23 | 15 | 21 | 24 | 13 | 16 | 22 | 12 | 17 | 22 | 18 | 17 | 18 |
| CL229 | 14 | 11 | 18 | 17 | 20 | 21 | 15 | 15 | 13 | 13 | 13 | 22 | 23 | 21 | 8 | 6 | 11 | 9 | 17 |
| CL231 | 17 | 22 | 13 | 18 | 16 | 10 | 9 | 13 | 8 | 8 | 35 | 16 | 12 | 11 | 9 | 13 | 24 | 14 | 12 |
| CL232 | 15 | 11 | 12 | 11 | 14 | 11 | 18 | 15 | 8 | 10 | 13 | 10 | 19 | 15 | 11 | 17 | 10 | 10 | 9 |
| CL234 | 9 | 6 | 17 | 6 | 20 | 6 | 10 | 13 | 14 | 13 | 9 | 13 | 7 | 11 | 12 | 11 | 10 | 16 | 7 |
| CL235 | 15 | 8 | 11 | 17 | 8 | 6 | 11 | 7 | 13 | 11 | 14 | 15 | 12 | 11 | 4 | 10 | 6 | 9 | 16 |
| CL236 | 14 | 7 | 14 | 6 | 12 | 7 | 8 | 16 | 6 | 9 | 8 | 13 | 15 | 5 | 8 | 6 | 5 | 7 | 10 |
| CL237 | 11 | 11 | 6 | 10 | 10 | 9 | 7 | 10 | 11 | 13 | 3 | 7 | 5 | 10 | 13 | 15 | 14 | 13 | 13 |
| CL239 | 10 | 5 | 12 | 10 | 12 | 11 | 8 | 16 | 8 | 12 | 8 | 7 | 11 | 9 | 10 | 11 | 9 | 14 | 9 |

| | A2_142 | A2_147 | A2_154 | A2_156 | A2_158 | A2_164 | A2_255 | A2_lin | A2_NC3 | A2_G42 | A2_G43 | A2_G40 | A2_G41 | A2_G38 | A2_G39 | A2_G36 | A2_G37 | A2_G56 | A2_G57 |
|---|---|---|---|---|---|---|---|---|---|---|---|---|---|---|---|---|---|---|---|
| CL002 | 4610 | 5158 | 4929 | 5070 | 4795 | 5061 | 4872 | 4469 | 4918 | 4502 | 4753 | 4951 | 4847 | 4709 | 4732 | 4797 | 4577 | 4750 | 4540 |
| CL003 | 2882 | 3102 | 3049 | 3073 | 2933 | 2896 | 3114 | 2757 | 2826 | 2881 | 2987 | 3133 | 3005 | 3078 | 3016 | 3104 | 3025 | 3065 | 2914 |
| CL004 | 2695 | 2933 | 2966 | 2793 | 2792 | 2756 | 2878 | 2555 | 2741 | 3062 | 2979 | 3099 | 3231 | 2903 | 3230 | 3044 | 3137 | 3113 | 3014 |
| CL005 | 2446 | 2582 | 2591 | 2674 | 2536 | 2552 | 2613 | 2332 | 2758 | 2533 | 2549 | 2445 | 2495 | 2604 | 2648 | 2574 | 2532 | 2603 | 2483 |
| CL008 | 2212 | 2371 | 2251 | 2195 | 2171 | 2196 | 2462 | 2113 | 2272 | 2470 | 2489 | 2637 | 2714 | 2485 | 2502 | 2476 | 2480 | 2512 | 2422 |
| CL009 | 2000 | 2187 | 2302 | 2163 | 2152 | 2152 | 2459 | 2058 | 2210 | 2332 | 2339 | 2442 | 2467 | 2318 | 2387 | 2344 | 2305 | 2337 | 2276 |
| CL010 | 2080 | 2370 | 2291 | 2280 | 2174 | 2268 | 1971 | 2003 | 2095 | 2071 | 2105 | 2169 | 2286 | 2129 | 2120 | 2208 | 2198 | 2183 | 2059 |
| CL011 | 2272 | 2347 | 2271 | 2299 | 2271 | 2199 | 3205 | 1964 | 2168 | 2060 | 2075 | 2073 | 2059 | 2070 | 2133 | 2064 | 2077 | 2016 | 1954 |
| CL012 | 1893 | 2043 | 1940 | 2094 | 2027 | 1997 | 1954 | 1839 | 2135 | 1984 | 2112 | 2040 | 2138 | 2157 | 2222 | 2212 | 2110 | 2164 | 2039 |
| CL013 | 1936 | 2094 | 2050 | 1952 | 1942 | 1968 | 2171 | 1833 | 2002 | 2116 | 2212 | 2335 | 2302 | 2265 | 2189 | 2331 | 2226 | 2216 | 2052 |
| CL014 | 1910 | 2187 | 2115 | 2074 | 2093 | 2142 | 2489 | 1935 | 2164 | 1970 | 1982 | 2037 | 2088 | 2000 | 2035 | 2007 | 1957 | 2024 | 1985 |
| CL015 | 1943 | 2222 | 2136 | 1887 | 2076 | 1990 | 2176 | 1774 | 1902 | 2000 | 2094 | 2158 | 2192 | 1977 | 2058 | 2144 | 2160 | 2095 | 2059 |
| CL016 | 1695 | 1545 | 1620 | 1643 | 1884 | 1687 | 1543 | 1697 | 1983 | 1766 | 1548 | 1526 | 1428 | 1807 | 1694 | 1784 | 1707 | 1749 | 1636 |
| CL017 | 1814 | 1972 | 2047 | 1901 | 1885 | 1843 | 1794 | 1773 | 1674 | 1704 | 1768 | 1831 | 1862 | 1814 | 1831 | 1828 | 1857 | 1950 | 1694 |
| CL020 | 1456 | 1570 | 1638 | 1560 | 1618 | 1565 | 1487 | 1525 | 1618 | 1646 | 1621 | 1709 | 1637 | 1743 | 1619 | 1702 | 1645 | 1632 | 1576 |
| CL021 | 1527 | 1675 | 1567 | 1630 | 1555 | 1532 | 1622 | 1455 | 1581 | 1601 | 1701 | 1666 | 1655 | 1626 | 1718 | 1663 | 1632 | 1646 | 1558 |
| CL023 | 1513 | 1582 | 1620 | 1596 | 1498 | 1578 | 1717 | 1448 | 1567 | 1511 | 1486 | 1514 | 1538 | 1542 | 1556 | 1531 | 1505 | 1582 | 1442 |
| CL024 | 1447 | 1596 | 1544 | 1496 | 1501 | 1505 | 1609 | 1420 | 1424 | 1593 | 1543 | 1609 | 1552 | 1506 | 1569 | 1611 | 1567 | 1599 | 1535 |
| CL025 | 1359 | 1350 | 1429 | 1425 | 1400 | 1473 | 1608 | 1331 | 1587 | 1446 | 1408 | 1427 | 1422 | 1417 | 1530 | 1527 | 1450 | 1434 | 1455 |
| CL026 | 1337 | 1357 | 1508 | 1370 | 1432 | 1453 | 1364 | 1328 | 1539 | 1593 | 1433 | 1430 | 1377 | 1465 | 1474 | 1537 | 1439 | 1420 | 1474 |
| CL027 | 1411 | 1470 | 1438 | 1440 | 1445 | 1388 | 1622 | 1386 | 1487 | 1462 | 1581 | 1529 | 1601 | 1505 | 1470 | 1497 | 1421 | 1456 | 1407 |
| CL030 | 1344 | 1377 | 1414 | 1421 | 1313 | 1347 | 1415 | 1271 | 1388 | 1350 | 1321 | 1357 | 1326 | 1383 | 1308 | 1373 | 1325 | 1299 | 1261 |
| CL032 | 1296 | 1353 | 1349 | 1335 | 1324 | 1298 | 1221 | 1251 | 1192 | 1247 | 1317 | 1291 | 1278 | 1298 | 1348 | 1315 | 1296 | 1297 | 1298 |
| CL033 | 1118 | 1180 | 1159 | 1177 | 1217 | 1208 | 1083 | 1094 | 1205 | 1178 | 1198 | 1212 | 1207 | 1168 | 1137 | 1268 | 1175 | 1211 | 1177 |
| CL034 | 1161 | 1354 | 1278 | 1311 | 1196 | 1261 | 1391 | 1075 | 1196 | 1174 | 1158 | 1254 | 1237 | 1175 | 1179 | 1137 | 1195 | 1148 | 1142 |
| CL037 | 1111 | 1104 | 1153 | 1054 | 1145 | 1111 | 1323 | 1044 | 1165 | 1063 | 1110 | 1152 | 1115 | 1117 | 1117 | 1201 | 1098 | 1081 | 1090 |
| CL038 | 1044 | 1085 | 1088 | 1010 | 1048 | 1083 | 1235 | 1007 | 1010 | 1104 | 1133 | 1166 | 1144 | 1166 | 1163 | 1166 | 1144 | 1163 | 1064 |
| CL040 | 998 | 1018 | 1043 | 1021 | 1023 | 983 | 1178 | 972 | 1036 | 1033 | 1100 | 1123 | 1107 | 1138 | 1144 | 1105 | 1082 | 1103 | 1095 |
| CL046 | 953 | 1097 | 1085 | 1050 | 1003 | 1016 | 1002 | 911 | 900 | 1074 | 1037 | 1133 | 1102 | 1008 | 1100 | 1085 | 1061 | 1078 | 1061 |
| CL047 | 992 | 1084 | 1047 | 1057 | 1046 | 1071 | 1068 | 977 | 1033 | 976 | 912 | 853 | 936 | 976 | 941 | 960 | 901 | 931 | 885 |
| CL050 | 827 | 876 | 960 | 936 | 832 | 870 | 857 | 835 | 790 | 803 | 751 | 863 | 846 | 819 | 813 | 781 | 862 | 806 | 802 |
| CL052 | 772 | 833 | 841 | 849 | 795 | 808 | 796 | 733 | 826 | 755 | 786 | 767 | 718 | 772 | 761 | 761 | 773 | 729 | 721 |
| CL053 | 727 | 743 | 795 | 744 | 774 | 764 | 797 | 739 | 858 | 735 | 749 | 786 | 759 | 784 | 759 | 863 | 841 | 777 | 788 |
| CL057 | 704 | 787 | 754 | 764 | 712 | 735 | 800 | 720 | 693 | 744 | 777 | 798 | 843 | 781 | 783 | 794 | 782 | 776 | 718 |
| CL058 | 698 | 786 | 784 | 772 | 827 | 742 | 858 | 640 | 753 | 695 | 720 | 769 | 744 | 732 | 739 | 702 | 673 | 697 | 666 |
| CL065 | 558 | 679 | 583 | 639 | 584 | 559 | 599 | 525 | 559 | 604 | 577 | 657 | 631 | 564 | 632 | 594 | 637 | 602 | 591 |
| CL066 | 589 | 595 | 612 | 610 | 562 | 600 | 705 | 556 | 638 | 639 | 627 | 603 | 641 | 596 | 626 | 605 | 673 | 643 | 585 |
| CL067 | 503 | 644 | 611 | 582 | 633 | 595 | 707 | 598 | 606 | 633 | 629 | 586 | 658 | 659 | 652 | 655 | 606 | 621 | 591 |
| CL072 | 523 | 610 | 581 | 538 | 565 | 560 | 627 | 587 | 555 | 552 | 564 | 628 | 569 | 579 | 540 | 622 | 568 | 610 | 611 |
| CL081 | 470 | 504 | 510 | 460 | 517 | 466 | 649 | 468 | 521 | 569 | 508 | 538 | 577 | 534 | 534 | 517 | 537 | 552 | 542 |
| CL093 | 469 | 412 | 396 | 411 | 475 | 466 | 292 | 512 | 514 | 367 | 278 | 361 | 318 | 427 | 322 | 371 | 367 | 343 | 315 |
| CL099 | 411 | 426 | 383 | 379 | 411 | 406 | 405 | 373 | 407 | 383 | 433 | 430 | 359 | 371 | 399 | 390 | 394 | 382 | 384 |

| | A2_142 | A2_147 | A2_154 | A2_156 | A2_158 | A2_164 | A2_255 | A2_lin | A2_NC3 | A2_G42 | A2_G43 | A2_G40 | A2_G41 | A2_G38 | A2_G39 | A2_G36 | A2_G37 | A2_G56 | A2_G57 |
|---|---|---|---|---|---|---|---|---|---|---|---|---|---|---|---|---|---|---|---|
| CL100 | 407 | 419 | 346 | 338 | 361 | 336 | 444 | 389 | 399 | 431 | 482 | 384 | 441 | 340 | 397 | 452 | 432 | 410 | 475 |
| CL101 | 368 | 413 | 382 | 398 | 360 | 379 | 373 | 394 | 388 | 396 | 420 | 392 | 394 | 418 | 378 | 413 | 403 | 370 | 378 |
| CL113 | 322 | 264 | 315 | 348 | 343 | 393 | 217 | 322 | 405 | 299 | 293 | 293 | 249 | 326 | 290 | 346 | 310 | 311 | 278 |
| CL118 | 285 | 302 | 297 | 314 | 288 | 347 | 351 | 306 | 309 | 323 | 260 | 282 | 275 | 308 | 301 | 326 | 300 | 329 | 271 |
| CL128 | 293 | 293 | 311 | 266 | 287 | 302 | 357 | 298 | 331 | 316 | 315 | 309 | 322 | 311 | 344 | 329 | 300 | 298 | 279 |
| CL133 | 233 | 211 | 212 | 232 | 257 | 274 | 257 | 202 | 306 | 281 | 240 | 276 | 231 | 262 | 288 | 264 | 274 | 277 | 255 |
| CL137 | 249 | 169 | 235 | 236 | 253 | 248 | 216 | 234 | 300 | 266 | 258 | 225 | 217 | 249 | 244 | 260 | 216 | 241 | 260 |
| CL146 | 208 | 220 | 219 | 216 | 217 | 241 | 226 | 189 | 246 | 226 | 205 | 230 | 213 | 218 | 240 | 245 | 222 | 229 | 221 |
| CL148 | 226 | 235 | 207 | 227 | 217 | 232 | 222 | 205 | 211 | 216 | 230 | 237 | 195 | 227 | 209 | 217 | 236 | 220 | 240 |
| CL149 | 178 | 185 | 185 | 191 | 194 | 225 | 104 | 166 | 234 | 210 | 192 | 188 | 189 | 198 | 259 | 236 | 187 | 226 | 209 |
| CL152 | 163 | 202 | 188 | 166 | 239 | 223 | 217 | 168 | 178 | 204 | 209 | 225 | 222 | 209 | 194 | 209 | 199 | 204 | 190 |
| CL155 | 159 | 170 | 152 | 153 | 175 | 185 | 133 | 160 | 197 | 185 | 182 | 171 | 149 | 192 | 184 | 185 | 167 | 193 | 181 |
| CL168 | 123 | 117 | 121 | 123 | 143 | 127 | 124 | 132 | 163 | 154 | 138 | 140 | 113 | 145 | 124 | 116 | 132 | 115 | 135 |
| CL170 | 109 | 151 | 147 | 125 | 132 | 147 | 116 | 134 | 126 | 130 | 118 | 143 | 114 | 147 | 132 | 147 | 136 | 156 | 111 |
| CL174 | 101 | 98 | 100 | 93 | 126 | 100 | 88 | 112 | 114 | 119 | 121 | 85 | 98 | 126 | 109 | 134 | 112 | 127 | 120 |
| CL176 | 92 | 98 | 105 | 98 | 98 | 101 | 99 | 89 | 97 | 98 | 96 | 108 | 85 | 117 | 98 | 98 | 98 | 123 | 94 |
| CL184 | 72 | 78 | 74 | 85 | 88 | 80 | 62 | 79 | 94 | 71 | 79 | 66 | 61 | 81 | 72 | 88 | 74 | 91 | 62 |
| CL196 | 39 | 51 | 41 | 38 | 41 | 38 | 59 | 41 | 51 | 47 | 51 | 47 | 47 | 41 | 36 | 57 | 45 | 45 | 45 |
| CL198 | 47 | 58 | 47 | 39 | 39 | 46 | 29 | 50 | 51 | 39 | 38 | 30 | 55 | 59 | 55 | 58 | 42 | 40 | 57 |
| CL210 | 32 | 36 | 30 | 33 | 34 | 30 | 46 | 32 | 32 | 29 | 32 | 30 | 36 | 31 | 39 | 41 | 36 | 33 | 26 |
| CL219 | 14 | 22 | 10 | 18 | 14 | 15 | 33 | 23 | 26 | 18 | 24 | 22 | 11 | 29 | 16 | 32 | 27 | 19 | 28 |
| CL238 | 6 | 7 | 7 | 10 | 2 | 5 | 13 | 9 | 11 | 15 | 9 | 13 | 11 | 15 | 14 | 12 | 16 | 14 | 6 |
| CL001 | 4738 | 5249 | 5153 | 4974 | 4871 | 5038 | 4818 | 4675 | 4687 | 4732 | 4774 | 4766 | 4948 | 4942 | 4820 | 4914 | 4692 | 4713 | 4772 |
| CL006 | 2487 | 2508 | 2628 | 2530 | 2422 | 2550 | 2536 | 2311 | 2637 | 2389 | 2394 | 2339 | 2383 | 2447 | 2447 | 2474 | 2377 | 2415 | 2455 |
| CL007 | 2119 | 2213 | 2330 | 2371 | 2300 | 2314 | 2197 | 1984 | 2474 | 2334 | 2255 | 2306 | 2283 | 2362 | 2328 | 2414 | 2355 | 2407 | 2274 |
| CL018 | 1614 | 1645 | 1811 | 1748 | 1724 | 1719 | 1411 | 1542 | 1761 | 1707 | 1706 | 1667 | 1582 | 1772 | 1750 | 1769 | 1705 | 1729 | 1731 |
| CL019 | 1547 | 1780 | 1649 | 1573 | 1676 | 1615 | 2033 | 1646 | 1757 | 1794 | 1749 | 1778 | 1787 | 1830 | 1841 | 1801 | 1785 | 1764 | 1666 |
| CL022 | 1496 | 1603 | 1598 | 1490 | 1435 | 1546 | 1754 | 1397 | 1574 | 1614 | 1530 | 1613 | 1624 | 1606 | 1580 | 1615 | 1673 | 1574 | 1575 |
| CL028 | 1337 | 1430 | 1568 | 1475 | 1379 | 1529 | 1509 | 1304 | 1472 | 1391 | 1364 | 1284 | 1391 | 1352 | 1380 | 1447 | 1334 | 1382 | 1306 |
| CL029 | 1250 | 1367 | 1336 | 1393 | 1317 | 1398 | 1353 | 1148 | 1477 | 1256 | 1293 | 1357 | 1298 | 1357 | 1394 | 1357 | 1348 | 1320 | 1326 |
| CL031 | 1232 | 1234 | 1155 | 1113 | 1277 | 1274 | 1453 | 1248 | 1302 | 1243 | 1250 | 1309 | 1318 | 1255 | 1322 | 1355 | 1293 | 1365 | 1262 |
| CL035 | 1080 | 1142 | 1220 | 1164 | 1118 | 1184 | 1206 | 1010 | 1247 | 1081 | 1128 | 1151 | 1111 | 1135 | 1172 | 1151 | 1155 | 1179 | 1124 |
| CL036 | 1054 | 1020 | 1147 | 1081 | 1133 | 1071 | 1149 | 1010 | 1195 | 1056 | 1070 | 1079 | 1059 | 1125 | 1173 | 1167 | 1097 | 1104 | 1091 |
| CL039 | 1043 | 1125 | 1069 | 1061 | 1051 | 1062 | 1061 | 1022 | 1145 | 1060 | 1103 | 1025 | 1033 | 1092 | 1105 | 1107 | 1052 | 1108 | 1053 |
| CL041 | 980 | 944 | 975 | 992 | 1037 | 1032 | 914 | 929 | 1105 | 994 | 1017 | 1046 | 992 | 1027 | 1128 | 1094 | 1032 | 1018 | 1038 |
| CL042 | 912 | 987 | 975 | 990 | 1014 | 1015 | 957 | 954 | 948 | 1030 | 1018 | 1059 | 1073 | 990 | 1041 | 1036 | 1028 | 994 | 974 |
| CL043 | 993 | 1072 | 985 | 1026 | 1027 | 1003 | 1105 | 978 | 1000 | 992 | 1005 | 1068 | 1104 | 1031 | 1037 | 1096 | 1048 | 1000 | 1023 |
| CL044 | 904 | 969 | 1000 | 947 | 950 | 949 | 996 | 930 | 950 | 985 | 919 | 983 | 961 | 1024 | 1012 | 1005 | 973 | 1022 | 932 |
| CL045 | 952 | 961 | 1019 | 968 | 928 | 1000 | 1088 | 960 | 1059 | 913 | 932 | 899 | 971 | 971 | 971 | 994 | 941 | 961 | 958 |
| CL048 | 917 | 975 | 967 | 915 | 975 | 925 | 1065 | 876 | 944 | 860 | 896 | 820 | 834 | 901 | 938 | 894 | 927 | 873 | 881 |
| CL049 | 833 | 957 | 944 | 964 | 921 | 900 | 900 | 843 | 988 | 937 | 902 | 877 | 917 | 937 | 1002 | 970 | 886 | 876 | 859 |
| CL051 | 850 | 880 | 919 | 845 | 893 | 835 | 935 | 802 | 906 | 788 | 852 | 787 | 830 | 847 | 868 | 863 | 843 | 879 | 807 |
| CL055 | 667 | 689 | 761 | 712 | 739 | 704 | 670 | 690 | 694 | 713 | 719 | 674 | 660 | 702 | 767 | 730 | 754 | 694 | 683 |
| CL056 | 697 | 682 | 704 | 720 | 748 | 730 | 769 | 687 | 816 | 783 | 747 | 736 | 712 | 735 | 731 | 782 | 705 | 713 | 738 |

| | A2_142 | A2_147 | A2_154 | A2_156 | A2_158 | A2_164 | A2_255 | A2_lin | A2_NC3 | A2_G42 | A2_G43 | A2_G40 | A2_G41 | A2_G38 | A2_G39 | A2_G36 | A2_G37 | A2_G56 | A2_G57 |
|---|---|---|---|---|---|---|---|---|---|---|---|---|---|---|---|---|---|---|---|
| CL059 | 737 | 750 | 716 | 669 | 680 | 677 | 792 | 684 | 735 | 672 | 716 | 679 | 679 | 689 | 746 | 744 | 705 | 694 | 651 |
| CL060 | 665 | 680 | 688 | 677 | 685 | 757 | 660 | 646 | 694 | 656 | 673 | 670 | 623 | 682 | 644 | 718 | 699 | 677 | 633 |
| CL061 | 696 | 639 | 697 | 623 | 695 | 720 | 685 | 620 | 783 | 735 | 667 | 748 | 666 | 744 | 743 | 748 | 709 | 720 | 728 |
| CL062 | 672 | 676 | 695 | 752 | 647 | 714 | 666 | 606 | 683 | 633 | 635 | 647 | 629 | 662 | 690 | 676 | 659 | 671 | 686 |
| CL063 | 669 | 545 | 580 | 619 | 710 | 686 | 560 | 597 | 712 | 755 | 621 | 713 | 565 | 722 | 755 | 784 | 691 | 745 | 686 |
| CL064 | 631 | 685 | 605 | 646 | 611 | 633 | 603 | 579 | 553 | 585 | 658 | 657 | 649 | 624 | 579 | 623 | 655 | 612 | 619 |
| CL068 | 602 | 607 | 564 | 577 | 584 | 599 | 650 | 532 | 596 | 602 | 624 | 642 | 597 | 558 | 596 | 582 | 598 | 629 | 602 |
| CL070 | 569 | 564 | 580 | 549 | 525 | 579 | 575 | 564 | 592 | 522 | 575 | 517 | 514 | 563 | 594 | 627 | 561 | 564 | 541 |
| CL071 | 566 | 513 | 513 | 554 | 609 | 529 | 587 | 512 | 580 | 520 | 535 | 581 | 511 | 560 | 553 | 546 | 544 | 561 | 522 |
| CL074 | 450 | 549 | 536 | 505 | 513 | 542 | 547 | 477 | 538 | 548 | 538 | 531 | 515 | 535 | 551 | 553 | 521 | 575 | 497 |
| CL075 | 518 | 561 | 559 | 526 | 525 | 563 | 504 | 497 | 546 | 516 | 558 | 606 | 579 | 536 | 551 | 567 | 579 | 534 | 511 |
| CL076 | 511 | 457 | 476 | 462 | 514 | 512 | 518 | 452 | 623 | 510 | 574 | 524 | 488 | 550 | 552 | 579 | 541 | 570 | 543 |
| CL078 | 502 | 520 | 516 | 477 | 493 | 484 | 578 | 509 | 551 | 545 | 516 | 524 | 515 | 542 | 561 | 540 | 568 | 545 | 498 |
| CL079 | 441 | 463 | 567 | 507 | 495 | 544 | 549 | 494 | 518 | 552 | 475 | 526 | 504 | 507 | 540 | 559 | 507 | 515 | 502 |
| CL080 | 507 | 488 | 475 | 459 | 498 | 462 | 576 | 478 | 548 | 489 | 491 | 493 | 478 | 495 | 506 | 533 | 524 | 495 | 492 |
| CL082 | 458 | 494 | 534 | 522 | 476 | 511 | 444 | 492 | 393 | 453 | 436 | 438 | 489 | 492 | 474 | 535 | 491 | 504 | 467 |
| CL084 | 450 | 502 | 547 | 489 | 466 | 469 | 418 | 479 | 481 | 448 | 462 | 416 | 456 | 502 | 520 | 501 | 463 | 474 | 503 |
| CL085 | 481 | 513 | 499 | 445 | 471 | 463 | 525 | 453 | 500 | 479 | 462 | 490 | 499 | 539 | 456 | 453 | 471 | 477 | 462 |
| CL091 | 407 | 417 | 454 | 423 | 413 | 464 | 501 | 442 | 446 | 476 | 455 | 496 | 493 | 438 | 476 | 452 | 470 | 448 | 434 |
| CL092 | 415 | 398 | 461 | 444 | 438 | 440 | 489 | 380 | 468 | 479 | 443 | 455 | 450 | 429 | 474 | 463 | 464 | 505 | 455 |
| CL094 | 377 | 399 | 424 | 397 | 426 | 380 | 359 | 393 | 471 | 456 | 410 | 452 | 394 | 473 | 489 | 437 | 440 | 437 | 447 |
| CL095 | 372 | 532 | 340 | 402 | 263 | 283 | 360 | 569 | 233 | 472 | 655 | 310 | 541 | 378 | 359 | 409 | 471 | 412 | 394 |
| CL102 | 353 | 348 | 419 | 331 | 364 | 387 | 349 | 363 | 397 | 431 | 387 | 378 | 421 | 419 | 422 | 421 | 447 | 395 | 402 |
| CL104 | 316 | 297 | 319 | 334 | 375 | 362 | 282 | 334 | 441 | 374 | 320 | 342 | 286 | 351 | 413 | 380 | 370 | 402 | 394 |
| CL106 | 360 | 308 | 354 | 355 | 393 | 344 | 380 | 309 | 420 | 330 | 362 | 323 | 313 | 378 | 344 | 370 | 346 | 330 | 332 |
| CL107 | 357 | 373 | 344 | 348 | 367 | 375 | 326 | 303 | 418 | 343 | 324 | 324 | 314 | 354 | 334 | 344 | 329 | 329 | 337 |
| CL109 | 322 | 323 | 336 | 312 | 365 | 331 | 391 | 328 | 380 | 381 | 366 | 385 | 360 | 372 | 333 | 366 | 391 | 334 | 343 |
| CL112 | 314 | 317 | 306 | 306 | 309 | 319 | 346 | 329 | 359 | 337 | 317 | 348 | 336 | 323 | 339 | 351 | 302 | 346 | 339 |
| CL117 | 329 | 296 | 297 | 304 | 291 | 303 | 357 | 289 | 330 | 351 | 302 | 337 | 347 | 332 | 329 | 359 | 316 | 325 | 304 |
| CL119 | 306 | 295 | 334 | 264 | 301 | 293 | 331 | 301 | 297 | 312 | 291 | 283 | 296 | 304 | 295 | 300 | 295 | 276 | 295 |
| CL120 | 302 | 311 | 344 | 316 | 286 | 328 | 344 | 299 | 311 | 272 | 288 | 319 | 320 | 261 | 317 | 341 | 307 | 309 | 315 |
| CL121 | 277 | 286 | 268 | 298 | 290 | 300 | 357 | 262 | 286 | 319 | 344 | 348 | 323 | 290 | 308 | 318 | 346 | 320 | 360 |
| CL122 | 274 | 242 | 261 | 269 | 320 | 312 | 237 | 265 | 342 | 339 | 278 | 341 | 292 | 352 | 348 | 316 | 340 | 356 | 345 |
| CL129 | 298 | 288 | 286 | 253 | 285 | 277 | 294 | 257 | 300 | 283 | 271 | 292 | 308 | 307 | 339 | 339 | 306 | 340 | 315 |
| CL130 | 264 | 231 | 272 | 222 | 257 | 244 | 277 | 259 | 275 | 266 | 232 | 266 | 322 | 287 | 265 | 314 | 338 | 233 | 284 |
| CL131 | 232 | 239 | 267 | 252 | 257 | 260 | 247 | 241 | 312 | 260 | 251 | 284 | 246 | 278 | 274 | 298 | 270 | 273 | 245 |
| CL139 | 236 | 234 | 276 | 233 | 270 | 270 | 257 | 243 | 278 | 228 | 250 | 252 | 246 | 251 | 251 | 264 | 261 | 263 | 246 |
| CL143 | 209 | 238 | 234 | 229 | 235 | 223 | 234 | 215 | 242 | 241 | 222 | 226 | 242 | 254 | 199 | 201 | 222 | 241 | 248 |
| CL147 | 191 | 219 | 200 | 176 | 239 | 198 | 232 | 183 | 230 | 209 | 236 | 211 | 224 | 213 | 227 | 208 | 196 | 211 | 209 |
| CL151 | 173 | 173 | 174 | 190 | 217 | 206 | 232 | 206 | 215 | 232 | 202 | 197 | 218 | 228 | 222 | 247 | 208 | 241 | 200 |
| CL157 | 156 | 135 | 144 | 168 | 180 | 183 | 137 | 168 | 175 | 196 | 189 | 182 | 167 | 188 | 195 | 189 | 155 | 176 | 195 |
| CL158 | 171 | 168 | 176 | 185 | 158 | 168 | 191 | 156 | 149 | 175 | 167 | 171 | 167 | 163 | 163 | 171 | 157 | 178 | 193 |
| CL159 | 156 | 163 | 161 | 195 | 157 | 155 | 144 | 167 | 182 | 160 | 155 | 116 | 139 | 160 | 163 | 187 | 142 | 158 | 154 |
| CL160 | 156 | 175 | 144 | 156 | 154 | 176 | 174 | 183 | 201 | 144 | 157 | 203 | 162 | 178 | 187 | 171 | 181 | 163 | 172 |

| | A2_142 | A2_147 | A2_154 | A2_156 | A2_158 | A2_164 | A2_255 | A2_lin | A2_NC3 | A2_G42 | A2_G43 | A2_G40 | A2_G41 | A2_G38 | A2_G39 | A2_G36 | A2_G37 | A2_G56 | A2_G57 |
|---|---|---|---|---|---|---|---|---|---|---|---|---|---|---|---|---|---|---|---|
| CL162 | 152 | 161 | 180 | 142 | 153 | 164 | 191 | 164 | 166 | 165 | 168 | 168 | 164 | 163 | 153 | 155 | 157 | 185 | 164 |
| CL163 | 148 | 151 | 122 | 140 | 152 | 175 | 211 | 153 | 179 | 142 | 137 | 158 | 141 | 181 | 204 | 142 | 158 | 149 | 166 |
| CL164 | 167 | 160 | 162 | 153 | 148 | 150 | 139 | 135 | 158 | 152 | 153 | 141 | 128 | 142 | 156 | 150 | 155 | 153 | 145 |
| CL166 | 142 | 120 | 121 | 141 | 138 | 151 | 105 | 142 | 167 | 156 | 157 | 129 | 132 | 160 | 156 | 159 | 148 | 164 | 146 |
| CL171 | 121 | 136 | 133 | 114 | 124 | 114 | 144 | 116 | 131 | 138 | 107 | 132 | 143 | 123 | 118 | 123 | 135 | 128 | 118 |
| CL173 | 107 | 109 | 112 | 124 | 94 | 121 | 117 | 99 | 145 | 101 | 97 | 92 | 104 | 133 | 135 | 128 | 109 | 141 | 118 |
| CL175 | 108 | 91 | 119 | 116 | 105 | 104 | 130 | 92 | 103 | 128 | 105 | 118 | 103 | 121 | 106 | 116 | 127 | 118 | 100 |
| CL177 | 116 | 96 | 79 | 97 | 103 | 96 | 69 | 92 | 91 | 96 | 98 | 119 | 118 | 110 | 114 | 116 | 106 | 103 | 104 |
| CL182 | 90 | 58 | 79 | 71 | 75 | 88 | 90 | 58 | 86 | 75 | 81 | 64 | 87 | 97 | 95 | 85 | 85 | 89 | 75 |
| CL186 | 79 | 75 | 72 | 70 | 72 | 53 | 83 | 74 | 79 | 89 | 67 | 83 | 65 | 78 | 85 | 93 | 78 | 83 | 68 |
| CL187 | 74 | 62 | 67 | 72 | 73 | 79 | 93 | 67 | 63 | 83 | 66 | 74 | 66 | 87 | 85 | 81 | 73 | 84 | 65 |
| CL189 | 59 | 59 | 59 | 67 | 67 | 66 | 57 | 63 | 53 | 57 | 65 | 76 | 58 | 58 | 76 | 54 | 72 | 70 | 70 |
| CL191 | 40 | 50 | 75 | 45 | 51 | 51 | 64 | 59 | 67 | 54 | 52 | 71 | 64 | 64 | 56 | 57 | 51 | 73 | 69 |
| CL194 | 54 | 50 | 41 | 39 | 51 | 44 | 58 | 48 | 52 | 52 | 59 | 41 | 56 | 62 | 60 | 59 | 48 | 47 | 40 |
| CL195 | 44 | 24 | 35 | 41 | 42 | 42 | 49 | 36 | 65 | 45 | 45 | 36 | 34 | 42 | 44 | 56 | 44 | 51 | 39 |
| CL197 | 36 | 51 | 40 | 35 | 37 | 47 | 39 | 46 | 36 | 56 | 46 | 37 | 52 | 38 | 36 | 53 | 43 | 45 | 34 |
| CL199 | 29 | 47 | 42 | 36 | 40 | 38 | 48 | 47 | 49 | 59 | 43 | 52 | 44 | 44 | 50 | 35 | 49 | 42 | 47 |
| CL200 | 38 | 27 | 43 | 34 | 51 | 32 | 48 | 37 | 40 | 41 | 23 | 29 | 43 | 41 | 34 | 33 | 41 | 40 | 36 |
| CL202 | 29 | 29 | 37 | 39 | 25 | 32 | 48 | 28 | 38 | 41 | 34 | 34 | 37 | 39 | 41 | 36 | 37 | 26 | 31 |
| CL203 | 24 | 22 | 28 | 29 | 44 | 48 | 29 | 21 | 29 | 38 | 31 | 31 | 29 | 49 | 20 | 44 | 34 | 36 | 49 |
| CL204 | 41 | 33 | 31 | 34 | 28 | 28 | 33 | 27 | 30 | 47 | 32 | 37 | 28 | 29 | 30 | 28 | 30 | 26 | 36 |
| CL205 | 23 | 31 | 26 | 34 | 38 | 28 | 28 | 33 | 47 | 32 | 24 | 20 | 29 | 25 | 34 | 24 | 40 | 33 | 28 |
| CL206 | 30 | 30 | 36 | 41 | 38 | 31 | 39 | 32 | 28 | 39 | 32 | 35 | 29 | 25 | 34 | 31 | 36 | 32 | 30 |
| CL209 | 28 | 32 | 27 | 32 | 22 | 34 | 28 | 19 | 29 | 39 | 23 | 29 | 24 | 23 | 26 | 31 | 23 | 26 | 23 |
| CL211 | 29 | 28 | 35 | 30 | 20 | 31 | 30 | 26 | 28 | 21 | 27 | 26 | 28 | 30 | 18 | 41 | 29 | 28 | 19 |
| CL212 | 27 | 32 | 35 | 24 | 28 | 24 | 22 | 18 | 31 | 26 | 20 | 28 | 28 | 20 | 48 | 15 | 22 | 28 | 27 |
| CL213 | 16 | 21 | 21 | 16 | 27 | 20 | 20 | 23 | 23 | 24 | 20 | 21 | 20 | 26 | 30 | 27 | 23 | 28 | 25 |
| CL214 | 17 | 21 | 20 | 22 | 33 | 31 | 18 | 27 | 34 | 21 | 30 | 18 | 17 | 28 | 22 | 23 | 19 | 31 | 20 |
| CL215 | 22 | 29 | 16 | 23 | 20 | 14 | 29 | 13 | 21 | 21 | 20 | 21 | 21 | 24 | 20 | 27 | 22 | 22 | 27 |
| CL216 | 14 | 15 | 25 | 17 | 24 | 23 | 21 | 19 | 20 | 28 | 28 | 21 | 31 | 33 | 18 | 22 | 17 | 19 | 27 |
| CL218 | 13 | 17 | 26 | 25 | 23 | 12 | 18 | 27 | 17 | 18 | 21 | 23 | 21 | 16 | 25 | 22 | 19 | 19 | 21 |
| CL220 | 24 | 19 | 19 | 18 | 16 | 25 | 12 | 13 | 19 | 23 | 22 | 31 | 19 | 22 | 24 | 19 | 21 | 29 | 23 |
| CL221 | 20 | 13 | 19 | 21 | 17 | 19 | 16 | 15 | 20 | 19 | 13 | 22 | 21 | 25 | 14 | 20 | 16 | 13 | 15 |
| CL222 | 18 | 20 | 24 | 27 | 23 | 15 | 19 | 14 | 23 | 21 | 31 | 12 | 19 | 27 | 21 | 14 | 23 | 14 | 13 |
| CL223 | 15 | 17 | 17 | 12 | 19 | 9 | 15 | 25 | 17 | 11 | 22 | 15 | 23 | 17 | 15 | 20 | 14 | 19 | 18 |
| CL226 | 16 | 16 | 16 | 23 | 20 | 26 | 21 | 11 | 16 | 20 | 14 | 24 | 23 | 19 | 11 | 19 | 21 | 21 | 19 |
| CL228 | 13 | 19 | 23 | 23 | 20 | 26 | 18 | 19 | 16 | 22 | 16 | 14 | 24 | 20 | 16 | 18 | 15 | 17 | 12 |
| CL229 | 17 | 11 | 20 | 11 | 21 | 19 | 17 | 21 | 15 | 25 | 25 | 14 | 14 | 19 | 20 | 20 | 19 | 13 | 19 |
| CL231 | 15 | 15 | 21 | 13 | 8 | 13 | 21 | 13 | 17 | 12 | 18 | 11 | 13 | 13 | 18 | 12 | 11 | 14 | 12 |
| CL232 | 9 | 15 | 11 | 10 | 14 | 13 | 18 | 7 | 14 | 19 | 11 | 12 | 8 | 15 | 9 | 12 | 12 | 7 | 17 |
| CL234 | 17 | 9 | 11 | 9 | 8 | 15 | 14 | 14 | 13 | 12 | 7 | 14 | 10 | 11 | 23 | 18 | 10 | 7 | 13 |
| CL235 | 10 | 15 | 6 | 6 | 10 | 12 | 11 | 17 | 13 | 18 | 16 | 9 | 13 | 7 | 17 | 14 | 10 | 7 | 11 |
| CL236 | 6 | 10 | 5 | 6 | 7 | 7 | 16 | 17 | 12 | 11 | 9 | 13 | 7 | 14 | 9 | 15 | 11 | 9 | 10 |
| CL237 | 11 | 3 | 12 | 6 | 12 | 10 | 8 | 11 | 16 | 13 | 14 | 9 | 11 | 7 | 14 | 14 | 8 | 13 | 13 |
| CL239 | 12 | 13 | 7 | 15 | 14 | 6 | 9 | 12 | 10 | 10 | 9 | 11 | 8 | 5 | 8 | 13 | 7 | 6 | 10 |

| | A2_G58 | A2_G59 | A2_G60 | A2_G61 | A2_G44 | A2_G45 | A2_G34 | A2_G35 | A2_G32 | A2_G33 | A2_G30 | A2_G31 | A2_G03 | A2_G04 | A2_G01 | A2_G02 | A2_G65 | A2_G66 |
|---|---|---|---|---|---|---|---|---|---|---|---|---|---|---|---|---|---|---|
| CL002 | 4407 | 4672 | 4644 | 4564 | 4712 | 4792 | 4689 | 4786 | 4647 | 4726 | 4853 | 4608 | 4754 | 4824 | 4522 | 4681 | 4711 | 4582 |
| CL003 | 2891 | 3001 | 2929 | 2961 | 3054 | 3038 | 2986 | 3058 | 3035 | 2953 | 3036 | 2989 | 2977 | 3043 | 3028 | 3063 | 3032 | 3016 |
| CL004 | 2814 | 3093 | 3016 | 2945 | 3009 | 3094 | 2778 | 2973 | 3000 | 2940 | 3201 | 2962 | 2975 | 3150 | 2851 | 3141 | 3058 | 2904 |
| CL005 | 2469 | 2470 | 2481 | 2620 | 2539 | 2640 | 2602 | 2558 | 2655 | 2614 | 2416 | 2465 | 2563 | 2598 | 2581 | 2595 | 2585 | 2508 |
| CL008 | 2276 | 2493 | 2394 | 2438 | 2561 | 2551 | 2184 | 2546 | 2527 | 2426 | 2687 | 2480 | 2462 | 2537 | 2429 | 2552 | 2565 | 2380 |
| CL009 | 2299 | 2363 | 2308 | 2271 | 2257 | 2404 | 2231 | 2330 | 2380 | 2378 | 2489 | 2306 | 2410 | 2334 | 2226 | 2383 | 2394 | 2235 |
| CL010 | 2092 | 2154 | 2085 | 2178 | 2203 | 2169 | 2057 | 2237 | 2175 | 2217 | 2187 | 2138 | 2241 | 2169 | 2075 | 2260 | 2181 | 2152 |
| CL011 | 1944 | 1951 | 1977 | 1968 | 2034 | 2034 | 2022 | 2029 | 2105 | 2059 | 2016 | 1957 | 2044 | 2080 | 2085 | 2073 | 2044 | 2061 |
| CL012 | 2042 | 2010 | 2017 | 2084 | 2131 | 2203 | 2012 | 2105 | 2119 | 2025 | 2058 | 2051 | 2013 | 2168 | 2086 | 2188 | 2091 | 2038 |
| CL013 | 2049 | 2232 | 2189 | 2096 | 2236 | 2145 | 2004 | 2252 | 2218 | 2143 | 2328 | 2174 | 2164 | 2246 | 2064 | 2257 | 2169 | 2086 |
| CL014 | 1849 | 1984 | 1920 | 2014 | 2066 | 2002 | 2056 | 1956 | 1943 | 2017 | 1942 | 1965 | 1866 | 2036 | 1995 | 2010 | 1977 | 1947 |
| CL015 | 1952 | 2086 | 1994 | 1948 | 1987 | 2070 | 2022 | 2006 | 2078 | 2037 | 2137 | 2077 | 2050 | 2075 | 2003 | 2077 | 2148 | 2068 |
| CL016 | 1655 | 1737 | 1741 | 1779 | 1659 | 1711 | 1745 | 1694 | 1805 | 1660 | 1613 | 1721 | 1667 | 1730 | 1701 | 1813 | 1804 | 1623 |
| CL017 | 1704 | 1808 | 1747 | 1745 | 1761 | 1811 | 1676 | 1729 | 1853 | 1748 | 1869 | 1777 | 1729 | 1865 | 1724 | 1798 | 1758 | 1778 |
| CL020 | 1623 | 1593 | 1594 | 1666 | 1750 | 1678 | 1582 | 1638 | 1689 | 1563 | 1675 | 1666 | 1582 | 1719 | 1614 | 1662 | 1737 | 1613 |
| CL021 | 1516 | 1555 | 1607 | 1607 | 1705 | 1695 | 1498 | 1616 | 1685 | 1690 | 1650 | 1635 | 1692 | 1680 | 1596 | 1700 | 1746 | 1649 |
| CL023 | 1440 | 1542 | 1508 | 1612 | 1559 | 1565 | 1545 | 1537 | 1437 | 1526 | 1493 | 1506 | 1490 | 1580 | 1457 | 1542 | 1616 | 1515 |
| CL024 | 1563 | 1550 | 1473 | 1500 | 1618 | 1531 | 1389 | 1516 | 1559 | 1570 | 1497 | 1574 | 1573 | 1554 | 1480 | 1498 | 1564 | 1447 |
| CL025 | 1385 | 1448 | 1416 | 1465 | 1418 | 1437 | 1532 | 1443 | 1455 | 1500 | 1481 | 1422 | 1475 | 1470 | 1464 | 1455 | 1471 | 1418 |
| CL026 | 1353 | 1458 | 1508 | 1536 | 1477 | 1507 | 1544 | 1483 | 1510 | 1451 | 1505 | 1334 | 1474 | 1488 | 1527 | 1492 | 1597 | 1342 |
| CL027 | 1393 | 1424 | 1454 | 1470 | 1469 | 1581 | 1345 | 1417 | 1528 | 1499 | 1409 | 1513 | 1419 | 1488 | 1347 | 1588 | 1475 | 1459 |
| CL030 | 1204 | 1313 | 1256 | 1256 | 1283 | 1324 | 1278 | 1321 | 1390 | 1365 | 1337 | 1236 | 1264 | 1302 | 1261 | 1374 | 1315 | 1246 |
| CL032 | 1263 | 1304 | 1332 | 1280 | 1366 | 1352 | 1180 | 1325 | 1415 | 1312 | 1255 | 1302 | 1274 | 1360 | 1234 | 1336 | 1369 | 1261 |
| CL033 | 1133 | 1235 | 1180 | 1210 | 1120 | 1174 | 1107 | 1194 | 1186 | 1139 | 1211 | 1218 | 1186 | 1244 | 1127 | 1236 | 1239 | 1165 |
| CL034 | 1102 | 1183 | 1146 | 1142 | 1137 | 1179 | 1097 | 1155 | 1146 | 1171 | 1182 | 1168 | 1171 | 1221 | 1131 | 1164 | 1221 | 1201 |
| CL037 | 1034 | 1067 | 1174 | 1071 | 1136 | 1176 | 1099 | 1107 | 1107 | 1108 | 1135 | 1082 | 1135 | 1190 | 1126 | 1151 | 1095 | 1016 |
| CL038 | 1021 | 1141 | 1171 | 1136 | 1120 | 1177 | 1116 | 1138 | 1172 | 1143 | 1087 | 1172 | 1104 | 1103 | 1116 | 1158 | 1137 | 1100 |
| CL040 | 1012 | 1052 | 1114 | 1095 | 1114 | 1106 | 1057 | 1147 | 1121 | 1103 | 1161 | 1062 | 1075 | 1109 | 1078 | 1066 | 1153 | 1081 |
| CL046 | 1057 | 1053 | 1096 | 1051 | 1132 | 1063 | 951 | 1121 | 1001 | 1114 | 1148 | 1085 | 1068 | 1083 | 1044 | 1127 | 1059 | 1062 |
| CL047 | 891 | 924 | 928 | 941 | 914 | 951 | 960 | 963 | 934 | 924 | 918 | 917 | 973 | 914 | 977 | 977 | 935 | 924 |
| CL050 | 802 | 789 | 800 | 808 | 773 | 822 | 823 | 791 | 776 | 776 | 812 | 761 | 784 | 811 | 802 | 798 | 800 | 811 |
| CL052 | 670 | 744 | 742 | 782 | 764 | 794 | 751 | 739 | 777 | 762 | 749 | 751 | 761 | 797 | 729 | 817 | 784 | 738 |
| CL053 | 733 | 754 | 783 | 784 | 725 | 761 | 826 | 702 | 854 | 769 | 790 | 791 | 764 | 796 | 809 | 783 | 819 | 699 |
| CL057 | 713 | 782 | 805 | 785 | 831 | 865 | 704 | 771 | 779 | 782 | 804 | 738 | 795 | 783 | 784 | 785 | 882 | 731 |
| CL058 | 722 | 746 | 717 | 688 | 766 | 742 | 660 | 723 | 765 | 659 | 709 | 716 | 723 | 686 | 638 | 754 | 710 | 676 |
| CL065 | 555 | 589 | 588 | 577 | 603 | 617 | 568 | 618 | 562 | 582 | 610 | 577 | 629 | 612 | 584 | 609 | 637 | 624 |
| CL066 | 563 | 611 | 609 | 626 | 656 | 635 | 685 | 620 | 631 | 564 | 610 | 560 | 632 | 651 | 587 | 708 | 646 | 635 |
| CL067 | 572 | 588 | 602 | 563 | 667 | 687 | 588 | 654 | 610 | 630 | 616 | 646 | 635 | 653 | 607 | 642 | 667 | 636 |
| CL072 | 579 | 568 | 561 | 575 | 593 | 548 | 517 | 572 | 565 | 573 | 515 | 547 | 561 | 576 | 540 | 614 | 589 | 524 |
| CL081 | 518 | 548 | 565 | 533 | 570 | 531 | 530 | 553 | 546 | 560 | 605 | 528 | 559 | 555 | 528 | 545 | 554 | 501 |
| CL093 | 319 | 346 | 359 | 336 | 340 | 359 | 438 | 363 | 382 | 367 | 334 | 332 | 287 | 335 | 366 | 363 | 336 | 378 |
| CL099 | 360 | 382 | 398 | 375 | 360 | 390 | 395 | 397 | 398 | 424 | 393 | 402 | 370 | 454 | 404 | 398 | 391 | 371 |

| | A2_G58 | A2_G59 | A2_G60 | A2_G61 | A2_G44 | A2_G45 | A2_G34 | A2_G35 | A2_G32 | A2_G33 | A2_G30 | A2_G31 | A2_G03 | A2_G04 | A2_G01 | A2_G02 | A2_G65 | A2_G66 |
|---|---|---|---|---|---|---|---|---|---|---|---|---|---|---|---|---|---|---|
| CL100 | 402 | 356 | 414 | 454 | 407 | 412 | 425 | 390 | 371 | 345 | 316 | 438 | 340 | 398 | 354 | 387 | 392 | 335 |
| CL101 | 397 | 382 | 390 | 379 | 369 | 380 | 351 | 362 | 352 | 338 | 374 | 372 | 389 | 377 | 389 | 407 | 372 | 369 |
| CL113 | 277 | 289 | 268 | 319 | 312 | 314 | 332 | 290 | 302 | 271 | 287 | 301 | 292 | 290 | 312 | 328 | 340 | 287 |
| CL118 | 289 | 289 | 289 | 281 | 318 | 294 | 311 | 280 | 328 | 297 | 326 | 264 | 304 | 368 | 280 | 310 | 330 | 299 |
| CL128 | 324 | 293 | 313 | 326 | 290 | 324 | 328 | 332 | 316 | 340 | 285 | 315 | 314 | 291 | 327 | 297 | 322 | 310 |
| CL133 | 230 | 231 | 281 | 300 | 282 | 284 | 294 | 286 | 284 | 277 | 250 | 280 | 244 | 290 | 241 | 292 | 277 | 259 |
| CL137 | 228 | 244 | 249 | 236 | 284 | 226 | 294 | 262 | 241 | 239 | 230 | 240 | 237 | 246 | 267 | 229 | 230 | 244 |
| CL146 | 207 | 221 | 211 | 214 | 254 | 246 | 237 | 228 | 241 | 215 | 214 | 257 | 235 | 220 | 220 | 222 | 272 | 221 |
| CL148 | 223 | 211 | 217 | 174 | 231 | 199 | 213 | 210 | 225 | 220 | 217 | 223 | 235 | 202 | 234 | 219 | 236 | 216 |
| CL149 | 190 | 187 | 207 | 220 | 224 | 214 | 212 | 210 | 234 | 172 | 218 | 179 | 214 | 230 | 208 | 240 | 206 | 207 |
| CL152 | 202 | 189 | 212 | 205 | 215 | 238 | 210 | 218 | 172 | 216 | 219 | 178 | 190 | 188 | 194 | 219 | 200 | 200 |
| CL155 | 191 | 192 | 160 | 177 | 167 | 162 | 223 | 178 | 182 | 151 | 151 | 190 | 172 | 170 | 183 | 164 | 172 | 184 |
| CL168 | 143 | 133 | 137 | 142 | 133 | 149 | 169 | 129 | 126 | 121 | 154 | 134 | 137 | 133 | 139 | 148 | 140 | 129 |
| CL170 | 133 | 145 | 137 | 133 | 162 | 146 | 136 | 114 | 146 | 128 | 129 | 125 | 165 | 146 | 134 | 144 | 142 | 154 |
| CL174 | 119 | 112 | 117 | 117 | 97 | 113 | 131 | 125 | 124 | 113 | 108 | 103 | 97 | 120 | 121 | 133 | 121 | 97 |
| CL176 | 97 | 105 | 103 | 101 | 120 | 104 | 115 | 98 | 87 | 101 | 93 | 90 | 99 | 93 | 111 | 105 | 100 | 91 |
| CL184 | 65 | 72 | 87 | 70 | 84 | 82 | 98 | 64 | 84 | 72 | 72 | 58 | 65 | 73 | 69 | 74 | 84 | 68 |
| CL196 | 41 | 54 | 59 | 37 | 59 | 43 | 47 | 46 | 46 | 42 | 46 | 40 | 40 | 41 | 45 | 47 | 48 | 47 |
| CL198 | 46 | 36 | 37 | 50 | 38 | 49 | 47 | 43 | 48 | 51 | 44 | 35 | 44 | 33 | 31 | 49 | 38 | 35 |
| CL210 | 38 | 33 | 32 | 36 | 33 | 21 | 25 | 35 | 32 | 31 | 30 | 23 | 43 | 26 | 27 | 25 | 22 | 23 |
| CL219 | 29 | 28 | 16 | 27 | 28 | 30 | 19 | 25 | 29 | 25 | 11 | 22 | 12 | 13 | 14 | 17 | 12 | 15 |
| CL238 | 10 | 19 | 12 | 14 | 15 | 15 | 8 | 16 | 11 | 17 | 11 | 9 | 9 | 13 | 7 | 7 | 15 | 10 |
| CL001 | 4449 | 4649 | 4716 | 4650 | 4668 | 4948 | 4439 | 4684 | 4780 | 4662 | 4775 | 4764 | 4917 | 4833 | 4544 | 4819 | 5112 | 4743 |
| CL006 | 2344 | 2271 | 2501 | 2440 | 2385 | 2386 | 2601 | 2437 | 2387 | 2401 | 2417 | 2388 | 2438 | 2494 | 2419 | 2417 | 2366 | 2272 |
| CL007 | 2164 | 2330 | 2413 | 2343 | 2308 | 2259 | 2460 | 2365 | 2403 | 2361 | 2334 | 2330 | 2311 | 2375 | 2325 | 2379 | 2341 | 2226 |
| CL018 | 1639 | 1677 | 1765 | 1762 | 1710 | 1787 | 1818 | 1720 | 1787 | 1720 | 1682 | 1705 | 1685 | 1710 | 1780 | 1749 | 1721 | 1664 |
| CL019 | 1670 | 1724 | 1791 | 1750 | 1762 | 1871 | 1717 | 1810 | 1758 | 1831 | 1859 | 1710 | 1776 | 1747 | 1675 | 1811 | 1817 | 1718 |
| CL022 | 1493 | 1589 | 1592 | 1582 | 1593 | 1610 | 1543 | 1591 | 1584 | 1543 | 1601 | 1544 | 1584 | 1564 | 1564 | 1615 | 1642 | 1630 |
| CL028 | 1348 | 1295 | 1295 | 1331 | 1376 | 1339 | 1395 | 1362 | 1378 | 1289 | 1343 | 1285 | 1366 | 1414 | 1392 | 1419 | 1382 | 1302 |
| CL029 | 1223 | 1328 | 1333 | 1347 | 1275 | 1357 | 1335 | 1388 | 1315 | 1307 | 1292 | 1281 | 1347 | 1341 | 1295 | 1370 | 1333 | 1273 |
| CL031 | 1252 | 1267 | 1320 | 1286 | 1276 | 1359 | 1264 | 1331 | 1362 | 1258 | 1325 | 1337 | 1306 | 1322 | 1215 | 1352 | 1350 | 1269 |
| CL035 | 1069 | 1114 | 1092 | 1189 | 1109 | 1155 | 1179 | 1114 | 1193 | 1142 | 1074 | 1117 | 1083 | 1164 | 1134 | 1186 | 1190 | 1156 |
| CL036 | 1045 | 1069 | 1098 | 1076 | 1064 | 1157 | 1175 | 1143 | 1187 | 1102 | 1089 | 1094 | 1096 | 1161 | 1146 | 1178 | 1094 | 1057 |
| CL039 | 1065 | 1004 | 1110 | 1039 | 1146 | 1122 | 1083 | 1096 | 1152 | 1110 | 1040 | 1107 | 1060 | 1107 | 1035 | 1042 | 1061 | 1085 |
| CL041 | 1059 | 1100 | 1058 | 1040 | 1065 | 1116 | 1104 | 1100 | 1044 | 1060 | 1048 | 1037 | 1050 | 1107 | 1040 | 1117 | 1030 | 1054 |
| CL042 | 994 | 1025 | 991 | 1040 | 998 | 1043 | 1057 | 1092 | 979 | 1009 | 1062 | 1022 | 1064 | 1030 | 987 | 1067 | 1007 | 985 |
| CL043 | 929 | 1004 | 944 | 1039 | 1001 | 1070 | 1023 | 1083 | 1030 | 981 | 1006 | 1030 | 1015 | 1054 | 955 | 1042 | 1065 | 1006 |
| CL044 | 969 | 1046 | 975 | 976 | 936 | 983 | 945 | 988 | 1029 | 985 | 976 | 1008 | 966 | 1001 | 985 | 980 | 995 | 925 |
| CL045 | 896 | 917 | 984 | 932 | 938 | 945 | 985 | 993 | 981 | 945 | 919 | 944 | 867 | 981 | 924 | 927 | 998 | 918 |
| CL048 | 861 | 873 | 898 | 890 | 949 | 811 | 912 | 840 | 942 | 893 | 870 | 839 | 887 | 917 | 868 | 889 | 859 | 834 |
| CL049 | 917 | 969 | 933 | 924 | 899 | 941 | 982 | 862 | 947 | 954 | 952 | 905 | 894 | 959 | 977 | 938 | 944 | 851 |
| CL051 | 814 | 909 | 903 | 809 | 856 | 874 | 847 | 880 | 858 | 857 | 828 | 861 | 818 | 915 | 879 | 884 | 877 | 836 |
| CL055 | 728 | 666 | 709 | 728 | 736 | 688 | 712 | 697 | 773 | 712 | 712 | 720 | 694 | 691 | 721 | 750 | 769 | 689 |
| CL056 | 698 | 731 | 742 | 708 | 713 | 712 | 760 | 720 | 750 | 735 | 756 | 712 | 706 | 731 | 720 | 724 | 765 | 714 |

| | A2_G58 | A2_G59 | A2_G60 | A2_G61 | A2_G44 | A2_G45 | A2_G34 | A2_G35 | A2_G32 | A2_G33 | A2_G30 | A2_G31 | A2_G03 | A2_G04 | A2_G01 | A2_G02 | A2_G65 | A2_G66 |
|---|---|---|---|---|---|---|---|---|---|---|---|---|---|---|---|---|---|---|
| CL059 | 657 | 679 | 666 | 698 | 735 | 696 | 700 | 673 | 740 | 665 | 706 | 688 | 694 | 686 | 680 | 726 | 658 | 703 |
| CL060 | 616 | 720 | 709 | 700 | 713 | 744 | 712 | 753 | 697 | 692 | 675 | 723 | 679 | 697 | 633 | 737 | 677 | 616 |
| CL061 | 683 | 755 | 729 | 672 | 673 | 711 | 767 | 748 | 808 | 670 | 720 | 718 | 745 | 711 | 727 | 736 | 732 | 713 |
| CL062 | 611 | 649 | 672 | 687 | 692 | 683 | 689 | 638 | 670 | 695 | 655 | 632 | 709 | 607 | 638 | 671 | 660 | 639 |
| CL063 | 670 | 707 | 733 | 727 | 709 | 717 | 726 | 744 | 721 | 657 | 693 | 754 | 648 | 748 | 730 | 767 | 734 | 647 |
| CL064 | 589 | 636 | 643 | 602 | 639 | 676 | 643 | 647 | 654 | 647 | 703 | 618 | 607 | 647 | 583 | 613 | 597 | 606 |
| CL068 | 576 | 605 | 571 | 583 | 544 | 603 | 535 | 597 | 655 | 648 | 605 | 587 | 584 | 635 | 546 | 636 | 617 | 532 |
| CL070 | 544 | 535 | 529 | 547 | 571 | 566 | 514 | 598 | 564 | 583 | 547 | 536 | 573 | 601 | 578 | 592 | 572 | 564 |
| CL071 | 495 | 563 | 569 | 515 | 547 | 544 | 578 | 575 | 598 | 514 | 564 | 566 | 532 | 590 | 543 | 567 | 532 | 522 |
| CL074 | 520 | 515 | 535 | 500 | 546 | 545 | 522 | 529 | 551 | 519 | 522 | 535 | 519 | 550 | 543 | 603 | 552 | 531 |
| CL075 | 491 | 571 | 527 | 538 | 557 | 553 | 473 | 568 | 604 | 572 | 589 | 479 | 558 | 502 | 548 | 569 | 534 | 557 |
| CL076 | 553 | 503 | 547 | 579 | 550 | 522 | 600 | 550 | 539 | 518 | 508 | 556 | 495 | 554 | 524 | 590 | 587 | 523 |
| CL078 | 542 | 530 | 557 | 544 | 567 | 562 | 518 | 539 | 560 | 511 | 531 | 563 | 576 | 522 | 530 | 575 | 571 | 537 |
| CL079 | 476 | 461 | 499 | 535 | 532 | 484 | 555 | 497 | 543 | 514 | 477 | 499 | 486 | 528 | 516 | 557 | 507 | 525 |
| CL080 | 427 | 467 | 496 | 511 | 493 | 476 | 526 | 518 | 511 | 489 | 495 | 511 | 498 | 527 | 485 | 529 | 495 | 499 |
| CL082 | 454 | 496 | 512 | 455 | 503 | 521 | 484 | 479 | 480 | 484 | 491 | 471 | 545 | 484 | 451 | 543 | 492 | 498 |
| CL084 | 472 | 502 | 521 | 503 | 461 | 496 | 519 | 522 | 489 | 485 | 502 | 506 | 502 | 498 | 492 | 534 | 539 | 459 |
| CL085 | 434 | 498 | 489 | 508 | 468 | 523 | 504 | 495 | 509 | 509 | 509 | 464 | 470 | 522 | 475 | 497 | 440 | 481 |
| CL091 | 477 | 440 | 515 | 452 | 467 | 489 | 493 | 476 | 488 | 476 | 497 | 446 | 437 | 472 | 473 | 501 | 482 | 475 |
| CL092 | 409 | 419 | 452 | 464 | 449 | 480 | 489 | 452 | 472 | 423 | 463 | 448 | 424 | 492 | 444 | 441 | 454 | 433 |
| CL094 | 420 | 477 | 463 | 438 | 401 | 447 | 431 | 447 | 457 | 455 | 424 | 408 | 442 | 467 | 481 | 470 | 475 | 421 |
| CL095 | 344 | 256 | 325 | 463 | 487 | 448 | 398 | 393 | 362 | 350 | 440 | 464 | 302 | 225 | 322 | 289 | 366 | 299 |
| CL102 | 392 | 417 | 432 | 438 | 355 | 417 | 406 | 383 | 444 | 406 | 366 | 408 | 426 | 430 | 403 | 394 | 415 | 400 |
| CL104 | 360 | 337 | 393 | 387 | 376 | 396 | 442 | 378 | 369 | 336 | 386 | 393 | 363 | 404 | 406 | 370 | 408 | 346 |
| CL106 | 315 | 330 | 313 | 326 | 344 | 363 | 392 | 382 | 360 | 370 | 361 | 369 | 366 | 380 | 344 | 367 | 349 | 363 |
| CL107 | 338 | 333 | 345 | 334 | 376 | 387 | 370 | 348 | 332 | 377 | 328 | 316 | 367 | 347 | 391 | 357 | 368 | 360 |
| CL109 | 337 | 350 | 366 | 356 | 344 | 360 | 393 | 365 | 393 | 340 | 335 | 366 | 339 | 389 | 322 | 377 | 358 | 332 |
| CL112 | 318 | 340 | 324 | 329 | 351 | 314 | 319 | 359 | 340 | 326 | 338 | 337 | 317 | 350 | 323 | 341 | 349 | 318 |
| CL117 | 330 | 331 | 318 | 347 | 323 | 340 | 298 | 341 | 326 | 347 | 326 | 284 | 332 | 307 | 306 | 342 | 359 | 342 |
| CL119 | 258 | 297 | 276 | 283 | 300 | 289 | 298 | 312 | 316 | 272 | 294 | 315 | 263 | 296 | 282 | 320 | 301 | 252 |
| CL120 | 263 | 310 | 283 | 294 | 293 | 289 | 310 | 342 | 293 | 290 | 328 | 314 | 313 | 273 | 290 | 325 | 304 | 291 |
| CL121 | 313 | 330 | 317 | 317 | 301 | 322 | 328 | 323 | 345 | 321 | 307 | 315 | 348 | 293 | 321 | 335 | 319 | 337 |
| CL122 | 342 | 352 | 345 | 333 | 344 | 332 | 342 | 336 | 313 | 328 | 311 | 325 | 316 | 320 | 294 | 355 | 360 | 327 |
| CL129 | 279 | 280 | 300 | 307 | 266 | 300 | 299 | 272 | 294 | 303 | 308 | 311 | 315 | 296 | 311 | 302 | 313 | 262 |
| CL130 | 245 | 230 | 298 | 248 | 228 | 283 | 277 | 283 | 261 | 272 | 293 | 283 | 264 | 243 | 321 | 292 | 271 | 340 |
| CL131 | 259 | 275 | 295 | 261 | 275 | 273 | 293 | 286 | 290 | 265 | 238 | 268 | 274 | 271 | 269 | 279 | 288 | 252 |
| CL139 | 265 | 247 | 247 | 256 | 237 | 246 | 267 | 245 | 265 | 263 | 222 | 235 | 217 | 254 | 256 | 281 | 253 | 221 |
| CL143 | 228 | 252 | 239 | 245 | 247 | 236 | 212 | 240 | 208 | 205 | 212 | 208 | 213 | 242 | 205 | 241 | 245 | 233 |
| CL147 | 181 | 201 | 201 | 223 | 198 | 228 | 207 | 202 | 218 | 209 | 207 | 207 | 239 | 205 | 199 | 215 | 211 | 215 |
| CL151 | 178 | 227 | 222 | 215 | 186 | 195 | 238 | 205 | 220 | 200 | 188 | 222 | 204 | 204 | 224 | 240 | 233 | 216 |
| CL157 | 176 | 178 | 172 | 182 | 163 | 193 | 189 | 183 | 205 | 184 | 163 | 193 | 206 | 185 | 178 | 188 | 192 | 180 |
| CL158 | 175 | 151 | 212 | 181 | 169 | 166 | 177 | 179 | 175 | 147 | 187 | 163 | 165 | 176 | 175 | 183 | 157 | 163 |
| CL159 | 173 | 148 | 155 | 161 | 177 | 150 | 180 | 161 | 152 | 129 | 175 | 143 | 158 | 174 | 167 | 185 | 153 | 148 |
| CL160 | 187 | 148 | 159 | 171 | 184 | 170 | 175 | 176 | 179 | 183 | 176 | 157 | 166 | 179 | 194 | 192 | 162 | 183 |

| | A2_G58 | A2_G59 | A2_G60 | A2_G61 | A2_G44 | A2_G45 | A2_G34 | A2_G35 | A2_G32 | A2_G33 | A2_G30 | A2_G31 | A2_G03 | A2_G04 | A2_G01 | A2_G02 | A2_G65 | A2_G66 |
|---|---|---|---|---|---|---|---|---|---|---|---|---|---|---|---|---|---|---|
| CL162 | 178 | 143 | 160 | 172 | 149 | 183 | 152 | 182 | 153 | 164 | 191 | 160 | 168 | 171 | 166 | 179 | 152 | 164 |
| CL163 | 173 | 170 | 173 | 154 | 176 | 152 | 181 | 178 | 170 | 162 | 161 | 153 | 150 | 159 | 181 | 169 | 150 | 148 |
| CL164 | 145 | 170 | 121 | 155 | 152 | 144 | 137 | 148 | 154 | 158 | 138 | 142 | 141 | 148 | 156 | 139 | 140 | 143 |
| CL166 | 145 | 154 | 138 | 157 | 160 | 173 | 179 | 161 | 162 | 155 | 176 | 158 | 153 | 155 | 171 | 180 | 151 | 146 |
| CL171 | 125 | 138 | 135 | 139 | 141 | 110 | 119 | 138 | 116 | 109 | 127 | 125 | 129 | 145 | 125 | 108 | 121 | 122 |
| CL173 | 93 | 124 | 115 | 107 | 125 | 123 | 144 | 107 | 115 | 103 | 133 | 116 | 102 | 126 | 112 | 126 | 127 | 119 |
| CL175 | 122 | 115 | 100 | 107 | 120 | 114 | 104 | 106 | 87 | 128 | 112 | 108 | 116 | 123 | 99 | 113 | 106 | 110 |
| CL177 | 100 | 99 | 102 | 117 | 102 | 111 | 90 | 98 | 121 | 95 | 116 | 103 | 102 | 118 | 81 | 110 | 104 | 103 |
| CL182 | 92 | 94 | 80 | 87 | 74 | 98 | 100 | 68 | 90 | 90 | 70 | 85 | 92 | 90 | 78 | 86 | 82 | 72 |
| CL186 | 74 | 65 | 79 | 76 | 74 | 62 | 80 | 73 | 83 | 80 | 77 | 71 | 77 | 79 | 64 | 70 | 91 | 64 |
| CL187 | 74 | 66 | 71 | 66 | 76 | 67 | 78 | 83 | 92 | 65 | 76 | 68 | 73 | 84 | 85 | 81 | 73 | 80 |
| CL189 | 67 | 58 | 63 | 71 | 74 | 57 | 65 | 72 | 55 | 65 | 72 | 59 | 55 | 57 | 63 | 69 | 53 | 81 |
| CL191 | 61 | 61 | 59 | 55 | 58 | 56 | 62 | 59 | 66 | 63 | 58 | 65 | 68 | 54 | 57 | 57 | 75 | 66 |
| CL194 | 54 | 47 | 51 | 57 | 57 | 56 | 59 | 52 | 53 | 51 | 60 | 55 | 61 | 64 | 55 | 57 | 44 | 52 |
| CL195 | 45 | 63 | 47 | 45 | 49 | 56 | 50 | 50 | 61 | 36 | 57 | 46 | 55 | 45 | 59 | 59 | 40 | 48 |
| CL197 | 48 | 46 | 51 | 39 | 43 | 43 | 42 | 41 | 41 | 45 | 42 | 37 | 46 | 47 | 47 | 46 | 39 | 45 |
| CL199 | 32 | 36 | 42 | 35 | 50 | 45 | 41 | 45 | 53 | 33 | 40 | 50 | 35 | 38 | 48 | 33 | 34 | 35 |
| CL200 | 37 | 32 | 40 | 43 | 38 | 39 | 47 | 37 | 32 | 35 | 49 | 43 | 47 | 31 | 46 | 40 | 34 | 37 |
| CL202 | 48 | 33 | 35 | 26 | 27 | 30 | 42 | 51 | 23 | 38 | 27 | 35 | 42 | 32 | 27 | 38 | 29 | 31 |
| CL203 | 26 | 42 | 43 | 37 | 35 | 39 | 29 | 40 | 28 | 38 | 37 | 41 | 38 | 41 | 33 | 33 | 41 | 40 |
| CL204 | 37 | 36 | 35 | 35 | 34 | 38 | 45 | 40 | 48 | 36 | 43 | 42 | 31 | 41 | 26 | 35 | 36 | 30 |
| CL205 | 37 | 37 | 34 | 34 | 35 | 39 | 41 | 33 | 36 | 21 | 30 | 35 | 28 | 31 | 36 | 34 | 34 | 30 |
| CL206 | 19 | 29 | 29 | 30 | 34 | 34 | 35 | 28 | 26 | 43 | 28 | 34 | 33 | 33 | 30 | 30 | 41 | 33 |
| CL209 | 29 | 29 | 31 | 35 | 25 | 36 | 30 | 20 | 23 | 31 | 28 | 33 | 39 | 30 | 23 | 29 | 20 | 27 |
| CL211 | 14 | 23 | 30 | 35 | 26 | 27 | 25 | 24 | 27 | 26 | 23 | 18 | 21 | 33 | 27 | 25 | 20 | 20 |
| CL212 | 17 | 34 | 30 | 25 | 17 | 25 | 37 | 31 | 22 | 26 | 31 | 27 | 31 | 36 | 40 | 23 | 47 | 29 |
| CL213 | 21 | 26 | 16 | 24 | 26 | 24 | 24 | 28 | 30 | 24 | 20 | 30 | 22 | 33 | 20 | 34 | 25 | 30 |
| CL214 | 16 | 24 | 22 | 27 | 26 | 18 | 23 | 23 | 26 | 25 | 24 | 22 | 20 | 20 | 20 | 23 | 30 | 15 |
| CL215 | 19 | 19 | 27 | 20 | 20 | 21 | 19 | 20 | 20 | 19 | 29 | 25 | 22 | 23 | 17 | 19 | 25 | 29 |
| CL216 | 28 | 23 | 16 | 21 | 23 | 22 | 24 | 17 | 22 | 26 | 27 | 29 | 18 | 30 | 24 | 23 | 23 | 27 |
| CL218 | 13 | 23 | 13 | 14 | 17 | 20 | 22 | 16 | 28 | 21 | 15 | 26 | 23 | 23 | 16 | 23 | 20 | 27 |
| CL220 | 19 | 13 | 14 | 23 | 22 | 27 | 18 | 21 | 28 | 24 | 15 | 18 | 18 | 15 | 17 | 10 | 20 | 25 |
| CL221 | 13 | 18 | 23 | 25 | 15 | 25 | 26 | 27 | 22 | 24 | 15 | 24 | 34 | 12 | 18 | 16 | 23 | 18 |
| CL222 | 25 | 20 | 18 | 21 | 13 | 22 | 33 | 20 | 17 | 17 | 19 | 11 | 17 | 23 | 20 | 16 | 18 | 17 |
| CL223 | 20 | 17 | 14 | 19 | 23 | 20 | 21 | 21 | 16 | 16 | 16 | 11 | 20 | 17 | 12 | 16 | 23 | 19 |
| CL226 | 19 | 21 | 14 | 23 | 17 | 20 | 16 | 16 | 23 | 21 | 14 | 25 | 21 | 17 | 15 | 24 | 18 | 20 |
| CL228 | 6 | 13 | 10 | 13 | 20 | 11 | 16 | 14 | 15 | 18 | 17 | 12 | 11 | 16 | 22 | 21 | 12 | 23 |
| CL229 | 11 | 22 | 21 | 20 | 13 | 23 | 16 | 17 | 18 | 24 | 10 | 15 | 20 | 22 | 17 | 15 | 18 | 18 |
| CL231 | 9 | 14 | 15 | 19 | 14 | 23 | 18 | 16 | 16 | 13 | 19 | 9 | 17 | 18 | 14 | 9 | 8 | 11 |
| CL232 | 18 | 11 | 19 | 8 | 17 | 14 | 12 | 12 | 16 | 16 | 11 | 15 | 14 | 14 | 14 | 11 | 10 | 9 |
| CL234 | 10 | 10 | 10 | 12 | 13 | 13 | 8 | 11 | 6 | 11 | 8 | 11 | 7 | 5 | 8 | 11 | 15 | 10 |
| CL235 | 8 | 10 | 7 | 21 | 8 | 24 | 11 | 12 | 15 | 12 | 12 | 12 | 12 | 9 | 16 | 12 | 10 | 16 |
| CL236 | 7 | 18 | 12 | 11 | 9 | 14 | 12 | 13 | 9 | 9 | 8 | 11 | 17 | 13 | 11 | 9 | 16 | 8 |
| CL237 | 9 | 5 | 12 | 11 | 11 | 10 | 8 | 11 | 14 | 13 | 13 | 13 | 13 | 9 | 14 | 13 | 16 | 7 |
| CL239 | 9 | 12 | 7 | 13 | 14 | 14 | 11 | 12 | 9 | 11 | 13 | 12 | 10 | 5 | 11 | 10 | 7 | 8 |

| | A2_G55 | A2_G54 | A2_G53 | A2_G52 | A2_G51 | A2_G50 | A2_G49 | A2_G48 | A2_G47 | A2_G46 | A2_G62 | A2_G63 | A2_G64 |
|---|---|---|---|---|---|---|---|---|---|---|---|---|---|
| CL002 | 4875 | 4881 | 4891 | 4518 | 4631 | 4713 | 4705 | 4613 | 4671 | 4560 | 4860 | 4585 | 4705 |
| CL003 | 2933 | 3028 | 2997 | 2967 | 2890 | 2895 | 2911 | 2887 | 2934 | 3061 | 3021 | 3000 | 2983 |
| CL004 | 3148 | 3072 | 2957 | 2833 | 3046 | 3046 | 3062 | 3120 | 3107 | 3034 | 3228 | 3005 | 2920 |
| CL005 | 2545 | 2691 | 2481 | 2676 | 2524 | 2485 | 2532 | 2450 | 2407 | 2608 | 2445 | 2559 | 2609 |
| CL008 | 2548 | 2530 | 2581 | 2313 | 2367 | 2475 | 2457 | 2564 | 2645 | 2478 | 2731 | 2503 | 2432 |
| CL009 | 2398 | 2332 | 2380 | 2297 | 2253 | 2426 | 2328 | 2378 | 2351 | 2461 | 2439 | 2321 | 2318 |
| CL010 | 2308 | 2278 | 2148 | 1997 | 2216 | 2144 | 2155 | 2399 | 2175 | 2198 | 2266 | 2162 | 2159 |
| CL011 | 2054 | 2059 | 2105 | 2008 | 2013 | 1941 | 2020 | 2003 | 1988 | 2011 | 1966 | 1997 | 2140 |
| CL012 | 2195 | 2090 | 2029 | 2141 | 2141 | 2024 | 2099 | 2025 | 2091 | 2091 | 2152 | 2180 | 2103 |
| CL013 | 2179 | 2190 | 2264 | 2051 | 2140 | 2196 | 2290 | 2247 | 2236 | 2291 | 2422 | 2157 | 2202 |
| CL014 | 2055 | 2029 | 2053 | 2012 | 2087 | 1988 | 1998 | 1981 | 2059 | 1996 | 1960 | 1994 | 1927 |
| CL015 | 2150 | 2088 | 2111 | 1856 | 2001 | 2083 | 2068 | 2037 | 2125 | 1994 | 2047 | 2020 | 2065 |
| CL016 | 1727 | 1735 | 1562 | 1884 | 1761 | 1675 | 1684 | 1511 | 1637 | 1636 | 1686 | 1703 | 1744 |
| CL017 | 1823 | 1860 | 1886 | 1764 | 1788 | 1707 | 1784 | 1812 | 1956 | 1824 | 1992 | 1741 | 1736 |
| CL020 | 1707 | 1662 | 1611 | 1652 | 1645 | 1687 | 1608 | 1671 | 1569 | 1679 | 1658 | 1634 | 1629 |
| CL021 | 1700 | 1649 | 1608 | 1564 | 1722 | 1614 | 1713 | 1756 | 1653 | 1647 | 1664 | 1640 | 1704 |
| CL023 | 1506 | 1560 | 1519 | 1481 | 1515 | 1540 | 1484 | 1548 | 1528 | 1540 | 1471 | 1478 | 1500 |
| CL024 | 1591 | 1557 | 1389 | 1449 | 1585 | 1460 | 1525 | 1575 | 1504 | 1558 | 1555 | 1539 | 1497 |
| CL025 | 1387 | 1560 | 1441 | 1597 | 1411 | 1338 | 1370 | 1320 | 1380 | 1376 | 1365 | 1431 | 1434 |
| CL026 | 1449 | 1512 | 1446 | 1528 | 1570 | 1462 | 1450 | 1431 | 1416 | 1503 | 1372 | 1504 | 1480 |
| CL027 | 1510 | 1421 | 1430 | 1354 | 1442 | 1406 | 1421 | 1546 | 1416 | 1434 | 1582 | 1544 | 1474 |
| CL030 | 1310 | 1345 | 1332 | 1255 | 1340 | 1247 | 1368 | 1298 | 1281 | 1234 | 1274 | 1319 | 1317 |
| CL032 | 1374 | 1370 | 1253 | 1260 | 1354 | 1323 | 1297 | 1383 | 1258 | 1272 | 1363 | 1386 | 1226 |
| CL033 | 1210 | 1209 | 1160 | 1129 | 1173 | 1196 | 1131 | 1132 | 1133 | 1195 | 1160 | 1186 | 1193 |
| CL034 | 1198 | 1190 | 1167 | 1077 | 1120 | 1102 | 1172 | 1207 | 1177 | 1155 | 1241 | 1137 | 1200 |
| CL037 | 1109 | 1181 | 1070 | 1128 | 1144 | 1105 | 1099 | 1156 | 1134 | 1100 | 1102 | 1066 | 1033 |
| CL038 | 1147 | 1223 | 1060 | 1006 | 1134 | 1161 | 1128 | 1105 | 1119 | 1140 | 1171 | 1127 | 1116 |
| CL040 | 1079 | 1049 | 1083 | 1115 | 1060 | 1077 | 1072 | 1116 | 1056 | 1071 | 1117 | 1143 | 1093 |
| CL046 | 1081 | 1086 | 1039 | 933 | 1055 | 1104 | 1112 | 1114 | 1166 | 1063 | 1149 | 1031 | 1100 |
| CL047 | 923 | 952 | 954 | 930 | 948 | 918 | 951 | 945 | 906 | 912 | 918 | 888 | 953 |
| CL050 | 820 | 810 | 809 | 879 | 819 | 807 | 853 | 781 | 801 | 799 | 775 | 789 | 793 |
| CL052 | 769 | 835 | 767 | 835 | 776 | 755 | 819 | 690 | 697 | 765 | 760 | 745 | 788 |
| CL053 | 730 | 816 | 779 | 877 | 753 | 780 | 786 | 776 | 802 | 743 | 726 | 788 | 782 |
| CL057 | 795 | 724 | 802 | 721 | 781 | 734 | 844 | 793 | 811 | 807 | 863 | 770 | 730 |
| CL058 | 670 | 713 | 753 | 728 | 730 | 703 | 708 | 731 | 703 | 744 | 734 | 686 | 723 |
| CL065 | 644 | 604 | 584 | 515 | 605 | 536 | 624 | 637 | 603 | 608 | 649 | 634 | 574 |
| CL066 | 644 | 585 | 611 | 675 | 629 | 602 | 614 | 568 | 631 | 609 | 669 | 652 | 558 |
| CL067 | 611 | 594 | 618 | 562 | 596 | 643 | 627 | 571 | 577 | 602 | 608 | 627 | 599 |
| CL072 | 556 | 586 | 550 | 559 | 568 | 591 | 578 | 560 | 568 | 556 | 583 | 558 | 552 |
| CL081 | 525 | 549 | 542 | 522 | 560 | 538 | 534 | 550 | 533 | 520 | 589 | 542 | 521 |
| CL093 | 337 | 347 | 309 | 358 | 377 | 371 | 325 | 335 | 366 | 344 | 368 | 339 | 346 |
| CL099 | 383 | 385 | 394 | 447 | 433 | 402 | 420 | 374 | 421 | 436 | 360 | 406 | 384 |

| | A2_G55 | A2_G54 | A2_G53 | A2_G52 | A2_G51 | A2_G50 | A2_G49 | A2_G48 | A2_G47 | A2_G46 | A2_G62 | A2_G63 | A2_G64 |
|---|---|---|---|---|---|---|---|---|---|---|---|---|---|
| CL100 | 359 | 354 | 418 | 378 | 454 | 384 | 372 | 335 | 445 | 371 | 384 | 402 | 409 |
| CL101 | 415 | 419 | 406 | 377 | 372 | 358 | 385 | 374 | 370 | 376 | 378 | 417 | 389 |
| CL113 | 308 | 287 | 249 | 356 | 321 | 297 | 286 | 275 | 256 | 298 | 234 | 291 | 301 |
| CL118 | 291 | 305 | 307 | 323 | 311 | 299 | 290 | 280 | 284 | 299 | 282 | 283 | 285 |
| CL128 | 322 | 302 | 297 | 345 | 332 | 287 | 348 | 291 | 298 | 312 | 342 | 318 | 328 |
| CL133 | 244 | 269 | 236 | 286 | 297 | 273 | 268 | 250 | 260 | 239 | 278 | 274 | 247 |
| CL137 | 243 | 264 | 224 | 312 | 264 | 237 | 239 | 223 | 240 | 248 | 232 | 224 | 231 |
| CL146 | 228 | 245 | 212 | 232 | 234 | 228 | 219 | 245 | 238 | 229 | 229 | 229 | 238 |
| CL148 | 211 | 224 | 246 | 224 | 209 | 233 | 208 | 192 | 191 | 207 | 241 | 181 | 206 |
| CL149 | 193 | 211 | 218 | 232 | 222 | 191 | 219 | 199 | 190 | 204 | 184 | 228 | 218 |
| CL152 | 219 | 196 | 209 | 192 | 202 | 211 | 206 | 203 | 191 | 198 | 240 | 176 | 192 |
| CL155 | 183 | 167 | 185 | 184 | 185 | 170 | 190 | 176 | 173 | 158 | 159 | 178 | 182 |
| CL168 | 149 | 145 | 123 | 155 | 144 | 153 | 133 | 139 | 110 | 147 | 141 | 139 | 145 |
| CL170 | 122 | 133 | 122 | 129 | 135 | 145 | 147 | 135 | 134 | 145 | 145 | 150 | 145 |
| CL174 | 113 | 105 | 115 | 138 | 124 | 109 | 117 | 101 | 124 | 103 | 95 | 117 | 112 |
| CL176 | 107 | 93 | 102 | 100 | 111 | 101 | 82 | 84 | 95 | 80 | 89 | 92 | 106 |
| CL184 | 65 | 68 | 63 | 80 | 85 | 82 | 70 | 69 | 87 | 70 | 67 | 78 | 69 |
| CL196 | 49 | 54 | 43 | 52 | 35 | 38 | 39 | 44 | 36 | 52 | 44 | 51 | 47 |
| CL198 | 39 | 49 | 38 | 44 | 40 | 48 | 42 | 46 | 32 | 27 | 54 | 50 | 64 |
| CL210 | 34 | 21 | 26 | 24 | 30 | 27 | 28 | 26 | 35 | 24 | 26 | 33 | 32 |
| CL219 | 30 | 40 | 20 | 27 | 22 | 19 | 27 | 24 | 21 | 41 | 26 | 20 | 24 |
| CL238 | 22 | 11 | 8 | 13 | 17 | 13 | 9 | 14 | 11 | 20 | 15 | 13 | 11 |
| CL001 | 4809 | 4801 | 4758 | 4509 | 4827 | 4645 | 4873 | 4823 | 4798 | 4692 | 4857 | 4851 | 4699 |
| CL006 | 2375 | 2534 | 2354 | 2574 | 2442 | 2463 | 2373 | 2288 | 2324 | 2369 | 2289 | 2425 | 2432 |
| CL007 | 2205 | 2318 | 2369 | 2490 | 2359 | 2282 | 2341 | 2298 | 2273 | 2341 | 2330 | 2326 | 2352 |
| CL018 | 1777 | 1757 | 1653 | 1828 | 1760 | 1740 | 1710 | 1686 | 1705 | 1770 | 1655 | 1698 | 1683 |
| CL019 | 1744 | 1758 | 1783 | 1762 | 1720 | 1728 | 1727 | 1782 | 1759 | 1739 | 1808 | 1791 | 1707 |
| CL022 | 1593 | 1567 | 1631 | 1634 | 1588 | 1568 | 1612 | 1577 | 1575 | 1608 | 1657 | 1595 | 1582 |
| CL028 | 1362 | 1379 | 1386 | 1411 | 1358 | 1363 | 1417 | 1251 | 1310 | 1327 | 1278 | 1333 | 1372 |
| CL029 | 1311 | 1359 | 1248 | 1350 | 1294 | 1351 | 1296 | 1342 | 1303 | 1271 | 1268 | 1335 | 1230 |
| CL031 | 1405 | 1323 | 1219 | 1242 | 1315 | 1301 | 1398 | 1341 | 1291 | 1349 | 1385 | 1315 | 1311 |
| CL035 | 1108 | 1181 | 1117 | 1205 | 1194 | 1154 | 1132 | 1128 | 1065 | 1174 | 1046 | 1110 | 1161 |
| CL036 | 1102 | 1137 | 1087 | 1194 | 1127 | 1063 | 1124 | 983 | 1016 | 1157 | 1054 | 1090 | 1114 |
| CL039 | 1048 | 1120 | 1037 | 1131 | 1132 | 1127 | 1084 | 1025 | 1043 | 1021 | 981 | 1031 | 1032 |
| CL041 | 1007 | 1088 | 996 | 1123 | 1043 | 1044 | 1052 | 956 | 995 | 1013 | 1033 | 1050 | 1057 |
| CL042 | 1025 | 1043 | 984 | 1066 | 1063 | 1064 | 1086 | 1030 | 1034 | 1026 | 1002 | 1008 | 957 |
| CL043 | 1017 | 989 | 991 | 924 | 1051 | 1016 | 1031 | 1023 | 1046 | 1072 | 1054 | 1038 | 989 |
| CL044 | 977 | 1098 | 997 | 984 | 921 | 1049 | 1016 | 1026 | 1019 | 1003 | 1028 | 975 | 1014 |
| CL045 | 959 | 943 | 914 | 1020 | 928 | 905 | 914 | 930 | 950 | 966 | 942 | 965 | 993 |
| CL048 | 855 | 852 | 883 | 865 | 944 | 911 | 855 | 852 | 906 | 872 | 838 | 898 | 889 |
| CL049 | 971 | 909 | 916 | 942 | 898 | 913 | 906 | 933 | 909 | 864 | 936 | 938 | 877 |
| CL051 | 839 | 871 | 842 | 916 | 928 | 863 | 890 | 851 | 870 | 896 | 882 | 880 | 929 |
| CL055 | 706 | 797 | 692 | 757 | 723 | 701 | 737 | 677 | 677 | 739 | 695 | 780 | 732 |
| CL056 | 758 | 758 | 706 | 762 | 774 | 726 | 695 | 742 | 681 | 791 | 693 | 731 | 764 |

| | A2_G55 | A2_G54 | A2_G53 | A2_G52 | A2_G51 | A2_G50 | A2_G49 | A2_G48 | A2_G47 | A2_G46 | A2_G62 | A2_G63 | A2_G64 |
|---|---|---|---|---|---|---|---|---|---|---|---|---|---|
| CL059 | 668 | 719 | 674 | 711 | 711 | 683 | 719 | 671 | 744 | 630 | 741 | 686 | 741 |
| CL060 | 706 | 712 | 672 | 742 | 713 | 711 | 713 | 665 | 670 | 678 | 633 | 647 | 625 |
| CL061 | 744 | 737 | 665 | 743 | 763 | 717 | 716 | 713 | 732 | 733 | 739 | 684 | 720 |
| CL062 | 628 | 669 | 674 | 653 | 662 | 673 | 662 | 613 | 647 | 656 | 646 | 685 | 667 |
| CL063 | 710 | 694 | 609 | 705 | 733 | 705 | 704 | 620 | 697 | 699 | 568 | 716 | 707 |
| CL064 | 660 | 653 | 619 | 581 | 623 | 611 | 631 | 697 | 627 | 642 | 660 | 677 | 634 |
| CL068 | 651 | 604 | 568 | 571 | 620 | 629 | 587 | 533 | 587 | 579 | 593 | 641 | 642 |
| CL070 | 605 | 603 | 553 | 542 | 569 | 573 | 585 | 570 | 571 | 576 | 493 | 596 | 582 |
| CL071 | 547 | 538 | 545 | 598 | 578 | 573 | 581 | 501 | 492 | 550 | 529 | 569 | 549 |
| CL074 | 535 | 539 | 475 | 578 | 527 | 528 | 546 | 500 | 487 | 581 | 513 | 527 | 538 |
| CL075 | 579 | 580 | 539 | 473 | 483 | 517 | 568 | 539 | 534 | 572 | 584 | 557 | 535 |
| CL076 | 497 | 564 | 591 | 648 | 574 | 552 | 491 | 506 | 499 | 559 | 510 | 545 | 605 |
| CL078 | 554 | 528 | 513 | 535 | 562 | 534 | 557 | 590 | 580 | 566 | 546 | 592 | 542 |
| CL079 | 533 | 503 | 513 | 576 | 548 | 536 | 529 | 504 | 476 | 501 | 522 | 517 | 532 |
| CL080 | 547 | 508 | 483 | 500 | 529 | 461 | 484 | 538 | 483 | 503 | 482 | 520 | 499 |
| CL082 | 471 | 501 | 490 | 462 | 524 | 502 | 499 | 487 | 487 | 533 | 535 | 525 | 497 |
| CL084 | 530 | 471 | 498 | 532 | 570 | 520 | 494 | 440 | 440 | 499 | 473 | 498 | 474 |
| CL085 | 499 | 480 | 462 | 506 | 496 | 511 | 506 | 482 | 500 | 496 | 448 | 511 | 513 |
| CL091 | 473 | 488 | 484 | 490 | 471 | 455 | 429 | 440 | 487 | 496 | 496 | 476 | 476 |
| CL092 | 452 | 448 | 449 | 477 | 468 | 439 | 398 | 453 | 467 | 432 | 450 | 443 | 468 |
| CL094 | 461 | 423 | 417 | 447 | 470 | 433 | 449 | 424 | 433 | 422 | 467 | 501 | 437 |
| CL095 | 317 | 289 | 437 | 216 | 494 | 356 | 249 | 358 | 548 | 284 | 490 | 271 | 196 |
| CL102 | 406 | 423 | 384 | 430 | 424 | 408 | 423 | 376 | 377 | 397 | 409 | 392 | 407 |
| CL104 | 371 | 373 | 370 | 402 | 375 | 391 | 402 | 327 | 366 | 363 | 359 | 345 | 370 |
| CL106 | 354 | 329 | 311 | 364 | 349 | 358 | 373 | 311 | 359 | 388 | 322 | 340 | 375 |
| CL107 | 343 | 376 | 345 | 393 | 421 | 342 | 355 | 336 | 329 | 345 | 345 | 366 | 372 |
| CL109 | 335 | 362 | 372 | 374 | 331 | 412 | 330 | 348 | 342 | 350 | 348 | 353 | 359 |
| CL112 | 324 | 312 | 304 | 372 | 330 | 348 | 290 | 307 | 348 | 356 | 316 | 291 | 345 |
| CL117 | 345 | 309 | 323 | 313 | 333 | 334 | 331 | 358 | 339 | 327 | 338 | 288 | 332 |
| CL119 | 297 | 332 | 319 | 315 | 301 | 303 | 279 | 304 | 283 | 301 | 270 | 293 | 289 |
| CL120 | 324 | 300 | 303 | 270 | 294 | 326 | 339 | 304 | 263 | 298 | 298 | 316 | 300 |
| CL121 | 372 | 334 | 333 | 315 | 332 | 330 | 331 | 330 | 317 | 274 | 337 | 358 | 302 |
| CL122 | 325 | 345 | 316 | 341 | 363 | 318 | 318 | 254 | 348 | 327 | 315 | 306 | 329 |
| CL129 | 304 | 301 | 267 | 286 | 309 | 285 | 324 | 275 | 325 | 279 | 277 | 328 | 287 |
| CL130 | 262 | 232 | 241 | 199 | 285 | 267 | 282 | 268 | 295 | 266 | 256 | 251 | 288 |
| CL131 | 238 | 266 | 257 | 289 | 251 | 286 | 257 | 283 | 246 | 289 | 243 | 276 | 276 |
| CL139 | 252 | 211 | 235 | 288 | 240 | 261 | 231 | 251 | 239 | 239 | 260 | 246 | 275 |
| CL143 | 218 | 250 | 252 | 208 | 247 | 229 | 257 | 226 | 211 | 218 | 225 | 228 | 229 |
| CL147 | 228 | 213 | 228 | 244 | 219 | 226 | 187 | 202 | 188 | 232 | 208 | 234 | 201 |
| CL151 | 223 | 205 | 196 | 235 | 200 | 218 | 196 | 193 | 209 | 195 | 211 | 209 | 208 |
| CL157 | 187 | 169 | 182 | 194 | 192 | 194 | 181 | 174 | 182 | 174 | 143 | 172 | 183 |
| CL158 | 172 | 193 | 152 | 157 | 196 | 183 | 174 | 172 | 160 | 174 | 150 | 174 | 181 |
| CL159 | 161 | 166 | 149 | 197 | 165 | 156 | 161 | 146 | 126 | 139 | 137 | 147 | 166 |
| CL160 | 171 | 168 | 160 | 186 | 178 | 194 | 191 | 161 | 177 | 185 | 152 | 155 | 173 |

| | A2_G55 | A2_G54 | A2_G53 | A2_G52 | A2_G51 | A2_G50 | A2_G49 | A2_G48 | A2_G47 | A2_G46 | A2_G62 | A2_G63 | A2_G64 |
|---|---|---|---|---|---|---|---|---|---|---|---|---|---|
| CL162 | 182 | 165 | 174 | 168 | 173 | 153 | 186 | 163 | 137 | 173 | 179 | 144 | 187 |
| CL163 | 168 | 156 | 155 | 194 | 189 | 177 | 156 | 150 | 150 | 127 | 162 | 171 | 145 |
| CL164 | 155 | 140 | 172 | 156 | 133 | 158 | 147 | 142 | 145 | 137 | 140 | 153 | 153 |
| CL166 | 131 | 169 | 131 | 176 | 171 | 145 | 158 | 127 | 154 | 159 | 157 | 139 | 144 |
| CL171 | 149 | 144 | 133 | 132 | 144 | 110 | 119 | 131 | 105 | 130 | 115 | 150 | 140 |
| CL173 | 116 | 132 | 97 | 124 | 115 | 117 | 131 | 107 | 132 | 149 | 101 | 82 | 136 |
| CL175 | 121 | 115 | 93 | 109 | 107 | 109 | 109 | 86 | 108 | 134 | 104 | 112 | 116 |
| CL177 | 86 | 100 | 102 | 79 | 111 | 98 | 102 | 85 | 97 | 98 | 107 | 104 | 97 |
| CL182 | 93 | 88 | 69 | 83 | 79 | 97 | 74 | 83 | 71 | 70 | 81 | 87 | 84 |
| CL186 | 93 | 76 | 76 | 82 | 83 | 83 | 75 | 73 | 66 | 73 | 81 | 93 | 79 |
| CL187 | 61 | 65 | 69 | 72 | 80 | 80 | 75 | 86 | 80 | 76 | 84 | 59 | 78 |
| CL189 | 77 | 56 | 61 | 59 | 52 | 77 | 52 | 68 | 60 | 65 | 84 | 63 | 57 |
| CL191 | 54 | 58 | 69 | 62 | 59 | 64 | 70 | 59 | 77 | 55 | 55 | 49 | 67 |
| CL194 | 55 | 42 | 51 | 52 | 51 | 66 | 40 | 73 | 56 | 46 | 62 | 50 | 49 |
| CL195 | 51 | 40 | 38 | 60 | 50 | 51 | 49 | 44 | 50 | 50 | 38 | 37 | 44 |
| CL197 | 32 | 47 | 47 | 45 | 41 | 34 | 46 | 45 | 37 | 48 | 46 | 48 | 47 |
| CL199 | 38 | 44 | 40 | 29 | 42 | 42 | 38 | 41 | 35 | 34 | 43 | 38 | 32 |
| CL200 | 34 | 34 | 41 | 57 | 43 | 46 | 46 | 29 | 37 | 36 | 41 | 39 | 41 |
| CL202 | 41 | 32 | 40 | 39 | 32 | 31 | 39 | 38 | 35 | 43 | 32 | 34 | 30 |
| CL203 | 37 | 50 | 27 | 40 | 34 | 43 | 43 | 36 | 32 | 35 | 35 | 29 | 52 |
| CL204 | 37 | 39 | 35 | 30 | 42 | 34 | 44 | 31 | 41 | 34 | 43 | 30 | 39 |
| CL205 | 47 | 30 | 31 | 30 | 32 | 33 | 29 | 33 | 27 | 27 | 31 | 36 | 36 |
| CL206 | 38 | 28 | 23 | 36 | 37 | 30 | 25 | 29 | 32 | 24 | 32 | 38 | 33 |
| CL209 | 17 | 25 | 33 | 28 | 17 | 35 | 35 | 35 | 30 | 33 | 27 | 23 | 34 |
| CL211 | 29 | 37 | 24 | 27 | 34 | 16 | 26 | 25 | 24 | 31 | 23 | 23 | 30 |
| CL212 | 29 | 33 | 24 | 37 | 23 | 28 | 20 | 20 | 30 | 30 | 27 | 23 | 31 |
| CL213 | 24 | 18 | 23 | 34 | 28 | 25 | 23 | 30 | 21 | 33 | 19 | 27 | 20 |
| CL214 | 27 | 21 | 22 | 32 | 20 | 10 | 26 | 15 | 17 | 21 | 26 | 24 | 26 |
| CL215 | 24 | 26 | 18 | 21 | 23 | 23 | 20 | 31 | 22 | 25 | 19 | 22 | 17 |
| CL216 | 25 | 22 | 26 | 24 | 12 | 36 | 29 | 21 | 24 | 18 | 33 | 30 | 24 |
| CL218 | 21 | 20 | 24 | 21 | 20 | 17 | 25 | 15 | 14 | 27 | 20 | 18 | 31 |
| CL220 | 21 | 16 | 20 | 19 | 21 | 26 | 23 | 22 | 19 | 19 | 26 | 19 | 16 |
| CL221 | 24 | 28 | 19 | 14 | 10 | 20 | 18 | 14 | 18 | 29 | 23 | 18 | 15 |
| CL222 | 18 | 35 | 24 | 14 | 10 | 20 | 19 | 19 | 21 | 17 | 22 | 35 | 17 |
| CL223 | 13 | 25 | 18 | 15 | 12 | 16 | 19 | 21 | 15 | 16 | 15 | 17 | 16 |
| CL226 | 28 | 21 | 12 | 17 | 24 | 18 | 13 | 15 | 9 | 18 | 24 | 17 | 16 |
| CL228 | 15 | 21 | 19 | 19 | 11 | 17 | 14 | 9 | 14 | 11 | 17 | 13 | 18 |
| CL229 | 12 | 27 | 22 | 20 | 16 | 11 | 15 | 19 | 16 | 16 | 12 | 19 | 18 |
| CL231 | 16 | 14 | 13 | 22 | 13 | 11 | 18 | 11 | 12 | 15 | 16 | 15 | 12 |
| CL232 | 12 | 13 | 14 | 5 | 14 | 6 | 23 | 13 | 11 | 15 | 11 | 14 | 18 |
| CL234 | 16 | 15 | 13 | 10 | 16 | 14 | 8 | 10 | 9 | 17 | 12 | 11 | 13 |
| CL235 | 16 | 9 | 9 | 8 | 11 | 13 | 11 | 11 | 8 | 7 | 6 | 4 | 11 |
| CL236 | 12 | 14 | 14 | 14 | 10 | 11 | 11 | 5 | 11 | 17 | 11 | 12 | 9 |
| CL237 | 11 | 16 | 8 | 9 | 14 | 12 | 13 | 10 | 13 | 10 | 16 | 11 | 11 |
| CL239 | 7 | 14 | 8 | 9 | 10 | 10 | 10 | 11 | 9 | 12 | 11 | 6 | 11 |

**Supplementary Table 10**: Correlation between cluster abundance and the number of clusters whose abundance is correlated (column 1).

| | CL002 | CL003 | CL004 | CL005 | CL006 | CL007 | CL008 | CL009 | CL010 | CL011 | CL012 | CL013 | CL014 | CL015 | CL016 | CL017 | CL018 | CL019 | CL020 | CL021 | CL022 | CL023 | CL024 | CL025 | CL026 | CL027 | CL028 | CL029 |
|---|---|---|---|---|---|---|---|---|---|---|---|---|---|---|---|---|---|---|---|---|---|---|---|---|---|---|---|---|
| CL001 | 0.30 | 0.29 | | 0.44 | 0.41 | 0.43 | | 0.26 | 0.48 | 0.36 | | | 0.39 | 0.25 | | 0.39 | | | 0.28 | | | 0.36 | 0.27 | | 0.25 | | 0.48 | 0.52 |
| CL002 | | 0.86 | 0.70 | 0.71 | | | 0.68 | 0.64 | 0.77 | 0.82 | 0.50 | 0.72 | 0.80 | 0.81 | -0.58 | 0.82 | | | 0.61 | 0.63 | 0.26 | 0.90 | 0.58 | -0.30 | -0.34 | 0.58 | | |
| CL003 | | | 0.84 | 0.57 | -0.25 | | 0.86 | 0.68 | 0.84 | 0.74 | 0.48 | 0.87 | 0.73 | 0.90 | -0.69 | 0.92 | | | 0.77 | 0.79 | 0.41 | 0.87 | 0.75 | -0.50 | -0.40 | 0.71 | | |
| CL004 | | | | 0.29 | -0.51 | -0.24 | 0.97 | 0.80 | 0.82 | 0.45 | 0.53 | 0.97 | 0.47 | 0.93 | -0.73 | 0.81 | -0.34 | 0.33 | 0.78 | 0.92 | 0.62 | 0.70 | 0.82 | -0.65 | -0.50 | 0.85 | -0.50 | |
| CL005 | | | | | 0.37 | 0.59 | 0.30 | 0.48 | 0.48 | 0.67 | 0.55 | 0.34 | 0.71 | 0.39 | | 0.46 | 0.39 | | 0.45 | 0.25 | | 0.67 | 0.26 | 0.25 | | | 0.39 | 0.50 |
| CL006 | | | | | | 0.75 | -0.43 | | | | -0.26 | -0.46 | 0.34 | -0.37 | 0.38 | -0.28 | 0.54 | -0.38 | -0.40 | -0.59 | -0.55 | | -0.50 | 0.68 | 0.72 | -0.55 | 0.93 | 0.72 |
| CL007 | | | | | | | | | | | 0.27 | | 0.28 | | 0.45 | | 0.76 | | | -0.29 | | | | 0.73 | 0.80 | -0.32 | 0.74 | 0.77 |
| CL008 | | | | | | | | 0.79 | 0.81 | 0.50 | 0.48 | 0.97 | 0.52 | 0.92 | -0.75 | 0.80 | -0.35 | 0.30 | 0.76 | 0.88 | 0.57 | 0.72 | 0.79 | -0.63 | -0.46 | 0.82 | -0.44 | |
| CL009 | | | | | | | | | 0.68 | 0.41 | 0.65 | 0.81 | 0.45 | 0.75 | -0.42 | 0.61 | | 0.50 | 0.75 | 0.74 | 0.64 | 0.62 | 0.71 | | | 0.70 | | |
| CL010 | | | | | | | | | | 0.58 | 0.40 | 0.80 | 0.60 | 0.87 | -0.66 | 0.88 | -0.25 | | 0.73 | 0.80 | 0.48 | 0.79 | 0.77 | -0.51 | -0.37 | 0.74 | | |
| CL011 | | | | | | | | | | | | 0.49 | 0.94 | 0.63 | -0.56 | 0.68 | | -0.25 | 0.29 | 0.37 | | 0.87 | 0.36 | | | 0.38 | 0.25 | |
| CL012 | | | | | | | | | | | | 0.54 | | 0.42 | | 0.36 | 0.34 | 0.37 | 0.70 | 0.54 | 0.48 | 0.32 | 0.49 | | | 0.42 | | |
| CL013 | | | | | | | | | | | | | 0.51 | 0.92 | -0.72 | 0.81 | -0.33 | 0.29 | 0.79 | 0.90 | 0.58 | 0.73 | 0.79 | -0.62 | -0.47 | 0.83 | -0.46 | |
| CL014 | | | | | | | | | | | | | | 0.62 | -0.52 | 0.67 | | -0.28 | 0.33 | 0.34 | | 0.90 | 0.32 | | | 0.33 | 0.32 | 0.30 |
| CL015 | | | | | | | | | | | | | | | -0.77 | 0.88 | -0.36 | | 0.72 | 0.87 | 0.55 | 0.81 | 0.79 | -0.60 | -0.51 | 0.82 | -0.35 | |
| CL016 | | | | | | | | | | | | | | | | -0.67 | 0.65 | | -0.37 | -0.62 | -0.28 | -0.66 | -0.55 | 0.72 | 0.72 | -0.65 | 0.40 | 0.29 |
| CL017 | | | | | | | | | | | | | | | | | -0.31 | | 0.71 | 0.77 | 0.41 | 0.83 | 0.75 | -0.57 | -0.46 | 0.71 | -0.25 | |
| CL018 | | | | | | | | | | | | | | | | | | | | -0.36 | | | -0.29 | 0.68 | 0.77 | -0.46 | 0.53 | 0.54 |
| CL019 | | | | | | | | | | | | | | | | | | | 0.41 | 0.46 | 0.77 | | 0.43 | | | 0.48 | -0.28 | |
| CL020 | | | | | | | | | | | | | | | | | | | | 0.81 | 0.66 | 0.55 | 0.80 | -0.39 | | 0.71 | -0.33 | |
| CL021 | | | | | | | | | | | | | | | | | | | | | 0.73 | 0.60 | 0.86 | -0.62 | -0.51 | 0.90 | -0.53 | -0.25 |
| CL022 | | | | | | | | | | | | | | | | | | | | | | | 0.68 | -0.28 | -0.26 | 0.73 | -0.42 | |
| CL023 | | | | | | | | | | | | | | | | | | | | | | | 0.55 | -0.34 | -0.32 | 0.57 | | |
| CL024 | | | | | | | | | | | | | | | | | | | | | | | | -0.53 | -0.40 | 0.81 | -0.44 | |
| CL025 | | | | | | | | | | | | | | | | | | | | | | | | | 0.77 | -0.58 | 0.71 | 0.67 |
| CL026 | | | | | | | | | | | | | | | | | | | | | | | | | | -0.55 | 0.69 | 0.61 |
| CL027 | | | | | | | | | | | | | | | | | | | | | | | | | | | -0.49 | |
| CL028 | | | | | | | | | | | | | | | | | | | | | | | | | | | | 0.74 |
| CL029 | | | | | | | | | | | | | | | | | | | | | | | | | | | | |
| CL030 | | | | | | | | | | | | | | | | | | | | | | | | | | | | |
| CL031 | | | | | | | | | | | | | | | | | | | | | | | | | | | | |
| CL032 | | | | | | | | | | | | | | | | | | | | | | | | | | | | |
| CL033 | | | | | | | | | | | | | | | | | | | | | | | | | | | | |
| CL034 | | | | | | | | | | | | | | | | | | | | | | | | | | | | |
| CL035 | | | | | | | | | | | | | | | | | | | | | | | | | | | | |
| CL036 | | | | | | | | | | | | | | | | | | | | | | | | | | | | |
| CL037 | | | | | | | | | | | | | | | | | | | | | | | | | | | | |
| CL038 | | | | | | | | | | | | | | | | | | | | | | | | | | | | |
| CL039 | | | | | | | | | | | | | | | | | | | | | | | | | | | | |
| CL040 | | | | | | | | | | | | | | | | | | | | | | | | | | | | |
| CL041 | | | | | | | | | | | | | | | | | | | | | | | | | | | | |
| CL042 | | | | | | | | | | | | | | | | | | | | | | | | | | | | |
| CL043 | | | | | | | | | | | | | | | | | | | | | | | | | | | | |
| CL044 | | | | | | | | | | | | | | | | | | | | | | | | | | | | |
| CL045 | | | | | | | | | | | | | | | | | | | | | | | | | | | | |
| CL046 | | | | | | | | | | | | | | | | | | | | | | | | | | | | |
| CL047 | | | | | | | | | | | | | | | | | | | | | | | | | | | | |
| CL048 | | | | | | | | | | | | | | | | | | | | | | | | | | | | |
| CL049 | | | | | | | | | | | | | | | | | | | | | | | | | | | | |
| CL050 | | | | | | | | | | | | | | | | | | | | | | | | | | | | |

| | CL030 | CL031 | CL032 | CL033 | CL034 | CL035 | CL036 | CL037 | CL038 | CL039 | CL040 | CL041 | CL042 | CL043 | CL044 | CL045 | CL046 | CL047 | CL048 | CL049 | CL050 | CL051 | CL052 | CL053 | CL054 | CL055 | CL056 | CL057 |
|---|---|---|---|---|---|---|---|---|---|---|---|---|---|---|---|---|---|---|---|---|---|---|---|---|---|---|---|---|
| CL001 | 0.78 | | 0.56 | 0.29 | 0.40 | 0.28 | | 0.28 | | 0.53 | | | 0.27 | 0.53 | 0.29 | | | 0.45 | 0.46 | 0.57 | 0.46 | | 0.55 | | | 0.32 | | |
| CL002 | 0.54 | | 0.66 | -0.25 | 0.90 | | -0.30 | | -0.31 | 0.30 | 0.45 | -0.31 | 0.26 | | 0.29 | | 0.68 | 0.73 | | | -0.49 | | | 0.60 | 0.78 | | -0.40 | 0.69 |
| CL003 | 0.48 | | 0.79 | -0.45 | 0.86 | | -0.31 | 0.32 | | 0.33 | 0.56 | -0.40 | 0.33 | 0.47 | 0.37 | | 0.85 | 0.61 | | | -0.60 | | -0.33 | 0.60 | 0.91 | | -0.35 | 0.83 |
| CL004 | | 0.33 | 0.73 | -0.42 | 0.68 | | -0.48 | | | 0.26 | 0.73 | -0.44 | 0.41 | 0.51 | 0.52 | -0.40 | 0.95 | 0.29 | -0.38 | | -0.66 | -0.27 | -0.52 | 0.38 | 0.83 | | -0.43 | 0.93 |
| CL005 | 0.69 | | 0.46 | | 0.63 | 0.53 | 0.24 | 0.28 | | 0.46 | 0.27 | | 0.27 | | | 0.32 | | 0.70 | 0.51 | 0.54 | | | 0.30 | 0.67 | 0.53 | 0.45 | | 0.24 |
| CL006 | 0.48 | -0.45 | | 0.32 | | 0.34 | 0.74 | | | | -0.42 | 0.39 | | -0.30 | -0.44 | 0.79 | -0.59 | 0.46 | 0.81 | 0.60 | 0.56 | 0.24 | 0.82 | 0.34 | -0.24 | 0.39 | 0.45 | -0.62 |
| CL007 | 0.54 | | | 0.53 | | 0.66 | 0.73 | 0.31 | | 0.38 | | 0.66 | 0.34 | | | 0.55 | -0.36 | 0.33 | 0.68 | 0.75 | 0.36 | 0.35 | 0.69 | 0.42 | | 0.64 | 0.51 | -0.35 |
| CL008 | 0.25 | 0.29 | 0.71 | -0.47 | 0.71 | | -0.41 | 0.26 | | | 0.70 | -0.46 | 0.37 | 0.51 | 0.47 | -0.31 | 0.93 | 0.34 | -0.35 | | -0.66 | -0.25 | -0.51 | 0.44 | 0.84 | -0.27 | -0.38 | 0.90 |
| CL009 | 0.38 | 0.42 | 0.63 | | 0.57 | | | 0.32 | 0.28 | 0.44 | 0.81 | | 0.60 | 0.48 | 0.59 | | 0.67 | 0.29 | | 0.27 | -0.36 | | | 0.42 | 0.68 | | | 0.69 |
| CL010 | 0.52 | | 0.86 | -0.29 | 0.79 | | -0.41 | 0.24 | | 0.39 | 0.56 | -0.43 | 0.37 | 0.57 | 0.47 | | 0.81 | 0.51 | | | -0.37 | -0.26 | | 0.36 | 0.78 | | -0.47 | 0.77 |
| CL011 | 0.62 | -0.27 | 0.47 | -0.38 | 0.89 | | | | -0.42 | | | -0.40 | | | | 0.31 | 0.44 | 0.89 | 0.43 | | -0.30 | | | 0.68 | 0.67 | | -0.25 | 0.43 |
| CL012 | | 0.34 | 0.45 | | 0.28 | 0.43 | | 0.35 | 0.29 | 0.48 | 0.65 | 0.35 | 0.55 | 0.28 | 0.43 | | 0.44 | | | 0.25 | -0.47 | | | 0.39 | 0.50 | 0.37 | | 0.51 |
| CL013 | 0.25 | 0.29 | 0.71 | -0.42 | 0.71 | | -0.42 | 0.25 | | 0.27 | 0.73 | -0.41 | 0.39 | 0.50 | 0.49 | -0.35 | 0.93 | 0.34 | -0.34 | | -0.68 | -0.25 | -0.50 | 0.44 | 0.84 | | -0.38 | 0.92 |
| CL014 | 0.67 | -0.31 | 0.50 | -0.34 | 0.88 | | | | -0.40 | | | -0.30 | | | | 0.35 | 0.43 | 0.92 | 0.46 | 0.33 | -0.28 | | | 0.75 | 0.67 | | | 0.41 |
| CL015 | 0.37 | 0.24 | 0.76 | -0.42 | 0.82 | | -0.48 | | | 0.28 | 0.66 | -0.51 | 0.33 | 0.51 | 0.46 | -0.29 | 0.90 | 0.48 | | | -0.57 | -0.29 | -0.39 | 0.43 | 0.85 | | -0.50 | 0.88 |
| CL016 | | | -0.46 | 0.64 | -0.69 | 0.43 | 0.62 | | 0.35 | | -0.32 | 0.78 | | -0.31 | | 0.24 | -0.75 | -0.39 | 0.30 | 0.31 | 0.51 | 0.47 | 0.49 | -0.26 | -0.62 | 0.53 | 0.59 | -0.72 |
| CL017 | 0.50 | | 0.84 | -0.43 | 0.85 | | -0.43 | 0.28 | | 0.31 | 0.50 | -0.50 | 0.24 | 0.51 | 0.39 | | 0.84 | 0.58 | | | -0.49 | -0.28 | -0.29 | 0.46 | 0.83 | | -0.50 | 0.81 |
| CL018 | | | | 0.58 | -0.25 | 0.64 | 0.71 | | | 0.28 | | 0.80 | 0.34 | -0.25 | | 0.41 | -0.43 | | 0.45 | 0.54 | 0.25 | 0.40 | 0.51 | 0.27 | | 0.69 | 0.59 | -0.41 |
| CL019 | | 0.82 | 0.25 | 0.27 | | | | 0.31 | 0.77 | 0.43 | 0.68 | | 0.51 | 0.56 | 0.61 | -0.31 | 0.24 | -0.41 | -0.30 | | | 0.28 | | | | | | 0.29 |
| CL020 | 0.36 | 0.44 | 0.81 | | 0.52 | | -0.24 | 0.42 | 0.29 | 0.55 | 0.73 | | 0.60 | 0.63 | 0.62 | -0.35 | 0.76 | | -0.25 | | -0.50 | | -0.34 | 0.38 | 0.77 | | | 0.77 |
| CL021 | | 0.49 | 0.76 | -0.33 | 0.59 | | -0.52 | 0.26 | | 0.39 | 0.76 | -0.42 | 0.41 | 0.65 | 0.60 | -0.48 | 0.90 | | -0.40 | | -0.56 | | -0.48 | | 0.78 | | -0.42 | 0.91 |
| CL022 | | 0.76 | 0.53 | | | | -0.37 | 0.29 | 0.54 | 0.46 | 0.77 | | 0.52 | 0.69 | 0.73 | -0.44 | 0.55 | | -0.37 | | | | -0.28 | | 0.43 | | | 0.60 |
| CL023 | 0.59 | | 0.67 | -0.37 | 0.94 | | -0.25 | | -0.32 | 0.24 | 0.41 | -0.42 | | 0.26 | | | 0.68 | 0.80 | | | -0.42 | -0.29 | | 0.63 | 0.80 | | -0.41 | 0.66 |
| CL024 | 0.30 | 0.48 | 0.78 | -0.26 | 0.57 | | -0.43 | 0.37 | 0.27 | 0.47 | 0.70 | -0.38 | 0.46 | 0.67 | 0.60 | -0.42 | 0.81 | | -0.31 | | -0.48 | | -0.43 | | 0.74 | | -0.31 | 0.80 |
| CL025 | | | -0.41 | 0.70 | -0.37 | 0.59 | 0.69 | | | | -0.27 | 0.73 | | -0.30 | | 0.53 | -0.75 | | 0.63 | 0.54 | 0.62 | 0.41 | 0.72 | | -0.50 | 0.62 | 0.55 | -0.71 |
| CL026 | 0.28 | | | 0.57 | -0.36 | 0.52 | 0.88 | 0.34 | 0.39 | 0.25 | | 0.76 | 0.26 | | | 0.50 | -0.59 | | 0.54 | 0.61 | 0.48 | 0.45 | 0.65 | | -0.33 | 0.60 | 0.73 | -0.60 |
| CL027 | | 0.50 | 0.69 | -0.29 | 0.59 | | -0.57 | | | 0.35 | 0.73 | -0.49 | 0.35 | 0.66 | 0.57 | -0.42 | 0.80 | | -0.35 | | -0.44 | | -0.43 | | 0.68 | | -0.48 | 0.83 |
| CL028 | 0.54 | -0.37 | | 0.40 | | 0.44 | 0.69 | | | | -0.38 | 0.38 | | | -0.33 | 0.74 | -0.58 | 0.45 | 0.85 | 0.64 | 0.61 | 0.30 | 0.86 | 0.28 | | 0.46 | 0.41 | -0.58 |
| CL029 | 0.51 | | | 0.53 | | 0.56 | 0.55 | | | 0.30 | | 0.45 | 0.24 | | | 0.54 | -0.36 | 0.34 | 0.68 | 0.68 | 0.50 | 0.27 | 0.75 | 0.26 | | 0.51 | 0.31 | -0.34 |
| CL030 | | | 0.57 | | 0.61 | 0.35 | 0.25 | 0.41 | | 0.50 | | | 0.26 | 0.37 | | 0.31 | | 0.69 | 0.57 | 0.66 | | | 0.48 | 0.50 | 0.41 | 0.33 | | |
| CL031 | | | 0.34 | | | | | 0.33 | 0.73 | 0.44 | 0.60 | | 0.46 | 0.64 | 0.65 | -0.38 | 0.27 | -0.43 | -0.38 | | | | | -0.29 | | | | 0.30 |
| CL032 | | | | | 0.66 | | -0.28 | 0.41 | | 0.54 | 0.59 | -0.26 | 0.45 | 0.67 | 0.49 | -0.25 | 0.73 | 0.41 | | 0.25 | -0.32 | | | 0.34 | 0.77 | | -0.34 | 0.71 |
| CL033 | | | | | -0.42 | 0.59 | 0.37 | | 0.43 | 0.29 | | 0.64 | 0.25 | | | | -0.53 | -0.26 | 0.32 | 0.41 | 0.62 | 0.31 | 0.57 | -0.26 | -0.46 | 0.62 | 0.35 | -0.48 |
| CL034 | | | | | | | -0.27 | | -0.35 | | 0.38 | -0.47 | | 0.28 | | | 0.67 | 0.83 | | | -0.41 | -0.25 | | 0.61 | 0.79 | | -0.40 | 0.66 |
| CL035 | | | | | | | 0.42 | 0.24 | 0.29 | 0.47 | | 0.60 | 0.38 | | | 0.28 | | | 0.50 | 0.54 | | 0.38 | 0.49 | | | 0.65 | 0.33 | |
| CL036 | | | | | | | | 0.41 | 0.25 | | -0.24 | 0.71 | | | -0.32 | 0.67 | -0.55 | | 0.51 | 0.51 | 0.32 | 0.53 | 0.59 | 0.36 | -0.25 | 0.47 | 0.81 | -0.56 |
| CL037 | | | | | | | | | 0.49 | 0.45 | 0.34 | | 0.39 | 0.44 | | | | | | 0.37 | | 0.43 | | 0.46 | 0.39 | | 0.43 | |
| CL038 | | | | | | | | | | 0.47 | 0.43 | 0.39 | 0.50 | 0.53 | 0.48 | | | -0.48 | | 0.26 | | 0.42 | | | | 0.25 | 0.36 | |
| CL039 | | | | | | | | | | | 0.51 | 0.24 | 0.51 | 0.50 | 0.51 | | | | | 0.47 | | 0.25 | | | 0.33 | 0.44 | | |
| CL040 | | | | | | | | | | | | | 0.52 | 0.57 | 0.64 | -0.35 | 0.64 | | -0.28 | | -0.37 | | -0.31 | | 0.58 | | | 0.67 |
| CL041 | | | | | | | | | | | | | 0.30 | | | 0.28 | -0.51 | -0.24 | 0.29 | 0.44 | 0.26 | 0.46 | 0.41 | | -0.36 | 0.65 | 0.65 | -0.48 |
| CL042 | | | | | | | | | | | | | | 0.44 | 0.47 | | 0.30 | | | 0.34 | | | | | 0.36 | 0.33 | | 0.31 |
| CL043 | | | | | | | | | | | | | | | 0.52 | -0.28 | 0.50 | | | | | | | | 0.42 | | | 0.50 |
| CL044 | | | | | | | | | | | | | | | | -0.40 | 0.46 | | -0.29 | | | | | | 0.36 | | | 0.46 |
| CL045 | | | | | | | | | | | | | | | | | -0.45 | 0.42 | 0.65 | 0.38 | 0.31 | 0.40 | 0.54 | 0.43 | | 0.25 | 0.52 | -0.44 |
| CL046 | | | | | | | | | | | | | | | | | | 0.27 | -0.46 | | -0.72 | -0.31 | -0.62 | 0.34 | 0.83 | -0.32 | -0.47 | 0.95 |
| CL047 | | | | | | | | | | | | | | | | | | | 0.55 | 0.36 | | | 0.29 | 0.68 | 0.54 | | | 0.25 |
| CL048 | | | | | | | | | | | | | | | | | | | | 0.60 | 0.47 | | 0.75 | 0.32 | | 0.48 | 0.24 | -0.44 |
| CL049 | | | | | | | | | | | | | | | | | | | | | 0.35 | 0.32 | 0.60 | 0.36 | | 0.52 | 0.33 | |
| CL050 | | | | | | | | | | | | | | | | | | | | | | | 0.76 | -0.45 | -0.63 | 0.32 | | -0.70 |

| | CL030 | CL031 | CL032 | CL033 | CL034 | CL035 | CL036 | CL037 | CL038 | CL039 | CL040 | CL041 | CL042 | CL043 | CL044 | CL045 | CL046 | CL047 | CL048 | CL049 | CL050 | CL051 | CL052 | CL053 | CL054 | CL055 | CL056 | CL057 |
|---|---|---|---|---|---|---|---|---|---|---|---|---|---|---|---|---|---|---|---|---|---|---|---|---|---|---|---|---|
| CL051 | | | | | | | | | | | | | | | | | | | | | | | 0.24 | | | 0.34 | 0.58 | -0.29 |
| CL052 | | | | | | | | | | | | | | | | | | | | | | | | | -0.33 | 0.50 | 0.30 | -0.61 |
| CL053 | | | | | | | | | | | | | | | | | | | | | | | | | 0.64 | | | 0.32 |
| CL054 | | | | | | | | | | | | | | | | | | | | | | | | | | | -0.24 | 0.80 |
| CL055 | | | | | | | | | | | | | | | | | | | | | | | | | | | 0.35 | -0.27 |
| CL056 | | | | | | | | | | | | | | | | | | | | | | | | | | | | -0.48 |
| CL057 | | | | | | | | | | | | | | | | | | | | | | | | | | | | |
| CL058 | | | | | | | | | | | | | | | | | | | | | | | | | | | | |
| CL059 | | | | | | | | | | | | | | | | | | | | | | | | | | | | |
| CL060 | | | | | | | | | | | | | | | | | | | | | | | | | | | | |
| CL061 | | | | | | | | | | | | | | | | | | | | | | | | | | | | |
| CL062 | | | | | | | | | | | | | | | | | | | | | | | | | | | | |
| CL063 | | | | | | | | | | | | | | | | | | | | | | | | | | | | |
| CL064 | | | | | | | | | | | | | | | | | | | | | | | | | | | | |
| CL065 | | | | | | | | | | | | | | | | | | | | | | | | | | | | |
| CL066 | | | | | | | | | | | | | | | | | | | | | | | | | | | | |
| CL067 | | | | | | | | | | | | | | | | | | | | | | | | | | | | |
| CL068 | | | | | | | | | | | | | | | | | | | | | | | | | | | | |
| CL069 | | | | | | | | | | | | | | | | | | | | | | | | | | | | |
| CL070 | | | | | | | | | | | | | | | | | | | | | | | | | | | | |
| CL071 | | | | | | | | | | | | | | | | | | | | | | | | | | | | |
| CL072 | | | | | | | | | | | | | | | | | | | | | | | | | | | | |
| CL073 | | | | | | | | | | | | | | | | | | | | | | | | | | | | |
| CL074 | | | | | | | | | | | | | | | | | | | | | | | | | | | | |
| CL075 | | | | | | | | | | | | | | | | | | | | | | | | | | | | |
| CL076 | | | | | | | | | | | | | | | | | | | | | | | | | | | | |
| CL077 | | | | | | | | | | | | | | | | | | | | | | | | | | | | |
| CL078 | | | | | | | | | | | | | | | | | | | | | | | | | | | | |
| CL079 | | | | | | | | | | | | | | | | | | | | | | | | | | | | |
| CL080 | | | | | | | | | | | | | | | | | | | | | | | | | | | | |
| CL081 | | | | | | | | | | | | | | | | | | | | | | | | | | | | |
| CL082 | | | | | | | | | | | | | | | | | | | | | | | | | | | | |
| CL083 | | | | | | | | | | | | | | | | | | | | | | | | | | | | |
| CL084 | | | | | | | | | | | | | | | | | | | | | | | | | | | | |
| CL085 | | | | | | | | | | | | | | | | | | | | | | | | | | | | |
| CL086 | | | | | | | | | | | | | | | | | | | | | | | | | | | | |
| CL087 | | | | | | | | | | | | | | | | | | | | | | | | | | | | |
| CL088 | | | | | | | | | | | | | | | | | | | | | | | | | | | | |
| CL089 | | | | | | | | | | | | | | | | | | | | | | | | | | | | |
| CL090 | | | | | | | | | | | | | | | | | | | | | | | | | | | | |
| CL091 | | | | | | | | | | | | | | | | | | | | | | | | | | | | |
| CL092 | | | | | | | | | | | | | | | | | | | | | | | | | | | | |
| CL093 | | | | | | | | | | | | | | | | | | | | | | | | | | | | |
| CL094 | | | | | | | | | | | | | | | | | | | | | | | | | | | | |
| CL095 | | | | | | | | | | | | | | | | | | | | | | | | | | | | |
| CL096 | | | | | | | | | | | | | | | | | | | | | | | | | | | | |
| CL097 | | | | | | | | | | | | | | | | | | | | | | | | | | | | |
| CL098 | | | | | | | | | | | | | | | | | | | | | | | | | | | | |
| CL099 | | | | | | | | | | | | | | | | | | | | | | | | | | | | |
| CL100 | | | | | | | | | | | | | | | | | | | | | | | | | | | | |
| CL101 | | | | | | | | | | | | | | | | | | | | | | | | | | | | |
| CL102 | | | | | | | | | | | | | | | | | | | | | | | | | | | | |
| CL103 | | | | | | | | | | | | | | | | | | | | | | | | | | | | |
| CL104 | | | | | | | | | | | | | | | | | | | | | | | | | | | | |
| CL105 | | | | | | | | | | | | | | | | | | | | | | | | | | | | |
| CL106 | | | | | | | | | | | | | | | | | | | | | | | | | | | | |
| CL107 | | | | | | | | | | | | | | | | | | | | | | | | | | | | |
| CL108 | | | | | | | | | | | | | | | | | | | | | | | | | | | | |
| CL109 | | | | | | | | | | | | | | | | | | | | | | | | | | | | |
| CL110 | | | | | | | | | | | | | | | | | | | | | | | | | | | | |

| | CL058 | CL059 | CL060 | CL061 | CL062 | CL063 | CL064 | CL065 | CL066 | CL067 | CL068 | CL069 | CL070 | CL071 | CL072 | CL073 | CL074 | CL075 | CL076 | CL077 | CL078 | CL079 | CL080 | CL081 | CL082 | CL083 | CL084 | CL085 |
|---|---|---|---|---|---|---|---|---|---|---|---|---|---|---|---|---|---|---|---|---|---|---|---|---|---|---|---|---|
| CL001 | 0.26 | 0.28 | 0.28 | | 0.60 | -0.30 | 0.41 | 0.33 | | | | | 0.26 | | | | | 0.39 | | -0.36 | | 0.38 | | | 0.68 | | | |
| CL002 | 0.82 | 0.28 | | | | -0.55 | | 0.37 | 0.50 | 0.74 | -0.31 | | | -0.39 | 0.81 | | | 0.29 | -0.52 | | | | -0.34 | 0.58 | 0.44 | | | |
| CL003 | 0.76 | 0.28 | -0.28 | | | -0.45 | 0.39 | 0.53 | 0.59 | 0.87 | | | | -0.38 | 0.88 | | | 0.45 | -0.48 | | | | -0.27 | 0.73 | 0.55 | | | |
| CL004 | 0.52 | 0.25 | -0.48 | | -0.38 | -0.44 | 0.53 | 0.68 | 0.52 | 0.85 | -0.26 | 0.33 | | -0.54 | 0.85 | | -0.30 | 0.59 | -0.52 | | | -0.35 | -0.30 | 0.73 | 0.48 | 0.29 | | |
| CL005 | 0.75 | 0.34 | 0.34 | | 0.42 | | | | 0.52 | 0.36 | | -0.26 | | | 0.51 | -0.31 | 0.38 | | | -0.36 | | 0.36 | | 0.45 | 0.29 | -0.26 | 0.42 | |
| CL006 | | | 0.73 | 0.25 | 0.80 | | -0.37 | -0.55 | | -0.49 | 0.48 | -0.40 | | 0.73 | -0.32 | -0.41 | 0.56 | -0.50 | 0.46 | -0.34 | -0.31 | 0.66 | 0.25 | -0.25 | | -0.43 | | |
| CL007 | 0.28 | | 0.79 | 0.54 | 0.72 | 0.34 | | | 0.24 | -0.29 | 0.52 | -0.58 | | 0.57 | | -0.67 | 0.70 | | 0.50 | -0.65 | | 0.75 | 0.35 | | | -0.60 | 0.47 | |
| CL008 | 0.56 | | -0.45 | | -0.34 | -0.42 | 0.50 | 0.61 | 0.53 | 0.84 | | 0.30 | | -0.46 | 0.84 | | -0.27 | 0.56 | -0.47 | | | -0.32 | -0.27 | 0.75 | 0.44 | 0.25 | | |
| CL009 | 0.52 | 0.33 | | | | -0.30 | 0.46 | 0.56 | 0.54 | 0.60 | | | 0.24 | -0.34 | 0.73 | -0.24 | | 0.52 | -0.28 | -0.33 | 0.36 | | | 0.71 | 0.47 | | 0.39 | |
| CL010 | 0.59 | 0.28 | -0.27 | | | -0.56 | 0.58 | 0.63 | 0.38 | 0.72 | -0.26 | | | -0.49 | 0.79 | | | 0.64 | -0.60 | | | | -0.32 | 0.56 | 0.70 | 0.25 | | |
| CL011 | 0.85 | | | -0.26 | | -0.54 | | | 0.37 | 0.56 | | | | | 0.62 | | | | -0.38 | | -0.33 | | | 0.43 | 0.32 | | | |
| CL012 | 0.40 | 0.42 | | 0.32 | | | 0.24 | 0.43 | 0.72 | 0.50 | | -0.25 | 0.35 | | 0.56 | -0.42 | 0.25 | 0.35 | | -0.46 | 0.51 | | | 0.64 | | -0.25 | 0.50 | |
| CL013 | 0.57 | 0.25 | -0.44 | | -0.35 | -0.41 | 0.49 | 0.63 | 0.55 | 0.85 | | 0.28 | | -0.49 | 0.86 | | -0.26 | 0.57 | -0.47 | | | -0.31 | -0.29 | 0.76 | 0.46 | 0.24 | | |
| CL014 | 0.86 | | | | 0.29 | -0.47 | | | 0.38 | 0.51 | | | | | 0.62 | | | | -0.29 | | -0.33 | | | 0.46 | 0.34 | | | |
| CL015 | 0.62 | 0.26 | -0.43 | -0.26 | -0.26 | -0.56 | 0.50 | 0.63 | 0.46 | 0.85 | -0.28 | 0.37 | | -0.52 | 0.88 | | -0.30 | 0.55 | -0.60 | | | -0.33 | -0.32 | 0.67 | 0.53 | 0.35 | | |
| CL016 | -0.45 | | 0.61 | 0.58 | 0.38 | 0.72 | -0.30 | -0.41 | | -0.68 | 0.44 | -0.59 | | 0.50 | -0.59 | -0.50 | 0.54 | -0.34 | 0.69 | -0.44 | 0.27 | 0.52 | 0.49 | -0.36 | -0.31 | -0.55 | | |
| CL017 | 0.67 | 0.27 | -0.35 | | | -0.54 | 0.49 | 0.57 | 0.46 | 0.80 | -0.30 | 0.26 | | -0.49 | 0.82 | | -0.30 | 0.54 | -0.60 | | | | -0.37 | 0.63 | 0.67 | 0.31 | | |
| CL018 | | | 0.75 | 0.64 | 0.53 | 0.58 | -0.27 | -0.31 | 0.24 | -0.36 | 0.49 | -0.60 | | 0.51 | | -0.66 | 0.67 | -0.28 | 0.61 | -0.62 | | 0.62 | 0.38 | | | -0.61 | 0.53 | |
| CL019 | | 0.42 | | | | | 0.55 | 0.57 | | | | | 0.41 | | | -0.36 | | 0.57 | | -0.45 | 0.70 | | 0.31 | 0.33 | | | | 0.32 |
| CL020 | 0.49 | 0.40 | | | | | 0.58 | 0.69 | 0.64 | 0.75 | | | 0.27 | -0.40 | 0.78 | -0.24 | | 0.63 | -0.29 | -0.36 | 0.45 | | | 0.75 | 0.54 | | 0.33 | |
| CL021 | 0.40 | 0.34 | -0.48 | | -0.35 | -0.42 | 0.64 | 0.79 | 0.48 | 0.83 | -0.28 | 0.30 | 0.28 | -0.62 | 0.79 | | -0.29 | 0.71 | -0.53 | | 0.34 | -0.35 | | 0.67 | 0.54 | 0.29 | | |
| CL022 | | 0.46 | -0.30 | | -0.28 | | 0.67 | 0.77 | 0.34 | 0.47 | | | 0.46 | -0.49 | 0.48 | | | 0.74 | -0.31 | -0.25 | 0.58 | | | 0.47 | 0.43 | | 0.24 | |
| CL023 | 0.82 | | | -0.29 | | -0.58 | | 0.31 | 0.41 | 0.70 | -0.28 | 0.27 | | -0.31 | 0.78 | | | 0.24 | -0.50 | | | | -0.36 | 0.55 | 0.47 | 0.25 | | |
| CL024 | 0.41 | 0.40 | -0.39 | | -0.25 | -0.33 | 0.64 | 0.76 | 0.48 | 0.76 | | | | -0.52 | 0.78 | | | 0.68 | -0.48 | | 0.36 | | | 0.65 | 0.56 | | | |
| CL025 | | | 0.73 | 0.43 | 0.60 | 0.43 | -0.37 | -0.45 | | -0.64 | 0.45 | -0.52 | | 0.57 | -0.49 | -0.51 | 0.66 | -0.37 | 0.57 | -0.46 | | 0.66 | 0.43 | -0.31 | -0.27 | -0.52 | 0.27 | |
| CL026 | | | 0.81 | 0.70 | 0.68 | 0.64 | | -0.36 | | -0.53 | 0.70 | -0.69 | | 0.68 | -0.36 | -0.77 | 0.70 | -0.30 | 0.74 | -0.70 | 0.26 | 0.77 | 0.51 | | | -0.73 | 0.40 | |
| CL027 | 0.35 | 0.36 | -0.51 | -0.30 | -0.37 | -0.53 | 0.60 | 0.74 | 0.34 | 0.73 | -0.30 | 0.40 | | -0.60 | 0.72 | | -0.30 | 0.68 | -0.59 | | 0.26 | -0.38 | | 0.54 | 0.49 | 0.40 | | |
| CL028 | | | 0.73 | 0.24 | 0.78 | | -0.33 | -0.44 | | -0.45 | 0.43 | -0.43 | | 0.65 | -0.29 | -0.40 | 0.61 | -0.39 | 0.39 | -0.36 | -0.26 | 0.69 | 0.25 | -0.25 | | -0.44 | | |
| CL029 | | | 0.69 | 0.25 | 0.69 | | | | | -0.31 | 0.35 | -0.33 | 0.24 | 0.47 | | -0.45 | 0.54 | | 0.25 | -0.46 | | 0.54 | 0.26 | | | -0.35 | 0.37 | |
| CL030 | 0.60 | 0.32 | 0.35 | | 0.59 | | 0.29 | | 0.29 | | | -0.24 | | | 0.40 | -0.32 | 0.27 | 0.26 | | -0.42 | | 0.46 | | 0.33 | 0.51 | | 0.27 | |
| CL031 | | 0.39 | | | | | 0.60 | 0.61 | | | | | 0.44 | -0.26 | | -0.29 | | 0.61 | | -0.41 | 0.70 | | 0.33 | 0.28 | 0.32 | | | |
| CL032 | 0.53 | 0.37 | | | | -0.35 | 0.65 | 0.66 | 0.49 | 0.70 | | | | -0.37 | 0.75 | | | 0.68 | -0.44 | -0.26 | 0.25 | | | 0.64 | 0.76 | | 0.27 | |
| CL033 | -0.31 | | 0.57 | 0.35 | 0.38 | 0.34 | | | | -0.56 | 0.28 | -0.39 | 0.35 | | -0.36 | -0.40 | 0.49 | | 0.33 | -0.43 | | 0.46 | 0.28 | -0.29 | | -0.37 | 0.41 | |
| CL034 | 0.83 | | | -0.32 | | -0.63 | 0.24 | 0.36 | 0.39 | 0.70 | -0.25 | 0.27 | | -0.28 | 0.76 | | | 0.27 | -0.53 | | | | -0.32 | 0.52 | 0.47 | 0.27 | | |
| CL035 | | 0.34 | 0.59 | 0.48 | 0.43 | 0.25 | | | 0.32 | | 0.33 | -0.52 | 0.39 | 0.24 | | -0.52 | 0.55 | | 0.33 | -0.55 | 0.31 | 0.54 | 0.29 | | | -0.50 | 0.54 | |
| CL036 | | | 0.78 | 0.67 | 0.63 | 0.64 | -0.36 | -0.44 | | -0.41 | 0.74 | -0.66 | | 0.80 | -0.32 | -0.77 | 0.69 | -0.44 | 0.82 | -0.67 | | 0.72 | 0.56 | | | -0.71 | 0.35 | 0.29 |
| CL037 | 0.35 | 0.39 | 0.27 | 0.53 | 0.25 | 0.28 | 0.32 | | 0.63 | 0.30 | 0.51 | -0.51 | | 0.24 | 0.35 | -0.66 | 0.25 | 0.27 | 0.28 | -0.68 | 0.48 | 0.37 | 0.48 | 0.61 | 0.24 | -0.50 | 0.26 | |
| CL038 | -0.30 | 0.36 | | 0.53 | | 0.43 | 0.51 | 0.37 | | | 0.48 | -0.43 | 0.35 | | | -0.62 | 0.28 | 0.47 | 0.31 | -0.67 | 0.75 | 0.35 | 0.47 | 0.26 | | -0.42 | 0.25 | 0.25 |
| CL039 | | 0.50 | 0.26 | 0.29 | 0.28 | | 0.43 | 0.39 | 0.36 | 0.24 | | -0.28 | 0.44 | | 0.35 | -0.46 | 0.27 | 0.43 | | -0.57 | 0.45 | 0.32 | | 0.36 | 0.45 | | 0.42 | |
| CL040 | 0.28 | 0.36 | | | | | 0.58 | 0.66 | 0.52 | 0.57 | | | 0.36 | -0.40 | 0.64 | | | 0.59 | -0.24 | -0.33 | 0.54 | | | 0.66 | 0.40 | | 0.33 | |
| CL041 | | | 0.70 | 0.72 | 0.39 | 0.71 | -0.24 | -0.28 | | -0.45 | 0.53 | -0.66 | | 0.50 | -0.34 | -0.70 | 0.66 | -0.24 | 0.73 | -0.65 | 0.37 | 0.58 | 0.45 | | -0.25 | -0.65 | 0.45 | |
| CL042 | | 0.37 | 0.25 | 0.35 | | | 0.39 | 0.44 | 0.47 | 0.25 | | -0.30 | 0.36 | | 0.39 | -0.51 | 0.33 | 0.43 | | -0.58 | 0.50 | 0.24 | | 0.47 | 0.34 | -0.30 | 0.48 | 0.24 |
| CL043 | | 0.37 | | | | | 0.73 | 0.76 | 0.28 | 0.45 | | | 0.26 | -0.32 | 0.42 | | | 0.75 | -0.31 | -0.35 | 0.42 | | | 0.43 | 0.65 | | | |
| CL044 | | 0.36 | | | | | 0.57 | 0.64 | | 0.37 | | | 0.44 | -0.48 | 0.44 | | | 0.65 | -0.36 | | 0.48 | | | 0.33 | 0.44 | | 0.25 | |
| CL045 | 0.26 | | 0.60 | | 0.56 | | -0.39 | -0.49 | | -0.28 | 0.46 | -0.30 | | 0.69 | | -0.37 | 0.46 | -0.50 | 0.46 | -0.29 | -0.25 | 0.51 | 0.35 | | | -0.35 | | 0.29 |
| CL046 | 0.51 | | -0.55 | | -0.42 | -0.42 | 0.53 | 0.67 | 0.50 | 0.89 | -0.32 | 0.35 | | -0.60 | 0.84 | | -0.40 | 0.59 | -0.56 | | | -0.42 | -0.36 | 0.71 | 0.48 | 0.33 | | |
| CL047 | 0.80 | | | | 0.37 | -0.45 | | | 0.26 | 0.38 | | | | | 0.47 | | | | -0.26 | | -0.44 | | | 0.29 | 0.33 | | | |
| CL048 | 0.35 | | 0.64 | | 0.70 | | -0.32 | -0.38 | | -0.31 | 0.26 | -0.28 | | 0.48 | | -0.27 | 0.51 | -0.37 | | -0.25 | -0.28 | 0.53 | | | | -0.29 | | |
| CL049 | 0.28 | 0.25 | 0.60 | 0.41 | 0.66 | | | | | | 0.41 | -0.45 | | 0.35 | | -0.58 | 0.53 | | 0.26 | -0.61 | | 0.64 | 0.29 | | | -0.44 | 0.44 | |
| CL050 | -0.48 | | 0.45 | | 0.56 | | | | -0.61 | -0.76 | | | | 0.34 | -0.64 | | 0.31 | | | | | 0.43 | | -0.68 | | | | |

| | CL058 | CL059 | CL060 | CL061 | CL062 | CL063 | CL064 | CL065 | CL066 | CL067 | CL068 | CL069 | CL070 | CL071 | CL072 | CL073 | CL074 | CL075 | CL076 | CL077 | CL078 | CL079 | CL080 | CL081 | CL082 | CL083 | CL084 | CL085 |
|---|---|---|---|---|---|---|---|---|---|---|---|---|---|---|---|---|---|---|---|---|---|---|---|---|---|---|---|---|
| CL051 | | 0.30 | 0.45 | 0.48 | 0.26 | 0.41 | | | | | 0.52 | -0.52 | 0.28 | 0.47 | | -0.56 | 0.45 | | 0.51 | -0.53 | 0.41 | 0.41 | 0.66 | | | -0.53 | 0.31 | 0.33 |
| CL052 | | | 0.70 | | 0.79 | | | -0.30 | -0.26 | -0.56 | 0.34 | -0.32 | | 0.52 | -0.37 | -0.31 | 0.55 | -0.27 | 0.30 | -0.32 | | 0.59 | 0.24 | -0.37 | | -0.30 | 0.24 | |
| CL053 | 0.81 | | 0.26 | 0.25 | 0.25 | | | | 0.66 | 0.46 | 0.24 | | | 0.25 | 0.55 | -0.37 | | | | -0.33 | | 0.28 | | 0.61 | | -0.27 | 0.27 | |
| CL054 | 0.73 | 0.26 | | | | -0.37 | 0.38 | 0.49 | 0.65 | 0.86 | | | | -0.30 | 0.89 | | | 0.41 | -0.37 | | | | | 0.77 | 0.48 | | | |
| CL055 | | 0.29 | 0.60 | 0.38 | 0.46 | 0.31 | | | | -0.26 | 0.28 | -0.47 | 0.35 | 0.25 | | -0.50 | 0.63 | | 0.36 | -0.52 | 0.24 | 0.57 | 0.30 | | | -0.43 | 0.53 | |
| CL056 | | | 0.62 | 0.68 | 0.36 | 0.72 | -0.25 | -0.31 | 0.24 | -0.36 | 0.74 | -0.65 | | 0.69 | -0.33 | -0.74 | 0.55 | -0.34 | 0.83 | -0.63 | 0.31 | 0.57 | 0.67 | | -0.36 | -0.71 | 0.25 | 0.31 |
| CL057 | 0.49 | | -0.54 | -0.25 | -0.44 | -0.45 | 0.51 | 0.69 | 0.51 | 0.89 | -0.37 | 0.38 | | -0.61 | 0.82 | | -0.38 | 0.59 | -0.57 | | | -0.43 | -0.35 | 0.69 | 0.47 | 0.36 | | |
| CL058 | | 0.24 | | | | -0.32 | | | 0.60 | 0.61 | | | | | 0.70 | | | | | | | | | 0.63 | 0.28 | | | |
| CL059 | | | | | | | 0.32 | 0.30 | 0.37 | 0.25 | | | 0.35 | | 0.34 | -0.30 | | 0.37 | | -0.39 | 0.39 | | | 0.36 | | | | |
| CL060 | | | | 0.53 | 0.68 | 0.47 | -0.25 | -0.37 | | -0.44 | 0.55 | -0.60 | | 0.65 | -0.28 | -0.62 | 0.66 | -0.35 | 0.60 | -0.58 | | 0.69 | 0.48 | | | -0.59 | 0.43 | |
| CL061 | | | | | 0.35 | 0.74 | | | 0.48 | | 0.67 | -0.72 | | 0.46 | | -0.80 | 0.49 | | 0.69 | -0.76 | 0.56 | 0.56 | 0.54 | 0.33 | | -0.72 | 0.37 | |
| CL062 | | | | | | | | | | -0.36 | 0.47 | -0.43 | | 0.53 | | -0.48 | 0.47 | | 0.34 | -0.50 | | 0.66 | 0.30 | | 0.28 | -0.40 | 0.31 | |
| CL063 | | | | | | | | -0.32 | | -0.38 | 0.61 | -0.62 | | 0.53 | -0.38 | -0.64 | 0.45 | -0.28 | 0.80 | -0.53 | 0.42 | 0.45 | 0.51 | | -0.36 | -0.63 | | |
| CL064 | | | | | | | | 0.72 | | 0.37 | | | | -0.36 | 0.40 | | | 0.79 | -0.36 | | 0.41 | | | 0.36 | 0.60 | | | |
| CL065 | | | | | | | | | 0.30 | 0.56 | | | 0.29 | -0.56 | 0.53 | | -0.26 | 0.76 | -0.49 | | 0.42 | -0.27 | | 0.46 | 0.57 | 0.24 | | |
| CL066 | | | | | | | | | | 0.61 | 0.28 | -0.34 | | | 0.64 | -0.50 | | | | -0.52 | 0.47 | | | 0.85 | | -0.37 | 0.39 | |
| CL067 | | | | | | | | | | | -0.26 | 0.25 | | -0.47 | 0.84 | | -0.32 | 0.45 | -0.48 | | | -0.35 | | 0.73 | 0.41 | 0.24 | | |
| CL068 | | | | | | | | | | | | -0.62 | | 0.68 | | -0.77 | 0.50 | | 0.69 | -0.70 | 0.36 | 0.58 | 0.57 | | | -0.67 | | 0.36 |
| CL069 | | | | | | | | | | | | | | -0.54 | | 0.67 | -0.55 | | -0.59 | 0.63 | -0.39 | -0.68 | -0.52 | | | 0.96 | -0.32 | |
| CL070 | | | | | | | | | | | | | | | | | | 0.32 | | -0.26 | 0.33 | | | | | | 0.33 | |
| CL071 | | | | | | | | | | | | | | | -0.42 | -0.54 | 0.54 | -0.50 | 0.70 | -0.43 | | 0.56 | 0.49 | | -0.29 | -0.59 | | 0.28 |
| CL072 | | | | | | | | | | | | | | | | | | 0.46 | -0.43 | | | | | 0.78 | 0.45 | | | |
| CL073 | | | | | | | | | | | | | | | | | -0.64 | | -0.70 | 0.96 | -0.56 | -0.68 | -0.62 | -0.38 | | 0.71 | -0.48 | -0.33 |
| CL074 | | | | | | | | | | | | | | | | | | | 0.57 | -0.58 | | 0.63 | 0.39 | | | -0.58 | 0.38 | 0.25 |
| CL075 | | | | | | | | | | | | | | | | | | | -0.47 | | 0.40 | | | 0.39 | 0.58 | | | |
| CL076 | | | | | | | | | | | | | | | | | | | | -0.57 | 0.27 | 0.58 | 0.57 | | -0.43 | -0.66 | | 0.24 |
| CL077 | | | | | | | | | | | | | | | | | | | | | -0.60 | -0.65 | -0.60 | -0.41 | | 0.63 | -0.49 | -0.30 |
| CL078 | | | | | | | | | | | | | | | | | | | | | | | 0.44 | 0.47 | | -0.38 | 0.31 | 0.26 |
| CL079 | | | | | | | | | | | | | | | | | | | | | | | 0.44 | | | -0.68 | 0.37 | |
| CL080 | | | | | | | | | | | | | | | | | | | | | | | | | | -0.52 | | 0.33 |
| CL081 | | | | | | | | | | | | | | | | | | | | | | | | | 0.30 | | 0.28 | |
| CL082 | | | | | | | | | | | | | | | | | | | | | | | | | | | 0.26 | |
| CL083 | | | | | | | | | | | | | | | | | | | | | | | | | | | -0.35 | |
| CL084 | | | | | | | | | | | | | | | | | | | | | | | | | | | | |
| CL085 | | | | | | | | | | | | | | | | | | | | | | | | | | | | |
| CL086 | | | | | | | | | | | | | | | | | | | | | | | | | | | | |
| CL087 | | | | | | | | | | | | | | | | | | | | | | | | | | | | |
| CL088 | | | | | | | | | | | | | | | | | | | | | | | | | | | | |
| CL089 | | | | | | | | | | | | | | | | | | | | | | | | | | | | |
| CL090 | | | | | | | | | | | | | | | | | | | | | | | | | | | | |
| CL091 | | | | | | | | | | | | | | | | | | | | | | | | | | | | |
| CL092 | | | | | | | | | | | | | | | | | | | | | | | | | | | | |
| CL093 | | | | | | | | | | | | | | | | | | | | | | | | | | | | |
| CL094 | | | | | | | | | | | | | | | | | | | | | | | | | | | | |
| CL095 | | | | | | | | | | | | | | | | | | | | | | | | | | | | |
| CL096 | | | | | | | | | | | | | | | | | | | | | | | | | | | | |
| CL097 | | | | | | | | | | | | | | | | | | | | | | | | | | | | |
| CL098 | | | | | | | | | | | | | | | | | | | | | | | | | | | | |
| CL099 | | | | | | | | | | | | | | | | | | | | | | | | | | | | |
| CL100 | | | | | | | | | | | | | | | | | | | | | | | | | | | | |
| CL101 | | | | | | | | | | | | | | | | | | | | | | | | | | | | |
| CL102 | | | | | | | | | | | | | | | | | | | | | | | | | | | | |
| CL103 | | | | | | | | | | | | | | | | | | | | | | | | | | | | |
| CL104 | | | | | | | | | | | | | | | | | | | | | | | | | | | | |
| CL105 | | | | | | | | | | | | | | | | | | | | | | | | | | | | |
| CL106 | | | | | | | | | | | | | | | | | | | | | | | | | | | | |
| CL107 | | | | | | | | | | | | | | | | | | | | | | | | | | | | |
| CL108 | | | | | | | | | | | | | | | | | | | | | | | | | | | | |
| CL109 | | | | | | | | | | | | | | | | | | | | | | | | | | | | |
| CL110 | | | | | | | | | | | | | | | | | | | | | | | | | | | | |

| | CL086 | CL087 | CL088 | CL089 | CL090 | CL091 | CL092 | CL093 | CL094 | CL095 | CL096 | CL097 | CL098 | CL099 | CL100 | CL101 | CL102 | CL103 | CL104 | CL105 | CL106 | CL107 | CL108 | CL109 | CL110 | CL111 | CL112 | CL113 |
|---|---|---|---|---|---|---|---|---|---|---|---|---|---|---|---|---|---|---|---|---|---|---|---|---|---|---|---|---|
| CL001 | -0.43 | -0.36 | -0.37 | -0.29 | -0.34 | | | | | | -0.28 | -0.33 | -0.32 | 0.27 | | 0.34 | | | -0.44 | | | | | | -0.41 | | | |
| CL002 | | | | | | | -0.37 | -0.47 | -0.32 | | | | | -0.39 | -0.32 | | -0.29 | | -0.53 | | -0.42 | -0.30 | | | | | -0.37 | -0.45 |
| CL003 | | | | | -0.29 | 0.27 | -0.33 | -0.68 | | | | -0.28 | | -0.45 | -0.27 | -0.37 | | -0.27 | -0.53 | | -0.44 | -0.36 | -0.28 | | | | -0.40 | -0.62 |
| CL004 | | | | | -0.25 | 0.50 | -0.38 | -0.79 | | 0.32 | | -0.26 | | -0.54 | | -0.54 | | -0.26 | -0.53 | | -0.55 | -0.50 | -0.29 | | | 0.32 | -0.53 | -0.74 |
| CL005 | -0.38 | -0.37 | -0.39 | -0.37 | -0.37 | | | -0.25 | | -0.30 | -0.35 | -0.38 | -0.33 | | -0.47 | 0.27 | | -0.32 | | -0.29 | | | -0.30 | | -0.38 | -0.28 | | |
| CL006 | -0.36 | -0.36 | -0.35 | -0.33 | -0.26 | -0.50 | 0.71 | 0.24 | | -0.46 | -0.31 | -0.26 | -0.33 | 0.68 | -0.32 | 0.79 | | -0.26 | 0.26 | -0.29 | 0.40 | 0.58 | | | -0.35 | -0.41 | 0.73 | 0.33 |
| CL007 | -0.60 | -0.66 | -0.65 | -0.62 | -0.51 | | 0.56 | | 0.35 | -0.60 | -0.58 | -0.52 | -0.64 | 0.62 | -0.33 | 0.67 | 0.35 | -0.48 | | -0.51 | 0.43 | 0.60 | -0.43 | 0.37 | -0.59 | -0.60 | 0.62 | 0.35 |
| CL008 | | | | | -0.27 | 0.46 | -0.30 | -0.84 | | 0.29 | | -0.27 | | -0.50 | | -0.51 | | -0.30 | -0.52 | -0.24 | -0.52 | -0.47 | -0.31 | | | 0.29 | -0.45 | -0.78 |
| CL009 | | -0.32 | -0.33 | -0.40 | -0.50 | 0.55 | | -0.63 | | | -0.44 | -0.50 | -0.32 | | | | | -0.48 | -0.45 | -0.43 | -0.29 | -0.29 | -0.50 | | | | -0.25 | -0.53 |
| CL010 | | | | | | 0.32 | -0.38 | -0.62 | | 0.26 | | | | -0.41 | | -0.32 | | | -0.67 | | -0.55 | -0.38 | | | | | -0.47 | -0.62 |
| CL011 | | | | | | | | -0.46 | -0.44 | | | | | | -0.32 | | -0.40 | | -0.48 | | -0.30 | | | | | | | -0.45 |
| CL012 | -0.31 | -0.46 | -0.49 | -0.49 | -0.46 | 0.44 | | -0.27 | 0.35 | -0.24 | -0.48 | -0.47 | -0.47 | | | | 0.24 | -0.44 | | -0.43 | | | -0.44 | 0.30 | -0.32 | -0.25 | | |
| CL013 | | | | | -0.27 | 0.47 | -0.36 | -0.79 | | 0.27 | | -0.27 | | -0.51 | | -0.49 | | -0.28 | -0.51 | | -0.50 | -0.47 | -0.30 | | | 0.27 | -0.48 | -0.74 |
| CL014 | | | | | -0.25 | | | -0.48 | -0.32 | | | -0.25 | | | -0.38 | | -0.31 | -0.26 | -0.39 | | -0.29 | | -0.24 | | | | | -0.43 |
| CL015 | | | | | -0.26 | 0.37 | -0.40 | -0.74 | | 0.36 | | -0.25 | | -0.51 | | -0.43 | | -0.24 | -0.63 | | -0.56 | -0.51 | -0.27 | | | 0.37 | -0.53 | -0.73 |
| CL016 | -0.51 | -0.45 | -0.43 | -0.34 | | | 0.38 | 0.79 | 0.54 | -0.55 | -0.29 | | -0.45 | 0.58 | | 0.46 | 0.47 | | 0.68 | | 0.68 | 0.62 | | 0.30 | -0.50 | -0.59 | 0.57 | 0.85 |
| CL017 | | | | | | 0.28 | -0.41 | -0.56 | -0.24 | 0.31 | | | | -0.46 | -0.24 | -0.38 | | | -0.59 | | -0.55 | -0.43 | | | | 0.27 | -0.49 | -0.58 |
| CL018 | -0.58 | -0.62 | -0.61 | -0.57 | -0.42 | | 0.49 | 0.37 | 0.50 | -0.61 | -0.53 | -0.42 | -0.61 | 0.56 | -0.32 | 0.52 | 0.40 | -0.41 | 0.53 | -0.48 | 0.58 | 0.61 | -0.37 | 0.37 | -0.57 | -0.62 | 0.59 | 0.60 |
| CL019 | | -0.42 | -0.42 | -0.44 | -0.49 | 0.77 | | | 0.35 | | -0.47 | -0.49 | -0.45 | | | -0.26 | 0.45 | -0.43 | | -0.39 | | | -0.45 | 0.42 | | | | |
| CL020 | | -0.36 | -0.37 | -0.38 | -0.42 | 0.58 | -0.34 | -0.52 | | | -0.38 | -0.44 | -0.35 | -0.32 | | -0.37 | | -0.37 | -0.36 | -0.32 | -0.26 | -0.24 | -0.40 | 0.26 | | | -0.35 | -0.40 |
| CL021 | | | | | -0.26 | 0.59 | -0.49 | -0.67 | | 0.33 | | -0.27 | | -0.53 | | -0.60 | | | -0.55 | | -0.47 | -0.47 | -0.27 | | | 0.29 | -0.59 | -0.66 |
| CL022 | | | -0.24 | -0.27 | -0.35 | 0.77 | -0.39 | -0.39 | | | -0.28 | -0.37 | | -0.30 | | -0.44 | 0.28 | -0.28 | -0.45 | | | -0.34 | -0.33 | 0.27 | | | -0.46 | -0.43 |
| CL023 | | | | | | | | -0.60 | -0.36 | | | | | -0.38 | -0.29 | | -0.31 | | -0.57 | | -0.46 | -0.29 | | | | 0.27 | -0.30 | -0.56 |
| CL024 | | | | | -0.34 | 0.55 | -0.42 | -0.61 | | | -0.26 | -0.33 | | -0.40 | | -0.52 | | -0.28 | -0.52 | | -0.41 | -0.40 | -0.32 | | | | -0.50 | -0.58 |
| CL025 | -0.46 | -0.46 | -0.46 | -0.41 | -0.28 | | 0.50 | 0.53 | 0.25 | -0.54 | -0.36 | -0.29 | -0.45 | 0.69 | | 0.71 | 0.28 | -0.26 | 0.40 | -0.29 | 0.58 | 0.58 | | | -0.44 | -0.52 | 0.63 | 0.62 |
| CL026 | -0.66 | -0.72 | -0.70 | -0.64 | -0.44 | | 0.68 | 0.37 | 0.59 | -0.69 | -0.58 | -0.45 | -0.71 | 0.79 | | 0.66 | 0.54 | -0.43 | 0.54 | -0.48 | 0.61 | 0.74 | -0.38 | 0.46 | -0.66 | -0.72 | 0.76 | 0.56 |
| CL027 | 0.24 | | | | -0.25 | 0.56 | -0.47 | -0.64 | | 0.43 | | -0.26 | | -0.53 | | -0.51 | | | -0.66 | | -0.49 | -0.54 | -0.27 | | 0.24 | 0.40 | -0.63 | -0.69 |
| CL028 | -0.40 | -0.37 | -0.37 | -0.34 | -0.27 | -0.42 | 0.62 | 0.29 | | -0.46 | -0.31 | -0.29 | -0.34 | 0.63 | -0.32 | 0.76 | | -0.25 | | -0.26 | 0.40 | 0.55 | | | -0.38 | -0.43 | 0.66 | 0.36 |
| CL029 | -0.38 | -0.46 | -0.47 | -0.44 | -0.41 | | 0.46 | | | -0.37 | -0.43 | -0.43 | -0.43 | 0.51 | | 0.66 | | -0.36 | | -0.36 | 0.29 | 0.44 | -0.37 | | -0.37 | -0.34 | 0.46 | |
| CL030 | -0.42 | -0.42 | -0.44 | -0.40 | -0.43 | | | | | | -0.38 | -0.44 | -0.39 | 0.26 | -0.41 | 0.35 | | -0.35 | -0.35 | -0.28 | | | -0.32 | | -0.40 | | | |
| CL031 | | -0.39 | -0.39 | -0.38 | -0.42 | 0.75 | -0.26 | | 0.34 | | -0.40 | -0.42 | -0.40 | | 0.26 | -0.35 | 0.47 | -0.33 | | -0.29 | | | -0.40 | 0.38 | | | -0.24 | |
| CL032 | -0.25 | -0.26 | -0.28 | -0.26 | -0.35 | 0.42 | -0.35 | -0.48 | | | -0.26 | -0.34 | -0.24 | -0.25 | -0.26 | -0.27 | | -0.25 | -0.54 | | -0.40 | -0.25 | -0.28 | | -0.25 | | -0.39 | -0.48 |
| CL033 | -0.43 | -0.42 | -0.42 | -0.35 | -0.25 | | | 0.48 | 0.30 | -0.35 | -0.32 | -0.26 | -0.41 | 0.48 | | 0.52 | 0.41 | | | | 0.39 | 0.31 | | 0.24 | -0.42 | -0.39 | 0.33 | 0.56 |
| CL034 | | | | | | | | -0.58 | -0.39 | | | | | -0.35 | -0.33 | | -0.34 | | -0.60 | | -0.45 | -0.31 | | | | 0.28 | -0.31 | -0.59 |
| CL035 | -0.56 | -0.55 | -0.56 | -0.49 | -0.42 | | | 0.29 | 0.31 | -0.52 | -0.44 | -0.43 | -0.52 | 0.40 | -0.25 | 0.38 | 0.36 | -0.33 | | -0.33 | 0.38 | 0.41 | -0.31 | 0.36 | -0.56 | -0.53 | 0.31 | 0.40 |
| CL036 | -0.55 | -0.69 | -0.67 | -0.66 | -0.49 | | 0.79 | 0.27 | 0.51 | -0.69 | -0.62 | -0.50 | -0.68 | 0.74 | -0.26 | 0.57 | 0.40 | -0.51 | 0.63 | -0.59 | 0.71 | 0.72 | -0.46 | 0.47 | -0.54 | -0.68 | 0.82 | 0.48 |
| CL037 | -0.59 | -0.70 | -0.68 | -0.64 | -0.53 | 0.38 | 0.29 | | 0.43 | -0.45 | -0.61 | -0.52 | -0.69 | 0.28 | -0.28 | | 0.43 | -0.49 | | -0.50 | 0.25 | 0.29 | -0.47 | 0.56 | -0.59 | -0.52 | 0.24 | |
| CL038 | -0.47 | -0.66 | -0.64 | -0.60 | -0.50 | 0.66 | | | 0.60 | -0.31 | -0.59 | -0.51 | -0.68 | 0.29 | | | 0.72 | -0.45 | | -0.44 | 0.27 | | -0.45 | 0.60 | -0.45 | -0.43 | | |
| CL039 | -0.54 | -0.57 | -0.57 | -0.52 | -0.49 | 0.45 | | | 0.28 | | -0.50 | -0.50 | -0.55 | | | | 0.46 | -0.38 | | -0.33 | | | -0.40 | 0.35 | -0.52 | -0.28 | | |
| CL040 | | -0.31 | -0.33 | -0.37 | -0.45 | 0.69 | -0.26 | -0.52 | | | -0.39 | -0.44 | -0.33 | -0.25 | | -0.35 | | -0.39 | -0.40 | -0.35 | | -0.27 | -0.42 | 0.32 | | | -0.36 | -0.47 |
| CL041 | -0.61 | -0.66 | -0.64 | -0.57 | -0.39 | | 0.42 | 0.47 | 0.62 | -0.65 | -0.52 | -0.39 | -0.66 | 0.57 | | 0.46 | 0.50 | -0.38 | 0.62 | -0.43 | 0.66 | 0.64 | -0.36 | 0.48 | -0.61 | -0.67 | 0.60 | 0.66 |
| CL042 | -0.36 | -0.58 | -0.57 | -0.57 | -0.55 | 0.53 | | | 0.36 | -0.25 | -0.57 | -0.56 | -0.56 | | | | 0.40 | -0.52 | | -0.51 | | | -0.54 | 0.38 | -0.36 | -0.30 | | |
| CL043 | | -0.34 | -0.36 | -0.34 | -0.41 | 0.59 | | -0.37 | | | -0.36 | -0.42 | -0.33 | | | -0.33 | 0.29 | -0.32 | -0.45 | | | -0.25 | -0.35 | 0.25 | | | -0.31 | -0.41 |
| CL044 | | | | | -0.25 | 0.62 | -0.42 | -0.29 | | | | -0.26 | | | | -0.32 | 0.29 | | -0.43 | | -0.27 | -0.36 | | | | | -0.41 | -0.32 |
| CL045 | -0.24 | -0.29 | -0.31 | -0.33 | -0.30 | -0.47 | 0.69 | | | -0.39 | -0.32 | -0.30 | -0.27 | 0.50 | -0.27 | 0.52 | | -0.31 | 0.27 | -0.35 | 0.41 | 0.45 | -0.29 | | -0.24 | -0.32 | 0.67 | |
| CL046 | | | | | | 0.46 | -0.47 | -0.73 | | 0.36 | | | | -0.61 | | -0.63 | | | -0.51 | | -0.56 | -0.52 | | | | 0.34 | -0.58 | -0.71 |
| CL047 | | | | | | -0.34 | | -0.31 | -0.35 | | | | | | -0.45 | | -0.35 | | -0.34 | | | | | -0.26 | | | | -0.28 |
| CL048 | -0.34 | -0.27 | -0.28 | -0.25 | | -0.47 | 0.41 | 0.30 | | -0.34 | | -0.25 | | 0.48 | -0.31 | 0.69 | | | | | 0.31 | 0.40 | | | -0.33 | -0.29 | 0.50 | 0.30 |
| CL049 | -0.57 | -0.62 | -0.62 | -0.58 | -0.53 | | 0.38 | | 0.32 | -0.43 | -0.55 | -0.55 | -0.60 | 0.51 | -0.37 | 0.55 | 0.37 | -0.47 | | -0.44 | 0.27 | 0.43 | -0.43 | 0.27 | -0.55 | -0.46 | 0.41 | |
| CL050 | | | | | | | 0.31 | 0.54 | | | | | | 0.52 | | 0.58 | | | | | 0.26 | 0.28 | | | | | 0.36 | 0.44 |

| | CL086 | CL087 | CL088 | CL089 | CL090 | CL091 | CL092 | CL093 | CL094 | CL095 | CL096 | CL097 | CL098 | CL099 | CL100 | CL101 | CL102 | CL103 | CL104 | CL105 | CL106 | CL107 | CL108 | CL109 | CL110 | CL111 | CL112 | CL113 |
|---|---|---|---|---|---|---|---|---|---|---|---|---|---|---|---|---|---|---|---|---|---|---|---|---|---|---|---|---|
| CL051 | -0.47 | -0.54 | -0.54 | -0.50 | -0.40 | | 0.38 | | 0.39 | -0.53 | -0.47 | -0.41 | -0.55 | 0.49 | | | 0.37 | -0.37 | 0.32 | -0.39 | 0.56 | 0.50 | -0.35 | 0.38 | -0.48 | -0.52 | 0.39 | 0.24 |
| CL052 | -0.38 | -0.32 | -0.31 | -0.27 | | -0.31 | 0.45 | 0.41 | | -0.31 | -0.25 | -0.24 | -0.28 | 0.60 | | 0.76 | | | | | 0.36 | 0.50 | | | -0.37 | -0.32 | 0.53 | 0.43 |
| CL053 | | -0.35 | -0.36 | -0.41 | -0.42 | | 0.30 | -0.45 | | -0.29 | -0.41 | -0.41 | -0.34 | | -0.49 | | | -0.45 | | -0.47 | | | -0.41 | | | | 0.25 | -0.24 |
| CL054 | | | | | -0.32 | 0.31 | -0.25 | -0.71 | | | | -0.31 | | -0.41 | -0.31 | -0.37 | | -0.30 | -0.44 | -0.26 | -0.36 | -0.25 | -0.32 | | | | -0.32 | -0.61 |
| CL055 | -0.53 | -0.53 | -0.53 | -0.47 | -0.38 | | | 0.32 | 0.33 | -0.44 | -0.42 | -0.39 | -0.52 | 0.43 | | 0.47 | 0.36 | -0.30 | | -0.31 | 0.36 | 0.39 | -0.31 | | -0.54 | -0.48 | 0.31 | 0.46 |
| CL056 | -0.52 | -0.64 | -0.62 | -0.61 | -0.42 | | 0.67 | 0.27 | 0.53 | -0.69 | -0.57 | -0.42 | -0.65 | 0.62 | | 0.34 | 0.40 | -0.46 | 0.67 | -0.52 | 0.73 | 0.68 | -0.40 | 0.55 | -0.52 | -0.67 | 0.70 | 0.48 |
| CL057 | | | | | | 0.46 | -0.51 | -0.69 | | 0.39 | | | | -0.62 | | -0.63 | | | -0.52 | | -0.56 | -0.53 | | | | 0.38 | -0.60 | -0.68 |
| CL058 | | | | | -0.29 | | | -0.48 | | | | -0.29 | | | -0.48 | | -0.24 | -0.29 | -0.31 | -0.24 | | | -0.27 | | | | | -0.38 |
| CL059 | -0.28 | -0.38 | -0.38 | -0.36 | -0.37 | 0.38 | | | | | -0.34 | -0.38 | -0.37 | | | | 0.24 | -0.29 | | | | | -0.31 | 0.26 | -0.29 | | | |
| CL060 | -0.61 | -0.60 | -0.58 | -0.52 | -0.39 | -0.25 | 0.56 | 0.39 | 0.30 | -0.59 | -0.48 | -0.40 | -0.57 | 0.66 | -0.25 | 0.65 | 0.34 | -0.36 | 0.42 | -0.40 | 0.57 | 0.67 | -0.34 | 0.34 | -0.59 | -0.60 | 0.68 | 0.55 |
| CL061 | -0.67 | -0.77 | -0.76 | -0.68 | -0.48 | | 0.43 | | 0.75 | -0.68 | -0.63 | -0.47 | -0.78 | 0.50 | -0.26 | | 0.65 | -0.47 | 0.57 | -0.51 | 0.59 | 0.57 | -0.43 | 0.63 | -0.67 | -0.73 | 0.54 | 0.43 |
| CL062 | -0.57 | -0.51 | -0.50 | -0.44 | -0.36 | -0.24 | 0.48 | 0.27 | | -0.38 | -0.41 | -0.35 | -0.47 | 0.58 | -0.24 | 0.63 | | -0.29 | | -0.29 | 0.31 | 0.53 | -0.26 | | -0.56 | -0.43 | 0.52 | 0.34 |
| CL063 | -0.45 | -0.55 | -0.53 | -0.50 | -0.27 | | 0.47 | 0.42 | 0.68 | -0.60 | -0.45 | -0.27 | -0.56 | 0.49 | | | 0.52 | -0.32 | 0.83 | -0.41 | 0.67 | 0.54 | -0.27 | 0.51 | -0.45 | -0.63 | 0.58 | 0.64 |
| CL064 | | | | | | 0.63 | -0.32 | -0.33 | | | | -0.26 | | | | -0.35 | | | -0.47 | | -0.30 | -0.28 | | | | | -0.42 | -0.41 |
| CL065 | | | | | -0.27 | 0.61 | -0.46 | -0.40 | | 0.29 | | -0.27 | | -0.36 | | -0.46 | | | -0.51 | | -0.38 | -0.39 | -0.24 | | | | -0.56 | -0.47 |
| CL066 | -0.43 | -0.53 | -0.53 | -0.53 | -0.46 | 0.37 | | -0.38 | 0.37 | -0.34 | -0.50 | -0.45 | -0.53 | | -0.35 | | 0.25 | -0.45 | | -0.45 | | | -0.43 | 0.47 | -0.44 | -0.36 | | |
| CL067 | | | | | | 0.35 | -0.44 | -0.66 | | 0.25 | | | | -0.55 | | -0.58 | | | -0.43 | | -0.41 | -0.41 | | | | 0.24 | -0.49 | -0.63 |
| CL068 | -0.57 | -0.71 | -0.70 | -0.68 | -0.52 | | 0.66 | | 0.57 | -0.62 | -0.65 | -0.51 | -0.72 | 0.69 | | 0.35 | 0.46 | -0.53 | 0.45 | -0.57 | 0.57 | 0.57 | -0.47 | 0.56 | -0.56 | -0.65 | 0.65 | 0.25 |
| CL069 | 0.73 | 0.65 | 0.63 | 0.51 | | | -0.44 | -0.29 | -0.58 | 0.95 | 0.42 | 0.26 | 0.65 | -0.60 | 0.55 | -0.37 | -0.52 | | -0.45 | 0.27 | -0.58 | -0.62 | | -0.50 | 0.73 | 0.99 | -0.53 | -0.43 |
| CL070 | | -0.25 | -0.27 | | | 0.29 | | | | | | -0.29 | -0.24 | | | | | | | | | | | | | | | |
| CL071 | -0.37 | -0.44 | -0.42 | -0.41 | -0.27 | -0.36 | 0.76 | 0.26 | 0.32 | -0.60 | -0.37 | -0.27 | -0.44 | 0.70 | -0.28 | 0.55 | | -0.30 | 0.59 | -0.38 | 0.63 | 0.67 | -0.25 | 0.35 | -0.36 | -0.55 | 0.78 | 0.39 |
| CL072 | | | | | -0.31 | 0.37 | -0.36 | -0.65 | | | | -0.30 | | -0.42 | -0.26 | -0.35 | | -0.28 | -0.48 | | -0.45 | -0.35 | -0.30 | | | | -0.38 | -0.59 |
| CL073 | 0.71 | 0.97 | 0.96 | 0.95 | 0.80 | -0.29 | -0.58 | | -0.70 | 0.66 | 0.91 | 0.80 | 0.97 | -0.58 | | -0.32 | -0.65 | 0.78 | -0.46 | 0.82 | -0.59 | -0.59 | 0.75 | -0.72 | 0.71 | 0.70 | -0.62 | -0.28 |
| CL074 | -0.47 | -0.59 | -0.57 | -0.58 | -0.46 | | 0.51 | | 0.38 | -0.58 | -0.55 | -0.49 | -0.58 | 0.53 | -0.25 | 0.52 | 0.29 | -0.46 | 0.38 | -0.50 | 0.54 | 0.55 | -0.44 | 0.32 | -0.47 | -0.57 | 0.61 | 0.42 |
| CL075 | | | | | | 0.66 | -0.46 | -0.35 | | | | | | -0.30 | | -0.39 | | | -0.53 | | -0.36 | -0.33 | | | | | -0.51 | -0.42 |
| CL076 | -0.46 | -0.58 | -0.56 | -0.57 | -0.36 | | 0.70 | 0.35 | 0.58 | -0.63 | -0.52 | -0.36 | -0.60 | 0.64 | | 0.37 | 0.46 | -0.42 | 0.81 | -0.51 | 0.73 | 0.68 | -0.36 | 0.53 | -0.46 | -0.61 | 0.77 | 0.57 |
| CL077 | 0.74 | 0.99 | 0.99 | 0.96 | 0.85 | -0.39 | -0.45 | | -0.67 | 0.57 | 0.93 | 0.84 | 0.99 | -0.52 | | -0.27 | -0.67 | 0.79 | -0.33 | 0.79 | -0.51 | -0.52 | 0.77 | -0.68 | 0.73 | 0.64 | -0.48 | |
| CL078 | -0.45 | -0.59 | -0.58 | -0.55 | -0.44 | 0.66 | | | 0.61 | -0.30 | -0.53 | -0.44 | -0.62 | | | | 0.64 | -0.40 | | -0.40 | 0.24 | | -0.41 | 0.58 | -0.45 | -0.40 | | |
| CL079 | -0.62 | -0.67 | -0.64 | -0.59 | -0.43 | | 0.58 | 0.24 | 0.44 | -0.65 | -0.53 | -0.45 | -0.64 | 0.66 | -0.32 | 0.55 | 0.52 | -0.39 | 0.32 | -0.43 | 0.48 | 0.57 | -0.35 | 0.44 | -0.62 | -0.69 | 0.64 | 0.39 |
| CL080 | -0.46 | -0.60 | -0.58 | -0.55 | -0.43 | | 0.45 | | 0.40 | -0.50 | -0.51 | -0.42 | -0.60 | 0.46 | | | 0.40 | -0.40 | 0.41 | -0.44 | 0.62 | 0.49 | -0.38 | 0.49 | -0.45 | -0.52 | 0.45 | 0.30 |
| CL081 | -0.27 | -0.42 | -0.42 | -0.44 | -0.43 | 0.47 | | -0.59 | 0.27 | | -0.45 | -0.42 | -0.43 | | -0.33 | -0.33 | | -0.43 | | -0.41 | | | -0.42 | 0.40 | -0.28 | | | -0.45 |
| CL082 | | | | | -0.26 | 0.34 | -0.29 | -0.27 | | | | -0.26 | | | | | | | -0.50 | | -0.36 | | | | | | -0.31 | -0.32 |
| CL083 | 0.66 | 0.66 | 0.63 | 0.55 | 0.28 | | -0.54 | | -0.60 | 0.98 | 0.47 | 0.31 | 0.66 | -0.61 | 0.53 | -0.38 | -0.50 | 0.29 | -0.48 | 0.37 | -0.57 | -0.65 | | -0.53 | 0.66 | 0.98 | -0.59 | -0.35 |
| CL084 | -0.38 | -0.50 | -0.50 | -0.49 | -0.45 | | | | 0.35 | -0.32 | -0.47 | -0.45 | -0.49 | | | | 0.36 | -0.41 | | -0.41 | | 0.26 | -0.42 | 0.28 | -0.39 | -0.33 | | |
| CL085 | | -0.28 | -0.29 | -0.35 | -0.36 | | 0.28 | | | | -0.37 | -0.39 | -0.29 | | | | | -0.40 | | -0.45 | 0.35 | | -0.42 | 0.37 | | | 0.24 | |
| CL086 | | 0.76 | 0.73 | 0.59 | 0.40 | | -0.25 | -0.26 | -0.56 | 0.62 | 0.52 | 0.38 | 0.74 | -0.57 | 0.30 | -0.32 | -0.53 | 0.31 | -0.27 | 0.30 | -0.40 | -0.53 | 0.27 | -0.52 | 0.99 | 0.73 | -0.43 | -0.40 |
| CL087 | | | 0.99 | 0.95 | 0.82 | -0.37 | -0.46 | | -0.68 | 0.60 | 0.91 | 0.82 | 0.99 | -0.53 | 0.24 | -0.28 | -0.67 | 0.77 | -0.34 | 0.78 | -0.52 | -0.54 | 0.75 | -0.69 | 0.75 | 0.67 | -0.51 | -0.24 |
| CL088 | | | | 0.96 | 0.86 | -0.36 | -0.44 | | -0.66 | 0.59 | 0.93 | 0.85 | 0.98 | -0.51 | 0.26 | -0.26 | -0.65 | 0.80 | -0.32 | 0.80 | -0.51 | -0.53 | 0.79 | -0.65 | 0.72 | 0.64 | -0.48 | |
| CL089 | | | | | 0.93 | -0.37 | -0.49 | | -0.60 | 0.50 | 0.98 | 0.92 | 0.95 | -0.45 | | | -0.59 | 0.91 | -0.33 | 0.91 | -0.47 | -0.46 | 0.88 | -0.65 | 0.59 | 0.53 | -0.48 | |
| CL090 | | | | | | -0.39 | -0.37 | | -0.42 | 0.24 | 0.95 | 0.98 | 0.82 | -0.29 | | | -0.45 | 0.96 | | 0.92 | -0.32 | -0.29 | 0.96 | -0.50 | 0.38 | 0.26 | -0.33 | |
| CL091 | | | | | | | | -0.32 | 0.34 | | -0.39 | -0.39 | -0.38 | | | -0.40 | 0.46 | -0.33 | | -0.31 | | | -0.37 | 0.44 | | | -0.27 | -0.32 |
| CL092 | | | | | | | | | 0.31 | -0.52 | -0.48 | -0.37 | -0.47 | 0.61 | | 0.48 | 0.24 | -0.44 | 0.51 | -0.52 | 0.51 | 0.54 | -0.38 | 0.36 | -0.25 | -0.47 | 0.81 | 0.24 |
| CL093 | | | | | | | | | | | | | | 0.33 | | 0.37 | | 0.24 | 0.48 | 0.24 | 0.45 | 0.36 | 0.26 | | -0.26 | -0.25 | 0.30 | 0.87 |
| CL094 | | | | | | | | | | -0.55 | -0.56 | -0.40 | -0.71 | 0.42 | | | 0.69 | -0.39 | 0.52 | -0.45 | 0.46 | 0.47 | -0.36 | 0.59 | -0.56 | -0.60 | 0.37 | 0.31 |
| CL095 | | | | | | | | | | | 0.42 | 0.27 | 0.61 | -0.59 | 0.58 | -0.40 | -0.44 | 0.25 | -0.48 | 0.33 | -0.60 | -0.66 | | -0.47 | 0.62 | 0.96 | -0.58 | -0.39 |
| CL096 | | | | | | | | | | | | 0.95 | 0.91 | -0.40 | | | -0.56 | 0.94 | -0.29 | 0.94 | -0.43 | -0.42 | 0.92 | -0.63 | 0.51 | 0.44 | -0.45 | |
| CL097 | | | | | | | | | | | | | 0.81 | -0.28 | | | -0.46 | 0.96 | | 0.93 | -0.32 | -0.29 | 0.96 | -0.52 | 0.37 | 0.28 | -0.32 | |
| CL098 | | | | | | | | | | | | | | -0.54 | | -0.26 | -0.68 | 0.77 | -0.36 | 0.78 | -0.51 | -0.53 | 0.75 | -0.69 | 0.73 | 0.67 | -0.50 | |
| CL099 | | | | | | | | | | | | | | | | 0.58 | 0.41 | -0.25 | 0.42 | -0.30 | 0.57 | 0.61 | | 0.34 | -0.55 | -0.61 | 0.67 | 0.41 |
| CL100 | | | | | | | | | | | | | | | | | | | | | | -0.28 | | | 0.28 | 0.55 | -0.26 | |
| CL101 | | | | | | | | | | | | | | | | | | | | | 0.30 | 0.51 | | | -0.32 | -0.38 | 0.55 | 0.43 |
| CL102 | | | | | | | | | | | | | | | | | | -0.41 | 0.33 | -0.43 | 0.32 | 0.32 | -0.40 | 0.58 | -0.54 | -0.53 | 0.25 | 0.26 |
| CL103 | | | | | | | | | | | | | | | | | | | | 0.96 | -0.34 | -0.30 | 0.96 | -0.50 | 0.30 | 0.24 | -0.37 | |
| CL104 | | | | | | | | | | | | | | | | | | | | -0.32 | 0.65 | 0.58 | | 0.32 | -0.28 | -0.47 | 0.60 | 0.67 |
| CL105 | | | | | | | | | | | | | | | | | | | | | -0.40 | -0.35 | 0.94 | -0.55 | 0.29 | 0.31 | -0.45 | |
| CL106 | | | | | | | | | | | | | | | | | | | | | | 0.63 | -0.31 | 0.40 | -0.40 | -0.57 | 0.61 | 0.60 |
| CL107 | | | | | | | | | | | | | | | | | | | | | | | -0.24 | 0.31 | -0.52 | -0.64 | 0.60 | 0.53 |
| CL108 | | | | | | | | | | | | | | | | | | | | | | | | -0.48 | 0.26 | | -0.32 | |
| CL109 | | | | | | | | | | | | | | | | | | | | | | | | | -0.52 | -0.52 | 0.39 | |
| CL110 | | | | | | | | | | | | | | | | | | | | | | | | | | 0.73 | -0.43 | -0.40 |

| | CL086 | CL087 | CL088 | CL089 | CL090 | CL091 | CL092 | CL093 | CL094 | CL095 | CL096 | CL097 | CL098 | CL099 | CL100 | CL101 | CL102 | CL103 | CL104 | CL105 | CL106 | CL107 | CL108 | CL109 | CL110 | CL111 | CL112 | CL113 |
|---|---|---|---|---|---|---|---|---|---|---|---|---|---|---|---|---|---|---|---|---|---|---|---|---|---|---|---|---|
| CL111 | | | | | | | | | | | | | | | | | | | | | | | | | | | -0.55 | -0.42 |
| CL112 | | | | | | | | | | | | | | | | | | | | | | | | | | | | 0.46 |
| CL113 | | | | | | | | | | | | | | | | | | | | | | | | | | | | |
| CL114 | | | | | | | | | | | | | | | | | | | | | | | | | | | | |
| CL115 | | | | | | | | | | | | | | | | | | | | | | | | | | | | |
| CL116 | | | | | | | | | | | | | | | | | | | | | | | | | | | | |
| CL117 | | | | | | | | | | | | | | | | | | | | | | | | | | | | |
| CL118 | | | | | | | | | | | | | | | | | | | | | | | | | | | | |
| CL119 | | | | | | | | | | | | | | | | | | | | | | | | | | | | |
| CL120 | | | | | | | | | | | | | | | | | | | | | | | | | | | | |
| CL121 | | | | | | | | | | | | | | | | | | | | | | | | | | | | |
| CL122 | | | | | | | | | | | | | | | | | | | | | | | | | | | | |
| CL123 | | | | | | | | | | | | | | | | | | | | | | | | | | | | |
| CL124 | | | | | | | | | | | | | | | | | | | | | | | | | | | | |
| CL125 | | | | | | | | | | | | | | | | | | | | | | | | | | | | |
| CL126 | | | | | | | | | | | | | | | | | | | | | | | | | | | | |
| CL127 | | | | | | | | | | | | | | | | | | | | | | | | | | | | |
| CL128 | | | | | | | | | | | | | | | | | | | | | | | | | | | | |
| CL129 | | | | | | | | | | | | | | | | | | | | | | | | | | | | |
| CL130 | | | | | | | | | | | | | | | | | | | | | | | | | | | | |
| CL131 | | | | | | | | | | | | | | | | | | | | | | | | | | | | |
| CL132 | | | | | | | | | | | | | | | | | | | | | | | | | | | | |
| CL133 | | | | | | | | | | | | | | | | | | | | | | | | | | | | |
| CL134 | | | | | | | | | | | | | | | | | | | | | | | | | | | | |
| CL135 | | | | | | | | | | | | | | | | | | | | | | | | | | | | |
| CL136 | | | | | | | | | | | | | | | | | | | | | | | | | | | | |
| CL137 | | | | | | | | | | | | | | | | | | | | | | | | | | | | |
| CL138 | | | | | | | | | | | | | | | | | | | | | | | | | | | | |
| CL139 | | | | | | | | | | | | | | | | | | | | | | | | | | | | |
| CL140 | | | | | | | | | | | | | | | | | | | | | | | | | | | | |
| CL141 | | | | | | | | | | | | | | | | | | | | | | | | | | | | |
| CL142 | | | | | | | | | | | | | | | | | | | | | | | | | | | | |
| CL143 | | | | | | | | | | | | | | | | | | | | | | | | | | | | |
| CL144 | | | | | | | | | | | | | | | | | | | | | | | | | | | | |
| CL145 | | | | | | | | | | | | | | | | | | | | | | | | | | | | |
| CL146 | | | | | | | | | | | | | | | | | | | | | | | | | | | | |
| CL147 | | | | | | | | | | | | | | | | | | | | | | | | | | | | |
| CL148 | | | | | | | | | | | | | | | | | | | | | | | | | | | | |
| CL149 | | | | | | | | | | | | | | | | | | | | | | | | | | | | |
| CL150 | | | | | | | | | | | | | | | | | | | | | | | | | | | | |
| CL151 | | | | | | | | | | | | | | | | | | | | | | | | | | | | |
| CL152 | | | | | | | | | | | | | | | | | | | | | | | | | | | | |
| CL153 | | | | | | | | | | | | | | | | | | | | | | | | | | | | |
| CL154 | | | | | | | | | | | | | | | | | | | | | | | | | | | | |
| CL155 | | | | | | | | | | | | | | | | | | | | | | | | | | | | |
| CL156 | | | | | | | | | | | | | | | | | | | | | | | | | | | | |
| CL157 | | | | | | | | | | | | | | | | | | | | | | | | | | | | |
| CL158 | | | | | | | | | | | | | | | | | | | | | | | | | | | | |
| CL159 | | | | | | | | | | | | | | | | | | | | | | | | | | | | |
| CL160 | | | | | | | | | | | | | | | | | | | | | | | | | | | | |
| CL161 | | | | | | | | | | | | | | | | | | | | | | | | | | | | |
| CL162 | | | | | | | | | | | | | | | | | | | | | | | | | | | | |
| CL163 | | | | | | | | | | | | | | | | | | | | | | | | | | | | |
| CL164 | | | | | | | | | | | | | | | | | | | | | | | | | | | | |
| CL165 | | | | | | | | | | | | | | | | | | | | | | | | | | | | |
| CL166 | | | | | | | | | | | | | | | | | | | | | | | | | | | | |
| CL167 | | | | | | | | | | | | | | | | | | | | | | | | | | | | |
| CL168 | | | | | | | | | | | | | | | | | | | | | | | | | | | | |
| CL169 | | | | | | | | | | | | | | | | | | | | | | | | | | | | |
| CL170 | | | | | | | | | | | | | | | | | | | | | | | | | | | | |

| | CL114 | CL115 | CL116 | CL117 | CL118 | CL119 | CL120 | CL121 | CL122 | CL123 | CL124 | CL125 | CL126 | CL127 | CL128 | CL129 | CL130 | CL131 | CL132 | CL133 | CL134 | CL135 | CL136 | CL137 | CL138 | CL139 | CL140 | CL141 |
|---|---|---|---|---|---|---|---|---|---|---|---|---|---|---|---|---|---|---|---|---|---|---|---|---|---|---|---|---|
| CL001 | -0.42 | -0.33 | | | | 0.28 | | | -0.34 | -0.30 | -0.32 | -0.32 | | | | | -0.51 | | -0.31 | -0.29 | -0.33 | -0.44 | -0.26 | | -0.25 | | -0.42 | |
| CL002 | | | | | -0.35 | | | | -0.50 | | | | | | 0.31 | -0.38 | | -0.46 | | -0.53 | | | | -0.43 | | -0.48 | | |
| CL003 | | | | | -0.41 | | | | -0.38 | | | | -0.29 | | 0.50 | | | -0.48 | | -0.53 | -0.25 | -0.25 | -0.27 | -0.54 | -0.29 | -0.43 | | -0.25 |
| CL004 | | | | | -0.62 | -0.43 | | | -0.26 | | | | -0.30 | | 0.51 | | | -0.59 | | -0.48 | | | -0.26 | -0.63 | -0.29 | -0.47 | | -0.26 |
| CL005 | -0.39 | -0.38 | -0.33 | | | | | | | -0.36 | -0.38 | -0.38 | -0.30 | -0.28 | 0.34 | | -0.43 | | -0.37 | | -0.35 | -0.40 | -0.34 | | -0.29 | | -0.36 | -0.29 |
| CL006 | -0.38 | -0.28 | -0.43 | | 0.70 | 0.75 | -0.37 | -0.26 | | -0.33 | -0.30 | -0.33 | -0.25 | -0.41 | | | -0.31 | 0.50 | -0.35 | 0.33 | -0.28 | -0.30 | -0.27 | 0.50 | | 0.43 | -0.36 | -0.25 |
| CL007 | -0.61 | -0.55 | -0.68 | 0.30 | 0.51 | 0.44 | -0.29 | | 0.29 | -0.59 | -0.58 | -0.61 | -0.41 | -0.67 | | | -0.52 | 0.51 | -0.60 | 0.42 | -0.51 | -0.56 | -0.49 | 0.53 | -0.43 | 0.43 | -0.59 | -0.47 |
| CL008 | | | | | -0.55 | -0.35 | | | -0.25 | | | | -0.33 | | 0.54 | | | -0.55 | | -0.46 | -0.25 | | -0.28 | -0.63 | -0.31 | -0.43 | | -0.28 |
| CL009 | | -0.45 | -0.27 | 0.36 | -0.42 | -0.26 | | 0.41 | | -0.43 | -0.47 | -0.42 | -0.49 | | 0.50 | | | -0.32 | -0.42 | -0.26 | -0.48 | -0.47 | -0.49 | -0.36 | -0.51 | -0.26 | | -0.47 |
| CL010 | | | | | -0.49 | | | | -0.46 | | | | | | 0.37 | -0.28 | | -0.59 | | -0.59 | | | | -0.62 | | -0.48 | | |
| CL011 | | | | | | 0.24 | | -0.34 | -0.56 | | | | | | 0.26 | -0.39 | | -0.32 | | -0.48 | | | | -0.36 | | -0.34 | | |
| CL012 | -0.29 | -0.49 | -0.42 | 0.31 | -0.26 | -0.38 | | 0.30 | | -0.46 | -0.48 | -0.47 | -0.38 | -0.39 | 0.50 | | -0.27 | | -0.44 | | -0.44 | -0.45 | -0.44 | | -0.42 | | -0.29 | -0.44 |
| CL013 | | | | | -0.57 | -0.40 | | | -0.25 | | | | -0.31 | | 0.53 | | | -0.54 | | -0.46 | -0.24 | | -0.27 | -0.60 | -0.30 | -0.45 | | -0.27 |
| CL014 | | | | | | 0.28 | | -0.31 | -0.44 | | | | -0.29 | | 0.30 | -0.32 | | -0.26 | | -0.40 | | | -0.26 | -0.27 | -0.27 | -0.28 | | |
| CL015 | | | | | -0.55 | -0.28 | | | -0.42 | | | | -0.29 | | 0.44 | | | -0.61 | | -0.61 | | | -0.25 | -0.67 | -0.28 | -0.53 | | |
| CL016 | -0.51 | -0.26 | -0.46 | | 0.54 | | | | 0.57 | -0.28 | -0.25 | -0.32 | | -0.51 | | 0.43 | -0.36 | 0.69 | -0.29 | 0.73 | | | | 0.75 | | 0.62 | -0.53 | |
| CL017 | | | | | -0.46 | | | | -0.48 | | | | | | 0.39 | -0.26 | | -0.58 | | -0.62 | | | | -0.61 | | -0.50 | | |
| CL018 | -0.58 | -0.51 | -0.67 | 0.24 | 0.47 | 0.28 | | | 0.50 | -0.54 | -0.50 | -0.54 | -0.34 | -0.66 | | 0.33 | -0.44 | 0.65 | -0.53 | 0.62 | -0.42 | -0.45 | -0.42 | 0.70 | -0.37 | 0.57 | -0.58 | -0.42 |
| CL019 | | -0.46 | -0.33 | 0.51 | | -0.37 | 0.32 | 0.72 | | -0.45 | -0.48 | -0.45 | -0.42 | -0.35 | 0.35 | 0.28 | | | -0.45 | | -0.49 | -0.48 | -0.44 | | -0.43 | | | -0.44 |
| CL020 | | -0.42 | | 0.26 | -0.45 | -0.39 | | 0.29 | | -0.37 | -0.42 | -0.39 | -0.36 | | 0.56 | | -0.29 | -0.35 | -0.35 | -0.26 | -0.40 | -0.43 | -0.39 | -0.37 | -0.38 | | | -0.35 |
| CL021 | | | | | -0.63 | -0.51 | 0.28 | 0.32 | -0.30 | | | | -0.27 | | 0.47 | | | -0.57 | | -0.48 | | | -0.24 | -0.64 | -0.27 | -0.47 | | |
| CL022 | | -0.32 | | 0.40 | -0.48 | -0.49 | 0.39 | 0.62 | | -0.28 | -0.31 | -0.28 | -0.30 | | 0.35 | | | -0.37 | -0.27 | -0.24 | -0.35 | -0.35 | -0.31 | -0.39 | -0.32 | -0.24 | | -0.28 |
| CL023 | | | | | -0.30 | | | | -0.51 | | | | | | 0.32 | -0.37 | | -0.47 | | -0.57 | | | | -0.47 | | -0.49 | | |
| CL024 | | -0.30 | | | -0.52 | -0.39 | 0.29 | 0.33 | -0.24 | | -0.29 | | -0.29 | | 0.45 | | | -0.49 | | -0.42 | -0.30 | -0.33 | -0.29 | -0.55 | -0.33 | -0.36 | | -0.26 |
| CL025 | -0.47 | -0.35 | -0.49 | | 0.63 | 0.46 | | | 0.32 | -0.38 | -0.36 | -0.40 | | -0.50 | | | -0.35 | 0.59 | -0.40 | 0.54 | -0.32 | -0.34 | -0.27 | 0.72 | | 0.48 | -0.45 | -0.26 |
| CL026 | -0.68 | -0.53 | -0.75 | 0.39 | 0.66 | 0.42 | | | 0.55 | -0.59 | -0.55 | -0.60 | -0.34 | -0.78 | | 0.44 | -0.50 | 0.74 | -0.59 | 0.68 | -0.47 | -0.50 | -0.45 | 0.69 | -0.39 | 0.68 | -0.67 | -0.43 |
| CL027 | 0.25 | | | | -0.64 | -0.42 | 0.28 | 0.32 | -0.41 | | | | -0.28 | | 0.35 | | | -0.63 | | -0.54 | -0.24 | | | -0.69 | -0.28 | -0.48 | 0.25 | |
| CL028 | -0.42 | -0.30 | -0.41 | | 0.68 | 0.70 | -0.30 | | | -0.32 | -0.31 | -0.34 | | -0.40 | | | -0.36 | 0.45 | -0.36 | 0.26 | -0.27 | -0.32 | -0.26 | 0.46 | -0.24 | 0.40 | -0.40 | |
| CL029 | -0.39 | -0.43 | -0.43 | | 0.38 | 0.46 | | | | -0.45 | -0.44 | -0.44 | -0.37 | -0.42 | | | -0.33 | 0.33 | -0.47 | | -0.44 | -0.48 | -0.41 | 0.34 | -0.35 | 0.27 | -0.38 | -0.36 |
| CL030 | -0.43 | -0.42 | -0.31 | | | 0.31 | | | -0.25 | -0.40 | -0.43 | -0.44 | -0.34 | -0.30 | 0.24 | | -0.49 | | -0.41 | | -0.42 | -0.50 | -0.36 | | -0.36 | | -0.42 | -0.30 |
| CL031 | | -0.43 | -0.25 | 0.48 | -0.26 | -0.35 | 0.40 | 0.70 | | -0.38 | -0.41 | -0.40 | -0.34 | -0.28 | 0.32 | 0.31 | | | -0.38 | | -0.43 | -0.44 | -0.37 | | -0.37 | | | -0.35 |
| CL032 | -0.24 | -0.33 | | | -0.38 | | | | -0.29 | -0.26 | -0.30 | -0.27 | -0.26 | | 0.45 | | -0.38 | -0.43 | -0.25 | -0.43 | -0.30 | -0.38 | -0.29 | -0.47 | -0.30 | -0.33 | -0.25 | |
| CL033 | -0.43 | -0.32 | -0.36 | 0.37 | 0.28 | | | 0.34 | 0.26 | -0.34 | -0.31 | -0.35 | | -0.39 | | | -0.36 | 0.36 | -0.33 | 0.37 | -0.28 | -0.34 | | 0.45 | | 0.32 | -0.43 | |
| CL034 | | | | | -0.30 | | | | -0.57 | | | | -0.24 | | 0.29 | -0.39 | | -0.50 | | -0.59 | | | | -0.53 | | -0.47 | | |
| CL035 | -0.55 | -0.48 | -0.49 | 0.30 | 0.28 | | | | | -0.47 | -0.45 | -0.49 | -0.28 | -0.50 | | | -0.50 | 0.37 | -0.49 | 0.32 | -0.40 | -0.48 | -0.36 | 0.42 | -0.32 | 0.32 | -0.57 | -0.31 |
| CL036 | -0.57 | -0.56 | -0.78 | 0.30 | 0.73 | 0.53 | | | 0.56 | -0.63 | -0.57 | -0.63 | -0.44 | -0.78 | | 0.45 | -0.37 | 0.80 | -0.63 | 0.70 | -0.51 | -0.51 | -0.51 | 0.69 | -0.45 | 0.74 | -0.55 | -0.52 |
| CL037 | -0.58 | -0.58 | -0.62 | 0.42 | | | | | 0.24 | -0.59 | -0.59 | -0.61 | -0.41 | -0.66 | 0.59 | 0.46 | -0.49 | 0.27 | -0.58 | | -0.51 | -0.57 | -0.49 | | -0.45 | 0.34 | -0.59 | -0.47 |
| CL038 | -0.46 | -0.57 | -0.55 | 0.58 | | | 0.28 | 0.72 | 0.45 | -0.56 | -0.56 | -0.58 | -0.40 | -0.62 | 0.33 | 0.52 | -0.35 | 0.25 | -0.56 | 0.38 | -0.53 | -0.54 | -0.47 | | -0.43 | 0.34 | -0.47 | -0.45 |
| CL039 | -0.53 | -0.54 | -0.39 | 0.29 | | | | 0.37 | | -0.48 | -0.53 | -0.52 | -0.33 | -0.42 | 0.34 | | -0.57 | | -0.49 | | -0.47 | -0.56 | -0.42 | | -0.39 | | -0.52 | -0.35 |
| CL040 | | -0.42 | | 0.36 | -0.50 | -0.43 | 0.25 | 0.51 | | -0.37 | -0.42 | -0.37 | -0.39 | | 0.50 | | | -0.31 | -0.37 | | -0.44 | -0.42 | -0.42 | -0.36 | -0.41 | -0.25 | | -0.39 |
| CL041 | -0.61 | -0.50 | -0.68 | 0.37 | 0.47 | | | | 0.65 | -0.53 | -0.49 | -0.54 | -0.28 | -0.70 | | 0.41 | -0.44 | 0.71 | -0.51 | 0.75 | -0.41 | -0.44 | -0.39 | 0.73 | -0.31 | 0.64 | -0.60 | -0.39 |
| CL042 | -0.35 | -0.55 | -0.49 | 0.40 | | | | 0.48 | | -0.56 | -0.58 | -0.56 | -0.48 | -0.47 | 0.41 | | -0.32 | | -0.53 | | -0.55 | -0.56 | -0.55 | | -0.53 | | -0.36 | -0.53 |
| CL043 | | -0.39 | | 0.36 | -0.31 | -0.27 | | 0.47 | | -0.34 | -0.38 | -0.35 | -0.33 | | 0.33 | | | -0.34 | -0.33 | -0.26 | -0.40 | -0.45 | -0.35 | -0.46 | -0.36 | | | -0.29 |
| CL044 | | | | 0.28 | -0.43 | -0.39 | 0.29 | 0.45 | | | -0.24 | | | | | | | -0.36 | | -0.30 | -0.24 | -0.27 | | -0.33 | | -0.28 | | |
| CL045 | -0.27 | -0.29 | -0.42 | | 0.65 | 0.72 | | -0.36 | | -0.34 | -0.29 | -0.33 | -0.31 | -0.39 | | | | 0.50 | -0.35 | 0.33 | -0.32 | -0.29 | -0.31 | 0.41 | -0.27 | 0.44 | -0.24 | -0.31 |
| CL046 | | | | | -0.64 | -0.48 | 0.25 | | -0.29 | | | | | | 0.50 | | | -0.61 | | -0.52 | | | | -0.68 | | -0.50 | | |
| CL047 | | | | | | 0.40 | | -0.42 | -0.46 | | | | | | | -0.37 | | | | -0.37 | | | | | | -0.24 | | |
| CL048 | -0.37 | -0.25 | -0.29 | | 0.52 | 0.62 | -0.33 | -0.27 | | -0.26 | -0.25 | -0.26 | | -0.26 | | | -0.32 | 0.30 | -0.29 | | | -0.31 | | 0.35 | | | -0.34 | |
| CL049 | -0.57 | -0.57 | -0.57 | 0.31 | 0.34 | 0.34 | | | | -0.55 | -0.58 | -0.59 | -0.42 | -0.56 | | | -0.54 | 0.30 | -0.58 | | -0.52 | -0.60 | -0.52 | 0.32 | -0.46 | 0.27 | -0.56 | -0.43 |
| CL050 | | | | | 0.46 | 0.45 | | | | | | | | | -0.48 | | | 0.26 | | | | | | 0.33 | | 0.24 | | |

| | CL114 | CL115 | CL116 | CL117 | CL118 | CL119 | CL120 | CL121 | CL122 | CL123 | CL124 | CL125 | CL126 | CL127 | CL128 | CL129 | CL130 | CL131 | CL132 | CL133 | CL134 | CL135 | CL136 | CL137 | CL138 | CL139 | CL140 | CL141 |
|---|---|---|---|---|---|---|---|---|---|---|---|---|---|---|---|---|---|---|---|---|---|---|---|---|---|---|---|---|
| CL051 | -0.49 | -0.44 | -0.56 | 0.40 | 0.37 | 0.24 | | | 0.36 | -0.48 | -0.44 | -0.48 | -0.31 | -0.56 | | 0.30 | -0.31 | 0.53 | -0.47 | 0.46 | -0.42 | -0.41 | -0.39 | 0.40 | -0.32 | 0.52 | -0.49 | -0.36 |
| CL052 | -0.39 | -0.25 | -0.31 | | 0.57 | 0.60 | -0.27 | | | -0.28 | -0.25 | -0.28 | | -0.31 | -0.25 | | -0.37 | 0.37 | -0.31 | | -0.24 | -0.31 | | 0.44 | | 0.30 | -0.39 | |
| CL053 | -0.24 | -0.40 | -0.43 | | | | | -0.25 | | -0.40 | -0.42 | -0.41 | -0.42 | -0.35 | 0.56 | | | | -0.40 | | -0.38 | -0.37 | -0.42 | | -0.41 | | | -0.45 |
| CL054 | | -0.27 | | | -0.37 | | | | -0.26 | | -0.27 | | -0.31 | | 0.57 | | | -0.38 | | -0.40 | -0.27 | -0.27 | -0.30 | -0.47 | -0.31 | -0.30 | | -0.28 |
| CL055 | -0.55 | -0.44 | -0.48 | | 0.25 | | | | | -0.45 | -0.42 | -0.47 | -0.25 | -0.49 | | | -0.51 | 0.34 | -0.43 | 0.33 | -0.38 | -0.45 | -0.32 | 0.42 | -0.29 | 0.30 | -0.55 | -0.30 |
| CL056 | -0.54 | -0.49 | -0.73 | 0.37 | 0.60 | 0.32 | | | 0.67 | -0.56 | -0.51 | -0.57 | -0.35 | -0.76 | 0.24 | 0.53 | -0.31 | 0.78 | -0.53 | 0.74 | -0.44 | -0.42 | -0.44 | 0.67 | -0.37 | 0.73 | -0.52 | -0.47 |
| CL057 | | | | | -0.64 | -0.49 | 0.24 | | -0.31 | | | | | | 0.48 | | | -0.61 | | -0.51 | | | | -0.66 | | -0.51 | | |
| CL058 | | -0.26 | | | | | | -0.27 | -0.28 | -0.24 | -0.26 | -0.25 | -0.29 | | 0.45 | -0.25 | | | | -0.29 | -0.26 | -0.27 | -0.29 | -0.24 | -0.28 | | | -0.27 |
| CL059 | -0.27 | -0.38 | -0.27 | 0.26 | | | | 0.33 | | -0.36 | -0.37 | -0.39 | -0.27 | -0.27 | 0.34 | | | | -0.35 | | -0.40 | -0.41 | -0.33 | | -0.30 | | -0.28 | -0.27 |
| CL060 | -0.62 | -0.46 | -0.62 | 0.24 | 0.62 | 0.53 | | | 0.35 | -0.50 | -0.46 | -0.51 | -0.30 | -0.62 | | | -0.50 | 0.68 | -0.49 | 0.56 | -0.39 | -0.45 | -0.38 | 0.61 | -0.32 | 0.56 | -0.61 | -0.36 |
| CL061 | -0.68 | -0.60 | -0.75 | 0.49 | 0.42 | | | 0.31 | 0.69 | -0.62 | -0.60 | -0.64 | -0.35 | -0.81 | 0.42 | 0.56 | -0.50 | 0.69 | -0.61 | 0.69 | -0.47 | -0.52 | -0.48 | 0.58 | -0.41 | 0.65 | -0.68 | -0.46 |
| CL062 | -0.57 | -0.41 | -0.44 | | 0.58 | 0.55 | -0.25 | | | -0.44 | -0.39 | -0.43 | -0.26 | -0.48 | | | -0.54 | 0.43 | -0.44 | 0.25 | -0.36 | -0.43 | -0.33 | 0.42 | -0.29 | 0.34 | -0.56 | -0.27 |
| CL063 | -0.46 | -0.38 | -0.61 | 0.29 | 0.49 | | | | 0.85 | -0.42 | -0.37 | -0.43 | | -0.66 | | 0.66 | -0.25 | 0.74 | -0.42 | 0.85 | -0.28 | -0.28 | -0.29 | 0.66 | -0.26 | 0.68 | -0.46 | -0.33 |
| CL064 | | | | | -0.43 | -0.36 | 0.24 | 0.52 | | | | | | | | | -0.24 | -0.38 | | -0.32 | | -0.29 | | -0.43 | | -0.25 | | |
| CL065 | | | | 0.24 | -0.53 | -0.49 | 0.31 | 0.45 | | | | | | | 0.30 | | | -0.48 | | -0.39 | -0.25 | -0.28 | | -0.57 | -0.24 | -0.38 | | |
| CL066 | -0.43 | -0.50 | -0.49 | 0.31 | | | | | | -0.49 | -0.50 | -0.51 | -0.36 | -0.49 | 0.72 | 0.25 | -0.39 | | -0.47 | | -0.43 | -0.44 | -0.43 | | -0.42 | | -0.43 | -0.45 |
| CL067 | | | | | -0.48 | -0.36 | | | -0.29 | | | | | | 0.53 | | | -0.47 | | -0.45 | | | | -0.57 | | -0.42 | | |
| CL068 | -0.57 | -0.58 | -0.75 | 0.43 | 0.56 | 0.33 | | 0.26 | 0.62 | -0.64 | -0.59 | -0.63 | -0.45 | -0.79 | 0.34 | 0.61 | -0.39 | 0.61 | -0.64 | 0.65 | -0.55 | -0.53 | -0.51 | 0.46 | -0.49 | 0.68 | -0.56 | -0.53 |
| CL069 | 0.71 | 0.37 | 0.62 | -0.36 | -0.45 | | | | -0.54 | 0.42 | 0.37 | 0.46 | | 0.70 | -0.26 | -0.45 | 0.59 | -0.63 | 0.39 | -0.58 | 0.25 | 0.31 | | -0.55 | | -0.60 | 0.73 | |
| CL070 | -0.24 | -0.27 | | 0.26 | | | 0.28 | 0.30 | | -0.24 | -0.25 | | | | | | | | -0.24 | | -0.26 | -0.31 | | | | | -0.25 | |
| CL071 | -0.38 | -0.31 | -0.56 | | 0.72 | 0.63 | | | 0.43 | -0.38 | -0.33 | -0.38 | -0.26 | -0.56 | | 0.32 | | 0.70 | -0.39 | 0.63 | -0.30 | -0.26 | -0.29 | 0.63 | | 0.67 | -0.37 | -0.32 |
| CL072 | | -0.26 | | | -0.45 | -0.27 | | | -0.26 | | -0.26 | | -0.31 | | 0.55 | | | -0.44 | | -0.43 | -0.27 | -0.27 | -0.29 | -0.48 | -0.30 | -0.34 | | -0.27 |
| CL073 | 0.72 | 0.87 | 0.97 | -0.55 | -0.45 | | | -0.38 | -0.64 | 0.90 | 0.88 | 0.92 | 0.69 | 0.99 | -0.52 | -0.59 | 0.52 | -0.63 | 0.90 | -0.66 | 0.80 | 0.82 | 0.78 | -0.51 | 0.72 | -0.69 | 0.71 | 0.78 |
| CL074 | -0.49 | -0.52 | -0.64 | 0.28 | 0.52 | 0.35 | | | 0.38 | -0.57 | -0.53 | -0.56 | -0.42 | -0.62 | | | -0.38 | 0.55 | -0.56 | 0.52 | -0.47 | -0.49 | -0.49 | 0.52 | -0.44 | 0.51 | -0.49 | -0.47 |
| CL075 | | | | | -0.51 | -0.47 | 0.30 | 0.50 | | | | | | | 0.27 | | | -0.45 | | -0.38 | | -0.25 | | -0.48 | | -0.32 | | |
| CL076 | -0.47 | -0.45 | -0.72 | 0.31 | 0.61 | 0.36 | | | 0.76 | -0.51 | -0.46 | -0.52 | -0.32 | -0.73 | | 0.52 | -0.25 | 0.83 | -0.51 | 0.86 | -0.38 | -0.35 | -0.39 | 0.72 | -0.34 | 0.81 | -0.47 | -0.43 |
| CL077 | 0.75 | 0.91 | 0.93 | -0.56 | -0.36 | | | -0.43 | -0.52 | 0.92 | 0.91 | 0.94 | 0.71 | 0.95 | -0.54 | -0.53 | 0.58 | -0.51 | 0.91 | -0.55 | 0.84 | 0.88 | 0.81 | -0.41 | 0.75 | -0.59 | 0.73 | 0.78 |
| CL078 | -0.43 | -0.52 | -0.50 | 0.52 | | -0.35 | | 0.62 | 0.44 | -0.50 | -0.50 | -0.53 | -0.34 | -0.56 | 0.46 | 0.49 | -0.33 | | -0.48 | 0.33 | -0.46 | -0.47 | -0.41 | | -0.39 | | -0.45 | -0.42 |
| CL079 | -0.64 | -0.51 | -0.66 | 0.33 | 0.56 | 0.42 | | | 0.38 | -0.53 | -0.52 | -0.58 | -0.32 | -0.68 | | 0.30 | -0.54 | 0.53 | -0.54 | 0.51 | -0.43 | -0.48 | -0.41 | 0.59 | -0.35 | 0.57 | -0.64 | -0.38 |
| CL080 | -0.47 | -0.48 | -0.59 | 0.40 | 0.40 | | | | 0.43 | -0.52 | -0.48 | -0.53 | -0.34 | -0.64 | | 0.48 | -0.27 | 0.57 | -0.49 | 0.57 | -0.46 | -0.45 | -0.39 | 0.46 | -0.34 | 0.56 | -0.47 | -0.43 |
| CL081 | -0.26 | -0.45 | -0.38 | 0.28 | -0.26 | -0.31 | | | | -0.42 | -0.46 | -0.43 | -0.38 | -0.36 | 0.74 | | -0.28 | | -0.41 | | -0.41 | -0.41 | -0.43 | -0.24 | -0.40 | | -0.26 | -0.42 |
| CL082 | | -0.24 | | | | | | | -0.35 | | | | | | | | -0.33 | -0.37 | | -0.44 | | -0.32 | | -0.42 | | -0.31 | | |
| CL083 | 0.65 | 0.40 | 0.68 | -0.42 | -0.47 | | | | -0.60 | 0.47 | 0.42 | 0.49 | | 0.74 | -0.31 | -0.47 | 0.51 | -0.64 | 0.45 | -0.62 | 0.30 | 0.33 | 0.29 | -0.57 | | -0.63 | 0.67 | 0.28 |
| CL084 | -0.38 | -0.48 | -0.48 | 0.28 | | | | | | -0.48 | -0.46 | -0.51 | -0.35 | -0.45 | 0.29 | | -0.37 | 0.25 | -0.46 | | -0.41 | -0.47 | -0.42 | 0.24 | -0.41 | | -0.40 | -0.39 |
| CL085 | | -0.35 | -0.37 | 0.31 | | | | | | -0.38 | -0.35 | -0.35 | -0.43 | -0.32 | 0.25 | | | | -0.40 | | -0.42 | -0.33 | -0.40 | | -0.40 | 0.28 | | -0.45 |
| CL086 | 0.98 | 0.52 | 0.63 | -0.36 | -0.38 | | | | -0.35 | 0.52 | 0.49 | 0.55 | | 0.71 | -0.29 | -0.36 | 0.91 | -0.49 | 0.51 | -0.40 | 0.39 | 0.49 | 0.33 | -0.43 | 0.25 | -0.49 | 0.98 | 0.30 |
| CL087 | 0.77 | 0.90 | 0.93 | -0.56 | -0.37 | | | -0.42 | -0.54 | 0.91 | 0.90 | 0.93 | 0.68 | 0.95 | -0.53 | -0.54 | 0.60 | -0.53 | 0.90 | -0.55 | 0.82 | 0.87 | 0.79 | -0.42 | 0.73 | -0.60 | 0.75 | 0.76 |
| CL088 | 0.74 | 0.92 | 0.93 | -0.56 | -0.37 | | | -0.41 | -0.53 | 0.93 | 0.92 | 0.95 | 0.72 | 0.95 | -0.53 | -0.53 | 0.57 | -0.52 | 0.92 | -0.54 | 0.85 | 0.89 | 0.82 | -0.40 | 0.76 | -0.58 | 0.72 | 0.79 |
| CL089 | 0.61 | 0.96 | 0.95 | -0.57 | -0.36 | | | -0.42 | -0.53 | 0.97 | 0.97 | 0.98 | 0.84 | 0.93 | -0.56 | -0.51 | 0.43 | -0.48 | 0.97 | -0.53 | 0.92 | 0.92 | 0.91 | -0.35 | 0.87 | -0.55 | 0.59 | 0.90 |
| CL090 | 0.43 | 0.97 | 0.82 | -0.48 | -0.24 | | | -0.38 | -0.33 | 0.95 | 0.97 | 0.95 | 0.94 | 0.77 | -0.55 | -0.36 | 0.28 | -0.29 | 0.95 | -0.35 | 0.97 | 0.96 | 0.96 | | 0.96 | -0.38 | 0.39 | 0.95 |
| CL091 | | -0.38 | -0.25 | 0.41 | -0.39 | -0.48 | 0.33 | 0.68 | | -0.36 | -0.40 | -0.37 | -0.31 | -0.28 | 0.43 | | | | -0.36 | | -0.39 | -0.38 | -0.35 | | -0.35 | | | -0.33 |
| CL092 | -0.27 | -0.39 | -0.62 | 0.25 | 0.65 | 0.54 | | | 0.48 | -0.46 | -0.44 | -0.46 | -0.40 | -0.60 | | 0.36 | | 0.63 | -0.48 | 0.62 | -0.39 | -0.35 | -0.42 | 0.49 | -0.37 | 0.65 | -0.26 | -0.45 |
| CL093 | -0.26 | | | | 0.41 | | | | | | | | 0.30 | | -0.49 | | | 0.45 | | 0.42 | | | | 0.54 | 0.24 | 0.33 | -0.28 | 0.25 |
| CL094 | -0.54 | -0.51 | -0.67 | 0.43 | 0.25 | | | 0.46 | 0.70 | -0.53 | -0.52 | -0.56 | -0.29 | -0.71 | 0.40 | 0.63 | -0.40 | 0.53 | -0.52 | 0.63 | -0.41 | -0.43 | -0.41 | 0.41 | -0.36 | 0.57 | -0.57 | -0.41 |
| CL095 | 0.61 | 0.36 | 0.64 | -0.33 | -0.47 | | | | -0.56 | 0.42 | 0.38 | 0.44 | | 0.69 | -0.26 | -0.43 | 0.47 | -0.64 | 0.40 | -0.60 | 0.26 | 0.29 | 0.25 | -0.58 | | -0.62 | 0.62 | 0.24 |
| CL096 | 0.54 | 0.97 | 0.92 | -0.55 | -0.32 | | | -0.42 | -0.49 | 0.97 | 0.98 | 0.97 | 0.89 | 0.89 | -0.56 | -0.48 | 0.37 | -0.42 | 0.98 | -0.50 | 0.94 | 0.94 | 0.94 | -0.29 | 0.91 | -0.51 | 0.51 | 0.93 |
| CL097 | 0.41 | 0.96 | 0.82 | -0.50 | | | | -0.39 | -0.32 | 0.94 | 0.96 | 0.94 | 0.94 | 0.76 | -0.54 | -0.34 | 0.26 | -0.26 | 0.95 | -0.34 | 0.97 | 0.96 | 0.96 | | 0.95 | -0.37 | 0.38 | 0.94 |
| CL098 | 0.74 | 0.89 | 0.94 | -0.57 | -0.36 | | | -0.45 | -0.57 | 0.91 | 0.90 | 0.93 | 0.68 | 0.96 | -0.56 | -0.57 | 0.57 | -0.53 | 0.89 | -0.57 | 0.82 | 0.85 | 0.79 | -0.41 | 0.72 | -0.60 | 0.73 | 0.77 |
| CL099 | -0.58 | -0.35 | -0.56 | 0.25 | 0.60 | 0.46 | | | 0.40 | -0.41 | -0.36 | -0.42 | | -0.60 | | 0.35 | -0.42 | 0.58 | -0.42 | 0.54 | -0.32 | -0.34 | -0.27 | 0.60 | | 0.58 | -0.57 | -0.26 |
| CL100 | 0.29 | | 0.26 | | | | | | | | | | | 0.26 | -0.27 | | 0.28 | | | | | | | | | | 0.29 | |
| CL101 | -0.33 | | -0.32 | | 0.52 | 0.59 | -0.40 | | | | | | | -0.31 | -0.24 | | -0.27 | 0.41 | | 0.26 | | | | 0.49 | | 0.29 | -0.33 | |
| CL102 | -0.53 | -0.54 | -0.60 | 0.47 | | | | 0.52 | 0.51 | -0.53 | -0.54 | -0.57 | -0.33 | -0.66 | 0.37 | 0.49 | -0.40 | 0.37 | -0.53 | 0.47 | -0.45 | -0.48 | -0.41 | 0.33 | -0.38 | 0.42 | -0.54 | -0.41 |
| CL103 | 0.34 | 0.93 | 0.83 | -0.48 | -0.25 | | | -0.34 | -0.41 | 0.93 | 0.94 | 0.92 | 0.96 | 0.76 | -0.53 | -0.38 | | -0.29 | 0.94 | -0.40 | 0.94 | 0.91 | 0.96 | | 0.96 | -0.40 | 0.30 | 0.97 |
| CL104 | -0.28 | | -0.48 | | 0.55 | | | | 0.74 | -0.28 | | -0.28 | | -0.49 | | 0.51 | | 0.75 | -0.26 | 0.82 | | | | 0.69 | | 0.66 | -0.29 | -0.25 |
| CL105 | 0.34 | 0.91 | 0.87 | -0.47 | -0.31 | | | -0.33 | -0.48 | 0.93 | 0.93 | 0.91 | 0.94 | 0.80 | -0.53 | -0.44 | | -0.39 | 0.93 | -0.49 | 0.91 | 0.86 | 0.94 | -0.27 | 0.94 | -0.47 | 0.30 | 0.97 |
| CL106 | -0.42 | -0.39 | -0.58 | | 0.56 | 0.27 | | | 0.58 | -0.43 | -0.39 | -0.45 | -0.26 | -0.60 | | 0.41 | | 0.74 | -0.43 | 0.73 | -0.34 | -0.33 | -0.33 | 0.70 | -0.30 | 0.69 | -0.41 | -0.34 |
| CL107 | -0.52 | -0.36 | -0.58 | | 0.59 | 0.34 | -0.25 | | 0.49 | -0.41 | -0.39 | -0.43 | | -0.61 | | 0.37 | -0.36 | 0.69 | -0.40 | 0.63 | -0.29 | -0.33 | -0.31 | 0.66 | -0.25 | 0.60 | -0.53 | -0.29 |
| CL108 | 0.31 | 0.93 | 0.79 | -0.47 | | | | -0.34 | -0.36 | 0.92 | 0.94 | 0.90 | 0.96 | 0.71 | -0.53 | -0.34 | | -0.26 | 0.93 | -0.35 | 0.95 | 0.91 | 0.96 | | 0.96 | -0.36 | 0.27 | 0.96 |
| CL109 | -0.52 | -0.57 | -0.70 | 0.44 | | | | 0.43 | 0.51 | -0.60 | -0.58 | -0.61 | -0.43 | -0.72 | 0.50 | 0.44 | -0.36 | 0.45 | -0.59 | 0.49 | -0.52 | -0.52 | -0.51 | 0.32 | -0.43 | 0.55 | -0.52 | -0.51 |
| CL110 | 0.98 | 0.50 | 0.63 | -0.36 | -0.36 | | | | -0.36 | 0.51 | 0.48 | 0.54 | | 0.71 | -0.30 | -0.36 | 0.90 | -0.49 | 0.50 | -0.40 | 0.38 | 0.48 | 0.32 | -0.43 | 0.24 | -0.48 | 0.99 | 0.28 |

| | CL114 | CL115 | CL116 | CL117 | CL118 | CL119 | CL120 | CL121 | CL122 | CL123 | CL124 | CL125 | CL126 | CL127 | CL128 | CL129 | CL130 | CL131 | CL132 | CL133 | CL134 | CL135 | CL136 | CL137 | CL138 | CL139 | CL140 | CL141 |
|---|---|---|---|---|---|---|---|---|---|---|---|---|---|---|---|---|---|---|---|---|---|---|---|---|---|---|---|---|
| CL111 | 0.72 | 0.39 | 0.65 | -0.38 | -0.46 | | | | -0.56 | 0.45 | 0.40 | 0.48 | | 0.73 | -0.28 | -0.46 | 0.59 | -0.63 | 0.42 | -0.60 | 0.27 | 0.33 | 0.25 | -0.56 | | -0.62 | 0.74 | |
| CL112 | -0.45 | -0.39 | -0.64 | 0.25 | 0.71 | 0.59 | | | 0.52 | -0.44 | -0.41 | -0.46 | -0.31 | -0.62 | | 0.36 | -0.28 | 0.70 | -0.45 | 0.66 | -0.34 | -0.33 | -0.37 | 0.66 | -0.31 | 0.73 | -0.43 | -0.37 |
| CL113 | -0.40 | | -0.25 | | 0.51 | | | | 0.42 | | | | | -0.30 | -0.30 | 0.25 | -0.30 | 0.62 | | 0.61 | | | | 0.67 | | 0.53 | -0.41 | |
| CL114 | | 0.54 | 0.65 | -0.35 | -0.39 | | | | -0.35 | 0.54 | 0.52 | 0.58 | | 0.71 | -0.30 | -0.34 | 0.90 | -0.50 | 0.54 | -0.41 | 0.42 | 0.52 | 0.37 | -0.44 | 0.29 | -0.50 | 0.98 | 0.33 |
| CL115 | | | 0.88 | -0.52 | -0.29 | | | -0.41 | -0.42 | 0.97 | 0.97 | 0.97 | 0.89 | 0.85 | -0.55 | -0.43 | 0.39 | -0.36 | 0.97 | -0.43 | 0.95 | 0.96 | 0.94 | -0.26 | 0.92 | -0.45 | 0.51 | 0.92 |
| CL116 | | | | -0.54 | -0.46 | -0.26 | | -0.34 | -0.64 | 0.91 | 0.89 | 0.92 | 0.75 | 0.97 | -0.54 | -0.56 | 0.44 | -0.62 | 0.91 | -0.67 | 0.82 | 0.81 | 0.82 | -0.50 | 0.77 | -0.69 | 0.63 | 0.83 |
| CL117 | | | | | | | | 0.54 | 0.36 | -0.56 | -0.53 | -0.56 | -0.42 | -0.54 | 0.34 | 0.28 | | 0.25 | -0.54 | 0.32 | -0.52 | -0.50 | -0.49 | | -0.45 | 0.34 | -0.37 | -0.46 |
| CL118 | | | | | | 0.62 | -0.26 | | 0.39 | -0.33 | -0.29 | -0.34 | | -0.47 | | 0.24 | -0.25 | 0.69 | -0.36 | 0.59 | -0.24 | -0.25 | -0.24 | 0.57 | | 0.56 | -0.39 | -0.26 |
| CL119 | | | | | | | | -0.32 | | | | | | | | | | 0.37 | -0.24 | | | | | 0.34 | | 0.30 | | |
| CL120 | | | | | | | | 0.32 | | | | | | | | | | | | | | | | | | | | |
| CL121 | | | | | | | | | 0.25 | -0.41 | -0.41 | -0.41 | -0.31 | -0.37 | 0.24 | 0.29 | | | -0.41 | | -0.38 | -0.40 | -0.38 | | -0.36 | | | -0.35 |
| CL122 | | | | | | | | | | -0.47 | -0.43 | -0.47 | -0.29 | -0.66 | 0.27 | 0.65 | | 0.65 | -0.45 | 0.82 | -0.34 | -0.29 | -0.35 | 0.59 | -0.32 | 0.64 | -0.36 | -0.43 |
| CL123 | | | | | | | | | | | 0.96 | 0.97 | 0.89 | 0.88 | -0.54 | -0.44 | 0.37 | -0.42 | 0.97 | -0.48 | 0.95 | 0.94 | 0.94 | -0.29 | 0.90 | -0.50 | 0.51 | 0.93 |
| CL124 | | | | | | | | | | | | 0.97 | 0.90 | 0.85 | -0.55 | -0.43 | 0.36 | -0.37 | 0.97 | -0.44 | 0.95 | 0.95 | 0.95 | -0.27 | 0.92 | -0.45 | 0.49 | 0.93 |
| CL125 | | | | | | | | | | | | | 0.86 | 0.89 | -0.56 | -0.46 | 0.41 | -0.42 | 0.96 | -0.49 | 0.94 | 0.94 | 0.92 | -0.32 | 0.89 | -0.51 | 0.55 | 0.91 |
| CL126 | | | | | | | | | | | | | | 0.66 | -0.49 | -0.29 | | | 0.90 | -0.31 | 0.93 | 0.88 | 0.95 | | 0.96 | -0.31 | | 0.96 |
| CL127 | | | | | | | | | | | | | | | -0.52 | -0.62 | 0.51 | -0.65 | 0.87 | -0.68 | 0.77 | 0.79 | 0.75 | -0.51 | 0.69 | -0.70 | 0.71 | 0.76 |
| CL128 | | | | | | | | | | | | | | | | 0.33 | -0.24 | | -0.54 | | -0.53 | -0.49 | -0.52 | | -0.50 | | -0.28 | -0.54 |
| CL129 | | | | | | | | | | | | | | | | | | 0.42 | -0.45 | 0.56 | -0.35 | -0.35 | -0.34 | 0.34 | -0.32 | 0.47 | -0.37 | -0.39 |
| CL130 | | | | | | | | | | | | | | | | | | -0.29 | 0.36 | | 0.26 | 0.38 | | -0.26 | | -0.30 | 0.90 | |
| CL131 | | | | | | | | | | | | | | | | | | | -0.41 | 0.79 | -0.29 | -0.28 | -0.28 | 0.72 | | 0.76 | -0.51 | -0.32 |
| CL132 | | | | | | | | | | | | | | | | | | | | -0.48 | 0.95 | 0.94 | 0.93 | -0.30 | 0.91 | -0.50 | 0.51 | 0.92 |
| CL133 | | | | | | | | | | | | | | | | | | | | | -0.36 | -0.33 | -0.38 | 0.72 | -0.34 | 0.75 | -0.41 | -0.42 |
| CL134 | | | | | | | | | | | | | | | | | | | | | | 0.94 | 0.95 | | 0.94 | -0.40 | 0.38 | 0.94 |
| CL135 | | | | | | | | | | | | | | | | | | | | | | | 0.92 | | 0.91 | -0.39 | 0.49 | 0.88 |
| CL136 | | | | | | | | | | | | | | | | | | | | | | | | | 0.95 | -0.39 | 0.33 | 0.95 |
| CL137 | | | | | | | | | | | | | | | | | | | | | | | | | | 0.58 | -0.44 | |
| CL138 | | | | | | | | | | | | | | | | | | | | | | | | | | -0.34 | 0.24 | 0.95 |
| CL139 | | | | | | | | | | | | | | | | | | | | | | | | | | | -0.50 | -0.40 |
| CL140 | | | | | | | | | | | | | | | | | | | | | | | | | | | | 0.29 |
| CL141 | | | | | | | | | | | | | | | | | | | | | | | | | | | | |
| CL142 | | | | | | | | | | | | | | | | | | | | | | | | | | | | |
| CL143 | | | | | | | | | | | | | | | | | | | | | | | | | | | | |
| CL144 | | | | | | | | | | | | | | | | | | | | | | | | | | | | |
| CL145 | | | | | | | | | | | | | | | | | | | | | | | | | | | | |
| CL146 | | | | | | | | | | | | | | | | | | | | | | | | | | | | |
| CL147 | | | | | | | | | | | | | | | | | | | | | | | | | | | | |
| CL148 | | | | | | | | | | | | | | | | | | | | | | | | | | | | |
| CL149 | | | | | | | | | | | | | | | | | | | | | | | | | | | | |
| CL150 | | | | | | | | | | | | | | | | | | | | | | | | | | | | |
| CL151 | | | | | | | | | | | | | | | | | | | | | | | | | | | | |
| CL152 | | | | | | | | | | | | | | | | | | | | | | | | | | | | |
| CL153 | | | | | | | | | | | | | | | | | | | | | | | | | | | | |
| CL154 | | | | | | | | | | | | | | | | | | | | | | | | | | | | |
| CL155 | | | | | | | | | | | | | | | | | | | | | | | | | | | | |
| CL156 | | | | | | | | | | | | | | | | | | | | | | | | | | | | |
| CL157 | | | | | | | | | | | | | | | | | | | | | | | | | | | | |
| CL158 | | | | | | | | | | | | | | | | | | | | | | | | | | | | |
| CL159 | | | | | | | | | | | | | | | | | | | | | | | | | | | | |
| CL160 | | | | | | | | | | | | | | | | | | | | | | | | | | | | |
| CL161 | | | | | | | | | | | | | | | | | | | | | | | | | | | | |
| CL162 | | | | | | | | | | | | | | | | | | | | | | | | | | | | |
| CL163 | | | | | | | | | | | | | | | | | | | | | | | | | | | | |
| CL164 | | | | | | | | | | | | | | | | | | | | | | | | | | | | |
| CL165 | | | | | | | | | | | | | | | | | | | | | | | | | | | | |
| CL166 | | | | | | | | | | | | | | | | | | | | | | | | | | | | |
| CL167 | | | | | | | | | | | | | | | | | | | | | | | | | | | | |
| CL168 | | | | | | | | | | | | | | | | | | | | | | | | | | | | |
| CL169 | | | | | | | | | | | | | | | | | | | | | | | | | | | | |
| CL170 | | | | | | | | | | | | | | | | | | | | | | | | | | | | |

| | CL142 | CL143 | CL144 | CL145 | CL146 | CL147 | CL148 | CL149 | CL150 | CL151 | CL152 | CL153 | CL154 | CL155 | CL156 | CL157 | CL158 | CL159 | CL160 | CL161 | CL162 | CL163 | CL164 | CL165 | CL166 | CL167 | CL168 | CL169 |
|---|---|---|---|---|---|---|---|---|---|---|---|---|---|---|---|---|---|---|---|---|---|---|---|---|---|---|---|---|
| CL001 | -0.36 | | | -0.27 | -0.29 | | 0.26 | | -0.27 | -0.32 | | -0.42 | -0.26 | | | | | 0.31 | | -0.43 | | -0.26 | 0.33 | -0.35 | -0.44 | -0.33 | -0.28 | -0.27 |
| CL002 | | 0.29 | 0.71 | | | -0.28 | 0.80 | | | -0.54 | 0.24 | | | -0.55 | | -0.41 | -0.27 | | | | -0.28 | -0.27 | | | -0.50 | | -0.62 | |
| CL003 | -0.30 | 0.35 | 0.76 | | | -0.26 | 0.77 | | | -0.43 | 0.46 | | -0.32 | -0.56 | -0.29 | -0.35 | | | | | | -0.25 | | | -0.46 | | -0.67 | -0.26 |
| CL004 | -0.27 | 0.49 | 0.76 | | | -0.39 | 0.51 | | | -0.40 | 0.55 | | -0.29 | -0.58 | -0.33 | -0.29 | -0.30 | -0.46 | 0.37 | | -0.26 | | -0.27 | | -0.36 | | -0.68 | -0.25 |
| CL005 | -0.36 | | 0.26 | -0.31 | | | 0.71 | | -0.32 | | | -0.40 | -0.37 | | -0.29 | | | 0.39 | | -0.37 | | | 0.45 | -0.37 | | -0.36 | -0.28 | -0.35 |
| CL006 | -0.25 | -0.43 | -0.38 | -0.34 | | 0.46 | | | -0.30 | 0.34 | -0.38 | -0.37 | -0.24 | 0.43 | | | 0.30 | 0.83 | -0.42 | -0.35 | 0.29 | | 0.51 | -0.33 | | -0.29 | 0.33 | -0.26 |
| CL007 | -0.47 | -0.26 | -0.35 | -0.61 | | 0.38 | 0.26 | | -0.58 | 0.30 | | -0.60 | -0.46 | 0.42 | -0.41 | 0.37 | 0.28 | 0.67 | | -0.58 | | | 0.51 | -0.57 | | -0.57 | 0.26 | -0.51 |
| CL008 | -0.28 | 0.45 | 0.72 | | | -0.33 | 0.56 | | | -0.34 | 0.56 | | -0.31 | -0.56 | -0.36 | -0.27 | -0.27 | -0.41 | 0.33 | | | | -0.25 | | -0.38 | | -0.68 | -0.27 |
| CL009 | -0.50 | 0.39 | 0.44 | -0.35 | | | 0.46 | | -0.38 | -0.28 | 0.50 | | -0.50 | -0.38 | -0.52 | | | | 0.34 | | | | | -0.41 | -0.25 | -0.46 | -0.49 | -0.49 |
| CL010 | -0.26 | 0.48 | 0.63 | | | -0.35 | 0.58 | | | -0.55 | 0.46 | | | -0.57 | | -0.41 | -0.33 | | | | | -0.39 | | | -0.53 | | -0.72 | |
| CL011 | | | 0.56 | | | | 0.82 | -0.53 | | -0.35 | | | | -0.45 | | -0.46 | | | | | | | | | -0.65 | | -0.54 | |
| CL012 | -0.44 | | 0.29 | -0.45 | 0.43 | | 0.33 | 0.39 | -0.46 | | 0.51 | -0.30 | -0.47 | | -0.42 | | | | 0.34 | -0.28 | | | | -0.41 | | -0.49 | | -0.45 |
| CL013 | -0.27 | 0.43 | 0.74 | | | -0.35 | 0.56 | | | -0.36 | 0.56 | | -0.30 | -0.55 | -0.34 | -0.29 | -0.29 | -0.41 | 0.36 | | -0.24 | | -0.26 | | -0.36 | | -0.68 | -0.26 |
| CL014 | -0.25 | | 0.55 | | | | 0.85 | -0.46 | | -0.27 | | | -0.30 | -0.35 | -0.28 | -0.36 | | 0.33 | | | | | | | -0.58 | | -0.49 | -0.24 |
| CL015 | -0.28 | 0.47 | 0.77 | | | -0.37 | 0.61 | -0.25 | | -0.50 | 0.46 | | -0.28 | -0.63 | -0.31 | -0.41 | -0.35 | -0.34 | 0.27 | | -0.27 | -0.31 | | | -0.54 | | -0.74 | -0.24 |
| CL016 | | -0.40 | -0.71 | -0.39 | 0.28 | 0.39 | -0.47 | 0.51 | -0.32 | 0.52 | | -0.52 | | 0.69 | | 0.63 | 0.36 | 0.38 | | -0.51 | | 0.38 | 0.28 | -0.30 | 0.65 | -0.25 | 0.72 | |
| CL017 | | 0.36 | 0.74 | | | -0.34 | 0.68 | | | -0.55 | 0.44 | | | -0.59 | | -0.41 | -0.38 | | | | -0.30 | -0.35 | | | -0.50 | | -0.69 | |
| CL018 | -0.39 | -0.40 | -0.37 | -0.58 | 0.40 | 0.38 | | 0.39 | -0.54 | 0.41 | | -0.60 | -0.36 | 0.57 | -0.32 | 0.62 | 0.44 | 0.53 | | -0.57 | | 0.27 | 0.44 | -0.50 | 0.47 | -0.50 | 0.50 | -0.45 |
| CL019 | -0.49 | 0.36 | | -0.44 | | | -0.27 | 0.25 | -0.45 | | 0.49 | | -0.42 | | -0.45 | | | -0.42 | 0.49 | | | | | -0.44 | | -0.46 | | -0.47 |
| CL020 | -0.44 | 0.42 | 0.50 | -0.34 | | | 0.46 | | -0.37 | -0.29 | 0.64 | | -0.40 | -0.36 | -0.36 | | | -0.30 | 0.45 | -0.24 | | | | -0.37 | | -0.41 | -0.42 | -0.40 |
| CL021 | -0.28 | 0.55 | 0.67 | | | -0.39 | 0.37 | | | -0.46 | 0.58 | | -0.28 | -0.59 | -0.29 | -0.25 | -0.31 | -0.51 | 0.48 | | -0.27 | | -0.28 | | -0.33 | | -0.66 | -0.25 |
| CL022 | -0.36 | 0.54 | | -0.25 | | -0.29 | | | -0.27 | -0.30 | 0.53 | | -0.32 | -0.40 | -0.33 | | | -0.54 | 0.56 | | | | | -0.29 | | -0.31 | -0.40 | -0.32 |
| CL023 | | 0.28 | 0.72 | | | -0.25 | 0.79 | -0.35 | | -0.48 | | | -0.25 | -0.57 | -0.24 | -0.45 | -0.29 | | | | -0.26 | -0.32 | | | -0.60 | | -0.67 | |
| CL024 | -0.35 | 0.48 | 0.56 | | | -0.32 | 0.38 | | | -0.40 | 0.59 | | -0.30 | -0.53 | -0.30 | | | -0.46 | 0.38 | | | | | -0.25 | -0.30 | -0.29 | -0.58 | -0.31 |
| CL025 | -0.26 | -0.39 | -0.65 | -0.42 | | 0.43 | | | -0.38 | 0.36 | -0.35 | -0.45 | -0.24 | 0.51 | | 0.41 | 0.34 | 0.59 | | -0.45 | 0.25 | 0.30 | 0.52 | -0.39 | 0.34 | -0.34 | 0.50 | -0.28 |
| CL026 | -0.43 | -0.41 | -0.60 | -0.67 | 0.37 | 0.52 | | 0.31 | -0.60 | 0.56 | | -0.67 | -0.36 | 0.67 | -0.32 | 0.61 | 0.44 | 0.59 | | -0.66 | 0.34 | 0.28 | 0.41 | -0.56 | 0.41 | -0.55 | 0.57 | -0.46 |
| CL027 | -0.27 | 0.57 | 0.62 | | | -0.39 | 0.29 | | | -0.48 | 0.49 | 0.26 | -0.28 | -0.62 | -0.31 | -0.38 | -0.39 | -0.54 | 0.40 | 0.24 | -0.26 | -0.25 | -0.29 | | -0.44 | | -0.70 | |
| CL028 | -0.26 | -0.37 | -0.39 | -0.34 | | 0.42 | | | -0.31 | 0.25 | -0.34 | -0.41 | -0.25 | 0.36 | | | 0.30 | 0.80 | -0.39 | -0.40 | 0.31 | | 0.52 | -0.33 | | -0.29 | 0.28 | -0.26 |
| CL029 | -0.40 | | -0.27 | -0.42 | | 0.32 | | | -0.41 | | | -0.38 | -0.39 | 0.26 | -0.36 | | | 0.58 | | -0.37 | | | 0.48 | -0.44 | | -0.42 | | -0.40 |
| CL030 | -0.44 | | | -0.37 | | | 0.58 | | -0.38 | | | -0.43 | -0.38 | | -0.29 | | | 0.44 | | -0.42 | | | 0.43 | -0.43 | -0.38 | -0.42 | -0.29 | -0.40 |
| CL031 | -0.45 | 0.43 | | -0.38 | | | -0.30 | 0.32 | -0.40 | | 0.50 | | -0.34 | | -0.38 | | | -0.48 | 0.54 | | | | -0.26 | -0.38 | | -0.43 | | -0.40 |
| CL032 | -0.37 | 0.41 | 0.52 | | | -0.24 | 0.54 | | -0.25 | -0.47 | 0.52 | -0.25 | -0.29 | -0.45 | -0.25 | | -0.25 | | 0.27 | -0.25 | -0.24 | -0.24 | | -0.31 | -0.35 | -0.30 | -0.56 | -0.31 |
| CL033 | -0.27 | | -0.57 | -0.35 | | | -0.37 | 0.42 | -0.33 | | | -0.41 | | 0.36 | | 0.34 | | 0.29 | | -0.42 | | | 0.36 | -0.33 | 0.29 | -0.29 | 0.35 | |
| CL034 | | 0.27 | 0.71 | | | | 0.81 | -0.41 | | -0.49 | | | -0.24 | -0.56 | | -0.49 | -0.27 | | | | | -0.31 | | | -0.64 | | -0.65 | |
| CL035 | -0.39 | | -0.26 | -0.48 | 0.24 | | | 0.35 | -0.46 | | | -0.56 | -0.34 | | -0.28 | 0.34 | | 0.37 | | -0.54 | | | 0.44 | -0.47 | | -0.45 | | -0.41 |
| CL036 | -0.46 | -0.52 | -0.48 | -0.68 | 0.45 | 0.60 | | | -0.64 | 0.69 | | -0.56 | -0.45 | 0.63 | -0.39 | 0.61 | 0.53 | 0.60 | | -0.55 | 0.46 | 0.40 | 0.34 | -0.57 | 0.41 | -0.59 | 0.56 | -0.51 |
| CL037 | -0.54 | | | -0.65 | 0.49 | 0.31 | 0.30 | | -0.63 | 0.25 | 0.58 | -0.59 | -0.46 | | -0.38 | 0.32 | 0.29 | | | -0.60 | 0.29 | 0.26 | | -0.58 | | -0.62 | | -0.53 |
| CL038 | -0.51 | | -0.34 | -0.64 | 0.33 | | -0.34 | 0.39 | -0.60 | 0.29 | 0.49 | -0.43 | -0.41 | | -0.38 | 0.49 | 0.24 | | 0.35 | -0.47 | | 0.26 | | -0.56 | 0.34 | -0.57 | | -0.50 |
| CL039 | -0.51 | | | -0.50 | | | | 0.34 | -0.47 | | 0.36 | -0.51 | -0.41 | | -0.34 | | | | 0.29 | -0.52 | | | | -0.51 | | -0.51 | | -0.47 |
| CL040 | -0.45 | 0.41 | 0.34 | -0.32 | | -0.24 | | | -0.35 | -0.26 | 0.58 | | -0.42 | -0.37 | -0.44 | | | -0.39 | 0.44 | | | | | -0.36 | | -0.40 | -0.47 | -0.44 |
| CL041 | -0.36 | -0.36 | -0.51 | -0.60 | 0.47 | 0.39 | | 0.56 | -0.56 | 0.49 | | -0.61 | -0.36 | 0.66 | -0.27 | 0.68 | 0.39 | 0.40 | | -0.59 | | 0.45 | 0.33 | -0.51 | 0.66 | -0.49 | 0.58 | -0.39 |
| CL042 | -0.54 | | | -0.56 | 0.26 | | | 0.27 | -0.56 | | 0.48 | -0.35 | -0.51 | | -0.52 | 0.32 | | | 0.34 | -0.35 | | | | -0.53 | | -0.58 | | -0.55 |
| CL043 | -0.42 | 0.47 | | -0.33 | | | | | -0.35 | -0.25 | 0.57 | | -0.38 | -0.31 | -0.33 | | | -0.32 | 0.47 | | | | -0.24 | -0.36 | | -0.40 | -0.41 | -0.38 |
| CL044 | -0.27 | 0.45 | | | | -0.25 | | | | -0.32 | 0.38 | | | -0.37 | | | | -0.45 | 0.42 | | | | | | | | -0.37 | -0.25 |
| CL045 | -0.29 | -0.36 | -0.26 | -0.33 | | 0.42 | | -0.30 | -0.32 | 0.38 | -0.28 | -0.26 | -0.31 | 0.35 | -0.27 | | 0.34 | 0.63 | -0.37 | | 0.35 | | 0.33 | -0.32 | | -0.30 | 0.26 | -0.29 |
| CL046 | | 0.47 | 0.79 | | | -0.42 | 0.51 | | | -0.45 | 0.58 | | | -0.61 | | -0.32 | -0.32 | -0.50 | 0.36 | | -0.29 | -0.25 | -0.32 | | -0.34 | | -0.67 | |
| CL047 | | | 0.46 | | | | 0.79 | -0.44 | | -0.26 | | | | -0.29 | | -0.34 | | 0.46 | -0.26 | | | | 0.30 | | -0.57 | | -0.39 | |
| CL048 | | -0.27 | | | | 0.29 | 0.30 | | | | -0.39 | -0.35 | -0.24 | | | | | 0.73 | -0.40 | -0.33 | | | 0.58 | -0.30 | | -0.24 | | |
| CL049 | -0.51 | | | -0.56 | | 0.30 | 0.26 | | -0.56 | | | -0.56 | -0.49 | 0.26 | -0.42 | 0.24 | | 0.53 | | -0.55 | | | 0.49 | -0.59 | | -0.56 | | -0.52 |
| CL050 | | | -0.66 | | -0.33 | | -0.48 | | | | -0.45 | | | 0.32 | | | | 0.40 | | | 0.24 | | 0.32 | | | | 0.31 | |

| | CL142 | CL143 | CL144 | CL145 | CL146 | CL147 | CL148 | CL149 | CL150 | CL151 | CL152 | CL153 | CL154 | CL155 | CL156 | CL157 | CL158 | CL159 | CL160 | CL161 | CL162 | CL163 | CL164 | CL165 | CL166 | CL167 | CL168 | CL169 |
|---|---|---|---|---|---|---|---|---|---|---|---|---|---|---|---|---|---|---|---|---|---|---|---|---|---|---|---|---|
| CL051 | -0.37 | | -0.40 | -0.53 | 0.42 | 0.38 | | | -0.50 | 0.38 | | -0.49 | -0.36 | 0.38 | -0.27 | 0.37 | 0.42 | | | -0.47 | 0.38 | 0.42 | | -0.43 | 0.34 | -0.45 | 0.32 | -0.38 |
| CL052 | | -0.25 | -0.46 | -0.27 | | 0.34 | | | -0.24 | | -0.38 | -0.38 | | 0.29 | | | | 0.71 | -0.28 | -0.38 | 0.24 | | 0.51 | -0.31 | | -0.24 | 0.31 | |
| CL053 | -0.40 | | 0.42 | -0.37 | 0.38 | | 0.82 | | -0.40 | | | -0.25 | -0.44 | | -0.42 | | | 0.31 | | | | | | -0.39 | | -0.40 | | -0.43 |
| CL054 | -0.32 | 0.34 | 0.69 | | | | 0.72 | | | -0.31 | 0.48 | | -0.33 | -0.47 | -0.32 | | | | 0.24 | | | | | -0.24 | -0.39 | -0.25 | -0.56 | -0.29 |
| CL055 | -0.38 | | -0.30 | -0.45 | | 0.25 | | 0.41 | -0.42 | | | -0.55 | -0.31 | 0.29 | -0.25 | 0.32 | | 0.42 | | -0.52 | | | 0.45 | -0.46 | 0.30 | -0.41 | 0.27 | -0.37 |
| CL056 | -0.39 | -0.41 | -0.52 | -0.64 | 0.61 | 0.60 | | | -0.59 | 0.73 | | -0.53 | -0.36 | 0.60 | -0.33 | 0.68 | 0.62 | 0.36 | | -0.52 | 0.42 | 0.51 | | -0.51 | 0.52 | -0.53 | 0.61 | -0.44 |
| CL057 | | 0.48 | 0.76 | | | -0.43 | 0.47 | | | -0.45 | 0.55 | | | -0.62 | | -0.34 | -0.34 | -0.52 | 0.37 | | -0.31 | | -0.31 | | -0.31 | | -0.65 | |
| CL058 | -0.28 | | 0.56 | | | | 0.88 | -0.31 | | | 0.27 | | -0.33 | -0.31 | -0.26 | | | | | | | | | -0.25 | -0.39 | -0.25 | -0.42 | -0.29 |
| CL059 | -0.38 | | | -0.35 | | | | | -0.35 | | 0.32 | -0.29 | -0.36 | | -0.27 | | | | 0.30 | -0.29 | | | | -0.39 | | -0.36 | | -0.34 |
| CL060 | -0.38 | -0.37 | -0.46 | -0.54 | 0.28 | 0.43 | | | -0.50 | 0.40 | | -0.61 | -0.32 | 0.54 | -0.27 | 0.46 | 0.43 | 0.65 | -0.26 | -0.59 | 0.32 | | 0.44 | -0.48 | 0.29 | -0.45 | 0.46 | -0.38 |
| CL061 | -0.47 | -0.31 | -0.34 | -0.73 | 0.62 | 0.38 | | 0.46 | -0.68 | 0.57 | 0.27 | -0.69 | -0.40 | 0.59 | -0.33 | 0.73 | 0.42 | | | -0.67 | 0.25 | 0.42 | | -0.59 | 0.60 | -0.62 | 0.47 | -0.51 |
| CL062 | -0.34 | -0.27 | -0.38 | -0.44 | | 0.41 | | | -0.39 | | | -0.59 | -0.27 | 0.32 | | 0.25 | | 0.69 | -0.26 | -0.55 | | | 0.51 | -0.46 | | -0.41 | | -0.34 |
| CL063 | | -0.40 | -0.45 | -0.54 | 0.60 | 0.40 | -0.30 | 0.49 | -0.48 | 0.73 | | -0.46 | | 0.70 | | 0.81 | 0.46 | | | -0.44 | 0.27 | 0.47 | | -0.38 | 0.77 | -0.39 | 0.66 | -0.31 |
| CL064 | -0.27 | 0.47 | | | | -0.24 | | | | -0.38 | 0.53 | | | -0.32 | | | | -0.39 | 0.41 | | | | | -0.24 | | | -0.37 | |
| CL065 | -0.28 | 0.52 | 0.37 | | | -0.31 | | | | -0.44 | 0.55 | | -0.24 | -0.49 | | | | -0.51 | 0.44 | | | | -0.28 | | | | -0.44 | |
| CL066 | -0.46 | | 0.32 | -0.50 | 0.56 | | 0.56 | | -0.51 | | 0.59 | -0.45 | -0.45 | | -0.36 | 0.24 | | | | -0.43 | | | | -0.47 | | -0.51 | | -0.46 |
| CL067 | | 0.39 | 0.76 | | | -0.27 | 0.59 | | | -0.35 | 0.55 | | | -0.58 | | -0.29 | | -0.40 | 0.30 | | -0.24 | | -0.24 | | -0.31 | | -0.57 | |
| CL068 | -0.50 | -0.28 | -0.43 | -0.71 | 0.46 | 0.52 | | | -0.66 | 0.60 | | -0.56 | -0.44 | 0.49 | -0.42 | 0.62 | 0.50 | 0.30 | | -0.55 | 0.44 | 0.37 | | -0.58 | 0.36 | -0.62 | 0.43 | -0.53 |
| CL069 | | 0.35 | 0.47 | 0.55 | -0.49 | -0.44 | | -0.34 | 0.47 | -0.47 | | 0.73 | | -0.50 | | -0.59 | -0.37 | -0.36 | | 0.71 | -0.27 | -0.35 | -0.25 | 0.40 | -0.43 | 0.39 | -0.47 | 0.25 |
| CL070 | -0.29 | | | | | | | | -0.24 | | | | | | | | | | 0.31 | | | | | -0.25 | | -0.26 | | |
| CL071 | | -0.49 | -0.50 | -0.45 | 0.33 | 0.57 | | | -0.40 | 0.63 | | -0.39 | -0.24 | 0.60 | | 0.47 | 0.50 | 0.58 | -0.29 | -0.36 | 0.38 | 0.38 | 0.24 | -0.34 | 0.28 | -0.32 | 0.55 | -0.27 |
| CL072 | -0.32 | 0.41 | 0.73 | | | -0.29 | 0.69 | | | -0.36 | 0.50 | | -0.31 | -0.53 | -0.31 | -0.24 | | -0.25 | 0.32 | | | | | | -0.37 | -0.25 | -0.62 | -0.30 |
| CL073 | 0.78 | | 0.34 | 0.96 | -0.60 | -0.43 | | -0.33 | 0.92 | -0.57 | -0.37 | 0.71 | 0.72 | -0.51 | 0.66 | -0.68 | -0.48 | -0.34 | | 0.70 | -0.38 | -0.47 | | 0.86 | -0.44 | 0.88 | -0.37 | 0.80 |
| CL074 | -0.44 | -0.27 | -0.35 | -0.57 | 0.29 | 0.43 | | | -0.54 | 0.42 | | -0.49 | -0.46 | 0.45 | -0.42 | 0.41 | 0.39 | 0.49 | | -0.47 | 0.29 | | 0.34 | -0.54 | 0.29 | -0.53 | 0.33 | -0.46 |
| CL075 | | 0.53 | 0.27 | | | -0.30 | | | | -0.44 | 0.57 | | | -0.41 | | | -0.33 | -0.46 | 0.50 | | | | | | | | -0.48 | |
| CL076 | -0.33 | -0.44 | -0.54 | -0.60 | 0.55 | 0.54 | | 0.30 | -0.55 | 0.79 | | -0.47 | -0.32 | 0.75 | -0.32 | 0.72 | 0.52 | 0.39 | | -0.45 | 0.42 | 0.51 | | -0.47 | 0.63 | -0.46 | 0.69 | -0.39 |
| CL077 | 0.83 | | 0.30 | 0.96 | -0.52 | -0.36 | | -0.38 | 0.93 | -0.42 | -0.44 | 0.74 | 0.75 | -0.40 | 0.68 | -0.60 | -0.41 | -0.28 | | 0.73 | -0.33 | -0.40 | | 0.89 | -0.37 | 0.91 | -0.27 | 0.83 |
| CL078 | -0.46 | | | -0.57 | 0.53 | | | 0.48 | -0.54 | | 0.49 | -0.43 | -0.35 | | -0.33 | 0.46 | | -0.33 | 0.40 | -0.44 | | 0.29 | | -0.48 | 0.42 | -0.53 | | -0.47 |
| CL079 | -0.43 | -0.33 | -0.48 | -0.61 | 0.28 | 0.45 | | 0.30 | -0.56 | 0.47 | | -0.63 | -0.35 | 0.49 | -0.31 | 0.45 | 0.36 | 0.57 | | -0.62 | 0.35 | 0.25 | 0.39 | -0.53 | 0.24 | -0.49 | 0.36 | -0.42 |
| CL080 | -0.41 | | -0.45 | -0.57 | 0.38 | 0.48 | | | -0.54 | 0.53 | | -0.47 | -0.36 | 0.41 | -0.32 | 0.49 | 0.48 | | | -0.46 | 0.42 | 0.43 | | -0.47 | 0.40 | -0.52 | 0.45 | -0.42 |
| CL081 | -0.43 | | 0.49 | -0.42 | 0.42 | | 0.61 | | -0.44 | | 0.69 | -0.27 | -0.43 | -0.26 | -0.38 | | | | 0.27 | -0.27 | | | | -0.41 | | -0.46 | -0.33 | -0.45 |
| CL082 | -0.29 | 0.36 | 0.28 | | | | 0.29 | | | -0.45 | 0.32 | | | -0.36 | | -0.24 | | | | | | -0.36 | | | -0.35 | | -0.43 | |
| CL083 | 0.26 | 0.33 | 0.45 | 0.59 | -0.52 | -0.46 | | -0.25 | 0.51 | -0.56 | | 0.66 | | -0.52 | | -0.62 | -0.42 | -0.37 | | 0.65 | -0.31 | -0.38 | | 0.43 | -0.41 | 0.44 | -0.47 | 0.31 |
| CL084 | -0.44 | | | -0.47 | 0.29 | | | 0.30 | -0.46 | | | -0.39 | -0.39 | | -0.34 | 0.30 | | 0.26 | | -0.36 | | | 0.27 | -0.46 | | -0.47 | | -0.41 |
| CL085 | -0.37 | | | -0.36 | | | | | -0.37 | | | | -0.42 | | -0.42 | | 0.25 | | | | 0.29 | 0.31 | | -0.34 | | -0.35 | | -0.40 |
| CL086 | 0.40 | | 0.41 | 0.62 | -0.39 | -0.37 | | -0.44 | 0.56 | | -0.26 | 0.98 | 0.27 | -0.39 | | -0.48 | -0.31 | -0.35 | | 0.98 | | | -0.34 | 0.59 | -0.32 | 0.52 | -0.29 | 0.36 |
| CL087 | 0.82 | | 0.30 | 0.96 | -0.52 | -0.38 | | -0.40 | 0.92 | -0.43 | -0.44 | 0.76 | 0.73 | -0.42 | 0.65 | -0.61 | -0.41 | -0.29 | | 0.75 | -0.33 | -0.40 | | 0.88 | -0.37 | 0.90 | -0.28 | 0.81 |
| CL088 | 0.84 | | 0.27 | 0.96 | -0.51 | -0.35 | | -0.37 | 0.94 | -0.41 | -0.43 | 0.74 | 0.76 | -0.40 | 0.69 | -0.58 | -0.39 | -0.29 | | 0.72 | -0.32 | -0.41 | | 0.89 | -0.35 | 0.92 | -0.26 | 0.84 |
| CL089 | 0.91 | | | 0.98 | -0.52 | -0.32 | | -0.28 | 0.96 | -0.44 | -0.41 | 0.60 | 0.86 | -0.37 | 0.82 | -0.56 | -0.37 | -0.27 | | 0.59 | -0.33 | -0.41 | | 0.93 | -0.31 | 0.96 | | 0.92 |
| CL090 | 0.97 | | | 0.90 | -0.37 | | -0.25 | | 0.93 | -0.27 | -0.40 | 0.41 | 0.93 | | 0.92 | -0.37 | -0.28 | | -0.24 | 0.40 | -0.26 | -0.33 | | 0.92 | | 0.94 | | 0.96 |
| CL091 | -0.39 | 0.37 | | -0.38 | | | | 0.32 | -0.37 | | 0.63 | | -0.35 | | -0.35 | | | -0.52 | 0.58 | | | | -0.31 | -0.37 | | -0.39 | | -0.38 |
| CL092 | -0.33 | -0.43 | -0.42 | -0.51 | 0.33 | 0.51 | | | -0.48 | 0.68 | | -0.27 | -0.38 | 0.57 | -0.38 | 0.45 | 0.48 | 0.50 | -0.27 | -0.26 | 0.41 | 0.30 | | -0.42 | | -0.43 | 0.43 | -0.40 |
| CL093 | | -0.36 | -0.56 | | | | -0.49 | 0.37 | | | -0.43 | -0.27 | 0.27 | 0.46 | 0.32 | 0.29 | | 0.32 | | -0.26 | | | | | 0.45 | | 0.63 | |
| CL094 | -0.40 | | -0.34 | -0.64 | 0.52 | 0.29 | | 0.56 | -0.59 | 0.53 | 0.36 | -0.56 | -0.33 | 0.49 | -0.31 | 0.67 | 0.32 | | | -0.56 | | 0.41 | | -0.50 | 0.54 | -0.53 | 0.38 | -0.43 |
| CL095 | | 0.33 | 0.41 | 0.53 | -0.49 | -0.44 | | -0.24 | 0.46 | -0.52 | | 0.63 | | -0.50 | | -0.57 | -0.42 | -0.40 | | 0.61 | -0.28 | -0.39 | -0.25 | 0.38 | -0.38 | 0.38 | -0.48 | 0.26 |
| CL096 | 0.93 | | | 0.96 | -0.48 | -0.29 | | | 0.96 | -0.42 | -0.43 | 0.53 | 0.89 | -0.30 | 0.87 | -0.52 | -0.37 | | | 0.52 | -0.29 | -0.40 | | 0.93 | -0.27 | 0.96 | | 0.95 |
| CL097 | 0.96 | | | 0.89 | -0.35 | | | | 0.92 | -0.26 | -0.41 | 0.39 | 0.94 | | 0.92 | -0.36 | -0.26 | | -0.26 | 0.38 | -0.26 | -0.31 | | 0.92 | | 0.94 | | 0.96 |
| CL098 | 0.81 | | 0.31 | 0.96 | -0.55 | -0.37 | | -0.39 | 0.92 | -0.45 | -0.45 | 0.74 | 0.72 | -0.43 | 0.65 | -0.62 | -0.41 | -0.27 | | 0.73 | -0.33 | -0.43 | | 0.87 | -0.39 | 0.90 | -0.28 | 0.81 |
| CL099 | -0.28 | -0.42 | -0.61 | -0.50 | | 0.45 | | | -0.44 | 0.45 | | -0.56 | | 0.55 | | 0.47 | 0.36 | 0.46 | | -0.56 | 0.32 | 0.28 | 0.45 | -0.39 | 0.29 | -0.38 | 0.48 | -0.28 |
| CL100 | | | | | | | -0.50 | | | | | 0.32 | | | | | | -0.32 | | 0.28 | | | | | | | | |
| CL101 | | -0.33 | -0.46 | | | 0.36 | | | | | -0.42 | -0.32 | | 0.43 | | | | 0.72 | -0.34 | -0.31 | 0.24 | | 0.50 | | | | 0.37 | |
| CL102 | -0.45 | | -0.34 | -0.63 | 0.36 | 0.24 | | 0.54 | -0.60 | 0.36 | 0.25 | -0.52 | -0.32 | 0.43 | -0.32 | 0.58 | | | | -0.54 | | | | -0.52 | 0.45 | -0.53 | 0.28 | -0.46 |
| CL103 | 0.93 | | | 0.87 | -0.40 | | -0.24 | | 0.90 | -0.36 | -0.36 | 0.32 | 0.94 | | 0.94 | -0.39 | -0.30 | | | 0.31 | -0.27 | -0.34 | | 0.89 | | 0.92 | | 0.95 |
| CL104 | | -0.50 | -0.43 | -0.37 | 0.53 | 0.38 | -0.25 | 0.32 | -0.31 | 0.73 | | -0.30 | | 0.71 | | 0.69 | 0.43 | 0.29 | | -0.27 | 0.25 | 0.47 | | -0.24 | 0.72 | | 0.74 | |
| CL105 | 0.89 | | | 0.88 | -0.46 | -0.24 | | | 0.90 | -0.45 | -0.34 | 0.32 | 0.91 | -0.27 | 0.93 | -0.47 | -0.38 | | | 0.31 | -0.31 | -0.41 | | 0.87 | -0.24 | 0.90 | | 0.93 |
| CL106 | -0.29 | -0.42 | -0.51 | -0.50 | 0.43 | 0.49 | | 0.27 | -0.47 | 0.61 | | -0.42 | -0.30 | 0.58 | -0.25 | 0.60 | 0.48 | 0.33 | | -0.40 | 0.36 | 0.49 | | -0.38 | 0.56 | -0.41 | 0.64 | -0.33 |
| CL107 | -0.26 | -0.36 | -0.50 | -0.48 | 0.39 | 0.50 | | 0.24 | -0.43 | 0.48 | | -0.54 | -0.26 | 0.59 | | 0.50 | 0.40 | 0.54 | | -0.50 | 0.27 | 0.39 | 0.26 | -0.38 | 0.39 | -0.39 | 0.56 | -0.29 |
| CL108 | 0.94 | | | 0.85 | -0.37 | | | | 0.89 | -0.30 | -0.38 | 0.29 | 0.94 | | 0.95 | -0.35 | -0.27 | | -0.25 | 0.28 | -0.26 | -0.33 | | 0.89 | | 0.91 | | 0.94 |
| CL109 | -0.51 | | -0.29 | -0.68 | 0.57 | 0.29 | | 0.29 | -0.65 | 0.46 | 0.46 | -0.51 | -0.44 | 0.30 | -0.43 | 0.57 | 0.32 | | | -0.53 | | 0.39 | | -0.60 | 0.40 | -0.57 | 0.26 | -0.55 |
| CL110 | 0.39 | | 0.40 | 0.61 | -0.41 | -0.36 | | -0.46 | 0.56 | | -0.25 | 0.98 | 0.26 | -0.40 | | -0.48 | -0.31 | -0.34 | | 0.97 | | -0.24 | -0.33 | 0.58 | -0.33 | 0.51 | -0.31 | 0.35 |

| | CL142 | CL143 | CL144 | CL145 | CL146 | CL147 | CL148 | CL149 | CL150 | CL151 | CL152 | CL153 | CL154 | CL155 | CL156 | CL157 | CL158 | CL159 | CL160 | CL161 | CL162 | CL163 | CL164 | CL165 | CL166 | CL167 | CL168 | CL169 |
|---|---|---|---|---|---|---|---|---|---|---|---|---|---|---|---|---|---|---|---|---|---|---|---|---|---|---|---|---|
| CL111 | 0.24 | 0.33 | 0.47 | 0.57 | -0.50 | -0.46 | | -0.34 | 0.49 | -0.49 | | 0.73 | | -0.52 | | -0.61 | -0.39 | -0.37 | | 0.72 | -0.29 | -0.34 | -0.25 | 0.43 | -0.43 | 0.41 | -0.48 | 0.27 |
| CL112 | -0.31 | -0.51 | -0.48 | -0.51 | 0.36 | 0.52 | | | -0.47 | 0.67 | | -0.45 | -0.31 | 0.59 | -0.28 | 0.54 | 0.49 | 0.59 | | -0.43 | 0.39 | 0.33 | 0.31 | -0.44 | 0.35 | -0.40 | 0.56 | -0.34 |
| CL113 | | -0.42 | -0.56 | | | 0.33 | -0.38 | 0.48 | | 0.40 | -0.35 | -0.42 | | 0.62 | | 0.51 | 0.27 | 0.44 | | -0.40 | | 0.25 | | | 0.59 | | 0.71 | |
| CL114 | 0.44 | | 0.39 | 0.64 | -0.39 | -0.35 | | -0.41 | 0.59 | | -0.24 | 0.98 | 0.30 | -0.40 | | -0.47 | -0.34 | -0.35 | | 0.98 | | -0.24 | -0.35 | 0.62 | -0.30 | 0.54 | -0.30 | 0.39 |
| CL115 | 0.95 | | | 0.94 | -0.44 | | | | 0.96 | -0.32 | -0.42 | 0.52 | 0.90 | -0.25 | 0.86 | -0.46 | -0.33 | | | 0.51 | -0.26 | -0.36 | | 0.94 | | 0.96 | | 0.94 |
| CL116 | 0.80 | | 0.30 | 0.95 | -0.58 | -0.43 | | -0.27 | 0.92 | -0.59 | -0.33 | 0.64 | 0.76 | -0.49 | 0.73 | -0.66 | -0.49 | -0.36 | | 0.63 | -0.41 | -0.48 | | 0.86 | -0.42 | 0.89 | -0.36 | 0.83 |
| CL117 | -0.50 | | -0.26 | -0.56 | 0.30 | | | | -0.54 | 0.28 | 0.31 | -0.35 | -0.47 | | -0.41 | 0.37 | | | 0.29 | -0.37 | 0.30 | | | -0.49 | | -0.57 | | -0.49 |
| CL118 | | -0.48 | -0.51 | -0.38 | | 0.50 | | | -0.34 | 0.56 | -0.29 | -0.40 | | 0.51 | | 0.38 | 0.39 | 0.60 | -0.31 | -0.37 | 0.35 | 0.32 | 0.30 | -0.33 | 0.26 | -0.28 | 0.48 | |
| CL119 | | | -0.27 | | | 0.35 | | -0.28 | | 0.26 | -0.41 | | | | | | 0.25 | 0.59 | -0.43 | | 0.30 | | 0.36 | | | | | |
| CL120 | | 0.24 | | | | | | | | | 0.27 | | | | | | | -0.42 | 0.24 | | | | | | | | | |
| CL121 | -0.39 | 0.25 | | -0.43 | | | -0.26 | 0.29 | -0.42 | | 0.42 | | -0.32 | | -0.36 | 0.30 | | -0.33 | 0.41 | | | | | -0.38 | | -0.42 | | -0.38 |
| CL122 | -0.30 | | -0.39 | -0.56 | 0.59 | 0.36 | | 0.41 | -0.52 | 0.73 | | -0.36 | -0.28 | 0.61 | -0.30 | 0.74 | 0.39 | | | -0.34 | 0.27 | 0.50 | | -0.41 | 0.72 | -0.43 | 0.59 | -0.38 |
| CL123 | 0.93 | | | 0.96 | -0.45 | -0.29 | | | 0.95 | -0.39 | -0.40 | 0.53 | 0.89 | -0.30 | 0.86 | -0.50 | -0.35 | -0.24 | | 0.52 | -0.31 | -0.37 | | 0.93 | -0.26 | 0.97 | | 0.94 |
| CL124 | 0.95 | | | 0.94 | -0.44 | -0.25 | | | 0.96 | -0.36 | -0.41 | 0.50 | 0.92 | -0.27 | 0.88 | -0.47 | -0.32 | | | 0.50 | -0.28 | -0.37 | | 0.94 | | 0.96 | | 0.95 |
| CL125 | 0.93 | | | 0.96 | -0.47 | -0.29 | | -0.25 | 0.96 | -0.40 | -0.40 | 0.56 | 0.88 | -0.33 | 0.83 | -0.51 | -0.37 | -0.26 | | 0.56 | -0.32 | -0.38 | | 0.93 | -0.28 | 0.96 | | 0.93 |
| CL126 | 0.92 | | | 0.80 | -0.31 | | -0.25 | | 0.84 | -0.28 | -0.32 | | 0.93 | | 0.96 | -0.30 | -0.25 | | | | -0.26 | -0.26 | | 0.84 | | 0.87 | | 0.94 |
| CL127 | 0.75 | | 0.38 | 0.94 | -0.60 | -0.47 | | -0.34 | 0.90 | -0.59 | -0.36 | 0.72 | 0.68 | -0.53 | 0.63 | -0.69 | -0.48 | -0.34 | | 0.70 | -0.40 | -0.48 | | 0.83 | -0.46 | 0.86 | -0.41 | 0.78 |
| CL128 | -0.54 | | 0.28 | -0.56 | 0.50 | | 0.43 | | -0.57 | | 0.63 | -0.30 | -0.49 | | -0.48 | 0.25 | | | 0.28 | -0.28 | | 0.25 | | -0.50 | | -0.55 | | -0.55 |
| CL129 | -0.35 | | -0.30 | -0.53 | 0.48 | 0.34 | -0.24 | 0.35 | -0.50 | 0.57 | 0.28 | -0.36 | -0.24 | 0.35 | -0.30 | 0.57 | 0.37 | | | -0.35 | 0.24 | 0.36 | | -0.39 | 0.52 | -0.46 | 0.38 | -0.39 |
| CL130 | 0.28 | | 0.24 | 0.46 | -0.24 | -0.24 | | -0.36 | 0.40 | | | 0.89 | | | | -0.32 | | -0.33 | | 0.89 | | | -0.36 | 0.48 | | 0.37 | | 0.24 |
| CL131 | -0.25 | -0.54 | -0.53 | -0.52 | 0.53 | 0.55 | | 0.25 | -0.47 | 0.70 | | -0.52 | | 0.72 | | 0.67 | 0.51 | 0.40 | | -0.47 | 0.35 | 0.47 | | -0.38 | 0.52 | -0.37 | 0.68 | -0.29 |
| CL132 | 0.93 | | | 0.95 | -0.43 | -0.29 | | | 0.95 | -0.38 | -0.37 | 0.52 | 0.90 | -0.29 | 0.87 | -0.47 | -0.36 | -0.26 | | 0.51 | -0.31 | -0.39 | | 0.93 | | 0.95 | | 0.94 |
| CL133 | -0.31 | -0.40 | -0.53 | -0.58 | 0.57 | 0.45 | -0.28 | 0.39 | -0.52 | 0.77 | | -0.40 | -0.30 | 0.74 | -0.31 | 0.74 | 0.52 | 0.28 | | -0.38 | 0.33 | 0.57 | | -0.42 | 0.73 | -0.43 | 0.70 | -0.37 |
| CL134 | 0.96 | | | 0.90 | -0.36 | | | | 0.92 | -0.29 | -0.39 | 0.39 | 0.93 | | 0.91 | -0.38 | -0.26 | | -0.24 | 0.39 | -0.26 | -0.31 | | 0.91 | | 0.94 | | 0.95 |
| CL135 | 0.96 | | | 0.90 | -0.33 | | | | 0.93 | | -0.40 | 0.50 | 0.90 | | 0.85 | -0.37 | -0.29 | | | 0.49 | -0.24 | -0.32 | | 0.92 | | 0.94 | | 0.93 |
| CL136 | 0.94 | | | 0.88 | -0.40 | | -0.25 | | 0.90 | -0.32 | -0.39 | 0.34 | 0.94 | | 0.93 | -0.38 | -0.29 | | -0.24 | 0.34 | -0.24 | -0.35 | | 0.90 | | 0.93 | | 0.95 |
| CL137 | | -0.48 | -0.55 | -0.39 | 0.37 | 0.45 | | 0.33 | -0.35 | 0.57 | | -0.44 | | 0.65 | | 0.61 | 0.42 | 0.51 | | -0.42 | 0.24 | 0.43 | 0.29 | -0.28 | 0.61 | -0.27 | 0.70 | |
| CL138 | 0.93 | | | 0.84 | -0.34 | | | | 0.87 | -0.28 | -0.35 | 0.27 | 0.93 | | 0.95 | -0.35 | -0.27 | | | 0.25 | | -0.30 | | 0.87 | | 0.90 | | 0.94 |
| CL139 | -0.35 | -0.44 | -0.52 | -0.59 | 0.47 | 0.49 | | | -0.55 | 0.75 | | -0.51 | -0.32 | 0.65 | -0.28 | 0.67 | 0.52 | 0.32 | | -0.49 | 0.41 | 0.48 | | -0.46 | 0.50 | -0.45 | 0.62 | -0.39 |
| CL140 | 0.40 | | 0.40 | 0.61 | -0.40 | -0.37 | | -0.44 | 0.57 | | -0.25 | 0.98 | 0.26 | -0.41 | | -0.48 | -0.30 | -0.35 | | 0.98 | | -0.24 | -0.32 | 0.58 | -0.32 | 0.51 | -0.33 | 0.35 |
| CL141 | 0.92 | | | 0.87 | -0.42 | | | | 0.89 | -0.38 | -0.37 | 0.31 | 0.92 | | 0.95 | -0.43 | -0.33 | | | 0.30 | -0.28 | -0.36 | | 0.89 | | 0.90 | | 0.94 |
| CL142 | | | | 0.87 | -0.34 | | -0.25 | | 0.91 | | -0.40 | 0.41 | 0.90 | | 0.89 | -0.32 | -0.26 | | -0.25 | 0.41 | -0.29 | -0.28 | | 0.90 | | 0.93 | | 0.94 |
| CL143 | | | | | -0.30 | -0.33 | | | | -0.44 | 0.25 | | | -0.41 | | -0.39 | -0.29 | -0.33 | 0.36 | | | | -0.26 | | -0.27 | | -0.44 | |
| CL144 | | | | | | -0.42 | 0.56 | -0.30 | | -0.43 | 0.24 | 0.40 | | -0.57 | | -0.40 | -0.34 | -0.29 | | 0.40 | -0.39 | -0.29 | -0.24 | | -0.40 | | -0.58 | |
| CL145 | | | | | -0.55 | -0.33 | | -0.29 | 0.96 | -0.48 | -0.45 | 0.63 | 0.82 | -0.41 | 0.78 | -0.62 | -0.41 | -0.24 | | 0.62 | -0.32 | -0.43 | | 0.91 | -0.36 | 0.94 | -0.25 | 0.89 |
| CL146 | | | | | | 0.30 | | 0.26 | -0.52 | 0.49 | 0.36 | -0.41 | -0.31 | 0.40 | -0.31 | 0.59 | 0.34 | | | -0.38 | | 0.42 | | -0.41 | 0.52 | -0.43 | 0.32 | -0.40 |
| CL147 | | | | | | | | | -0.30 | 0.54 | | -0.38 | | 0.37 | | 0.43 | 0.33 | 0.33 | | -0.36 | 0.40 | 0.27 | | -0.26 | | | 0.46 | |
| CL148 | | | | | | | | -0.32 | | | | | -0.26 | -0.27 | | | | 0.25 | | | | | | | -0.38 | | -0.40 | -0.25 |
| CL149 | | | | | | | | | -0.24 | | | -0.44 | | 0.38 | | 0.44 | | | 0.24 | -0.42 | | | | | 0.61 | | 0.32 | |
| CL150 | | | | | | | | | | -0.43 | -0.44 | 0.57 | 0.86 | -0.34 | 0.82 | -0.55 | -0.38 | | | 0.57 | -0.32 | -0.42 | | 0.92 | -0.29 | 0.94 | | 0.90 |
| CL151 | | | | | | | | | | | | -0.24 | -0.25 | 0.63 | -0.28 | 0.64 | 0.48 | 0.26 | | | 0.37 | 0.50 | | -0.31 | 0.52 | -0.34 | 0.59 | -0.30 |
| CL152 | | | | | | | | | | | | -0.24 | -0.36 | | -0.35 | | | -0.38 | 0.41 | -0.25 | | | | -0.37 | | -0.42 | | -0.39 |
| CL153 | | | | | | | | | | | | | 0.28 | -0.41 | | -0.50 | -0.33 | -0.37 | | 0.97 | | -0.25 | -0.34 | 0.60 | -0.32 | 0.53 | -0.32 | 0.37 |
| CL154 | | | | | | | | | | | | | | | 0.92 | -0.31 | -0.27 | | -0.25 | 0.28 | | -0.32 | | 0.87 | | 0.89 | | 0.91 |
| CL155 | | | | | | | | | | | | | | | | 0.63 | 0.33 | 0.38 | | -0.38 | 0.28 | 0.35 | | -0.29 | 0.56 | -0.28 | 0.66 | |
| CL156 | | | | | | | | | | | | | | | | -0.29 | -0.24 | | -0.26 | | | -0.28 | | 0.80 | | 0.85 | | 0.92 |
| CL157 | | | | | | | | | | | | | | | | | 0.44 | | | -0.47 | | 0.43 | | -0.47 | 0.67 | -0.49 | 0.57 | -0.41 |
| CL158 | | | | | | | | | | | | | | | | | | | | -0.33 | 0.38 | 0.38 | | -0.31 | 0.32 | -0.34 | 0.43 | -0.28 |
| CL159 | | | | | | | | | | | | | | | | | | | -0.37 | -0.33 | | | 0.42 | -0.26 | | | 0.35 | |
| CL160 | | | | | | | | | | | | | | | | | | | | | | | -0.27 | -0.24 | | -0.26 | | |
| CL161 | | | | | | | | | | | | | | | | | | | | | | | -0.32 | 0.59 | -0.30 | 0.52 | -0.31 | 0.36 |
| CL162 | | | | | | | | | | | | | | | | | | | | | | | | | | -0.29 | 0.35 | -0.24 |
| CL163 | | | | | | | | | | | | | | | | | | | | | | | | -0.37 | 0.43 | -0.35 | 0.42 | -0.28 |
| CL164 | | | | | | | | | | | | | | | | | | | | | | | | | | | | |
| CL165 | | | | | | | | | | | | | | | | | | | | | | | | | | 0.91 | | 0.89 |
| CL166 | | | | | | | | | | | | | | | | | | | | | | | | | | | 0.64 | |
| CL167 | | | | | | | | | | | | | | | | | | | | | | | | | | | | 0.94 |
| CL168 | | | | | | | | | | | | | | | | | | | | | | | | | | | | |
| CL169 | | | | | | | | | | | | | | | | | | | | | | | | | | | | |
| CL170 | | | | | | | | | | | | | | | | | | | | | | | | | | | | |

| | CL170 | CL171 | CL172 | CL173 | CL174 | CL175 | CL176 | CL177 | CL178 | CL179 | CL180 | CL181 | CL182 | CL183 | CL184 | CL185 | CL186 | CL187 | CL188 | CL189 | CL190 | CL191 | CL192 | CL193 | CL194 | CL195 | CL196 | CL197 |
|---|---|---|---|---|---|---|---|---|---|---|---|---|---|---|---|---|---|---|---|---|---|---|---|---|---|---|---|---|
| CL001 | | | -0.24 | | -0.36 | | 0.28 | | -0.40 | -0.36 | -0.26 | | -0.38 | -0.26 | | | | | -0.39 | | -0.30 | | -0.43 | -0.40 | -0.30 | -0.33 | | |
| CL002 | 0.42 | | | -0.33 | -0.68 | -0.33 | -0.46 | | | | | -0.27 | -0.57 | -0.45 | -0.53 | 0.69 | -0.45 | -0.38 | | | -0.29 | | | | | -0.37 | | |
| CL003 | 0.50 | | | -0.27 | -0.67 | | -0.56 | | | -0.24 | | | -0.53 | -0.58 | -0.58 | 0.67 | -0.44 | -0.30 | | | -0.38 | | | | | -0.35 | 0.27 | |
| CL004 | 0.46 | 0.26 | | -0.29 | -0.64 | | -0.64 | | | | | | -0.42 | -0.64 | -0.62 | 0.53 | -0.40 | -0.29 | 0.26 | | -0.42 | | | | | -0.36 | 0.25 | |
| CL005 | 0.24 | | -0.30 | | -0.29 | | | | -0.38 | -0.41 | -0.37 | | -0.25 | | | 0.53 | | | -0.35 | | -0.30 | | -0.35 | -0.41 | | | | |
| CL006 | -0.38 | | -0.30 | 0.28 | 0.25 | | 0.69 | | -0.38 | -0.34 | -0.33 | | | 0.26 | 0.29 | | 0.30 | 0.26 | -0.35 | | | | -0.34 | -0.36 | -0.24 | 0.28 | | 0.27 |
| CL007 | | | -0.50 | 0.36 | 0.26 | | 0.52 | | -0.59 | -0.58 | -0.61 | 0.28 | | 0.28 | | | 0.31 | 0.24 | -0.57 | 0.36 | -0.32 | | -0.55 | -0.61 | | 0.34 | | 0.37 |
| CL008 | 0.41 | | | -0.26 | -0.65 | | -0.60 | | | | | | -0.41 | -0.66 | -0.64 | 0.53 | -0.37 | -0.25 | | | -0.44 | | | | | -0.31 | 0.24 | |
| CL009 | 0.32 | 0.27 | -0.43 | | -0.49 | | -0.33 | | | -0.44 | -0.31 | | -0.27 | -0.47 | -0.52 | 0.45 | | | | | -0.57 | | | | | | | |
| CL010 | 0.41 | 0.27 | | -0.30 | -0.74 | -0.27 | -0.45 | | | | | -0.28 | -0.57 | -0.61 | -0.61 | 0.59 | -0.52 | -0.31 | | | -0.35 | | | | | -0.49 | | |
| CL011 | | | | -0.28 | -0.60 | -0.30 | | -0.39 | | | | | -0.47 | -0.45 | -0.46 | 0.68 | -0.32 | | | | -0.28 | | | | | -0.29 | | |
| CL012 | 0.49 | | -0.42 | | | | -0.35 | 0.28 | -0.29 | -0.45 | -0.49 | | | | | 0.30 | | | -0.26 | 0.24 | -0.40 | | -0.27 | -0.33 | 0.30 | | 0.27 | |
| CL013 | 0.49 | 0.24 | | -0.27 | -0.62 | | -0.63 | | | | | -0.24 | -0.43 | -0.61 | -0.62 | 0.55 | -0.36 | -0.26 | | | -0.42 | | | | | -0.33 | 0.27 | |
| CL014 | | | | | -0.54 | -0.33 | | -0.38 | | -0.24 | | | -0.44 | -0.42 | -0.40 | 0.68 | -0.30 | | | | -0.34 | | | | | | | |
| CL015 | 0.40 | 0.30 | | -0.35 | -0.73 | | -0.58 | | | | | -0.26 | -0.53 | -0.67 | -0.65 | 0.61 | -0.47 | -0.34 | 0.24 | | -0.42 | | | | | -0.44 | | |
| CL016 | | | | 0.51 | 0.75 | 0.28 | 0.55 | 0.36 | -0.52 | | -0.39 | 0.37 | 0.57 | 0.76 | 0.73 | -0.45 | 0.57 | 0.33 | -0.52 | 0.24 | | | -0.49 | -0.47 | | 0.54 | | 0.29 |
| CL017 | 0.47 | | | -0.35 | -0.72 | -0.24 | -0.56 | | | | | -0.32 | -0.62 | -0.58 | -0.54 | 0.66 | -0.52 | -0.35 | | | -0.26 | | | | | -0.47 | | -0.26 |
| CL018 | | | -0.44 | 0.46 | 0.51 | | 0.43 | | -0.58 | -0.53 | -0.59 | 0.36 | 0.39 | 0.49 | 0.46 | | 0.42 | | -0.55 | 0.39 | | | -0.53 | -0.55 | | 0.51 | | 0.40 |
| CL019 | | | -0.41 | | | 0.25 | | 0.34 | | -0.39 | -0.38 | | | | | | | | | | -0.47 | | | | 0.39 | | | |
| CL020 | 0.57 | | -0.34 | | -0.43 | | -0.47 | 0.30 | | -0.38 | -0.32 | | -0.36 | -0.37 | -0.40 | 0.49 | -0.25 | | | | -0.42 | | | | 0.29 | | | |
| CL021 | 0.47 | 0.29 | | | -0.63 | | -0.61 | | | | | | -0.46 | -0.60 | -0.55 | 0.47 | -0.36 | -0.30 | | | -0.40 | | | | | -0.39 | 0.26 | |
| CL022 | 0.28 | 0.30 | -0.26 | | -0.40 | | -0.32 | 0.25 | | -0.27 | | | -0.27 | -0.38 | -0.39 | | | | | | -0.41 | | | | | -0.28 | 0.27 | |
| CL023 | 0.30 | 0.24 | | -0.32 | -0.72 | -0.32 | -0.37 | -0.31 | | | | -0.28 | -0.56 | -0.54 | -0.57 | 0.69 | -0.47 | -0.27 | | | -0.34 | | | | | -0.39 | | |
| CL024 | 0.44 | 0.28 | -0.25 | | -0.57 | | -0.55 | | | -0.25 | | | -0.42 | -0.53 | -0.50 | 0.44 | -0.35 | | | | -0.39 | | | | | -0.38 | 0.29 | |
| CL025 | -0.41 | | -0.31 | 0.40 | 0.47 | | 0.68 | | -0.47 | -0.37 | -0.44 | 0.36 | 0.38 | 0.55 | 0.41 | -0.31 | 0.45 | 0.26 | -0.45 | 0.30 | | | -0.43 | -0.45 | | 0.41 | | 0.39 |
| CL026 | | | -0.48 | 0.52 | 0.57 | | 0.64 | | -0.67 | -0.54 | -0.65 | 0.47 | 0.48 | 0.47 | 0.52 | | 0.59 | 0.37 | -0.64 | 0.39 | | | -0.62 | -0.64 | | 0.55 | | 0.43 |
| CL027 | 0.32 | 0.36 | | -0.36 | -0.67 | | -0.53 | | 0.26 | | | -0.28 | -0.48 | -0.62 | -0.62 | 0.41 | -0.42 | -0.29 | 0.28 | | -0.41 | -0.25 | | | | -0.47 | | -0.25 |
| CL028 | -0.32 | | -0.28 | 0.26 | | | 0.69 | | -0.40 | -0.36 | -0.34 | | | | 0.26 | | | 0.29 | -0.38 | | | | -0.39 | -0.39 | -0.27 | | | 0.29 |
| CL029 | -0.26 | | -0.43 | | | | 0.56 | | -0.37 | -0.46 | -0.42 | | | | | | | | -0.33 | 0.25 | -0.32 | | -0.34 | -0.41 | | | | |
| CL030 | | | -0.33 | | -0.34 | | | | -0.40 | -0.44 | -0.36 | | -0.31 | | | 0.42 | | | -0.38 | | -0.35 | | -0.42 | -0.43 | | | | |
| CL031 | | | -0.34 | | | | | 0.39 | | -0.34 | -0.34 | | | | | | | | | | -0.36 | | | | 0.33 | | | |
| CL032 | 0.46 | | -0.24 | | -0.59 | | -0.39 | | | -0.32 | | | -0.48 | -0.51 | -0.43 | 0.57 | -0.37 | -0.27 | | | -0.34 | -0.24 | -0.25 | | | -0.36 | | |
| CL033 | -0.26 | | | | 0.34 | | 0.51 | 0.30 | -0.41 | -0.29 | -0.35 | | | 0.42 | 0.26 | -0.33 | 0.25 | | -0.40 | 0.28 | | | -0.40 | -0.41 | | | | 0.30 |
| CL034 | 0.30 | 0.26 | | -0.35 | -0.73 | -0.33 | -0.40 | -0.30 | | | | -0.30 | -0.57 | -0.56 | -0.58 | 0.71 | -0.48 | -0.26 | | | -0.34 | | | | | -0.42 | | |
| CL035 | | | -0.37 | 0.30 | | | 0.27 | | -0.53 | -0.46 | -0.50 | 0.25 | | 0.28 | | | | | -0.54 | | | | -0.51 | -0.54 | | | | 0.26 |
| CL036 | | | -0.56 | 0.57 | 0.59 | | 0.53 | | -0.56 | -0.58 | -0.68 | 0.51 | 0.55 | 0.47 | 0.54 | | 0.63 | 0.42 | -0.51 | 0.38 | -0.26 | 0.33 | -0.50 | -0.52 | | 0.64 | | 0.40 |
| CL037 | 0.28 | | -0.53 | 0.32 | | | | 0.27 | -0.59 | -0.54 | -0.65 | 0.33 | | | | | | | -0.52 | | -0.39 | | -0.55 | -0.58 | 0.34 | 0.26 | 0.32 | |
| CL038 | | | -0.47 | 0.36 | | 0.43 | | 0.51 | -0.42 | -0.47 | -0.59 | 0.31 | 0.31 | | | -0.27 | 0.33 | 0.27 | -0.42 | 0.30 | -0.32 | | -0.45 | -0.40 | 0.36 | | | |
| CL039 | | | -0.39 | | | | | 0.29 | -0.50 | -0.49 | -0.46 | | | | | | | | -0.50 | | -0.40 | | -0.52 | -0.50 | | | | |
| CL040 | 0.35 | 0.24 | -0.36 | | -0.39 | | -0.38 | | | -0.36 | -0.30 | | | -0.43 | -0.41 | 0.32 | | | | | -0.49 | | | | 0.32 | | 0.24 | |
| CL041 | | -0.25 | -0.43 | 0.52 | 0.63 | 0.26 | 0.42 | 0.36 | -0.60 | -0.48 | -0.62 | 0.47 | 0.49 | 0.62 | 0.54 | -0.26 | 0.51 | 0.24 | -0.57 | 0.42 | | | -0.56 | -0.59 | | 0.60 | | 0.38 |
| CL042 | | | -0.53 | | | | | 0.28 | -0.33 | -0.53 | -0.50 | | | | | | | | -0.31 | 0.25 | -0.48 | | -0.35 | -0.37 | 0.31 | | | |
| CL043 | | 0.25 | -0.31 | | -0.39 | | | 0.38 | | -0.33 | -0.28 | | -0.33 | -0.44 | -0.31 | | | | | | -0.43 | | | | | -0.27 | | |
| CL044 | 0.26 | 0.24 | | | -0.35 | | -0.26 | | | | | | -0.28 | -0.32 | -0.41 | | | | | | -0.28 | | | | | -0.34 | | |
| CL045 | -0.28 | | -0.36 | 0.30 | 0.26 | | 0.45 | -0.28 | -0.28 | -0.37 | -0.33 | 0.26 | | | 0.26 | | 0.32 | 0.29 | | | -0.25 | 0.26 | | | | 0.30 | | |
| CL046 | 0.51 | | | -0.31 | -0.64 | | -0.73 | | | | | -0.25 | -0.46 | -0.63 | -0.59 | 0.55 | -0.44 | -0.33 | 0.24 | | -0.30 | | | | | -0.38 | | -0.25 |
| CL047 | | | | | -0.47 | -0.36 | | -0.41 | | | | | -0.41 | -0.32 | -0.31 | 0.60 | -0.32 | | | | | | | | -0.26 | | | |
| CL048 | | | -0.24 | | | | 0.54 | | -0.36 | -0.31 | -0.25 | | | | | | | | -0.33 | | | | -0.33 | -0.36 | -0.27 | | | |
| CL049 | | | -0.49 | 0.28 | | | 0.41 | | -0.56 | -0.56 | -0.55 | 0.26 | | | | | 0.24 | | -0.51 | | -0.39 | | -0.55 | -0.59 | | | | 0.32 |
| CL050 | -0.55 | | | | 0.24 | | 0.79 | | | | | | | 0.26 | 0.26 | -0.44 | | | | | | | | | -0.35 | | -0.27 | |

| | CL170 | CL171 | CL172 | CL173 | CL174 | CL175 | CL176 | CL177 | CL178 | CL179 | CL180 | CL181 | CL182 | CL183 | CL184 | CL185 | CL186 | CL187 | CL188 | CL189 | CL190 | CL191 | CL192 | CL193 | CL194 | CL195 | CL196 | CL197 |
|---|---|---|---|---|---|---|---|---|---|---|---|---|---|---|---|---|---|---|---|---|---|---|---|---|---|---|---|---|
| CL051 | | | -0.42 | 0.48 | 0.39 | 0.28 | 0.27 | | -0.48 | -0.43 | -0.54 | 0.38 | 0.38 | 0.30 | 0.39 | | 0.47 | 0.30 | -0.46 | | -0.27 | | -0.44 | -0.45 | | 0.41 | | 0.34 |
| CL052 | -0.38 | | | | | | 0.72 | | -0.38 | -0.31 | -0.25 | | | 0.26 | 0.25 | | | | -0.37 | | | | -0.38 | -0.37 | -0.33 | | | |
| CL053 | 0.29 | | -0.43 | | | | | | | -0.41 | -0.38 | | | | | 0.57 | | | | | -0.40 | | | -0.25 | | | | |
| CL054 | 0.50 | | -0.27 | | -0.58 | | -0.56 | | | -0.26 | | | -0.41 | -0.56 | -0.51 | 0.65 | -0.32 | -0.26 | | | -0.43 | | | | | -0.24 | 0.33 | |
| CL055 | | | -0.34 | 0.25 | 0.27 | | 0.33 | | -0.53 | -0.45 | -0.46 | 0.24 | | 0.37 | | | | | -0.52 | 0.28 | | | -0.51 | -0.50 | | | | 0.26 |
| CL056 | | -0.27 | -0.49 | 0.60 | 0.65 | 0.36 | 0.31 | | -0.53 | -0.45 | -0.64 | 0.50 | 0.60 | 0.49 | 0.56 | | 0.69 | 0.43 | -0.49 | 0.40 | | 0.33 | -0.45 | -0.49 | 0.32 | 0.62 | 0.25 | 0.37 |
| CL057 | 0.54 | | | -0.32 | -0.63 | | -0.69 | | 0.24 | | | -0.31 | -0.47 | -0.59 | -0.57 | 0.54 | -0.44 | -0.35 | 0.25 | | -0.31 | | | | | -0.39 | 0.25 | -0.27 |
| CL058 | 0.34 | | -0.27 | | -0.47 | -0.24 | -0.31 | | | -0.30 | | | -0.33 | -0.34 | -0.33 | 0.74 | | | | | -0.33 | | | | | | | |
| CL059 | | | -0.31 | | | | | 0.28 | -0.27 | -0.36 | -0.35 | | | | | | | | -0.24 | | -0.32 | | -0.32 | -0.28 | | | | |
| CL060 | | | -0.42 | 0.44 | 0.44 | | 0.57 | | -0.60 | -0.48 | -0.54 | 0.38 | 0.33 | 0.43 | 0.42 | | 0.40 | 0.29 | -0.58 | 0.34 | | | -0.56 | -0.58 | | 0.46 | | 0.37 |
| CL061 | | -0.29 | -0.50 | 0.58 | 0.51 | 0.25 | | 0.46 | -0.66 | -0.54 | -0.74 | 0.50 | 0.48 | 0.43 | 0.51 | | 0.58 | 0.33 | -0.61 | 0.33 | | | -0.61 | -0.63 | 0.36 | 0.63 | | 0.39 |
| CL062 | | | -0.35 | 0.28 | | | 0.60 | | -0.56 | -0.45 | -0.43 | | | | 0.30 | | 0.25 | | -0.55 | | | | -0.56 | -0.54 | | | | 0.29 |
| CL063 | | -0.32 | -0.34 | 0.60 | 0.79 | 0.39 | 0.26 | 0.42 | -0.45 | -0.31 | -0.53 | 0.45 | 0.68 | 0.60 | 0.68 | -0.40 | 0.63 | 0.35 | -0.44 | 0.29 | | | -0.40 | -0.41 | 0.35 | 0.72 | | 0.30 |
| CL064 | | 0.25 | | | -0.42 | | -0.26 | 0.33 | | | | | -0.28 | -0.46 | -0.29 | | -0.26 | | | | -0.28 | -0.26 | | | | -0.38 | | |
| CL065 | 0.34 | 0.29 | | -0.25 | -0.49 | | -0.41 | 0.31 | | | | | -0.46 | -0.47 | -0.44 | 0.27 | -0.27 | -0.25 | | | -0.34 | | | | | -0.42 | | |
| CL066 | 0.55 | | -0.46 | | | | -0.42 | | -0.43 | -0.47 | -0.56 | | | | | 0.48 | | | -0.37 | 0.26 | -0.35 | | -0.39 | -0.45 | 0.39 | | 0.34 | |
| CL067 | 0.57 | | | -0.25 | -0.57 | | -0.71 | | | | | | -0.42 | -0.51 | -0.48 | 0.58 | -0.36 | -0.33 | | | -0.32 | | | | | -0.31 | 0.33 | |
| CL068 | | | -0.56 | 0.47 | 0.49 | 0.32 | 0.35 | 0.24 | -0.57 | -0.57 | -0.69 | 0.42 | 0.54 | 0.27 | 0.42 | | 0.62 | 0.34 | -0.52 | 0.38 | -0.28 | | -0.51 | -0.51 | 0.25 | 0.50 | | 0.35 |
| CL069 | | | 0.27 | -0.48 | -0.43 | -0.33 | -0.26 | -0.35 | 0.71 | 0.33 | 0.59 | -0.46 | -0.46 | -0.41 | -0.43 | | -0.41 | -0.35 | 0.72 | -0.47 | | | 0.66 | 0.66 | | -0.44 | | -0.43 |
| CL070 | | | | | | | | | | -0.27 | | | | | | | | | | | | | | | | | | |
| CL071 | -0.27 | | -0.35 | 0.48 | 0.55 | | 0.54 | | -0.38 | -0.32 | -0.46 | 0.41 | 0.52 | 0.41 | 0.51 | | 0.52 | 0.46 | -0.36 | 0.26 | | 0.25 | -0.33 | -0.33 | | 0.55 | | 0.25 |
| CL072 | 0.52 | | -0.25 | -0.26 | -0.59 | | -0.58 | | | -0.24 | | | -0.43 | -0.53 | -0.50 | 0.64 | -0.34 | -0.29 | | | -0.38 | | | | | -0.32 | 0.25 | |
| CL073 | | | 0.82 | -0.57 | -0.45 | -0.28 | -0.30 | -0.35 | 0.70 | 0.83 | 0.93 | -0.54 | -0.47 | -0.29 | -0.40 | | -0.54 | -0.35 | 0.64 | -0.41 | 0.54 | | 0.67 | 0.67 | -0.39 | -0.53 | | -0.40 |
| CL074 | | | -0.50 | 0.49 | 0.35 | | 0.47 | | -0.48 | -0.55 | -0.59 | 0.24 | 0.31 | 0.37 | 0.28 | | 0.38 | 0.34 | -0.46 | 0.32 | -0.29 | | -0.48 | -0.46 | | 0.44 | | 0.38 |
| CL075 | 0.31 | | | | -0.50 | | -0.36 | 0.37 | | | | -0.24 | -0.40 | -0.42 | -0.39 | | -0.36 | -0.24 | | | -0.26 | | | | | -0.37 | | |
| CL076 | | -0.29 | -0.46 | 0.62 | 0.78 | 0.29 | 0.43 | | -0.47 | -0.40 | -0.59 | 0.54 | 0.71 | 0.59 | 0.67 | -0.33 | 0.68 | 0.42 | -0.43 | 0.34 | | 0.35 | -0.41 | -0.43 | 0.31 | 0.69 | | 0.39 |
| CL077 | | | 0.83 | -0.51 | -0.31 | -0.27 | -0.25 | -0.42 | 0.72 | 0.87 | 0.93 | -0.48 | -0.35 | | -0.31 | | -0.45 | -0.29 | 0.66 | -0.38 | 0.58 | | 0.70 | 0.69 | -0.36 | -0.42 | | -0.37 |
| CL078 | 0.28 | | -0.43 | 0.30 | | 0.38 | | 0.50 | -0.41 | -0.45 | -0.54 | 0.34 | | | | | 0.34 | | -0.41 | | -0.30 | | -0.42 | -0.39 | 0.50 | 0.25 | | |
| CL079 | | | -0.45 | 0.40 | 0.33 | | 0.53 | | -0.62 | -0.50 | -0.60 | 0.33 | 0.30 | 0.33 | 0.33 | | 0.40 | 0.38 | -0.60 | 0.37 | -0.24 | | -0.59 | -0.58 | | 0.36 | | 0.39 |
| CL080 | | | -0.48 | 0.47 | 0.48 | 0.26 | 0.28 | | -0.46 | -0.43 | -0.55 | 0.45 | 0.47 | 0.38 | 0.44 | | 0.58 | 0.44 | -0.43 | 0.26 | -0.24 | | -0.42 | -0.42 | 0.36 | 0.46 | | 0.33 |
| CL081 | 0.51 | | -0.41 | | -0.32 | | -0.51 | | -0.25 | -0.42 | -0.44 | | | -0.35 | -0.27 | 0.54 | | | | | -0.41 | | -0.25 | -0.28 | 0.39 | | 0.34 | |
| CL082 | 0.34 | 0.25 | | | -0.50 | | | | | -0.25 | | -0.27 | -0.47 | -0.45 | -0.35 | 0.37 | -0.39 | | | | -0.29 | | -0.24 | | | -0.40 | | |
| CL083 | | | 0.34 | -0.48 | -0.45 | -0.31 | -0.27 | -0.27 | 0.65 | 0.36 | 0.62 | -0.50 | -0.52 | -0.36 | -0.40 | | -0.47 | -0.36 | 0.64 | -0.50 | | | 0.59 | 0.61 | | -0.49 | | -0.45 |
| CL084 | | | -0.42 | | | | | | -0.35 | -0.51 | -0.47 | | | | | | | | -0.33 | | -0.33 | | -0.36 | -0.38 | | | | |
| CL085 | | | -0.40 | 0.30 | | | | | | -0.35 | -0.31 | | 0.24 | | | | 0.26 | | | | -0.41 | | | | 0.35 | 0.24 | | |
| CL086 | | | 0.39 | -0.39 | -0.27 | | -0.25 | -0.44 | 0.97 | 0.49 | 0.65 | -0.38 | | -0.29 | -0.37 | | -0.34 | | 0.96 | -0.34 | | | 0.95 | 0.95 | | -0.29 | | -0.35 |
| CL087 | | | 0.81 | -0.51 | -0.32 | -0.27 | -0.26 | -0.43 | 0.75 | 0.85 | 0.93 | -0.48 | -0.35 | | -0.32 | | -0.45 | -0.30 | 0.69 | -0.37 | 0.57 | | 0.72 | 0.72 | -0.34 | -0.44 | | -0.37 |
| CL088 | | | 0.83 | -0.51 | -0.31 | -0.25 | -0.24 | -0.41 | 0.71 | 0.87 | 0.94 | -0.46 | -0.32 | | -0.30 | | -0.44 | -0.26 | 0.65 | -0.36 | 0.60 | | 0.69 | 0.68 | -0.32 | -0.42 | | -0.35 |
| CL089 | | | 0.92 | -0.49 | -0.29 | | | -0.31 | 0.58 | 0.91 | 0.93 | -0.45 | -0.35 | | -0.24 | | -0.43 | -0.28 | 0.51 | -0.33 | 0.71 | | 0.56 | 0.55 | -0.38 | -0.42 | | -0.32 |
| CL090 | | | 0.94 | -0.38 | | | | | 0.39 | 0.93 | 0.82 | -0.32 | | | | | -0.30 | | 0.31 | | 0.84 | | 0.39 | 0.37 | -0.32 | -0.29 | | |
| CL091 | | | -0.30 | | | | -0.26 | 0.35 | | -0.33 | -0.32 | | | -0.31 | | | | | | | -0.37 | | | | 0.35 | | | |
| CL092 | -0.34 | | -0.47 | 0.46 | 0.48 | | 0.54 | | -0.30 | -0.39 | -0.47 | 0.41 | 0.55 | 0.25 | 0.45 | | 0.53 | 0.47 | | 0.29 | -0.28 | 0.29 | | -0.24 | | 0.52 | | 0.29 |
| CL093 | | | | 0.25 | 0.56 | | 0.42 | 0.26 | -0.27 | | | | 0.29 | 0.71 | 0.65 | -0.42 | 0.27 | | -0.31 | | 0.43 | | -0.27 | | | 0.27 | | |
| CL094 | | -0.24 | -0.43 | 0.46 | 0.48 | 0.31 | | 0.48 | -0.56 | -0.46 | -0.66 | 0.43 | 0.50 | 0.33 | 0.41 | | 0.52 | | -0.52 | 0.30 | | | -0.52 | -0.53 | 0.32 | 0.51 | | 0.30 |
| CL095 | | | 0.30 | -0.47 | -0.45 | -0.28 | -0.26 | | 0.61 | 0.32 | 0.57 | -0.47 | -0.50 | -0.40 | -0.41 | | -0.44 | -0.34 | 0.60 | -0.46 | | | 0.55 | 0.58 | | -0.48 | | -0.42 |
| CL096 | | | 0.94 | -0.45 | -0.24 | | | -0.29 | 0.52 | 0.92 | 0.90 | -0.42 | -0.31 | | | | -0.41 | -0.25 | 0.43 | -0.30 | 0.76 | | 0.50 | 0.49 | -0.41 | -0.40 | | -0.29 |
| CL097 | | | 0.94 | -0.37 | | | | | 0.37 | 0.93 | 0.82 | -0.31 | | | | | -0.29 | | 0.30 | | 0.85 | | 0.37 | 0.35 | -0.34 | -0.27 | | |
| CL098 | | | 0.81 | -0.52 | -0.34 | -0.29 | | -0.43 | 0.72 | 0.84 | 0.93 | -0.50 | -0.38 | | -0.32 | | -0.48 | -0.29 | 0.66 | -0.38 | 0.56 | | 0.69 | 0.69 | -0.36 | -0.45 | | -0.39 |
| CL099 | -0.34 | | -0.31 | 0.39 | 0.50 | | 0.59 | | -0.58 | -0.36 | -0.49 | 0.39 | 0.41 | 0.40 | 0.46 | -0.24 | 0.50 | 0.29 | -0.55 | 0.32 | | | -0.51 | -0.53 | | 0.40 | | 0.40 |
| CL100 | -0.29 | | | | | | | | 0.29 | | 0.24 | | | | | -0.30 | | | 0.28 | | | | 0.26 | 0.25 | | | | |
| CL101 | -0.37 | | | | 0.29 | | 0.64 | | -0.33 | | | | | 0.31 | 0.28 | | 0.28 | | -0.31 | | | | -0.32 | -0.35 | -0.24 | 0.24 | | 0.33 |
| CL102 | | | -0.43 | 0.36 | 0.38 | 0.35 | | 0.56 | -0.52 | -0.44 | -0.58 | 0.36 | 0.38 | 0.28 | 0.31 | | 0.36 | | -0.50 | | | | -0.50 | -0.48 | 0.26 | 0.31 | | 0.35 |
| CL103 | | | 0.95 | -0.38 | | | | | 0.30 | 0.90 | 0.80 | -0.32 | -0.26 | | | | -0.33 | | | | 0.83 | | 0.30 | 0.29 | -0.40 | -0.35 | | |
| CL104 | | -0.34 | -0.26 | 0.57 | 0.82 | 0.28 | 0.26 | | -0.29 | | -0.38 | 0.44 | 0.70 | 0.68 | 0.73 | -0.37 | 0.58 | 0.32 | -0.27 | | | 0.31 | -0.25 | | 0.28 | 0.69 | | 0.24 |
| CL105 | | | 0.94 | -0.43 | -0.25 | | | | 0.30 | 0.87 | 0.81 | -0.36 | -0.35 | | | | -0.38 | -0.26 | | -0.25 | 0.81 | | 0.29 | 0.29 | -0.43 | -0.41 | | |
| CL106 | | | -0.37 | 0.61 | 0.67 | 0.34 | 0.34 | | -0.42 | -0.36 | -0.52 | 0.39 | 0.59 | 0.63 | 0.59 | -0.31 | 0.55 | 0.37 | -0.38 | | | | -0.36 | -0.34 | | 0.66 | | 0.30 |
| CL107 | | | -0.33 | 0.50 | 0.55 | | 0.44 | | -0.52 | -0.36 | -0.51 | 0.42 | 0.46 | 0.52 | 0.53 | -0.26 | 0.43 | 0.33 | -0.53 | 0.26 | | | -0.47 | -0.51 | | 0.52 | | 0.33 |
| CL108 | | | 0.94 | -0.37 | | | | | 0.27 | 0.91 | 0.78 | -0.30 | | | | | -0.28 | | | | 0.86 | | 0.28 | 0.25 | -0.37 | -0.31 | | |
| CL109 | | | -0.54 | 0.37 | 0.32 | 0.26 | | 0.38 | -0.51 | -0.51 | -0.64 | 0.49 | 0.37 | | 0.27 | | 0.43 | 0.34 | -0.48 | 0.33 | -0.36 | | -0.50 | -0.47 | 0.51 | 0.34 | 0.25 | 0.31 |
| CL110 | | | 0.39 | -0.38 | -0.28 | | | -0.45 | 0.97 | 0.48 | 0.65 | -0.37 | | -0.29 | -0.37 | | -0.34 | | 0.96 | -0.32 | | | 0.94 | 0.94 | | -0.29 | | -0.35 |

| | CL170 | CL171 | CL172 | CL173 | CL174 | CL175 | CL176 | CL177 | CL178 | CL179 | CL180 | CL181 | CL182 | CL183 | CL184 | CL185 | CL186 | CL187 | CL188 | CL189 | CL190 | CL191 | CL192 | CL193 | CL194 | CL195 | CL196 | CL197 |
|---|---|---|---|---|---|---|---|---|---|---|---|---|---|---|---|---|---|---|---|---|---|---|---|---|---|---|---|---|
| CL111 | | | 0.30 | -0.49 | -0.44 | -0.33 | -0.27 | -0.34 | 0.72 | 0.35 | 0.61 | -0.48 | -0.47 | -0.39 | -0.43 | | -0.44 | -0.33 | 0.72 | -0.48 | | | 0.66 | 0.67 | | -0.46 | | -0.43 |
| CL112 | -0.25 | | -0.43 | 0.54 | 0.56 | | 0.58 | | -0.47 | -0.38 | -0.49 | 0.41 | 0.53 | 0.46 | 0.53 | | 0.52 | 0.45 | -0.43 | 0.30 | | 0.29 | -0.39 | -0.40 | | 0.58 | | 0.33 |
| CL113 | | -0.28 | | 0.39 | 0.69 | | 0.43 | 0.25 | -0.41 | | | | 0.45 | 0.80 | 0.70 | -0.38 | 0.39 | | -0.42 | | 0.30 | | -0.38 | -0.37 | | 0.48 | | |
| CL114 | | | 0.43 | -0.40 | -0.29 | -0.24 | -0.26 | -0.40 | 0.97 | 0.52 | 0.66 | -0.37 | | -0.29 | -0.36 | | -0.35 | | 0.94 | -0.35 | | | 0.94 | 0.94 | | -0.29 | | -0.34 |
| CL115 | | | 0.93 | -0.44 | | | | -0.30 | 0.50 | 0.94 | 0.88 | -0.38 | | | | | -0.35 | | 0.42 | -0.28 | 0.78 | | 0.50 | 0.48 | -0.34 | -0.33 | | -0.26 |
| CL116 | | | 0.85 | -0.56 | -0.44 | | -0.29 | -0.27 | 0.63 | 0.83 | 0.91 | -0.53 | -0.51 | -0.27 | -0.37 | | -0.54 | -0.35 | 0.56 | -0.38 | 0.60 | -0.24 | 0.59 | 0.60 | -0.39 | -0.56 | | -0.41 |
| CL117 | | | -0.49 | 0.26 | | | | 0.31 | -0.35 | -0.46 | -0.53 | 0.34 | | | | | 0.35 | 0.25 | -0.29 | 0.35 | -0.42 | | -0.32 | -0.34 | 0.33 | 0.25 | | 0.25 |
| CL118 | -0.25 | | -0.29 | 0.50 | 0.52 | | 0.58 | | -0.40 | -0.31 | -0.41 | 0.34 | 0.42 | 0.45 | 0.56 | -0.27 | 0.53 | 0.36 | -0.37 | | | 0.26 | -0.38 | -0.32 | | 0.46 | | 0.34 |
| CL119 | -0.41 | | | | | | 0.51 | -0.29 | | | | | | | | | | | | | | | | | -0.24 | | | |
| CL120 | | | | | | | -0.25 | | | | | | | | | | | | | | | | | | | | | |
| CL121 | | | -0.35 | | | | | 0.38 | | -0.32 | -0.36 | | | | | | | | | | -0.32 | | | | 0.26 | | | |
| CL122 | | -0.26 | -0.40 | 0.57 | 0.71 | 0.37 | | 0.38 | -0.35 | -0.34 | -0.55 | 0.44 | 0.68 | 0.49 | 0.55 | -0.37 | 0.59 | 0.26 | -0.32 | 0.35 | | 0.28 | -0.29 | -0.32 | 0.36 | 0.64 | | 0.31 |
| CL123 | | | 0.94 | -0.45 | | | | -0.27 | 0.51 | 0.94 | 0.90 | -0.39 | -0.30 | | | | -0.39 | -0.24 | 0.43 | -0.32 | 0.76 | | 0.49 | 0.49 | -0.36 | -0.39 | | -0.28 |
| CL124 | | | 0.94 | -0.43 | | | | -0.27 | 0.49 | 0.93 | 0.89 | -0.39 | -0.25 | | | | -0.36 | | 0.41 | -0.28 | 0.80 | | 0.48 | 0.47 | -0.37 | -0.36 | | -0.28 |
| CL125 | | | 0.92 | -0.47 | -0.25 | | | -0.29 | 0.54 | 0.92 | 0.91 | -0.42 | -0.30 | | | | -0.40 | -0.25 | 0.47 | -0.30 | 0.74 | | 0.53 | 0.51 | -0.35 | -0.38 | | -0.29 |
| CL126 | | -0.24 | 0.92 | -0.31 | | | | | | 0.87 | 0.71 | -0.26 | | | | | -0.26 | | | | 0.87 | | | | -0.35 | -0.26 | | |
| CL127 | | | 0.79 | -0.58 | -0.48 | -0.29 | -0.30 | -0.36 | 0.70 | 0.80 | 0.92 | -0.54 | -0.50 | -0.29 | -0.42 | | -0.56 | -0.37 | 0.64 | -0.41 | 0.51 | -0.24 | 0.66 | 0.66 | -0.40 | -0.55 | | -0.42 |
| CL128 | 0.33 | | -0.52 | | | | -0.33 | | -0.28 | -0.50 | -0.54 | 0.30 | | | | 0.36 | | | | | -0.46 | | | -0.27 | 0.45 | | 0.26 | |
| CL129 | | -0.27 | -0.38 | 0.48 | 0.51 | 0.32 | | 0.39 | -0.36 | -0.35 | -0.49 | 0.36 | 0.55 | 0.31 | 0.48 | -0.28 | 0.50 | 0.29 | -0.31 | | | | -0.32 | -0.33 | 0.36 | 0.46 | | |
| CL130 | | | 0.27 | | | | | -0.36 | 0.89 | 0.38 | 0.48 | | | | | | | | 0.88 | -0.29 | | | 0.89 | 0.87 | | | | |
| CL131 | | -0.30 | -0.35 | 0.59 | 0.71 | 0.30 | 0.41 | | -0.50 | -0.36 | -0.54 | 0.53 | 0.61 | 0.60 | 0.68 | -0.31 | 0.69 | 0.37 | -0.47 | 0.29 | | 0.34 | -0.45 | -0.45 | | 0.66 | | 0.39 |
| CL132 | | | 0.94 | -0.46 | | | -0.25 | -0.27 | 0.50 | 0.93 | 0.90 | -0.40 | -0.29 | | | | -0.40 | | 0.42 | -0.30 | 0.77 | | 0.49 | 0.49 | -0.35 | -0.38 | | -0.29 |
| CL133 | | -0.27 | -0.44 | 0.60 | 0.79 | 0.31 | 0.36 | 0.32 | -0.41 | -0.38 | -0.57 | 0.47 | 0.75 | 0.67 | 0.67 | -0.39 | 0.66 | 0.38 | -0.39 | 0.33 | | 0.27 | -0.35 | -0.36 | 0.34 | 0.69 | | 0.32 |
| CL134 | | | 0.94 | -0.38 | | | | | 0.38 | 0.93 | 0.82 | -0.32 | | | | | -0.32 | | 0.31 | -0.25 | 0.83 | | 0.39 | 0.36 | -0.35 | -0.29 | | |
| CL135 | | | 0.91 | -0.35 | | | | -0.27 | 0.48 | 0.93 | 0.83 | -0.31 | | | | | -0.27 | | 0.40 | | 0.79 | | 0.49 | 0.46 | -0.30 | -0.26 | | |
| CL136 | | | 0.95 | -0.38 | | | | | 0.33 | 0.91 | 0.81 | -0.34 | | | | | -0.32 | | 0.25 | | 0.83 | | 0.33 | 0.32 | -0.38 | -0.32 | | |
| CL137 | | | | 0.47 | 0.67 | 0.26 | 0.49 | | -0.44 | -0.24 | -0.38 | 0.39 | 0.53 | 0.68 | 0.61 | -0.33 | 0.50 | 0.34 | -0.43 | 0.26 | | | -0.38 | -0.42 | | 0.61 | | 0.36 |
| CL138 | | | 0.93 | -0.32 | | | | | 0.25 | 0.89 | 0.75 | -0.24 | | | | | -0.26 | | | | 0.85 | | 0.26 | | -0.35 | -0.29 | | |
| CL139 | | | -0.47 | 0.54 | 0.68 | | 0.40 | | -0.51 | -0.41 | -0.57 | 0.51 | 0.65 | 0.49 | 0.64 | -0.29 | 0.59 | 0.36 | -0.45 | 0.33 | | 0.26 | -0.46 | -0.45 | 0.35 | 0.61 | | 0.34 |
| CL140 | | | 0.39 | -0.39 | -0.29 | -0.24 | -0.26 | -0.44 | 0.97 | 0.48 | 0.65 | -0.37 | | -0.30 | -0.39 | | -0.36 | | 0.95 | -0.34 | | | 0.95 | 0.94 | | -0.30 | | -0.34 |
| CL141 | | | 0.95 | -0.39 | | | | | 0.29 | 0.88 | 0.79 | -0.31 | -0.27 | | | | -0.35 | | | | 0.84 | | 0.29 | 0.28 | -0.40 | -0.37 | | |
| CL142 | | | 0.93 | -0.35 | | | | | 0.40 | 0.93 | 0.79 | -0.31 | | | | | -0.26 | | 0.31 | | 0.84 | | 0.40 | 0.37 | -0.33 | | | |
| CL143 | | 0.36 | | | -0.38 | | -0.25 | | | | | -0.32 | -0.33 | -0.41 | -0.40 | | -0.36 | -0.25 | | -0.24 | | | | | | -0.43 | | |
| CL144 | 0.37 | | | -0.35 | -0.60 | -0.24 | -0.63 | | 0.42 | | 0.26 | -0.32 | -0.47 | -0.50 | -0.48 | 0.59 | -0.45 | -0.39 | 0.42 | -0.25 | | | 0.38 | 0.35 | | -0.32 | | -0.31 |
| CL145 | | | 0.89 | -0.52 | -0.32 | -0.25 | -0.25 | -0.35 | 0.61 | 0.90 | 0.93 | -0.47 | -0.38 | | -0.30 | | -0.48 | -0.31 | 0.54 | -0.34 | 0.66 | | 0.59 | 0.58 | -0.39 | -0.46 | | -0.33 |
| CL146 | 0.32 | -0.26 | -0.45 | 0.44 | 0.36 | 0.30 | | 0.30 | -0.40 | -0.40 | -0.56 | 0.50 | 0.46 | | 0.35 | | 0.49 | | -0.34 | 0.26 | | 0.27 | -0.34 | -0.35 | 0.47 | 0.53 | | 0.26 |
| CL147 | | | -0.24 | 0.30 | 0.43 | | 0.27 | | -0.38 | | -0.36 | 0.27 | 0.35 | 0.34 | 0.37 | | 0.45 | | -0.34 | | | 0.29 | -0.31 | -0.35 | | 0.42 | | |
| CL148 | 0.37 | | | | -0.45 | -0.28 | -0.29 | | | -0.24 | | | -0.32 | -0.36 | -0.37 | 0.68 | | | | | -0.25 | | | | | | | |
| CL149 | | | | 0.32 | 0.36 | | | 0.60 | -0.41 | | -0.31 | | | 0.40 | 0.34 | -0.28 | | | -0.44 | | | | -0.42 | -0.40 | | 0.27 | | |
| CL150 | | | 0.91 | -0.48 | -0.27 | | | -0.31 | 0.56 | 0.91 | 0.90 | -0.45 | -0.32 | | -0.26 | | -0.42 | -0.25 | 0.48 | -0.29 | 0.72 | | 0.54 | 0.53 | -0.38 | -0.43 | | -0.31 |
| CL151 | | -0.28 | -0.38 | 0.54 | 0.72 | | 0.32 | | | -0.26 | -0.49 | 0.42 | 0.72 | 0.45 | 0.58 | -0.30 | 0.63 | 0.46 | | 0.27 | | 0.34 | | | 0.37 | 0.68 | | 0.30 |
| CL152 | 0.42 | | -0.35 | | | | -0.39 | 0.29 | | -0.36 | -0.42 | | | -0.30 | | 0.25 | | | | | -0.38 | | -0.26 | -0.25 | 0.43 | | 0.24 | |
| CL153 | | | 0.40 | -0.40 | -0.29 | | -0.25 | -0.42 | 0.97 | 0.50 | 0.67 | -0.37 | | -0.28 | -0.40 | | -0.33 | | 0.94 | -0.32 | | | 0.95 | 0.94 | | -0.33 | | -0.36 |
| CL154 | | | 0.92 | -0.31 | | | | | 0.26 | 0.88 | 0.78 | -0.27 | | | | | -0.24 | | | | 0.86 | | 0.27 | 0.26 | -0.33 | -0.28 | | |
| CL155 | | -0.27 | -0.26 | 0.51 | 0.74 | | 0.50 | | -0.40 | | -0.41 | 0.35 | 0.61 | 0.57 | 0.61 | -0.42 | 0.56 | 0.29 | -0.39 | | | | -0.36 | -0.37 | | 0.59 | | 0.25 |
| CL156 | | | 0.90 | -0.31 | | | | | | 0.84 | 0.69 | -0.24 | | | | | -0.24 | | | | 0.87 | | | | -0.36 | -0.28 | | |
| CL157 | | -0.32 | -0.45 | 0.52 | 0.63 | 0.33 | | 0.41 | -0.47 | -0.39 | -0.56 | 0.42 | 0.60 | 0.44 | 0.59 | -0.31 | 0.51 | 0.32 | -0.46 | 0.28 | | 0.29 | -0.45 | -0.43 | 0.42 | 0.67 | | 0.33 |
| CL158 | | | -0.34 | 0.42 | 0.44 | | | | -0.30 | -0.32 | -0.42 | 0.25 | 0.41 | 0.28 | 0.36 | | 0.42 | 0.35 | -0.26 | 0.27 | | | -0.31 | -0.29 | | 0.43 | | 0.28 |
| CL159 | | | | | 0.28 | | 0.52 | | -0.35 | -0.28 | -0.25 | | | 0.31 | 0.32 | | | 0.24 | -0.36 | | | | -0.35 | -0.38 | | 0.35 | | |
| CL160 | | | | | | | | 0.26 | | | | | | | | | | | | | -0.27 | | | | | | | |
| CL161 | | | 0.40 | -0.38 | -0.27 | | -0.24 | -0.43 | 0.96 | 0.49 | 0.65 | -0.36 | | -0.28 | -0.36 | | -0.33 | | 0.93 | -0.36 | | | 0.95 | 0.94 | | -0.29 | | -0.34 |
| CL162 | | | -0.26 | 0.27 | 0.31 | | 0.27 | | | -0.29 | -0.30 | | 0.33 | | | | 0.30 | 0.29 | | | | 0.28 | | | | 0.29 | | |
| CL163 | | | -0.39 | 0.35 | 0.47 | | | | | -0.32 | -0.45 | 0.48 | 0.44 | 0.33 | 0.42 | -0.25 | 0.45 | 0.25 | | 0.27 | | | | | 0.36 | 0.46 | | |
| CL164 | | | | | | | 0.42 | | -0.36 | | | | | | | | | | -0.33 | 0.24 | | | -0.35 | -0.28 | -0.26 | | | |
| CL165 | | | 0.91 | -0.43 | | | | -0.29 | 0.58 | 0.90 | 0.86 | -0.38 | -0.24 | | | | -0.37 | | 0.51 | -0.27 | 0.73 | | 0.59 | 0.56 | -0.33 | -0.35 | -0.26 | |
| CL166 | | -0.37 | | 0.52 | 0.72 | 0.32 | | 0.43 | -0.32 | | -0.36 | 0.28 | 0.59 | 0.65 | 0.62 | -0.42 | 0.44 | 0.27 | -0.32 | | | | -0.29 | -0.27 | 0.35 | 0.62 | | 0.27 |
| CL167 | | | 0.93 | -0.42 | | | | -0.28 | 0.51 | 0.93 | 0.88 | -0.39 | -0.28 | | | | -0.37 | -0.24 | 0.43 | -0.27 | 0.76 | | 0.50 | 0.49 | -0.36 | -0.36 | | -0.27 |
| CL168 | | -0.27 | | 0.41 | 0.77 | 0.25 | 0.40 | | -0.30 | | | 0.30 | 0.59 | 0.67 | 0.67 | -0.43 | 0.49 | 0.26 | -0.31 | | | | -0.27 | -0.26 | | 0.59 | | |
| CL169 | | | 0.93 | -0.37 | | | | | 0.36 | 0.91 | 0.80 | -0.34 | | | | | -0.31 | | 0.28 | | 0.82 | | 0.34 | 0.33 | -0.37 | -0.32 | | |
| CL170 | | | | | | | -0.46 | 0.30 | | | | | | | | 0.33 | | | | | | | | | | | | |

| | CL170 | CL171 | CL172 | CL173 | CL174 | CL175 | CL176 | CL177 | CL178 | CL179 | CL180 | CL181 | CL182 | CL183 | CL184 | CL185 | CL186 | CL187 | CL188 | CL189 | CL190 | CL191 | CL192 | CL193 | CL194 | CL195 | CL196 | CL197 |
|---|---|---|---|---|---|---|---|---|---|---|---|---|---|---|---|---|---|---|---|---|---|---|---|---|---|---|---|---|
| CL171 | | | | | -0.34 | | | | | | | | | | -0.29 | | | | | | -0.30 | -0.27 | | | | -0.30 | | |
| CL172 | | | | -0.39 | | | | | 0.39 | 0.92 | 0.82 | -0.39 | -0.29 | | | | -0.36 | | 0.30 | -0.28 | 0.81 | | 0.40 | 0.38 | -0.41 | -0.36 | | |
| CL173 | | | | | 0.48 | 0.28 | 0.28 | | -0.39 | -0.38 | -0.50 | 0.29 | 0.52 | 0.38 | 0.53 | -0.29 | 0.43 | 0.34 | -0.34 | | | 0.27 | -0.36 | -0.36 | | 0.54 | | |
| CL174 | | | | | | 0.26 | 0.37 | | -0.30 | | -0.32 | 0.34 | 0.70 | 0.69 | 0.74 | -0.50 | 0.60 | 0.34 | -0.29 | | | 0.25 | -0.24 | -0.26 | 0.27 | 0.61 | | 0.28 |
| CL175 | | | | | | | | | | | -0.27 | | 0.32 | | | | 0.30 | 0.24 | -0.25 | | | | | | | | | |
| CL176 | | | | | | | | | -0.25 | | | | 0.30 | 0.27 | 0.37 | -0.42 | 0.33 | 0.33 | -0.25 | | | | -0.26 | | | | | 0.26 |
| CL177 | | | | | | | | | -0.42 | | -0.35 | | | | 0.26 | | | | -0.43 | | | | -0.42 | -0.39 | | | | |
| CL178 | | | | | | | | | | 0.47 | 0.64 | -0.38 | -0.25 | -0.32 | -0.38 | | -0.36 | | 0.93 | -0.33 | | | 0.93 | 0.93 | | -0.31 | | -0.34 |
| CL179 | | | | | | | | | | | 0.84 | -0.36 | | | | | -0.35 | | 0.42 | -0.28 | 0.75 | | 0.49 | 0.45 | -0.27 | -0.32 | | |
| CL180 | | | | | | | | | | | | -0.49 | -0.35 | | -0.30 | | -0.48 | -0.28 | 0.57 | -0.37 | 0.58 | | 0.61 | 0.60 | -0.31 | -0.45 | | -0.38 |
| CL181 | | | | | | | | | | | | | 0.45 | 0.26 | 0.35 | | 0.54 | 0.27 | -0.35 | 0.31 | | 0.25 | -0.32 | -0.34 | 0.25 | 0.38 | | 0.31 |
| CL182 | | | | | | | | | | | | | | 0.53 | 0.57 | -0.40 | 0.60 | 0.39 | | | | | | | 0.35 | 0.61 | | |
| CL183 | | | | | | | | | | | | | | | 0.64 | -0.36 | 0.43 | | -0.31 | | 0.26 | | -0.26 | -0.24 | | 0.46 | | |
| CL184 | | | | | | | | | | | | | | | | -0.40 | 0.54 | 0.32 | -0.36 | | | | -0.36 | -0.36 | | 0.57 | | |
| CL185 | | | | | | | | | | | | | | | | | | | | | | | | | | | | |
| CL186 | | | | | | | | | | | | | | | | | | 0.31 | -0.32 | 0.26 | | 0.25 | -0.31 | -0.31 | 0.29 | 0.53 | | 0.35 |
| CL187 | | | | | | | | | | | | | | | | | | | | | | | | | 0.29 | 0.35 | | |
| CL188 | | | | | | | | | | | | | | | | | | | | -0.32 | | | 0.91 | 0.90 | | -0.27 | | -0.34 |
| CL189 | | | | | | | | | | | | | | | | | | | | | | | -0.33 | -0.30 | | | | |
| CL190 | | | | | | | | | | | | | | | | | | | | | | | | | -0.28 | | | |
| CL191 | | | | | | | | | | | | | | | | | | | | | | | | | | | | |
| CL192 | | | | | | | | | | | | | | | | | | | | | | | | 0.90 | | -0.27 | | -0.38 |
| CL193 | | | | | | | | | | | | | | | | | | | | | | | | | | -0.28 | | -0.33 |
| CL194 | | | | | | | | | | | | | | | | | | | | | | | | | | 0.33 | | |
| CL195 | | | | | | | | | | | | | | | | | | | | | | | | | | | | 0.29 |
| CL196 | | | | | | | | | | | | | | | | | | | | | | | | | | | | |
| CL197 | | | | | | | | | | | | | | | | | | | | | | | | | | | | |
| CL198 | | | | | | | | | | | | | | | | | | | | | | | | | | | | |
| CL199 | | | | | | | | | | | | | | | | | | | | | | | | | | | | |
| CL200 | | | | | | | | | | | | | | | | | | | | | | | | | | | | |
| CL201 | | | | | | | | | | | | | | | | | | | | | | | | | | | | |
| CL202 | | | | | | | | | | | | | | | | | | | | | | | | | | | | |
| CL203 | | | | | | | | | | | | | | | | | | | | | | | | | | | | |
| CL204 | | | | | | | | | | | | | | | | | | | | | | | | | | | | |
| CL205 | | | | | | | | | | | | | | | | | | | | | | | | | | | | |
| CL206 | | | | | | | | | | | | | | | | | | | | | | | | | | | | |
| CL207 | | | | | | | | | | | | | | | | | | | | | | | | | | | | |
| CL208 | | | | | | | | | | | | | | | | | | | | | | | | | | | | |
| CL209 | | | | | | | | | | | | | | | | | | | | | | | | | | | | |
| CL210 | | | | | | | | | | | | | | | | | | | | | | | | | | | | |
| CL211 | | | | | | | | | | | | | | | | | | | | | | | | | | | | |
| CL212 | | | | | | | | | | | | | | | | | | | | | | | | | | | | |
| CL213 | | | | | | | | | | | | | | | | | | | | | | | | | | | | |
| CL214 | | | | | | | | | | | | | | | | | | | | | | | | | | | | |
| CL215 | | | | | | | | | | | | | | | | | | | | | | | | | | | | |
| CL216 | | | | | | | | | | | | | | | | | | | | | | | | | | | | |
| CL217 | | | | | | | | | | | | | | | | | | | | | | | | | | | | |
| CL218 | | | | | | | | | | | | | | | | | | | | | | | | | | | | |
| CL219 | | | | | | | | | | | | | | | | | | | | | | | | | | | | |
| CL220 | | | | | | | | | | | | | | | | | | | | | | | | | | | | |
| CL221 | | | | | | | | | | | | | | | | | | | | | | | | | | | | |
| CL222 | | | | | | | | | | | | | | | | | | | | | | | | | | | | |
| CL223 | | | | | | | | | | | | | | | | | | | | | | | | | | | | |
| CL224 | | | | | | | | | | | | | | | | | | | | | | | | | | | | |
| CL225 | | | | | | | | | | | | | | | | | | | | | | | | | | | | |
| CL226 | | | | | | | | | | | | | | | | | | | | | | | | | | | | |
| CL227 | | | | | | | | | | | | | | | | | | | | | | | | | | | | |
| CL228 | | | | | | | | | | | | | | | | | | | | | | | | | | | | |
| CL229 | | | | | | | | | | | | | | | | | | | | | | | | | | | | |
| CL230 | | | | | | | | | | | | | | | | | | | | | | | | | | | | |

| | CL198 | CL199 | CL200 | CL201 | CL202 | CL203 | CL204 | CL205 | CL206 | CL207 | CL208 | CL209 | CL210 | CL211 | CL212 | CL213 | CL214 | CL215 | CL216 | CL217 | CL218 | CL219 | CL220 | CL221 | CL222 | CL223 | CL224 | CL225 |
|---|---|---|---|---|---|---|---|---|---|---|---|---|---|---|---|---|---|---|---|---|---|---|---|---|---|---|---|---|
| CL001 | | | | | | -0.24 | | | | | | | | | | -0.24 | | | | -0.29 | -0.25 | | | | | | | -0.28 |
| CL002 | 0.26 | | | -0.28 | | -0.26 | -0.27 | | | -0.28 | | | 0.45 | | | -0.36 | | | | -0.29 | | | | | | | | -0.28 |
| CL003 | 0.37 | | | -0.40 | | | -0.24 | | | -0.40 | | | 0.60 | | | -0.28 | -0.30 | | | -0.42 | | | | | | | | |
| CL004 | 0.26 | | | -0.45 | | | | | | -0.46 | | -0.24 | 0.50 | -0.25 | | -0.31 | -0.45 | | | -0.55 | | | | | | | | |
| CL005 | | | | -0.30 | | | | | | | | | 0.27 | | | | | | | | | | | | | | | |
| CL006 | | | 0.30 | | | | | | | | | 0.40 | | 0.31 | | | 0.58 | | | 0.24 | | | | | | 0.25 | | |
| CL007 | | | | -0.31 | | | 0.24 | | | | | 0.38 | | 0.30 | | | 0.40 | | | | | | | | | | | |
| CL008 | 0.31 | | | -0.46 | | | | | | -0.48 | | | 0.53 | -0.25 | | -0.29 | -0.40 | | | -0.57 | | | | | | | | |
| CL009 | | | | -0.57 | | | | | | -0.29 | | | 0.33 | | | -0.26 | -0.34 | | | -0.48 | | 0.27 | | | | | | |
| CL010 | 0.27 | | | -0.33 | | -0.25 | -0.27 | | | -0.42 | | | 0.40 | | | -0.40 | -0.34 | | | -0.53 | | | | | | | | -0.25 |
| CL011 | 0.29 | | | -0.29 | | -0.35 | -0.25 | -0.24 | | -0.38 | | | 0.48 | | -0.28 | -0.29 | | | | -0.26 | | | | | | | | -0.27 |
| CL012 | | | | -0.39 | | | | | | | | | 0.31 | | | | | | | | | 0.25 | | | | | | |
| CL013 | 0.30 | | | -0.45 | | | | | | -0.43 | | | 0.53 | | | -0.29 | -0.39 | | | -0.53 | | | | | | | | |
| CL014 | 0.36 | | | -0.34 | | -0.27 | | | | -0.36 | | | 0.45 | | -0.29 | | | | | -0.25 | | | | | | | | |
| CL015 | 0.26 | | | -0.43 | | -0.25 | -0.24 | | | -0.50 | | -0.25 | 0.49 | -0.28 | | -0.40 | -0.42 | | | -0.57 | | | | | | | | |
| CL016 | -0.25 | | | | | 0.41 | 0.36 | 0.34 | | 0.67 | | | -0.43 | 0.33 | | 0.42 | 0.45 | | | 0.59 | | | | | | | | |
| CL017 | 0.30 | | | -0.26 | | | -0.30 | | | -0.37 | | -0.25 | 0.52 | -0.24 | | -0.33 | -0.31 | | | -0.42 | | | | | | | | -0.29 |
| CL018 | | | | | | 0.34 | 0.34 | 0.25 | | 0.49 | | 0.25 | | 0.40 | | 0.31 | 0.40 | | | 0.41 | | | | | | | | |
| CL019 | | 0.27 | | -0.43 | 0.28 | | | | | | | | | | 0.31 | | -0.31 | 0.32 | | -0.32 | | | | | | | | |
| CL020 | | | | -0.44 | | | | | | | | | 0.42 | | | | -0.36 | | | -0.38 | | | | | | | | |
| CL021 | | | | -0.40 | | | | | | -0.36 | | -0.27 | 0.45 | -0.26 | | -0.36 | -0.51 | | | -0.52 | | | | | | | | |
| CL022 | | | | -0.40 | 0.24 | | | | | | | | | | 0.27 | -0.28 | -0.52 | 0.24 | | -0.49 | | | | | | | | |
| CL023 | 0.34 | | | -0.32 | | -0.31 | -0.30 | -0.26 | | -0.40 | | | 0.46 | | | -0.33 | | | | -0.36 | | | | | | | | -0.27 |
| CL024 | | | | -0.41 | | | | | | -0.32 | | -0.26 | 0.53 | | | -0.32 | -0.45 | | | -0.45 | | | | | | | | |
| CL025 | | | | | | | | | | 0.39 | | 0.28 | -0.40 | 0.29 | | 0.24 | 0.42 | | | 0.37 | | | | | | | | |
| CL026 | | | | | | 0.35 | 0.44 | 0.28 | | 0.39 | | 0.42 | | 0.43 | | 0.34 | 0.46 | | 0.24 | 0.32 | | | | | | | | |
| CL027 | | | -0.24 | -0.43 | | -0.27 | -0.26 | | | -0.38 | | -0.27 | 0.34 | -0.28 | | -0.41 | -0.53 | | | -0.56 | | | | | | | | |
| CL028 | | | 0.25 | | | | | | | | | 0.35 | | 0.31 | | | 0.49 | | | | | | | | | 0.28 | | |
| CL029 | | | | -0.30 | | | | | | | | 0.26 | | 0.25 | | | 0.25 | | | | | | | | | | | |
| CL030 | | | | -0.33 | | | | | | | | | 0.26 | | | | | | | | | | | | | | | -0.24 |
| CL031 | | 0.25 | | -0.35 | | | | | | | | | | | 0.30 | | -0.35 | 0.32 | | -0.32 | | | | | | | | |
| CL032 | 0.24 | | | -0.34 | | | | | | -0.24 | | | 0.42 | | | -0.30 | -0.36 | | | -0.42 | | | | | | | | -0.25 |
| CL033 | -0.40 | | | | | | | | | 0.41 | | | -0.49 | | | | | 0.33 | | | | | | | | | | |
| CL034 | 0.33 | | | -0.35 | | -0.33 | -0.28 | -0.25 | | -0.41 | | | 0.48 | | | -0.36 | | | | -0.41 | | | | | | | | -0.25 |
| CL035 | | | | | | | | | | 0.29 | | | | 0.28 | | | | 0.32 | | | | | | | | | | |
| CL036 | | | 0.31 | -0.30 | | 0.37 | 0.41 | 0.30 | | 0.35 | | 0.39 | | 0.48 | | 0.41 | 0.54 | | 0.24 | 0.37 | | | | | | | | 0.29 |
| CL037 | | | | -0.40 | 0.25 | 0.25 | | | | | | | 0.46 | 0.37 | | | | | | | | | | | | | 0.24 | |
| CL038 | | 0.38 | | -0.35 | | 0.36 | 0.27 | | | | | | | | 0.34 | | | 0.27 | 0.26 | | | | 0.26 | 0.24 | | | | |
| CL039 | | | | -0.32 | | | | | | | | | | | | | | 0.26 | | | | | | | | | | |
| CL040 | | | | -0.47 | | | | | | | | | 0.29 | | | | -0.39 | 0.31 | | -0.46 | | | | | | | | |
| CL041 | | | | | | 0.45 | 0.36 | 0.28 | | 0.53 | | 0.31 | | 0.35 | 0.30 | 0.35 | 0.30 | | 0.28 | 0.45 | | | | | | | | 0.27 |
| CL042 | | | | -0.46 | 0.29 | | | | | | | | | | | | | | | | | | | | | | | |
| CL043 | | | | -0.38 | | | | | | | | | 0.28 | | | -0.27 | -0.43 | | | -0.49 | | | | | | | | |
| CL044 | | | | -0.28 | | | | | -0.27 | | | | | | 0.29 | | -0.41 | 0.24 | | -0.40 | | | | | | | | |
| CL045 | | | 0.34 | | | | | | | | | | | 0.37 | | | 0.54 | | | | | | | | | 0.27 | | |
| CL046 | 0.29 | | | -0.32 | | | | | | -0.43 | | -0.27 | 0.55 | -0.28 | | -0.30 | -0.47 | | | -0.50 | | | | | | | | |
| CL047 | 0.38 | | | | | -0.29 | -0.24 | | | -0.25 | | | 0.33 | | -0.30 | | | | | | | | | | | | | |
| CL048 | | | | | | | | | | | -0.30 | 0.26 | | | | | 0.37 | | | | | | | | | | | |
| CL049 | | | | -0.35 | | | | | | | | 0.28 | | | | | | | | | | | | | | | | |
| CL050 | -0.37 | | | | | | | | | | -0.31 | 0.28 | -0.62 | | | | 0.24 | | | | | -0.25 | | | | 0.28 | | |

| | CL198 | CL199 | CL200 | CL201 | CL202 | CL203 | CL204 | CL205 | CL206 | CL207 | CL208 | CL209 | CL210 | CL211 | CL212 | CL213 | CL214 | CL215 | CL216 | CL217 | CL218 | CL219 | CL220 | CL221 | CL222 | CL223 | CL224 | CL225 |
|---|---|---|---|---|---|---|---|---|---|---|---|---|---|---|---|---|---|---|---|---|---|---|---|---|---|---|---|---|
| CL051 | | | 0.28 | -0.26 | | 0.26 | 0.34 | | | 0.29 | | | | 0.44 | 0.25 | | 0.28 | | | | | | | | | | | 0.37 |
| CL052 | | | | | | | | | | | | 0.36 | -0.40 | | | | 0.43 | | | | | | | | | 0.26 | | |
| CL053 | 0.41 | | | -0.44 | | | | | | | | | 0.56 | | | | | | | | | 0.28 | | | | | | |
| CL054 | 0.39 | | | -0.43 | | | | | | -0.39 | | | 0.62 | | | -0.24 | -0.24 | | | -0.39 | | | | | | | | |
| CL055 | | | | | | | | | | 0.41 | | | -0.25 | 0.27 | | | | | | | | | | | | | | |
| CL056 | | | | | | 0.45 | 0.43 | 0.32 | 0.25 | 0.36 | | 0.27 | | 0.47 | | 0.42 | 0.43 | | 0.31 | 0.37 | | | | | | | | 0.43 |
| CL057 | | | | -0.34 | | | | | | -0.38 | | -0.31 | 0.50 | -0.25 | | -0.32 | -0.47 | | | -0.47 | | | | | | | | |
| CL058 | 0.38 | | | -0.34 | | | | | | | | | 0.57 | | | | | | | -0.24 | | 0.26 | | | | | | |
| CL059 | | | | -0.29 | | | | | | | | | | | | -0.25 | | | | | | | | | | | | |
| CL060 | | | | | | 0.24 | 0.35 | | | 0.38 | | 0.33 | | 0.41 | | 0.27 | 0.42 | | | 0.38 | | | | | | 0.30 | | |
| CL061 | | | | -0.25 | | 0.54 | 0.45 | 0.36 | | 0.43 | | 0.26 | | 0.34 | 0.33 | 0.31 | 0.25 | | 0.30 | 0.28 | | | 0.27 | 0.24 | | | | 0.27 |
| CL062 | | | | | | | | | | | | 0.35 | | | | | 0.40 | | | | | | | | | 0.28 | | |
| CL063 | | | | | | 0.62 | 0.47 | 0.44 | | 0.53 | | 0.25 | | 0.38 | 0.28 | 0.54 | 0.35 | | 0.28 | 0.50 | 0.24 | | | | | | | 0.34 |
| CL064 | | | -0.24 | -0.25 | | | | | | | | | | | | -0.28 | -0.35 | | | -0.47 | | | | | | | | |
| CL065 | | | -0.28 | -0.34 | | | | | | -0.27 | | | 0.24 | -0.25 | | -0.29 | -0.56 | | | -0.49 | | | | | | | | |
| CL066 | 0.26 | | | -0.40 | | | | | | | | | 0.60 | | | | | | | | | 0.29 | | | | | | |
| CL067 | 0.32 | | | -0.32 | | | | | | -0.37 | | -0.31 | 0.64 | | | | -0.34 | | | -0.38 | | | | | | | | |
| CL068 | | | | -0.33 | | 0.36 | 0.52 | 0.30 | 0.28 | | | 0.30 | | 0.38 | | 0.33 | 0.37 | | 0.35 | | | | | | | | | 0.30 |
| CL069 | | | | | | -0.39 | -0.37 | -0.27 | | -0.35 | | -0.26 | | -0.38 | | -0.34 | -0.34 | | -0.26 | -0.33 | | | | | | | | -0.25 |
| CL070 | | | | | | | | | -0.27 | | | | | | | | | 0.26 | | | | | | | | | | |
| CL071 | | | 0.44 | | | | 0.46 | | 0.29 | | | 0.25 | | 0.38 | | 0.38 | 0.62 | | | 0.29 | | | | | | | | 0.32 |
| CL072 | 0.30 | | | -0.40 | | | | | | -0.31 | | -0.26 | 0.57 | | | -0.26 | -0.27 | | | -0.41 | | 0.24 | | | | | | |
| CL073 | | -0.30 | | 0.56 | | -0.43 | -0.45 | -0.31 | | -0.28 | | -0.26 | | -0.39 | -0.26 | -0.25 | -0.27 | | -0.35 | | | -0.25 | | | | | | -0.32 |
| CL074 | | | | -0.29 | | | 0.25 | | | 0.27 | | 0.35 | | 0.30 | | 0.33 | 0.39 | | | 0.27 | | | | | | | | |
| CL075 | | | -0.26 | | | | | | | | | | | | | -0.34 | -0.44 | | | -0.47 | | | | | | | | |
| CL076 | | | | | | 0.53 | 0.47 | 0.38 | 0.24 | 0.46 | | 0.36 | | 0.46 | | 0.48 | 0.48 | | 0.28 | 0.44 | | | | | | | | 0.41 |
| CL077 | | -0.31 | | 0.60 | | -0.36 | -0.38 | -0.27 | | -0.26 | | | | -0.31 | -0.28 | | | -0.26 | -0.29 | | | | | | | | | -0.24 |
| CL078 | | 0.25 | | -0.33 | 0.25 | 0.31 | 0.27 | 0.31 | | | | | | | 0.36 | | | 0.34 | | | | | | 0.26 | | | | 0.27 |
| CL079 | | | | | | 0.26 | 0.26 | | | 0.31 | | 0.37 | | 0.35 | | 0.24 | 0.39 | | | | | | | | | | | |
| CL080 | | | | -0.26 | 0.27 | | 0.35 | 0.30 | | | | | | 0.45 | | 0.28 | | | | | | | | | | | | 0.33 |
| CL081 | 0.31 | | | -0.45 | | | | | | | 0.28 | | 0.67 | | | | -0.24 | | | -0.27 | | 0.28 | | | | | | |
| CL082 | | | | | | | | | | | | | | | | -0.25 | -0.28 | | | -0.41 | | | | | | | | -0.29 |
| CL083 | | | | | | -0.43 | -0.42 | -0.28 | | -0.29 | | -0.29 | | -0.42 | | -0.37 | -0.37 | | -0.29 | -0.28 | | | | | | | | -0.31 |
| CL084 | | | | -0.31 | | | | | | | | | | | 0.29 | | | | | | | | | | | | | |
| CL085 | | | | -0.41 | | | | | | | | | | | | | | | | | | | | | | | | 0.29 |
| CL086 | | | | | | -0.29 | -0.26 | -0.24 | | -0.35 | | | | -0.26 | | | -0.26 | -0.24 | | | | | | | | | | |
| CL087 | | -0.29 | | 0.57 | | -0.39 | -0.38 | -0.28 | | -0.26 | | | | -0.32 | -0.27 | | | -0.24 | -0.30 | | | | | | | | | -0.25 |
| CL088 | | -0.31 | | 0.60 | | -0.36 | -0.37 | -0.29 | | -0.26 | | | | -0.31 | -0.27 | | | -0.25 | -0.30 | | | | | | | | | -0.24 |
| CL089 | | -0.28 | | 0.73 | -0.24 | -0.34 | -0.37 | -0.27 | | | | | | -0.31 | -0.26 | | | | -0.32 | | | | | | | | | -0.27 |
| CL090 | | -0.27 | | 0.84 | -0.24 | | -0.26 | | | | | | | | | | | | | | | | | | | | | |
| CL091 | | | | -0.37 | 0.25 | | | | | | | | | | 0.34 | | -0.41 | | | -0.36 | | | | | | | | |
| CL092 | | | 0.38 | -0.30 | | 0.28 | 0.42 | | | | | 0.39 | | 0.45 | | 0.31 | 0.48 | | 0.25 | | | | | | | | | 0.37 |
| CL093 | -0.31 | | | 0.45 | | | | | | 0.62 | | | -0.52 | | | 0.25 | 0.29 | | | 0.63 | | | | | | | | |
| CL094 | | | | | | 0.52 | 0.41 | 0.36 | | 0.41 | | | | | 0.40 | 0.33 | | | 0.36 | | | | 0.29 | | | | | 0.31 |
| CL095 | | | | | | -0.39 | -0.39 | -0.26 | | -0.30 | | -0.27 | | -0.43 | | -0.35 | -0.39 | | -0.26 | -0.34 | | | | | | | | -0.31 |
| CL096 | | -0.26 | | 0.78 | -0.24 | -0.34 | -0.33 | -0.25 | | | | | | -0.28 | -0.25 | | | | -0.31 | | | | | | | | | -0.26 |
| CL097 | | -0.26 | | 0.86 | | | -0.25 | | | | | | | | | | | | | | | | | | | | | |
| CL098 | | -0.30 | | 0.57 | | -0.39 | -0.40 | -0.29 | | -0.27 | | | | -0.32 | -0.29 | | | -0.26 | -0.33 | | | | | | | | | -0.27 |
| CL099 | | | | | | | 0.34 | | | 0.32 | | 0.33 | | 0.40 | | 0.31 | 0.45 | | | 0.26 | | | | | | | | 0.24 |
| CL100 | -0.29 | | | | | | | | | | | | | | | | -0.25 | | | | | | | | | | | |
| CL101 | | | | | | | | | | | -0.27 | 0.31 | -0.30 | 0.33 | | | 0.47 | | | 0.28 | | | | | | | | |
| CL102 | | 0.32 | | -0.29 | | 0.46 | 0.30 | 0.28 | | 0.32 | | | | 0.26 | 0.39 | | | | 0.27 | | | | 0.28 | 0.31 | | | | |
| CL103 | | | | 0.87 | | -0.25 | -0.30 | | | | | | | | | | | | -0.26 | | | | | | | | | |
| CL104 | | | | | | 0.52 | 0.44 | 0.48 | 0.25 | 0.52 | | | | 0.33 | | 0.53 | 0.48 | | | 0.64 | 0.25 | | | | | | | 0.33 |
| CL105 | | | | 0.84 | -0.27 | -0.31 | -0.34 | | | | | | | -0.28 | | | | | -0.30 | | | | | | -0.24 | | | -0.30 |
| CL106 | | | 0.30 | | | 0.43 | 0.40 | 0.26 | | 0.44 | | 0.31 | | 0.44 | | 0.38 | 0.35 | | 0.26 | 0.47 | | | | | | | | 0.31 |
| CL107 | | | | | | 0.29 | 0.36 | 0.28 | 0.26 | 0.36 | | 0.26 | | 0.40 | | 0.32 | 0.50 | | | 0.41 | | | | | | | | 0.29 |
| CL108 | | -0.24 | | 0.88 | -0.24 | | -0.27 | | | | | | | | | | | | -0.24 | | | | | | | | | |
| CL109 | | 0.29 | | -0.39 | | 0.38 | 0.37 | | | | 0.27 | | | 0.24 | 0.30 | | | | 0.35 | | | | | | | | | 0.36 |
| CL110 | | | | | | -0.30 | -0.26 | -0.24 | | -0.34 | | | | -0.27 | | | -0.24 | | | | | | | | | | | |

| | CL198 | CL199 | CL200 | CL201 | CL202 | CL203 | CL204 | CL205 | CL206 | CL207 | CL208 | CL209 | CL210 | CL211 | CL212 | CL213 | CL214 | CL215 | CL216 | CL217 | CL218 | CL219 | CL220 | CL221 | CL222 | CL223 | CL224 | CL225 |
|---|---|---|---|---|---|---|---|---|---|---|---|---|---|---|---|---|---|---|---|---|---|---|---|---|---|---|---|---|
| CL111 | | | | | | -0.41 | -0.41 | -0.28 | | -0.34 | | -0.29 | | -0.38 | | -0.36 | -0.36 | | -0.27 | -0.32 | | | | | | | | -0.24 |
| CL112 | | | 0.26 | | | 0.34 | 0.41 | | | 0.27 | | 0.37 | | 0.44 | | 0.41 | 0.57 | | | 0.34 | | | | | | | | 0.25 |
| CL113 | | | | 0.31 | | 0.35 | | 0.24 | | 0.67 | | | -0.42 | 0.30 | | 0.37 | 0.43 | | | 0.69 | | | | | | | | |
| CL114 | | | | | | -0.31 | -0.28 | -0.25 | | -0.35 | | | | -0.27 | | | -0.27 | | | | | | | | | | | |
| CL115 | | -0.28 | | 0.79 | | -0.30 | -0.28 | -0.24 | | | | | | | | | | | -0.26 | | | | | | | | | |
| CL116 | | -0.26 | | 0.61 | | -0.43 | -0.46 | -0.31 | | -0.26 | | -0.28 | | -0.41 | | -0.25 | -0.33 | | -0.35 | | | -0.29 | | | | | | -0.37 |
| CL117 | | | | -0.40 | 0.25 | 0.25 | | | | | | | | 0.29 | | | | 0.27 | | | | | 0.30 | | | | | |
| CL118 | | | 0.29 | | | | 0.24 | | 0.24 | 0.32 | | 0.34 | | 0.32 | | 0.39 | 0.57 | | | 0.42 | | | | | | | | |
| CL119 | | | 0.26 | | | | | | | | -0.27 | | | | | | 0.47 | | | | | | | | | 0.26 | | |
| CL120 | | | | | | | | | | | | | | | | | | | | | | | | | | | | |
| CL121 | | 0.26 | | -0.34 | | | | | | | | | | | | | | 0.30 | | -0.33 | | | | | | | | |
| CL122 | | | | | | 0.61 | 0.48 | 0.38 | | 0.40 | | | | 0.30 | 0.30 | 0.48 | 0.29 | | 0.32 | 0.31 | 0.25 | | 0.30 | | | | | 0.36 |
| CL123 | | | | 0.77 | -0.24 | -0.30 | -0.35 | -0.25 | | | | | | -0.27 | | | | | -0.29 | | | | | -0.24 | | | | |
| CL124 | | -0.30 | | 0.80 | | -0.28 | -0.32 | | | | | | | | | | | | -0.28 | | | | | | | | | -0.24 |
| CL125 | | -0.31 | | 0.75 | | -0.31 | -0.34 | -0.25 | | | | | | -0.30 | | | | | -0.28 | | | | | | | | | |
| CL126 | | | | 0.89 | | | | | | | | | | | | | | | | | | | | | | | | |
| CL127 | | -0.29 | | 0.53 | | -0.43 | -0.45 | -0.34 | | -0.29 | | -0.28 | | -0.39 | -0.29 | -0.28 | -0.28 | | -0.37 | | | | -0.24 | | | | | -0.34 |
| CL128 | 0.27 | | | -0.53 | 0.25 | | | | | | 0.26 | | 0.55 | | 0.24 | | | | | | | 0.30 | | | | | | |
| CL129 | | | | | | 0.40 | 0.38 | 0.31 | | 0.30 | | | | 0.27 | 0.29 | 0.34 | | | 0.32 | 0.24 | | | 0.29 | | | | | 0.34 |
| CL130 | | | | | | | | | | -0.28 | | | | | | | | | | | | | | | | | | |
| CL131 | | | | | | 0.44 | 0.39 | 0.32 | | 0.50 | | 0.34 | | 0.39 | | 0.48 | 0.46 | | | 0.51 | | | | | | | | 0.39 |
| CL132 | | -0.25 | | 0.79 | -0.24 | -0.28 | -0.32 | | | | | | | -0.28 | | | | | -0.29 | | | | | | | | | |
| CL133 | | | | | | 0.56 | 0.55 | 0.40 | 0.29 | 0.53 | | 0.33 | | 0.44 | | 0.51 | 0.43 | | 0.33 | 0.50 | | | 0.25 | | | | | 0.40 |
| CL134 | | -0.27 | | 0.85 | | | -0.28 | | | | | | | | | | | | -0.24 | | | | | | | | | |
| CL135 | | -0.30 | | 0.80 | | | | | | | | | | | | | | | | | | | | | | | | |
| CL136 | | -0.24 | | 0.86 | | -0.24 | -0.29 | | | | | | | | | | | | | | | | | | | | | |
| CL137 | | | | | | 0.41 | 0.31 | 0.28 | | 0.48 | | | -0.25 | 0.38 | | 0.39 | 0.51 | | | 0.54 | | | | | | | | 0.25 |
| CL138 | | -0.24 | | 0.87 | | | -0.25 | | | | | | | | | | | | | | | | | | | | | |
| CL139 | | | | | | 0.41 | 0.41 | 0.31 | | 0.36 | | 0.27 | | 0.47 | | 0.35 | 0.45 | | 0.33 | 0.34 | | | | | | | | 0.38 |
| CL140 | | | | | | -0.30 | -0.26 | -0.25 | | -0.36 | | | | -0.28 | | | -0.27 | | | | | | | | | | | |
| CL141 | | | | 0.85 | -0.24 | -0.26 | -0.30 | | | | | | | | | | | | -0.26 | | | -0.25 | | | | | | -0.26 |
| CL142 | | -0.28 | | 0.84 | | | -0.24 | | | | | | | | | | | | -0.24 | | | | | | | | | |
| CL143 | | | | | | | | | | -0.36 | | | | | | -0.37 | -0.38 | | | -0.48 | | | | | | | | |
| CL144 | 0.37 | | | | | | -0.30 | | | -0.31 | | -0.24 | 0.50 | | | -0.24 | -0.34 | | | -0.27 | | | | | | | | |
| CL145 | | -0.30 | | 0.70 | | -0.40 | -0.40 | -0.28 | | -0.24 | | | | -0.33 | -0.25 | | | | -0.33 | | | | | | | | | -0.28 |
| CL146 | | | | -0.28 | | 0.54 | 0.43 | 0.30 | 0.27 | 0.33 | | | 0.28 | 0.35 | 0.29 | 0.32 | | | | | 0.25 | 0.27 | 0.26 | | | | | 0.35 |
| CL147 | | | | | | | | | 0.27 | 0.25 | | 0.28 | | 0.24 | | 0.28 | 0.37 | | | 0.27 | | | | | | | | 0.28 |
| CL148 | 0.39 | | | -0.28 | | | | | | -0.24 | | | 0.53 | | | | | | | | | | | | | | | |
| CL149 | | | | | | 0.38 | | | | 0.47 | | | | | 0.36 | | | | | 0.24 | | | | | | | | |
| CL150 | | -0.30 | | 0.74 | | -0.38 | -0.34 | -0.26 | | | | | | -0.32 | | | | | -0.32 | | | -0.28 | | | | | | -0.25 |
| CL151 | | | 0.26 | | | 0.49 | 0.38 | 0.33 | 0.29 | 0.29 | | 0.26 | | 0.41 | | 0.44 | 0.48 | | 0.34 | 0.34 | 0.26 | | | | | | | 0.44 |
| CL152 | | | | -0.35 | | 0.25 | | | | | 0.26 | | 0.46 | | | | -0.27 | | 0.24 | -0.32 | | | | | | | | |
| CL153 | | | | | | -0.30 | -0.25 | -0.25 | | -0.35 | | | | -0.27 | | | -0.28 | | | | | | | | | | | |
| CL154 | | | | 0.86 | | | | | | | | | | | | | | | | | | | | | | | | |
| CL155 | | | | | | 0.43 | 0.40 | 0.35 | | 0.44 | | 0.35 | -0.25 | 0.35 | | 0.41 | 0.47 | | 0.31 | 0.40 | | | | | | | | 0.29 |
| CL156 | | | | 0.87 | | | -0.25 | | | | | | | | | | | | | | | | | | | | | |
| CL157 | | | | | | 0.56 | 0.45 | 0.34 | | 0.45 | | 0.25 | | 0.28 | 0.26 | 0.42 | 0.26 | | 0.31 | 0.38 | 0.24 | | 0.25 | | | | | 0.40 |
| CL158 | | | | | | 0.33 | 0.41 | | | | | | | 0.42 | | | | | | | | | | | | | | 0.37 |
| CL159 | | | | | | | | | | | | 0.30 | | 0.27 | | | 0.51 | | | 0.33 | | | | | | 0.24 | | |
| CL160 | | | | | | | | | | | | | | | | | -0.28 | | | | | | | | | | | |
| CL161 | | -0.24 | | | | -0.29 | -0.25 | | | -0.34 | | | | -0.28 | | | -0.24 | -0.25 | | | | | | | | | | |
| CL162 | | | 0.28 | | | | | | | | | | | 0.33 | | | 0.25 | | | | | | | | | | | |
| CL163 | | | 0.26 | | 0.30 | 0.29 | 0.25 | 0.27 | | | | | | | | | | | | 0.24 | | | | | | | | 0.40 |
| CL164 | | | | | -0.25 | | | | | | | | | | | | 0.27 | | | | | | | | | | | |
| CL165 | | -0.29 | | 0.73 | | -0.26 | -0.30 | | | | | | | | | | | | -0.26 | | | | | | | | | |
| CL166 | | | | | | 0.53 | 0.42 | 0.38 | | 0.52 | | | | 0.29 | 0.31 | 0.45 | | | | 0.48 | | | 0.24 | | | | | 0.24 |
| CL167 | | -0.26 | | 0.77 | -0.27 | -0.28 | -0.32 | -0.26 | | | | | | -0.25 | | | | | -0.26 | | | | | -0.25 | | | | |
| CL168 | | | | | | 0.43 | 0.37 | 0.35 | | 0.46 | | | -0.31 | 0.36 | | 0.44 | 0.43 | | | 0.58 | | | | | | | | 0.30 |
| CL169 | | -0.27 | | 0.84 | | | -0.30 | | | | | | | | | | | | -0.24 | | | | | | | | | |
| CL170 | | | | | | | | | | | | | 0.33 | | | | | | | | | | | | | | | |

| | CL198 | CL199 | CL200 | CL201 | CL202 | CL203 | CL204 | CL205 | CL206 | CL207 | CL208 | CL209 | CL210 | CL211 | CL212 | CL213 | CL214 | CL215 | CL216 | CL217 | CL218 | CL219 | CL220 | CL221 | CL222 | CL223 | CL224 | CL225 |
|---|---|---|---|---|---|---|---|---|---|---|---|---|---|---|---|---|---|---|---|---|---|---|---|---|---|---|---|---|
| CL171 | | | | -0.29 | | | | | | -0.28 | | | | | | -0.28 | | | | -0.29 | | | | | | | | |
| CL172 | | | | 0.82 | -0.26 | | -0.31 | | | | | | | | | | | | -0.24 | | | | | | | | | |
| CL173 | | | | | | 0.38 | 0.38 | 0.24 | | 0.33 | | 0.27 | | 0.31 | 0.31 | 0.27 | 0.29 | | | 0.32 | | | | | | | | |
| CL174 | | | | | | 0.45 | 0.42 | 0.36 | | 0.48 | | | -0.26 | 0.43 | | 0.51 | 0.46 | | 0.30 | 0.55 | | | | | | | | 0.35 |
| CL175 | | | | | | 0.34 | | | | | | | | | | | | | | | | | | | | | | |
| CL176 | | | | | | | | | | | | 0.30 | -0.48 | | | | 0.40 | | | | | | | | | 0.26 | | |
| CL177 | | | | | | 0.38 | | | | 0.38 | | | | | 0.25 | | | | | | | | 0.29 | | | | | |
| CL178 | | | | | | -0.31 | -0.25 | -0.27 | | -0.37 | | | | -0.29 | | | -0.30 | | | -0.24 | | | | | | | | |
| CL179 | | | | 0.79 | | | -0.32 | | | | | | | | | | | | | | | | | | | | | |
| CL180 | | -0.29 | | 0.61 | | -0.35 | -0.38 | -0.28 | | -0.25 | | | | -0.31 | -0.27 | | | -0.29 | -0.26 | | | -0.26 | | | | | | -0.27 |
| CL181 | | | | | | | 0.30 | 0.36 | | | | | | | | | | | | | | 0.25 | | | | | | 0.40 |
| CL182 | | | | | | 0.45 | 0.50 | 0.41 | 0.29 | 0.38 | | 0.24 | | 0.41 | | 0.45 | 0.38 | | 0.27 | 0.34 | | | | | 0.24 | | | 0.41 |
| CL183 | | | | | | 0.34 | 0.29 | 0.26 | | 0.64 | | | -0.32 | 0.32 | | 0.42 | 0.41 | | | 0.66 | | | | | | | | |
| CL184 | | | | | | 0.39 | 0.27 | 0.36 | | 0.50 | | | | 0.43 | | 0.31 | 0.42 | | | 0.60 | | | | | | | | 0.26 |
| CL185 | 0.28 | | | | | | | | | | | | 0.44 | | | | | | | | | | | | | | | |
| CL186 | | | | | | 0.35 | 0.46 | 0.32 | | 0.37 | | 0.24 | | 0.33 | | 0.34 | 0.30 | | | 0.25 | | | | | | | | 0.33 |
| CL187 | | | | | | | 0.31 | | | | | | | | | 0.27 | | | | | | | | | | | | 0.28 |
| CL188 | | | | | | -0.27 | -0.25 | -0.25 | | -0.33 | | | | -0.24 | | | -0.26 | | | -0.26 | | | | | | | | |
| CL189 | | | | | | | | | | | | | | | | 0.24 | | | | | | | | | | | | |
| CL190 | | | | 0.85 | | | | | | | | | -0.26 | | | | | | | 0.32 | | | | | | | | |
| CL191 | | | | | | | | | 0.25 | | | | | | | | | | | | | | | | | | | |
| CL192 | | | | | | -0.25 | | | | -0.33 | | | | | | | | | | | | | | | | | | |
| CL193 | | | | | | -0.26 | | | | -0.28 | | | | -0.30 | | | -0.26 | | | | | | | | | | | |
| CL194 | | | | -0.35 | 0.30 | | 0.26 | | | | 0.28 | | 0.28 | | | | | | 0.36 | | | | | | 0.27 | | | 0.35 |
| CL195 | | | | | | 0.45 | 0.45 | 0.39 | | 0.36 | | 0.31 | | 0.40 | | 0.39 | 0.30 | | | 0.46 | | | | | | | | 0.36 |
| CL196 | | | | | | | | | | -0.25 | | | 0.31 | | | | | | | | | | | | | | | |
| CL197 | | | | | | 0.25 | | | | | | | | 0.27 | | | | | | | | | | | | | | |
| CL198 | | | | | | | | | | | | | 0.45 | | | | | | | | | | | | | | 0.26 | |
| CL199 | | | | | | | | | | | | | | | | | | | | | | | 0.24 | | | | | |
| CL200 | | | | | | | | | | | | | | | | | 0.26 | | | | | | | | | | | |
| CL201 | | | | | -0.24 | | | | | | | | -0.30 | | | | | | | 0.30 | | | | -0.24 | | | | |
| CL202 | | | | | | | | | | | | | | | | | | | | | | | | | | | | |
| CL203 | | | | | | | 0.25 | | | 0.37 | | 0.27 | | 0.35 | | 0.27 | | | 0.32 | 0.29 | | 0.27 | 0.24 | | | | | |
| CL204 | | | | | | | | 0.25 | | | | | | | | 0.31 | | | | | | | | | | | | 0.37 |
| CL205 | | | | | | | | | | 0.30 | | | | | | 0.40 | | | | | | | | | | | -0.26 | |
| CL206 | | | | | | | | | | | | | | | | | | | | | | | | | | | | |
| CL207 | | | | | | | | | | | | | -0.32 | | | 0.36 | 0.28 | | | 0.57 | | | | | | | | |
| CL208 | | | | | | | | | | | | | | | | | | -0.25 | | | | | | | | | | |
| CL209 | | | | | | | | | | | | | | | | | | | 0.26 | | | | | | | | | |
| CL210 | | | | | | | | | | | | | | | | | | | | | | | | | | | | |
| CL211 | | | | | | | | | | | | | | | | | 0.30 | | | | | | | | | | | 0.30 |
| CL212 | | | | | | | | | | | | | | | | | | | | | | | | | | | | |
| CL213 | | | | | | | | | | | | | | | | | 0.36 | | | 0.29 | | | | | | | | |
| CL214 | | | | | | | | | | | | | | | | | | | | 0.35 | | | | | | | | |
| CL215 | | | | | | | | | | | | | | | | | | | | | | | | | | | | |
| CL216 | | | | | | | | | | | | | | | | | | | | | | | | | 0.28 | | | |
| CL217 | | | | | | | | | | | | | | | | | | | | | | | | | | | | |
| CL218 | | | | | | | | | | | | | | | | | | | | | | | | | | | | |
| CL219 | | | | | | | | | | | | | | | | | | | | | | | | | | | | |
| CL220 | | | | | | | | | | | | | | | | | | | | | | | | | | | | |
| CL221 | | | | | | | | | | | | | | | | | | | | | | | | | | | | |
| CL222 | | | | | | | | | | | | | | | | | | | | | | | | | | | | |
| CL223 | | | | | | | | | | | | | | | | | | | | | | | | | | | | |
| CL224 | | | | | | | | | | | | | | | | | | | | | | | | | | | | |
| CL225 | | | | | | | | | | | | | | | | | | | | | | | | | | | | |
| CL226 | | | | | | | | | | | | | | | | | | | | | | | | | | | | |
| CL227 | | | | | | | | | | | | | | | | | | | | | | | | | | | | |
| CL228 | | | | | | | | | | | | | | | | | | | | | | | | | | | | |
| CL229 | | | | | | | | | | | | | | | | | | | | | | | | | | | | |
| CL230 | | | | | | | | | | | | | | | | | | | | | | | | | | | | |

| | CL226 | CL227 | CL228 | CL229 | CL230 | CL231 | CL232 | CL233 | CL234 | CL235 | CL236 | CL237 | CL238 | CL239 | CL240 | Number of correlated clusters |
|---|---|---|---|---|---|---|---|---|---|---|---|---|---|---|---|---|
| CL001 | | | | -0.27 | | 0.27 | | -0.32 | | | | | | | | 111 |
| CL002 | | | | | | | -0.33 | | -0.27 | -0.28 | -0.39 | | | | | 112 |
| CL003 | | | | -0.24 | | | -0.30 | | | -0.26 | -0.30 | | 0.31 | -0.28 | -0.28 | 128 |
| CL004 | | | | | | | -0.37 | 0.27 | | -0.27 | -0.32 | | 0.32 | -0.31 | | 144 |
| CL005 | | | | | | | | -0.30 | | | -0.24 | | | | | 118 |
| CL006 | | | | | | 0.48 | | -0.32 | | | | | | | | 155 |
| CL007 | | 0.29 | | | | 0.33 | | -0.56 | | | | | | | | 145 |
| CL008 | | | | | | | -0.35 | 0.25 | | -0.25 | -0.26 | | 0.35 | -0.33 | | 138 |
| CL009 | | | | | | | -0.25 | | | | | | 0.35 | | | 127 |
| CL010 | | | | -0.32 | | | -0.35 | | | | -0.44 | | 0.26 | -0.29 | -0.24 | 116 |
| CL011 | | | | | | | | | | | -0.32 | | | | | 85 |
| CL012 | | | | | | | | | | | | 0.24 | 0.31 | | | 124 |
| CL013 | | | | | | | -0.34 | | -0.24 | -0.32 | -0.27 | | 0.35 | -0.29 | | 135 |
| CL014 | | | | | | 0.25 | -0.26 | | | | -0.27 | | | | | 93 |
| CL015 | | | | -0.26 | | | -0.36 | 0.27 | -0.30 | -0.25 | -0.38 | | | -0.31 | -0.25 | 137 |
| CL016 | | | | 0.38 | | | 0.38 | -0.52 | 0.24 | 0.33 | 0.39 | | | 0.27 | 0.24 | 154 |
| CL017 | | | | -0.26 | | | -0.34 | | -0.26 | -0.27 | -0.35 | | | -0.28 | -0.27 | 118 |
| CL018 | | 0.28 | | 0.32 | | | | -0.54 | | | | | | | | 152 |
| CL019 | | | | | | | | | | | | 0.31 | | | | 96 |
| CL020 | | | | | | | | | | | | | 0.34 | | | 122 |
| CL021 | | | | -0.27 | | | -0.30 | | | -0.24 | -0.29 | | 0.30 | -0.28 | | 124 |
| CL022 | | | | -0.25 | | | | | | | | 0.30 | | -0.25 | | 116 |
| CL023 | | | | | | | -0.37 | | -0.26 | -0.26 | -0.39 | | | | | 102 |
| CL024 | | | | | | | | | | | -0.24 | | 0.37 | | | 115 |
| CL025 | | | | 0.25 | | 0.27 | 0.25 | -0.45 | | | | | | | | 145 |
| CL026 | | 0.30 | | 0.41 | | 0.33 | 0.35 | -0.62 | | 0.30 | 0.31 | | | | | 156 |
| CL027 | | | | -0.32 | | | -0.26 | 0.31 | | | -0.31 | | 0.25 | -0.30 | | 130 |
| CL028 | | | | | | 0.47 | | -0.33 | | | | | | | | 127 |
| CL029 | | | | | | 0.30 | | -0.35 | | | | | | | | 108 |
| CL030 | | | | | | 0.31 | | -0.31 | | | | | | | | 98 |
| CL031 | | | | | | | | | | | | 0.40 | | | | 92 |
| CL032 | | | | | | | -0.24 | | | | -0.30 | | 0.29 | -0.24 | -0.26 | 126 |
| CL033 | | | | | | | | -0.41 | | | | | | | | 116 |
| CL034 | | | | | | | -0.30 | | -0.30 | -0.27 | -0.40 | | | | | 92 |
| CL035 | | | | | | | | -0.53 | | | | | | | | 111 |
| CL036 | | 0.36 | | 0.40 | | 0.37 | 0.31 | -0.51 | | 0.24 | 0.33 | | | | | 155 |
| CL037 | | | | | | | | -0.54 | | | | | 0.32 | | | 123 |
| CL038 | 0.29 | | | | | | | -0.43 | 0.27 | | 0.25 | 0.32 | | | | 130 |
| CL039 | | | | | | | | -0.43 | | | | | 0.27 | | | 91 |
| CL040 | | | | | | | | | | | | 0.25 | 0.32 | | | 103 |
| CL041 | | 0.31 | | 0.42 | | | 0.28 | -0.58 | 0.24 | 0.27 | 0.27 | | | 0.28 | | 152 |
| CL042 | | | | | | | | -0.32 | | | | | | | | 94 |
| CL043 | | | | -0.33 | | | | | | | | | 0.25 | | | 90 |
| CL044 | | | | -0.24 | | | | | | | | 0.32 | | -0.27 | | 70 |
| CL045 | | | | | | 0.44 | | | | | | | | | | 118 |
| CL046 | | | | | | | -0.32 | 0.26 | -0.24 | -0.27 | -0.29 | | 0.29 | -0.30 | | 98 |
| CL047 | | | | | | 0.25 | | | | | -0.28 | | | | | 58 |
| CL048 | | | | | | 0.37 | | -0.26 | | | | | | | | 84 |
| CL049 | | | | | | 0.32 | | -0.46 | | | | | | | | 101 |
| CL050 | | | | | | 0.33 | | | | | | | -0.29 | | | 55 |

| | CL226 | CL227 | CL228 | CL229 | CL230 | CL231 | CL232 | CL233 | CL234 | CL235 | CL236 | CL237 | CL238 | CL239 | CL240 | Number of correlated clusters |
|---|---|---|---|---|---|---|---|---|---|---|---|---|---|---|---|---|
| CL051 | | | | | | | | -0.41 | | | | | | | | 123 |
| CL052 | | | | | | 0.43 | | -0.35 | | | | | | | | 94 |
| CL053 | | | | | | | | | | | | | 0.24 | | | 74 |
| CL054 | | | | | | | -0.28 | | | | -0.25 | | 0.32 | -0.24 | | 86 |
| CL055 | | | | | | | | -0.48 | | | | | | | | 102 |
| CL056 | 0.24 | 0.34 | | 0.38 | | | 0.30 | -0.48 | 0.32 | | 0.41 | | | 0.26 | | 144 |
| CL057 | | | | | | | -0.35 | 0.29 | | -0.30 | -0.27 | | 0.32 | -0.28 | | 88 |
| CL058 | | | | | | | | | | | -0.27 | | | | | 63 |
| CL059 | | | | | | | | -0.24 | | | | | | | | 67 |
| CL060 | | | | 0.27 | | 0.34 | 0.25 | -0.56 | | 0.26 | | | | 0.25 | | 129 |
| CL061 | 0.27 | 0.35 | | 0.37 | | | 0.30 | -0.63 | 0.26 | | 0.35 | | | | | 135 |
| CL062 | | | | | | 0.44 | | -0.52 | | | | | | | | 94 |
| CL063 | | 0.35 | | 0.45 | | | 0.36 | -0.47 | 0.27 | 0.31 | 0.46 | | | | | 129 |
| CL064 | | | | -0.30 | | | | | | | | | | | | 56 |
| CL065 | | | | -0.32 | | | | | | | | | 0.24 | | | 76 |
| CL066 | | | | | | | | -0.36 | | | | | 0.36 | | | 84 |
| CL067 | | | | | | | -0.26 | | | | | | 0.30 | -0.28 | | 68 |
| CL068 | 0.36 | 0.26 | | 0.30 | | 0.26 | 0.27 | -0.47 | | 0.29 | 0.29 | | | | | 128 |
| CL069 | | | | -0.32 | | | | 0.76 | -0.24 | | -0.26 | | | | | 113 |
| CL070 | | -0.30 | | | | | | | | | | 0.31 | | | | 30 |
| CL071 | | 0.25 | | 0.33 | | 0.33 | 0.26 | -0.39 | | 0.27 | 0.33 | | | | | 120 |
| CL072 | | | | | | | -0.29 | | | | -0.27 | | 0.27 | -0.28 | | 80 |
| CL073 | -0.29 | -0.38 | | -0.29 | | | -0.28 | 0.61 | | -0.24 | -0.31 | | | | | 130 |
| CL074 | | | | | | | 0.25 | -0.43 | | | | | | | | 110 |
| CL075 | | | | -0.30 | | | | | | | | | | -0.27 | -0.27 | 53 |
| CL076 | | 0.38 | | 0.47 | | | 0.31 | -0.44 | 0.29 | 0.27 | 0.41 | | | 0.24 | | 126 |
| CL077 | -0.28 | -0.29 | | | | | -0.26 | 0.63 | | | | -0.25 | | | | 120 |
| CL078 | | | | | | | | -0.38 | | | | 0.37 | | | | 93 |
| CL079 | | 0.30 | | | | 0.35 | | -0.58 | | | | | | | | 110 |
| CL080 | 0.29 | | | | | | 0.26 | -0.47 | 0.30 | 0.28 | 0.31 | | | | | 106 |
| CL081 | | | | | | | | | | | | | 0.40 | | | 82 |
| CL082 | | | | -0.24 | | | | | | | | | | | | 47 |
| CL083 | -0.24 | -0.31 | | -0.38 | | | | 0.67 | -0.24 | | -0.30 | | | | | 109 |
| CL084 | | | | | | | | -0.33 | | | | 0.26 | | | | 65 |
| CL085 | | | | | | | | | | | | | | | | 52 |
| CL086 | | | | | | | | 0.86 | | | | | | | | 97 |
| CL087 | -0.28 | -0.31 | | | | | -0.26 | 0.66 | | -0.24 | | -0.24 | | | | 114 |
| CL088 | -0.27 | -0.30 | | | | | -0.26 | 0.62 | | | | -0.24 | | | | 111 |
| CL089 | -0.26 | -0.32 | | | | | | 0.48 | | | | | -0.24 | | | 102 |
| CL090 | | -0.25 | | | | | | 0.26 | | | | | | | | 81 |
| CL091 | | | | | | | | | | | | 0.28 | | | | 60 |
| CL092 | | 0.38 | | 0.30 | | 0.34 | | -0.25 | | 0.24 | 0.27 | | | | 0.29 | 103 |
| CL093 | | | | | | | 0.28 | -0.30 | | 0.25 | | | -0.31 | 0.30 | | 63 |
| CL094 | | 0.35 | | 0.35 | | | 0.29 | -0.52 | | 0.24 | 0.39 | | | | | 101 |
| CL095 | | -0.28 | | -0.37 | | | | 0.65 | | | -0.27 | | | | | 97 |
| CL096 | -0.25 | -0.31 | | | | | | 0.39 | | | | -0.25 | | | | 93 |
| CL097 | | | | | | | | 0.24 | | | | | | | | 71 |
| CL098 | -0.27 | -0.31 | | | | | -0.25 | 0.64 | | | -0.26 | -0.25 | | | | 101 |
| CL099 | | | | 0.28 | | 0.33 | 0.27 | -0.53 | | | 0.26 | | | | | 90 |
| CL100 | | | | | | | | 0.29 | | | | | | | | 24 |
| CL101 | | | | | | 0.32 | | -0.29 | | | | | | | | 52 |
| CL102 | 0.25 | 0.28 | | 0.29 | | | | -0.52 | | | 0.26 | 0.30 | | | | 92 |
| CL103 | | -0.28 | | | | | | | | | | | | | | 68 |
| CL104 | | 0.34 | | 0.49 | | | 0.35 | -0.31 | | 0.32 | 0.48 | | | 0.24 | | 82 |
| CL105 | | -0.34 | | | | | | | | | | | | | | 77 |
| CL106 | | | | 0.28 | | | 0.37 | -0.43 | 0.28 | 0.28 | 0.38 | | | 0.29 | | 93 |
| CL107 | | 0.26 | | 0.37 | | 0.28 | 0.34 | -0.51 | | 0.32 | 0.25 | | | 0.27 | | 93 |
| CL108 | | | | | | | | | | | | | | | | 61 |
| CL109 | 0.29 | 0.33 | | 0.28 | | | | -0.51 | | | 0.28 | | | | | 86 |
| CL110 | | | | | | | -0.24 | 0.86 | | | | | | | | 74 |

| | CL226 | CL227 | CL228 | CL229 | CL230 | CL231 | CL232 | CL233 | CL234 | CL235 | CL236 | CL237 | CL238 | CL239 | CL240 | Number of correlated clusters |
|---|---|---|---|---|---|---|---|---|---|---|---|---|---|---|---|---|
| CL111 | | -0.27 | | -0.34 | | | -0.25 | 0.75 | | | -0.27 | | | | | 84 |
| CL112 | | 0.35 | | 0.34 | | 0.33 | | -0.39 | | 0.29 | 0.39 | | | 0.25 | | 88 |
| CL113 | | | | 0.32 | | | 0.31 | -0.41 | | 0.33 | 0.31 | | -0.25 | 0.29 | | 61 |
| CL114 | | | | | | | -0.27 | 0.85 | | | | | | | | 72 |
| CL115 | -0.24 | -0.27 | | | | | | 0.38 | | | | -0.25 | | | | 68 |
| CL116 | -0.29 | -0.40 | | -0.31 | | -0.24 | -0.24 | 0.53 | | | -0.31 | | | | | 91 |
| CL117 | 0.25 | | | | | | | -0.29 | 0.26 | | | 0.29 | 0.29 | | | 64 |
| CL118 | | | | 0.26 | | 0.38 | 0.25 | -0.35 | | | 0.34 | | | | | 75 |
| CL119 | | | | | | 0.37 | | | | | | | | | | 24 |
| CL120 | | | | | | | | | | | | | | | | 6 |
| CL121 | 0.25 | | | | | | | | | | | | | | | 40 |
| CL122 | 0.32 | 0.42 | | 0.37 | | | 0.25 | -0.36 | 0.27 | | 0.46 | | | | | 84 |
| CL123 | -0.26 | -0.28 | | | | | | 0.40 | | | | | | | | 68 |
| CL124 | | -0.28 | | | | | | 0.37 | | | | -0.26 | | | | 62 |
| CL125 | -0.24 | -0.27 | | | | | | 0.43 | | | | | | | | 67 |
| CL126 | | | | | | -0.24 | | | | | | | | | | 39 |
| CL127 | -0.29 | -0.38 | | -0.28 | | | -0.29 | 0.63 | | -0.25 | -0.33 | | | | | 83 |
| CL128 | | | | | | | | | | | | | 0.33 | | | 46 |
| CL129 | | 0.25 | | 0.27 | | | 0.27 | -0.35 | 0.25 | 0.24 | 0.39 | | | | | 74 |
| CL130 | | | | | | | | 0.77 | | | | | | | | 33 |
| CL131 | 0.24 | 0.31 | | 0.43 | | | 0.36 | -0.51 | 0.26 | 0.26 | 0.40 | | | 0.30 | | 73 |
| CL132 | | -0.26 | | | | -0.25 | | 0.38 | | | | | | | | 57 |
| CL133 | 0.31 | 0.39 | | 0.45 | | | 0.32 | -0.45 | | 0.34 | 0.44 | | | | 0.25 | 79 |
| CL134 | | | | | | | | 0.25 | | | | -0.24 | | | | 44 |
| CL135 | | | | | | | | 0.36 | | | | | | | | 38 |
| CL136 | | -0.27 | | | | | | | | | | | | | | 41 |
| CL137 | | 0.30 | | 0.36 | | | 0.26 | -0.49 | | | 0.29 | | | 0.30 | | 59 |
| CL138 | | | | | | | | | | | | | | | | 33 |
| CL139 | 0.29 | 0.36 | | 0.35 | | | 0.32 | -0.45 | 0.31 | 0.32 | 0.37 | | | | | 66 |
| CL140 | | | | | | | -0.24 | 0.86 | | | | | | | | 50 |
| CL141 | | -0.27 | | | | | | | | | | | | | | 38 |
| CL142 | | | | | | | | 0.26 | | | | -0.29 | | | | 35 |
| CL143 | | | | -0.32 | | | | | | | -0.24 | | | -0.30 | | 30 |
| CL144 | | | | | | | -0.36 | 0.49 | -0.35 | -0.30 | -0.33 | | | | | 49 |
| CL145 | -0.27 | -0.33 | | | | | -0.25 | 0.50 | | | -0.25 | -0.24 | | | | 59 |
| CL146 | 0.25 | 0.36 | | 0.41 | | | | -0.36 | | | 0.32 | 0.29 | 0.29 | | | 61 |
| CL147 | 0.28 | | | | | 0.30 | | -0.36 | 0.25 | 0.26 | 0.27 | | | | | 42 |
| CL148 | | | | | | | | | | | | | | | | 21 |
| CL149 | | | | | | | | -0.44 | | | | | | | | 25 |
| CL150 | -0.29 | -0.32 | | | | | | 0.44 | | | | -0.24 | | | | 48 |
| CL151 | 0.30 | 0.44 | | 0.38 | | | 0.24 | | | 0.30 | 0.51 | | | | 0.29 | 54 |
| CL152 | | | | | | | | | | | | | 0.30 | | | 30 |
| CL153 | | | | | | | -0.24 | 0.86 | | | | | | | | 39 |
| CL154 | | | | | | | | | | | | | | | | 22 |
| CL155 | | 0.29 | | 0.40 | | | 0.30 | -0.40 | | 0.32 | 0.34 | | | 0.25 | | 48 |
| CL156 | | | | | | | | | | | | | | | | 18 |
| CL157 | 0.24 | 0.36 | | 0.45 | | | 0.24 | -0.46 | | 0.39 | 0.39 | | | | | 53 |
| CL158 | | | | | | | 0.33 | -0.30 | | | 0.25 | | | 0.24 | | 34 |
| CL159 | | | | | | 0.44 | | -0.31 | | | | | | 0.25 | | 25 |
| CL160 | | -0.28 | | -0.24 | | | | | | | | | | | | 8 |
| CL161 | | | | | | | -0.24 | 0.86 | | | | | | | | 33 |
| CL162 | 0.24 | | | | | 0.36 | | | | 0.25 | | | | | | 20 |
| CL163 | | | | | | | | -0.26 | 0.28 | | 0.31 | | | | | 30 |
| CL164 | | | | | | | | -0.29 | | | | | | | | 10 |
| CL165 | | -0.24 | | | | | | 0.44 | | | | | | | | 26 |
| CL166 | | 0.30 | | 0.37 | | | 0.26 | -0.36 | | | 0.37 | | | | | 36 |
| CL167 | | -0.27 | | | | | | 0.40 | | | | | | | | 30 |
| CL168 | | | | 0.37 | | | 0.40 | -0.36 | 0.24 | 0.32 | 0.43 | | | 0.37 | | 34 |
| CL169 | | -0.28 | | | | | | 0.25 | | | | -0.24 | | | | 20 |
| CL170 | | | | | | | | | | | | | | | | 4 |

| | CL226 | CL227 | CL228 | CL229 | CL230 | CL231 | CL232 | CL233 | CL234 | CL235 | CL236 | CL237 | CL238 | CL239 | CL240 | Number of correlated clusters |
|---|---|---|---|---|---|---|---|---|---|---|---|---|---|---|---|---|
| CL171 | | | | -0.26 | | | | | | | | | | | | 10 |
| CL172 | | -0.29 | | | | | | 0.25 | | | | | | | | 20 |
| CL173 | | 0.26 | | | | | 0.25 | -0.38 | 0.25 | 0.25 | 0.30 | | | | | 34 |
| CL174 | | 0.29 | | 0.47 | | | 0.33 | -0.31 | | 0.31 | 0.45 | | | 0.25 | | 36 |
| CL175 | 0.24 | | | | | | | -0.26 | | | | | | | | 8 |
| CL176 | | | | | | 0.33 | | -0.24 | | | | | | | | 16 |
| CL177 | | | | | | | | -0.39 | | | | | | | | 11 |
| CL178 | | | | -0.25 | | | | 0.84 | | | | | | | | 22 |
| CL179 | | | | | | | | 0.34 | | | | -0.25 | | | | 14 |
| CL180 | -0.29 | -0.28 | | | | | -0.26 | 0.55 | | | -0.24 | | -0.24 | | | 31 |
| CL181 | | | | 0.33 | | | | -0.35 | 0.24 | | 0.28 | 0.28 | | | | 22 |
| CL182 | 0.28 | 0.33 | | 0.43 | | | | -0.24 | | 0.31 | 0.43 | | | | 0.31 | 27 |
| CL183 | | | | 0.37 | | | 0.28 | -0.35 | 0.25 | | 0.28 | | | | 0.24 | 23 |
| CL184 | | | | 0.36 | | | 0.28 | -0.38 | 0.25 | 0.31 | 0.36 | | | 0.28 | 0.29 | 24 |
| CL185 | | | | | | | | | | -0.25 | -0.27 | | | | | 4 |
| CL186 | | | | 0.32 | | | | -0.24 | 0.33 | | 0.41 | | | | 0.31 | 24 |
| CL187 | | | | | | | | | | | | | | | | 5 |
| CL188 | | | | | | | | 0.84 | | | | | | | | 13 |
| CL189 | | | | | | | | -0.33 | | | | | | | | 4 |
| CL190 | | | | | | | | | | | | | | | | 4 |
| CL191 | | | | | | | | | | | | | | | | 1 |
| CL192 | | | | | | | -0.24 | 0.80 | | | | | | | | 7 |
| CL193 | | | | | | | | 0.82 | | | | | | | | 7 |
| CL194 | | | | | | | | | | | | | | | | 9 |
| CL195 | | 0.30 | | 0.31 | | | 0.27 | -0.29 | | 0.35 | 0.42 | | | | | 17 |
| CL196 | | | | | | | | | | | | | 0.24 | | | 3 |
| CL197 | | 0.26 | | | | | | -0.32 | | | | | | | | 4 |
| CL198 | | | | | | | | | | | | | | | | 2 |
| CL199 | | | | | | | | | | 0.33 | | | | | | 2 |
| CL200 | | | | | | | | | | | | | | | | 1 |
| CL201 | | | | | | | | | | | | | | | | 4 |
| CL202 | | | | | | | | | | | | | | | | 0 |
| CL203 | | 0.40 | | 0.42 | | | | -0.31 | | | 0.31 | | | | | 13 |
| CL204 | | 0.28 | | 0.31 | 0.26 | | 0.31 | -0.28 | | 0.25 | 0.25 | | | | | 10 |
| CL205 | | | | | | | | -0.26 | | 0.27 | | | | | | 5 |
| CL206 | 0.30 | | | 0.24 | | | | | | | | | | | | 2 |
| CL207 | | | | 0.28 | | | | -0.32 | | | 0.28 | | | | | 7 |
| CL208 | | | | | | | | | | | | | | | | 1 |
| CL209 | | | | | | | | -0.24 | | | | | | | | 2 |
| CL210 | | | | | | | | | | | | | 0.28 | | | 1 |
| CL211 | | | | 0.39 | | 0.24 | | -0.27 | | | 0.27 | | | | 0.29 | 7 |
| CL212 | | | | | | | | -0.24 | | | | | | | | 1 |
| CL213 | | 0.26 | | 0.35 | | | | | | | 0.40 | | | | | 5 |
| CL214 | | | | 0.32 | | 0.32 | | -0.25 | | | 0.25 | | | | | 5 |
| CL215 | | | | | | | | | | | | | | | | 0 |
| CL216 | | | | | | | | | | | | | | | | 1 |
| CL217 | | | | 0.32 | | | 0.27 | -0.26 | | 0.27 | | | | | | 4 |
| CL218 | | | | | | | | | | | | | | | | 0 |
| CL219 | | | | 0.26 | | | | | | | | | 0.35 | | | 2 |
| CL220 | | | | | | | | | | | | | | | | 0 |
| CL221 | | | | | | | | | | | | | | | | 0 |
| CL222 | | | | | | | | | | | | | | | | 0 |
| CL223 | | | | | | | | | | | | | | | | 0 |
| CL224 | | | | | | | | | | | | | | | | 0 |
| CL225 | | | | | | | | | | | | | | | | 0 |
| CL226 | | | | | | | | | | | | | | | | 0 |
| CL227 | | | | 0.26 | | | | | | | | | | | | 1 |
| CL228 | | | | | | | | | | | | | | | | 0 |
| CL229 | | | | | | | | | | | 0.32 | | | | | 1 |
| CL230 | | | | | | | | | | | | | | | | 0 |

| | CL226 | CL227 | CL228 | CL229 | CL230 | CL231 | CL232 | CL233 | CL234 | CL235 | CL236 | CL237 | CL238 | CL239 | CL240 | Number of correlated clusters |
|---|---|---|---|---|---|---|---|---|---|---|---|---|---|---|---|---|
| CL231 | | | | | | | | | | | | | | | | 0 |
| CL232 | | | | | | | | -0.25 | | | | | | | | 1 |
| CL233 | | | | | | | | | | | | | | | | 0 |
| CL234 | | | | | | | | | | | | | | | | 0 |
| CL235 | | | | | | | | | | | | | | | | 0 |
| CL236 | | | | | | | | | | | | | | | 0.26 | 1 |
| CL237 | | | | | | | | | | | | | | | | 0 |
| CL238 | | | | | | | | | | | | | | | | 0 |
| CL239 | | | | | | | | | | | | | | | 0.26 | 1 |
| CL240 | | | | | | | | | | | | | | | | 0 |

Supplementary Table 11. Summary statistics for all estimates of Ks between *G. herbaceum* (A1) and *G. arboreum* (A2) among 112,831 pairwise permutations of all A1 and A2 individuals.

| **Filter Level** | **Mean** | **Median** | **Mode*** | **Range** | **Standard Deviation** | **95% CIs**** | **Mean (outliers excluded)***** | **Median (outliers excluded)** | **Mode (outliers excluded)** | **Range (outliers excluded)** | **Standard Deviation (outliers excluded)** | **95% CIs (outliers excluded)** | **Divergence time ******* |
|---|---|---|---|---|---|---|---|---|---|---|---|---|---|
| Low Filter***** | 0.0086 | 0.0074 | 0.0067 | 0.0002 - 0.1835 | 0.006 | 0.0016 - 0.0240 | 0.0082 | 0.0074 | 0.0068 | 0.0002 - 0.1835 | 0.0045 | 0.0021 - 0.0197 | 899 kya (230 kya - 2.2 mya) |
| Medium Filter****** | 0.0069 | 0.0058 | 0.0056 | 0.0002 - 0.3201 | 0.005 | 0.0010 - 0.0223 | 0.0062 | 0.0059 | 0.0056 | 0.0002 - 0.3201 | 0.0033 | 0.0015 - 0.0139 | 680 kya (164 kya - 1.5 mya) |
| High Filter******* | 0.0068 | 0.0058 | 0.0061 | 0.0002 - 0.3900 | 0.005 | 0.0010 - 0.0223 | 0.0061 | 0.0058 | 0.0061 | 0.0002 - 0.3900 | 0.0032 | 0.0016 - 0.0138 | 669 kya (175 kya - 1.5 mya) |
| No Filter | 0.0088 | 0.0074 | 0.0059 | 0.0002 - 1.0150 | 0.006 | 0.0016 - 0.0246 | 0.0084 | 0.0074 | 0.0059 | 0.0002 - 1.0150 | 0.0050 | 0.0020 - 0.0207 | 921 kya (219 kya - 2.3 mya) |

* Modes were estimated using the R density function, which produces a Kernal Density Estimate for a given dataset, from which the maximum density was identified.
** 95% confidence intervals were estimated by sorting the pairwise permuted dS values, using the 2.5 %ile and the 97.5%ile as the lower and upper bounds for CIs, respectively.
*** Three outlier indivdiuals were identified by persistently elevated dS values in all pairwise comparisons: two from *G. herbaceum* (A1_037 and A1_148) and one from *G. arboreum* (A2_038). These were also identified via PCA
**** Divergence time is calculated via T=dS/2r, where dS is represented by the dS median and CI for each filter level
***** The "Low Filter" removed genes from consideration if ≥ 70% of the nucleotides in the alignment were ambiguous Ns.
****** The "Medium Filter" removed genes from consideration if either ≥ 70% of the nucleotides in the alignment were ambiguous Ns and/or >4 sequences contained stop codons.
****** The "High Filter" removed genes from consideration if either ≥ 70% of the nucleotides in the alignment were ambiguous Ns and/or any sequences contained stop codons.

Supplementary Table 12. Details regarding the 50-gene windows where the mean dS is outside of the 95% CI for the genome-wide dS. Chromosomes and positions are based off of the *G. longicalyx* genome. The window number for each position is given, along with the total number of 50-gene windows for that chromosome (562 gene windows total).

| Chromosome | Position | dS | Window Position | Total Windows on chromosome |
|---|---|---|---|---|
| F01 | --- | --- | --- | 40 |
| F02 | --- | --- | --- | 31 |
| F03 | --- | --- | --- | 40 |
| F04 | --- | --- | --- | 33 |
| F05 | 70.61 | 0.0252 | 60 | 65 |
| | 73.47 | 0.0221 | 62 | |
| F06 | --- | --- | --- | 35 |
| F07 | 69.73 | 0.0243 | 32 | 41 |
| | 72.89 | 0.0207 | 33 | |
| F08 | 6.57 | 0.0229 | 9 | 43 |
| F09 | 3.89 | 0.0211 | 4 | 42 |
| F10 | 1.72 | 0.0029 | 4 | 45 |
| | 18.1 | 0.0209 | 18 | |
| | 89.53 | 0.0205 | 39 | |
| | 91.95 | 0.0206 | 41 | |
| F11 | 64.32 | 0.0267 | 38 | 59 |
| | 90.59 | 0.0219 | 50 | |
| | 91.8 | 0.0226 | 51 | |
| | 95.42 | 0.0228 | 54 | |
| F12 | --- | --- | --- | 46 |
| F13 | --- | --- | --- | 42 |
| **Total** | | | | **647** |

**Supplementary Table 13**. Details regarding the accessions and genomic windows that fall outside of the genome-wide 95% CI for *G. herbaceum* (A1) versus *G. arboreum* (A2).

Accessions with dS outside of 95% CI (0.0031 - 0.0198)

| **Accession** | **Chr** | **Position** | **dS** | |
|---|---|---|---|---|
| | | | 0.0006 | **Minimum** |
| | | | 0.0370 | **Maximum** |
| A2_004 | 1 | 2.04 | 0.0030 | |
| A2_SRR8283970_Gar43 | 1 | 4.54 | 0.0205 | |
| A2_021 | 1 | 5.46 | 0.0206 | |
| A2_026 | 1 | 5.46 | 0.0218 | |
| A2_029 | 1 | 5.46 | 0.0214 | |
| A2_034 | 1 | 5.46 | 0.0213 | |
| A2_041 | 1 | 5.46 | 0.0226 | |
| A2_043 | 1 | 5.46 | 0.0219 | |
| A2_044 | 1 | 5.46 | 0.0221 | |
| A2_046 | 1 | 5.46 | 0.0224 | |
| A2_047 | 1 | 5.46 | 0.0219 | |
| A2_057 | 1 | 5.46 | 0.0226 | |
| A2_061 | 1 | 5.46 | 0.0219 | |
| A2_073 | 1 | 5.46 | 0.0203 | |
| A2_074 | 1 | 5.46 | 0.0220 | |
| A2_076 | 1 | 5.46 | 0.0219 | |
| A2_087 | 1 | 5.46 | 0.0220 | |
| A2_113 | 1 | 5.46 | 0.0217 | |
| A2_119 | 1 | 5.46 | 0.0221 | |
| A2_123 | 1 | 5.46 | 0.0213 | |
| A2_154 | 1 | 5.46 | 0.0211 | |
| A2_156 | 1 | 5.46 | 0.0218 | |
| A2_158 | 1 | 5.46 | 0.0225 | |
| A2_164 | 1 | 5.46 | 0.0227 | |
| A2_255 | 1 | 5.46 | 0.0215 | |
| A2_lintless | 1 | 5.46 | 0.0223 | |
| A2_SRR8283970_Gar43 | 1 | 5.46 | 0.0205 | |
| A2_SRR8283973_Gar38 | 1 | 5.46 | 0.0222 | |
| A2_SRR8283974_Gar39 | 1 | 5.46 | 0.0201 | |
| A2_SRR8283980_Gar59 | 1 | 5.46 | 0.0218 | |
| A2_SRR8283986_Gar35 | 1 | 5.46 | 0.0199 | |
| A2_SRR8283989_Gar30 | 1 | 5.46 | 0.0223 | |
| A2_SRR8283992_Gar04 | 1 | 5.46 | 0.0202 | |
| A2_SRR8283999_Gar55 | 1 | 5.46 | 0.0216 | |
| A2_SRR8284000_Gar54 | 1 | 5.46 | 0.0207 | |
| A2_SRR8284004_Gar50 | 1 | 5.46 | 0.0215 | |
| A2_SRR8284005_Gar49 | 1 | 5.46 | 0.0212 | |
| A2_SRR8284006_Gar48 | 1 | 5.46 | 0.0211 | |
| A2_SRR8284010_Gar63 | 1 | 5.46 | 0.0210 | |
| A1_125 | 1 | 18.77 | 0.0232 | |
| A1_SRR8283996_Ghe01 | 1 | 18.77 | 0.0235 | |
| A1_SRR8283998_Ghe02 | 1 | 18.77 | 0.0251 | |
| A2_SRR8283969_Gar42 | 1 | 18.77 | 0.0204 | |
| A2_SRR8283972_Gar41 | 1 | 18.77 | 0.0200 | |
| A2_SRR8283973_Gar38 | 1 | 18.77 | 0.0207 | |
| A2_SRR8283974_Gar39 | 1 | 18.77 | 0.0204 | |
| A2_SRR8283976_Gar37 | 1 | 18.77 | 0.0204 | |
| A2_SRR8283978_Gar57 | 1 | 18.77 | 0.0201 | |
| A2_SRR8283979_Gar58 | 1 | 18.77 | 0.0211 | |
| A2_SRR8283982_Gar61 | 1 | 18.77 | 0.0202 | |
| A2_SRR8283983_Gar44 | 1 | 18.77 | 0.0217 | |
| A2_SRR8283984_Gar45 | 1 | 18.77 | 0.0210 | |
| A2_SRR8283989_Gar30 | 1 | 18.77 | 0.0201 | |
| A2_SRR8283990_Gar31 | 1 | 18.77 | 0.0243 | |
| A2_SRR8283991_Gar03 | 1 | 18.77 | 0.0199 | |
| A2_SRR8283992_Gar04 | 1 | 18.77 | 0.0236 | |
| A2_SRR8283993_Gar01 | 1 | 18.77 | 0.0210 | |
| A2_SRR8283995_Gar65 | 1 | 18.77 | 0.0234 | |
| A2_SRR8284003_Gar51 | 1 | 18.77 | 0.0215 | |
| A2_SRR8284004_Gar50 | 1 | 18.77 | 0.0201 | |
| A2_SRR8284007_Gar47 | 1 | 18.77 | 0.0201 | |
| A2_SRR8284008_Gar46 | 1 | 18.77 | 0.0200 | |
| A2_SRR8284010_Gar63 | 1 | 18.77 | 0.0199 | |
| A2_SRR8284011_Gar64 | 1 | 18.77 | 0.0213 | |
| A1_155 | 1 | 91.33 | 0.0026 | |
| A2_124 | 1 | 92.64 | 0.0030 | |
| A2_SRR8284001_Gar53 | 1 | 92.64 | 0.0027 | |
| A1_155 | 1 | 93.91 | 0.0024 | |
| A2_021 | 1 | 98.44 | 0.0022 | |
| A2_040 | 1 | 98.44 | 0.0026 | |
| A2_043 | 1 | 98.44 | 0.0021 | |
| A2_044 | 1 | 98.44 | 0.0020 | |
| A2_047 | 1 | 98.44 | 0.0021 | |
| A2_057 | 1 | 98.44 | 0.0022 | |
| A2_059 | 1 | 98.44 | 0.0020 | |
| A2_060 | 1 | 98.44 | 0.0021 | |
| A2_061 | 1 | 98.44 | 0.0025 | |
| A2_062 | 1 | 98.44 | 0.0022 | |
| A2_064 | 1 | 98.44 | 0.0024 | |
| A2_065 | 1 | 98.44 | 0.0026 | |
| A2_076 | 1 | 98.44 | 0.0029 | |
| A2_100 | 1 | 98.44 | 0.0026 | |

Number of windows each accession is below/above 95% CI

| **Accession** | **total** | **under dS** | **over dS** | |
|---|---|---|---|---|
| | **13** | **0** | **9** | **Minimum** |
| | **44** | **34** | **18** | **Maximum** |
| | **18** | **5** | **13** | **Mean** |
| | **17** | **5** | **13** | **Median** |
| | **120** | **116** | **120** | **>0% windows** |
| | **120** | **16** | **120** | **>10% windows** |
| A1_019 | 22 | 10 | 12 | |
| A1_028 | 24 | 12 | 12 | |
| A1_030 | 22 | 9 | 13 | |
| A1_051 | 29 | 16 | 13 | |
| A1_054 | 26 | 14 | 12 | |
| A1_073 | 16 | 0 | 16 | |
| A1_074 | 13 | 0 | 13 | |
| A1_077 | 23 | 10 | 13 | |
| A1_079 | 18 | 7 | 11 | |
| A1_084 | 22 | 9 | 13 | |
| A1_088 | 22 | 10 | 12 | |
| A1_097 | 20 | 8 | 12 | |
| A1_108 | 19 | 8 | 11 | |
| A1_125 | 25 | 12 | 13 | |
| A1_132 | 27 | 16 | 11 | |
| A1_133 | 30 | 18 | 12 | |
| A1_155 | 44 | 34 | 10 | |
| A1_Af | 16 | 0 | 16 | |
| A1_Nisa | 16 | 0 | 16 | |
| A1_SRR8283996_Ghe01 | 16 | 1 | 15 | |
| A1_SRR8283998_Ghe02 | 16 | 6 | 10 | |
| A2_001 | 15 | 4 | 11 | |
| A2_004 | 14 | 5 | 9 | |
| A2_008 | 14 | 3 | 11 | |
| A2_016 | 15 | 4 | 11 | |
| A2_019 | 14 | 3 | 11 | |
| A2_021 | 17 | 5 | 12 | |
| A2_026 | 17 | 3 | 14 | |
| A2_029 | 17 | 5 | 12 | |
| A2_030 | 18 | 6 | 12 | |
| A2_031 | 13 | 2 | 11 | |
| A2_034 | 16 | 4 | 12 | |
| A2_040 | 18 | 5 | 13 | |
| A2_041 | 17 | 4 | 13 | |
| A2_043 | 20 | 9 | 11 | |
| A2_044 | 18 | 5 | 13 | |
| A2_045 | 17 | 7 | 10 | |
| A2_046 | 17 | 5 | 12 | |
| A2_047 | 18 | 5 | 13 | |
| A2_056 | 19 | 7 | 12 | |
| A2_057 | 19 | 6 | 13 | |
| A2_059 | 15 | 3 | 12 | |
| A2_060 | 17 | 6 | 11 | |
| A2_061 | 20 | 9 | 11 | |
| A2_062 | 21 | 10 | 11 | |
| A2_064 | 17 | 5 | 12 | |
| A2_065 | 16 | 6 | 10 | |
| A2_066 | 16 | 4 | 12 | |
| A2_067 | 17 | 5 | 12 | |
| A2_069 | 13 | 1 | 12 | |
| A2_073 | 15 | 5 | 10 | |
| A2_074 | 17 | 4 | 13 | |
| A2_076 | 20 | 7 | 13 | |
| A2_079 | 17 | 5 | 12 | |
| A2_080 | 16 | 5 | 11 | |
| A2_084 | 15 | 3 | 12 | |
| A2_085 | 15 | 6 | 9 | |
| A2_087 | 16 | 2 | 14 | |
| A2_091 | 20 | 7 | 13 | |
| A2_096 | 16 | 4 | 12 | |
| A2_100 | 17 | 5 | 12 | |
| A2_101 | 16 | 5 | 11 | |
| A2_113 | 17 | 4 | 13 | |
| A2_117 | 18 | 6 | 12 | |
| A2_118 | 16 | 5 | 11 | |
| A2_119 | 18 | 5 | 13 | |
| A2_123 | 14 | 3 | 11 | |
| A2_124 | 17 | 5 | 12 | |
| A2_131 | 14 | 2 | 12 | |
| A2_141 | 15 | 4 | 11 | |
| A2_142 | 16 | 4 | 12 | |
| A2_147 | 19 | 6 | 13 | |
| A2_154 | 15 | 4 | 11 | |
| A2_156 | 18 | 5 | 13 | |
| A2_158 | 16 | 5 | 11 | |
| A2_164 | 15 | 4 | 11 | |
| A2_255 | 16 | 5 | 11 | |
| A2_lintless | 14 | 2 | 12 | |

Number of accessions outside 95% CI per genome window

| **Chr** | **Position** | **total** | **under dS** | **over dS** | |
|---|---|---|---|---|---|
| | | **1** | **1** | **1** | **Minimum** |
| | | **120** | **96** | **120** | **Maximum** |
| | | **17** | **7** | **38** | **Mean** |
| | | **2** | **2** | **5** | **Median** |
| | | **15** | **2** | **13** | **>50% accessions** |
| | | **11** | **1** | **10** | **>75% accessions** |
| 1 | 2.04 | 1 | 1 | --- | |
| 1 | 4.54 | 1 | --- | 1 | |
| 1 | 5.46 | 37 | --- | 37 | |
| 1 | 18.77 | 25 | --- | 25 | |
| 1 | 91.33 | 1 | 1 | --- | |
| 1 | 92.64 | 2 | 2 | --- | |
| 1 | 93.91 | 1 | 1 | --- | |
| 1 | 98.44 | 26 | 26 | --- | |
| 2 | 0.16 | 1 | 1 | --- | |
| 2 | 17.21 | 1 | --- | 1 | |
| 2 | 66.73 | 2 | 2 | --- | |
| 3 | 0.22 | 4 | 4 | --- | |
| 3 | 0.80 | 4 | --- | 4 | |
| 3 | 2.17 | 34 | --- | 34 | |
| 3 | 3.46 | 7 | --- | 7 | |
| 3 | 67.14 | 1 | 1 | --- | |
| 3 | 109.80 | 4 | 4 | --- | |
| 4 | 68.49 | 2 | --- | 2 | |
| 4 | 72.18 | 1 | --- | 1 | |
| 5 | 4.45 | 1 | 1 | --- | |
| 5 | 4.83 | 1 | 1 | --- | |
| 5 | 5.23 | 1 | 1 | --- | |
| 5 | 5.65 | 6 | 6 | --- | |
| 5 | 6.16 | 1 | 1 | --- | |
| 5 | 6.67 | 1 | 1 | --- | |
| 5 | 7.11 | 1 | 1 | --- | |
| 5 | 7.50 | 1 | 1 | --- | |
| 5 | 8.43 | 6 | 6 | --- | |
| 5 | 8.98 | 1 | 1 | --- | |
| 5 | 10.16 | 8 | 8 | --- | |
| 5 | 12.34 | 2 | 2 | --- | |
| 5 | 12.87 | 6 | 6 | --- | |
| 5 | 13.47 | 2 | 2 | --- | |
| 5 | 14.01 | 1 | 1 | --- | |
| 5 | 14.97 | 1 | --- | 1 | |
| 5 | 51.55 | 2 | --- | 2 | |
| 5 | 68.18 | 1 | --- | 1 | |
| 5 | 70.61 | 114 | --- | 114 | |
| 5 | 73.47 | 112 | --- | 112 | |
| 5 | 75.43 | 1 | --- | 1 | |
| 6 | 0.98 | 4 | --- | 4 | |
| 6 | 28.83 | 1 | 1 | --- | |
| 7 | 1.60 | 5 | 5 | --- | |
| 7 | 3.31 | 18 | 18 | --- | |
| 7 | 3.92 | 47 | 47 | --- | |
| 7 | 4.68 | 23 | 23 | --- | |
| 7 | 5.36 | 48 | 48 | --- | |
| 7 | 11.85 | 2 | --- | 2 | |
| 7 | 14.27 | 2 | --- | 2 | |
| 7 | 24.59 | 1 | 1 | --- | |
| 7 | 26.61 | 1 | 1 | --- | |
| 7 | 29.39 | 5 | 5 | --- | |
| 7 | 41.78 | 1 | 1 | --- | |
| 7 | 48.64 | 5 | --- | 5 | |
| 7 | 69.73 | 115 | --- | 115 | |
| 7 | 72.89 | 88 | --- | 88 | |
| 7 | 77.30 | 5 | 5 | --- | |
| 7 | 78.32 | 1 | 1 | --- | |
| 7 | 79.31 | 1 | 1 | --- | |
| 7 | 80.08 | 1 | 1 | --- | |
| 8 | 6.57 | 116 | --- | 116 | |
| 8 | 76.89 | 5 | 5 | --- | |
| 8 | 80.33 | 4 | 4 | --- | |
| 8 | 83.55 | 3 | 3 | --- | |
| 8 | 88.96 | 3 | 3 | --- | |
| 8 | 92.03 | 1 | 1 | --- | |
| 8 | 96.46 | 1 | 1 | --- | |
| 8 | 99.32 | 3 | 3 | --- | |
| 8 | 100.18 | 1 | --- | 1 | |
| 8 | 102.41 | 1 | 1 | --- | |
| 8 | 103.08 | 1 | 1 | --- | |
| 9 | 0.38 | 1 | 1 | --- | |
| 9 | 1.41 | 1 | 1 | --- | |
| 9 | 3.89 | 101 | --- | 101 | |
| 9 | 16.72 | 36 | --- | 36 | |
| 9 | 47.05 | 15 | 15 | --- | |
| 9 | 50.13 | 16 | 16 | --- | |
| 9 | 55.41 | 4 | 4 | --- | |

Accessions with dS outside of 95% CI (0.0031 - 0.0198)

| Accession | Chr | Position | dS |
|---|---|---|---|
| A2_113 | 1 | 98.44 | 0.0021 |
| A2_117 | 1 | 98.44 | 0.0022 |
| A2_141 | 1 | 98.44 | 0.0022 |
| A2_142 | 1 | 98.44 | 0.0022 |
| A2_154 | 1 | 98.44 | 0.0021 |
| A2_156 | 1 | 98.44 | 0.0024 |
| A2_158 | 1 | 98.44 | 0.0023 |
| A2_164 | 1 | 98.44 | 0.0022 |
| A2_SRR8283978_Gar57 | 1 | 98.44 | 0.0030 |
| A2_SRR8283990_Gar31 | 1 | 98.44 | 0.0028 |
| A2_SRR8283991_Gar03 | 1 | 98.44 | 0.0025 |
| A2_SRR8283993_Gar01 | 1 | 98.44 | 0.0028 |
| A1_155 | 2 | 0.16 | 0.0017 |
| A1_SRR8283996_Ghe01 | 2 | 17.21 | 0.0199 |
| A1_028 | 2 | 66.73 | 0.0030 |
| A1_088 | 2 | 66.73 | 0.0029 |
| A1_088 | 3 | 0.22 | 0.0030 |
| A1_125 | 3 | 0.22 | 0.0027 |
| A2_SRR8284006_Gar48 | 3 | 0.22 | 0.0028 |
| A2_SRR8284011_Gar64 | 3 | 0.22 | 0.0030 |
| A1_073 | 3 | 0.80 | 0.0210 |
| A1_074 | 3 | 0.80 | 0.0205 |
| A1_Af | 3 | 0.80 | 0.0216 |
| A1_Nisa | 3 | 0.80 | 0.0213 |
| A1_054 | 3 | 2.17 | 0.0199 |
| A1_073 | 3 | 2.17 | 0.0220 |
| A1_084 | 3 | 2.17 | 0.0202 |
| A1_Af | 3 | 2.17 | 0.0205 |
| A1_Nisa | 3 | 2.17 | 0.0209 |
| A1_SRR8283996_Ghe01 | 3 | 2.17 | 0.0204 |
| A2_021 | 3 | 2.17 | 0.0200 |
| A2_045 | 3 | 2.17 | 0.0206 |
| A2_SRR8283969_Gar42 | 3 | 2.17 | 0.0212 |
| A2_SRR8283971_Gar40 | 3 | 2.17 | 0.0200 |
| A2_SRR8283972_Gar41 | 3 | 2.17 | 0.0215 |
| A2_SRR8283973_Gar38 | 3 | 2.17 | 0.0199 |
| A2_SRR8283974_Gar39 | 3 | 2.17 | 0.0203 |
| A2_SRR8283976_Gar37 | 3 | 2.17 | 0.0205 |
| A2_SRR8283978_Gar57 | 3 | 2.17 | 0.0235 |
| A2_SRR8283980_Gar59 | 3 | 2.17 | 0.0225 |
| A2_SRR8283981_Gar60 | 3 | 2.17 | 0.0209 |
| A2_SRR8283984_Gar45 | 3 | 2.17 | 0.0206 |
| A2_SRR8283987_Gar32 | 3 | 2.17 | 0.0202 |
| A2_SRR8283988_Gar33 | 3 | 2.17 | 0.0202 |
| A2_SRR8283989_Gar30 | 3 | 2.17 | 0.0202 |
| A2_SRR8283990_Gar31 | 3 | 2.17 | 0.0219 |
| A2_SRR8283993_Gar01 | 3 | 2.17 | 0.0205 |
| A2_SRR8283994_Gar02 | 3 | 2.17 | 0.0227 |
| A2_SRR8283995_Gar65 | 3 | 2.17 | 0.0220 |
| A2_SRR8283999_Gar55 | 3 | 2.17 | 0.0206 |
| A2_SRR8284001_Gar53 | 3 | 2.17 | 0.0200 |
| A2_SRR8284002_Gar52 | 3 | 2.17 | 0.0219 |
| A2_SRR8284003_Gar51 | 3 | 2.17 | 0.0209 |
| A2_SRR8284005_Gar49 | 3 | 2.17 | 0.0199 |
| A2_SRR8284006_Gar48 | 3 | 2.17 | 0.0203 |
| A2_SRR8284007_Gar47 | 3 | 2.17 | 0.0210 |
| A2_SRR8284010_Gar63 | 3 | 2.17 | 0.0222 |
| A2_SRR8284011_Gar64 | 3 | 2.17 | 0.0200 |
| A2_SRR8283969_Gar42 | 3 | 3.46 | 0.0207 |
| A2_SRR8283978_Gar57 | 3 | 3.46 | 0.0206 |
| A2_SRR8283979_Gar58 | 3 | 3.46 | 0.0207 |
| A2_SRR8283989_Gar30 | 3 | 3.46 | 0.0199 |
| A2_SRR8283993_Gar01 | 3 | 3.46 | 0.0206 |
| A2_SRR8283994_Gar02 | 3 | 3.46 | 0.0200 |
| A2_SRR8284011_Gar64 | 3 | 3.46 | 0.0200 |
| A2_SRR8283993_Gar01 | 3 | 67.14 | 0.0030 |
| A1_019 | 3 | 109.80 | 0.0030 |
| A1_028 | 3 | 109.80 | 0.0023 |
| A1_077 | 3 | 109.80 | 0.0028 |
| A1_088 | 3 | 109.80 | 0.0029 |
| A2_SRR8283983_Gar44 | 4 | 68.49 | 0.0202 |
| A2_SRR8283999_Gar55 | 4 | 68.49 | 0.0210 |
| A2_SRR8284008_Gar46 | 4 | 72.18 | 0.0230 |
| A1_155 | 5 | 4.45 | 0.0030 |
| A1_155 | 5 | 4.83 | 0.0013 |
| A1_155 | 5 | 5.23 | 0.0017 |
| A1_028 | 5 | 5.65 | 0.0027 |
| A1_030 | 5 | 5.65 | 0.0028 |
| A1_077 | 5 | 5.65 | 0.0028 |
| A1_108 | 5 | 5.65 | 0.0030 |
| A1_155 | 5 | 5.65 | 0.0016 |
| A2_084 | 5 | 5.65 | 0.0026 |
| A1_155 | 5 | 6.16 | 0.0030 |
| A1_155 | 5 | 6.67 | 0.0024 |
| A1_155 | 5 | 7.11 | 0.0006 |
| A1_155 | 5 | 7.50 | 0.0020 |

Number of windows each accession is below/above 95% CI

| Accession | total | under dS | over dS |
|---|---|---|---|
| A2_NC503 | 14 | 3 | 11 |
| A2_SRR8283969_Gar42 | 19 | 4 | 15 |
| A2_SRR8283970_Gar43 | 15 | 3 | 12 |
| A2_SRR8283971_Gar40 | 15 | 1 | 14 |
| A2_SRR8283972_Gar41 | 19 | 2 | 17 |
| A2_SRR8283973_Gar38 | 18 | 3 | 15 |
| A2_SRR8283974_Gar39 | 19 | 4 | 15 |
| A2_SRR8283975_Gar36 | 16 | 2 | 14 |
| A2_SRR8283976_Gar37 | 24 | 7 | 17 |
| A2_SRR8283977_Gar56 | 19 | 4 | 15 |
| A2_SRR8283978_Gar57 | 22 | 5 | 17 |
| A2_SRR8283979_Gar58 | 19 | 5 | 14 |
| A2_SRR8283980_Gar59 | 19 | 4 | 15 |
| A2_SRR8283981_Gar60 | 13 | 1 | 12 |
| A2_SRR8283982_Gar61 | 17 | 4 | 13 |
| A2_SRR8283983_Gar44 | 17 | 3 | 14 |
| A2_SRR8283984_Gar45 | 16 | 2 | 14 |
| A2_SRR8283985_Gar34 | 20 | 9 | 11 |
| A2_SRR8283986_Gar35 | 19 | 6 | 13 |
| A2_SRR8283987_Gar32 | 15 | 2 | 13 |
| A2_SRR8283988_Gar33 | 17 | 4 | 13 |
| A2_SRR8283989_Gar30 | 19 | 1 | 18 |
| A2_SRR8283990_Gar31 | 19 | 4 | 15 |
| A2_SRR8283991_Gar03 | 16 | 4 | 12 |
| A2_SRR8283992_Gar04 | 19 | 6 | 13 |
| A2_SRR8283993_Gar01 | 20 | 6 | 14 |
| A2_SRR8283994_Gar02 | 20 | 6 | 14 |
| A2_SRR8283995_Gar65 | 18 | 3 | 15 |
| A2_SRR8283997_Gar66 | 16 | 4 | 12 |
| A2_SRR8283999_Gar55 | 22 | 4 | 18 |
| A2_SRR8284000_Gar54 | 16 | 1 | 15 |
| A2_SRR8284001_Gar53 | 17 | 3 | 14 |
| A2_SRR8284002_Gar52 | 17 | 3 | 14 |
| A2_SRR8284003_Gar51 | 17 | 3 | 14 |
| A2_SRR8284004_Gar50 | 18 | 5 | 13 |
| A2_SRR8284005_Gar49 | 16 | 2 | 14 |
| A2_SRR8284006_Gar48 | 18 | 2 | 16 |
| A2_SRR8284007_Gar47 | 22 | 5 | 17 |
| A2_SRR8284008_Gar46 | 18 | 2 | 16 |
| A2_SRR8284009_Gar62 | 17 | 1 | 16 |
| A2_SRR8284010_Gar63 | 22 | 4 | 18 |
| A2_SRR8284011_Gar64 | 24 | 7 | 17 |

Number of accessions outside 95% CI per genome window

| Chr | Position | total | under dS | over dS |
|---|---|---|---|---|
| 9 | 58.52 | 9 | 9 | --- |
| 9 | 59.57 | 1 | 1 | --- |
| 9 | 66.69 | 1 | 1 | --- |
| 9 | 67.69 | 4 | 4 | --- |
| 9 | 70.53 | 4 | 4 | --- |
| 10 | 1.72 | 96 | 96 | --- |
| 10 | 2.31 | 11 | 11 | --- |
| 10 | 2.82 | 5 | 5 | --- |
| 10 | 3.41 | 2 | 2 | --- |
| 10 | 7.72 | 1 | 1 | --- |
| 10 | 8.99 | 3 | 3 | --- |
| 10 | 18.10 | 64 | --- | 64 |
| 10 | 33.12 | 3 | 3 | --- |
| 10 | 36.28 | 1 | 1 | --- |
| 10 | 79.32 | 3 | 3 | --- |
| 10 | 83.98 | 18 | --- | 18 |
| 10 | 89.53 | 72 | --- | 72 |
| 10 | 90.98 | 2 | --- | 2 |
| 10 | 91.95 | 95 | --- | 95 |
| 10 | 93.58 | 20 | --- | 20 |
| 10 | 95.02 | 2 | 2 | --- |
| 11 | 1.73 | 1 | 1 | --- |
| 11 | 2.15 | 1 | 1 | --- |
| 11 | 2.62 | 1 | 1 | --- |
| 11 | 4.10 | 1 | 1 | --- |
| 11 | 7.60 | 1 | 1 | --- |
| 11 | 18.49 | 28 | 28 | --- |
| 11 | 45.55 | 1 | 1 | --- |
| 11 | 64.32 | 120 | --- | 120 |
| 11 | 85.43 | 1 | --- | 1 |
| 11 | 89.98 | 1 | --- | 1 |
| 11 | 90.59 | 103 | --- | 103 |
| 11 | 91.80 | 113 | --- | 113 |
| 11 | 93.14 | 1 | --- | 1 |
| 11 | 95.42 | 115 | --- | 115 |
| 11 | 98.35 | 1 | --- | 1 |
| 11 | 99.47 | 61 | 61 | --- |
| 12 | 0.26 | 3 | 3 | --- |
| 12 | 79.63 | 37 | 37 | --- |
| 12 | 86.32 | 2 | 2 | --- |
| 12 | 86.96 | 31 | 31 | --- |
| 12 | 88.44 | 1 | 1 | --- |
| 12 | 91.02 | 7 | 7 | --- |
| 13 | 85.68 | 1 | --- | 1 |
| 13 | 93.06 | 5 | --- | 5 |
| 13 | 95.64 | 1 | 1 | --- |
| 13 | 95.82 | 5 | 5 | --- |

Accessions with dS outside of 95% CI (0.0031 - 0.0198)

| Accession | Chr | Position | dS |
|---|---|---|---|
| A1_028 | 5 | 8.43 | 0.0028 |
| A1_051 | 5 | 8.43 | 0.0021 |
| A1_054 | 5 | 8.43 | 0.0016 |
| A1_132 | 5 | 8.43 | 0.0022 |
| A1_133 | 5 | 8.43 | 0.0021 |
| A1_155 | 5 | 8.43 | 0.0009 |
| A1_155 | 5 | 8.98 | 0.0021 |
| A2_021 | 5 | 10.16 | 0.0030 |
| A2_043 | 5 | 10.16 | 0.0027 |
| A2_046 | 5 | 10.16 | 0.0029 |
| A2_057 | 5 | 10.16 | 0.0030 |
| A2_066 | 5 | 10.16 | 0.0030 |
| A2_073 | 5 | 10.16 | 0.0030 |
| A2_101 | 5 | 10.16 | 0.0030 |
| A2_SRR8283984_Gar45 | 5 | 10.16 | 0.0028 |
| A1_132 | 5 | 12.34 | 0.0029 |
| A1_155 | 5 | 12.34 | 0.0024 |
| A1_019 | 5 | 12.87 | 0.0030 |
| A1_028 | 5 | 12.87 | 0.0013 |
| A1_030 | 5 | 12.87 | 0.0030 |
| A1_054 | 5 | 12.87 | 0.0027 |
| A1_155 | 5 | 12.87 | 0.0006 |
| A2_SRR8283993_Gar01 | 5 | 12.87 | 0.0030 |
| A1_054 | 5 | 13.47 | 0.0011 |
| A1_155 | 5 | 13.47 | 0.0010 |
| A1_155 | 5 | 14.01 | 0.0026 |
| A2_SRR8283999_Gar55 | 5 | 14.97 | 0.0199 |
| A1_SRR8283996_Ghe01 | 5 | 51.55 | 0.0208 |
| A2_SRR8283995_Gar65 | 5 | 51.55 | 0.0205 |
| A2_031 | 5 | 68.18 | 0.0201 |
| A1_019 | 5 | 70.61 | 0.0291 |
| A1_028 | 5 | 70.61 | 0.0278 |
| A1_030 | 5 | 70.61 | 0.0289 |
| A1_051 | 5 | 70.61 | 0.0260 |
| A1_054 | 5 | 70.61 | 0.0252 |
| A1_073 | 5 | 70.61 | 0.0250 |
| A1_074 | 5 | 70.61 | 0.0253 |
| A1_077 | 5 | 70.61 | 0.0274 |
| A1_079 | 5 | 70.61 | 0.0296 |
| A1_084 | 5 | 70.61 | 0.0249 |
| A1_088 | 5 | 70.61 | 0.0260 |
| A1_097 | 5 | 70.61 | 0.0290 |
| A1_108 | 5 | 70.61 | 0.0281 |
| A1_125 | 5 | 70.61 | 0.0219 |
| A1_132 | 5 | 70.61 | 0.0252 |
| A1_133 | 5 | 70.61 | 0.0248 |
| A1_Af | 5 | 70.61 | 0.0290 |
| A1_Nisa | 5 | 70.61 | 0.0285 |
| A1_SRR8283996_Ghe01 | 5 | 70.61 | 0.0210 |
| A2_001 | 5 | 70.61 | 0.0287 |
| A2_004 | 5 | 70.61 | 0.0279 |
| A2_008 | 5 | 70.61 | 0.0261 |
| A2_016 | 5 | 70.61 | 0.0257 |
| A2_019 | 5 | 70.61 | 0.0270 |
| A2_021 | 5 | 70.61 | 0.0252 |
| A2_026 | 5 | 70.61 | 0.0263 |
| A2_029 | 5 | 70.61 | 0.0258 |
| A2_030 | 5 | 70.61 | 0.0264 |
| A2_031 | 5 | 70.61 | 0.0279 |
| A2_034 | 5 | 70.61 | 0.0259 |
| A2_040 | 5 | 70.61 | 0.0259 |
| A2_041 | 5 | 70.61 | 0.0269 |
| A2_043 | 5 | 70.61 | 0.0249 |
| A2_044 | 5 | 70.61 | 0.0261 |
| A2_045 | 5 | 70.61 | 0.0222 |
| A2_046 | 5 | 70.61 | 0.0268 |
| A2_047 | 5 | 70.61 | 0.0265 |
| A2_056 | 5 | 70.61 | 0.0261 |
| A2_057 | 5 | 70.61 | 0.0261 |
| A2_059 | 5 | 70.61 | 0.0261 |
| A2_060 | 5 | 70.61 | 0.0257 |
| A2_061 | 5 | 70.61 | 0.0250 |
| A2_062 | 5 | 70.61 | 0.0252 |
| A2_064 | 5 | 70.61 | 0.0263 |
| A2_065 | 5 | 70.61 | 0.0288 |
| A2_066 | 5 | 70.61 | 0.0263 |
| A2_067 | 5 | 70.61 | 0.0281 |
| A2_069 | 5 | 70.61 | 0.0279 |
| A2_073 | 5 | 70.61 | 0.0246 |
| A2_074 | 5 | 70.61 | 0.0262 |
| A2_076 | 5 | 70.61 | 0.0256 |
| A2_079 | 5 | 70.61 | 0.0284 |
| A2_080 | 5 | 70.61 | 0.0257 |
| A2_084 | 5 | 70.61 | 0.0262 |
| A2_087 | 5 | 70.61 | 0.0281 |
| A2_091 | 5 | 70.61 | 0.0280 |

| Accession | Chr | Position | dS |
|---|---|---|---|
| A2_096 | 5 | 70.61 | 0.0263 |
| A2_100 | 5 | 70.61 | 0.0254 |
| A2_101 | 5 | 70.61 | 0.0255 |
| A2_113 | 5 | 70.61 | 0.0266 |
| A2_117 | 5 | 70.61 | 0.0259 |
| A2_118 | 5 | 70.61 | 0.0287 |
| A2_119 | 5 | 70.61 | 0.0274 |
| A2_123 | 5 | 70.61 | 0.0257 |
| A2_124 | 5 | 70.61 | 0.0283 |
| A2_131 | 5 | 70.61 | 0.0261 |
| A2_141 | 5 | 70.61 | 0.0259 |
| A2_142 | 5 | 70.61 | 0.0262 |
| A2_147 | 5 | 70.61 | 0.0274 |
| A2_154 | 5 | 70.61 | 0.0269 |
| A2_156 | 5 | 70.61 | 0.0262 |
| A2_158 | 5 | 70.61 | 0.0263 |
| A2_164 | 5 | 70.61 | 0.0258 |
| A2_255 | 5 | 70.61 | 0.0259 |
| A2_lintless | 5 | 70.61 | 0.0279 |
| A2_NC503 | 5 | 70.61 | 0.0274 |
| A2_SRR8283969_Gar42 | 5 | 70.61 | 0.0233 |
| A2_SRR8283971_Gar40 | 5 | 70.61 | 0.0271 |
| A2_SRR8283972_Gar41 | 5 | 70.61 | 0.0276 |
| A2_SRR8283973_Gar38 | 5 | 70.61 | 0.0242 |
| A2_SRR8283974_Gar39 | 5 | 70.61 | 0.0262 |
| A2_SRR8283975_Gar36 | 5 | 70.61 | 0.0262 |
| A2_SRR8283976_Gar37 | 5 | 70.61 | 0.0234 |
| A2_SRR8283977_Gar56 | 5 | 70.61 | 0.0219 |
| A2_SRR8283978_Gar57 | 5 | 70.61 | 0.0244 |
| A2_SRR8283979_Gar58 | 5 | 70.61 | 0.0237 |
| A2_SRR8283980_Gar59 | 5 | 70.61 | 0.0255 |
| A2_SRR8283981_Gar60 | 5 | 70.61 | 0.0211 |
| A2_SRR8283982_Gar61 | 5 | 70.61 | 0.0280 |
| A2_SRR8283983_Gar44 | 5 | 70.61 | 0.0224 |
| A2_SRR8283984_Gar45 | 5 | 70.61 | 0.0238 |
| A2_SRR8283985_Gar34 | 5 | 70.61 | 0.0208 |
| A2_SRR8283986_Gar35 | 5 | 70.61 | 0.0288 |
| A2_SRR8283987_Gar32 | 5 | 70.61 | 0.0253 |
| A2_SRR8283988_Gar33 | 5 | 70.61 | 0.0265 |
| A2_SRR8283989_Gar30 | 5 | 70.61 | 0.0235 |
| A2_SRR8283990_Gar31 | 5 | 70.61 | 0.0286 |
| A2_SRR8283991_Gar03 | 5 | 70.61 | 0.0257 |
| A2_SRR8283992_Gar04 | 5 | 70.61 | 0.0223 |
| A2_SRR8283994_Gar02 | 5 | 70.61 | 0.0252 |
| A2_SRR8283995_Gar65 | 5 | 70.61 | 0.0213 |
| A2_SRR8283999_Gar55 | 5 | 70.61 | 0.0261 |
| A2_SRR8284000_Gar54 | 5 | 70.61 | 0.0211 |
| A2_SRR8284001_Gar53 | 5 | 70.61 | 0.0261 |
| A2_SRR8284002_Gar52 | 5 | 70.61 | 0.0221 |
| A2_SRR8284003_Gar51 | 5 | 70.61 | 0.0252 |
| A2_SRR8284004_Gar50 | 5 | 70.61 | 0.0213 |
| A2_SRR8284005_Gar49 | 5 | 70.61 | 0.0210 |
| A2_SRR8284006_Gar48 | 5 | 70.61 | 0.0282 |
| A2_SRR8284007_Gar47 | 5 | 70.61 | 0.0243 |
| A2_SRR8284008_Gar46 | 5 | 70.61 | 0.0241 |
| A2_SRR8284009_Gar62 | 5 | 70.61 | 0.0246 |
| A2_SRR8284010_Gar63 | 5 | 70.61 | 0.0223 |
| A2_SRR8284011_Gar64 | 5 | 70.61 | 0.0237 |
| A1_019 | 5 | 73.47 | 0.0226 |
| A1_028 | 5 | 73.47 | 0.0228 |
| A1_030 | 5 | 73.47 | 0.0223 |
| A1_051 | 5 | 73.47 | 0.0220 |
| A1_054 | 5 | 73.47 | 0.0237 |
| A1_073 | 5 | 73.47 | 0.0214 |
| A1_074 | 5 | 73.47 | 0.0215 |
| A1_077 | 5 | 73.47 | 0.0225 |
| A1_079 | 5 | 73.47 | 0.0217 |
| A1_084 | 5 | 73.47 | 0.0225 |
| A1_088 | 5 | 73.47 | 0.0219 |
| A1_097 | 5 | 73.47 | 0.0226 |
| A1_108 | 5 | 73.47 | 0.0218 |
| A1_125 | 5 | 73.47 | 0.0213 |
| A1_132 | 5 | 73.47 | 0.0218 |
| A1_133 | 5 | 73.47 | 0.0229 |
| A1_155 | 5 | 73.47 | 0.0214 |
| A1_Af | 5 | 73.47 | 0.0222 |
| A1_Nisa | 5 | 73.47 | 0.0222 |
| A1_SRR8283996_Ghe01 | 5 | 73.47 | 0.0223 |
| A2_001 | 5 | 73.47 | 0.0218 |
| A2_004 | 5 | 73.47 | 0.0227 |
| A2_008 | 5 | 73.47 | 0.0235 |
| A2_016 | 5 | 73.47 | 0.0217 |
| A2_019 | 5 | 73.47 | 0.0224 |
| A2_021 | 5 | 73.47 | 0.0230 |
| A2_026 | 5 | 73.47 | 0.0247 |
| A2_029 | 5 | 73.47 | 0.0228 |

| Accession | Chr | Position | dS |
|---|---|---|---|
| A2_030 | 5 | 73.47 | 0.0227 |
| A2_031 | 5 | 73.47 | 0.0239 |
| A2_034 | 5 | 73.47 | 0.0220 |
| A2_040 | 5 | 73.47 | 0.0233 |
| A2_041 | 5 | 73.47 | 0.0237 |
| A2_043 | 5 | 73.47 | 0.0216 |
| A2_044 | 5 | 73.47 | 0.0227 |
| A2_046 | 5 | 73.47 | 0.0228 |
| A2_047 | 5 | 73.47 | 0.0238 |
| A2_056 | 5 | 73.47 | 0.0227 |
| A2_057 | 5 | 73.47 | 0.0240 |
| A2_059 | 5 | 73.47 | 0.0233 |
| A2_060 | 5 | 73.47 | 0.0231 |
| A2_061 | 5 | 73.47 | 0.0214 |
| A2_062 | 5 | 73.47 | 0.0216 |
| A2_064 | 5 | 73.47 | 0.0224 |
| A2_065 | 5 | 73.47 | 0.0231 |
| A2_066 | 5 | 73.47 | 0.0224 |
| A2_067 | 5 | 73.47 | 0.0245 |
| A2_069 | 5 | 73.47 | 0.0232 |
| A2_073 | 5 | 73.47 | 0.0210 |
| A2_074 | 5 | 73.47 | 0.0231 |
| A2_076 | 5 | 73.47 | 0.0213 |
| A2_079 | 5 | 73.47 | 0.0236 |
| A2_080 | 5 | 73.47 | 0.0217 |
| A2_084 | 5 | 73.47 | 0.0238 |
| A2_085 | 5 | 73.47 | 0.0228 |
| A2_087 | 5 | 73.47 | 0.0205 |
| A2_091 | 5 | 73.47 | 0.0230 |
| A2_096 | 5 | 73.47 | 0.0200 |
| A2_100 | 5 | 73.47 | 0.0210 |
| A2_101 | 5 | 73.47 | 0.0205 |
| A2_113 | 5 | 73.47 | 0.0212 |
| A2_117 | 5 | 73.47 | 0.0221 |
| A2_118 | 5 | 73.47 | 0.0216 |
| A2_119 | 5 | 73.47 | 0.0213 |
| A2_123 | 5 | 73.47 | 0.0233 |
| A2_124 | 5 | 73.47 | 0.0228 |
| A2_131 | 5 | 73.47 | 0.0216 |
| A2_141 | 5 | 73.47 | 0.0211 |
| A2_142 | 5 | 73.47 | 0.0234 |
| A2_147 | 5 | 73.47 | 0.0199 |
| A2_154 | 5 | 73.47 | 0.0209 |
| A2_156 | 5 | 73.47 | 0.0212 |
| A2_158 | 5 | 73.47 | 0.0199 |
| A2_255 | 5 | 73.47 | 0.0217 |
| A2_lintless | 5 | 73.47 | 0.0246 |
| A2_NC503 | 5 | 73.47 | 0.0219 |
| A2_SRR8283969_Gar4 | 5 | 73.47 | 0.0221 |
| A2_SRR8283970_Gar4 | 5 | 73.47 | 0.0207 |
| A2_SRR8283971_Gar4 | 5 | 73.47 | 0.0239 |
| A2_SRR8283972_Gar4 | 5 | 73.47 | 0.0218 |
| A2_SRR8283973_Gar3 | 5 | 73.47 | 0.0254 |
| A2_SRR8283974_Gar3 | 5 | 73.47 | 0.0230 |
| A2_SRR8283975_Gar3 | 5 | 73.47 | 0.0213 |
| A2_SRR8283976_Gar3 | 5 | 73.47 | 0.0223 |
| A2_SRR8283977_Gar5 | 5 | 73.47 | 0.0219 |
| A2_SRR8283978_Gar5 | 5 | 73.47 | 0.0228 |
| A2_SRR8283979_Gar5 | 5 | 73.47 | 0.0217 |
| A2_SRR8283980_Gar5 | 5 | 73.47 | 0.0221 |
| A2_SRR8283981_Gar6 | 5 | 73.47 | 0.0236 |
| A2_SRR8283982_Gar6 | 5 | 73.47 | 0.0202 |
| A2_SRR8283983_Gar4 | 5 | 73.47 | 0.0229 |
| A2_SRR8283984_Gar4 | 5 | 73.47 | 0.0207 |
| A2_SRR8283985_Gar3 | 5 | 73.47 | 0.0206 |
| A2_SRR8283987_Gar3 | 5 | 73.47 | 0.0199 |
| A2_SRR8283988_Gar3 | 5 | 73.47 | 0.0221 |
| A2_SRR8283989_Gar3 | 5 | 73.47 | 0.0214 |
| A2_SRR8283991_Gar0 | 5 | 73.47 | 0.0207 |
| A2_SRR8283992_Gar0 | 5 | 73.47 | 0.0202 |
| A2_SRR8283994_Gar0 | 5 | 73.47 | 0.0235 |
| A2_SRR8283997_Gar6 | 5 | 73.47 | 0.0228 |
| A2_SRR8283999_Gar5 | 5 | 73.47 | 0.0230 |
| A2_SRR8284000_Gar5 | 5 | 73.47 | 0.0220 |
| A2_SRR8284001_Gar5 | 5 | 73.47 | 0.0204 |
| A2_SRR8284002_Gar5 | 5 | 73.47 | 0.0264 |
| A2_SRR8284003_Gar5 | 5 | 73.47 | 0.0199 |
| A2_SRR8284004_Gar5 | 5 | 73.47 | 0.0234 |
| A2_SRR8284005_Gar4 | 5 | 73.47 | 0.0230 |
| A2_SRR8284006_Gar4 | 5 | 73.47 | 0.0275 |
| A2_SRR8284007_Gar4 | 5 | 73.47 | 0.0207 |
| A2_SRR8284008_Gar4 | 5 | 73.47 | 0.0264 |
| A2_SRR8284009_Gar6 | 5 | 73.47 | 0.0225 |
| A2_SRR8284011_Gar6 | 5 | 73.47 | 0.0203 |
| A2_SRR8284007_Gar4 | 5 | 75.43 | 0.0202 |
| A1_SRR8283998_Ghe0 | 6 | 0.98 | 0.0212 |

Accessions with dS outside of 95% CI (0.0031 - 0.0198)

| Accession | Chr | Position | dS | Accession | Chr | Position | dS | Accession | Chr | Position | dS |
|---|---|---|---|---|---|---|---|---|---|---|---|
| A2_045 | 6 | 0.98 | 0.0212 | A2_091 | 7 | 4.68 | 0.0025 | A1_088 | 7 | 69.73 | 0.0249 |
| A2_SRR8283979_Gar58 | 6 | 0.98 | 0.0199 | A2_117 | 7 | 4.68 | 0.0029 | A1_097 | 7 | 69.73 | 0.0257 |
| A2_SRR8284002_Gar52 | 6 | 0.98 | 0.0238 | A2_124 | 7 | 4.68 | 0.0026 | A1_108 | 7 | 69.73 | 0.0258 |
| A2_062 | 6 | 28.83 | 0.0027 | A2_147 | 7 | 4.68 | 0.0028 | A1_125 | 7 | 69.73 | 0.0201 |
| A1_051 | 7 | 1.60 | 0.0026 | A2_255 | 7 | 4.68 | 0.0028 | A1_132 | 7 | 69.73 | 0.0249 |
| A1_054 | 7 | 1.60 | 0.0026 | A2_lintless | 7 | 4.68 | 0.0030 | A1_133 | 7 | 69.73 | 0.0259 |
| A1_132 | 7 | 1.60 | 0.0025 | A2_SRR8283977_Gar56 | 7 | 4.68 | 0.0026 | A1_Af | 7 | 69.73 | 0.0261 |
| A1_133 | 7 | 1.60 | 0.0026 | A2_SRR8283988_Gar33 | 7 | 4.68 | 0.0029 | A1_Nisa | 7 | 69.73 | 0.0258 |
| A2_SRR8283993_Gar01 | 7 | 1.60 | 0.0029 | A2_SRR8283990_Gar31 | 7 | 4.68 | 0.0028 | A1_SRR8283998_Ghe0 | 7 | 69.73 | 0.0199 |
| A1_125 | 7 | 3.31 | 0.0028 | A2_SRR8283994_Gar02 | 7 | 4.68 | 0.0022 | A2_001 | 7 | 69.73 | 0.0258 |
| A1_SRR8283996_Ghe01 | 7 | 3.31 | 0.0028 | A2_SRR8284007_Gar47 | 7 | 4.68 | 0.0028 | A2_004 | 7 | 69.73 | 0.0233 |
| A2_016 | 7 | 3.31 | 0.0028 | A1_019 | 7 | 5.36 | 0.0028 | A2_008 | 7 | 69.73 | 0.0258 |
| A2_019 | 7 | 3.31 | 0.0027 | A1_028 | 7 | 5.36 | 0.0029 | A2_016 | 7 | 69.73 | 0.0252 |
| A2_031 | 7 | 3.31 | 0.0030 | A1_030 | 7 | 5.36 | 0.0028 | A2_019 | 7 | 69.73 | 0.0219 |
| A2_040 | 7 | 3.31 | 0.0025 | A1_051 | 7 | 5.36 | 0.0029 | A2_021 | 7 | 69.73 | 0.0240 |
| A2_044 | 7 | 3.31 | 0.0029 | A1_054 | 7 | 5.36 | 0.0027 | A2_026 | 7 | 69.73 | 0.0263 |
| A2_059 | 7 | 3.31 | 0.0030 | A1_077 | 7 | 5.36 | 0.0030 | A2_029 | 7 | 69.73 | 0.0249 |
| A2_060 | 7 | 3.31 | 0.0026 | A1_079 | 7 | 5.36 | 0.0027 | A2_030 | 7 | 69.73 | 0.0246 |
| A2_061 | 7 | 3.31 | 0.0022 | A1_084 | 7 | 5.36 | 0.0028 | A2_031 | 7 | 69.73 | 0.0249 |
| A2_062 | 7 | 3.31 | 0.0026 | A1_088 | 7 | 5.36 | 0.0024 | A2_034 | 7 | 69.73 | 0.0238 |
| A2_064 | 7 | 3.31 | 0.0028 | A1_097 | 7 | 5.36 | 0.0028 | A2_040 | 7 | 69.73 | 0.0252 |
| A2_100 | 7 | 3.31 | 0.0030 | A1_108 | 7 | 5.36 | 0.0027 | A2_041 | 7 | 69.73 | 0.0256 |
| A2_158 | 7 | 3.31 | 0.0028 | A1_132 | 7 | 5.36 | 0.0028 | A2_043 | 7 | 69.73 | 0.0233 |
| A2_164 | 7 | 3.31 | 0.0028 | A1_133 | 7 | 5.36 | 0.0027 | A2_044 | 7 | 69.73 | 0.0251 |
| A2_SRR8283975_Gar36 | 7 | 3.31 | 0.0027 | A2_001 | 7 | 5.36 | 0.0030 | A2_046 | 7 | 69.73 | 0.0256 |
| A2_SRR8283976_Gar37 | 7 | 3.31 | 0.0030 | A2_004 | 7 | 5.36 | 0.0027 | A2_047 | 7 | 69.73 | 0.0257 |
| A2_SRR8283979_Gar58 | 7 | 3.31 | 0.0028 | A2_019 | 7 | 5.36 | 0.0030 | A2_056 | 7 | 69.73 | 0.0243 |
| A1_019 | 7 | 3.92 | 0.0030 | A2_029 | 7 | 5.36 | 0.0030 | A2_057 | 7 | 69.73 | 0.0246 |
| A1_028 | 7 | 3.92 | 0.0026 | A2_030 | 7 | 5.36 | 0.0026 | A2_059 | 7 | 69.73 | 0.0247 |
| A1_030 | 7 | 3.92 | 0.0027 | A2_034 | 7 | 5.36 | 0.0027 | A2_060 | 7 | 69.73 | 0.0255 |
| A1_051 | 7 | 3.92 | 0.0027 | A2_040 | 7 | 5.36 | 0.0030 | A2_061 | 7 | 69.73 | 0.0232 |
| A1_077 | 7 | 3.92 | 0.0026 | A2_045 | 7 | 5.36 | 0.0030 | A2_062 | 7 | 69.73 | 0.0227 |
| A1_079 | 7 | 3.92 | 0.0025 | A2_046 | 7 | 5.36 | 0.0030 | A2_064 | 7 | 69.73 | 0.0248 |
| A1_084 | 7 | 3.92 | 0.0027 | A2_056 | 7 | 5.36 | 0.0027 | A2_065 | 7 | 69.73 | 0.0235 |
| A1_088 | 7 | 3.92 | 0.0028 | A2_062 | 7 | 5.36 | 0.0030 | A2_066 | 7 | 69.73 | 0.0261 |
| A1_097 | 7 | 3.92 | 0.0028 | A2_066 | 7 | 5.36 | 0.0030 | A2_067 | 7 | 69.73 | 0.0250 |
| A1_108 | 7 | 3.92 | 0.0025 | A2_080 | 7 | 5.36 | 0.0026 | A2_069 | 7 | 69.73 | 0.0259 |
| A1_125 | 7 | 3.92 | 0.0030 | A2_085 | 7 | 5.36 | 0.0026 | A2_073 | 7 | 69.73 | 0.0228 |
| A2_001 | 7 | 3.92 | 0.0029 | A2_091 | 7 | 5.36 | 0.0029 | A2_074 | 7 | 69.73 | 0.0248 |
| A2_026 | 7 | 3.92 | 0.0028 | A2_096 | 7 | 5.36 | 0.0028 | A2_076 | 7 | 69.73 | 0.0263 |
| A2_030 | 7 | 3.92 | 0.0023 | A2_117 | 7 | 5.36 | 0.0026 | A2_079 | 7 | 69.73 | 0.0252 |
| A2_034 | 7 | 3.92 | 0.0024 | A2_118 | 7 | 5.36 | 0.0029 | A2_080 | 7 | 69.73 | 0.0244 |
| A2_043 | 7 | 3.92 | 0.0030 | A2_119 | 7 | 5.36 | 0.0026 | A2_084 | 7 | 69.73 | 0.0253 |
| A2_056 | 7 | 3.92 | 0.0026 | A2_124 | 7 | 5.36 | 0.0027 | A2_085 | 7 | 69.73 | 0.0251 |
| A2_061 | 7 | 3.92 | 0.0019 | A2_147 | 7 | 5.36 | 0.0026 | A2_087 | 7 | 69.73 | 0.0231 |
| A2_062 | 7 | 3.92 | 0.0028 | A2_255 | 7 | 5.36 | 0.0029 | A2_091 | 7 | 69.73 | 0.0249 |
| A2_065 | 7 | 3.92 | 0.0030 | A2_NC503 | 7 | 5.36 | 0.0026 | A2_096 | 7 | 69.73 | 0.0252 |
| A2_067 | 7 | 3.92 | 0.0026 | A2_SRR8283969_Gar42 | 7 | 5.36 | 0.0030 | A2_100 | 7 | 69.73 | 0.0239 |
| A2_079 | 7 | 3.92 | 0.0026 | A2_SRR8283974_Gar39 | 7 | 5.36 | 0.0029 | A2_101 | 7 | 69.73 | 0.0233 |
| A2_080 | 7 | 3.92 | 0.0030 | A2_SRR8283976_Gar37 | 7 | 5.36 | 0.0029 | A2_113 | 7 | 69.73 | 0.0250 |
| A2_085 | 7 | 3.92 | 0.0023 | A2_SRR8283977_Gar56 | 7 | 5.36 | 0.0027 | A2_117 | 7 | 69.73 | 0.0255 |
| A2_091 | 7 | 3.92 | 0.0024 | A2_SRR8283978_Gar57 | 7 | 5.36 | 0.0029 | A2_118 | 7 | 69.73 | 0.0245 |
| A2_096 | 7 | 3.92 | 0.0026 | A2_SRR8283985_Gar34 | 7 | 5.36 | 0.0030 | A2_119 | 7 | 69.73 | 0.0240 |
| A2_101 | 7 | 3.92 | 0.0026 | A2_SRR8283986_Gar35 | 7 | 5.36 | 0.0028 | A2_123 | 7 | 69.73 | 0.0253 |
| A2_113 | 7 | 3.92 | 0.0023 | A2_SRR8283988_Gar33 | 7 | 5.36 | 0.0028 | A2_124 | 7 | 69.73 | 0.0253 |
| A2_117 | 7 | 3.92 | 0.0028 | A2_SRR8283993_Gar01 | 7 | 5.36 | 0.0029 | A2_131 | 7 | 69.73 | 0.0253 |
| A2_118 | 7 | 3.92 | 0.0024 | A2_SRR8283995_Gar65 | 7 | 5.36 | 0.0030 | A2_141 | 7 | 69.73 | 0.0253 |
| A2_119 | 7 | 3.92 | 0.0025 | A2_SRR8284007_Gar47 | 7 | 5.36 | 0.0030 | A2_142 | 7 | 69.73 | 0.0257 |
| A2_123 | 7 | 3.92 | 0.0024 | A2_SRR8284010_Gar63 | 7 | 5.36 | 0.0030 | A2_147 | 7 | 69.73 | 0.0252 |
| A2_124 | 7 | 3.92 | 0.0030 | A2_SRR8283991_Gar03 | 7 | 11.85 | 0.0202 | A2_154 | 7 | 69.73 | 0.0255 |
| A2_131 | 7 | 3.92 | 0.0025 | A2_SRR8284010_Gar63 | 7 | 11.85 | 0.0200 | A2_156 | 7 | 69.73 | 0.0240 |
| A2_147 | 7 | 3.92 | 0.0030 | A2_045 | 7 | 14.27 | 0.0212 | A2_158 | 7 | 69.73 | 0.0255 |
| A2_NC503 | 7 | 3.92 | 0.0023 | A2_SRR8283993_Gar01 | 7 | 14.27 | 0.0202 | A2_164 | 7 | 69.73 | 0.0253 |
| A2_SRR8283971_Gar40 | 7 | 3.92 | 0.0028 | A1_155 | 7 | 24.59 | 0.0019 | A2_255 | 7 | 69.73 | 0.0230 |
| A2_SRR8283977_Gar56 | 7 | 3.92 | 0.0024 | A1_155 | 7 | 26.61 | 0.0030 | A2_lintless | 7 | 69.73 | 0.0251 |
| A2_SRR8283983_Gar44 | 7 | 3.92 | 0.0027 | A2_030 | 7 | 29.39 | 0.0024 | A2_NC503 | 7 | 69.73 | 0.0233 |
| A2_SRR8283985_Gar34 | 7 | 3.92 | 0.0030 | A2_047 | 7 | 29.39 | 0.0028 | A2_SRR8283969_Gar4 | 7 | 69.73 | 0.0247 |
| A2_SRR8283986_Gar35 | 7 | 3.92 | 0.0027 | A2_076 | 7 | 29.39 | 0.0030 | A2_SRR8283970_Gar4 | 7 | 69.73 | 0.0200 |
| A2_SRR8283988_Gar33 | 7 | 3.92 | 0.0030 | A2_091 | 7 | 29.39 | 0.0027 | A2_SRR8283971_Gar4 | 7 | 69.73 | 0.0256 |
| A2_SRR8283994_Gar02 | 7 | 3.92 | 0.0027 | A2_156 | 7 | 29.39 | 0.0018 | A2_SRR8283972_Gar4 | 7 | 69.73 | 0.0257 |
| A2_SRR8284001_Gar53 | 7 | 3.92 | 0.0026 | A1_155 | 7 | 41.78 | 0.0013 | A2_SRR8283973_Gar3 | 7 | 69.73 | 0.0230 |
| A2_SRR8284005_Gar49 | 7 | 3.92 | 0.0030 | A1_SRR8283998_Ghe02 | 7 | 48.64 | 0.0248 | A2_SRR8283974_Gar3 | 7 | 69.73 | 0.0231 |
| A2_SRR8284010_Gar63 | 7 | 3.92 | 0.0029 | A2_SRR8283995_Gar65 | 7 | 48.64 | 0.0203 | A2_SRR8283975_Gar3 | 7 | 69.73 | 0.0252 |
| A2_SRR8284011_Gar64 | 7 | 3.92 | 0.0027 | A2_SRR8283999_Gar55 | 7 | 48.64 | 0.0204 | A2_SRR8283976_Gar3 | 7 | 69.73 | 0.0302 |
| A1_125 | 7 | 4.68 | 0.0029 | A2_SRR8284000_Gar54 | 7 | 48.64 | 0.0203 | A2_SRR8283977_Gar5 | 7 | 69.73 | 0.0263 |
| A2_008 | 7 | 4.68 | 0.0030 | A2_SRR8284002_Gar52 | 7 | 48.64 | 0.0218 | A2_SRR8283978_Gar5 | 7 | 69.73 | 0.0205 |
| A2_029 | 7 | 4.68 | 0.0027 | A1_019 | 7 | 69.73 | 0.0264 | A2_SRR8283979_Gar5 | 7 | 69.73 | 0.0226 |
| A2_043 | 7 | 4.68 | 0.0022 | A1_028 | 7 | 69.73 | 0.0260 | A2_SRR8283980_Gar5 | 7 | 69.73 | 0.0272 |
| A2_056 | 7 | 4.68 | 0.0026 | A1_030 | 7 | 69.73 | 0.0267 | A2_SRR8283981_Gar6 | 7 | 69.73 | 0.0203 |
| A2_057 | 7 | 4.68 | 0.0022 | A1_051 | 7 | 69.73 | 0.0254 | A2_SRR8283982_Gar6 | 7 | 69.73 | 0.0248 |
| A2_062 | 7 | 4.68 | 0.0025 | A1_054 | 7 | 69.73 | 0.0262 | A2_SRR8283983_Gar4 | 7 | 69.73 | 0.0223 |
| A2_065 | 7 | 4.68 | 0.0027 | A1_073 | 7 | 69.73 | 0.0243 | A2_SRR8283984_Gar4 | 7 | 69.73 | 0.0252 |
| A2_066 | 7 | 4.68 | 0.0028 | A1_074 | 7 | 69.73 | 0.0231 | A2_SRR8283985_Gar3 | 7 | 69.73 | 0.0208 |
| A2_067 | 7 | 4.68 | 0.0029 | A1_077 | 7 | 69.73 | 0.0259 | A2_SRR8283986_Gar3 | 7 | 69.73 | 0.0297 |
| A2_069 | 7 | 4.68 | 0.0030 | A1_079 | 7 | 69.73 | 0.0255 | A2_SRR8283987_Gar3 | 7 | 69.73 | 0.0277 |
| A2_079 | 7 | 4.68 | 0.0027 | A1_084 | 7 | 69.73 | 0.0251 | A2_SRR8283988_Gar3 | 7 | 69.73 | 0.0235 |

Accessions with dS outside of 95% CI (0.0031 - 0.0198)

| Accession | Chr | Position | dS | Accession | Chr | Position | dS | Accession | Chr | Position | dS |
|---|---|---|---|---|---|---|---|---|---|---|---|
| A2_SRR8283989_Gar30 | 7 | 69.73 | 0.0251 | A2_SRR8283977_Gar56 | 7 | 72.89 | 0.0217 | A2_100 | 8 | 6.57 | 0.0220 |
| A2_SRR8283990_Gar31 | 7 | 69.73 | 0.0215 | A2_SRR8283978_Gar57 | 7 | 72.89 | 0.0239 | A2_101 | 8 | 6.57 | 0.0206 |
| A2_SRR8283991_Gar03 | 7 | 69.73 | 0.0228 | A2_SRR8283979_Gar58 | 7 | 72.89 | 0.0202 | A2_113 | 8 | 6.57 | 0.0221 |
| A2_SRR8283992_Gar04 | 7 | 69.73 | 0.0264 | A2_SRR8283981_Gar60 | 7 | 72.89 | 0.0226 | A2_117 | 8 | 6.57 | 0.0218 |
| A2_SRR8283994_Gar02 | 7 | 69.73 | 0.0236 | A2_SRR8283982_Gar61 | 7 | 72.89 | 0.0218 | A2_118 | 8 | 6.57 | 0.0215 |
| A2_SRR8283995_Gar65 | 7 | 69.73 | 0.0240 | A2_SRR8283986_Gar35 | 7 | 72.89 | 0.0237 | A2_119 | 8 | 6.57 | 0.0218 |
| A2_SRR8283997_Gar66 | 7 | 69.73 | 0.0265 | A2_SRR8283988_Gar33 | 7 | 72.89 | 0.0238 | A2_123 | 8 | 6.57 | 0.0216 |
| A2_SRR8283999_Gar55 | 7 | 69.73 | 0.0260 | A2_SRR8283989_Gar30 | 7 | 72.89 | 0.0231 | A2_124 | 8 | 6.57 | 0.0219 |
| A2_SRR8284000_Gar54 | 7 | 69.73 | 0.0228 | A2_SRR8283990_Gar31 | 7 | 72.89 | 0.0228 | A2_131 | 8 | 6.57 | 0.0219 |
| A2_SRR8284001_Gar53 | 7 | 69.73 | 0.0267 | A2_SRR8283993_Gar01 | 7 | 72.89 | 0.0201 | A2_141 | 8 | 6.57 | 0.0221 |
| A2_SRR8284002_Gar52 | 7 | 69.73 | 0.0239 | A2_SRR8283995_Gar65 | 7 | 72.89 | 0.0227 | A2_142 | 8 | 6.57 | 0.0222 |
| A2_SRR8284004_Gar50 | 7 | 69.73 | 0.0244 | A2_SRR8283999_Gar55 | 7 | 72.89 | 0.0221 | A2_147 | 8 | 6.57 | 0.0223 |
| A2_SRR8284005_Gar49 | 7 | 69.73 | 0.0206 | A2_SRR8284000_Gar54 | 7 | 72.89 | 0.0209 | A2_154 | 8 | 6.57 | 0.0241 |
| A2_SRR8284006_Gar48 | 7 | 69.73 | 0.0239 | A2_SRR8284001_Gar53 | 7 | 72.89 | 0.0230 | A2_156 | 8 | 6.57 | 0.0224 |
| A2_SRR8284007_Gar47 | 7 | 69.73 | 0.0210 | A2_SRR8284005_Gar49 | 7 | 72.89 | 0.0205 | A2_158 | 8 | 6.57 | 0.0215 |
| A2_SRR8284008_Gar46 | 7 | 69.73 | 0.0286 | A2_SRR8284006_Gar48 | 7 | 72.89 | 0.0221 | A2_164 | 8 | 6.57 | 0.0220 |
| A2_SRR8284009_Gar62 | 7 | 69.73 | 0.0208 | A2_SRR8284007_Gar47 | 7 | 72.89 | 0.0238 | A2_255 | 8 | 6.57 | 0.0202 |
| A2_SRR8284010_Gar63 | 7 | 69.73 | 0.0237 | A2_SRR8284008_Gar46 | 7 | 72.89 | 0.0226 | A2_lintless | 8 | 6.57 | 0.0224 |
| A2_SRR8284011_Gar64 | 7 | 69.73 | 0.0253 | A2_SRR8284009_Gar62 | 7 | 72.89 | 0.0251 | A2_NC503 | 8 | 6.57 | 0.0202 |
| A1_019 | 7 | 72.89 | 0.0222 | A2_SRR8284010_Gar63 | 7 | 72.89 | 0.0205 | A2_SRR8283969_Gar4 | 8 | 6.57 | 0.0202 |
| A1_028 | 7 | 72.89 | 0.0228 | A2_SRR8284011_Gar64 | 7 | 72.89 | 0.0199 | A2_SRR8283970_Gar4 | 8 | 6.57 | 0.0247 |
| A1_030 | 7 | 72.89 | 0.0224 | A1_125 | 7 | 77.30 | 0.0029 | A2_SRR8283971_Gar4 | 8 | 6.57 | 0.0230 |
| A1_051 | 7 | 72.89 | 0.0220 | A1_155 | 7 | 77.30 | 0.0015 | A2_SRR8283972_Gar4 | 8 | 6.57 | 0.0265 |
| A1_054 | 7 | 72.89 | 0.0221 | A1_SRR8283998_Ghe02 | 7 | 77.30 | 0.0029 | A2_SRR8283973_Gar3 | 8 | 6.57 | 0.0266 |
| A1_073 | 7 | 72.89 | 0.0204 | A2_SRR8283997_Gar66 | 7 | 77.30 | 0.0027 | A2_SRR8283974_Gar3 | 8 | 6.57 | 0.0220 |
| A1_077 | 7 | 72.89 | 0.0220 | A2_SRR8284008_Gar46 | 7 | 77.30 | 0.0029 | A2_SRR8283975_Gar3 | 8 | 6.57 | 0.0229 |
| A1_079 | 7 | 72.89 | 0.0223 | A1_155 | 7 | 78.32 | 0.0028 | A2_SRR8283976_Gar3 | 8 | 6.57 | 0.0226 |
| A1_084 | 7 | 72.89 | 0.0226 | A1_155 | 7 | 79.31 | 0.0027 | A2_SRR8283977_Gar5 | 8 | 6.57 | 0.0221 |
| A1_088 | 7 | 72.89 | 0.0212 | A1_155 | 7 | 80.08 | 0.0030 | A2_SRR8283978_Gar5 | 8 | 6.57 | 0.0237 |
| A1_097 | 7 | 72.89 | 0.0215 | A1_019 | 8 | 6.57 | 0.0236 | A2_SRR8283979_Gar5 | 8 | 6.57 | 0.0283 |
| A1_108 | 7 | 72.89 | 0.0222 | A1_028 | 8 | 6.57 | 0.0243 | A2_SRR8283980_Gar5 | 8 | 6.57 | 0.0241 |
| A1_125 | 7 | 72.89 | 0.0206 | A1_030 | 8 | 6.57 | 0.0232 | A2_SRR8283981_Gar6 | 8 | 6.57 | 0.0251 |
| A1_132 | 7 | 72.89 | 0.0220 | A1_051 | 8 | 6.57 | 0.0243 | A2_SRR8283982_Gar6 | 8 | 6.57 | 0.0220 |
| A1_133 | 7 | 72.89 | 0.0224 | A1_054 | 8 | 6.57 | 0.0234 | A2_SRR8283983_Gar4 | 8 | 6.57 | 0.0252 |
| A1_Af | 7 | 72.89 | 0.0221 | A1_073 | 8 | 6.57 | 0.0221 | A2_SRR8283984_Gar4 | 8 | 6.57 | 0.0248 |
| A1_Nisa | 7 | 72.89 | 0.0215 | A1_074 | 8 | 6.57 | 0.0244 | A2_SRR8283985_Gar3 | 8 | 6.57 | 0.0309 |
| A2_001 | 7 | 72.89 | 0.0210 | A1_077 | 8 | 6.57 | 0.0235 | A2_SRR8283986_Gar3 | 8 | 6.57 | 0.0234 |
| A2_008 | 7 | 72.89 | 0.0211 | A1_079 | 8 | 6.57 | 0.0228 | A2_SRR8283987_Gar3 | 8 | 6.57 | 0.0245 |
| A2_016 | 7 | 72.89 | 0.0210 | A1_084 | 8 | 6.57 | 0.0237 | A2_SRR8283988_Gar3 | 8 | 6.57 | 0.0265 |
| A2_021 | 7 | 72.89 | 0.0212 | A1_088 | 8 | 6.57 | 0.0229 | A2_SRR8283989_Gar3 | 8 | 6.57 | 0.0282 |
| A2_026 | 7 | 72.89 | 0.0217 | A1_097 | 8 | 6.57 | 0.0236 | A2_SRR8283990_Gar3 | 8 | 6.57 | 0.0231 |
| A2_029 | 7 | 72.89 | 0.0207 | A1_108 | 8 | 6.57 | 0.0233 | A2_SRR8283991_Gar0 | 8 | 6.57 | 0.0223 |
| A2_030 | 7 | 72.89 | 0.0208 | A1_125 | 8 | 6.57 | 0.0233 | A2_SRR8283992_Gar0 | 8 | 6.57 | 0.0266 |
| A2_031 | 7 | 72.89 | 0.0206 | A1_132 | 8 | 6.57 | 0.0238 | A2_SRR8283994_Gar0 | 8 | 6.57 | 0.0255 |
| A2_040 | 7 | 72.89 | 0.0213 | A1_133 | 8 | 6.57 | 0.0237 | A2_SRR8283995_Gar6 | 8 | 6.57 | 0.0211 |
| A2_041 | 7 | 72.89 | 0.0212 | A1_155 | 8 | 6.57 | 0.0221 | A2_SRR8283997_Gar6 | 8 | 6.57 | 0.0262 |
| A2_044 | 7 | 72.89 | 0.0208 | A1_Af | 8 | 6.57 | 0.0237 | A2_SRR8283999_Gar5 | 8 | 6.57 | 0.0238 |
| A2_046 | 7 | 72.89 | 0.0210 | A1_Nisa | 8 | 6.57 | 0.0237 | A2_SRR8284000_Gar5 | 8 | 6.57 | 0.0229 |
| A2_047 | 7 | 72.89 | 0.0203 | A1_SRR8283996_Ghe01 | 8 | 6.57 | 0.0205 | A2_SRR8284001_Gar5 | 8 | 6.57 | 0.0256 |
| A2_056 | 7 | 72.89 | 0.0225 | A2_001 | 8 | 6.57 | 0.0220 | A2_SRR8284002_Gar5 | 8 | 6.57 | 0.0280 |
| A2_057 | 7 | 72.89 | 0.0218 | A2_004 | 8 | 6.57 | 0.0202 | A2_SRR8284003_Gar5 | 8 | 6.57 | 0.0230 |
| A2_059 | 7 | 72.89 | 0.0206 | A2_008 | 8 | 6.57 | 0.0219 | A2_SRR8284004_Gar5 | 8 | 6.57 | 0.0227 |
| A2_060 | 7 | 72.89 | 0.0210 | A2_016 | 8 | 6.57 | 0.0224 | A2_SRR8284005_Gar4 | 8 | 6.57 | 0.0248 |
| A2_062 | 7 | 72.89 | 0.0218 | A2_019 | 8 | 6.57 | 0.0211 | A2_SRR8284006_Gar4 | 8 | 6.57 | 0.0244 |
| A2_064 | 7 | 72.89 | 0.0211 | A2_021 | 8 | 6.57 | 0.0214 | A2_SRR8284007_Gar4 | 8 | 6.57 | 0.0243 |
| A2_065 | 7 | 72.89 | 0.0213 | A2_026 | 8 | 6.57 | 0.0225 | A2_SRR8284008_Gar4 | 8 | 6.57 | 0.0290 |
| A2_066 | 7 | 72.89 | 0.0217 | A2_029 | 8 | 6.57 | 0.0220 | A2_SRR8284009_Gar6 | 8 | 6.57 | 0.0232 |
| A2_067 | 7 | 72.89 | 0.0206 | A2_030 | 8 | 6.57 | 0.0213 | A2_SRR8284010_Gar6 | 8 | 6.57 | 0.0224 |
| A2_069 | 7 | 72.89 | 0.0202 | A2_031 | 8 | 6.57 | 0.0221 | A2_SRR8284011_Gar6 | 8 | 6.57 | 0.0223 |
| A2_074 | 7 | 72.89 | 0.0213 | A2_034 | 8 | 6.57 | 0.0205 | A1_051 | 8 | 76.89 | 0.0028 |
| A2_076 | 7 | 72.89 | 0.0219 | A2_040 | 8 | 6.57 | 0.0228 | A1_054 | 8 | 76.89 | 0.0029 |
| A2_079 | 7 | 72.89 | 0.0206 | A2_041 | 8 | 6.57 | 0.0223 | A1_077 | 8 | 76.89 | 0.0029 |
| A2_080 | 7 | 72.89 | 0.0207 | A2_043 | 8 | 6.57 | 0.0209 | A1_132 | 8 | 76.89 | 0.0028 |
| A2_084 | 7 | 72.89 | 0.0216 | A2_044 | 8 | 6.57 | 0.0225 | A1_133 | 8 | 76.89 | 0.0027 |
| A2_085 | 7 | 72.89 | 0.0213 | A2_046 | 8 | 6.57 | 0.0221 | A1_051 | 8 | 80.33 | 0.0028 |
| A2_087 | 7 | 72.89 | 0.0215 | A2_047 | 8 | 6.57 | 0.0223 | A1_077 | 8 | 80.33 | 0.0030 |
| A2_091 | 7 | 72.89 | 0.0212 | A2_056 | 8 | 6.57 | 0.0217 | A1_132 | 8 | 80.33 | 0.0028 |
| A2_096 | 7 | 72.89 | 0.0211 | A2_057 | 8 | 6.57 | 0.0218 | A1_133 | 8 | 80.33 | 0.0028 |
| A2_100 | 7 | 72.89 | 0.0205 | A2_059 | 8 | 6.57 | 0.0225 | A1_051 | 8 | 83.55 | 0.0028 |
| A2_113 | 7 | 72.89 | 0.0214 | A2_060 | 8 | 6.57 | 0.0218 | A1_077 | 8 | 83.55 | 0.0028 |
| A2_117 | 7 | 72.89 | 0.0210 | A2_062 | 8 | 6.57 | 0.0210 | A1_133 | 8 | 83.55 | 0.0029 |
| A2_118 | 7 | 72.89 | 0.0208 | A2_064 | 8 | 6.57 | 0.0237 | A1_155 | 8 | 88.96 | 0.0009 |
| A2_119 | 7 | 72.89 | 0.0200 | A2_065 | 8 | 6.57 | 0.0222 | A2_062 | 8 | 88.96 | 0.0010 |
| A2_123 | 7 | 72.89 | 0.0212 | A2_066 | 8 | 6.57 | 0.0221 | A2_SRR8283994_Gar0 | 8 | 88.96 | 0.0019 |
| A2_124 | 7 | 72.89 | 0.0208 | A2_067 | 8 | 6.57 | 0.0227 | A1_155 | 8 | 92.03 | 0.0026 |
| A2_131 | 7 | 72.89 | 0.0208 | A2_069 | 8 | 6.57 | 0.0213 | A1_155 | 8 | 96.46 | 0.0022 |
| A2_142 | 7 | 72.89 | 0.0207 | A2_073 | 8 | 6.57 | 0.0221 | A1_155 | 8 | 99.32 | 0.0020 |
| A2_147 | 7 | 72.89 | 0.0206 | A2_074 | 8 | 6.57 | 0.0222 | A2_041 | 8 | 99.32 | 0.0028 |
| A2_156 | 7 | 72.89 | 0.0204 | A2_076 | 8 | 6.57 | 0.0241 | A2_SRR8283992_Gar0 | 8 | 99.32 | 0.0029 |
| A2_158 | 7 | 72.89 | 0.0206 | A2_079 | 8 | 6.57 | 0.0221 | A2_SRR8284008_Gar4 | 8 | 100.18 | 0.0227 |
| A2_164 | 7 | 72.89 | 0.0213 | A2_080 | 8 | 6.57 | 0.0222 | A1_155 | 8 | 102.41 | 0.0023 |
| A2_lintless | 7 | 72.89 | 0.0212 | A2_084 | 8 | 6.57 | 0.0222 | A1_155 | 8 | 103.08 | 0.0020 |
| A2_NC503 | 7 | 72.89 | 0.0201 | A2_085 | 8 | 6.57 | 0.0232 | A2_043 | 9 | 0.38 | 0.0030 |
| A2_SRR8283969_Gar42 | 7 | 72.89 | 0.0206 | A2_087 | 8 | 6.57 | 0.0218 | A2_043 | 9 | 1.41 | 0.0030 |
| A2_SRR8283971_Gar40 | 7 | 72.89 | 0.0241 | A2_091 | 8 | 6.57 | 0.0222 | A1_019 | 9 | 3.89 | 0.0222 |
| A2_SRR8283972_Gar41 | 7 | 72.89 | 0.0259 | A2_096 | 8 | 6.57 | 0.0232 | A1_028 | 9 | 3.89 | 0.0199 |

Accessions with dS outside of 95% CI (0.0031 - 0.0198)

| Accession | Chr | Position | dS |
|---|---|---|---|
| A1_030 | 9 | 3.89 | 0.0213 |
| A1_051 | 9 | 3.89 | 0.0206 |
| A1_054 | 9 | 3.89 | 0.0215 |
| A1_073 | 9 | 3.89 | 0.0205 |
| A1_077 | 9 | 3.89 | 0.0218 |
| A1_079 | 9 | 3.89 | 0.0208 |
| A1_084 | 9 | 3.89 | 0.0208 |
| A1_088 | 9 | 3.89 | 0.0209 |
| A1_097 | 9 | 3.89 | 0.0214 |
| A1_108 | 9 | 3.89 | 0.0216 |
| A1_125 | 9 | 3.89 | 0.0204 |
| A1_132 | 9 | 3.89 | 0.0216 |
| A1_133 | 9 | 3.89 | 0.0220 |
| A1_155 | 9 | 3.89 | 0.0226 |
| A1_Af | 9 | 3.89 | 0.0208 |
| A1_Nisa | 9 | 3.89 | 0.0208 |
| A1_SRR8283996_Ghe01 | 9 | 3.89 | 0.0206 |
| A1_SRR8283998_Ghe02 | 9 | 3.89 | 0.0217 |
| A2_001 | 9 | 3.89 | 0.0216 |
| A2_008 | 9 | 3.89 | 0.0215 |
| A2_016 | 9 | 3.89 | 0.0207 |
| A2_019 | 9 | 3.89 | 0.0205 |
| A2_021 | 9 | 3.89 | 0.0208 |
| A2_026 | 9 | 3.89 | 0.0202 |
| A2_030 | 9 | 3.89 | 0.0200 |
| A2_034 | 9 | 3.89 | 0.0201 |
| A2_040 | 9 | 3.89 | 0.0211 |
| A2_041 | 9 | 3.89 | 0.0208 |
| A2_043 | 9 | 3.89 | 0.0201 |
| A2_044 | 9 | 3.89 | 0.0203 |
| A2_046 | 9 | 3.89 | 0.0223 |
| A2_047 | 9 | 3.89 | 0.0217 |
| A2_056 | 9 | 3.89 | 0.0215 |
| A2_057 | 9 | 3.89 | 0.0214 |
| A2_059 | 9 | 3.89 | 0.0216 |
| A2_060 | 9 | 3.89 | 0.0203 |
| A2_061 | 9 | 3.89 | 0.0204 |
| A2_064 | 9 | 3.89 | 0.0208 |
| A2_065 | 9 | 3.89 | 0.0199 |
| A2_066 | 9 | 3.89 | 0.0212 |
| A2_067 | 9 | 3.89 | 0.0218 |
| A2_069 | 9 | 3.89 | 0.0228 |
| A2_074 | 9 | 3.89 | 0.0214 |
| A2_076 | 9 | 3.89 | 0.0214 |
| A2_079 | 9 | 3.89 | 0.0211 |
| A2_080 | 9 | 3.89 | 0.0217 |
| A2_084 | 9 | 3.89 | 0.0213 |
| A2_087 | 9 | 3.89 | 0.0216 |
| A2_091 | 9 | 3.89 | 0.0228 |
| A2_096 | 9 | 3.89 | 0.0209 |
| A2_100 | 9 | 3.89 | 0.0208 |
| A2_101 | 9 | 3.89 | 0.0208 |
| A2_113 | 9 | 3.89 | 0.0217 |
| A2_117 | 9 | 3.89 | 0.0222 |
| A2_118 | 9 | 3.89 | 0.0206 |
| A2_119 | 9 | 3.89 | 0.0206 |
| A2_124 | 9 | 3.89 | 0.0205 |
| A2_131 | 9 | 3.89 | 0.0215 |
| A2_141 | 9 | 3.89 | 0.0219 |
| A2_142 | 9 | 3.89 | 0.0217 |
| A2_147 | 9 | 3.89 | 0.0218 |
| A2_154 | 9 | 3.89 | 0.0223 |
| A2_156 | 9 | 3.89 | 0.0209 |
| A2_158 | 9 | 3.89 | 0.0216 |
| A2_164 | 9 | 3.89 | 0.0215 |
| A2_SRR8283969_Gar42 | 9 | 3.89 | 0.0225 |
| A2_SRR8283971_Gar40 | 9 | 3.89 | 0.0223 |
| A2_SRR8283972_Gar41 | 9 | 3.89 | 0.0217 |
| A2_SRR8283973_Gar38 | 9 | 3.89 | 0.0212 |
| A2_SRR8283974_Gar39 | 9 | 3.89 | 0.0210 |
| A2_SRR8283975_Gar36 | 9 | 3.89 | 0.0224 |
| A2_SRR8283976_Gar37 | 9 | 3.89 | 0.0231 |
| A2_SRR8283977_Gar56 | 9 | 3.89 | 0.0209 |
| A2_SRR8283978_Gar57 | 9 | 3.89 | 0.0236 |
| A2_SRR8283980_Gar59 | 9 | 3.89 | 0.0213 |
| A2_SRR8283981_Gar60 | 9 | 3.89 | 0.0219 |
| A2_SRR8283982_Gar61 | 9 | 3.89 | 0.0240 |
| A2_SRR8283984_Gar45 | 9 | 3.89 | 0.0200 |
| A2_SRR8283985_Gar34 | 9 | 3.89 | 0.0223 |
| A2_SRR8283986_Gar35 | 9 | 3.89 | 0.0214 |
| A2_SRR8283987_Gar32 | 9 | 3.89 | 0.0234 |
| A2_SRR8283988_Gar33 | 9 | 3.89 | 0.0204 |
| A2_SRR8283989_Gar30 | 9 | 3.89 | 0.0252 |
| A2_SRR8283990_Gar31 | 9 | 3.89 | 0.0230 |
| A2_SRR8283991_Gar03 | 9 | 3.89 | 0.0219 |
| A2_SRR8283992_Gar04 | 9 | 3.89 | 0.0236 |

| Accession | Chr | Position | dS |
|---|---|---|---|
| A2_SRR8283993_Gar01 | 9 | 3.89 | 0.0228 |
| A2_SRR8283997_Gar66 | 9 | 3.89 | 0.0217 |
| A2_SRR8284000_Gar54 | 9 | 3.89 | 0.0209 |
| A2_SRR8284001_Gar53 | 9 | 3.89 | 0.0225 |
| A2_SRR8284002_Gar52 | 9 | 3.89 | 0.0205 |
| A2_SRR8284003_Gar51 | 9 | 3.89 | 0.0210 |
| A2_SRR8284005_Gar49 | 9 | 3.89 | 0.0217 |
| A2_SRR8284006_Gar48 | 9 | 3.89 | 0.0210 |
| A2_SRR8284007_Gar47 | 9 | 3.89 | 0.0237 |
| A2_SRR8284008_Gar46 | 9 | 3.89 | 0.0203 |
| A2_SRR8284009_Gar62 | 9 | 3.89 | 0.0238 |
| A2_SRR8284010_Gar63 | 9 | 3.89 | 0.0237 |
| A2_SRR8284011_Gar64 | 9 | 3.89 | 0.0220 |
| A1_125 | 9 | 16.72 | 0.0201 |
| A1_SRR8283996_Ghe01 | 9 | 16.72 | 0.0286 |
| A1_SRR8283998_Ghe02 | 9 | 16.72 | 0.0238 |
| A2_045 | 9 | 16.72 | 0.0239 |
| A2_091 | 9 | 16.72 | 0.0200 |
| A2_124 | 9 | 16.72 | 0.0199 |
| A2_SRR8283969_Gar42 | 9 | 16.72 | 0.0292 |
| A2_SRR8283970_Gar43 | 9 | 16.72 | 0.0209 |
| A2_SRR8283971_Gar40 | 9 | 16.72 | 0.0254 |
| A2_SRR8283972_Gar41 | 9 | 16.72 | 0.0199 |
| A2_SRR8283973_Gar38 | 9 | 16.72 | 0.0222 |
| A2_SRR8283974_Gar39 | 9 | 16.72 | 0.0225 |
| A2_SRR8283975_Gar36 | 9 | 16.72 | 0.0207 |
| A2_SRR8283976_Gar37 | 9 | 16.72 | 0.0202 |
| A2_SRR8283977_Gar56 | 9 | 16.72 | 0.0236 |
| A2_SRR8283978_Gar57 | 9 | 16.72 | 0.0237 |
| A2_SRR8283979_Gar58 | 9 | 16.72 | 0.0222 |
| A2_SRR8283980_Gar59 | 9 | 16.72 | 0.0215 |
| A2_SRR8283983_Gar44 | 9 | 16.72 | 0.0202 |
| A2_SRR8283984_Gar45 | 9 | 16.72 | 0.0223 |
| A2_SRR8283985_Gar34 | 9 | 16.72 | 0.0214 |
| A2_SRR8283986_Gar35 | 9 | 16.72 | 0.0218 |
| A2_SRR8283987_Gar32 | 9 | 16.72 | 0.0199 |
| A2_SRR8283990_Gar31 | 9 | 16.72 | 0.0280 |
| A2_SRR8283992_Gar04 | 9 | 16.72 | 0.0213 |
| A2_SRR8283993_Gar01 | 9 | 16.72 | 0.0236 |
| A2_SRR8283995_Gar65 | 9 | 16.72 | 0.0224 |
| A2_SRR8283997_Gar66 | 9 | 16.72 | 0.0247 |
| A2_SRR8283999_Gar55 | 9 | 16.72 | 0.0243 |
| A2_SRR8284000_Gar54 | 9 | 16.72 | 0.0215 |
| A2_SRR8284001_Gar53 | 9 | 16.72 | 0.0221 |
| A2_SRR8284002_Gar52 | 9 | 16.72 | 0.0247 |
| A2_SRR8284003_Gar51 | 9 | 16.72 | 0.0214 |
| A2_SRR8284009_Gar62 | 9 | 16.72 | 0.0206 |
| A2_SRR8284010_Gar63 | 9 | 16.72 | 0.0209 |
| A2_SRR8284011_Gar64 | 9 | 16.72 | 0.0201 |
| A1_019 | 9 | 47.05 | 0.0028 |
| A1_028 | 9 | 47.05 | 0.0028 |
| A1_030 | 9 | 47.05 | 0.0030 |
| A1_051 | 9 | 47.05 | 0.0028 |
| A1_054 | 9 | 47.05 | 0.0026 |
| A1_077 | 9 | 47.05 | 0.0029 |
| A1_079 | 9 | 47.05 | 0.0029 |
| A1_084 | 9 | 47.05 | 0.0027 |
| A1_088 | 9 | 47.05 | 0.0024 |
| A1_097 | 9 | 47.05 | 0.0028 |
| A1_108 | 9 | 47.05 | 0.0026 |
| A1_125 | 9 | 47.05 | 0.0023 |
| A1_132 | 9 | 47.05 | 0.0027 |
| A1_133 | 9 | 47.05 | 0.0028 |
| A2_SRR8283976_Gar37 | 9 | 47.05 | 0.0030 |
| A1_019 | 9 | 50.13 | 0.0022 |
| A1_028 | 9 | 50.13 | 0.0024 |
| A1_030 | 9 | 50.13 | 0.0024 |
| A1_051 | 9 | 50.13 | 0.0026 |
| A1_054 | 9 | 50.13 | 0.0025 |
| A1_077 | 9 | 50.13 | 0.0024 |
| A1_079 | 9 | 50.13 | 0.0022 |
| A1_084 | 9 | 50.13 | 0.0023 |
| A1_088 | 9 | 50.13 | 0.0024 |
| A1_097 | 9 | 50.13 | 0.0022 |
| A1_108 | 9 | 50.13 | 0.0020 |
| A1_125 | 9 | 50.13 | 0.0024 |
| A1_132 | 9 | 50.13 | 0.0024 |
| A1_133 | 9 | 50.13 | 0.0024 |
| A2_045 | 9 | 50.13 | 0.0028 |
| A2_061 | 9 | 50.13 | 0.0030 |
| A1_051 | 9 | 55.41 | 0.0027 |
| A1_054 | 9 | 55.41 | 0.0028 |
| A1_132 | 9 | 55.41 | 0.0025 |
| A1_133 | 9 | 55.41 | 0.0027 |
| A2_029 | 9 | 58.52 | 0.0011 |
| A2_067 | 9 | 58.52 | 0.0023 |

| Accession | Chr | Position | dS |
|---|---|---|---|
| A2_079 | 9 | 58.52 | 0.0023 |
| A2_091 | 9 | 58.52 | 0.0017 |
| A2_147 | 9 | 58.52 | 0.0023 |
| A2_255 | 9 | 58.52 | 0.0009 |
| A2_SRR8283986_Gar35 | 9 | 58.52 | 0.0029 |
| A2_SRR8283990_Gar31 | 9 | 58.52 | 0.0030 |
| A2_SRR8283995_Gar65 | 9 | 58.52 | 0.0022 |
| A2_004 | 9 | 59.57 | 0.0022 |
| A2_074 | 9 | 66.69 | 0.0029 |
| A2_045 | 9 | 67.69 | 0.0023 |
| A2_SRR8283980_Gar59 | 9 | 67.69 | 0.0030 |
| A2_SRR8283999_Gar55 | 9 | 67.69 | 0.0026 |
| A2_SRR8284006_Gar48 | 9 | 67.69 | 0.0021 |
| A2_074 | 9 | 70.53 | 0.0008 |
| A2_SRR8283979_Gar58 | 9 | 70.53 | 0.0030 |
| A2_SRR8283985_Gar34 | 9 | 70.53 | 0.0026 |
| A2_SRR8284002_Gar52 | 9 | 70.53 | 0.0025 |
| A1_019 | 10 | 1.72 | 0.0011 |
| A1_028 | 10 | 1.72 | 0.0009 |
| A1_030 | 10 | 1.72 | 0.0011 |
| A1_051 | 10 | 1.72 | 0.0008 |
| A1_054 | 10 | 1.72 | 0.0007 |
| A1_079 | 10 | 1.72 | 0.0011 |
| A1_084 | 10 | 1.72 | 0.0009 |
| A1_088 | 10 | 1.72 | 0.0009 |
| A1_097 | 10 | 1.72 | 0.0007 |
| A1_108 | 10 | 1.72 | 0.0011 |
| A1_125 | 10 | 1.72 | 0.0016 |
| A1_132 | 10 | 1.72 | 0.0009 |
| A1_133 | 10 | 1.72 | 0.0009 |
| A1_SRR8283998_Ghe02 | 10 | 1.72 | 0.0011 |
| A2_001 | 10 | 1.72 | 0.0028 |
| A2_004 | 10 | 1.72 | 0.0030 |
| A2_008 | 10 | 1.72 | 0.0029 |
| A2_016 | 10 | 1.72 | 0.0026 |
| A2_019 | 10 | 1.72 | 0.0029 |
| A2_021 | 10 | 1.72 | 0.0029 |
| A2_026 | 10 | 1.72 | 0.0029 |
| A2_029 | 10 | 1.72 | 0.0029 |
| A2_030 | 10 | 1.72 | 0.0029 |
| A2_031 | 10 | 1.72 | 0.0029 |
| A2_034 | 10 | 1.72 | 0.0028 |
| A2_040 | 10 | 1.72 | 0.0028 |
| A2_041 | 10 | 1.72 | 0.0029 |
| A2_043 | 10 | 1.72 | 0.0028 |
| A2_044 | 10 | 1.72 | 0.0030 |
| A2_045 | 10 | 1.72 | 0.0026 |
| A2_046 | 10 | 1.72 | 0.0030 |
| A2_047 | 10 | 1.72 | 0.0029 |
| A2_056 | 10 | 1.72 | 0.0026 |
| A2_057 | 10 | 1.72 | 0.0028 |
| A2_060 | 10 | 1.72 | 0.0029 |
| A2_061 | 10 | 1.72 | 0.0029 |
| A2_062 | 10 | 1.72 | 0.0029 |
| A2_064 | 10 | 1.72 | 0.0030 |
| A2_065 | 10 | 1.72 | 0.0030 |
| A2_066 | 10 | 1.72 | 0.0030 |
| A2_067 | 10 | 1.72 | 0.0030 |
| A2_073 | 10 | 1.72 | 0.0028 |
| A2_074 | 10 | 1.72 | 0.0029 |
| A2_076 | 10 | 1.72 | 0.0029 |
| A2_079 | 10 | 1.72 | 0.0028 |
| A2_080 | 10 | 1.72 | 0.0029 |
| A2_084 | 10 | 1.72 | 0.0028 |
| A2_085 | 10 | 1.72 | 0.0028 |
| A2_087 | 10 | 1.72 | 0.0029 |
| A2_091 | 10 | 1.72 | 0.0029 |
| A2_096 | 10 | 1.72 | 0.0029 |
| A2_100 | 10 | 1.72 | 0.0029 |
| A2_101 | 10 | 1.72 | 0.0029 |
| A2_113 | 10 | 1.72 | 0.0029 |
| A2_117 | 10 | 1.72 | 0.0030 |
| A2_118 | 10 | 1.72 | 0.0028 |
| A2_119 | 10 | 1.72 | 0.0028 |
| A2_123 | 10 | 1.72 | 0.0029 |
| A2_124 | 10 | 1.72 | 0.0029 |
| A2_131 | 10 | 1.72 | 0.0029 |
| A2_141 | 10 | 1.72 | 0.0029 |
| A2_142 | 10 | 1.72 | 0.0030 |
| A2_147 | 10 | 1.72 | 0.0029 |
| A2_154 | 10 | 1.72 | 0.0030 |
| A2_156 | 10 | 1.72 | 0.0029 |
| A2_158 | 10 | 1.72 | 0.0029 |
| A2_164 | 10 | 1.72 | 0.0029 |
| A2_255 | 10 | 1.72 | 0.0029 |
| A2_lintless | 10 | 1.72 | 0.0029 |

Accessions with dS outside of 95% CI (0.0031 - 0.0198)

| Accession | Chr | Position | dS |
|---|---|---|---|
| A2_NC503 | 10 | 1.72 | 0.0027 |
| A2_SRR8283969_Gar42 | 10 | 1.72 | 0.0030 |
| A2_SRR8283970_Gar43 | 10 | 1.72 | 0.0028 |
| A2_SRR8283973_Gar38 | 10 | 1.72 | 0.0029 |
| A2_SRR8283974_Gar39 | 10 | 1.72 | 0.0028 |
| A2_SRR8283975_Gar36 | 10 | 1.72 | 0.0029 |
| A2_SRR8283976_Gar37 | 10 | 1.72 | 0.0025 |
| A2_SRR8283978_Gar57 | 10 | 1.72 | 0.0029 |
| A2_SRR8283979_Gar58 | 10 | 1.72 | 0.0027 |
| A2_SRR8283980_Gar59 | 10 | 1.72 | 0.0028 |
| A2_SRR8283982_Gar61 | 10 | 1.72 | 0.0030 |
| A2_SRR8283983_Gar44 | 10 | 1.72 | 0.0028 |
| A2_SRR8283986_Gar35 | 10 | 1.72 | 0.0029 |
| A2_SRR8283987_Gar32 | 10 | 1.72 | 0.0027 |
| A2_SRR8283988_Gar33 | 10 | 1.72 | 0.0028 |
| A2_SRR8283991_Gar03 | 10 | 1.72 | 0.0030 |
| A2_SRR8283997_Gar66 | 10 | 1.72 | 0.0030 |
| A2_SRR8283999_Gar55 | 10 | 1.72 | 0.0029 |
| A2_SRR8284000_Gar54 | 10 | 1.72 | 0.0029 |
| A2_SRR8284002_Gar52 | 10 | 1.72 | 0.0026 |
| A2_SRR8284003_Gar51 | 10 | 1.72 | 0.0023 |
| A2_SRR8284004_Gar50 | 10 | 1.72 | 0.0027 |
| A2_SRR8284007_Gar47 | 10 | 1.72 | 0.0030 |
| A2_SRR8284008_Gar46 | 10 | 1.72 | 0.0030 |
| A2_SRR8284009_Gar62 | 10 | 1.72 | 0.0030 |
| A2_SRR8284010_Gar63 | 10 | 1.72 | 0.0026 |
| A2_SRR8284011_Gar64 | 10 | 1.72 | 0.0026 |
| A1_019 | 10 | 2.31 | 0.0020 |
| A1_028 | 10 | 2.31 | 0.0018 |
| A1_051 | 10 | 2.31 | 0.0021 |
| A1_054 | 10 | 2.31 | 0.0020 |
| A1_084 | 10 | 2.31 | 0.0011 |
| A1_097 | 10 | 2.31 | 0.0020 |
| A1_125 | 10 | 2.31 | 0.0027 |
| A1_132 | 10 | 2.31 | 0.0018 |
| A1_133 | 10 | 2.31 | 0.0020 |
| A1_SRR8283998_Ghe02 | 10 | 2.31 | 0.0024 |
| A2_045 | 10 | 2.31 | 0.0028 |
| A1_051 | 10 | 2.82 | 0.0018 |
| A1_054 | 10 | 2.82 | 0.0020 |
| A1_084 | 10 | 2.82 | 0.0022 |
| A1_132 | 10 | 2.82 | 0.0017 |
| A1_133 | 10 | 2.82 | 0.0019 |
| A2_041 | 10 | 3.41 | 0.0027 |
| A2_SRR8283992_Gar04 | 10 | 3.41 | 0.0022 |
| A1_133 | 10 | 7.72 | 0.0030 |
| A2_047 | 10 | 8.99 | 0.0029 |
| A2_061 | 10 | 8.99 | 0.0030 |
| A2_158 | 10 | 8.99 | 0.0027 |
| A1_019 | 10 | 18.10 | 0.0223 |
| A1_028 | 10 | 18.10 | 0.0219 |
| A1_030 | 10 | 18.10 | 0.0220 |
| A1_051 | 10 | 18.10 | 0.0219 |
| A1_054 | 10 | 18.10 | 0.0216 |
| A1_073 | 10 | 18.10 | 0.0207 |
| A1_077 | 10 | 18.10 | 0.0214 |
| A1_079 | 10 | 18.10 | 0.0223 |
| A1_084 | 10 | 18.10 | 0.0216 |
| A1_088 | 10 | 18.10 | 0.0222 |
| A1_097 | 10 | 18.10 | 0.0217 |
| A1_108 | 10 | 18.10 | 0.0217 |
| A1_132 | 10 | 18.10 | 0.0215 |
| A1_133 | 10 | 18.10 | 0.0219 |
| A1_Af | 10 | 18.10 | 0.0213 |
| A1_Nisa | 10 | 18.10 | 0.0214 |
| A1_SRR8283996_Ghe01 | 10 | 18.10 | 0.0206 |
| A2_026 | 10 | 18.10 | 0.0199 |
| A2_040 | 10 | 18.10 | 0.0200 |
| A2_045 | 10 | 18.10 | 0.0232 |
| A2_076 | 10 | 18.10 | 0.0205 |
| A2_087 | 10 | 18.10 | 0.0203 |
| A2_147 | 10 | 18.10 | 0.0205 |
| A2_SRR8283969_Gar42 | 10 | 18.10 | 0.0213 |
| A2_SRR8283970_Gar43 | 10 | 18.10 | 0.0215 |
| A2_SRR8283971_Gar40 | 10 | 18.10 | 0.0218 |
| A2_SRR8283972_Gar41 | 10 | 18.10 | 0.0211 |
| A2_SRR8283973_Gar38 | 10 | 18.10 | 0.0227 |
| A2_SRR8283974_Gar39 | 10 | 18.10 | 0.0262 |
| A2_SRR8283975_Gar36 | 10 | 18.10 | 0.0229 |
| A2_SRR8283976_Gar37 | 10 | 18.10 | 0.0212 |
| A2_SRR8283977_Gar56 | 10 | 18.10 | 0.0261 |
| A2_SRR8283978_Gar57 | 10 | 18.10 | 0.0228 |
| A2_SRR8283979_Gar58 | 10 | 18.10 | 0.0255 |
| A2_SRR8283980_Gar59 | 10 | 18.10 | 0.0252 |
| A2_SRR8283981_Gar60 | 10 | 18.10 | 0.0218 |
| A2_SRR8283982_Gar61 | 10 | 18.10 | 0.0199 |
| A2_SRR8283983_Gar44 | 10 | 18.10 | 0.0218 |
| A2_SRR8283984_Gar45 | 10 | 18.10 | 0.0260 |
| A2_SRR8283985_Gar34 | 10 | 18.10 | 0.0255 |
| A2_SRR8283986_Gar35 | 10 | 18.10 | 0.0233 |
| A2_SRR8283987_Gar32 | 10 | 18.10 | 0.0224 |
| A2_SRR8283988_Gar33 | 10 | 18.10 | 0.0217 |
| A2_SRR8283989_Gar30 | 10 | 18.10 | 0.0211 |
| A2_SRR8283990_Gar31 | 10 | 18.10 | 0.0260 |
| A2_SRR8283991_Gar03 | 10 | 18.10 | 0.0216 |
| A2_SRR8283992_Gar04 | 10 | 18.10 | 0.0221 |
| A2_SRR8283993_Gar01 | 10 | 18.10 | 0.0243 |
| A2_SRR8283994_Gar02 | 10 | 18.10 | 0.0220 |
| A2_SRR8283995_Gar65 | 10 | 18.10 | 0.0243 |
| A2_SRR8283997_Gar66 | 10 | 18.10 | 0.0245 |
| A2_SRR8283999_Gar55 | 10 | 18.10 | 0.0264 |
| A2_SRR8284000_Gar54 | 10 | 18.10 | 0.0203 |
| A2_SRR8284001_Gar53 | 10 | 18.10 | 0.0248 |
| A2_SRR8284002_Gar52 | 10 | 18.10 | 0.0255 |
| A2_SRR8284003_Gar51 | 10 | 18.10 | 0.0238 |
| A2_SRR8284004_Gar50 | 10 | 18.10 | 0.0218 |
| A2_SRR8284005_Gar49 | 10 | 18.10 | 0.0244 |
| A2_SRR8284006_Gar48 | 10 | 18.10 | 0.0226 |
| A2_SRR8284007_Gar47 | 10 | 18.10 | 0.0230 |
| A2_SRR8284008_Gar46 | 10 | 18.10 | 0.0216 |
| A2_SRR8284009_Gar62 | 10 | 18.10 | 0.0206 |
| A2_SRR8284010_Gar63 | 10 | 18.10 | 0.0250 |
| A2_SRR8284011_Gar64 | 10 | 18.10 | 0.0229 |
| A2_067 | 10 | 33.12 | 0.0027 |
| A2_079 | 10 | 33.12 | 0.0025 |
| A2_SRR8283982_Gar61 | 10 | 33.12 | 0.0025 |
| A2_046 | 10 | 36.28 | 0.0028 |
| A2_045 | 10 | 79.32 | 0.0027 |
| A2_SRR8283985_Gar34 | 10 | 79.32 | 0.0028 |
| A2_SRR8283992_Gar04 | 10 | 79.32 | 0.0019 |
| A1_074 | 10 | 83.98 | 0.0206 |
| A1_125 | 10 | 83.98 | 0.0202 |
| A2_045 | 10 | 83.98 | 0.0201 |
| A2_SRR8283971_Gar40 | 10 | 83.98 | 0.0221 |
| A2_SRR8283972_Gar41 | 10 | 83.98 | 0.0203 |
| A2_SRR8283975_Gar36 | 10 | 83.98 | 0.0208 |
| A2_SRR8283976_Gar37 | 10 | 83.98 | 0.0220 |
| A2_SRR8283977_Gar56 | 10 | 83.98 | 0.0208 |
| A2_SRR8283982_Gar61 | 10 | 83.98 | 0.0215 |
| A2_SRR8283984_Gar45 | 10 | 83.98 | 0.0201 |
| A2_SRR8283986_Gar35 | 10 | 83.98 | 0.0199 |
| A2_SRR8283989_Gar30 | 10 | 83.98 | 0.0213 |
| A2_SRR8284002_Gar52 | 10 | 83.98 | 0.0206 |
| A2_SRR8284003_Gar51 | 10 | 83.98 | 0.0203 |
| A2_SRR8284006_Gar48 | 10 | 83.98 | 0.0200 |
| A2_SRR8284007_Gar47 | 10 | 83.98 | 0.0225 |
| A2_SRR8284009_Gar62 | 10 | 83.98 | 0.0200 |
| A2_SRR8284011_Gar64 | 10 | 83.98 | 0.0199 |
| A1_051 | 10 | 89.53 | 0.0248 |
| A1_054 | 10 | 89.53 | 0.0211 |
| A1_073 | 10 | 89.53 | 0.0240 |
| A1_074 | 10 | 89.53 | 0.0239 |
| A1_077 | 10 | 89.53 | 0.0213 |
| A1_132 | 10 | 89.53 | 0.0234 |
| A1_133 | 10 | 89.53 | 0.0247 |
| A1_155 | 10 | 89.53 | 0.0262 |
| A1_Af | 10 | 89.53 | 0.0254 |
| A1_Nisa | 10 | 89.53 | 0.0254 |
| A2_004 | 10 | 89.53 | 0.0231 |
| A2_016 | 10 | 89.53 | 0.0212 |
| A2_019 | 10 | 89.53 | 0.0217 |
| A2_021 | 10 | 89.53 | 0.0234 |
| A2_026 | 10 | 89.53 | 0.0247 |
| A2_029 | 10 | 89.53 | 0.0241 |
| A2_030 | 10 | 89.53 | 0.0207 |
| A2_031 | 10 | 89.53 | 0.0238 |
| A2_034 | 10 | 89.53 | 0.0231 |
| A2_040 | 10 | 89.53 | 0.0212 |
| A2_041 | 10 | 89.53 | 0.0233 |
| A2_043 | 10 | 89.53 | 0.0215 |
| A2_044 | 10 | 89.53 | 0.0209 |
| A2_046 | 10 | 89.53 | 0.0200 |
| A2_047 | 10 | 89.53 | 0.0212 |
| A2_056 | 10 | 89.53 | 0.0239 |
| A2_057 | 10 | 89.53 | 0.0227 |
| A2_059 | 10 | 89.53 | 0.0236 |
| A2_060 | 10 | 89.53 | 0.0242 |
| A2_061 | 10 | 89.53 | 0.0205 |
| A2_062 | 10 | 89.53 | 0.0202 |
| A2_064 | 10 | 89.53 | 0.0206 |
| A2_066 | 10 | 89.53 | 0.0202 |
| A2_067 | 10 | 89.53 | 0.0203 |
| A2_069 | 10 | 89.53 | 0.0207 |
| A2_074 | 10 | 89.53 | 0.0207 |
| A2_076 | 10 | 89.53 | 0.0215 |
| A2_079 | 10 | 89.53 | 0.0206 |
| A2_084 | 10 | 89.53 | 0.0234 |
| A2_087 | 10 | 89.53 | 0.0211 |
| A2_091 | 10 | 89.53 | 0.0214 |
| A2_096 | 10 | 89.53 | 0.0253 |
| A2_100 | 10 | 89.53 | 0.0218 |
| A2_101 | 10 | 89.53 | 0.0245 |
| A2_113 | 10 | 89.53 | 0.0210 |
| A2_117 | 10 | 89.53 | 0.0238 |
| A2_119 | 10 | 89.53 | 0.0207 |
| A2_141 | 10 | 89.53 | 0.0209 |
| A2_142 | 10 | 89.53 | 0.0237 |
| A2_147 | 10 | 89.53 | 0.0213 |
| A2_156 | 10 | 89.53 | 0.0201 |
| A2_158 | 10 | 89.53 | 0.0208 |
| A2_164 | 10 | 89.53 | 0.0213 |
| A2_255 | 10 | 89.53 | 0.0227 |
| A2_lintless | 10 | 89.53 | 0.0220 |
| A2_NC503 | 10 | 89.53 | 0.0245 |
| A2_SRR8283975_Gar36 | 10 | 89.53 | 0.0220 |
| A2_SRR8283976_Gar37 | 10 | 89.53 | 0.0216 |
| A2_SRR8283977_Gar56 | 10 | 89.53 | 0.0217 |
| A2_SRR8283979_Gar58 | 10 | 89.53 | 0.0205 |
| A2_SRR8283980_Gar59 | 10 | 89.53 | 0.0222 |
| A2_SRR8283989_Gar30 | 10 | 89.53 | 0.0205 |
| A2_SRR8283990_Gar31 | 10 | 89.53 | 0.0210 |
| A2_SRR8283994_Gar02 | 10 | 89.53 | 0.0217 |
| A2_SRR8283997_Gar66 | 10 | 89.53 | 0.0200 |
| A2_SRR8284001_Gar53 | 10 | 89.53 | 0.0238 |
| A2_SRR8284004_Gar50 | 10 | 89.53 | 0.0201 |
| A2_SRR8284006_Gar48 | 10 | 89.53 | 0.0201 |
| A2_SRR8284007_Gar47 | 10 | 89.53 | 0.0239 |
| A2_SRR8284008_Gar46 | 10 | 89.53 | 0.0208 |
| A2_SRR8284009_Gar62 | 10 | 89.53 | 0.0202 |
| A2_SRR8284010_Gar63 | 10 | 89.53 | 0.0212 |
| A2_SRR8283990_Gar31 | 10 | 90.98 | 0.0205 |
| A2_SRR8283994_Gar02 | 10 | 90.98 | 0.0205 |
| A1_019 | 10 | 91.95 | 0.0219 |
| A1_028 | 10 | 91.95 | 0.0224 |
| A1_030 | 10 | 91.95 | 0.0221 |
| A1_051 | 10 | 91.95 | 0.0203 |
| A1_073 | 10 | 91.95 | 0.0209 |
| A1_074 | 10 | 91.95 | 0.0201 |
| A1_079 | 10 | 91.95 | 0.0226 |
| A1_084 | 10 | 91.95 | 0.0221 |
| A1_088 | 10 | 91.95 | 0.0222 |
| A1_097 | 10 | 91.95 | 0.0219 |
| A1_108 | 10 | 91.95 | 0.0218 |
| A1_133 | 10 | 91.95 | 0.0201 |
| A1_155 | 10 | 91.95 | 0.0224 |
| A1_Af | 10 | 91.95 | 0.0224 |
| A1_Nisa | 10 | 91.95 | 0.0214 |
| A2_001 | 10 | 91.95 | 0.0203 |
| A2_008 | 10 | 91.95 | 0.0207 |
| A2_019 | 10 | 91.95 | 0.0202 |
| A2_026 | 10 | 91.95 | 0.0221 |
| A2_029 | 10 | 91.95 | 0.0216 |
| A2_030 | 10 | 91.95 | 0.0215 |
| A2_031 | 10 | 91.95 | 0.0205 |
| A2_034 | 10 | 91.95 | 0.0206 |
| A2_040 | 10 | 91.95 | 0.0210 |
| A2_041 | 10 | 91.95 | 0.0203 |
| A2_044 | 10 | 91.95 | 0.0201 |
| A2_047 | 10 | 91.95 | 0.0199 |
| A2_056 | 10 | 91.95 | 0.0209 |
| A2_057 | 10 | 91.95 | 0.0202 |
| A2_059 | 10 | 91.95 | 0.0202 |
| A2_061 | 10 | 91.95 | 0.0208 |
| A2_062 | 10 | 91.95 | 0.0202 |
| A2_064 | 10 | 91.95 | 0.0200 |
| A2_066 | 10 | 91.95 | 0.0200 |
| A2_067 | 10 | 91.95 | 0.0214 |
| A2_069 | 10 | 91.95 | 0.0212 |
| A2_073 | 10 | 91.95 | 0.0202 |
| A2_074 | 10 | 91.95 | 0.0204 |
| A2_079 | 10 | 91.95 | 0.0209 |
| A2_080 | 10 | 91.95 | 0.0216 |
| A2_084 | 10 | 91.95 | 0.0211 |
| A2_085 | 10 | 91.95 | 0.0206 |
| A2_087 | 10 | 91.95 | 0.0207 |
| A2_091 | 10 | 91.95 | 0.0214 |
| A2_096 | 10 | 91.95 | 0.0205 |
| A2_100 | 10 | 91.95 | 0.0201 |

Accessions with dS outside of 95% CI (0.0031 - 0.0198)

| Accession | Chr | Position | dS |
|---|---|---|---|
| A2_101 | 10 | 91.95 | 0.0207 |
| A2_113 | 10 | 91.95 | 0.0204 |
| A2_117 | 10 | 91.95 | 0.0210 |
| A2_118 | 10 | 91.95 | 0.0208 |
| A2_119 | 10 | 91.95 | 0.0207 |
| A2_123 | 10 | 91.95 | 0.0203 |
| A2_124 | 10 | 91.95 | 0.0206 |
| A2_131 | 10 | 91.95 | 0.0211 |
| A2_141 | 10 | 91.95 | 0.0201 |
| A2_142 | 10 | 91.95 | 0.0205 |
| A2_147 | 10 | 91.95 | 0.0217 |
| A2_156 | 10 | 91.95 | 0.0207 |
| A2_164 | 10 | 91.95 | 0.0202 |
| A2_255 | 10 | 91.95 | 0.0206 |
| A2_lintless | 10 | 91.95 | 0.0208 |
| A2_NC503 | 10 | 91.95 | 0.0206 |
| A2_SRR8283970_Gar43 | 10 | 91.95 | 0.0212 |
| A2_SRR8283972_Gar41 | 10 | 91.95 | 0.0202 |
| A2_SRR8283973_Gar38 | 10 | 91.95 | 0.0222 |
| A2_SRR8283974_Gar39 | 10 | 91.95 | 0.0212 |
| A2_SRR8283975_Gar36 | 10 | 91.95 | 0.0210 |
| A2_SRR8283976_Gar37 | 10 | 91.95 | 0.0223 |
| A2_SRR8283977_Gar56 | 10 | 91.95 | 0.0199 |
| A2_SRR8283978_Gar57 | 10 | 91.95 | 0.0211 |
| A2_SRR8283980_Gar59 | 10 | 91.95 | 0.0200 |
| A2_SRR8283981_Gar60 | 10 | 91.95 | 0.0200 |
| A2_SRR8283982_Gar61 | 10 | 91.95 | 0.0201 |
| A2_SRR8283983_Gar44 | 10 | 91.95 | 0.0226 |
| A2_SRR8283984_Gar45 | 10 | 91.95 | 0.0215 |
| A2_SRR8283987_Gar32 | 10 | 91.95 | 0.0241 |
| A2_SRR8283989_Gar30 | 10 | 91.95 | 0.0226 |
| A2_SRR8283990_Gar31 | 10 | 91.95 | 0.0203 |
| A2_SRR8283991_Gar03 | 10 | 91.95 | 0.0202 |
| A2_SRR8283993_Gar01 | 10 | 91.95 | 0.0201 |
| A2_SRR8283994_Gar02 | 10 | 91.95 | 0.0211 |
| A2_SRR8283995_Gar65 | 10 | 91.95 | 0.0224 |
| A2_SRR8283997_Gar66 | 10 | 91.95 | 0.0222 |
| A2_SRR8283999_Gar55 | 10 | 91.95 | 0.0199 |
| A2_SRR8284000_Gar54 | 10 | 91.95 | 0.0218 |
| A2_SRR8284001_Gar53 | 10 | 91.95 | 0.0217 |
| A2_SRR8284003_Gar51 | 10 | 91.95 | 0.0204 |
| A2_SRR8284004_Gar50 | 10 | 91.95 | 0.0221 |
| A2_SRR8284005_Gar49 | 10 | 91.95 | 0.0211 |
| A2_SRR8284006_Gar48 | 10 | 91.95 | 0.0203 |
| A2_SRR8284007_Gar47 | 10 | 91.95 | 0.0226 |
| A2_SRR8284008_Gar46 | 10 | 91.95 | 0.0213 |
| A2_SRR8284009_Gar62 | 10 | 91.95 | 0.0221 |
| A2_SRR8284010_Gar63 | 10 | 91.95 | 0.0212 |
| A2_SRR8284011_Gar64 | 10 | 91.95 | 0.0208 |
| A1_030 | 10 | 93.58 | 0.0204 |
| A1_073 | 10 | 93.58 | 0.0221 |
| A1_074 | 10 | 93.58 | 0.0202 |
| A1_088 | 10 | 93.58 | 0.0205 |
| A1_155 | 10 | 93.58 | 0.0233 |
| A1_Af | 10 | 93.58 | 0.0241 |
| A1_Nisa | 10 | 93.58 | 0.0234 |
| A1_SRR8283996_Ghe01 | 10 | 93.58 | 0.0200 |
| A2_131 | 10 | 93.58 | 0.0206 |
| A2_154 | 10 | 93.58 | 0.0209 |
| A2_156 | 10 | 93.58 | 0.0204 |
| A2_SRR8283972_Gar41 | 10 | 93.58 | 0.0199 |
| A2_SRR8283976_Gar37 | 10 | 93.58 | 0.0211 |
| A2_SRR8283983_Gar44 | 10 | 93.58 | 0.0201 |
| A2_SRR8283988_Gar33 | 10 | 93.58 | 0.0200 |
| A2_SRR8283993_Gar01 | 10 | 93.58 | 0.0207 |
| A2_SRR8283999_Gar55 | 10 | 93.58 | 0.0200 |
| A2_SRR8284003_Gar51 | 10 | 93.58 | 0.0229 |
| A2_SRR8284009_Gar62 | 10 | 93.58 | 0.0205 |
| A2_SRR8284010_Gar63 | 10 | 93.58 | 0.0208 |
| A1_132 | 10 | 95.02 | 0.0024 |
| A1_133 | 10 | 95.02 | 0.0025 |
| A1_155 | 11 | 1.73 | 0.0009 |
| A1_155 | 11 | 2.15 | 0.0008 |
| A1_155 | 11 | 2.62 | 0.0018 |
| A2_SRR8283974_Gar39 | 11 | 4.10 | 0.0026 |
| A2_SRR8284004_Gar50 | 11 | 7.60 | 0.0029 |
| A1_051 | 11 | 18.49 | 0.0028 |
| A1_125 | 11 | 18.49 | 0.0030 |
| A1_132 | 11 | 18.49 | 0.0029 |
| A1_133 | 11 | 18.49 | 0.0028 |
| A2_016 | 11 | 18.49 | 0.0026 |
| A2_021 | 11 | 18.49 | 0.0026 |
| A2_040 | 11 | 18.49 | 0.0026 |
| A2_044 | 11 | 18.49 | 0.0026 |
| A2_047 | 11 | 18.49 | 0.0026 |
| A2_060 | 11 | 18.49 | 0.0026 |

| Accession | Chr | Position | dS |
|---|---|---|---|
| A2_062 | 11 | 18.49 | 0.0023 |
| A2_064 | 11 | 18.49 | 0.0026 |
| A2_073 | 11 | 18.49 | 0.0028 |
| A2_076 | 11 | 18.49 | 0.0030 |
| A2_100 | 11 | 18.49 | 0.0028 |
| A2_141 | 11 | 18.49 | 0.0028 |
| A2_142 | 11 | 18.49 | 0.0026 |
| A2_154 | 11 | 18.49 | 0.0028 |
| A2_156 | 11 | 18.49 | 0.0024 |
| A2_SRR8283969_Gar42 | 11 | 18.49 | 0.0026 |
| A2_SRR8283977_Gar56 | 11 | 18.49 | 0.0030 |
| A2_SRR8283982_Gar61 | 11 | 18.49 | 0.0030 |
| A2_SRR8283985_Gar34 | 11 | 18.49 | 0.0027 |
| A2_SRR8283986_Gar35 | 11 | 18.49 | 0.0030 |
| A2_SRR8283991_Gar03 | 11 | 18.49 | 0.0024 |
| A2_SRR8283992_Gar04 | 11 | 18.49 | 0.0028 |
| A2_SRR8283994_Gar02 | 11 | 18.49 | 0.0027 |
| A2_SRR8284011_Gar64 | 11 | 18.49 | 0.0029 |
| A2_085 | 11 | 45.55 | 0.0030 |
| A1_019 | 11 | 64.32 | 0.0264 |
| A1_028 | 11 | 64.32 | 0.0261 |
| A1_030 | 11 | 64.32 | 0.0259 |
| A1_051 | 11 | 64.32 | 0.0265 |
| A1_054 | 11 | 64.32 | 0.0257 |
| A1_073 | 11 | 64.32 | 0.0254 |
| A1_074 | 11 | 64.32 | 0.0268 |
| A1_077 | 11 | 64.32 | 0.0262 |
| A1_079 | 11 | 64.32 | 0.0270 |
| A1_084 | 11 | 64.32 | 0.0260 |
| A1_088 | 11 | 64.32 | 0.0274 |
| A1_097 | 11 | 64.32 | 0.0255 |
| A1_108 | 11 | 64.32 | 0.0259 |
| A1_125 | 11 | 64.32 | 0.0298 |
| A1_132 | 11 | 64.32 | 0.0262 |
| A1_133 | 11 | 64.32 | 0.0257 |
| A1_155 | 11 | 64.32 | 0.0255 |
| A1_Af | 11 | 64.32 | 0.0262 |
| A1_Nisa | 11 | 64.32 | 0.0262 |
| A1_SRR8283996_Ghe01 | 11 | 64.32 | 0.0313 |
| A1_SRR8283998_Ghe02 | 11 | 64.32 | 0.0297 |
| A2_001 | 11 | 64.32 | 0.0249 |
| A2_004 | 11 | 64.32 | 0.0228 |
| A2_008 | 11 | 64.32 | 0.0240 |
| A2_016 | 11 | 64.32 | 0.0234 |
| A2_019 | 11 | 64.32 | 0.0232 |
| A2_021 | 11 | 64.32 | 0.0235 |
| A2_026 | 11 | 64.32 | 0.0238 |
| A2_029 | 11 | 64.32 | 0.0242 |
| A2_030 | 11 | 64.32 | 0.0243 |
| A2_031 | 11 | 64.32 | 0.0243 |
| A2_034 | 11 | 64.32 | 0.0229 |
| A2_040 | 11 | 64.32 | 0.0232 |
| A2_041 | 11 | 64.32 | 0.0231 |
| A2_043 | 11 | 64.32 | 0.0228 |
| A2_044 | 11 | 64.32 | 0.0232 |
| A2_045 | 11 | 64.32 | 0.0301 |
| A2_046 | 11 | 64.32 | 0.0235 |
| A2_047 | 11 | 64.32 | 0.0237 |
| A2_056 | 11 | 64.32 | 0.0249 |
| A2_057 | 11 | 64.32 | 0.0240 |
| A2_059 | 11 | 64.32 | 0.0228 |
| A2_060 | 11 | 64.32 | 0.0231 |
| A2_061 | 11 | 64.32 | 0.0248 |
| A2_062 | 11 | 64.32 | 0.0264 |
| A2_064 | 11 | 64.32 | 0.0233 |
| A2_065 | 11 | 64.32 | 0.0235 |
| A2_066 | 11 | 64.32 | 0.0229 |
| A2_067 | 11 | 64.32 | 0.0239 |
| A2_069 | 11 | 64.32 | 0.0238 |
| A2_073 | 11 | 64.32 | 0.0222 |
| A2_074 | 11 | 64.32 | 0.0232 |
| A2_076 | 11 | 64.32 | 0.0241 |
| A2_079 | 11 | 64.32 | 0.0237 |
| A2_080 | 11 | 64.32 | 0.0242 |
| A2_084 | 11 | 64.32 | 0.0232 |
| A2_085 | 11 | 64.32 | 0.0257 |
| A2_087 | 11 | 64.32 | 0.0228 |
| A2_091 | 11 | 64.32 | 0.0259 |
| A2_096 | 11 | 64.32 | 0.0244 |
| A2_100 | 11 | 64.32 | 0.0233 |
| A2_101 | 11 | 64.32 | 0.0221 |
| A2_113 | 11 | 64.32 | 0.0229 |
| A2_117 | 11 | 64.32 | 0.0239 |
| A2_118 | 11 | 64.32 | 0.0237 |
| A2_119 | 11 | 64.32 | 0.0225 |
| A2_123 | 11 | 64.32 | 0.0241 |

| Accession | Chr | Position | dS |
|---|---|---|---|
| A2_124 | 11 | 64.32 | 0.0247 |
| A2_131 | 11 | 64.32 | 0.0232 |
| A2_141 | 11 | 64.32 | 0.0235 |
| A2_142 | 11 | 64.32 | 0.0235 |
| A2_147 | 11 | 64.32 | 0.0232 |
| A2_154 | 11 | 64.32 | 0.0242 |
| A2_156 | 11 | 64.32 | 0.0238 |
| A2_158 | 11 | 64.32 | 0.0231 |
| A2_164 | 11 | 64.32 | 0.0249 |
| A2_255 | 11 | 64.32 | 0.0225 |
| A2_lintless | 11 | 64.32 | 0.0242 |
| A2_NC503 | 11 | 64.32 | 0.0227 |
| A2_SRR8283969_Gar4 | 11 | 64.32 | 0.0317 |
| A2_SRR8283970_Gar4 | 11 | 64.32 | 0.0308 |
| A2_SRR8283971_Gar4 | 11 | 64.32 | 0.0303 |
| A2_SRR8283972_Gar4 | 11 | 64.32 | 0.0299 |
| A2_SRR8283973_Gar3 | 11 | 64.32 | 0.0293 |
| A2_SRR8283974_Gar3 | 11 | 64.32 | 0.0310 |
| A2_SRR8283975_Gar3 | 11 | 64.32 | 0.0312 |
| A2_SRR8283976_Gar3 | 11 | 64.32 | 0.0304 |
| A2_SRR8283977_Gar5 | 11 | 64.32 | 0.0294 |
| A2_SRR8283978_Gar5 | 11 | 64.32 | 0.0334 |
| A2_SRR8283979_Gar5 | 11 | 64.32 | 0.0314 |
| A2_SRR8283980_Gar5 | 11 | 64.32 | 0.0339 |
| A2_SRR8283981_Gar6 | 11 | 64.32 | 0.0293 |
| A2_SRR8283982_Gar61 | 11 | 64.32 | 0.0294 |
| A2_SRR8283983_Gar4 | 11 | 64.32 | 0.0296 |
| A2_SRR8283984_Gar4 | 11 | 64.32 | 0.0312 |
| A2_SRR8283985_Gar3 | 11 | 64.32 | 0.0315 |
| A2_SRR8283986_Gar3 | 11 | 64.32 | 0.0269 |
| A2_SRR8283987_Gar3 | 11 | 64.32 | 0.0300 |
| A2_SRR8283988_Gar3 | 11 | 64.32 | 0.0306 |
| A2_SRR8283989_Gar3 | 11 | 64.32 | 0.0293 |
| A2_SRR8283990_Gar31 | 11 | 64.32 | 0.0329 |
| A2_SRR8283991_Gar0 | 11 | 64.32 | 0.0281 |
| A2_SRR8283992_Gar0 | 11 | 64.32 | 0.0288 |
| A2_SRR8283993_Gar01 | 11 | 64.32 | 0.0340 |
| A2_SRR8283994_Gar0 | 11 | 64.32 | 0.0330 |
| A2_SRR8283995_Gar6 | 11 | 64.32 | 0.0366 |
| A2_SRR8283997_Gar6 | 11 | 64.32 | 0.0291 |
| A2_SRR8283999_Gar5 | 11 | 64.32 | 0.0317 |
| A2_SRR8284000_Gar5 | 11 | 64.32 | 0.0370 |
| A2_SRR8284001_Gar5 | 11 | 64.32 | 0.0278 |
| A2_SRR8284002_Gar5 | 11 | 64.32 | 0.0345 |
| A2_SRR8284003_Gar51 | 11 | 64.32 | 0.0319 |
| A2_SRR8284004_Gar5 | 11 | 64.32 | 0.0280 |
| A2_SRR8284005_Gar4 | 11 | 64.32 | 0.0303 |
| A2_SRR8284006_Gar4 | 11 | 64.32 | 0.0305 |
| A2_SRR8284007_Gar4 | 11 | 64.32 | 0.0301 |
| A2_SRR8284008_Gar4 | 11 | 64.32 | 0.0287 |
| A2_SRR8284009_Gar6 | 11 | 64.32 | 0.0304 |
| A2_SRR8284010_Gar6 | 11 | 64.32 | 0.0325 |
| A2_SRR8284011_Gar6 | 11 | 64.32 | 0.0294 |
| A1_SRR8283998_Ghe0 | 11 | 85.43 | 0.0201 |
| A1_077 | 11 | 89.98 | 0.0199 |
| A1_019 | 11 | 90.59 | 0.0246 |
| A1_028 | 11 | 90.59 | 0.0210 |
| A1_030 | 11 | 90.59 | 0.0224 |
| A1_051 | 11 | 90.59 | 0.0204 |
| A1_054 | 11 | 90.59 | 0.0231 |
| A1_073 | 11 | 90.59 | 0.0221 |
| A1_074 | 11 | 90.59 | 0.0219 |
| A1_077 | 11 | 90.59 | 0.0229 |
| A1_079 | 11 | 90.59 | 0.0226 |
| A1_084 | 11 | 90.59 | 0.0210 |
| A1_088 | 11 | 90.59 | 0.0215 |
| A1_097 | 11 | 90.59 | 0.0222 |
| A1_108 | 11 | 90.59 | 0.0223 |
| A1_125 | 11 | 90.59 | 0.0214 |
| A1_132 | 11 | 90.59 | 0.0208 |
| A1_133 | 11 | 90.59 | 0.0209 |
| A1_155 | 11 | 90.59 | 0.0239 |
| A1_Af | 11 | 90.59 | 0.0227 |
| A1_Nisa | 11 | 90.59 | 0.0223 |
| A1_SRR8283996_Ghe0 | 11 | 90.59 | 0.0212 |
| A2_001 | 11 | 90.59 | 0.0200 |
| A2_004 | 11 | 90.59 | 0.0268 |
| A2_008 | 11 | 90.59 | 0.0211 |
| A2_016 | 11 | 90.59 | 0.0218 |
| A2_019 | 11 | 90.59 | 0.0212 |
| A2_026 | 11 | 90.59 | 0.0216 |
| A2_029 | 11 | 90.59 | 0.0208 |
| A2_030 | 11 | 90.59 | 0.0211 |
| A2_034 | 11 | 90.59 | 0.0211 |
| A2_040 | 11 | 90.59 | 0.0213 |
| A2_041 | 11 | 90.59 | 0.0276 |

Accessions with dS outside of 95% CI (0.0031 - 0.0198)

| Accession | Chr | Position | dS | Accession | Chr | Position | dS | Accession | Chr | Position | dS |
|---|---|---|---|---|---|---|---|---|---|---|---|
| A2_043 | 11 | 90.59 | 0.0202 | A1_Af | 11 | 91.80 | 0.0255 | A2_SRR8283997_Gar6 | 11 | 91.80 | 0.0235 |
| A2_044 | 11 | 90.59 | 0.0213 | A1_Nisa | 11 | 91.80 | 0.0251 | A2_SRR8283999_Gar5 | 11 | 91.80 | 0.0235 |
| A2_046 | 11 | 90.59 | 0.0219 | A2_001 | 11 | 91.80 | 0.0236 | A2_SRR8284000_Gar5 | 11 | 91.80 | 0.0240 |
| A2_047 | 11 | 90.59 | 0.0214 | A2_004 | 11 | 91.80 | 0.0231 | A2_SRR8284001_Gar5 | 11 | 91.80 | 0.0232 |
| A2_056 | 11 | 90.59 | 0.0275 | A2_008 | 11 | 91.80 | 0.0218 | A2_SRR8284003_Gar51 | 11 | 91.80 | 0.0226 |
| A2_057 | 11 | 90.59 | 0.0214 | A2_016 | 11 | 91.80 | 0.0227 | A2_SRR8284004_Gar5 | 11 | 91.80 | 0.0233 |
| A2_059 | 11 | 90.59 | 0.0230 | A2_019 | 11 | 91.80 | 0.0220 | A2_SRR8284005_Gar4 | 11 | 91.80 | 0.0218 |
| A2_060 | 11 | 90.59 | 0.0216 | A2_021 | 11 | 91.80 | 0.0218 | A2_SRR8284006_Gar4 | 11 | 91.80 | 0.0232 |
| A2_061 | 11 | 90.59 | 0.0206 | A2_026 | 11 | 91.80 | 0.0235 | A2_SRR8284007_Gar4 | 11 | 91.80 | 0.0218 |
| A2_062 | 11 | 90.59 | 0.0203 | A2_029 | 11 | 91.80 | 0.0229 | A2_SRR8284008_Gar4 | 11 | 91.80 | 0.0223 |
| A2_064 | 11 | 90.59 | 0.0205 | A2_030 | 11 | 91.80 | 0.0231 | A2_SRR8284009_Gar6 | 11 | 91.80 | 0.0231 |
| A2_065 | 11 | 90.59 | 0.0229 | A2_031 | 11 | 91.80 | 0.0215 | A2_SRR8284010_Gar6 | 11 | 91.80 | 0.0243 |
| A2_066 | 11 | 90.59 | 0.0218 | A2_034 | 11 | 91.80 | 0.0228 | A2_SRR8284011_Gar6 | 11 | 91.80 | 0.0229 |
| A2_067 | 11 | 90.59 | 0.0280 | A2_040 | 11 | 91.80 | 0.0227 | A1_SRR8283998_Ghe0 | 11 | 93.14 | 0.0205 |
| A2_069 | 11 | 90.59 | 0.0274 | A2_041 | 11 | 91.80 | 0.0220 | A1_019 | 11 | 95.42 | 0.0233 |
| A2_073 | 11 | 90.59 | 0.0207 | A2_043 | 11 | 91.80 | 0.0215 | A1_028 | 11 | 95.42 | 0.0228 |
| A2_074 | 11 | 90.59 | 0.0218 | A2_044 | 11 | 91.80 | 0.0231 | A1_030 | 11 | 95.42 | 0.0226 |
| A2_076 | 11 | 90.59 | 0.0210 | A2_045 | 11 | 91.80 | 0.0231 | A1_051 | 11 | 95.42 | 0.0201 |
| A2_079 | 11 | 90.59 | 0.0282 | A2_046 | 11 | 91.80 | 0.0242 | A1_073 | 11 | 95.42 | 0.0264 |
| A2_080 | 11 | 90.59 | 0.0270 | A2_047 | 11 | 91.80 | 0.0234 | A1_074 | 11 | 95.42 | 0.0264 |
| A2_084 | 11 | 90.59 | 0.0248 | A2_056 | 11 | 91.80 | 0.0231 | A1_077 | 11 | 95.42 | 0.0284 |
| A2_085 | 11 | 90.59 | 0.0228 | A2_057 | 11 | 91.80 | 0.0231 | A1_079 | 11 | 95.42 | 0.0233 |
| A2_087 | 11 | 90.59 | 0.0207 | A2_059 | 11 | 91.80 | 0.0235 | A1_084 | 11 | 95.42 | 0.0231 |
| A2_091 | 11 | 90.59 | 0.0281 | A2_060 | 11 | 91.80 | 0.0229 | A1_088 | 11 | 95.42 | 0.0216 |
| A2_096 | 11 | 90.59 | 0.0214 | A2_061 | 11 | 91.80 | 0.0210 | A1_097 | 11 | 95.42 | 0.0228 |
| A2_100 | 11 | 90.59 | 0.0214 | A2_062 | 11 | 91.80 | 0.0210 | A1_108 | 11 | 95.42 | 0.0232 |
| A2_101 | 11 | 90.59 | 0.0208 | A2_064 | 11 | 91.80 | 0.0226 | A1_125 | 11 | 95.42 | 0.0210 |
| A2_113 | 11 | 90.59 | 0.0207 | A2_065 | 11 | 91.80 | 0.0221 | A1_155 | 11 | 95.42 | 0.0283 |
| A2_117 | 11 | 90.59 | 0.0213 | A2_066 | 11 | 91.80 | 0.0233 | A1_Af | 11 | 95.42 | 0.0283 |
| A2_118 | 11 | 90.59 | 0.0201 | A2_067 | 11 | 91.80 | 0.0233 | A1_Nisa | 11 | 95.42 | 0.0281 |
| A2_119 | 11 | 90.59 | 0.0212 | A2_069 | 11 | 91.80 | 0.0234 | A1_SRR8283996_Ghe0 | 11 | 95.42 | 0.0230 |
| A2_123 | 11 | 90.59 | 0.0219 | A2_073 | 11 | 91.80 | 0.0224 | A2_001 | 11 | 95.42 | 0.0220 |
| A2_124 | 11 | 90.59 | 0.0280 | A2_074 | 11 | 91.80 | 0.0230 | A2_004 | 11 | 95.42 | 0.0244 |
| A2_141 | 11 | 90.59 | 0.0214 | A2_076 | 11 | 91.80 | 0.0211 | A2_008 | 11 | 95.42 | 0.0228 |
| A2_142 | 11 | 90.59 | 0.0224 | A2_079 | 11 | 91.80 | 0.0234 | A2_016 | 11 | 95.42 | 0.0223 |
| A2_147 | 11 | 90.59 | 0.0210 | A2_080 | 11 | 91.80 | 0.0228 | A2_019 | 11 | 95.42 | 0.0218 |
| A2_154 | 11 | 90.59 | 0.0217 | A2_084 | 11 | 91.80 | 0.0232 | A2_021 | 11 | 95.42 | 0.0219 |
| A2_255 | 11 | 90.59 | 0.0204 | A2_085 | 11 | 91.80 | 0.0224 | A2_026 | 11 | 95.42 | 0.0230 |
| A2_lintless | 11 | 90.59 | 0.0281 | A2_087 | 11 | 91.80 | 0.0230 | A2_029 | 11 | 95.42 | 0.0213 |
| A2_NC503 | 11 | 90.59 | 0.0261 | A2_091 | 11 | 91.80 | 0.0235 | A2_030 | 11 | 95.42 | 0.0212 |
| A2_SRR8283969_Gar42 | 11 | 90.59 | 0.0216 | A2_096 | 11 | 91.80 | 0.0226 | A2_031 | 11 | 95.42 | 0.0222 |
| A2_SRR8283970_Gar43 | 11 | 90.59 | 0.0219 | A2_100 | 11 | 91.80 | 0.0226 | A2_034 | 11 | 95.42 | 0.0238 |
| A2_SRR8283971_Gar40 | 11 | 90.59 | 0.0222 | A2_101 | 11 | 91.80 | 0.0227 | A2_040 | 11 | 95.42 | 0.0216 |
| A2_SRR8283972_Gar41 | 11 | 90.59 | 0.0207 | A2_113 | 11 | 91.80 | 0.0235 | A2_041 | 11 | 95.42 | 0.0227 |
| A2_SRR8283973_Gar38 | 11 | 90.59 | 0.0201 | A2_117 | 11 | 91.80 | 0.0235 | A2_043 | 11 | 95.42 | 0.0211 |
| A2_SRR8283974_Gar39 | 11 | 90.59 | 0.0206 | A2_118 | 11 | 91.80 | 0.0229 | A2_044 | 11 | 95.42 | 0.0223 |
| A2_SRR8283975_Gar36 | 11 | 90.59 | 0.0240 | A2_119 | 11 | 91.80 | 0.0231 | A2_045 | 11 | 95.42 | 0.0264 |
| A2_SRR8283976_Gar37 | 11 | 90.59 | 0.0214 | A2_123 | 11 | 91.80 | 0.0223 | A2_046 | 11 | 95.42 | 0.0224 |
| A2_SRR8283977_Gar56 | 11 | 90.59 | 0.0267 | A2_124 | 11 | 91.80 | 0.0226 | A2_047 | 11 | 95.42 | 0.0219 |
| A2_SRR8283978_Gar57 | 11 | 90.59 | 0.0225 | A2_131 | 11 | 91.80 | 0.0232 | A2_056 | 11 | 95.42 | 0.0220 |
| A2_SRR8283980_Gar59 | 11 | 90.59 | 0.0211 | A2_141 | 11 | 91.80 | 0.0225 | A2_057 | 11 | 95.42 | 0.0222 |
| A2_SRR8283983_Gar44 | 11 | 90.59 | 0.0261 | A2_142 | 11 | 91.80 | 0.0232 | A2_059 | 11 | 95.42 | 0.0224 |
| A2_SRR8283985_Gar34 | 11 | 90.59 | 0.0204 | A2_147 | 11 | 91.80 | 0.0231 | A2_060 | 11 | 95.42 | 0.0218 |
| A2_SRR8283986_Gar35 | 11 | 90.59 | 0.0203 | A2_154 | 11 | 91.80 | 0.0216 | A2_061 | 11 | 95.42 | 0.0255 |
| A2_SRR8283987_Gar32 | 11 | 90.59 | 0.0214 | A2_156 | 11 | 91.80 | 0.0215 | A2_062 | 11 | 95.42 | 0.0217 |
| A2_SRR8283988_Gar33 | 11 | 90.59 | 0.0199 | A2_158 | 11 | 91.80 | 0.0215 | A2_064 | 11 | 95.42 | 0.0222 |
| A2_SRR8283989_Gar30 | 11 | 90.59 | 0.0210 | A2_164 | 11 | 91.80 | 0.0222 | A2_065 | 11 | 95.42 | 0.0212 |
| A2_SRR8283992_Gar04 | 11 | 90.59 | 0.0229 | A2_255 | 11 | 91.80 | 0.0223 | A2_066 | 11 | 95.42 | 0.0222 |
| A2_SRR8283994_Gar02 | 11 | 90.59 | 0.0222 | A2_lintless | 11 | 91.80 | 0.0234 | A2_067 | 11 | 95.42 | 0.0223 |
| A2_SRR8283995_Gar65 | 11 | 90.59 | 0.0202 | A2_NC503 | 11 | 91.80 | 0.0220 | A2_069 | 11 | 95.42 | 0.0224 |
| A2_SRR8283997_Gar66 | 11 | 90.59 | 0.0296 | A2_SRR8283969_Gar42 | 11 | 91.80 | 0.0234 | A2_073 | 11 | 95.42 | 0.0223 |
| A2_SRR8283999_Gar55 | 11 | 90.59 | 0.0220 | A2_SRR8283971_Gar40 | 11 | 91.80 | 0.0231 | A2_074 | 11 | 95.42 | 0.0222 |
| A2_SRR8284000_Gar54 | 11 | 90.59 | 0.0237 | A2_SRR8283972_Gar41 | 11 | 91.80 | 0.0248 | A2_076 | 11 | 95.42 | 0.0205 |
| A2_SRR8284002_Gar52 | 11 | 90.59 | 0.0202 | A2_SRR8283973_Gar38 | 11 | 91.80 | 0.0248 | A2_079 | 11 | 95.42 | 0.0222 |
| A2_SRR8284004_Gar50 | 11 | 90.59 | 0.0280 | A2_SRR8283974_Gar39 | 11 | 91.80 | 0.0207 | A2_080 | 11 | 95.42 | 0.0223 |
| A2_SRR8284005_Gar49 | 11 | 90.59 | 0.0205 | A2_SRR8283975_Gar36 | 11 | 91.80 | 0.0221 | A2_084 | 11 | 95.42 | 0.0213 |
| A2_SRR8284006_Gar48 | 11 | 90.59 | 0.0224 | A2_SRR8283976_Gar37 | 11 | 91.80 | 0.0214 | A2_085 | 11 | 95.42 | 0.0233 |
| A2_SRR8284007_Gar47 | 11 | 90.59 | 0.0255 | A2_SRR8283977_Gar56 | 11 | 91.80 | 0.0225 | A2_087 | 11 | 95.42 | 0.0217 |
| A2_SRR8284008_Gar46 | 11 | 90.59 | 0.0267 | A2_SRR8283978_Gar57 | 11 | 91.80 | 0.0217 | A2_091 | 11 | 95.42 | 0.0219 |
| A2_SRR8284009_Gar62 | 11 | 90.59 | 0.0227 | A2_SRR8283979_Gar58 | 11 | 91.80 | 0.0209 | A2_096 | 11 | 95.42 | 0.0248 |
| A2_SRR8284010_Gar63 | 11 | 90.59 | 0.0244 | A2_SRR8283980_Gar59 | 11 | 91.80 | 0.0233 | A2_100 | 11 | 95.42 | 0.0224 |
| A2_SRR8284011_Gar64 | 11 | 90.59 | 0.0208 | A2_SRR8283981_Gar60 | 11 | 91.80 | 0.0215 | A2_101 | 11 | 95.42 | 0.0210 |
| A1_019 | 11 | 91.80 | 0.0220 | A2_SRR8283982_Gar61 | 11 | 91.80 | 0.0237 | A2_113 | 11 | 95.42 | 0.0254 |
| A1_028 | 11 | 91.80 | 0.0236 | A2_SRR8283983_Gar44 | 11 | 91.80 | 0.0216 | A2_117 | 11 | 95.42 | 0.0221 |
| A1_030 | 11 | 91.80 | 0.0237 | A2_SRR8283984_Gar45 | 11 | 91.80 | 0.0211 | A2_118 | 11 | 95.42 | 0.0213 |
| A1_051 | 11 | 91.80 | 0.0233 | A2_SRR8283985_Gar34 | 11 | 91.80 | 0.0202 | A2_119 | 11 | 95.42 | 0.0217 |
| A1_054 | 11 | 91.80 | 0.0233 | A2_SRR8283986_Gar35 | 11 | 91.80 | 0.0225 | A2_123 | 11 | 95.42 | 0.0217 |
| A1_073 | 11 | 91.80 | 0.0277 | A2_SRR8283987_Gar32 | 11 | 91.80 | 0.0236 | A2_124 | 11 | 95.42 | 0.0229 |
| A1_074 | 11 | 91.80 | 0.0249 | A2_SRR8283988_Gar33 | 11 | 91.80 | 0.0230 | A2_131 | 11 | 95.42 | 0.0243 |
| A1_077 | 11 | 91.80 | 0.0284 | A2_SRR8283989_Gar30 | 11 | 91.80 | 0.0240 | A2_141 | 11 | 95.42 | 0.0226 |
| A1_084 | 11 | 91.80 | 0.0243 | A2_SRR8283990_Gar31 | 11 | 91.80 | 0.0221 | A2_142 | 11 | 95.42 | 0.0221 |
| A1_097 | 11 | 91.80 | 0.0206 | A2_SRR8283991_Gar03 | 11 | 91.80 | 0.0238 | A2_147 | 11 | 95.42 | 0.0221 |
| A1_125 | 11 | 91.80 | 0.0209 | A2_SRR8283992_Gar04 | 11 | 91.80 | 0.0205 | A2_154 | 11 | 95.42 | 0.0216 |
| A1_132 | 11 | 91.80 | 0.0224 | A2_SRR8283993_Gar01 | 11 | 91.80 | 0.0212 | A2_156 | 11 | 95.42 | 0.0228 |
| A1_133 | 11 | 91.80 | 0.0241 | A2_SRR8283994_Gar02 | 11 | 91.80 | 0.0222 | A2_158 | 11 | 95.42 | 0.0213 |
| A1_155 | 11 | 91.80 | 0.0257 | A2_SRR8283995_Gar65 | 11 | 91.80 | 0.0223 | A2_164 | 11 | 95.42 | 0.0221 |

Accessions with dS outside of 95% CI (0.0031 - 0.0198)

| Accession | Chr | Position | dS |
|---|---|---|---|
| A2_255 | 11 | 95.42 | 0.0217 |
| A2_lintless | 11 | 95.42 | 0.0233 |
| A2_NC503 | 11 | 95.42 | 0.0208 |
| A2_SRR8283969_Gar42 | 11 | 95.42 | 0.0234 |
| A2_SRR8283970_Gar43 | 11 | 95.42 | 0.0221 |
| A2_SRR8283971_Gar40 | 11 | 95.42 | 0.0222 |
| A2_SRR8283972_Gar41 | 11 | 95.42 | 0.0236 |
| A2_SRR8283973_Gar38 | 11 | 95.42 | 0.0237 |
| A2_SRR8283974_Gar39 | 11 | 95.42 | 0.0236 |
| A2_SRR8283975_Gar36 | 11 | 95.42 | 0.0230 |
| A2_SRR8283976_Gar37 | 11 | 95.42 | 0.0235 |
| A2_SRR8283977_Gar56 | 11 | 95.42 | 0.0231 |
| A2_SRR8283978_Gar57 | 11 | 95.42 | 0.0230 |
| A2_SRR8283979_Gar58 | 11 | 95.42 | 0.0237 |
| A2_SRR8283980_Gar59 | 11 | 95.42 | 0.0260 |
| A2_SRR8283981_Gar60 | 11 | 95.42 | 0.0237 |
| A2_SRR8283982_Gar61 | 11 | 95.42 | 0.0229 |
| A2_SRR8283983_Gar44 | 11 | 95.42 | 0.0231 |
| A2_SRR8283984_Gar45 | 11 | 95.42 | 0.0233 |
| A2_SRR8283985_Gar34 | 11 | 95.42 | 0.0228 |
| A2_SRR8283986_Gar35 | 11 | 95.42 | 0.0221 |
| A2_SRR8283987_Gar32 | 11 | 95.42 | 0.0242 |
| A2_SRR8283988_Gar33 | 11 | 95.42 | 0.0234 |
| A2_SRR8283989_Gar30 | 11 | 95.42 | 0.0233 |
| A2_SRR8283990_Gar31 | 11 | 95.42 | 0.0225 |
| A2_SRR8283991_Gar03 | 11 | 95.42 | 0.0227 |
| A2_SRR8283992_Gar04 | 11 | 95.42 | 0.0211 |
| A2_SRR8283993_Gar01 | 11 | 95.42 | 0.0224 |
| A2_SRR8283994_Gar02 | 11 | 95.42 | 0.0240 |
| A2_SRR8283995_Gar65 | 11 | 95.42 | 0.0226 |
| A2_SRR8283997_Gar66 | 11 | 95.42 | 0.0238 |
| A2_SRR8283999_Gar55 | 11 | 95.42 | 0.0249 |
| A2_SRR8284000_Gar54 | 11 | 95.42 | 0.0263 |
| A2_SRR8284002_Gar52 | 11 | 95.42 | 0.0243 |
| A2_SRR8284003_Gar51 | 11 | 95.42 | 0.0270 |
| A2_SRR8284004_Gar50 | 11 | 95.42 | 0.0264 |
| A2_SRR8284005_Gar49 | 11 | 95.42 | 0.0240 |
| A2_SRR8284006_Gar48 | 11 | 95.42 | 0.0250 |
| A2_SRR8284007_Gar47 | 11 | 95.42 | 0.0229 |
| A2_SRR8284008_Gar46 | 11 | 95.42 | 0.0221 |
| A2_SRR8284009_Gar62 | 11 | 95.42 | 0.0236 |
| A2_SRR8284010_Gar63 | 11 | 95.42 | 0.0250 |
| A2_SRR8284011_Gar64 | 11 | 95.42 | 0.0215 |
| A2_131 | 11 | 98.35 | 0.0199 |
| A1_019 | 11 | 99.47 | 0.0028 |
| A1_030 | 11 | 99.47 | 0.0027 |
| A1_051 | 11 | 99.47 | 0.0028 |
| A1_054 | 11 | 99.47 | 0.0028 |
| A1_079 | 11 | 99.47 | 0.0027 |
| A1_084 | 11 | 99.47 | 0.0026 |
| A1_088 | 11 | 99.47 | 0.0029 |
| A1_097 | 11 | 99.47 | 0.0029 |
| A1_108 | 11 | 99.47 | 0.0030 |
| A1_125 | 11 | 99.47 | 0.0030 |
| A1_132 | 11 | 99.47 | 0.0030 |
| A1_133 | 11 | 99.47 | 0.0028 |
| A2_001 | 11 | 99.47 | 0.0027 |
| A2_004 | 11 | 99.47 | 0.0013 |
| A2_008 | 11 | 99.47 | 0.0027 |
| A2_016 | 11 | 99.47 | 0.0030 |
| A2_030 | 11 | 99.47 | 0.0014 |
| A2_041 | 11 | 99.47 | 0.0009 |
| A2_044 | 11 | 99.47 | 0.0025 |
| A2_061 | 11 | 99.47 | 0.0023 |
| A2_064 | 11 | 99.47 | 0.0029 |
| A2_065 | 11 | 99.47 | 0.0019 |
| A2_073 | 11 | 99.47 | 0.0030 |
| A2_076 | 11 | 99.47 | 0.0030 |
| A2_080 | 11 | 99.47 | 0.0029 |
| A2_085 | 11 | 99.47 | 0.0030 |
| A2_087 | 11 | 99.47 | 0.0027 |
| A2_091 | 11 | 99.47 | 0.0030 |
| A2_096 | 11 | 99.47 | 0.0016 |
| A2_100 | 11 | 99.47 | 0.0029 |
| A2_101 | 11 | 99.47 | 0.0025 |
| A2_113 | 11 | 99.47 | 0.0026 |
| A2_117 | 11 | 99.47 | 0.0023 |
| A2_118 | 11 | 99.47 | 0.0030 |
| A2_119 | 11 | 99.47 | 0.0025 |
| A2_123 | 11 | 99.47 | 0.0026 |
| A2_141 | 11 | 99.47 | 0.0026 |
| A2_142 | 11 | 99.47 | 0.0019 |
| A2_156 | 11 | 99.47 | 0.0015 |
| A2_158 | 11 | 99.47 | 0.0012 |
| A2_164 | 11 | 99.47 | 0.0011 |
| A2_SRR8283972_Gar41 | 11 | 99.47 | 0.0027 |
| A2_SRR8283974_Gar39 | 11 | 99.47 | 0.0030 |
| A2_SRR8283976_Gar37 | 11 | 99.47 | 0.0022 |
| A2_SRR8283978_Gar57 | 11 | 99.47 | 0.0029 |
| A2_SRR8283979_Gar58 | 11 | 99.47 | 0.0021 |
| A2_SRR8283980_Gar59 | 11 | 99.47 | 0.0023 |
| A2_SRR8283981_Gar60 | 11 | 99.47 | 0.0019 |
| A2_SRR8283982_Gar61 | 11 | 99.47 | 0.0024 |
| A2_SRR8283985_Gar34 | 11 | 99.47 | 0.0019 |
| A2_SRR8283986_Gar35 | 11 | 99.47 | 0.0021 |
| A2_SRR8283990_Gar31 | 11 | 99.47 | 0.0016 |
| A2_SRR8283992_Gar04 | 11 | 99.47 | 0.0008 |
| A2_SRR8283997_Gar66 | 11 | 99.47 | 0.0027 |
| A2_SRR8283999_Gar55 | 11 | 99.47 | 0.0029 |
| A2_SRR8284003_Gar51 | 11 | 99.47 | 0.0027 |
| A2_SRR8284004_Gar50 | 11 | 99.47 | 0.0029 |
| A2_SRR8284005_Gar49 | 11 | 99.47 | 0.0028 |
| A2_SRR8284007_Gar47 | 11 | 99.47 | 0.0029 |
| A2_SRR8284010_Gar63 | 11 | 99.47 | 0.0027 |
| A2_SRR8284011_Gar64 | 11 | 99.47 | 0.0030 |
| A2_SRR8283976_Gar37 | 12 | 0.26 | 0.0030 |
| A2_SRR8283985_Gar34 | 12 | 0.26 | 0.0029 |
| A2_SRR8283995_Gar65 | 12 | 0.26 | 0.0025 |
| A2_026 | 12 | 79.63 | 0.0023 |
| A2_029 | 12 | 79.63 | 0.0023 |
| A2_030 | 12 | 79.63 | 0.0024 |
| A2_034 | 12 | 79.63 | 0.0024 |
| A2_043 | 12 | 79.63 | 0.0026 |
| A2_045 | 12 | 79.63 | 0.0011 |
| A2_046 | 12 | 79.63 | 0.0025 |
| A2_056 | 12 | 79.63 | 0.0023 |
| A2_057 | 12 | 79.63 | 0.0026 |
| A2_059 | 12 | 79.63 | 0.0025 |
| A2_060 | 12 | 79.63 | 0.0024 |
| A2_061 | 12 | 79.63 | 0.0024 |
| A2_062 | 12 | 79.63 | 0.0020 |
| A2_065 | 12 | 79.63 | 0.0024 |
| A2_073 | 12 | 79.63 | 0.0022 |
| A2_076 | 12 | 79.63 | 0.0024 |
| A2_080 | 12 | 79.63 | 0.0026 |
| A2_084 | 12 | 79.63 | 0.0030 |
| A2_101 | 12 | 79.63 | 0.0025 |
| A2_118 | 12 | 79.63 | 0.0028 |
| A2_119 | 12 | 79.63 | 0.0026 |
| A2_147 | 12 | 79.63 | 0.0030 |
| A2_154 | 12 | 79.63 | 0.0026 |
| A2_255 | 12 | 79.63 | 0.0024 |
| A2_SRR8283970_Gar43 | 12 | 79.63 | 0.0022 |
| A2_SRR8283972_Gar41 | 12 | 79.63 | 0.0030 |
| A2_SRR8283973_Gar38 | 12 | 79.63 | 0.0028 |
| A2_SRR8283976_Gar37 | 12 | 79.63 | 0.0020 |
| A2_SRR8283983_Gar44 | 12 | 79.63 | 0.0026 |
| A2_SRR8283985_Gar34 | 12 | 79.63 | 0.0029 |
| A2_SRR8283987_Gar32 | 12 | 79.63 | 0.0023 |
| A2_SRR8283989_Gar30 | 12 | 79.63 | 0.0019 |
| A2_SRR8283994_Gar02 | 12 | 79.63 | 0.0021 |
| A2_SRR8284001_Gar53 | 12 | 79.63 | 0.0020 |
| A2_SRR8284004_Gar50 | 12 | 79.63 | 0.0030 |
| A2_SRR8284007_Gar47 | 12 | 79.63 | 0.0025 |
| A2_SRR8284011_Gar64 | 12 | 79.63 | 0.0028 |
| A1_SRR8283998_Ghe02 | 12 | 86.32 | 0.0029 |
| A2_056 | 12 | 86.32 | 0.0030 |
| A1_019 | 12 | 86.96 | 0.0021 |
| A1_028 | 12 | 86.96 | 0.0022 |
| A1_030 | 12 | 86.96 | 0.0021 |
| A1_051 | 12 | 86.96 | 0.0015 |
| A1_054 | 12 | 86.96 | 0.0021 |
| A1_077 | 12 | 86.96 | 0.0020 |
| A1_079 | 12 | 86.96 | 0.0021 |
| A1_084 | 12 | 86.96 | 0.0021 |
| A1_088 | 12 | 86.96 | 0.0019 |
| A1_097 | 12 | 86.96 | 0.0021 |
| A1_108 | 12 | 86.96 | 0.0020 |
| A1_125 | 12 | 86.96 | 0.0016 |
| A1_132 | 12 | 86.96 | 0.0016 |
| A1_133 | 12 | 86.96 | 0.0015 |
| A1_SRR8283998_Ghe02 | 12 | 86.96 | 0.0018 |
| A2_056 | 12 | 86.96 | 0.0030 |
| A2_061 | 12 | 86.96 | 0.0028 |
| A2_SRR8283970_Gar43 | 12 | 86.96 | 0.0028 |
| A2_SRR8283973_Gar38 | 12 | 86.96 | 0.0024 |
| A2_SRR8283978_Gar57 | 12 | 86.96 | 0.0027 |
| A2_SRR8283980_Gar59 | 12 | 86.96 | 0.0028 |
| A2_SRR8283984_Gar45 | 12 | 86.96 | 0.0029 |
| A2_SRR8283985_Gar34 | 12 | 86.96 | 0.0022 |
| A2_SRR8283991_Gar03 | 12 | 86.96 | 0.0028 |
| A2_SRR8283992_Gar04 | 12 | 86.96 | 0.0029 |
| A2_SRR8283994_Gar02 | 12 | 86.96 | 0.0025 |
| A2_SRR8283997_Gar66 | 12 | 86.96 | 0.0026 |
| A2_SRR8283999_Gar55 | 12 | 86.96 | 0.0030 |
| A2_SRR8284003_Gar51 | 12 | 86.96 | 0.0030 |
| A2_SRR8284004_Gar50 | 12 | 86.96 | 0.0027 |
| A2_SRR8284011_Gar64 | 12 | 86.96 | 0.0030 |
| A1_133 | 12 | 88.44 | 0.0030 |
| A2_021 | 12 | 91.02 | 0.0016 |
| A2_043 | 12 | 91.02 | 0.0015 |
| A2_057 | 12 | 91.02 | 0.0017 |
| A2_060 | 12 | 91.02 | 0.0016 |
| A2_074 | 12 | 91.02 | 0.0019 |
| A2_076 | 12 | 91.02 | 0.0019 |
| A2_085 | 12 | 91.02 | 0.0030 |
| A2_SRR8283993_Gar01 | 13 | 85.68 | 0.0209 |
| A1_SRR8283996_Ghe0 | 13 | 93.06 | 0.0210 |
| A1_SRR8283998_Ghe02 | 13 | 93.06 | 0.0228 |
| A2_SRR8283970_Gar43 | 13 | 93.06 | 0.0206 |
| A2_SRR8283978_Gar57 | 13 | 93.06 | 0.0204 |
| A2_SRR8284001_Gar53 | 13 | 93.06 | 0.0201 |
| A1_155 | 13 | 95.64 | 0.0019 |
| A1_SRR8283998_Ghe02 | 13 | 95.82 | 0.0023 |
| A2_SRR8283969_Gar42 | 13 | 95.82 | 0.0021 |
| A2_SRR8283979_Gar58 | 13 | 95.82 | 0.0030 |
| A2_SRR8283993_Gar01 | 13 | 95.82 | 0.0018 |
| A2_SRR8284002_Gar52 | 13 | 95.82 | 0.0026 |
